# Molecular profiling of XPO1 inhibitor and gemcitabine-nab-paclitaxel combination in cellular and LSL-Kras G12D/+; Trp53 fl/+; Pdx1-Cre (KPC) pancreatic cancer model

**DOI:** 10.1101/2022.03.08.481108

**Authors:** Md. Hafiz Uddin, Amro Aboukameel, Yiwei Li, Husain Yar Khan, Rachel E. Sexton, Sahar Bannoura, Gregory Dyson, Mohammad Najeeb Al-Hallak, Yosef Mzannar, Amr Mohamed, Yosef Landesman, Steve Kim, Rafic Beydoun, Ramzi M. Mohammad, Anthony F. Shields, Asfar S. Azmi

**Author notes:** Address of correspondence ^1^Asfar S. Azmi, Department of Oncology, Ph.D., Karmanos Cancer Institute, Wayne State University School of Medicine, 4100 John R, Detroit MI 48201. **Author contribution statement:** ASA designed the study, interpreted the results, analyzed the data, wrote, and edited the paper. MHU, HYK, RES, SB performed *in vitro* experiments, wrote and edited the paper. AA performed KCP mice experiment. YL performed human tissue analysis. GD performed bioinformatics and statistics. MNA, AM, SK, RB and AFS provided the patient derived tumors, wrote and edited the manuscript. RMM analyzed the animal studies, wrote and edited the manuscript. All authors provided significant scientific input in this study, wrote and edited the paper. All authors are aware of the content in this paper.

## Abstract

The majority of pancreatic ductal adenocarcinoma (PDAC) patients experience disease progression while on treatment with gemcitabine and nab-Paclitaxel (GemPac) treatment indicating the need for more effective combinations for this recalcitrant disease.

Earlier we showed that nuclear exporter protein exportin 1 (XPO1) is a valid therapeutic target in PDAC and the selective inhibitor of nuclear export (SINE) selinexor (Sel), synergistically enhances the efficacy of GemPac in pancreatic cancer cells, spheroids, patient derived tumors and had promising activity in a phase I study in patients with PDAC. Here we investigated the mechanisms of synergy by molecular profiling of Sel or Sel-GemPac treated PDAC cells, *in vitro* and by utilizing genetically modified LSL-Kras G12D/+; Trp53 fl/+; Pdx1-Cre (KPC) mouse model.

In KPC model, Sel given with GemPac at a sub-MTD dose enhanced the survival compared to controls (*p* < 0.05). Molecular analysis of residual KPC tumors showed re-organization of tumor stromal architecture, suppression of proliferation and nuclear retention of tumor suppressors. Single cell nuclear RNA sequencing (snRNAseq) revealed significant loss of cellular clusters in the Sel-GemPac treated mice including CD44 stem cell population. RNA-seq, Gene Ontology (GO) and Gene Set Enrichment Analysis (GSEA) analysis showed inhibition of several tumor promoting molecules.

Prioritized RNA-seq identified molecules were validated in *in vitro* or in the PDAC patient samples through siRNA mediated silencing, quantitative gene expression, cytotoxicity assays and confirmed their role in observed synergy. Sel or Sel-GemPac caused broad penetration in PDAC supporting signaling networks.

## Introduction

Pancreatic ductal adenocarcinoma (PDAC) is one of the leading causes of cancer associated deaths [1]. The survival rate of PDAC is dreadfully low [2]. Due to frequent late presentation of the disease, surgery cannot be considered as a treatment option in majority of patients with PDAC. For patients with better performance status, FOLFIRINOX is used in front line. However, the median overall survival (OS) of metastatic PDAC patients remains below 1 year, even with this highly potent and poorly tolerated chemotherapy combination [3]. For patients with poor performance status, gemcitabine (Gem) along with nanoparticle albumin-bound (nab) paclitaxel (GemPac) has been considered as a standard of care first-line therapy for following the MPACT phase III trial published in 2013, where median OS improved to 8.7 months for GemPac compared to 6.6 months for Gem alone (*p* = 0.001) [4]. A large fraction of patients with advanced PDAC receive GemPac based therapy [5] however, most PDAC patients experience disease progression suggesting an urgent need for novel therapeutic strategies.

Nuclear export protein exportin 1 or XPO1 is the major export protein of the cells [6] and frequently overexpressed in PDAC tumor tissue [7, 8]. XPO1 expression was also shown to be associated with tumor size, lymph node invasion, liver metastasis, and can act as a prognostic marker [9]. The overexpressed XPO1 protein exports many tumor suppressor proteins (TSPs) from the nucleus to the cytoplasm leading to their functional inactivation [6]. Therefore, simultaneous targeting of PDAC with GemPac and nuclear export inhibitors is considered a reasonable approach. We have demonstrated in our earlier studies that Selective Inhibitor of Nuclear Export (SINE) compounds (Selinexor-Sel and analogs) synergize with Gem or GemPac and suppress the growth of PDAC cells, spheroids and patient derived tumors. In a Phase I study, durable responses were observed in patients administered Sel-GemPac [9, 10].

In the present study, we have tested the combination in genetically modified LSL-Kras G12D/+; Trp53 fl/+; Pdx1-Cre (KPC) pancreatic cancer mouse model and assessed tumor microenvironment utilizing single nuclear RNA sequencing approach. We also investigated the broader implications of Sel-GemPac on PDAC signaling using high throughput approaches.

## Materials and methods

### Cell lines and reagents

The human pancreatic cancer cell line MiaPaCa-2 (Cat # CRL-1420; RRID: CVCL_0428) was purchased from the American Type Culture Collection (Rockville, MD, USA). Cells were cultured in DMEM media. The medium was supplemented with 10% fetal bovine serum (FBS) and 1% penicillin and streptomycin, and maintained at 37 °C in a humidified atmosphere of 5% CO_2_ (Stericycle CO_2_ incubator, Thermo Scientific, Waltham, MA, USA). The cell line was authenticated (STR profiling) and tested negative for *Mycoplasma* spp. by polymerase chain reaction (PCR). Ki-67 (Catalog no. M7240) and CD4 (Catalog no. M731029-2) antibodies were purchased from Dako (CA, USA). FOXO3a (Catalog no. 12829) antibody was purchased from cell signaling technology (CA, USA). For HRP conjugated secondary antibody Pollink-2 Plus HRP Broad for DAB Kit (Catalog no. D41-110) was used (GBI Labs, WA, USA).

### KPC mouse model

Here we investigated the survival and pathological changes utilizing genetically modified LSL-Kras G12D/+; Trp53 fl/+; Pdx1-Cre (KPC) mouse model of spontaneous pancreatic ductal adenocarcinoma (PDAC). The animal protocol used in the study was approved by Wayne State University Institutional Animal Care and Use Committee (IACUC# 20-07-2492). These mice express activated Kras and mutated Trp53 in the developing pancreas conditionally and spontaneously develop sequential ductal lesions from pancreatic intraepithelial neoplasias (PanINs) to metastatic PDAC. This model almost accurately mimics the human disease from initiation to progression. The mice were crossed and bred in the animal facility of Wayne State University. Mouse DNA was extracted from the ear using the Wizard® Genomic DNA Purification Kit from Promega (Madison, WI, USA), as per the manufacturer’s instructions. The ear tissue was digested overnight with continuous agitation and genomic DNA was extracted. To confirm the pdx-Cre introduction and mutations of KRAS and p53, conventional PCR was performed. Applying rolling bases, 4-5-week-old genotyped and confirmed KPC mice were enrolled in two groups. Mice were randomly divided in to two groups: control (n = 13) and treatment (n = 14). The control group received the vehicle and treated group received Sel-GemPac combination whereas Sel was given orally at 15 mg/kg twice a week for 4 weeks and gemcitabine and nab-paclitaxel were given intravenously (i.v.) at 30 mg/kg twice a week for 2 weeks. During treatments, mice were followed daily for any signs of distress from either treatment or illness due to tumor growth. After euthanasia, gross pathological examination was performed. The survival curve was generated using GraphPad Prism (GraphPad software, https://www.graphpad.com).

### Ultrasonography of KPC mice pancreas

Mice were also examined for the development of tumors in pancreas using high frequency ultrasonography. One day before imaging, a small portion of hair (about 1-2 cm in diameter) on the left abdominal flank was shaved using clipper. Animals were anesthetized with isoflurane regulated by a precision vaporizer anesthesia machine. Initially mice will be placed in a jar supplied with isoflurane for a very short period, once they are immobile, they be removed and put on a table connected to cone that had a continuous regulated isoflurane. Eye lubricate (Puralube Vet, Dechra, Northwich, UK) was applied to both eyes. Animals were checked to make sure they are under general anesthesia by pinching the foot and look for a reflex, once no reflex is noted, a small amount of warm Aquasonic ultrasound gel (warmed using Thermosonic Gel Warmer) was placed on the shaved site. Using a single element probe (PB506e; 30-60 MHz) directly on the gel, organ/tumor images were acquired (Scintica, TX, USA). The whole procedure took no more than 5 minutes, after which the mice were removed from anesthesia, placed in a dry cage without bedding on top of a circulating hot water blanket (on half of the cage to allow mice to move to the unheated half of cage if mice feel too warm). Once animals regained mobilization, they were put back in their cage.

### Patient samples

Fresh PDAC tissue was obtained under an approved Institutional Review Board (IRB) protocol. All samples were obtained with patient consent and approval from our Institutional Ethics Committee (protocol # 1603014771). The PDAC origin was confirmed by pathologist on site and put on ice. Tissues were processed immediately for the extraction of RNA.

### Single nuclear RNA sequencing

Randomly selected tumors from KPC mice with or without treatment were harvested and Single nuclear RNA sequencing was performed in a blinded fashion (Singulomics Corporation, NY, USA). Available at: http://singulomics.com/ (accessed on September 7, 2021). In brief, tumor cell nuclei were isolated from frozen mice pancreatic cancer tissue samples and 3’ single cell gene expression libraries (Next GEM v3.1) were constructed using the 10x Genomics Chromium system. The libraries were sequenced with ~400 million PE150 reads per sample on Illumina NovaSeq. After sequencing, clean reads were analyzed with mouse reference genome mm10 using Cell Ranger v6.0.1 (https://support.10xgenomics.com/single-cell-geneexpression/software/pipelines/latest/what-is-cell-ranger). Introns were included in the analysis and aggregation of the samples were also performed. Principal Component Analysis (PCA) was performed to reduce the number of feature dimensions, t-distributed Stochastic Neighbor Embedding (t-SNE) and Uniform Manifold Approximation and Projection for Dimension Reduction (UMAP) to visualize cells in a 2-D space, clusterings of cells that have similar expression profiles.

### Growth inhibition assay

The growth inhibition was determined by 3-(4,5-dimethylthiazol-2-yl)-2,5-diphenyltetrazolium bromide (MTT) assay as described previously [11]. In brief, MiaPaCa-2 cells was plated onto 96-well plates at a density of 4,000 cells per well. The cells were cultured for 72 hrs in the presence of drugs dissolved in dimethyl sulfoxide (DMSO). After incubation, 10 μl of MTT solution (5 mg/ml) was added to the media for 2 hrs at 37 °C. The media was removed, and the formazan crystals were solubilized in 100 μl DMSO for 10 mins in a shaker. The optical density was measured at 570 nm.

### RNA-seq analysis

MiaPaCa-2 cells were treated with vehicle DMSO control (Con) or 1 μM (μmol/L) selinexor (Sel) or 1 μM Sel plus 300 nM (nmol/L) Gemcitabine and 3 nM (nmol/L) nab-paclitaxel (Sel-GemPac) for 24 hrs. Total RNA was extracted using RNeasy mini kit (QIAGEN, Valencia, CA, USA) and submitted to LC Sciences on dry ice for poly(A)RNA-seq analysis (https://lcsciences.com/services). In brief, the quantity and purity were measured with Bioanalyzer 2100 and RNA 6000 Nano LabChip Kit (Agilent, CA, USA) with RIN number >7.0. Approximately 10 μg of total RNA was subjected to isolate Poly (A) mRNA with poly-T oligoattached magnetic beads (Invitrogen). The poly(A)-or poly(A)+ RNA fractions is fragmented into small pieces using divalent cations under elevated temperature. Then the cleaved RNA fragments were reverse-transcribed to create the final cDNA library (Illumina, San Diego, USA) followed by paired-end sequencing on an Illumina Hiseq 4000. The raw data was cleaned by removing low quality reads and adapters. Mapping of genome was performed using HISAT2 software followed by transcripts assembly by StringTie. All assemblies were merged to identify differentially expressed genes via mRNA expression profiling. StringTie [12] was used to perform expression level for mRNAs by calculating FPKM. The differentially expressed mRNAs were selected with log2 (fold change) >1 or log2 (fold change) <−1 and with statistical significance (p value < 0.05) by R package Ballgown [13].

### Real-time RT-qPCR

Total RNAs were extracted using the RNeasy Mini Kit (QIAGEN, Valencia, CA, USA) following manufacturer’s protocol. Complementary DNA was prepared using High Capacity cDNA Reverse Transcription Kit (Applied Biosystems, Waltham, MA, USA). Gene expression was detected by SYBR Green using StepOnePlus real-time PCR system (Applied Biosystems, Waltham, MA, USA) [14].

### Immunohistochemical analysis

Immunohistochemical analysis (IHC) of KPC residual tumor tissues for Ki-67, FOXO3a and CD4 were performed in the histopathology core facility of Karmanos Cancer Institute, Wayne State University in a double blinded fashion. For IHC analysis, all the tumor tissues from KPC mice were stored in 10% neutral-buffered formalin (v/v). Standard procedures were followed for IHC. Briefly, paraffin sections were de-waxed and rehydrated in a xylene-ethanol series. Endogenous peroxides were removed by a methanol/1.2% hydrogen peroxide incubation at room temperature for 25 minutes. Heat induced epitope retrieval (HIER) was performed with a pH 6.0 citrate buffer (ki67), pH 9.0 EDTA buffer (CD4 and Foxo3a) using the BIOCARE Decloaking Chamber. A 40-minute blocking step with Super Block Blocking buffer (Thermo Scientific) was performed prior to adding the primary antibody. The primary antibody dilutions were 1:50, 1:100 and 1:200 for CD4, Ki-67 and FOXO3a respectively. Detection was obtained using GBI Labs DAB chromogen kit and counterstained with Mayer’s Hematoxylin. Sections were then de-hydrated through a series of ethanol to xylene washes and cover slipped with Permount. Routine H&E staining also performed to observe tissue histology. For Picrosirius staining, Picro Sirius Red stain Kit (Catalog no. ab150681) from Abcam (Cambridge, UK) was used according to manufacturer instructions to visualize collagen distribution. Images were captured at 100x or 200x magnification.

### Colony formation assay

After control or target siRNA transfections for 48 hrs, a total of 500 cells were seeded in 24 well culture plates in triplicate and allowed to grow for 7 days. After the incubation period, cells were washed in PBS and fixed in 100% methanol. Colonies were stained with 0.5% crystal violet for 20 mins and washed carefully in tap water. Colonies are air-dried before scanning.

### siRNA mediated silencing

MiaPaCa-2 cells were transfected with *PFN* siRNA (siPFN), *HMGA* siRNA (siHMGA), *YBX* siRNA (siYBX) or negative control siRNA (siC) using lipofectamine 3000 (Invitrogen, San Diego, CA) according to manufacturer’s instruction. Cells were seeded at 1.5× 10^5^ cells/well in a 6-well plate in DMEM with 10% FBS. After 6–8 h of siRNA transfection, media were replaced. After 2-4 days cells were collected for experiments.

### STRING analysis

STRING analysis was performed on differentially overexpressed or underexpressed gene associated proteins to generate protein-protein interaction networks (STRING version 11.5; https://string-db.org/). All the filters were set to default.

### Gene set enrichment analysis (GSEA)

GSEA was performed to determine differential expression levels of a predefined gene set in two different treatment groups. Genome-wide expression profiles from Con vs Sel. Or Con vs Sel-GemPac were used to rank all genes in the data set, and the ranking list was then used to calculate enrichment score (ES). The steps involve acquiring the gene-ranking list, calculating an ES, estimating the level of significance etc. The parameter “Metric for ranking genes” is set to “Singal2Noise,” while other parameters are set to their default values. The GSEA packages was downloaded from Broad Institute, Inc. website (http://www.gsea-msigdb.org/gsea/index.jsp).

## Results

### Selinexor-Gemcitabine-nab-Paclitaxel treatment enhanced the survival of genetically modified KPC mouse model

All KPC mice have been genetically confirmed to carry *KRAS* and *p53* mutations by PCR (Fig. 1A and 1B) and randomly assigned to control and treated groups. Histopathological examination has confirmed the tumors as PDAC (Fig. 1C). Poorly differentiated adenocarcinoma is evident in the histopathological section. An ultrasonographic examination was performed to visualize suspected tumor in pancreas (Fig. 1D; Media file included in the supplementary information). A total of 13 KPC mice in control group and 14 KPC mice in the treatment group were used for the survival analysis. We have observed an increase in median survival by a month in the Selinexor-Gemcitabine-nab-Paclitaxel treatment (Sel-GemPac) treated mice (median survival 105 and 136.5 days for control and treated groups respectively). About 80% animal survived in the Sel-GemPac treatment group after 100 days, whereas in the control group, it was 54% (Fig. 1E). After 150 days survival rates were 15% and 43% in control and treated group respectively. The overall survival between control and Sel-GemPac treated group was statistically significant (*p* = 0.047, Log-Rank (Mantel-Cox) test). Immunohistochemical analysis revealed lower expression of proliferation marker Ki-67 along with nuclear accumulation of FOXO3a tumor suppressors in the Sel-GemPac treated KPC tumors. In the tumor microenvironment, picrosirius staining demonstrated a lower level of collagen fibers in the treated mice (Fig. 2A). To explore the effect of Sel-GemPAC on tumor microenvironment, we have performed single nuclear RNA sequencing (snRNAseq) both for untreated and treated murine PDAC KPC tumors (Fig. 2B). After quality control of single nuclear transcriptome, we have observed a reduced number of clusters in Sel-GemPAC treated KPC tumors (Fig. 2B and supplementary Fig. S1). The composition of tumors was determined by applying two-dimensional t-distributed stochastic neighbor embedding (t-SNE) analysis, and a total of 17 and 22 clusters were observed in treatment naïve KPC tumors whereas it has been reduced to 4 and 7 clusters in Sel-GemPac treated KPC tumors. A combined color-coded cluster distribution of untreated (control) and treated KPC tumors is shown in Fig. S2. Among different cell types, reduced expression of CD44 stem cells, PECAM1 positive endothelial cells are noted in the drug treatment group (Fig. S3). These results indicate a pronounced effect of triple combo (Sel-GemPac) in the murine PDAC KPC tumors.

**Figure 1.**
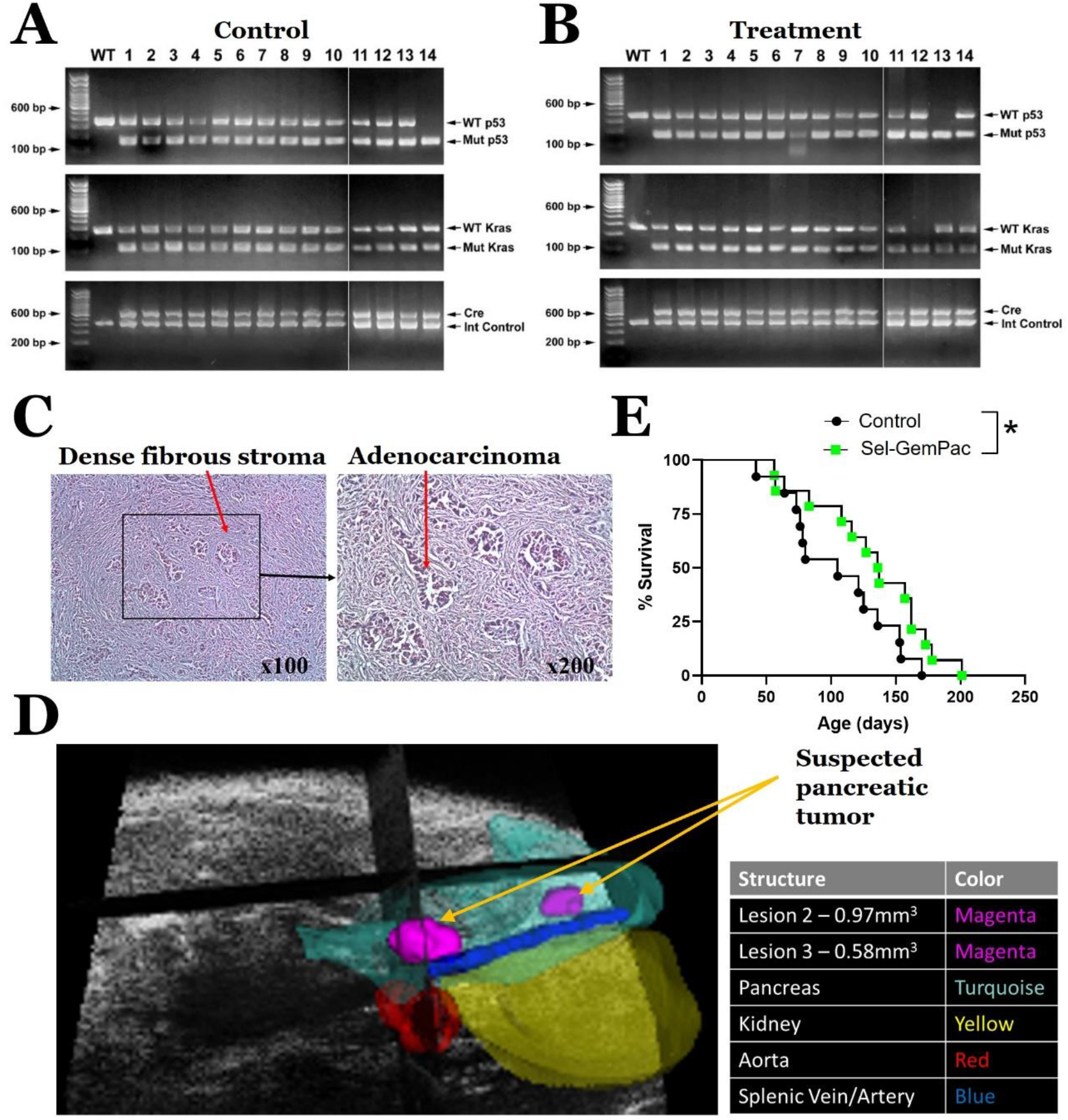
Survival of Sel-GemPac treated KPC mice with tumors. A-B. Confirmation of pdx-Cre introduction and mutations of KRAS and p53 in the KPC mice both in control and Sel-GemPac treated group using conventional PCR and subsequent 2% agarose gel electrophoresis. DNA was isolated from the ear of mice (collected using ear punch) using commercially available kit. C. Hematoxylin and Eosin (H & E) staining of pancreatic tumor tissue showing desmoplastic changes (indicated with red arrow in the left) confirming pancreatic ductal adenocarcinoma (PDAC, indicated with red arrow) in representative mouse. Specific area highlighting adenocarcinoma shown in 200x magnification on the right. D. Ultrasonographic evaluation of suspected tumors in the KPC. Suspected tumors are shown as spheres of magenta color. The pancreas, kidney, aorta, and splenic circulation also depicted in the picture. E. Kaplan-Meier Survival plot for control (n = 13) and Sel-GemPac (n = 14) treated KPC mice. Sel was given at a dose of 15 mg/kg orally (2x/week x4). Gem and Pac were given at a dose of 30 mg/kg i.v. twice a week for 2 weeks. Sel, Selinexor and GemPac, gemcitabine-nab-paclitaxel. *, *p*<0.05.

**Figure 2.**
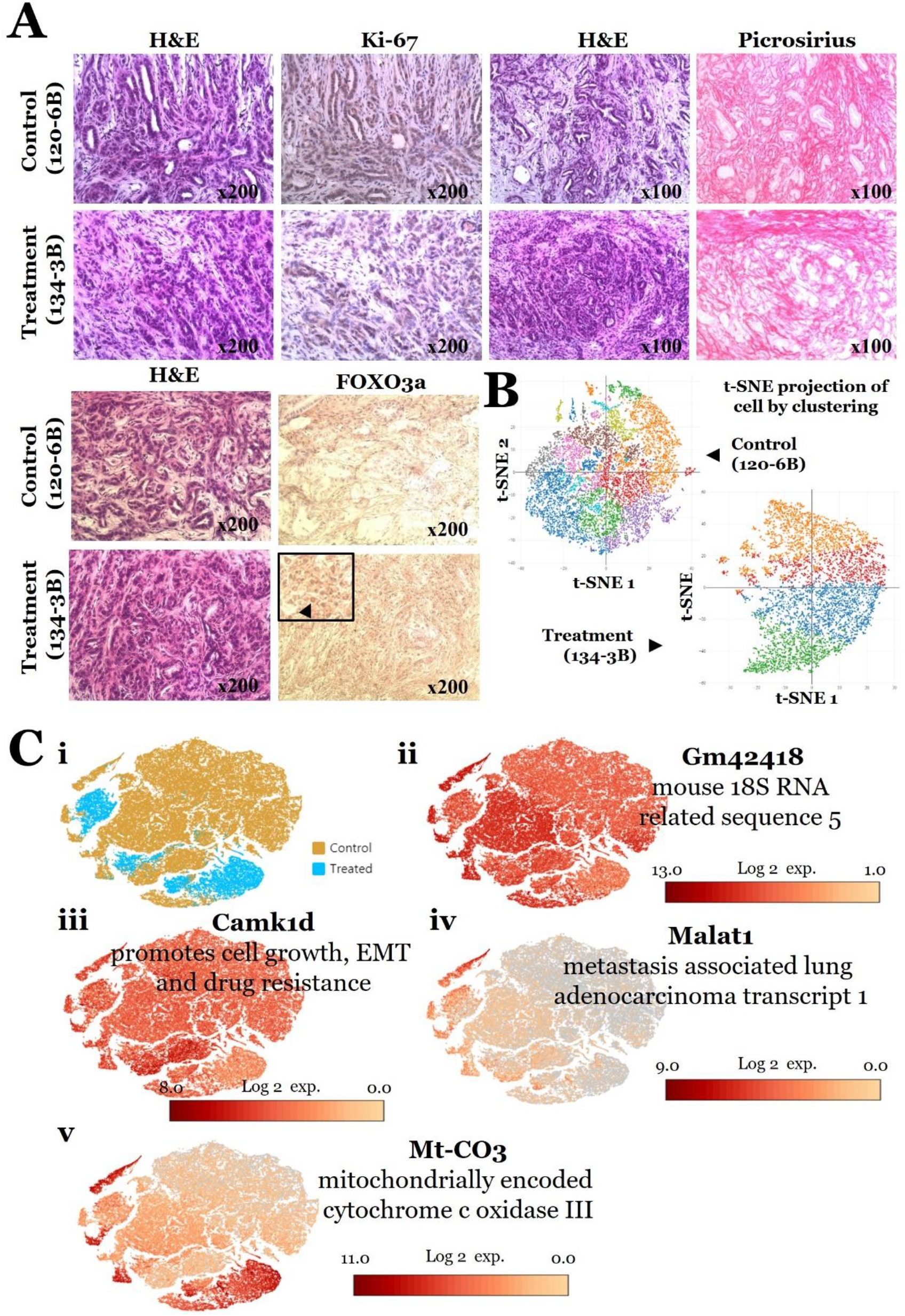
Immunohistochemical and two-dimensional t-distributed stochastic neighbor embedding (t-SNE) combined analysis of single nuclear RNA sequences from untreated (control) and treated KPC tumor cells. A. Immunohistochemical staining of KPC tumors from untreated and treated mice along with H&E staining. Nuclear accumulation of FOXO3a is enlarged and marked with arrowhead (lower panel). Original magnifications are shown on each histopathological image. B. Two-dimensional t-SNE analysis of single nuclear RNA sequences from a control (120-6B) and treated (134-3B) KPC mice tumor cells. Different cluster of cells are color-coded. C i). Merged clusters of control and treated KPC tumor cells. Clusters containing positive expression of *Gm42418* (C ii), *Camk1d* (C iii), *Malat1* (C iv), *Mt-CO3* (C v).

Further molecular analysis showed that Sel-GemPac treatment caused suppression of *Camk1d*, and long non-coding RNA (lncRNA) *Gm42418* and *Malat1* in the clusters of treated KPC tumor cells. On the other hand, the treatment caused upregulation of mitochondrially encoded cytochrome c oxidase III (Mt-CO3) (Fig. 2C). The top upregulated and downregulated genes are listed in Table S1 and S2 respectively. These changes involve several clusters most importantly cluster 1, 5, 28 (Table S3-S5).

### RNA-seq analysis identified differential expressed genes in Selinexor or Selinexor-Gemcitabine-nab-Paclitaxel treated PDAC cells

To analyze transcriptional alterations in PDAC, we performed a Poly(A) RNA-sequencing of MiaPaCa-2 cells that were treated with either DMSO (Con) or Sel or Sel-GemPac (Fig. 3A). A total of 8690 transcripts were increased in Sel and Sel-GemPac treated cells (Supplementary Table S6). Whereas a total of 2860 transcripts were decreased in the treated groups (Supplementary Table S7). Heatmap presented in the figure is showing top 20 differentially expressed genes that are common in both Sel and Sel-GemPac treated groups (Fig. 3B). Detailed differentially expressed genes between Con vs Sel, Con vs Sel-GemPac and Sel vs Sel-GemPac has shown in the supplementary figure (Fig. S4). We have analyzed the top upregulated and downregulated genes (shown in Fig. 3B) in STRING database. Interestingly, genes that downregulated by Sel or Sel-GemPac treatment showed strong interaction at the protein level (protein-protein interaction) compared to the genes that are upregulated (Fig. 3C). The existing known interactions and predicted interactions among proteins encoded by these genes suggest that Sel or Sel-GemPac can target different well-connected network that can be involved in PDAC progression. We have selected 6 genes from the downregulated genes list and evaluated further for their association in growth suppression *in vitro* and in patient samples (Fig. 3B).

**Figure 3.**
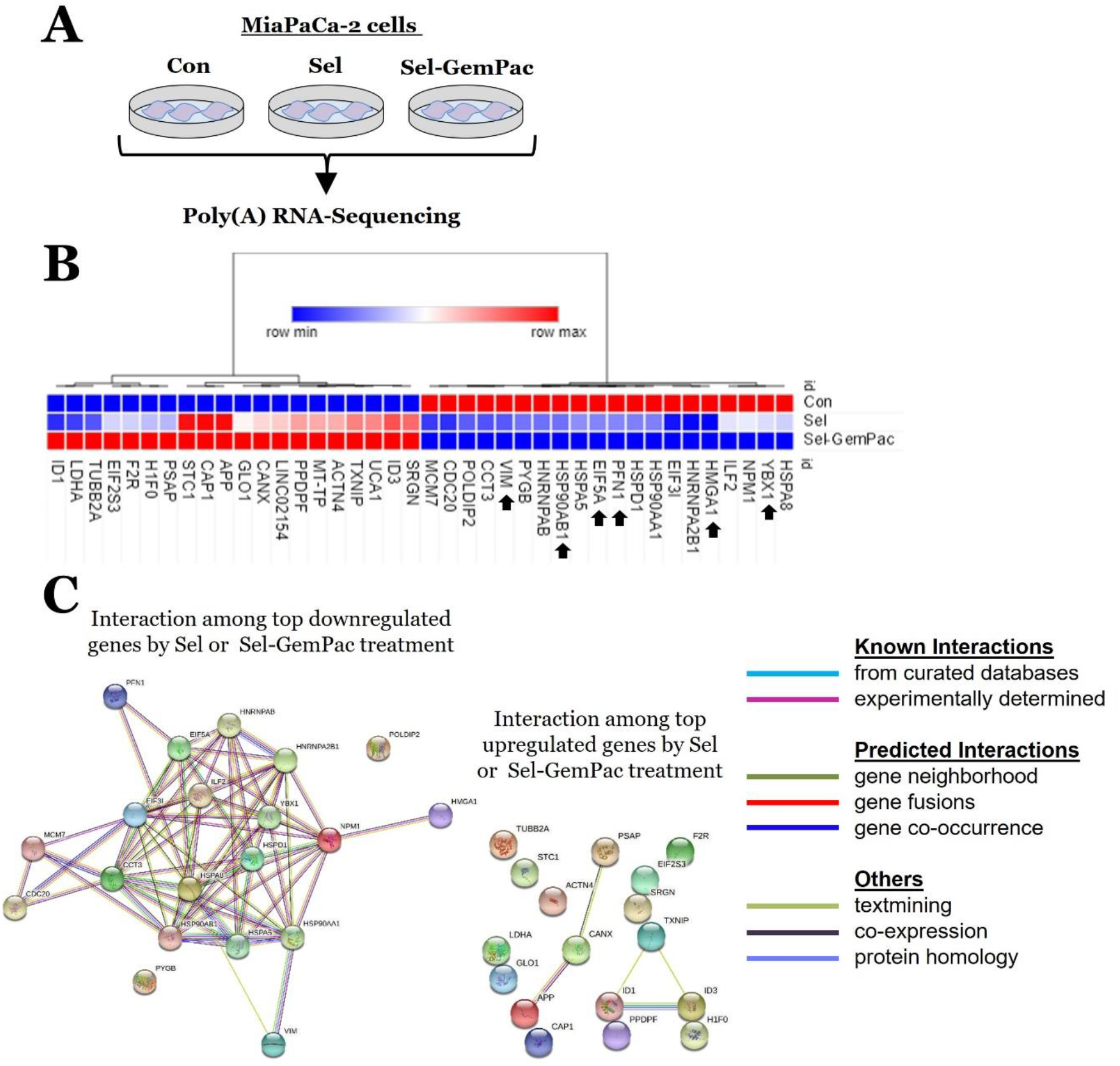
Differential expression of genes in Sel and Sel-GemPac treated MiaPcCa-2 cells. A. Experimental scheme. MiaPaCa-2 pancreatic cancer cells were treated with vehicle DMSO (Con), or selinexor or Sel. in combination with GemPac. B. Heatmap of top 20 differentially expressed genes in Con, Sel and Sel-GemPac treated groups. Heatmap was created using online interactive visualization and analysis tools in Morpheus from the Broad Institute (https://software.broadinstitute.org/morpheus/configuration.html) C. String analysis of topmost upregulated and downregulated gene encoding proteins. The protein-protein interaction map was created from STRING website (https://string-db.org/). The interactions among proteins were depicted with color coded lines. Con, control; Sel, Selinexor and GemPac, gemcitabine-nab-paclitaxel.

### Gene Ontology (GO) analysis predicted several pathways involved Selinexor or Selinexor-Gemcitabine-nab-Paclitaxel treated PDAC cells

The treatment with either Sel or Sel-GemPac were predicted to be most frequently involved in ‘DNA templated regulation of transcription’ or ‘transcription’ (Fig. 4A-4B). For example, Sel treatment caused alterations in 76 genes participating in ‘regulation of transcription’ process which is increased to 135 participated genes when cells were treated with Sel-GemPac combination. Among them, there were 55 differentially expressed genes that are common in both Sel and Sel-GemPac groups and there were 21 and 80 genes in Sel and Sel-GemPac groups respectively that are differentially expressed in each group (gene names along with the Venn diagram (Fig. 4C)). A similar trend was observed in ‘transport’, ‘signal transduction’, ‘cell differentiation’, ‘negative regulation of cell proliferation’ etc. pathways. The number of genes participating in cellular compartments and molecular functions were also significantly increased in the combination treatment (Supplementary Figure S5 and S6). Sel caused highest alteration of genes in the nuclear compartment, whereas combination treatment caused in the genes associated with membrane. In molecular functions, ‘protein binding’ was the topmost modulated function followed by ‘metal ion binding’, ‘DNA binding’, as well as ‘hydrolase activity’, ‘ATP binding’, ‘RNA binding’ etc. These findings suggest a potentiated inhibitory effects by combination treatment.

**Figure 4.**
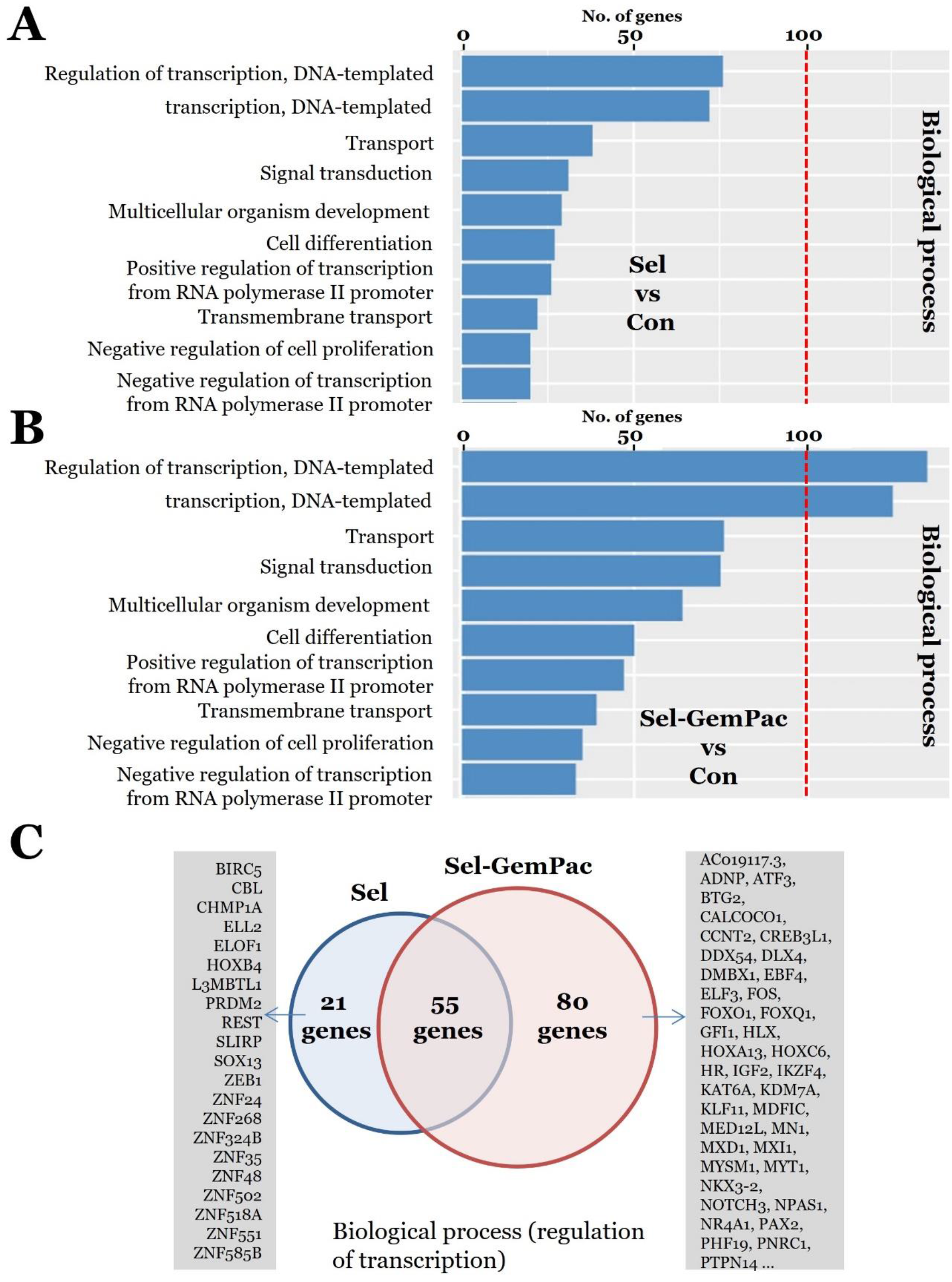
Genes involved in the biological process and genes interactions in Sel and Sel-GemPac treated MiaPaCa-2 cells compared to control. A-B. Top ten pathways modulation in relation to the number of genes from Gene Ontology (GO) enrichment analysis. A. Number of genes participated in different biological processes when treated with Sel relative to control. B. Number of genes participated in different biological processes when treated with Sel-GemPac relative to control. C. Venn-diagram showing common and differential gene lists induced by either Sel or Sel-GemPac treatment. Con, control; Sel, selinexor and GemPac, gemcitabine-nab-paclitaxel.

### Gene Set Enrichment Analysis (GSEA) predicted several gene sets differentially regulated by Selinexor or Selinexor-Gemcitabine-nab-Paclitaxel treatment

We have investigated the expression of different gene sets in Sel or Sel-GemPac treated groups by GSEA. GSEA was performed using C2 curated gene set, C2 KEGG gene set, C5 GO gene set and C3 TFT gene set databases. Total 10449 gene sets were used in the analysis while comparing Sel or Sel-GemPac treatment with control. In Sel vs Con group, top upregulated gene sets include small nucleolar RNA *SNORA44*, calcium voltage-gated channel auxiliary subunit gamma 8 (*CACNG8*), protease serine 35 (*PRSS35*) etc. whereas. top downregulated gene sets include microRNA 1224, adenosylhomocysteinase pseudogene 5, microRNA 675 etc. On the other hand, in Sel-GemPac vs Con group, top upregulated gene sets include protocadherin gamma subfamily A8, *CACNG8*, *PRSS35*. Whereas the top downregulated gene sets include coiled-coil-helix domain containing 4 pseudogene 3, leucine rich repeat containing 38, heterogeneous nuclear RNP U pseudogene 1 (Top 50 differentially expressed gene sets for Sel vs Con and Sel-GemPac vs Con are presented in Table S3 and S4). The common downregulated gene sets in both Sel and Sel-GemPac treated groups were DNA replication, protein export, RNP biogenesis, ribosome, spliceosome, proteasome etc (Fig. 5A and Fig. S7). The upregulated gene sets involve SATO silenced by deacetylation in pancreatic cancer and MEF2-02 (Fig. 5B).

**Figure 5.**
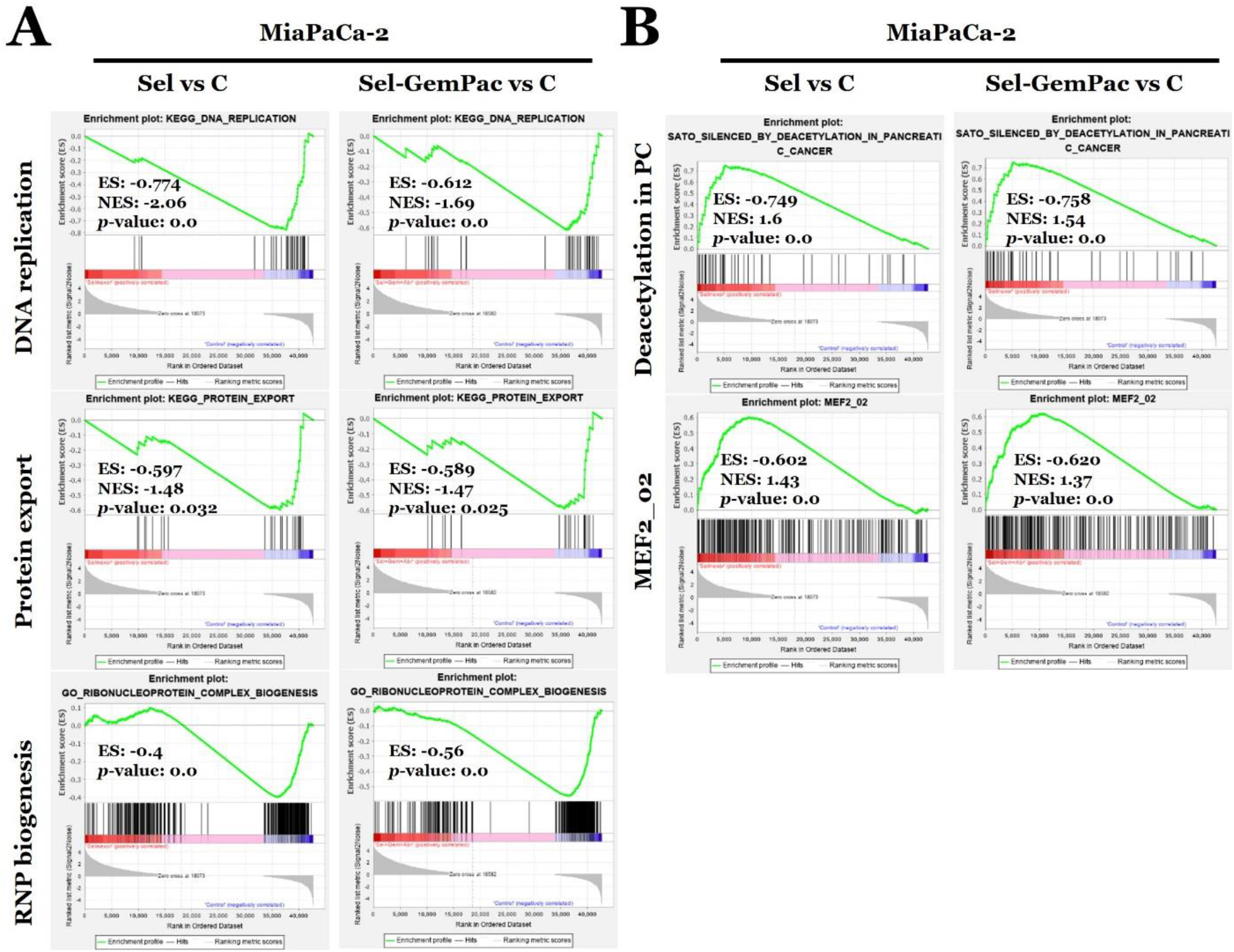
Gene Set Enrichment Analysis (GSEA) enrichment score curves for Sel and Sel-GemPac treated MiaPcCa-2 cells. A. Selected gene sets with ranked, non-redundant, and down-regulated (low enrichment scores) upon both Sel and Sel-GemPac treatment compared to Con group. B. Selected gene sets with ranked, non-redundant, and up-regulated (high enrichment scores) upon both Sel and Sel-GemPac treatment compared to control. In each GSEA set, probes on the far left (red) correlated with the most up-regulated probes and probes on the far right (blue) correlated with the most down-regulated probes. The vertical lines indicate the position of each of the probes of the studied probe set in the ordered, non-redundant data set. The green curve denotes the ES (enrichment score) curve, the running sum of the weighted enrichment score in GSEA. Con, control; Sel, Selinexor; GemPac, gemcitabine-nab-paclitaxel; ES, enrichment score; and NES, normalized enrichment score. *p*-values are shown on each gene set analysis.

### Selinexor and Selinexor-Gemcitabine-nab-Paclitaxel causes downregulation of genes that are overexpressed in PDAC

We next checked the gene expression of topmost downregulated genes observed in RNA-seq analysis in both Sel or Sel-GemPac groups including *HSP90AB1, YBX1, PFN1, HMGA1, EIF5A,* and *VIM* in PDAC patient samples. The tissue samples were obtained from baseline biopsies of patients enrolled in Phase Ib/II clinical trials who underwent Sel-GemPac treatment regimen. The expression of above genes was normalized utilizing normal patient pancreatic tissues. We observed higher baseline expressions for all the genes compared to normal tissues (Fig. 6A) suggesting their role in the progression of disease. The HSP90AB1 or high-mobility group is a chromosomal protein that frequently overexpressed in cancer promote proliferation, metastasis, and malignancy [15][16]. The Y-box transcription factor or nuclease-sensitive element-binding protein 1 (YBX1) is a protein that is known to promote metastasis [17]. Whereas PFN (profilin1) is a ubiquitous actin monomer-binding protein can cause drug resistance and progression via regulation of actin polymerization [18]. To evaluate the findings, we treated MiaPaCa-2 cells with either selinexor or Sel-GemPac combination and assess the expression of above genes. All the gene expressions were significantly suppressed by Sel alone except *EIF5A*. The Sel-GemPac combination caused significant downregulation of all genes to a much superior extent including *EIF5A* (Fig. 6B). These findings validate the results obtained from the RNA-seq data as well as the data obtained from the patient samples.

**Figure 6.**
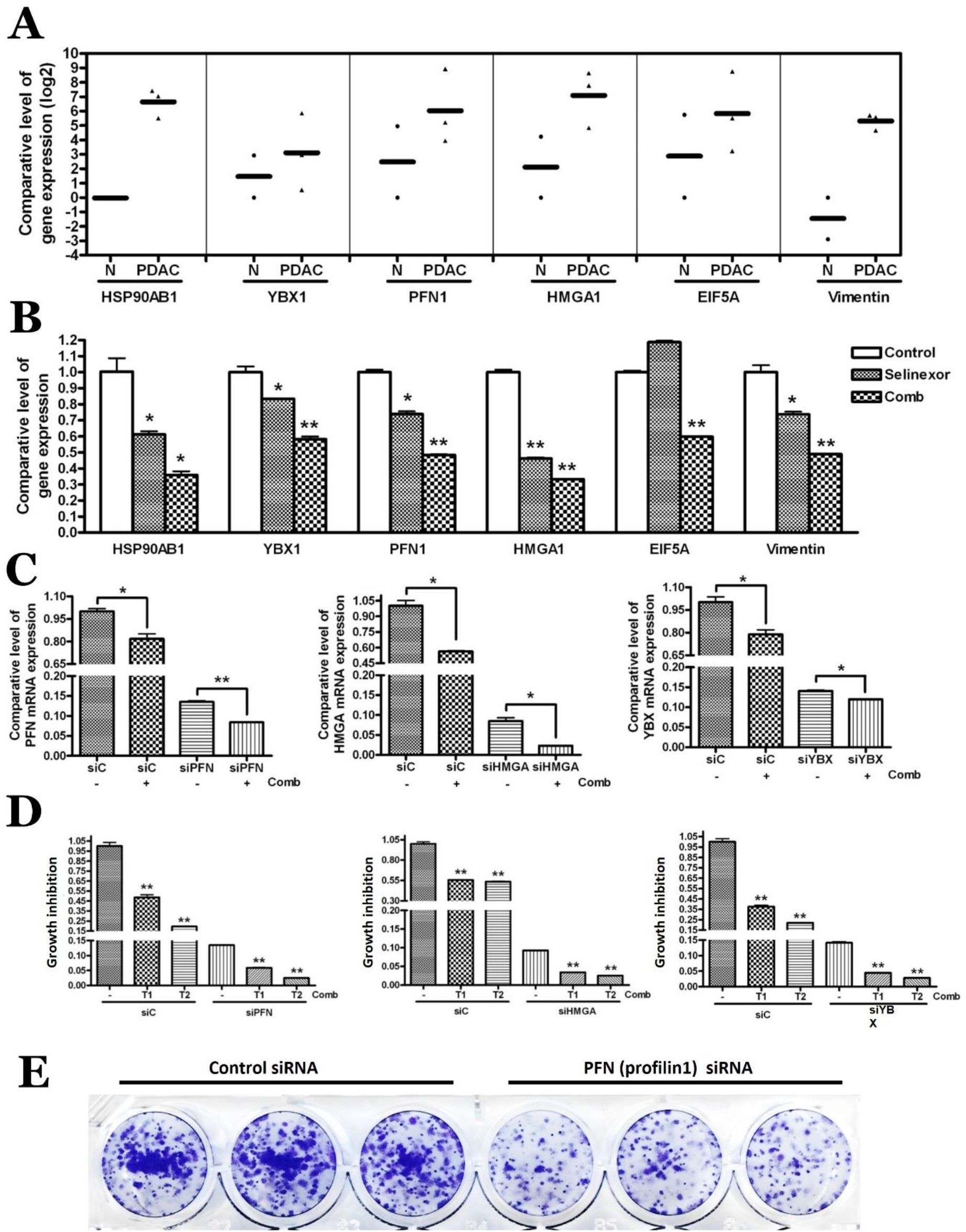
Validation of transcriptomic data *in vitro* and in patient samples. A. Validation of RNA-seq identified genes in pancreatic ductal adenocarcinoma (PDAC) patient samples. RT-PCR on PDAC patient tissue RNA from baseline biopsies of patients on the Phase Ib/II clinical study involving Sel-GemPac and compared it to normal pancreas tissue. The samples were probed for prioritized down regulated markers shown in Fig. 1. (i.e. RNA-seq identified targets post Sel-GemPac treatment in cellular models). B. Biological validation of RNA-seq and patient tumor identified targets. PDAC cells (data for MiaPaCa-2 cell line shown here) were grown in six well plates and exposed to either vehicle or 1 μM Sel alone or 1 μM Sel in combination with 300 nM Gem and 3 nM Pac for 24 hrs. At the end of the treatment period, RNA was isolated, and RT-PCR performed for selected set of markers (that were identified and prioritized in the RNA-seq experiments). C. MiaPaCa-2 cells were transfected with *PFN* siRNA (siPFN), *HMGA* siRNA (siHMGA), *YBX* siRNA (siYBX) or negative control siRNA (siC) using lipofectamine for 48 hrs and, then, the cells were treated with DMSO as control or 1μM Sel, or combination of 1μM Sel, 300 nM Gem and 3 nM Pac (Comb) for 24 hrs. Total RNA from each sample was extracted and the expression of various mRNA was measured by real-time RT-qPCR. D. Growth inhibition in siRNA silenced MiaPaCa-2 cells after transfected with siPFN, siHMGA, siYBX or siC for respective targets. Growth inhibition was determined using MTT assay after DMSO or combination treatments for 72 hrs. T1, treatment 1 (1μM Sel); T2, treatment 2 (1μM Sel, 300 nM Gem and 3 nM Pac) E. Colony formation in siRNA silenced MiaPaCa-2 cells after transfected with siPFN or siC for respective targets. Cells were allowed to grow for additional 7 days after siC or siPFN transfection for 48 hrs. Colony formation was observed using crystal violet staining in triplicates. Con, control; Sel, Selinexor and GemPac, gemcitabine-nab-paclitaxel. *, *p*<0.05; **, *p*<0.01.

### Silencing of *PFN*, *HMGA* or *YBX* sensitize PDAC cells to Selinexor or Selinexor-Gemcitabine-nab-Paclitaxel treatment

We tested the role of *PFN, HMGA* and *YBX* gene on synergy using siRNA approach. The Sel-GemPac treatment caused the downregulation of all these genes either transfected with control siRNA or target siRNA (Fig. 6C). Then, we checked the effect of silencing of these genes on the growth inhibition of MiaPaCa-2 cells. Silencing of these target genes itself caused pronounced inhibition of growth in PDAC cells. More significantly treatment with Sel (T1) or Sel in combination with GemPac (T2) caused further suppression of growth in these silenced cells (Fig. 6D). The growth inhibition was supported by colony formation ability of PDAC cells. When *PFN* gene was silenced, there was a lower colony formation efficiency (Fig. 6E), suggesting an association of *PFN, HMGA* and *YBX* genes with the vulnerability of PDAC cells for growth suppression.

## Discussion

Novel therapeutic strategies for pancreatic ductal adenocarcinoma (PDAC) are a necessity for many patients experiencing modest benefits from GemPac therapy. Selective inhibitor of nuclear export (SINE) compound Sel can work synergistically with GemPac, however, the mechanisms for such synergy are currently not known. Here we presented clues to understand molecular synergy between Sel and GemPac through bioinformatic analysis of snRNA-seq and RNA-seq data from KPC tumors and PDAC cells respectively. The snRNA-seq analysis demonstrated a significant reduction of cellular clusters in the Sel-GemPac treated tumors. Whereas RNA-seq analysis revealed several genes that are under expressed when PDAC cells are treated with Sel or Sel-GemPac and were verified using RNAi for their role in the molecular synergy. The siRNA mediated silencing of *PFN1, HMGA1, YBX1* and their growth suppressive role further support this mechanistic association.

Several studies have reported cellular heterogeneity utilizing single cell RNA-seq earlier focusing on untreated KPC tumors only [19–21]. Such studies observed normal or tumor ductal cell types, fibroblasts, different immune cells infiltrations [19, 20]. In our study, we have observed the presence of CD44 positive stem cells in the untreated KPC tumors which is greatly reduced in the treated tumors. The CD44 positive stem cells are associated with Gem or chemotherapy resistance in PDAC [22, 23]. A limited infiltration of immune cells (Fig. S3) was observed in the current study, both in untreated and treated KPC tumors. This might be due to the harvesting of late-stage KPC tumors as observed by other independent groups study [21]. Though CD4 expressing cells are rarely detected both in untreated and treated KPC tumor’s snRNA-seq, we have observed CD4 cells in untreated tumors using IHC staining. Importantly, the expression was decreased in the treated tumor tissues (Fig. S8). It is possible we might have loss some cells during the isolation of single cells via multiple digestion cycle [21].

Our snRNA-seq analysis showed that Sel-GemPac treatment was significantly effective in reducing the expression of *Camk1d* and long non-coding RNA *Gm42418* and *Malat1*. The calcium/calmodulin‐dependent protein kinase (CAMK) has been shown to regulate a diverse range of cancer‐related functions in different tumor types [24]. In 2021, the role of Camk1 has been demonstrated in pancreatic cancer [25]. The study found higher expression Camk1 in pancreatic cancer from bioinformatics as well as TMA‐IHC analyses. The protein‐protein interactions network predicted an association of Camk1 with a number of genes including *CALM1, CALM3, CREB1, CALM2, SYN1, NOS3, ATF1, GAPDH, PPM1F* and *FBXL12*. Most of these genes are involved in aldosterone synthesis and secretion, oxytocin signaling pathway and upregulated in pancreatic cancer. The role of *Malat1* in pancreatic cancer aggressiveness are well documented [26–28] due to its involvement in *KRAS* expression regulation [29]. These observations indicates and important role of *Camk1d, Mt-CO3* genes and *Gm42418, Malat1* lncRNAs in the regulation of tumor growth in KPC genetic mouse model. Our analysis has identified a novel less recognized lncRNA. The role of *Gm42418* not yet demonstrated in pancreatic cancer and can be subjected to further investigation.

We have identified several differential expressed genes in Sel or Sel-GemPac treated PDAC cells. We also verified such gene expressions in the baseline tissue samples of PDAC patients on the Phase Ib/II clinical study involving Sel-GemPac treatment. Among them, *HMGA1* was shown to be associated with advance tumor grade, lymph node metastasis and decreased survival in PDAC [30, 31]. Overexpression of *YBX1* shown to promote the growth of PDAC by modulation cell cycle regulatory machinery such as *GSK3B, Cyclin D1* and *Cyclin E1* [32]. Both Sel and Sel-GemPac treatment caused the suppression of these genes and helps in sensitizing PDAC cells. We have observed *PFN1* suppression which is opposite of the published literature in PDAC so far and its exact role remains elusive [33, 34]. We also observed suppression of *PFN1* with Sel and Sel-GemPac treatments and its silencing sensitized cells to the treatments. In fact, in some human and rat gastric cancer *PFN1* shown to be overexpressed [35]. It has been concluded that PFN1 expression has spatial and temporal specificity in cancer [36] and we are the first to show its opposing effect in PDAC. Remarkably, the top downregulated genes upon treatment were all well connected as revealed from STRING analysis. YBX1 act as a central molecule in the network. It is connected with HSP90AB1, NPM1, HSP90AA1, HNRNPA2B1, ILF2, HSPD1 etc. which is experimentally determined in cancer. Sel treatment disrupts such network that may play a role leading to synergy between Sel and GemPac.

Moreover, a range of GO terms related to biological processes, including cell differentiation and negative regulation of cell proliferation were found significantly enriched in Sel or Sel-GemPac treated PDAC cells. Expectedly, Sel treatment affected the genes of nuclear compartment, whereas in combination with GemPac it affects membrane mostly. In molecular function, protein, metal ion and DNA binding largely affected by Sel or Sel-GemPac treatment. GSEA can precisely and consistently identify gene sets with biological significance [37]. Our GSEA analysis revealed robust disruption in the gene set enrichment of DNA replication, protein export, RNP biogenesis, ribosome, spliceosome and proteasome machinery both in Sel or Sel-GemPac treated cells, whereas “deacetylation in pancreatic cancer” gene set get enriched suggesting the biological relevance of the findings. Sel alone or in combination with GemPac upregulates protease serine 35 (PRSS35). There are limited studies performed on PRSS35. It belongs to the trypsin class of serine proteases [38]. Though originally identified as a novel mouse ovary-selective gene [38, 39] it is involved in collagen remodeling [40]. PRSS35 ablation in mouse models of chemically induced tumorigenesis demonstrated aberrant collagen composition in the extra cellular matrix (ECM) and increased tumor incidence [40]. MicroRNA *1224* and adenosylhomocysteinase pseudogene 5 gene sets are downregulated by Sel only. However, in the combination treatment gene sets coiled-coil-helix domain containing 4 pseudogene 3 and leucine rich repeat containing 38 were found to be suppressed. The association of diverse gene sets with Sel or Sel-GemPac might constitute a rationale for a combined treatment regimen.

Collectively, we have identified a series of key genes including *Camk1d, PFN1, HMGA1, YBX1* and lncRNA *Malat1* that are associated with PDAC growth and plays an important role in the Sel-GemPac mediated growth suppression. These findings provide supports for the understanding of molecular mechanism of Sel and GemPac mediated synergy in pancreatic cancer.

## Supplementary figure legends

**Supplementary figure S1.** Two-dimensional t-distributed stochastic neighbor embedding (t-SNE) analysis of single nuclear RNA sequences from an untreated (left, 117-6B) and a treated (right, 136-6B) KPC tumor cells. Different cluster of cells are color-coded and shown on the right of each t-SNE image.

**Supplementary figure S2.** Two-dimensional t-distributed stochastic neighbor embedding (t-SNE) combined analysis of single nuclear RNA sequences from all untreated (top; 117-6B, 120-6B) and treated (bottom; 134-6B, 136-6B) KPC tumor cells. Different cluster of cells are color-coded and shown on the right of the image.

**Supplementary figure S3.** Two-dimensional t-distributed stochastic neighbor embedding (t-SNE) combined analysis of single nuclear RNA sequences from all untreated (control) and treated KPC tumor cells. A. Merged clusters of control and treated KPC tumor cells. B. Clusters containing positive CD44 stem cells. The log2 expressions of CD44 stem cell marker is shown in color gradients. C. Clusters containing positive markers for other cell types including endothelial, fibroblast and immune cells.

**Supplementary figure S4.** Differential expression of genes in Sel and Sel-GemPac treated MiaPcCa-2 cells. MiaPaCa-2 pancreatic cancer cells were treated with vehicle DMSO (Con), or Sel or Sel-GemPac. A. Heatmap of differentially expressed genes in Sel treated group compared to Con. B. Heatmap of differentially expressed genes in Sel-GemPac treated group compared to Con. C. Heatmap of differentially expressed genes in Sel-GemPac treated group compared to Sel. Con, control; Sel, selinexor and GemPac, gemcitabine-nab-paclitaxel.

**Supplementary figure S5.** Genes involved with different cellular compartments in Sel and Sel-GemPac treated MiaPaCa-2 cells compared to control. A-B. Top ten pathways modulation in relation to the number of genes from Gene Ontology (GO) enrichment analysis. A. Number of genes participated in different cellular compartments when treated with Sel relative to control. B. Number of genes participated in different cellular compartments when treated with Sel-GemPac relative to control. Sel, selinexor and GemPac, gemcitabine-nab-paclitaxel.

**Supplementary figure S6.** Genes involved in the molecular functions in Sel and Sel-GemPac treated MiaPaCa-2 cells compared to control. A-B. Top ten pathways modulation in relation to the number of genes from Gene Ontology (GO) enrichment analysis. A. Number of genes participated in the molecular functions when treated with Sel relative to control. B. Number of genes participated in the molecular functions when treated with Sel-GemPac relative to control. Sel, selinexor and GemPac, gemcitabine-nab-paclitaxel.

**Supplementary figure S7.** Gene Set Enrichment Analysis (GSEA) enrichment score curves for Sel and Sel-GemPac treated MiaPcCa-2 cells. Selected gene sets with ranked, non-redundant, and down-regulated (low enrichment scores) upon both Sel and Sel-GemPac treatment compared to control. Sel, selinexor and GemPac, gemcitabine-nab-paclitaxel.

**Supplementary figure S8.** Immunohistochemical staining of KPC tumors for CD4 from untreated and treated mice along with H&E staining. Original magnifications are shown on each histopathological image.

**Supplementary table S1.** Top downregulated genes in the treated KPC mouse compared to control. *, *p*<0.05; **, *p*<0.01; ***, *p*<0.001.

**Supplementary table S2.** Top upregulated genes in the treated KPC mouse compared to control. *, *p*<0.05; ***, *p*<0.001.

**Supplementary table S3.** Log2 fold change in cluster 1 between control and treated KPC mouse tumor. Adjusted *p-*values are in the last column.

**Supplementary table S4.** Log2 fold change in cluster 5 between control and treated KPC mouse tumor. Adjusted *p-*values are in the last column.

**Supplementary table S5.** Log2 fold change in cluster 28 between control and treated KPC mouse tumor. Adjusted *p-*values are in the last column.

**Supplementary table S6.** List of upregulated genes upon Sel and Sel-GemPac treatment in MiaPaCa-2 cells obtained from RNA-seq data analysis. Sel, selinexor and GemPac, gemcitabine-nab-paclitaxel.

**Supplementary table S7.** List of downregulated genes upon Sel and Sel-GemPac treatment in MiaPaCa-2 cells obtained from RNA-seq data analysis. Sel, selinexor and GemPac, gemcitabine-nab-paclitaxel.

