## Supplementary Figure S1-S8 and Table S1-S2 for "Molecular profiling of XPO1 inhibitor and gemcitabine-nab-paclitaxel combination in cellular and LSL-Kras G12D/+; Trp53 fl/+; Pdx1-Cre (KPC) pancreatic cancer model"

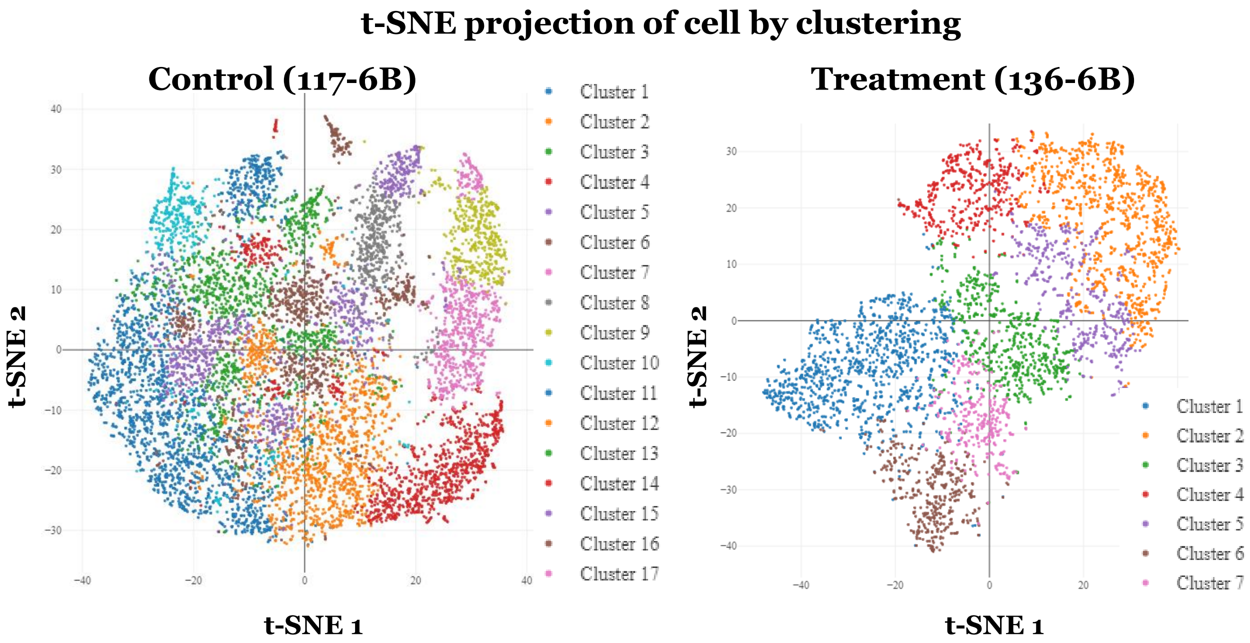

### Supplementary figure S2

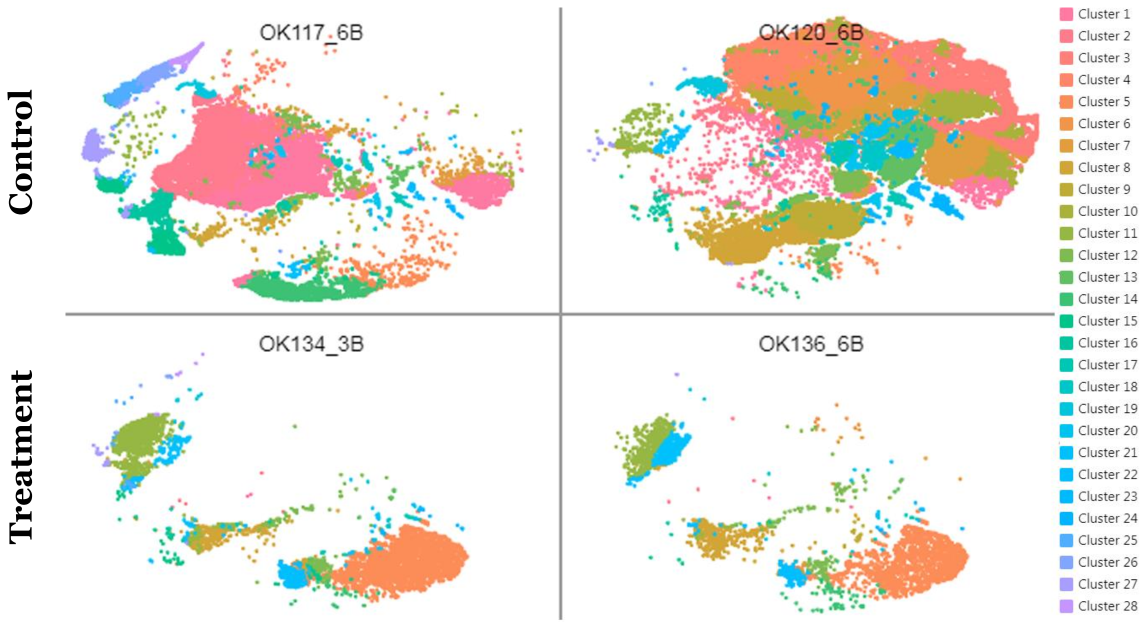

### Supplementary figure S3

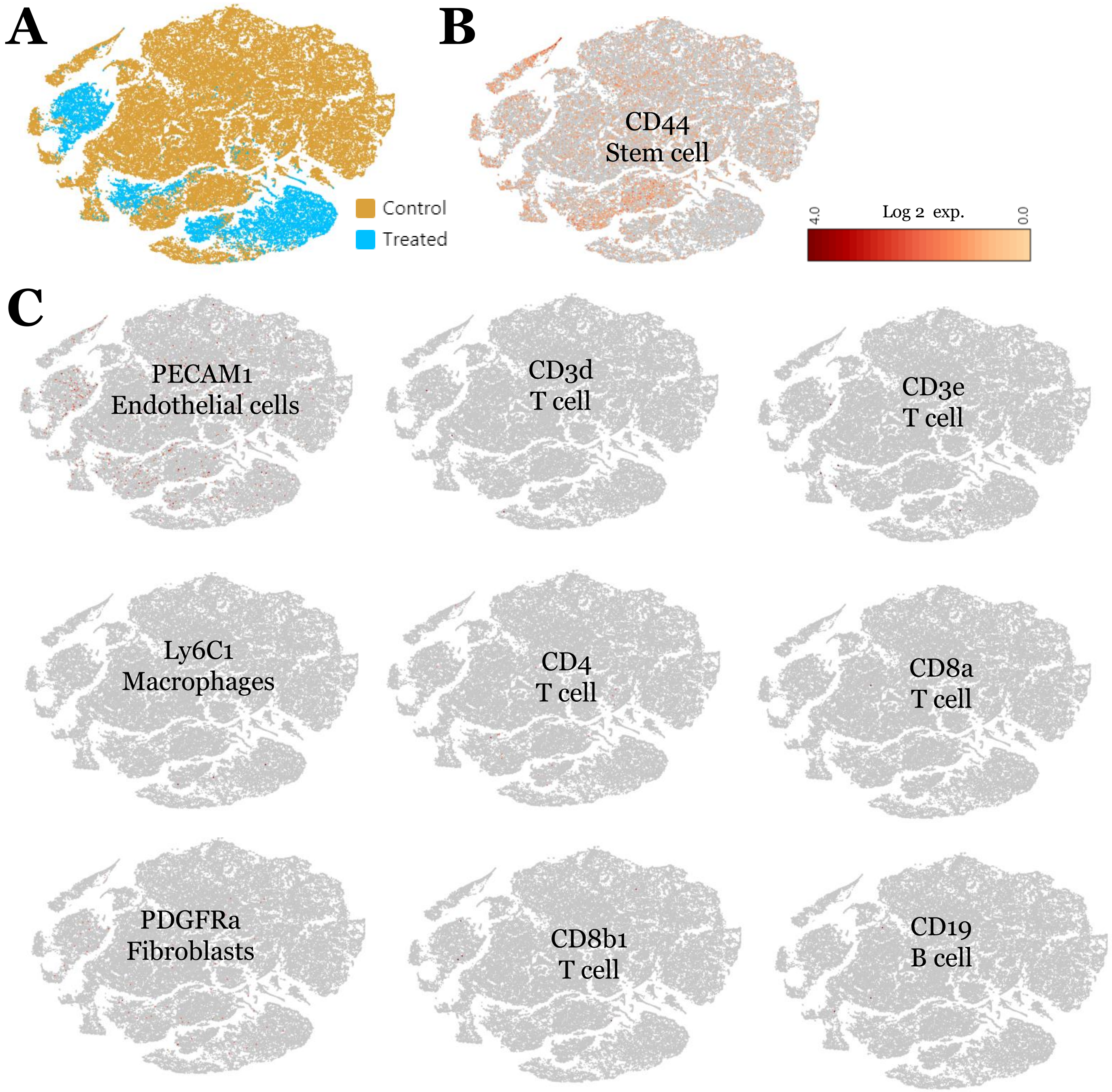

### Supplementary figure S4

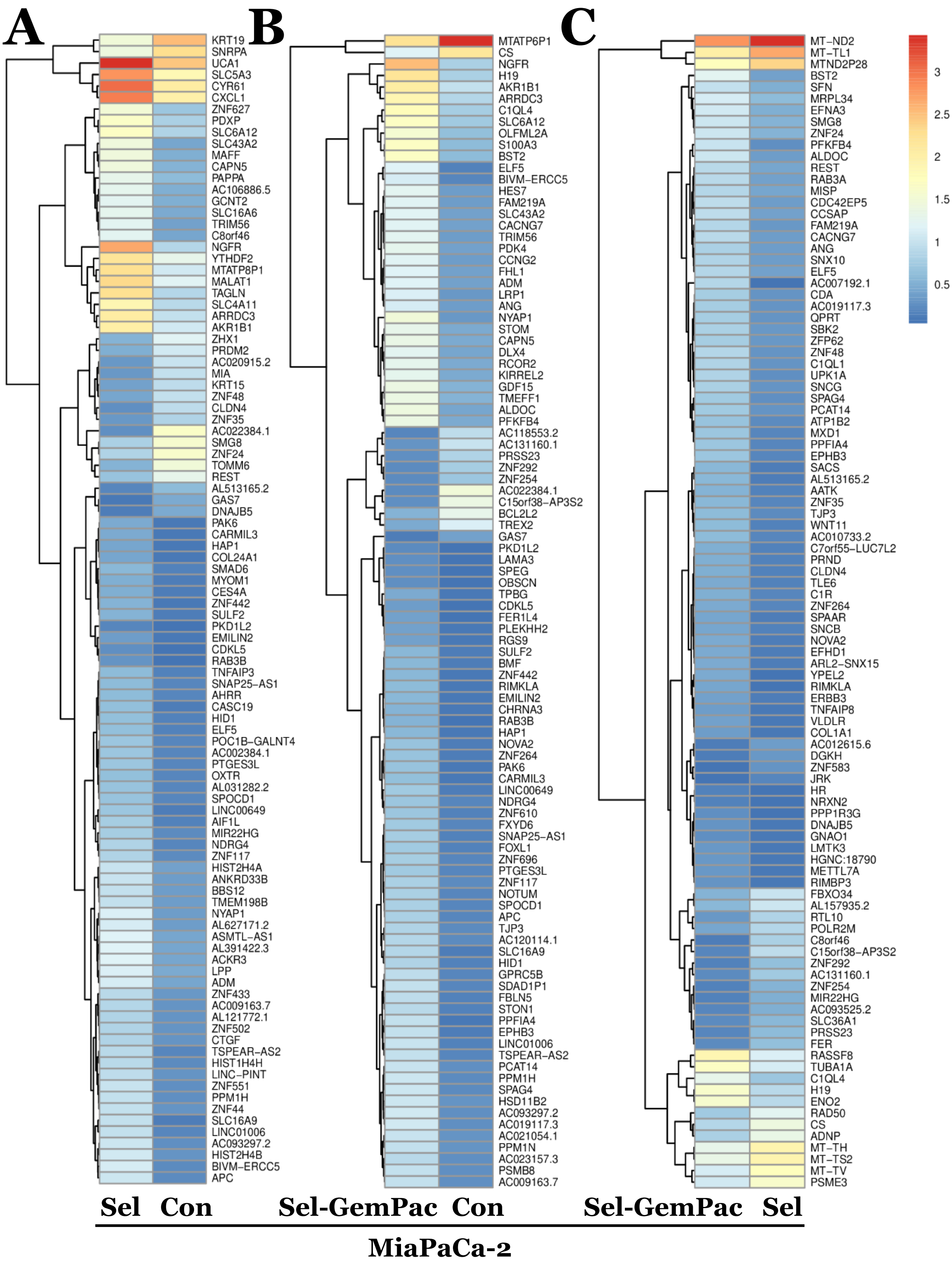

### Supplementary figure S5

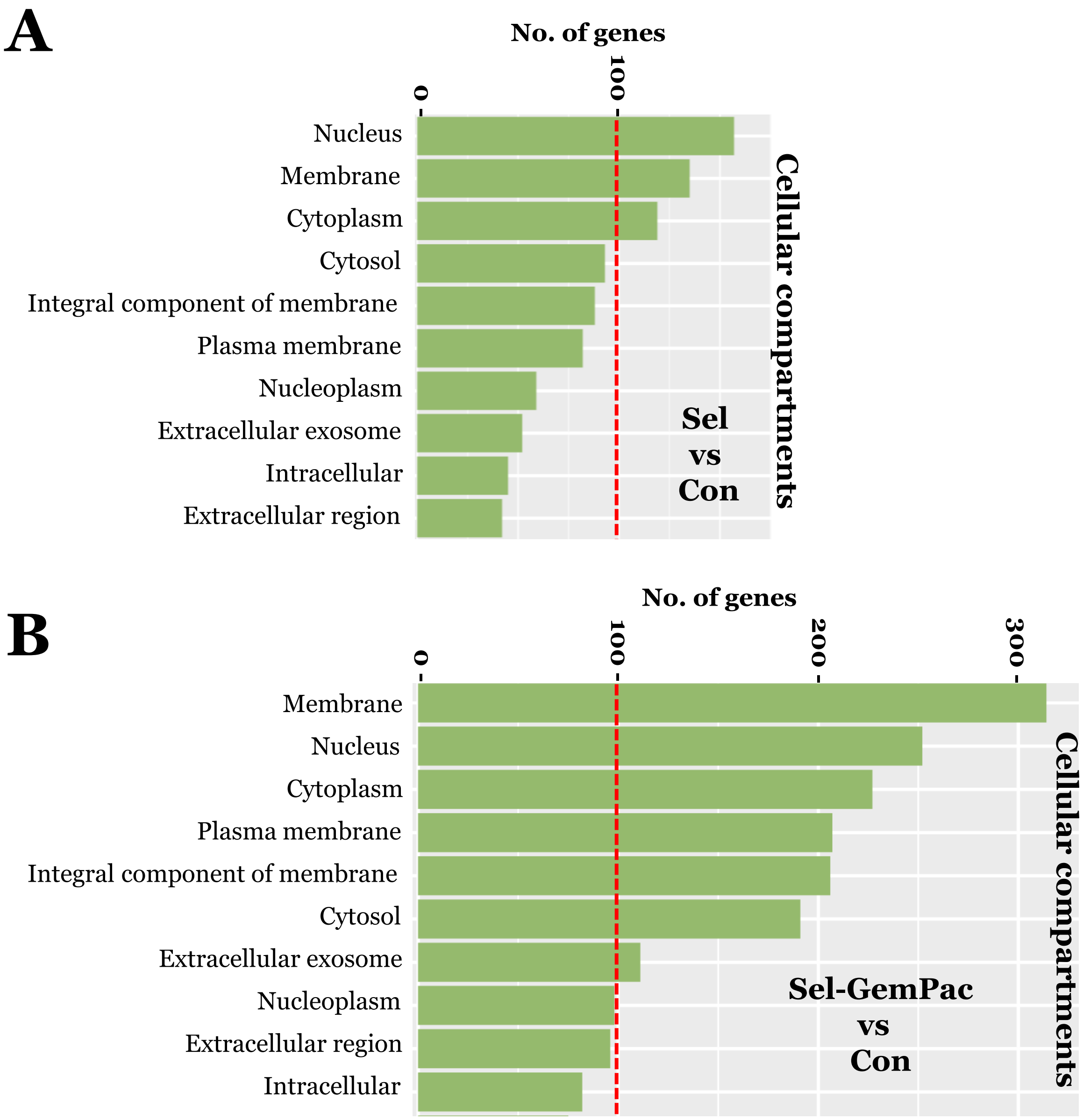

### Supplementary figure S6

A

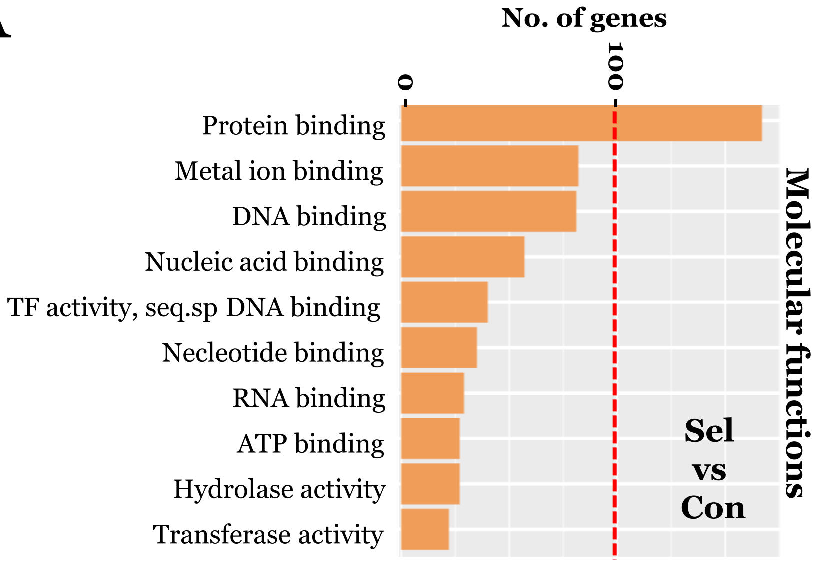


B

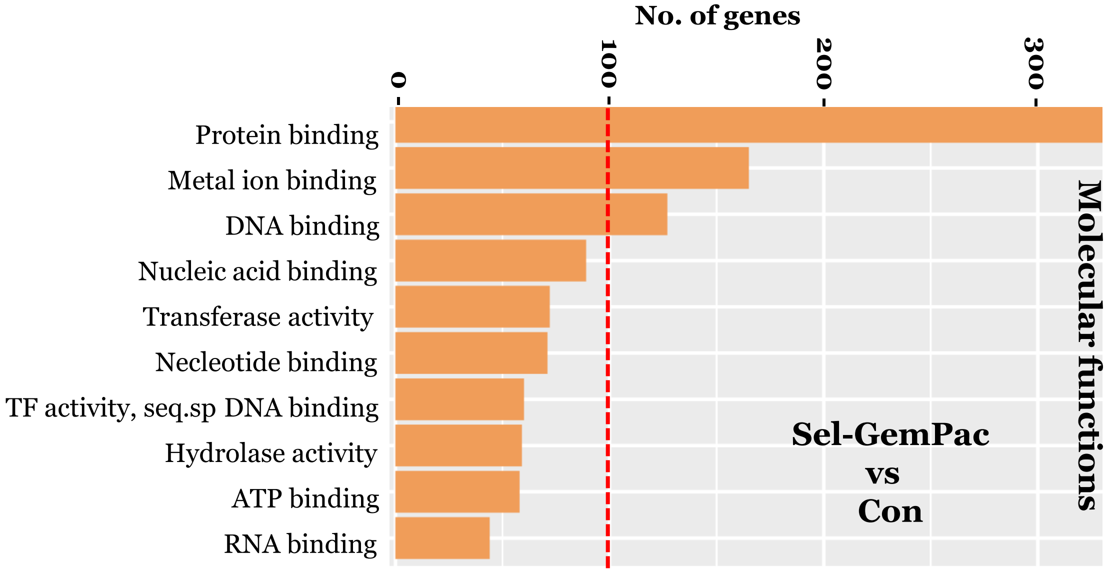

### Supplementary figure S7

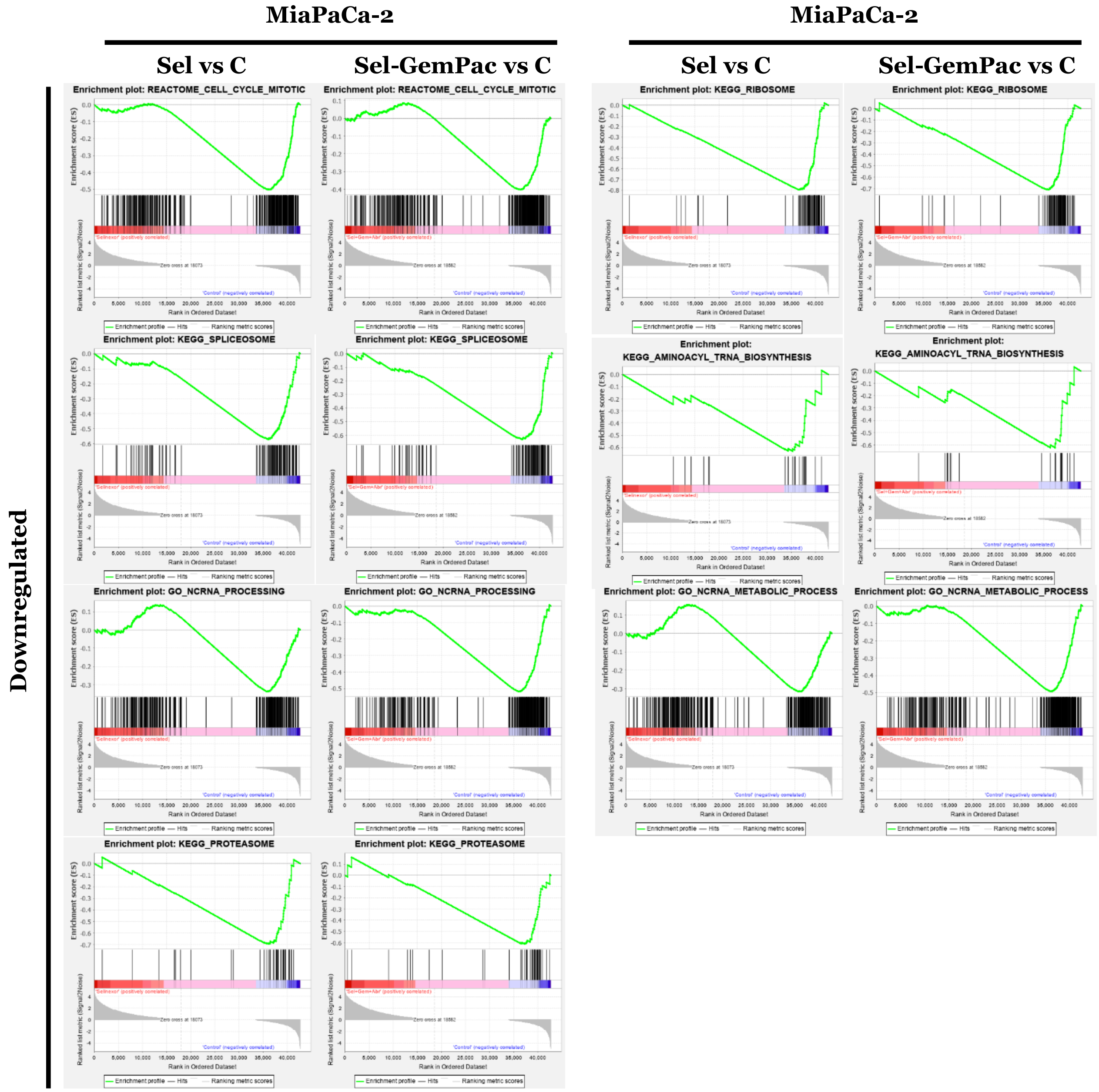

### Supplementary figure S8

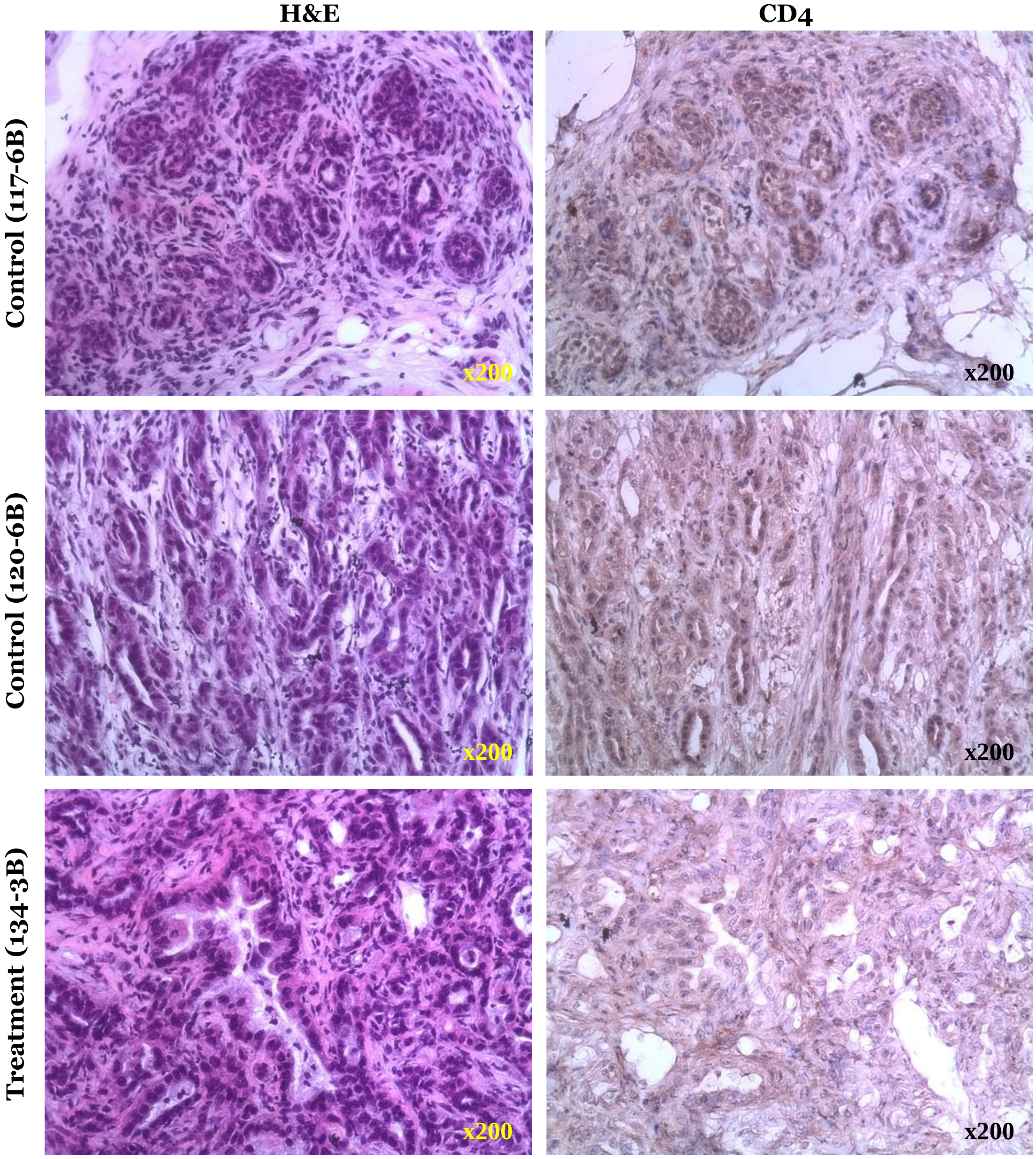

### Supplementary table S1

Table S1: Top downregulated genes in the treated KPC mouse compared to control

| FeatureID | Name | Control Ave | Control Log2 FC | Control P-Value | Treated Ave | Treated Log2 FC | Treated P-Value | |
|---|---|---|---|---|---|---|---|---|
| ENSMUSG00000098178 | Gm42418 | 165.03731 | 1.828185652 | 3.43E-06 | 46.478875 | -1.828185652 | 3.43E-06 | *** |
| ENSMUSG00000039145 | Camk1d | 9.2905329 | 1.353242243 | 0.000663036 | 3.636491 | -1.353242243 | 0.000663036 | *** |
| ENSMUSG00000035202 | Lars2 | 9.0061998 | 1.095564973 | 0.006994068 | 4.2145644 | -1.095564973 | 0.006994068 | ** |
| ENSMUSG00000047454 | Gphn | 3.1987594 | 0.935692563 | 0.023855471 | 1.6722989 | -0.935692563 | 0.023855471 | * |
| ENSMUSG00000029635 | Cdk8 | 1.4207631 | 0.789638986 | 0.06272897 | 0.8218864 | -0.789638986 | 0.06272897 | |
| ENSMUSG00000022748 | Cmss1 | 8.6494701 | 0.240609552 | 0.66128517 | 7.3209853 | -0.240609552 | 0.66128517 | |

### Supplementary table S2

Table S2: Top upregulated genes in the treated KPC mouse compared to control

| FeatureID | Name | Control Ave | Control Log2 FC | Control P-Value | Treated Ave | Treated Log2 FC | Treated P-Value | |
|---|---|---|---|---|---|---|---|---|
| ENSMUSG00000064370 | mt-Cytb | 1.074255 | -2.903516006 | 6.26E-27 | 8.03835311 | 2.903516006 | 6.26E-27 | *** |
| ENSMUSG00000064357 | mt-Atp6 | 3.1930501 | -2.594939496 | 3.42E-21 | 19.2918251 | 2.594939496 | 3.42E-21 | *** |
| ENSMUSG00000064356 | mt-Atp8 | 0.2701692 | -2.519410338 | 7.18E-20 | 1.54906961 | 2.519410338 | 7.18E-20 | *** |
| ENSMUSG00000064351 | mt-Co1 | 5.5953424 | -2.480489126 | 3.12E-19 | 31.2277741 | 2.480489126 | 3.12E-19 | *** |
| ENSMUSG00000064363 | mt-Nd4 | 1.5647203 | -2.479949398 | 3.15E-19 | 8.72949535 | 2.479949398 | 3.15E-19 | *** |
| ENSMUSG00000064354 | mt-Co2 | 4.5641682 | -2.439500397 | 1.47E-18 | 24.7592334 | 2.439500397 | 1.47E-18 | *** |
| ENSMUSG00000064341 | mt-Nd1 | 1.7904313 | -2.342258665 | 5.74E-17 | 9.07947574 | 2.342258665 | 5.74E-17 | *** |
| ENSMUSG00000064345 | mt-Nd2 | 0.7693457 | -2.306191834 | 2.07E-16 | 3.80511647 | 2.306191834 | 2.07E-16 | *** |
| ENSMUSG00000064360 | mt-Nd3 | 0.5271624 | -2.26497665 | 9.67E-16 | 2.53386871 | 2.26497665 | 9.67E-16 | *** |
| ENSMUSG00000064367 | mt-Nd5 | 0.598685 | -2.119518067 | 1.36E-13 | 2.60165632 | 2.119518067 | 1.36E-13 | *** |
| ENSMUSG00000064358 | mt-Co3 | 7.7249006 | -1.974798493 | 9.93E-12 | 30.3653992 | 1.974798493 | 9.93E-12 | *** |
| ENSMUSG00000105361 | AY036118 | 2.6699631 | -1.762674776 | 3.16E-09 | 9.06015167 | 1.762674776 | 3.16E-09 | *** |
| ENSMUSG00000097971 | Gm26917 | 0.835596 | -0.849136622 | 0.016209626 | 1.50528372 | 0.849136622 | 0.016209626 | * |
