## Supplementary Table S4 for "Molecular profiling of XPO1 inhibitor and gemcitabine-nab-paclitaxel combination in cellular and LSL-Kras G12D/+; Trp53 fl/+; Pdx1-Cre (KPC) pancreatic cancer model"

| Feature ID | Feature Name | Cluster 5 Mean Counts | Cluster 5 Log2 fold change | Cluster 5 Adjusted p value |
|---|---|---|---|---|
| ENSMUSG00000064358 | mt-Co3 | 41.5372318 | 2.460506155 | 5.11E-15 |
| ENSMUSG00000064351 | mt-Co1 | 41.29662304 | 2.830753146 | 7.91E-21 |
| ENSMUSG00000064354 | mt-Co2 | 32.87033389 | 2.805855145 | 1.72E-20 |
| ENSMUSG00000098178 | Gm42418 | 29.62111915 | -2.42741158 | 2.38E-06 |
| ENSMUSG00000064357 | mt-Atp6 | 25.46361466 | 2.926756821 | 1.84E-22 |
| ENSMUSG00000064341 | mt-Nd1 | 12.20267291 | 2.754329583 | 1.31E-19 |
| ENSMUSG00000064363 | mt-Nd4 | 11.58692266 | 2.841675624 | 5.91E-21 |
| ENSMUSG00000064370 | mt-Cytb | 10.42606569 | 3.135332221 | 1.88E-26 |
| ENSMUSG00000105361 | AY036118 | 5.406499781 | 0.54523018 | 0.359213366 |
| ENSMUSG00000064345 | mt-Nd2 | 5.02089463 | 2.667246509 | 3.40E-18 |
| ENSMUSG00000064367 | mt-Nd5 | 3.488937686 | 2.542105136 | 3.11E-16 |
| ENSMUSG00000022748 | Cmss1 | 3.460591693 | -1.406096301 | 0.006078728 |
| ENSMUSG00000064360 | mt-Nd3 | 3.453117652 | 2.723940715 | 4.24E-19 |
| ENSMUSG00000035202 | Lars2 | 2.484924849 | -1.84756098 | 0.000274457 |
| ENSMUSG00000064356 | mt-Atp8 | 2.024136383 | 2.82845058 | 8.46E-21 |
| ENSMUSG00000039145 | Camk1d | 1.915790471 | -2.258168993 | 1.02E-05 |
| ENSMUSG00000065947 | mt-Nd4l | 1.324953692 | 2.545563306 | 3.11E-16 |
| ENSMUSG00000047454 | Gphn | 0.787044195 | -2.037815705 | 6.27E-05 |
| ENSMUSG00000029635 | Cdk8 | 0.432165658 | -1.736813863 | 0.000646062 |
| ENSMUSG00000113136 | Gm19951 | 0.412511698 | -0.912345276 | 0.109343152 |
| ENSMUSG00000097971 | Gm26917 | 0.214089751 | -2.346249038 | 5.11E-06 |
| ENSMUSG00000064368 | mt-Nd6 | 0.199086306 | 1.021842799 | 0.014746263 |
| ENSMUSG00000001025 | S100a6 | 0.135307823 | 2.011532078 | 9.51E-09 |
| ENSMUSG00000060126 | Tpt1 | 0.110394353 | 1.188418149 | 0.003112598 |
| ENSMUSG00000024032 | Tff1 | 0.09373001 | 3.291574383 | 5.73E-27 |
| ENSMUSG00000092341 | Malat1 | 0.084373618 | -2.906719135 | 2.12E-08 |
| ENSMUSG00000029304 | Spp1 | 0.067653912 | 0.249513517 | 0.88176882 |
| ENSMUSG00000117465 | Gm49980 | 0.064221389 | 1.097820383 | 0.008444043 |
| ENSMUSG00000022037 | Clu | 0.061951495 | 1.184727446 | 0.003630595 |
| ENSMUSG00000037742 | Eef1a1 | 0.056082989 | 1.471027812 | 0.0001414 |
| ENSMUSG00000047675 | Rps8 | 0.054975724 | 1.414878378 | 0.0003174 |
| ENSMUSG00000038274 | Fau | 0.05204147 | 1.281928071 | 0.001438867 |
| ENSMUSG00000049775 | Tmsb4x | 0.050657389 | 1.658316604 | 1.23E-05 |
| ENSMUSG00000061983 | Rps12 | 0.04999303 | 1.627450166 | 1.82E-05 |
| ENSMUSG00000025314 | Ptprj | 0.049384034 | -1.264502854 | 0.02307354 |
| ENSMUSG00000032518 | Rpsa | 0.047557046 | 1.104131646 | 0.00592361 |
| ENSMUSG00000007892 | Rplp1 | 0.046449781 | 0.823437563 | 0.062039544 |
| ENSMUSG00000025290 | Rps24 | 0.045287152 | 1.16039211 | 0.003437576 |
| ENSMUSG00000086054 | Hnf1aos1 | 0.044954972 | -2.156616832 | 5.72E-05 |
| ENSMUSG00000093674 | Rpl41 | 0.044899609 | 1.479700419 | 9.26E-05 |
| ENSMUSG00000079523 | Tmsb10 | 0.044622793 | 1.346707699 | 0.000519924 |
| ENSMUSG00000111056 | Gm39317 | 0.04135636 | -2.409980986 | 6.69E-06 |
| ENSMUSG00000040952 | Rps19 | 0.041134907 | 1.779664622 | 1.04E-06 |
| ENSMUSG00000034892 | Rps29 | 0.04102418 | 1.47761491 | 0.00010639 |
| ENSMUSG00000046330 | Rpl37a | 0.040802727 | 1.801273126 | 8.12E-07 |
| ENSMUSG00000000740 | Rpl13 | 0.040138368 | 1.29390319 | 0.000965898 |
| ENSMUSG00000028081 | Rps3a1 | 0.039695462 | 1.166569695 | 0.003330075 |
| ENSMUSG00000067288 | Rps28 | 0.03880965 | 1.687365244 | 5.74E-06 |
| ENSMUSG00000044533 | Rps2 | 0.037702384 | 1.41051487 | 0.000229249 |
| ENSMUSG00000029580 | Actb | 0.036871935 | 1.366820398 | 0.000428474 |

| | | | | |
|---|---|---|---|---|
| ENSMUSG00000039001 | Rps21 | 0.036761209 | 1.645585017 | 1.02E-05 |
| ENSMUSG00000028234 | Rps20 | 0.036761209 | 1.384897484 | 0.00033621 |
| ENSMUSG00000041841 | Rpl37 | 0.036539756 | 1.931205788 | 7.10E-08 |
| ENSMUSG00000045128 | Rpl18a | 0.036484393 | 1.766044313 | 1.53E-06 |
| ENSMUSG00000074129 | Rpl13a | 0.036262939 | 1.326907457 | 0.00064047 |
| ENSMUSG00000006333 | Rps9 | 0.036152213 | 1.060934771 | 0.009232869 |
| ENSMUSG00000025508 | Rplp2 | 0.03593076 | 1.180151988 | 0.003021141 |
| ENSMUSG00000075015 | Gm10801 | 0.03559858 | 1.350395809 | 0.000581282 |
| ENSMUSG00000024661 | Fth1 | 0.035377127 | 1.914125931 | 1.19E-07 |
| ENSMUSG00000069670 | Nkain2 | 0.035211037 | -1.660696128 | 0.001939074 |
| ENSMUSG00000067274 | Rplp0 | 0.034934221 | 1.004979362 | 0.015417072 |
| ENSMUSG00000057841 | Rpl32 | 0.034823495 | 1.484519197 | 0.000106037 |
| ENSMUSG00000090733 | Rps27 | 0.034325225 | 1.580771276 | 2.65E-05 |
| ENSMUSG00000071415 | Rpl23 | 0.033826956 | 1.210995103 | 0.002335057 |
| ENSMUSG00000060636 | Rpl35a | 0.03321796 | 1.493116487 | 0.000105546 |
| ENSMUSG00000020460 | Rps27a | 0.03271969 | 1.578678336 | 2.82E-05 |
| ENSMUSG00000052146 | Rps10 | 0.032332148 | 1.710557803 | 4.27E-06 |
| ENSMUSG00000037563 | Rps16 | 0.031889241 | 1.563875217 | 3.47E-05 |
| ENSMUSG00000046364 | Rpl27a | 0.03067125 | 1.705835052 | 5.16E-06 |
| ENSMUSG00000025362 | Rps26 | 0.030615886 | 1.299053344 | 0.001014164 |
| ENSMUSG00000057322 | Rpl38 | 0.03050516 | 1.477749427 | 0.000124522 |
| ENSMUSG00000041453 | Rpl21 | 0.030062254 | 1.85824748 | 3.69E-07 |
| ENSMUSG00000050708 | Ftl1 | 0.029674711 | 1.643472367 | 1.16E-05 |
| ENSMUSG00000030432 | Rpl28 | 0.029231805 | 1.619143867 | 1.86E-05 |
| ENSMUSG00000024608 | Rps14 | 0.028512082 | 1.323556247 | 0.000760735 |
| ENSMUSG00000063457 | Rps15 | 0.028124539 | 1.137525593 | 0.005062549 |
| ENSMUSG00000047215 | Rpl9 | 0.02795845 | 1.629014111 | 1.54E-05 |
| ENSMUSG00000062328 | Rpl17 | 0.027128001 | 1.250792864 | 0.001641759 |
| ENSMUSG00000043336 | Filip1l | 0.026574368 | 0.412796489 | 0.530172915 |
| ENSMUSG00000062006 | Rpl34 | 0.026519005 | 1.647333526 | 1.33E-05 |
| ENSMUSG00000079641 | Rpl39 | 0.025799282 | 1.615247483 | 2.01E-05 |
| ENSMUSG00000061477 | Rps7 | 0.025411739 | 1.21936513 | 0.002319335 |
| ENSMUSG00000003970 | Rpl8 | 0.025301013 | 1.382130831 | 0.00043944 |
| ENSMUSG00000038900 | Rpl12 | 0.025134923 | 1.391095445 | 0.000366437 |
| ENSMUSG00000049517 | Rps23 | 0.025024196 | 1.39005375 | 0.000366437 |
| ENSMUSG00000008668 | Rps18 | 0.02491347 | 1.457390553 | 0.000161297 |
| ENSMUSG00000057666 | Gapdh | 0.024027658 | 2.021389719 | 2.28E-08 |
| ENSMUSG00000031320 | Rps4x | 0.023972294 | 1.593403161 | 2.85E-05 |
| ENSMUSG00000058600 | Rpl30 | 0.023584751 | 1.481540806 | 0.000136945 |
| ENSMUSG00000023043 | Krt18 | 0.023529388 | 0.890690551 | 0.041441321 |
| ENSMUSG00000012848 | Rps5 | 0.023307935 | 1.556153549 | 5.09E-05 |
| ENSMUSG00000022283 | Pabpc1 | 0.022754302 | 1.021319375 | 0.018196199 |
| ENSMUSG00000030050 | Gkn1 | 0.022422123 | 3.513271569 | 8.29E-28 |
| ENSMUSG00000043716 | Rpl7 | 0.02220067 | 1.397512128 | 0.000368819 |
| ENSMUSG00000075014 | Gm10800 | 0.022145307 | 1.326188087 | 0.000983199 |
| ENSMUSG00000030049 | Gkn2 | 0.021923853 | 4.162889676 | 1.64E-38 |
| ENSMUSG00000008683 | Rps15a | 0.021647037 | 1.48382604 | 0.000134016 |
| ENSMUSG00000027907 | S100a11 | 0.020982678 | 1.465474411 | 0.000173266 |
| ENSMUSG00000030744 | Rps3 | 0.020705862 | 1.403542728 | 0.000365509 |
| ENSMUSG00000060938 | Rpl26 | 0.020318319 | 1.336347152 | 0.000876027 |
| ENSMUSG00000057863 | Rpl36 | 0.01915569 | 1.659267231 | 1.47E-05 |

| | | | | |
|---|---|---|---|---|
| ENSMUSG00000026238 | Ptma | 0.018878874 | 0.259333144 | 0.800428907 |
| ENSMUSG00000059291 | Rpl11 | 0.018768147 | 1.423415437 | 0.000355339 |
| ENSMUSG00000023944 | Hsp90ab1 | 0.018768147 | 0.135642085 | 1 |
| ENSMUSG00000058558 | Rpl5 | 0.018214515 | 1.439618519 | 0.000255136 |
| ENSMUSG00000003429 | Rps11 | 0.018159151 | 1.662051171 | 1.61E-05 |
| ENSMUSG00000079435 | Rpl36a | 0.017993062 | 1.744898273 | 4.96E-06 |
| ENSMUSG00000059070 | Rpl18 | 0.017882335 | 1.717222526 | 8.60E-06 |
| ENSMUSG00000090862 | Rps13 | 0.017716245 | 1.503589824 | 0.000142784 |
| ENSMUSG00000037805 | Rpl10a | 0.017716245 | 1.095681789 | 0.009429212 |
| ENSMUSG00000029614 | Rpl6 | 0.017439429 | 1.158200433 | 0.005513373 |
| ENSMUSG00000047907 | Tshz2 | 0.017217976 | -1.730431724 | 0.001396809 |
| ENSMUSG00000017404 | Rpl19 | 0.016885796 | 1.19541696 | 0.003613093 |
| ENSMUSG00000031765 | Mt1 | 0.016830433 | 2.170181953 | 2.83E-09 |
| ENSMUSG00000062825 | Actg1 | 0.016332164 | 1.278972979 | 0.001648667 |
| ENSMUSG00000008682 | Rpl10 | 0.016332164 | 1.394764544 | 0.000530015 |
| ENSMUSG00000049382 | Krt8 | 0.016055347 | 2.129807321 | 5.54E-09 |
| ENSMUSG00000025794 | Rpl14 | 0.015501715 | 1.56167475 | 8.32E-05 |
| ENSMUSG00000004637 | Wwox | 0.015390988 | -1.706065484 | 0.001608894 |
| ENSMUSG00000067851 | Arfgef1 | 0.015280262 | -1.209325029 | 0.034492686 |
| ENSMUSG00000039199 | Zdhhc1 | 0.015224898 | -1.09906552 | 0.063572047 |
| ENSMUSG00000028936 | Rpl22 | 0.015169535 | 1.338495431 | 0.001278866 |
| ENSMUSG00000028495 | Rps6 | 0.015058808 | 1.103156778 | 0.008872754 |
| ENSMUSG00000003617 | Cp | 0.014671266 | -1.251168659 | 0.028914368 |
| ENSMUSG00000028639 | Ybx1 | 0.014615902 | 0.791793505 | 0.096008389 |
| ENSMUSG00000020372 | Rack1 | 0.014505176 | 1.033118512 | 0.016311236 |
| ENSMUSG00000098274 | Rpl24 | 0.014505176 | 0.846976617 | 0.070609852 |
| ENSMUSG00000032231 | Anxa2 | 0.014449813 | -0.54415364 | 0.578640165 |
| ENSMUSG00000027684 | Mecom | 0.014394449 | -1.852933014 | 0.000593259 |
| ENSMUSG00000045991 | Onecut2 | 0.014228359 | -1.868635477 | 0.000533458 |
| ENSMUSG00000071866 | Ppia | 0.014006906 | 1.518579165 | 0.000136945 |
| ENSMUSG00000012405 | Rpl15 | 0.013951543 | 1.355704721 | 0.000887428 |
| ENSMUSG00000010608 | Rbm25 | 0.013674727 | -2.377286256 | 1.05E-05 |
| ENSMUSG00000060036 | Rpl3 | 0.013674727 | 1.189403933 | 0.004294426 |
| ENSMUSG00000074305 | Peak1 | 0.013564 | -0.877782963 | 0.182638688 |
| ENSMUSG00000025967 | Eef1b2 | 0.012955004 | 1.203424403 | 0.004015829 |
| ENSMUSG00000052397 | Ezr | 0.012899641 | 0.30488115 | 0.748188375 |
| ENSMUSG00000028691 | Prdx1 | 0.012678188 | 1.024683785 | 0.01904133 |
| ENSMUSG00000034994 | Eef2 | 0.012567461 | 1.399576524 | 0.000600167 |
| ENSMUSG00000057113 | Npm1 | 0.012567461 | 0.823514083 | 0.082076736 |
| ENSMUSG00000058546 | Rpl23a | 0.012512098 | 1.518281997 | 0.000158234 |
| ENSMUSG00000046876 | Atxn1 | 0.012401372 | -1.340707176 | 0.016961017 |
| ENSMUSG00000031626 | Sorbs2 | 0.011792376 | -1.648273565 | 0.002585743 |
| ENSMUSG00000006360 | Crip1 | 0.011792376 | 2.081963508 | 4.53E-08 |
| ENSMUSG00000022938 | Fam3b | 0.011681649 | -0.113091954 | 1 |
| ENSMUSG00000031841 | Cdh13 | 0.01134947 | -0.781007883 | 0.278325364 |
| ENSMUSG00000056608 | Chd9 | 0.011238743 | 0.682824979 | 0.194179714 |
| ENSMUSG00000017723 | Wfdc2 | 0.01101729 | 1.900374318 | 1.28E-06 |
| ENSMUSG00000062647 | Rpl7a | 0.010961927 | 1.708031344 | 1.82E-05 |
| ENSMUSG00000023039 | Krt7 | 0.010961927 | -0.135552684 | 1 |
| ENSMUSG00000009927 | Rps25 | 0.010906563 | 1.392495798 | 0.000899695 |
| ENSMUSG00000046727 | Cystm1 | 0.01068511 | 0.723227585 | 0.156554801 |

| | | | | |
|---|---|---|---|---|
| ENSMUSG00000086550 | Pla2g10os | 0.010352931 | 0.467022472 | 0.47966612 |
| ENSMUSG00000027620 | Rbm39 | 0.010076114 | -1.927561025 | 0.000385719 |
| ENSMUSG00000031906 | Smpd3 | 0.010076114 | -1.201240246 | 0.037959551 |
| ENSMUSG00000022892 | App | 0.010076114 | -2.211861361 | 4.70E-05 |
| ENSMUSG00000061787 | Rps17 | 0.009910025 | 1.718623882 | 2.16E-05 |
| ENSMUSG00000110386 | Gm42031 | 0.009910025 | 1.63686291 | 5.24E-05 |
| ENSMUSG00000069833 | Ahnak | 0.009910025 | -2.328444372 | 1.76E-05 |
| ENSMUSG00000073702 | Rpl31 | 0.009743935 | 1.728423816 | 2.27E-05 |
| ENSMUSG00000069601 | Ank3 | 0.009522482 | -1.629922736 | 0.003112598 |
| ENSMUSG00000061436 | Hipk2 | 0.009467119 | -1.292860403 | 0.023669966 |
| ENSMUSG00000029381 | Shroom3 | 0.009411755 | -0.936126949 | 0.146698911 |
| ENSMUSG00000062997 | Rpl35 | 0.009301029 | 1.124902484 | 0.012147019 |
| ENSMUSG00000005087 | Cd44 | 0.009245665 | -2.514634428 | 4.44E-06 |
| ENSMUSG00000032289 | Thsd4 | 0.009245665 | -1.63964606 | 0.002779402 |
| ENSMUSG00000090386 | Mir99ahg | 0.009190302 | -1.287696601 | 0.025318334 |
| ENSMUSG00000092274 | Neat1 | 0.009079576 | -2.734711662 | 6.21E-07 |
| ENSMUSG00000003847 | Nfat5 | 0.008858123 | -1.936500269 | 0.000373042 |
| ENSMUSG00000061665 | Cd2ap | 0.008747396 | -1.277283251 | 0.025925336 |
| ENSMUSG00000086324 | Gm15564 | 0.008692033 | -2.634223488 | 1.59E-06 |
| ENSMUSG00000038244 | Mical2 | 0.008581306 | -1.651663679 | 0.00277148 |
| ENSMUSG00000025006 | Sorbs1 | 0.008581306 | -1.368972179 | 0.016701832 |
| ENSMUSG00000032473 | Cldn18 | 0.008525943 | -0.113959033 | 1 |
| ENSMUSG00000019820 | Utrn | 0.008525943 | -1.454959861 | 0.009213836 |
| ENSMUSG00000056612 | Ppp1r14b | 0.008525943 | 1.511332028 | 0.000305057 |
| ENSMUSG00000024985 | Tcf7l2 | 0.00847058 | -2.014006191 | 0.000217681 |
| ENSMUSG00000030695 | Aldoa | 0.008415216 | 1.818760553 | 8.21E-06 |
| ENSMUSG00000058056 | Palld | 0.008415216 | -1.287917845 | 0.028938539 |
| ENSMUSG00000015932 | Dstn | 0.008359853 | 1.199795539 | 0.006637019 |
| ENSMUSG00000070493 | Chchd2 | 0.008359853 | 0.899583584 | 0.062231591 |
| ENSMUSG00000001175 | Calm1 | 0.008359853 | 1.06704033 | 0.017974172 |
| ENSMUSG00000029322 | Plac8 | 0.00830449 | 0.405440859 | 0.596503438 |
| ENSMUSG00000031818 | Cox4i1 | 0.00830449 | 1.747688938 | 2.16E-05 |
| ENSMUSG00000053332 | Gas5 | 0.008249127 | 0.632101536 | 0.26331908 |
| ENSMUSG00000038732 | Mboat1 | 0.008249127 | -1.714802795 | 0.00181069 |
| ENSMUSG00000090841 | Myl6 | 0.008193763 | 0.868004072 | 0.078516003 |
| ENSMUSG00000063316 | Rpl27 | 0.0081384 | 1.250958603 | 0.005089043 |
| ENSMUSG00000021215 | Net1 | 0.0081384 | 0.429928744 | 0.557051363 |
| ENSMUSG00000021699 | Pde4d | 0.008083037 | -1.912212051 | 0.000461905 |
| ENSMUSG00000068220 | Lgals1 | 0.008083037 | 1.558489049 | 0.000206806 |
| ENSMUSG00000004552 | Ctse | 0.008027674 | 2.097193124 | 1.74E-07 |
| ENSMUSG00000028333 | Anp32b | 0.00797231 | 0.208524816 | 0.923036689 |
| ENSMUSG00000073408 | Mucl3 | 0.00797231 | 1.552620236 | 0.000227496 |
| ENSMUSG00000032216 | Nedd4 | 0.007916947 | -1.106051951 | 0.067274027 |
| ENSMUSG00000017778 | Cox7c | 0.007916947 | 1.310965913 | 0.002322357 |
| ENSMUSG00000035530 | Eif1 | 0.007861584 | 1.063873051 | 0.019528034 |
| ENSMUSG00000030067 | Foxp1 | 0.007750857 | -1.408829789 | 0.012632774 |
| ENSMUSG00000090137 | Uba52 | 0.007750857 | 1.648835551 | 0.000108795 |
| ENSMUSG00000064052 | Gm5089 | 0.007750857 | -1.443434275 | 0.010226213 |
| ENSMUSG00000039607 | Rbms3 | 0.007640131 | -1.398386458 | 0.01400891 |
| ENSMUSG00000021488 | Nsd1 | 0.007640131 | -1.695236789 | 0.002060158 |
| ENSMUSG00000056201 | Cfl1 | 0.007640131 | 1.534156947 | 0.000303106 |

| | | | | |
|---|---|---|---|---|
| ENSMUSG00000062070 | Pgk1 | 0.007640131 | 0.496313596 | 0.465296823 |
| ENSMUSG00000063229 | Ldha | 0.007529404 | 1.571941952 | 0.000241256 |
| ENSMUSG00000026234 | Ncl | 0.007474041 | 0.181709541 | 0.976386142 |
| ENSMUSG00000029994 | Anxa4 | 0.007474041 | -1.032712928 | 0.097987606 |
| ENSMUSG00000105703 | Gm43305 | 0.007474041 | 1.566373391 | 0.000229249 |
| ENSMUSG00000031075 | Ano1 | 0.007418678 | -1.697464762 | 0.002074662 |
| ENSMUSG00000032399 | Rpl4 | 0.007418678 | 0.915079514 | 0.05622212 |
| ENSMUSG00000023175 | Bsg | 0.007418678 | -0.164613129 | 1 |
| ENSMUSG00000015656 | Hspa8 | 0.007363314 | 0.760728432 | 0.141423134 |
| ENSMUSG00000023393 | Slc17a9 | 0.007307951 | -2.883659529 | 1.74E-07 |
| ENSMUSG00000000538 | Tom1l2 | 0.007307951 | -1.023802468 | 0.105099811 |
| ENSMUSG00000020911 | Krt19 | 0.007197225 | 1.311571146 | 0.002594038 |
| ENSMUSG00000035885 | Cox8a | 0.007197225 | 1.578970745 | 0.000216481 |
| ENSMUSG00000039529 | Atp8b1 | 0.007141861 | -0.61766271 | 0.47966612 |
| ENSMUSG00000059005 | Hnrnpa3 | 0.007086498 | 0.194496402 | 0.955892779 |
| ENSMUSG00000039221 | Rpl22l1 | 0.007086498 | 1.117030987 | 0.01607245 |
| ENSMUSG00000076432 | Ywhaq | 0.007086498 | -1.087291023 | 0.076862095 |
| ENSMUSG00000033307 | Mif | 0.006975772 | 1.817341276 | 1.52E-05 |
| ENSMUSG00000037965 | Zc3h7a | 0.006920408 | -1.508310845 | 0.007505624 |
| ENSMUSG00000060477 | Irak2 | 0.006865045 | -1.920169375 | 0.000501136 |
| ENSMUSG00000045962 | Wnk1 | 0.006865045 | -1.317594916 | 0.022359957 |
| ENSMUSG00000039087 | Rreb1 | 0.006865045 | -1.597268458 | 0.00403405 |
| ENSMUSG00000026796 | Fam129b | 0.006809682 | -1.82689788 | 0.000949133 |
| ENSMUSG00000059456 | Ptk2b | 0.006809682 | -1.444773692 | 0.01067424 |
| ENSMUSG00000035569 | Ankrd11 | 0.006754319 | -2.061046805 | 0.0001721 |
| ENSMUSG00000018412 | Kansl1 | 0.006698955 | -1.600336201 | 0.004015829 |
| ENSMUSG00000063011 | Msln | 0.006698955 | -1.911277988 | 0.000516191 |
| ENSMUSG00000024998 | Plce1 | 0.006698955 | -1.048872651 | 0.095021946 |
| ENSMUSG00000074813 | Morrbid | 0.006643592 | -1.581413202 | 0.004581608 |
| ENSMUSG00000068523 | Gng5 | 0.006643592 | 1.205755015 | 0.00757325 |
| ENSMUSG00000030122 | Ptms | 0.006643592 | 0.254815833 | 0.858831093 |
| ENSMUSG00000028649 | Macf1 | 0.006588229 | -1.483192842 | 0.008383243 |
| ENSMUSG00000000303 | Cdh1 | 0.006588229 | -0.491397765 | 0.66791649 |
| ENSMUSG00000049751 | Rpl36al | 0.006588229 | 1.084050933 | 0.021923154 |
| ENSMUSG00000033149 | Phldb2 | 0.006588229 | -2.260057397 | 3.68E-05 |
| ENSMUSG00000071644 | Eef1g | 0.006588229 | 0.758064621 | 0.155047762 |
| ENSMUSG00000024991 | Eif3a | 0.006588229 | -1.068155343 | 0.084413034 |
| ENSMUSG00000069662 | Marcks | 0.006532865 | 0.437744806 | 0.549953572 |
| ENSMUSG00000007041 | Clic1 | 0.006532865 | 0.798819477 | 0.124712353 |
| ENSMUSG00000024811 | Tnks2 | 0.006532865 | -2.150704572 | 9.04E-05 |
| ENSMUSG00000027940 | Tpm3 | 0.006477502 | -0.922820693 | 0.166504886 |
| ENSMUSG00000029761 | Cald1 | 0.006477502 | -0.868333359 | 0.213441582 |
| ENSMUSG00000047517 | Dmbt1 | 0.006477502 | 0.407357289 | 0.606852834 |
| ENSMUSG00000075706 | Gpx4 | 0.006477502 | -0.198142143 | 1 |
| ENSMUSG00000019505 | Ubb | 0.006477502 | 1.366571674 | 0.001946259 |
| ENSMUSG00000020863 | Luc7l3 | 0.006477502 | -2.330383739 | 2.15E-05 |
| ENSMUSG00000061315 | Naca | 0.006422139 | 0.930485635 | 0.054667757 |
| ENSMUSG00000056758 | Hmga2 | 0.006256049 | -1.827182238 | 0.001022622 |
| ENSMUSG00000036241 | Ube2r2 | 0.006200686 | -1.217486013 | 0.039995951 |
| ENSMUSG00000031762 | Mt2 | 0.006200686 | 2.202227973 | 1.06E-07 |
| ENSMUSG00000054889 | Dsp | 0.006200686 | -1.449848723 | 0.010685899 |

| | | | | |
|---|---|---|---|---|
| ENSMUSG00000043501 | Lgals2 | 0.006200686 | 2.017803402 | 1.92E-06 |
| ENSMUSG00000025019 | Lcor | 0.006200686 | -1.769390225 | 0.001412356 |
| ENSMUSG00000020184 | Mdm2 | 0.006145323 | -1.278519147 | 0.029702344 |
| ENSMUSG00000022285 | Ywhaz | 0.006145323 | 0.610700423 | 0.316909787 |
| ENSMUSG00000026193 | Fn1 | 0.006089959 | -0.548270428 | 0.600946203 |
| ENSMUSG00000033377 | Palmd | 0.006089959 | -1.807639118 | 0.001112545 |
| ENSMUSG00000029064 | Gnb1 | 0.006089959 | -0.67698646 | 0.433829004 |
| ENSMUSG00000063856 | Gpx1 | 0.006089959 | 1.566734471 | 0.000319528 |
| ENSMUSG00000034165 | Ccnd3 | 0.006089959 | -1.712831658 | 0.002044714 |
| ENSMUSG00000025340 | Rabgef1 | 0.006034596 | -1.602539264 | 0.004220296 |
| ENSMUSG00000021253 | Tgfb3 | 0.005979233 | -2.145699551 | 0.000110213 |
| ENSMUSG00000037896 | Rcor1 | 0.005979233 | -1.898129686 | 0.000600167 |
| ENSMUSG00000026547 | Tagln2 | 0.00592387 | 1.521080004 | 0.000557602 |
| ENSMUSG00000026970 | Rbms1 | 0.00592387 | -1.901768739 | 0.000615213 |
| ENSMUSG00000027797 | Dclk1 | 0.00592387 | -1.557002941 | 0.00592361 |
| ENSMUSG00000038872 | Zfhx3 | 0.00592387 | -0.975655313 | 0.131906476 |
| ENSMUSG00000027356 | Fermt1 | 0.005868506 | -2.251749155 | 4.59E-05 |
| ENSMUSG00000035299 | Mid1 | 0.005813143 | -3.399677026 | 7.33E-10 |
| ENSMUSG00000074884 | Serf2 | 0.005702416 | 1.436385933 | 0.001314883 |
| ENSMUSG00000049470 | Aff4 | 0.005702416 | -1.328444372 | 0.022386331 |
| ENSMUSG00000024028 | Tff2 | 0.005702416 | 1.80238081 | 3.21E-05 |
| ENSMUSG00000027993 | Trim2 | 0.005647053 | -0.887458824 | 0.20161331 |
| ENSMUSG00000034981 | Parm1 | 0.005647053 | -0.956985554 | 0.146744515 |
| ENSMUSG00000030587 | 2200002D01Rik | 0.005647053 | 1.346651392 | 0.002693122 |
| ENSMUSG00000027523 | Gnas | 0.00559169 | 0.999332799 | 0.042237573 |
| ENSMUSG00000039286 | Fndc3b | 0.00559169 | -2.023589791 | 0.000251527 |
| ENSMUSG00000068823 | Csde1 | 0.00559169 | -0.561382156 | 0.586721871 |
| ENSMUSG00000029622 | Arpc1b | 0.00559169 | 1.233798052 | 0.006937952 |
| ENSMUSG00000034127 | Tspan8 | 0.00559169 | -0.034090203 | 1 |
| ENSMUSG00000021270 | Hsp90aa1 | 0.00559169 | -1.009655985 | 0.116637579 |
| ENSMUSG00000022306 | Zfpm2 | 0.00559169 | -2.021369405 | 0.000302289 |
| ENSMUSG00000032348 | Gsta4 | 0.005536327 | 0.872418807 | 0.092468958 |
| ENSMUSG00000025068 | Gsto1 | 0.005536327 | 1.152228925 | 0.014140013 |
| ENSMUSG00000039159 | Ube2h | 0.005480963 | -1.598332449 | 0.0044369 |
| ENSMUSG00000021870 | Slmap | 0.005480963 | -1.696702759 | 0.002408232 |
| ENSMUSG00000033014 | Trim33 | 0.0054256 | -1.302066396 | 0.025915431 |
| ENSMUSG00000004980 | Hnrnpa2b1 | 0.0054256 | -1.79661829 | 0.001293124 |
| ENSMUSG00000001794 | Capns1 | 0.0054256 | 0.480530646 | 0.511928788 |
| ENSMUSG00000003316 | Glg1 | 0.0054256 | -1.204615785 | 0.043751983 |
| ENSMUSG00000039419 | Cntnap2 | 0.005370237 | -0.668669356 | 0.466464457 |
| ENSMUSG00000045095 | Magi1 | 0.005370237 | -2.712660695 | 1.05E-06 |
| ENSMUSG00000032220 | Myo1e | 0.005370237 | -1.719448179 | 0.002083251 |
| ENSMUSG00000032425 | Zfp949 | 0.005370237 | -0.822590829 | 0.262284476 |
| ENSMUSG00000024969 | Mark2 | 0.005370237 | -1.821262992 | 0.001097448 |
| ENSMUSG00000006304 | Arpc2 | 0.005314874 | 0.093930285 | 1 |
| ENSMUSG00000038518 | Jarid2 | 0.005314874 | -1.347441363 | 0.021623593 |
| ENSMUSG00000038170 | Pde4dip | 0.00525951 | -1.413660374 | 0.014077762 |
| ENSMUSG00000029088 | Kcnip4 | 0.00525951 | -1.538516522 | 0.006565796 |
| ENSMUSG00000029484 | Anxa3 | 0.00525951 | -1.085874069 | 0.083337667 |
| ENSMUSG00000037974 | Muc5ac | 0.00525951 | 1.432898152 | 0.001656499 |
| ENSMUSG00000070348 | Ccnd1 | 0.00525951 | 0.369611908 | 0.675561045 |

| | | | | |
|---|---|---|---|---|
| ENSMUSG00000064373 | Selenop | 0.00525951 | 1.948395833 | 8.70E-06 |
| ENSMUSG00000061751 | Kalrn | 0.00525951 | -0.529414315 | 0.643790066 |
| ENSMUSG00000005371 | Fbxo11 | 0.00525951 | -1.357724294 | 0.019633115 |
| ENSMUSG00000041959 | S100a10 | 0.005204147 | 1.385801146 | 0.002408232 |
| ENSMUSG00000028664 | Ephb2 | 0.005204147 | -1.945587279 | 0.000537421 |
| ENSMUSG00000066551 | Hmgb1 | 0.005204147 | -0.30290928 | 0.974161742 |
| ENSMUSG00000057278 | Snrpg | 0.005204147 | 1.914910412 | 1.87E-05 |
| ENSMUSG00000001506 | Col1a1 | 0.005204147 | -0.213850741 | 1 |
| ENSMUSG00000030342 | Cd9 | 0.005148784 | 1.203424403 | 0.010326707 |
| ENSMUSG00000033545 | Znrf1 | 0.005148784 | -1.012108597 | 0.116475189 |
| ENSMUSG00000032435 | Dync1li1 | 0.005148784 | -1.90117435 | 0.000683341 |
| ENSMUSG00000031167 | Rbm3 | 0.005148784 | 1.079303091 | 0.02567871 |
| ENSMUSG00000027800 | Tm4sf1 | 0.005093421 | 1.515569021 | 0.000882856 |
| ENSMUSG00000025809 | Itgb1 | 0.005093421 | -0.882126582 | 0.212080558 |
| ENSMUSG00000013663 | Pten | 0.005093421 | -1.575218004 | 0.005298129 |
| ENSMUSG00000025016 | Tm9sf3 | 0.005093421 | -0.688522584 | 0.424933774 |
| ENSMUSG00000028082 | Sh3d19 | 0.005038057 | -1.829714269 | 0.001064964 |
| ENSMUSG00000032294 | Pkm | 0.005038057 | 0.130738356 | 1 |
| ENSMUSG00000020745 | Pafah1b1 | 0.005038057 | -1.617950989 | 0.004133756 |
| ENSMUSG00000001525 | Tubb5 | 0.005038057 | 1.229883002 | 0.008646994 |
| ENSMUSG00000026728 | Vim | 0.004982694 | 0.131490629 | 1 |
| ENSMUSG00000058835 | Abi1 | 0.004982694 | -0.008276735 | 1 |
| ENSMUSG00000023456 | Tpi1 | 0.004982694 | 1.011971949 | 0.039390861 |
| ENSMUSG00000021171 | Esyt2 | 0.004982694 | -1.721420721 | 0.002119258 |
| ENSMUSG00000022201 | Zfr | 0.004982694 | -1.80047185 | 0.001305524 |
| ENSMUSG00000060743 | H3f3a | 0.004927331 | 0.761310281 | 0.174915761 |
| ENSMUSG00000039852 | Rere | 0.004927331 | -2.193584638 | 7.90E-05 |
| ENSMUSG00000029632 | Ndufa4 | 0.004927331 | 2.085406212 | 4.67E-06 |
| ENSMUSG00000024248 | Cox7a2l | 0.004927331 | 0.423012225 | 0.611716824 |
| ENSMUSG00000095041 | AC149090.1 | 0.004927331 | -3.042636577 | 4.53E-08 |
| ENSMUSG00000028416 | Bag1 | 0.004871967 | 0.719427519 | 0.217655393 |
| ENSMUSG00000030228 | Pik3c2g | 0.004871967 | -1.466018068 | 0.01067424 |
| ENSMUSG00000047139 | Cd24a | 0.004871967 | 0.400055129 | 0.661861928 |
| ENSMUSG00000025730 | Rab40c | 0.004871967 | -1.461332467 | 0.011088332 |
| ENSMUSG00000024501 | Dpysl3 | 0.004871967 | -1.605901155 | 0.00502152 |
| ENSMUSG00000024812 | Tjp2 | 0.004871967 | -1.81721226 | 0.001200854 |
| ENSMUSG00000006699 | Cdc42 | 0.004816604 | -1.038937755 | 0.109827081 |
| ENSMUSG00000028780 | Sema3c | 0.004816604 | -2.742310377 | 9.25E-07 |
| ENSMUSG00000092570 | Plut | 0.004816604 | -0.987129853 | 0.13195254 |
| ENSMUSG00000000440 | Pparg | 0.004816604 | -0.734083173 | 0.375416882 |
| ENSMUSG00000031584 | Gsr | 0.004816604 | 0.244081025 | 0.896076486 |
| ENSMUSG00000037852 | Cpe | 0.004816604 | 0.195377834 | 0.971854959 |
| ENSMUSG00000037646 | Vps13b | 0.004816604 | -1.407432668 | 0.014921593 |
| ENSMUSG00000048249 | Crebrf | 0.004816604 | -1.21027564 | 0.045984312 |
| ENSMUSG00000005656 | Snx6 | 0.004761241 | -1.474274991 | 0.010251963 |
| ENSMUSG00000060147 | Serpinb6a | 0.004761241 | 0.916853365 | 0.078555425 |
| ENSMUSG00000021660 | Btf3 | 0.004761241 | 0.636451827 | 0.307900693 |
| ENSMUSG00000025278 | Flnb | 0.004761241 | -1.705071748 | 0.002457575 |
| ENSMUSG00000053617 | Sh3pxd2a | 0.004761241 | 0.058356046 | 1 |
| ENSMUSG00000032966 | Fkbp1a | 0.004705878 | 1.025251757 | 0.04321855 |
| ENSMUSG00000028063 | Lmna | 0.004705878 | -0.623604361 | 0.497214025 |

| | | | | |
|---|---|---|---|---|
| ENSMUSG00000060512 | 0610040J01Rik | 0.004705878 | -0.813642943 | 0.27459352 |
| ENSMUSG00000032479 | Map4 | 0.004705878 | -1.219520443 | 0.043231237 |
| ENSMUSG00000054452 | Tle5 | 0.004705878 | 0.567305666 | 0.405392586 |
| ENSMUSG00000016559 | H3f3b | 0.004705878 | 0.316877091 | 0.792441218 |
| ENSMUSG00000064138 | Fam172a | 0.004705878 | -1.676443753 | 0.00302914 |
| ENSMUSG00000024201 | Kdm4b | 0.004705878 | -1.360569955 | 0.02007424 |
| ENSMUSG00000029661 | Col1a2 | 0.004650514 | -0.131598071 | 1 |
| ENSMUSG00000053964 | Lgals4 | 0.004650514 | 2.473263987 | 1.59E-08 |
| ENSMUSG00000020048 | Hsp90b1 | 0.004650514 | -0.073630473 | 1 |
| ENSMUSG00000021939 | Ctsb | 0.004650514 | 0.566373391 | 0.415501353 |
| ENSMUSG00000030064 | Frmd4b | 0.004595151 | -0.819933932 | 0.274319469 |
| ENSMUSG00000059474 | Mbtd1 | 0.004595151 | -1.78618973 | 0.001506566 |
| ENSMUSG00000018042 | Cyb5r3 | 0.004595151 | 1.279601742 | 0.006478925 |
| ENSMUSG00000062210 | Tnfaip8 | 0.004595151 | -1.727858347 | 0.002106549 |
| ENSMUSG00000031229 | Atrx | 0.004595151 | -2.253467775 | 5.24E-05 |
| ENSMUSG00000039202 | Abhd2 | 0.004539788 | -1.60821143 | 0.004586201 |
| ENSMUSG00000019889 | Ptprk | 0.004539788 | -2.271893952 | 4.65E-05 |
| ENSMUSG00000038127 | Ccdc50 | 0.004539788 | -1.668740519 | 0.003195324 |
| ENSMUSG00000038859 | Baiap2l1 | 0.004484425 | -1.530830844 | 0.007505624 |
| ENSMUSG00000018293 | Pfn1 | 0.004484425 | 0.644443782 | 0.313943937 |
| ENSMUSG00000021111 | Papola | 0.004484425 | -1.153483781 | 0.061337104 |
| ENSMUSG00000017843 | Ppp2r5c | 0.004484425 | -0.651823927 | 0.466464457 |
| ENSMUSG00000040225 | Prrc2c | 0.004429061 | -2.38309109 | 1.87E-05 |
| ENSMUSG00000042228 | Lyn | 0.004429061 | -1.326464008 | 0.025976209 |
| ENSMUSG00000036617 | Etl4 | 0.004373698 | -1.794936841 | 0.00146822 |
| ENSMUSG00000042548 | Asxl1 | 0.004373698 | -1.853399563 | 0.001014164 |
| ENSMUSG00000028521 | Slc35d1 | 0.004373698 | -1.363509639 | 0.022438532 |
| ENSMUSG00000039542 | Ncam1 | 0.004373698 | -1.069526075 | 0.096412134 |
| ENSMUSG00000020857 | Nme2 | 0.004373698 | 1.022396415 | 0.053612203 |
| ENSMUSG00000071655 | Ubxn1 | 0.004373698 | -0.490715801 | 0.688085317 |
| ENSMUSG00000027210 | Meis2 | 0.004318335 | -1.798090687 | 0.001456709 |
| ENSMUSG00000047843 | Bri3 | 0.004318335 | 0.662597453 | 0.309146024 |
| ENSMUSG00000063870 | Chd4 | 0.004318335 | -1.461011629 | 0.011662663 |
| ENSMUSG00000074227 | Spint2 | 0.004318335 | 0.241809866 | 0.886442531 |
| ENSMUSG00000014313 | Cox6c | 0.004318335 | 2.010101794 | 2.05E-05 |
| ENSMUSG00000061080 | Lsamp | 0.004318335 | 0.480941085 | 0.530793253 |
| ENSMUSG00000040732 | Erg | 0.004318335 | -2.418071107 | 1.72E-05 |
| ENSMUSG00000037062 | Sh3glb1 | 0.004262972 | -0.082570517 | 1 |
| ENSMUSG00000002985 | Apoe | 0.004262972 | 1.063873051 | 0.041272021 |
| ENSMUSG00000030688 | Stard10 | 0.004262972 | 0.830949962 | 0.139155838 |
| ENSMUSG00000035242 | Oaz1 | 0.004262972 | 1.499609543 | 0.001294434 |
| ENSMUSG00000001441 | Npepps | 0.004262972 | -1.18080588 | 0.054186866 |
| ENSMUSG00000017493 | Igfbp4 | 0.004262972 | 1.015374533 | 0.045091139 |
| ENSMUSG00000020580 | Rock2 | 0.004262972 | -1.826482177 | 0.001228827 |
| ENSMUSG00000024222 | Fkbp5 | 0.004262972 | -0.877939535 | 0.219470377 |
| ENSMUSG00000016252 | Atp5e | 0.004207608 | 1.667694579 | 0.000433557 |
| ENSMUSG00000028692 | Akr1a1 | 0.004207608 | 0.672448109 | 0.274319469 |
| ENSMUSG00000038690 | Atp5j2 | 0.004207608 | 1.639497687 | 0.00039298 |
| ENSMUSG00000036371 | Serbp1 | 0.004207608 | 0.137544963 | 1 |
| ENSMUSG00000030849 | Fgfr2 | 0.004207608 | -0.383585926 | 0.874366423 |
| ENSMUSG00000004383 | Large1 | 0.004207608 | -1.719188958 | 0.002459077 |

| | | | | |
|---|---|---|---|---|
| ENSMUSG00000038717 | Atp5l | 0.004207608 | 1.140835033 | 0.02307354 |
| ENSMUSG00000021676 | Iqgap2 | 0.004207608 | -2.06594519 | 0.000227247 |
| ENSMUSG00000056124 | B4galt6 | 0.004207608 | -0.76561632 | 0.344436715 |
| ENSMUSG00000038400 | Pmepa1 | 0.004152245 | -0.512627871 | 0.66416768 |
| ENSMUSG00000028399 | Ptprd | 0.004152245 | -0.836252257 | 0.267681437 |
| ENSMUSG00000028788 | Ptp4a2 | 0.004152245 | 0.03878207 | 1 |
| ENSMUSG00000068250 | Amn1 | 0.004152245 | -1.316341155 | 0.028062645 |
| ENSMUSG00000057092 | Fxyd3 | 0.004152245 | 1.919483141 | 2.57E-05 |
| ENSMUSG00000062683 | Atp5g2 | 0.004152245 | 1.189403933 | 0.014846268 |
| ENSMUSG00000022885 | St6gal1 | 0.004152245 | -1.475013579 | 0.011006052 |
| ENSMUSG00000022808 | Snx4 | 0.004152245 | -1.67214309 | 0.00335241 |
| ENSMUSG00000040681 | Hmgn1 | 0.004152245 | -0.188045323 | 1 |
| ENSMUSG00000039218 | Srrm2 | 0.004152245 | -2.059613092 | 0.000239964 |
| ENSMUSG00000024981 | Acsl5 | 0.004152245 | -1.352350388 | 0.022357178 |
| ENSMUSG00000049252 | Lrp1b | 0.004096882 | -0.92162738 | 0.198212447 |
| ENSMUSG00000007815 | Rhoa | 0.004096882 | -0.845153289 | 0.254490204 |
| ENSMUSG00000019997 | Ccn2 | 0.004096882 | 0.173750668 | 1 |
| ENSMUSG00000113586 | Gm48606 | 0.004096882 | -2.545986953 | 6.69E-06 |
| ENSMUSG00000058152 | Chsy3 | 0.004096882 | -2.209808599 | 8.22E-05 |
| ENSMUSG00000062995 | Ica1 | 0.004041518 | -1.513487727 | 0.009240275 |
| ENSMUSG00000031750 | Il34 | 0.004041518 | -2.235788769 | 7.10E-05 |
| ENSMUSG00000020821 | Kif1c | 0.004041518 | 0.386613703 | 0.668835613 |
| ENSMUSG00000025085 | Ablim1 | 0.004041518 | -1.133232929 | 0.071310956 |
| ENSMUSG00000027422 | Rrbp1 | 0.003986155 | -0.523283663 | 0.66416768 |
| ENSMUSG00000042784 | Muc1 | 0.003986155 | 0.821851823 | 0.14317358 |
| ENSMUSG00000063882 | Uqcrh | 0.003986155 | 1.238747268 | 0.016567866 |
| ENSMUSG00000034042 | Gpbp1l1 | 0.003986155 | -1.04245385 | 0.112433946 |
| ENSMUSG00000037443 | Cep85 | 0.003986155 | 0.063873051 | 1 |
| ENSMUSG00000032582 | Rbm6 | 0.003986155 | -1.666158877 | 0.003460332 |
| ENSMUSG00000078812 | Eif5a | 0.003986155 | 1.472337896 | 0.001871203 |
| ENSMUSG00000045763 | Basp1 | 0.003986155 | -0.238155487 | 1 |
| ENSMUSG00000024843 | Chka | 0.003986155 | -3.66913453 | 5.56E-11 |
| ENSMUSG00000025246 | Tbl1x | 0.003986155 | -1.430466569 | 0.014373157 |
| ENSMUSG00000026031 | Cflar | 0.003930792 | -1.892906421 | 0.000825737 |
| ENSMUSG00000008475 | Arpc5 | 0.003930792 | 0.801952265 | 0.167904846 |
| ENSMUSG00000026743 | Mllt10 | 0.003930792 | -1.912558478 | 0.000727108 |
| ENSMUSG00000068267 | Cenpb | 0.003930792 | 0.847216999 | 0.130342151 |
| ENSMUSG00000028771 | Ptpn12 | 0.003930792 | -2.289110361 | 4.52E-05 |
| ENSMUSG00000050732 | Vamp8 | 0.003930792 | -0.500911568 | 0.680536135 |
| ENSMUSG00000031029 | Eif3f | 0.003930792 | 1.496832458 | 0.001713818 |
| ENSMUSG00000005161 | Prdx2 | 0.003930792 | 2.090840098 | 5.53E-06 |
| ENSMUSG00000020152 | Actr2 | 0.003930792 | -0.343630776 | 0.915692354 |
| ENSMUSG00000056342 | Usp34 | 0.003930792 | -1.486161117 | 0.01067424 |
| ENSMUSG00000048264 | Dip2c | 0.003930792 | -2.344260087 | 2.87E-05 |
| ENSMUSG00000023022 | Lima1 | 0.003930792 | -1.389473805 | 0.018271716 |
| ENSMUSG00000034868 | Myl12b | 0.003930792 | 0.779841564 | 0.180821216 |
| ENSMUSG00000024758 | Rtn3 | 0.003930792 | -1.169924134 | 0.060407117 |
| ENSMUSG00000048612 | Myof | 0.003930792 | -1.83945093 | 0.00118021 |
| ENSMUSG00000025060 | Slk | 0.003930792 | -1.399197145 | 0.017291146 |
| ENSMUSG00000026991 | Pkp4 | 0.003875429 | -0.825298819 | 0.279592421 |
| ENSMUSG00000027276 | Jag1 | 0.003875429 | -2.536249257 | 7.74E-06 |

| | | | | |
|---|---|---|---|---|
| ENSMUSG00000028367 | Txn1 | 0.003875429 | 1.070662216 | 0.035450567 |
| ENSMUSG00000040003 | Magi2 | 0.003875429 | -1.237590942 | 0.045084086 |
| ENSMUSG00000000184 | Ccnd2 | 0.003875429 | 0.050389821 | 1 |
| ENSMUSG00000045659 | Plekha7 | 0.003875429 | -1.589434614 | 0.005674039 |
| ENSMUSG00000031586 | Rbpms | 0.003875429 | -1.856053358 | 0.001052851 |
| ENSMUSG00000020315 | Sptbn1 | 0.003875429 | -1.351164449 | 0.023148235 |
| ENSMUSG00000039168 | Dap | 0.003875429 | 0.347371559 | 0.757955408 |
| ENSMUSG00000022450 | Ndufa6 | 0.003875429 | 2.275020715 | 1.04E-06 |
| ENSMUSG00000031010 | Usp9x | 0.003875429 | -1.352433868 | 0.023300198 |
| ENSMUSG00000062590 | Armc9 | 0.003820065 | -1.219060913 | 0.049367015 |
| ENSMUSG00000027506 | Tpd52 | 0.003820065 | 0.163408724 | 1 |
| ENSMUSG00000005610 | Eif4g2 | 0.003820065 | 0.298338304 | 0.829198173 |
| ENSMUSG00000031561 | Tenm3 | 0.003820065 | -0.502652202 | 0.677195462 |
| ENSMUSG00000069769 | Msi2 | 0.003820065 | -1.355023644 | 0.022940117 |
| ENSMUSG00000034801 | Sos2 | 0.003820065 | -2.701661696 | 1.92E-06 |
| ENSMUSG00000058317 | Ube2e2 | 0.003820065 | -2.503231947 | 8.71E-06 |
| ENSMUSG00000003279 | Dlgap1 | 0.003820065 | -0.125763277 | 1 |
| ENSMUSG00000028028 | Alpk1 | 0.003764702 | -2.444151336 | 1.67E-05 |
| ENSMUSG00000028452 | Vcp | 0.003764702 | -1.010824317 | 0.128972787 |
| ENSMUSG00000004849 | Ap1s1 | 0.003764702 | -1.204089686 | 0.05230171 |
| ENSMUSG00000093904 | Tomm20 | 0.003764702 | 1.225978548 | 0.013337086 |
| ENSMUSG00000056851 | Pcbp2 | 0.003764702 | -0.45938585 | 0.74886907 |
| ENSMUSG00000054428 | Atpif1 | 0.003709339 | 0.968114068 | 0.082214652 |
| ENSMUSG00000036751 | Cox6b1 | 0.003709339 | 1.659482796 | 0.000755174 |
| ENSMUSG00000078974 | Sec61g | 0.003709339 | 1.566373391 | 0.00119626 |
| ENSMUSG00000006169 | Clint1 | 0.003709339 | -1.209145444 | 0.051295394 |
| ENSMUSG00000022108 | Itm2b | 0.003709339 | 0.502678716 | 0.507847339 |
| ENSMUSG00000055839 | Elob | 0.003709339 | 1.782102082 | 0.000391161 |
| ENSMUSG00000024498 | Tcerg1 | 0.003709339 | -1.715942848 | 0.003095952 |
| ENSMUSG00000058624 | Gda | 0.003709339 | -0.041778737 | 1 |
| ENSMUSG00000054766 | Set | 0.003653976 | 0.247319192 | 0.909998684 |
| ENSMUSG00000036256 | Igfbp7 | 0.003653976 | -1.359216721 | 0.02280603 |
| ENSMUSG00000020422 | Tns3 | 0.003653976 | -1.479832113 | 0.011297904 |
| ENSMUSG00000020267 | Hint1 | 0.003653976 | 1.466997228 | 0.003705514 |
| ENSMUSG00000018593 | Sparc | 0.003653976 | -0.09243018 | 1 |
| ENSMUSG00000018567 | Gabarap | 0.003653976 | 1.152682318 | 0.022357178 |
| ENSMUSG00000021838 | Samd4 | 0.003653976 | -1.517496185 | 0.009262075 |
| ENSMUSG00000022914 | Brwd1 | 0.003653976 | -2.506058109 | 8.88E-06 |
| ENSMUSG00000036438 | Calm2 | 0.003653976 | 0.243249528 | 0.911026545 |
| ENSMUSG00000026131 | Dst | 0.003598612 | -1.962301052 | 0.000557602 |
| ENSMUSG00000026034 | Clk1 | 0.003598612 | -1.722512097 | 0.00263959 |
| ENSMUSG00000067336 | Bmpr2 | 0.003598612 | -1.263872371 | 0.038173389 |
| ENSMUSG00000026425 | Srgap2 | 0.003598612 | -1.278313883 | 0.034685647 |
| ENSMUSG00000017897 | Eya2 | 0.003598612 | -1.380911792 | 0.020162889 |
| ENSMUSG00000027712 | Anxa5 | 0.003598612 | 0.73903336 | 0.234177923 |
| ENSMUSG00000029705 | Cux1 | 0.003598612 | -1.101186196 | 0.086523965 |
| ENSMUSG00000032562 | Gnai2 | 0.003598612 | 0.835248676 | 0.151190638 |
| ENSMUSG00000061904 | Slc25a3 | 0.003598612 | 0.999742713 | 0.061348023 |
| ENSMUSG00000018707 | Dync1h1 | 0.003598612 | -1.651066237 | 0.004015829 |
| ENSMUSG00000050335 | Lgals3 | 0.003598612 | 1.209413894 | 0.01628893 |
| ENSMUSG00000097589 | Dleu2 | 0.003598612 | -1.698013919 | 0.003089995 |

| | | | | |
|---|---|---|---|---|
| ENSMUSG00000022312 | Eif3h | 0.003598612 | 0.450055687 | 0.614552622 |
| ENSMUSG00000052920 | Prkg1 | 0.003598612 | -0.66651989 | 0.466464457 |
| ENSMUSG00000071528 | Atp5md | 0.003598612 | 1.704544984 | 0.001014665 |
| ENSMUSG00000004897 | Hdgf | 0.003543249 | 0.060178567 | 1 |
| ENSMUSG00000033161 | Atp1a1 | 0.003543249 | -0.004753668 | 1 |
| ENSMUSG00000066043 | Phactr4 | 0.003543249 | -1.389492567 | 0.019471235 |
| ENSMUSG00000070733 | Fryl | 0.003543249 | -0.934276154 | 0.184013112 |
| ENSMUSG00000020163 | Uqcr11 | 0.003543249 | 1.776385601 | 0.000305057 |
| ENSMUSG00000071454 | Dtnb | 0.003543249 | -1.704922025 | 0.003094221 |
| ENSMUSG00000021794 | Glud1 | 0.003543249 | -1.681943461 | 0.00337143 |
| ENSMUSG00000006740 | Kif5b | 0.003543249 | -0.132927657 | 1 |
| ENSMUSG00000046138 | 9930021J03Rik | 0.003543249 | -1.08701985 | 0.096053423 |
| ENSMUSG00000025261 | Huwe1 | 0.003543249 | -1.750861402 | 0.002182685 |
| ENSMUSG00000063972 | Nr6a1 | 0.003487886 | -1.596717155 | 0.005814889 |
| ENSMUSG00000040506 | Ambra1 | 0.003487886 | -1.466208099 | 0.01274526 |
| ENSMUSG00000054280 | Prr14l | 0.003487886 | -1.494547663 | 0.01086068 |
| ENSMUSG00000050856 | Atp5k | 0.003487886 | 1.400908038 | 0.007127824 |
| ENSMUSG00000029686 | Cul1 | 0.003487886 | -1.7185792 | 0.002747766 |
| ENSMUSG00000031523 | Dlc1 | 0.003487886 | -1.198221795 | 0.060572541 |
| ENSMUSG00000032402 | Smad3 | 0.003487886 | -2.264053785 | 6.28E-05 |
| ENSMUSG00000096054 | Syne1 | 0.003487886 | -0.858959089 | 0.25645617 |
| ENSMUSG00000017781 | Pitpna | 0.003487886 | -1.02888409 | 0.123033833 |
| ENSMUSG00000035954 | Dock4 | 0.003487886 | -2.42597491 | 1.74E-05 |
| ENSMUSG00000069729 | Arid1b | 0.003487886 | -1.168148072 | 0.062517662 |
| ENSMUSG00000037815 | Ctnna1 | 0.003487886 | -1.491994219 | 0.011088332 |
| ENSMUSG00000034154 | Ino80 | 0.003432523 | -1.010509146 | 0.135445236 |
| ENSMUSG00000102758 | Naaladl2 | 0.003432523 | -1.07605721 | 0.103463818 |
| ENSMUSG00000028053 | Ash1l | 0.003432523 | -1.770489306 | 0.001987967 |
| ENSMUSG00000042312 | S100a13 | 0.003432523 | 0.713226139 | 0.271484429 |
| ENSMUSG00000078578 | Ube2d3 | 0.003432523 | 0.592692473 | 0.418506646 |
| ENSMUSG00000034040 | Galnt17 | 0.003432523 | -0.698065072 | 0.463241392 |
| ENSMUSG00000019943 | Atp2b1 | 0.003432523 | -1.33243724 | 0.02789416 |
| ENSMUSG00000034566 | Atp5h | 0.003432523 | 1.517591018 | 0.002606179 |
| ENSMUSG00000021957 | Tkt | 0.003432523 | 0.281819567 | 0.855776926 |
| ENSMUSG00000022000 | Zc3h13 | 0.003432523 | -1.687901229 | 0.003399554 |
| ENSMUSG00000058655 | Eif4b | 0.003432523 | -0.128772027 | 1 |
| ENSMUSG00000023048 | Prr13 | 0.003432523 | 0.522827667 | 0.496010946 |
| ENSMUSG00000078247 | Airn | 0.003432523 | -2.224071363 | 8.67E-05 |
| ENSMUSG00000034382 | AI661453 | 0.003432523 | 0.392495798 | 0.692227143 |
| ENSMUSG00000014294 | Ndufa2 | 0.003432523 | 1.818760553 | 0.000178388 |
| ENSMUSG00000031207 | Msn | 0.003432523 | -0.973797367 | 0.168437012 |
| ENSMUSG00000002107 | Celf2 | 0.003377159 | -1.20271201 | 0.054462494 |
| ENSMUSG00000036893 | Ehmt1 | 0.003377159 | -1.777429203 | 0.001952189 |
| ENSMUSG00000029713 | Gnb2 | 0.003377159 | 1.504999779 | 0.002322357 |
| ENSMUSG00000036427 | Gpi1 | 0.003377159 | 0.536942671 | 0.47966612 |
| ENSMUSG00000033685 | Ucp2 | 0.003377159 | 1.15182075 | 0.028163619 |
| ENSMUSG00000004207 | Psap | 0.003377159 | 0.553183312 | 0.466464457 |
| ENSMUSG00000022707 | Gbe1 | 0.003377159 | -0.70758092 | 0.453142191 |
| ENSMUSG00000044477 | Zfand3 | 0.003377159 | -2.175866614 | 0.00012563 |
| ENSMUSG00000024002 | Brd4 | 0.003377159 | -1.697320699 | 0.003195324 |
| ENSMUSG00000041607 | Mbp | 0.003377159 | -1.207539773 | 0.053533761 |

| | | | | |
|---|---|---|---|---|
| ENSMUSG00000053333 | Dis3l2 | 0.003321796 | -1.293871224 | 0.034141187 |
| ENSMUSG00000061859 | Patj | 0.003321796 | -1.295408459 | 0.032981099 |
| ENSMUSG00000097462 | 9530026P05Rik | 0.003321796 | -1.014751715 | 0.140524002 |
| ENSMUSG00000050271 | Prag1 | 0.003321796 | -1.733310066 | 0.002627996 |
| ENSMUSG00000003814 | Calr | 0.003321796 | 0.886085932 | 0.126855892 |
| ENSMUSG00000025812 | Pard3 | 0.003321796 | -1.943989067 | 0.000662625 |
| ENSMUSG00000015501 | Hivep2 | 0.003321796 | -1.414780548 | 0.017802109 |
| ENSMUSG00000050965 | Prkca | 0.003321796 | -1.361474717 | 0.023487243 |
| ENSMUSG00000020993 | Trappc6b | 0.003321796 | -0.936126949 | 0.183791735 |
| ENSMUSG00000021477 | Ctsl | 0.003321796 | 0.928521198 | 0.108201045 |
| ENSMUSG00000047888 | Tnrc6b | 0.003321796 | -1.351164449 | 0.025088292 |
| ENSMUSG00000023010 | Tmbim6 | 0.003321796 | 0.063873051 | 1 |
| ENSMUSG00000022770 | Dlg1 | 0.003321796 | -1.684869711 | 0.003477777 |
| ENSMUSG00000022636 | Alcam | 0.003321796 | -2.651648292 | 3.43E-06 |
| ENSMUSG00000044393 | Dsg2 | 0.003321796 | -0.677814954 | 0.463312474 |
| ENSMUSG00000054843 | Atrnl1 | 0.003321796 | -1.34968552 | 0.024890795 |
| ENSMUSG00000026074 | Map4k4 | 0.003266433 | -1.068155343 | 0.107391034 |
| ENSMUSG00000036155 | Mgat5 | 0.003266433 | -1.402826568 | 0.018905476 |
| ENSMUSG00000026509 | Capn2 | 0.003266433 | -1.886180557 | 0.001014164 |
| ENSMUSG00000068205 | Macrod2 | 0.003266433 | -0.311552593 | 0.980745941 |
| ENSMUSG00000061143 | Maml3 | 0.003266433 | -1.816050497 | 0.001550379 |
| ENSMUSG00000037894 | H2afz | 0.003266433 | 1.279601742 | 0.01344034 |
| ENSMUSG00000042541 | Sem1 | 0.003266433 | 1.261105398 | 0.022626543 |
| ENSMUSG00000030447 | Cyfip1 | 0.003266433 | -2.382383179 | 2.75E-05 |
| ENSMUSG00000052572 | Dlg2 | 0.003266433 | 0.125273595 | 1 |
| ENSMUSG00000031715 | Smarca5 | 0.003266433 | -0.668275527 | 0.466464457 |
| ENSMUSG00000032290 | Ptpn9 | 0.003266433 | -1.424413431 | 0.017132257 |
| ENSMUSG00000032238 | Rora | 0.003266433 | -1.237064147 | 0.051295394 |
| ENSMUSG00000039835 | Nhsl1 | 0.003266433 | -2.097118826 | 0.000238811 |
| ENSMUSG00000033499 | Larp4b | 0.003266433 | -1.331494468 | 0.028456883 |
| ENSMUSG00000021665 | Hexb | 0.003266433 | -2.520421381 | 9.63E-06 |
| ENSMUSG00000024290 | Rock1 | 0.003266433 | -0.668275527 | 0.466464457 |
| ENSMUSG00000024659 | Anxa1 | 0.003266433 | 0.067886113 | 1 |
| ENSMUSG00000038393 | Txnip | 0.003211069 | 0.887622411 | 0.124372705 |
| ENSMUSG00000045083 | Lingo2 | 0.003211069 | -0.240010869 | 1 |
| ENSMUSG00000028843 | Sh3bgrl3 | 0.003211069 | 1.844978074 | 0.000148372 |
| ENSMUSG00000029648 | Flt1 | 0.003211069 | -0.539983762 | 0.66416768 |
| ENSMUSG00000029767 | Calu | 0.003211069 | 0.011842348 | 1 |
| ENSMUSG00000052512 | Nav2 | 0.003211069 | -0.772302147 | 0.356416343 |
| ENSMUSG00000030536 | Iqgap1 | 0.003211069 | -1.31714434 | 0.030430985 |
| ENSMUSG00000034390 | Cmip | 0.003211069 | -2.240835973 | 8.05E-05 |
| ENSMUSG00000006932 | Ctnnb1 | 0.003211069 | -0.115979826 | 1 |
| ENSMUSG00000003746 | Man1a | 0.003211069 | -0.974332953 | 0.171582618 |
| ENSMUSG00000034163 | Zfc3h1 | 0.003211069 | -1.893737302 | 0.000984963 |
| ENSMUSG00000025393 | Atp5b | 0.003211069 | 1.371607264 | 0.006326993 |
| ENSMUSG00000034621 | Gpatch8 | 0.003211069 | -1.851145426 | 0.00128813 |
| ENSMUSG00000035133 | Arhgap5 | 0.003211069 | -1.196441854 | 0.058052325 |
| ENSMUSG00000021546 | Hnrnpk | 0.003211069 | 0.165156387 | 1 |
| ENSMUSG00000055065 | Ddx17 | 0.003211069 | -2.001851132 | 0.000451458 |
| ENSMUSG00000024983 | Vti1a | 0.003211069 | -2.033623478 | 0.000366437 |
| ENSMUSG00000042133 | Ppig | 0.003155706 | -1.3697004 | 0.023472221 |

| | | | | |
|---|---|---|---|---|
| ENSMUSG00000027365 | Trpm7 | 0.003155706 | -0.757626054 | 0.37105914 |
| ENSMUSG00000028249 | Sdcbp | 0.003155706 | 0.855764855 | 0.152182498 |
| ENSMUSG00000037822 | Smim14 | 0.003155706 | -0.769307859 | 0.370945697 |
| ENSMUSG00000029311 | Hsd17b11 | 0.003155706 | -1.60562296 | 0.00592361 |
| ENSMUSG00000005534 | Insr | 0.003155706 | -1.176178037 | 0.064562465 |
| ENSMUSG00000037940 | Inpp4b | 0.003155706 | -1.14064188 | 0.084475788 |
| ENSMUSG00000019978 | Epb41l2 | 0.003155706 | -1.879045854 | 0.001061285 |
| ENSMUSG00000113902 | Ndufb1-ps | 0.003155706 | 1.76534956 | 0.000358192 |
| ENSMUSG00000034751 | Mast4 | 0.003155706 | -1.973973655 | 0.000557602 |
| ENSMUSG00000021745 | Ptprg | 0.003155706 | -2.0696678 | 0.000287323 |
| ENSMUSG00000063458 | Lrmda | 0.003155706 | -2.23549289 | 8.67E-05 |
| ENSMUSG00000060373 | Hnrnpc | 0.003155706 | -1.341806394 | 0.026752204 |
| ENSMUSG00000021996 | Esd | 0.003155706 | 1.023000769 | 0.058090234 |
| ENSMUSG00000036461 | Elf1 | 0.003155706 | -1.575332495 | 0.007109019 |
| ENSMUSG00000033306 | Lpp | 0.003155706 | -1.736357437 | 0.002693122 |
| ENSMUSG00000025428 | Atp5a1 | 0.003155706 | 0.535272993 | 0.485776178 |
| ENSMUSG00000025283 | Sat1 | 0.003155706 | -0.556578535 | 0.625377538 |
| ENSMUSG00000057914 | Cacnb2 | 0.003100343 | -1.691951571 | 0.003568342 |
| ENSMUSG00000043940 | Wdfy3 | 0.003100343 | -1.518274434 | 0.010399855 |
| ENSMUSG00000009013 | Dynll1 | 0.003100343 | 0.989872469 | 0.083474887 |
| ENSMUSG00000036199 | Ndufa13 | 0.003100343 | 2.437331446 | 6.21E-07 |
| ENSMUSG00000031799 | Tpm4 | 0.003100343 | 1.030514454 | 0.059512215 |
| ENSMUSG00000036550 | Cnot1 | 0.003100343 | -1.609709821 | 0.005847415 |
| ENSMUSG00000062270 | Morf4l1 | 0.003100343 | -0.655265546 | 0.474032441 |
| ENSMUSG00000018474 | Chd3 | 0.003100343 | -1.929785422 | 0.000785466 |
| ENSMUSG00000021686 | Ap3b1 | 0.003100343 | -1.172910463 | 0.064852501 |
| ENSMUSG00000022443 | Myh9 | 0.003100343 | -1.688199436 | 0.003613491 |
| ENSMUSG00000071984 | Fndc1 | 0.003100343 | -0.50979628 | 0.739569616 |
| ENSMUSG00000041881 | Ndufa7 | 0.003100343 | 1.493040879 | 0.002793064 |
| ENSMUSG00000031328 | Flna | 0.003100343 | -1.555821388 | 0.00820443 |
| ENSMUSG00000104960 | Snhg8 | 0.00304498 | 0.701302971 | 0.284190911 |
| ENSMUSG00000028337 | Coro2a | 0.00304498 | -0.662750599 | 0.466464457 |
| ENSMUSG00000030189 | Ybx3 | 0.00304498 | 0.783765132 | 0.210994085 |
| ENSMUSG00000006519 | Cyba | 0.00304498 | 1.43382266 | 0.004666096 |
| ENSMUSG00000025351 | Cd63 | 0.00304498 | 1.932628517 | 0.00010639 |
| ENSMUSG00000036644 | Tbc1d9b | 0.00304498 | -2.239037818 | 8.85E-05 |
| ENSMUSG00000021218 | Gdi2 | 0.00304498 | -1.051604167 | 0.117709307 |
| ENSMUSG00000041112 | Elmo1 | 0.00304498 | -0.867990073 | 0.252352914 |
| ENSMUSG00000021792 | Prxl2a | 0.00304498 | -0.46862203 | 0.751934995 |
| ENSMUSG00000022565 | Plec | 0.00304498 | -1.747157529 | 0.002544624 |
| ENSMUSG00000001289 | Pfdn5 | 0.00304498 | 0.996758855 | 0.078980382 |
| ENSMUSG00000022500 | Litaf | 0.00304498 | -1.235553046 | 0.048996826 |
| ENSMUSG00000022708 | Zbtb20 | 0.00304498 | -2.280211458 | 6.36E-05 |
| ENSMUSG00000032624 | Eml4 | 0.00304498 | -1.675666487 | 0.003929137 |
| ENSMUSG00000026721 | Rabgap1l | 0.002989616 | -1.133954225 | 0.080282494 |
| ENSMUSG00000039630 | Hnrnpu | 0.002989616 | -1.836591276 | 0.001501658 |
| ENSMUSG00000026864 | Hspa5 | 0.002989616 | -0.065659762 | 1 |
| ENSMUSG00000027562 | Car2 | 0.002989616 | 2.407827452 | 1.04E-06 |
| ENSMUSG00000054091 | 1810037I17Rik | 0.002989616 | 1.475998955 | 0.005555646 |
| ENSMUSG00000028809 | Srrm1 | 0.002989616 | -1.130137569 | 0.082214652 |
| ENSMUSG00000029004 | Kmt2e | 0.002989616 | -1.544219325 | 0.008989808 |

| | | | | |
|---|---|---|---|---|
| ENSMUSG00000038056 | Kmt2c | 0.002989616 | -1.326361058 | 0.030355209 |
| ENSMUSG00000001098 | Kctd10 | 0.002989616 | 0.922400625 | 0.110800285 |
| ENSMUSG00000029642 | Polr1d | 0.002989616 | 0.648835551 | 0.372782494 |
| ENSMUSG00000055320 | Tead1 | 0.002989616 | -1.3723211 | 0.023913246 |
| ENSMUSG00000031925 | Maml2 | 0.002989616 | -3.077103859 | 6.76E-08 |
| ENSMUSG00000041408 | Wapl | 0.002989616 | -1.699087752 | 0.00345597 |
| ENSMUSG00000075602 | Ly6a | 0.002989616 | 2.220741899 | 6.69E-06 |
| ENSMUSG00000022952 | Runx1 | 0.002989616 | -1.929554295 | 0.000833813 |
| ENSMUSG00000024730 | Ms4a8a | 0.002989616 | -2.159455882 | 0.00024492 |
| ENSMUSG00000026740 | Dnajc1 | 0.002934253 | -1.220158436 | 0.054044548 |
| ENSMUSG00000027248 | Pdia3 | 0.002934253 | -0.286274918 | 1 |
| ENSMUSG00000032959 | Pebp1 | 0.002934253 | 1.759866864 | 0.000516965 |
| ENSMUSG00000020440 | Arf5 | 0.002934253 | 1.127598648 | 0.039730049 |
| ENSMUSG00000064080 | Fbln2 | 0.002934253 | -0.890897695 | 0.235305538 |
| ENSMUSG00000066406 | Akap13 | 0.002934253 | -2.287147956 | 6.27E-05 |
| ENSMUSG00000038371 | Sbf2 | 0.002934253 | -1.43390488 | 0.017201466 |
| ENSMUSG00000004936 | Map2k1 | 0.002934253 | -1.380911792 | 0.022940117 |
| ENSMUSG00000009647 | Mcu | 0.002934253 | -1.440197998 | 0.016202833 |
| ENSMUSG00000020074 | Ccar1 | 0.002934253 | -1.735828299 | 0.002850297 |
| ENSMUSG00000020368 | Canx | 0.002934253 | -0.441766997 | 0.792441218 |
| ENSMUSG00000025132 | Arhgdia | 0.002934253 | 0.233798052 | 0.968189706 |
| ENSMUSG00000076431 | Sox4 | 0.002934253 | -0.303636078 | 1 |
| ENSMUSG00000050377 | Il31ra | 0.002934253 | -1.757408829 | 0.002647379 |
| ENSMUSG00000033060 | Lmo7 | 0.002934253 | -2.237425544 | 9.04E-05 |
| ENSMUSG00000036501 | Fam13b | 0.002934253 | -1.948320372 | 0.000731778 |
| ENSMUSG00000024646 | Cyb5a | 0.002934253 | 0.703283335 | 0.306626259 |
| ENSMUSG00000016150 | Tenm1 | 0.002934253 | -0.837068279 | 0.279592421 |
| ENSMUSG00000073676 | Hspe1 | 0.00287889 | 1.257814933 | 0.024470179 |
| ENSMUSG00000070565 | Rasal2 | 0.00287889 | -2.176873288 | 0.000144292 |
| ENSMUSG00000014077 | Chp1 | 0.00287889 | -0.856863671 | 0.26465685 |
| ENSMUSG00000002233 | Rhoc | 0.00287889 | 1.46986541 | 0.003974539 |
| ENSMUSG00000039967 | Zfp292 | 0.00287889 | -1.850859937 | 0.001389409 |
| ENSMUSG00000058440 | Nrf1 | 0.00287889 | -1.340920417 | 0.028938329 |
| ENSMUSG00000041540 | Sox5 | 0.00287889 | -0.539123373 | 0.66416768 |
| ENSMUSG00000041571 | Selenow | 0.00287889 | 0.310666816 | 0.822358126 |
| ENSMUSG00000030616 | Sytl2 | 0.00287889 | 0.299940409 | 0.854285611 |
| ENSMUSG00000007891 | Ctsd | 0.00287889 | 1.010433792 | 0.088250241 |
| ENSMUSG00000071369 | Map3k5 | 0.00287889 | -1.328876381 | 0.03126869 |
| ENSMUSG00000021290 | Atp5mpl | 0.00287889 | 1.595396292 | 0.001718437 |
| ENSMUSG00000021360 | Gcnt2 | 0.00287889 | -1.780590929 | 0.002145048 |
| ENSMUSG00000021375 | Kif13a | 0.00287889 | -0.705393035 | 0.441319775 |
| ENSMUSG00000021973 | Micu2 | 0.00287889 | -1.488977265 | 0.01253654 |
| ENSMUSG00000022598 | Psca | 0.00287889 | 3.85319405 | 2.01E-14 |
| ENSMUSG00000022533 | Atp13a3 | 0.00287889 | -1.89353932 | 0.00107162 |
| ENSMUSG00000024360 | Etf1 | 0.00287889 | -0.609085931 | 0.544908989 |
| ENSMUSG00000031430 | Vsig1 | 0.00287889 | 1.621868504 | 0.00152573 |
| ENSMUSG00000034088 | Hdlbp | 0.002823527 | -0.931495501 | 0.210550914 |
| ENSMUSG00000039349 | C130074G19Rik | 0.002823527 | 0.267126228 | 0.907125767 |
| ENSMUSG00000005625 | Psmd4 | 0.002823527 | -0.025764162 | 1 |
| ENSMUSG00000029203 | Ube2k | 0.002823527 | -1.276433573 | 0.04054889 |
| ENSMUSG00000029621 | Arpc1a | 0.002823527 | -0.154550468 | 1 |

| | | | | |
|---|---|---|---|---|
| ENSMUSG00000038156 | Spon1 | 0.002823527 | -1.097807957 | 0.11066193 |
| ENSMUSG00000006589 | Aprt | 0.002823527 | 1.857422173 | 0.000239964 |
| ENSMUSG00000047409 | Ctdspl | 0.002823527 | -1.344211688 | 0.028297253 |
| ENSMUSG00000020846 | Rflnb | 0.002823527 | 1.406760764 | 0.006811186 |
| ENSMUSG00000020530 | Ggnbp2 | 0.002823527 | -2.072789497 | 0.000312002 |
| ENSMUSG00000091803 | Cox16 | 0.002823527 | -2.204354024 | 0.000120117 |
| ENSMUSG00000021877 | Arf4 | 0.002823527 | -0.412275963 | 0.827142217 |
| ENSMUSG00000022680 | Pdxdc1 | 0.002823527 | -1.70057328 | 0.003603181 |
| ENSMUSG00000025612 | Bach1 | 0.002823527 | -1.023589791 | 0.134206505 |
| ENSMUSG00000024359 | Hspa9 | 0.002823527 | 0.235533104 | 0.951660573 |
| ENSMUSG00000024576 | Csnk1a1 | 0.002823527 | -1.091593898 | 0.099626479 |
| ENSMUSG00000024566 | Atp9b | 0.002823527 | -1.030103097 | 0.131195826 |
| ENSMUSG00000038344 | Txlng | 0.002823527 | -1.51800347 | 0.010847767 |
| ENSMUSG00000051177 | Plcb1 | 0.002768163 | -1.814448393 | 0.00194589 |
| ENSMUSG00000038729 | Pakap | 0.002768163 | -0.86609118 | 0.262284476 |
| ENSMUSG00000008348 | Ubc | 0.002768163 | 0.233798052 | 0.971854959 |
| ENSMUSG00000001847 | Rac1 | 0.002768163 | 0.789879433 | 0.215910103 |
| ENSMUSG00000025477 | Inpp5a | 0.002768163 | -0.221770927 | 1 |
| ENSMUSG00000060924 | Csmd1 | 0.002768163 | -0.800272624 | 0.377307523 |
| ENSMUSG00000038708 | Golga4 | 0.002768163 | -1.223703539 | 0.054350902 |
| ENSMUSG00000039031 | Arhgap18 | 0.002768163 | -1.387822919 | 0.022755145 |
| ENSMUSG00000020390 | Ube2b | 0.002768163 | -1.746173467 | 0.002803749 |
| ENSMUSG00000021242 | Npc2 | 0.002768163 | 0.607015376 | 0.423814296 |
| ENSMUSG00000060739 | Nsa2 | 0.002768163 | 0.911869957 | 0.139039734 |
| ENSMUSG00000009470 | Tnpo1 | 0.002768163 | -1.612429762 | 0.006207271 |
| ENSMUSG00000033278 | Ptprm | 0.002768163 | -2.167583453 | 0.000199484 |
| ENSMUSG00000038773 | Kdm3b | 0.002768163 | -0.980235302 | 0.167024802 |
| ENSMUSG00000025862 | Stag2 | 0.002768163 | -1.000667201 | 0.152459667 |
| ENSMUSG00000031231 | Cox7b | 0.002768163 | 0.870049782 | 0.148656704 |
| ENSMUSG00000034480 | Diaph2 | 0.002768163 | -2.057033963 | 0.000365509 |
| ENSMUSG00000043241 | Upf2 | 0.0027128 | -1.354766685 | 0.027444373 |
| ENSMUSG00000027309 | 4930402H24Rik | 0.0027128 | -1.047158262 | 0.128972787 |
| ENSMUSG00000029202 | Pds5a | 0.0027128 | -1.033737746 | 0.133665336 |
| ENSMUSG00000041697 | Cox6a1 | 0.0027128 | 2.007289522 | 7.37E-05 |
| ENSMUSG00000087380 | 2210408F21Rik | 0.0027128 | -1.629149196 | 0.005979622 |
| ENSMUSG00000004347 | Pde1c | 0.0027128 | -0.537030994 | 0.66416768 |
| ENSMUSG00000029992 | Gfpt1 | 0.0027128 | -0.011089007 | 1 |
| ENSMUSG00000071076 | Jund | 0.0027128 | 0.287067192 | 0.886203497 |
| ENSMUSG00000000088 | Cox5a | 0.0027128 | 1.866426987 | 0.000271858 |
| ENSMUSG00000032249 | Anp32a | 0.0027128 | 0.866426987 | 0.157935867 |
| ENSMUSG00000037286 | Stag1 | 0.0027128 | -1.52108945 | 0.010987762 |
| ENSMUSG00000038240 | Pdss2 | 0.0027128 | -1.112449722 | 0.092636114 |
| ENSMUSG00000042331 | Specc1 | 0.0027128 | -1.43862729 | 0.017810489 |
| ENSMUSG00000020738 | Sumo2 | 0.0027128 | -0.458852318 | 0.769402862 |
| ENSMUSG00000021282 | Eif5 | 0.0027128 | -0.732447821 | 0.422790236 |
| ENSMUSG00000038014 | Fam120a | 0.0027128 | -0.571494403 | 0.618995574 |
| ENSMUSG00000021725 | Parp8 | 0.0027128 | -1.790787586 | 0.002145048 |
| ENSMUSG00000055421 | Pcdh9 | 0.0027128 | -0.746227248 | 0.40935944 |
| ENSMUSG00000022607 | Ptk2 | 0.0027128 | -1.569945616 | 0.008054668 |
| ENSMUSG00000041852 | Tcf20 | 0.0027128 | -1.678851813 | 0.004176784 |
| ENSMUSG00000015575 | Atp6v0e | 0.0027128 | 0.557982121 | 0.494808857 |

| | | | | |
|---|---|---|---|---|
| ENSMUSG00000061232 | H2-K1 | 0.0027128 | -0.781449722 | 0.367810373 |
| ENSMUSG00000041482 | Piezo2 | 0.0027128 | -1.500098553 | 0.013476078 |
| ENSMUSG00000026043 | Col3a1 | 0.002657437 | 0.029925719 | 1 |
| ENSMUSG00000026596 | Abl2 | 0.002657437 | -1.489671354 | 0.013366234 |
| ENSMUSG00000034101 | Ctnnd1 | 0.002657437 | -1.846285259 | 0.001521072 |
| ENSMUSG00000028161 | Ppp3ca | 0.002657437 | -1.649343941 | 0.005298129 |
| ENSMUSG00000028998 | Tomm7 | 0.002657437 | 1.837280641 | 0.000357676 |
| ENSMUSG00000079426 | Arpc4 | 0.002657437 | -0.098398378 | 1 |
| ENSMUSG00000030231 | Plekha5 | 0.002657437 | -1.469469926 | 0.015349166 |
| ENSMUSG00000031502 | Col4a1 | 0.002657437 | -0.166907156 | 1 |
| ENSMUSG00000031633 | Slc25a4 | 0.002657437 | 1.368727632 | 0.012969842 |
| ENSMUSG00000032502 | Stac | 0.002657437 | -2.427325614 | 2.86E-05 |
| ENSMUSG00000059439 | Bcas3 | 0.002657437 | -1.267836065 | 0.044042118 |
| ENSMUSG00000020719 | Ddx5 | 0.002657437 | -1.915119455 | 0.001014665 |
| ENSMUSG00000037486 | Asxl2 | 0.002657437 | -1.537354504 | 0.010114343 |
| ENSMUSG00000020594 | Pum2 | 0.002657437 | -1.650839737 | 0.005097082 |
| ENSMUSG00000052632 | Asap2 | 0.002657437 | -1.842363113 | 0.001608894 |
| ENSMUSG00000053754 | Chd8 | 0.002657437 | -1.466075348 | 0.015090026 |
| ENSMUSG00000115186 | Gm49417 | 0.002657437 | -2.150082533 | 0.000192324 |
| ENSMUSG00000023991 | Foxp4 | 0.002657437 | -1.481288442 | 0.014140013 |
| ENSMUSG00000037058 | Paip2 | 0.002657437 | -0.799849686 | 0.352215149 |
| ENSMUSG00000024695 | Zfp91 | 0.002657437 | -0.474546864 | 0.759927903 |
| ENSMUSG00000026303 | Mlph | 0.002602074 | -0.410058138 | 0.850654272 |
| ENSMUSG00000026576 | Atp1b1 | 0.002602074 | 0.582746361 | 0.466464457 |
| ENSMUSG00000028559 | Osbpl9 | 0.002602074 | -1.08135151 | 0.112433946 |
| ENSMUSG00000028868 | Wasf2 | 0.002602074 | -1.106051951 | 0.099734556 |
| ENSMUSG00000029201 | Ugdh | 0.002602074 | -0.145580315 | 1 |
| ENSMUSG00000071424 | Grid2 | 0.002602074 | -0.128145722 | 1 |
| ENSMUSG00000066232 | Ipo7 | 0.002602074 | -0.673092543 | 0.495337286 |
| ENSMUSG00000030852 | Tacc2 | 0.002602074 | -0.928593277 | 0.208051765 |
| ENSMUSG00000005054 | Cstb | 0.002602074 | 1.465613727 | 0.007127824 |
| ENSMUSG00000036019 | Tmtc2 | 0.002602074 | -1.436858197 | 0.019442592 |
| ENSMUSG00000045671 | Spred2 | 0.002602074 | -2.560617814 | 8.97E-06 |
| ENSMUSG00000035992 | Fnip1 | 0.002602074 | -0.946111038 | 0.208883511 |
| ENSMUSG00000061981 | Flot2 | 0.002602074 | -0.794107944 | 0.344723315 |
| ENSMUSG00000021895 | Arhgef3 | 0.002602074 | -1.889029943 | 0.001206416 |
| ENSMUSG00000022051 | Bnip3l | 0.002602074 | 0.758064621 | 0.267492155 |
| ENSMUSG00000041736 | Tspo | 0.002602074 | 1.338980289 | 0.013235138 |
| ENSMUSG00000000346 | Dazap2 | 0.002602074 | 1.726003412 | 0.001046072 |
| ENSMUSG00000002365 | Snx9 | 0.002602074 | -1.395558568 | 0.023745467 |
| ENSMUSG00000033475 | Tomm6 | 0.002602074 | 1.388308001 | 0.012717738 |
| ENSMUSG00000026341 | Actr3 | 0.00254671 | -0.247786709 | 1 |
| ENSMUSG00000090553 | Snrpe | 0.00254671 | 1.489178885 | 0.006863256 |
| ENSMUSG00000040423 | Rc3h1 | 0.00254671 | -1.171615028 | 0.074493339 |
| ENSMUSG00000075284 | Wipf1 | 0.00254671 | -1.286423557 | 0.040498642 |
| ENSMUSG00000007659 | Bcl2l1 | 0.00254671 | -1.756577529 | 0.002779402 |
| ENSMUSG00000027777 | Schip1 | 0.00254671 | -2.138650265 | 0.000217681 |
| ENSMUSG00000015697 | Setdb1 | 0.00254671 | -0.931001917 | 0.21397135 |
| ENSMUSG00000010517 | Faf1 | 0.00254671 | -1.433200217 | 0.018719769 |
| ENSMUSG00000028811 | Yars | 0.00254671 | -0.666940316 | 0.47966612 |
| ENSMUSG00000028760 | Eif4g3 | 0.00254671 | -0.928432557 | 0.213040163 |

| | | | | |
|---|---|---|---|---|
| ENSMUSG00000062604 | Srpk2 | 0.00254671 | -1.075025055 | 0.116121883 |
| ENSMUSG00000037210 | Fam193a | 0.00254671 | -1.890653283 | 0.00125629 |
| ENSMUSG00000060961 | Slc4a4 | 0.00254671 | -1.931001917 | 0.000952239 |
| ENSMUSG00000063015 | Ccni | 0.00254671 | 0.596094089 | 0.463009681 |
| ENSMUSG00000032754 | Slc8b1 | 0.00254671 | -1.692908001 | 0.004863274 |
| ENSMUSG00000029455 | Aldh2 | 0.00254671 | 0.121275362 | 1 |
| ENSMUSG00000029673 | Auts2 | 0.00254671 | -1.104914627 | 0.103463818 |
| ENSMUSG00000062980 | Cped1 | 0.00254671 | -1.068038625 | 0.13144915 |
| ENSMUSG00000030077 | Chl1 | 0.00254671 | -0.26009997 | 1 |
| ENSMUSG00000030583 | Sipa1l3 | 0.00254671 | -1.389032634 | 0.023955428 |
| ENSMUSG00000032405 | Pias1 | 0.00254671 | -1.390900201 | 0.024689002 |
| ENSMUSG00000059534 | Uqcr10 | 0.00254671 | 2.094899946 | 9.25E-05 |
| ENSMUSG00000020849 | Ywhae | 0.00254671 | -0.816166325 | 0.322665395 |
| ENSMUSG00000020647 | Ncoa1 | 0.00254671 | -1.460133701 | 0.016559383 |
| ENSMUSG00000020961 | Ston2 | 0.00254671 | -1.405754335 | 0.0224005 |
| ENSMUSG00000021585 | Cast | 0.00254671 | -1.20171706 | 0.063585454 |
| ENSMUSG00000062397 | Zfp706 | 0.00254671 | 0.872507525 | 0.165729006 |
| ENSMUSG00000022452 | Smdt1 | 0.00254671 | 2.475503949 | 1.75E-06 |
| ENSMUSG00000022811 | Zfp148 | 0.00254671 | -1.850501478 | 0.001604302 |
| ENSMUSG00000023826 | Prkn | 0.00254671 | -1.015744117 | 0.145964731 |
| ENSMUSG00000025203 | Scd2 | 0.00254671 | -1.276355861 | 0.045933679 |
| ENSMUSG00000026083 | Eif5b | 0.002491347 | -1.600741699 | 0.007426793 |
| ENSMUSG00000026335 | Pam | 0.002491347 | -0.977349612 | 0.172404208 |
| ENSMUSG00000026479 | Lamc2 | 0.002491347 | -0.362877869 | 0.904300125 |
| ENSMUSG00000041235 | Chd7 | 0.002491347 | -2.118773192 | 0.000266578 |
| ENSMUSG00000028565 | Nfia | 0.002491347 | -1.69023985 | 0.004210624 |
| ENSMUSG00000042446 | Zmym4 | 0.002491347 | -1.693335763 | 0.004101636 |
| ENSMUSG00000061288 | Taok3 | 0.002491347 | -1.597440336 | 0.007644399 |
| ENSMUSG00000052144 | Ppp4r2 | 0.002491347 | -1.650969733 | 0.005523435 |
| ENSMUSG00000030770 | Parva | 0.002491347 | -1.178541674 | 0.071751408 |
| ENSMUSG00000030660 | Pik3c2a | 0.002491347 | -1.820540829 | 0.00194589 |
| ENSMUSG00000031503 | Col4a2 | 0.002491347 | -1.393656531 | 0.023303677 |
| ENSMUSG00000003848 | Nob1 | 0.002491347 | -0.962028813 | 0.18412711 |
| ENSMUSG00000032330 | Cox7a2 | 0.002491347 | 2.150029694 | 3.30E-05 |
| ENSMUSG00000046402 | Rbp1 | 0.002491347 | 1.780080085 | 0.00105249 |
| ENSMUSG00000045518 | Onecut3 | 0.002491347 | -1.694881232 | 0.004201222 |
| ENSMUSG00000004044 | Cavin1 | 0.002491347 | -0.575795342 | 0.624974096 |
| ENSMUSG00000017119 | Nbr1 | 0.002491347 | -0.558921524 | 0.642722979 |
| ENSMUSG00000020700 | Map3k3 | 0.002491347 | -1.226346184 | 0.055795486 |
| ENSMUSG00000004698 | Hdac9 | 0.002491347 | -1.925634575 | 0.001052647 |
| ENSMUSG00000015143 | Actn1 | 0.002491347 | -2.249667259 | 0.000100573 |
| ENSMUSG00000052727 | Map1b | 0.002491347 | -2.101398092 | 0.000301165 |
| ENSMUSG00000022185 | Acin1 | 0.002491347 | -2.063616684 | 0.000384507 |
| ENSMUSG00000022982 | Sod1 | 0.002491347 | 1.265506912 | 0.021544088 |
| ENSMUSG00000024511 | Rab27b | 0.002491347 | -1.24756267 | 0.051440957 |
| ENSMUSG00000040385 | Ppp1ca | 0.002491347 | 1.032846155 | 0.082879941 |
| ENSMUSG00000054611 | Kdm2a | 0.002491347 | -2.405975184 | 3.04E-05 |
| ENSMUSG00000024740 | Ddb1 | 0.002491347 | 0.410846275 | 0.687112677 |
| ENSMUSG00000011752 | Pgam1 | 0.002491347 | 0.982572949 | 0.107391034 |
| ENSMUSG00000026305 | Lrrfip1 | 0.002435984 | -1.806217627 | 0.002108876 |
| ENSMUSG00000026511 | Srp9 | 0.002435984 | 1.454188121 | 0.007443294 |

| | | | | |
|---|---|---|---|---|
| ENSMUSG00000035168 | Tanc1 | 0.002435984 | -1.253774047 | 0.04883823 |
| ENSMUSG00000018770 | Atp5g3 | 0.002435984 | 1.555726147 | 0.00431473 |
| ENSMUSG00000047187 | Rab2a | 0.002435984 | -0.488667972 | 0.736456705 |
| ENSMUSG00000028382 | Ptbp3 | 0.002435984 | -0.382873308 | 0.886203497 |
| ENSMUSG00000029313 | Aff1 | 0.002435984 | -1.711060394 | 0.003902624 |
| ENSMUSG00000031447 | Lamp1 | 0.002435984 | 0.756984355 | 0.27201021 |
| ENSMUSG00000096914 | Galntl6 | 0.002435984 | 0.931235282 | 0.137228254 |
| ENSMUSG00000031635 | Anxa10 | 0.002435984 | 0.765649216 | 0.268502936 |
| ENSMUSG00000058799 | Nap1l1 | 0.002435984 | -0.264452815 | 1 |
| ENSMUSG00000020456 | Ogdh | 0.002435984 | -1.373038948 | 0.028201696 |
| ENSMUSG00000020758 | Itgb4 | 0.002435984 | -1.881679165 | 0.001343478 |
| ENSMUSG00000020592 | Sdc1 | 0.002435984 | -0.107238866 | 1 |
| ENSMUSG00000044912 | Syt16 | 0.002435984 | -0.55975107 | 0.651901451 |
| ENSMUSG00000021051 | Ppp2r5e | 0.002435984 | -2.07605721 | 0.00036427 |
| ENSMUSG00000066392 | Nrxn3 | 0.002435984 | 0.42644313 | 0.686078534 |
| ENSMUSG00000021486 | Prelid1 | 0.002435984 | 1.282707653 | 0.030456171 |
| ENSMUSG00000010406 | Mrpl52 | 0.002435984 | 1.001137295 | 0.094701574 |
| ENSMUSG00000021969 | Zdhhc20 | 0.002435984 | -1.251628775 | 0.051152519 |
| ENSMUSG00000022272 | Myo10 | 0.002435984 | -1.82365222 | 0.001894736 |
| ENSMUSG00000046516 | Cox17 | 0.002435984 | 1.714423893 | 0.001116263 |
| ENSMUSG00000024121 | Atp6v0c | 0.002435984 | 1.333333726 | 0.014800323 |
| ENSMUSG00000073411 | H2-D1 | 0.002435984 | -0.903705472 | 0.234018719 |
| ENSMUSG00000024544 | Ldlrad4 | 0.002435984 | -1.720398258 | 0.003714408 |
| ENSMUSG00000040565 | Btaf1 | 0.002435984 | -1.728133907 | 0.003584399 |
| ENSMUSG00000045103 | Dmd | 0.002435984 | -0.314638573 | 0.993830483 |
| ENSMUSG00000026029 | Casp8 | 0.00238062 | -1.11090135 | 0.101578009 |
| ENSMUSG00000026879 | Gsn | 0.00238062 | 0.523304669 | 0.563788155 |
| ENSMUSG00000036053 | Fmnl2 | 0.00238062 | -2.696670502 | 3.54E-06 |
| ENSMUSG00000005506 | Celf1 | 0.00238062 | -0.831312044 | 0.306535101 |
| ENSMUSG00000045078 | Rnf216 | 0.00238062 | -1.95782202 | 0.000873396 |
| ENSMUSG00000051331 | Cacna1c | 0.00238062 | -1.081557389 | 0.119414767 |
| ENSMUSG00000054808 | Actn4 | 0.00238062 | -0.209145444 | 1 |
| ENSMUSG00000041203 | Trir | 0.00238062 | 0.508354328 | 0.572095301 |
| ENSMUSG00000001472 | Tcf25 | 0.00238062 | -0.919638826 | 0.225039093 |
| ENSMUSG00000007656 | Arpp19 | 0.00238062 | 0.87944848 | 0.167654895 |
| ENSMUSG00000006673 | Qrich1 | 0.00238062 | -1.76209755 | 0.002869506 |
| ENSMUSG00000039497 | Dse | 0.00238062 | -2.929279784 | 6.31E-07 |
| ENSMUSG00000044894 | Uqcrq | 0.00238062 | 2.04217798 | 0.000216481 |
| ENSMUSG00000018501 | Ncor1 | 0.00238062 | -1.837176666 | 0.001791632 |
| ENSMUSG00000020671 | Rab10 | 0.00238062 | -1.508281674 | 0.013250594 |
| ENSMUSG00000032621 | Srek1 | 0.00238062 | -2.889617958 | 6.46E-07 |
| ENSMUSG00000058571 | Gpc6 | 0.00238062 | -0.900161452 | 0.252412956 |
| ENSMUSG00000023027 | Atf1 | 0.00238062 | -0.938856742 | 0.212931091 |
| ENSMUSG00000038965 | Ube2l3 | 0.00238062 | -0.499063144 | 0.727874612 |
| ENSMUSG00000012114 | Med15 | 0.00238062 | -1.581730801 | 0.008348692 |
| ENSMUSG00000036572 | Upf3b | 0.00238062 | -2.20007186 | 0.00015723 |
| ENSMUSG00000026385 | Dbi | 0.002325257 | 1.320212804 | 0.020482717 |
| ENSMUSG00000075376 | Rc3h2 | 0.002325257 | -0.607894278 | 0.57206629 |
| ENSMUSG00000038467 | Chmp4b | 0.002325257 | -0.182287537 | 1 |
| ENSMUSG00000027801 | Tm4sf4 | 0.002325257 | 0.482643269 | 0.602791066 |
| ENSMUSG00000043962 | Thrap3 | 0.002325257 | -1.37198292 | 0.027990713 |

| | | | | |
|---|---|---|---|---|
| ENSMUSG00000001089 | Luzp1 | 0.002325257 | -0.792178434 | 0.363322631 |
| ENSMUSG00000106918 | Mrpl33 | 0.002325257 | 1.33363332 | 0.017241035 |
| ENSMUSG00000029823 | Luc7l2 | 0.002325257 | -1.720939303 | 0.003742303 |
| ENSMUSG00000052040 | Klf13 | 0.002325257 | -1.365768861 | 0.029638079 |
| ENSMUSG00000053877 | Srcap | 0.002325257 | -2.194026373 | 0.000165037 |
| ENSMUSG00000040459 | Arglu1 | 0.002325257 | -2.684787877 | 3.96E-06 |
| ENSMUSG00000065954 | Tacc1 | 0.002325257 | -1.275838875 | 0.05041615 |
| ENSMUSG00000052837 | Junb | 0.002325257 | -0.269195602 | 1 |
| ENSMUSG00000031924 | Cyb5b | 0.002325257 | -0.561524314 | 0.642722979 |
| ENSMUSG00000031822 | Gse1 | 0.002325257 | -1.214918541 | 0.062517662 |
| ENSMUSG00000058587 | Tmod3 | 0.002325257 | -1.114724253 | 0.10245709 |
| ENSMUSG00000020170 | Frs2 | 0.002325257 | -1.586677792 | 0.008644519 |
| ENSMUSG00000020458 | Rtn4 | 0.002325257 | -1.353259867 | 0.030336396 |
| ENSMUSG00000018171 | Vmp1 | 0.002325257 | -1.262521596 | 0.050837911 |
| ENSMUSG00000047126 | Cltc | 0.002325257 | -1.289034349 | 0.043003741 |
| ENSMUSG00000041685 | Fcho2 | 0.002325257 | -1.91642154 | 0.001132566 |
| ENSMUSG00000025555 | Farp1 | 0.002325257 | -2.298852349 | 7.90E-05 |
| ENSMUSG00000072501 | Phf20l1 | 0.002325257 | -1.905039271 | 0.001206416 |
| ENSMUSG00000022587 | Ly6e | 0.002325257 | 0.075100306 | 1 |
| ENSMUSG00000039903 | Eva1c | 0.002325257 | -1.581324557 | 0.008890736 |
| ENSMUSG00000068039 | Tcp1 | 0.002325257 | 0.691396014 | 0.361622075 |
| ENSMUSG00000071659 | Hnrnpul2 | 0.002325257 | -1.077184536 | 0.120140246 |
| ENSMUSG00000025204 | Ndufb8 | 0.002325257 | 1.648835551 | 0.00224524 |
| ENSMUSG00000073678 | Pgap1 | 0.002269894 | -0.756508552 | 0.411460922 |
| ENSMUSG00000078193 | Gm2000 | 0.002269894 | 1.533358334 | 0.006168268 |
| ENSMUSG00000043535 | Setx | 0.002269894 | -1.503811459 | 0.013768928 |
| ENSMUSG00000027184 | Caprin1 | 0.002269894 | -0.52108945 | 0.68046121 |
| ENSMUSG00000000563 | Atp5f1 | 0.002269894 | 1.146335211 | 0.048701631 |
| ENSMUSG00000039865 | Slc44a3 | 0.002269894 | -1.650590545 | 0.005982282 |
| ENSMUSG00000050608 | Micos10 | 0.002269894 | 0.881281637 | 0.185619905 |
| ENSMUSG00000028745 | Capzb | 0.002269894 | 0.152409725 | 1 |
| ENSMUSG00000041301 | Cftr | 0.002269894 | -2.048959445 | 0.000516191 |
| ENSMUSG00000038538 | Ubn2 | 0.002269894 | -1.876219105 | 0.001491796 |
| ENSMUSG00000030208 | Emp1 | 0.002269894 | -0.397638878 | 0.857508859 |
| ENSMUSG00000063838 | Cdc42ep5 | 0.002269894 | -0.187665716 | 1 |
| ENSMUSG00000078671 | Chd2 | 0.002269894 | -2.563400255 | 1.05E-05 |
| ENSMUSG00000036180 | Gatad2a | 0.002269894 | -1.494122402 | 0.014606422 |
| ENSMUSG00000032420 | Nt5e | 0.002269894 | -1.022242107 | 0.150952178 |
| ENSMUSG00000079227 | Ccr5 | 0.002269894 | 0.471297366 | 0.672182136 |
| ENSMUSG00000056427 | Slit3 | 0.002269894 | -0.927513819 | 0.228120793 |
| ENSMUSG00000048076 | Arf1 | 0.002269894 | 0.841480629 | 0.201723858 |
| ENSMUSG00000005417 | Mprip | 0.002269894 | -2.16951837 | 0.000206806 |
| ENSMUSG00000038290 | Smg6 | 0.002269894 | -1.753262892 | 0.003112598 |
| ENSMUSG00000018363 | Smurf2 | 0.002269894 | -1.255763233 | 0.05110605 |
| ENSMUSG00000021248 | Tmed10 | 0.002269894 | -0.32516924 | 0.971210385 |
| ENSMUSG00000021520 | Uqcrb | 0.002269894 | 1.871227973 | 0.000456223 |
| ENSMUSG00000060579 | Fhit | 0.002269894 | -1.054244049 | 0.143740392 |
| ENSMUSG00000022010 | Tsc22d1 | 0.002269894 | -1.27625964 | 0.046298134 |
| ENSMUSG00000064210 | Ano6 | 0.002269894 | -1.420326473 | 0.022466238 |
| ENSMUSG00000022772 | Senp5 | 0.002269894 | -1.133772708 | 0.092711052 |
| ENSMUSG00000053477 | Tcf4 | 0.002269894 | -1.329534433 | 0.034384865 |

| | | | | |
|---|---|---|---|---|
| ENSMUSG00000031169 | Porcn | 0.002269894 | -1.436605292 | 0.02045337 |
| ENSMUSG00000006005 | Tpr | 0.002214531 | -1.685355963 | 0.004970293 |
| ENSMUSG00000026956 | Uap1l1 | 0.002214531 | 0.179839069 | 1 |
| ENSMUSG00000027087 | Itgav | 0.002214531 | -1.048894879 | 0.14097975 |
| ENSMUSG00000027808 | Serp1 | 0.002214531 | 0.787219036 | 0.261873336 |
| ENSMUSG00000074457 | S100a16 | 0.002214531 | 0.866836204 | 0.179916358 |
| ENSMUSG00000037072 | Selenof | 0.002214531 | 1.59699662 | 0.003424593 |
| ENSMUSG00000008575 | Nfib | 0.002214531 | -1.657170548 | 0.005872899 |
| ENSMUSG00000014592 | Camta1 | 0.002214531 | -0.493458331 | 0.740075579 |
| ENSMUSG00000029062 | Cdk11b | 0.002214531 | -1.446883078 | 0.020146995 |
| ENSMUSG00000014956 | Ppp1cb | 0.002214531 | -0.081075285 | 1 |
| ENSMUSG00000070780 | Rbm47 | 0.002214531 | -1.337908352 | 0.034182785 |
| ENSMUSG00000056737 | Capg | 0.002214531 | 0.836462555 | 0.218817775 |
| ENSMUSG00000035199 | Arl6ip5 | 0.002214531 | -1.424064996 | 0.023815668 |
| ENSMUSG00000030314 | Atg7 | 0.002214531 | -1.038006563 | 0.149130287 |
| ENSMUSG00000030313 | Dennd5b | 0.002214531 | -0.39449188 | 0.871850653 |
| ENSMUSG00000054342 | Kcnn4 | 0.002214531 | 0.436531948 | 0.687112677 |
| ENSMUSG00000066568 | Lsm14a | 0.002214531 | -0.962279237 | 0.196917094 |
| ENSMUSG00000025503 | Taldo1 | 0.002214531 | 0.4828256 | 0.610828132 |
| ENSMUSG00000084786 | Ubl5 | 0.002214531 | -0.054308376 | 1 |
| ENSMUSG00000032423 | Syncrip | 0.002214531 | -1.587937048 | 0.008961687 |
| ENSMUSG00000032410 | Xrn1 | 0.002214531 | -1.455091891 | 0.020809074 |
| ENSMUSG00000036478 | Btg1 | 0.002214531 | 0.167183768 | 1 |
| ENSMUSG00000057522 | Spop | 0.002214531 | -1.950714486 | 0.000980834 |
| ENSMUSG00000025130 | P4hb | 0.002214531 | -0.5898022 | 0.616879894 |
| ENSMUSG00000052609 | Plekhg3 | 0.002214531 | -1.587937048 | 0.008961687 |
| ENSMUSG00000114304 | Gm48099 | 0.002214531 | -3.294536935 | 1.59E-08 |
| ENSMUSG00000007817 | Zmiz1 | 0.002214531 | -1.6247895 | 0.007081075 |
| ENSMUSG00000021910 | Nisch | 0.002214531 | -2.784775332 | 1.79E-06 |
| ENSMUSG00000021796 | Bmpr1a | 0.002214531 | -1.326767794 | 0.036614542 |
| ENSMUSG00000010376 | Nedd8 | 0.002214531 | -0.178487787 | 1 |
| ENSMUSG00000032849 | Abcc4 | 0.002214531 | -1.78802831 | 0.002606179 |
| ENSMUSG00000042292 | Mrtfa | 0.002214531 | -1.274383214 | 0.047323433 |
| ENSMUSG00000009575 | Cbx5 | 0.002214531 | -1.083724864 | 0.117170911 |
| ENSMUSG00000014769 | Psmb1 | 0.002214531 | 0.546955937 | 0.543691231 |
| ENSMUSG00000034647 | Ankrd12 | 0.002214531 | -1.569151691 | 0.00958188 |
| ENSMUSG00000091896 | Ube2d2a | 0.002214531 | -1.062390722 | 0.129326112 |
| ENSMUSG00000056763 | Cspp1 | 0.002159167 | -1.955532348 | 0.000976139 |
| ENSMUSG00000025980 | Hspd1 | 0.002159167 | -0.286624196 | 1 |
| ENSMUSG00000042699 | Dhx9 | 0.002159167 | -1.344385916 | 0.033658021 |
| ENSMUSG00000026473 | Glul | 0.002159167 | 0.741944956 | 0.32389073 |
| ENSMUSG00000027630 | Tbl1xr1 | 0.002159167 | -0.939129437 | 0.213598989 |
| ENSMUSG00000027737 | Slc7a11 | 0.002159167 | 0.082020397 | 1 |
| ENSMUSG00000005779 | Psmb4 | 0.002159167 | 1.356053802 | 0.016225033 |
| ENSMUSG00000038612 | Mcl1 | 0.002159167 | -0.777429203 | 0.395118486 |
| ENSMUSG00000027881 | Prpf38b | 0.002159167 | -3.108721137 | 8.95E-08 |
| ENSMUSG00000028478 | Clta | 0.002159167 | 0.771091301 | 0.279592421 |
| ENSMUSG00000066113 | Adamtsl1 | 0.002159167 | -0.382383179 | 0.915313016 |
| ENSMUSG00000061298 | Agbl4 | 0.002159167 | 0.595724215 | 0.497790175 |
| ENSMUSG00000018965 | Ywhah | 0.002159167 | -0.19414328 | 1 |
| ENSMUSG00000057406 | Nsd2 | 0.002159167 | -1.369086357 | 0.029826887 |

| | | | | |
|---|---|---|---|---|
| ENSMUSG00000029207 | Apbb2 | 0.002159167 | -1.660413409 | 0.005923922 |
| ENSMUSG00000035851 | Ythdc1 | 0.002159167 | -2.443921589 | 2.81E-05 |
| ENSMUSG00000029467 | Atp2a2 | 0.002159167 | -1.858959089 | 0.001729929 |
| ENSMUSG00000038095 | Sbno1 | 0.002159167 | -1.214111697 | 0.066698557 |
| ENSMUSG00000029649 | Pomp | 0.002159167 | 1.298338304 | 0.045774348 |
| ENSMUSG00000063063 | Ctnna2 | 0.002159167 | -0.084518789 | 1 |
| ENSMUSG00000033769 | Exoc6b | 0.002159167 | -2.148177426 | 0.000253444 |
| ENSMUSG00000079477 | Rab7 | 0.002159167 | -1.267641093 | 0.054186866 |
| ENSMUSG00000030717 | Nupr1 | 0.002159167 | 0.408521222 | 0.757517141 |
| ENSMUSG00000031644 | Nek1 | 0.002159167 | -0.614198854 | 0.572809984 |
| ENSMUSG00000014329 | Bicc1 | 0.002159167 | -1.333017102 | 0.036113193 |
| ENSMUSG00000032714 | Syde1 | 0.002159167 | -1.148177426 | 0.095188999 |
| ENSMUSG00000041126 | H2afv | 0.002159167 | 0.519552534 | 0.587484274 |
| ENSMUSG00000053519 | Kcnip1 | 0.002159167 | -2.545921303 | 1.59E-05 |
| ENSMUSG00000059796 | Eif4a1 | 0.002159167 | -0.209145444 | 1 |
| ENSMUSG00000035441 | Myo1d | 0.002159167 | -1.787459568 | 0.002768072 |
| ENSMUSG00000052812 | Atad2b | 0.002159167 | -1.810596067 | 0.00233013 |
| ENSMUSG00000039114 | Nrn1 | 0.002159167 | -0.034860903 | 1 |
| ENSMUSG00000047497 | Adamts12 | 0.002159167 | -1.054375916 | 0.149611214 |
| ENSMUSG00000059586 | Nsmce2 | 0.002159167 | -1.562566086 | 0.010649327 |
| ENSMUSG00000023004 | Tuba1b | 0.002159167 | 0.771091301 | 0.28540778 |
| ENSMUSG00000046434 | Hnrnpa1 | 0.002159167 | -0.643946198 | 0.529519243 |
| ENSMUSG00000033581 | Igf2bp2 | 0.002159167 | -1.629149196 | 0.007817567 |
| ENSMUSG00000056692 | Ilrun | 0.002159167 | -0.046044641 | 1 |
| ENSMUSG00000048915 | Efna5 | 0.002159167 | -2.877859294 | 8.77E-07 |
| ENSMUSG00000024346 | Pfdn1 | 0.002159167 | 0.144215159 | 1 |
| ENSMUSG00000024425 | Ndfip1 | 0.002159167 | -0.951077291 | 0.207356329 |
| ENSMUSG00000024855 | Pacs1 | 0.002159167 | -1.654945197 | 0.006082255 |
| ENSMUSG00000036372 | Tmem258 | 0.002159167 | 2.385801146 | 1.82E-05 |
| ENSMUSG00000043760 | Pkhd1 | 0.002103804 | -2.192466703 | 0.000197475 |
| ENSMUSG00000073616 | Cops9 | 0.002103804 | 2.003500433 | 0.000244583 |
| ENSMUSG00000027660 | Skil | 0.002103804 | -1.460224924 | 0.018671925 |
| ENSMUSG00000027828 | Ssr3 | 0.002103804 | 0.334324928 | 0.846569175 |
| ENSMUSG00000050359 | Sprr1a | 0.002103804 | 2.379648919 | 1.50E-05 |
| ENSMUSG00000015711 | Prune1 | 0.002103804 | -0.29458092 | 1 |
| ENSMUSG00000070372 | Capza1 | 0.002103804 | -0.698757427 | 0.466464457 |
| ENSMUSG00000028248 | Pnisr | 0.002103804 | -2.3207908 | 7.82E-05 |
| ENSMUSG00000050390 | C77080 | 0.002103804 | -1.453779515 | 0.019209067 |
| ENSMUSG00000056076 | Eif3b | 0.002103804 | -0.620351081 | 0.575389694 |
| ENSMUSG00000019370 | Calm3 | 0.002103804 | 1.379648919 | 0.016178743 |
| ENSMUSG00000009687 | Fxyd5 | 0.002103804 | 0.956957847 | 0.129344751 |
| ENSMUSG00000095115 | Itpripl2 | 0.002103804 | 0.19952815 | 1 |
| ENSMUSG00000006276 | Eps15l1 | 0.002103804 | -1.66381473 | 0.006004198 |
| ENSMUSG00000032383 | Ppib | 0.002103804 | 1.804954753 | 0.001191908 |
| ENSMUSG00000048758 | Rpl29 | 0.002103804 | -0.008276735 | 1 |
| ENSMUSG00000091537 | Tma7 | 0.002103804 | 0.418537932 | 0.705679946 |
| ENSMUSG00000019866 | Crybg1 | 0.002103804 | -1.39891758 | 0.027444373 |
| ENSMUSG00000037876 | Jmjd1c | 0.002103804 | -1.142577827 | 0.094179373 |
| ENSMUSG00000020457 | Drg1 | 0.002103804 | -1.481503998 | 0.017800854 |
| ENSMUSG00000018585 | Atox1 | 0.002103804 | 1.126882848 | 0.075389977 |
| ENSMUSG00000060073 | Psma3 | 0.002103804 | 0.283186079 | 0.930100175 |

| | | | | |
|---|---|---|---|---|
| ENSMUSG00000038372 | Gmds | 0.002103804 | -1.253114303 | 0.055310547 |
| ENSMUSG00000021771 | Vdac2 | 0.002103804 | 0.364382162 | 0.792441218 |
| ENSMUSG00000061244 | Exoc5 | 0.002103804 | -2.496214781 | 2.43E-05 |
| ENSMUSG00000060143 | Gm10076 | 0.002103804 | 0.559198339 | 0.504738484 |
| ENSMUSG00000022498 | Txndc11 | 0.002103804 | -0.783438653 | 0.387239085 |
| ENSMUSG00000022890 | Atp5j | 0.002103804 | 1.166053446 | 0.063407823 |
| ENSMUSG00000034377 | Tulp4 | 0.002103804 | -1.110156349 | 0.108089961 |
| ENSMUSG00000033863 | Klf9 | 0.002103804 | -0.436449637 | 0.80085216 |
| ENSMUSG00000031012 | Cask | 0.002103804 | -0.879543421 | 0.268394649 |
| ENSMUSG00000052062 | Pard3b | 0.002048441 | -1.177378398 | 0.082214652 |
| ENSMUSG00000055013 | Agap1 | 0.002048441 | -1.671193011 | 0.005873683 |
| ENSMUSG00000052534 | Pbx1 | 0.002048441 | -3.684437517 | 2.10E-10 |
| ENSMUSG00000027175 | Tcp11l1 | 0.002048441 | 0.045014023 | 1 |
| ENSMUSG00000028108 | Ecm1 | 0.002048441 | 0.304306028 | 0.877127085 |
| ENSMUSG00000053317 | Sec61b | 0.002048441 | 0.908078378 | 0.17471932 |
| ENSMUSG00000086320 | Gm12840 | 0.002048441 | 1.629976524 | 0.00400581 |
| ENSMUSG00000040274 | Cdk6 | 0.002048441 | -1.814043369 | 0.002408232 |
| ENSMUSG00000015653 | Steap2 | 0.002048441 | -0.891139495 | 0.268597079 |
| ENSMUSG00000031558 | Slit2 | 0.002048441 | -2.335657862 | 7.37E-05 |
| ENSMUSG00000029247 | Paics | 0.002048441 | 0.089408143 | 1 |
| ENSMUSG00000037605 | Adgrl3 | 0.002048441 | -2.301681977 | 9.25E-05 |
| ENSMUSG00000060261 | Gtf2i | 0.002048441 | -1.451965089 | 0.02007424 |
| ENSMUSG00000034021 | Pds5b | 0.002048441 | -1.49125422 | 0.016801443 |
| ENSMUSG00000029840 | Mtpn | 0.002048441 | -0.247895178 | 1 |
| ENSMUSG00000060032 | H2afj | 0.002048441 | 1.747015945 | 0.001709467 |
| ENSMUSG00000110424 | 1700012D14Rik | 0.002048441 | -1.348789641 | 0.039339084 |
| ENSMUSG00000030655 | Smg1 | 0.002048441 | -2.133904071 | 0.000303106 |
| ENSMUSG00000030738 | Eif3c | 0.002048441 | -1.174699298 | 0.082076736 |
| ENSMUSG00000032336 | Nptn | 0.002048441 | -2.216979101 | 0.000188836 |
| ENSMUSG00000032278 | Paqr5 | 0.002048441 | -1.734413991 | 0.004015829 |
| ENSMUSG00000032232 | Cgnl1 | 0.002048441 | -0.958105318 | 0.209966431 |
| ENSMUSG00000019795 | Pcmt1 | 0.002048441 | -1.544106103 | 0.012019151 |
| ENSMUSG00000004798 | Ulk2 | 0.002048441 | -1.063238867 | 0.133783875 |
| ENSMUSG00000086841 | 2410006H16Rik | 0.002048441 | 0.874395252 | 0.209230199 |
| ENSMUSG00000020859 | Spag9 | 0.002048441 | -1.756081908 | 0.003427316 |
| ENSMUSG00000040481 | Bptf | 0.002048441 | -1.591076969 | 0.009507975 |
| ENSMUSG00000063142 | Kcnma1 | 0.002048441 | -1.310251255 | 0.050925387 |
| ENSMUSG00000021948 | Prkcd | 0.002048441 | -1.327238609 | 0.038867277 |
| ENSMUSG00000022387 | Brd1 | 0.002048441 | -1.777429203 | 0.002970317 |
| ENSMUSG00000022521 | Crebbp | 0.002048441 | -1.690076723 | 0.00517567 |
| ENSMUSG00000062203 | Gspt1 | 0.002048441 | -0.650131395 | 0.517196103 |
| ENSMUSG00000022822 | Abcc5 | 0.002048441 | -2.246620149 | 0.000134111 |
| ENSMUSG00000022812 | Gsk3b | 0.002048441 | -1.219580896 | 0.065665243 |
| ENSMUSG00000022883 | Robo1 | 0.002048441 | -0.583017199 | 0.652136175 |
| ENSMUSG00000019432 | Ddx39b | 0.002048441 | -2.18937316 | 0.000206806 |
| ENSMUSG00000023951 | Vegfa | 0.002048441 | -1.122827663 | 0.105567237 |
| ENSMUSG00000024251 | Thada | 0.002048441 | -1.471743397 | 0.018323811 |
| ENSMUSG00000045394 | Epcam | 0.002048441 | -0.153085485 | 1 |
| ENSMUSG00000041915 | Ammecr1l | 0.002048441 | -1.650131395 | 0.006684444 |
| ENSMUSG00000024589 | Nedd4l | 0.002048441 | -1.886235144 | 0.001556178 |
| ENSMUSG00000024914 | Drap1 | 0.002048441 | 1.373201109 | 0.0180259 |

| | | | | |
|---|---|---|---|---|
| ENSMUSG00000024712 | Rfk | 0.002048441 | 1.767480048 | 0.001554857 |
| ENSMUSG00000051223 | Bzw1 | 0.001993078 | 0.334726961 | 0.859724344 |
| ENSMUSG00000026219 | Trip12 | 0.001993078 | -2.190197957 | 0.00021533 |
| ENSMUSG00000049690 | Nckap5 | 0.001993078 | -0.118991006 | 1 |
| ENSMUSG00000041782 | Lad1 | 0.001993078 | -0.450050113 | 0.806496094 |
| ENSMUSG00000013698 | Pea15a | 0.001993078 | 1.273326416 | 0.033421986 |
| ENSMUSG00000026618 | Iars2 | 0.001993078 | -1.678929483 | 0.005743689 |
| ENSMUSG00000025781 | Atp5c1 | 0.001993078 | 1.334726961 | 0.0224005 |
| ENSMUSG00000057738 | Sptan1 | 0.001993078 | -1.60319053 | 0.009112331 |
| ENSMUSG00000054580 | Pla2r1 | 0.001993078 | -1.561671261 | 0.011248594 |
| ENSMUSG00000027288 | Zfp106 | 0.001993078 | -0.30661801 | 1 |
| ENSMUSG00000027395 | Polr1b | 0.001993078 | -1.231823503 | 0.071100693 |
| ENSMUSG00000028060 | Khdc4 | 0.001993078 | -2.044086197 | 0.00058535 |
| ENSMUSG00000068739 | Sars | 0.001993078 | -0.244998891 | 1 |
| ENSMUSG00000029086 | Prom1 | 0.001993078 | 1.382555486 | 0.019304523 |
| ENSMUSG00000059518 | Znhit1 | 0.001993078 | -0.555339012 | 0.685568829 |
| ENSMUSG00000036402 | Gng12 | 0.001993078 | 0.214432727 | 1 |
| ENSMUSG00000030172 | Erc1 | 0.001993078 | -0.849070215 | 0.30480191 |
| ENSMUSG00000040824 | Snrpd2 | 0.001993078 | 1.243579073 | 0.036215483 |
| ENSMUSG00000041308 | Sntb2 | 0.001993078 | -1.548978847 | 0.011888285 |
| ENSMUSG00000031954 | Cfdp1 | 0.001993078 | -1.210489361 | 0.070886768 |
| ENSMUSG00000033538 | Casp4 | 0.001993078 | -1.06057432 | 0.154554575 |
| ENSMUSG00000033577 | Myo6 | 0.001993078 | -1.516750514 | 0.014575482 |
| ENSMUSG00000037608 | Bclaf1 | 0.001993078 | -1.942610983 | 0.001121749 |
| ENSMUSG00000060843 | Ctnna3 | 0.001993078 | -0.147335632 | 1 |
| ENSMUSG00000019944 | Rhobtb1 | 0.001993078 | -0.113254637 | 1 |
| ENSMUSG00000043822 | Adamtsl5 | 0.001993078 | -0.726673584 | 0.455131546 |
| ENSMUSG00000034024 | Cct2 | 0.001993078 | 0.000307922 | 1 |
| ENSMUSG00000020402 | Vdac1 | 0.001993078 | 0.090104592 | 1 |
| ENSMUSG00000020704 | Asic2 | 0.001993078 | 0.465971494 | 0.677791694 |
| ENSMUSG00000059060 | Rad51b | 0.001993078 | -2.147335632 | 0.000290024 |
| ENSMUSG00000042043 | Tbca | 0.001993078 | 0.200077434 | 1 |
| ENSMUSG00000022112 | Gpc5 | 0.001993078 | 0.536360822 | 0.587484274 |
| ENSMUSG00000022139 | Mbnl2 | 0.001993078 | -1.881491693 | 0.001645173 |
| ENSMUSG00000022240 | Ctnnd2 | 0.001993078 | 0.26583188 | 0.970970089 |
| ENSMUSG00000055114 | Anxa13 | 0.001993078 | -0.629551117 | 0.563068654 |
| ENSMUSG00000033020 | Polr2f | 0.001993078 | 0.975645868 | 0.124712353 |
| ENSMUSG00000022710 | Usp7 | 0.001993078 | -0.394376515 | 0.877117624 |
| ENSMUSG00000000385 | Tmprss2 | 0.001993078 | -1.574252983 | 0.01067424 |
| ENSMUSG00000024238 | Zeb1 | 0.001993078 | -2.121850661 | 0.00033621 |
| ENSMUSG00000039616 | Mocos | 0.001993078 | -2.028931698 | 0.000804371 |
| ENSMUSG00000024953 | Prdx5 | 0.001993078 | 1.448897981 | 0.01181109 |
| ENSMUSG00000040913 | Fbxw4 | 0.001993078 | -0.486006991 | 0.748235869 |
| ENSMUSG00000039901 | Armh3 | 0.001993078 | -1.090078315 | 0.123023696 |
| ENSMUSG00000031059 | Ndufb11 | 0.001993078 | 1.951398321 | 0.000448859 |
| ENSMUSG00000016534 | Lamp2 | 0.001993078 | -0.118991006 | 1 |
| ENSMUSG00000032666 | 1700025G04Rik | 0.001937714 | -0.577844228 | 0.66416768 |
| ENSMUSG00000025782 | Taf3 | 0.001937714 | -1.412460628 | 0.026168672 |
| ENSMUSG00000102805 | Gm37240 | 0.001937714 | -0.27927153 | 1 |
| ENSMUSG00000028080 | Lrba | 0.001937714 | -1.624182943 | 0.008054668 |
| ENSMUSG00000052539 | Magi3 | 0.001937714 | -2.374764808 | 5.71E-05 |

| | | | | |
|---|---|---|---|---|
| ENSMUSG00000068240 | Gm11808 | 0.001937714 | 1.551974012 | 0.006670473 |
| ENSMUSG00000044303 | Cdkn2a | 0.001937714 | -0.443334292 | 0.825446518 |
| ENSMUSG00000028763 | Hspg2 | 0.001937714 | -1.111976785 | 0.109430726 |
| ENSMUSG00000028926 | Cdk14 | 0.001937714 | -0.832291138 | 0.330090423 |
| ENSMUSG00000063531 | Sema3e | 0.001937714 | -1.452702475 | 0.021072216 |
| ENSMUSG00000029110 | Rnf4 | 0.001937714 | -1.106051951 | 0.114102131 |
| ENSMUSG00000029153 | Ociad2 | 0.001937714 | 1.024344687 | 0.119907873 |
| ENSMUSG00000048578 | Mlec | 0.001937714 | -0.825095637 | 0.354971784 |
| ENSMUSG00000019054 | Fis1 | 0.001937714 | 0.461208548 | 0.687112677 |
| ENSMUSG00000029655 | N4bp2l2 | 0.001937714 | -1.100102684 | 0.119351941 |
| ENSMUSG00000029629 | Phf14 | 0.001937714 | -1.663038879 | 0.00657428 |
| ENSMUSG00000015733 | Capza2 | 0.001937714 | 0.570833039 | 0.530469833 |
| ENSMUSG00000030697 | Ppp4c | 0.001937714 | -0.885143021 | 0.271138872 |
| ENSMUSG00000058325 | Dock1 | 0.001937714 | -1.383363271 | 0.030770816 |
| ENSMUSG00000031557 | Plekha2 | 0.001937714 | -1.85719648 | 0.002194569 |
| ENSMUSG00000031585 | Gtf2e2 | 0.001937714 | -1.114930102 | 0.110207286 |
| ENSMUSG00000036990 | Otud4 | 0.001937714 | -0.9057533 | 0.253672958 |
| ENSMUSG00000055835 | Zfp1 | 0.001937714 | -1.167081384 | 0.088605926 |
| ENSMUSG00000032228 | Tcf12 | 0.001937714 | -1.784865792 | 0.003038683 |
| ENSMUSG00000044976 | Wdr72 | 0.001937714 | -0.919331707 | 0.244907767 |
| ENSMUSG00000019920 | Lims1 | 0.001937714 | -1.724271267 | 0.004497171 |
| ENSMUSG00000019947 | Arid5b | 0.001937714 | -1.855431715 | 0.002104835 |
| ENSMUSG00000017291 | Taok1 | 0.001937714 | -1.642718894 | 0.007428355 |
| ENSMUSG00000000631 | Myo18a | 0.001937714 | -1.648844997 | 0.006823284 |
| ENSMUSG00000038366 | Lasp1 | 0.001937714 | 0.011405631 | 1 |
| ENSMUSG00000019132 | BC005537 | 0.001937714 | -1.405241121 | 0.026412566 |
| ENSMUSG00000021553 | Slc28a3 | 0.001937714 | -0.922706434 | 0.246055697 |
| ENSMUSG00000074867 | Zfp808 | 0.001937714 | -2.702839887 | 4.64E-06 |
| ENSMUSG00000042323 | Pbrm1 | 0.001937714 | -1.584251471 | 0.010114343 |
| ENSMUSG00000033004 | Mycbp2 | 0.001937714 | -2.725237897 | 3.96E-06 |
| ENSMUSG00000055737 | Ghr | 0.001937714 | -1.741572281 | 0.003968764 |
| ENSMUSG00000037487 | Ubr5 | 0.001937714 | -1.988594369 | 0.00087662 |
| ENSMUSG00000022336 | Eif3e | 0.001937714 | 0.050576228 | 1 |
| ENSMUSG00000016664 | Pacsin2 | 0.001937714 | -1.263388488 | 0.056547843 |
| ENSMUSG00000008658 | Rbfox1 | 0.001937714 | -0.35616513 | 0.961379127 |
| ENSMUSG00000022781 | Pak2 | 0.001937714 | -1.209145444 | 0.074103722 |
| ENSMUSG00000024006 | Stk38 | 0.001937714 | -2.292375117 | 0.000105658 |
| ENSMUSG00000071172 | Srsf3 | 0.001937714 | -1.088130043 | 0.123403868 |
| ENSMUSG00000024073 | Birc6 | 0.001937714 | -1.756778694 | 0.003603181 |
| ENSMUSG00000035473 | Galm | 0.001937714 | -0.135435758 | 1 |
| ENSMUSG00000024750 | Zfand5 | 0.001937714 | -0.30018052 | 1 |
| ENSMUSG00000016319 | Slc25a5 | 0.001937714 | 2.42644313 | 1.74E-05 |
| ENSMUSG00000025860 | Xiap | 0.001937714 | -1.114930102 | 0.110207286 |
| ENSMUSG00000033813 | Tcea1 | 0.001882351 | -0.52566218 | 0.699902136 |
| ENSMUSG00000025920 | Stau2 | 0.001882351 | -1.656509659 | 0.006672082 |
| ENSMUSG00000009418 | Nav1 | 0.001882351 | -1.705697209 | 0.005091765 |
| ENSMUSG00000026688 | Mgst3 | 0.001882351 | 2.193156068 | 0.000111427 |
| ENSMUSG00000096929 | A330023F24Rik | 0.001882351 | -1.073630473 | 0.140206858 |
| ENSMUSG00000026839 | Upp2 | 0.001882351 | -2.048429919 | 0.00061775 |
| ENSMUSG00000074582 | Arfgef2 | 0.001882351 | -0.993370901 | 0.181903548 |
| ENSMUSG00000033400 | Agl | 0.001882351 | -1.230310054 | 0.068439514 |

| | | | | |
|---|---|---|---|---|
| ENSMUSG00000033386 | Frrs1 | 0.001882351 | -1.658592974 | 0.006756729 |
| ENSMUSG00000049122 | Frmd3 | 0.001882351 | -1.741517684 | 0.004064718 |
| ENSMUSG00000053510 | Nrd1 | 0.001882351 | -2.52566218 | 1.87E-05 |
| ENSMUSG00000063524 | Eno1 | 0.001882351 | 1.119907086 | 0.073372104 |
| ENSMUSG00000038342 | Mlxip | 0.001882351 | -2.008151059 | 0.000812252 |
| ENSMUSG00000029390 | Tmed2 | 0.001882351 | 0.983702702 | 0.138139802 |
| ENSMUSG00000029430 | Ran | 0.001882351 | 0.755750755 | 0.341912702 |
| ENSMUSG00000066621 | Tecpr1 | 0.001882351 | -1.119726888 | 0.117170911 |
| ENSMUSG00000029772 | Ahcyl2 | 0.001882351 | -0.579433436 | 0.633493255 |
| ENSMUSG00000038784 | Cnot4 | 0.001882351 | -1.518797638 | 0.01485218 |
| ENSMUSG00000030180 | Kdm5a | 0.001882351 | -1.666896277 | 0.006654031 |
| ENSMUSG00000003452 | Bicd1 | 0.001882351 | -0.976768934 | 0.19917628 |
| ENSMUSG00000053565 | Eif3k | 0.001882351 | 0.970763646 | 0.141839788 |
| ENSMUSG00000030417 | Pdcd5 | 0.001882351 | 2.163408724 | 0.000139104 |
| ENSMUSG00000040212 | Emp3 | 0.001882351 | 1.871227973 | 0.002074662 |
| ENSMUSG00000004508 | Gab2 | 0.001882351 | -1.032453066 | 0.158846729 |
| ENSMUSG00000031642 | Sh3rf1 | 0.001882351 | -2.766845864 | 3.54E-06 |
| ENSMUSG00000003812 | Dnase2a | 0.001882351 | -0.818071188 | 0.364159489 |
| ENSMUSG00000031731 | Ap1g1 | 0.001882351 | -1.164395937 | 0.093228366 |
| ENSMUSG00000032038 | St3gal4 | 0.001882351 | -4.941270253 | 7.01E-17 |
| ENSMUSG00000032366 | Tpm1 | 0.001882351 | -0.963348418 | 0.20542113 |
| ENSMUSG00000111709 | Gm3776 | 0.001882351 | 4.31868695 | 2.25E-12 |
| ENSMUSG00000003226 | Ranbp2 | 0.001882351 | -1.78793547 | 0.003136686 |
| ENSMUSG00000052613 | Pcdh15 | 0.001882351 | -1.057931787 | 0.154954309 |
| ENSMUSG00000020265 | Sumo3 | 0.001882351 | -0.41170599 | 0.868661374 |
| ENSMUSG00000020358 | Hnrnpab | 0.001882351 | 0.084631611 | 1 |
| ENSMUSG00000037926 | Ssh2 | 0.001882351 | -1.951502175 | 0.001267248 |
| ENSMUSG00000014195 | Dnajc7 | 0.001882351 | -0.936126949 | 0.231737538 |
| ENSMUSG00000036333 | Kidins220 | 0.001882351 | -2.330405888 | 8.38E-05 |
| ENSMUSG00000020644 | Id2 | 0.001882351 | 1.148761948 | 0.06813757 |
| ENSMUSG00000063450 | Syne2 | 0.001882351 | -2.54268198 | 1.67E-05 |
| ENSMUSG00000021482 | Prxl2c | 0.001882351 | -1.555036782 | 0.0133372 |
| ENSMUSG00000021944 | Gata4 | 0.001882351 | -0.904876015 | 0.262029351 |
| ENSMUSG00000022329 | Stk3 | 0.001882351 | -1.546061978 | 0.013235138 |
| ENSMUSG00000036158 | Prickle1 | 0.001882351 | -1.41170599 | 0.027328142 |
| ENSMUSG00000022855 | Senp2 | 0.001882351 | -1.703680863 | 0.005645141 |
| ENSMUSG00000003955 | Fam162a | 0.001882351 | 1.134262379 | 0.070717344 |
| ENSMUSG00000023800 | Tiam2 | 0.001882351 | -2.062659355 | 0.000548215 |
| ENSMUSG00000024091 | Vapa | 0.001882351 | -0.244249245 | 1 |
| ENSMUSG00000043384 | Gprasp1 | 0.001882351 | -1.032453066 | 0.155002026 |
| ENSMUSG00000031438 | Rnf128 | 0.001882351 | -0.249787428 | 1 |
| ENSMUSG00000026153 | Fam135a | 0.001826988 | -1.670969372 | 0.00638313 |
| ENSMUSG00000061518 | Cox5b | 0.001826988 | 1.853655343 | 0.001942582 |
| ENSMUSG00000036815 | Dpp10 | 0.001826988 | -0.458458462 | 0.815591899 |
| ENSMUSG00000026434 | Nucks1 | 0.001826988 | -1.149397981 | 0.100676805 |
| ENSMUSG00000026701 | Prdx6 | 0.001826988 | 1.035858675 | 0.114102131 |
| ENSMUSG00000052533 | Nup188 | 0.001826988 | -1.031885932 | 0.160065028 |
| ENSMUSG00000027189 | Trim44 | 0.001826988 | -2.115450649 | 0.00040506 |
| ENSMUSG00000086141 | 9030622O22Rik | 0.001826988 | -2.27911666 | 0.000124429 |
| ENSMUSG00000053141 | Ptprt | 0.001826988 | -2.703532491 | 7.54E-06 |
| ENSMUSG00000034544 | Rsrc1 | 0.001826988 | -1.562909626 | 0.012197062 |

| | | | | |
|---|---|---|---|---|
| ENSMUSG00000053819 | Camk2d | 0.001826988 | -1.874726405 | 0.001888585 |
| ENSMUSG00000038764 | Ptpn3 | 0.001826988 | -1.272130229 | 0.05514173 |
| ENSMUSG00000041120 | Nbl1 | 0.001826988 | 1.106941773 | 0.082076736 |
| ENSMUSG00000003161 | Sri | 0.001826988 | 0.793783887 | 0.305378889 |
| ENSMUSG00000097726 | 9530036O11Rik | 0.001826988 | -0.038488667 | 1 |
| ENSMUSG00000029394 | Cdk2ap1 | 0.001826988 | -0.212068839 | 1 |
| ENSMUSG00000040731 | Eif4h | 0.001826988 | -0.121682602 | 1 |
| ENSMUSG00000016128 | Stard13 | 0.001826988 | -1.220803649 | 0.06973296 |
| ENSMUSG00000030126 | Tmcc1 | 0.001826988 | -2.262292037 | 0.000139738 |
| ENSMUSG00000042079 | Hnrnpf | 0.001826988 | -0.217897918 | 1 |
| ENSMUSG00000025790 | Slco3a1 | 0.001826988 | -2.386529602 | 6.14E-05 |
| ENSMUSG00000062991 | Nrg1 | 0.001826988 | -0.630023822 | 0.589669066 |
| ENSMUSG00000031812 | Map1lc3b | 0.001826988 | 0.968114068 | 0.166514237 |
| ENSMUSG00000049577 | Zfpm1 | 0.001826988 | -1.636566667 | 0.008184643 |
| ENSMUSG00000037287 | Tbcel | 0.001826988 | -1.035191077 | 0.161655655 |
| ENSMUSG00000032418 | Me1 | 0.001826988 | -0.651718893 | 0.533955593 |
| ENSMUSG00000032356 | Rasgrf1 | 0.001826988 | -1.410906532 | 0.027328142 |
| ENSMUSG00000038774 | Ascc3 | 0.001826988 | -1.614640789 | 0.010795678 |
| ENSMUSG00000058589 | Anks1b | 0.001826988 | 0.128968079 | 1 |
| ENSMUSG00000020716 | Nf1 | 0.001826988 | -1.487703282 | 0.018532615 |
| ENSMUSG00000000275 | Trim25 | 0.001826988 | -1.182564845 | 0.088250241 |
| ENSMUSG00000020883 | Fbxl20 | 0.001826988 | -0.743481871 | 0.441676749 |
| ENSMUSG00000017466 | Timp2 | 0.001826988 | 0.293354897 | 0.913408058 |
| ENSMUSG00000020640 | Itsn2 | 0.001826988 | -1.953699579 | 0.001209043 |
| ENSMUSG00000035021 | Baz1a | 0.001826988 | -2.616848433 | 1.02E-05 |
| ENSMUSG00000033102 | Cdc14b | 0.001826988 | -0.833557216 | 0.339660071 |
| ENSMUSG00000021709 | Erbin | 0.001826988 | -1.313550157 | 0.046054041 |
| ENSMUSG00000022311 | Csmd3 | 0.001826988 | 0.301670165 | 0.92947099 |
| ENSMUSG00000022403 | St13 | 0.001826988 | -0.351164449 | 0.954530998 |
| ENSMUSG00000022400 | Rbx1 | 0.001826988 | 1.212736437 | 0.051203786 |
| ENSMUSG00000045983 | Eif4g1 | 0.001826988 | -1.551413987 | 0.013154731 |
| ENSMUSG00000022884 | Eif4a2 | 0.001826988 | -2.013571035 | 0.000845808 |
| ENSMUSG00000025613 | Cct8 | 0.001826988 | -0.258055044 | 1 |
| ENSMUSG00000022983 | Scaf4 | 0.001826988 | -1.839240854 | 0.002435902 |
| ENSMUSG00000046201 | Scaf8 | 0.001826988 | -1.747517385 | 0.004083284 |
| ENSMUSG00000042644 | Itpr3 | 0.001826988 | -1.343186391 | 0.039583356 |
| ENSMUSG00000024456 | Diaph1 | 0.001826988 | -0.567482355 | 0.66416768 |
| ENSMUSG00000024597 | Slc12a2 | 0.001826988 | -1.914753298 | 0.001557955 |
| ENSMUSG00000024516 | Sec11c | 0.001826988 | 1.260564962 | 0.036215483 |
| ENSMUSG00000054453 | Sytl5 | 0.001826988 | -1.953699579 | 0.001209043 |
| ENSMUSG00000046873 | Mbtps2 | 0.001826988 | -1.395558568 | 0.029059832 |
| ENSMUSG00000025907 | Rb1cc1 | 0.001771625 | -1.818522983 | 0.002869506 |
| ENSMUSG00000028354 | Fmn2 | 0.001771625 | -1.818522983 | 0.003521444 |
| ENSMUSG00000037499 | Nenf | 0.001771625 | 1.063873051 | 0.116992527 |
| ENSMUSG00000036792 | Mbd5 | 0.001771625 | -1.934304214 | 0.001356788 |
| ENSMUSG00000063714 | Sp3os | 0.001771625 | -0.982727362 | 0.200147186 |
| ENSMUSG00000060802 | B2m | 0.001771625 | 1.250286175 | 0.041296798 |
| ENSMUSG00000074656 | Eif2s2 | 0.001771625 | -0.737222881 | 0.458119266 |
| ENSMUSG00000018326 | Ywhab | 0.001771625 | 0.435841828 | 0.721281222 |
| ENSMUSG00000105265 | Sox2ot | 0.001771625 | 0.445302157 | 0.703048872 |
| ENSMUSG00000027864 | Ptgfrn | 0.001771625 | 0.138640819 | 1 |

| | | | | |
|---|---|---|---|---|
| ENSMUSG00000055761 | Nkain3 | 0.001771625 | -1.417905999 | 0.032072771 |
| ENSMUSG00000078713 | Tomm5 | 0.001771625 | 1.563946654 | 0.00864105 |
| ENSMUSG00000066036 | Ubr4 | 0.001771625 | -1.668714103 | 0.008307884 |
| ENSMUSG00000058183 | Mmel1 | 0.001771625 | -0.624182943 | 0.621016316 |
| ENSMUSG00000061535 | C1qtnf7 | 0.001771625 | -0.094672889 | 1 |
| ENSMUSG00000034573 | Ptpn13 | 0.001771625 | -1.741398557 | 0.004317489 |
| ENSMUSG00000029616 | Erp29 | 0.001771625 | 0.013749571 | 1 |
| ENSMUSG00000029447 | Cct6a | 0.001771625 | -0.711911792 | 0.466464457 |
| ENSMUSG00000009376 | Met | 0.001771625 | -0.814564969 | 0.36612346 |
| ENSMUSG00000106874 | Gm20186 | 0.001771625 | -0.101186196 | 1 |
| ENSMUSG00000029763 | Exoc4 | 0.001771625 | -1.901094933 | 0.001737647 |
| ENSMUSG00000029822 | Osbpl3 | 0.001771625 | -2.139660343 | 0.000355547 |
| ENSMUSG00000051695 | Pcbp1 | 0.001771625 | 0.978984153 | 0.156554801 |
| ENSMUSG00000030120 | Mlf2 | 0.001771625 | 1.078519827 | 0.094849255 |
| ENSMUSG00000048078 | Tenm4 | 0.001771625 | -0.910396674 | 0.265312568 |
| ENSMUSG00000031563 | Wwc2 | 0.001771625 | -1.714038093 | 0.005140767 |
| ENSMUSG00000031608 | Galnt7 | 0.001771625 | -1.696939285 | 0.005670902 |
| ENSMUSG00000004319 | Clcn3 | 0.001771625 | -1.174049069 | 0.088605926 |
| ENSMUSG00000032187 | Smarca4 | 0.001771625 | -0.972106246 | 0.20807227 |
| ENSMUSG00000059495 | Arhgef12 | 0.001771625 | -1.174049069 | 0.088605926 |
| ENSMUSG00000034135 | Sik3 | 0.001771625 | -1.323578617 | 0.04310517 |
| ENSMUSG00000003072 | Atp5d | 0.001771625 | 1.483776305 | 0.018228581 |
| ENSMUSG00000036112 | Metap2 | 0.001771625 | -1.003737855 | 0.183490997 |
| ENSMUSG00000020817 | Rabep1 | 0.001771625 | -1.240460984 | 0.067856911 |
| ENSMUSG00000020717 | Pecam1 | 0.001771625 | -0.711911792 | 0.497214025 |
| ENSMUSG00000066643 | Wdr35 | 0.001771625 | -2.005474996 | 0.001052647 |
| ENSMUSG00000035247 | Hectd1 | 0.001771625 | -1.716161265 | 0.005197445 |
| ENSMUSG00000021112 | Mpp5 | 0.001771625 | -1.893610117 | 0.001752947 |
| ENSMUSG00000033713 | Foxn3 | 0.001771625 | -1.081557389 | 0.130876638 |
| ENSMUSG00000007411 | Mark3 | 0.001771625 | -1.694787615 | 0.005982282 |
| ENSMUSG00000039219 | Arid4b | 0.001771625 | -2.717221681 | 4.72E-06 |
| ENSMUSG00000021635 | Rad17 | 0.001771625 | -1.407432668 | 0.02878527 |
| ENSMUSG00000095280 | Gm21738 | 0.001771625 | 0.493557326 | 0.66416768 |
| ENSMUSG00000021807 | Rtraf | 0.001771625 | 0.62714048 | 0.466464457 |
| ENSMUSG00000021767 | Kat6b | 0.001771625 | -1.603686536 | 0.009638741 |
| ENSMUSG00000042682 | Selenok | 0.001771625 | 1.10826717 | 0.102317499 |
| ENSMUSG00000037580 | Gch1 | 0.001771625 | 0.553678318 | 0.579226907 |
| ENSMUSG00000115432 | D130009I18Rik | 0.001771625 | -0.989764913 | 0.21778593 |
| ENSMUSG00000042961 | Egflam | 0.001771625 | -0.95782202 | 0.219707865 |
| ENSMUSG00000005125 | Ndrg1 | 0.001771625 | 0.60576683 | 0.505435723 |
| ENSMUSG00000055762 | Eef1d | 0.001771625 | -0.456517449 | 0.812026069 |
| ENSMUSG00000033565 | Rbfox2 | 0.001771625 | -2.781490276 | 2.96E-06 |
| ENSMUSG00000000420 | Galnt1 | 0.001771625 | -1.516223695 | 0.015927 |
| ENSMUSG00000060534 | Dcc | 0.001771625 | -0.501527184 | 0.771834758 |
| ENSMUSG00000052942 | Glis3 | 0.001771625 | -2.874726405 | 1.26E-06 |
| ENSMUSG00000042712 | Tceal9 | 0.001771625 | 0.911869957 | 0.231101892 |
| ENSMUSG00000031299 | Pdha1 | 0.001771625 | -0.306770329 | 1 |
| ENSMUSG00000038235 | F11r | 0.001716261 | -1.715299104 | 0.005435019 |
| ENSMUSG00000015092 | Edf1 | 0.001716261 | 1.694639241 | 0.004168617 |
| ENSMUSG00000026917 | Wdr5 | 0.001716261 | -0.909585162 | 0.262284476 |
| ENSMUSG00000044627 | Swi5 | 0.001716261 | 0.815945537 | 0.29714275 |

| | | | | |
|---|---|---|---|---|
| ENSMUSG00000026872 | Zeb2 | 0.001716261 | -1.051604167 | 0.155026807 |
| ENSMUSG00000026987 | Baz2b | 0.001716261 | -2.027121482 | 0.00081463 |
| ENSMUSG00000027111 | Itga6 | 0.001716261 | -1.339849135 | 0.040750047 |
| ENSMUSG00000027104 | Atf2 | 0.001716261 | -1.398288361 | 0.031559235 |
| ENSMUSG00000008200 | Fnbp4 | 0.001716261 | -1.909585162 | 0.001700368 |
| ENSMUSG00000017707 | Serinc3 | 0.001716261 | -0.89032326 | 0.28748987 |
| ENSMUSG00000044080 | S100a1 | 0.001716261 | 1.363433333 | 0.049126005 |
| ENSMUSG00000038712 | Mindy1 | 0.001716261 | -1.668577062 | 0.007061791 |
| ENSMUSG00000028580 | Pum1 | 0.001716261 | -2.004009422 | 0.000930142 |
| ENSMUSG00000028854 | Slc9a1 | 0.001716261 | -1.348343929 | 0.039339084 |
| ENSMUSG00000046447 | Camk2n1 | 0.001716261 | 0.828656589 | 0.261022536 |
| ENSMUSG00000028964 | Park7 | 0.001716261 | 2.034125707 | 0.000708492 |
| ENSMUSG00000028883 | Sema3a | 0.001716261 | -0.245982212 | 1 |
| ENSMUSG00000028945 | Rheb | 0.001716261 | -0.677593936 | 0.500515172 |
| ENSMUSG00000029104 | Htt | 0.001716261 | -0.822839663 | 0.346568875 |
| ENSMUSG00000018076 | Med13l | 0.001716261 | -2.009375931 | 0.000909854 |
| ENSMUSG00000029465 | Arpc3 | 0.001716261 | 1.034125707 | 0.133437803 |
| ENSMUSG00000051391 | Ywhag | 0.001716261 | -1.119348773 | 0.12034531 |
| ENSMUSG00000038780 | Smurf1 | 0.001716261 | -1.622627476 | 0.009432168 |
| ENSMUSG00000001424 | Snd1 | 0.001716261 | -2.249009905 | 0.000197224 |
| ENSMUSG00000033420 | Antxr1 | 0.001716261 | -1.290743663 | 0.054350902 |
| ENSMUSG00000041560 | Nop53 | 0.001716261 | 1.156982455 | 0.070555673 |
| ENSMUSG00000066306 | Numa1 | 0.001716261 | -1.704311274 | 0.005847048 |
| ENSMUSG00000031555 | Adam9 | 0.001716261 | -0.511035785 | 0.744828569 |
| ENSMUSG00000031902 | Nfatc3 | 0.001716261 | -1.739181734 | 0.004732823 |
| ENSMUSG00000095547 | Gm10719 | 0.001716261 | 1.648835551 | 0.006113379 |
| ENSMUSG00000062257 | Opcml | 0.001716261 | 0.519552534 | 0.641107521 |
| ENSMUSG00000032076 | Cadm1 | 0.001716261 | -1.978698333 | 0.001196704 |
| ENSMUSG00000038412 | Higd1a | 0.001716261 | 1.867475838 | 0.001408958 |
| ENSMUSG00000020154 | Ptprb | 0.001716261 | -0.659503479 | 0.547533991 |
| ENSMUSG00000037533 | Rapgef6 | 0.001716261 | -2.224608561 | 0.00021489 |
| ENSMUSG00000010277 | 2610507B11Rik | 0.001716261 | -0.909585162 | 0.262075002 |
| ENSMUSG00000025583 | Rptor | 0.001716261 | -1.594338432 | 0.011044864 |
| ENSMUSG00000036095 | Dgkb | 0.001716261 | -1.249009905 | 0.077471852 |
| ENSMUSG00000048118 | Arid4a | 0.001716261 | -1.996822881 | 0.00098929 |
| ENSMUSG00000021279 | Cdc42bpb | 0.001716261 | -1.985975499 | 0.001024813 |
| ENSMUSG00000039197 | Adk | 0.001716261 | -1.985975499 | 0.001024813 |
| ENSMUSG00000061731 | Ext1 | 0.001716261 | -2.247496852 | 0.000173266 |
| ENSMUSG00000034243 | Golgb1 | 0.001716261 | -2.233807498 | 0.00019453 |
| ENSMUSG00000064115 | Cadm2 | 0.001716261 | -0.464906615 | 0.810591941 |
| ENSMUSG00000022956 | Atp5o | 0.001716261 | 0.8033455 | 0.31726506 |
| ENSMUSG00000054640 | Slc8a1 | 0.001716261 | -1.233807498 | 0.076640437 |
| ENSMUSG00000024500 | Ppp2r2b | 0.001716261 | -1.381831585 | 0.033909435 |
| ENSMUSG00000071856 | Mcc | 0.001716261 | -1.675344995 | 0.006838229 |
| ENSMUSG00000024639 | Gnaq | 0.001716261 | -2.38458745 | 6.48E-05 |
| ENSMUSG00000026670 | Uap1 | 0.001660898 | -1.291785901 | 0.05392641 |
| ENSMUSG00000026643 | Nmt2 | 0.001660898 | -1.618555259 | 0.010082056 |
| ENSMUSG00000000194 | Gpr107 | 0.001660898 | -2.056964031 | 0.000692658 |
| ENSMUSG00000040549 | Ckap5 | 0.001660898 | -1.596640483 | 0.011081818 |
| ENSMUSG00000050587 | Lrrc4c | 0.001660898 | -0.138435125 | 1 |
| ENSMUSG00000074578 | Zfas1 | 0.001660898 | 1.281103767 | 0.044759969 |

| | | | | |
|---|---|---|---|---|
| ENSMUSG00000027852 | Nras | 0.001660898 | -0.055179621 | 1 |
| ENSMUSG00000032826 | Ank2 | 0.001660898 | 0.025603034 | 1 |
| ENSMUSG00000040270 | Bach2 | 0.001660898 | -0.682370357 | 0.506030384 |
| ENSMUSG00000037583 | Nr0b2 | 0.001660898 | 0.143600243 | 1 |
| ENSMUSG00000028967 | Errfi1 | 0.001660898 | -0.839911634 | 0.364834983 |
| ENSMUSG00000055302 | Mrfap1 | 0.001660898 | 1.374213171 | 0.026976566 |
| ENSMUSG00000042605 | Atxn2 | 0.001660898 | -1.647266556 | 0.008203873 |
| ENSMUSG00000002748 | Baz1b | 0.001660898 | -2.773093527 | 3.54E-06 |
| ENSMUSG00000004285 | Atp6v1f | 0.001660898 | 1.393578496 | 0.025329352 |
| ENSMUSG00000004535 | Tax1bp1 | 0.001660898 | -0.441362258 | 0.846566779 |
| ENSMUSG00000053119 | Chmp3 | 0.001660898 | -0.56689314 | 0.66416768 |
| ENSMUSG00000030062 | Rpn1 | 0.001660898 | 1.04844301 | 0.118021173 |
| ENSMUSG00000064293 | Cntn4 | 0.001660898 | 0.056137402 | 1 |
| ENSMUSG00000015766 | Eps8 | 0.001660898 | -1.581843481 | 0.01181598 |
| ENSMUSG00000030287 | Itpr2 | 0.001660898 | -1.239318482 | 0.069702155 |
| ENSMUSG00000030403 | Vasp | 0.001660898 | -0.273485085 | 1 |
| ENSMUSG00000030516 | Tjp1 | 0.001660898 | -1.124888593 | 0.114587779 |
| ENSMUSG00000060591 | Ifitm2 | 0.001660898 | 1.625751938 | 0.006459516 |
| ENSMUSG00000019428 | Fkbp8 | 0.001660898 | 0.84814436 | 0.267492155 |
| ENSMUSG00000031807 | Pgls | 0.001660898 | -0.291785901 | 1 |
| ENSMUSG00000031729 | Ist1 | 0.001660898 | -1.327705476 | 0.046003064 |
| ENSMUSG00000033335 | Dnm2 | 0.001660898 | -2.079962722 | 0.000597711 |
| ENSMUSG00000019804 | Snx3 | 0.001660898 | -0.184870698 | 1 |
| ENSMUSG00000036955 | Kif1bp | 0.001660898 | -1.251836521 | 0.067360743 |
| ENSMUSG00000020151 | Ptprr | 0.001660898 | -0.223516625 | 1 |
| ENSMUSG00000018849 | Wwc1 | 0.001660898 | -2.138435125 | 0.00039298 |
| ENSMUSG00000049336 | Tenm2 | 0.001660898 | 0.033176253 | 1 |
| ENSMUSG00000035152 | Ap2b1 | 0.001660898 | -0.515909211 | 0.716109464 |
| ENSMUSG00000020483 | Dynll2 | 0.001660898 | 0.888786344 | 0.250869635 |
| ENSMUSG00000018378 | Cuedc1 | 0.001660898 | -1.058746236 | 0.155026807 |
| ENSMUSG00000058070 | Eml1 | 0.001660898 | -2.51461124 | 2.46E-05 |
| ENSMUSG00000021266 | Wars | 0.001660898 | -0.611287259 | 0.618469788 |
| ENSMUSG00000021277 | Traf3 | 0.001660898 | -1.924445144 | 0.00161209 |
| ENSMUSG00000021556 | Golm1 | 0.001660898 | -0.47911718 | 0.772933445 |
| ENSMUSG00000068758 | Il3ra | 0.001660898 | -1.569395647 | 0.012849063 |
| ENSMUSG00000056234 | Ncoa4 | 0.001660898 | 0.708214099 | 0.408564937 |
| ENSMUSG00000021843 | Ktn1 | 0.001660898 | -2.4221077 | 5.09E-05 |
| ENSMUSG00000005148 | Klf5 | 0.001660898 | -1.062304055 | 0.146115505 |
| ENSMUSG00000022141 | Nipbl | 0.001660898 | -2.217147101 | 0.000229249 |
| ENSMUSG00000022353 | Mtss1 | 0.001660898 | -1.682370357 | 0.006613366 |
| ENSMUSG00000022378 | Fam49b | 0.001660898 | -1.01535364 | 0.185053751 |
| ENSMUSG00000009585 | Apobec3 | 0.001660898 | -0.309857474 | 1 |
| ENSMUSG00000024208 | Uqcc2 | 0.001660898 | 1.959175672 | 0.002861483 |
| ENSMUSG00000056515 | Rab31 | 0.001660898 | -0.52108945 | 0.717944159 |
| ENSMUSG00000024095 | Hnrnpll | 0.001660898 | -0.963022176 | 0.217655393 |
| ENSMUSG00000045215 | Asxl3 | 0.001660898 | -1.135060398 | 0.121170765 |
| ENSMUSG00000024483 | Ankhd1 | 0.001660898 | -1.700748886 | 0.006004198 |
| ENSMUSG00000040929 | Rfx3 | 0.001660898 | -2.09219643 | 0.000716101 |
| ENSMUSG00000055044 | Pdlim1 | 0.001660898 | 0.326907457 | 0.874962283 |
| ENSMUSG00000025066 | Sfr1 | 0.001660898 | 0.202152426 | 1 |
| ENSMUSG00000031357 | Syap1 | 0.001660898 | -0.99688098 | 0.193977729 |

| | | | | |
|---|---|---|---|---|
| ENSMUSG00000066842 | Hmcn1 | 0.001605535 | -1.372051815 | 0.070297939 |
| ENSMUSG00000062963 | Ufc1 | 0.001605535 | 1.44720169 | 0.018507952 |
| ENSMUSG00000026727 | Rsu1 | 0.001605535 | -1.175592884 | 0.093356919 |
| ENSMUSG00000027624 | Epb41l1 | 0.001605535 | -0.983432664 | 0.20428579 |
| ENSMUSG00000027566 | Psma7 | 0.001605535 | -0.444273853 | 0.834451374 |
| ENSMUSG00000015647 | Lama5 | 0.001605535 | -2.212458178 | 0.000254311 |
| ENSMUSG00000063273 | Naa15 | 0.001605535 | -2.39992376 | 6.34E-05 |
| ENSMUSG00000027827 | Kcnab1 | 0.001605535 | -1.080898473 | 0.173990867 |
| ENSMUSG00000040998 | Npnt | 0.001605535 | -0.793002007 | 0.394449161 |
| ENSMUSG00000028410 | Dnaja1 | 0.001605535 | 0.511332028 | 0.63765603 |
| ENSMUSG00000090053 | Pakap | 0.001605535 | -1.601620788 | 0.014369421 |
| ENSMUSG00000028656 | Cap1 | 0.001605535 | 0.500443712 | 0.663838662 |
| ENSMUSG00000020220 | Vps13d | 0.001605535 | -1.515736216 | 0.018125621 |
| ENSMUSG00000029014 | Dnajc2 | 0.001605535 | 0.722836133 | 0.385094015 |
| ENSMUSG00000038803 | Ost4 | 0.001605535 | 1.883300805 | 0.001641385 |
| ENSMUSG00000037235 | Mxd4 | 0.001605535 | -0.36313709 | 0.947393387 |
| ENSMUSG00000029470 | P2rx4 | 0.001605535 | -1.588932096 | 0.011662663 |
| ENSMUSG00000005514 | Por | 0.001605535 | -1.148177426 | 0.106668556 |
| ENSMUSG00000055239 | Kcmf1 | 0.001605535 | -1.345140661 | 0.043715437 |
| ENSMUSG00000097048 | 1600020E01Rik | 0.001605535 | -1.202497068 | 0.087253032 |
| ENSMUSG00000004264 | Phb2 | 0.001605535 | 1.215876144 | 0.066709581 |
| ENSMUSG00000006599 | Gtf2h1 | 0.001605535 | -1.444273853 | 0.025925336 |
| ENSMUSG00000005621 | Zfp592 | 0.001605535 | -0.629149196 | 0.577550711 |
| ENSMUSG00000043866 | Taf10 | 0.001605535 | 1.163408724 | 0.106668556 |
| ENSMUSG00000030759 | Far1 | 0.001605535 | -1.251628775 | 0.065872489 |
| ENSMUSG00000031078 | Cttn | 0.001605535 | -0.866338619 | 0.31603319 |
| ENSMUSG00000037148 | Arhgap10 | 0.001605535 | -0.499556288 | 0.757651781 |
| ENSMUSG00000025810 | Nrp1 | 0.001605535 | -1.404275785 | 0.03467347 |
| ENSMUSG00000036781 | Rps27l | 0.001605535 | 0.955813305 | 0.222101622 |
| ENSMUSG00000032212 | Sltm | 0.001605535 | -2.733139927 | 5.18E-06 |
| ENSMUSG00000032369 | Plscr1 | 0.001605535 | 0.589941862 | 0.551643788 |
| ENSMUSG00000020023 | Tmcc3 | 0.001605535 | -1.480447466 | 0.024804914 |
| ENSMUSG00000033066 | Gas7 | 0.001605535 | -0.36313709 | 0.951469479 |
| ENSMUSG00000020903 | Stx8 | 0.001605535 | -1.123753952 | 0.123404777 |
| ENSMUSG00000020532 | Acaca | 0.001605535 | -1.080898473 | 0.144776283 |
| ENSMUSG00000001440 | Kpnb1 | 0.001605535 | -1.258055044 | 0.067114333 |
| ENSMUSG00000020585 | Laptm4a | 0.001605535 | 0.42644313 | 0.745725374 |
| ENSMUSG00000035181 | Heatr5a | 0.001605535 | -1.836591276 | 0.002872939 |
| ENSMUSG00000021103 | Mnat1 | 0.001605535 | -1.680288045 | 0.006936048 |
| ENSMUSG00000021036 | Sptlc2 | 0.001605535 | -1.853664789 | 0.002627996 |
| ENSMUSG00000015671 | Psma2 | 0.001605535 | 1.096294528 | 0.114304227 |
| ENSMUSG00000021500 | Ddx46 | 0.001605535 | -2.275581097 | 0.000157524 |
| ENSMUSG00000021715 | Cwc27 | 0.001605535 | -1.866338619 | 0.002325047 |
| ENSMUSG00000022076 | Klhl1 | 0.001605535 | -0.185740839 | 1 |
| ENSMUSG00000022064 | Pibf1 | 0.001605535 | -1.663442373 | 0.007765102 |
| ENSMUSG00000022280 | Rnf19a | 0.001605535 | -1.155080287 | 0.10147754 |
| ENSMUSG00000051359 | Ncald | 0.001605535 | -2.329970226 | 0.000125764 |
| ENSMUSG00000011960 | Ccnt1 | 0.001605535 | -1.314638573 | 0.04883823 |
| ENSMUSG00000001280 | Sp1 | 0.001605535 | -1.141241379 | 0.112238591 |
| ENSMUSG00000063358 | Mapk1 | 0.001605535 | -1.55279831 | 0.015017644 |
| ENSMUSG00000055540 | Epha6 | 0.001605535 | -0.006516277 | 1 |

| | | | | |
|---|---|---|---|---|
| ENSMUSG00000087590 | Epb41l4aos | 0.001605535 | 0.511332028 | 0.63765603 |
| ENSMUSG00000025027 | Xpnpep1 | 0.001605535 | -1.165373042 | 0.09684834 |
| ENSMUSG00000031431 | Tsc22d3 | 0.001605535 | 0.233798052 | 1 |
| ENSMUSG00000041133 | Smc1a | 0.001605535 | -1.106051951 | 0.126801989 |
| ENSMUSG00000031355 | Arhgap6 | 0.001605535 | -1.757156808 | 0.005156146 |
| ENSMUSG00000041144 | Dnah7b | 0.001550171 | -2.107893298 | 0.000782952 |
| ENSMUSG00000025950 | Idh1 | 0.001550171 | 0.541032262 | 0.603636648 |
| ENSMUSG00000062209 | Erbb4 | 0.001550171 | 0.419353705 | 0.792441218 |
| ENSMUSG00000033257 | Ttll4 | 0.001550171 | -1.846330279 | 0.002869506 |
| ENSMUSG00000026343 | Gpr39 | 0.001550171 | -3.207428971 | 7.46E-08 |
| ENSMUSG00000026393 | Nek7 | 0.001550171 | -0.447379764 | 0.811581337 |
| ENSMUSG00000053286 | Trmt1l | 0.001550171 | -1.606925619 | 0.011271367 |
| ENSMUSG00000026594 | Ralgps2 | 0.001550171 | -1.612124526 | 0.010795678 |
| ENSMUSG00000027012 | Dync1i2 | 0.001550171 | -0.931975306 | 0.252106712 |
| ENSMUSG00000027167 | Elp4 | 0.001550171 | -1.297314475 | 0.058171406 |
| ENSMUSG00000027680 | Fxr1 | 0.001550171 | -0.86387086 | 0.331314809 |
| ENSMUSG00000054199 | Gon4l | 0.001550171 | -1.960789004 | 0.001397255 |
| ENSMUSG00000028136 | Snx27 | 0.001550171 | -0.86387086 | 0.331314809 |
| ENSMUSG00000053931 | Cnn3 | 0.001550171 | -1.412046691 | 0.031754639 |
| ENSMUSG00000028164 | Manba | 0.001550171 | -0.764646481 | 0.451099851 |
| ENSMUSG00000028197 | Col24a1 | 0.001550171 | 0.158088392 | 1 |
| ENSMUSG00000003032 | Klf4 | 0.001550171 | 1.131777115 | 0.124712353 |
| ENSMUSG00000039105 | Atp6v1g1 | 0.001550171 | 1.184888452 | 0.087502216 |
| ENSMUSG00000028804 | Csmd2 | 0.001550171 | 0.212195798 | 1 |
| ENSMUSG00000028790 | Khdrbs1 | 0.001550171 | -1.248070956 | 0.068316354 |
| ENSMUSG00000029505 | Ep400 | 0.001550171 | -1.066830641 | 0.152663609 |
| ENSMUSG00000049327 | Kmt5a | 0.001550171 | 0.09742561 | 1 |
| ENSMUSG00000019179 | Mdh2 | 0.001550171 | 0.952227695 | 0.19917628 |
| ENSMUSG00000029647 | Pan3 | 0.001550171 | -1.861689915 | 0.002585723 |
| ENSMUSG00000029836 | Cbx3 | 0.001550171 | -0.388001217 | 0.889654654 |
| ENSMUSG00000052337 | Immt | 0.001550171 | -1.580646295 | 0.013392012 |
| ENSMUSG00000030007 | Cct7 | 0.001550171 | 1.114499124 | 0.100910102 |
| ENSMUSG00000030082 | Sec61a1 | 0.001550171 | -1.529357066 | 0.02280887 |
| ENSMUSG00000030357 | Fkbp4 | 0.001550171 | 0.37753353 | 0.840804483 |
| ENSMUSG00000030275 | Etnk1 | 0.001550171 | -1.137039643 | 0.119081751 |
| ENSMUSG00000030306 | Tmtc1 | 0.001550171 | -1.126178651 | 0.184013112 |
| ENSMUSG00000040811 | Eml2 | 0.001550171 | -0.948510674 | 0.242159739 |
| ENSMUSG00000015165 | Hnrnpl | 0.001550171 | -0.627609774 | 0.584388918 |
| ENSMUSG00000071064 | Zfp827 | 0.001550171 | -1.577991841 | 0.013182548 |
| ENSMUSG00000039067 | Psmd7 | 0.001550171 | -1.300538375 | 0.055368915 |
| ENSMUSG00000051495 | Irf2bp2 | 0.001550171 | 0.203035798 | 1 |
| ENSMUSG00000032376 | Usp3 | 0.001550171 | -1.787804202 | 0.00403405 |
| ENSMUSG00000033590 | Myo5c | 0.001550171 | -0.632734806 | 0.586600655 |
| ENSMUSG00000034252 | Senp6 | 0.001550171 | -1.794679648 | 0.003944628 |
| ENSMUSG00000032612 | Usp4 | 0.001550171 | -0.227893074 | 1 |
| ENSMUSG00000020219 | Timm13 | 0.001550171 | 0.725456833 | 0.446811748 |
| ENSMUSG00000019888 | Mgat4c | 0.001550171 | -0.158519371 | 1 |
| ENSMUSG00000020121 | Srgap1 | 0.001550171 | -1.670602991 | 0.008075158 |
| ENSMUSG00000078429 | Ctdsp2 | 0.001550171 | 0.277997856 | 0.958555629 |
| ENSMUSG00000020454 | Eif4enif1 | 0.001550171 | -1.794679648 | 0.003944628 |
| ENSMUSG00000001891 | Ugp2 | 0.001550171 | -0.40607279 | 0.871508522 |

| | | | | |
|---|---|---|---|---|
| ENSMUSG00000020354 | Sgcd | 0.001550171 | -0.024564914 | 1 |
| ENSMUSG00000037601 | Nme1 | 0.001550171 | 0.599925951 | 0.532882217 |
| ENSMUSG00000034120 | Srsf2 | 0.001550171 | -2.392540376 | 7.63E-05 |
| ENSMUSG00000056899 | Immp2l | 0.001550171 | -2.341806394 | 0.000107956 |
| ENSMUSG00000021027 | Ralgapa1 | 0.001550171 | -2.006911049 | 0.001061285 |
| ENSMUSG00000021000 | Mia2 | 0.001550171 | -1.409062832 | 0.031456064 |
| ENSMUSG00000021224 | Numb | 0.001550171 | -1.7832023 | 0.00394563 |
| ENSMUSG00000041702 | Btbd7 | 0.001550171 | -2.306964645 | 0.000136945 |
| ENSMUSG00000000078 | Klf6 | 0.001550171 | -0.326073468 | 0.992876904 |
| ENSMUSG00000000957 | Mmp14 | 0.001550171 | -0.98903848 | 0.212848515 |
| ENSMUSG00000058997 | Vwa8 | 0.001550171 | -1.281086013 | 0.060525223 |
| ENSMUSG00000039801 | Cplane1 | 0.001550171 | -1.244727513 | 0.072832619 |
| ENSMUSG00000043556 | Fbxl7 | 0.001550171 | -1.207428971 | 0.097034518 |
| ENSMUSG00000022255 | Mtdh | 0.001550171 | -1.778585672 | 0.004091555 |
| ENSMUSG00000024479 | Mal2 | 0.001550171 | 1.806376828 | 0.00281843 |
| ENSMUSG00000005732 | Ranbp1 | 0.001550171 | 0.76534956 | 0.344671683 |
| ENSMUSG00000068036 | Afdn | 0.001550171 | -2.349608982 | 0.000104671 |
| ENSMUSG00000059208 | Hnrnpm | 0.001550171 | -1.588580476 | 0.012887418 |
| ENSMUSG00000057440 | Mpp7 | 0.001550171 | -0.881200739 | 0.297155035 |
| ENSMUSG00000024620 | Pdgfrb | 0.001550171 | -0.98903848 | 0.219470377 |
| ENSMUSG00000024588 | Fech | 0.001550171 | -2.083770503 | 0.000852612 |
| ENSMUSG00000024845 | Tmem134 | 0.001550171 | 1.484448734 | 0.017880447 |
| ENSMUSG00000024853 | Sf3b2 | 0.001550171 | -1.553879385 | 0.014575482 |
| ENSMUSG00000024940 | Ltbp3 | 0.001550171 | -1.833033456 | 0.0030443 |
| ENSMUSG00000024725 | Ostf1 | 0.001550171 | 0.047384928 | 1 |
| ENSMUSG00000052387 | Trpm3 | 0.001550171 | -0.23465044 | 1 |
| ENSMUSG00000024899 | Papss2 | 0.001550171 | 1.184888452 | 0.070328284 |
| ENSMUSG00000013662 | Atad1 | 0.001550171 | -1.074384035 | 0.144776283 |
| ENSMUSG00000044948 | Cfap43 | 0.001550171 | -1.815111548 | 0.004154208 |
| ENSMUSG00000031103 | Elf4 | 0.001550171 | -1.750571296 | 0.005002256 |
| ENSMUSG00000008136 | Fhl2 | 0.001494808 | -0.503811459 | 0.806496094 |
| ENSMUSG00000048000 | Gigyf2 | 0.001494808 | -1.733634085 | 0.00592361 |
| ENSMUSG00000026309 | Ilkap | 0.001494808 | -1.230310054 | 0.079998381 |
| ENSMUSG00000041757 | Plekha6 | 0.001494808 | -1.838430275 | 0.003170271 |
| ENSMUSG00000041570 | Camsap2 | 0.001494808 | -1.459688905 | 0.024804914 |
| ENSMUSG00000038473 | Nos1ap | 0.001494808 | -0.647097335 | 0.587454349 |
| ENSMUSG00000026604 | Ptpn14 | 0.001494808 | -1.910131741 | 0.001994407 |
| ENSMUSG00000026657 | Frmd4a | 0.001494808 | -0.543809527 | 0.697198109 |
| ENSMUSG00000026915 | Strbp | 0.001494808 | -0.698627636 | 0.496010946 |
| ENSMUSG00000027130 | Slc12a6 | 0.001494808 | -1.370358014 | 0.039998622 |
| ENSMUSG00000027479 | Mapre1 | 0.001494808 | 0.585825754 | 0.58453522 |
| ENSMUSG00000027646 | Src | 0.001494808 | -2.003241145 | 0.00128813 |
| ENSMUSG00000017009 | Sdc4 | 0.001494808 | -0.738566381 | 0.466464457 |
| ENSMUSG00000039671 | Zmynd8 | 0.001494808 | -1.94682155 | 0.001584919 |
| ENSMUSG00000027692 | Tnik | 0.001494808 | -1.420326473 | 0.03461849 |
| ENSMUSG00000027677 | Ttc14 | 0.001494808 | -1.465650464 | 0.0254129 |
| ENSMUSG00000040412 | 5330417C22Rik | 0.001494808 | -0.462672764 | 0.811740881 |
| ENSMUSG00000028121 | Bcar3 | 0.001494808 | -0.944688963 | 0.249952095 |
| ENSMUSG00000028252 | Ccnc | 0.001494808 | -1.0073339 | 0.198290676 |
| ENSMUSG00000028464 | Tpm2 | 0.001494808 | 1.525453136 | 0.017823676 |
| ENSMUSG00000028618 | Tmem59 | 0.001494808 | -0.748380671 | 0.463850347 |

| | | | | |
|---|---|---|---|---|
| ENSMUSG00000040659 | Efhd2 | 0.001494808 | 1.098638469 | 0.113700853 |
| ENSMUSG00000040407 | Akap9 | 0.001494808 | -1.817605126 | 0.003432497 |
| ENSMUSG00000057315 | Arhgap24 | 0.001494808 | -1.879199881 | 0.003064878 |
| ENSMUSG00000033209 | Ttc28 | 0.001494808 | -0.363988489 | 0.955253611 |
| ENSMUSG00000042744 | Hectd4 | 0.001494808 | -1.176804724 | 0.099113432 |
| ENSMUSG00000001687 | Ubl3 | 0.001494808 | -1.051604167 | 0.162298752 |
| ENSMUSG00000056602 | Fry | 0.001494808 | -0.657551693 | 0.57597659 |
| ENSMUSG00000029775 | Klhdc10 | 0.001494808 | -1.338225393 | 0.048635862 |
| ENSMUSG00000039578 | Ccser1 | 0.001494808 | 0.170788255 | 1 |
| ENSMUSG00000073002 | Vamp5 | 0.001494808 | -0.462672764 | 0.796840156 |
| ENSMUSG00000057230 | Aak1 | 0.001494808 | -1.441654983 | 0.0289692 |
| ENSMUSG00000030057 | Cnbp | 0.001494808 | 0.306443354 | 0.948737557 |
| ENSMUSG00000056755 | Grm7 | 0.001494808 | -0.237296484 | 1 |
| ENSMUSG00000035203 | Epn1 | 0.001494808 | 0.029925719 | 1 |
| ENSMUSG00000002981 | Clptm1 | 0.001494808 | -0.486324032 | 0.789622116 |
| ENSMUSG00000063511 | Snrnp70 | 0.001494808 | -2.370358014 | 9.35E-05 |
| ENSMUSG00000005533 | Igf1r | 0.001494808 | -2.174986582 | 0.00036427 |
| ENSMUSG00000047721 | Bola2 | 0.001494808 | 1.134262379 | 0.109815511 |
| ENSMUSG00000086742 | Gm16201 | 0.001494808 | -0.128772027 | 1 |
| ENSMUSG00000025487 | Psmd13 | 0.001494808 | -0.647097335 | 0.57597659 |
| ENSMUSG00000031701 | Dnaja2 | 0.001494808 | -0.209145444 | 1 |
| ENSMUSG00000031930 | Wwp2 | 0.001494808 | -0.888105434 | 0.295355925 |
| ENSMUSG00000053110 | Yap1 | 0.001494808 | -1.194861218 | 0.089980814 |
| ENSMUSG00000059974 | Ntm | 0.001494808 | -1.106051951 | 0.153381132 |
| ENSMUSG00000032036 | Kirrel3 | 0.001494808 | -2.516789313 | 3.04E-05 |
| ENSMUSG00000040111 | Gramd1b | 0.001494808 | -0.285276513 | 1 |
| ENSMUSG00000074415 | Mir100hg | 0.001494808 | -3.210033529 | 8.21E-08 |
| ENSMUSG00000040524 | Zfp609 | 0.001494808 | -1.721229064 | 0.006380589 |
| ENSMUSG00000044791 | Setd2 | 0.001494808 | -1.596377577 | 0.012790012 |
| ENSMUSG00000032525 | Nktr | 0.001494808 | -1.955320515 | 0.001567357 |
| ENSMUSG00000038481 | Cdk19 | 0.001494808 | -1.535331373 | 0.017425186 |
| ENSMUSG00000020159 | Gabrp | 0.001494808 | -1.46862203 | 0.024960134 |
| ENSMUSG00000020349 | Ppp2ca | 0.001494808 | -0.577232528 | 0.66416768 |
| ENSMUSG00000001123 | Lgals9 | 0.001494808 | 1.029925719 | 0.1562152 |
| ENSMUSG00000075595 | Zfp652 | 0.001494808 | -1.147435871 | 0.123797963 |
| ENSMUSG00000069744 | Psmb3 | 0.001494808 | 1.368727632 | 0.046387048 |
| ENSMUSG00000008859 | Rala | 0.001494808 | -0.641841609 | 0.56360526 |
| ENSMUSG00000085936 | 2610307P16Rik | 0.001494808 | -2.889214773 | 1.62E-06 |
| ENSMUSG00000006191 | Cdkal1 | 0.001494808 | -1.549434076 | 0.015997866 |
| ENSMUSG00000021474 | Sfxn1 | 0.001494808 | -0.209145444 | 1 |
| ENSMUSG00000007836 | Hnrnpa0 | 0.001494808 | 0.598209478 | 0.559918909 |
| ENSMUSG00000044258 | Ctla2a | 0.001494808 | -0.292002376 | 1 |
| ENSMUSG00000022234 | Cct5 | 0.001494808 | 0.116340471 | 1 |
| ENSMUSG00000022295 | Atp6v1c1 | 0.001494808 | -0.15113984 | 1 |
| ENSMUSG00000022681 | Ntan1 | 0.001494808 | -1.230310054 | 0.082935667 |
| ENSMUSG00000022865 | Cxadr | 0.001494808 | -1.223289626 | 0.083958642 |
| ENSMUSG00000022961 | Son | 0.001494808 | -2.798838096 | 3.54E-06 |
| ENSMUSG00000022957 | Itsn1 | 0.001494808 | -1.953200463 | 0.001550379 |
| ENSMUSG00000038762 | Abcf1 | 0.001494808 | -1.636566667 | 0.009900358 |
| ENSMUSG00000023923 | Tbc1d5 | 0.001494808 | -1.392432467 | 0.039833998 |
| ENSMUSG00000034320 | Slc26a2 | 0.001494808 | -0.843017545 | 0.361297544 |

| | | | | |
|---|---|---|---|---|
| ENSMUSG00000024856 | Cdk2ap2 | 0.001494808 | 1.501994163 | 0.019039678 |
| ENSMUSG00000046230 | Vps13a | 0.001494808 | -2.106051951 | 0.000573507 |
| ENSMUSG00000025034 | Trim8 | 0.001494808 | -0.995020638 | 0.207356329 |
| ENSMUSG00000031146 | Plp2 | 0.001494808 | 1.152409725 | 0.084483671 |
| ENSMUSG00000002014 | Ssr4 | 0.001494808 | 1.841480629 | 0.002661694 |
| ENSMUSG00000052372 | Il1rapl1 | 0.001494808 | 0.623300459 | 0.510771494 |
| ENSMUSG00000031314 | Taf1 | 0.001494808 | -1.43862729 | 0.028681695 |
| ENSMUSG00000031246 | Sh3bgrl | 0.001494808 | 0.046799537 | 1 |
| ENSMUSG00000025332 | Kdm5c | 0.001494808 | -2.041661363 | 0.000929748 |
| ENSMUSG00000049176 | Frmpd4 | 0.001494808 | 0.623300459 | 0.513117327 |
| ENSMUSG00000033671 | Cep350 | 0.001439445 | -1.469721059 | 0.025616174 |
| ENSMUSG00000009549 | Srp14 | 0.001439445 | 1.703283335 | 0.007698034 |
| ENSMUSG00000087056 | Gm14004 | 0.001439445 | -2.42758419 | 8.32E-05 |
| ENSMUSG00000040209 | Zfp704 | 0.001439445 | -1.733268058 | 0.006207271 |
| ENSMUSG00000027763 | Mbnl1 | 0.001439445 | -2.135435758 | 0.000524026 |
| ENSMUSG00000046519 | Golph3l | 0.001439445 | -0.984294232 | 0.224125257 |
| ENSMUSG00000033308 | Dpyd | 0.001439445 | -0.005667883 | 1 |
| ENSMUSG00000028494 | Plin2 | 0.001439445 | 0.305690971 | 0.92282267 |
| ENSMUSG00000063077 | Kif1b | 0.001439445 | -1.781152289 | 0.00439578 |
| ENSMUSG00000029552 | Tes | 0.001439445 | -0.527014284 | 0.756790446 |
| ENSMUSG00000008036 | Ap2s1 | 0.001439445 | 0.648835551 | 0.515310426 |
| ENSMUSG00000066571 | 4931406P16Rik | 0.001439445 | -0.810596067 | 0.395118486 |
| ENSMUSG00000015133 | Lrrk1 | 0.001439445 | -1.895484965 | 0.00236773 |
| ENSMUSG00000038650 | Rnh1 | 0.001439445 | -0.303636078 | 1 |
| ENSMUSG00000054823 | Nsd3 | 0.001439445 | -1.254488429 | 0.07173713 |
| ENSMUSG00000031885 | Cbfb | 0.001439445 | -0.57355687 | 0.66416768 |
| ENSMUSG00000031753 | Cog4 | 0.001439445 | -1.478919996 | 0.025596136 |
| ENSMUSG00000033596 | Rfwd3 | 0.001439445 | -0.791033801 | 0.406526815 |
| ENSMUSG00000039016 | Timm8b | 0.001439445 | 1.521080004 | 0.020631722 |
| ENSMUSG00000055334 | Snupn | 0.001439445 | -1.275757046 | 0.070858113 |
| ENSMUSG00000032226 | Gcnt3 | 0.001439445 | 1.911869957 | 0.0020979 |
| ENSMUSG00000034858 | Fam214a | 0.001439445 | -0.997156383 | 0.215670938 |
| ENSMUSG00000032554 | Trf | 0.001439445 | -0.005667883 | 1 |
| ENSMUSG00000019873 | Reep3 | 0.001439445 | -1.406848581 | 0.036737509 |
| ENSMUSG00000020250 | Txnrd1 | 0.001439445 | -0.282777474 | 1 |
| ENSMUSG00000025404 | R3hdm2 | 0.001439445 | -0.953828951 | 0.245842568 |
| ENSMUSG00000094483 | Purb | 0.001439445 | -0.047488058 | 1 |
| ENSMUSG00000044949 | Ubtd2 | 0.001439445 | -1.79840077 | 0.004191433 |
| ENSMUSG00000017631 | Abr | 0.001439445 | -1.52108945 | 0.021034129 |
| ENSMUSG00000018425 | Dhx40 | 0.001439445 | 0.020018761 | 1 |
| ENSMUSG00000003119 | Cdk12 | 0.001439445 | -1.927193825 | 0.001902483 |
| ENSMUSG00000037361 | Sf3b6 | 0.001439445 | 0.545742058 | 0.613550092 |
| ENSMUSG00000020576 | Nbas | 0.001439445 | -1.047488058 | 0.176201216 |
| ENSMUSG00000011179 | Odc1 | 0.001439445 | 0.42644313 | 0.792441218 |
| ENSMUSG00000052593 | Adam17 | 0.001439445 | -1.080092724 | 0.150952178 |
| ENSMUSG00000020564 | Atxn7l1 | 0.001439445 | -2.354500161 | 0.000112953 |
| ENSMUSG00000021096 | Ppm1a | 0.001439445 | -0.710019112 | 0.476074732 |
| ENSMUSG00000042808 | Gpx2 | 0.001439445 | 0.184554533 | 1 |
| ENSMUSG00000021065 | Fut8 | 0.001439445 | -2.037146114 | 0.001014164 |
| ENSMUSG00000090935 | Synj2bp | 0.001439445 | -1.791033801 | 0.004320639 |
| ENSMUSG00000079036 | Alkbh1 | 0.001439445 | -1.310522464 | 0.057484546 |

| | | | | |
|---|---|---|---|---|
| ENSMUSG00000034928 | Rnf44 | 0.001439445 | 0.818760553 | 0.321073309 |
| ENSMUSG00000021493 | Pdlim7 | 0.001439445 | -1.196189789 | 0.096100696 |
| ENSMUSG00000021693 | Kif2a | 0.001439445 | -0.556278878 | 0.680154972 |
| ENSMUSG00000015533 | Itga2 | 0.001439445 | -2.611691999 | 1.61E-05 |
| ENSMUSG00000040651 | Tasor | 0.001439445 | -2.30018052 | 0.00016102 |
| ENSMUSG00000041028 | Ghitm | 0.001439445 | 0.774366434 | 0.3939206 |
| ENSMUSG00000022174 | Dad1 | 0.001439445 | 1.213898495 | 0.079721397 |
| ENSMUSG00000021983 | Atp8a2 | 0.001439445 | -0.410058138 | 0.881940904 |
| ENSMUSG00000022314 | Rad21 | 0.001439445 | -0.927193825 | 0.268379364 |
| ENSMUSG00000102018 | Iqank1 | 0.001439445 | -1.181239447 | 0.099755203 |
| ENSMUSG00000022791 | Tnk2 | 0.001439445 | -1.139308767 | 0.120696884 |
| ENSMUSG00000002250 | Ppard | 0.001439445 | -1.3207908 | 0.051203786 |
| ENSMUSG00000024096 | Ralbp1 | 0.001439445 | -0.261612864 | 1 |
| ENSMUSG00000055471 | Alk | 0.001439445 | -0.503167542 | 0.77466292 |
| ENSMUSG00000045817 | Zfp36l2 | 0.001439445 | -0.303636078 | 1 |
| ENSMUSG00000036103 | Colec12 | 0.001439445 | -0.478919996 | 0.792441218 |
| ENSMUSG00000048410 | Zfp407 | 0.001439445 | -1.825095637 | 0.003399554 |
| ENSMUSG00000048720 | Tbc1d12 | 0.001439445 | -1.397176846 | 0.035651073 |
| ENSMUSG00000036097 | Slf2 | 0.001439445 | -1.846575364 | 0.003112598 |
| ENSMUSG00000024974 | Smc3 | 0.001439445 | -0.550473257 | 0.677800556 |
| ENSMUSG00000024976 | Shoc2 | 0.001439445 | -2.296716665 | 0.000166828 |
| ENSMUSG00000074733 | Zfp950 | 0.001439445 | -2.216496772 | 0.000296362 |
| ENSMUSG00000046032 | Snx12 | 0.001439445 | -1.018341713 | 0.193977729 |
| ENSMUSG00000047045 | Tmem164 | 0.001439445 | -1.846575364 | 0.003112598 |
| ENSMUSG00000067194 | Eif1ax | 0.001439445 | 1.381355241 | 0.031497833 |
| ENSMUSG00000028033 | Kcnq5 | 0.001384082 | -1.030103097 | 0.188834491 |
| ENSMUSG00000026021 | Sumo1 | 0.001384082 | -0.922187758 | 0.277974519 |
| ENSMUSG00000026204 | Ptprn | 0.001384082 | -0.893898714 | 0.305378889 |
| ENSMUSG00000034066 | Farp2 | 0.001384082 | -0.70057328 | 0.501646057 |
| ENSMUSG00000026278 | Bok | 0.001384082 | 1.045494521 | 0.14991131 |
| ENSMUSG00000064302 | Clasp1 | 0.001384082 | -1.106051951 | 0.139919923 |
| ENSMUSG00000026457 | Adipor1 | 0.001384082 | 0.857422173 | 0.285257732 |
| ENSMUSG00000056220 | Pla2g4a | 0.001384082 | -1.80047185 | 0.005053277 |
| ENSMUSG00000026504 | Sdccag8 | 0.001384082 | -1.850397075 | 0.003399678 |
| ENSMUSG00000026640 | Plxna2 | 0.001384082 | -1.290969667 | 0.061096399 |
| ENSMUSG00000026655 | Fam107b | 0.001384082 | -1.106051951 | 0.139919923 |
| ENSMUSG00000026842 | Abl1 | 0.001384082 | -1.578502693 | 0.014476156 |
| ENSMUSG00000026867 | Gapvd1 | 0.001384082 | -1.675864293 | 0.008958881 |
| ENSMUSG00000036202 | Rif1 | 0.001384082 | -1.803009573 | 0.004102547 |
| ENSMUSG00000001768 | Rin2 | 0.001384082 | -0.855295875 | 0.364063958 |
| ENSMUSG00000027613 | Eif6 | 0.001384082 | -1.209145444 | 0.10105371 |
| ENSMUSG00000069094 | Pde7a | 0.001384082 | -1.445140597 | 0.02937793 |
| ENSMUSG00000027810 | Eif2a | 0.001384082 | -1.162477384 | 0.110207286 |
| ENSMUSG00000075700 | Selenot | 0.001384082 | -0.759249187 | 0.466464457 |
| ENSMUSG00000027782 | Kpna4 | 0.001384082 | -1.979399658 | 0.00146822 |
| ENSMUSG00000042747 | Krtcap2 | 0.001384082 | 0.734565425 | 0.434836739 |
| ENSMUSG00000027939 | Nup210l | 0.001384082 | -1.138564764 | 0.121084639 |
| ENSMUSG00000028106 | Rprd2 | 0.001384082 | -1.305360759 | 0.05909562 |
| ENSMUSG00000000001 | Gnai3 | 0.001384082 | -0.527241677 | 0.720623338 |
| ENSMUSG00000028266 | Lmo4 | 0.001384082 | 0.082488729 | 1 |
| ENSMUSG00000028413 | B4galt1 | 0.001384082 | -1.4222142 | 0.034492686 |

| | | | | |
|---|---|---|---|---|
| ENSMUSG00000028412 | Slc44a1 | 0.001384082 | 0.621354815 | 0.56359992 |
| ENSMUSG00000011257 | Pabpc4 | 0.001384082 | -0.764466896 | 0.449024837 |
| ENSMUSG00000040928 | S100pbp | 0.001384082 | -1.480447466 | 0.024497997 |
| ENSMUSG00000028879 | Stx12 | 0.001384082 | -1.060115667 | 0.179411863 |
| ENSMUSG00000028937 | Acot7 | 0.001384082 | -0.265434575 | 1 |
| ENSMUSG00000039410 | Prdm16 | 0.001384082 | -1.653539746 | 0.010186111 |
| ENSMUSG00000029038 | Ssu72 | 0.001384082 | -0.235687231 | 1 |
| ENSMUSG00000061576 | Dpp6 | 0.001384082 | 0.326907457 | 0.921320733 |
| ENSMUSG00000037685 | Atp8a1 | 0.001384082 | -1.92452033 | 0.002079494 |
| ENSMUSG00000036435 | Exoc1 | 0.001384082 | -0.959063761 | 0.240510494 |
| ENSMUSG00000049550 | Clip1 | 0.001384082 | -2.043042153 | 0.000981983 |
| ENSMUSG00000053768 | Chchd3 | 0.001384082 | -1.283719928 | 0.064921475 |
| ENSMUSG00000063694 | Cycs | 0.001384082 | 1.676849928 | 0.013025276 |
| ENSMUSG00000034269 | Setd5 | 0.001384082 | -1.979399658 | 0.00146822 |
| ENSMUSG00000041372 | B4galnt3 | 0.001384082 | -1.542508434 | 0.020439604 |
| ENSMUSG00000019210 | Atp6v1e1 | 0.001384082 | -0.780007747 | 0.426429627 |
| ENSMUSG00000072772 | Grcc10 | 0.001384082 | 0.149602925 | 1 |
| ENSMUSG00000030591 | Psmd8 | 0.001384082 | 0.676849928 | 0.480188651 |
| ENSMUSG00000055978 | Fut2 | 0.001384082 | -0.33020483 | 0.997253937 |
| ENSMUSG00000048897 | Zfp710 | 0.001384082 | -1.619391524 | 0.01181598 |
| ENSMUSG00000055723 | Rras2 | 0.001384082 | -1.483614745 | 0.024821097 |
| ENSMUSG00000030795 | Fus | 0.001384082 | -1.385434351 | 0.040680007 |
| ENSMUSG00000030847 | Bag3 | 0.001384082 | -1.785151051 | 0.004662801 |
| ENSMUSG00000037541 | Shank2 | 0.001384082 | -1.70057328 | 0.009157516 |
| ENSMUSG00000070056 | Mfhas1 | 0.001384082 | -1.347692257 | 0.049315587 |
| ENSMUSG00000030465 | Psd3 | 0.001384082 | -1.572565667 | 0.015221042 |
| ENSMUSG00000034656 | Cacna1a | 0.001384082 | -0.53336778 | 0.723208723 |
| ENSMUSG00000041949 | Tango6 | 0.001384082 | -1.337225258 | 0.051667273 |
| ENSMUSG00000032239 | Rp9 | 0.001384082 | -0.790276083 | 0.430746607 |
| ENSMUSG00000001833 | 7-Sep | 0.001384082 | -0.53336778 | 0.723208723 |
| ENSMUSG00000032384 | Csnk1g1 | 0.001384082 | -1.418909055 | 0.033421986 |
| ENSMUSG00000032217 | Rnf111 | 0.001384082 | -1.545542494 | 0.017175873 |
| ENSMUSG00000032580 | Rbm5 | 0.001384082 | -1.917511271 | 0.002131249 |
| ENSMUSG00000039952 | Dag1 | 0.001384082 | -1.40226879 | 0.036798276 |
| ENSMUSG00000020019 | Ntn4 | 0.001384082 | -0.65634928 | 0.584485366 |
| ENSMUSG00000036499 | Eea1 | 0.001384082 | -1.415596321 | 0.032981099 |
| ENSMUSG00000071072 | Ptges3 | 0.001384082 | -0.954505479 | 0.256504754 |
| ENSMUSG00000040054 | Baz2a | 0.001384082 | -2.019231192 | 0.00117591 |
| ENSMUSG00000020122 | Egfr | 0.001384082 | -0.769665803 | 0.451565197 |
| ENSMUSG00000020361 | Hspa4 | 0.001384082 | -1.189883542 | 0.101297285 |
| ENSMUSG00000001036 | Epn2 | 0.001384082 | -2.212967155 | 0.000320842 |
| ENSMUSG00000070394 | Tmem256 | 0.001384082 | 1.567915556 | 0.014575482 |
| ENSMUSG00000017774 | Myo1c | 0.001384082 | -1.182106191 | 0.09943433 |
| ENSMUSG00000017776 | Crk | 0.001384082 | -1.4222142 | 0.034053737 |
| ENSMUSG00000020680 | Taf15 | 0.001384082 | -1.527241677 | 0.020146995 |
| ENSMUSG00000034297 | Med13 | 0.001384082 | -1.805542839 | 0.004168617 |
| ENSMUSG00000035775 | Krt20 | 0.001384082 | 0.70541908 | 0.450971315 |
| ENSMUSG00000020695 | Mrc2 | 0.001384082 | -1.458079652 | 0.029677057 |
| ENSMUSG00000053580 | Tanc2 | 0.001384082 | -0.599091962 | 0.66416768 |
| ENSMUSG00000047798 | Cd300lf | 0.001384082 | -0.684147732 | 0.551482001 |
| ENSMUSG00000020609 | Apob | 0.001384082 | 0.749362427 | 0.430214327 |

| | | | | |
|---|---|---|---|---|
| ENSMUSG00000021458 | Aopep | 0.001384082 | -1.943046363 | 0.001758822 |
| ENSMUSG00000021606 | Ndufs6 | 0.001384082 | 0.47891055 | 0.708275224 |
| ENSMUSG00000013701 | Timm23 | 0.001384082 | 0.889843651 | 0.274176098 |
| ENSMUSG00000060012 | Kif13b | 0.001384082 | -1.792831788 | 0.004497171 |
| ENSMUSG00000034731 | Dgkh | 0.001384082 | -1.008276735 | 0.214662905 |
| ENSMUSG00000034333 | Zbed4 | 0.001384082 | -0.527241677 | 0.720623338 |
| ENSMUSG00000041974 | Spidr | 0.001384082 | -1.886738922 | 0.00257107 |
| ENSMUSG00000006998 | Psmd2 | 0.001384082 | -0.051604167 | 1 |
| ENSMUSG00000052384 | Nrros | 0.001384082 | -2.036587131 | 0.001319925 |
| ENSMUSG00000034158 | Lrrc58 | 0.001384082 | -0.060115667 | 1 |
| ENSMUSG00000002845 | Tmem39a | 0.001384082 | -1.838076804 | 0.003437912 |
| ENSMUSG00000032855 | Pkd1 | 0.001384082 | -1.333719314 | 0.051152519 |
| ENSMUSG00000061613 | U2af1 | 0.001384082 | -0.70057328 | 0.501646057 |
| ENSMUSG00000023965 | Fbxl17 | 0.001384082 | -1.508705726 | 0.021921798 |
| ENSMUSG00000035051 | Dhx57 | 0.001384082 | -2.085352958 | 0.000752229 |
| ENSMUSG00000043991 | Pura | 0.001384082 | -0.445140597 | 0.836964118 |
| ENSMUSG00000024782 | Ak3 | 0.001384082 | 0.764312769 | 0.373232174 |
| ENSMUSG00000000787 | Ddx3x | 0.001384082 | -0.489928519 | 0.788248107 |
| ENSMUSG00000067567 | Hdac8 | 0.001384082 | -1.548570186 | 0.017369611 |
| ENSMUSG00000025909 | Sntg1 | 0.001328718 | 0.315411818 | 0.961451902 |
| ENSMUSG00000053153 | Spag16 | 0.001328718 | 0.410048692 | 0.833666942 |
| ENSMUSG00000055322 | Tns1 | 0.001328718 | -0.857055378 | 0.355090064 |
| ENSMUSG00000026814 | Eng | 0.001328718 | -0.841734579 | 0.408970699 |
| ENSMUSG00000068882 | Ssb | 0.001328718 | -0.386788358 | 0.918330572 |
| ENSMUSG00000044042 | Fmn1 | 0.001328718 | -0.643946198 | 0.586774596 |
| ENSMUSG00000048911 | Rnf24 | 0.001328718 | -1.120936188 | 0.143506057 |
| ENSMUSG00000027598 | Itch | 0.001328718 | -1.690301833 | 0.008534388 |
| ENSMUSG00000047459 | Dynlrb1 | 0.001328718 | 1.485336819 | 0.042972006 |
| ENSMUSG00000027673 | Ndufb5 | 0.001328718 | 1.2052289 | 0.080855102 |
| ENSMUSG00000037174 | Elf2 | 0.001328718 | -1.562176642 | 0.016410626 |
| ENSMUSG00000034109 | Golim4 | 0.001328718 | -1.754432171 | 0.006130108 |
| ENSMUSG00000042306 | S100a14 | 0.001328718 | 1.769129785 | 0.005872899 |
| ENSMUSG00000045576 | St7l | 0.001328718 | -2.327528084 | 0.000151735 |
| ENSMUSG00000068749 | Psma5 | 0.001328718 | 1.385801146 | 0.038042319 |
| ENSMUSG00000040151 | Hs2st1 | 0.001328718 | -1.468859491 | 0.027444373 |
| ENSMUSG00000028188 | Spata1 | 0.001328718 | -1.673092543 | 0.009584046 |
| ENSMUSG00000028696 | Ipp | 0.001328718 | -1.051604167 | 0.17273405 |
| ENSMUSG00000028648 | Ndufs5 | 0.001328718 | 1.73810289 | 0.018305209 |
| ENSMUSG00000028837 | Psmb2 | 0.001328718 | 1.385801146 | 0.038042319 |
| ENSMUSG00000036940 | Kdm1a | 0.001328718 | -1.602126022 | 0.014575482 |
| ENSMUSG00000028975 | Pex14 | 0.001328718 | -1.215102899 | 0.091791571 |
| ENSMUSG00000029131 | Dnajb6 | 0.001328718 | -0.940927936 | 0.262029351 |
| ENSMUSG00000052139 | Babam2 | 0.001328718 | -1.069252033 | 0.166514237 |
| ENSMUSG00000005103 | Wdr1 | 0.001328718 | 0.025905201 | 1 |
| ENSMUSG00000029179 | Zcchc4 | 0.001328718 | -1.174913809 | 0.113235376 |
| ENSMUSG00000055204 | Ankrd17 | 0.001328718 | -2.77070334 | 5.80E-06 |
| ENSMUSG00000029287 | Tgfbr3 | 0.001328718 | -1.069252033 | 0.173532034 |
| ENSMUSG00000051339 | 2900026A02Rik | 0.001328718 | -1.51143928 | 0.021725703 |
| ENSMUSG00000029550 | Sppl3 | 0.001328718 | -1.629149196 | 0.011642972 |
| ENSMUSG00000041740 | Rnf10 | 0.001328718 | -0.191124036 | 1 |
| ENSMUSG00000039477 | Tnrc18 | 0.001328718 | -1.721328729 | 0.007010781 |

| | | | | |
|---|---|---|---|---|
| ENSMUSG00000029623 | Pdap1 | 0.001328718 | -0.322018103 | 1 |
| ENSMUSG00000004633 | Chn2 | 0.001328718 | -1.974094799 | 0.001554188 |
| ENSMUSG00000056091 | St3gal5 | 0.001328718 | -0.735214255 | 0.466464457 |
| ENSMUSG00000030168 | Adipor2 | 0.001328718 | -0.365519742 | 0.966942564 |
| ENSMUSG00000004328 | Hif3a | 0.001328718 | -0.684588182 | 0.530543686 |
| ENSMUSG00000002603 | Tgfb1 | 0.001328718 | -0.116699195 | 1 |
| ENSMUSG00000055116 | Arntl | 0.001328718 | -0.307221101 | 1 |
| ENSMUSG00000063065 | Mapk3 | 0.001328718 | 0.620266399 | 0.538110273 |
| ENSMUSG00000078566 | Bnip3 | 0.001328718 | 1.2052289 | 0.101869423 |
| ENSMUSG00000004626 | Stxbp2 | 0.001328718 | -0.038225137 | 1 |
| ENSMUSG00000037234 | Hook3 | 0.001328718 | -0.661504569 | 0.557051363 |
| ENSMUSG00000070047 | Fat1 | 0.001328718 | -2.267641093 | 0.000241299 |
| ENSMUSG00000054717 | Hmgb2 | 0.001328718 | 0.194659658 | 1 |
| ENSMUSG00000031703 | Itfg1 | 0.001328718 | -1.372644176 | 0.043242687 |
| ENSMUSG00000034593 | Myo5a | 0.001328718 | -0.997327106 | 0.213374766 |
| ENSMUSG00000032481 | Smarcc1 | 0.001328718 | -1.914322579 | 0.002194711 |
| ENSMUSG00000040675 | Mthfd1l | 0.001328718 | -1.029236354 | 0.200079844 |
| ENSMUSG00000019996 | Map7 | 0.001328718 | -0.421553776 | 0.885240416 |
| ENSMUSG00000019990 | Pde7b | 0.001328718 | -0.729676072 | 0.496209758 |
| ENSMUSG00000057098 | Ebf1 | 0.001328718 | -0.983432664 | 0.223409161 |
| ENSMUSG00000036309 | Skp1a | 0.001328718 | -0.038225137 | 1 |
| ENSMUSG00000001552 | Jup | 0.001328718 | -0.421553776 | 0.885240416 |
| ENSMUSG00000020661 | Dnmt3a | 0.001328718 | -1.76800419 | 0.005638811 |
| ENSMUSG00000021133 | Susd6 | 0.001328718 | -1.945712998 | 0.001942582 |
| ENSMUSG00000021188 | Trip11 | 0.001328718 | -2.020191214 | 0.001184321 |
| ENSMUSG00000021413 | Prpf4b | 0.001328718 | -2.652752095 | 1.46E-05 |
| ENSMUSG00000004562 | Arhgef40 | 0.001328718 | -1.504969785 | 0.022883134 |
| ENSMUSG00000035726 | Supt16 | 0.001328718 | -2.056036413 | 0.000987149 |
| ENSMUSG00000022021 | Diaph3 | 0.001328718 | -1.504969785 | 0.022883134 |
| ENSMUSG00000025545 | Clybl | 0.001328718 | -0.632120762 | 0.611716824 |
| ENSMUSG00000036698 | Ago2 | 0.001328718 | -0.902065113 | 0.302586904 |
| ENSMUSG00000022451 | Twf1 | 0.001328718 | 1.083238376 | 0.154807284 |
| ENSMUSG00000050783 | Htr1f | 0.001328718 | -0.199161355 | 1 |
| ENSMUSG00000024012 | Mtch1 | 0.001328718 | 0.184167284 | 1 |
| ENSMUSG00000023122 | Sult1c2 | 0.001328718 | 1.063873051 | 0.159648194 |
| ENSMUSG00000000127 | Fer | 0.001328718 | -1.133573013 | 0.128972787 |
| ENSMUSG00000024268 | Celf4 | 0.001328718 | -1.112449722 | 0.153658826 |
| ENSMUSG00000005873 | Reep5 | 0.001328718 | 0.524507416 | 0.673152643 |
| ENSMUSG00000001383 | Zmat2 | 0.001328718 | -0.495210818 | 0.767977795 |
| ENSMUSG00000024603 | Dctn4 | 0.001328718 | -1.468859491 | 0.027444373 |
| ENSMUSG00000040560 | Wdr7 | 0.001328718 | -0.596051508 | 0.66416768 |
| ENSMUSG00000024942 | Capn1 | 0.001328718 | -0.695992946 | 0.503623269 |
| ENSMUSG00000024759 | Atl3 | 0.001328718 | -1.555931199 | 0.017363741 |
| ENSMUSG00000036098 | Myrf | 0.001328718 | -0.735214255 | 0.466464457 |
| ENSMUSG00000024713 | Pcsk5 | 0.001328718 | -0.955235772 | 0.286730633 |
| ENSMUSG00000025220 | Oga | 0.001328718 | -1.844299371 | 0.003354396 |
| ENSMUSG00000025026 | Add3 | 0.001328718 | -1.170832632 | 0.112238591 |
| ENSMUSG00000024975 | Pdcd4 | 0.001328718 | -0.386788358 | 0.911880553 |
| ENSMUSG00000031392 | Irak1 | 0.001328718 | 1.025905201 | 0.170846453 |
| ENSMUSG00000056537 | Rlim | 0.001328718 | -1.318333056 | 0.05923335 |
| ENSMUSG00000031298 | Adgrg2 | 0.001328718 | -0.778770621 | 0.452471727 |

| | | | | |
|---|---|---|---|---|
| ENSMUSG00000026042 | Col5a2 | 0.001273355 | -0.816050497 | 0.409269374 |
| ENSMUSG00000026032 | Ndufb3 | 0.001273355 | 1.807533297 | 0.011305389 |
| ENSMUSG00000026277 | Stk25 | 0.001273355 | -1.301477324 | 0.063949329 |
| ENSMUSG00000073557 | Ppp1r12b | 0.001273355 | -0.895484965 | 0.315589992 |
| ENSMUSG00000026361 | Cdc73 | 0.001273355 | -1.699892603 | 0.008534388 |
| ENSMUSG00000052748 | Swt1 | 0.001273355 | -1.645785197 | 0.011816249 |
| ENSMUSG00000026469 | Xpr1 | 0.001273355 | -1.188266714 | 0.128972787 |
| ENSMUSG00000014867 | Surf4 | 0.001273355 | 0.505877598 | 0.720623338 |
| ENSMUSG00000023224 | Serping1 | 0.001273355 | 0.710236096 | 0.47966612 |
| ENSMUSG00000058318 | Phf21a | 0.001273355 | -1.908309005 | 0.002440951 |
| ENSMUSG00000085438 | Oip5os1 | 0.001273355 | -0.52775318 | 0.751723078 |
| ENSMUSG00000027375 | Mal | 0.001273355 | 0.967011511 | 0.209790273 |
| ENSMUSG00000038116 | Phf20 | 0.001273355 | -2.429760052 | 8.67E-05 |
| ENSMUSG00000027642 | Rpn2 | 0.001273355 | 0.452438339 | 0.792441218 |
| ENSMUSG00000017950 | Hnf4a | 0.001273355 | -1.406446884 | 0.038546493 |
| ENSMUSG00000000827 | Tpd52l2 | 0.001273355 | -0.895484965 | 0.318264659 |
| ENSMUSG00000002428 | Hltf | 0.001273355 | -2.083614562 | 0.000876027 |
| ENSMUSG00000027706 | Sec62 | 0.001273355 | -1.69397991 | 0.009018945 |
| ENSMUSG00000037152 | Ndufc1 | 0.001273355 | 1.125273595 | 0.137540713 |
| ENSMUSG00000061175 | Fnip2 | 0.001273355 | -1.651898321 | 0.011191687 |
| ENSMUSG00000028019 | Pdgfc | 0.001273355 | -2.419046921 | 9.04E-05 |
| ENSMUSG00000041734 | Kirrel | 0.001273355 | -2.645785197 | 1.61E-05 |
| ENSMUSG00000008958 | Vps72 | 0.001273355 | -1.273996588 | 0.078768438 |
| ENSMUSG00000054312 | Mrps21 | 0.001273355 | 1.400908038 | 0.040750047 |
| ENSMUSG00000053965 | Pde5a | 0.001273355 | -1.427980046 | 0.037136816 |
| ENSMUSG00000004591 | Pkn2 | 0.001273355 | -1.167081384 | 0.120203977 |
| ENSMUSG00000039988 | Ankrd13c | 0.001273355 | -0.885143021 | 0.336079046 |
| ENSMUSG00000028496 | Mllt3 | 0.001273355 | -2.624182943 | 1.82E-05 |
| ENSMUSG00000028759 | Hp1bp3 | 0.001273355 | -0.749195523 | 0.467537667 |
| ENSMUSG00000041459 | Tardbp | 0.001273355 | -1.943621486 | 0.002052334 |
| ENSMUSG00000029063 | Nadk | 0.001273355 | 0.063873051 | 1 |
| ENSMUSG00000014932 | Yes1 | 0.001273355 | -1.258055044 | 0.079721397 |
| ENSMUSG00000029106 | Add1 | 0.001273355 | -2.119348773 | 0.000723318 |
| ENSMUSG00000029092 | D5Ertd615e | 0.001273355 | -1.705781162 | 0.008383243 |
| ENSMUSG00000029169 | Dhx15 | 0.001273355 | -1.560617814 | 0.018532615 |
| ENSMUSG00000039191 | Rbpj | 0.001273355 | -2.058523581 | 0.001052851 |
| ENSMUSG00000029238 | Clock | 0.001273355 | -0.642718894 | 0.590424306 |
| ENSMUSG00000029260 | Ugt2b34 | 0.001273355 | 1.004979362 | 0.195746456 |
| ENSMUSG00000044221 | Grsf1 | 0.001273355 | -1.229726321 | 0.086523965 |
| ENSMUSG00000029472 | Anapc5 | 0.001273355 | -0.514394797 | 0.747239297 |
| ENSMUSG00000029674 | Limk1 | 0.001273355 | 1.648835551 | 0.014219736 |
| ENSMUSG00000052581 | Lrrtm4 | 0.001273355 | 0.303060715 | 0.950123751 |
| ENSMUSG00000030096 | Slc6a6 | 0.001273355 | -0.910860191 | 0.299063909 |
| ENSMUSG00000000441 | Raf1 | 0.001273355 | -1.200830175 | 0.102424142 |
| ENSMUSG00000044709 | Gemin7 | 0.001273355 | -0.679091284 | 0.536465189 |
| ENSMUSG00000003380 | Rabac1 | 0.001273355 | 1.876246048 | 0.004005818 |
| ENSMUSG00000003123 | Lipe | 0.001273355 | -1.069982696 | 0.169739731 |
| ENSMUSG00000003762 | Coq8b | 0.001273355 | -0.799624949 | 0.431388315 |
| ENSMUSG00000030990 | Pgap2 | 0.001273355 | -0.58638091 | 0.66416768 |
| ENSMUSG00000054555 | Adam12 | 0.001273355 | -1.285838201 | 0.066614198 |
| ENSMUSG00000045969 | Ing1 | 0.001273355 | 1.400908038 | 0.040750047 |

| | | | | |
|---|---|---|---|---|
| ENSMUSG00000031537 | Ikbkb | 0.001273355 | -0.636566667 | 0.623042427 |
| ENSMUSG00000070044 | Fam149a | 0.001273355 | 0.841480629 | 0.343481345 |
| ENSMUSG00000002395 | Use1 | 0.001273355 | 1.948395833 | 0.006062433 |
| ENSMUSG00000031754 | Nudt21 | 0.001273355 | -0.636566667 | 0.634616388 |
| ENSMUSG00000035770 | Dync1li2 | 0.001273355 | -0.480447466 | 0.80085216 |
| ENSMUSG00000032097 | Ddx6 | 0.001273355 | -1.074540978 | 0.171185497 |
| ENSMUSG00000034007 | Scaper | 0.001273355 | -1.673092543 | 0.009613871 |
| ENSMUSG00000025234 | Arih1 | 0.001273355 | -1.402826568 | 0.038042319 |
| ENSMUSG00000032548 | Slco2a1 | 0.001273355 | -0.970773092 | 0.245924683 |
| ENSMUSG00000033392 | Clasp2 | 0.001273355 | -1.281901786 | 0.072757287 |
| ENSMUSG00000052336 | Cx3cr1 | 0.001273355 | -0.605405736 | 0.66416768 |
| ENSMUSG00000061032 | Rrp1 | 0.001273355 | 0.492331066 | 0.726066583 |
| ENSMUSG00000035781 | R3hdm4 | 0.001273355 | 0.619088208 | 0.578855097 |
| ENSMUSG00000019979 | Apaf1 | 0.001273355 | -1.72620388 | 0.007061791 |
| ENSMUSG00000020205 | Phlda1 | 0.001273355 | -0.760555385 | 0.466464457 |
| ENSMUSG00000034813 | Grip1 | 0.001273355 | -0.500911568 | 0.792441218 |
| ENSMUSG00000025432 | Avil | 0.001273355 | -0.52108945 | 0.749195699 |
| ENSMUSG00000020484 | Xbp1 | 0.001273355 | 1.104515035 | 0.132703831 |
| ENSMUSG00000007739 | Cct4 | 0.001273355 | 0.56137271 | 0.66416768 |
| ENSMUSG00000007850 | Hnrnph1 | 0.001273355 | -2.273996588 | 0.000241036 |
| ENSMUSG00000018920 | Cxcl16 | 0.001273355 | -0.805120937 | 0.405663956 |
| ENSMUSG00000020803 | Txndc17 | 0.001273355 | 1.351155003 | 0.048434208 |
| ENSMUSG00000020774 | Aspa | 0.001273355 | 0.400908038 | 0.823212269 |
| ENSMUSG00000037857 | Nufip2 | 0.001273355 | -1.534386273 | 0.021455695 |
| ENSMUSG00000020612 | Prkar1a | 0.001273355 | 0.256518129 | 1 |
| ENSMUSG00000087259 | 2610035D17Rik | 0.001273355 | -2.343630776 | 0.000148372 |
| ENSMUSG00000035933 | Cog5 | 0.001273355 | -1.167081384 | 0.120203977 |
| ENSMUSG00000061603 | Akap6 | 0.001273355 | 1.256518129 | 0.094160212 |
| ENSMUSG00000034912 | Mdga2 | 0.001273355 | 0.575586569 | 0.631007587 |
| ENSMUSG00000021178 | Psmc1 | 0.001273355 | -0.743481871 | 0.466464457 |
| ENSMUSG00000001542 | Ell2 | 0.001273355 | -1.511035785 | 0.02298647 |
| ENSMUSG00000021708 | Rasgrf2 | 0.001273355 | -0.931108875 | 0.286643284 |
| ENSMUSG00000069089 | Cdk7 | 0.001273355 | -1.874726405 | 0.00294545 |
| ENSMUSG00000015759 | Cnih1 | 0.001273355 | 0.363433333 | 0.871107617 |
| ENSMUSG00000022194 | Pabpn1 | 0.001273355 | -3.704311274 | 7.33E-10 |
| ENSMUSG00000022052 | Ppp2r2a | 0.001273355 | -1.258055044 | 0.079721397 |
| ENSMUSG00000022263 | Trio | 0.001273355 | -2.570332969 | 2.79E-05 |
| ENSMUSG00000022354 | Ndufb9 | 0.001273355 | 1.911869957 | 0.00335241 |
| ENSMUSG00000022377 | Asap1 | 0.001273355 | -2.209145444 | 0.000365509 |
| ENSMUSG00000033228 | Scaf11 | 0.001273355 | -2.021230518 | 0.00128813 |
| ENSMUSG00000022698 | Naa50 | 0.001273355 | 0.043973493 | 1 |
| ENSMUSG00000023904 | Hcfc1r1 | 0.001273355 | 1.648835551 | 0.014219736 |
| ENSMUSG00000004865 | Srpk1 | 0.001273355 | -1.664047404 | 0.0100807 |
| ENSMUSG00000039153 | Runx2 | 0.001273355 | -1.324622661 | 0.057823895 |
| ENSMUSG00000045038 | Prkce | 0.001273355 | -0.749195523 | 0.467537667 |
| ENSMUSG00000005871 | Apc | 0.001273355 | -1.277954602 | 0.070876897 |
| ENSMUSG00000024480 | Ap3s1 | 0.001273355 | -0.014129461 | 1 |
| ENSMUSG00000073563 | Csnk1g3 | 0.001273355 | -1.167081384 | 0.120203977 |
| ENSMUSG00000024513 | Mbd2 | 0.001273355 | -1.106051951 | 0.143740392 |
| ENSMUSG00000036908 | Unc93b1 | 0.001273355 | -0.766201948 | 0.466464457 |
| ENSMUSG00000035735 | Dagla | 0.001273355 | 0.291283547 | 0.9980272 |

| | | | | |
|---|---|---|---|---|
| ENSMUSG00000040451 | Sgms1 | 0.001273355 | -2.309233768 | 0.000197786 |
| ENSMUSG00000023307 | 5-Mar | 0.001273355 | -0.951077291 | 0.25774053 |
| ENSMUSG00000025224 | Gbf1 | 0.001273355 | -1.424413431 | 0.03461849 |
| ENSMUSG00000001127 | Araf | 0.001273355 | -1.864234031 | 0.003263045 |
| ENSMUSG00000016308 | Ube2a | 0.001273355 | -0.654945197 | 0.595281189 |
| ENSMUSG00000037475 | Thoc2 | 0.001273355 | -2.544279077 | 3.32E-05 |
| ENSMUSG00000031375 | Bgn | 0.001273355 | 0.967011511 | 0.238797341 |
| ENSMUSG00000025151 | Maged1 | 0.001273355 | -0.402826568 | 0.894898658 |
| ENSMUSG00000005886 | Ncoa2 | 0.001217992 | -1.471458682 | 0.029649257 |
| ENSMUSG00000079658 | Eloc | 0.001217992 | 0.48589698 | 0.706050453 |
| ENSMUSG00000073643 | Wdfy1 | 0.001217992 | -1.061222169 | 0.185472119 |
| ENSMUSG00000036707 | Cab39 | 0.001217992 | -0.569069479 | 0.733451889 |
| ENSMUSG00000049866 | Arl4c | 0.001217992 | -0.245455007 | 1 |
| ENSMUSG00000026353 | Ubxn4 | 0.001217992 | -1.625264018 | 0.013045269 |
| ENSMUSG00000026663 | Atf6 | 0.001217992 | -1.517600464 | 0.02307354 |
| ENSMUSG00000062169 | Cnih4 | 0.001217992 | 0.391037794 | 0.847265357 |
| ENSMUSG00000039128 | Cdc123 | 0.001217992 | -0.26223072 | 1 |
| ENSMUSG00000026781 | Acbd5 | 0.001217992 | -0.075530006 | 1 |
| ENSMUSG00000038860 | Garnl3 | 0.001217992 | -1.176330647 | 0.120903112 |
| ENSMUSG00000044308 | Ubr3 | 0.001217992 | -1.43862729 | 0.033836695 |
| ENSMUSG00000040591 | 1110051M20Rik | 0.001217992 | -1.475060919 | 0.03005022 |
| ENSMUSG00000012350 | Ehf | 0.001217992 | -1.315442526 | 0.064252863 |
| ENSMUSG00000027133 | Nop10 | 0.001217992 | 1.680544411 | 0.023472221 |
| ENSMUSG00000033943 | Mga | 0.001217992 | -1.719386196 | 0.007880333 |
| ENSMUSG00000074793 | Hspa12b | 0.001217992 | -0.202641924 | 1 |
| ENSMUSG00000074698 | Csnk2a1 | 0.001217992 | -0.821955929 | 0.408970699 |
| ENSMUSG00000067847 | Romo1 | 0.001217992 | 1.022650388 | 0.224000282 |
| ENSMUSG00000102869 | 2900097C17Rik | 0.001217992 | 0.106308317 | 1 |
| ENSMUSG00000006800 | Sulf2 | 0.001217992 | -0.849970306 | 0.367040718 |
| ENSMUSG00000027546 | Atp9a | 0.001217992 | -1.232743956 | 0.093377644 |
| ENSMUSG00000039717 | Ralyl | 0.001217992 | 0.277579744 | 0.999122005 |
| ENSMUSG00000063600 | Egfem1 | 0.001217992 | -0.849970306 | 0.424188363 |
| ENSMUSG00000027799 | Nbea | 0.001217992 | -1.431228837 | 0.035917176 |
| ENSMUSG00000027803 | Wwtr1 | 0.001217992 | -1.524570019 | 0.023673997 |
| ENSMUSG00000056071 | S100a9 | 0.001217992 | 1.780080085 | 0.009479588 |
| ENSMUSG00000005968 | Tuft1 | 0.001217992 | -1.821955929 | 0.004552202 |
| ENSMUSG00000038374 | Rbm8a | 0.001217992 | 0.339507493 | 0.912904617 |
| ENSMUSG00000058135 | Gstm1 | 0.001217992 | 2.365042585 | 0.000429764 |
| ENSMUSG00000028180 | Zranb2 | 0.001217992 | -1.386023206 | 0.043688957 |
| ENSMUSG00000035696 | Rnf38 | 0.001217992 | -1.81344443 | 0.004732823 |
| ENSMUSG00000044813 | Shb | 0.001217992 | -1.149530587 | 0.126185633 |
| ENSMUSG00000028556 | Dock7 | 0.001217992 | -1.946543565 | 0.002182685 |
| ENSMUSG00000035305 | Ror1 | 0.001217992 | -1.94913601 | 0.002052334 |
| ENSMUSG00000028603 | Scp2 | 0.001217992 | 0.471957789 | 0.789829535 |
| ENSMUSG00000028538 | St3gal3 | 0.001217992 | -1.48938059 | 0.028740185 |
| ENSMUSG00000028633 | Ctps | 0.001217992 | -1.370634313 | 0.049367015 |
| ENSMUSG00000028820 | Sfpq | 0.001217992 | -1.917714912 | 0.002547455 |
| ENSMUSG00000028796 | Phc2 | 0.001217992 | -1.524570019 | 0.023673997 |
| ENSMUSG00000037426 | Depdc5 | 0.001217992 | -0.500027834 | 0.792441218 |
| ENSMUSG00000029108 | Pcdh7 | 0.001217992 | -0.219919915 | 1 |
| ENSMUSG00000029178 | Klf3 | 0.001217992 | -0.634957415 | 0.633493255 |

| | | | | |
|---|---|---|---|---|
| ENSMUSG00000058013 | 11-Sep | 0.001217992 | -1.108373263 | 0.145809751 |
| ENSMUSG00000011831 | Evi5 | 0.001217992 | -1.405031321 | 0.042735074 |
| ENSMUSG00000029578 | Wipi2 | 0.001217992 | -0.972260735 | 0.257440232 |
| ENSMUSG00000029826 | Zc3hav1 | 0.001217992 | -0.956885509 | 0.2689688 |
| ENSMUSG00000059187 | Tafa1 | 0.001217992 | 0.962944142 | 0.256384946 |
| ENSMUSG00000030232 | Aebp2 | 0.001217992 | -0.679351534 | 0.538463052 |
| ENSMUSG00000032712 | Resf1 | 0.001217992 | -2.041921613 | 0.001278866 |
| ENSMUSG00000035674 | Ndufa3 | 0.001217992 | 0.183712821 | 1 |
| ENSMUSG00000005575 | Ube2m | 0.001217992 | 0.850469413 | 0.315648377 |
| ENSMUSG00000052833 | Sae1 | 0.001217992 | -1.15403198 | 0.127386871 |
| ENSMUSG00000005447 | Pafah1b3 | 0.001217992 | 1.499972166 | 0.027306242 |
| ENSMUSG00000040725 | Hnrnpul1 | 0.001217992 | -0.007511583 | 1 |
| ENSMUSG00000038406 | Scaf1 | 0.001217992 | -1.528042211 | 0.022226859 |
| ENSMUSG00000002778 | Kdelr1 | 0.001217992 | 0.982572949 | 0.221100123 |
| ENSMUSG00000030605 | Mfge8 | 0.001217992 | 1.417510005 | 0.044392129 |
| ENSMUSG00000029461 | Fam168a | 0.001217992 | -1.291126866 | 0.07322814 |
| ENSMUSG00000052707 | Tnrc6a | 0.001217992 | -2.211306785 | 0.000391183 |
| ENSMUSG00000054065 | Pkp3 | 0.001217992 | -0.602389552 | 0.66416768 |
| ENSMUSG00000037706 | Cd81 | 0.001217992 | 1.471957789 | 0.039344003 |
| ENSMUSG00000037519 | Ppfia1 | 0.001217992 | -0.666806281 | 0.573944243 |
| ENSMUSG00000039539 | Sgcz | 0.001217992 | 0.084934666 | 1 |
| ENSMUSG00000031591 | Asah1 | 0.001217992 | -0.351164449 | 0.964337983 |
| ENSMUSG00000036469 | 1-Mar | 0.001217992 | -0.534961625 | 0.748964803 |
| ENSMUSG00000047747 | Rnf150 | 0.001217992 | -0.319455589 | 1 |
| ENSMUSG00000033938 | Ndufb7 | 0.001217992 | 1.106308317 | 0.153647653 |
| ENSMUSG00000013160 | Atp6v0d1 | 0.001217992 | -0.471458682 | 0.82558911 |
| ENSMUSG00000095941 | Tle7 | 0.001217992 | -1.679351534 | 0.012519972 |
| ENSMUSG00000035941 | Ibtk | 0.001217992 | -1.219919915 | 0.097987606 |
| ENSMUSG00000032504 | Pdcd6ip | 0.001217992 | -1.228481929 | 0.092468958 |
| ENSMUSG00000032536 | Trak1 | 0.001217992 | -1.569069479 | 0.022200985 |
| ENSMUSG00000019806 | Aig1 | 0.001217992 | -0.781798803 | 0.452280643 |
| ENSMUSG00000087177 | E130307A14Rik | 0.001217992 | -1.638174127 | 0.012879784 |
| ENSMUSG00000020091 | Eif4ebp2 | 0.001217992 | -0.7284693 | 0.491653643 |
| ENSMUSG00000034612 | Chst11 | 0.001217992 | -1.282929713 | 0.076432558 |
| ENSMUSG00000074781 | Ube2n | 0.001217992 | -0.628502392 | 0.630163115 |
| ENSMUSG00000019929 | Dcn | 0.001217992 | 0.982572949 | 0.221100123 |
| ENSMUSG00000019966 | Kitl | 0.001217992 | -1.755380455 | 0.008611176 |
| ENSMUSG00000025795 | Rassf3 | 0.001217992 | 0.022650388 | 1 |
| ENSMUSG00000005683 | Cs | 0.001217992 | -0.787604425 | 0.454819697 |
| ENSMUSG00000025369 | Smarcc2 | 0.001217992 | -1.198289899 | 0.109935582 |
| ENSMUSG00000042302 | Ehbp1 | 0.001217992 | -1.989993821 | 0.001752947 |
| ENSMUSG00000020463 | Ppp4r3b | 0.001217992 | 0.032846155 | 1 |
| ENSMUSG00000020305 | Asb3 | 0.001217992 | -1.631733514 | 0.013437516 |
| ENSMUSG00000055670 | Zzef1 | 0.001217992 | -1.74646573 | 0.007068303 |
| ENSMUSG00000020844 | Nxn | 0.001217992 | -1.274685719 | 0.07443787 |
| ENSMUSG00000037750 | Fam222b | 0.001217992 | -1.176330647 | 0.113235376 |
| ENSMUSG00000040430 | Pitpnc1 | 0.001217992 | -2.784704534 | 6.03E-06 |
| ENSMUSG00000020571 | Pdia6 | 0.001217992 | 0.012526171 | 1 |
| ENSMUSG00000021219 | Rgs6 | 0.001217992 | -1.113004711 | 0.177165715 |
| ENSMUSG00000001729 | Akt1 | 0.001217992 | 0.277579744 | 0.987157952 |
| ENSMUSG00000072812 | Ahnak2 | 0.001217992 | 0.022650388 | 1 |

| | | | | |
|---|---|---|---|---|
| ENSMUSG00000056553 | Ptprn2 | 0.001217992 | -0.374496952 | 0.985672229 |
| ENSMUSG00000055137 | Sugct | 0.001217992 | -1.416317128 | 0.037122674 |
| ENSMUSG00000038415 | Foxq1 | 0.001217992 | 0.229883002 | 1 |
| ENSMUSG00000021420 | Fars2 | 0.001217992 | -1.650969733 | 0.01127169 |
| ENSMUSG00000021431 | Snrnp48 | 0.001217992 | -2.245455007 | 0.000328058 |
| ENSMUSG00000025873 | Faf2 | 0.001217992 | -1.641383684 | 0.012887418 |
| ENSMUSG00000021496 | Pcbd2 | 0.001217992 | -0.287034111 | 1 |
| ENSMUSG00000074865 | Zfp934 | 0.001217992 | -2.584158815 | 2.86E-05 |
| ENSMUSG00000021596 | Mctp1 | 0.001217992 | 0.195117584 | 1 |
| ENSMUSG00000021537 | Cetn3 | 0.001217992 | 1.022650388 | 0.25647791 |
| ENSMUSG00000021814 | Anxa7 | 0.001217992 | 0.265506912 | 0.989737947 |
| ENSMUSG00000021770 | Samd8 | 0.001217992 | -1.027274837 | 0.205339793 |
| ENSMUSG00000021917 | Spcs1 | 0.001217992 | 1.814845503 | 0.006295279 |
| ENSMUSG00000014496 | Ankrd28 | 0.001217992 | -2.836031115 | 4.27E-06 |
| ENSMUSG00000021840 | Mapk1ip1l | 0.001217992 | -0.893691683 | 0.33333983 |
| ENSMUSG00000022226 | Mcpt2 | 0.001217992 | 2.084934666 | 0.001737647 |
| ENSMUSG00000047911 | Npm2 | 0.001217992 | -0.185154497 | 1 |
| ENSMUSG00000022205 | Sub1 | 0.001217992 | 0.868616759 | 0.306626259 |
| ENSMUSG00000040420 | Cdh18 | 0.001217992 | 0.043114491 | 1 |
| ENSMUSG00000039007 | Cpq | 0.001217992 | -1.534961625 | 0.020939914 |
| ENSMUSG00000022307 | Oxr1 | 0.001217992 | -2.824781973 | 4.67E-06 |
| ENSMUSG00000022419 | Deptor | 0.001217992 | -0.641383684 | 0.598253125 |
| ENSMUSG00000022427 | Tomm22 | 0.001217992 | 1.924469994 | 0.004183188 |
| ENSMUSG00000042428 | Mgat3 | 0.001217992 | -1.343302331 | 0.053843381 |
| ENSMUSG00000035891 | Cerk | 0.001217992 | -1.176330647 | 0.115171587 |
| ENSMUSG00000022515 | Anks3 | 0.001217992 | -1.089697338 | 0.166936301 |
| ENSMUSG00000022807 | Osbpl11 | 0.001217992 | -1.012477835 | 0.217194106 |
| ENSMUSG00000022799 | Arhgap31 | 0.001217992 | -0.647781455 | 0.608204428 |
| ENSMUSG00000022935 | Grik1 | 0.001217992 | -1.09906552 | 0.176575961 |
| ENSMUSG00000040785 | Ttc3 | 0.001217992 | -2.660492507 | 1.61E-05 |
| ENSMUSG00000024130 | Abca3 | 0.001217992 | -0.54184801 | 0.736456705 |
| ENSMUSG00000015120 | Ube2i | 0.001217992 | -0.319455589 | 1 |
| ENSMUSG00000002279 | Lmf1 | 0.001217992 | -1.909751534 | 0.00337143 |
| ENSMUSG00000062202 | Btbd9 | 0.001217992 | -1.888298424 | 0.00293439 |
| ENSMUSG00000024038 | Ndufv3 | 0.001217992 | -0.236993429 | 1 |
| ENSMUSG00000024387 | Csnk2b | 0.001217992 | 1.195117584 | 0.127257642 |
| ENSMUSG00000023932 | Cdc5l | 0.001217992 | -1.228481929 | 0.092468958 |
| ENSMUSG00000036568 | Bicral | 0.001217992 | -1.962028813 | 0.001932744 |
| ENSMUSG00000002379 | Ndufa11 | 0.001217992 | 1.648835551 | 0.031609741 |
| ENSMUSG00000039770 | Ypel5 | 0.001217992 | -1.066007232 | 0.172064687 |
| ENSMUSG00000033382 | Trappc8 | 0.001217992 | -0.992509419 | 0.226567468 |
| ENSMUSG00000033032 | Afap1l1 | 0.001217992 | -0.915065334 | 0.295355925 |
| ENSMUSG00000086128 | St8sia3os | 0.001217992 | -2.339355146 | 0.000161297 |
| ENSMUSG00000024587 | Nars | 0.001217992 | 0.002472506 | 1 |
| ENSMUSG00000024539 | Ptpn2 | 0.001217992 | -1.456959113 | 0.030588995 |
| ENSMUSG00000024908 | Ppp6r3 | 0.001217992 | -2.082631062 | 0.000987149 |
| ENSMUSG00000045098 | Kmt5b | 0.001217992 | -2.395558568 | 0.000110956 |
| ENSMUSG00000024955 | Esrra | 0.001217992 | -1.682470876 | 0.010154662 |
| ENSMUSG00000024943 | Smc5 | 0.001217992 | -1.189546266 | 0.107795092 |
| ENSMUSG00000024789 | Jak2 | 0.001217992 | -1.287034111 | 0.070562421 |
| ENSMUSG00000041362 | Shtn1 | 0.001217992 | -0.412564993 | 0.886203497 |

| | | | | |
|---|---|---|---|---|
| ENSMUSG00000031109 | Enox2 | 0.001217992 | -2.029726316 | 0.001569694 |
| ENSMUSG00000059203 | Il1rapl2 | 0.001217992 | 0.002472506 | 1 |
| ENSMUSG00000068457 | Uty | 0.001217992 | -2.536686305 | 4.79E-05 |
| ENSMUSG00000025921 | Rdh10 | 0.001162629 | -0.743481871 | 0.467800117 |
| ENSMUSG00000038349 | Plcl1 | 0.001162629 | -1.041479949 | 0.20542113 |
| ENSMUSG00000026028 | Trak2 | 0.001162629 | 0.340082845 | 0.89270014 |
| ENSMUSG00000067028 | Cntnap5b | 0.001162629 | -0.139660343 | 1 |
| ENSMUSG00000042207 | Kdm5b | 0.001162629 | -1.919638826 | 0.002739666 |
| ENSMUSG00000026483 | Fam129a | 0.001162629 | -0.491645672 | 0.817758884 |
| ENSMUSG00000019699 | Akt3 | 0.001162629 | -1.743481871 | 0.007372949 |
| ENSMUSG00000026494 | Kif26b | 0.001162629 | -1.081557389 | 0.167528217 |
| ENSMUSG00000026843 | Fubp3 | 0.001162629 | -1.34706005 | 0.056352204 |
| ENSMUSG00000061136 | Prpf40a | 0.001162629 | -1.427008207 | 0.039723841 |
| ENSMUSG00000042410 | Agps | 0.001162629 | -1.383585926 | 0.050877179 |
| ENSMUSG00000044033 | Ccdc141 | 0.001162629 | -1.531977766 | 0.023822655 |
| ENSMUSG00000033955 | Tnks1bp1 | 0.001162629 | -0.743481871 | 0.467800117 |
| ENSMUSG00000027170 | Eif3m | 0.001162629 | 0.584705214 | 0.605573874 |
| ENSMUSG00000057147 | Dph6 | 0.001162629 | -1.578233357 | 0.018507952 |
| ENSMUSG00000046229 | Scand1 | 0.001162629 | 1.616414074 | 0.021198462 |
| ENSMUSG00000070544 | Top1 | 0.001162629 | -1.046550939 | 0.202018463 |
| ENSMUSG00000056501 | Cebpb | 0.001162629 | 0.074844169 | 1 |
| ENSMUSG00000033502 | Cdc14a | 0.001162629 | -0.886086267 | 0.350554273 |
| ENSMUSG00000028128 | F3 | 0.001162629 | 1.250286175 | 0.098930652 |
| ENSMUSG00000028124 | Gclm | 0.001162629 | 0.450055687 | 0.811247775 |
| ENSMUSG00000039735 | Fnbp1l | 0.001162629 | -1.724622844 | 0.007982873 |
| ENSMUSG00000050931 | Sgms2 | 0.001162629 | -1.679635389 | 0.010987762 |
| ENSMUSG00000041272 | Tox | 0.001162629 | -1.395558568 | 0.046054041 |
| ENSMUSG00000040720 | Virma | 0.001162629 | -1.12055152 | 0.151868261 |
| ENSMUSG00000028274 | Rngtt | 0.001162629 | -1.177135049 | 0.123736816 |
| ENSMUSG00000028409 | Smu1 | 0.001162629 | -1.465380017 | 0.032028637 |
| ENSMUSG00000028456 | Unc13b | 0.001162629 | -2.294744853 | 0.000251527 |
| ENSMUSG00000028309 | Rnf20 | 0.001162629 | -2.373532262 | 0.000146025 |
| ENSMUSG00000050812 | Ecpas | 0.001162629 | -1.076608173 | 0.179855359 |
| ENSMUSG00000028373 | Astn2 | 0.001162629 | -0.021015847 | 1 |
| ENSMUSG00000028798 | Eif3i | 0.001162629 | 1.553678318 | 0.0315747 |
| ENSMUSG00000040945 | Rcc2 | 0.001162629 | 0.10826717 | 1 |
| ENSMUSG00000009863 | Sdhb | 0.001162629 | -0.021015847 | 1 |
| ENSMUSG00000005907 | Pex1 | 0.001162629 | -1.091405175 | 0.170180554 |
| ENSMUSG00000013622 | Atraid | 0.001162629 | -1.190940848 | 0.116637579 |
| ENSMUSG00000036323 | Srp72 | 0.001162629 | -1.036391073 | 0.19917628 |
| ENSMUSG00000029338 | Antxr2 | 0.001162629 | -1.502757627 | 0.02789416 |
| ENSMUSG00000057816 | Cfap299 | 0.001162629 | -0.919638826 | 0.346711083 |
| ENSMUSG00000054720 | Lrrc8c | 0.001162629 | -3.581730801 | 4.09E-09 |
| ENSMUSG00000029359 | Tesc | 0.001162629 | 2.10826717 | 0.005551659 |
| ENSMUSG00000066735 | Vkorc1l1 | 0.001162629 | -1.62982509 | 0.013611347 |
| ENSMUSG00000039959 | Hip1 | 0.001162629 | -1.695863851 | 0.011633761 |
| ENSMUSG00000039683 | Sdk1 | 0.001162629 | -0.391578717 | 0.982068793 |
| ENSMUSG00000079111 | Kdelr2 | 0.001162629 | 1.978984153 | 0.003427316 |
| ENSMUSG00000029640 | Usp12 | 0.001162629 | -1.564158172 | 0.01904133 |
| ENSMUSG00000053470 | Kdm3a | 0.001162629 | -1.971217614 | 0.001942582 |
| ENSMUSG00000043131 | Mob1a | 0.001162629 | -0.010673903 | 1 |

| | | | | |
|---|---|---|---|---|
| ENSMUSG00000051586 | Mical3 | 0.001162629 | -2.132524162 | 0.00078463 |
| ENSMUSG00000030213 | Atf7ip | 0.001162629 | -1.851734762 | 0.00425152 |
| ENSMUSG00000030223 | Ptpro | 0.001162629 | -2.041479949 | 0.001743806 |
| ENSMUSG00000060860 | Ube2s | 0.001162629 | 0.87944848 | 0.341720553 |
| ENSMUSG00000003435 | Supt5 | 0.001162629 | -1.585219787 | 0.017420371 |
| ENSMUSG00000031068 | Glrx3 | 0.001162629 | 0.201376574 | 1 |
| ENSMUSG00000101609 | Kcnq1ot1 | 0.001162629 | -1.430891641 | 0.036215483 |
| ENSMUSG00000063626 | Unc5d | 0.001162629 | 0.733227739 | 0.466464457 |
| ENSMUSG00000003039 | Fam32a | 0.001162629 | 0.394021652 | 0.832472219 |
| ENSMUSG00000060098 | Prmt7 | 0.001162629 | -0.43862729 | 0.877036949 |
| ENSMUSG00000031950 | Gabarapl2 | 0.001162629 | 0.978984153 | 0.245298386 |
| ENSMUSG00000052557 | Gan | 0.001162629 | -1.041479949 | 0.200799422 |
| ENSMUSG00000031938 | 4931406C07Rik | 0.001162629 | -0.542784521 | 0.707525726 |
| ENSMUSG00000042138 | Msantd2 | 0.001162629 | -0.936126949 | 0.295347805 |
| ENSMUSG00000032332 | Col12a1 | 0.001162629 | -0.186353579 | 1 |
| ENSMUSG00000015755 | Tab2 | 0.001162629 | -0.430891641 | 0.875352165 |
| ENSMUSG00000060487 | Samd5 | 0.001162629 | -2.57648145 | 3.32E-05 |
| ENSMUSG00000020009 | Ifngr1 | 0.001162629 | -0.186353579 | 1 |
| ENSMUSG00000019802 | Sec63 | 0.001162629 | -1.158519371 | 0.128972787 |
| ENSMUSG00000020076 | Ddx50 | 0.001162629 | -0.941581379 | 0.27550706 |
| ENSMUSG00000003068 | Stk11 | 0.001162629 | -1.262420237 | 0.083056159 |
| ENSMUSG00000020018 | Snrpf | 0.001162629 | 1.786339075 | 0.008282606 |
| ENSMUSG00000020427 | Igfbp3 | 0.001162629 | 1.786339075 | 0.008282606 |
| ENSMUSG00000018583 | G3bp1 | 0.001162629 | -1.676367675 | 0.010838314 |
| ENSMUSG00000038188 | Scarf1 | 0.001162629 | -0.952428762 | 0.279592421 |
| ENSMUSG00000019173 | Rab5c | 0.001162629 | -0.571212929 | 0.675561045 |
| ENSMUSG00000034708 | Grn | 0.001162629 | 0.493557326 | 0.75496808 |
| ENSMUSG00000025571 | Tnrc6c | 0.001162629 | -1.411369083 | 0.040634685 |
| ENSMUSG00000020570 | Sypl | 0.001162629 | 0.288088208 | 0.971854959 |
| ENSMUSG00000020952 | Scfd1 | 0.001162629 | -1.279750115 | 0.080437551 |
| ENSMUSG00000021010 | Npas3 | 0.001162629 | 0.262777119 | 1 |
| ENSMUSG00000020994 | Pnn | 0.001162629 | -1.592172548 | 0.016410626 |
| ENSMUSG00000021109 | Hif1a | 0.001162629 | -0.963195192 | 0.26676534 |
| ENSMUSG00000021124 | Vti1b | 0.001162629 | 0.898813804 | 0.33333983 |
| ENSMUSG00000021134 | Srsf5 | 0.001162629 | -2.969883638 | 1.59E-06 |
| ENSMUSG00000041645 | Ddx24 | 0.001162629 | -1.430891641 | 0.036215483 |
| ENSMUSG00000021281 | Tnfaip2 | 0.001162629 | -2.279750115 | 0.000281729 |
| ENSMUSG00000006356 | Crip2 | 0.001162629 | 1.353379668 | 0.059512215 |
| ENSMUSG00000039233 | Tbce | 0.001162629 | -0.857517115 | 0.365386077 |
| ENSMUSG00000038175 | Mylip | 0.001162629 | -0.469161658 | 0.866111192 |
| ENSMUSG00000025878 | Uimc1 | 0.001162629 | -1.524728027 | 0.023162612 |
| ENSMUSG00000021501 | Caml | 0.001162629 | -1.679635389 | 0.010987762 |
| ENSMUSG00000035248 | Tut7 | 0.001162629 | -1.746601213 | 0.007463152 |
| ENSMUSG00000021559 | Dapk1 | 0.001162629 | -1.560617814 | 0.021021848 |
| ENSMUSG00000021576 | Pdcd6 | 0.001162629 | 0.682002415 | 0.604880543 |
| ENSMUSG00000007617 | Homer1 | 0.001162629 | -1.43476465 | 0.045356901 |
| ENSMUSG00000021687 | Scamp1 | 0.001162629 | -1.096303974 | 0.183728346 |
| ENSMUSG00000021662 | Arhgef28 | 0.001162629 | -2.134906813 | 0.000732354 |
| ENSMUSG00000021754 | Map3k1 | 0.001162629 | -2.084025644 | 0.001046072 |
| ENSMUSG00000040640 | Erc2 | 0.001162629 | -1.553510928 | 0.020028214 |
| ENSMUSG00000022100 | Xpo7 | 0.001162629 | -1.922399966 | 0.00256978 |

| | | | | |
|---|---|---|---|---|
| ENSMUSG00000115704 | Gm49171 | 0.001162629 | 0.353379668 | 0.909076441 |
| ENSMUSG00000022200 | Golph3 | 0.001162629 | -0.101186196 | 1 |
| ENSMUSG00000042406 | Atf4 | 0.001162629 | -0.804622166 | 0.433763754 |
| ENSMUSG00000023020 | Cox14 | 0.001162629 | 1.523304669 | 0.05095448 |
| ENSMUSG00000039457 | Ppl | 0.001162629 | -0.919638826 | 0.311028128 |
| ENSMUSG00000009569 | Mrtfb | 0.001162629 | -0.52108945 | 0.74230625 |
| ENSMUSG00000022816 | Fstl1 | 0.001162629 | 0.020804329 | 1 |
| ENSMUSG00000004843 | Chmp2b | 0.001162629 | 0.380346716 | 0.897739091 |
| ENSMUSG00000001576 | Ergic1 | 0.001162629 | -0.249284835 | 1 |
| ENSMUSG00000002625 | Akap8l | 0.001162629 | -2.091405175 | 0.00094815 |
| ENSMUSG00000038954 | Supt3 | 0.001162629 | -0.880417517 | 0.347684915 |
| ENSMUSG00000000708 | Kat2b | 0.001162629 | -1.391578717 | 0.045609829 |
| ENSMUSG00000049760 | Micos13 | 0.001162629 | 1.523304669 | 0.034892018 |
| ENSMUSG00000024242 | Map4k3 | 0.001162629 | -0.89173283 | 0.325200788 |
| ENSMUSG00000055817 | Mta3 | 0.001162629 | -1.659917155 | 0.012749115 |
| ENSMUSG00000071037 | Camkmt | 0.001162629 | -0.874726405 | 0.345705585 |
| ENSMUSG00000024236 | Svil | 0.001162629 | -1.86614742 | 0.003580054 |
| ENSMUSG00000097222 | 1010001N08Rik | 0.001162629 | -1.046550939 | 0.202018463 |
| ENSMUSG00000034295 | Fhod3 | 0.001162629 | -1.149120673 | 0.136825431 |
| ENSMUSG00000118257 | 9630014M24Rik | 0.001162629 | -1.86614742 | 0.003580054 |
| ENSMUSG00000037416 | Dmxl1 | 0.001162629 | -2.049079765 | 0.001228072 |
| ENSMUSG00000044646 | Zbtb7c | 0.001162629 | -0.35933838 | 0.989834251 |
| ENSMUSG00000024913 | Lrp5 | 0.001162629 | -1.305360759 | 0.07188116 |
| ENSMUSG00000024867 | Pip5k1b | 0.001162629 | -0.968548427 | 0.274319469 |
| ENSMUSG00000024803 | Ankrd1 | 0.001162629 | -2.689394356 | 1.47E-05 |
| ENSMUSG00000047731 | Wbp1l | 0.001162629 | -1.371513094 | 0.051203786 |
| ENSMUSG00000057421 | Las1l | 0.001162629 | -1.419209836 | 0.041758382 |
| ENSMUSG00000034160 | Ogt | 0.001162629 | -2.170182288 | 0.000581282 |
| ENSMUSG00000031353 | Rbbp7 | 0.001162629 | -0.585219787 | 0.680611375 |
| ENSMUSG00000096768 | Gm47283 | 0.001162629 | -2.132524162 | 0.00078463 |
| ENSMUSG00000051951 | Xkr4 | 0.001107265 | -0.981214839 | 0.262284476 |
| ENSMUSG00000026123 | Plekhb2 | 0.001107265 | 0.631762038 | 0.574003246 |
| ENSMUSG00000025982 | Sf3b1 | 0.001107265 | -1.720398258 | 0.009262075 |
| ENSMUSG00000025969 | Nrp2 | 0.001107265 | -2.703680863 | 1.50E-05 |
| ENSMUSG00000070695 | Cntnap5a | 0.001107265 | 0.246737108 | 1 |
| ENSMUSG00000026466 | Tor1aip1 | 0.001107265 | -1.602703216 | 0.016342424 |
| ENSMUSG00000038733 | Wdr26 | 0.001107265 | -1.89842624 | 0.00335241 |
| ENSMUSG00000026623 | Lpgat1 | 0.001107265 | -0.865737621 | 0.363322631 |
| ENSMUSG00000015094 | Npdc1 | 0.001107265 | 1.286265472 | 0.081548655 |
| ENSMUSG00000026878 | Rab14 | 0.001107265 | -0.262627774 | 1 |
| ENSMUSG00000035392 | Dennd1a | 0.001107265 | -1.978437754 | 0.002124365 |
| ENSMUSG00000027363 | Usp8 | 0.001107265 | -1.717070241 | 0.009112331 |
| ENSMUSG00000068290 | Ddrgk1 | 0.001107265 | 0.975063784 | 0.303468298 |
| ENSMUSG00000027593 | Raly | 0.001107265 | 2.041152974 | 0.003136686 |
| ENSMUSG00000038914 | Dido1 | 0.001107265 | -1.547561661 | 0.022755145 |
| ENSMUSG00000016344 | Ppdpf | 0.001107265 | 1.648835551 | 0.066465211 |
| ENSMUSG00000027550 | Lrrcc1 | 0.001107265 | -1.410058138 | 0.045308174 |
| ENSMUSG00000041977 | Arhgef11 | 0.001107265 | -1.779025988 | 0.006613418 |
| ENSMUSG00000001416 | Cct3 | 0.001107265 | 0.146335211 | 1 |
| ENSMUSG00000027944 | Hax1 | 0.001107265 | -1.285276513 | 0.07932676 |
| ENSMUSG00000042520 | Ubap2l | 0.001107265 | -2.014129461 | 0.001729963 |

| | | | | |
|---|---|---|---|---|
| ENSMUSG00000042390 | Gatad2b | 0.001107265 | -2.678235847 | 1.67E-05 |
| ENSMUSG00000032913 | Lrig2 | 0.001107265 | -0.693556646 | 0.557948937 |
| ENSMUSG00000036745 | Ttll7 | 0.001107265 | -0.766201948 | 0.466464457 |
| ENSMUSG00000062908 | Acadm | 0.001107265 | 0.029925719 | 1 |
| ENSMUSG00000028207 | Asph | 0.001107265 | -0.551304063 | 0.730132058 |
| ENSMUSG00000028557 | Rnf11 | 0.001107265 | 0.831699609 | 0.383578797 |
| ENSMUSG00000028710 | Atpaf1 | 0.001107265 | -0.727031351 | 0.510984604 |
| ENSMUSG00000028830 | AU040320 | 0.001107265 | -0.829211745 | 0.405913817 |
| ENSMUSG00000040859 | Bsdc1 | 0.001107265 | -0.700314012 | 0.521275193 |
| ENSMUSG00000028961 | Pgd | 0.001107265 | -0.014129461 | 1 |
| ENSMUSG00000061601 | Pclo | 0.001107265 | -0.859713834 | 0.399109893 |
| ENSMUSG00000028944 | Prkag2 | 0.001107265 | -2.090703986 | 0.001343992 |
| ENSMUSG00000029135 | Fosl2 | 0.001107265 | -0.759746926 | 0.47563319 |
| ENSMUSG00000029093 | Sorcs2 | 0.001107265 | -1.524901064 | 0.024821097 |
| ENSMUSG00000061755 | Bod1l | 0.001107265 | -2.000506177 | 0.00188145 |
| ENSMUSG00000037736 | Limch1 | 0.001107265 | -0.746749585 | 0.533752558 |
| ENSMUSG00000029212 | Gabrb1 | 0.001107265 | 0.272968649 | 1 |
| ENSMUSG00000029229 | Chic2 | 0.001107265 | -0.992270027 | 0.245842568 |
| ENSMUSG00000029291 | Rufy3 | 0.001107265 | -0.727031351 | 0.496209758 |
| ENSMUSG00000029422 | Rsrc2 | 0.001107265 | -1.372474955 | 0.0530054 |
| ENSMUSG00000060882 | Kcnd2 | 0.001107265 | 0.195662923 | 1 |
| ENSMUSG00000025609 | Mkln1 | 0.001107265 | -1.89842624 | 0.00335241 |
| ENSMUSG00000091625 | Lsm5 | 0.001107265 | 1.286265472 | 0.154807284 |
| ENSMUSG00000030335 | Mrpl51 | 0.001107265 | 0.812334284 | 0.396214287 |
| ENSMUSG00000030203 | Dusp16 | 0.001107265 | -1.772628217 | 0.00701405 |
| ENSMUSG00000046733 | Gprc5a | 0.001107265 | -0.609898717 | 0.647835155 |
| ENSMUSG00000030265 | Kras | 0.001107265 | -1.31639903 | 0.071480332 |
| ENSMUSG00000002984 | Tomm40 | 0.001107265 | -0.216234868 | 1 |
| ENSMUSG00000109324 | Prmt1 | 0.001107265 | 1.259793261 | 0.110579724 |
| ENSMUSG00000025326 | Ube3a | 0.001107265 | -1.303142934 | 0.075935796 |
| ENSMUSG00000108532 | Gm32647 | 0.001107265 | 0.31323252 | 0.969214425 |
| ENSMUSG00000035623 | Rsf1 | 0.001107265 | -2.220941871 | 0.000440079 |
| ENSMUSG00000052889 | Prkcb | 0.001107265 | -0.216234868 | 1 |
| ENSMUSG00000030779 | Rbbp6 | 0.001107265 | -1.883659529 | 0.00364093 |
| ENSMUSG00000030805 | Stx4a | 0.001107265 | 0.299685988 | 0.958639125 |
| ENSMUSG00000038611 | Phrf1 | 0.001107265 | -1.158519371 | 0.128972787 |
| ENSMUSG00000039057 | Myo16 | 0.001107265 | 1.134262379 | 0.172061305 |
| ENSMUSG00000006273 | Atp6v1b2 | 0.001107265 | -0.342551318 | 1 |
| ENSMUSG00000031622 | Sin3b | 0.001107265 | 0.233798052 | 1 |
| ENSMUSG00000019362 | D8Ertd738e | 0.001107265 | 1.134262379 | 0.139375619 |
| ENSMUSG00000031652 | N4bp1 | 0.001107265 | -1.133772708 | 0.153743585 |
| ENSMUSG00000036510 | Cdh8 | 0.001107265 | 0.737372226 | 0.47966612 |
| ENSMUSG00000096188 | Cmtm4 | 0.001107265 | -0.38511178 | 0.935904351 |
| ENSMUSG00000031827 | Cotl1 | 0.001107265 | 1.368727632 | 0.065226458 |
| ENSMUSG00000039488 | Cntn5 | 0.001107265 | -0.118718363 | 1 |
| ENSMUSG00000074505 | Fat3 | 0.001107265 | -0.513435877 | 0.774790843 |
| ENSMUSG00000004771 | Rab11a | 0.001107265 | -0.253467775 | 1 |
| ENSMUSG00000032413 | Rasa2 | 0.001107265 | -1.458692913 | 0.033547406 |
| ENSMUSG00000019846 | Lama4 | 0.001107265 | -0.753262892 | 0.472949383 |
| ENSMUSG00000111368 | Gm47889 | 0.001107265 | -0.197251765 | 1 |
| ENSMUSG00000004665 | Cnn2 | 0.001107265 | 0.471297366 | 0.769719461 |

| | | | | |
|---|---|---|---|---|
| ENSMUSG00000069565 | Dazap1 | 0.001107265 | -1.359726462 | 0.055580646 |
| ENSMUSG00000052681 | Rap1b | 0.001107265 | -0.727031351 | 0.496209758 |
| ENSMUSG00000020224 | Llph | 0.001107265 | -0.271729981 | 1 |
| ENSMUSG00000034602 | Mon2 | 0.001107265 | -1.540047608 | 0.023955525 |
| ENSMUSG00000040043 | Rbms2 | 0.001107265 | -1.289763904 | 0.080293673 |
| ENSMUSG00000041961 | Znrf3 | 0.001107265 | -1.148671585 | 0.13735263 |
| ENSMUSG00000015837 | Sqstm1 | 0.001107265 | 0.146335211 | 1 |
| ENSMUSG00000037331 | Larp1 | 0.001107265 | -1.108594145 | 0.165537404 |
| ENSMUSG00000020794 | Ube2g1 | 0.001107265 | -1.972867496 | 0.002193744 |
| ENSMUSG00000017615 | Tnfaip1 | 0.001107265 | 0.549299878 | 0.705338822 |
| ENSMUSG00000020696 | Rffl | 0.001107265 | -1.280775121 | 0.081728936 |
| ENSMUSG00000020627 | Klhl29 | 0.001107265 | 0.368727632 | 0.884935781 |
| ENSMUSG00000054302 | Eapp | 0.001107265 | -1.019542957 | 0.217448938 |
| ENSMUSG00000021127 | Zfp36l1 | 0.001107265 | -0.234971431 | 1 |
| ENSMUSG00000049106 | Dcaf5 | 0.001107265 | -1.056879109 | 0.195690131 |
| ENSMUSG00000020962 | Gtf2a1 | 0.001107265 | -0.536275854 | 0.724721153 |
| ENSMUSG00000041236 | Vps41 | 0.001107265 | -1.798050814 | 0.006159437 |
| ENSMUSG00000071252 | 2210408I21Rik | 0.001107265 | -1.303142934 | 0.075935796 |
| ENSMUSG00000012422 | Tmem167 | 0.001107265 | 0.220974012 | 1 |
| ENSMUSG00000049658 | Bdp1 | 0.001107265 | -2.715403348 | 1.40E-05 |
| ENSMUSG00000041014 | Nrg3 | 0.001107265 | -0.056879109 | 1 |
| ENSMUSG00000022193 | Psmb5 | 0.001107265 | 1.456190473 | 0.039995951 |
| ENSMUSG00000044447 | Dock5 | 0.001107265 | -2.225633566 | 0.000457984 |
| ENSMUSG00000068015 | Lrch1 | 0.001107265 | -1.652333983 | 0.012887418 |
| ENSMUSG00000022024 | Sugt1 | 0.001107265 | -0.740206739 | 0.499942088 |
| ENSMUSG00000033083 | Tbc1d4 | 0.001107265 | -0.992270027 | 0.257354963 |
| ENSMUSG00000022246 | Rai14 | 0.001107265 | -1.844543399 | 0.004559973 |
| ENSMUSG00000040452 | Cdh12 | 0.001107265 | 0.07536869 | 1 |
| ENSMUSG00000047921 | Trappc9 | 0.001107265 | -0.964471575 | 0.27550706 |
| ENSMUSG00000033047 | Eif3l | 0.001107265 | 0.029925719 | 1 |
| ENSMUSG00000022383 | Ppara | 0.001107265 | -1.014129461 | 0.238808981 |
| ENSMUSG00000016028 | Celsr1 | 0.001107265 | -0.52108945 | 0.765086656 |
| ENSMUSG00000099083 | Atf7 | 0.001107265 | -2.303142934 | 0.000251527 |
| ENSMUSG00000005846 | Rsl1d1 | 0.001107265 | -1.267186056 | 0.082214652 |
| ENSMUSG00000022528 | Hes1 | 0.001107265 | -0.652333983 | 0.617771696 |
| ENSMUSG00000022946 | Dop1b | 0.001107265 | -1.720398258 | 0.009262075 |
| ENSMUSG00000050272 | Dscam | 0.001107265 | -0.128772027 | 1 |
| ENSMUSG00000037098 | Rab11fip3 | 0.001107265 | -0.458692913 | 0.82350809 |
| ENSMUSG00000023067 | Cdkn1a | 0.001107265 | 1.793225461 | 0.009461327 |
| ENSMUSG00000024300 | Myo1f | 0.001107265 | -0.187665716 | 1 |
| ENSMUSG00000024393 | Prrc2a | 0.001107265 | -1.389299577 | 0.046343118 |
| ENSMUSG00000023915 | Tnfrsf21 | 0.001107265 | -0.418278644 | 0.886746766 |
| ENSMUSG00000024099 | Ndufv2 | 0.001107265 | -0.30757518 | 1 |
| ENSMUSG00000001870 | Ltbp1 | 0.001107265 | -1.558759868 | 0.021725703 |
| ENSMUSG00000024109 | Nrxn1 | 0.001107265 | -0.38511178 | 0.943363727 |
| ENSMUSG00000024294 | Mib1 | 0.001107265 | -0.631272368 | 0.656787419 |
| ENSMUSG00000024404 | Riok3 | 0.001107265 | -1.225633566 | 0.09684834 |
| ENSMUSG00000024357 | Sil1 | 0.001107265 | -1.655814552 | 0.013104109 |
| ENSMUSG00000032656 | 3-Mar | 0.001107265 | -1.43862729 | 0.040575148 |
| ENSMUSG00000044176 | Spink10 | 0.001107265 | -0.93039058 | 0.30756351 |
| ENSMUSG00000041258 | Zfp236 | 0.001107265 | -1.478483279 | 0.032780298 |

| | | | | |
|---|---|---|---|---|
| ENSMUSG00000024921 | Smarca2 | 0.001107265 | -0.997766015 | 0.249496428 |
| ENSMUSG00000025025 | Mxi1 | 0.001107265 | -0.953200463 | 0.271909314 |
| ENSMUSG00000031153 | Gripap1 | 0.001107265 | -1.638327125 | 0.014476156 |
| ENSMUSG00000031309 | Rps6ka3 | 0.001107265 | -1.128772027 | 0.144078128 |
| ENSMUSG00000033569 | Adgrb3 | 0.001051902 | 0.544498892 | 0.697198109 |
| ENSMUSG00000026070 | Il18r1 | 0.001051902 | -2.849415316 | 7.09E-06 |
| ENSMUSG00000025979 | Mob4 | 0.001051902 | -0.708716453 | 0.529111943 |
| ENSMUSG00000073664 | Nbeal1 | 0.001051902 | -2.335298043 | 0.000239934 |
| ENSMUSG00000026421 | Csrp1 | 0.001051902 | 0.202579322 | 1 |
| ENSMUSG00000026667 | Uhmk1 | 0.001051902 | 0.125273595 | 1 |
| ENSMUSG00000055067 | Smyd3 | 0.001051902 | -2.310007124 | 0.000266937 |
| ENSMUSG00000022995 | Enah | 0.001051902 | -1.911879403 | 0.003299476 |
| ENSMUSG00000073481 | 2-Mar | 0.001051902 | -1.13776081 | 0.147740585 |
| ENSMUSG00000036591 | Arhgap21 | 0.001051902 | -2.01792104 | 0.001807071 |
| ENSMUSG00000015087 | Rabl6 | 0.001051902 | -0.576130814 | 0.706929043 |
| ENSMUSG00000039844 | Rapgef1 | 0.001051902 | -1.204162036 | 0.119732322 |
| ENSMUSG00000026851 | BC005624 | 0.001051902 | -1.173894597 | 0.138589466 |
| ENSMUSG00000027204 | Fbn1 | 0.001051902 | -0.488667972 | 0.802072052 |
| ENSMUSG00000027404 | Snrpb | 0.001051902 | 1.040026309 | 0.224151767 |
| ENSMUSG00000019188 | H13 | 0.001051902 | -1.084518789 | 0.175506002 |
| ENSMUSG00000027536 | Chmp4c | 0.001051902 | -0.116699195 | 1 |
| ENSMUSG00000036834 | Plch1 | 0.001051902 | -2.316948733 | 0.000259495 |
| ENSMUSG00000028062 | Lamtor2 | 0.001051902 | 1.189403933 | 0.148130557 |
| ENSMUSG00000028059 | Arhgef2 | 0.001051902 | -1.817138913 | 0.00592361 |
| ENSMUSG00000038766 | Gabpb2 | 0.001051902 | -1.219060913 | 0.113023865 |
| ENSMUSG00000046722 | Cdc42se1 | 0.001051902 | 0.163408724 | 1 |
| ENSMUSG00000028030 | Tbck | 0.001051902 | -1.369086357 | 0.054245239 |
| ENSMUSG00000040128 | Pnrc1 | 0.001051902 | -0.062659355 | 1 |
| ENSMUSG00000028437 | Ubap1 | 0.001051902 | -1.736595486 | 0.011841136 |
| ENSMUSG00000028519 | Dab1 | 0.001051902 | -0.095325544 | 1 |
| ENSMUSG00000034557 | Zfyve9 | 0.001051902 | -0.722723311 | 0.534119278 |
| ENSMUSG00000028849 | Map7d1 | 0.001051902 | -1.243555474 | 0.099922289 |
| ENSMUSG00000057236 | Rbbp4 | 0.001051902 | -1.560617814 | 0.02307354 |
| ENSMUSG00000047945 | Marcksl1 | 0.001051902 | 0.86223919 | 0.365716212 |
| ENSMUSG00000028899 | Taf12 | 0.001051902 | -0.743481871 | 0.51221187 |
| ENSMUSG00000028851 | Nudc | 0.001051902 | 0.326907457 | 0.943363727 |
| ENSMUSG00000028822 | Tmem50a | 0.001051902 | -1.238689719 | 0.098608858 |
| ENSMUSG00000040761 | Spen | 0.001051902 | -1.753750207 | 0.008115089 |
| ENSMUSG00000049606 | Zfp644 | 0.001051902 | -1.583825205 | 0.021072979 |
| ENSMUSG00000029364 | Wsb2 | 0.001051902 | 0.088120597 | 1 |
| ENSMUSG00000079215 | Zfp664 | 0.001051902 | -1.286624196 | 0.084124052 |
| ENSMUSG00000053094 | Tmem248 | 0.001051902 | -0.13776081 | 1 |
| ENSMUSG00000006143 | Upk3bl | 0.001051902 | 0.312552164 | 0.958408446 |
| ENSMUSG00000029727 | Cyp3a13 | 0.001051902 | -0.351164449 | 1 |
| ENSMUSG00000041298 | Katnal1 | 0.001051902 | -0.77070334 | 0.466464457 |
| ENSMUSG00000044471 | Lncpint | 0.001051902 | -2.291331199 | 0.000323891 |
| ENSMUSG00000073155 | 1810058I24Rik | 0.001051902 | 0.841480629 | 0.416069222 |
| ENSMUSG00000002416 | Ndufb2 | 0.001051902 | 1.761310281 | 0.014353091 |
| ENSMUSG00000002413 | Braf | 0.001051902 | -1.089932285 | 0.185619905 |
| ENSMUSG00000053012 | Krcc1 | 0.001051902 | -1.948099591 | 0.002599819 |
| ENSMUSG00000030000 | Add2 | 0.001051902 | -1.106051951 | 0.176593224 |

| | | | | |
|---|---|---|---|---|
| ENSMUSG00000109336 | Samd4b | 0.001051902 | 0.075945883 | 1 |
| ENSMUSG00000001247 | Lsr | 0.001051902 | 0.821016527 | 0.424249514 |
| ENSMUSG00000015134 | Aldh1a3 | 0.001051902 | 0.416174795 | 0.849516228 |
| ENSMUSG00000039361 | Picalm | 0.001051902 | -1.199161355 | 0.118076677 |
| ENSMUSG00000030774 | Pak1 | 0.001051902 | -1.525091381 | 0.026888478 |
| ENSMUSG00000032812 | Arap1 | 0.001051902 | -0.855784841 | 0.398040691 |
| ENSMUSG00000030786 | Itgam | 0.001051902 | -1.52108945 | 0.032495347 |
| ENSMUSG00000040268 | Plekha1 | 0.001051902 | -0.954048857 | 0.294699981 |
| ENSMUSG00000031004 | Mki67 | 0.001051902 | -1.484563574 | 0.033469112 |
| ENSMUSG00000004568 | Arhgef18 | 0.001051902 | -1.463864581 | 0.034221489 |
| ENSMUSG00000031451 | Gas6 | 0.001051902 | 0.56137271 | 0.703945114 |
| ENSMUSG00000008892 | Vdac3 | 0.001051902 | 0.511332028 | 0.769512189 |
| ENSMUSG00000045636 | Mtus1 | 0.001051902 | -0.314638573 | 1 |
| ENSMUSG00000031684 | Slc10a7 | 0.001051902 | -1.463864581 | 0.034221489 |
| ENSMUSG00000031708 | Tecr | 0.001051902 | 0.17634778 | 1 |
| ENSMUSG00000001911 | Nfix | 0.001051902 | -1.023589791 | 0.22623675 |
| ENSMUSG00000031877 | Ces2g | 0.001051902 | 2.883300805 | 8.29E-05 |
| ENSMUSG00000013150 | Gfod2 | 0.001051902 | -1.18910769 | 0.127308076 |
| ENSMUSG00000037415 | Ranbp10 | 0.001051902 | -1.662231551 | 0.013677655 |
| ENSMUSG00000015023 | Ddx19a | 0.001051902 | -0.529082241 | 0.737638245 |
| ENSMUSG00000050730 | Arhgap42 | 0.001051902 | -3.069526075 | 8.77E-07 |
| ENSMUSG00000035382 | Pcsk7 | 0.001051902 | -0.687447836 | 0.561560978 |
| ENSMUSG00000039217 | Il18 | 0.001051902 | 0.595724215 | 0.616280812 |
| ENSMUSG00000039585 | Myo9a | 0.001051902 | -1.488667972 | 0.031064467 |
| ENSMUSG00000042444 | Mindy2 | 0.001051902 | -0.948099591 | 0.289810004 |
| ENSMUSG00000032407 | U2surp | 0.001051902 | -1.774070191 | 0.007443294 |
| ENSMUSG00000032458 | Copb2 | 0.001051902 | 0.648835551 | 0.610511243 |
| ENSMUSG00000015305 | Sash1 | 0.001051902 | -1.62542611 | 0.015437976 |
| ENSMUSG00000019998 | Stx7 | 0.001051902 | -0.314638573 | 1 |
| ENSMUSG00000019791 | Hint3 | 0.001051902 | 0.341407026 | 0.929782276 |
| ENSMUSG00000063760 | Rnf217 | 0.001051902 | -1.492760727 | 0.031456064 |
| ENSMUSG00000020089 | Ppa1 | 0.001051902 | 1.242843192 | 0.139039734 |
| ENSMUSG00000006498 | Ptbp1 | 0.001051902 | -0.228908699 | 1 |
| ENSMUSG00000008398 | Elk3 | 0.001051902 | -1.673092543 | 0.015397826 |
| ENSMUSG00000020021 | Fgd6 | 0.001051902 | -1.525091381 | 0.030588995 |
| ENSMUSG00000020451 | Limk2 | 0.001051902 | -1.073630473 | 0.189457792 |
| ENSMUSG00000009079 | Ewsr1 | 0.001051902 | -1.417253639 | 0.044662814 |
| ENSMUSG00000020149 | Rab1a | 0.001051902 | -0.583825205 | 0.724721153 |
| ENSMUSG00000040711 | Sh3pxd2b | 0.001051902 | -1.219060913 | 0.113023865 |
| ENSMUSG00000055134 | 9130017K11Rik | 0.001051902 | -2.029236354 | 0.001634767 |
| ENSMUSG00000018569 | Cldn7 | 0.001051902 | 2.722836133 | 0.000175432 |
| ENSMUSG00000017286 | Glod4 | 0.001051902 | -0.817138913 | 0.436887165 |
| ENSMUSG00000020709 | Adap2 | 0.001051902 | -0.817138913 | 0.454419454 |
| ENSMUSG00000017428 | Psmd11 | 0.001051902 | -0.47217985 | 0.847265357 |
| ENSMUSG00000047040 | Prr15l | 0.001051902 | -0.757156808 | 0.475884921 |
| ENSMUSG00000049807 | Arhgap23 | 0.001051902 | -0.810596067 | 0.426815929 |
| ENSMUSG00000020737 | Jpt1 | 0.001051902 | -0.095325544 | 1 |
| ENSMUSG00000020777 | Acox1 | 0.001051902 | -1.046044641 | 0.215910103 |
| ENSMUSG00000042700 | Sipa1l1 | 0.001051902 | -1.484563574 | 0.033469112 |
| ENSMUSG00000021264 | Yy1 | 0.001051902 | -1.647621856 | 0.014140013 |
| ENSMUSG00000052551 | Adarb2 | 0.001051902 | 0.431604835 | 0.868661374 |

| | | | | |
|---|---|---|---|---|
| ENSMUSG00000046159 | Chrm3 | 0.001051902 | -2.166227466 | 0.001013348 |
| ENSMUSG00000015937 | H2afy | 0.001051902 | -1.19414328 | 0.118554709 |
| ENSMUSG00000021595 | Nsun2 | 0.001051902 | -0.708716453 | 0.529111943 |
| ENSMUSG00000042348 | Arl15 | 0.001051902 | -1.45130112 | 0.039785293 |
| ENSMUSG00000062014 | Gmfb | 0.001051902 | -1.595290392 | 0.018165507 |
| ENSMUSG00000040123 | Zmym5 | 0.001051902 | -2.009375931 | 0.001871203 |
| ENSMUSG00000021945 | Zmym2 | 0.001051902 | -2.369086357 | 0.000177656 |
| ENSMUSG00000034205 | Loxl2 | 0.001051902 | 0.202579322 | 1 |
| ENSMUSG00000025558 | Dock9 | 0.001051902 | -2.003650943 | 0.001961053 |
| ENSMUSG00000039828 | Wdr70 | 0.001051902 | -1.560617814 | 0.02307354 |
| ENSMUSG00000039385 | Cdh6 | 0.001051902 | -0.729676072 | 0.499921258 |
| ENSMUSG00000055024 | Ep300 | 0.001051902 | -2.01792104 | 0.001807071 |
| ENSMUSG00000036046 | 5031439G07Rik | 0.001051902 | 0.004979362 | 1 |
| ENSMUSG00000016541 | Atxn10 | 0.001051902 | -1.647621856 | 0.014140013 |
| ENSMUSG00000116305 | Lncppara | 0.001051902 | -1.800725823 | 0.00653297 |
| ENSMUSG00000008690 | Ncaph2 | 0.001051902 | -0.062659355 | 1 |
| ENSMUSG00000063296 | Tmem117 | 0.001051902 | -1.248404874 | 0.092468958 |
| ENSMUSG00000003360 | Ddx23 | 0.001051902 | -1.346648967 | 0.061463568 |
| ENSMUSG00000051853 | Arf3 | 0.001051902 | -1.029236354 | 0.228120793 |
| ENSMUSG00000022858 | Tra2b | 0.001051902 | -1.820399242 | 0.005483665 |
| ENSMUSG00000022800 | Fyttd1 | 0.001051902 | -0.43011579 | 0.881661525 |
| ENSMUSG00000055447 | Cd47 | 0.001051902 | -1.272410337 | 0.089176403 |
| ENSMUSG00000062078 | Qk | 0.001051902 | -1.986338395 | 0.00252798 |
| ENSMUSG00000038002 | Cramp1l | 0.001051902 | -1.45130112 | 0.039785293 |
| ENSMUSG00000038048 | Cntnap5c | 0.001051902 | 0.44720169 | 0.792441218 |
| ENSMUSG00000052105 | Mtcl1 | 0.001051902 | -1.82689788 | 0.005164338 |
| ENSMUSG00000024048 | Myl12a | 0.001051902 | 0.298338304 | 1 |
| ENSMUSG00000005836 | Gata6 | 0.001051902 | -0.658592974 | 0.590875107 |
| ENSMUSG00000024529 | Lox | 0.001051902 | 1.578446224 | 0.031497833 |
| ENSMUSG00000024563 | Smad2 | 0.001051902 | -2.763945974 | 1.04E-05 |
| ENSMUSG00000024548 | Setbp1 | 0.001051902 | -1.977603585 | 0.002153076 |
| ENSMUSG00000052085 | Dock8 | 0.001051902 | -0.777429203 | 0.489129831 |
| ENSMUSG00000024805 | Pcgf5 | 0.001051902 | -1.568395165 | 0.021820859 |
| ENSMUSG00000074852 | Hpse2 | 0.001051902 | -0.084518789 | 1 |
| ENSMUSG00000025231 | Sufu | 0.001051902 | -1.386788358 | 0.052213587 |
| ENSMUSG00000031171 | Ftsj1 | 0.001051902 | -1.572268174 | 0.022189564 |
| ENSMUSG00000031176 | Dynlt3 | 0.001051902 | 0.189403933 | 1 |
| ENSMUSG00000035150 | Eif2s3x | 0.001051902 | -0.395558568 | 0.91587537 |
| ENSMUSG00000031311 | Nono | 0.001051902 | -0.855784841 | 0.384195668 |
| ENSMUSG00000031232 | Magt1 | 0.001051902 | -1.94212569 | 0.0027444 |
| ENSMUSG00000031278 | Acsl4 | 0.001051902 | -0.377964508 | 0.968284003 |
| ENSMUSG00000025260 | Hsd17b10 | 0.001051902 | 1.47891055 | 0.101504196 |
| ENSMUSG00000026478 | Lamc1 | 0.000996539 | -2.010127531 | 0.002090644 |
| ENSMUSG00000052428 | Tmco1 | 0.000996539 | 0.141875563 | 1 |
| ENSMUSG00000006412 | Pfdn2 | 0.000996539 | -0.010127531 | 1 |
| ENSMUSG00000026499 | Acbd3 | 0.000996539 | -2.975142173 | 2.37E-06 |
| ENSMUSG00000026609 | Ush2a | 0.000996539 | 0.282053221 | 1 |
| ENSMUSG00000063275 | Hacd1 | 0.000996539 | 0.224337723 | 1 |
| ENSMUSG00000026788 | Zbtb43 | 0.000996539 | -1.684437517 | 0.013392251 |
| ENSMUSG00000034780 | B3galt1 | 0.000996539 | 0.629976524 | 0.66215991 |
| ENSMUSG00000027002 | Nckap1 | 0.000996539 | -1.630713941 | 0.016395322 |

| | | | | |
|---|---|---|---|---|
| ENSMUSG00000027236 | Eif3j1 | 0.000996539 | -2.057433246 | 0.001521072 |
| ENSMUSG00000027433 | Xrn2 | 0.000996539 | -0.942440723 | 0.295523053 |
| ENSMUSG00000033059 | Pygb | 0.000996539 | -0.929785422 | 0.316909787 |
| ENSMUSG00000027456 | Sdcbp2 | 0.000996539 | 1.063873051 | 0.244874301 |
| ENSMUSG00000038324 | Trpc4ap | 0.000996539 | -1.429666422 | 0.051312064 |
| ENSMUSG00000037820 | Tgm2 | 0.000996539 | 0.238551582 | 1 |
| ENSMUSG00000039536 | Stau1 | 0.000996539 | -2.028049439 | 0.001871203 |
| ENSMUSG00000013523 | Bcas1 | 0.000996539 | 0.16884261 | 1 |
| ENSMUSG00000038848 | Ythdf1 | 0.000996539 | -1.728945778 | 0.0105575 |
| ENSMUSG00000000826 | Dnajc5 | 0.000996539 | -1.634618396 | 0.016684459 |
| ENSMUSG00000063887 | Nlgn1 | 0.000996539 | -0.252984055 | 1 |
| ENSMUSG00000027695 | Pld1 | 0.000996539 | -2.158519371 | 0.000876027 |
| ENSMUSG00000042626 | Shc1 | 0.000996539 | -0.618936773 | 0.66416768 |
| ENSMUSG00000027878 | Notch2 | 0.000996539 | -1.858124437 | 0.004683913 |
| ENSMUSG00000068798 | Rap1a | 0.000996539 | -0.680665763 | 0.567714406 |
| ENSMUSG00000040322 | Slc25a24 | 0.000996539 | -1.443086938 | 0.043272281 |
| ENSMUSG00000028132 | Tmem56 | 0.000996539 | -1.447532843 | 0.051829371 |
| ENSMUSG00000074212 | Dnajb14 | 0.000996539 | -0.929785422 | 0.316909787 |
| ENSMUSG00000039131 | Gipc2 | 0.000996539 | -0.125604748 | 1 |
| ENSMUSG00000087402 | Gm11867 | 0.000996539 | -0.546180431 | 0.76647565 |
| ENSMUSG00000035649 | Zcchc7 | 0.000996539 | -2.222178008 | 0.000558159 |
| ENSMUSG00000028343 | Erp44 | 0.000996539 | -1.36533178 | 0.061620281 |
| ENSMUSG00000025791 | Pgm1 | 0.000996539 | 0.155296078 | 1 |
| ENSMUSG00000028530 | Jak1 | 0.000996539 | -1.293061494 | 0.082954581 |
| ENSMUSG00000061887 | Ssbp3 | 0.000996539 | -1.973601655 | 0.002431629 |
| ENSMUSG00000032998 | Foxj3 | 0.000996539 | -1.355902368 | 0.060182632 |
| ENSMUSG00000032726 | Bmp8a | 0.000996539 | -0.844703922 | 0.415055165 |
| ENSMUSG00000073147 | 5031425E22Rik | 0.000996539 | -1.516864864 | 0.030947694 |
| ENSMUSG00000029189 | Sel1l3 | 0.000996539 | -0.851429785 | 0.406329712 |
| ENSMUSG00000006262 | Mob1b | 0.000996539 | -0.96121793 | 0.27866723 |
| ENSMUSG00000058427 | Cxcl2 | 0.000996539 | 2.311800564 | 0.00153265 |
| ENSMUSG00000034724 | Cnot6l | 0.000996539 | -0.397857684 | 0.952096994 |
| ENSMUSG00000000568 | Hnrnpd | 0.000996539 | -1.537865163 | 0.027621844 |
| ENSMUSG00000029538 | Srsf9 | 0.000996539 | 0.53921106 | 0.755370971 |
| ENSMUSG00000029528 | Pxn | 0.000996539 | -1.016126271 | 0.232693869 |
| ENSMUSG00000061578 | Ksr2 | 0.000996539 | 0.53921106 | 0.775197196 |
| ENSMUSG00000034310 | Tmem132d | 0.000996539 | 0.12857874 | 1 |
| ENSMUSG00000053293 | Pom121 | 0.000996539 | -0.837946555 | 0.434290508 |
| ENSMUSG00000029918 | Mrps33 | 0.000996539 | 0.629976524 | 0.618148714 |
| ENSMUSG00000036390 | Gadd45a | 0.000996539 | -1.341641674 | 0.070644516 |
| ENSMUSG00000056429 | Tgoln1 | 0.000996539 | -0.341641674 | 0.98880775 |
| ENSMUSG00000034832 | Tet3 | 0.000996539 | -1.942440723 | 0.00292746 |
| ENSMUSG00000030103 | Bhlhe40 | 0.000996539 | -0.360624778 | 1 |
| ENSMUSG00000030218 | Mgp | 0.000996539 | -0.125604748 | 1 |
| ENSMUSG00000053898 | Ech1 | 0.000996539 | 2.089408143 | 0.004581608 |
| ENSMUSG00000030451 | Herc2 | 0.000996539 | -1.473924342 | 0.03552266 |
| ENSMUSG00000030538 | Cib1 | 0.000996539 | 1.373201109 | 0.100453001 |
| ENSMUSG00000030691 | Fchsd2 | 0.000996539 | -1.022100172 | 0.23504169 |
| ENSMUSG00000030654 | Arl6ip1 | 0.000996539 | 0.747015945 | 0.466464457 |
| ENSMUSG00000037772 | Mrpl23 | 0.000996539 | 2.252906875 | 0.00194589 |
| ENSMUSG00000002948 | Map2k7 | 0.000996539 | 0.224337723 | 1 |

| | | | | |
|---|---|---|---|---|
| ENSMUSG00000039842 | Mcph1 | 0.000996539 | -1.185385976 | 0.126339952 |
| ENSMUSG00000031538 | Plat | 0.000996539 | -0.147631054 | 1 |
| ENSMUSG00000058833 | Rex1bd | 0.000996539 | 1.437331446 | 0.054859373 |
| ENSMUSG00000004677 | Myo9b | 0.000996539 | -1.486941228 | 0.033673469 |
| ENSMUSG00000031683 | Lsm6 | 0.000996539 | 1.063873051 | 0.244874301 |
| ENSMUSG00000031776 | Arl2bp | 0.000996539 | -0.775662277 | 0.466464457 |
| ENSMUSG00000031823 | Zdhhc7 | 0.000996539 | -0.942440723 | 0.295523053 |
| ENSMUSG00000031939 | Taf1d | 0.000996539 | -2.247895178 | 0.000471573 |
| ENSMUSG00000111059 | Gm33838 | 0.000996539 | 1.224337723 | 0.128972787 |
| ENSMUSG00000002028 | Kmt2a | 0.000996539 | -2.613011939 | 3.65E-05 |
| ENSMUSG00000003131 | Pafah1b2 | 0.000996539 | -0.673092543 | 0.621590003 |
| ENSMUSG00000032047 | Acat1 | 0.000996539 | -0.96742308 | 0.290284345 |
| ENSMUSG00000054693 | Adam10 | 0.000996539 | -1.273161937 | 0.087198749 |
| ENSMUSG00000032199 | Polr2m | 0.000996539 | 0.437331446 | 0.826843374 |
| ENSMUSG00000062151 | Unc13c | 0.000996539 | -0.578970366 | 0.737916568 |
| ENSMUSG00000040661 | Rad54l2 | 0.000996539 | -1.43862729 | 0.042735074 |
| ENSMUSG00000032594 | Ip6k1 | 0.000996539 | -1.665479359 | 0.014189547 |
| ENSMUSG00000038775 | Vill | 0.000996539 | -1.595090031 | 0.02111872 |
| ENSMUSG00000019899 | Lama2 | 0.000996539 | 0.063873051 | 1 |
| ENSMUSG00000019837 | Gtf3c6 | 0.000996539 | 0.326907457 | 0.980745941 |
| ENSMUSG00000049422 | Chchd10 | 0.000996539 | 3.726838063 | 8.97E-06 |
| ENSMUSG00000003346 | Abhd17a | 0.000996539 | 1.404909969 | 0.060951656 |
| ENSMUSG00000035027 | Map2k2 | 0.000996539 | -0.103236935 | 1 |
| ENSMUSG00000035873 | Pawr | 0.000996539 | -1.268143862 | 0.094775592 |
| ENSMUSG00000034613 | Ppm1h | 0.000996539 | -0.948726986 | 0.316397507 |
| ENSMUSG00000000378 | Ccm2 | 0.000996539 | -1.37936134 | 0.058171406 |
| ENSMUSG00000020321 | Mdh1 | 0.000996539 | 0.989872469 | 0.266705895 |
| ENSMUSG00000020290 | Xpo1 | 0.000996539 | -2.001082391 | 0.002159501 |
| ENSMUSG00000020362 | Cnot6 | 0.000996539 | -0.252984055 | 1 |
| ENSMUSG00000002064 | Sdf2 | 0.000996539 | -1.142155924 | 0.155026807 |
| ENSMUSG00000018481 | Appbp2 | 0.000996539 | -0.858124437 | 0.424188363 |
| ENSMUSG00000056158 | Car10 | 0.000996539 | 0.326907457 | 0.998731799 |
| ENSMUSG00000078676 | Casc3 | 0.000996539 | -1.52950169 | 0.027940391 |
| ENSMUSG00000018677 | Slc25a39 | 0.000996539 | 0.747015945 | 0.466464457 |
| ENSMUSG00000040548 | Tex2 | 0.000996539 | -1.43862729 | 0.044042118 |
| ENSMUSG00000017132 | Cyth1 | 0.000996539 | -1.554448047 | 0.024213692 |
| ENSMUSG00000020590 | Snx13 | 0.000996539 | -1.793234907 | 0.007207413 |
| ENSMUSG00000020547 | Bzw2 | 0.000996539 | -0.642395746 | 0.63765603 |
| ENSMUSG00000050103 | Agmo | 0.000996539 | -0.878023995 | 0.388385939 |
| ENSMUSG00000001627 | Ifrd1 | 0.000996539 | -1.566761308 | 0.022940117 |
| ENSMUSG00000044408 | Sptssa | 0.000996539 | 0.57483497 | 0.66416768 |
| ENSMUSG00000021250 | Fos | 0.000996539 | -0.114464191 | 1 |
| ENSMUSG00000021102 | Glrx5 | 0.000996539 | 0.57483497 | 0.66416768 |
| ENSMUSG00000056770 | Setd3 | 0.000996539 | -1.691951571 | 0.011858726 |
| ENSMUSG00000037957 | Wdr20 | 0.000996539 | -1.495554358 | 0.031497833 |
| ENSMUSG00000021427 | Ssr1 | 0.000996539 | 0.311800564 | 0.992895409 |
| ENSMUSG00000047547 | Cltb | 0.000996539 | 0.210262538 | 1 |
| ENSMUSG00000021495 | Fam193b | 0.000996539 | -1.546180431 | 0.025692018 |
| ENSMUSG00000021610 | Clptm1l | 0.000996539 | 0.155296078 | 1 |
| ENSMUSG00000021823 | Vcl | 0.000996539 | -2.06902122 | 0.001501658 |
| ENSMUSG00000021831 | Ero1l | 0.000996539 | 0.296850223 | 0.98880775 |

| | | | | |
|---|---|---|---|---|
| ENSMUSG00000021981 | Cab39l | 0.000996539 | -0.703149777 | 0.531301099 |
| ENSMUSG00000075486 | Commd6 | 0.000996539 | 2.437331446 | 0.000909854 |
| ENSMUSG00000041650 | Pcca | 0.000996539 | -1.41611989 | 0.048881736 |
| ENSMUSG00000022186 | Oxct1 | 0.000996539 | -1.039874874 | 0.229243081 |
| ENSMUSG00000022197 | Pdzd2 | 0.000996539 | -0.52108945 | 0.81042937 |
| ENSMUSG00000037119 | Fam91a1 | 0.000996539 | -0.443086938 | 0.866287757 |
| ENSMUSG00000042303 | Sgsm3 | 0.000996539 | -1.057433246 | 0.207969581 |
| ENSMUSG00000043091 | Tuba1c | 0.000996539 | 0.53921106 | 0.735367024 |
| ENSMUSG00000071669 | Snx29 | 0.000996539 | -0.917018126 | 0.342462615 |
| ENSMUSG00000060657 | Marf1 | 0.000996539 | -1.237663255 | 0.1126701 |
| ENSMUSG00000041215 | Yeats2 | 0.000996539 | -2.051604167 | 0.001634767 |
| ENSMUSG00000022841 | Ap2m1 | 0.000996539 | 0.141875563 | 1 |
| ENSMUSG00000022797 | Tfrc | 0.000996539 | -1.258055044 | 0.092707526 |
| ENSMUSG00000075254 | Heg1 | 0.000996539 | -0.478276366 | 0.829198173 |
| ENSMUSG00000022641 | Bbx | 0.000996539 | -0.851429785 | 0.406329712 |
| ENSMUSG00000071533 | Pcnp | 0.000996539 | -0.991980184 | 0.266705895 |
| ENSMUSG00000022897 | Dyrk1a | 0.000996539 | -2.074780489 | 0.001424242 |
| ENSMUSG00000023868 | Pde10a | 0.000996539 | -1.258055044 | 0.112238591 |
| ENSMUSG00000023905 | Tnfrsf12a | 0.000996539 | -0.022100172 | 1 |
| ENSMUSG00000024026 | Glo1 | 0.000996539 | 0.629976524 | 0.653235135 |
| ENSMUSG00000079557 | 2-Mar | 0.000996539 | -0.923415898 | 0.314709743 |
| ENSMUSG00000024335 | Brd2 | 0.000996539 | -0.83115739 | 0.432072793 |
| ENSMUSG00000013787 | Ehmt2 | 0.000996539 | -1.384007705 | 0.053533761 |
| ENSMUSG00000002833 | Hdgfl2 | 0.000996539 | -1.810596067 | 0.006613418 |
| ENSMUSG00000047407 | Tgif1 | 0.000996539 | 0.404909969 | 0.850873693 |
| ENSMUSG00000058704 | Memo1 | 0.000996539 | -1.216979101 | 0.111961696 |
| ENSMUSG00000024074 | Crim1 | 0.000996539 | -2.182721719 | 0.000725712 |
| ENSMUSG00000024081 | Cebpz | 0.000996539 | -1.237663255 | 0.106586836 |
| ENSMUSG00000024283 | Wac | 0.000996539 | -1.091921622 | 0.183057083 |
| ENSMUSG00000014504 | Srp19 | 0.000996539 | 0.809300224 | 0.462134007 |
| ENSMUSG00000024583 | Txnl1 | 0.000996539 | 0.115403351 | 1 |
| ENSMUSG00000035765 | Dym | 0.000996539 | -1.103236935 | 0.170135426 |
| ENSMUSG00000052928 | Ctif | 0.000996539 | -1.566761308 | 0.024804715 |
| ENSMUSG00000090000 | Ier3ip1 | 0.000996539 | 0.611360846 | 0.661446724 |
| ENSMUSG00000025423 | Pias2 | 0.000996539 | -1.96742308 | 0.002579787 |
| ENSMUSG00000061451 | Tmem151a | 0.000996539 | 0.852368946 | 0.382191952 |
| ENSMUSG00000024960 | Plcb3 | 0.000996539 | -0.87142126 | 0.384884537 |
| ENSMUSG00000108414 | Snhg1 | 0.000996539 | -1.611031575 | 0.018410436 |
| ENSMUSG00000024817 | Uhrf2 | 0.000996539 | -1.650131395 | 0.014605485 |
| ENSMUSG00000024781 | Lipa | 0.000996539 | 1.504445642 | 0.045084086 |
| ENSMUSG00000025190 | Got1 | 0.000996539 | 0.989872469 | 0.266705895 |
| ENSMUSG00000063434 | Sorcs3 | 0.000996539 | 0.076584102 | 1 |
| ENSMUSG00000001924 | Uba1 | 0.000996539 | 0.706938506 | 0.543824129 |
| ENSMUSG00000001131 | Timp1 | 0.000996539 | 0.373201109 | 0.90563095 |
| ENSMUSG00000006423 | C330007P06Rik | 0.000996539 | -0.293061494 | 1 |
| ENSMUSG00000086370 | Ftx | 0.000996539 | -1.174699298 | 0.133860161 |
| ENSMUSG00000031422 | Morf4l2 | 0.000996539 | -0.954985977 | 0.300304685 |
| ENSMUSG00000042225 | Ammecr1 | 0.000996539 | -0.942440723 | 0.295523053 |
| ENSMUSG00000040990 | Sh3kbp1 | 0.000996539 | -0.796723893 | 0.455131546 |
| ENSMUSG00000048960 | Prex2 | 0.000941176 | 0.204050709 | 1 |
| ENSMUSG00000041763 | Tpp2 | 0.000941176 | -1.014129461 | 0.242941974 |

| | | | | |
|---|---|---|---|---|
| ENSMUSG00000002881 | Nab1 | 0.000941176 | -0.648844997 | 0.653542789 |
| ENSMUSG00000026229 | Psmd1 | 0.000941176 | -0.803291267 | 0.454181515 |
| ENSMUSG00000004451 | Ralb | 0.000941176 | 0.264171701 | 1 |
| ENSMUSG00000036104 | Rab3gap1 | 0.000941176 | -2.27927153 | 0.000443732 |
| ENSMUSG00000016528 | Mapkapk2 | 0.000941176 | -0.995020638 | 0.26104641 |
| ENSMUSG00000056268 | Dennd1b | 0.000941176 | -1.656972878 | 0.015417887 |
| ENSMUSG00000035131 | Brinp3 | 0.000941176 | -0.83945093 | 0.466464457 |
| ENSMUSG00000042772 | Smg7 | 0.000941176 | -1.689034087 | 0.013145489 |
| ENSMUSG00000033684 | Qsox1 | 0.000941176 | -0.07605721 | 1 |
| ENSMUSG00000058076 | Sdhc | 0.000941176 | 0.461208548 | 0.792441218 |
| ENSMUSG00000026490 | Cdc42bpa | 0.000941176 | -2.263388488 | 0.000468588 |
| ENSMUSG00000026748 | Plxdc2 | 0.000941176 | -1.754886635 | 0.011284174 |
| ENSMUSG00000026790 | Odf2 | 0.000941176 | -0.640671066 | 0.65079351 |
| ENSMUSG00000079484 | Phyhd1 | 0.000941176 | -1.871237419 | 0.005404702 |
| ENSMUSG00000038696 | Mapkap1 | 0.000941176 | -1.480447466 | 0.036400837 |
| ENSMUSG00000035437 | Rabgap1 | 0.000941176 | -2.007787934 | 0.002865534 |
| ENSMUSG00000036890 | Gtdc1 | 0.000941176 | -1.498652061 | 0.041758382 |
| ENSMUSG00000026971 | Itgb6 | 0.000941176 | 0.024344687 | 1 |
| ENSMUSG00000015839 | Nfe2l2 | 0.000941176 | -1.603304213 | 0.020814694 |
| ENSMUSG00000002102 | Psmc3 | 0.000941176 | 0.609307187 | 0.630903907 |
| ENSMUSG00000040479 | Dgkz | 0.000941176 | -1.164233022 | 0.145085302 |
| ENSMUSG00000027238 | Frmd5 | 0.000941176 | -0.225633566 | 1 |
| ENSMUSG00000027423 | Snx5 | 0.000941176 | -1.192466703 | 0.128531628 |
| ENSMUSG00000027447 | Cst3 | 0.000941176 | 1.609307187 | 0.059151955 |
| ENSMUSG00000032046 | Abhd12 | 0.000941176 | -0.607504202 | 0.66416768 |
| ENSMUSG00000038533 | Cbfa2t2 | 0.000941176 | -1.462010217 | 0.042309732 |
| ENSMUSG00000078923 | Ube2v1 | 0.000941176 | 0.42644313 | 0.875465725 |
| ENSMUSG00000015627 | Gata5 | 0.000941176 | -0.236521883 | 1 |
| ENSMUSG00000037270 | 4932438A13Rik | 0.000941176 | -1.594867376 | 0.022357178 |
| ENSMUSG00000008730 | Hipk1 | 0.000941176 | 0.011405631 | 1 |
| ENSMUSG00000027962 | Vcam1 | 0.000941176 | -1.020443235 | 0.245298386 |
| ENSMUSG00000028031 | Dkk2 | 0.000941176 | -1.429166961 | 0.053643891 |
| ENSMUSG00000028029 | Aimp1 | 0.000941176 | -0.390692813 | 0.929361739 |
| ENSMUSG00000028149 | Rap1gds1 | 0.000941176 | -2.244634528 | 0.000541951 |
| ENSMUSG00000052430 | Bmpr1b | 0.000941176 | 0.752671362 | 0.51581176 |
| ENSMUSG00000038975 | Rabggtb | 0.000941176 | 0.118320835 | 1 |
| ENSMUSG00000040037 | Negr1 | 0.000941176 | 0.174904363 | 1 |
| ENSMUSG00000028173 | Wls | 0.000941176 | -1.656972878 | 0.015417887 |
| ENSMUSG00000028465 | Tln1 | 0.000941176 | -1.543183221 | 0.027568457 |
| ENSMUSG00000028434 | Epb41l4b | 0.000941176 | -0.895484965 | 0.355761423 |
| ENSMUSG00000028488 | Sh3gl2 | 0.000941176 | 0.628935994 | 0.663838662 |
| ENSMUSG00000028552 | Eps15 | 0.000941176 | -1.380911792 | 0.060940556 |
| ENSMUSG00000086290 | Snhg12 | 0.000941176 | -0.475860196 | 0.858030174 |
| ENSMUSG00000041351 | Rap1gap | 0.000941176 | -1.00141841 | 0.262071024 |
| ENSMUSG00000028920 | Fbxo42 | 0.000941176 | -1.590630383 | 0.0224005 |
| ENSMUSG00000028960 | Ube4b | 0.000941176 | -1.507668934 | 0.032495347 |
| ENSMUSG00000029068 | Ccnl2 | 0.000941176 | -2.161379025 | 0.000917711 |
| ENSMUSG00000029154 | Cwh43 | 0.000941176 | 0.864564242 | 0.410540658 |
| ENSMUSG00000082361 | Btc | 0.000941176 | -0.380911792 | 0.975624465 |
| ENSMUSG00000029405 | G3bp2 | 0.000941176 | -1.220158436 | 0.131969432 |
| ENSMUSG00000033623 | Pcgf3 | 0.000941176 | -0.512156325 | 0.778722129 |

| | | | | |
|---|---|---|---|---|
| ENSMUSG00000004530 | Coro1c | 0.000941176 | 0.174904363 | 1 |
| ENSMUSG00000011884 | Gltp | 0.000941176 | 1.233798052 | 0.171055101 |
| ENSMUSG00000029478 | Ncor2 | 0.000941176 | -2.079084903 | 0.001448268 |
| ENSMUSG00000050552 | Lamtor4 | 0.000941176 | 1.960779558 | 0.00814171 |
| ENSMUSG00000056413 | Adap1 | 0.000941176 | -0.400407968 | 0.930842647 |
| ENSMUSG00000018001 | Cyth3 | 0.000941176 | -1.175592884 | 0.134836815 |
| ENSMUSG00000016520 | Lnx2 | 0.000941176 | -1.045425592 | 0.231596497 |
| ENSMUSG00000042460 | C1galt1 | 0.000941176 | -0.247328638 | 1 |
| ENSMUSG00000024211 | Grm8 | 0.000941176 | -0.192466703 | 1 |
| ENSMUSG00000029920 | Smarcad1 | 0.000941176 | -1.268702288 | 0.093015331 |
| ENSMUSG00000030016 | Zfp638 | 0.000941176 | -2.390692813 | 0.000197826 |
| ENSMUSG00000005893 | Nr2c2 | 0.000941176 | -2.060822697 | 0.001699321 |
| ENSMUSG00000033940 | Brk1 | 0.000941176 | -0.380911792 | 0.975624465 |
| ENSMUSG00000004268 | Emg1 | 0.000941176 | 0.375817057 | 0.886203497 |
| ENSMUSG00000030353 | Tead4 | 0.000941176 | -0.063882496 | 1 |
| ENSMUSG00000033916 | Chmp2a | 0.000941176 | 0.774366434 | 0.488035387 |
| ENSMUSG00000030413 | Pglyrp1 | 0.000941176 | 2.063873051 | 0.005443828 |
| ENSMUSG00000099481 | Xndc1 | 0.000941176 | -1.164233022 | 0.165798952 |
| ENSMUSG00000030987 | Stim1 | 0.000941176 | -1.828697874 | 0.006369168 |
| ENSMUSG00000030879 | Mrpl17 | 0.000941176 | 1.818760553 | 0.028837933 |
| ENSMUSG00000030754 | Copb1 | 0.000941176 | 0.132260026 | 1 |
| ENSMUSG00000035064 | Eef2k | 0.000941176 | -1.198047735 | 0.128972787 |
| ENSMUSG00000040028 | Elavl1 | 0.000941176 | -0.410058138 | 0.933335341 |
| ENSMUSG00000001300 | Efnb2 | 0.000941176 | -0.400407968 | 0.930842647 |
| ENSMUSG00000079157 | Fam155a | 0.000941176 | 0.589941862 | 0.700484316 |
| ENSMUSG00000047495 | Dlgap2 | 0.000941176 | 0.710236096 | 0.54606373 |
| ENSMUSG00000037656 | Slc20a2 | 0.000941176 | -1.111976785 | 0.176201216 |
| ENSMUSG00000031529 | Tnks | 0.000941176 | -1.36611479 | 0.064891891 |
| ENSMUSG00000015568 | Lpl | 0.000941176 | -1.07605721 | 0.232106082 |
| ENSMUSG00000036810 | Cnep1r1 | 0.000941176 | -1.020443235 | 0.245298386 |
| ENSMUSG00000031667 | Aktip | 0.000941176 | 0.409369617 | 0.87923507 |
| ENSMUSG00000050079 | Rspry1 | 0.000941176 | -0.795949291 | 0.466464457 |
| ENSMUSG00000048310 | Pskh1 | 0.000941176 | -1.247328638 | 0.112515907 |
| ENSMUSG00000015027 | Galns | 0.000941176 | 0.174904363 | 1 |
| ENSMUSG00000039960 | Rhou | 0.000941176 | 0.461208548 | 0.792441218 |
| ENSMUSG00000095186 | Gm10718 | 0.000941176 | 0.514979805 | 0.792441218 |
| ENSMUSG00000019471 | Cdc37 | 0.000941176 | -0.825095637 | 0.426482132 |
| ENSMUSG00000057193 | Slc44a2 | 0.000941176 | -1.777429203 | 0.008644519 |
| ENSMUSG00000031996 | Aplp2 | 0.000941176 | -0.803291267 | 0.47563319 |
| ENSMUSG00000042496 | Prdm10 | 0.000941176 | -1.147023732 | 0.154859761 |
| ENSMUSG00000034342 | Cbl | 0.000941176 | -1.494122402 | 0.034375132 |
| ENSMUSG00000048537 | Phldb1 | 0.000941176 | -1.07605721 | 0.201910942 |
| ENSMUSG00000037674 | Rfx7 | 0.000941176 | -2.004606687 | 0.002309101 |
| ENSMUSG00000032412 | Atp1b3 | 0.000941176 | -0.410058138 | 0.940012028 |
| ENSMUSG00000062867 | Impdh2 | 0.000941176 | 0.960779558 | 0.306313838 |
| ENSMUSG00000050621 | Rps27rt | 0.000941176 | 1.233798052 | 0.128972787 |
| ENSMUSG00000055493 | Epm2a | 0.000941176 | -2.032988489 | 0.002746919 |
| ENSMUSG00000039697 | Ncoa7 | 0.000941176 | -0.781152289 | 0.466464457 |
| ENSMUSG00000019841 | Rev3l | 0.000941176 | -1.485020195 | 0.036927154 |
| ENSMUSG00000020064 | Herc4 | 0.000941176 | -1.507668934 | 0.032495347 |
| ENSMUSG00000001435 | Col18a1 | 0.000941176 | -0.853664789 | 0.403955877 |

| | | | | |
|---|---|---|---|---|
| ENSMUSG00000020190 | Mknk2 | 0.000941176 | 0.752671362 | 0.496010946 |
| ENSMUSG00000053825 | Ppfia2 | 0.000941176 | 1.174904363 | 0.218817775 |
| ENSMUSG00000035864 | Syt1 | 0.000941176 | 0.146335211 | 1 |
| ENSMUSG00000006728 | Cdk4 | 0.000941176 | 0.132260026 | 1 |
| ENSMUSG00000040249 | Lrp1 | 0.000941176 | -1.158519371 | 0.150993311 |
| ENSMUSG00000025357 | Dgka | 0.000941176 | -0.258055044 | 1 |
| ENSMUSG00000020134 | Peli1 | 0.000941176 | -0.781152289 | 0.466464457 |
| ENSMUSG00000097887 | Gm26542 | 0.000941176 | -0.758668275 | 0.543347452 |
| ENSMUSG00000001056 | Nhp2 | 0.000941176 | 1.037400839 | 0.257432522 |
| ENSMUSG00000017548 | Suz12 | 0.000941176 | -1.915949067 | 0.004015829 |
| ENSMUSG00000014349 | Ube2z | 0.000941176 | -1.129606679 | 0.171496486 |
| ENSMUSG00000035007 | Rundc1 | 0.000941176 | -0.902338636 | 0.358457412 |
| ENSMUSG00000040838 | Gm11639 | 0.000941176 | 0.533358334 | 0.731681545 |
| ENSMUSG00000059923 | Grb2 | 0.000941176 | -1.516629802 | 0.030563094 |
| ENSMUSG00000039640 | Mrpl12 | 0.000941176 | 0.936117504 | 0.316952476 |
| ENSMUSG00000020634 | Ubxn2a | 0.000941176 | -0.743481871 | 0.509878973 |
| ENSMUSG00000071379 | Hpcal1 | 0.000941176 | -2.540989007 | 7.37E-05 |
| ENSMUSG00000020659 | Cbll1 | 0.000941176 | -0.982139347 | 0.276168778 |
| ENSMUSG00000020954 | Strn3 | 0.000941176 | -2.704801403 | 2.00E-05 |
| ENSMUSG00000045690 | Wdr89 | 0.000941176 | 0.551974012 | 0.701892102 |
| ENSMUSG00000016756 | Cmah | 0.000941176 | -0.640671066 | 0.655144705 |
| ENSMUSG00000037960 | Card19 | 0.000941176 | 0.936117504 | 0.316952476 |
| ENSMUSG00000021395 | Spin1 | 0.000941176 | -0.494122402 | 0.802584425 |
| ENSMUSG00000034488 | Edil3 | 0.000941176 | 0.669013433 | 0.605745417 |
| ENSMUSG00000021615 | Xrcc4 | 0.000941176 | -1.43862729 | 0.043463239 |
| ENSMUSG00000021638 | Ocln | 0.000941176 | -1.20360726 | 0.129397681 |
| ENSMUSG00000040760 | Appl1 | 0.000941176 | -1.480447466 | 0.036400837 |
| ENSMUSG00000021991 | Cacna2d3 | 0.000941176 | 0.443721122 | 0.817764435 |
| ENSMUSG00000054509 | Parp4 | 0.000941176 | -1.656972878 | 0.015417887 |
| ENSMUSG00000022098 | Bmp1 | 0.000941176 | -0.922706434 | 0.335454037 |
| ENSMUSG00000042104 | Uggt2 | 0.000941176 | -2.395558568 | 0.000189328 |
| ENSMUSG00000115520 | Gm41335 | 0.000941176 | 0.47891055 | 0.795570493 |
| ENSMUSG00000063480 | Snu13 | 0.000941176 | 1.146335211 | 0.228008347 |
| ENSMUSG00000036606 | Plxnb2 | 0.000941176 | -0.696939285 | 0.580988658 |
| ENSMUSG00000036561 | Ppp6r2 | 0.000941176 | -0.624182943 | 0.66416768 |
| ENSMUSG00000023026 | Dip2b | 0.000941176 | -1.485020195 | 0.036927154 |
| ENSMUSG00000036966 | Spryd3 | 0.000941176 | -0.867739974 | 0.408535739 |
| ENSMUSG00000022507 | 1810013L24Rik | 0.000941176 | -0.615867675 | 0.66416768 |
| ENSMUSG00000006134 | Crkl | 0.000941176 | -1.07605721 | 0.201910942 |
| ENSMUSG00000047714 | Ppp1r2 | 0.000941176 | -0.640671066 | 0.65079351 |
| ENSMUSG00000005615 | Pcyt1a | 0.000941176 | -0.371064006 | 0.974915472 |
| ENSMUSG00000035629 | Rubcn | 0.000941176 | -1.448025988 | 0.044761838 |
| ENSMUSG00000022817 | Itgb5 | 0.000941176 | -0.982139347 | 0.276168778 |
| ENSMUSG00000024122 | Pdpk1 | 0.000941176 | -1.516629802 | 0.030563094 |
| ENSMUSG00000024182 | Axin1 | 0.000941176 | -1.258055044 | 0.099922289 |
| ENSMUSG00000090083 | Rnf8 | 0.000941176 | -1.051604167 | 0.213570906 |
| ENSMUSG00000003534 | Ddr1 | 0.000941176 | -1.268702288 | 0.093015331 |
| ENSMUSG00000036026 | Tmem63b | 0.000941176 | -1.494122402 | 0.034375132 |
| ENSMUSG00000013236 | Ptprs | 0.000941176 | -1.405241121 | 0.053398165 |
| ENSMUSG00000024068 | Spast | 0.000941176 | -0.773696484 | 0.491466936 |
| ENSMUSG00000033632 | AW554918 | 0.000941176 | -1.878206973 | 0.004848505 |

| | | | | |
|---|---|---|---|---|
| ENSMUSG00000024598 | Fbn2 | 0.000941176 | -0.915949067 | 0.347971352 |
| ENSMUSG00000024970 | Spindoc | 0.000941176 | -1.551926854 | 0.025594044 |
| ENSMUSG00000024810 | Il33 | 0.000941176 | -0.83945093 | 0.432023774 |
| ENSMUSG00000025041 | Nt5c2 | 0.000941176 | -1.636566667 | 0.017420371 |
| ENSMUSG00000074746 | Pdzd8 | 0.000941176 | -1.475860196 | 0.039827167 |
| ENSMUSG00000016427 | Ndufa1 | 0.000941176 | 3.011405631 | 0.00010315 |
| ENSMUSG00000062184 | Hs6st2 | 0.000941176 | -1.225633566 | 0.112824885 |
| ENSMUSG00000025630 | Hprt | 0.000941176 | -0.158519371 | 1 |
| ENSMUSG00000031189 | Aff2 | 0.000941176 | -1.181239447 | 0.147628491 |
| ENSMUSG00000016382 | Pls3 | 0.000941176 | -1.56494374 | 0.024213692 |
| ENSMUSG00000033845 | Mrpl15 | 0.000885812 | 0.035858675 | 1 |
| ENSMUSG00000045210 | Vcpip1 | 0.000885812 | -1.44857695 | 0.045635603 |
| ENSMUSG00000025937 | Lactb2 | 0.000885812 | -0.825944032 | 0.456648093 |
| ENSMUSG00000026110 | Mgat4a | 0.000885812 | -1.433626609 | 0.048635862 |
| ENSMUSG00000018417 | Myo1b | 0.000885812 | -1.829755645 | 0.007127824 |
| ENSMUSG00000026020 | Nop58 | 0.000885812 | -2.077482799 | 0.001709467 |
| ENSMUSG00000026223 | Itm2c | 0.000885812 | 1.181709541 | 0.162595422 |
| ENSMUSG00000003051 | Elf3 | 0.000885812 | -0.638741039 | 0.66416768 |
| ENSMUSG00000040265 | Dnm3 | 0.000885812 | 0.488370879 | 0.790198002 |
| ENSMUSG00000026558 | Uck2 | 0.000885812 | -1.714912719 | 0.013362945 |
| ENSMUSG00000026554 | Dcaf8 | 0.000885812 | -1.759556634 | 0.00958188 |
| ENSMUSG00000053702 | Nebl | 0.000885812 | -2.640912138 | 3.90E-05 |
| ENSMUSG00000039515 | Ptpa | 0.000885812 | -0.096591621 | 1 |
| ENSMUSG00000026914 | Psmd14 | 0.000885812 | -1.217897918 | 0.121939966 |
| ENSMUSG00000050199 | Lgr4 | 0.000885812 | -0.28606942 | 1 |
| ENSMUSG00000027304 | Rtf1 | 0.000885812 | -1.246695182 | 0.116637579 |
| ENSMUSG00000037514 | Pank2 | 0.000885812 | -1.433626609 | 0.048635862 |
| ENSMUSG00000037110 | Ralgapa2 | 0.000885812 | -2.747517385 | 1.82E-05 |
| ENSMUSG00000037025 | Foxa2 | 0.000885812 | 0.648835551 | 0.66416768 |
| ENSMUSG00000027652 | Ralgapb | 0.000885812 | -1.625645381 | 0.019307676 |
| ENSMUSG00000035268 | Pkig | 0.000885812 | -0.240981531 | 1 |
| ENSMUSG00000017929 | B4galt5 | 0.000885812 | -0.482870128 | 0.83220417 |
| ENSMUSG00000054455 | Vapb | 0.000885812 | 0.022052875 | 1 |
| ENSMUSG00000025762 | Larp1b | 0.000885812 | -1.340517204 | 0.079479908 |
| ENSMUSG00000028086 | Fbxw7 | 0.000885812 | -1.863614449 | 0.005659765 |
| ENSMUSG00000001419 | Mef2d | 0.000885812 | -1.625645381 | 0.019307676 |
| ENSMUSG00000052310 | Slc39a1 | 0.000885812 | 0.526845027 | 0.766729124 |
| ENSMUSG00000027957 | Slc35a3 | 0.000885812 | -0.950202134 | 0.306471805 |
| ENSMUSG00000106245 | Gm43824 | 0.000885812 | 1.063873051 | 0.26104641 |
| ENSMUSG00000028156 | Eif4e | 0.000885812 | -0.576584563 | 0.712758955 |
| ENSMUSG00000028211 | Trp53inp1 | 0.000885812 | -0.473154973 | 0.829198173 |
| ENSMUSG00000045205 | Dpy19l4 | 0.000885812 | -1.134066327 | 0.170180652 |
| ENSMUSG00000028238 | Atp6v0d2 | 0.000885812 | -0.463373952 | 0.887847724 |
| ENSMUSG00000028282 | Casp8ap2 | 0.000885812 | -1.492520298 | 0.037978522 |
| ENSMUSG00000028433 | Ubap2 | 0.000885812 | -1.867327952 | 0.00579486 |
| ENSMUSG00000055296 | Tmem245 | 0.000885812 | -1.428608534 | 0.053198683 |
| ENSMUSG00000052407 | Ccdc171 | 0.000885812 | -0.991622062 | 0.271380007 |
| ENSMUSG00000034610 | Tut4 | 0.000885812 | -1.229485892 | 0.124965758 |
| ENSMUSG00000003810 | Mast2 | 0.000885812 | -2.051604167 | 0.00194589 |
| ENSMUSG00000028677 | Rnf220 | 0.000885812 | -0.005168594 | 1 |
| ENSMUSG00000028797 | Tmem234 | 0.000885812 | 0.310033638 | 1 |

| | | | | |
|---|---|---|---|---|
| ENSMUSG00000040025 | Ythdf2 | 0.000885812 | -0.863614449 | 0.402406531 |
| ENSMUSG00000037242 | Clic4 | 0.000885812 | -0.706645103 | 0.589936698 |
| ENSMUSG00000040964 | Arhgef10l | 0.000885812 | -1.280509895 | 0.093233636 |
| ENSMUSG00000006219 | Fblim1 | 0.000885812 | -0.392984624 | 0.968136327 |
| ENSMUSG00000028948 | Nol9 | 0.000885812 | -0.621253612 | 0.676854064 |
| ENSMUSG00000029030 | Tprgl | 0.000885812 | 1.450896174 | 0.088742706 |
| ENSMUSG00000007415 | Gatad1 | 0.000885812 | 0.450896174 | 0.85084719 |
| ENSMUSG00000028995 | Fam126a | 0.000885812 | -0.170592203 | 1 |
| ENSMUSG00000028954 | Nub1 | 0.000885812 | -1.096591621 | 0.196126878 |
| ENSMUSG00000047881 | Rell1 | 0.000885812 | -0.182564845 | 1 |
| ENSMUSG00000029245 | Epha5 | 0.000885812 | -1.706645103 | 0.013149543 |
| ENSMUSG00000029371 | Cxcl5 | 0.000885812 | -0.351164449 | 1 |
| ENSMUSG00000032867 | Fbxw8 | 0.000885812 | -0.795083068 | 0.466464457 |
| ENSMUSG00000043733 | Ptpn11 | 0.000885812 | -1.530488148 | 0.031456064 |
| ENSMUSG00000004455 | Ppp1cc | 0.000885812 | -0.630023822 | 0.661861928 |
| ENSMUSG00000029432 | Nipsnap2 | 0.000885812 | -1.127887752 | 0.168779114 |
| ENSMUSG00000029729 | Zkscan1 | 0.000885812 | -0.998411228 | 0.271680772 |
| ENSMUSG00000029554 | Mad1l1 | 0.000885812 | -1.258055044 | 0.10831708 |
| ENSMUSG00000029534 | St7 | 0.000885812 | -0.698329835 | 0.588477963 |
| ENSMUSG00000029672 | Fam3c | 0.000885812 | 0.691904273 | 0.584901683 |
| ENSMUSG00000023089 | Ndufa5 | 0.000885812 | 1.035858675 | 0.257440232 |
| ENSMUSG00000029684 | Wasl | 0.000885812 | -1.235245161 | 0.113922494 |
| ENSMUSG00000043421 | Hilpda | 0.000885812 | 0.035858675 | 1 |
| ENSMUSG00000033542 | Arhgef5 | 0.000885812 | -0.943181707 | 0.334416634 |
| ENSMUSG00000038301 | Snx10 | 0.000885812 | -1.280509895 | 0.093233636 |
| ENSMUSG00000059182 | Skap2 | 0.000885812 | -0.914753298 | 0.366731284 |
| ENSMUSG00000044162 | Tnip3 | 0.000885812 | -0.984800796 | 0.29714275 |
| ENSMUSG00000000693 | Loxl3 | 0.000885812 | -1.848664108 | 0.007343761 |
| ENSMUSG00000004263 | Atn1 | 0.000885812 | -0.706645103 | 0.589936698 |
| ENSMUSG00000039985 | Sinhcaf | 0.000885812 | -0.998411228 | 0.27201021 |
| ENSMUSG00000086784 | Isoc2a | 0.000885812 | -1.907557797 | 0.005200942 |
| ENSMUSG00000058709 | Egln2 | 0.000885812 | 1.181709541 | 0.162595422 |
| ENSMUSG00000004610 | Etfb | 0.000885812 | 0.691904273 | 0.655144705 |
| ENSMUSG00000003865 | Gys1 | 0.000885812 | -0.802860418 | 0.466464457 |
| ENSMUSG00000030513 | Pcsk6 | 0.000885812 | -1.44857695 | 0.045635603 |
| ENSMUSG00000030629 | Zfand6 | 0.000885812 | -0.621253612 | 0.676854064 |
| ENSMUSG00000039428 | Tmem135 | 0.000885812 | -1.492520298 | 0.037978522 |
| ENSMUSG00000030982 | Vps35l | 0.000885812 | -1.146344657 | 0.157417169 |
| ENSMUSG00000030869 | Ndufab1 | 0.000885812 | -0.182564845 | 1 |
| ENSMUSG00000002957 | Ap2a2 | 0.000885812 | -1.058117474 | 0.222835955 |
| ENSMUSG00000087687 | Pet100 | 0.000885812 | 1.310033638 | 0.167167286 |
| ENSMUSG00000009630 | Ppp2cb | 0.000885812 | -0.977947125 | 0.289833436 |
| ENSMUSG00000031601 | Cnot7 | 0.000885812 | -0.530488148 | 0.792441218 |
| ENSMUSG00000031605 | Klhl2 | 0.000885812 | -0.594618485 | 0.669116588 |
| ENSMUSG00000070002 | Ell | 0.000885812 | -1.297124609 | 0.087270847 |
| ENSMUSG00000036779 | Tent4b | 0.000885812 | -1.735376822 | 0.01181855 |
| ENSMUSG00000032000 | Birc3 | 0.000885812 | -1.918337636 | 0.0044369 |
| ENSMUSG00000037808 | Fam76b | 0.000885812 | -0.971060739 | 0.287049594 |
| ENSMUSG00000011114 | Tbrg1 | 0.000885812 | 0.71393058 | 0.559529464 |
| ENSMUSG00000032096 | Arcn1 | 0.000885812 | -0.747517385 | 0.515072602 |
| ENSMUSG00000000171 | Sdhd | 0.000885812 | 2.414370298 | 0.001737647 |

| | | | | |
|---|---|---|---|---|
| ENSMUSG00000037112 | Sik2 | 0.000885812 | -1.856158645 | 0.00604468 |
| ENSMUSG00000032397 | Tipin | 0.000885812 | -1.206216113 | 0.130094223 |
| ENSMUSG00000032350 | Gclc | 0.000885812 | -1.616848433 | 0.020850515 |
| ENSMUSG00000032560 | Dnajc13 | 0.000885812 | -1.345850649 | 0.072014244 |
| ENSMUSG00000032440 | Tgfbr2 | 0.000885812 | -0.723133226 | 0.555706947 |
| ENSMUSG00000025786 | Zdhhc3 | 0.000885812 | -0.096591621 | 1 |
| ENSMUSG00000039116 | Adgrg6 | 0.000885812 | -1.44857695 | 0.045635603 |
| ENSMUSG00000039089 | L3mbtl3 | 0.000885812 | -1.291607604 | 0.095375956 |
| ENSMUSG00000019779 | Frk | 0.000885812 | -1.511629121 | 0.036400837 |
| ENSMUSG00000037012 | Hk1 | 0.000885812 | -0.630023822 | 0.661861928 |
| ENSMUSG00000009681 | Bcr | 0.000885812 | -1.418519716 | 0.051667273 |
| ENSMUSG00000020180 | Snrpd3 | 0.000885812 | 0.244445296 | 1 |
| ENSMUSG00000035852 | Misp | 0.000885812 | 0.244445296 | 1 |
| ENSMUSG00000003345 | Csnk1g2 | 0.000885812 | -0.567482355 | 0.7096926 |
| ENSMUSG00000035011 | Zbtb7a | 0.000885812 | -0.567482355 | 0.7096926 |
| ENSMUSG00000020042 | Btbd11 | 0.000885812 | -1.170592203 | 0.148225054 |
| ENSMUSG00000020181 | Nav3 | 0.000885812 | -2.188514111 | 0.000857868 |
| ENSMUSG00000020189 | Osbpl8 | 0.000885812 | -1.235245161 | 0.113922494 |
| ENSMUSG00000040462 | Os9 | 0.000885812 | -1.43862729 | 0.05819568 |
| ENSMUSG00000025381 | Cnpy2 | 0.000885812 | 1.414370298 | 0.078395375 |
| ENSMUSG00000053838 | Nudcd3 | 0.000885812 | -0.60355161 | 0.672254461 |
| ENSMUSG00000020160 | Meis1 | 0.000885812 | -0.971060739 | 0.287049594 |
| ENSMUSG00000032985 | 5730522E02Rik | 0.000885812 | -0.423572944 | 0.937368223 |
| ENSMUSG00000020271 | Fbxw11 | 0.000885812 | -1.967605181 | 0.003263045 |
| ENSMUSG00000018339 | Gpx3 | 0.000885812 | 0.648835551 | 0.616879894 |
| ENSMUSG00000040158 | Tax1bp3 | 0.000885812 | 1.736298393 | 0.051152519 |
| ENSMUSG00000017376 | Nlk | 0.000885812 | -1.66884307 | 0.016754406 |
| ENSMUSG00000017421 | Zfp207 | 0.000885812 | -1.090250095 | 0.194209701 |
| ENSMUSG00000047181 | Samd14 | 0.000885812 | -0.848664108 | 0.431803825 |
| ENSMUSG00000038909 | Kat7 | 0.000885812 | -0.848664108 | 0.448770968 |
| ENSMUSG00000006057 | Atp5g1 | 0.000885812 | 0.878317398 | 0.420286951 |
| ENSMUSG00000002580 | Mien1 | 0.000885812 | 2.035858675 | 0.021623593 |
| ENSMUSG00000038013 | Wipf2 | 0.000885812 | -1.590131094 | 0.023140688 |
| ENSMUSG00000004040 | Stat3 | 0.000885812 | -1.280509895 | 0.106312068 |
| ENSMUSG00000020923 | Ubtf | 0.000885812 | -0.739435038 | 0.51221187 |
| ENSMUSG00000020705 | Ddx42 | 0.000885812 | -0.779401446 | 0.487205201 |
| ENSMUSG00000041895 | Wipi1 | 0.000885812 | -0.08388057 | 1 |
| ENSMUSG00000034714 | Ttyh2 | 0.000885812 | 0.151335892 | 1 |
| ENSMUSG00000033880 | Lgals3bp | 0.000885812 | 2.212736437 | 0.003808841 |
| ENSMUSG00000048445 | Ccdc57 | 0.000885812 | -0.731307157 | 0.568338892 |
| ENSMUSG00000054364 | Rhob | 0.000885812 | 0.022052875 | 1 |
| ENSMUSG00000020669 | Sh3yl1 | 0.000885812 | -0.511629121 | 0.792441218 |
| ENSMUSG00000035472 | Slc25a21 | 0.000885812 | -0.755554704 | 0.532882217 |
| ENSMUSG00000041324 | Inhba | 0.000885812 | 0.310033638 | 1 |
| ENSMUSG00000054728 | Phactr1 | 0.000885812 | 0.432517645 | 0.868770472 |
| ENSMUSG00000037946 | Fgd3 | 0.000885812 | -0.6474059 | 0.66416768 |
| ENSMUSG00000025868 | Higd2a | 0.000885812 | 1.526845027 | 0.071100693 |
| ENSMUSG00000034675 | Dbn1 | 0.000885812 | -0.329790798 | 1 |
| ENSMUSG00000058569 | Tmed9 | 0.000885812 | -0.194438945 | 1 |
| ENSMUSG00000054423 | Cadps | 0.000885812 | 0.805561055 | 0.471405909 |
| ENSMUSG00000021810 | Ecd | 0.000885812 | -1.011894457 | 0.254677217 |

| | | | | |
|---|---|---|---|---|
| ENSMUSG00000063787 | Chchd1 | 0.000885812 | -0.833557216 | 0.767883539 |
| ENSMUSG00000021820 | Camk2g | 0.000885812 | -1.673092543 | 0.019037404 |
| ENSMUSG00000021832 | Psmc6 | 0.000885812 | -0.841130435 | 0.429114808 |
| ENSMUSG00000021952 | Xpo4 | 0.000885812 | -2.465825431 | 0.000143457 |
| ENSMUSG00000114797 | Gm49336 | 0.000885812 | -0.681554122 | 0.584353605 |
| ENSMUSG00000022003 | Slc25a30 | 0.000885812 | -0.217897918 | 1 |
| ENSMUSG00000063410 | Stk24 | 0.000885812 | -2.188514111 | 0.000857868 |
| ENSMUSG00000041765 | Ubac2 | 0.000885812 | -1.535164635 | 0.031835944 |
| ENSMUSG00000039100 | 6-Mar | 0.000885812 | -1.78333786 | 0.00864105 |
| ENSMUSG00000058656 | Samd12 | 0.000885812 | -0.511629121 | 0.792441218 |
| ENSMUSG00000016554 | Eif3d | 0.000885812 | 0.378746388 | 0.94237544 |
| ENSMUSG00000068099 | 1500009C09Rik | 0.000885812 | 0.736298393 | 0.549239067 |
| ENSMUSG00000044250 | Pced1b | 0.000885812 | -0.121682602 | 1 |
| ENSMUSG00000022475 | Hdac7 | 0.000885812 | -2.188514111 | 0.000857868 |
| ENSMUSG00000023033 | Scn8a | 0.000885812 | 0.151335892 | 1 |
| ENSMUSG00000004070 | Hmox2 | 0.000885812 | -0.463373952 | 0.874192437 |
| ENSMUSG00000008393 | Carhsp1 | 0.000885812 | 0.928943471 | 0.395118486 |
| ENSMUSG00000022523 | Fgf12 | 0.000885812 | -0.52108945 | 0.814261366 |
| ENSMUSG00000107002 | 0610012G03Rik | 0.000885812 | 1.566373391 | 0.053533761 |
| ENSMUSG00000022665 | Ccdc80 | 0.000885812 | 0.310033638 | 1 |
| ENSMUSG00000052504 | Epha3 | 0.000885812 | -0.423572944 | 0.930842647 |
| ENSMUSG00000023852 | Chd1 | 0.000885812 | -1.313550157 | 0.082076736 |
| ENSMUSG00000024163 | Mapk8ip3 | 0.000885812 | -1.607997515 | 0.022405467 |
| ENSMUSG00000024187 | Fam234a | 0.000885812 | -1.413448727 | 0.056280922 |
| ENSMUSG00000024213 | Nudt3 | 0.000885812 | -0.274928863 | 1 |
| ENSMUSG00000024219 | Anks1 | 0.000885812 | -1.971060739 | 0.002944921 |
| ENSMUSG00000053436 | Mapk14 | 0.000885812 | -1.423572944 | 0.05392641 |
| ENSMUSG00000024036 | Slc37a1 | 0.000885812 | -1.200339546 | 0.128972787 |
| ENSMUSG00000092203 | 1110038B12Rik | 0.000885812 | 0.71393058 | 0.559529464 |
| ENSMUSG00000054951 | 9130008F23Rik | 0.000885812 | -0.731307157 | 0.554265357 |
| ENSMUSG00000002768 | Mea1 | 0.000885812 | 1.782102082 | 0.022226859 |
| ENSMUSG00000002372 | Ranbp3 | 0.000885812 | -1.953699579 | 0.003274704 |
| ENSMUSG00000061950 | Ppp4r1 | 0.000885812 | -1.43862729 | 0.049278849 |
| ENSMUSG00000041225 | Arhgap12 | 0.000885812 | -1.398127928 | 0.060182632 |
| ENSMUSG00000002477 | Snrpd1 | 0.000885812 | -0.005168594 | 1 |
| ENSMUSG00000024491 | Rbm27 | 0.000885812 | -1.925479705 | 0.004187446 |
| ENSMUSG00000025880 | Smad7 | 0.000885812 | -2.263701607 | 0.000554426 |
| ENSMUSG00000024892 | Pcx | 0.000885812 | -0.755554704 | 0.518463617 |
| ENSMUSG00000024769 | Cdc42bpg | 0.000885812 | -0.217897918 | 1 |
| ENSMUSG00000024949 | Sf1 | 0.000885812 | -1.562909626 | 0.028853812 |
| ENSMUSG00000075010 | AW112010 | 0.000885812 | 2.928943471 | 0.000219056 |
| ENSMUSG00000039126 | Prune2 | 0.000885812 | -0.096591621 | 1 |
| ENSMUSG00000053279 | Aldh1a1 | 0.000885812 | 0.670209202 | 0.591397544 |
| ENSMUSG00000025212 | Sfxn3 | 0.000885812 | 0.151335892 | 1 |
| ENSMUSG00000015176 | Nolc1 | 0.000885812 | -0.943181707 | 0.334416634 |
| ENSMUSG00000079480 | Pin4 | 0.000885812 | 1.244445296 | 0.208395894 |
| ENSMUSG00000033965 | Slc16a2 | 0.000885812 | -1.274928863 | 0.101718411 |
| ENSMUSG00000042501 | Cpa6 | 0.000830449 | -0.171343411 | 1 |
| ENSMUSG00000016918 | Sulf1 | 0.000830449 | -1.367972736 | 0.075331067 |
| ENSMUSG00000026058 | Khdrbs2 | 0.000830449 | 0.56137271 | 0.74382594 |
| ENSMUSG00000010453 | Kansl3 | 0.000830449 | -1.485590769 | 0.041727945 |

| | | | | |
|---|---|---|---|---|
| ENSMUSG00000100580 | 4933436I20Rik | 0.000830449 | 1.439382186 | 0.101935624 |
| ENSMUSG00000070738 | Dgkd | 0.000830449 | -1.184054463 | 0.153855756 |
| ENSMUSG00000044340 | Phlpp1 | 0.000830449 | -2.284855104 | 0.00056346 |
| ENSMUSG00000026374 | Tsn | 0.000830449 | -0.570332969 | 0.733448846 |
| ENSMUSG00000042305 | Tmem183a | 0.000830449 | -0.490715801 | 0.800251178 |
| ENSMUSG00000041879 | Ipo9 | 0.000830449 | -1.209145444 | 0.130860671 |
| ENSMUSG00000015829 | Tnr | 0.000830449 | 0.741944956 | 0.548202245 |
| ENSMUSG00000026568 | Mpc2 | 0.000830449 | -0.079084903 | 1 |
| ENSMUSG00000042901 | Aida | 0.000830449 | 0.604441432 | 0.710103834 |
| ENSMUSG00000026918 | Brd3 | 0.000830449 | -1.184054463 | 0.153855756 |
| ENSMUSG00000026883 | Dab2ip | 0.000830449 | -1.092631435 | 0.201080744 |
| ENSMUSG00000014959 | Gorasp2 | 0.000830449 | 0.256518129 | 1 |
| ENSMUSG00000041997 | Tlk1 | 0.000830449 | -1.47528576 | 0.045180838 |
| ENSMUSG00000041219 | Arhgap11a | 0.000830449 | 0.0942467 | 1 |
| ENSMUSG00000040152 | Thbs1 | 0.000830449 | -1.830944713 | 0.009099911 |
| ENSMUSG00000027287 | Snap23 | 0.000830449 | -1.603829881 | 0.023828756 |
| ENSMUSG00000033411 | Ctdspl2 | 0.000830449 | -1.82689788 | 0.008046368 |
| ENSMUSG00000027200 | Sema6d | 0.000830449 | -0.270027686 | 1 |
| ENSMUSG00000027405 | Nop56 | 0.000830449 | -2.620291128 | 5.72E-05 |
| ENSMUSG00000039033 | Tasp1 | 0.000830449 | -1.449196532 | 0.05217028 |
| ENSMUSG00000027618 | Nfs1 | 0.000830449 | -2.00221614 | 0.003299476 |
| ENSMUSG00000027615 | Hps3 | 0.000830449 | -1.275976952 | 0.110821201 |
| ENSMUSG00000027806 | Tsc22d2 | 0.000830449 | -1.928593277 | 0.004218263 |
| ENSMUSG00000043461 | Sptssb | 0.000830449 | 1.326907457 | 0.144078128 |
| ENSMUSG00000062232 | Rapgef2 | 0.000830449 | -1.969549951 | 0.003447696 |
| ENSMUSG00000068876 | Cgn | 0.000830449 | -2.020049433 | 0.002745698 |
| ENSMUSG00000028126 | Pip5k1a | 0.000830449 | -2.119348773 | 0.001489683 |
| ENSMUSG00000008763 | Man1a2 | 0.000830449 | -1.311166381 | 0.087270847 |
| ENSMUSG00000044098 | Rsbn1 | 0.000830449 | -2.142327337 | 0.001280305 |
| ENSMUSG00000027883 | Gpsm2 | 0.000830449 | -2.016500366 | 0.002690223 |
| ENSMUSG00000027882 | Stxbp3 | 0.000830449 | -1.495822691 | 0.038511752 |
| ENSMUSG00000059921 | Unc5c | 0.000830449 | -0.965874293 | 0.301813761 |
| ENSMUSG00000028273 | Pdlim5 | 0.000830449 | -2.645785197 | 4.59E-05 |
| ENSMUSG00000028184 | Adgrl2 | 0.000830449 | -1.704311274 | 0.01433775 |
| ENSMUSG00000039058 | Ak5 | 0.000830449 | 0.10967674 | 1 |
| ENSMUSG00000052544 | St6galnac3 | 0.000830449 | 0.048922709 | 1 |
| ENSMUSG00000028439 | Fam219a | 0.000830449 | -1.401012998 | 0.063228528 |
| ENSMUSG00000039813 | Tbc1d2 | 0.000830449 | -1.132524162 | 0.173210682 |
| ENSMUSG00000028431 | Elp1 | 0.000830449 | -1.878641455 | 0.005647183 |
| ENSMUSG00000066149 | Cdc26 | 0.000830449 | -0.943621486 | 0.324049807 |
| ENSMUSG00000028397 | Kdm4c | 0.000830449 | -1.743481871 | 0.012204918 |
| ENSMUSG00000028483 | Snapc3 | 0.000830449 | -1.708716453 | 0.014575482 |
| ENSMUSG00000038368 | Focad | 0.000830449 | -1.287802388 | 0.106828177 |
| ENSMUSG00000070867 | Trabd2b | 0.000830449 | 0.291283547 | 1 |
| ENSMUSG00000028719 | Cmpk1 | 0.000830449 | -0.293678954 | 1 |
| ENSMUSG00000028842 | Ago3 | 0.000830449 | -0.898058908 | 0.374573031 |
| ENSMUSG00000037692 | Ahdc1 | 0.000830449 | -1.401012998 | 0.066307071 |
| ENSMUSG00000037266 | Rsrp1 | 0.000830449 | -2.606193018 | 5.98E-05 |
| ENSMUSG00000028962 | Slc4a2 | 0.000830449 | -0.449196532 | 0.889426255 |
| ENSMUSG00000062761 | Zfp512 | 0.000830449 | -1.822839663 | 0.008096296 |
| ENSMUSG00000029221 | Slc30a9 | 0.000830449 | -0.785792676 | 0.491653643 |

| | | | | |
|---|---|---|---|---|
| ENSMUSG00000029249 | Rest | 0.000830449 | -1.427980046 | 0.054462494 |
| ENSMUSG00000089809 | Rasgef1b | 0.000830449 | -1.112715681 | 0.186482365 |
| ENSMUSG00000063406 | Tmed5 | 0.000830449 | -0.490715801 | 0.800251178 |
| ENSMUSG00000066900 | Suds3 | 0.000830449 | 0.189403933 | 1 |
| ENSMUSG00000029427 | Zcchc8 | 0.000830449 | -1.270027686 | 0.109343152 |
| ENSMUSG00000000915 | Hip1r | 0.000830449 | -1.085874069 | 0.219023377 |
| ENSMUSG00000044017 | Adgrd1 | 0.000830449 | 0.56137271 | 0.727265277 |
| ENSMUSG00000025337 | Sbds | 0.000830449 | -1.132524162 | 0.17471932 |
| ENSMUSG00000029651 | Mtus2 | 0.000830449 | 0.400908038 | 0.943363727 |
| ENSMUSG00000029762 | Akr1b8 | 0.000830449 | 0.519552534 | 0.806496094 |
| ENSMUSG00000042599 | Kdm7a | 0.000830449 | -1.636566667 | 0.021279439 |
| ENSMUSG00000029911 | Ssbp1 | 0.000830449 | -0.316948733 | 1 |
| ENSMUSG00000030105 | Arl8b | 0.000830449 | -0.281901786 | 1 |
| ENSMUSG00000024104 | Washc2 | 0.000830449 | -2.085874069 | 0.001793217 |
| ENSMUSG00000030199 | Etv6 | 0.000830449 | -2.565483569 | 7.79E-05 |
| ENSMUSG00000030282 | Cmas | 0.000830449 | -0.184054463 | 1 |
| ENSMUSG00000046456 | Tmem150b | 0.000830449 | -0.065409966 | 1 |
| ENSMUSG00000030435 | U2af2 | 0.000830449 | -0.221529168 | 1 |
| ENSMUSG00000059273 | Zc3h4 | 0.000830449 | -1.345517885 | 0.076797763 |
| ENSMUSG00000040703 | Cyp2s1 | 0.000830449 | 0.648835551 | 0.636289884 |
| ENSMUSG00000036835 | Psenen | 0.000830449 | 1.256518129 | 0.220013747 |
| ENSMUSG00000052997 | Uba2 | 0.000830449 | -0.673092543 | 0.6352512 |
| ENSMUSG00000030421 | Uri1 | 0.000830449 | -1.589569188 | 0.027621844 |
| ENSMUSG00000014402 | Tsg101 | 0.000830449 | -0.316948733 | 1 |
| ENSMUSG00000055026 | Gabrg3 | 0.000830449 | 0.671555628 | 0.66416768 |
| ENSMUSG00000030557 | Mef2a | 0.000830449 | -2.245982212 | 0.000715304 |
| ENSMUSG00000025724 | Sec11a | 0.000830449 | -0.708716453 | 0.603815648 |
| ENSMUSG00000030647 | Ndufc2 | 0.000830449 | 1.47891055 | 0.129397681 |
| ENSMUSG00000030704 | Rab6a | 0.000830449 | -0.417253639 | 0.950123751 |
| ENSMUSG00000030663 | 1110004F10Rik | 0.000830449 | -0.106051951 | 1 |
| ENSMUSG00000030657 | Xylt1 | 0.000830449 | -1.058523581 | 0.232213601 |
| ENSMUSG00000030884 | Uqcrc2 | 0.000830449 | 0.459010993 | 0.829270208 |
| ENSMUSG00000030682 | Cdipt | 0.000830449 | 0.671555628 | 0.624848895 |
| ENSMUSG00000030681 | Mvp | 0.000830449 | 0.048922709 | 1 |
| ENSMUSG00000025499 | Hras | 0.000830449 | 1.790854556 | 0.023162612 |
| ENSMUSG00000025504 | Eps8l2 | 0.000830449 | -1.037664976 | 0.250264692 |
| ENSMUSG00000031098 | Syt8 | 0.000830449 | -0.654945197 | 0.628168638 |
| ENSMUSG00000031505 | Naxd | 0.000830449 | -0.691014451 | 0.59254712 |
| ENSMUSG00000031540 | Kat6a | 0.000830449 | -1.673092543 | 0.017542038 |
| ENSMUSG00000031592 | Pcm1 | 0.000830449 | -1.839004482 | 0.007081075 |
| ENSMUSG00000038143 | Stox2 | 0.000830449 | -0.958494762 | 0.352215149 |
| ENSMUSG00000036356 | Csgalnact1 | 0.000830449 | -0.351164449 | 0.992008082 |
| ENSMUSG00000031858 | Mau2 | 0.000830449 | -0.743481871 | 0.517510782 |
| ENSMUSG00000034518 | Hmgxb4 | 0.000830449 | -0.645785197 | 0.66416768 |
| ENSMUSG00000052926 | Rnaseh2a | 0.000830449 | -0.599091962 | 0.688771131 |
| ENSMUSG00000052566 | Hook2 | 0.000830449 | -0.079084903 | 1 |
| ENSMUSG00000031696 | Vps35 | 0.000830449 | -0.316948733 | 1 |
| ENSMUSG00000036879 | Phkb | 0.000830449 | -1.673092543 | 0.017542038 |
| ENSMUSG00000031751 | Amfr | 0.000830449 | -0.449196532 | 0.889426255 |
| ENSMUSG00000000738 | Spg7 | 0.000830449 | -0.734868741 | 0.561631281 |
| ENSMUSG00000001482 | Def8 | 0.000830449 | 0.0942467 | 1 |

| | | | | |
|---|---|---|---|---|
| ENSMUSG00000031985 | Gnpat | 0.000830449 | -0.362391704 | 0.993747522 |
| ENSMUSG00000041624 | Gucy1a2 | 0.000830449 | -0.023589791 | 1 |
| ENSMUSG00000045519 | Zfp560 | 0.000830449 | 0.125273595 | 1 |
| ENSMUSG00000036611 | Eepd1 | 0.000830449 | -1.058523581 | 0.240746297 |
| ENSMUSG00000041737 | Tmem45b | 0.000830449 | -1.699892603 | 0.015705148 |
| ENSMUSG00000032314 | Etfa | 0.000830449 | -0.52108945 | 0.808536712 |
| ENSMUSG00000034485 | Uaca | 0.000830449 | -0.752043885 | 0.521275193 |
| ENSMUSG00000034563 | Ccpg1 | 0.000830449 | -0.079084903 | 1 |
| ENSMUSG00000042688 | Mapk6 | 0.000830449 | -1.227681395 | 0.125315747 |
| ENSMUSG00000009828 | Ick | 0.000830449 | -1.305360759 | 0.095779728 |
| ENSMUSG00000032411 | Tfdp2 | 0.000830449 | -1.579983139 | 0.026700028 |
| ENSMUSG00000035048 | Anapc13 | 0.000830449 | 1.948395833 | 0.040680007 |
| ENSMUSG00000040021 | Lats1 | 0.000830449 | -1.798247675 | 0.008996849 |
| ENSMUSG00000049420 | Tmem200a | 0.000830449 | -0.459688905 | 0.85084719 |
| ENSMUSG00000044499 | Hs3st5 | 0.000830449 | 1.125273595 | 0.25710887 |
| ENSMUSG00000019818 | Cd164 | 0.000830449 | 1.156982455 | 0.197460275 |
| ENSMUSG00000038122 | Tbc1d32 | 0.000830449 | -1.132524162 | 0.219707865 |
| ENSMUSG00000020077 | Srgn | 0.000830449 | 0.718098214 | 0.615418676 |
| ENSMUSG00000004667 | Polr2e | 0.000830449 | 0.867475838 | 0.422124485 |
| ENSMUSG00000004937 | Sgta | 0.000830449 | 0.078979943 | 1 |
| ENSMUSG00000020232 | Hmg20b | 0.000830449 | 0.156982455 | 1 |
| ENSMUSG00000019961 | Tmpo | 0.000830449 | -1.079084903 | 0.217570959 |
| ENSMUSG00000020029 | Nudt4 | 0.000830449 | 0.363433333 | 0.951660573 |
| ENSMUSG00000019907 | Ppp1r12a | 0.000830449 | -2.296608285 | 0.000503874 |
| ENSMUSG00000002129 | Sf3a1 | 0.000830449 | -0.009375931 | 1 |
| ENSMUSG00000009073 | Nf2 | 0.000830449 | -1.119348773 | 0.188776548 |
| ENSMUSG00000051355 | Commd1 | 0.000830449 | -0.184054463 | 1 |
| ENSMUSG00000011256 | Adam19 | 0.000830449 | -1.43331349 | 0.055240506 |
| ENSMUSG00000047804 | Akap10 | 0.000830449 | -1.367972736 | 0.072487717 |
| ENSMUSG00000018736 | Ndel1 | 0.000830449 | -2.379070445 | 0.000427357 |
| ENSMUSG00000018476 | Kdm6b | 0.000830449 | -0.245982212 | 1 |
| ENSMUSG00000005198 | Polr2a | 0.000830449 | -1.599091962 | 0.025522804 |
| ENSMUSG00000040463 | Mybbp1a | 0.000830449 | -0.89032326 | 0.40611166 |
| ENSMUSG00000045287 | Rtn4rl1 | 0.000830449 | -0.459688905 | 0.85084719 |
| ENSMUSG00000017677 | Wsb1 | 0.000830449 | -1.550836793 | 0.029931867 |
| ENSMUSG00000001507 | Itga3 | 0.000830449 | -1.575166123 | 0.02922066 |
| ENSMUSG00000057058 | Skap1 | 0.000830449 | -0.82689788 | 0.466464457 |
| ENSMUSG00000020922 | Lsm12 | 0.000830449 | -0.132524162 | 1 |
| ENSMUSG00000059248 | 9-Sep | 0.000830449 | -1.80237556 | 0.009173922 |
| ENSMUSG00000102098 | 2310016D03Rik | 0.000830449 | -2.435972836 | 0.000274457 |
| ENSMUSG00000020955 | Ap4s1 | 0.000830449 | -0.943621486 | 0.324049807 |
| ENSMUSG00000062929 | Cfl2 | 0.000830449 | -0.531073539 | 0.77665738 |
| ENSMUSG00000021067 | Sav1 | 0.000830449 | -1.516071375 | 0.036737509 |
| ENSMUSG00000046768 | Rhoj | 0.000830449 | -1.505982558 | 0.039855326 |
| ENSMUSG00000021116 | Eif2s1 | 0.000830449 | -1.233807498 | 0.125263762 |
| ENSMUSG00000021118 | Plek2 | 0.000830449 | 0.27379612 | 1 |
| ENSMUSG00000041846 | Ppp4r3a | 0.000830449 | -2.353979464 | 0.000355547 |
| ENSMUSG00000021189 | Atxn3 | 0.000830449 | -1.987789069 | 0.003112598 |
| ENSMUSG00000021288 | Klc1 | 0.000830449 | -1.954790794 | 0.003974539 |
| ENSMUSG00000072825 | Cep170b | 0.000830449 | -0.589569188 | 0.685568829 |
| ENSMUSG00000041992 | Rapgef5 | 0.000830449 | -0.490715801 | 0.800251178 |

| | | | | |
|---|---|---|---|---|
| ENSMUSG00000097709 | 2810429I04Rik | 0.000830449 | -2.311166381 | 0.000668725 |
| ENSMUSG00000021149 | Gtpbp4 | 0.000830449 | -1.976873292 | 0.003263045 |
| ENSMUSG00000018102 | Hist1h2bc | 0.000830449 | -1.009375931 | 0.263004591 |
| ENSMUSG00000039109 | F13a1 | 0.000830449 | -0.092631435 | 1 |
| ENSMUSG00000021484 | Lman2 | 0.000830449 | -0.608552291 | 0.689413368 |
| ENSMUSG00000048376 | F2r | 0.000830449 | -0.898058908 | 0.374573031 |
| ENSMUSG00000021669 | Col4a3bp | 0.000830449 | -1.991409385 | 0.003189069 |
| ENSMUSG00000021749 | Oit1 | 0.000830449 | 0.363433333 | 0.923036689 |
| ENSMUSG00000056296 | Synpr | 0.000830449 | -1.334158023 | 0.084226426 |
| ENSMUSG00000036339 | Tmem260 | 0.000830449 | -1.668577062 | 0.017255096 |
| ENSMUSG00000079197 | Psme2 | 0.000830449 | 0.867475838 | 0.422124485 |
| ENSMUSG00000021929 | Kpna3 | 0.000830449 | -1.334158023 | 0.082735457 |
| ENSMUSG00000021972 | Hmbox1 | 0.000830449 | -1.627288854 | 0.022883134 |
| ENSMUSG00000022035 | Ccdc25 | 0.000830449 | -2.184054463 | 0.001021658 |
| ENSMUSG00000022048 | Dpysl2 | 0.000830449 | -1.379070445 | 0.066698557 |
| ENSMUSG00000033487 | Fndc3a | 0.000830449 | -1.555735593 | 0.030394611 |
| ENSMUSG00000022146 | Osmr | 0.000830449 | -0.80237556 | 0.466464457 |
| ENSMUSG00000022324 | Matn2 | 0.000830449 | -0.065409966 | 1 |
| ENSMUSG00000022338 | Eny2 | 0.000830449 | 0.156982455 | 1 |
| ENSMUSG00000038725 | Pkhd1l1 | 0.000830449 | -0.645785197 | 0.66416768 |
| ENSMUSG00000037106 | Fer1l6 | 0.000830449 | 0.308985549 | 1 |
| ENSMUSG00000033039 | Micall1 | 0.000830449 | -0.760555385 | 0.542568449 |
| ENSMUSG00000022477 | Aco2 | 0.000830449 | -1.464906615 | 0.048929027 |
| ENSMUSG00000035900 | Gramd4 | 0.000830449 | -1.526090131 | 0.034053737 |
| ENSMUSG00000023025 | Larp4 | 0.000830449 | -1.682081327 | 0.016292718 |
| ENSMUSG00000023031 | Cela1 | 0.000830449 | 1.56137271 | 0.064891891 |
| ENSMUSG00000060992 | Copz1 | 0.000830449 | -1.00221614 | 0.290284345 |
| ENSMUSG00000022788 | Fgd4 | 0.000830449 | -1.764792378 | 0.010795678 |
| ENSMUSG00000022771 | Ppil2 | 0.000830449 | -1.258055044 | 0.120203977 |
| ENSMUSG00000035356 | Nfkbiz | 0.000830449 | -2.348343929 | 0.00033621 |
| ENSMUSG00000048490 | Nrip1 | 0.000830449 | -2.511035785 | 0.000120114 |
| ENSMUSG00000052712 | BC004004 | 0.000830449 | -0.654945197 | 0.628168638 |
| ENSMUSG00000015597 | Zfp318 | 0.000830449 | -0.921020057 | 0.349567117 |
| ENSMUSG00000039316 | Rftn1 | 0.000830449 | -0.80237556 | 0.476063048 |
| ENSMUSG00000024197 | Plin3 | 0.000830449 | -0.037664976 | 1 |
| ENSMUSG00000007670 | Khsrp | 0.000830449 | 0.400908038 | 0.912694223 |
| ENSMUSG00000041057 | Wdr43 | 0.000830449 | -2.545921303 | 9.31E-05 |
| ENSMUSG00000024150 | Mcfd2 | 0.000830449 | 1.063873051 | 0.274225983 |
| ENSMUSG00000036918 | Ttc7 | 0.000830449 | -0.158519371 | 1 |
| ENSMUSG00000024240 | Epc1 | 0.000830449 | -0.958494762 | 0.330721885 |
| ENSMUSG00000033628 | Pik3c3 | 0.000830449 | -0.184054463 | 1 |
| ENSMUSG00000024400 | Wdr33 | 0.000830449 | -2.337006386 | 0.000373042 |
| ENSMUSG00000024376 | Epb41l4a | 0.000830449 | -1.227681395 | 0.123869634 |
| ENSMUSG00000024552 | Slc14a2 | 0.000830449 | 0.308985549 | 1 |
| ENSMUSG00000082848 | Gm16066 | 0.000830449 | -0.52108945 | 0.808536712 |
| ENSMUSG00000024858 | Grk2 | 0.000830449 | -0.794107944 | 0.466464457 |
| ENSMUSG00000024966 | Stip1 | 0.000830449 | -1.470105521 | 0.045358829 |
| ENSMUSG00000024687 | Osbp | 0.000830449 | -1.125951508 | 0.191296504 |
| ENSMUSG00000047368 | Abhd17b | 0.000830449 | -1.806491669 | 0.008348479 |
| ENSMUSG00000033207 | Mamdc2 | 0.000830449 | 0.0942467 | 1 |
| ENSMUSG00000024827 | Gldc | 0.000830449 | -0.673092543 | 0.6352512 |

| | | | | |
|---|---|---|---|---|
| ENSMUSG00000025195 | Dnmbp | 0.000830449 | -1.016500366 | 0.274319469 |
| ENSMUSG00000025217 | Btrc | 0.000830449 | -1.608552291 | 0.023669966 |
| ENSMUSG00000043531 | Sorcs1 | 0.000830449 | 0.063873051 | 1 |
| ENSMUSG00000062949 | Atp11c | 0.000830449 | -1.589569188 | 0.027621844 |
| ENSMUSG00000031393 | Mecp2 | 0.000830449 | -1.239907697 | 0.127124729 |
| ENSMUSG00000019087 | Atp6ap1 | 0.000830449 | -0.051604167 | 1 |
| ENSMUSG00000047230 | Cldn2 | 0.000830449 | 0.239444615 | 1 |
| ENSMUSG00000031274 | Col4a5 | 0.000830449 | -1.227681395 | 0.124445804 |
| ENSMUSG00000031370 | Zrsr2 | 0.000830449 | -0.851010336 | 0.428669128 |
| ENSMUSG00000057715 | A830018L16Rik | 0.000775086 | 0.365901588 | 0.986676554 |
| ENSMUSG00000026141 | Col19a1 | 0.000775086 | 0.748371225 | 0.589936698 |
| ENSMUSG00000026116 | Tmem131 | 0.000775086 | -1.165373042 | 0.168491196 |
| ENSMUSG00000026103 | Gls | 0.000775086 | -1.251628775 | 0.128972787 |
| ENSMUSG00000006301 | Tmbim1 | 0.000775086 | -0.073630473 | 1 |
| ENSMUSG00000038608 | Dock10 | 0.000775086 | -1.449898402 | 0.061096399 |
| ENSMUSG00000026159 | Agfg1 | 0.000775086 | -2.058983697 | 0.002322357 |
| ENSMUSG00000026288 | Inpp5d | 0.000775086 | 0.047931507 | 1 |
| ENSMUSG00000026281 | Dtymk | 0.000775086 | -0.499556288 | 0.836374502 |
| ENSMUSG00000040648 | Ppip5k2 | 0.000775086 | -1.715736881 | 0.015208289 |
| ENSMUSG00000056211 | R3hdm1 | 0.000775086 | -1.588932096 | 0.029638079 |
| ENSMUSG00000026417 | Pigr | 0.000775086 | -1.537030994 | 0.052775647 |
| ENSMUSG00000054387 | Mdm4 | 0.000775086 | -2.336057556 | 0.000462341 |
| ENSMUSG00000062580 | Timm17a | 0.000775086 | 1.001137295 | 0.333922922 |
| ENSMUSG00000041605 | Inava | 0.000775086 | -1.624182943 | 0.023815668 |
| ENSMUSG00000026600 | Soat1 | 0.000775086 | -1.095325544 | 0.208124838 |
| ENSMUSG00000026626 | Ppp2r5a | 0.000775086 | -1.212458178 | 0.146698911 |
| ENSMUSG00000026775 | Yme1l1 | 0.000775086 | -1.296022894 | 0.101644331 |
| ENSMUSG00000026965 | Anapc2 | 0.000775086 | -0.158519371 | 1 |
| ENSMUSG00000039157 | Fam102a | 0.000775086 | 0.251945399 | 1 |
| ENSMUSG00000026749 | Nek6 | 0.000775086 | -1.029236354 | 0.260229808 |
| ENSMUSG00000049744 | Arhgap15 | 0.000775086 | -1.427267428 | 0.077471852 |
| ENSMUSG00000086353 | Gm13481 | 0.000775086 | 0.047931507 | 1 |
| ENSMUSG00000027030 | Stk39 | 0.000775086 | -1.10960977 | 0.231596497 |
| ENSMUSG00000041777 | Cir1 | 0.000775086 | -1.205825085 | 0.144776283 |
| ENSMUSG00000027006 | Dnajc10 | 0.000775086 | -0.836591276 | 0.461930846 |
| ENSMUSG00000027361 | Gabpb1 | 0.000775086 | -0.499556288 | 0.836374502 |
| ENSMUSG00000027366 | Sppl2a | 0.000775086 | -0.784123856 | 0.491653643 |
| ENSMUSG00000045624 | Esf1 | 0.000775086 | -1.878902081 | 0.006474516 |
| ENSMUSG00000006418 | Rnf114 | 0.000775086 | -0.653727219 | 0.642722979 |
| ENSMUSG00000016256 | Ctsz | 0.000775086 | 0.533358334 | 0.792441218 |
| ENSMUSG00000040269 | Mrps28 | 0.000775086 | -1.588932096 | 0.029638079 |
| ENSMUSG00000048332 | Lhfp | 0.000775086 | -0.952068493 | 0.351766416 |
| ENSMUSG00000036885 | Arhgef26 | 0.000775086 | -1.151632985 | 0.183392342 |
| ENSMUSG00000027822 | Slc33a1 | 0.000775086 | -0.903705472 | 0.405392586 |
| ENSMUSG00000034098 | Fstl5 | 0.000775086 | 0.063873051 | 1 |
| ENSMUSG00000027804 | Ppid | 0.000775086 | 0.270323928 | 1 |
| ENSMUSG00000068917 | Clk2 | 0.000775086 | -2.205825085 | 0.00098929 |
| ENSMUSG00000042572 | Ube2q1 | 0.000775086 | -0.466641666 | 0.878987102 |
| ENSMUSG00000027935 | Rab13 | 0.000775086 | -0.044186695 | 1 |
| ENSMUSG00000001020 | S100a4 | 0.000775086 | 1.911869957 | 0.01884098 |
| ENSMUSG00000015750 | Aph1a | 0.000775086 | 0.941016303 | 0.394341469 |

| | | | | |
|---|---|---|---|---|
| ENSMUSG00000028099 | Polr3c | 0.000775086 | -1.43295854 | 0.057125834 |
| ENSMUSG00000053398 | Phgdh | 0.000775086 | 1.855286429 | 0.022883134 |
| ENSMUSG00000040774 | Cept1 | 0.000775086 | -1.251628775 | 0.128972787 |
| ENSMUSG00000058897 | Col25a1 | 0.000775086 | 0.511332028 | 0.792441218 |
| ENSMUSG00000028194 | Ddah1 | 0.000775086 | 0.624988809 | 0.679269439 |
| ENSMUSG00000028360 | Slc44a5 | 0.000775086 | 1.096294528 | 0.290722159 |
| ENSMUSG00000073910 | Mob3b | 0.000775086 | -1.494122402 | 0.04321855 |
| ENSMUSG00000028419 | Chmp5 | 0.000775086 | -0.766201948 | 0.559988061 |
| ENSMUSG00000073889 | Il11ra1 | 0.000775086 | -1.302254848 | 0.102734511 |
| ENSMUSG00000028426 | Rad23b | 0.000775086 | -1.245173753 | 0.127356309 |
| ENSMUSG00000028370 | Pappa | 0.000775086 | -0.766201948 | 0.53074305 |
| ENSMUSG00000035212 | Leprot | 0.000775086 | 0.941016303 | 0.438297617 |
| ENSMUSG00000028514 | Usp24 | 0.000775086 | -2.18910769 | 0.001078058 |
| ENSMUSG00000078598 | Skint5 | 0.000775086 | 0.800838645 | 0.495054752 |
| ENSMUSG00000033295 | Ptprf | 0.000775086 | -0.375011191 | 0.97378787 |
| ENSMUSG00000043333 | Rhbdl2 | 0.000775086 | -0.643946198 | 0.664426832 |
| ENSMUSG00000032643 | Fhl3 | 0.000775086 | -0.775190731 | 0.488986361 |
| ENSMUSG00000023232 | Serinc2 | 0.000775086 | 0.096294528 | 1 |
| ENSMUSG00000028906 | Epb41 | 0.000775086 | -1.849415316 | 0.007771978 |
| ENSMUSG00000028886 | Eya3 | 0.000775086 | -1.351164449 | 0.09086825 |
| ENSMUSG00000037295 | Ldlrap1 | 0.000775086 | -1.308459998 | 0.093638582 |
| ENSMUSG00000028757 | Ddost | 0.000775086 | 0.385801146 | 0.975624465 |
| ENSMUSG00000028741 | Mrto4 | 0.000775086 | 0.251945399 | 1 |
| ENSMUSG00000028583 | Pdpn | 0.000775086 | 0.233798052 | 1 |
| ENSMUSG00000028991 | Mtor | 0.000775086 | -0.810596067 | 0.466464457 |
| ENSMUSG00000039768 | Dnajc11 | 0.000775086 | -0.720398258 | 0.564487875 |
| ENSMUSG00000092094 | Zfp804b | 0.000775086 | 0.346272781 | 0.987410771 |
| ENSMUSG00000023353 | Agap3 | 0.000775086 | -1.302254848 | 0.102734511 |
| ENSMUSG00000048271 | Rbm33 | 0.000775086 | -1.797420678 | 0.009680621 |
| ENSMUSG00000106864 | Gtf3c2 | 0.000775086 | -1.283477641 | 0.110389286 |
| ENSMUSG00000079562 | Maea | 0.000775086 | -0.692201366 | 0.601112074 |
| ENSMUSG00000054814 | Usp46 | 0.000775086 | -0.784123856 | 0.491653643 |
| ENSMUSG00000097516 | 4930467D21Rik | 0.000775086 | 0.180686716 | 1 |
| ENSMUSG00000035325 | Sec31a | 0.000775086 | -1.314638573 | 0.094802366 |
| ENSMUSG00000041609 | Bicdl1 | 0.000775086 | -0.542305936 | 0.792441218 |
| ENSMUSG00000023106 | Denr | 0.000775086 | 0.233798052 | 1 |
| ENSMUSG00000004951 | Hspb1 | 0.000775086 | 2.555726147 | 0.001945531 |
| ENSMUSG00000036817 | Sun1 | 0.000775086 | -0.073630473 | 1 |
| ENSMUSG00000056493 | Foxk1 | 0.000775086 | 0.346272781 | 0.991930303 |
| ENSMUSG00000029617 | Ccz1 | 0.000775086 | -1.102485336 | 0.210550914 |
| ENSMUSG00000038722 | Bud31 | 0.000775086 | 1.163408724 | 0.215910103 |
| ENSMUSG00000015112 | Slc25a13 | 0.000775086 | -0.199161355 | 1 |
| ENSMUSG00000089862 | Umad1 | 0.000775086 | -0.983432664 | 0.29603254 |
| ENSMUSG00000000058 | Cav2 | 0.000775086 | 1.346272781 | 0.200079844 |
| ENSMUSG00000025607 | Copg2 | 0.000775086 | -0.810596067 | 0.466464457 |
| ENSMUSG00000029833 | Trim24 | 0.000775086 | -0.701661696 | 0.604317526 |
| ENSMUSG00000025823 | Pdia4 | 0.000775086 | -0.116699195 | 1 |
| ENSMUSG00000029810 | Tmem176b | 0.000775086 | -0.212458178 | 1 |
| ENSMUSG00000038388 | Mpp6 | 0.000775086 | -0.775190731 | 0.488986361 |
| ENSMUSG00000053007 | Creb5 | 0.000775086 | -0.701661696 | 0.615909658 |
| ENSMUSG00000030002 | Dusp11 | 0.000775086 | -0.701661696 | 0.604317526 |

| | | | | |
|---|---|---|---|---|
| ENSMUSG00000061838 | Suclg2 | 0.000775086 | -0.853664789 | 0.462504539 |
| ENSMUSG00000030075 | Cntn3 | 0.000775086 | 0.288939606 | 1 |
| ENSMUSG00000056952 | Tatdn2 | 0.000775086 | -1.165373042 | 0.168491196 |
| ENSMUSG00000023191 | P3h3 | 0.000775086 | 0.533358334 | 0.792441218 |
| ENSMUSG00000072704 | Smim10l1 | 0.000775086 | 0.601529837 | 0.710918288 |
| ENSMUSG00000030209 | Grin2b | 0.000775086 | 0.648835551 | 0.695372548 |
| ENSMUSG00000001630 | Stk38l | 0.000775086 | -1.483192842 | 0.04673182 |
| ENSMUSG00000052296 | Ppp6r1 | 0.000775086 | 0.648835551 | 0.665027112 |
| ENSMUSG00000051527 | Usp29 | 0.000775086 | 0.112782651 | 1 |
| ENSMUSG00000006095 | Tbcb | 0.000775086 | -0.088130043 | 1 |
| ENSMUSG00000063931 | Pepd | 0.000775086 | -0.044186695 | 1 |
| ENSMUSG00000003873 | Bax | 0.000775086 | 0.555726147 | 0.790186865 |
| ENSMUSG00000030530 | Furin | 0.000775086 | -0.421553776 | 0.924442362 |
| ENSMUSG00000049583 | Grm5 | 0.000775086 | 0.385801146 | 0.911880553 |
| ENSMUSG00000030748 | Il4ra | 0.000775086 | -0.936126949 | 0.345739288 |
| ENSMUSG00000030965 | Abraxas2 | 0.000775086 | -0.701661696 | 0.624732991 |
| ENSMUSG00000025505 | Tmem80 | 0.000775086 | -1.029236354 | 0.260229808 |
| ENSMUSG00000031543 | Ank1 | 0.000775086 | -0.073630473 | 1 |
| ENSMUSG00000037492 | Zmat4 | 0.000775086 | 0.326907457 | 1 |
| ENSMUSG00000031490 | Eif4ebp1 | 0.000775086 | 0.883300805 | 0.47622561 |
| ENSMUSG00000031516 | Dctn6 | 0.000775086 | 0.468263306 | 0.879677127 |
| ENSMUSG00000038225 | Primpol | 0.000775086 | -1.205825085 | 0.144776283 |
| ENSMUSG00000031628 | Casp3 | 0.000775086 | -1.073630473 | 0.223172237 |
| ENSMUSG00000095026 | Gm3336 | 0.000775086 | 1.855286429 | 0.022883134 |
| ENSMUSG00000007721 | Ccdc124 | 0.000775086 | 0.941016303 | 0.438297617 |
| ENSMUSG00000002393 | Nr2f6 | 0.000775086 | 0.270323928 | 1 |
| ENSMUSG00000031813 | Mvb12a | 0.000775086 | 0.601529837 | 0.683838517 |
| ENSMUSG00000003037 | Rab8a | 0.000775086 | 2.032164191 | 0.012525552 |
| ENSMUSG00000059355 | Wdr83os | 0.000775086 | 1.233798052 | 0.192431778 |
| ENSMUSG00000031783 | Polr2c | 0.000775086 | 1.307798634 | 0.173382478 |
| ENSMUSG00000061048 | Cdh3 | 0.000775086 | -0.594020972 | 0.703415131 |
| ENSMUSG00000017478 | Zc3h18 | 0.000775086 | -2.123753952 | 0.001641385 |
| ENSMUSG00000000743 | Chmp1a | 0.000775086 | 0.307798634 | 1 |
| ENSMUSG00000031984 | 2810004N23Rik | 0.000775086 | -1.668275527 | 0.020028214 |
| ENSMUSG00000057060 | Slc35f3 | 0.000775086 | 0.601529837 | 0.711282364 |
| ENSMUSG00000095891 | Gm10717 | 0.000775086 | 1.42644313 | 0.110207286 |
| ENSMUSG00000025892 | Gria4 | 0.000775086 | -0.199161355 | 1 |
| ENSMUSG00000031995 | St14 | 0.000775086 | 0.405979028 | 0.886203497 |
| ENSMUSG00000032116 | Stt3a | 0.000775086 | -1.302254848 | 0.10822118 |
| ENSMUSG00000042790 | Rnf214 | 0.000775086 | -1.52108945 | 0.037122674 |
| ENSMUSG00000032078 | Zpr1 | 0.000775086 | -1.333017102 | 0.093042393 |
| ENSMUSG00000032329 | Hmg20a | 0.000775086 | -1.289763904 | 0.112194197 |
| ENSMUSG00000032243 | Itga11 | 0.000775086 | -0.339091616 | 1 |
| ENSMUSG00000032816 | Igdcc4 | 0.000775086 | -0.952068493 | 0.351766416 |
| ENSMUSG00000007908 | Hmgcll1 | 0.000775086 | -1.526422894 | 0.037800765 |
| ENSMUSG00000040433 | Zbtb38 | 0.000775086 | -2.720398258 | 3.04E-05 |
| ENSMUSG00000032454 | Rbp2 | 0.000775086 | 1.468263306 | 0.173382478 |
| ENSMUSG00000032564 | Cpne4 | 0.000775086 | 0.079992716 | 1 |
| ENSMUSG00000032601 | Prkar2a | 0.000775086 | -0.583825205 | 0.70039169 |
| ENSMUSG00000040875 | Osbpl10 | 0.000775086 | -0.653727219 | 0.642722979 |
| ENSMUSG00000070280 | Slc22a14 | 0.000775086 | -0.895484965 | 0.402012541 |

| | | | | |
|---|---|---|---|---|
| ENSMUSG00000019775 | Rgs17 | 0.000775086 | -2.375011191 | 0.000425958 |
| ENSMUSG00000019828 | Grm1 | 0.000775086 | 0.307798634 | 1 |
| ENSMUSG00000019987 | Arg1 | 0.000775086 | 1.468263306 | 0.110928515 |
| ENSMUSG00000019782 | Rwdd1 | 0.000775086 | 0.941016303 | 0.438297617 |
| ENSMUSG00000019843 | Fyn | 0.000775086 | -0.55279831 | 0.797063201 |
| ENSMUSG00000019891 | Dcbld1 | 0.000775086 | -1.779664208 | 0.011271367 |
| ENSMUSG00000038039 | Gcc2 | 0.000775086 | -1.793002007 | 0.010721159 |
| ENSMUSG00000019916 | P4ha1 | 0.000775086 | -0.351164449 | 1 |
| ENSMUSG00000020111 | Micu1 | 0.000775086 | -0.887217349 | 0.430285568 |
| ENSMUSG00000009291 | Pttg1ip | 0.000775086 | -0.144713571 | 1 |
| ENSMUSG00000020277 | Pfkl | 0.000775086 | 0.555726147 | 0.790186865 |
| ENSMUSG00000059602 | Syn3 | 0.000775086 | -0.466641666 | 0.886203497 |
| ENSMUSG00000019960 | Dusp6 | 0.000775086 | 0.215876144 | 1 |
| ENSMUSG00000025410 | Dctn2 | 0.000775086 | 0.001137295 | 1 |
| ENSMUSG00000002147 | Stat6 | 0.000775086 | -0.421553776 | 0.926564113 |
| ENSMUSG00000025364 | Pa2g4 | 0.000775086 | -1.066325672 | 0.245975468 |
| ENSMUSG00000032740 | Ccdc88a | 0.000775086 | -2.091732279 | 0.002048947 |
| ENSMUSG00000040850 | Psme4 | 0.000775086 | -0.862126368 | 0.427868543 |
| ENSMUSG00000050567 | Maml1 | 0.000775086 | -1.82365222 | 0.009984592 |
| ENSMUSG00000036391 | Sec24a | 0.000775086 | -1.455501108 | 0.054268544 |
| ENSMUSG00000056752 | Dnah9 | 0.000775086 | 0.800838645 | 0.482421209 |
| ENSMUSG00000053930 | Shisa6 | 0.000775086 | -1.43295854 | 0.057125834 |
| ENSMUSG00000001323 | Srr | 0.000775086 | -0.878902081 | 0.433732559 |
| ENSMUSG00000002052 | Supt6 | 0.000775086 | -1.351164449 | 0.086523965 |
| ENSMUSG00000018651 | Tada2a | 0.000775086 | -2.002694566 | 0.003250596 |
| ENSMUSG00000038615 | Nfe2l1 | 0.000775086 | -0.663442373 | 0.645149417 |
| ENSMUSG00000038485 | Socs7 | 0.000775086 | -0.911879403 | 0.371691552 |
| ENSMUSG00000017607 | Tns4 | 0.000775086 | -1.936126949 | 0.00493417 |
| ENSMUSG00000019302 | Atp6v0a1 | 0.000775086 | -2.720398258 | 3.04E-05 |
| ENSMUSG00000034993 | Vat1 | 0.000775086 | 1.970763646 | 0.015383621 |
| ENSMUSG00000020687 | Cdc27 | 0.000775086 | -1.258055044 | 0.116513075 |
| ENSMUSG00000041629 | Fam104a | 0.000775086 | 0.270323928 | 1 |
| ENSMUSG00000020607 | Lratd1 | 0.000775086 | 0.827805692 | 0.59863828 |
| ENSMUSG00000019256 | Ahr | 0.000775086 | -1.073630473 | 0.223172237 |
| ENSMUSG00000021024 | Psma6 | 0.000775086 | -0.43295854 | 0.927267615 |
| ENSMUSG00000021113 | Snapc1 | 0.000775086 | -0.60414519 | 0.705975399 |
| ENSMUSG00000042523 | Dnal1 | 0.000775086 | -1.60414519 | 0.028201696 |
| ENSMUSG00000042507 | Elmsan1 | 0.000775086 | -1.653727219 | 0.021277364 |
| ENSMUSG00000034290 | Nek9 | 0.000775086 | -1.663442373 | 0.019611962 |
| ENSMUSG00000051166 | Eml5 | 0.000775086 | -1.634098412 | 0.022226859 |
| ENSMUSG00000057963 | Itpk1 | 0.000775086 | -0.614198854 | 0.708141045 |
| ENSMUSG00000021147 | Wdr37 | 0.000775086 | -1.531736694 | 0.038471909 |
| ENSMUSG00000021313 | Ryr2 | 0.000775086 | 0.326907457 | 0.9980272 |
| ENSMUSG00000038462 | Uqcrfs1 | 0.000775086 | 1.032164191 | 0.319437035 |
| ENSMUSG00000044734 | Serpinb1a | 0.000775086 | 1.748371225 | 0.032072771 |
| ENSMUSG00000021365 | Nedd9 | 0.000775086 | -1.43295854 | 0.060841459 |
| ENSMUSG00000063200 | Nol7 | 0.000775086 | 0.405979028 | 0.905499991 |
| ENSMUSG00000038546 | Ranbp9 | 0.000775086 | -2.062659355 | 0.002378744 |
| ENSMUSG00000021385 | Ippk | 0.000775086 | -1.461082141 | 0.049333764 |
| ENSMUSG00000021461 | Fancc | 0.000775086 | -0.766201948 | 0.529126167 |
| ENSMUSG00000021567 | Nkd2 | 0.000775086 | -1.044186695 | 0.262284476 |

| | | | | |
|---|---|---|---|---|
| ENSMUSG00000021694 | Ercc8 | 0.000775086 | -0.952068493 | 0.36612346 |
| ENSMUSG00000021756 | Il6st | 0.000775086 | -0.967835809 | 0.32188193 |
| ENSMUSG00000016018 | Mtrex | 0.000775086 | -1.55279831 | 0.032655306 |
| ENSMUSG00000053453 | Thoc7 | 0.000775086 | 0.001137295 | 1 |
| ENSMUSG00000022159 | Rab2b | 0.000775086 | -1.788569761 | 0.010521644 |
| ENSMUSG00000016831 | Tox4 | 0.000775086 | -2.333017102 | 0.000452645 |
| ENSMUSG00000022216 | Psme1 | 0.000775086 | 0.326907457 | 0.995687918 |
| ENSMUSG00000021938 | Pspc1 | 0.000775086 | -1.682678592 | 0.01883138 |
| ENSMUSG00000054733 | Msra | 0.000775086 | 0.063873051 | 1 |
| ENSMUSG00000014725 | Adam28 | 0.000775086 | 0.385801146 | 0.975624465 |
| ENSMUSG00000067995 | Gtf2f2 | 0.000775086 | -0.94411974 | 0.348467824 |
| ENSMUSG00000072294 | Klf12 | 0.000775086 | -1.238689719 | 0.138856843 |
| ENSMUSG00000025544 | Tm9sf2 | 0.000775086 | -1.277163867 | 0.110026241 |
| ENSMUSG00000022191 | Drosha | 0.000775086 | -0.594020972 | 0.703415131 |
| ENSMUSG00000022270 | Retreg1 | 0.000775086 | -0.870538608 | 0.432443282 |
| ENSMUSG00000045996 | Polr2k | 0.000775086 | 0.911869957 | 0.455131546 |
| ENSMUSG00000022286 | Grhl2 | 0.000775086 | -0.36313709 | 1 |
| ENSMUSG00000038679 | Trps1 | 0.000775086 | -0.144713571 | 1 |
| ENSMUSG00000097039 | Pvt1 | 0.000775086 | -2.415817407 | 0.000251527 |
| ENSMUSG00000013846 | St3gal1 | 0.000775086 | -1.326916902 | 0.109343152 |
| ENSMUSG00000005354 | Txn2 | 0.000775086 | 0.774366434 | 0.554322179 |
| ENSMUSG00000022391 | Rangap1 | 0.000775086 | -1.386788358 | 0.071808577 |
| ENSMUSG00000022437 | Samm50 | 0.000775086 | 0.533358334 | 0.792441218 |
| ENSMUSG00000022536 | Glyr1 | 0.000775086 | -1.073630473 | 0.223172237 |
| ENSMUSG00000038084 | Opa1 | 0.000775086 | -1.806217627 | 0.011416719 |
| ENSMUSG00000022836 | Mylk | 0.000775086 | -1.158519371 | 0.167904846 |
| ENSMUSG00000022906 | Parp9 | 0.000775086 | -1.766201948 | 0.011908213 |
| ENSMUSG00000022601 | Zbtb11 | 0.000775086 | -1.857901782 | 0.007243475 |
| ENSMUSG00000116976 | 4930500H12Rik | 0.000775086 | -0.158519371 | 1 |
| ENSMUSG00000023805 | Synj2 | 0.000775086 | -1.158519371 | 0.167024802 |
| ENSMUSG00000034345 | Gtf2h5 | 0.000775086 | 1.129461392 | 0.296122408 |
| ENSMUSG00000023845 | Lnpep | 0.000775086 | -1.421553776 | 0.062237474 |
| ENSMUSG00000040048 | Ndufb10 | 0.000775086 | 1.270323928 | 0.256372267 |
| ENSMUSG00000024181 | Mrpl28 | 0.000775086 | -0.634098412 | 0.66416768 |
| ENSMUSG00000004864 | Mapk13 | 0.000775086 | 0.146335211 | 1 |
| ENSMUSG00000097305 | Gm17276 | 0.000775086 | -0.836591276 | 0.460623385 |
| ENSMUSG00000024308 | Tapbp | 0.000775086 | -0.314638573 | 1 |
| ENSMUSG00000015478 | Rnf5 | 0.000775086 | 2.648835551 | 0.001412437 |
| ENSMUSG00000003541 | Ier3 | 0.000775086 | -0.251628775 | 1 |
| ENSMUSG00000034595 | Ppp1r18 | 0.000775086 | -0.088130043 | 1 |
| ENSMUSG00000023939 | Mrpl14 | 0.000775086 | 0.511332028 | 0.792441218 |
| ENSMUSG00000017831 | Rab5a | 0.000775086 | -0.277163867 | 1 |
| ENSMUSG00000019578 | Ubxn6 | 0.000775086 | 0.800838645 | 0.545242312 |
| ENSMUSG00000019579 | Mydgf | 0.000775086 | 0.326907457 | 1 |
| ENSMUSG00000044279 | Crb3 | 0.000775086 | 1.346272781 | 0.121949489 |
| ENSMUSG00000024044 | Epb41l3 | 0.000775086 | -0.444273853 | 0.952107111 |
| ENSMUSG00000024054 | Smchd1 | 0.000775086 | -1.903705472 | 0.005873683 |
| ENSMUSG00000024063 | Lbh | 0.000775086 | 0.063873051 | 1 |
| ENSMUSG00000059811 | Atl2 | 0.000775086 | -0.757156808 | 0.538441519 |
| ENSMUSG00000061130 | Ppm1b | 0.000775086 | -1.682678592 | 0.01883138 |
| ENSMUSG00000063889 | Crem | 0.000775086 | -0.998862705 | 0.302165678 |

| | | | | |
|---|---|---|---|---|
| ENSMUSG00000024286 | Ccny | 0.000775086 | -0.959973691 | 0.318877315 |
| ENSMUSG00000047879 | Usp14 | 0.000775086 | -1.058983697 | 0.24342539 |
| ENSMUSG00000049411 | Tmem241 | 0.000775086 | -1.444273853 | 0.052792081 |
| ENSMUSG00000041923 | Nol4 | 0.000775086 | 1.063873051 | 0.346237121 |
| ENSMUSG00000040446 | Rprd1a | 0.000775086 | -1.77070334 | 0.012187597 |
| ENSMUSG00000024474 | Ik | 0.000775086 | -1.658592974 | 0.021725703 |
| ENSMUSG00000024431 | Nr3c1 | 0.000775086 | -1.836591276 | 0.008209564 |
| ENSMUSG00000001700 | Gramd3 | 0.000775086 | -0.903705472 | 0.405392586 |
| ENSMUSG00000024515 | Smad4 | 0.000775086 | -1.302254848 | 0.102734511 |
| ENSMUSG00000056153 | Socs6 | 0.000775086 | 0.129461392 | 1 |
| ENSMUSG00000001750 | Tcirg1 | 0.000775086 | -2.407169858 | 0.000290138 |
| ENSMUSG00000080268 | Brms1 | 0.000775086 | -0.614198854 | 0.736456705 |
| ENSMUSG00000036278 | Macrod1 | 0.000775086 | 0.047931507 | 1 |
| ENSMUSG00000114469 | C730002L08Rik | 0.000775086 | -0.212458178 | 1 |
| ENSMUSG00000024749 | Tmc1 | 0.000775086 | 0.673083098 | 0.649282271 |
| ENSMUSG00000032702 | Kank1 | 0.000775086 | -1.27082234 | 0.119907873 |
| ENSMUSG00000034321 | Exosc1 | 0.000775086 | -1.144713571 | 0.180821216 |
| ENSMUSG00000003228 | Grk5 | 0.000775086 | -1.296022894 | 0.101811693 |
| ENSMUSG00000060090 | Rp2 | 0.000775086 | -1.212458178 | 0.146698911 |
| ENSMUSG00000063785 | Utp14a | 0.000775086 | -0.911879403 | 0.371691552 |
| ENSMUSG00000085396 | Firre | 0.000775086 | -2.507668934 | 0.000148372 |
| ENSMUSG00000001962 | Fam50a | 0.000775086 | -0.116699195 | 1 |
| ENSMUSG00000004221 | Ikbkg | 0.000775086 | 0.016567336 | 1 |
| ENSMUSG00000031402 | Mpp1 | 0.000775086 | -1.27082234 | 0.119907873 |
| ENSMUSG00000046532 | Ar | 0.000775086 | -0.289763904 | 1 |
| ENSMUSG00000031333 | Abcb7 | 0.000775086 | -1.599091962 | 0.029460939 |
| ENSMUSG00000042473 | Tbc1d8b | 0.000775086 | -1.192466703 | 0.158324873 |
| ENSMUSG00000031373 | Car5b | 0.000775086 | -1.619199535 | 0.031171876 |
| ENSMUSG00000002459 | Rgs20 | 0.000719722 | 0.525453136 | 0.847265357 |
| ENSMUSG00000101337 | Dnah7c | 0.000719722 | 0.47891055 | 0.885348816 |
| ENSMUSG00000006299 | Aamp | 0.000719722 | -0.067371483 | 1 |
| ENSMUSG00000026269 | Rnpepl1 | 0.000719722 | -1.509593811 | 0.045609829 |
| ENSMUSG00000026384 | Ptpn4 | 0.000719722 | -1.555036782 | 0.036384144 |
| ENSMUSG00000058665 | En1 | 0.000719722 | -0.43862729 | 0.89889079 |
| ENSMUSG00000026414 | Tnnt2 | 0.000719722 | 0.841480629 | 0.495337286 |
| ENSMUSG00000098243 | Gm4258 | 0.000719722 | -1.538162963 | 0.04799268 |
| ENSMUSG00000090394 | 4930523C07Rik | 0.000719722 | -2.0073339 | 0.003517094 |
| ENSMUSG00000040848 | Sft2d2 | 0.000719722 | -0.113665135 | 1 |
| ENSMUSG00000040843 | Tiprl | 0.000719722 | 0.525453136 | 0.847265357 |
| ENSMUSG00000026566 | Mpzl1 | 0.000719722 | -0.856692482 | 0.466464457 |
| ENSMUSG00000058715 | Fcer1g | 0.000719722 | -1.027625304 | 0.288242813 |
| ENSMUSG00000039046 | Usp6nl | 0.000719722 | -1.593658076 | 0.029147762 |
| ENSMUSG00000026972 | Arrdc1 | 0.000719722 | -0.450700122 | 0.901829791 |
| ENSMUSG00000026837 | Col5a1 | 0.000719722 | -1.338225393 | 0.096412134 |
| ENSMUSG00000075415 | Fnbp1 | 0.000719722 | -1.426452576 | 0.069204997 |
| ENSMUSG00000039205 | Ciz1 | 0.000719722 | -1.56617734 | 0.033469112 |
| ENSMUSG00000068966 | Zbtb34 | 0.000719722 | -1.090703986 | 0.239078271 |
| ENSMUSG00000026885 | Ttll11 | 0.000719722 | -0.927513819 | 0.368467521 |
| ENSMUSG00000026750 | Psmb7 | 0.000719722 | -0.829211745 | 0.494229899 |
| ENSMUSG00000035236 | Scai | 0.000719722 | -1.113665135 | 0.232220731 |
| ENSMUSG00000026753 | Ppp6c | 0.000719722 | -1.090703986 | 0.239078271 |

| | | | | |
|---|---|---|---|---|
| ENSMUSG00000027035 | Cers6 | 0.000719722 | -1.738566381 | 0.015378262 |
| ENSMUSG00000059173 | Pde1a | 0.000719722 | -0.733634085 | 0.587484274 |
| ENSMUSG00000027067 | Ssrp1 | 0.000719722 | -0.474546864 | 0.857786543 |
| ENSMUSG00000074968 | Ano3 | 0.000719722 | 0.701302971 | 0.66416768 |
| ENSMUSG00000057378 | Ryr3 | 0.000719722 | 0.47891055 | 0.868869354 |
| ENSMUSG00000027293 | Ehd4 | 0.000719722 | -0.376699541 | 1 |
| ENSMUSG00000027245 | Hypk | 0.000719722 | 0.871227973 | 0.479831833 |
| ENSMUSG00000027203 | Dut | 0.000719722 | 1.456190473 | 0.138667568 |
| ENSMUSG00000014355 | Anapc1 | 0.000719722 | -1.901361531 | 0.006613366 |
| ENSMUSG00000027339 | Rassf2 | 0.000719722 | -1.292002376 | 0.124712353 |
| ENSMUSG00000027340 | Slc23a2 | 0.000719722 | -1.344709426 | 0.087198749 |
| ENSMUSG00000038844 | Kif16b | 0.000719722 | -1.538162963 | 0.038938939 |
| ENSMUSG00000027466 | Rbck1 | 0.000719722 | -0.113665135 | 1 |
| ENSMUSG00000061411 | Nol4l | 0.000719722 | -0.733634085 | 0.57597659 |
| ENSMUSG00000027602 | Map1lc3a | 0.000719722 | 1.134262379 | 0.303632843 |
| ENSMUSG00000038241 | Cep250 | 0.000719722 | -1.599091962 | 0.029612256 |
| ENSMUSG00000013338 | Fer1l4 | 0.000719722 | 0.573547424 | 0.751236211 |
| ENSMUSG00000016921 | Srsf6 | 0.000719722 | -0.52108945 | 0.80515297 |
| ENSMUSG00000039463 | Slc9a8 | 0.000719722 | -0.498005837 | 0.861315678 |
| ENSMUSG00000090213 | Tmem189 | 0.000719722 | -0.723718617 | 0.573177912 |
| ENSMUSG00000051149 | Adnp | 0.000719722 | -2.176804724 | 0.001437097 |
| ENSMUSG00000027551 | Zfp64 | 0.000719722 | -1.059509365 | 0.255976738 |
| ENSMUSG00000037400 | Atp11b | 0.000719722 | -0.703680863 | 0.613701629 |
| ENSMUSG00000027722 | Spata5 | 0.000719722 | -2.039664553 | 0.003965984 |
| ENSMUSG00000037111 | Setd7 | 0.000719722 | -1.003241145 | 0.290854149 |
| ENSMUSG00000098207 | Arl14 | 0.000719722 | -0.970074281 | 0.347431871 |
| ENSMUSG00000103428 | Gm20754 | 0.000719722 | 0.47891055 | 0.878272211 |
| ENSMUSG00000027835 | Pdcd10 | 0.000719722 | -0.311993851 | 1 |
| ENSMUSG00000001415 | Smg5 | 0.000719722 | -0.662750599 | 0.661446724 |
| ENSMUSG00000005687 | Bcas2 | 0.000719722 | -0.311993851 | 1 |
| ENSMUSG00000027893 | Ahcyl1 | 0.000719722 | -1.223289626 | 0.150322605 |
| ENSMUSG00000039701 | Usp53 | 0.000719722 | -1.805904372 | 0.010987762 |
| ENSMUSG00000027971 | Ndst4 | 0.000719722 | 0.368727632 | 0.984944183 |
| ENSMUSG00000001052 | Sec24b | 0.000719722 | -1.805904372 | 0.010987762 |
| ENSMUSG00000066324 | Impad1 | 0.000719722 | -0.298697029 | 1 |
| ENSMUSG00000049119 | Fam110b | 0.000719722 | 0.20826296 | 1 |
| ENSMUSG00000028217 | Cdh17 | 0.000719722 | 1.501994163 | 0.103956898 |
| ENSMUSG00000040044 | Orc3 | 0.000719722 | -1.237296484 | 0.138021363 |
| ENSMUSG00000039853 | Trim14 | 0.000719722 | -0.631272368 | 0.676381993 |
| ENSMUSG00000028369 | Svep1 | 0.000719722 | 1.170788255 | 0.242941974 |
| ENSMUSG00000038047 | Haus6 | 0.000719722 | -2.278519147 | 0.00076725 |
| ENSMUSG00000038024 | Dennd4c | 0.000719722 | -2.194861218 | 0.001293124 |
| ENSMUSG00000028577 | Plaa | 0.000719722 | -1.298697029 | 0.111578305 |
| ENSMUSG00000028576 | Ift74 | 0.000719722 | -0.79173704 | 0.495592345 |
| ENSMUSG00000028517 | Plpp3 | 0.000719722 | -0.509593811 | 0.801989789 |
| ENSMUSG00000034636 | Zyg11b | 0.000719722 | -0.588203646 | 0.72276051 |
| ENSMUSG00000028567 | Txndc12 | 0.000719722 | -1.128772027 | 0.202977999 |
| ENSMUSG00000033423 | Eri3 | 0.000719722 | -1.216234868 | 0.148136321 |
| ENSMUSG00000054405 | Dnajc8 | 0.000719722 | -0.003241145 | 1 |
| ENSMUSG00000028673 | Fuca1 | 0.000719722 | -0.450700122 | 0.910105983 |
| ENSMUSG00000078515 | Ddi2 | 0.000719722 | -1.027625304 | 0.271138872 |

| | | | | |
|---|---|---|---|---|
| ENSMUSG00000028977 | Casz1 | 0.000719722 | -0.426452576 | 0.896193257 |
| ENSMUSG00000029050 | Ski | 0.000719722 | -1.927513819 | 0.007572946 |
| ENSMUSG00000040351 | Ankib1 | 0.000719722 | -1.833828373 | 0.009782658 |
| ENSMUSG00000001467 | Cyp51 | 0.000719722 | 0.063873051 | 1 |
| ENSMUSG00000038525 | Armc10 | 0.000719722 | 0.081151042 | 1 |
| ENSMUSG00000025745 | Hadha | 0.000719722 | -0.216234868 | 1 |
| ENSMUSG00000029146 | Snx17 | 0.000719722 | 0.411796354 | 0.956910276 |
| ENSMUSG00000039706 | Ldb2 | 0.000719722 | 0.098638469 | 1 |
| ENSMUSG00000037999 | Arap2 | 0.000719722 | -1.106051951 | 0.218817775 |
| ENSMUSG00000029171 | Pgm2 | 0.000719722 | 0.623300459 | 0.736456705 |
| ENSMUSG00000029185 | Fam114a1 | 0.000719722 | -1.582728516 | 0.031456064 |
| ENSMUSG00000029191 | Rfc1 | 0.000719722 | -1.143722369 | 0.185472119 |
| ENSMUSG00000054598 | 9130230L23Rik | 0.000719722 | -0.713734528 | 0.616891292 |
| ENSMUSG00000029426 | Scarb2 | 0.000719722 | -0.187665716 | 1 |
| ENSMUSG00000062234 | Gak | 0.000719722 | -1.582728516 | 0.031456064 |
| ENSMUSG00000039886 | Tmem120a | 0.000719722 | -1.011415077 | 0.294131497 |
| ENSMUSG00000037411 | Serpine1 | 0.000719722 | 0.152409725 | 1 |
| ENSMUSG00000037364 | Srrt | 0.000719722 | -1.892537681 | 0.007109019 |
| ENSMUSG00000032625 | Thsd7a | 0.000719722 | -1.86122214 | 0.008080818 |
| ENSMUSG00000038507 | Parp12 | 0.000719722 | -1.338225393 | 0.096412134 |
| ENSMUSG00000025821 | Zfp282 | 0.000719722 | -1.549434076 | 0.035802687 |
| ENSMUSG00000029776 | Hibadh | 0.000719722 | -0.662750599 | 0.661446724 |
| ENSMUSG00000029780 | Nt5c3 | 0.000719722 | -0.298697029 | 1 |
| ENSMUSG00000049553 | Polr1a | 0.000719722 | -1.543809527 | 0.039720134 |
| ENSMUSG00000053907 | Mat2a | 0.000719722 | -0.936126949 | 0.371413601 |
| ENSMUSG00000033720 | Sfxn5 | 0.000719722 | -2.03164956 | 0.003399554 |
| ENSMUSG00000034312 | Iqsec1 | 0.000719722 | -2.202021009 | 0.001215361 |
| ENSMUSG00000030059 | Tmf1 | 0.000719722 | -1.067371483 | 0.258513752 |
| ENSMUSG00000030315 | Vgll4 | 0.000719722 | -2.2477131 | 0.000967034 |
| ENSMUSG00000025702 | 8-Mar | 0.000719722 | -0.113665135 | 1 |
| ENSMUSG00000007458 | M6pr | 0.000719722 | 0.306443354 | 1 |
| ENSMUSG00000040242 | Fgfr1op2 | 0.000719722 | -0.599091962 | 0.726119591 |
| ENSMUSG00000016487 | Ppfibp1 | 0.000719722 | -2.244249245 | 0.001052851 |
| ENSMUSG00000005566 | Trim28 | 0.000719722 | -0.143722369 | 1 |
| ENSMUSG00000062300 | Nectin2 | 0.000719722 | -0.143722369 | 1 |
| ENSMUSG00000046223 | Plaur | 0.000719722 | -0.56617734 | 0.777063553 |
| ENSMUSG00000008496 | Pou2f2 | 0.000719722 | 0.755750755 | 0.559611683 |
| ENSMUSG00000004056 | Akt2 | 0.000719722 | -0.462672764 | 0.85551189 |
| ENSMUSG00000092416 | Zfp141 | 0.000719722 | -1.351164449 | 0.088488623 |
| ENSMUSG00000038502 | Ptov1 | 0.000719722 | 0.623300459 | 0.736456705 |
| ENSMUSG00000055409 | Nell1 | 0.000719722 | 0.189403933 | 1 |
| ENSMUSG00000030450 | Oca2 | 0.000719722 | 1.20826296 | 0.246055697 |
| ENSMUSG00000101970 | 1810026B05Rik | 0.000719722 | -1.264908715 | 0.129397681 |
| ENSMUSG00000025754 | Agbl1 | 0.000719722 | -0.609898717 | 0.73719709 |
| ENSMUSG00000025103 | Btbd1 | 0.000719722 | -0.187665716 | 1 |
| ENSMUSG00000038459 | Abhd17c | 0.000719722 | -0.782214266 | 0.492327909 |
| ENSMUSG00000025439 | Clns1a | 0.000719722 | -1.237296484 | 0.138021363 |
| ENSMUSG00000070436 | Serpinh1 | 0.000719722 | 0.246737108 | 1 |
| ENSMUSG00000030842 | Lamtor1 | 0.000719722 | 0.623300459 | 0.736456705 |
| ENSMUSG00000089847 | Timm10b | 0.000719722 | 1.098638469 | 0.269992279 |
| ENSMUSG00000030751 | Psma1 | 0.000719722 | -0.285276513 | 1 |

| | | | | |
|---|---|---|---|---|
| ENSMUSG00000030876 | Mettl9 | 0.000719722 | -0.035662623 | 1 |
| ENSMUSG00000025465 | Echs1 | 0.000719722 | -0.035662623 | 1 |
| ENSMUSG00000038497 | Tmco3 | 0.000719722 | -0.113665135 | 1 |
| ENSMUSG00000031556 | Tm2d2 | 0.000719722 | 2.871227973 | 0.000987149 |
| ENSMUSG00000031488 | Rab11fip1 | 0.000719722 | -1.223289626 | 0.150322605 |
| ENSMUSG00000031578 | Mak16 | 0.000719722 | 0.67483076 | 0.65280571 |
| ENSMUSG00000031513 | Leprotl1 | 0.000719722 | 0.901601622 | 0.472949383 |
| ENSMUSG00000031532 | Saraf | 0.000719722 | -1.444676334 | 0.061141158 |
| ENSMUSG00000031627 | Irf2 | 0.000719722 | -1.743481871 | 0.015312842 |
| ENSMUSG00000039375 | Wdr17 | 0.000719722 | 1.063873051 | 0.384228838 |
| ENSMUSG00000031864 | Ints10 | 0.000719722 | -0.986753022 | 0.345505232 |
| ENSMUSG00000031833 | Mast3 | 0.000719722 | 0.871227973 | 0.481689361 |
| ENSMUSG00000110485 | Gm45740 | 0.000719722 | 0.456190473 | 0.875685162 |
| ENSMUSG00000074235 | Gm10649 | 0.000719722 | 1.411796354 | 0.126701228 |
| ENSMUSG00000055932 | Fto | 0.000719722 | -2.176804724 | 0.001437097 |
| ENSMUSG00000031948 | Kars | 0.000719722 | 0.227371783 | 1 |
| ENSMUSG00000032006 | Pdgfd | 0.000719722 | -2.308681117 | 0.000816307 |
| ENSMUSG00000058443 | Rpl10-ps3 | 0.000719722 | 1.098638469 | 0.282985014 |
| ENSMUSG00000053289 | Ddx10 | 0.000719722 | -1.662750599 | 0.023030501 |
| ENSMUSG00000036466 | Megf11 | 0.000719722 | -1.901361531 | 0.00828411 |
| ENSMUSG00000032386 | Trip4 | 0.000719722 | -1.713734528 | 0.017305126 |
| ENSMUSG00000032263 | Bckdhb | 0.000719722 | -1.462672764 | 0.074055496 |
| ENSMUSG00000032547 | Ryk | 0.000719722 | -0.113665135 | 1 |
| ENSMUSG00000032803 | Cdv3 | 0.000719722 | 0.029925719 | 1 |
| ENSMUSG00000032575 | Manf | 0.000719722 | 0.046799537 | 1 |
| ENSMUSG00000019659 | Ccdc12 | 0.000719722 | -0.631272368 | 0.676381993 |
| ENSMUSG00000039285 | Azi2 | 0.000719722 | -0.271729981 | 1 |
| ENSMUSG00000039115 | Itga9 | 0.000719722 | -0.79173704 | 0.495592345 |
| ENSMUSG00000025239 | Limd1 | 0.000719722 | -0.32516924 | 1 |
| ENSMUSG00000038587 | Akap12 | 0.000719722 | -0.298697029 | 1 |
| ENSMUSG00000019970 | Sgk1 | 0.000719722 | 0.598209478 | 0.792441218 |
| ENSMUSG00000019787 | Trdn | 0.000719722 | 0.456190473 | 0.876895022 |
| ENSMUSG00000019777 | Hdac2 | 0.000719722 | 0.390101283 | 0.962548966 |
| ENSMUSG00000038332 | Sesn1 | 0.000719722 | -0.79173704 | 0.495592345 |
| ENSMUSG00000020097 | Sgpl1 | 0.000719722 | -0.187665716 | 1 |
| ENSMUSG00000037151 | Lrrc20 | 0.000719722 | -1.098398378 | 0.262284476 |
| ENSMUSG00000037031 | Tspan15 | 0.000719722 | -0.874726405 | 0.426815929 |
| ENSMUSG00000035673 | Sbno2 | 0.000719722 | -2.043655414 | 0.00335241 |
| ENSMUSG00000020198 | Ap3d1 | 0.000719722 | -1.331712086 | 0.096412134 |
| ENSMUSG00000034781 | Gna11 | 0.000719722 | -0.067371483 | 1 |
| ENSMUSG00000020044 | Timp3 | 0.000719722 | -0.158519371 | 1 |
| ENSMUSG00000019951 | Uhrf1bp1l | 0.000719722 | -1.914496933 | 0.006295279 |
| ENSMUSG00000035916 | Ptprq | 0.000719722 | 0.266365915 | 1 |
| ENSMUSG00000020186 | Csrp2 | 0.000719722 | -0.019542957 | 1 |
| ENSMUSG00000074748 | Atxn7l3b | 0.000719722 | 1.134262379 | 0.257413002 |
| ENSMUSG00000075702 | Selenom | 0.000719722 | 1.755750755 | 0.03830656 |
| ENSMUSG00000034354 | Mtmr3 | 0.000719722 | -1.838430275 | 0.010114343 |
| ENSMUSG00000020412 | Ascc2 | 0.000719722 | -0.953200463 | 0.341777725 |
| ENSMUSG00000020476 | Dbnl | 0.000719722 | 0.841480629 | 0.491886316 |
| ENSMUSG00000049659 | Aftph | 0.000719722 | -1.599091962 | 0.029612256 |
| ENSMUSG00000044950 | Pwwp2a | 0.000719722 | -1.896956352 | 0.006613418 |

| | | | | |
|---|---|---|---|---|
| ENSMUSG00000036545 | Adamts2 | 0.000719722 | -1.762978047 | 0.01465972 |
| ENSMUSG00000005237 | Dnah2 | 0.000719722 | 0.871227973 | 0.479831833 |
| ENSMUSG00000018765 | Fxr2 | 0.000719722 | -1.708716453 | 0.016962438 |
| ENSMUSG00000038807 | Rap1gap2 | 0.000719722 | -1.027625304 | 0.271138872 |
| ENSMUSG00000018809 | Smyd4 | 0.000719722 | -1.223289626 | 0.152825523 |
| ENSMUSG00000020841 | Cpd | 0.000719722 | -0.555036782 | 0.774854805 |
| ENSMUSG00000037791 | Phf12 | 0.000719722 | -1.923187894 | 0.005872899 |
| ENSMUSG00000000804 | Usp32 | 0.000719722 | -1.667930838 | 0.020915629 |
| ENSMUSG00000058756 | Thra | 0.000719722 | -0.652333983 | 0.657805628 |
| ENSMUSG00000020694 | Tlk2 | 0.000719722 | -1.901361531 | 0.006613366 |
| ENSMUSG00000039230 | Tbcd | 0.000719722 | -1.318596586 | 0.10343288 |
| ENSMUSG00000044573 | Acp1 | 0.000719722 | 0.67483076 | 0.65280571 |
| ENSMUSG00000002688 | Prkd1 | 0.000719722 | 0.411796354 | 0.963349652 |
| ENSMUSG00000035142 | Nubpl | 0.000719722 | -0.961662041 | 0.344436715 |
| ENSMUSG00000035105 | Egln3 | 0.000719722 | -1.067371483 | 0.258513752 |
| ENSMUSG00000021025 | Nfkbia | 0.000719722 | -0.910131741 | 0.423814296 |
| ENSMUSG00000020982 | Nemf | 0.000719722 | -2.755697821 | 2.81E-05 |
| ENSMUSG00000032705 | Exd2 | 0.000719722 | -1.395558568 | 0.078253055 |
| ENSMUSG00000012609 | Ttll5 | 0.000719722 | -1.059509365 | 0.255976738 |
| ENSMUSG00000021009 | Ptpn21 | 0.000719722 | -1.782214266 | 0.012668216 |
| ENSMUSG00000037679 | Inf2 | 0.000719722 | -1.662750599 | 0.023030501 |
| ENSMUSG00000036181 | Hist1h1c | 0.000719722 | 1.812334284 | 0.032028637 |
| ENSMUSG00000021366 | Hivep1 | 0.000719722 | -1.370358014 | 0.095188999 |
| ENSMUSG00000021377 | Dek | 0.000719722 | -1.075190987 | 0.233527036 |
| ENSMUSG00000025871 | 4833439L19Rik | 0.000719722 | -0.258055044 | 1 |
| ENSMUSG00000021456 | Fbp2 | 0.000719722 | 2.063873051 | 0.014428788 |
| ENSMUSG00000021643 | Serf1 | 0.000719722 | 1.098638469 | 0.356350819 |
| ENSMUSG00000021716 | Srek1ip1 | 0.000719722 | -1.509593811 | 0.045609829 |
| ENSMUSG00000025451 | Paip1 | 0.000719722 | -1.414174246 | 0.072273892 |
| ENSMUSG00000093930 | Hmgcs1 | 0.000719722 | 0.227371783 | 1 |
| ENSMUSG00000033885 | Pxk | 0.000719722 | -1.158519371 | 0.190366478 |
| ENSMUSG00000021738 | Atxn7 | 0.000719722 | -1.56617734 | 0.033469112 |
| ENSMUSG00000017485 | Top2b | 0.000719722 | -1.113665135 | 0.220806928 |
| ENSMUSG00000040818 | Dennd6a | 0.000719722 | -1.560617814 | 0.037009644 |
| ENSMUSG00000015968 | Cacna1d | 0.000719722 | -0.082968338 | 1 |
| ENSMUSG00000006527 | Sfmbt1 | 0.000719722 | -0.298697029 | 1 |
| ENSMUSG00000021891 | Mettl6 | 0.000719722 | -2.0073339 | 0.003517094 |
| ENSMUSG00000041078 | Grid1 | 0.000719722 | 0.456190473 | 0.875072755 |
| ENSMUSG00000040997 | Abhd4 | 0.000719722 | 0.701302971 | 0.66416768 |
| ENSMUSG00000079184 | Mphosph8 | 0.000719722 | -2.711227672 | 3.93E-05 |
| ENSMUSG00000040040 | Ift88 | 0.000719722 | -1.426452576 | 0.066116266 |
| ENSMUSG00000022012 | Enox1 | 0.000719722 | 0.134262379 | 1 |
| ENSMUSG00000053253 | Ndfip2 | 0.000719722 | 0.152409725 | 1 |
| ENSMUSG00000025551 | Fgf14 | 0.000719722 | -0.743481871 | 0.611716824 |
| ENSMUSG00000050310 | Rictor | 0.000719722 | -1.815272554 | 0.010154662 |
| ENSMUSG00000022231 | Sema5a | 0.000719722 | -1.376699541 | 0.09026005 |
| ENSMUSG00000038879 | Nipal2 | 0.000719722 | -0.782214266 | 0.495276006 |
| ENSMUSG00000022309 | Angpt1 | 0.000719722 | -0.883659529 | 0.430332878 |
| ENSMUSG00000062373 | Tmem65 | 0.000719722 | -0.810596067 | 0.466464457 |
| ENSMUSG00000036661 | Dennd3 | 0.000719722 | 0.20826296 | 1 |
| ENSMUSG00000096210 | H1f0 | 0.000719722 | 1.029925719 | 0.36612346 |

| | | | | |
|---|---|---|---|---|
| ENSMUSG00000022433 | Csnk1e | 0.000719722 | -0.944688963 | 0.374672325 |
| ENSMUSG00000089715 | Cbx6 | 0.000719722 | 0.525453136 | 0.847265357 |
| ENSMUSG00000022404 | Slc25a17 | 0.000719722 | -0.311993851 | 1 |
| ENSMUSG00000052560 | Cpne8 | 0.000719722 | -1.609898717 | 0.027444373 |
| ENSMUSG00000021018 | Polr2h | 0.000719722 | 0.525453136 | 0.847265357 |
| ENSMUSG00000053774 | Ubxn7 | 0.000719722 | -1.982601379 | 0.004372181 |
| ENSMUSG00000022801 | Lrch3 | 0.000719722 | -2.36079315 | 0.000457984 |
| ENSMUSG00000022905 | Kpna1 | 0.000719722 | -0.532494213 | 0.808235359 |
| ENSMUSG00000049436 | Upk1b | 0.000719722 | -0.311993851 | 1 |
| ENSMUSG00000068196 | Col8a1 | 0.000719722 | -1.136266564 | 0.217243313 |
| ENSMUSG00000022876 | Samsn1 | 0.000719722 | 0.783765132 | 0.555706947 |
| ENSMUSG00000022867 | Usp25 | 0.000719722 | -1.113665135 | 0.220806928 |
| ENSMUSG00000022762 | Ncam2 | 0.000719722 | -0.641841609 | 0.709183387 |
| ENSMUSG00000022969 | Il10rb | 0.000719722 | -0.230310054 | 1 |
| ENSMUSG00000006818 | Sod2 | 0.000719722 | 0.623300459 | 0.736456705 |
| ENSMUSG00000039615 | Stub1 | 0.000719722 | -1.059509365 | 0.255976738 |
| ENSMUSG00000025733 | Rhot2 | 0.000719722 | 0.368727632 | 0.968854578 |
| ENSMUSG00000037326 | Capn15 | 0.000719722 | -0.995020638 | 0.319437035 |
| ENSMUSG00000024188 | Luc7l | 0.000719722 | -2.601801249 | 8.67E-05 |
| ENSMUSG00000077450 | Rab11b | 0.000719722 | 0.246737108 | 1 |
| ENSMUSG00000007029 | Vars | 0.000719722 | 0.996758855 | 0.37936848 |
| ENSMUSG00000023977 | Ubr2 | 0.000719722 | -1.599091962 | 0.029612256 |
| ENSMUSG00000042625 | Safb2 | 0.000719722 | -2.45370261 | 0.000237464 |
| ENSMUSG00000019487 | Trip10 | 0.000719722 | -0.588203646 | 0.72276051 |
| ENSMUSG00000024052 | Lpin2 | 0.000719722 | -0.723718617 | 0.573177912 |
| ENSMUSG00000024120 | Lrpprc | 0.000719722 | -0.683360879 | 0.607698939 |
| ENSMUSG00000073639 | Rab18 | 0.000719722 | 0.189403933 | 1 |
| ENSMUSG00000024306 | Ccdc178 | 0.000719722 | 0.549299878 | 0.775410248 |
| ENSMUSG00000024384 | Iws1 | 0.000719722 | -1.879199881 | 0.007443294 |
| ENSMUSG00000001380 | Hars | 0.000719722 | -1.311993851 | 0.104437622 |
| ENSMUSG00000052713 | Zfp608 | 0.000719722 | -1.625958568 | 0.025562997 |
| ENSMUSG00000036299 | BC031181 | 0.000719722 | 1.246737108 | 0.2224613 |
| ENSMUSG00000034006 | Pqlc1 | 0.000719722 | -0.244249245 | 1 |
| ENSMUSG00000024883 | Rin1 | 0.000719722 | 0.549299878 | 0.76262822 |
| ENSMUSG00000024844 | Banf1 | 0.000719722 | 0.996758855 | 0.37936848 |
| ENSMUSG00000024925 | Rnaseh2c | 0.000719722 | 1.501994163 | 0.1150226 |
| ENSMUSG00000024927 | Rela | 0.000719722 | -1.708716453 | 0.016962438 |
| ENSMUSG00000024767 | Otub1 | 0.000719722 | 0.170788255 | 1 |
| ENSMUSG00000024642 | Tle4 | 0.000719722 | -1.194861218 | 0.163620065 |
| ENSMUSG00000041360 | Pum3 | 0.000719722 | -0.743481871 | 0.580047121 |
| ENSMUSG00000039652 | Cpeb3 | 0.000719722 | -0.051604167 | 1 |
| ENSMUSG00000056999 | Ide | 0.000719722 | -0.838430275 | 0.455131546 |
| ENSMUSG00000053799 | Exoc6 | 0.000719722 | -1.824580297 | 0.009364429 |
| ENSMUSG00000025199 | Chuk | 0.000719722 | -0.838430275 | 0.455131546 |
| ENSMUSG00000055491 | Pprc1 | 0.000719722 | -1.395558568 | 0.078253055 |
| ENSMUSG00000062376 | Borcs7 | 0.000719722 | -0.450700122 | 1 |
| ENSMUSG00000031007 | Atp6ap2 | 0.000719722 | -0.414174246 | 0.966591578 |
| ENSMUSG00000067873 | Htatsf1 | 0.000719722 | -0.762978047 | 0.531673093 |
| ENSMUSG00000031201 | Brcc3 | 0.000719722 | -0.051604167 | 1 |
| ENSMUSG00000031245 | Hmgn5 | 0.000719722 | 0.116340471 | 1 |
| ENSMUSG00000055109 | Gm15155 | 0.000719722 | 0.812334284 | 0.50324127 |

| | | | | |
|---|---|---|---|---|
| ENSMUSG00000102045 | Gm21294 | 0.000719722 | -1.351164449 | 0.341107651 |
| ENSMUSG00000025935 | Tram1 | 0.000664359 | -0.142577827 | 1 |
| ENSMUSG00000026245 | Farsb | 0.000664359 | -0.025764162 | 1 |
| ENSMUSG00000026289 | Atg16l1 | 0.000664359 | -1.344211688 | 0.093904647 |
| ENSMUSG00000056536 | Pign | 0.000664359 | -2.212967155 | 0.001340467 |
| ENSMUSG00000026383 | Epb41l5 | 0.000664359 | -1.678630727 | 0.022755145 |
| ENSMUSG00000026356 | Dars | 0.000664359 | -1.337225258 | 0.103956898 |
| ENSMUSG00000042581 | Thsd7b | 0.000664359 | -0.110156349 | 1 |
| ENSMUSG00000018196 | Glrx2 | 0.000664359 | -0.265434575 | 1 |
| ENSMUSG00000004110 | Cacna1e | 0.000664359 | 0.567915556 | 0.832061506 |
| ENSMUSG00000026470 | Stx6 | 0.000664359 | -1.33020483 | 0.102521481 |
| ENSMUSG00000033557 | Fam20b | 0.000664359 | -0.470903693 | 0.886203497 |
| ENSMUSG00000013973 | Dedd | 0.000664359 | -0.593239236 | 0.743457936 |
| ENSMUSG00000026553 | Copa | 0.000664359 | -1.514910875 | 0.046387376 |
| ENSMUSG00000026527 | Rgs7 | 0.000664359 | 0.261812428 | 1 |
| ENSMUSG00000055214 | Pld5 | 0.000664359 | 0.326907457 | 1 |
| ENSMUSG00000026502 | Desi2 | 0.000664359 | -1.385434351 | 0.081057143 |
| ENSMUSG00000026617 | Bpnt1 | 0.000664359 | 1.139821904 | 0.298812586 |
| ENSMUSG00000037375 | Hhat | 0.000664359 | -0.174286686 | 1 |
| ENSMUSG00000026662 | Sephs1 | 0.000664359 | -0.80047185 | 0.499189033 |
| ENSMUSG00000037197 | Rbm17 | 0.000664359 | -0.337225258 | 1 |
| ENSMUSG00000026932 | Nacc2 | 0.000664359 | -0.205313582 | 1 |
| ENSMUSG00000039678 | Tbc1d13 | 0.000664359 | -0.458079652 | 0.884935781 |
| ENSMUSG00000050737 | Ptges | 0.000664359 | -0.706007166 | 0.627920558 |
| ENSMUSG00000026854 | Usp20 | 0.000664359 | -0.790276083 | 0.496427464 |
| ENSMUSG00000026797 | Stxbp1 | 0.000664359 | -0.392191717 | 0.998731799 |
| ENSMUSG00000026766 | Mmadhc | 0.000664359 | 2.179350268 | 0.013220053 |
| ENSMUSG00000026977 | 7-Mar | 0.000664359 | -1.990574733 | 0.004421576 |
| ENSMUSG00000026893 | Gca | 0.000664359 | 1.825713314 | 0.048635862 |
| ENSMUSG00000034903 | Cobll1 | 0.000664359 | -0.860178096 | 0.543824129 |
| ENSMUSG00000027108 | Ola1 | 0.000664359 | -0.405612233 | 0.938647404 |
| ENSMUSG00000075270 | Pde11a | 0.000664359 | 0.120456579 | 1 |
| ENSMUSG00000027011 | Ube2e3 | 0.000664359 | -1.189883542 | 0.181408286 |
| ENSMUSG00000027247 | Arhgap1 | 0.000664359 | 0.491294274 | 0.832031027 |
| ENSMUSG00000027187 | Cat | 0.000664359 | 1.491294274 | 0.155026807 |
| ENSMUSG00000027160 | Ccdc34 | 0.000664359 | -1.110156349 | 0.24200918 |
| ENSMUSG00000000308 | Ckmt1 | 0.000664359 | 1.442384674 | 0.136432253 |
| ENSMUSG00000027227 | Sord | 0.000664359 | 1.34927527 | 0.1616851 |
| ENSMUSG00000033256 | Shf | 0.000664359 | 0.491294274 | 0.83395241 |
| ENSMUSG00000033213 | AA467197 | 0.000664359 | 1.648835551 | 0.066238753 |
| ENSMUSG00000027364 | Usp50 | 0.000664359 | 1.648835551 | 0.068940102 |
| ENSMUSG00000042745 | Id1 | 0.000664359 | -0.076989485 | 1 |
| ENSMUSG00000027649 | Ctnnbl1 | 0.000664359 | 0.101347756 | 1 |
| ENSMUSG00000057133 | Chd6 | 0.000664359 | -2.097807957 | 0.002693122 |
| ENSMUSG00000047213 | Ythdf3 | 0.000664359 | -2.418909055 | 0.000362341 |
| ENSMUSG00000037643 | Prkci | 0.000664359 | -0.945345479 | 0.364693089 |
| ENSMUSG00000044864 | Ankrd50 | 0.000664359 | -1.458079652 | 0.066238753 |
| ENSMUSG00000027784 | Ppm1l | 0.000664359 | -1.126458161 | 0.209444759 |
| ENSMUSG00000027787 | Nmd3 | 0.000664359 | -0.936126949 | 0.401140948 |
| ENSMUSG00000028073 | Pear1 | 0.000664359 | -1.912819576 | 0.007127824 |
| ENSMUSG00000042737 | Dpm3 | 0.000664359 | 1.70541908 | 0.054654277 |

| | | | | |
|---|---|---|---|---|
| ENSMUSG00000049128 | Ivl | 0.000664359 | 1.889843651 | 0.038870212 |
| ENSMUSG00000038495 | Otud7b | 0.000664359 | -1.83063382 | 0.010464728 |
| ENSMUSG00000049288 | Lix1l | 0.000664359 | 0.326907457 | 1 |
| ENSMUSG00000027867 | Spag17 | 0.000664359 | 0.648835551 | 0.758506969 |
| ENSMUSG00000087260 | Lamtor5 | 0.000664359 | 1.825713314 | 0.058022225 |
| ENSMUSG00000039887 | Alg14 | 0.000664359 | -1.051604167 | 0.263425186 |
| ENSMUSG00000039756 | Dnttip2 | 0.000664359 | -0.093668226 | 1 |
| ENSMUSG00000041084 | Ostc | 0.000664359 | -0.820649732 | 0.466464457 |
| ENSMUSG00000028034 | Fubp1 | 0.000664359 | -2.572565667 | 0.000139738 |
| ENSMUSG00000039068 | Zzz3 | 0.000664359 | -2.146579757 | 0.001982991 |
| ENSMUSG00000028176 | Lrrc7 | 0.000664359 | -0.093668226 | 1 |
| ENSMUSG00000028228 | Cpne3 | 0.000664359 | -1.464505922 | 0.062686275 |
| ENSMUSG00000085632 | Gm12381 | 0.000664359 | 0.70541908 | 0.706929043 |
| ENSMUSG00000045672 | Col27a1 | 0.000664359 | -3.45162463 | 1.23E-07 |
| ENSMUSG00000028484 | Psip1 | 0.000664359 | -0.810596067 | 0.501921673 |
| ENSMUSG00000062937 | Mtap | 0.000664359 | -0.850397075 | 0.455131546 |
| ENSMUSG00000028573 | Fggy | 0.000664359 | -0.545542494 | 0.792441218 |
| ENSMUSG00000028532 | Cachd1 | 0.000664359 | -1.780007747 | 0.013903775 |
| ENSMUSG00000028522 | Mier1 | 0.000664359 | -0.879543421 | 0.463155604 |
| ENSMUSG00000028619 | Tceanc2 | 0.000664359 | -0.684147732 | 0.718012086 |
| ENSMUSG00000087194 | Skint6 | 0.000664359 | 0.991723265 | 0.430332878 |
| ENSMUSG00000028706 | Nsun4 | 0.000664359 | -0.445140597 | 0.882296038 |
| ENSMUSG00000000085 | Scmh1 | 0.000664359 | -1.893898714 | 0.007372949 |
| ENSMUSG00000028779 | Pef1 | 0.000664359 | -0.684147732 | 0.618995574 |
| ENSMUSG00000007880 | Arid1a | 0.000664359 | -2.209145444 | 0.001305524 |
| ENSMUSG00000012117 | Dhdds | 0.000664359 | -1.43862729 | 0.080836646 |
| ENSMUSG00000043924 | Ncmap | 0.000664359 | -0.265434575 | 1 |
| ENSMUSG00000001604 | Tcea3 | 0.000664359 | 0.794686418 | 0.570144904 |
| ENSMUSG00000040842 | Szrd1 | 0.000664359 | -0.706007166 | 0.62591081 |
| ENSMUSG00000110067 | 6330411D24Rik | 0.000664359 | -0.250637573 | 1 |
| ENSMUSG00000057637 | Prdm2 | 0.000664359 | -1.874726405 | 0.008996849 |
| ENSMUSG00000066026 | Dhrs3 | 0.000664359 | -1.52108945 | 0.047343204 |
| ENSMUSG00000028980 | H6pd | 0.000664359 | 0.082488729 | 1 |
| ENSMUSG00000029053 | Prkcz | 0.000664359 | -0.727540327 | 0.604880543 |
| ENSMUSG00000023286 | Ube2j2 | 0.000664359 | -0.323150072 | 1 |
| ENSMUSG00000041936 | Agrn | 0.000664359 | -2.860178096 | 1.82E-05 |
| ENSMUSG00000040473 | Cfap69 | 0.000664359 | -1.716813921 | 0.02037731 |
| ENSMUSG00000042508 | Dmtf1 | 0.000664359 | -0.83063382 | 0.466464457 |
| ENSMUSG00000039968 | Rsbn1l | 0.000664359 | -1.29458092 | 0.121570223 |
| ENSMUSG00000039000 | Ube3c | 0.000664359 | -1.820649732 | 0.011278518 |
| ENSMUSG00000045302 | Preb | 0.000664359 | -0.093668226 | 1 |
| ENSMUSG00000039753 | Fbxl5 | 0.000664359 | -2.922187758 | 1.14E-05 |
| ENSMUSG00000029186 | Pi4k2b | 0.000664359 | -0.716813921 | 0.630177036 |
| ENSMUSG00000029227 | Fip1l1 | 0.000664359 | -1.205313582 | 0.165970855 |
| ENSMUSG00000036377 | C530008M17Rik | 0.000664359 | 0.857422173 | 0.489324213 |
| ENSMUSG00000029376 | Mthfd2l | 0.000664359 | -1.545542494 | 0.045008199 |
| ENSMUSG00000029328 | Hnrnpdl | 0.000664359 | -1.748756813 | 0.015726022 |
| ENSMUSG00000029361 | Nos1 | 0.000664359 | -1.80047185 | 0.017371408 |
| ENSMUSG00000029442 | Wdr66 | 0.000664359 | -0.076989485 | 1 |
| ENSMUSG00000038126 | Mphosph9 | 0.000664359 | -1.684147732 | 0.02307354 |
| ENSMUSG00000056310 | Tyw1 | 0.000664359 | -0.496214781 | 0.835787719 |

| | | | | |
|---|---|---|---|---|
| ENSMUSG00000063146 | Clip2 | 0.000664359 | -1.034429023 | 0.288021971 |
| ENSMUSG00000032827 | Ppp1r9a | 0.000664359 | -2.774846042 | 3.96E-05 |
| ENSMUSG00000044155 | Lsm8 | 0.000664359 | 0.857422173 | 0.482937109 |
| ENSMUSG00000017978 | Cadps2 | 0.000664359 | -0.557615326 | 0.80694623 |
| ENSMUSG00000012535 | Tnpo3 | 0.000664359 | -1.633718305 | 0.028214665 |
| ENSMUSG00000038648 | Creb3l2 | 0.000664359 | -1.785151051 | 0.014189547 |
| ENSMUSG00000029922 | Mkrn1 | 0.000664359 | 0.027347175 | 1 |
| ENSMUSG00000029687 | Ezh2 | 0.000664359 | -1.205313582 | 0.165970855 |
| ENSMUSG00000029817 | Tra2a | 0.000664359 | -3.263593228 | 8.13E-07 |
| ENSMUSG00000029802 | Abcg2 | 0.000664359 | -0.043042153 | 1 |
| ENSMUSG00000055850 | Rnf181 | 0.000664359 | 0.34927527 | 1 |
| ENSMUSG00000053460 | Ggcx | 0.000664359 | -1.931495501 | 0.006162498 |
| ENSMUSG00000014748 | Tex261 | 0.000664359 | 0.30488115 | 1 |
| ENSMUSG00000001158 | Snrnp27 | 0.000664359 | -1.008276735 | 0.423814296 |
| ENSMUSG00000030058 | Copg1 | 0.000664359 | -0.351164449 | 1 |
| ENSMUSG00000035357 | Pdzrn3 | 0.000664359 | -1.110156349 | 0.228120793 |
| ENSMUSG00000030102 | Itpr1 | 0.000664359 | -1.489928519 | 0.056306476 |
| ENSMUSG00000030264 | Thumpd3 | 0.000664359 | -0.80047185 | 0.499189033 |
| ENSMUSG00000009112 | Bcl2l13 | 0.000664359 | -1.228153558 | 0.152182101 |
| ENSMUSG00000030116 | Mfap5 | 0.000664359 | 0.676849928 | 0.704871626 |
| ENSMUSG00000038521 | C1s1 | 0.000664359 | 2.179350268 | 0.013220053 |
| ENSMUSG00000030330 | Ing4 | 0.000664359 | -1.371823919 | 0.088327259 |
| ENSMUSG00000030226 | Lmo3 | 0.000664359 | -1.563614067 | 0.053198683 |
| ENSMUSG00000035632 | Cnot3 | 0.000664359 | -1.514910875 | 0.046387376 |
| ENSMUSG00000058230 | Arhgap35 | 0.000664359 | -1.795382973 | 0.013105536 |
| ENSMUSG00000057177 | Gsk3a | 0.000664359 | 1.594387767 | 0.076341241 |
| ENSMUSG00000074221 | Zfp568 | 0.000664359 | -2.201471516 | 0.001402697 |
| ENSMUSG00000030494 | Rhpn2 | 0.000664359 | -0.936126949 | 0.401140948 |
| ENSMUSG00000021217 | Tshz3 | 0.000664359 | -1.514910875 | 0.052841181 |
| ENSMUSG00000063297 | Luzp2 | 0.000664359 | 0.923010515 | 0.475407382 |
| ENSMUSG00000025324 | Atp10a | 0.000664359 | -1.235687231 | 0.154350241 |
| ENSMUSG00000032640 | Chsy1 | 0.000664359 | -0.954505479 | 0.367736443 |
| ENSMUSG00000030753 | Thap12 | 0.000664359 | 0.825713314 | 0.557051363 |
| ENSMUSG00000035227 | Spcs2 | 0.000664359 | 1.027347175 | 0.440944074 |
| ENSMUSG00000058761 | Rnf169 | 0.000664359 | -1.477273218 | 0.053988377 |
| ENSMUSG00000030921 | Trim30a | 0.000664359 | -0.963607686 | 0.410658253 |
| ENSMUSG00000101585 | 1600010M07Rik | 0.000664359 | 0.621354815 | 0.712988472 |
| ENSMUSG00000059263 | Usp47 | 0.000664359 | -2.034429023 | 0.003427316 |
| ENSMUSG00000051910 | Sox6 | 0.000664359 | -0.661951986 | 0.668261021 |
| ENSMUSG00000030652 | Coq7 | 0.000664359 | 2.179350268 | 0.013220053 |
| ENSMUSG00000032743 | D430042O09Rik | 0.000664359 | -1.738187572 | 0.019160749 |
| ENSMUSG00000032637 | Atxn2l | 0.000664359 | -0.496214781 | 0.835787719 |
| ENSMUSG00000059981 | Taok2 | 0.000664359 | -1.489928519 | 0.055578762 |
| ENSMUSG00000048170 | Mcmbp | 0.000664359 | -1.385434351 | 0.081057143 |
| ENSMUSG00000009545 | Kcnq1 | 0.000664359 | -1.945345479 | 0.005851747 |
| ENSMUSG00000031446 | Cul4a | 0.000664359 | -1.392191717 | 0.082076736 |
| ENSMUSG00000031545 | Gpat4 | 0.000664359 | -1.639409417 | 0.028807627 |
| ENSMUSG00000056313 | Tcim | 0.000664359 | 0.261812428 | 1 |
| ENSMUSG00000031570 | Plpp5 | 0.000664359 | 0.676849928 | 0.686078534 |
| ENSMUSG00000031590 | Frg1 | 0.000664359 | -0.093668226 | 1 |
| ENSMUSG00000054162 | Spock3 | 0.000664359 | -0.759249187 | 0.629310788 |

| | | | | |
|---|---|---|---|---|
| ENSMUSG00000031838 | Ifi30 | 0.000664359 | 1.219992253 | 0.262536718 |
| ENSMUSG00000031820 | Babam1 | 0.000664359 | 1.34927527 | 0.1616851 |
| ENSMUSG00000031618 | Nr3c2 | 0.000664359 | -1.287349351 | 0.119771064 |
| ENSMUSG00000053560 | Ier2 | 0.000664359 | 0.139821904 | 1 |
| ENSMUSG00000031691 | Tnpo2 | 0.000664359 | -1.458079652 | 0.066238753 |
| ENSMUSG00000074194 | Zfp791 | 0.000664359 | -0.008276735 | 1 |
| ENSMUSG00000036934 | 4921524J17Rik | 0.000664359 | -0.158519371 | 1 |
| ENSMUSG00000031770 | Herpud1 | 0.000664359 | 0.395078959 | 0.95295629 |
| ENSMUSG00000036534 | Slc38a7 | 0.000664359 | -1.470903693 | 0.060182632 |
| ENSMUSG00000012519 | Mlkl | 0.000664359 | -0.759249187 | 0.535651684 |
| ENSMUSG00000031772 | Cntnap4 | 0.000664359 | 0.956957847 | 0.458392812 |
| ENSMUSG00000046844 | Vat1l | 0.000664359 | 1.541920348 | 0.158846729 |
| ENSMUSG00000055435 | Maf | 0.000664359 | -0.557615326 | 0.796840156 |
| ENSMUSG00000025316 | Banp | 0.000664359 | -2.382043761 | 0.000480462 |
| ENSMUSG00000015013 | Trappc2l | 0.000664359 | 1.261812428 | 0.19382896 |
| ENSMUSG00000010154 | Spire2 | 0.000664359 | -0.470903693 | 0.886203497 |
| ENSMUSG00000031967 | Afg3l1 | 0.000664359 | -0.581462068 | 0.872990655 |
| ENSMUSG00000089704 | Galnt2 | 0.000664359 | -0.65072473 | 0.66416768 |
| ENSMUSG00000031987 | Egln1 | 0.000664359 | -0.850397075 | 0.455131546 |
| ENSMUSG00000031963 | Bmper | 0.000664359 | -0.235687231 | 1 |
| ENSMUSG00000032020 | Ubash3b | 0.000664359 | -1.557615326 | 0.043588231 |
| ENSMUSG00000032017 | Grik4 | 0.000664359 | -0.458079652 | 0.884935781 |
| ENSMUSG00000032026 | Rexo2 | 0.000664359 | -1.068577245 | 0.279917418 |
| ENSMUSG00000032244 | Fem1b | 0.000664359 | -1.706007166 | 0.019471235 |
| ENSMUSG00000058444 | Map2k5 | 0.000664359 | -0.850397075 | 0.455131546 |
| ENSMUSG00000032381 | Ciao2a | 0.000664359 | 0.34927527 | 1 |
| ENSMUSG00000032373 | Car12 | 0.000664359 | 0.491294274 | 0.8327465 |
| ENSMUSG00000066415 | Msl2 | 0.000664359 | -1.418909055 | 0.077478519 |
| ENSMUSG00000043154 | Ppp2r3a | 0.000664359 | -0.917511271 | 0.395118486 |
| ENSMUSG00000032786 | Alas1 | 0.000664359 | -0.569587968 | 0.737668143 |
| ENSMUSG00000032497 | Lrrfip2 | 0.000664359 | -1.017046945 | 0.314661273 |
| ENSMUSG00000039163 | Cmc1 | 0.000664359 | -0.142577827 | 1 |
| ENSMUSG00000006941 | Eif1b | 0.000664359 | 1.70541908 | 0.054654277 |
| ENSMUSG00000019768 | Esr1 | 0.000664359 | -1.45162463 | 0.080446524 |
| ENSMUSG00000047712 | Ust | 0.000664359 | 0.30488115 | 1 |
| ENSMUSG00000019790 | Stxbp5 | 0.000664359 | -0.810596067 | 0.515426845 |
| ENSMUSG00000010461 | Eya4 | 0.000664359 | -0.265434575 | 1 |
| ENSMUSG00000035621 | Midn | 0.000664359 | -0.025764162 | 1 |
| ENSMUSG00000060002 | Chpt1 | 0.000664359 | 1.027347175 | 0.418771429 |
| ENSMUSG00000062981 | Mrpl42 | 0.000664359 | 0.794686418 | 0.570144904 |
| ENSMUSG00000020140 | Lgr5 | 0.000664359 | -1.174286686 | 0.184487376 |
| ENSMUSG00000078427 | Sarnp | 0.000664359 | -1.258055044 | 0.150480064 |
| ENSMUSG00000023764 | Sfi1 | 0.000664359 | -0.80047185 | 0.499189033 |
| ENSMUSG00000020448 | Rnf185 | 0.000664359 | -0.879543421 | 0.463155604 |
| ENSMUSG00000009090 | Ap1b1 | 0.000664359 | -1.025764162 | 0.284707509 |
| ENSMUSG00000020477 | Mrps24 | 0.000664359 | 2.101347756 | 0.056595096 |
| ENSMUSG00000000581 | C1d | 0.000664359 | 1.101347756 | 0.303101829 |
| ENSMUSG00000049800 | Sertad2 | 0.000664359 | -0.898652244 | 0.429419964 |
| ENSMUSG00000042208 | 0610010F05Rik | 0.000664359 | -1.265434575 | 0.128972787 |
| ENSMUSG00000020280 | Pus10 | 0.000664359 | -0.28008135 | 1 |
| ENSMUSG00000036620 | Mgat4b | 0.000664359 | 0.261812428 | 1 |

| | | | | |
|---|---|---|---|---|
| ENSMUSG00000020400 | Tnip1 | 0.000664359 | -1.358083862 | 0.096620181 |
| ENSMUSG00000042650 | Alkbh5 | 0.000664359 | -0.060115667 | 1 |
| ENSMUSG00000059278 | Naa38 | 0.000664359 | 2.764312769 | 0.002052334 |
| ENSMUSG00000018919 | Tm4sf5 | 0.000664359 | 2.34927527 | 0.007624707 |
| ENSMUSG00000001100 | Poldip2 | 0.000664359 | -1.051604167 | 0.318970047 |
| ENSMUSG00000018334 | Ksr1 | 0.000664359 | -1.912819576 | 0.007127824 |
| ENSMUSG00000018927 | Ccl6 | 0.000664359 | 1.261812428 | 0.233878333 |
| ENSMUSG00000020865 | Abcc3 | 0.000664359 | -0.418909055 | 0.941335619 |
| ENSMUSG00000017417 | Plxdc1 | 0.000664359 | -0.483614745 | 0.886203497 |
| ENSMUSG00000020917 | Acly | 0.000664359 | -0.337225258 | 1 |
| ENSMUSG00000020611 | Gna13 | 0.000664359 | -0.29458092 | 1 |
| ENSMUSG00000041592 | Sdk2 | 0.000664359 | -1.917511271 | 0.006505658 |
| ENSMUSG00000045980 | Tmem104 | 0.000664359 | -0.790276083 | 0.496427464 |
| ENSMUSG00000020736 | Nt5c | 0.000664359 | 0.923010515 | 0.455131546 |
| ENSMUSG00000034341 | Wbp2 | 0.000664359 | 0.923010515 | 0.455131546 |
| ENSMUSG00000052949 | Rnf157 | 0.000664359 | -0.673092543 | 0.66448336 |
| ENSMUSG00000039908 | Slc26a11 | 0.000664359 | -0.569587968 | 0.737668143 |
| ENSMUSG00000078572 | Ndufaf8 | 0.000664359 | 1.261812428 | 0.276154954 |
| ENSMUSG00000061111 | Mcrip1 | 0.000664359 | -1.879543421 | 0.008872754 |
| ENSMUSG00000061911 | Myt1l | 0.000664359 | 0.594387767 | 0.736456705 |
| ENSMUSG00000104861 | 3110039M20Rik | 0.000664359 | 0.045494521 | 1 |
| ENSMUSG00000021076 | Actr10 | 0.000664359 | 0.045494521 | 1 |
| ENSMUSG00000034574 | Daam1 | 0.000664359 | -1.483614745 | 0.054654277 |
| ENSMUSG00000021108 | Prkch | 0.000664359 | -1.043042153 | 0.319801491 |
| ENSMUSG00000033454 | Zbtb1 | 0.000664359 | -1.684147732 | 0.02307354 |
| ENSMUSG00000066440 | Zfyve26 | 0.000664359 | -2.126458161 | 0.002793064 |
| ENSMUSG00000042734 | Ttc9 | 0.000664359 | -0.780007747 | 0.542583607 |
| ENSMUSG00000021240 | Abcd4 | 0.000664359 | -1.80047185 | 0.013359473 |
| ENSMUSG00000007867 | Ift43 | 0.000664359 | -1.025764162 | 0.284707509 |
| ENSMUSG00000020964 | Sel1l | 0.000664359 | -0.860178096 | 0.456406193 |
| ENSMUSG00000021180 | Rps6ka5 | 0.000664359 | -1.110156349 | 0.228120793 |
| ENSMUSG00000021285 | Ppp1r13b | 0.000664359 | -2.448386256 | 0.000324247 |
| ENSMUSG00000021143 | Pacs2 | 0.000664359 | -0.879543421 | 0.463155604 |
| ENSMUSG00000021156 | Zmynd11 | 0.000664359 | -1.316060648 | 0.112544287 |
| ENSMUSG00000042842 | Serpinb6b | 0.000664359 | 0.764312769 | 0.583067882 |
| ENSMUSG00000086693 | A730081D07Rik | 0.000664359 | -1.158519371 | 0.195601357 |
| ENSMUSG00000113074 | Gm3772 | 0.000664359 | -1.076989485 | 0.308966189 |
| ENSMUSG00000087143 | A830082K12Rik | 0.000664359 | -2.545542494 | 0.000256348 |
| ENSMUSG00000069170 | Adgrv1 | 0.000664359 | -0.639409417 | 0.709183387 |
| ENSMUSG00000021549 | Rasa1 | 0.000664359 | -2.154550468 | 0.001866323 |
| ENSMUSG00000021666 | Gfm2 | 0.000664359 | -2.250637573 | 0.001078058 |
| ENSMUSG00000078945 | Naip2 | 0.000664359 | -0.926849136 | 0.397039162 |
| ENSMUSG00000071180 | Smim15 | 0.000664359 | -0.684147732 | 0.665635898 |
| ENSMUSG00000021764 | Ndufs4 | 0.000664359 | -1.27277655 | 0.129885117 |
| ENSMUSG00000034235 | Usp54 | 0.000664359 | -0.628004654 | 0.694502633 |
| ENSMUSG00000021866 | Anxa11 | 0.000664359 | -0.820649732 | 0.466464457 |
| ENSMUSG00000037722 | Gnpnat1 | 0.000664359 | 1.179350268 | 0.400280857 |
| ENSMUSG00000098682 | Otx2os1 | 0.000664359 | 0.70541908 | 0.699192777 |
| ENSMUSG00000021963 | Sap18 | 0.000664359 | 0.923010515 | 0.455131546 |
| ENSMUSG00000021273 | Fdft1 | 0.000664359 | -0.639409417 | 0.70310512 |
| ENSMUSG00000022094 | Slc39a14 | 0.000664359 | -0.80047185 | 0.499189033 |

| | | | | |
|---|---|---|---|---|
| ENSMUSG00000022110 | Sucla2 | 0.000664359 | -1.017046945 | 0.314661273 |
| ENSMUSG00000030662 | Ipo5 | 0.000664359 | -0.545542494 | 0.792441218 |
| ENSMUSG00000022148 | Fyb | 0.000664359 | 0.159450711 | 1 |
| ENSMUSG00000037458 | Azin1 | 0.000664359 | 0.045494521 | 1 |
| ENSMUSG00000022358 | Fbxo32 | 0.000664359 | 0.442384674 | 0.92370707 |
| ENSMUSG00000022350 | Washc5 | 0.000664359 | -0.458079652 | 0.991293659 |
| ENSMUSG00000022575 | Gsdmd | 0.000664359 | -0.769665803 | 0.538110273 |
| ENSMUSG00000022558 | Mroh1 | 0.000664359 | -1.344211688 | 0.093904647 |
| ENSMUSG00000042622 | Maff | 0.000664359 | 0.676849928 | 0.686078534 |
| ENSMUSG00000036298 | Slc2a13 | 0.000664359 | -0.189883542 | 1 |
| ENSMUSG00000022635 | Zcrb1 | 0.000664359 | 0.179350268 | 1 |
| ENSMUSG00000033237 | Arid2 | 0.000664359 | -2.536421104 | 0.000175945 |
| ENSMUSG00000022479 | Vdr | 0.000664359 | -2.220580339 | 0.001510358 |
| ENSMUSG00000067713 | Prkag1 | 0.000664359 | -0.990574733 | 0.343295555 |
| ENSMUSG00000072235 | Tuba1a | 0.000664359 | 1.30488115 | 0.175780319 |
| ENSMUSG00000005981 | Trap1 | 0.000664359 | -0.628004654 | 0.698110157 |
| ENSMUSG00000022505 | Emp2 | 0.000664359 | -0.810596067 | 0.543691231 |
| ENSMUSG00000022789 | Dnm1l | 0.000664359 | -1.569587968 | 0.037394962 |
| ENSMUSG00000052459 | Atp6v1a | 0.000664359 | -0.483614745 | 0.886203497 |
| ENSMUSG00000022752 | Tomm70a | 0.000664359 | -1.017046945 | 0.314661273 |
| ENSMUSG00000052516 | Robo2 | 0.000664359 | -0.470903693 | 0.899603262 |
| ENSMUSG00000000159 | Igsf5 | 0.000664359 | -1.893898714 | 0.007372949 |
| ENSMUSG00000004945 | Tmem242 | 0.000664359 | 1.70541908 | 0.054654277 |
| ENSMUSG00000024194 | Cuta | 0.000664359 | 0.648835551 | 0.705975399 |
| ENSMUSG00000063952 | Brpf3 | 0.000664359 | -1.051604167 | 0.326043364 |
| ENSMUSG00000005936 | Kctd20 | 0.000664359 | -1.711420662 | 0.019931772 |
| ENSMUSG00000024019 | Cmtr1 | 0.000664359 | -0.990574733 | 0.3498121 |
| ENSMUSG00000041354 | Rgl2 | 0.000664359 | -0.569587968 | 0.737668143 |
| ENSMUSG00000024327 | Slc39a7 | 0.000664359 | -1.118330281 | 0.23119629 |
| ENSMUSG00000059714 | Flot1 | 0.000664359 | -0.616509015 | 0.691084416 |
| ENSMUSG00000002763 | Pex6 | 0.000664359 | -1.126458161 | 0.208883511 |
| ENSMUSG00000064043 | Trerf1 | 0.000664359 | -2.106051951 | 0.002515482 |
| ENSMUSG00000034729 | Mrps10 | 0.000664359 | 0.30488115 | 1 |
| ENSMUSG00000024164 | C3 | 0.000664359 | -0.337225258 | 1 |
| ENSMUSG00000043705 | Capn13 | 0.000664359 | 1.34927527 | 0.1616851 |
| ENSMUSG00000066057 | Gm1976 | 0.000664359 | -1.874726405 | 0.00864105 |
| ENSMUSG00000024271 | Elp2 | 0.000664359 | -1.949932749 | 0.007817567 |
| ENSMUSG00000038128 | Camk4 | 0.000664359 | 0.764312769 | 0.653235135 |
| ENSMUSG00000043079 | Synpo | 0.000664359 | -1.412275963 | 0.076166309 |
| ENSMUSG00000024622 | Hmgxb3 | 0.000664359 | -0.748756813 | 0.613567373 |
| ENSMUSG00000033871 | Ppargc1b | 0.000664359 | -0.025764162 | 1 |
| ENSMUSG00000024580 | Grpel2 | 0.000664359 | 0.889843651 | 0.466464457 |
| ENSMUSG00000079614 | Seh1l | 0.000664359 | -1.085352958 | 0.248128495 |
| ENSMUSG00000025885 | Myo5b | 0.000664359 | -2.134540508 | 0.002069296 |
| ENSMUSG00000036880 | Acaa2 | 0.000664359 | 0.240750813 | 1 |
| ENSMUSG00000024644 | Cndp2 | 0.000664359 | 0.923010515 | 0.455131546 |
| ENSMUSG00000073514 | Dok6 | 0.000664359 | -0.378645185 | 1 |
| ENSMUSG00000024900 | Cpt1a | 0.000664359 | -0.727540327 | 0.581736296 |
| ENSMUSG00000024937 | Ehbp1l1 | 0.000664359 | -1.727540327 | 0.018625118 |
| ENSMUSG00000024773 | Atg2a | 0.000664359 | -1.835600073 | 0.01067424 |
| ENSMUSG00000038812 | Trmt112 | 0.000664359 | 0.857422173 | 0.544908989 |

| | | | | |
|---|---|---|---|---|
| ENSMUSG00000024664 | Fads3 | 0.000664359 | -0.126458161 | 1 |
| ENSMUSG00000024668 | Sdhaf2 | 0.000664359 | 0.326907457 | 1 |
| ENSMUSG00000024727 | Trpm6 | 0.000664359 | -0.850397075 | 0.455131546 |
| ENSMUSG00000041731 | Pgm5 | 0.000664359 | 0.240750813 | 1 |
| ENSMUSG00000035342 | Lzts2 | 0.000664359 | -1.093668226 | 0.25111499 |
| ENSMUSG00000025228 | Actr1a | 0.000664359 | -0.126458161 | 1 |
| ENSMUSG00000042694 | Stn1 | 0.000664359 | -0.52108945 | 0.841346887 |
| ENSMUSG00000040363 | Bcor | 0.000664359 | -2.378645185 | 0.000467353 |
| ENSMUSG00000006373 | Pgrmc1 | 0.000664359 | 1.027347175 | 0.418771429 |
| ENSMUSG00000031095 | Cul4b | 0.000664359 | -0.126458161 | 1 |
| ENSMUSG00000006678 | Pola1 | 0.000664359 | -1.358083862 | 0.096620181 |
| ENSMUSG00000043929 | Klhl15 | 0.000664359 | -0.917511271 | 0.395118486 |
| ENSMUSG00000047797 | Gjb1 | 0.000664359 | 0.418537932 | 0.930842647 |
| ENSMUSG00000079481 | Nhsl2 | 0.000664359 | -1.008276735 | 0.322665395 |
| ENSMUSG00000097571 | Jpx | 0.000664359 | -0.557615326 | 0.792441218 |
| ENSMUSG00000067369 | Trmt2b | 0.000664359 | -0.639409417 | 0.697198109 |
| ENSMUSG00000048007 | Timm8a1 | 0.000664359 | 2.541920348 | 0.004143395 |
| ENSMUSG00000025262 | Fam120c | 0.000664359 | -1.458079652 | 0.066238753 |
| ENSMUSG00000056673 | Kdm5d | 0.000664359 | -2.008276735 | 0.004293776 |
| ENSMUSG00000025903 | Lypla1 | 0.000608996 | -1.57355687 | 0.042622948 |
| ENSMUSG00000025925 | Terf1 | 0.000608996 | -0.777429203 | 0.545310874 |
| ENSMUSG00000025939 | Ube2w | 0.000608996 | -1.042326353 | 0.307900693 |
| ENSMUSG00000048874 | Phf3 | 0.000608996 | -2.723303989 | 6.36E-05 |
| ENSMUSG00000058407 | Txndc9 | 0.000608996 | 0.452438339 | 0.912904617 |
| ENSMUSG00000037138 | Aff3 | 0.000608996 | 0.167708862 | 1 |
| ENSMUSG00000026095 | Asnsd1 | 0.000608996 | 0.948395833 | 0.471767627 |
| ENSMUSG00000042807 | Hecw2 | 0.000608996 | -0.289763904 | 1 |
| ENSMUSG00000041040 | Fam117b | 0.000608996 | -1.069982696 | 0.284008359 |
| ENSMUSG00000084799 | Ino80dos | 0.000608996 | -0.956026507 | 0.398007836 |
| ENSMUSG00000101599 | Gm20342 | 0.000608996 | -0.507668934 | 0.866342478 |
| ENSMUSG00000025959 | Klf7 | 0.000608996 | -1.560617814 | 0.046596585 |
| ENSMUSG00000026174 | Cnot9 | 0.000608996 | 0.024344687 | 1 |
| ENSMUSG00000026202 | Tuba4a | 0.000608996 | -0.225633566 | 1 |
| ENSMUSG00000045658 | Pid1 | 0.000608996 | -1.258055044 | 0.156455592 |
| ENSMUSG00000073633 | Fbxo36 | 0.000608996 | -0.788569761 | 0.549953572 |
| ENSMUSG00000026260 | Ndufa10 | 0.000608996 | -0.305360759 | 1 |
| ENSMUSG00000034107 | Ano7 | 0.000608996 | 0.279601742 | 1 |
| ENSMUSG00000026319 | Relch | 0.000608996 | -1.679091284 | 0.027328142 |
| ENSMUSG00000026339 | Ccdc93 | 0.000608996 | 0.063873051 | 1 |
| ENSMUSG00000026349 | Ccnt2 | 0.000608996 | -1.879944114 | 0.009405028 |
| ENSMUSG00000042268 | Slc26a9 | 0.000608996 | -1.192466703 | 0.185619905 |
| ENSMUSG00000026455 | Klhl12 | 0.000608996 | -1.250017725 | 0.158846729 |
| ENSMUSG00000057335 | Cep170 | 0.000608996 | -1.816050497 | 0.013903775 |
| ENSMUSG00000026491 | Ahctf1 | 0.000608996 | -2.110497856 | 0.00292746 |
| ENSMUSG00000026608 | Kctd3 | 0.000608996 | -0.305360759 | 1 |
| ENSMUSG00000026637 | Traf5 | 0.000608996 | 1.146335211 | 0.331463239 |
| ENSMUSG00000016194 | Hsd11b1 | 0.000608996 | -1.114930102 | 0.262284476 |
| ENSMUSG00000026718 | Stam | 0.000608996 | -2.119348773 | 0.002745698 |
| ENSMUSG00000075520 | Malrd1 | 0.000608996 | -0.032988489 | 1 |
| ENSMUSG00000036752 | Tubb4b | 0.000608996 | 0.56137271 | 0.792441218 |
| ENSMUSG00000015095 | Fbxw5 | 0.000608996 | -0.088130043 | 1 |

| | | | | |
|---|---|---|---|---|
| ENSMUSG00000026924 | Sec16a | 0.000608996 | -1.52108945 | 0.054277566 |
| ENSMUSG00000026860 | Sh3glb2 | 0.000608996 | -0.351164449 | 1 |
| ENSMUSG00000009555 | Cdk9 | 0.000608996 | -1.079084903 | 0.25896442 |
| ENSMUSG00000038831 | Ralgps1 | 0.000608996 | -1.250017725 | 0.158846729 |
| ENSMUSG00000026895 | Ndufa8 | 0.000608996 | 2.233798052 | 0.015201129 |
| ENSMUSG00000026755 | Arpc5l | 0.000608996 | 1.063873051 | 0.364909386 |
| ENSMUSG00000026754 | Golga1 | 0.000608996 | -1.343630776 | 0.115171587 |
| ENSMUSG00000036093 | Arl5a | 0.000608996 | -0.810596067 | 0.529993091 |
| ENSMUSG00000026960 | Arl6ip6 | 0.000608996 | -1.431537865 | 0.078395375 |
| ENSMUSG00000060988 | Galnt13 | 0.000608996 | -0.799624949 | 0.598570743 |
| ENSMUSG00000026827 | Gpd2 | 0.000608996 | -0.58638091 | 0.771834758 |
| ENSMUSG00000027015 | Cybrd1 | 0.000608996 | 0.452438339 | 0.891922215 |
| ENSMUSG00000027007 | Itprid2 | 0.000608996 | -1.097118826 | 0.262284476 |
| ENSMUSG00000005510 | Ndufs3 | 0.000608996 | -1.158519371 | 0.200799422 |
| ENSMUSG00000027195 | Hsd17b12 | 0.000608996 | -0.995020638 | 0.342524494 |
| ENSMUSG00000048058 | Ldlrad3 | 0.000608996 | -1.946111038 | 0.007344049 |
| ENSMUSG00000074994 | Qser1 | 0.000608996 | -1.611691999 | 0.036508071 |
| ENSMUSG00000003660 | Snrnp200 | 0.000608996 | -1.731986233 | 0.022085266 |
| ENSMUSG00000051319 | Mtln | 0.000608996 | 1.841480629 | 0.046343118 |
| ENSMUSG00000039943 | Plcb4 | 0.000608996 | -0.534386273 | 0.81635018 |
| ENSMUSG00000085071 | Gm14066 | 0.000608996 | 0.42644313 | 1 |
| ENSMUSG00000027637 | Rab5if | 0.000608996 | 0.589941862 | 0.796840156 |
| ENSMUSG00000067787 | Blcap | 0.000608996 | -0.410058138 | 0.975624465 |
| ENSMUSG00000016024 | Lbp | 0.000608996 | 1.104515035 | 0.347515171 |
| ENSMUSG00000017734 | Dbndd2 | 0.000608996 | 1.985870539 | 0.030983864 |
| ENSMUSG00000017760 | Ctsa | 0.000608996 | 0.56137271 | 0.792441218 |
| ENSMUSG00000039621 | Prex1 | 0.000608996 | 0.375817057 | 1 |
| ENSMUSG00000027509 | Rae1 | 0.000608996 | 0.841480629 | 0.513117327 |
| ENSMUSG00000039050 | Osbpl2 | 0.000608996 | -1.097118826 | 0.262284476 |
| ENSMUSG00000027583 | Zbtb46 | 0.000608996 | -0.051604167 | 1 |
| ENSMUSG00000039335 | Spata16 | 0.000608996 | 0.533358334 | 0.802702877 |
| ENSMUSG00000027698 | Nceh1 | 0.000608996 | -1.014129461 | 0.33727435 |
| ENSMUSG00000037225 | Fgf2 | 0.000608996 | -0.466641666 | 0.922546053 |
| ENSMUSG00000025757 | Hspa4l | 0.000608996 | -0.410058138 | 0.975624465 |
| ENSMUSG00000025764 | Jade1 | 0.000608996 | 0.400908038 | 1 |
| ENSMUSG00000028069 | Gpatch4 | 0.000608996 | -1.313096408 | 0.123478546 |
| ENSMUSG00000027954 | Efna1 | 0.000608996 | -1.395558568 | 0.088078849 |
| ENSMUSG00000028044 | Cks1b | 0.000608996 | 0.741944956 | 0.66416768 |
| ENSMUSG00000001017 | Chtop | 0.000608996 | 0.125273595 | 1 |
| ENSMUSG00000028089 | Chd1l | 0.000608996 | -0.810596067 | 0.503004381 |
| ENSMUSG00000027994 | Mcub | 0.000608996 | -0.946111038 | 0.395118486 |
| ENSMUSG00000028032 | Papss1 | 0.000608996 | -0.351164449 | 1 |
| ENSMUSG00000040943 | Tet2 | 0.000608996 | -1.985370468 | 0.005474255 |
| ENSMUSG00000097911 | Gm26691 | 0.000608996 | 0.505877598 | 0.887209732 |
| ENSMUSG00000104710 | Gm31243 | 0.000608996 | 0.774366434 | 0.663534823 |
| ENSMUSG00000028138 | Adh5 | 0.000608996 | 0.876246048 | 0.498537716 |
| ENSMUSG00000106053 | Gm20752 | 0.000608996 | 0.648835551 | 0.792441218 |
| ENSMUSG00000041058 | Wwp1 | 0.000608996 | 0.043973493 | 1 |
| ENSMUSG00000073987 | Ggh | 0.000608996 | 1.104515035 | 0.370799436 |
| ENSMUSG00000028423 | Nfx1 | 0.000608996 | -1.648844997 | 0.031609741 |
| ENSMUSG00000028447 | Dctn3 | 0.000608996 | 1.104515035 | 0.347515171 |

| | | | | |
|---|---|---|---|---|
| ENSMUSG00000061455 | Stx17 | 0.000608996 | -0.241935379 | 1 |
| ENSMUSG00000060206 | Zfp462 | 0.000608996 | -1.702839887 | 0.02280603 |
| ENSMUSG00000028393 | Alad | 0.000608996 | 1.589941862 | 0.09266756 |
| ENSMUSG00000028364 | Tnc | 0.000608996 | -1.014129461 | 0.354502808 |
| ENSMUSG00000084939 | Gm830 | 0.000608996 | -2.424413431 | 0.000736032 |
| ENSMUSG00000028525 | Pde4b | 0.000608996 | 0.189403933 | 1 |
| ENSMUSG00000057375 | Yipf1 | 0.000608996 | -0.305360759 | 1 |
| ENSMUSG00000028643 | Svbp | 0.000608996 | 0.084050933 | 1 |
| ENSMUSG00000028634 | Hivep3 | 0.000608996 | -0.685065185 | 0.682854035 |
| ENSMUSG00000028793 | Rnf19b | 0.000608996 | -0.088130043 | 1 |
| ENSMUSG00000046671 | Mtfr1l | 0.000608996 | -0.032988489 | 1 |
| ENSMUSG00000028826 | Maco1 | 0.000608996 | -1.417253639 | 0.081728936 |
| ENSMUSG00000037139 | Myom3 | 0.000608996 | 0.167708862 | 1 |
| ENSMUSG00000043411 | Usp48 | 0.000608996 | -1.507668934 | 0.052676838 |
| ENSMUSG00000057530 | Ece1 | 0.000608996 | -0.832291138 | 0.466464457 |
| ENSMUSG00000028599 | Tnfrsf1b | 0.000608996 | -1.250017725 | 0.158846729 |
| ENSMUSG00000059423 | Zfp933 | 0.000608996 | -1.931108875 | 0.007360617 |
| ENSMUSG00000073684 | Faap20 | 0.000608996 | -0.273996588 | 1 |
| ENSMUSG00000040118 | Cacna2d1 | 0.000608996 | -2.119348773 | 0.002713623 |
| ENSMUSG00000029166 | Mapre3 | 0.000608996 | -1.358658985 | 0.104688592 |
| ENSMUSG00000037373 | Ctbp1 | 0.000608996 | -0.273996588 | 1 |
| ENSMUSG00000036693 | Nop14 | 0.000608996 | -1.720398258 | 0.021277364 |
| ENSMUSG00000015882 | Lcorl | 0.000608996 | -1.720398258 | 0.021277364 |
| ENSMUSG00000105320 | Gm43016 | 0.000608996 | 0.375817057 | 1 |
| ENSMUSG00000036087 | Slain2 | 0.000608996 | -0.995020638 | 0.330355416 |
| ENSMUSG00000029334 | Prkg2 | 0.000608996 | -0.380911792 | 0.971854959 |
| ENSMUSG00000029559 | 2210016L21Rik | 0.000608996 | -0.3207908 | 1 |
| ENSMUSG00000043913 | Ccdc60 | 0.000608996 | -0.158519371 | 1 |
| ENSMUSG00000029420 | Rimbp2 | 0.000608996 | 0.948395833 | 0.503644124 |
| ENSMUSG00000043279 | Trim56 | 0.000608996 | -0.069982696 | 1 |
| ENSMUSG00000029718 | Pcolce | 0.000608996 | 1.42644313 | 0.168602819 |
| ENSMUSG00000029723 | Tsc22d4 | 0.000608996 | -0.507668934 | 0.866342478 |
| ENSMUSG00000037007 | Zfp113 | 0.000608996 | -0.975655313 | 0.363492638 |
| ENSMUSG00000087484 | 2900089D17Rik | 0.000608996 | -0.743481871 | 0.591705455 |
| ENSMUSG00000047735 | Samd9l | 0.000608996 | -1.810596067 | 0.013547701 |
| ENSMUSG00000052419 | 2610001J05Rik | 0.000608996 | 2.233798052 | 0.015201129 |
| ENSMUSG00000029563 | Foxp2 | 0.000608996 | -0.661019711 | 0.695372548 |
| ENSMUSG00000085264 | Gm15581 | 0.000608996 | 1.911869957 | 0.052791116 |
| ENSMUSG00000029765 | Plxna4 | 0.000608996 | -1.258055044 | 0.147041569 |
| ENSMUSG00000071477 | Zfp777 | 0.000608996 | 0.589941862 | 0.786367309 |
| ENSMUSG00000002797 | Ggct | 0.000608996 | -0.766201948 | 0.542568449 |
| ENSMUSG00000029781 | Fkbp9 | 0.000608996 | -0.258055044 | 1 |
| ENSMUSG00000031668 | Eif2ak3 | 0.000608996 | -1.766201948 | 0.018507952 |
| ENSMUSG00000002222 | Rmnd5a | 0.000608996 | -1.088130043 | 0.279592421 |
| ENSMUSG00000030042 | Pole4 | 0.000608996 | 0.303060715 | 1 |
| ENSMUSG00000045160 | Bola3 | 0.000608996 | 1.648835551 | 0.079734846 |
| ENSMUSG00000051343 | Rab11fip5 | 0.000608996 | -0.192466703 | 1 |
| ENSMUSG00000034192 | Lsm3 | 0.000608996 | 1.024344687 | 0.410681194 |
| ENSMUSG00000014550 | Rbsn | 0.000608996 | 0.189403933 | 1 |
| ENSMUSG00000030286 | Emc3 | 0.000608996 | -0.051604167 | 1 |
| ENSMUSG00000030302 | Atp2b2 | 0.000608996 | 0.948395833 | 0.508263132 |

| | | | | |
|---|---|---|---|---|
| ENSMUSG00000030123 | Plxnd1 | 0.000608996 | -1.184054463 | 0.183490997 |
| ENSMUSG00000038429 | Usp5 | 0.000608996 | 0.375817057 | 1 |
| ENSMUSG00000035585 | Tsen34 | 0.000608996 | 0.256518129 | 1 |
| ENSMUSG00000056394 | Lig1 | 0.000608996 | 0.589941862 | 0.786367309 |
| ENSMUSG00000002983 | Relb | 0.000608996 | -1.088130043 | 0.261873336 |
| ENSMUSG00000053175 | Bcl3 | 0.000608996 | -0.351164449 | 1 |
| ENSMUSG00000040940 | Arhgef1 | 0.000608996 | -1.336057556 | 0.113461807 |
| ENSMUSG00000030796 | Tead2 | 0.000608996 | -0.810596067 | 0.523075307 |
| ENSMUSG00000033676 | Gabrb3 | 0.000608996 | -0.014129461 | 1 |
| ENSMUSG00000075701 | Selenos | 0.000608996 | 0.6792092 | 0.664933061 |
| ENSMUSG00000109532 | Gm42397 | 0.000608996 | -0.141241379 | 1 |
| ENSMUSG00000030541 | Idh2 | 0.000608996 | 0.024344687 | 1 |
| ENSMUSG00000030630 | Fah | 0.000608996 | -1.500911568 | 0.071137013 |
| ENSMUSG00000030760 | Acer3 | 0.000608996 | -1.895484965 | 0.010987762 |
| ENSMUSG00000030978 | Rrm1 | 0.000608996 | 0.42644313 | 1 |
| ENSMUSG00000031024 | St5 | 0.000608996 | -0.673092543 | 0.664309971 |
| ENSMUSG00000030678 | Maz | 0.000608996 | -0.3207908 | 1 |
| ENSMUSG00000055319 | Sec23ip | 0.000608996 | -0.599091962 | 0.774790843 |
| ENSMUSG00000096938 | 9530052E02Rik | 0.000608996 | -0.410058138 | 0.976198489 |
| ENSMUSG00000000759 | Tubgcp3 | 0.000608996 | -0.547561661 | 0.806496094 |
| ENSMUSG00000055681 | Cope | 0.000608996 | 0.876246048 | 0.521929444 |
| ENSMUSG00000034880 | Mrpl34 | 0.000608996 | 2.063873051 | 0.024825015 |
| ENSMUSG00000045248 | Med26 | 0.000608996 | -1.258055044 | 0.142268504 |
| ENSMUSG00000044600 | Smim7 | 0.000608996 | -0.106051951 | 1 |
| ENSMUSG00000031714 | Gab1 | 0.000608996 | -1.445682047 | 0.076754389 |
| ENSMUSG00000005481 | Ddx39 | 0.000608996 | 0.146335211 | 1 |
| ENSMUSG00000052456 | Asna1 | 0.000608996 | 1.710236096 | 0.066942815 |
| ENSMUSG00000036712 | Cyld | 0.000608996 | -0.832291138 | 0.466464457 |
| ENSMUSG00000031788 | Kifc3 | 0.000608996 | -0.799624949 | 0.499921258 |
| ENSMUSG00000052479 | A330008L17Rik | 0.000608996 | 0.619088208 | 0.792441218 |
| ENSMUSG00000031871 | Cdh5 | 0.000608996 | -0.560617814 | 0.850505377 |
| ENSMUSG00000031839 | Hsbp1 | 0.000608996 | 0.125273595 | 1 |
| ENSMUSG00000001995 | Sipa1l2 | 0.000608996 | -1.097118826 | 0.262284476 |
| ENSMUSG00000033931 | Rbm34 | 0.000608996 | -0.624182943 | 0.712431478 |
| ENSMUSG00000096519 | Gm10721 | 0.000608996 | 2.063873051 | 0.024825015 |
| ENSMUSG00000070319 | Eif3g | 0.000608996 | 0.167708862 | 1 |
| ENSMUSG00000032177 | Pde4a | 0.000608996 | -0.273996588 | 1 |
| ENSMUSG00000003402 | Prkcsh | 0.000608996 | 0.104515035 | 1 |
| ENSMUSG00000087047 | 1700110K17Rik | 0.000608996 | -0.611691999 | 0.708791588 |
| ENSMUSG00000034584 | Exph5 | 0.000608996 | -1.250017725 | 0.158846729 |
| ENSMUSG00000061559 | Wdr61 | 0.000608996 | 0.351155003 | 1 |
| ENSMUSG00000025232 | Hexa | 0.000608996 | 0.589941862 | 0.786367309 |
| ENSMUSG00000032382 | Snx1 | 0.000608996 | -2.092631435 | 0.003012992 |
| ENSMUSG00000011958 | Bnip2 | 0.000608996 | -0.696939285 | 0.637116417 |
| ENSMUSG00000032357 | Tinag | 0.000608996 | -0.534386273 | 0.803386053 |
| ENSMUSG00000032328 | Tmem30a | 0.000608996 | -0.731986233 | 0.588684589 |
| ENSMUSG00000032261 | Sh3bgrl2 | 0.000608996 | -0.507668934 | 0.874366423 |
| ENSMUSG00000032359 | Ctsh | 0.000608996 | -0.3207908 | 1 |
| ENSMUSG00000032452 | Clstn2 | 0.000608996 | 0.063873051 | 1 |
| ENSMUSG00000045594 | Glb1 | 0.000608996 | -1.21741306 | 0.170415155 |
| ENSMUSG00000035498 | Cdcp1 | 0.000608996 | -1.642718894 | 0.031105605 |

| | | | | |
|---|---|---|---|---|
| ENSMUSG00000093803 | Ppp2r3d | 0.000608996 | -1.014129461 | 0.393268759 |
| ENSMUSG00000040624 | Plekhg1 | 0.000608996 | -1.297583408 | 0.143159082 |
| ENSMUSG00000078453 | Abracl | 0.000608996 | -0.014129461 | 1 |
| ENSMUSG00000019850 | Tnfaip3 | 0.000608996 | -0.192466703 | 1 |
| ENSMUSG00000019986 | Ahi1 | 0.000608996 | -1.209145444 | 0.217859532 |
| ENSMUSG00000038822 | Hace1 | 0.000608996 | 0.256518129 | 1 |
| ENSMUSG00000056073 | Grik2 | 0.000608996 | -0.192466703 | 1 |
| ENSMUSG00000044475 | Ascc1 | 0.000608996 | -0.624182943 | 0.742128326 |
| ENSMUSG00000020075 | Ddx21 | 0.000608996 | -1.667068756 | 0.02738711 |
| ENSMUSG00000020153 | Ndufs7 | 0.000608996 | 1.189403933 | 0.316598354 |
| ENSMUSG00000035311 | Gnptab | 0.000608996 | -0.843017545 | 0.466464457 |
| ENSMUSG00000036099 | Vezt | 0.000608996 | -1.547561661 | 0.045084086 |
| ENSMUSG00000020022 | Ndufa12 | 0.000608996 | 0.619088208 | 0.789767653 |
| ENSMUSG00000020027 | Socs2 | 0.000608996 | -0.141241379 | 1 |
| ENSMUSG00000019952 | Poc1b | 0.000608996 | -0.380911792 | 0.971854959 |
| ENSMUSG00000020132 | Rab21 | 0.000608996 | -0.380911792 | 1 |
| ENSMUSG00000020114 | Cand1 | 0.000608996 | -0.636566667 | 0.716896651 |
| ENSMUSG00000051236 | Msrb3 | 0.000608996 | -0.821484383 | 0.526094357 |
| ENSMUSG00000020124 | Usp15 | 0.000608996 | -1.685065185 | 0.02425942 |
| ENSMUSG00000020467 | Efemp1 | 0.000608996 | -1.788569761 | 0.016178743 |
| ENSMUSG00000018848 | Rars | 0.000608996 | -0.864234031 | 0.455131546 |
| ENSMUSG00000055415 | Atp10b | 0.000608996 | -0.480447466 | 0.955073065 |
| ENSMUSG00000036275 | 9530068E07Rik | 0.000608996 | 0.42644313 | 1 |
| ENSMUSG00000018239 | Zcchc10 | 0.000608996 | -1.167081384 | 0.20277243 |
| ENSMUSG00000020523 | Fam114a2 | 0.000608996 | -0.52108945 | 0.800428907 |
| ENSMUSG00000062115 | Rai1 | 0.000608996 | -2.2049938 | 0.001674533 |
| ENSMUSG00000018572 | Phf23 | 0.000608996 | -0.799624949 | 0.505918528 |
| ENSMUSG00000060216 | Arrb2 | 0.000608996 | 0.774366434 | 0.641976873 |
| ENSMUSG00000018442 | Derl2 | 0.000608996 | -1.380911792 | 0.096782144 |
| ENSMUSG00000085208 | Brip1os | 0.000608996 | -0.766201948 | 0.542568449 |
| ENSMUSG00000000278 | Scpep1 | 0.000608996 | -1.225633566 | 0.174296547 |
| ENSMUSG00000020869 | Lrrc59 | 0.000608996 | -1.069982696 | 0.295355925 |
| ENSMUSG00000018666 | Cbx1 | 0.000608996 | -0.995020638 | 0.330355416 |
| ENSMUSG00000017747 | Ghdc | 0.000608996 | -0.624182943 | 0.712431478 |
| ENSMUSG00000006920 | Ezh1 | 0.000608996 | -1.023589791 | 0.317852689 |
| ENSMUSG00000017724 | Etv4 | 0.000608996 | -1.534386273 | 0.049518535 |
| ENSMUSG00000034255 | Arhgap27 | 0.000608996 | 0.303060715 | 1 |
| ENSMUSG00000000567 | Sox9 | 0.000608996 | -1.648844997 | 0.031609741 |
| ENSMUSG00000041654 | Slc39a11 | 0.000608996 | -1.642718894 | 0.031105605 |
| ENSMUSG00000020778 | Ten1 | 0.000608996 | -0.209145444 | 1 |
| ENSMUSG00000025135 | Anapc11 | 0.000608996 | -1.167081384 | 0.205339793 |
| ENSMUSG00000020608 | Smc6 | 0.000608996 | -1.642718894 | 0.031105605 |
| ENSMUSG00000002997 | Prkar2b | 0.000608996 | 0.911869957 | 0.482785805 |
| ENSMUSG00000020561 | Twistnb | 0.000608996 | -1.175592884 | 0.205339793 |
| ENSMUSG00000043153 | Crppa | 0.000608996 | -1.57355687 | 0.042622948 |
| ENSMUSG00000036257 | Pnpla8 | 0.000608996 | -0.494122402 | 0.863905162 |
| ENSMUSG00000035329 | Fbxo33 | 0.000608996 | -0.175592884 | 1 |
| ENSMUSG00000035597 | Prpf39 | 0.000608996 | -2.72620388 | 5.96E-05 |
| ENSMUSG00000020988 | L2hgdh | 0.000608996 | -1.200830175 | 0.188750351 |
| ENSMUSG00000021068 | Nin | 0.000608996 | -1.114930102 | 0.240233158 |
| ENSMUSG00000033373 | Fntb | 0.000608996 | -0.3207908 | 1 |

| | | | | |
|---|---|---|---|---|
| ENSMUSG00000021244 | Ylpm1 | 0.000608996 | -3.420837976 | 2.38E-07 |
| ENSMUSG00000021012 | Zc3h14 | 0.000608996 | -1.507668934 | 0.052676838 |
| ENSMUSG00000021187 | Tc2n | 0.000608996 | -1.611691999 | 0.036508071 |
| ENSMUSG00000021192 | Golga5 | 0.000608996 | -1.592750435 | 0.039390861 |
| ENSMUSG00000078984 | Gm11027 | 0.000608996 | -1.106051951 | 0.23731931 |
| ENSMUSG00000019726 | Lyst | 0.000608996 | -1.192466703 | 0.185619905 |
| ENSMUSG00000021338 | Carmil1 | 0.000608996 | -2.557364836 | 0.000199484 |
| ENSMUSG00000045827 | Serpinb9 | 0.000608996 | -0.175592884 | 1 |
| ENSMUSG00000062248 | Cks2 | 0.000608996 | 0.807533297 | 0.629092968 |
| ENSMUSG00000045767 | B230219D22Rik | 0.000608996 | -0.57355687 | 0.767831378 |
| ENSMUSG00000021497 | Txndc15 | 0.000608996 | 0.375817057 | 1 |
| ENSMUSG00000056222 | Spock1 | 0.000608996 | -1.241935379 | 0.187153246 |
| ENSMUSG00000021536 | Adcy2 | 0.000608996 | 0.56137271 | 0.792441218 |
| ENSMUSG00000034525 | Ice1 | 0.000608996 | -2.369828293 | 0.000642327 |
| ENSMUSG00000021577 | Sdha | 0.000608996 | -0.43862729 | 0.912904617 |
| ENSMUSG00000062382 | Ftl1-ps1 | 0.000608996 | 0.619088208 | 0.775789276 |
| ENSMUSG00000021629 | Slc30a5 | 0.000608996 | -1.380911792 | 0.096782144 |
| ENSMUSG00000021774 | Ube2e1 | 0.000608996 | -0.480447466 | 0.860520775 |
| ENSMUSG00000039367 | Sec24c | 0.000608996 | -1.459688905 | 0.069190439 |
| ENSMUSG00000021905 | Dph3 | 0.000608996 | 0.505877598 | 0.887052657 |
| ENSMUSG00000048379 | Socs4 | 0.000608996 | -0.36611479 | 1 |
| ENSMUSG00000035161 | Ints6 | 0.000608996 | -2.167081384 | 0.002153076 |
| ENSMUSG00000021975 | Ints9 | 0.000608996 | -1.004606687 | 0.33333983 |
| ENSMUSG00000022023 | Wbp4 | 0.000608996 | -0.480447466 | 0.860520775 |
| ENSMUSG00000033186 | Mzt1 | 0.000608996 | -0.708716453 | 0.641730377 |
| ENSMUSG00000005360 | Slc1a3 | 0.000608996 | -0.995020638 | 0.404341805 |
| ENSMUSG00000046034 | Otulin | 0.000608996 | -1.424413431 | 0.082594755 |
| ENSMUSG00000015002 | Efr3a | 0.000608996 | -0.534386273 | 0.803386053 |
| ENSMUSG00000022551 | Cyc1 | 0.000608996 | 1.279601742 | 0.215910103 |
| ENSMUSG00000033170 | Card10 | 0.000608996 | -1.402826568 | 0.089464519 |
| ENSMUSG00000022472 | Desi1 | 0.000608996 | -0.648844997 | 0.740459298 |
| ENSMUSG00000022463 | Srebf2 | 0.000608996 | -1.788569761 | 0.016178743 |
| ENSMUSG00000054277 | Arfgap3 | 0.000608996 | -0.141241379 | 1 |
| ENSMUSG00000051864 | Tbc1d22a | 0.000608996 | -0.946111038 | 0.395118486 |
| ENSMUSG00000036218 | Pdzrn4 | 0.000608996 | 0.189403933 | 1 |
| ENSMUSG00000022999 | Lmbr1l | 0.000608996 | -1.654945197 | 0.028651998 |
| ENSMUSG00000051934 | Spats2 | 0.000608996 | -1.965874293 | 0.006432995 |
| ENSMUSG00000023009 | Nckap5l | 0.000608996 | -0.209145444 | 1 |
| ENSMUSG00000009733 | Tfcp2 | 0.000608996 | -1.794107944 | 0.01448405 |
| ENSMUSG00000004069 | Dnaja3 | 0.000608996 | -1.14990624 | 0.2224613 |
| ENSMUSG00000022674 | Ube2v2 | 0.000608996 | -1.507668934 | 0.052676838 |
| ENSMUSG00000022702 | Hira | 0.000608996 | -1.885143021 | 0.010114343 |
| ENSMUSG00000043391 | 2510009E07Rik | 0.000608996 | -1.788569761 | 0.016178743 |
| ENSMUSG00000023791 | Pigx | 0.000608996 | -0.43862729 | 0.915313016 |
| ENSMUSG00000052917 | Senp7 | 0.000608996 | -1.858959089 | 0.011192226 |
| ENSMUSG00000054604 | Cggbp1 | 0.000608996 | -0.088130043 | 1 |
| ENSMUSG00000041134 | Cyyr1 | 0.000608996 | -0.336057556 | 1 |
| ENSMUSG00000023861 | Mpc1 | 0.000608996 | 0.807533297 | 0.629092968 |
| ENSMUSG00000080316 | Spaca6 | 0.000608996 | -2.494122402 | 0.000303424 |
| ENSMUSG00000016946 | Kctd5 | 0.000608996 | -1.184054463 | 0.196126878 |
| ENSMUSG00000079605 | Zbtb9 | 0.000608996 | -0.853664789 | 0.469317816 |

| | | | | |
|---|---|---|---|---|
| ENSMUSG00000024217 | Snrpc | 0.000608996 | 0.256518129 | 1 |
| ENSMUSG00000043557 | Mdga1 | 0.000608996 | 0.104515035 | 1 |
| ENSMUSG00000024309 | Pfdn6 | 0.000608996 | 0.841480629 | 0.513117327 |
| ENSMUSG00000024422 | Dhx16 | 0.000608996 | -1.466641666 | 0.070555673 |
| ENSMUSG00000067148 | Polr1c | 0.000608996 | 1.146335211 | 0.36612346 |
| ENSMUSG00000062619 | 2310039H08Rik | 0.000608996 | 2.42644313 | 0.014195545 |
| ENSMUSG00000023990 | Tfeb | 0.000608996 | -0.258055044 | 1 |
| ENSMUSG00000024085 | Man2a1 | 0.000608996 | -1.079084903 | 0.25896442 |
| ENSMUSG00000045036 | Tmem232 | 0.000608996 | 1.279601742 | 0.215910103 |
| ENSMUSG00000024043 | Arhgap28 | 0.000608996 | -0.624182943 | 0.741428753 |
| ENSMUSG00000041565 | L3mbtl4 | 0.000608996 | 0.211430239 | 1 |
| ENSMUSG00000056121 | Fez2 | 0.000608996 | -0.3207908 | 1 |
| ENSMUSG00000033960 | Jcad | 0.000608996 | 0.619088208 | 0.775789276 |
| ENSMUSG00000042680 | Garem1 | 0.000608996 | -1.720398258 | 0.022226859 |
| ENSMUSG00000090523 | Gypc | 0.000608996 | -1.281901786 | 0.140206858 |
| ENSMUSG00000057561 | Eif1a | 0.000608996 | -0.936126949 | 0.401637143 |
| ENSMUSG00000024535 | Snx24 | 0.000608996 | -2.621070331 | 0.000121771 |
| ENSMUSG00000024526 | Cidea | 0.000608996 | 0.6792092 | 0.66416768 |
| ENSMUSG00000037253 | Mex3c | 0.000608996 | -0.624182943 | 0.712431478 |
| ENSMUSG00000024571 | Gm16286 | 0.000608996 | -0.480447466 | 0.860520775 |
| ENSMUSG00000041845 | Rhod | 0.000608996 | -1.209145444 | 0.168602819 |
| ENSMUSG00000024902 | Mrpl11 | 0.000608996 | 1.985870539 | 0.030983864 |
| ENSMUSG00000056629 | Fkbp2 | 0.000608996 | 1.710236096 | 0.066942815 |
| ENSMUSG00000010110 | Stx5a | 0.000608996 | -1.281901786 | 0.130342151 |
| ENSMUSG00000071654 | Uqcc3 | 0.000608996 | 0.948395833 | 0.471767627 |
| ENSMUSG00000024735 | Prpf19 | 0.000608996 | 1.063873051 | 0.364909386 |
| ENSMUSG00000024691 | Fam111a | 0.000608996 | 0.004979362 | 1 |
| ENSMUSG00000038843 | Gcnt1 | 0.000608996 | -0.305360759 | 1 |
| ENSMUSG00000024785 | Rcl1 | 0.000608996 | -1.617950989 | 0.032620065 |
| ENSMUSG00000046324 | Ermp1 | 0.000608996 | -0.123753952 | 1 |
| ENSMUSG00000062456 | Rpl9-ps6 | 0.000608996 | 2.233798052 | 0.015201129 |
| ENSMUSG00000025159 | Mms19 | 0.000608996 | -1.673092543 | 0.026690314 |
| ENSMUSG00000025223 | Ldb1 | 0.000608996 | -1.258055044 | 0.142268504 |
| ENSMUSG00000035173 | Ccdc186 | 0.000608996 | -1.410058138 | 0.080463993 |
| ENSMUSG00000031154 | Otud5 | 0.000608996 | -2.645785197 | 0.000107956 |
| ENSMUSG00000036769 | Wdr44 | 0.000608996 | -1.192466703 | 0.185619905 |
| ENSMUSG00000001986 | Gria3 | 0.000608996 | 0.084050933 | 1 |
| ENSMUSG00000001173 | Ocrl | 0.000608996 | -1.466641666 | 0.070555673 |
| ENSMUSG00000035776 | Cd99l2 | 0.000608996 | 0.351155003 | 1 |
| ENSMUSG00000002015 | Bcap31 | 0.000608996 | -0.754886635 | 0.596197889 |
| ENSMUSG00000086629 | 2810403D21Rik | 0.000608996 | -0.985370468 | 0.367009511 |
| ENSMUSG00000031284 | Pak3 | 0.000608996 | -0.788569761 | 0.549953572 |
| ENSMUSG00000045180 | Shroom2 | 0.000608996 | -1.079084903 | 0.25936473 |
| ENSMUSG00000031360 | Ctps2 | 0.000608996 | -1.032988489 | 0.305378889 |
| ENSMUSG00000098234 | Snhg6 | 0.000553633 | 0.978984153 | 0.455131546 |
| ENSMUSG00000025917 | Cops5 | 0.000553633 | 0.250286175 | 1 |
| ENSMUSG00000026112 | Coa5 | 0.000553633 | 1.10826717 | 0.400280857 |
| ENSMUSG00000026078 | Pdcl3 | 0.000553633 | 0.616414074 | 0.840804483 |
| ENSMUSG00000026072 | Il1r1 | 0.000553633 | -1.091405175 | 0.274319469 |
| ENSMUSG00000066877 | Nck2 | 0.000553633 | -2.309585345 | 0.001215098 |
| ENSMUSG00000025995 | Wdr75 | 0.000553633 | -1.11090135 | 0.251393677 |

| | | | | |
|---|---|---|---|---|
| ENSMUSG00000026109 | Tmeff2 | 0.000553633 | -0.081557389 | 1 |
| ENSMUSG00000026035 | Ppil3 | 0.000553633 | 0.275377156 | 1 |
| ENSMUSG00000026014 | Raph1 | 0.000553633 | -1.711911792 | 0.024405815 |
| ENSMUSG00000025958 | Creb1 | 0.000553633 | -1.724622844 | 0.025490595 |
| ENSMUSG00000026003 | Acadl | 0.000553633 | -1.07164192 | 0.309680928 |
| ENSMUSG00000026179 | Pnkd | 0.000553633 | 0.523304669 | 0.858831093 |
| ENSMUSG00000099706 | Gm29536 | 0.000553633 | -0.724622844 | 0.668608522 |
| ENSMUSG00000004364 | Cul3 | 0.000553633 | -1.828370769 | 0.014189547 |
| ENSMUSG00000026150 | Mff | 0.000553633 | 0.523304669 | 0.858831093 |
| ENSMUSG00000026254 | Eif4e2 | 0.000553633 | -0.857517115 | 0.467537667 |
| ENSMUSG00000026259 | Ngef | 0.000553633 | -2.031284182 | 0.005091765 |
| ENSMUSG00000086825 | Gm15675 | 0.000553633 | -1.542784521 | 0.051667273 |
| ENSMUSG00000041926 | Rnpep | 0.000553633 | -1.973881871 | 0.006637019 |
| ENSMUSG00000052726 | Kcnt2 | 0.000553633 | 0.860339657 | 0.597803097 |
| ENSMUSG00000026482 | Rgl1 | 0.000553633 | -0.686148696 | 0.699192777 |
| ENSMUSG00000014226 | Cacybp | 0.000553633 | 0.584705214 | 0.846566779 |
| ENSMUSG00000026715 | Serpinc1 | 0.000553633 | 0.020804329 | 1 |
| ENSMUSG00000026674 | Ddr2 | 0.000553633 | 0.584705214 | 0.854285611 |
| ENSMUSG00000038949 | Cnst | 0.000553633 | -0.231582833 | 1 |
| ENSMUSG00000038855 | Itpkb | 0.000553633 | -0.383585926 | 1 |
| ENSMUSG00000026610 | Esrrg | 0.000553633 | -0.383585926 | 1 |
| ENSMUSG00000026605 | Cenpf | 0.000553633 | -2.35933838 | 0.000904756 |
| ENSMUSG00000026639 | Lamb3 | 0.000553633 | -1.051604167 | 0.295355925 |
| ENSMUSG00000048186 | Bend7 | 0.000553633 | -1.231582833 | 0.175666076 |
| ENSMUSG00000001855 | Nup214 | 0.000553633 | -0.880417517 | 0.455131546 |
| ENSMUSG00000039262 | Prrc2b | 0.000553633 | -2.560617814 | 0.000244583 |
| ENSMUSG00000026880 | Stom | 0.000553633 | 0.648835551 | 0.733004386 |
| ENSMUSG00000026836 | Acvr1 | 0.000553633 | -1.453975254 | 0.082076736 |
| ENSMUSG00000027022 | Xirp2 | 0.000553633 | 0.648835551 | 0.784038142 |
| ENSMUSG00000002731 | Prkra | 0.000553633 | 0.10826717 | 1 |
| ENSMUSG00000085976 | Gm13816 | 0.000553633 | -1.42311429 | 0.087613188 |
| ENSMUSG00000027180 | Fbxo3 | 0.000553633 | 0.250286175 | 1 |
| ENSMUSG00000027351 | Spred1 | 0.000553633 | -1.07164192 | 0.301591206 |
| ENSMUSG00000043909 | Trp53bp1 | 0.000553633 | -2.000257287 | 0.00657865 |
| ENSMUSG00000033486 | Catsper2 | 0.000553633 | -1.167857235 | 0.208892302 |
| ENSMUSG00000033396 | Spg11 | 0.000553633 | -1.724622844 | 0.033084874 |
| ENSMUSG00000005803 | Sqor | 0.000553633 | 0.085899357 | 1 |
| ENSMUSG00000001998 | Ap4e1 | 0.000553633 | -0.461588438 | 0.886203497 |
| ENSMUSG00000027381 | Bcl2l11 | 0.000553633 | -0.549944313 | 0.790752525 |
| ENSMUSG00000014361 | Mertk | 0.000553633 | -1.484189867 | 0.07666551 |
| ENSMUSG00000027333 | Smox | 0.000553633 | 0.978984153 | 0.455131546 |
| ENSMUSG00000018322 | Tomm34 | 0.000553633 | -0.605978348 | 0.733672617 |
| ENSMUSG00000018209 | Stk4 | 0.000553633 | -1.39952747 | 0.095278337 |
| ENSMUSG00000017721 | Pigt | 0.000553633 | 0.353379668 | 1 |
| ENSMUSG00000027519 | Rab22a | 0.000553633 | -1.081557389 | 0.271444635 |
| ENSMUSG00000000305 | Cdh4 | 0.000553633 | -0.249284835 | 1 |
| ENSMUSG00000027575 | Arfgap1 | 0.000553633 | -1.585219787 | 0.043272281 |
| ENSMUSG00000038628 | Polr3k | 0.000553633 | 1.201376574 | 0.281836872 |
| ENSMUSG00000078784 | Rbis | 0.000553633 | 0.225624121 | 1 |
| ENSMUSG00000039519 | Cyp7b1 | 0.000553633 | -1.564158172 | 0.046981468 |
| ENSMUSG00000037610 | Kcnmb2 | 0.000553633 | 0.786339075 | 0.612434853 |

| | | | | |
|---|---|---|---|---|
| ENSMUSG00000027668 | Mfn1 | 0.000553633 | -0.415294786 | 0.955878311 |
| ENSMUSG00000074604 | Mgst2 | 0.000553633 | -0.301123766 | 1 |
| ENSMUSG00000056476 | Med12l | 0.000553633 | 0.407827452 | 1 |
| ENSMUSG00000027823 | Gmps | 0.000553633 | -1.351164449 | 0.191885686 |
| ENSMUSG00000034349 | Smc4 | 0.000553633 | -1.851734762 | 0.013508949 |
| ENSMUSG00000019710 | Mrpl24 | 0.000553633 | -0.673092543 | 0.681660809 |
| ENSMUSG00000041355 | Ssr2 | 0.000553633 | 0.750715165 | 0.696370246 |
| ENSMUSG00000028049 | Scamp3 | 0.000553633 | 1.020804329 | 0.43456208 |
| ENSMUSG00000015714 | Cers2 | 0.000553633 | -0.686148696 | 0.649601793 |
| ENSMUSG00000028098 | Rnf115 | 0.000553633 | -1.549944313 | 0.05188053 |
| ENSMUSG00000051076 | Vtcn1 | 0.000553633 | -1.249284835 | 0.158846729 |
| ENSMUSG00000007379 | Dennd2c | 0.000553633 | -0.659917155 | 0.677768314 |
| ENSMUSG00000062127 | Cttnbp2nl | 0.000553633 | -2.472933412 | 0.000429764 |
| ENSMUSG00000040339 | Fam102b | 0.000553633 | -0.334676326 | 1 |
| ENSMUSG00000059857 | Ntng1 | 0.000553633 | 0.353379668 | 1 |
| ENSMUSG00000028007 | Snx7 | 0.000553633 | -0.284050253 | 1 |
| ENSMUSG00000028127 | Abcd3 | 0.000553633 | -0.84592914 | 0.466464457 |
| ENSMUSG00000039234 | Sec24d | 0.000553633 | -0.902960085 | 0.460248995 |
| ENSMUSG00000027977 | Ndst3 | 0.000553633 | -1.774375879 | 0.025181726 |
| ENSMUSG00000027968 | Larp7 | 0.000553633 | -1.592172548 | 0.044171611 |
| ENSMUSG00000050549 | Fam241a | 0.000553633 | -0.936126949 | 0.425555641 |
| ENSMUSG00000027984 | Hadh | 0.000553633 | -0.041479949 | 1 |
| ENSMUSG00000045328 | Cenpe | 0.000553633 | -2.106051951 | 0.003319543 |
| ENSMUSG00000028152 | Tspan5 | 0.000553633 | -1.091405175 | 0.274319469 |
| ENSMUSG00000028256 | Odf2l | 0.000553633 | -0.810596067 | 0.503807982 |
| ENSMUSG00000028191 | Bcl10 | 0.000553633 | 0.063873051 | 1 |
| ENSMUSG00000025437 | Usp33 | 0.000553633 | -2.602539264 | 0.000189248 |
| ENSMUSG00000039047 | Pigk | 0.000553633 | -0.231582833 | 1 |
| ENSMUSG00000055436 | Srsf11 | 0.000553633 | -2.204615785 | 0.001994407 |
| ENSMUSG00000086080 | 1700123O12Rik | 0.000553633 | 0.553678318 | 0.873400121 |
| ENSMUSG00000058006 | Mdn1 | 0.000553633 | -1.804622166 | 0.017062925 |
| ENSMUSG00000071014 | Ndufb6 | 0.000553633 | 1.523304669 | 0.129242108 |
| ENSMUSG00000028436 | Dcaf12 | 0.000553633 | -1.091405175 | 0.274319469 |
| ENSMUSG00000035495 | Tstd2 | 0.000553633 | -0.081557389 | 1 |
| ENSMUSG00000070985 | Acnat1 | 0.000553633 | -1.158519371 | 0.233562839 |
| ENSMUSG00000044018 | Mrpl50 | 0.000553633 | 0.020804329 | 1 |
| ENSMUSG00000038816 | Ctnnal1 | 0.000553633 | -1.914100643 | 0.010114343 |
| ENSMUSG00000028378 | Ptgr1 | 0.000553633 | 1.648835551 | 0.096834041 |
| ENSMUSG00000038607 | Gng10 | 0.000553633 | 1.860339657 | 0.080733669 |
| ENSMUSG00000028392 | Bspry | 0.000553633 | 2.407827452 | 0.01643282 |
| ENSMUSG00000028609 | Magoh | 0.000553633 | 0.898813804 | 0.553980644 |
| ENSMUSG00000028568 | Btf3l4 | 0.000553633 | -0.158519371 | 1 |
| ENSMUSG00000028703 | Lrrc41 | 0.000553633 | 0.275377156 | 1 |
| ENSMUSG00000028683 | Eif2b3 | 0.000553633 | -0.914100643 | 0.463850347 |
| ENSMUSG00000028861 | Mrps15 | 0.000553633 | 0.435841828 | 0.973694431 |
| ENSMUSG00000053841 | Txlna | 0.000553633 | -1.653284062 | 0.03467347 |
| ENSMUSG00000003731 | Kpna6 | 0.000553633 | -0.633199816 | 0.736456705 |
| ENSMUSG00000085241 | Snhg3 | 0.000553633 | 1.938342169 | 0.067122487 |
| ENSMUSG00000028832 | Stmn1 | 0.000553633 | -0.195513578 | 1 |
| ENSMUSG00000028821 | Syf2 | 0.000553633 | 0.020804329 | 1 |
| ENSMUSG00000029048 | Rer1 | 0.000553633 | 0.130987247 | 1 |

| | | | | |
|---|---|---|---|---|
| ENSMUSG00000029076 | Sdf4 | 0.000553633 | -0.351164449 | 1 |
| ENSMUSG00000095567 | Noc2l | 0.000553633 | -1.11090135 | 0.250264692 |
| ENSMUSG00000040584 | Abcb1a | 0.000553633 | 1.300912248 | 0.243566181 |
| ENSMUSG00000010721 | Lmbr1 | 0.000553633 | -1.979195671 | 0.006797116 |
| ENSMUSG00000029148 | Nrbp1 | 0.000553633 | -0.39952747 | 0.954021409 |
| ENSMUSG00000029123 | Stk32b | 0.000553633 | 1.15407086 | 0.387414024 |
| ENSMUSG00000051022 | Hs3st1 | 0.000553633 | -1.653284062 | 0.034991708 |
| ENSMUSG00000039682 | Lap3 | 0.000553633 | -0.101186196 | 1 |
| ENSMUSG00000029213 | Commd8 | 0.000553633 | 1.250286175 | 0.308966189 |
| ENSMUSG00000029234 | Tmem165 | 0.000553633 | -0.231582833 | 1 |
| ENSMUSG00000029246 | Ppat | 0.000553633 | -0.673092543 | 0.681660809 |
| ENSMUSG00000029407 | Uso1 | 0.000553633 | -1.874726405 | 0.011271367 |
| ENSMUSG00000034826 | Nup54 | 0.000553633 | -0.619653284 | 0.736456705 |
| ENSMUSG00000046709 | Mapk10 | 0.000553633 | 0.225624121 | 1 |
| ENSMUSG00000029290 | Zfp326 | 0.000553633 | -0.834247335 | 0.510743339 |
| ENSMUSG00000029267 | Mtf2 | 0.000553633 | -1.564158172 | 0.046808743 |
| ENSMUSG00000029265 | Dr1 | 0.000553633 | 0.523304669 | 0.865117569 |
| ENSMUSG00000041870 | Ankrd13a | 0.000553633 | 0.250286175 | 1 |
| ENSMUSG00000087292 | Gm13832 | 0.000553633 | -0.673092543 | 0.693613791 |
| ENSMUSG00000041638 | Gcn1 | 0.000553633 | -1.041479949 | 0.330909214 |
| ENSMUSG00000087516 | Tbx3os1 | 0.000553633 | 0.225624121 | 1 |
| ENSMUSG00000029599 | Ddx54 | 0.000553633 | -0.89173283 | 0.456446681 |
| ENSMUSG00000029468 | P2rx7 | 0.000553633 | -1.249284835 | 0.221100123 |
| ENSMUSG00000049686 | Orai1 | 0.000553633 | -1.407432668 | 0.096782144 |
| ENSMUSG00000029434 | Vps33a | 0.000553633 | -0.76209755 | 0.605764786 |
| ENSMUSG00000029401 | Rilpl2 | 0.000553633 | -1.578233357 | 0.048860005 |
| ENSMUSG00000070498 | Tmem132b | 0.000553633 | 0.898813804 | 0.588477963 |
| ENSMUSG00000060371 | Caln1 | 0.000553633 | -0.177135049 | 1 |
| ENSMUSG00000015942 | Gtf2ird2 | 0.000553633 | -1.469161658 | 0.081714616 |
| ENSMUSG00000023079 | Gtf2ird1 | 0.000553633 | -2.096303974 | 0.003569729 |
| ENSMUSG00000039771 | Polr2j | 0.000553633 | 0.750715165 | 0.727318386 |
| ENSMUSG00000045482 | Trrap | 0.000553633 | -2.061657831 | 0.004467147 |
| ENSMUSG00000029627 | Zkscan14 | 0.000553633 | 0.201376574 | 1 |
| ENSMUSG00000042742 | Bmt2 | 0.000553633 | -0.979195671 | 0.411579031 |
| ENSMUSG00000025608 | Podxl | 0.000553633 | -0.798623426 | 0.603605773 |
| ENSMUSG00000001642 | Akr1b3 | 0.000553633 | 0.46441098 | 0.971854959 |
| ENSMUSG00000038665 | Dgki | 0.000553633 | 0.15407086 | 1 |
| ENSMUSG00000023367 | Tmem176a | 0.000553633 | -0.000257287 | 1 |
| ENSMUSG00000000628 | Hk2 | 0.000553633 | -2.010673903 | 0.006061218 |
| ENSMUSG00000030035 | Wbp1 | 0.000553633 | 1.786339075 | 0.098608858 |
| ENSMUSG00000031865 | Dctn1 | 0.000553633 | -1.686148696 | 0.029588339 |
| ENSMUSG00000033216 | Eefsec | 0.000553633 | -1.081557389 | 0.271444635 |
| ENSMUSG00000033174 | Mgll | 0.000553633 | -1.612832019 | 0.045630079 |
| ENSMUSG00000030086 | Chchd6 | 0.000553633 | -0.549944313 | 0.851250572 |
| ENSMUSG00000030341 | Tnfrsf1a | 0.000553633 | 0.978984153 | 0.455131546 |
| ENSMUSG00000061414 | Cracr2a | 0.000553633 | -1.139660343 | 0.231236006 |
| ENSMUSG00000032899 | Styk1 | 0.000553633 | -1.768249776 | 0.019655136 |
| ENSMUSG00000030222 | Rerg | 0.000553633 | -0.53558902 | 0.843284619 |
| ENSMUSG00000040029 | Ipo8 | 0.000553633 | -1.139660343 | 0.227871652 |
| ENSMUSG00000070808 | Bicra | 0.000553633 | -1.222649708 | 0.173500492 |
| ENSMUSG00000019158 | Tmem160 | 0.000553633 | 1.715949747 | 0.106312068 |

| | | | | |
|---|---|---|---|---|
| ENSMUSG00000045039 | Megf8 | 0.000553633 | -0.592172548 | 0.792441218 |
| ENSMUSG00000053714 | 4732471J01Rik | 0.000553633 | -0.84592914 | 0.466464457 |
| ENSMUSG00000046865 | Fbl | 0.000553633 | 0.225624121 | 1 |
| ENSMUSG00000003444 | Med29 | 0.000553633 | 0.353379668 | 1 |
| ENSMUSG00000056216 | Cebpg | 0.000553633 | 0.682002415 | 0.718266608 |
| ENSMUSG00000043456 | Zfp536 | 0.000553633 | 0.225624121 | 1 |
| ENSMUSG00000060279 | Ap2a1 | 0.000553633 | -1.317997585 | 0.13735263 |
| ENSMUSG00000087138 | Gm15545 | 0.000553633 | -1.646620332 | 0.046003064 |
| ENSMUSG00000030607 | Acan | 0.000553633 | 0.063873051 | 1 |
| ENSMUSG00000039176 | Polg | 0.000553633 | 0.04217798 | 1 |
| ENSMUSG00000070469 | Adamtsl3 | 0.000553633 | -0.213660925 | 1 |
| ENSMUSG00000030560 | Ctsc | 0.000553633 | -1.66651989 | 0.031582307 |
| ENSMUSG00000035547 | Capn5 | 0.000553633 | -1.041479949 | 0.330909214 |
| ENSMUSG00000035314 | Gdpd5 | 0.000553633 | 0.085899357 | 1 |
| ENSMUSG00000030718 | Ppme1 | 0.000553633 | -1.484189867 | 0.066698557 |
| ENSMUSG00000001829 | Clpb | 0.000553633 | -0.52108945 | 0.846566779 |
| ENSMUSG00000057143 | Trim12c | 0.000553633 | -1.139660343 | 0.227871652 |
| ENSMUSG00000037060 | Cavin3 | 0.000553633 | 1.938342169 | 0.055187251 |
| ENSMUSG00000061079 | Zfp143 | 0.000553633 | -1.07164192 | 0.301591206 |
| ENSMUSG00000042246 | Tmc7 | 0.000553633 | -0.446321682 | 0.961918768 |
| ENSMUSG00000030922 | Lyrm1 | 0.000553633 | -0.925155831 | 0.42263302 |
| ENSMUSG00000042462 | Dctpp1 | 0.000553633 | 2.407827452 | 0.013182548 |
| ENSMUSG00000030822 | Prr14 | 0.000553633 | -1.52835745 | 0.05604143 |
| ENSMUSG00000030815 | Phkg2 | 0.000553633 | -1.139660343 | 0.227871652 |
| ENSMUSG00000039990 | Edrf1 | 0.000553633 | -1.249284835 | 0.158846729 |
| ENSMUSG00000060260 | Pwwp2b | 0.000553633 | 0.201376574 | 1 |
| ENSMUSG00000038489 | Polr2l | 0.000553633 | 1.938342169 | 0.082076736 |
| ENSMUSG00000025511 | Tspan4 | 0.000553633 | -0.39952747 | 0.95526509 |
| ENSMUSG00000059119 | Nap1l4 | 0.000553633 | -0.914100643 | 0.528505953 |
| ENSMUSG00000056228 | Cars2 | 0.000553633 | -0.936126949 | 0.425555641 |
| ENSMUSG00000015341 | Golga7 | 0.000553633 | -0.000257287 | 1 |
| ENSMUSG00000031575 | Ash2l | 0.000553633 | -0.902960085 | 0.460248995 |
| ENSMUSG00000031576 | Kcnu1 | 0.000553633 | 0.250286175 | 1 |
| ENSMUSG00000031483 | Erlin2 | 0.000553633 | 0.130987247 | 1 |
| ENSMUSG00000031583 | Wrn | 0.000553633 | -0.774375879 | 0.549953572 |
| ENSMUSG00000038291 | Snx25 | 0.000553633 | -0.737222881 | 0.601474804 |
| ENSMUSG00000053626 | Tll1 | 0.000553633 | -1.834247335 | 0.015701711 |
| ENSMUSG00000051147 | Nat2 | 0.000553633 | 2.648835551 | 0.006727538 |
| ENSMUSG00000036054 | Sugp2 | 0.000553633 | -1.484189867 | 0.066698557 |
| ENSMUSG00000031848 | Lsm4 | 0.000553633 | 1.300912248 | 0.262284476 |
| ENSMUSG00000034807 | Colgalt1 | 0.000553633 | -1.334676326 | 0.125161377 |
| ENSMUSG00000057672 | Pkn1 | 0.000553633 | -0.989764913 | 0.354935311 |
| ENSMUSG00000005469 | Prkaca | 0.000553633 | 1.15407086 | 0.387414024 |
| ENSMUSG00000036686 | Cc2d1a | 0.000553633 | 0.407827452 | 0.980050262 |
| ENSMUSG00000031666 | Rbl2 | 0.000553633 | -0.902960085 | 0.460248995 |
| ENSMUSG00000031774 | Fam192a | 0.000553633 | -1.139660343 | 0.227871652 |
| ENSMUSG00000031792 | Usb1 | 0.000553633 | -1.342943942 | 0.139855038 |
| ENSMUSG00000031672 | Got2 | 0.000553633 | -0.549944313 | 0.790752525 |
| ENSMUSG00000031673 | Cdh11 | 0.000553633 | -0.195513578 | 1 |
| ENSMUSG00000005698 | Ctcf | 0.000553633 | -1.383585926 | 0.105255472 |
| ENSMUSG00000014444 | Piezo1 | 0.000553633 | -0.249284835 | 1 |

| | | | | |
|---|---|---|---|---|
| ENSMUSG00000034796 | Cpne7 | 0.000553633 | -0.491645672 | 0.896076486 |
| ENSMUSG00000110697 | Gm31718 | 0.000553633 | -0.158519371 | 1 |
| ENSMUSG00000091028 | Gm10722 | 0.000553633 | 1.201376574 | 0.281836872 |
| ENSMUSG00000047193 | Dync2h1 | 0.000553633 | -1.149120673 | 0.230787384 |
| ENSMUSG00000032198 | Dock6 | 0.000553633 | -0.317997585 | 1 |
| ENSMUSG00000047412 | Zbtb44 | 0.000553633 | -1.326361058 | 0.123403868 |
| ENSMUSG00000032041 | Tirap | 0.000553633 | -0.195513578 | 1 |
| ENSMUSG00000032042 | Srpr | 0.000553633 | -0.774375879 | 0.549953572 |
| ENSMUSG00000032024 | Clmp | 0.000553633 | 0.063873051 | 1 |
| ENSMUSG00000063382 | Bcl9l | 0.000553633 | -0.798623426 | 0.559434132 |
| ENSMUSG00000035164 | Zc3h12c | 0.000553633 | -0.646620332 | 0.674297981 |
| ENSMUSG00000032301 | Psma4 | 0.000553633 | -0.619653284 | 0.736456705 |
| ENSMUSG00000032324 | Tspan3 | 0.000553633 | 1.020804329 | 0.43456208 |
| ENSMUSG00000036986 | Pml | 0.000553633 | -1.284050253 | 0.146946082 |
| ENSMUSG00000053641 | Dennd4a | 0.000553633 | -2.581730801 | 0.000366437 |
| ENSMUSG00000033629 | Hacd3 | 0.000553633 | -0.84592914 | 0.466464457 |
| ENSMUSG00000052698 | Tln2 | 0.000553633 | -0.383585926 | 1 |
| ENSMUSG00000041361 | Myzap | 0.000553633 | -0.549944313 | 0.790752525 |
| ENSMUSG00000032215 | Rsl24d1 | 0.000553633 | -1.204615785 | 0.210550914 |
| ENSMUSG00000032352 | Lrrc1 | 0.000553633 | -0.659917155 | 0.677768314 |
| ENSMUSG00000034898 | Filip1 | 0.000553633 | 0.353379668 | 1 |
| ENSMUSG00000032253 | Phip | 0.000553633 | -2.819510792 | 4.25E-05 |
| ENSMUSG00000034973 | Dop1a | 0.000553633 | -0.737222881 | 0.599135879 |
| ENSMUSG00000032480 | Dhx30 | 0.000553633 | -0.902960085 | 0.460248995 |
| ENSMUSG00000090112 | Shprh | 0.000553633 | -1.653284062 | 0.03467347 |
| ENSMUSG00000062866 | Phactr2 | 0.000553633 | -0.549944313 | 0.847320319 |
| ENSMUSG00000019977 | Hbs1l | 0.000553633 | -0.786550593 | 0.554308261 |
| ENSMUSG00000037370 | Enpp1 | 0.000553633 | -0.53558902 | 0.843284619 |
| ENSMUSG00000038446 | Cdc40 | 0.000553633 | -0.857517115 | 0.467537667 |
| ENSMUSG00000075225 | Ccdc162 | 0.000553633 | -0.301123766 | 1 |
| ENSMUSG00000019797 | 1700021F05Rik | 0.000553633 | 0.898813804 | 0.553980644 |
| ENSMUSG00000019877 | Serinc1 | 0.000553633 | 0.353379668 | 1 |
| ENSMUSG00000020098 | Pcbd1 | 0.000553633 | 1.648835551 | 0.096834041 |
| ENSMUSG00000020078 | Vps26a | 0.000553633 | -1.605978348 | 0.046344122 |
| ENSMUSG00000019923 | Zwint | 0.000553633 | -1.810596067 | 0.015201129 |
| ENSMUSG00000020329 | Polrmt | 0.000553633 | -0.869012753 | 0.471767627 |
| ENSMUSG00000035478 | Mbd3 | 0.000553633 | 0.130987247 | 1 |
| ENSMUSG00000034674 | Tdg | 0.000553633 | -0.564158172 | 0.792441218 |
| ENSMUSG00000046841 | Ckap4 | 0.000553633 | 0.493557326 | 0.868550295 |
| ENSMUSG00000045867 | Cradd | 0.000553633 | -2.11090135 | 0.003386784 |
| ENSMUSG00000112290 | Gm33782 | 0.000553633 | -0.334676326 | 1 |
| ENSMUSG00000074776 | Gm10754 | 0.000553633 | 0.978984153 | 0.455131546 |
| ENSMUSG00000019892 | Lrriq1 | 0.000553633 | 1.648835551 | 0.096834041 |
| ENSMUSG00000020171 | Yeats4 | 0.000553633 | 0.130987247 | 1 |
| ENSMUSG00000020228 | Helb | 0.000553633 | -0.446321682 | 0.961918768 |
| ENSMUSG00000018166 | Erbb3 | 0.000553633 | 0.15407086 | 1 |
| ENSMUSG00000000711 | Rab5b | 0.000553633 | -0.195513578 | 1 |
| ENSMUSG00000084941 | Gm11944 | 0.000553633 | -0.53558902 | 0.843284619 |
| ENSMUSG00000048834 | Vstm2a | 0.000553633 | 0.523304669 | 0.865413877 |
| ENSMUSG00000033953 | Ppp3r1 | 0.000553633 | 0.04217798 | 1 |
| ENSMUSG00000044072 | Eml6 | 0.000553633 | -0.491645672 | 0.89542114 |

| | | | | |
|---|---|---|---|---|
| ENSMUSG00000060923 | Acyp2 | 0.000553633 | -1.130137569 | 0.256718203 |
| ENSMUSG00000020300 | Cpeb4 | 0.000553633 | -0.430891641 | 0.958408446 |
| ENSMUSG00000020289 | Nprl3 | 0.000553633 | 0.201376574 | 1 |
| ENSMUSG00000020143 | Dock2 | 0.000553633 | -0.284050253 | 1 |
| ENSMUSG00000069911 | Insyn2b | 0.000553633 | -2.031284182 | 0.007201453 |
| ENSMUSG00000020328 | Nudcd2 | 0.000553633 | 0.407827452 | 0.99651149 |
| ENSMUSG00000020326 | Ccng1 | 0.000553633 | 1.10826717 | 0.400280857 |
| ENSMUSG00000020376 | Rnf130 | 0.000553633 | -0.415294786 | 0.971854959 |
| ENSMUSG00000020385 | Clk4 | 0.000553633 | -1.874726405 | 0.011271367 |
| ENSMUSG00000010025 | Aldh3a2 | 0.000553633 | 0.715949747 | 0.706154373 |
| ENSMUSG00000059552 | Trp53 | 0.000553633 | -1.66651989 | 0.031582307 |
| ENSMUSG00000018286 | Psmb6 | 0.000553633 | 0.715949747 | 0.77388575 |
| ENSMUSG00000020790 | Ankfy1 | 0.000553633 | -1.679635389 | 0.028983805 |
| ENSMUSG00000020741 | Cluh | 0.000553633 | -1.491645672 | 0.06783677 |
| ENSMUSG00000038351 | Sgsm2 | 0.000553633 | -1.857517115 | 0.012088308 |
| ENSMUSG00000020679 | Hnf1b | 0.000553633 | -2.021015847 | 0.005551659 |
| ENSMUSG00000018548 | Trim37 | 0.000553633 | -1.275437122 | 0.144856744 |
| ENSMUSG00000010080 | Epn3 | 0.000553633 | -0.430891641 | 0.958408446 |
| ENSMUSG00000076435 | Acsf2 | 0.000553633 | -2.930651819 | 2.58E-05 |
| ENSMUSG00000006058 | Snf8 | 0.000553633 | -0.266772261 | 1 |
| ENSMUSG00000020941 | Map3k14 | 0.000553633 | -1.186353579 | 0.217655393 |
| ENSMUSG00000034187 | Nsf | 0.000553633 | -1.53558902 | 0.061620805 |
| ENSMUSG00000020708 | Psmc5 | 0.000553633 | 0.938342169 | 0.538334238 |
| ENSMUSG00000020715 | Ern1 | 0.000553633 | -2.419209836 | 0.000632836 |
| ENSMUSG00000020623 | Map2k6 | 0.000553633 | -1.222649708 | 0.173500492 |
| ENSMUSG00000075420 | Smim6 | 0.000553633 | 2.300912248 | 0.017593853 |
| ENSMUSG00000070327 | Rnf213 | 0.000553633 | -1.407432668 | 0.096782144 |
| ENSMUSG00000078249 | Hmga1b | 0.000553633 | 1.46441098 | 0.209790273 |
| ENSMUSG00000037669 | Ldah | 0.000553633 | -2.484189867 | 0.000413098 |
| ENSMUSG00000037149 | Ddx1 | 0.000553633 | -1.101186196 | 0.276938641 |
| ENSMUSG00000055917 | Zfp277 | 0.000553633 | -0.564158172 | 0.792441218 |
| ENSMUSG00000113105 | Gm47371 | 0.000553633 | 0.275377156 | 1 |
| ENSMUSG00000047022 | Mipol1 | 0.000553633 | -1.718281317 | 0.024881876 |
| ENSMUSG00000090258 | Churc1 | 0.000553633 | 1.020804329 | 0.452280643 |
| ENSMUSG00000042628 | Zfyve1 | 0.000553633 | -2.213660925 | 0.001866323 |
| ENSMUSG00000021236 | Entpd5 | 0.000553633 | -1.542784521 | 0.051152519 |
| ENSMUSG00000021243 | Fcf1 | 0.000553633 | -1.391578717 | 0.093904647 |
| ENSMUSG00000004788 | Eif2b2 | 0.000553633 | -1.301123766 | 0.132846599 |
| ENSMUSG00000021245 | Mlh3 | 0.000553633 | -1.461588438 | 0.072659903 |
| ENSMUSG00000021255 | Esrrb | 0.000553633 | 0.938342169 | 0.564419844 |
| ENSMUSG00000021258 | Ccnk | 0.000553633 | -0.53558902 | 0.843284619 |
| ENSMUSG00000021262 | Evl | 0.000553633 | -1.792599638 | 0.018724441 |
| ENSMUSG00000033799 | Tasor2 | 0.000553633 | -2.296874293 | 0.001278866 |
| ENSMUSG00000021213 | Akr1c13 | 0.000553633 | 1.584705214 | 0.113023865 |
| ENSMUSG00000021301 | Hecw1 | 0.000553633 | 0.225624121 | 1 |
| ENSMUSG00000037966 | Ninj1 | 0.000553633 | 0.682002415 | 0.786367309 |
| ENSMUSG00000025867 | Cplx2 | 0.000553633 | 0.15407086 | 1 |
| ENSMUSG00000035493 | Tgfbi | 0.000553633 | -2.415294786 | 0.00098929 |
| ENSMUSG00000005312 | Ubqln1 | 0.000553633 | -0.699087752 | 0.653235135 |
| ENSMUSG00000058093 | Zfp729b | 0.000553633 | -0.774375879 | 0.549953572 |
| ENSMUSG00000021598 | Med10 | 0.000553633 | 0.786339075 | 0.59945267 |

| | | | | |
|---|---|---|---|---|
| ENSMUSG00000042082 | Arsb | 0.000553633 | -0.89173283 | 0.466464457 |
| ENSMUSG00000041773 | Enc1 | 0.000553633 | -1.62644245 | 0.036798276 |
| ENSMUSG00000110235 | Gm5086 | 0.000553633 | -2.330524683 | 0.001554857 |
| ENSMUSG00000078941 | Ak6 | 0.000553633 | 0.250286175 | 1 |
| ENSMUSG00000021747 | 4930452B06Rik | 0.000553633 | -0.231582833 | 1 |
| ENSMUSG00000021737 | Psmd6 | 0.000553633 | -0.301123766 | 1 |
| ENSMUSG00000017491 | Rarb | 0.000553633 | -1.258055044 | 0.170610013 |
| ENSMUSG00000021779 | Thrb | 0.000553633 | -1.646620332 | 0.034108291 |
| ENSMUSG00000021775 | Nr1d2 | 0.000553633 | -1.130137569 | 0.256497687 |
| ENSMUSG00000042354 | Gnl3 | 0.000553633 | -1.407432668 | 0.096782144 |
| ENSMUSG00000021892 | Sh3bp5 | 0.000553633 | -0.231582833 | 1 |
| ENSMUSG00000063506 | Arhgap22 | 0.000553633 | -1.592172548 | 0.044171611 |
| ENSMUSG00000021950 | Anxa8 | 0.000553633 | -0.673092543 | 0.681660809 |
| ENSMUSG00000037824 | Tspan14 | 0.000553633 | -1.599091962 | 0.054186866 |
| ENSMUSG00000040618 | Pck2 | 0.000553633 | -1.513784649 | 0.062237474 |
| ENSMUSG00000047153 | Khnyn | 0.000553633 | -1.653284062 | 0.03467347 |
| ENSMUSG00000060548 | Tnfrsf19 | 0.000553633 | -1.453975254 | 0.096834041 |
| ENSMUSG00000022085 | Pebp4 | 0.000553633 | 0.353379668 | 1 |
| ENSMUSG00000053465 | Hs6st3 | 0.000553633 | -0.659917155 | 0.705089874 |
| ENSMUSG00000022261 | Sdc2 | 0.000553633 | -0.902960085 | 0.460248995 |
| ENSMUSG00000037386 | Rims2 | 0.000553633 | -1.925155831 | 0.009194696 |
| ENSMUSG00000038622 | Med30 | 0.000553633 | 1.250286175 | 0.262284476 |
| ENSMUSG00000036800 | Fam135b | 0.000553633 | 1.063873051 | 0.43934811 |
| ENSMUSG00000022596 | Slurp1 | 0.000553633 | 6.10826717 | 3.54E-06 |
| ENSMUSG00000053929 | Cyhr1 | 0.000553633 | -0.158519371 | 1 |
| ENSMUSG00000049521 | Cdc42ep1 | 0.000553633 | -0.564158172 | 0.792441218 |
| ENSMUSG00000000489 | Pdgfb | 0.000553633 | -1.612832019 | 0.040680007 |
| ENSMUSG00000016619 | Nup50 | 0.000553633 | -0.446321682 | 0.961918768 |
| ENSMUSG00000054863 | Tafa5 | 0.000553633 | -0.12055152 | 1 |
| ENSMUSG00000036167 | Pphln1 | 0.000553633 | -1.375548607 | 0.103463818 |
| ENSMUSG00000022462 | Slc38a2 | 0.000553633 | -1.446321682 | 0.081057143 |
| ENSMUSG00000023007 | Prpf40b | 0.000553633 | -1.542784521 | 0.069190439 |
| ENSMUSG00000009739 | Pou6f1 | 0.000553633 | -1.499063144 | 0.069323271 |
| ENSMUSG00000001288 | Rarg | 0.000553633 | -1.139660343 | 0.227871652 |
| ENSMUSG00000000555 | Itga5 | 0.000553633 | -1.886086267 | 0.01181855 |
| ENSMUSG00000005580 | Adcy9 | 0.000553633 | -1.351164449 | 0.11286528 |
| ENSMUSG00000096215 | Smim22 | 0.000553633 | -0.367466261 | 1 |
| ENSMUSG00000039473 | Ubn1 | 0.000553633 | -2.780476079 | 5.24E-05 |
| ENSMUSG00000022711 | Pmm2 | 0.000553633 | -0.351164449 | 1 |
| ENSMUSG00000059003 | Grin2a | 0.000553633 | 0.063873051 | 1 |
| ENSMUSG00000065979 | Cpped1 | 0.000553633 | -0.633199816 | 0.736456705 |
| ENSMUSG00000022685 | Parn | 0.000553633 | -1.249284835 | 0.158846729 |
| ENSMUSG00000041720 | Pi4ka | 0.000553633 | -1.686148696 | 0.032878985 |
| ENSMUSG00000000326 | Comt | 0.000553633 | -1.605978348 | 0.039883588 |
| ENSMUSG00000071632 | 2510002D24Rik | 0.000553633 | -0.231582833 | 1 |
| ENSMUSG00000062901 | Klhl24 | 0.000553633 | -0.914100643 | 0.463850347 |
| ENSMUSG00000033653 | Vps8 | 0.000553633 | -1.158519371 | 0.233562839 |
| ENSMUSG00000044626 | Liph | 0.000553633 | -1.231582833 | 0.177165715 |
| ENSMUSG00000034422 | Parp14 | 0.000553633 | -0.711911792 | 0.65747066 |
| ENSMUSG00000022701 | Ccdc191 | 0.000553633 | -2.317997585 | 0.00112891 |
| ENSMUSG00000022863 | Btg3 | 0.000553633 | -1.851734762 | 0.013508949 |

| | | | | |
|---|---|---|---|---|
| ENSMUSG00000022974 | Paxbp1 | 0.000553633 | -1.851734762 | 0.013508949 |
| ENSMUSG00000022971 | Ifnar2 | 0.000553633 | -0.869012753 | 0.483322383 |
| ENSMUSG00000014039 | Prdm15 | 0.000553633 | -0.822470167 | 0.507610106 |
| ENSMUSG00000116858 | Gm49797 | 0.000553633 | -1.89173283 | 0.01060834 |
| ENSMUSG00000014426 | Map3k4 | 0.000553633 | -0.686148696 | 0.649601793 |
| ENSMUSG00000014763 | Fam120b | 0.000553633 | -1.266772261 | 0.163935241 |
| ENSMUSG00000116564 | Riok2 | 0.000553633 | -0.76209755 | 0.605764786 |
| ENSMUSG00000007564 | Ppp2r1a | 0.000553633 | 0.201376574 | 1 |
| ENSMUSG00000024014 | Pim1 | 0.000553633 | -0.947015265 | 0.430269377 |
| ENSMUSG00000033826 | Dnah8 | 0.000553633 | 0.275377156 | 1 |
| ENSMUSG00000024397 | Aif1 | 0.000553633 | -0.430891641 | 0.958408446 |
| ENSMUSG00000071074 | Yipf3 | 0.000553633 | 0.380346716 | 0.98846816 |
| ENSMUSG00000023999 | Kif6 | 0.000553633 | 0.648835551 | 0.756090103 |
| ENSMUSG00000003199 | Mpnd | 0.000553633 | -0.081557389 | 1 |
| ENSMUSG00000071054 | Safb | 0.000553633 | -2.843017545 | 3.32E-05 |
| ENSMUSG00000024098 | Twsg1 | 0.000553633 | 0.407827452 | 0.996793385 |
| ENSMUSG00000024077 | Strn | 0.000553633 | -0.711911792 | 0.66416768 |
| ENSMUSG00000024146 | Cript | 0.000553633 | 0.353379668 | 1 |
| ENSMUSG00000034998 | Foxn2 | 0.000553633 | -1.792599638 | 0.018820073 |
| ENSMUSG00000024287 | Thoc1 | 0.000553633 | -0.822470167 | 0.507610106 |
| ENSMUSG00000024304 | Cdh2 | 0.000553633 | -1.653284062 | 0.03467347 |
| ENSMUSG00000024486 | Hbegf | 0.000553633 | -2.1725036 | 0.002459077 |
| ENSMUSG00000036452 | Arhgap26 | 0.000553633 | -1.149120673 | 0.240589043 |
| ENSMUSG00000057719 | Sh3rf2 | 0.000553633 | 0.648835551 | 0.733004386 |
| ENSMUSG00000024493 | Lars | 0.000553633 | -2.491645672 | 0.000384507 |
| ENSMUSG00000034484 | Snx2 | 0.000553633 | -1.679635389 | 0.031586725 |
| ENSMUSG00000024538 | Ppic | 0.000553633 | 0.860339657 | 0.575389694 |
| ENSMUSG00000024590 | Lmnb1 | 0.000553633 | -0.711911792 | 0.65747066 |
| ENSMUSG00000032688 | Malt1 | 0.000553633 | -0.659917155 | 0.694202668 |
| ENSMUSG00000046318 | Ccbe1 | 0.000553633 | -2.305360759 | 0.001184321 |
| ENSMUSG00000024556 | Me2 | 0.000553633 | 0.682002415 | 0.718266608 |
| ENSMUSG00000025429 | Pstpip2 | 0.000553633 | -0.699087752 | 0.66416768 |
| ENSMUSG00000023066 | Rttn | 0.000553633 | -0.869012753 | 0.471767627 |
| ENSMUSG00000024870 | Rab1b | 0.000553633 | 0.553678318 | 0.880201363 |
| ENSMUSG00000095098 | Ccdc85b | 0.000553633 | -0.101186196 | 1 |
| ENSMUSG00000024911 | Fibp | 0.000553633 | -0.301123766 | 1 |
| ENSMUSG00000024944 | Arl2 | 0.000553633 | -0.39952747 | 0.954021409 |
| ENSMUSG00000010095 | Slc3a2 | 0.000553633 | -1.499063144 | 0.069323271 |
| ENSMUSG00000071660 | Ttc9c | 0.000553633 | 0.407827452 | 0.980050262 |
| ENSMUSG00000041488 | Stx3 | 0.000553633 | -1.091405175 | 0.29131171 |
| ENSMUSG00000024726 | Carnmt1 | 0.000553633 | -0.592172548 | 0.792441218 |
| ENSMUSG00000024897 | Apba1 | 0.000553633 | -0.249284835 | 1 |
| ENSMUSG00000024780 | Cdc37l1 | 0.000553633 | -0.476695331 | 0.886203497 |
| ENSMUSG00000024990 | Rbp4 | 0.000553633 | 1.10826717 | 0.400280857 |
| ENSMUSG00000025020 | Slit1 | 0.000553633 | -1.633199816 | 0.0478898 |
| ENSMUSG00000025184 | R3hcc1l | 0.000553633 | -1.461588438 | 0.072659903 |
| ENSMUSG00000037071 | Scd1 | 0.000553633 | 2.300912248 | 0.017593853 |
| ENSMUSG00000025047 | Pdcd11 | 0.000553633 | -1.686148696 | 0.029588339 |
| ENSMUSG00000025076 | Casp7 | 0.000553633 | 1.201376574 | 0.281836872 |
| ENSMUSG00000025089 | Gfra1 | 0.000553633 | 0.822864951 | 0.608747763 |
| ENSMUSG00000033417 | Cacul1 | 0.000553633 | -1.130137569 | 0.256497687 |

| | | | | |
|---|---|---|---|---|
| ENSMUSG00000000134 | Tfe3 | 0.000553633 | 0.063873051 | 1 |
| ENSMUSG00000064127 | Med14 | 0.000553633 | -1.863276384 | 0.01239886 |
| ENSMUSG00000036985 | Zdhhc9 | 0.000553633 | 0.435841828 | 0.973694431 |
| ENSMUSG00000031134 | Rbmx | 0.000553633 | -1.231582833 | 0.175666076 |
| ENSMUSG00000031198 | Fundc2 | 0.000553633 | 0.435841828 | 0.973694431 |
| ENSMUSG00000047996 | Prrg1 | 0.000553633 | -2.222649708 | 0.001952189 |
| ENSMUSG00000031214 | Ophn1 | 0.000553633 | -0.822470167 | 0.507610106 |
| ENSMUSG00000034055 | Phka1 | 0.000553633 | -1.240460984 | 0.178410618 |
| ENSMUSG00000031226 | Pbdc1 | 0.000553633 | -0.605978348 | 0.733672617 |
| ENSMUSG00000025269 | Apex2 | 0.000553633 | -1.53558902 | 0.057177697 |
| ENSMUSG00000025266 | Gnl3l | 0.000553633 | -0.383585926 | 1 |
| ENSMUSG00000051285 | Pcmtd1 | 0.000498269 | -1.305360759 | 0.15064762 |
| ENSMUSG00000092083 | Kcnb2 | 0.000498269 | -0.757156808 | 0.645880837 |
| ENSMUSG00000073725 | Lmbrd1 | 0.000498269 | -1.228908699 | 0.204365753 |
| ENSMUSG00000045216 | Hs6st1 | 0.000498269 | -0.504969785 | 0.867965264 |
| ENSMUSG00000060771 | Tsga10 | 0.000498269 | -0.296022894 | 1 |
| ENSMUSG00000026087 | Mrpl30 | 0.000498269 | -0.006516277 | 1 |
| ENSMUSG00000041439 | Mfsd6 | 0.000498269 | -0.568395165 | 0.803386053 |
| ENSMUSG00000063558 | Aox1 | 0.000498269 | -1.729676072 | 0.027328142 |
| ENSMUSG00000026037 | Orc2 | 0.000498269 | -1.544936192 | 0.059512215 |
| ENSMUSG00000040865 | Ino80d | 0.000498269 | -2.034860903 | 0.006565796 |
| ENSMUSG00000025968 | Ndufs1 | 0.000498269 | -0.757156808 | 0.611716824 |
| ENSMUSG00000033364 | Usp37 | 0.000498269 | -2.148177426 | 0.003630036 |
| ENSMUSG00000026135 | Zfp142 | 0.000498269 | -1.983432664 | 0.007932154 |
| ENSMUSG00000049339 | Retreg2 | 0.000498269 | -0.199161355 | 1 |
| ENSMUSG00000006576 | Slc4a3 | 0.000498269 | -1.360153232 | 0.127175897 |
| ENSMUSG00000054702 | Ap1s3 | 0.000498269 | -0.899601073 | 0.455131546 |
| ENSMUSG00000036766 | Dner | 0.000498269 | 0.040026309 | 1 |
| ENSMUSG00000049608 | Gpr55 | 0.000498269 | -0.673092543 | 0.716109464 |
| ENSMUSG00000034343 | Ube2f | 0.000498269 | -0.178983473 | 1 |
| ENSMUSG00000044768 | D1Ertd622e | 0.000498269 | -0.899601073 | 0.455131546 |
| ENSMUSG00000067006 | Serpinb5 | 0.000498269 | 1.215876144 | 0.318970047 |
| ENSMUSG00000026377 | Nifk | 0.000498269 | 0.215876144 | 1 |
| ENSMUSG00000048402 | Gli2 | 0.000498269 | 0.685361427 | 0.792441218 |
| ENSMUSG00000026387 | Sctr | 0.000498269 | 0.063873051 | 1 |
| ENSMUSG00000003721 | Insig2 | 0.000498269 | -0.333017102 | 1 |
| ENSMUSG00000001674 | Ddx18 | 0.000498269 | -1.369086357 | 0.128972787 |
| ENSMUSG00000059149 | Mfsd4a | 0.000498269 | -1.106051951 | 0.294131497 |
| ENSMUSG00000079330 | Lemd1 | 0.000498269 | -1.599091962 | 0.051204769 |
| ENSMUSG00000026395 | Ptprc | 0.000498269 | -0.614198854 | 0.767831378 |
| ENSMUSG00000079283 | 2310009B15Rik | 0.000498269 | 1.970763646 | 0.064562465 |
| ENSMUSG00000026365 | Cfh | 0.000498269 | 0.016567336 | 1 |
| ENSMUSG00000023150 | Ivns1abp | 0.000498269 | -1.568395165 | 0.054363019 |
| ENSMUSG00000033701 | Acbd6 | 0.000498269 | -1.651288173 | 0.037959551 |
| ENSMUSG00000058267 | Mrps14 | 0.000498269 | 0.926369527 | 0.503644124 |
| ENSMUSG00000026586 | Prrx1 | 0.000498269 | 0.326907457 | 1 |
| ENSMUSG00000026675 | Hsd17b7 | 0.000498269 | 3.163408724 | 0.002869506 |
| ENSMUSG00000013593 | Ndufs2 | 0.000498269 | 0.137873632 | 1 |
| ENSMUSG00000007097 | Atp1a2 | 0.000498269 | -0.258055044 | 1 |
| ENSMUSG00000047539 | Fbxo28 | 0.000498269 | -1.750335542 | 0.026833613 |
| ENSMUSG00000046836 | Brox | 0.000498269 | 0.685361427 | 0.792441218 |

| | | | | |
|---|---|---|---|---|
| ENSMUSG00000026615 | Eprs | 0.000498269 | -2.310007124 | 0.001540198 |
| ENSMUSG00000097325 | Gm16897 | 0.000498269 | -1.463864581 | 0.101935624 |
| ENSMUSG00000050530 | Fam171a1 | 0.000498269 | -0.277163867 | 1 |
| ENSMUSG00000026664 | Phyh | 0.000498269 | 0.883300805 | 0.637094164 |
| ENSMUSG00000026672 | Optn | 0.000498269 | -1.643946198 | 0.04321855 |
| ENSMUSG00000026712 | Mrc1 | 0.000498269 | -0.404275785 | 1 |
| ENSMUSG00000025716 | Myo3a | 0.000498269 | 0.761310281 | 0.66416768 |
| ENSMUSG00000026786 | Apbb1ip | 0.000498269 | 0.511332028 | 0.946435944 |
| ENSMUSG00000047617 | Paxx | 0.000498269 | 2.163408724 | 0.034789439 |
| ENSMUSG00000009621 | Vav2 | 0.000498269 | -2.14297852 | 0.003533684 |
| ENSMUSG00000026799 | Med27 | 0.000498269 | -0.874726405 | 0.469666084 |
| ENSMUSG00000070953 | Rabepk | 0.000498269 | -1.342119309 | 0.140153028 |
| ENSMUSG00000026869 | Psmd5 | 0.000498269 | 0.761310281 | 0.66416768 |
| ENSMUSG00000069495 | Epc2 | 0.000498269 | -1.148177426 | 0.240589043 |
| ENSMUSG00000017144 | Rnd3 | 0.000498269 | -0.911879403 | 0.455680108 |
| ENSMUSG00000060703 | Cd302 | 0.000498269 | -1.576130814 | 0.063798339 |
| ENSMUSG00000059742 | Kcnh7 | 0.000498269 | -0.455501108 | 0.938986057 |
| ENSMUSG00000042359 | Osbpl6 | 0.000498269 | -0.47217985 | 0.932615392 |
| ENSMUSG00000042272 | Sestd1 | 0.000498269 | -1.599091962 | 0.051204769 |
| ENSMUSG00000027091 | Zc3h15 | 0.000498269 | -1.480447466 | 0.080165108 |
| ENSMUSG00000002105 | Slc39a13 | 0.000498269 | 0.44720169 | 0.952530143 |
| ENSMUSG00000032841 | Prr5l | 0.000498269 | -0.369086357 | 0.997721814 |
| ENSMUSG00000027176 | Cstf3 | 0.000498269 | -2.121993495 | 0.007651296 |
| ENSMUSG00000034278 | Dnajc17 | 0.000498269 | 0.511332028 | 0.936966169 |
| ENSMUSG00000027306 | Nusap1 | 0.000498269 | -1.395558568 | 0.117548226 |
| ENSMUSG00000023572 | Ccndbp1 | 0.000498269 | 0.544498892 | 0.874652376 |
| ENSMUSG00000068394 | Cep152 | 0.000498269 | -0.887217349 | 0.473330531 |
| ENSMUSG00000027377 | Mall | 0.000498269 | -0.614198854 | 0.758211246 |
| ENSMUSG00000042851 | Zc3h6 | 0.000498269 | -1.757156808 | 0.025616174 |
| ENSMUSG00000027303 | Ptpra | 0.000498269 | -2.223993208 | 0.002392684 |
| ENSMUSG00000027312 | Atrn | 0.000498269 | -2.18910769 | 0.002606179 |
| ENSMUSG00000027342 | Pcna | 0.000498269 | 0.416174795 | 1 |
| ENSMUSG00000027358 | Bmp2 | 0.000498269 | -0.238689719 | 1 |
| ENSMUSG00000027429 | Sec23b | 0.000498269 | -0.369086357 | 0.994894633 |
| ENSMUSG00000063873 | Slc24a3 | 0.000498269 | -0.658592974 | 0.711796933 |
| ENSMUSG00000027596 | a | 0.000498269 | 0.270323928 | 1 |
| ENSMUSG00000038383 | Pigu | 0.000498269 | -2.774070191 | 0.000148372 |
| ENSMUSG00000038375 | Trp53inp2 | 0.000498269 | 1.385801146 | 0.261780615 |
| ENSMUSG00000027605 | Acss2 | 0.000498269 | 0.511332028 | 0.943363727 |
| ENSMUSG00000027610 | Gss | 0.000498269 | -0.006516277 | 1 |
| ENSMUSG00000005881 | Ergic3 | 0.000498269 | 0.613211642 | 0.798829579 |
| ENSMUSG00000027634 | Ndrg3 | 0.000498269 | 0.215876144 | 1 |
| ENSMUSG00000027639 | Samhd1 | 0.000498269 | -1.073630473 | 0.32653955 |
| ENSMUSG00000027651 | Rprd1b | 0.000498269 | -1.106051951 | 0.294131497 |
| ENSMUSG00000045503 | Sys1 | 0.000498269 | 0.063873051 | 1 |
| ENSMUSG00000053166 | Cdh22 | 0.000498269 | 1.215876144 | 0.358457412 |
| ENSMUSG00000002455 | Prpf6 | 0.000498269 | -1.797420678 | 0.019039678 |
| ENSMUSG00000043542 | Zc2hc1a | 0.000498269 | -1.52108945 | 0.065235331 |
| ENSMUSG00000027533 | Fabp5 | 0.000498269 | 1.326907457 | 0.27685776 |
| ENSMUSG00000027708 | Dcun1d1 | 0.000498269 | -0.599091962 | 0.755498536 |
| ENSMUSG00000037325 | Bbs7 | 0.000498269 | -1.736595486 | 0.04249798 |

| | | | | |
|---|---|---|---|---|
| ENSMUSG00000027751 | Supt20 | 0.000498269 | -2.26285603 | 0.001980192 |
| ENSMUSG00000047557 | Lxn | 0.000498269 | 2.511332028 | 0.014759624 |
| ENSMUSG00000027778 | Ift80 | 0.000498269 | -0.687447836 | 0.66416768 |
| ENSMUSG00000048039 | Isg20l2 | 0.000498269 | -0.47217985 | 0.932615392 |
| ENSMUSG00000068921 | Dap3 | 0.000498269 | -1.18910769 | 0.220677407 |
| ENSMUSG00000027947 | Il6ra | 0.000498269 | -0.614198854 | 0.763475704 |
| ENSMUSG00000056054 | S100a8 | 0.000498269 | 1.385801146 | 0.304871131 |
| ENSMUSG00000028140 | Mrpl9 | 0.000498269 | -0.277163867 | 1 |
| ENSMUSG00000044313 | Mab21l3 | 0.000498269 | 0.215876144 | 1 |
| ENSMUSG00000040896 | Kcnd3 | 0.000498269 | 0.416174795 | 1 |
| ENSMUSG00000000561 | Wdr77 | 0.000498269 | -0.095325544 | 1 |
| ENSMUSG00000000340 | Dbt | 0.000498269 | -0.959973691 | 0.423592395 |
| ENSMUSG00000028010 | Gar1 | 0.000498269 | -0.051604167 | 1 |
| ENSMUSG00000027999 | Pla2g12a | 0.000498269 | 0.648835551 | 0.792441218 |
| ENSMUSG00000047940 | Stpg2 | 0.000498269 | 0.47891055 | 0.945126684 |
| ENSMUSG00000005034 | Prkacb | 0.000498269 | -1.537030994 | 0.06783677 |
| ENSMUSG00000087098 | Gm11884 | 0.000498269 | 1.722836133 | 0.101604496 |
| ENSMUSG00000028463 | Car9 | 0.000498269 | -0.158519371 | 1 |
| ENSMUSG00000035517 | Tdrd7 | 0.000498269 | -0.757156808 | 0.611716824 |
| ENSMUSG00000038668 | Lpar1 | 0.000498269 | -0.810596067 | 0.56836173 |
| ENSMUSG00000028381 | Ugcg | 0.000498269 | -1.116699195 | 0.262284476 |
| ENSMUSG00000066151 | Fkbp15 | 0.000498269 | -0.614198854 | 0.758211246 |
| ENSMUSG00000059810 | Rgs3 | 0.000498269 | -1.314638573 | 0.153414334 |
| ENSMUSG00000028358 | Zfp618 | 0.000498269 | -1.983432664 | 0.007932154 |
| ENSMUSG00000039270 | Megf9 | 0.000498269 | -0.862126368 | 0.466464457 |
| ENSMUSG00000059049 | Frem1 | 0.000498269 | 0.215876144 | 1 |
| ENSMUSG00000028487 | Bnc2 | 0.000498269 | 0.137873632 | 1 |
| ENSMUSG00000038070 | Cntln | 0.000498269 | -0.13776081 | 1 |
| ENSMUSG00000085183 | Wincr1 | 0.000498269 | -0.715736881 | 0.66416768 |
| ENSMUSG00000028578 | Caap1 | 0.000498269 | -2.073630473 | 0.005140767 |
| ENSMUSG00000034210 | Efcab14 | 0.000498269 | -0.658592974 | 0.708723873 |
| ENSMUSG00000028657 | Ppt1 | 0.000498269 | 0.016567336 | 1 |
| ENSMUSG00000028907 | Utp11 | 0.000498269 | -0.238689719 | 1 |
| ENSMUSG00000028869 | Gnl2 | 0.000498269 | -1.722723311 | 0.037122674 |
| ENSMUSG00000040690 | Col16a1 | 0.000498269 | -0.936126949 | 0.463850347 |
| ENSMUSG00000028898 | Trnau1ap | 0.000498269 | -1.743481871 | 0.024497997 |
| ENSMUSG00000028878 | Fam76a | 0.000498269 | -0.199161355 | 1 |
| ENSMUSG00000037622 | Wdtc1 | 0.000498269 | -1.006516277 | 0.394013503 |
| ENSMUSG00000003644 | Rps6ka1 | 0.000498269 | -1.55279831 | 0.060700655 |
| ENSMUSG00000037306 | Man1c1 | 0.000498269 | -1.178983473 | 0.218611747 |
| ENSMUSG00000028676 | Srsf10 | 0.000498269 | -1.797420678 | 0.019039678 |
| ENSMUSG00000028670 | Lypla2 | 0.000498269 | 0.326907457 | 1 |
| ENSMUSG00000097995 | Gm24362 | 0.000498269 | -1.377964508 | 0.113700853 |
| ENSMUSG00000006445 | Epha2 | 0.000498269 | -2.111385395 | 0.004015829 |
| ENSMUSG00000028917 | Plekhm2 | 0.000498269 | -0.959973691 | 0.423592395 |
| ENSMUSG00000051435 | Fhad1 | 0.000498269 | 0.063873051 | 1 |
| ENSMUSG00000040616 | Tmem51 | 0.000498269 | -0.568395165 | 0.803386053 |
| ENSMUSG00000029020 | Mfn2 | 0.000498269 | -0.52108945 | 0.870739619 |
| ENSMUSG00000029074 | Ttll10 | 0.000498269 | -1.296022894 | 0.148292407 |
| ENSMUSG00000105068 | Gm30835 | 0.000498269 | 0.47891055 | 0.951890397 |
| ENSMUSG00000028932 | Psmc2 | 0.000498269 | 0.416174795 | 1 |

| | | | | |
|---|---|---|---|---|
| ENSMUSG00000097207 | 6030443J06Rik | 0.000498269 | -1.084518789 | 0.365611667 |
| ENSMUSG00000059447 | Hadhb | 0.000498269 | 0.063873051 | 1 |
| ENSMUSG00000038828 | Tmem214 | 0.000498269 | -0.178983473 | 1 |
| ENSMUSG00000029190 | D5Ertd579e | 0.000498269 | -1.868440142 | 0.013814241 |
| ENSMUSG00000005107 | Slc2a9 | 0.000498269 | 0.137873632 | 1 |
| ENSMUSG00000015804 | Med28 | 0.000498269 | 0.326907457 | 1 |
| ENSMUSG00000039156 | Stim2 | 0.000498269 | -2.029236354 | 0.006400827 |
| ENSMUSG00000029199 | Lias | 0.000498269 | -1.209145444 | 0.267421113 |
| ENSMUSG00000037720 | Tmem33 | 0.000498269 | 1.215876144 | 0.318970047 |
| ENSMUSG00000062110 | Scfd2 | 0.000498269 | -1.395558568 | 0.117548226 |
| ENSMUSG00000029253 | Cenpc1 | 0.000498269 | -1.55279831 | 0.060700655 |
| ENSMUSG00000035898 | Uba6 | 0.000498269 | -0.862126368 | 0.466464457 |
| ENSMUSG00000035310 | Lin54 | 0.000498269 | -1.504969785 | 0.077278951 |
| ENSMUSG00000029309 | Sparcl1 | 0.000498269 | 0.385801146 | 1 |
| ENSMUSG00000079065 | BC005561 | 0.000498269 | -2.715736881 | 0.000120117 |
| ENSMUSG00000070639 | Lrrc8b | 0.000498269 | 0.016567336 | 1 |
| ENSMUSG00000046079 | Lrrc8d | 0.000498269 | -1.936126949 | 0.010114343 |
| ENSMUSG00000042121 | Ssh1 | 0.000498269 | -0.836591276 | 0.511600603 |
| ENSMUSG00000029545 | Acads | 0.000498269 | -0.314638573 | 1 |
| ENSMUSG00000029469 | Ift81 | 0.000498269 | -0.673092543 | 0.70513931 |
| ENSMUSG00000029440 | Psmd9 | 0.000498269 | -0.199161355 | 1 |
| ENSMUSG00000015944 | Castor2 | 0.000498269 | -0.158519371 | 1 |
| ENSMUSG00000023104 | Rfc2 | 0.000498269 | -0.849415316 | 0.539270944 |
| ENSMUSG00000086220 | Gm16599 | 0.000498269 | 0.578446224 | 0.857460146 |
| ENSMUSG00000029726 | Mepce | 0.000498269 | 0.416174795 | 1 |
| ENSMUSG00000029560 | Snx8 | 0.000498269 | -0.504969785 | 0.867965264 |
| ENSMUSG00000036526 | Card11 | 0.000498269 | 0.578446224 | 0.853268347 |
| ENSMUSG00000029581 | Fscn1 | 0.000498269 | -0.238689719 | 1 |
| ENSMUSG00000038970 | Lmtk2 | 0.000498269 | -1.621693391 | 0.067319757 |
| ENSMUSG00000029659 | Medag | 0.000498269 | 0.088120597 | 1 |
| ENSMUSG00000041147 | Brca2 | 0.000498269 | -0.258055044 | 1 |
| ENSMUSG00000038759 | Nup205 | 0.000498269 | -1.874726405 | 0.014157897 |
| ENSMUSG00000029925 | Tbxas1 | 0.000498269 | -0.77070334 | 0.61612409 |
| ENSMUSG00000038456 | Dennd2a | 0.000498269 | 0.47891055 | 0.945126684 |
| ENSMUSG00000029735 | Tpk1 | 0.000498269 | -2.111385395 | 0.004015829 |
| ENSMUSG00000029814 | Igf2bp3 | 0.000498269 | -0.258055044 | 1 |
| ENSMUSG00000029777 | Gars | 0.000498269 | -1.238689719 | 0.179916358 |
| ENSMUSG00000052861 | Dnah6 | 0.000498269 | -0.199161355 | 1 |
| ENSMUSG00000052738 | Suclg1 | 0.000498269 | -0.116699195 | 1 |
| ENSMUSG00000068328 | Aup1 | 0.000498269 | 0.970763646 | 0.483322383 |
| ENSMUSG00000071337 | Tia1 | 0.000498269 | -1.722723311 | 0.030603822 |
| ENSMUSG00000001157 | Gmcl1 | 0.000498269 | 0.215876144 | 1 |
| ENSMUSG00000030098 | Grip2 | 0.000498269 | -0.051604167 | 1 |
| ENSMUSG00000072872 | Rybp | 0.000498269 | -1.544936192 | 0.059512215 |
| ENSMUSG00000035378 | Shq1 | 0.000498269 | -1.351164449 | 0.126339952 |
| ENSMUSG00000030104 | Edem1 | 0.000498269 | -1.277163867 | 0.17909875 |
| ENSMUSG00000042042 | Csgalnact2 | 0.000498269 | -0.504969785 | 0.977096893 |
| ENSMUSG00000030138 | Bms1 | 0.000498269 | -1.694572271 | 0.032072771 |
| ENSMUSG00000000399 | Ndufa9 | 0.000498269 | 0.883300805 | 0.66416768 |
| ENSMUSG00000030352 | Tspan9 | 0.000498269 | -1.369086357 | 0.128972787 |
| ENSMUSG00000030161 | Gabarapl1 | 0.000498269 | 0.326907457 | 1 |

| | | | | |
|---|---|---|---|---|
| ENSMUSG00000030201 | Lrp6 | 0.000498269 | -1.995020638 | 0.007243475 |
| ENSMUSG00000041741 | Pde3a | 0.000498269 | -0.095325544 | 1 |
| ENSMUSG00000030279 | C2cd5 | 0.000498269 | -1.560617814 | 0.053583845 |
| ENSMUSG00000006024 | Napa | 0.000498269 | 0.189403933 | 1 |
| ENSMUSG00000030374 | Strn4 | 0.000498269 | -0.333017102 | 1 |
| ENSMUSG00000030602 | Pak4 | 0.000498269 | 1.970763646 | 0.056021813 |
| ENSMUSG00000015149 | Sirt2 | 0.000498269 | 0.761310281 | 0.66416768 |
| ENSMUSG00000011427 | Zfp790 | 0.000498269 | -0.55279831 | 0.800135369 |
| ENSMUSG00000006307 | Kmt2b | 0.000498269 | -2.55279831 | 0.000323891 |
| ENSMUSG00000006315 | Tmem147 | 0.000498269 | 0.841480629 | 0.651935656 |
| ENSMUSG00000063808 | Gpatch1 | 0.000498269 | -0.219060913 | 1 |
| ENSMUSG00000101643 | Gm28307 | 0.000498269 | 0.016567336 | 1 |
| ENSMUSG00000008140 | Emc10 | 0.000498269 | -0.238689719 | 1 |
| ENSMUSG00000003269 | Cyth2 | 0.000498269 | 0.578446224 | 0.813594433 |
| ENSMUSG00000047037 | Nipa1 | 0.000498269 | -0.537030994 | 0.796840156 |
| ENSMUSG00000033510 | Otud7a | 0.000498269 | -0.77070334 | 0.637094164 |
| ENSMUSG00000030518 | Fam189a1 | 0.000498269 | -0.258055044 | 1 |
| ENSMUSG00000030539 | Sema4b | 0.000498269 | -1.01792104 | 0.349040761 |
| ENSMUSG00000030527 | Crtc3 | 0.000498269 | -0.924054117 | 0.460059915 |
| ENSMUSG00000025813 | Homer2 | 0.000498269 | -0.887217349 | 0.478060318 |
| ENSMUSG00000062797 | Hikeshi | 0.000498269 | -1.116699195 | 0.262284476 |
| ENSMUSG00000018995 | Nars2 | 0.000498269 | -0.277163867 | 1 |
| ENSMUSG00000030980 | Knop1 | 0.000498269 | -1.971750859 | 0.008702421 |
| ENSMUSG00000030870 | Ubfd1 | 0.000498269 | 1.215876144 | 0.356350819 |
| ENSMUSG00000078591 | Hs3st4 | 0.000498269 | 0.926369527 | 0.544908989 |
| ENSMUSG00000000131 | Xpo6 | 0.000498269 | -1.629149196 | 0.041312073 |
| ENSMUSG00000030967 | Zranb1 | 0.000498269 | -2.412940646 | 0.000873102 |
| ENSMUSG00000030970 | Ctbp2 | 0.000498269 | 0.163408724 | 1 |
| ENSMUSG00000041836 | Ptpre | 0.000498269 | -1.708716453 | 0.029403796 |
| ENSMUSG00000025466 | Fuom | 0.000498269 | 1.800838645 | 0.101924945 |
| ENSMUSG00000004565 | Pnpla6 | 0.000498269 | -0.995020638 | 0.388755846 |
| ENSMUSG00000110286 | Gm7706 | 0.000498269 | 0.883300805 | 0.637094164 |
| ENSMUSG00000031479 | Vps36 | 0.000498269 | -1.480447466 | 0.080165108 |
| ENSMUSG00000031549 | Ido2 | 0.000498269 | -0.258055044 | 1 |
| ENSMUSG00000015994 | Fnta | 0.000498269 | 0.242843192 | 1 |
| ENSMUSG00000025044 | Msr1 | 0.000498269 | -0.369086357 | 1 |
| ENSMUSG00000084954 | Gm16351 | 0.000498269 | -1.13776081 | 0.268597079 |
| ENSMUSG00000031634 | Ufsp2 | 0.000498269 | 0.016567336 | 1 |
| ENSMUSG00000054408 | Spcs3 | 0.000498269 | -0.13776081 | 1 |
| ENSMUSG00000031613 | Hpgd | 0.000498269 | 2.385801146 | 0.031809473 |
| ENSMUSG00000031834 | Pik3r2 | 0.000498269 | 0.040026309 | 1 |
| ENSMUSG00000042870 | Tom1 | 0.000498269 | -1.560617814 | 0.053583845 |
| ENSMUSG00000031620 | Iqcm | 0.000498269 | -0.959973691 | 0.463155604 |
| ENSMUSG00000086451 | 4933431K23Rik | 0.000498269 | 0.761310281 | 0.675561045 |
| ENSMUSG00000031616 | Ednra | 0.000498269 | 0.416174795 | 0.971720917 |
| ENSMUSG00000036977 | Anapc10 | 0.000498269 | -0.47217985 | 0.932615392 |
| ENSMUSG00000063253 | Scoc | 0.000498269 | 1.44720169 | 0.274922737 |
| ENSMUSG00000031700 | Gpt2 | 0.000498269 | -0.258055044 | 1 |
| ENSMUSG00000047866 | Lonp2 | 0.000498269 | -1.148177426 | 0.240589043 |
| ENSMUSG00000031660 | Brd7 | 0.000498269 | -1.488667972 | 0.081728936 |
| ENSMUSG00000074151 | Nlrc5 | 0.000498269 | -0.073630473 | 1 |

| | | | | |
|---|---|---|---|---|
| ENSMUSG00000031775 | Pllp | 0.000498269 | 0.613211642 | 0.798829579 |
| ENSMUSG00000031878 | Nae1 | 0.000498269 | -0.296022894 | 1 |
| ENSMUSG00000031879 | Ciao2b | 0.000498269 | 2.385801146 | 0.020391041 |
| ENSMUSG00000031904 | Slc7a6 | 0.000498269 | -2.106051951 | 0.004486593 |
| ENSMUSG00000041438 | Utp4 | 0.000498269 | -1.665860974 | 0.034221489 |
| ENSMUSG00000033732 | Sf3b3 | 0.000498269 | -1.606665182 | 0.045091139 |
| ENSMUSG00000110455 | Gm45904 | 0.000498269 | 1.063873051 | 0.470296964 |
| ENSMUSG00000031758 | Cdyl2 | 0.000498269 | -1.314638573 | 0.153414334 |
| ENSMUSG00000037300 | Ttc13 | 0.000498269 | -0.784123856 | 0.559529464 |
| ENSMUSG00000032002 | Dcun1d5 | 0.000498269 | -0.948099591 | 0.41958714 |
| ENSMUSG00000004096 | Cwc15 | 0.000498269 | -0.277163867 | 1 |
| ENSMUSG00000013076 | Amotl1 | 0.000498269 | -1.006516277 | 0.401140948 |
| ENSMUSG00000032194 | Kank2 | 0.000498269 | -0.095325544 | 1 |
| ENSMUSG00000035919 | Bbs9 | 0.000498269 | -1.377964508 | 0.113922494 |
| ENSMUSG00000032035 | Ets1 | 0.000498269 | -1.084518789 | 0.287486356 |
| ENSMUSG00000050471 | Fam118b | 0.000498269 | -1.178983473 | 0.291563202 |
| ENSMUSG00000002031 | Ift46 | 0.000498269 | -0.673092543 | 0.70029785 |
| ENSMUSG00000059890 | Ube4a | 0.000498269 | -1.463864581 | 0.088605926 |
| ENSMUSG00000043987 | Cep164 | 0.000498269 | -1.743481871 | 0.024497997 |
| ENSMUSG00000034908 | Sidt2 | 0.000498269 | -1.209145444 | 0.19917628 |
| ENSMUSG00000042396 | Rbm7 | 0.000498269 | 0.761310281 | 0.66416768 |
| ENSMUSG00000059820 | Nkapd1 | 0.000498269 | -0.614198854 | 0.758211246 |
| ENSMUSG00000032281 | Acsbg1 | 0.000498269 | -0.687447836 | 0.66416768 |
| ENSMUSG00000035284 | Vps13c | 0.000498269 | -0.959973691 | 0.423592395 |
| ENSMUSG00000031129 | Slc9a9 | 0.000498269 | -0.238689719 | 1 |
| ENSMUSG00000049493 | Pls1 | 0.000498269 | -1.168787706 | 0.246604415 |
| ENSMUSG00000032537 | Ephb1 | 0.000498269 | -0.178983473 | 1 |
| ENSMUSG00000064145 | Arih2 | 0.000498269 | -1.43862729 | 0.096782144 |
| ENSMUSG00000025651 | Uqcrc1 | 0.000498269 | 1.112782651 | 0.462023498 |
| ENSMUSG00000097736 | 9530059O14Rik | 0.000498269 | 0.040026309 | 1 |
| ENSMUSG00000036768 | Kif15 | 0.000498269 | -0.073630473 | 1 |
| ENSMUSG00000015757 | Ppil4 | 0.000498269 | -0.971750859 | 0.427581535 |
| ENSMUSG00000039166 | Akap7 | 0.000498269 | -0.757156808 | 0.611716824 |
| ENSMUSG00000038302 | Afg1l | 0.000498269 | -0.862126368 | 0.53834393 |
| ENSMUSG00000038280 | Ostm1 | 0.000498269 | 1.163408724 | 0.408970699 |
| ENSMUSG00000019861 | Gopc | 0.000498269 | -0.658592974 | 0.697198109 |
| ENSMUSG00000038602 | Slc35f1 | 0.000498269 | -0.006516277 | 1 |
| ENSMUSG00000020107 | Anapc16 | 0.000498269 | -1.040463609 | 0.358146077 |
| ENSMUSG00000048701 | Ccdc6 | 0.000498269 | -1.43862729 | 0.096782144 |
| ENSMUSG00000060733 | Ipmk | 0.000498269 | 0.685361427 | 0.792441218 |
| ENSMUSG00000020196 | Cabin1 | 0.000498269 | -0.887217349 | 0.482965166 |
| ENSMUSG00000001151 | Pcnt | 0.000498269 | -1.936126949 | 0.010114343 |
| ENSMUSG00000020260 | Pofut2 | 0.000498269 | 1.063873051 | 0.466464457 |
| ENSMUSG00000052151 | Plpp2 | 0.000498269 | 0.722836133 | 0.748991588 |
| ENSMUSG00000045193 | Cirbp | 0.000498269 | 0.416174795 | 1 |
| ENSMUSG00000035278 | Plekhj1 | 0.000498269 | 0.613211642 | 0.798829579 |
| ENSMUSG00000055053 | Nfic | 0.000498269 | -1.305360759 | 0.15064762 |
| ENSMUSG00000020255 | D10Wsu102e | 0.000498269 | -0.658592974 | 0.697198109 |
| ENSMUSG00000020263 | Appl2 | 0.000498269 | -1.199161355 | 0.223631971 |
| ENSMUSG00000020038 | Cry1 | 0.000498269 | -0.537030994 | 0.796840156 |
| ENSMUSG00000047638 | Nr1h4 | 0.000498269 | 0.215876144 | 1 |

| | | | | |
|---|---|---|---|---|
| ENSMUSG00000019975 | Ikbip | 0.000498269 | -1.342119309 | 0.140072303 |
| ENSMUSG00000020015 | Cdk17 | 0.000498269 | -1.736595486 | 0.027791338 |
| ENSMUSG00000050663 | Trhde | 0.000498269 | 0.385801146 | 1 |
| ENSMUSG00000020130 | Tbc1d15 | 0.000498269 | -0.983432664 | 0.424249514 |
| ENSMUSG00000069520 | Tmem19 | 0.000498269 | -0.404275785 | 0.998731799 |
| ENSMUSG00000069516 | Lyz2 | 0.000498269 | 0.511332028 | 0.95254727 |
| ENSMUSG00000006731 | B4galnt1 | 0.000498269 | -0.757156808 | 0.611716824 |
| ENSMUSG00000025373 | Rnf41 | 0.000498269 | -1.836591276 | 0.016369484 |
| ENSMUSG00000020182 | Ddc | 0.000498269 | -0.757156808 | 0.611716824 |
| ENSMUSG00000020128 | Vps54 | 0.000498269 | -1.708716453 | 0.029403796 |
| ENSMUSG00000020319 | Wdpcp | 0.000498269 | -1.116699195 | 0.262284476 |
| ENSMUSG00000052298 | Cdc42se2 | 0.000498269 | -1.480447466 | 0.080165108 |
| ENSMUSG00000086020 | Gm12239 | 0.000498269 | 0.761310281 | 0.66416768 |
| ENSMUSG00000020515 | Cnot8 | 0.000498269 | -0.687447836 | 0.66416768 |
| ENSMUSG00000014245 | Pigl | 0.000498269 | 0.040026309 | 1 |
| ENSMUSG00000051790 | Nlgn2 | 0.000498269 | -1.544936192 | 0.059512215 |
| ENSMUSG00000020827 | Mink1 | 0.000498269 | -1.158519371 | 0.243247585 |
| ENSMUSG00000072753 | 9230020A06Rik | 0.000498269 | 0.137873632 | 1 |
| ENSMUSG00000020783 | Ncbp3 | 0.000498269 | -1.386788358 | 0.115348677 |
| ENSMUSG00000047260 | Emc6 | 0.000498269 | 0.511332028 | 0.916315568 |
| ENSMUSG00000017550 | Atad5 | 0.000498269 | -0.455501108 | 0.929541218 |
| ENSMUSG00000072620 | Slfn2 | 0.000498269 | -1.52108945 | 0.112111296 |
| ENSMUSG00000072582 | Ptrh2 | 0.000498269 | 0.298338304 | 1 |
| ENSMUSG00000010342 | Tex14 | 0.000498269 | -0.095325544 | 1 |
| ENSMUSG00000086193 | Gm11508 | 0.000498269 | -0.369086357 | 0.994894633 |
| ENSMUSG00000018379 | Srsf1 | 0.000498269 | -1.52108945 | 0.065235331 |
| ENSMUSG00000018428 | Akap1 | 0.000498269 | -0.959973691 | 0.430332878 |
| ENSMUSG00000047773 | Ankfn1 | 0.000498269 | -0.404275785 | 1 |
| ENSMUSG00000020864 | Ankrd40 | 0.000498269 | -1.421553776 | 0.093694723 |
| ENSMUSG00000017188 | Coa3 | 0.000498269 | 1.270323928 | 0.296122408 |
| ENSMUSG00000035086 | Becn1 | 0.000498269 | -1.421553776 | 0.093694723 |
| ENSMUSG00000034931 | Dhx8 | 0.000498269 | -1.178983473 | 0.218611747 |
| ENSMUSG00000003518 | Dusp3 | 0.000498269 | 0.761310281 | 0.66416768 |
| ENSMUSG00000034520 | Gjc1 | 0.000498269 | -0.82365222 | 0.508263132 |
| ENSMUSG00000018411 | Mapt | 0.000498269 | 0.385801146 | 1 |
| ENSMUSG00000019590 | Cyb561 | 0.000498269 | 0.270323928 | 1 |
| ENSMUSG00000020706 | Ftsj3 | 0.000498269 | -0.658592974 | 0.826148082 |
| ENSMUSG00000020721 | Helz | 0.000498269 | -2.046044641 | 0.006012657 |
| ENSMUSG00000018661 | Cog1 | 0.000498269 | -0.52108945 | 0.870739619 |
| ENSMUSG00000020770 | Unk | 0.000498269 | -1.843017545 | 0.016760227 |
| ENSMUSG00000048277 | Syngr2 | 0.000498269 | -0.029236354 | 1 |
| ENSMUSG00000025372 | Baiap2 | 0.000498269 | -0.687447836 | 0.66416768 |
| ENSMUSG00000025793 | Hgs | 0.000498269 | -0.743481871 | 0.608155934 |
| ENSMUSG00000020668 | Kif3c | 0.000498269 | -0.614198854 | 0.758211246 |
| ENSMUSG00000071456 | 1110002L01Rik | 0.000498269 | -2.351164449 | 0.00116846 |
| ENSMUSG00000002900 | Lamb1 | 0.000498269 | -1.199161355 | 0.223631971 |
| ENSMUSG00000090946 | Ccdc71l | 0.000498269 | 1.800838645 | 0.084746177 |
| ENSMUSG00000047446 | Arl4a | 0.000498269 | 1.063873051 | 0.466464457 |
| ENSMUSG00000035293 | G2e3 | 0.000498269 | -2.040463609 | 0.005873683 |
| ENSMUSG00000020978 | Klhdc2 | 0.000498269 | -0.43862729 | 0.926303366 |
| ENSMUSG00000085793 | Lin52 | 0.000498269 | -1.13776081 | 0.268597079 |

| | | | | |
|---|---|---|---|---|
| ENSMUSG00000021039 | Snw1 | 0.000498269 | -1.219060913 | 0.20161331 |
| ENSMUSG00000061533 | Cep128 | 0.000498269 | -1.708716453 | 0.029521649 |
| ENSMUSG00000021007 | Spata7 | 0.000498269 | -0.369086357 | 1 |
| ENSMUSG00000021190 | Lgmn | 0.000498269 | -1.095325544 | 0.298470864 |
| ENSMUSG00000021275 | Tecpr2 | 0.000498269 | -0.924054117 | 0.460059915 |
| ENSMUSG00000021144 | Mta1 | 0.000498269 | -1.258055044 | 0.185619905 |
| ENSMUSG00000018581 | Dnah11 | 0.000498269 | 0.016567336 | 1 |
| ENSMUSG00000041297 | Cdk13 | 0.000498269 | -3.186583289 | 4.33E-06 |
| ENSMUSG00000051627 | Hist1h1e | 0.000498269 | 0.112782651 | 1 |
| ENSMUSG00000045136 | Tubb2b | 0.000498269 | 0.544498892 | 0.858831093 |
| ENSMUSG00000038267 | Slc22a23 | 0.000498269 | -2.111385395 | 0.004015829 |
| ENSMUSG00000021417 | Eci2 | 0.000498269 | -2.111385395 | 0.004154208 |
| ENSMUSG00000059288 | Cdyl | 0.000498269 | -1.296022894 | 0.148292407 |
| ENSMUSG00000114828 | AI463229 | 0.000498269 | -0.314638573 | 1 |
| ENSMUSG00000051335 | Gfod1 | 0.000498269 | -1.314638573 | 0.153414334 |
| ENSMUSG00000037933 | Bicd2 | 0.000498269 | -0.643946198 | 0.69466012 |
| ENSMUSG00000021391 | Cenpp | 0.000498269 | -1.286624196 | 0.169030494 |
| ENSMUSG00000021460 | Auh | 0.000498269 | -0.936126949 | 0.508263132 |
| ENSMUSG00000021468 | Sptlc1 | 0.000498269 | -0.743481871 | 0.608155934 |
| ENSMUSG00000021555 | Naa35 | 0.000498269 | -1.887217349 | 0.012887418 |
| ENSMUSG00000059839 | Zfp874b | 0.000498269 | -1.219060913 | 0.20161331 |
| ENSMUSG00000021594 | Srd5a1 | 0.000498269 | -0.504969785 | 0.867965264 |
| ENSMUSG00000005583 | Mef2c | 0.000498269 | -0.810596067 | 0.56836173 |
| ENSMUSG00000021668 | Polk | 0.000498269 | -1.614198854 | 0.045933679 |
| ENSMUSG00000021639 | Gtf2h2 | 0.000498269 | -0.073630473 | 1 |
| ENSMUSG00000021720 | Rnf180 | 0.000498269 | 0.578446224 | 0.825797108 |
| ENSMUSG00000042590 | Ipo11 | 0.000498269 | -0.936126949 | 0.463850347 |
| ENSMUSG00000032846 | Zswim6 | 0.000498269 | -2.777429203 | 7.72E-05 |
| ENSMUSG00000062822 | 4833420G17Rik | 0.000498269 | -1.843017545 | 0.016760227 |
| ENSMUSG00000021733 | Slc4a7 | 0.000498269 | -2.377964508 | 0.000937453 |
| ENSMUSG00000049960 | Mrps16 | 0.000498269 | 0.648835551 | 0.792441218 |
| ENSMUSG00000040717 | Il17rd | 0.000498269 | -1.643946198 | 0.045943268 |
| ENSMUSG00000006526 | Stimate | 0.000498269 | -0.568395165 | 0.803386053 |
| ENSMUSG00000021916 | Glt8d1 | 0.000498269 | -0.784123856 | 0.559529464 |
| ENSMUSG00000021884 | Hacl1 | 0.000498269 | -1.529082241 | 0.066532957 |
| ENSMUSG00000058690 | Ccser2 | 0.000498269 | -1.447088869 | 0.085527234 |
| ENSMUSG00000037536 | Fbxo34 | 0.000498269 | -1.899601073 | 0.011649133 |
| ENSMUSG00000035953 | Pip4p1 | 0.000498269 | 1.511332028 | 0.160607547 |
| ENSMUSG00000002320 | Tm9sf1 | 0.000498269 | -1.148177426 | 0.240589043 |
| ENSMUSG00000021947 | Cryl1 | 0.000498269 | -1.209145444 | 0.200616383 |
| ENSMUSG00000035469 | Rcbtb1 | 0.000498269 | -2.447088869 | 0.000661954 |
| ENSMUSG00000021990 | Spata13 | 0.000498269 | -2.680288045 | 0.000146944 |
| ENSMUSG00000014547 | Wdfy2 | 0.000498269 | -1.544936192 | 0.059512215 |
| ENSMUSG00000021978 | Extl3 | 0.000498269 | -1.797420678 | 0.019039678 |
| ENSMUSG00000022040 | Ephx2 | 0.000498269 | 0.215876144 | 1 |
| ENSMUSG00000022089 | Bin3 | 0.000498269 | -1.158519371 | 0.243247585 |
| ENSMUSG00000034795 | Ccdc122 | 0.000498269 | -1.930103162 | 0.012835386 |
| ENSMUSG00000022019 | Tdrd3 | 0.000498269 | -2.228908699 | 0.002124365 |
| ENSMUSG00000022265 | Ank | 0.000498269 | -1.01792104 | 0.349040761 |
| ENSMUSG00000022370 | Mrpl13 | 0.000498269 | -0.277163867 | 1 |
| ENSMUSG00000060429 | Sntb1 | 0.000498269 | 0.112782651 | 1 |

| | | | | |
|---|---|---|---|---|
| ENSMUSG00000071757 | Zhx2 | 0.000498269 | -1.636566667 | 0.042309732 |
| ENSMUSG00000115987 | Vps28 | 0.000498269 | 1.722836133 | 0.101604496 |
| ENSMUSG00000033819 | Ppp1r16a | 0.000498269 | -1.537030994 | 0.06783677 |
| ENSMUSG00000036529 | Sbf1 | 0.000498269 | -0.583825205 | 0.807749549 |
| ENSMUSG00000081534 | Slc48a1 | 0.000498269 | 1.163408724 | 0.346237121 |
| ENSMUSG00000022992 | Kansl2 | 0.000498269 | -1.209145444 | 0.19917628 |
| ENSMUSG00000023008 | Fmnl3 | 0.000498269 | -1.168787706 | 0.246604415 |
| ENSMUSG00000054619 | Mettl7a1 | 0.000498269 | 0.578446224 | 0.813594433 |
| ENSMUSG00000053559 | Smagp | 0.000498269 | 1.511332028 | 0.160607547 |
| ENSMUSG00000046897 | Zfp740 | 0.000498269 | -1.323857103 | 0.135445236 |
| ENSMUSG00000001285 | Myg1 | 0.000498269 | -0.421553776 | 1 |
| ENSMUSG00000068663 | Clec16a | 0.000498269 | -1.219060913 | 0.20161331 |
| ENSMUSG00000022672 | Prkdc | 0.000498269 | -1.777429203 | 0.030355209 |
| ENSMUSG00000003166 | Dgcr2 | 0.000498269 | -1.701661696 | 0.033863211 |
| ENSMUSG00000003235 | Eif2b5 | 0.000498269 | -0.55279831 | 0.800135369 |
| ENSMUSG00000013089 | Etv5 | 0.000498269 | -1.095325544 | 0.307566094 |
| ENSMUSG00000052133 | Sema5b | 0.000498269 | 0.242843192 | 1 |
| ENSMUSG00000022757 | Tfg | 0.000498269 | -0.936126949 | 0.466464457 |
| ENSMUSG00000022754 | Tmem45a | 0.000498269 | -0.924054117 | 0.460059915 |
| ENSMUSG00000022747 | St3gal6 | 0.000498269 | -1.333017102 | 0.137862055 |
| ENSMUSG00000022912 | Pros1 | 0.000498269 | -1.238689719 | 0.194823812 |
| ENSMUSG00000053414 | Hunk | 0.000498269 | -1.286624196 | 0.200147186 |
| ENSMUSG00000051989 | Smim11 | 0.000498269 | 1.648835551 | 0.173260859 |
| ENSMUSG00000116673 | A630089N07Rik | 0.000498269 | -1.537030994 | 0.06783677 |
| ENSMUSG00000057388 | Mrpl18 | 0.000498269 | 1.270323928 | 0.29603254 |
| ENSMUSG00000071267 | Zfp942 | 0.000498269 | -0.504969785 | 0.867965264 |
| ENSMUSG00000040097 | Flywch1 | 0.000498269 | -1.463864581 | 0.088605926 |
| ENSMUSG00000034681 | Rnps1 | 0.000498269 | -0.55279831 | 0.800135369 |
| ENSMUSG00000071230 | Npw | 0.000498269 | 2.648835551 | 0.080225907 |
| ENSMUSG00000075705 | Msrb1 | 0.000498269 | 1.44720169 | 0.245298386 |
| ENSMUSG00000025731 | Mettl26 | 0.000498269 | 0.356053802 | 1 |
| ENSMUSG00000097217 | Gm26549 | 0.000498269 | -1.116699195 | 0.351477434 |
| ENSMUSG00000090231 | Cfb | 0.000498269 | 0.385801146 | 1 |
| ENSMUSG00000024446 | Rpp21 | 0.000498269 | 0.613211642 | 0.828352648 |
| ENSMUSG00000042256 | Ptchd4 | 0.000498269 | 1.016567336 | 0.471767627 |
| ENSMUSG00000039601 | Rcan2 | 0.000498269 | -0.258055044 | 1 |
| ENSMUSG00000059409 | Ppp2r5d | 0.000498269 | -0.219060913 | 1 |
| ENSMUSG00000038910 | Plcl2 | 0.000498269 | -2.089932285 | 0.004794066 |
| ENSMUSG00000035580 | Kcnh8 | 0.000498269 | -1.899601073 | 0.019989375 |
| ENSMUSG00000047123 | Ticam1 | 0.000498269 | -0.116699195 | 1 |
| ENSMUSG00000041168 | Lonp1 | 0.000498269 | -0.006516277 | 1 |
| ENSMUSG00000002660 | Clpp | 0.000498269 | 0.063873051 | 1 |
| ENSMUSG00000024067 | Dpy30 | 0.000498269 | 0.137873632 | 1 |
| ENSMUSG00000039193 | Nlrc4 | 0.000498269 | 0.648835551 | 0.792441218 |
| ENSMUSG00000024143 | Rhoq | 0.000498269 | 0.242843192 | 1 |
| ENSMUSG00000061802 | Armc4 | 0.000498269 | -0.258055044 | 1 |
| ENSMUSG00000042942 | Greb1l | 0.000498269 | -0.862126368 | 0.466464457 |
| ENSMUSG00000024293 | Esco1 | 0.000498269 | -0.369086357 | 0.994894633 |
| ENSMUSG00000041238 | Rbbp8 | 0.000498269 | -0.296022894 | 1 |
| ENSMUSG00000024421 | Lama3 | 0.000498269 | -0.784123856 | 0.559529464 |
| ENSMUSG00000047161 | Chst9 | 0.000498269 | -0.421553776 | 1 |

| | | | | |
|---|---|---|---|---|
| ENSMUSG00000024277 | Mapre2 | 0.000498269 | -0.47217985 | 0.932615392 |
| ENSMUSG00000057455 | Rit2 | 0.000498269 | 1.511332028 | 0.224626635 |
| ENSMUSG00000024381 | Bin1 | 0.000498269 | -1.029236354 | 0.352215149 |
| ENSMUSG00000006050 | Sra1 | 0.000498269 | -0.219060913 | 1 |
| ENSMUSG00000060450 | Rnf14 | 0.000498269 | -0.537030994 | 0.886766194 |
| ENSMUSG00000033319 | Fem1c | 0.000498269 | -1.544936192 | 0.059512215 |
| ENSMUSG00000042705 | Commd10 | 0.000498269 | -1.694572271 | 0.032072771 |
| ENSMUSG00000024507 | Hsd17b4 | 0.000498269 | 0.040026309 | 1 |
| ENSMUSG00000053644 | Aldh7a1 | 0.000498269 | 0.088120597 | 1 |
| ENSMUSG00000008301 | Phax | 0.000498269 | -0.43862729 | 0.926583891 |
| ENSMUSG00000024594 | Prrc1 | 0.000498269 | 0.47891055 | 0.95254727 |
| ENSMUSG00000042211 | Fbxo38 | 0.000498269 | -1.219060913 | 0.20161331 |
| ENSMUSG00000001473 | Tubb6 | 0.000498269 | -0.43862729 | 1 |
| ENSMUSG00000024561 | Mbd1 | 0.000498269 | -1.029236354 | 0.352215149 |
| ENSMUSG00000047466 | 8030462N17Rik | 0.000498269 | -1.899601073 | 0.011649133 |
| ENSMUSG00000058881 | Zfp516 | 0.000498269 | -1.777429203 | 0.025747509 |
| ENSMUSG00000118094 | AC142100.1 | 0.000498269 | 2.270323928 | 0.029612256 |
| ENSMUSG00000024847 | Aip | 0.000498269 | 0.648835551 | 0.827142217 |
| ENSMUSG00000040663 | Clcf1 | 0.000498269 | -0.924054117 | 0.460059915 |
| ENSMUSG00000006456 | Rbm14 | 0.000498269 | -1.463864581 | 0.088605926 |
| ENSMUSG00000045664 | Cdc42ep2 | 0.000498269 | 0.926369527 | 0.503644124 |
| ENSMUSG00000053080 | 2700081O15Rik | 0.000498269 | -0.82365222 | 0.508263132 |
| ENSMUSG00000035171 | 1110059E24Rik | 0.000498269 | -1.395558568 | 0.128972787 |
| ENSMUSG00000025192 | Entpd7 | 0.000498269 | -0.568395165 | 0.803386053 |
| ENSMUSG00000025035 | Arl3 | 0.000498269 | -0.614198854 | 0.758211246 |
| ENSMUSG00000025050 | Pcgf6 | 0.000498269 | -0.959973691 | 0.423592395 |
| ENSMUSG00000084957 | Bbip1 | 0.000498269 | -1.051604167 | 0.315589992 |
| ENSMUSG00000044148 | 1810030O07Rik | 0.000498269 | -1.178983473 | 0.218611747 |
| ENSMUSG00000031093 | Dock11 | 0.000498269 | -2.654945197 | 0.000175432 |
| ENSMUSG00000000355 | Mcts1 | 0.000498269 | 0.137873632 | 1 |
| ENSMUSG00000035847 | Ids | 0.000498269 | 1.215876144 | 0.318970047 |
| ENSMUSG00000031337 | Mtm1 | 0.000498269 | -0.971750859 | 0.446038896 |
| ENSMUSG00000015217 | Hmgb3 | 0.000498269 | 0.685361427 | 0.792441218 |
| ENSMUSG00000073131 | Vma21 | 0.000498269 | 1.578446224 | 0.138717949 |
| ENSMUSG00000031347 | Cetn2 | 0.000498269 | 1.326907457 | 0.27685776 |
| ENSMUSG00000031425 | Plp1 | 0.000498269 | -1.715736881 | 0.029965083 |
| ENSMUSG00000042595 | Fam199x | 0.000498269 | -1.095325544 | 0.290284345 |
| ENSMUSG00000087403 | Kantr | 0.000498269 | -1.277163867 | 0.166923355 |
| ENSMUSG00000033740 | St18 | 0.000442906 | -0.158519371 | 1 |
| ENSMUSG00000101640 | Gm28376 | 0.000442906 | 0.609307187 | 0.886203497 |
| ENSMUSG00000041670 | Rims1 | 0.000442906 | -0.225633566 | 1 |
| ENSMUSG00000101227 | Gm29506 | 0.000442906 | 1.648835551 | 0.1477368 |
| ENSMUSG00000026155 | Smap1 | 0.000442906 | -1.648844997 | 0.049126005 |
| ENSMUSG00000037351 | Actr1b | 0.000442906 | 0.146335211 | 1 |
| ENSMUSG00000026090 | 2010300C02Rik | 0.000442906 | -0.689034087 | 0.66416768 |
| ENSMUSG00000026082 | Rev1 | 0.000442906 | -1.147023732 | 0.284698396 |
| ENSMUSG00000070939 | Tgfbrap1 | 0.000442906 | 0.42644313 | 1 |
| ENSMUSG00000056870 | Gulp1 | 0.000442906 | -0.088130043 | 1 |
| ENSMUSG00000052331 | Ankrd44 | 0.000442906 | -0.673092543 | 0.723846946 |
| ENSMUSG00000038331 | Satb2 | 0.000442906 | -0.83945093 | 0.528505953 |
| ENSMUSG00000087022 | 9130024F11Rik | 0.000442906 | 2.496832458 | 0.040680007 |

| | | | | |
|---|---|---|---|---|
| ENSMUSG00000038174 | Fam126b | 0.000442906 | -0.909159902 | 0.475407382 |
| ENSMUSG00000026024 | Als2 | 0.000442906 | -0.936126949 | 0.455131546 |
| ENSMUSG00000049439 | Cyp20a1 | 0.000442906 | -0.825095637 | 0.57709448 |
| ENSMUSG00000026782 | Abi2 | 0.000442906 | 0.496832458 | 0.924536002 |
| ENSMUSG00000026249 | Serpine2 | 0.000442906 | 0.461208548 | 1 |
| ENSMUSG00000026163 | Sphkap | 0.000442906 | 0.648835551 | 0.795877354 |
| ENSMUSG00000052477 | C130026I21Rik | 0.000442906 | 0.774366434 | 0.740864041 |
| ENSMUSG00000026222 | Sp100 | 0.000442906 | -1.258055044 | 0.187579947 |
| ENSMUSG00000005501 | Usp40 | 0.000442906 | -0.881679165 | 0.467537667 |
| ENSMUSG00000101214 | Gm28403 | 0.000442906 | -0.810596067 | 0.573149109 |
| ENSMUSG00000057329 | Bcl2 | 0.000442906 | -2.385810591 | 0.001172078 |
| ENSMUSG00000026380 | Tfcp2l1 | 0.000442906 | 0.037400839 | 1 |
| ENSMUSG00000026342 | Slc35f5 | 0.000442906 | -0.310522464 | 1 |
| ENSMUSG00000026347 | Tmem163 | 0.000442906 | -0.624182943 | 0.806201078 |
| ENSMUSG00000026437 | Cdk18 | 0.000442906 | -0.111976785 | 1 |
| ENSMUSG00000042229 | Rabif | 0.000442906 | -1.111976785 | 0.314582476 |
| ENSMUSG00000087230 | Mroh3 | 0.000442906 | -2.43390488 | 0.000905629 |
| ENSMUSG00000042684 | Npl | 0.000442906 | 0.118320835 | 1 |
| ENSMUSG00000050565 | Tor1aip2 | 0.000442906 | -1.538791452 | 0.07003776 |
| ENSMUSG00000026500 | Cox20 | 0.000442906 | 0.570833039 | 0.886203497 |
| ENSMUSG00000056050 | Mia3 | 0.000442906 | -1.615867675 | 0.052676838 |
| ENSMUSG00000039239 | Tgfb2 | 0.000442906 | -0.088130043 | 1 |
| ENSMUSG00000026611 | Spata17 | 0.000442906 | -0.135435758 | 1 |
| ENSMUSG00000058248 | Kcnh1 | 0.000442906 | -0.371064006 | 1 |
| ENSMUSG00000039449 | Prpf18 | 0.000442906 | -0.689034087 | 0.66416768 |
| ENSMUSG00000037262 | Kin | 0.000442906 | -1.922706434 | 0.013677655 |
| ENSMUSG00000026773 | Pfkfb3 | 0.000442906 | -0.962599161 | 0.466464457 |
| ENSMUSG00000026730 | Pter | 0.000442906 | 0.496832458 | 0.915242623 |
| ENSMUSG00000026739 | Bmi1 | 0.000442906 | 0.960779558 | 0.559918909 |
| ENSMUSG00000036646 | Man1b1 | 0.000442906 | -1.743481871 | 0.029744952 |
| ENSMUSG00000075467 | Dnlz | 0.000442906 | 1.648835551 | 0.182139012 |
| ENSMUSG00000026807 | Ak8 | 0.000442906 | 0.648835551 | 0.792441218 |
| ENSMUSG00000039660 | Spout1 | 0.000442906 | 0.609307187 | 0.886203497 |
| ENSMUSG00000007476 | Lrrc8a | 0.000442906 | -1.214662449 | 0.235934029 |
| ENSMUSG00000026858 | Miga2 | 0.000442906 | -0.485020195 | 0.901829791 |
| ENSMUSG00000055632 | Hmcn2 | 0.000442906 | 0.648835551 | 0.792441218 |
| ENSMUSG00000039195 | 1110008P14Rik | 0.000442906 | 1.295198597 | 0.409864206 |
| ENSMUSG00000035949 | Fbxw2 | 0.000442906 | -1.341110784 | 0.157935867 |
| ENSMUSG00000047250 | Ptgs1 | 0.000442906 | -0.689034087 | 0.734875632 |
| ENSMUSG00000055371 | Stam2 | 0.000442906 | -0.795949291 | 0.568577582 |
| ENSMUSG00000006494 | Pdk1 | 0.000442906 | -0.20360726 | 1 |
| ENSMUSG00000004085 | Map3k20 | 0.000442906 | -1.665055224 | 0.050877179 |
| ENSMUSG00000027016 | Zfp385b | 0.000442906 | 0.264171701 | 1 |
| ENSMUSG00000034075 | Zdhhc5 | 0.000442906 | -0.810596067 | 0.573149109 |
| ENSMUSG00000068373 | D430041D05Rik | 0.000442906 | 0.960779558 | 0.620520753 |
| ENSMUSG00000042670 | Immp1l | 0.000442906 | -0.949423772 | 0.466464457 |
| ENSMUSG00000057234 | Mettl15 | 0.000442906 | -0.590630383 | 0.792441218 |
| ENSMUSG00000027132 | Katnbl1 | 0.000442906 | -0.909159902 | 0.475407382 |
| ENSMUSG00000055943 | Emc7 | 0.000442906 | 0.570833039 | 0.886203497 |
| ENSMUSG00000101448 | Gm28494 | 0.000442906 | 0.011405631 | 1 |
| ENSMUSG00000046804 | Phgr1 | 0.000442906 | 2.818760553 | 0.011159735 |

| | | | | |
|---|---|---|---|---|
| ENSMUSG00000034032 | Rpap1 | 0.000442906 | -1.039220442 | 0.387414024 |
| ENSMUSG00000027201 | Myef2 | 0.000442906 | -1.429166961 | 0.102454279 |
| ENSMUSG00000027380 | Acoxl | 0.000442906 | 0.090840098 | 1 |
| ENSMUSG00000014353 | Tmem87b | 0.000442906 | -1.56494374 | 0.063366891 |
| ENSMUSG00000027406 | Idh3b | 0.000442906 | 0.648835551 | 0.877529363 |
| ENSMUSG00000039913 | Pak7 | 0.000442906 | 0.359328934 | 1 |
| ENSMUSG00000027425 | Kat14 | 0.000442906 | -1.147023732 | 0.284698396 |
| ENSMUSG00000001767 | Crnkl1 | 0.000442906 | 0.118320835 | 1 |
| ENSMUSG00000027454 | Gins1 | 0.000442906 | -1.247328638 | 0.226567468 |
| ENSMUSG00000038369 | Ncoa6 | 0.000442906 | -2.063882496 | 0.007127824 |
| ENSMUSG00000074643 | Cpne1 | 0.000442906 | -1.795949291 | 0.025711081 |
| ENSMUSG00000039804 | Ncoa5 | 0.000442906 | -1.648844997 | 0.049126005 |
| ENSMUSG00000027544 | Nfatc2 | 0.000442906 | -1.158519371 | 0.254892207 |
| ENSMUSG00000027525 | Phactr3 | 0.000442906 | 1.011405631 | 0.551482001 |
| ENSMUSG00000039069 | Mtg2 | 0.000442906 | -0.111976785 | 1 |
| ENSMUSG00000017688 | Hnf4g | 0.000442906 | -0.039220442 | 1 |
| ENSMUSG00000027531 | Impa1 | 0.000442906 | -0.673092543 | 0.723846946 |
| ENSMUSG00000027601 | Mtfr1 | 0.000442906 | -0.268702288 | 1 |
| ENSMUSG00000019528 | Gyg | 0.000442906 | 2.496832458 | 0.040680007 |
| ENSMUSG00000027665 | Pik3ca | 0.000442906 | -2.026729498 | 0.008224754 |
| ENSMUSG00000027667 | Zfp639 | 0.000442906 | -0.656972878 | 0.719591951 |
| ENSMUSG00000027669 | Gnb4 | 0.000442906 | -0.881679165 | 0.467537667 |
| ENSMUSG00000027679 | Dnajc19 | 0.000442906 | 1.648835551 | 0.1477368 |
| ENSMUSG00000037818 | Abhd18 | 0.000442906 | -1.361148537 | 0.153802689 |
| ENSMUSG00000027746 | Ufm1 | 0.000442906 | -0.014129461 | 1 |
| ENSMUSG00000027750 | Postn | 0.000442906 | 0.496832458 | 0.909161999 |
| ENSMUSG00000036503 | Rnf13 | 0.000442906 | -1.988594369 | 0.009568987 |
| ENSMUSG00000027829 | Ccnl1 | 0.000442906 | -3.195259917 | 5.68E-06 |
| ENSMUSG00000103181 | A330069K06Rik | 0.000442906 | 1.648835551 | 0.1477368 |
| ENSMUSG00000102692 | Dchs2 | 0.000442906 | 0.295198597 | 1 |
| ENSMUSG00000033767 | Tmem131l | 0.000442906 | -1.795949291 | 0.025711081 |
| ENSMUSG00000001418 | Glmp | 0.000442906 | -0.640671066 | 0.801225115 |
| ENSMUSG00000008604 | Ubqln4 | 0.000442906 | -0.371064006 | 1 |
| ENSMUSG00000059743 | Fdps | 0.000442906 | 1.011405631 | 0.555109572 |
| ENSMUSG00000042642 | Flad1 | 0.000442906 | 1.174904363 | 0.395423178 |
| ENSMUSG00000027952 | Pmvk | 0.000442906 | -0.063882496 | 1 |
| ENSMUSG00000000794 | Kcnn3 | 0.000442906 | -0.825095637 | 0.597124264 |
| ENSMUSG00000038902 | Pogz | 0.000442906 | -1.874726405 | 0.018221288 |
| ENSMUSG00000028115 | Bnipl | 0.000442906 | -0.247328638 | 1 |
| ENSMUSG00000038642 | Ctss | 0.000442906 | -0.57355687 | 0.862751913 |
| ENSMUSG00000028107 | Tars2 | 0.000442906 | -0.704801403 | 0.66416768 |
| ENSMUSG00000015749 | Anp32e | 0.000442906 | -1.247328638 | 0.213853707 |
| ENSMUSG00000050064 | Zfp697 | 0.000442906 | -2.039220442 | 0.007443294 |
| ENSMUSG00000033721 | Vav3 | 0.000442906 | -0.825095637 | 0.588078615 |
| ENSMUSG00000089911 | Mfsd14a | 0.000442906 | -1.111976785 | 0.314582476 |
| ENSMUSG00000028134 | Ptbp2 | 0.000442906 | -1.615867675 | 0.052676838 |
| ENSMUSG00000028165 | Cisd2 | 0.000442906 | 0.118320835 | 1 |
| ENSMUSG00000028163 | Nfkb1 | 0.000442906 | -2.100102684 | 0.00542227 |
| ENSMUSG00000037922 | Bank1 | 0.000442906 | 0.533358334 | 0.894464956 |
| ENSMUSG00000054942 | Miga1 | 0.000442906 | -0.936126949 | 0.455131546 |
| ENSMUSG00000040550 | Otud6b | 0.000442906 | -0.181239447 | 1 |

| | | | | |
|---|---|---|---|---|
| ENSMUSG00000028223 | Decr1 | 0.000442906 | 0.174904363 | 1 |
| ENSMUSG00000056494 | Cngb3 | 0.000442906 | -0.158519371 | 1 |
| ENSMUSG00000040359 | Ufl1 | 0.000442906 | 0.774366434 | 0.732187634 |
| ENSMUSG00000028284 | Map3k7 | 0.000442906 | -0.975655313 | 0.415257602 |
| ENSMUSG00000028405 | Aco1 | 0.000442906 | -0.466641666 | 0.889426255 |
| ENSMUSG00000045973 | Slc25a51 | 0.000442906 | -0.20360726 | 1 |
| ENSMUSG00000039774 | Galnt12 | 0.000442906 | -1.123753952 | 0.278293619 |
| ENSMUSG00000008305 | Tle1 | 0.000442906 | -1.225633566 | 0.207756141 |
| ENSMUSG00000038172 | Ttc39b | 0.000442906 | -1.735828299 | 0.033649319 |
| ENSMUSG00000038007 | Acer2 | 0.000442906 | -0.656972878 | 0.719591951 |
| ENSMUSG00000028528 | Dnajc6 | 0.000442906 | -0.538791452 | 0.840804483 |
| ENSMUSG00000028524 | Sgip1 | 0.000442906 | 0.533358334 | 0.930100175 |
| ENSMUSG00000028617 | Lrrc42 | 0.000442906 | -0.881679165 | 0.541865293 |
| ENSMUSG00000063172 | Hspb11 | 0.000442906 | 1.174904363 | 0.395423178 |
| ENSMUSG00000028693 | Nasp | 0.000442906 | -1.795949291 | 0.025711081 |
| ENSMUSG00000033379 | Atp6v0b | 0.000442906 | -1.673092543 | 0.04496186 |
| ENSMUSG00000032897 | Nfyc | 0.000442906 | -1.735828299 | 0.033649319 |
| ENSMUSG00000032870 | Smap2 | 0.000442906 | -0.781152289 | 0.630177036 |
| ENSMUSG00000023075 | Akirin1 | 0.000442906 | 0.118320835 | 1 |
| ENSMUSG00000028646 | Rragc | 0.000442906 | -0.289763904 | 1 |
| ENSMUSG00000028863 | Meaf6 | 0.000442906 | 0.063873051 | 1 |
| ENSMUSG00000042608 | Stk40 | 0.000442906 | -1.247328638 | 0.213853707 |
| ENSMUSG00000042408 | Zmym6 | 0.000442906 | -1.039220442 | 0.387414024 |
| ENSMUSG00000046623 | Gjb4 | 0.000442906 | 0.911869957 | 0.66416768 |
| ENSMUSG00000028772 | Zcchc17 | 0.000442906 | -0.922706434 | 0.455038142 |
| ENSMUSG00000028911 | Srsf4 | 0.000442906 | -1.712620908 | 0.036637611 |
| ENSMUSG00000036896 | C1qc | 0.000442906 | 0.731297712 | 0.773746672 |
| ENSMUSG00000028923 | Necap2 | 0.000442906 | 0.570833039 | 0.886203497 |
| ENSMUSG00000039953 | Clstn1 | 0.000442906 | 0.359328934 | 1 |
| ENSMUSG00000065990 | Aurkaip1 | 0.000442906 | 1.118320835 | 0.47966612 |
| ENSMUSG00000058503 | Fam133b | 0.000442906 | -2.07605721 | 0.006539077 |
| ENSMUSG00000094230 | Gm21847 | 0.000442906 | 0.461208548 | 1 |
| ENSMUSG00000057614 | Gnai1 | 0.000442906 | 0.533358334 | 0.894464956 |
| ENSMUSG00000106379 | Lhfpl3 | 0.000442906 | -0.795949291 | 0.66416768 |
| ENSMUSG00000028999 | Rint1 | 0.000442906 | -0.135435758 | 1 |
| ENSMUSG00000001569 | Nom1 | 0.000442906 | -1.111976785 | 0.314582476 |
| ENSMUSG00000029147 | Ppm1g | 0.000442906 | -0.735828299 | 0.664426832 |
| ENSMUSG00000023452 | Pisd | 0.000442906 | -0.83945093 | 0.529886422 |
| ENSMUSG00000029103 | Lrpap1 | 0.000442906 | 0.774366434 | 0.732187634 |
| ENSMUSG00000029095 | Ablim2 | 0.000442906 | 0.118320835 | 1 |
| ENSMUSG00000029198 | Grpel1 | 0.000442906 | 0.911869957 | 0.601220428 |
| ENSMUSG00000106741 | 4930513D17Rik | 0.000442906 | 0.818760553 | 0.66416768 |
| ENSMUSG00000061461 | Smim20 | 0.000442906 | 0.295198597 | 1 |
| ENSMUSG00000029174 | Tbc1d1 | 0.000442906 | -2.051604167 | 0.00679769 |
| ENSMUSG00000037795 | N4bp2 | 0.000442906 | -2.063882496 | 0.007127824 |
| ENSMUSG00000029217 | Tec | 0.000442906 | -1.547561661 | 0.071310956 |
| ENSMUSG00000029152 | Ociad1 | 0.000442906 | -0.330986567 | 1 |
| ENSMUSG00000106943 | Dancr | 0.000442906 | -0.52108945 | 0.837985936 |
| ENSMUSG00000029231 | Pdgfra | 0.000442906 | 0.264171701 | 1 |
| ENSMUSG00000070697 | Utp3 | 0.000442906 | -0.735828299 | 0.664426832 |
| ENSMUSG00000029377 | Ereg | 0.000442906 | -0.466641666 | 0.889426255 |

| | | | | |
|---|---|---|---|---|
| ENSMUSG00000091034 | Gm17660 | 0.000442906 | -0.225633566 | 1 |
| ENSMUSG00000029270 | Dipk1a | 0.000442906 | -1.94279068 | 0.013074066 |
| ENSMUSG00000014668 | Chfr | 0.000442906 | -2.289763904 | 0.002052334 |
| ENSMUSG00000043510 | Hscb | 0.000442906 | 0.570833039 | 0.886203497 |
| ENSMUSG00000058153 | Sez6l | 0.000442906 | 0.063873051 | 1 |
| ENSMUSG00000018974 | Sart3 | 0.000442906 | -1.419644186 | 0.116436313 |
| ENSMUSG00000025825 | Iscu | 0.000442906 | 0.392495798 | 1 |
| ENSMUSG00000029577 | Ube3b | 0.000442906 | -0.795949291 | 0.568577582 |
| ENSMUSG00000029524 | Sirt4 | 0.000442906 | -1.704801403 | 0.035862436 |
| ENSMUSG00000029518 | Rab35 | 0.000442906 | -0.181239447 | 1 |
| ENSMUSG00000029513 | Prkab1 | 0.000442906 | -0.289763904 | 1 |
| ENSMUSG00000042726 | Trafd1 | 0.000442906 | 0.731297712 | 0.741090984 |
| ENSMUSG00000029458 | Brap | 0.000442906 | -1.599091962 | 0.061096399 |
| ENSMUSG00000038023 | Atp6v0a2 | 0.000442906 | -1.781152289 | 0.024569482 |
| ENSMUSG00000079173 | Zan | 0.000442906 | -0.590630383 | 0.786092784 |
| ENSMUSG00000029714 | Gigyf1 | 0.000442906 | -2.867739974 | 6.32E-05 |
| ENSMUSG00000037017 | Zscan21 | 0.000442906 | -1.982139347 | 0.010947317 |
| ENSMUSG00000019494 | Cops6 | 0.000442906 | -0.310522464 | 1 |
| ENSMUSG00000029730 | Mcm7 | 0.000442906 | -1.410058138 | 0.114388824 |
| ENSMUSG00000045438 | Cox19 | 0.000442906 | 0.42644313 | 1 |
| ENSMUSG00000053581 | Zfand2a | 0.000442906 | -0.20360726 | 1 |
| ENSMUSG00000029547 | Ints1 | 0.000442906 | -0.909159902 | 0.475407382 |
| ENSMUSG00000018143 | Mafk | 0.000442906 | -0.390692813 | 1 |
| ENSMUSG00000016503 | Gtf3a | 0.000442906 | 1.174904363 | 0.395423178 |
| ENSMUSG00000041313 | Slc7a1 | 0.000442906 | -2.039220442 | 0.007443294 |
| ENSMUSG00000029658 | Wdr95 | 0.000442906 | 1.42644313 | 0.349040761 |
| ENSMUSG00000001376 | Vps50 | 0.000442906 | -1.929432297 | 0.014115255 |
| ENSMUSG00000032757 | Bet1 | 0.000442906 | 1.359328934 | 0.415055165 |
| ENSMUSG00000042505 | Sdhaf3 | 0.000442906 | -1.547561661 | 0.071310956 |
| ENSMUSG00000000416 | Cttnbp2 | 0.000442906 | 1.118320835 | 0.435791313 |
| ENSMUSG00000029669 | Tspan12 | 0.000442906 | -0.825095637 | 0.57709448 |
| ENSMUSG00000029701 | Rbm28 | 0.000442906 | -1.881679165 | 0.016000576 |
| ENSMUSG00000029924 | Slc37a3 | 0.000442906 | -1.547561661 | 0.071310956 |
| ENSMUSG00000073067 | 9130019P16Rik | 0.000442906 | 1.496832458 | 0.243691584 |
| ENSMUSG00000038022 | Mindy4 | 0.000442906 | -1.135435758 | 0.281150145 |
| ENSMUSG00000076609 | Igkc | 0.000442906 | -0.268702288 | 1 |
| ENSMUSG00000068335 | Dok1 | 0.000442906 | 1.011405631 | 0.538882935 |
| ENSMUSG00000034744 | Nagk | 0.000442906 | -0.014129461 | 1 |
| ENSMUSG00000033735 | Spr | 0.000442906 | 0.689477536 | 0.771569221 |
| ENSMUSG00000029993 | Nfu1 | 0.000442906 | -0.014129461 | 1 |
| ENSMUSG00000030020 | Prickle2 | 0.000442906 | -0.181239447 | 1 |
| ENSMUSG00000045100 | Slc25a26 | 0.000442906 | 0.011405631 | 1 |
| ENSMUSG00000030074 | Gxylt2 | 0.000442906 | -0.310522464 | 1 |
| ENSMUSG00000030254 | Rad18 | 0.000442906 | -0.52108945 | 0.836964118 |
| ENSMUSG00000030170 | Wnt5b | 0.000442906 | 0.037400839 | 1 |
| ENSMUSG00000041377 | Ninj2 | 0.000442906 | -1.063882496 | 0.361093181 |
| ENSMUSG00000030177 | Ccdc77 | 0.000442906 | -1.582118883 | 0.066307071 |
| ENSMUSG00000001518 | Itfg2 | 0.000442906 | -0.410058138 | 0.971854959 |
| ENSMUSG00000032652 | Crebl2 | 0.000442906 | -1.123753952 | 0.278293619 |
| ENSMUSG00000030220 | Arhgdib | 0.000442906 | -0.867739974 | 0.584850095 |
| ENSMUSG00000030224 | Strap | 0.000442906 | -1.330986567 | 0.154833135 |

| | | | | |
|---|---|---|---|---|
| ENSMUSG00000008540 | Mgst1 | 0.000442906 | 2.359328934 | 0.031557722 |
| ENSMUSG00000030291 | Med21 | 0.000442906 | 0.264171701 | 1 |
| ENSMUSG00000068566 | Myadm | 0.000442906 | 1.174904363 | 0.466464457 |
| ENSMUSG00000006154 | Eps8l1 | 0.000442906 | -0.247328638 | 1 |
| ENSMUSG00000030431 | Tmem238 | 0.000442906 | 0.911869957 | 0.582339791 |
| ENSMUSG00000044876 | Zfp444 | 0.000442906 | -2.106051951 | 0.005551659 |
| ENSMUSG00000003099 | Ppp5c | 0.000442906 | 0.533358334 | 0.894464956 |
| ENSMUSG00000060188 | Cxcl17 | 0.000442906 | 0.570833039 | 0.886203497 |
| ENSMUSG00000061479 | Snrpa | 0.000442906 | -0.825095637 | 0.599135879 |
| ENSMUSG00000060791 | Gmfg | 0.000442906 | 0.960779558 | 0.559918909 |
| ENSMUSG00000030598 | Fbxo17 | 0.000442906 | 0.648835551 | 0.792441218 |
| ENSMUSG00000045948 | Mrps12 | 0.000442906 | 0.359328934 | 1 |
| ENSMUSG00000049421 | Zfp260 | 0.000442906 | -1.123753952 | 0.278293619 |
| ENSMUSG00000058239 | Usf2 | 0.000442906 | 0.011405631 | 1 |
| ENSMUSG00000044452 | Zfp507 | 0.000442906 | -0.810596067 | 0.573149109 |
| ENSMUSG00000030739 | Myh14 | 0.000442906 | -0.247328638 | 1 |
| ENSMUSG00000002205 | Vrk3 | 0.000442906 | -1.720398258 | 0.032072771 |
| ENSMUSG00000038520 | Tbc1d17 | 0.000442906 | -1.225633566 | 0.207756141 |
| ENSMUSG00000011096 | Akt1s1 | 0.000442906 | -1.039220442 | 0.387414024 |
| ENSMUSG00000038387 | Rras | 0.000442906 | 0.774366434 | 0.69613433 |
| ENSMUSG00000003423 | Pih1d1 | 0.000442906 | 0.063873051 | 1 |
| ENSMUSG00000030839 | Sergef | 0.000442906 | -0.781152289 | 0.663124861 |
| ENSMUSG00000040026 | Saa3 | 0.000442906 | 1.118320835 | 0.423364274 |
| ENSMUSG00000100005 | B130024G19Rik | 0.000442906 | 1.174904363 | 0.42263302 |
| ENSMUSG00000038646 | Ramac | 0.000442906 | 1.063873051 | 0.523007753 |
| ENSMUSG00000041328 | Pcf11 | 0.000442906 | -2.36611479 | 0.001371562 |
| ENSMUSG00000035713 | Usp35 | 0.000442906 | -1.529967601 | 0.080542848 |
| ENSMUSG00000051727 | Kctd14 | 0.000442906 | -0.181239447 | 1 |
| ENSMUSG00000035354 | Uvrag | 0.000442906 | -2.039220442 | 0.007443294 |
| ENSMUSG00000030706 | Mrpl48 | 0.000442906 | -0.330986567 | 1 |
| ENSMUSG00000030888 | Rrp8 | 0.000442906 | -0.795949291 | 0.568577582 |
| ENSMUSG00000031016 | Wee1 | 0.000442906 | -0.371064006 | 1 |
| ENSMUSG00000030877 | Mfsd13b | 0.000442906 | -0.063882496 | 1 |
| ENSMUSG00000042492 | Tbc1d10b | 0.000442906 | -1.247328638 | 0.213853707 |
| ENSMUSG00000042105 | Inpp5f | 0.000442906 | -0.962599161 | 0.463155604 |
| ENSMUSG00000054612 | Mgmt | 0.000442906 | 0.174904363 | 1 |
| ENSMUSG00000047264 | Zfp358 | 0.000442906 | 1.174904363 | 0.395423178 |
| ENSMUSG00000038482 | Tfdp1 | 0.000442906 | -1.56494374 | 0.063366891 |
| ENSMUSG00000031546 | Gins4 | 0.000442906 | 1.063873051 | 0.523007753 |
| ENSMUSG00000037260 | Hgsnat | 0.000442906 | -0.735828299 | 0.664426832 |
| ENSMUSG00000031486 | Adgra2 | 0.000442906 | -1.225633566 | 0.220900379 |
| ENSMUSG00000039530 | Tusc3 | 0.000442906 | -0.949423772 | 0.458976782 |
| ENSMUSG00000109907 | Gm45321 | 0.000442906 | 1.496832458 | 0.20161331 |
| ENSMUSG00000038206 | Fbxo8 | 0.000442906 | -0.014129461 | 1 |
| ENSMUSG00000110406 | 4930470O06Rik | 0.000442906 | 2.496832458 | 0.031456064 |
| ENSMUSG00000053886 | Sh2d4a | 0.000442906 | -1.656972878 | 0.042972006 |
| ENSMUSG00000070003 | Ssbp4 | 0.000442906 | 0.146335211 | 1 |
| ENSMUSG00000052794 | 1700030K09Rik | 0.000442906 | -0.673092543 | 0.723846946 |
| ENSMUSG00000031709 | Tbc1d9 | 0.000442906 | 0.911869957 | 0.586600655 |
| ENSMUSG00000005483 | Dnajb1 | 0.000442906 | 0.392495798 | 1 |
| ENSMUSG00000002885 | Adgre5 | 0.000442906 | -0.853664789 | 0.516400387 |

| | | | | |
|---|---|---|---|---|
| ENSMUSG00000003813 | Rad23a | 0.000442906 | 0.461208548 | 1 |
| ENSMUSG00000005142 | Man2b1 | 0.000442906 | 0.42644313 | 1 |
| ENSMUSG00000036902 | Neto2 | 0.000442906 | 1.063873051 | 0.523007753 |
| ENSMUSG00000031740 | Mmp2 | 0.000442906 | 0.960779558 | 0.559918909 |
| ENSMUSG00000032939 | Nup93 | 0.000442906 | -1.158519371 | 0.254892207 |
| ENSMUSG00000031785 | Adgrg1 | 0.000442906 | -0.063882496 | 1 |
| ENSMUSG00000046707 | Csnk2a2 | 0.000442906 | -0.720398258 | 0.66416768 |
| ENSMUSG00000014856 | Tmem208 | 0.000442906 | 0.204050709 | 1 |
| ENSMUSG00000031913 | Vps4a | 0.000442906 | 0.648835551 | 0.877529363 |
| ENSMUSG00000031917 | Nip7 | 0.000442906 | 0.42644313 | 1 |
| ENSMUSG00000097393 | D030068K23Rik | 0.000442906 | 0.609307187 | 0.886203497 |
| ENSMUSG00000059854 | Hydin | 0.000442906 | -0.390692813 | 1 |
| ENSMUSG00000033658 | Ddx19b | 0.000442906 | -0.624182943 | 0.792441218 |
| ENSMUSG00000031959 | Wdr59 | 0.000442906 | -1.310522464 | 0.173990867 |
| ENSMUSG00000031835 | Mbtps1 | 0.000442906 | -1.07605721 | 0.349674462 |
| ENSMUSG00000001065 | Zfp276 | 0.000442906 | -0.429166961 | 0.971854959 |
| ENSMUSG00000056820 | Tsnax | 0.000442906 | 0.864564242 | 0.600946203 |
| ENSMUSG00000025898 | Cwf19l2 | 0.000442906 | -1.351164449 | 0.13833117 |
| ENSMUSG00000049723 | Mmp12 | 0.000442906 | 0.911869957 | 0.623042427 |
| ENSMUSG00000002820 | Atg4d | 0.000442906 | 0.063873051 | 1 |
| ENSMUSG00000032178 | Ilf3 | 0.000442906 | -1.582118883 | 0.066307071 |
| ENSMUSG00000008429 | Herpud2 | 0.000442906 | -1.214662449 | 0.235934029 |
| ENSMUSG00000016087 | Fli1 | 0.000442906 | -0.83945093 | 0.606655093 |
| ENSMUSG00000035934 | Pknox2 | 0.000442906 | 0.392495798 | 1 |
| ENSMUSG00000001942 | Siae | 0.000442906 | 0.233798052 | 1 |
| ENSMUSG00000070305 | Mpzl3 | 0.000442906 | 0.174904363 | 1 |
| ENSMUSG00000111669 | Gm31698 | 0.000442906 | -1.788569761 | 0.034011079 |
| ENSMUSG00000044229 | Nxpe4 | 0.000442906 | -1.766201948 | 0.031171876 |
| ENSMUSG00000032050 | Rdx | 0.000442906 | -1.512156325 | 0.113922494 |
| ENSMUSG00000032320 | Rcn2 | 0.000442906 | 0.146335211 | 1 |
| ENSMUSG00000040722 | Scamp5 | 0.000442906 | -0.390692813 | 1 |
| ENSMUSG00000040188 | Scamp2 | 0.000442906 | -1.214662449 | 0.244858026 |
| ENSMUSG00000032334 | Loxl1 | 0.000442906 | -0.20360726 | 1 |
| ENSMUSG00000032280 | Tle3 | 0.000442906 | -2.082106255 | 0.006684444 |
| ENSMUSG00000032254 | Kif23 | 0.000442906 | -1.351164449 | 0.13833117 |
| ENSMUSG00000040652 | Oaz2 | 0.000442906 | -0.225633566 | 1 |
| ENSMUSG00000038664 | Herc1 | 0.000442906 | -2.129606679 | 0.004533725 |
| ENSMUSG00000038535 | Zfp280d | 0.000442906 | -2.952728917 | 3.32E-05 |
| ENSMUSG00000032377 | Plscr4 | 0.000442906 | -1.247328638 | 0.219707865 |
| ENSMUSG00000041440 | Gk5 | 0.000442906 | -0.810596067 | 0.573149109 |
| ENSMUSG00000051234 | Rnf7 | 0.000442906 | 0.648835551 | 0.792441218 |
| ENSMUSG00000037784 | Dzip1l | 0.000442906 | -2.543183221 | 0.000519924 |
| ENSMUSG00000040325 | Dcaf1 | 0.000442906 | -2.117877386 | 0.005052992 |
| ENSMUSG00000010067 | Rassf1 | 0.000442906 | -0.538791452 | 0.855885826 |
| ENSMUSG00000010054 | Tusc2 | 0.000442906 | 1.063873051 | 0.472949383 |
| ENSMUSG00000032436 | Cmtm7 | 0.000442906 | -0.181239447 | 1 |
| ENSMUSG00000036737 | Oxsr1 | 0.000442906 | -0.988594369 | 0.420065063 |
| ENSMUSG00000040936 | Ulk4 | 0.000442906 | -0.225633566 | 1 |
| ENSMUSG00000032530 | Lyzl4 | 0.000442906 | -1.429166961 | 0.13351417 |
| ENSMUSG00000038145 | Snrk | 0.000442906 | -0.57355687 | 0.845086935 |
| ENSMUSG00000037949 | Ano10 | 0.000442906 | -1.341110784 | 0.181169872 |

| | | | | |
|---|---|---|---|---|
| ENSMUSG00000047036 | Zfp445 | 0.000442906 | -1.825095637 | 0.020996898 |
| ENSMUSG00000025240 | Sacm1l | 0.000442906 | -0.867739974 | 0.521275193 |
| ENSMUSG00000019851 | Perp | 0.000442906 | 0.911869957 | 0.582339791 |
| ENSMUSG00000037455 | Slc18b1 | 0.000442906 | -0.088130043 | 1 |
| ENSMUSG00000075232 | Amd1 | 0.000442906 | -0.720398258 | 0.66416768 |
| ENSMUSG00000019826 | Zbtb24 | 0.000442906 | -0.975655313 | 0.415257602 |
| ENSMUSG00000071324 | Armc2 | 0.000442906 | -0.689034087 | 0.736456705 |
| ENSMUSG00000044770 | Scml4 | 0.000442906 | -1.448025988 | 0.106203553 |
| ENSMUSG00000058298 | Mcm9 | 0.000442906 | -0.975655313 | 0.415257602 |
| ENSMUSG00000012819 | Cdh23 | 0.000442906 | -0.371064006 | 1 |
| ENSMUSG00000020088 | Sar1a | 0.000442906 | -0.640671066 | 0.716109464 |
| ENSMUSG00000033318 | Gstt2 | 0.000442906 | 0.731297712 | 0.781012813 |
| ENSMUSG00000009293 | Ube2g2 | 0.000442906 | -0.088130043 | 1 |
| ENSMUSG00000000374 | Trappc10 | 0.000442906 | -0.704801403 | 0.66416768 |
| ENSMUSG00000001211 | Agpat3 | 0.000442906 | -1.590630383 | 0.059481058 |
| ENSMUSG00000020156 | Pwwp3a | 0.000442906 | -1.410058138 | 0.114388824 |
| ENSMUSG00000048696 | Mex3d | 0.000442906 | 1.42644313 | 0.304871131 |
| ENSMUSG00000020167 | Tcf3 | 0.000442906 | -1.100102684 | 0.310034885 |
| ENSMUSG00000035215 | Lsm7 | 0.000442906 | -0.371064006 | 1 |
| ENSMUSG00000004934 | Pias4 | 0.000442906 | -0.795949291 | 0.568577582 |
| ENSMUSG00000020248 | Nfyb | 0.000442906 | 0.570833039 | 0.886203497 |
| ENSMUSG00000020053 | Igf1 | 0.000442906 | -0.088130043 | 1 |
| ENSMUSG00000004356 | Utp20 | 0.000442906 | -1.720398258 | 0.033147592 |
| ENSMUSG00000036446 | Lum | 0.000442906 | 0.359328934 | 1 |
| ENSMUSG00000019971 | Cep290 | 0.000442906 | -1.766201948 | 0.027469977 |
| ENSMUSG00000063334 | Krr1 | 0.000442906 | -0.975655313 | 0.415257602 |
| ENSMUSG00000020166 | Cnot2 | 0.000442906 | -1.94279068 | 0.012749115 |
| ENSMUSG00000064181 | Rab3ip | 0.000442906 | -1.83945093 | 0.022106374 |
| ENSMUSG00000025353 | Ormdl2 | 0.000442906 | 1.496832458 | 0.298197303 |
| ENSMUSG00000034543 | Morc2a | 0.000442906 | -1.529967601 | 0.080542848 |
| ENSMUSG00000020432 | Tcn2 | 0.000442906 | -0.014129461 | 1 |
| ENSMUSG00000034493 | 4930556J24Rik | 0.000442906 | 1.731297712 | 0.145553211 |
| ENSMUSG00000051650 | B3gnt2 | 0.000442906 | -1.52108945 | 0.078980179 |
| ENSMUSG00000020459 | Mtif2 | 0.000442906 | -1.158519371 | 0.254892207 |
| ENSMUSG00000069917 | Hba-a2 | 0.000442906 | -1.982139347 | 0.180537169 |
| ENSMUSG00000069892 | 9930111J21Rik2 | 0.000442906 | 0.011405631 | 1 |
| ENSMUSG00000036264 | Fstl4 | 0.000442906 | -0.52108945 | 0.949822782 |
| ENSMUSG00000049755 | Zfp672 | 0.000442906 | -1.43862729 | 0.103956898 |
| ENSMUSG00000020444 | Guk1 | 0.000442906 | 0.960779558 | 0.559918909 |
| ENSMUSG00000002812 | Flii | 0.000442906 | 0.392495798 | 1 |
| ENSMUSG00000042569 | Dhrs7b | 0.000442906 | -1.158519371 | 0.257907731 |
| ENSMUSG00000046731 | Kctd11 | 0.000442906 | 1.295198597 | 0.373169426 |
| ENSMUSG00000018559 | Ctdnep1 | 0.000442906 | 0.359328934 | 1 |
| ENSMUSG00000093989 | Rnasek | 0.000442906 | 0.648835551 | 0.787439548 |
| ENSMUSG00000018287 | Spag7 | 0.000442906 | 0.496832458 | 0.920245613 |
| ENSMUSG00000040712 | Camta2 | 0.000442906 | -1.341110784 | 0.175382301 |
| ENSMUSG00000040667 | Nup88 | 0.000442906 | -0.039220442 | 1 |
| ENSMUSG00000005947 | Itgae | 0.000442906 | 1.011405631 | 0.587484274 |
| ENSMUSG00000037958 | Nsrp1 | 0.000442906 | -1.681085334 | 0.039214372 |
| ENSMUSG00000011877 | Git1 | 0.000442906 | 0.533358334 | 0.894464956 |
| ENSMUSG00000017386 | Traf4 | 0.000442906 | -1.063882496 | 0.345490666 |

| | | | | |
|---|---|---|---|---|
| ENSMUSG00000034940 | Synrg | 0.000442906 | -0.83945093 | 0.523546561 |
| ENSMUSG00000018697 | Aatf | 0.000442906 | -1.607504202 | 0.051667273 |
| ENSMUSG00000034329 | Brip1 | 0.000442906 | -3.132524162 | 1.60E-05 |
| ENSMUSG00000018377 | Vezf1 | 0.000442906 | -0.556278878 | 0.866287757 |
| ENSMUSG00000033983 | Coil | 0.000442906 | -1.330986567 | 0.154833135 |
| ENSMUSG00000020541 | Tom1l1 | 0.000442906 | -1.475860196 | 0.096607172 |
| ENSMUSG00000038845 | Phb | 0.000442906 | 0.264171701 | 1 |
| ENSMUSG00000018669 | Cdk5rap3 | 0.000442906 | 0.295198597 | 1 |
| ENSMUSG00000017221 | Psmd3 | 0.000442906 | 2.011405631 | 0.082214652 |
| ENSMUSG00000037992 | Rara | 0.000442906 | -1.810596067 | 0.023147462 |
| ENSMUSG00000020914 | Top2a | 0.000442906 | -1.503167542 | 0.092421376 |
| ENSMUSG00000037935 | Smarce1 | 0.000442906 | -0.781152289 | 0.630177036 |
| ENSMUSG00000020919 | Stat5b | 0.000442906 | -0.704801403 | 0.66416768 |
| ENSMUSG00000020740 | Gga3 | 0.000442906 | -0.825095637 | 0.57709448 |
| ENSMUSG00000020752 | Recql5 | 0.000442906 | -0.225633566 | 1 |
| ENSMUSG00000025580 | Eif4a3 | 0.000442906 | -0.039220442 | 1 |
| ENSMUSG00000025137 | Pcyt2 | 0.000442906 | -1.07605721 | 0.349674462 |
| ENSMUSG00000025142 | Aspscr1 | 0.000442906 | 0.233798052 | 1 |
| ENSMUSG00000025162 | Csnk1d | 0.000442906 | -0.853664789 | 0.516400387 |
| ENSMUSG00000020589 | Fam49a | 0.000442906 | 0.609307187 | 0.886203497 |
| ENSMUSG00000036613 | Eipr1 | 0.000442906 | -0.624182943 | 0.791749236 |
| ENSMUSG00000020572 | Nampt | 0.000442906 | -0.624182943 | 0.791749236 |
| ENSMUSG00000020598 | Nrcam | 0.000442906 | -0.158519371 | 1 |
| ENSMUSG00000046314 | Stxbp6 | 0.000442906 | -2.792264244 | 0.000106037 |
| ENSMUSG00000043998 | Mgat2 | 0.000442906 | -0.039220442 | 1 |
| ENSMUSG00000021072 | Tmx1 | 0.000442906 | -0.135435758 | 1 |
| ENSMUSG00000021140 | Pcnx | 0.000442906 | -1.599091962 | 0.058882736 |
| ENSMUSG00000021222 | Dcaf4 | 0.000442906 | -1.039220442 | 0.387414024 |
| ENSMUSG00000019969 | Psen1 | 0.000442906 | -1.503167542 | 0.088040549 |
| ENSMUSG00000004789 | Dlst | 0.000442906 | -1.289763904 | 0.169119671 |
| ENSMUSG00000021038 | Vipas39 | 0.000442906 | -1.147023732 | 0.284698396 |
| ENSMUSG00000041781 | Cpsf2 | 0.000442906 | -0.895484965 | 0.471405909 |
| ENSMUSG00000041771 | Slc24a4 | 0.000442906 | 1.731297712 | 0.172546568 |
| ENSMUSG00000021260 | Hhipl1 | 0.000442906 | -0.962599161 | 0.463155604 |
| ENSMUSG00000037787 | Apopt1 | 0.000442906 | -1.088130043 | 0.451099851 |
| ENSMUSG00000011158 | Brf1 | 0.000442906 | -2.452702475 | 0.000887428 |
| ENSMUSG00000071551 | Akr1c19 | 0.000442906 | -0.158519371 | 1 |
| ENSMUSG00000036006 | Ripor2 | 0.000442906 | -1.599091962 | 0.078980382 |
| ENSMUSG00000113061 | Gm11361 | 0.000442906 | 1.496832458 | 0.20161331 |
| ENSMUSG00000016477 | E2f3 | 0.000442906 | -1.915949067 | 0.013369592 |
| ENSMUSG00000021373 | Cap2 | 0.000442906 | 0.864564242 | 0.618078673 |
| ENSMUSG00000038025 | Phf2 | 0.000442906 | -0.962599161 | 0.463155604 |
| ENSMUSG00000021481 | Zfp346 | 0.000442906 | -0.371064006 | 1 |
| ENSMUSG00000021470 | Ercc6l2 | 0.000442906 | -1.969142007 | 0.010394537 |
| ENSMUSG00000017756 | Slc12a7 | 0.000442906 | -1.485020195 | 0.084487163 |
| ENSMUSG00000034152 | Exoc3 | 0.000442906 | -0.795949291 | 0.568577582 |
| ENSMUSG00000021597 | Slf1 | 0.000442906 | -1.100102684 | 0.310034885 |
| ENSMUSG00000021614 | Vcan | 0.000442906 | -1.881679165 | 0.016000576 |
| ENSMUSG00000021619 | Atg10 | 0.000442906 | -1.380911792 | 0.126838325 |
| ENSMUSG00000003992 | Ssbp2 | 0.000442906 | -1.147023732 | 0.284698396 |
| ENSMUSG00000042015 | Wdr41 | 0.000442906 | -0.135435758 | 1 |

| | | | | |
|---|---|---|---|---|
| ENSMUSG00000046169 | Adamts6 | 0.000442906 | -1.902338636 | 0.014749215 |
| ENSMUSG00000114564 | Gm10739 | 0.000442906 | 1.818760553 | 0.109855104 |
| ENSMUSG00000021759 | Plpp1 | 0.000442906 | -1.147023732 | 0.284698396 |
| ENSMUSG00000074634 | Tmem267 | 0.000442906 | 0.011405631 | 1 |
| ENSMUSG00000021782 | Dlg5 | 0.000442906 | -2.129606679 | 0.004533725 |
| ENSMUSG00000042286 | Stab1 | 0.000442906 | 0.496832458 | 0.909161999 |
| ENSMUSG00000021911 | Parg | 0.000442906 | -2.336057556 | 0.001528947 |
| ENSMUSG00000054051 | Ercc6 | 0.000442906 | -1.860719547 | 0.017363741 |
| ENSMUSG00000037697 | Ddhd1 | 0.000442906 | -2.147023732 | 0.004232796 |
| ENSMUSG00000000958 | Slc7a7 | 0.000442906 | -1.624182943 | 0.053730218 |
| ENSMUSG00000022204 | Ngdn | 0.000442906 | -0.014129461 | 1 |
| ENSMUSG00000022210 | Dhrs4 | 0.000442906 | 0.392495798 | 1 |
| ENSMUSG00000022214 | Dcaf11 | 0.000442906 | -1.169924134 | 0.257688304 |
| ENSMUSG00000021987 | Mtmr6 | 0.000442906 | -1.330986567 | 0.154833135 |
| ENSMUSG00000034463 | Scara3 | 0.000442906 | -2.330986567 | 0.001701831 |
| ENSMUSG00000048922 | Cdca2 | 0.000442906 | -2.466641666 | 0.000849494 |
| ENSMUSG00000034248 | Slc25a37 | 0.000442906 | -4.211906583 | 4.50E-10 |
| ENSMUSG00000022092 | Ppp3cc | 0.000442906 | -0.673092543 | 0.723846946 |
| ENSMUSG00000022105 | Rb1 | 0.000442906 | -1.30018052 | 0.171582618 |
| ENSMUSG00000022014 | Epsti1 | 0.000442906 | 0.233798052 | 1 |
| ENSMUSG00000022016 | Akap11 | 0.000442906 | -2.147023732 | 0.004232796 |
| ENSMUSG00000022020 | Naa16 | 0.000442906 | -2.27927153 | 0.002230174 |
| ENSMUSG00000033166 | Dis3 | 0.000442906 | -1.147023732 | 0.284698396 |
| ENSMUSG00000022136 | Dnajc3 | 0.000442906 | -1.135435758 | 0.281150145 |
| ENSMUSG00000000197 | Nalcn | 0.000442906 | -0.225633566 | 1 |
| ENSMUSG00000032925 | Itgbl1 | 0.000442906 | -0.949423772 | 0.466464457 |
| ENSMUSG00000048424 | Ranbp3l | 0.000442906 | 0.864564242 | 0.608155934 |
| ENSMUSG00000039458 | Mtmr12 | 0.000442906 | -1.00141841 | 0.424188363 |
| ENSMUSG00000052253 | Zfp622 | 0.000442906 | 0.533358334 | 0.896319431 |
| ENSMUSG00000022269 | 11-Mar | 0.000442906 | 1.731297712 | 0.162298752 |
| ENSMUSG00000022323 | Rida | 0.000442906 | -1.100102684 | 0.310034885 |
| ENSMUSG00000002524 | Puf60 | 0.000442906 | 0.818760553 | 0.717242831 |
| ENSMUSG00000071711 | Mpst | 0.000442906 | 3.818760553 | 0.001289676 |
| ENSMUSG00000033128 | Gga1 | 0.000442906 | -1.088130043 | 0.306626259 |
| ENSMUSG00000033088 | Triobp | 0.000442906 | -0.867739974 | 0.521275193 |
| ENSMUSG00000033055 | Ankrd54 | 0.000442906 | 0.090840098 | 1 |
| ENSMUSG00000022414 | Tab1 | 0.000442906 | -0.640671066 | 0.716109464 |
| ENSMUSG00000022390 | Zc3h7b | 0.000442906 | -1.30018052 | 0.171582618 |
| ENSMUSG00000041815 | Poldip3 | 0.000442906 | -2.494122402 | 0.000684267 |
| ENSMUSG00000035828 | Pim3 | 0.000442906 | 0.392495798 | 1 |
| ENSMUSG00000048175 | Asb8 | 0.000442906 | -0.607504202 | 0.789505358 |
| ENSMUSG00000022994 | Adcy6 | 0.000442906 | -3.010962182 | 2.16E-05 |
| ENSMUSG00000023021 | Cers5 | 0.000442906 | -1.720398258 | 0.036113193 |
| ENSMUSG00000014232 | Cluap1 | 0.000442906 | -2.198047735 | 0.003491159 |
| ENSMUSG00000039738 | Slx4 | 0.000442906 | -0.448025988 | 1 |
| ENSMUSG00000014301 | Pam16 | 0.000442906 | 1.118320835 | 0.454909192 |
| ENSMUSG00000039637 | Coro7 | 0.000442906 | -1.100102684 | 0.310034885 |
| ENSMUSG00000022679 | Mpv17l | 0.000442906 | -0.503167542 | 0.896653623 |
| ENSMUSG00000022677 | Fopnl | 0.000442906 | -0.330986567 | 1 |
| ENSMUSG00000071637 | Cebpd | 0.000442906 | 1.174904363 | 0.395423178 |
| ENSMUSG00000003528 | Slc25a1 | 0.000442906 | 0.392495798 | 1 |

| | | | | |
|---|---|---|---|---|
| ENSMUSG00000004460 | Dnajb11 | 0.000442906 | 0.609307187 | 0.886203497 |
| ENSMUSG00000022828 | Gtf2e1 | 0.000442906 | -0.881679165 | 0.467537667 |
| ENSMUSG00000022820 | Ndufb4 | 0.000442906 | -0.181239447 | 1 |
| ENSMUSG00000022637 | Cblb | 0.000442906 | -2.057756394 | 0.006984748 |
| ENSMUSG00000022965 | Ifngr2 | 0.000442906 | 0.090840098 | 1 |
| ENSMUSG00000022962 | Gart | 0.000442906 | -1.268702288 | 0.198085003 |
| ENSMUSG00000058240 | Cryzl1 | 0.000442906 | -1.781152289 | 0.024569482 |
| ENSMUSG00000039680 | Mrps6 | 0.000442906 | -0.795949291 | 0.568577582 |
| ENSMUSG00000034265 | Zdhhc14 | 0.000442906 | -0.225633566 | 1 |
| ENSMUSG00000023830 | Igf2r | 0.000442906 | -0.704801403 | 0.66416768 |
| ENSMUSG00000014773 | Dll1 | 0.000442906 | 0.911869957 | 0.582339791 |
| ENSMUSG00000014771 | Pdcd2 | 0.000442906 | 1.648835551 | 0.1477368 |
| ENSMUSG00000036820 | Amdhd2 | 0.000442906 | -0.310522464 | 1 |
| ENSMUSG00000024137 | E4f1 | 0.000442906 | -0.766201948 | 0.624849798 |
| ENSMUSG00000002504 | Slc9a3r2 | 0.000442906 | -0.158519371 | 1 |
| ENSMUSG00000002257 | Def6 | 0.000442906 | -0.590630383 | 0.786092784 |
| ENSMUSG00000034673 | Pbx2 | 0.000442906 | -0.225633566 | 1 |
| ENSMUSG00000024392 | Bag6 | 0.000442906 | -0.268702288 | 1 |
| ENSMUSG00000042419 | Nfkbil1 | 0.000442906 | 0.204050709 | 1 |
| ENSMUSG00000024457 | Trim26 | 0.000442906 | -1.100102684 | 0.310034885 |
| ENSMUSG00000036398 | Ppp1r11 | 0.000442906 | 1.295198597 | 0.34594864 |
| ENSMUSG00000023967 | Mrps18a | 0.000442906 | 0.359328934 | 1 |
| ENSMUSG00000023972 | Ptk7 | 0.000442906 | -1.341110784 | 0.162257313 |
| ENSMUSG00000040771 | Oard1 | 0.000442906 | 0.731297712 | 0.778426679 |
| ENSMUSG00000002661 | Alkbh7 | 0.000442906 | 3.233798052 | 0.004321381 |
| ENSMUSG00000023460 | Rab12 | 0.000442906 | -1.457363852 | 0.093377644 |
| ENSMUSG00000024064 | Galnt14 | 0.000442906 | 0.204050709 | 1 |
| ENSMUSG00000024066 | Xdh | 0.000442906 | -3.225633566 | 4.64E-06 |
| ENSMUSG00000024078 | Ttc27 | 0.000442906 | -1.485020195 | 0.084487163 |
| ENSMUSG00000024097 | Srsf7 | 0.000442906 | -1.361148537 | 0.140513734 |
| ENSMUSG00000024270 | Slc39a6 | 0.000442906 | 0.037400839 | 1 |
| ENSMUSG00000024383 | Map3k2 | 0.000442906 | -2.414859124 | 0.00105249 |
| ENSMUSG00000003778 | Brd8 | 0.000442906 | -1.773696484 | 0.028062645 |
| ENSMUSG00000046668 | Cxxc5 | 0.000442906 | 0.063873051 | 1 |
| ENSMUSG00000024477 | Pggt1b | 0.000442906 | -1.026729498 | 0.383513703 |
| ENSMUSG00000038059 | Smim3 | 0.000442906 | -0.014129461 | 1 |
| ENSMUSG00000054008 | Ndst1 | 0.000442906 | -1.07605721 | 0.349674462 |
| ENSMUSG00000042439 | Zfp532 | 0.000442906 | -2.410058138 | 0.001021934 |
| ENSMUSG00000041891 | Lman1 | 0.000442906 | -0.766201948 | 0.624849798 |
| ENSMUSG00000024521 | Pmaip1 | 0.000442906 | 0.204050709 | 1 |
| ENSMUSG00000057130 | Txnl4a | 0.000442906 | 0.090840098 | 1 |
| ENSMUSG00000033016 | Nfatc1 | 0.000442906 | -1.225633566 | 0.207756141 |
| ENSMUSG00000024614 | Tmx3 | 0.000442906 | -1.758668275 | 0.031083987 |
| ENSMUSG00000100969 | 1700030N03Rik | 0.000442906 | 1.174904363 | 0.396731698 |
| ENSMUSG00000024829 | Mrpl21 | 0.000442906 | -1.07605721 | 0.37296511 |
| ENSMUSG00000035372 | 1810055G02Rik | 0.000442906 | -1.371064006 | 0.14342984 |
| ENSMUSG00000034616 | Ssh3 | 0.000442906 | 0.204050709 | 1 |
| ENSMUSG00000034108 | Ccs | 0.000442906 | -1.704801403 | 0.04031668 |
| ENSMUSG00000024826 | Dpf2 | 0.000442906 | -2.07605721 | 0.006539077 |
| ENSMUSG00000071662 | Polr2g | 0.000442906 | 0.063873051 | 1 |
| ENSMUSG00000071657 | Bscl2 | 0.000442906 | 0.864564242 | 0.600946203 |

| | | | | |
|---|---|---|---|---|
| ENSMUSG00000071650 | Ganab | 0.000442906 | -0.975655313 | 0.415257602 |
| ENSMUSG00000024660 | Incenp | 0.000442906 | -1.766201948 | 0.027469977 |
| ENSMUSG00000036192 | Rorb | 0.000442906 | 0.174904363 | 1 |
| ENSMUSG00000038658 | Ric1 | 0.000442906 | -1.27927153 | 0.201326391 |
| ENSMUSG00000018821 | Avpi1 | 0.000442906 | 0.818760553 | 0.717242831 |
| ENSMUSG00000044345 | Marveld1 | 0.000442906 | -1.371064006 | 0.14342984 |
| ENSMUSG00000025207 | Sema4g | 0.000442906 | 0.118320835 | 1 |
| ENSMUSG00000041035 | Gm17018 | 0.000442906 | 0.648835551 | 0.792441218 |
| ENSMUSG00000025225 | Nfkb2 | 0.000442906 | -1.158519371 | 0.254892207 |
| ENSMUSG00000036748 | Cuedc2 | 0.000442906 | -0.735828299 | 0.664426832 |
| ENSMUSG00000043639 | Rbm20 | 0.000442906 | -0.310522464 | 1 |
| ENSMUSG00000033478 | Fam160b1 | 0.000442906 | -1.051604167 | 0.35315595 |
| ENSMUSG00000058254 | Tspan7 | 0.000442906 | -0.039220442 | 1 |
| ENSMUSG00000008035 | Mid1ip1 | 0.000442906 | 0.864564242 | 0.686735271 |
| ENSMUSG00000048970 | C1galt1c1 | 0.000442906 | -0.640671066 | 0.716109464 |
| ENSMUSG00000036916 | Zfp280c | 0.000442906 | -1.43862729 | 0.103956898 |
| ENSMUSG00000050029 | Rap2c | 0.000442906 | -1.111976785 | 0.314582476 |
| ENSMUSG00000031119 | Gpc4 | 0.000442906 | -1.27927153 | 0.194109095 |
| ENSMUSG00000002010 | Idh3g | 0.000442906 | 0.118320835 | 1 |
| ENSMUSG00000031388 | Naa10 | 0.000442906 | 1.42644313 | 0.304871131 |
| ENSMUSG00000031403 | Dkc1 | 0.000442906 | -1.100102684 | 0.310034885 |
| ENSMUSG00000031196 | F8 | 0.000442906 | 0.326907457 | 1 |
| ENSMUSG00000031221 | Igbp1 | 0.000442906 | -0.135435758 | 1 |
| ENSMUSG00000045427 | Hnrnph2 | 0.000442906 | -0.063882496 | 1 |
| ENSMUSG00000046432 | Bex3 | 0.000442906 | 1.42644313 | 0.304871131 |
| ENSMUSG00000041710 | Trpc5 | 0.000442906 | 1.359328934 | 0.392708123 |
| ENSMUSG00000025265 | Fgd1 | 0.000442906 | -0.039220442 | 1 |
| ENSMUSG00000031292 | Cdkl5 | 0.000442906 | -0.83945093 | 0.512602663 |
| ENSMUSG00000059493 | Nhs | 0.000442906 | -1.111976785 | 0.34594864 |
| ENSMUSG00000040855 | Reps2 | 0.000442906 | -1.640671066 | 0.048860005 |
| ENSMUSG00000079316 | Rab9 | 0.000442906 | 1.063873051 | 0.523007753 |
| ENSMUSG00000025915 | Sgk3 | 0.000387543 | -0.459688905 | 0.946435944 |
| ENSMUSG00000025938 | Slco5a1 | 0.000387543 | -1.106051951 | 0.383025294 |
| ENSMUSG00000086727 | 4931428L18Rik | 0.000387543 | 0.948395833 | 0.653235135 |
| ENSMUSG00000026134 | Prim2 | 0.000387543 | -1.1966545 | 0.275000936 |
| ENSMUSG00000001143 | Lman2l | 0.000387543 | -1.339849135 | 0.180637196 |
| ENSMUSG00000087589 | D430040D24Rik | 0.000387543 | 2.47891055 | 0.036895541 |
| ENSMUSG00000026077 | Npas2 | 0.000387543 | -0.395558568 | 1 |
| ENSMUSG00000048234 | Rnf149 | 0.000387543 | 0.604441432 | 0.857399533 |
| ENSMUSG00000025986 | Slc39a10 | 0.000387543 | -0.89032326 | 0.526306254 |
| ENSMUSG00000096141 | Dnah7a | 0.000387543 | 1.189403933 | 0.466464457 |
| ENSMUSG00000026023 | Cdk15 | 0.000387543 | 0.0942467 | 1 |
| ENSMUSG00000025964 | Adam23 | 0.000387543 | 0.790854556 | 0.699112874 |
| ENSMUSG00000039372 | 4-Mar | 0.000387543 | 1.004979362 | 0.663984692 |
| ENSMUSG00000026185 | Igfbp5 | 0.000387543 | 0.291283547 | 1 |
| ENSMUSG00000026142 | Rhbdd1 | 0.000387543 | -0.654945197 | 0.76683182 |
| ENSMUSG00000026239 | Pde6d | 0.000387543 | -1.245982212 | 0.217655393 |
| ENSMUSG00000034432 | Cops8 | 0.000387543 | 0.222570797 | 1 |
| ENSMUSG00000026280 | Atg4b | 0.000387543 | -0.995020638 | 0.466464457 |
| ENSMUSG00000009907 | Vps4b | 0.000387543 | -1.664047404 | 0.049569528 |
| ENSMUSG00000026436 | Elk4 | 0.000387543 | -1.734868741 | 0.042660478 |

| | | | | |
|---|---|---|---|---|
| ENSMUSG00000026426 | Arl8a | 0.000387543 | -0.540989007 | 0.8730922 |
| ENSMUSG00000041498 | Kif14 | 0.000387543 | 0.56137271 | 0.98880775 |
| ENSMUSG00000018199 | Ro60 | 0.000387543 | 0.741944956 | 0.840804483 |
| ENSMUSG00000018189 | Uchl5 | 0.000387543 | -0.184054463 | 1 |
| ENSMUSG00000026484 | Rnf2 | 0.000387543 | -0.560617814 | 0.875465725 |
| ENSMUSG00000026578 | Ccdc181 | 0.000387543 | 1.125273595 | 0.466464457 |
| ENSMUSG00000026575 | Nme7 | 0.000387543 | -0.579983139 | 0.806496094 |
| ENSMUSG00000038463 | Olfml2b | 0.000387543 | 1.004979362 | 0.633493255 |
| ENSMUSG00000052423 | B4galt3 | 0.000387543 | -1.233807498 | 0.249952095 |
| ENSMUSG00000038034 | Igsf8 | 0.000387543 | -0.654945197 | 0.741406232 |
| ENSMUSG00000026526 | Fh1 | 0.000387543 | 0.519552534 | 0.996628004 |
| ENSMUSG00000063659 | Zbtb18 | 0.000387543 | -1.171343411 | 0.267806521 |
| ENSMUSG00000015961 | Adss | 0.000387543 | -1.874726405 | 0.020746096 |
| ENSMUSG00000058729 | Lin9 | 0.000387543 | -0.43862729 | 1 |
| ENSMUSG00000026516 | Nvl | 0.000387543 | -1.654945197 | 0.05650534 |
| ENSMUSG00000072258 | Taf1a | 0.000387543 | -1.145580315 | 0.300304685 |
| ENSMUSG00000089872 | Rps6kc1 | 0.000387543 | -1.943621486 | 0.020043687 |
| ENSMUSG00000016481 | Cr1l | 0.000387543 | -1.184054463 | 0.271553405 |
| ENSMUSG00000109865 | Hspa14 | 0.000387543 | -0.023589791 | 1 |
| ENSMUSG00000045967 | Gpr158 | 0.000387543 | -0.051604167 | 1 |
| ENSMUSG00000004113 | Cacna1b | 0.000387543 | 1.189403933 | 0.497329546 |
| ENSMUSG00000036833 | Pnpla7 | 0.000387543 | -1.158519371 | 0.320528813 |
| ENSMUSG00000015488 | Cacfd1 | 0.000387543 | -0.233807498 | 1 |
| ENSMUSG00000026785 | Pkn3 | 0.000387543 | -0.858959089 | 0.51724245 |
| ENSMUSG00000039476 | Prrx2 | 0.000387543 | -0.051604167 | 1 |
| ENSMUSG00000038740 | Mvb12b | 0.000387543 | -0.209145444 | 1 |
| ENSMUSG00000038718 | Pbx3 | 0.000387543 | -1.866864287 | 0.023669966 |
| ENSMUSG00000036641 | Ccdc148 | 0.000387543 | -0.89032326 | 0.553628878 |
| ENSMUSG00000034848 | Ttc21b | 0.000387543 | -0.89032326 | 0.526094357 |
| ENSMUSG00000027010 | Slc25a12 | 0.000387543 | -0.560617814 | 0.916860107 |
| ENSMUSG00000051747 | Ttn | 0.000387543 | -0.328444372 | 1 |
| ENSMUSG00000034683 | Ppp1r1c | 0.000387543 | 1.326907457 | 0.44168045 |
| ENSMUSG00000059588 | Calcrl | 0.000387543 | -0.373532262 | 1 |
| ENSMUSG00000050043 | Tmx2 | 0.000387543 | -0.106051951 | 1 |
| ENSMUSG00000002111 | Spi1 | 0.000387543 | 1.125273595 | 0.466464457 |
| ENSMUSG00000027215 | Cd82 | 0.000387543 | -0.023589791 | 1 |
| ENSMUSG00000027198 | Ext2 | 0.000387543 | -1.065409966 | 0.391016627 |
| ENSMUSG00000027194 | Ttc17 | 0.000387543 | -1.145580315 | 0.300304685 |
| ENSMUSG00000027188 | Pamr1 | 0.000387543 | -1.777429203 | 0.034288602 |
| ENSMUSG00000027177 | Hipk3 | 0.000387543 | -1.065409966 | 0.383551613 |
| ENSMUSG00000087203 | Gm13986 | 0.000387543 | 0.0942467 | 1 |
| ENSMUSG00000027314 | Dll4 | 0.000387543 | -0.579983139 | 0.793880199 |
| ENSMUSG00000027272 | Ubr1 | 0.000387543 | -2.030644548 | 0.010934137 |
| ENSMUSG00000005802 | Slc30a4 | 0.000387543 | 0.893948049 | 0.733004386 |
| ENSMUSG00000005804 | Bloc1s6 | 0.000387543 | 0.841480629 | 0.664309971 |
| ENSMUSG00000091337 | Eid1 | 0.000387543 | 0.326907457 | 1 |
| ENSMUSG00000027207 | Galk2 | 0.000387543 | -0.794107944 | 0.646349374 |
| ENSMUSG00000027398 | Il1b | 0.000387543 | 4.063873051 | 0.001705734 |
| ENSMUSG00000027408 | Cpxm1 | 0.000387543 | 0.189403933 | 1 |
| ENSMUSG00000074796 | Slc4a11 | 0.000387543 | 0.893948049 | 0.66416768 |
| ENSMUSG00000027318 | Adam33 | 0.000387543 | -0.9057533 | 0.544908989 |

| | | | | |
|---|---|---|---|---|
| ENSMUSG00000062098 | Btbd3 | 0.000387543 | -1.171343411 | 0.267806521 |
| ENSMUSG00000002728 | Naa20 | 0.000387543 | -0.858959089 | 0.51724245 |
| ENSMUSG00000027455 | Nsfl1c | 0.000387543 | 0.0942467 | 1 |
| ENSMUSG00000027475 | Kif3b | 0.000387543 | -1.80237556 | 0.031456064 |
| ENSMUSG00000056941 | Commd7 | 0.000387543 | -0.305360759 | 1 |
| ENSMUSG00000027488 | Snta1 | 0.000387543 | -0.810596067 | 0.587484274 |
| ENSMUSG00000000876 | Pxmp4 | 0.000387543 | -0.351164449 | 1 |
| ENSMUSG00000061689 | Dlgap4 | 0.000387543 | -1.293678954 | 0.198966953 |
| ENSMUSG00000074625 | Arhgap40 | 0.000387543 | -1.459688905 | 0.13735263 |
| ENSMUSG00000017299 | Dnttip1 | 0.000387543 | -1.293678954 | 0.200520458 |
| ENSMUSG00000045822 | Zswim3 | 0.000387543 | -0.72620388 | 0.678664944 |
| ENSMUSG00000027678 | Ncoa3 | 0.000387543 | -2.221529168 | 0.003846186 |
| ENSMUSG00000039501 | Znfx1 | 0.000387543 | -0.540989007 | 0.8730922 |
| ENSMUSG00000039263 | Npepl1 | 0.000387543 | 0.790854556 | 0.69941771 |
| ENSMUSG00000074555 | Gm10714 | 0.000387543 | 1.648835551 | 0.189783899 |
| ENSMUSG00000039041 | Adrm1 | 0.000387543 | 0.948395833 | 0.653235135 |
| ENSMUSG00000049401 | Ogfr | 0.000387543 | 0.004979362 | 1 |
| ENSMUSG00000038705 | Gmeb2 | 0.000387543 | -1.158519371 | 0.305264898 |
| ENSMUSG00000038685 | Rtel1 | 0.000387543 | -0.636566667 | 0.737916568 |
| ENSMUSG00000027589 | Pcmtd2 | 0.000387543 | -0.579983139 | 0.793880199 |
| ENSMUSG00000027671 | Actl6a | 0.000387543 | -1.065409966 | 0.374573031 |
| ENSMUSG00000027674 | Pex5l | 0.000387543 | 0.400908038 | 1 |
| ENSMUSG00000027770 | Dhx36 | 0.000387543 | -1.810596067 | 0.027621844 |
| ENSMUSG00000104093 | A330015K06Rik | 0.000387543 | 0.363433333 | 1 |
| ENSMUSG00000027956 | Tmem144 | 0.000387543 | 0.56137271 | 0.886203497 |
| ENSMUSG00000041842 | Fhdc1 | 0.000387543 | 0.841480629 | 0.684284975 |
| ENSMUSG00000098912 | 1500004A13Rik | 0.000387543 | -0.328444372 | 1 |
| ENSMUSG00000015748 | Prpf3 | 0.000387543 | -2.119348773 | 0.006319812 |
| ENSMUSG00000038256 | Bcl9 | 0.000387543 | -1.427980046 | 0.128972787 |
| ENSMUSG00000027879 | Sec22b | 0.000387543 | -0.023589791 | 1 |
| ENSMUSG00000004233 | Wars2 | 0.000387543 | -0.89032326 | 0.54950665 |
| ENSMUSG00000104543 | Gm6602 | 0.000387543 | 0.604441432 | 0.876040999 |
| ENSMUSG00000033439 | Trmt13 | 0.000387543 | -1.52108945 | 0.096053423 |
| ENSMUSG00000050315 | Synpo2 | 0.000387543 | 0.125273595 | 1 |
| ENSMUSG00000074238 | Ap1ar | 0.000387543 | -1.834980226 | 0.025141936 |
| ENSMUSG00000105816 | D030025E07Rik | 0.000387543 | 1.189403933 | 0.466464457 |
| ENSMUSG00000105331 | Gm29865 | 0.000387543 | -0.500911568 | 0.971854959 |
| ENSMUSG00000028013 | Ppa2 | 0.000387543 | -0.480447466 | 0.949822782 |
| ENSMUSG00000028271 | Gtf2b | 0.000387543 | -0.023589791 | 1 |
| ENSMUSG00000028195 | Ccn1 | 0.000387543 | -0.106051951 | 1 |
| ENSMUSG00000040086 | Tnni3k | 0.000387543 | -0.305360759 | 1 |
| ENSMUSG00000040016 | Ptger3 | 0.000387543 | 0.47891055 | 1 |
| ENSMUSG00000063052 | Lrrc40 | 0.000387543 | 0.156982455 | 1 |
| ENSMUSG00000040738 | Ints8 | 0.000387543 | -2.152064348 | 0.006222976 |
| ENSMUSG00000040536 | Necab1 | 0.000387543 | 1.004979362 | 0.633493255 |
| ENSMUSG00000028226 | Mmp16 | 0.000387543 | 0.741944956 | 0.850854802 |
| ENSMUSG00000073991 | Cnbd1 | 0.000387543 | 1.125273595 | 0.466464457 |
| ENSMUSG00000040455 | Usp45 | 0.000387543 | -0.760555385 | 0.66416768 |
| ENSMUSG00000085885 | Gm11906 | 0.000387543 | 0.222570797 | 1 |
| ENSMUSG00000055373 | Fut9 | 0.000387543 | 0.189403933 | 1 |
| ENSMUSG00000028277 | Ube2j1 | 0.000387543 | -0.209145444 | 1 |

| | | | | |
|---|---|---|---|---|
| ENSMUSG00000028291 | Akirin2 | 0.000387543 | -1.158519371 | 0.305264898 |
| ENSMUSG00000036822 | Topors | 0.000387543 | -1.743481871 | 0.036756237 |
| ENSMUSG00000028461 | Ccdc107 | 0.000387543 | 1.189403933 | 0.455131546 |
| ENSMUSG00000028479 | Gne | 0.000387543 | -0.305360759 | 1 |
| ENSMUSG00000035601 | Trmt10b | 0.000387543 | 0.893948049 | 0.66416768 |
| ENSMUSG00000028334 | Nans | 0.000387543 | 0.56137271 | 0.98880775 |
| ENSMUSG00000028339 | Col15a1 | 0.000387543 | -0.760555385 | 0.636683133 |
| ENSMUSG00000028344 | Invs | 0.000387543 | -1.43862729 | 0.113452361 |
| ENSMUSG00000028345 | Tex10 | 0.000387543 | -2.221529168 | 0.003846186 |
| ENSMUSG00000028347 | Tmeff1 | 0.000387543 | -0.579983139 | 0.911642089 |
| ENSMUSG00000028420 | Tmem38b | 0.000387543 | -0.636566667 | 0.752038064 |
| ENSMUSG00000045071 | E130308A19Rik | 0.000387543 | -1.579983139 | 0.079144658 |
| ENSMUSG00000028385 | Snx30 | 0.000387543 | -0.843017545 | 0.587454349 |
| ENSMUSG00000066148 | Prpf4 | 0.000387543 | -0.72620388 | 0.678664944 |
| ENSMUSG00000048706 | Lurap1l | 0.000387543 | -0.995020638 | 0.462504539 |
| ENSMUSG00000070934 | Rraga | 0.000387543 | 0.291283547 | 1 |
| ENSMUSG00000037996 | Slc24a2 | 0.000387543 | 0.841480629 | 0.667778757 |
| ENSMUSG00000073802 | Cdkn2b | 0.000387543 | -0.599091962 | 0.796840156 |
| ENSMUSG00000086035 | Gm12610 | 0.000387543 | 0.741944956 | 0.821038733 |
| ENSMUSG00000008489 | Elavl2 | 0.000387543 | 2.063873051 | 0.086523965 |
| ENSMUSG00000084833 | Gm12649 | 0.000387543 | 0.694639241 | 0.847265357 |
| ENSMUSG00000085931 | Gm12648 | 0.000387543 | -0.810596067 | 0.577550711 |
| ENSMUSG00000028563 | Tm2d1 | 0.000387543 | 0.948395833 | 0.665635898 |
| ENSMUSG00000034926 | Dhcr24 | 0.000387543 | 0.256518129 | 1 |
| ENSMUSG00000028622 | Mrpl37 | 0.000387543 | -0.281901786 | 1 |
| ENSMUSG00000028607 | Cpt2 | 0.000387543 | 0.004979362 | 1 |
| ENSMUSG00000033326 | Kdm4a | 0.000387543 | -2.184054463 | 0.005214857 |
| ENSMUSG00000049878 | Rlf | 0.000387543 | -2.270027686 | 0.003050114 |
| ENSMUSG00000028902 | Sf3a3 | 0.000387543 | -1.80237556 | 0.031456064 |
| ENSMUSG00000001985 | Grik3 | 0.000387543 | 0.841480629 | 0.673811773 |
| ENSMUSG00000042500 | Ago4 | 0.000387543 | -0.72620388 | 0.687112677 |
| ENSMUSG00000028776 | Tinagl1 | 0.000387543 | -0.480447466 | 0.969079264 |
| ENSMUSG00000028901 | Gmeb1 | 0.000387543 | -1.752043885 | 0.037652954 |
| ENSMUSG00000037553 | Zdhhc18 | 0.000387543 | -1.362391704 | 0.159425617 |
| ENSMUSG00000028841 | Cnksr1 | 0.000387543 | 0.222570797 | 1 |
| ENSMUSG00000037348 | Paqr7 | 0.000387543 | -0.417253639 | 1 |
| ENSMUSG00000028668 | Eloa | 0.000387543 | -1.636566667 | 0.054245239 |
| ENSMUSG00000066037 | Hnrnpr | 0.000387543 | -1.973216268 | 0.012887418 |
| ENSMUSG00000036887 | C1qa | 0.000387543 | -0.560617814 | 0.886203497 |
| ENSMUSG00000028755 | Cda | 0.000387543 | 0.439382186 | 1 |
| ENSMUSG00000041241 | Mul1 | 0.000387543 | 1.326907457 | 0.406329712 |
| ENSMUSG00000028743 | Akr7a5 | 0.000387543 | 0.063873051 | 1 |
| ENSMUSG00000040972 | Igsf21 | 0.000387543 | 0.291283547 | 1 |
| ENSMUSG00000019055 | Plod1 | 0.000387543 | 0.893948049 | 0.66416768 |
| ENSMUSG00000039759 | Thap3 | 0.000387543 | -0.794107944 | 0.646349374 |
| ENSMUSG00000057751 | Megf6 | 0.000387543 | 0.893948049 | 0.66416768 |
| ENSMUSG00000029071 | Dvl1 | 0.000387543 | -0.980521069 | 0.466464457 |
| ENSMUSG00000106588 | Gm17590 | 0.000387543 | -1.995020638 | 0.016962438 |
| ENSMUSG00000029012 | Orc5 | 0.000387543 | -0.579983139 | 0.793880199 |
| ENSMUSG00000057541 | Pus7 | 0.000387543 | -1.80237556 | 0.031456064 |
| ENSMUSG00000085234 | Gm15614 | 0.000387543 | 0.326907457 | 1 |

| | | | | |
|---|---|---|---|---|
| ENSMUSG00000039315 | Clnk | 0.000387543 | 0.291283547 | 1 |
| ENSMUSG00000029176 | Anapc4 | 0.000387543 | -1.233807498 | 0.249952095 |
| ENSMUSG00000000560 | Gabra2 | 0.000387543 | 0.47891055 | 1 |
| ENSMUSG00000062960 | Kdr | 0.000387543 | 1.063873051 | 0.480093937 |
| ENSMUSG00000029248 | Thegl | 0.000387543 | 0.291283547 | 1 |
| ENSMUSG00000029250 | Polr2b | 0.000387543 | -1.092631435 | 0.332159278 |
| ENSMUSG00000029385 | Ccng2 | 0.000387543 | -0.328444372 | 1 |
| ENSMUSG00000034687 | Fras1 | 0.000387543 | 0.519552534 | 1 |
| ENSMUSG00000016833 | Mrps18c | 0.000387543 | 0.189403933 | 1 |
| ENSMUSG00000029314 | Gpat3 | 0.000387543 | -0.72620388 | 0.678664944 |
| ENSMUSG00000104956 | 4930429D17Rik | 0.000387543 | 1.948395833 | 0.113284116 |
| ENSMUSG00000029310 | Nudt9 | 0.000387543 | 1.189403933 | 0.455131546 |
| ENSMUSG00000029501 | Ankle2 | 0.000387543 | -1.851010336 | 0.02269114 |
| ENSMUSG00000007080 | Pole | 0.000387543 | -0.654945197 | 0.741406232 |
| ENSMUSG00000072720 | Myo18b | 0.000387543 | 1.004979362 | 0.66416768 |
| ENSMUSG00000041733 | Coq5 | 0.000387543 | -0.079084903 | 1 |
| ENSMUSG00000029516 | Cit | 0.000387543 | -1.023589791 | 0.414942392 |
| ENSMUSG00000066894 | Vsig10 | 0.000387543 | -0.43862729 | 0.943572777 |
| ENSMUSG00000029428 | Stx2 | 0.000387543 | -0.184054463 | 1 |
| ENSMUSG00000029439 | Sfswap | 0.000387543 | -2.287802388 | 0.002852874 |
| ENSMUSG00000029681 | Bcl7b | 0.000387543 | 0.439382186 | 1 |
| ENSMUSG00000039737 | Prkrip1 | 0.000387543 | -1.037664976 | 0.42039982 |
| ENSMUSG00000004415 | Col26a1 | 0.000387543 | 1.189403933 | 0.484692065 |
| ENSMUSG00000029710 | Ephb4 | 0.000387543 | -1.106051951 | 0.336079046 |
| ENSMUSG00000029722 | Agfg2 | 0.000387543 | -0.9057533 | 0.46833724 |
| ENSMUSG00000025856 | Pdgfa | 0.000387543 | -0.209145444 | 1 |
| ENSMUSG00000036565 | Ttyh3 | 0.000387543 | 0.222570797 | 1 |
| ENSMUSG00000036555 | Iqce | 0.000387543 | -1.777429203 | 0.034288602 |
| ENSMUSG00000041264 | Uspl1 | 0.000387543 | -1.570332969 | 0.077364646 |
| ENSMUSG00000051950 | B3glct | 0.000387543 | -0.89032326 | 0.526094357 |
| ENSMUSG00000068794 | Col28a1 | 0.000387543 | 0.439382186 | 1 |
| ENSMUSG00000041390 | Mdfic | 0.000387543 | -1.079084903 | 0.37949434 |
| ENSMUSG00000007655 | Cav1 | 0.000387543 | -0.079084903 | 1 |
| ENSMUSG00000049939 | Lrrc4 | 0.000387543 | 0.326907457 | 1 |
| ENSMUSG00000003500 | Impdh1 | 0.000387543 | 1.063873051 | 0.480093937 |
| ENSMUSG00000038600 | Atp6v0a4 | 0.000387543 | -0.743481871 | 0.68463297 |
| ENSMUSG00000029860 | Zyx | 0.000387543 | 0.189403933 | 1 |
| ENSMUSG00000029815 | Malsu1 | 0.000387543 | 0.063873051 | 1 |
| ENSMUSG00000037788 | Vopp1 | 0.000387543 | 0.256518129 | 1 |
| ENSMUSG00000097603 | A430010J10Rik | 0.000387543 | 0.222570797 | 1 |
| ENSMUSG00000052852 | Reep1 | 0.000387543 | -0.023589791 | 1 |
| ENSMUSG00000063884 | Ptcd3 | 0.000387543 | -0.951077291 | 0.452335407 |
| ENSMUSG00000030037 | Mrpl53 | 0.000387543 | 1.125273595 | 0.472831736 |
| ENSMUSG00000063810 | Alms1 | 0.000387543 | -1.258055044 | 0.220022719 |
| ENSMUSG00000029999 | Tgfa | 0.000387543 | -1.936126949 | 0.017363741 |
| ENSMUSG00000030051 | Aplf | 0.000387543 | -1.092631435 | 0.332159278 |
| ENSMUSG00000055403 | 4933427D06Rik | 0.000387543 | 0.291283547 | 1 |
| ENSMUSG00000030094 | Xpc | 0.000387543 | -0.599091962 | 0.796840156 |
| ENSMUSG00000030022 | Adamts9 | 0.000387543 | -0.936126949 | 0.502158856 |
| ENSMUSG00000035158 | Mitf | 0.000387543 | -0.936126949 | 0.49126082 |
| ENSMUSG00000030257 | Srgap3 | 0.000387543 | -2.356788998 | 0.001915848 |

| | | | | |
|---|---|---|---|---|
| ENSMUSG00000068011 | Mkrn2os | 0.000387543 | 1.189403933 | 0.472180092 |
| ENSMUSG00000025701 | Alox5 | 0.000387543 | -1.158519371 | 0.342598917 |
| ENSMUSG00000059878 | Zfp422 | 0.000387543 | 0.741944956 | 0.840264756 |
| ENSMUSG00000003154 | Foxj2 | 0.000387543 | -1.209145444 | 0.342316353 |
| ENSMUSG00000004270 | Lpcat3 | 0.000387543 | -1.362391704 | 0.159425617 |
| ENSMUSG00000030127 | Cops7a | 0.000387543 | 0.156982455 | 1 |
| ENSMUSG00000030339 | Ltbr | 0.000387543 | -0.281901786 | 1 |
| ENSMUSG00000001521 | Tulp3 | 0.000387543 | -1.560617814 | 0.075810288 |
| ENSMUSG00000042770 | Hebp1 | 0.000387543 | -0.258055044 | 1 |
| ENSMUSG00000054966 | Lmntd1 | 0.000387543 | 0.841480629 | 0.704951233 |
| ENSMUSG00000030301 | Ccdc91 | 0.000387543 | -1.270027686 | 0.306074679 |
| ENSMUSG00000063802 | Hspbp1 | 0.000387543 | 0.004979362 | 1 |
| ENSMUSG00000000605 | Clcn4 | 0.000387543 | -0.258055044 | 1 |
| ENSMUSG00000074364 | Ehd2 | 0.000387543 | -0.43862729 | 0.943572777 |
| ENSMUSG00000040891 | Foxa3 | 0.000387543 | 0.56137271 | 0.98880775 |
| ENSMUSG00000052214 | Opa3 | 0.000387543 | -0.810596067 | 0.578855097 |
| ENSMUSG00000003549 | Ercc1 | 0.000387543 | 0.604441432 | 0.857399533 |
| ENSMUSG00000002980 | Bcam | 0.000387543 | -0.72620388 | 0.678664944 |
| ENSMUSG00000053291 | Rab4b | 0.000387543 | -0.328444372 | 1 |
| ENSMUSG00000003363 | Pld3 | 0.000387543 | -0.023589791 | 1 |
| ENSMUSG00000003438 | Timm50 | 0.000387543 | -0.258055044 | 1 |
| ENSMUSG00000059975 | Zfp74 | 0.000387543 | 0.741944956 | 0.821038733 |
| ENSMUSG00000001248 | Gramd1a | 0.000387543 | -0.500911568 | 0.868661374 |
| ENSMUSG00000034875 | Nudt19 | 0.000387543 | 1.841480629 | 0.130996353 |
| ENSMUSG00000034867 | Ankrd27 | 0.000387543 | -1.339849135 | 0.180637196 |
| ENSMUSG00000108754 | Gm44748 | 0.000387543 | 0.47891055 | 1 |
| ENSMUSG00000069727 | Zfp975 | 0.000387543 | -1.874726405 | 0.020746096 |
| ENSMUSG00000038670 | Mybpc2 | 0.000387543 | 3.063873051 | 0.011284174 |
| ENSMUSG00000007837 | Prrg2 | 0.000387543 | 0.034125707 | 1 |
| ENSMUSG00000003271 | Sult2b1 | 0.000387543 | -0.417253639 | 1 |
| ENSMUSG00000030505 | Prmt3 | 0.000387543 | -0.52108945 | 0.871107617 |
| ENSMUSG00000030452 | Nipa2 | 0.000387543 | -2.085874069 | 0.007450793 |
| ENSMUSG00000100826 | Snhg14 | 0.000387543 | 1.004979362 | 0.66416768 |
| ENSMUSG00000030519 | Apba2 | 0.000387543 | 0.604441432 | 0.864675985 |
| ENSMUSG00000030512 | Snrpa1 | 0.000387543 | 0.400908038 | 1 |
| ENSMUSG00000109097 | Gm29683 | 0.000387543 | 0.648835551 | 0.846566779 |
| ENSMUSG00000030551 | Nr2f2 | 0.000387543 | -0.480447466 | 0.971854959 |
| ENSMUSG00000059146 | Ntrk3 | 0.000387543 | 0.326907457 | 1 |
| ENSMUSG00000045795 | Whamm | 0.000387543 | -0.777429203 | 0.641730377 |
| ENSMUSG00000038563 | Efl1 | 0.000387543 | -1.43862729 | 0.113452361 |
| ENSMUSG00000070462 | Tlnrd1 | 0.000387543 | 2.063873051 | 0.086523965 |
| ENSMUSG00000038503 | Mesd | 0.000387543 | 0.222570797 | 1 |
| ENSMUSG00000061119 | Prcp | 0.000387543 | 0.604441432 | 0.916365786 |
| ENSMUSG00000035401 | Emsy | 0.000387543 | -2.390083438 | 0.001701831 |
| ENSMUSG00000018909 | Arrb1 | 0.000387543 | -1.051604167 | 0.374171924 |
| ENSMUSG00000035901 | Dennd5a | 0.000387543 | -2.079084903 | 0.008534388 |
| ENSMUSG00000030788 | Rnf141 | 0.000387543 | -0.258055044 | 1 |
| ENSMUSG00000005611 | Mrvi1 | 0.000387543 | 0.256518129 | 1 |
| ENSMUSG00000038296 | Galnt18 | 0.000387543 | -1.617950989 | 0.061133438 |
| ENSMUSG00000030659 | Nucb2 | 0.000387543 | -0.079084903 | 1 |
| ENSMUSG00000030889 | Vwa3a | 0.000387543 | 0.222570797 | 1 |

| | | | | |
|---|---|---|---|---|
| ENSMUSG00000046182 | Gsg1l | 0.000387543 | 1.256518129 | 0.455131546 |
| ENSMUSG00000030727 | Rabep2 | 0.000387543 | 0.648835551 | 0.846566779 |
| ENSMUSG00000086502 | B130055M24Rik | 0.000387543 | -1.951077291 | 0.016410626 |
| ENSMUSG00000030814 | Bcl7c | 0.000387543 | 0.063873051 | 1 |
| ENSMUSG00000096145 | Vkorc1 | 0.000387543 | 1.256518129 | 0.430499017 |
| ENSMUSG00000030802 | Bckdk | 0.000387543 | 1.004979362 | 0.633493255 |
| ENSMUSG00000030846 | Tial1 | 0.000387543 | -2.470105521 | 0.001112545 |
| ENSMUSG00000063179 | Pstk | 0.000387543 | 1.400908038 | 0.423814296 |
| ENSMUSG00000025464 | Paox | 0.000387543 | -0.52108945 | 0.874945517 |
| ENSMUSG00000038618 | Rassf7 | 0.000387543 | 1.47891055 | 0.254875603 |
| ENSMUSG00000058886 | Deaf1 | 0.000387543 | -1.38458745 | 0.14097975 |
| ENSMUSG00000031511 | Arhgef7 | 0.000387543 | -0.980521069 | 0.456648093 |
| ENSMUSG00000010435 | Spaca7 | 0.000387543 | -2.995020638 | 3.97E-05 |
| ENSMUSG00000038416 | Cdc16 | 0.000387543 | -0.654945197 | 0.741406232 |
| ENSMUSG00000026317 | Cln8 | 0.000387543 | 2.326907457 | 0.050877179 |
| ENSMUSG00000031533 | Mrps31 | 0.000387543 | 1.004979362 | 0.633493255 |
| ENSMUSG00000031565 | Fgfr1 | 0.000387543 | -0.328444372 | 1 |
| ENSMUSG00000031530 | Dusp4 | 0.000387543 | 0.256518129 | 1 |
| ENSMUSG00000025521 | Tmem192 | 0.000387543 | 0.156982455 | 1 |
| ENSMUSG00000043090 | Zfp866 | 0.000387543 | -0.328444372 | 1 |
| ENSMUSG00000011306 | Sugp1 | 0.000387543 | -1.500911568 | 0.123190608 |
| ENSMUSG00000035559 | Mpv17l2 | 0.000387543 | 1.841480629 | 0.130996353 |
| ENSMUSG00000034799 | Unc13a | 0.000387543 | 0.063873051 | 1 |
| ENSMUSG00000031803 | B3gnt3 | 0.000387543 | -1.065409966 | 0.374573031 |
| ENSMUSG00000003033 | Ap1m1 | 0.000387543 | -0.636566667 | 0.737916568 |
| ENSMUSG00000079011 | Gm11033 | 0.000387543 | 1.063873051 | 0.517439702 |
| ENSMUSG00000037101 | Ttc29 | 0.000387543 | 0.694639241 | 0.841229868 |
| ENSMUSG00000031681 | Smad1 | 0.000387543 | -1.305360759 | 0.201837433 |
| ENSMUSG00000045333 | Zfp423 | 0.000387543 | -0.184054463 | 1 |
| ENSMUSG00000031657 | Heatr3 | 0.000387543 | -2.328444372 | 0.002197062 |
| ENSMUSG00000031781 | Ciapin1 | 0.000387543 | 0.648835551 | 0.854285611 |
| ENSMUSG00000005705 | Agrp | 0.000387543 | -1.119348773 | 0.341720553 |
| ENSMUSG00000036270 | Edc4 | 0.000387543 | -2.516071375 | 0.000825737 |
| ENSMUSG00000031901 | Dus2 | 0.000387543 | -1.560617814 | 0.084812586 |
| ENSMUSG00000031916 | Cog8 | 0.000387543 | -0.258055044 | 1 |
| ENSMUSG00000031921 | Terf2 | 0.000387543 | -1.184054463 | 0.271553405 |
| ENSMUSG00000033579 | Fa2h | 0.000387543 | 1.125273595 | 0.466464457 |
| ENSMUSG00000031955 | Bcar1 | 0.000387543 | -0.980521069 | 0.456648093 |
| ENSMUSG00000040263 | Klhdc4 | 0.000387543 | -1.362391704 | 0.16368308 |
| ENSMUSG00000033862 | Cdk10 | 0.000387543 | -2.119348773 | 0.006319812 |
| ENSMUSG00000062380 | Tubb3 | 0.000387543 | -0.417253639 | 1 |
| ENSMUSG00000019478 | Rab4a | 0.000387543 | 1.648835551 | 0.188603791 |
| ENSMUSG00000060212 | Pcnx2 | 0.000387543 | 0.519552534 | 1 |
| ENSMUSG00000033998 | Kcnk1 | 0.000387543 | 1.189403933 | 0.455131546 |
| ENSMUSG00000037419 | Endod1 | 0.000387543 | 0.741944956 | 0.821038733 |
| ENSMUSG00000032171 | Pin1 | 0.000387543 | 1.256518129 | 0.466464457 |
| ENSMUSG00000086016 | 1700084C06Rik | 0.000387543 | -0.43862729 | 0.943572777 |
| ENSMUSG00000035047 | Kri1 | 0.000387543 | -0.72620388 | 0.68439554 |
| ENSMUSG00000040146 | Rgl3 | 0.000387543 | -1.132524162 | 0.29614584 |
| ENSMUSG00000066829 | Zfp810 | 0.000387543 | -1.037664976 | 0.430332878 |
| ENSMUSG00000041444 | Arhgap32 | 0.000387543 | -1.209145444 | 0.242837209 |

| | | | | |
|---|---|---|---|---|
| ENSMUSG00000074452 | Pate2 | 0.000387543 | 1.47891055 | 0.259053361 |
| ENSMUSG00000032092 | Mpzl2 | 0.000387543 | -0.351164449 | 1 |
| ENSMUSG00000048534 | Jaml | 0.000387543 | -0.132524162 | 1 |
| ENSMUSG00000032087 | Dscaml1 | 0.000387543 | -0.209145444 | 1 |
| ENSMUSG00000032059 | Alg9 | 0.000387543 | -0.636566667 | 0.737916568 |
| ENSMUSG00000034218 | Atm | 0.000387543 | -1.119348773 | 0.341720553 |
| ENSMUSG00000032293 | Ireb2 | 0.000387543 | -1.810596067 | 0.027621844 |
| ENSMUSG00000032307 | Ube2q2 | 0.000387543 | -1.636566667 | 0.054245239 |
| ENSMUSG00000032288 | Imp3 | 0.000387543 | 1.326907457 | 0.406329712 |
| ENSMUSG00000032316 | Clk3 | 0.000387543 | -0.72620388 | 0.678664944 |
| ENSMUSG00000035914 | Cd276 | 0.000387543 | -0.691014451 | 0.672182136 |
| ENSMUSG00000025237 | Parp6 | 0.000387543 | -1.106051951 | 0.336079046 |
| ENSMUSG00000032246 | Calml4 | 0.000387543 | 0.948395833 | 0.66416768 |
| ENSMUSG00000032393 | Dpp8 | 0.000387543 | -2.449196532 | 0.001138977 |
| ENSMUSG00000032380 | Dapk2 | 0.000387543 | 0.648835551 | 0.857093467 |
| ENSMUSG00000032360 | Hcrtr2 | 0.000387543 | 1.841480629 | 0.130996353 |
| ENSMUSG00000097655 | Gm26713 | 0.000387543 | 0.363433333 | 1 |
| ENSMUSG00000032374 | Plod2 | 0.000387543 | 0.256518129 | 1 |
| ENSMUSG00000032409 | Atr | 0.000387543 | 0.034125707 | 1 |
| ENSMUSG00000100807 | Gm29521 | 0.000387543 | 1.56137271 | 0.262284476 |
| ENSMUSG00000032549 | Rab6b | 0.000387543 | 0.256518129 | 1 |
| ENSMUSG00000032553 | Srprb | 0.000387543 | -0.480447466 | 0.949822782 |
| ENSMUSG00000035032 | Nek11 | 0.000387543 | 1.004979362 | 0.66416768 |
| ENSMUSG00000032570 | Atp2c1 | 0.000387543 | -1.794107944 | 0.030770816 |
| ENSMUSG00000032577 | Mapkapk3 | 0.000387543 | -2.406446884 | 0.00165757 |
| ENSMUSG00000034684 | Sema3f | 0.000387543 | -1.281901786 | 0.196126878 |
| ENSMUSG00000032599 | Ip6k2 | 0.000387543 | -1.184054463 | 0.271553405 |
| ENSMUSG00000025647 | Shisa5 | 0.000387543 | 0.034125707 | 1 |
| ENSMUSG00000032431 | Crtap | 0.000387543 | 0.694639241 | 0.833666942 |
| ENSMUSG00000032437 | Stt3b | 0.000387543 | -1.734868741 | 0.042660478 |
| ENSMUSG00000036138 | Acaa1a | 0.000387543 | -0.106051951 | 1 |
| ENSMUSG00000025785 | Exosc7 | 0.000387543 | -0.52108945 | 0.871107617 |
| ENSMUSG00000061759 | Armt1 | 0.000387543 | -1.500911568 | 0.10343288 |
| ENSMUSG00000019767 | Ccdc170 | 0.000387543 | 0.648835551 | 0.866342478 |
| ENSMUSG00000000766 | Oprm1 | 0.000387543 | 0.291283547 | 1 |
| ENSMUSG00000101621 | Gm28905 | 0.000387543 | 0.326907457 | 1 |
| ENSMUSG00000112929 | Gm47729 | 0.000387543 | 2.063873051 | 0.107857035 |
| ENSMUSG00000112629 | Gm47771 | 0.000387543 | 1.189403933 | 0.455131546 |
| ENSMUSG00000039531 | Zup1 | 0.000387543 | -0.540989007 | 0.8730922 |
| ENSMUSG00000019838 | Slc16a10 | 0.000387543 | -1.079084903 | 0.391578102 |
| ENSMUSG00000038160 | Atg5 | 0.000387543 | -1.233807498 | 0.249952095 |
| ENSMUSG00000019849 | Prep | 0.000387543 | -0.395558568 | 1 |
| ENSMUSG00000023068 | Nus1 | 0.000387543 | -0.980521069 | 0.479878818 |
| ENSMUSG00000019872 | Smpdl3a | 0.000387543 | 2.189403933 | 0.067154141 |
| ENSMUSG00000037990 | Sh3rf3 | 0.000387543 | 0.400908038 | 1 |
| ENSMUSG00000020108 | Ddit4 | 0.000387543 | 0.790854556 | 0.689981618 |
| ENSMUSG00000058806 | Col13a1 | 0.000387543 | 1.326907457 | 0.466464457 |
| ENSMUSG00000020080 | Hkdc1 | 0.000387543 | -0.395558568 | 1 |
| ENSMUSG00000020069 | Hnrnph3 | 0.000387543 | -0.480447466 | 0.949822782 |
| ENSMUSG00000047417 | Rexo1 | 0.000387543 | -0.579983139 | 0.793880199 |
| ENSMUSG00000034974 | Dapk3 | 0.000387543 | -0.980521069 | 0.456648093 |

| | | | | |
|---|---|---|---|---|
| ENSMUSG00000034932 | Mrpl54 | 0.000387543 | 0.790854556 | 0.697234589 |
| ENSMUSG00000047129 | 1700113H08Rik | 0.000387543 | 1.189403933 | 0.466464457 |
| ENSMUSG00000035798 | Zdhhc17 | 0.000387543 | -1.579983139 | 0.079144658 |
| ENSMUSG00000112430 | Gm30624 | 0.000387543 | -1.958494762 | 0.014316904 |
| ENSMUSG00000028630 | Dyrk2 | 0.000387543 | -0.351164449 | 1 |
| ENSMUSG00000112716 | 4930471E19Rik | 0.000387543 | 1.47891055 | 0.279041806 |
| ENSMUSG00000034707 | Gns | 0.000387543 | 0.156982455 | 1 |
| ENSMUSG00000112527 | Gm35696 | 0.000387543 | 0.034125707 | 1 |
| ENSMUSG00000044071 | Tafa2 | 0.000387543 | -0.079084903 | 1 |
| ENSMUSG00000025354 | Dnajc14 | 0.000387543 | -0.936126949 | 0.47622561 |
| ENSMUSG00000034614 | Pik3ip1 | 0.000387543 | -0.106051951 | 1 |
| ENSMUSG00000049721 | Gal3st1 | 0.000387543 | 0.291283547 | 1 |
| ENSMUSG00000004394 | Tmed4 | 0.000387543 | 0.893948049 | 0.66416768 |
| ENSMUSG00000004668 | Abca13 | 0.000387543 | 0.326907457 | 1 |
| ENSMUSG00000020176 | Grb10 | 0.000387543 | 0.400908038 | 1 |
| ENSMUSG00000020120 | Plek | 0.000387543 | 1.004979362 | 0.65280571 |
| ENSMUSG00000087085 | Gm12068 | 0.000387543 | 1.948395833 | 0.108201045 |
| ENSMUSG00000069919 | Hba-a1 | 0.000387543 | -0.951077291 | 1 |
| ENSMUSG00000018846 | Pank3 | 0.000387543 | -0.636566667 | 0.83628546 |
| ENSMUSG00000020436 | Gabrg2 | 0.000387543 | 1.948395833 | 0.13735263 |
| ENSMUSG00000040365 | Trim41 | 0.000387543 | -0.794107944 | 0.646349374 |
| ENSMUSG00000018395 | Kif3a | 0.000387543 | -2.152064348 | 0.006222976 |
| ENSMUSG00000018906 | P4ha2 | 0.000387543 | 0.326907457 | 1 |
| ENSMUSG00000000594 | Gm2a | 0.000387543 | 1.47891055 | 0.254875603 |
| ENSMUSG00000020524 | Gria1 | 0.000387543 | -0.480447466 | 0.971854959 |
| ENSMUSG00000020496 | Rnf187 | 0.000387543 | 0.256518129 | 1 |
| ENSMUSG00000019102 | Aldh3a1 | 0.000387543 | 1.648835551 | 0.188603791 |
| ENSMUSG00000042298 | Ttc19 | 0.000387543 | -1.092631435 | 0.332159278 |
| ENSMUSG00000046417 | Lrrc75a | 0.000387543 | -1.777429203 | 0.041457324 |
| ENSMUSG00000048497 | Mmgt2 | 0.000387543 | -0.106051951 | 1 |
| ENSMUSG00000018761 | Mpdu1 | 0.000387543 | -0.617950989 | 0.801225115 |
| ENSMUSG00000005204 | Senp3 | 0.000387543 | 0.034125707 | 1 |
| ENSMUSG00000047284 | Neurl4 | 0.000387543 | -2.281901786 | 0.002768072 |
| ENSMUSG00000014606 | Slc25a11 | 0.000387543 | 0.893948049 | 0.680536135 |
| ENSMUSG00000018446 | C1qbp | 0.000387543 | -0.673092543 | 0.765052883 |
| ENSMUSG00000020788 | Atp2a3 | 0.000387543 | -2.367972736 | 0.002465504 |
| ENSMUSG00000017288 | Vps53 | 0.000387543 | -0.52108945 | 0.871107617 |
| ENSMUSG00000020840 | Blmh | 0.000387543 | -1.531073539 | 0.083271608 |
| ENSMUSG00000050944 | Efcab5 | 0.000387543 | -0.599091962 | 0.814575544 |
| ENSMUSG00000035413 | Tmem98 | 0.000387543 | -0.617950989 | 0.801225115 |
| ENSMUSG00000018841 | Rad51d | 0.000387543 | 0.063873051 | 1 |
| ENSMUSG00000019122 | Ccl9 | 0.000387543 | 1.948395833 | 0.115288807 |
| ENSMUSG00000069792 | Wfdc17 | 0.000387543 | 2.47891055 | 0.036895541 |
| ENSMUSG00000020677 | Ddx52 | 0.000387543 | -0.500911568 | 0.971854959 |
| ENSMUSG00000018427 | Ypel2 | 0.000387543 | -3.125951508 | 1.61E-05 |
| ENSMUSG00000034177 | Rnf43 | 0.000387543 | -2.526090131 | 0.000752229 |
| ENSMUSG00000013418 | B4galnt2 | 0.000387543 | 0.948395833 | 0.662054817 |
| ENSMUSG00000018547 | Pip4k2b | 0.000387543 | -1.132524162 | 0.30756351 |
| ENSMUSG00000018160 | Med1 | 0.000387543 | -1.579983139 | 0.081618814 |
| ENSMUSG00000052915 | Msl1 | 0.000387543 | -0.72620388 | 0.699496878 |
| ENSMUSG00000006931 | P3h4 | 0.000387543 | 0.326907457 | 1 |

| | | | | |
|---|---|---|---|---|
| ENSMUSG00000078652 | Psme3 | 0.000387543 | 0.56137271 | 0.98880775 |
| ENSMUSG00000059995 | Atxn7l3 | 0.000387543 | -0.617950989 | 0.801225115 |
| ENSMUSG00000040699 | Limd2 | 0.000387543 | 1.004979362 | 0.633493255 |
| ENSMUSG00000020720 | Psmd12 | 0.000387543 | -0.43862729 | 1 |
| ENSMUSG00000051043 | Gprc5c | 0.000387543 | 2.648835551 | 0.041296623 |
| ENSMUSG00000034427 | Myo15b | 0.000387543 | 0.741944956 | 0.821038733 |
| ENSMUSG00000050628 | Ubald2 | 0.000387543 | 0.0942467 | 1 |
| ENSMUSG00000020818 | Mfsd11 | 0.000387543 | -1.38458745 | 0.14097975 |
| ENSMUSG00000061306 | Slc38a10 | 0.000387543 | -1.106051951 | 0.336079046 |
| ENSMUSG00000046605 | B3gntl1 | 0.000387543 | -0.708716453 | 0.674297981 |
| ENSMUSG00000111971 | Gm48678 | 0.000387543 | 0.326907457 | 1 |
| ENSMUSG00000061458 | Nol10 | 0.000387543 | -1.608552291 | 0.071137013 |
| ENSMUSG00000062054 | Iah1 | 0.000387543 | 0.56137271 | 0.98880775 |
| ENSMUSG00000002996 | Hbp1 | 0.000387543 | -0.82689788 | 0.587484274 |
| ENSMUSG00000035799 | Twist1 | 0.000387543 | 1.189403933 | 0.469131246 |
| ENSMUSG00000071342 | Lsmem1 | 0.000387543 | 2.063873051 | 0.201578467 |
| ENSMUSG00000020950 | Foxg1 | 0.000387543 | -0.777429203 | 0.641730377 |
| ENSMUSG00000021023 | Prorp | 0.000387543 | -1.52108945 | 0.096053423 |
| ENSMUSG00000020986 | Sec23a | 0.000387543 | -2.47528576 | 0.000993739 |
| ENSMUSG00000020992 | 4930512B01Rik | 0.000387543 | 0.47891055 | 1 |
| ENSMUSG00000034761 | Map4k5 | 0.000387543 | -1.500911568 | 0.092707526 |
| ENSMUSG00000021086 | Ccdc175 | 0.000387543 | 2.063873051 | 0.108364916 |
| ENSMUSG00000021114 | Atp6v1d | 0.000387543 | 0.004979362 | 1 |
| ENSMUSG00000060716 | Plekhh1 | 0.000387543 | -1.406446884 | 0.145704891 |
| ENSMUSG00000021131 | Erh | 0.000387543 | -0.43862729 | 0.943572777 |
| ENSMUSG00000066438 | Plekhd1 | 0.000387543 | -1.023589791 | 0.414942392 |
| ENSMUSG00000021044 | Adck1 | 0.000387543 | 0.063873051 | 1 |
| ENSMUSG00000047014 | Catsperb | 0.000387543 | 0.56137271 | 0.885240416 |
| ENSMUSG00000021208 | Ifi27l2b | 0.000387543 | 1.256518129 | 0.430499017 |
| ENSMUSG00000021091 | Serpina3n | 0.000387543 | 1.948395833 | 0.108201045 |
| ENSMUSG00000021115 | Vrk1 | 0.000387543 | -0.936126949 | 0.47622561 |
| ENSMUSG00000040877 | Wdr25 | 0.000387543 | -1.645785197 | 0.055187251 |
| ENSMUSG00000064326 | Siva1 | 0.000387543 | 1.125273595 | 0.466464457 |
| ENSMUSG00000002804 | Nudt14 | 0.000387543 | 1.004979362 | 0.633493255 |
| ENSMUSG00000042050 | Wdr60 | 0.000387543 | -2.112715681 | 0.007127824 |
| ENSMUSG00000033781 | Asb13 | 0.000387543 | -0.158519371 | 1 |
| ENSMUSG00000097009 | Gm26861 | 0.000387543 | -1.682081327 | 0.067084058 |
| ENSMUSG00000039242 | B3galnt2 | 0.000387543 | -1.965874293 | 0.01465972 |
| ENSMUSG00000015672 | Mrpl32 | 0.000387543 | 0.400908038 | 1 |
| ENSMUSG00000075054 | Yae1d1 | 0.000387543 | 0.519552534 | 0.996793385 |
| ENSMUSG00000003062 | Stard3nl | 0.000387543 | -0.72620388 | 0.678664944 |
| ENSMUSG00000022228 | Zscan26 | 0.000387543 | -2.171343411 | 0.004911267 |
| ENSMUSG00000006715 | Gmnn | 0.000387543 | -0.079084903 | 1 |
| ENSMUSG00000035910 | Dcdc2a | 0.000387543 | -0.810596067 | 0.577550711 |
| ENSMUSG00000038408 | 1700018A04Rik | 0.000387543 | 1.004979362 | 0.660989068 |
| ENSMUSG00000114838 | Gm48703 | 0.000387543 | 1.400908038 | 0.415257602 |
| ENSMUSG00000114407 | Gm48765 | 0.000387543 | 1.741944956 | 0.167110871 |
| ENSMUSG00000038683 | Pak1ip1 | 0.000387543 | -1.328444372 | 0.177083054 |
| ENSMUSG00000021464 | Ror2 | 0.000387543 | -0.540989007 | 0.882342583 |
| ENSMUSG00000021504 | B4galt7 | 0.000387543 | 0.604441432 | 0.857399533 |
| ENSMUSG00000021552 | Gkap1 | 0.000387543 | -0.743481871 | 0.68463297 |

| | | | | |
|---|---|---|---|---|
| ENSMUSG00000044792 | Isca1 | 0.000387543 | 0.948395833 | 0.653235135 |
| ENSMUSG00000038212 | Mfsd14b | 0.000387543 | -0.106051951 | 1 |
| ENSMUSG00000114616 | Gm26754 | 0.000387543 | -0.560617814 | 0.875465725 |
| ENSMUSG00000021607 | Mrpl36 | 0.000387543 | 1.648835551 | 0.196126878 |
| ENSMUSG00000057649 | Brd9 | 0.000387543 | -2.112715681 | 0.007127824 |
| ENSMUSG00000089940 | Gm4117 | 0.000387543 | -2.406446884 | 0.001608894 |
| ENSMUSG00000051111 | Sv2c | 0.000387543 | 0.948395833 | 0.653235135 |
| ENSMUSG00000021645 | Smn1 | 0.000387543 | 0.034125707 | 1 |
| ENSMUSG00000021710 | Nln | 0.000387543 | -1.965874293 | 0.01465972 |
| ENSMUSG00000021712 | Trim23 | 0.000387543 | -0.636566667 | 0.737916568 |
| ENSMUSG00000021696 | Elovl7 | 0.000387543 | -1.589569188 | 0.067856911 |
| ENSMUSG00000021765 | Fst | 0.000387543 | -1.245982212 | 0.217655393 |
| ENSMUSG00000015536 | Mocs2 | 0.000387543 | 0.063873051 | 1 |
| ENSMUSG00000095930 | Nim1k | 0.000387543 | 0.363433333 | 1 |
| ENSMUSG00000114786 | Gm48594 | 0.000387543 | 1.125273595 | 0.467242784 |
| ENSMUSG00000021785 | Ngly1 | 0.000387543 | -2.112715681 | 0.007127824 |
| ENSMUSG00000055538 | Zcchc24 | 0.000387543 | -1.540989007 | 0.101048675 |
| ENSMUSG00000021936 | Mapk8 | 0.000387543 | -0.636566667 | 0.737916568 |
| ENSMUSG00000041471 | Shld2 | 0.000387543 | -1.459688905 | 0.117170911 |
| ENSMUSG00000049092 | Gpr137c | 0.000387543 | 1.326907457 | 0.409502039 |
| ENSMUSG00000068399 | Gm7247 | 0.000387543 | 0.47891055 | 1 |
| ENSMUSG00000021967 | Mrpl57 | 0.000387543 | 1.189403933 | 0.455131546 |
| ENSMUSG00000021930 | Spryd7 | 0.000387543 | 0.694639241 | 0.833666942 |
| ENSMUSG00000034522 | Zfp395 | 0.000387543 | -1.936126949 | 0.015523381 |
| ENSMUSG00000022053 | Ebf2 | 0.000387543 | 0.604441432 | 0.871494441 |
| ENSMUSG00000022075 | Rhobtb2 | 0.000387543 | -1.037664976 | 0.42039982 |
| ENSMUSG00000022124 | Fbxl3 | 0.000387543 | -1.500911568 | 0.092707526 |
| ENSMUSG00000054263 | Lifr | 0.000387543 | -1.328444372 | 0.184885592 |
| ENSMUSG00000044224 | Dnajc21 | 0.000387543 | -1.373532262 | 0.162595422 |
| ENSMUSG00000115773 | Gm31458 | 0.000387543 | 1.004979362 | 0.66416768 |
| ENSMUSG00000022237 | Ankrd33b | 0.000387543 | 0.519552534 | 1 |
| ENSMUSG00000022299 | Slc25a32 | 0.000387543 | 0.0942467 | 1 |
| ENSMUSG00000086541 | Has2os | 0.000387543 | 0.841480629 | 0.709111889 |
| ENSMUSG00000022376 | Adcy8 | 0.000387543 | -0.43862729 | 1 |
| ENSMUSG00000075600 | Zc3h3 | 0.000387543 | -2.373532262 | 0.00180223 |
| ENSMUSG00000022564 | Grina | 0.000387543 | -1.233807498 | 0.249952095 |
| ENSMUSG00000075555 | Gm10863 | 0.000387543 | 1.326907457 | 0.466464457 |
| ENSMUSG00000022474 | Pmm1 | 0.000387543 | 1.063873051 | 0.480093937 |
| ENSMUSG00000022449 | Adamts20 | 0.000387543 | 1.400908038 | 0.317364054 |
| ENSMUSG00000022454 | Nell2 | 0.000387543 | -0.9057533 | 0.581366187 |
| ENSMUSG00000033075 | Senp1 | 0.000387543 | -1.717486663 | 0.040535668 |
| ENSMUSG00000003352 | Cacnb3 | 0.000387543 | 0.790854556 | 0.687112677 |
| ENSMUSG00000037991 | Rmi2 | 0.000387543 | -3.664047404 | 2.16E-07 |
| ENSMUSG00000018830 | Myh11 | 0.000387543 | 0.189403933 | 1 |
| ENSMUSG00000023088 | Abcc1 | 0.000387543 | -2.495822691 | 0.001138977 |
| ENSMUSG00000041957 | Pkp2 | 0.000387543 | -0.708716453 | 0.674297981 |
| ENSMUSG00000033918 | Parl | 0.000387543 | -1.023589791 | 0.414942392 |
| ENSMUSG00000097290 | 1300002E11Rik | 0.000387543 | 1.004979362 | 0.66416768 |
| ENSMUSG00000022514 | Il1rap | 0.000387543 | 0.56137271 | 0.98880775 |
| ENSMUSG00000035764 | Fbxo45 | 0.000387543 | -0.72620388 | 0.694502633 |
| ENSMUSG00000022787 | Wdr53 | 0.000387543 | -0.395558568 | 1 |

| | | | | |
|---|---|---|---|---|
| ENSMUSG00000014074 | Rnf168 | 0.000387543 | -2.099357298 | 0.007916911 |
| ENSMUSG00000078489 | Gm17106 | 0.000387543 | 0.648835551 | 0.857969607 |
| ENSMUSG00000049502 | Dtx3l | 0.000387543 | -0.708716453 | 0.674297981 |
| ENSMUSG00000075229 | Ccdc58 | 0.000387543 | -1.106051951 | 0.361093181 |
| ENSMUSG00000022829 | Stxbp5l | 0.000387543 | 1.063873051 | 0.520356359 |
| ENSMUSG00000075070 | 4932412D23Rik | 0.000387543 | 0.741944956 | 0.794761217 |
| ENSMUSG00000116689 | 4930405D01Rik | 0.000387543 | 0.948395833 | 0.668575194 |
| ENSMUSG00000075033 | Nxpe3 | 0.000387543 | -2.401012998 | 0.001556178 |
| ENSMUSG00000022604 | Cep97 | 0.000387543 | 0.790854556 | 0.706069688 |
| ENSMUSG00000025616 | Usp16 | 0.000387543 | -1.664047404 | 0.049287094 |
| ENSMUSG00000022973 | Synj1 | 0.000387543 | -0.777429203 | 0.641730377 |
| ENSMUSG00000022951 | Rcan1 | 0.000387543 | -0.72620388 | 0.708161645 |
| ENSMUSG00000051483 | Cbr1 | 0.000387543 | 1.326907457 | 0.406329712 |
| ENSMUSG00000022947 | Cbr3 | 0.000387543 | 3.063873051 | 0.011284174 |
| ENSMUSG00000039456 | Morc3 | 0.000387543 | -2.362391704 | 0.001963455 |
| ENSMUSG00000022895 | Ets2 | 0.000387543 | -0.579983139 | 0.793880199 |
| ENSMUSG00000095687 | Rnaset2a | 0.000387543 | 0.034125707 | 1 |
| ENSMUSG00000060475 | Wtap | 0.000387543 | -2.00221614 | 0.012155548 |
| ENSMUSG00000047786 | Lix1 | 0.000387543 | -2.316948733 | 0.002642488 |
| ENSMUSG00000041130 | Zfp598 | 0.000387543 | -1.540989007 | 0.084875472 |
| ENSMUSG00000024165 | Jpt2 | 0.000387543 | 0.291283547 | 1 |
| ENSMUSG00000002274 | Metrn | 0.000387543 | 0.694639241 | 0.833666942 |
| ENSMUSG00000078838 | Gm17382 | 0.000387543 | -0.777429203 | 0.641730377 |
| ENSMUSG00000057789 | Bak1 | 0.000387543 | 0.790854556 | 0.687112677 |
| ENSMUSG00000024225 | Clps | 0.000387543 | 2.47891055 | 0.036895541 |
| ENSMUSG00000024045 | Akap8 | 0.000387543 | -1.80237556 | 0.031456064 |
| ENSMUSG00000067430 | Zfp763 | 0.000387543 | -0.82689788 | 0.581736296 |
| ENSMUSG00000067424 | Zfp563 | 0.000387543 | -1.417253639 | 0.128531628 |
| ENSMUSG00000090877 | Hspa1b | 0.000387543 | 1.326907457 | 0.406329712 |
| ENSMUSG00000091971 | Hspa1a | 0.000387543 | 3.063873051 | 0.011284174 |
| ENSMUSG00000007050 | Lsm2 | 0.000387543 | 1.256518129 | 0.430499017 |
| ENSMUSG00000038500 | Prr3 | 0.000387543 | -1.293678954 | 0.198966953 |
| ENSMUSG00000117210 | Gm49857 | 0.000387543 | -2.93236503 | 0.000105658 |
| ENSMUSG00000023952 | Gtpbp2 | 0.000387543 | -1.132524162 | 0.29614584 |
| ENSMUSG00000063888 | Rpl7l1 | 0.000387543 | -0.858959089 | 0.51724245 |
| ENSMUSG00000040260 | Daam2 | 0.000387543 | 0.189403933 | 1 |
| ENSMUSG00000001229 | Dpp9 | 0.000387543 | -0.023589791 | 1 |
| ENSMUSG00000002658 | Gtf2f1 | 0.000387543 | -0.52108945 | 0.871107617 |
| ENSMUSG00000005823 | Gpr108 | 0.000387543 | -1.221529168 | 0.245842568 |
| ENSMUSG00000117191 | Gm49853 | 0.000387543 | -1.339849135 | 0.217194106 |
| ENSMUSG00000024083 | Pja2 | 0.000387543 | -1.051604167 | 0.370945697 |
| ENSMUSG00000024069 | Slc30a6 | 0.000387543 | -0.158519371 | 1 |
| ENSMUSG00000024072 | Yipf4 | 0.000387543 | -0.459688905 | 0.946435944 |
| ENSMUSG00000071042 | Rasgrp3 | 0.000387543 | 0.0942467 | 1 |
| ENSMUSG00000002017 | Fam98a | 0.000387543 | 0.326907457 | 1 |
| ENSMUSG00000024076 | Vit | 0.000387543 | -0.233807498 | 1 |
| ENSMUSG00000005370 | Msh6 | 0.000387543 | 0.0942467 | 1 |
| ENSMUSG00000033855 | Ston1 | 0.000387543 | -0.921020057 | 0.472222633 |
| ENSMUSG00000024231 | Cul2 | 0.000387543 | -1.065409966 | 0.395650934 |
| ENSMUSG00000040957 | Cables1 | 0.000387543 | 0.439382186 | 1 |
| ENSMUSG00000036225 | Kctd1 | 0.000387543 | -0.233807498 | 1 |

| | | | | |
|---|---|---|---|---|
| ENSMUSG00000042514 | Klhl14 | 0.000387543 | -0.281901786 | 1 |
| ENSMUSG00000024378 | Stard4 | 0.000387543 | 1.326907457 | 0.406329712 |
| ENSMUSG00000037236 | Matr3 | 0.000387543 | -2.258055044 | 0.00335241 |
| ENSMUSG00000024350 | Dnajc18 | 0.000387543 | -0.951077291 | 0.496313716 |
| ENSMUSG00000060275 | Nrg2 | 0.000387543 | 0.56137271 | 0.994391499 |
| ENSMUSG00000024472 | Dcp2 | 0.000387543 | -0.184054463 | 1 |
| ENSMUSG00000033184 | Tmed7 | 0.000387543 | -0.459688905 | 0.946435944 |
| ENSMUSG00000024505 | Dtwd2 | 0.000387543 | 0.948395833 | 0.653235135 |
| ENSMUSG00000024613 | Tcof1 | 0.000387543 | -1.293678954 | 0.198966953 |
| ENSMUSG00000109901 | Chmp1b | 0.000387543 | 0.0942467 | 1 |
| ENSMUSG00000024525 | Impa2 | 0.000387543 | 1.326907457 | 0.406329712 |
| ENSMUSG00000024558 | Mapk4 | 0.000387543 | -0.417253639 | 1 |
| ENSMUSG00000024560 | Cxxc1 | 0.000387543 | 0.125273595 | 1 |
| ENSMUSG00000025421 | Hdhd2 | 0.000387543 | 0.125273595 | 1 |
| ENSMUSG00000059336 | Slc14a1 | 0.000387543 | -0.106051951 | 1 |
| ENSMUSG00000059734 | Ndufs8 | 0.000387543 | 1.125273595 | 0.466464457 |
| ENSMUSG00000024851 | Pitpnm1 | 0.000387543 | -0.691014451 | 0.672182136 |
| ENSMUSG00000024854 | Pold4 | 0.000387543 | 0.741944956 | 0.821038733 |
| ENSMUSG00000024889 | Rce1 | 0.000387543 | 0.741944956 | 0.850634638 |
| ENSMUSG00000024909 | Efemp2 | 0.000387543 | -0.351164449 | 1 |
| ENSMUSG00000024816 | Frmd8 | 0.000387543 | -2.422626811 | 0.001306782 |
| ENSMUSG00000024959 | Bad | 0.000387543 | -0.617950989 | 0.801225115 |
| ENSMUSG00000071647 | Eml3 | 0.000387543 | -0.52108945 | 0.871107617 |
| ENSMUSG00000034820 | Cpsf7 | 0.000387543 | -2.939879084 | 5.72E-05 |
| ENSMUSG00000024673 | Ms4a1 | 0.000387543 | -0.233807498 | 1 |
| ENSMUSG00000039982 | Dtx4 | 0.000387543 | -0.52108945 | 0.871107617 |
| ENSMUSG00000024754 | Cemip2 | 0.000387543 | -2.41186038 | 0.001437097 |
| ENSMUSG00000024878 | Cbwd1 | 0.000387543 | -0.951077291 | 0.452335407 |
| ENSMUSG00000016495 | Plgrkt | 0.000387543 | -0.395558568 | 1 |
| ENSMUSG00000024887 | Asah2 | 0.000387543 | -0.72620388 | 0.705274189 |
| ENSMUSG00000025188 | Hps1 | 0.000387543 | -0.079084903 | 1 |
| ENSMUSG00000040414 | Slc25a28 | 0.000387543 | -0.500911568 | 0.868661374 |
| ENSMUSG00000057506 | Bloc1s2 | 0.000387543 | -0.417253639 | 1 |
| ENSMUSG00000036450 | Hif1an | 0.000387543 | 0.222570797 | 1 |
| ENSMUSG00000056209 | Npm3 | 0.000387543 | 0.156982455 | 1 |
| ENSMUSG00000099988 | 1700054A03Rik | 0.000387543 | -1.480447466 | 0.132703831 |
| ENSMUSG00000040022 | Rab11fip2 | 0.000387543 | -0.691014451 | 0.672182136 |
| ENSMUSG00000057858 | Fam204a | 0.000387543 | -0.43862729 | 0.943572777 |
| ENSMUSG00000024993 | Fam45a | 0.000387543 | -1.987789069 | 0.013518154 |
| ENSMUSG00000031157 | Pqbp1 | 0.000387543 | 0.256518129 | 1 |
| ENSMUSG00000031168 | Ebp | 0.000387543 | 0.694639241 | 0.833666942 |
| ENSMUSG00000101233 | Gm29242 | 0.000387543 | 1.256518129 | 0.466464457 |
| ENSMUSG00000036959 | Bcorl1 | 0.000387543 | -1.866864287 | 0.023669966 |
| ENSMUSG00000031197 | Vbp1 | 0.000387543 | -0.500911568 | 0.868661374 |
| ENSMUSG00000035725 | Prkx | 0.000387543 | 0.519552534 | 0.996628004 |
| ENSMUSG00000059327 | Eda | 0.000387543 | -0.691014451 | 0.672182136 |
| ENSMUSG00000031266 | Gla | 0.000387543 | 2.189403933 | 0.067154141 |
| ENSMUSG00000018595 | Glra4 | 0.000387543 | -1.540989007 | 0.087399301 |
| ENSMUSG00000031429 | Psmd10 | 0.000387543 | 2.189403933 | 0.093649769 |
| ENSMUSG00000031273 | Col4a6 | 0.000387543 | -1.645785197 | 0.068476472 |
| ENSMUSG00000041096 | Tspyl2 | 0.000387543 | -1.119348773 | 0.341720553 |

| | | | | |
|---|---|---|---|---|
| ENSMUSG00000086316 | Nbdy | 0.000387543 | 0.694639241 | 0.833666942 |
| ENSMUSG00000031379 | Pir | 0.000387543 | -0.158519371 | 1 |
| ENSMUSG00000079671 | 2610203C22Rik | 0.00033218 | 1.368727632 | 0.466958516 |
| ENSMUSG00000042686 | Jph1 | 0.00033218 | 1.063873051 | 0.559988061 |
| ENSMUSG00000109887 | Gm28756 | 0.00033218 | 1.549299878 | 0.278293619 |
| ENSMUSG00000102336 | Gm37233 | 0.00033218 | -0.351164449 | 1 |
| ENSMUSG00000026062 | Slc9a2 | 0.00033218 | -0.883659529 | 0.662198438 |
| ENSMUSG00000045954 | Cavin2 | 0.00033218 | -2.121238355 | 0.009680621 |
| ENSMUSG00000026094 | Stk17b | 0.00033218 | 0.411796354 | 1 |
| ENSMUSG00000025978 | Rftn2 | 0.00033218 | -0.271729981 | 1 |
| ENSMUSG00000054770 | Kctd18 | 0.00033218 | 0.134262379 | 1 |
| ENSMUSG00000026017 | Carf | 0.00033218 | -0.450700122 | 0.9980272 |
| ENSMUSG00000025997 | Ikzf2 | 0.00033218 | -1.865737621 | 0.027697231 |
| ENSMUSG00000026192 | Atic | 0.00033218 | -0.986753022 | 0.45060963 |
| ENSMUSG00000100534 | Gm816 | 0.00033218 | 2.286265472 | 0.117709307 |
| ENSMUSG00000032883 | Acsl3 | 0.00033218 | -0.426452576 | 0.994894633 |
| ENSMUSG00000079470 | Utp14b | 0.00033218 | -1.733634085 | 0.046362955 |
| ENSMUSG00000036206 | Sh3bp4 | 0.00033218 | -2.216234868 | 0.006130108 |
| ENSMUSG00000044337 | Ackr3 | 0.00033218 | -1.498005837 | 0.115347233 |
| ENSMUSG00000034292 | Traf3ip1 | 0.00033218 | -1.32516924 | 0.207356329 |
| ENSMUSG00000026275 | Ppp1r7 | 0.00033218 | -0.936126949 | 0.467957538 |
| ENSMUSG00000026279 | Thap4 | 0.00033218 | -1.462672764 | 0.139842794 |
| ENSMUSG00000073602 | Serpinb3b | 0.00033218 | 1.134262379 | 0.575521514 |
| ENSMUSG00000026389 | Steap3 | 0.00033218 | 0.246737108 | 1 |
| ENSMUSG00000042510 | AA986860 | 0.00033218 | -1.158519371 | 0.324379889 |
| ENSMUSG00000042349 | Ikbke | 0.00033218 | -2.128772027 | 0.008348479 |
| ENSMUSG00000026463 | Atp2b4 | 0.00033218 | -1.56617734 | 0.095375956 |
| ENSMUSG00000026452 | Syt2 | 0.00033218 | 0.701302971 | 0.82150906 |
| ENSMUSG00000032649 | Colgalt2 | 0.00033218 | 1.286265472 | 0.484833447 |
| ENSMUSG00000100627 | A830008E24Rik | 0.00033218 | -2.538162963 | 0.001608894 |
| ENSMUSG00000056708 | Ier5 | 0.00033218 | 0.501994163 | 0.971854959 |
| ENSMUSG00000026587 | Astn1 | 0.00033218 | -0.901361531 | 0.57206629 |
| ENSMUSG00000073530 | Pappa2 | 0.00033218 | 0.648835551 | 0.969214425 |
| ENSMUSG00000040782 | Cop1 | 0.00033218 | -2.46862203 | 0.001502231 |
| ENSMUSG00000104302 | Gm36975 | 0.00033218 | 0.598209478 | 0.970970089 |
| ENSMUSG00000040297 | Suco | 0.00033218 | -1.52108945 | 0.101869423 |
| ENSMUSG00000026577 | Blzf1 | 0.00033218 | -0.810596067 | 0.66416768 |
| ENSMUSG00000026571 | Dcaf6 | 0.00033218 | -3.295353586 | 8.21E-06 |
| ENSMUSG00000040723 | Rcsd1 | 0.00033218 | 0.812334284 | 0.771834758 |
| ENSMUSG00000060568 | Fam78b | 0.00033218 | -1.620625124 | 0.073315436 |
| ENSMUSG00000003458 | Ncstn | 0.00033218 | -0.351164449 | 1 |
| ENSMUSG00000091476 | Catspere2 | 0.00033218 | -1.271729981 | 0.228247077 |
| ENSMUSG00000038806 | Sde2 | 0.00033218 | -0.588203646 | 0.840804483 |
| ENSMUSG00000047369 | Dnah14 | 0.00033218 | 0.598209478 | 0.953305458 |
| ENSMUSG00000004880 | Lbr | 0.00033218 | -1.970074281 | 0.019037404 |
| ENSMUSG00000037461 | Ints7 | 0.00033218 | -1.426452576 | 0.143740392 |
| ENSMUSG00000026648 | Dclre1c | 0.00033218 | -0.609898717 | 0.844591606 |
| ENSMUSG00000058594 | Fbh1 | 0.00033218 | -1.035662623 | 0.4594611 |
| ENSMUSG00000086358 | Gm13270 | 0.00033218 | 2.456190473 | 0.090628403 |
| ENSMUSG00000051154 | Commd3 | 0.00033218 | 0.812334284 | 0.771834758 |
| ENSMUSG00000037683 | Armc3 | 0.00033218 | 1.063873051 | 0.59863828 |

| | | | | |
|---|---|---|---|---|
| ENSMUSG00000026779 | Mastl | 0.00033218 | -0.158519371 | 1 |
| ENSMUSG00000036850 | Mrpl41 | 0.00033218 | 1.648835551 | 0.279592421 |
| ENSMUSG00000026959 | Grin1 | 0.00033218 | -0.067371483 | 1 |
| ENSMUSG00000015787 | Abo | 0.00033218 | 3.871227973 | 0.00517567 |
| ENSMUSG00000026803 | Ttf1 | 0.00033218 | -1.414174246 | 0.167654895 |
| ENSMUSG00000035513 | Ntng2 | 0.00033218 | -0.035662623 | 1 |
| ENSMUSG00000026857 | Ntmt1 | 0.00033218 | -1.051604167 | 0.405981552 |
| ENSMUSG00000079442 | St6galnac4 | 0.00033218 | -0.376699541 | 1 |
| ENSMUSG00000009566 | Fpgs | 0.00033218 | 0.411796354 | 1 |
| ENSMUSG00000057110 | Cntrl | 0.00033218 | -1.847590275 | 0.031101228 |
| ENSMUSG00000052155 | Acvr2a | 0.00033218 | -1.202021009 | 0.289810004 |
| ENSMUSG00000000392 | Fap | 0.00033218 | -0.953200463 | 0.49132419 |
| ENSMUSG00000026888 | Grb14 | 0.00033218 | 0.648835551 | 0.941472227 |
| ENSMUSG00000075316 | Scn9a | 0.00033218 | 0.755750755 | 0.791515963 |
| ENSMUSG00000042064 | Myo3b | 0.00033218 | -1.202021009 | 0.331463239 |
| ENSMUSG00000041921 | Metap1d | 0.00033218 | -0.847590275 | 0.588477963 |
| ENSMUSG00000070866 | Zfp804a | 0.00033218 | 0.326907457 | 1 |
| ENSMUSG00000051329 | Nup160 | 0.00033218 | -1.723718617 | 0.054117915 |
| ENSMUSG00000063235 | Ptpmt1 | 0.00033218 | 0.501994163 | 0.971854959 |
| ENSMUSG00000027244 | Atg13 | 0.00033218 | -0.810596067 | 0.66416768 |
| ENSMUSG00000027217 | Tspan18 | 0.00033218 | -1.067371483 | 0.431388315 |
| ENSMUSG00000032724 | Abtb2 | 0.00033218 | -1.244249245 | 0.261609949 |
| ENSMUSG00000040383 | Aqr | 0.00033218 | -0.936126949 | 0.467957538 |
| ENSMUSG00000040282 | BC052040 | 0.00033218 | -1.555036782 | 0.091420641 |
| ENSMUSG00000033808 | Tmem87a | 0.00033218 | -2.173166147 | 0.007109019 |
| ENSMUSG00000033705 | Stard9 | 0.00033218 | -0.401790522 | 1 |
| ENSMUSG00000033526 | Ppip5k1 | 0.00033218 | 0.134262379 | 1 |
| ENSMUSG00000035093 | Secisbp2l | 0.00033218 | -0.609898717 | 0.844591606 |
| ENSMUSG00000027206 | Cops2 | 0.00033218 | -0.474546864 | 0.901106111 |
| ENSMUSG00000098024 | Gm27003 | 0.00033218 | -1.113665135 | 0.394507393 |
| ENSMUSG00000001999 | Blvra | 0.00033218 | 0.20826296 | 1 |
| ENSMUSG00000027367 | Stard7 | 0.00033218 | -1.426452576 | 0.176830814 |
| ENSMUSG00000086831 | Gm14280 | 0.00033218 | -0.351164449 | 1 |
| ENSMUSG00000087377 | AV099323 | 0.00033218 | 1.286265472 | 0.485776178 |
| ENSMUSG00000027346 | Gpcpd1 | 0.00033218 | -1.143722369 | 0.324016775 |
| ENSMUSG00000027384 | Ndufaf5 | 0.00033218 | -1.051604167 | 0.405214457 |
| ENSMUSG00000074764 | Sel1l2 | 0.00033218 | 0.134262379 | 1 |
| ENSMUSG00000027419 | Pcsk2 | 0.00033218 | -0.79173704 | 0.667778757 |
| ENSMUSG00000074749 | Kiz | 0.00033218 | -1.244249245 | 0.261609949 |
| ENSMUSG00000033068 | Entpd6 | 0.00033218 | -0.713734528 | 0.706929043 |
| ENSMUSG00000068115 | Ninl | 0.00033218 | -1.426452576 | 0.143740392 |
| ENSMUSG00000027469 | Tpx2 | 0.00033218 | -1.874726405 | 0.028429451 |
| ENSMUSG00000059842 | Zfp341 | 0.00033218 | -0.673092543 | 0.792441218 |
| ENSMUSG00000027612 | Mmp24 | 0.00033218 | 0.326907457 | 1 |
| ENSMUSG00000074649 | BC029722 | 0.00033218 | 0.812334284 | 0.792441218 |
| ENSMUSG00000055485 | Soga1 | 0.00033218 | -1.414174246 | 0.167654895 |
| ENSMUSG00000035877 | Zhx3 | 0.00033218 | -1.143722369 | 0.319437035 |
| ENSMUSG00000017817 | Jph2 | 0.00033218 | 0.326907457 | 1 |
| ENSMUSG00000002718 | Cse1l | 0.00033218 | -0.351164449 | 1 |
| ENSMUSG00000085596 | Gm11476 | 0.00033218 | -0.035662623 | 1 |
| ENSMUSG00000027540 | Ptpn1 | 0.00033218 | -1.067371483 | 0.410337547 |

| | | | | |
|---|---|---|---|---|
| ENSMUSG00000044641 | Pard6b | 0.00033218 | -1.035662623 | 0.463954265 |
| ENSMUSG00000078919 | Dpm1 | 0.00033218 | -1.244249245 | 0.262284476 |
| ENSMUSG00000052033 | Pfdn4 | 0.00033218 | 0.134262379 | 1 |
| ENSMUSG00000016257 | Prelid3b | 0.00033218 | 0.932628517 | 0.736456705 |
| ENSMUSG00000039117 | Taf4 | 0.00033218 | -1.723718617 | 0.054117915 |
| ENSMUSG00000025255 | Zfhx4 | 0.00033218 | 0.134262379 | 1 |
| ENSMUSG00000027508 | Pag1 | 0.00033218 | -0.187665716 | 1 |
| ENSMUSG00000027530 | Fabp12 | 0.00033218 | 0.932628517 | 0.736456705 |
| ENSMUSG00000102886 | Gm30667 | 0.00033218 | 0.812334284 | 0.796840156 |
| ENSMUSG00000027699 | Ect2 | 0.00033218 | -1.414174246 | 0.167654895 |
| ENSMUSG00000069072 | Slc7a14 | 0.00033218 | 0.098638469 | 1 |
| ENSMUSG00000049940 | Pgrmc2 | 0.00033218 | 0.501994163 | 0.971854959 |
| ENSMUSG00000023087 | Noct | 0.00033218 | 0.755750755 | 0.791515963 |
| ENSMUSG00000027739 | Rab33b | 0.00033218 | -0.216234868 | 1 |
| ENSMUSG00000049504 | Proser1 | 0.00033218 | -0.810596067 | 0.66416768 |
| ENSMUSG00000036894 | Rap2b | 0.00033218 | 0.063873051 | 1 |
| ENSMUSG00000027998 | Plrg1 | 0.00033218 | 0.286265472 | 1 |
| ENSMUSG00000003382 | Etv3 | 0.00033218 | -0.401790522 | 0.992008082 |
| ENSMUSG00000028066 | Pmf1 | 0.00033218 | -0.098398378 | 1 |
| ENSMUSG00000106218 | Gm43713 | 0.00033218 | 0.368727632 | 1 |
| ENSMUSG00000042766 | Trim46 | 0.00033218 | 0.063873051 | 1 |
| ENSMUSG00000028042 | Zbtb7b | 0.00033218 | -0.753262892 | 0.737916568 |
| ENSMUSG00000042613 | Pbxip1 | 0.00033218 | -1.555036782 | 0.091420641 |
| ENSMUSG00000027933 | Ints3 | 0.00033218 | -1.32516924 | 0.207356329 |
| ENSMUSG00000068874 | Selenbp1 | 0.00033218 | 0.996758855 | 0.591388707 |
| ENSMUSG00000038861 | Pi4kb | 0.00033218 | -2.223289626 | 0.00530821 |
| ENSMUSG00000092607 | Scnm1 | 0.00033218 | -0.652333983 | 0.774790843 |
| ENSMUSG00000053192 | Mllt11 | 0.00033218 | -1.216234868 | 0.293900105 |
| ENSMUSG00000015522 | Arnt | 0.00033218 | -1.927513819 | 0.023487243 |
| ENSMUSG00000068856 | Sf3b4 | 0.00033218 | -0.067371483 | 1 |
| ENSMUSG00000057123 | Gja5 | 0.00033218 | 0.246737108 | 1 |
| ENSMUSG00000027858 | Tspan2 | 0.00033218 | -0.631272368 | 0.771834758 |
| ENSMUSG00000002227 | Mov10 | 0.00033218 | -1.389299577 | 0.170147907 |
| ENSMUSG00000040809 | Chil3 | 0.00033218 | -1.098398378 | 0.466464457 |
| ENSMUSG00000027900 | Dram2 | 0.00033218 | -1.883659529 | 0.029198945 |
| ENSMUSG00000056260 | Lrif1 | 0.00033218 | -1.216234868 | 0.293900105 |
| ENSMUSG00000048109 | Rbm15 | 0.00033218 | -1.995020638 | 0.017369611 |
| ENSMUSG00000068740 | Celsr2 | 0.00033218 | -0.543809527 | 0.912904617 |
| ENSMUSG00000054414 | Slc30a7 | 0.00033218 | -1.486324032 | 0.113617082 |
| ENSMUSG00000097852 | 4933405D12Rik | 0.00033218 | 1.368727632 | 0.393605552 |
| ENSMUSG00000051278 | Zgrf1 | 0.00033218 | -0.986753022 | 0.45060963 |
| ENSMUSG00000105085 | Gm42567 | 0.00033218 | 0.812334284 | 0.771834758 |
| ENSMUSG00000028159 | Dapp1 | 0.00033218 | -1.298697029 | 0.236225921 |
| ENSMUSG00000074182 | Znhit6 | 0.00033218 | -2.098398378 | 0.010622787 |
| ENSMUSG00000039167 | Adgrl4 | 0.00033218 | -0.187665716 | 1 |
| ENSMUSG00000039037 | St6galnac5 | 0.00033218 | 0.368727632 | 1 |
| ENSMUSG00000028199 | Cryz | 0.00033218 | 0.368727632 | 1 |
| ENSMUSG00000040728 | Esrp1 | 0.00033218 | -1.202021009 | 0.303461557 |
| ENSMUSG00000028218 | Fam92a | 0.00033218 | 0.246737108 | 1 |
| ENSMUSG00000028221 | Pip4p2 | 0.00033218 | -2.278519147 | 0.004062275 |
| ENSMUSG00000085522 | 4930556G01Rik | 0.00033218 | 0.456190473 | 1 |

| | | | | |
|---|---|---|---|---|
| ENSMUSG00000040183 | Ankrd6 | 0.00033218 | 0.170788255 | 1 |
| ENSMUSG00000028292 | Rars2 | 0.00033218 | -1.098398378 | 0.369032695 |
| ENSMUSG00000028293 | Slc35a1 | 0.00033218 | 0.598209478 | 0.960947216 |
| ENSMUSG00000028455 | Stoml2 | 0.00033218 | -0.035662623 | 1 |
| ENSMUSG00000036002 | Fam214b | 0.00033218 | -1.311993851 | 0.203883351 |
| ENSMUSG00000039809 | Gabbr2 | 0.00033218 | -0.187665716 | 1 |
| ENSMUSG00000007613 | Tgfbr1 | 0.00033218 | -2.662750599 | 0.000519924 |
| ENSMUSG00000063446 | Plppr1 | 0.00033218 | -0.003241145 | 1 |
| ENSMUSG00000015247 | Nipsnap3b | 0.00033218 | 1.549299878 | 0.278293619 |
| ENSMUSG00000015243 | Abca1 | 0.00033218 | -1.693556646 | 0.060182632 |
| ENSMUSG00000028403 | Zdhhc21 | 0.00033218 | -1.401790522 | 0.172973155 |
| ENSMUSG00000070923 | Klhl9 | 0.00033218 | -0.450700122 | 0.9980272 |
| ENSMUSG00000052684 | Jun | 0.00033218 | -1.258055044 | 0.277974519 |
| ENSMUSG00000028549 | Itgb3bp | 0.00033218 | -1.577232528 | 0.095045546 |
| ENSMUSG00000057722 | Lepr | 0.00033218 | -0.693556646 | 0.815616279 |
| ENSMUSG00000028708 | Mknk1 | 0.00033218 | 0.098638469 | 1 |
| ENSMUSG00000028698 | Pik3r3 | 0.00033218 | -0.035662623 | 1 |
| ENSMUSG00000028729 | Ebna1bp2 | 0.00033218 | -0.32516924 | 1 |
| ENSMUSG00000028645 | Slc2a1 | 0.00033218 | -0.351164449 | 1 |
| ENSMUSG00000028647 | Mycbp | 0.00033218 | 0.029925719 | 1 |
| ENSMUSG00000028894 | Inpp5b | 0.00033218 | -0.693556646 | 0.69244122 |
| ENSMUSG00000028890 | Mtf1 | 0.00033218 | -0.098398378 | 1 |
| ENSMUSG00000028847 | Trappc3 | 0.00033218 | -0.847590275 | 0.588477963 |
| ENSMUSG00000028800 | Hdac1 | 0.00033218 | -1.641841609 | 0.077204065 |
| ENSMUSG00000028874 | Fgr | 0.00033218 | 0.134262379 | 1 |
| ENSMUSG00000047281 | Sfn | 0.00033218 | -0.376699541 | 1 |
| ENSMUSG00000037366 | Pafah2 | 0.00033218 | -1.035662623 | 0.466464457 |
| ENSMUSG00000028803 | Nipal3 | 0.00033218 | -0.772628217 | 0.66416768 |
| ENSMUSG00000036905 | C1qb | 0.00033218 | 1.368727632 | 0.466464457 |
| ENSMUSG00000041143 | Tmco4 | 0.00033218 | -0.158519371 | 1 |
| ENSMUSG00000078517 | Emc1 | 0.00033218 | -1.003241145 | 0.452335407 |
| ENSMUSG00000040697 | Dnajc16 | 0.00033218 | 0.326907457 | 1 |
| ENSMUSG00000029007 | Agtrap | 0.00033218 | 0.812334284 | 0.771834758 |
| ENSMUSG00000029003 | Mad2l2 | 0.00033218 | -0.543809527 | 0.919998879 |
| ENSMUSG00000017264 | Exosc10 | 0.00033218 | -1.762978047 | 0.044637458 |
| ENSMUSG00000028955 | Vamp3 | 0.00033218 | 0.701302971 | 0.929603462 |
| ENSMUSG00000028943 | Espn | 0.00033218 | 0.456190473 | 1 |
| ENSMUSG00000029032 | Arhgef16 | 0.00033218 | 0.286265472 | 1 |
| ENSMUSG00000042202 | Slc35e2 | 0.00033218 | -0.865737621 | 0.593002749 |
| ENSMUSG00000029036 | Atad3a | 0.00033218 | -1.474546864 | 0.130342151 |
| ENSMUSG00000029066 | Mrpl20 | 0.00033218 | -0.216234868 | 1 |
| ENSMUSG00000040570 | Rundc3b | 0.00033218 | -0.271729981 | 1 |
| ENSMUSG00000042476 | Abcb4 | 0.00033218 | 1.368727632 | 0.409502039 |
| ENSMUSG00000003974 | Grm3 | 0.00033218 | -0.098398378 | 1 |
| ENSMUSG00000048520 | Fbxl13 | 0.00033218 | 1.134262379 | 0.571758706 |
| ENSMUSG00000029017 | Pmpcb | 0.00033218 | -0.158519371 | 1 |
| ENSMUSG00000042453 | Reln | 0.00033218 | -0.298697029 | 1 |
| ENSMUSG00000075703 | Selenoi | 0.00033218 | -2.474546864 | 0.001550379 |
| ENSMUSG00000029177 | Cenpa | 0.00033218 | -0.56617734 | 0.83628546 |
| ENSMUSG00000037339 | Fam53a | 0.00033218 | -1.426452576 | 0.143740392 |
| ENSMUSG00000029094 | Afap1 | 0.00033218 | -2.019542957 | 0.015900288 |

| | | | | |
|---|---|---|---|---|
| ENSMUSG00000029192 | Tbc1d14 | 0.00033218 | -0.953200463 | 0.472222633 |
| ENSMUSG00000054920 | Klhl5 | 0.00033218 | -0.829211745 | 0.587454349 |
| ENSMUSG00000005220 | Corin | 0.00033218 | -1.363988489 | 0.222382174 |
| ENSMUSG00000035540 | Gc | 0.00033218 | -0.543809527 | 0.951660573 |
| ENSMUSG00000029415 | Sdad1 | 0.00033218 | -1.693556646 | 0.060182632 |
| ENSMUSG00000033773 | Rpap2 | 0.00033218 | -0.79173704 | 0.659031063 |
| ENSMUSG00000056531 | Ccdc18 | 0.00033218 | -1.035662623 | 0.4594611 |
| ENSMUSG00000029502 | Golga3 | 0.00033218 | -1.599091962 | 0.083247511 |
| ENSMUSG00000043323 | Fbrsl1 | 0.00033218 | -1.52108945 | 0.101869423 |
| ENSMUSG00000029512 | Ulk1 | 0.00033218 | -0.772628217 | 0.66416768 |
| ENSMUSG00000050017 | Pitpnb | 0.00033218 | -1.216234868 | 0.293900105 |
| ENSMUSG00000041930 | Fam222a | 0.00033218 | 1.871227973 | 0.167904846 |
| ENSMUSG00000038582 | Pptc7 | 0.00033218 | -1.474546864 | 0.139039734 |
| ENSMUSG00000029463 | Fam216a | 0.00033218 | -0.865737621 | 0.593002749 |
| ENSMUSG00000029466 | Anapc7 | 0.00033218 | -0.376699541 | 1 |
| ENSMUSG00000029471 | Camkk2 | 0.00033218 | -1.733634085 | 0.046362955 |
| ENSMUSG00000029475 | Kdm2b | 0.00033218 | -1.673092543 | 0.094535947 |
| ENSMUSG00000054434 | Tmem120b | 0.00033218 | -0.829211745 | 0.587454349 |
| ENSMUSG00000029387 | Gtf2h3 | 0.00033218 | 0.501994163 | 0.971854959 |
| ENSMUSG00000034324 | Tmem132c | 0.00033218 | -0.450700122 | 1 |
| ENSMUSG00000025533 | Asl | 0.00033218 | 0.411796354 | 1 |
| ENSMUSG00000034118 | Tpst1 | 0.00033218 | -0.351164449 | 1 |
| ENSMUSG00000004947 | Dtx2 | 0.00033218 | -0.298697029 | 1 |
| ENSMUSG00000058291 | Zfp68 | 0.00033218 | 0.456190473 | 1 |
| ENSMUSG00000025858 | Get4 | 0.00033218 | 0.170788255 | 1 |
| ENSMUSG00000029576 | Radil | 0.00033218 | 1.368727632 | 0.398502279 |
| ENSMUSG00000029634 | Rnf6 | 0.00033218 | -0.216234868 | 1 |
| ENSMUSG00000033970 | Rfc3 | 0.00033218 | 0.932628517 | 0.736456705 |
| ENSMUSG00000032667 | Pon2 | 0.00033218 | -0.376699541 | 1 |
| ENSMUSG00000029638 | Glcci1 | 0.00033218 | 0.871227973 | 0.748991588 |
| ENSMUSG00000029571 | Tmem106b | 0.00033218 | -0.271729981 | 1 |
| ENSMUSG00000039130 | Zc3hc1 | 0.00033218 | -0.158519371 | 1 |
| ENSMUSG00000087379 | Gm13849 | 0.00033218 | 1.549299878 | 0.318970047 |
| ENSMUSG00000038871 | Bpgm | 0.00033218 | 0.063873051 | 1 |
| ENSMUSG00000005225 | Plekha8 | 0.00033218 | 0.598209478 | 0.95815809 |
| ENSMUSG00000037973 | Itprid1 | 0.00033218 | 1.456190473 | 0.33333983 |
| ENSMUSG00000029787 | Avl9 | 0.00033218 | -0.498005837 | 1 |
| ENSMUSG00000029913 | Prdm5 | 0.00033218 | -0.829211745 | 0.599865044 |
| ENSMUSG00000056698 | Elmod3 | 0.00033218 | 0.996758855 | 0.639438208 |
| ENSMUSG00000071356 | Reg3b | 0.00033218 | 1.456190473 | 0.350255143 |
| ENSMUSG00000030045 | Mrpl19 | 0.00033218 | 0.812334284 | 0.771834758 |
| ENSMUSG00000034930 | Rtkn | 0.00033218 | -0.32516924 | 1 |
| ENSMUSG00000046679 | C87436 | 0.00033218 | -1.901361531 | 0.037941996 |
| ENSMUSG00000030048 | Gkn3 | 0.00033218 | 0.368727632 | 1 |
| ENSMUSG00000030047 | Arhgap25 | 0.00033218 | 1.286265472 | 0.486197134 |
| ENSMUSG00000030055 | Rab43 | 0.00033218 | -2.609898717 | 0.000715085 |
| ENSMUSG00000030093 | Wnt7a | 0.00033218 | -1.158519371 | 0.349674462 |
| ENSMUSG00000030061 | Uba3 | 0.00033218 | -1.143722369 | 0.452335407 |
| ENSMUSG00000030092 | Cntn6 | 0.00033218 | 0.098638469 | 1 |
| ENSMUSG00000030278 | Cidec | 0.00033218 | 0.411796354 | 1 |
| ENSMUSG00000033933 | Vhl | 0.00033218 | -0.376699541 | 1 |

| | | | | |
|---|---|---|---|---|
| ENSMUSG00000030322 | Mbd4 | 0.00033218 | -0.298697029 | 1 |
| ENSMUSG00000059689 | Zfp637 | 0.00033218 | 0.871227973 | 0.748991588 |
| ENSMUSG00000041460 | Cacna2d4 | 0.00033218 | 1.549299878 | 0.334416634 |
| ENSMUSG00000107736 | Gm44148 | 0.00033218 | 0.456190473 | 1 |
| ENSMUSG00000098318 | Lockd | 0.00033218 | -0.035662623 | 1 |
| ENSMUSG00000030245 | Golt1b | 0.00033218 | 0.932628517 | 0.66416768 |
| ENSMUSG00000030259 | Rassf8 | 0.00033218 | -0.986753022 | 0.45060963 |
| ENSMUSG00000030255 | Sspn | 0.00033218 | -0.986753022 | 0.499088689 |
| ENSMUSG00000040112 | Mrps35 | 0.00033218 | -0.158519371 | 1 |
| ENSMUSG00000030304 | Ergic2 | 0.00033218 | -1.351164449 | 0.180821216 |
| ENSMUSG00000039958 | Etfbkmt | 0.00033218 | -0.158519371 | 1 |
| ENSMUSG00000019734 | Tmc4 | 0.00033218 | -0.673092543 | 0.778722129 |
| ENSMUSG00000008435 | Rdh13 | 0.00033218 | 0.996758855 | 0.591388707 |
| ENSMUSG00000001918 | Slc1a5 | 0.00033218 | -1.019542957 | 0.455131546 |
| ENSMUSG00000002043 | Trappc6a | 0.00033218 | -1.244249245 | 0.261609949 |
| ENSMUSG00000002992 | Apoc2 | 0.00033218 | 2.871227973 | 0.0289692 |
| ENSMUSG00000040511 | Pvr | 0.00033218 | -0.713734528 | 0.733672617 |
| ENSMUSG00000045693 | Nlrp4e | 0.00033218 | 2.286265472 | 0.082902688 |
| ENSMUSG00000040857 | Erf | 0.00033218 | 0.648835551 | 0.863905162 |
| ENSMUSG00000061286 | Exosc5 | 0.00033218 | 0.20826296 | 1 |
| ENSMUSG00000002602 | Axl | 0.00033218 | -1.426452576 | 0.143740392 |
| ENSMUSG00000040488 | Ltbp4 | 0.00033218 | -1.338225393 | 0.210892072 |
| ENSMUSG00000040466 | Blvrb | 0.00033218 | -0.035662623 | 1 |
| ENSMUSG00000055200 | Sertad3 | 0.00033218 | 1.134262379 | 0.528505953 |
| ENSMUSG00000049643 | 2310022A10Rik | 0.00033218 | -0.401790522 | 0.99227179 |
| ENSMUSG00000037640 | Zfp60 | 0.00033218 | -0.187665716 | 1 |
| ENSMUSG00000030592 | Ryr1 | 0.00033218 | 0.501994163 | 0.97378787 |
| ENSMUSG00000078768 | Zfp566 | 0.00033218 | -1.202021009 | 0.296122408 |
| ENSMUSG00000036459 | Wtip | 0.00033218 | -1.338225393 | 0.217655393 |
| ENSMUSG00000023072 | Cep89 | 0.00033218 | -1.641841609 | 0.077204065 |
| ENSMUSG00000074158 | Zfp976 | 0.00033218 | -0.588203646 | 0.840804483 |
| ENSMUSG00000038539 | Atf5 | 0.00033218 | 0.411796354 | 1 |
| ENSMUSG00000002963 | Pnkp | 0.00033218 | 1.134262379 | 0.528505953 |
| ENSMUSG00000003421 | Nosip | 0.00033218 | 0.20826296 | 1 |
| ENSMUSG00000019539 | Rcn3 | 0.00033218 | 1.755750755 | 0.211237702 |
| ENSMUSG00000030825 | Hsd17b14 | 0.00033218 | -0.003241145 | 1 |
| ENSMUSG00000054161 | Fam83e | 0.00033218 | 0.648835551 | 0.941472227 |
| ENSMUSG00000039745 | Htatip2 | 0.00033218 | -0.003241145 | 1 |
| ENSMUSG00000030498 | Gas2 | 0.00033218 | -0.216234868 | 1 |
| ENSMUSG00000033429 | Mcee | 0.00033218 | 1.134262379 | 0.528505953 |
| ENSMUSG00000030515 | Tarsl2 | 0.00033218 | -0.474546864 | 0.901106111 |
| ENSMUSG00000058145 | Adamts17 | 0.00033218 | -0.298697029 | 1 |
| ENSMUSG00000039043 | Arpin | 0.00033218 | 1.286265472 | 0.477132639 |
| ENSMUSG00000038943 | Prc1 | 0.00033218 | -1.631272368 | 0.075237108 |
| ENSMUSG00000025586 | Cpeb1 | 0.00033218 | -0.588203646 | 0.840804483 |
| ENSMUSG00000030649 | Anapc15 | 0.00033218 | 0.701302971 | 0.810473388 |
| ENSMUSG00000070426 | Rnf121 | 0.00033218 | -1.082968338 | 0.413835764 |
| ENSMUSG00000063550 | Nup98 | 0.00033218 | -1.652333983 | 0.067853661 |
| ENSMUSG00000044465 | Fam160a2 | 0.00033218 | -0.733634085 | 0.699192777 |
| ENSMUSG00000036989 | Trim3 | 0.00033218 | -0.32516924 | 1 |
| ENSMUSG00000030881 | Arfip2 | 0.00033218 | 0.063873051 | 1 |

| | | | | |
|---|---|---|---|---|
| ENSMUSG00000030894 | Tpp1 | 0.00033218 | 1.20826296 | 0.499921258 |
| ENSMUSG00000036111 | Lmo1 | 0.00033218 | 1.648835551 | 0.262284476 |
| ENSMUSG00000031015 | Swap70 | 0.00033218 | -1.098398378 | 0.361297544 |
| ENSMUSG00000005609 | Ctr9 | 0.00033218 | -1.450700122 | 0.127679449 |
| ENSMUSG00000087475 | 4933406I18Rik | 0.00033218 | -1.961662041 | 0.018507952 |
| ENSMUSG00000048787 | Dcun1d3 | 0.00033218 | -0.543809527 | 0.912904617 |
| ENSMUSG00000063317 | Usp31 | 0.00033218 | -0.829211745 | 0.587454349 |
| ENSMUSG00000030868 | Dctn5 | 0.00033218 | 0.246737108 | 1 |
| ENSMUSG00000030750 | Nsmce1 | 0.00033218 | -0.128772027 | 1 |
| ENSMUSG00000032777 | Gtf3c1 | 0.00033218 | -1.772628217 | 0.042972006 |
| ENSMUSG00000042675 | Ypel3 | 0.00033218 | 1.063873051 | 0.559988061 |
| ENSMUSG00000060538 | Tmem219 | 0.00033218 | -0.128772027 | 1 |
| ENSMUSG00000030811 | Fbxl19 | 0.00033218 | -1.673092543 | 0.057375443 |
| ENSMUSG00000030844 | Rgs10 | 0.00033218 | -1.230310054 | 0.257518409 |
| ENSMUSG00000066979 | Bub3 | 0.00033218 | 0.063873051 | 1 |
| ENSMUSG00000025467 | Prap1 | 0.00033218 | 2.134262379 | 0.120274235 |
| ENSMUSG00000025492 | Ifitm3 | 0.00033218 | 1.20826296 | 0.499921258 |
| ENSMUSG00000051007 | Gatd1 | 0.00033218 | -0.098398378 | 1 |
| ENSMUSG00000045777 | Ifitm10 | 0.00033218 | -1.753262892 | 0.052441913 |
| ENSMUSG00000018819 | Lsp1 | 0.00033218 | -0.753262892 | 0.705338822 |
| ENSMUSG00000101360 | 4933417O13Rik | 0.00033218 | 2.456190473 | 0.060841459 |
| ENSMUSG00000000154 | Slc22a18 | 0.00033218 | -0.829211745 | 0.587454349 |
| ENSMUSG00000037613 | Tnfrsf23 | 0.00033218 | -1.733634085 | 0.046362955 |
| ENSMUSG00000115783 | Bc1 | 0.00033218 | -1.230310054 | 0.257518409 |
| ENSMUSG00000048644 | Ctxn1 | 0.00033218 | 1.368727632 | 0.466464457 |
| ENSMUSG00000002949 | Timm44 | 0.00033218 | -0.543809527 | 0.912904617 |
| ENSMUSG00000023235 | Ccl25 | 0.00033218 | -1.244249245 | 0.261609949 |
| ENSMUSG00000109127 | Gm31135 | 0.00033218 | -0.158519371 | 1 |
| ENSMUSG00000031441 | Atp11a | 0.00033218 | 0.246737108 | 1 |
| ENSMUSG00000038398 | Upf3a | 0.00033218 | -0.865737621 | 0.63765603 |
| ENSMUSG00000047710 | Champ1 | 0.00033218 | 0.063873051 | 1 |
| ENSMUSG00000055675 | Kbtbd11 | 0.00033218 | 0.20826296 | 1 |
| ENSMUSG00000031534 | Smim19 | 0.00033218 | 0.170788255 | 1 |
| ENSMUSG00000031548 | Sfrp1 | 0.00033218 | -0.035662623 | 1 |
| ENSMUSG00000100335 | 2310008N11Rik | 0.00033218 | 1.20826296 | 0.548293945 |
| ENSMUSG00000009628 | Tex15 | 0.00033218 | 1.549299878 | 0.278293619 |
| ENSMUSG00000031596 | Slc7a2 | 0.00033218 | -1.56617734 | 0.093233636 |
| ENSMUSG00000097534 | Gm16675 | 0.00033218 | -0.376699541 | 1 |
| ENSMUSG00000063049 | Ing2 | 0.00033218 | -0.829211745 | 0.587454349 |
| ENSMUSG00000031562 | Dctd | 0.00033218 | -1.703680863 | 0.051793654 |
| ENSMUSG00000031519 | Asb5 | 0.00033218 | -0.244249245 | 1 |
| ENSMUSG00000038257 | Glra3 | 0.00033218 | 0.701302971 | 0.810565466 |
| ENSMUSG00000038005 | Hpf1 | 0.00033218 | -0.753262892 | 0.705338822 |
| ENSMUSG00000002345 | Borcs8 | 0.00033218 | 1.648835551 | 0.376851667 |
| ENSMUSG00000057788 | Ddx49 | 0.00033218 | -1.847590275 | 0.031101228 |
| ENSMUSG00000058301 | Upf1 | 0.00033218 | -1.258055044 | 0.262410478 |
| ENSMUSG00000055707 | Klhl26 | 0.00033218 | -1.311993851 | 0.203883351 |
| ENSMUSG00000031842 | Pde4c | 0.00033218 | 0.170788255 | 1 |
| ENSMUSG00000002910 | Arrdc2 | 0.00033218 | -0.883659529 | 0.521297728 |
| ENSMUSG00000019261 | Map1s | 0.00033218 | -0.401790522 | 0.999962098 |
| ENSMUSG00000074247 | Dda1 | 0.00033218 | 0.701302971 | 0.929603462 |

| | | | | |
|---|---|---|---|---|
| ENSMUSG00000019731 | Slc35e1 | 0.00033218 | 0.996758855 | 0.591388707 |
| ENSMUSG00000005413 | Hmox1 | 0.00033218 | -0.376699541 | 1 |
| ENSMUSG00000058355 | Abce1 | 0.00033218 | -1.158519371 | 0.324379889 |
| ENSMUSG00000019433 | Gipc1 | 0.00033218 | 0.286265472 | 1 |
| ENSMUSG00000043668 | Tox3 | 0.00033218 | 0.501994163 | 1 |
| ENSMUSG00000031748 | Gnao1 | 0.00033218 | 0.286265472 | 1 |
| ENSMUSG00000031778 | Cx3cl1 | 0.00033218 | 0.932628517 | 0.775989718 |
| ENSMUSG00000014776 | Nol3 | 0.00033218 | 0.812334284 | 0.771834758 |
| ENSMUSG00000038604 | Ripor1 | 0.00033218 | -0.772628217 | 0.653247672 |
| ENSMUSG00000053687 | Dpep2 | 0.00033218 | 0.246737108 | 1 |
| ENSMUSG00000033106 | Slc7a6os | 0.00033218 | -1.901361531 | 0.025726109 |
| ENSMUSG00000001672 | Marveld3 | 0.00033218 | -0.918848958 | 0.530172915 |
| ENSMUSG00000010936 | Vac14 | 0.00033218 | -0.652333983 | 0.774790843 |
| ENSMUSG00000031957 | Ctrb1 | 0.00033218 | 2.134262379 | 0.114997164 |
| ENSMUSG00000031952 | Chst5 | 0.00033218 | -1.801197369 | 0.050463874 |
| ENSMUSG00000031845 | Bco1 | 0.00033218 | -0.474546864 | 1 |
| ENSMUSG00000031844 | Hsd17b2 | 0.00033218 | 2.648835551 | 0.046003064 |
| ENSMUSG00000061410 | Zcchc14 | 0.00033218 | -1.801197369 | 0.038715462 |
| ENSMUSG00000019278 | Dpep1 | 0.00033218 | 0.812334284 | 0.778722129 |
| ENSMUSG00000031983 | 2310022B05Rik | 0.00033218 | 0.549299878 | 0.903303381 |
| ENSMUSG00000031986 | Sprtn | 0.00033218 | -1.271729981 | 0.228247077 |
| ENSMUSG00000043051 | Disc1 | 0.00033218 | 0.286265472 | 1 |
| ENSMUSG00000025808 | Ccdc7a | 0.00033218 | 0.368727632 | 1 |
| ENSMUSG00000096385 | Gm11168 | 0.00033218 | 0.598209478 | 0.971854959 |
| ENSMUSG00000025899 | Alkbh8 | 0.00033218 | -1.098398378 | 0.361256382 |
| ENSMUSG00000041124 | Msantd4 | 0.00033218 | -0.733634085 | 0.699192777 |
| ENSMUSG00000031997 | Trpc6 | 0.00033218 | 0.246737108 | 1 |
| ENSMUSG00000046111 | Cep295 | 0.00033218 | -1.555036782 | 0.091420641 |
| ENSMUSG00000050901 | Mtnr1b | 0.00033218 | 1.871227973 | 0.177177527 |
| ENSMUSG00000060510 | Zfp266 | 0.00033218 | -1.753262892 | 0.048881736 |
| ENSMUSG00000032175 | Tyk2 | 0.00033218 | -0.067371483 | 1 |
| ENSMUSG00000057191 | AB124611 | 0.00033218 | 1.063873051 | 0.582735301 |
| ENSMUSG00000040883 | Tmem205 | 0.00033218 | 1.549299878 | 0.278293619 |
| ENSMUSG00000043659 | Npsr1 | 0.00033218 | -0.298697029 | 1 |
| ENSMUSG00000085576 | Dpy19l2 | 0.00033218 | 1.063873051 | 0.562149835 |
| ENSMUSG00000032040 | Dcps | 0.00033218 | -0.401790522 | 0.992008082 |
| ENSMUSG00000049313 | Sorl1 | 0.00033218 | -0.035662623 | 1 |
| ENSMUSG00000032015 | Pou2f3 | 0.00033218 | 1.549299878 | 0.325968639 |
| ENSMUSG00000047880 | Cxcr5 | 0.00033218 | 1.755750755 | 0.20807227 |
| ENSMUSG00000066687 | Zbtb16 | 0.00033218 | -0.918848958 | 0.573970806 |
| ENSMUSG00000000168 | Dlat | 0.00033218 | -0.098398378 | 1 |
| ENSMUSG00000037971 | 1110032A03Rik | 0.00033218 | 1.368727632 | 0.466464457 |
| ENSMUSG00000032051 | Fdx1 | 0.00033218 | 1.20826296 | 0.499921258 |
| ENSMUSG00000111520 | Gm33125 | 0.00033218 | 1.996758855 | 0.157767958 |
| ENSMUSG00000074284 | Gm10658 | 0.00033218 | 0.368727632 | 1 |
| ENSMUSG00000042557 | Sin3a | 0.00033218 | -1.936126949 | 0.020391041 |
| ENSMUSG00000032300 | 1700017B05Rik | 0.00033218 | -1.474546864 | 0.167024802 |
| ENSMUSG00000032312 | Csk | 0.00033218 | 0.134262379 | 1 |
| ENSMUSG00000055720 | Ubl7 | 0.00033218 | 0.098638469 | 1 |
| ENSMUSG00000097651 | 4930461G14Rik | 0.00033218 | 1.755750755 | 0.203485308 |
| ENSMUSG00000025235 | Bbs4 | 0.00033218 | 0.648835551 | 0.941472227 |

| | | | | |
|---|---|---|---|---|
| ENSMUSG00000036867 | Smad6 | 0.00033218 | -1.43862729 | 0.145704891 |
| ENSMUSG00000032394 | Igdcc3 | 0.00033218 | 0.648835551 | 0.951890397 |
| ENSMUSG00000015357 | Clpx | 0.00033218 | -1.819933932 | 0.034141187 |
| ENSMUSG00000036943 | Rab8b | 0.00033218 | -0.865737621 | 0.622315002 |
| ENSMUSG00000032202 | Rab27a | 0.00033218 | 0.326907457 | 1 |
| ENSMUSG00000097836 | Gm26903 | 0.00033218 | 0.170788255 | 1 |
| ENSMUSG00000046186 | Cd109 | 0.00033218 | 0.871227973 | 0.757651781 |
| ENSMUSG00000079427 | Mthfsl | 0.00033218 | 1.063873051 | 0.559988061 |
| ENSMUSG00000032353 | Tmed3 | 0.00033218 | 1.286265472 | 0.477132639 |
| ENSMUSG00000037410 | Tbc1d2b | 0.00033218 | -0.733634085 | 0.700503166 |
| ENSMUSG00000032462 | Pik3cb | 0.00033218 | -1.713734528 | 0.052841181 |
| ENSMUSG00000032534 | Cep63 | 0.00033218 | -1.829211745 | 0.034899857 |
| ENSMUSG00000032531 | Amotl2 | 0.00033218 | -1.051604167 | 0.405214457 |
| ENSMUSG00000039716 | Dock3 | 0.00033218 | -0.035662623 | 1 |
| ENSMUSG00000032590 | Apeh | 0.00033218 | 0.598209478 | 0.951660573 |
| ENSMUSG00000032589 | Bsn | 0.00033218 | -0.035662623 | 1 |
| ENSMUSG00000036057 | Ptpn23 | 0.00033218 | -1.509593811 | 0.117709307 |
| ENSMUSG00000087038 | 2900079G21Rik | 0.00033218 | 1.755750755 | 0.222246903 |
| ENSMUSG00000041012 | Cmtm8 | 0.00033218 | -2.753262892 | 0.000323891 |
| ENSMUSG00000111360 | Gm38642 | 0.00033218 | 1.286265472 | 0.5249062 |
| ENSMUSG00000032540 | Abhd5 | 0.00033218 | 0.501994163 | 0.971854959 |
| ENSMUSG00000032551 | 1110059G10Rik | 0.00033218 | 2.648835551 | 0.042972006 |
| ENSMUSG00000019774 | Mtrf1l | 0.00033218 | -1.723718617 | 0.054921419 |
| ENSMUSG00000019794 | Katna1 | 0.00033218 | -1.363988489 | 0.184141965 |
| ENSMUSG00000050994 | Adgb | 0.00033218 | -0.79173704 | 0.66416768 |
| ENSMUSG00000112089 | Gm47712 | 0.00033218 | 0.701302971 | 0.797939559 |
| ENSMUSG00000112800 | Gm40604 | 0.00033218 | 2.286265472 | 0.089456479 |
| ENSMUSG00000039879 | Heca | 0.00033218 | -0.986753022 | 0.45060963 |
| ENSMUSG00000020003 | Pex7 | 0.00033218 | -0.376699541 | 1 |
| ENSMUSG00000071359 | Tbpl1 | 0.00033218 | 1.755750755 | 0.206412457 |
| ENSMUSG00000037440 | Vnn1 | 0.00033218 | 1.20826296 | 0.499921258 |
| ENSMUSG00000038510 | Rpf2 | 0.00033218 | -1.067371483 | 0.408535739 |
| ENSMUSG00000019813 | Cep57l1 | 0.00033218 | -0.128772027 | 1 |
| ENSMUSG00000048756 | Foxo3 | 0.00033218 | -1.486324032 | 0.113617082 |
| ENSMUSG00000075184 | F930017D23Rik | 0.00033218 | 0.20826296 | 1 |
| ENSMUSG00000112148 | Lilrb4a | 0.00033218 | 0.871227973 | 0.775410248 |
| ENSMUSG00000050953 | Gja1 | 0.00033218 | 0.170788255 | 1 |
| ENSMUSG00000019876 | Pkib | 0.00033218 | 0.411796354 | 1 |
| ENSMUSG00000009646 | Pla2g12b | 0.00033218 | 1.996758855 | 0.135334273 |
| ENSMUSG00000020109 | Dnajb12 | 0.00033218 | -0.847590275 | 0.588477963 |
| ENSMUSG00000020063 | Sirt1 | 0.00033218 | -0.450700122 | 0.9980272 |
| ENSMUSG00000112189 | C730027H18Rik | 0.00033218 | 2.286265472 | 0.082902688 |
| ENSMUSG00000019927 | Ube2d1 | 0.00033218 | -0.128772027 | 1 |
| ENSMUSG00000040009 | Gnaz | 0.00033218 | -1.271729981 | 0.324016775 |
| ENSMUSG00000033416 | Gucd1 | 0.00033218 | -0.918848958 | 0.570195249 |
| ENSMUSG00000001119 | Col6a1 | 0.00033218 | -1.285276513 | 0.231596497 |
| ENSMUSG00000035041 | Creb3l3 | 0.00033218 | 1.456190473 | 0.318057572 |
| ENSMUSG00000034591 | Slc41a2 | 0.00033218 | -0.216234868 | 1 |
| ENSMUSG00000020061 | Mybpc1 | 0.00033218 | 0.549299878 | 0.969187697 |
| ENSMUSG00000019935 | Slc17a8 | 0.00033218 | 1.871227973 | 0.167904846 |
| ENSMUSG00000020024 | Cep83 | 0.00033218 | -1.577232528 | 0.095045546 |

| | | | | |
|---|---|---|---|---|
| ENSMUSG00000074785 | Plxnc1 | 0.00033218 | 0.598209478 | 0.964262694 |
| ENSMUSG00000036676 | Tmtc3 | 0.00033218 | -0.498005837 | 0.912904617 |
| ENSMUSG00000055531 | Cpsf6 | 0.00033218 | -2.128772027 | 0.008348479 |
| ENSMUSG00000052798 | Nup107 | 0.00033218 | -1.113665135 | 0.365716212 |
| ENSMUSG00000020225 | Tmbim4 | 0.00033218 | -0.426452576 | 0.999171521 |
| ENSMUSG00000020115 | Tbk1 | 0.00033218 | -1.143722369 | 0.354250365 |
| ENSMUSG00000034667 | Xpot | 0.00033218 | -1.019542957 | 0.455131546 |
| ENSMUSG00000047642 | D930020B18Rik | 0.00033218 | 0.549299878 | 0.964003521 |
| ENSMUSG00000112873 | Gm48877 | 0.00033218 | 1.20826296 | 0.554625414 |
| ENSMUSG00000025409 | Mbd6 | 0.00033218 | -0.652333983 | 0.774790843 |
| ENSMUSG00000020430 | Pes1 | 0.00033218 | -0.351164449 | 1 |
| ENSMUSG00000009076 | Zmat5 | 0.00033218 | -0.810596067 | 0.66416768 |
| ENSMUSG00000020473 | Aebp1 | 0.00033218 | -0.271729981 | 1 |
| ENSMUSG00000002741 | Ykt6 | 0.00033218 | 0.029925719 | 1 |
| ENSMUSG00000046634 | Pkd1l1 | 0.00033218 | 2.134262379 | 0.108201045 |
| ENSMUSG00000020116 | Pno1 | 0.00033218 | 0.063873051 | 1 |
| ENSMUSG00000020288 | Ahsa2 | 0.00033218 | -0.693556646 | 0.69244122 |
| ENSMUSG00000064090 | Vrk2 | 0.00033218 | -1.285276513 | 0.231596497 |
| ENSMUSG00000020272 | Stk10 | 0.00033218 | -1.801197369 | 0.038715462 |
| ENSMUSG00000085301 | Gm12132 | 0.00033218 | -1.019542957 | 0.472764596 |
| ENSMUSG00000042032 | Mat2b | 0.00033218 | -0.003241145 | 1 |
| ENSMUSG00000087306 | A230004M16Rik | 0.00033218 | -0.753262892 | 0.741652762 |
| ENSMUSG00000063564 | Col23a1 | 0.00033218 | -1.271729981 | 0.24525903 |
| ENSMUSG00000020392 | Cdkn2aipnl | 0.00033218 | 0.598209478 | 0.951660573 |
| ENSMUSG00000000263 | Glra1 | 0.00033218 | 0.501994163 | 0.97950322 |
| ENSMUSG00000020519 | Sap30l | 0.00033218 | -1.143722369 | 0.319437035 |
| ENSMUSG00000037145 | 2210407C18Rik | 0.00033218 | 2.871227973 | 0.0289692 |
| ENSMUSG00000036860 | Mrpl55 | 0.00033218 | 1.456190473 | 0.404572446 |
| ENSMUSG00000019373 | Cops3 | 0.00033218 | -0.79173704 | 0.659031063 |
| ENSMUSG00000020528 | Prpsap2 | 0.00033218 | 0.246737108 | 1 |
| ENSMUSG00000047821 | Trim16 | 0.00033218 | -0.376699541 | 1 |
| ENSMUSG00000020900 | Myh10 | 0.00033218 | -1.003241145 | 0.452335407 |
| ENSMUSG00000041189 | Chrnb1 | 0.00033218 | -0.067371483 | 1 |
| ENSMUSG00000019461 | Plscr3 | 0.00033218 | -0.035662623 | 1 |
| ENSMUSG00000000282 | Mnt | 0.00033218 | -0.450700122 | 0.9980272 |
| ENSMUSG00000020850 | Prpf8 | 0.00033218 | -1.019542957 | 0.455131546 |
| ENSMUSG00000001105 | Ift20 | 0.00033218 | 0.286265472 | 1 |
| ENSMUSG00000017686 | Rhot1 | 0.00033218 | -1.56617734 | 0.093233636 |
| ENSMUSG00000020485 | Supt4a | 0.00033218 | 0.098638469 | 1 |
| ENSMUSG00000039110 | Mycbpap | 0.00033218 | 0.871227973 | 0.748991588 |
| ENSMUSG00000020873 | Slc35b1 | 0.00033218 | 0.871227973 | 0.792441218 |
| ENSMUSG00000038692 | Hoxb4 | 0.00033218 | -1.051604167 | 0.405214457 |
| ENSMUSG00000078695 | Cisd3 | 0.00033218 | 0.871227973 | 0.785504436 |
| ENSMUSG00000018167 | Stard3 | 0.00033218 | -0.52108945 | 0.909293502 |
| ENSMUSG00000006777 | Krt23 | 0.00033218 | -1.113665135 | 0.403635177 |
| ENSMUSG00000001555 | Fkbp10 | 0.00033218 | -1.376699541 | 0.188182248 |
| ENSMUSG00000017801 | Mlx | 0.00033218 | 1.996758855 | 0.154554575 |
| ENSMUSG00000020929 | Eftud2 | 0.00033218 | -1.143722369 | 0.319437035 |
| ENSMUSG00000020936 | Nmt1 | 0.00033218 | -0.474546864 | 0.901106111 |
| ENSMUSG00000020937 | Plcd3 | 0.00033218 | -1.158519371 | 0.324379889 |
| ENSMUSG00000062421 | Arf2 | 0.00033218 | -1.32516924 | 0.207356329 |

| | | | | |
|---|---|---|---|---|
| ENSMUSG00000001983 | Taco1 | 0.00033218 | -2.318596586 | 0.003488509 |
| ENSMUSG00000020723 | Cacng4 | 0.00033218 | 0.932628517 | 0.736456705 |
| ENSMUSG00000020728 | Cep112 | 0.00033218 | -1.883659529 | 0.029198945 |
| ENSMUSG00000041623 | D11Wsu47e | 0.00033218 | -0.067371483 | 1 |
| ENSMUSG00000020733 | Slc9a3r1 | 0.00033218 | -0.271729981 | 1 |
| ENSMUSG00000015542 | Nat9 | 0.00033218 | -0.271729981 | 1 |
| ENSMUSG00000015869 | Prpsap1 | 0.00033218 | -1.143722369 | 0.368445326 |
| ENSMUSG00000017715 | Pgs1 | 0.00033218 | -0.401790522 | 0.992008082 |
| ENSMUSG00000033987 | Dnah17 | 0.00033218 | 0.648835551 | 0.952530143 |
| ENSMUSG00000025138 | Sirt7 | 0.00033218 | -0.52108945 | 0.909293502 |
| ENSMUSG00000025156 | Gps1 | 0.00033218 | 2.134262379 | 0.108201045 |
| ENSMUSG00000039275 | Foxk2 | 0.00033218 | -1.703680863 | 0.051793654 |
| ENSMUSG00000025173 | Wdr45b | 0.00033218 | -2.432552775 | 0.001980192 |
| ENSMUSG00000020654 | Adcy3 | 0.00033218 | -0.128772027 | 1 |
| ENSMUSG00000054459 | Vsnl1 | 0.00033218 | 0.411796354 | 1 |
| ENSMUSG00000020593 | Lpin1 | 0.00033218 | -0.067371483 | 1 |
| ENSMUSG00000116736 | Gm49692 | 0.00033218 | -0.128772027 | 1 |
| ENSMUSG00000020649 | Rrm2 | 0.00033218 | -0.401790522 | 0.992008082 |
| ENSMUSG00000020633 | Dcdc2c | 0.00033218 | 1.456190473 | 0.363008484 |
| ENSMUSG00000020674 | Pxdn | 0.00033218 | -1.713734528 | 0.052841181 |
| ENSMUSG00000020672 | Sntg2 | 0.00033218 | -0.003241145 | 1 |
| ENSMUSG00000036188 | Ankmy2 | 0.00033218 | -1.79173704 | 0.037924279 |
| ENSMUSG00000062198 | 2700097O09Rik | 0.00033218 | -1.389299577 | 0.161349026 |
| ENSMUSG00000035614 | Togaram1 | 0.00033218 | -1.986753022 | 0.016885859 |
| ENSMUSG00000044147 | Arf6 | 0.00033218 | -0.298697029 | 1 |
| ENSMUSG00000034601 | 2700049A03Rik | 0.00033218 | -2.549434076 | 0.001072379 |
| ENSMUSG00000087700 | Gm15283 | 0.00033218 | -1.389299577 | 0.161349026 |
| ENSMUSG00000021061 | Sptb | 0.00033218 | 0.501994163 | 0.980745941 |
| ENSMUSG00000021130 | Galnt16 | 0.00033218 | -0.498005837 | 0.923530731 |
| ENSMUSG00000002679 | Med6 | 0.00033218 | -1.743481871 | 0.04760001 |
| ENSMUSG00000072946 | Ptgr2 | 0.00033218 | -0.883659529 | 0.521297728 |
| ENSMUSG00000042472 | Zfp410 | 0.00033218 | -0.652333983 | 0.774790843 |
| ENSMUSG00000047414 | Flrt2 | 0.00033218 | -0.901361531 | 0.534735391 |
| ENSMUSG00000033530 | Ttc7b | 0.00033218 | -0.609898717 | 0.844591606 |
| ENSMUSG00000041712 | Ubr7 | 0.00033218 | -0.450700122 | 0.9980272 |
| ENSMUSG00000021097 | Clmn | 0.00033218 | -2.258055044 | 0.004365699 |
| ENSMUSG00000084883 | Ccdc85c | 0.00033218 | -1.723718617 | 0.054117915 |
| ENSMUSG00000021263 | Degs2 | 0.00033218 | -0.035662623 | 1 |
| ENSMUSG00000021268 | Meg3 | 0.00033218 | -0.865737621 | 0.593002749 |
| ENSMUSG00000095079 | Igha | 0.00033218 | 0.755750755 | 0.791515963 |
| ENSMUSG00000025323 | Sp4 | 0.00033218 | -0.56617734 | 0.83628546 |
| ENSMUSG00000021211 | Akr1c12 | 0.00033218 | 1.286265472 | 0.477132639 |
| ENSMUSG00000113017 | Gm46401 | 0.00033218 | -0.035662623 | 1 |
| ENSMUSG00000021311 | Mtr | 0.00033218 | -1.772628217 | 0.042972006 |
| ENSMUSG00000057554 | Lgals8 | 0.00033218 | -1.082968338 | 0.413835764 |
| ENSMUSG00000005397 | Nid1 | 0.00033218 | 0.134262379 | 1 |
| ENSMUSG00000021327 | Zkscan3 | 0.00033218 | -1.713734528 | 0.052841181 |
| ENSMUSG00000112963 | Gm6093 | 0.00033218 | 0.812334284 | 0.771834758 |
| ENSMUSG00000021423 | Ly86 | 0.00033218 | -0.772628217 | 0.66416768 |
| ENSMUSG00000114767 | Gm46392 | 0.00033218 | 1.134262379 | 0.566058593 |
| ENSMUSG00000021361 | Tmem14c | 0.00033218 | 0.411796354 | 1 |

| | | | | |
|---|---|---|---|---|
| ENSMUSG00000021371 | Mcur1 | 0.00033218 | -0.450700122 | 0.9980272 |
| ENSMUSG00000021374 | Nup153 | 0.00033218 | -2.098398378 | 0.010622787 |
| ENSMUSG00000021392 | Nol8 | 0.00033218 | -2.003241145 | 0.014974159 |
| ENSMUSG00000021453 | Gadd45g | 0.00033218 | 1.134262379 | 0.528505953 |
| ENSMUSG00000021457 | Syk | 0.00033218 | -1.363988489 | 0.19020889 |
| ENSMUSG00000025869 | Nop16 | 0.00033218 | -0.216234868 | 1 |
| ENSMUSG00000021541 | Trpc7 | 0.00033218 | 0.598209478 | 0.974170342 |
| ENSMUSG00000060176 | Kif27 | 0.00033218 | 1.648835551 | 0.241308529 |
| ENSMUSG00000114384 | 4930455J16Rik | 0.00033218 | 0.932628517 | 0.747121178 |
| ENSMUSG00000021557 | Agtpbp1 | 0.00033218 | -1.772628217 | 0.042972006 |
| ENSMUSG00000074863 | Platr25 | 0.00033218 | -1.32516924 | 0.207356329 |
| ENSMUSG00000097101 | 1810034E14Rik | 0.00033218 | 0.871227973 | 0.748991588 |
| ENSMUSG00000021518 | Ptdss1 | 0.00033218 | -0.56617734 | 0.83628546 |
| ENSMUSG00000021578 | Ccdc127 | 0.00033218 | -1.389299577 | 0.161349026 |
| ENSMUSG00000021583 | Erap1 | 0.00033218 | 0.286265472 | 1 |
| ENSMUSG00000097098 | 9330111N05Rik | 0.00033218 | 0.701302971 | 0.929603462 |
| ENSMUSG00000035762 | Tmem161b | 0.00033218 | -1.311993851 | 0.203883351 |
| ENSMUSG00000021621 | Zcchc9 | 0.00033218 | 0.326907457 | 1 |
| ENSMUSG00000021707 | Dhfr | 0.00033218 | -1.035662623 | 0.466464457 |
| ENSMUSG00000021703 | Serinc5 | 0.00033218 | -0.401790522 | 0.992008082 |
| ENSMUSG00000021702 | Thbs4 | 0.00033218 | 0.063873051 | 1 |
| ENSMUSG00000042167 | Tent2 | 0.00033218 | -1.801197369 | 0.038715462 |
| ENSMUSG00000045312 | Lhfpl2 | 0.00033218 | 0.170788255 | 1 |
| ENSMUSG00000114926 | Gm2379 | 0.00033218 | -1.128772027 | 0.37105914 |
| ENSMUSG00000021636 | Marveld2 | 0.00033218 | -1.202021009 | 0.366575239 |
| ENSMUSG00000048924 | Ccdc125 | 0.00033218 | -1.874726405 | 0.028429451 |
| ENSMUSG00000061474 | Mrps36 | 0.00033218 | 0.20826296 | 1 |
| ENSMUSG00000041417 | Pik3r1 | 0.00033218 | -2.683360879 | 0.000459869 |
| ENSMUSG00000021700 | Rab3c | 0.00033218 | 0.20826296 | 1 |
| ENSMUSG00000021701 | Plk2 | 0.00033218 | -0.609898717 | 0.844591606 |
| ENSMUSG00000032745 | Gpbp1 | 0.00033218 | -3.132524162 | 2.27E-05 |
| ENSMUSG00000032727 | Mier3 | 0.00033218 | -1.509593811 | 0.117709307 |
| ENSMUSG00000085204 | Gm15327 | 0.00033218 | -1.733634085 | 0.051667273 |
| ENSMUSG00000021730 | Hcn1 | 0.00033218 | -1.098398378 | 0.405913817 |
| ENSMUSG00000051729 | Gm5087 | 0.00033218 | 1.134262379 | 0.528505953 |
| ENSMUSG00000043004 | Gng2 | 0.00033218 | -0.003241145 | 1 |
| ENSMUSG00000015971 | Actr8 | 0.00033218 | -0.474546864 | 0.901106111 |
| ENSMUSG00000058351 | Smim4 | 0.00033218 | -0.543809527 | 0.914537994 |
| ENSMUSG00000021798 | Ldb3 | 0.00033218 | 1.368727632 | 0.411460922 |
| ENSMUSG00000049235 | Gm7324 | 0.00033218 | 1.755750755 | 0.252495441 |
| ENSMUSG00000085866 | Gm15218 | 0.00033218 | 0.456190473 | 1 |
| ENSMUSG00000021846 | Peli2 | 0.00033218 | -1.173166147 | 0.342524494 |
| ENSMUSG00000115338 | Pnp | 0.00033218 | 0.812334284 | 0.792441218 |
| ENSMUSG00000004558 | Ndrg2 | 0.00033218 | 0.996758855 | 0.591388707 |
| ENSMUSG00000049295 | Zfp219 | 0.00033218 | -1.588203646 | 0.082076736 |
| ENSMUSG00000049532 | Sall2 | 0.00033218 | 2.648835551 | 0.042972006 |
| ENSMUSG00000022175 | Lrp10 | 0.00033218 | -0.035662623 | 1 |
| ENSMUSG00000023110 | Prmt5 | 0.00033218 | -0.810596067 | 0.66416768 |
| ENSMUSG00000002332 | Dhrs1 | 0.00033218 | 0.456190473 | 1 |
| ENSMUSG00000021951 | Eef1akmt1 | 0.00033218 | -0.271729981 | 1 |
| ENSMUSG00000021982 | Cdadc1 | 0.00033218 | -1.673092543 | 0.057375443 |

| | | | | |
|---|---|---|---|---|
| ENSMUSG00000021993 | Mipep | 0.00033218 | 0.411796354 | 1 |
| ENSMUSG00000022043 | Trim35 | 0.00033218 | -1.52108945 | 0.101869423 |
| ENSMUSG00000022095 | Fam160b2 | 0.00033218 | -1.376699541 | 0.225938083 |
| ENSMUSG00000021998 | Lcp1 | 0.00033218 | 0.286265472 | 1 |
| ENSMUSG00000022009 | Nufip1 | 0.00033218 | -1.577232528 | 0.095045546 |
| ENSMUSG00000055639 | Dach1 | 0.00033218 | -0.772628217 | 0.66416768 |
| ENSMUSG00000091509 | Gm17066 | 0.00033218 | -1.838430275 | 0.030430985 |
| ENSMUSG00000041594 | Tmtc4 | 0.00033218 | -1.474546864 | 0.130342151 |
| ENSMUSG00000022151 | Ttc33 | 0.00033218 | -0.918848958 | 0.581736296 |
| ENSMUSG00000022150 | Dab2 | 0.00033218 | -0.772628217 | 0.66416768 |
| ENSMUSG00000072663 | Spef2 | 0.00033218 | 0.932628517 | 0.66416768 |
| ENSMUSG00000022247 | Brix1 | 0.00033218 | -0.693556646 | 0.789505358 |
| ENSMUSG00000022249 | Ttc23l | 0.00033218 | 0.134262379 | 1 |
| ENSMUSG00000025370 | Cdh9 | 0.00033218 | 1.755750755 | 0.203485308 |
| ENSMUSG00000115204 | Gm36642 | 0.00033218 | 1.456190473 | 0.318057572 |
| ENSMUSG00000115405 | Gm41307 | 0.00033218 | 1.456190473 | 0.318057572 |
| ENSMUSG00000115009 | G930009F23Rik | 0.00033218 | -0.652333983 | 0.775410248 |
| ENSMUSG00000022297 | Fzd6 | 0.00033218 | -0.693556646 | 0.69244122 |
| ENSMUSG00000022425 | Enpp2 | 0.00033218 | 0.134262379 | 1 |
| ENSMUSG00000022371 | Col14a1 | 0.00033218 | 0.368727632 | 1 |
| ENSMUSG00000022360 | Atad2 | 0.00033218 | -2.414174246 | 0.002096839 |
| ENSMUSG00000022372 | Sla | 0.00033218 | 0.411796354 | 1 |
| ENSMUSG00000091412 | Gm2895 | 0.00033218 | 0.20826296 | 1 |
| ENSMUSG00000116068 | Gm49457 | 0.00033218 | 1.755750755 | 0.207969581 |
| ENSMUSG00000022570 | Tsta3 | 0.00033218 | 0.549299878 | 0.971854959 |
| ENSMUSG00000022556 | Hsf1 | 0.00033218 | 0.098638469 | 1 |
| ENSMUSG00000033728 | Lrrc14 | 0.00033218 | -0.376699541 | 1 |
| ENSMUSG00000033697 | Arhgap39 | 0.00033218 | -1.486324032 | 0.113617082 |
| ENSMUSG00000042632 | Pla2g6 | 0.00033218 | -0.829211745 | 0.587454349 |
| ENSMUSG00000009035 | Tmem184b | 0.00033218 | -1.173166147 | 0.343998736 |
| ENSMUSG00000053411 | Cbx7 | 0.00033218 | -0.187665716 | 1 |
| ENSMUSG00000042351 | Grap2 | 0.00033218 | -1.363988489 | 0.193049754 |
| ENSMUSG00000022401 | Xpnpep3 | 0.00033218 | -1.703680863 | 0.059369924 |
| ENSMUSG00000035845 | Alg12 | 0.00033218 | -0.810596067 | 0.66416768 |
| ENSMUSG00000015363 | Trabd | 0.00033218 | -1.128772027 | 0.37105914 |
| ENSMUSG00000055022 | Cntn1 | 0.00033218 | 1.063873051 | 0.559988061 |
| ENSMUSG00000036197 | Gxylt1 | 0.00033218 | -1.801197369 | 0.038715462 |
| ENSMUSG00000022469 | Rapgef3 | 0.00033218 | -1.414174246 | 0.189783899 |
| ENSMUSG00000023018 | Smarcd1 | 0.00033218 | 0.871227973 | 0.748991588 |
| ENSMUSG00000014303 | Glis2 | 0.00033218 | -1.599091962 | 0.083247511 |
| ENSMUSG00000022517 | Mgrn1 | 0.00033218 | -1.311993851 | 0.203883351 |
| ENSMUSG00000022503 | Nubp1 | 0.00033218 | 0.932628517 | 0.775870935 |
| ENSMUSG00000097537 | 2610020C07Rik | 0.00033218 | -1.32516924 | 0.212709851 |
| ENSMUSG00000022684 | Bfar | 0.00033218 | -1.202021009 | 0.289810004 |
| ENSMUSG00000012017 | Scarf2 | 0.00033218 | -0.986753022 | 0.494188684 |
| ENSMUSG00000003531 | Dgcr6 | 0.00033218 | 0.598209478 | 0.951660573 |
| ENSMUSG00000003526 | Prodh | 0.00033218 | -0.401790522 | 0.992008082 |
| ENSMUSG00000022706 | Mrpl40 | 0.00033218 | 0.932628517 | 0.775870935 |
| ENSMUSG00000022856 | Tmem41a | 0.00033218 | 0.755750755 | 0.791515963 |
| ENSMUSG00000022861 | Dgkg | 0.00033218 | -0.376699541 | 1 |
| ENSMUSG00000038168 | P3h2 | 0.00033218 | 0.549299878 | 0.971854959 |

| | | | | |
|---|---|---|---|---|
| ENSMUSG00000068428 | Gmnc | 0.00033218 | 2.648835551 | 0.042972006 |
| ENSMUSG00000022774 | Ncbp2 | 0.00033218 | -0.631272368 | 0.875465725 |
| ENSMUSG00000034473 | Sec22a | 0.00033218 | -2.829211745 | 0.000193862 |
| ENSMUSG00000022849 | Hspbap1 | 0.00033218 | -0.543809527 | 0.913930369 |
| ENSMUSG00000022901 | Cd86 | 0.00033218 | 1.648835551 | 0.241308529 |
| ENSMUSG00000022837 | Iqcb1 | 0.00033218 | -1.082968338 | 0.420799142 |
| ENSMUSG00000022831 | Hcls1 | 0.00033218 | 0.932628517 | 0.739073334 |
| ENSMUSG00000034206 | Polq | 0.00033218 | -0.067371483 | 1 |
| ENSMUSG00000036292 | Gramd1c | 0.00033218 | 0.648835551 | 0.941472227 |
| ENSMUSG00000053062 | Jam2 | 0.00033218 | 0.098638469 | 1 |
| ENSMUSG00000022893 | Adamts1 | 0.00033218 | 0.20826296 | 1 |
| ENSMUSG00000116555 | Gm38477 | 0.00033218 | 1.368727632 | 0.466464457 |
| ENSMUSG00000052299 | Ltn1 | 0.00033218 | -1.865737621 | 0.027697231 |
| ENSMUSG00000022967 | Ifnar1 | 0.00033218 | -0.901361531 | 0.525722649 |
| ENSMUSG00000040605 | Bace2 | 0.00033218 | -0.450700122 | 0.998731799 |
| ENSMUSG00000005251 | Ripk4 | 0.00033218 | -1.216234868 | 0.293900105 |
| ENSMUSG00000045975 | C2cd2 | 0.00033218 | -0.426452576 | 0.994894633 |
| ENSMUSG00000080717 | B230307C23Rik | 0.00033218 | -1.910131741 | 0.02307354 |
| ENSMUSG00000023809 | Rps6ka2 | 0.00033218 | -0.035662623 | 1 |
| ENSMUSG00000037196 | Pacrg | 0.00033218 | -1.003241145 | 0.466464457 |
| ENSMUSG00000023827 | Agpat4 | 0.00033218 | -1.498005837 | 0.115347233 |
| ENSMUSG00000023883 | Phf10 | 0.00033218 | -0.543809527 | 0.912904617 |
| ENSMUSG00000014767 | Tbp | 0.00033218 | 0.246737108 | 1 |
| ENSMUSG00000048027 | Rgmb | 0.00033218 | 0.063873051 | 1 |
| ENSMUSG00000041319 | Thoc6 | 0.00033218 | -0.351164449 | 1 |
| ENSMUSG00000002496 | Tsc2 | 0.00033218 | -1.856692482 | 0.031796625 |
| ENSMUSG00000090101 | Snhg9 | 0.00033218 | 0.411796354 | 1 |
| ENSMUSG00000039183 | Nubp2 | 0.00033218 | 1.549299878 | 0.278293619 |
| ENSMUSG00000036636 | Clcn7 | 0.00033218 | 0.411796354 | 1 |
| ENSMUSG00000057411 | Fam173a | 0.00033218 | 0.598209478 | 0.951660573 |
| ENSMUSG00000025732 | Mcrip2 | 0.00033218 | 1.871227973 | 0.176214126 |
| ENSMUSG00000025728 | Pigq | 0.00033218 | -0.673092543 | 0.930393555 |
| ENSMUSG00000024018 | Ccdc167 | 0.00033218 | 0.063873051 | 1 |
| ENSMUSG00000024030 | Abcg1 | 0.00033218 | -0.32516924 | 1 |
| ENSMUSG00000024050 | Wiz | 0.00033218 | -2.498005837 | 0.001288623 |
| ENSMUSG00000073418 | C4b | 0.00033218 | -0.098398378 | 1 |
| ENSMUSG00000024369 | Nelfe | 0.00033218 | -0.128772027 | 1 |
| ENSMUSG00000023961 | Enpp4 | 0.00033218 | 1.456190473 | 0.318057572 |
| ENSMUSG00000003546 | Klc4 | 0.00033218 | -0.098398378 | 1 |
| ENSMUSG00000002767 | Mrpl2 | 0.00033218 | 0.20826296 | 1 |
| ENSMUSG00000063576 | Klhdc3 | 0.00033218 | -0.098398378 | 1 |
| ENSMUSG00000117222 | Gm49906 | 0.00033218 | 0.063873051 | 1 |
| ENSMUSG00000054723 | Vmac | 0.00033218 | 3.871227973 | 0.00517567 |
| ENSMUSG00000024212 | Mllt1 | 0.00033218 | 0.286265472 | 1 |
| ENSMUSG00000024101 | Washc1 | 0.00033218 | -0.52108945 | 0.909293502 |
| ENSMUSG00000117220 | Gm49932 | 0.00033218 | 1.456190473 | 0.318057572 |
| ENSMUSG00000117358 | Gm31645 | 0.00033218 | 0.812334284 | 0.771834758 |
| ENSMUSG00000024070 | Prkd3 | 0.00033218 | -0.829211745 | 0.587454349 |
| ENSMUSG00000024140 | Epas1 | 0.00033218 | -0.271729981 | 1 |
| ENSMUSG00000037104 | Socs5 | 0.00033218 | -2.251168659 | 0.005080271 |
| ENSMUSG00000024420 | Zfp521 | 0.00033218 | -0.158519371 | 1 |

| | | | | |
|---|---|---|---|---|
| ENSMUSG00000024331 | Dsc2 | 0.00033218 | -0.003241145 | 1 |
| ENSMUSG00000038299 | Wdr36 | 0.00033218 | 0.170788255 | 1 |
| ENSMUSG00000042834 | Nrep | 0.00033218 | 0.871227973 | 0.748991588 |
| ENSMUSG00000118380 | Gm36037 | 0.00033218 | -0.067371483 | 1 |
| ENSMUSG00000051439 | Cd14 | 0.00033218 | 0.411796354 | 1 |
| ENSMUSG00000024454 | Hdac3 | 0.00033218 | -0.652333983 | 0.774790843 |
| ENSMUSG00000036585 | Fgf1 | 0.00033218 | -0.498005837 | 0.904743019 |
| ENSMUSG00000054477 | Kcnn2 | 0.00033218 | 0.134262379 | 1 |
| ENSMUSG00000098284 | A330093E20Rik | 0.00033218 | -0.158519371 | 1 |
| ENSMUSG00000024534 | Sncaip | 0.00033218 | 0.326907457 | 1 |
| ENSMUSG00000058925 | Ccdc192 | 0.00033218 | 0.063873051 | 1 |
| ENSMUSG00000118391 | Gm19519 | 0.00033218 | 0.648835551 | 0.847265357 |
| ENSMUSG00000053441 | Adamts19 | 0.00033218 | 1.063873051 | 0.602791066 |
| ENSMUSG00000024610 | Cd74 | 0.00033218 | 0.063873051 | 1 |
| ENSMUSG00000024621 | Csf1r | 0.00033218 | 1.996758855 | 0.136198393 |
| ENSMUSG00000024575 | Pde6a | 0.00033218 | -1.733634085 | 0.064788899 |
| ENSMUSG00000032735 | Ablim3 | 0.00033218 | -1.298697029 | 0.238301367 |
| ENSMUSG00000118134 | 4930549G23Rik | 0.00033218 | -0.543809527 | 0.912904617 |
| ENSMUSG00000118012 | Gm46620 | 0.00033218 | 2.286265472 | 0.082902688 |
| ENSMUSG00000058216 | Gstp3 | 0.00033218 | 2.871227973 | 0.0289692 |
| ENSMUSG00000024757 | Slc22a19 | 0.00033218 | 1.871227973 | 0.192076162 |
| ENSMUSG00000071649 | B3gat3 | 0.00033218 | -1.298697029 | 0.236225921 |
| ENSMUSG00000059363 | Fxn | 0.00033218 | -0.79173704 | 0.659031063 |
| ENSMUSG00000012443 | Kif11 | 0.00033218 | -1.693556646 | 0.060182632 |
| ENSMUSG00000018820 | Zfyve27 | 0.00033218 | -1.019542957 | 0.455131546 |
| ENSMUSG00000117703 | Gm50306 | 0.00033218 | -1.389299577 | 0.199564169 |
| ENSMUSG00000064105 | Cnnm2 | 0.00033218 | -1.555036782 | 0.091420641 |
| ENSMUSG00000025024 | Smndc1 | 0.00033218 | -1.43862729 | 0.145704891 |
| ENSMUSG00000024982 | Zdhhc6 | 0.00033218 | -1.376699541 | 0.188182248 |
| ENSMUSG00000025091 | Pnliprp2 | 0.00033218 | 2.134262379 | 0.108201045 |
| ENSMUSG00000025092 | Hspa12a | 0.00033218 | 0.648835551 | 0.969915348 |
| ENSMUSG00000024997 | Prdx3 | 0.00033218 | 0.326907457 | 1 |
| ENSMUSG00000031166 | Wdr13 | 0.00033218 | -0.609898717 | 0.844591606 |
| ENSMUSG00000039201 | Tbc1d25 | 0.00033218 | -1.43862729 | 0.171021969 |
| ENSMUSG00000025040 | Fundc1 | 0.00033218 | 0.368727632 | 1 |
| ENSMUSG00000037369 | Kdm6a | 0.00033218 | -2.503811459 | 0.00132167 |
| ENSMUSG00000031065 | Cdk16 | 0.00033218 | -0.244249245 | 1 |
| ENSMUSG00000017057 | Il13ra1 | 0.00033218 | -0.713734528 | 0.695372548 |
| ENSMUSG00000086157 | Gm14643 | 0.00033218 | 0.326907457 | 1 |
| ENSMUSG00000031133 | Arhgef6 | 0.00033218 | 1.134262379 | 0.565425077 |
| ENSMUSG00000019088 | Dnase1l1 | 0.00033218 | -0.003241145 | 1 |
| ENSMUSG00000015290 | Ubl4a | 0.00033218 | 1.648835551 | 0.241308529 |
| ENSMUSG00000035476 | Tab3 | 0.00033218 | -0.244249245 | 1 |
| ENSMUSG00000079509 | Zfx | 0.00033218 | -2.158519371 | 0.007920214 |
| ENSMUSG00000047694 | Yipf6 | 0.00033218 | 0.996758855 | 0.591388707 |
| ENSMUSG00000034403 | Pja1 | 0.00033218 | -0.498005837 | 0.904743019 |
| ENSMUSG00000034311 | Kif4 | 0.00033218 | -0.631272368 | 0.771834758 |
| ENSMUSG00000046449 | Nexmif | 0.00033218 | -2.082968338 | 0.011858726 |
| ENSMUSG00000034551 | Hdx | 0.00033218 | 0.648835551 | 0.950457641 |
| ENSMUSG00000109156 | Gm45194 | 0.00033218 | 1.996758855 | 0.143740392 |
| ENSMUSG00000025525 | Apool | 0.00033218 | 1.286265472 | 0.477132639 |

| | | | | |
|---|---|---|---|---|
| ENSMUSG00000031432 | Prps1 | 0.00033218 | -0.883659529 | 0.521297728 |
| ENSMUSG00000000266 | Mid2 | 0.00033218 | -1.098398378 | 0.361256382 |
| ENSMUSG00000041688 | Amot | 0.00033218 | 0.549299878 | 0.964003521 |
| ENSMUSG00000084771 | A230072E10Rik | 0.00033218 | 0.063873051 | 1 |
| ENSMUSG00000025287 | Acot9 | 0.00033218 | -0.351164449 | 1 |
| ENSMUSG00000031342 | Gpm6b | 0.00033218 | -2.492176758 | 0.001462538 |
| ENSMUSG00000031358 | Msl3 | 0.00033218 | -0.970074281 | 0.476638318 |
| ENSMUSG00000025900 | Rp1 | 0.000276816 | -0.289763904 | 1 |
| ENSMUSG00000097797 | Gm26901 | 0.000276816 | 0.146335211 | 1 |
| ENSMUSG00000061024 | Rrs1 | 0.000276816 | -0.3207908 | 1 |
| ENSMUSG00000025934 | Gsta3 | 0.000276816 | 2.911869957 | 0.048102214 |
| ENSMUSG00000042197 | Zfp451 | 0.000276816 | -2.417253639 | 0.003399554 |
| ENSMUSG00000037503 | Fam168b | 0.000276816 | -1.175592884 | 0.354502808 |
| ENSMUSG00000037470 | Uggt1 | 0.000276816 | -1.685065185 | 0.083095907 |
| ENSMUSG00000047180 | Neurl3 | 0.000276816 | -0.258055044 | 1 |
| ENSMUSG00000050122 | Vwa3b | 0.000276816 | 1.533358334 | 0.394013503 |
| ENSMUSG00000026088 | Mitd1 | 0.000276816 | -0.466641666 | 0.96367904 |
| ENSMUSG00000026065 | Slc9a4 | 0.000276816 | -0.014129461 | 1 |
| ENSMUSG00000010290 | AI597479 | 0.000276816 | 0.710236096 | 0.886203497 |
| ENSMUSG00000026048 | Ercc5 | 0.000276816 | -0.547561661 | 0.871850653 |
| ENSMUSG00000101674 | 4930444A19Rik | 0.000276816 | 1.774366434 | 0.278415007 |
| ENSMUSG00000025981 | Coq10b | 0.000276816 | -0.258055044 | 1 |
| ENSMUSG00000025971 | Maip1 | 0.000276816 | 1.146335211 | 0.66416768 |
| ENSMUSG00000038305 | Spats2l | 0.000276816 | -0.743481871 | 0.736456705 |
| ENSMUSG00000047361 | Gm973 | 0.000276816 | 0.589941862 | 1 |
| ENSMUSG00000101156 | Gm29114 | 0.000276816 | 1.648835551 | 0.30756351 |
| ENSMUSG00000026189 | Pecr | 0.000276816 | 0.42644313 | 1 |
| ENSMUSG00000039354 | Smarcal1 | 0.000276816 | -1.273996588 | 0.27550706 |
| ENSMUSG00000026173 | Plcd4 | 0.000276816 | -0.466641666 | 0.970917467 |
| ENSMUSG00000033159 | Cnppd1 | 0.000276816 | 0.279601742 | 1 |
| ENSMUSG00000026207 | Speg | 0.000276816 | 1.146335211 | 0.653235135 |
| ENSMUSG00000044783 | Hjurp | 0.000276816 | -2.060822697 | 0.015769506 |
| ENSMUSG00000048126 | Col6a3 | 0.000276816 | -0.289763904 | 1 |
| ENSMUSG00000034353 | Ramp1 | 0.000276816 | -0.380911792 | 1 |
| ENSMUSG00000026313 | Hdac4 | 0.000276816 | -2.708716453 | 0.001150334 |
| ENSMUSG00000026283 | Ing5 | 0.000276816 | -1.636566667 | 0.091420641 |
| ENSMUSG00000051185 | Fam174a | 0.000276816 | 1.146335211 | 0.653751605 |
| ENSMUSG00000040710 | St8sia4 | 0.000276816 | -0.351164449 | 1 |
| ENSMUSG00000047496 | Rnf152 | 0.000276816 | -1.3207908 | 0.245794249 |
| ENSMUSG00000026321 | Tnfrsf11a | 0.000276816 | -1.088130043 | 0.455131546 |
| ENSMUSG00000107722 | 2900060B14Rik | 0.000276816 | -1.611691999 | 0.110753494 |
| ENSMUSG00000026355 | Mcm6 | 0.000276816 | -1.258055044 | 0.27201021 |
| ENSMUSG00000026427 | Eif2d | 0.000276816 | -1.014129461 | 0.47563319 |
| ENSMUSG00000026447 | Pik3c2b | 0.000276816 | -1.648844997 | 0.093610056 |
| ENSMUSG00000020423 | Btg2 | 0.000276816 | 0.42644313 | 1 |
| ENSMUSG00000026456 | Cyb5r1 | 0.000276816 | -0.258055044 | 1 |
| ENSMUSG00000063681 | Crb1 | 0.000276816 | 0.710236096 | 0.894256887 |
| ENSMUSG00000042751 | Nmnat2 | 0.000276816 | -0.410058138 | 1 |
| ENSMUSG00000066800 | Rnasel | 0.000276816 | -0.158519371 | 1 |
| ENSMUSG00000060519 | Tor3a | 0.000276816 | 0.104515035 | 1 |
| ENSMUSG00000033488 | Cryzl2 | 0.000276816 | 0.104515035 | 1 |

| | | | | |
|---|---|---|---|---|
| ENSMUSG00000026589 | Sec16b | 0.000276816 | -0.466641666 | 0.96367904 |
| ENSMUSG00000040124 | Gorab | 0.000276816 | -0.192466703 | 1 |
| ENSMUSG00000040596 | Pogk | 0.000276816 | -0.975655313 | 0.466464457 |
| ENSMUSG00000059089 | Fcgr4 | 0.000276816 | 0.533358334 | 1 |
| ENSMUSG00000026649 | Cfap126 | 0.000276816 | -0.874726405 | 0.597898961 |
| ENSMUSG00000013997 | Nit1 | 0.000276816 | -0.874726405 | 0.597898961 |
| ENSMUSG00000048865 | Arhgap30 | 0.000276816 | 0.189403933 | 1 |
| ENSMUSG00000073489 | Ifi204 | 0.000276816 | -0.743481871 | 0.732124826 |
| ENSMUSG00000102483 | Catspere1 | 0.000276816 | 0.841480629 | 0.870739619 |
| ENSMUSG00000026492 | Tfb2m | 0.000276816 | 0.279601742 | 1 |
| ENSMUSG00000038633 | Degs1 | 0.000276816 | 0.063873051 | 1 |
| ENSMUSG00000026510 | Trp53bp2 | 0.000276816 | -1.336057556 | 0.250559418 |
| ENSMUSG00000030768 | Disp1 | 0.000276816 | -1.843017545 | 0.045731953 |
| ENSMUSG00000039210 | Gpatch2 | 0.000276816 | -2.83766431 | 0.00033828 |
| ENSMUSG00000037568 | Vash2 | 0.000276816 | -0.088130043 | 1 |
| ENSMUSG00000026628 | Atf3 | 0.000276816 | 0.841480629 | 0.886203497 |
| ENSMUSG00000037395 | Rcor3 | 0.000276816 | -1.58638091 | 0.100676805 |
| ENSMUSG00000016200 | Syt14 | 0.000276816 | 2.233798052 | 0.134836815 |
| ENSMUSG00000025816 | Sec61a2 | 0.000276816 | -1.410058138 | 0.191885686 |
| ENSMUSG00000061186 | Sfmbt2 | 0.000276816 | 0.648835551 | 0.948737557 |
| ENSMUSG00000026778 | Prkcq | 0.000276816 | -0.123753952 | 1 |
| ENSMUSG00000026767 | Mindy3 | 0.000276816 | -0.43862729 | 0.96070793 |
| ENSMUSG00000003418 | St8sia6 | 0.000276816 | -1.810596067 | 0.058667604 |
| ENSMUSG00000085057 | Gm13415 | 0.000276816 | 0.911869957 | 0.847265357 |
| ENSMUSG00000006476 | Nsmf | 0.000276816 | -0.696939285 | 0.716109464 |
| ENSMUSG00000026966 | Ssna1 | 0.000276816 | 0.42644313 | 1 |
| ENSMUSG00000026933 | Camsap1 | 0.000276816 | -1.599091962 | 0.102454279 |
| ENSMUSG00000036352 | Ubac1 | 0.000276816 | -0.57355687 | 0.874192437 |
| ENSMUSG00000087679 | Tmem250-ps | 0.000276816 | 0.911869957 | 0.847265357 |
| ENSMUSG00000026927 | Entr1 | 0.000276816 | -1.480447466 | 0.145553211 |
| ENSMUSG00000039686 | Zer1 | 0.000276816 | -0.832291138 | 0.66416768 |
| ENSMUSG00000026848 | Tor1b | 0.000276816 | -0.599091962 | 0.877036949 |
| ENSMUSG00000062661 | Ncs1 | 0.000276816 | -0.225633566 | 1 |
| ENSMUSG00000026840 | Lamc3 | 0.000276816 | 1.063873051 | 0.66416768 |
| ENSMUSG00000002546 | Golga2 | 0.000276816 | -2.004606687 | 0.023140688 |
| ENSMUSG00000026819 | Slc25a25 | 0.000276816 | -0.975655313 | 0.560430577 |
| ENSMUSG00000038765 | Lmx1b | 0.000276816 | -0.599091962 | 0.877036949 |
| ENSMUSG00000026764 | Kif5c | 0.000276816 | -1.58638091 | 0.100676805 |
| ENSMUSG00000026950 | Neb | 0.000276816 | 0.375817057 | 1 |
| ENSMUSG00000017412 | Cacnb4 | 0.000276816 | -0.351164449 | 1 |
| ENSMUSG00000079353 | Gm11099 | 0.000276816 | 2.063873051 | 0.210550914 |
| ENSMUSG00000026904 | Slc4a10 | 0.000276816 | -1.225633566 | 0.358945041 |
| ENSMUSG00000026994 | Galnt3 | 0.000276816 | -1.720398258 | 0.061086889 |
| ENSMUSG00000027086 | Fastkd1 | 0.000276816 | 0.47891055 | 1 |
| ENSMUSG00000051730 | Mettl5 | 0.000276816 | -0.52108945 | 0.971210385 |
| ENSMUSG00000041975 | Mettl8 | 0.000276816 | -1.241935379 | 0.316344908 |
| ENSMUSG00000049044 | Rapgef4 | 0.000276816 | -2.21741306 | 0.012879784 |
| ENSMUSG00000100303 | 2600014E21Rik | 0.000276816 | 0.375817057 | 1 |
| ENSMUSG00000076437 | Selenoh | 0.000276816 | 0.279601742 | 1 |
| ENSMUSG00000027080 | Med19 | 0.000276816 | -0.051604167 | 1 |
| ENSMUSG00000027282 | Mtch2 | 0.000276816 | -0.258055044 | 1 |

| | | | | |
|---|---|---|---|---|
| ENSMUSG00000027230 | Creb3l1 | 0.000276816 | -0.494122402 | 0.967976676 |
| ENSMUSG00000087197 | Gm13780 | 0.000276816 | 0.104515035 | 1 |
| ENSMUSG00000068742 | Cry2 | 0.000276816 | 0.533358334 | 1 |
| ENSMUSG00000075020 | Mir670hg | 0.000276816 | -0.192466703 | 1 |
| ENSMUSG00000027193 | Api5 | 0.000276816 | -0.43862729 | 1 |
| ENSMUSG00000032698 | Lmo2 | 0.000276816 | 1.911869957 | 0.224329705 |
| ENSMUSG00000045106 | Ccdc73 | 0.000276816 | -0.192466703 | 1 |
| ENSMUSG00000016458 | Wt1 | 0.000276816 | 1.063873051 | 0.66416768 |
| ENSMUSG00000027162 | Lin7c | 0.000276816 | -0.766201948 | 0.766975255 |
| ENSMUSG00000040084 | Bub1b | 0.000276816 | 0.233798052 | 1 |
| ENSMUSG00000040035 | Disp2 | 0.000276816 | 1.648835551 | 0.334416634 |
| ENSMUSG00000027326 | Knl1 | 0.000276816 | -1.410058138 | 0.191885686 |
| ENSMUSG00000070730 | Rmdn3 | 0.000276816 | 1.42644313 | 0.408970699 |
| ENSMUSG00000079110 | Capn3 | 0.000276816 | 0.47891055 | 1 |
| ENSMUSG00000027199 | Gatm | 0.000276816 | -1.123753952 | 0.400891089 |
| ENSMUSG00000027209 | Fam227b | 0.000276816 | 0.911869957 | 0.727318386 |
| ENSMUSG00000034850 | Tmem127 | 0.000276816 | -1.032988489 | 0.443153295 |
| ENSMUSG00000027394 | Ttl | 0.000276816 | 0.233798052 | 1 |
| ENSMUSG00000037902 | Sirpa | 0.000276816 | 0.104515035 | 1 |
| ENSMUSG00000027327 | 1700037H04Rik | 0.000276816 | 0.841480629 | 0.858831093 |
| ENSMUSG00000027330 | Cdc25b | 0.000276816 | 0.589941862 | 1 |
| ENSMUSG00000086756 | Gm14051 | 0.000276816 | 0.774366434 | 0.872990655 |
| ENSMUSG00000027341 | Tmem230 | 0.000276816 | 0.326907457 | 1 |
| ENSMUSG00000058793 | Cds2 | 0.000276816 | -0.956026507 | 0.534443935 |
| ENSMUSG00000085631 | 9630028H03Rik | 0.000276816 | -0.051604167 | 1 |
| ENSMUSG00000074766 | Ism1 | 0.000276816 | 0.710236096 | 0.899066862 |
| ENSMUSG00000074758 | Gm5535 | 0.000276816 | 1.911869957 | 0.225663244 |
| ENSMUSG00000027428 | Rbbp9 | 0.000276816 | 1.648835551 | 0.330777885 |
| ENSMUSG00000037143 | Cfap61 | 0.000276816 | -0.853664789 | 0.714585823 |
| ENSMUSG00000074736 | Syndig1 | 0.000276816 | 0.589941862 | 1 |
| ENSMUSG00000027465 | Tbc1d20 | 0.000276816 | 0.189403933 | 1 |
| ENSMUSG00000068062 | Gm14164 | 0.000276816 | 0.533358334 | 1 |
| ENSMUSG00000068040 | Tm9sf4 | 0.000276816 | -1.534386273 | 0.13144915 |
| ENSMUSG00000046020 | Pofut1 | 0.000276816 | -0.158519371 | 1 |
| ENSMUSG00000038312 | Edem2 | 0.000276816 | -2.032988489 | 0.02580662 |
| ENSMUSG00000005882 | Uqcc1 | 0.000276816 | -1.123753952 | 0.436069664 |
| ENSMUSG00000038085 | Cnbd2 | 0.000276816 | -0.995020638 | 0.470839676 |
| ENSMUSG00000027628 | Aar2 | 0.000276816 | -1.648844997 | 0.094160212 |
| ENSMUSG00000085741 | 5430405H02Rik | 0.000276816 | -2.514394797 | 0.003094221 |
| ENSMUSG00000027641 | Rbl1 | 0.000276816 | -0.599091962 | 0.877036949 |
| ENSMUSG00000037754 | Ppp1r16b | 0.000276816 | 0.42644313 | 1 |
| ENSMUSG00000074622 | Mafb | 0.000276816 | -0.351164449 | 1 |
| ENSMUSG00000017858 | Ift52 | 0.000276816 | -1.273996588 | 0.289497193 |
| ENSMUSG00000017697 | Ada | 0.000276816 | 1.648835551 | 0.30756351 |
| ENSMUSG00000050373 | Snx21 | 0.000276816 | 0.189403933 | 1 |
| ENSMUSG00000086166 | Gm14342 | 0.000276816 | 2.063873051 | 0.173532034 |
| ENSMUSG00000038671 | Arfrp1 | 0.000276816 | -0.158519371 | 1 |
| ENSMUSG00000027582 | Zgpat | 0.000276816 | -0.810596067 | 0.66416768 |
| ENSMUSG00000089917 | Uckl1 | 0.000276816 | 0.841480629 | 0.886203497 |
| ENSMUSG00000086224 | 2700069I18Rik | 0.000276816 | -1.158519371 | 0.370278211 |
| ENSMUSG00000040374 | Pex2 | 0.000276816 | -0.720398258 | 0.719591951 |

| | | | | |
|---|---|---|---|---|
| ENSMUSG00000062515 | Fabp4 | 0.000276816 | 1.326907457 | 0.466464457 |
| ENSMUSG00000027599 | Armc1 | 0.000276816 | -0.088130043 | 1 |
| ENSMUSG00000037661 | Gpr160 | 0.000276816 | 0.774366434 | 0.872990655 |
| ENSMUSG00000037652 | Phc3 | 0.000276816 | -1.915949067 | 0.031334679 |
| ENSMUSG00000027663 | Zmat3 | 0.000276816 | -0.599091962 | 0.917185102 |
| ENSMUSG00000056900 | Usp13 | 0.000276816 | 0.774366434 | 0.872990655 |
| ENSMUSG00000027709 | Mccc1 | 0.000276816 | -0.410058138 | 1 |
| ENSMUSG00000037211 | Spry1 | 0.000276816 | 1.063873051 | 0.66416768 |
| ENSMUSG00000059834 | Sclt1 | 0.000276816 | -1.336057556 | 0.250559418 |
| ENSMUSG00000036513 | Commd2 | 0.000276816 | 0.47891055 | 1 |
| ENSMUSG00000097725 | 4930593A02Rik | 0.000276816 | 0.104515035 | 1 |
| ENSMUSG00000027820 | Mme | 0.000276816 | 0.279601742 | 1 |
| ENSMUSG00000027774 | Gfm1 | 0.000276816 | -1.014129461 | 0.47563319 |
| ENSMUSG00000027775 | Mfsd1 | 0.000276816 | -0.624182943 | 0.792441218 |
| ENSMUSG00000104301 | Wdr49 | 0.000276816 | -0.380911792 | 1 |
| ENSMUSG00000033752 | Mnd1 | 0.000276816 | 0.911869957 | 0.699192777 |
| ENSMUSG00000028078 | Dclk2 | 0.000276816 | 0.42644313 | 1 |
| ENSMUSG00000004895 | Prcc | 0.000276816 | -1.534386273 | 0.13144915 |
| ENSMUSG00000028064 | Sema4a | 0.000276816 | -0.43862729 | 0.96070793 |
| ENSMUSG00000068923 | Syt11 | 0.000276816 | -1.258055044 | 0.290284345 |
| ENSMUSG00000041263 | Rusc1 | 0.000276816 | -0.936126949 | 0.529126167 |
| ENSMUSG00000064068 | Mtx1 | 0.000276816 | 2.063873051 | 0.173532034 |
| ENSMUSG00000028040 | Efna4 | 0.000276816 | 0.279601742 | 1 |
| ENSMUSG00000028041 | Adam15 | 0.000276816 | -1.965874293 | 0.024825015 |
| ENSMUSG00000047824 | Pygo2 | 0.000276816 | -2.069982696 | 0.016215729 |
| ENSMUSG00000027942 | 4933434E20Rik | 0.000276816 | -1.225633566 | 0.310683162 |
| ENSMUSG00000001018 | Snapin | 0.000276816 | 1.326907457 | 0.466464457 |
| ENSMUSG00000052415 | Tchh | 0.000276816 | -1.956026507 | 0.02926227 |
| ENSMUSG00000028150 | Rorc | 0.000276816 | -1.424413431 | 0.196126878 |
| ENSMUSG00000041912 | Tdrkh | 0.000276816 | 0.326907457 | 1 |
| ENSMUSG00000105734 | 4930558C23Rik | 0.000276816 | 1.146335211 | 0.587484274 |
| ENSMUSG00000028111 | Ctsk | 0.000276816 | 0.774366434 | 0.872990655 |
| ENSMUSG00000038619 | Ensa | 0.000276816 | -0.494122402 | 0.96881158 |
| ENSMUSG00000080727 | C920021L13Rik | 0.000276816 | 0.710236096 | 0.886203497 |
| ENSMUSG00000028104 | Polr3gl | 0.000276816 | -1.466641666 | 0.143740392 |
| ENSMUSG00000090210 | Itga10 | 0.000276816 | -0.088130043 | 1 |
| ENSMUSG00000042035 | Igsf3 | 0.000276816 | -2.737745502 | 0.000533458 |
| ENSMUSG00000032902 | Slc16a1 | 0.000276816 | -0.956026507 | 0.534443935 |
| ENSMUSG00000074340 | Ovgp1 | 0.000276816 | 0.189403933 | 1 |
| ENSMUSG00000010136 | Pifo | 0.000276816 | 3.233798052 | 0.029538332 |
| ENSMUSG00000040747 | Cd53 | 0.000276816 | 0.774366434 | 0.88422754 |
| ENSMUSG00000046213 | Cym | 0.000276816 | 1.648835551 | 0.30756351 |
| ENSMUSG00000058952 | Cfi | 0.000276816 | -0.936126949 | 0.529126167 |
| ENSMUSG00000050150 | Slc9b1 | 0.000276816 | 2.42644313 | 0.102043984 |
| ENSMUSG00000091512 | Lamtor3 | 0.000276816 | -0.088130043 | 1 |
| ENSMUSG00000028187 | Rpf1 | 0.000276816 | -0.289763904 | 1 |
| ENSMUSG00000048652 | Samd13 | 0.000276816 | 1.533358334 | 0.395268819 |
| ENSMUSG00000041261 | Car8 | 0.000276816 | 0.774366434 | 0.886203497 |
| ENSMUSG00000023151 | Lrrc69 | 0.000276816 | 0.774366434 | 0.886203497 |
| ENSMUSG00000028247 | Coq3 | 0.000276816 | -1.599091962 | 0.102454279 |
| ENSMUSG00000040387 | Klhl32 | 0.000276816 | -0.810596067 | 0.689704239 |

| | | | | |
|---|---|---|---|---|
| ENSMUSG00000115420 | Rmrp | 0.000276816 | -2.36611479 | 0.004176784 |
| ENSMUSG00000028480 | Glipr2 | 0.000276816 | -0.624182943 | 0.792683407 |
| ENSMUSG00000035615 | Frmpd1 | 0.000276816 | -1.885143021 | 0.034614 |
| ENSMUSG00000066191 | Anks6 | 0.000276816 | 0.774366434 | 0.872990655 |
| ENSMUSG00000039579 | Grin3a | 0.000276816 | 0.146335211 | 1 |
| ENSMUSG00000057280 | Musk | 0.000276816 | 1.326907457 | 0.580988658 |
| ENSMUSG00000066150 | Slc31a1 | 0.000276816 | -0.410058138 | 1 |
| ENSMUSG00000028398 | Dmac1 | 0.000276816 | 0.911869957 | 0.847265357 |
| ENSMUSG00000028497 | Hacd4 | 0.000276816 | 1.326907457 | 0.580988658 |
| ENSMUSG00000006386 | Tek | 0.000276816 | -0.599091962 | 0.897772995 |
| ENSMUSG00000062627 | Mysm1 | 0.000276816 | -2.266047835 | 0.006295279 |
| ENSMUSG00000028572 | Hook1 | 0.000276816 | -0.995020638 | 0.470839676 |
| ENSMUSG00000048747 | E130114P18Rik | 0.000276816 | 0.233798052 | 1 |
| ENSMUSG00000035407 | Kank4 | 0.000276816 | 0.710236096 | 0.899615503 |
| ENSMUSG00000035069 | Oma1 | 0.000276816 | -1.158519371 | 0.3498121 |
| ENSMUSG00000028518 | Prkaa2 | 0.000276816 | 0.533358334 | 1 |
| ENSMUSG00000028582 | Cc2d1b | 0.000276816 | -1.175592884 | 0.354502808 |
| ENSMUSG00000003411 | Rab3b | 0.000276816 | -0.289763904 | 1 |
| ENSMUSG00000028716 | Pdzk1ip1 | 0.000276816 | 1.911869957 | 0.395423178 |
| ENSMUSG00000033948 | Zswim5 | 0.000276816 | 0.710236096 | 0.886203497 |
| ENSMUSG00000048772 | Tmem53 | 0.000276816 | 1.533358334 | 0.359933138 |
| ENSMUSG00000033191 | Tie1 | 0.000276816 | 0.648835551 | 0.896076486 |
| ENSMUSG00000060288 | Ppih | 0.000276816 | -1.424413431 | 0.211844791 |
| ENSMUSG00000028636 | Ppcs | 0.000276816 | 0.589941862 | 1 |
| ENSMUSG00000043207 | Zmpste24 | 0.000276816 | -1.225633566 | 0.310683162 |
| ENSMUSG00000028651 | Ppie | 0.000276816 | -0.874726405 | 0.597898961 |
| ENSMUSG00000084757 | 1700057H15Rik | 0.000276816 | 1.233798052 | 0.651252873 |
| ENSMUSG00000098760 | Gm2164 | 0.000276816 | 0.146335211 | 1 |
| ENSMUSG00000028833 | Ncdn | 0.000276816 | -0.225633566 | 1 |
| ENSMUSG00000028799 | Zfp362 | 0.000276816 | -0.088130043 | 1 |
| ENSMUSG00000028789 | Azin2 | 0.000276816 | -1.305360759 | 0.24200918 |
| ENSMUSG00000028792 | Ak2 | 0.000276816 | -0.380911792 | 1 |
| ENSMUSG00000025743 | Sdc3 | 0.000276816 | 0.985870539 | 0.695372548 |
| ENSMUSG00000028581 | Laptm5 | 0.000276816 | -0.258055044 | 1 |
| ENSMUSG00000003038 | Hmgn2 | 0.000276816 | 1.326907457 | 0.580988658 |
| ENSMUSG00000062157 | Ifnlr1 | 0.000276816 | 0.47891055 | 1 |
| ENSMUSG00000028756 | Pink1 | 0.000276816 | 1.146335211 | 0.627920558 |
| ENSMUSG00000036622 | Atp13a2 | 0.000276816 | 0.841480629 | 0.858831093 |
| ENSMUSG00000040606 | Kazn | 0.000276816 | -0.547561661 | 0.998731799 |
| ENSMUSG00000006442 | Srm | 0.000276816 | 2.063873051 | 0.173532034 |
| ENSMUSG00000028974 | Dffa | 0.000276816 | -1.534386273 | 0.13144915 |
| ENSMUSG00000039936 | Pik3cd | 0.000276816 | -0.351164449 | 1 |
| ENSMUSG00000073700 | Klhl21 | 0.000276816 | 0.911869957 | 0.732581709 |
| ENSMUSG00000028931 | Kcnab2 | 0.000276816 | 1.233798052 | 0.642722979 |
| ENSMUSG00000003623 | Crot | 0.000276816 | -1.424413431 | 0.196126878 |
| ENSMUSG00000039934 | Gsap | 0.000276816 | -2.21741306 | 0.010112645 |
| ENSMUSG00000105987 | AI506816 | 0.000276816 | 0.47891055 | 1 |
| ENSMUSG00000056367 | Actr3b | 0.000276816 | 2.648835551 | 0.072659903 |
| ENSMUSG00000045294 | Insig1 | 0.000276816 | 1.774366434 | 0.262284476 |
| ENSMUSG00000029165 | Agbl5 | 0.000276816 | -0.915949067 | 0.524670309 |
| ENSMUSG00000029162 | Khk | 0.000276816 | 1.326907457 | 0.580988658 |

| | | | | |
|---|---|---|---|---|
| ENSMUSG00000029145 | Eif2b4 | 0.000276816 | -0.599091962 | 0.877036949 |
| ENSMUSG00000029149 | Krtcap3 | 0.000276816 | 0.648835551 | 0.89889079 |
| ENSMUSG00000038564 | Ift172 | 0.000276816 | 0.233798052 | 1 |
| ENSMUSG00000029136 | Rbks | 0.000276816 | 0.024344687 | 1 |
| ENSMUSG00000005299 | Letm1 | 0.000276816 | 0.146335211 | 1 |
| ENSMUSG00000067365 | Tmem128 | 0.000276816 | 0.42644313 | 1 |
| ENSMUSG00000029128 | Rab28 | 0.000276816 | -1.225633566 | 0.33333983 |
| ENSMUSG00000015806 | Qdpr | 0.000276816 | 0.841480629 | 0.858831093 |
| ENSMUSG00000045790 | Ccdc149 | 0.000276816 | -1.289763904 | 0.279592421 |
| ENSMUSG00000039178 | Tbc1d19 | 0.000276816 | -0.743481871 | 0.725185838 |
| ENSMUSG00000097640 | Gm20033 | 0.000276816 | -2.132524162 | 0.011441934 |
| ENSMUSG00000068082 | Grxcr1 | 0.000276816 | 1.911869957 | 0.236879967 |
| ENSMUSG00000072889 | Nfxl1 | 0.000276816 | -1.258055044 | 0.27201021 |
| ENSMUSG00000067219 | Nipal1 | 0.000276816 | 0.146335211 | 1 |
| ENSMUSG00000036285 | Noa1 | 0.000276816 | -0.995020638 | 0.470839676 |
| ENSMUSG00000029370 | Rassf6 | 0.000276816 | -1.088130043 | 0.455131546 |
| ENSMUSG00000029504 | Ddx51 | 0.000276816 | 0.911869957 | 0.847265357 |
| ENSMUSG00000029521 | Chek2 | 0.000276816 | -1.106051951 | 0.395423178 |
| ENSMUSG00000042249 | Grk3 | 0.000276816 | -1.424413431 | 0.228871882 |
| ENSMUSG00000042096 | Dao | 0.000276816 | 2.42644313 | 0.108201045 |
| ENSMUSG00000042078 | Svop | 0.000276816 | 0.985870539 | 0.669958667 |
| ENSMUSG00000072694 | 1500011B03Rik | 0.000276816 | 1.326907457 | 0.580988658 |
| ENSMUSG00000041827 | Oasl1 | 0.000276816 | 0.911869957 | 0.847265357 |
| ENSMUSG00000079278 | Tmem233 | 0.000276816 | 1.326907457 | 0.580988658 |
| ENSMUSG00000106909 | Gm43122 | 0.000276816 | 0.375817057 | 1 |
| ENSMUSG00000032741 | Tpcn1 | 0.000276816 | -0.975655313 | 0.466464457 |
| ENSMUSG00000042719 | Naa25 | 0.000276816 | -1.995020638 | 0.022629778 |
| ENSMUSG00000029462 | Vps29 | 0.000276816 | 0.233798052 | 1 |
| ENSMUSG00000029449 | Rhof | 0.000276816 | 0.326907457 | 1 |
| ENSMUSG00000090086 | AI480526 | 0.000276816 | -0.014129461 | 1 |
| ENSMUSG00000038384 | Setd1b | 0.000276816 | -1.380911792 | 0.220566815 |
| ENSMUSG00000066278 | Vps37b | 0.000276816 | -1.766201948 | 0.055368915 |
| ENSMUSG00000029404 | Arl6ip4 | 0.000276816 | -1.014129461 | 0.47563319 |
| ENSMUSG00000029386 | Tctn2 | 0.000276816 | -1.466641666 | 0.143740392 |
| ENSMUSG00000037979 | Ccdc92 | 0.000276816 | -0.936126949 | 0.571688481 |
| ENSMUSG00000037905 | Bri3bp | 0.000276816 | -0.599091962 | 0.877036949 |
| ENSMUSG00000034211 | Mrps17 | 0.000276816 | 0.063873051 | 1 |
| ENSMUSG00000025534 | Gusb | 0.000276816 | -1.241935379 | 0.316344908 |
| ENSMUSG00000061979 | Rcc1l | 0.000276816 | -0.975655313 | 0.466464457 |
| ENSMUSG00000029675 | Eln | 0.000276816 | -0.720398258 | 0.719591951 |
| ENSMUSG00000039860 | Srrm3 | 0.000276816 | 0.47891055 | 1 |
| ENSMUSG00000093445 | Lrch4 | 0.000276816 | 0.146335211 | 1 |
| ENSMUSG00000000149 | Gna12 | 0.000276816 | -0.673092543 | 0.797063201 |
| ENSMUSG00000091351 | Gm17135 | 0.000276816 | 1.911869957 | 0.245794249 |
| ENSMUSG00000029625 | Cpsf4 | 0.000276816 | -1.141241379 | 0.438808325 |
| ENSMUSG00000007812 | Zfp655 | 0.000276816 | -1.258055044 | 0.27201021 |
| ENSMUSG00000029636 | Wasf3 | 0.000276816 | 2.648835551 | 0.09197283 |
| ENSMUSG00000029657 | Hsph1 | 0.000276816 | -1.636566667 | 0.091420641 |
| ENSMUSG00000044156 | Hepacam2 | 0.000276816 | 0.648835551 | 0.899737888 |
| ENSMUSG00000029759 | Pon3 | 0.000276816 | -0.766201948 | 0.732525536 |
| ENSMUSG00000012483 | Rpa3 | 0.000276816 | 0.47891055 | 1 |

| | | | | |
|---|---|---|---|---|
| ENSMUSG00000046178 | Nxph1 | 0.000276816 | 0.233798052 | 1 |
| ENSMUSG00000107847 | Gm35822 | 0.000276816 | 1.911869957 | 0.250559418 |
| ENSMUSG00000079679 | Vwde | 0.000276816 | 0.911869957 | 0.728745035 |
| ENSMUSG00000039841 | Zfp800 | 0.000276816 | -2.417253639 | 0.003399554 |
| ENSMUSG00000052131 | Akr1b7 | 0.000276816 | 3.648835551 | 0.0159954 |
| ENSMUSG00000073154 | 9330158H04Rik | 0.000276816 | 1.063873051 | 0.675933929 |
| ENSMUSG00000019689 | Fmc1 | 0.000276816 | 0.648835551 | 0.896811934 |
| ENSMUSG00000037172 | E330009J07Rik | 0.000276816 | -0.289763904 | 1 |
| ENSMUSG00000062519 | Zfp398 | 0.000276816 | -0.915949067 | 0.524670309 |
| ENSMUSG00000029832 | Nfe2l3 | 0.000276816 | -1.069982696 | 0.490769731 |
| ENSMUSG00000063568 | Jazf1 | 0.000276816 | -0.351164449 | 1 |
| ENSMUSG00000086040 | Wipf3 | 0.000276816 | 1.326907457 | 0.466464457 |
| ENSMUSG00000025889 | Snca | 0.000276816 | 1.648835551 | 0.349567117 |
| ENSMUSG00000055027 | Smyd1 | 0.000276816 | 1.233798052 | 0.624497001 |
| ENSMUSG00000052656 | Rnf103 | 0.000276816 | -0.088130043 | 1 |
| ENSMUSG00000030043 | Tacr1 | 0.000276816 | 1.063873051 | 0.66416768 |
| ENSMUSG00000030041 | M1ap | 0.000276816 | 0.985870539 | 0.700914788 |
| ENSMUSG00000068323 | Slc4a5 | 0.000276816 | 0.841480629 | 0.874192437 |
| ENSMUSG00000029998 | Pcyox1 | 0.000276816 | -0.351164449 | 1 |
| ENSMUSG00000030060 | Hmces | 0.000276816 | -0.874726405 | 0.597898961 |
| ENSMUSG00000030079 | Ruvbl1 | 0.000276816 | -0.192466703 | 1 |
| ENSMUSG00000030084 | Plxna1 | 0.000276816 | -1.106051951 | 0.395423178 |
| ENSMUSG00000034203 | Chchd4 | 0.000276816 | 0.42644313 | 1 |
| ENSMUSG00000014551 | Mrps25 | 0.000276816 | 0.774366434 | 0.872990655 |
| ENSMUSG00000030029 | Lrig1 | 0.000276816 | -1.424413431 | 0.196126878 |
| ENSMUSG00000030101 | Sumf1 | 0.000276816 | -0.853664789 | 0.592845519 |
| ENSMUSG00000086429 | Gt(ROSA)26Sor | 0.000276816 | -0.088130043 | 1 |
| ENSMUSG00000030269 | Mtmr14 | 0.000276816 | -1.507668934 | 0.128531628 |
| ENSMUSG00000043088 | Il17re | 0.000276816 | 0.279601742 | 1 |
| ENSMUSG00000030284 | Creld1 | 0.000276816 | -1.258055044 | 0.275781942 |
| ENSMUSG00000009394 | Syn2 | 0.000276816 | 1.233798052 | 0.615418676 |
| ENSMUSG00000042389 | Tsen2 | 0.000276816 | -1.032988489 | 0.443153295 |
| ENSMUSG00000048489 | Depp1 | 0.000276816 | 1.911869957 | 0.217655393 |
| ENSMUSG00000030145 | Zfp248 | 0.000276816 | 0.710236096 | 0.898335435 |
| ENSMUSG00000072623 | Zfp9 | 0.000276816 | -0.624182943 | 0.886203497 |
| ENSMUSG00000004446 | Bid | 0.000276816 | -0.853664789 | 0.592845519 |
| ENSMUSG00000023505 | Cdca3 | 0.000276816 | 0.911869957 | 0.847265357 |
| ENSMUSG00000030337 | Vamp1 | 0.000276816 | 0.326907457 | 1 |
| ENSMUSG00000030344 | Akap3 | 0.000276816 | 2.648835551 | 0.072659903 |
| ENSMUSG00000030200 | Bcl2l14 | 0.000276816 | 0.42644313 | 1 |
| ENSMUSG00000003031 | Cdkn1b | 0.000276816 | -0.743481871 | 0.725185838 |
| ENSMUSG00000030205 | Gprc5d | 0.000276816 | 4.233798052 | 0.007061791 |
| ENSMUSG00000030216 | Wbp11 | 0.000276816 | -1.995020638 | 0.022629778 |
| ENSMUSG00000084927 | Gm5724 | 0.000276816 | 0.146335211 | 1 |
| ENSMUSG00000030249 | Abcc9 | 0.000276816 | -0.466641666 | 0.970917467 |
| ENSMUSG00000030283 | St8sia1 | 0.000276816 | 0.985870539 | 0.66416768 |
| ENSMUSG00000030256 | Bhlhe41 | 0.000276816 | 0.911869957 | 0.847265357 |
| ENSMUSG00000061374 | Fiz1 | 0.000276816 | 0.063873051 | 1 |
| ENSMUSG00000046792 | Zfp787 | 0.000276816 | 0.024344687 | 1 |
| ENSMUSG00000002265 | Peg3 | 0.000276816 | 0.710236096 | 0.89270014 |
| ENSMUSG00000040987 | Mill2 | 0.000276816 | -1.209145444 | 0.306535101 |

| | | | | |
|---|---|---|---|---|
| ENSMUSG00000044030 | Irf2bp1 | 0.000276816 | -1.088130043 | 0.455131546 |
| ENSMUSG00000047649 | Cd3eap | 0.000276816 | 0.233798052 | 1 |
| ENSMUSG00000040525 | Cblc | 0.000276816 | -1.424413431 | 0.197227651 |
| ENSMUSG00000002210 | Smg9 | 0.000276816 | -0.3207908 | 1 |
| ENSMUSG00000064254 | Ethe1 | 0.000276816 | -0.547561661 | 0.871850653 |
| ENSMUSG00000005442 | Cic | 0.000276816 | -1.225633566 | 0.310683162 |
| ENSMUSG00000060376 | Bckdha | 0.000276816 | -1.175592884 | 0.391578102 |
| ENSMUSG00000003752 | Itpkc | 0.000276816 | -0.936126949 | 0.533072206 |
| ENSMUSG00000040390 | Map3k10 | 0.000276816 | -0.599091962 | 0.877036949 |
| ENSMUSG00000098022 | Zfp82 | 0.000276816 | -2.273996588 | 0.006478925 |
| ENSMUSG00000037020 | Wdr62 | 0.000276816 | 0.533358334 | 0.971854959 |
| ENSMUSG00000046056 | Sbsn | 0.000276816 | 0.063873051 | 1 |
| ENSMUSG00000030468 | Siglecg | 0.000276816 | 2.648835551 | 0.072659903 |
| ENSMUSG00000097178 | 2310002F09Rik | 0.000276816 | -0.051604167 | 1 |
| ENSMUSG00000038695 | Josd2 | 0.000276816 | 1.326907457 | 0.580988658 |
| ENSMUSG00000046574 | Prr12 | 0.000276816 | -2.52108945 | 0.001982991 |
| ENSMUSG00000003868 | Ruvbl2 | 0.000276816 | 0.911869957 | 0.847265357 |
| ENSMUSG00000030824 | Nucb1 | 0.000276816 | -0.673092543 | 0.812486317 |
| ENSMUSG00000030826 | Bcat2 | 0.000276816 | -1.480447466 | 0.146050741 |
| ENSMUSG00000043262 | Uevld | 0.000276816 | -1.424413431 | 0.203201152 |
| ENSMUSG00000109394 | A230057D06Rik | 0.000276816 | -0.289763904 | 1 |
| ENSMUSG00000030522 | Mtmr10 | 0.000276816 | -1.069982696 | 0.452189485 |
| ENSMUSG00000030555 | Ttc23 | 0.000276816 | -1.351164449 | 0.236225921 |
| ENSMUSG00000042659 | Arrdc4 | 0.000276816 | 0.47891055 | 1 |
| ENSMUSG00000039236 | Isg20 | 0.000276816 | 0.911869957 | 0.847265357 |
| ENSMUSG00000015709 | Arnt2 | 0.000276816 | 0.42644313 | 1 |
| ENSMUSG00000030562 | Nox4 | 0.000276816 | -0.788569761 | 0.66416768 |
| ENSMUSG00000030621 | Me3 | 0.000276816 | -0.351164449 | 1 |
| ENSMUSG00000051451 | Crebzf | 0.000276816 | -2.358658985 | 0.004443316 |
| ENSMUSG00000030643 | Rab30 | 0.000276816 | -0.648844997 | 0.792441218 |
| ENSMUSG00000055407 | Map6 | 0.000276816 | -1.452702475 | 0.168813981 |
| ENSMUSG00000030729 | Pgm2l1 | 0.000276816 | -0.696939285 | 0.836964118 |
| ENSMUSG00000047248 | C2cd3 | 0.000276816 | -1.088130043 | 0.466464457 |
| ENSMUSG00000032875 | Arhgef17 | 0.000276816 | -1.106051951 | 0.395423178 |
| ENSMUSG00000110195 | Pde2a | 0.000276816 | 0.146335211 | 1 |
| ENSMUSG00000032737 | Inppl1 | 0.000276816 | -0.466641666 | 0.96367904 |
| ENSMUSG00000030890 | Ilk | 0.000276816 | 0.648835551 | 0.896076486 |
| ENSMUSG00000036862 | Dchs1 | 0.000276816 | 2.233798052 | 0.134836815 |
| ENSMUSG00000047554 | Tmem41b | 0.000276816 | -0.494122402 | 0.967976676 |
| ENSMUSG00000030772 | Dkk3 | 0.000276816 | 1.326907457 | 0.466464457 |
| ENSMUSG00000030671 | Pde3b | 0.000276816 | -1.507668934 | 0.128531628 |
| ENSMUSG00000073856 | Iqck | 0.000276816 | -0.956026507 | 0.557068455 |
| ENSMUSG00000030942 | Thumpd1 | 0.000276816 | 0.233798052 | 1 |
| ENSMUSG00000051900 | Abca16 | 0.000276816 | 1.146335211 | 0.642722979 |
| ENSMUSG00000046321 | Hs3st2 | 0.000276816 | 0.47891055 | 1 |
| ENSMUSG00000034951 | Cog7 | 0.000276816 | -0.192466703 | 1 |
| ENSMUSG00000030871 | Ears2 | 0.000276816 | -0.351164449 | 1 |
| ENSMUSG00000030689 | Ino80e | 0.000276816 | -0.648844997 | 0.792441218 |
| ENSMUSG00000042423 | Fbrs | 0.000276816 | -1.241935379 | 0.316344908 |
| ENSMUSG00000042308 | Setd1a | 0.000276816 | -1.685065185 | 0.083095907 |
| ENSMUSG00000070369 | Itgad | 0.000276816 | 2.648835551 | 0.093649769 |

| | | | | |
|---|---|---|---|---|
| ENSMUSG00000040298 | Btbd16 | 0.000276816 | -0.810596067 | 0.66416768 |
| ENSMUSG00000006205 | Htra1 | 0.000276816 | 0.985870539 | 0.66416768 |
| ENSMUSG00000030861 | Acadsb | 0.000276816 | -0.466641666 | 0.98880775 |
| ENSMUSG00000030934 | Oat | 0.000276816 | 1.233798052 | 0.601786238 |
| ENSMUSG00000030983 | Bccip | 0.000276816 | -0.599091962 | 0.877036949 |
| ENSMUSG00000030994 | D7Ertd443e | 0.000276816 | 0.279601742 | 1 |
| ENSMUSG00000091002 | Tcerg1l | 0.000276816 | 1.233798052 | 0.600946203 |
| ENSMUSG00000025510 | Cd151 | 0.000276816 | 1.42644313 | 0.408970699 |
| ENSMUSG00000025139 | Tollip | 0.000276816 | 0.189403933 | 1 |
| ENSMUSG00000000244 | Tspan32 | 0.000276816 | 1.42644313 | 0.433763754 |
| ENSMUSG00000044433 | Camsap3 | 0.000276816 | -0.547561661 | 0.871850653 |
| ENSMUSG00000019470 | Xab2 | 0.000276816 | 1.146335211 | 0.627920558 |
| ENSMUSG00000109260 | Gm30954 | 0.000276816 | 0.42644313 | 1 |
| ENSMUSG00000031442 | Mcf2l | 0.000276816 | 0.589941862 | 1 |
| ENSMUSG00000038542 | Pcid2 | 0.000276816 | -1.754886635 | 0.054264589 |
| ENSMUSG00000031453 | Rasa3 | 0.000276816 | -2.567101848 | 0.001828826 |
| ENSMUSG00000051978 | Erich1 | 0.000276816 | -0.494122402 | 0.967976676 |
| ENSMUSG00000110217 | Gm45408 | 0.000276816 | 2.063873051 | 0.173532034 |
| ENSMUSG00000031467 | Agpat5 | 0.000276816 | -0.258055044 | 1 |
| ENSMUSG00000031478 | Nek3 | 0.000276816 | 0.985870539 | 0.66416768 |
| ENSMUSG00000031552 | Adam18 | 0.000276816 | 1.533358334 | 0.359933138 |
| ENSMUSG00000031554 | Adam5 | 0.000276816 | 2.233798052 | 0.169181054 |
| ENSMUSG00000013878 | Rnf170 | 0.000276816 | 0.104515035 | 1 |
| ENSMUSG00000063932 | Poteg | 0.000276816 | 1.774366434 | 0.274319469 |
| ENSMUSG00000039328 | Rnf122 | 0.000276816 | -0.158519371 | 1 |
| ENSMUSG00000046152 | Fut10 | 0.000276816 | 0.375817057 | 1 |
| ENSMUSG00000052906 | Ubxn8 | 0.000276816 | 0.146335211 | 1 |
| ENSMUSG00000031527 | Eri1 | 0.000276816 | -0.936126949 | 0.529126167 |
| ENSMUSG00000109764 | Klkb1 | 0.000276816 | -1.52108945 | 0.166514237 |
| ENSMUSG00000031639 | Tlr3 | 0.000276816 | -0.57355687 | 0.886203497 |
| ENSMUSG00000031631 | Cfap97 | 0.000276816 | -0.43862729 | 1 |
| ENSMUSG00000031568 | Rwdd4a | 0.000276816 | -0.123753952 | 1 |
| ENSMUSG00000110246 | C130073E24Rik | 0.000276816 | 0.146335211 | 1 |
| ENSMUSG00000031604 | Msmo1 | 0.000276816 | -1.599091962 | 0.102454279 |
| ENSMUSG00000025591 | Tma16 | 0.000276816 | -0.351164449 | 1 |
| ENSMUSG00000055553 | Kxd1 | 0.000276816 | -0.494122402 | 0.971854959 |
| ENSMUSG00000031808 | Slc27a1 | 0.000276816 | -0.696939285 | 0.886203497 |
| ENSMUSG00000109583 | Gm39204 | 0.000276816 | 1.146335211 | 0.627920558 |
| ENSMUSG00000064325 | Hhip | 0.000276816 | 1.42644313 | 0.458610057 |
| ENSMUSG00000031711 | Zfp330 | 0.000276816 | -1.192466703 | 0.360150586 |
| ENSMUSG00000008129 | 4930432K21Rik | 0.000276816 | 1.063873051 | 0.66416768 |
| ENSMUSG00000004996 | Mri1 | 0.000276816 | -0.673092543 | 0.792441218 |
| ENSMUSG00000001910 | Nacc1 | 0.000276816 | -0.289763904 | 1 |
| ENSMUSG00000003824 | Syce2 | 0.000276816 | 0.911869957 | 0.847265357 |
| ENSMUSG00000005150 | Wdr83 | 0.000276816 | 0.47891055 | 1 |
| ENSMUSG00000031662 | Snx20 | 0.000276816 | 1.42644313 | 0.439898472 |
| ENSMUSG00000031789 | Cngb1 | 0.000276816 | 0.589941862 | 1 |
| ENSMUSG00000031796 | Cfap20 | 0.000276816 | -0.648844997 | 0.792441218 |
| ENSMUSG00000110483 | Gm45702 | 0.000276816 | 0.024344687 | 1 |
| ENSMUSG00000036564 | Ndrg4 | 0.000276816 | 1.648835551 | 0.374672325 |
| ENSMUSG00000031889 | D230025D16Rik | 0.000276816 | -1.088130043 | 0.455131546 |

| | | | | |
|---|---|---|---|---|
| ENSMUSG00000031887 | Tradd | 0.000276816 | 1.42644313 | 0.408970699 |
| ENSMUSG00000038000 | Acd | 0.000276816 | 1.911869957 | 0.217655393 |
| ENSMUSG00000017765 | Slc12a4 | 0.000276816 | -1.106051951 | 0.395423178 |
| ENSMUSG00000084128 | Esrp2 | 0.000276816 | -0.466641666 | 0.96367904 |
| ENSMUSG00000074113 | Gm10629 | 0.000276816 | 1.063873051 | 0.662993938 |
| ENSMUSG00000031907 | Zfp90 | 0.000276816 | -1.410058138 | 0.191885686 |
| ENSMUSG00000037993 | Dhx38 | 0.000276816 | -1.106051951 | 0.395423178 |
| ENSMUSG00000031749 | St3gal2 | 0.000276816 | -1.241935379 | 0.316344908 |
| ENSMUSG00000034330 | Plcg2 | 0.000276816 | -0.995020638 | 0.51581176 |
| ENSMUSG00000074064 | Mlycd | 0.000276816 | 1.648835551 | 0.30756351 |
| ENSMUSG00000031826 | Usp10 | 0.000276816 | -2.452702475 | 0.00313074 |
| ENSMUSG00000043687 | 1190005I06Rik | 0.000276816 | 0.326907457 | 1 |
| ENSMUSG00000097384 | Gm26815 | 0.000276816 | 2.233798052 | 0.149519292 |
| ENSMUSG00000038697 | Taf5l | 0.000276816 | -0.766201948 | 0.749204141 |
| ENSMUSG00000031851 | Ntpcr | 0.000276816 | 1.233798052 | 0.600946203 |
| ENSMUSG00000079084 | Ccdc82 | 0.000276816 | -1.985370468 | 0.022085266 |
| ENSMUSG00000032009 | Sesn3 | 0.000276816 | -1.305360759 | 0.245924683 |
| ENSMUSG00000063108 | Zfp26 | 0.000276816 | -2.313096408 | 0.006780324 |
| ENSMUSG00000004098 | Col5a3 | 0.000276816 | -0.258055044 | 1 |
| ENSMUSG00000079677 | Fdx2 | 0.000276816 | -1.209145444 | 0.306535101 |
| ENSMUSG00000003308 | Keap1 | 0.000276816 | -0.52108945 | 0.971210385 |
| ENSMUSG00000032193 | Ldlr | 0.000276816 | -1.788569761 | 0.048635862 |
| ENSMUSG00000019066 | Rab3d | 0.000276816 | 0.648835551 | 0.896076486 |
| ENSMUSG00000010607 | Pigyl | 0.000276816 | 2.233798052 | 0.134836815 |
| ENSMUSG00000036777 | Anln | 0.000276816 | -1.624182943 | 0.089456479 |
| ENSMUSG00000074465 | Gm10701 | 0.000276816 | -0.52108945 | 0.979191062 |
| ENSMUSG00000035443 | Thyn1 | 0.000276816 | 1.326907457 | 0.466464457 |
| ENSMUSG00000038119 | Cdon | 0.000276816 | -1.936126949 | 0.030355209 |
| ENSMUSG00000025602 | Zfp202 | 0.000276816 | -2.636566667 | 0.001267919 |
| ENSMUSG00000032119 | Hinfp | 0.000276816 | -1.241935379 | 0.316344908 |
| ENSMUSG00000032115 | Hyou1 | 0.000276816 | -0.995020638 | 0.66416768 |
| ENSMUSG00000032091 | Tmprss4 | 0.000276816 | 1.146335211 | 0.627920558 |
| ENSMUSG00000040219 | Ttc12 | 0.000276816 | -0.192466703 | 1 |
| ENSMUSG00000055069 | Rab39 | 0.000276816 | 1.063873051 | 0.662993938 |
| ENSMUSG00000032311 | Nrg4 | 0.000276816 | -1.57355687 | 0.136125028 |
| ENSMUSG00000032313 | Tmem266 | 0.000276816 | 0.104515035 | 1 |
| ENSMUSG00000091908 | Gm17231 | 0.000276816 | 1.774366434 | 0.306950973 |
| ENSMUSG00000032323 | Cyp11a1 | 0.000276816 | -0.158519371 | 1 |
| ENSMUSG00000032340 | Neo1 | 0.000276816 | -2.004606687 | 0.023140688 |
| ENSMUSG00000074259 | Gramd2 | 0.000276816 | -1.088130043 | 0.455131546 |
| ENSMUSG00000032403 | 2300009A05Rik | 0.000276816 | 1.326907457 | 0.580988658 |
| ENSMUSG00000086539 | Gm16759 | 0.000276816 | -0.289763904 | 1 |
| ENSMUSG00000032392 | Parp16 | 0.000276816 | 0.589941862 | 1 |
| ENSMUSG00000032388 | Spg21 | 0.000276816 | -0.088130043 | 1 |
| ENSMUSG00000032370 | Lactb | 0.000276816 | -0.624182943 | 0.792441218 |
| ENSMUSG00000033543 | Gtf2a2 | 0.000276816 | -0.192466703 | 1 |
| ENSMUSG00000032207 | Lipc | 0.000276816 | -0.380911792 | 1 |
| ENSMUSG00000036030 | Prtg | 0.000276816 | -0.088130043 | 1 |
| ENSMUSG00000032358 | Fam83b | 0.000276816 | -0.696939285 | 0.716109464 |
| ENSMUSG00000032349 | Elovl5 | 0.000276816 | -1.241935379 | 0.316344908 |
| ENSMUSG00000001366 | Fbxo9 | 0.000276816 | -0.258055044 | 1 |

| | | | | |
|---|---|---|---|---|
| ENSMUSG00000032342 | Mto1 | 0.000276816 | -2.3207908 | 0.005496087 |
| ENSMUSG00000049624 | Slc17a5 | 0.000276816 | -2.097118826 | 0.014679132 |
| ENSMUSG00000043289 | Mei4 | 0.000276816 | -0.3207908 | 1 |
| ENSMUSG00000032872 | Cyb5r4 | 0.000276816 | -1.032988489 | 0.443153295 |
| ENSMUSG00000043587 | Pxylp1 | 0.000276816 | 0.233798052 | 1 |
| ENSMUSG00000032449 | Slc25a36 | 0.000276816 | -1.258055044 | 0.274319469 |
| ENSMUSG00000042757 | Tmem108 | 0.000276816 | -0.624182943 | 0.886203497 |
| ENSMUSG00000020257 | Wdr82 | 0.000276816 | -0.088130043 | 1 |
| ENSMUSG00000023277 | Twf2 | 0.000276816 | -0.014129461 | 1 |
| ENSMUSG00000118396 | Iqcf3 | 0.000276816 | 0.533358334 | 1 |
| ENSMUSG00000040813 | Tex264 | 0.000276816 | 0.146335211 | 1 |
| ENSMUSG00000074102 | Rbm15b | 0.000276816 | 1.326907457 | 0.580988658 |
| ENSMUSG00000010066 | Cacna2d2 | 0.000276816 | 0.375817057 | 1 |
| ENSMUSG00000032606 | Nicn1 | 0.000276816 | 1.326907457 | 0.580988658 |
| ENSMUSG00000053646 | Plxnb1 | 0.000276816 | -1.452702475 | 0.168813981 |
| ENSMUSG00000032489 | Kif9 | 0.000276816 | 0.774366434 | 0.872990655 |
| ENSMUSG00000056724 | Nbeal2 | 0.000276816 | -1.560617814 | 0.115171587 |
| ENSMUSG00000044037 | Als2cl | 0.000276816 | -1.494122402 | 0.148292407 |
| ENSMUSG00000046785 | Epm2aip1 | 0.000276816 | -2.599091962 | 0.001292431 |
| ENSMUSG00000032503 | Arpp21 | 0.000276816 | 0.375817057 | 1 |
| ENSMUSG00000009741 | Ubp1 | 0.000276816 | -2.266047835 | 0.006295279 |
| ENSMUSG00000056167 | Cnot10 | 0.000276816 | -1.708716453 | 0.072273892 |
| ENSMUSG00000032443 | Zcwpw2 | 0.000276816 | -0.192466703 | 1 |
| ENSMUSG00000032511 | Scn5a | 0.000276816 | -0.123753952 | 1 |
| ENSMUSG00000041794 | Myrip | 0.000276816 | -0.52108945 | 0.987137113 |
| ENSMUSG00000046603 | Tcaim | 0.000276816 | 0.841480629 | 0.858831093 |
| ENSMUSG00000075327 | Zbtb2 | 0.000276816 | -1.547561661 | 0.113322984 |
| ENSMUSG00000078348 | Sf3b5 | 0.000276816 | 0.985870539 | 0.686078534 |
| ENSMUSG00000019868 | Vta1 | 0.000276816 | -1.351164449 | 0.24901188 |
| ENSMUSG00000112444 | 4930401C15Rik | 0.000276816 | 2.42644313 | 0.102043984 |
| ENSMUSG00000038876 | Rnf146 | 0.000276816 | -1.410058138 | 0.191885686 |
| ENSMUSG00000019785 | Clvs2 | 0.000276816 | 1.326907457 | 0.466464457 |
| ENSMUSG00000039480 | Nt5dc1 | 0.000276816 | -1.069982696 | 0.524144449 |
| ENSMUSG00000091415 | Ak9 | 0.000276816 | 0.985870539 | 0.66416768 |
| ENSMUSG00000020100 | Slc29a3 | 0.000276816 | -0.051604167 | 1 |
| ENSMUSG00000047146 | Tet1 | 0.000276816 | -0.225633566 | 1 |
| ENSMUSG00000042846 | Lrrtm3 | 0.000276816 | 1.911869957 | 0.236944647 |
| ENSMUSG00000033444 | Specc1l | 0.000276816 | -1.885143021 | 0.034614 |
| ENSMUSG00000033208 | S100b | 0.000276816 | 1.146335211 | 0.66416768 |
| ENSMUSG00000001120 | Pcbp3 | 0.000276816 | -1.192466703 | 0.360150586 |
| ENSMUSG00000035890 | Rnf126 | 0.000276816 | 0.774366434 | 0.872990655 |
| ENSMUSG00000035863 | Palm | 0.000276816 | 0.063873051 | 1 |
| ENSMUSG00000020133 | 2310011J03Rik | 0.000276816 | -0.624182943 | 0.792441218 |
| ENSMUSG00000003344 | Btbd2 | 0.000276816 | 0.533358334 | 1 |
| ENSMUSG00000048240 | Gng7 | 0.000276816 | 0.42644313 | 1 |
| ENSMUSG00000034949 | Zfr2 | 0.000276816 | 0.146335211 | 1 |
| ENSMUSG00000020037 | Rfx4 | 0.000276816 | 0.326907457 | 1 |
| ENSMUSG00000060935 | Tmem263 | 0.000276816 | -0.52108945 | 0.971210385 |
| ENSMUSG00000035529 | Prdm4 | 0.000276816 | -0.225633566 | 1 |
| ENSMUSG00000001783 | Rtcb | 0.000276816 | -0.766201948 | 0.73806297 |
| ENSMUSG00000050108 | Bpifc | 0.000276816 | 2.648835551 | 0.072659903 |

| | | | | |
|---|---|---|---|---|
| ENSMUSG00000054027 | Nt5dc3 | 0.000276816 | -0.853664789 | 0.592845519 |
| ENSMUSG00000035459 | Stab2 | 0.000276816 | -0.696939285 | 0.803482361 |
| ENSMUSG00000060904 | Arl1 | 0.000276816 | -1.175592884 | 0.354502808 |
| ENSMUSG00000098040 | Gm20757 | 0.000276816 | 0.47891055 | 1 |
| ENSMUSG00000015889 | Lta4h | 0.000276816 | -0.52108945 | 1 |
| ENSMUSG00000091455 | Otogl | 0.000276816 | 1.648835551 | 0.34594864 |
| ENSMUSG00000048661 | Lemd3 | 0.000276816 | -1.810596067 | 0.051203786 |
| ENSMUSG00000034620 | Rxylt1 | 0.000276816 | -1.106051951 | 0.395423178 |
| ENSMUSG00000020105 | Lrig3 | 0.000276816 | -1.946111038 | 0.028429451 |
| ENSMUSG00000040033 | Stat2 | 0.000276816 | -1.380911792 | 0.220566815 |
| ENSMUSG00000025374 | Nabp2 | 0.000276816 | -0.547561661 | 0.886203497 |
| ENSMUSG00000064030 | Pym1 | 0.000276816 | -0.158519371 | 1 |
| ENSMUSG00000020453 | Patz1 | 0.000276816 | -0.599091962 | 0.877036949 |
| ENSMUSG00000056579 | Tug1 | 0.000276816 | -1.123753952 | 0.400891089 |
| ENSMUSG00000034285 | Nipsnap1 | 0.000276816 | -1.141241379 | 0.405981552 |
| ENSMUSG00000034201 | Gas2l1 | 0.000276816 | 0.589941862 | 1 |
| ENSMUSG00000020393 | Kremen1 | 0.000276816 | -0.648844997 | 0.81042937 |
| ENSMUSG00000049680 | Urgcp | 0.000276816 | -1.424413431 | 0.196126878 |
| ENSMUSG00000020471 | Pold2 | 0.000276816 | 2.063873051 | 0.239441732 |
| ENSMUSG00000041798 | Gck | 0.000276816 | 1.774366434 | 0.311026798 |
| ENSMUSG00000085156 | Snhg15 | 0.000276816 | 0.985870539 | 0.707847898 |
| ENSMUSG00000020193 | Zpbp | 0.000276816 | -2.141241379 | 0.017363741 |
| ENSMUSG00000020173 | Cobl | 0.000276816 | -1.289763904 | 0.279592421 |
| ENSMUSG00000020283 | Pex13 | 0.000276816 | 0.233798052 | 1 |
| ENSMUSG00000000861 | Bcl11a | 0.000276816 | 1.233798052 | 0.621279144 |
| ENSMUSG00000020309 | Chac2 | 0.000276816 | 0.42644313 | 1 |
| ENSMUSG00000007653 | Gabrb2 | 0.000276816 | 1.063873051 | 0.673152643 |
| ENSMUSG00000020409 | Slu7 | 0.000276816 | -0.466641666 | 0.972350276 |
| ENSMUSG00000044707 | Ccnjl | 0.000276816 | -0.014129461 | 1 |
| ENSMUSG00000019189 | Rnf145 | 0.000276816 | -0.743481871 | 0.792441218 |
| ENSMUSG00000020340 | Cyfip2 | 0.000276816 | 0.648835551 | 0.897366006 |
| ENSMUSG00000020359 | Phykpl | 0.000276816 | -0.874726405 | 0.597898961 |
| ENSMUSG00000081769 | Gm12216 | 0.000276816 | -1.673092543 | 0.107857035 |
| ENSMUSG00000018340 | Anxa6 | 0.000276816 | -0.52108945 | 0.972192372 |
| ENSMUSG00000009894 | Snap47 | 0.000276816 | -1.175592884 | 0.354502808 |
| ENSMUSG00000072893 | 4933439C10Rik | 0.000276816 | -1.424413431 | 0.196126878 |
| ENSMUSG00000018415 | Gid4 | 0.000276816 | -0.915949067 | 0.524670309 |
| ENSMUSG00000042506 | Usp22 | 0.000276816 | -1.069982696 | 0.47966612 |
| ENSMUSG00000018509 | Cenpv | 0.000276816 | -0.743481871 | 0.725185838 |
| ENSMUSG00000042148 | Cox10 | 0.000276816 | -0.766201948 | 0.732525536 |
| ENSMUSG00000033389 | Arhgap44 | 0.000276816 | -0.57355687 | 0.887975703 |
| ENSMUSG00000084967 | Gm12296 | 0.000276816 | 1.063873051 | 0.667581691 |
| ENSMUSG00000018750 | Zbtb4 | 0.000276816 | -1.158519371 | 0.3498121 |
| ENSMUSG00000043602 | Zfp3 | 0.000276816 | -1.014129461 | 0.47563319 |
| ENSMUSG00000010554 | Mettl16 | 0.000276816 | -1.685065185 | 0.083095907 |
| ENSMUSG00000037907 | Ankrd13b | 0.000276816 | 0.189403933 | 1 |
| ENSMUSG00000041958 | Pigs | 0.000276816 | 0.024344687 | 1 |
| ENSMUSG00000017561 | Crlf3 | 0.000276816 | -1.636566667 | 0.093377644 |
| ENSMUSG00000020516 | Rps6kb1 | 0.000276816 | -1.946111038 | 0.028429451 |
| ENSMUSG00000023723 | Mrps23 | 0.000276816 | 0.533358334 | 0.971854959 |
| ENSMUSG00000092399 | 4930556N13Rik | 0.000276816 | 0.841480629 | 0.867171841 |

| | | | | |
|---|---|---|---|---|
| ENSMUSG00000020546 | Stxbp4 | 0.000276816 | -1.209145444 | 0.306535101 |
| ENSMUSG00000038976 | Ppp1r9b | 0.000276816 | -0.673092543 | 0.798829579 |
| ENSMUSG00000038534 | Osbpl7 | 0.000276816 | -0.547561661 | 0.871850653 |
| ENSMUSG00000085586 | Gm11613 | 0.000276816 | -1.141241379 | 0.405981552 |
| ENSMUSG00000038437 | Mllt6 | 0.000276816 | -1.273996588 | 0.27550706 |
| ENSMUSG00000010362 | Rdm1 | 0.000276816 | -0.289763904 | 1 |
| ENSMUSG00000008855 | Hdac5 | 0.000276816 | -0.466641666 | 0.981852873 |
| ENSMUSG00000034247 | Plekhm1 | 0.000276816 | -0.853664789 | 0.592845519 |
| ENSMUSG00000020946 | Gosr2 | 0.000276816 | 0.911869957 | 0.847265357 |
| ENSMUSG00000049354 | Dcaf7 | 0.000276816 | -0.351164449 | 1 |
| ENSMUSG00000018433 | Nol11 | 0.000276816 | -1.395558568 | 0.188182248 |
| ENSMUSG00000020610 | Amz2 | 0.000276816 | -0.43862729 | 0.96070793 |
| ENSMUSG00000020604 | Arsg | 0.000276816 | -0.410058138 | 1 |
| ENSMUSG00000087139 | Gm11683 | 0.000276816 | 0.841480629 | 0.792441218 |
| ENSMUSG00000020739 | Nup85 | 0.000276816 | -0.351164449 | 1 |
| ENSMUSG00000020747 | Tmem94 | 0.000276816 | -0.673092543 | 0.792441218 |
| ENSMUSG00000020782 | Llgl2 | 0.000276816 | -1.069982696 | 0.452189485 |
| ENSMUSG00000020780 | Srp68 | 0.000276816 | -1.507668934 | 0.128531628 |
| ENSMUSG00000020792 | Exoc7 | 0.000276816 | 0.189403933 | 1 |
| ENSMUSG00000020802 | Ube2o | 0.000276816 | -1.123753952 | 0.400891089 |
| ENSMUSG00000020810 | Cygb | 0.000276816 | 0.47891055 | 1 |
| ENSMUSG00000017716 | Birc5 | 0.000276816 | -0.466641666 | 0.96367904 |
| ENSMUSG00000025575 | Cant1 | 0.000276816 | -0.57355687 | 0.874192437 |
| ENSMUSG00000025371 | Chmp6 | 0.000276816 | -1.225633566 | 0.367167007 |
| ENSMUSG00000039741 | Bahcc1 | 0.000276816 | -1.141241379 | 0.405981552 |
| ENSMUSG00000039703 | Nploc4 | 0.000276816 | -2.141241379 | 0.013239496 |
| ENSMUSG00000025134 | Alyref | 0.000276816 | 0.47891055 | 1 |
| ENSMUSG00000025155 | Dus1l | 0.000276816 | 0.911869957 | 0.847265357 |
| ENSMUSG00000025153 | Fasn | 0.000276816 | -0.52108945 | 0.992008082 |
| ENSMUSG00000096199 | Ptrhd1 | 0.000276816 | 1.063873051 | 0.66416768 |
| ENSMUSG00000020629 | Adi1 | 0.000276816 | -0.547561661 | 0.871850653 |
| ENSMUSG00000020650 | Bcap29 | 0.000276816 | -0.696939285 | 0.716109464 |
| ENSMUSG00000085664 | Atxn7l1os2 | 0.000276816 | 2.648835551 | 0.072659903 |
| ENSMUSG00000014905 | Dnajb9 | 0.000276816 | 0.233798052 | 1 |
| ENSMUSG00000035653 | Lrfn5 | 0.000276816 | 0.146335211 | 1 |
| ENSMUSG00000020949 | Fkbp3 | 0.000276816 | -0.051604167 | 1 |
| ENSMUSG00000021071 | Trim9 | 0.000276816 | 1.42644313 | 0.451785624 |
| ENSMUSG00000048285 | Frmd6 | 0.000276816 | -2.158519371 | 0.010489786 |
| ENSMUSG00000021079 | Timm9 | 0.000276816 | -0.410058138 | 1 |
| ENSMUSG00000021098 | 4930447C04Rik | 0.000276816 | 1.648835551 | 0.313757032 |
| ENSMUSG00000112459 | Gm47689 | 0.000276816 | -2.946111038 | 0.000227334 |
| ENSMUSG00000059436 | Max | 0.000276816 | -1.175592884 | 0.354502808 |
| ENSMUSG00000021136 | Smoc1 | 0.000276816 | 0.326907457 | 1 |
| ENSMUSG00000042724 | Map3k9 | 0.000276816 | -1.754886635 | 0.054264589 |
| ENSMUSG00000021221 | Dpf3 | 0.000276816 | -0.051604167 | 1 |
| ENSMUSG00000021252 | Erg28 | 0.000276816 | 0.42644313 | 1 |
| ENSMUSG00000021254 | Gpatch2l | 0.000276816 | -1.241935379 | 0.316344908 |
| ENSMUSG00000021033 | Gstz1 | 0.000276816 | -0.289763904 | 1 |
| ENSMUSG00000021037 | Ahsa1 | 0.000276816 | -0.051604167 | 1 |
| ENSMUSG00000097071 | 4930544I03Rik | 0.000276816 | 0.104515035 | 1 |
| ENSMUSG00000020963 | Tshr | 0.000276816 | 0.47891055 | 1 |

| | | | | |
|---|---|---|---|---|
| ENSMUSG00000021003 | Galc | 0.000276816 | -1.273996588 | 0.27550706 |
| ENSMUSG00000044456 | Rin3 | 0.000276816 | -2.184054463 | 0.009479588 |
| ENSMUSG00000041323 | Ak7 | 0.000276816 | 0.104515035 | 1 |
| ENSMUSG00000042063 | Zfp386 | 0.000276816 | -0.810596067 | 0.66416768 |
| ENSMUSG00000072791 | Abcb5 | 0.000276816 | 0.911869957 | 0.720612886 |
| ENSMUSG00000114098 | Gm40660 | 0.000276816 | 1.326907457 | 0.587484274 |
| ENSMUSG00000021196 | Pfkp | 0.000276816 | -1.380911792 | 0.220566815 |
| ENSMUSG00000050244 | Heatr1 | 0.000276816 | -1.624182943 | 0.089456479 |
| ENSMUSG00000021318 | Gli3 | 0.000276816 | -1.685065185 | 0.109343152 |
| ENSMUSG00000009734 | Pou6f2 | 0.000276816 | -0.225633566 | 1 |
| ENSMUSG00000021322 | Aoah | 0.000276816 | -0.225633566 | 1 |
| ENSMUSG00000084136 | Vmn1r207-ps | 0.000276816 | 2.911869957 | 0.095375956 |
| ENSMUSG00000021336 | Slc17a4 | 0.000276816 | -0.766201948 | 0.775967531 |
| ENSMUSG00000035936 | Aldh5a1 | 0.000276816 | 1.326907457 | 0.47966612 |
| ENSMUSG00000071451 | Psmg4 | 0.000276816 | -0.43862729 | 0.96070793 |
| ENSMUSG00000038991 | Txndc5 | 0.000276816 | -0.599091962 | 0.877036949 |
| ENSMUSG00000001707 | Eef1e1 | 0.000276816 | -0.624182943 | 0.792441218 |
| ENSMUSG00000047094 | Ofcc1 | 0.000276816 | 0.063873051 | 1 |
| ENSMUSG00000057531 | Dtnbp1 | 0.000276816 | -1.192466703 | 0.360150586 |
| ENSMUSG00000069237 | Fam8a1 | 0.000276816 | -0.494122402 | 0.967976676 |
| ENSMUSG00000038080 | Kdm1b | 0.000276816 | 0.146335211 | 1 |
| ENSMUSG00000101136 | Gm8739 | 0.000276816 | 1.42644313 | 0.408970699 |
| ENSMUSG00000034987 | Hrh2 | 0.000276816 | 1.533358334 | 0.364784642 |
| ENSMUSG00000025872 | Thoc3 | 0.000276816 | -0.466641666 | 0.96367904 |
| ENSMUSG00000043183 | Simc1 | 0.000276816 | -1.158519371 | 0.3498121 |
| ENSMUSG00000055254 | Ntrk2 | 0.000276816 | -0.547561661 | 0.998731799 |
| ENSMUSG00000021476 | Habp4 | 0.000276816 | -1.106051951 | 0.410540658 |
| ENSMUSG00000074832 | 2410141K09Rik | 0.000276816 | 2.063873051 | 0.241724181 |
| ENSMUSG00000095432 | Zfp748 | 0.000276816 | -0.466641666 | 0.96367904 |
| ENSMUSG00000021510 | Zfp729a | 0.000276816 | -1.051604167 | 0.448359534 |
| ENSMUSG00000048280 | Zfp738 | 0.000276816 | -1.708716453 | 0.072273892 |
| ENSMUSG00000114038 | Gm40999 | 0.000276816 | 0.985870539 | 0.681660809 |
| ENSMUSG00000021608 | Lpcat1 | 0.000276816 | 0.063873051 | 1 |
| ENSMUSG00000021591 | Glrx | 0.000276816 | 0.146335211 | 1 |
| ENSMUSG00000033991 | Ttc37 | 0.000276816 | -2.209145444 | 0.008626067 |
| ENSMUSG00000114401 | Gm38604 | 0.000276816 | 1.648835551 | 0.30756351 |
| ENSMUSG00000100168 | Gm28526 | 0.000276816 | -0.380911792 | 1 |
| ENSMUSG00000074794 | Arrdc3 | 0.000276816 | -1.241935379 | 0.316344908 |
| ENSMUSG00000021706 | Zfyve16 | 0.000276816 | -2.47356108 | 0.003246102 |
| ENSMUSG00000021684 | Pde8b | 0.000276816 | 0.42644313 | 1 |
| ENSMUSG00000021678 | F2rl1 | 0.000276816 | -1.175592884 | 0.354502808 |
| ENSMUSG00000057762 | Gm6169 | 0.000276816 | 2.42644313 | 0.102043984 |
| ENSMUSG00000041632 | Mrps27 | 0.000276816 | -0.975655313 | 0.466464457 |
| ENSMUSG00000100733 | 4932411K12Rik | 0.000276816 | -0.351164449 | 1 |
| ENSMUSG00000114206 | Gm30551 | 0.000276816 | 0.841480629 | 0.885240416 |
| ENSMUSG00000021713 | Ppwd1 | 0.000276816 | -1.777429203 | 0.056971848 |
| ENSMUSG00000068184 | Ndufaf2 | 0.000276816 | -1.032988489 | 0.503117077 |
| ENSMUSG00000049985 | Ankrd55 | 0.000276816 | 0.533358334 | 1 |
| ENSMUSG00000021728 | Emb | 0.000276816 | 0.774366434 | 0.887369997 |
| ENSMUSG00000094870 | Zfp131 | 0.000276816 | -1.351164449 | 0.213441582 |
| ENSMUSG00000033111 | 3830406C13Rik | 0.000276816 | -0.410058138 | 1 |

| | | | | |
|---|---|---|---|---|
| ENSMUSG00000042567 | Nek10 | 0.000276816 | -0.956026507 | 0.559918909 |
| ENSMUSG00000045107 | Saysd1 | 0.000276816 | -0.088130043 | 1 |
| ENSMUSG00000021816 | Ppp3cb | 0.000276816 | -2.494122402 | 0.002048947 |
| ENSMUSG00000114774 | Gm35281 | 0.000276816 | 2.42644313 | 0.102043984 |
| ENSMUSG00000091898 | Tnnc1 | 0.000276816 | -0.995020638 | 0.470839676 |
| ENSMUSG00000021890 | Eaf1 | 0.000276816 | -0.192466703 | 1 |
| ENSMUSG00000037572 | Wdhd1 | 0.000276816 | -1.936126949 | 0.027678678 |
| ENSMUSG00000022160 | Mettl3 | 0.000276816 | 0.42644313 | 1 |
| ENSMUSG00000002329 | Mdp1 | 0.000276816 | 0.326907457 | 1 |
| ENSMUSG00000022157 | Mcpt8 | 0.000276816 | -0.088130043 | 1 |
| ENSMUSG00000048279 | Sacs | 0.000276816 | -1.673092543 | 0.081728936 |
| ENSMUSG00000021928 | Ebpl | 0.000276816 | -0.258055044 | 1 |
| ENSMUSG00000035067 | Xkr6 | 0.000276816 | -1.799624949 | 0.061725817 |
| ENSMUSG00000034327 | Kctd9 | 0.000276816 | -1.696939285 | 0.070585664 |
| ENSMUSG00000034190 | Chmp7 | 0.000276816 | 0.279601742 | 1 |
| ENSMUSG00000115289 | Gm49164 | 0.000276816 | -0.853664789 | 0.620556057 |
| ENSMUSG00000115479 | Gm33203 | 0.000276816 | 1.774366434 | 0.273353405 |
| ENSMUSG00000022130 | Tgds | 0.000276816 | -0.289763904 | 1 |
| ENSMUSG00000022132 | Cldn10 | 0.000276816 | 1.326907457 | 0.587484274 |
| ENSMUSG00000041625 | Ggact | 0.000276816 | -1.3207908 | 0.262879729 |
| ENSMUSG00000041935 | AW549877 | 0.000276816 | -2.52775318 | 0.001729963 |
| ENSMUSG00000049148 | Plcxd3 | 0.000276816 | 0.104515035 | 1 |
| ENSMUSG00000022155 | Mroh2b | 0.000276816 | 1.42644313 | 0.408970699 |
| ENSMUSG00000022253 | Nadk2 | 0.000276816 | 0.326907457 | 1 |
| ENSMUSG00000039704 | Lmbrd2 | 0.000276816 | -0.874726405 | 0.597898961 |
| ENSMUSG00000098150 | 4921515E04Rik | 0.000276816 | 0.326907457 | 1 |
| ENSMUSG00000022262 | Dnah5 | 0.000276816 | 0.589941862 | 0.932255214 |
| ENSMUSG00000022257 | Laptm4b | 0.000276816 | -1.289763904 | 0.279592421 |
| ENSMUSG00000048307 | Ankrd46 | 0.000276816 | 0.533358334 | 1 |
| ENSMUSG00000115172 | Gm2582 | 0.000276816 | 1.063873051 | 0.662993938 |
| ENSMUSG00000022365 | Derl1 | 0.000276816 | -0.088130043 | 1 |
| ENSMUSG00000101892 | 9130401M01Rik | 0.000276816 | 0.47891055 | 1 |
| ENSMUSG00000022351 | Sqle | 0.000276816 | 0.063873051 | 1 |
| ENSMUSG00000116498 | Gm38563 | 0.000276816 | 0.589941862 | 1 |
| ENSMUSG00000116031 | D030024E09Rik | 0.000276816 | -0.351164449 | 1 |
| ENSMUSG00000022346 | Myc | 0.000276816 | -3.136889355 | 5.09E-05 |
| ENSMUSG00000036944 | Tmem71 | 0.000276816 | 1.911869957 | 0.217655393 |
| ENSMUSG00000093387 | Gm20732 | 0.000276816 | -1.175592884 | 0.354502808 |
| ENSMUSG00000116483 | Gm19782 | 0.000276816 | 1.648835551 | 0.354200337 |
| ENSMUSG00000079022 | Col22a1 | 0.000276816 | 1.326907457 | 0.466464457 |
| ENSMUSG00000059895 | Ptp4a3 | 0.000276816 | 0.533358334 | 1 |
| ENSMUSG00000034429 | Zfp707 | 0.000276816 | 1.146335211 | 0.627920558 |
| ENSMUSG00000022561 | Gpaa1 | 0.000276816 | -0.673092543 | 0.792441218 |
| ENSMUSG00000034161 | Scx | 0.000276816 | 1.648835551 | 0.30756351 |
| ENSMUSG00000063354 | Slc39a4 | 0.000276816 | 0.911869957 | 0.847265357 |
| ENSMUSG00000044986 | Tst | 0.000276816 | 1.533358334 | 0.359933138 |
| ENSMUSG00000006378 | Gcat | 0.000276816 | 1.233798052 | 0.53074305 |
| ENSMUSG00000022426 | Josd1 | 0.000276816 | -0.853664789 | 0.592845519 |
| ENSMUSG00000022415 | Syngr1 | 0.000276816 | 0.533358334 | 1 |
| ENSMUSG00000022389 | Tef | 0.000276816 | -1.051604167 | 0.448359534 |
| ENSMUSG00000022476 | Polr3h | 0.000276816 | -1.788569761 | 0.048635862 |

| | | | | |
|---|---|---|---|---|
| ENSMUSG00000016757 | Ttll12 | 0.000276816 | -0.051604167 | 1 |
| ENSMUSG00000062760 | Shisal1 | 0.000276816 | 1.146335211 | 0.66416768 |
| ENSMUSG00000078954 | Arhgap8 | 0.000276816 | -1.52108945 | 0.128972787 |
| ENSMUSG00000006369 | Fbln1 | 0.000276816 | 0.774366434 | 0.872990655 |
| ENSMUSG00000022386 | Trmu | 0.000276816 | 0.146335211 | 1 |
| ENSMUSG00000036273 | Lrrk2 | 0.000276816 | -0.051604167 | 1 |
| ENSMUSG00000048154 | Kmt2d | 0.000276816 | -3.52775318 | 3.46E-06 |
| ENSMUSG00000116518 | Gm49494 | 0.000276816 | -0.599091962 | 0.877036949 |
| ENSMUSG00000037280 | Galnt6 | 0.000276816 | -0.192466703 | 1 |
| ENSMUSG00000023044 | Csad | 0.000276816 | -0.832291138 | 0.669438508 |
| ENSMUSG00000036678 | Aaas | 0.000276816 | -1.209145444 | 0.306535101 |
| ENSMUSG00000022783 | Spag6l | 0.000276816 | 0.710236096 | 0.919998879 |
| ENSMUSG00000005262 | Ufd1 | 0.000276816 | -1.560617814 | 0.115171587 |
| ENSMUSG00000003233 | Dvl3 | 0.000276816 | 0.648835551 | 0.896076486 |
| ENSMUSG00000022887 | Masp1 | 0.000276816 | -0.547561661 | 0.886203497 |
| ENSMUSG00000033355 | Rtp4 | 0.000276816 | 2.42644313 | 0.102043984 |
| ENSMUSG00000051065 | Mb21d2 | 0.000276816 | -1.051604167 | 0.448359534 |
| ENSMUSG00000022538 | Lsg1 | 0.000276816 | -2.106051951 | 0.015201129 |
| ENSMUSG00000047434 | Xxylt1 | 0.000276816 | -0.52108945 | 0.971210385 |
| ENSMUSG00000014075 | Tctex1d2 | 0.000276816 | 1.146335211 | 0.66416768 |
| ENSMUSG00000022802 | Lmln | 0.000276816 | -0.225633566 | 1 |
| ENSMUSG00000022833 | Ccdc14 | 0.000276816 | -0.895484965 | 0.602434911 |
| ENSMUSG00000022840 | Adcy5 | 0.000276816 | 0.42644313 | 1 |
| ENSMUSG00000075122 | Cd80 | 0.000276816 | 1.063873051 | 0.674297981 |
| ENSMUSG00000034064 | Poglut1 | 0.000276816 | -2.023589791 | 0.02040298 |
| ENSMUSG00000022663 | Atg3 | 0.000276816 | -0.380911792 | 1 |
| ENSMUSG00000022656 | Nectin3 | 0.000276816 | -1.636566667 | 0.101935624 |
| ENSMUSG00000022651 | Retnlg | 0.000276816 | 3.648835551 | 0.0159954 |
| ENSMUSG00000035258 | Abi3bp | 0.000276816 | 0.233798052 | 1 |
| ENSMUSG00000084999 | Gm15839 | 0.000276816 | 1.42644313 | 0.442955829 |
| ENSMUSG00000022749 | Tbc1d23 | 0.000276816 | -1.289763904 | 0.279592421 |
| ENSMUSG00000035107 | Dcbld2 | 0.000276816 | -1.175592884 | 0.354502808 |
| ENSMUSG00000022723 | Crybg3 | 0.000276816 | -0.57355687 | 0.874192437 |
| ENSMUSG00000022911 | Arl13b | 0.000276816 | -1.673092543 | 0.081728936 |
| ENSMUSG00000068167 | Csnka2ip | 0.000276816 | 0.911869957 | 0.722638283 |
| ENSMUSG00000008976 | Gabpa | 0.000276816 | -1.720398258 | 0.061086889 |
| ENSMUSG00000022894 | Adamts5 | 0.000276816 | 0.375817057 | 1 |
| ENSMUSG00000044442 | N6amt1 | 0.000276816 | 1.233798052 | 0.600946203 |
| ENSMUSG00000099968 | 1700029J03Rik | 0.000276816 | 1.326907457 | 0.580988658 |
| ENSMUSG00000062713 | Sim2 | 0.000276816 | -1.424413431 | 0.196126878 |
| ENSMUSG00000040820 | Hlcs | 0.000276816 | -2.023589791 | 0.02040298 |
| ENSMUSG00000043301 | Kcnj6 | 0.000276816 | -1.696939285 | 0.092468958 |
| ENSMUSG00000023806 | Rsph3b | 0.000276816 | -0.895484965 | 0.602434911 |
| ENSMUSG00000069135 | Fgfr1op | 0.000276816 | -0.696939285 | 0.716109464 |
| ENSMUSG00000117143 | Gm49807 | 0.000276816 | 2.063873051 | 0.210550914 |
| ENSMUSG00000117042 | 2700054A10Rik | 0.000276816 | -2.032988489 | 0.021103588 |
| ENSMUSG00000051341 | Zfp52 | 0.000276816 | -0.720398258 | 0.719591951 |
| ENSMUSG00000061544 | Zfp229 | 0.000276816 | -0.720398258 | 0.719591951 |
| ENSMUSG00000085185 | BC028777 | 0.000276816 | 3.648835551 | 0.0159954 |
| ENSMUSG00000038880 | Mrps34 | 0.000276816 | 0.104515035 | 1 |
| ENSMUSG00000024180 | Tmem8 | 0.000276816 | 1.533358334 | 0.359933138 |

| | | | | |
|---|---|---|---|---|
| ENSMUSG00000062753 | AI413582 | 0.000276816 | 1.326907457 | 0.580988658 |
| ENSMUSG00000007570 | Fance | 0.000276816 | 0.710236096 | 0.886203497 |
| ENSMUSG00000006705 | Pknox1 | 0.000276816 | -0.975655313 | 0.466464457 |
| ENSMUSG00000024299 | Adamts10 | 0.000276816 | -0.788569761 | 0.66416768 |
| ENSMUSG00000002308 | Cd320 | 0.000276816 | 1.911869957 | 0.217655393 |
| ENSMUSG00000039656 | Rxrb | 0.000276816 | -1.289763904 | 0.279592421 |
| ENSMUSG00000034254 | Agpat1 | 0.000276816 | -0.225633566 | 1 |
| ENSMUSG00000024371 | C2 | 0.000276816 | -0.123753952 | 1 |
| ENSMUSG00000007036 | Abhd16a | 0.000276816 | 0.911869957 | 0.847265357 |
| ENSMUSG00000039220 | Ppp1r10 | 0.000276816 | -1.57355687 | 0.117709307 |
| ENSMUSG00000067212 | H2-T23 | 0.000276816 | 1.774366434 | 0.262284476 |
| ENSMUSG00000024462 | Gabbr1 | 0.000276816 | -2.599091962 | 0.001292431 |
| ENSMUSG00000025431 | Crisp1 | 0.000276816 | 1.42644313 | 0.450590406 |
| ENSMUSG00000041293 | Adgrf1 | 0.000276816 | 1.774366434 | 0.262284476 |
| ENSMUSG00000056492 | Adgrf5 | 0.000276816 | -0.52108945 | 0.98374804 |
| ENSMUSG00000023959 | Clic5 | 0.000276816 | -0.225633566 | 1 |
| ENSMUSG00000023947 | Nfkbie | 0.000276816 | -0.158519371 | 1 |
| ENSMUSG00000023953 | Polh | 0.000276816 | -1.3207908 | 0.255869172 |
| ENSMUSG00000040327 | Cul9 | 0.000276816 | -0.743481871 | 0.725185838 |
| ENSMUSG00000038545 | Cul7 | 0.000276816 | -1.720398258 | 0.061086889 |
| ENSMUSG00000023973 | Cnpy3 | 0.000276816 | -1.088130043 | 0.455131546 |
| ENSMUSG00000092558 | Med20 | 0.000276816 | -2.051604167 | 0.021899967 |
| ENSMUSG00000041754 | Trem3 | 0.000276816 | 1.233798052 | 0.640346301 |
| ENSMUSG00000043592 | Unc5cl | 0.000276816 | 0.189403933 | 1 |
| ENSMUSG00000100150 | Gm19585 | 0.000276816 | -0.088130043 | 1 |
| ENSMUSG00000024174 | Pot1b | 0.000276816 | -2.679091284 | 0.000746706 |
| ENSMUSG00000001228 | Uhrf1 | 0.000276816 | 0.063873051 | 1 |
| ENSMUSG00000034116 | Vav1 | 0.000276816 | 1.533358334 | 0.360150586 |
| ENSMUSG00000117243 | Gm49944 | 0.000276816 | 0.279601742 | 1 |
| ENSMUSG00000117231 | Gm41609 | 0.000276816 | -1.336057556 | 0.250559418 |
| ENSMUSG00000039414 | Heatr5b | 0.000276816 | -0.936126949 | 0.529126167 |
| ENSMUSG00000062691 | Cebpzos | 0.000276816 | 1.42644313 | 0.45354452 |
| ENSMUSG00000040852 | Plekhh2 | 0.000276816 | -0.258055044 | 1 |
| ENSMUSG00000024135 | Srbd1 | 0.000276816 | -2.702839887 | 0.000727108 |
| ENSMUSG00000024151 | Msh2 | 0.000276816 | -1.158519371 | 0.475407382 |
| ENSMUSG00000032937 | Fshr | 0.000276816 | -0.289763904 | 1 |
| ENSMUSG00000063087 | Gm10125 | 0.000276816 | -1.241935379 | 0.316344908 |
| ENSMUSG00000024413 | Npc1 | 0.000276816 | -1.720398258 | 0.061086889 |
| ENSMUSG00000057766 | Ankrd29 | 0.000276816 | 1.233798052 | 0.612182491 |
| ENSMUSG00000024424 | Ttc39c | 0.000276816 | 0.189403933 | 1 |
| ENSMUSG00000024423 | Impact | 0.000276816 | -1.351164449 | 0.213441582 |
| ENSMUSG00000054321 | Taf4b | 0.000276816 | -1.43862729 | 0.166121613 |
| ENSMUSG00000047989 | Ino80c | 0.000276816 | 0.533358334 | 1 |
| ENSMUSG00000024273 | 2700062C07Rik | 0.000276816 | 0.42644313 | 1 |
| ENSMUSG00000117604 | Gm33228 | 0.000276816 | -0.014129461 | 1 |
| ENSMUSG00000024388 | Myo7b | 0.000276816 | 1.911869957 | 0.217655393 |
| ENSMUSG00000110185 | Igip | 0.000276816 | 0.589941862 | 1 |
| ENSMUSG00000033272 | Slc35a4 | 0.000276816 | 0.533358334 | 1 |
| ENSMUSG00000051401 | Kctd16 | 0.000276816 | -0.599091962 | 0.891195418 |
| ENSMUSG00000039954 | Stk32a | 0.000276816 | 0.146335211 | 1 |
| ENSMUSG00000019647 | Sema6a | 0.000276816 | -2.547561661 | 0.001620452 |

| | | | | |
|---|---|---|---|---|
| ENSMUSG00000118259 | Gm5095 | 0.000276816 | 0.375817057 | 1 |
| ENSMUSG00000073565 | Prr16 | 0.000276816 | -1.766201948 | 0.064033463 |
| ENSMUSG00000118362 | Gm19466 | 0.000276816 | 1.063873051 | 0.66416768 |
| ENSMUSG00000096948 | Gm4221 | 0.000276816 | -1.106051951 | 0.395423178 |
| ENSMUSG00000056812 | St8sia3 | 0.000276816 | 1.063873051 | 0.66416768 |
| ENSMUSG00000032845 | Alpk2 | 0.000276816 | 0.589941862 | 1 |
| ENSMUSG00000118376 | Gm9926 | 0.000276816 | -2.023589791 | 0.02040298 |
| ENSMUSG00000024527 | Afg3l2 | 0.000276816 | -2.821484383 | 0.000336123 |
| ENSMUSG00000024533 | Spire1 | 0.000276816 | -1.661019711 | 0.079998381 |
| ENSMUSG00000024542 | Cep192 | 0.000276816 | -2.200830175 | 0.008348692 |
| ENSMUSG00000038903 | Ccdc68 | 0.000276816 | -1.560617814 | 0.115171587 |
| ENSMUSG00000025425 | St8sia5 | 0.000276816 | 1.911869957 | 0.262288269 |
| ENSMUSG00000039840 | Epg5 | 0.000276816 | -1.507668934 | 0.128531628 |
| ENSMUSG00000049090 | Zadh2 | 0.000276816 | -0.410058138 | 1 |
| ENSMUSG00000037916 | Ndufv1 | 0.000276816 | -0.43862729 | 0.96070793 |
| ENSMUSG00000049303 | Syt12 | 0.000276816 | 0.710236096 | 0.886203497 |
| ENSMUSG00000063904 | Dpp3 | 0.000276816 | -1.273996588 | 0.27550706 |
| ENSMUSG00000024841 | Eif1ad | 0.000276816 | -1.088130043 | 0.455131546 |
| ENSMUSG00000039148 | Sart1 | 0.000276816 | -1.885143021 | 0.034614 |
| ENSMUSG00000024939 | Fam89b | 0.000276816 | -0.410058138 | 1 |
| ENSMUSG00000024941 | Scyl1 | 0.000276816 | -1.225633566 | 0.310683162 |
| ENSMUSG00000024958 | Gpr137 | 0.000276816 | 0.063873051 | 1 |
| ENSMUSG00000071646 | Mta2 | 0.000276816 | -1.175592884 | 0.354502808 |
| ENSMUSG00000024654 | Asrgl1 | 0.000276816 | 0.279601742 | 1 |
| ENSMUSG00000024658 | Gm9750 | 0.000276816 | 1.146335211 | 0.627920558 |
| ENSMUSG00000034371 | Tkfc | 0.000276816 | 1.42644313 | 0.408970699 |
| ENSMUSG00000048832 | Vps37c | 0.000276816 | -1.788569761 | 0.048635862 |
| ENSMUSG00000046139 | Patl1 | 0.000276816 | -1.534386273 | 0.13144915 |
| ENSMUSG00000046805 | Mpeg1 | 0.000276816 | 1.533358334 | 0.359933138 |
| ENSMUSG00000094732 | 1500015L24Rik | 0.000276816 | -0.874726405 | 0.639754062 |
| ENSMUSG00000074925 | Ptar1 | 0.000276816 | -1.225633566 | 0.310683162 |
| ENSMUSG00000071604 | Fam189a2 | 0.000276816 | -0.258055044 | 1 |
| ENSMUSG00000024924 | Vldlr | 0.000276816 | -1.3207908 | 0.245794249 |
| ENSMUSG00000064202 | 4430402I18Rik | 0.000276816 | 0.589941862 | 1 |
| ENSMUSG00000024766 | Lipo3 | 0.000276816 | 0.648835551 | 0.913923844 |
| ENSMUSG00000024795 | Kif20b | 0.000276816 | -2.305360759 | 0.005137215 |
| ENSMUSG00000118212 | Gm41836 | 0.000276816 | 2.063873051 | 0.173532034 |
| ENSMUSG00000024986 | Hhex | 0.000276816 | -0.832291138 | 0.66416768 |
| ENSMUSG00000067242 | Lgi1 | 0.000276816 | 1.648835551 | 0.342187905 |
| ENSMUSG00000061132 | Blnk | 0.000276816 | 0.024344687 | 1 |
| ENSMUSG00000025171 | Ubtd1 | 0.000276816 | -1.534386273 | 0.13144915 |
| ENSMUSG00000049670 | Morn4 | 0.000276816 | 2.42644313 | 0.10330121 |
| ENSMUSG00000044220 | Nkx2-3 | 0.000276816 | 0.024344687 | 1 |
| ENSMUSG00000025202 | Scd3 | 0.000276816 | 1.233798052 | 0.611036796 |
| ENSMUSG00000025209 | Twnk | 0.000276816 | -0.624182943 | 0.792441218 |
| ENSMUSG00000033033 | Calhm2 | 0.000276816 | -1.708716453 | 0.094521215 |
| ENSMUSG00000004317 | Clcn5 | 0.000276816 | -2.673092543 | 0.000857868 |
| ENSMUSG00000031143 | Ccdc22 | 0.000276816 | -0.673092543 | 0.792441218 |
| ENSMUSG00000031150 | Ccdc120 | 0.000276816 | -0.014129461 | 1 |
| ENSMUSG00000031158 | Timm17b | 0.000276816 | -0.52108945 | 0.982652955 |
| ENSMUSG00000036782 | Klhl13 | 0.000276816 | 1.146335211 | 0.627920558 |

| | | | | |
|---|---|---|---|---|
| ENSMUSG00000031139 | Mcf2 | 0.000276816 | 2.648835551 | 0.072659903 |
| ENSMUSG00000000838 | Fmr1 | 0.000276816 | -1.410058138 | 0.191885686 |
| ENSMUSG00000058328 | Xlr5a | 0.000276816 | -2.395558568 | 0.006222976 |
| ENSMUSG00000015291 | Gdi1 | 0.000276816 | -1.351164449 | 0.213441582 |
| ENSMUSG00000032750 | Gab3 | 0.000276816 | -0.289763904 | 1 |
| ENSMUSG00000031217 | Efnb1 | 0.000276816 | -0.696939285 | 0.716109464 |
| ENSMUSG00000000881 | Dlg3 | 0.000276816 | 0.104515035 | 1 |
| ENSMUSG00000031327 | Chic1 | 0.000276816 | -1.611691999 | 0.142157135 |
| ENSMUSG00000073016 | Uprt | 0.000276816 | -0.853664789 | 0.592845519 |
| ENSMUSG00000031242 | 2610002M06Rik | 0.000276816 | 0.841480629 | 0.880492974 |
| ENSMUSG00000034755 | Pcdh11x | 0.000276816 | -0.743481871 | 0.74230625 |
| ENSMUSG00000086807 | Platr21 | 0.000276816 | 1.326907457 | 0.580988658 |
| ENSMUSG00000031433 | Rbm41 | 0.000276816 | -0.410058138 | 1 |
| ENSMUSG00000041718 | Alg13 | 0.000276816 | -1.336057556 | 0.250559418 |
| ENSMUSG00000044150 | Bclaf3 | 0.000276816 | -2.097118826 | 0.014679132 |
| ENSMUSG00000031380 | Vegfd | 0.000276816 | 1.146335211 | 0.627920558 |
| ENSMUSG00000079317 | Trappc2 | 0.000276816 | 1.774366434 | 0.262284476 |
| ENSMUSG00000069045 | Ddx3y | 0.000276816 | 0.104515035 | 1 |
| ENSMUSG00000033793 | Atp6v1h | 0.000221453 | -1.836591276 | 0.058667604 |
| ENSMUSG00000025927 | Tfap2b | 0.000221453 | 0.215876144 | 1 |
| ENSMUSG00000042215 | Bag2 | 0.000221453 | -1.797420678 | 0.067360743 |
| ENSMUSG00000026126 | Ptpn18 | 0.000221453 | -0.351164449 | 1 |
| ENSMUSG00000037509 | Arhgef4 | 0.000221453 | 0.970763646 | 0.792441218 |
| ENSMUSG00000037447 | Arid5a | 0.000221453 | -0.673092543 | 0.833666942 |
| ENSMUSG00000046337 | Fam178b | 0.000221453 | 1.063873051 | 0.75757939 |
| ENSMUSG00000026073 | Il1r2 | 0.000221453 | 1.163408724 | 0.759570628 |
| ENSMUSG00000100832 | Gm29260 | 0.000221453 | -0.386788358 | 1 |
| ENSMUSG00000060679 | Mrps9 | 0.000221453 | -1.948099591 | 0.040663018 |
| ENSMUSG00000057363 | Uxs1 | 0.000221453 | -1.77070334 | 0.080452994 |
| ENSMUSG00000041303 | Gtf3c3 | 0.000221453 | -1.199161355 | 0.391578102 |
| ENSMUSG00000038242 | Aox4 | 0.000221453 | 1.648835551 | 0.452956236 |
| ENSMUSG00000099752 | Gm29018 | 0.000221453 | 1.511332028 | 0.463009681 |
| ENSMUSG00000069085 | Dytn | 0.000221453 | -0.810596067 | 0.670365203 |
| ENSMUSG00000025949 | Pikfyve | 0.000221453 | -2.258055044 | 0.010947317 |
| ENSMUSG00000026004 | Kansl1l | 0.000221453 | -2.228908699 | 0.012088308 |
| ENSMUSG00000045648 | Vwc2l | 0.000221453 | -0.455501108 | 1 |
| ENSMUSG00000026187 | Xrcc5 | 0.000221453 | -2.277163867 | 0.009507975 |
| ENSMUSG00000100301 | 6030407O03Rik | 0.000221453 | 0.44720169 | 1 |
| ENSMUSG00000026172 | Bcs1l | 0.000221453 | 1.063873051 | 0.75757939 |
| ENSMUSG00000026200 | Glb1l | 0.000221453 | -1.073630473 | 0.472298934 |
| ENSMUSG00000026203 | Dnajb2 | 0.000221453 | -1.095325544 | 0.433313155 |
| ENSMUSG00000026209 | Dnpep | 0.000221453 | 0.215876144 | 1 |
| ENSMUSG00000026235 | Epha4 | 0.000221453 | -1.701661696 | 0.103047269 |
| ENSMUSG00000067158 | Col4a4 | 0.000221453 | -0.073630473 | 1 |
| ENSMUSG00000026149 | Tm4sf20 | 0.000221453 | 2.648835551 | 0.124712353 |
| ENSMUSG00000070031 | Sp140 | 0.000221453 | -0.199161355 | 1 |
| ENSMUSG00000036251 | Trpm8 | 0.000221453 | 0.44720169 | 1 |
| ENSMUSG00000067071 | Hes6 | 0.000221453 | 0.112782651 | 1 |
| ENSMUSG00000026274 | Pask | 0.000221453 | -0.029236354 | 1 |
| ENSMUSG00000009905 | Kdsr | 0.000221453 | -0.673092543 | 0.833666942 |
| ENSMUSG00000026312 | Cdh7 | 0.000221453 | 1.385801146 | 0.491653643 |

| | | | | |
|---|---|---|---|---|
| ENSMUSG00000051590 | Map3k19 | 0.000221453 | 0.722836133 | 1 |
| ENSMUSG00000036086 | Zranb3 | 0.000221453 | -0.729676072 | 0.754374521 |
| ENSMUSG00000013275 | Slc41a1 | 0.000221453 | -0.029236354 | 1 |
| ENSMUSG00000026439 | Rbbp5 | 0.000221453 | -1.687447836 | 0.100676805 |
| ENSMUSG00000046062 | Ppp1r15b | 0.000221453 | -1.386788358 | 0.219470377 |
| ENSMUSG00000026398 | Nr5a2 | 0.000221453 | 0.215876144 | 1 |
| ENSMUSG00000026360 | Rgs2 | 0.000221453 | -0.729676072 | 0.744267071 |
| ENSMUSG00000099964 | Gm29514 | 0.000221453 | 1.648835551 | 0.405392586 |
| ENSMUSG00000032487 | Ptgs2 | 0.000221453 | 0.800838645 | 0.998731799 |
| ENSMUSG00000006010 | Odr4 | 0.000221453 | -1.404275785 | 0.2224613 |
| ENSMUSG00000093650 | Gm20631 | 0.000221453 | 0.800838645 | 0.844591606 |
| ENSMUSG00000073538 | E330020D12Rik | 0.000221453 | 3.385801146 | 0.048720096 |
| ENSMUSG00000045968 | Teddm2 | 0.000221453 | 0.016567336 | 1 |
| ENSMUSG00000015484 | Fam163a | 0.000221453 | -0.701661696 | 0.873294632 |
| ENSMUSG00000043467 | Zbtb37 | 0.000221453 | -1.47217985 | 0.19917628 |
| ENSMUSG00000026709 | Dars2 | 0.000221453 | -0.643946198 | 0.828559607 |
| ENSMUSG00000026696 | Vamp4 | 0.000221453 | -1.238689719 | 0.333941363 |
| ENSMUSG00000102302 | Gm38190 | 0.000221453 | -2.395558568 | 0.005551659 |
| ENSMUSG00000040713 | Creg1 | 0.000221453 | 0.270323928 | 1 |
| ENSMUSG00000093538 | Gm20711 | 0.000221453 | 1.270323928 | 0.57709448 |
| ENSMUSG00000026678 | Rgs5 | 0.000221453 | 0.326907457 | 1 |
| ENSMUSG00000026641 | Usf1 | 0.000221453 | -0.887217349 | 0.600946203 |
| ENSMUSG00000091993 | B930036N10Rik | 0.000221453 | -0.614198854 | 0.829657084 |
| ENSMUSG00000003464 | Pex19 | 0.000221453 | -0.277163867 | 1 |
| ENSMUSG00000005338 | Cadm3 | 0.000221453 | 2.385801146 | 0.171970178 |
| ENSMUSG00000037860 | Aim2 | 0.000221453 | 0.270323928 | 1 |
| ENSMUSG00000039783 | Kmo | 0.000221453 | 2.648835551 | 0.124712353 |
| ENSMUSG00000026523 | Wdr64 | 0.000221453 | 0.112782651 | 1 |
| ENSMUSG00000026519 | Tmem63a | 0.000221453 | -2.158519371 | 0.017601978 |
| ENSMUSG00000102682 | Gm37223 | 0.000221453 | 1.063873051 | 0.772933445 |
| ENSMUSG00000103961 | Gm36388 | 0.000221453 | 1.970763646 | 0.341667562 |
| ENSMUSG00000026603 | Smyd2 | 0.000221453 | -2.006516277 | 0.031339065 |
| ENSMUSG00000016493 | Cd46 | 0.000221453 | -1.836591276 | 0.060407117 |
| ENSMUSG00000025815 | Dhtkd1 | 0.000221453 | -1.051604167 | 0.466464457 |
| ENSMUSG00000087125 | A230108P19Rik | 0.000221453 | -1.006516277 | 0.584777539 |
| ENSMUSG00000026768 | Itga8 | 0.000221453 | 0.112782651 | 1 |
| ENSMUSG00000026726 | Cubn | 0.000221453 | 0.511332028 | 1 |
| ENSMUSG00000026737 | Pip4k2a | 0.000221453 | -2.52108945 | 0.004015829 |
| ENSMUSG00000026981 | Il1rn | 0.000221453 | 0.016567336 | 1 |
| ENSMUSG00000026974 | Zmynd19 | 0.000221453 | -1.369086357 | 0.263753194 |
| ENSMUSG00000026975 | Dph7 | 0.000221453 | -0.277163867 | 1 |
| ENSMUSG00000013465 | Nelfb | 0.000221453 | -1.199161355 | 0.391578102 |
| ENSMUSG00000036731 | Cysrt1 | 0.000221453 | 2.163408724 | 0.226617702 |
| ENSMUSG00000044628 | Rnf208 | 0.000221453 | 2.648835551 | 0.124712353 |
| ENSMUSG00000049086 | Bmyc | 0.000221453 | 1.511332028 | 0.466464457 |
| ENSMUSG00000036327 | Qsox2 | 0.000221453 | 0.800838645 | 0.998731799 |
| ENSMUSG00000015790 | Surf1 | 0.000221453 | 0.163408724 | 1 |
| ENSMUSG00000009614 | Sardh | 0.000221453 | -1.178983473 | 0.384884537 |
| ENSMUSG00000086425 | F730016J06Rik | 0.000221453 | 3.385801146 | 0.048720096 |
| ENSMUSG00000026812 | Tsc1 | 0.000221453 | -1.488667972 | 0.167024802 |
| ENSMUSG00000026806 | Ddx31 | 0.000221453 | -1.687447836 | 0.100676805 |

| | | | | |
|---|---|---|---|---|
| ENSMUSG00000069020 | Urm1 | 0.000221453 | -0.583825205 | 0.921320733 |
| ENSMUSG00000039715 | Wdr34 | 0.000221453 | -0.784123856 | 0.775410248 |
| ENSMUSG00000015335 | Zdhhc12 | 0.000221453 | 1.385801146 | 0.491653643 |
| ENSMUSG00000039648 | Kyat1 | 0.000221453 | -1.404275785 | 0.2224613 |
| ENSMUSG00000087269 | D330023K18Rik | 0.000221453 | -0.936126949 | 0.611141945 |
| ENSMUSG00000076441 | Ass1 | 0.000221453 | 0.578446224 | 1 |
| ENSMUSG00000053746 | Ptrh1 | 0.000221453 | 1.648835551 | 0.405392586 |
| ENSMUSG00000026870 | Cutal | 0.000221453 | 0.722836133 | 1 |
| ENSMUSG00000026875 | Traf1 | 0.000221453 | -0.757156808 | 0.77665738 |
| ENSMUSG00000026874 | Hc | 0.000221453 | 1.270323928 | 0.571786485 |
| ENSMUSG00000026887 | Mrrf | 0.000221453 | 0.648835551 | 1 |
| ENSMUSG00000009030 | Pdcl | 0.000221453 | 0.578446224 | 1 |
| ENSMUSG00000026834 | Acvr1c | 0.000221453 | 1.648835551 | 0.405392586 |
| ENSMUSG00000075324 | Fign | 0.000221453 | -0.52108945 | 0.920236372 |
| ENSMUSG00000064329 | Scn1a | 0.000221453 | 0.270323928 | 1 |
| ENSMUSG00000034738 | Nostrin | 0.000221453 | -0.158519371 | 1 |
| ENSMUSG00000063145 | Bbs5 | 0.000221453 | 0.215876144 | 1 |
| ENSMUSG00000070883 | Ccdc173 | 0.000221453 | -0.421553776 | 1 |
| ENSMUSG00000027109 | Sp3 | 0.000221453 | -2.386788358 | 0.006613418 |
| ENSMUSG00000085405 | 4930441J16Rik | 0.000221453 | 0.063873051 | 1 |
| ENSMUSG00000026999 | Nup35 | 0.000221453 | -0.836591276 | 0.673811773 |
| ENSMUSG00000086495 | Gm13778 | 0.000221453 | -0.673092543 | 0.833666942 |
| ENSMUSG00000040687 | Madd | 0.000221453 | -2.323857103 | 0.007505624 |
| ENSMUSG00000002109 | Ddb2 | 0.000221453 | -1.874726405 | 0.05622212 |
| ENSMUSG00000040174 | Alkbh3 | 0.000221453 | -0.116699195 | 1 |
| ENSMUSG00000027164 | Traf6 | 0.000221453 | -2.377964508 | 0.006295279 |
| ENSMUSG00000027163 | Commd9 | 0.000221453 | 0.883300805 | 0.796840156 |
| ENSMUSG00000005089 | Slc1a2 | 0.000221453 | -0.277163867 | 1 |
| ENSMUSG00000027186 | Elf5 | 0.000221453 | 1.648835551 | 0.410761383 |
| ENSMUSG00000027168 | Pax6 | 0.000221453 | 2.648835551 | 0.127356309 |
| ENSMUSG00000027166 | Dnajc24 | 0.000221453 | -1.258055044 | 0.340118707 |
| ENSMUSG00000016386 | Mpped2 | 0.000221453 | -0.701661696 | 0.740459298 |
| ENSMUSG00000027122 | Arl14ep | 0.000221453 | -1.849415316 | 0.060372549 |
| ENSMUSG00000086758 | Gm13912 | 0.000221453 | 0.800838645 | 0.847265357 |
| ENSMUSG00000027157 | 4930430A15Rik | 0.000221453 | 1.800838645 | 0.344436715 |
| ENSMUSG00000027134 | Lpcat4 | 0.000221453 | -0.314638573 | 1 |
| ENSMUSG00000003604 | Aven | 0.000221453 | -1.568395165 | 0.150632966 |
| ENSMUSG00000086308 | G630016G05Rik | 0.000221453 | -1.47217985 | 0.19917628 |
| ENSMUSG00000027347 | Rasgrp1 | 0.000221453 | 0.883300805 | 0.796840156 |
| ENSMUSG00000005102 | Eif2ak4 | 0.000221453 | -2.701661696 | 0.001400623 |
| ENSMUSG00000040093 | Bmf | 0.000221453 | 1.163408724 | 0.73806297 |
| ENSMUSG00000039983 | Ccdc32 | 0.000221453 | 0.800838645 | 0.998731799 |
| ENSMUSG00000068580 | Zfyve19 | 0.000221453 | -0.277163867 | 1 |
| ENSMUSG00000062646 | Ganc | 0.000221453 | -0.643946198 | 0.828559607 |
| ENSMUSG00000027285 | Haus2 | 0.000221453 | 0.883300805 | 0.796840156 |
| ENSMUSG00000027259 | Adal | 0.000221453 | -1.314638573 | 0.322556056 |
| ENSMUSG00000068479 | Mfap1a | 0.000221453 | 0.800838645 | 0.998731799 |
| ENSMUSG00000060131 | Atp8b4 | 0.000221453 | 0.016567336 | 1 |
| ENSMUSG00000037740 | Mrps26 | 0.000221453 | 0.970763646 | 0.789767653 |
| ENSMUSG00000027357 | Crls1 | 0.000221453 | -0.488667972 | 1 |
| ENSMUSG00000085640 | Gm14064 | 0.000221453 | 0.800838645 | 0.998731799 |

| | | | | |
|---|---|---|---|---|
| ENSMUSG00000027611 | Procr | 0.000221453 | 0.722836133 | 1 |
| ENSMUSG00000063019 | Manbal | 0.000221453 | -0.836591276 | 0.667778757 |
| ENSMUSG00000078956 | Gm14221 | 0.000221453 | 0.970763646 | 0.789767653 |
| ENSMUSG00000016933 | Plcg1 | 0.000221453 | -1.47217985 | 0.19917628 |
| ENSMUSG00000001403 | Ube2c | 0.000221453 | 0.800838645 | 0.998731799 |
| ENSMUSG00000017664 | Slc35c2 | 0.000221453 | -0.073630473 | 1 |
| ENSMUSG00000017670 | Elmo2 | 0.000221453 | 0.44720169 | 1 |
| ENSMUSG00000018459 | Slc13a3 | 0.000221453 | 1.648835551 | 0.430269377 |
| ENSMUSG00000017999 | Ddx27 | 0.000221453 | -1.55279831 | 0.147260869 |
| ENSMUSG00000050556 | Kcnb1 | 0.000221453 | -0.386788358 | 1 |
| ENSMUSG00000027560 | Dok5 | 0.000221453 | 0.326907457 | 1 |
| ENSMUSG00000074570 | Cass4 | 0.000221453 | 0.326907457 | 1 |
| ENSMUSG00000027502 | Rtf2 | 0.000221453 | -1.006516277 | 0.537139526 |
| ENSMUSG00000078899 | Gm4631 | 0.000221453 | 0.970763646 | 0.789767653 |
| ENSMUSG00000039108 | Lsm14b | 0.000221453 | -0.199161355 | 1 |
| ENSMUSG00000097804 | Gm16685 | 0.000221453 | 0.722836133 | 1 |
| ENSMUSG00000087108 | Gm16093 | 0.000221453 | -1.959973691 | 0.034141187 |
| ENSMUSG00000037531 | Mrpl47 | 0.000221453 | -0.887217349 | 0.600946203 |
| ENSMUSG00000114832 | Gm21388 | 0.000221453 | 2.385801146 | 0.205095549 |
| ENSMUSG00000027710 | Acad9 | 0.000221453 | -0.073630473 | 1 |
| ENSMUSG00000044167 | Foxo1 | 0.000221453 | -2.568395165 | 0.002484871 |
| ENSMUSG00000036580 | Spg20 | 0.000221453 | -1.797420678 | 0.066785139 |
| ENSMUSG00000036353 | P2ry12 | 0.000221453 | 1.385801146 | 0.51581176 |
| ENSMUSG00000036334 | Igsf10 | 0.000221453 | 1.511332028 | 0.463009681 |
| ENSMUSG00000062200 | Vmn2r7 | 0.000221453 | 1.800838645 | 0.344436715 |
| ENSMUSG00000034640 | Tiparp | 0.000221453 | -0.911879403 | 0.605887572 |
| ENSMUSG00000094763 | Gm1647 | 0.000221453 | 0.648835551 | 1 |
| ENSMUSG00000102362 | 4930509J09Rik | 0.000221453 | 0.970763646 | 0.790129914 |
| ENSMUSG00000027809 | Etfdh | 0.000221453 | -0.673092543 | 0.833666942 |
| ENSMUSG00000033910 | Gucy1a1 | 0.000221453 | 1.970763646 | 0.282889199 |
| ENSMUSG00000074513 | Arfip1 | 0.000221453 | -0.643946198 | 0.828559607 |
| ENSMUSG00000004896 | Rrnad1 | 0.000221453 | -0.836591276 | 0.66416768 |
| ENSMUSG00000028068 | Iqgap3 | 0.000221453 | -0.158519371 | 1 |
| ENSMUSG00000060671 | Atp8b2 | 0.000221453 | -0.810596067 | 0.66416768 |
| ENSMUSG00000027937 | Jtb | 0.000221453 | -1.178983473 | 0.384884537 |
| ENSMUSG00000001016 | Ilf2 | 0.000221453 | -2.219060913 | 0.014337901 |
| ENSMUSG00000104791 | 4930511M18Rik | 0.000221453 | 2.385801146 | 0.181341877 |
| ENSMUSG00000074445 | Sprr2a3 | 0.000221453 | 3.970763646 | 0.035887064 |
| ENSMUSG00000102476 | Gm37245 | 0.000221453 | -0.421553776 | 1 |
| ENSMUSG00000019338 | Zfp687 | 0.000221453 | -0.073630473 | 1 |
| ENSMUSG00000093424 | 6330562C20Rik | 0.000221453 | -0.116699195 | 1 |
| ENSMUSG00000063689 | Hist2h2ab | 0.000221453 | -1.568395165 | 0.150632966 |
| ENSMUSG00000028096 | Gpr89 | 0.000221453 | -0.983432664 | 0.530172915 |
| ENSMUSG00000044468 | Tent5c | 0.000221453 | 1.648835551 | 0.405392586 |
| ENSMUSG00000027860 | Vangl1 | 0.000221453 | -0.488667972 | 1 |
| ENSMUSG00000105891 | A230001M10Rik | 0.000221453 | 0.270323928 | 1 |
| ENSMUSG00000027854 | Sike1 | 0.000221453 | -0.729676072 | 0.744267071 |
| ENSMUSG00000027848 | Olfml3 | 0.000221453 | 0.270323928 | 1 |
| ENSMUSG00000062778 | Chia1 | 0.000221453 | 2.970763646 | 0.09957688 |
| ENSMUSG00000040600 | Eps8l3 | 0.000221453 | -0.238689719 | 1 |
| ENSMUSG00000004032 | Gstm5 | 0.000221453 | 0.44720169 | 1 |

| | | | | |
|---|---|---|---|---|
| ENSMUSG00000085318 | Gm12524 | 0.000221453 | 1.800838645 | 0.451785624 |
| ENSMUSG00000068732 | Tmem167b | 0.000221453 | -0.116699195 | 1 |
| ENSMUSG00000027965 | Olfm3 | 0.000221453 | 0.270323928 | 1 |
| ENSMUSG00000033554 | Dph5 | 0.000221453 | -1.537030994 | 0.144776283 |
| ENSMUSG00000000339 | Rtca | 0.000221453 | -0.643946198 | 0.855178417 |
| ENSMUSG00000098097 | 6530403H02Rik | 0.000221453 | -0.238689719 | 1 |
| ENSMUSG00000028133 | Rwdd3 | 0.000221453 | 0.163408724 | 1 |
| ENSMUSG00000028114 | Mettl14 | 0.000221453 | -0.488667972 | 1 |
| ENSMUSG00000097031 | Gm26820 | 0.000221453 | 0.326907457 | 1 |
| ENSMUSG00000028167 | Bdh2 | 0.000221453 | 1.648835551 | 0.405392586 |
| ENSMUSG00000043020 | Wdr63 | 0.000221453 | -0.116699195 | 1 |
| ENSMUSG00000028189 | Ctbs | 0.000221453 | 1.511332028 | 0.463009681 |
| ENSMUSG00000104785 | Gm31121 | 0.000221453 | 0.722836133 | 0.877887538 |
| ENSMUSG00000085996 | A830012C17Rik | 0.000221453 | 1.970763646 | 0.281836872 |
| ENSMUSG00000028245 | Nsmaf | 0.000221453 | -2.455501108 | 0.004768037 |
| ENSMUSG00000041216 | Clvs1 | 0.000221453 | 0.326907457 | 1 |
| ENSMUSG00000006586 | Runx1t1 | 0.000221453 | 0.511332028 | 1 |
| ENSMUSG00000028224 | Nbn | 0.000221453 | -0.810596067 | 0.66416768 |
| ENSMUSG00000028261 | Ndufaf4 | 0.000221453 | 0.800838645 | 0.998731799 |
| ENSMUSG00000028407 | Smim27 | 0.000221453 | 2.385801146 | 0.171970178 |
| ENSMUSG00000028411 | Aptx | 0.000221453 | 0.215876144 | 1 |
| ENSMUSG00000028444 | Cntfr | 0.000221453 | -1.995020638 | 0.034978902 |
| ENSMUSG00000036078 | Sigmar1 | 0.000221453 | -0.351164449 | 1 |
| ENSMUSG00000073888 | Ccl27a | 0.000221453 | 1.648835551 | 0.405392586 |
| ENSMUSG00000028457 | Atp8b5 | 0.000221453 | 0.270323928 | 1 |
| ENSMUSG00000035969 | Rusc2 | 0.000221453 | -2.544936192 | 0.002693122 |
| ENSMUSG00000028458 | Tesk1 | 0.000221453 | 0.970763646 | 0.816561732 |
| ENSMUSG00000028470 | Hint2 | 0.000221453 | 0.511332028 | 1 |
| ENSMUSG00000035683 | Melk | 0.000221453 | -0.386788358 | 1 |
| ENSMUSG00000035572 | Dcaf10 | 0.000221453 | -1.995020638 | 0.030563094 |
| ENSMUSG00000038544 | Inip | 0.000221453 | 0.883300805 | 0.796840156 |
| ENSMUSG00000066152 | Slc31a2 | 0.000221453 | 0.016567336 | 1 |
| ENSMUSG00000028394 | Pole3 | 0.000221453 | 0.270323928 | 1 |
| ENSMUSG00000039005 | Tlr4 | 0.000221453 | 0.063873051 | 1 |
| ENSMUSG00000043003 | Rasef | 0.000221453 | 0.722836133 | 1 |
| ENSMUSG00000028402 | Mpdz | 0.000221453 | -0.836591276 | 0.674297981 |
| ENSMUSG00000117123 | Gm49890 | 0.000221453 | -3.500911568 | 8.86E-06 |
| ENSMUSG00000085569 | Gm12602 | 0.000221453 | -2.158519371 | 0.017601978 |
| ENSMUSG00000085147 | Gm12609 | 0.000221453 | -0.351164449 | 1 |
| ENSMUSG00000087366 | Junos | 0.000221453 | -0.757156808 | 0.777158385 |
| ENSMUSG00000028550 | Atg4c | 0.000221453 | -0.386788358 | 1 |
| ENSMUSG00000069733 | Ube2u | 0.000221453 | 0.970763646 | 0.792441218 |
| ENSMUSG00000087361 | 0610043K17Rik | 0.000221453 | -0.116699195 | 1 |
| ENSMUSG00000028527 | Ak4 | 0.000221453 | -0.55279831 | 0.916104407 |
| ENSMUSG00000028520 | 4921539E11Rik | 0.000221453 | 1.385801146 | 0.520824233 |
| ENSMUSG00000028614 | Ndc1 | 0.000221453 | -2.01792104 | 0.032072771 |
| ENSMUSG00000085549 | 1700047F07Rik | 0.000221453 | 1.163408724 | 0.761315087 |
| ENSMUSG00000028601 | Echdc2 | 0.000221453 | -0.52108945 | 0.912904617 |
| ENSMUSG00000048351 | Coa7 | 0.000221453 | -0.386788358 | 1 |
| ENSMUSG00000059816 | Shisal2a | 0.000221453 | 1.800838645 | 0.344436715 |
| ENSMUSG00000028555 | Ttc39a | 0.000221453 | -0.351164449 | 1 |

| | | | | |
|---|---|---|---|---|
| ENSMUSG00000087641 | Gm12811 | 0.000221453 | 0.578446224 | 1 |
| ENSMUSG00000034401 | Spata6 | 0.000221453 | -1.333017102 | 0.300304685 |
| ENSMUSG00000046861 | Hectd3 | 0.000221453 | 0.970763646 | 0.789767653 |
| ENSMUSG00000028542 | Slc6a9 | 0.000221453 | -0.158519371 | 1 |
| ENSMUSG00000006392 | Med8 | 0.000221453 | -0.614198854 | 0.971854959 |
| ENSMUSG00000006389 | Mpl | 0.000221453 | -0.810596067 | 0.687925796 |
| ENSMUSG00000078584 | AU022252 | 0.000221453 | 0.385801146 | 1 |
| ENSMUSG00000028876 | Epha10 | 0.000221453 | 1.511332028 | 0.466464457 |
| ENSMUSG00000042677 | Zc3h12a | 0.000221453 | -0.421553776 | 1 |
| ENSMUSG00000042489 | Clspn | 0.000221453 | -2.658592974 | 0.00153265 |
| ENSMUSG00000073755 | 5730409E04Rik | 0.000221453 | 1.970763646 | 0.281836872 |
| ENSMUSG00000043872 | Zmym1 | 0.000221453 | -1.82365222 | 0.057125834 |
| ENSMUSG00000057572 | Zbtb8os | 0.000221453 | 0.722836133 | 1 |
| ENSMUSG00000028807 | Zbtb8a | 0.000221453 | -0.583825205 | 0.921320733 |
| ENSMUSG00000074088 | Snrnp40 | 0.000221453 | -0.701661696 | 0.740459298 |
| ENSMUSG00000078532 | Nkain1 | 0.000221453 | -0.643946198 | 0.846742525 |
| ENSMUSG00000050511 | Oprd1 | 0.000221453 | 1.063873051 | 0.774790843 |
| ENSMUSG00000028896 | Rcc1 | 0.000221453 | 0.016567336 | 1 |
| ENSMUSG00000028893 | Sesn2 | 0.000221453 | 0.163408724 | 1 |
| ENSMUSG00000048003 | Catsper4 | 0.000221453 | 0.578446224 | 1 |
| ENSMUSG00000028825 | Rhd | 0.000221453 | 1.648835551 | 0.411939273 |
| ENSMUSG00000037157 | Il22ra1 | 0.000221453 | 1.063873051 | 0.75757939 |
| ENSMUSG00000028675 | Pnrc2 | 0.000221453 | -0.386788358 | 1 |
| ENSMUSG00000028669 | Pithd1 | 0.000221453 | -2.323857103 | 0.008555928 |
| ENSMUSG00000060862 | Zbtb40 | 0.000221453 | -1.296022894 | 0.290284345 |
| ENSMUSG00000041193 | Pla2g5 | 0.000221453 | 2.385801146 | 0.171970178 |
| ENSMUSG00000041025 | Iffo2 | 0.000221453 | -1.55279831 | 0.147260869 |
| ENSMUSG00000028927 | Padi2 | 0.000221453 | -0.199161355 | 1 |
| ENSMUSG00000040860 | Crocc | 0.000221453 | -1.386788358 | 0.219470377 |
| ENSMUSG00000028914 | Casp9 | 0.000221453 | 1.270323928 | 0.706154373 |
| ENSMUSG00000029009 | Mthfr | 0.000221453 | -0.55279831 | 0.916104407 |
| ENSMUSG00000044700 | Tmem201 | 0.000221453 | 0.326907457 | 1 |
| ENSMUSG00000039911 | Spsb1 | 0.000221453 | -1.095325544 | 0.433313155 |
| ENSMUSG00000028965 | Tnfrsf9 | 0.000221453 | 0.722836133 | 1 |
| ENSMUSG00000028963 | Uts2 | 0.000221453 | 2.648835551 | 0.124712353 |
| ENSMUSG00000039523 | Cep104 | 0.000221453 | -0.959973691 | 0.526094357 |
| ENSMUSG00000029026 | Trp73 | 0.000221453 | 1.063873051 | 0.759652247 |
| ENSMUSG00000029049 | Morn1 | 0.000221453 | -0.199161355 | 1 |
| ENSMUSG00000000600 | Krit1 | 0.000221453 | -1.537030994 | 0.144776283 |
| ENSMUSG00000044674 | Fzd1 | 0.000221453 | 0.648835551 | 1 |
| ENSMUSG00000046798 | Cldn12 | 0.000221453 | -0.386788358 | 1 |
| ENSMUSG00000040537 | Adam22 | 0.000221453 | -0.277163867 | 1 |
| ENSMUSG00000039987 | Phtf2 | 0.000221453 | -1.629149196 | 0.112238591 |
| ENSMUSG00000029015 | Slc26a5 | 0.000221453 | 1.648835551 | 0.405392586 |
| ENSMUSG00000028969 | Cdk5 | 0.000221453 | -0.386788358 | 1 |
| ENSMUSG00000002221 | Paxip1 | 0.000221453 | -0.983432664 | 0.674472331 |
| ENSMUSG00000025747 | Tyms | 0.000221453 | -0.199161355 | 1 |
| ENSMUSG00000053194 | Cib4 | 0.000221453 | 0.722836133 | 1 |
| ENSMUSG00000029168 | Dpysl5 | 0.000221453 | 1.163408724 | 0.733004386 |
| ENSMUSG00000029151 | Slc30a3 | 0.000221453 | 3.970763646 | 0.023955525 |
| ENSMUSG00000107283 | Mpv17 | 0.000221453 | 1.800838645 | 0.344436715 |

| | | | | |
|---|---|---|---|---|
| ENSMUSG00000029141 | Slc4a1ap | 0.000221453 | -1.715736881 | 0.086523965 |
| ENSMUSG00000029134 | Plb1 | 0.000221453 | 0.648835551 | 1 |
| ENSMUSG00000037313 | Tacc3 | 0.000221453 | -0.455501108 | 1 |
| ENSMUSG00000037224 | Zfyve28 | 0.000221453 | -0.116699195 | 1 |
| ENSMUSG00000059866 | Tnip2 | 0.000221453 | -1.404275785 | 0.2224613 |
| ENSMUSG00000029119 | Man2b2 | 0.000221453 | 0.385801146 | 1 |
| ENSMUSG00000063646 | Jakmip1 | 0.000221453 | 1.511332028 | 0.463009681 |
| ENSMUSG00000039765 | Cc2d2a | 0.000221453 | -1.296022894 | 0.290284345 |
| ENSMUSG00000029082 | Bst1 | 0.000221453 | 0.016567336 | 1 |
| ENSMUSG00000029089 | 5730480H06Rik | 0.000221453 | -0.421553776 | 1 |
| ENSMUSG00000029090 | Adgra3 | 0.000221453 | -1.488667972 | 0.167024802 |
| ENSMUSG00000097216 | 4932441J04Rik | 0.000221453 | 1.648835551 | 0.405392586 |
| ENSMUSG00000090061 | Nwd2 | 0.000221453 | -0.673092543 | 0.847265357 |
| ENSMUSG00000037890 | Wdr19 | 0.000221453 | -0.614198854 | 0.825311622 |
| ENSMUSG00000087261 | Gm15477 | 0.000221453 | 1.648835551 | 0.421384409 |
| ENSMUSG00000005672 | Kit | 0.000221453 | 2.163408724 | 0.226617702 |
| ENSMUSG00000036403 | Cep135 | 0.000221453 | -1.729676072 | 0.088605926 |
| ENSMUSG00000029254 | Stap1 | 0.000221453 | 0.063873051 | 1 |
| ENSMUSG00000029273 | Sult1d1 | 0.000221453 | 1.970763646 | 0.281836872 |
| ENSMUSG00000029397 | Rchy1 | 0.000221453 | -0.757156808 | 0.749778237 |
| ENSMUSG00000034842 | Art3 | 0.000221453 | -0.029236354 | 1 |
| ENSMUSG00000050050 | Ccdc158 | 0.000221453 | 0.44720169 | 1 |
| ENSMUSG00000029486 | Mrpl1 | 0.000221453 | -1.333017102 | 0.463850347 |
| ENSMUSG00000035297 | Cops4 | 0.000221453 | -1.029236354 | 0.466464457 |
| ENSMUSG00000034462 | Pkd2 | 0.000221453 | -2.817138913 | 0.000716101 |
| ENSMUSG00000029299 | Abcg3 | 0.000221453 | 0.44720169 | 1 |
| ENSMUSG00000111375 | Btbd8 | 0.000221453 | -0.643946198 | 0.828559607 |
| ENSMUSG00000029276 | Glmn | 0.000221453 | -0.936126949 | 0.611141945 |
| ENSMUSG00000033434 | Gtpbp6 | 0.000221453 | -0.314638573 | 1 |
| ENSMUSG00000029500 | Pgam5 | 0.000221453 | -0.238689719 | 1 |
| ENSMUSG00000033294 | Noc4l | 0.000221453 | -0.386788358 | 1 |
| ENSMUSG00000042328 | Hps4 | 0.000221453 | -1.687447836 | 0.100676805 |
| ENSMUSG00000042216 | Sgsm1 | 0.000221453 | 0.578446224 | 1 |
| ENSMUSG00000054675 | Tmem119 | 0.000221453 | 3.385801146 | 0.087209077 |
| ENSMUSG00000044339 | Alkbh2 | 0.000221453 | 1.648835551 | 0.405392586 |
| ENSMUSG00000041890 | Git2 | 0.000221453 | -3.084518789 | 0.000139388 |
| ENSMUSG00000086122 | Gm14507 | 0.000221453 | 1.385801146 | 0.521275193 |
| ENSMUSG00000063919 | Srrm4 | 0.000221453 | -0.351164449 | 1 |
| ENSMUSG00000032898 | Fbxo21 | 0.000221453 | -0.314638573 | 1 |
| ENSMUSG00000029596 | Sdsl | 0.000221453 | -1.333017102 | 0.300304685 |
| ENSMUSG00000029602 | Rasal1 | 0.000221453 | 0.648835551 | 1 |
| ENSMUSG00000029474 | Rnf34 | 0.000221453 | -0.887217349 | 0.606122355 |
| ENSMUSG00000029392 | Rilpl1 | 0.000221453 | 0.270323928 | 1 |
| ENSMUSG00000029389 | Ddx55 | 0.000221453 | -1.386788358 | 0.219470377 |
| ENSMUSG00000105039 | Gm32585 | 0.000221453 | 0.326907457 | 1 |
| ENSMUSG00000029423 | Piwil1 | 0.000221453 | 1.648835551 | 0.405392586 |
| ENSMUSG00000029446 | Psph | 0.000221453 | 1.063873051 | 0.75757939 |
| ENSMUSG00000007207 | Stx1a | 0.000221453 | -2.360153232 | 0.007109019 |
| ENSMUSG00000005378 | Bud23 | 0.000221453 | 0.511332028 | 1 |
| ENSMUSG00000005374 | Tbl2 | 0.000221453 | -0.199161355 | 1 |
| ENSMUSG00000004952 | Rasa4 | 0.000221453 | -0.314638573 | 1 |

| | | | | |
|---|---|---|---|---|
| ENSMUSG00000004846 | Plod3 | 0.000221453 | -0.836591276 | 0.66416768 |
| ENSMUSG00000079165 | Sap25 | 0.000221453 | -0.029236354 | 1 |
| ENSMUSG00000025854 | Fam20c | 0.000221453 | -1.351164449 | 0.259656769 |
| ENSMUSG00000025855 | Prkar1b | 0.000221453 | -0.862126368 | 0.700227059 |
| ENSMUSG00000025857 | Dnaaf5 | 0.000221453 | -1.599091962 | 0.128972787 |
| ENSMUSG00000053553 | 3110082I17Rik | 0.000221453 | -1.743481871 | 0.107393453 |
| ENSMUSG00000036718 | Micall2 | 0.000221453 | -1.095325544 | 0.491872181 |
| ENSMUSG00000000148 | Brat1 | 0.000221453 | 0.326907457 | 1 |
| ENSMUSG00000075569 | Rsph10b | 0.000221453 | -0.116699195 | 1 |
| ENSMUSG00000107132 | Gm15997 | 0.000221453 | 1.385801146 | 0.500596163 |
| ENSMUSG00000029757 | Dync1i1 | 0.000221453 | -0.643946198 | 0.861533036 |
| ENSMUSG00000029752 | Asns | 0.000221453 | 1.648835551 | 0.457077215 |
| ENSMUSG00000068748 | Ptprz1 | 0.000221453 | -0.455501108 | 1 |
| ENSMUSG00000046192 | Iqub | 0.000221453 | 1.511332028 | 0.466464457 |
| ENSMUSG00000084861 | 1700095J07Rik | 0.000221453 | 0.648835551 | 1 |
| ENSMUSG00000058486 | Wdr91 | 0.000221453 | 0.511332028 | 1 |
| ENSMUSG00000029838 | Ptn | 0.000221453 | 0.722836133 | 1 |
| ENSMUSG00000063455 | D630045J12Rik | 0.000221453 | -0.455501108 | 1 |
| ENSMUSG00000064262 | Gimap8 | 0.000221453 | 1.648835551 | 0.405392586 |
| ENSMUSG00000058446 | Znrf2 | 0.000221453 | -1.948099591 | 0.040663018 |
| ENSMUSG00000038058 | Nod1 | 0.000221453 | -0.199161355 | 1 |
| ENSMUSG00000003476 | Crhr2 | 0.000221453 | 1.511332028 | 0.476074732 |
| ENSMUSG00000059486 | Kbtbd2 | 0.000221453 | -1.277163867 | 0.346101473 |
| ENSMUSG00000029804 | Herc3 | 0.000221453 | -0.55279831 | 0.916104407 |
| ENSMUSG00000107799 | Gm38825 | 0.000221453 | -1.599091962 | 0.147574014 |
| ENSMUSG00000053604 | Rpia | 0.000221453 | -0.862126368 | 0.675262063 |
| ENSMUSG00000052962 | Mrpl35 | 0.000221453 | 0.970763646 | 0.789767653 |
| ENSMUSG00000072476 | Gm9008 | 0.000221453 | 2.648835551 | 0.173253468 |
| ENSMUSG00000059654 | Reg1 | 0.000221453 | 0.883300805 | 0.867531944 |
| ENSMUSG00000035125 | Gcfc2 | 0.000221453 | -0.614198854 | 0.825311622 |
| ENSMUSG00000005667 | Mthfd2 | 0.000221453 | -1.073630473 | 0.472298934 |
| ENSMUSG00000059430 | Actg2 | 0.000221453 | 0.722836133 | 1 |
| ENSMUSG00000006906 | Stambp | 0.000221453 | -2.116699195 | 0.019446635 |
| ENSMUSG00000033788 | Dysf | 0.000221453 | -1.314638573 | 0.302168577 |
| ENSMUSG00000057497 | Fam136a | 0.000221453 | 0.016567336 | 1 |
| ENSMUSG00000001156 | Mxd1 | 0.000221453 | 0.016567336 | 1 |
| ENSMUSG00000030056 | Isy1 | 0.000221453 | -1.258055044 | 0.355090064 |
| ENSMUSG00000030095 | Tmem43 | 0.000221453 | -0.614198854 | 0.825311622 |
| ENSMUSG00000034083 | Ccdc174 | 0.000221453 | -1.55279831 | 0.147260869 |
| ENSMUSG00000034037 | Fgd5 | 0.000221453 | -1.178983473 | 0.384884537 |
| ENSMUSG00000090667 | Gm765 | 0.000221453 | -0.810596067 | 0.792441218 |
| ENSMUSG00000107614 | Gm44196 | 0.000221453 | 0.44720169 | 1 |
| ENSMUSG00000005362 | Crbn | 0.000221453 | -0.455501108 | 1 |
| ENSMUSG00000057604 | Lmcd1 | 0.000221453 | -0.784123856 | 0.775504385 |
| ENSMUSG00000001632 | Brpf1 | 0.000221453 | -1.568395165 | 0.158915235 |
| ENSMUSG00000030272 | Camk1 | 0.000221453 | -0.158519371 | 1 |
| ENSMUSG00000042213 | Zfand4 | 0.000221453 | -0.583825205 | 0.921320733 |
| ENSMUSG00000099869 | 1700030F04Rik | 0.000221453 | 1.163408724 | 0.736456705 |
| ENSMUSG00000030134 | Rasgef1a | 0.000221453 | -0.314638573 | 1 |
| ENSMUSG00000041477 | Dcp1b | 0.000221453 | -1.568395165 | 0.150632966 |
| ENSMUSG00000030019 | Fbxl14 | 0.000221453 | 1.385801146 | 0.491653643 |

| | | | | |
|---|---|---|---|---|
| ENSMUSG00000030144 | Clec4d | 0.000221453 | 2.648835551 | 0.124712353 |
| ENSMUSG00000055172 | C1ra | 0.000221453 | 1.163408724 | 0.771834758 |
| ENSMUSG00000038346 | Zfp384 | 0.000221453 | -2.199161355 | 0.013392012 |
| ENSMUSG00000038279 | Nop2 | 0.000221453 | -1.219060913 | 0.330277691 |
| ENSMUSG00000038252 | Ncapd2 | 0.000221453 | -1.277163867 | 0.346101473 |
| ENSMUSG00000030336 | Cd27 | 0.000221453 | 1.385801146 | 0.508263132 |
| ENSMUSG00000001930 | Vwf | 0.000221453 | 0.163408724 | 1 |
| ENSMUSG00000030347 | D6Wsu163e | 0.000221453 | -0.386788358 | 1 |
| ENSMUSG00000097166 | 9330179D12Rik | 0.000221453 | 1.270323928 | 0.718266608 |
| ENSMUSG00000059659 | Gm10069 | 0.000221453 | 0.385801146 | 1 |
| ENSMUSG00000079852 | Klra4 | 0.000221453 | -0.983432664 | 0.530172915 |
| ENSMUSG00000094113 | 5430401F13Rik | 0.000221453 | 1.648835551 | 0.427459603 |
| ENSMUSG00000042992 | Borcs5 | 0.000221453 | -0.836591276 | 0.66416768 |
| ENSMUSG00000030204 | Ddx47 | 0.000221453 | -1.386788358 | 0.219470377 |
| ENSMUSG00000042638 | Gucy2c | 0.000221453 | 0.970763646 | 0.796840156 |
| ENSMUSG00000030225 | Dera | 0.000221453 | -1.537030994 | 0.144776283 |
| ENSMUSG00000089957 | A830011K09Rik | 0.000221453 | 2.163408724 | 0.226617702 |
| ENSMUSG00000040250 | Ints13 | 0.000221453 | -0.455501108 | 1 |
| ENSMUSG00000030303 | Far2 | 0.000221453 | 0.578446224 | 1 |
| ENSMUSG00000035596 | Mboat7 | 0.000221453 | -1.729676072 | 0.088605926 |
| ENSMUSG00000086155 | 9430041J12Rik | 0.000221453 | -0.614198854 | 0.825311622 |
| ENSMUSG00000035545 | Leng8 | 0.000221453 | -2.01792104 | 0.032072771 |
| ENSMUSG00000019254 | Ppp1r12c | 0.000221453 | -0.351164449 | 1 |
| ENSMUSG00000085272 | Sbk3 | 0.000221453 | 1.511332028 | 0.474258566 |
| ENSMUSG00000117863 | Vmn1r61 | 0.000221453 | 3.970763646 | 0.023955525 |
| ENSMUSG00000056696 | Olfr1350 | 0.000221453 | 3.385801146 | 0.051069295 |
| ENSMUSG00000047031 | Vmn1r68 | 0.000221453 | 2.385801146 | 0.171970178 |
| ENSMUSG00000030385 | 2900092C05Rik | 0.000221453 | 1.163408724 | 0.786364013 |
| ENSMUSG00000054310 | Obox1 | 0.000221453 | 1.063873051 | 0.75757939 |
| ENSMUSG00000023118 | Sympk | 0.000221453 | -1.899601073 | 0.044759969 |
| ENSMUSG00000030407 | Qpctl | 0.000221453 | 1.511332028 | 0.466464457 |
| ENSMUSG00000040734 | Ppp1r13l | 0.000221453 | -0.238689719 | 1 |
| ENSMUSG00000051403 | Ppp1r37 | 0.000221453 | -1.874726405 | 0.05188053 |
| ENSMUSG00000040498 | Igsf23 | 0.000221453 | 1.270323928 | 0.56359992 |
| ENSMUSG00000041037 | Irgq | 0.000221453 | -1.47217985 | 0.19917628 |
| ENSMUSG00000051768 | Xrcc1 | 0.000221453 | -0.199161355 | 1 |
| ENSMUSG00000003378 | Grik5 | 0.000221453 | -1.687447836 | 0.100676805 |
| ENSMUSG00000054499 | Dedd2 | 0.000221453 | -0.614198854 | 0.947833588 |
| ENSMUSG00000057229 | Dmac2 | 0.000221453 | 0.883300805 | 0.796840156 |
| ENSMUSG00000063160 | Numbl | 0.000221453 | -0.277163867 | 1 |
| ENSMUSG00000089832 | Shkbp1 | 0.000221453 | 0.44720169 | 1 |
| ENSMUSG00000008384 | Sertad1 | 0.000221453 | 0.385801146 | 1 |
| ENSMUSG00000063047 | Zfp780b | 0.000221453 | -0.455501108 | 1 |
| ENSMUSG00000096916 | Zfp850 | 0.000221453 | 1.970763646 | 0.281836872 |
| ENSMUSG00000030603 | Psmc4 | 0.000221453 | -0.757156808 | 0.750362092 |
| ENSMUSG00000002409 | Dyrk1b | 0.000221453 | 0.648835551 | 1 |
| ENSMUSG00000046185 | Zfp84 | 0.000221453 | -0.757156808 | 0.749778237 |
| ENSMUSG00000042831 | Alkbh6 | 0.000221453 | 1.800838645 | 0.344436715 |
| ENSMUSG00000030579 | Tyrobp | 0.000221453 | 1.800838645 | 0.356350819 |
| ENSMUSG00000006649 | Nphs1 | 0.000221453 | 1.800838645 | 0.382095022 |
| ENSMUSG00000036733 | Rbm42 | 0.000221453 | 0.112782651 | 1 |

| | | | | |
|---|---|---|---|---|
| ENSMUSG00000019194 | Scn1b | 0.000221453 | 0.722836133 | 1 |
| ENSMUSG00000085124 | Gm12766 | 0.000221453 | 1.163408724 | 0.733004386 |
| ENSMUSG00000108332 | D530033B14Rik | 0.000221453 | 1.970763646 | 0.281836872 |
| ENSMUSG00000064194 | Zfp936 | 0.000221453 | 1.063873051 | 0.75757939 |
| ENSMUSG00000045411 | 2410002F23Rik | 0.000221453 | -1.095325544 | 0.433313155 |
| ENSMUSG00000109491 | Gm15396 | 0.000221453 | 1.970763646 | 0.307184021 |
| ENSMUSG00000003420 | Fcgrt | 0.000221453 | 0.578446224 | 1 |
| ENSMUSG00000038260 | Trpm4 | 0.000221453 | -1.314638573 | 0.301813761 |
| ENSMUSG00000040435 | Ppp1r15a | 0.000221453 | 0.578446224 | 1 |
| ENSMUSG00000030835 | Nomo1 | 0.000221453 | -0.583825205 | 0.921320733 |
| ENSMUSG00000040136 | Abcc8 | 0.000221453 | 0.970763646 | 0.797728148 |
| ENSMUSG00000068151 | A230006K03Rik | 0.000221453 | 0.648835551 | 1 |
| ENSMUSG00000108986 | Gm32061 | 0.000221453 | 0.648835551 | 1 |
| ENSMUSG00000030525 | Chrna7 | 0.000221453 | 1.800838645 | 0.344436715 |
| ENSMUSG00000078681 | Tm2d3 | 0.000221453 | -0.029236354 | 1 |
| ENSMUSG00000030556 | Lrrc28 | 0.000221453 | -0.199161355 | 1 |
| ENSMUSG00000032776 | Mctp2 | 0.000221453 | -0.701661696 | 0.866565928 |
| ENSMUSG00000025789 | St8sia2 | 0.000221453 | 1.800838645 | 0.378412439 |
| ENSMUSG00000053025 | Sv2b | 0.000221453 | -0.701661696 | 0.864965452 |
| ENSMUSG00000097415 | AU020206 | 0.000221453 | -0.729676072 | 0.766882369 |
| ENSMUSG00000030611 | Mrps11 | 0.000221453 | 1.648835551 | 0.435901584 |
| ENSMUSG00000039187 | Fanci | 0.000221453 | 0.578446224 | 1 |
| ENSMUSG00000063801 | Ap3s2 | 0.000221453 | -1.258055044 | 0.340118707 |
| ENSMUSG00000025584 | Pde8a | 0.000221453 | -0.784123856 | 0.768296548 |
| ENSMUSG00000038623 | Tm6sf1 | 0.000221453 | 2.970763646 | 0.088786705 |
| ENSMUSG00000057706 | Mex3b | 0.000221453 | 2.385801146 | 0.171970178 |
| ENSMUSG00000091205 | Vmn2r71 | 0.000221453 | 1.385801146 | 0.51067985 |
| ENSMUSG00000109278 | Gm45144 | 0.000221453 | 0.970763646 | 0.792441218 |
| ENSMUSG00000030619 | Eed | 0.000221453 | -1.77070334 | 0.078628118 |
| ENSMUSG00000097644 | Gm26862 | 0.000221453 | 1.163408724 | 0.633928457 |
| ENSMUSG00000035582 | Gdpd4 | 0.000221453 | 1.063873051 | 0.75757939 |
| ENSMUSG00000030726 | Pold3 | 0.000221453 | -1.386788358 | 0.219470377 |
| ENSMUSG00000047767 | Atg16l2 | 0.000221453 | 1.163408724 | 0.733004386 |
| ENSMUSG00000073982 | Rhog | 0.000221453 | 0.385801146 | 1 |
| ENSMUSG00000052305 | Hbb-bs | 0.000221453 | -2.480447466 | 1 |
| ENSMUSG00000072244 | Trim6 | 0.000221453 | -0.238689719 | 1 |
| ENSMUSG00000036528 | Ppfibp2 | 0.000221453 | -1.455501108 | 0.216846462 |
| ENSMUSG00000005686 | Ampd3 | 0.000221453 | -0.238689719 | 1 |
| ENSMUSG00000058420 | Syt17 | 0.000221453 | -0.862126368 | 0.668575194 |
| ENSMUSG00000030650 | Tmc5 | 0.000221453 | -0.784123856 | 0.75757939 |
| ENSMUSG00000052273 | Dnah3 | 0.000221453 | -0.488667972 | 1 |
| ENSMUSG00000046096 | Mosmo | 0.000221453 | -1.277163867 | 0.394034904 |
| ENSMUSG00000030878 | Cdr2 | 0.000221453 | 0.270323928 | 1 |
| ENSMUSG00000066189 | Cacng3 | 0.000221453 | 0.883300805 | 0.815645145 |
| ENSMUSG00000030766 | Arhgap17 | 0.000221453 | -1.948099591 | 0.05882722 |
| ENSMUSG00000030763 | Lcmt1 | 0.000221453 | -0.784123856 | 0.75757939 |
| ENSMUSG00000030752 | Kdm8 | 0.000221453 | -0.810596067 | 0.66416768 |
| ENSMUSG00000030733 | Sh2b1 | 0.000221453 | -0.238689719 | 1 |
| ENSMUSG00000030720 | Cln3 | 0.000221453 | -1.178983473 | 0.384884537 |
| ENSMUSG00000030830 | Itgal | 0.000221453 | 0.578446224 | 1 |
| ENSMUSG00000108600 | Gm32884 | 0.000221453 | 0.578446224 | 1 |

| | | | | |
|---|---|---|---|---|
| ENSMUSG00000030850 | Ate1 | 0.000221453 | 0.722836133 | 1 |
| ENSMUSG00000040331 | Nsmce4a | 0.000221453 | 0.163408724 | 1 |
| ENSMUSG00000006204 | 5430419D17Rik | 0.000221453 | 0.722836133 | 1 |
| ENSMUSG00000041769 | Ppp2r2d | 0.000221453 | -0.757156808 | 0.800428907 |
| ENSMUSG00000025509 | Pnpla2 | 0.000221453 | 0.44720169 | 1 |
| ENSMUSG00000010751 | Tnfrsf22 | 0.000221453 | -0.158519371 | 1 |
| ENSMUSG00000037606 | Osbpl5 | 0.000221453 | -1.333017102 | 0.300304685 |
| ENSMUSG00000031072 | LTO1 | 0.000221453 | 0.722836133 | 1 |
| ENSMUSG00000040236 | Trappc5 | 0.000221453 | 0.326907457 | 1 |
| ENSMUSG00000065987 | Cd209b | 0.000221453 | 2.648835551 | 0.124712353 |
| ENSMUSG00000011832 | Evi5l | 0.000221453 | -0.959973691 | 0.526094357 |
| ENSMUSG00000008206 | Cers4 | 0.000221453 | 0.970763646 | 0.789767653 |
| ENSMUSG00000031508 | Ankrd10 | 0.000221453 | -1.643946198 | 0.114102131 |
| ENSMUSG00000038515 | Grtp1 | 0.000221453 | -0.116699195 | 1 |
| ENSMUSG00000037738 | Nek5 | 0.000221453 | 0.578446224 | 1 |
| ENSMUSG00000043794 | D830025C05Rik | 0.000221453 | 1.800838645 | 0.372408878 |
| ENSMUSG00000037296 | Lsm1 | 0.000221453 | -0.614198854 | 0.8327465 |
| ENSMUSG00000039620 | Trmt9b | 0.000221453 | 1.385801146 | 0.498919678 |
| ENSMUSG00000039478 | Micu3 | 0.000221453 | -1.051604167 | 0.466464457 |
| ENSMUSG00000039470 | Zdhhc2 | 0.000221453 | -0.52108945 | 0.912904617 |
| ENSMUSG00000039431 | Mtmr7 | 0.000221453 | -1.13776081 | 0.447196769 |
| ENSMUSG00000102046 | 2810404M03Rik | 0.000221453 | 1.385801146 | 0.503644124 |
| ENSMUSG00000038069 | Cdkn2aip | 0.000221453 | -0.810596067 | 0.66416768 |
| ENSMUSG00000031521 | Aga | 0.000221453 | 0.883300805 | 0.809009233 |
| ENSMUSG00000031520 | Vegfc | 0.000221453 | -0.936126949 | 0.611141945 |
| ENSMUSG00000031517 | Gpm6a | 0.000221453 | 0.270323928 | 1 |
| ENSMUSG00000031609 | Sap30 | 0.000221453 | 0.44720169 | 1 |
| ENSMUSG00000086681 | Gm16178 | 0.000221453 | 0.063873051 | 1 |
| ENSMUSG00000037921 | Ddx60 | 0.000221453 | -0.386788358 | 1 |
| ENSMUSG00000109994 | Gm45459 | 0.000221453 | -0.643946198 | 0.847265357 |
| ENSMUSG00000054648 | Zfp869 | 0.000221453 | -1.687447836 | 0.100676805 |
| ENSMUSG00000002342 | Tmem161a | 0.000221453 | -0.673092543 | 0.833666942 |
| ENSMUSG00000003575 | Crtc1 | 0.000221453 | -1.455501108 | 0.195601357 |
| ENSMUSG00000007888 | Crlf1 | 0.000221453 | 0.270323928 | 1 |
| ENSMUSG00000056204 | Pgpep1 | 0.000221453 | -0.029236354 | 1 |
| ENSMUSG00000002908 | Kcnn1 | 0.000221453 | 1.511332028 | 0.463009681 |
| ENSMUSG00000007950 | Abhd8 | 0.000221453 | 1.800838645 | 0.344436715 |
| ENSMUSG00000034863 | Ano8 | 0.000221453 | -1.13776081 | 0.4443026 |
| ENSMUSG00000070000 | Fcho1 | 0.000221453 | 1.648835551 | 0.405392586 |
| ENSMUSG00000089857 | Zfp882 | 0.000221453 | 1.648835551 | 0.405392586 |
| ENSMUSG00000052446 | Zfp961 | 0.000221453 | -0.158519371 | 1 |
| ENSMUSG00000062007 | Hsh2d | 0.000221453 | 1.163408724 | 0.733004386 |
| ENSMUSG00000031791 | Tmem38a | 0.000221453 | -0.116699195 | 1 |
| ENSMUSG00000037134 | Prmt9 | 0.000221453 | -1.887217349 | 0.055795486 |
| ENSMUSG00000031617 | Tmem184c | 0.000221453 | -0.116699195 | 1 |
| ENSMUSG00000097174 | Gm4890 | 0.000221453 | 2.970763646 | 0.086513796 |
| ENSMUSG00000038250 | Usp38 | 0.000221453 | -2.127268437 | 0.019633115 |
| ENSMUSG00000005470 | Asf1b | 0.000221453 | -0.583825205 | 0.921320733 |
| ENSMUSG00000079003 | Samd1 | 0.000221453 | -1.277163867 | 0.346101473 |
| ENSMUSG00000001909 | Trmt1 | 0.000221453 | -1.784123856 | 0.065287234 |
| ENSMUSG00000053226 | Dand5 | 0.000221453 | -0.614198854 | 0.825311622 |

| | | | | |
|---|---|---|---|---|
| ENSMUSG00000033751 | Gadd45gip1 | 0.000221453 | 1.163408724 | 0.755370971 |
| ENSMUSG00000008167 | Fbxw9 | 0.000221453 | -0.158519371 | 1 |
| ENSMUSG00000060038 | Dhps | 0.000221453 | 0.970763646 | 0.789767653 |
| ENSMUSG00000098257 | Gm27169 | 0.000221453 | 2.163408724 | 0.254575024 |
| ENSMUSG00000031736 | Crnde | 0.000221453 | 0.578446224 | 1 |
| ENSMUSG00000031782 | Coq9 | 0.000221453 | -1.051604167 | 0.47329858 |
| ENSMUSG00000031875 | Cmtm3 | 0.000221453 | 1.800838645 | 0.344436715 |
| ENSMUSG00000055730 | Ces2a | 0.000221453 | 3.385801146 | 0.048720096 |
| ENSMUSG00000061825 | Ces2c | 0.000221453 | 2.970763646 | 0.082902688 |
| ENSMUSG00000099465 | Gm3830 | 0.000221453 | 0.722836133 | 1 |
| ENSMUSG00000054320 | Lrrc36 | 0.000221453 | 1.163408724 | 0.733004386 |
| ENSMUSG00000104168 | Gm38250 | 0.000221453 | -0.073630473 | 1 |
| ENSMUSG00000031893 | Tsnaxip1 | 0.000221453 | 0.648835551 | 1 |
| ENSMUSG00000048827 | Pkd1l3 | 0.000221453 | 1.511332028 | 0.466464457 |
| ENSMUSG00000069895 | Atxn1l | 0.000221453 | 0.800838645 | 0.998731799 |
| ENSMUSG00000003657 | Calb2 | 0.000221453 | 2.970763646 | 0.082902688 |
| ENSMUSG00000031949 | Adat1 | 0.000221453 | -1.095325544 | 0.433313155 |
| ENSMUSG00000053399 | Adamts18 | 0.000221453 | -0.029236354 | 1 |
| ENSMUSG00000014633 | Cmc2 | 0.000221453 | -1.568395165 | 0.185619905 |
| ENSMUSG00000006517 | Mvd | 0.000221453 | 0.648835551 | 1 |
| ENSMUSG00000014470 | Rnf166 | 0.000221453 | 0.970763646 | 0.789767653 |
| ENSMUSG00000032815 | Fanca | 0.000221453 | 0.215876144 | 1 |
| ENSMUSG00000031979 | Cog2 | 0.000221453 | -0.421553776 | 1 |
| ENSMUSG00000031853 | Map3k21 | 0.000221453 | -2.168787706 | 0.01485218 |
| ENSMUSG00000051671 | Coa6 | 0.000221453 | 1.970763646 | 0.342560092 |
| ENSMUSG00000057367 | Birc2 | 0.000221453 | -0.983432664 | 0.530172915 |
| ENSMUSG00000031918 | Mtmr2 | 0.000221453 | -1.073630473 | 0.472298934 |
| ENSMUSG00000031922 | Cep57 | 0.000221453 | -0.701661696 | 0.740459298 |
| ENSMUSG00000031935 | Med17 | 0.000221453 | -1.715736881 | 0.100628726 |
| ENSMUSG00000043943 | Naalad2 | 0.000221453 | 0.800838645 | 0.84781195 |
| ENSMUSG00000059475 | Zfp426 | 0.000221453 | -0.073630473 | 1 |
| ENSMUSG00000066892 | Fbxl12 | 0.000221453 | -1.971750859 | 0.053242359 |
| ENSMUSG00000037405 | Icam1 | 0.000221453 | -1.614198854 | 0.13195254 |
| ENSMUSG00000001348 | Acp5 | 0.000221453 | 0.511332028 | 1 |
| ENSMUSG00000031969 | Acad8 | 0.000221453 | -0.116699195 | 1 |
| ENSMUSG00000035024 | Ncapd3 | 0.000221453 | -0.862126368 | 0.668575194 |
| ENSMUSG00000031993 | Snx19 | 0.000221453 | -0.757156808 | 0.749778237 |
| ENSMUSG00000111673 | Gm29724 | 0.000221453 | -0.983432664 | 0.572501893 |
| ENSMUSG00000032121 | Tmem218 | 0.000221453 | 0.722836133 | 1 |
| ENSMUSG00000032122 | Slc37a2 | 0.000221453 | 2.648835551 | 0.124712353 |
| ENSMUSG00000110747 | Gm35657 | 0.000221453 | 1.970763646 | 0.31693419 |
| ENSMUSG00000032018 | Sc5d | 0.000221453 | 0.215876144 | 1 |
| ENSMUSG00000095385 | D630033O11Rik | 0.000221453 | 1.648835551 | 0.405392586 |
| ENSMUSG00000032126 | Hmbs | 0.000221453 | 1.270323928 | 0.706154373 |
| ENSMUSG00000032114 | Slc37a4 | 0.000221453 | 0.44720169 | 1 |
| ENSMUSG00000032112 | Trappc4 | 0.000221453 | -0.784123856 | 0.766729124 |
| ENSMUSG00000066705 | Fxyd6 | 0.000221453 | 0.970763646 | 0.792441218 |
| ENSMUSG00000032085 | Tagln | 0.000221453 | -0.488667972 | 1 |
| ENSMUSG00000032077 | Bud13 | 0.000221453 | 0.063873051 | 1 |
| ENSMUSG00000032267 | Usp28 | 0.000221453 | -1.43862729 | 0.192076162 |
| ENSMUSG00000032064 | Dixdc1 | 0.000221453 | -0.701661696 | 0.740459298 |

| | | | | |
|---|---|---|---|---|
| ENSMUSG00000111020 | Gm1715 | 0.000221453 | 0.722836133 | 1 |
| ENSMUSG00000032030 | Cul5 | 0.000221453 | -2.051604167 | 0.023745467 |
| ENSMUSG00000049556 | Lingo1 | 0.000221453 | 0.578446224 | 1 |
| ENSMUSG00000063849 | Ppcdc | 0.000221453 | -0.277163867 | 1 |
| ENSMUSG00000032308 | Ulk3 | 0.000221453 | 0.722836133 | 1 |
| ENSMUSG00000038957 | Edc3 | 0.000221453 | -0.55279831 | 0.916104407 |
| ENSMUSG00000111761 | 9230112J17Rik | 0.000221453 | 1.063873051 | 0.75757939 |
| ENSMUSG00000032245 | Cln6 | 0.000221453 | 0.800838645 | 0.998731799 |
| ENSMUSG00000032400 | Zwilch | 0.000221453 | -0.959973691 | 0.526094357 |
| ENSMUSG00000111722 | Gm47270 | 0.000221453 | 1.970763646 | 0.295874115 |
| ENSMUSG00000032235 | Ice2 | 0.000221453 | -0.729676072 | 0.744267071 |
| ENSMUSG00000032224 | Fam81a | 0.000221453 | 0.511332028 | 1 |
| ENSMUSG00000032218 | Ccnb2 | 0.000221453 | 0.511332028 | 1 |
| ENSMUSG00000032192 | Gnb5 | 0.000221453 | -0.386788358 | 1 |
| ENSMUSG00000110839 | Gm47800 | 0.000221453 | 2.163408724 | 0.226617702 |
| ENSMUSG00000032343 | Impg1 | 0.000221453 | 0.970763646 | 0.789767653 |
| ENSMUSG00000066456 | Hmgn3 | 0.000221453 | 1.385801146 | 0.491653643 |
| ENSMUSG00000099724 | 4930554C24Rik | 0.000221453 | 2.385801146 | 0.204071921 |
| ENSMUSG00000032415 | Ube2cbp | 0.000221453 | -1.614198854 | 0.134289966 |
| ENSMUSG00000056131 | Pgm3 | 0.000221453 | -0.673092543 | 0.833666942 |
| ENSMUSG00000056919 | Cep162 | 0.000221453 | -2.576130814 | 0.002585743 |
| ENSMUSG00000032422 | Snx14 | 0.000221453 | -1.948099591 | 0.040663018 |
| ENSMUSG00000097177 | 9330159M07Rik | 0.000221453 | 0.215876144 | 1 |
| ENSMUSG00000066442 | Mthfs | 0.000221453 | -0.729676072 | 0.744267071 |
| ENSMUSG00000056267 | Cep70 | 0.000221453 | -0.959973691 | 0.526094357 |
| ENSMUSG00000043719 | Col6a6 | 0.000221453 | -0.073630473 | 1 |
| ENSMUSG00000042073 | Abhd14b | 0.000221453 | 0.883300805 | 0.796840156 |
| ENSMUSG00000006676 | Usp19 | 0.000221453 | -2.084518789 | 0.021377252 |
| ENSMUSG00000032604 | Qars | 0.000221453 | -0.836591276 | 0.66416768 |
| ENSMUSG00000006675 | P4htm | 0.000221453 | 1.270323928 | 0.71704651 |
| ENSMUSG00000032477 | Cdc25a | 0.000221453 | -1.386788358 | 0.219470377 |
| ENSMUSG00000032485 | Scap | 0.000221453 | -0.784123856 | 0.75757939 |
| ENSMUSG00000110876 | 4933411E02Rik | 0.000221453 | 2.648835551 | 0.124712353 |
| ENSMUSG00000032434 | Cmtm6 | 0.000221453 | 0.163408724 | 1 |
| ENSMUSG00000050627 | Gpd1l | 0.000221453 | -0.029236354 | 1 |
| ENSMUSG00000111627 | Gm48749 | 0.000221453 | 0.883300805 | 0.832285197 |
| ENSMUSG00000019810 | Fuca2 | 0.000221453 | 1.970763646 | 0.281836872 |
| ENSMUSG00000019808 | Adat2 | 0.000221453 | 2.385801146 | 0.171970178 |
| ENSMUSG00000071392 | Ect2l | 0.000221453 | 1.648835551 | 0.423814296 |
| ENSMUSG00000111200 | Gm33728 | 0.000221453 | 1.800838645 | 0.409864206 |
| ENSMUSG00000097327 | E030030I06Rik | 0.000221453 | -0.583825205 | 0.954649416 |
| ENSMUSG00000020000 | Moxd1 | 0.000221453 | -0.199161355 | 1 |
| ENSMUSG00000019883 | Echdc1 | 0.000221453 | -0.784123856 | 0.75757939 |
| ENSMUSG00000000295 | Hddc2 | 0.000221453 | -0.488667972 | 1 |
| ENSMUSG00000086428 | Gm15199 | 0.000221453 | 1.970763646 | 0.283819349 |
| ENSMUSG00000038151 | Prdm1 | 0.000221453 | 0.016567336 | 1 |
| ENSMUSG00000019848 | Popdc3 | 0.000221453 | 0.970763646 | 0.789767653 |
| ENSMUSG00000063804 | Lin28b | 0.000221453 | -0.757156808 | 0.792441218 |
| ENSMUSG00000019893 | Ros1 | 0.000221453 | -0.277163867 | 1 |
| ENSMUSG00000019917 | 10-Sep | 0.000221453 | -1.583825205 | 0.145505537 |
| ENSMUSG00000071253 | Slc25a16 | 0.000221453 | 0.016567336 | 1 |

| | | | | |
|---|---|---|---|---|
| ENSMUSG00000112314 | Gm49454 | 0.000221453 | 1.063873051 | 0.774790843 |
| ENSMUSG00000047692 | 4930533K18Rik | 0.000221453 | 2.163408724 | 0.261873336 |
| ENSMUSG00000112126 | Gm34544 | 0.000221453 | 0.578446224 | 1 |
| ENSMUSG00000085456 | Gm15398 | 0.000221453 | 1.063873051 | 0.792441218 |
| ENSMUSG00000000902 | Smarcb1 | 0.000221453 | -1.488667972 | 0.167024802 |
| ENSMUSG00000033255 | Gm5134 | 0.000221453 | 1.270323928 | 0.706154373 |
| ENSMUSG00000020226 | Slc5a4b | 0.000221453 | 1.270323928 | 0.564663656 |
| ENSMUSG00000020231 | Dip2a | 0.000221453 | -1.743481871 | 0.091285738 |
| ENSMUSG00000020262 | Adarb1 | 0.000221453 | -2.314638573 | 0.01450148 |
| ENSMUSG00000032788 | Pdxk | 0.000221453 | 0.215876144 | 1 |
| ENSMUSG00000109776 | Gm30400 | 0.000221453 | 0.063873051 | 1 |
| ENSMUSG00000020307 | Cdc34 | 0.000221453 | -1.116699195 | 0.437750651 |
| ENSMUSG00000013833 | Med16 | 0.000221453 | -0.936126949 | 0.617074698 |
| ENSMUSG00000013858 | Tmem259 | 0.000221453 | -1.82365222 | 0.057125834 |
| ENSMUSG00000035697 | Arhgap45 | 0.000221453 | -2.178983473 | 0.01534632 |
| ENSMUSG00000035504 | Reep6 | 0.000221453 | 1.970763646 | 0.281836872 |
| ENSMUSG00000015312 | Gadd45b | 0.000221453 | 1.511332028 | 0.463009681 |
| ENSMUSG00000034854 | Mfsd12 | 0.000221453 | 1.385801146 | 0.491653643 |
| ENSMUSG00000020235 | Fzr1 | 0.000221453 | -0.784123856 | 0.75757939 |
| ENSMUSG00000034560 | Washc4 | 0.000221453 | -1.658592974 | 0.116962967 |
| ENSMUSG00000111062 | Gm47253 | 0.000221453 | 1.970763646 | 0.443904862 |
| ENSMUSG00000020034 | Tcp11l2 | 0.000221453 | -0.073630473 | 1 |
| ENSMUSG00000034453 | Polr3b | 0.000221453 | -1.504969785 | 0.169370438 |
| ENSMUSG00000020057 | Dram1 | 0.000221453 | -0.614198854 | 0.953669611 |
| ENSMUSG00000035189 | Ano4 | 0.000221453 | -1.029236354 | 0.568923354 |
| ENSMUSG00000019948 | Actr6 | 0.000221453 | -0.277163867 | 1 |
| ENSMUSG00000112117 | Rmst | 0.000221453 | 0.578446224 | 1 |
| ENSMUSG00000020014 | Cfap54 | 0.000221453 | -0.421553776 | 1 |
| ENSMUSG00000112874 | Gm34297 | 0.000221453 | -0.887217349 | 0.725732014 |
| ENSMUSG00000090035 | Galnt4 | 0.000221453 | 1.163408724 | 0.733004386 |
| ENSMUSG00000044633 | B530045E10Rik | 0.000221453 | 0.270323928 | 1 |
| ENSMUSG00000035948 | Acss3 | 0.000221453 | -0.673092543 | 0.851986518 |
| ENSMUSG00000006764 | Tph2 | 0.000221453 | 1.163408724 | 0.751288778 |
| ENSMUSG00000052291 | 5330438D12Rik | 0.000221453 | 0.016567336 | 1 |
| ENSMUSG00000112230 | Ifngas1 | 0.000221453 | -0.029236354 | 1 |
| ENSMUSG00000020227 | Irak3 | 0.000221453 | 0.800838645 | 0.853723469 |
| ENSMUSG00000006732 | Mettl1 | 0.000221453 | -1.051604167 | 0.466464457 |
| ENSMUSG00000025417 | Pip4k2c | 0.000221453 | 0.44720169 | 1 |
| ENSMUSG00000090247 | Bloc1s1 | 0.000221453 | 1.511332028 | 0.466464457 |
| ENSMUSG00000034833 | Tespa1 | 0.000221453 | 0.800838645 | 0.998731799 |
| ENSMUSG00000034587 | 8430429K09Rik | 0.000221453 | 0.44720169 | 1 |
| ENSMUSG00000051427 | Ccdc157 | 0.000221453 | -0.862126368 | 0.668575194 |
| ENSMUSG00000016984 | Etaa1 | 0.000221453 | -0.386788358 | 1 |
| ENSMUSG00000042363 | Lgalsl | 0.000221453 | 1.648835551 | 0.405392586 |
| ENSMUSG00000044068 | Zrsr1 | 0.000221453 | 0.063873051 | 1 |
| ENSMUSG00000020273 | Papolg | 0.000221453 | -0.983432664 | 0.530172915 |
| ENSMUSG00000004018 | Fancl | 0.000221453 | 0.063873051 | 1 |
| ENSMUSG00000020311 | Erlec1 | 0.000221453 | -0.55279831 | 0.916104407 |
| ENSMUSG00000044502 | Bod1 | 0.000221453 | 0.063873051 | 1 |
| ENSMUSG00000040594 | Ranbp17 | 0.000221453 | -1.629149196 | 0.135560909 |
| ENSMUSG00000002699 | Lcp2 | 0.000221453 | 0.883300805 | 0.833666942 |

| | | | | |
|---|---|---|---|---|
| ENSMUSG00000069899 | Gm12166 | 0.000221453 | 2.970763646 | 0.082902688 |
| ENSMUSG00000040328 | Olfr56 | 0.000221453 | 1.063873051 | 0.786902265 |
| ENSMUSG00000001054 | Rmnd5b | 0.000221453 | -0.911879403 | 0.611716824 |
| ENSMUSG00000020386 | Sar1b | 0.000221453 | -1.701661696 | 0.103047269 |
| ENSMUSG00000020389 | Cdkl3 | 0.000221453 | -1.629149196 | 0.117043635 |
| ENSMUSG00000000782 | Tcf7 | 0.000221453 | -1.455501108 | 0.195601357 |
| ENSMUSG00000018398 | 8-Sep | 0.000221453 | -0.784123856 | 0.75757939 |
| ENSMUSG00000020380 | Rad50 | 0.000221453 | -2.333017102 | 0.007765102 |
| ENSMUSG00000020334 | Slc22a4 | 0.000221453 | -0.488667972 | 1 |
| ENSMUSG00000020388 | Pdlim4 | 0.000221453 | 0.326907457 | 1 |
| ENSMUSG00000086962 | Gm12248 | 0.000221453 | 0.648835551 | 1 |
| ENSMUSG00000037275 | Gemin5 | 0.000221453 | -1.404275785 | 0.261932394 |
| ENSMUSG00000032615 | Nt5m | 0.000221453 | -1.333017102 | 0.330109505 |
| ENSMUSG00000061650 | Med9 | 0.000221453 | 2.648835551 | 0.124712353 |
| ENSMUSG00000020537 | Drg2 | 0.000221453 | -0.277163867 | 1 |
| ENSMUSG00000002814 | Top3a | 0.000221453 | -1.219060913 | 0.330777885 |
| ENSMUSG00000043284 | Tmem11 | 0.000221453 | -0.158519371 | 1 |
| ENSMUSG00000047342 | Zfp286 | 0.000221453 | 0.215876144 | 1 |
| ENSMUSG00000070407 | Hs3st3b1 | 0.000221453 | 1.800838645 | 0.360664474 |
| ENSMUSG00000085162 | Gm12295 | 0.000221453 | 1.511332028 | 0.463954265 |
| ENSMUSG00000033352 | Map2k4 | 0.000221453 | -2.228908699 | 0.012088308 |
| ENSMUSG00000052921 | Arhgef15 | 0.000221453 | 1.270323928 | 0.706154373 |
| ENSMUSG00000020893 | Per1 | 0.000221453 | -0.673092543 | 0.833666942 |
| ENSMUSG00000001583 | Tnk1 | 0.000221453 | 0.163408724 | 1 |
| ENSMUSG00000020886 | Dlg4 | 0.000221453 | -2.687447836 | 0.002083251 |
| ENSMUSG00000018921 | Pelp1 | 0.000221453 | -1.995020638 | 0.030563094 |
| ENSMUSG00000020828 | Pld2 | 0.000221453 | -1.386788358 | 0.219470377 |
| ENSMUSG00000018449 | Rpain | 0.000221453 | 0.063873051 | 1 |
| ENSMUSG00000020798 | Spns3 | 0.000221453 | 1.511332028 | 0.463009681 |
| ENSMUSG00000038335 | Tsr1 | 0.000221453 | -1.52108945 | 0.172614487 |
| ENSMUSG00000038178 | Slc43a2 | 0.000221453 | -2.148177426 | 0.017095924 |
| ENSMUSG00000020847 | Rph3al | 0.000221453 | -1.238689719 | 0.333941363 |
| ENSMUSG00000020843 | Timm22 | 0.000221453 | 0.648835551 | 1 |
| ENSMUSG00000002059 | Rab34 | 0.000221453 | 0.578446224 | 1 |
| ENSMUSG00000001095 | Slc13a2 | 0.000221453 | 1.800838645 | 0.344436715 |
| ENSMUSG00000051232 | Tmem199 | 0.000221453 | 0.648835551 | 1 |
| ENSMUSG00000020826 | Nos2 | 0.000221453 | 1.511332028 | 0.466464457 |
| ENSMUSG00000035575 | Utp6 | 0.000221453 | -1.995020638 | 0.030563094 |
| ENSMUSG00000086507 | Adap2os | 0.000221453 | 1.511332028 | 0.463009681 |
| ENSMUSG00000057181 | 5730455P16Rik | 0.000221453 | -1.314638573 | 0.32758856 |
| ENSMUSG00000109297 | Gm31522 | 0.000221453 | 0.883300805 | 0.832285197 |
| ENSMUSG00000020702 | Ccl1 | 0.000221453 | 0.800838645 | 0.853723469 |
| ENSMUSG00000020698 | Cct6b | 0.000221453 | 0.800838645 | 0.849773563 |
| ENSMUSG00000020682 | Mmp28 | 0.000221453 | -0.729676072 | 0.744267071 |
| ENSMUSG00000000976 | Heatr6 | 0.000221453 | -1.158519371 | 0.3798128 |
| ENSMUSG00000020521 | Rnft1 | 0.000221453 | -0.810596067 | 0.66416768 |
| ENSMUSG00000061666 | Gdpd1 | 0.000221453 | -0.277163867 | 1 |
| ENSMUSG00000046442 | Ppm1e | 0.000221453 | 1.511332028 | 0.463009681 |
| ENSMUSG00000003948 | Mmd | 0.000221453 | -0.936126949 | 0.618991694 |
| ENSMUSG00000086191 | Zfp652os | 0.000221453 | 1.511332028 | 0.463009681 |
| ENSMUSG00000038811 | Gngt2 | 0.000221453 | 2.385801146 | 0.171970178 |

| | | | | |
|---|---|---|---|---|
| ENSMUSG00000018882 | Mrpl45 | 0.000221453 | -0.836591276 | 0.66416768 |
| ENSMUSG00000062312 | Erbb2 | 0.000221453 | -1.874726405 | 0.05188053 |
| ENSMUSG00000019312 | Grb7 | 0.000221453 | -0.158519371 | 1 |
| ENSMUSG00000017210 | Med24 | 0.000221453 | -1.13776081 | 0.4443026 |
| ENSMUSG00000017802 | Retreg3 | 0.000221453 | -0.614198854 | 0.838350879 |
| ENSMUSG00000010358 | Ifi35 | 0.000221453 | -1.599091962 | 0.128972787 |
| ENSMUSG00000034685 | Fam171a2 | 0.000221453 | -0.073630473 | 1 |
| ENSMUSG00000020925 | Ccdc43 | 0.000221453 | -1.158519371 | 0.466464457 |
| ENSMUSG00000020689 | Itgb3 | 0.000221453 | 0.385801146 | 1 |
| ENSMUSG00000069631 | Strada | 0.000221453 | -0.158519371 | 1 |
| ENSMUSG00000020617 | 1700012B07Rik | 0.000221453 | 1.511332028 | 0.466464457 |
| ENSMUSG00000041797 | Abca9 | 0.000221453 | -0.314638573 | 1 |
| ENSMUSG00000018800 | Abca5 | 0.000221453 | -1.887217349 | 0.05350651 |
| ENSMUSG00000020732 | Rab37 | 0.000221453 | 0.578446224 | 1 |
| ENSMUSG00000020773 | Trim47 | 0.000221453 | -0.158519371 | 1 |
| ENSMUSG00000020823 | Sec14l1 | 0.000221453 | -1.296022894 | 0.290284345 |
| ENSMUSG00000025576 | Rbfox3 | 0.000221453 | -0.614198854 | 0.94561855 |
| ENSMUSG00000039989 | Cbx4 | 0.000221453 | 0.163408724 | 1 |
| ENSMUSG00000039850 | Endov | 0.000221453 | -0.073630473 | 1 |
| ENSMUSG00000025384 | Faap100 | 0.000221453 | 0.648835551 | 1 |
| ENSMUSG00000025792 | Slc25a10 | 0.000221453 | -0.029236354 | 1 |
| ENSMUSG00000051510 | Mafg | 0.000221453 | -1.643946198 | 0.114102131 |
| ENSMUSG00000025170 | Rab40b | 0.000221453 | 0.722836133 | 1 |
| ENSMUSG00000020658 | Efr3b | 0.000221453 | -0.421553776 | 1 |
| ENSMUSG00000051721 | Wdcp | 0.000221453 | -0.614198854 | 0.825311622 |
| ENSMUSG00000112454 | Gm48619 | 0.000221453 | 1.648835551 | 0.405392586 |
| ENSMUSG00000112323 | Gm32828 | 0.000221453 | 0.215876144 | 1 |
| ENSMUSG00000112694 | Gm33037 | 0.000221453 | 0.578446224 | 1 |
| ENSMUSG00000113157 | Gm48693 | 0.000221453 | 0.883300805 | 0.796840156 |
| ENSMUSG00000020656 | Grhl1 | 0.000221453 | 0.44720169 | 1 |
| ENSMUSG00000020653 | Klf11 | 0.000221453 | -1.862126368 | 0.050837911 |
| ENSMUSG00000020646 | Mboat2 | 0.000221453 | -1.959973691 | 0.034141187 |
| ENSMUSG00000113669 | Gm36723 | 0.000221453 | 0.44720169 | 1 |
| ENSMUSG00000113550 | Gm29687 | 0.000221453 | 2.385801146 | 0.188834491 |
| ENSMUSG00000020642 | Rnf144a | 0.000221453 | 1.270323928 | 0.706154373 |
| ENSMUSG00000113387 | Gm31508 | 0.000221453 | 0.800838645 | 0.998731799 |
| ENSMUSG00000020664 | Dld | 0.000221453 | -0.729676072 | 0.747958595 |
| ENSMUSG00000112407 | Gm33340 | 0.000221453 | 1.270323928 | 0.706154373 |
| ENSMUSG00000112372 | Gm48779 | 0.000221453 | 0.016567336 | 1 |
| ENSMUSG00000111986 | 1700008C04Rik | 0.000221453 | 1.270323928 | 0.565001618 |
| ENSMUSG00000112437 | Gm35188 | 0.000221453 | 0.326907457 | 1 |
| ENSMUSG00000073079 | Srp54a | 0.000221453 | -0.887217349 | 0.600946203 |
| ENSMUSG00000021022 | Ppp2r3c | 0.000221453 | -1.277163867 | 0.346101473 |
| ENSMUSG00000046782 | Ttc6 | 0.000221453 | 0.511332028 | 1 |
| ENSMUSG00000060121 | Gemin2 | 0.000221453 | -1.006516277 | 0.537139526 |
| ENSMUSG00000020974 | Pole2 | 0.000221453 | -0.614198854 | 0.825311622 |
| ENSMUSG00000051890 | Klhdc1 | 0.000221453 | -1.810596067 | 0.097376298 |
| ENSMUSG00000113900 | Gm40437 | 0.000221453 | 0.511332028 | 1 |
| ENSMUSG00000113441 | Gm40438 | 0.000221453 | -0.314638573 | 1 |
| ENSMUSG00000114028 | 9630002D21Rik | 0.000221453 | 0.578446224 | 1 |
| ENSMUSG00000005078 | Jkamp | 0.000221453 | -1.455501108 | 0.195601357 |

| | | | | |
|---|---|---|---|---|
| ENSMUSG00000044712 | Slc38a6 | 0.000221453 | -2.47217985 | 0.005272109 |
| ENSMUSG00000021054 | Sgpp1 | 0.000221453 | 0.326907457 | 1 |
| ENSMUSG00000112784 | Gm34868 | 0.000221453 | 1.970763646 | 0.281836872 |
| ENSMUSG00000079055 | Slc8a3 | 0.000221453 | -0.238689719 | 1 |
| ENSMUSG00000002020 | Ltbp2 | 0.000221453 | -2.062659355 | 0.028030037 |
| ENSMUSG00000050671 | Ism2 | 0.000221453 | 1.648835551 | 0.405392586 |
| ENSMUSG00000097157 | Gm26512 | 0.000221453 | -1.77070334 | 0.098210219 |
| ENSMUSG00000021176 | Efcab11 | 0.000221453 | -0.158519371 | 1 |
| ENSMUSG00000021185 | Dglucy | 0.000221453 | -1.643946198 | 0.114102131 |
| ENSMUSG00000021209 | Ppp4r4 | 0.000221453 | 1.800838645 | 0.344436715 |
| ENSMUSG00000041341 | Atg2b | 0.000221453 | -2.18910769 | 0.015900288 |
| ENSMUSG00000044715 | Gskip | 0.000221453 | 0.163408724 | 1 |
| ENSMUSG00000054003 | Tdrd9 | 0.000221453 | 1.163408724 | 0.733004386 |
| ENSMUSG00000021294 | Kif26a | 0.000221453 | 2.385801146 | 0.171970178 |
| ENSMUSG00000002799 | Jag2 | 0.000221453 | -1.421553776 | 0.226813511 |
| ENSMUSG00000002803 | Btbd6 | 0.000221453 | 2.970763646 | 0.082902688 |
| ENSMUSG00000021175 | Cdca7l | 0.000221453 | -1.073630473 | 0.472298934 |
| ENSMUSG00000041886 | Macc1 | 0.000221453 | 0.44720169 | 1 |
| ENSMUSG00000048004 | Tmem196 | 0.000221453 | 0.800838645 | 0.998731799 |
| ENSMUSG00000113759 | Gm47695 | 0.000221453 | 1.800838645 | 0.344436715 |
| ENSMUSG00000021148 | Gm9745 | 0.000221453 | -2.404275785 | 0.008989808 |
| ENSMUSG00000021306 | Gpr137b | 0.000221453 | -0.643946198 | 0.828559607 |
| ENSMUSG00000053101 | Gpr141 | 0.000221453 | -1.583825205 | 0.19773933 |
| ENSMUSG00000060981 | Hist1h4h | 0.000221453 | 1.970763646 | 0.281836872 |
| ENSMUSG00000006717 | Acot13 | 0.000221453 | 0.648835551 | 1 |
| ENSMUSG00000035958 | Tdp2 | 0.000221453 | -1.488667972 | 0.167024802 |
| ENSMUSG00000021400 | Wrnip1 | 0.000221453 | -0.029236354 | 1 |
| ENSMUSG00000021408 | Ripk1 | 0.000221453 | -2.804023413 | 0.000795775 |
| ENSMUSG00000113457 | Gm47152 | 0.000221453 | 0.578446224 | 1 |
| ENSMUSG00000021428 | Riok1 | 0.000221453 | -2.636566667 | 0.001641759 |
| ENSMUSG00000051606 | 2010001K21Rik | 0.000221453 | -1.029236354 | 0.466464457 |
| ENSMUSG00000038068 | Rnf144b | 0.000221453 | 1.511332028 | 0.463009681 |
| ENSMUSG00000114959 | G630093K05Rik | 0.000221453 | 3.970763646 | 0.023955525 |
| ENSMUSG00000037851 | Iars | 0.000221453 | -2.583825205 | 0.002193744 |
| ENSMUSG00000021396 | Nxnl2 | 0.000221453 | 1.270323928 | 0.706154373 |
| ENSMUSG00000035139 | Secisbp2 | 0.000221453 | -2.106051951 | 0.022743266 |
| ENSMUSG00000021451 | Sema4d | 0.000221453 | -1.614198854 | 0.13195254 |
| ENSMUSG00000034918 | Cdhr2 | 0.000221453 | 0.578446224 | 1 |
| ENSMUSG00000025875 | Tspan17 | 0.000221453 | 1.163408724 | 0.733004386 |
| ENSMUSG00000021494 | Ddx41 | 0.000221453 | -1.351164449 | 0.256718203 |
| ENSMUSG00000114493 | Gm47071 | 0.000221453 | 0.511332028 | 1 |
| ENSMUSG00000050002 | Idnk | 0.000221453 | -0.52108945 | 0.912904617 |
| ENSMUSG00000035367 | Rmi1 | 0.000221453 | -0.983432664 | 0.530172915 |
| ENSMUSG00000021466 | Ptch1 | 0.000221453 | -2.219060913 | 0.022711572 |
| ENSMUSG00000033114 | Slc35d2 | 0.000221453 | -1.199161355 | 0.391578102 |
| ENSMUSG00000097360 | 9430065F17Rik | 0.000221453 | 1.063873051 | 0.75757939 |
| ENSMUSG00000021589 | Rhobtb3 | 0.000221453 | 0.016567336 | 1 |
| ENSMUSG00000043190 | Rfesd | 0.000221453 | 0.385801146 | 1 |
| ENSMUSG00000109228 | Fam81b | 0.000221453 | 0.511332028 | 1 |
| ENSMUSG00000035834 | Polr3g | 0.000221453 | -0.199161355 | 1 |
| ENSMUSG00000114461 | Gm48155 | 0.000221453 | 0.578446224 | 1 |

| | | | | |
|---|---|---|---|---|
| ENSMUSG00000041995 | Zbed3 | 0.000221453 | -0.073630473 | 1 |
| ENSMUSG00000069117 | Gm10260 | 0.000221453 | -0.351164449 | 1 |
| ENSMUSG00000114598 | D130062J10Rik | 0.000221453 | -0.757156808 | 0.749778237 |
| ENSMUSG00000050919 | Zfp366 | 0.000221453 | -1.488667972 | 0.256389682 |
| ENSMUSG00000021650 | Ptcd2 | 0.000221453 | 0.578446224 | 1 |
| ENSMUSG00000042655 | Shisal2b | 0.000221453 | 1.970763646 | 0.475407382 |
| ENSMUSG00000085634 | Gm15290 | 0.000221453 | 1.511332028 | 0.463009681 |
| ENSMUSG00000114615 | Gm48370 | 0.000221453 | 4.970763646 | 0.008075158 |
| ENSMUSG00000114794 | Gm47782 | 0.000221453 | 0.44720169 | 1 |
| ENSMUSG00000039599 | Fam149b | 0.000221453 | -1.629149196 | 0.112238591 |
| ENSMUSG00000021811 | Dnajc9 | 0.000221453 | 1.970763646 | 0.281836872 |
| ENSMUSG00000021824 | Ap3m1 | 0.000221453 | -0.52108945 | 0.912904617 |
| ENSMUSG00000114875 | Gm30363 | 0.000221453 | 0.112782651 | 1 |
| ENSMUSG00000021773 | Comtd1 | 0.000221453 | 1.270323928 | 0.706154373 |
| ENSMUSG00000087535 | Zmiz1os1 | 0.000221453 | -1.199161355 | 0.402770121 |
| ENSMUSG00000021879 | Dnah12 | 0.000221453 | 0.063873051 | 1 |
| ENSMUSG00000046753 | Ccdc66 | 0.000221453 | -1.296022894 | 0.290284345 |
| ENSMUSG00000052395 | Rft1 | 0.000221453 | 0.163408724 | 1 |
| ENSMUSG00000021918 | Nek4 | 0.000221453 | -1.757156808 | 0.076538465 |
| ENSMUSG00000021893 | Capn7 | 0.000221453 | -1.874726405 | 0.05188053 |
| ENSMUSG00000021906 | Oxnad1 | 0.000221453 | -1.421553776 | 0.226813511 |
| ENSMUSG00000050666 | Vstm4 | 0.000221453 | 0.385801146 | 1 |
| ENSMUSG00000051506 | Wdfy4 | 0.000221453 | -1.643946198 | 0.139351384 |
| ENSMUSG00000036242 | Armh4 | 0.000221453 | 1.163408724 | 0.733004386 |
| ENSMUSG00000071470 | Ccnb1ip1 | 0.000221453 | -1.983432664 | 0.089404923 |
| ENSMUSG00000021876 | Rnase4 | 0.000221453 | 1.063873051 | 0.75757939 |
| ENSMUSG00000057132 | Rpgrip1 | 0.000221453 | -0.351164449 | 1 |
| ENSMUSG00000040701 | Ap1g2 | 0.000221453 | -1.073630473 | 0.472298934 |
| ENSMUSG00000000365 | Rnf17 | 0.000221453 | 0.970763646 | 0.792441218 |
| ENSMUSG00000021959 | Lats2 | 0.000221453 | -1.715736881 | 0.086523965 |
| ENSMUSG00000115627 | Gm41168 | 0.000221453 | 0.270323928 | 1 |
| ENSMUSG00000035296 | Sgcg | 0.000221453 | 1.970763646 | 0.281836872 |
| ENSMUSG00000035184 | Fam124a | 0.000221453 | -0.277163867 | 1 |
| ENSMUSG00000021958 | Pinx1 | 0.000221453 | -1.643946198 | 0.114102131 |
| ENSMUSG00000022033 | Pbk | 0.000221453 | 2.648835551 | 0.207356329 |
| ENSMUSG00000045875 | Adra1a | 0.000221453 | 0.578446224 | 1 |
| ENSMUSG00000034194 | R3hcc1 | 0.000221453 | -1.351164449 | 0.256718203 |
| ENSMUSG00000022008 | Gpalpp1 | 0.000221453 | -0.351164449 | 1 |
| ENSMUSG00000022013 | Dnajc15 | 0.000221453 | -0.158519371 | 1 |
| ENSMUSG00000060377 | Rpl36a-ps1 | 0.000221453 | 2.970763646 | 0.082902688 |
| ENSMUSG00000022111 | Uchl3 | 0.000221453 | -1.455501108 | 0.195601357 |
| ENSMUSG00000022123 | Scel | 0.000221453 | -1.936126949 | 0.039541007 |
| ENSMUSG00000022119 | Rbm26 | 0.000221453 | -3.480447466 | 9.75E-06 |
| ENSMUSG00000022114 | Spry2 | 0.000221453 | -2.01792104 | 0.032072771 |
| ENSMUSG00000115754 | Gm48972 | 0.000221453 | -1.029236354 | 0.592845519 |
| ENSMUSG00000115146 | Gm4681 | 0.000221453 | 0.163408724 | 1 |
| ENSMUSG00000022184 | Fbxo4 | 0.000221453 | -0.029236354 | 1 |
| ENSMUSG00000041849 | Card6 | 0.000221453 | 0.270323928 | 1 |
| ENSMUSG00000022149 | C9 | 0.000221453 | 1.163408724 | 0.755314751 |
| ENSMUSG00000115480 | Gm49249 | 0.000221453 | -1.051604167 | 0.471773113 |
| ENSMUSG00000099349 | 1700047G03Rik | 0.000221453 | 3.970763646 | 0.023955525 |

| | | | | |
|---|---|---|---|---|
| ENSMUSG00000022321 | Cdh10 | 0.000221453 | 0.112782651 | 1 |
| ENSMUSG00000114971 | Gm49162 | 0.000221453 | 1.970763646 | 0.281836872 |
| ENSMUSG00000115261 | Gm35769 | 0.000221453 | 0.215876144 | 1 |
| ENSMUSG00000051237 | 9230109A22Rik | 0.000221453 | 0.883300805 | 0.821997796 |
| ENSMUSG00000115127 | Gm46515 | 0.000221453 | 1.063873051 | 0.790501196 |
| ENSMUSG00000022300 | Dcaf13 | 0.000221453 | -0.836591276 | 0.708502317 |
| ENSMUSG00000022304 | Dpys | 0.000221453 | 1.385801146 | 0.493584846 |
| ENSMUSG00000022339 | Ebag9 | 0.000221453 | -0.073630473 | 1 |
| ENSMUSG00000022366 | Slc22a22 | 0.000221453 | 0.800838645 | 0.851814318 |
| ENSMUSG00000086801 | Gm15943 | 0.000221453 | -0.029236354 | 1 |
| ENSMUSG00000050891 | Tatdn1 | 0.000221453 | -1.537030994 | 0.153855756 |
| ENSMUSG00000115681 | Gm2682 | 0.000221453 | -1.629149196 | 0.114102131 |
| ENSMUSG00000072568 | Lratd2 | 0.000221453 | -0.277163867 | 1 |
| ENSMUSG00000116000 | Gm49497 | 0.000221453 | -0.029236354 | 1 |
| ENSMUSG00000056258 | Kcnq3 | 0.000221453 | -0.614198854 | 0.945126684 |
| ENSMUSG00000068391 | Chrac1 | 0.000221453 | -1.314638573 | 0.324228447 |
| ENSMUSG00000022568 | Scrib | 0.000221453 | -0.887217349 | 0.600946203 |
| ENSMUSG00000034259 | Exosc4 | 0.000221453 | 1.270323928 | 0.706154373 |
| ENSMUSG00000022553 | Maf1 | 0.000221453 | 0.163408724 | 1 |
| ENSMUSG00000022546 | Gpt | 0.000221453 | -0.887217349 | 0.600946203 |
| ENSMUSG00000033669 | Zfp7 | 0.000221453 | -0.238689719 | 1 |
| ENSMUSG00000010601 | Apol7a | 0.000221453 | 0.800838645 | 0.886203497 |
| ENSMUSG00000019146 | Cacng2 | 0.000221453 | -0.116699195 | 1 |
| ENSMUSG00000016942 | Tmprss6 | 0.000221453 | 1.163408724 | 0.775421797 |
| ENSMUSG00000018008 | Cyth4 | 0.000221453 | 1.163408724 | 0.747902755 |
| ENSMUSG00000033099 | Nol12 | 0.000221453 | -0.314638573 | 1 |
| ENSMUSG00000010830 | Kdelr3 | 0.000221453 | -1.455501108 | 0.195601357 |
| ENSMUSG00000022407 | Adsl | 0.000221453 | -0.757156808 | 0.749778237 |
| ENSMUSG00000022471 | Xrcc6 | 0.000221453 | -1.421553776 | 0.226813511 |
| ENSMUSG00000058099 | Nfam1 | 0.000221453 | 0.215876144 | 1 |
| ENSMUSG00000018040 | Rrp7a | 0.000221453 | -0.314638573 | 1 |
| ENSMUSG00000022438 | Parvb | 0.000221453 | -2.47217985 | 0.006565796 |
| ENSMUSG00000036106 | Prr5 | 0.000221453 | 0.800838645 | 0.998731799 |
| ENSMUSG00000022382 | Wnt7b | 0.000221453 | -1.421553776 | 0.263599559 |
| ENSMUSG00000051786 | Tubgcp6 | 0.000221453 | -1.537030994 | 0.144776283 |
| ENSMUSG00000053137 | Mapk11 | 0.000221453 | -1.219060913 | 0.330277691 |
| ENSMUSG00000022614 | Lmf2 | 0.000221453 | 0.44720169 | 1 |
| ENSMUSG00000022620 | Arsa | 0.000221453 | 1.385801146 | 0.491653643 |
| ENSMUSG00000022623 | Shank3 | 0.000221453 | 0.44720169 | 1 |
| ENSMUSG00000022634 | Yaf2 | 0.000221453 | -2.158519371 | 0.017601978 |
| ENSMUSG00000023169 | Slc38a1 | 0.000221453 | -0.836591276 | 0.689152163 |
| ENSMUSG00000022464 | Slc38a4 | 0.000221453 | 1.800838645 | 0.381685516 |
| ENSMUSG00000022466 | Rpap3 | 0.000221453 | -1.333017102 | 0.300304685 |
| ENSMUSG00000052369 | Tmem106c | 0.000221453 | 0.215876144 | 1 |
| ENSMUSG00000037525 | Bcdin3d | 0.000221453 | 1.511332028 | 0.463009681 |
| ENSMUSG00000000532 | Acvr1b | 0.000221453 | -1.029236354 | 0.466464457 |
| ENSMUSG00000000531 | Grasp | 0.000221453 | -0.911879403 | 0.605887572 |
| ENSMUSG00000061397 | Krt79 | 0.000221453 | 2.385801146 | 0.171970178 |
| ENSMUSG00000022529 | Zfp263 | 0.000221453 | -1.701661696 | 0.103047269 |
| ENSMUSG00000004071 | Cdip1 | 0.000221453 | -1.715736881 | 0.086523965 |
| ENSMUSG00000057880 | Abat | 0.000221453 | -1.55279831 | 0.171611512 |

| | | | | |
|---|---|---|---|---|
| ENSMUSG00000022494 | Shisa9 | 0.000221453 | 0.112782651 | 1 |
| ENSMUSG00000087526 | Gm15738 | 0.000221453 | 0.722836133 | 1 |
| ENSMUSG00000022682 | Rrn3 | 0.000221453 | -1.836591276 | 0.058667604 |
| ENSMUSG00000044117 | Bmerb1 | 0.000221453 | 0.883300805 | 0.796840156 |
| ENSMUSG00000022678 | Nde1 | 0.000221453 | -0.488667972 | 1 |
| ENSMUSG00000022779 | Top3b | 0.000221453 | -0.757156808 | 0.749778237 |
| ENSMUSG00000022765 | Snap29 | 0.000221453 | 0.016567336 | 1 |
| ENSMUSG00000022763 | Aifm3 | 0.000221453 | -0.029236354 | 1 |
| ENSMUSG00000022761 | Lztr1 | 0.000221453 | -0.836591276 | 0.66416768 |
| ENSMUSG00000022758 | P2rx6 | 0.000221453 | 2.970763646 | 0.082902688 |
| ENSMUSG00000041378 | Cldn5 | 0.000221453 | -0.488667972 | 1 |
| ENSMUSG00000050821 | Fam131a | 0.000221453 | -0.55279831 | 0.931049705 |
| ENSMUSG00000022843 | Clcn2 | 0.000221453 | -0.386788358 | 1 |
| ENSMUSG00000022881 | Rfc4 | 0.000221453 | -0.351164449 | 1 |
| ENSMUSG00000022510 | Trp63 | 0.000221453 | 0.648835551 | 1 |
| ENSMUSG00000116660 | 9030404E10Rik | 0.000221453 | -0.277163867 | 1 |
| ENSMUSG00000049076 | Acap2 | 0.000221453 | -3.214111697 | 6.28E-05 |
| ENSMUSG00000022848 | Slc49a4 | 0.000221453 | -1.643946198 | 0.144267323 |
| ENSMUSG00000051980 | Casr | 0.000221453 | 0.215876144 | 1 |
| ENSMUSG00000022790 | Igsf11 | 0.000221453 | -0.936126949 | 0.611141945 |
| ENSMUSG00000047261 | Gap43 | 0.000221453 | 1.511332028 | 0.466464457 |
| ENSMUSG00000068284 | Usf3 | 0.000221453 | -2.333017102 | 0.00862911 |
| ENSMUSG00000022696 | Sidt1 | 0.000221453 | -0.199161355 | 1 |
| ENSMUSG00000022687 | Boc | 0.000221453 | 0.016567336 | 1 |
| ENSMUSG00000033031 | Cip2a | 0.000221453 | -0.073630473 | 1 |
| ENSMUSG00000064310 | Zpld1 | 0.000221453 | 1.163408724 | 0.733004386 |
| ENSMUSG00000022724 | Riox2 | 0.000221453 | -0.073630473 | 1 |
| ENSMUSG00000059920 | 4930453N24Rik | 0.000221453 | 0.016567336 | 1 |
| ENSMUSG00000116800 | Gm29823 | 0.000221453 | 2.648835551 | 0.159425617 |
| ENSMUSG00000109338 | Gm45030 | 0.000221453 | 0.883300805 | 0.826074164 |
| ENSMUSG00000109841 | E330011O21Rik | 0.000221453 | 0.326907457 | 1 |
| ENSMUSG00000047109 | Cldn14 | 0.000221453 | 0.270323928 | 1 |
| ENSMUSG00000022898 | Vps26c | 0.000221453 | -1.47217985 | 0.19917628 |
| ENSMUSG00000116720 | Gm31641 | 0.000221453 | 0.270323928 | 1 |
| ENSMUSG00000022913 | Psmg1 | 0.000221453 | -0.614198854 | 0.828331908 |
| ENSMUSG00000023147 | Wrb | 0.000221453 | -0.488667972 | 1 |
| ENSMUSG00000094724 | Rnaset2b | 0.000221453 | 2.648835551 | 0.16368308 |
| ENSMUSG00000090069 | E430024P14Rik | 0.000221453 | 0.326907457 | 1 |
| ENSMUSG00000023828 | Slc22a3 | 0.000221453 | 0.44720169 | 1 |
| ENSMUSG00000023886 | Smoc2 | 0.000221453 | 0.385801146 | 1 |
| ENSMUSG00000060149 | BC002059 | 0.000221453 | -1.13776081 | 0.466464457 |
| ENSMUSG00000116802 | Gm5165 | 0.000221453 | 0.511332028 | 1 |
| ENSMUSG00000097781 | 9330136K24Rik | 0.000221453 | -0.614198854 | 0.825311622 |
| ENSMUSG00000053347 | Zfp943 | 0.000221453 | -1.687447836 | 0.100676805 |
| ENSMUSG00000096433 | Zfp994 | 0.000221453 | -0.158519371 | 1 |
| ENSMUSG00000036473 | Tbc1d24 | 0.000221453 | -1.296022894 | 0.290284345 |
| ENSMUSG00000043445 | Pgp | 0.000221453 | -0.729676072 | 0.744267071 |
| ENSMUSG00000052752 | Traf7 | 0.000221453 | -0.52108945 | 0.912904617 |
| ENSMUSG00000024158 | Hagh | 0.000221453 | -1.199161355 | 0.452280643 |
| ENSMUSG00000045316 | Fahd1 | 0.000221453 | -0.199161355 | 1 |
| ENSMUSG00000046070 | Igfals | 0.000221453 | 1.163408724 | 0.737668143 |

| | | | | |
|---|---|---|---|---|
| ENSMUSG00000024169 | Ift140 | 0.000221453 | -1.504969785 | 0.231596497 |
| ENSMUSG00000024170 | Telo2 | 0.000221453 | 0.800838645 | 0.846566779 |
| ENSMUSG00000024186 | Rgs11 | 0.000221453 | 0.800838645 | 0.998731799 |
| ENSMUSG00000024190 | Dusp1 | 0.000221453 | 0.112782651 | 1 |
| ENSMUSG00000024191 | Bnip1 | 0.000221453 | -1.095325544 | 0.433313155 |
| ENSMUSG00000118147 | Gm41555 | 0.000221453 | 1.063873051 | 0.75757939 |
| ENSMUSG00000116657 | Gm49774 | 0.000221453 | -0.314638573 | 1 |
| ENSMUSG00000024215 | Spdef | 0.000221453 | -1.583825205 | 0.130094223 |
| ENSMUSG00000024218 | Taf11 | 0.000221453 | -0.784123856 | 0.75757939 |
| ENSMUSG00000117786 | Gm10503 | 0.000221453 | 2.385801146 | 0.211477132 |
| ENSMUSG00000041119 | Pde9a | 0.000221453 | -0.116699195 | 1 |
| ENSMUSG00000024042 | Sik1 | 0.000221453 | -0.673092543 | 0.833666942 |
| ENSMUSG00000058392 | Rrp1b | 0.000221453 | -1.488667972 | 0.167024802 |
| ENSMUSG00000117872 | A530088E08Rik | 0.000221453 | 0.722836133 | 1 |
| ENSMUSG00000091586 | Cyp4f17 | 0.000221453 | 0.326907457 | 1 |
| ENSMUSG00000024298 | Zfp871 | 0.000221453 | -2.029236354 | 0.027444373 |
| ENSMUSG00000095253 | Zfp799 | 0.000221453 | -1.52108945 | 0.172614487 |
| ENSMUSG00000003929 | Zfp81 | 0.000221453 | 0.112782651 | 1 |
| ENSMUSG00000002289 | Angptl4 | 0.000221453 | 0.883300805 | 0.796840156 |
| ENSMUSG00000002307 | Daxx | 0.000221453 | -0.386788358 | 1 |
| ENSMUSG00000007034 | Slc44a4 | 0.000221453 | -0.614198854 | 0.825311622 |
| ENSMUSG00000007039 | Ddah2 | 0.000221453 | 0.112782651 | 1 |
| ENSMUSG00000040312 | Cchcr1 | 0.000221453 | -2.029236354 | 0.027444373 |
| ENSMUSG00000024436 | Mrps18b | 0.000221453 | 0.270323928 | 1 |
| ENSMUSG00000024429 | Gnl1 | 0.000221453 | -1.673092543 | 0.098436701 |
| ENSMUSG00000076439 | Mog | 0.000221453 | 2.648835551 | 0.124712353 |
| ENSMUSG00000023913 | Pla2g7 | 0.000221453 | -0.55279831 | 1 |
| ENSMUSG00000067150 | Xpo5 | 0.000221453 | -1.911879403 | 0.052733229 |
| ENSMUSG00000023971 | Rrp36 | 0.000221453 | -0.029236354 | 1 |
| ENSMUSG00000040490 | Lrfn2 | 0.000221453 | 0.648835551 | 1 |
| ENSMUSG00000032915 | Adgre4 | 0.000221453 | 1.163408724 | 0.780759125 |
| ENSMUSG00000003198 | Zfp959 | 0.000221453 | 1.511332028 | 0.463009681 |
| ENSMUSG00000002835 | Chaf1a | 0.000221453 | -2.248404874 | 0.010622787 |
| ENSMUSG00000046329 | Slc25a23 | 0.000221453 | -1.051604167 | 0.46833724 |
| ENSMUSG00000062591 | Tubb4a | 0.000221453 | 0.215876144 | 1 |
| ENSMUSG00000117100 | A330072L02Rik | 0.000221453 | 0.883300805 | 0.800143934 |
| ENSMUSG00000024053 | Emilin2 | 0.000221453 | 0.44720169 | 1 |
| ENSMUSG00000085492 | Trmt61b | 0.000221453 | -1.421553776 | 0.226813511 |
| ENSMUSG00000054469 | Lclat1 | 0.000221453 | -0.701661696 | 0.740459298 |
| ENSMUSG00000024082 | Ndufaf7 | 0.000221453 | 0.578446224 | 1 |
| ENSMUSG00000024241 | Sos1 | 0.000221453 | -2.13776081 | 0.020391041 |
| ENSMUSG00000050945 | Zfp438 | 0.000221453 | -1.258055044 | 0.340118707 |
| ENSMUSG00000002475 | Abhd3 | 0.000221453 | 1.163408724 | 0.77665738 |
| ENSMUSG00000117633 | 8430422H06Rik | 0.000221453 | -1.629149196 | 0.173040376 |
| ENSMUSG00000024317 | Rnf138 | 0.000221453 | -1.849415316 | 0.060372549 |
| ENSMUSG00000024302 | Dtna | 0.000221453 | -1.537030994 | 0.144776283 |
| ENSMUSG00000024276 | Zfp397 | 0.000221453 | -1.849415316 | 0.060372549 |
| ENSMUSG00000117618 | 4930578E11Rik | 0.000221453 | 0.883300805 | 0.829166895 |
| ENSMUSG00000024259 | Slc25a46 | 0.000221453 | -1.199161355 | 0.391578102 |
| ENSMUSG00000024258 | Polr2d | 0.000221453 | 0.215876144 | 1 |
| ENSMUSG00000024386 | Proc | 0.000221453 | 0.163408724 | 1 |

| | | | | |
|---|---|---|---|---|
| ENSMUSG00000035984 | Nme5 | 0.000221453 | 1.063873051 | 0.75757939 |
| ENSMUSG00000003779 | Kif20a | 0.000221453 | 0.722836133 | 1 |
| ENSMUSG00000038418 | Egr1 | 0.000221453 | -0.029236354 | 1 |
| ENSMUSG00000073599 | Ecscr | 0.000221453 | 0.163408724 | 1 |
| ENSMUSG00000090264 | Eif4ebp3 | 0.000221453 | 1.063873051 | 0.665635898 |
| ENSMUSG00000042660 | Wdr55 | 0.000221453 | 0.112782651 | 1 |
| ENSMUSG00000097330 | Gm26672 | 0.000221453 | 0.648835551 | 1 |
| ENSMUSG00000118056 | Gm30093 | 0.000221453 | 0.44720169 | 1 |
| ENSMUSG00000056671 | Prelid2 | 0.000221453 | 0.112782651 | 1 |
| ENSMUSG00000043424 | Eif3j2 | 0.000221453 | 0.883300805 | 0.796840156 |
| ENSMUSG00000024502 | Jakmip2 | 0.000221453 | -0.52108945 | 1 |
| ENSMUSG00000032905 | Atg12 | 0.000221453 | 0.326907457 | 1 |
| ENSMUSG00000069378 | Prdm6 | 0.000221453 | 0.578446224 | 1 |
| ENSMUSG00000048799 | Cep120 | 0.000221453 | -2.01792104 | 0.032072771 |
| ENSMUSG00000024593 | Megf10 | 0.000221453 | 0.270323928 | 1 |
| ENSMUSG00000024604 | Rbm22 | 0.000221453 | -1.488667972 | 0.173426677 |
| ENSMUSG00000024579 | Pcyox1l | 0.000221453 | -1.238689719 | 0.333941363 |
| ENSMUSG00000024537 | Psmg2 | 0.000221453 | 0.215876144 | 1 |
| ENSMUSG00000038121 | Fam210a | 0.000221453 | -1.743481871 | 0.091285738 |
| ENSMUSG00000009535 | Rnmt | 0.000221453 | -1.874726405 | 0.05188053 |
| ENSMUSG00000079608 | Stard6 | 0.000221453 | 1.063873051 | 0.772125261 |
| ENSMUSG00000041840 | Haus1 | 0.000221453 | -0.386788358 | 1 |
| ENSMUSG00000033323 | Ctdp1 | 0.000221453 | -0.583825205 | 0.921320733 |
| ENSMUSG00000118352 | Gm50401 | 0.000221453 | 1.970763646 | 0.281836872 |
| ENSMUSG00000034028 | Cd226 | 0.000221453 | 1.385801146 | 0.511693804 |
| ENSMUSG00000024831 | Ighmbp2 | 0.000221453 | -1.743481871 | 0.091285738 |
| ENSMUSG00000033760 | Rbm4b | 0.000221453 | -0.614198854 | 0.825311622 |
| ENSMUSG00000047423 | AI837181 | 0.000221453 | 0.326907457 | 1 |
| ENSMUSG00000024912 | Fosl1 | 0.000221453 | -0.455501108 | 1 |
| ENSMUSG00000024906 | Mus81 | 0.000221453 | -0.351164449 | 1 |
| ENSMUSG00000056185 | Snx32 | 0.000221453 | -1.178983473 | 0.384884537 |
| ENSMUSG00000024807 | Syvn1 | 0.000221453 | -0.983432664 | 0.530172915 |
| ENSMUSG00000007338 | Mrpl49 | 0.000221453 | 0.163408724 | 1 |
| ENSMUSG00000024790 | Sac3d1 | 0.000221453 | 3.970763646 | 0.023955525 |
| ENSMUSG00000024777 | Ppp2r5b | 0.000221453 | -0.158519371 | 1 |
| ENSMUSG00000024772 | Ehd1 | 0.000221453 | -1.43862729 | 0.192076162 |
| ENSMUSG00000024962 | Vegfb | 0.000221453 | 0.385801146 | 1 |
| ENSMUSG00000024963 | Dnajc4 | 0.000221453 | 1.648835551 | 0.405392586 |
| ENSMUSG00000060675 | Plaat3 | 0.000221453 | 0.163408724 | 1 |
| ENSMUSG00000032773 | Chrm1 | 0.000221453 | 0.578446224 | 1 |
| ENSMUSG00000042729 | Wdr74 | 0.000221453 | -1.836591276 | 0.058667604 |
| ENSMUSG00000024743 | Syt7 | 0.000221453 | 0.163408724 | 1 |
| ENSMUSG00000024669 | Cd5 | 0.000221453 | 1.970763646 | 0.295355925 |
| ENSMUSG00000047044 | D030056L22Rik | 0.000221453 | -1.116699195 | 0.437750651 |
| ENSMUSG00000097930 | C330002G04Rik | 0.000221453 | 0.063873051 | 1 |
| ENSMUSG00000118020 | Gm35438 | 0.000221453 | 1.970763646 | 0.358457412 |
| ENSMUSG00000117752 | Gm35554 | 0.000221453 | 0.800838645 | 0.847265357 |
| ENSMUSG00000097855 | A930007I19Rik | 0.000221453 | 0.270323928 | 1 |
| ENSMUSG00000074909 | Ranbp6 | 0.000221453 | -0.455501108 | 1 |
| ENSMUSG00000024776 | Stambpl1 | 0.000221453 | -2.158519371 | 0.017601978 |
| ENSMUSG00000035783 | Acta2 | 0.000221453 | 0.112782651 | 1 |

| | | | | |
|---|---|---|---|---|
| ENSMUSG00000117865 | Gm50137 | 0.000221453 | 1.800838645 | 0.344436715 |
| ENSMUSG00000054237 | Fra10ac1 | 0.000221453 | -0.073630473 | 1 |
| ENSMUSG00000049164 | Zfp518a | 0.000221453 | -1.43862729 | 0.253473638 |
| ENSMUSG00000025017 | Pik3ap1 | 0.000221453 | -2.073630473 | 0.025562997 |
| ENSMUSG00000025176 | Hoga1 | 0.000221453 | -0.52108945 | 0.929973967 |
| ENSMUSG00000025178 | Pi4k2a | 0.000221453 | -1.369086357 | 0.261609949 |
| ENSMUSG00000042401 | Crtac1 | 0.000221453 | 0.270323928 | 1 |
| ENSMUSG00000025194 | Abcc2 | 0.000221453 | 0.44720169 | 1 |
| ENSMUSG00000025198 | Erlin1 | 0.000221453 | -0.983432664 | 0.530172915 |
| ENSMUSG00000004231 | Pax2 | 0.000221453 | -0.314638573 | 1 |
| ENSMUSG00000025069 | Gsto2 | 0.000221453 | 1.385801146 | 0.491653643 |
| ENSMUSG00000046585 | Cfap58 | 0.000221453 | 1.648835551 | 0.423990399 |
| ENSMUSG00000097446 | Gm26629 | 0.000221453 | 1.063873051 | 0.791874736 |
| ENSMUSG00000097636 | Mirt1 | 0.000221453 | -1.051604167 | 0.466464457 |
| ENSMUSG00000034765 | Dusp5 | 0.000221453 | -2.959973691 | 0.000279265 |
| ENSMUSG00000071497 | Nutf2-ps1 | 0.000221453 | 1.511332028 | 0.463009681 |
| ENSMUSG00000025075 | Habp2 | 0.000221453 | 1.063873051 | 0.774601541 |
| ENSMUSG00000025083 | Afap1l2 | 0.000221453 | -0.277163867 | 1 |
| ENSMUSG00000025086 | Trub1 | 0.000221453 | 0.270323928 | 1 |
| ENSMUSG00000048029 | Eno4 | 0.000221453 | 2.385801146 | 0.193580702 |
| ENSMUSG00000039231 | Suv39h1 | 0.000221453 | -2.127268437 | 0.027697231 |
| ENSMUSG00000031174 | Rpgr | 0.000221453 | -0.614198854 | 0.825311622 |
| ENSMUSG00000037341 | Slc9a7 | 0.000221453 | -0.810596067 | 0.68463297 |
| ENSMUSG00000037315 | Jade3 | 0.000221453 | -1.643946198 | 0.114102131 |
| ENSMUSG00000044149 | Nkrf | 0.000221453 | 0.722836133 | 1 |
| ENSMUSG00000036932 | Aifm1 | 0.000221453 | -1.404275785 | 0.2224613 |
| ENSMUSG00000023074 | Mospd1 | 0.000221453 | 1.063873051 | 0.776081332 |
| ENSMUSG00000031137 | Fgf13 | 0.000221453 | -1.369086357 | 0.277473977 |
| ENSMUSG00000031387 | Renbp | 0.000221453 | -1.178983473 | 0.405354468 |
| ENSMUSG00000009995 | Taz | 0.000221453 | -1.504969785 | 0.169370438 |
| ENSMUSG00000015289 | Lage3 | 0.000221453 | 0.722836133 | 1 |
| ENSMUSG00000087640 | Gm14798 | 0.000221453 | 0.722836133 | 1 |
| ENSMUSG00000045802 | Hsf3 | 0.000221453 | -0.314638573 | 1 |
| ENSMUSG00000042903 | Foxo4 | 0.000221453 | 0.970763646 | 0.789767653 |
| ENSMUSG00000033792 | Atp7a | 0.000221453 | -2.158519371 | 0.017601978 |
| ENSMUSG00000031239 | Itm2a | 0.000221453 | 1.385801146 | 0.512458849 |
| ENSMUSG00000063663 | Brwd3 | 0.000221453 | -1.911879403 | 0.045883682 |
| ENSMUSG00000025531 | Chm | 0.000221453 | -2.116699195 | 0.019039678 |
| ENSMUSG00000025592 | Dach2 | 0.000221453 | -0.729676072 | 0.775504385 |
| ENSMUSG00000031256 | Cstf2 | 0.000221453 | -0.757156808 | 0.749778237 |
| ENSMUSG00000052854 | Nrk | 0.000221453 | -1.158519371 | 0.47966612 |
| ENSMUSG00000031434 | Morc4 | 0.000221453 | -1.673092543 | 0.098436701 |
| ENSMUSG00000042282 | Gucy2f | 0.000221453 | 1.970763646 | 0.335256906 |
| ENSMUSG00000031283 | Chrdl1 | 0.000221453 | 2.163408724 | 0.226617702 |
| ENSMUSG00000041380 | Htr2c | 0.000221453 | -2.01792104 | 0.045774348 |
| ENSMUSG00000031290 | Lrch2 | 0.000221453 | 0.270323928 | 1 |
| ENSMUSG00000041353 | Tmem29 | 0.000221453 | -0.073630473 | 1 |
| ENSMUSG00000041115 | Iqsec2 | 0.000221453 | -1.701661696 | 0.103047269 |
| ENSMUSG00000025289 | Prdx4 | 0.000221453 | 1.270323928 | 0.706154373 |
| ENSMUSG00000071708 | Sms | 0.000221453 | -0.729676072 | 0.744267071 |
| ENSMUSG00000025658 | Cnksr2 | 0.000221453 | -2.006516277 | 0.031339065 |

| | | | | |
|---|---|---|---|---|
| ENSMUSG00000041020 | Map7d2 | 0.000221453 | 1.800838645 | 0.344436715 |
| ENSMUSG00000031295 | Phka2 | 0.000221453 | -1.073630473 | 0.472298934 |
| ENSMUSG00000040808 | S100g | 0.000221453 | 2.648835551 | 0.124712353 |
| ENSMUSG00000079800 | AC125149.3 | 0.000221453 | 0.326907457 | 1 |
| ENSMUSG00000025940 | Tmem70 | 0.00016609 | 0.741944956 | 0.970970089 |
| ENSMUSG00000042596 | Tfap2d | 0.00016609 | 0.948395833 | 0.912904617 |
| ENSMUSG00000025931 | Paqr8 | 0.00016609 | -1.636566667 | 0.144776283 |
| ENSMUSG00000026154 | Sdhaf4 | 0.00016609 | 0.47891055 | 1 |
| ENSMUSG00000026147 | Col9a1 | 0.00016609 | 1.648835551 | 0.536343234 |
| ENSMUSG00000042182 | Bend6 | 0.00016609 | 0.948395833 | 0.912904617 |
| ENSMUSG00000026127 | Imp4 | 0.00016609 | -0.43862729 | 1 |
| ENSMUSG00000037408 | Cnnm4 | 0.00016609 | -0.480447466 | 0.972350276 |
| ENSMUSG00000026121 | Sema4c | 0.00016609 | -1.305360759 | 0.371776814 |
| ENSMUSG00000003135 | Cnot11 | 0.00016609 | 0.256518129 | 1 |
| ENSMUSG00000060424 | Pantr1 | 0.00016609 | 0.400908038 | 1 |
| ENSMUSG00000026049 | Tex30 | 0.00016609 | 0.400908038 | 1 |
| ENSMUSG00000041684 | Bivm | 0.00016609 | -1.158519371 | 0.466464457 |
| ENSMUSG00000100120 | Gm553 | 0.00016609 | -0.43862729 | 1 |
| ENSMUSG00000026102 | Inpp1 | 0.00016609 | -1.417253639 | 0.269874327 |
| ENSMUSG00000026097 | Ormdl1 | 0.00016609 | -0.258055044 | 1 |
| ENSMUSG00000025983 | Ccdc150 | 0.00016609 | 0.400908038 | 1 |
| ENSMUSG00000097649 | Gm10561 | 0.00016609 | -0.051604167 | 1 |
| ENSMUSG00000048495 | Tyw5 | 0.00016609 | -1.023589791 | 0.526815951 |
| ENSMUSG00000079554 | Aox2 | 0.00016609 | -0.051604167 | 1 |
| ENSMUSG00000026036 | Nif3l1 | 0.00016609 | -1.636566667 | 0.217655393 |
| ENSMUSG00000038079 | Tmem237 | 0.00016609 | 0.47891055 | 1 |
| ENSMUSG00000101895 | Gm28981 | 0.00016609 | 0.741944956 | 0.972350276 |
| ENSMUSG00000051344 | Plekhm3 | 0.00016609 | -1.540989007 | 0.201837433 |
| ENSMUSG00000055567 | Unc80 | 0.00016609 | 0.56137271 | 1 |
| ENSMUSG00000026005 | Rpe | 0.00016609 | 0.189403933 | 1 |
| ENSMUSG00000086204 | Gm15668 | 0.00016609 | 2.648835551 | 0.218817775 |
| ENSMUSG00000101112 | Gm29185 | 0.00016609 | 1.063873051 | 0.886203497 |
| ENSMUSG00000026162 | Nhej1 | 0.00016609 | -1.373532262 | 0.317734263 |
| ENSMUSG00000026199 | Ankzf1 | 0.00016609 | -1.843017545 | 0.082214652 |
| ENSMUSG00000026201 | Stk16 | 0.00016609 | -0.258055044 | 1 |
| ENSMUSG00000026208 | Des | 0.00016609 | 2.063873051 | 0.413835764 |
| ENSMUSG00000033021 | Gmppa | 0.00016609 | -0.43862729 | 1 |
| ENSMUSG00000032908 | Sgpp2 | 0.00016609 | -2.500911568 | 0.006838229 |
| ENSMUSG00000099364 | 5730419F03Rik | 0.00016609 | -0.209145444 | 1 |
| ENSMUSG00000054976 | Nyap2 | 0.00016609 | -0.106051951 | 1 |
| ENSMUSG00000055980 | Irs1 | 0.00016609 | -2.132524162 | 0.031023691 |
| ENSMUSG00000026220 | Slc16a14 | 0.00016609 | 0.56137271 | 1 |
| ENSMUSG00000090252 | Gm16028 | 0.00016609 | 1.648835551 | 0.502227476 |
| ENSMUSG00000026237 | Nmur1 | 0.00016609 | 1.47891055 | 0.582159468 |
| ENSMUSG00000026240 | Cops7b | 0.00016609 | -1.579983139 | 0.185472119 |
| ENSMUSG00000026255 | Efhd1 | 0.00016609 | 1.063873051 | 0.904300125 |
| ENSMUSG00000056055 | Sag | 0.00016609 | -0.560617814 | 0.994229741 |
| ENSMUSG00000089943 | Ugt1a5 | 0.00016609 | -0.209145444 | 1 |
| ENSMUSG00000062310 | Glrp1 | 0.00016609 | 1.326907457 | 0.642722979 |
| ENSMUSG00000026307 | Scly | 0.00016609 | -1.184054463 | 0.419360593 |
| ENSMUSG00000034220 | Gpc1 | 0.00016609 | -0.52108945 | 0.97950322 |

| | | | | |
|---|---|---|---|---|
| ENSMUSG00000014602 | Kif1a | 0.00016609 | 0.47891055 | 1 |
| ENSMUSG00000047793 | Sned1 | 0.00016609 | -2.1966545 | 0.027444373 |
| ENSMUSG00000026333 | Gin1 | 0.00016609 | -1.233807498 | 0.430332878 |
| ENSMUSG00000100684 | Gm20268 | 0.00016609 | 3.063873051 | 0.177083054 |
| ENSMUSG00000099375 | Gm28187 | 0.00016609 | 1.648835551 | 0.499921258 |
| ENSMUSG00000038866 | Zcchc2 | 0.00016609 | -2.009375931 | 0.045701951 |
| ENSMUSG00000101253 | Gm29088 | 0.00016609 | 1.648835551 | 0.508434024 |
| ENSMUSG00000097648 | 9330185C12Rik | 0.00016609 | -1.43862729 | 0.313649744 |
| ENSMUSG00000036975 | Tmem177 | 0.00016609 | 2.326907457 | 0.295355925 |
| ENSMUSG00000026388 | 3110009E18Rik | 0.00016609 | 0.741944956 | 0.971854959 |
| ENSMUSG00000099957 | 2610027F03Rik | 0.00016609 | -0.708716453 | 0.775410248 |
| ENSMUSG00000097881 | Celrr | 0.00016609 | 0.189403933 | 1 |
| ENSMUSG00000085526 | Gm16083 | 0.00016609 | 1.063873051 | 0.886203497 |
| ENSMUSG00000042554 | Zp3r | 0.00016609 | 2.063873051 | 0.406526815 |
| ENSMUSG00000009772 | Nuak2 | 0.00016609 | 0.125273595 | 1 |
| ENSMUSG00000026442 | Nfasc | 0.00016609 | -0.708716453 | 0.886252908 |
| ENSMUSG00000115958 | Gm28040 | 0.00016609 | 1.063873051 | 0.886203497 |
| ENSMUSG00000070643 | Sox13 | 0.00016609 | -1.52108945 | 0.198212447 |
| ENSMUSG00000042793 | Lgr6 | 0.00016609 | -1.708716453 | 0.128972787 |
| ENSMUSG00000031506 | Ptpn7 | 0.00016609 | 0.125273595 | 1 |
| ENSMUSG00000026411 | Tmem9 | 0.00016609 | 0.948395833 | 0.792441218 |
| ENSMUSG00000041642 | Kif21b | 0.00016609 | -1.9057533 | 0.096053423 |
| ENSMUSG00000026404 | Ddx59 | 0.00016609 | -0.810596067 | 0.805794066 |
| ENSMUSG00000033964 | Zbtb41 | 0.00016609 | -1.395558568 | 0.26547175 |
| ENSMUSG00000033952 | Aspm | 0.00016609 | -1.579983139 | 0.169119671 |
| ENSMUSG00000057037 | Cfhr1 | 0.00016609 | 2.648835551 | 0.218817775 |
| ENSMUSG00000026480 | Ncf2 | 0.00016609 | -0.843017545 | 0.697198109 |
| ENSMUSG00000026700 | Tnfsf4 | 0.00016609 | -0.305360759 | 1 |
| ENSMUSG00000103393 | Gm38304 | 0.00016609 | -2.608552291 | 0.004410478 |
| ENSMUSG00000040170 | Fmo2 | 0.00016609 | 0.841480629 | 0.945922569 |
| ENSMUSG00000026585 | Kifap3 | 0.00016609 | -1.636566667 | 0.144776283 |
| ENSMUSG00000026584 | Scyl3 | 0.00016609 | -1.023589791 | 0.524144449 |
| ENSMUSG00000102897 | Gm37853 | 0.00016609 | 2.063873051 | 0.404717745 |
| ENSMUSG00000103659 | Gm32569 | 0.00016609 | 1.326907457 | 0.661891221 |
| ENSMUSG00000104117 | Gm20743 | 0.00016609 | 0.948395833 | 0.929961175 |
| ENSMUSG00000026567 | Adcy10 | 0.00016609 | 0.125273595 | 1 |
| ENSMUSG00000026687 | Aldh9a1 | 0.00016609 | -0.480447466 | 0.987137113 |
| ENSMUSG00000026686 | Lmx1a | 0.00016609 | 0.125273595 | 1 |
| ENSMUSG00000026656 | Fcgr2b | 0.00016609 | -0.258055044 | 1 |
| ENSMUSG00000005674 | Tomm40l | 0.00016609 | -0.965874293 | 0.611716824 |
| ENSMUSG00000037995 | Igsf9 | 0.00016609 | -1.540989007 | 0.201326391 |
| ENSMUSG00000026546 | Cfap45 | 0.00016609 | 0.841480629 | 0.943363727 |
| ENSMUSG00000073490 | Ifi207 | 0.00016609 | -0.158519371 | 1 |
| ENSMUSG00000039997 | Ifi203 | 0.00016609 | -0.810596067 | 0.791730654 |
| ENSMUSG00000026535 | Ifi202b | 0.00016609 | 0.741944956 | 0.970970089 |
| ENSMUSG00000104021 | Gm36536 | 0.00016609 | 1.326907457 | 0.642722979 |
| ENSMUSG00000097892 | Gm26801 | 0.00016609 | 0.948395833 | 0.938564865 |
| ENSMUSG00000038936 | Sccpdh | 0.00016609 | -0.051604167 | 1 |
| ENSMUSG00000038776 | Ephx1 | 0.00016609 | -0.258055044 | 1 |
| ENSMUSG00000038576 | Susd4 | 0.00016609 | -0.051604167 | 1 |
| ENSMUSG00000103305 | Gm34068 | 0.00016609 | 1.841480629 | 0.455131546 |

| | | | | |
|---|---|---|---|---|
| ENSMUSG00000039318 | Rab3gap2 | 0.00016609 | -2.834980226 | 0.001300227 |
| ENSMUSG00000103449 | Gm37134 | 0.00016609 | 2.063873051 | 0.383513703 |
| ENSMUSG00000001305 | Rrp15 | 0.00016609 | -0.936126949 | 0.606655093 |
| ENSMUSG00000104245 | Gm38155 | 0.00016609 | 1.326907457 | 0.653235135 |
| ENSMUSG00000037624 | Kcnk2 | 0.00016609 | 0.004979362 | 1 |
| ENSMUSG00000103510 | Gm30725 | 0.00016609 | 1.47891055 | 0.57597659 |
| ENSMUSG00000016181 | Utp25 | 0.00016609 | -0.9057533 | 0.687112677 |
| ENSMUSG00000026638 | Irf6 | 0.00016609 | -0.480447466 | 0.975624465 |
| ENSMUSG00000026645 | Olah | 0.00016609 | 1.189403933 | 0.874796537 |
| ENSMUSG00000025817 | Nudt5 | 0.00016609 | -1.051604167 | 0.529126167 |
| ENSMUSG00000045319 | Proser2 | 0.00016609 | -2.258055044 | 0.01835797 |
| ENSMUSG00000015619 | Gata3 | 0.00016609 | 0.56137271 | 1 |
| ENSMUSG00000086748 | Gm13261 | 0.00016609 | 1.47891055 | 0.576114773 |
| ENSMUSG00000023206 | Il15ra | 0.00016609 | 0.56137271 | 1 |
| ENSMUSG00000026723 | Trdmt1 | 0.00016609 | 1.47891055 | 0.600248432 |
| ENSMUSG00000084901 | Gm13266 | 0.00016609 | 2.063873051 | 0.395118486 |
| ENSMUSG00000026707 | Nsun6 | 0.00016609 | -0.673092543 | 0.872990655 |
| ENSMUSG00000026771 | Spopl | 0.00016609 | -2.209145444 | 0.024328557 |
| ENSMUSG00000087028 | Gm13387 | 0.00016609 | -1.89032326 | 0.07134433 |
| ENSMUSG00000048707 | Tprn | 0.00016609 | 0.648835551 | 1 |
| ENSMUSG00000026961 | Lrrc26 | 0.00016609 | 3.648835551 | 0.082076736 |
| ENSMUSG00000026944 | Abca2 | 0.00016609 | 0.125273595 | 1 |
| ENSMUSG00000036281 | Snapc4 | 0.00016609 | -0.743481871 | 0.778722129 |
| ENSMUSG00000026926 | Pmpca | 0.00016609 | -1.654945197 | 0.1477368 |
| ENSMUSG00000036160 | Surf6 | 0.00016609 | -1.281901786 | 0.386539267 |
| ENSMUSG00000091192 | Sardhos | 0.00016609 | 1.47891055 | 0.578548252 |
| ENSMUSG00000015846 | Rxra | 0.00016609 | -1.051604167 | 0.529126167 |
| ENSMUSG00000063611 | Gm10134 | 0.00016609 | 1.063873051 | 0.912067357 |
| ENSMUSG00000026821 | Ralgds | 0.00016609 | -1.654945197 | 0.16251927 |
| ENSMUSG00000079502 | Cfap77 | 0.00016609 | -0.599091962 | 1 |
| ENSMUSG00000086994 | Gm13402 | 0.00016609 | 0.648835551 | 1 |
| ENSMUSG00000084892 | Gm14471 | 0.00016609 | 0.47891055 | 1 |
| ENSMUSG00000086405 | 9330198N18Rik | 0.00016609 | 0.648835551 | 0.997917736 |
| ENSMUSG00000026841 | Fibcd1 | 0.00016609 | 3.648835551 | 0.082076736 |
| ENSMUSG00000026822 | Lcn2 | 0.00016609 | 0.125273595 | 1 |
| ENSMUSG00000026820 | Ptges2 | 0.00016609 | -0.599091962 | 0.865542683 |
| ENSMUSG00000026811 | St6galnac6 | 0.00016609 | 0.948395833 | 0.912904617 |
| ENSMUSG00000059013 | Sh2d3c | 0.00016609 | -0.708716453 | 0.915766329 |
| ENSMUSG00000026792 | Lrsam1 | 0.00016609 | -1.373532262 | 0.395118486 |
| ENSMUSG00000087461 | C230014O12Rik | 0.00016609 | 1.326907457 | 0.655049556 |
| ENSMUSG00000035778 | Ggta1 | 0.00016609 | -1.858959089 | 0.084357602 |
| ENSMUSG00000085201 | Nr6a1os | 0.00016609 | -2.579983139 | 0.005834122 |
| ENSMUSG00000050447 | Lypd6 | 0.00016609 | 0.741944956 | 0.970970089 |
| ENSMUSG00000026824 | Kcnj3 | 0.00016609 | 0.948395833 | 0.938505839 |
| ENSMUSG00000026832 | Cytip | 0.00016609 | 0.326907457 | 1 |
| ENSMUSG00000026988 | Wdsub1 | 0.00016609 | -2.065409966 | 0.041260008 |
| ENSMUSG00000026980 | Ly75 | 0.00016609 | -0.43862729 | 1 |
| ENSMUSG00000064289 | Tank | 0.00016609 | -1.023589791 | 0.524144449 |
| ENSMUSG00000075318 | Scn2a | 0.00016609 | -0.258055044 | 1 |
| ENSMUSG00000034810 | Scn7a | 0.00016609 | 0.326907457 | 1 |
| ENSMUSG00000005233 | Spc25 | 0.00016609 | 0.841480629 | 0.943363727 |

| | | | | |
|---|---|---|---|---|
| ENSMUSG00000042155 | Klhl23 | 0.00016609 | 3.063873051 | 0.144078128 |
| ENSMUSG00000075302 | Erich2 | 0.00016609 | -0.965874293 | 0.611716824 |
| ENSMUSG00000070880 | Gad1 | 0.00016609 | 1.47891055 | 0.587484274 |
| ENSMUSG00000027018 | Hat1 | 0.00016609 | -1.52108945 | 0.198212447 |
| ENSMUSG00000056486 | Chn1 | 0.00016609 | -1.636566667 | 0.144776283 |
| ENSMUSG00000009207 | Lnpk | 0.00016609 | -0.673092543 | 0.872990655 |
| ENSMUSG00000027099 | Mtx2 | 0.00016609 | -0.777429203 | 0.792441218 |
| ENSMUSG00000042369 | Rbm45 | 0.00016609 | -0.843017545 | 0.674297981 |
| ENSMUSG00000027014 | Cwc22 | 0.00016609 | -1.43862729 | 0.27459352 |
| ENSMUSG00000054510 | Gm14461 | 0.00016609 | 2.326907457 | 0.295355925 |
| ENSMUSG00000027009 | Itga4 | 0.00016609 | -0.599091962 | 1 |
| ENSMUSG00000027076 | Timm10 | 0.00016609 | 1.841480629 | 0.479831536 |
| ENSMUSG00000002103 | Acp2 | 0.00016609 | -2.490715801 | 0.006613366 |
| ENSMUSG00000075040 | Zfp408 | 0.00016609 | 0.256518129 | 1 |
| ENSMUSG00000027222 | Pex16 | 0.00016609 | 1.47891055 | 0.57597659 |
| ENSMUSG00000049922 | Slc35c1 | 0.00016609 | 0.326907457 | 1 |
| ENSMUSG00000027221 | Chst1 | 0.00016609 | 1.47891055 | 0.57597659 |
| ENSMUSG00000027220 | Syt13 | 0.00016609 | -0.599091962 | 0.871850653 |
| ENSMUSG00000075028 | Prdm11 | 0.00016609 | -1.051604167 | 0.529126167 |
| ENSMUSG00000027165 | B230118H07Rik | 0.00016609 | -0.209145444 | 1 |
| ENSMUSG00000074987 | Wt1os | 0.00016609 | -0.810596067 | 0.791730654 |
| ENSMUSG00000055312 | Them7 | 0.00016609 | -0.051604167 | 1 |
| ENSMUSG00000005973 | Rcn1 | 0.00016609 | 0.648835551 | 1 |
| ENSMUSG00000027131 | Emc4 | 0.00016609 | 0.256518129 | 1 |
| ENSMUSG00000023236 | Scg5 | 0.00016609 | -0.106051951 | 1 |
| ENSMUSG00000027355 | Tmco5 | 0.00016609 | 2.648835551 | 0.218817775 |
| ENSMUSG00000027344 | Fsip1 | 0.00016609 | 0.063873051 | 1 |
| ENSMUSG00000040133 | Gpr176 | 0.00016609 | -0.106051951 | 1 |
| ENSMUSG00000074923 | Pak6 | 0.00016609 | -0.351164449 | 1 |
| ENSMUSG00000027332 | Ivd | 0.00016609 | -0.777429203 | 0.785942057 |
| ENSMUSG00000033902 | Mapkbp1 | 0.00016609 | 0.189403933 | 1 |
| ENSMUSG00000027263 | Tubgcp4 | 0.00016609 | -0.480447466 | 0.972350276 |
| ENSMUSG00000044145 | 1810024B03Rik | 0.00016609 | -0.936126949 | 0.606655093 |
| ENSMUSG00000027397 | Slc20a1 | 0.00016609 | -2.106051951 | 0.036215483 |
| ENSMUSG00000027399 | Il1a | 0.00016609 | 1.063873051 | 0.919428997 |
| ENSMUSG00000074797 | Itpa | 0.00016609 | -0.258055044 | 1 |
| ENSMUSG00000079037 | Prnp | 0.00016609 | 0.56137271 | 1 |
| ENSMUSG00000050558 | Prokr2 | 0.00016609 | 1.326907457 | 0.642722979 |
| ENSMUSG00000086379 | 1700026D11Rik | 0.00016609 | 1.47891055 | 0.57597659 |
| ENSMUSG00000037376 | Trmt6 | 0.00016609 | -1.158519371 | 0.466464457 |
| ENSMUSG00000051379 | Flrt3 | 0.00016609 | 0.256518129 | 1 |
| ENSMUSG00000008333 | Snrpb2 | 0.00016609 | -0.43862729 | 1 |
| ENSMUSG00000027427 | Polr3f | 0.00016609 | -2.351164449 | 0.015579272 |
| ENSMUSG00000086509 | Nkx2-2os | 0.00016609 | 0.189403933 | 1 |
| ENSMUSG00000027435 | Cd93 | 0.00016609 | -0.305360759 | 1 |
| ENSMUSG00000027439 | Gzf1 | 0.00016609 | -1.281901786 | 0.397083363 |
| ENSMUSG00000095315 | Gm10130 | 0.00016609 | 0.063873051 | 1 |
| ENSMUSG00000003283 | Hck | 0.00016609 | -0.673092543 | 0.891817216 |
| ENSMUSG00000037761 | Actr5 | 0.00016609 | -0.395558568 | 1 |
| ENSMUSG00000085198 | 4933416E03Rik | 0.00016609 | 0.47891055 | 1 |
| ENSMUSG00000086653 | 9130015L21Rik | 0.00016609 | 1.648835551 | 0.53393869 |

| | | | | |
|---|---|---|---|---|
| ENSMUSG00000017861 | Mybl2 | 0.00016609 | -0.636566667 | 0.868661374 |
| ENSMUSG00000017679 | Ttpal | 0.00016609 | -0.106051951 | 1 |
| ENSMUSG00000039873 | Neurl2 | 0.00016609 | 0.56137271 | 1 |
| ENSMUSG00000039849 | Pcif1 | 0.00016609 | 0.125273595 | 1 |
| ENSMUSG00000039834 | Zfp335 | 0.00016609 | 0.063873051 | 1 |
| ENSMUSG00000017737 | Mmp9 | 0.00016609 | 1.326907457 | 0.643065264 |
| ENSMUSG00000017969 | Ptgis | 0.00016609 | 0.47891055 | 1 |
| ENSMUSG00000052056 | Zfp217 | 0.00016609 | -2.171343411 | 0.027621844 |
| ENSMUSG00000085905 | Gm14273 | 0.00016609 | 0.56137271 | 1 |
| ENSMUSG00000008999 | Bmp7 | 0.00016609 | 1.841480629 | 0.455131546 |
| ENSMUSG00000078887 | Gm6710 | 0.00016609 | 3.648835551 | 0.127356309 |
| ENSMUSG00000095199 | Zfp967 | 0.00016609 | 0.189403933 | 1 |
| ENSMUSG00000078862 | Gm14326 | 0.00016609 | 0.47891055 | 1 |
| ENSMUSG00000084843 | B230312C02Rik | 0.00016609 | 2.063873051 | 0.472764596 |
| ENSMUSG00000027569 | Mrgbp | 0.00016609 | -0.106051951 | 1 |
| ENSMUSG00000000823 | Zfp512b | 0.00016609 | -1.305360759 | 0.371776814 |
| ENSMUSG00000038605 | Samd10 | 0.00016609 | 0.56137271 | 1 |
| ENSMUSG00000027500 | Stmn2 | 0.00016609 | 1.063873051 | 0.901934379 |
| ENSMUSG00000039795 | Zfand1 | 0.00016609 | -1.079084903 | 0.552193432 |
| ENSMUSG00000027552 | E2f5 | 0.00016609 | -0.743481871 | 0.787908961 |
| ENSMUSG00000011463 | Cpb1 | 0.00016609 | 0.948395833 | 0.96070793 |
| ENSMUSG00000105662 | Gm6639 | 0.00016609 | 1.648835551 | 0.499921258 |
| ENSMUSG00000085655 | Gm15952 | 0.00016609 | -1.560617814 | 0.250264692 |
| ENSMUSG00000086392 | Mccc1os | 0.00016609 | 0.948395833 | 0.926077759 |
| ENSMUSG00000047696 | Ccdc144b | 0.00016609 | 1.648835551 | 0.501319486 |
| ENSMUSG00000027715 | Ccna2 | 0.00016609 | 0.841480629 | 0.943363727 |
| ENSMUSG00000106461 | Gm20755 | 0.00016609 | -0.52108945 | 0.992008082 |
| ENSMUSG00000060798 | Intu | 0.00016609 | -0.43862729 | 1 |
| ENSMUSG00000097040 | 2610316D01Rik | 0.00016609 | 0.004979362 | 1 |
| ENSMUSG00000027744 | Stoml3 | 0.00016609 | 0.741944956 | 0.986163943 |
| ENSMUSG00000037016 | Frem2 | 0.00016609 | 0.841480629 | 0.967730476 |
| ENSMUSG00000027752 | Exosc8 | 0.00016609 | 0.125273595 | 1 |
| ENSMUSG00000036615 | Rfxap | 0.00016609 | -0.106051951 | 1 |
| ENSMUSG00000027805 | Pfn2 | 0.00016609 | 1.47891055 | 0.57597659 |
| ENSMUSG00000048581 | E130311K13Rik | 0.00016609 | 0.189403933 | 1 |
| ENSMUSG00000092049 | Vmn2r4 | 0.00016609 | 3.063873051 | 0.144078128 |
| ENSMUSG00000074579 | Lekr1 | 0.00016609 | -1.158519371 | 0.466464457 |
| ENSMUSG00000034151 | Zbbx | 0.00016609 | 1.189403933 | 0.691044471 |
| ENSMUSG00000104448 | Gm37256 | 0.00016609 | 0.741944956 | 0.971854959 |
| ENSMUSG00000033900 | Map9 | 0.00016609 | 0.648835551 | 1 |
| ENSMUSG00000086273 | Rbm46os | 0.00016609 | 2.063873051 | 0.37936848 |
| ENSMUSG00000104004 | 4930565D16Rik | 0.00016609 | 1.326907457 | 0.642722979 |
| ENSMUSG00000015852 | Fcrls | 0.00016609 | 3.063873051 | 0.144078128 |
| ENSMUSG00000048031 | Fcrl5 | 0.00016609 | 0.948395833 | 0.928159492 |
| ENSMUSG00000004891 | Nes | 0.00016609 | -2.184054463 | 0.030219786 |
| ENSMUSG00000028070 | Naxe | 0.00016609 | 0.400908038 | 1 |
| ENSMUSG00000074480 | Mex3a | 0.00016609 | -0.106051951 | 1 |
| ENSMUSG00000068922 | Msto1 | 0.00016609 | 0.189403933 | 1 |
| ENSMUSG00000041237 | Pklr | 0.00016609 | 0.948395833 | 0.912904617 |
| ENSMUSG00000027953 | Slc50a1 | 0.00016609 | -0.480447466 | 0.972350276 |
| ENSMUSG00000074466 | Gm15417 | 0.00016609 | 0.004979362 | 1 |

| | | | | |
|---|---|---|---|---|
| ENSMUSG00000027936 | Crtc2 | 0.00016609 | -0.351164449 | 1 |
| ENSMUSG00000042244 | Pglyrp3 | 0.00016609 | 4.648835551 | 0.031809473 |
| ENSMUSG00000005774 | Rfx5 | 0.00016609 | -0.351164449 | 1 |
| ENSMUSG00000015702 | Anxa9 | 0.00016609 | 0.841480629 | 0.943363727 |
| ENSMUSG00000045934 | Mtmr11 | 0.00016609 | -0.305360759 | 1 |
| ENSMUSG00000038354 | Ankrd35 | 0.00016609 | -0.051604167 | 1 |
| ENSMUSG00000028093 | Acp6 | 0.00016609 | -0.743481871 | 0.778722129 |
| ENSMUSG00000033285 | Wdr3 | 0.00016609 | -2.1966545 | 0.023519181 |
| ENSMUSG00000048938 | Nr1h5 | 0.00016609 | 1.47891055 | 0.57597659 |
| ENSMUSG00000044165 | Bcl2l15 | 0.00016609 | 1.326907457 | 0.651476428 |
| ENSMUSG00000027843 | Ptpn22 | 0.00016609 | -1.654945197 | 0.150607708 |
| ENSMUSG00000058388 | Phtf1 | 0.00016609 | -1.417253639 | 0.269874327 |
| ENSMUSG00000027901 | Dennd2d | 0.00016609 | -2.305360759 | 0.016521187 |
| ENSMUSG00000098108 | Gm27008 | 0.00016609 | 0.47891055 | 1 |
| ENSMUSG00000014601 | Strip1 | 0.00016609 | -0.351164449 | 1 |
| ENSMUSG00000068747 | Sort1 | 0.00016609 | -1.373532262 | 0.332003096 |
| ENSMUSG00000048100 | Taf13 | 0.00016609 | -0.051604167 | 1 |
| ENSMUSG00000040389 | Wdr47 | 0.00016609 | -0.743481871 | 0.788332288 |
| ENSMUSG00000049565 | Aknad1 | 0.00016609 | 1.063873051 | 0.886203497 |
| ENSMUSG00000027981 | Rnpc3 | 0.00016609 | -2.145580315 | 0.031872898 |
| ENSMUSG00000027966 | Col11a1 | 0.00016609 | -1.951077291 | 0.077471852 |
| ENSMUSG00000044667 | Plppr4 | 0.00016609 | -0.708716453 | 0.775410248 |
| ENSMUSG00000028125 | Abca4 | 0.00016609 | -0.743481871 | 0.796920709 |
| ENSMUSG00000027973 | 1700006A11Rik | 0.00016609 | 2.326907457 | 0.33333983 |
| ENSMUSG00000046561 | Arsj | 0.00016609 | 1.063873051 | 0.886203497 |
| ENSMUSG00000028024 | Enpep | 0.00016609 | 1.189403933 | 0.697198109 |
| ENSMUSG00000028017 | Egf | 0.00016609 | 0.004979362 | 1 |
| ENSMUSG00000027997 | Casp6 | 0.00016609 | -0.209145444 | 1 |
| ENSMUSG00000019232 | Etnppl | 0.00016609 | 3.648835551 | 0.082076736 |
| ENSMUSG00000106095 | 2010110G14Rik | 0.00016609 | 0.063873051 | 1 |
| ENSMUSG00000027985 | Lef1 | 0.00016609 | -1.051604167 | 0.557141493 |
| ENSMUSG00000028018 | Gstcd | 0.00016609 | -3.058523581 | 0.000412599 |
| ENSMUSG00000040969 | Arhgef38 | 0.00016609 | -0.52108945 | 0.979007222 |
| ENSMUSG00000028172 | Tacr3 | 0.00016609 | 1.189403933 | 0.874796537 |
| ENSMUSG00000053897 | Slc39a8 | 0.00016609 | -0.051604167 | 1 |
| ENSMUSG00000054690 | Emcn | 0.00016609 | -0.52108945 | 1 |
| ENSMUSG00000004127 | Trmt10a | 0.00016609 | 0.189403933 | 1 |
| ENSMUSG00000055301 | Adh7 | 0.00016609 | 2.063873051 | 0.37936848 |
| ENSMUSG00000005813 | Metap1 | 0.00016609 | -2.560617814 | 0.004533725 |
| ENSMUSG00000105279 | Gm6260 | 0.00016609 | 1.47891055 | 0.57597659 |
| ENSMUSG00000105811 | Gm42707 | 0.00016609 | 0.948395833 | 0.931397339 |
| ENSMUSG00000036832 | Lpar3 | 0.00016609 | 1.47891055 | 0.588078615 |
| ENSMUSG00000036825 | Ssx2ip | 0.00016609 | -1.480447466 | 0.279592421 |
| ENSMUSG00000028186 | Uox | 0.00016609 | 1.063873051 | 0.925873597 |
| ENSMUSG00000039103 | Nexn | 0.00016609 | -1.874726405 | 0.069729442 |
| ENSMUSG00000028182 | Lrriq3 | 0.00016609 | 1.063873051 | 0.912283435 |
| ENSMUSG00000028232 | Tmem68 | 0.00016609 | -0.106051951 | 1 |
| ENSMUSG00000028229 | Rmdn1 | 0.00016609 | -1.843017545 | 0.082214652 |
| ENSMUSG00000095139 | Pou3f2 | 0.00016609 | 2.326907457 | 0.353346469 |
| ENSMUSG00000045751 | Mms22l | 0.00016609 | -2.009375931 | 0.045701951 |
| ENSMUSG00000086257 | 4930548K13Rik | 0.00016609 | 1.47891055 | 0.589485743 |

| | | | | |
|---|---|---|---|---|
| ENSMUSG00000028289 | Epha7 | 0.00016609 | -2.184054463 | 0.030336396 |
| ENSMUSG00000087180 | Gm11924 | 0.00016609 | 0.189403933 | 1 |
| ENSMUSG00000040296 | Ddx58 | 0.00016609 | -0.743481871 | 0.778722129 |
| ENSMUSG00000028443 | Nudt2 | 0.00016609 | 0.56137271 | 1 |
| ENSMUSG00000028454 | Pigo | 0.00016609 | -0.158519371 | 1 |
| ENSMUSG00000087326 | Gm12503 | 0.00016609 | 2.326907457 | 0.295355925 |
| ENSMUSG00000014030 | Pax5 | 0.00016609 | 0.063873051 | 1 |
| ENSMUSG00000035539 | Ccdc180 | 0.00016609 | 0.948395833 | 0.941335619 |
| ENSMUSG00000087383 | Gm12446 | 0.00016609 | 1.47891055 | 0.592845519 |
| ENSMUSG00000039693 | Msantd3 | 0.00016609 | -0.743481871 | 0.778722129 |
| ENSMUSG00000039611 | Tmem246 | 0.00016609 | 1.47891055 | 0.57597659 |
| ENSMUSG00000054752 | Fsd1l | 0.00016609 | -0.43862729 | 1 |
| ENSMUSG00000087614 | Gm12514 | 0.00016609 | 0.189403933 | 1 |
| ENSMUSG00000086346 | Gm12510 | 0.00016609 | 3.063873051 | 0.169342649 |
| ENSMUSG00000066176 | Gm12511 | 0.00016609 | 0.326907457 | 1 |
| ENSMUSG00000070972 | Dnajc25 | 0.00016609 | -1.051604167 | 0.539935316 |
| ENSMUSG00000038598 | Shoc1 | 0.00016609 | 0.063873051 | 1 |
| ENSMUSG00000038578 | Susd1 | 0.00016609 | -1.599091962 | 0.231596497 |
| ENSMUSG00000028383 | Hsdl2 | 0.00016609 | 0.400908038 | 1 |
| ENSMUSG00000050395 | Tnfsf15 | 0.00016609 | -0.158519371 | 1 |
| ENSMUSG00000028362 | Tnfsf8 | 0.00016609 | -1.691014451 | 0.2022061 |
| ENSMUSG00000028351 | Brinp1 | 0.00016609 | 0.256518129 | 1 |
| ENSMUSG00000028492 | Saxo1 | 0.00016609 | 1.326907457 | 0.642722979 |
| ENSMUSG00000054351 | 4930553M12Rik | 0.00016609 | -1.636566667 | 0.224818264 |
| ENSMUSG00000052914 | Cyp2j6 | 0.00016609 | 1.189403933 | 0.874796537 |
| ENSMUSG00000028560 | Usp1 | 0.00016609 | -0.965874293 | 0.611716824 |
| ENSMUSG00000086592 | Gm12682 | 0.00016609 | 1.648835551 | 0.508263132 |
| ENSMUSG00000073792 | Alg6 | 0.00016609 | -0.158519371 | 1 |
| ENSMUSG00000085721 | Gm12796 | 0.00016609 | 0.741944956 | 0.971854959 |
| ENSMUSG00000034853 | Acot11 | 0.00016609 | -0.258055044 | 1 |
| ENSMUSG00000034762 | Glis1 | 0.00016609 | 0.741944956 | 0.971854959 |
| ENSMUSG00000028608 | Czib | 0.00016609 | 0.56137271 | 1 |
| ENSMUSG00000094958 | 3110021N24Rik | 0.00016609 | 0.648835551 | 1 |
| ENSMUSG00000085399 | Foxd2os | 0.00016609 | 0.741944956 | 0.970970089 |
| ENSMUSG00000028718 | Stil | 0.00016609 | -1.351164449 | 0.312100107 |
| ENSMUSG00000028700 | Pomgnt1 | 0.00016609 | -0.106051951 | 1 |
| ENSMUSG00000055900 | Tmem69 | 0.00016609 | -0.843017545 | 0.707643264 |
| ENSMUSG00000028690 | Mmachc | 0.00016609 | 3.648835551 | 0.082076736 |
| ENSMUSG00000060268 | Armh1 | 0.00016609 | 0.326907457 | 1 |
| ENSMUSG00000033365 | Ipo13 | 0.00016609 | -0.43862729 | 1 |
| ENSMUSG00000086554 | 9530034E10Rik | 0.00016609 | 2.648835551 | 0.262284476 |
| ENSMUSG00000033253 | Szt2 | 0.00016609 | -1.209145444 | 0.423814296 |
| ENSMUSG00000087238 | Gm12865 | 0.00016609 | 1.47891055 | 0.57597659 |
| ENSMUSG00000028637 | Ccdc30 | 0.00016609 | -0.209145444 | 1 |
| ENSMUSG00000085875 | Gm12905 | 0.00016609 | 1.326907457 | 0.642722979 |
| ENSMUSG00000078570 | 1110065P20Rik | 0.00016609 | 1.326907457 | 0.66416768 |
| ENSMUSG00000050188 | Lsm10 | 0.00016609 | 1.648835551 | 0.52650018 |
| ENSMUSG00000050212 | Eva1b | 0.00016609 | 0.741944956 | 0.970970089 |
| ENSMUSG00000073758 | Sh3d21 | 0.00016609 | -0.158519371 | 1 |
| ENSMUSG00000056174 | Col8a2 | 0.00016609 | 0.256518129 | 1 |
| ENSMUSG00000041530 | Ago1 | 0.00016609 | -0.599091962 | 0.865542683 |

| | | | | |
|---|---|---|---|---|
| ENSMUSG00000042380 | Smim12 | 0.00016609 | 1.063873051 | 0.747182608 |
| ENSMUSG00000028910 | Mecr | 0.00016609 | -0.52108945 | 0.975624465 |
| ENSMUSG00000100813 | Gm28874 | 0.00016609 | 1.841480629 | 0.466329251 |
| ENSMUSG00000028882 | Ppp1r8 | 0.00016609 | 0.326907457 | 1 |
| ENSMUSG00000028860 | Sytl1 | 0.00016609 | 1.326907457 | 0.642722979 |
| ENSMUSG00000028857 | Tmem222 | 0.00016609 | 0.256518129 | 1 |
| ENSMUSG00000028848 | Gpn2 | 0.00016609 | 0.400908038 | 1 |
| ENSMUSG00000043257 | Pigv | 0.00016609 | 0.841480629 | 0.955646176 |
| ENSMUSG00000000682 | Cd52 | 0.00016609 | 1.063873051 | 0.886203497 |
| ENSMUSG00000028840 | Zfp593 | 0.00016609 | 1.47891055 | 0.593915351 |
| ENSMUSG00000050989 | Selenon | 0.00016609 | 0.189403933 | 1 |
| ENSMUSG00000070691 | Runx3 | 0.00016609 | 0.741944956 | 0.976813659 |
| ENSMUSG00000028801 | Stpg1 | 0.00016609 | 0.400908038 | 1 |
| ENSMUSG00000062585 | Cnr2 | 0.00016609 | 1.189403933 | 0.686735271 |
| ENSMUSG00000028672 | Hmgcl | 0.00016609 | -1.936126949 | 0.062531119 |
| ENSMUSG00000070687 | Htr1d | 0.00016609 | 1.47891055 | 0.57597659 |
| ENSMUSG00000036856 | Wnt4 | 0.00016609 | -0.843017545 | 0.701794743 |
| ENSMUSG00000028753 | Vwa5b1 | 0.00016609 | 1.47891055 | 0.587484274 |
| ENSMUSG00000041161 | Otud3 | 0.00016609 | -0.209145444 | 1 |
| ENSMUSG00000028744 | Pqlc2 | 0.00016609 | 0.841480629 | 0.950123751 |
| ENSMUSG00000028736 | Pax7 | 0.00016609 | 0.063873051 | 1 |
| ENSMUSG00000025330 | Padi4 | 0.00016609 | 0.400908038 | 1 |
| ENSMUSG00000060572 | Mfap2 | 0.00016609 | 3.648835551 | 0.082076736 |
| ENSMUSG00000062478 | Ctrc | 0.00016609 | 1.063873051 | 0.745688239 |
| ENSMUSG00000085816 | Gm13052 | 0.00016609 | 0.47891055 | 1 |
| ENSMUSG00000028602 | Tnfrsf8 | 0.00016609 | 0.741944956 | 0.970970089 |
| ENSMUSG00000056300 | Zfp981 | 0.00016609 | 0.47891055 | 1 |
| ENSMUSG00000044496 | 2510039O18Rik | 0.00016609 | -0.106051951 | 1 |
| ENSMUSG00000029016 | Clcn6 | 0.00016609 | -1.82689788 | 0.100320414 |
| ENSMUSG00000055401 | Fbxo6 | 0.00016609 | 1.47891055 | 0.590885473 |
| ENSMUSG00000041544 | Disp3 | 0.00016609 | 1.841480629 | 0.455131546 |
| ENSMUSG00000073705 | Cenps | 0.00016609 | 1.189403933 | 0.705223486 |
| ENSMUSG00000085819 | Ube4bos1 | 0.00016609 | 2.063873051 | 0.37936848 |
| ENSMUSG00000028982 | Slc25a33 | 0.00016609 | -0.480447466 | 0.972350276 |
| ENSMUSG00000039662 | Icmt | 0.00016609 | 1.326907457 | 0.642722979 |
| ENSMUSG00000039577 | Nphp4 | 0.00016609 | 0.841480629 | 0.975624465 |
| ENSMUSG00000029055 | Plch2 | 0.00016609 | 0.841480629 | 0.943363727 |
| ENSMUSG00000084803 | 5830444B04Rik | 0.00016609 | 0.841480629 | 0.955892779 |
| ENSMUSG00000078490 | Cfap74 | 0.00016609 | -0.560617814 | 0.980745941 |
| ENSMUSG00000029061 | Mmp23 | 0.00016609 | -1.079084903 | 0.536465189 |
| ENSMUSG00000035692 | Isg15 | 0.00016609 | 3.063873051 | 0.144078128 |
| ENSMUSG00000015652 | Steap1 | 0.00016609 | -0.52108945 | 0.975624465 |
| ENSMUSG00000012428 | Steap4 | 0.00016609 | 2.326907457 | 0.310311603 |
| ENSMUSG00000002297 | Dbf4 | 0.00016609 | -1.281901786 | 0.365716212 |
| ENSMUSG00000105157 | Gm31831 | 0.00016609 | 0.948395833 | 0.926141556 |
| ENSMUSG00000040254 | Sema3d | 0.00016609 | -0.673092543 | 0.885435607 |
| ENSMUSG00000028986 | Klhl7 | 0.00016609 | -2.540989007 | 0.005200942 |
| ENSMUSG00000038295 | Atg9b | 0.00016609 | -1.373532262 | 0.317734263 |
| ENSMUSG00000028953 | Abcf2 | 0.00016609 | -1.560617814 | 0.20542113 |
| ENSMUSG00000038181 | Chpf2 | 0.00016609 | -0.599091962 | 0.865542683 |
| ENSMUSG00000028949 | Smarcd3 | 0.00016609 | -1.500911568 | 0.23866117 |

| | | | | |
|---|---|---|---|---|
| ENSMUSG00000038072 | Galnt11 | 0.00016609 | -1.921020057 | 0.060210487 |
| ENSMUSG00000029044 | 4930584F24Rik | 0.00016609 | 1.189403933 | 0.874796537 |
| ENSMUSG00000002633 | Shh | 0.00016609 | 1.841480629 | 0.455131546 |
| ENSMUSG00000053214 | Gm9899 | 0.00016609 | 0.256518129 | 1 |
| ENSMUSG00000107078 | Gm36840 | 0.00016609 | 1.841480629 | 0.466464457 |
| ENSMUSG00000004642 | Slbp | 0.00016609 | -1.258055044 | 0.361093181 |
| ENSMUSG00000029098 | Acox3 | 0.00016609 | -1.708716453 | 0.128972787 |
| ENSMUSG00000029121 | Crmp1 | 0.00016609 | 3.648835551 | 0.104546659 |
| ENSMUSG00000029125 | Stx18 | 0.00016609 | -1.209145444 | 0.423814296 |
| ENSMUSG00000067367 | Lyar | 0.00016609 | -0.936126949 | 0.928107834 |
| ENSMUSG00000029084 | Cd38 | 0.00016609 | -1.184054463 | 0.428944094 |
| ENSMUSG00000015879 | Fam184b | 0.00016609 | 0.841480629 | 0.957062426 |
| ENSMUSG00000029167 | Ppargc1a | 0.00016609 | -0.209145444 | 1 |
| ENSMUSG00000097145 | 9230114K14Rik | 0.00016609 | -0.209145444 | 1 |
| ENSMUSG00000029205 | Chrna9 | 0.00016609 | 0.004979362 | 1 |
| ENSMUSG00000029206 | Nsun7 | 0.00016609 | 0.189403933 | 1 |
| ENSMUSG00000087601 | Uchl1os | 0.00016609 | 1.063873051 | 0.886203497 |
| ENSMUSG00000029223 | Uchl1 | 0.00016609 | -0.965874293 | 0.66416768 |
| ENSMUSG00000037653 | Kctd8 | 0.00016609 | -0.106051951 | 1 |
| ENSMUSG00000029209 | Gnpda2 | 0.00016609 | 1.063873051 | 0.895427854 |
| ENSMUSG00000001260 | Gabrg1 | 0.00016609 | 2.326907457 | 0.33333983 |
| ENSMUSG00000051674 | Dcun1d4 | 0.00016609 | -2.009375931 | 0.045701951 |
| ENSMUSG00000067206 | Lrrc66 | 0.00016609 | -0.351164449 | 1 |
| ENSMUSG00000099605 | 1700066N21Rik | 0.00016609 | 1.063873051 | 0.887369997 |
| ENSMUSG00000070690 | Albfm1 | 0.00016609 | 2.063873051 | 0.37936848 |
| ENSMUSG00000029403 | Cdkl2 | 0.00016609 | -1.079084903 | 0.536465189 |
| ENSMUSG00000029413 | Naaa | 0.00016609 | 0.741944956 | 0.970970089 |
| ENSMUSG00000029335 | Bmp3 | 0.00016609 | 2.326907457 | 0.296030202 |
| ENSMUSG00000035273 | Hpse | 0.00016609 | 0.063873051 | 1 |
| ENSMUSG00000029321 | Slc10a6 | 0.00016609 | 3.063873051 | 0.144078128 |
| ENSMUSG00000013495 | Tmem175 | 0.00016609 | -0.43862729 | 1 |
| ENSMUSG00000066613 | Zfp932 | 0.00016609 | 0.648835551 | 1 |
| ENSMUSG00000109771 | Gm35315 | 0.00016609 | 0.004979362 | 1 |
| ENSMUSG00000033316 | Galnt9 | 0.00016609 | 1.47891055 | 0.57597659 |
| ENSMUSG00000063430 | Wscd2 | 0.00016609 | 1.189403933 | 0.689413368 |
| ENSMUSG00000042184 | 1700069L16Rik | 0.00016609 | 1.648835551 | 0.605997429 |
| ENSMUSG00000042010 | Acacb | 0.00016609 | -0.051604167 | 1 |
| ENSMUSG00000002486 | Tchp | 0.00016609 | -0.995020638 | 0.618995574 |
| ENSMUSG00000029561 | Oasl2 | 0.00016609 | -0.810596067 | 0.792441218 |
| ENSMUSG00000041548 | Hspb8 | 0.00016609 | 1.841480629 | 0.455131546 |
| ENSMUSG00000032850 | Rnft2 | 0.00016609 | -0.743481871 | 0.778722129 |
| ENSMUSG00000032840 | 2410131K14Rik | 0.00016609 | 0.948395833 | 0.912904617 |
| ENSMUSG00000029598 | Plbd2 | 0.00016609 | -0.560617814 | 1 |
| ENSMUSG00000029601 | Iqcd | 0.00016609 | 1.841480629 | 0.466464457 |
| ENSMUSG00000032661 | Oas3 | 0.00016609 | 2.063873051 | 0.466464457 |
| ENSMUSG00000042594 | Sh2b3 | 0.00016609 | 0.125273595 | 1 |
| ENSMUSG00000038569 | Rad9b | 0.00016609 | 0.948395833 | 0.923527923 |
| ENSMUSG00000029464 | Gpn3 | 0.00016609 | 0.004979362 | 1 |
| ENSMUSG00000029433 | Diablo | 0.00016609 | -0.258055044 | 1 |
| ENSMUSG00000029414 | Kntc1 | 0.00016609 | -0.810596067 | 0.791730654 |
| ENSMUSG00000023707 | Ogfod2 | 0.00016609 | -0.395558568 | 1 |

| | | | | |
|---|---|---|---|---|
| ENSMUSG00000029388 | Eif2b1 | 0.00016609 | 1.47891055 | 0.59022739 |
| ENSMUSG00000029482 | Aacs | 0.00016609 | -0.636566667 | 0.868661374 |
| ENSMUSG00000105673 | 1700081B01Rik | 0.00016609 | 1.47891055 | 0.587484274 |
| ENSMUSG00000029416 | Slc15a4 | 0.00016609 | -0.480447466 | 0.972350276 |
| ENSMUSG00000049971 | Glt1d1 | 0.00016609 | -0.395558568 | 1 |
| ENSMUSG00000025538 | Sumf2 | 0.00016609 | -1.305360759 | 0.371776814 |
| ENSMUSG00000005373 | Mlxipl | 0.00016609 | 0.841480629 | 0.946435944 |
| ENSMUSG00000039747 | Orai2 | 0.00016609 | -0.560617814 | 0.9980272 |
| ENSMUSG00000005057 | Sh2b2 | 0.00016609 | -0.599091962 | 1 |
| ENSMUSG00000097908 | 4933404O12Rik | 0.00016609 | 2.063873051 | 0.37936848 |
| ENSMUSG00000037390 | Muc3 | 0.00016609 | 1.47891055 | 0.598570743 |
| ENSMUSG00000029715 | Pop7 | 0.00016609 | 2.648835551 | 0.218817775 |
| ENSMUSG00000037221 | Mospd3 | 0.00016609 | 0.400908038 | 1 |
| ENSMUSG00000029725 | Ppp1r35 | 0.00016609 | 1.189403933 | 0.874796537 |
| ENSMUSG00000037108 | Zcwpw1 | 0.00016609 | -0.43862729 | 1 |
| ENSMUSG00000036980 | Taf6 | 0.00016609 | -0.43862729 | 1 |
| ENSMUSG00000056014 | A430033K04Rik | 0.00016609 | 0.648835551 | 1 |
| ENSMUSG00000036687 | Tmem184a | 0.00016609 | 0.841480629 | 0.957779577 |
| ENSMUSG00000036639 | Nudt1 | 0.00016609 | 0.47891055 | 1 |
| ENSMUSG00000036599 | Chst12 | 0.00016609 | -0.305360759 | 1 |
| ENSMUSG00000050822 | Slc29a4 | 0.00016609 | 1.648835551 | 0.50315098 |
| ENSMUSG00000029587 | Zfp12 | 0.00016609 | -0.305360759 | 1 |
| ENSMUSG00000093720 | Gm20635 | 0.00016609 | 1.063873051 | 0.886203497 |
| ENSMUSG00000051306 | Usp42 | 0.00016609 | -0.395558568 | 1 |
| ENSMUSG00000043388 | Tmem130 | 0.00016609 | 2.648835551 | 0.278293619 |
| ENSMUSG00000055991 | Zkscan5 | 0.00016609 | 0.256518129 | 1 |
| ENSMUSG00000106719 | Gm42556 | 0.00016609 | 1.189403933 | 0.69286889 |
| ENSMUSG00000015189 | Casd1 | 0.00016609 | -1.951077291 | 0.063804743 |
| ENSMUSG00000093482 | Gm20619 | 0.00016609 | -0.051604167 | 1 |
| ENSMUSG00000042447 | Mios | 0.00016609 | -1.395558568 | 0.26547175 |
| ENSMUSG00000029569 | Tmem168 | 0.00016609 | -0.52108945 | 0.975681674 |
| ENSMUSG00000010797 | Wnt2 | 0.00016609 | -2.092631435 | 0.034881114 |
| ENSMUSG00000010796 | Asz1 | 0.00016609 | 0.189403933 | 1 |
| ENSMUSG00000029670 | Ing3 | 0.00016609 | 0.648835551 | 0.998731799 |
| ENSMUSG00000029695 | Aass | 0.00016609 | 1.648835551 | 0.499921258 |
| ENSMUSG00000029700 | Slc13a1 | 0.00016609 | 1.47891055 | 0.57597659 |
| ENSMUSG00000029708 | Gcc1 | 0.00016609 | -1.132524162 | 0.466464457 |
| ENSMUSG00000029790 | Cep41 | 0.00016609 | 0.56137271 | 1 |
| ENSMUSG00000056215 | Lrguk | 0.00016609 | 0.56137271 | 1 |
| ENSMUSG00000045613 | Chrm2 | 0.00016609 | 0.648835551 | 0.9980272 |
| ENSMUSG00000046947 | Adck2 | 0.00016609 | 0.47891055 | 1 |
| ENSMUSG00000057716 | Tmem178b | 0.00016609 | -0.995020638 | 0.66416768 |
| ENSMUSG00000076490 | Trbc1 | 0.00016609 | 1.841480629 | 0.466464457 |
| ENSMUSG00000029864 | Gstk1 | 0.00016609 | 3.063873051 | 0.144078128 |
| ENSMUSG00000029863 | Casp2 | 0.00016609 | -0.777429203 | 0.785942057 |
| ENSMUSG00000029851 | Tcaf2 | 0.00016609 | -0.051604167 | 1 |
| ENSMUSG00000009281 | Rarres2 | 0.00016609 | 0.326907457 | 1 |
| ENSMUSG00000023403 | Stk31 | 0.00016609 | -0.051604167 | 1 |
| ENSMUSG00000029819 | Npy | 0.00016609 | 0.648835551 | 1 |
| ENSMUSG00000108071 | Gm3455 | 0.00016609 | 1.648835551 | 0.508263132 |
| ENSMUSG00000029828 | 4921507P07Rik | 0.00016609 | 0.189403933 | 1 |

| | | | | |
|---|---|---|---|---|
| ENSMUSG00000038065 | Mturn | 0.00016609 | -0.673092543 | 0.908885177 |
| ENSMUSG00000037709 | Fam13a | 0.00016609 | 0.063873051 | 1 |
| ENSMUSG00000073045 | Gm5570 | 0.00016609 | 0.400908038 | 1 |
| ENSMUSG00000018341 | Il12rb2 | 0.00016609 | 0.841480629 | 0.956368165 |
| ENSMUSG00000054474 | Thnsl2 | 0.00016609 | 1.841480629 | 0.455131546 |
| ENSMUSG00000056305 | Usp39 | 0.00016609 | -1.72620388 | 0.131190036 |
| ENSMUSG00000058706 | 0610030E20Rik | 0.00016609 | -0.52108945 | 0.975624465 |
| ENSMUSG00000055912 | Tmem150a | 0.00016609 | -0.158519371 | 1 |
| ENSMUSG00000096887 | Gm20594 | 0.00016609 | 0.841480629 | 0.943363727 |
| ENSMUSG00000107419 | Gm38839 | 0.00016609 | 1.648835551 | 0.499921258 |
| ENSMUSG00000014554 | Dguok | 0.00016609 | -1.158519371 | 0.475407382 |
| ENSMUSG00000108660 | Gm21284 | 0.00016609 | 2.648835551 | 0.259407688 |
| ENSMUSG00000045896 | Paip2b | 0.00016609 | -0.965874293 | 0.611716824 |
| ENSMUSG00000033706 | Smyd5 | 0.00016609 | -0.351164449 | 1 |
| ENSMUSG00000068263 | Efcc1 | 0.00016609 | 0.948395833 | 0.929563033 |
| ENSMUSG00000002870 | Mcm2 | 0.00016609 | 0.189403933 | 1 |
| ENSMUSG00000034430 | Zxdc | 0.00016609 | -1.023589791 | 0.524144449 |
| ENSMUSG00000030088 | Aldh1l1 | 0.00016609 | -1.965874293 | 0.075551022 |
| ENSMUSG00000034245 | Hdac11 | 0.00016609 | 1.189403933 | 0.874796537 |
| ENSMUSG00000068180 | AY512915 | 0.00016609 | 0.948395833 | 0.936016139 |
| ENSMUSG00000093661 | Eif4e3 | 0.00016609 | -0.673092543 | 0.886203497 |
| ENSMUSG00000042873 | Lhfpl4 | 0.00016609 | 1.648835551 | 0.51221187 |
| ENSMUSG00000048930 | Tada3 | 0.00016609 | -0.636566667 | 0.872788504 |
| ENSMUSG00000087083 | Gm15083 | 0.00016609 | 1.648835551 | 0.526094357 |
| ENSMUSG00000053004 | Hrh1 | 0.00016609 | -0.743481871 | 0.778722129 |
| ENSMUSG00000030316 | Tamm41 | 0.00016609 | -0.965874293 | 0.611716824 |
| ENSMUSG00000002897 | Il17ra | 0.00016609 | -0.52108945 | 0.975624465 |
| ENSMUSG00000108171 | Gm43915 | 0.00016609 | 0.648835551 | 1 |
| ENSMUSG00000040669 | Phc1 | 0.00016609 | 0.56137271 | 1 |
| ENSMUSG00000003153 | Slc2a3 | 0.00016609 | -0.351164449 | 1 |
| ENSMUSG00000040552 | C3ar1 | 0.00016609 | 1.189403933 | 0.692233739 |
| ENSMUSG00000030327 | Necap1 | 0.00016609 | -1.106051951 | 0.466464457 |
| ENSMUSG00000005069 | Pex5 | 0.00016609 | -0.843017545 | 0.674297981 |
| ENSMUSG00000072770 | Acrbp | 0.00016609 | -1.281901786 | 0.365716212 |
| ENSMUSG00000038167 | Plekhg6 | 0.00016609 | 1.326907457 | 0.642722979 |
| ENSMUSG00000038115 | Ano2 | 0.00016609 | -1.560617814 | 0.246745962 |
| ENSMUSG00000038077 | Kcna6 | 0.00016609 | 1.326907457 | 0.645058099 |
| ENSMUSG00000030346 | Rad51ap1 | 0.00016609 | -1.743481871 | 0.109343152 |
| ENSMUSG00000107153 | Gm38404 | 0.00016609 | 2.648835551 | 0.218817775 |
| ENSMUSG00000037997 | Parp11 | 0.00016609 | -0.52108945 | 0.975624465 |
| ENSMUSG00000030351 | Tspan11 | 0.00016609 | 0.56137271 | 1 |
| ENSMUSG00000048668 | Rhno1 | 0.00016609 | -1.810596067 | 0.106876393 |
| ENSMUSG00000030325 | Klrb1c | 0.00016609 | 1.47891055 | 0.588078615 |
| ENSMUSG00000067591 | Klra3 | 0.00016609 | 4.648835551 | 0.089729229 |
| ENSMUSG00000079853 | Klra1 | 0.00016609 | 1.648835551 | 0.499921258 |
| ENSMUSG00000107666 | 4933406J09Rik | 0.00016609 | 1.063873051 | 0.886203497 |
| ENSMUSG00000030207 | Fam234b | 0.00016609 | -0.258055044 | 1 |
| ENSMUSG00000107768 | Gm44275 | 0.00016609 | -1.843017545 | 0.082214652 |
| ENSMUSG00000030214 | Plbd1 | 0.00016609 | 0.326907457 | 1 |
| ENSMUSG00000100707 | Gm28523 | 0.00016609 | 1.841480629 | 0.455131546 |
| ENSMUSG00000030237 | Slco1a4 | 0.00016609 | 0.56137271 | 1 |

| | | | | |
|---|---|---|---|---|
| ENSMUSG00000092500 | Gm20400 | 0.00016609 | 3.648835551 | 0.082076736 |
| ENSMUSG00000041671 | Pyroxd1 | 0.00016609 | -0.995020638 | 0.618995574 |
| ENSMUSG00000030244 | Gys2 | 0.00016609 | 1.326907457 | 0.650180218 |
| ENSMUSG00000030263 | Lrmp | 0.00016609 | -1.794107944 | 0.094802366 |
| ENSMUSG00000086013 | Gm15706 | 0.00016609 | 3.063873051 | 0.189855518 |
| ENSMUSG00000072660 | Gm6288 | 0.00016609 | 1.063873051 | 0.886203497 |
| ENSMUSG00000030309 | Caprin2 | 0.00016609 | -1.328444372 | 0.307566094 |
| ENSMUSG00000035390 | Brsk1 | 0.00016609 | -0.305360759 | 1 |
| ENSMUSG00000074405 | Zfp865 | 0.00016609 | -0.743481871 | 0.778722129 |
| ENSMUSG00000034690 | Nlrp4c | 0.00016609 | 3.063873051 | 0.210757491 |
| ENSMUSG00000006019 | Dhx34 | 0.00016609 | -0.874726405 | 0.680536135 |
| ENSMUSG00000041375 | Ccdc9 | 0.00016609 | -2.943621486 | 0.000714323 |
| ENSMUSG00000074358 | Ccdc61 | 0.00016609 | -2.037664976 | 0.038867277 |
| ENSMUSG00000040866 | Rsph6a | 0.00016609 | 1.841480629 | 0.455131546 |
| ENSMUSG00000030409 | Dmpk | 0.00016609 | 0.004979362 | 1 |
| ENSMUSG00000060508 | Nlrp9b | 0.00016609 | 1.47891055 | 0.591388707 |
| ENSMUSG00000015721 | Nlrp5 | 0.00016609 | 2.326907457 | 0.295355925 |
| ENSMUSG00000052675 | Zfp112 | 0.00016609 | 1.063873051 | 0.886203497 |
| ENSMUSG00000040907 | Atp1a3 | 0.00016609 | 0.326907457 | 1 |
| ENSMUSG00000054385 | Ceacam2 | 0.00016609 | 1.326907457 | 0.66416768 |
| ENSMUSG00000002608 | Ccdc97 | 0.00016609 | 1.063873051 | 0.886203497 |
| ENSMUSG00000011751 | Sptbn4 | 0.00016609 | -0.051604167 | 1 |
| ENSMUSG00000020420 | Zfp607a | 0.00016609 | -0.106051951 | 1 |
| ENSMUSG00000070709 | Zfp974 | 0.00016609 | -0.560617814 | 0.980745941 |
| ENSMUSG00000046058 | Eid2 | 0.00016609 | 1.841480629 | 0.466464457 |
| ENSMUSG00000037552 | Plekhg2 | 0.00016609 | -1.373532262 | 0.317734263 |
| ENSMUSG00000109455 | Gm44710 | 0.00016609 | 0.648835551 | 1 |
| ENSMUSG00000003437 | Paf1 | 0.00016609 | -1.132524162 | 0.47966612 |
| ENSMUSG00000108905 | Gm44618 | 0.00016609 | 1.47891055 | 0.590958085 |
| ENSMUSG00000051735 | Rinl | 0.00016609 | 1.648835551 | 0.499921258 |
| ENSMUSG00000030584 | Dpf1 | 0.00016609 | 1.47891055 | 0.57597659 |
| ENSMUSG00000050855 | Zfp940 | 0.00016609 | 1.648835551 | 0.499921258 |
| ENSMUSG00000036957 | Lrfn3 | 0.00016609 | 1.841480629 | 0.455131546 |
| ENSMUSG00000006651 | Aplp1 | 0.00016609 | 1.648835551 | 0.499921258 |
| ENSMUSG00000006313 | Upk1a | 0.00016609 | 0.400908038 | 1 |
| ENSMUSG00000043671 | Dpy19l3 | 0.00016609 | -0.480447466 | 0.972350276 |
| ENSMUSG00000054676 | 1600014C10Rik | 0.00016609 | 0.648835551 | 1 |
| ENSMUSG00000108621 | Gm37494 | 0.00016609 | -2.245982212 | 0.022024798 |
| ENSMUSG00000072259 | Gm5592 | 0.00016609 | -0.209145444 | 1 |
| ENSMUSG00000030469 | Zfp719 | 0.00016609 | 0.189403933 | 1 |
| ENSMUSG00000038644 | Pold1 | 0.00016609 | -0.158519371 | 1 |
| ENSMUSG00000060601 | Nr1h2 | 0.00016609 | -0.673092543 | 0.896076486 |
| ENSMUSG00000109511 | Nup62 | 0.00016609 | 0.400908038 | 1 |
| ENSMUSG00000002968 | Med25 | 0.00016609 | -0.43862729 | 1 |
| ENSMUSG00000003190 | Bcl2l12 | 0.00016609 | 0.841480629 | 0.943363727 |
| ENSMUSG00000003184 | Irf3 | 0.00016609 | -0.480447466 | 0.972350276 |
| ENSMUSG00000053801 | Grwd1 | 0.00016609 | 0.326907457 | 1 |
| ENSMUSG00000002781 | Tmem143 | 0.00016609 | 1.47891055 | 0.587484274 |
| ENSMUSG00000110240 | Gm45486 | 0.00016609 | 0.063873051 | 1 |
| ENSMUSG00000030851 | Ldhc | 0.00016609 | 1.648835551 | 0.520134349 |
| ENSMUSG00000085265 | Nell1os | 0.00016609 | 1.063873051 | 0.886203497 |

| | | | | |
|---|---|---|---|---|
| ENSMUSG00000074093 | Svip | 0.00016609 | -1.106051951 | 0.625388802 |
| ENSMUSG00000033790 | Tubgcp5 | 0.00016609 | -0.995020638 | 0.618995574 |
| ENSMUSG00000087178 | A230056P14Rik | 0.00016609 | -0.560617814 | 0.980745941 |
| ENSMUSG00000102252 | Snrpn | 0.00016609 | 0.256518129 | 1 |
| ENSMUSG00000097682 | 4930554H23Rik | 0.00016609 | 0.948395833 | 0.928063181 |
| ENSMUSG00000092187 | Gm20457 | 0.00016609 | -1.158519371 | 0.466464457 |
| ENSMUSG00000097634 | Gm26827 | 0.00016609 | 1.063873051 | 0.897629756 |
| ENSMUSG00000030609 | Aen | 0.00016609 | -0.936126949 | 0.606655093 |
| ENSMUSG00000047084 | Ngrn | 0.00016609 | -0.106051951 | 1 |
| ENSMUSG00000038886 | Man2a2 | 0.00016609 | -1.617950989 | 0.204073384 |
| ENSMUSG00000030528 | Blm | 0.00016609 | -1.258055044 | 0.361093181 |
| ENSMUSG00000038763 | Alpk3 | 0.00016609 | 0.841480629 | 0.943363727 |
| ENSMUSG00000030638 | Sh3gl3 | 0.00016609 | -0.708716453 | 0.886203497 |
| ENSMUSG00000038540 | Tmc3 | 0.00016609 | 3.648835551 | 0.082076736 |
| ENSMUSG00000046027 | Stard5 | 0.00016609 | 0.189403933 | 1 |
| ENSMUSG00000090362 | Vmn2r79 | 0.00016609 | 1.326907457 | 0.642722979 |
| ENSMUSG00000039405 | Prss23 | 0.00016609 | -0.777429203 | 0.792441218 |
| ENSMUSG00000030615 | Tmem126a | 0.00016609 | -0.258055044 | 1 |
| ENSMUSG00000041343 | Ankrd42 | 0.00016609 | 0.400908038 | 1 |
| ENSMUSG00000025133 | Ints4 | 0.00016609 | -2.051604167 | 0.039883588 |
| ENSMUSG00000086433 | Rsf1os1 | 0.00016609 | 0.741944956 | 0.971854959 |
| ENSMUSG00000042797 | Aqp11 | 0.00016609 | 1.648835551 | 0.53074305 |
| ENSMUSG00000030761 | Myo7a | 0.00016609 | -0.777429203 | 0.785942057 |
| ENSMUSG00000030747 | Dgat2 | 0.00016609 | 0.063873051 | 1 |
| ENSMUSG00000030737 | Slco2b1 | 0.00016609 | 0.741944956 | 0.970970089 |
| ENSMUSG00000109186 | Gm34821 | 0.00016609 | 3.648835551 | 0.082076736 |
| ENSMUSG00000035165 | Kcne3 | 0.00016609 | 0.125273595 | 1 |
| ENSMUSG00000051048 | P4ha3 | 0.00016609 | 0.326907457 | 1 |
| ENSMUSG00000064307 | Lrrc51 | 0.00016609 | -1.417253639 | 0.287921264 |
| ENSMUSG00000078616 | Trim30c | 0.00016609 | -0.051604167 | 1 |
| ENSMUSG00000037049 | Smpd1 | 0.00016609 | 1.648835551 | 0.499921258 |
| ENSMUSG00000045868 | Gvin1 | 0.00016609 | -0.209145444 | 1 |
| ENSMUSG00000057215 | Platr28 | 0.00016609 | 1.47891055 | 0.57597659 |
| ENSMUSG00000062542 | Syt9 | 0.00016609 | 0.189403933 | 1 |
| ENSMUSG00000031028 | Tub | 0.00016609 | 1.648835551 | 0.527226787 |
| ENSMUSG00000031027 | Stk33 | 0.00016609 | -0.158519371 | 1 |
| ENSMUSG00000085945 | 2310014F06Rik | 0.00016609 | -1.079084903 | 0.536465189 |
| ENSMUSG00000048782 | Insc | 0.00016609 | 1.326907457 | 0.66416768 |
| ENSMUSG00000030909 | Anks4b | 0.00016609 | 0.004979362 | 1 |
| ENSMUSG00000109113 | Gm32916 | 0.00016609 | 1.841480629 | 0.455131546 |
| ENSMUSG00000030880 | Polr3e | 0.00016609 | -0.305360759 | 1 |
| ENSMUSG00000030872 | Gga2 | 0.00016609 | -0.965874293 | 0.65079351 |
| ENSMUSG00000109814 | Gm45847 | 0.00016609 | 1.063873051 | 0.912067357 |
| ENSMUSG00000045928 | 4933440M02Rik | 0.00016609 | 1.326907457 | 0.66416768 |
| ENSMUSG00000044701 | Il27 | 0.00016609 | -2.480447466 | 0.013074066 |
| ENSMUSG00000030703 | Gdpd3 | 0.00016609 | 0.256518129 | 1 |
| ENSMUSG00000030685 | Kctd13 | 0.00016609 | 1.063873051 | 0.886203497 |
| ENSMUSG00000000486 | 1-Sep | 0.00016609 | -0.810596067 | 0.791730654 |
| ENSMUSG00000047371 | Zfp768 | 0.00016609 | 1.47891055 | 0.57597659 |
| ENSMUSG00000030816 | Rnf40 | 0.00016609 | -1.560617814 | 0.20542113 |
| ENSMUSG00000042178 | Armc5 | 0.00016609 | 0.56137271 | 1 |

| | | | | |
|---|---|---|---|---|
| ENSMUSG00000040167 | Ikzf5 | 0.00016609 | 0.47891055 | 1 |
| ENSMUSG00000030946 | Lhpp | 0.00016609 | 1.47891055 | 0.57597659 |
| ENSMUSG00000030956 | Fam53b | 0.00016609 | 2.063873051 | 0.406929086 |
| ENSMUSG00000109917 | Gm45671 | 0.00016609 | 2.326907457 | 0.314582476 |
| ENSMUSG00000030986 | Dhx32 | 0.00016609 | -1.540989007 | 0.201326391 |
| ENSMUSG00000059136 | Olfr539 | 0.00016609 | 3.063873051 | 0.144078128 |
| ENSMUSG00000025481 | Urah | 0.00016609 | 0.648835551 | 1 |
| ENSMUSG00000025494 | Sigirr | 0.00016609 | -0.106051951 | 1 |
| ENSMUSG00000025498 | Irf7 | 0.00016609 | 3.063873051 | 0.144078128 |
| ENSMUSG00000025147 | Mob2 | 0.00016609 | -2.490715801 | 0.006613366 |
| ENSMUSG00000031085 | Acte1 | 0.00016609 | 1.841480629 | 0.466464457 |
| ENSMUSG00000065952 | C330021F23Rik | 0.00016609 | 2.326907457 | 0.329244596 |
| ENSMUSG00000086805 | 4932443L11Rik | 0.00016609 | 1.47891055 | 0.580529935 |
| ENSMUSG00000040396 | Abhd13 | 0.00016609 | -0.051604167 | 1 |
| ENSMUSG00000031497 | Tnfsf13b | 0.00016609 | 1.648835551 | 0.536876159 |
| ENSMUSG00000031504 | Rab20 | 0.00016609 | -0.560617814 | 0.984666001 |
| ENSMUSG00000031509 | 1700016D06Rik | 0.00016609 | 1.063873051 | 0.886945503 |
| ENSMUSG00000038365 | Fbxo25 | 0.00016609 | -0.599091962 | 1 |
| ENSMUSG00000071176 | Arhgef10 | 0.00016609 | -2.500911568 | 0.007443294 |
| ENSMUSG00000031461 | Myom2 | 0.00016609 | 1.648835551 | 0.507847339 |
| ENSMUSG00000031465 | Angpt2 | 0.00016609 | 0.56137271 | 1 |
| ENSMUSG00000031536 | Polb | 0.00016609 | -1.599091962 | 0.172061305 |
| ENSMUSG00000031553 | Adam3 | 0.00016609 | 1.841480629 | 0.466464457 |
| ENSMUSG00000037316 | Bag4 | 0.00016609 | -1.051604167 | 0.529126167 |
| ENSMUSG00000031574 | Star | 0.00016609 | 2.063873051 | 0.37936848 |
| ENSMUSG00000085795 | Zfp703 | 0.00016609 | 0.326907457 | 1 |
| ENSMUSG00000049184 | Purg | 0.00016609 | -0.480447466 | 0.972350276 |
| ENSMUSG00000109372 | Gm19410 | 0.00016609 | 0.56137271 | 1 |
| ENSMUSG00000031600 | Vps37a | 0.00016609 | -2.092631435 | 0.042735074 |
| ENSMUSG00000031595 | Pdgfrl | 0.00016609 | -1.395558568 | 0.266000629 |
| ENSMUSG00000085672 | Gm16193 | 0.00016609 | 2.326907457 | 0.307184021 |
| ENSMUSG00000110260 | Gm30504 | 0.00016609 | 1.648835551 | 0.523881086 |
| ENSMUSG00000038102 | Trappc11 | 0.00016609 | -1.184054463 | 0.419360593 |
| ENSMUSG00000109823 | Gm45388 | 0.00016609 | 3.648835551 | 0.082076736 |
| ENSMUSG00000039396 | Neil3 | 0.00016609 | -0.258055044 | 1 |
| ENSMUSG00000109629 | Gm34099 | 0.00016609 | 1.841480629 | 0.474032441 |
| ENSMUSG00000096917 | 2500002B13Rik | 0.00016609 | -1.079084903 | 0.536465189 |
| ENSMUSG00000109966 | Gm34623 | 0.00016609 | 0.326907457 | 1 |
| ENSMUSG00000074300 | Smim31 | 0.00016609 | -1.184054463 | 0.544024047 |
| ENSMUSG00000059897 | Zfp930 | 0.00016609 | 0.648835551 | 1 |
| ENSMUSG00000031862 | Atp13a1 | 0.00016609 | -0.995020638 | 0.618995574 |
| ENSMUSG00000036246 | Gmip | 0.00016609 | -2.470105521 | 0.00753629 |
| ENSMUSG00000071078 | Nr2c2ap | 0.00016609 | 0.063873051 | 1 |
| ENSMUSG00000003573 | Homer3 | 0.00016609 | -0.708716453 | 0.775410248 |
| ENSMUSG00000019139 | Isyna1 | 0.00016609 | 0.948395833 | 0.933335341 |
| ENSMUSG00000002396 | Ocel1 | 0.00016609 | 1.648835551 | 0.499921258 |
| ENSMUSG00000046718 | Bst2 | 0.00016609 | 0.326907457 | 1 |
| ENSMUSG00000087502 | Gm16091 | 0.00016609 | 0.004979362 | 1 |
| ENSMUSG00000055148 | Klf2 | 0.00016609 | -0.305360759 | 1 |
| ENSMUSG00000048148 | Nwd1 | 0.00016609 | 0.648835551 | 0.997931184 |
| ENSMUSG00000035671 | Zswim4 | 0.00016609 | -1.305360759 | 0.407870155 |

| | | | | |
|---|---|---|---|---|
| ENSMUSG00000003808 | Farsa | 0.00016609 | -2.589569188 | 0.004083284 |
| ENSMUSG00000053693 | Mast1 | 0.00016609 | 0.948395833 | 0.934418125 |
| ENSMUSG00000031697 | Orc6 | 0.00016609 | -0.936126949 | 0.606655093 |
| ENSMUSG00000031698 | Mylk3 | 0.00016609 | 0.56137271 | 1 |
| ENSMUSG00000069971 | 4933402J07Rik | 0.00016609 | 1.326907457 | 0.642722979 |
| ENSMUSG00000110534 | Gm45708 | 0.00016609 | 2.648835551 | 0.27550706 |
| ENSMUSG00000110559 | Gm26843 | 0.00016609 | 1.47891055 | 0.57597659 |
| ENSMUSG00000040631 | Dok4 | 0.00016609 | 0.189403933 | 1 |
| ENSMUSG00000063605 | Ccdc102a | 0.00016609 | -0.560617814 | 0.99942319 |
| ENSMUSG00000060470 | Adgrg3 | 0.00016609 | 1.841480629 | 0.455131546 |
| ENSMUSG00000031790 | Mmp15 | 0.00016609 | 0.741944956 | 0.970970089 |
| ENSMUSG00000110623 | Gm32005 | 0.00016609 | 1.326907457 | 0.642722979 |
| ENSMUSG00000090116 | Gm15680 | 0.00016609 | 1.063873051 | 0.886203497 |
| ENSMUSG00000031872 | Bean1 | 0.00016609 | 0.189403933 | 1 |
| ENSMUSG00000054400 | Cklf | 0.00016609 | -0.43862729 | 1 |
| ENSMUSG00000052616 | Terb1 | 0.00016609 | -0.874726405 | 0.680536135 |
| ENSMUSG00000023737 | 1700082M22Rik | 0.00016609 | 2.326907457 | 0.327521427 |
| ENSMUSG00000014859 | E2f4 | 0.00016609 | -1.023589791 | 0.524144449 |
| ENSMUSG00000014778 | Fhod1 | 0.00016609 | -2.395558568 | 0.011305389 |
| ENSMUSG00000014786 | Slc9a5 | 0.00016609 | -0.480447466 | 0.972350276 |
| ENSMUSG00000036442 | Thap11 | 0.00016609 | 1.648835551 | 0.499921258 |
| ENSMUSG00000031728 | Zfp821 | 0.00016609 | -1.459688905 | 0.229335266 |
| ENSMUSG00000031953 | Tmem170 | 0.00016609 | -0.051604167 | 1 |
| ENSMUSG00000034416 | Pkd1l2 | 0.00016609 | -0.43862729 | 1 |
| ENSMUSG00000110494 | Gm31774 | 0.00016609 | -0.9057533 | 0.736456705 |
| ENSMUSG00000031843 | Mphosph6 | 0.00016609 | -0.995020638 | 0.624094312 |
| ENSMUSG00000034112 | Atp2c2 | 0.00016609 | -0.158519371 | 1 |
| ENSMUSG00000031825 | Crispld2 | 0.00016609 | 0.189403933 | 1 |
| ENSMUSG00000031819 | Emc8 | 0.00016609 | -0.936126949 | 0.606655093 |
| ENSMUSG00000040010 | Slc7a5 | 0.00016609 | -0.673092543 | 0.872990655 |
| ENSMUSG00000025317 | Car5a | 0.00016609 | 0.400908038 | 1 |
| ENSMUSG00000109957 | Gm45353 | 0.00016609 | 1.648835551 | 0.499921258 |
| ENSMUSG00000015016 | Acsf3 | 0.00016609 | -0.258055044 | 1 |
| ENSMUSG00000099881 | 2810013P06Rik | 0.00016609 | -0.395558568 | 1 |
| ENSMUSG00000110537 | Gm4316 | 0.00016609 | 0.948395833 | 0.926955707 |
| ENSMUSG00000048478 | Spata33 | 0.00016609 | 2.063873051 | 0.37936848 |
| ENSMUSG00000040220 | Gas8 | 0.00016609 | -1.281901786 | 0.365716212 |
| ENSMUSG00000050751 | Pgbd5 | 0.00016609 | 2.063873051 | 0.37936848 |
| ENSMUSG00000074030 | Exoc8 | 0.00016609 | 0.841480629 | 0.943363727 |
| ENSMUSG00000090290 | Tarbp1 | 0.00016609 | -1.184054463 | 0.419360593 |
| ENSMUSG00000110626 | Gm45805 | 0.00016609 | 4.648835551 | 0.031809473 |
| ENSMUSG00000110498 | A630001O12Rik | 0.00016609 | 1.189403933 | 0.874796537 |
| ENSMUSG00000042360 | 4930433N12Rik | 0.00016609 | 0.741944956 | 0.970970089 |
| ENSMUSG00000025894 | Aasdhppt | 0.00016609 | -0.158519371 | 1 |
| ENSMUSG00000050578 | Mmp13 | 0.00016609 | 0.948395833 | 0.921679287 |
| ENSMUSG00000111602 | Gm32014 | 0.00016609 | 1.063873051 | 0.893759439 |
| ENSMUSG00000079083 | Jrkl | 0.00016609 | -0.777429203 | 0.785942057 |
| ENSMUSG00000031931 | Ankrd49 | 0.00016609 | -1.708716453 | 0.128972787 |
| ENSMUSG00000031928 | Mre11a | 0.00016609 | -1.810596067 | 0.09684834 |
| ENSMUSG00000031934 | Panx1 | 0.00016609 | 0.189403933 | 1 |
| ENSMUSG00000001774 | Chordc1 | 0.00016609 | -1.654945197 | 0.1477368 |

| | | | | |
|---|---|---|---|---|
| ENSMUSG00000004100 | Ppan | 0.00016609 | -0.673092543 | 0.872990655 |
| ENSMUSG00000004099 | Dnmt1 | 0.00016609 | -1.82689788 | 0.100320414 |
| ENSMUSG00000003299 | Mrpl4 | 0.00016609 | 0.741944956 | 0.970970089 |
| ENSMUSG00000003309 | Ap1m2 | 0.00016609 | 0.125273595 | 1 |
| ENSMUSG00000002825 | Qtrt1 | 0.00016609 | -1.051604167 | 0.529126167 |
| ENSMUSG00000032185 | Carm1 | 0.00016609 | -1.673092543 | 0.153341465 |
| ENSMUSG00000040563 | Plppr2 | 0.00016609 | 0.741944956 | 0.970970089 |
| ENSMUSG00000038895 | Zfp653 | 0.00016609 | -0.209145444 | 1 |
| ENSMUSG00000096981 | Gm16845 | 0.00016609 | -0.158519371 | 1 |
| ENSMUSG00000001349 | Cnn1 | 0.00016609 | 1.063873051 | 0.74230625 |
| ENSMUSG00000057982 | Zfp809 | 0.00016609 | -1.023589791 | 0.524144449 |
| ENSMUSG00000043067 | Dpy19l1 | 0.00016609 | -0.965874293 | 0.611716824 |
| ENSMUSG00000031988 | Vps26b | 0.00016609 | -0.777429203 | 0.785942057 |
| ENSMUSG00000070315 | 4930581F22Rik | 0.00016609 | -0.965874293 | 0.611716824 |
| ENSMUSG00000062762 | Ei24 | 0.00016609 | -0.480447466 | 0.972350276 |
| ENSMUSG00000111593 | Gm48742 | 0.00016609 | 0.004979362 | 1 |
| ENSMUSG00000111511 | AC160966.1 | 0.00016609 | 2.326907457 | 0.341869836 |
| ENSMUSG00000032012 | Nectin1 | 0.00016609 | -0.560617814 | 0.980745941 |
| ENSMUSG00000097617 | Gm10687 | 0.00016609 | 0.47891055 | 1 |
| ENSMUSG00000032086 | Bace1 | 0.00016609 | -2.258055044 | 0.017911707 |
| ENSMUSG00000110864 | Gm31374 | 0.00016609 | 0.648835551 | 1 |
| ENSMUSG00000109844 | Gm31816 | 0.00016609 | -0.106051951 | 1 |
| ENSMUSG00000032264 | Zw10 | 0.00016609 | -0.480447466 | 0.972350276 |
| ENSMUSG00000032068 | Plet1 | 0.00016609 | 2.326907457 | 0.336336902 |
| ENSMUSG00000032060 | Cryab | 0.00016609 | 0.648835551 | 1 |
| ENSMUSG00000060594 | Layn | 0.00016609 | -0.480447466 | 0.972350276 |
| ENSMUSG00000005131 | 4930550C14Rik | 0.00016609 | 0.648835551 | 1 |
| ENSMUSG00000033054 | Npat | 0.00016609 | -1.980521069 | 0.053865131 |
| ENSMUSG00000074345 | Tnfaip8l3 | 0.00016609 | 1.326907457 | 0.66416768 |
| ENSMUSG00000032274 | Cyp19a1 | 0.00016609 | 0.841480629 | 0.953305458 |
| ENSMUSG00000032295 | Man2c1 | 0.00016609 | -0.395558568 | 1 |
| ENSMUSG00000032305 | Fam219b | 0.00016609 | 0.841480629 | 0.943363727 |
| ENSMUSG00000037206 | Islr | 0.00016609 | 1.841480629 | 0.455131546 |
| ENSMUSG00000032338 | Hcn4 | 0.00016609 | 3.648835551 | 0.082076736 |
| ENSMUSG00000049526 | Tmem202 | 0.00016609 | 1.648835551 | 0.521275193 |
| ENSMUSG00000110803 | Gm20275 | 0.00016609 | -0.708716453 | 0.886203497 |
| ENSMUSG00000041729 | Coro2b | 0.00016609 | -1.051604167 | 0.580988658 |
| ENSMUSG00000037257 | Aagab | 0.00016609 | -1.079084903 | 0.536465189 |
| ENSMUSG00000032401 | Lctl | 0.00016609 | 1.47891055 | 0.587484274 |
| ENSMUSG00000035337 | Uchl4 | 0.00016609 | 3.648835551 | 0.126162788 |
| ENSMUSG00000034452 | Slc24a1 | 0.00016609 | 2.063873051 | 0.37936848 |
| ENSMUSG00000041837 | Pdcd7 | 0.00016609 | -1.43862729 | 0.27459352 |
| ENSMUSG00000111818 | Gm17749 | 0.00016609 | 2.648835551 | 0.264734265 |
| ENSMUSG00000050721 | Plekho2 | 0.00016609 | -0.636566667 | 0.868661374 |
| ENSMUSG00000040204 | Pclaf | 0.00016609 | -0.258055044 | 1 |
| ENSMUSG00000111556 | Gm19299 | 0.00016609 | 0.741944956 | 0.971854959 |
| ENSMUSG00000090626 | Tex9 | 0.00016609 | -0.599091962 | 1 |
| ENSMUSG00000034910 | Pygo1 | 0.00016609 | 1.841480629 | 0.455131546 |
| ENSMUSG00000098332 | 2310009A05Rik | 0.00016609 | 1.841480629 | 0.455131546 |
| ENSMUSG00000110824 | Gm48677 | 0.00016609 | 3.063873051 | 0.144078128 |
| ENSMUSG00000032355 | Mlip | 0.00016609 | 0.004979362 | 1 |

| | | | | |
|---|---|---|---|---|
| ENSMUSG00000099718 | Gm2087 | 0.00016609 | 2.063873051 | 0.399109893 |
| ENSMUSG00000038379 | Ttk | 0.00016609 | 0.004979362 | 1 |
| ENSMUSG00000042761 | Mrap2 | 0.00016609 | 2.648835551 | 0.218817775 |
| ENSMUSG00000032363 | Adamts7 | 0.00016609 | 0.47891055 | 1 |
| ENSMUSG00000015354 | Pcolce2 | 0.00016609 | -0.560617814 | 0.980745941 |
| ENSMUSG00000101581 | C430002N11Rik | 0.00016609 | -1.106051951 | 0.466464457 |
| ENSMUSG00000032459 | Mrps22 | 0.00016609 | 0.741944956 | 0.970970089 |
| ENSMUSG00000102013 | E330023G01Rik | 0.00016609 | 0.004979362 | 1 |
| ENSMUSG00000032463 | Faim | 0.00016609 | -0.106051951 | 1 |
| ENSMUSG00000032470 | Mras | 0.00016609 | -0.9057533 | 0.687112677 |
| ENSMUSG00000032468 | Armc8 | 0.00016609 | -2.281901786 | 0.015417072 |
| ENSMUSG00000032469 | Dbr1 | 0.00016609 | 0.063873051 | 1 |
| ENSMUSG00000035606 | Ky | 0.00016609 | 0.400908038 | 1 |
| ENSMUSG00000033688 | 1300017J02Rik | 0.00016609 | 0.256518129 | 1 |
| ENSMUSG00000032555 | Topbp1 | 0.00016609 | -2.866864287 | 0.001269736 |
| ENSMUSG00000032557 | Uba5 | 0.00016609 | -0.708716453 | 0.775410248 |
| ENSMUSG00000090150 | Acad11 | 0.00016609 | 0.326907457 | 1 |
| ENSMUSG00000053716 | Dusp7 | 0.00016609 | 0.326907457 | 1 |
| ENSMUSG00000042210 | Abhd14a | 0.00016609 | -0.209145444 | 1 |
| ENSMUSG00000041506 | Rrp9 | 0.00016609 | 0.004979362 | 1 |
| ENSMUSG00000010045 | Tmem115 | 0.00016609 | 2.326907457 | 0.295355925 |
| ENSMUSG00000079334 | Naa80 | 0.00016609 | -0.599091962 | 0.865542683 |
| ENSMUSG00000032583 | Mon1a | 0.00016609 | -1.459688905 | 0.229335266 |
| ENSMUSG00000052911 | Lamb2 | 0.00016609 | -1.858959089 | 0.084357602 |
| ENSMUSG00000032602 | Slc25a20 | 0.00016609 | -1.43862729 | 0.27459352 |
| ENSMUSG00000104614 | Fbxw27 | 0.00016609 | 2.648835551 | 0.218817775 |
| ENSMUSG00000032492 | Pth1r | 0.00016609 | 1.189403933 | 0.684924577 |
| ENSMUSG00000062296 | Trank1 | 0.00016609 | 1.47891055 | 0.592845519 |
| ENSMUSG00000032507 | Fbxl2 | 0.00016609 | -0.560617814 | 0.980745941 |
| ENSMUSG00000061393 | Acvr2b | 0.00016609 | -1.843017545 | 0.082214652 |
| ENSMUSG00000042787 | Exog | 0.00016609 | -0.305360759 | 1 |
| ENSMUSG00000032526 | Ss18l2 | 0.00016609 | 2.326907457 | 0.295355925 |
| ENSMUSG00000013419 | Zfp651 | 0.00016609 | -0.743481871 | 0.778722129 |
| ENSMUSG00000025241 | Fyco1 | 0.00016609 | -1.258055044 | 0.361093181 |
| ENSMUSG00000064065 | Ipcef1 | 0.00016609 | -0.305360759 | 1 |
| ENSMUSG00000079685 | Ulbp1 | 0.00016609 | -1.708716453 | 0.128972787 |
| ENSMUSG00000040006 | Ginm1 | 0.00016609 | 0.400908038 | 1 |
| ENSMUSG00000097007 | 4930432B10Rik | 0.00016609 | 2.326907457 | 0.295355925 |
| ENSMUSG00000039232 | Stx11 | 0.00016609 | -1.233807498 | 0.430332878 |
| ENSMUSG00000039910 | Cited2 | 0.00016609 | -1.459688905 | 0.319014843 |
| ENSMUSG00000019854 | Reps1 | 0.00016609 | -2.43862729 | 0.008383243 |
| ENSMUSG00000053219 | Raet1e | 0.00016609 | 0.256518129 | 1 |
| ENSMUSG00000112868 | 4930520K02Rik | 0.00016609 | 0.326907457 | 1 |
| ENSMUSG00000019984 | Med23 | 0.00016609 | -1.673092543 | 0.151721823 |
| ENSMUSG00000101537 | Gm29571 | 0.00016609 | 1.47891055 | 0.590105758 |
| ENSMUSG00000051354 | Samd3 | 0.00016609 | 1.326907457 | 0.662009956 |
| ENSMUSG00000112206 | Gm30228 | 0.00016609 | 1.47891055 | 0.57597659 |
| ENSMUSG00000110831 | Gm48159 | 0.00016609 | 1.063873051 | 0.916969874 |
| ENSMUSG00000000296 | Tpd52l1 | 0.00016609 | -1.89032326 | 0.256785562 |
| ENSMUSG00000019842 | Traf3ip2 | 0.00016609 | -0.743481871 | 0.778722129 |
| ENSMUSG00000038214 | Bend3 | 0.00016609 | 0.004979362 | 1 |

| | | | | |
|---|---|---|---|---|
| ENSMUSG00000110893 | Gm47392 | 0.00016609 | 0.56137271 | 1 |
| ENSMUSG00000071317 | Bves | 0.00016609 | 1.189403933 | 0.691084416 |
| ENSMUSG00000112659 | Gm40646 | 0.00016609 | 0.648835551 | 1 |
| ENSMUSG00000038594 | Cep85l | 0.00016609 | -1.106051951 | 0.466464457 |
| ENSMUSG00000019856 | Fam184a | 0.00016609 | -0.305360759 | 1 |
| ENSMUSG00000019878 | Hsf2 | 0.00016609 | -0.743481871 | 0.778722129 |
| ENSMUSG00000098188 | Sowahc | 0.00016609 | 0.256518129 | 1 |
| ENSMUSG00000020101 | Vsir | 0.00016609 | -0.480447466 | 0.996647518 |
| ENSMUSG00000020099 | Unc5b | 0.00016609 | -2.760555385 | 0.002006197 |
| ENSMUSG00000020090 | Npffr1 | 0.00016609 | -0.395558568 | 1 |
| ENSMUSG00000020085 | Aifm2 | 0.00016609 | -0.636566667 | 0.886203497 |
| ENSMUSG00000020083 | Fam241b | 0.00016609 | 2.063873051 | 0.37936848 |
| ENSMUSG00000085347 | Hk1os | 0.00016609 | 0.063873051 | 1 |
| ENSMUSG00000020067 | Mypn | 0.00016609 | -0.051604167 | 1 |
| ENSMUSG00000112425 | Gm47896 | 0.00016609 | 1.063873051 | 0.909693468 |
| ENSMUSG00000037868 | Egr2 | 0.00016609 | 3.648835551 | 0.082076736 |
| ENSMUSG00000116971 | Gm49766 | 0.00016609 | 2.063873051 | 0.37936848 |
| ENSMUSG00000019942 | Cdk1 | 0.00016609 | 0.189403933 | 1 |
| ENSMUSG00000043259 | Fam13c | 0.00016609 | -0.636566667 | 0.868661374 |
| ENSMUSG00000112414 | Gm34609 | 0.00016609 | 1.189403933 | 0.874796537 |
| ENSMUSG00000111994 | A330049N07Rik | 0.00016609 | 1.326907457 | 0.661861928 |
| ENSMUSG00000020230 | Prmt2 | 0.00016609 | -0.936126949 | 0.792441218 |
| ENSMUSG00000020241 | Col6a2 | 0.00016609 | -0.560617814 | 0.990994686 |
| ENSMUSG00000032977 | Fam207a | 0.00016609 | -0.708716453 | 0.775410248 |
| ENSMUSG00000112433 | Gm30122 | 0.00016609 | 3.063873051 | 0.144078128 |
| ENSMUSG00000019564 | Arid3a | 0.00016609 | -1.305360759 | 0.371776814 |
| ENSMUSG00000035595 | 1600002K03Rik | 0.00016609 | 1.648835551 | 0.499921258 |
| ENSMUSG00000035397 | Klf16 | 0.00016609 | 2.063873051 | 0.41958714 |
| ENSMUSG00000020211 | Sf3a2 | 0.00016609 | -0.158519371 | 1 |
| ENSMUSG00000004929 | Thop1 | 0.00016609 | 0.648835551 | 1 |
| ENSMUSG00000004931 | Apba3 | 0.00016609 | -0.051604167 | 1 |
| ENSMUSG00000034902 | Pip5k1c | 0.00016609 | -0.673092543 | 0.872990655 |
| ENSMUSG00000034889 | Cactin | 0.00016609 | -0.777429203 | 0.785942057 |
| ENSMUSG00000020256 | Aldh1l2 | 0.00016609 | -1.52108945 | 0.35608845 |
| ENSMUSG00000035620 | Ric8b | 0.00016609 | -1.654945197 | 0.157469272 |
| ENSMUSG00000069539 | Scyl2 | 0.00016609 | -1.743481871 | 0.109343152 |
| ENSMUSG00000019988 | Nedd1 | 0.00016609 | -1.43862729 | 0.27459352 |
| ENSMUSG00000015890 | Amdhd1 | 0.00016609 | 1.648835551 | 0.499921258 |
| ENSMUSG00000020020 | Usp44 | 0.00016609 | 3.063873051 | 0.254892207 |
| ENSMUSG00000112354 | Gm33843 | 0.00016609 | 0.56137271 | 1 |
| ENSMUSG00000111962 | Gm20110 | 0.00016609 | 1.648835551 | 0.499921258 |
| ENSMUSG00000112104 | Gm35722 | 0.00016609 | 1.47891055 | 0.618995574 |
| ENSMUSG00000036009 | Mettl25 | 0.00016609 | -0.843017545 | 0.674297981 |
| ENSMUSG00000019906 | Lin7a | 0.00016609 | -0.874726405 | 0.73705235 |
| ENSMUSG00000112910 | Gm29685 | 0.00016609 | 1.326907457 | 0.654432257 |
| ENSMUSG00000112398 | 9230102K24Rik | 0.00016609 | 1.189403933 | 0.690401213 |
| ENSMUSG00000035681 | Kcnc2 | 0.00016609 | 0.400908038 | 1 |
| ENSMUSG00000112513 | 4930422I22Rik | 0.00016609 | -1.617950989 | 0.175890234 |
| ENSMUSG00000034057 | Myrfl | 0.00016609 | 0.56137271 | 1 |
| ENSMUSG00000020183 | Cpm | 0.00016609 | 0.189403933 | 1 |
| ENSMUSG00000060181 | Slc35e3 | 0.00016609 | -0.636566667 | 0.868661374 |

| | | | | |
|---|---|---|---|---|
| ENSMUSG00000112155 | 4930477N07Rik | 0.00016609 | 3.063873051 | 0.144078128 |
| ENSMUSG00000006736 | Tspan31 | 0.00016609 | -0.258055044 | 1 |
| ENSMUSG00000040415 | Dtx3 | 0.00016609 | -0.351164449 | 1 |
| ENSMUSG00000040354 | Mars | 0.00016609 | 0.004979362 | 1 |
| ENSMUSG00000025402 | Nab2 | 0.00016609 | 0.326907457 | 1 |
| ENSMUSG00000040195 | Nemp1 | 0.00016609 | -1.106051951 | 0.466464457 |
| ENSMUSG00000014498 | Ankrd52 | 0.00016609 | -1.305360759 | 0.371776814 |
| ENSMUSG00000025366 | Esyt1 | 0.00016609 | -0.673092543 | 0.872990655 |
| ENSMUSG00000002578 | Ikzf4 | 0.00016609 | -1.132524162 | 0.466464457 |
| ENSMUSG00000025358 | Cdk2 | 0.00016609 | -1.258055044 | 0.361093181 |
| ENSMUSG00000053263 | Gm12592 | 0.00016609 | 0.400908038 | 1 |
| ENSMUSG00000020435 | Osbp2 | 0.00016609 | 0.189403933 | 1 |
| ENSMUSG00000048807 | Slc35e4 | 0.00016609 | 0.189403933 | 1 |
| ENSMUSG00000034412 | Tbc1d10a | 0.00016609 | 0.004979362 | 1 |
| ENSMUSG00000020482 | Ccdc117 | 0.00016609 | -0.305360759 | 1 |
| ENSMUSG00000057897 | Camk2b | 0.00016609 | -0.258055044 | 1 |
| ENSMUSG00000041164 | Zmiz2 | 0.00016609 | -1.708716453 | 0.128972787 |
| ENSMUSG00000000384 | Tbrg4 | 0.00016609 | -1.281901786 | 0.365716212 |
| ENSMUSG00000020407 | Upp1 | 0.00016609 | 1.063873051 | 0.886203497 |
| ENSMUSG00000078970 | Wdr92 | 0.00016609 | 0.948395833 | 0.937074365 |
| ENSMUSG00000032878 | Ccdc85a | 0.00016609 | -1.051604167 | 0.548358353 |
| ENSMUSG00000020464 | Pnpt1 | 0.00016609 | -0.52108945 | 0.975624465 |
| ENSMUSG00000020303 | Stc2 | 0.00016609 | 1.189403933 | 0.874796537 |
| ENSMUSG00000050541 | Adra1b | 0.00016609 | -0.777429203 | 0.821051673 |
| ENSMUSG00000087172 | Gm12153 | 0.00016609 | 0.47891055 | 1 |
| ENSMUSG00000046311 | Zfp62 | 0.00016609 | -2.184054463 | 0.022940117 |
| ENSMUSG00000020375 | Rufy1 | 0.00016609 | -1.540989007 | 0.202907954 |
| ENSMUSG00000007777 | 0610009B22Rik | 0.00016609 | 0.948395833 | 0.951890397 |
| ENSMUSG00000020387 | Jade2 | 0.00016609 | -2.500911568 | 0.006838229 |
| ENSMUSG00000018387 | Shroom1 | 0.00016609 | -1.233807498 | 0.430332878 |
| ENSMUSG00000018899 | Irf1 | 0.00016609 | -1.051604167 | 0.529126167 |
| ENSMUSG00000020522 | Mfap3 | 0.00016609 | 0.189403933 | 1 |
| ENSMUSG00000020520 | Galnt10 | 0.00016609 | -3.009375931 | 0.000558159 |
| ENSMUSG00000020514 | Mrpl22 | 0.00016609 | 0.189403933 | 1 |
| ENSMUSG00000020455 | Trim11 | 0.00016609 | -0.599091962 | 0.865542683 |
| ENSMUSG00000032633 | Flcn | 0.00016609 | -1.351164449 | 0.317734263 |
| ENSMUSG00000020538 | Srebf1 | 0.00016609 | -1.459688905 | 0.229335266 |
| ENSMUSG00000056598 | Drc3 | 0.00016609 | -0.106051951 | 1 |
| ENSMUSG00000020536 | Llgl1 | 0.00016609 | -0.708716453 | 0.775410248 |
| ENSMUSG00000020534 | Shmt1 | 0.00016609 | 0.47891055 | 1 |
| ENSMUSG00000001034 | Mapk7 | 0.00016609 | -0.599091962 | 0.865542683 |
| ENSMUSG00000042377 | Fam83g | 0.00016609 | 0.063873051 | 1 |
| ENSMUSG00000047759 | Hs3st3a1 | 0.00016609 | 1.47891055 | 0.590703035 |
| ENSMUSG00000020542 | Myocd | 0.00016609 | -2.258055044 | 0.02307354 |
| ENSMUSG00000020902 | Ntn1 | 0.00016609 | -0.351164449 | 1 |
| ENSMUSG00000032892 | Rangrf | 0.00016609 | 1.47891055 | 0.57597659 |
| ENSMUSG00000020899 | Pfas | 0.00016609 | -0.051604167 | 1 |
| ENSMUSG00000049299 | Trappc1 | 0.00016609 | 0.256518129 | 1 |
| ENSMUSG00000097328 | Tnfsf12 | 0.00016609 | 0.841480629 | 0.943363727 |
| ENSMUSG00000023170 | Gps2 | 0.00016609 | -2.785792676 | 0.001866828 |
| ENSMUSG00000018574 | Acadvl | 0.00016609 | 0.47891055 | 1 |

| | | | | |
|---|---|---|---|---|
| ENSMUSG00000020831 | 0610010K14Rik | 0.00016609 | 1.841480629 | 0.466464457 |
| ENSMUSG00000040746 | Rnf167 | 0.00016609 | -0.874726405 | 0.705067264 |
| ENSMUSG00000057054 | Inca1 | 0.00016609 | -0.158519371 | 1 |
| ENSMUSG00000070390 | Nlrp1b | 0.00016609 | -1.184054463 | 0.51115569 |
| ENSMUSG00000085148 | Mir22hg | 0.00016609 | -0.9057533 | 0.687112677 |
| ENSMUSG00000006127 | Inpp5k | 0.00016609 | -2.037664976 | 0.038867277 |
| ENSMUSG00000069808 | Fam57a | 0.00016609 | -1.373532262 | 0.317734263 |
| ENSMUSG00000010392 | Gosr1 | 0.00016609 | -0.395558568 | 1 |
| ENSMUSG00000000632 | Sez6 | 0.00016609 | 1.648835551 | 0.523151006 |
| ENSMUSG00000002058 | Unc119 | 0.00016609 | 1.063873051 | 0.89270014 |
| ENSMUSG00000017639 | Rab11fip4 | 0.00016609 | -0.395558568 | 1 |
| ENSMUSG00000020707 | Rnf135 | 0.00016609 | -1.051604167 | 0.529126167 |
| ENSMUSG00000017692 | Rhbdl3 | 0.00016609 | -0.874726405 | 0.680536135 |
| ENSMUSG00000020697 | Lig3 | 0.00016609 | -0.599091962 | 0.865542683 |
| ENSMUSG00000000982 | Ccl3 | 0.00016609 | 1.841480629 | 0.455131546 |
| ENSMUSG00000018930 | Ccl4 | 0.00016609 | 2.648835551 | 0.218817775 |
| ENSMUSG00000000983 | Wfdc18 | 0.00016609 | 3.063873051 | 0.268335928 |
| ENSMUSG00000051748 | Wfdc21 | 0.00016609 | 4.648835551 | 0.031809473 |
| ENSMUSG00000020527 | Myo19 | 0.00016609 | -0.743481871 | 0.778722129 |
| ENSMUSG00000018068 | Ints2 | 0.00016609 | -0.810596067 | 0.791730654 |
| ENSMUSG00000020513 | Tubd1 | 0.00016609 | -1.132524162 | 0.506177813 |
| ENSMUSG00000000276 | Dgke | 0.00016609 | -1.132524162 | 0.476457341 |
| ENSMUSG00000037573 | Tob1 | 0.00016609 | -0.743481871 | 0.783218548 |
| ENSMUSG00000020866 | Cacna1g | 0.00016609 | 0.948395833 | 0.918572342 |
| ENSMUSG00000024414 | Mrpl27 | 0.00016609 | 0.47891055 | 1 |
| ENSMUSG00000020868 | Xylt2 | 0.00016609 | 0.948395833 | 0.912904617 |
| ENSMUSG00000000120 | Ngfr | 0.00016609 | 2.648835551 | 0.257440232 |
| ENSMUSG00000078706 | Gm53 | 0.00016609 | 3.063873051 | 0.144078128 |
| ENSMUSG00000070336 | Fbxo47 | 0.00016609 | -1.158519371 | 0.466464457 |
| ENSMUSG00000055937 | Krt28 | 0.00016609 | -0.777429203 | 0.804682567 |
| ENSMUSG00000006784 | Ttc25 | 0.00016609 | 1.47891055 | 0.57597659 |
| ENSMUSG00000085604 | Dhx58os | 0.00016609 | 1.648835551 | 0.499921258 |
| ENSMUSG00000001755 | Coasy | 0.00016609 | -0.777429203 | 0.785942057 |
| ENSMUSG00000075528 | Aarsd1 | 0.00016609 | 0.189403933 | 1 |
| ENSMUSG00000051378 | Kif18b | 0.00016609 | 0.400908038 | 1 |
| ENSMUSG00000020935 | Dcakd | 0.00016609 | -1.209145444 | 0.423814296 |
| ENSMUSG00000055805 | Fmnl1 | 0.00016609 | -1.579983139 | 0.169119671 |
| ENSMUSG00000078632 | Lrrc37a | 0.00016609 | 1.189403933 | 0.687112677 |
| ENSMUSG00000061086 | Myl4 | 0.00016609 | 1.841480629 | 0.455131546 |
| ENSMUSG00000099681 | 1700052K11Rik | 0.00016609 | 1.063873051 | 0.886203497 |
| ENSMUSG00000040528 | Milr1 | 0.00016609 | 1.326907457 | 0.66416768 |
| ENSMUSG00000000142 | Axin2 | 0.00016609 | -0.636566667 | 0.868661374 |
| ENSMUSG00000041828 | Abca8a | 0.00016609 | 1.648835551 | 0.5249062 |
| ENSMUSG00000041674 | BC006965 | 0.00016609 | -0.599091962 | 0.878725021 |
| ENSMUSG00000034706 | Dnaic2 | 0.00016609 | 1.648835551 | 0.517601878 |
| ENSMUSG00000034586 | Hid1 | 0.00016609 | -0.560617814 | 0.980745941 |
| ENSMUSG00000057219 | Armc7 | 0.00016609 | -0.874726405 | 0.680536135 |
| ENSMUSG00000020743 | Mif4gd | 0.00016609 | 0.004979362 | 1 |
| ENSMUSG00000020744 | Slc25a19 | 0.00016609 | -0.560617814 | 0.980745941 |
| ENSMUSG00000020766 | Galk1 | 0.00016609 | 0.400908038 | 1 |
| ENSMUSG00000020776 | Fbf1 | 0.00016609 | -1.636566667 | 0.144776283 |

| | | | | |
|---|---|---|---|---|
| ENSMUSG00000034282 | Evpl | 0.00016609 | -1.258055044 | 0.361093181 |
| ENSMUSG00000020806 | Rhbdf2 | 0.00016609 | -1.233807498 | 0.430332878 |
| ENSMUSG00000020814 | Mxra7 | 0.00016609 | -0.743481871 | 0.783356215 |
| ENSMUSG00000090266 | Mettl23 | 0.00016609 | 1.47891055 | 0.57597659 |
| ENSMUSG00000025572 | Tmc6 | 0.00016609 | -1.459688905 | 0.229335266 |
| ENSMUSG00000025574 | Tk1 | 0.00016609 | 0.47891055 | 1 |
| ENSMUSG00000053113 | Socs3 | 0.00016609 | -1.023589791 | 0.524144449 |
| ENSMUSG00000039976 | Tbc1d16 | 0.00016609 | -0.52108945 | 1 |
| ENSMUSG00000025579 | Gaa | 0.00016609 | 0.648835551 | 1 |
| ENSMUSG00000085081 | Rptoros | 0.00016609 | 3.648835551 | 0.082076736 |
| ENSMUSG00000075389 | 2810410L24Rik | 0.00016609 | 0.326907457 | 1 |
| ENSMUSG00000025161 | Slc16a3 | 0.00016609 | -0.52108945 | 0.975624465 |
| ENSMUSG00000000056 | Narf | 0.00016609 | -0.874726405 | 0.705975399 |
| ENSMUSG00000020657 | Dnajc27 | 0.00016609 | 0.326907457 | 1 |
| ENSMUSG00000112819 | Gm46332 | 0.00016609 | 2.063873051 | 0.466464457 |
| ENSMUSG00000112276 | 5033421B08Rik | 0.00016609 | 0.400908038 | 1 |
| ENSMUSG00000020600 | Slc7a15 | 0.00016609 | 0.400908038 | 1 |
| ENSMUSG00000066637 | Ttc32 | 0.00016609 | -0.305360759 | 1 |
| ENSMUSG00000113522 | Gm48479 | 0.00016609 | 1.47891055 | 0.587484274 |
| ENSMUSG00000036523 | Greb1 | 0.00016609 | -0.051604167 | 1 |
| ENSMUSG00000045679 | Pqlc3 | 0.00016609 | 2.326907457 | 0.295355925 |
| ENSMUSG00000073197 | 5730507C01Rik | 0.00016609 | 0.648835551 | 1 |
| ENSMUSG00000054309 | Cpsf3 | 0.00016609 | -1.459688905 | 0.229335266 |
| ENSMUSG00000062563 | Cys1 | 0.00016609 | -1.843017545 | 0.082214652 |
| ENSMUSG00000113621 | 2900060N12Rik | 0.00016609 | 0.004979362 | 1 |
| ENSMUSG00000113986 | Gm31333 | 0.00016609 | 0.400908038 | 1 |
| ENSMUSG00000036136 | Fam110c | 0.00016609 | -1.281901786 | 0.365716212 |
| ENSMUSG00000113741 | Gm48236 | 0.00016609 | 0.189403933 | 1 |
| ENSMUSG00000020956 | Dtd2 | 0.00016609 | 0.56137271 | 1 |
| ENSMUSG00000012076 | Brms1l | 0.00016609 | -1.500911568 | 0.23866117 |
| ENSMUSG00000112343 | Sfta3-ps | 0.00016609 | 2.648835551 | 0.218817775 |
| ENSMUSG00000001497 | Pax9 | 0.00016609 | 0.741944956 | 0.971210385 |
| ENSMUSG00000112704 | Gm46329 | 0.00016609 | 0.648835551 | 0.997248381 |
| ENSMUSG00000114112 | Gm48301 | 0.00016609 | -0.9057533 | 0.73806297 |
| ENSMUSG00000047534 | Mis18bp1 | 0.00016609 | -1.858959089 | 0.084357602 |
| ENSMUSG00000113158 | Gm31447 | 0.00016609 | 3.648835551 | 0.082076736 |
| ENSMUSG00000054894 | Dmac2l | 0.00016609 | -0.965874293 | 0.803345306 |
| ENSMUSG00000085622 | 3110056K07Rik | 0.00016609 | -2.258055044 | 0.017911707 |
| ENSMUSG00000056359 | D830013O20Rik | 0.00016609 | 1.648835551 | 0.513921295 |
| ENSMUSG00000034402 | Kcnh5 | 0.00016609 | -1.82689788 | 0.100320414 |
| ENSMUSG00000021056 | Tex21 | 0.00016609 | 1.189403933 | 0.688461306 |
| ENSMUSG00000021048 | Mthfd1 | 0.00016609 | -1.936126949 | 0.061652631 |
| ENSMUSG00000056459 | Zbtb25 | 0.00016609 | -1.560617814 | 0.20542113 |
| ENSMUSG00000048833 | Slc39a9 | 0.00016609 | -1.351164449 | 0.312100107 |
| ENSMUSG00000090843 | Heatr4 | 0.00016609 | 0.56137271 | 1 |
| ENSMUSG00000042320 | Prox2 | 0.00016609 | 0.948395833 | 0.912904617 |
| ENSMUSG00000004791 | Pgf | 0.00016609 | 0.648835551 | 1 |
| ENSMUSG00000034258 | Flvcr2 | 0.00016609 | 0.841480629 | 0.943363727 |
| ENSMUSG00000021256 | Vash1 | 0.00016609 | 1.063873051 | 0.886203497 |
| ENSMUSG00000034168 | Irf2bpl | 0.00016609 | -1.395558568 | 0.26547175 |
| ENSMUSG00000034157 | Cipc | 0.00016609 | -0.874726405 | 0.680536135 |

| | | | | |
|---|---|---|---|---|
| ENSMUSG00000034111 | Tmed8 | 0.00016609 | 0.004979362 | 1 |
| ENSMUSG00000113001 | Gm40538 | 0.00016609 | 2.648835551 | 0.218817775 |
| ENSMUSG00000097917 | Gm26839 | 0.00016609 | 2.063873051 | 0.37936848 |
| ENSMUSG00000021182 | Ccdc88c | 0.00016609 | -3.917218487 | 1.52E-06 |
| ENSMUSG00000046675 | Tmem251 | 0.00016609 | 1.648835551 | 0.50472994 |
| ENSMUSG00000021198 | Unc79 | 0.00016609 | -0.351164449 | 1 |
| ENSMUSG00000064215 | Ifi27 | 0.00016609 | -0.810596067 | 0.791730654 |
| ENSMUSG00000041415 | Dicer1 | 0.00016609 | -3.264053785 | 0.000110956 |
| ENSMUSG00000113063 | Gm34667 | 0.00016609 | 0.326907457 | 1 |
| ENSMUSG00000113585 | Gm19605 | 0.00016609 | -0.258055044 | 1 |
| ENSMUSG00000021271 | Zfp839 | 0.00016609 | -2.158519371 | 0.027510407 |
| ENSMUSG00000113056 | 6030440G07Rik | 0.00016609 | 1.063873051 | 0.894433906 |
| ENSMUSG00000001270 | Ckb | 0.00016609 | -0.843017545 | 0.674297981 |
| ENSMUSG00000011148 | Adssl1 | 0.00016609 | -0.560617814 | 0.980745941 |
| ENSMUSG00000037638 | Zbtb42 | 0.00016609 | 0.741944956 | 0.970970089 |
| ENSMUSG00000037594 | Clba1 | 0.00016609 | 2.063873051 | 0.37936848 |
| ENSMUSG00000114053 | Gm35330 | 0.00016609 | -0.9057533 | 0.732525536 |
| ENSMUSG00000113757 | Gm47507 | 0.00016609 | -0.936126949 | 0.606655093 |
| ENSMUSG00000113543 | Gm36264 | 0.00016609 | 0.256518129 | 1 |
| ENSMUSG00000052374 | Actn2 | 0.00016609 | 0.841480629 | 0.957843399 |
| ENSMUSG00000095105 | Edaradd | 0.00016609 | 0.648835551 | 1 |
| ENSMUSG00000021302 | Ggps1 | 0.00016609 | -0.43862729 | 1 |
| ENSMUSG00000021326 | Trim27 | 0.00016609 | -2.221529168 | 0.020322338 |
| ENSMUSG00000036721 | Zscan12 | 0.00016609 | 0.326907457 | 1 |
| ENSMUSG00000036376 | Abt1 | 0.00016609 | -0.051604167 | 1 |
| ENSMUSG00000087173 | Gm11337 | 0.00016609 | 0.189403933 | 1 |
| ENSMUSG00000006711 | D130043K22Rik | 0.00016609 | 0.648835551 | 1 |
| ENSMUSG00000113216 | Gm40841 | 0.00016609 | -0.9057533 | 0.687112677 |
| ENSMUSG00000021357 | Exoc2 | 0.00016609 | -2.38458745 | 0.010976115 |
| ENSMUSG00000044951 | Mylk4 | 0.00016609 | 1.189403933 | 0.684326407 |
| ENSMUSG00000038286 | Bphl | 0.00016609 | -1.43862729 | 0.27459352 |
| ENSMUSG00000058672 | Tubb2a | 0.00016609 | 0.841480629 | 0.949402664 |
| ENSMUSG00000021411 | Pxdc1 | 0.00016609 | -1.106051951 | 0.466464457 |
| ENSMUSG00000114784 | Gm47754 | 0.00016609 | -0.874726405 | 0.70936601 |
| ENSMUSG00000038982 | Bloc1s5 | 0.00016609 | 0.004979362 | 1 |
| ENSMUSG00000114841 | Gm47352 | 0.00016609 | -0.158519371 | 1 |
| ENSMUSG00000114662 | Gm31683 | 0.00016609 | -0.636566667 | 0.868661374 |
| ENSMUSG00000091264 | Smim13 | 0.00016609 | 0.56137271 | 1 |
| ENSMUSG00000021367 | Edn1 | 0.00016609 | 0.648835551 | 1 |
| ENSMUSG00000044164 | Rnf182 | 0.00016609 | 0.189403933 | 1 |
| ENSMUSG00000098144 | 1700029N11Rik | 0.00016609 | 3.648835551 | 0.082076736 |
| ENSMUSG00000097622 | A330033J07Rik | 0.00016609 | 1.648835551 | 0.5118121 |
| ENSMUSG00000021448 | Shc3 | 0.00016609 | 0.741944956 | 0.971854959 |
| ENSMUSG00000113834 | Gm34022 | 0.00016609 | 0.648835551 | 0.998731799 |
| ENSMUSG00000035711 | Dok3 | 0.00016609 | 0.400908038 | 1 |
| ENSMUSG00000021499 | Catsper3 | 0.00016609 | -0.158519371 | 1 |
| ENSMUSG00000097399 | Gm26555 | 0.00016609 | 1.841480629 | 0.466464457 |
| ENSMUSG00000113295 | Gm49357 | 0.00016609 | 0.56137271 | 1 |
| ENSMUSG00000114036 | Gm48228 | 0.00016609 | 0.47891055 | 1 |
| ENSMUSG00000113566 | Gm40977 | 0.00016609 | 3.648835551 | 0.110500179 |
| ENSMUSG00000113825 | 1700024I08Rik | 0.00016609 | 1.47891055 | 0.592845519 |

| | | | | |
|---|---|---|---|---|
| ENSMUSG00000033063 | Cntnap3 | 0.00016609 | -0.209145444 | 1 |
| ENSMUSG00000113618 | Gm48166 | 0.00016609 | -0.636566667 | 0.878615048 |
| ENSMUSG00000069206 | Zfp874a | 0.00016609 | 2.648835551 | 0.218817775 |
| ENSMUSG00000071291 | Zfp58 | 0.00016609 | -1.158519371 | 0.499942088 |
| ENSMUSG00000049538 | Adamts16 | 0.00016609 | 0.004979362 | 1 |
| ENSMUSG00000112964 | E430024I08Rik | 0.00016609 | -2.921020057 | 0.000937453 |
| ENSMUSG00000035840 | Lysmd3 | 0.00016609 | -0.258055044 | 1 |
| ENSMUSG00000051098 | Mblac2 | 0.00016609 | 0.56137271 | 1 |
| ENSMUSG00000050334 | C130071C03Rik | 0.00016609 | 0.400908038 | 1 |
| ENSMUSG00000042857 | Gm9776 | 0.00016609 | 0.004979362 | 1 |
| ENSMUSG00000021681 | Aggf1 | 0.00016609 | -2.051604167 | 0.039883588 |
| ENSMUSG00000021671 | Poc5 | 0.00016609 | -0.351164449 | 1 |
| ENSMUSG00000047117 | Ankdd1b | 0.00016609 | -0.743481871 | 0.778722129 |
| ENSMUSG00000021670 | Hmgcr | 0.00016609 | -1.351164449 | 0.312100107 |
| ENSMUSG00000078952 | Lncenc1 | 0.00016609 | 0.063873051 | 1 |
| ENSMUSG00000021640 | Naip1 | 0.00016609 | 1.648835551 | 0.504119351 |
| ENSMUSG00000021714 | Cenpk | 0.00016609 | 0.189403933 | 1 |
| ENSMUSG00000021719 | Rgs7bp | 0.00016609 | 0.189403933 | 1 |
| ENSMUSG00000021692 | Dimt1 | 0.00016609 | 0.400908038 | 1 |
| ENSMUSG00000021697 | Depdc1b | 0.00016609 | 0.004979362 | 1 |
| ENSMUSG00000042284 | Itga1 | 0.00016609 | -2.245982212 | 0.023955525 |
| ENSMUSG00000021732 | Fgf10 | 0.00016609 | -0.395558568 | 1 |
| ENSMUSG00000025279 | Dnase1l3 | 0.00016609 | 1.841480629 | 0.466464457 |
| ENSMUSG00000023156 | Rpp14 | 0.00016609 | -1.417253639 | 0.269874327 |
| ENSMUSG00000021748 | Pdhb | 0.00016609 | -0.874726405 | 0.734120426 |
| ENSMUSG00000097148 | Gm3839 | 0.00016609 | 2.326907457 | 0.295355925 |
| ENSMUSG00000045201 | Lrrc3b | 0.00016609 | 0.47891055 | 1 |
| ENSMUSG00000021786 | Oxsm | 0.00016609 | 1.47891055 | 0.57597659 |
| ENSMUSG00000114504 | Gm48321 | 0.00016609 | 0.648835551 | 0.9980272 |
| ENSMUSG00000068697 | Myoz1 | 0.00016609 | 2.648835551 | 0.218817775 |
| ENSMUSG00000039357 | Fut11 | 0.00016609 | 0.256518129 | 1 |
| ENSMUSG00000021819 | Zswim8 | 0.00016609 | -2.339849135 | 0.014721679 |
| ENSMUSG00000063821 | Dupd1 | 0.00016609 | 1.47891055 | 0.57597659 |
| ENSMUSG00000021962 | Dcp1a | 0.00016609 | -2.560617814 | 0.004497171 |
| ENSMUSG00000037833 | Sh2d4b | 0.00016609 | -1.158519371 | 0.469530167 |
| ENSMUSG00000115272 | Gm49965 | 0.00016609 | 1.063873051 | 0.889311043 |
| ENSMUSG00000115735 | Gm49180 | 0.00016609 | 2.063873051 | 0.394676564 |
| ENSMUSG00000053205 | Styx | 0.00016609 | -2.406446884 | 0.009429212 |
| ENSMUSG00000061510 | Gm10101 | 0.00016609 | 3.063873051 | 0.144078128 |
| ENSMUSG00000115669 | Gm49151 | 0.00016609 | -1.691014451 | 0.159544618 |
| ENSMUSG00000036291 | Ap5m1 | 0.00016609 | -0.843017545 | 0.674297981 |
| ENSMUSG00000021852 | Slc35f4 | 0.00016609 | 0.400908038 | 1 |
| ENSMUSG00000068431 | Olfr742 | 0.00016609 | 4.648835551 | 0.031809473 |
| ENSMUSG00000006288 | Ttc5 | 0.00016609 | -2.106051951 | 0.036215483 |
| ENSMUSG00000006281 | Tep1 | 0.00016609 | -2.682081327 | 0.002606179 |
| ENSMUSG00000090799 | Klhl33 | 0.00016609 | 0.741944956 | 0.979457724 |
| ENSMUSG00000035960 | Apex1 | 0.00016609 | -0.673092543 | 0.872990655 |
| ENSMUSG00000000959 | Oxa1l | 0.00016609 | -0.305360759 | 1 |
| ENSMUSG00000022178 | Ajuba | 0.00016609 | -2.636566667 | 0.003263045 |
| ENSMUSG00000022179 | 4931414P19Rik | 0.00016609 | 0.004979362 | 1 |
| ENSMUSG00000040721 | Zfhx2 | 0.00016609 | -1.936126949 | 0.061652631 |

| | | | | |
|---|---|---|---|---|
| ENSMUSG00000021965 | Ska3 | 0.00016609 | -0.305360759 | 1 |
| ENSMUSG00000071350 | Setdb2 | 0.00016609 | -1.106051951 | 0.466464457 |
| ENSMUSG00000101797 | Gm29266 | 0.00016609 | 0.256518129 | 1 |
| ENSMUSG00000084951 | Mipepos | 0.00016609 | 2.063873051 | 0.402021228 |
| ENSMUSG00000035078 | Mtmr9 | 0.00016609 | -1.209145444 | 0.423814296 |
| ENSMUSG00000061356 | Nuggc | 0.00016609 | 2.063873051 | 0.37936848 |
| ENSMUSG00000022032 | Scara5 | 0.00016609 | -1.579983139 | 0.263599559 |
| ENSMUSG00000022039 | Adam2 | 0.00016609 | -0.209145444 | 1 |
| ENSMUSG00000114697 | Gm30806 | 0.00016609 | 0.400908038 | 1 |
| ENSMUSG00000093472 | 4930438E09Rik | 0.00016609 | 0.841480629 | 0.956396131 |
| ENSMUSG00000014813 | Stc1 | 0.00016609 | 0.125273595 | 1 |
| ENSMUSG00000022074 | Tnfrsf10b | 0.00016609 | -1.051604167 | 0.534963044 |
| ENSMUSG00000114446 | Gm33524 | 0.00016609 | 0.400908038 | 1 |
| ENSMUSG00000044551 | 9930012K11Rik | 0.00016609 | 1.648835551 | 0.499921258 |
| ENSMUSG00000033644 | Piwil2 | 0.00016609 | 0.063873051 | 1 |
| ENSMUSG00000115304 | 4930434J06Rik | 0.00016609 | 0.326907457 | 1 |
| ENSMUSG00000091722 | Siah3 | 0.00016609 | 0.648835551 | 0.999122005 |
| ENSMUSG00000044350 | Lacc1 | 0.00016609 | -0.743481871 | 0.778722129 |
| ENSMUSG00000022015 | Tnfsf11 | 0.00016609 | 0.256518129 | 1 |
| ENSMUSG00000035566 | Pcdh17 | 0.00016609 | 0.47891055 | 1 |
| ENSMUSG00000115623 | Gm41225 | 0.00016609 | 1.47891055 | 0.590312322 |
| ENSMUSG00000115001 | 4930474H20Rik | 0.00016609 | -0.158519371 | 1 |
| ENSMUSG00000022070 | Bora | 0.00016609 | -1.995020638 | 0.055374754 |
| ENSMUSG00000098557 | Kctd12 | 0.00016609 | 0.948395833 | 0.912904617 |
| ENSMUSG00000115762 | Gm34907 | 0.00016609 | 0.400908038 | 1 |
| ENSMUSG00000115424 | D130079A08Rik | 0.00016609 | -0.158519371 | 1 |
| ENSMUSG00000022120 | Rnf219 | 0.00016609 | -1.079084903 | 0.536465189 |
| ENSMUSG00000022129 | Dct | 0.00016609 | 0.841480629 | 0.953305458 |
| ENSMUSG00000022131 | Gpr180 | 0.00016609 | -0.351164449 | 1 |
| ENSMUSG00000051615 | Rap2a | 0.00016609 | -0.52108945 | 0.992511838 |
| ENSMUSG00000115016 | Gm33906 | 0.00016609 | 0.004979362 | 1 |
| ENSMUSG00000022144 | Gdnf | 0.00016609 | 0.841480629 | 0.943363727 |
| ENSMUSG00000022142 | Nup155 | 0.00016609 | -2.328444372 | 0.014195545 |
| ENSMUSG00000003882 | Il7r | 0.00016609 | 0.841480629 | 0.964654104 |
| ENSMUSG00000022244 | Amacr | 0.00016609 | 1.841480629 | 0.463009681 |
| ENSMUSG00000022241 | Tars | 0.00016609 | -1.209145444 | 0.437556558 |
| ENSMUSG00000022235 | Cmbl | 0.00016609 | 0.648835551 | 1 |
| ENSMUSG00000039065 | Atpsckmt | 0.00016609 | -0.106051951 | 1 |
| ENSMUSG00000037627 | Rgs22 | 0.00016609 | -0.599091962 | 1 |
| ENSMUSG00000037617 | Spag1 | 0.00016609 | -1.209145444 | 0.423814296 |
| ENSMUSG00000037465 | Klf10 | 0.00016609 | -1.560617814 | 0.231741911 |
| ENSMUSG00000115547 | 1100001I12Rik | 0.00016609 | 1.841480629 | 0.455131546 |
| ENSMUSG00000054196 | Cthrc1 | 0.00016609 | 0.841480629 | 0.943363727 |
| ENSMUSG00000051920 | Rspo2 | 0.00016609 | -0.395558568 | 1 |
| ENSMUSG00000022337 | Emc2 | 0.00016609 | -0.599091962 | 0.865542683 |
| ENSMUSG00000038760 | Trhr | 0.00016609 | 0.125273595 | 1 |
| ENSMUSG00000038736 | Nudcd1 | 0.00016609 | -0.708716453 | 0.775410248 |
| ENSMUSG00000022340 | Sybu | 0.00016609 | 0.256518129 | 1 |
| ENSMUSG00000115654 | Gm19303 | 0.00016609 | 0.125273595 | 1 |
| ENSMUSG00000022367 | Has2 | 0.00016609 | -0.351164449 | 1 |
| ENSMUSG00000022362 | Gm29394 | 0.00016609 | 0.841480629 | 0.943363727 |

| | | | | |
|---|---|---|---|---|
| ENSMUSG00000022359 | Wdyhv1 | 0.00016609 | -1.673092543 | 0.20161331 |
| ENSMUSG00000032501 | Trib1 | 0.00016609 | -1.132524162 | 0.466464457 |
| ENSMUSG00000115862 | Gm41333 | 0.00016609 | 0.47891055 | 1 |
| ENSMUSG00000115164 | Gm30159 | 0.00016609 | 1.841480629 | 0.455131546 |
| ENSMUSG00000115369 | Gm49014 | 0.00016609 | 3.648835551 | 0.082076736 |
| ENSMUSG00000115169 | Gm30563 | 0.00016609 | 2.063873051 | 0.37936848 |
| ENSMUSG00000022335 | Zfat | 0.00016609 | -1.579983139 | 0.169119671 |
| ENSMUSG00000022332 | Khdrbs3 | 0.00016609 | 0.400908038 | 1 |
| ENSMUSG00000046761 | Fam83h | 0.00016609 | -0.258055044 | 1 |
| ENSMUSG00000063268 | Parp10 | 0.00016609 | -1.43862729 | 0.297381526 |
| ENSMUSG00000109179 | Gm35339 | 0.00016609 | -0.351164449 | 1 |
| ENSMUSG00000016637 | Ift27 | 0.00016609 | 1.648835551 | 0.517004328 |
| ENSMUSG00000022440 | C1qtnf6 | 0.00016609 | -0.305360759 | 1 |
| ENSMUSG00000068206 | Pick1 | 0.00016609 | -0.9057533 | 0.687112677 |
| ENSMUSG00000042564 | Fam227a | 0.00016609 | -1.579983139 | 0.169119671 |
| ENSMUSG00000022412 | Mief1 | 0.00016609 | -2.258055044 | 0.017911707 |
| ENSMUSG00000022416 | Cacna1i | 0.00016609 | 0.326907457 | 1 |
| ENSMUSG00000050439 | Enthd1 | 0.00016609 | -0.305360759 | 1 |
| ENSMUSG00000022408 | Fam83f | 0.00016609 | -1.233807498 | 0.430332878 |
| ENSMUSG00000115970 | 8430426J06Rik | 0.00016609 | -0.258055044 | 1 |
| ENSMUSG00000048546 | Tob2 | 0.00016609 | -2.245982212 | 0.021820859 |
| ENSMUSG00000068114 | Ccdc134 | 0.00016609 | 0.004979362 | 1 |
| ENSMUSG00000116130 | Gm49431 | 0.00016609 | 1.326907457 | 0.642722979 |
| ENSMUSG00000022442 | Ttll1 | 0.00016609 | -0.106051951 | 1 |
| ENSMUSG00000048755 | Mcat | 0.00016609 | 0.326907457 | 1 |
| ENSMUSG00000016763 | Scube1 | 0.00016609 | -1.023589791 | 0.66416768 |
| ENSMUSG00000016624 | Phf21b | 0.00016609 | 0.189403933 | 1 |
| ENSMUSG00000022432 | Smc1b | 0.00016609 | 0.741944956 | 0.971650633 |
| ENSMUSG00000115918 | A930001M01Rik | 0.00016609 | 0.948395833 | 0.938117612 |
| ENSMUSG00000116171 | 4930543I03Rik | 0.00016609 | 1.326907457 | 0.642722979 |
| ENSMUSG00000023272 | Creld2 | 0.00016609 | 0.741944956 | 0.970970089 |
| ENSMUSG00000015365 | Mov10l1 | 0.00016609 | 1.841480629 | 0.455131546 |
| ENSMUSG00000022629 | Kif21a | 0.00016609 | -1.540989007 | 0.207756141 |
| ENSMUSG00000059883 | Irak4 | 0.00016609 | -0.843017545 | 0.674297981 |
| ENSMUSG00000022993 | 4930415O20Rik | 0.00016609 | 0.741944956 | 0.970970089 |
| ENSMUSG00000037570 | Mcrs1 | 0.00016609 | 0.004979362 | 1 |
| ENSMUSG00000023019 | Gpd1 | 0.00016609 | 0.948395833 | 0.912904617 |
| ENSMUSG00000090500 | Gm17057 | 0.00016609 | 2.648835551 | 0.221090891 |
| ENSMUSG00000045350 | Fam186a | 0.00016609 | 1.326907457 | 0.642722979 |
| ENSMUSG00000023030 | Slc11a2 | 0.00016609 | 0.189403933 | 1 |
| ENSMUSG00000098112 | Bin2 | 0.00016609 | 0.741944956 | 0.971854959 |
| ENSMUSG00000023032 | Slc4a8 | 0.00016609 | 0.948395833 | 0.925937889 |
| ENSMUSG00000000530 | Acvrl1 | 0.00016609 | 0.841480629 | 0.951907087 |
| ENSMUSG00000037003 | Tns2 | 0.00016609 | -0.480447466 | 0.981266767 |
| ENSMUSG00000023046 | Igfbp6 | 0.00016609 | 3.063873051 | 0.190974831 |
| ENSMUSG00000045665 | Mfsd5 | 0.00016609 | 1.648835551 | 0.499921258 |
| ENSMUSG00000058290 | Espl1 | 0.00016609 | -0.480447466 | 0.972350276 |
| ENSMUSG00000036061 | Smug1 | 0.00016609 | -0.209145444 | 1 |
| ENSMUSG00000000552 | Zfp385a | 0.00016609 | 0.948395833 | 0.912904617 |
| ENSMUSG00000022489 | Pde1b | 0.00016609 | 1.648835551 | 0.515378764 |
| ENSMUSG00000039789 | Zfp597 | 0.00016609 | -0.673092543 | 0.872990655 |

| | | | | |
|---|---|---|---|---|
| ENSMUSG00000022519 | Srl | 0.00016609 | -1.794107944 | 0.094802366 |
| ENSMUSG00000039646 | Vasn | 0.00016609 | 0.326907457 | 1 |
| ENSMUSG00000022516 | Nudt16l1 | 0.00016609 | -0.209145444 | 1 |
| ENSMUSG00000116029 | Gm41414 | 0.00016609 | -0.636566667 | 0.875728762 |
| ENSMUSG00000022504 | Ciita | 0.00016609 | -2.1966545 | 0.023519181 |
| ENSMUSG00000055015 | Gm9961 | 0.00016609 | 2.648835551 | 0.218817775 |
| ENSMUSG00000022545 | Ercc4 | 0.00016609 | -1.500911568 | 0.23866117 |
| ENSMUSG00000079737 | 3110001I22Rik | 0.00016609 | -0.106051951 | 1 |
| ENSMUSG00000116324 | Gm33696 | 0.00016609 | 1.326907457 | 0.658243753 |
| ENSMUSG00000041774 | Ydjc | 0.00016609 | 1.063873051 | 0.886203497 |
| ENSMUSG00000005899 | Smpd4 | 0.00016609 | -1.079084903 | 0.548942955 |
| ENSMUSG00000022750 | Klhl22 | 0.00016609 | -0.636566667 | 0.885228764 |
| ENSMUSG00000003527 | Ess2 | 0.00016609 | -0.708716453 | 0.775410248 |
| ENSMUSG00000000325 | Arvcf | 0.00016609 | -0.936126949 | 0.624094312 |
| ENSMUSG00000003234 | Abcf3 | 0.00016609 | -0.810596067 | 0.791730654 |
| ENSMUSG00000004462 | Tbccd1 | 0.00016609 | -0.777429203 | 0.785942057 |
| ENSMUSG00000022508 | Bcl6 | 0.00016609 | -1.417253639 | 0.282916407 |
| ENSMUSG00000038094 | Atp13a4 | 0.00016609 | 0.189403933 | 1 |
| ENSMUSG00000097313 | Gm26569 | 0.00016609 | 0.841480629 | 0.943363727 |
| ENSMUSG00000022537 | Tmem44 | 0.00016609 | 0.400908038 | 1 |
| ENSMUSG00000046546 | Fam43a | 0.00016609 | -1.500911568 | 0.23866117 |
| ENSMUSG00000046598 | Bdh1 | 0.00016609 | -1.233807498 | 0.495065067 |
| ENSMUSG00000035790 | Cep19 | 0.00016609 | 0.063873051 | 1 |
| ENSMUSG00000035699 | Slc51a | 0.00016609 | 3.063873051 | 0.144078128 |
| ENSMUSG00000052363 | Zdhhc19 | 0.00016609 | 0.256518129 | 1 |
| ENSMUSG00000079620 | Muc4 | 0.00016609 | -0.743481871 | 0.778722129 |
| ENSMUSG00000035506 | Slc12a8 | 0.00016609 | -0.995020638 | 0.66416768 |
| ENSMUSG00000022824 | Muc13 | 0.00016609 | 1.648835551 | 0.515310426 |
| ENSMUSG00000022844 | Pdia5 | 0.00016609 | -0.810596067 | 0.792441218 |
| ENSMUSG00000079594 | Cstdc6 | 0.00016609 | 4.648835551 | 0.031809473 |
| ENSMUSG00000022899 | Slc15a2 | 0.00016609 | 1.841480629 | 0.455131546 |
| ENSMUSG00000046961 | Gpr156 | 0.00016609 | 0.400908038 | 1 |
| ENSMUSG00000002847 | Pla1a | 0.00016609 | -0.305360759 | 1 |
| ENSMUSG00000002846 | Timmdc1 | 0.00016609 | -1.351164449 | 0.312100107 |
| ENSMUSG00000022704 | Qtrt2 | 0.00016609 | -2.051604167 | 0.039883588 |
| ENSMUSG00000036208 | Nepro | 0.00016609 | 0.648835551 | 1 |
| ENSMUSG00000033210 | Slc9c1 | 0.00016609 | 0.189403933 | 1 |
| ENSMUSG00000116755 | 6430553K19Rik | 0.00016609 | 1.326907457 | 0.642722979 |
| ENSMUSG00000030775 | Trat1 | 0.00016609 | 2.648835551 | 0.218817775 |
| ENSMUSG00000092009 | Myh15 | 0.00016609 | 1.189403933 | 0.685793194 |
| ENSMUSG00000032965 | Ift57 | 0.00016609 | -0.258055044 | 1 |
| ENSMUSG00000022639 | Dubr | 0.00016609 | 0.741944956 | 0.970970089 |
| ENSMUSG00000116920 | Gm49575 | 0.00016609 | 0.741944956 | 0.970970089 |
| ENSMUSG00000091243 | Vgll3 | 0.00016609 | 0.400908038 | 1 |
| ENSMUSG00000032932 | Hspa13 | 0.00016609 | 0.326907457 | 1 |
| ENSMUSG00000100857 | 1700041M19Rik | 0.00016609 | -0.560617814 | 0.997956586 |
| ENSMUSG00000022864 | D16Ertd472e | 0.00016609 | -1.023589791 | 0.524144449 |
| ENSMUSG00000025610 | Map3k7cl | 0.00016609 | 1.841480629 | 0.466464457 |
| ENSMUSG00000002489 | Tiam1 | 0.00016609 | -1.760555385 | 0.128972787 |
| ENSMUSG00000085169 | Gm10785 | 0.00016609 | 0.004979362 | 1 |
| ENSMUSG00000116747 | Gm41496 | 0.00016609 | 0.648835551 | 0.997212359 |

| | | | | |
|---|---|---|---|---|
| ENSMUSG00000022949 | Clic6 | 0.00016609 | -1.777429203 | 0.114811919 |
| ENSMUSG00000100277 | 1810053B23Rik | 0.00016609 | 1.47891055 | 0.592797015 |
| ENSMUSG00000117232 | 1600002D24Rik | 0.00016609 | -0.965874293 | 0.660275527 |
| ENSMUSG00000117270 | Gm49953 | 0.00016609 | 1.841480629 | 0.455131546 |
| ENSMUSG00000045275 | Lca5l | 0.00016609 | -0.743481871 | 0.919691163 |
| ENSMUSG00000040666 | Sh3bgr | 0.00016609 | 1.189403933 | 0.874796537 |
| ENSMUSG00000023802 | Nox3 | 0.00016609 | 1.326907457 | 0.661591638 |
| ENSMUSG00000073471 | Rsph3a | 0.00016609 | -1.051604167 | 0.529126167 |
| ENSMUSG00000085705 | Gm16046 | 0.00016609 | -2.051604167 | 0.055993444 |
| ENSMUSG00000038347 | Tcte2 | 0.00016609 | -0.936126949 | 0.606655093 |
| ENSMUSG00000116590 | AC154200.1 | 0.00016609 | 0.004979362 | 1 |
| ENSMUSG00000090417 | Vmn2r94 | 0.00016609 | 2.326907457 | 0.295355925 |
| ENSMUSG00000095961 | Vmn2r102 | 0.00016609 | 2.326907457 | 0.295355925 |
| ENSMUSG00000091151 | Vmn1r224 | 0.00016609 | 1.648835551 | 0.50741962 |
| ENSMUSG00000116974 | Gm49761 | 0.00016609 | 2.326907457 | 0.302168577 |
| ENSMUSG00000067942 | Zfp160 | 0.00016609 | -1.810596067 | 0.09684834 |
| ENSMUSG00000057409 | Zfp53 | 0.00016609 | -2.145580315 | 0.031872898 |
| ENSMUSG00000078546 | Zfp995 | 0.00016609 | -0.560617814 | 0.980745941 |
| ENSMUSG00000059142 | Zfp945 | 0.00016609 | 0.841480629 | 0.943363727 |
| ENSMUSG00000043782 | Bicdl2 | 0.00016609 | -0.395558568 | 1 |
| ENSMUSG00000023906 | Cldn6 | 0.00016609 | 0.326907457 | 1 |
| ENSMUSG00000023911 | Flywch2 | 0.00016609 | 0.648835551 | 1 |
| ENSMUSG00000035435 | Abca17 | 0.00016609 | 0.47891055 | 1 |
| ENSMUSG00000040888 | Gfer | 0.00016609 | 1.841480629 | 0.455131546 |
| ENSMUSG00000117379 | Gm50022 | 0.00016609 | 3.648835551 | 0.082076736 |
| ENSMUSG00000002280 | Ciao3 | 0.00016609 | -1.500911568 | 0.23866117 |
| ENSMUSG00000036775 | Decr2 | 0.00016609 | -1.43862729 | 0.287872755 |
| ENSMUSG00000024301 | Kifc5b | 0.00016609 | -0.843017545 | 0.674297981 |
| ENSMUSG00000044857 | Lemd2 | 0.00016609 | -0.874726405 | 0.680536135 |
| ENSMUSG00000002249 | Tead3 | 0.00016609 | -0.599091962 | 0.865542683 |
| ENSMUSG00000037446 | Tulp1 | 0.00016609 | 0.948395833 | 0.792441218 |
| ENSMUSG00000086154 | Gm16196 | 0.00016609 | 0.648835551 | 1 |
| ENSMUSG00000042203 | Tbc1d22b | 0.00016609 | -1.132524162 | 0.466464457 |
| ENSMUSG00000024037 | Wdr4 | 0.00016609 | -1.52108945 | 0.198212447 |
| ENSMUSG00000079553 | Kifc1 | 0.00016609 | 0.125273595 | 1 |
| ENSMUSG00000024319 | Vps52 | 0.00016609 | -0.258055044 | 1 |
| ENSMUSG00000073422 | H2-Ke6 | 0.00016609 | 0.56137271 | 1 |
| ENSMUSG00000015468 | Notch4 | 0.00016609 | 0.648835551 | 1 |
| ENSMUSG00000061207 | Stk19 | 0.00016609 | -0.936126949 | 0.606655093 |
| ENSMUSG00000040482 | Dxo | 0.00016609 | 0.841480629 | 0.943363727 |
| ENSMUSG00000040356 | Skiv2l | 0.00016609 | -1.328444372 | 0.307566094 |
| ENSMUSG00000024426 | Atat1 | 0.00016609 | 0.326907457 | 1 |
| ENSMUSG00000092368 | A930015D03Rik | 0.00016609 | -1.051604167 | 0.529126167 |
| ENSMUSG00000092243 | Gm7030 | 0.00016609 | 1.189403933 | 0.874796537 |
| ENSMUSG00000045409 | Trim39 | 0.00016609 | -1.459688905 | 0.229335266 |
| ENSMUSG00000097727 | F630040K05Rik | 0.00016609 | -1.328444372 | 0.438467012 |
| ENSMUSG00000034509 | Mad2l1bp | 0.00016609 | 0.648835551 | 1 |
| ENSMUSG00000032842 | Abcc10 | 0.00016609 | 0.189403933 | 1 |
| ENSMUSG00000047150 | 1700001C19Rik | 0.00016609 | -1.184054463 | 0.534323592 |
| ENSMUSG00000032717 | Mdfi | 0.00016609 | -0.52108945 | 0.975624465 |
| ENSMUSG00000110276 | Gm45330 | 0.00016609 | 1.326907457 | 0.642722979 |

| | | | | |
|---|---|---|---|---|
| ENSMUSG00000024172 | St6gal2 | 0.00016609 | 2.063873051 | 0.408535739 |
| ENSMUSG00000038781 | Stap2 | 0.00016609 | -0.051604167 | 1 |
| ENSMUSG00000003200 | Sh3gl1 | 0.00016609 | -1.079084903 | 0.536465189 |
| ENSMUSG00000007603 | Dus3l | 0.00016609 | -0.708716453 | 0.886203497 |
| ENSMUSG00000024206 | Rfx2 | 0.00016609 | 0.326907457 | 1 |
| ENSMUSG00000079414 | Gm11110 | 0.00016609 | -1.233807498 | 0.466329251 |
| ENSMUSG00000032796 | Lama1 | 0.00016609 | -1.305360759 | 0.409269374 |
| ENSMUSG00000091636 | Akain1 | 0.00016609 | 1.841480629 | 0.455131546 |
| ENSMUSG00000097154 | Gm26510 | 0.00016609 | 0.256518129 | 1 |
| ENSMUSG00000097546 | Gm26749 | 0.00016609 | 0.741944956 | 0.971178138 |
| ENSMUSG00000024079 | Eif2ak2 | 0.00016609 | -2.106051951 | 0.036215483 |
| ENSMUSG00000117509 | Gm46606 | 0.00016609 | 1.326907457 | 0.659434776 |
| ENSMUSG00000000673 | Haao | 0.00016609 | 1.326907457 | 0.642722979 |
| ENSMUSG00000117647 | Gm49969 | 0.00016609 | 0.326907457 | 1 |
| ENSMUSG00000024254 | Abcg8 | 0.00016609 | 0.47891055 | 1 |
| ENSMUSG00000024145 | Pigf | 0.00016609 | 1.063873051 | 0.886203497 |
| ENSMUSG00000117351 | Gm31615 | 0.00016609 | -0.52108945 | 0.996150047 |
| ENSMUSG00000034709 | Ppp1r21 | 0.00016609 | -2.743481871 | 0.002276816 |
| ENSMUSG00000024107 | Lhcgr | 0.00016609 | 0.56137271 | 1 |
| ENSMUSG00000117511 | Gm50061 | 0.00016609 | 2.326907457 | 0.295355925 |
| ENSMUSG00000087425 | Gm15404 | 0.00016609 | 1.189403933 | 0.690819967 |
| ENSMUSG00000117433 | Gm41654 | 0.00016609 | 1.47891055 | 0.620556057 |
| ENSMUSG00000050996 | Cetn1 | 0.00016609 | 4.648835551 | 0.031809473 |
| ENSMUSG00000037346 | Hrh4 | 0.00016609 | 2.063873051 | 0.37936848 |
| ENSMUSG00000117404 | Gm50035 | 0.00016609 | 3.063873051 | 0.179296195 |
| ENSMUSG00000117574 | Gm50084 | 0.00016609 | 0.400908038 | 1 |
| ENSMUSG00000044322 | Dsc1 | 0.00016609 | 3.063873051 | 0.144078128 |
| ENSMUSG00000034300 | Fam53c | 0.00016609 | -1.184054463 | 0.419360593 |
| ENSMUSG00000117869 | Snhg4 | 0.00016609 | -1.023589791 | 0.53393869 |
| ENSMUSG00000024485 | Slc4a9 | 0.00016609 | 0.56137271 | 1 |
| ENSMUSG00000051375 | Pcdh1 | 0.00016609 | -2.119348773 | 0.030152024 |
| ENSMUSG00000117994 | Gm5820 | 0.00016609 | -0.874726405 | 0.719591951 |
| ENSMUSG00000024487 | Yipf5 | 0.00016609 | -1.743481871 | 0.109343152 |
| ENSMUSG00000033022 | Cdo1 | 0.00016609 | 2.648835551 | 0.230984459 |
| ENSMUSG00000118365 | Gm41724 | 0.00016609 | 2.326907457 | 0.295355925 |
| ENSMUSG00000086607 | 4930511M06Rik | 0.00016609 | -0.743481871 | 0.792441218 |
| ENSMUSG00000024600 | Slc27a6 | 0.00016609 | 0.189403933 | 1 |
| ENSMUSG00000024601 | Isoc1 | 0.00016609 | 0.47891055 | 1 |
| ENSMUSG00000049173 | Myoz3 | 0.00016609 | -0.305360759 | 1 |
| ENSMUSG00000024617 | Camk2a | 0.00016609 | 0.004979362 | 1 |
| ENSMUSG00000045094 | Arhgef37 | 0.00016609 | 0.400908038 | 1 |
| ENSMUSG00000098702 | 1500015A07Rik | 0.00016609 | -1.52108945 | 0.198212447 |
| ENSMUSG00000026322 | Htr4 | 0.00016609 | -0.395558568 | 1 |
| ENSMUSG00000118170 | Gm30094 | 0.00016609 | 0.948395833 | 0.939582412 |
| ENSMUSG00000024524 | Gnal | 0.00016609 | -1.051604167 | 0.581923692 |
| ENSMUSG00000035394 | Cfap53 | 0.00016609 | 0.063873051 | 1 |
| ENSMUSG00000025420 | Katnal2 | 0.00016609 | 1.326907457 | 0.642722979 |
| ENSMUSG00000117717 | Gm30192 | 0.00016609 | 1.326907457 | 0.66162259 |
| ENSMUSG00000118227 | Gm30288 | 0.00016609 | 2.063873051 | 0.37936848 |
| ENSMUSG00000100430 | 1700095A13Rik | 0.00016609 | -2.079084903 | 0.034053737 |
| ENSMUSG00000046982 | Tshz1 | 0.00016609 | -1.921020057 | 0.060210487 |

| | | | | |
|---|---|---|---|---|
| ENSMUSG00000050321 | Neto1 | 0.00016609 | 1.841480629 | 0.455131546 |
| ENSMUSG00000037263 | Aldh3b3 | 0.00016609 | 3.063873051 | 0.144078128 |
| ENSMUSG00000060803 | Gstp1 | 0.00016609 | 0.841480629 | 0.946764296 |
| ENSMUSG00000054874 | Pcnx3 | 0.00016609 | -1.281901786 | 0.365716212 |
| ENSMUSG00000004054 | Map3k11 | 0.00016609 | -1.43862729 | 0.296607321 |
| ENSMUSG00000079478 | Znrd2 | 0.00016609 | -1.258055044 | 0.363644968 |
| ENSMUSG00000075227 | Znhit2 | 0.00016609 | 0.47891055 | 1 |
| ENSMUSG00000024799 | Tm7sf2 | 0.00016609 | 0.948395833 | 0.912904617 |
| ENSMUSG00000024797 | Vps51 | 0.00016609 | -0.599091962 | 0.865542683 |
| ENSMUSG00000039699 | Batf2 | 0.00016609 | 1.189403933 | 0.685371374 |
| ENSMUSG00000024947 | Men1 | 0.00016609 | -0.106051951 | 1 |
| ENSMUSG00000024948 | Map4k2 | 0.00016609 | -1.079084903 | 0.599135879 |
| ENSMUSG00000010097 | Nxf1 | 0.00016609 | -2.717486663 | 0.002484871 |
| ENSMUSG00000024663 | Rab3il1 | 0.00016609 | 1.189403933 | 0.688085317 |
| ENSMUSG00000024742 | Fen1 | 0.00016609 | 0.841480629 | 0.943363727 |
| ENSMUSG00000024667 | Tmem216 | 0.00016609 | -0.560617814 | 0.980745941 |
| ENSMUSG00000034445 | Cyb561a3 | 0.00016609 | -1.373532262 | 0.317734263 |
| ENSMUSG00000067577 | A430093F15Rik | 0.00016609 | 1.648835551 | 0.499921258 |
| ENSMUSG00000024670 | Cd6 | 0.00016609 | 1.47891055 | 0.587484274 |
| ENSMUSG00000024736 | Tmem132a | 0.00016609 | 0.004979362 | 1 |
| ENSMUSG00000097091 | 4930526L06Rik | 0.00016609 | 1.326907457 | 0.652166789 |
| ENSMUSG00000024683 | Mrpl16 | 0.00016609 | 0.841480629 | 0.943363727 |
| ENSMUSG00000086213 | A330040F15Rik | 0.00016609 | -0.051604167 | 1 |
| ENSMUSG00000024640 | Psat1 | 0.00016609 | -0.106051951 | 1 |
| ENSMUSG00000041491 | Cep78 | 0.00016609 | -0.43862729 | 1 |
| ENSMUSG00000085003 | Pip5k1bos | 0.00016609 | 1.841480629 | 0.455131546 |
| ENSMUSG00000118005 | Gm50377 | 0.00016609 | 2.063873051 | 0.413417657 |
| ENSMUSG00000024896 | Minpp1 | 0.00016609 | -1.209145444 | 0.423814296 |
| ENSMUSG00000117998 | Gm32081 | 0.00016609 | 1.841480629 | 0.475598944 |
| ENSMUSG00000054200 | Ffar4 | 0.00016609 | 0.125273595 | 1 |
| ENSMUSG00000024999 | Noc3l | 0.00016609 | -2.682081327 | 0.002606179 |
| ENSMUSG00000067231 | Cyp2c65 | 0.00016609 | 3.063873051 | 0.197743955 |
| ENSMUSG00000025007 | Aldh18a1 | 0.00016609 | -0.43862729 | 1 |
| ENSMUSG00000025008 | Tctn3 | 0.00016609 | 0.189403933 | 1 |
| ENSMUSG00000035049 | Rrp12 | 0.00016609 | -0.874726405 | 0.680536135 |
| ENSMUSG00000090117 | Gm16541 | 0.00016609 | 0.841480629 | 0.955815331 |
| ENSMUSG00000040018 | Cox15 | 0.00016609 | 0.47891055 | 1 |
| ENSMUSG00000036961 | Wnt8b | 0.00016609 | 1.841480629 | 0.473176854 |
| ENSMUSG00000117911 | Gm35460 | 0.00016609 | 1.189403933 | 0.874796537 |
| ENSMUSG00000025208 | Mrpl43 | 0.00016609 | -0.480447466 | 0.972350276 |
| ENSMUSG00000025227 | Mfsd13a | 0.00016609 | -0.43862729 | 1 |
| ENSMUSG00000025036 | Sfxn2 | 0.00016609 | 0.125273595 | 1 |
| ENSMUSG00000003559 | As3mt | 0.00016609 | 0.063873051 | 1 |
| ENSMUSG00000117981 | 4930484I04Rik | 0.00016609 | 0.326907457 | 1 |
| ENSMUSG00000025078 | Nhlrc2 | 0.00016609 | -3.112715681 | 0.000324247 |
| ENSMUSG00000025082 | Vwa2 | 0.00016609 | 1.189403933 | 0.688956689 |
| ENSMUSG00000117655 | 6720468P15Rik | 0.00016609 | 2.326907457 | 0.295355925 |
| ENSMUSG00000046008 | Pnlip | 0.00016609 | 1.648835551 | 0.525136458 |
| ENSMUSG00000087095 | Emx2os | 0.00016609 | -0.051604167 | 1 |
| ENSMUSG00000059326 | Csf2ra | 0.00016609 | 1.326907457 | 0.642927607 |
| ENSMUSG00000068270 | Shroom4 | 0.00016609 | -1.673092543 | 0.168724646 |

| | | | | |
|---|---|---|---|---|
| ENSMUSG00000063018 | 2010204K13Rik | 0.00016609 | 2.063873051 | 0.37936848 |
| ENSMUSG00000031144 | Syp | 0.00016609 | 0.400908038 | 1 |
| ENSMUSG00000039382 | Wdr45 | 0.00016609 | -1.395558568 | 0.26547175 |
| ENSMUSG00000009731 | Kcnd1 | 0.00016609 | -1.158519371 | 0.466464457 |
| ENSMUSG00000031161 | Hdac6 | 0.00016609 | -1.106051951 | 0.466464457 |
| ENSMUSG00000031163 | Glod5 | 0.00016609 | -0.158519371 | 1 |
| ENSMUSG00000047344 | Lancl3 | 0.00016609 | -0.106051951 | 1 |
| ENSMUSG00000015340 | Cybb | 0.00016609 | 0.256518129 | 1 |
| ENSMUSG00000040147 | Maob | 0.00016609 | 1.189403933 | 0.693613791 |
| ENSMUSG00000001134 | Uxt | 0.00016609 | 1.47891055 | 0.57597659 |
| ENSMUSG00000059708 | Akap17b | 0.00016609 | 0.256518129 | 1 |
| ENSMUSG00000050379 | 6-Sep | 0.00016609 | -0.560617814 | 0.980745941 |
| ENSMUSG00000016409 | Nkap | 0.00016609 | -1.579983139 | 0.174915761 |
| ENSMUSG00000061082 | Plac1 | 0.00016609 | 0.648835551 | 0.998731799 |
| ENSMUSG00000067835 | Gm14661 | 0.00016609 | 0.47891055 | 1 |
| ENSMUSG00000059401 | Mamld1 | 0.00016609 | -0.480447466 | 0.972350276 |
| ENSMUSG00000015214 | Mtmr1 | 0.00016609 | 0.063873051 | 1 |
| ENSMUSG00000031349 | Nsdhl | 0.00016609 | 0.741944956 | 0.970970089 |
| ENSMUSG00000031371 | Haus7 | 0.00016609 | -0.158519371 | 1 |
| ENSMUSG00000031378 | Abcd1 | 0.00016609 | -0.599091962 | 0.874796537 |
| ENSMUSG00000002006 | Pdzd4 | 0.00016609 | -0.636566667 | 0.870078045 |
| ENSMUSG00000031386 | Hcfc1 | 0.00016609 | -1.794107944 | 0.094802366 |
| ENSMUSG00000087396 | 4933407K13Rik | 0.00016609 | -1.395558568 | 0.26547175 |
| ENSMUSG00000073077 | Cfap47 | 0.00016609 | 0.56137271 | 1 |
| ENSMUSG00000035232 | Pdk3 | 0.00016609 | 0.841480629 | 0.954844735 |
| ENSMUSG00000079508 | Apoo | 0.00016609 | 0.841480629 | 0.943363727 |
| ENSMUSG00000031216 | Stard8 | 0.00016609 | 0.741944956 | 0.970970089 |
| ENSMUSG00000031310 | Zmym3 | 0.00016609 | -2.023589791 | 0.046981468 |
| ENSMUSG00000049191 | Rtl5 | 0.00016609 | 0.948395833 | 0.912904617 |
| ENSMUSG00000109088 | Gm44593 | 0.00016609 | 0.948395833 | 0.952108693 |
| ENSMUSG00000034607 | Pof1b | 0.00016609 | 0.125273595 | 1 |
| ENSMUSG00000079450 | Cldn34c1 | 0.00016609 | -0.051604167 | 1 |
| ENSMUSG00000031264 | Btk | 0.00016609 | -0.777429203 | 0.792441218 |
| ENSMUSG00000051579 | Tceal8 | 0.00016609 | 1.326907457 | 0.66416768 |
| ENSMUSG00000042425 | Frmpd3 | 0.00016609 | 0.063873051 | 1 |
| ENSMUSG00000071679 | Rtl4 | 0.00016609 | 0.648835551 | 0.999962098 |
| ENSMUSG00000079387 | Luzp4 | 0.00016609 | 0.648835551 | 0.998814138 |
| ENSMUSG00000095082 | Gm15091 | 0.00016609 | 1.326907457 | 0.65747066 |
| ENSMUSG00000025268 | Maged2 | 0.00016609 | 1.326907457 | 0.642722979 |
| ENSMUSG00000062168 | Ppef1 | 0.00016609 | 1.326907457 | 0.66416768 |
| ENSMUSG00000061778 | Mospd2 | 0.00016609 | -2.092631435 | 0.034881114 |
| ENSMUSG00000093806 | Asmt | 0.00016609 | 2.063873051 | 0.403758949 |
| ENSMUSG00000063897 | CAAA01118383.1 | 0.00016609 | -1.874726405 | 0.081714616 |
| ENSMUSG00000051412 | Vamp7 | 0.00016609 | 0.400908038 | 1 |
| ENSMUSG00000089699 | Gm1992 | 0.000110727 | 1.42644313 | 0.736456705 |
| ENSMUSG00000046101 | Mcmdc2 | 0.000110727 | 0.533358334 | 1 |
| ENSMUSG00000025916 | Ppp1r42 | 0.000110727 | 1.648835551 | 0.66416768 |
| ENSMUSG00000101476 | Gm29570 | 0.000110727 | 2.648835551 | 0.395118486 |
| ENSMUSG00000054493 | Gm9947 | 0.000110727 | 1.233798052 | 0.792441218 |
| ENSMUSG00000067795 | 4930444P10Rik | 0.000110727 | 0.774366434 | 1 |
| ENSMUSG00000025779 | Ly96 | 0.000110727 | -0.014129461 | 1 |

| | | | | |
|---|---|---|---|---|
| ENSMUSG00000067773 | Defb41 | 0.000110727 | 3.233798052 | 0.262284476 |
| ENSMUSG00000026156 | B3gat2 | 0.000110727 | 0.146335211 | 1 |
| ENSMUSG00000037432 | Fer1l5 | 0.000110727 | 1.42644313 | 0.736456705 |
| ENSMUSG00000001138 | Cnnm3 | 0.000110727 | -1.720398258 | 0.169030494 |
| ENSMUSG00000026113 | Inpp4a | 0.000110727 | -2.788569761 | 0.00418953 |
| ENSMUSG00000103270 | Gm37915 | 0.000110727 | 3.233798052 | 0.262284476 |
| ENSMUSG00000079588 | Tmem182 | 0.000110727 | 1.233798052 | 0.792441218 |
| ENSMUSG00000093436 | Gm20646 | 0.000110727 | 0.648835551 | 1 |
| ENSMUSG00000091844 | Gm8251 | 0.000110727 | 0.233798052 | 1 |
| ENSMUSG00000026047 | Poglut2 | 0.000110727 | -0.895484965 | 0.691084416 |
| ENSMUSG00000051616 | C230029F24Rik | 0.000110727 | -0.014129461 | 1 |
| ENSMUSG00000101799 | 9330175M20Rik | 0.000110727 | -0.410058138 | 1 |
| ENSMUSG00000043015 | Nemp2 | 0.000110727 | 0.233798052 | 1 |
| ENSMUSG00000043629 | 1700019D03Rik | 0.000110727 | 1.42644313 | 0.736456705 |
| ENSMUSG00000026098 | Pms1 | 0.000110727 | -1.158519371 | 0.521275193 |
| ENSMUSG00000025977 | Boll | 0.000110727 | 0.42644313 | 1 |
| ENSMUSG00000026039 | Sgo2a | 0.000110727 | -0.52108945 | 1 |
| ENSMUSG00000072295 | C2cd6 | 0.000110727 | 0.911869957 | 1 |
| ENSMUSG00000041075 | Fzd7 | 0.000110727 | 2.233798052 | 0.47966612 |
| ENSMUSG00000026012 | Cd28 | 0.000110727 | 0.774366434 | 1 |
| ENSMUSG00000026011 | Ctla4 | 0.000110727 | 0.648835551 | 1 |
| ENSMUSG00000085842 | Pard3bos1 | 0.000110727 | 1.648835551 | 0.66416768 |
| ENSMUSG00000046856 | Gpr1 | 0.000110727 | 1.648835551 | 0.66416768 |
| ENSMUSG00000087561 | Gm11608 | 0.000110727 | 2.648835551 | 0.431443405 |
| ENSMUSG00000101832 | Gm28982 | 0.000110727 | 0.774366434 | 1 |
| ENSMUSG00000045005 | Fzd5 | 0.000110727 | -0.225633566 | 1 |
| ENSMUSG00000015222 | Map2 | 0.000110727 | -1.351164449 | 0.45354452 |
| ENSMUSG00000025991 | Cps1 | 0.000110727 | 1.063873051 | 0.850527636 |
| ENSMUSG00000050296 | Abca12 | 0.000110727 | 0.233798052 | 1 |
| ENSMUSG00000039395 | Mreg | 0.000110727 | -0.088130043 | 1 |
| ENSMUSG00000039342 | Ankar | 0.000110727 | 0.911869957 | 1 |
| ENSMUSG00000061815 | Rufy4 | 0.000110727 | -0.673092543 | 0.929961175 |
| ENSMUSG00000026177 | Slc11a1 | 0.000110727 | 3.233798052 | 0.318839109 |
| ENSMUSG00000026176 | Ctdsp1 | 0.000110727 | -0.810596067 | 0.796840156 |
| ENSMUSG00000006542 | Prkag3 | 0.000110727 | 0.911869957 | 1 |
| ENSMUSG00000026197 | Zfand2b | 0.000110727 | 0.774366434 | 1 |
| ENSMUSG00000033124 | Atg9a | 0.000110727 | -1.380911792 | 0.408535739 |
| ENSMUSG00000101840 | Gm28294 | 0.000110727 | 1.233798052 | 0.792441218 |
| ENSMUSG00000100827 | Gm29069 | 0.000110727 | 0.146335211 | 1 |
| ENSMUSG00000101483 | 1700016L21Rik | 0.000110727 | 4.233798052 | 0.128972787 |
| ENSMUSG00000079465 | Col4a3 | 0.000110727 | -1.088130043 | 0.66416768 |
| ENSMUSG00000053161 | Daw1 | 0.000110727 | 1.42644313 | 0.736456705 |
| ENSMUSG00000073628 | Gm10552 | 0.000110727 | 0.063873051 | 1 |
| ENSMUSG00000073627 | C130036L24Rik | 0.000110727 | -0.720398258 | 0.792441218 |
| ENSMUSG00000079429 | Mroh2a | 0.000110727 | 1.233798052 | 0.792441218 |
| ENSMUSG00000100798 | Gm19589 | 0.000110727 | 0.533358334 | 1 |
| ENSMUSG00000102009 | 4933400F21Rik | 0.000110727 | 2.648835551 | 0.411460922 |
| ENSMUSG00000026301 | Iqca | 0.000110727 | -0.225633566 | 1 |
| ENSMUSG00000090322 | Gm17090 | 0.000110727 | 4.233798052 | 0.128972787 |
| ENSMUSG00000026270 | Capn10 | 0.000110727 | 0.42644313 | 1 |
| ENSMUSG00000084989 | Crocc2 | 0.000110727 | 1.648835551 | 0.66416768 |

| | | | | |
|---|---|---|---|---|
| ENSMUSG00000026276 | 2-Sep | 0.000110727 | -2.380911792 | 0.02142809 |
| ENSMUSG00000101952 | Gm10550 | 0.000110727 | -0.52108945 | 1 |
| ENSMUSG00000040693 | Slco4c1 | 0.000110727 | 1.648835551 | 0.66416768 |
| ENSMUSG00000026336 | Slco6d1 | 0.000110727 | 1.063873051 | 1 |
| ENSMUSG00000050840 | Cdh20 | 0.000110727 | -0.936126949 | 0.736456705 |
| ENSMUSG00000099843 | Gm7160 | 0.000110727 | 2.648835551 | 0.395118486 |
| ENSMUSG00000101583 | Gm20753 | 0.000110727 | 1.911869957 | 0.587484274 |
| ENSMUSG00000062345 | Serpinb2 | 0.000110727 | 1.648835551 | 0.66416768 |
| ENSMUSG00000100457 | D830032E09Rik | 0.000110727 | 0.42644313 | 1 |
| ENSMUSG00000047216 | Cdh19 | 0.000110727 | -0.289763904 | 1 |
| ENSMUSG00000037035 | Inhbb | 0.000110727 | -0.466641666 | 1 |
| ENSMUSG00000036962 | Cfap221 | 0.000110727 | 0.774366434 | 1 |
| ENSMUSG00000050777 | Tmem37 | 0.000110727 | 1.911869957 | 0.587484274 |
| ENSMUSG00000026390 | Marco | 0.000110727 | -1.014129461 | 0.608426921 |
| ENSMUSG00000101791 | 2210011K15Rik | 0.000110727 | 4.233798052 | 0.15358091 |
| ENSMUSG00000099960 | Gm28590 | 0.000110727 | 1.42644313 | 0.736456705 |
| ENSMUSG00000026399 | Cd55 | 0.000110727 | -1.014129461 | 0.608426921 |
| ENSMUSG00000026409 | Pfkfb2 | 0.000110727 | -1.696939285 | 0.210550914 |
| ENSMUSG00000026435 | Slc45a3 | 0.000110727 | -0.720398258 | 0.792441218 |
| ENSMUSG00000048174 | Tmem81 | 0.000110727 | 1.42644313 | 0.736456705 |
| ENSMUSG00000101693 | Gm19461 | 0.000110727 | 2.233798052 | 0.508263132 |
| ENSMUSG00000051998 | Lax1 | 0.000110727 | 2.648835551 | 0.395118486 |
| ENSMUSG00000026450 | Chit1 | 0.000110727 | 1.648835551 | 0.66416768 |
| ENSMUSG00000026458 | Ppfia4 | 0.000110727 | 1.063873051 | 1 |
| ENSMUSG00000048096 | Lmod1 | 0.000110727 | 0.146335211 | 1 |
| ENSMUSG00000102717 | Gm37759 | 0.000110727 | 4.233798052 | 0.197130662 |
| ENSMUSG00000104414 | Gm37799 | 0.000110727 | 0.911869957 | 1 |
| ENSMUSG00000041483 | Zfp281 | 0.000110727 | -0.766201948 | 0.792441218 |
| ENSMUSG00000097113 | Gm19705 | 0.000110727 | 0.146335211 | 1 |
| ENSMUSG00000097448 | Platr22 | 0.000110727 | -1.599091962 | 0.242941974 |
| ENSMUSG00000099510 | Gm28501 | 0.000110727 | 1.233798052 | 0.792441218 |
| ENSMUSG00000051079 | Rgs13 | 0.000110727 | 0.911869957 | 1 |
| ENSMUSG00000026357 | Rgs18 | 0.000110727 | 1.42644313 | 0.736456705 |
| ENSMUSG00000085822 | Gm15584 | 0.000110727 | 0.774366434 | 1 |
| ENSMUSG00000043019 | Edem3 | 0.000110727 | -0.936126949 | 0.706929043 |
| ENSMUSG00000042671 | Rgs8 | 0.000110727 | 1.063873051 | 1 |
| ENSMUSG00000026475 | Rgs16 | 0.000110727 | 0.533358334 | 1 |
| ENSMUSG00000042641 | Rgsl1 | 0.000110727 | -0.158519371 | 1 |
| ENSMUSG00000026471 | Mr1 | 0.000110727 | 0.42644313 | 1 |
| ENSMUSG00000026468 | Lhx4 | 0.000110727 | 1.648835551 | 0.66416768 |
| ENSMUSG00000026601 | Axdnd1 | 0.000110727 | 0.233798052 | 1 |
| ENSMUSG00000101768 | Gm28694 | 0.000110727 | -0.014129461 | 1 |
| ENSMUSG00000087093 | Gm15486 | 0.000110727 | 0.774366434 | 1 |
| ENSMUSG00000004031 | Brinp2 | 0.000110727 | -0.088130043 | 1 |
| ENSMUSG00000103971 | Gm37679 | 0.000110727 | 0.533358334 | 1 |
| ENSMUSG00000026705 | Klhl20 | 0.000110727 | -1.648844997 | 0.201326391 |
| ENSMUSG00000086277 | 4930558K02Rik | 0.000110727 | -0.673092543 | 0.951660573 |
| ENSMUSG00000026698 | Pigc | 0.000110727 | -0.52108945 | 1 |
| ENSMUSG00000026694 | Eef1aknmt | 0.000110727 | -1.158519371 | 0.521275193 |
| ENSMUSG00000026692 | Fmo4 | 0.000110727 | 0.911869957 | 1 |
| ENSMUSG00000095576 | Fmo6 | 0.000110727 | 2.648835551 | 0.395118486 |

| | | | | |
|---|---|---|---|---|
| ENSMUSG00000103288 | Gm37273 | 0.000110727 | -0.351164449 | 1 |
| ENSMUSG00000041396 | Mettl18 | 0.000110727 | 0.146335211 | 1 |
| ENSMUSG00000026580 | Selp | 0.000110727 | 0.533358334 | 1 |
| ENSMUSG00000026574 | Dpt | 0.000110727 | 2.233798052 | 0.512602663 |
| ENSMUSG00000005763 | Cd247 | 0.000110727 | 0.774366434 | 1 |
| ENSMUSG00000026565 | Pou2f1 | 0.000110727 | -2.946111038 | 0.002012957 |
| ENSMUSG00000026564 | Dusp27 | 0.000110727 | 1.911869957 | 0.604880543 |
| ENSMUSG00000026563 | Tada1 | 0.000110727 | -0.351164449 | 1 |
| ENSMUSG00000026683 | Nuf2 | 0.000110727 | 0.063873051 | 1 |
| ENSMUSG00000053925 | Gm9929 | 0.000110727 | 1.648835551 | 0.66416768 |
| ENSMUSG00000038370 | Pcp4l1 | 0.000110727 | -0.014129461 | 1 |
| ENSMUSG00000038209 | Itln1 | 0.000110727 | 1.648835551 | 0.66416768 |
| ENSMUSG00000015314 | Slamf6 | 0.000110727 | 1.42644313 | 0.736456705 |
| ENSMUSG00000092085 | Gm17224 | 0.000110727 | 4.233798052 | 0.128972787 |
| ENSMUSG00000044708 | Kcnj10 | 0.000110727 | 1.42644313 | 0.736456705 |
| ENSMUSG00000050229 | Pigm | 0.000110727 | -0.52108945 | 1 |
| ENSMUSG00000026544 | Dusp23 | 0.000110727 | 1.648835551 | 0.665635898 |
| ENSMUSG00000037849 | Ifi206 | 0.000110727 | -0.410058138 | 1 |
| ENSMUSG00000073491 | Ifi213 | 0.000110727 | 2.648835551 | 0.395118486 |
| ENSMUSG00000026532 | Spta1 | 0.000110727 | 2.648835551 | 0.395118486 |
| ENSMUSG00000078185 | Chml | 0.000110727 | -0.014129461 | 1 |
| ENSMUSG00000039748 | Exo1 | 0.000110727 | -1.43862729 | 0.361297544 |
| ENSMUSG00000087236 | Kif28 | 0.000110727 | 0.233798052 | 1 |
| ENSMUSG00000026489 | Coq8a | 0.000110727 | 0.774366434 | 1 |
| ENSMUSG00000026496 | Parp1 | 0.000110727 | -1.466641666 | 0.346237121 |
| ENSMUSG00000103649 | Gm37768 | 0.000110727 | 0.648835551 | 1 |
| ENSMUSG00000026520 | Pycr2 | 0.000110727 | -0.351164449 | 1 |
| ENSMUSG00000026514 | Cnih3 | 0.000110727 | 0.648835551 | 1 |
| ENSMUSG00000038599 | Capn8 | 0.000110727 | 0.911869957 | 1 |
| ENSMUSG00000039384 | Dusp10 | 0.000110727 | -2.410058138 | 0.023019447 |
| ENSMUSG00000101574 | 1700112H15Rik | 0.000110727 | 2.233798052 | 0.498350473 |
| ENSMUSG00000103308 | Gm37800 | 0.000110727 | 1.648835551 | 0.66416768 |
| ENSMUSG00000102158 | Gm34342 | 0.000110727 | 1.42644313 | 0.736456705 |
| ENSMUSG00000079045 | Prox1os | 0.000110727 | -0.624182943 | 0.952107111 |
| ENSMUSG00000026634 | Angel2 | 0.000110727 | -1.225633566 | 0.463954265 |
| ENSMUSG00000026627 | Pacc1 | 0.000110727 | 0.146335211 | 1 |
| ENSMUSG00000103114 | Gm32200 | 0.000110727 | 1.911869957 | 0.587484274 |
| ENSMUSG00000037434 | Slc30a1 | 0.000110727 | 1.063873051 | 1 |
| ENSMUSG00000079144 | A130010J15Rik | 0.000110727 | 2.648835551 | 0.395118486 |
| ENSMUSG00000102887 | Gm10857 | 0.000110727 | 2.233798052 | 0.47966612 |
| ENSMUSG00000025780 | Itih5 | 0.000110727 | -0.975655313 | 0.706929043 |
| ENSMUSG00000085257 | Gm13264 | 0.000110727 | 0.774366434 | 1 |
| ENSMUSG00000086006 | Gm13293 | 0.000110727 | 0.233798052 | 1 |
| ENSMUSG00000054057 | A930004D18Rik | 0.000110727 | -0.088130043 | 1 |
| ENSMUSG00000086340 | 1810059C17Rik | 0.000110727 | 4.233798052 | 0.128972787 |
| ENSMUSG00000023094 | Msrb2 | 0.000110727 | 0.911869957 | 0.880201363 |
| ENSMUSG00000026784 | Pdss1 | 0.000110727 | -1.599091962 | 0.242941974 |
| ENSMUSG00000026774 | 4931423N10Rik | 0.000110727 | 0.911869957 | 1 |
| ENSMUSG00000026979 | Psd4 | 0.000110727 | -0.673092543 | 0.919998879 |
| ENSMUSG00000026976 | Pax8 | 0.000110727 | 0.326907457 | 1 |
| ENSMUSG00000006471 | Ndor1 | 0.000110727 | -0.853664789 | 0.806496094 |

| | | | | |
|---|---|---|---|---|
| ENSMUSG00000036504 | Phpt1 | 0.000110727 | 1.233798052 | 0.792441218 |
| ENSMUSG00000052403 | Fcnaos | 0.000110727 | 1.911869957 | 0.601156657 |
| ENSMUSG00000026939 | Tmem141 | 0.000110727 | 1.648835551 | 0.66416768 |
| ENSMUSG00000026938 | Fcna | 0.000110727 | 2.648835551 | 0.425537134 |
| ENSMUSG00000026925 | Inpp5e | 0.000110727 | -0.975655313 | 0.705975399 |
| ENSMUSG00000026921 | Egfl7 | 0.000110727 | -1.225633566 | 0.463954265 |
| ENSMUSG00000015776 | Med22 | 0.000110727 | 0.326907457 | 1 |
| ENSMUSG00000052406 | Rexo4 | 0.000110727 | -1.915949067 | 0.103463818 |
| ENSMUSG00000009214 | Mymk | 0.000110727 | 3.233798052 | 0.336482224 |
| ENSMUSG00000026816 | Gtf3c5 | 0.000110727 | -0.57355687 | 0.910050347 |
| ENSMUSG00000035666 | Gtf3c4 | 0.000110727 | -0.895484965 | 0.691084416 |
| ENSMUSG00000089762 | Ier5l | 0.000110727 | 0.233798052 | 1 |
| ENSMUSG00000026849 | Tor1a | 0.000110727 | -1.123753952 | 0.514974685 |
| ENSMUSG00000002550 | Uck1 | 0.000110727 | 0.233798052 | 1 |
| ENSMUSG00000026825 | Dnm1 | 0.000110727 | -1.43862729 | 0.358945041 |
| ENSMUSG00000026817 | Ak1 | 0.000110727 | 0.911869957 | 1 |
| ENSMUSG00000009563 | Tor2a | 0.000110727 | 1.42644313 | 0.736456705 |
| ENSMUSG00000026889 | Rbm18 | 0.000110727 | -0.766201948 | 0.792441218 |
| ENSMUSG00000085141 | Gm13429 | 0.000110727 | 1.648835551 | 0.66416768 |
| ENSMUSG00000059429 | Olfr365 | 0.000110727 | 0.326907457 | 1 |
| ENSMUSG00000066798 | Zbtb6 | 0.000110727 | -0.810596067 | 0.827142217 |
| ENSMUSG00000085950 | Gm13589 | 0.000110727 | 1.063873051 | 0.849896406 |
| ENSMUSG00000046618 | Olfml2a | 0.000110727 | 0.063873051 | 1 |
| ENSMUSG00000086644 | Gm13470 | 0.000110727 | -0.289763904 | 1 |
| ENSMUSG00000085862 | Gm13483 | 0.000110727 | -0.52108945 | 1 |
| ENSMUSG00000085014 | Gm13490 | 0.000110727 | 0.774366434 | 1 |
| ENSMUSG00000026946 | Nmi | 0.000110727 | -0.410058138 | 1 |
| ENSMUSG00000084938 | BB557941 | 0.000110727 | 0.911869957 | 1 |
| ENSMUSG00000086196 | Gm13571 | 0.000110727 | 2.648835551 | 0.430269377 |
| ENSMUSG00000044647 | Csrnp3 | 0.000110727 | 0.233798052 | 1 |
| ENSMUSG00000100197 | Gm28638 | 0.000110727 | -0.810596067 | 0.855313747 |
| ENSMUSG00000087301 | Gm13629 | 0.000110727 | 1.648835551 | 0.66416768 |
| ENSMUSG00000053896 | 4933409G03Rik | 0.000110727 | 2.648835551 | 0.433234003 |
| ENSMUSG00000109667 | Gm36231 | 0.000110727 | 1.233798052 | 0.792441218 |
| ENSMUSG00000027048 | Abcb11 | 0.000110727 | 1.063873051 | 1 |
| ENSMUSG00000027068 | Dhrs9 | 0.000110727 | 0.911869957 | 1 |
| ENSMUSG00000027070 | Lrp2 | 0.000110727 | -0.466641666 | 1 |
| ENSMUSG00000027088 | Phospho2 | 0.000110727 | -0.810596067 | 0.796840156 |
| ENSMUSG00000087264 | Gad1os | 0.000110727 | 1.233798052 | 0.792441218 |
| ENSMUSG00000041966 | Dcaf17 | 0.000110727 | -1.289763904 | 0.466464457 |
| ENSMUSG00000085439 | Rapgef4os1 | 0.000110727 | 1.233798052 | 0.792441218 |
| ENSMUSG00000055612 | Cdca7 | 0.000110727 | 0.774366434 | 1 |
| ENSMUSG00000085427 | 6430710C18Rik | 0.000110727 | 0.326907457 | 1 |
| ENSMUSG00000041762 | Gpr155 | 0.000110727 | 0.233798052 | 1 |
| ENSMUSG00000086544 | Chn1os3 | 0.000110727 | 2.648835551 | 0.395118486 |
| ENSMUSG00000075267 | Pjvk | 0.000110727 | 2.648835551 | 0.435791313 |
| ENSMUSG00000002732 | Fkbp7 | 0.000110727 | 0.911869957 | 1 |
| ENSMUSG00000085431 | 4930440I19Rik | 0.000110727 | 0.648835551 | 1 |
| ENSMUSG00000048388 | Fam171b | 0.000110727 | 1.911869957 | 0.597246405 |
| ENSMUSG00000087362 | Gm13710 | 0.000110727 | -1.380911792 | 0.411787226 |
| ENSMUSG00000097187 | Gm19426 | 0.000110727 | 1.233798052 | 0.792441218 |

| | | | | |
|---|---|---|---|---|
| ENSMUSG00000027075 | Slc43a1 | 0.000110727 | 0.911869957 | 1 |
| ENSMUSG00000027071 | P2rx3 | 0.000110727 | 0.42644313 | 1 |
| ENSMUSG00000075224 | Lrrc55 | 0.000110727 | -0.225633566 | 1 |
| ENSMUSG00000033850 | Olfr1015 | 0.000110727 | 4.233798052 | 0.128972787 |
| ENSMUSG00000075143 | Olfr1157 | 0.000110727 | 4.233798052 | 0.128972787 |
| ENSMUSG00000075107 | Olfr1216 | 0.000110727 | 4.233798052 | 0.208124838 |
| ENSMUSG00000027257 | Pacsin3 | 0.000110727 | -0.720398258 | 1 |
| ENSMUSG00000027223 | Mapk8ip1 | 0.000110727 | 0.774366434 | 1 |
| ENSMUSG00000085261 | Gm13814 | 0.000110727 | 0.774366434 | 1 |
| ENSMUSG00000010914 | Pdhx | 0.000110727 | -0.158519371 | 1 |
| ENSMUSG00000027185 | Nat10 | 0.000110727 | -1.720398258 | 0.169030494 |
| ENSMUSG00000046085 | 4931422A03Rik | 0.000110727 | 2.233798052 | 0.495023111 |
| ENSMUSG00000027173 | Depdc7 | 0.000110727 | -0.810596067 | 0.796840156 |
| ENSMUSG00000027171 | Prrg4 | 0.000110727 | -0.766201948 | 0.972192372 |
| ENSMUSG00000074934 | Grem1 | 0.000110727 | 0.42644313 | 1 |
| ENSMUSG00000040321 | Zfp770 | 0.000110727 | 1.063873051 | 1 |
| ENSMUSG00000027349 | Fam98b | 0.000110727 | -0.466641666 | 1 |
| ENSMUSG00000040061 | Plcb2 | 0.000110727 | 0.42644313 | 1 |
| ENSMUSG00000045838 | Ccdc9b | 0.000110727 | -2.032988489 | 0.076038495 |
| ENSMUSG00000027323 | Rad51 | 0.000110727 | -0.466641666 | 1 |
| ENSMUSG00000046814 | Gchfr | 0.000110727 | 3.233798052 | 0.262284476 |
| ENSMUSG00000034216 | Vps18 | 0.000110727 | -0.466641666 | 1 |
| ENSMUSG00000027305 | Ndufaf1 | 0.000110727 | -1.57355687 | 0.23866117 |
| ENSMUSG00000090100 | Ttbk2 | 0.000110727 | -0.088130043 | 1 |
| ENSMUSG00000048222 | Mfap1b | 0.000110727 | -0.014129461 | 1 |
| ENSMUSG00000027233 | Patl2 | 0.000110727 | 0.648835551 | 1 |
| ENSMUSG00000079071 | Gm14085 | 0.000110727 | 1.911869957 | 0.602510455 |
| ENSMUSG00000027202 | Slc12a1 | 0.000110727 | 0.774366434 | 0.924487348 |
| ENSMUSG00000027208 | Fgf7 | 0.000110727 | 0.533358334 | 1 |
| ENSMUSG00000023330 | Dtwd1 | 0.000110727 | 1.648835551 | 0.66416768 |
| ENSMUSG00000074825 | Itpripl1 | 0.000110727 | 0.911869957 | 1 |
| ENSMUSG00000027374 | Mrps5 | 0.000110727 | -0.158519371 | 1 |
| ENSMUSG00000027386 | Fbln7 | 0.000110727 | 1.063873051 | 0.847265357 |
| ENSMUSG00000087528 | 9830144P21Rik | 0.000110727 | 1.233798052 | 0.792441218 |
| ENSMUSG00000048327 | Ckap2l | 0.000110727 | -1.051604167 | 0.614718876 |
| ENSMUSG00000037885 | Stk35 | 0.000110727 | -0.895484965 | 0.691084416 |
| ENSMUSG00000053552 | Ebf4 | 0.000110727 | 0.42644313 | 1 |
| ENSMUSG00000027409 | 1700020A23Rik | 0.000110727 | 4.233798052 | 0.211844791 |
| ENSMUSG00000027411 | Vps16 | 0.000110727 | -3.051604167 | 0.001471155 |
| ENSMUSG00000027329 | Spef1 | 0.000110727 | -0.720398258 | 0.938724818 |
| ENSMUSG00000068264 | Ap5s1 | 0.000110727 | -0.158519371 | 1 |
| ENSMUSG00000027345 | 4921508D12Rik | 0.000110727 | 2.233798052 | 0.47966612 |
| ENSMUSG00000027353 | Mcm8 | 0.000110727 | -0.351164449 | 1 |
| ENSMUSG00000086650 | A430048G15Rik | 0.000110727 | 1.648835551 | 0.66416768 |
| ENSMUSG00000027261 | Hao1 | 0.000110727 | 1.648835551 | 0.664933061 |
| ENSMUSG00000027273 | Snap25 | 0.000110727 | -0.158519371 | 1 |
| ENSMUSG00000027281 | Slx4ip | 0.000110727 | -2.241935379 | 0.049826098 |
| ENSMUSG00000039092 | Sptlc3 | 0.000110727 | -0.52108945 | 1 |
| ENSMUSG00000086450 | Macrod2os1 | 0.000110727 | 0.326907457 | 1 |
| ENSMUSG00000086384 | Banf2os | 0.000110727 | 3.233798052 | 0.327136963 |
| ENSMUSG00000037279 | Ovol2 | 0.000110727 | -0.466641666 | 1 |

| | | | | |
|---|---|---|---|---|
| ENSMUSG00000037259 | Dzank1 | 0.000110727 | 0.326907457 | 1 |
| ENSMUSG00000074754 | Smim26 | 0.000110727 | 1.648835551 | 0.66416768 |
| ENSMUSG00000027430 | Dtd1 | 0.000110727 | -1.895484965 | 0.128972787 |
| ENSMUSG00000036992 | Nxt1 | 0.000110727 | 0.648835551 | 1 |
| ENSMUSG00000027438 | Napb | 0.000110727 | 1.42644313 | 0.736456705 |
| ENSMUSG00000087029 | Gm14133 | 0.000110727 | 1.063873051 | 0.847320319 |
| ENSMUSG00000033096 | Apmap | 0.000110727 | 0.063873051 | 1 |
| ENSMUSG00000027452 | Acss1 | 0.000110727 | -1.289763904 | 0.466464457 |
| ENSMUSG00000032802 | Srxn1 | 0.000110727 | 0.648835551 | 1 |
| ENSMUSG00000032715 | Trib3 | 0.000110727 | -0.936126949 | 0.697198109 |
| ENSMUSG00000051817 | Sox12 | 0.000110727 | 1.233798052 | 0.792441218 |
| ENSMUSG00000074681 | Defb23 | 0.000110727 | 3.233798052 | 0.305407814 |
| ENSMUSG00000009876 | Cox4i2 | 0.000110727 | 1.911869957 | 0.587484274 |
| ENSMUSG00000027472 | Pdrg1 | 0.000110727 | 0.533358334 | 1 |
| ENSMUSG00000027474 | Ccm2l | 0.000110727 | -0.52108945 | 1 |
| ENSMUSG00000073236 | 2500004C02Rik | 0.000110727 | 2.233798052 | 0.47966612 |
| ENSMUSG00000044083 | Efcab8 | 0.000110727 | 1.42644313 | 0.736456705 |
| ENSMUSG00000027487 | Cdk5rap1 | 0.000110727 | -2.57355687 | 0.012336561 |
| ENSMUSG00000027490 | E2f1 | 0.000110727 | -1.192466703 | 0.600618836 |
| ENSMUSG00000074627 | Mroh8 | 0.000110727 | -1.123753952 | 0.514974685 |
| ENSMUSG00000027650 | Tti1 | 0.000110727 | -0.766201948 | 0.792441218 |
| ENSMUSG00000085385 | Snhg17 | 0.000110727 | -1.599091962 | 0.242941974 |
| ENSMUSG00000086496 | Gm14204 | 0.000110727 | 2.648835551 | 0.395118486 |
| ENSMUSG00000027412 | Lpin3 | 0.000110727 | 0.233798052 | 1 |
| ENSMUSG00000035399 | Oser1 | 0.000110727 | -0.853664789 | 0.806496094 |
| ENSMUSG00000078949 | R3hdml | 0.000110727 | 1.648835551 | 0.66416768 |
| ENSMUSG00000017754 | Pltp | 0.000110727 | 0.648835551 | 1 |
| ENSMUSG00000085436 | Zfp335os | 0.000110727 | 0.648835551 | 1 |
| ENSMUSG00000039725 | Trp53rka | 0.000110727 | 0.326907457 | 1 |
| ENSMUSG00000085914 | Gm14320 | 0.000110727 | 1.233798052 | 0.792441218 |
| ENSMUSG00000085495 | Gm16796 | 0.000110727 | 0.774366434 | 1 |
| ENSMUSG00000043583 | 4930470P17Rik | 0.000110727 | 1.42644313 | 0.736456705 |
| ENSMUSG00000027495 | Fam210b | 0.000110727 | 0.648835551 | 1 |
| ENSMUSG00000027510 | Rbm38 | 0.000110727 | 0.648835551 | 0.916431043 |
| ENSMUSG00000087382 | Ctcflos | 0.000110727 | -0.57355687 | 1 |
| ENSMUSG00000027522 | Stx16 | 0.000110727 | -2.995020638 | 0.001619835 |
| ENSMUSG00000086537 | Gnasas1 | 0.000110727 | -0.52108945 | 1 |
| ENSMUSG00000095362 | Gm14325 | 0.000110727 | -0.410058138 | 1 |
| ENSMUSG00000039086 | Ss18l1 | 0.000110727 | -0.289763904 | 1 |
| ENSMUSG00000038990 | Cables2 | 0.000110727 | -0.766201948 | 0.792441218 |
| ENSMUSG00000027573 | Gid8 | 0.000110727 | -0.720398258 | 0.792441218 |
| ENSMUSG00000027580 | Helz2 | 0.000110727 | -0.624182943 | 0.912904617 |
| ENSMUSG00000046470 | Sox18 | 0.000110727 | 2.648835551 | 0.395118486 |
| ENSMUSG00000103034 | Gm8797 | 0.000110727 | 1.648835551 | 0.66416768 |
| ENSMUSG00000027499 | Pkia | 0.000110727 | -0.936126949 | 0.756492825 |
| ENSMUSG00000027555 | Car13 | 0.000110727 | -0.088130043 | 1 |
| ENSMUSG00000106019 | Gm43672 | 0.000110727 | 0.42644313 | 1 |
| ENSMUSG00000037730 | Mynn | 0.000110727 | -1.410058138 | 0.455131546 |
| ENSMUSG00000027703 | Lrriq4 | 0.000110727 | 2.648835551 | 0.395118486 |
| ENSMUSG00000103998 | Gm38025 | 0.000110727 | 4.233798052 | 0.128972787 |
| ENSMUSG00000092196 | Gm38505 | 0.000110727 | 0.774366434 | 1 |

| | | | | |
|---|---|---|---|---|
| ENSMUSG00000106525 | Gm42695 | 0.000110727 | 2.648835551 | 0.395118486 |
| ENSMUSG00000033883 | Zfp267 | 0.000110727 | -1.014129461 | 0.608426921 |
| ENSMUSG00000085320 | Gm11548 | 0.000110727 | 2.233798052 | 0.506177813 |
| ENSMUSG00000027714 | Exosc9 | 0.000110727 | 0.063873051 | 1 |
| ENSMUSG00000027719 | Adad1 | 0.000110727 | 1.648835551 | 0.66416768 |
| ENSMUSG00000027720 | Il2 | 0.000110727 | 1.233798052 | 0.792441218 |
| ENSMUSG00000106441 | Gm42921 | 0.000110727 | 2.233798052 | 0.5015528 |
| ENSMUSG00000025766 | D3Ertd751e | 0.000110727 | -0.810596067 | 0.796840156 |
| ENSMUSG00000103965 | Gm30173 | 0.000110727 | 1.911869957 | 0.597898961 |
| ENSMUSG00000027742 | Cog6 | 0.000110727 | -1.123753952 | 0.514974685 |
| ENSMUSG00000042997 | Nhlrc3 | 0.000110727 | -0.289763904 | 1 |
| ENSMUSG00000027796 | Smad9 | 0.000110727 | 0.911869957 | 1 |
| ENSMUSG00000056306 | Sertm1 | 0.000110727 | 0.326907457 | 1 |
| ENSMUSG00000105975 | Gm42609 | 0.000110727 | 3.233798052 | 0.307566094 |
| ENSMUSG00000036381 | P2ry14 | 0.000110727 | -0.289763904 | 1 |
| ENSMUSG00000027832 | Ptx3 | 0.000110727 | 1.063873051 | 0.843995018 |
| ENSMUSG00000048416 | Mlf1 | 0.000110727 | 0.326907457 | 1 |
| ENSMUSG00000034317 | Trim59 | 0.000110727 | -2.624182943 | 0.007725935 |
| ENSMUSG00000027790 | Sis | 0.000110727 | 1.063873051 | 1 |
| ENSMUSG00000027834 | Serpini1 | 0.000110727 | -0.766201948 | 0.801811393 |
| ENSMUSG00000104585 | 4921511C10Rik | 0.000110727 | 0.326907457 | 1 |
| ENSMUSG00000034009 | Rxfp1 | 0.000110727 | -0.673092543 | 0.919998879 |
| ENSMUSG00000102191 | Gm36569 | 0.000110727 | 2.233798052 | 0.485615738 |
| ENSMUSG00000027955 | Gask1b | 0.000110727 | 0.648835551 | 1 |
| ENSMUSG00000028020 | Glrb | 0.000110727 | -2.241935379 | 0.060315801 |
| ENSMUSG00000056498 | Tmem154 | 0.000110727 | 0.063873051 | 1 |
| ENSMUSG00000028085 | Gatb | 0.000110727 | -1.788569761 | 0.142787052 |
| ENSMUSG00000028076 | Cd1d1 | 0.000110727 | 1.42644313 | 0.736456705 |
| ENSMUSG00000015854 | Cd5l | 0.000110727 | 0.911869957 | 1 |
| ENSMUSG00000008601 | Rab25 | 0.000110727 | 0.42644313 | 1 |
| ENSMUSG00000032657 | Fam189b | 0.000110727 | -0.225633566 | 1 |
| ENSMUSG00000109293 | Dcst2 | 0.000110727 | 1.063873051 | 1 |
| ENSMUSG00000027938 | Creb3l4 | 0.000110727 | 2.648835551 | 0.395118486 |
| ENSMUSG00000049133 | Flg2 | 0.000110727 | 1.911869957 | 0.602510455 |
| ENSMUSG00000013707 | Tnfaip8l2 | 0.000110727 | 2.233798052 | 0.509597331 |
| ENSMUSG00000015850 | Adamtsl4 | 0.000110727 | -1.410058138 | 0.342524494 |
| ENSMUSG00000038543 | BC028528 | 0.000110727 | 1.911869957 | 0.590972065 |
| ENSMUSG00000015747 | Vps45 | 0.000110727 | -2.158519371 | 0.044761838 |
| ENSMUSG00000015943 | Bola1 | 0.000110727 | 1.233798052 | 0.792441218 |
| ENSMUSG00000118406 | AC121792.1 | 0.000110727 | 0.911869957 | 1 |
| ENSMUSG00000028100 | Nudt17 | 0.000110727 | 2.233798052 | 0.504119351 |
| ENSMUSG00000038304 | Cd160 | 0.000110727 | 1.911869957 | 0.587484274 |
| ENSMUSG00000038298 | Pdzk1 | 0.000110727 | 2.648835551 | 0.395118486 |
| ENSMUSG00000028088 | Fmo5 | 0.000110727 | 0.533358334 | 1 |
| ENSMUSG00000038205 | Prkab2 | 0.000110727 | -1.014129461 | 0.608426921 |
| ENSMUSG00000027868 | Tbx15 | 0.000110727 | 0.063873051 | 1 |
| ENSMUSG00000033222 | Ttf2 | 0.000110727 | -1.956026507 | 0.086523965 |
| ENSMUSG00000086564 | Cd101 | 0.000110727 | 1.063873051 | 1 |
| ENSMUSG00000033147 | Slc22a15 | 0.000110727 | -1.624182943 | 0.24944595 |
| ENSMUSG00000106296 | 4632404M16Rik | 0.000110727 | 1.911869957 | 0.587484274 |
| ENSMUSG00000027859 | Ngf | 0.000110727 | -1.380911792 | 0.435791313 |

| | | | | |
|---|---|---|---|---|
| ENSMUSG00000027855 | Sycp1 | 0.000110727 | 0.648835551 | 1 |
| ENSMUSG00000027849 | Syt6 | 0.000110727 | 0.774366434 | 1 |
| ENSMUSG00000104721 | Gm42696 | 0.000110727 | 1.648835551 | 0.66416768 |
| ENSMUSG00000027840 | Wnt2b | 0.000110727 | 2.233798052 | 0.493461798 |
| ENSMUSG00000074344 | Tmigd3 | 0.000110727 | 1.911869957 | 0.597898961 |
| ENSMUSG00000043873 | Chil5 | 0.000110727 | 3.233798052 | 0.262284476 |
| ENSMUSG00000027902 | Chil6 | 0.000110727 | 4.233798052 | 0.128972787 |
| ENSMUSG00000094613 | A630076J17Rik | 0.000110727 | 1.911869957 | 0.587484274 |
| ENSMUSG00000027896 | Slc16a4 | 0.000110727 | 1.911869957 | 0.587484274 |
| ENSMUSG00000040562 | Gstm2 | 0.000110727 | 0.911869957 | 1 |
| ENSMUSG00000027886 | 1700013F07Rik | 0.000110727 | 1.911869957 | 0.587484274 |
| ENSMUSG00000045326 | Fndc7 | 0.000110727 | 0.42644313 | 1 |
| ENSMUSG00000039831 | Arhgap29 | 0.000110727 | -2.141241379 | 0.054954924 |
| ENSMUSG00000057359 | Gm17494 | 0.000110727 | -0.895484965 | 0.736456705 |
| ENSMUSG00000023057 | Fabp2 | 0.000110727 | 3.233798052 | 0.262284476 |
| ENSMUSG00000027978 | Prss12 | 0.000110727 | -1.832291138 | 0.182205252 |
| ENSMUSG00000104763 | 9830132P13Rik | 0.000110727 | -1.624182943 | 0.295890024 |
| ENSMUSG00000041220 | Elovl6 | 0.000110727 | -2.3207908 | 0.029965083 |
| ENSMUSG00000105789 | Gm42997 | 0.000110727 | 0.233798052 | 1 |
| ENSMUSG00000104556 | Gm43192 | 0.000110727 | 1.233798052 | 0.792441218 |
| ENSMUSG00000046818 | Ddit4l | 0.000110727 | -0.351164449 | 1 |
| ENSMUSG00000105427 | Gm31678 | 0.000110727 | 3.233798052 | 0.274319469 |
| ENSMUSG00000090066 | 1110002E22Rik | 0.000110727 | 1.911869957 | 0.587484274 |
| ENSMUSG00000089635 | Gm16559 | 0.000110727 | 4.233798052 | 0.128972787 |
| ENSMUSG00000105504 | Gbp5 | 0.000110727 | 0.326907457 | 1 |
| ENSMUSG00000040253 | Gbp7 | 0.000110727 | 0.911869957 | 1 |
| ENSMUSG00000028268 | Gbp3 | 0.000110727 | 1.42644313 | 0.736456705 |
| ENSMUSG00000105511 | Gm33758 | 0.000110727 | 3.233798052 | 0.262284476 |
| ENSMUSG00000105324 | Gm42705 | 0.000110727 | 0.911869957 | 1 |
| ENSMUSG00000028262 | Clca3a2 | 0.000110727 | 2.648835551 | 0.409925097 |
| ENSMUSG00000036853 | Mcoln3 | 0.000110727 | 2.233798052 | 0.47966612 |
| ENSMUSG00000103349 | Gm36888 | 0.000110727 | 3.233798052 | 0.290854149 |
| ENSMUSG00000104786 | Gm43573 | 0.000110727 | 0.774366434 | 1 |
| ENSMUSG00000106515 | Gm30382 | 0.000110727 | -1.192466703 | 0.580988658 |
| ENSMUSG00000105797 | Gm5149 | 0.000110727 | 0.774366434 | 1 |
| ENSMUSG00000028037 | Ifi44 | 0.000110727 | 1.648835551 | 0.66416768 |
| ENSMUSG00000028035 | Dnajb4 | 0.000110727 | -1.599091962 | 0.242941974 |
| ENSMUSG00000005493 | Msh4 | 0.000110727 | -0.57355687 | 0.923648502 |
| ENSMUSG00000053870 | Fpgt | 0.000110727 | 0.911869957 | 1 |
| ENSMUSG00000028233 | Tgs1 | 0.000110727 | -2.547561661 | 0.012733015 |
| ENSMUSG00000042198 | Chchd7 | 0.000110727 | -0.766201948 | 0.792441218 |
| ENSMUSG00000050323 | Ndufaf6 | 0.000110727 | -0.57355687 | 0.910050347 |
| ENSMUSG00000078773 | Rad54b | 0.000110727 | -1.088130043 | 0.624094312 |
| ENSMUSG00000049225 | Pdp1 | 0.000110727 | 0.648835551 | 1 |
| ENSMUSG00000052137 | Rbm12b2 | 0.000110727 | -2.754886635 | 0.004683913 |
| ENSMUSG00000055963 | Triqk | 0.000110727 | 1.233798052 | 0.792441218 |
| ENSMUSG00000086927 | Gm11823 | 0.000110727 | 0.774366434 | 1 |
| ENSMUSG00000040569 | Slc26a7 | 0.000110727 | 3.233798052 | 0.262284476 |
| ENSMUSG00000085112 | A530072M11Rik | 0.000110727 | -0.810596067 | 0.796840156 |
| ENSMUSG00000086072 | Gm11861 | 0.000110727 | 1.063873051 | 0.848685201 |
| ENSMUSG00000078772 | Gm12353 | 0.000110727 | 0.42644313 | 1 |

| | | | | |
|---|---|---|---|---|
| ENSMUSG00000040478 | Prdm13 | 0.000110727 | 3.233798052 | 0.264129704 |
| ENSMUSG00000028278 | Rragd | 0.000110727 | 0.648835551 | 1 |
| ENSMUSG00000085037 | 4933421O10Rik | 0.000110727 | 4.233798052 | 0.128972787 |
| ENSMUSG00000084828 | Gm12367 | 0.000110727 | 0.42644313 | 1 |
| ENSMUSG00000028430 | Nol6 | 0.000110727 | -0.936126949 | 0.697198109 |
| ENSMUSG00000028438 | Kif24 | 0.000110727 | -1.788569761 | 0.142787052 |
| ENSMUSG00000036114 | Rpp25l | 0.000110727 | 0.233798052 | 1 |
| ENSMUSG00000028466 | Creb3 | 0.000110727 | -0.289763904 | 1 |
| ENSMUSG00000028468 | Rgp1 | 0.000110727 | 0.326907457 | 1 |
| ENSMUSG00000086043 | Gm12473 | 0.000110727 | 3.233798052 | 0.310582691 |
| ENSMUSG00000028469 | Npr2 | 0.000110727 | -0.766201948 | 0.792441218 |
| ENSMUSG00000078716 | Tmem8b | 0.000110727 | -1.696939285 | 0.210550914 |
| ENSMUSG00000028476 | Reck | 0.000110727 | 0.063873051 | 1 |
| ENSMUSG00000086284 | Frmpd1os | 0.000110727 | -0.225633566 | 1 |
| ENSMUSG00000028322 | Exosc3 | 0.000110727 | 0.146335211 | 1 |
| ENSMUSG00000028330 | Ncbp1 | 0.000110727 | -1.915949067 | 0.103463818 |
| ENSMUSG00000028329 | Xpa | 0.000110727 | -2.720398258 | 0.005298129 |
| ENSMUSG00000028348 | Cavin4 | 0.000110727 | 0.911869957 | 1 |
| ENSMUSG00000028312 | Smc2 | 0.000110727 | -1.956026507 | 0.086523965 |
| ENSMUSG00000028414 | Fktn | 0.000110727 | -1.766201948 | 0.177083054 |
| ENSMUSG00000087044 | 1700042G15Rik | 0.000110727 | 0.648835551 | 1 |
| ENSMUSG00000073846 | Gm12526 | 0.000110727 | 1.42644313 | 0.736456705 |
| ENSMUSG00000028389 | Zfp37 | 0.000110727 | 1.233798052 | 0.792441218 |
| ENSMUSG00000038422 | Hdhd3 | 0.000110727 | 2.648835551 | 0.455131546 |
| ENSMUSG00000039158 | Akna | 0.000110727 | -0.088130043 | 1 |
| ENSMUSG00000039137 | Whrn | 0.000110727 | 0.233798052 | 1 |
| ENSMUSG00000058935 | Tex48 | 0.000110727 | -2.273996588 | 0.043231237 |
| ENSMUSG00000039298 | Cdk5rap2 | 0.000110727 | -3.250017725 | 0.000428474 |
| ENSMUSG00000084937 | Gm11228 | 0.000110727 | 1.42644313 | 0.736456705 |
| ENSMUSG00000087110 | Gm11264 | 0.000110727 | 2.648835551 | 0.395118486 |
| ENSMUSG00000086074 | Gm13274 | 0.000110727 | 1.648835551 | 0.66416768 |
| ENSMUSG00000086251 | Gm12694 | 0.000110727 | 0.911869957 | 1 |
| ENSMUSG00000028571 | Cyp2j13 | 0.000110727 | 0.42644313 | 1 |
| ENSMUSG00000086249 | Gm12724 | 0.000110727 | 1.063873051 | 0.85551189 |
| ENSMUSG00000085933 | Tmem61 | 0.000110727 | 1.648835551 | 0.66416768 |
| ENSMUSG00000054362 | Lexm | 0.000110727 | 0.774366434 | 1 |
| ENSMUSG00000028613 | Lrp8 | 0.000110727 | -1.351164449 | 0.401963028 |
| ENSMUSG00000008932 | Slc1a7 | 0.000110727 | 1.063873051 | 1 |
| ENSMUSG00000084968 | Gm12743 | 0.000110727 | 0.648835551 | 1 |
| ENSMUSG00000086483 | 8030443G20Rik | 0.000110727 | 1.42644313 | 0.736456705 |
| ENSMUSG00000028546 | Elavl4 | 0.000110727 | -0.52108945 | 1 |
| ENSMUSG00000049214 | Skint7 | 0.000110727 | 0.911869957 | 0.886203497 |
| ENSMUSG00000028709 | Mob3c | 0.000110727 | 0.42644313 | 1 |
| ENSMUSG00000028699 | Tspan1 | 0.000110727 | 4.233798052 | 0.128972787 |
| ENSMUSG00000033985 | Tesk2 | 0.000110727 | -2.946111038 | 0.002012957 |
| ENSMUSG00000028688 | Toe1 | 0.000110727 | -0.853664789 | 0.847265357 |
| ENSMUSG00000028678 | Kif2c | 0.000110727 | -0.895484965 | 0.691084416 |
| ENSMUSG00000006390 | Elovl1 | 0.000110727 | -0.52108945 | 1 |
| ENSMUSG00000006398 | Cdc20 | 0.000110727 | 0.911869957 | 1 |
| ENSMUSG00000085895 | Gm12861 | 0.000110727 | 1.648835551 | 0.66416768 |
| ENSMUSG00000070806 | Zmynd12 | 0.000110727 | 1.42644313 | 0.736456705 |

| | | | | |
|---|---|---|---|---|
| ENSMUSG00000091297 | Gm8439 | 0.000110727 | -0.410058138 | 1 |
| ENSMUSG00000028631 | Kcnq4 | 0.000110727 | 0.648835551 | 1 |
| ENSMUSG00000032890 | Rims3 | 0.000110727 | 2.648835551 | 0.395118486 |
| ENSMUSG00000073764 | Gm12888 | 0.000110727 | 4.233798052 | 0.128972787 |
| ENSMUSG00000086955 | 1700021L23Rik | 0.000110727 | 3.233798052 | 0.262284476 |
| ENSMUSG00000086589 | Gm12915 | 0.000110727 | 0.326907457 | 1 |
| ENSMUSG00000050213 | Snip1 | 0.000110727 | -0.57355687 | 0.910050347 |
| ENSMUSG00000085562 | 2610028E06Rik | 0.000110727 | 4.233798052 | 0.128972787 |
| ENSMUSG00000028859 | Csf3r | 0.000110727 | -2.032988489 | 0.08660337 |
| ENSMUSG00000042616 | Oscp1 | 0.000110727 | -0.225633566 | 1 |
| ENSMUSG00000042388 | Dlgap3 | 0.000110727 | 0.233798052 | 1 |
| ENSMUSG00000061894 | Zscan20 | 0.000110727 | -0.410058138 | 1 |
| ENSMUSG00000001333 | Sync | 0.000110727 | 0.326907457 | 1 |
| ENSMUSG00000040795 | Iqcc | 0.000110727 | -0.52108945 | 1 |
| ENSMUSG00000053730 | Tmem39b | 0.000110727 | -0.088130043 | 1 |
| ENSMUSG00000028784 | Spocd1 | 0.000110727 | 0.533358334 | 1 |
| ENSMUSG00000028782 | Adgrb2 | 0.000110727 | 1.233798052 | 0.792441218 |
| ENSMUSG00000028909 | Ptpru | 0.000110727 | 0.533358334 | 1 |
| ENSMUSG00000028862 | Map3k6 | 0.000110727 | 1.911869957 | 0.587484274 |
| ENSMUSG00000059713 | Rcan3 | 0.000110727 | -1.192466703 | 0.529126167 |
| ENSMUSG00000037188 | Grhl3 | 0.000110727 | 0.911869957 | 1 |
| ENSMUSG00000028671 | Gale | 0.000110727 | 0.774366434 | 1 |
| ENSMUSG00000007872 | Id3 | 0.000110727 | 1.648835551 | 0.66416768 |
| ENSMUSG00000036995 | Asap3 | 0.000110727 | -1.895484965 | 0.128972787 |
| ENSMUSG00000051351 | Zfp46 | 0.000110727 | 0.533358334 | 1 |
| ENSMUSG00000087476 | Rap1gapos | 0.000110727 | 1.233798052 | 0.792441218 |
| ENSMUSG00000028766 | Alpl | 0.000110727 | -0.52108945 | 1 |
| ENSMUSG00000045349 | Sh2d5 | 0.000110727 | 2.648835551 | 0.42263302 |
| ENSMUSG00000028758 | Kif17 | 0.000110727 | -0.410058138 | 1 |
| ENSMUSG00000070661 | Rnf186 | 0.000110727 | 3.233798052 | 0.262284476 |
| ENSMUSG00000028747 | Htr6 | 0.000110727 | 2.233798052 | 0.498076455 |
| ENSMUSG00000085928 | 4933427I22Rik | 0.000110727 | 1.063873051 | 0.854285611 |
| ENSMUSG00000086960 | Gm13017 | 0.000110727 | 2.648835551 | 0.433732559 |
| ENSMUSG00000086159 | Gm13025 | 0.000110727 | 1.42644313 | 0.736456705 |
| ENSMUSG00000025329 | Padi1 | 0.000110727 | 0.326907457 | 1 |
| ENSMUSG00000028919 | Arhgef19 | 0.000110727 | -0.225633566 | 1 |
| ENSMUSG00000085395 | Gm13056 | 0.000110727 | 1.648835551 | 0.66416768 |
| ENSMUSG00000006218 | Fam131c | 0.000110727 | 2.233798052 | 0.481842598 |
| ENSMUSG00000046862 | Pramef8 | 0.000110727 | 1.42644313 | 0.736456705 |
| ENSMUSG00000086046 | 1700095A21Rik | 0.000110727 | 0.233798052 | 1 |
| ENSMUSG00000078495 | Zfp984 | 0.000110727 | 1.063873051 | 1 |
| ENSMUSG00000041556 | Fbxo2 | 0.000110727 | 3.233798052 | 0.262284476 |
| ENSMUSG00000028989 | Angptl7 | 0.000110727 | 2.648835551 | 0.432730061 |
| ENSMUSG00000086740 | Gm17029 | 0.000110727 | 2.233798052 | 0.47966612 |
| ENSMUSG00000028992 | Nmnat1 | 0.000110727 | -0.088130043 | 1 |
| ENSMUSG00000028990 | Lzic | 0.000110727 | 0.063873051 | 1 |
| ENSMUSG00000028957 | Per3 | 0.000110727 | -0.52108945 | 1 |
| ENSMUSG00000028952 | Zbtb48 | 0.000110727 | -0.351164449 | 1 |
| ENSMUSG00000039713 | Plekhg5 | 0.000110727 | -1.289763904 | 0.466464457 |
| ENSMUSG00000058498 | Rnf207 | 0.000110727 | 2.648835551 | 0.421384409 |
| ENSMUSG00000078350 | Smim1 | 0.000110727 | 1.42644313 | 0.736456705 |

| | | | | |
|---|---|---|---|---|
| ENSMUSG00000029056 | Pank4 | 0.000110727 | -0.720398258 | 0.792441218 |
| ENSMUSG00000029047 | Pex10 | 0.000110727 | 1.648835551 | 0.66416768 |
| ENSMUSG00000073682 | Gm10563 | 0.000110727 | 0.911869957 | 1 |
| ENSMUSG00000074738 | Fndc10 | 0.000110727 | 4.233798052 | 0.128972787 |
| ENSMUSG00000029070 | Mxra8 | 0.000110727 | -0.088130043 | 1 |
| ENSMUSG00000029033 | Acap3 | 0.000110727 | -0.673092543 | 0.919998879 |
| ENSMUSG00000087381 | Gm16008 | 0.000110727 | -1.648844997 | 0.201326391 |
| ENSMUSG00000078485 | Plekhn1 | 0.000110727 | -1.123753952 | 0.514974685 |
| ENSMUSG00000040302 | Rbm48 | 0.000110727 | -1.289763904 | 0.466464457 |
| ENSMUSG00000040429 | Mterf1a | 0.000110727 | 0.42644313 | 1 |
| ENSMUSG00000086528 | Gm15731 | 0.000110727 | 0.774366434 | 1 |
| ENSMUSG00000028970 | Abcb1b | 0.000110727 | -2.106051951 | 0.100510303 |
| ENSMUSG00000079659 | Tmem243 | 0.000110727 | -0.466641666 | 1 |
| ENSMUSG00000056004 | 9330182L06Rik | 0.000110727 | 0.774366434 | 1 |
| ENSMUSG00000106004 | Gm43391 | 0.000110727 | 1.063873051 | 1 |
| ENSMUSG00000028864 | Hgf | 0.000110727 | 1.42644313 | 0.736456705 |
| ENSMUSG00000109903 | Gm28710 | 0.000110727 | 0.911869957 | 0.886203497 |
| ENSMUSG00000002944 | Cd36 | 0.000110727 | 0.533358334 | 1 |
| ENSMUSG00000045435 | Tmem60 | 0.000110727 | 0.063873051 | 1 |
| ENSMUSG00000101013 | A630072M18Rik | 0.000110727 | -0.225633566 | 1 |
| ENSMUSG00000064280 | Ccdc146 | 0.000110727 | 0.648835551 | 1 |
| ENSMUSG00000039899 | Fgl2 | 0.000110727 | 1.648835551 | 0.66416768 |
| ENSMUSG00000044968 | Napepld | 0.000110727 | 0.146335211 | 1 |
| ENSMUSG00000028973 | Abcb8 | 0.000110727 | -2.810596067 | 0.003605503 |
| ENSMUSG00000028959 | Fastk | 0.000110727 | -0.624182943 | 0.912904617 |
| ENSMUSG00000067724 | Gbx1 | 0.000110727 | 1.063873051 | 0.853473606 |
| ENSMUSG00000101856 | 1700096K18Rik | 0.000110727 | 1.42644313 | 0.736456705 |
| ENSMUSG00000059645 | Gm7361 | 0.000110727 | 4.233798052 | 0.128972787 |
| ENSMUSG00000073102 | Drc1 | 0.000110727 | -0.014129461 | 1 |
| ENSMUSG00000062372 | Otof | 0.000110727 | -0.351164449 | 1 |
| ENSMUSG00000029175 | Slc35f6 | 0.000110727 | 0.533358334 | 1 |
| ENSMUSG00000013629 | Cad | 0.000110727 | -0.289763904 | 1 |
| ENSMUSG00000053134 | Supt7l | 0.000110727 | 0.533358334 | 1 |
| ENSMUSG00000054252 | Fgfr3 | 0.000110727 | 0.648835551 | 1 |
| ENSMUSG00000029111 | Nelfa | 0.000110727 | -0.410058138 | 1 |
| ENSMUSG00000045102 | Poln | 0.000110727 | -0.810596067 | 0.825653816 |
| ENSMUSG00000052783 | Grk4 | 0.000110727 | -1.696939285 | 0.210550914 |
| ENSMUSG00000029102 | Hgfac | 0.000110727 | -0.466641666 | 1 |
| ENSMUSG00000029097 | Trmt44 | 0.000110727 | -0.810596067 | 0.796840156 |
| ENSMUSG00000036553 | Sh3tc1 | 0.000110727 | -1.466641666 | 0.346237121 |
| ENSMUSG00000029196 | Tada2b | 0.000110727 | 0.42644313 | 1 |
| ENSMUSG00000060708 | Bloc1s4 | 0.000110727 | 0.774366434 | 1 |
| ENSMUSG00000029122 | Evc | 0.000110727 | -0.52108945 | 1 |
| ENSMUSG00000050248 | Evc2 | 0.000110727 | -1.258055044 | 0.466464457 |
| ENSMUSG00000029126 | Nsg1 | 0.000110727 | 2.233798052 | 0.47966612 |
| ENSMUSG00000106794 | Gm40293 | 0.000110727 | -1.810596067 | 0.145553211 |
| ENSMUSG00000067285 | Gm16223 | 0.000110727 | 0.326907457 | 1 |
| ENSMUSG00000105423 | Gm36814 | 0.000110727 | 0.326907457 | 1 |
| ENSMUSG00000029173 | Sepsecs | 0.000110727 | -0.975655313 | 0.705975399 |
| ENSMUSG00000106352 | 5033403H07Rik | 0.000110727 | -1.766201948 | 0.250869635 |
| ENSMUSG00000106013 | 4933402J10Rik | 0.000110727 | 2.233798052 | 0.47966612 |

| | | | | |
|---|---|---|---|---|
| ENSMUSG00000090326 | Dthd1 | 0.000110727 | 0.533358334 | 1 |
| ENSMUSG00000106219 | 5830416I19Rik | 0.000110727 | -1.832291138 | 0.201230294 |
| ENSMUSG00000105105 | Gm31363 | 0.000110727 | 0.533358334 | 1 |
| ENSMUSG00000105402 | Gm3716 | 0.000110727 | -1.088130043 | 0.624094312 |
| ENSMUSG00000037913 | Tmem156 | 0.000110727 | 1.233798052 | 0.792441218 |
| ENSMUSG00000029195 | Klb | 0.000110727 | -0.158519371 | 1 |
| ENSMUSG00000029204 | Rhoh | 0.000110727 | 0.911869957 | 1 |
| ENSMUSG00000097092 | Gm26725 | 0.000110727 | 0.774366434 | 1 |
| ENSMUSG00000049387 | Cox7b2 | 0.000110727 | 0.774366434 | 1 |
| ENSMUSG00000105489 | Gm38562 | 0.000110727 | 1.911869957 | 0.587484274 |
| ENSMUSG00000029233 | Srd5a3 | 0.000110727 | -0.088130043 | 1 |
| ENSMUSG00000061184 | Tmprss11c | 0.000110727 | 0.648835551 | 1 |
| ENSMUSG00000048764 | Tmprss11f | 0.000110727 | -0.225633566 | 1 |
| ENSMUSG00000029269 | Sult1b1 | 0.000110727 | 1.648835551 | 0.66416768 |
| ENSMUSG00000029272 | Sult1e1 | 0.000110727 | 1.911869957 | 0.596304264 |
| ENSMUSG00000029288 | Ambn | 0.000110727 | 2.648835551 | 0.395118486 |
| ENSMUSG00000035528 | Npffr2 | 0.000110727 | 1.648835551 | 0.66416768 |
| ENSMUSG00000106863 | Gm42109 | 0.000110727 | 0.911869957 | 1 |
| ENSMUSG00000045314 | Sowahb | 0.000110727 | 0.648835551 | 1 |
| ENSMUSG00000106357 | Gm8013 | 0.000110727 | 1.648835551 | 0.66416768 |
| ENSMUSG00000034663 | Bmp2k | 0.000110727 | -2.123753952 | 0.053533761 |
| ENSMUSG00000105850 | Gm2861 | 0.000110727 | 2.233798052 | 0.47966612 |
| ENSMUSG00000105579 | Gm43251 | 0.000110727 | -0.088130043 | 1 |
| ENSMUSG00000089743 | Gm16226 | 0.000110727 | 0.774366434 | 1 |
| ENSMUSG00000105218 | Gm38413 | 0.000110727 | 2.233798052 | 0.509878973 |
| ENSMUSG00000105078 | Vamp9 | 0.000110727 | 1.063873051 | 1 |
| ENSMUSG00000029326 | Enoph1 | 0.000110727 | -0.014129461 | 1 |
| ENSMUSG00000105615 | Gm31048 | 0.000110727 | 3.233798052 | 0.262284476 |
| ENSMUSG00000097392 | D930016D06Rik | 0.000110727 | -2.051604167 | 0.078768438 |
| ENSMUSG00000072774 | Zfp951 | 0.000110727 | -1.123753952 | 0.514974685 |
| ENSMUSG00000034438 | Gbp8 | 0.000110727 | -0.52108945 | 1 |
| ENSMUSG00000105531 | 4930458A03Rik | 0.000110727 | 1.911869957 | 0.588199275 |
| ENSMUSG00000090558 | Gm17202 | 0.000110727 | 2.233798052 | 0.47966612 |
| ENSMUSG00000090030 | A430072P03Rik | 0.000110727 | 4.233798052 | 0.128972787 |
| ENSMUSG00000029491 | Pde6b | 0.000110727 | 0.42644313 | 1 |
| ENSMUSG00000004815 | Dgkq | 0.000110727 | -1.051604167 | 0.661861928 |
| ENSMUSG00000091635 | Vmn2r13 | 0.000110727 | 4.233798052 | 0.128972787 |
| ENSMUSG00000090963 | Gm17655 | 0.000110727 | -1.743481871 | 0.172404208 |
| ENSMUSG00000086401 | Gm15559 | 0.000110727 | -0.088130043 | 1 |
| ENSMUSG00000029507 | Pus1 | 0.000110727 | -1.3207908 | 0.395686633 |
| ENSMUSG00000052848 | C130026L21Rik | 0.000110727 | 1.911869957 | 0.587484274 |
| ENSMUSG00000029344 | Tpst2 | 0.000110727 | -2.225633566 | 0.039583356 |
| ENSMUSG00000029591 | Ung | 0.000110727 | 1.42644313 | 0.736456705 |
| ENSMUSG00000066952 | Myo1h | 0.000110727 | -1.3207908 | 0.50809997 |
| ENSMUSG00000087477 | Gm13822 | 0.000110727 | 1.911869957 | 0.587484274 |
| ENSMUSG00000060152 | Pop5 | 0.000110727 | 0.648835551 | 1 |
| ENSMUSG00000086368 | Gm13830 | 0.000110727 | -1.225633566 | 0.463954265 |
| ENSMUSG00000029536 | Gatc | 0.000110727 | 1.233798052 | 0.792441218 |
| ENSMUSG00000029535 | Triap1 | 0.000110727 | 1.063873051 | 0.847265357 |
| ENSMUSG00000054256 | Msi1 | 0.000110727 | -0.014129461 | 1 |
| ENSMUSG00000046607 | Hrk | 0.000110727 | 1.233798052 | 0.792441218 |

| | | | | |
|---|---|---|---|---|
| ENSMUSG00000018604 | Tbx3 | 0.000110727 | 1.063873051 | 1 |
| ENSMUSG00000001166 | Oas1c | 0.000110727 | 2.233798052 | 0.488204097 |
| ENSMUSG00000029452 | Tmem116 | 0.000110727 | -1.624182943 | 0.27804835 |
| ENSMUSG00000042647 | Acad12 | 0.000110727 | -0.158519371 | 1 |
| ENSMUSG00000029456 | Acad10 | 0.000110727 | -0.895484965 | 0.697312717 |
| ENSMUSG00000064267 | Hvcn1 | 0.000110727 | 0.533358334 | 1 |
| ENSMUSG00000038593 | Tctn1 | 0.000110727 | -0.225633566 | 1 |
| ENSMUSG00000086700 | Gm15747 | 0.000110727 | 0.648835551 | 1 |
| ENSMUSG00000106261 | Gm34086 | 0.000110727 | 0.42644313 | 1 |
| ENSMUSG00000104818 | Gm43661 | 0.000110727 | -0.158519371 | 1 |
| ENSMUSG00000029408 | Abcb9 | 0.000110727 | -0.158519371 | 1 |
| ENSMUSG00000029406 | Pitpnm2 | 0.000110727 | -1.547561661 | 0.290722159 |
| ENSMUSG00000047635 | 2810006K23Rik | 0.000110727 | 0.326907457 | 1 |
| ENSMUSG00000029402 | Snrnp35 | 0.000110727 | -0.225633566 | 1 |
| ENSMUSG00000038011 | Dnah10 | 0.000110727 | -0.766201948 | 0.83580699 |
| ENSMUSG00000104932 | Gm40323 | 0.000110727 | 1.233798052 | 0.792441218 |
| ENSMUSG00000029480 | Dhx37 | 0.000110727 | -1.051604167 | 0.614718876 |
| ENSMUSG00000106644 | Gm40331 | 0.000110727 | 2.233798052 | 0.509878973 |
| ENSMUSG00000105490 | 4933438B17Rik | 0.000110727 | 0.911869957 | 1 |
| ENSMUSG00000034173 | Zbed5 | 0.000110727 | 0.533358334 | 1 |
| ENSMUSG00000106395 | Gm42439 | 0.000110727 | 3.233798052 | 0.262284476 |
| ENSMUSG00000105227 | Gm36667 | 0.000110727 | 1.648835551 | 0.66416768 |
| ENSMUSG00000015950 | Ncf1 | 0.000110727 | 0.42644313 | 1 |
| ENSMUSG00000107090 | Gm42884 | 0.000110727 | 2.648835551 | 0.395118486 |
| ENSMUSG00000107318 | Gm30003 | 0.000110727 | 0.063873051 | 1 |
| ENSMUSG00000047501 | Cldn4 | 0.000110727 | -0.673092543 | 0.954617295 |
| ENSMUSG00000070473 | Cldn3 | 0.000110727 | 2.233798052 | 0.47966612 |
| ENSMUSG00000039917 | Rhbdd2 | 0.000110727 | -0.810596067 | 0.796840156 |
| ENSMUSG00000004948 | Zp3 | 0.000110727 | 3.233798052 | 0.262284476 |
| ENSMUSG00000051502 | Ufsp1 | 0.000110727 | 1.911869957 | 0.587484274 |
| ENSMUSG00000037344 | Slc12a9 | 0.000110727 | -1.696939285 | 0.210550914 |
| ENSMUSG00000089984 | Fbxo24 | 0.000110727 | 1.063873051 | 0.849985619 |
| ENSMUSG00000045348 | Nyap1 | 0.000110727 | 0.063873051 | 1 |
| ENSMUSG00000019518 | Ap4m1 | 0.000110727 | -0.624182943 | 0.912904617 |
| ENSMUSG00000036968 | Cnpy4 | 0.000110727 | 0.233798052 | 1 |
| ENSMUSG00000044197 | Gpr146 | 0.000110727 | -1.57355687 | 0.245475583 |
| ENSMUSG00000029551 | Psmg3 | 0.000110727 | 0.233798052 | 1 |
| ENSMUSG00000066640 | Fbxl18 | 0.000110727 | -1.788569761 | 0.144034706 |
| ENSMUSG00000046658 | Zfp316 | 0.000110727 | -1.696939285 | 0.210550914 |
| ENSMUSG00000039206 | Daglb | 0.000110727 | -1.051604167 | 0.614718876 |
| ENSMUSG00000059991 | Nptx2 | 0.000110727 | 2.233798052 | 0.47966612 |
| ENSMUSG00000038770 | Kpna7 | 0.000110727 | -1.895484965 | 0.167024802 |
| ENSMUSG00000056035 | Cyp3a11 | 0.000110727 | 1.42644313 | 0.736456705 |
| ENSMUSG00000029644 | Pdx1 | 0.000110727 | -1.743481871 | 0.172404208 |
| ENSMUSG00000042817 | Flt3 | 0.000110727 | 0.146335211 | 1 |
| ENSMUSG00000108207 | 1810059H22Rik | 0.000110727 | -0.975655313 | 0.705975399 |
| ENSMUSG00000107384 | Gm42557 | 0.000110727 | 1.42644313 | 0.736456705 |
| ENSMUSG00000056586 | Zar1l | 0.000110727 | 2.648835551 | 0.420597765 |
| ENSMUSG00000106734 | Gm20559 | 0.000110727 | -1.466641666 | 0.346237121 |
| ENSMUSG00000032766 | Gng11 | 0.000110727 | -0.014129461 | 1 |
| ENSMUSG00000087305 | A430035B10Rik | 0.000110727 | 2.648835551 | 0.395118486 |

| | | | | |
|---|---|---|---|---|
| ENSMUSG00000048216 | Gpr85 | 0.000110727 | 1.911869957 | 0.587484274 |
| ENSMUSG00000097616 | 1110019D14Rik | 0.000110727 | -1.599091962 | 0.242941974 |
| ENSMUSG00000108025 | Gm36503 | 0.000110727 | 1.911869957 | 0.598778059 |
| ENSMUSG00000029553 | Tfec | 0.000110727 | -3.417253639 | 0.000393279 |
| ENSMUSG00000054556 | Gm4876 | 0.000110727 | -0.225633566 | 1 |
| ENSMUSG00000087389 | Gm15592 | 0.000110727 | 1.911869957 | 0.600855395 |
| ENSMUSG00000029676 | Pot1a | 0.000110727 | -1.088130043 | 0.624094312 |
| ENSMUSG00000043340 | 6530409C15Rik | 0.000110727 | 1.911869957 | 0.607698939 |
| ENSMUSG00000054446 | Cpa1 | 0.000110727 | 1.911869957 | 0.595281189 |
| ENSMUSG00000039032 | Tsga13 | 0.000110727 | 1.063873051 | 1 |
| ENSMUSG00000061758 | Akr1b10 | 0.000110727 | 0.063873051 | 1 |
| ENSMUSG00000038836 | Agbl3 | 0.000110727 | -2.258055044 | 0.033084874 |
| ENSMUSG00000046806 | Cyren | 0.000110727 | -1.014129461 | 0.608426921 |
| ENSMUSG00000029847 | Slc23a4 | 0.000110727 | 0.326907457 | 1 |
| ENSMUSG00000115026 | Gm49041 | 0.000110727 | 2.648835551 | 0.395118486 |
| ENSMUSG00000037159 | Wee2 | 0.000110727 | 1.911869957 | 0.600497859 |
| ENSMUSG00000057163 | Prss2 | 0.000110727 | 3.233798052 | 0.262284476 |
| ENSMUSG00000036667 | Tcaf1 | 0.000110727 | -1.599091962 | 0.242941974 |
| ENSMUSG00000052763 | Zfp212 | 0.000110727 | -1.3207908 | 0.509528214 |
| ENSMUSG00000045466 | Zfp956 | 0.000110727 | 1.063873051 | 1 |
| ENSMUSG00000029797 | Sspo | 0.000110727 | 0.911869957 | 1 |
| ENSMUSG00000051124 | Gimap9 | 0.000110727 | 0.326907457 | 1 |
| ENSMUSG00000050786 | Ccdc126 | 0.000110727 | 0.326907457 | 1 |
| ENSMUSG00000107993 | G930045G22Rik | 0.000110727 | 2.233798052 | 0.47966612 |
| ENSMUSG00000107660 | Gm6559 | 0.000110727 | 1.233798052 | 0.792441218 |
| ENSMUSG00000038227 | Hoxa9 | 0.000110727 | 2.648835551 | 0.459931813 |
| ENSMUSG00000108062 | Gm44028 | 0.000110727 | 3.233798052 | 0.262284476 |
| ENSMUSG00000108175 | Gm16499 | 0.000110727 | 0.911869957 | 0.886203497 |
| ENSMUSG00000086946 | Gm15527 | 0.000110727 | 3.233798052 | 0.299296681 |
| ENSMUSG00000045725 | Prr15 | 0.000110727 | 0.146335211 | 1 |
| ENSMUSG00000004655 | Aqp1 | 0.000110727 | -0.289763904 | 1 |
| ENSMUSG00000071893 | Vmn1r4 | 0.000110727 | 1.233798052 | 0.792441218 |
| ENSMUSG00000115792 | Vmn1r16 | 0.000110727 | 3.233798052 | 0.262284476 |
| ENSMUSG00000029798 | Herc6 | 0.000110727 | -0.014129461 | 1 |
| ENSMUSG00000062190 | Lancl2 | 0.000110727 | 1.911869957 | 0.587484274 |
| ENSMUSG00000107960 | Gm35077 | 0.000110727 | 2.233798052 | 0.508022164 |
| ENSMUSG00000045441 | Gprin3 | 0.000110727 | -0.466641666 | 1 |
| ENSMUSG00000054641 | Mmrn1 | 0.000110727 | -0.088130043 | 1 |
| ENSMUSG00000029917 | Qrfprl | 0.000110727 | 0.42644313 | 1 |
| ENSMUSG00000049001 | Ndnf | 0.000110727 | -1.192466703 | 0.529126167 |
| ENSMUSG00000060699 | Vmn1r35 | 0.000110727 | 3.233798052 | 0.295705877 |
| ENSMUSG00000049093 | Il23r | 0.000110727 | 0.774366434 | 1 |
| ENSMUSG00000056370 | Sftpb | 0.000110727 | 2.233798052 | 0.487205201 |
| ENSMUSG00000108435 | Gm45051 | 0.000110727 | 1.911869957 | 0.600132691 |
| ENSMUSG00000055799 | Tcf7l1 | 0.000110727 | -2.985370468 | 0.001902483 |
| ENSMUSG00000107408 | 1700065L07Rik | 0.000110727 | 2.233798052 | 0.508841397 |
| ENSMUSG00000086739 | Gm4409 | 0.000110727 | 0.911869957 | 1 |
| ENSMUSG00000107499 | Ccdc142 | 0.000110727 | 0.911869957 | 1 |
| ENSMUSG00000030034 | Ino80b | 0.000110727 | -0.624182943 | 0.912904617 |
| ENSMUSG00000047013 | Fbxo41 | 0.000110727 | 1.063873051 | 1 |
| ENSMUSG00000054226 | Tprkb | 0.000110727 | -0.673092543 | 1 |

| | | | | |
|---|---|---|---|---|
| ENSMUSG00000107509 | Gm44941 | 0.000110727 | 4.233798052 | 0.128972787 |
| ENSMUSG00000048206 | Dnajb8 | 0.000110727 | 1.233798052 | 0.792441218 |
| ENSMUSG00000033182 | Kbtbd12 | 0.000110727 | 0.911869957 | 1 |
| ENSMUSG00000030083 | Abtb1 | 0.000110727 | -1.088130043 | 0.624094312 |
| ENSMUSG00000100680 | 1810044D09Rik | 0.000110727 | -0.225633566 | 1 |
| ENSMUSG00000034063 | 4930590J08Rik | 0.000110727 | 1.911869957 | 0.603006227 |
| ENSMUSG00000048636 | A730049H05Rik | 0.000110727 | 1.42644313 | 0.736456705 |
| ENSMUSG00000046500 | Tafa4 | 0.000110727 | -0.673092543 | 0.934902134 |
| ENSMUSG00000035245 | Eogt | 0.000110727 | -0.673092543 | 0.919998879 |
| ENSMUSG00000108151 | Gm33201 | 0.000110727 | 1.42644313 | 0.736456705 |
| ENSMUSG00000107427 | Gm43948 | 0.000110727 | 0.774366434 | 1 |
| ENSMUSG00000005364 | Il5ra | 0.000110727 | 0.42644313 | 1 |
| ENSMUSG00000107714 | Gm34933 | 0.000110727 | 1.233798052 | 0.792441218 |
| ENSMUSG00000087341 | 0610040F04Rik | 0.000110727 | 0.774366434 | 1 |
| ENSMUSG00000030276 | Ttll3 | 0.000110727 | -1.599091962 | 0.242941974 |
| ENSMUSG00000051256 | Jagn1 | 0.000110727 | -1.123753952 | 0.530172915 |
| ENSMUSG00000030281 | Il17rc | 0.000110727 | -0.936126949 | 1 |
| ENSMUSG00000045009 | Prrt3 | 0.000110727 | 1.648835551 | 0.66416768 |
| ENSMUSG00000034023 | Fancd2 | 0.000110727 | -0.766201948 | 0.992008082 |
| ENSMUSG00000030298 | Sec13 | 0.000110727 | -0.624182943 | 0.952530143 |
| ENSMUSG00000059900 | Tmem40 | 0.000110727 | -0.466641666 | 1 |
| ENSMUSG00000030323 | Ift122 | 0.000110727 | -1.466641666 | 0.346237121 |
| ENSMUSG00000108030 | 9530062K07Rik | 0.000110727 | 1.233798052 | 0.792441218 |
| ENSMUSG00000061353 | Cxcl12 | 0.000110727 | 0.326907457 | 1 |
| ENSMUSG00000094626 | Tmem121b | 0.000110727 | 3.233798052 | 0.262284476 |
| ENSMUSG00000058979 | Hdhd5 | 0.000110727 | -0.853664789 | 0.806496094 |
| ENSMUSG00000071226 | Cecr2 | 0.000110727 | -0.351164449 | 1 |
| ENSMUSG00000067825 | Pex26 | 0.000110727 | -0.624182943 | 0.912904617 |
| ENSMUSG00000040797 | Iqsec3 | 0.000110727 | 0.774366434 | 1 |
| ENSMUSG00000059908 | Mug1 | 0.000110727 | 1.648835551 | 0.66416768 |
| ENSMUSG00000040613 | Apobec1 | 0.000110727 | -0.225633566 | 1 |
| ENSMUSG00000049037 | Clec4a1 | 0.000110727 | 2.233798052 | 0.47966612 |
| ENSMUSG00000030148 | Clec4a2 | 0.000110727 | 2.648835551 | 0.408970699 |
| ENSMUSG00000030142 | Clec4e | 0.000110727 | 2.648835551 | 0.395118486 |
| ENSMUSG00000008153 | Clstn3 | 0.000110727 | 2.648835551 | 0.395118486 |
| ENSMUSG00000038527 | C1rl | 0.000110727 | 1.063873051 | 0.840238065 |
| ENSMUSG00000098470 | C1rb | 0.000110727 | 1.648835551 | 0.66416768 |
| ENSMUSG00000004266 | Ptpn6 | 0.000110727 | 0.42644313 | 1 |
| ENSMUSG00000030125 | Lrrc23 | 0.000110727 | 1.233798052 | 0.792441218 |
| ENSMUSG00000055818 | A230083G16Rik | 0.000110727 | 0.063873051 | 1 |
| ENSMUSG00000108129 | 4930417O13Rik | 0.000110727 | 2.233798052 | 0.47966612 |
| ENSMUSG00000030345 | Dyrk4 | 0.000110727 | 1.648835551 | 0.66416768 |
| ENSMUSG00000038028 | Tigar | 0.000110727 | -0.014129461 | 1 |
| ENSMUSG00000097525 | Platr31 | 0.000110727 | 0.911869957 | 1 |
| ENSMUSG00000085813 | Gm15870 | 0.000110727 | 0.911869957 | 1 |
| ENSMUSG00000030361 | Klrb1a | 0.000110727 | 1.648835551 | 0.665635898 |
| ENSMUSG00000000248 | Clec2g | 0.000110727 | 0.533358334 | 1 |
| ENSMUSG00000030157 | Clec2d | 0.000110727 | -1.52108945 | 0.301375981 |
| ENSMUSG00000097354 | 2310001H17Rik | 0.000110727 | -0.410058138 | 1 |
| ENSMUSG00000030167 | Klrc1 | 0.000110727 | 1.42644313 | 0.736456705 |
| ENSMUSG00000030188 | Magohb | 0.000110727 | -0.158519371 | 1 |

| | | | | |
|---|---|---|---|---|
| ENSMUSG00000095577 | Gm6619 | 0.000110727 | 2.648835551 | 0.395118486 |
| ENSMUSG00000058295 | Prp2 | 0.000110727 | 3.233798052 | 0.266705895 |
| ENSMUSG00000108276 | Gm36640 | 0.000110727 | -2.123753952 | 0.053533761 |
| ENSMUSG00000047104 | Pbp2 | 0.000110727 | 0.063873051 | 1 |
| ENSMUSG00000045291 | E330021D16Rik | 0.000110727 | 1.648835551 | 0.66416768 |
| ENSMUSG00000030219 | Erp27 | 0.000110727 | 2.648835551 | 0.395118486 |
| ENSMUSG00000044378 | Slc15a5 | 0.000110727 | 0.233798052 | 1 |
| ENSMUSG00000079262 | Slco1a6 | 0.000110727 | 0.233798052 | 1 |
| ENSMUSG00000107802 | 1700126G02Rik | 0.000110727 | -0.936126949 | 0.73915793 |
| ENSMUSG00000086887 | 1700060C16Rik | 0.000110727 | 0.533358334 | 1 |
| ENSMUSG00000030268 | Bcat1 | 0.000110727 | -0.088130043 | 1 |
| ENSMUSG00000040370 | Etfrf1 | 0.000110727 | 0.648835551 | 1 |
| ENSMUSG00000040187 | Arntl2 | 0.000110727 | -1.57355687 | 0.23866117 |
| ENSMUSG00000085355 | 3010003L21Rik | 0.000110727 | 0.774366434 | 1 |
| ENSMUSG00000054753 | AU018091 | 0.000110727 | 3.233798052 | 0.344436715 |
| ENSMUSG00000078817 | Nlrp12 | 0.000110727 | 0.42644313 | 1 |
| ENSMUSG00000078816 | Prkcg | 0.000110727 | -1.289763904 | 0.466464457 |
| ENSMUSG00000107985 | Gm35037 | 0.000110727 | 2.233798052 | 0.497095023 |
| ENSMUSG00000006335 | Tfpt | 0.000110727 | -0.466641666 | 1 |
| ENSMUSG00000078813 | Leng1 | 0.000110727 | -1.57355687 | 0.23866117 |
| ENSMUSG00000058818 | Pirb | 0.000110727 | 0.326907457 | 1 |
| ENSMUSG00000089942 | Pira2 | 0.000110727 | 1.42644313 | 0.736456705 |
| ENSMUSG00000078810 | Gp6 | 0.000110727 | 2.648835551 | 0.395118486 |
| ENSMUSG00000091177 | Gm15494 | 0.000110727 | 1.063873051 | 1 |
| ENSMUSG00000055809 | Dnaaf3 | 0.000110727 | 1.42644313 | 0.736456705 |
| ENSMUSG00000035429 | Ptprh | 0.000110727 | 1.648835551 | 0.66416768 |
| ENSMUSG00000059851 | Kmt5c | 0.000110727 | -1.936126949 | 0.106828177 |
| ENSMUSG00000092518 | Fam71e2 | 0.000110727 | 0.326907457 | 1 |
| ENSMUSG00000062861 | Zfp28 | 0.000110727 | 1.063873051 | 0.85551189 |
| ENSMUSG00000054938 | Olfr1346 | 0.000110727 | 3.233798052 | 0.305609173 |
| ENSMUSG00000034538 | Zfp418 | 0.000110727 | 1.648835551 | 0.66416768 |
| ENSMUSG00000091651 | Vmn2r36 | 0.000110727 | 3.233798052 | 0.262284476 |
| ENSMUSG00000057894 | Zfp329 | 0.000110727 | -0.766201948 | 0.792441218 |
| ENSMUSG00000033847 | Pla2g4c | 0.000110727 | 0.648835551 | 1 |
| ENSMUSG00000074375 | Sult2a3 | 0.000110727 | 3.233798052 | 0.262284476 |
| ENSMUSG00000094156 | Sult2a7 | 0.000110727 | 2.648835551 | 0.395118486 |
| ENSMUSG00000066772 | Obox3 | 0.000110727 | 1.063873051 | 1 |
| ENSMUSG00000006021 | Kptn | 0.000110727 | -0.289763904 | 1 |
| ENSMUSG00000030376 | Slc8a2 | 0.000110727 | 1.911869957 | 0.607010878 |
| ENSMUSG00000091811 | Inafm1 | 0.000110727 | 0.911869957 | 1 |
| ENSMUSG00000041187 | Prkd2 | 0.000110727 | -1.289763904 | 0.466464457 |
| ENSMUSG00000078794 | Dact3 | 0.000110727 | 1.063873051 | 1 |
| ENSMUSG00000030406 | Gipr | 0.000110727 | 0.648835551 | 1 |
| ENSMUSG00000030399 | Ckm | 0.000110727 | 1.911869957 | 0.587484274 |
| ENSMUSG00000030397 | Mark4 | 0.000110727 | -1.410058138 | 0.33333983 |
| ENSMUSG00000030486 | Zfp108 | 0.000110727 | 1.42644313 | 0.736456705 |
| ENSMUSG00000054793 | Cadm4 | 0.000110727 | -1.192466703 | 0.529126167 |
| ENSMUSG00000074277 | Phldb3 | 0.000110727 | -1.466641666 | 0.346237121 |
| ENSMUSG00000074272 | Ceacam1 | 0.000110727 | 1.063873051 | 1 |
| ENSMUSG00000040660 | Cyp2b9 | 0.000110727 | 4.233798052 | 0.128972787 |
| ENSMUSG00000078786 | BC024978 | 0.000110727 | -0.289763904 | 1 |

| | | | | |
|---|---|---|---|---|
| ENSMUSG00000053198 | Prx | 0.000110727 | -1.380911792 | 0.425992858 |
| ENSMUSG00000044786 | Zfp36 | 0.000110727 | -1.410058138 | 0.33333983 |
| ENSMUSG00000037469 | Acp7 | 0.000110727 | 0.233798052 | 1 |
| ENSMUSG00000030590 | Fam98c | 0.000110727 | -0.52108945 | 1 |
| ENSMUSG00000049676 | Catsperg1 | 0.000110727 | -0.014129461 | 1 |
| ENSMUSG00000062040 | Zfp27 | 0.000110727 | 0.146335211 | 1 |
| ENSMUSG00000019738 | Polr2i | 0.000110727 | -0.410058138 | 1 |
| ENSMUSG00000013928 | Thap8 | 0.000110727 | 3.233798052 | 0.262284476 |
| ENSMUSG00000064109 | Hcst | 0.000110727 | 2.648835551 | 0.395118486 |
| ENSMUSG00000036931 | Nfkbid | 0.000110727 | 4.233798052 | 0.128972787 |
| ENSMUSG00000036864 | Proser3 | 0.000110727 | -0.673092543 | 0.923527923 |
| ENSMUSG00000013083 | 2200002J24Rik | 0.000110727 | 4.233798052 | 0.128972787 |
| ENSMUSG00000060962 | Dmkn | 0.000110727 | 1.911869957 | 0.587484274 |
| ENSMUSG00000030577 | Cd22 | 0.000110727 | -0.466641666 | 1 |
| ENSMUSG00000001249 | Hpn | 0.000110727 | -0.351164449 | 1 |
| ENSMUSG00000086631 | Gm12784 | 0.000110727 | 1.911869957 | 0.587484274 |
| ENSMUSG00000060402 | Chst8 | 0.000110727 | -0.289763904 | 1 |
| ENSMUSG00000034957 | Cebpa | 0.000110727 | 2.233798052 | 0.47966612 |
| ENSMUSG00000044997 | E130304I02Rik | 0.000110727 | 1.648835551 | 0.66416768 |
| ENSMUSG00000056383 | AI987944 | 0.000110727 | -0.088130043 | 1 |
| ENSMUSG00000090744 | Gm6871 | 0.000110727 | 3.233798052 | 0.262284476 |
| ENSMUSG00000091474 | 2610021A01Rik | 0.000110727 | -0.810596067 | 0.801720351 |
| ENSMUSG00000090619 | Vmn2r60 | 0.000110727 | 1.42644313 | 0.736456705 |
| ENSMUSG00000108695 | Gm2511 | 0.000110727 | 4.233798052 | 0.128972787 |
| ENSMUSG00000056592 | Zfp658 | 0.000110727 | -0.014129461 | 1 |
| ENSMUSG00000002204 | Napsa | 0.000110727 | 1.233798052 | 0.792441218 |
| ENSMUSG00000059891 | Tsks | 0.000110727 | 1.063873051 | 0.85084719 |
| ENSMUSG00000044562 | Rasip1 | 0.000110727 | 0.774366434 | 1 |
| ENSMUSG00000057342 | Sphk2 | 0.000110727 | -0.766201948 | 0.792441218 |
| ENSMUSG00000109602 | Gm31597 | 0.000110727 | -1.158519371 | 0.556893363 |
| ENSMUSG00000002771 | Grin2d | 0.000110727 | -1.3207908 | 0.395686633 |
| ENSMUSG00000009487 | Otog | 0.000110727 | 0.911869957 | 1 |
| ENSMUSG00000058975 | Kcnc1 | 0.000110727 | 1.233798052 | 0.792441218 |
| ENSMUSG00000040046 | Tph1 | 0.000110727 | 1.911869957 | 0.59021625 |
| ENSMUSG00000006763 | Saal1 | 0.000110727 | -0.895484965 | 0.691084416 |
| ENSMUSG00000014418 | Hps5 | 0.000110727 | -2.648844997 | 0.008348692 |
| ENSMUSG00000049516 | Spty2d1 | 0.000110727 | -2.141241379 | 0.054954924 |
| ENSMUSG00000096719 | Mrgpra2b | 0.000110727 | 2.648835551 | 0.395118486 |
| ENSMUSG00000109006 | B230209E15Rik | 0.000110727 | -1.014129461 | 0.774790843 |
| ENSMUSG00000108393 | Gm32633 | 0.000110727 | 1.42644313 | 0.736456705 |
| ENSMUSG00000030523 | Trpm1 | 0.000110727 | 0.911869957 | 1 |
| ENSMUSG00000030521 | Mphosph10 | 0.000110727 | -2.648844997 | 0.008348692 |
| ENSMUSG00000078677 | Gm10974 | 0.000110727 | -0.351164449 | 1 |
| ENSMUSG00000030509 | Asb7 | 0.000110727 | -1.696939285 | 0.210550914 |
| ENSMUSG00000030510 | Cers3 | 0.000110727 | -0.158519371 | 1 |
| ENSMUSG00000043831 | Lysmd4 | 0.000110727 | -0.936126949 | 0.697198109 |
| ENSMUSG00000109193 | Gm39027 | 0.000110727 | 4.233798052 | 0.128972787 |
| ENSMUSG00000030553 | Pgpep1l | 0.000110727 | 0.774366434 | 1 |
| ENSMUSG00000110028 | Gm34838 | 0.000110727 | 1.648835551 | 0.66416768 |
| ENSMUSG00000109161 | Gm39033 | 0.000110727 | 4.233798052 | 0.128972787 |
| ENSMUSG00000108800 | 1700011C11Rik | 0.000110727 | 0.533358334 | 1 |

| | | | | |
|---|---|---|---|---|
| ENSMUSG00000097756 | A730056A06Rik | 0.000110727 | 0.233798052 | 1 |
| ENSMUSG00000078670 | Fam174b | 0.000110727 | 2.233798052 | 0.498537716 |
| ENSMUSG00000097247 | 1500012K07Rik | 0.000110727 | 1.911869957 | 0.605780185 |
| ENSMUSG00000055652 | Klhl25 | 0.000110727 | -0.289763904 | 1 |
| ENSMUSG00000030612 | Mrpl46 | 0.000110727 | 0.648835551 | 1 |
| ENSMUSG00000039099 | Wdr93 | 0.000110727 | 0.774366434 | 1 |
| ENSMUSG00000039062 | Anpep | 0.000110727 | 0.533358334 | 1 |
| ENSMUSG00000038930 | Rccd1 | 0.000110727 | -0.351164449 | 1 |
| ENSMUSG00000084821 | Gm15880 | 0.000110727 | 2.648835551 | 0.395118486 |
| ENSMUSG00000025722 | Wdr73 | 0.000110727 | -0.936126949 | 0.697198109 |
| ENSMUSG00000109008 | Gm44899 | 0.000110727 | -0.720398258 | 0.792441218 |
| ENSMUSG00000025726 | Slc28a1 | 0.000110727 | 0.063873051 | 1 |
| ENSMUSG00000025102 | 3110040N11Rik | 0.000110727 | -0.57355687 | 0.910050347 |
| ENSMUSG00000025104 | Hdgfl3 | 0.000110727 | 0.533358334 | 1 |
| ENSMUSG00000087088 | Gm16638 | 0.000110727 | 1.911869957 | 0.587484274 |
| ENSMUSG00000011154 | Cfap161 | 0.000110727 | 2.648835551 | 0.395118486 |
| ENSMUSG00000109248 | Gm44993 | 0.000110727 | 1.648835551 | 0.664426832 |
| ENSMUSG00000097789 | Gm2115 | 0.000110727 | 1.911869957 | 0.597803097 |
| ENSMUSG00000066372 | Vmn2r65 | 0.000110727 | 0.533358334 | 1 |
| ENSMUSG00000091239 | Vmn2r76 | 0.000110727 | 1.42644313 | 0.736456705 |
| ENSMUSG00000004651 | Tyr | 0.000110727 | 1.911869957 | 0.589394798 |
| ENSMUSG00000097162 | 2310010J17Rik | 0.000110727 | 1.233798052 | 0.792441218 |
| ENSMUSG00000030614 | Tmem126b | 0.000110727 | 1.911869957 | 0.587484274 |
| ENSMUSG00000030641 | Ddias | 0.000110727 | 1.911869957 | 0.587484274 |
| ENSMUSG00000085560 | C230038L03Rik | 0.000110727 | 1.648835551 | 0.66416768 |
| ENSMUSG00000074006 | Omp | 0.000110727 | 0.146335211 | 1 |
| ENSMUSG00000099137 | Gm10603 | 0.000110727 | 1.911869957 | 0.587484274 |
| ENSMUSG00000030701 | Plekhb1 | 0.000110727 | 1.648835551 | 0.66416768 |
| ENSMUSG00000048779 | P2ry6 | 0.000110727 | -1.43862729 | 0.349763156 |
| ENSMUSG00000043073 | Usp17le | 0.000110727 | 1.911869957 | 0.588477963 |
| ENSMUSG00000030882 | Dnhd1 | 0.000110727 | -1.051604167 | 0.66416768 |
| ENSMUSG00000078606 | Gm4070 | 0.000110727 | -0.410058138 | 1 |
| ENSMUSG00000045581 | Olfr710 | 0.000110727 | 2.233798052 | 0.47966612 |
| ENSMUSG00000031021 | Tmem9b | 0.000110727 | 0.648835551 | 1 |
| ENSMUSG00000084984 | Far1os | 0.000110727 | -0.624182943 | 0.912904617 |
| ENSMUSG00000108722 | Gm34225 | 0.000110727 | 1.42644313 | 0.736456705 |
| ENSMUSG00000033917 | Gde1 | 0.000110727 | -1.088130043 | 0.628253603 |
| ENSMUSG00000008734 | Gprc5b | 0.000110727 | -1.088130043 | 0.66416768 |
| ENSMUSG00000030924 | Rexo5 | 0.000110727 | -0.810596067 | 0.796840156 |
| ENSMUSG00000062017 | Abca14 | 0.000110727 | 0.42644313 | 1 |
| ENSMUSG00000054746 | Abca15 | 0.000110727 | 0.911869957 | 1 |
| ENSMUSG00000035004 | Igsf6 | 0.000110727 | 1.42644313 | 0.736456705 |
| ENSMUSG00000108351 | Gm36736 | 0.000110727 | 1.42644313 | 0.736456705 |
| ENSMUSG00000030867 | Plk1 | 0.000110727 | -0.351164449 | 1 |
| ENSMUSG00000030745 | Il21r | 0.000110727 | 0.911869957 | 1 |
| ENSMUSG00000042978 | Sbk1 | 0.000110727 | 1.42644313 | 0.736456705 |
| ENSMUSG00000073838 | Tufm | 0.000110727 | 0.146335211 | 1 |
| ENSMUSG00000030707 | Coro1a | 0.000110727 | 0.648835551 | 1 |
| ENSMUSG00000058966 | Fam57b | 0.000110727 | 1.063873051 | 0.848260747 |
| ENSMUSG00000049350 | Zg16 | 0.000110727 | -0.088130043 | 1 |
| ENSMUSG00000042502 | Cd2bp2 | 0.000110727 | -0.57355687 | 1 |

| | | | | |
|---|---|---|---|---|
| ENSMUSG00000054716 | Zfp771 | 0.000110727 | -1.351164449 | 0.421384409 |
| ENSMUSG00000049091 | Sephs2 | 0.000110727 | 0.233798052 | 1 |
| ENSMUSG00000054381 | Zfp747 | 0.000110727 | 0.146335211 | 1 |
| ENSMUSG00000078580 | E430018J23Rik | 0.000110727 | 1.42644313 | 0.736456705 |
| ENSMUSG00000101225 | 1700008J07Rik | 0.000110727 | 1.063873051 | 1 |
| ENSMUSG00000106715 | Tmem265 | 0.000110727 | -1.547561661 | 0.290722159 |
| ENSMUSG00000045639 | Zfp629 | 0.000110727 | -1.258055044 | 0.466464457 |
| ENSMUSG00000042340 | Ctf1 | 0.000110727 | 2.233798052 | 0.47966612 |
| ENSMUSG00000049739 | Zfp646 | 0.000110727 | -1.088130043 | 0.639764279 |
| ENSMUSG00000108354 | 4931431B13Rik | 0.000110727 | 2.648835551 | 0.42417782 |
| ENSMUSG00000030782 | Tgfb1i1 | 0.000110727 | 0.533358334 | 1 |
| ENSMUSG00000042055 | Wdr11 | 0.000110727 | -1.051604167 | 0.614718876 |
| ENSMUSG00000108580 | Gm39094 | 0.000110727 | 0.648835551 | 1 |
| ENSMUSG00000030960 | Eef1akmt2 | 0.000110727 | 1.063873051 | 1 |
| ENSMUSG00000110101 | Gm6249 | 0.000110727 | 0.326907457 | 1 |
| ENSMUSG00000040139 | 9430038I01Rik | 0.000110727 | -0.624182943 | 0.921320733 |
| ENSMUSG00000109881 | Gm45507 | 0.000110727 | 0.146335211 | 1 |
| ENSMUSG00000025475 | Adgra1 | 0.000110727 | 0.42644313 | 1 |
| ENSMUSG00000066129 | Kndc1 | 0.000110727 | 1.911869957 | 0.607693164 |
| ENSMUSG00000025473 | Adam8 | 0.000110727 | -0.410058138 | 1 |
| ENSMUSG00000025484 | Bet1l | 0.000110727 | 0.326907457 | 1 |
| ENSMUSG00000025485 | Ric8a | 0.000110727 | -0.410058138 | 1 |
| ENSMUSG00000025486 | Sirt3 | 0.000110727 | -0.225633566 | 1 |
| ENSMUSG00000055629 | B4galnt4 | 0.000110727 | 0.911869957 | 1 |
| ENSMUSG00000025495 | Ptdss2 | 0.000110727 | -0.624182943 | 0.912904617 |
| ENSMUSG00000038637 | Lrrc56 | 0.000110727 | -0.088130043 | 1 |
| ENSMUSG00000048200 | Cracr2b | 0.000110727 | -0.014129461 | 1 |
| ENSMUSG00000031097 | Tnni2 | 0.000110727 | -0.720398258 | 0.792441218 |
| ENSMUSG00000110125 | Gm33148 | 0.000110727 | 0.648835551 | 1 |
| ENSMUSG00000009246 | Trpm5 | 0.000110727 | 2.233798052 | 0.47966612 |
| ENSMUSG00000073778 | Faddos | 0.000110727 | 2.648835551 | 0.40935944 |
| ENSMUSG00000031077 | Fadd | 0.000110727 | 1.233798052 | 0.792441218 |
| ENSMUSG00000048677 | Tpcn2 | 0.000110727 | -1.289763904 | 0.466464457 |
| ENSMUSG00000005540 | Fcer2a | 0.000110727 | 1.063873051 | 0.851260516 |
| ENSMUSG00000089956 | Gm16553 | 0.000110727 | 1.648835551 | 0.66416768 |
| ENSMUSG00000046589 | Lrrc8e | 0.000110727 | 1.063873051 | 1 |
| ENSMUSG00000011837 | Snapc2 | 0.000110727 | 1.911869957 | 0.587484274 |
| ENSMUSG00000022322 | Shcbp1 | 0.000110727 | -0.853664789 | 0.825405528 |
| ENSMUSG00000109371 | Gm44626 | 0.000110727 | 0.533358334 | 1 |
| ENSMUSG00000038894 | Irs2 | 0.000110727 | -1.648844997 | 0.201326391 |
| ENSMUSG00000047935 | Sox1ot | 0.000110727 | 1.233798052 | 0.792441218 |
| ENSMUSG00000054910 | 4931415C17Rik | 0.000110727 | 2.233798052 | 0.501301436 |
| ENSMUSG00000031444 | F10 | 0.000110727 | -1.258055044 | 0.49552919 |
| ENSMUSG00000038506 | Dcun1d2 | 0.000110727 | -1.380911792 | 0.408535739 |
| ENSMUSG00000090336 | Cfap97d2 | 0.000110727 | 0.911869957 | 1 |
| ENSMUSG00000037725 | Ckap2 | 0.000110727 | -0.810596067 | 0.796840156 |
| ENSMUSG00000037363 | Letm2 | 0.000110727 | 0.063873051 | 1 |
| ENSMUSG00000091514 | Gm17484 | 0.000110727 | -0.289763904 | 1 |
| ENSMUSG00000037214 | Thap1 | 0.000110727 | 0.063873051 | 1 |
| ENSMUSG00000109682 | Gm45307 | 0.000110727 | 0.911869957 | 0.886203497 |
| ENSMUSG00000110345 | Gm45572 | 0.000110727 | 2.648835551 | 0.395118486 |

| | | | | |
|---|---|---|---|---|
| ENSMUSG00000031485 | Plpbp | 0.000110727 | -0.720398258 | 0.792441218 |
| ENSMUSG00000063412 | Gm10131 | 0.000110727 | 2.648835551 | 0.423814296 |
| ENSMUSG00000110335 | 1700015I17Rik | 0.000110727 | 4.233798052 | 0.128972787 |
| ENSMUSG00000109827 | Gm6213 | 0.000110727 | 0.911869957 | 1 |
| ENSMUSG00000084960 | B430010I23Rik | 0.000110727 | 1.911869957 | 0.587484274 |
| ENSMUSG00000079058 | Adam34 | 0.000110727 | 0.648835551 | 1 |
| ENSMUSG00000079057 | Cyp4v3 | 0.000110727 | 1.233798052 | 0.792441218 |
| ENSMUSG00000085440 | Sorbs2os | 0.000110727 | 0.533358334 | 1 |
| ENSMUSG00000050914 | Ankrd37 | 0.000110727 | 0.063873051 | 1 |
| ENSMUSG00000031637 | Lrp2bp | 0.000110727 | -0.225633566 | 1 |
| ENSMUSG00000018796 | Acsl1 | 0.000110727 | -0.936126949 | 0.733004386 |
| ENSMUSG00000038173 | Enpp6 | 0.000110727 | -0.158519371 | 1 |
| ENSMUSG00000074357 | AA386476 | 0.000110727 | -0.975655313 | 0.705975399 |
| ENSMUSG00000110464 | 1700061N14Rik | 0.000110727 | -0.673092543 | 0.930886129 |
| ENSMUSG00000100706 | Gm19744 | 0.000110727 | 1.063873051 | 0.847265357 |
| ENSMUSG00000109838 | Gm45341 | 0.000110727 | 0.774366434 | 1 |
| ENSMUSG00000110013 | 1190028D05Rik | 0.000110727 | 1.648835551 | 0.66416768 |
| ENSMUSG00000110362 | Gm45554 | 0.000110727 | 0.911869957 | 1 |
| ENSMUSG00000110000 | Gm33501 | 0.000110727 | 1.063873051 | 1 |
| ENSMUSG00000110381 | Gm34370 | 0.000110727 | 1.648835551 | 0.66416768 |
| ENSMUSG00000057228 | Aadat | 0.000110727 | 0.533358334 | 1 |
| ENSMUSG00000014907 | Naf1 | 0.000110727 | -1.014129461 | 0.627323375 |
| ENSMUSG00000110468 | Gm39185 | 0.000110727 | 3.233798052 | 0.262284476 |
| ENSMUSG00000056426 | Nat3 | 0.000110727 | 2.233798052 | 0.47966612 |
| ENSMUSG00000036306 | Lzts1 | 0.000110727 | 2.648835551 | 0.397697717 |
| ENSMUSG00000079038 | D130040H23Rik | 0.000110727 | -1.696939285 | 0.210550914 |
| ENSMUSG00000048967 | Yjefn3 | 0.000110727 | 1.911869957 | 0.588014228 |
| ENSMUSG00000036151 | Tm6sf2 | 0.000110727 | 3.233798052 | 0.262284476 |
| ENSMUSG00000036120 | Rfxank | 0.000110727 | 0.648835551 | 1 |
| ENSMUSG00000049988 | Lrrc25 | 0.000110727 | -0.466641666 | 1 |
| ENSMUSG00000101854 | 1700026F02Rik | 0.000110727 | -0.466641666 | 1 |
| ENSMUSG00000110554 | Gm35256 | 0.000110727 | 0.774366434 | 1 |
| ENSMUSG00000034845 | Plvap | 0.000110727 | -0.158519371 | 1 |
| ENSMUSG00000108900 | Ccdc194 | 0.000110727 | 2.233798052 | 0.47966612 |
| ENSMUSG00000031805 | Jak3 | 0.000110727 | -1.874726405 | 0.125315747 |
| ENSMUSG00000110462 | Gm35572 | 0.000110727 | 0.774366434 | 1 |
| ENSMUSG00000003484 | Cyp4f18 | 0.000110727 | 1.063873051 | 0.850654272 |
| ENSMUSG00000019732 | Calr3 | 0.000110727 | 0.533358334 | 1 |
| ENSMUSG00000052488 | Cherp | 0.000110727 | -1.648844997 | 0.201326391 |
| ENSMUSG00000097000 | Gm17435 | 0.000110727 | 0.42644313 | 1 |
| ENSMUSG00000110094 | Gm45326 | 0.000110727 | -1.52108945 | 0.284607766 |
| ENSMUSG00000042353 | Frem3 | 0.000110727 | 0.233798052 | 1 |
| ENSMUSG00000037982 | Gm9725 | 0.000110727 | 0.326907457 | 1 |
| ENSMUSG00000031712 | Il15 | 0.000110727 | -1.494122402 | 0.279592421 |
| ENSMUSG00000109643 | Gm31545 | 0.000110727 | 3.233798052 | 0.262284476 |
| ENSMUSG00000002190 | Clgn | 0.000110727 | 1.233798052 | 0.792441218 |
| ENSMUSG00000013033 | Adgrl1 | 0.000110727 | -0.52108945 | 1 |
| ENSMUSG00000031706 | Rfx1 | 0.000110727 | -0.52108945 | 1 |
| ENSMUSG00000012889 | Podnl1 | 0.000110727 | 3.233798052 | 0.262284476 |
| ENSMUSG00000056155 | Nanos3 | 0.000110727 | -0.014129461 | 1 |
| ENSMUSG00000097296 | Gm26532 | 0.000110727 | 2.648835551 | 0.395118486 |

| | | | | |
|---|---|---|---|---|
| ENSMUSG00000074203 | G430095P16Rik | 0.000110727 | 1.233798052 | 0.792441218 |
| ENSMUSG00000036872 | Abcc12 | 0.000110727 | 1.233798052 | 0.792441218 |
| ENSMUSG00000098854 | Gm5118 | 0.000110727 | 3.233798052 | 0.262284476 |
| ENSMUSG00000097248 | Gm2694 | 0.000110727 | 1.063873051 | 0.847320319 |
| ENSMUSG00000033282 | Rpgrip1l | 0.000110727 | -2.158519371 | 0.044761838 |
| ENSMUSG00000109696 | Gm36670 | 0.000110727 | 3.233798052 | 0.329075396 |
| ENSMUSG00000110134 | Gm45334 | 0.000110727 | 2.648835551 | 0.39591051 |
| ENSMUSG00000055368 | Slc6a2 | 0.000110727 | 2.233798052 | 0.47966612 |
| ENSMUSG00000061959 | Ces1e | 0.000110727 | 0.648835551 | 1 |
| ENSMUSG00000058019 | Ces5a | 0.000110727 | 1.233798052 | 0.792441218 |
| ENSMUSG00000033009 | Ogfod1 | 0.000110727 | -2.305360759 | 0.028985062 |
| ENSMUSG00000034361 | Cpne2 | 0.000110727 | -1.123753952 | 0.514974685 |
| ENSMUSG00000031787 | Katnb1 | 0.000110727 | 0.533358334 | 1 |
| ENSMUSG00000031669 | Gins3 | 0.000110727 | 1.233798052 | 0.792441218 |
| ENSMUSG00000031671 | Setd6 | 0.000110727 | 1.063873051 | 1 |
| ENSMUSG00000109701 | Gm32531 | 0.000110727 | -0.158519371 | 1 |
| ENSMUSG00000110336 | 4933400L20Rik | 0.000110727 | 0.42644313 | 1 |
| ENSMUSG00000110053 | Gm29682 | 0.000110727 | 3.233798052 | 0.262284476 |
| ENSMUSG00000035824 | Tk2 | 0.000110727 | -1.051604167 | 0.614718876 |
| ENSMUSG00000014837 | 4931428F04Rik | 0.000110727 | -0.351164449 | 1 |
| ENSMUSG00000031891 | Hsd11b2 | 0.000110727 | 2.648835551 | 0.428685419 |
| ENSMUSG00000036672 | Cenpt | 0.000110727 | -1.766201948 | 0.177083054 |
| ENSMUSG00000008450 | Nutf2 | 0.000110727 | -2.424413431 | 0.018617059 |
| ENSMUSG00000031897 | Psmb10 | 0.000110727 | -0.466641666 | 1 |
| ENSMUSG00000110613 | Lncbate1 | 0.000110727 | 2.233798052 | 0.47966612 |
| ENSMUSG00000031722 | Hp | 0.000110727 | -0.351164449 | 1 |
| ENSMUSG00000031732 | Phlpp2 | 0.000110727 | -1.975655313 | 0.088605926 |
| ENSMUSG00000044676 | Zfp612 | 0.000110727 | -0.766201948 | 0.792441218 |
| ENSMUSG00000046441 | Cmtr2 | 0.000110727 | 1.233798052 | 0.792441218 |
| ENSMUSG00000033703 | Fcsk | 0.000110727 | 0.063873051 | 1 |
| ENSMUSG00000031960 | Aars | 0.000110727 | -1.874726405 | 0.125315747 |
| ENSMUSG00000033633 | Clec18a | 0.000110727 | 0.648835551 | 1 |
| ENSMUSG00000084808 | 9430091E24Rik | 0.000110727 | -1.123753952 | 0.514974685 |
| ENSMUSG00000031958 | Ldhd | 0.000110727 | -0.351164449 | 1 |
| ENSMUSG00000008874 | Clec3a | 0.000110727 | 1.911869957 | 0.603006227 |
| ENSMUSG00000047388 | Atmin | 0.000110727 | -0.088130043 | 1 |
| ENSMUSG00000034424 | Gcsh | 0.000110727 | -0.936126949 | 0.743814404 |
| ENSMUSG00000034105 | Meak7 | 0.000110727 | -0.088130043 | 1 |
| ENSMUSG00000031821 | Gins2 | 0.000110727 | 0.233798052 | 1 |
| ENSMUSG00000097336 | Fendrr | 0.000110727 | 0.911869957 | 1 |
| ENSMUSG00000031816 | Mthfsd | 0.000110727 | -2.141241379 | 0.054954924 |
| ENSMUSG00000097424 | 1700030M09Rik | 0.000110727 | 1.648835551 | 0.66416768 |
| ENSMUSG00000052934 | Fbxo31 | 0.000110727 | -1.648844997 | 0.201326391 |
| ENSMUSG00000097812 | Gm26812 | 0.000110727 | 0.774366434 | 1 |
| ENSMUSG00000006362 | Cbfa2t3 | 0.000110727 | -0.225633566 | 1 |
| ENSMUSG00000110523 | C230057M02Rik | 0.000110727 | 1.42644313 | 0.736456705 |
| ENSMUSG00000110440 | Gm45894 | 0.000110727 | 1.42644313 | 0.736456705 |
| ENSMUSG00000074037 | Mc1r | 0.000110727 | -2.158519371 | 0.084002784 |
| ENSMUSG00000097203 | 4732419C18Rik | 0.000110727 | 0.533358334 | 1 |
| ENSMUSG00000031974 | Abcb10 | 0.000110727 | -0.810596067 | 0.796840156 |
| ENSMUSG00000031980 | Agt | 0.000110727 | 0.911869957 | 1 |

| | | | | |
|---|---|---|---|---|
| ENSMUSG00000110629 | 4930567H12Rik | 0.000110727 | 1.063873051 | 0.849591317 |
| ENSMUSG00000056018 | Ccdc7b | 0.000110727 | -0.088130043 | 1 |
| ENSMUSG00000071302 | 2610044O15Rik8 | 0.000110727 | -1.158519371 | 0.521275193 |
| ENSMUSG00000110605 | Gm32856 | 0.000110727 | 0.326907457 | 1 |
| ENSMUSG00000025893 | Kbtbd3 | 0.000110727 | -0.720398258 | 0.964262694 |
| ENSMUSG00000070323 | Mmp27 | 0.000110727 | 2.648835551 | 0.395118486 |
| ENSMUSG00000050912 | Tmem123 | 0.000110727 | -0.225633566 | 1 |
| ENSMUSG00000053070 | Cfap300 | 0.000110727 | 0.533358334 | 1 |
| ENSMUSG00000085253 | Gm16833 | 0.000110727 | 1.233798052 | 0.792441218 |
| ENSMUSG00000111061 | Gm46102 | 0.000110727 | -0.720398258 | 0.949822782 |
| ENSMUSG00000111691 | Gm47002 | 0.000110727 | 1.911869957 | 0.606122355 |
| ENSMUSG00000049307 | Fut4 | 0.000110727 | 0.911869957 | 1 |
| ENSMUSG00000031927 | 1700012B09Rik | 0.000110727 | 1.063873051 | 1 |
| ENSMUSG00000039977 | Deup1 | 0.000110727 | 0.774366434 | 1 |
| ENSMUSG00000043885 | Slc36a4 | 0.000110727 | 0.146335211 | 1 |
| ENSMUSG00000109564 | Muc16 | 0.000110727 | -0.014129461 | 1 |
| ENSMUSG00000057551 | Zfp317 | 0.000110727 | -1.088130043 | 0.624094312 |
| ENSMUSG00000058192 | Zfp846 | 0.000110727 | -1.3207908 | 0.430269377 |
| ENSMUSG00000062470 | Fbxl12os | 0.000110727 | 1.648835551 | 0.66416768 |
| ENSMUSG00000032172 | Olfm2 | 0.000110727 | 0.533358334 | 1 |
| ENSMUSG00000043895 | S1pr2 | 0.000110727 | -0.088130043 | 1 |
| ENSMUSG00000045087 | S1pr5 | 0.000110727 | 1.42644313 | 0.736456705 |
| ENSMUSG00000096472 | Cdkn2d | 0.000110727 | 1.911869957 | 0.600492565 |
| ENSMUSG00000032182 | Yipf2 | 0.000110727 | -0.014129461 | 1 |
| ENSMUSG00000003410 | Elavl3 | 0.000110727 | 0.326907457 | 1 |
| ENSMUSG00000066839 | Ecsit | 0.000110727 | -1.289763904 | 0.466464457 |
| ENSMUSG00000045994 | B3gat1 | 0.000110727 | 1.648835551 | 0.66416768 |
| ENSMUSG00000031966 | Glb1l3 | 0.000110727 | 2.233798052 | 0.487308813 |
| ENSMUSG00000079644 | Gm1110 | 0.000110727 | 2.648835551 | 0.395118486 |
| ENSMUSG00000031990 | Jam3 | 0.000110727 | -0.766201948 | 0.799253004 |
| ENSMUSG00000033453 | Adamts15 | 0.000110727 | -1.289763904 | 0.508263132 |
| ENSMUSG00000111771 | Gm47465 | 0.000110727 | 0.648835551 | 1 |
| ENSMUSG00000099025 | Gm27162 | 0.000110727 | 1.648835551 | 0.66416768 |
| ENSMUSG00000111645 | Gm32171 | 0.000110727 | 0.648835551 | 1 |
| ENSMUSG00000111467 | Gm47775 | 0.000110727 | 3.233798052 | 0.278054278 |
| ENSMUSG00000098358 | Gm27164 | 0.000110727 | 0.774366434 | 1 |
| ENSMUSG00000032103 | Pus3 | 0.000110727 | -1.547561661 | 0.290722159 |
| ENSMUSG00000098304 | 1700027I24Rik | 0.000110727 | 2.648835551 | 0.395118486 |
| ENSMUSG00000032118 | Fez1 | 0.000110727 | 0.911869957 | 1 |
| ENSMUSG00000034303 | Ccdc15 | 0.000110727 | -0.57355687 | 0.910050347 |
| ENSMUSG00000001943 | Vsig2 | 0.000110727 | 0.911869957 | 1 |
| ENSMUSG00000023186 | Vwa5a | 0.000110727 | -0.810596067 | 0.840804483 |
| ENSMUSG00000094745 | Olfr954 | 0.000110727 | 3.233798052 | 0.262676162 |
| ENSMUSG00000111504 | Gm48284 | 0.000110727 | 3.233798052 | 0.262284476 |
| ENSMUSG00000110862 | Gm35835 | 0.000110727 | 0.648835551 | 0.946081192 |
| ENSMUSG00000111767 | 4930546K05Rik | 0.000110727 | 0.911869957 | 1 |
| ENSMUSG00000087456 | Gm16322 | 0.000110727 | 2.648835551 | 0.418061772 |
| ENSMUSG00000111409 | Gm49380 | 0.000110727 | 1.911869957 | 0.589659371 |
| ENSMUSG00000032123 | Dpagt1 | 0.000110727 | 0.326907457 | 1 |
| ENSMUSG00000049932 | H2afx | 0.000110727 | 0.533358334 | 1 |
| ENSMUSG00000032127 | Vps11 | 0.000110727 | -2.069982696 | 0.072192623 |

| | | | | |
|---|---|---|---|---|
| ENSMUSG00000043923 | Ccdc84 | 0.000110727 | 0.533358334 | 1 |
| ENSMUSG00000060380 | C030014I23Rik | 0.000110727 | 1.233798052 | 0.792441218 |
| ENSMUSG00000111378 | Gm47230 | 0.000110727 | 2.233798052 | 0.47966612 |
| ENSMUSG00000032098 | Treh | 0.000110727 | 2.648835551 | 0.423592395 |
| ENSMUSG00000111482 | Gm47153 | 0.000110727 | 2.648835551 | 0.395118486 |
| ENSMUSG00000110924 | Gm40518 | 0.000110727 | 4.233798052 | 0.128972787 |
| ENSMUSG00000032028 | Nxpe2 | 0.000110727 | 1.063873051 | 1 |
| ENSMUSG00000042293 | Gm5617 | 0.000110727 | 3.233798052 | 0.295890024 |
| ENSMUSG00000032269 | Htr3a | 0.000110727 | 1.648835551 | 0.66416768 |
| ENSMUSG00000032062 | 2310030G06Rik | 0.000110727 | 1.233798052 | 0.792441218 |
| ENSMUSG00000032058 | Ppp2r1b | 0.000110727 | -1.158519371 | 0.559529464 |
| ENSMUSG00000032057 | 4833427G06Rik | 0.000110727 | 0.533358334 | 1 |
| ENSMUSG00000110266 | Gm32742 | 0.000110727 | 1.063873051 | 0.854285611 |
| ENSMUSG00000053199 | Arhgap20 | 0.000110727 | 0.146335211 | 1 |
| ENSMUSG00000074354 | Arhgap20os | 0.000110727 | 3.233798052 | 0.262284476 |
| ENSMUSG00000034487 | Poglut3 | 0.000110727 | -0.853664789 | 0.806496094 |
| ENSMUSG00000086914 | Gm16124 | 0.000110727 | -1.123753952 | 0.514974685 |
| ENSMUSG00000041986 | Elmod1 | 0.000110727 | 0.533358334 | 1 |
| ENSMUSG00000041268 | Dmxl2 | 0.000110727 | -1.014129461 | 0.608426921 |
| ENSMUSG00000035594 | Chrna5 | 0.000110727 | 2.648835551 | 0.395118486 |
| ENSMUSG00000032322 | Pstpip1 | 0.000110727 | -1.123753952 | 0.522267426 |
| ENSMUSG00000032911 | Cspg4 | 0.000110727 | -0.351164449 | 1 |
| ENSMUSG00000032299 | Commd4 | 0.000110727 | -0.088130043 | 1 |
| ENSMUSG00000070298 | Trcg1 | 0.000110727 | 2.233798052 | 0.47966612 |
| ENSMUSG00000032333 | Stoml1 | 0.000110727 | 0.233798052 | 1 |
| ENSMUSG00000111846 | Gm47271 | 0.000110727 | 0.911869957 | 1 |
| ENSMUSG00000032292 | Nr2e3 | 0.000110727 | 2.233798052 | 0.47966612 |
| ENSMUSG00000047766 | Lrrc49 | 0.000110727 | -1.3207908 | 0.497790175 |
| ENSMUSG00000037801 | Iqch | 0.000110727 | -0.351164449 | 1 |
| ENSMUSG00000032398 | Snapc5 | 0.000110727 | 0.233798052 | 1 |
| ENSMUSG00000111712 | Gm39363 | 0.000110727 | 0.774366434 | 1 |
| ENSMUSG00000041696 | Rasl12 | 0.000110727 | -2.032988489 | 0.085297803 |
| ENSMUSG00000059183 | Mtfmt | 0.000110727 | -0.936126949 | 0.697198109 |
| ENSMUSG00000050503 | Fbxl22 | 0.000110727 | 1.648835551 | 0.66416768 |
| ENSMUSG00000032375 | Aph1b | 0.000110727 | 2.648835551 | 0.451785624 |
| ENSMUSG00000053040 | Aph1c | 0.000110727 | 1.063873051 | 0.847265357 |
| ENSMUSG00000032204 | Aqp9 | 0.000110727 | 0.233798052 | 1 |
| ENSMUSG00000097211 | BC065403 | 0.000110727 | 0.533358334 | 1 |
| ENSMUSG00000092192 | Dnaaf4 | 0.000110727 | 1.063873051 | 1 |
| ENSMUSG00000079469 | Pigb | 0.000110727 | -0.014129461 | 1 |
| ENSMUSG00000042487 | Leo1 | 0.000110727 | -2.123753952 | 0.07443787 |
| ENSMUSG00000032186 | Tmod2 | 0.000110727 | 1.42644313 | 0.736456705 |
| ENSMUSG00000032184 | Lysmd2 | 0.000110727 | 0.774366434 | 1 |
| ENSMUSG00000098685 | Gm27224 | 0.000110727 | 0.326907457 | 1 |
| ENSMUSG00000111839 | Gm39383 | 0.000110727 | 1.233798052 | 0.792441218 |
| ENSMUSG00000032258 | Lca5 | 0.000110727 | -0.810596067 | 0.814645943 |
| ENSMUSG00000032262 | Elovl4 | 0.000110727 | 2.233798052 | 0.47966612 |
| ENSMUSG00000032265 | Tent5a | 0.000110727 | -0.810596067 | 0.872990655 |
| ENSMUSG00000110956 | Gm48834 | 0.000110727 | 1.233798052 | 0.792441218 |
| ENSMUSG00000035274 | Tpbg | 0.000110727 | 1.42644313 | 0.736456705 |
| ENSMUSG00000097195 | Snhg5 | 0.000110727 | -2.661019711 | 0.006892135 |

| | | | | |
|---|---|---|---|---|
| ENSMUSG00000099930 | Gm2396 | 0.000110727 | 2.233798052 | 0.47966612 |
| ENSMUSG00000098627 | 4930524O08Rik | 0.000110727 | 1.911869957 | 0.601514099 |
| ENSMUSG00000074139 | 1700057G04Rik | 0.000110727 | 1.648835551 | 0.66416768 |
| ENSMUSG00000032839 | Trpc1 | 0.000110727 | 0.533358334 | 1 |
| ENSMUSG00000100931 | Gm28085 | 0.000110727 | 4.233798052 | 0.152304756 |
| ENSMUSG00000099564 | Gm28729 | 0.000110727 | 1.063873051 | 1 |
| ENSMUSG00000046997 | Spsb4 | 0.000110727 | 0.774366434 | 1 |
| ENSMUSG00000087167 | Gm15891 | 0.000110727 | 2.648835551 | 0.44064783 |
| ENSMUSG00000032456 | Nmnat3 | 0.000110727 | -3.388253767 | 0.000225399 |
| ENSMUSG00000037681 | Esyt3 | 0.000110727 | 2.233798052 | 0.49106711 |
| ENSMUSG00000089655 | Gm16004 | 0.000110727 | -2.777429203 | 0.004046078 |
| ENSMUSG00000032475 | Nck1 | 0.000110727 | -2.141241379 | 0.054954924 |
| ENSMUSG00000032556 | Bfsp2 | 0.000110727 | 1.233798052 | 0.792441218 |
| ENSMUSG00000079355 | Ackr4 | 0.000110727 | 1.233798052 | 0.792441218 |
| ENSMUSG00000099373 | Gm19667 | 0.000110727 | 3.233798052 | 0.276544772 |
| ENSMUSG00000091345 | Col6a5 | 0.000110727 | 1.648835551 | 0.66416768 |
| ENSMUSG00000023495 | Pcbp4 | 0.000110727 | -0.014129461 | 1 |
| ENSMUSG00000103835 | Gm37612 | 0.000110727 | 0.911869957 | 1 |
| ENSMUSG00000010047 | Hyal2 | 0.000110727 | -0.014129461 | 1 |
| ENSMUSG00000036091 | Hyal3 | 0.000110727 | 1.42644313 | 0.736456705 |
| ENSMUSG00000102224 | 4930447F24Rik | 0.000110727 | 1.063873051 | 0.853996752 |
| ENSMUSG00000039461 | Tcta | 0.000110727 | 0.648835551 | 1 |
| ENSMUSG00000047220 | Ccdc36 | 0.000110727 | 0.326907457 | 1 |
| ENSMUSG00000032609 | Klhdc8b | 0.000110727 | -1.956026507 | 0.086523965 |
| ENSMUSG00000032598 | Nckipsd | 0.000110727 | 0.233798052 | 1 |
| ENSMUSG00000025650 | Col7a1 | 0.000110727 | -2.088130043 | 0.066402406 |
| ENSMUSG00000025648 | Pfkfb4 | 0.000110727 | -0.410058138 | 1 |
| ENSMUSG00000025645 | Ccdc51 | 0.000110727 | -0.57355687 | 1 |
| ENSMUSG00000054836 | Elp6 | 0.000110727 | 1.233798052 | 0.792441218 |
| ENSMUSG00000054792 | Klhl18 | 0.000110727 | -2.731986233 | 0.006138621 |
| ENSMUSG00000049719 | Prss46 | 0.000110727 | 0.533358334 | 1 |
| ENSMUSG00000110784 | Gm39436 | 0.000110727 | 1.42644313 | 0.736456705 |
| ENSMUSG00000109890 | AU023762 | 0.000110727 | 0.326907457 | 1 |
| ENSMUSG00000056880 | Gadl1 | 0.000110727 | -1.832291138 | 0.198212447 |
| ENSMUSG00000010660 | Plcd1 | 0.000110727 | -0.810596067 | 0.796840156 |
| ENSMUSG00000038060 | Dlec1 | 0.000110727 | 2.233798052 | 0.47966612 |
| ENSMUSG00000032528 | Vipr1 | 0.000110727 | 0.911869957 | 1 |
| ENSMUSG00000061536 | Sec22c | 0.000110727 | -0.351164449 | 1 |
| ENSMUSG00000038233 | Gask1a | 0.000110727 | 3.233798052 | 0.262284476 |
| ENSMUSG00000025787 | Tgm4 | 0.000110727 | 1.063873051 | 1 |
| ENSMUSG00000029530 | Ccr9 | 0.000110727 | 0.233798052 | 1 |
| ENSMUSG00000025245 | Lztfl1 | 0.000110727 | -0.975655313 | 0.718012086 |
| ENSMUSG00000060509 | Xcr1 | 0.000110727 | 1.911869957 | 0.595785668 |
| ENSMUSG00000025804 | Ccr1 | 0.000110727 | 1.233798052 | 0.792441218 |
| ENSMUSG00000110834 | Gm39469 | 0.000110727 | -0.225633566 | 1 |
| ENSMUSG00000019763 | Rmnd1 | 0.000110727 | -1.410058138 | 0.357595703 |
| ENSMUSG00000085687 | Gm16153 | 0.000110727 | -1.258055044 | 0.466464457 |
| ENSMUSG00000015202 | Cnksr3 | 0.000110727 | -1.088130043 | 0.647074599 |
| ENSMUSG00000092190 | Gm20470 | 0.000110727 | 1.648835551 | 0.66416768 |
| ENSMUSG00000109160 | 4930598N05Rik | 0.000110727 | 1.911869957 | 0.611036796 |
| ENSMUSG00000019817 | Plagl1 | 0.000110727 | 0.146335211 | 1 |

| | | | | |
|---|---|---|---|---|
| ENSMUSG00000019814 | Ltv1 | 0.000110727 | -0.351164449 | 1 |
| ENSMUSG00000019809 | Pex3 | 0.000110727 | -0.895484965 | 0.863673952 |
| ENSMUSG00000019865 | Nmbr | 0.000110727 | 2.648835551 | 0.395118486 |
| ENSMUSG00000111927 | Gm47770 | 0.000110727 | 2.648835551 | 0.395118486 |
| ENSMUSG00000059554 | Ccdc28a | 0.000110727 | 0.146335211 | 1 |
| ENSMUSG00000096977 | Gm10827 | 0.000110727 | -0.289763904 | 1 |
| ENSMUSG00000111052 | Gm20139 | 0.000110727 | 0.533358334 | 1 |
| ENSMUSG00000111447 | Gm48249 | 0.000110727 | 1.911869957 | 0.595281189 |
| ENSMUSG00000111091 | Gm48651 | 0.000110727 | -0.351164449 | 1 |
| ENSMUSG00000097705 | Gm26740 | 0.000110727 | -1.874726405 | 0.133676174 |
| ENSMUSG00000019989 | Enpp3 | 0.000110727 | -0.158519371 | 1 |
| ENSMUSG00000112610 | Gm30165 | 0.000110727 | 0.233798052 | 1 |
| ENSMUSG00000049109 | Themis | 0.000110727 | -1.123753952 | 0.547344776 |
| ENSMUSG00000038916 | Soga3 | 0.000110727 | -0.014129461 | 1 |
| ENSMUSG00000004360 | 9330159F19Rik | 0.000110727 | 0.533358334 | 1 |
| ENSMUSG00000075266 | Cenpw | 0.000110727 | 2.648835551 | 0.395118486 |
| ENSMUSG00000019792 | Trmt11 | 0.000110727 | -1.832291138 | 0.150263833 |
| ENSMUSG00000111778 | Gm30676 | 0.000110727 | 0.533358334 | 1 |
| ENSMUSG00000112707 | D830005E20Rik | 0.000110727 | 3.233798052 | 0.295355925 |
| ENSMUSG00000090298 | Sult3a2 | 0.000110727 | 2.648835551 | 0.411579031 |
| ENSMUSG00000071340 | Trappc3l | 0.000110727 | 0.774366434 | 1 |
| ENSMUSG00000047514 | Tspyl1 | 0.000110727 | -0.895484965 | 0.691084416 |
| ENSMUSG00000086915 | Gm16364 | 0.000110727 | -0.52108945 | 1 |
| ENSMUSG00000038522 | Mfsd4b1 | 0.000110727 | -2.673092543 | 0.012650612 |
| ENSMUSG00000038417 | Fig4 | 0.000110727 | -1.3207908 | 0.395686633 |
| ENSMUSG00000019822 | Smpd2 | 0.000110727 | 1.42644313 | 0.736456705 |
| ENSMUSG00000102114 | BC048559 | 0.000110727 | 1.063873051 | 0.853587691 |
| ENSMUSG00000038248 | Sobp | 0.000110727 | -0.895484965 | 0.700503166 |
| ENSMUSG00000111535 | Gm35154 | 0.000110727 | 0.233798052 | 1 |
| ENSMUSG00000111182 | Gm35430 | 0.000110727 | 0.911869957 | 1 |
| ENSMUSG00000110843 | Gm35552 | 0.000110727 | 1.648835551 | 0.66416768 |
| ENSMUSG00000112023 | Lilr4b | 0.000110727 | 0.233798052 | 1 |
| ENSMUSG00000038624 | Nepn | 0.000110727 | 2.233798052 | 0.47966612 |
| ENSMUSG00000019857 | Asf1a | 0.000110727 | -0.936126949 | 0.697198109 |
| ENSMUSG00000020092 | Pald1 | 0.000110727 | -1.832291138 | 0.150263833 |
| ENSMUSG00000101792 | Gm28447 | 0.000110727 | -1.466641666 | 0.409912463 |
| ENSMUSG00000020086 | H2afy2 | 0.000110727 | -0.158519371 | 1 |
| ENSMUSG00000036923 | Stox1 | 0.000110727 | 1.063873051 | 1 |
| ENSMUSG00000020070 | Rufy2 | 0.000110727 | -1.673092543 | 0.205249116 |
| ENSMUSG00000020072 | Pbld2 | 0.000110727 | -0.158519371 | 1 |
| ENSMUSG00000085372 | Gm16145 | 0.000110727 | 2.648835551 | 0.395118486 |
| ENSMUSG00000100000 | 1700023F02Rik | 0.000110727 | 1.063873051 | 0.85084719 |
| ENSMUSG00000112210 | 4930407I19Rik | 0.000110727 | 2.233798052 | 0.491789642 |
| ENSMUSG00000075000 | Nrbf2 | 0.000110727 | -0.351164449 | 1 |
| ENSMUSG00000037846 | Rtkn2 | 0.000110727 | 1.063873051 | 0.853996752 |
| ENSMUSG00000019933 | Mrln | 0.000110727 | 0.533358334 | 1 |
| ENSMUSG00000037747 | Phyhipl | 0.000110727 | 0.146335211 | 1 |
| ENSMUSG00000112870 | Gm40689 | 0.000110727 | 0.42644313 | 1 |
| ENSMUSG00000112125 | Gm48024 | 0.000110727 | 2.233798052 | 0.499921258 |
| ENSMUSG00000112366 | C730027H18Rik | 0.000110727 | -1.351164449 | 0.401963028 |
| ENSMUSG00000003923 | Tfam | 0.000110727 | -1.014129461 | 0.641920893 |

| | | | | |
|---|---|---|---|---|
| ENSMUSG00000020178 | Adora2a | 0.000110727 | 0.533358334 | 1 |
| ENSMUSG00000006344 | Ggt5 | 0.000110727 | 0.648835551 | 1 |
| ENSMUSG00000001665 | Gstt3 | 0.000110727 | 1.063873051 | 0.855776926 |
| ENSMUSG00000001663 | Gstt1 | 0.000110727 | 1.063873051 | 0.847265357 |
| ENSMUSG00000033126 | Ybey | 0.000110727 | 0.326907457 | 1 |
| ENSMUSG00000001150 | Mcm3ap | 0.000110727 | -1.853664789 | 0.121970859 |
| ENSMUSG00000069581 | Tspear | 0.000110727 | -0.289763904 | 1 |
| ENSMUSG00000020284 | Cfap410 | 0.000110727 | 0.533358334 | 1 |
| ENSMUSG00000060301 | 2610008E11Rik | 0.000110727 | -1.696939285 | 0.210550914 |
| ENSMUSG00000091888 | Vmn2r80 | 0.000110727 | 2.233798052 | 0.47966612 |
| ENSMUSG00000020308 | Tpgs1 | 0.000110727 | 1.42644313 | 0.736456705 |
| ENSMUSG00000035835 | Plppr3 | 0.000110727 | 1.233798052 | 0.792441218 |
| ENSMUSG00000035722 | Abca7 | 0.000110727 | -2.696939285 | 0.006179697 |
| ENSMUSG00000003348 | Mob3a | 0.000110727 | -1.43862729 | 0.392877738 |
| ENSMUSG00000061589 | Dot1l | 0.000110727 | -2.410058138 | 0.023019447 |
| ENSMUSG00000062075 | Lmnb2 | 0.000110727 | -0.351164449 | 1 |
| ENSMUSG00000034958 | Atcay | 0.000110727 | 1.42644313 | 0.736456705 |
| ENSMUSG00000004933 | Matk | 0.000110727 | 1.233798052 | 0.792441218 |
| ENSMUSG00000034917 | Tjp3 | 0.000110727 | -1.289763904 | 0.466464457 |
| ENSMUSG00000078440 | Dohh | 0.000110727 | 0.648835551 | 1 |
| ENSMUSG00000020238 | Ncln | 0.000110727 | -1.088130043 | 0.624094312 |
| ENSMUSG00000034792 | Gna15 | 0.000110727 | 0.911869957 | 1 |
| ENSMUSG00000096856 | Gm10778 | 0.000110727 | -0.466641666 | 1 |
| ENSMUSG00000063320 | 1190007I07Rik | 0.000110727 | 1.42644313 | 0.736456705 |
| ENSMUSG00000112375 | Gm38560 | 0.000110727 | 0.774366434 | 1 |
| ENSMUSG00000109864 | Eid3 | 0.000110727 | -2.069982696 | 0.062490345 |
| ENSMUSG00000020032 | Nuak1 | 0.000110727 | -0.936126949 | 0.737239571 |
| ENSMUSG00000089959 | Gm16268 | 0.000110727 | 1.42644313 | 0.736456705 |
| ENSMUSG00000112213 | 4930555G07Rik | 0.000110727 | 1.42644313 | 0.736456705 |
| ENSMUSG00000058934 | Igf1os | 0.000110727 | 0.911869957 | 1 |
| ENSMUSG00000020056 | Washc3 | 0.000110727 | -0.673092543 | 0.919998879 |
| ENSMUSG00000074802 | Gas2l3 | 0.000110727 | -0.088130043 | 1 |
| ENSMUSG00000110647 | Gm17745 | 0.000110727 | 1.648835551 | 0.66416768 |
| ENSMUSG00000036168 | Ccdc38 | 0.000110727 | 0.533358334 | 1 |
| ENSMUSG00000086269 | Tmcc3os | 0.000110727 | 0.326907457 | 1 |
| ENSMUSG00000112477 | Gm48657 | 0.000110727 | 1.063873051 | 0.850873693 |
| ENSMUSG00000101517 | 4732465J04Rik | 0.000110727 | 1.233798052 | 0.792441218 |
| ENSMUSG00000111912 | Gm48521 | 0.000110727 | 0.774366434 | 1 |
| ENSMUSG00000112488 | Gm33981 | 0.000110727 | 0.774366434 | 1 |
| ENSMUSG00000111924 | 4930556N09Rik | 0.000110727 | 0.648835551 | 1 |
| ENSMUSG00000112805 | Gm34574 | 0.000110727 | 1.911869957 | 0.587484274 |
| ENSMUSG00000112384 | Gm34921 | 0.000110727 | 2.233798052 | 0.47966612 |
| ENSMUSG00000056912 | 1700017N19Rik | 0.000110727 | 2.648835551 | 0.428469537 |
| ENSMUSG00000036602 | Alx1 | 0.000110727 | 1.648835551 | 0.66416768 |
| ENSMUSG00000019897 | Ccdc59 | 0.000110727 | -2.158519371 | 0.044761838 |
| ENSMUSG00000112201 | 4930532I03Rik | 0.000110727 | 2.233798052 | 0.504544056 |
| ENSMUSG00000101801 | 1700020G17Rik | 0.000110727 | 0.233798052 | 1 |
| ENSMUSG00000056888 | Glipr1 | 0.000110727 | -1.158519371 | 0.521275193 |
| ENSMUSG00000020137 | Thap2 | 0.000110727 | -1.832291138 | 0.150263833 |
| ENSMUSG00000052302 | Tbc1d30 | 0.000110727 | -0.225633566 | 1 |
| ENSMUSG00000080115 | Eef1akmt3 | 0.000110727 | 1.233798052 | 0.792441218 |

| | | | | |
|---|---|---|---|---|
| ENSMUSG00000025407 | Gli1 | 0.000110727 | 1.648835551 | 0.66416768 |
| ENSMUSG00000025405 | Inhbc | 0.000110727 | 1.648835551 | 0.66416768 |
| ENSMUSG00000040280 | Ndufa4l2 | 0.000110727 | 2.233798052 | 0.47966612 |
| ENSMUSG00000025403 | Shmt2 | 0.000110727 | 0.42644313 | 1 |
| ENSMUSG00000039994 | Timeless | 0.000110727 | -1.743481871 | 0.181684975 |
| ENSMUSG00000025350 | Rdh5 | 0.000110727 | 1.911869957 | 0.596991088 |
| ENSMUSG00000025348 | Itga7 | 0.000110727 | 1.42644313 | 0.736456705 |
| ENSMUSG00000112742 | Gm31793 | 0.000110727 | -1.014129461 | 0.6352512 |
| ENSMUSG00000020439 | Smtn | 0.000110727 | -0.895484965 | 0.716899444 |
| ENSMUSG00000047205 | Dusp18 | 0.000110727 | 1.911869957 | 0.587484274 |
| ENSMUSG00000034274 | Thoc5 | 0.000110727 | -0.57355687 | 0.914654891 |
| ENSMUSG00000034175 | Rhbdd3 | 0.000110727 | 0.648835551 | 1 |
| ENSMUSG00000004393 | Ddx56 | 0.000110727 | -2.088130043 | 0.064551115 |
| ENSMUSG00000086808 | Gm11981 | 0.000110727 | 2.648835551 | 0.395118486 |
| ENSMUSG00000020431 | Adcy1 | 0.000110727 | 1.063873051 | 0.855066236 |
| ENSMUSG00000020191 | Spata48 | 0.000110727 | 0.42644313 | 1 |
| ENSMUSG00000035455 | Fignl1 | 0.000110727 | 1.911869957 | 0.606122355 |
| ENSMUSG00000085806 | Gm12023 | 0.000110727 | 0.533358334 | 1 |
| ENSMUSG00000020142 | Slc1a4 | 0.000110727 | -0.158519371 | 1 |
| ENSMUSG00000033015 | 4933427E13Rik | 0.000110727 | 0.063873051 | 1 |
| ENSMUSG00000085510 | Gm12082 | 0.000110727 | 1.063873051 | 1 |
| ENSMUSG00000020462 | Cfap36 | 0.000110727 | -1.547561661 | 0.290722159 |
| ENSMUSG00000032673 | Prorsd1 | 0.000110727 | 0.774366434 | 1 |
| ENSMUSG00000020297 | Nsg2 | 0.000110727 | 0.326907457 | 1 |
| ENSMUSG00000020287 | Mpg | 0.000110727 | 0.533358334 | 1 |
| ENSMUSG00000020270 | Smim23 | 0.000110727 | 1.063873051 | 1 |
| ENSMUSG00000085402 | Gm12111 | 0.000110727 | 1.648835551 | 0.66416768 |
| ENSMUSG00000057967 | Fgf18 | 0.000110727 | 1.648835551 | 0.66416768 |
| ENSMUSG00000085951 | Gm12145 | 0.000110727 | 2.648835551 | 0.395118486 |
| ENSMUSG00000086791 | Gm12147 | 0.000110727 | 1.233798052 | 0.792441218 |
| ENSMUSG00000020415 | Pttg1 | 0.000110727 | 0.533358334 | 1 |
| ENSMUSG00000041278 | Ttc1 | 0.000110727 | -1.743481871 | 0.185619905 |
| ENSMUSG00000041231 | Ublcp1 | 0.000110727 | 0.533358334 | 1 |
| ENSMUSG00000087089 | Gm12160 | 0.000110727 | 0.774366434 | 1 |
| ENSMUSG00000020411 | Nipal4 | 0.000110727 | 4.233798052 | 0.128972787 |
| ENSMUSG00000046974 | BC053393 | 0.000110727 | 0.533358334 | 1 |
| ENSMUSG00000064010 | Dppa1 | 0.000110727 | 2.648835551 | 0.415055165 |
| ENSMUSG00000040405 | Havcr1 | 0.000110727 | -0.853664789 | 0.847265357 |
| ENSMUSG00000020346 | Mgat1 | 0.000110727 | -1.380911792 | 0.408535739 |
| ENSMUSG00000020357 | Flt4 | 0.000110727 | -0.810596067 | 0.843284619 |
| ENSMUSG00000020374 | Rasgef1c | 0.000110727 | 1.42644313 | 0.736456705 |
| ENSMUSG00000044807 | Zfp354c | 0.000110727 | -1.051604167 | 0.626822479 |
| ENSMUSG00000052563 | D930048N14Rik | 0.000110727 | -0.225633566 | 1 |
| ENSMUSG00000049491 | Slc36a3 | 0.000110727 | 0.911869957 | 1 |
| ENSMUSG00000069874 | Irgm2 | 0.000110727 | 1.063873051 | 1 |
| ENSMUSG00000061462 | Obscn | 0.000110727 | 0.774366434 | 1 |
| ENSMUSG00000020441 | 2310033P09Rik | 0.000110727 | -1.158519371 | 0.521275193 |
| ENSMUSG00000000301 | Pemt | 0.000110727 | 0.648835551 | 1 |
| ENSMUSG00000018931 | Natd1 | 0.000110727 | -0.410058138 | 1 |
| ENSMUSG00000018932 | Map2k3 | 0.000110727 | -1.853664789 | 0.121970859 |
| ENSMUSG00000069855 | Slc47a2 | 0.000110727 | 0.233798052 | 1 |

| | | | | |
|---|---|---|---|---|
| ENSMUSG00000010122 | Slc47a1 | 0.000110727 | 0.911869957 | 1 |
| ENSMUSG00000018507 | Trpv2 | 0.000110727 | -0.624182943 | 0.943363727 |
| ENSMUSG00000014177 | Tvp23b | 0.000110727 | -0.158519371 | 1 |
| ENSMUSG00000042200 | Cdrt4 | 0.000110727 | 0.533358334 | 1 |
| ENSMUSG00000042189 | Tekt3 | 0.000110727 | 3.233798052 | 0.262284476 |
| ENSMUSG00000085683 | Tmem238l | 0.000110727 | 2.233798052 | 0.494808857 |
| ENSMUSG00000020910 | Adprm | 0.000110727 | -0.289763904 | 1 |
| ENSMUSG00000056328 | Myh1 | 0.000110727 | 3.233798052 | 0.274319469 |
| ENSMUSG00000020905 | Usp43 | 0.000110727 | -1.43862729 | 0.339627968 |
| ENSMUSG00000032782 | Cntrob | 0.000110727 | -0.52108945 | 1 |
| ENSMUSG00000040938 | Slc16a11 | 0.000110727 | 1.648835551 | 0.66416768 |
| ENSMUSG00000040829 | Zmynd15 | 0.000110727 | -0.088130043 | 1 |
| ENSMUSG00000040620 | Dhx33 | 0.000110727 | -0.766201948 | 0.792441218 |
| ENSMUSG00000040599 | Mis12 | 0.000110727 | -1.494122402 | 0.279592421 |
| ENSMUSG00000020811 | Wscd1 | 0.000110727 | 2.648835551 | 0.448108027 |
| ENSMUSG00000020801 | Med31 | 0.000110727 | 0.648835551 | 1 |
| ENSMUSG00000072708 | Olfr381 | 0.000110727 | 3.233798052 | 0.262284476 |
| ENSMUSG00000000751 | Rpa1 | 0.000110727 | -1.810596067 | 0.145553211 |
| ENSMUSG00000000753 | Serpinf1 | 0.000110727 | -0.088130043 | 1 |
| ENSMUSG00000087059 | Gm12339 | 0.000110727 | 2.233798052 | 0.510207151 |
| ENSMUSG00000002057 | Foxn1 | 0.000110727 | 1.42644313 | 0.736456705 |
| ENSMUSG00000049489 | Ccnq | 0.000110727 | 4.233798052 | 0.21769511 |
| ENSMUSG00000035385 | Ccl2 | 0.000110727 | 1.648835551 | 0.66416768 |
| ENSMUSG00000018845 | Unc45b | 0.000110727 | 1.42644313 | 0.736456705 |
| ENSMUSG00000054404 | Slfn5 | 0.000110727 | -0.52108945 | 1 |
| ENSMUSG00000069793 | Slfn9 | 0.000110727 | 0.326907457 | 1 |
| ENSMUSG00000051832 | E230016K23Rik | 0.000110727 | 0.146335211 | 1 |
| ENSMUSG00000020526 | Znhit3 | 0.000110727 | 0.911869957 | 1 |
| ENSMUSG00000020525 | Ppm1d | 0.000110727 | -1.088130043 | 0.624094312 |
| ENSMUSG00000086199 | Bcas3os1 | 0.000110727 | 2.233798052 | 0.503644124 |
| ENSMUSG00000018401 | Mtmr4 | 0.000110727 | -0.936126949 | 0.762251949 |
| ENSMUSG00000070345 | Hsf5 | 0.000110727 | -0.351164449 | 1 |
| ENSMUSG00000087574 | C030037D09Rik | 0.000110727 | -0.895484965 | 0.691084416 |
| ENSMUSG00000087338 | Scpep1os | 0.000110727 | -0.014129461 | 1 |
| ENSMUSG00000085940 | 4930405D11Rik | 0.000110727 | 1.42644313 | 0.736456705 |
| ENSMUSG00000039096 | Rsad1 | 0.000110727 | -0.225633566 | 1 |
| ENSMUSG00000038967 | Pdk2 | 0.000110727 | 2.233798052 | 0.47966612 |
| ENSMUSG00000038893 | Fam117a | 0.000110727 | -2.032988489 | 0.076038495 |
| ENSMUSG00000018381 | Abi3 | 0.000110727 | 1.233798052 | 0.792441218 |
| ENSMUSG00000013415 | Igf2bp1 | 0.000110727 | -0.624182943 | 0.912904617 |
| ENSMUSG00000000690 | Hoxb6 | 0.000110727 | 0.911869957 | 1 |
| ENSMUSG00000085860 | 2410003L11Rik | 0.000110727 | 1.42644313 | 0.736456705 |
| ENSMUSG00000018541 | Cwc25 | 0.000110727 | -2.708716453 | 0.006432995 |
| ENSMUSG00000086917 | Gm11630 | 0.000110727 | 0.146335211 | 1 |
| ENSMUSG00000020889 | Nr1d1 | 0.000110727 | 1.233798052 | 0.792441218 |
| ENSMUSG00000038020 | Rapgefl1 | 0.000110727 | 0.774366434 | 1 |
| ENSMUSG00000043485 | Krt34 | 0.000110727 | 4.233798052 | 0.128972787 |
| ENSMUSG00000017176 | Nt5c3b | 0.000110727 | -1.380911792 | 0.408535739 |
| ENSMUSG00000017837 | Nkiras2 | 0.000110727 | 1.648835551 | 0.66416768 |
| ENSMUSG00000020918 | Kat2a | 0.000110727 | -0.289763904 | 1 |
| ENSMUSG00000001751 | Naglu | 0.000110727 | 3.233798052 | 0.262284476 |

| | | | | |
|---|---|---|---|---|
| ENSMUSG00000034947 | Tmem106a | 0.000110727 | -0.810596067 | 0.854989385 |
| ENSMUSG00000034768 | Asb16 | 0.000110727 | 1.911869957 | 0.596503438 |
| ENSMUSG00000034757 | Tmub2 | 0.000110727 | -0.014129461 | 1 |
| ENSMUSG00000034664 | Itga2b | 0.000110727 | 0.326907457 | 1 |
| ENSMUSG00000056938 | Acbd4 | 0.000110727 | -0.014129461 | 1 |
| ENSMUSG00000048878 | Hexim1 | 0.000110727 | -1.258055044 | 0.70936601 |
| ENSMUSG00000000125 | Wnt3 | 0.000110727 | 2.233798052 | 0.503644124 |
| ENSMUSG00000018634 | Crhr1 | 0.000110727 | 1.42644313 | 0.736456705 |
| ENSMUSG00000020691 | Mettl2 | 0.000110727 | -1.380911792 | 0.408535739 |
| ENSMUSG00000078627 | 10-Mar | 0.000110727 | 0.326907457 | 1 |
| ENSMUSG00000078622 | Ccdc47 | 0.000110727 | -2.636566667 | 0.008032984 |
| ENSMUSG00000078619 | Smarcd2 | 0.000110727 | -1.599091962 | 0.242941974 |
| ENSMUSG00000018372 | Cep95 | 0.000110727 | -2.241935379 | 0.041054791 |
| ENSMUSG00000018362 | Kpna2 | 0.000110727 | -0.810596067 | 0.796840156 |
| ENSMUSG00000087113 | Gm11714 | 0.000110727 | 1.233798052 | 0.792441218 |
| ENSMUSG00000085864 | C330019F10Rik | 0.000110727 | 0.326907457 | 1 |
| ENSMUSG00000020722 | Cacng1 | 0.000110727 | 2.233798052 | 0.504738484 |
| ENSMUSG00000020614 | Fam20a | 0.000110727 | -2.3207908 | 0.029965083 |
| ENSMUSG00000020620 | Abca8b | 0.000110727 | -1.410058138 | 0.33333983 |
| ENSMUSG00000051497 | Kcnj16 | 0.000110727 | 0.533358334 | 1 |
| ENSMUSG00000010021 | Kif19a | 0.000110727 | 1.911869957 | 0.606122355 |
| ENSMUSG00000063193 | Cd300lb | 0.000110727 | -0.158519371 | 1 |
| ENSMUSG00000034641 | Cd300ld | 0.000110727 | 2.233798052 | 0.47966612 |
| ENSMUSG00000020734 | Grin2c | 0.000110727 | 1.911869957 | 0.587484274 |
| ENSMUSG00000018858 | Mrpl58 | 0.000110727 | 0.146335211 | 1 |
| ENSMUSG00000016940 | Kctd2 | 0.000110727 | -1.088130043 | 0.624094312 |
| ENSMUSG00000087072 | Recql5os1 | 0.000110727 | 4.233798052 | 0.224109065 |
| ENSMUSG00000020755 | Sap30bp | 0.000110727 | -1.810596067 | 0.145553211 |
| ENSMUSG00000054517 | Trim65 | 0.000110727 | 0.533358334 | 1 |
| ENSMUSG00000009588 | St6galnac1 | 0.000110727 | 2.648835551 | 0.423592395 |
| ENSMUSG00000017718 | Afmid | 0.000110727 | -0.810596067 | 0.796840156 |
| ENSMUSG00000033909 | Usp36 | 0.000110727 | -2.336057556 | 0.024405815 |
| ENSMUSG00000025577 | Cbx2 | 0.000110727 | 1.063873051 | 1 |
| ENSMUSG00000013483 | Card14 | 0.000110727 | 0.146335211 | 1 |
| ENSMUSG00000025140 | Pycr1 | 0.000110727 | -1.599091962 | 0.242941974 |
| ENSMUSG00000025144 | Cenpx | 0.000110727 | 1.063873051 | 1 |
| ENSMUSG00000039450 | Dcxr | 0.000110727 | 2.233798052 | 0.47966612 |
| ENSMUSG00000039364 | Sectm1b | 0.000110727 | 2.648835551 | 0.395118486 |
| ENSMUSG00000039294 | Cybc1 | 0.000110727 | 1.233798052 | 0.792441218 |
| ENSMUSG00000079179 | Rab10os | 0.000110727 | -0.014129461 | 1 |
| ENSMUSG00000112308 | Gm48512 | 0.000110727 | -0.088130043 | 1 |
| ENSMUSG00000020652 | Cenpo | 0.000110727 | -1.380911792 | 0.408535739 |
| ENSMUSG00000111916 | 4921501I09Rik | 0.000110727 | 4.233798052 | 0.128972787 |
| ENSMUSG00000118449 | AC241534.1 | 0.000110727 | 2.233798052 | 0.47966612 |
| ENSMUSG00000037735 | 2810032G03Rik | 0.000110727 | 1.911869957 | 0.587484274 |
| ENSMUSG00000020583 | Matn3 | 0.000110727 | 1.063873051 | 0.837805191 |
| ENSMUSG00000113913 | Gm46323 | 0.000110727 | 0.063873051 | 1 |
| ENSMUSG00000020621 | Rdh14 | 0.000110727 | 0.146335211 | 1 |
| ENSMUSG00000112792 | Gm38407 | 0.000110727 | 0.42644313 | 1 |
| ENSMUSG00000051235 | Gen1 | 0.000110727 | -0.766201948 | 0.792441218 |
| ENSMUSG00000097580 | Gm5432 | 0.000110727 | 0.42644313 | 1 |

| | | | | |
|---|---|---|---|---|
| ENSMUSG00000057469 | E2f6 | 0.000110727 | -1.123753952 | 0.514974685 |
| ENSMUSG00000113447 | Gm48538 | 0.000110727 | 2.648835551 | 0.395118486 |
| ENSMUSG00000020566 | Atp6v1c2 | 0.000110727 | 0.233798052 | 1 |
| ENSMUSG00000062352 | Itgb1bp1 | 0.000110727 | -1.051604167 | 0.614718876 |
| ENSMUSG00000073158 | 9030624G23Rik | 0.000110727 | 0.533358334 | 1 |
| ENSMUSG00000059669 | Taf1b | 0.000110727 | -2.685065185 | 0.007443294 |
| ENSMUSG00000113813 | Gm29968 | 0.000110727 | 1.911869957 | 0.587484274 |
| ENSMUSG00000020641 | Rsad2 | 0.000110727 | 0.774366434 | 1 |
| ENSMUSG00000063632 | Sox11 | 0.000110727 | -2.611691999 | 0.043715437 |
| ENSMUSG00000020636 | Allc | 0.000110727 | 0.911869957 | 1 |
| ENSMUSG00000020630 | Rnaseh1 | 0.000110727 | 0.146335211 | 1 |
| ENSMUSG00000020673 | Tpo | 0.000110727 | 0.233798052 | 1 |
| ENSMUSG00000112876 | Gm32443 | 0.000110727 | 2.648835551 | 0.451565197 |
| ENSMUSG00000035860 | Cdhr3 | 0.000110727 | 1.233798052 | 0.792441218 |
| ENSMUSG00000113548 | Gm34047 | 0.000110727 | -0.52108945 | 1 |
| ENSMUSG00000020577 | Tspan13 | 0.000110727 | -0.289763904 | 1 |
| ENSMUSG00000113989 | Gm47013 | 0.000110727 | 1.063873051 | 1 |
| ENSMUSG00000112258 | Gm10165 | 0.000110727 | 1.911869957 | 0.587484274 |
| ENSMUSG00000112547 | Gm47096 | 0.000110727 | 0.911869957 | 1 |
| ENSMUSG00000021047 | Nova1 | 0.000110727 | -0.624182943 | 0.939194134 |
| ENSMUSG00000079108 | Srp54c | 0.000110727 | -0.088130043 | 1 |
| ENSMUSG00000020948 | Klhl28 | 0.000110727 | -2.946111038 | 0.002012957 |
| ENSMUSG00000113233 | Gm31063 | 0.000110727 | 4.233798052 | 0.128972787 |
| ENSMUSG00000113320 | Gm47515 | 0.000110727 | 1.648835551 | 0.664883298 |
| ENSMUSG00000020990 | Cdkl1 | 0.000110727 | 0.326907457 | 1 |
| ENSMUSG00000021069 | Pygl | 0.000110727 | -1.351164449 | 0.444576901 |
| ENSMUSG00000113694 | Gm48747 | 0.000110727 | 2.233798052 | 0.485878728 |
| ENSMUSG00000114005 | Gm24474 | 0.000110727 | 0.648835551 | 1 |
| ENSMUSG00000021087 | Rtn1 | 0.000110727 | -1.52108945 | 0.296157668 |
| ENSMUSG00000034501 | Pcnx4 | 0.000110727 | -0.975655313 | 0.705975399 |
| ENSMUSG00000109482 | Gm4756 | 0.000110727 | 0.911869957 | 1 |
| ENSMUSG00000034435 | Tmem30b | 0.000110727 | 2.648835551 | 0.395118486 |
| ENSMUSG00000021057 | Akap5 | 0.000110727 | -1.599091962 | 0.242941974 |
| ENSMUSG00000059970 | Hspa2 | 0.000110727 | 1.42644313 | 0.736456705 |
| ENSMUSG00000021062 | Rab15 | 0.000110727 | 0.233798052 | 1 |
| ENSMUSG00000112857 | Gm35041 | 0.000110727 | -0.088130043 | 1 |
| ENSMUSG00000021120 | Pigh | 0.000110727 | 0.233798052 | 1 |
| ENSMUSG00000021125 | Arg2 | 0.000110727 | 1.063873051 | 1 |
| ENSMUSG00000066441 | Rdh11 | 0.000110727 | 0.146335211 | 1 |
| ENSMUSG00000044062 | Plekhd1os | 0.000110727 | 0.774366434 | 1 |
| ENSMUSG00000096953 | Gm26571 | 0.000110727 | 0.648835551 | 1 |
| ENSMUSG00000021226 | Acot2 | 0.000110727 | 1.42644313 | 0.736456705 |
| ENSMUSG00000072949 | Acot1 | 0.000110727 | 1.648835551 | 0.66416768 |
| ENSMUSG00000057265 | Bbof1 | 0.000110727 | -0.088130043 | 1 |
| ENSMUSG00000021238 | Aldh6a1 | 0.000110727 | -0.225633566 | 1 |
| ENSMUSG00000042350 | Arel1 | 0.000110727 | -3.592750435 | 6.34E-05 |
| ENSMUSG00000019235 | Rps6kl1 | 0.000110727 | -0.975655313 | 0.705975399 |
| ENSMUSG00000008822 | Acyp1 | 0.000110727 | 1.063873051 | 1 |
| ENSMUSG00000034126 | Pomt2 | 0.000110727 | -1.258055044 | 0.466464457 |
| ENSMUSG00000072919 | Noxred1 | 0.000110727 | -0.720398258 | 0.950123751 |
| ENSMUSG00000021040 | Slirp | 0.000110727 | -0.466641666 | 1 |

| | | | | |
|---|---|---|---|---|
| ENSMUSG00000007682 | Dio2 | 0.000110727 | 2.648835551 | 0.420414118 |
| ENSMUSG00000021177 | Tdp1 | 0.000110727 | -1.380911792 | 0.408535739 |
| ENSMUSG00000045404 | Kcnk13 | 0.000110727 | -0.014129461 | 1 |
| ENSMUSG00000113679 | Gm10432 | 0.000110727 | 1.233798052 | 0.792441218 |
| ENSMUSG00000021186 | Fbln5 | 0.000110727 | -0.351164449 | 1 |
| ENSMUSG00000096458 | Moap1 | 0.000110727 | 1.233798052 | 0.792441218 |
| ENSMUSG00000054150 | Syne3 | 0.000110727 | -1.014129461 | 0.6155493 |
| ENSMUSG00000021070 | Bdkrb2 | 0.000110727 | -1.410058138 | 0.368337985 |
| ENSMUSG00000114149 | Gm47646 | 0.000110727 | 1.42644313 | 0.736456705 |
| ENSMUSG00000085120 | Gm16086 | 0.000110727 | 4.233798052 | 0.128972787 |
| ENSMUSG00000048856 | Slc25a47 | 0.000110727 | -1.43862729 | 0.339627968 |
| ENSMUSG00000097391 | Mirg | 0.000110727 | -0.853664789 | 0.806496094 |
| ENSMUSG00000113868 | Gm34719 | 0.000110727 | 0.774366434 | 1 |
| ENSMUSG00000113581 | Dio3os | 0.000110727 | 0.233798052 | 1 |
| ENSMUSG00000097381 | A230087F16Rik | 0.000110727 | 1.063873051 | 1 |
| ENSMUSG00000021280 | Exoc3l4 | 0.000110727 | 0.533358334 | 1 |
| ENSMUSG00000060950 | Trmt61a | 0.000110727 | -0.088130043 | 1 |
| ENSMUSG00000054013 | Tmem179 | 0.000110727 | 1.911869957 | 0.587484274 |
| ENSMUSG00000012211 | Tex22 | 0.000110727 | 0.063873051 | 1 |
| ENSMUSG00000042029 | Ncapg2 | 0.000110727 | -1.158519371 | 0.521275193 |
| ENSMUSG00000113875 | D230030E09Rik | 0.000110727 | 2.648835551 | 0.403518669 |
| ENSMUSG00000045410 | Akr1e1 | 0.000110727 | -0.895484965 | 0.691084416 |
| ENSMUSG00000058258 | Idi1 | 0.000110727 | -0.225633566 | 1 |
| ENSMUSG00000057069 | Ero1lb | 0.000110727 | -0.088130043 | 1 |
| ENSMUSG00000056457 | Prl2c3 | 0.000110727 | 1.233798052 | 0.792441218 |
| ENSMUSG00000113338 | A530046M15Rik | 0.000110727 | 0.774366434 | 1 |
| ENSMUSG00000012429 | Mplkip | 0.000110727 | -1.3207908 | 0.395686633 |
| ENSMUSG00000021314 | Amph | 0.000110727 | -0.975655313 | 0.745189742 |
| ENSMUSG00000041138 | Nme8 | 0.000110727 | 0.42644313 | 1 |
| ENSMUSG00000079941 | Gm11273 | 0.000110727 | 1.911869957 | 0.598226668 |
| ENSMUSG00000087097 | 4930586N03Rik | 0.000110727 | 0.533358334 | 1 |
| ENSMUSG00000069272 | Hist1h2ae | 0.000110727 | 1.42644313 | 0.736456705 |
| ENSMUSG00000084788 | Gm11342 | 0.000110727 | -0.014129461 | 1 |
| ENSMUSG00000086531 | Gm11351 | 0.000110727 | 0.42644313 | 1 |
| ENSMUSG00000045835 | Hdgfl1 | 0.000110727 | 2.648835551 | 0.395118486 |
| ENSMUSG00000114646 | Gm47888 | 0.000110727 | 0.774366434 | 1 |
| ENSMUSG00000049115 | Agtr1a | 0.000110727 | 1.911869957 | 0.596454325 |
| ENSMUSG00000113515 | Gm29675 | 0.000110727 | 2.648835551 | 0.395118486 |
| ENSMUSG00000085620 | G630018N14Rik | 0.000110727 | 1.233798052 | 0.792441218 |
| ENSMUSG00000085457 | 1110046J04Rik | 0.000110727 | -0.410058138 | 1 |
| ENSMUSG00000046949 | Nqo2 | 0.000110727 | -1.192466703 | 0.538933945 |
| ENSMUSG00000113615 | Gm47150 | 0.000110727 | 4.233798052 | 0.128972787 |
| ENSMUSG00000113484 | Gm47157 | 0.000110727 | 1.911869957 | 0.587484274 |
| ENSMUSG00000114647 | Gm30127 | 0.000110727 | 0.774366434 | 0.912904617 |
| ENSMUSG00000114732 | Gm40918 | 0.000110727 | -0.52108945 | 1 |
| ENSMUSG00000114866 | Gm40922 | 0.000110727 | 2.648835551 | 0.421384409 |
| ENSMUSG00000114319 | Gm47509 | 0.000110727 | -0.289763904 | 1 |
| ENSMUSG00000021432 | Slc35b3 | 0.000110727 | -2.014129461 | 0.073735838 |
| ENSMUSG00000114491 | A230103O09Rik | 0.000110727 | 1.233798052 | 0.792441218 |
| ENSMUSG00000021363 | Mak | 0.000110727 | 0.326907457 | 1 |
| ENSMUSG00000038651 | Sycp2l | 0.000110727 | 0.774366434 | 1 |

| | | | | |
|---|---|---|---|---|
| ENSMUSG00000089710 | Gm15810 | 0.000110727 | 3.233798052 | 0.262284476 |
| ENSMUSG00000113780 | Gm33195 | 0.000110727 | 1.233798052 | 0.792441218 |
| ENSMUSG00000113977 | Gm34276 | 0.000110727 | 1.063873051 | 0.847265357 |
| ENSMUSG00000114961 | A930002C04Rik | 0.000110727 | 3.233798052 | 0.262284476 |
| ENSMUSG00000114843 | Gm35733 | 0.000110727 | 1.648835551 | 0.66416768 |
| ENSMUSG00000047324 | 4931429P17Rik | 0.000110727 | 0.063873051 | 1 |
| ENSMUSG00000050954 | Zfp169 | 0.000110727 | -1.410058138 | 0.33333983 |
| ENSMUSG00000037989 | Wnk2 | 0.000110727 | -0.624182943 | 0.949989126 |
| ENSMUSG00000021384 | Susd3 | 0.000110727 | 4.233798052 | 0.128972787 |
| ENSMUSG00000112959 | Gm32834 | 0.000110727 | 1.233798052 | 0.792441218 |
| ENSMUSG00000067586 | S1pr3 | 0.000110727 | 2.233798052 | 0.503644124 |
| ENSMUSG00000113090 | Gm48190 | 0.000110727 | 2.648835551 | 0.433732559 |
| ENSMUSG00000047842 | Diras2 | 0.000110727 | 1.911869957 | 0.605948267 |
| ENSMUSG00000025877 | Hk3 | 0.000110727 | 2.648835551 | 0.395118486 |
| ENSMUSG00000034789 | Rab24 | 0.000110727 | -1.766201948 | 0.177083054 |
| ENSMUSG00000021492 | F12 | 0.000110727 | 0.911869957 | 1 |
| ENSMUSG00000074886 | Grk6 | 0.000110727 | -2.051604167 | 0.078768438 |
| ENSMUSG00000097361 | 4930550C17Rik | 0.000110727 | 3.233798052 | 0.262284476 |
| ENSMUSG00000021508 | Cxcl14 | 0.000110727 | 0.648835551 | 1 |
| ENSMUSG00000021540 | Smad5 | 0.000110727 | -2.395558568 | 0.022222922 |
| ENSMUSG00000021550 | 2210016F16Rik | 0.000110727 | -0.810596067 | 0.796840156 |
| ENSMUSG00000114231 | Gm40968 | 0.000110727 | 2.648835551 | 0.423639929 |
| ENSMUSG00000114854 | Gm34354 | 0.000110727 | 1.911869957 | 0.589666002 |
| ENSMUSG00000114354 | Gm34961 | 0.000110727 | 2.648835551 | 0.404997369 |
| ENSMUSG00000074874 | Ctla2b | 0.000110727 | 0.42644313 | 1 |
| ENSMUSG00000113450 | Gm49359 | 0.000110727 | -2.336057556 | 0.024405815 |
| ENSMUSG00000069805 | Fbp1 | 0.000110727 | 1.42644313 | 0.736456705 |
| ENSMUSG00000044934 | Zfp367 | 0.000110727 | -1.52108945 | 0.284607766 |
| ENSMUSG00000113244 | Gm31218 | 0.000110727 | 2.233798052 | 0.47966612 |
| ENSMUSG00000021519 | Mterf3 | 0.000110727 | -0.766201948 | 0.792441218 |
| ENSMUSG00000071281 | Zfp65 | 0.000110727 | -1.192466703 | 0.606122355 |
| ENSMUSG00000052981 | Ube2ql1 | 0.000110727 | 1.648835551 | 0.66416768 |
| ENSMUSG00000113794 | Gm36607 | 0.000110727 | 2.648835551 | 0.404713153 |
| ENSMUSG00000021565 | Slc6a19 | 0.000110727 | 3.233798052 | 0.262284476 |
| ENSMUSG00000021569 | Trip13 | 0.000110727 | -1.158519371 | 0.521275193 |
| ENSMUSG00000069184 | Zfp72 | 0.000110727 | -0.624182943 | 0.912904617 |
| ENSMUSG00000021592 | Arsk | 0.000110727 | -0.673092543 | 0.919998879 |
| ENSMUSG00000114522 | Gm32067 | 0.000110727 | 0.911869957 | 1 |
| ENSMUSG00000114594 | Gm48402 | 0.000110727 | 4.233798052 | 0.128972787 |
| ENSMUSG00000114372 | 2310067P03Rik | 0.000110727 | 0.774366434 | 1 |
| ENSMUSG00000098087 | Gm17750 | 0.000110727 | -0.975655313 | 0.705975399 |
| ENSMUSG00000021613 | Hapln1 | 0.000110727 | -0.52108945 | 1 |
| ENSMUSG00000014850 | Msh3 | 0.000110727 | -3.14990624 | 0.000752229 |
| ENSMUSG00000021690 | Jmy | 0.000110727 | -3.480447466 | 0.00011788 |
| ENSMUSG00000109561 | Ankrd31 | 0.000110727 | 0.063873051 | 1 |
| ENSMUSG00000041817 | Fam169a | 0.000110727 | -0.895484965 | 0.856504447 |
| ENSMUSG00000114415 | Gm41030 | 0.000110727 | -0.014129461 | 1 |
| ENSMUSG00000021661 | Ankra2 | 0.000110727 | -0.158519371 | 1 |
| ENSMUSG00000096330 | Gm21976 | 0.000110727 | 0.911869957 | 1 |
| ENSMUSG00000021646 | Mccc2 | 0.000110727 | -2.975655313 | 0.003846186 |
| ENSMUSG00000045273 | Cenph | 0.000110727 | -1.648844997 | 0.201326391 |

| | | | | |
|---|---|---|---|---|
| ENSMUSG00000021711 | Trappc13 | 0.000110727 | -2.051604167 | 0.078768438 |
| ENSMUSG00000114534 | Gm32004 | 0.000110727 | 2.648835551 | 0.426063356 |
| ENSMUSG00000086861 | Gm15325 | 0.000110727 | 4.233798052 | 0.128972787 |
| ENSMUSG00000086402 | Gm15322 | 0.000110727 | 3.233798052 | 0.295355925 |
| ENSMUSG00000021758 | Ddx4 | 0.000110727 | 2.648835551 | 0.395519648 |
| ENSMUSG00000047789 | Slc38a9 | 0.000110727 | -2.799624949 | 0.00438405 |
| ENSMUSG00000042426 | Dhx29 | 0.000110727 | -0.466641666 | 1 |
| ENSMUSG00000021760 | Gpx8 | 0.000110727 | 1.063873051 | 1 |
| ENSMUSG00000114333 | 1700084D21Rik | 0.000110727 | 3.233798052 | 0.262284476 |
| ENSMUSG00000042385 | Gzmk | 0.000110727 | -0.351164449 | 1 |
| ENSMUSG00000114715 | Gm34471 | 0.000110727 | 0.774366434 | 1 |
| ENSMUSG00000114935 | Gm47040 | 0.000110727 | 2.233798052 | 0.47966612 |
| ENSMUSG00000042275 | Pelo | 0.000110727 | 1.233798052 | 0.792441218 |
| ENSMUSG00000114731 | Gm47914 | 0.000110727 | 0.911869957 | 1 |
| ENSMUSG00000114613 | Gm10732 | 0.000110727 | 2.648835551 | 0.395118486 |
| ENSMUSG00000086709 | Gm16263 | 0.000110727 | 0.648835551 | 1 |
| ENSMUSG00000074715 | Ccl28 | 0.000110727 | 1.233798052 | 0.792441218 |
| ENSMUSG00000114369 | Gm41077 | 0.000110727 | 1.648835551 | 0.66416768 |
| ENSMUSG00000095609 | Gm21188 | 0.000110727 | 1.063873051 | 1 |
| ENSMUSG00000097278 | Gm26650 | 0.000110727 | 3.233798052 | 0.262284476 |
| ENSMUSG00000090585 | 4933406F09Rik | 0.000110727 | 2.233798052 | 0.47966612 |
| ENSMUSG00000114532 | 2610318M16Rik | 0.000110727 | 2.233798052 | 0.47966612 |
| ENSMUSG00000084902 | Gm281 | 0.000110727 | 0.911869957 | 1 |
| ENSMUSG00000114883 | Gm48860 | 0.000110727 | -0.57355687 | 0.910050347 |
| ENSMUSG00000097547 | B230110C06Rik | 0.000110727 | 2.648835551 | 0.395118486 |
| ENSMUSG00000114243 | Gm48163 | 0.000110727 | -0.810596067 | 0.796840156 |
| ENSMUSG00000084925 | 1810062O18Rik | 0.000110727 | -0.853664789 | 0.806496094 |
| ENSMUSG00000090564 | A430057M04Rik | 0.000110727 | 3.233798052 | 0.279041806 |
| ENSMUSG00000039081 | Zfp503 | 0.000110727 | -1.380911792 | 0.408535739 |
| ENSMUSG00000068680 | Gm10248 | 0.000110727 | 1.911869957 | 0.607010878 |
| ENSMUSG00000118519 | AC163638.2 | 0.000110727 | 1.233798052 | 0.792441218 |
| ENSMUSG00000021994 | Wnt5a | 0.000110727 | -1.088130043 | 0.624094312 |
| ENSMUSG00000114704 | Gm48003 | 0.000110727 | 1.063873051 | 0.847390568 |
| ENSMUSG00000114553 | Gm7644 | 0.000110727 | 1.42644313 | 0.736456705 |
| ENSMUSG00000021904 | Sema3g | 0.000110727 | 0.911869957 | 1 |
| ENSMUSG00000019027 | Dnah1 | 0.000110727 | 1.42644313 | 0.736456705 |
| ENSMUSG00000115311 | Gm35823 | 0.000110727 | 2.648835551 | 0.395118486 |
| ENSMUSG00000057606 | Colq | 0.000110727 | 0.063873051 | 1 |
| ENSMUSG00000021900 | Btd | 0.000110727 | -0.57355687 | 0.910050347 |
| ENSMUSG00000021903 | Galnt15 | 0.000110727 | -0.088130043 | 1 |
| ENSMUSG00000021913 | Ogdhl | 0.000110727 | 0.774366434 | 1 |
| ENSMUSG00000041707 | Tmem273 | 0.000110727 | 1.063873051 | 1 |
| ENSMUSG00000108841 | Frmpd2 | 0.000110727 | 0.774366434 | 1 |
| ENSMUSG00000115726 | Gm30083 | 0.000110727 | 4.233798052 | 0.128972787 |
| ENSMUSG00000095493 | A630023A22Rik | 0.000110727 | 0.326907457 | 1 |
| ENSMUSG00000043681 | Fam25c | 0.000110727 | 4.233798052 | 0.128972787 |
| ENSMUSG00000041445 | Mmrn2 | 0.000110727 | 0.233798052 | 1 |
| ENSMUSG00000093383 | Gm20642 | 0.000110727 | 0.533358334 | 1 |
| ENSMUSG00000021795 | Sftpd | 0.000110727 | 2.233798052 | 0.47966612 |
| ENSMUSG00000093926 | Gm17027 | 0.000110727 | 2.233798052 | 0.503513099 |
| ENSMUSG00000096446 | Gm8104 | 0.000110727 | 4.233798052 | 0.128972787 |

| | | | | |
|---|---|---|---|---|
| ENSMUSG00000115050 | Gm21154 | 0.000110727 | 1.648835551 | 0.66416768 |
| ENSMUSG00000115801 | AC160336.1 | 0.000110727 | 0.42644313 | 1 |
| ENSMUSG00000115324 | 4930527F14Rik | 0.000110727 | 1.648835551 | 0.66416768 |
| ENSMUSG00000115000 | Gm49192 | 0.000110727 | 1.648835551 | 0.66416768 |
| ENSMUSG00000037628 | Cdkn3 | 0.000110727 | 0.533358334 | 1 |
| ENSMUSG00000055128 | Cgrrf1 | 0.000110727 | 0.063873051 | 1 |
| ENSMUSG00000072589 | Gm10371 | 0.000110727 | 0.42644313 | 1 |
| ENSMUSG00000037526 | Atg14 | 0.000110727 | -1.599091962 | 0.242941974 |
| ENSMUSG00000115215 | Gm41148 | 0.000110727 | 0.911869957 | 1 |
| ENSMUSG00000095917 | Olfr740 | 0.000110727 | 2.648835551 | 0.424957368 |
| ENSMUSG00000095765 | Olfr741 | 0.000110727 | 4.233798052 | 0.213902962 |
| ENSMUSG00000036023 | Parp2 | 0.000110727 | -1.192466703 | 0.529126167 |
| ENSMUSG00000115540 | 4930597G03Rik | 0.000110727 | 4.233798052 | 0.128972787 |
| ENSMUSG00000072572 | Slc39a2 | 0.000110727 | -0.289763904 | 1 |
| ENSMUSG00000105963 | Gm43647 | 0.000110727 | 1.233798052 | 0.792441218 |
| ENSMUSG00000022177 | Haus4 | 0.000110727 | -1.014129461 | 0.608426921 |
| ENSMUSG00000040822 | 1700123O20Rik | 0.000110727 | -0.673092543 | 0.919998879 |
| ENSMUSG00000072494 | Ppp1r3e | 0.000110727 | 0.648835551 | 1 |
| ENSMUSG00000089682 | Bcl2l2 | 0.000110727 | -0.57355687 | 0.910050347 |
| ENSMUSG00000115610 | Gm31251 | 0.000110727 | 3.233798052 | 0.325969989 |
| ENSMUSG00000002325 | Irf9 | 0.000110727 | -1.52108945 | 0.284607766 |
| ENSMUSG00000002319 | Ipo4 | 0.000110727 | -1.410058138 | 0.33333983 |
| ENSMUSG00000040472 | Rabggta | 0.000110727 | -0.52108945 | 1 |
| ENSMUSG00000022220 | Adcy4 | 0.000110727 | 0.648835551 | 1 |
| ENSMUSG00000023411 | Nfatc4 | 0.000110727 | -1.225633566 | 0.463954265 |
| ENSMUSG00000075592 | Nynrin | 0.000110727 | 0.233798052 | 1 |
| ENSMUSG00000040380 | Cbln3 | 0.000110727 | 3.233798052 | 0.262284476 |
| ENSMUSG00000061068 | Mcpt4 | 0.000110727 | 3.233798052 | 0.262284476 |
| ENSMUSG00000114690 | Gm33321 | 0.000110727 | 1.648835551 | 0.66416768 |
| ENSMUSG00000114720 | Gm41162 | 0.000110727 | 1.648835551 | 0.66416768 |
| ENSMUSG00000071347 | C1qtnf9 | 0.000110727 | 1.42644313 | 0.736456705 |
| ENSMUSG00000115531 | Gm35419 | 0.000110727 | 1.648835551 | 0.66416768 |
| ENSMUSG00000021932 | Rnaseh2b | 0.000110727 | -0.895484965 | 0.691084416 |
| ENSMUSG00000035121 | Neil2 | 0.000110727 | -0.225633566 | 1 |
| ENSMUSG00000021961 | 4930578I06Rik | 0.000110727 | 4.233798052 | 0.128972787 |
| ENSMUSG00000007989 | Fzd3 | 0.000110727 | -1.936126949 | 0.106828177 |
| ENSMUSG00000114209 | Gm48433 | 0.000110727 | 1.233798052 | 0.792441218 |
| ENSMUSG00000022031 | Elp3 | 0.000110727 | -1.853664789 | 0.121970859 |
| ENSMUSG00000022034 | Esco2 | 0.000110727 | -0.57355687 | 0.910050347 |
| ENSMUSG00000034450 | Gulo | 0.000110727 | 1.911869957 | 0.587484274 |
| ENSMUSG00000093386 | 1700120O09Rik | 0.000110727 | 1.648835551 | 0.66416768 |
| ENSMUSG00000022057 | Adamdec1 | 0.000110727 | 1.648835551 | 0.664426832 |
| ENSMUSG00000114833 | Gm41192 | 0.000110727 | 0.533358334 | 1 |
| ENSMUSG00000033712 | Ccar2 | 0.000110727 | -2.843017545 | 0.003274704 |
| ENSMUSG00000022090 | Pdlim2 | 0.000110727 | 0.063873051 | 1 |
| ENSMUSG00000000776 | Polr3d | 0.000110727 | -2.547561661 | 0.012733015 |
| ENSMUSG00000033589 | Reep4 | 0.000110727 | -0.289763904 | 1 |
| ENSMUSG00000022099 | Dmtn | 0.000110727 | 0.911869957 | 1 |
| ENSMUSG00000099528 | 2410012E07Rik | 0.000110727 | 2.648835551 | 0.4474981 |
| ENSMUSG00000022103 | Gfra2 | 0.000110727 | 0.533358334 | 1 |
| ENSMUSG00000115532 | Gm41206 | 0.000110727 | 1.648835551 | 0.66416768 |

| | | | | |
|---|---|---|---|---|
| ENSMUSG00000034959 | Rubcnl | 0.000110727 | -0.289763904 | 1 |
| ENSMUSG00000021997 | Lrrc63 | 0.000110727 | 0.911869957 | 1 |
| ENSMUSG00000021999 | Cpb2 | 0.000110727 | 0.648835551 | 1 |
| ENSMUSG00000115615 | Gm49200 | 0.000110727 | 3.233798052 | 0.262284476 |
| ENSMUSG00000034893 | Cog3 | 0.000110727 | -3.273996588 | 0.00039298 |
| ENSMUSG00000052584 | Serp2 | 0.000110727 | -0.410058138 | 1 |
| ENSMUSG00000115363 | Gm49003 | 0.000110727 | -0.014129461 | 1 |
| ENSMUSG00000115257 | Gm30467 | 0.000110727 | 2.648835551 | 0.434488598 |
| ENSMUSG00000022018 | Rgcc | 0.000110727 | 2.648835551 | 0.395118486 |
| ENSMUSG00000097927 | Gm6999 | 0.000110727 | 0.774366434 | 1 |
| ENSMUSG00000115542 | Gm49046 | 0.000110727 | 2.233798052 | 0.47966612 |
| ENSMUSG00000115271 | 4930529K09Rik | 0.000110727 | 0.911869957 | 1 |
| ENSMUSG00000115664 | Gm32913 | 0.000110727 | 2.648835551 | 0.455131546 |
| ENSMUSG00000115601 | Gm49007 | 0.000110727 | 1.911869957 | 0.60764178 |
| ENSMUSG00000110273 | Gm41231 | 0.000110727 | 2.648835551 | 0.395118486 |
| ENSMUSG00000115696 | Gm49292 | 0.000110727 | 1.648835551 | 0.66416768 |
| ENSMUSG00000115099 | 1700087I21Rik | 0.000110727 | 0.326907457 | 1 |
| ENSMUSG00000055717 | Slain1 | 0.000110727 | 1.063873051 | 0.85566943 |
| ENSMUSG00000115548 | 5430440P10Rik | 0.000110727 | 2.648835551 | 0.426815929 |
| ENSMUSG00000115149 | 9330188P03Rik | 0.000110727 | 0.648835551 | 1 |
| ENSMUSG00000058126 | Tpm3-rs7 | 0.000110727 | 0.533358334 | 1 |
| ENSMUSG00000089726 | Mir17hg | 0.000110727 | -0.720398258 | 0.792441218 |
| ENSMUSG00000093551 | 4930505G20Rik | 0.000110727 | 1.911869957 | 0.601314378 |
| ENSMUSG00000090534 | Gm4675 | 0.000110727 | 1.233798052 | 0.792441218 |
| ENSMUSG00000115258 | Gm32093 | 0.000110727 | 4.233798052 | 0.128972787 |
| ENSMUSG00000050350 | Gpr18 | 0.000110727 | 2.233798052 | 0.47966612 |
| ENSMUSG00000075463 | 4930594M22Rik | 0.000110727 | -0.158519371 | 1 |
| ENSMUSG00000022181 | C6 | 0.000110727 | 0.648835551 | 1 |
| ENSMUSG00000050697 | Prkaa1 | 0.000110727 | -2.43862729 | 0.019307676 |
| ENSMUSG00000039942 | Ptger4 | 0.000110727 | 0.42644313 | 1 |
| ENSMUSG00000115591 | Gm31282 | 0.000110727 | 0.063873051 | 1 |
| ENSMUSG00000054115 | Skp2 | 0.000110727 | 0.648835551 | 1 |
| ENSMUSG00000039676 | Capsl | 0.000110727 | 3.233798052 | 0.262284476 |
| ENSMUSG00000005268 | Prlr | 0.000110727 | -0.088130043 | 1 |
| ENSMUSG00000089678 | Agxt2 | 0.000110727 | 1.063873051 | 1 |
| ENSMUSG00000096971 | 4930556M19Rik | 0.000110727 | -0.810596067 | 0.851260516 |
| ENSMUSG00000115151 | Gm19276 | 0.000110727 | 1.911869957 | 0.587484274 |
| ENSMUSG00000022206 | Npr3 | 0.000110727 | 1.063873051 | 0.848985747 |
| ENSMUSG00000115485 | Gm49126 | 0.000110727 | 0.774366434 | 1 |
| ENSMUSG00000115198 | Gm49729 | 0.000110727 | 0.648835551 | 1 |
| ENSMUSG00000097452 | Gm2824 | 0.000110727 | 1.648835551 | 0.66416768 |
| ENSMUSG00000115138 | Gm36899 | 0.000110727 | 3.233798052 | 0.262284476 |
| ENSMUSG00000056069 | Otulinl | 0.000110727 | -0.624182943 | 0.936637573 |
| ENSMUSG00000115474 | 4930430F21Rik | 0.000110727 | 0.911869957 | 1 |
| ENSMUSG00000096956 | Snhg18 | 0.000110727 | 0.533358334 | 1 |
| ENSMUSG00000115749 | Gm41289 | 0.000110727 | 2.233798052 | 0.47966612 |
| ENSMUSG00000044726 | Erich5 | 0.000110727 | -1.158519371 | 0.613899307 |
| ENSMUSG00000097259 | Gm26766 | 0.000110727 | 0.648835551 | 1 |
| ENSMUSG00000022288 | 4930447A16Rik | 0.000110727 | 2.648835551 | 0.395118486 |
| ENSMUSG00000022292 | Rrm2b | 0.000110727 | -0.766201948 | 0.806008864 |
| ENSMUSG00000102106 | 2310043O21Rik | 0.000110727 | -0.158519371 | 1 |

| | | | | |
|---|---|---|---|---|
| ENSMUSG00000115620 | Gm45924 | 0.000110727 | 1.911869957 | 0.587484274 |
| ENSMUSG00000022303 | Dcstamp | 0.000110727 | 1.911869957 | 0.588477963 |
| ENSMUSG00000022305 | Lrp12 | 0.000110727 | -0.624182943 | 0.912904617 |
| ENSMUSG00000072592 | Gm10373 | 0.000110727 | 0.911869957 | 1 |
| ENSMUSG00000115478 | Gm49198 | 0.000110727 | 1.233798052 | 0.792441218 |
| ENSMUSG00000022313 | Utp23 | 0.000110727 | -1.351164449 | 0.401963028 |
| ENSMUSG00000037362 | Ccn3 | 0.000110727 | -0.225633566 | 1 |
| ENSMUSG00000022369 | Mtbp | 0.000110727 | -0.57355687 | 0.928798266 |
| ENSMUSG00000022364 | Tbc1d31 | 0.000110727 | -0.720398258 | 1 |
| ENSMUSG00000051225 | Fam83a | 0.000110727 | 1.42644313 | 0.736456705 |
| ENSMUSG00000037085 | Trmt12 | 0.000110727 | 1.648835551 | 0.66416768 |
| ENSMUSG00000037075 | Rnf139 | 0.000110727 | -0.766201948 | 0.796840156 |
| ENSMUSG00000115896 | Gm36677 | 0.000110727 | 3.233798052 | 0.262284476 |
| ENSMUSG00000097493 | 9930014A18Rik | 0.000110727 | 2.233798052 | 0.47966612 |
| ENSMUSG00000116299 | Gm29904 | 0.000110727 | 0.146335211 | 1 |
| ENSMUSG00000094296 | Gm21798 | 0.000110727 | 0.326907457 | 1 |
| ENSMUSG00000079024 | Gm21961 | 0.000110727 | -0.351164449 | 1 |
| ENSMUSG00000022375 | Lrrc6 | 0.000110727 | 1.063873051 | 1 |
| ENSMUSG00000069996 | Gm7125 | 0.000110727 | 0.533358334 | 1 |
| ENSMUSG00000022603 | Mroh4 | 0.000110727 | -2.926073285 | 0.004210624 |
| ENSMUSG00000046380 | Jrk | 0.000110727 | 1.233798052 | 0.792441218 |
| ENSMUSG00000022594 | Lynx1 | 0.000110727 | 0.774366434 | 1 |
| ENSMUSG00000079018 | Ly6c1 | 0.000110727 | 3.233798052 | 0.262284476 |
| ENSMUSG00000047728 | Ly6g2 | 0.000110727 | 2.648835551 | 0.423814296 |
| ENSMUSG00000000934 | Top1mt | 0.000110727 | 0.233798052 | 1 |
| ENSMUSG00000022571 | Pycrl | 0.000110727 | 1.42644313 | 0.736456705 |
| ENSMUSG00000050846 | Zfp623 | 0.000110727 | -0.088130043 | 1 |
| ENSMUSG00000075590 | Nrbp2 | 0.000110727 | -0.410058138 | 1 |
| ENSMUSG00000115388 | Eppk1 | 0.000110727 | -0.624182943 | 0.912904617 |
| ENSMUSG00000022562 | Oplah | 0.000110727 | 0.063873051 | 1 |
| ENSMUSG00000022559 | Fbxl6 | 0.000110727 | 0.648835551 | 1 |
| ENSMUSG00000022550 | Adck5 | 0.000110727 | -1.158519371 | 0.521275193 |
| ENSMUSG00000034022 | Cpsf1 | 0.000110727 | -1.051604167 | 0.614718876 |
| ENSMUSG00000054967 | Zfp647 | 0.000110727 | 0.911869957 | 1 |
| ENSMUSG00000018893 | Mb | 0.000110727 | 0.146335211 | 1 |
| ENSMUSG00000071713 | Csf2rb | 0.000110727 | 1.233798052 | 0.792441218 |
| ENSMUSG00000033287 | Kctd17 | 0.000110727 | 1.233798052 | 0.792441218 |
| ENSMUSG00000116097 | Gm36738 | 0.000110727 | 0.648835551 | 1 |
| ENSMUSG00000043460 | Elfn2 | 0.000110727 | 1.648835551 | 0.66416768 |
| ENSMUSG00000018126 | Baiap2l2 | 0.000110727 | -0.936126949 | 0.697198109 |
| ENSMUSG00000022429 | Dmc1 | 0.000110727 | 0.648835551 | 1 |
| ENSMUSG00000022428 | Cby1 | 0.000110727 | 0.233798052 | 1 |
| ENSMUSG00000042535 | Gtpbp1 | 0.000110727 | -0.014129461 | 1 |
| ENSMUSG00000116125 | Gm41361 | 0.000110727 | -1.051604167 | 0.614718876 |
| ENSMUSG00000051518 | Rps19bp1 | 0.000110727 | 0.146335211 | 1 |
| ENSMUSG00000022394 | L3mbtl2 | 0.000110727 | -1.766201948 | 0.177083054 |
| ENSMUSG00000063765 | Chadl | 0.000110727 | 1.42644313 | 0.736456705 |
| ENSMUSG00000042109 | Csdc2 | 0.000110727 | 0.42644313 | 1 |
| ENSMUSG00000022453 | Naga | 0.000110727 | -0.289763904 | 1 |
| ENSMUSG00000058586 | Serhl | 0.000110727 | -0.853664789 | 0.806496094 |
| ENSMUSG00000047878 | A4galt | 0.000110727 | -0.410058138 | 1 |

| | | | | |
|---|---|---|---|---|
| ENSMUSG00000041708 | Mpped1 | 0.000110727 | 1.063873051 | 1 |
| ENSMUSG00000022441 | Efcab6 | 0.000110727 | -0.466641666 | 1 |
| ENSMUSG00000022434 | Fam118a | 0.000110727 | -0.673092543 | 0.925807344 |
| ENSMUSG00000022385 | Gtse1 | 0.000110727 | 1.648835551 | 0.66416768 |
| ENSMUSG00000022388 | Ttll8 | 0.000110727 | 1.233798052 | 0.792441218 |
| ENSMUSG00000022610 | Mapk12 | 0.000110727 | -0.466641666 | 1 |
| ENSMUSG00000015377 | Dennd6b | 0.000110727 | -1.158519371 | 0.521275193 |
| ENSMUSG00000091780 | Sco2 | 0.000110727 | 1.911869957 | 0.587484274 |
| ENSMUSG00000063260 | Syt10 | 0.000110727 | 1.911869957 | 0.590105758 |
| ENSMUSG00000116428 | Gm36618 | 0.000110727 | 2.233798052 | 0.47966612 |
| ENSMUSG00000055782 | Abcd2 | 0.000110727 | 2.233798052 | 0.504119351 |
| ENSMUSG00000044021 | Muc19 | 0.000110727 | 1.42644313 | 0.736456705 |
| ENSMUSG00000115569 | Gm49169 | 0.000110727 | 3.233798052 | 0.262284476 |
| ENSMUSG00000097536 | 2610037D02Rik | 0.000110727 | -3.167081384 | 0.000661954 |
| ENSMUSG00000022468 | Endou | 0.000110727 | 2.233798052 | 0.492903817 |
| ENSMUSG00000054036 | Olfr279 | 0.000110727 | 0.911869957 | 1 |
| ENSMUSG00000003355 | Fkbp11 | 0.000110727 | 0.146335211 | 1 |
| ENSMUSG00000038009 | Dnajc22 | 0.000110727 | -0.466641666 | 1 |
| ENSMUSG00000078907 | Fam186b | 0.000110727 | 0.42644313 | 1 |
| ENSMUSG00000023015 | Racgap1 | 0.000110727 | -1.975655313 | 0.088605926 |
| ENSMUSG00000090406 | Gm17058 | 0.000110727 | 2.233798052 | 0.496010946 |
| ENSMUSG00000037353 | Letmd1 | 0.000110727 | -0.014129461 | 1 |
| ENSMUSG00000116180 | Gm49492 | 0.000110727 | 2.233798052 | 0.47966612 |
| ENSMUSG00000023034 | Nr4a1 | 0.000110727 | -0.289763904 | 1 |
| ENSMUSG00000037185 | Krt80 | 0.000110727 | -0.853664789 | 0.806496094 |
| ENSMUSG00000023050 | Map3k12 | 0.000110727 | -0.289763904 | 1 |
| ENSMUSG00000023051 | Tarbp2 | 0.000110727 | -2.175592884 | 0.045933679 |
| ENSMUSG00000023055 | Calcoco1 | 0.000110727 | -1.225633566 | 0.463954265 |
| ENSMUSG00000115961 | Gm41409 | 0.000110727 | -0.895484965 | 0.709951754 |
| ENSMUSG00000005982 | Naa60 | 0.000110727 | -0.673092543 | 0.919998879 |
| ENSMUSG00000039568 | Ubald1 | 0.000110727 | 0.326907457 | 1 |
| ENSMUSG00000023143 | Nagpa | 0.000110727 | 0.648835551 | 1 |
| ENSMUSG00000087168 | Gm15983 | 0.000110727 | 1.233798052 | 0.792441218 |
| ENSMUSG00000116525 | Gm30371 | 0.000110727 | 2.233798052 | 0.508263132 |
| ENSMUSG00000039200 | Atf7ip2 | 0.000110727 | 0.533358334 | 1 |
| ENSMUSG00000038055 | Dexi | 0.000110727 | -1.915949067 | 0.103463818 |
| ENSMUSG00000096921 | Gm26822 | 0.000110727 | 0.911869957 | 1 |
| ENSMUSG00000116520 | 4930414F18Rik | 0.000110727 | -0.975655313 | 0.743702746 |
| ENSMUSG00000022683 | Pla2g10 | 0.000110727 | 1.063873051 | 1 |
| ENSMUSG00000065968 | Ifitm7 | 0.000110727 | 3.233798052 | 0.262284476 |
| ENSMUSG00000075395 | A630010A05Rik | 0.000110727 | 0.648835551 | 1 |
| ENSMUSG00000022673 | Mcm4 | 0.000110727 | -0.853664789 | 0.806496094 |
| ENSMUSG00000022773 | Ypel1 | 0.000110727 | -0.52108945 | 1 |
| ENSMUSG00000022760 | Thap7 | 0.000110727 | 0.42644313 | 1 |
| ENSMUSG00000060166 | Zdhhc8 | 0.000110727 | -0.410058138 | 1 |
| ENSMUSG00000013539 | Tango2 | 0.000110727 | -2.088130043 | 0.064551115 |
| ENSMUSG00000075335 | 4930588K23Rik | 0.000110727 | 0.533358334 | 1 |
| ENSMUSG00000046613 | Vwa5b2 | 0.000110727 | 1.911869957 | 0.587484274 |
| ENSMUSG00000115219 | Eef1akmt4 | 0.000110727 | -1.051604167 | 0.614718876 |
| ENSMUSG00000033618 | Map3k13 | 0.000110727 | -1.648844997 | 0.207356329 |
| ENSMUSG00000022871 | Fetub | 0.000110727 | 1.911869957 | 0.591702939 |

| | | | | |
|---|---|---|---|---|
| ENSMUSG00000116290 | Gm49518 | 0.000110727 | 1.648835551 | 0.66416768 |
| ENSMUSG00000115869 | Gm31814 | 0.000110727 | 0.146335211 | 1 |
| ENSMUSG00000048399 | Tprg | 0.000110727 | -0.351164449 | 1 |
| ENSMUSG00000092545 | Gm20319 | 0.000110727 | 1.648835551 | 0.66416768 |
| ENSMUSG00000116654 | Gm10823 | 0.000110727 | 0.063873051 | 1 |
| ENSMUSG00000048939 | Atp13a5 | 0.000110727 | 0.648835551 | 1 |
| ENSMUSG00000075286 | Gm1968 | 0.000110727 | 0.533358334 | 1 |
| ENSMUSG00000052316 | Lrrc15 | 0.000110727 | 2.648835551 | 0.424188363 |
| ENSMUSG00000116581 | Gm46565 | 0.000110727 | 1.42644313 | 0.736456705 |
| ENSMUSG00000116597 | Gm536 | 0.000110727 | 0.774366434 | 1 |
| ENSMUSG00000022780 | Meltf | 0.000110727 | 1.42644313 | 0.736456705 |
| ENSMUSG00000022814 | Umps | 0.000110727 | -0.351164449 | 1 |
| ENSMUSG00000035376 | Hacd2 | 0.000110727 | -2.305360759 | 0.028985062 |
| ENSMUSG00000079597 | Cstdc4 | 0.000110727 | 2.233798052 | 0.47966612 |
| ENSMUSG00000022838 | Eaf2 | 0.000110727 | -1.788569761 | 0.142787052 |
| ENSMUSG00000022821 | Hgd | 0.000110727 | 0.42644313 | 1 |
| ENSMUSG00000022805 | Maats1 | 0.000110727 | -0.014129461 | 1 |
| ENSMUSG00000022798 | Tex55 | 0.000110727 | 0.911869957 | 1 |
| ENSMUSG00000086175 | Gm15802 | 0.000110727 | 0.326907457 | 1 |
| ENSMUSG00000116573 | Gm36903 | 0.000110727 | 0.533358334 | 1 |
| ENSMUSG00000101463 | Gm28750 | 0.000110727 | 2.233798052 | 0.49224519 |
| ENSMUSG00000022705 | Drd3 | 0.000110727 | 0.146335211 | 1 |
| ENSMUSG00000022667 | Cd200r1 | 0.000110727 | 1.42644313 | 0.736456705 |
| ENSMUSG00000036172 | Cd200r3 | 0.000110727 | 2.233798052 | 0.50047911 |
| ENSMUSG00000022664 | Slc35a5 | 0.000110727 | 0.774366434 | 1 |
| ENSMUSG00000022657 | Cd96 | 0.000110727 | 0.774366434 | 1 |
| ENSMUSG00000022652 | Morc1 | 0.000110727 | -0.673092543 | 0.94896181 |
| ENSMUSG00000064061 | Dzip3 | 0.000110727 | -2.032988489 | 0.082475901 |
| ENSMUSG00000116848 | 4930542D17Rik | 0.000110727 | 1.063873051 | 0.854811373 |
| ENSMUSG00000035270 | Impg2 | 0.000110727 | -1.289763904 | 0.466464457 |
| ENSMUSG00000022742 | Cpox | 0.000110727 | -0.014129461 | 1 |
| ENSMUSG00000022744 | Cldnd1 | 0.000110727 | -0.158519371 | 1 |
| ENSMUSG00000050312 | Nsun3 | 0.000110727 | -0.936126949 | 0.778568111 |
| ENSMUSG00000047141 | Zfp654 | 0.000110727 | -2.258055044 | 0.033084874 |
| ENSMUSG00000108903 | Gm45029 | 0.000110727 | 1.911869957 | 0.599135879 |
| ENSMUSG00000022889 | Mrpl39 | 0.000110727 | -0.289763904 | 1 |
| ENSMUSG00000116556 | 1700007H22Rik | 0.000110727 | 1.42644313 | 0.736456705 |
| ENSMUSG00000116656 | Gm49708 | 0.000110727 | -1.123753952 | 0.66416768 |
| ENSMUSG00000039929 | Urb1 | 0.000110727 | -1.158519371 | 0.521275193 |
| ENSMUSG00000022972 | Cfap298 | 0.000110727 | -0.351164449 | 1 |
| ENSMUSG00000085865 | Gm15966 | 0.000110727 | -0.225633566 | 1 |
| ENSMUSG00000116987 | Gm49625 | 0.000110727 | -0.57355687 | 1 |
| ENSMUSG00000022964 | Tmem50b | 0.000110727 | 0.146335211 | 1 |
| ENSMUSG00000089774 | Slc5a3 | 0.000110727 | -1.258055044 | 0.466464457 |
| ENSMUSG00000000157 | Itgb2l | 0.000110727 | 2.233798052 | 0.47966612 |
| ENSMUSG00000117179 | Gm49907 | 0.000110727 | 3.233798052 | 0.262284476 |
| ENSMUSG00000046962 | Zbtb21 | 0.000110727 | -0.624182943 | 0.912904617 |
| ENSMUSG00000015659 | Serac1 | 0.000110727 | -0.673092543 | 0.919998879 |
| ENSMUSG00000058159 | T2 | 0.000110727 | 0.063873051 | 1 |
| ENSMUSG00000117026 | C87487 | 0.000110727 | 1.648835551 | 0.66416768 |
| ENSMUSG00000089633 | A230009B12Rik | 0.000110727 | 2.648835551 | 0.395118486 |

| | | | | |
|---|---|---|---|---|
| ENSMUSG00000059481 | Plg | 0.000110727 | 0.533358334 | 1 |
| ENSMUSG00000117030 | Gm49959 | 0.000110727 | 3.233798052 | 0.262284476 |
| ENSMUSG00000073460 | Pnldc1 | 0.000110727 | 0.774366434 | 1 |
| ENSMUSG00000117278 | Gm36684 | 0.000110727 | 4.233798052 | 0.128972787 |
| ENSMUSG00000052469 | Tcp10c | 0.000110727 | 0.774366434 | 1 |
| ENSMUSG00000023885 | Thbs2 | 0.000110727 | -0.52108945 | 1 |
| ENSMUSG00000046991 | Wdr27 | 0.000110727 | 0.533358334 | 1 |
| ENSMUSG00000050088 | 1600012H06Rik | 0.000110727 | -0.895484965 | 0.718667768 |
| ENSMUSG00000036552 | Ermard | 0.000110727 | 0.326907457 | 1 |
| ENSMUSG00000116851 | Gm35576 | 0.000110727 | 0.648835551 | 1 |
| ENSMUSG00000096696 | Zfp960 | 0.000110727 | 4.233798052 | 0.128972787 |
| ENSMUSG00000090304 | Vmn2r99 | 0.000110727 | 0.911869957 | 1 |
| ENSMUSG00000116984 | 4930549P19Rik | 0.000110727 | 1.911869957 | 0.587484274 |
| ENSMUSG00000067931 | Zfp948 | 0.000110727 | -1.494122402 | 0.279592421 |
| ENSMUSG00000117070 | Gm41541 | 0.000110727 | 0.911869957 | 1 |
| ENSMUSG00000117284 | Gm7072 | 0.000110727 | -1.936126949 | 0.106828177 |
| ENSMUSG00000044501 | Zfp758 | 0.000110727 | -0.158519371 | 1 |
| ENSMUSG00000091945 | Vmn2r114 | 0.000110727 | 2.233798052 | 0.47966612 |
| ENSMUSG00000024142 | Mlst8 | 0.000110727 | 2.233798052 | 0.47966612 |
| ENSMUSG00000117423 | Gm33508 | 0.000110727 | 1.648835551 | 0.66416768 |
| ENSMUSG00000041429 | Nthl1 | 0.000110727 | 1.233798052 | 0.792441218 |
| ENSMUSG00000024155 | Meiob | 0.000110727 | 0.911869957 | 1 |
| ENSMUSG00000073436 | Eme2 | 0.000110727 | 1.233798052 | 0.792441218 |
| ENSMUSG00000073435 | Nme3 | 0.000110727 | 3.233798052 | 0.262284476 |
| ENSMUSG00000067722 | BC003965 | 0.000110727 | -0.810596067 | 0.796840156 |
| ENSMUSG00000015127 | Unkl | 0.000110727 | -2.069982696 | 0.062490345 |
| ENSMUSG00000061046 | Haghl | 0.000110727 | -1.766201948 | 0.177083054 |
| ENSMUSG00000071202 | Ccdc78 | 0.000110727 | 1.42644313 | 0.736456705 |
| ENSMUSG00000073434 | Wdr90 | 0.000110727 | 0.911869957 | 1 |
| ENSMUSG00000118107 | Gm34455 | 0.000110727 | 2.233798052 | 0.47966612 |
| ENSMUSG00000024193 | Phf1 | 0.000110727 | -0.720398258 | 0.792441218 |
| ENSMUSG00000067629 | Syngap1 | 0.000110727 | -1.410058138 | 0.33333983 |
| ENSMUSG00000097090 | Gm26724 | 0.000110727 | 1.063873051 | 1 |
| ENSMUSG00000063239 | Grm4 | 0.000110727 | 0.146335211 | 1 |
| ENSMUSG00000046711 | Hmga1 | 0.000110727 | -0.410058138 | 1 |
| ENSMUSG00000040276 | Pacsin1 | 0.000110727 | 0.146335211 | 1 |
| ENSMUSG00000039512 | Uhrf1bp1 | 0.000110727 | -0.57355687 | 0.910050347 |
| ENSMUSG00000024220 | Zfp523 | 0.000110727 | -1.743481871 | 0.172404208 |
| ENSMUSG00000117084 | Gm22146 | 0.000110727 | 1.42644313 | 0.736456705 |
| ENSMUSG00000043286 | Pnpla1 | 0.000110727 | 2.233798052 | 0.47966612 |
| ENSMUSG00000117080 | 1700063J08Rik | 0.000110727 | 2.233798052 | 0.488986361 |
| ENSMUSG00000117007 | Gm41556 | 0.000110727 | 1.233798052 | 0.792441218 |
| ENSMUSG00000064147 | Rab44 | 0.000110727 | 0.146335211 | 1 |
| ENSMUSG00000024011 | Pi16 | 0.000110727 | -0.853664789 | 0.828908937 |
| ENSMUSG00000024013 | Fgd2 | 0.000110727 | 0.533358334 | 1 |
| ENSMUSG00000097125 | Gm26885 | 0.000110727 | -1.624182943 | 0.306526877 |
| ENSMUSG00000079563 | Pglyrp2 | 0.000110727 | 3.233798052 | 0.262284476 |
| ENSMUSG00000073423 | Zfp414 | 0.000110727 | 1.911869957 | 0.587484274 |
| ENSMUSG00000051390 | Zbtb22 | 0.000110727 | 0.648835551 | 1 |
| ENSMUSG00000024312 | Wdr46 | 0.000110727 | -0.720398258 | 0.792441218 |
| ENSMUSG00000079547 | H2-DMb1 | 0.000110727 | -1.088130043 | 0.66416768 |

| | | | | |
|---|---|---|---|---|
| ENSMUSG00000024338 | Psmb8 | 0.000110727 | 1.42644313 | 0.736456705 |
| ENSMUSG00000041538 | H2-Ob | 0.000110727 | 1.233798052 | 0.792441218 |
| ENSMUSG00000036594 | H2-Aa | 0.000110727 | -0.158519371 | 1 |
| ENSMUSG00000092415 | Gm20513 | 0.000110727 | 1.911869957 | 0.587484274 |
| ENSMUSG00000033739 | Fkbpl | 0.000110727 | 1.233798052 | 0.792441218 |
| ENSMUSG00000024365 | Cyp21a1 | 0.000110727 | 0.326907457 | 1 |
| ENSMUSG00000049823 | Zbtb12 | 0.000110727 | -0.410058138 | 1 |
| ENSMUSG00000007038 | Neu1 | 0.000110727 | -1.410058138 | 0.33333983 |
| ENSMUSG00000007035 | Msh5 | 0.000110727 | -0.766201948 | 0.844591606 |
| ENSMUSG00000092417 | Gpank1 | 0.000110727 | -0.088130043 | 1 |
| ENSMUSG00000024399 | Ltb | 0.000110727 | 1.233798052 | 0.792441218 |
| ENSMUSG00000035929 | H2-Q4 | 0.000110727 | -0.014129461 | 1 |
| ENSMUSG00000090509 | Sfta2 | 0.000110727 | 0.233798052 | 1 |
| ENSMUSG00000061607 | Mdc1 | 0.000110727 | -2.507668934 | 0.01433775 |
| ENSMUSG00000087443 | Ppp1r18os | 0.000110727 | 1.063873051 | 1 |
| ENSMUSG00000050705 | 2310061I04Rik | 0.000110727 | 1.42644313 | 0.736456705 |
| ENSMUSG00000056116 | H2-T22 | 0.000110727 | 1.911869957 | 0.587484274 |
| ENSMUSG00000073399 | Trim40 | 0.000110727 | 3.233798052 | 0.262284476 |
| ENSMUSG00000036214 | Znrd1as | 0.000110727 | 0.233798052 | 1 |
| ENSMUSG00000024459 | H2-M5 | 0.000110727 | 0.146335211 | 1 |
| ENSMUSG00000059964 | Olfr119 | 0.000110727 | 3.233798052 | 0.262284476 |
| ENSMUSG00000059030 | Olfr128 | 0.000110727 | 2.648835551 | 0.395118486 |
| ENSMUSG00000095540 | Esp4 | 0.000110727 | 4.233798052 | 0.128972787 |
| ENSMUSG00000023930 | Crisp2 | 0.000110727 | 1.911869957 | 0.587484274 |
| ENSMUSG00000023926 | Rhag | 0.000110727 | 2.233798052 | 0.47966612 |
| ENSMUSG00000043972 | Opn5 | 0.000110727 | 1.911869957 | 0.60637478 |
| ENSMUSG00000117011 | E130008D07Rik | 0.000110727 | -0.014129461 | 1 |
| ENSMUSG00000096140 | Ankrd66 | 0.000110727 | 4.233798052 | 0.128972787 |
| ENSMUSG00000023912 | Slc25a27 | 0.000110727 | -1.158519371 | 0.521275193 |
| ENSMUSG00000023960 | Enpp5 | 0.000110727 | 3.233798052 | 0.262284476 |
| ENSMUSG00000037089 | Slc35b2 | 0.000110727 | -0.975655313 | 0.705975399 |
| ENSMUSG00000023942 | Slc29a1 | 0.000110727 | -0.673092543 | 0.919998879 |
| ENSMUSG00000012296 | Tjap1 | 0.000110727 | -3.023589791 | 0.002579787 |
| ENSMUSG00000015605 | Srf | 0.000110727 | 0.648835551 | 1 |
| ENSMUSG00000023982 | Guca1a | 0.000110727 | 3.233798052 | 0.274319469 |
| ENSMUSG00000023980 | Taf8 | 0.000110727 | -0.895484965 | 0.73806297 |
| ENSMUSG00000023988 | Bysl | 0.000110727 | -0.351164449 | 1 |
| ENSMUSG00000090115 | Usp49 | 0.000110727 | -0.766201948 | 0.792441218 |
| ENSMUSG00000051682 | Treml4 | 0.000110727 | 3.233798052 | 0.262284476 |
| ENSMUSG00000023927 | Satb1 | 0.000110727 | -0.158519371 | 1 |
| ENSMUSG00000023931 | Efhb | 0.000110727 | -0.624182943 | 0.912904617 |
| ENSMUSG00000044957 | Pp2d1 | 0.000110727 | 1.648835551 | 0.66416768 |
| ENSMUSG00000099747 | 1700025K24Rik | 0.000110727 | 2.233798052 | 0.519011117 |
| ENSMUSG00000003206 | Ebi3 | 0.000110727 | 1.911869957 | 0.599865044 |
| ENSMUSG00000003208 | Yju2 | 0.000110727 | 0.533358334 | 1 |
| ENSMUSG00000037095 | Lrg1 | 0.000110727 | 0.911869957 | 0.886203497 |
| ENSMUSG00000001227 | Sema6b | 0.000110727 | 0.146335211 | 1 |
| ENSMUSG00000024207 | Acsbg2 | 0.000110727 | -3.336057556 | 0.000471117 |
| ENSMUSG00000045019 | Acer1 | 0.000110727 | -0.466641666 | 1 |
| ENSMUSG00000002668 | Dennd1c | 0.000110727 | 1.648835551 | 0.66416768 |
| ENSMUSG00000035678 | Tnfsf9 | 0.000110727 | 1.063873051 | 0.85551189 |

| | | | | |
|---|---|---|---|---|
| ENSMUSG00000024227 | Pdzph1 | 0.000110727 | 0.648835551 | 1 |
| ENSMUSG00000101606 | Gm29051 | 0.000110727 | 0.774366434 | 1 |
| ENSMUSG00000073377 | AU016765 | 0.000110727 | 2.648835551 | 0.395118486 |
| ENSMUSG00000049672 | Zbtb14 | 0.000110727 | 0.063873051 | 1 |
| ENSMUSG00000024056 | Ndc80 | 0.000110727 | -0.673092543 | 0.919998879 |
| ENSMUSG00000024059 | Clip4 | 0.000110727 | 0.42644313 | 1 |
| ENSMUSG00000117560 | Gm31759 | 0.000110727 | 2.648835551 | 0.452335407 |
| ENSMUSG00000117539 | Gm4710 | 0.000110727 | 2.233798052 | 0.47966612 |
| ENSMUSG00000024084 | Qpct | 0.000110727 | 1.648835551 | 0.664426832 |
| ENSMUSG00000024087 | Cyp1b1 | 0.000110727 | 0.42644313 | 1 |
| ENSMUSG00000117616 | Gm49984 | 0.000110727 | 3.233798052 | 0.284008359 |
| ENSMUSG00000055760 | Gemin6 | 0.000110727 | 1.648835551 | 0.66416768 |
| ENSMUSG00000054901 | Arhgef33 | 0.000110727 | -0.289763904 | 1 |
| ENSMUSG00000024245 | Tmem178 | 0.000110727 | -0.289763904 | 1 |
| ENSMUSG00000024246 | Thumpd2 | 0.000110727 | -0.289763904 | 1 |
| ENSMUSG00000117399 | Gm19689 | 0.000110727 | 2.233798052 | 0.487205201 |
| ENSMUSG00000097254 | C430042M11Rik | 0.000110727 | 4.233798052 | 0.128972787 |
| ENSMUSG00000024253 | Dync2li1 | 0.000110727 | -0.895484965 | 0.707410086 |
| ENSMUSG00000097047 | 1110020A21Rik | 0.000110727 | 1.063873051 | 0.852547761 |
| ENSMUSG00000024127 | Prepl | 0.000110727 | -1.014129461 | 0.633928457 |
| ENSMUSG00000085287 | 4833418N02Rik | 0.000110727 | -0.52108945 | 1 |
| ENSMUSG00000036557 | Stpg4 | 0.000110727 | 1.063873051 | 0.847265357 |
| ENSMUSG00000055660 | Mettl4 | 0.000110727 | -1.051604167 | 0.614718876 |
| ENSMUSG00000095193 | Gm20939 | 0.000110727 | 0.233798052 | 1 |
| ENSMUSG00000097746 | Gm6225 | 0.000110727 | 2.233798052 | 0.47966612 |
| ENSMUSG00000117521 | Gm50024 | 0.000110727 | 2.233798052 | 0.47966612 |
| ENSMUSG00000117519 | Gm41664 | 0.000110727 | 0.648835551 | 1 |
| ENSMUSG00000061013 | Mkx | 0.000110727 | 0.42644313 | 1 |
| ENSMUSG00000097440 | Gm6277 | 0.000110727 | 1.233798052 | 0.792441218 |
| ENSMUSG00000037013 | Ss18 | 0.000110727 | -2.480447466 | 0.016974693 |
| ENSMUSG00000036743 | Psma8 | 0.000110727 | 1.911869957 | 0.602789838 |
| ENSMUSG00000024411 | Aqp4 | 0.000110727 | 1.911869957 | 0.587484274 |
| ENSMUSG00000034774 | Dsg1c | 0.000110727 | 2.233798052 | 0.49612097 |
| ENSMUSG00000056632 | Dsg3 | 0.000110727 | 0.774366434 | 1 |
| ENSMUSG00000061808 | Ttr | 0.000110727 | 1.063873051 | 0.850318498 |
| ENSMUSG00000063281 | Zfp35 | 0.000110727 | -1.43862729 | 0.339627968 |
| ENSMUSG00000024269 | Tpgs2 | 0.000110727 | 1.233798052 | 0.792441218 |
| ENSMUSG00000117456 | Gm49976 | 0.000110727 | 4.233798052 | 0.128972787 |
| ENSMUSG00000117473 | Gm33948 | 0.000110727 | 0.774366434 | 1 |
| ENSMUSG00000024382 | Ercc3 | 0.000110727 | -1.3207908 | 0.395686633 |
| ENSMUSG00000109232 | Gm44577 | 0.000110727 | 2.233798052 | 0.47966612 |
| ENSMUSG00000049357 | 4933408B17Rik | 0.000110727 | -0.014129461 | 1 |
| ENSMUSG00000024366 | Gfra3 | 0.000110727 | 0.648835551 | 1 |
| ENSMUSG00000117694 | Snhg4 | 0.000110727 | 4.233798052 | 0.128972787 |
| ENSMUSG00000117840 | Gm50399 | 0.000110727 | 1.233798052 | 0.792441218 |
| ENSMUSG00000006850 | Tmco6 | 0.000110727 | -0.088130043 | 1 |
| ENSMUSG00000019143 | Hars2 | 0.000110727 | -1.766201948 | 0.177083054 |
| ENSMUSG00000051242 | Pcdhb9 | 0.000110727 | 3.233798052 | 0.262284476 |
| ENSMUSG00000024451 | Arap3 | 0.000110727 | -1.289763904 | 0.466464457 |
| ENSMUSG00000086312 | Gm15336 | 0.000110727 | 1.42644313 | 0.736456705 |
| ENSMUSG00000117922 | Gm50397 | 0.000110727 | -4.705781162 | 1.28E-08 |

| | | | | |
|---|---|---|---|---|
| ENSMUSG00000117745 | Gm50410 | 0.000110727 | 0.42644313 | 1 |
| ENSMUSG00000117662 | Gm4013 | 0.000110727 | 1.648835551 | 0.66416768 |
| ENSMUSG00000117792 | 4930588A03Rik | 0.000110727 | -0.225633566 | 1 |
| ENSMUSG00000034653 | Ythdc2 | 0.000110727 | -2.141241379 | 0.054954924 |
| ENSMUSG00000071855 | Ccdc112 | 0.000110727 | -0.720398258 | 0.792441218 |
| ENSMUSG00000118267 | G630055G22Rik | 0.000110727 | -0.351164449 | 1 |
| ENSMUSG00000059040 | Eno1b | 0.000110727 | 1.42644313 | 0.736456705 |
| ENSMUSG00000024528 | Srfbp1 | 0.000110727 | -1.52108945 | 0.284607766 |
| ENSMUSG00000087597 | Gm15345 | 0.000110727 | 2.233798052 | 0.48396292 |
| ENSMUSG00000024619 | Cdx1 | 0.000110727 | 1.648835551 | 0.66416768 |
| ENSMUSG00000045629 | Sh3tc2 | 0.000110727 | 0.326907457 | 1 |
| ENSMUSG00000024581 | Napg | 0.000110727 | -2.051604167 | 0.078768438 |
| ENSMUSG00000046610 | Oacyl | 0.000110727 | 0.42644313 | 1 |
| ENSMUSG00000024518 | Rax | 0.000110727 | 1.911869957 | 0.587484274 |
| ENSMUSG00000085055 | Gm15958 | 0.000110727 | 0.42644313 | 1 |
| ENSMUSG00000117968 | Gm50162 | 0.000110727 | 0.648835551 | 1 |
| ENSMUSG00000118216 | Gm19784 | 0.000110727 | 3.233798052 | 0.262284476 |
| ENSMUSG00000073542 | Cep76 | 0.000110727 | -0.52108945 | 1 |
| ENSMUSG00000045569 | Mc2r | 0.000110727 | 2.648835551 | 0.395118486 |
| ENSMUSG00000024546 | 4930546C10Rik | 0.000110727 | 0.146335211 | 1 |
| ENSMUSG00000084948 | 1700061H18Rik | 0.000110727 | -0.158519371 | 1 |
| ENSMUSG00000036941 | Elac1 | 0.000110727 | 0.233798052 | 1 |
| ENSMUSG00000118004 | BC026513 | 0.000110727 | 0.146335211 | 1 |
| ENSMUSG00000118108 | Gm50284 | 0.000110727 | 2.233798052 | 0.51115569 |
| ENSMUSG00000032818 | Loxhd1 | 0.000110727 | 0.146335211 | 1 |
| ENSMUSG00000118373 | Gm41787 | 0.000110727 | 1.233798052 | 0.792441218 |
| ENSMUSG00000059852 | Kcng2 | 0.000110727 | 0.648835551 | 1 |
| ENSMUSG00000097181 | 4930594M17Rik | 0.000110727 | 1.648835551 | 0.664971056 |
| ENSMUSG00000117676 | Gm50314 | 0.000110727 | 2.648835551 | 0.449026583 |
| ENSMUSG00000047992 | Dipk1c | 0.000110727 | 0.533358334 | 1 |
| ENSMUSG00000110277 | Gm45871 | 0.000110727 | 1.063873051 | 1 |
| ENSMUSG00000024905 | Tesmin | 0.000110727 | 0.146335211 | 1 |
| ENSMUSG00000118249 | Gm36608 | 0.000110727 | 0.911869957 | 1 |
| ENSMUSG00000110949 | Nudt8 | 0.000110727 | 0.774366434 | 1 |
| ENSMUSG00000024871 | Doc2g | 0.000110727 | 0.42644313 | 1 |
| ENSMUSG00000024830 | Rps6kb2 | 0.000110727 | -1.192466703 | 0.529126167 |
| ENSMUSG00000097233 | Gm17552 | 0.000110727 | 2.648835551 | 0.395118486 |
| ENSMUSG00000005986 | Ankrd13d | 0.000110727 | 0.146335211 | 1 |
| ENSMUSG00000067889 | Sptbn2 | 0.000110727 | -0.810596067 | 0.843995018 |
| ENSMUSG00000094936 | Rbm4 | 0.000110727 | -2.624182943 | 0.007725935 |
| ENSMUSG00000006457 | Actn3 | 0.000110727 | 2.233798052 | 0.47966612 |
| ENSMUSG00000006464 | Bbs1 | 0.000110727 | -0.853664789 | 0.806496094 |
| ENSMUSG00000024891 | Slc29a2 | 0.000110727 | -0.52108945 | 1 |
| ENSMUSG00000047379 | B4gat1 | 0.000110727 | 0.533358334 | 1 |
| ENSMUSG00000024875 | Yif1a | 0.000110727 | -0.014129461 | 1 |
| ENSMUSG00000086938 | 4930481A15Rik | 0.000110727 | -0.014129461 | 1 |
| ENSMUSG00000100937 | 1700020D05Rik | 0.000110727 | 0.774366434 | 1 |
| ENSMUSG00000049562 | Ap5b1 | 0.000110727 | 2.648835551 | 0.444576901 |
| ENSMUSG00000024792 | Zfpl1 | 0.000110727 | -0.225633566 | 1 |
| ENSMUSG00000079467 | Gm14966 | 0.000110727 | -0.158519371 | 1 |
| ENSMUSG00000084825 | 2410152P15Rik | 0.000110727 | 0.911869957 | 0.886203497 |

| | | | | |
|---|---|---|---|---|
| ENSMUSG00000024965 | Fermt3 | 0.000110727 | -1.766201948 | 0.218040004 |
| ENSMUSG00000047787 | Flrt1 | 0.000110727 | 2.233798052 | 0.47966612 |
| ENSMUSG00000067656 | Slc22a27 | 0.000110727 | 1.063873051 | 1 |
| ENSMUSG00000117924 | Tmem223 | 0.000110727 | 0.063873051 | 1 |
| ENSMUSG00000118346 | Tmem179b | 0.000110727 | 0.42644313 | 1 |
| ENSMUSG00000003680 | Taf6l | 0.000110727 | -2.141241379 | 0.054954924 |
| ENSMUSG00000024665 | Fads2 | 0.000110727 | 0.774366434 | 1 |
| ENSMUSG00000010663 | Fads1 | 0.000110727 | 0.533358334 | 1 |
| ENSMUSG00000034659 | Tmem109 | 0.000110727 | -1.915949067 | 0.103463818 |
| ENSMUSG00000067571 | Ms4a15 | 0.000110727 | 2.648835551 | 0.395118486 |
| ENSMUSG00000057240 | Ms4a13 | 0.000110727 | 2.233798052 | 0.497541978 |
| ENSMUSG00000101031 | Ms4a12 | 0.000110727 | 2.233798052 | 0.47966612 |
| ENSMUSG00000024677 | Ms4a6b | 0.000110727 | 1.42644313 | 0.736456705 |
| ENSMUSG00000024679 | Ms4a6d | 0.000110727 | 1.42644313 | 0.736456705 |
| ENSMUSG00000048292 | Olfr1417 | 0.000110727 | 2.648835551 | 0.395118486 |
| ENSMUSG00000060878 | Olfr1420 | 0.000110727 | 1.063873051 | 0.852297462 |
| ENSMUSG00000067524 | Olfr1428 | 0.000110727 | 4.233798052 | 0.206083156 |
| ENSMUSG00000024694 | Keg1 | 0.000110727 | 1.233798052 | 0.792441218 |
| ENSMUSG00000024696 | Lpxn | 0.000110727 | -0.225633566 | 1 |
| ENSMUSG00000061387 | Olfr1490 | 0.000110727 | 3.233798052 | 0.262284476 |
| ENSMUSG00000109520 | Olfr1493-ps1 | 0.000110727 | 4.233798052 | 0.128972787 |
| ENSMUSG00000118079 | Gm50181 | 0.000110727 | 2.648835551 | 0.44267062 |
| ENSMUSG00000037847 | Nmrk1 | 0.000110727 | -0.720398258 | 0.792441218 |
| ENSMUSG00000117909 | Gm50212 | 0.000110727 | 2.648835551 | 0.404997369 |
| ENSMUSG00000118150 | Gm3728 | 0.000110727 | -1.743481871 | 0.215910103 |
| ENSMUSG00000024747 | Aldh1a7 | 0.000110727 | 1.648835551 | 0.66416768 |
| ENSMUSG00000118086 | AC106834.1 | 0.000110727 | -0.289763904 | 1 |
| ENSMUSG00000118273 | Gm41819 | 0.000110727 | -0.720398258 | 0.971854959 |
| ENSMUSG00000033053 | 1700028P14Rik | 0.000110727 | -0.289763904 | 1 |
| ENSMUSG00000093772 | 4931403E22Rik | 0.000110727 | -0.466641666 | 1 |
| ENSMUSG00000050957 | Insl6 | 0.000110727 | 3.233798052 | 0.262284476 |
| ENSMUSG00000016498 | Pdcd1lg2 | 0.000110727 | 0.533358334 | 1 |
| ENSMUSG00000053536 | Cstf2t | 0.000110727 | -0.014129461 | 1 |
| ENSMUSG00000117664 | Gm36860 | 0.000110727 | 2.233798052 | 0.47966612 |
| ENSMUSG00000024771 | Lipk | 0.000110727 | 1.063873051 | 1 |
| ENSMUSG00000024778 | Fas | 0.000110727 | -0.810596067 | 0.796840156 |
| ENSMUSG00000067297 | Ifit1bl2 | 0.000110727 | 1.911869957 | 0.595720535 |
| ENSMUSG00000009378 | Slc16a12 | 0.000110727 | -0.289763904 | 1 |
| ENSMUSG00000094707 | A830019P07Rik | 0.000110727 | 0.42644313 | 1 |
| ENSMUSG00000024800 | Rpp30 | 0.000110727 | -0.895484965 | 0.707410086 |
| ENSMUSG00000118032 | Gm50147 | 0.000110727 | 4.233798052 | 0.128972787 |
| ENSMUSG00000087579 | Hectd2os | 0.000110727 | -0.014129461 | 1 |
| ENSMUSG00000024989 | Cep55 | 0.000110727 | -0.975655313 | 0.705975399 |
| ENSMUSG00000024992 | Pde6c | 0.000110727 | 0.911869957 | 1 |
| ENSMUSG00000025001 | Hells | 0.000110727 | -2.273996588 | 0.034182785 |
| ENSMUSG00000067229 | Cyp2c66 | 0.000110727 | 3.233798052 | 0.262284476 |
| ENSMUSG00000003053 | Cyp2c29 | 0.000110727 | 1.911869957 | 0.587484274 |
| ENSMUSG00000025003 | Cyp2c39 | 0.000110727 | 1.42644313 | 0.736456705 |
| ENSMUSG00000092008 | Cyp2c69 | 0.000110727 | 4.233798052 | 0.128972787 |
| ENSMUSG00000048120 | Entpd1 | 0.000110727 | -0.52108945 | 1 |
| ENSMUSG00000108929 | Cc2d2b | 0.000110727 | 1.233798052 | 0.792441218 |

| | | | | |
|---|---|---|---|---|
| ENSMUSG00000025010 | Ccnj | 0.000110727 | 0.774366434 | 1 |
| ENSMUSG00000025013 | Tll2 | 0.000110727 | 0.146335211 | 1 |
| ENSMUSG00000025154 | Arhgap19 | 0.000110727 | -0.936126949 | 0.697198109 |
| ENSMUSG00000047604 | Frat2 | 0.000110727 | 0.533358334 | 1 |
| ENSMUSG00000025157 | Zdhhc16 | 0.000110727 | -0.466641666 | 1 |
| ENSMUSG00000117880 | Gm34517 | 0.000110727 | 0.533358334 | 1 |
| ENSMUSG00000025189 | Cnnm1 | 0.000110727 | 0.774366434 | 1 |
| ENSMUSG00000025193 | Cutc | 0.000110727 | -0.895484965 | 0.855990803 |
| ENSMUSG00000025197 | Cyp2c23 | 0.000110727 | 1.233798052 | 0.792441218 |
| ENSMUSG00000025200 | Cwf19l1 | 0.000110727 | -1.258055044 | 0.466510427 |
| ENSMUSG00000051984 | Sec31b | 0.000110727 | 0.326907457 | 1 |
| ENSMUSG00000117848 | Gm35867 | 0.000110727 | -0.014129461 | 1 |
| ENSMUSG00000100417 | Gm28578 | 0.000110727 | 0.326907457 | 1 |
| ENSMUSG00000025218 | Poll | 0.000110727 | -0.014129461 | 1 |
| ENSMUSG00000025221 | Kcnip2 | 0.000110727 | -0.52108945 | 1 |
| ENSMUSG00000025226 | Fbxl15 | 0.000110727 | 1.42644313 | 0.736456705 |
| ENSMUSG00000079258 | Calhm1 | 0.000110727 | 2.233798052 | 0.47966612 |
| ENSMUSG00000118127 | Gm36693 | 0.000110727 | 2.648835551 | 0.395118486 |
| ENSMUSG00000117690 | Gm50430 | 0.000110727 | 1.42644313 | 0.736456705 |
| ENSMUSG00000117896 | AA387883 | 0.000110727 | 2.648835551 | 0.395118486 |
| ENSMUSG00000055523 | Gucy2g | 0.000110727 | 0.911869957 | 0.886203497 |
| ENSMUSG00000117695 | Gm31912 | 0.000110727 | 2.233798052 | 0.503853932 |
| ENSMUSG00000035818 | Plekhs1 | 0.000110727 | -1.673092543 | 0.205249116 |
| ENSMUSG00000097785 | B230217O12Rik | 0.000110727 | -1.547561661 | 0.290722159 |
| ENSMUSG00000042179 | Pnliprp1 | 0.000110727 | 1.063873051 | 0.847265357 |
| ENSMUSG00000043969 | Emx2 | 0.000110727 | 1.42644313 | 0.736456705 |
| ENSMUSG00000117790 | Gm46652 | 0.000110727 | 3.233798052 | 0.262284476 |
| ENSMUSG00000062393 | Dgkk | 0.000110727 | 0.063873051 | 1 |
| ENSMUSG00000031145 | Prickle3 | 0.000110727 | -0.088130043 | 1 |
| ENSMUSG00000031148 | Gpkow | 0.000110727 | -0.624182943 | 0.912904617 |
| ENSMUSG00000031155 | Pim2 | 0.000110727 | 1.648835551 | 0.66416768 |
| ENSMUSG00000087591 | Gm14635 | 0.000110727 | 2.648835551 | 0.397039162 |
| ENSMUSG00000073274 | Gm14636 | 0.000110727 | -1.225633566 | 0.463954265 |
| ENSMUSG00000031060 | Rbm10 | 0.000110727 | -2.624182943 | 0.007725935 |
| ENSMUSG00000037217 | Syn1 | 0.000110727 | -2.032988489 | 0.076038495 |
| ENSMUSG00000009406 | Elk1 | 0.000110727 | 0.42644313 | 1 |
| ENSMUSG00000054737 | Zfp182 | 0.000110727 | -1.624182943 | 0.24944595 |
| ENSMUSG00000087358 | 4930453H23Rik | 0.000110727 | 1.648835551 | 0.66416768 |
| ENSMUSG00000064137 | Rhox8 | 0.000110727 | -1.380911792 | 0.460958452 |
| ENSMUSG00000036502 | Tmem255a | 0.000110727 | 0.063873051 | 1 |
| ENSMUSG00000055653 | Gpc3 | 0.000110727 | -0.810596067 | 0.846622462 |
| ENSMUSG00000025626 | Phf6 | 0.000110727 | -2.534386273 | 0.014126898 |
| ENSMUSG00000067925 | Rtl8a | 0.000110727 | 2.233798052 | 0.495747121 |
| ENSMUSG00000054727 | 1700013H16Rik | 0.000110727 | 2.233798052 | 0.47966612 |
| ENSMUSG00000054626 | Xlr | 0.000110727 | 3.233798052 | 0.262284476 |
| ENSMUSG00000035967 | Ints6l | 0.000110727 | 0.063873051 | 1 |
| ENSMUSG00000060681 | Slc9a6 | 0.000110727 | -0.088130043 | 1 |
| ENSMUSG00000045237 | 1110012L19Rik | 0.000110727 | 0.911869957 | 0.85084719 |
| ENSMUSG00000031343 | Gabra3 | 0.000110727 | 0.146335211 | 1 |
| ENSMUSG00000031351 | Zfp185 | 0.000110727 | 1.233798052 | 0.792441218 |
| ENSMUSG00000019558 | Slc6a8 | 0.000110727 | 0.42644313 | 1 |

| | | | | |
|---|---|---|---|---|
| ENSMUSG00000031400 | G6pdx | 0.000110727 | -0.766201948 | 0.792441218 |
| ENSMUSG00000035491 | Fthl17a | 0.000110727 | 2.648835551 | 0.395118486 |
| ENSMUSG00000025059 | Gk | 0.000110727 | -0.014129461 | 1 |
| ENSMUSG00000067649 | Mageb18 | 0.000110727 | -0.673092543 | 0.95813886 |
| ENSMUSG00000035246 | Pcyt1b | 0.000110727 | -0.720398258 | 0.961379127 |
| ENSMUSG00000073062 | Zxdb | 0.000110727 | 0.774366434 | 1 |
| ENSMUSG00000025656 | Arhgef9 | 0.000110727 | 0.648835551 | 1 |
| ENSMUSG00000035045 | Zc3h12b | 0.000110727 | -2.051604167 | 0.084475788 |
| ENSMUSG00000015668 | Pdzd11 | 0.000110727 | 0.326907457 | 1 |
| ENSMUSG00000009670 | Tex11 | 0.000110727 | 0.42644313 | 1 |
| ENSMUSG00000031312 | Itgb1bp2 | 0.000110727 | 1.42644313 | 0.736456705 |
| ENSMUSG00000051220 | Ercc6l | 0.000110727 | 0.326907457 | 1 |
| ENSMUSG00000032894 | 1700031F05Rik | 0.000110727 | 1.063873051 | 1 |
| ENSMUSG00000033906 | Zdhhc15 | 0.000110727 | -2.956026507 | 0.002779402 |
| ENSMUSG00000087174 | 5530601H04Rik | 0.000110727 | -1.088130043 | 0.624094312 |
| ENSMUSG00000047686 | Rtl3 | 0.000110727 | 4.233798052 | 0.128972787 |
| ENSMUSG00000073007 | Tent5d | 0.000110727 | 0.774366434 | 1 |
| ENSMUSG00000025665 | Rps6ka6 | 0.000110727 | 1.063873051 | 1 |
| ENSMUSG00000062791 | Gm382 | 0.000110727 | 4.233798052 | 0.128972787 |
| ENSMUSG00000031257 | Nox1 | 0.000110727 | -0.466641666 | 1 |
| ENSMUSG00000049047 | Armcx3 | 0.000110727 | -0.088130043 | 1 |
| ENSMUSG00000042498 | Radx | 0.000110727 | 1.42644313 | 0.736456705 |
| ENSMUSG00000079418 | Atg4a | 0.000110727 | -0.810596067 | 0.796840156 |
| ENSMUSG00000042271 | Nxt2 | 0.000110727 | 1.233798052 | 0.792441218 |
| ENSMUSG00000041229 | Phf8 | 0.000110727 | -2.673092543 | 0.007127824 |
| ENSMUSG00000056679 | Gpr173 | 0.000110727 | 0.146335211 | 1 |
| ENSMUSG00000050148 | Ubqln2 | 0.000110727 | -0.351164449 | 1 |
| ENSMUSG00000031303 | Map3k15 | 0.000110727 | -0.088130043 | 1 |
| ENSMUSG00000018589 | Glra2 | 0.000110727 | 0.648835551 | 1 |
| ENSMUSG00000040621 | Gemin8 | 0.000110727 | -1.014129461 | 0.608426921 |
| ENSMUSG00000000402 | Egfl6 | 0.000110727 | 1.648835551 | 0.66416768 |
| ENSMUSG00000031352 | Hccs | 0.000110727 | -1.158519371 | 0.521275193 |
| ENSMUSG00000095562 | Gm21887 | 0.000110727 | -0.466641666 | 1 |
| ENSMUSG00000053211 | Zfy1 | 0.000110727 | 2.648835551 | 0.416069222 |
| ENSMUSG00000069049 | Eif2s3y | 0.000110727 | -1.915949067 | 0.103463818 |
| ENSMUSG00000079834 | Tmlhe | 0.000110727 | -3.042326353 | 0.001300227 |
| ENSMUSG00000102331 | Gm19938 | 5.54E-05 | 3.648835551 | 0.466464457 |
| ENSMUSG00000102343 | Gm37381 | 5.54E-05 | 0.47891055 | 1 |
| ENSMUSG00000025902 | Sox17 | 5.54E-05 | 2.648835551 | 0.66416768 |
| ENSMUSG00000090031 | 4732440D04Rik | 5.54E-05 | -1.673092543 | 0.290284345 |
| ENSMUSG00000025911 | Adhfe1 | 5.54E-05 | -0.158519371 | 1 |
| ENSMUSG00000025912 | Mybl1 | 5.54E-05 | -1.158519371 | 0.599135879 |
| ENSMUSG00000097893 | 1700034P13Rik | 5.54E-05 | 0.841480629 | 1 |
| ENSMUSG00000032769 | Trpa1 | 5.54E-05 | 1.648835551 | 0.886203497 |
| ENSMUSG00000100868 | Gm28095 | 5.54E-05 | 2.063873051 | 0.792441218 |
| ENSMUSG00000025777 | Gdap1 | 5.54E-05 | 2.063873051 | 0.792441218 |
| ENSMUSG00000099899 | Gm28784 | 5.54E-05 | 2.648835551 | 0.66416768 |
| ENSMUSG00000099895 | Gm28153 | 5.54E-05 | -0.351164449 | 1 |
| ENSMUSG00000089787 | Gm15825 | 5.54E-05 | 1.326907457 | 0.950123751 |
| ENSMUSG00000099906 | Gm28653 | 5.54E-05 | -0.351164449 | 1 |
| ENSMUSG00000041859 | Mcm3 | 5.54E-05 | -1.209145444 | 0.835382858 |

| | | | | |
|---|---|---|---|---|
| ENSMUSG00000041779 | Tram2 | 5.54E-05 | -1.995020638 | 0.167528217 |
| ENSMUSG00000025933 | Tmem14a | 5.54E-05 | 2.648835551 | 0.66416768 |
| ENSMUSG00000099503 | Gm28822 | 5.54E-05 | 1.326907457 | 0.953462889 |
| ENSMUSG00000048411 | Gm597 | 5.54E-05 | 3.648835551 | 0.466464457 |
| ENSMUSG00000117310 | Ptp4a1 | 5.54E-05 | 2.648835551 | 0.66416768 |
| ENSMUSG00000004768 | Rab23 | 5.54E-05 | -1.599091962 | 0.363644968 |
| ENSMUSG00000042111 | Ccdc115 | 5.54E-05 | -0.051604167 | 1 |
| ENSMUSG00000045174 | Amer3 | 5.54E-05 | 3.648835551 | 0.466464457 |
| ENSMUSG00000104417 | Gm37068 | 5.54E-05 | 2.063873051 | 0.790752525 |
| ENSMUSG00000103708 | Gm33222 | 5.54E-05 | 1.648835551 | 0.886203497 |
| ENSMUSG00000079610 | Ankrd39 | 5.54E-05 | -0.051604167 | 1 |
| ENSMUSG00000102416 | 4933424G06Rik | 5.54E-05 | -0.743481871 | 1 |
| ENSMUSG00000026114 | Cnga3 | 5.54E-05 | 0.47891055 | 1 |
| ENSMUSG00000102716 | Gm5099 | 5.54E-05 | 1.648835551 | 0.884688852 |
| ENSMUSG00000048814 | Lonrf2 | 5.54E-05 | 1.648835551 | 0.867965264 |
| ENSMUSG00000003134 | Tbc1d8 | 5.54E-05 | -2.43862729 | 0.044042118 |
| ENSMUSG00000050967 | Creg2 | 5.54E-05 | 0.648835551 | 1 |
| ENSMUSG00000057173 | Rfx8 | 5.54E-05 | -0.051604167 | 1 |
| ENSMUSG00000070942 | Il1rl2 | 5.54E-05 | 0.189403933 | 1 |
| ENSMUSG00000026069 | Il1rl1 | 5.54E-05 | 1.326907457 | 0.949822639 |
| ENSMUSG00000026068 | Il18rap | 5.54E-05 | -2.328444372 | 0.062688794 |
| ENSMUSG00000041945 | Mfsd9 | 5.54E-05 | 0.326907457 | 1 |
| ENSMUSG00000045515 | Pou3f3 | 5.54E-05 | 2.648835551 | 0.66416768 |
| ENSMUSG00000101701 | 4930521E06Rik | 5.54E-05 | 3.648835551 | 0.466464457 |
| ENSMUSG00000025993 | Slc40a1 | 5.54E-05 | -0.874726405 | 0.841317093 |
| ENSMUSG00000101447 | Gm28826 | 5.54E-05 | 1.648835551 | 0.886203497 |
| ENSMUSG00000099924 | Gm28320 | 5.54E-05 | 1.648835551 | 0.859013091 |
| ENSMUSG00000026107 | Nabp1 | 5.54E-05 | -1.52108945 | 0.452184898 |
| ENSMUSG00000099413 | Gm17767 | 5.54E-05 | 0.47891055 | 1 |
| ENSMUSG00000101741 | Gm28055 | 5.54E-05 | 0.189403933 | 1 |
| ENSMUSG00000062939 | Stat4 | 5.54E-05 | -0.874726405 | 0.866303883 |
| ENSMUSG00000041426 | Hibch | 5.54E-05 | -0.599091962 | 0.980745941 |
| ENSMUSG00000060715 | 1700019A02Rik | 5.54E-05 | 1.326907457 | 0.971854959 |
| ENSMUSG00000026096 | Osgepl1 | 5.54E-05 | 0.189403933 | 1 |
| ENSMUSG00000101429 | BC055402 | 5.54E-05 | 3.648835551 | 0.466464457 |
| ENSMUSG00000064294 | Aox3 | 5.54E-05 | -0.743481871 | 1 |
| ENSMUSG00000099811 | Gm28411 | 5.54E-05 | 2.648835551 | 0.66416768 |
| ENSMUSG00000026019 | Wdr12 | 5.54E-05 | -2.540989007 | 0.029897056 |
| ENSMUSG00000026009 | Icos | 5.54E-05 | 1.063873051 | 1 |
| ENSMUSG00000100893 | Gm28083 | 5.54E-05 | 2.648835551 | 0.66416768 |
| ENSMUSG00000100811 | Gm29083 | 5.54E-05 | 3.648835551 | 0.466464457 |
| ENSMUSG00000044689 | Gm13749 | 5.54E-05 | 3.648835551 | 0.466464457 |
| ENSMUSG00000025956 | Mettl21a | 5.54E-05 | 0.189403933 | 1 |
| ENSMUSG00000100437 | Gm29152 | 5.54E-05 | 3.648835551 | 0.466464457 |
| ENSMUSG00000073656 | Gm10558 | 5.54E-05 | 0.326907457 | 1 |
| ENSMUSG00000026196 | Bard1 | 5.54E-05 | -1.305360759 | 0.491653643 |
| ENSMUSG00000073652 | Apol7d | 5.54E-05 | 1.648835551 | 0.886203497 |
| ENSMUSG00000039323 | Igfbp2 | 5.54E-05 | 0.326907457 | 1 |
| ENSMUSG00000099877 | Pinc | 5.54E-05 | 2.648835551 | 0.66416768 |
| ENSMUSG00000100429 | Gm29186 | 5.54E-05 | 2.063873051 | 0.792441218 |
| ENSMUSG00000026180 | Cxcr2 | 5.54E-05 | 0.841480629 | 1 |

| | | | | |
|---|---|---|---|---|
| ENSMUSG00000064272 | Gpbar1 | 5.54E-05 | 2.648835551 | 0.66416768 |
| ENSMUSG00000026175 | Vil1 | 5.54E-05 | -3.52108945 | 0.000717333 |
| ENSMUSG00000026170 | Cyp27a1 | 5.54E-05 | -0.051604167 | 1 |
| ENSMUSG00000033227 | Wnt6 | 5.54E-05 | -1.52108945 | 0.466464457 |
| ENSMUSG00000047021 | Cfap65 | 5.54E-05 | 0.189403933 | 1 |
| ENSMUSG00000096094 | A630095N17Rik | 5.54E-05 | 2.063873051 | 0.792441218 |
| ENSMUSG00000085653 | Gm15179 | 5.54E-05 | 1.648835551 | 0.868661374 |
| ENSMUSG00000086053 | Gm15178 | 5.54E-05 | 2.648835551 | 0.66416768 |
| ENSMUSG00000032997 | Chpf | 5.54E-05 | -0.351164449 | 1 |
| ENSMUSG00000032968 | Inha | 5.54E-05 | 1.648835551 | 0.886203497 |
| ENSMUSG00000026213 | Stk11ip | 5.54E-05 | -1.843017545 | 0.255869172 |
| ENSMUSG00000102052 | Gm28387 | 5.54E-05 | 1.648835551 | 0.886203497 |
| ENSMUSG00000004872 | Pax3 | 5.54E-05 | -0.351164449 | 1 |
| ENSMUSG00000099561 | Gm28410 | 5.54E-05 | 1.326907457 | 0.958408446 |
| ENSMUSG00000012187 | Mogat1 | 5.54E-05 | 1.648835551 | 0.886203497 |
| ENSMUSG00000026166 | Ccl20 | 5.54E-05 | 1.063873051 | 1 |
| ENSMUSG00000070034 | Sp110 | 5.54E-05 | -0.43862729 | 1 |
| ENSMUSG00000090246 | Gm17017 | 5.54E-05 | 1.648835551 | 0.886203497 |
| ENSMUSG00000026224 | 4933407L21Rik | 5.54E-05 | 1.326907457 | 0.971854959 |
| ENSMUSG00000100108 | Gm28100 | 5.54E-05 | 2.063873051 | 0.792441218 |
| ENSMUSG00000079445 | B3gnt7 | 5.54E-05 | 1.648835551 | 0.865542683 |
| ENSMUSG00000099401 | Gm28375 | 5.54E-05 | -1.43862729 | 0.460248995 |
| ENSMUSG00000026258 | Snorc | 5.54E-05 | 2.063873051 | 0.792441218 |
| ENSMUSG00000079434 | Neu2 | 5.54E-05 | -0.599091962 | 1 |
| ENSMUSG00000090165 | Ugt1a10 | 5.54E-05 | 2.648835551 | 0.66416768 |
| ENSMUSG00000092220 | Gm20528 | 5.54E-05 | -1.051604167 | 0.706929043 |
| ENSMUSG00000090145 | Ugt1a6b | 5.54E-05 | 2.063873051 | 0.792441218 |
| ENSMUSG00000054545 | Ugt1a6a | 5.54E-05 | 0.841480629 | 1 |
| ENSMUSG00000090171 | Ugt1a2 | 5.54E-05 | 1.063873051 | 1 |
| ENSMUSG00000101288 | Gm28342 | 5.54E-05 | 1.648835551 | 0.886203497 |
| ENSMUSG00000067081 | Asb18 | 5.54E-05 | -0.051604167 | 1 |
| ENSMUSG00000090550 | Prlh | 5.54E-05 | 3.648835551 | 0.466464457 |
| ENSMUSG00000026304 | Rab17 | 5.54E-05 | 0.841480629 | 1 |
| ENSMUSG00000100791 | Gm28199 | 5.54E-05 | 2.648835551 | 0.66416768 |
| ENSMUSG00000070732 | Rbm44 | 5.54E-05 | 2.648835551 | 0.66416768 |
| ENSMUSG00000055866 | Per2 | 5.54E-05 | -2.52108945 | 0.037394962 |
| ENSMUSG00000100781 | Gm29481 | 5.54E-05 | 0.841480629 | 1 |
| ENSMUSG00000034212 | Ankmy1 | 5.54E-05 | 1.326907457 | 0.971854959 |
| ENSMUSG00000047067 | Dusp28 | 5.54E-05 | 0.47891055 | 1 |
| ENSMUSG00000100632 | 9430060I03Rik | 5.54E-05 | 0.47891055 | 1 |
| ENSMUSG00000026271 | Gpr35 | 5.54E-05 | 0.326907457 | 1 |
| ENSMUSG00000101378 | Gm28086 | 5.54E-05 | 3.648835551 | 0.466464457 |
| ENSMUSG00000026273 | Mterf4 | 5.54E-05 | -0.599091962 | 0.991046228 |
| ENSMUSG00000026285 | Pdcd1 | 5.54E-05 | 3.648835551 | 0.466464457 |
| ENSMUSG00000100336 | Gm29601 | 5.54E-05 | 2.063873051 | 0.792441218 |
| ENSMUSG00000100589 | 4930533P14Rik | 5.54E-05 | 0.841480629 | 1 |
| ENSMUSG00000100182 | 1810006J02Rik | 5.54E-05 | 3.648835551 | 0.466464457 |
| ENSMUSG00000100711 | Gm29461 | 5.54E-05 | 2.648835551 | 0.66416768 |
| ENSMUSG00000054006 | D630008O14Rik | 5.54E-05 | 0.841480629 | 1 |
| ENSMUSG00000067001 | Serpinb7 | 5.54E-05 | 2.063873051 | 0.792441218 |
| ENSMUSG00000100265 | Gm28363 | 5.54E-05 | 2.648835551 | 0.66416768 |

| | | | | |
|---|---|---|---|---|
| ENSMUSG00000101179 | Gm29455 | 5.54E-05 | -0.351164449 | 1 |
| ENSMUSG00000098447 | Gm27184 | 5.54E-05 | 2.648835551 | 0.66416768 |
| ENSMUSG00000101922 | Gm29345 | 5.54E-05 | 2.063873051 | 0.792441218 |
| ENSMUSG00000099336 | Gm29348 | 5.54E-05 | 2.648835551 | 0.66416768 |
| ENSMUSG00000026344 | Lypd1 | 5.54E-05 | 0.841480629 | 1 |
| ENSMUSG00000026348 | Acmsd | 5.54E-05 | 0.841480629 | 1 |
| ENSMUSG00000102014 | 2900009J06Rik | 5.54E-05 | 0.189403933 | 1 |
| ENSMUSG00000026354 | Lct | 5.54E-05 | 0.189403933 | 1 |
| ENSMUSG00000097283 | Gm26686 | 5.54E-05 | 1.648835551 | 0.8730922 |
| ENSMUSG00000046404 | Yod1 | 5.54E-05 | 0.063873051 | 1 |
| ENSMUSG00000100106 | Gm28856 | 5.54E-05 | -1.843017545 | 0.295355925 |
| ENSMUSG00000026415 | Fcamr | 5.54E-05 | 3.648835551 | 0.466464457 |
| ENSMUSG00000086706 | Gm15848 | 5.54E-05 | 2.063873051 | 0.792441218 |
| ENSMUSG00000016524 | Il19 | 5.54E-05 | 2.648835551 | 0.66416768 |
| ENSMUSG00000016526 | Dyrk3 | 5.54E-05 | -0.673092543 | 1 |
| ENSMUSG00000026430 | Rassf5 | 5.54E-05 | -0.995020638 | 0.720253965 |
| ENSMUSG00000052688 | Rab7b | 5.54E-05 | -0.351164449 | 1 |
| ENSMUSG00000042251 | Pm20d1 | 5.54E-05 | 0.47891055 | 1 |
| ENSMUSG00000100658 | F730311O21Rik | 5.54E-05 | 2.063873051 | 0.792441218 |
| ENSMUSG00000042066 | Tmcc2 | 5.54E-05 | 0.063873051 | 1 |
| ENSMUSG00000070644 | Etnk2 | 5.54E-05 | -0.351164449 | 1 |
| ENSMUSG00000102976 | Zc3h11a | 5.54E-05 | 0.063873051 | 1 |
| ENSMUSG00000094410 | Gm38394 | 5.54E-05 | -0.599091962 | 0.980745941 |
| ENSMUSG00000064246 | Chil1 | 5.54E-05 | 0.47891055 | 1 |
| ENSMUSG00000101061 | Platr1 | 5.54E-05 | 3.648835551 | 0.466464457 |
| ENSMUSG00000026429 | Ube2t | 5.54E-05 | 0.47891055 | 1 |
| ENSMUSG00000041889 | Shisa4 | 5.54E-05 | 1.326907457 | 0.971854959 |
| ENSMUSG00000041801 | Phlda3 | 5.54E-05 | 0.648835551 | 1 |
| ENSMUSG00000026418 | Tnni1 | 5.54E-05 | -0.351164449 | 1 |
| ENSMUSG00000051985 | Igfn1 | 5.54E-05 | 2.063873051 | 0.792441218 |
| ENSMUSG00000026407 | Cacna1s | 5.54E-05 | 0.841480629 | 1 |
| ENSMUSG00000103594 | A430034D21Rik | 5.54E-05 | 1.648835551 | 0.886203497 |
| ENSMUSG00000097229 | Platr23 | 5.54E-05 | 1.063873051 | 1 |
| ENSMUSG00000019230 | Lhx9 | 5.54E-05 | -1.106051951 | 0.743748632 |
| ENSMUSG00000026368 | F13b | 5.54E-05 | 1.326907457 | 0.971854959 |
| ENSMUSG00000026358 | Rgs1 | 5.54E-05 | 1.063873051 | 1 |
| ENSMUSG00000102074 | Gm29020 | 5.54E-05 | 1.648835551 | 0.886203497 |
| ENSMUSG00000097028 | Ptgs2os | 5.54E-05 | 0.47891055 | 1 |
| ENSMUSG00000097754 | Ptgs2os2 | 5.54E-05 | 2.648835551 | 0.66416768 |
| ENSMUSG00000100954 | Gm10138 | 5.54E-05 | -0.258055044 | 1 |
| ENSMUSG00000100685 | Gm28610 | 5.54E-05 | 0.326907457 | 1 |
| ENSMUSG00000099969 | Gm28960 | 5.54E-05 | 0.841480629 | 1 |
| ENSMUSG00000099933 | 4930452N14Rik | 5.54E-05 | 2.648835551 | 0.66416768 |
| ENSMUSG00000073535 | Gm5532 | 5.54E-05 | 0.841480629 | 1 |
| ENSMUSG00000026592 | Tex35 | 5.54E-05 | 1.648835551 | 0.886203497 |
| ENSMUSG00000026708 | Cenpl | 5.54E-05 | 0.326907457 | 1 |
| ENSMUSG00000078190 | Dnm3os | 5.54E-05 | 0.648835551 | 1 |
| ENSMUSG00000040181 | Fmo1 | 5.54E-05 | 0.326907457 | 1 |
| ENSMUSG00000026691 | Fmo3 | 5.54E-05 | 0.841480629 | 1 |
| ENSMUSG00000071890 | Mroh9 | 5.54E-05 | 0.063873051 | 1 |
| ENSMUSG00000041406 | BC055324 | 5.54E-05 | -0.743481871 | 0.827142217 |

| | | | | |
|---|---|---|---|---|
| ENSMUSG00000097281 | Gm26685 | 5.54E-05 | 2.063873051 | 0.792441218 |
| ENSMUSG00000102761 | Gm37184 | 5.54E-05 | 0.326907457 | 1 |
| ENSMUSG00000089829 | Gm16565 | 5.54E-05 | 2.063873051 | 0.792441218 |
| ENSMUSG00000000544 | Gpa33 | 5.54E-05 | -0.743481871 | 1 |
| ENSMUSG00000040629 | Mael | 5.54E-05 | 2.648835551 | 0.66416768 |
| ENSMUSG00000040612 | Ildr2 | 5.54E-05 | -1.209145444 | 0.606122355 |
| ENSMUSG00000097823 | Gm16701 | 5.54E-05 | 2.063873051 | 0.792441218 |
| ENSMUSG00000103139 | Gm37982 | 5.54E-05 | 0.326907457 | 1 |
| ENSMUSG00000070532 | Ccdc190 | 5.54E-05 | 0.47891055 | 1 |
| ENSMUSG00000102418 | Sh2d1b1 | 5.54E-05 | 2.063873051 | 0.792441218 |
| ENSMUSG00000102752 | Gm7694 | 5.54E-05 | 1.063873051 | 1 |
| ENSMUSG00000026659 | Dusp12 | 5.54E-05 | -1.106051951 | 0.719591951 |
| ENSMUSG00000059498 | Fcgr3 | 5.54E-05 | -0.673092543 | 1 |
| ENSMUSG00000056569 | Mpz | 5.54E-05 | 1.326907457 | 0.971854959 |
| ENSMUSG00000006403 | Adamts4 | 5.54E-05 | -0.158519371 | 1 |
| ENSMUSG00000053483 | Usp21 | 5.54E-05 | -0.051604167 | 1 |
| ENSMUSG00000006411 | Nectin4 | 5.54E-05 | -0.599091962 | 1 |
| ENSMUSG00000004709 | Cd244a | 5.54E-05 | -0.52108945 | 1 |
| ENSMUSG00000038179 | Slamf7 | 5.54E-05 | 2.063873051 | 0.785343077 |
| ENSMUSG00000015355 | Cd48 | 5.54E-05 | 1.648835551 | 0.886203497 |
| ENSMUSG00000015316 | Slamf1 | 5.54E-05 | 1.063873051 | 1 |
| ENSMUSG00000038147 | Cd84 | 5.54E-05 | -1.560617814 | 0.371776814 |
| ENSMUSG00000026556 | Vangl2 | 5.54E-05 | -2.691014451 | 0.018000019 |
| ENSMUSG00000085184 | 4933439K11Rik | 5.54E-05 | 0.841480629 | 1 |
| ENSMUSG00000049598 | Vsig8 | 5.54E-05 | 3.648835551 | 0.466464457 |
| ENSMUSG00000053318 | Slamf8 | 5.54E-05 | 1.063873051 | 1 |
| ENSMUSG00000037942 | Crp | 5.54E-05 | -1.395558568 | 0.551049466 |
| ENSMUSG00000062527 | Olfr1408 | 5.54E-05 | 3.648835551 | 0.466464457 |
| ENSMUSG00000070501 | Ifi214 | 5.54E-05 | 2.063873051 | 0.778568111 |
| ENSMUSG00000026536 | Ifi211 | 5.54E-05 | -1.673092543 | 0.290284345 |
| ENSMUSG00000045381 | Olfr433 | 5.54E-05 | 3.648835551 | 0.466464457 |
| ENSMUSG00000049528 | Olfr429 | 5.54E-05 | 2.648835551 | 0.66416768 |
| ENSMUSG00000046486 | Olfr231 | 5.54E-05 | 3.648835551 | 0.466464457 |
| ENSMUSG00000055033 | Olfr420 | 5.54E-05 | 2.648835551 | 0.66416768 |
| ENSMUSG00000050069 | Grem2 | 5.54E-05 | -0.158519371 | 1 |
| ENSMUSG00000015962 | 1700016C15Rik | 5.54E-05 | 1.326907457 | 0.971854959 |
| ENSMUSG00000010609 | Psen2 | 5.54E-05 | -1.708716453 | 0.355320248 |
| ENSMUSG00000102923 | Gm31728 | 5.54E-05 | 1.326907457 | 0.962548966 |
| ENSMUSG00000053963 | Stum | 5.54E-05 | 3.648835551 | 0.466464457 |
| ENSMUSG00000102608 | Gm37267 | 5.54E-05 | 2.063873051 | 0.789767653 |
| ENSMUSG00000056699 | Gm5533 | 5.54E-05 | 3.648835551 | 0.466464457 |
| ENSMUSG00000044854 | 1700056E22Rik | 5.54E-05 | 1.326907457 | 0.971854959 |
| ENSMUSG00000102784 | Gm38251 | 5.54E-05 | 2.063873051 | 0.792441218 |
| ENSMUSG00000026614 | Slc30a10 | 5.54E-05 | 2.648835551 | 0.66416768 |
| ENSMUSG00000039246 | Lyplal1 | 5.54E-05 | -0.158519371 | 1 |
| ENSMUSG00000103669 | 2900042K21Rik | 5.54E-05 | 2.063873051 | 0.792441218 |
| ENSMUSG00000099404 | Gm28172 | 5.54E-05 | 3.648835551 | 0.466464457 |
| ENSMUSG00000026630 | Batf3 | 5.54E-05 | 2.063873051 | 0.792441218 |
| ENSMUSG00000073478 | D730003I15Rik | 5.54E-05 | 1.648835551 | 0.886203497 |
| ENSMUSG00000037474 | Dtl | 5.54E-05 | -1.43862729 | 0.433313155 |
| ENSMUSG00000026622 | Nek2 | 5.54E-05 | 1.326907457 | 0.971854959 |

| | | | | |
|---|---|---|---|---|
| ENSMUSG00000055833 | 1700034H15Rik | 5.54E-05 | 1.063873051 | 1 |
| ENSMUSG00000049353 | Rd3 | 5.54E-05 | 2.063873051 | 0.792441218 |
| ENSMUSG00000016262 | Sertad4 | 5.54E-05 | 0.063873051 | 1 |
| ENSMUSG00000009633 | G0s2 | 5.54E-05 | 2.063873051 | 0.792441218 |
| ENSMUSG00000016494 | Cd34 | 5.54E-05 | -0.158519371 | 1 |
| ENSMUSG00000026616 | Cr2 | 5.54E-05 | 0.648835551 | 1 |
| ENSMUSG00000049950 | Rpp38 | 5.54E-05 | 1.063873051 | 1 |
| ENSMUSG00000026650 | Meig1 | 5.54E-05 | 1.326907457 | 0.949907407 |
| ENSMUSG00000051396 | Gm45902 | 5.54E-05 | 0.841480629 | 1 |
| ENSMUSG00000026669 | Mcm10 | 5.54E-05 | -0.051604167 | 1 |
| ENSMUSG00000039063 | Echdc3 | 5.54E-05 | -0.936126949 | 0.695372548 |
| ENSMUSG00000087485 | Gm13383 | 5.54E-05 | 3.648835551 | 0.466464457 |
| ENSMUSG00000086109 | Gm13391 | 5.54E-05 | 3.648835551 | 0.466464457 |
| ENSMUSG00000085070 | Gm13211 | 5.54E-05 | -0.52108945 | 0.980050262 |
| ENSMUSG00000079604 | Gm13219 | 5.54E-05 | -0.351164449 | 1 |
| ENSMUSG00000037254 | Itih2 | 5.54E-05 | 1.063873051 | 0.992259159 |
| ENSMUSG00000102674 | 8030442B05Rik | 5.54E-05 | -1.43862729 | 0.444743375 |
| ENSMUSG00000086657 | Stamos | 5.54E-05 | 2.648835551 | 0.66416768 |
| ENSMUSG00000036949 | Slc39a12 | 5.54E-05 | 0.47891055 | 1 |
| ENSMUSG00000087086 | Gm13269 | 5.54E-05 | 1.326907457 | 0.948624873 |
| ENSMUSG00000017418 | Arl5b | 5.54E-05 | -1.351164449 | 0.499088689 |
| ENSMUSG00000037708 | Spag6 | 5.54E-05 | -1.209145444 | 0.642722979 |
| ENSMUSG00000026734 | 4921504E06Rik | 5.54E-05 | -0.351164449 | 1 |
| ENSMUSG00000043415 | Otud1 | 5.54E-05 | 2.063873051 | 0.792441218 |
| ENSMUSG00000026787 | Gad2 | 5.54E-05 | -0.158519371 | 1 |
| ENSMUSG00000026986 | Hnmt | 5.54E-05 | 0.189403933 | 1 |
| ENSMUSG00000026983 | Il1f5 | 5.54E-05 | 2.648835551 | 0.66416768 |
| ENSMUSG00000006469 | Slc34a3 | 5.54E-05 | 3.648835551 | 0.466464457 |
| ENSMUSG00000078201 | Tmem203 | 5.54E-05 | 2.648835551 | 0.66416768 |
| ENSMUSG00000026958 | Dpp7 | 5.54E-05 | 0.326907457 | 1 |
| ENSMUSG00000015085 | Entpd2 | 5.54E-05 | 0.841480629 | 1 |
| ENSMUSG00000015090 | Ptgds | 5.54E-05 | 2.648835551 | 0.66416768 |
| ENSMUSG00000026942 | Traf2 | 5.54E-05 | -2.82689788 | 0.011086379 |
| ENSMUSG00000026940 | Ccdc183 | 5.54E-05 | 2.063873051 | 0.792441218 |
| ENSMUSG00000036401 | Glt6d1 | 5.54E-05 | 1.648835551 | 0.871107617 |
| ENSMUSG00000023210 | Lcn9 | 5.54E-05 | 3.648835551 | 0.466464457 |
| ENSMUSG00000026930 | Gpsm1 | 5.54E-05 | -0.52108945 | 0.977096893 |
| ENSMUSG00000026928 | Card9 | 5.54E-05 | 1.063873051 | 1 |
| ENSMUSG00000026923 | Notch1 | 5.54E-05 | -1.708716453 | 0.296030202 |
| ENSMUSG00000026922 | Agpat2 | 5.54E-05 | -0.995020638 | 0.699192777 |
| ENSMUSG00000009216 | Fam163b | 5.54E-05 | 1.063873051 | 1 |
| ENSMUSG00000000889 | Dbh | 5.54E-05 | 2.063873051 | 0.792441218 |
| ENSMUSG00000026835 | Fcnb | 5.54E-05 | 2.063873051 | 0.790186865 |
| ENSMUSG00000026831 | 1700007K13Rik | 5.54E-05 | 3.648835551 | 0.466464457 |
| ENSMUSG00000035772 | Mrps2 | 5.54E-05 | 0.648835551 | 1 |
| ENSMUSG00000026809 | Spaca9 | 5.54E-05 | 2.648835551 | 0.66416768 |
| ENSMUSG00000039826 | Trub2 | 5.54E-05 | -1.636566667 | 0.433295324 |
| ENSMUSG00000026798 | Coq4 | 5.54E-05 | -0.936126949 | 0.695372548 |
| ENSMUSG00000059316 | Slc27a4 | 5.54E-05 | -2.023589791 | 0.130342151 |
| ENSMUSG00000019715 | Gle1 | 5.54E-05 | -3.691014451 | 0.000251527 |
| ENSMUSG00000075416 | Gm14488 | 5.54E-05 | 2.063873051 | 0.788941408 |

| | | | | |
|---|---|---|---|---|
| ENSMUSG00000044320 | 1700001O22Rik | 5.54E-05 | 3.648835551 | 0.466464457 |
| ENSMUSG00000039483 | Asb6 | 5.54E-05 | -0.743481871 | 0.827142217 |
| ENSMUSG00000087549 | Gm13427 | 5.54E-05 | 0.648835551 | 1 |
| ENSMUSG00000050592 | Fam78a | 5.54E-05 | -1.480447466 | 0.466464457 |
| ENSMUSG00000051373 | Plpp7 | 5.54E-05 | 1.648835551 | 0.868661374 |
| ENSMUSG00000086216 | Gm13610 | 5.54E-05 | 2.648835551 | 0.66416768 |
| ENSMUSG00000039254 | Pomt1 | 5.54E-05 | -0.743481871 | 0.827142217 |
| ENSMUSG00000026810 | Dpm2 | 5.54E-05 | 0.648835551 | 1 |
| ENSMUSG00000039021 | Ttc16 | 5.54E-05 | 2.063873051 | 0.792441218 |
| ENSMUSG00000038987 | Cfap157 | 5.54E-05 | 0.841480629 | 1 |
| ENSMUSG00000087496 | Gm13523 | 5.54E-05 | 3.648835551 | 0.466464457 |
| ENSMUSG00000085087 | Gm13528 | 5.54E-05 | 1.326907457 | 0.957537943 |
| ENSMUSG00000084908 | C79798 | 5.54E-05 | 0.326907457 | 1 |
| ENSMUSG00000035875 | AI182371 | 5.54E-05 | 0.648835551 | 1 |
| ENSMUSG00000079421 | 4930402F06Rik | 5.54E-05 | 2.648835551 | 0.66416768 |
| ENSMUSG00000086449 | 9030204H09Rik | 5.54E-05 | 3.648835551 | 0.466464457 |
| ENSMUSG00000050714 | Zbtb26 | 5.54E-05 | -0.52108945 | 0.975624465 |
| ENSMUSG00000087062 | Gm13587 | 5.54E-05 | 3.648835551 | 0.466464457 |
| ENSMUSG00000026866 | Kynu | 5.54E-05 | -0.874726405 | 0.860952303 |
| ENSMUSG00000085089 | Arhgap15os | 5.54E-05 | 2.648835551 | 0.66416768 |
| ENSMUSG00000100135 | Gm28643 | 5.54E-05 | 2.648835551 | 0.66416768 |
| ENSMUSG00000085591 | Gm13479 | 5.54E-05 | 0.648835551 | 1 |
| ENSMUSG00000026761 | Orc4 | 5.54E-05 | -2.233807498 | 0.09684834 |
| ENSMUSG00000026765 | Lypd6b | 5.54E-05 | -2.480447466 | 0.050877179 |
| ENSMUSG00000086523 | Gm13499 | 5.54E-05 | 3.648835551 | 0.466464457 |
| ENSMUSG00000053475 | Tnfaip6 | 5.54E-05 | 0.189403933 | 1 |
| ENSMUSG00000086447 | Gm13522 | 5.54E-05 | 1.648835551 | 0.886203497 |
| ENSMUSG00000085315 | A430018G15Rik | 5.54E-05 | 2.648835551 | 0.66416768 |
| ENSMUSG00000026826 | Nr4a2 | 5.54E-05 | -0.673092543 | 0.993144405 |
| ENSMUSG00000110205 | Gm33594 | 5.54E-05 | 3.648835551 | 0.466464457 |
| ENSMUSG00000026828 | Galnt5 | 5.54E-05 | 1.326907457 | 0.971854959 |
| ENSMUSG00000026989 | Dapl1 | 5.54E-05 | 1.326907457 | 0.950123751 |
| ENSMUSG00000087283 | Gm13580 | 5.54E-05 | 3.648835551 | 0.466464457 |
| ENSMUSG00000035000 | Dpp4 | 5.54E-05 | -0.673092543 | 0.9980272 |
| ENSMUSG00000087518 | Gm13561 | 5.54E-05 | 0.189403933 | 1 |
| ENSMUSG00000087681 | Gm13565 | 5.54E-05 | 1.648835551 | 0.877701541 |
| ENSMUSG00000026896 | Ifih1 | 5.54E-05 | -1.305360759 | 0.491653643 |
| ENSMUSG00000061171 | Slc38a11 | 5.54E-05 | 0.326907457 | 1 |
| ENSMUSG00000085160 | Gm13617 | 5.54E-05 | 2.648835551 | 0.66416768 |
| ENSMUSG00000086908 | Gm13598 | 5.54E-05 | 1.326907457 | 0.971854959 |
| ENSMUSG00000087043 | Gm13599 | 5.54E-05 | 0.648835551 | 1 |
| ENSMUSG00000087467 | Gm13601 | 5.54E-05 | 2.063873051 | 0.792441218 |
| ENSMUSG00000079324 | 4932414N04Rik | 5.54E-05 | -0.351164449 | 1 |
| ENSMUSG00000086819 | Gm13613 | 5.54E-05 | 1.648835551 | 0.886203497 |
| ENSMUSG00000008226 | Scrn3 | 5.54E-05 | -0.995020638 | 0.699192777 |
| ENSMUSG00000087176 | D230022J07Rik | 5.54E-05 | 0.326907457 | 1 |
| ENSMUSG00000086069 | Gm13708 | 5.54E-05 | 1.326907457 | 0.958555629 |
| ENSMUSG00000084791 | Gm13667 | 5.54E-05 | 2.648835551 | 0.66416768 |
| ENSMUSG00000001819 | Hoxd13 | 5.54E-05 | 3.648835551 | 0.466464457 |
| ENSMUSG00000043342 | Hoxd9 | 5.54E-05 | 3.648835551 | 0.466464457 |
| ENSMUSG00000052371 | Hoxd3os1 | 5.54E-05 | 0.648835551 | 1 |

| | | | | |
|---|---|---|---|---|
| ENSMUSG00000087333 | Gm13652 | 5.54E-05 | 2.063873051 | 0.792441218 |
| ENSMUSG00000002733 | Plekha3 | 5.54E-05 | -1.874726405 | 0.198601369 |
| ENSMUSG00000085845 | Gm13944 | 5.54E-05 | 1.326907457 | 0.971854959 |
| ENSMUSG00000050114 | Prdx6b | 5.54E-05 | 3.648835551 | 0.466464457 |
| ENSMUSG00000027004 | Frzb | 5.54E-05 | -1.52108945 | 0.455131546 |
| ENSMUSG00000087061 | A130030D18Rik | 5.54E-05 | 1.326907457 | 0.967564517 |
| ENSMUSG00000075249 | Fsip2 | 5.54E-05 | 0.47891055 | 1 |
| ENSMUSG00000102082 | Gm28675 | 5.54E-05 | 0.47891055 | 1 |
| ENSMUSG00000085335 | Gm13684 | 5.54E-05 | -0.673092543 | 0.994783189 |
| ENSMUSG00000027082 | Tfpi | 5.54E-05 | -0.43862729 | 1 |
| ENSMUSG00000027079 | Clp1 | 5.54E-05 | 0.189403933 | 1 |
| ENSMUSG00000034059 | Ypel4 | 5.54E-05 | 1.326907457 | 0.971854959 |
| ENSMUSG00000027078 | Ube2l6 | 5.54E-05 | 0.648835551 | 1 |
| ENSMUSG00000027072 | Prg3 | 5.54E-05 | 2.648835551 | 0.66416768 |
| ENSMUSG00000085135 | Gm13713 | 5.54E-05 | 1.326907457 | 0.971854959 |
| ENSMUSG00000075202 | Olfr1042 | 5.54E-05 | 2.648835551 | 0.66416768 |
| ENSMUSG00000075175 | Olfr1086 | 5.54E-05 | 3.648835551 | 0.466464457 |
| ENSMUSG00000049843 | Olfr1102 | 5.54E-05 | 3.648835551 | 0.466464457 |
| ENSMUSG00000025839 | Pramel7 | 5.54E-05 | 0.47891055 | 1 |
| ENSMUSG00000061520 | Olfr153 | 5.54E-05 | 3.648835551 | 0.466464457 |
| ENSMUSG00000075115 | Olfr1200 | 5.54E-05 | 2.063873051 | 0.792441218 |
| ENSMUSG00000075104 | Olfr1219 | 5.54E-05 | 2.648835551 | 0.66416768 |
| ENSMUSG00000049149 | Olfr1258 | 5.54E-05 | 2.648835551 | 0.66416768 |
| ENSMUSG00000082980 | Olfr1274-ps | 5.54E-05 | 2.648835551 | 0.66416768 |
| ENSMUSG00000040812 | Agbl2 | 5.54E-05 | 0.189403933 | 1 |
| ENSMUSG00000040794 | C1qtnf4 | 5.54E-05 | 2.648835551 | 0.66416768 |
| ENSMUSG00000002100 | Mybpc3 | 5.54E-05 | 2.648835551 | 0.66416768 |
| ENSMUSG00000002108 | Nr1h3 | 5.54E-05 | -0.995020638 | 0.699192777 |
| ENSMUSG00000027255 | Arfgap2 | 5.54E-05 | -2.184054463 | 0.094802366 |
| ENSMUSG00000027253 | Lrp4 | 5.54E-05 | 0.47891055 | 1 |
| ENSMUSG00000027243 | Harbi1 | 5.54E-05 | -0.743481871 | 0.829413358 |
| ENSMUSG00000027239 | Mdk | 5.54E-05 | 0.47891055 | 1 |
| ENSMUSG00000085293 | Gm13817 | 5.54E-05 | 2.063873051 | 0.792441218 |
| ENSMUSG00000091061 | 4930445B16Rik | 5.54E-05 | -0.43862729 | 1 |
| ENSMUSG00000032864 | Rag2 | 5.54E-05 | 3.648835551 | 0.466464457 |
| ENSMUSG00000085417 | Gm13919 | 5.54E-05 | 0.648835551 | 1 |
| ENSMUSG00000075012 | Fjx1 | 5.54E-05 | -0.673092543 | 0.98880775 |
| ENSMUSG00000010911 | Apip | 5.54E-05 | 0.47891055 | 1 |
| ENSMUSG00000070794 | BC016548 | 5.54E-05 | -2.417253639 | 0.079721397 |
| ENSMUSG00000032679 | Cd59a | 5.54E-05 | -1.43862729 | 0.630177036 |
| ENSMUSG00000087226 | Gm14015 | 5.54E-05 | 0.47891055 | 1 |
| ENSMUSG00000027115 | Kif18a | 5.54E-05 | -1.52108945 | 0.452184898 |
| ENSMUSG00000048482 | Bdnf | 5.54E-05 | -1.936126949 | 0.158846729 |
| ENSMUSG00000085444 | Gm13936 | 5.54E-05 | 0.326907457 | 1 |
| ENSMUSG00000074971 | Fibin | 5.54E-05 | 3.648835551 | 0.466464457 |
| ENSMUSG00000079169 | Gm15130 | 5.54E-05 | 3.648835551 | 0.466464457 |
| ENSMUSG00000094747 | Olfr1307 | 5.54E-05 | 3.648835551 | 0.466464457 |
| ENSMUSG00000041255 | Tmco5b | 5.54E-05 | 3.648835551 | 0.466464457 |
| ENSMUSG00000085448 | Gm13963 | 5.54E-05 | -0.43862729 | 1 |
| ENSMUSG00000084785 | Gm13972 | 5.54E-05 | 1.648835551 | 0.886203497 |
| ENSMUSG00000086197 | Gm13976 | 5.54E-05 | 3.648835551 | 0.466464457 |

| | | | | |
|---|---|---|---|---|
| ENSMUSG00000086221 | 4930412B13Rik | 5.54E-05 | -0.810596067 | 0.847265357 |
| ENSMUSG00000027331 | Knstrn | 5.54E-05 | -1.051604167 | 0.713570913 |
| ENSMUSG00000086421 | Gm14091 | 5.54E-05 | 1.648835551 | 0.886203497 |
| ENSMUSG00000040007 | Bahd1 | 5.54E-05 | -1.874726405 | 0.232414399 |
| ENSMUSG00000055926 | Gm14137 | 5.54E-05 | 0.47891055 | 1 |
| ENSMUSG00000027317 | Ppp1r14d | 5.54E-05 | 1.326907457 | 0.971854959 |
| ENSMUSG00000027315 | Spint1 | 5.54E-05 | -0.874726405 | 0.886203497 |
| ENSMUSG00000072980 | Oip5 | 5.54E-05 | 0.063873051 | 1 |
| ENSMUSG00000027298 | Tyro3 | 5.54E-05 | 0.189403933 | 1 |
| ENSMUSG00000098789 | Jmjd7 | 5.54E-05 | 0.189403933 | 1 |
| ENSMUSG00000027291 | Vps39 | 5.54E-05 | -3.305360759 | 0.001866323 |
| ENSMUSG00000087519 | AV039307 | 5.54E-05 | 0.326907457 | 1 |
| ENSMUSG00000054484 | Tmem62 | 5.54E-05 | -1.52108945 | 0.452184898 |
| ENSMUSG00000053675 | Tgm5 | 5.54E-05 | 0.648835551 | 1 |
| ENSMUSG00000074890 | Lcmt2 | 5.54E-05 | -0.158519371 | 1 |
| ENSMUSG00000027254 | Map1a | 5.54E-05 | -1.051604167 | 0.744267071 |
| ENSMUSG00000027246 | Ell3 | 5.54E-05 | 2.648835551 | 0.66416768 |
| ENSMUSG00000060227 | Casc4 | 5.54E-05 | -1.305360759 | 0.491653643 |
| ENSMUSG00000033368 | Trim69 | 5.54E-05 | 0.47891055 | 1 |
| ENSMUSG00000027229 | Terb2 | 5.54E-05 | 1.063873051 | 1 |
| ENSMUSG00000068452 | Duox2 | 5.54E-05 | -0.43862729 | 1 |
| ENSMUSG00000027225 | Duoxa2 | 5.54E-05 | 0.648835551 | 1 |
| ENSMUSG00000033268 | Duox1 | 5.54E-05 | 1.326907457 | 0.946342172 |
| ENSMUSG00000035183 | Slc24a5 | 5.54E-05 | 2.648835551 | 0.66416768 |
| ENSMUSG00000035109 | Shc4 | 5.54E-05 | -0.810596067 | 0.844410019 |
| ENSMUSG00000034906 | Ncaph | 5.54E-05 | -0.258055044 | 1 |
| ENSMUSG00000003662 | Ciao1 | 5.54E-05 | 0.326907457 | 1 |
| ENSMUSG00000027371 | Fahd2a | 5.54E-05 | 1.063873051 | 1 |
| ENSMUSG00000079056 | Kcnip3 | 5.54E-05 | 0.063873051 | 1 |
| ENSMUSG00000027376 | Prom2 | 5.54E-05 | 3.648835551 | 0.466464457 |
| ENSMUSG00000034800 | Zfp661 | 5.54E-05 | 0.841480629 | 1 |
| ENSMUSG00000087162 | Gm14244 | 5.54E-05 | 1.063873051 | 1 |
| ENSMUSG00000027378 | Nphp1 | 5.54E-05 | -1.43862729 | 0.433313155 |
| ENSMUSG00000085750 | Gm14009 | 5.54E-05 | -0.051604167 | 1 |
| ENSMUSG00000109931 | Gm39929 | 5.54E-05 | 1.648835551 | 0.876040999 |
| ENSMUSG00000087151 | Gm14022 | 5.54E-05 | 1.063873051 | 0.971854959 |
| ENSMUSG00000037938 | Chchd5 | 5.54E-05 | -0.874726405 | 0.840804483 |
| ENSMUSG00000085485 | Gm14024 | 5.54E-05 | 1.648835551 | 0.884935781 |
| ENSMUSG00000027403 | Tgm6 | 5.54E-05 | 0.326907457 | 1 |
| ENSMUSG00000060332 | Tmc2 | 5.54E-05 | -0.810596067 | 1 |
| ENSMUSG00000037773 | Pced1a | 5.54E-05 | -1.158519371 | 0.599135879 |
| ENSMUSG00000060029 | 4930473A02Rik | 5.54E-05 | 1.326907457 | 0.971854959 |
| ENSMUSG00000027300 | Ubox5 | 5.54E-05 | -1.52108945 | 0.673811773 |
| ENSMUSG00000085566 | A730017L22Rik | 5.54E-05 | 0.189403933 | 1 |
| ENSMUSG00000027316 | Gfra4 | 5.54E-05 | 3.648835551 | 0.466464457 |
| ENSMUSG00000087405 | Gm14232 | 5.54E-05 | 1.648835551 | 0.86916119 |
| ENSMUSG00000037523 | Mavs | 5.54E-05 | 0.648835551 | 1 |
| ENSMUSG00000086822 | 5330413P13Rik | 5.54E-05 | 2.648835551 | 0.66416768 |
| ENSMUSG00000074783 | AU019990 | 5.54E-05 | 0.47891055 | 1 |
| ENSMUSG00000034723 | Tmx4 | 5.54E-05 | -1.52108945 | 0.452184898 |
| ENSMUSG00000027274 | Mkks | 5.54E-05 | 0.063873051 | 1 |

| | | | | |
|---|---|---|---|---|
| ENSMUSG00000085504 | Gm14055 | 5.54E-05 | 0.841480629 | 1 |
| ENSMUSG00000085911 | Gm14061 | 5.54E-05 | 1.326907457 | 0.950709758 |
| ENSMUSG00000078197 | Gm17374 | 5.54E-05 | 2.648835551 | 0.66416768 |
| ENSMUSG00000027420 | Bfsp1 | 5.54E-05 | -0.351164449 | 1 |
| ENSMUSG00000037307 | Banf2 | 5.54E-05 | 0.326907457 | 1 |
| ENSMUSG00000027424 | Mgme1 | 5.54E-05 | -1.636566667 | 0.373232174 |
| ENSMUSG00000068154 | Insm1 | 5.54E-05 | 3.648835551 | 0.466464457 |
| ENSMUSG00000027434 | Nkx2-2 | 5.54E-05 | -1.209145444 | 0.663030707 |
| ENSMUSG00000068134 | Zfp120 | 5.54E-05 | -0.52108945 | 0.975624465 |
| ENSMUSG00000079009 | Gm14139 | 5.54E-05 | 3.648835551 | 0.466464457 |
| ENSMUSG00000063364 | 3300002I08Rik | 5.54E-05 | -1.209145444 | 0.606122355 |
| ENSMUSG00000086407 | Gm14123 | 5.54E-05 | 3.648835551 | 0.466464457 |
| ENSMUSG00000033080 | Vsx1 | 5.54E-05 | 3.648835551 | 0.466464457 |
| ENSMUSG00000053916 | Nanp | 5.54E-05 | 0.326907457 | 1 |
| ENSMUSG00000087433 | Gm14167 | 5.54E-05 | 1.326907457 | 0.919691163 |
| ENSMUSG00000027457 | Snph | 5.54E-05 | 0.648835551 | 1 |
| ENSMUSG00000027460 | Angpt4 | 5.54E-05 | 0.648835551 | 1 |
| ENSMUSG00000027459 | Fam110a | 5.54E-05 | 0.189403933 | 1 |
| ENSMUSG00000027463 | Slc52a3 | 5.54E-05 | 1.326907457 | 0.971854959 |
| ENSMUSG00000032680 | 6820408C15Rik | 5.54E-05 | 1.326907457 | 0.971854959 |
| ENSMUSG00000090485 | Gm17416 | 5.54E-05 | 3.648835551 | 0.466464457 |
| ENSMUSG00000042814 | Mcts2 | 5.54E-05 | -1.395558568 | 0.508263132 |
| ENSMUSG00000027478 | Dnmt3b | 5.54E-05 | -0.673092543 | 0.98880775 |
| ENSMUSG00000027481 | Bpifb2 | 5.54E-05 | 1.648835551 | 0.886203497 |
| ENSMUSG00000042459 | Bpifa2 | 5.54E-05 | 3.648835551 | 0.466464457 |
| ENSMUSG00000027485 | Bpifb1 | 5.54E-05 | 1.326907457 | 0.950472516 |
| ENSMUSG00000027603 | Ggt7 | 5.54E-05 | 0.326907457 | 1 |
| ENSMUSG00000085010 | Gssos1 | 5.54E-05 | 3.648835551 | 0.466464457 |
| ENSMUSG00000074652 | Myh7b | 5.54E-05 | 2.063873051 | 0.792441218 |
| ENSMUSG00000086754 | Gm16098 | 5.54E-05 | 2.063873051 | 0.792441218 |
| ENSMUSG00000074646 | 6430550D23Rik | 5.54E-05 | -0.43862729 | 1 |
| ENSMUSG00000085546 | Gm14252 | 5.54E-05 | 2.063873051 | 0.792441218 |
| ENSMUSG00000086118 | Gm14169 | 5.54E-05 | 0.47891055 | 1 |
| ENSMUSG00000067818 | Myl9 | 5.54E-05 | -0.743481871 | 1 |
| ENSMUSG00000027636 | Sla2 | 5.54E-05 | 0.841480629 | 1 |
| ENSMUSG00000067786 | Nnat | 5.54E-05 | 1.063873051 | 1 |
| ENSMUSG00000087547 | Platr27 | 5.54E-05 | 2.648835551 | 0.66416768 |
| ENSMUSG00000037813 | D630003M21Rik | 5.54E-05 | -3.132524162 | 0.00337143 |
| ENSMUSG00000027655 | Dhx35 | 5.54E-05 | -0.810596067 | 0.840804483 |
| ENSMUSG00000035576 | L3mbtl1 | 5.54E-05 | -0.051604167 | 1 |
| ENSMUSG00000074607 | Tox2 | 5.54E-05 | -0.351164449 | 1 |
| ENSMUSG00000035226 | Rims4 | 5.54E-05 | 0.063873051 | 1 |
| ENSMUSG00000040154 | Wfdc5 | 5.54E-05 | 3.648835551 | 0.466464457 |
| ENSMUSG00000050383 | Svs3b | 5.54E-05 | 2.648835551 | 0.66416768 |
| ENSMUSG00000017002 | Slpi | 5.54E-05 | -0.051604167 | 1 |
| ENSMUSG00000074596 | Spint3 | 5.54E-05 | 0.841480629 | 1 |
| ENSMUSG00000086453 | Gm11457 | 5.54E-05 | 0.189403933 | 1 |
| ENSMUSG00000017307 | Acot8 | 5.54E-05 | -2.184054463 | 0.132948501 |
| ENSMUSG00000017740 | Slc12a5 | 5.54E-05 | -0.258055044 | 1 |
| ENSMUSG00000017667 | Zfp334 | 5.54E-05 | -0.673092543 | 0.98880775 |
| ENSMUSG00000085151 | 1110018N20Rik | 5.54E-05 | 3.648835551 | 0.466464457 |

| | | | | |
|---|---|---|---|---|
| ENSMUSG00000006462 | A530013C23Rik | 5.54E-05 | -0.810596067 | 1 |
| ENSMUSG00000087648 | E130018N17Rik | 5.54E-05 | 3.648835551 | 0.466464457 |
| ENSMUSG00000085350 | 4930529I22Rik | 5.54E-05 | 3.648835551 | 0.466464457 |
| ENSMUSG00000087074 | Gm14249 | 5.54E-05 | -0.43862729 | 1 |
| ENSMUSG00000087409 | Gm14271 | 5.54E-05 | 3.648835551 | 0.466464457 |
| ENSMUSG00000085949 | Gm14275 | 5.54E-05 | 3.648835551 | 0.466464457 |
| ENSMUSG00000074569 | Gcnt7 | 5.54E-05 | 2.063873051 | 0.792441218 |
| ENSMUSG00000016255 | Tubb1 | 5.54E-05 | 2.648835551 | 0.66416768 |
| ENSMUSG00000078897 | Gm4724 | 5.54E-05 | 3.648835551 | 0.466464457 |
| ENSMUSG00000078877 | Gm14295 | 5.54E-05 | 0.841480629 | 1 |
| ENSMUSG00000074527 | Gm14296 | 5.54E-05 | 0.648835551 | 1 |
| ENSMUSG00000094786 | Gm14403 | 5.54E-05 | 3.648835551 | 0.466464457 |
| ENSMUSG00000074519 | Zfp971 | 5.54E-05 | -2.233807498 | 0.09684834 |
| ENSMUSG00000049999 | Ppp1r3d | 5.54E-05 | 3.648835551 | 0.466464457 |
| ENSMUSG00000070476 | Fam217b | 5.54E-05 | 0.648835551 | 1 |
| ENSMUSG00000039155 | Cdh26 | 5.54E-05 | 0.063873051 | 1 |
| ENSMUSG00000085704 | 4921531C22Rik | 5.54E-05 | 1.326907457 | 0.899950578 |
| ENSMUSG00000038963 | Slco4a1 | 5.54E-05 | 0.189403933 | 1 |
| ENSMUSG00000027568 | Ntsr1 | 5.54E-05 | 0.47891055 | 1 |
| ENSMUSG00000070461 | 9230112E08Rik | 5.54E-05 | 1.648835551 | 0.886203497 |
| ENSMUSG00000016356 | Col20a1 | 5.54E-05 | -0.258055044 | 1 |
| ENSMUSG00000016346 | Kcnq2 | 5.54E-05 | 0.326907457 | 1 |
| ENSMUSG00000027579 | Srms | 5.54E-05 | 1.063873051 | 1 |
| ENSMUSG00000040289 | Hey1 | 5.54E-05 | 2.648835551 | 0.66416768 |
| ENSMUSG00000103116 | 4930539M17Rik | 5.54E-05 | 3.648835551 | 0.466464457 |
| ENSMUSG00000069114 | Zbtb10 | 5.54E-05 | -1.395558568 | 0.508263132 |
| ENSMUSG00000089794 | Gm16337 | 5.54E-05 | 2.063873051 | 0.792441218 |
| ENSMUSG00000058921 | Slc10a5 | 5.54E-05 | 3.648835551 | 0.466464457 |
| ENSMUSG00000027534 | Snx16 | 5.54E-05 | -0.673092543 | 0.98880775 |
| ENSMUSG00000104015 | Gm10745 | 5.54E-05 | 1.648835551 | 0.86953619 |
| ENSMUSG00000027556 | Car1 | 5.54E-05 | 0.063873051 | 1 |
| ENSMUSG00000098008 | A930001A20Rik | 5.54E-05 | 1.326907457 | 0.941472227 |
| ENSMUSG00000078780 | Gm5150 | 5.54E-05 | 0.063873051 | 1 |
| ENSMUSG00000027606 | Dnajc5b | 5.54E-05 | -0.743481871 | 0.827142217 |
| ENSMUSG00000082766 | 1700064H15Rik | 5.54E-05 | 1.648835551 | 0.886203497 |
| ENSMUSG00000049796 | Crh | 5.54E-05 | 3.648835551 | 0.466464457 |
| ENSMUSG00000001865 | Cpa3 | 5.54E-05 | 0.648835551 | 1 |
| ENSMUSG00000102203 | 4930412E21Rik | 5.54E-05 | 2.063873051 | 0.773756651 |
| ENSMUSG00000039304 | Tnfsf10 | 5.54E-05 | 0.47891055 | 1 |
| ENSMUSG00000043164 | Tmem212 | 5.54E-05 | -0.43862729 | 1 |
| ENSMUSG00000109716 | Gm42196 | 5.54E-05 | 1.063873051 | 1 |
| ENSMUSG00000050192 | Eif5a2 | 5.54E-05 | 2.063873051 | 0.775870935 |
| ENSMUSG00000102574 | Gm33206 | 5.54E-05 | 0.326907457 | 1 |
| ENSMUSG00000037737 | Actrt3 | 5.54E-05 | 2.063873051 | 0.792441218 |
| ENSMUSG00000074653 | Lrrc31 | 5.54E-05 | 0.063873051 | 1 |
| ENSMUSG00000051860 | Samd7 | 5.54E-05 | 0.47891055 | 1 |
| ENSMUSG00000037625 | Cldn11 | 5.54E-05 | 1.648835551 | 0.870819142 |
| ENSMUSG00000104283 | Gm37459 | 5.54E-05 | 0.189403933 | 1 |
| ENSMUSG00000104588 | Gm43666 | 5.54E-05 | 1.326907457 | 0.970970089 |
| ENSMUSG00000027676 | Ccdc39 | 5.54E-05 | 1.648835551 | 0.886203497 |
| ENSMUSG00000105475 | 4930431L21Rik | 5.54E-05 | 1.648835551 | 0.886203497 |

| | | | | |
|---|---|---|---|---|
| ENSMUSG00000074637 | Sox2 | 5.54E-05 | 3.648835551 | 0.466464457 |
| ENSMUSG00000105153 | Gm3143 | 5.54E-05 | 0.648835551 | 1 |
| ENSMUSG00000106519 | Gm38509 | 5.54E-05 | 2.648835551 | 0.66416768 |
| ENSMUSG00000105651 | 1700017M07Rik | 5.54E-05 | 3.648835551 | 0.466464457 |
| ENSMUSG00000058400 | Qrfpr | 5.54E-05 | 1.326907457 | 0.971854959 |
| ENSMUSG00000085007 | Gm11549 | 5.54E-05 | 2.648835551 | 0.66416768 |
| ENSMUSG00000027716 | Trpc3 | 5.54E-05 | 0.189403933 | 1 |
| ENSMUSG00000045031 | Cetn4 | 5.54E-05 | 1.063873051 | 1 |
| ENSMUSG00000054779 | Fgf2os | 5.54E-05 | 1.648835551 | 0.886203497 |
| ENSMUSG00000050174 | Nudt6 | 5.54E-05 | -0.051604167 | 1 |
| ENSMUSG00000058174 | Gm5148 | 5.54E-05 | -1.209145444 | 0.606122355 |
| ENSMUSG00000105219 | Gm43821 | 5.54E-05 | 1.063873051 | 1 |
| ENSMUSG00000046743 | Fat4 | 5.54E-05 | -0.43862729 | 1 |
| ENSMUSG00000069041 | Slc25a31 | 5.54E-05 | -0.936126949 | 0.868661374 |
| ENSMUSG00000025759 | Mfsd8 | 5.54E-05 | -0.743481871 | 0.827142217 |
| ENSMUSG00000105297 | 1700052H01Rik | 5.54E-05 | 0.326907457 | 1 |
| ENSMUSG00000102679 | Gm20557 | 5.54E-05 | 2.063873051 | 0.792441218 |
| ENSMUSG00000102892 | Gm37854 | 5.54E-05 | 1.326907457 | 0.971854959 |
| ENSMUSG00000102715 | Gm6209 | 5.54E-05 | -1.673092543 | 0.290284345 |
| ENSMUSG00000102544 | Gm5103 | 5.54E-05 | 3.648835551 | 0.466464457 |
| ENSMUSG00000037161 | Mgarp | 5.54E-05 | 1.648835551 | 0.886203497 |
| ENSMUSG00000102539 | Gm37229 | 5.54E-05 | 2.648835551 | 0.66416768 |
| ENSMUSG00000104161 | Gm20089 | 5.54E-05 | 3.648835551 | 0.466464457 |
| ENSMUSG00000103655 | Gm37202 | 5.54E-05 | 0.189403933 | 1 |
| ENSMUSG00000102785 | Gm2447 | 5.54E-05 | 0.648835551 | 1 |
| ENSMUSG00000106577 | Gm31026 | 5.54E-05 | 2.063873051 | 0.788226642 |
| ENSMUSG00000106589 | 4931419H13Rik | 5.54E-05 | 1.648835551 | 0.886203497 |
| ENSMUSG00000104677 | Gm43376 | 5.54E-05 | 1.648835551 | 0.886203497 |
| ENSMUSG00000074591 | Ankub1 | 5.54E-05 | 0.648835551 | 1 |
| ENSMUSG00000104586 | 4921539H07Rik | 5.54E-05 | 1.326907457 | 0.971854959 |
| ENSMUSG00000100666 | 1700007F19Rik | 5.54E-05 | 2.648835551 | 0.66416768 |
| ENSMUSG00000070471 | Erich6 | 5.54E-05 | 0.47891055 | 1 |
| ENSMUSG00000036432 | Siah2 | 5.54E-05 | -0.051604167 | 1 |
| ENSMUSG00000027761 | Aadac | 5.54E-05 | 2.063873051 | 0.792441218 |
| ENSMUSG00000027765 | P2ry1 | 5.54E-05 | -1.106051951 | 0.719591951 |
| ENSMUSG00000104336 | Gm34240 | 5.54E-05 | 1.648835551 | 0.874658224 |
| ENSMUSG00000103620 | Gm37359 | 5.54E-05 | 2.063873051 | 0.792441218 |
| ENSMUSG00000027831 | Veph1 | 5.54E-05 | -2.184054463 | 0.115010622 |
| ENSMUSG00000027833 | Shox2 | 5.54E-05 | 1.063873051 | 1 |
| ENSMUSG00000027792 | Bche | 5.54E-05 | -0.673092543 | 1 |
| ENSMUSG00000034139 | Serpini2 | 5.54E-05 | 1.326907457 | 0.954524296 |
| ENSMUSG00000099370 | Platr10 | 5.54E-05 | 2.063873051 | 0.792441218 |
| ENSMUSG00000100962 | Gm29133 | 5.54E-05 | 0.648835551 | 1 |
| ENSMUSG00000091685 | Gm17359 | 5.54E-05 | -1.258055044 | 0.66416768 |
| ENSMUSG00000033981 | Gria2 | 5.54E-05 | -0.351164449 | 1 |
| ENSMUSG00000105092 | Gm42468 | 5.54E-05 | 2.648835551 | 0.66416768 |
| ENSMUSG00000028011 | Tdo2 | 5.54E-05 | 2.648835551 | 0.66416768 |
| ENSMUSG00000028005 | Gucy1b1 | 5.54E-05 | 0.47891055 | 1 |
| ENSMUSG00000104898 | Gm30043 | 5.54E-05 | 3.648835551 | 0.466464457 |
| ENSMUSG00000105034 | Gm43348 | 5.54E-05 | 3.648835551 | 0.466464457 |
| ENSMUSG00000102595 | Gm30097 | 5.54E-05 | 1.326907457 | 0.971854959 |

| | | | | |
|---|---|---|---|---|
| ENSMUSG00000033860 | Fgg | 5.54E-05 | 2.648835551 | 0.66416768 |
| ENSMUSG00000101334 | 1700028M03Rik | 5.54E-05 | 1.326907457 | 0.971854959 |
| ENSMUSG00000027996 | Sfrp2 | 5.54E-05 | 1.326907457 | 0.948823388 |
| ENSMUSG00000097250 | Gm26771 | 5.54E-05 | 0.841480629 | 1 |
| ENSMUSG00000085527 | Gm15535 | 5.54E-05 | 0.841480629 | 0.956297469 |
| ENSMUSG00000051000 | Fam160a1 | 5.54E-05 | -0.43862729 | 1 |
| ENSMUSG00000049013 | Prss48 | 5.54E-05 | 1.648835551 | 0.886203497 |
| ENSMUSG00000028071 | Sh2d2a | 5.54E-05 | 1.326907457 | 0.939707052 |
| ENSMUSG00000004892 | Bcan | 5.54E-05 | 1.648835551 | 0.886203497 |
| ENSMUSG00000097428 | AW047730 | 5.54E-05 | 3.648835551 | 0.466464457 |
| ENSMUSG00000097156 | Gm3764 | 5.54E-05 | 0.648835551 | 1 |
| ENSMUSG00000010538 | Tsacc | 5.54E-05 | -2.708716453 | 0.018639833 |
| ENSMUSG00000001420 | Tmem79 | 5.54E-05 | -0.995020638 | 0.699192777 |
| ENSMUSG00000074489 | Bglap3 | 5.54E-05 | 3.648835551 | 0.466464457 |
| ENSMUSG00000050144 | Slc25a44 | 5.54E-05 | -0.158519371 | 1 |
| ENSMUSG00000103115 | Gm37584 | 5.54E-05 | 1.648835551 | 0.886203497 |
| ENSMUSG00000028057 | Rit1 | 5.54E-05 | -2.184054463 | 0.091007367 |
| ENSMUSG00000028051 | Hcn3 | 5.54E-05 | 2.063873051 | 0.792441218 |
| ENSMUSG00000028048 | Gba | 5.54E-05 | -1.395558568 | 0.508263132 |
| ENSMUSG00000086389 | Gm15998 | 5.54E-05 | 1.648835551 | 0.886203497 |
| ENSMUSG00000027950 | Chrnb2 | 5.54E-05 | 2.648835551 | 0.66416768 |
| ENSMUSG00000046280 | She | 5.54E-05 | 0.47891055 | 1 |
| ENSMUSG00000042404 | Dennd4b | 5.54E-05 | -0.743481871 | 0.827142217 |
| ENSMUSG00000027932 | Slc27a3 | 5.54E-05 | 0.841480629 | 1 |
| ENSMUSG00000001021 | S100a3 | 5.54E-05 | 0.47891055 | 0.999962098 |
| ENSMUSG00000068893 | Sprr2a2 | 5.54E-05 | 2.648835551 | 0.66416768 |
| ENSMUSG00000050635 | Sprr2f | 5.54E-05 | 3.648835551 | 0.466464457 |
| ENSMUSG00000042165 | Gm9774 | 5.54E-05 | 0.841480629 | 1 |
| ENSMUSG00000042031 | Lce3b | 5.54E-05 | 3.648835551 | 0.466464457 |
| ENSMUSG00000102439 | Flg | 5.54E-05 | 2.648835551 | 0.66416768 |
| ENSMUSG00000074424 | Gm10696 | 5.54E-05 | 3.648835551 | 0.466464457 |
| ENSMUSG00000028141 | Oaz3 | 5.54E-05 | 2.648835551 | 0.66416768 |
| ENSMUSG00000028139 | Riiad1 | 5.54E-05 | -1.9057533 | 0.203406755 |
| ENSMUSG00000105412 | Gm43768 | 5.54E-05 | 1.063873051 | 1 |
| ENSMUSG00000084846 | A730011C13Rik | 5.54E-05 | 1.063873051 | 1 |
| ENSMUSG00000086288 | Gm15265 | 5.54E-05 | -1.158519371 | 0.599135879 |
| ENSMUSG00000053769 | Lysmd1 | 5.54E-05 | -1.051604167 | 0.706929043 |
| ENSMUSG00000097167 | Gm16740 | 5.54E-05 | 0.841480629 | 1 |
| ENSMUSG00000028109 | Hormad1 | 5.54E-05 | 1.326907457 | 0.971854959 |
| ENSMUSG00000097189 | Gm26594 | 5.54E-05 | 1.326907457 | 0.971854959 |
| ENSMUSG00000015745 | Plekho1 | 5.54E-05 | -1.52108945 | 0.453994425 |
| ENSMUSG00000093507 | Gm20627 | 5.54E-05 | 2.648835551 | 0.66416768 |
| ENSMUSG00000046317 | BC107364 | 5.54E-05 | 1.648835551 | 0.886203497 |
| ENSMUSG00000074398 | Gm15441 | 5.54E-05 | 1.648835551 | 0.886203497 |
| ENSMUSG00000049097 | Ankrd34a | 5.54E-05 | 2.648835551 | 0.66416768 |
| ENSMUSG00000028102 | Pex11b | 5.54E-05 | -0.258055044 | 1 |
| ENSMUSG00000028101 | Pias3 | 5.54E-05 | -1.43862729 | 0.433313155 |
| ENSMUSG00000104766 | 4930573H18Rik | 5.54E-05 | 0.841480629 | 1 |
| ENSMUSG00000086557 | Gm15999 | 5.54E-05 | 1.648835551 | 0.872990655 |
| ENSMUSG00000027876 | Reg4 | 5.54E-05 | 0.47891055 | 1 |
| ENSMUSG00000027875 | Hmgcs2 | 5.54E-05 | -0.051604167 | 1 |

| | | | | |
|---|---|---|---|---|
| ENSMUSG00000095388 | Gm10681 | 5.54E-05 | 1.326907457 | 0.971854959 |
| ENSMUSG00000063730 | Hsd3b2 | 5.54E-05 | 0.841480629 | 1 |
| ENSMUSG00000062410 | Hsd3b3 | 5.54E-05 | 1.063873051 | 1 |
| ENSMUSG00000105689 | Gm43120 | 5.54E-05 | 3.648835551 | 0.466464457 |
| ENSMUSG00000104639 | Gm42939 | 5.54E-05 | 0.648835551 | 1 |
| ENSMUSG00000106070 | Gm43135 | 5.54E-05 | 3.648835551 | 0.466464457 |
| ENSMUSG00000027861 | Casq2 | 5.54E-05 | 0.648835551 | 1 |
| ENSMUSG00000104925 | Gm43061 | 5.54E-05 | -1.636566667 | 0.373232174 |
| ENSMUSG00000085754 | Gm15886 | 5.54E-05 | 1.063873051 | 1 |
| ENSMUSG00000027845 | Dclre1b | 5.54E-05 | -1.52108945 | 0.466464457 |
| ENSMUSG00000089998 | Phtf1os | 5.54E-05 | 2.063873051 | 0.792441218 |
| ENSMUSG00000105083 | Gm42699 | 5.54E-05 | -1.843017545 | 0.255869172 |
| ENSMUSG00000105933 | Gm43260 | 5.54E-05 | 1.326907457 | 0.971854959 |
| ENSMUSG00000022187 | Gm5546 | 5.54E-05 | 1.063873051 | 1 |
| ENSMUSG00000100188 | Gm29561 | 5.54E-05 | 3.648835551 | 0.466464457 |
| ENSMUSG00000048458 | Inka2 | 5.54E-05 | 2.063873051 | 0.786367309 |
| ENSMUSG00000106317 | Gm43848 | 5.54E-05 | 3.648835551 | 0.466464457 |
| ENSMUSG00000104910 | Gm43331 | 5.54E-05 | 2.063873051 | 0.792441218 |
| ENSMUSG00000106093 | Gm42722 | 5.54E-05 | 0.841480629 | 1 |
| ENSMUSG00000063779 | Chil4 | 5.54E-05 | 2.063873051 | 0.792441218 |
| ENSMUSG00000105102 | Gm35507 | 5.54E-05 | -1.258055044 | 0.616879894 |
| ENSMUSG00000027895 | Kcnc4 | 5.54E-05 | 1.063873051 | 1 |
| ENSMUSG00000106426 | Gm36211 | 5.54E-05 | 2.063873051 | 0.772099175 |
| ENSMUSG00000087357 | Gm12498 | 5.54E-05 | 2.648835551 | 0.66416768 |
| ENSMUSG00000048997 | Atxn7l2 | 5.54E-05 | -0.599091962 | 0.980745941 |
| ENSMUSG00000105961 | Gm40123 | 5.54E-05 | 1.648835551 | 0.871433572 |
| ENSMUSG00000027884 | Clcc1 | 5.54E-05 | -2.184054463 | 0.091007367 |
| ENSMUSG00000027880 | Slc25a54 | 5.54E-05 | 1.326907457 | 0.971854959 |
| ENSMUSG00000045092 | S1pr1 | 5.54E-05 | 0.326907457 | 1 |
| ENSMUSG00000054426 | A930005H10Rik | 5.54E-05 | -0.258055044 | 1 |
| ENSMUSG00000027961 | Lrrc39 | 5.54E-05 | 0.841480629 | 1 |
| ENSMUSG00000027959 | Sass6 | 5.54E-05 | -2.417253639 | 0.054636426 |
| ENSMUSG00000033342 | Plppr5 | 5.54E-05 | 0.47891055 | 1 |
| ENSMUSG00000097311 | Gm26871 | 5.54E-05 | 0.326907457 | 1 |
| ENSMUSG00000104554 | Gm4610 | 5.54E-05 | 2.063873051 | 0.792441218 |
| ENSMUSG00000106583 | 4930447N08Rik | 5.54E-05 | 0.648835551 | 1 |
| ENSMUSG00000106423 | Gm42825 | 5.54E-05 | 0.648835551 | 1 |
| ENSMUSG00000032854 | Ugt8a | 5.54E-05 | -0.158519371 | 1 |
| ENSMUSG00000046688 | Tifa | 5.54E-05 | -0.673092543 | 0.98880775 |
| ENSMUSG00000105527 | 4933424H11Rik | 5.54E-05 | 3.648835551 | 0.466464457 |
| ENSMUSG00000028023 | Pitx2 | 5.54E-05 | 2.063873051 | 0.792441218 |
| ENSMUSG00000105837 | Gm35986 | 5.54E-05 | 1.326907457 | 0.971854959 |
| ENSMUSG00000106120 | Gm42697 | 5.54E-05 | 3.648835551 | 0.466464457 |
| ENSMUSG00000105596 | Gm42650 | 5.54E-05 | 3.648835551 | 0.466464457 |
| ENSMUSG00000106086 | Lef1os1 | 5.54E-05 | 3.648835551 | 0.466464457 |
| ENSMUSG00000091721 | Gimd1 | 5.54E-05 | -1.810596067 | 0.246776726 |
| ENSMUSG00000028016 | Ints12 | 5.54E-05 | -1.395558568 | 0.508263132 |
| ENSMUSG00000104645 | Gm30484 | 5.54E-05 | 3.648835551 | 0.466464457 |
| ENSMUSG00000106139 | Gm30648 | 5.54E-05 | 2.648835551 | 0.66416768 |
| ENSMUSG00000044365 | Cxxc4 | 5.54E-05 | -0.673092543 | 0.98880775 |
| ENSMUSG00000037994 | Slc9b2 | 5.54E-05 | -0.874726405 | 0.870548115 |

| | | | | |
|---|---|---|---|---|
| ENSMUSG00000099508 | 1700030L20Rik | 5.54E-05 | 2.648835551 | 0.66416768 |
| ENSMUSG00000104610 | Gm567 | 5.54E-05 | 2.648835551 | 0.66416768 |
| ENSMUSG00000049349 | Gm5105 | 5.54E-05 | 2.063873051 | 0.792441218 |
| ENSMUSG00000028158 | Mttp | 5.54E-05 | -1.52108945 | 0.452184898 |
| ENSMUSG00000012042 | 4930579F01Rik | 5.54E-05 | 0.47891055 | 1 |
| ENSMUSG00000105777 | Gm43049 | 5.54E-05 | 2.063873051 | 0.780656988 |
| ENSMUSG00000106550 | Gm43646 | 5.54E-05 | 2.648835551 | 0.66416768 |
| ENSMUSG00000056025 | Clca3a1 | 5.54E-05 | 0.189403933 | 1 |
| ENSMUSG00000037033 | Clca3b | 5.54E-05 | 2.648835551 | 0.66416768 |
| ENSMUSG00000028255 | Clca1 | 5.54E-05 | 0.326907457 | 1 |
| ENSMUSG00000097183 | Gm17501 | 5.54E-05 | -0.351164449 | 1 |
| ENSMUSG00000036873 | 2410004B18Rik | 5.54E-05 | -1.051604167 | 0.706929043 |
| ENSMUSG00000036863 | Syde2 | 5.54E-05 | -1.158519371 | 0.791730654 |
| ENSMUSG00000011008 | Mcoln2 | 5.54E-05 | 0.063873051 | 1 |
| ENSMUSG00000105602 | Gm10636 | 5.54E-05 | 1.648835551 | 0.868869354 |
| ENSMUSG00000106023 | Gm36831 | 5.54E-05 | 1.326907457 | 0.966795599 |
| ENSMUSG00000028036 | Ptgfr | 5.54E-05 | 0.841480629 | 1 |
| ENSMUSG00000104909 | Gm43617 | 5.54E-05 | 3.648835551 | 0.466464457 |
| ENSMUSG00000047583 | Tyw3 | 5.54E-05 | 0.063873051 | 1 |
| ENSMUSG00000078161 | Erich3 | 5.54E-05 | -0.051604167 | 1 |
| ENSMUSG00000085084 | 4930570G19Rik | 5.54E-05 | 2.648835551 | 0.66416768 |
| ENSMUSG00000086708 | Gm15577 | 5.54E-05 | 2.063873051 | 0.789505358 |
| ENSMUSG00000028179 | Cth | 5.54E-05 | -0.158519371 | 1 |
| ENSMUSG00000028177 | 1810013D15Rik | 5.54E-05 | 1.326907457 | 0.971854959 |
| ENSMUSG00000003282 | Plag1 | 5.54E-05 | -1.708716453 | 0.296030202 |
| ENSMUSG00000045573 | Penk | 5.54E-05 | -1.673092543 | 0.466464457 |
| ENSMUSG00000086066 | 4930423M02Rik | 5.54E-05 | 1.648835551 | 0.886203497 |
| ENSMUSG00000028243 | Ubxn2b | 5.54E-05 | -0.743481871 | 0.827142217 |
| ENSMUSG00000087564 | Gm11817 | 5.54E-05 | 0.841480629 | 1 |
| ENSMUSG00000085558 | 4930412C18Rik | 5.54E-05 | -0.351164449 | 1 |
| ENSMUSG00000059482 | 2610301B20Rik | 5.54E-05 | 0.841480629 | 1 |
| ENSMUSG00000049969 | Plekhf2 | 5.54E-05 | -2.079084903 | 0.13833117 |
| ENSMUSG00000028212 | Ccne2 | 5.54E-05 | 0.47891055 | 1 |
| ENSMUSG00000085027 | Gm11840 | 5.54E-05 | 2.648835551 | 0.66416768 |
| ENSMUSG00000043252 | Tmem64 | 5.54E-05 | -0.810596067 | 0.844591606 |
| ENSMUSG00000041135 | Ripk2 | 5.54E-05 | -1.743481871 | 0.305378889 |
| ENSMUSG00000041052 | Slc7a13 | 5.54E-05 | 1.063873051 | 1 |
| ENSMUSG00000073988 | Ttpa | 5.54E-05 | 1.326907457 | 0.958189678 |
| ENSMUSG00000028251 | Tstd3 | 5.54E-05 | 0.648835551 | 1 |
| ENSMUSG00000028246 | Faxc | 5.54E-05 | -1.051604167 | 0.736456705 |
| ENSMUSG00000040410 | Fbxl4 | 5.54E-05 | -2.743481871 | 0.015417072 |
| ENSMUSG00000046219 | C230012O17Rik | 5.54E-05 | 3.648835551 | 0.466464457 |
| ENSMUSG00000028259 | Fhl5 | 5.54E-05 | 0.648835551 | 1 |
| ENSMUSG00000040520 | Manea | 5.54E-05 | -0.52108945 | 0.975624465 |
| ENSMUSG00000086150 | Bach2os | 5.54E-05 | -0.43862729 | 1 |
| ENSMUSG00000045854 | Lyrm2 | 5.54E-05 | -0.810596067 | 0.832285197 |
| ENSMUSG00000028280 | Gabrr1 | 5.54E-05 | 3.648835551 | 0.466464457 |
| ENSMUSG00000028294 | Cfap206 | 5.54E-05 | 1.326907457 | 0.956389871 |
| ENSMUSG00000071015 | Gm136 | 5.54E-05 | 2.648835551 | 0.66416768 |
| ENSMUSG00000028295 | Smim8 | 5.54E-05 | 0.841480629 | 1 |
| ENSMUSG00000084898 | Gm12371 | 5.54E-05 | 1.326907457 | 0.971854959 |

| | | | | |
|---|---|---|---|---|
| ENSMUSG00000086026 | Gm12374 | 5.54E-05 | 1.326907457 | 0.971854959 |
| ENSMUSG00000090839 | Gm17094 | 5.54E-05 | 1.326907457 | 0.971854959 |
| ENSMUSG00000028441 | 1110017D15Rik | 5.54E-05 | 0.326907457 | 1 |
| ENSMUSG00000061322 | Dnaic1 | 5.54E-05 | 0.063873051 | 1 |
| ENSMUSG00000028445 | Enho | 5.54E-05 | 1.326907457 | 0.948148009 |
| ENSMUSG00000036073 | Galt | 5.54E-05 | 0.189403933 | 1 |
| ENSMUSG00000036062 | Phf24 | 5.54E-05 | 0.648835551 | 1 |
| ENSMUSG00000028451 | 1700022I11Rik | 5.54E-05 | -1.209145444 | 0.66280018 |
| ENSMUSG00000042788 | Fam166b | 5.54E-05 | 2.648835551 | 0.66416768 |
| ENSMUSG00000087142 | Gm12454 | 5.54E-05 | 1.326907457 | 0.946281031 |
| ENSMUSG00000028475 | Spaar | 5.54E-05 | 1.326907457 | 0.971854959 |
| ENSMUSG00000086211 | Gm12462 | 5.54E-05 | 0.648835551 | 1 |
| ENSMUSG00000035637 | Grhpr | 5.54E-05 | -0.158519371 | 1 |
| ENSMUSG00000070997 | 1700055D18Rik | 5.54E-05 | 1.326907457 | 0.971854959 |
| ENSMUSG00000028318 | Polr1e | 5.54E-05 | -0.258055044 | 1 |
| ENSMUSG00000048232 | Fbxo10 | 5.54E-05 | -0.52108945 | 0.975624465 |
| ENSMUSG00000028327 | Stra6l | 5.54E-05 | -0.158519371 | 1 |
| ENSMUSG00000028331 | Trmo | 5.54E-05 | -0.52108945 | 0.975624465 |
| ENSMUSG00000028332 | Hemgn | 5.54E-05 | 0.063873051 | 1 |
| ENSMUSG00000039740 | Alg2 | 5.54E-05 | -0.874726405 | 0.840804483 |
| ENSMUSG00000085660 | Gm12434 | 5.54E-05 | 2.648835551 | 0.66416768 |
| ENSMUSG00000060317 | Acnat2 | 5.54E-05 | 0.189403933 | 1 |
| ENSMUSG00000039634 | Zfp189 | 5.54E-05 | 0.47891055 | 1 |
| ENSMUSG00000085515 | C630028M04Rik | 5.54E-05 | 0.326907457 | 1 |
| ENSMUSG00000087192 | 4930412L05Rik | 5.54E-05 | 2.063873051 | 0.792441218 |
| ENSMUSG00000085375 | Gm12506 | 5.54E-05 | -0.258055044 | 1 |
| ENSMUSG00000045589 | Frrs1l | 5.54E-05 | 0.326907457 | 1 |
| ENSMUSG00000085654 | Gm12536 | 5.54E-05 | -0.995020638 | 0.699192777 |
| ENSMUSG00000086952 | Gm12596 | 5.54E-05 | 1.326907457 | 0.951907087 |
| ENSMUSG00000078688 | Mup2 | 5.54E-05 | -0.743481871 | 0.827142217 |
| ENSMUSG00000028391 | Wdr31 | 5.54E-05 | -1.158519371 | 0.599135879 |
| ENSMUSG00000028356 | Ambp | 5.54E-05 | 1.648835551 | 0.886203497 |
| ENSMUSG00000073821 | 8030451A03Rik | 5.54E-05 | 0.189403933 | 1 |
| ENSMUSG00000085032 | Gm11217 | 5.54E-05 | 0.841480629 | 1 |
| ENSMUSG00000051675 | Trim32 | 5.54E-05 | 0.189403933 | 1 |
| ENSMUSG00000086364 | Gm11751 | 5.54E-05 | -0.43862729 | 1 |
| ENSMUSG00000086696 | Gm11234 | 5.54E-05 | 2.648835551 | 0.66416768 |
| ENSMUSG00000084910 | C630043F03Rik | 5.54E-05 | 0.648835551 | 1 |
| ENSMUSG00000028396 | 2310002L09Rik | 5.54E-05 | 1.063873051 | 1 |
| ENSMUSG00000085022 | Gm5860 | 5.54E-05 | 1.648835551 | 0.875465725 |
| ENSMUSG00000087413 | Gm11266 | 5.54E-05 | -1.209145444 | 0.606122355 |
| ENSMUSG00000117251 | Gm49891 | 5.54E-05 | 0.063873051 | 1 |
| ENSMUSG00000087659 | Gm12606 | 5.54E-05 | 1.063873051 | 1 |
| ENSMUSG00000085209 | Gm12637 | 5.54E-05 | 2.648835551 | 0.66416768 |
| ENSMUSG00000054000 | Tusc1 | 5.54E-05 | 0.47891055 | 1 |
| ENSMUSG00000049799 | Lrrc19 | 5.54E-05 | 1.648835551 | 0.886203497 |
| ENSMUSG00000081362 | Cyp2j7 | 5.54E-05 | 0.841480629 | 1 |
| ENSMUSG00000082932 | Cyp2j8 | 5.54E-05 | -0.158519371 | 1 |
| ENSMUSG00000052520 | Cyp2j5 | 5.54E-05 | 0.841480629 | 1 |
| ENSMUSG00000078639 | Gm12695 | 5.54E-05 | -0.52108945 | 1 |
| ENSMUSG00000070891 | Gm12689 | 5.54E-05 | -0.351164449 | 1 |

| | | | | |
|---|---|---|---|---|
| ENSMUSG00000073791 | Efcab7 | 5.54E-05 | -0.995020638 | 0.699192777 |
| ENSMUSG00000035275 | Raver2 | 5.54E-05 | 0.648835551 | 1 |
| ENSMUSG00000086782 | E130102H24Rik | 5.54E-05 | -0.936126949 | 0.695372548 |
| ENSMUSG00000035201 | B020004J07Rik | 5.54E-05 | -0.599091962 | 0.989737947 |
| ENSMUSG00000046133 | C130073F10Rik | 5.54E-05 | 1.648835551 | 0.886203497 |
| ENSMUSG00000078626 | Gm12790 | 5.54E-05 | 2.648835551 | 0.66416768 |
| ENSMUSG00000028523 | Tctex1d1 | 5.54E-05 | 1.063873051 | 1 |
| ENSMUSG00000035126 | Wdr78 | 5.54E-05 | -1.480447466 | 0.441576406 |
| ENSMUSG00000078618 | 4930456L15Rik | 5.54E-05 | 1.648835551 | 0.886203497 |
| ENSMUSG00000095386 | Gm17662 | 5.54E-05 | 0.648835551 | 1 |
| ENSMUSG00000078612 | Fyb2 | 5.54E-05 | -2.373532262 | 0.07714516 |
| ENSMUSG00000044254 | Pcsk9 | 5.54E-05 | 2.648835551 | 0.66416768 |
| ENSMUSG00000047502 | Mroh7 | 5.54E-05 | -0.051604167 | 1 |
| ENSMUSG00000047636 | Cdcp2 | 5.54E-05 | 2.063873051 | 0.792441218 |
| ENSMUSG00000085304 | Gm12802 | 5.54E-05 | 1.648835551 | 0.886203497 |
| ENSMUSG00000087200 | Lrp8os3 | 5.54E-05 | -0.158519371 | 1 |
| ENSMUSG00000028600 | Podn | 5.54E-05 | 0.189403933 | 1 |
| ENSMUSG00000034645 | Zyg11a | 5.54E-05 | 0.326907457 | 1 |
| ENSMUSG00000028597 | Gpx7 | 5.54E-05 | 1.063873051 | 1 |
| ENSMUSG00000063800 | Prpf38a | 5.54E-05 | -1.209145444 | 0.606122355 |
| ENSMUSG00000073775 | Kti12 | 5.54E-05 | -0.258055044 | 1 |
| ENSMUSG00000060491 | 4930522H14Rik | 5.54E-05 | 2.063873051 | 0.792441218 |
| ENSMUSG00000028551 | Cdkn2c | 5.54E-05 | -0.810596067 | 0.832285197 |
| ENSMUSG00000028544 | Slc5a9 | 5.54E-05 | -0.158519371 | 1 |
| ENSMUSG00000089773 | Skint1 | 5.54E-05 | 2.648835551 | 0.66416768 |
| ENSMUSG00000057977 | Skint11 | 5.54E-05 | 1.063873051 | 1 |
| ENSMUSG00000055198 | Gm12830 | 5.54E-05 | 1.063873051 | 1 |
| ENSMUSG00000047155 | Cyp4x1 | 5.54E-05 | -0.258055044 | 1 |
| ENSMUSG00000066071 | Cyp4a12a | 5.54E-05 | 1.326907457 | 0.95526509 |
| ENSMUSG00000028713 | Cyp4b1 | 5.54E-05 | 0.648835551 | 1 |
| ENSMUSG00000044556 | Tex38 | 5.54E-05 | 3.648835551 | 0.466464457 |
| ENSMUSG00000034171 | Faah | 5.54E-05 | -1.258055044 | 0.616879894 |
| ENSMUSG00000028702 | Rad54l | 5.54E-05 | -1.351164449 | 0.499088689 |
| ENSMUSG00000028689 | Ccdc163 | 5.54E-05 | -1.395558568 | 0.508263132 |
| ENSMUSG00000028684 | Urod | 5.54E-05 | -0.258055044 | 1 |
| ENSMUSG00000028681 | Ptch2 | 5.54E-05 | 0.648835551 | 1 |
| ENSMUSG00000028680 | Plk3 | 5.54E-05 | -0.158519371 | 1 |
| ENSMUSG00000086876 | Gm12828 | 5.54E-05 | 2.648835551 | 0.66416768 |
| ENSMUSG00000009640 | Dmap1 | 5.54E-05 | -0.743481871 | 0.827142217 |
| ENSMUSG00000091237 | Gm17114 | 5.54E-05 | 1.063873051 | 0.986886956 |
| ENSMUSG00000028540 | Dph2 | 5.54E-05 | 0.326907457 | 1 |
| ENSMUSG00000050854 | Tmem125 | 5.54E-05 | 2.063873051 | 0.792441218 |
| ENSMUSG00000070821 | Olfr1340 | 5.54E-05 | 3.648835551 | 0.466464457 |
| ENSMUSG00000073769 | Olfr1331 | 5.54E-05 | 2.063873051 | 0.792441218 |
| ENSMUSG00000066060 | Gm12866 | 5.54E-05 | -1.158519371 | 0.599135879 |
| ENSMUSG00000045268 | Zfp691 | 5.54E-05 | 0.326907457 | 1 |
| ENSMUSG00000086187 | Gm12860 | 5.54E-05 | -0.810596067 | 0.854682219 |
| ENSMUSG00000070803 | Cited4 | 5.54E-05 | 1.648835551 | 0.860176358 |
| ENSMUSG00000028629 | Exo5 | 5.54E-05 | -1.209145444 | 0.606122355 |
| ENSMUSG00000064141 | Zfp69 | 5.54E-05 | -0.743481871 | 0.827142217 |
| ENSMUSG00000078576 | Gm12886 | 5.54E-05 | 3.648835551 | 0.466464457 |

| | | | | |
|---|---|---|---|---|
| ENSMUSG00000078575 | Gm12887 | 5.54E-05 | 3.648835551 | 0.466464457 |
| ENSMUSG00000028655 | Mfsd2a | 5.54E-05 | 0.47891055 | 1 |
| ENSMUSG00000028653 | Trit1 | 5.54E-05 | -2.281901786 | 0.078768438 |
| ENSMUSG00000002384 | Bmp8b | 5.54E-05 | -0.43862729 | 1 |
| ENSMUSG00000076438 | Oxct2b | 5.54E-05 | 3.648835551 | 0.466464457 |
| ENSMUSG00000046093 | Hpcal4 | 5.54E-05 | 0.063873051 | 1 |
| ENSMUSG00000086275 | 1700121C08Rik | 5.54E-05 | 1.648835551 | 0.868559276 |
| ENSMUSG00000073761 | 4933427I04Rik | 5.54E-05 | 3.648835551 | 0.466464457 |
| ENSMUSG00000086978 | 4933435F18Rik | 5.54E-05 | 3.648835551 | 0.466464457 |
| ENSMUSG00000073759 | 4933407E24Rik | 5.54E-05 | -0.810596067 | 0.847265357 |
| ENSMUSG00000028889 | Yrdc | 5.54E-05 | -0.43862729 | 1 |
| ENSMUSG00000085263 | 1700125G02Rik | 5.54E-05 | 2.648835551 | 0.66416768 |
| ENSMUSG00000044730 | 9930104L06Rik | 5.54E-05 | 0.063873051 | 1 |
| ENSMUSG00000087011 | Gm12930 | 5.54E-05 | 0.47891055 | 1 |
| ENSMUSG00000085334 | Gm12940 | 5.54E-05 | -2.184054463 | 0.091007367 |
| ENSMUSG00000070737 | Tmem35b | 5.54E-05 | 1.648835551 | 0.886203497 |
| ENSMUSG00000042367 | Gjb3 | 5.54E-05 | -0.051604167 | 1 |
| ENSMUSG00000042357 | Gjb5 | 5.54E-05 | 1.063873051 | 1 |
| ENSMUSG00000087470 | A630031M04Rik | 5.54E-05 | 2.063873051 | 0.792441218 |
| ENSMUSG00000028794 | A3galt2 | 5.54E-05 | -0.810596067 | 0.832285197 |
| ENSMUSG00000085701 | Gm12968 | 5.54E-05 | 0.47891055 | 1 |
| ENSMUSG00000085961 | Gm15904 | 5.54E-05 | 2.063873051 | 0.792441218 |
| ENSMUSG00000087575 | Gm12976 | 5.54E-05 | 0.841480629 | 1 |
| ENSMUSG00000078554 | Fam229a | 5.54E-05 | 3.648835551 | 0.466464457 |
| ENSMUSG00000086336 | Gm12979 | 5.54E-05 | 0.326907457 | 1 |
| ENSMUSG00000000409 | Lck | 5.54E-05 | 0.648835551 | 1 |
| ENSMUSG00000028795 | Ccdc28b | 5.54E-05 | 0.189403933 | 1 |
| ENSMUSG00000085517 | Gm12963 | 5.54E-05 | 0.841480629 | 1 |
| ENSMUSG00000028773 | Fabp3 | 5.54E-05 | 1.648835551 | 0.886203497 |
| ENSMUSG00000078531 | Gm831 | 5.54E-05 | 2.063873051 | 0.792441218 |
| ENSMUSG00000087171 | Gm12962 | 5.54E-05 | 1.648835551 | 0.878674585 |
| ENSMUSG00000056529 | Ptafr | 5.54E-05 | -0.43862729 | 1 |
| ENSMUSG00000028885 | Smpdl3b | 5.54E-05 | -0.43862729 | 1 |
| ENSMUSG00000028884 | Rpa2 | 5.54E-05 | -1.351164449 | 0.514824729 |
| ENSMUSG00000037731 | Themis2 | 5.54E-05 | -0.52108945 | 1 |
| ENSMUSG00000046694 | Tent5b | 5.54E-05 | 1.648835551 | 0.886203497 |
| ENSMUSG00000056596 | Trnp1 | 5.54E-05 | -0.874726405 | 0.840804483 |
| ENSMUSG00000028850 | Gpatch3 | 5.54E-05 | -1.743481871 | 0.305378889 |
| ENSMUSG00000012123 | Crybg2 | 5.54E-05 | 2.063873051 | 0.792441218 |
| ENSMUSG00000028838 | Extl1 | 5.54E-05 | 0.47891055 | 1 |
| ENSMUSG00000087343 | 1700021N21Rik | 5.54E-05 | 3.648835551 | 0.466464457 |
| ENSMUSG00000078521 | Aunip | 5.54E-05 | -0.351164449 | 1 |
| ENSMUSG00000094035 | Ldlrad2 | 5.54E-05 | 0.326907457 | 1 |
| ENSMUSG00000046109 | AB041806 | 5.54E-05 | 2.648835551 | 0.66416768 |
| ENSMUSG00000028737 | Aldh4a1 | 5.54E-05 | -0.599091962 | 1 |
| ENSMUSG00000078234 | Klhdc7a | 5.54E-05 | 1.326907457 | 0.954844735 |
| ENSMUSG00000040935 | Padi6 | 5.54E-05 | 3.648835551 | 0.466464457 |
| ENSMUSG00000033770 | Clcnka | 5.54E-05 | 2.648835551 | 0.66416768 |
| ENSMUSG00000006216 | Clcnkb | 5.54E-05 | 3.648835551 | 0.466464457 |
| ENSMUSG00000006215 | Zbtb17 | 5.54E-05 | -2.184054463 | 0.091007367 |
| ENSMUSG00000085922 | Fhad1os2 | 5.54E-05 | 1.648835551 | 0.872893507 |

| | | | | |
|---|---|---|---|---|
| ENSMUSG00000073721 | Pramef20 | 5.54E-05 | 2.063873051 | 0.792441218 |
| ENSMUSG00000028591 | Pramef12 | 5.54E-05 | 2.063873051 | 0.792441218 |
| ENSMUSG00000028589 | 1700012P22Rik | 5.54E-05 | 1.063873051 | 1 |
| ENSMUSG00000095213 | Gm9944 | 5.54E-05 | 3.648835551 | 0.466464457 |
| ENSMUSG00000041735 | Gm13178 | 5.54E-05 | 2.063873051 | 0.792441218 |
| ENSMUSG00000078504 | Gm438 | 5.54E-05 | 3.648835551 | 0.466464457 |
| ENSMUSG00000085101 | Platr16 | 5.54E-05 | 0.841480629 | 1 |
| ENSMUSG00000078500 | Zfp986 | 5.54E-05 | 3.648835551 | 0.466464457 |
| ENSMUSG00000066009 | Zfp987 | 5.54E-05 | 2.648835551 | 0.66416768 |
| ENSMUSG00000085525 | Gm13166 | 5.54E-05 | 1.326907457 | 0.949989126 |
| ENSMUSG00000070605 | Zfp992 | 5.54E-05 | -0.351164449 | 1 |
| ENSMUSG00000086147 | Zfp989 | 5.54E-05 | 2.648835551 | 0.66416768 |
| ENSMUSG00000093617 | 4933438K21Rik | 5.54E-05 | -0.673092543 | 0.98880775 |
| ENSMUSG00000067916 | Zfp991 | 5.54E-05 | 1.648835551 | 0.886203497 |
| ENSMUSG00000078498 | Zfp988 | 5.54E-05 | 1.648835551 | 0.886203497 |
| ENSMUSG00000078496 | Zfp982 | 5.54E-05 | 1.063873051 | 1 |
| ENSMUSG00000093594 | Gm20707 | 5.54E-05 | 3.648835551 | 0.466464457 |
| ENSMUSG00000029022 | Miip | 5.54E-05 | -0.743481871 | 1 |
| ENSMUSG00000029001 | Fbxo44 | 5.54E-05 | 0.189403933 | 1 |
| ENSMUSG00000028988 | Ctnnbip1 | 5.54E-05 | -1.9057533 | 0.203406755 |
| ENSMUSG00000047875 | Gpr157 | 5.54E-05 | 2.063873051 | 0.792441218 |
| ENSMUSG00000028976 | Slc2a5 | 5.54E-05 | 0.648835551 | 1 |
| ENSMUSG00000028972 | Car6 | 5.54E-05 | 2.063873051 | 0.792441218 |
| ENSMUSG00000085593 | 4930589P08Rik | 5.54E-05 | 3.648835551 | 0.466464457 |
| ENSMUSG00000047777 | Phf13 | 5.54E-05 | -0.743481871 | 0.827142217 |
| ENSMUSG00000042804 | Gpr153 | 5.54E-05 | -1.480447466 | 0.466464457 |
| ENSMUSG00000005045 | Chd5 | 5.54E-05 | 0.063873051 | 1 |
| ENSMUSG00000029027 | Dffb | 5.54E-05 | 0.47891055 | 1 |
| ENSMUSG00000029028 | Lrrc47 | 5.54E-05 | -1.43862729 | 0.433313155 |
| ENSMUSG00000087071 | Trp73os | 5.54E-05 | 2.648835551 | 0.66416768 |
| ENSMUSG00000029029 | Wrap73 | 5.54E-05 | -0.673092543 | 1 |
| ENSMUSG00000097871 | B230104I21Rik | 5.54E-05 | 2.648835551 | 0.66416768 |
| ENSMUSG00000086810 | Gm13110 | 5.54E-05 | 2.648835551 | 0.66416768 |
| ENSMUSG00000046637 | Ttc34 | 5.54E-05 | 2.648835551 | 0.66416768 |
| ENSMUSG00000029059 | Prxl2b | 5.54E-05 | 0.47891055 | 1 |
| ENSMUSG00000029060 | Mib2 | 5.54E-05 | -1.480447466 | 0.441576406 |
| ENSMUSG00000096221 | 1500002C15Rik | 5.54E-05 | 3.648835551 | 0.466464457 |
| ENSMUSG00000029073 | Cptp | 5.54E-05 | 0.841480629 | 1 |
| ENSMUSG00000029034 | Ints11 | 5.54E-05 | -2.051604167 | 0.143740392 |
| ENSMUSG00000023571 | C1qtnf12 | 5.54E-05 | 2.648835551 | 0.66416768 |
| ENSMUSG00000050796 | B3galt6 | 5.54E-05 | 0.47891055 | 1 |
| ENSMUSG00000059939 | 9430015G10Rik | 5.54E-05 | -2.158519371 | 0.115633409 |
| ENSMUSG00000078349 | AW011738 | 5.54E-05 | 0.189403933 | 1 |
| ENSMUSG00000014529 | Tmbim7 | 5.54E-05 | 2.648835551 | 0.66416768 |
| ENSMUSG00000040367 | Lrrd1 | 5.54E-05 | 1.063873051 | 1 |
| ENSMUSG00000106535 | Gm43031 | 5.54E-05 | 2.648835551 | 0.66416768 |
| ENSMUSG00000106472 | Gm43111 | 5.54E-05 | 2.063873051 | 0.792441218 |
| ENSMUSG00000040464 | Gtpbp10 | 5.54E-05 | -0.874726405 | 0.840804483 |
| ENSMUSG00000054099 | Slc25a40 | 5.54E-05 | -2.051604167 | 0.133549019 |
| ENSMUSG00000073226 | Gm10482 | 5.54E-05 | 1.648835551 | 0.874192437 |
| ENSMUSG00000104789 | Gm43640 | 5.54E-05 | 3.648835551 | 0.466464457 |

| | | | | |
|---|---|---|---|---|
| ENSMUSG00000099516 | Gm29066 | 5.54E-05 | 3.648835551 | 0.466464457 |
| ENSMUSG00000085352 | Gm4128 | 5.54E-05 | 1.648835551 | 0.872990655 |
| ENSMUSG00000097976 | Gm26918 | 5.54E-05 | 2.063873051 | 0.792441218 |
| ENSMUSG00000098233 | Gm26954 | 5.54E-05 | 1.063873051 | 1 |
| ENSMUSG00000089871 | Speer4cos | 5.54E-05 | -0.351164449 | 1 |
| ENSMUSG00000047221 | Fam185a | 5.54E-05 | -3.427980046 | 0.00094815 |
| ENSMUSG00000039883 | Lrrc17 | 5.54E-05 | 0.841480629 | 1 |
| ENSMUSG00000073152 | Gm10475 | 5.54E-05 | 2.648835551 | 0.66416768 |
| ENSMUSG00000105106 | Gm43657 | 5.54E-05 | 2.648835551 | 0.66416768 |
| ENSMUSG00000104866 | Gm42948 | 5.54E-05 | 2.063873051 | 0.792441218 |
| ENSMUSG00000105254 | Gm42951 | 5.54E-05 | 2.648835551 | 0.66416768 |
| ENSMUSG00000038319 | Kcnh2 | 5.54E-05 | 0.326907457 | 1 |
| ENSMUSG00000086769 | Gm15587 | 5.54E-05 | 1.648835551 | 0.882650692 |
| ENSMUSG00000055235 | Wdr86 | 5.54E-05 | -0.936126949 | 0.925002796 |
| ENSMUSG00000059631 | 1500035N22Rik | 5.54E-05 | 1.326907457 | 0.971854959 |
| ENSMUSG00000043833 | 2900005J15Rik | 5.54E-05 | 0.648835551 | 1 |
| ENSMUSG00000028938 | Galntl5 | 5.54E-05 | 0.47891055 | 1 |
| ENSMUSG00000059244 | Gm10062 | 5.54E-05 | -2.132524162 | 0.146946082 |
| ENSMUSG00000028933 | Xrcc2 | 5.54E-05 | 1.063873051 | 1 |
| ENSMUSG00000029130 | Rnf32 | 5.54E-05 | -2.305360759 | 0.082076736 |
| ENSMUSG00000067642 | Adgrf3 | 5.54E-05 | 1.648835551 | 0.886203497 |
| ENSMUSG00000029163 | Emilin1 | 5.54E-05 | -0.351164449 | 1 |
| ENSMUSG00000006642 | Tcf23 | 5.54E-05 | 2.063873051 | 0.792441218 |
| ENSMUSG00000053856 | Dnajc5g | 5.54E-05 | 0.648835551 | 1 |
| ENSMUSG00000062077 | Trim54 | 5.54E-05 | 2.648835551 | 0.66416768 |
| ENSMUSG00000043059 | Zfp513 | 5.54E-05 | -2.051604167 | 0.133549019 |
| ENSMUSG00000059434 | Gckr | 5.54E-05 | -0.673092543 | 0.98880775 |
| ENSMUSG00000064037 | Gpn1 | 5.54E-05 | -1.43862729 | 0.433313155 |
| ENSMUSG00000107352 | Gm43660 | 5.54E-05 | 2.063873051 | 0.790752525 |
| ENSMUSG00000011034 | Slc5a1 | 5.54E-05 | 0.47891055 | 1 |
| ENSMUSG00000037379 | Spon2 | 5.54E-05 | 1.063873051 | 1 |
| ENSMUSG00000092275 | Gm20465 | 5.54E-05 | 2.648835551 | 0.66416768 |
| ENSMUSG00000037355 | Uvssa | 5.54E-05 | -2.305360759 | 0.138139802 |
| ENSMUSG00000070858 | Gm1673 | 5.54E-05 | 2.063873051 | 0.792441218 |
| ENSMUSG00000106902 | 4930557J02Rik | 5.54E-05 | -0.936126949 | 0.882179532 |
| ENSMUSG00000079555 | Haus3 | 5.54E-05 | -2.373532262 | 0.051203786 |
| ENSMUSG00000109572 | Cfap99 | 5.54E-05 | 0.841480629 | 1 |
| ENSMUSG00000107287 | Gm43458 | 5.54E-05 | 1.326907457 | 0.948823388 |
| ENSMUSG00000001082 | Mfsd10 | 5.54E-05 | 0.063873051 | 1 |
| ENSMUSG00000029101 | Rgs12 | 5.54E-05 | -2.579983139 | 0.032789124 |
| ENSMUSG00000036596 | Cpz | 5.54E-05 | 0.47891055 | 1 |
| ENSMUSG00000039563 | 2210406O10Rik | 5.54E-05 | 0.47891055 | 1 |
| ENSMUSG00000039474 | Wfs1 | 5.54E-05 | -0.995020638 | 0.699192777 |
| ENSMUSG00000117286 | Gm1043 | 5.54E-05 | 1.648835551 | 0.886203497 |
| ENSMUSG00000048450 | Msx1 | 5.54E-05 | 1.648835551 | 0.886203497 |
| ENSMUSG00000046572 | Zfp518b | 5.54E-05 | -0.351164449 | 1 |
| ENSMUSG00000107193 | Gm43700 | 5.54E-05 | 2.648835551 | 0.66416768 |
| ENSMUSG00000106501 | Gm42711 | 5.54E-05 | 1.648835551 | 0.866187285 |
| ENSMUSG00000085720 | Gm7854 | 5.54E-05 | -0.673092543 | 1 |
| ENSMUSG00000039782 | Cpeb2 | 5.54E-05 | -2.051604167 | 0.133549019 |
| ENSMUSG00000104691 | Gm43021 | 5.54E-05 | 3.648835551 | 0.466464457 |

| | | | | |
|---|---|---|---|---|
| ENSMUSG00000087290 | Gm15866 | 5.54E-05 | 2.648835551 | 0.66416768 |
| ENSMUSG00000048373 | Fgfbp1 | 5.54E-05 | 2.648835551 | 0.66416768 |
| ENSMUSG00000105264 | Gm42716 | 5.54E-05 | 2.648835551 | 0.66416768 |
| ENSMUSG00000046985 | Tapt1 | 5.54E-05 | -2.82689788 | 0.011086379 |
| ENSMUSG00000104800 | Gm36017 | 5.54E-05 | 2.063873051 | 0.792441218 |
| ENSMUSG00000104894 | Gm43507 | 5.54E-05 | 1.648835551 | 0.886203497 |
| ENSMUSG00000105224 | Gm3364 | 5.54E-05 | 1.648835551 | 0.886203497 |
| ENSMUSG00000105255 | Gm42413 | 5.54E-05 | 1.063873051 | 1 |
| ENSMUSG00000015880 | Ncapg | 5.54E-05 | -0.673092543 | 0.98880775 |
| ENSMUSG00000072941 | Sod3 | 5.54E-05 | 0.841480629 | 1 |
| ENSMUSG00000106538 | Gm30301 | 5.54E-05 | 0.189403933 | 1 |
| ENSMUSG00000029188 | Slc34a2 | 5.54E-05 | 0.648835551 | 1 |
| ENSMUSG00000106981 | Gm45495 | 5.54E-05 | 1.326907457 | 0.941335619 |
| ENSMUSG00000105912 | Gm10440 | 5.54E-05 | 2.063873051 | 0.792441218 |
| ENSMUSG00000106978 | N4bp2os | 5.54E-05 | 0.648835551 | 1 |
| ENSMUSG00000012520 | Phox2b | 5.54E-05 | 3.648835551 | 0.466464457 |
| ENSMUSG00000085363 | Gm15478 | 5.54E-05 | 2.648835551 | 0.66416768 |
| ENSMUSG00000029211 | Gabra4 | 5.54E-05 | 1.326907457 | 0.971854959 |
| ENSMUSG00000093400 | Gm20647 | 5.54E-05 | 0.47891055 | 1 |
| ENSMUSG00000054892 | Txk | 5.54E-05 | -0.936126949 | 0.847320319 |
| ENSMUSG00000063656 | Gm10135 | 5.54E-05 | 1.648835551 | 0.879358566 |
| ENSMUSG00000100175 | 1700025M24Rik | 5.54E-05 | 2.648835551 | 0.66416768 |
| ENSMUSG00000087250 | Usp46os1 | 5.54E-05 | 1.063873051 | 1 |
| ENSMUSG00000086234 | Usp46os2 | 5.54E-05 | 1.326907457 | 0.946040281 |
| ENSMUSG00000029228 | Lnx1 | 5.54E-05 | -0.743481871 | 0.840624557 |
| ENSMUSG00000107192 | Gm19583 | 5.54E-05 | -0.810596067 | 0.847265357 |
| ENSMUSG00000106754 | Gm43101 | 5.54E-05 | 1.648835551 | 0.882179532 |
| ENSMUSG00000106756 | Gm32780 | 5.54E-05 | 2.063873051 | 0.792441218 |
| ENSMUSG00000087037 | 4930432L08Rik | 5.54E-05 | 3.648835551 | 0.466464457 |
| ENSMUSG00000055923 | Aasdh | 5.54E-05 | -2.305360759 | 0.082076736 |
| ENSMUSG00000085858 | Gm15831 | 5.54E-05 | 2.648835551 | 0.66416768 |
| ENSMUSG00000106985 | Gm42761 | 5.54E-05 | 2.063873051 | 0.792441218 |
| ENSMUSG00000106598 | Gm43532 | 5.54E-05 | 3.648835551 | 0.466464457 |
| ENSMUSG00000061259 | Tmprss11d | 5.54E-05 | -0.52108945 | 0.975624465 |
| ENSMUSG00000035861 | Tmprss11b | 5.54E-05 | 2.063873051 | 0.792441218 |
| ENSMUSG00000054537 | Tmprss11e | 5.54E-05 | 1.326907457 | 0.971854959 |
| ENSMUSG00000035811 | Ugt2b35 | 5.54E-05 | 1.648835551 | 0.886203497 |
| ENSMUSG00000070704 | Ugt2b36 | 5.54E-05 | 0.47891055 | 1 |
| ENSMUSG00000054630 | Ugt2b5 | 5.54E-05 | 2.063873051 | 0.792441218 |
| ENSMUSG00000061937 | Csn1s2a | 5.54E-05 | 2.063873051 | 0.792441218 |
| ENSMUSG00000029281 | Smr2 | 5.54E-05 | 1.648835551 | 0.878817358 |
| ENSMUSG00000064156 | Prol1 | 5.54E-05 | 3.648835551 | 0.466464457 |
| ENSMUSG00000067149 | Jchain | 5.54E-05 | 0.063873051 | 1 |
| ENSMUSG00000043635 | Adamts3 | 5.54E-05 | -1.051604167 | 0.74155121 |
| ENSMUSG00000054945 | Gm9958 | 5.54E-05 | 0.648835551 | 1 |
| ENSMUSG00000029368 | Alb | 5.54E-05 | -0.43862729 | 1 |
| ENSMUSG00000106788 | Gm43449 | 5.54E-05 | 2.648835551 | 0.66416768 |
| ENSMUSG00000029373 | Pf4 | 5.54E-05 | 0.189403933 | 1 |
| ENSMUSG00000029379 | Cxcl3 | 5.54E-05 | 0.841480629 | 1 |
| ENSMUSG00000029375 | Cxcl15 | 5.54E-05 | 1.648835551 | 0.873283731 |
| ENSMUSG00000029378 | Areg | 5.54E-05 | -1.52108945 | 0.452184898 |

| | | | | |
|---|---|---|---|---|
| ENSMUSG00000029410 | Ppef2 | 5.54E-05 | 0.648835551 | 1 |
| ENSMUSG00000096044 | Gm16427 | 5.54E-05 | 2.063873051 | 0.792441218 |
| ENSMUSG00000107392 | Gm7792 | 5.54E-05 | 3.648835551 | 0.466464457 |
| ENSMUSG00000072822 | BC061212 | 5.54E-05 | 3.648835551 | 0.466464457 |
| ENSMUSG00000073497 | AA792892 | 5.54E-05 | 2.648835551 | 0.66416768 |
| ENSMUSG00000023078 | Cxcl13 | 5.54E-05 | 2.648835551 | 0.66416768 |
| ENSMUSG00000046000 | Naa11 | 5.54E-05 | 2.063873051 | 0.792441218 |
| ENSMUSG00000067101 | 1700010H22Rik | 5.54E-05 | 2.063873051 | 0.792441218 |
| ENSMUSG00000050640 | Tmem150c | 5.54E-05 | -2.560617814 | 0.030770816 |
| ENSMUSG00000029319 | Coq2 | 5.54E-05 | -2.209145444 | 0.093610056 |
| ENSMUSG00000085269 | Gm15777 | 5.54E-05 | 2.063873051 | 0.792441218 |
| ENSMUSG00000035266 | Helq | 5.54E-05 | -1.995020638 | 0.167528217 |
| ENSMUSG00000035234 | Abraxas1 | 5.54E-05 | 0.063873051 | 1 |
| ENSMUSG00000087611 | 4930458D05Rik | 5.54E-05 | 2.063873051 | 0.792441218 |
| ENSMUSG00000035187 | Nkx6-1 | 5.54E-05 | 2.063873051 | 0.79083603 |
| ENSMUSG00000029330 | Cds1 | 5.54E-05 | -0.43862729 | 1 |
| ENSMUSG00000092354 | Gm20548 | 5.54E-05 | 3.648835551 | 0.466464457 |
| ENSMUSG00000106586 | 1700013M08Rik | 5.54E-05 | 3.648835551 | 0.466464457 |
| ENSMUSG00000104580 | Gm42620 | 5.54E-05 | 0.189403933 | 1 |
| ENSMUSG00000029312 | Klhl8 | 5.54E-05 | 0.063873051 | 1 |
| ENSMUSG00000034528 | Hsd17b13 | 5.54E-05 | 2.063873051 | 0.792441218 |
| ENSMUSG00000097877 | Gm26703 | 5.54E-05 | 2.648835551 | 0.66416768 |
| ENSMUSG00000053268 | Dspp | 5.54E-05 | 2.648835551 | 0.66416768 |
| ENSMUSG00000106438 | Gm32051 | 5.54E-05 | 2.648835551 | 0.66416768 |
| ENSMUSG00000099061 | Gm28050 | 5.54E-05 | -1.051604167 | 0.737916568 |
| ENSMUSG00000029283 | Cdc7 | 5.54E-05 | -1.52108945 | 0.452184898 |
| ENSMUSG00000029279 | Brdt | 5.54E-05 | -0.810596067 | 0.832285197 |
| ENSMUSG00000033805 | Ephx4 | 5.54E-05 | -0.52108945 | 0.975624465 |
| ENSMUSG00000089798 | 1700028K03Rik | 5.54E-05 | 1.648835551 | 0.886203497 |
| ENSMUSG00000097035 | Gm26692 | 5.54E-05 | 1.326907457 | 0.929973967 |
| ENSMUSG00000106333 | 4930428O21Rik | 5.54E-05 | 2.648835551 | 0.66416768 |
| ENSMUSG00000029263 | Pigg | 5.54E-05 | -1.258055044 | 0.616879894 |
| ENSMUSG00000072769 | Gm10419 | 5.54E-05 | 2.648835551 | 0.66416768 |
| ENSMUSG00000008090 | Fgfrl1 | 5.54E-05 | 0.189403933 | 1 |
| ENSMUSG00000090961 | Vmn2r8 | 5.54E-05 | 2.063873051 | 0.792441218 |
| ENSMUSG00000090688 | Vmn2r12 | 5.54E-05 | 3.648835551 | 0.466464457 |
| ENSMUSG00000092080 | Vmn2r16 | 5.54E-05 | 2.648835551 | 0.66416768 |
| ENSMUSG00000033467 | Crlf2 | 5.54E-05 | 1.326907457 | 0.971854959 |
| ENSMUSG00000072763 | 5430403G16Rik | 5.54E-05 | 0.648835551 | 1 |
| ENSMUSG00000086247 | Gm15787 | 5.54E-05 | -1.051604167 | 0.706929043 |
| ENSMUSG00000029499 | Pxmp2 | 5.54E-05 | -1.209145444 | 0.6523678 |
| ENSMUSG00000048215 | A630023P12Rik | 5.54E-05 | -0.051604167 | 1 |
| ENSMUSG00000104559 | Gm43118 | 5.54E-05 | 3.648835551 | 0.466464457 |
| ENSMUSG00000070576 | Mn1 | 5.54E-05 | 0.47891055 | 1 |
| ENSMUSG00000085271 | E130006D01Rik | 5.54E-05 | 1.326907457 | 0.971854959 |
| ENSMUSG00000105376 | Gm36535 | 5.54E-05 | 1.648835551 | 0.886203497 |
| ENSMUSG00000029343 | Crybb1 | 5.54E-05 | 3.648835551 | 0.466464457 |
| ENSMUSG00000029345 | Tfip11 | 5.54E-05 | 0.47891055 | 1 |
| ENSMUSG00000029346 | Srrd | 5.54E-05 | 0.47891055 | 1 |
| ENSMUSG00000029348 | Asphd2 | 5.54E-05 | 2.648835551 | 0.66416768 |
| ENSMUSG00000106659 | Gm42161 | 5.54E-05 | 1.063873051 | 1 |

| | | | | |
|---|---|---|---|---|
| ENSMUSG00000066964 | Tmem211 | 5.54E-05 | 2.063873051 | 0.792441218 |
| ENSMUSG00000029592 | Usp30 | 5.54E-05 | -1.810596067 | 0.246776726 |
| ENSMUSG00000014158 | Trpv4 | 5.54E-05 | -0.810596067 | 1 |
| ENSMUSG00000029564 | 4930519G04Rik | 5.54E-05 | 2.063873051 | 0.792441218 |
| ENSMUSG00000029522 | Pla2g1b | 5.54E-05 | 1.063873051 | 1 |
| ENSMUSG00000086715 | B230112J18Rik | 5.54E-05 | 2.063873051 | 0.792441218 |
| ENSMUSG00000029363 | Rfc5 | 5.54E-05 | 0.326907457 | 1 |
| ENSMUSG00000107144 | 4930413E15Rik | 5.54E-05 | 1.063873051 | 1 |
| ENSMUSG00000085708 | Gm16063 | 5.54E-05 | 2.648835551 | 0.66416768 |
| ENSMUSG00000085345 | Gm16064 | 5.54E-05 | 0.47891055 | 1 |
| ENSMUSG00000086847 | Tbx3os2 | 5.54E-05 | 0.841480629 | 1 |
| ENSMUSG00000018263 | Tbx5 | 5.54E-05 | -0.43862729 | 1 |
| ENSMUSG00000029594 | Rbm19 | 5.54E-05 | -1.395558568 | 0.508263132 |
| ENSMUSG00000029595 | Lhx5 | 5.54E-05 | 0.326907457 | 1 |
| ENSMUSG00000029600 | Rita1 | 5.54E-05 | -0.936126949 | 0.695372548 |
| ENSMUSG00000001168 | Oas1h | 5.54E-05 | 3.648835551 | 0.466464457 |
| ENSMUSG00000052776 | Oas1a | 5.54E-05 | 3.648835551 | 0.466464457 |
| ENSMUSG00000032623 | Oas1d | 5.54E-05 | 2.648835551 | 0.66416768 |
| ENSMUSG00000029608 | Rph3a | 5.54E-05 | 0.189403933 | 1 |
| ENSMUSG00000029454 | Mapkapk5 | 5.54E-05 | 1.063873051 | 1 |
| ENSMUSG00000089729 | Gm16552 | 5.54E-05 | 1.326907457 | 0.914092674 |
| ENSMUSG00000044134 | Pheta1 | 5.54E-05 | -1.599091962 | 0.384748897 |
| ENSMUSG00000042589 | Cux2 | 5.54E-05 | -1.395558568 | 0.553404132 |
| ENSMUSG00000043036 | Ccdc63 | 5.54E-05 | 0.841480629 | 1 |
| ENSMUSG00000106555 | Gm2479 | 5.54E-05 | 1.063873051 | 1 |
| ENSMUSG00000087336 | Gm15860 | 5.54E-05 | 0.189403933 | 1 |
| ENSMUSG00000029438 | Bcl7a | 5.54E-05 | -1.351164449 | 0.499088689 |
| ENSMUSG00000106636 | Gm43813 | 5.54E-05 | -2.654945197 | 0.025465458 |
| ENSMUSG00000061882 | Ccdc62 | 5.54E-05 | -1.560617814 | 0.375505958 |
| ENSMUSG00000037936 | Scarb1 | 5.54E-05 | -1.395558568 | 0.508263132 |
| ENSMUSG00000105274 | 1700048F04Rik | 5.54E-05 | 2.063873051 | 0.775410248 |
| ENSMUSG00000105593 | Gm42952 | 5.54E-05 | 3.648835551 | 0.466464457 |
| ENSMUSG00000106275 | Gm42495 | 5.54E-05 | 1.648835551 | 0.886203497 |
| ENSMUSG00000029436 | Mmp17 | 5.54E-05 | 1.326907457 | 0.971854959 |
| ENSMUSG00000086916 | Gm15903 | 5.54E-05 | 2.063873051 | 0.792441218 |
| ENSMUSG00000025537 | Phkg1 | 5.54E-05 | -1.480447466 | 0.508841397 |
| ENSMUSG00000025532 | Crcp | 5.54E-05 | -1.395558568 | 0.508263132 |
| ENSMUSG00000105387 | Gm29681 | 5.54E-05 | 2.063873051 | 0.792441218 |
| ENSMUSG00000105773 | 4930563F08Rik | 5.54E-05 | 3.648835551 | 0.466464457 |
| ENSMUSG00000040751 | Lat2 | 5.54E-05 | -0.051604167 | 1 |
| ENSMUSG00000040532 | Abhd11 | 5.54E-05 | 0.063873051 | 1 |
| ENSMUSG00000053388 | Trim50 | 5.54E-05 | 1.326907457 | 0.945880581 |
| ENSMUSG00000004814 | Ccl24 | 5.54E-05 | 3.648835551 | 0.466464457 |
| ENSMUSG00000029699 | Ssc4d | 5.54E-05 | 0.063873051 | 1 |
| ENSMUSG00000042985 | Upk3b | 5.54E-05 | 3.648835551 | 0.466464457 |
| ENSMUSG00000029703 | Lrwd1 | 5.54E-05 | -1.560617814 | 0.356350819 |
| ENSMUSG00000039754 | Alkbh4 | 5.54E-05 | -0.52108945 | 0.975624465 |
| ENSMUSG00000007987 | Ift22 | 5.54E-05 | 0.063873051 | 1 |
| ENSMUSG00000037428 | Vgf | 5.54E-05 | 3.648835551 | 0.466464457 |
| ENSMUSG00000106466 | Gm40348 | 5.54E-05 | 2.063873051 | 0.790186865 |
| ENSMUSG00000023348 | Trip6 | 5.54E-05 | -0.52108945 | 1 |

| | | | | |
|---|---|---|---|---|
| ENSMUSG00000029711 | Epo | 5.54E-05 | 0.326907457 | 1 |
| ENSMUSG00000090873 | Gm17112 | 5.54E-05 | 2.063873051 | 0.792441218 |
| ENSMUSG00000047182 | Irs3 | 5.54E-05 | 2.648835551 | 0.66416768 |
| ENSMUSG00000046245 | Pilra | 5.54E-05 | -0.351164449 | 1 |
| ENSMUSG00000066682 | Pilrb2 | 5.54E-05 | -0.810596067 | 0.832285197 |
| ENSMUSG00000056966 | Gjc3 | 5.54E-05 | 1.326907457 | 0.969214425 |
| ENSMUSG00000106338 | Gm43645 | 5.54E-05 | 3.648835551 | 0.466464457 |
| ENSMUSG00000047592 | Nxpe5 | 5.54E-05 | 0.841480629 | 1 |
| ENSMUSG00000085227 | 6330418K02Rik | 5.54E-05 | -1.209145444 | 0.642722979 |
| ENSMUSG00000086873 | Gm15672 | 5.54E-05 | 1.063873051 | 1 |
| ENSMUSG00000044092 | C130050O18Rik | 5.54E-05 | -1.43862729 | 0.590885473 |
| ENSMUSG00000106056 | 4930500L23Rik | 5.54E-05 | 1.063873051 | 1 |
| ENSMUSG00000105335 | Gm42423 | 5.54E-05 | 2.063873051 | 0.792441218 |
| ENSMUSG00000085673 | Gm16122 | 5.54E-05 | 0.841480629 | 1 |
| ENSMUSG00000106350 | Gm4869 | 5.54E-05 | -1.158519371 | 0.599135879 |
| ENSMUSG00000029570 | Lfng | 5.54E-05 | -3.079084903 | 0.004720588 |
| ENSMUSG00000050022 | Amz1 | 5.54E-05 | -0.599091962 | 1 |
| ENSMUSG00000084934 | Gm16035 | 5.54E-05 | 3.648835551 | 0.466464457 |
| ENSMUSG00000039623 | Ap5z1 | 5.54E-05 | -1.480447466 | 0.441576406 |
| ENSMUSG00000039533 | Mmd2 | 5.54E-05 | -0.43862729 | 1 |
| ENSMUSG00000097559 | D430018E03Rik | 5.54E-05 | 1.648835551 | 0.875465725 |
| ENSMUSG00000061898 | Rbak | 5.54E-05 | -0.936126949 | 0.695372548 |
| ENSMUSG00000039244 | E130309D02Rik | 5.54E-05 | -0.43862729 | 1 |
| ENSMUSG00000010825 | Grid2ip | 5.54E-05 | 0.47891055 | 1 |
| ENSMUSG00000118332 | Fam220a | 5.54E-05 | 1.648835551 | 0.886203497 |
| ENSMUSG00000029613 | Eif2ak1 | 5.54E-05 | -2.52108945 | 0.037394962 |
| ENSMUSG00000029610 | Aimp2 | 5.54E-05 | -0.995020638 | 0.699192777 |
| ENSMUSG00000079109 | Pms2 | 5.54E-05 | -1.305360759 | 0.5074242 |
| ENSMUSG00000052271 | Bhlha15 | 5.54E-05 | 2.063873051 | 0.792441218 |
| ENSMUSG00000029624 | Ptcd1 | 5.54E-05 | -1.995020638 | 0.167528217 |
| ENSMUSG00000070419 | Cyp3a57 | 5.54E-05 | 0.189403933 | 1 |
| ENSMUSG00000061292 | Cyp3a59 | 5.54E-05 | 0.841480629 | 1 |
| ENSMUSG00000106547 | B230303O12Rik | 5.54E-05 | -0.158519371 | 1 |
| ENSMUSG00000105636 | Gm43625 | 5.54E-05 | -0.051604167 | 1 |
| ENSMUSG00000029641 | Rasl11a | 5.54E-05 | 1.326907457 | 0.971854959 |
| ENSMUSG00000016510 | Mtif3 | 5.54E-05 | -1.560617814 | 0.356350819 |
| ENSMUSG00000106845 | Gm43156 | 5.54E-05 | 3.648835551 | 0.466464457 |
| ENSMUSG00000106892 | Gm42791 | 5.54E-05 | 0.648835551 | 1 |
| ENSMUSG00000100465 | Gm29264 | 5.54E-05 | 1.648835551 | 0.886203497 |
| ENSMUSG00000060063 | Alox5ap | 5.54E-05 | -0.258055044 | 1 |
| ENSMUSG00000053368 | Rxfp2 | 5.54E-05 | 0.189403933 | 1 |
| ENSMUSG00000107178 | Gm42531 | 5.54E-05 | 2.648835551 | 0.66416768 |
| ENSMUSG00000041132 | N4bp2l1 | 5.54E-05 | -0.673092543 | 0.98880775 |
| ENSMUSG00000107096 | Gm43597 | 5.54E-05 | 0.326907457 | 1 |
| ENSMUSG00000106692 | Gm36447 | 5.54E-05 | 1.326907457 | 0.947742944 |
| ENSMUSG00000107047 | Gm43298 | 5.54E-05 | 1.326907457 | 0.971854959 |
| ENSMUSG00000058488 | Kl | 5.54E-05 | 0.648835551 | 1 |
| ENSMUSG00000107387 | 5430435K18Rik | 5.54E-05 | 2.063873051 | 0.792441218 |
| ENSMUSG00000097321 | 1700028E10Rik | 5.54E-05 | -1.395558568 | 0.574741398 |
| ENSMUSG00000091794 | Vmn2r18 | 5.54E-05 | 2.063873051 | 0.792441218 |
| ENSMUSG00000106779 | Gm8579 | 5.54E-05 | 2.063873051 | 0.792441218 |

| | | | | |
|---|---|---|---|---|
| ENSMUSG00000023964 | Calcr | 5.54E-05 | 1.648835551 | 0.871107617 |
| ENSMUSG00000004631 | Sgce | 5.54E-05 | -0.52108945 | 1 |
| ENSMUSG00000042607 | Asb4 | 5.54E-05 | 1.326907457 | 0.971854959 |
| ENSMUSG00000019577 | Pdk4 | 5.54E-05 | 0.326907457 | 1 |
| ENSMUSG00000093658 | Gm20617 | 5.54E-05 | 0.326907457 | 1 |
| ENSMUSG00000098501 | Gm19272 | 5.54E-05 | 2.063873051 | 0.780444312 |
| ENSMUSG00000061762 | Tac1 | 5.54E-05 | 2.648835551 | 0.66416768 |
| ENSMUSG00000029685 | Asb15 | 5.54E-05 | -0.158519371 | 1 |
| ENSMUSG00000079652 | Fam71f2 | 5.54E-05 | -0.258055044 | 1 |
| ENSMUSG00000068699 | Flnc | 5.54E-05 | -2.480447466 | 0.046272089 |
| ENSMUSG00000001761 | Smo | 5.54E-05 | -2.89032326 | 0.011194612 |
| ENSMUSG00000039629 | Strip2 | 5.54E-05 | 0.841480629 | 1 |
| ENSMUSG00000029782 | Tmem209 | 5.54E-05 | -2.560617814 | 0.030770816 |
| ENSMUSG00000071553 | Cpa2 | 5.54E-05 | 1.648835551 | 0.886203497 |
| ENSMUSG00000029788 | Cpa5 | 5.54E-05 | 2.648835551 | 0.66416768 |
| ENSMUSG00000051855 | Mest | 5.54E-05 | 0.326907457 | 1 |
| ENSMUSG00000085321 | 4930412F09Rik | 5.54E-05 | 2.063873051 | 0.790129914 |
| ENSMUSG00000086212 | Mkln1os | 5.54E-05 | -0.43862729 | 1 |
| ENSMUSG00000086855 | Gm13844 | 5.54E-05 | 1.326907457 | 0.971854959 |
| ENSMUSG00000086804 | Gm43154 | 5.54E-05 | -1.051604167 | 0.718012086 |
| ENSMUSG00000084973 | Gm13848 | 5.54E-05 | 0.841480629 | 1 |
| ENSMUSG00000107010 | Gm43293 | 5.54E-05 | 0.063873051 | 1 |
| ENSMUSG00000085847 | Gm14540 | 5.54E-05 | 2.648835551 | 0.66416768 |
| ENSMUSG00000018999 | Slc35b4 | 5.54E-05 | -1.673092543 | 0.290284345 |
| ENSMUSG00000086762 | Gm14546 | 5.54E-05 | 1.648835551 | 0.871122629 |
| ENSMUSG00000057137 | Tmem140 | 5.54E-05 | -0.351164449 | 1 |
| ENSMUSG00000029843 | Slc13a4 | 5.54E-05 | 0.648835551 | 1 |
| ENSMUSG00000038641 | Akr1d1 | 5.54E-05 | 2.648835551 | 0.66416768 |
| ENSMUSG00000029830 | Svopl | 5.54E-05 | -0.351164449 | 1 |
| ENSMUSG00000047749 | Zc3hav1l | 5.54E-05 | 0.648835551 | 1 |
| ENSMUSG00000056832 | Ttc26 | 5.54E-05 | -1.258055044 | 0.626515359 |
| ENSMUSG00000071537 | Klrg2 | 5.54E-05 | 0.47891055 | 1 |
| ENSMUSG00000100121 | 1700025N23Rik | 5.54E-05 | 2.648835551 | 0.66416768 |
| ENSMUSG00000029923 | Rab19 | 5.54E-05 | -0.158519371 | 1 |
| ENSMUSG00000085058 | 8030453O22Rik | 5.54E-05 | 0.189403933 | 1 |
| ENSMUSG00000029916 | Agk | 5.54E-05 | -1.52108945 | 0.466464457 |
| ENSMUSG00000068259 | Olfr461 | 5.54E-05 | 3.648835551 | 0.466464457 |
| ENSMUSG00000102802 | Mgam2-ps | 5.54E-05 | 0.326907457 | 1 |
| ENSMUSG00000029883 | 1700074P13Rik | 5.54E-05 | 2.063873051 | 0.792441218 |
| ENSMUSG00000036938 | Try5 | 5.54E-05 | 2.648835551 | 0.66416768 |
| ENSMUSG00000071521 | Try10 | 5.54E-05 | 3.648835551 | 0.466464457 |
| ENSMUSG00000076498 | Trbc2 | 5.54E-05 | 1.326907457 | 0.971854959 |
| ENSMUSG00000029869 | Ephb6 | 5.54E-05 | 0.47891055 | 1 |
| ENSMUSG00000018656 | Tcaf3 | 5.54E-05 | 1.648835551 | 0.886203497 |
| ENSMUSG00000054431 | Olfr450 | 5.54E-05 | 2.648835551 | 0.66416768 |
| ENSMUSG00000071481 | Olfr437 | 5.54E-05 | 3.648835551 | 0.466464457 |
| ENSMUSG00000051499 | Zfp786 | 5.54E-05 | 0.648835551 | 1 |
| ENSMUSG00000057691 | Zfp746 | 5.54E-05 | -1.843017545 | 0.255869172 |
| ENSMUSG00000097434 | Gm16630 | 5.54E-05 | 0.189403933 | 1 |
| ENSMUSG00000042810 | Krba1 | 5.54E-05 | 0.189403933 | 1 |
| ENSMUSG00000068551 | Zfp467 | 5.54E-05 | -1.43862729 | 0.433313155 |

| | | | | |
|---|---|---|---|---|
| ENSMUSG00000007216 | Zfp775 | 5.54E-05 | -1.158519371 | 0.60072014 |
| ENSMUSG00000054435 | Gimap4 | 5.54E-05 | 0.47891055 | 1 |
| ENSMUSG00000047867 | Gimap6 | 5.54E-05 | 1.063873051 | 1 |
| ENSMUSG00000043505 | Gimap5 | 5.54E-05 | 0.648835551 | 1 |
| ENSMUSG00000107481 | 4833403J16Rik | 5.54E-05 | 3.648835551 | 0.466464457 |
| ENSMUSG00000029816 | Gpnmb | 5.54E-05 | -0.258055044 | 1 |
| ENSMUSG00000029821 | Gsdme | 5.54E-05 | -1.9057533 | 0.203406755 |
| ENSMUSG00000085776 | 5430402O13Rik | 5.54E-05 | -0.158519371 | 1 |
| ENSMUSG00000107667 | C530044C16Rik | 5.54E-05 | 0.063873051 | 1 |
| ENSMUSG00000108145 | Gm38811 | 5.54E-05 | 1.063873051 | 1 |
| ENSMUSG00000085412 | Halr1 | 5.54E-05 | -0.52108945 | 0.986300373 |
| ENSMUSG00000087626 | Gm15050 | 5.54E-05 | 2.648835551 | 0.66416768 |
| ENSMUSG00000085696 | Hoxaas3 | 5.54E-05 | 1.063873051 | 1 |
| ENSMUSG00000000942 | Hoxa4 | 5.54E-05 | 2.063873051 | 0.792441218 |
| ENSMUSG00000086427 | Hoxa11os | 5.54E-05 | 1.648835551 | 0.886203497 |
| ENSMUSG00000055408 | Hottip | 5.54E-05 | 2.648835551 | 0.66416768 |
| ENSMUSG00000038074 | Fkbp14 | 5.54E-05 | -0.995020638 | 0.699192777 |
| ENSMUSG00000108285 | Gm34358 | 5.54E-05 | 3.648835551 | 0.466464457 |
| ENSMUSG00000004654 | Ghrhr | 5.54E-05 | 3.648835551 | 0.466464457 |
| ENSMUSG00000108228 | 6430584L05Rik | 5.54E-05 | 3.648835551 | 0.466464457 |
| ENSMUSG00000057981 | Vmn1r12 | 5.54E-05 | 2.648835551 | 0.66416768 |
| ENSMUSG00000064259 | Vmn1r13 | 5.54E-05 | -0.810596067 | 0.865350008 |
| ENSMUSG00000091382 | Vmn1r18 | 5.54E-05 | 2.063873051 | 0.792441218 |
| ENSMUSG00000093535 | 4930533I22Rik | 5.54E-05 | 0.47891055 | 1 |
| ENSMUSG00000037826 | Ppm1k | 5.54E-05 | -1.258055044 | 0.620160408 |
| ENSMUSG00000115091 | Vmn1r22 | 5.54E-05 | 2.063873051 | 0.783231522 |
| ENSMUSG00000092323 | BB365896 | 5.54E-05 | 2.063873051 | 0.792441218 |
| ENSMUSG00000046764 | A530053G22Rik | 5.54E-05 | -0.673092543 | 1 |
| ENSMUSG00000029910 | Mad2l1 | 5.54E-05 | -0.051604167 | 1 |
| ENSMUSG00000059375 | Vmn1r33 | 5.54E-05 | 3.648835551 | 0.466464457 |
| ENSMUSG00000087231 | E230016M11Rik | 5.54E-05 | -1.777429203 | 0.241307011 |
| ENSMUSG00000106494 | Igkv2-107 | 5.54E-05 | 3.648835551 | 0.466464457 |
| ENSMUSG00000076573 | Igkv1-35 | 5.54E-05 | 3.648835551 | 0.466464457 |
| ENSMUSG00000105646 | Gm30211 | 5.54E-05 | 1.648835551 | 0.886203497 |
| ENSMUSG00000108289 | Gm44174 | 5.54E-05 | 2.063873051 | 0.774790843 |
| ENSMUSG00000108213 | Gm44172 | 5.54E-05 | 2.063873051 | 0.792441218 |
| ENSMUSG00000108356 | 9130221F21Rik | 5.54E-05 | 2.063873051 | 0.771834758 |
| ENSMUSG00000037621 | Atoh8 | 5.54E-05 | -0.810596067 | 0.832285197 |
| ENSMUSG00000052631 | Sh2d6 | 5.54E-05 | 1.648835551 | 0.886203497 |
| ENSMUSG00000097569 | Gm26640 | 5.54E-05 | 3.648835551 | 0.466464457 |
| ENSMUSG00000023140 | Reg2 | 5.54E-05 | 2.063873051 | 0.792441218 |
| ENSMUSG00000108027 | Gm44271 | 5.54E-05 | 1.648835551 | 0.886203497 |
| ENSMUSG00000035104 | Eva1a | 5.54E-05 | -0.52108945 | 0.991306982 |
| ENSMUSG00000108015 | Gm32591 | 5.54E-05 | 0.326907457 | 1 |
| ENSMUSG00000000627 | Sema4f | 5.54E-05 | -2.89032326 | 0.01005046 |
| ENSMUSG00000009145 | Dqx1 | 5.54E-05 | -0.936126949 | 0.695372548 |
| ENSMUSG00000030036 | Mogs | 5.54E-05 | -1.599091962 | 0.575497924 |
| ENSMUSG00000030030 | 1700003E16Rik | 5.54E-05 | 1.326907457 | 0.971854959 |
| ENSMUSG00000034777 | Vax2 | 5.54E-05 | 1.326907457 | 0.94561855 |
| ENSMUSG00000107797 | Gm10445 | 5.54E-05 | 1.063873051 | 1 |
| ENSMUSG00000063415 | Cyp26b1 | 5.54E-05 | 0.326907457 | 1 |

| | | | | |
|---|---|---|---|---|
| ENSMUSG00000072952 | Gm5878 | 5.54E-05 | 2.648835551 | 0.66416768 |
| ENSMUSG00000030008 | Pradc1 | 5.54E-05 | 1.648835551 | 0.886203497 |
| ENSMUSG00000051262 | Nat8f3 | 5.54E-05 | -0.051604167 | 1 |
| ENSMUSG00000107689 | Gm44386 | 5.54E-05 | 0.648835551 | 1 |
| ENSMUSG00000100164 | 2610306M01Rik | 5.54E-05 | 0.841480629 | 1 |
| ENSMUSG00000090891 | D6Ertd527e | 5.54E-05 | 3.648835551 | 0.466464457 |
| ENSMUSG00000049409 | Prokr1 | 5.54E-05 | 2.648835551 | 0.66416768 |
| ENSMUSG00000089997 | 1810020O05Rik | 5.54E-05 | 0.648835551 | 1 |
| ENSMUSG00000030054 | Gp9 | 5.54E-05 | 2.648835551 | 0.662198438 |
| ENSMUSG00000084950 | Gm5577 | 5.54E-05 | -0.051604167 | 1 |
| ENSMUSG00000085828 | Gm15612 | 5.54E-05 | 3.648835551 | 0.466464457 |
| ENSMUSG00000002871 | Tpra1 | 5.54E-05 | -0.995020638 | 0.699192777 |
| ENSMUSG00000000811 | Txnrd3 | 5.54E-05 | -0.051604167 | 1 |
| ENSMUSG00000068232 | Vmn1r42 | 5.54E-05 | 0.063873051 | 1 |
| ENSMUSG00000061653 | Vmn1r46 | 5.54E-05 | 3.648835551 | 0.466464457 |
| ENSMUSG00000094553 | Vmn1r50 | 5.54E-05 | 2.063873051 | 0.792441218 |
| ENSMUSG00000047203 | Vmn1r54 | 5.54E-05 | 1.063873051 | 1 |
| ENSMUSG00000034456 | Uroc1 | 5.54E-05 | 1.063873051 | 1 |
| ENSMUSG00000030091 | Nup210 | 5.54E-05 | -1.305360759 | 0.491653643 |
| ENSMUSG00000107882 | Gm44278 | 5.54E-05 | 3.648835551 | 0.466464457 |
| ENSMUSG00000107528 | Gm44442 | 5.54E-05 | 2.648835551 | 0.66416768 |
| ENSMUSG00000101553 | 1700049E22Rik | 5.54E-05 | 2.648835551 | 0.66416768 |
| ENSMUSG00000097834 | Gm26911 | 5.54E-05 | 1.648835551 | 0.874796537 |
| ENSMUSG00000107563 | 4930587E11Rik | 5.54E-05 | 0.841480629 | 1 |
| ENSMUSG00000013736 | Trnt1 | 5.54E-05 | -3.233807498 | 0.00236773 |
| ENSMUSG00000034387 | Ssu2 | 5.54E-05 | 1.063873051 | 1 |
| ENSMUSG00000062694 | Cav3 | 5.54E-05 | 2.648835551 | 0.66416768 |
| ENSMUSG00000049112 | Oxtr | 5.54E-05 | 2.063873051 | 0.792441218 |
| ENSMUSG00000030271 | Ogg1 | 5.54E-05 | 0.326907457 | 1 |
| ENSMUSG00000097994 | Gm26982 | 5.54E-05 | 1.648835551 | 0.886203497 |
| ENSMUSG00000030317 | Timp4 | 5.54E-05 | 1.326907457 | 0.971854959 |
| ENSMUSG00000000439 | Mkrn2 | 5.54E-05 | -1.258055044 | 0.616879894 |
| ENSMUSG00000030319 | Cand2 | 5.54E-05 | 1.326907457 | 0.971854959 |
| ENSMUSG00000030321 | Efcab12 | 5.54E-05 | 0.189403933 | 1 |
| ENSMUSG00000042279 | H1foo | 5.54E-05 | 3.648835551 | 0.466464457 |
| ENSMUSG00000108020 | 6820426E19Rik | 5.54E-05 | 1.648835551 | 0.886203497 |
| ENSMUSG00000104222 | Gm7292 | 5.54E-05 | 2.648835551 | 0.66416768 |
| ENSMUSG00000004988 | Fxyd4 | 5.54E-05 | 2.648835551 | 0.66416768 |
| ENSMUSG00000007827 | Ankrd26 | 5.54E-05 | -3.158519371 | 0.002869506 |
| ENSMUSG00000004902 | Slc25a18 | 5.54E-05 | 0.063873051 | 1 |
| ENSMUSG00000030108 | Slc6a13 | 5.54E-05 | 2.063873051 | 0.792441218 |
| ENSMUSG00000030111 | A2m | 5.54E-05 | 0.841480629 | 1 |
| ENSMUSG00000108022 | Gm7298 | 5.54E-05 | 0.841480629 | 1 |
| ENSMUSG00000040649 | Rimklb | 5.54E-05 | -0.599091962 | 0.98833461 |
| ENSMUSG00000012396 | Nanog | 5.54E-05 | 0.841480629 | 1 |
| ENSMUSG00000097635 | Gm26826 | 5.54E-05 | 2.063873051 | 0.792441218 |
| ENSMUSG00000043832 | Clec4a3 | 5.54E-05 | 3.648835551 | 0.466464457 |
| ENSMUSG00000067767 | Clec4b2 | 5.54E-05 | 1.648835551 | 0.886203497 |
| ENSMUSG00000023349 | Clec4n | 5.54E-05 | 0.189403933 | 1 |
| ENSMUSG00000094145 | Vmn2r20 | 5.54E-05 | 1.326907457 | 0.94250747 |
| ENSMUSG00000095486 | Vmn2r22 | 5.54E-05 | 2.648835551 | 0.66416768 |

| | | | | |
|---|---|---|---|---|
| ENSMUSG00000072780 | Vmn2r24 | 5.54E-05 | 1.326907457 | 0.937917675 |
| ENSMUSG00000038451 | Spsb2 | 5.54E-05 | 1.326907457 | 0.954524296 |
| ENSMUSG00000023274 | Cd4 | 5.54E-05 | -0.351164449 | 1 |
| ENSMUSG00000030329 | Pianp | 5.54E-05 | 2.648835551 | 0.66416768 |
| ENSMUSG00000108034 | 4930557K07Rik | 5.54E-05 | 0.648835551 | 1 |
| ENSMUSG00000038271 | Iffo1 | 5.54E-05 | 1.063873051 | 1 |
| ENSMUSG00000038213 | Tapbpl | 5.54E-05 | 0.063873051 | 1 |
| ENSMUSG00000030340 | Scnn1a | 5.54E-05 | -0.258055044 | 1 |
| ENSMUSG00000101162 | Gm26728 | 5.54E-05 | 0.189403933 | 1 |
| ENSMUSG00000049107 | Ntf3 | 5.54E-05 | 0.841480629 | 1 |
| ENSMUSG00000107623 | Gm10415 | 5.54E-05 | 1.063873051 | 1 |
| ENSMUSG00000000182 | Fgf23 | 5.54E-05 | 0.47891055 | 1 |
| ENSMUSG00000030350 | Prmt8 | 5.54E-05 | -0.051604167 | 1 |
| ENSMUSG00000079304 | Tex52 | 5.54E-05 | 1.063873051 | 1 |
| ENSMUSG00000079299 | Klrb1 | 5.54E-05 | 1.648835551 | 0.886203497 |
| ENSMUSG00000090164 | BC035044 | 5.54E-05 | 0.189403933 | 1 |
| ENSMUSG00000030155 | Clec2e | 5.54E-05 | 2.648835551 | 0.66416768 |
| ENSMUSG00000047720 | 4922502D21Rik | 5.54E-05 | 1.648835551 | 0.886203497 |
| ENSMUSG00000030162 | Olr1 | 5.54E-05 | 0.063873051 | 1 |
| ENSMUSG00000030149 | Klrk1 | 5.54E-05 | 0.47891055 | 1 |
| ENSMUSG00000014543 | Klra17 | 5.54E-05 | 1.063873051 | 1 |
| ENSMUSG00000067599 | Klra7 | 5.54E-05 | 2.063873051 | 0.787641865 |
| ENSMUSG00000030187 | Klra2 | 5.54E-05 | -0.936126949 | 0.85084719 |
| ENSMUSG00000067541 | A630073D07Rik | 5.54E-05 | 1.326907457 | 0.951890397 |
| ENSMUSG00000032758 | Kap | 5.54E-05 | 3.648835551 | 0.466464457 |
| ENSMUSG00000092134 | Gm17089 | 5.54E-05 | 2.648835551 | 0.66416768 |
| ENSMUSG00000032718 | Mansc1 | 5.54E-05 | -0.351164449 | 1 |
| ENSMUSG00000032641 | Gpr19 | 5.54E-05 | -0.158519371 | 1 |
| ENSMUSG00000100876 | 2810454H06Rik | 5.54E-05 | 0.841480629 | 1 |
| ENSMUSG00000107730 | Gm44238 | 5.54E-05 | 2.648835551 | 0.66416768 |
| ENSMUSG00000090698 | Apold1 | 5.54E-05 | 1.326907457 | 0.971854959 |
| ENSMUSG00000030206 | Gsg1 | 5.54E-05 | -0.158519371 | 1 |
| ENSMUSG00000107396 | 4930425L21Rik | 5.54E-05 | 1.648835551 | 0.874796537 |
| ENSMUSG00000086239 | Gm14329 | 5.54E-05 | 2.063873051 | 0.792441218 |
| ENSMUSG00000047515 | BC049715 | 5.54E-05 | 0.326907457 | 1 |
| ENSMUSG00000064330 | Pde6h | 5.54E-05 | -1.480447466 | 0.456626102 |
| ENSMUSG00000030230 | Plcz1 | 5.54E-05 | 0.648835551 | 1 |
| ENSMUSG00000079264 | Gm11077 | 5.54E-05 | 1.326907457 | 0.949432525 |
| ENSMUSG00000030235 | Slco1c1 | 5.54E-05 | 0.189403933 | 1 |
| ENSMUSG00000030236 | Slco1b2 | 5.54E-05 | -0.158519371 | 1 |
| ENSMUSG00000063975 | Slco1a5 | 5.54E-05 | 0.648835551 | 1 |
| ENSMUSG00000030243 | Recql | 5.54E-05 | -0.936126949 | 0.695372548 |
| ENSMUSG00000071112 | Spx | 5.54E-05 | 1.326907457 | 0.971854959 |
| ENSMUSG00000108094 | Gm31108 | 5.54E-05 | -0.810596067 | 0.874796537 |
| ENSMUSG00000072679 | D6Ertd474e | 5.54E-05 | 0.841480629 | 1 |
| ENSMUSG00000087157 | Sox5os2 | 5.54E-05 | 1.648835551 | 0.875261697 |
| ENSMUSG00000086282 | Sox5os4 | 5.54E-05 | 0.189403933 | 1 |
| ENSMUSG00000086863 | Gm15543 | 5.54E-05 | -1.305360759 | 0.66416768 |
| ENSMUSG00000040234 | Tm7sf3 | 5.54E-05 | -1.560617814 | 0.391016627 |
| ENSMUSG00000040121 | Rep15 | 5.54E-05 | -0.810596067 | 0.832285197 |
| ENSMUSG00000040102 | Klhl42 | 5.54E-05 | -1.874726405 | 0.198601369 |

| | | | | |
|---|---|---|---|---|
| ENSMUSG00000085600 | E330012B07Rik | 5.54E-05 | 1.326907457 | 0.971854959 |
| ENSMUSG00000096299 | Gm21814 | 5.54E-05 | 1.648835551 | 0.886203497 |
| ENSMUSG00000053395 | Cacng8 | 5.54E-05 | -0.351164449 | 1 |
| ENSMUSG00000053338 | Tarm1 | 5.54E-05 | -0.936126949 | 0.858831093 |
| ENSMUSG00000054594 | Oscar | 5.54E-05 | 0.841480629 | 1 |
| ENSMUSG00000008373 | Prpf31 | 5.54E-05 | -1.480447466 | 0.45696969 |
| ENSMUSG00000043432 | Leng9 | 5.54E-05 | 0.841480629 | 0.955333935 |
| ENSMUSG00000064179 | Tnnt1 | 5.54E-05 | 1.063873051 | 1 |
| ENSMUSG00000074406 | Zfp628 | 5.54E-05 | 1.326907457 | 0.971854959 |
| ENSMUSG00000051184 | Zfp524 | 5.54E-05 | 0.189403933 | 1 |
| ENSMUSG00000091652 | Vmn1r57 | 5.54E-05 | 2.648835551 | 0.66416768 |
| ENSMUSG00000035177 | Nlrp2 | 5.54E-05 | 3.648835551 | 0.466464457 |
| ENSMUSG00000078808 | Vmn1r58 | 5.54E-05 | 1.063873051 | 1 |
| ENSMUSG00000118298 | Vmn1r60 | 5.54E-05 | 0.326907457 | 1 |
| ENSMUSG00000058028 | Zscan5b | 5.54E-05 | 1.648835551 | 0.886203497 |
| ENSMUSG00000053367 | Epp13 | 5.54E-05 | 1.648835551 | 0.871850653 |
| ENSMUSG00000030443 | Zfp583 | 5.54E-05 | 2.063873051 | 0.792441218 |
| ENSMUSG00000055150 | Zfp78 | 5.54E-05 | 0.189403933 | 1 |
| ENSMUSG00000096228 | Olfr5 | 5.54E-05 | 2.648835551 | 0.66416768 |
| ENSMUSG00000034583 | Olfr1347 | 5.54E-05 | 3.648835551 | 0.466464457 |
| ENSMUSG00000002266 | Zim1 | 5.54E-05 | 0.841480629 | 1 |
| ENSMUSG00000108043 | Zim3 | 5.54E-05 | 0.189403933 | 1 |
| ENSMUSG00000062116 | Zfp954 | 5.54E-05 | 0.326907457 | 1 |
| ENSMUSG00000095730 | Vmn2r29 | 5.54E-05 | 1.648835551 | 0.886203497 |
| ENSMUSG00000070847 | Vmn2r30 | 5.54E-05 | 0.648835551 | 1 |
| ENSMUSG00000096399 | Vmn2r35 | 5.54E-05 | 3.648835551 | 0.466464457 |
| ENSMUSG00000034087 | Nlrp4b | 5.54E-05 | 1.326907457 | 0.971117005 |
| ENSMUSG00000115267 | Vmn1r76 | 5.54E-05 | 0.841480629 | 1 |
| ENSMUSG00000066803 | Vmn1r84 | 5.54E-05 | 3.648835551 | 0.466464457 |
| ENSMUSG00000030386 | Zfp606 | 5.54E-05 | -0.936126949 | 0.695372548 |
| ENSMUSG00000091045 | Vmn2r55 | 5.54E-05 | 2.063873051 | 0.792441218 |
| ENSMUSG00000070822 | Zscan18 | 5.54E-05 | -0.673092543 | 0.98880775 |
| ENSMUSG00000058638 | Zfp110 | 5.54E-05 | -0.936126949 | 0.695372548 |
| ENSMUSG00000097115 | 1810019N24Rik | 5.54E-05 | -0.52108945 | 0.975624465 |
| ENSMUSG00000033961 | Zfp446 | 5.54E-05 | 0.326907457 | 1 |
| ENSMUSG00000049600 | Zbtb45 | 5.54E-05 | 0.648835551 | 1 |
| ENSMUSG00000070817 | Vmn1r85 | 5.54E-05 | 0.189403933 | 1 |
| ENSMUSG00000070814 | Zswim9 | 5.54E-05 | -0.351164449 | 1 |
| ENSMUSG00000074378 | Bsph1 | 5.54E-05 | 2.648835551 | 0.66416768 |
| ENSMUSG00000078800 | Bsph2 | 5.54E-05 | 3.648835551 | 0.466464457 |
| ENSMUSG00000070811 | Sult2a2 | 5.54E-05 | 0.189403933 | 1 |
| ENSMUSG00000078798 | Sult2a1 | 5.54E-05 | 2.648835551 | 0.66416768 |
| ENSMUSG00000074377 | Sult2a4 | 5.54E-05 | 0.063873051 | 1 |
| ENSMUSG00000074365 | Crxos | 5.54E-05 | -0.258055044 | 1 |
| ENSMUSG00000078796 | Zfp541 | 5.54E-05 | 1.063873051 | 1 |
| ENSMUSG00000074361 | C5ar2 | 5.54E-05 | 2.648835551 | 0.66416768 |
| ENSMUSG00000049130 | C5ar1 | 5.54E-05 | 0.47891055 | 1 |
| ENSMUSG00000002083 | Bbc3 | 5.54E-05 | 0.47891055 | 1 |
| ENSMUSG00000001988 | Npas1 | 5.54E-05 | 1.326907457 | 0.971854959 |
| ENSMUSG00000100583 | Gm29443 | 5.54E-05 | 0.841480629 | 1 |
| ENSMUSG00000097842 | 9330104G04Rik | 5.54E-05 | 3.648835551 | 0.466464457 |

| | | | | |
|---|---|---|---|---|
| ENSMUSG00000043017 | Ptgir | 5.54E-05 | 2.648835551 | 0.66416768 |
| ENSMUSG00000041117 | Ccdc8 | 5.54E-05 | 1.063873051 | 1 |
| ENSMUSG00000023159 | Psg29 | 5.54E-05 | 1.648835551 | 0.886203497 |
| ENSMUSG00000108441 | Gm32846 | 5.54E-05 | 2.063873051 | 0.792441218 |
| ENSMUSG00000078793 | Gm5155 | 5.54E-05 | 1.326907457 | 0.971854959 |
| ENSMUSG00000054005 | Mill1 | 5.54E-05 | 3.648835551 | 0.466464457 |
| ENSMUSG00000003505 | Psg18 | 5.54E-05 | 3.648835551 | 0.466464457 |
| ENSMUSG00000063305 | Psg20 | 5.54E-05 | 3.648835551 | 0.466464457 |
| ENSMUSG00000048481 | Mypop | 5.54E-05 | 1.326907457 | 0.971854959 |
| ENSMUSG00000030410 | Dmwd | 5.54E-05 | -1.395558568 | 0.508263132 |
| ENSMUSG00000040841 | Six5 | 5.54E-05 | -0.43862729 | 1 |
| ENSMUSG00000089636 | 1700058P15Rik | 5.54E-05 | 2.648835551 | 0.66416768 |
| ENSMUSG00000003545 | Fosb | 5.54E-05 | 0.063873051 | 1 |
| ENSMUSG00000108854 | D830036C21Rik | 5.54E-05 | 0.648835551 | 1 |
| ENSMUSG00000030400 | Ercc2 | 5.54E-05 | -1.305360759 | 0.491653643 |
| ENSMUSG00000040714 | Klc3 | 5.54E-05 | -1.351164449 | 0.499088689 |
| ENSMUSG00000040705 | A930016O22Rik | 5.54E-05 | -0.995020638 | 0.749195699 |
| ENSMUSG00000011263 | Exoc3l2 | 5.54E-05 | -1.158519371 | 0.631419854 |
| ENSMUSG00000057667 | Bloc1s3 | 5.54E-05 | 1.326907457 | 0.971854959 |
| ENSMUSG00000061028 | Clasrp | 5.54E-05 | -2.965874293 | 0.007243475 |
| ENSMUSG00000040564 | Apoc1 | 5.54E-05 | 2.063873051 | 0.792441218 |
| ENSMUSG00000049848 | Ceacam19 | 5.54E-05 | 1.648835551 | 0.877036949 |
| ENSMUSG00000070777 | Ceacam20 | 5.54E-05 | 2.063873051 | 0.792441218 |
| ENSMUSG00000090715 | Vmn1r167 | 5.54E-05 | 3.648835551 | 0.466464457 |
| ENSMUSG00000115021 | Vmn1r173 | 5.54E-05 | 3.648835551 | 0.466464457 |
| ENSMUSG00000090411 | Vmn1r174 | 5.54E-05 | 3.648835551 | 0.466464457 |
| ENSMUSG00000095632 | Vmn1r175 | 5.54E-05 | 3.648835551 | 0.466464457 |
| ENSMUSG00000096859 | Vmn1r176 | 5.54E-05 | 3.648835551 | 0.466464457 |
| ENSMUSG00000097425 | Vmn1r181 | 5.54E-05 | 2.648835551 | 0.66416768 |
| ENSMUSG00000108613 | Gm36694 | 5.54E-05 | 1.326907457 | 0.950123751 |
| ENSMUSG00000047603 | Zfp235 | 5.54E-05 | -0.599091962 | 0.980745941 |
| ENSMUSG00000055305 | Zfp93 | 5.54E-05 | 1.326907457 | 0.971854959 |
| ENSMUSG00000050605 | Zfp61 | 5.54E-05 | 0.189403933 | 1 |
| ENSMUSG00000074282 | Zfp94 | 5.54E-05 | -0.258055044 | 1 |
| ENSMUSG00000062028 | Irgc1 | 5.54E-05 | 1.326907457 | 0.954410036 |
| ENSMUSG00000108360 | Gm36159 | 5.54E-05 | 2.648835551 | 0.66416768 |
| ENSMUSG00000064264 | Zfp428 | 5.54E-05 | 0.648835551 | 1 |
| ENSMUSG00000066721 | Zfp575 | 5.54E-05 | 2.063873051 | 0.792441218 |
| ENSMUSG00000052212 | Cd177 | 5.54E-05 | 1.648835551 | 0.877036949 |
| ENSMUSG00000011350 | Gm5893 | 5.54E-05 | -1.106051951 | 0.719591951 |
| ENSMUSG00000062732 | Lypd4 | 5.54E-05 | 2.063873051 | 0.792441218 |
| ENSMUSG00000011349 | Dmrtc2 | 5.54E-05 | 2.648835551 | 0.66416768 |
| ENSMUSG00000003379 | Cd79a | 5.54E-05 | 2.063873051 | 0.792441218 |
| ENSMUSG00000108367 | Gm4881 | 5.54E-05 | 0.326907457 | 1 |
| ENSMUSG00000045252 | Zfp574 | 5.54E-05 | -0.743481871 | 0.827142217 |
| ENSMUSG00000043843 | Tmem145 | 5.54E-05 | 2.063873051 | 0.792441218 |
| ENSMUSG00000061702 | Tmem91 | 5.54E-05 | 0.841480629 | 1 |
| ENSMUSG00000063439 | B9d2 | 5.54E-05 | 2.063873051 | 0.792441218 |
| ENSMUSG00000040614 | Nlrp9c | 5.54E-05 | 1.648835551 | 0.886203497 |
| ENSMUSG00000040601 | Nlrp4a | 5.54E-05 | 1.326907457 | 0.970970089 |
| ENSMUSG00000054102 | Nlrp9a | 5.54E-05 | 1.648835551 | 0.886203497 |

| | | | | |
|---|---|---|---|---|
| ENSMUSG00000066704 | Cyp2b19 | 5.54E-05 | 1.648835551 | 0.886203497 |
| ENSMUSG00000049685 | Cyp2g1 | 5.54E-05 | 2.648835551 | 0.66416768 |
| ENSMUSG00000005547 | Cyp2a5 | 5.54E-05 | 3.648835551 | 0.466464457 |
| ENSMUSG00000060407 | Cyp2a12 | 5.54E-05 | 3.648835551 | 0.466464457 |
| ENSMUSG00000052974 | Cyp2f2 | 5.54E-05 | 1.326907457 | 0.970970089 |
| ENSMUSG00000089661 | Mia | 5.54E-05 | 1.326907457 | 0.948626544 |
| ENSMUSG00000057093 | Zfp607b | 5.54E-05 | -0.43862729 | 1 |
| ENSMUSG00000030604 | Zfp626 | 5.54E-05 | -1.051604167 | 0.706929043 |
| ENSMUSG00000030600 | Lrfn1 | 5.54E-05 | -0.158519371 | 1 |
| ENSMUSG00000047586 | Nccrp1 | 5.54E-05 | 1.648835551 | 0.886203497 |
| ENSMUSG00000030595 | Nfkbib | 5.54E-05 | -0.258055044 | 1 |
| ENSMUSG00000053522 | Lgals7 | 5.54E-05 | 1.063873051 | 1 |
| ENSMUSG00000030589 | Rasgrp4 | 5.54E-05 | 0.841480629 | 1 |
| ENSMUSG00000037239 | Spred3 | 5.54E-05 | -2.921020057 | 0.008282606 |
| ENSMUSG00000046410 | Kcnk6 | 5.54E-05 | 0.648835551 | 1 |
| ENSMUSG00000030588 | Yif1b | 5.54E-05 | -0.599091962 | 1 |
| ENSMUSG00000097809 | Gm26554 | 5.54E-05 | 3.648835551 | 0.466464457 |
| ENSMUSG00000058402 | Zfp420 | 5.54E-05 | -0.351164449 | 1 |
| ENSMUSG00000053985 | Zfp14 | 5.54E-05 | -1.965874293 | 0.167024802 |
| ENSMUSG00000013921 | Clip3 | 5.54E-05 | 0.47891055 | 1 |
| ENSMUSG00000074211 | Sdhaf1 | 5.54E-05 | 0.648835551 | 1 |
| ENSMUSG00000036892 | Prodh2 | 5.54E-05 | 1.326907457 | 0.943363727 |
| ENSMUSG00000006311 | Etv2 | 5.54E-05 | 2.648835551 | 0.66416768 |
| ENSMUSG00000005553 | Atp4a | 5.54E-05 | 1.648835551 | 0.886203497 |
| ENSMUSG00000051314 | Ffar2 | 5.54E-05 | 1.326907457 | 0.971854959 |
| ENSMUSG00000036634 | Mag | 5.54E-05 | -0.158519371 | 1 |
| ENSMUSG00000046826 | Fam187b | 5.54E-05 | -0.258055044 | 1 |
| ENSMUSG00000002635 | Pdcd2l | 5.54E-05 | -1.43862729 | 0.434488598 |
| ENSMUSG00000085105 | Gm12758 | 5.54E-05 | 1.326907457 | 0.971854959 |
| ENSMUSG00000086184 | Gm12764 | 5.54E-05 | -0.051604167 | 1 |
| ENSMUSG00000030499 | Kctd15 | 5.54E-05 | -0.051604167 | 1 |
| ENSMUSG00000085923 | Gm12781 | 5.54E-05 | 1.648835551 | 0.886203497 |
| ENSMUSG00000001802 | Lrp3 | 5.54E-05 | 2.063873051 | 0.792441218 |
| ENSMUSG00000030493 | Faap24 | 5.54E-05 | 0.648835551 | 1 |
| ENSMUSG00000030492 | Slc7a9 | 5.54E-05 | 0.47891055 | 1 |
| ENSMUSG00000030491 | Tdrd12 | 5.54E-05 | 0.189403933 | 1 |
| ENSMUSG00000108749 | Gm44767 | 5.54E-05 | 1.063873051 | 1 |
| ENSMUSG00000068959 | Zfp619 | 5.54E-05 | 0.47891055 | 1 |
| ENSMUSG00000100650 | Gm28807 | 5.54E-05 | -0.158519371 | 1 |
| ENSMUSG00000102004 | 4933402C06Rik | 5.54E-05 | 0.326907457 | 1 |
| ENSMUSG00000108490 | 1700110I07Rik | 5.54E-05 | 2.648835551 | 0.66416768 |
| ENSMUSG00000074165 | Zfp788 | 5.54E-05 | -1.305360759 | 0.491653643 |
| ENSMUSG00000092032 | Vmn2r59 | 5.54E-05 | 1.063873051 | 1 |
| ENSMUSG00000090967 | Vmn2r61 | 5.54E-05 | 1.648835551 | 0.886203497 |
| ENSMUSG00000108446 | Gm44997 | 5.54E-05 | 0.326907457 | 1 |
| ENSMUSG00000091594 | Gm17067 | 5.54E-05 | 2.063873051 | 0.792441218 |
| ENSMUSG00000090751 | Vmn2r63 | 5.54E-05 | 0.47891055 | 1 |
| ENSMUSG00000074887 | EU599041 | 5.54E-05 | 2.648835551 | 0.66416768 |
| ENSMUSG00000013353 | 4931406B18Rik | 5.54E-05 | 3.648835551 | 0.466464457 |
| ENSMUSG00000030472 | Ceacam18 | 5.54E-05 | 1.063873051 | 1 |
| ENSMUSG00000038888 | Ctu1 | 5.54E-05 | 0.063873051 | 1 |

| | | | | |
|---|---|---|---|---|
| ENSMUSG00000030693 | Klk10 | 5.54E-05 | 2.648835551 | 0.66416768 |
| ENSMUSG00000108432 | Gm44756 | 5.54E-05 | 0.841480629 | 1 |
| ENSMUSG00000038738 | Shank1 | 5.54E-05 | -0.351164449 | 1 |
| ENSMUSG00000008028 | 1700008O03Rik | 5.54E-05 | 0.648835551 | 1 |
| ENSMUSG00000030731 | Syt3 | 5.54E-05 | -0.936126949 | 0.86218124 |
| ENSMUSG00000047085 | Lrrc4b | 5.54E-05 | 0.648835551 | 1 |
| ENSMUSG00000066500 | Izumo2 | 5.54E-05 | 2.063873051 | 0.792441218 |
| ENSMUSG00000048012 | Zfp473 | 5.54E-05 | 1.648835551 | 0.87923507 |
| ENSMUSG00000109947 | Gm45669 | 5.54E-05 | 0.063873051 | 1 |
| ENSMUSG00000007833 | Aldh16a1 | 5.54E-05 | -0.351164449 | 1 |
| ENSMUSG00000038292 | Ccdc155 | 5.54E-05 | 1.326907457 | 0.971854959 |
| ENSMUSG00000030798 | Cd37 | 5.54E-05 | 0.326907457 | 1 |
| ENSMUSG00000038201 | Kcna7 | 5.54E-05 | 1.648835551 | 0.877701541 |
| ENSMUSG00000011382 | Dhdh | 5.54E-05 | -1.209145444 | 0.622539808 |
| ENSMUSG00000023467 | Tulp2 | 5.54E-05 | -0.158519371 | 1 |
| ENSMUSG00000110357 | A030001D20Rik | 5.54E-05 | 3.648835551 | 0.466464457 |
| ENSMUSG00000042918 | Mamstr | 5.54E-05 | 0.326907457 | 1 |
| ENSMUSG00000003273 | Car11 | 5.54E-05 | 3.648835551 | 0.466464457 |
| ENSMUSG00000059824 | Dbp | 5.54E-05 | 0.841480629 | 1 |
| ENSMUSG00000040189 | Ccdc114 | 5.54E-05 | -2.636566667 | 0.027656333 |
| ENSMUSG00000030834 | Abcc6 | 5.54E-05 | 0.189403933 | 1 |
| ENSMUSG00000096146 | Kcnj11 | 5.54E-05 | 0.47891055 | 1 |
| ENSMUSG00000110316 | Gm45311 | 5.54E-05 | 1.648835551 | 0.886203497 |
| ENSMUSG00000052303 | Mrgpra6 | 5.54E-05 | 2.648835551 | 0.66416768 |
| ENSMUSG00000070552 | Mrgprx1 | 5.54E-05 | -1.209145444 | 0.607677014 |
| ENSMUSG00000108950 | 9130015G15Rik | 5.54E-05 | 2.063873051 | 0.787908961 |
| ENSMUSG00000039728 | Slc6a5 | 5.54E-05 | 0.648835551 | 1 |
| ENSMUSG00000055489 | Ano5 | 5.54E-05 | -0.599091962 | 1 |
| ENSMUSG00000092118 | Fancf | 5.54E-05 | 1.326907457 | 0.971854959 |
| ENSMUSG00000108659 | Gm34121 | 5.54E-05 | 3.648835551 | 0.466464457 |
| ENSMUSG00000055048 | Gm9962 | 5.54E-05 | -0.051604167 | 1 |
| ENSMUSG00000109321 | A330076H08Rik | 5.54E-05 | 1.063873051 | 1 |
| ENSMUSG00000045813 | Gm9801 | 5.54E-05 | 0.326907457 | 1 |
| ENSMUSG00000033458 | Fan1 | 5.54E-05 | -0.673092543 | 1 |
| ENSMUSG00000070520 | Nsmce3 | 5.54E-05 | 0.47891055 | 1 |
| ENSMUSG00000086628 | Gm16157 | 5.54E-05 | 1.063873051 | 1 |
| ENSMUSG00000074071 | Fam169b | 5.54E-05 | -0.52108945 | 1 |
| ENSMUSG00000086410 | Gm16158 | 5.54E-05 | 0.47891055 | 1 |
| ENSMUSG00000108828 | Gm34549 | 5.54E-05 | 2.063873051 | 0.792441218 |
| ENSMUSG00000109407 | Gm34664 | 5.54E-05 | -0.051604167 | 1 |
| ENSMUSG00000109021 | Gm44812 | 5.54E-05 | 2.648835551 | 0.66416768 |
| ENSMUSG00000099853 | Gm29328 | 5.54E-05 | -0.52108945 | 1 |
| ENSMUSG00000108669 | 4930441H08Rik | 5.54E-05 | 1.326907457 | 0.912904617 |
| ENSMUSG00000108778 | Gm20083 | 5.54E-05 | 3.648835551 | 0.466464457 |
| ENSMUSG00000070509 | Rgma | 5.54E-05 | -1.051604167 | 0.730578151 |
| ENSMUSG00000108652 | 0610006L08Rik | 5.54E-05 | 0.648835551 | 1 |
| ENSMUSG00000097441 | Gm26633 | 5.54E-05 | 0.648835551 | 1 |
| ENSMUSG00000097023 | Mir9-3hg | 5.54E-05 | 1.063873051 | 1 |
| ENSMUSG00000030546 | Plin1 | 5.54E-05 | 1.648835551 | 0.864413622 |
| ENSMUSG00000108425 | Gm44706 | 5.54E-05 | 3.648835551 | 0.466464457 |
| ENSMUSG00000050973 | Gdpgp1 | 5.54E-05 | -0.52108945 | 0.975624465 |

| | | | | |
|---|---|---|---|---|
| ENSMUSG00000045467 | Ttll13 | 5.54E-05 | 1.063873051 | 1 |
| ENSMUSG00000030534 | Vps33b | 5.54E-05 | -0.743481871 | 0.827142217 |
| ENSMUSG00000030533 | Unc45a | 5.54E-05 | -1.051604167 | 0.724721153 |
| ENSMUSG00000053158 | Fes | 5.54E-05 | -1.480447466 | 0.471015773 |
| ENSMUSG00000038797 | Zscan2 | 5.54E-05 | -1.673092543 | 0.290284345 |
| ENSMUSG00000062444 | Ap3b2 | 5.54E-05 | 1.326907457 | 0.971650633 |
| ENSMUSG00000038663 | Fsd2 | 5.54E-05 | 0.841480629 | 1 |
| ENSMUSG00000025105 | Bnc1 | 5.54E-05 | 2.063873051 | 0.792441218 |
| ENSMUSG00000086690 | 4933406J10Rik | 5.54E-05 | 3.648835551 | 0.466464457 |
| ENSMUSG00000053528 | A530021J07Rik | 5.54E-05 | -0.351164449 | 1 |
| ENSMUSG00000001741 | Il16 | 5.54E-05 | -0.673092543 | 0.98880775 |
| ENSMUSG00000097016 | Gm26708 | 5.54E-05 | -0.874726405 | 0.853905061 |
| ENSMUSG00000109091 | Gm44827 | 5.54E-05 | 2.063873051 | 0.789767653 |
| ENSMUSG00000085236 | 2610206C17Rik | 5.54E-05 | -0.351164449 | 1 |
| ENSMUSG00000091006 | Vmn2r69 | 5.54E-05 | 2.063873051 | 0.792441218 |
| ENSMUSG00000090806 | Vmn2r70 | 5.54E-05 | 3.648835551 | 0.466464457 |
| ENSMUSG00000054054 | Olfr309 | 5.54E-05 | 3.648835551 | 0.466464457 |
| ENSMUSG00000055610 | Olfr307 | 5.54E-05 | 3.648835551 | 0.466464457 |
| ENSMUSG00000059319 | Olfr295 | 5.54E-05 | 2.648835551 | 0.66416768 |
| ENSMUSG00000001773 | Folh1 | 5.54E-05 | 0.841480629 | 1 |
| ENSMUSG00000090949 | Vmn2r77 | 5.54E-05 | 2.648835551 | 0.66416768 |
| ENSMUSG00000091962 | Vmn2r78 | 5.54E-05 | 2.648835551 | 0.66416768 |
| ENSMUSG00000109244 | Gm44751 | 5.54E-05 | 1.063873051 | 1 |
| ENSMUSG00000049791 | Fzd4 | 5.54E-05 | 1.326907457 | 0.971854959 |
| ENSMUSG00000109311 | AI314278 | 5.54E-05 | 2.063873051 | 0.792441218 |
| ENSMUSG00000097585 | E230029C05Rik | 5.54E-05 | -1.395558568 | 0.559373995 |
| ENSMUSG00000030617 | Ccdc83 | 5.54E-05 | 0.47891055 | 1 |
| ENSMUSG00000109382 | 4930567K12Rik | 5.54E-05 | -2.023589791 | 0.13144915 |
| ENSMUSG00000030613 | Ccdc90b | 5.54E-05 | -0.936126949 | 0.695372548 |
| ENSMUSG00000098041 | Gm26981 | 5.54E-05 | 0.47891055 | 1 |
| ENSMUSG00000054061 | Gm9934 | 5.54E-05 | 1.326907457 | 0.971854959 |
| ENSMUSG00000109175 | Gm44947 | 5.54E-05 | 0.648835551 | 1 |
| ENSMUSG00000044952 | Kctd21 | 5.54E-05 | 0.063873051 | 1 |
| ENSMUSG00000035704 | Alg8 | 5.54E-05 | -1.351164449 | 0.499088689 |
| ENSMUSG00000035642 | Aamdc | 5.54E-05 | -2.328444372 | 0.062688794 |
| ENSMUSG00000074004 | B3gnt6 | 5.54E-05 | 1.326907457 | 0.971854959 |
| ENSMUSG00000049580 | Tsku | 5.54E-05 | 0.063873051 | 1 |
| ENSMUSG00000052396 | Mogat2 | 5.54E-05 | 0.841480629 | 1 |
| ENSMUSG00000035211 | Xrra1 | 5.54E-05 | 0.841480629 | 1 |
| ENSMUSG00000030725 | Lipt2 | 5.54E-05 | 0.841480629 | 1 |
| ENSMUSG00000030708 | Dnajb13 | 5.54E-05 | 1.326907457 | 0.971854959 |
| ENSMUSG00000044881 | Coa4 | 5.54E-05 | 0.648835551 | 1 |
| ENSMUSG00000008318 | Relt | 5.54E-05 | 1.648835551 | 0.886203497 |
| ENSMUSG00000110301 | Gm35363 | 5.54E-05 | 1.326907457 | 0.950046029 |
| ENSMUSG00000073985 | Gm10602 | 5.54E-05 | 0.841480629 | 1 |
| ENSMUSG00000001827 | Folr1 | 5.54E-05 | 2.063873051 | 0.788062228 |
| ENSMUSG00000030996 | Art1 | 5.54E-05 | 2.648835551 | 0.66416768 |
| ENSMUSG00000066279 | Chrna10 | 5.54E-05 | 3.648835551 | 0.466464457 |
| ENSMUSG00000043925 | Olfr544 | 5.54E-05 | 1.648835551 | 0.886203497 |
| ENSMUSG00000073967 | Olfr557 | 5.54E-05 | 1.648835551 | 0.847265357 |
| ENSMUSG00000066273 | Olfr33 | 5.54E-05 | 2.648835551 | 0.66416768 |

| | | | | |
|---|---|---|---|---|
| ENSMUSG00000084262 | Olfr588-ps1 | 5.54E-05 | 2.063873051 | 0.792441218 |
| ENSMUSG00000058976 | Usp17lc | 5.54E-05 | 1.326907457 | 0.971854959 |
| ENSMUSG00000045584 | Olfr610 | 5.54E-05 | 2.648835551 | 0.66416768 |
| ENSMUSG00000073945 | Olfr618 | 5.54E-05 | 1.648835551 | 0.886203497 |
| ENSMUSG00000094520 | Olfr635 | 5.54E-05 | 2.063873051 | 0.792441218 |
| ENSMUSG00000066262 | Olfr640 | 5.54E-05 | 2.648835551 | 0.66416768 |
| ENSMUSG00000073932 | Olfr641 | 5.54E-05 | 2.648835551 | 0.66416768 |
| ENSMUSG00000052749 | Trim30b | 5.54E-05 | 3.648835551 | 0.466464457 |
| ENSMUSG00000044705 | Olfr670 | 5.54E-05 | 3.648835551 | 0.466464457 |
| ENSMUSG00000094531 | Olfr671 | 5.54E-05 | 2.648835551 | 0.66416768 |
| ENSMUSG00000096773 | Olfr675 | 5.54E-05 | 1.063873051 | 1 |
| ENSMUSG00000093378 | Gm20663 | 5.54E-05 | 1.063873051 | 1 |
| ENSMUSG00000036744 | Olfr701 | 5.54E-05 | 3.648835551 | 0.466464457 |
| ENSMUSG00000056863 | Olfr702 | 5.54E-05 | 2.648835551 | 0.66416768 |
| ENSMUSG00000036647 | Olfr6 | 5.54E-05 | 3.648835551 | 0.466464457 |
| ENSMUSG00000070417 | Olfr2 | 5.54E-05 | 2.648835551 | 0.66416768 |
| ENSMUSG00000062987 | Olfr715b | 5.54E-05 | 2.648835551 | 0.66416768 |
| ENSMUSG00000016626 | Nlrp14 | 5.54E-05 | 2.648835551 | 0.66416768 |
| ENSMUSG00000051041 | Olfml1 | 5.54E-05 | -1.480447466 | 0.443153295 |
| ENSMUSG00000048236 | Ovch2 | 5.54E-05 | 1.326907457 | 0.971854959 |
| ENSMUSG00000043855 | Olfr479 | 5.54E-05 | 2.648835551 | 0.66416768 |
| ENSMUSG00000063120 | Olfr480 | 5.54E-05 | 3.648835551 | 0.466464457 |
| ENSMUSG00000046431 | Olfr518 | 5.54E-05 | 3.648835551 | 0.466464457 |
| ENSMUSG00000049709 | Nlrp10 | 5.54E-05 | 1.063873051 | 1 |
| ENSMUSG00000031026 | Trim66 | 5.54E-05 | 1.063873051 | 1 |
| ENSMUSG00000031023 | Akip1 | 5.54E-05 | 0.841480629 | 1 |
| ENSMUSG00000031022 | BC051019 | 5.54E-05 | 3.648835551 | 0.466464457 |
| ENSMUSG00000034825 | Nrip3 | 5.54E-05 | 2.063873051 | 0.792441218 |
| ENSMUSG00000099803 | Gm28863 | 5.54E-05 | -0.258055044 | 1 |
| ENSMUSG00000030790 | Adm | 5.54E-05 | -0.258055044 | 1 |
| ENSMUSG00000109700 | Gm21123 | 5.54E-05 | 1.648835551 | 0.871433572 |
| ENSMUSG00000030787 | Lyve1 | 5.54E-05 | 1.326907457 | 0.971854959 |
| ENSMUSG00000110207 | Gm33586 | 5.54E-05 | 2.063873051 | 0.792441218 |
| ENSMUSG00000038187 | Btbd10 | 5.54E-05 | -2.52108945 | 0.037394962 |
| ENSMUSG00000073862 | Gm5600 | 5.54E-05 | 2.648835551 | 0.66416768 |
| ENSMUSG00000108489 | Gm45025 | 5.54E-05 | 2.648835551 | 0.66416768 |
| ENSMUSG00000087621 | 1700003G18Rik | 5.54E-05 | 2.648835551 | 0.66416768 |
| ENSMUSG00000108461 | AV356131 | 5.54E-05 | 0.841480629 | 1 |
| ENSMUSG00000108303 | Gm39075 | 5.54E-05 | 1.326907457 | 0.971854959 |
| ENSMUSG00000033904 | Ccp110 | 5.54E-05 | -0.810596067 | 0.841517001 |
| ENSMUSG00000109517 | Gm44763 | 5.54E-05 | 2.648835551 | 0.66416768 |
| ENSMUSG00000066197 | Gpr139 | 5.54E-05 | 1.326907457 | 0.958408446 |
| ENSMUSG00000030945 | Acsm2 | 5.54E-05 | 2.063873051 | 0.792441218 |
| ENSMUSG00000047026 | Acsm4 | 5.54E-05 | 0.841480629 | 1 |
| ENSMUSG00000030935 | Acsm3 | 5.54E-05 | 0.648835551 | 1 |
| ENSMUSG00000030917 | Tmem159 | 5.54E-05 | 0.841480629 | 1 |
| ENSMUSG00000030905 | Crym | 5.54E-05 | 1.063873051 | 1 |
| ENSMUSG00000034990 | Otoa | 5.54E-05 | -0.673092543 | 1 |
| ENSMUSG00000108811 | 4933432K03Rik | 5.54E-05 | 0.648835551 | 1 |
| ENSMUSG00000000216 | Scnn1g | 5.54E-05 | 2.648835551 | 0.66416768 |
| ENSMUSG00000044702 | Palb2 | 5.54E-05 | -1.351164449 | 0.499088689 |

| | | | | |
|---|---|---|---|---|
| ENSMUSG00000086942 | Gm15489 | 5.54E-05 | 0.47891055 | 1 |
| ENSMUSG00000030866 | Ern2 | 5.54E-05 | -0.936126949 | 0.695372548 |
| ENSMUSG00000030865 | Chp2 | 5.54E-05 | 3.648835551 | 0.466464457 |
| ENSMUSG00000108551 | Gm20274 | 5.54E-05 | 1.648835551 | 0.886203497 |
| ENSMUSG00000030769 | Slc5a11 | 5.54E-05 | 0.47891055 | 1 |
| ENSMUSG00000086254 | 3100003L05Rik | 5.54E-05 | -0.158519371 | 1 |
| ENSMUSG00000108676 | 1700123J17Rik | 5.54E-05 | -0.258055044 | 1 |
| ENSMUSG00000030741 | Spns1 | 5.54E-05 | -0.43862729 | 1 |
| ENSMUSG00000030722 | Nfatc2ip | 5.54E-05 | -0.874726405 | 0.840804483 |
| ENSMUSG00000042759 | Apobr | 5.54E-05 | 2.648835551 | 0.66416768 |
| ENSMUSG00000059772 | Slx1b | 5.54E-05 | -1.560617814 | 0.405849979 |
| ENSMUSG00000093768 | Gm20650 | 5.54E-05 | 2.063873051 | 0.792441218 |
| ENSMUSG00000030683 | Sez6l2 | 5.54E-05 | -0.258055044 | 1 |
| ENSMUSG00000030677 | Kif22 | 5.54E-05 | -1.480447466 | 0.441576406 |
| ENSMUSG00000045165 | AI467606 | 5.54E-05 | 0.47891055 | 1 |
| ENSMUSG00000045598 | Zfp553 | 5.54E-05 | -0.743481871 | 1 |
| ENSMUSG00000045757 | Zfp764 | 5.54E-05 | 1.063873051 | 0.970970089 |
| ENSMUSG00000045251 | Zfp688 | 5.54E-05 | 1.326907457 | 0.971854959 |
| ENSMUSG00000048921 | Zfp689 | 5.54E-05 | 0.648835551 | 1 |
| ENSMUSG00000101845 | Gm28198 | 5.54E-05 | 1.648835551 | 0.886203497 |
| ENSMUSG00000060034 | Ctf2 | 5.54E-05 | 3.648835551 | 0.466464457 |
| ENSMUSG00000043964 | Orai3 | 5.54E-05 | 1.063873051 | 1 |
| ENSMUSG00000042289 | Hsd3b7 | 5.54E-05 | -1.158519371 | 0.599135879 |
| ENSMUSG00000030806 | Stx1b | 5.54E-05 | -0.351164449 | 1 |
| ENSMUSG00000049728 | Zfp668 | 5.54E-05 | -1.051604167 | 0.706929043 |
| ENSMUSG00000030801 | Kat8 | 5.54E-05 | -1.051604167 | 0.706929043 |
| ENSMUSG00000030800 | Prss8 | 5.54E-05 | 0.326907457 | 1 |
| ENSMUSG00000070371 | Prss36 | 5.54E-05 | -0.52108945 | 0.975624465 |
| ENSMUSG00000030789 | Itgax | 5.54E-05 | 1.326907457 | 0.971854959 |
| ENSMUSG00000030780 | BC017158 | 5.54E-05 | -1.395558568 | 0.542200603 |
| ENSMUSG00000108668 | Gm32816 | 5.54E-05 | 0.47891055 | 1 |
| ENSMUSG00000070366 | Plpp4 | 5.54E-05 | -1.158519371 | 0.612182491 |
| ENSMUSG00000108504 | Gm4265 | 5.54E-05 | 0.841480629 | 1 |
| ENSMUSG00000109014 | Gm44546 | 5.54E-05 | 0.841480629 | 1 |
| ENSMUSG00000108870 | Gm44893 | 5.54E-05 | -1.52108945 | 0.452184898 |
| ENSMUSG00000087348 | Gm15582 | 5.54E-05 | 3.648835551 | 0.466464457 |
| ENSMUSG00000109816 | 4930483O08Rik | 5.54E-05 | -1.43862729 | 0.466464457 |
| ENSMUSG00000030981 | Mmp21 | 5.54E-05 | 2.063873051 | 0.792441218 |
| ENSMUSG00000030979 | Uros | 5.54E-05 | -1.480447466 | 0.441576406 |
| ENSMUSG00000053111 | Fank1 | 5.54E-05 | -1.106051951 | 0.743798867 |
| ENSMUSG00000073805 | Insyn2a | 5.54E-05 | 1.063873051 | 1 |
| ENSMUSG00000109891 | Gm45339 | 5.54E-05 | 0.47891055 | 1 |
| ENSMUSG00000050866 | Clrn3 | 5.54E-05 | 2.063873051 | 0.792441218 |
| ENSMUSG00000109831 | Gm36431 | 5.54E-05 | 0.063873051 | 1 |
| ENSMUSG00000110027 | C030029H02Rik | 5.54E-05 | 1.063873051 | 1 |
| ENSMUSG00000110037 | 4930543N07Rik | 5.54E-05 | 0.063873051 | 1 |
| ENSMUSG00000041775 | Mapk1ip1 | 5.54E-05 | -0.810596067 | 0.847265357 |
| ENSMUSG00000025478 | Dpysl4 | 5.54E-05 | 2.648835551 | 0.66416768 |
| ENSMUSG00000015980 | Lrrc27 | 5.54E-05 | 0.648835551 | 1 |
| ENSMUSG00000025474 | Tubgcp2 | 5.54E-05 | -1.106051951 | 0.719591951 |
| ENSMUSG00000025470 | Zfp511 | 5.54E-05 | -0.258055044 | 1 |

| | | | | |
|---|---|---|---|---|
| ENSMUSG00000025469 | Msx3 | 5.54E-05 | 3.648835551 | 0.466464457 |
| ENSMUSG00000054672 | 5830411N06Rik | 5.54E-05 | 2.648835551 | 0.66416768 |
| ENSMUSG00000060112 | Olfr60 | 5.54E-05 | 2.063873051 | 0.786367309 |
| ENSMUSG00000093942 | Olfr46 | 5.54E-05 | 1.648835551 | 0.886203497 |
| ENSMUSG00000054662 | Ano9 | 5.54E-05 | 0.189403933 | 1 |
| ENSMUSG00000025512 | Chid1 | 5.54E-05 | -1.051604167 | 0.706929043 |
| ENSMUSG00000048191 | Muc6 | 5.54E-05 | -0.43862729 | 1 |
| ENSMUSG00000109936 | Gm45889 | 5.54E-05 | 3.648835551 | 0.466464457 |
| ENSMUSG00000043795 | Prr33 | 5.54E-05 | 2.648835551 | 0.66416768 |
| ENSMUSG00000087090 | Nctc1 | 5.54E-05 | 2.063873051 | 0.792441218 |
| ENSMUSG00000010755 | Cars | 5.54E-05 | -2.794107944 | 0.013337086 |
| ENSMUSG00000045362 | Tnfrsf26 | 5.54E-05 | 0.326907457 | 1 |
| ENSMUSG00000109251 | E230032D23Rik | 5.54E-05 | -1.305360759 | 0.668075609 |
| ENSMUSG00000048965 | Mrgpre | 5.54E-05 | 1.326907457 | 0.971854959 |
| ENSMUSG00000085745 | Gm14372 | 5.54E-05 | 0.841480629 | 1 |
| ENSMUSG00000108977 | Gm39119 | 5.54E-05 | 3.648835551 | 0.466464457 |
| ENSMUSG00000051207 | Mrgprd | 5.54E-05 | 1.648835551 | 0.886203497 |
| ENSMUSG00000004567 | Mcoln1 | 5.54E-05 | -1.777429203 | 0.241307011 |
| ENSMUSG00000012705 | Retn | 5.54E-05 | 3.648835551 | 0.466464457 |
| ENSMUSG00000089665 | Fcor | 5.54E-05 | 1.063873051 | 0.954021409 |
| ENSMUSG00000031495 | Cd209d | 5.54E-05 | 3.648835551 | 0.466464457 |
| ENSMUSG00000089736 | Tgfbr3l | 5.54E-05 | -0.051604167 | 1 |
| ENSMUSG00000058748 | Zfp958 | 5.54E-05 | -1.52108945 | 0.452184898 |
| ENSMUSG00000023073 | Slc10a2 | 5.54E-05 | -0.995020638 | 0.699192777 |
| ENSMUSG00000092092 | Gm17215 | 5.54E-05 | 0.47891055 | 1 |
| ENSMUSG00000109436 | Gm45075 | 5.54E-05 | -0.051604167 | 1 |
| ENSMUSG00000109346 | Gm44515 | 5.54E-05 | 2.063873051 | 0.792441218 |
| ENSMUSG00000067627 | Gm10217 | 5.54E-05 | 3.648835551 | 0.466464457 |
| ENSMUSG00000109545 | Gm44788 | 5.54E-05 | 0.47891055 | 1 |
| ENSMUSG00000109029 | Gm44717 | 5.54E-05 | 2.063873051 | 0.792441218 |
| ENSMUSG00000085573 | Gm15418 | 5.54E-05 | 0.189403933 | 1 |
| ENSMUSG00000031512 | Tex29 | 5.54E-05 | 1.326907457 | 0.94640403 |
| ENSMUSG00000074469 | Gm15348 | 5.54E-05 | 2.063873051 | 0.791976577 |
| ENSMUSG00000031443 | F7 | 5.54E-05 | 1.063873051 | 1 |
| ENSMUSG00000091443 | Gm17023 | 5.54E-05 | 3.648835551 | 0.466464457 |
| ENSMUSG00000099338 | 2810030D12Rik | 5.54E-05 | 0.47891055 | 1 |
| ENSMUSG00000110278 | Gm5608 | 5.54E-05 | 0.841480629 | 1 |
| ENSMUSG00000110058 | Gm5907 | 5.54E-05 | 3.648835551 | 0.466464457 |
| ENSMUSG00000098322 | 5830468F06Rik | 5.54E-05 | 3.648835551 | 0.466464457 |
| ENSMUSG00000087300 | Gm16725 | 5.54E-05 | 1.648835551 | 0.886203497 |
| ENSMUSG00000110530 | Gm35934 | 5.54E-05 | 0.841480629 | 1 |
| ENSMUSG00000061847 | Defb39 | 5.54E-05 | 2.063873051 | 0.792441218 |
| ENSMUSG00000110542 | Gm39139 | 5.54E-05 | 2.063873051 | 0.792441218 |
| ENSMUSG00000037790 | Defb7 | 5.54E-05 | 0.47891055 | 1 |
| ENSMUSG00000110463 | Gm35998 | 5.54E-05 | -0.051604167 | 1 |
| ENSMUSG00000071165 | Gm6040 | 5.54E-05 | 2.648835551 | 0.66416768 |
| ENSMUSG00000110616 | Gm36879 | 5.54E-05 | 0.841480629 | 1 |
| ENSMUSG00000063362 | Alg11 | 5.54E-05 | -0.874726405 | 0.840804483 |
| ENSMUSG00000031481 | Tpte | 5.54E-05 | -1.395558568 | 0.554032151 |
| ENSMUSG00000031482 | Slc25a15 | 5.54E-05 | -1.158519371 | 0.633493255 |
| ENSMUSG00000097042 | Gm17491 | 5.54E-05 | -1.743481871 | 0.305378889 |

| | | | | |
|---|---|---|---|---|
| ENSMUSG00000031539 | Ap3m2 | 5.54E-05 | -1.708716453 | 0.296030202 |
| ENSMUSG00000054822 | 1700041G16Rik | 5.54E-05 | 3.648835551 | 0.466464457 |
| ENSMUSG00000053979 | A730045E13Rik | 5.54E-05 | 3.648835551 | 0.466464457 |
| ENSMUSG00000097654 | Gm26714 | 5.54E-05 | 1.326907457 | 0.952530143 |
| ENSMUSG00000037437 | Adam32 | 5.54E-05 | -3.305360759 | 0.001866323 |
| ENSMUSG00000037406 | Htra4 | 5.54E-05 | 1.648835551 | 0.886203497 |
| ENSMUSG00000110148 | 5830408C22Rik | 5.54E-05 | -0.351164449 | 1 |
| ENSMUSG00000110063 | Gm31045 | 5.54E-05 | -0.158519371 | 1 |
| ENSMUSG00000061313 | Ddhd2 | 5.54E-05 | -1.936126949 | 0.158846729 |
| ENSMUSG00000109961 | Gm31983 | 5.54E-05 | 1.063873051 | 0.998731799 |
| ENSMUSG00000109794 | Gm45571 | 5.54E-05 | 3.648835551 | 0.466464457 |
| ENSMUSG00000110223 | Gm45489 | 5.54E-05 | 2.648835551 | 0.66416768 |
| ENSMUSG00000079070 | Gm3985 | 5.54E-05 | 0.326907457 | 1 |
| ENSMUSG00000110214 | Gm45587 | 5.54E-05 | 0.47891055 | 1 |
| ENSMUSG00000094500 | Smim18 | 5.54E-05 | 1.648835551 | 0.886203497 |
| ENSMUSG00000109964 | Gm45350 | 5.54E-05 | 2.063873051 | 0.792441218 |
| ENSMUSG00000110369 | Gm45349 | 5.54E-05 | 0.326907457 | 1 |
| ENSMUSG00000109880 | Gm34096 | 5.54E-05 | 3.648835551 | 0.466464457 |
| ENSMUSG00000110130 | Gm34419 | 5.54E-05 | 3.648835551 | 0.466464457 |
| ENSMUSG00000109607 | Gm34533 | 5.54E-05 | 3.648835551 | 0.466464457 |
| ENSMUSG00000109698 | Gm45469 | 5.54E-05 | 1.063873051 | 1 |
| ENSMUSG00000046794 | Ppp1r3b | 5.54E-05 | 0.47891055 | 1 |
| ENSMUSG00000097357 | Gm16793 | 5.54E-05 | -1.708716453 | 0.341869836 |
| ENSMUSG00000109761 | 5430403N17Rik | 5.54E-05 | 1.326907457 | 0.950123751 |
| ENSMUSG00000039633 | Lonrf1 | 5.54E-05 | -1.106051951 | 0.792441218 |
| ENSMUSG00000110285 | Gm45524 | 5.54E-05 | 0.189403933 | 1 |
| ENSMUSG00000031603 | Fgf20 | 5.54E-05 | 2.648835551 | 0.662993938 |
| ENSMUSG00000110168 | Gm45488 | 5.54E-05 | 0.326907457 | 1 |
| ENSMUSG00000110409 | Gm9908 | 5.54E-05 | 1.063873051 | 1 |
| ENSMUSG00000054764 | Mtnr1a | 5.54E-05 | 0.841480629 | 1 |
| ENSMUSG00000110231 | Gm6329 | 5.54E-05 | 2.648835551 | 0.66416768 |
| ENSMUSG00000071104 | Ccdc110 | 5.54E-05 | 1.648835551 | 0.886203497 |
| ENSMUSG00000071103 | 1700029J07Rik | 5.54E-05 | 0.063873051 | 1 |
| ENSMUSG00000110179 | Gm39168 | 5.54E-05 | 0.47891055 | 1 |
| ENSMUSG00000110264 | E330018M18Rik | 5.54E-05 | 1.326907457 | 0.971854959 |
| ENSMUSG00000110363 | Gm45576 | 5.54E-05 | 2.063873051 | 0.792441218 |
| ENSMUSG00000110539 | Gm45864 | 5.54E-05 | 3.648835551 | 0.466464457 |
| ENSMUSG00000031559 | 4930555F03Rik | 5.54E-05 | -1.209145444 | 0.66416768 |
| ENSMUSG00000110375 | Gm45323 | 5.54E-05 | 1.326907457 | 0.950506499 |
| ENSMUSG00000110310 | 4930518J21Rik | 5.54E-05 | 1.648835551 | 0.88176882 |
| ENSMUSG00000038215 | Cep44 | 5.54E-05 | -1.965874293 | 0.162298752 |
| ENSMUSG00000100510 | Hand2os1 | 5.54E-05 | 0.47891055 | 1 |
| ENSMUSG00000038193 | Hand2 | 5.54E-05 | 2.648835551 | 0.66416768 |
| ENSMUSG00000110162 | Gm45356 | 5.54E-05 | 1.326907457 | 0.971854959 |
| ENSMUSG00000049946 | BC030500 | 5.54E-05 | 3.648835551 | 0.466464457 |
| ENSMUSG00000070023 | Gm10283 | 5.54E-05 | 0.063873051 | 1 |
| ENSMUSG00000031647 | Mfap3l | 5.54E-05 | 1.648835551 | 0.880252773 |
| ENSMUSG00000031641 | Cbr4 | 5.54E-05 | 1.648835551 | 0.886203497 |
| ENSMUSG00000110368 | Gm45518 | 5.54E-05 | 2.648835551 | 0.66416768 |
| ENSMUSG00000109854 | Gm45345 | 5.54E-05 | 2.648835551 | 0.66416768 |
| ENSMUSG00000079042 | Apela | 5.54E-05 | 2.648835551 | 0.66416768 |

| | | | | |
|---|---|---|---|---|
| ENSMUSG00000087266 | Gm15991 | 5.54E-05 | 1.648835551 | 0.872990655 |
| ENSMUSG00000036330 | Slc18a1 | 5.54E-05 | -1.305360759 | 0.491653643 |
| ENSMUSG00000092260 | Zfp963 | 5.54E-05 | -0.051604167 | 1 |
| ENSMUSG00000031860 | Pbx4 | 5.54E-05 | -1.051604167 | 0.740075579 |
| ENSMUSG00000002341 | Ncan | 5.54E-05 | 1.326907457 | 0.968182137 |
| ENSMUSG00000079033 | Mef2b | 5.54E-05 | -0.258055044 | 1 |
| ENSMUSG00000002346 | Slc25a42 | 5.54E-05 | -2.777429203 | 0.02111872 |
| ENSMUSG00000002343 | Armc6 | 5.54E-05 | 0.063873051 | 1 |
| ENSMUSG00000038508 | Gdf15 | 5.54E-05 | 2.063873051 | 0.792441218 |
| ENSMUSG00000034911 | Ushbp1 | 5.54E-05 | 0.841480629 | 1 |
| ENSMUSG00000046295 | Ankle1 | 5.54E-05 | 2.648835551 | 0.66416768 |
| ENSMUSG00000007610 | Gtpbp3 | 5.54E-05 | -0.743481871 | 1 |
| ENSMUSG00000066880 | Zfp617 | 5.54E-05 | -2.184054463 | 0.091007367 |
| ENSMUSG00000061561 | Olfr373 | 5.54E-05 | 1.648835551 | 0.86556459 |
| ENSMUSG00000097472 | Gm26586 | 5.54E-05 | 3.648835551 | 0.466464457 |
| ENSMUSG00000031621 | Isx | 5.54E-05 | 1.648835551 | 0.886203497 |
| ENSMUSG00000110593 | Gm45822 | 5.54E-05 | 3.648835551 | 0.466464457 |
| ENSMUSG00000005410 | Mcm5 | 5.54E-05 | 0.189403933 | 1 |
| ENSMUSG00000110410 | 4933421D24Rik | 5.54E-05 | 2.648835551 | 0.66416768 |
| ENSMUSG00000110382 | Gm29895 | 5.54E-05 | 3.648835551 | 0.466464457 |
| ENSMUSG00000037070 | Rbmxl1 | 5.54E-05 | -0.158519371 | 1 |
| ENSMUSG00000031682 | 1700011L22Rik | 5.54E-05 | 1.648835551 | 0.886203497 |
| ENSMUSG00000109945 | Gm45601 | 5.54E-05 | 2.648835551 | 0.66416768 |
| ENSMUSG00000051839 | Gypa | 5.54E-05 | 2.648835551 | 0.66416768 |
| ENSMUSG00000109980 | Gm45538 | 5.54E-05 | 2.648835551 | 0.66416768 |
| ENSMUSG00000031710 | Ucp1 | 5.54E-05 | 2.648835551 | 0.66416768 |
| ENSMUSG00000035151 | Elmod2 | 5.54E-05 | -0.599091962 | 0.98880775 |
| ENSMUSG00000035057 | Mgat4d | 5.54E-05 | 2.063873051 | 0.792441218 |
| ENSMUSG00000110580 | D830024N08Rik | 5.54E-05 | -0.258055044 | 1 |
| ENSMUSG00000046408 | 1700067K01Rik | 5.54E-05 | 2.063873051 | 0.792441218 |
| ENSMUSG00000074217 | 2210011C24Rik | 5.54E-05 | 3.648835551 | 0.466464457 |
| ENSMUSG00000037103 | Dcaf15 | 5.54E-05 | -0.351164449 | 1 |
| ENSMUSG00000097120 | Gm26887 | 5.54E-05 | 1.063873051 | 1 |
| ENSMUSG00000004994 | Ccdc130 | 5.54E-05 | -0.43862729 | 1 |
| ENSMUSG00000086067 | Gm16183 | 5.54E-05 | 3.648835551 | 0.466464457 |
| ENSMUSG00000034041 | Lyl1 | 5.54E-05 | 1.063873051 | 1 |
| ENSMUSG00000003809 | Gcdh | 5.54E-05 | 0.063873051 | 1 |
| ENSMUSG00000087026 | A230103J11Rik | 5.54E-05 | 3.648835551 | 0.466464457 |
| ENSMUSG00000109754 | Gm39214 | 5.54E-05 | 0.47891055 | 1 |
| ENSMUSG00000036840 | Siah1a | 5.54E-05 | -2.158519371 | 0.115633409 |
| ENSMUSG00000098768 | Gm27207 | 5.54E-05 | 3.648835551 | 0.466464457 |
| ENSMUSG00000031661 | Nkd1 | 5.54E-05 | -0.158519371 | 1 |
| ENSMUSG00000031665 | Sall1 | 5.54E-05 | 1.648835551 | 0.886203497 |
| ENSMUSG00000109672 | Gm45639 | 5.54E-05 | 1.326907457 | 0.964003521 |
| ENSMUSG00000110089 | Gm36243 | 5.54E-05 | 2.648835551 | 0.66416768 |
| ENSMUSG00000110138 | 4831440D22Rik | 5.54E-05 | 0.648835551 | 1 |
| ENSMUSG00000031734 | Irx3 | 5.54E-05 | -1.158519371 | 0.599135879 |
| ENSMUSG00000046413 | Irx3os | 5.54E-05 | 1.063873051 | 1 |
| ENSMUSG00000101613 | Gm28516 | 5.54E-05 | 3.648835551 | 0.466464457 |
| ENSMUSG00000110380 | Gm45332 | 5.54E-05 | 2.648835551 | 0.66416768 |
| ENSMUSG00000031737 | Irx5 | 5.54E-05 | 0.326907457 | 1 |

| | | | | |
|---|---|---|---|---|
| ENSMUSG00000078964 | Ces1b | 5.54E-05 | 3.648835551 | 0.466464457 |
| ENSMUSG00000110718 | Gm45765 | 5.54E-05 | 2.648835551 | 0.66416768 |
| ENSMUSG00000031755 | Bbs2 | 5.54E-05 | -0.351164449 | 1 |
| ENSMUSG00000110524 | Gm39228 | 5.54E-05 | 2.648835551 | 0.66416768 |
| ENSMUSG00000110517 | Gm31224 | 5.54E-05 | 3.648835551 | 0.466464457 |
| ENSMUSG00000046556 | Zfp319 | 5.54E-05 | 0.648835551 | 1 |
| ENSMUSG00000110447 | Gm31518 | 5.54E-05 | 1.063873051 | 1 |
| ENSMUSG00000110489 | Gm31659 | 5.54E-05 | -0.158519371 | 1 |
| ENSMUSG00000061104 | Sap18b | 5.54E-05 | 1.063873051 | 0.975624465 |
| ENSMUSG00000110634 | Gm45895 | 5.54E-05 | 2.063873051 | 0.792441218 |
| ENSMUSG00000110633 | Gm32122 | 5.54E-05 | -0.051604167 | 1 |
| ENSMUSG00000089819 | Gm15679 | 5.54E-05 | 2.063873051 | 0.792441218 |
| ENSMUSG00000109769 | Gm39232 | 5.54E-05 | 0.189403933 | 1 |
| ENSMUSG00000109848 | Gm45391 | 5.54E-05 | 1.063873051 | 1 |
| ENSMUSG00000109699 | Gm45277 | 5.54E-05 | 1.648835551 | 0.863493415 |
| ENSMUSG00000110497 | 9230110F11Rik | 5.54E-05 | 2.648835551 | 0.66416768 |
| ENSMUSG00000069922 | Ces3a | 5.54E-05 | 1.326907457 | 0.971854959 |
| ENSMUSG00000062181 | Ces3b | 5.54E-05 | 1.648835551 | 0.886203497 |
| ENSMUSG00000060560 | Ces4a | 5.54E-05 | 1.063873051 | 1 |
| ENSMUSG00000043251 | Exoc3l | 5.54E-05 | 1.648835551 | 0.886203497 |
| ENSMUSG00000014791 | Elmo3 | 5.54E-05 | -0.43862729 | 1 |
| ENSMUSG00000041679 | Lrrc29 | 5.54E-05 | -1.52108945 | 0.452184898 |
| ENSMUSG00000051648 | Kctd19 | 5.54E-05 | 1.648835551 | 0.886203497 |
| ENSMUSG00000013158 | 4933405L10Rik | 5.54E-05 | 3.648835551 | 0.466464457 |
| ENSMUSG00000087436 | Gm16156 | 5.54E-05 | 1.326907457 | 0.971854959 |
| ENSMUSG00000110457 | Gm45752 | 5.54E-05 | 1.063873051 | 1 |
| ENSMUSG00000115067 | Dpep2 | 5.54E-05 | 0.326907457 | 1 |
| ENSMUSG00000115768 | Dpep2nb | 5.54E-05 | 2.063873051 | 0.791749236 |
| ENSMUSG00000086390 | 1810019D21Rik | 5.54E-05 | -0.995020638 | 0.699192777 |
| ENSMUSG00000031903 | Pla2g15 | 5.54E-05 | -0.810596067 | 0.844591606 |
| ENSMUSG00000003849 | Nqo1 | 5.54E-05 | 0.189403933 | 1 |
| ENSMUSG00000031727 | Pmfbp1 | 5.54E-05 | 0.648835551 | 1 |
| ENSMUSG00000031723 | Txnl4b | 5.54E-05 | 0.47891055 | 1 |
| ENSMUSG00000031730 | Dhodh | 5.54E-05 | 0.841480629 | 1 |
| ENSMUSG00000110507 | 8030455M16Rik | 5.54E-05 | 2.063873051 | 0.792441218 |
| ENSMUSG00000091041 | Gm17720 | 5.54E-05 | -0.351164449 | 1 |
| ENSMUSG00000033763 | Mtss2 | 5.54E-05 | 0.063873051 | 1 |
| ENSMUSG00000031951 | Tmem231 | 5.54E-05 | -0.810596067 | 0.873717735 |
| ENSMUSG00000033430 | Terf2ip | 5.54E-05 | -1.52108945 | 0.452184898 |
| ENSMUSG00000110541 | Gm10280 | 5.54E-05 | 2.648835551 | 0.66416768 |
| ENSMUSG00000078908 | Mon1b | 5.54E-05 | 0.189403933 | 1 |
| ENSMUSG00000031767 | Nudt7 | 5.54E-05 | -1.106051951 | 0.758130694 |
| ENSMUSG00000034467 | Dynlrb2 | 5.54E-05 | 0.326907457 | 1 |
| ENSMUSG00000031756 | Cenpn | 5.54E-05 | 0.841480629 | 1 |
| ENSMUSG00000031847 | 1700030J22Rik | 5.54E-05 | 0.47891055 | 1 |
| ENSMUSG00000110151 | Gm38416 | 5.54E-05 | -0.599091962 | 1 |
| ENSMUSG00000031837 | Necab2 | 5.54E-05 | 0.648835551 | 1 |
| ENSMUSG00000031831 | Dnaaf1 | 5.54E-05 | 1.326907457 | 0.971854959 |
| ENSMUSG00000031832 | Taf1c | 5.54E-05 | -2.023589791 | 0.130342151 |
| ENSMUSG00000024266 | Adad2 | 5.54E-05 | 3.648835551 | 0.466464457 |
| ENSMUSG00000023336 | Wfdc1 | 5.54E-05 | 2.063873051 | 0.792441218 |

| | | | | |
|---|---|---|---|---|
| ENSMUSG00000031828 | Klhl36 | 5.54E-05 | -0.995020638 | 0.718161064 |
| ENSMUSG00000041515 | Irf8 | 5.54E-05 | -0.258055044 | 1 |
| ENSMUSG00000097438 | Gm26784 | 5.54E-05 | 3.648835551 | 0.466464457 |
| ENSMUSG00000110631 | Gm42047 | 5.54E-05 | 0.841480629 | 1 |
| ENSMUSG00000043903 | Zfp469 | 5.54E-05 | -0.051604167 | 1 |
| ENSMUSG00000049482 | Ctu2 | 5.54E-05 | 0.063873051 | 1 |
| ENSMUSG00000006585 | Cdt1 | 5.54E-05 | 0.063873051 | 1 |
| ENSMUSG00000031962 | Cdh15 | 5.54E-05 | 2.063873051 | 0.790186865 |
| ENSMUSG00000000739 | Sult5a1 | 5.54E-05 | -0.351164449 | 1 |
| ENSMUSG00000031970 | Dbndd1 | 5.54E-05 | 2.063873051 | 0.792441218 |
| ENSMUSG00000031971 | Ccsap | 5.54E-05 | 0.47891055 | 1 |
| ENSMUSG00000031976 | Urb2 | 5.54E-05 | -1.395558568 | 0.508263132 |
| ENSMUSG00000031981 | Capn9 | 5.54E-05 | 2.063873051 | 0.792441218 |
| ENSMUSG00000036913 | Trim67 | 5.54E-05 | 1.063873051 | 1 |
| ENSMUSG00000110706 | Gm45709 | 5.54E-05 | 1.648835551 | 0.886203497 |
| ENSMUSG00000100812 | A730098A19Rik | 5.54E-05 | 1.063873051 | 1 |
| ENSMUSG00000097060 | Gm26759 | 5.54E-05 | 0.063873051 | 1 |
| ENSMUSG00000074564 | Gm10720 | 5.54E-05 | 1.063873051 | 1 |
| ENSMUSG00000096201 | Gm10715 | 5.54E-05 | 1.648835551 | 0.886203497 |
| ENSMUSG00000110491 | Gm31138 | 5.54E-05 | 2.063873051 | 0.792441218 |
| ENSMUSG00000025887 | Casp12 | 5.54E-05 | -0.158519371 | 1 |
| ENSMUSG00000018620 | Mmp20 | 5.54E-05 | 0.841480629 | 1 |
| ENSMUSG00000018623 | Mmp7 | 5.54E-05 | -1.560617814 | 0.401963028 |
| ENSMUSG00000040729 | Cep126 | 5.54E-05 | 0.648835551 | 1 |
| ENSMUSG00000111871 | Gm47557 | 5.54E-05 | -0.351164449 | 1 |
| ENSMUSG00000111555 | Gm47326 | 5.54E-05 | 2.063873051 | 0.784017887 |
| ENSMUSG00000031937 | Vstm5 | 5.54E-05 | -3.43862729 | 0.00098929 |
| ENSMUSG00000058173 | Smco4 | 5.54E-05 | -0.673092543 | 1 |
| ENSMUSG00000038691 | Mbd3l1 | 5.54E-05 | 3.648835551 | 0.466464457 |
| ENSMUSG00000054141 | Olfr24 | 5.54E-05 | 2.648835551 | 0.66416768 |
| ENSMUSG00000051118 | Olfr77 | 5.54E-05 | 2.648835551 | 0.66416768 |
| ENSMUSG00000044106 | Olfr868 | 5.54E-05 | 3.648835551 | 0.466464457 |
| ENSMUSG00000049028 | Olfr873 | 5.54E-05 | 0.841480629 | 1 |
| ENSMUSG00000038884 | Shfl | 5.54E-05 | -0.158519371 | 1 |
| ENSMUSG00000038742 | Angptl6 | 5.54E-05 | 1.063873051 | 1 |
| ENSMUSG00000097665 | Gm16853 | 5.54E-05 | 0.648835551 | 1 |
| ENSMUSG00000074476 | Spc24 | 5.54E-05 | -0.810596067 | 0.832285197 |
| ENSMUSG00000013822 | Elof1 | 5.54E-05 | -0.351164449 | 1 |
| ENSMUSG00000090925 | 1810064F22Rik | 5.54E-05 | 0.063873051 | 1 |
| ENSMUSG00000111205 | Gm39307 | 5.54E-05 | 2.648835551 | 0.66416768 |
| ENSMUSG00000036411 | 9530077C05Rik | 5.54E-05 | 0.189403933 | 1 |
| ENSMUSG00000091159 | Gm17545 | 5.54E-05 | 0.841480629 | 1 |
| ENSMUSG00000031965 | Tbx20 | 5.54E-05 | 0.47891055 | 1 |
| ENSMUSG00000111866 | E130101E03Rik | 5.54E-05 | 1.648835551 | 0.85551189 |
| ENSMUSG00000034275 | Igsf9b | 5.54E-05 | 0.47891055 | 1 |
| ENSMUSG00000110741 | Gm48091 | 5.54E-05 | -0.351164449 | 1 |
| ENSMUSG00000111109 | Gm30933 | 5.54E-05 | 0.841480629 | 1 |
| ENSMUSG00000111867 | Gm31013 | 5.54E-05 | 0.841480629 | 1 |
| ENSMUSG00000111655 | 4930434F21Rik | 5.54E-05 | 1.326907457 | 0.971854959 |
| ENSMUSG00000032033 | Barx2 | 5.54E-05 | -0.351164449 | 1 |
| ENSMUSG00000098645 | Gm27201 | 5.54E-05 | 2.063873051 | 0.789538366 |

| | | | | |
|---|---|---|---|---|
| ENSMUSG00000098519 | Gm27166 | 5.54E-05 | 0.648835551 | 1 |
| ENSMUSG00000098758 | 7630403G23Rik | 5.54E-05 | 1.326907457 | 0.926171157 |
| ENSMUSG00000039048 | Foxred1 | 5.54E-05 | 0.47891055 | 1 |
| ENSMUSG00000032044 | Rpusd4 | 5.54E-05 | -0.43862729 | 1 |
| ENSMUSG00000032101 | Ddx25 | 5.54E-05 | 1.648835551 | 0.867915716 |
| ENSMUSG00000032108 | Pate6 | 5.54E-05 | 3.648835551 | 0.466464457 |
| ENSMUSG00000111748 | Gm34885 | 5.54E-05 | 2.063873051 | 0.774601541 |
| ENSMUSG00000046240 | Hepacam | 5.54E-05 | 1.648835551 | 0.886203497 |
| ENSMUSG00000001946 | Esam | 5.54E-05 | 0.47891055 | 1 |
| ENSMUSG00000094778 | Olfr143 | 5.54E-05 | 2.648835551 | 0.66416768 |
| ENSMUSG00000096794 | Olfr905 | 5.54E-05 | 1.063873051 | 1 |
| ENSMUSG00000047050 | Olfr914 | 5.54E-05 | 3.648835551 | 0.466464457 |
| ENSMUSG00000059106 | Olfr961 | 5.54E-05 | -0.051604167 | 1 |
| ENSMUSG00000094353 | Olfr150 | 5.54E-05 | 2.063873051 | 0.792441218 |
| ENSMUSG00000095903 | Olfr968 | 5.54E-05 | 3.648835551 | 0.466464457 |
| ENSMUSG00000046678 | Olfr981 | 5.54E-05 | 1.063873051 | 1 |
| ENSMUSG00000032023 | Jhy | 5.54E-05 | -0.258055044 | 1 |
| ENSMUSG00000111854 | Gm48740 | 5.54E-05 | 2.063873051 | 0.792441218 |
| ENSMUSG00000110727 | Gm48784 | 5.54E-05 | 3.648835551 | 0.466464457 |
| ENSMUSG00000111726 | Gm40513 | 5.54E-05 | 3.648835551 | 0.466464457 |
| ENSMUSG00000032014 | Oaf | 5.54E-05 | -1.743481871 | 0.305378889 |
| ENSMUSG00000032013 | Trim29 | 5.54E-05 | 3.648835551 | 0.466464457 |
| ENSMUSG00000110823 | Gm36799 | 5.54E-05 | 0.47891055 | 1 |
| ENSMUSG00000111110 | Gm36855 | 5.54E-05 | 2.063873051 | 0.774790843 |
| ENSMUSG00000110988 | Gm29909 | 5.54E-05 | 3.648835551 | 0.466464457 |
| ENSMUSG00000111334 | Gm30015 | 5.54E-05 | 0.841480629 | 1 |
| ENSMUSG00000111758 | Gm30373 | 5.54E-05 | -0.874726405 | 0.840804483 |
| ENSMUSG00000111229 | Gm39323 | 5.54E-05 | 0.189403933 | 1 |
| ENSMUSG00000032011 | Thy1 | 5.54E-05 | -0.52108945 | 1 |
| ENSMUSG00000032010 | Usp2 | 5.54E-05 | 0.47891055 | 1 |
| ENSMUSG00000097467 | Gm26737 | 5.54E-05 | 1.326907457 | 0.971854959 |
| ENSMUSG00000032135 | Mcam | 5.54E-05 | -0.43862729 | 1 |
| ENSMUSG00000070306 | Ccdc153 | 5.54E-05 | 2.648835551 | 0.66416768 |
| ENSMUSG00000032120 | C2cd2l | 5.54E-05 | -0.158519371 | 1 |
| ENSMUSG00000110773 | Gm39326 | 5.54E-05 | 1.648835551 | 0.886203497 |
| ENSMUSG00000002033 | Cd3g | 5.54E-05 | 1.326907457 | 0.971854959 |
| ENSMUSG00000032093 | Cd3e | 5.54E-05 | 1.326907457 | 0.971854959 |
| ENSMUSG00000111203 | Gm48719 | 5.54E-05 | 2.063873051 | 0.792441218 |
| ENSMUSG00000070304 | Scn2b | 5.54E-05 | 2.648835551 | 0.66416768 |
| ENSMUSG00000091996 | BC049352 | 5.54E-05 | 0.326907457 | 1 |
| ENSMUSG00000037129 | Tmprss13 | 5.54E-05 | 0.841480629 | 1 |
| ENSMUSG00000032081 | Apoc3 | 5.54E-05 | 3.648835551 | 0.466464457 |
| ENSMUSG00000111115 | Gm47144 | 5.54E-05 | 0.189403933 | 1 |
| ENSMUSG00000032271 | Nnmt | 5.54E-05 | 2.063873051 | 0.792441218 |
| ENSMUSG00000032268 | Tmprss5 | 5.54E-05 | -0.258055044 | 1 |
| ENSMUSG00000032257 | Ankk1 | 5.54E-05 | 0.326907457 | 1 |
| ENSMUSG00000101304 | Plet1os | 5.54E-05 | 0.189403933 | 1 |
| ENSMUSG00000032067 | Pts | 5.54E-05 | -0.936126949 | 0.695372548 |
| ENSMUSG00000032066 | Bco2 | 5.54E-05 | 1.648835551 | 0.886203497 |
| ENSMUSG00000038086 | Hspb2 | 5.54E-05 | 3.648835551 | 0.466464457 |
| ENSMUSG00000037845 | Fdxacb1 | 5.54E-05 | -1.051604167 | 0.706929043 |

| | | | | |
|---|---|---|---|---|
| ENSMUSG00000044694 | 2010007H06Rik | 5.54E-05 | 0.326907457 | 1 |
| ENSMUSG00000036027 | 1810046K07Rik | 5.54E-05 | 1.648835551 | 0.886203497 |
| ENSMUSG00000078308 | Gm47854 | 5.54E-05 | 2.648835551 | 0.66416768 |
| ENSMUSG00000111836 | 4933407I05Rik | 5.54E-05 | 3.648835551 | 0.466464457 |
| ENSMUSG00000032048 | 4930510E17Rik | 5.54E-05 | 2.063873051 | 0.792441218 |
| ENSMUSG00000042195 | Slc35f2 | 5.54E-05 | 0.648835551 | 1 |
| ENSMUSG00000074343 | Gldnos | 5.54E-05 | 0.648835551 | 1 |
| ENSMUSG00000046167 | Gldn | 5.54E-05 | 2.063873051 | 0.786367309 |
| ENSMUSG00000046460 | Sh2d7 | 5.54E-05 | 2.648835551 | 0.66416768 |
| ENSMUSG00000032279 | Idh3a | 5.54E-05 | -1.051604167 | 0.747340115 |
| ENSMUSG00000032291 | Crabp1 | 5.54E-05 | 2.648835551 | 0.66416768 |
| ENSMUSG00000035878 | Hykk | 5.54E-05 | -0.874726405 | 0.840804483 |
| ENSMUSG00000032309 | Fbxo22 | 5.54E-05 | -1.673092543 | 0.290284345 |
| ENSMUSG00000032733 | Snx33 | 5.54E-05 | -1.636566667 | 0.373232174 |
| ENSMUSG00000062309 | Rpp25 | 5.54E-05 | 2.648835551 | 0.66416768 |
| ENSMUSG00000056271 | Lman1l | 5.54E-05 | 2.648835551 | 0.66416768 |
| ENSMUSG00000032315 | Cyp1a1 | 5.54E-05 | 2.648835551 | 0.663388579 |
| ENSMUSG00000004661 | Arid3b | 5.54E-05 | -0.599091962 | 1 |
| ENSMUSG00000037716 | Ccdc33 | 5.54E-05 | -0.43862729 | 1 |
| ENSMUSG00000032327 | Stra6 | 5.54E-05 | -1.599091962 | 0.409269374 |
| ENSMUSG00000025236 | Adpgk | 5.54E-05 | -1.743481871 | 0.305378889 |
| ENSMUSG00000051705 | Senp8 | 5.54E-05 | -1.051604167 | 0.706929043 |
| ENSMUSG00000111809 | Gm49334 | 5.54E-05 | 1.648835551 | 0.88176882 |
| ENSMUSG00000111290 | Gm5122 | 5.54E-05 | 2.648835551 | 0.66416768 |
| ENSMUSG00000111147 | Gm33699 | 5.54E-05 | 0.326907457 | 1 |
| ENSMUSG00000111354 | Gm33914 | 5.54E-05 | 0.189403933 | 1 |
| ENSMUSG00000032252 | Glce | 5.54E-05 | -1.351164449 | 0.499088689 |
| ENSMUSG00000111193 | Gm35288 | 5.54E-05 | -0.874726405 | 0.853473606 |
| ENSMUSG00000022245 | Skor1 | 5.54E-05 | 2.063873051 | 0.792441218 |
| ENSMUSG00000111160 | Gm48855 | 5.54E-05 | 2.648835551 | 0.66416768 |
| ENSMUSG00000034263 | Ints14 | 5.54E-05 | -2.351164449 | 0.072659903 |
| ENSMUSG00000111471 | Gm47350 | 5.54E-05 | 1.326907457 | 0.929361739 |
| ENSMUSG00000042254 | Cilp | 5.54E-05 | 1.326907457 | 0.910865313 |
| ENSMUSG00000086228 | Ubap1l | 5.54E-05 | 1.326907457 | 0.971854959 |
| ENSMUSG00000053862 | Slc51b | 5.54E-05 | 2.648835551 | 0.66416768 |
| ENSMUSG00000066510 | Ankdd1a | 5.54E-05 | 0.47891055 | 1 |
| ENSMUSG00000041064 | Pif1 | 5.54E-05 | 1.063873051 | 1 |
| ENSMUSG00000086387 | Gm15563 | 5.54E-05 | 0.326907457 | 1 |
| ENSMUSG00000111303 | 4930502A04Rik | 5.54E-05 | 0.648835551 | 1 |
| ENSMUSG00000086648 | Gm15511 | 5.54E-05 | 2.648835551 | 0.66416768 |
| ENSMUSG00000097623 | B230323A14Rik | 5.54E-05 | -1.9057533 | 0.261700715 |
| ENSMUSG00000013584 | Aldh1a2 | 5.54E-05 | -1.158519371 | 0.653235135 |
| ENSMUSG00000032221 | Mns1 | 5.54E-05 | -0.258055044 | 1 |
| ENSMUSG00000099051 | Gm27241 | 5.54E-05 | 0.841480629 | 1 |
| ENSMUSG00000092622 | Khdc3 | 5.54E-05 | 2.648835551 | 0.66416768 |
| ENSMUSG00000043013 | Onecut1 | 5.54E-05 | 0.063873051 | 1 |
| ENSMUSG00000032181 | Scg3 | 5.54E-05 | 0.841480629 | 1 |
| ENSMUSG00000098370 | Gm19541 | 5.54E-05 | 3.648835551 | 0.466464457 |
| ENSMUSG00000099170 | 5730403I07Rik | 5.54E-05 | 0.326907457 | 1 |
| ENSMUSG00000044938 | Klhl31 | 5.54E-05 | 1.648835551 | 0.886203497 |
| ENSMUSG00000111225 | Gm47790 | 5.54E-05 | 1.063873051 | 1 |

| | | | | |
|---|---|---|---|---|
| ENSMUSG00000111335 | Gm39375 | 5.54E-05 | 2.648835551 | 0.66416768 |
| ENSMUSG00000074183 | Gsta1 | 5.54E-05 | 0.063873051 | 1 |
| ENSMUSG00000074179 | Gm10639 | 5.54E-05 | 3.648835551 | 0.466464457 |
| ENSMUSG00000110923 | Gm47499 | 5.54E-05 | 1.326907457 | 0.964003521 |
| ENSMUSG00000111638 | Gm47501 | 5.54E-05 | 0.326907457 | 1 |
| ENSMUSG00000111765 | Gm10635 | 5.54E-05 | 0.648835551 | 1 |
| ENSMUSG00000086918 | 4930429F24Rik | 5.54E-05 | 2.063873051 | 0.775097267 |
| ENSMUSG00000097241 | Gm26907 | 5.54E-05 | -0.051604167 | 1 |
| ENSMUSG00000111697 | Gm35548 | 5.54E-05 | 1.326907457 | 0.954524296 |
| ENSMUSG00000032251 | Irak1bp1 | 5.54E-05 | 3.648835551 | 0.466464457 |
| ENSMUSG00000101144 | Gm29054 | 5.54E-05 | 2.063873051 | 0.792441218 |
| ENSMUSG00000111820 | Gm36120 | 5.54E-05 | 2.063873051 | 0.792441218 |
| ENSMUSG00000111769 | Gm36278 | 5.54E-05 | 3.648835551 | 0.466464457 |
| ENSMUSG00000111281 | Gm48830 | 5.54E-05 | 2.063873051 | 0.792441218 |
| ENSMUSG00000033419 | Snap91 | 5.54E-05 | -0.52108945 | 1 |
| ENSMUSG00000089929 | Bcl2a1b | 5.54E-05 | 0.189403933 | 1 |
| ENSMUSG00000111806 | Gm47403 | 5.54E-05 | 3.648835551 | 0.466464457 |
| ENSMUSG00000047606 | Ankrd34c | 5.54E-05 | 3.648835551 | 0.466464457 |
| ENSMUSG00000036972 | Zic4 | 5.54E-05 | 0.648835551 | 1 |
| ENSMUSG00000100341 | Gm29478 | 5.54E-05 | -0.158519371 | 1 |
| ENSMUSG00000032372 | Plscr2 | 5.54E-05 | -0.874726405 | 0.886203497 |
| ENSMUSG00000101734 | 4933400C23Rik | 5.54E-05 | 0.063873051 | 1 |
| ENSMUSG00000045414 | Dipk2a | 5.54E-05 | -0.051604167 | 1 |
| ENSMUSG00000033350 | Chst2 | 5.54E-05 | 0.648835551 | 1 |
| ENSMUSG00000032451 | Trim42 | 5.54E-05 | 1.063873051 | 1 |
| ENSMUSG00000090255 | 4921534H16Rik | 5.54E-05 | 2.648835551 | 0.66416768 |
| ENSMUSG00000044860 | Gm1123 | 5.54E-05 | 0.841480629 | 1 |
| ENSMUSG00000037953 | A4gnt | 5.54E-05 | 2.648835551 | 0.66416768 |
| ENSMUSG00000044244 | Il20rb | 5.54E-05 | -0.599091962 | 1 |
| ENSMUSG00000032527 | Pccb | 5.54E-05 | -1.305360759 | 0.521297728 |
| ENSMUSG00000057710 | 9630041A04Rik | 5.54E-05 | 0.326907457 | 1 |
| ENSMUSG00000111399 | Gm47328 | 5.54E-05 | 1.326907457 | 0.971854959 |
| ENSMUSG00000086236 | 5830418P13Rik | 5.54E-05 | 1.063873051 | 1 |
| ENSMUSG00000085168 | Gm16252 | 5.54E-05 | 3.648835551 | 0.466464457 |
| ENSMUSG00000111278 | Gm32743 | 5.54E-05 | 2.063873051 | 0.792441218 |
| ENSMUSG00000032558 | Nphp3 | 5.54E-05 | -0.051604167 | 1 |
| ENSMUSG00000032561 | Acpp | 5.54E-05 | -0.351164449 | 1 |
| ENSMUSG00000032563 | Mrpl3 | 5.54E-05 | -2.158519371 | 0.115633409 |
| ENSMUSG00000086296 | D030055H07Rik | 5.54E-05 | 3.648835551 | 0.466464457 |
| ENSMUSG00000020253 | Ppm1m | 5.54E-05 | 0.326907457 | 1 |
| ENSMUSG00000023345 | Poc1a | 5.54E-05 | -2.184054463 | 0.091007367 |
| ENSMUSG00000023192 | Grm2 | 5.54E-05 | 3.648835551 | 0.466464457 |
| ENSMUSG00000032578 | Cish | 5.54E-05 | -0.599091962 | 0.980745941 |
| ENSMUSG00000037190 | Cyb561d2 | 5.54E-05 | 2.648835551 | 0.66416768 |
| ENSMUSG00000010051 | Hyal1 | 5.54E-05 | 1.063873051 | 1 |
| ENSMUSG00000010048 | Ifrd2 | 5.54E-05 | -0.874726405 | 0.840804483 |
| ENSMUSG00000057969 | Sema3b | 5.54E-05 | 0.47891055 | 1 |
| ENSMUSG00000032584 | Mst1r | 5.54E-05 | -2.106051951 | 0.109343152 |
| ENSMUSG00000032586 | Traip | 5.54E-05 | 0.47891055 | 1 |
| ENSMUSG00000032596 | Uba7 | 5.54E-05 | -1.258055044 | 0.616879894 |
| ENSMUSG00000070284 | Gmppb | 5.54E-05 | -0.051604167 | 1 |

| | | | | |
|---|---|---|---|---|
| ENSMUSG00000049305 | Ccdc71 | 5.54E-05 | 0.47891055 | 1 |
| ENSMUSG00000103103 | 4833445I07Rik | 5.54E-05 | 1.063873051 | 1 |
| ENSMUSG00000019039 | Dalrd3 | 5.54E-05 | 0.326907457 | 1 |
| ENSMUSG00000025646 | Atrip | 5.54E-05 | -0.258055044 | 1 |
| ENSMUSG00000047237 | Fbxw21 | 5.54E-05 | 0.841480629 | 1 |
| ENSMUSG00000105589 | Fbxw14 | 5.54E-05 | 2.063873051 | 0.792441218 |
| ENSMUSG00000070324 | Fbxw22 | 5.54E-05 | 1.326907457 | 0.971854959 |
| ENSMUSG00000032478 | Nme6 | 5.54E-05 | -0.599091962 | 1 |
| ENSMUSG00000032482 | Cspg5 | 5.54E-05 | 1.063873051 | 1 |
| ENSMUSG00000032491 | Nradd | 5.54E-05 | -1.43862729 | 0.433313155 |
| ENSMUSG00000059741 | Myl3 | 5.54E-05 | 0.648835551 | 1 |
| ENSMUSG00000058398 | Prss43 | 5.54E-05 | 0.326907457 | 1 |
| ENSMUSG00000032496 | Ltf | 5.54E-05 | -0.599091962 | 1 |
| ENSMUSG00000032498 | Mlh1 | 5.54E-05 | -1.874726405 | 0.198601369 |
| ENSMUSG00000032500 | Dclk3 | 5.54E-05 | 1.326907457 | 0.971854959 |
| ENSMUSG00000111022 | Gm36367 | 5.54E-05 | 2.648835551 | 0.66416768 |
| ENSMUSG00000086596 | Susd5 | 5.54E-05 | 2.648835551 | 0.66416768 |
| ENSMUSG00000111843 | 4933432G23Rik | 5.54E-05 | 3.648835551 | 0.466464457 |
| ENSMUSG00000032446 | Eomes | 5.54E-05 | 2.648835551 | 0.66416768 |
| ENSMUSG00000086119 | Gm2415 | 5.54E-05 | 0.47891055 | 1 |
| ENSMUSG00000010651 | Acaa1b | 5.54E-05 | 1.063873051 | 1 |
| ENSMUSG00000074028 | Slc22a13 | 5.54E-05 | 3.648835551 | 0.466464457 |
| ENSMUSG00000032508 | Myd88 | 5.54E-05 | -0.258055044 | 1 |
| ENSMUSG00000035769 | Xylb | 5.54E-05 | -0.936126949 | 0.695372548 |
| ENSMUSG00000034533 | Scn10a | 5.54E-05 | -0.258055044 | 1 |
| ENSMUSG00000034115 | Scn11a | 5.54E-05 | 0.063873051 | 1 |
| ENSMUSG00000032514 | Ttc21a | 5.54E-05 | 0.189403933 | 1 |
| ENSMUSG00000032515 | Csrnp1 | 5.54E-05 | 1.648835551 | 0.886203497 |
| ENSMUSG00000032517 | Mobp | 5.54E-05 | 0.841480629 | 1 |
| ENSMUSG00000111831 | Gm47064 | 5.54E-05 | 2.063873051 | 0.766975255 |
| ENSMUSG00000111528 | Gm39460 | 5.54E-05 | 1.326907457 | 0.909998684 |
| ENSMUSG00000032532 | Cck | 5.54E-05 | 2.648835551 | 0.66416768 |
| ENSMUSG00000084995 | Lyzl4os | 5.54E-05 | 2.648835551 | 0.66416768 |
| ENSMUSG00000079235 | Ccdc13 | 5.54E-05 | -0.051604167 | 1 |
| ENSMUSG00000092042 | Gm17163 | 5.54E-05 | 3.648835551 | 0.466464457 |
| ENSMUSG00000044534 | Ackr2 | 5.54E-05 | 2.648835551 | 0.66416768 |
| ENSMUSG00000111873 | Gm39463 | 5.54E-05 | 2.063873051 | 0.792441218 |
| ENSMUSG00000097572 | Gm26797 | 5.54E-05 | 2.063873051 | 0.792441218 |
| ENSMUSG00000093598 | A730085K08Rik | 5.54E-05 | 3.648835551 | 0.466464457 |
| ENSMUSG00000111068 | Gm35501 | 5.54E-05 | 2.063873051 | 0.792441218 |
| ENSMUSG00000111577 | Gm47136 | 5.54E-05 | 2.063873051 | 0.790550166 |
| ENSMUSG00000094985 | Topaz1 | 5.54E-05 | 0.841480629 | 1 |
| ENSMUSG00000063488 | Zkscan7 | 5.54E-05 | -1.209145444 | 0.606122355 |
| ENSMUSG00000057895 | Zfp105 | 5.54E-05 | 0.47891055 | 1 |
| ENSMUSG00000066233 | Tmem42 | 5.54E-05 | 2.063873051 | 0.792441218 |
| ENSMUSG00000025243 | Slc6a20b | 5.54E-05 | -1.051604167 | 0.737638245 |
| ENSMUSG00000091661 | Gm17021 | 5.54E-05 | 1.648835551 | 0.886203497 |
| ENSMUSG00000040653 | Ppp1r14c | 5.54E-05 | -0.874726405 | 0.875465725 |
| ENSMUSG00000046916 | Myct1 | 5.54E-05 | 0.648835551 | 1 |
| ENSMUSG00000111215 | Gm48654 | 5.54E-05 | 2.648835551 | 0.66416768 |
| ENSMUSG00000111731 | Gm48727 | 5.54E-05 | 0.326907457 | 1 |

| | | | | |
|---|---|---|---|---|
| ENSMUSG00000112303 | Gm48759 | 5.54E-05 | 1.648835551 | 0.878539089 |
| ENSMUSG00000112146 | Gm46210 | 5.54E-05 | 2.648835551 | 0.66416768 |
| ENSMUSG00000019832 | Rab32 | 5.54E-05 | 1.326907457 | 0.971854959 |
| ENSMUSG00000112730 | 4930567K20Rik | 5.54E-05 | 2.063873051 | 0.792441218 |
| ENSMUSG00000047648 | Fbxo30 | 5.54E-05 | -1.843017545 | 0.255869172 |
| ENSMUSG00000112172 | Gm32172 | 5.54E-05 | 2.648835551 | 0.66416768 |
| ENSMUSG00000112225 | Gm48854 | 5.54E-05 | 2.063873051 | 0.792441218 |
| ENSMUSG00000111888 | Gm46212 | 5.54E-05 | -3.293678954 | 0.001758822 |
| ENSMUSG00000112928 | Gm47710 | 5.54E-05 | 0.189403933 | 1 |
| ENSMUSG00000101969 | Gm20125 | 5.54E-05 | -0.810596067 | 0.856287335 |
| ENSMUSG00000111922 | Gm47760 | 5.54E-05 | 2.063873051 | 0.792441218 |
| ENSMUSG00000093672 | Gm20655 | 5.54E-05 | 3.648835551 | 0.466464457 |
| ENSMUSG00000019852 | Arfgef3 | 5.54E-05 | -0.936126949 | 0.872990655 |
| ENSMUSG00000111474 | Gm32926 | 5.54E-05 | 0.47891055 | 1 |
| ENSMUSG00000110917 | Gm33056 | 5.54E-05 | 1.063873051 | 1 |
| ENSMUSG00000019992 | Mtfr2 | 5.54E-05 | -0.43862729 | 1 |
| ENSMUSG00000019982 | Myb | 5.54E-05 | 1.648835551 | 0.863572577 |
| ENSMUSG00000050844 | 1700020N01Rik | 5.54E-05 | 2.063873051 | 0.777257911 |
| ENSMUSG00000056716 | Gm5420 | 5.54E-05 | 1.648835551 | 0.886203497 |
| ENSMUSG00000097107 | Platr6 | 5.54E-05 | 2.063873051 | 0.792441218 |
| ENSMUSG00000069712 | 4930444G20Rik | 5.54E-05 | 3.648835551 | 0.466464457 |
| ENSMUSG00000075297 | H60b | 5.54E-05 | 3.648835551 | 0.466464457 |
| ENSMUSG00000037490 | Slc2a12 | 5.54E-05 | 0.841480629 | 1 |
| ENSMUSG00000111894 | Gm47843 | 5.54E-05 | 1.063873051 | 1 |
| ENSMUSG00000085451 | Gm40614 | 5.54E-05 | 0.841480629 | 1 |
| ENSMUSG00000111970 | Gm47715 | 5.54E-05 | 0.841480629 | 1 |
| ENSMUSG00000112630 | Gm36908 | 5.54E-05 | 3.648835551 | 0.466464457 |
| ENSMUSG00000111950 | Gm48893 | 5.54E-05 | 1.063873051 | 1 |
| ENSMUSG00000112751 | Gm48084 | 5.54E-05 | 0.063873051 | 1 |
| ENSMUSG00000111080 | Gm20300 | 5.54E-05 | 0.326907457 | 1 |
| ENSMUSG00000069668 | Sult3a1 | 5.54E-05 | 2.648835551 | 0.66416768 |
| ENSMUSG00000112280 | A830082N09Rik | 5.54E-05 | -0.995020638 | 0.699192777 |
| ENSMUSG00000039485 | Tspyl4 | 5.54E-05 | -0.743481871 | 0.827142217 |
| ENSMUSG00000051736 | Fam229b | 5.54E-05 | 0.841480629 | 1 |
| ENSMUSG00000096687 | Mfsd4b4 | 5.54E-05 | -0.158519371 | 1 |
| ENSMUSG00000019834 | Slc22a16 | 5.54E-05 | 1.063873051 | 1 |
| ENSMUSG00000063428 | Ddo | 5.54E-05 | -1.480447466 | 0.466464457 |
| ENSMUSG00000045555 | Mettl24 | 5.54E-05 | -0.051604167 | 1 |
| ENSMUSG00000019831 | Wasf1 | 5.54E-05 | -0.52108945 | 1 |
| ENSMUSG00000111576 | Gm32284 | 5.54E-05 | 1.648835551 | 0.858831093 |
| ENSMUSG00000111587 | Gm32338 | 5.54E-05 | 2.648835551 | 0.66416768 |
| ENSMUSG00000078451 | Ppil6 | 5.54E-05 | 1.648835551 | 0.867965264 |
| ENSMUSG00000045008 | 9030612E09Rik | 5.54E-05 | 2.063873051 | 0.792441218 |
| ENSMUSG00000045886 | Gm9803 | 5.54E-05 | 2.648835551 | 0.66416768 |
| ENSMUSG00000111749 | Gm47927 | 5.54E-05 | 3.648835551 | 0.466464457 |
| ENSMUSG00000019863 | Qrsl1 | 5.54E-05 | -0.051604167 | 1 |
| ENSMUSG00000019864 | Rtn4ip1 | 5.54E-05 | -0.599091962 | 0.980745941 |
| ENSMUSG00000110874 | D030045P18Rik | 5.54E-05 | 1.648835551 | 0.886203497 |
| ENSMUSG00000019913 | Sim1 | 5.54E-05 | 0.189403933 | 1 |
| ENSMUSG00000049641 | Vgll2 | 5.54E-05 | 3.648835551 | 0.466464457 |
| ENSMUSG00000112731 | Gm47739 | 5.54E-05 | 3.648835551 | 0.466464457 |

| | | | | |
|---|---|---|---|---|
| ENSMUSG00000112809 | Gm47915 | 5.54E-05 | 3.648835551 | 0.466464457 |
| ENSMUSG00000112837 | Gm48053 | 5.54E-05 | 2.648835551 | 0.66416768 |
| ENSMUSG00000059901 | Adamts14 | 5.54E-05 | -0.158519371 | 1 |
| ENSMUSG00000037202 | Prf1 | 5.54E-05 | 2.648835551 | 0.66416768 |
| ENSMUSG00000112080 | Gm47595 | 5.54E-05 | 3.648835551 | 0.466464457 |
| ENSMUSG00000112581 | 1700022H01Rik | 5.54E-05 | 3.648835551 | 0.466464457 |
| ENSMUSG00000036875 | Dna2 | 5.54E-05 | -0.673092543 | 1 |
| ENSMUSG00000112129 | Pbld1 | 5.54E-05 | -0.051604167 | 1 |
| ENSMUSG00000112383 | Gm47895 | 5.54E-05 | 3.648835551 | 0.466464457 |
| ENSMUSG00000112267 | Gm47903 | 5.54E-05 | 2.648835551 | 0.66416768 |
| ENSMUSG00000097921 | Gm26576 | 5.54E-05 | 1.648835551 | 0.886203497 |
| ENSMUSG00000112925 | Gm31763 | 5.54E-05 | 0.063873051 | 1 |
| ENSMUSG00000057134 | Ado | 5.54E-05 | 0.189403933 | 1 |
| ENSMUSG00000037762 | Slc16a9 | 5.54E-05 | -0.258055044 | 1 |
| ENSMUSG00000112320 | Gm48022 | 5.54E-05 | 3.648835551 | 0.466464457 |
| ENSMUSG00000112813 | Gm34376 | 5.54E-05 | 2.063873051 | 0.7724237 |
| ENSMUSG00000037710 | Cisd1 | 5.54E-05 | -0.051604167 | 1 |
| ENSMUSG00000020175 | Rab36 | 5.54E-05 | 2.063873051 | 0.792441218 |
| ENSMUSG00000006345 | Ggt1 | 5.54E-05 | 0.841480629 | 1 |
| ENSMUSG00000001666 | Ddt | 5.54E-05 | -0.43862729 | 1 |
| ENSMUSG00000009093 | Gstt4 | 5.54E-05 | 3.648835551 | 0.466464457 |
| ENSMUSG00000000901 | Mmp11 | 5.54E-05 | 0.648835551 | 1 |
| ENSMUSG00000049764 | Zfp280b | 5.54E-05 | 0.189403933 | 1 |
| ENSMUSG00000020229 | Slc5a4a | 5.54E-05 | 0.841480629 | 1 |
| ENSMUSG00000033105 | Lss | 5.54E-05 | 0.189403933 | 1 |
| ENSMUSG00000112773 | Gm3137 | 5.54E-05 | 3.648835551 | 0.466464457 |
| ENSMUSG00000001436 | Slc19a1 | 5.54E-05 | 0.326907457 | 1 |
| ENSMUSG00000078444 | Gm10941 | 5.54E-05 | 1.063873051 | 1 |
| ENSMUSG00000112818 | Gm35920 | 5.54E-05 | 1.326907457 | 0.948737557 |
| ENSMUSG00000000290 | Itgb2 | 5.54E-05 | -1.051604167 | 0.729168925 |
| ENSMUSG00000009292 | Trpm2 | 5.54E-05 | 0.648835551 | 1 |
| ENSMUSG00000000730 | Dnmt3l | 5.54E-05 | 2.648835551 | 0.661145551 |
| ENSMUSG00000000732 | Icosl | 5.54E-05 | 1.063873051 | 1 |
| ENSMUSG00000053329 | Gatd3a | 5.54E-05 | -0.351164449 | 1 |
| ENSMUSG00000032834 | Pwp2 | 5.54E-05 | -0.258055044 | 1 |
| ENSMUSG00000112732 | Gm47343 | 5.54E-05 | 0.326907457 | 1 |
| ENSMUSG00000055515 | Vmn2r81 | 5.54E-05 | 1.063873051 | 1 |
| ENSMUSG00000042570 | Mier2 | 5.54E-05 | -0.743481871 | 0.827142217 |
| ENSMUSG00000045912 | C2cd4c | 5.54E-05 | 0.189403933 | 1 |
| ENSMUSG00000020312 | Shc2 | 5.54E-05 | -1.209145444 | 0.606122355 |
| ENSMUSG00000020310 | Madcam1 | 5.54E-05 | 2.648835551 | 0.66416768 |
| ENSMUSG00000020325 | Fstl3 | 5.54E-05 | 0.189403933 | 1 |
| ENSMUSG00000020323 | Prss57 | 5.54E-05 | 2.063873051 | 0.790198002 |
| ENSMUSG00000035640 | Cbarp | 5.54E-05 | -1.673092543 | 0.292736724 |
| ENSMUSG00000003070 | Efna2 | 5.54E-05 | 0.47891055 | 1 |
| ENSMUSG00000020131 | Pcsk4 | 5.54E-05 | 0.841480629 | 1 |
| ENSMUSG00000055862 | Izumo4 | 5.54E-05 | 0.326907457 | 1 |
| ENSMUSG00000075558 | Gm17151 | 5.54E-05 | 3.648835551 | 0.466464457 |
| ENSMUSG00000059406 | Tmprss9 | 5.54E-05 | 0.648835551 | 1 |
| ENSMUSG00000046822 | Slc39a3 | 5.54E-05 | 0.063873051 | 1 |
| ENSMUSG00000085779 | Atcayos | 5.54E-05 | 2.648835551 | 0.66416768 |

| | | | | |
|---|---|---|---|---|
| ENSMUSG00000034881 | Tbxa2r | 5.54E-05 | 0.648835551 | 1 |
| ENSMUSG00000034818 | Celf5 | 5.54E-05 | -1.051604167 | 0.706929043 |
| ENSMUSG00000074862 | BC025920 | 5.54E-05 | 0.841480629 | 1 |
| ENSMUSG00000034748 | Sirt6 | 5.54E-05 | -1.599091962 | 0.363644968 |
| ENSMUSG00000054708 | Ankrd24 | 5.54E-05 | 0.189403933 | 1 |
| ENSMUSG00000096795 | Zfp433 | 5.54E-05 | 1.326907457 | 0.971854959 |
| ENSMUSG00000112160 | BC024063 | 5.54E-05 | -0.258055044 | 1 |
| ENSMUSG00000062931 | Zfp938 | 5.54E-05 | -0.051604167 | 1 |
| ENSMUSG00000020251 | Glt8d2 | 5.54E-05 | 0.47891055 | 1 |
| ENSMUSG00000112286 | Gm47561 | 5.54E-05 | 1.063873051 | 1 |
| ENSMUSG00000087651 | 1500009L16Rik | 5.54E-05 | 1.648835551 | 0.886203497 |
| ENSMUSG00000111699 | C230099D08Rik | 5.54E-05 | 0.648835551 | 1 |
| ENSMUSG00000111049 | Gm34184 | 5.54E-05 | -0.158519371 | 1 |
| ENSMUSG00000111583 | Gm47962 | 5.54E-05 | -0.258055044 | 1 |
| ENSMUSG00000020033 | 4930463O16Rik | 5.54E-05 | 2.648835551 | 0.66416768 |
| ENSMUSG00000097309 | Gm17249 | 5.54E-05 | 3.648835551 | 0.466464457 |
| ENSMUSG00000049038 | Mterf2 | 5.54E-05 | 0.326907457 | 1 |
| ENSMUSG00000111752 | Gm38575 | 5.54E-05 | 0.841480629 | 1 |
| ENSMUSG00000001785 | Pwp1 | 5.54E-05 | -1.106051951 | 0.719591951 |
| ENSMUSG00000085111 | Ascl4 | 5.54E-05 | -0.051604167 | 1 |
| ENSMUSG00000001786 | Fbxo7 | 5.54E-05 | -1.158519371 | 0.599135879 |
| ENSMUSG00000089656 | Gm16271 | 5.54E-05 | 2.063873051 | 0.792441218 |
| ENSMUSG00000035365 | Parpbp | 5.54E-05 | -1.158519371 | 0.599135879 |
| ENSMUSG00000020059 | Sycp3 | 5.54E-05 | 0.063873051 | 1 |
| ENSMUSG00000020062 | Slc5a8 | 5.54E-05 | 0.841480629 | 1 |
| ENSMUSG00000111657 | Gm31592 | 5.54E-05 | 0.189403933 | 1 |
| ENSMUSG00000111470 | Gm31822 | 5.54E-05 | 2.063873051 | 0.792441218 |
| ENSMUSG00000111013 | Gm32468 | 5.54E-05 | 2.063873051 | 0.792441218 |
| ENSMUSG00000020017 | Hal | 5.54E-05 | -0.158519371 | 1 |
| ENSMUSG00000111366 | Gm47156 | 5.54E-05 | 0.47891055 | 1 |
| ENSMUSG00000085723 | Gm15915 | 5.54E-05 | 2.648835551 | 0.66416768 |
| ENSMUSG00000005897 | Nr2c1 | 5.54E-05 | -2.132524162 | 0.112236795 |
| ENSMUSG00000053420 | Gm4792 | 5.54E-05 | 2.648835551 | 0.66416768 |
| ENSMUSG00000112669 | Gm48689 | 5.54E-05 | 3.648835551 | 0.466464457 |
| ENSMUSG00000112261 | Gm47599 | 5.54E-05 | 1.063873051 | 1 |
| ENSMUSG00000112762 | 4930459C07Rik | 5.54E-05 | 0.063873051 | 1 |
| ENSMUSG00000112749 | Gm48428 | 5.54E-05 | 3.648835551 | 0.466464457 |
| ENSMUSG00000112497 | 4930525C09Rik | 5.54E-05 | 0.841480629 | 1 |
| ENSMUSG00000112896 | Gm8633 | 5.54E-05 | 1.648835551 | 0.872990655 |
| ENSMUSG00000113032 | Gm47578 | 5.54E-05 | -0.599091962 | 1 |
| ENSMUSG00000019890 | Nts | 5.54E-05 | 2.648835551 | 0.66416768 |
| ENSMUSG00000044921 | Rassf9 | 5.54E-05 | -0.673092543 | 1 |
| ENSMUSG00000035759 | Bbs10 | 5.54E-05 | 1.326907457 | 0.971854959 |
| ENSMUSG00000097082 | 4933440J02Rik | 5.54E-05 | 3.648835551 | 0.466464457 |
| ENSMUSG00000112249 | Gm30262 | 5.54E-05 | 0.47891055 | 1 |
| ENSMUSG00000020214 | Glipr1l2 | 5.54E-05 | 1.326907457 | 0.971854959 |
| ENSMUSG00000112611 | Glipr1l3 | 5.54E-05 | 2.648835551 | 0.66416768 |
| ENSMUSG00000035694 | Caps2 | 5.54E-05 | 1.648835551 | 0.886203497 |
| ENSMUSG00000100314 | 1700010J16Rik | 5.54E-05 | 0.063873051 | 1 |
| ENSMUSG00000112767 | Gm47532 | 5.54E-05 | 2.648835551 | 0.66416768 |
| ENSMUSG00000092210 | A930009A15Rik | 5.54E-05 | 1.063873051 | 1 |

| | | | | |
|---|---|---|---|---|
| ENSMUSG00000100198 | 1700030O20Rik | 5.54E-05 | 1.648835551 | 0.886203497 |
| ENSMUSG00000112924 | 4930579P08Rik | 5.54E-05 | 2.648835551 | 0.66416768 |
| ENSMUSG00000069518 | Gm10271 | 5.54E-05 | 3.648835551 | 0.466464457 |
| ENSMUSG00000112728 | Gm48903 | 5.54E-05 | 2.063873051 | 0.792441218 |
| ENSMUSG00000112796 | Gm40770 | 5.54E-05 | 2.063873051 | 0.792441218 |
| ENSMUSG00000112609 | Gm47480 | 5.54E-05 | 2.648835551 | 0.66416768 |
| ENSMUSG00000100558 | 1700025F24Rik | 5.54E-05 | 2.648835551 | 0.66416768 |
| ENSMUSG00000085418 | Grip1os1 | 5.54E-05 | 2.648835551 | 0.66416768 |
| ENSMUSG00000034764 | 1700006J14Rik | 5.54E-05 | 2.648835551 | 0.66416768 |
| ENSMUSG00000044544 | 4921513I03Rik | 5.54E-05 | 1.063873051 | 1 |
| ENSMUSG00000086522 | Gm4473 | 5.54E-05 | 3.648835551 | 0.466464457 |
| ENSMUSG00000020218 | Wif1 | 5.54E-05 | 0.326907457 | 1 |
| ENSMUSG00000112526 | Gm35865 | 5.54E-05 | 3.648835551 | 0.466464457 |
| ENSMUSG00000053684 | BC048403 | 5.54E-05 | 1.648835551 | 0.859604755 |
| ENSMUSG00000112095 | A130077B15Rik | 5.54E-05 | 3.648835551 | 0.466464457 |
| ENSMUSG00000112294 | Gm4129 | 5.54E-05 | 3.648835551 | 0.466464457 |
| ENSMUSG00000020102 | Slc16a7 | 5.54E-05 | -0.258055044 | 1 |
| ENSMUSG00000025436 | Atp23 | 5.54E-05 | 0.841480629 | 1 |
| ENSMUSG00000040521 | Tsfm | 5.54E-05 | -0.874726405 | 0.840804483 |
| ENSMUSG00000025422 | Agap2 | 5.54E-05 | 0.189403933 | 1 |
| ENSMUSG00000040441 | Slc26a10 | 5.54E-05 | -0.158519371 | 1 |
| ENSMUSG00000019467 | Arhgef25 | 5.54E-05 | -0.936126949 | 0.695372548 |
| ENSMUSG00000025408 | Ddit3 | 5.54E-05 | -1.106051951 | 0.719591951 |
| ENSMUSG00000087163 | Gm16230 | 5.54E-05 | 3.648835551 | 0.466464457 |
| ENSMUSG00000044617 | Zbtb39 | 5.54E-05 | -0.43862729 | 1 |
| ENSMUSG00000069456 | Rdh16 | 5.54E-05 | 1.063873051 | 1 |
| ENSMUSG00000074639 | Rdh16f2 | 5.54E-05 | 2.648835551 | 0.66416768 |
| ENSMUSG00000044005 | Gls2 | 5.54E-05 | 0.648835551 | 1 |
| ENSMUSG00000051346 | Spryd4 | 5.54E-05 | 1.326907457 | 0.95254727 |
| ENSMUSG00000025383 | Il23a | 5.54E-05 | -0.995020638 | 0.747182608 |
| ENSMUSG00000005682 | Pan2 | 5.54E-05 | -2.106051951 | 0.109343152 |
| ENSMUSG00000039914 | Coq10a | 5.54E-05 | 0.47891055 | 1 |
| ENSMUSG00000039810 | Zc3h10 | 5.54E-05 | 2.063873051 | 0.792441218 |
| ENSMUSG00000049858 | Suox | 5.54E-05 | -0.051604167 | 1 |
| ENSMUSG00000025355 | Mmp19 | 5.54E-05 | -0.258055044 | 1 |
| ENSMUSG00000047090 | Tmem198b | 5.54E-05 | 1.063873051 | 1 |
| ENSMUSG00000097691 | 9030616G12Rik | 5.54E-05 | 0.841480629 | 1 |
| ENSMUSG00000047868 | Olfr770 | 5.54E-05 | 2.648835551 | 0.66416768 |
| ENSMUSG00000095138 | Olfr781 | 5.54E-05 | 3.648835551 | 0.466464457 |
| ENSMUSG00000058513 | Olfr801 | 5.54E-05 | 2.648835551 | 0.66416768 |
| ENSMUSG00000049052 | Olfr812 | 5.54E-05 | 3.648835551 | 0.466464457 |
| ENSMUSG00000070601 | Vmn2r84 | 5.54E-05 | 1.648835551 | 0.886203497 |
| ENSMUSG00000092048 | Vmn2r85 | 5.54E-05 | 2.648835551 | 0.66416768 |
| ENSMUSG00000020424 | Castor1 | 5.54E-05 | 1.648835551 | 0.886203497 |
| ENSMUSG00000034394 | Lif | 5.54E-05 | -0.599091962 | 0.984748965 |
| ENSMUSG00000020419 | Hormad2 | 5.54E-05 | -0.051604167 | 1 |
| ENSMUSG00000034209 | Rasl10a | 5.54E-05 | 3.648835551 | 0.466464457 |
| ENSMUSG00000085189 | Gm11963 | 5.54E-05 | 0.189403933 | 1 |
| ENSMUSG00000020481 | Ankrd36 | 5.54E-05 | -0.43862729 | 1 |
| ENSMUSG00000086839 | Gm11973 | 5.54E-05 | 0.648835551 | 1 |
| ENSMUSG00000020437 | Myo1g | 5.54E-05 | 1.063873051 | 0.998731799 |

| | | | | |
|---|---|---|---|---|
| ENSMUSG00000087637 | Gm11986 | 5.54E-05 | 1.648835551 | 0.873400121 |
| ENSMUSG00000040985 | Sun3 | 5.54E-05 | 2.648835551 | 0.66416768 |
| ENSMUSG00000050830 | Vwc2 | 5.54E-05 | 0.47891055 | 1 |
| ENSMUSG00000087452 | Gm11998 | 5.54E-05 | 0.063873051 | 1 |
| ENSMUSG00000062511 | 4930512M02Rik | 5.54E-05 | 1.648835551 | 0.886203497 |
| ENSMUSG00000085652 | Gm11999 | 5.54E-05 | 1.063873051 | 1 |
| ENSMUSG00000018654 | Ikzf1 | 5.54E-05 | -1.52108945 | 0.499921258 |
| ENSMUSG00000085452 | 4930554G24Rik | 5.54E-05 | 0.326907457 | 1 |
| ENSMUSG00000044629 | Cnrip1 | 5.54E-05 | -0.43862729 | 1 |
| ENSMUSG00000085675 | Gm12022 | 5.54E-05 | 2.063873051 | 0.772933445 |
| ENSMUSG00000085743 | 8430419K02Rik | 5.54E-05 | -0.995020638 | 0.699192777 |
| ENSMUSG00000049811 | Fam161a | 5.54E-05 | -0.995020638 | 0.699192777 |
| ENSMUSG00000020286 | 1700093K21Rik | 5.54E-05 | 2.063873051 | 0.792441218 |
| ENSMUSG00000020275 | Rel | 5.54E-05 | -0.43862729 | 1 |
| ENSMUSG00000086612 | Gm12064 | 5.54E-05 | 0.189403933 | 1 |
| ENSMUSG00000085165 | Gm12089 | 5.54E-05 | 1.326907457 | 0.971854959 |
| ENSMUSG00000040919 | 4930505A04Rik | 5.54E-05 | 0.648835551 | 1 |
| ENSMUSG00000040767 | Snrnp25 | 5.54E-05 | -0.43862729 | 1 |
| ENSMUSG00000020282 | Rhbdf1 | 5.54E-05 | -1.480447466 | 0.441576406 |
| ENSMUSG00000044056 | Efcab9 | 5.54E-05 | 2.063873051 | 0.757762987 |
| ENSMUSG00000087318 | Gm12114 | 5.54E-05 | 1.326907457 | 0.950457641 |
| ENSMUSG00000085684 | 4930469K13Rik | 5.54E-05 | -0.673092543 | 1 |
| ENSMUSG00000069910 | Spdl1 | 5.54E-05 | -1.051604167 | 0.747121178 |
| ENSMUSG00000087296 | Gm12130 | 5.54E-05 | 0.841480629 | 1 |
| ENSMUSG00000020330 | Hmmr | 5.54E-05 | -1.395558568 | 0.508263132 |
| ENSMUSG00000010803 | Gabra1 | 5.54E-05 | 1.063873051 | 1 |
| ENSMUSG00000086686 | F630206G17Rik | 5.54E-05 | -0.673092543 | 0.98880775 |
| ENSMUSG00000044847 | Lsm11 | 5.54E-05 | 0.189403933 | 1 |
| ENSMUSG00000040489 | Sox30 | 5.54E-05 | 1.326907457 | 0.968182137 |
| ENSMUSG00000085853 | Gm12167 | 5.54E-05 | 3.648835551 | 0.466464457 |
| ENSMUSG00000020397 | Med7 | 5.54E-05 | -1.560617814 | 0.356350819 |
| ENSMUSG00000020399 | Havcr2 | 5.54E-05 | -1.258055044 | 0.616879894 |
| ENSMUSG00000078154 | Gm12184 | 5.54E-05 | 2.063873051 | 0.792441218 |
| ENSMUSG00000040350 | Trim7 | 5.54E-05 | 0.47891055 | 1 |
| ENSMUSG00000046879 | Irgm1 | 5.54E-05 | -0.158519371 | 1 |
| ENSMUSG00000048852 | Gm12185 | 5.54E-05 | -2.079084903 | 0.13833117 |
| ENSMUSG00000078153 | Psme2b | 5.54E-05 | 1.063873051 | 1 |
| ENSMUSG00000078921 | Tgtp2 | 5.54E-05 | 2.648835551 | 0.66416768 |
| ENSMUSG00000101750 | Olfr1392 | 5.54E-05 | 2.648835551 | 0.66416768 |
| ENSMUSG00000063386 | Olfr1387 | 5.54E-05 | 2.648835551 | 0.66416768 |
| ENSMUSG00000059729 | Olfr1385 | 5.54E-05 | 2.648835551 | 0.66416768 |
| ENSMUSG00000107417 | Olfr1383 | 5.54E-05 | 0.326907457 | 1 |
| ENSMUSG00000020366 | Mapk9 | 5.54E-05 | -0.995020638 | 0.699192777 |
| ENSMUSG00000020381 | Mrnip | 5.54E-05 | -0.936126949 | 0.695372548 |
| ENSMUSG00000020377 | Ltc4s | 5.54E-05 | 1.648835551 | 0.886203497 |
| ENSMUSG00000085564 | Gm12198 | 5.54E-05 | -0.051604167 | 1 |
| ENSMUSG00000049321 | Zfp2 | 5.54E-05 | 0.841480629 | 1 |
| ENSMUSG00000050343 | Olfr1378 | 5.54E-05 | 2.648835551 | 0.66416768 |
| ENSMUSG00000060918 | Olfr51 | 5.54E-05 | 2.648835551 | 0.66416768 |
| ENSMUSG00000072982 | 9230009I02Rik | 5.54E-05 | 2.063873051 | 0.792441218 |
| ENSMUSG00000001053 | N4bp3 | 5.54E-05 | -0.051604167 | 1 |

| | | | | |
|---|---|---|---|---|
| ENSMUSG00000018238 | Gdf9 | 5.54E-05 | 1.648835551 | 0.886203497 |
| ENSMUSG00000086209 | 4933405E24Rik | 5.54E-05 | 0.47891055 | 1 |
| ENSMUSG00000018916 | Csf2 | 5.54E-05 | 2.648835551 | 0.66162286 |
| ENSMUSG00000020333 | Acsl6 | 5.54E-05 | -0.051604167 | 1 |
| ENSMUSG00000020332 | Meikin | 5.54E-05 | 0.841480629 | 1 |
| ENSMUSG00000055333 | Fat2 | 5.54E-05 | -0.351164449 | 1 |
| ENSMUSG00000037393 | Nmur2 | 5.54E-05 | 1.326907457 | 0.951890397 |
| ENSMUSG00000087165 | 2010001A14Rik | 5.54E-05 | -0.52108945 | 0.975624465 |
| ENSMUSG00000087629 | Gm12246 | 5.54E-05 | 2.648835551 | 0.66416768 |
| ENSMUSG00000069873 | 4930438A08Rik | 5.54E-05 | 0.648835551 | 1 |
| ENSMUSG00000013646 | Sh3bp5l | 5.54E-05 | -0.158519371 | 1 |
| ENSMUSG00000056564 | Olfr324 | 5.54E-05 | 3.648835551 | 0.466464457 |
| ENSMUSG00000037124 | Trim58 | 5.54E-05 | 1.648835551 | 0.886203497 |
| ENSMUSG00000060030 | Olfr317 | 5.54E-05 | 2.063873051 | 0.792441218 |
| ENSMUSG00000096806 | Olfr312 | 5.54E-05 | 3.648835551 | 0.466464457 |
| ENSMUSG00000020491 | 2810021J22Rik | 5.54E-05 | -0.351164449 | 1 |
| ENSMUSG00000037001 | Zfp39 | 5.54E-05 | 0.841480629 | 1 |
| ENSMUSG00000085379 | 2310058D17Rik | 5.54E-05 | -0.351164449 | 1 |
| ENSMUSG00000049287 | Iba57 | 5.54E-05 | 1.648835551 | 0.886203497 |
| ENSMUSG00000000126 | Wnt9a | 5.54E-05 | 1.326907457 | 0.971854959 |
| ENSMUSG00000036819 | Jmjd4 | 5.54E-05 | 1.063873051 | 1 |
| ENSMUSG00000020472 | Zkscan17 | 5.54E-05 | -1.874726405 | 0.254892207 |
| ENSMUSG00000086330 | 1700007J10Rik | 5.54E-05 | 3.648835551 | 0.466464457 |
| ENSMUSG00000084766 | Gm12714 | 5.54E-05 | 0.841480629 | 1 |
| ENSMUSG00000042678 | Myo15 | 5.54E-05 | 0.063873051 | 1 |
| ENSMUSG00000049323 | Smcr8 | 5.54E-05 | -1.673092543 | 0.290284345 |
| ENSMUSG00000042549 | Map2k3os | 5.54E-05 | -1.106051951 | 0.771834758 |
| ENSMUSG00000042529 | Kcnj12 | 5.54E-05 | 0.841480629 | 1 |
| ENSMUSG00000042436 | Mfap4 | 5.54E-05 | -0.051604167 | 1 |
| ENSMUSG00000004837 | Grap | 5.54E-05 | 2.648835551 | 0.66416768 |
| ENSMUSG00000072847 | A530017D24Rik | 5.54E-05 | 0.063873051 | 1 |
| ENSMUSG00000018500 | Adora2b | 5.54E-05 | 1.648835551 | 0.886203497 |
| ENSMUSG00000014243 | Zswim7 | 5.54E-05 | 0.841480629 | 1 |
| ENSMUSG00000087604 | Cdrt4os1 | 5.54E-05 | 2.063873051 | 0.792441218 |
| ENSMUSG00000018217 | Pmp22 | 5.54E-05 | 3.648835551 | 0.466464457 |
| ENSMUSG00000085532 | B430319H21Rik | 5.54E-05 | 1.648835551 | 0.886203497 |
| ENSMUSG00000087497 | 2810001G20Rik | 5.54E-05 | -0.743481871 | 0.827142217 |
| ENSMUSG00000018347 | Zkscan6 | 5.54E-05 | -0.995020638 | 0.721329891 |
| ENSMUSG00000048070 | Pirt | 5.54E-05 | 0.189403933 | 1 |
| ENSMUSG00000050270 | Tmem220 | 5.54E-05 | 0.189403933 | 1 |
| ENSMUSG00000069844 | Sco1 | 5.54E-05 | -0.051604167 | 1 |
| ENSMUSG00000033196 | Myh2 | 5.54E-05 | 2.063873051 | 0.792441218 |
| ENSMUSG00000085348 | Myhas | 5.54E-05 | 1.648835551 | 0.886203497 |
| ENSMUSG00000060180 | Myh13 | 5.54E-05 | 0.841480629 | 1 |
| ENSMUSG00000049928 | Glp2r | 5.54E-05 | 0.47891055 | 1 |
| ENSMUSG00000020904 | Cfap52 | 5.54E-05 | -0.351164449 | 1 |
| ENSMUSG00000084993 | Gm12305 | 5.54E-05 | 2.648835551 | 0.66416768 |
| ENSMUSG00000020901 | Pik3r5 | 5.54E-05 | -0.810596067 | 0.844591606 |
| ENSMUSG00000046207 | Pik3r6 | 5.54E-05 | -1.636566667 | 0.451785624 |
| ENSMUSG00000072809 | 9330160F10Rik | 5.54E-05 | -0.158519371 | 1 |
| ENSMUSG00000020892 | Aloxe3 | 5.54E-05 | 1.326907457 | 0.952108693 |

| | | | | |
|---|---|---|---|---|
| ENSMUSG00000032807 | Alox12b | 5.54E-05 | 2.648835551 | 0.66416768 |
| ENSMUSG00000043419 | Rnf227 | 5.54E-05 | 0.648835551 | 1 |
| ENSMUSG00000044795 | Cyb5d1 | 5.54E-05 | 1.648835551 | 0.886203497 |
| ENSMUSG00000069835 | Sat2 | 5.54E-05 | 2.648835551 | 0.66416768 |
| ENSMUSG00000018774 | Cd68 | 5.54E-05 | 1.326907457 | 0.971854959 |
| ENSMUSG00000042826 | Fgf11 | 5.54E-05 | 2.063873051 | 0.792441218 |
| ENSMUSG00000001588 | Acap1 | 5.54E-05 | 0.189403933 | 1 |
| ENSMUSG00000018565 | Elp5 | 5.54E-05 | -2.395558568 | 0.067721718 |
| ENSMUSG00000040950 | Mgl2 | 5.54E-05 | 2.648835551 | 0.66416768 |
| ENSMUSG00000044367 | Slc16a13 | 5.54E-05 | 0.326907457 | 1 |
| ENSMUSG00000000320 | Alox12 | 5.54E-05 | 1.063873051 | 1 |
| ENSMUSG00000018923 | Med11 | 5.54E-05 | 0.189403933 | 1 |
| ENSMUSG00000085952 | Gm12315 | 5.54E-05 | 2.648835551 | 0.66416768 |
| ENSMUSG00000060600 | Eno3 | 5.54E-05 | 3.648835551 | 0.466464457 |
| ENSMUSG00000085502 | Gm12320 | 5.54E-05 | 0.841480629 | 1 |
| ENSMUSG00000057135 | Scimp | 5.54E-05 | 0.841480629 | 1 |
| ENSMUSG00000087116 | Gm12324 | 5.54E-05 | 1.063873051 | 0.9980272 |
| ENSMUSG00000040543 | Pitpnm3 | 5.54E-05 | -1.810596067 | 0.246776726 |
| ENSMUSG00000040483 | Xaf1 | 5.54E-05 | 0.47891055 | 1 |
| ENSMUSG00000040447 | Spns2 | 5.54E-05 | -0.810596067 | 0.868661374 |
| ENSMUSG00000020787 | P2rx1 | 5.54E-05 | 2.063873051 | 0.792441218 |
| ENSMUSG00000005949 | Ctns | 5.54E-05 | -0.995020638 | 0.699192777 |
| ENSMUSG00000005951 | Shpk | 5.54E-05 | 0.326907457 | 1 |
| ENSMUSG00000112920 | Spata22 | 5.54E-05 | 3.648835551 | 0.466464457 |
| ENSMUSG00000069818 | Olfr390 | 5.54E-05 | 2.063873051 | 0.792441218 |
| ENSMUSG00000069814 | Ccdc92b | 5.54E-05 | 0.648835551 | 1 |
| ENSMUSG00000085609 | 1700016P03Rik | 5.54E-05 | -0.258055044 | 1 |
| ENSMUSG00000038268 | Ovca2 | 5.54E-05 | 0.47891055 | 1 |
| ENSMUSG00000045374 | Wdr81 | 5.54E-05 | -1.743481871 | 0.305378889 |
| ENSMUSG00000038217 | Tlcd2 | 5.54E-05 | 2.648835551 | 0.66416768 |
| ENSMUSG00000020848 | Doc2b | 5.54E-05 | -0.258055044 | 1 |
| ENSMUSG00000053783 | 1700016K19Rik | 5.54E-05 | 0.063873051 | 1 |
| ENSMUSG00000020838 | Slc6a4 | 5.54E-05 | 0.648835551 | 1 |
| ENSMUSG00000000686 | Abhd15 | 5.54E-05 | 0.648835551 | 1 |
| ENSMUSG00000017453 | Pipox | 5.54E-05 | 2.648835551 | 0.66416768 |
| ENSMUSG00000020834 | Dhrs13 | 5.54E-05 | 1.063873051 | 1 |
| ENSMUSG00000017405 | Nek8 | 5.54E-05 | -0.258055044 | 1 |
| ENSMUSG00000019437 | Tlcd1 | 5.54E-05 | 0.841480629 | 0.941282226 |
| ENSMUSG00000044122 | Proca1 | 5.54E-05 | -0.051604167 | 1 |
| ENSMUSG00000037593 | Rskr | 5.54E-05 | 0.841480629 | 1 |
| ENSMUSG00000002055 | Spag5 | 5.54E-05 | -2.209145444 | 0.093610056 |
| ENSMUSG00000020829 | Slc46a1 | 5.54E-05 | 2.648835551 | 0.66416768 |
| ENSMUSG00000108822 | Gm44787 | 5.54E-05 | -0.051604167 | 1 |
| ENSMUSG00000086569 | Rab11fip4os2 | 5.54E-05 | 2.648835551 | 0.66416768 |
| ENSMUSG00000084969 | Gm11205 | 5.54E-05 | 1.648835551 | 0.886203497 |
| ENSMUSG00000085130 | Gm11417 | 5.54E-05 | 0.648835551 | 1 |
| ENSMUSG00000092157 | Gm17268 | 5.54E-05 | 1.326907457 | 0.971854959 |
| ENSMUSG00000035352 | Ccl12 | 5.54E-05 | 2.063873051 | 0.792441218 |
| ENSMUSG00000020701 | Tmem132e | 5.54E-05 | 0.47891055 | 1 |
| ENSMUSG00000018844 | Fndc8 | 5.54E-05 | 1.063873051 | 0.971854959 |
| ENSMUSG00000020692 | Nle1 | 5.54E-05 | 0.47891055 | 1 |

| | | | | |
|---|---|---|---|---|
| ENSMUSG00000035085 | 1700020L24Rik | 5.54E-05 | 0.189403933 | 1 |
| ENSMUSG00000085235 | Gm12576 | 5.54E-05 | -2.560617814 | 0.037615804 |
| ENSMUSG00000018648 | Dusp14 | 5.54E-05 | 2.063873051 | 0.792441218 |
| ENSMUSG00000018698 | Lhx1 | 5.54E-05 | 2.063873051 | 0.782183909 |
| ENSMUSG00000034449 | Dhrs11 | 5.54E-05 | -0.743481871 | 1 |
| ENSMUSG00000045140 | Pigw | 5.54E-05 | -0.52108945 | 1 |
| ENSMUSG00000085628 | Appbp2os | 5.54E-05 | 0.841480629 | 1 |
| ENSMUSG00000085514 | Bcas3os2 | 5.54E-05 | 3.648835551 | 0.466464457 |
| ENSMUSG00000000093 | Tbx2 | 5.54E-05 | 0.326907457 | 1 |
| ENSMUSG00000000094 | Tbx4 | 5.54E-05 | 1.063873051 | 1 |
| ENSMUSG00000020493 | Prr11 | 5.54E-05 | -0.874726405 | 0.840804483 |
| ENSMUSG00000085310 | Gm11491 | 5.54E-05 | 1.063873051 | 1 |
| ENSMUSG00000007646 | Rad51c | 5.54E-05 | -0.258055044 | 1 |
| ENSMUSG00000020486 | 4-Sep | 5.54E-05 | 0.063873051 | 1 |
| ENSMUSG00000009350 | Mpo | 5.54E-05 | 1.326907457 | 0.943363727 |
| ENSMUSG00000009356 | Lpo | 5.54E-05 | 0.47891055 | 1 |
| ENSMUSG00000087373 | Gm15892 | 5.54E-05 | -0.743481871 | 0.827142217 |
| ENSMUSG00000110344 | Gm45716 | 5.54E-05 | 0.648835551 | 1 |
| ENSMUSG00000003949 | Hlf | 5.54E-05 | 0.189403933 | 1 |
| ENSMUSG00000054079 | Utp18 | 5.54E-05 | -1.810596067 | 0.246776726 |
| ENSMUSG00000085051 | Gm11542 | 5.54E-05 | 0.841480629 | 1 |
| ENSMUSG00000056008 | Gm11541 | 5.54E-05 | 0.326907457 | 1 |
| ENSMUSG00000086552 | Dlx4os | 5.54E-05 | -1.305360759 | 0.491653643 |
| ENSMUSG00000085143 | Gm11520 | 5.54E-05 | 0.063873051 | 1 |
| ENSMUSG00000101258 | Gm29477 | 5.54E-05 | 2.648835551 | 0.66416768 |
| ENSMUSG00000006056 | Calcoco2 | 5.54E-05 | 1.063873051 | 1 |
| ENSMUSG00000085645 | Hoxb5os | 5.54E-05 | 1.648835551 | 0.886203497 |
| ENSMUSG00000084844 | Hoxb3os | 5.54E-05 | 1.648835551 | 0.886203497 |
| ENSMUSG00000020876 | Snx11 | 5.54E-05 | -0.936126949 | 0.695372548 |
| ENSMUSG00000087117 | Gm11523 | 5.54E-05 | 2.063873051 | 0.792441218 |
| ENSMUSG00000078700 | D030028A08Rik | 5.54E-05 | -2.560617814 | 0.030770816 |
| ENSMUSG00000018659 | Pnpo | 5.54E-05 | -0.43862729 | 1 |
| ENSMUSG00000038560 | Sp6 | 5.54E-05 | -0.43862729 | 1 |
| ENSMUSG00000001445 | Mrpl10 | 5.54E-05 | -1.158519371 | 0.605389401 |
| ENSMUSG00000038517 | Tbkbp1 | 5.54E-05 | 1.326907457 | 0.971854959 |
| ENSMUSG00000070337 | Gpr179 | 5.54E-05 | -0.351164449 | 1 |
| ENSMUSG00000038453 | Srcin1 | 5.54E-05 | 0.841480629 | 1 |
| ENSMUSG00000038352 | Arl5c | 5.54E-05 | -0.258055044 | 1 |
| ENSMUSG00000061718 | Ppp1r1b | 5.54E-05 | 1.326907457 | 0.971854959 |
| ENSMUSG00000017204 | Gsdma | 5.54E-05 | 0.47891055 | 1 |
| ENSMUSG00000038067 | Csf3 | 5.54E-05 | 0.841480629 | 1 |
| ENSMUSG00000017499 | Cdc6 | 5.54E-05 | -0.936126949 | 0.695372548 |
| ENSMUSG00000035849 | Krt222 | 5.54E-05 | 1.648835551 | 0.886203497 |
| ENSMUSG00000019761 | Krt10 | 5.54E-05 | 1.063873051 | 1 |
| ENSMUSG00000054146 | Krt15 | 5.54E-05 | 3.648835551 | 0.466464457 |
| ENSMUSG00000006782 | Cnp | 5.54E-05 | 0.189403933 | 1 |
| ENSMUSG00000035355 | Kcnh4 | 5.54E-05 | 0.648835551 | 1 |
| ENSMUSG00000035112 | Wnk4 | 5.54E-05 | -1.43862729 | 0.433313155 |
| ENSMUSG00000019326 | Aoc3 | 5.54E-05 | -0.874726405 | 0.840804483 |
| ENSMUSG00000001493 | Meox1 | 5.54E-05 | 0.189403933 | 1 |
| ENSMUSG00000010841 | Cfap97d1 | 5.54E-05 | 3.648835551 | 0.466464457 |

| | | | | |
|---|---|---|---|---|
| ENSMUSG00000017309 | Cd300lg | 5.54E-05 | 2.648835551 | 0.66416768 |
| ENSMUSG00000020921 | Tmem101 | 5.54E-05 | 3.648835551 | 0.466464457 |
| ENSMUSG00000034793 | G6pc3 | 5.54E-05 | -1.209145444 | 0.606122355 |
| ENSMUSG00000034773 | BC030867 | 5.54E-05 | 0.841480629 | 1 |
| ENSMUSG00000006575 | Rundc3a | 5.54E-05 | 2.063873051 | 0.792441218 |
| ENSMUSG00000050288 | Fzd2 | 5.54E-05 | 0.841480629 | 1 |
| ENSMUSG00000020926 | Adam11 | 5.54E-05 | 2.648835551 | 0.66416768 |
| ENSMUSG00000045532 | C1ql1 | 5.54E-05 | 3.648835551 | 0.466464457 |
| ENSMUSG00000020940 | 1700023F06Rik | 5.54E-05 | 0.189403933 | 1 |
| ENSMUSG00000085887 | Arhgap27os3 | 5.54E-05 | 3.648835551 | 0.466464457 |
| ENSMUSG00000085360 | Arhgap27os2 | 5.54E-05 | 0.648835551 | 1 |
| ENSMUSG00000086932 | Gm22000 | 5.54E-05 | 2.063873051 | 0.792441218 |
| ENSMUSG00000020681 | Ace | 5.54E-05 | -0.051604167 | 1 |
| ENSMUSG00000020712 | Tcam1 | 5.54E-05 | 0.648835551 | 1 |
| ENSMUSG00000020718 | Polg2 | 5.54E-05 | -2.132524162 | 0.14929927 |
| ENSMUSG00000087683 | Gm11712 | 5.54E-05 | 2.063873051 | 0.792441218 |
| ENSMUSG00000040373 | Cacng5 | 5.54E-05 | 1.648835551 | 0.871508522 |
| ENSMUSG00000000049 | Apoh | 5.54E-05 | -0.051604167 | 1 |
| ENSMUSG00000085811 | Cep112it | 5.54E-05 | -0.351164449 | 1 |
| ENSMUSG00000085324 | Gm11680 | 5.54E-05 | 2.063873051 | 0.792441218 |
| ENSMUSG00000020624 | 4933434M16Rik | 5.54E-05 | 1.326907457 | 0.953117731 |
| ENSMUSG00000085421 | 4732490B19Rik | 5.54E-05 | 0.648835551 | 1 |
| ENSMUSG00000018727 | Cpsf4l | 5.54E-05 | 1.326907457 | 0.907125767 |
| ENSMUSG00000041598 | Cdc42ep4 | 5.54E-05 | -1.209145444 | 0.606122355 |
| ENSMUSG00000000202 | Btbd17 | 5.54E-05 | 3.648835551 | 0.466464457 |
| ENSMUSG00000034652 | Cd300a | 5.54E-05 | -1.599091962 | 0.426429205 |
| ENSMUSG00000048498 | Cd300e | 5.54E-05 | 1.063873051 | 1 |
| ENSMUSG00000044788 | Fads6 | 5.54E-05 | -0.158519371 | 1 |
| ENSMUSG00000045288 | Ush1g | 5.54E-05 | 2.063873051 | 0.792441218 |
| ENSMUSG00000018862 | Otop3 | 5.54E-05 | -1.708716453 | 0.296030202 |
| ENSMUSG00000045775 | Slc16a5 | 5.54E-05 | 0.841480629 | 1 |
| ENSMUSG00000034471 | Caskin2 | 5.54E-05 | 0.063873051 | 1 |
| ENSMUSG00000020781 | Tsen54 | 5.54E-05 | -0.743481871 | 1 |
| ENSMUSG00000020775 | Mrpl38 | 5.54E-05 | -2.184054463 | 0.110207286 |
| ENSMUSG00000034227 | Foxj1 | 5.54E-05 | 3.648835551 | 0.466464457 |
| ENSMUSG00000075410 | Prcd | 5.54E-05 | 1.326907457 | 0.925873597 |
| ENSMUSG00000056962 | Jmjd6 | 5.54E-05 | -1.995020638 | 0.167528217 |
| ENSMUSG00000043857 | Mgat5b | 5.54E-05 | 0.47891055 | 1 |
| ENSMUSG00000085104 | Gm11731 | 5.54E-05 | 2.063873051 | 0.792441218 |
| ENSMUSG00000086859 | Snhg20 | 5.54E-05 | -1.158519371 | 0.599135879 |
| ENSMUSG00000087057 | Gm11730 | 5.54E-05 | 3.648835551 | 0.466464457 |
| ENSMUSG00000087669 | Gm11724 | 5.54E-05 | 2.063873051 | 0.792441218 |
| ENSMUSG00000076433 | Cep295nl | 5.54E-05 | -1.743481871 | 0.305378889 |
| ENSMUSG00000033857 | Engase | 5.54E-05 | -0.351164449 | 1 |
| ENSMUSG00000039963 | Ccdc40 | 5.54E-05 | 0.189403933 | 1 |
| ENSMUSG00000109408 | A930037H05Rik | 5.54E-05 | -0.158519371 | 1 |
| ENSMUSG00000025375 | Aatk | 5.54E-05 | -0.258055044 | 1 |
| ENSMUSG00000086750 | Gm11770 | 5.54E-05 | 3.648835551 | 0.466464457 |
| ENSMUSG00000043644 | 0610009L18Rik | 5.54E-05 | 1.648835551 | 0.886203497 |
| ENSMUSG00000025386 | Pde6g | 5.54E-05 | 0.063873051 | 1 |
| ENSMUSG00000049957 | Ccdc137 | 5.54E-05 | -1.599091962 | 0.375375289 |

| | | | | |
|---|---|---|---|---|
| ENSMUSG00000053613 | Notumos | 5.54E-05 | 3.648835551 | 0.466464457 |
| ENSMUSG00000018012 | Rac3 | 5.54E-05 | 1.326907457 | 0.967976676 |
| ENSMUSG00000086835 | Gm11775 | 5.54E-05 | -1.051604167 | 0.786691619 |
| ENSMUSG00000025169 | Ogfod3 | 5.54E-05 | 0.648835551 | 1 |
| ENSMUSG00000039307 | Hexdc | 5.54E-05 | -1.305360759 | 0.667778757 |
| ENSMUSG00000092286 | Dtnbos | 5.54E-05 | 0.189403933 | 1 |
| ENSMUSG00000112422 | Gm31938 | 5.54E-05 | 3.648835551 | 0.466464457 |
| ENSMUSG00000079177 | Fam228a | 5.54E-05 | -0.051604167 | 1 |
| ENSMUSG00000020639 | Pfn4 | 5.54E-05 | 3.648835551 | 0.466464457 |
| ENSMUSG00000096940 | Gm4755 | 5.54E-05 | 2.648835551 | 0.66416768 |
| ENSMUSG00000111925 | Gm48791 | 5.54E-05 | 0.189403933 | 1 |
| ENSMUSG00000020622 | Nt5c1b | 5.54E-05 | 2.648835551 | 0.66416768 |
| ENSMUSG00000112041 | 9530020I12Rik | 5.54E-05 | 2.063873051 | 0.792441218 |
| ENSMUSG00000114136 | 4921511I17Rik | 5.54E-05 | 1.648835551 | 0.886203497 |
| ENSMUSG00000097627 | 4930519A11Rik | 5.54E-05 | 1.648835551 | 0.886203497 |
| ENSMUSG00000113928 | Gm48209 | 5.54E-05 | 2.648835551 | 0.66416768 |
| ENSMUSG00000113666 | Gm35725 | 5.54E-05 | 0.47891055 | 1 |
| ENSMUSG00000113735 | Gm48607 | 5.54E-05 | 2.063873051 | 0.792441218 |
| ENSMUSG00000114049 | Gm48140 | 5.54E-05 | 2.648835551 | 0.662993938 |
| ENSMUSG00000114018 | Gm36495 | 5.54E-05 | 2.648835551 | 0.66416768 |
| ENSMUSG00000114024 | Gm46336 | 5.54E-05 | 2.063873051 | 0.792441218 |
| ENSMUSG00000073184 | Gm49957 | 5.54E-05 | 3.648835551 | 0.466464457 |
| ENSMUSG00000092373 | Gm20535 | 5.54E-05 | 2.648835551 | 0.66416768 |
| ENSMUSG00000093846 | Gm4425 | 5.54E-05 | 1.326907457 | 0.958492465 |
| ENSMUSG00000073164 | 2410018L13Rik | 5.54E-05 | 1.648835551 | 0.885194626 |
| ENSMUSG00000113587 | Gm36287 | 5.54E-05 | -0.43862729 | 1 |
| ENSMUSG00000113628 | 1700020D12Rik | 5.54E-05 | -0.52108945 | 0.975624465 |
| ENSMUSG00000094002 | Gm9866 | 5.54E-05 | -0.051604167 | 1 |
| ENSMUSG00000113616 | Gm45941 | 5.54E-05 | 1.648835551 | 0.877036949 |
| ENSMUSG00000112663 | C630031E19Rik | 5.54E-05 | 3.648835551 | 0.466464457 |
| ENSMUSG00000001225 | Slc26a3 | 5.54E-05 | -0.351164449 | 1 |
| ENSMUSG00000020648 | Dus4l | 5.54E-05 | 0.189403933 | 1 |
| ENSMUSG00000020573 | Pik3cg | 5.54E-05 | 0.648835551 | 1 |
| ENSMUSG00000097494 | 4933406C10Rik | 5.54E-05 | 0.326907457 | 1 |
| ENSMUSG00000052125 | F730043M19Rik | 5.54E-05 | -2.500911568 | 0.036096973 |
| ENSMUSG00000079143 | Atxn7l1os1 | 5.54E-05 | 0.47891055 | 1 |
| ENSMUSG00000114082 | Gm40383 | 5.54E-05 | 1.326907457 | 0.946281031 |
| ENSMUSG00000113985 | 9130015A21Rik | 5.54E-05 | 1.326907457 | 0.971854959 |
| ENSMUSG00000020581 | Agr2 | 5.54E-05 | 0.47891055 | 1 |
| ENSMUSG00000113507 | Gm48613 | 5.54E-05 | 2.063873051 | 0.792441218 |
| ENSMUSG00000114052 | Gm48616 | 5.54E-05 | 0.841480629 | 1 |
| ENSMUSG00000036144 | Meox2 | 5.54E-05 | 0.648835551 | 1 |
| ENSMUSG00000113005 | Gm34408 | 5.54E-05 | 2.648835551 | 0.66416768 |
| ENSMUSG00000113608 | Gm48715 | 5.54E-05 | 2.063873051 | 0.790550166 |
| ENSMUSG00000004151 | Etv1 | 5.54E-05 | -0.810596067 | 0.871107617 |
| ENSMUSG00000035983 | Gm7008 | 5.54E-05 | 2.648835551 | 0.66416768 |
| ENSMUSG00000113086 | Gm47376 | 5.54E-05 | 2.648835551 | 0.66416768 |
| ENSMUSG00000036295 | Lrrn3 | 5.54E-05 | 1.648835551 | 0.886203497 |
| ENSMUSG00000113370 | Gm46348 | 5.54E-05 | 0.326907457 | 1 |
| ENSMUSG00000112898 | Gm47020 | 5.54E-05 | 1.648835551 | 0.875465725 |
| ENSMUSG00000092404 | Gm9921 | 5.54E-05 | 3.648835551 | 0.466464457 |

| | | | | |
|---|---|---|---|---|
| ENSMUSG00000112054 | Gm33680 | 5.54E-05 | 0.326907457 | 1 |
| ENSMUSG00000112685 | Gm40421 | 5.54E-05 | 2.648835551 | 0.66416768 |
| ENSMUSG00000020953 | Coch | 5.54E-05 | 2.063873051 | 0.792441218 |
| ENSMUSG00000112139 | Gm47431 | 5.54E-05 | -1.106051951 | 0.792441218 |
| ENSMUSG00000112211 | Gm47682 | 5.54E-05 | 2.063873051 | 0.775870935 |
| ENSMUSG00000021028 | Mbip | 5.54E-05 | -3.936126949 | 6.15E-05 |
| ENSMUSG00000001496 | Nkx2-1 | 5.54E-05 | 3.648835551 | 0.466464457 |
| ENSMUSG00000058669 | Nkx2-9 | 5.54E-05 | 3.648835551 | 0.466464457 |
| ENSMUSG00000089894 | Gm15524 | 5.54E-05 | 2.648835551 | 0.66416768 |
| ENSMUSG00000079104 | Prps1l3 | 5.54E-05 | -1.351164449 | 0.499088689 |
| ENSMUSG00000085851 | 4921518K17Rik | 5.54E-05 | 0.326907457 | 1 |
| ENSMUSG00000097443 | Gm17529 | 5.54E-05 | 0.841480629 | 1 |
| ENSMUSG00000113208 | Gm48421 | 5.54E-05 | 0.326907457 | 1 |
| ENSMUSG00000055884 | Fancm | 5.54E-05 | -2.560617814 | 0.030770816 |
| ENSMUSG00000113228 | 4930471E15Rik | 5.54E-05 | 1.648835551 | 0.886203497 |
| ENSMUSG00000114093 | Gm47979 | 5.54E-05 | 1.326907457 | 0.971854959 |
| ENSMUSG00000113526 | Gm47453 | 5.54E-05 | 2.648835551 | 0.66416768 |
| ENSMUSG00000113433 | Gm47489 | 5.54E-05 | 2.648835551 | 0.66416768 |
| ENSMUSG00000113098 | Gm2912 | 5.54E-05 | 0.648835551 | 1 |
| ENSMUSG00000113190 | Gm47545 | 5.54E-05 | 1.648835551 | 0.880201363 |
| ENSMUSG00000049882 | Vcpkmt | 5.54E-05 | -1.351164449 | 0.587566484 |
| ENSMUSG00000021066 | Atl1 | 5.54E-05 | -0.673092543 | 0.98880775 |
| ENSMUSG00000090121 | Abhd12b | 5.54E-05 | 3.648835551 | 0.466464457 |
| ENSMUSG00000085298 | F730035M05Rik | 5.54E-05 | 1.326907457 | 0.971854959 |
| ENSMUSG00000113302 | Gm32219 | 5.54E-05 | 2.063873051 | 0.783520679 |
| ENSMUSG00000113630 | 4930404H11Rik | 5.54E-05 | 1.326907457 | 0.971854959 |
| ENSMUSG00000019718 | L3hypdh | 5.54E-05 | -1.52108945 | 0.498867935 |
| ENSMUSG00000021090 | Lrrc9 | 5.54E-05 | -0.874726405 | 0.862828211 |
| ENSMUSG00000113779 | Gm33785 | 5.54E-05 | 3.648835551 | 0.466464457 |
| ENSMUSG00000034442 | Trmt5 | 5.54E-05 | -0.351164449 | 1 |
| ENSMUSG00000113099 | Gm34016 | 5.54E-05 | 2.648835551 | 0.66416768 |
| ENSMUSG00000113699 | Gm48823 | 5.54E-05 | 3.648835551 | 0.466464457 |
| ENSMUSG00000052221 | Ppp1r36 | 5.54E-05 | -0.051604167 | 1 |
| ENSMUSG00000046157 | Tmem229b | 5.54E-05 | 0.326907457 | 1 |
| ENSMUSG00000112489 | 9230116L04Rik | 5.54E-05 | 1.063873051 | 1 |
| ENSMUSG00000097729 | 2310015A10Rik | 5.54E-05 | 0.326907457 | 1 |
| ENSMUSG00000112278 | Gm30025 | 5.54E-05 | 0.841480629 | 1 |
| ENSMUSG00000112093 | Gm47941 | 5.54E-05 | 2.648835551 | 0.66416768 |
| ENSMUSG00000112880 | Gm20337 | 5.54E-05 | 0.841480629 | 1 |
| ENSMUSG00000072972 | Adam4 | 5.54E-05 | -1.995020638 | 0.167528217 |
| ENSMUSG00000091947 | 6530401F13Rik | 5.54E-05 | -0.051604167 | 1 |
| ENSMUSG00000090244 | Gm16572 | 5.54E-05 | 2.063873051 | 0.792441218 |
| ENSMUSG00000102029 | Gm29530 | 5.54E-05 | 0.648835551 | 1 |
| ENSMUSG00000099708 | Gm29361 | 5.54E-05 | 3.648835551 | 0.466464457 |
| ENSMUSG00000021223 | Papln | 5.54E-05 | 1.648835551 | 0.886203497 |
| ENSMUSG00000046791 | Riox1 | 5.54E-05 | 1.063873051 | 1 |
| ENSMUSG00000021241 | Isca2 | 5.54E-05 | -1.106051951 | 0.762809221 |
| ENSMUSG00000113940 | Gm40477 | 5.54E-05 | 0.189403933 | 1 |
| ENSMUSG00000034271 | Jdp2 | 5.54E-05 | -1.106051951 | 0.763475704 |
| ENSMUSG00000097543 | Gm805 | 5.54E-05 | 0.063873051 | 1 |
| ENSMUSG00000021257 | Angel1 | 5.54E-05 | 1.326907457 | 0.971854959 |

| | | | | |
|---|---|---|---|---|
| ENSMUSG00000059114 | Lrrc74a | 5.54E-05 | 0.47891055 | 1 |
| ENSMUSG00000034145 | Tmem63c | 5.54E-05 | 0.47891055 | 1 |
| ENSMUSG00000113087 | 4930473H19Rik | 5.54E-05 | 2.648835551 | 0.66416768 |
| ENSMUSG00000085004 | 5430427M07Rik | 5.54E-05 | 1.648835551 | 0.886203497 |
| ENSMUSG00000113164 | 4930559C10Rik | 5.54E-05 | 2.063873051 | 0.792441218 |
| ENSMUSG00000112812 | Gm47415 | 5.54E-05 | 1.063873051 | 1 |
| ENSMUSG00000021013 | Ttc8 | 5.54E-05 | -1.158519371 | 0.599135879 |
| ENSMUSG00000113634 | Gm36756 | 5.54E-05 | 1.063873051 | 1 |
| ENSMUSG00000095174 | Gm20069 | 5.54E-05 | 3.648835551 | 0.466464457 |
| ENSMUSG00000113907 | Gm47167 | 5.54E-05 | -0.051604167 | 1 |
| ENSMUSG00000041669 | Prima1 | 5.54E-05 | 1.063873051 | 1 |
| ENSMUSG00000096753 | Fam181a | 5.54E-05 | 3.648835551 | 0.466464457 |
| ENSMUSG00000071178 | Serpina1b | 5.54E-05 | 0.189403933 | 1 |
| ENSMUSG00000072849 | Serpina1e | 5.54E-05 | 2.063873051 | 0.792441218 |
| ENSMUSG00000058260 | Serpina9 | 5.54E-05 | -0.258055044 | 1 |
| ENSMUSG00000041567 | Serpina12 | 5.54E-05 | 3.648835551 | 0.466464457 |
| ENSMUSG00000079013 | Serpina3j | 5.54E-05 | 0.841480629 | 1 |
| ENSMUSG00000114826 | Gm10000 | 5.54E-05 | 0.648835551 | 1 |
| ENSMUSG00000021101 | 4930408O17Rik | 5.54E-05 | 1.326907457 | 0.970970089 |
| ENSMUSG00000113722 | Snhg10 | 5.54E-05 | 0.648835551 | 1 |
| ENSMUSG00000000701 | Tcl1b5 | 5.54E-05 | 2.648835551 | 0.66416768 |
| ENSMUSG00000091280 | Gm17032 | 5.54E-05 | 3.648835551 | 0.466464457 |
| ENSMUSG00000085801 | 1700013N06Rik | 5.54E-05 | 2.063873051 | 0.786367309 |
| ENSMUSG00000113415 | Gm3234 | 5.54E-05 | 1.648835551 | 0.886203497 |
| ENSMUSG00000021259 | Cyp46a1 | 5.54E-05 | -0.051604167 | 1 |
| ENSMUSG00000040867 | Begain | 5.54E-05 | 1.648835551 | 0.874192437 |
| ENSMUSG00000112977 | Gm34081 | 5.54E-05 | 2.648835551 | 0.66416768 |
| ENSMUSG00000097451 | Rian | 5.54E-05 | -1.965874293 | 0.162298752 |
| ENSMUSG00000098202 | B830012L14Rik | 5.54E-05 | 0.189403933 | 1 |
| ENSMUSG00000113331 | 3110009F21Rik | 5.54E-05 | 2.063873051 | 0.792441218 |
| ENSMUSG00000113193 | Gm40576 | 5.54E-05 | 2.648835551 | 0.66416768 |
| ENSMUSG00000056458 | Mok | 5.54E-05 | -1.209145444 | 0.606122355 |
| ENSMUSG00000097650 | 4921507G05Rik | 5.54E-05 | 2.648835551 | 0.66416768 |
| ENSMUSG00000049792 | Bag5 | 5.54E-05 | -0.52108945 | 0.975624465 |
| ENSMUSG00000113120 | E330035G20Rik | 5.54E-05 | 3.648835551 | 0.466464457 |
| ENSMUSG00000047832 | Cdca4 | 5.54E-05 | -0.673092543 | 0.98880775 |
| ENSMUSG00000113519 | Gm47541 | 5.54E-05 | 0.648835551 | 1 |
| ENSMUSG00000076614 | Ighg1 | 5.54E-05 | -0.258055044 | 1 |
| ENSMUSG00000076617 | Ighm | 5.54E-05 | 0.47891055 | 1 |
| ENSMUSG00000103168 | Gm30948 | 5.54E-05 | 3.648835551 | 0.466464457 |
| ENSMUSG00000101930 | Gm5441 | 5.54E-05 | 0.189403933 | 1 |
| ENSMUSG00000113831 | Gm49602 | 5.54E-05 | 0.47891055 | 1 |
| ENSMUSG00000025321 | Itgb8 | 5.54E-05 | -0.810596067 | 0.865413877 |
| ENSMUSG00000114091 | Gm40655 | 5.54E-05 | 3.648835551 | 0.466464457 |
| ENSMUSG00000033715 | Akr1c14 | 5.54E-05 | 2.063873051 | 0.792441218 |
| ENSMUSG00000054757 | Akr1c20 | 5.54E-05 | 1.648835551 | 0.886203497 |
| ENSMUSG00000021207 | Akr1c21 | 5.54E-05 | 2.648835551 | 0.66416768 |
| ENSMUSG00000114017 | Gm40662 | 5.54E-05 | -0.52108945 | 0.992252446 |
| ENSMUSG00000113241 | Gm40663 | 5.54E-05 | 1.326907457 | 0.971854959 |
| ENSMUSG00000033520 | Idi2 | 5.54E-05 | 0.189403933 | 1 |
| ENSMUSG00000113380 | Gm48682 | 5.54E-05 | 2.063873051 | 0.792441218 |

| | | | | |
|---|---|---|---|---|
| ENSMUSG00000113811 | Gm47882 | 5.54E-05 | 0.648835551 | 1 |
| ENSMUSG00000039182 | AW209491 | 5.54E-05 | 0.326907457 | 1 |
| ENSMUSG00000113851 | Gm48503 | 5.54E-05 | 1.063873051 | 1 |
| ENSMUSG00000114189 | 9330199G10Rik | 5.54E-05 | 0.648835551 | 1 |
| ENSMUSG00000113313 | Gm32036 | 5.54E-05 | 1.648835551 | 0.874192437 |
| ENSMUSG00000060404 | Olfr1369-ps1 | 5.54E-05 | 2.063873051 | 0.792441218 |
| ENSMUSG00000059395 | Nkapl | 5.54E-05 | -0.258055044 | 1 |
| ENSMUSG00000063894 | Zkscan8 | 5.54E-05 | -1.480447466 | 0.441576406 |
| ENSMUSG00000054890 | Olfr1535 | 5.54E-05 | 2.648835551 | 0.66416768 |
| ENSMUSG00000063021 | Hist1h2ak | 5.54E-05 | 2.648835551 | 0.66416768 |
| ENSMUSG00000095217 | Hist1h2bn | 5.54E-05 | 2.648835551 | 0.66416768 |
| ENSMUSG00000058773 | Hist1h1b | 5.54E-05 | -0.351164449 | 1 |
| ENSMUSG00000006720 | Zfp184 | 5.54E-05 | -0.051604167 | 1 |
| ENSMUSG00000095125 | Vmn1r198 | 5.54E-05 | 1.648835551 | 0.886203497 |
| ENSMUSG00000115020 | Vmn1r218 | 5.54E-05 | 1.063873051 | 1 |
| ENSMUSG00000082316 | Vmn1r221 | 5.54E-05 | 3.648835551 | 0.466464457 |
| ENSMUSG00000085488 | 4930557F10Rik | 5.54E-05 | 0.841480629 | 1 |
| ENSMUSG00000114456 | Hist1h2bh | 5.54E-05 | 3.648835551 | 0.466464457 |
| ENSMUSG00000069274 | Hist1h4f | 5.54E-05 | 3.648835551 | 0.466464457 |
| ENSMUSG00000052565 | Hist1h1d | 5.54E-05 | -0.874726405 | 0.840804483 |
| ENSMUSG00000061482 | Hist1h4d | 5.54E-05 | 0.648835551 | 1 |
| ENSMUSG00000047246 | Hist1h2be | 5.54E-05 | -1.305360759 | 0.491653643 |
| ENSMUSG00000061615 | Hist1h2ab | 5.54E-05 | 2.063873051 | 0.792441218 |
| ENSMUSG00000087603 | Gm11339 | 5.54E-05 | 3.648835551 | 0.466464457 |
| ENSMUSG00000086796 | 4932702P03Rik | 5.54E-05 | 3.648835551 | 0.466464457 |
| ENSMUSG00000021339 | Mrs2 | 5.54E-05 | -1.636566667 | 0.373232174 |
| ENSMUSG00000062737 | Prl3d2 | 5.54E-05 | 3.648835551 | 0.466464457 |
| ENSMUSG00000069259 | Prl6a1 | 5.54E-05 | 3.648835551 | 0.466464457 |
| ENSMUSG00000021346 | Prl8a8 | 5.54E-05 | 3.648835551 | 0.466464457 |
| ENSMUSG00000060738 | Prl7c1 | 5.54E-05 | 3.648835551 | 0.466464457 |
| ENSMUSG00000017064 | Prl5a1 | 5.54E-05 | 1.326907457 | 0.971854959 |
| ENSMUSG00000086363 | A330102I10Rik | 5.54E-05 | -0.673092543 | 1 |
| ENSMUSG00000069255 | Dusp22 | 5.54E-05 | -0.874726405 | 0.865542683 |
| ENSMUSG00000038402 | Foxf2 | 5.54E-05 | 1.063873051 | 1 |
| ENSMUSG00000051029 | Serpinb1b | 5.54E-05 | 1.648835551 | 0.867965264 |
| ENSMUSG00000054266 | Serpinb9d | 5.54E-05 | 3.648835551 | 0.466464457 |
| ENSMUSG00000057726 | Serpinb9g | 5.54E-05 | 2.648835551 | 0.66416768 |
| ENSMUSG00000052180 | Serpinb6c | 5.54E-05 | 1.326907457 | 0.971854959 |
| ENSMUSG00000021415 | 4933417A18Rik | 5.54E-05 | 2.063873051 | 0.792441218 |
| ENSMUSG00000114661 | Gm48707 | 5.54E-05 | 2.648835551 | 0.66416768 |
| ENSMUSG00000050423 | Ppp1r3g | 5.54E-05 | 2.648835551 | 0.66416768 |
| ENSMUSG00000046573 | Lyrm4 | 5.54E-05 | -0.43862729 | 1 |
| ENSMUSG00000114665 | Lncbate6 | 5.54E-05 | 0.326907457 | 1 |
| ENSMUSG00000114404 | Gm40915 | 5.54E-05 | 1.063873051 | 1 |
| ENSMUSG00000039004 | Bmp6 | 5.54E-05 | -2.184054463 | 0.091007367 |
| ENSMUSG00000114895 | 4930579J19Rik | 5.54E-05 | 2.648835551 | 0.66416768 |
| ENSMUSG00000114830 | Gm47400 | 5.54E-05 | 2.063873051 | 0.773495198 |
| ENSMUSG00000114198 | Gm47039 | 5.54E-05 | 0.841480629 | 1 |
| ENSMUSG00000021359 | Tfap2a | 5.54E-05 | 0.841480629 | 1 |
| ENSMUSG00000021364 | Elovl2 | 5.54E-05 | 2.648835551 | 0.66416768 |
| ENSMUSG00000114914 | Gm48571 | 5.54E-05 | 3.648835551 | 0.466464457 |

| | | | | |
|---|---|---|---|---|
| ENSMUSG00000058022 | Adtrp | 5.54E-05 | 0.063873051 | 1 |
| ENSMUSG00000113856 | Gm10790 | 5.54E-05 | 0.47891055 | 1 |
| ENSMUSG00000100071 | Gm28707 | 5.54E-05 | 2.063873051 | 0.792441218 |
| ENSMUSG00000054021 | Sirt5 | 5.54E-05 | -1.258055044 | 0.645167926 |
| ENSMUSG00000113891 | Gm47683 | 5.54E-05 | 2.648835551 | 0.66416768 |
| ENSMUSG00000113186 | A330076C08Rik | 5.54E-05 | 2.063873051 | 0.792441218 |
| ENSMUSG00000098107 | Gm27007 | 5.54E-05 | 1.326907457 | 0.958492465 |
| ENSMUSG00000113010 | Gm34084 | 5.54E-05 | 1.063873051 | 1 |
| ENSMUSG00000113730 | Gm47805 | 5.54E-05 | 2.648835551 | 0.66416768 |
| ENSMUSG00000113919 | Gm34466 | 5.54E-05 | 2.063873051 | 0.792441218 |
| ENSMUSG00000113987 | 4930453C13Rik | 5.54E-05 | 3.648835551 | 0.466464457 |
| ENSMUSG00000038152 | 5033430I15Rik | 5.54E-05 | 2.063873051 | 0.789767653 |
| ENSMUSG00000063529 | Stmnd1 | 5.54E-05 | 1.648835551 | 0.886203497 |
| ENSMUSG00000114596 | Gm47523 | 5.54E-05 | 3.648835551 | 0.466464457 |
| ENSMUSG00000114810 | Gm36101 | 5.54E-05 | 3.648835551 | 0.466464457 |
| ENSMUSG00000097375 | 6720427I07Rik | 5.54E-05 | -2.395558568 | 0.052731507 |
| ENSMUSG00000038042 | Ptpdc1 | 5.54E-05 | -2.132524162 | 0.112236795 |
| ENSMUSG00000113255 | Tes3-ps | 5.54E-05 | 2.648835551 | 0.66416768 |
| ENSMUSG00000043631 | Ecm2 | 5.54E-05 | 0.648835551 | 1 |
| ENSMUSG00000114035 | Gm31126 | 5.54E-05 | 3.648835551 | 0.466464457 |
| ENSMUSG00000037816 | Fbxw17 | 5.54E-05 | -1.351164449 | 0.499088689 |
| ENSMUSG00000113737 | BB123696 | 5.54E-05 | 1.648835551 | 0.886203497 |
| ENSMUSG00000113982 | Gm34557 | 5.54E-05 | 2.648835551 | 0.66416768 |
| ENSMUSG00000034891 | Sncb | 5.54E-05 | 3.648835551 | 0.466464457 |
| ENSMUSG00000005320 | Fgfr4 | 5.54E-05 | -2.843017545 | 0.011487643 |
| ENSMUSG00000052087 | Rgs14 | 5.54E-05 | 1.326907457 | 0.971854959 |
| ENSMUSG00000014164 | Klhl3 | 5.54E-05 | -1.305360759 | 0.491653643 |
| ENSMUSG00000114309 | Gm48295 | 5.54E-05 | 2.648835551 | 0.66416768 |
| ENSMUSG00000114808 | Gm47947 | 5.54E-05 | 1.326907457 | 0.971854959 |
| ENSMUSG00000114218 | Gm47423 | 5.54E-05 | 3.648835551 | 0.466464457 |
| ENSMUSG00000114437 | 4930415C11Rik | 5.54E-05 | 1.326907457 | 0.968284003 |
| ENSMUSG00000114743 | Gm34672 | 5.54E-05 | 0.648835551 | 1 |
| ENSMUSG00000114623 | Gm34788 | 5.54E-05 | 2.648835551 | 0.66416768 |
| ENSMUSG00000114531 | A530065N20Rik | 5.54E-05 | 0.063873051 | 1 |
| ENSMUSG00000114566 | Gm48500 | 5.54E-05 | 2.648835551 | 0.66416768 |
| ENSMUSG00000114452 | A530001N23Rik | 5.54E-05 | 1.063873051 | 1 |
| ENSMUSG00000062705 | Tpbpb | 5.54E-05 | 3.648835551 | 0.466464457 |
| ENSMUSG00000074871 | Ctsm | 5.54E-05 | 3.648835551 | 0.466464457 |
| ENSMUSG00000113683 | Gm47123 | 5.54E-05 | 1.326907457 | 0.971854959 |
| ENSMUSG00000032999 | Nlrp4f | 5.54E-05 | 1.648835551 | 0.886203497 |
| ENSMUSG00000113500 | Gm47249 | 5.54E-05 | 0.326907457 | 1 |
| ENSMUSG00000113135 | Gm48170 | 5.54E-05 | 0.47891055 | 1 |
| ENSMUSG00000114362 | 4933433G19Rik | 5.54E-05 | 0.47891055 | 1 |
| ENSMUSG00000114516 | Gm46440 | 5.54E-05 | 0.063873051 | 1 |
| ENSMUSG00000058900 | Rsl1 | 5.54E-05 | 0.326907457 | 1 |
| ENSMUSG00000055341 | Zfp457 | 5.54E-05 | 0.326907457 | 1 |
| ENSMUSG00000057842 | Zfp595 | 5.54E-05 | -0.810596067 | 0.832285197 |
| ENSMUSG00000078995 | Zfp456 | 5.54E-05 | 1.063873051 | 1 |
| ENSMUSG00000078994 | Zfp429 | 5.54E-05 | -0.599091962 | 0.980745941 |
| ENSMUSG00000097333 | Zfp87 | 5.54E-05 | -0.936126949 | 0.695372548 |
| ENSMUSG00000114264 | Gm48095 | 5.54E-05 | 3.648835551 | 0.466464457 |

| | | | | |
|---|---|---|---|---|
| ENSMUSG00000097276 | 4930525G20Rik | 5.54E-05 | -0.051604167 | 1 |
| ENSMUSG00000034617 | Mtrr | 5.54E-05 | -1.743481871 | 0.305378889 |
| ENSMUSG00000021532 | Fastkd3 | 5.54E-05 | -0.158519371 | 1 |
| ENSMUSG00000021534 | 1700001L19Rik | 5.54E-05 | 0.189403933 | 1 |
| ENSMUSG00000113121 | Gm35161 | 5.54E-05 | 2.063873051 | 0.792441218 |
| ENSMUSG00000113358 | 1700112M02Rik | 5.54E-05 | -0.673092543 | 1 |
| ENSMUSG00000021611 | Tert | 5.54E-05 | -1.743481871 | 0.305378889 |
| ENSMUSG00000069208 | Zfp825 | 5.54E-05 | -1.43862729 | 0.433313155 |
| ENSMUSG00000021587 | Pcsk1 | 5.54E-05 | 1.063873051 | 1 |
| ENSMUSG00000021590 | Spata9 | 5.54E-05 | 1.326907457 | 0.971854959 |
| ENSMUSG00000114242 | Gm31219 | 5.54E-05 | 1.648835551 | 0.886203497 |
| ENSMUSG00000093668 | Pou5f2 | 5.54E-05 | -0.52108945 | 0.975624465 |
| ENSMUSG00000114554 | 5430425K12Rik | 5.54E-05 | -0.673092543 | 0.98880775 |
| ENSMUSG00000097146 | Gm4211 | 5.54E-05 | 1.063873051 | 1 |
| ENSMUSG00000097793 | Gm17259 | 5.54E-05 | 0.648835551 | 1 |
| ENSMUSG00000021548 | Ccnh | 5.54E-05 | -1.560617814 | 0.356350819 |
| ENSMUSG00000114273 | Gm34585 | 5.54E-05 | -0.673092543 | 0.98880775 |
| ENSMUSG00000114541 | Gm29680 | 5.54E-05 | -2.351164449 | 0.065287234 |
| ENSMUSG00000114204 | Gm47520 | 5.54E-05 | 1.648835551 | 0.886203497 |
| ENSMUSG00000114688 | Gm47381 | 5.54E-05 | 2.648835551 | 0.66416768 |
| ENSMUSG00000074782 | 4833422C13Rik | 5.54E-05 | -0.43862729 | 1 |
| ENSMUSG00000021622 | Ckmt2 | 5.54E-05 | -0.351164449 | 1 |
| ENSMUSG00000021704 | Mtx3 | 5.54E-05 | -0.43862729 | 1 |
| ENSMUSG00000047419 | Cmya5 | 5.54E-05 | -0.258055044 | 1 |
| ENSMUSG00000074768 | Bhmt | 5.54E-05 | 1.063873051 | 1 |
| ENSMUSG00000042118 | Bhmt2 | 5.54E-05 | 2.648835551 | 0.66416768 |
| ENSMUSG00000113791 | Gm47216 | 5.54E-05 | 3.648835551 | 0.466464457 |
| ENSMUSG00000113494 | Gm48102 | 5.54E-05 | 1.648835551 | 0.886203497 |
| ENSMUSG00000021680 | Crhbp | 5.54E-05 | 1.648835551 | 0.886203497 |
| ENSMUSG00000114133 | Gm20075 | 5.54E-05 | 2.063873051 | 0.792441218 |
| ENSMUSG00000091387 | Gcnt4 | 5.54E-05 | 0.063873051 | 1 |
| ENSMUSG00000097346 | Gm26619 | 5.54E-05 | 2.063873051 | 0.792441218 |
| ENSMUSG00000041747 | Utp15 | 5.54E-05 | -2.459688905 | 0.044392129 |
| ENSMUSG00000114458 | Gm47551 | 5.54E-05 | 1.326907457 | 0.969860855 |
| ENSMUSG00000114414 | A930014D07Rik | 5.54E-05 | 1.063873051 | 1 |
| ENSMUSG00000097828 | 6430562O15Rik | 5.54E-05 | 1.648835551 | 0.886203497 |
| ENSMUSG00000041431 | Ccnb1 | 5.54E-05 | -0.673092543 | 0.98880775 |
| ENSMUSG00000100596 | Gm29502 | 5.54E-05 | 0.648835551 | 1 |
| ENSMUSG00000114590 | 5930438M14Rik | 5.54E-05 | 3.648835551 | 0.466464457 |
| ENSMUSG00000021624 | Cd180 | 5.54E-05 | 0.326907457 | 1 |
| ENSMUSG00000114637 | Gm47851 | 5.54E-05 | 2.648835551 | 0.66416768 |
| ENSMUSG00000118537 | Shld3 | 5.54E-05 | 0.47891055 | 1 |
| ENSMUSG00000021718 | 4933425L06Rik | 5.54E-05 | 2.648835551 | 0.66416768 |
| ENSMUSG00000071181 | 3830408C21Rik | 5.54E-05 | 0.189403933 | 1 |
| ENSMUSG00000114311 | Gm31452 | 5.54E-05 | 2.648835551 | 0.66416768 |
| ENSMUSG00000114780 | AI197445 | 5.54E-05 | 2.648835551 | 0.66416768 |
| ENSMUSG00000109853 | Gm33045 | 5.54E-05 | 2.648835551 | 0.66416768 |
| ENSMUSG00000055194 | Actbl2 | 5.54E-05 | 3.648835551 | 0.466464457 |
| ENSMUSG00000086870 | Gm15288 | 5.54E-05 | 1.326907457 | 0.971854959 |
| ENSMUSG00000086649 | Gm15286 | 5.54E-05 | -0.258055044 | 1 |
| ENSMUSG00000114745 | Gm48802 | 5.54E-05 | 2.063873051 | 0.774790843 |

| | | | | |
|---|---|---|---|---|
| ENSMUSG00000023132 | Gzma | 5.54E-05 | 3.648835551 | 0.466464457 |
| ENSMUSG00000021763 | BC067074 | 5.54E-05 | 1.063873051 | 1 |
| ENSMUSG00000114790 | 4921509O07Rik | 5.54E-05 | 0.326907457 | 1 |
| ENSMUSG00000051758 | 4930544M13Rik | 5.54E-05 | -0.051604167 | 1 |
| ENSMUSG00000021731 | Mrps30 | 5.54E-05 | -1.395558568 | 0.508263132 |
| ENSMUSG00000091849 | Gm17188 | 5.54E-05 | 0.326907457 | 1 |
| ENSMUSG00000095195 | Gm3005 | 5.54E-05 | 2.648835551 | 0.66416768 |
| ENSMUSG00000096793 | Gm3002 | 5.54E-05 | 2.063873051 | 0.792441218 |
| ENSMUSG00000097127 | Gm26886 | 5.54E-05 | 1.326907457 | 0.950123751 |
| ENSMUSG00000096003 | Gm3500 | 5.54E-05 | 3.648835551 | 0.466464457 |
| ENSMUSG00000091568 | Gm8206 | 5.54E-05 | 1.648835551 | 0.877701541 |
| ENSMUSG00000097447 | Gm26630 | 5.54E-05 | 2.063873051 | 0.789264184 |
| ENSMUSG00000091814 | Gm8050 | 5.54E-05 | 2.648835551 | 0.66416768 |
| ENSMUSG00000084775 | Gm16741 | 5.54E-05 | 0.063873051 | 1 |
| ENSMUSG00000025277 | Abhd6 | 5.54E-05 | -2.209145444 | 0.093610056 |
| ENSMUSG00000021751 | Acox2 | 5.54E-05 | -0.051604167 | 1 |
| ENSMUSG00000021750 | Fam107a | 5.54E-05 | 0.841480629 | 1 |
| ENSMUSG00000114298 | Gm48371 | 5.54E-05 | 2.648835551 | 0.66416768 |
| ENSMUSG00000089833 | Fhitos | 5.54E-05 | 0.648835551 | 1 |
| ENSMUSG00000085883 | Gm3848 | 5.54E-05 | -1.106051951 | 0.770341817 |
| ENSMUSG00000114230 | Gm48239 | 5.54E-05 | 2.648835551 | 0.66416768 |
| ENSMUSG00000044772 | Sntn | 5.54E-05 | 3.648835551 | 0.466464457 |
| ENSMUSG00000052417 | Olfr720 | 5.54E-05 | 1.063873051 | 1 |
| ENSMUSG00000114390 | 1700062C10Rik | 5.54E-05 | 0.47891055 | 1 |
| ENSMUSG00000114630 | Gm48105 | 5.54E-05 | 2.648835551 | 0.66416768 |
| ENSMUSG00000114214 | Gm41102 | 5.54E-05 | 3.648835551 | 0.466464457 |
| ENSMUSG00000021806 | Nid2 | 5.54E-05 | -0.599091962 | 0.980745941 |
| ENSMUSG00000114436 | Gm48283 | 5.54E-05 | 1.326907457 | 0.971854959 |
| ENSMUSG00000021809 | Nudt13 | 5.54E-05 | -1.560617814 | 0.356350819 |
| ENSMUSG00000039543 | Cfap70 | 5.54E-05 | 0.648835551 | 1 |
| ENSMUSG00000021815 | Mss51 | 5.54E-05 | -2.132524162 | 0.112236795 |
| ENSMUSG00000099966 | 2810402E24Rik | 5.54E-05 | -1.9057533 | 0.203406755 |
| ENSMUSG00000097577 | 6230400D17Rik | 5.54E-05 | -1.258055044 | 0.616879894 |
| ENSMUSG00000039308 | Ndst2 | 5.54E-05 | -2.132524162 | 0.112236795 |
| ENSMUSG00000114352 | 4931407E12Rik | 5.54E-05 | 1.648835551 | 0.886203497 |
| ENSMUSG00000114205 | Gm7480 | 5.54E-05 | 3.648835551 | 0.466464457 |
| ENSMUSG00000114722 | Gm31392 | 5.54E-05 | 0.326907457 | 1 |
| ENSMUSG00000114575 | Gm47814 | 5.54E-05 | 0.841480629 | 1 |
| ENSMUSG00000025280 | Polr3a | 5.54E-05 | -3.52108945 | 0.000717333 |
| ENSMUSG00000089887 | 4930428N03Rik | 5.54E-05 | 0.326907457 | 1 |
| ENSMUSG00000114709 | Gm47920 | 5.54E-05 | -0.43862729 | 1 |
| ENSMUSG00000021868 | Ppif | 5.54E-05 | -0.158519371 | 1 |
| ENSMUSG00000090861 | Arf4os | 5.54E-05 | 0.841480629 | 1 |
| ENSMUSG00000021898 | Asb14 | 5.54E-05 | 1.648835551 | 0.882296038 |
| ENSMUSG00000040726 | Hesx1 | 5.54E-05 | 1.648835551 | 0.886203497 |
| ENSMUSG00000109849 | Gm45645 | 5.54E-05 | 1.648835551 | 0.886203497 |
| ENSMUSG00000114760 | Gm48006 | 5.54E-05 | 2.648835551 | 0.66416768 |
| ENSMUSG00000021902 | Phf7 | 5.54E-05 | -1.305360759 | 0.491653643 |
| ENSMUSG00000021901 | Bap1 | 5.54E-05 | -1.351164449 | 0.499088689 |
| ENSMUSG00000115120 | 4930425P05Rik | 5.54E-05 | 2.063873051 | 0.792441218 |
| ENSMUSG00000115691 | Gm48914 | 5.54E-05 | 1.326907457 | 0.949873008 |

| | | | | |
|---|---|---|---|---|
| ENSMUSG00000072473 | 1700024G13Rik | 5.54E-05 | 2.063873051 | 0.791131141 |
| ENSMUSG00000021919 | Chat | 5.54E-05 | 0.47891055 | 1 |
| ENSMUSG00000041730 | Prrxl1 | 5.54E-05 | 0.47891055 | 1 |
| ENSMUSG00000021940 | Ptpn20 | 5.54E-05 | 1.648835551 | 0.886203497 |
| ENSMUSG00000021943 | Gdf10 | 5.54E-05 | 0.841480629 | 1 |
| ENSMUSG00000041534 | Rbp3 | 5.54E-05 | 2.648835551 | 0.66416768 |
| ENSMUSG00000096405 | 4930474N05Rik | 5.54E-05 | 2.648835551 | 0.66416768 |
| ENSMUSG00000021803 | Cdhr1 | 5.54E-05 | 2.648835551 | 0.66416768 |
| ENSMUSG00000096001 | 2610528A11Rik | 5.54E-05 | 1.326907457 | 0.971854959 |
| ENSMUSG00000093528 | Nrg3os | 5.54E-05 | 0.841480629 | 1 |
| ENSMUSG00000114196 | Gm47547 | 5.54E-05 | -0.52108945 | 0.990195911 |
| ENSMUSG00000021790 | Dydc1 | 5.54E-05 | 1.326907457 | 0.949424147 |
| ENSMUSG00000052334 | 1700024B05Rik | 5.54E-05 | 0.841480629 | 1 |
| ENSMUSG00000095384 | 1700001F09Rik | 5.54E-05 | 3.648835551 | 0.466464457 |
| ENSMUSG00000053961 | Ang5 | 5.54E-05 | 1.326907457 | 0.952096994 |
| ENSMUSG00000072596 | Ear2 | 5.54E-05 | 0.47891055 | 1 |
| ENSMUSG00000090709 | Gm17173 | 5.54E-05 | 0.326907457 | 1 |
| ENSMUSG00000037759 | Ptger2 | 5.54E-05 | 1.326907457 | 0.971854959 |
| ENSMUSG00000021830 | Txndc16 | 5.54E-05 | -0.351164449 | 1 |
| ENSMUSG00000115373 | Gm49194 | 5.54E-05 | 3.648835551 | 0.466464457 |
| ENSMUSG00000037712 | Fermt2 | 5.54E-05 | -2.540989007 | 0.029897056 |
| ENSMUSG00000021835 | Bmp4 | 5.54E-05 | -2.794107944 | 0.013337086 |
| ENSMUSG00000115168 | Gm48910 | 5.54E-05 | 3.648835551 | 0.466464457 |
| ENSMUSG00000085380 | E130120K24Rik | 5.54E-05 | 2.063873051 | 0.792441218 |
| ENSMUSG00000085123 | Rubie | 5.54E-05 | 0.841480629 | 1 |
| ENSMUSG00000115665 | 4933425B07Rik | 5.54E-05 | 1.326907457 | 0.947791455 |
| ENSMUSG00000115015 | D330046F09Rik | 5.54E-05 | 0.841480629 | 1 |
| ENSMUSG00000115417 | Gm48949 | 5.54E-05 | 1.648835551 | 0.886203497 |
| ENSMUSG00000037544 | Dlgap5 | 5.54E-05 | -1.743481871 | 0.305378889 |
| ENSMUSG00000115617 | Gm48975 | 5.54E-05 | 2.648835551 | 0.66416768 |
| ENSMUSG00000115609 | 4930447J18Rik | 5.54E-05 | 1.326907457 | 0.951890397 |
| ENSMUSG00000021848 | Otx2 | 5.54E-05 | 0.47891055 | 1 |
| ENSMUSG00000115398 | Gm534 | 5.54E-05 | 1.648835551 | 0.865542683 |
| ENSMUSG00000036282 | Naa30 | 5.54E-05 | -2.158519371 | 0.115633409 |
| ENSMUSG00000021850 | ccdc198 | 5.54E-05 | 0.47891055 | 1 |
| ENSMUSG00000115787 | Gm36146 | 5.54E-05 | 1.648835551 | 0.858831093 |
| ENSMUSG00000091873 | Olfr732 | 5.54E-05 | 2.063873051 | 0.792441218 |
| ENSMUSG00000045306 | Olfr734 | 5.54E-05 | 3.648835551 | 0.466464457 |
| ENSMUSG00000050028 | Olfr745 | 5.54E-05 | 2.648835551 | 0.66416768 |
| ENSMUSG00000060084 | Olfr748 | 5.54E-05 | 3.648835551 | 0.466464457 |
| ENSMUSG00000059069 | Olfr749 | 5.54E-05 | 1.648835551 | 0.886203497 |
| ENSMUSG00000097431 | Gm26782 | 5.54E-05 | -0.936126949 | 0.695372548 |
| ENSMUSG00000060615 | Ang4 | 5.54E-05 | 0.841480629 | 1 |
| ENSMUSG00000062148 | Ear6 | 5.54E-05 | 2.063873051 | 0.792441218 |
| ENSMUSG00000097533 | Gm26590 | 5.54E-05 | 2.063873051 | 0.792441218 |
| ENSMUSG00000095030 | Olfr1513 | 5.54E-05 | 2.648835551 | 0.66416768 |
| ENSMUSG00000094140 | Olfr1512 | 5.54E-05 | 2.063873051 | 0.792441218 |
| ENSMUSG00000063867 | Olfr1511 | 5.54E-05 | 1.648835551 | 0.875072755 |
| ENSMUSG00000106259 | Trav11n | 5.54E-05 | 3.648835551 | 0.466464457 |
| ENSMUSG00000095607 | Trav5-4 | 5.54E-05 | 3.648835551 | 0.466464457 |
| ENSMUSG00000076867 | Trdv4 | 5.54E-05 | 2.063873051 | 0.792441218 |

| | | | | |
|---|---|---|---|---|
| ENSMUSG00000022176 | Rem2 | 5.54E-05 | 0.189403933 | 1 |
| ENSMUSG00000072423 | Psmb11 | 5.54E-05 | 0.47891055 | 1 |
| ENSMUSG00000022180 | Slc7a8 | 5.54E-05 | -1.52108945 | 0.466464457 |
| ENSMUSG00000040759 | Cmtm5 | 5.54E-05 | 2.063873051 | 0.792441218 |
| ENSMUSG00000115234 | Gm49130 | 5.54E-05 | 2.648835551 | 0.66416768 |
| ENSMUSG00000040752 | Myh6 | 5.54E-05 | 2.648835551 | 0.66416768 |
| ENSMUSG00000093452 | Zfhx2os | 5.54E-05 | 2.648835551 | 0.66416768 |
| ENSMUSG00000022211 | Carmil3 | 5.54E-05 | 1.063873051 | 1 |
| ENSMUSG00000115778 | Gm10876 | 5.54E-05 | 3.648835551 | 0.466464457 |
| ENSMUSG00000047098 | Rnf31 | 5.54E-05 | -2.52108945 | 0.037394962 |
| ENSMUSG00000002324 | Rec8 | 5.54E-05 | 1.326907457 | 0.971854959 |
| ENSMUSG00000007591 | Tssk4 | 5.54E-05 | 1.326907457 | 0.971854959 |
| ENSMUSG00000115154 | Gm49067 | 5.54E-05 | -0.351164449 | 1 |
| ENSMUSG00000022218 | Tgm1 | 5.54E-05 | 1.063873051 | 1 |
| ENSMUSG00000019297 | Nop9 | 5.54E-05 | -2.023589791 | 0.130342151 |
| ENSMUSG00000046908 | Ltb4r1 | 5.54E-05 | 0.063873051 | 1 |
| ENSMUSG00000022221 | Ripk3 | 5.54E-05 | 0.47891055 | 1 |
| ENSMUSG00000068289 | Cma2 | 5.54E-05 | 2.648835551 | 0.66416768 |
| ENSMUSG00000022156 | Gzme | 5.54E-05 | 3.648835551 | 0.466464457 |
| ENSMUSG00000114303 | Gm4022 | 5.54E-05 | 0.841480629 | 1 |
| ENSMUSG00000022229 | Atp12a | 5.54E-05 | 3.648835551 | 0.466464457 |
| ENSMUSG00000064128 | Cenpj | 5.54E-05 | -1.636566667 | 0.373232174 |
| ENSMUSG00000021974 | Fgf9 | 5.54E-05 | -0.258055044 | 1 |
| ENSMUSG00000091649 | Phf11b | 5.54E-05 | 2.648835551 | 0.66416768 |
| ENSMUSG00000086112 | 4930563I02Rik | 5.54E-05 | 2.063873051 | 0.792441218 |
| ENSMUSG00000021986 | Amer2 | 5.54E-05 | 1.063873051 | 1 |
| ENSMUSG00000097997 | Gm27017 | 5.54E-05 | 0.648835551 | 1 |
| ENSMUSG00000048281 | Dleu7 | 5.54E-05 | 2.648835551 | 0.66416768 |
| ENSMUSG00000100486 | Gm4131 | 5.54E-05 | -0.351164449 | 1 |
| ENSMUSG00000091155 | Serpine3 | 5.54E-05 | 2.063873051 | 0.789022126 |
| ENSMUSG00000075572 | Defb43 | 5.54E-05 | 3.648835551 | 0.466464457 |
| ENSMUSG00000054763 | Defb42 | 5.54E-05 | 1.063873051 | 1 |
| ENSMUSG00000014453 | Blk | 5.54E-05 | 0.326907457 | 1 |
| ENSMUSG00000034623 | Prss55 | 5.54E-05 | 3.648835551 | 0.466464457 |
| ENSMUSG00000114693 | Gm41177 | 5.54E-05 | 2.648835551 | 0.66416768 |
| ENSMUSG00000034532 | Fbxo16 | 5.54E-05 | -0.673092543 | 1 |
| ENSMUSG00000114744 | Gm38409 | 5.54E-05 | 2.063873051 | 0.792441218 |
| ENSMUSG00000003469 | Phyhip | 5.54E-05 | 3.648835551 | 0.466464457 |
| ENSMUSG00000022096 | Hr | 5.54E-05 | 0.189403933 | 1 |
| ENSMUSG00000022102 | Dok2 | 5.54E-05 | -1.43862729 | 0.433313155 |
| ENSMUSG00000090216 | Gm16549 | 5.54E-05 | 2.648835551 | 0.66416768 |
| ENSMUSG00000033470 | Cysltr2 | 5.54E-05 | 2.063873051 | 0.792441218 |
| ENSMUSG00000022106 | Rcbtb2 | 5.54E-05 | -2.777429203 | 0.016866436 |
| ENSMUSG00000033446 | Lpar6 | 5.54E-05 | -1.673092543 | 0.290284345 |
| ENSMUSG00000022109 | Med4 | 5.54E-05 | -2.233807498 | 0.09684834 |
| ENSMUSG00000033405 | Nudt15 | 5.54E-05 | 1.648835551 | 0.886203497 |
| ENSMUSG00000034997 | Htr2a | 5.54E-05 | -3.589569188 | 0.000600167 |
| ENSMUSG00000022002 | Erich6b | 5.54E-05 | 0.648835551 | 1 |
| ENSMUSG00000067973 | Gm6994 | 5.54E-05 | 2.063873051 | 0.792441218 |
| ENSMUSG00000092272 | Gm19301 | 5.54E-05 | 2.063873051 | 0.773808512 |
| ENSMUSG00000071262 | Zfp957 | 5.54E-05 | 0.063873051 | 1 |

| | | | | |
|---|---|---|---|---|
| ENSMUSG00000022025 | Cnmd | 5.54E-05 | 1.648835551 | 0.886203497 |
| ENSMUSG00000115368 | Gm48942 | 5.54E-05 | 2.648835551 | 0.66416768 |
| ENSMUSG00000115229 | Gm33525 | 5.54E-05 | 1.648835551 | 0.864133138 |
| ENSMUSG00000115242 | Gm49018 | 5.54E-05 | 2.063873051 | 0.786830029 |
| ENSMUSG00000053821 | Gm9922 | 5.54E-05 | 3.648835551 | 0.466464457 |
| ENSMUSG00000114980 | 4933432I03Rik | 5.54E-05 | 0.189403933 | 1 |
| ENSMUSG00000115855 | Gm34643 | 5.54E-05 | 2.063873051 | 0.791887357 |
| ENSMUSG00000089707 | Slain1os | 5.54E-05 | 2.648835551 | 0.66416768 |
| ENSMUSG00000022122 | Ednrb | 5.54E-05 | 0.326907457 | 1 |
| ENSMUSG00000115727 | 4930432J09Rik | 5.54E-05 | 0.326907457 | 1 |
| ENSMUSG00000048349 | Pou4f1 | 5.54E-05 | 2.063873051 | 0.792441218 |
| ENSMUSG00000115386 | Gm35909 | 5.54E-05 | 2.063873051 | 0.792441218 |
| ENSMUSG00000022113 | Trim52 | 5.54E-05 | 2.648835551 | 0.66416768 |
| ENSMUSG00000099794 | 1700128A07Rik | 5.54E-05 | 2.063873051 | 0.790501196 |
| ENSMUSG00000115562 | Gm48977 | 5.54E-05 | 1.326907457 | 0.971854959 |
| ENSMUSG00000093497 | Gm20713 | 5.54E-05 | -0.051604167 | 1 |
| ENSMUSG00000099911 | 1700044C05Rik | 5.54E-05 | 3.648835551 | 0.466464457 |
| ENSMUSG00000042156 | Dzip1 | 5.54E-05 | -0.051604167 | 1 |
| ENSMUSG00000115718 | Gm49179 | 5.54E-05 | 2.648835551 | 0.66416768 |
| ENSMUSG00000087479 | Gm16835 | 5.54E-05 | 1.648835551 | 0.886203497 |
| ENSMUSG00000115121 | Gm49236 | 5.54E-05 | -0.258055044 | 1 |
| ENSMUSG00000100585 | 1700108J01Rik | 5.54E-05 | 1.648835551 | 0.886203497 |
| ENSMUSG00000041703 | Zic5 | 5.54E-05 | -0.158519371 | 1 |
| ENSMUSG00000085555 | 2610035F20Rik | 5.54E-05 | 2.063873051 | 0.792441218 |
| ENSMUSG00000097662 | Gm2093 | 5.54E-05 | 3.648835551 | 0.466464457 |
| ENSMUSG00000079105 | C7 | 5.54E-05 | -0.051604167 | 1 |
| ENSMUSG00000089647 | Gm2245 | 5.54E-05 | -1.351164449 | 0.499088689 |
| ENSMUSG00000072664 | Ugt3a1 | 5.54E-05 | 1.063873051 | 1 |
| ENSMUSG00000049152 | Ugt3a2 | 5.54E-05 | 2.648835551 | 0.66416768 |
| ENSMUSG00000022248 | Rad1 | 5.54E-05 | -1.051604167 | 0.708749425 |
| ENSMUSG00000022243 | Slc45a2 | 5.54E-05 | 2.648835551 | 0.66416768 |
| ENSMUSG00000115441 | Gm34759 | 5.54E-05 | 1.648835551 | 0.866736772 |
| ENSMUSG00000115266 | 4930557F08Rik | 5.54E-05 | 1.326907457 | 0.971854959 |
| ENSMUSG00000115492 | Gm41277 | 5.54E-05 | 1.326907457 | 0.971854959 |
| ENSMUSG00000084873 | C030047K22Rik | 5.54E-05 | 0.841480629 | 1 |
| ENSMUSG00000115803 | Gm35496 | 5.54E-05 | 1.648835551 | 0.886203497 |
| ENSMUSG00000115355 | 4930445E18Rik | 5.54E-05 | -0.158519371 | 1 |
| ENSMUSG00000115729 | Gm48957 | 5.54E-05 | 1.326907457 | 0.971854959 |
| ENSMUSG00000115433 | Gm49267 | 5.54E-05 | 0.063873051 | 1 |
| ENSMUSG00000115337 | 4930570B17Rik | 5.54E-05 | 1.326907457 | 0.971854959 |
| ENSMUSG00000115317 | Gm32618 | 5.54E-05 | -1.305360759 | 0.491653643 |
| ENSMUSG00000115140 | Gm32764 | 5.54E-05 | 0.47891055 | 1 |
| ENSMUSG00000115779 | 4930592A05Rik | 5.54E-05 | 1.063873051 | 1 |
| ENSMUSG00000115428 | Gm41290 | 5.54E-05 | 3.648835551 | 0.466464457 |
| ENSMUSG00000094447 | 9430069I07Rik | 5.54E-05 | 0.648835551 | 1 |
| ENSMUSG00000115715 | Gm33936 | 5.54E-05 | 2.648835551 | 0.66416768 |
| ENSMUSG00000114981 | Gm34093 | 5.54E-05 | 1.648835551 | 0.886203497 |
| ENSMUSG00000115395 | Gm49224 | 5.54E-05 | 3.648835551 | 0.466464457 |
| ENSMUSG00000115333 | Gm34590 | 5.54E-05 | 0.326907457 | 1 |
| ENSMUSG00000086125 | Gm15942 | 5.54E-05 | 1.648835551 | 0.886203497 |
| ENSMUSG00000115326 | Gm35019 | 5.54E-05 | -0.673092543 | 0.98880775 |

| | | | | |
|---|---|---|---|---|
| ENSMUSG00000115307 | Gm49313 | 5.54E-05 | 1.326907457 | 0.971854959 |
| ENSMUSG00000022296 | Baalc | 5.54E-05 | 2.063873051 | 0.792441218 |
| ENSMUSG00000115820 | Gm41311 | 5.54E-05 | 0.648835551 | 1 |
| ENSMUSG00000042895 | Abra | 5.54E-05 | 3.648835551 | 0.466464457 |
| ENSMUSG00000115986 | Gm49439 | 5.54E-05 | 1.326907457 | 0.971854959 |
| ENSMUSG00000099751 | 1700022A22Rik | 5.54E-05 | 0.326907457 | 1 |
| ENSMUSG00000051198 | 4930548G14Rik | 5.54E-05 | 0.841480629 | 1 |
| ENSMUSG00000063727 | Tnfrsf11b | 5.54E-05 | 0.063873051 | 1 |
| ENSMUSG00000037343 | Taf2 | 5.54E-05 | -3.270027686 | 0.002319335 |
| ENSMUSG00000115657 | Gm41325 | 5.54E-05 | 1.648835551 | 0.886203497 |
| ENSMUSG00000022361 | Zhx1 | 5.54E-05 | -2.158519371 | 0.115633409 |
| ENSMUSG00000093677 | Gm20712 | 5.54E-05 | 3.648835551 | 0.466464457 |
| ENSMUSG00000115062 | 4930544F09Rik | 5.54E-05 | 2.063873051 | 0.756795954 |
| ENSMUSG00000115118 | Gm36617 | 5.54E-05 | 3.648835551 | 0.466464457 |
| ENSMUSG00000115504 | Gm48946 | 5.54E-05 | 3.648835551 | 0.466464457 |
| ENSMUSG00000115231 | 4930402D18Rik | 5.54E-05 | 0.648835551 | 1 |
| ENSMUSG00000079025 | Gsdmc | 5.54E-05 | 0.648835551 | 1 |
| ENSMUSG00000056293 | Gsdmc2 | 5.54E-05 | 2.648835551 | 0.652166789 |
| ENSMUSG00000055748 | Gsdmc4 | 5.54E-05 | 2.063873051 | 0.792441218 |
| ENSMUSG00000115328 | Gm30691 | 5.54E-05 | 0.648835551 | 1 |
| ENSMUSG00000015001 | Oc90 | 5.54E-05 | 2.648835551 | 0.66416768 |
| ENSMUSG00000053469 | Tg | 5.54E-05 | -0.051604167 | 1 |
| ENSMUSG00000036760 | Kcnk9 | 5.54E-05 | 1.648835551 | 0.886203497 |
| ENSMUSG00000106847 | Peg13 | 5.54E-05 | -0.351164449 | 1 |
| ENSMUSG00000079020 | Slc45a4 | 5.54E-05 | -2.328444372 | 0.062688794 |
| ENSMUSG00000072487 | Mroh5 | 5.54E-05 | -0.051604167 | 1 |
| ENSMUSG00000034730 | Adgrb1 | 5.54E-05 | -0.874726405 | 0.885891259 |
| ENSMUSG00000022595 | Lypd2 | 5.54E-05 | 3.648835551 | 0.466464457 |
| ENSMUSG00000091709 | Gm17189 | 5.54E-05 | 0.841480629 | 0.971854959 |
| ENSMUSG00000116380 | Gm39556 | 5.54E-05 | -2.373532262 | 0.051203786 |
| ENSMUSG00000100287 | Gm28068 | 5.54E-05 | -0.43862729 | 1 |
| ENSMUSG00000022577 | Ly6h | 5.54E-05 | 2.648835551 | 0.66416768 |
| ENSMUSG00000022579 | Gpihbp1 | 5.54E-05 | 0.841480629 | 1 |
| ENSMUSG00000085523 | Gm15945 | 5.54E-05 | 3.648835551 | 0.466464457 |
| ENSMUSG00000022574 | Naprt | 5.54E-05 | 0.326907457 | 1 |
| ENSMUSG00000103906 | Tigd5 | 5.54E-05 | 1.063873051 | 1 |
| ENSMUSG00000098176 | Ccdc166 | 5.54E-05 | 2.063873051 | 0.792441218 |
| ENSMUSG00000115946 | Mirt2 | 5.54E-05 | 2.648835551 | 0.66416768 |
| ENSMUSG00000022552 | Sharpin | 5.54E-05 | -0.351164449 | 1 |
| ENSMUSG00000022557 | Bop1 | 5.54E-05 | -1.560617814 | 0.624094312 |
| ENSMUSG00000022555 | Dgat1 | 5.54E-05 | -1.43862729 | 0.433313155 |
| ENSMUSG00000022560 | Slc52a2 | 5.54E-05 | 0.648835551 | 1 |
| ENSMUSG00000059323 | Tonsl | 5.54E-05 | -1.395558568 | 0.589666002 |
| ENSMUSG00000116138 | C030006K11Rik | 5.54E-05 | 0.841480629 | 1 |
| ENSMUSG00000116024 | Gm49527 | 5.54E-05 | -1.560617814 | 0.356350819 |
| ENSMUSG00000063236 | 1110038F14Rik | 5.54E-05 | -1.9057533 | 0.203406755 |
| ENSMUSG00000071716 | Apol7e | 5.54E-05 | 2.063873051 | 0.792441218 |
| ENSMUSG00000092375 | A730060N03Rik | 5.54E-05 | 0.648835551 | 1 |
| ENSMUSG00000071715 | Ncf4 | 5.54E-05 | 0.063873051 | 1 |
| ENSMUSG00000115854 | Gm36644 | 5.54E-05 | 0.47891055 | 1 |
| ENSMUSG00000068227 | Il2rb | 5.54E-05 | 2.648835551 | 0.66416768 |

| | | | | |
|---|---|---|---|---|
| ENSMUSG00000033220 | Rac2 | 5.54E-05 | 0.326907457 | 1 |
| ENSMUSG00000022436 | Sh3bp1 | 5.54E-05 | -0.351164449 | 1 |
| ENSMUSG00000084864 | 1700027A07Rik | 5.54E-05 | 3.648835551 | 0.466464457 |
| ENSMUSG00000116165 | Pdxp | 5.54E-05 | 3.648835551 | 0.466464457 |
| ENSMUSG00000033029 | 1700088E04Rik | 5.54E-05 | 0.189403933 | 1 |
| ENSMUSG00000033006 | Sox10 | 5.54E-05 | 2.648835551 | 0.66416768 |
| ENSMUSG00000022420 | Dnal4 | 5.54E-05 | 0.47891055 | 1 |
| ENSMUSG00000115902 | AC113595.1 | 5.54E-05 | 2.648835551 | 0.66416768 |
| ENSMUSG00000050164 | Mchr1 | 5.54E-05 | 3.648835551 | 0.466464457 |
| ENSMUSG00000061360 | Phf5a | 5.54E-05 | -1.52108945 | 0.452184898 |
| ENSMUSG00000075524 | 4930407I10Rik | 5.54E-05 | 2.648835551 | 0.66416768 |
| ENSMUSG00000068117 | Mei1 | 5.54E-05 | 0.063873051 | 1 |
| ENSMUSG00000068101 | Cenpm | 5.54E-05 | 0.189403933 | 1 |
| ENSMUSG00000022455 | Wbp2nl | 5.54E-05 | 2.063873051 | 0.792441218 |
| ENSMUSG00000090964 | Gm17206 | 5.54E-05 | 2.648835551 | 0.66416768 |
| ENSMUSG00000116508 | Gm49463 | 5.54E-05 | 2.063873051 | 0.792441218 |
| ENSMUSG00000115924 | Gm19277 | 5.54E-05 | 3.648835551 | 0.466464457 |
| ENSMUSG00000096607 | 7530416G11Rik | 5.54E-05 | 2.063873051 | 0.792441218 |
| ENSMUSG00000097209 | AU022754 | 5.54E-05 | 0.648835551 | 1 |
| ENSMUSG00000064284 | Cdpf1 | 5.54E-05 | 0.648835551 | 1 |
| ENSMUSG00000035944 | Ttc38 | 5.54E-05 | -0.258055044 | 1 |
| ENSMUSG00000115880 | 4930445N06Rik | 5.54E-05 | 0.841480629 | 1 |
| ENSMUSG00000093394 | Gm20621 | 5.54E-05 | 2.648835551 | 0.66416768 |
| ENSMUSG00000035805 | Mlc1 | 5.54E-05 | 1.648835551 | 0.868550295 |
| ENSMUSG00000087331 | 1810021B22Rik | 5.54E-05 | 1.326907457 | 0.971854959 |
| ENSMUSG00000047394 | Odf3b | 5.54E-05 | 3.648835551 | 0.466464457 |
| ENSMUSG00000022617 | Chkb | 5.54E-05 | -1.258055044 | 0.651354034 |
| ENSMUSG00000084915 | C230037L18Rik | 5.54E-05 | -0.351164449 | 1 |
| ENSMUSG00000022621 | Rabl2 | 5.54E-05 | -0.743481871 | 1 |
| ENSMUSG00000075470 | Alg10b | 5.54E-05 | -1.209145444 | 0.606122355 |
| ENSMUSG00000078932 | CN725425 | 5.54E-05 | 1.326907457 | 0.948861429 |
| ENSMUSG00000097068 | Gm26760 | 5.54E-05 | 1.063873051 | 1 |
| ENSMUSG00000115858 | 4930588J15Rik | 5.54E-05 | 2.648835551 | 0.66416768 |
| ENSMUSG00000116190 | Gm41386 | 5.54E-05 | 3.648835551 | 0.466464457 |
| ENSMUSG00000116493 | Gm49445 | 5.54E-05 | 1.648835551 | 0.886203497 |
| ENSMUSG00000115965 | Gm30564 | 5.54E-05 | 2.648835551 | 0.66416768 |
| ENSMUSG00000033356 | Pus7l | 5.54E-05 | -0.52108945 | 0.993621492 |
| ENSMUSG00000115049 | Gm49172 | 5.54E-05 | 2.063873051 | 0.792441218 |
| ENSMUSG00000033065 | Pfkm | 5.54E-05 | -1.158519371 | 0.599135879 |
| ENSMUSG00000075427 | Olfr288 | 5.54E-05 | 1.063873051 | 0.99523757 |
| ENSMUSG00000063780 | Olfr282 | 5.54E-05 | 0.648835551 | 1 |
| ENSMUSG00000054855 | Rnd1 | 5.54E-05 | -0.810596067 | 0.861097337 |
| ENSMUSG00000003354 | Ccdc65 | 5.54E-05 | 0.189403933 | 1 |
| ENSMUSG00000022997 | Wnt1 | 5.54E-05 | 2.063873051 | 0.792441218 |
| ENSMUSG00000023755 | Rhebl1 | 5.54E-05 | -1.158519371 | 0.599135879 |
| ENSMUSG00000116223 | Gm41396 | 5.54E-05 | 1.648835551 | 0.886203497 |
| ENSMUSG00000032783 | Troap | 5.54E-05 | -0.743481871 | 0.827142217 |
| ENSMUSG00000037579 | Kcnh3 | 5.54E-05 | 0.326907457 | 1 |
| ENSMUSG00000100599 | 1700120C14Rik | 5.54E-05 | -0.158519371 | 1 |
| ENSMUSG00000023011 | Faim2 | 5.54E-05 | 0.648835551 | 1 |
| ENSMUSG00000044217 | Aqp5 | 5.54E-05 | -0.599091962 | 1 |

| | | | | |
|---|---|---|---|---|
| ENSMUSG00000023017 | Asic1 | 5.54E-05 | 2.648835551 | 0.66416768 |
| ENSMUSG00000044636 | Csrnp2 | 5.54E-05 | -0.673092543 | 1 |
| ENSMUSG00000095440 | Fignl2 | 5.54E-05 | -0.599091962 | 0.994229741 |
| ENSMUSG00000037204 | Atg101 | 5.54E-05 | -1.599091962 | 0.363644968 |
| ENSMUSG00000116116 | Gm671 | 5.54E-05 | -1.874726405 | 0.198601369 |
| ENSMUSG00000063971 | Krt88 | 5.54E-05 | 1.326907457 | 0.971854959 |
| ENSMUSG00000044294 | Krt84 | 5.54E-05 | 2.063873051 | 0.792441218 |
| ENSMUSG00000051879 | Krt71 | 5.54E-05 | 2.648835551 | 0.66416768 |
| ENSMUSG00000063661 | Krt73 | 5.54E-05 | 3.648835551 | 0.466464457 |
| ENSMUSG00000059668 | Krt4 | 5.54E-05 | 2.063873051 | 0.792441218 |
| ENSMUSG00000023045 | Soat2 | 5.54E-05 | -0.599091962 | 0.980745941 |
| ENSMUSG00000023047 | Amhr2 | 5.54E-05 | 1.648835551 | 0.886203497 |
| ENSMUSG00000023052 | Npff | 5.54E-05 | -0.810596067 | 0.832285197 |
| ENSMUSG00000097003 | D930007P13Rik | 5.54E-05 | -0.258055044 | 1 |
| ENSMUSG00000022487 | Gtsf1 | 5.54E-05 | 2.063873051 | 0.792441218 |
| ENSMUSG00000053508 | Gtsf2 | 5.54E-05 | 2.648835551 | 0.66416768 |
| ENSMUSG00000022488 | Nckap1l | 5.54E-05 | -0.874726405 | 0.840804483 |
| ENSMUSG00000022490 | Ppp1r1a | 5.54E-05 | 3.648835551 | 0.466464457 |
| ENSMUSG00000060311 | Mucl1 | 5.54E-05 | 3.648835551 | 0.466464457 |
| ENSMUSG00000059043 | Olfr15 | 5.54E-05 | 1.063873051 | 1 |
| ENSMUSG00000054939 | Zfp174 | 5.54E-05 | 1.326907457 | 0.971854959 |
| ENSMUSG00000049871 | Nlrc3 | 5.54E-05 | 0.189403933 | 1 |
| ENSMUSG00000022518 | 4930562C15Rik | 5.54E-05 | 1.063873051 | 1 |
| ENSMUSG00000022542 | 12-Sep | 5.54E-05 | 1.326907457 | 0.971854959 |
| ENSMUSG00000106967 | Gm42477 | 5.54E-05 | 0.326907457 | 1 |
| ENSMUSG00000022540 | Rogdi | 5.54E-05 | -0.43862729 | 1 |
| ENSMUSG00000091712 | Sec14l5 | 5.54E-05 | -0.599091962 | 1 |
| ENSMUSG00000022544 | Eef2kmt | 5.54E-05 | -1.258055044 | 0.616879894 |
| ENSMUSG00000039345 | Mettl22 | 5.54E-05 | 0.189403933 | 1 |
| ENSMUSG00000116469 | Gm49448 | 5.54E-05 | 2.648835551 | 0.66416768 |
| ENSMUSG00000087384 | Gm15558 | 5.54E-05 | 2.063873051 | 0.792441218 |
| ENSMUSG00000095369 | Gm21859 | 5.54E-05 | 0.189403933 | 1 |
| ENSMUSG00000115936 | 4833415N18Rik | 5.54E-05 | 2.063873051 | 0.790550166 |
| ENSMUSG00000068617 | Efcab1 | 5.54E-05 | 0.47891055 | 1 |
| ENSMUSG00000022671 | Mzt2 | 5.54E-05 | -0.52108945 | 0.975624465 |
| ENSMUSG00000022792 | Yars2 | 5.54E-05 | 0.841480629 | 1 |
| ENSMUSG00000026181 | Ppm1f | 5.54E-05 | -1.258055044 | 0.616879894 |
| ENSMUSG00000116645 | Gm49584 | 5.54E-05 | 1.063873051 | 1 |
| ENSMUSG00000022768 | Ccdc116 | 5.54E-05 | 0.648835551 | 1 |
| ENSMUSG00000050240 | Hic2 | 5.54E-05 | -2.209145444 | 0.093610056 |
| ENSMUSG00000055692 | Tmem191c | 5.54E-05 | 2.063873051 | 0.792441218 |
| ENSMUSG00000116898 | Gm49785 | 5.54E-05 | 1.326907457 | 0.935056557 |
| ENSMUSG00000090236 | Car15 | 5.54E-05 | 1.648835551 | 0.886203497 |
| ENSMUSG00000043811 | Rtn4r | 5.54E-05 | 1.063873051 | 1 |
| ENSMUSG00000022718 | Dgcr8 | 5.54E-05 | -4.560617814 | 1.03E-06 |
| ENSMUSG00000075704 | Txnrd2 | 5.54E-05 | -1.43862729 | 0.433950174 |
| ENSMUSG00000000884 | Gnb1l | 5.54E-05 | -0.936126949 | 0.695372548 |
| ENSMUSG00000000028 | Cdc45 | 5.54E-05 | -1.810596067 | 0.246776726 |
| ENSMUSG00000106567 | 2010309G21Rik | 5.54E-05 | 1.326907457 | 0.939400132 |
| ENSMUSG00000068535 | Olfr169 | 5.54E-05 | 2.648835551 | 0.66416768 |
| ENSMUSG00000041247 | Lamp3 | 5.54E-05 | 1.326907457 | 0.971854959 |

| | | | | |
|---|---|---|---|---|
| ENSMUSG00000043008 | Klhl6 | 5.54E-05 | -1.106051951 | 0.792441218 |
| ENSMUSG00000022818 | Cyp2ab1 | 5.54E-05 | -1.051604167 | 0.74583986 |
| ENSMUSG00000051146 | Camk2n2 | 5.54E-05 | 1.063873051 | 1 |
| ENSMUSG00000022842 | Ece2 | 5.54E-05 | 1.648835551 | 0.886203497 |
| ENSMUSG00000116850 | Gm49742 | 5.54E-05 | 2.648835551 | 0.66416768 |
| ENSMUSG00000096918 | Gm16863 | 5.54E-05 | 0.063873051 | 1 |
| ENSMUSG00000060459 | Kng2 | 5.54E-05 | -0.051604167 | 1 |
| ENSMUSG00000022875 | Kng1 | 5.54E-05 | 1.326907457 | 0.971854959 |
| ENSMUSG00000068463 | B630019A10Rik | 5.54E-05 | -0.051604167 | 1 |
| ENSMUSG00000045231 | BC106179 | 5.54E-05 | -0.051604167 | 1 |
| ENSMUSG00000047531 | Rtp2 | 5.54E-05 | 2.063873051 | 0.792441218 |
| ENSMUSG00000116173 | Gm32126 | 5.54E-05 | 3.648835551 | 0.466464457 |
| ENSMUSG00000038148 | Cldn16 | 5.54E-05 | 0.326907457 | 1 |
| ENSMUSG00000052276 | Ostn | 5.54E-05 | 0.648835551 | 1 |
| ENSMUSG00000097307 | Gm26834 | 5.54E-05 | 0.841480629 | 1 |
| ENSMUSG00000116617 | Gm49767 | 5.54E-05 | -1.051604167 | 0.771834758 |
| ENSMUSG00000090214 | Gm15657 | 5.54E-05 | 1.326907457 | 0.951890397 |
| ENSMUSG00000022832 | Ropn1 | 5.54E-05 | 0.063873051 | 1 |
| ENSMUSG00000116665 | E130310I04Rik | 5.54E-05 | 1.326907457 | 0.956923047 |
| ENSMUSG00000116961 | Gm49662 | 5.54E-05 | -0.051604167 | 1 |
| ENSMUSG00000022900 | Ildr1 | 5.54E-05 | 0.47891055 | 1 |
| ENSMUSG00000084834 | 4930565N06Rik | 5.54E-05 | 2.063873051 | 0.792441218 |
| ENSMUSG00000022809 | Nr1i2 | 5.54E-05 | 1.063873051 | 1 |
| ENSMUSG00000022793 | B4galt4 | 5.54E-05 | -0.874726405 | 0.840804483 |
| ENSMUSG00000116684 | Gm49735 | 5.54E-05 | 2.648835551 | 0.66416768 |
| ENSMUSG00000036304 | Zdhhc23 | 5.54E-05 | 2.648835551 | 0.66416768 |
| ENSMUSG00000071550 | Cfap44 | 5.54E-05 | -0.599091962 | 1 |
| ENSMUSG00000022668 | Gtpbp8 | 5.54E-05 | -0.351164449 | 1 |
| ENSMUSG00000090176 | Cd200r2 | 5.54E-05 | 2.648835551 | 0.66416768 |
| ENSMUSG00000022661 | Cd200 | 5.54E-05 | -1.258055044 | 0.616879894 |
| ENSMUSG00000033187 | BC016579 | 5.54E-05 | -1.43862729 | 0.433313155 |
| ENSMUSG00000022658 | Tagln3 | 5.54E-05 | 2.063873051 | 0.792441218 |
| ENSMUSG00000033157 | Abhd10 | 5.54E-05 | -0.743481871 | 0.827142217 |
| ENSMUSG00000085781 | Gm15640 | 5.54E-05 | 2.648835551 | 0.66416768 |
| ENSMUSG00000085826 | Gm15638 | 5.54E-05 | 0.648835551 | 1 |
| ENSMUSG00000048087 | Gm4737 | 5.54E-05 | 3.648835551 | 0.466464457 |
| ENSMUSG00000072419 | Dppa2 | 5.54E-05 | -0.743481871 | 1 |
| ENSMUSG00000110573 | Gm5485 | 5.54E-05 | 2.648835551 | 0.66416768 |
| ENSMUSG00000073976 | 1700026J12Rik | 5.54E-05 | 0.063873051 | 1 |
| ENSMUSG00000097694 | G730013B05Rik | 5.54E-05 | 2.063873051 | 0.792441218 |
| ENSMUSG00000116725 | Gm29686 | 5.54E-05 | -0.351164449 | 1 |
| ENSMUSG00000044763 | Trmt10c | 5.54E-05 | -1.43862729 | 0.433313155 |
| ENSMUSG00000022755 | Adgrg7 | 5.54E-05 | 0.47891055 | 1 |
| ENSMUSG00000022751 | Nit2 | 5.54E-05 | -0.874726405 | 0.840804483 |
| ENSMUSG00000097876 | Gm16892 | 5.54E-05 | 3.648835551 | 0.466464457 |
| ENSMUSG00000022753 | Tmem30c | 5.54E-05 | 1.648835551 | 0.886203497 |
| ENSMUSG00000116770 | Gm49576 | 5.54E-05 | 1.326907457 | 0.971854959 |
| ENSMUSG00000116717 | Gm49574 | 5.54E-05 | 2.063873051 | 0.792441218 |
| ENSMUSG00000063137 | Olfr177 | 5.54E-05 | 3.648835551 | 0.466464457 |
| ENSMUSG00000062105 | Olfr183 | 5.54E-05 | 3.648835551 | 0.466464457 |
| ENSMUSG00000064006 | Olfr190 | 5.54E-05 | 2.063873051 | 0.792441218 |

| | | | | |
|---|---|---|---|---|
| ENSMUSG00000060057 | Olfr193 | 5.54E-05 | 3.648835551 | 0.466464457 |
| ENSMUSG00000048810 | Olfr202 | 5.54E-05 | 2.648835551 | 0.66416768 |
| ENSMUSG00000116894 | 4930547E14Rik | 5.54E-05 | 2.063873051 | 0.792441218 |
| ENSMUSG00000116574 | Gm18870 | 5.54E-05 | 0.47891055 | 1 |
| ENSMUSG00000116871 | Gm33912 | 5.54E-05 | 3.648835551 | 0.466464457 |
| ENSMUSG00000045178 | 4933411O13Rik | 5.54E-05 | 1.648835551 | 0.821876923 |
| ENSMUSG00000116575 | Gm49648 | 5.54E-05 | 1.648835551 | 0.886203497 |
| ENSMUSG00000116588 | Gm49655 | 5.54E-05 | 0.189403933 | 1 |
| ENSMUSG00000116610 | Gm49678 | 5.54E-05 | 0.326907457 | 1 |
| ENSMUSG00000087214 | Gm15555 | 5.54E-05 | 2.648835551 | 0.66416768 |
| ENSMUSG00000116968 | Gm49587 | 5.54E-05 | 2.063873051 | 0.792441218 |
| ENSMUSG00000109333 | Gm30790 | 5.54E-05 | 1.063873051 | 1 |
| ENSMUSG00000116786 | 4930570E03Rik | 5.54E-05 | 1.648835551 | 0.864728805 |
| ENSMUSG00000022857 | Tmprss15 | 5.54E-05 | -0.52108945 | 1 |
| ENSMUSG00000116865 | Gm49560 | 5.54E-05 | 0.47891055 | 1 |
| ENSMUSG00000116937 | Gm41480 | 5.54E-05 | 3.648835551 | 0.466464457 |
| ENSMUSG00000116674 | 4930551I23Rik | 5.54E-05 | 2.063873051 | 0.792441218 |
| ENSMUSG00000116810 | Gm32624 | 5.54E-05 | 0.47891055 | 1 |
| ENSMUSG00000093606 | B130034C11Rik | 5.54E-05 | -0.599091962 | 0.999962098 |
| ENSMUSG00000116885 | 4930420G21Rik | 5.54E-05 | 1.063873051 | 1 |
| ENSMUSG00000116616 | Gm49688 | 5.54E-05 | 0.47891055 | 1 |
| ENSMUSG00000116849 | Gm49710 | 5.54E-05 | 1.648835551 | 0.874796537 |
| ENSMUSG00000022978 | Mis18a | 5.54E-05 | -1.52108945 | 0.518511033 |
| ENSMUSG00000116543 | Gm36363 | 5.54E-05 | 3.648835551 | 0.466464457 |
| ENSMUSG00000085398 | 4931406G06Rik | 5.54E-05 | 2.063873051 | 0.791749236 |
| ENSMUSG00000039763 | Dnajc28 | 5.54E-05 | 0.648835551 | 1 |
| ENSMUSG00000039672 | Kcne2 | 5.54E-05 | 1.326907457 | 0.949432525 |
| ENSMUSG00000116693 | Gm29805 | 5.54E-05 | 3.648835551 | 0.466464457 |
| ENSMUSG00000116964 | Gm46555 | 5.54E-05 | 2.648835551 | 0.66416768 |
| ENSMUSG00000022948 | Setd4 | 5.54E-05 | -1.560617814 | 0.356350819 |
| ENSMUSG00000097768 | 2310043M15Rik | 5.54E-05 | 3.648835551 | 0.466464457 |
| ENSMUSG00000022945 | Chaf1b | 5.54E-05 | -2.051604167 | 0.133549019 |
| ENSMUSG00000022941 | Ripply3 | 5.54E-05 | -0.258055044 | 1 |
| ENSMUSG00000022940 | Pigp | 5.54E-05 | -0.936126949 | 0.847390568 |
| ENSMUSG00000116589 | Gm31323 | 5.54E-05 | 1.648835551 | 0.886203497 |
| ENSMUSG00000116641 | Gm41505 | 5.54E-05 | 2.063873051 | 0.792441218 |
| ENSMUSG00000090223 | Pcp4 | 5.54E-05 | 1.063873051 | 1 |
| ENSMUSG00000117000 | Gm32509 | 5.54E-05 | 0.648835551 | 1 |
| ENSMUSG00000117324 | Gm32432 | 5.54E-05 | 3.648835551 | 0.466464457 |
| ENSMUSG00000116813 | 4930579D07Rik | 5.54E-05 | -0.874726405 | 0.858831093 |
| ENSMUSG00000116551 | Gm41517 | 5.54E-05 | 3.648835551 | 0.466464457 |
| ENSMUSG00000101133 | Gm29050 | 5.54E-05 | 1.648835551 | 0.886203497 |
| ENSMUSG00000101959 | Ldhal6b | 5.54E-05 | 1.063873051 | 1 |
| ENSMUSG00000038141 | Tmem181a | 5.54E-05 | -1.636566667 | 0.437556558 |
| ENSMUSG00000095677 | Dynlt1f | 5.54E-05 | 1.326907457 | 0.971854959 |
| ENSMUSG00000041831 | Sytl3 | 5.54E-05 | 1.063873051 | 1 |
| ENSMUSG00000052031 | Tagap1 | 5.54E-05 | -1.209145444 | 0.606122355 |
| ENSMUSG00000116955 | Gm49632 | 5.54E-05 | 3.648835551 | 0.466464457 |
| ENSMUSG00000087478 | 4930506C21Rik | 5.54E-05 | -0.258055044 | 1 |
| ENSMUSG00000073468 | Sft2d1 | 5.54E-05 | -1.158519371 | 0.599135879 |
| ENSMUSG00000023873 | 1700010I14Rik | 5.54E-05 | 1.063873051 | 1 |

| | | | | |
|---|---|---|---|---|
| ENSMUSG00000097049 | 6530411M01Rik | 5.54E-05 | 1.326907457 | 0.971854959 |
| ENSMUSG00000089907 | Gm16169 | 5.54E-05 | 2.648835551 | 0.66416768 |
| ENSMUSG00000100402 | Gm28505 | 5.54E-05 | 1.648835551 | 0.875465725 |
| ENSMUSG00000097350 | 4732491K20Rik | 5.54E-05 | -0.158519371 | 1 |
| ENSMUSG00000062480 | Acat3 | 5.54E-05 | -0.158519371 | 1 |
| ENSMUSG00000023832 | Acat2 | 5.54E-05 | -0.874726405 | 0.873717735 |
| ENSMUSG00000073458 | Smok2a | 5.54E-05 | 1.648835551 | 0.886203497 |
| ENSMUSG00000117185 | Gm34510 | 5.54E-05 | 3.648835551 | 0.466464457 |
| ENSMUSG00000117328 | 4930474M22Rik | 5.54E-05 | 2.648835551 | 0.66416768 |
| ENSMUSG00000116780 | Gm3417 | 5.54E-05 | 1.326907457 | 0.935784062 |
| ENSMUSG00000116953 | 9030025P20Rik | 5.54E-05 | 0.063873051 | 1 |
| ENSMUSG00000079710 | Gm3448 | 5.54E-05 | -0.351164449 | 1 |
| ENSMUSG00000097379 | Gm26873 | 5.54E-05 | 2.063873051 | 0.792441218 |
| ENSMUSG00000092120 | Vmn2r90 | 5.54E-05 | 2.648835551 | 0.66416768 |
| ENSMUSG00000003665 | Has1 | 5.54E-05 | 0.841480629 | 1 |
| ENSMUSG00000045551 | Fpr1 | 5.54E-05 | 2.063873051 | 0.781454058 |
| ENSMUSG00000079700 | Fpr3 | 5.54E-05 | -0.158519371 | 1 |
| ENSMUSG00000091631 | Vmn2r95 | 5.54E-05 | 0.841480629 | 1 |
| ENSMUSG00000091679 | Vmn2r96 | 5.54E-05 | 1.063873051 | 1 |
| ENSMUSG00000096717 | Vmn2r98 | 5.54E-05 | 3.648835551 | 0.466464457 |
| ENSMUSG00000116587 | Gm49753 | 5.54E-05 | 3.648835551 | 0.466464457 |
| ENSMUSG00000091771 | Vmn2r103 | 5.54E-05 | 0.841480629 | 1 |
| ENSMUSG00000090315 | Vmn2r104 | 5.54E-05 | 2.648835551 | 0.66416768 |
| ENSMUSG00000056910 | Vmn2r107 | 5.54E-05 | 1.063873051 | 1 |
| ENSMUSG00000043537 | Vmn1r225 | 5.54E-05 | 1.648835551 | 0.876061538 |
| ENSMUSG00000050102 | Vmn1r235 | 5.54E-05 | 3.648835551 | 0.466464457 |
| ENSMUSG00000116771 | Gm49691 | 5.54E-05 | 2.063873051 | 0.792441218 |
| ENSMUSG00000023882 | Zfp54 | 5.54E-05 | -0.52108945 | 0.975624465 |
| ENSMUSG00000023892 | Zfp51 | 5.54E-05 | 0.47891055 | 1 |
| ENSMUSG00000067928 | Zfp760 | 5.54E-05 | -0.258055044 | 1 |
| ENSMUSG00000071266 | Zfp946 | 5.54E-05 | -0.158519371 | 1 |
| ENSMUSG00000095093 | Vmn2r111 | 5.54E-05 | 1.063873051 | 1 |
| ENSMUSG00000095658 | Vmn2r-ps130 | 5.54E-05 | 1.326907457 | 0.971854959 |
| ENSMUSG00000002617 | Zfp40 | 5.54E-05 | -0.874726405 | 0.840804483 |
| ENSMUSG00000091407 | Vmn2r117 | 5.54E-05 | 2.063873051 | 0.770704476 |
| ENSMUSG00000023903 | Mmp25 | 5.54E-05 | 2.648835551 | 0.66416768 |
| ENSMUSG00000023909 | Paqr4 | 5.54E-05 | 0.648835551 | 1 |
| ENSMUSG00000040680 | Kremen2 | 5.54E-05 | 2.063873051 | 0.792441218 |
| ENSMUSG00000048992 | Prss32 | 5.54E-05 | 0.47891055 | 1 |
| ENSMUSG00000024124 | Prss30 | 5.54E-05 | 1.326907457 | 0.971854959 |
| ENSMUSG00000045027 | Prss22 | 5.54E-05 | -0.52108945 | 0.975624465 |
| ENSMUSG00000050762 | Prss27 | 5.54E-05 | 0.841480629 | 1 |
| ENSMUSG00000024118 | Tedc2 | 5.54E-05 | 0.648835551 | 1 |
| ENSMUSG00000024132 | Eci1 | 5.54E-05 | -1.965874293 | 0.162298752 |
| ENSMUSG00000093565 | Rab26os | 5.54E-05 | 1.326907457 | 0.971854959 |
| ENSMUSG00000007021 | Syngr3 | 5.54E-05 | -0.599091962 | 1 |
| ENSMUSG00000040688 | Tbl3 | 5.54E-05 | -0.673092543 | 0.98880775 |
| ENSMUSG00000024160 | Spsb3 | 5.54E-05 | 0.648835551 | 1 |
| ENSMUSG00000035521 | Gnptg | 5.54E-05 | 1.326907457 | 0.968182137 |
| ENSMUSG00000024112 | Cacna1h | 5.54E-05 | 0.326907457 | 1 |
| ENSMUSG00000024176 | Sox8 | 5.54E-05 | 1.648835551 | 0.886203497 |

| | | | | |
|---|---|---|---|---|
| ENSMUSG00000041199 | Rpusd1 | 5.54E-05 | -0.158519371 | 1 |
| ENSMUSG00000041062 | Mslnl | 5.54E-05 | -0.158519371 | 1 |
| ENSMUSG00000025737 | Wdr24 | 5.54E-05 | -0.051604167 | 1 |
| ENSMUSG00000025735 | Rhbdl1 | 5.54E-05 | 0.648835551 | 1 |
| ENSMUSG00000084862 | Gm16278 | 5.54E-05 | 0.648835551 | 1 |
| ENSMUSG00000097405 | D630044L22Rik | 5.54E-05 | 3.648835551 | 0.466464457 |
| ENSMUSG00000073433 | Arhgdig | 5.54E-05 | -0.051604167 | 1 |
| ENSMUSG00000117596 | Gm50013 | 5.54E-05 | 0.47891055 | 1 |
| ENSMUSG00000091078 | Gm17218 | 5.54E-05 | 3.648835551 | 0.466464457 |
| ENSMUSG00000048731 | Ggnbp1 | 5.54E-05 | -0.52108945 | 0.975624465 |
| ENSMUSG00000073430 | Gm10505 | 5.54E-05 | 2.063873051 | 0.792441218 |
| ENSMUSG00000087387 | Gm15420 | 5.54E-05 | 1.326907457 | 0.971854959 |
| ENSMUSG00000062859 | Tcp11 | 5.54E-05 | 0.189403933 | 1 |
| ENSMUSG00000038677 | Scube3 | 5.54E-05 | 1.648835551 | 0.886203497 |
| ENSMUSG00000117171 | C230013L11Rik | 5.54E-05 | 2.648835551 | 0.66416768 |
| ENSMUSG00000036196 | Slc26a8 | 5.54E-05 | -0.874726405 | 0.840804483 |
| ENSMUSG00000087615 | Pnpla1os | 5.54E-05 | 2.648835551 | 0.66416768 |
| ENSMUSG00000087693 | Gm16191 | 5.54E-05 | 3.648835551 | 0.466464457 |
| ENSMUSG00000085912 | Trp53cor1 | 5.54E-05 | -0.051604167 | 1 |
| ENSMUSG00000024008 | Cpne5 | 5.54E-05 | -0.158519371 | 1 |
| ENSMUSG00000024007 | Ppil1 | 5.54E-05 | -1.52108945 | 0.452403317 |
| ENSMUSG00000117427 | Gm36199 | 5.54E-05 | 0.47891055 | 1 |
| ENSMUSG00000079580 | Tmem217 | 5.54E-05 | 3.648835551 | 0.466464457 |
| ENSMUSG00000117489 | Gm50016 | 5.54E-05 | 2.648835551 | 0.66416768 |
| ENSMUSG00000099474 | 1700097N02Rik | 5.54E-05 | 3.648835551 | 0.466464457 |
| ENSMUSG00000024027 | Glp1r | 5.54E-05 | -1.52108945 | 0.493440512 |
| ENSMUSG00000054134 | Umodl1 | 5.54E-05 | 0.841480629 | 1 |
| ENSMUSG00000024029 | Tff3 | 5.54E-05 | 2.648835551 | 0.66416768 |
| ENSMUSG00000042345 | Ubash3a | 5.54E-05 | 0.063873051 | 1 |
| ENSMUSG00000117899 | Gm50218 | 5.54E-05 | 0.47891055 | 1 |
| ENSMUSG00000024033 | Rsph1 | 5.54E-05 | 0.648835551 | 1 |
| ENSMUSG00000118081 | Gm50138 | 5.54E-05 | 0.841480629 | 1 |
| ENSMUSG00000118071 | AC166172.1 | 5.54E-05 | 0.326907457 | 1 |
| ENSMUSG00000118239 | Gm50107 | 5.54E-05 | 3.648835551 | 0.466464457 |
| ENSMUSG00000024039 | Cbs | 5.54E-05 | -0.158519371 | 1 |
| ENSMUSG00000002076 | Hsf2bp | 5.54E-05 | -0.52108945 | 0.975624465 |
| ENSMUSG00000038146 | Notch3 | 5.54E-05 | -1.158519371 | 0.637223959 |
| ENSMUSG00000062464 | Cyp4f37 | 5.54E-05 | 3.648835551 | 0.466464457 |
| ENSMUSG00000090700 | Cyp4f40 | 5.54E-05 | 3.648835551 | 0.466464457 |
| ENSMUSG00000117443 | Gm50046 | 5.54E-05 | 1.648835551 | 0.886203497 |
| ENSMUSG00000117654 | Gm50050 | 5.54E-05 | -0.52108945 | 1 |
| ENSMUSG00000095325 | Zfp870 | 5.54E-05 | 0.326907457 | 1 |
| ENSMUSG00000024055 | Cyp4f13 | 5.54E-05 | -1.636566667 | 0.373232174 |
| ENSMUSG00000048602 | Morc2b | 5.54E-05 | 2.063873051 | 0.792441218 |
| ENSMUSG00000096169 | Olfr1564 | 5.54E-05 | 2.648835551 | 0.66416768 |
| ENSMUSG00000055240 | Zfp101 | 5.54E-05 | -1.708716453 | 0.296030202 |
| ENSMUSG00000090952 | Gm17251 | 5.54E-05 | 1.326907457 | 0.971854959 |
| ENSMUSG00000042099 | Kank3 | 5.54E-05 | -0.743481871 | 1 |
| ENSMUSG00000117772 | Gm50333 | 5.54E-05 | 1.063873051 | 1 |
| ENSMUSG00000096727 | Psmb9 | 5.54E-05 | 2.063873051 | 0.785799033 |
| ENSMUSG00000024339 | Tap2 | 5.54E-05 | -0.158519371 | 1 |

| | | | | |
|---|---|---|---|---|
| ENSMUSG00000073421 | H2-Ab1 | 5.54E-05 | -0.599091962 | 0.980745941 |
| ENSMUSG00000057246 | BC051142 | 5.54E-05 | 0.189403933 | 1 |
| ENSMUSG00000058435 | Btnl4 | 5.54E-05 | 3.648835551 | 0.466464457 |
| ENSMUSG00000034786 | Gpsm3 | 5.54E-05 | -0.051604167 | 1 |
| ENSMUSG00000015474 | Ppt2 | 5.54E-05 | -0.158519371 | 1 |
| ENSMUSG00000033327 | Tnxb | 5.54E-05 | -1.158519371 | 0.599135879 |
| ENSMUSG00000007033 | Hspa1l | 5.54E-05 | 2.063873051 | 0.792441218 |
| ENSMUSG00000034923 | Ly6g6f | 5.54E-05 | 1.648835551 | 0.876018808 |
| ENSMUSG00000073412 | Lst1 | 5.54E-05 | -1.258055044 | 0.647390088 |
| ENSMUSG00000024401 | Tnf | 5.54E-05 | -0.743481871 | 0.827142217 |
| ENSMUSG00000050410 | Tcf19 | 5.54E-05 | 0.47891055 | 1 |
| ENSMUSG00000001524 | Gtf2h4 | 5.54E-05 | 0.326907457 | 1 |
| ENSMUSG00000092600 | Gm20442 | 5.54E-05 | -0.810596067 | 0.866303883 |
| ENSMUSG00000092171 | 4833427F10Rik | 5.54E-05 | 0.47891055 | 1 |
| ENSMUSG00000073407 | Gm6034 | 5.54E-05 | 2.063873051 | 0.792441218 |
| ENSMUSG00000092365 | BC023719 | 5.54E-05 | 0.648835551 | 1 |
| ENSMUSG00000036315 | Znrd1 | 5.54E-05 | -0.051604167 | 1 |
| ENSMUSG00000058802 | Olfr97 | 5.54E-05 | 3.648835551 | 0.466464457 |
| ENSMUSG00000063660 | Olfr98 | 5.54E-05 | 2.648835551 | 0.66416768 |
| ENSMUSG00000092077 | Olfr101 | 5.54E-05 | 2.063873051 | 0.787383909 |
| ENSMUSG00000091531 | Olfr102 | 5.54E-05 | 3.648835551 | 0.466464457 |
| ENSMUSG00000016283 | H2-M2 | 5.54E-05 | 2.648835551 | 0.66416768 |
| ENSMUSG00000090675 | Olfr111 | 5.54E-05 | 2.063873051 | 0.792441218 |
| ENSMUSG00000062629 | Olfr114 | 5.54E-05 | 3.648835551 | 0.466464457 |
| ENSMUSG00000063994 | Olfr120 | 5.54E-05 | 2.648835551 | 0.66416768 |
| ENSMUSG00000092342 | Esp31 | 5.54E-05 | 2.648835551 | 0.66416768 |
| ENSMUSG00000090747 | Esp8 | 5.54E-05 | 2.063873051 | 0.792441218 |
| ENSMUSG00000023919 | Cenpq | 5.54E-05 | -0.351164449 | 1 |
| ENSMUSG00000023921 | Mmut | 5.54E-05 | -0.52108945 | 1 |
| ENSMUSG00000023918 | Adgrf4 | 5.54E-05 | 0.063873051 | 1 |
| ENSMUSG00000023963 | Cyp39a1 | 5.54E-05 | -2.82689788 | 0.011086379 |
| ENSMUSG00000073394 | Runx2os1 | 5.54E-05 | 1.326907457 | 0.943363727 |
| ENSMUSG00000023935 | Spats1 | 5.54E-05 | 0.648835551 | 1 |
| ENSMUSG00000096847 | Tmem151b | 5.54E-05 | 2.063873051 | 0.792441218 |
| ENSMUSG00000058626 | Capn11 | 5.54E-05 | 0.47891055 | 1 |
| ENSMUSG00000117139 | Gm35692 | 5.54E-05 | 1.326907457 | 0.948861429 |
| ENSMUSG00000117040 | Gm49805 | 5.54E-05 | 3.648835551 | 0.466464457 |
| ENSMUSG00000117287 | Gm36200 | 5.54E-05 | 2.648835551 | 0.66416768 |
| ENSMUSG00000067144 | Slc22a7 | 5.54E-05 | 2.648835551 | 0.66416768 |
| ENSMUSG00000015599 | Ttbk1 | 5.54E-05 | 0.189403933 | 1 |
| ENSMUSG00000036430 | Tbcc | 5.54E-05 | 0.189403933 | 1 |
| ENSMUSG00000085932 | Gm15556 | 5.54E-05 | 2.648835551 | 0.66416768 |
| ENSMUSG00000117334 | A730006G06Rik | 5.54E-05 | 2.063873051 | 0.792441218 |
| ENSMUSG00000042265 | Trem1 | 5.54E-05 | 2.063873051 | 0.771569813 |
| ENSMUSG00000071068 | Treml2 | 5.54E-05 | 1.063873051 | 1 |
| ENSMUSG00000023992 | Trem2 | 5.54E-05 | 3.648835551 | 0.466464457 |
| ENSMUSG00000023994 | Nfya | 5.54E-05 | -1.258055044 | 0.616879894 |
| ENSMUSG00000117019 | Gm49896 | 5.54E-05 | 2.063873051 | 0.792441218 |
| ENSMUSG00000064120 | Mocs1 | 5.54E-05 | -1.158519371 | 0.784626844 |
| ENSMUSG00000023940 | Sgo1 | 5.54E-05 | -0.43862729 | 1 |
| ENSMUSG00000023945 | Slc5a7 | 5.54E-05 | 2.648835551 | 0.66416768 |

| | | | | |
|---|---|---|---|---|
| ENSMUSG00000002831 | Plin4 | 5.54E-05 | 2.063873051 | 0.792441218 |
| ENSMUSG00000090273 | Prr22 | 5.54E-05 | 2.063873051 | 0.792441218 |
| ENSMUSG00000024209 | 1700061G19Rik | 5.54E-05 | 1.063873051 | 1 |
| ENSMUSG00000109739 | Gm17949 | 5.54E-05 | 0.063873051 | 1 |
| ENSMUSG00000091390 | Gm17168 | 5.54E-05 | 3.648835551 | 0.466464457 |
| ENSMUSG00000004730 | Adgre1 | 5.54E-05 | -1.480447466 | 0.466464457 |
| ENSMUSG00000117036 | Gm33890 | 5.54E-05 | 2.063873051 | 0.792441218 |
| ENSMUSG00000024105 | Themis3 | 5.54E-05 | -0.158519371 | 1 |
| ENSMUSG00000117193 | Gm36252 | 5.54E-05 | 2.648835551 | 0.66416768 |
| ENSMUSG00000096988 | A930029G22Rik | 5.54E-05 | 1.648835551 | 0.871476881 |
| ENSMUSG00000073374 | C030034I22Rik | 5.54E-05 | 1.648835551 | 0.886203497 |
| ENSMUSG00000117124 | Gm49895 | 5.54E-05 | 3.648835551 | 0.466464457 |
| ENSMUSG00000024049 | Myom1 | 5.54E-05 | -1.708716453 | 0.398502279 |
| ENSMUSG00000117315 | 1600022D10Rik | 5.54E-05 | 0.841480629 | 1 |
| ENSMUSG00000045761 | Togaram2 | 5.54E-05 | 0.47891055 | 1 |
| ENSMUSG00000117023 | Gm49876 | 5.54E-05 | 1.648835551 | 0.886203497 |
| ENSMUSG00000036368 | Rmdn2 | 5.54E-05 | -1.43862729 | 0.465580427 |
| ENSMUSG00000046196 | Ttc39d | 5.54E-05 | 1.063873051 | 1 |
| ENSMUSG00000033966 | Cdkl4 | 5.54E-05 | -0.351164449 | 1 |
| ENSMUSG00000024247 | Pkdcc | 5.54E-05 | -0.351164449 | 1 |
| ENSMUSG00000045053 | Kcng3 | 5.54E-05 | -1.051604167 | 0.771834758 |
| ENSMUSG00000117446 | Gm35551 | 5.54E-05 | 1.648835551 | 0.858623775 |
| ENSMUSG00000117393 | Gm36279 | 5.54E-05 | 0.47891055 | 1 |
| ENSMUSG00000093460 | Six3os1 | 5.54E-05 | 1.063873051 | 1 |
| ENSMUSG00000099635 | Gm28528 | 5.54E-05 | 3.648835551 | 0.466464457 |
| ENSMUSG00000117457 | Gm50011 | 5.54E-05 | 0.47891055 | 1 |
| ENSMUSG00000117545 | Gm30794 | 5.54E-05 | 2.063873051 | 0.77580133 |
| ENSMUSG00000117486 | Gm30938 | 5.54E-05 | 0.841480629 | 1 |
| ENSMUSG00000097676 | Gm26612 | 5.54E-05 | 0.189403933 | 1 |
| ENSMUSG00000024154 | Gtf2a1l | 5.54E-05 | 2.063873051 | 0.789767653 |
| ENSMUSG00000091539 | Vmn1r238 | 5.54E-05 | 3.648835551 | 0.466464457 |
| ENSMUSG00000024233 | Lyzl1 | 5.54E-05 | 2.063873051 | 0.792441218 |
| ENSMUSG00000024234 | Mtpap | 5.54E-05 | -2.351164449 | 0.065287234 |
| ENSMUSG00000097043 | Gm10556 | 5.54E-05 | 2.648835551 | 0.66416768 |
| ENSMUSG00000117520 | Gm50064 | 5.54E-05 | 1.063873051 | 1 |
| ENSMUSG00000117508 | Gm34804 | 5.54E-05 | 0.841480629 | 1 |
| ENSMUSG00000101320 | Gm28529 | 5.54E-05 | 0.326907457 | 1 |
| ENSMUSG00000117652 | Gm2350 | 5.54E-05 | 2.648835551 | 0.66416768 |
| ENSMUSG00000117634 | Gm50069 | 5.54E-05 | 1.326907457 | 0.971854959 |
| ENSMUSG00000024410 | Rmc1 | 5.54E-05 | -1.708716453 | 0.296030202 |
| ENSMUSG00000086935 | Gm15956 | 5.54E-05 | 2.063873051 | 0.792441218 |
| ENSMUSG00000117365 | Gm29992 | 5.54E-05 | 2.063873051 | 0.780696989 |
| ENSMUSG00000117450 | 4933424G05Rik | 5.54E-05 | -0.810596067 | 0.858831093 |
| ENSMUSG00000117541 | Gm50048 | 5.54E-05 | 2.648835551 | 0.66416768 |
| ENSMUSG00000087170 | 1700001G01Rik | 5.54E-05 | 0.189403933 | 1 |
| ENSMUSG00000117378 | Gm31086 | 5.54E-05 | 1.063873051 | 1 |
| ENSMUSG00000069441 | Dsg1a | 5.54E-05 | -0.051604167 | 1 |
| ENSMUSG00000001804 | Dsg4 | 5.54E-05 | -0.158519371 | 1 |
| ENSMUSG00000085595 | Gm16090 | 5.54E-05 | 0.841480629 | 1 |
| ENSMUSG00000117570 | Gm38410 | 5.54E-05 | 1.648835551 | 0.868869354 |
| ENSMUSG00000114511 | 0710001A04Rik | 5.54E-05 | 1.648835551 | 0.886203497 |

| | | | | |
|---|---|---|---|---|
| ENSMUSG00000117410 | Gm50304 | 5.54E-05 | 2.063873051 | 0.792441218 |
| ENSMUSG00000117529 | Gm49978 | 5.54E-05 | 3.648835551 | 0.466464457 |
| ENSMUSG00000117554 | 4930527G23Rik | 5.54E-05 | 2.063873051 | 0.782089013 |
| ENSMUSG00000096934 | Gm26658 | 5.54E-05 | 3.648835551 | 0.466464457 |
| ENSMUSG00000024260 | Sap130 | 5.54E-05 | -2.305360759 | 0.082076736 |
| ENSMUSG00000024395 | Lims2 | 5.54E-05 | -0.158519371 | 1 |
| ENSMUSG00000097413 | A830052D11Rik | 5.54E-05 | 1.326907457 | 0.971854959 |
| ENSMUSG00000014503 | Pkd2l2 | 5.54E-05 | 0.648835551 | 1 |
| ENSMUSG00000024370 | Cdc23 | 5.54E-05 | -1.708716453 | 0.296030202 |
| ENSMUSG00000078240 | Gm3550 | 5.54E-05 | 0.47891055 | 1 |
| ENSMUSG00000097755 | 2010110K18Rik | 5.54E-05 | 2.063873051 | 0.792441218 |
| ENSMUSG00000118030 | Gm50163 | 5.54E-05 | 0.841480629 | 1 |
| ENSMUSG00000073598 | 1700066B19Rik | 5.54E-05 | 0.326907457 | 1 |
| ENSMUSG00000024347 | Psd2 | 5.54E-05 | 0.47891055 | 1 |
| ENSMUSG00000073594 | Gm10545 | 5.54E-05 | 2.063873051 | 0.792441218 |
| ENSMUSG00000097430 | Gm10544 | 5.54E-05 | 0.189403933 | 1 |
| ENSMUSG00000043313 | Pcdhb19 | 5.54E-05 | 2.648835551 | 0.66416768 |
| ENSMUSG00000073590 | 3222401L13Rik | 5.54E-05 | 0.841480629 | 1 |
| ENSMUSG00000103144 | Pcdhga1 | 5.54E-05 | 0.648835551 | 1 |
| ENSMUSG00000103793 | Pcdhga6 | 5.54E-05 | -0.43862729 | 1 |
| ENSMUSG00000103472 | Pcdhga7 | 5.54E-05 | -0.43862729 | 1 |
| ENSMUSG00000104063 | Pcdhgb7 | 5.54E-05 | -1.777429203 | 0.241307011 |
| ENSMUSG00000102918 | Pcdhgc3 | 5.54E-05 | 0.063873051 | 1 |
| ENSMUSG00000052102 | Gnpda1 | 5.54E-05 | 1.648835551 | 0.886203497 |
| ENSMUSG00000024427 | Spry4 | 5.54E-05 | -0.810596067 | 0.832285197 |
| ENSMUSG00000118042 | Gm50395 | 5.54E-05 | 1.648835551 | 0.886203497 |
| ENSMUSG00000059455 | Plac8l1 | 5.54E-05 | 1.063873051 | 1 |
| ENSMUSG00000117719 | Gm50214 | 5.54E-05 | 0.47891055 | 1 |
| ENSMUSG00000118018 | AC156546.1 | 5.54E-05 | 3.648835551 | 0.466464457 |
| ENSMUSG00000071858 | Gm94 | 5.54E-05 | 3.648835551 | 0.466464457 |
| ENSMUSG00000118006 | Gm50207 | 5.54E-05 | 0.841480629 | 1 |
| ENSMUSG00000053729 | Spinkl | 5.54E-05 | 0.47891055 | 1 |
| ENSMUSG00000118002 | Gm4107 | 5.54E-05 | -1.106051951 | 0.72614554 |
| ENSMUSG00000117621 | Hspe1-rs1 | 5.54E-05 | 3.648835551 | 0.466464457 |
| ENSMUSG00000117350 | Gm49975 | 5.54E-05 | 2.648835551 | 0.66416768 |
| ENSMUSG00000118113 | Gm4146 | 5.54E-05 | 1.648835551 | 0.886203497 |
| ENSMUSG00000117885 | Gm50229 | 5.54E-05 | 3.648835551 | 0.466464457 |
| ENSMUSG00000100134 | 1700065O20Rik | 5.54E-05 | 0.841480629 | 1 |
| ENSMUSG00000046886 | Zfp474 | 5.54E-05 | 1.326907457 | 0.971854959 |
| ENSMUSG00000024532 | 1700034E13Rik | 5.54E-05 | 1.326907457 | 0.966751942 |
| ENSMUSG00000096982 | Redrum | 5.54E-05 | 1.648835551 | 0.886203497 |
| ENSMUSG00000118120 | Gm33732 | 5.54E-05 | 1.063873051 | 1 |
| ENSMUSG00000118141 | Gm4230 | 5.54E-05 | 2.648835551 | 0.66416768 |
| ENSMUSG00000118153 | Gm50288 | 5.54E-05 | 3.648835551 | 0.466464457 |
| ENSMUSG00000117864 | Gm19500 | 5.54E-05 | 1.063873051 | 1 |
| ENSMUSG00000118322 | 2210409D07Rik | 5.54E-05 | 2.063873051 | 0.792441218 |
| ENSMUSG00000118337 | Gm50227 | 5.54E-05 | 2.648835551 | 0.66416768 |
| ENSMUSG00000050875 | Minar2 | 5.54E-05 | 1.326907457 | 0.954021409 |
| ENSMUSG00000073555 | Gm4951 | 5.54E-05 | 0.841480629 | 1 |
| ENSMUSG00000068606 | Gm4841 | 5.54E-05 | 2.648835551 | 0.66416768 |
| ENSMUSG00000090942 | F830016B08Rik | 5.54E-05 | 0.648835551 | 1 |

| | | | | |
|---|---|---|---|---|
| ENSMUSG00000036412 | Arsi | 5.54E-05 | 1.326907457 | 0.934418125 |
| ENSMUSG00000098098 | Bvht | 5.54E-05 | 3.648835551 | 0.466464457 |
| ENSMUSG00000045730 | Adrb2 | 5.54E-05 | -0.351164449 | 1 |
| ENSMUSG00000073551 | Spink13 | 5.54E-05 | -0.673092543 | 0.98880775 |
| ENSMUSG00000117671 | Gm50128 | 5.54E-05 | 1.648835551 | 0.886203497 |
| ENSMUSG00000071847 | Apcdd1 | 5.54E-05 | 0.326907457 | 1 |
| ENSMUSG00000117895 | Gm50368 | 5.54E-05 | 0.648835551 | 1 |
| ENSMUSG00000024519 | Cplx4 | 5.54E-05 | 1.063873051 | 1 |
| ENSMUSG00000117750 | Gm30190 | 5.54E-05 | 0.47891055 | 1 |
| ENSMUSG00000062526 | Mppe1 | 5.54E-05 | -1.777429203 | 0.241307011 |
| ENSMUSG00000007480 | Mc5r | 5.54E-05 | 1.648835551 | 0.886203497 |
| ENSMUSG00000086824 | 4930448D08Rik | 5.54E-05 | 2.648835551 | 0.66416768 |
| ENSMUSG00000118497 | AC124764.1 | 5.54E-05 | 2.648835551 | 0.66416768 |
| ENSMUSG00000044906 | 4930503L19Rik | 5.54E-05 | -2.500911568 | 0.036096973 |
| ENSMUSG00000038425 | Poli | 5.54E-05 | -1.209145444 | 0.874796537 |
| ENSMUSG00000118309 | Gm50261 | 5.54E-05 | 2.063873051 | 0.792441218 |
| ENSMUSG00000117984 | Gm31819 | 5.54E-05 | 3.648835551 | 0.466464457 |
| ENSMUSG00000117985 | Gm32548 | 5.54E-05 | 2.648835551 | 0.66416768 |
| ENSMUSG00000064036 | Mro | 5.54E-05 | -0.051604167 | 1 |
| ENSMUSG00000086333 | 1700120E14Rik | 5.54E-05 | 2.063873051 | 0.792441218 |
| ENSMUSG00000097712 | Gm10532 | 5.54E-05 | 2.648835551 | 0.66416768 |
| ENSMUSG00000118311 | 1700003O11Rik | 5.54E-05 | 3.648835551 | 0.466464457 |
| ENSMUSG00000091519 | Skor2 | 5.54E-05 | 1.063873051 | 1 |
| ENSMUSG00000025427 | Rnf165 | 5.54E-05 | -0.158519371 | 1 |
| ENSMUSG00000073529 | F830208F22Rik | 5.54E-05 | 3.648835551 | 0.466464457 |
| ENSMUSG00000056214 | Pard6g | 5.54E-05 | 0.841480629 | 1 |
| ENSMUSG00000053950 | Adnp2 | 5.54E-05 | -1.843017545 | 0.255869172 |
| ENSMUSG00000118061 | Rbfaos | 5.54E-05 | 0.326907457 | 1 |
| ENSMUSG00000024570 | Rbfa | 5.54E-05 | -1.051604167 | 0.706929043 |
| ENSMUSG00000024565 | Sall3 | 5.54E-05 | 1.648835551 | 0.886203497 |
| ENSMUSG00000118254 | Gm46614 | 5.54E-05 | -0.051604167 | 1 |
| ENSMUSG00000118057 | B020010K11Rik | 5.54E-05 | 1.648835551 | 0.886203497 |
| ENSMUSG00000118339 | 4930592I03Rik | 5.54E-05 | 2.648835551 | 0.66416768 |
| ENSMUSG00000117715 | Gm50413 | 5.54E-05 | 1.063873051 | 1 |
| ENSMUSG00000118387 | Gm50415 | 5.54E-05 | 2.648835551 | 0.66416768 |
| ENSMUSG00000056162 | Cndp1 | 5.54E-05 | 0.47891055 | 1 |
| ENSMUSG00000117781 | Gm17266 | 5.54E-05 | 2.063873051 | 0.792441218 |
| ENSMUSG00000034391 | Fbxo15 | 5.54E-05 | 2.063873051 | 0.792441218 |
| ENSMUSG00000024645 | Timm21 | 5.54E-05 | -0.351164449 | 1 |
| ENSMUSG00000117736 | Gm50386 | 5.54E-05 | 1.326907457 | 0.953930683 |
| ENSMUSG00000117805 | Gm18280 | 5.54E-05 | 2.063873051 | 0.792441218 |
| ENSMUSG00000118132 | 4833408A19Rik | 5.54E-05 | 2.063873051 | 0.792441218 |
| ENSMUSG00000024835 | Coro1b | 5.54E-05 | -1.158519371 | 0.599135879 |
| ENSMUSG00000024824 | Rad9a | 5.54E-05 | -0.52108945 | 0.975624465 |
| ENSMUSG00000087132 | A930001C03Rik | 5.54E-05 | 0.648835551 | 1 |
| ENSMUSG00000045045 | Lrfn4 | 5.54E-05 | -0.995020638 | 0.699192777 |
| ENSMUSG00000024901 | Peli3 | 5.54E-05 | 2.063873051 | 0.792441218 |
| ENSMUSG00000024862 | Klc2 | 5.54E-05 | -0.936126949 | 0.695372548 |
| ENSMUSG00000047658 | Gal3st3 | 5.54E-05 | 0.648835551 | 1 |
| ENSMUSG00000024910 | Ctsw | 5.54E-05 | 3.648835551 | 0.466464457 |
| ENSMUSG00000024926 | Kat5 | 5.54E-05 | -2.82689788 | 0.011086379 |

| | | | | |
|---|---|---|---|---|
| ENSMUSG00000056917 | Sipa1 | 5.54E-05 | -2.184054463 | 0.102196437 |
| ENSMUSG00000043488 | Frmd8os | 5.54E-05 | 1.648835551 | 0.886203497 |
| ENSMUSG00000024818 | Slc25a45 | 5.54E-05 | -0.351164449 | 1 |
| ENSMUSG00000024791 | Cdca5 | 5.54E-05 | -0.258055044 | 1 |
| ENSMUSG00000024787 | Snx15 | 5.54E-05 | -1.965874293 | 0.162298752 |
| ENSMUSG00000024786 | Majin | 5.54E-05 | 0.648835551 | 1 |
| ENSMUSG00000085196 | Gm14963 | 5.54E-05 | -0.874726405 | 0.840804483 |
| ENSMUSG00000032648 | Pygm | 5.54E-05 | -0.051604167 | 1 |
| ENSMUSG00000033768 | Nrxn2 | 5.54E-05 | -0.52108945 | 0.975624465 |
| ENSMUSG00000118181 | CAAA01066123.1 | 5.54E-05 | 1.648835551 | 0.886203497 |
| ENSMUSG00000047810 | Ccdc88b | 5.54E-05 | -1.52108945 | 0.452184898 |
| ENSMUSG00000024957 | Kcnk4 | 5.54E-05 | 2.648835551 | 0.66416768 |
| ENSMUSG00000037349 | Nudt22 | 5.54E-05 | 0.47891055 | 1 |
| ENSMUSG00000024764 | Naa40 | 5.54E-05 | -1.843017545 | 0.255869172 |
| ENSMUSG00000024968 | Rcor2 | 5.54E-05 | 0.841480629 | 1 |
| ENSMUSG00000024972 | Lgals12 | 5.54E-05 | 0.326907457 | 1 |
| ENSMUSG00000063590 | Slc22a28 | 5.54E-05 | 0.841480629 | 1 |
| ENSMUSG00000075044 | Slc22a29 | 5.54E-05 | 3.648835551 | 0.466464457 |
| ENSMUSG00000052562 | Slc22a30 | 5.54E-05 | 2.063873051 | 0.792441218 |
| ENSMUSG00000063796 | Slc22a8 | 5.54E-05 | 0.841480629 | 1 |
| ENSMUSG00000071656 | Lrrn4cl | 5.54E-05 | 1.326907457 | 0.971854959 |
| ENSMUSG00000096740 | Lbhd1 | 5.54E-05 | 1.648835551 | 0.860483184 |
| ENSMUSG00000071653 | 1810009A15Rik | 5.54E-05 | 1.326907457 | 0.964262694 |
| ENSMUSG00000071652 | Ints5 | 5.54E-05 | -0.351164449 | 1 |
| ENSMUSG00000114220 | Gm10353 | 5.54E-05 | -0.43862729 | 1 |
| ENSMUSG00000037418 | Best1 | 5.54E-05 | 0.47891055 | 1 |
| ENSMUSG00000024666 | Tmem138 | 5.54E-05 | -1.395558568 | 0.508263132 |
| ENSMUSG00000118392 | 4930524O05Rik | 5.54E-05 | 2.648835551 | 0.66416768 |
| ENSMUSG00000024737 | Slc15a3 | 5.54E-05 | 0.648835551 | 1 |
| ENSMUSG00000024732 | Ccdc86 | 5.54E-05 | -1.209145444 | 0.606122355 |
| ENSMUSG00000024731 | Ms4a10 | 5.54E-05 | 1.648835551 | 0.882296038 |
| ENSMUSG00000024729 | 1700017D01Rik | 5.54E-05 | -0.743481871 | 1 |
| ENSMUSG00000024728 | 1700025F22Rik | 5.54E-05 | 0.189403933 | 1 |
| ENSMUSG00000054523 | Ms4a5 | 5.54E-05 | 3.648835551 | 0.466464457 |
| ENSMUSG00000101389 | Ms4a4a | 5.54E-05 | 1.326907457 | 0.971854959 |
| ENSMUSG00000024675 | Ms4a4c | 5.54E-05 | 1.326907457 | 0.971854959 |
| ENSMUSG00000079419 | Ms4a6c | 5.54E-05 | 0.648835551 | 1 |
| ENSMUSG00000099757 | BE692007 | 5.54E-05 | 0.189403933 | 1 |
| ENSMUSG00000058470 | Gm8369 | 5.54E-05 | -0.158519371 | 1 |
| ENSMUSG00000024678 | Ms4a4d | 5.54E-05 | -0.351164449 | 1 |
| ENSMUSG00000041857 | Oosp1 | 5.54E-05 | 2.648835551 | 0.66416768 |
| ENSMUSG00000024682 | Cblif | 5.54E-05 | 2.063873051 | 0.792441218 |
| ENSMUSG00000067522 | Olfr76 | 5.54E-05 | 2.648835551 | 0.66416768 |
| ENSMUSG00000045030 | Olfr1443 | 5.54E-05 | 3.648835551 | 0.466464457 |
| ENSMUSG00000079415 | Cntf | 5.54E-05 | 1.326907457 | 0.971854959 |
| ENSMUSG00000046913 | Olfr1451 | 5.54E-05 | 3.648835551 | 0.466464457 |
| ENSMUSG00000051156 | Olfr1491 | 5.54E-05 | 3.648835551 | 0.466464457 |
| ENSMUSG00000047207 | Olfr1495 | 5.54E-05 | 3.648835551 | 0.466464457 |
| ENSMUSG00000062314 | Olfr1505 | 5.54E-05 | 3.648835551 | 0.466464457 |
| ENSMUSG00000024697 | Gna14 | 5.54E-05 | -2.673092543 | 0.031064467 |
| ENSMUSG00000118067 | Gm32629 | 5.54E-05 | 2.063873051 | 0.792441218 |

| | | | | |
|---|---|---|---|---|
| ENSMUSG00000118293 | Gm32750 | 5.54E-05 | 1.648835551 | 0.874796537 |
| ENSMUSG00000118047 | 4930554I06Rik | 5.54E-05 | 2.648835551 | 0.66416768 |
| ENSMUSG00000074922 | Fam122a | 5.54E-05 | -0.52108945 | 0.975624465 |
| ENSMUSG00000024837 | Dmrt1 | 5.54E-05 | -0.743481871 | 1 |
| ENSMUSG00000118016 | 1700048O14Rik | 5.54E-05 | 3.648835551 | 0.466464457 |
| ENSMUSG00000093628 | Gm20616 | 5.54E-05 | 2.648835551 | 0.66416768 |
| ENSMUSG00000089952 | 4933413C19Rik | 5.54E-05 | 2.648835551 | 0.66416768 |
| ENSMUSG00000040105 | Plpp6 | 5.54E-05 | -0.52108945 | 1 |
| ENSMUSG00000118038 | Gm9895 | 5.54E-05 | 2.648835551 | 0.66416768 |
| ENSMUSG00000024806 | Mlana | 5.54E-05 | 3.648835551 | 0.466464457 |
| ENSMUSG00000074896 | Ifit3 | 5.54E-05 | 1.326907457 | 0.971854959 |
| ENSMUSG00000034459 | Ifit1 | 5.54E-05 | -0.936126949 | 0.695372548 |
| ENSMUSG00000033610 | Pank1 | 5.54E-05 | -0.52108945 | 1 |
| ENSMUSG00000032808 | Cyp2c38 | 5.54E-05 | 1.063873051 | 1 |
| ENSMUSG00000025004 | Cyp2c40 | 5.54E-05 | 0.841480629 | 1 |
| ENSMUSG00000054827 | Cyp2c50 | 5.54E-05 | 3.648835551 | 0.466464457 |
| ENSMUSG00000060613 | Cyp2c70 | 5.54E-05 | 0.648835551 | 1 |
| ENSMUSG00000025014 | Dntt | 5.54E-05 | 2.063873051 | 0.792441218 |
| ENSMUSG00000025172 | Ankrd2 | 5.54E-05 | 0.648835551 | 1 |
| ENSMUSG00000018822 | Sfrp5 | 5.54E-05 | 3.648835551 | 0.466464457 |
| ENSMUSG00000042532 | Golga7b | 5.54E-05 | 1.063873051 | 1 |
| ENSMUSG00000117728 | Gm34299 | 5.54E-05 | 1.648835551 | 0.886203497 |
| ENSMUSG00000118221 | Gm50194 | 5.54E-05 | -0.673092543 | 0.98880775 |
| ENSMUSG00000025196 | Cpn1 | 5.54E-05 | 0.841480629 | 1 |
| ENSMUSG00000050195 | Scd4 | 5.54E-05 | -0.158519371 | 1 |
| ENSMUSG00000118076 | Gm50337 | 5.54E-05 | 1.648835551 | 0.866342478 |
| ENSMUSG00000117796 | Gm35406 | 5.54E-05 | 2.648835551 | 0.66416768 |
| ENSMUSG00000118204 | 4930414N06Rik | 5.54E-05 | -0.52108945 | 1 |
| ENSMUSG00000074818 | Pdzd7 | 5.54E-05 | 0.189403933 | 1 |
| ENSMUSG00000092346 | Tlx1os | 5.54E-05 | 1.648835551 | 0.886203497 |
| ENSMUSG00000097690 | 4930505N22Rik | 5.54E-05 | -0.599091962 | 0.980745941 |
| ENSMUSG00000118087 | 4833438C02Rik | 5.54E-05 | -0.258055044 | 1 |
| ENSMUSG00000037126 | Psd | 5.54E-05 | 1.326907457 | 0.971854959 |
| ENSMUSG00000094219 | Calhm3 | 5.54E-05 | 1.648835551 | 0.886203497 |
| ENSMUSG00000025064 | Col17a1 | 5.54E-05 | 0.189403933 | 1 |
| ENSMUSG00000117975 | Itprip | 5.54E-05 | 1.326907457 | 0.971854959 |
| ENSMUSG00000118111 | Gm50442 | 5.54E-05 | 2.648835551 | 0.66416768 |
| ENSMUSG00000035804 | Ins1 | 5.54E-05 | 1.648835551 | 0.886203497 |
| ENSMUSG00000118369 | Gm30541 | 5.54E-05 | -1.965874293 | 0.162298752 |
| ENSMUSG00000033717 | Adra2a | 5.54E-05 | 3.648835551 | 0.466464457 |
| ENSMUSG00000024978 | Gpam | 5.54E-05 | -1.810596067 | 0.246776726 |
| ENSMUSG00000118099 | Gm31356 | 5.54E-05 | -1.158519371 | 0.599135879 |
| ENSMUSG00000024979 | Tectb | 5.54E-05 | 0.648835551 | 1 |
| ENSMUSG00000099746 | Ppnr | 5.54E-05 | -2.480447466 | 0.046272089 |
| ENSMUSG00000049134 | Nrap | 5.54E-05 | 0.326907457 | 1 |
| ENSMUSG00000117890 | Gm32441 | 5.54E-05 | 1.648835551 | 0.865542683 |
| ENSMUSG00000035283 | Adrb1 | 5.54E-05 | 1.326907457 | 0.94880235 |
| ENSMUSG00000025081 | Tdrd1 | 5.54E-05 | 0.326907457 | 1 |
| ENSMUSG00000117927 | Gm50271 | 5.54E-05 | 3.648835551 | 0.466464457 |
| ENSMUSG00000118178 | Gm32932 | 5.54E-05 | 2.063873051 | 0.792441218 |
| ENSMUSG00000117871 | Gm50287 | 5.54E-05 | 2.648835551 | 0.66416768 |

| | | | | |
|---|---|---|---|---|
| ENSMUSG00000025090 | Ccdc172 | 5.54E-05 | 1.648835551 | 0.886203497 |
| ENSMUSG00000025094 | Slc18a2 | 5.54E-05 | 0.326907457 | 1 |
| ENSMUSG00000100119 | 2700089I24Rik | 5.54E-05 | -0.258055044 | 1 |
| ENSMUSG00000072437 | Nanos1 | 5.54E-05 | 1.326907457 | 0.971854959 |
| ENSMUSG00000097342 | Gm26618 | 5.54E-05 | 3.648835551 | 0.466464457 |
| ENSMUSG00000031149 | Praf2 | 5.54E-05 | 0.326907457 | 1 |
| ENSMUSG00000031156 | Slc35a2 | 5.54E-05 | -1.305360759 | 0.491653643 |
| ENSMUSG00000031162 | Gata1 | 5.54E-05 | 1.648835551 | 0.847265357 |
| ENSMUSG00000087193 | Gm14820 | 5.54E-05 | 3.648835551 | 0.466464457 |
| ENSMUSG00000068218 | Ssxb9 | 5.54E-05 | 3.648835551 | 0.466464457 |
| ENSMUSG00000015342 | Xk | 5.54E-05 | -0.743481871 | 0.827142217 |
| ENSMUSG00000040456 | Hypm | 5.54E-05 | 2.648835551 | 0.66416768 |
| ENSMUSG00000031173 | Otc | 5.54E-05 | 0.841480629 | 1 |
| ENSMUSG00000085891 | Gm14634 | 5.54E-05 | -1.395558568 | 0.508263132 |
| ENSMUSG00000087424 | 5730405O15Rik | 5.54E-05 | 3.648835551 | 0.466464457 |
| ENSMUSG00000051228 | Nyx | 5.54E-05 | 0.841480629 | 1 |
| ENSMUSG00000084907 | Gm14505 | 5.54E-05 | 2.648835551 | 0.66416768 |
| ENSMUSG00000086048 | Drr1 | 5.54E-05 | 2.648835551 | 0.66416768 |
| ENSMUSG00000025038 | Efhc2 | 5.54E-05 | -0.43862729 | 1 |
| ENSMUSG00000001128 | Cfp | 5.54E-05 | -1.106051951 | 0.719591951 |
| ENSMUSG00000086877 | A230072C01Rik | 5.54E-05 | -0.351164449 | 1 |
| ENSMUSG00000037167 | Spaca5 | 5.54E-05 | 3.648835551 | 0.466464457 |
| ENSMUSG00000094945 | Rhox4a2 | 5.54E-05 | 3.648835551 | 0.466464457 |
| ENSMUSG00000079635 | Rhox4c | 5.54E-05 | 3.648835551 | 0.466464457 |
| ENSMUSG00000048047 | Zbtb33 | 5.54E-05 | 0.063873051 | 1 |
| ENSMUSG00000016327 | Atp1b4 | 5.54E-05 | 2.648835551 | 0.66416768 |
| ENSMUSG00000086948 | Gm14697 | 5.54E-05 | 1.063873051 | 1 |
| ENSMUSG00000056281 | Olfr1322 | 5.54E-05 | 3.648835551 | 0.466464457 |
| ENSMUSG00000031112 | Stk26 | 5.54E-05 | 0.648835551 | 1 |
| ENSMUSG00000036131 | Frmd7 | 5.54E-05 | 2.063873051 | 0.792441218 |
| ENSMUSG00000086704 | Gm14582 | 5.54E-05 | -1.258055044 | 0.66416768 |
| ENSMUSG00000055110 | A630012P03Rik | 5.54E-05 | 1.648835551 | 0.886203497 |
| ENSMUSG00000036022 | Fam122b | 5.54E-05 | 0.063873051 | 1 |
| ENSMUSG00000036013 | Fam122c | 5.54E-05 | 0.841480629 | 1 |
| ENSMUSG00000067924 | Rtl8b | 5.54E-05 | 2.063873051 | 0.789767653 |
| ENSMUSG00000055555 | 4930502E18Rik | 5.54E-05 | 2.648835551 | 0.66416768 |
| ENSMUSG00000073176 | Zfp449 | 5.54E-05 | 0.648835551 | 1 |
| ENSMUSG00000061273 | Mmgt1 | 5.54E-05 | -0.158519371 | 1 |
| ENSMUSG00000067878 | Map7d3 | 5.54E-05 | 0.063873051 | 1 |
| ENSMUSG00000053852 | Adgrg4 | 5.54E-05 | -0.051604167 | 1 |
| ENSMUSG00000031131 | Vgll1 | 5.54E-05 | 3.648835551 | 0.466464457 |
| ENSMUSG00000085808 | Gm14718 | 5.54E-05 | 1.326907457 | 0.971854959 |
| ENSMUSG00000079584 | Gm364 | 5.54E-05 | 2.648835551 | 0.66416768 |
| ENSMUSG00000031138 | F9 | 5.54E-05 | 1.648835551 | 0.886203497 |
| ENSMUSG00000071753 | Cdr1os | 5.54E-05 | 0.841480629 | 1 |
| ENSMUSG00000100511 | 1700111N16Rik | 5.54E-05 | 0.189403933 | 1 |
| ENSMUSG00000073130 | Gm1141 | 5.54E-05 | 2.648835551 | 0.66416768 |
| ENSMUSG00000053593 | Fate1 | 5.54E-05 | 1.648835551 | 0.886203497 |
| ENSMUSG00000031340 | Gabre | 5.54E-05 | -3.171343411 | 0.003007618 |
| ENSMUSG00000067768 | Xlr4b | 5.54E-05 | 2.063873051 | 0.792441218 |
| ENSMUSG00000031365 | Zfp275 | 5.54E-05 | 0.063873051 | 1 |

| | | | | |
|---|---|---|---|---|
| ENSMUSG00000097110 | Gm26726 | 5.54E-05 | 3.648835551 | 0.466464457 |
| ENSMUSG00000031376 | Atp2b3 | 5.54E-05 | 0.326907457 | 1 |
| ENSMUSG00000031383 | Dusp9 | 5.54E-05 | 2.063873051 | 0.792441218 |
| ENSMUSG00000002012 | Pnck | 5.54E-05 | 2.648835551 | 0.66416768 |
| ENSMUSG00000031389 | Arhgap4 | 5.54E-05 | -0.810596067 | 0.868661374 |
| ENSMUSG00000001964 | Emd | 5.54E-05 | -1.395558568 | 0.508263132 |
| ENSMUSG00000031398 | Plxna3 | 5.54E-05 | -1.43862729 | 0.433313155 |
| ENSMUSG00000086207 | Gm14717 | 5.54E-05 | 1.326907457 | 0.954089321 |
| ENSMUSG00000079525 | Cldn34d | 5.54E-05 | 2.648835551 | 0.66416768 |
| ENSMUSG00000046942 | Mageb16 | 5.54E-05 | 1.648835551 | 0.886203497 |
| ENSMUSG00000025058 | 5430427O19Rik | 5.54E-05 | 1.648835551 | 0.886203497 |
| ENSMUSG00000035387 | 4930415L06Rik | 5.54E-05 | 3.648835551 | 0.466464457 |
| ENSMUSG00000050332 | Amer1 | 5.54E-05 | -2.874726405 | 0.009566281 |
| ENSMUSG00000031204 | Asb12 | 5.54E-05 | 0.326907457 | 1 |
| ENSMUSG00000044206 | Vsig4 | 5.54E-05 | 2.063873051 | 0.792441218 |
| ENSMUSG00000031209 | Heph | 5.54E-05 | -0.258055044 | 1 |
| ENSMUSG00000071719 | Tmem28 | 5.54E-05 | 2.648835551 | 0.66416768 |
| ENSMUSG00000067597 | Dgat2l6 | 5.54E-05 | 2.063873051 | 0.792441218 |
| ENSMUSG00000060890 | Arr3 | 5.54E-05 | 2.063873051 | 0.792441218 |
| ENSMUSG00000031304 | Il2rg | 5.54E-05 | 2.063873051 | 0.771569813 |
| ENSMUSG00000085568 | Gm14858 | 5.54E-05 | 1.326907457 | 0.955146698 |
| ENSMUSG00000086503 | Xist | 5.54E-05 | 2.063873051 | 0.792441218 |
| ENSMUSG00000100364 | 1700121L16Rik | 5.54E-05 | 3.648835551 | 0.466464457 |
| ENSMUSG00000033777 | Tlr13 | 5.54E-05 | 1.326907457 | 0.928159492 |
| ENSMUSG00000073010 | Gm5127 | 5.54E-05 | -0.43862729 | 1 |
| ENSMUSG00000049929 | Lpar4 | 5.54E-05 | 2.063873051 | 0.792441218 |
| ENSMUSG00000056854 | Pou3f4 | 5.54E-05 | 3.648835551 | 0.466464457 |
| ENSMUSG00000118415 | BX088531.1 | 5.54E-05 | 3.648835551 | 0.466464457 |
| ENSMUSG00000072049 | Vmn2r121 | 5.54E-05 | 2.648835551 | 0.66416768 |
| ENSMUSG00000051323 | Pcdh19 | 5.54E-05 | -0.351164449 | 1 |
| ENSMUSG00000031253 | Srpx2 | 5.54E-05 | 0.326907457 | 1 |
| ENSMUSG00000052676 | Zmat1 | 5.54E-05 | -1.9057533 | 0.203406755 |
| ENSMUSG00000089759 | 3632454L22Rik | 5.54E-05 | 3.648835551 | 0.466464457 |
| ENSMUSG00000031410 | Nxf7 | 5.54E-05 | 3.648835551 | 0.466464457 |
| ENSMUSG00000072969 | Armcx5 | 5.54E-05 | -0.158519371 | 1 |
| ENSMUSG00000072964 | Bhlhb9 | 5.54E-05 | 1.063873051 | 1 |
| ENSMUSG00000072966 | Gprasp2 | 5.54E-05 | 2.648835551 | 0.66416768 |
| ENSMUSG00000057000 | Nxf3 | 5.54E-05 | 3.648835551 | 0.466464457 |
| ENSMUSG00000054034 | Tceal5 | 5.54E-05 | 3.648835551 | 0.466464457 |
| ENSMUSG00000057439 | Kir3dl2 | 5.54E-05 | 0.47891055 | 1 |
| ENSMUSG00000048155 | H2bfm | 5.54E-05 | 2.648835551 | 0.66416768 |
| ENSMUSG00000023443 | Esx1 | 5.54E-05 | 2.063873051 | 0.775470096 |
| ENSMUSG00000031271 | Serpina7 | 5.54E-05 | 3.648835551 | 0.466464457 |
| ENSMUSG00000067276 | Capn6 | 5.54E-05 | 2.063873051 | 0.792441218 |
| ENSMUSG00000031289 | Il13ra2 | 5.54E-05 | 1.063873051 | 1 |
| ENSMUSG00000095872 | Gm15128 | 5.54E-05 | 0.326907457 | 1 |
| ENSMUSG00000096834 | Gm15127 | 5.54E-05 | 2.063873051 | 0.792441218 |
| ENSMUSG00000025270 | Alas2 | 5.54E-05 | 3.648835551 | 0.466464457 |
| ENSMUSG00000025271 | Pfkfb1 | 5.54E-05 | -0.43862729 | 1 |
| ENSMUSG00000025264 | Tsr2 | 5.54E-05 | -2.158519371 | 0.115633409 |
| ENSMUSG00000025333 | Gpr143 | 5.54E-05 | 2.063873051 | 0.792441218 |

| | | | | |
|---|---|---|---|---|
| ENSMUSG00000041649 | Klf8 | 5.54E-05 | -1.52108945 | 0.452184898 |
| ENSMUSG00000057457 | Phex | 5.54E-05 | -0.810596067 | 0.832285197 |
| ENSMUSG00000086188 | Gm15169 | 5.54E-05 | -1.395558568 | 0.508263132 |
| ENSMUSG00000000037 | Scml2 | 5.54E-05 | 1.326907457 | 0.971854959 |
| ENSMUSG00000043518 | Rai2 | 5.54E-05 | 0.648835551 | 1 |
| ENSMUSG00000040749 | Siah1b | 5.54E-05 | 1.063873051 | 1 |
| ENSMUSG00000015401 | Cltrn | 5.54E-05 | 1.648835551 | 0.879677127 |
| ENSMUSG00000015405 | Ace2 | 5.54E-05 | 1.648835551 | 0.886203497 |
| ENSMUSG00000031377 | Bmx | 5.54E-05 | -0.673092543 | 1 |
| ENSMUSG00000031381 | Piga | 5.54E-05 | -1.560617814 | 0.356350819 |
| ENSMUSG00000031384 | Asb9 | 5.54E-05 | 2.648835551 | 0.66416768 |
| ENSMUSG00000040586 | Ofd1 | 5.54E-05 | -3.1966545 | 0.002570155 |
| ENSMUSG00000025742 | Prps2 | 5.54E-05 | -0.158519371 | 1 |
| ENSMUSG00000087201 | Gm15261 | 5.54E-05 | -0.258055044 | 1 |
| ENSMUSG00000072844 | G530011O06Rik | 5.54E-05 | 1.648835551 | 0.886203497 |
| ENSMUSG00000000103 | Zfy2 | 5.54E-05 | 1.648835551 | 0.886203497 |
| ENSMUSG00000093987 | Gm21704 | 5.54E-05 | 0.326907457 | 1 |
| ENSMUSG00000096520 | Gm3376 | 5.54E-05 | 3.648835551 | 0.466464457 |
| ENSMUSG00000101081 | Gm29193 | 5.54E-05 | 3.648835551 | 0.466464457 |
| ENSMUSG00000094556 | Gm20821 | 5.54E-05 | 3.648835551 | 0.466464457 |
| ENSMUSG00000096650 | Gm20896 | 5.54E-05 | 2.063873051 | 0.792441218 |
| ENSMUSG00000100045 | Gm20929 | 5.54E-05 | 2.063873051 | 0.792441218 |
| ENSMUSG00000094052 | Gm21118 | 5.54E-05 | 3.648835551 | 0.466464457 |
| ENSMUSG00000102111 | Gm29566 | 5.54E-05 | 3.648835551 | 0.466464457 |
| ENSMUSG00000101286 | Gm21317 | 5.54E-05 | 0.648835551 | 1 |
| ENSMUSG00000101915 | Gm28102 | 5.54E-05 | 1.648835551 | 0.886203497 |
| ENSMUSG00000100608 | Gm21996 | 5.54E-05 | 3.648835551 | 0.466464457 |
| ENSMUSG00000095552 | 4933409K07Rik | 5.54E-05 | 2.648835551 | 0.66416768 |
| ENSMUSG00000094855 | AC133095.1 | 5.54E-05 | 3.648835551 | 0.466464457 |
| ENSMUSG00000104328 | Gm37323 | 0 | 1.648835551 | 1 |
| ENSMUSG00000033774 | Npbwr1 | 0 | 1.648835551 | 1 |
| ENSMUSG00000087247 | Alkal1 | 0 | 1.648835551 | 1 |
| ENSMUSG00000103067 | Gm30414 | 0 | 0.648835551 | 1 |
| ENSMUSG00000067879 | Vxn | 0 | 1.063873051 | 1 |
| ENSMUSG00000099032 | Tcf24 | 0 | 1.648835551 | 1 |
| ENSMUSG00000099498 | Gm29283 | 0 | -0.810596067 | 1 |
| ENSMUSG00000042414 | Prdm14 | 0 | 1.648835551 | 1 |
| ENSMUSG00000067813 | Xkr9 | 0 | 0.326907457 | 1 |
| ENSMUSG00000025932 | Eya1 | 0 | -1.051604167 | 0.887698387 |
| ENSMUSG00000025930 | Msc | 0 | 1.063873051 | 1 |
| ENSMUSG00000032719 | Sbspon | 0 | -1.258055044 | 0.673811773 |
| ENSMUSG00000097744 | D030040B21Rik | 0 | 1.063873051 | 1 |
| ENSMUSG00000067780 | Pi15 | 0 | 0.063873051 | 1 |
| ENSMUSG00000085125 | Gm16070 | 0 | -0.52108945 | 1 |
| ENSMUSG00000025776 | Crispld1 | 0 | -1.158519371 | 0.954404283 |
| ENSMUSG00000025774 | Crisp4 | 0 | 0.063873051 | 1 |
| ENSMUSG00000089914 | 4930486I03Rik | 0 | 0.063873051 | 1 |
| ENSMUSG00000041872 | Il17f | 0 | 1.648835551 | 1 |
| ENSMUSG00000097934 | 6720483E21Rik | 0 | 1.648835551 | 1 |
| ENSMUSG00000041809 | Efhc1 | 0 | -0.52108945 | 1 |
| ENSMUSG00000099971 | Gm28287 | 0 | 1.648835551 | 1 |

| | | | | |
|---|---|---|---|---|
| ENSMUSG00000099343 | Gm28836 | 0 | 0.648835551 | 1 |
| ENSMUSG00000097970 | Gm27028 | 0 | 1.648835551 | 1 |
| ENSMUSG00000100775 | Gm29107 | 0 | 0.326907457 | 1 |
| ENSMUSG00000026158 | Ogfrl1 | 0 | -1.874726405 | 0.377958192 |
| ENSMUSG00000073722 | 4931408C20Rik | 0 | 0.648835551 | 1 |
| ENSMUSG00000101514 | Gm5524 | 0 | 1.648835551 | 1 |
| ENSMUSG00000026064 | Ptp4a1 | 0 | 0.326907457 | 1 |
| ENSMUSG00000100910 | Gm29128 | 0 | 1.648835551 | 1 |
| ENSMUSG00000050217 | Lgsn | 0 | 1.648835551 | 1 |
| ENSMUSG00000026124 | Cfc1 | 0 | 1.648835551 | 1 |
| ENSMUSG00000037529 | Prss40 | 0 | 1.063873051 | 1 |
| ENSMUSG00000104002 | Gm38336 | 0 | 1.648835551 | 1 |
| ENSMUSG00000101714 | Gm28415 | 0 | 1.648835551 | 1 |
| ENSMUSG00000102531 | 1110002O04Rik | 0 | 1.063873051 | 1 |
| ENSMUSG00000102845 | Gm38033 | 0 | 1.648835551 | 1 |
| ENSMUSG00000067653 | Ankrd23 | 0 | -2.305360759 | 0.231596497 |
| ENSMUSG00000026117 | Zap70 | 0 | 0.648835551 | 1 |
| ENSMUSG00000102930 | Gm38115 | 0 | -0.52108945 | 1 |
| ENSMUSG00000103603 | Gm37020 | 0 | 1.063873051 | 1 |
| ENSMUSG00000086064 | 4930439A04Rik | 0 | 1.648835551 | 1 |
| ENSMUSG00000026111 | Unc50 | 0 | -1.743481871 | 0.455131546 |
| ENSMUSG00000097268 | Gm26805 | 0 | 0.648835551 | 1 |
| ENSMUSG00000037216 | Lipt1 | 0 | 0.326907457 | 1 |
| ENSMUSG00000061584 | Lyg2 | 0 | 0.063873051 | 1 |
| ENSMUSG00000026085 | Lyg1 | 0 | 1.648835551 | 1 |
| ENSMUSG00000085391 | Gm16150 | 0 | 1.648835551 | 1 |
| ENSMUSG00000087131 | Gm16152 | 0 | 0.648835551 | 1 |
| ENSMUSG00000026080 | Chst10 | 0 | 0.326907457 | 1 |
| ENSMUSG00000067604 | Nms | 0 | 1.063873051 | 1 |
| ENSMUSG00000085894 | Gm15832 | 0 | -0.351164449 | 1 |
| ENSMUSG00000101634 | 1700066B17Rik | 0 | 0.326907457 | 1 |
| ENSMUSG00000097899 | Gm16894 | 0 | 0.326907457 | 1 |
| ENSMUSG00000100185 | Gm28634 | 0 | 1.648835551 | 1 |
| ENSMUSG00000053640 | Gm9915 | 0 | 1.648835551 | 1 |
| ENSMUSG00000101299 | Gm28175 | 0 | 0.326907457 | 1 |
| ENSMUSG00000100180 | Gm28140 | 0 | 0.648835551 | 1 |
| ENSMUSG00000097063 | Pantr2 | 0 | 1.648835551 | 1 |
| ENSMUSG00000099784 | Dalir | 0 | 1.648835551 | 1 |
| ENSMUSG00000041907 | Gpr45 | 0 | -0.351164449 | 1 |
| ENSMUSG00000099576 | Gm29040 | 0 | 0.326907457 | 1 |
| ENSMUSG00000026051 | Ecrg4 | 0 | 0.326907457 | 1 |
| ENSMUSG00000047343 | Mettl21c | 0 | 1.648835551 | 1 |
| ENSMUSG00000098549 | Gm5269 | 0 | 1.648835551 | 1 |
| ENSMUSG00000100213 | Gm28151 | 0 | 0.648835551 | 1 |
| ENSMUSG00000099625 | Gm29325 | 0 | 0.648835551 | 1 |
| ENSMUSG00000109599 | Gm31812 | 0 | 1.648835551 | 1 |
| ENSMUSG00000026104 | Stat1 | 0 | -2.258055044 | 0.222101622 |
| ENSMUSG00000099878 | Gm28178 | 0 | 1.063873051 | 1 |
| ENSMUSG00000109809 | Gm34066 | 0 | 1.063873051 | 1 |
| ENSMUSG00000046994 | Mars2 | 0 | -1.810596067 | 0.465580427 |
| ENSMUSG00000099950 | 9130227L01Rik | 0 | 1.063873051 | 1 |

| | | | | |
|---|---|---|---|---|
| ENSMUSG00000097519 | 4930558J18Rik | 0 | 1.063873051 | 1 |
| ENSMUSG00000038323 | 1700066M21Rik | 0 | 0.326907457 | 1 |
| ENSMUSG00000091283 | Gm17234 | 0 | 1.648835551 | 1 |
| ENSMUSG00000089822 | Gm15759 | 0 | 0.326907457 | 1 |
| ENSMUSG00000085054 | Gm15834 | 0 | -1.258055044 | 0.673811773 |
| ENSMUSG00000047528 | Flacc1 | 0 | -1.051604167 | 0.898863889 |
| ENSMUSG00000026027 | Stradb | 0 | -1.43862729 | 0.538110273 |
| ENSMUSG00000079550 | Mpp4 | 0 | 0.063873051 | 1 |
| ENSMUSG00000101586 | Gm29017 | 0 | 1.648835551 | 1 |
| ENSMUSG00000099922 | Gm29016 | 0 | 1.648835551 | 1 |
| ENSMUSG00000026018 | Ica1l | 0 | -0.351164449 | 1 |
| ENSMUSG00000085965 | 2310016D23Rik | 0 | 0.326907457 | 1 |
| ENSMUSG00000085447 | 9530026F06Rik | 0 | 0.648835551 | 1 |
| ENSMUSG00000086408 | Gm11588 | 0 | 1.063873051 | 1 |
| ENSMUSG00000086009 | 4930587A21Rik | 0 | 0.648835551 | 1 |
| ENSMUSG00000085019 | Pard3bos2 | 0 | 1.648835551 | 1 |
| ENSMUSG00000085400 | Pard3bos3 | 0 | 0.648835551 | 1 |
| ENSMUSG00000100426 | Gm4208 | 0 | 1.648835551 | 1 |
| ENSMUSG00000027520 | Zdbf2 | 0 | -0.158519371 | 1 |
| ENSMUSG00000115378 | Gm39653 | 0 | 1.648835551 | 1 |
| ENSMUSG00000025963 | Mdh1b | 0 | 0.063873051 | 1 |
| ENSMUSG00000025962 | Fastkd2 | 0 | -2.636566667 | 0.112813335 |
| ENSMUSG00000025961 | 4933402D24Rik | 0 | 0.063873051 | 1 |
| ENSMUSG00000086836 | Gm13748 | 0 | 1.648835551 | 1 |
| ENSMUSG00000087213 | 2810408I11Rik | 0 | -0.158519371 | 1 |
| ENSMUSG00000070871 | Ccnyl1 | 0 | -3.079084903 | 0.042197017 |
| ENSMUSG00000025955 | Akr1cl | 0 | 0.326907457 | 1 |
| ENSMUSG00000044816 | D630023F18Rik | 0 | -0.810596067 | 0.847265357 |
| ENSMUSG00000025946 | Pth2r | 0 | -1.43862729 | 0.538110273 |
| ENSMUSG00000061816 | Myl1 | 0 | -1.995020638 | 0.416154643 |
| ENSMUSG00000026000 | Lancl1 | 0 | -3.858959089 | 0.002420614 |
| ENSMUSG00000085075 | Gm15669 | 0 | 1.063873051 | 1 |
| ENSMUSG00000100815 | Gm29112 | 0 | 1.648835551 | 1 |
| ENSMUSG00000099954 | Gm28112 | 0 | 1.648835551 | 1 |
| ENSMUSG00000099707 | Gm8883 | 0 | 1.648835551 | 1 |
| ENSMUSG00000101211 | Gm28818 | 0 | 0.063873051 | 1 |
| ENSMUSG00000101848 | 4933417E11Rik | 0 | 0.648835551 | 1 |
| ENSMUSG00000101764 | 4930556G22Rik | 0 | 1.063873051 | 1 |
| ENSMUSG00000102123 | Gm4319 | 0 | 1.063873051 | 1 |
| ENSMUSG00000026188 | Tmem169 | 0 | 0.063873051 | 1 |
| ENSMUSG00000101968 | 1700027A15Rik | 0 | 1.063873051 | 1 |
| ENSMUSG00000101812 | D530049I02Rik | 0 | 0.648835551 | 1 |
| ENSMUSG00000099931 | Gm29358 | 0 | 1.063873051 | 1 |
| ENSMUSG00000073650 | Catip | 0 | 1.063873051 | 1 |
| ENSMUSG00000026171 | Rnf25 | 0 | -2.209145444 | 0.217194106 |
| ENSMUSG00000033276 | Stk36 | 0 | -2.258055044 | 0.290922985 |
| ENSMUSG00000026167 | Wnt10a | 0 | -0.351164449 | 1 |
| ENSMUSG00000089964 | Gm16582 | 0 | 0.063873051 | 1 |
| ENSMUSG00000090071 | Cdk5r2 | 0 | 1.063873051 | 1 |
| ENSMUSG00000055197 | Fev | 0 | 1.648835551 | 1 |
| ENSMUSG00000006538 | Ihh | 0 | 0.063873051 | 1 |

| | | | | |
|---|---|---|---|---|
| ENSMUSG00000026205 | Slc23a3 | 0 | 0.063873051 | 1 |
| ENSMUSG00000026198 | Abcb6 | 0 | -1.051604167 | 1 |
| ENSMUSG00000033061 | Resp18 | 0 | 1.063873051 | 1 |
| ENSMUSG00000100556 | Gm28902 | 0 | 1.648835551 | 1 |
| ENSMUSG00000033007 | Asic4 | 0 | -0.351164449 | 1 |
| ENSMUSG00000026211 | Obsl1 | 0 | -0.936126949 | 0.858831093 |
| ENSMUSG00000051703 | Tmem198 | 0 | 1.648835551 | 1 |
| ENSMUSG00000090486 | BC035947 | 0 | 0.063873051 | 1 |
| ENSMUSG00000100562 | Gm29611 | 0 | 1.063873051 | 1 |
| ENSMUSG00000047330 | Kcne4 | 0 | 0.326907457 | 1 |
| ENSMUSG00000101324 | Gm29187 | 0 | 1.648835551 | 1 |
| ENSMUSG00000026248 | Mrpl44 | 0 | -1.810596067 | 0.466464457 |
| ENSMUSG00000043230 | Fam124b | 0 | 0.648835551 | 1 |
| ENSMUSG00000022591 | Gm9747 | 0 | 1.648835551 | 1 |
| ENSMUSG00000100786 | Gm28940 | 0 | 1.063873051 | 1 |
| ENSMUSG00000097564 | C430014B12Rik | 0 | -0.351164449 | 1 |
| ENSMUSG00000113880 | A030005L19Rik | 0 | 1.063873051 | 1 |
| ENSMUSG00000114011 | Gm47955 | 0 | 1.648835551 | 1 |
| ENSMUSG00000038496 | Slc19a3 | 0 | -0.158519371 | 1 |
| ENSMUSG00000113267 | Gm47969 | 0 | 0.648835551 | 1 |
| ENSMUSG00000089844 | A530032D15Rik | 0 | 1.063873051 | 1 |
| ENSMUSG00000073631 | Gm10553 | 0 | 1.648835551 | 1 |
| ENSMUSG00000072109 | A530040E14Rik | 0 | 0.063873051 | 1 |
| ENSMUSG00000089652 | Gm16025 | 0 | 1.063873051 | 1 |
| ENSMUSG00000089714 | Gm16092 | 0 | 1.648835551 | 1 |
| ENSMUSG00000094127 | G530012D18Rik | 0 | 1.648835551 | 1 |
| ENSMUSG00000052760 | A630001G21Rik | 0 | -1.351164449 | 0.702214947 |
| ENSMUSG00000100217 | 9930111H07Rik | 0 | 1.648835551 | 1 |
| ENSMUSG00000026226 | Spata3 | 0 | 1.648835551 | 1 |
| ENSMUSG00000026227 | 2810459M11Rik | 0 | -0.936126949 | 0.847265357 |
| ENSMUSG00000026228 | Htr2b | 0 | 0.326907457 | 1 |
| ENSMUSG00000087538 | Gm16341 | 0 | -0.810596067 | 0.844591606 |
| ENSMUSG00000100768 | Gm29055 | 0 | -1.673092543 | 0.451785624 |
| ENSMUSG00000101625 | Gm29371 | 0 | 1.063873051 | 1 |
| ENSMUSG00000026251 | Chrnd | 0 | 1.648835551 | 1 |
| ENSMUSG00000026253 | Chrng | 0 | 1.063873051 | 1 |
| ENSMUSG00000079436 | Kcnj13 | 0 | 0.063873051 | 1 |
| ENSMUSG00000090124 | Ugt1a7c | 0 | -0.810596067 | 0.844591606 |
| ENSMUSG00000081984 | Dnajb3 | 0 | 1.648835551 | 1 |
| ENSMUSG00000089960 | Ugt1a1 | 0 | -0.936126949 | 0.847265357 |
| ENSMUSG00000099553 | Gm29538 | 0 | 0.063873051 | 1 |
| ENSMUSG00000100908 | Gm28888 | 0 | -0.810596067 | 0.844591606 |
| ENSMUSG00000100367 | Gm29336 | 0 | 1.648835551 | 1 |
| ENSMUSG00000099848 | Gm29337 | 0 | 0.648835551 | 1 |
| ENSMUSG00000099938 | C030007H22Rik | 0 | -1.258055044 | 0.673811773 |
| ENSMUSG00000034486 | Gbx2 | 0 | 1.648835551 | 1 |
| ENSMUSG00000092627 | D130058E05Rik | 0 | 1.063873051 | 1 |
| ENSMUSG00000100850 | 4930434B07Rik | 0 | 1.648835551 | 1 |
| ENSMUSG00000101951 | Gm28721 | 0 | 1.648835551 | 1 |
| ENSMUSG00000049515 | Espnl | 0 | 0.648835551 | 1 |
| ENSMUSG00000026308 | Klhl30 | 0 | 1.063873051 | 1 |

| | | | | |
|---|---|---|---|---|
| ENSMUSG00000047443 | Erfe | 0 | 0.326907457 | 1 |
| ENSMUSG00000026311 | Asb1 | 0 | -1.258055044 | 0.673811773 |
| ENSMUSG00000101705 | Gm28380 | 0 | 1.648835551 | 1 |
| ENSMUSG00000100980 | Gm29100 | 0 | 1.648835551 | 1 |
| ENSMUSG00000007805 | Twist2 | 0 | -0.351164449 | 1 |
| ENSMUSG00000101414 | Gm29101 | 0 | -0.673092543 | 1 |
| ENSMUSG00000101265 | Gm29099 | 0 | 1.648835551 | 1 |
| ENSMUSG00000067064 | Olfr1416 | 0 | 1.648835551 | 1 |
| ENSMUSG00000042849 | Olfr1414 | 0 | 1.063873051 | 1 |
| ENSMUSG00000101995 | Gm29480 | 0 | 1.063873051 | 1 |
| ENSMUSG00000045091 | Aqp12 | 0 | 1.648835551 | 1 |
| ENSMUSG00000026272 | Agxt | 0 | 1.063873051 | 1 |
| ENSMUSG00000034159 | Mab21l4 | 0 | -2.395558568 | 0.174915761 |
| ENSMUSG00000101172 | Gm28535 | 0 | 1.063873051 | 1 |
| ENSMUSG00000073609 | D2hgdh | 0 | -1.599091962 | 0.559918909 |
| ENSMUSG00000094651 | Gal3st2 | 0 | 0.326907457 | 1 |
| ENSMUSG00000093805 | Gal3st2b | 0 | -2.209145444 | 0.217194106 |
| ENSMUSG00000073608 | Gal3st2c | 0 | -0.351164449 | 1 |
| ENSMUSG00000100777 | 4930598F16Rik | 0 | 0.648835551 | 1 |
| ENSMUSG00000100062 | 1700063A18Rik | 0 | -0.52108945 | 1 |
| ENSMUSG00000097814 | Panct2 | 0 | 0.063873051 | 1 |
| ENSMUSG00000100454 | Gm28901 | 0 | 1.648835551 | 1 |
| ENSMUSG00000026331 | Slco6c1 | 0 | -1.158519371 | 0.886203497 |
| ENSMUSG00000089706 | B230216N24Rik | 0 | -0.351164449 | 1 |
| ENSMUSG00000100009 | Gm7967 | 0 | 1.063873051 | 1 |
| ENSMUSG00000089866 | Gm15699 | 0 | 1.648835551 | 1 |
| ENSMUSG00000097482 | Gm17634 | 0 | -0.351164449 | 1 |
| ENSMUSG00000100247 | A530053M12Rik | 0 | 1.648835551 | 1 |
| ENSMUSG00000059956 | Serpinb12 | 0 | 0.326907457 | 1 |
| ENSMUSG00000048775 | Serpinb13 | 0 | 1.063873051 | 1 |
| ENSMUSG00000044594 | Serpinb3a | 0 | 1.063873051 | 1 |
| ENSMUSG00000087210 | Gm15391 | 0 | 1.648835551 | 1 |
| ENSMUSG00000026315 | Serpinb8 | 0 | 0.648835551 | 1 |
| ENSMUSG00000038702 | Dsel | 0 | 1.648835551 | 1 |
| ENSMUSG00000094429 | Gm19965 | 0 | 1.648835551 | 1 |
| ENSMUSG00000100617 | Gm7145 | 0 | 1.063873051 | 1 |
| ENSMUSG00000102030 | Gm29106 | 0 | 1.648835551 | 1 |
| ENSMUSG00000099936 | Gm28867 | 0 | 1.063873051 | 1 |
| ENSMUSG00000097408 | Gm26831 | 0 | -0.673092543 | 1 |
| ENSMUSG00000099860 | Gm29456 | 0 | 0.326907457 | 1 |
| ENSMUSG00000098923 | Tmem185b | 0 | -1.936126949 | 0.358945041 |
| ENSMUSG00000101731 | 3830432H09Rik | 0 | 1.648835551 | 1 |
| ENSMUSG00000101224 | Gm28209 | 0 | 0.648835551 | 1 |
| ENSMUSG00000101386 | Gm29346 | 0 | 1.648835551 | 1 |
| ENSMUSG00000101700 | Gm29359 | 0 | 1.648835551 | 1 |
| ENSMUSG00000050534 | Htr5b | 0 | 1.648835551 | 1 |
| ENSMUSG00000099574 | 1700121L03Rik | 0 | 1.648835551 | 1 |
| ENSMUSG00000045382 | Cxcr4 | 0 | 0.063873051 | 1 |
| ENSMUSG00000026401 | Cd55b | 0 | 0.648835551 | 1 |
| ENSMUSG00000087691 | Gm15674 | 0 | 1.063873051 | 1 |
| ENSMUSG00000026405 | C4bp | 0 | 0.063873051 | 1 |

| | | | | |
|---|---|---|---|---|
| ENSMUSG00000042474 | Fcmr | 0 | 1.063873051 | 1 |
| ENSMUSG00000016529 | Il10 | 0 | 1.648835551 | 1 |
| ENSMUSG00000102055 | Gm28913 | 0 | 1.648835551 | 1 |
| ENSMUSG00000101778 | Gm29488 | 0 | 0.648835551 | 1 |
| ENSMUSG00000055184 | Fam72a | 0 | -0.52108945 | 1 |
| ENSMUSG00000026433 | Rab29 | 0 | -2.051604167 | 0.315094216 |
| ENSMUSG00000118219 | Gm29695 | 0 | 1.648835551 | 1 |
| ENSMUSG00000101549 | Gm29630 | 0 | 1.063873051 | 1 |
| ENSMUSG00000042115 | Klhdc8a | 0 | 1.648835551 | 1 |
| ENSMUSG00000042046 | Dstyk | 0 | -3.777429203 | 0.00334584 |
| ENSMUSG00000097760 | 6030442K20Rik | 0 | -2.305360759 | 0.262075002 |
| ENSMUSG00000053024 | Cntn2 | 0 | -0.810596067 | 0.844591606 |
| ENSMUSG00000101126 | Gm10538 | 0 | 1.648835551 | 1 |
| ENSMUSG00000026443 | Lrrn2 | 0 | -0.673092543 | 1 |
| ENSMUSG00000101102 | Gm28609 | 0 | -0.351164449 | 1 |
| ENSMUSG00000100830 | Gm28441 | 0 | 1.648835551 | 1 |
| ENSMUSG00000041577 | Prelp | 0 | -0.158519371 | 1 |
| ENSMUSG00000100987 | Gm1627 | 0 | 1.648835551 | 1 |
| ENSMUSG00000041559 | Fmod | 0 | 0.063873051 | 1 |
| ENSMUSG00000042429 | Adora1 | 0 | -0.52108945 | 1 |
| ENSMUSG00000046367 | Mgat4e | 0 | -0.673092543 | 1 |
| ENSMUSG00000097435 | Gm26783 | 0 | 1.063873051 | 1 |
| ENSMUSG00000101168 | Gm28892 | 0 | 1.063873051 | 1 |
| ENSMUSG00000097680 | Gm26642 | 0 | 0.648835551 | 1 |
| ENSMUSG00000054412 | Gm4793 | 0 | 1.063873051 | 1 |
| ENSMUSG00000026413 | Pkp1 | 0 | -1.43862729 | 0.736456705 |
| ENSMUSG00000086264 | Gm15850 | 0 | 1.648835551 | 1 |
| ENSMUSG00000097315 | Gm26568 | 0 | 1.648835551 | 1 |
| ENSMUSG00000097433 | Gm26781 | 0 | 1.063873051 | 1 |
| ENSMUSG00000097403 | 9230116N13Rik | 0 | -0.158519371 | 1 |
| ENSMUSG00000098045 | Gm26979 | 0 | 1.648835551 | 1 |
| ENSMUSG00000098145 | Gm26936 | 0 | 1.648835551 | 1 |
| ENSMUSG00000098206 | A430106G13Rik | 0 | 1.648835551 | 1 |
| ENSMUSG00000026394 | Atp6v1g3 | 0 | 1.648835551 | 1 |
| ENSMUSG00000100459 | Gm5833 | 0 | -0.52108945 | 1 |
| ENSMUSG00000104900 | 4930596I21Rik | 0 | 1.648835551 | 1 |
| ENSMUSG00000103844 | 4933436E23Rik | 0 | 0.326907457 | 1 |
| ENSMUSG00000070594 | Gm4788 | 0 | 0.063873051 | 1 |
| ENSMUSG00000033898 | Cfhr2 | 0 | 0.326907457 | 1 |
| ENSMUSG00000102368 | 4930590L20Rik | 0 | -1.158519371 | 0.886203497 |
| ENSMUSG00000033849 | B3galt2 | 0 | 1.648835551 | 1 |
| ENSMUSG00000100455 | Gm29170 | 0 | -0.158519371 | 1 |
| ENSMUSG00000098509 | Rgs21 | 0 | 1.063873051 | 1 |
| ENSMUSG00000099629 | Gm29515 | 0 | 1.648835551 | 1 |
| ENSMUSG00000101011 | Gm29398 | 0 | -0.351164449 | 1 |
| ENSMUSG00000006007 | Pdc | 0 | 1.648835551 | 1 |
| ENSMUSG00000099447 | 2310030A07Rik | 0 | 1.063873051 | 1 |
| ENSMUSG00000006014 | Prg4 | 0 | 0.648835551 | 1 |
| ENSMUSG00000100594 | 2810414N06Rik | 0 | 1.648835551 | 1 |
| ENSMUSG00000014980 | Tsen15 | 0 | -1.051604167 | 0.917930288 |
| ENSMUSG00000055547 | Apobec4 | 0 | -0.673092543 | 1 |

| | | | | |
|---|---|---|---|---|
| ENSMUSG00000100394 | Gm28791 | 0 | -2.209145444 | 0.217194106 |
| ENSMUSG00000042708 | Shcbp1l | 0 | 1.063873051 | 1 |
| ENSMUSG00000100945 | Gm29529 | 0 | 1.648835551 | 1 |
| ENSMUSG00000100334 | C230024C17Rik | 0 | 0.648835551 | 1 |
| ENSMUSG00000100183 | Gm28512 | 0 | 1.648835551 | 1 |
| ENSMUSG00000100502 | Gm28286 | 0 | 1.063873051 | 1 |
| ENSMUSG00000066797 | Zfp648 | 0 | 1.063873051 | 1 |
| ENSMUSG00000100221 | 4930532M18Rik | 0 | 1.648835551 | 1 |
| ENSMUSG00000101880 | Gm29282 | 0 | -0.52108945 | 1 |
| ENSMUSG00000096950 | Gm9530 | 0 | -0.810596067 | 1 |
| ENSMUSG00000033722 | BC034090 | 0 | -0.673092543 | 1 |
| ENSMUSG00000102558 | Gm37571 | 0 | 0.063873051 | 1 |
| ENSMUSG00000060985 | Tdrd5 | 0 | -0.52108945 | 1 |
| ENSMUSG00000103705 | 4930518J20Rik | 0 | 1.648835551 | 1 |
| ENSMUSG00000026602 | Nphs2 | 0 | 1.648835551 | 1 |
| ENSMUSG00000033544 | Angptl1 | 0 | 1.648835551 | 1 |
| ENSMUSG00000104014 | 4930562F17Rik | 0 | 0.326907457 | 1 |
| ENSMUSG00000026725 | Tnn | 0 | -0.52108945 | 1 |
| ENSMUSG00000103693 | Gm37529 | 0 | -2.305360759 | 0.231596497 |
| ENSMUSG00000118401 | Gpr52 | 0 | 1.648835551 | 1 |
| ENSMUSG00000102852 | Gm37083 | 0 | 1.063873051 | 1 |
| ENSMUSG00000102864 | Gm37634 | 0 | 1.648835551 | 1 |
| ENSMUSG00000044835 | Ankrd45 | 0 | 0.326907457 | 1 |
| ENSMUSG00000026697 | Myoc | 0 | 0.063873051 | 1 |
| ENSMUSG00000091060 | Myocos | 0 | 0.648835551 | 1 |
| ENSMUSG00000101869 | Gm29500 | 0 | 1.648835551 | 1 |
| ENSMUSG00000040113 | Mettl11b | 0 | 0.648835551 | 1 |
| ENSMUSG00000026581 | Sell | 0 | -0.351164449 | 1 |
| ENSMUSG00000086593 | Gm16548 | 0 | 1.648835551 | 1 |
| ENSMUSG00000026579 | F5 | 0 | -2.258055044 | 0.262284476 |
| ENSMUSG00000040918 | Slc19a2 | 0 | -0.936126949 | 0.847265357 |
| ENSMUSG00000104011 | Gm32391 | 0 | 1.648835551 | 1 |
| ENSMUSG00000103383 | Gm37754 | 0 | 0.648835551 | 1 |
| ENSMUSG00000102163 | Gm36945 | 0 | 1.063873051 | 1 |
| ENSMUSG00000026573 | Xcl1 | 0 | 1.648835551 | 1 |
| ENSMUSG00000026572 | Tbx19 | 0 | 0.326907457 | 1 |
| ENSMUSG00000040836 | Gpr161 | 0 | -2.636566667 | 0.112813335 |
| ENSMUSG00000102401 | Gm37864 | 0 | 0.326907457 | 1 |
| ENSMUSG00000103686 | Gm37073 | 0 | 1.063873051 | 1 |
| ENSMUSG00000104306 | Gm36972 | 0 | 1.648835551 | 1 |
| ENSMUSG00000097196 | Gm26665 | 0 | -1.599091962 | 0.637223959 |
| ENSMUSG00000091393 | 5330438I03Rik | 0 | 1.648835551 | 1 |
| ENSMUSG00000103400 | Gm15853 | 0 | 1.063873051 | 1 |
| ENSMUSG00000086056 | Gm4846 | 0 | -0.158519371 | 1 |
| ENSMUSG00000051081 | Gm4847 | 0 | 0.326907457 | 1 |
| ENSMUSG00000026560 | Fmo9 | 0 | 0.648835551 | 1 |
| ENSMUSG00000102548 | Gm16701 | 0 | -2.258055044 | 0.290722159 |
| ENSMUSG00000102909 | Gm2453 | 0 | 0.648835551 | 1 |
| ENSMUSG00000040485 | Lrrc52 | 0 | 1.063873051 | 1 |
| ENSMUSG00000015843 | Rxrg | 0 | -0.351164449 | 1 |
| ENSMUSG00000103530 | Gm37839 | 0 | 1.063873051 | 1 |

| | | | | |
|---|---|---|---|---|
| ENSMUSG00000038530 | Rgs4 | 0 | 1.648835551 | 1 |
| ENSMUSG00000103908 | Gm37502 | 0 | 1.648835551 | 1 |
| ENSMUSG00000073494 | Sh2d1b2 | 0 | 1.063873051 | 1 |
| ENSMUSG00000096968 | Gm26620 | 0 | 1.648835551 | 1 |
| ENSMUSG00000070524 | Fcrlb | 0 | 1.063873051 | 1 |
| ENSMUSG00000038421 | Fcrla | 0 | 0.648835551 | 1 |
| ENSMUSG00000103676 | Gm38204 | 0 | 1.648835551 | 1 |
| ENSMUSG00000104340 | Gm10522 | 0 | 1.648835551 | 1 |
| ENSMUSG00000005677 | Nr1i3 | 0 | 0.648835551 | 1 |
| ENSMUSG00000062729 | Ppox | 0 | -3.417253639 | 0.010795678 |
| ENSMUSG00000103983 | Gm20045 | 0 | -2.351164449 | 0.169846604 |
| ENSMUSG00000045259 | Klhdc9 | 0 | 1.648835551 | 1 |
| ENSMUSG00000103711 | Tstd1 | 0 | 1.063873051 | 1 |
| ENSMUSG00000060244 | Alyref2 | 0 | 1.648835551 | 1 |
| ENSMUSG00000004707 | Ly9 | 0 | 0.648835551 | 1 |
| ENSMUSG00000104413 | Gm37065 | 0 | 0.648835551 | 1 |
| ENSMUSG00000103380 | Gm37756 | 0 | 1.648835551 | 1 |
| ENSMUSG00000007122 | Casq1 | 0 | 0.063873051 | 1 |
| ENSMUSG00000007107 | Atp1a4 | 0 | 0.326907457 | 1 |
| ENSMUSG00000038026 | Kcnj9 | 0 | 0.648835551 | 1 |
| ENSMUSG00000026548 | Slamf9 | 0 | 1.648835551 | 1 |
| ENSMUSG00000043263 | Ifi209 | 0 | -0.158519371 | 1 |
| ENSMUSG00000066677 | Ifi208 | 0 | 0.326907457 | 1 |
| ENSMUSG00000090272 | Mndal | 0 | -2.158519371 | 0.290284345 |
| ENSMUSG00000054203 | Ifi205 | 0 | 0.326907457 | 1 |
| ENSMUSG00000059371 | Olfr427 | 0 | 1.648835551 | 1 |
| ENSMUSG00000066672 | Olfr417 | 0 | 1.648835551 | 1 |
| ENSMUSG00000051509 | Olfr414 | 0 | 1.648835551 | 1 |
| ENSMUSG00000026525 | Opn3 | 0 | 0.063873051 | 1 |
| ENSMUSG00000104157 | Gm38101 | 0 | 1.648835551 | 1 |
| ENSMUSG00000103043 | Gm37306 | 0 | 1.063873051 | 1 |
| ENSMUSG00000086350 | B230369F24Rik | 0 | 1.063873051 | 1 |
| ENSMUSG00000026495 | Efcab2 | 0 | -2.560617814 | 0.110800285 |
| ENSMUSG00000104520 | Gm37336 | 0 | 1.648835551 | 1 |
| ENSMUSG00000026497 | Mixl1 | 0 | 1.648835551 | 1 |
| ENSMUSG00000103156 | Gm38293 | 0 | -0.351164449 | 1 |
| ENSMUSG00000073485 | H3f3aos | 0 | 0.648835551 | 1 |
| ENSMUSG00000066652 | Lefty2 | 0 | 0.648835551 | 1 |
| ENSMUSG00000038793 | Lefty1 | 0 | -0.158519371 | 1 |
| ENSMUSG00000085563 | 2210411M09Rik | 0 | 0.063873051 | 1 |
| ENSMUSG00000038768 | 9130409I23Rik | 0 | 0.648835551 | 1 |
| ENSMUSG00000091013 | Vmn1r1 | 0 | 1.063873051 | 1 |
| ENSMUSG00000100147 | 1700047M11Rik | 0 | 0.326907457 | 1 |
| ENSMUSG00000079164 | Tlr5 | 0 | -1.158519371 | 0.909161999 |
| ENSMUSG00000104212 | Gm37986 | 0 | 0.326907457 | 1 |
| ENSMUSG00000053461 | Hhipl2 | 0 | 1.063873051 | 1 |
| ENSMUSG00000092578 | 4930488B22Rik | 0 | 1.063873051 | 1 |
| ENSMUSG00000039377 | Hlx | 0 | 0.063873051 | 1 |
| ENSMUSG00000026621 | 1-Mar | 0 | 0.326907457 | 1 |
| ENSMUSG00000026620 | Mark1 | 0 | -2.395558568 | 0.191663663 |
| ENSMUSG00000103613 | Gm34882 | 0 | 1.648835551 | 1 |

| | | | | |
|---|---|---|---|---|
| ENSMUSG00000104368 | 2010103J01Rik | 0 | 1.648835551 | 1 |
| ENSMUSG00000097261 | 4930532G15Rik | 0 | 0.326907457 | 1 |
| ENSMUSG00000051295 | 9630028B13Rik | 0 | 1.063873051 | 1 |
| ENSMUSG00000097615 | Gm2061 | 0 | 1.648835551 | 1 |
| ENSMUSG00000103799 | Gm38093 | 0 | 1.648835551 | 1 |
| ENSMUSG00000103690 | A730004F24Rik | 0 | 1.648835551 | 1 |
| ENSMUSG00000100393 | Gm29491 | 0 | 1.648835551 | 1 |
| ENSMUSG00000097784 | A430105J06Rik | 0 | -2.209145444 | 0.217194106 |
| ENSMUSG00000103503 | 1700023G09Rik | 0 | 1.648835551 | 1 |
| ENSMUSG00000089730 | 1700007P06Rik | 0 | 1.648835551 | 1 |
| ENSMUSG00000103923 | Gm37896 | 0 | 1.063873051 | 1 |
| ENSMUSG00000104482 | Gm38064 | 0 | 1.648835551 | 1 |
| ENSMUSG00000097925 | Gm26574 | 0 | 1.063873051 | 1 |
| ENSMUSG00000104435 | Gm37422 | 0 | 1.063873051 | 1 |
| ENSMUSG00000010175 | Prox1 | 0 | -1.810596067 | 0.465580427 |
| ENSMUSG00000103644 | Gm29678 | 0 | 0.648835551 | 1 |
| ENSMUSG00000066595 | Flvcr1 | 0 | -1.258055044 | 0.673811773 |
| ENSMUSG00000097845 | A230020J21Rik | 0 | 1.648835551 | 1 |
| ENSMUSG00000026632 | Tatdn3 | 0 | -0.936126949 | 0.847265357 |
| ENSMUSG00000062510 | Nsl1 | 0 | -1.874726405 | 0.352215149 |
| ENSMUSG00000091017 | Fam71a | 0 | 1.648835551 | 1 |
| ENSMUSG00000103174 | Gm37168 | 0 | 0.648835551 | 1 |
| ENSMUSG00000103517 | Gm37432 | 0 | 1.648835551 | 1 |
| ENSMUSG00000102840 | Gm38037 | 0 | 1.648835551 | 1 |
| ENSMUSG00000097416 | Gm26670 | 0 | 0.063873051 | 1 |
| ENSMUSG00000097089 | Gm26879 | 0 | 1.648835551 | 1 |
| ENSMUSG00000097316 | Gm10516 | 0 | -3.106051951 | 0.028853812 |
| ENSMUSG00000089812 | Gm15867 | 0 | 1.648835551 | 1 |
| ENSMUSG00000037318 | Traf3ip3 | 0 | -1.258055044 | 0.736043208 |
| ENSMUSG00000090080 | Gm15872 | 0 | 1.648835551 | 1 |
| ENSMUSG00000089683 | 4930570N18Rik | 0 | 1.648835551 | 1 |
| ENSMUSG00000016179 | Camk1g | 0 | 0.648835551 | 1 |
| ENSMUSG00000103692 | 4930503O07Rik | 0 | 1.063873051 | 1 |
| ENSMUSG00000104206 | Gm32250 | 0 | -0.52108945 | 1 |
| ENSMUSG00000026646 | Suv39h2 | 0 | 1.063873051 | 1 |
| ENSMUSG00000039496 | Cdnf | 0 | 1.063873051 | 1 |
| ENSMUSG00000085205 | Gm13185 | 0 | 1.063873051 | 1 |
| ENSMUSG00000085656 | Gm13180 | 0 | 1.063873051 | 1 |
| ENSMUSG00000087355 | Gm13187 | 0 | 0.063873051 | 1 |
| ENSMUSG00000102494 | Gm36988 | 0 | 1.063873051 | 1 |
| ENSMUSG00000026668 | Ucma | 0 | 0.648835551 | 1 |
| ENSMUSG00000086430 | 4930551O13Rik | 0 | 0.648835551 | 1 |
| ENSMUSG00000026676 | Ccdc3 | 0 | -1.874726405 | 0.370799436 |
| ENSMUSG00000085818 | Gm13267 | 0 | 1.063873051 | 1 |
| ENSMUSG00000085580 | 1700061F12Rik | 0 | 0.326907457 | 1 |
| ENSMUSG00000079603 | Gm13218 | 0 | -0.351164449 | 1 |
| ENSMUSG00000086618 | Gm13256 | 0 | 1.648835551 | 1 |
| ENSMUSG00000025783 | 4930412O13Rik | 0 | 1.648835551 | 1 |
| ENSMUSG00000075534 | Gm13262 | 0 | 1.648835551 | 1 |
| ENSMUSG00000102329 | Gm10851 | 0 | -0.351164449 | 1 |
| ENSMUSG00000026770 | Il2ra | 0 | -0.673092543 | 1 |

| | | | | |
|---|---|---|---|---|
| ENSMUSG00000047909 | Ankrd16 | 0 | -2.52108945 | 0.139351384 |
| ENSMUSG00000086843 | E030013I19Rik | 0 | 1.648835551 | 1 |
| ENSMUSG00000049630 | C1ql3 | 0 | 0.648835551 | 1 |
| ENSMUSG00000104354 | Gm37126 | 0 | 1.648835551 | 1 |
| ENSMUSG00000061531 | Tmem236 | 0 | -0.158519371 | 1 |
| ENSMUSG00000104106 | Gm10848 | 0 | -0.158519371 | 1 |
| ENSMUSG00000085319 | Gm13330 | 0 | 1.648835551 | 1 |
| ENSMUSG00000054074 | Skida1 | 0 | 0.326907457 | 1 |
| ENSMUSG00000092353 | Gm20539 | 0 | -0.52108945 | 1 |
| ENSMUSG00000087674 | 4930447M23Rik | 0 | 1.648835551 | 1 |
| ENSMUSG00000087103 | Gm13344 | 0 | 1.063873051 | 1 |
| ENSMUSG00000056329 | 1810010K12Rik | 0 | 1.648835551 | 1 |
| ENSMUSG00000087246 | Gm13346 | 0 | 0.326907457 | 1 |
| ENSMUSG00000091457 | Gm17171 | 0 | 1.063873051 | 1 |
| ENSMUSG00000085477 | Gm13335 | 0 | 1.063873051 | 1 |
| ENSMUSG00000085206 | Gm13380 | 0 | 0.648835551 | 1 |
| ENSMUSG00000075514 | Gm13375 | 0 | 1.648835551 | 1 |
| ENSMUSG00000087219 | Gm13376 | 0 | 1.648835551 | 1 |
| ENSMUSG00000026679 | Enkur | 0 | -1.258055044 | 0.703415131 |
| ENSMUSG00000048550 | Thnsl1 | 0 | -1.258055044 | 0.673811773 |
| ENSMUSG00000085353 | 1700092C17Rik | 0 | 1.063873051 | 1 |
| ENSMUSG00000069132 | Nxph2 | 0 | 1.063873051 | 1 |
| ENSMUSG00000061809 | Tbpl2 | 0 | 0.063873051 | 1 |
| ENSMUSG00000044103 | Il1f9 | 0 | 1.063873051 | 1 |
| ENSMUSG00000026984 | Il1f6 | 0 | 1.648835551 | 1 |
| ENSMUSG00000046845 | Il1f10 | 0 | 1.648835551 | 1 |
| ENSMUSG00000085330 | Gm13417 | 0 | 0.648835551 | 1 |
| ENSMUSG00000036813 | Entpd8 | 0 | 1.648835551 | 1 |
| ENSMUSG00000036805 | Noxa1 | 0 | 0.063873051 | 1 |
| ENSMUSG00000078202 | Nrarp | 0 | 0.063873051 | 1 |
| ENSMUSG00000059555 | Tor4a | 0 | 1.063873051 | 1 |
| ENSMUSG00000085830 | Grin1os | 0 | 0.648835551 | 1 |
| ENSMUSG00000026955 | Sapcd2 | 0 | 0.063873051 | 1 |
| ENSMUSG00000015093 | Clic3 | 0 | -0.810596067 | 0.844591606 |
| ENSMUSG00000015083 | C8g | 0 | 1.648835551 | 1 |
| ENSMUSG00000026941 | Mamdc4 | 0 | -0.351164449 | 1 |
| ENSMUSG00000029419 | Ajm1 | 0 | 0.648835551 | 1 |
| ENSMUSG00000062061 | Obp2a | 0 | 1.648835551 | 1 |
| ENSMUSG00000058740 | Kcnt1 | 0 | -0.810596067 | 0.844591606 |
| ENSMUSG00000026934 | Lhx3 | 0 | 1.648835551 | 1 |
| ENSMUSG00000048038 | Ccdc187 | 0 | 0.326907457 | 1 |
| ENSMUSG00000086779 | Gm13562 | 0 | 1.063873051 | 1 |
| ENSMUSG00000085767 | Gm13563 | 0 | -1.051604167 | 0.858831093 |
| ENSMUSG00000086714 | 0610009E02Rik | 0 | -0.936126949 | 0.847265357 |
| ENSMUSG00000036186 | Dipk1b | 0 | 1.063873051 | 1 |
| ENSMUSG00000014873 | Surf2 | 0 | -1.051604167 | 0.858831093 |
| ENSMUSG00000014852 | Adamts13 | 0 | -0.158519371 | 1 |
| ENSMUSG00000036067 | Slc2a6 | 0 | 1.648835551 | 1 |
| ENSMUSG00000036040 | Adamtsl2 | 0 | 0.326907457 | 1 |
| ENSMUSG00000085008 | Dbhos | 0 | 1.648835551 | 1 |
| ENSMUSG00000109946 | Brd3os | 0 | 1.063873051 | 1 |

| | | | | |
|---|---|---|---|---|
| ENSMUSG00000085929 | Gm13421 | 0 | -0.810596067 | 0.844591606 |
| ENSMUSG00000026833 | Olfm1 | 0 | -1.258055044 | 0.957378217 |
| ENSMUSG00000035829 | Ppp1r26 | 0 | 0.063873051 | 1 |
| ENSMUSG00000026818 | Cel | 0 | 1.063873051 | 1 |
| ENSMUSG00000087667 | Gm13381 | 0 | 0.063873051 | 1 |
| ENSMUSG00000026805 | Barhl1 | 0 | 1.648835551 | 1 |
| ENSMUSG00000079499 | 6530402F18Rik | 0 | 1.648835551 | 1 |
| ENSMUSG00000079497 | Gm13420 | 0 | 1.648835551 | 1 |
| ENSMUSG00000075425 | Gm13547 | 0 | 1.648835551 | 1 |
| ENSMUSG00000039798 | 2600006K01Rik | 0 | 1.063873051 | 1 |
| ENSMUSG00000039787 | Cercam | 0 | 0.063873051 | 1 |
| ENSMUSG00000015337 | Endog | 0 | 0.063873051 | 1 |
| ENSMUSG00000100747 | 1700084E18Rik | 0 | 1.063873051 | 1 |
| ENSMUSG00000075419 | Dolk | 0 | 1.648835551 | 1 |
| ENSMUSG00000026856 | Dolpp1 | 0 | -0.810596067 | 0.844591606 |
| ENSMUSG00000026853 | Crat | 0 | -2.209145444 | 0.417827589 |
| ENSMUSG00000085275 | Gm14487 | 0 | 1.648835551 | 1 |
| ENSMUSG00000047363 | Cstad | 0 | 0.063873051 | 1 |
| ENSMUSG00000087268 | Gm14486 | 0 | 1.063873051 | 1 |
| ENSMUSG00000085491 | 4930527E20Rik | 0 | 1.648835551 | 1 |
| ENSMUSG00000085224 | Gm13425 | 0 | 1.648835551 | 1 |
| ENSMUSG00000079466 | Prdm12 | 0 | 1.648835551 | 1 |
| ENSMUSG00000039356 | Exosc2 | 0 | -2.305360759 | 0.231596497 |
| ENSMUSG00000043102 | Qrfp | 0 | 1.063873051 | 1 |
| ENSMUSG00000086356 | Gm13441 | 0 | 1.648835551 | 1 |
| ENSMUSG00000001864 | Aif1l | 0 | -1.874726405 | 0.505481295 |
| ENSMUSG00000039164 | Naif1 | 0 | 1.063873051 | 1 |
| ENSMUSG00000085017 | Gm13412 | 0 | 0.648835551 | 1 |
| ENSMUSG00000046854 | Pip5kl1 | 0 | 1.063873051 | 1 |
| ENSMUSG00000009551 | 6330409D20Rik | 0 | 1.063873051 | 1 |
| ENSMUSG00000085397 | Gm13524 | 0 | 1.063873051 | 1 |
| ENSMUSG00000026791 | Slc2a8 | 0 | -0.158519371 | 1 |
| ENSMUSG00000004105 | Angptl2 | 0 | -2.305360759 | 0.231596497 |
| ENSMUSG00000085742 | Gm13536 | 0 | 1.063873051 | 1 |
| ENSMUSG00000085463 | 9430024E24Rik | 0 | 1.648835551 | 1 |
| ENSMUSG00000026873 | Phf19 | 0 | -0.158519371 | 1 |
| ENSMUSG00000026882 | 4930568D16Rik | 0 | 1.648835551 | 1 |
| ENSMUSG00000086555 | Gm13446 | 0 | 1.648835551 | 1 |
| ENSMUSG00000026894 | Morn5 | 0 | 0.326907457 | 1 |
| ENSMUSG00000026890 | Lhx6 | 0 | 1.063873051 | 1 |
| ENSMUSG00000087187 | Gm13431 | 0 | 1.648835551 | 1 |
| ENSMUSG00000075387 | Olfr341 | 0 | 1.648835551 | 1 |
| ENSMUSG00000111869 | Olfr342 | 0 | 1.648835551 | 1 |
| ENSMUSG00000111021 | Olfr50 | 0 | 1.063873051 | 1 |
| ENSMUSG00000075384 | Olfr3 | 0 | 1.648835551 | 1 |
| ENSMUSG00000050015 | Olfr350 | 0 | 1.063873051 | 1 |
| ENSMUSG00000075383 | Olfr351 | 0 | 1.648835551 | 1 |
| ENSMUSG00000053146 | Olfr352 | 0 | 1.648835551 | 1 |
| ENSMUSG00000055838 | Olfr357 | 0 | 1.648835551 | 1 |
| ENSMUSG00000075379 | Olfr358 | 0 | 1.648835551 | 1 |
| ENSMUSG00000078198 | Olfr364-ps1 | 0 | 1.063873051 | 1 |

| | | | | |
|---|---|---|---|---|
| ENSMUSG00000068947 | Olfr366 | 0 | -0.52108945 | 1 |
| ENSMUSG00000053164 | Gpr21 | 0 | -0.52108945 | 1 |
| ENSMUSG00000035403 | Crb2 | 0 | 0.063873051 | 1 |
| ENSMUSG00000000247 | Lhx2 | 0 | -0.351164449 | 1 |
| ENSMUSG00000087539 | Gm13584 | 0 | 1.648835551 | 1 |
| ENSMUSG00000087164 | Nr5a1os | 0 | 1.648835551 | 1 |
| ENSMUSG00000035295 | Wdr38 | 0 | -0.673092543 | 1 |
| ENSMUSG00000052248 | Zeb2os | 0 | -0.351164449 | 1 |
| ENSMUSG00000086349 | Gm13497 | 0 | 1.648835551 | 1 |
| ENSMUSG00000086007 | 1700057H21Rik | 0 | 1.648835551 | 1 |
| ENSMUSG00000036249 | Rbm43 | 0 | -0.673092543 | 1 |
| ENSMUSG00000087040 | Gm14033 | 0 | 1.648835551 | 1 |
| ENSMUSG00000086396 | 4930555B11Rik | 0 | -0.351164449 | 1 |
| ENSMUSG00000087455 | 5330411J11Rik | 0 | 0.648835551 | 1 |
| ENSMUSG00000086578 | Gm13583 | 0 | 0.326907457 | 1 |
| ENSMUSG00000075325 | Gm13582 | 0 | 1.648835551 | 1 |
| ENSMUSG00000035033 | Tbr1 | 0 | 0.648835551 | 1 |
| ENSMUSG00000087702 | Gm13575 | 0 | 1.063873051 | 1 |
| ENSMUSG00000086881 | Gm13594 | 0 | 1.063873051 | 1 |
| ENSMUSG00000057182 | Scn3a | 0 | -1.743481871 | 0.488578195 |
| ENSMUSG00000085386 | Gm13630 | 0 | 1.648835551 | 1 |
| ENSMUSG00000005232 | G6pc2 | 0 | 1.063873051 | 1 |
| ENSMUSG00000075307 | Klhl41 | 0 | 0.326907457 | 1 |
| ENSMUSG00000085735 | Mettl5os | 0 | -2.743481871 | 0.128647221 |
| ENSMUSG00000075304 | Sp5 | 0 | 1.648835551 | 1 |
| ENSMUSG00000085868 | Gm13632 | 0 | 1.063873051 | 1 |
| ENSMUSG00000084946 | Dlx1as | 0 | 0.648835551 | 1 |
| ENSMUSG00000041911 | Dlx1 | 0 | 1.063873051 | 1 |
| ENSMUSG00000064263 | Platr26 | 0 | 1.648835551 | 1 |
| ENSMUSG00000087570 | Gm13647 | 0 | 1.063873051 | 1 |
| ENSMUSG00000085001 | Rapgef4os2 | 0 | 1.648835551 | 1 |
| ENSMUSG00000086231 | Rapgef4os3 | 0 | 1.063873051 | 1 |
| ENSMUSG00000085252 | 8430437L04Rik | 0 | 0.648835551 | 1 |
| ENSMUSG00000079286 | Gm11084 | 0 | -0.52108945 | 1 |
| ENSMUSG00000068859 | Sp9 | 0 | 1.648835551 | 1 |
| ENSMUSG00000085519 | Gm13703 | 0 | 0.326907457 | 1 |
| ENSMUSG00000086546 | Gm13709 | 0 | 1.648835551 | 1 |
| ENSMUSG00000085768 | Gm13707 | 0 | 1.063873051 | 1 |
| ENSMUSG00000027107 | Chrna1 | 0 | 0.063873051 | 1 |
| ENSMUSG00000086382 | Chrna1os | 0 | 1.063873051 | 1 |
| ENSMUSG00000001823 | Hoxd12 | 0 | 1.648835551 | 1 |
| ENSMUSG00000027102 | Hoxd8 | 0 | 1.648835551 | 1 |
| ENSMUSG00000079277 | Hoxd3 | 0 | 0.063873051 | 1 |
| ENSMUSG00000075277 | Haglr | 0 | 1.648835551 | 1 |
| ENSMUSG00000086044 | A330043C09Rik | 0 | 0.063873051 | 1 |
| ENSMUSG00000086576 | Gm13660 | 0 | 1.648835551 | 1 |
| ENSMUSG00000087222 | E030042O20Rik | 0 | 1.648835551 | 1 |
| ENSMUSG00000086813 | Gm13657 | 0 | 0.648835551 | 1 |
| ENSMUSG00000075273 | Ttc30b | 0 | 0.326907457 | 1 |
| ENSMUSG00000075271 | Ttc30a1 | 0 | 1.063873051 | 1 |
| ENSMUSG00000075256 | Cerkl | 0 | -3.395558568 | 0.017810489 |

| | | | | |
|---|---|---|---|---|
| ENSMUSG00000034701 | Neurod1 | 0 | 1.648835551 | 1 |
| ENSMUSG00000027001 | Dusp19 | 0 | 1.648835551 | 1 |
| ENSMUSG00000034552 | Zswim2 | 0 | 1.063873051 | 1 |
| ENSMUSG00000086598 | Btbd18 | 0 | -0.158519371 | 1 |
| ENSMUSG00000027077 | Smtnl1 | 0 | 1.648835551 | 1 |
| ENSMUSG00000027074 | Slc43a3 | 0 | -1.43862729 | 0.740694319 |
| ENSMUSG00000027073 | Prg2 | 0 | 1.063873051 | 1 |
| ENSMUSG00000086969 | 4930443O20Rik | 0 | 1.648835551 | 1 |
| ENSMUSG00000044338 | Aplnr | 0 | -0.158519371 | 1 |
| ENSMUSG00000085025 | Gm13715 | 0 | 1.063873051 | 1 |
| ENSMUSG00000075223 | Olfr987 | 0 | 1.648835551 | 1 |
| ENSMUSG00000075217 | 4833423E24Rik | 0 | 0.063873051 | 1 |
| ENSMUSG00000075214 | Olfr1002 | 0 | 1.648835551 | 1 |
| ENSMUSG00000075211 | Olfr1006 | 0 | 1.648835551 | 1 |
| ENSMUSG00000075208 | Olfr1019 | 0 | 1.648835551 | 1 |
| ENSMUSG00000046975 | Olfr1020 | 0 | 1.648835551 | 1 |
| ENSMUSG00000042796 | Olfr1032 | 0 | 1.063873051 | 1 |
| ENSMUSG00000045392 | Olfr1033 | 0 | -2.351164449 | 0.169846604 |
| ENSMUSG00000102091 | Olfr1034 | 0 | 1.648835551 | 1 |
| ENSMUSG00000075205 | Olfr1037 | 0 | 1.063873051 | 1 |
| ENSMUSG00000075204 | Olfr1039 | 0 | 1.648835551 | 1 |
| ENSMUSG00000075201 | Olfr1043 | 0 | 1.648835551 | 1 |
| ENSMUSG00000075200 | Olfr1044 | 0 | 1.063873051 | 1 |
| ENSMUSG00000075199 | Olfr52 | 0 | 0.648835551 | 1 |
| ENSMUSG00000075195 | Olfr1048 | 0 | 1.648835551 | 1 |
| ENSMUSG00000075194 | Olfr1049 | 0 | 1.648835551 | 1 |
| ENSMUSG00000075188 | Olfr1056 | 0 | 1.063873051 | 1 |
| ENSMUSG00000090059 | Olfr1062 | 0 | 1.063873051 | 1 |
| ENSMUSG00000075179 | Olfr1079 | 0 | 1.648835551 | 1 |
| ENSMUSG00000111689 | Olfr1082 | 0 | 1.063873051 | 1 |
| ENSMUSG00000075176 | Olfr1085 | 0 | 1.648835551 | 1 |
| ENSMUSG00000075164 | Olfr1106 | 0 | 1.648835551 | 1 |
| ENSMUSG00000075160 | Olfr259 | 0 | -0.158519371 | 1 |
| ENSMUSG00000070856 | Olfr1115 | 0 | 1.648835551 | 1 |
| ENSMUSG00000070855 | Olfr1116 | 0 | 1.063873051 | 1 |
| ENSMUSG00000083706 | Olfr1118 | 0 | 1.648835551 | 1 |
| ENSMUSG00000070852 | Olfr1121 | 0 | 1.648835551 | 1 |
| ENSMUSG00000047594 | Olfr1122 | 0 | 1.648835551 | 1 |
| ENSMUSG00000050772 | Olfr1124 | 0 | 0.063873051 | 1 |
| ENSMUSG00000058194 | Olfr1126 | 0 | 1.648835551 | 1 |
| ENSMUSG00000068818 | Olfr1132 | 0 | 1.063873051 | 1 |
| ENSMUSG00000075150 | Olfr1137 | 0 | 1.648835551 | 1 |
| ENSMUSG00000075146 | Olfr1154 | 0 | 1.648835551 | 1 |
| ENSMUSG00000075144 | Olfr1156 | 0 | 1.648835551 | 1 |
| ENSMUSG00000075141 | Olfr1160 | 0 | 1.648835551 | 1 |
| ENSMUSG00000075140 | Olfr73 | 0 | 1.648835551 | 1 |
| ENSMUSG00000061342 | Olfr1168 | 0 | 1.648835551 | 1 |
| ENSMUSG00000083855 | Olfr1175-ps | 0 | 0.063873051 | 1 |
| ENSMUSG00000056995 | Olfr1178 | 0 | 1.648835551 | 1 |
| ENSMUSG00000075127 | Olfr1179 | 0 | -0.936126949 | 0.847265357 |
| ENSMUSG00000049372 | Olfr1183 | 0 | 1.648835551 | 1 |

| | | | | |
|---|---|---|---|---|
| ENSMUSG00000082882 | Olfr1186 | 0 | 1.648835551 | 1 |
| ENSMUSG00000068808 | Olfr1189 | 0 | 1.648835551 | 1 |
| ENSMUSG00000075121 | Olfr1195 | 0 | 1.648835551 | 1 |
| ENSMUSG00000075119 | Olfr1197 | 0 | 1.648835551 | 1 |
| ENSMUSG00000089892 | Olfr1199 | 0 | 1.648835551 | 1 |
| ENSMUSG00000075114 | Olfr1208 | 0 | 1.648835551 | 1 |
| ENSMUSG00000075112 | Olfr1211 | 0 | 1.648835551 | 1 |
| ENSMUSG00000101391 | Olfr1217 | 0 | 1.063873051 | 1 |
| ENSMUSG00000075100 | Olfr1223 | 0 | 1.648835551 | 1 |
| ENSMUSG00000075099 | Olfr1224-ps1 | 0 | 1.648835551 | 1 |
| ENSMUSG00000075097 | Olfr1226 | 0 | 1.063873051 | 1 |
| ENSMUSG00000075095 | Olfr1229 | 0 | 1.648835551 | 1 |
| ENSMUSG00000111517 | Olfr1238 | 0 | -0.351164449 | 1 |
| ENSMUSG00000075084 | Olfr1243 | 0 | 1.648835551 | 1 |
| ENSMUSG00000075081 | Olfr1247 | 0 | 1.063873051 | 1 |
| ENSMUSG00000075075 | Olfr1253 | 0 | 1.648835551 | 1 |
| ENSMUSG00000075073 | Olfr1256 | 0 | 1.648835551 | 1 |
| ENSMUSG00000042894 | Olfr1260 | 0 | 1.648835551 | 1 |
| ENSMUSG00000051313 | Olfr1262 | 0 | 0.648835551 | 1 |
| ENSMUSG00000059112 | Olfr1263 | 0 | 1.063873051 | 1 |
| ENSMUSG00000075061 | Olfr1272 | 0 | 1.648835551 | 1 |
| ENSMUSG00000085471 | 4933423P22Rik | 0 | 1.648835551 | 1 |
| ENSMUSG00000005505 | Kbtbd4 | 0 | -1.351164449 | 0.687112677 |
| ENSMUSG00000002104 | Rapsn | 0 | 0.063873051 | 1 |
| ENSMUSG00000027249 | F2 | 0 | 1.648835551 | 1 |
| ENSMUSG00000087018 | 2900072N19Rik | 0 | 1.648835551 | 1 |
| ENSMUSG00000040434 | Large2 | 0 | 0.648835551 | 1 |
| ENSMUSG00000044916 | 1700029I15Rik | 0 | 1.648835551 | 1 |
| ENSMUSG00000084781 | D930015M05Rik | 0 | 0.648835551 | 1 |
| ENSMUSG00000086980 | Gm13791 | 0 | 0.326907457 | 1 |
| ENSMUSG00000075027 | 4631405J19Rik | 0 | 1.648835551 | 1 |
| ENSMUSG00000068735 | Trp53i11 | 0 | -1.158519371 | 0.889462874 |
| ENSMUSG00000085852 | Gm13807 | 0 | 1.648835551 | 1 |
| ENSMUSG00000075025 | Gm10804 | 0 | 1.648835551 | 1 |
| ENSMUSG00000040310 | Alx4 | 0 | 0.648835551 | 1 |
| ENSMUSG00000040272 | Accs | 0 | 0.063873051 | 1 |
| ENSMUSG00000087006 | Gm13889 | 0 | 1.063873051 | 1 |
| ENSMUSG00000045464 | 2810002D19Rik | 0 | 1.063873051 | 1 |
| ENSMUSG00000084906 | Gm13794 | 0 | 0.326907457 | 1 |
| ENSMUSG00000087465 | Gm13905 | 0 | 1.648835551 | 1 |
| ENSMUSG00000087278 | A930006I01Rik | 0 | 1.648835551 | 1 |
| ENSMUSG00000068686 | Cd59b | 0 | -0.810596067 | 1 |
| ENSMUSG00000087473 | Gm13883 | 0 | 1.648835551 | 1 |
| ENSMUSG00000086029 | Pax6os1 | 0 | -1.52108945 | 0.624011826 |
| ENSMUSG00000100156 | Gm29053 | 0 | 1.648835551 | 1 |
| ENSMUSG00000063751 | Platr8 | 0 | 0.063873051 | 1 |
| ENSMUSG00000086186 | Ccdc34os | 0 | 0.326907457 | 1 |
| ENSMUSG00000041660 | Bbox1 | 0 | -0.673092543 | 1 |
| ENSMUSG00000041644 | Slc5a12 | 0 | 0.648835551 | 1 |
| ENSMUSG00000050808 | Muc15 | 0 | 1.648835551 | 1 |
| ENSMUSG00000079171 | Gm13942 | 0 | 1.648835551 | 1 |

| | | | | |
|---|---|---|---|---|
| ENSMUSG00000079170 | Gm13941 | 0 | 1.648835551 | 1 |
| ENSMUSG00000074965 | Olfr1277 | 0 | 1.648835551 | 1 |
| ENSMUSG00000068647 | Olfr1278 | 0 | 1.648835551 | 1 |
| ENSMUSG00000109449 | Olfr1280 | 0 | 1.648835551 | 1 |
| ENSMUSG00000095156 | Olfr1281 | 0 | 1.648835551 | 1 |
| ENSMUSG00000096554 | Olfr1282 | 0 | 1.648835551 | 1 |
| ENSMUSG00000109322 | Olfr1283 | 0 | 1.648835551 | 1 |
| ENSMUSG00000108931 | Olfr1284 | 0 | 1.648835551 | 1 |
| ENSMUSG00000096703 | Olfr1286 | 0 | 0.648835551 | 1 |
| ENSMUSG00000108891 | Olfr1291-ps1 | 0 | 1.063873051 | 1 |
| ENSMUSG00000108908 | Olfr1293-ps | 0 | 1.648835551 | 1 |
| ENSMUSG00000108919 | Olfr1295 | 0 | 1.648835551 | 1 |
| ENSMUSG00000094858 | Olfr1297 | 0 | 1.648835551 | 1 |
| ENSMUSG00000108827 | Olfr1310 | 0 | 1.648835551 | 1 |
| ENSMUSG00000074946 | Olfr1313 | 0 | 1.648835551 | 1 |
| ENSMUSG00000041358 | Nutm1 | 0 | 1.648835551 | 1 |
| ENSMUSG00000084798 | 4930533B01Rik | 0 | 1.063873051 | 1 |
| ENSMUSG00000068614 | Actc1 | 0 | 1.063873051 | 1 |
| ENSMUSG00000092397 | C130080G10Rik | 0 | 1.648835551 | 1 |
| ENSMUSG00000085222 | Gm13974 | 0 | -1.995020638 | 0.398502279 |
| ENSMUSG00000085582 | 3110099E03Rik | 0 | 0.326907457 | 1 |
| ENSMUSG00000086735 | Gm13977 | 0 | 0.648835551 | 1 |
| ENSMUSG00000086462 | Gm13990 | 0 | 1.648835551 | 1 |
| ENSMUSG00000085316 | D330050G23Rik | 0 | 1.648835551 | 1 |
| ENSMUSG00000086203 | Gm13985 | 0 | 1.648835551 | 1 |
| ENSMUSG00000045339 | A930104D05Rik | 0 | 0.648835551 | 1 |
| ENSMUSG00000085889 | 1700054M17Rik | 0 | 1.648835551 | 1 |
| ENSMUSG00000074918 | Inafm2 | 0 | -0.158519371 | 1 |
| ENSMUSG00000084802 | Gm14089 | 0 | 1.648835551 | 1 |
| ENSMUSG00000027324 | Rpusd2 | 0 | 1.063873051 | 1 |
| ENSMUSG00000034226 | Rhov | 0 | 1.063873051 | 1 |
| ENSMUSG00000074912 | Gm14207 | 0 | 0.326907457 | 1 |
| ENSMUSG00000027313 | Chac1 | 0 | 0.648835551 | 1 |
| ENSMUSG00000048647 | Exd1 | 0 | -1.995020638 | 0.395118486 |
| ENSMUSG00000027296 | Itpka | 0 | 1.063873051 | 1 |
| ENSMUSG00000027297 | Ltk | 0 | -0.158519371 | 1 |
| ENSMUSG00000087458 | Gm13999 | 0 | 1.648835551 | 1 |
| ENSMUSG00000098488 | Pla2g4b | 0 | -1.351164449 | 0.687112677 |
| ENSMUSG00000074899 | Sptbn5 | 0 | 0.063873051 | 1 |
| ENSMUSG00000050211 | Pla2g4e | 0 | -0.936126949 | 0.847265357 |
| ENSMUSG00000070719 | Pla2g4d | 0 | 0.648835551 | 1 |
| ENSMUSG00000046971 | Pla2g4f | 0 | -0.158519371 | 1 |
| ENSMUSG00000027286 | Lrrc57 | 0 | -1.258055044 | 0.930842647 |
| ENSMUSG00000027284 | Cdan1 | 0 | -1.673092543 | 0.451785624 |
| ENSMUSG00000050619 | Zscan29 | 0 | -2.936126949 | 0.042735074 |
| ENSMUSG00000046110 | Serinc4 | 0 | -0.351164449 | 1 |
| ENSMUSG00000027242 | Wdr76 | 0 | -4.079084903 | 0.001016474 |
| ENSMUSG00000074881 | Mageb3 | 0 | 1.648835551 | 1 |
| ENSMUSG00000027224 | Duoxa1 | 0 | 0.648835551 | 1 |
| ENSMUSG00000027219 | Slc28a2 | 0 | -0.158519371 | 1 |
| ENSMUSG00000074876 | Spata5l1 | 0 | 0.063873051 | 1 |

| | | | | |
|---|---|---|---|---|
| ENSMUSG00000085633 | Bloc1s6os | 0 | 1.648835551 | 1 |
| ENSMUSG00000074872 | Ctxn2 | 0 | 1.063873051 | 1 |
| ENSMUSG00000086416 | Gm14002 | 0 | 1.063873051 | 1 |
| ENSMUSG00000053615 | Gm9913 | 0 | 1.648835551 | 1 |
| ENSMUSG00000027359 | Slc27a2 | 0 | 1.648835551 | 1 |
| ENSMUSG00000027360 | Hdc | 0 | -0.351164449 | 1 |
| ENSMUSG00000027368 | Dusp2 | 0 | 0.063873051 | 1 |
| ENSMUSG00000050468 | Astl | 0 | 1.648835551 | 1 |
| ENSMUSG00000027379 | Bub1 | 0 | -2.158519371 | 0.290284345 |
| ENSMUSG00000114963 | Gm29010 | 0 | 0.063873051 | 1 |
| ENSMUSG00000027387 | Zc3h8 | 0 | -1.258055044 | 0.739346465 |
| ENSMUSG00000087267 | 4933427J07Rik | 0 | 1.063873051 | 1 |
| ENSMUSG00000074807 | Gm10762 | 0 | 0.648835551 | 1 |
| ENSMUSG00000090248 | Gm14027 | 0 | 1.063873051 | 1 |
| ENSMUSG00000079051 | Vinac1 | 0 | 0.648835551 | 1 |
| ENSMUSG00000084826 | AI847159 | 0 | -0.158519371 | 1 |
| ENSMUSG00000086652 | Gm14029 | 0 | 1.648835551 | 1 |
| ENSMUSG00000056738 | A730036I17Rik | 0 | 0.648835551 | 1 |
| ENSMUSG00000085498 | Gm14023 | 0 | 1.063873051 | 1 |
| ENSMUSG00000074805 | Il1bos | 0 | 1.648835551 | 1 |
| ENSMUSG00000043727 | F830045P16Rik | 0 | -0.158519371 | 1 |
| ENSMUSG00000027400 | Pdyn | 0 | 1.063873051 | 1 |
| ENSMUSG00000100255 | Gm28196 | 0 | 1.648835551 | 1 |
| ENSMUSG00000027401 | Tgm3 | 0 | 0.063873051 | 1 |
| ENSMUSG00000027301 | Oxt | 0 | -0.158519371 | 1 |
| ENSMUSG00000079043 | Fastkd5 | 0 | -0.158519371 | 1 |
| ENSMUSG00000037703 | Lzts3 | 0 | -2.158519371 | 0.290284345 |
| ENSMUSG00000027322 | Siglec1 | 0 | 0.326907457 | 1 |
| ENSMUSG00000027335 | Adra1d | 0 | 1.648835551 | 1 |
| ENSMUSG00000027338 | Prnd | 0 | 1.063873051 | 1 |
| ENSMUSG00000044991 | Shld1 | 0 | -1.52108945 | 0.590312322 |
| ENSMUSG00000084839 | Gm14097 | 0 | 1.648835551 | 1 |
| ENSMUSG00000043110 | Lrrn4 | 0 | 1.648835551 | 1 |
| ENSMUSG00000086478 | Gm14102 | 0 | 1.063873051 | 1 |
| ENSMUSG00000085247 | 4930545L23Rik | 0 | -1.673092543 | 0.6198239 |
| ENSMUSG00000027270 | Lamp5 | 0 | 1.063873051 | 1 |
| ENSMUSG00000074771 | Ankef1 | 0 | 0.326907457 | 1 |
| ENSMUSG00000086214 | Gm14062 | 0 | 1.648835551 | 1 |
| ENSMUSG00000086504 | Kif16bos | 0 | 1.648835551 | 1 |
| ENSMUSG00000086171 | Pcsk2os1 | 0 | 1.063873051 | 1 |
| ENSMUSG00000087115 | Pcsk2os2 | 0 | 1.063873051 | 1 |
| ENSMUSG00000085127 | 4930444E06Rik | 0 | 1.063873051 | 1 |
| ENSMUSG00000085958 | A930019D19Rik | 0 | 1.063873051 | 1 |
| ENSMUSG00000087521 | AI646519 | 0 | 1.648835551 | 1 |
| ENSMUSG00000037034 | Pax1 | 0 | 0.648835551 | 1 |
| ENSMUSG00000051986 | A530006G24Rik | 0 | 1.648835551 | 1 |
| ENSMUSG00000074743 | Thbd | 0 | 0.648835551 | 1 |
| ENSMUSG00000087678 | Gm14120 | 0 | 1.648835551 | 1 |
| ENSMUSG00000027446 | Cstdc2 | 0 | -0.158519371 | 1 |
| ENSMUSG00000086963 | Gm14135 | 0 | 1.648835551 | 1 |
| ENSMUSG00000060336 | Zfp937 | 0 | 0.063873051 | 1 |

| | | | | |
|---|---|---|---|---|
| ENSMUSG00000068130 | Zfp442 | 0 | -0.52108945 | 1 |
| ENSMUSG00000074731 | Zfp345 | 0 | 1.648835551 | 1 |
| ENSMUSG00000067352 | Gm14149 | 0 | 0.648835551 | 1 |
| ENSMUSG00000087495 | Gm14152 | 0 | 0.648835551 | 1 |
| ENSMUSG00000086659 | Gm14146 | 0 | 1.648835551 | 1 |
| ENSMUSG00000074704 | Rad21l | 0 | 0.648835551 | 1 |
| ENSMUSG00000087035 | Tmem74bos | 0 | 1.063873051 | 1 |
| ENSMUSG00000032869 | Psmf1 | 0 | -1.52108945 | 0.560378012 |
| ENSMUSG00000032852 | Rspo4 | 0 | 1.063873051 | 1 |
| ENSMUSG00000087004 | Gm14154 | 0 | 1.648835551 | 1 |
| ENSMUSG00000059361 | Nrsn2 | 0 | 1.063873051 | 1 |
| ENSMUSG00000074682 | Zcchc3 | 0 | 0.063873051 | 1 |
| ENSMUSG00000049560 | Defb20 | 0 | 1.648835551 | 1 |
| ENSMUSG00000074680 | Defb26 | 0 | 1.648835551 | 1 |
| ENSMUSG00000084935 | Gm14161 | 0 | 1.648835551 | 1 |
| ENSMUSG00000050645 | Defb19 | 0 | 1.648835551 | 1 |
| ENSMUSG00000000359 | Rem1 | 0 | 0.326907457 | 1 |
| ENSMUSG00000074676 | Foxs1 | 0 | 1.063873051 | 1 |
| ENSMUSG00000074673 | Ttll9 | 0 | -0.158519371 | 1 |
| ENSMUSG00000042631 | Xkr7 | 0 | -0.351164449 | 1 |
| ENSMUSG00000051413 | Plagl2 | 0 | -2.051604167 | 0.305609173 |
| ENSMUSG00000079000 | Dnmt3bos | 0 | 1.648835551 | 1 |
| ENSMUSG00000082079 | Dnmt3c | 0 | 1.063873051 | 1 |
| ENSMUSG00000027480 | Sun5 | 0 | 1.063873051 | 1 |
| ENSMUSG00000074665 | Bpifb4 | 0 | -0.158519371 | 1 |
| ENSMUSG00000078998 | Bpifa6 | 0 | 0.648835551 | 1 |
| ENSMUSG00000027482 | Bpifa3 | 0 | 1.648835551 | 1 |
| ENSMUSG00000027484 | Bpifa5 | 0 | 1.063873051 | 1 |
| ENSMUSG00000038572 | Bpifb5 | 0 | 0.063873051 | 1 |
| ENSMUSG00000067996 | Bpifb9b | 0 | 1.063873051 | 1 |
| ENSMUSG00000027489 | Necab3 | 0 | 1.648835551 | 1 |
| ENSMUSG00000038523 | 1700003F12Rik | 0 | 1.648835551 | 1 |
| ENSMUSG00000078129 | Actl10 | 0 | 1.648835551 | 1 |
| ENSMUSG00000097602 | 4930519P11Rik | 0 | 1.648835551 | 1 |
| ENSMUSG00000085483 | Gm14198 | 0 | 1.648835551 | 1 |
| ENSMUSG00000085691 | Gm14216 | 0 | 0.326907457 | 1 |
| ENSMUSG00000084897 | Gm14226 | 0 | 1.648835551 | 1 |
| ENSMUSG00000027597 | Ahcy | 0 | -0.673092543 | 1 |
| ENSMUSG00000109877 | Gm45609 | 0 | 0.063873051 | 1 |
| ENSMUSG00000085317 | Gssos2 | 0 | 1.648835551 | 1 |
| ENSMUSG00000038259 | Gdf5 | 0 | 1.648835551 | 1 |
| ENSMUSG00000038180 | Spag4 | 0 | 0.648835551 | 1 |
| ENSMUSG00000085524 | Gm14224 | 0 | 1.063873051 | 1 |
| ENSMUSG00000055494 | Gm14168 | 0 | -0.158519371 | 1 |
| ENSMUSG00000086638 | 4930405A21Rik | 0 | -0.158519371 | 1 |
| ENSMUSG00000087179 | Gm14230 | 0 | 0.648835551 | 1 |
| ENSMUSG00000062175 | Tgif2 | 0 | 0.326907457 | 1 |
| ENSMUSG00000027635 | Dsn1 | 0 | -1.743481871 | 0.455131546 |
| ENSMUSG00000074628 | Tldc2 | 0 | 1.648835551 | 1 |
| ENSMUSG00000087445 | Gm14286 | 0 | -1.051604167 | 0.858831093 |
| ENSMUSG00000037843 | Vstm2l | 0 | 1.063873051 | 1 |

| | | | | |
|---|---|---|---|---|
| ENSMUSG00000100860 | 2010009K17Rik | 0 | 0.648835551 | 1 |
| ENSMUSG00000078957 | 1700060C20Rik | 0 | 1.648835551 | 1 |
| ENSMUSG00000052922 | Bpi | 0 | -0.936126949 | 0.847265357 |
| ENSMUSG00000044349 | Snhg11 | 0 | -0.158519371 | 1 |
| ENSMUSG00000027654 | Fam83d | 0 | 0.326907457 | 1 |
| ENSMUSG00000086558 | Gm14223 | 0 | 0.648835551 | 1 |
| ENSMUSG00000078955 | Gm14222 | 0 | 1.648835551 | 1 |
| ENSMUSG00000085303 | Gm16751 | 0 | 0.648835551 | 1 |
| ENSMUSG00000087147 | Gm14228 | 0 | 1.648835551 | 1 |
| ENSMUSG00000109780 | Gm45447 | 0 | 0.648835551 | 1 |
| ENSMUSG00000067745 | Gm14243 | 0 | 1.648835551 | 1 |
| ENSMUSG00000017868 | Sgk2 | 0 | 0.326907457 | 1 |
| ENSMUSG00000086152 | Gm11454 | 0 | 1.063873051 | 1 |
| ENSMUSG00000099632 | 2900093K20Rik | 0 | -1.351164449 | 0.687112677 |
| ENSMUSG00000017943 | Gdap1l1 | 0 | 0.648835551 | 1 |
| ENSMUSG00000048486 | Fitm2 | 0 | -0.351164449 | 1 |
| ENSMUSG00000046840 | Hnf4aos | 0 | -0.52108945 | 1 |
| ENSMUSG00000087129 | Gm16316 | 0 | 1.648835551 | 1 |
| ENSMUSG00000027656 | Ccn5 | 0 | 0.063873051 | 1 |
| ENSMUSG00000054582 | Pabpc1l | 0 | -0.936126949 | 0.884678648 |
| ENSMUSG00000085538 | Gm11455 | 0 | 1.648835551 | 1 |
| ENSMUSG00000018211 | Wfdc15b | 0 | 1.648835551 | 1 |
| ENSMUSG00000016998 | Svs4 | 0 | 1.063873051 | 1 |
| ENSMUSG00000016995 | Matn4 | 0 | -0.52108945 | 1 |
| ENSMUSG00000017007 | Rbpjl | 0 | 0.648835551 | 1 |
| ENSMUSG00000086179 | Gm14317 | 0 | 0.063873051 | 1 |
| ENSMUSG00000074595 | Wfdc6a | 0 | 1.648835551 | 1 |
| ENSMUSG00000070533 | Wfdc8 | 0 | 0.648835551 | 1 |
| ENSMUSG00000070531 | Wfdc6b | 0 | 1.063873051 | 1 |
| ENSMUSG00000078940 | Wfdc11 | 0 | 1.648835551 | 1 |
| ENSMUSG00000076434 | Wfdc3 | 0 | 1.063873051 | 1 |
| ENSMUSG00000017764 | Zswim1 | 0 | 1.063873051 | 1 |
| ENSMUSG00000017652 | Cd40 | 0 | 0.326907457 | 1 |
| ENSMUSG00000056824 | Zfp663 | 0 | 1.063873051 | 1 |
| ENSMUSG00000027670 | Ocstamp | 0 | -0.673092543 | 1 |
| ENSMUSG00000085358 | Gm14437 | 0 | 1.648835551 | 1 |
| ENSMUSG00000027661 | Slc2a10 | 0 | 0.648835551 | 1 |
| ENSMUSG00000100796 | Gm28163 | 0 | 1.648835551 | 1 |
| ENSMUSG00000085274 | BC046401 | 0 | 0.648835551 | 1 |
| ENSMUSG00000086820 | Gm11465 | 0 | 1.648835551 | 1 |
| ENSMUSG00000084816 | Platr29 | 0 | -0.351164449 | 1 |
| ENSMUSG00000087351 | Gm11464 | 0 | 1.063873051 | 1 |
| ENSMUSG00000084840 | Gm11468 | 0 | -0.158519371 | 1 |
| ENSMUSG00000042854 | Trp53rkb | 0 | -0.673092543 | 1 |
| ENSMUSG00000047030 | Spata2 | 0 | -1.52108945 | 1 |
| ENSMUSG00000042821 | Snai1 | 0 | 1.063873051 | 1 |
| ENSMUSG00000084797 | Gm14321 | 0 | 1.063873051 | 1 |
| ENSMUSG00000087624 | 9230111E07Rik | 0 | 1.063873051 | 1 |
| ENSMUSG00000087684 | 1200007C13Rik | 0 | 0.063873051 | 1 |
| ENSMUSG00000085411 | Gm14319 | 0 | 1.648835551 | 1 |
| ENSMUSG00000074577 | Ripor3 | 0 | -0.673092543 | 1 |

| | | | | |
|---|---|---|---|---|
| ENSMUSG00000085267 | Gm14235 | 0 | 1.648835551 | 1 |
| ENSMUSG00000074576 | Mocs3 | 0 | 1.648835551 | 1 |
| ENSMUSG00000074575 | Kcng1 | 0 | -0.936126949 | 0.886203497 |
| ENSMUSG00000027547 | Sall4 | 0 | -0.810596067 | 1 |
| ENSMUSG00000085322 | Gm14261 | 0 | 1.648835551 | 1 |
| ENSMUSG00000052368 | Gm9873 | 0 | 1.063873051 | 1 |
| ENSMUSG00000086840 | Gm14250 | 0 | 0.326907457 | 1 |
| ENSMUSG00000078912 | Gm11011 | 0 | 1.648835551 | 1 |
| ENSMUSG00000086999 | Bcas1os2 | 0 | -0.351164449 | 1 |
| ENSMUSG00000086970 | Bcas1os1 | 0 | 1.648835551 | 1 |
| ENSMUSG00000038567 | Cyp24a1 | 0 | 1.648835551 | 1 |
| ENSMUSG00000085225 | Gm14637 | 0 | 1.063873051 | 1 |
| ENSMUSG00000084829 | Gm14266 | 0 | 0.063873051 | 1 |
| ENSMUSG00000067578 | Cbln4 | 0 | 1.063873051 | 1 |
| ENSMUSG00000027496 | Aurka | 0 | -0.351164449 | 1 |
| ENSMUSG00000027498 | Cstf1 | 0 | -1.351164449 | 0.687112677 |
| ENSMUSG00000087633 | Gm14455 | 0 | 1.063873051 | 1 |
| ENSMUSG00000027505 | Fam209 | 0 | 1.648835551 | 1 |
| ENSMUSG00000028640 | Tfap2c | 0 | 1.063873051 | 1 |
| ENSMUSG00000005883 | Spo11 | 0 | 0.648835551 | 1 |
| ENSMUSG00000087442 | Gm14453 | 0 | 1.648835551 | 1 |
| ENSMUSG00000070495 | Ctcfl | 0 | 1.063873051 | 1 |
| ENSMUSG00000027514 | Zbp1 | 0 | 0.648835551 | 1 |
| ENSMUSG00000087600 | Pmepa1os | 0 | 0.326907457 | 1 |
| ENSMUSG00000057047 | 1700010B08Rik | 0 | 1.648835551 | 1 |
| ENSMUSG00000090625 | Gm20721 | 0 | 1.063873051 | 1 |
| ENSMUSG00000016253 | Nelfcd | 0 | -1.52108945 | 0.544908989 |
| ENSMUSG00000050600 | Zfp831 | 0 | -1.743481871 | 0.466464457 |
| ENSMUSG00000027524 | Edn3 | 0 | 1.648835551 | 1 |
| ENSMUSG00000087215 | Gm14618 | 0 | 0.648835551 | 1 |
| ENSMUSG00000086177 | Gm14616 | 0 | 1.648835551 | 1 |
| ENSMUSG00000090093 | Gm14399 | 0 | 1.063873051 | 1 |
| ENSMUSG00000078902 | Gm14443 | 0 | 1.063873051 | 1 |
| ENSMUSG00000078903 | Gm14391 | 0 | 1.063873051 | 1 |
| ENSMUSG00000078898 | Zfp968 | 0 | 1.648835551 | 1 |
| ENSMUSG00000089756 | Zfp966 | 0 | 0.063873051 | 1 |
| ENSMUSG00000089951 | Gm14435 | 0 | 1.648835551 | 1 |
| ENSMUSG00000078879 | Zfp973 | 0 | 1.648835551 | 1 |
| ENSMUSG00000078878 | Gm14305 | 0 | 1.648835551 | 1 |
| ENSMUSG00000078872 | Gm14401 | 0 | -0.52108945 | 1 |
| ENSMUSG00000078870 | Gm14410 | 0 | 1.063873051 | 1 |
| ENSMUSG00000078868 | Gm14412 | 0 | 1.648835551 | 1 |
| ENSMUSG00000078867 | Gm14418 | 0 | 1.648835551 | 1 |
| ENSMUSG00000078866 | Zfp970 | 0 | -0.810596067 | 0.844591606 |
| ENSMUSG00000074521 | Gm14327 | 0 | -0.351164449 | 1 |
| ENSMUSG00000074529 | Zfp972 | 0 | -2.051604167 | 0.274319469 |
| ENSMUSG00000078861 | Zfp931 | 0 | -2.480447466 | 0.175065564 |
| ENSMUSG00000060445 | Sycp2 | 0 | -0.351164449 | 1 |
| ENSMUSG00000039059 | Hrh3 | 0 | 1.648835551 | 1 |
| ENSMUSG00000038980 | Rbbp8nl | 0 | 1.648835551 | 1 |
| ENSMUSG00000056061 | Gata5os | 0 | 0.648835551 | 1 |

| | | | | |
|---|---|---|---|---|
| ENSMUSG00000027570 | Col9a3 | 0 | 1.648835551 | 1 |
| ENSMUSG00000087550 | Gm14343 | 0 | 1.063873051 | 1 |
| ENSMUSG00000085428 | Gm14344 | 0 | 1.648835551 | 1 |
| ENSMUSG00000027574 | Nkain4 | 0 | -0.158519371 | 1 |
| ENSMUSG00000027577 | Chrna4 | 0 | 1.063873051 | 1 |
| ENSMUSG00000016349 | Eef1a2 | 0 | 1.063873051 | 1 |
| ENSMUSG00000038751 | Ptk6 | 0 | 1.063873051 | 1 |
| ENSMUSG00000027581 | Stmn3 | 0 | 0.648835551 | 1 |
| ENSMUSG00000055882 | Abhd16b | 0 | 1.648835551 | 1 |
| ENSMUSG00000010492 | Uckl1os | 0 | 1.063873051 | 1 |
| ENSMUSG00000002458 | Rgs19 | 0 | -0.52108945 | 1 |
| ENSMUSG00000027584 | Oprl1 | 0 | 1.648835551 | 1 |
| ENSMUSG00000010505 | Myt1 | 0 | -0.810596067 | 1 |
| ENSMUSG00000098659 | 1110015O18Rik | 0 | 1.648835551 | 1 |
| ENSMUSG00000040329 | Il7 | 0 | 0.326907457 | 1 |
| ENSMUSG00000100010 | 1700010I02Rik | 0 | 1.063873051 | 1 |
| ENSMUSG00000104102 | Gm32496 | 0 | 1.648835551 | 1 |
| ENSMUSG00000097365 | C030034L19Rik | 0 | 1.063873051 | 1 |
| ENSMUSG00000049230 | Gm9833 | 0 | -0.351164449 | 1 |
| ENSMUSG00000052468 | Pmp2 | 0 | 1.648835551 | 1 |
| ENSMUSG00000039710 | Slc7a12 | 0 | 1.063873051 | 1 |
| ENSMUSG00000027559 | Car3 | 0 | 0.648835551 | 1 |
| ENSMUSG00000103912 | Gm33819 | 0 | 0.326907457 | 1 |
| ENSMUSG00000078783 | Gm9733 | 0 | 1.063873051 | 1 |
| ENSMUSG00000095788 | Sirpb1a | 0 | -1.051604167 | 0.867965264 |
| ENSMUSG00000095028 | Sirpb1b | 0 | -0.351164449 | 1 |
| ENSMUSG00000074677 | Sirpb1c | 0 | -0.351164449 | 1 |
| ENSMUSG00000104411 | Gm30340 | 0 | 1.648835551 | 1 |
| ENSMUSG00000100252 | Mir124-2hg | 0 | 0.063873051 | 1 |
| ENSMUSG00000102699 | 4930433B08Rik | 0 | 1.648835551 | 1 |
| ENSMUSG00000060913 | Trim55 | 0 | 0.648835551 | 1 |
| ENSMUSG00000054988 | Agtr1b | 0 | 0.326907457 | 1 |
| ENSMUSG00000106321 | Gm43674 | 0 | 1.063873051 | 1 |
| ENSMUSG00000074664 | A830092H15Rik | 0 | 1.648835551 | 1 |
| ENSMUSG00000027690 | Slc2a2 | 0 | -1.158519371 | 0.667778757 |
| ENSMUSG00000091329 | 1700112D23Rik | 0 | -1.258055044 | 0.706929043 |
| ENSMUSG00000103827 | Gm32950 | 0 | 1.063873051 | 1 |
| ENSMUSG00000074655 | Gm1527 | 0 | 0.648835551 | 1 |
| ENSMUSG00000102590 | Mannr | 0 | 1.063873051 | 1 |
| ENSMUSG00000085702 | Mecomos | 0 | -0.351164449 | 1 |
| ENSMUSG00000027702 | Lrrc34 | 0 | 1.648835551 | 1 |
| ENSMUSG00000085953 | Gm15496 | 0 | 0.326907457 | 1 |
| ENSMUSG00000091091 | Kcnmb3 | 0 | 0.326907457 | 1 |
| ENSMUSG00000102862 | Gm37640 | 0 | 1.063873051 | 1 |
| ENSMUSG00000087625 | 4930419G24Rik | 0 | -0.936126949 | 0.847265357 |
| ENSMUSG00000105286 | Gm43140 | 0 | 1.648835551 | 1 |
| ENSMUSG00000105535 | Gm43077 | 0 | 1.648835551 | 1 |
| ENSMUSG00000092242 | Gm20515 | 0 | 1.063873051 | 1 |
| ENSMUSG00000027713 | 1810062G17Rik | 0 | 1.063873051 | 1 |
| ENSMUSG00000086090 | Gm12531 | 0 | 0.648835551 | 1 |
| ENSMUSG00000027718 | Il21 | 0 | 1.648835551 | 1 |

| | | | | |
|---|---|---|---|---|
| ENSMUSG00000051444 | Bbs12 | 0 | -1.743481871 | 0.455131546 |
| ENSMUSG00000106002 | 5430434I15Rik | 0 | 1.648835551 | 1 |
| ENSMUSG00000105516 | Gm36823 | 0 | 0.648835551 | 1 |
| ENSMUSG00000086181 | C230034O21Rik | 0 | 1.063873051 | 1 |
| ENSMUSG00000037884 | 1700017G19Rik | 0 | 1.063873051 | 1 |
| ENSMUSG00000025758 | Plk4 | 0 | -2.305360759 | 0.231596497 |
| ENSMUSG00000097583 | 6430590A07Rik | 0 | -2.158519371 | 0.290284345 |
| ENSMUSG00000074619 | 1700034I23Rik | 0 | 1.648835551 | 1 |
| ENSMUSG00000105258 | Gm40038 | 0 | 0.326907457 | 1 |
| ENSMUSG00000106000 | Gm16508 | 0 | 0.648835551 | 1 |
| ENSMUSG00000105260 | Gm40040 | 0 | 1.648835551 | 1 |
| ENSMUSG00000097639 | Platr4 | 0 | 1.063873051 | 1 |
| ENSMUSG00000049100 | Pcdh10 | 0 | -0.936126949 | 0.88337113 |
| ENSMUSG00000100799 | 1700027H10Rik | 0 | 1.063873051 | 1 |
| ENSMUSG00000090919 | Pabpc4l | 0 | 1.063873051 | 1 |
| ENSMUSG00000102263 | Gm31415 | 0 | 0.326907457 | 1 |
| ENSMUSG00000037892 | Pcdh18 | 0 | 1.063873051 | 1 |
| ENSMUSG00000102067 | Gm29230 | 0 | -0.158519371 | 1 |
| ENSMUSG00000097885 | 5031434O11Rik | 0 | -0.673092543 | 1 |
| ENSMUSG00000103726 | Gm30074 | 0 | 1.648835551 | 1 |
| ENSMUSG00000104401 | Gm20750 | 0 | 1.063873051 | 1 |
| ENSMUSG00000105168 | Gm30735 | 0 | 1.063873051 | 1 |
| ENSMUSG00000106270 | C820005J03Rik | 0 | 1.648835551 | 1 |
| ENSMUSG00000027748 | Trpc4 | 0 | -1.051604167 | 0.880033711 |
| ENSMUSG00000036632 | Alg5 | 0 | -2.673092543 | 0.084161134 |
| ENSMUSG00000027793 | Ccna1 | 0 | 1.063873051 | 1 |
| ENSMUSG00000048655 | Ccdc169 | 0 | 0.648835551 | 1 |
| ENSMUSG00000027794 | Sohlh2 | 0 | 1.648835551 | 1 |
| ENSMUSG00000106334 | Gm43549 | 0 | 1.648835551 | 1 |
| ENSMUSG00000097339 | Gm26671 | 0 | -0.52108945 | 1 |
| ENSMUSG00000101860 | Mindy4b-ps | 0 | -0.52108945 | 1 |
| ENSMUSG00000043850 | Clrn1 | 0 | -0.158519371 | 1 |
| ENSMUSG00000050075 | Gpr171 | 0 | 1.063873051 | 1 |
| ENSMUSG00000051431 | Gpr87 | 0 | 1.063873051 | 1 |
| ENSMUSG00000036362 | P2ry13 | 0 | 1.063873051 | 1 |
| ENSMUSG00000103668 | Gm38186 | 0 | 1.648835551 | 1 |
| ENSMUSG00000090527 | Gm5538 | 0 | -0.52108945 | 1 |
| ENSMUSG00000095522 | Gm8298 | 0 | 0.648835551 | 1 |
| ENSMUSG00000036951 | C130079G13Rik | 0 | 1.648835551 | 1 |
| ENSMUSG00000091376 | Aadacl2 | 0 | 1.648835551 | 1 |
| ENSMUSG00000027762 | Sucnr1 | 0 | 1.648835551 | 1 |
| ENSMUSG00000103587 | Gm38326 | 0 | 1.648835551 | 1 |
| ENSMUSG00000102564 | Gm37035 | 0 | 1.648835551 | 1 |
| ENSMUSG00000053706 | B430305J03Rik | 0 | 1.063873051 | 1 |
| ENSMUSG00000043441 | Gpr149 | 0 | 0.063873051 | 1 |
| ENSMUSG00000097724 | Gm26850 | 0 | 1.648835551 | 1 |
| ENSMUSG00000027824 | Vmn2r1 | 0 | 1.648835551 | 1 |
| ENSMUSG00000043897 | Vmn2r2 | 0 | 0.648835551 | 1 |
| ENSMUSG00000091572 | Vmn2r3 | 0 | -0.673092543 | 1 |
| ENSMUSG00000068999 | Vmn2r5 | 0 | 0.326907457 | 1 |
| ENSMUSG00000090581 | Vmn2r6 | 0 | 1.063873051 | 1 |

| | | | | |
|---|---|---|---|---|
| ENSMUSG00000090120 | A730090N16Rik | 0 | -1.43862729 | 0.538110273 |
| ENSMUSG00000103041 | Gm37305 | 0 | -1.258055044 | 0.673811773 |
| ENSMUSG00000090408 | Gm17402 | 0 | -0.351164449 | 1 |
| ENSMUSG00000049404 | Rarres1 | 0 | 0.326907457 | 1 |
| ENSMUSG00000027776 | Il12a | 0 | 1.063873051 | 1 |
| ENSMUSG00000043300 | B3galnt1 | 0 | 0.648835551 | 1 |
| ENSMUSG00000027788 | Otol1 | 0 | 1.648835551 | 1 |
| ENSMUSG00000048304 | Slitrk3 | 0 | 1.063873051 | 1 |
| ENSMUSG00000102545 | 6430573P05Rik | 0 | 0.063873051 | 1 |
| ENSMUSG00000068957 | 4930589L23Rik | 0 | 1.063873051 | 1 |
| ENSMUSG00000027811 | 4930579G24Rik | 0 | 1.063873051 | 1 |
| ENSMUSG00000103929 | Gm37892 | 0 | 1.063873051 | 1 |
| ENSMUSG00000102379 | Gm37971 | 0 | -0.52108945 | 1 |
| ENSMUSG00000085106 | Gm16000 | 0 | 1.648835551 | 1 |
| ENSMUSG00000056023 | Gm9989 | 0 | 1.648835551 | 1 |
| ENSMUSG00000028015 | Ctso | 0 | -1.051604167 | 0.858831093 |
| ENSMUSG00000028008 | Asic5 | 0 | -0.158519371 | 1 |
| ENSMUSG00000028004 | Npy2r | 0 | 1.648835551 | 1 |
| ENSMUSG00000028003 | Lrat | 0 | 1.648835551 | 1 |
| ENSMUSG00000027995 | Tlr2 | 0 | -0.52108945 | 1 |
| ENSMUSG00000047819 | Tigd4 | 0 | -0.52108945 | 1 |
| ENSMUSG00000031286 | Glt28d2 | 0 | 0.063873051 | 1 |
| ENSMUSG00000103408 | Gm37933 | 0 | 1.648835551 | 1 |
| ENSMUSG00000104108 | Gm37876 | 0 | 1.648835551 | 1 |
| ENSMUSG00000102893 | Gm37855 | 0 | 0.648835551 | 1 |
| ENSMUSG00000059994 | Fcrl1 | 0 | -0.351164449 | 1 |
| ENSMUSG00000023084 | Lrrc71 | 0 | 0.063873051 | 1 |
| ENSMUSG00000005640 | Insrr | 0 | -1.258055044 | 0.673811773 |
| ENSMUSG00000004885 | Crabp2 | 0 | 0.648835551 | 1 |
| ENSMUSG00000004894 | Hapln2 | 0 | 1.648835551 | 1 |
| ENSMUSG00000051036 | Ttc24 | 0 | 1.648835551 | 1 |
| ENSMUSG00000097280 | AI849053 | 0 | 0.326907457 | 1 |
| ENSMUSG00000041423 | Paqr6 | 0 | 1.063873051 | 1 |
| ENSMUSG00000093390 | Gm20652 | 0 | -0.158519371 | 1 |
| ENSMUSG00000105512 | Gm43714 | 0 | 1.063873051 | 1 |
| ENSMUSG00000086718 | Gm16069 | 0 | 1.063873051 | 1 |
| ENSMUSG00000028047 | Thbs3 | 0 | -2.051604167 | 0.274319469 |
| ENSMUSG00000028039 | Efna3 | 0 | 1.063873051 | 1 |
| ENSMUSG00000097004 | 4731419I09Rik | 0 | 0.063873051 | 1 |
| ENSMUSG00000042672 | Dcst1 | 0 | 1.063873051 | 1 |
| ENSMUSG00000027951 | Adar | 0 | -3.777429203 | 0.00334584 |
| ENSMUSG00000042579 | 4632404H12Rik | 0 | -0.351164449 | 1 |
| ENSMUSG00000105552 | Gm19710 | 0 | -0.158519371 | 1 |
| ENSMUSG00000090005 | Gm16540 | 0 | 0.063873051 | 1 |
| ENSMUSG00000118506 | 1700094D03Rik | 0 | -0.52108945 | 1 |
| ENSMUSG00000105350 | 4930537H20Rik | 0 | 0.326907457 | 1 |
| ENSMUSG00000027931 | Npr1 | 0 | 0.063873051 | 1 |
| ENSMUSG00000087411 | Gm16048 | 0 | 1.063873051 | 1 |
| ENSMUSG00000094018 | S100a2 | 0 | 1.063873051 | 1 |
| ENSMUSG00000001023 | S100a5 | 0 | 1.063873051 | 1 |
| ENSMUSG00000063767 | S100a7a | 0 | -0.351164449 | 1 |

| | | | | |
|---|---|---|---|---|
| ENSMUSG00000042250 | Pglyrp4 | 0 | 0.326907457 | 1 |
| ENSMUSG00000043165 | Lor | 0 | -0.158519371 | 1 |
| ENSMUSG00000106143 | Gm43735 | 0 | 1.648835551 | 1 |
| ENSMUSG00000105476 | Gm35439 | 0 | 1.648835551 | 1 |
| ENSMUSG00000042212 | Sprr2d | 0 | 1.648835551 | 1 |
| ENSMUSG00000055030 | Sprr2e | 0 | 1.648835551 | 1 |
| ENSMUSG00000046259 | Sprr2h | 0 | 1.648835551 | 1 |
| ENSMUSG00000027923 | Lce1b | 0 | 1.648835551 | 1 |
| ENSMUSG00000041984 | Rptn | 0 | 1.648835551 | 1 |
| ENSMUSG00000094084 | Tdpoz1 | 0 | 1.648835551 | 1 |
| ENSMUSG00000094328 | Gm9125 | 0 | 1.648835551 | 1 |
| ENSMUSG00000028145 | Them4 | 0 | -2.599091962 | 0.108201045 |
| ENSMUSG00000028148 | Them5 | 0 | 1.648835551 | 1 |
| ENSMUSG00000105708 | Gm36070 | 0 | 1.063873051 | 1 |
| ENSMUSG00000105391 | Gm43401 | 0 | 1.648835551 | 1 |
| ENSMUSG00000044505 | Lingo4 | 0 | 1.063873051 | 1 |
| ENSMUSG00000106032 | Gm42463 | 0 | 0.063873051 | 1 |
| ENSMUSG00000028137 | Celf3 | 0 | 0.648835551 | 1 |
| ENSMUSG00000106061 | Gm43773 | 0 | 1.648835551 | 1 |
| ENSMUSG00000104631 | Gm43774 | 0 | -0.351164449 | 1 |
| ENSMUSG00000085250 | Gm15234 | 0 | 1.648835551 | 1 |
| ENSMUSG00000068877 | Selenbp2 | 0 | -2.560617814 | 0.105255472 |
| ENSMUSG00000087660 | B230398E01Rik | 0 | 0.648835551 | 1 |
| ENSMUSG00000085956 | 4930481B07Rik | 0 | 0.648835551 | 1 |
| ENSMUSG00000005628 | Tmod4 | 0 | 1.063873051 | 1 |
| ENSMUSG00000038777 | Sema6c | 0 | -1.051604167 | 0.858831093 |
| ENSMUSG00000038550 | Ciart | 0 | 0.326907457 | 1 |
| ENSMUSG00000038526 | Car14 | 0 | 1.063873051 | 1 |
| ENSMUSG00000097204 | Gm17690 | 0 | -1.52108945 | 0.544908989 |
| ENSMUSG00000038486 | Sv2a | 0 | 0.648835551 | 1 |
| ENSMUSG00000086556 | Gm15444 | 0 | 0.326907457 | 1 |
| ENSMUSG00000068855 | Hist2h2ac | 0 | -2.051604167 | 0.274319469 |
| ENSMUSG00000068854 | Hist2h2be | 0 | 1.648835551 | 1 |
| ENSMUSG00000081058 | Hist2h3c2 | 0 | 1.648835551 | 1 |
| ENSMUSG00000093553 | Gm20633 | 0 | 1.648835551 | 1 |
| ENSMUSG00000064220 | Hist2h2aa1 | 0 | 1.063873051 | 1 |
| ENSMUSG00000093769 | Hist2h3c1 | 0 | 1.648835551 | 1 |
| ENSMUSG00000074403 | Hist2h3b | 0 | 1.648835551 | 1 |
| ENSMUSG00000050936 | Gm42743 | 0 | 1.648835551 | 1 |
| ENSMUSG00000015947 | Fcgr1 | 0 | -0.52108945 | 1 |
| ENSMUSG00000097807 | Gm26553 | 0 | 0.648835551 | 1 |
| ENSMUSG00000106429 | Gm10685 | 0 | 1.063873051 | 1 |
| ENSMUSG00000105245 | Gm31305 | 0 | 1.648835551 | 1 |
| ENSMUSG00000074388 | Gm5544 | 0 | -0.351164449 | 1 |
| ENSMUSG00000043468 | Adam30 | 0 | 1.648835551 | 1 |
| ENSMUSG00000038092 | Hsd3b5 | 0 | 0.648835551 | 1 |
| ENSMUSG00000027870 | Hao2 | 0 | 0.648835551 | 1 |
| ENSMUSG00000105472 | Gm43024 | 0 | 0.063873051 | 1 |
| ENSMUSG00000106197 | Gm42717 | 0 | 1.648835551 | 1 |
| ENSMUSG00000027865 | Gdap2 | 0 | -2.305360759 | 0.231596497 |
| ENSMUSG00000053957 | Gm12474 | 0 | -0.673092543 | 1 |

| | | | | |
|---|---|---|---|---|
| ENSMUSG00000104943 | Gm42868 | 0 | 1.648835551 | 1 |
| ENSMUSG00000105824 | Gm43466 | 0 | -0.158519371 | 1 |
| ENSMUSG00000105137 | Gm42869 | 0 | 1.648835551 | 1 |
| ENSMUSG00000105561 | Gm43462 | 0 | -0.52108945 | 1 |
| ENSMUSG00000106184 | Gm43463 | 0 | 0.326907457 | 1 |
| ENSMUSG00000106227 | Gm43464 | 0 | 1.648835551 | 1 |
| ENSMUSG00000033233 | Trim45 | 0 | -0.673092543 | 1 |
| ENSMUSG00000105717 | Gm43465 | 0 | 1.063873051 | 1 |
| ENSMUSG00000027863 | Cd2 | 0 | 0.648835551 | 1 |
| ENSMUSG00000105611 | Gm42938 | 0 | 0.648835551 | 1 |
| ENSMUSG00000104737 | Gm42937 | 0 | 0.063873051 | 1 |
| ENSMUSG00000104693 | Gm42941 | 0 | 0.326907457 | 1 |
| ENSMUSG00000105642 | Gm43134 | 0 | 1.063873051 | 1 |
| ENSMUSG00000105342 | Gm43244 | 0 | 1.648835551 | 1 |
| ENSMUSG00000089916 | Gm16160 | 0 | 1.648835551 | 1 |
| ENSMUSG00000104619 | Gm42678 | 0 | 1.648835551 | 1 |
| ENSMUSG00000086745 | Tspan2os | 0 | 1.063873051 | 1 |
| ENSMUSG00000104671 | Gm43062 | 0 | 1.648835551 | 1 |
| ENSMUSG00000070385 | Ampd1 | 0 | 0.648835551 | 1 |
| ENSMUSG00000105731 | Gm43063 | 0 | 0.326907457 | 1 |
| ENSMUSG00000104764 | Gm43066 | 0 | 1.648835551 | 1 |
| ENSMUSG00000087119 | Atg4a-ps | 0 | 1.648835551 | 1 |
| ENSMUSG00000105229 | Gm43149 | 0 | 1.063873051 | 1 |
| ENSMUSG00000032952 | Ap4b1 | 0 | -1.673092543 | 0.451785624 |
| ENSMUSG00000084760 | Gm15472 | 0 | 1.648835551 | 1 |
| ENSMUSG00000106371 | 9530097N15Rik | 0 | 0.648835551 | 1 |
| ENSMUSG00000105148 | Gm42700 | 0 | 0.326907457 | 1 |
| ENSMUSG00000106443 | Gm42702 | 0 | 0.063873051 | 1 |
| ENSMUSG00000104583 | Gm42701 | 0 | 0.063873051 | 1 |
| ENSMUSG00000105846 | Gm43258 | 0 | 1.063873051 | 1 |
| ENSMUSG00000106063 | C030032O16Rik | 0 | 0.326907457 | 1 |
| ENSMUSG00000106432 | Gm43259 | 0 | -0.158519371 | 1 |
| ENSMUSG00000105818 | Gm43256 | 0 | -1.258055044 | 0.673811773 |
| ENSMUSG00000104768 | Gm38412 | 0 | 0.326907457 | 1 |
| ENSMUSG00000105304 | Gm43696 | 0 | -0.52108945 | 1 |
| ENSMUSG00000104973 | A530041M06Rik | 0 | 1.063873051 | 1 |
| ENSMUSG00000105659 | 4930509H03Rik | 0 | 0.326907457 | 1 |
| ENSMUSG00000105199 | Gm43581 | 0 | 1.648835551 | 1 |
| ENSMUSG00000055865 | Tafa3 | 0 | 0.326907457 | 1 |
| ENSMUSG00000002228 | Ppm1j | 0 | -1.258055044 | 0.673811773 |
| ENSMUSG00000106251 | Gm42658 | 0 | 1.648835551 | 1 |
| ENSMUSG00000105655 | Gm42659 | 0 | -0.673092543 | 1 |
| ENSMUSG00000105925 | Gm43846 | 0 | 0.063873051 | 1 |
| ENSMUSG00000105355 | Gm40117 | 0 | 1.648835551 | 1 |
| ENSMUSG00000051788 | 4930564D02Rik | 0 | 0.648835551 | 1 |
| ENSMUSG00000027905 | Ddx20 | 0 | -1.995020638 | 0.370945697 |
| ENSMUSG00000105677 | Gm43328 | 0 | 1.648835551 | 1 |
| ENSMUSG00000105842 | Gm43329 | 0 | 0.648835551 | 1 |
| ENSMUSG00000106341 | Gm43330 | 0 | 0.063873051 | 1 |
| ENSMUSG00000074342 | I830077J02Rik | 0 | 0.648835551 | 1 |
| ENSMUSG00000104592 | Gm42721 | 0 | 1.063873051 | 1 |

| | | | | |
|---|---|---|---|---|
| ENSMUSG00000091575 | 2010016I18Rik | 0 | -0.673092543 | 1 |
| ENSMUSG00000043529 | Olfr266 | 0 | 0.063873051 | 1 |
| ENSMUSG00000056145 | AI504432 | 0 | 0.063873051 | 1 |
| ENSMUSG00000040724 | Kcna2 | 0 | -1.051604167 | 0.871107617 |
| ENSMUSG00000042861 | Kcna10 | 0 | 1.648835551 | 1 |
| ENSMUSG00000027894 | Slc6a17 | 0 | -0.158519371 | 1 |
| ENSMUSG00000014603 | Alx3 | 0 | 1.648835551 | 1 |
| ENSMUSG00000014599 | Csf1 | 0 | -1.673092543 | 0.451785624 |
| ENSMUSG00000086968 | 4933431E20Rik | 0 | 0.326907457 | 1 |
| ENSMUSG00000004035 | Gstm7 | 0 | 0.648835551 | 1 |
| ENSMUSG00000027890 | Gstm4 | 0 | 1.063873051 | 1 |
| ENSMUSG00000027889 | Ampd2 | 0 | -2.209145444 | 0.217194106 |
| ENSMUSG00000009108 | Gnat2 | 0 | 1.648835551 | 1 |
| ENSMUSG00000050947 | Amigo1 | 0 | -0.351164449 | 1 |
| ENSMUSG00000048796 | Cyb561d1 | 0 | 0.648835551 | 1 |
| ENSMUSG00000027887 | Sypl2 | 0 | 1.063873051 | 1 |
| ENSMUSG00000085102 | 1700010K24Rik | 0 | -1.995020638 | 0.370945697 |
| ENSMUSG00000068745 | Mybphl | 0 | 1.063873051 | 1 |
| ENSMUSG00000074281 | Gm12522 | 0 | 0.648835551 | 1 |
| ENSMUSG00000068744 | Psrc1 | 0 | 1.063873051 | 1 |
| ENSMUSG00000045662 | Henmt1 | 0 | 0.326907457 | 1 |
| ENSMUSG00000105139 | Gm19391 | 0 | -1.258055044 | 0.673811773 |
| ENSMUSG00000049300 | Prmt6 | 0 | -1.351164449 | 0.687112677 |
| ENSMUSG00000105509 | C130013H08Rik | 0 | 1.648835551 | 1 |
| ENSMUSG00000074264 | Amy1 | 0 | -2.051604167 | 0.316908951 |
| ENSMUSG00000027963 | Extl2 | 0 | -1.599091962 | 0.559918909 |
| ENSMUSG00000104653 | Gm43109 | 0 | 1.648835551 | 1 |
| ENSMUSG00000097362 | Gm26544 | 0 | -0.158519371 | 1 |
| ENSMUSG00000106005 | Gm29151 | 0 | 1.648835551 | 1 |
| ENSMUSG00000106415 | Gm42893 | 0 | 1.648835551 | 1 |
| ENSMUSG00000080907 | 4930455H04Rik | 0 | 0.063873051 | 1 |
| ENSMUSG00000028009 | 1700061I17Rik | 0 | -1.52108945 | 0.544908989 |
| ENSMUSG00000106099 | Gm42664 | 0 | -0.158519371 | 1 |
| ENSMUSG00000097124 | A530020G20Rik | 0 | -2.106051951 | 0.279592421 |
| ENSMUSG00000044522 | A730020M07Rik | 0 | 1.648835551 | 1 |
| ENSMUSG00000074248 | 4930432M17Rik | 0 | 1.063873051 | 1 |
| ENSMUSG00000106036 | Gm43608 | 0 | 1.063873051 | 1 |
| ENSMUSG00000104574 | Gm42836 | 0 | -0.673092543 | 1 |
| ENSMUSG00000106194 | Gm43569 | 0 | 0.063873051 | 1 |
| ENSMUSG00000106365 | Gm43568 | 0 | 1.648835551 | 1 |
| ENSMUSG00000028116 | Myoz2 | 0 | 0.326907457 | 1 |
| ENSMUSG00000104531 | Gm43731 | 0 | -0.673092543 | 1 |
| ENSMUSG00000044528 | Tram1l1 | 0 | 0.648835551 | 1 |
| ENSMUSG00000039174 | 1700003H04Rik | 0 | 1.063873051 | 1 |
| ENSMUSG00000090427 | Gm17225 | 0 | 1.648835551 | 1 |
| ENSMUSG00000105104 | Gm43729 | 0 | 1.648835551 | 1 |
| ENSMUSG00000105447 | Gm43653 | 0 | 1.063873051 | 1 |
| ENSMUSG00000105550 | Gm35585 | 0 | 1.648835551 | 1 |
| ENSMUSG00000093865 | Lrit3 | 0 | 0.326907457 | 1 |
| ENSMUSG00000028012 | Rrh | 0 | 0.648835551 | 1 |
| ENSMUSG00000027983 | Cyp2u1 | 0 | 1.648835551 | 1 |

| | | | | |
|---|---|---|---|---|
| ENSMUSG00000106230 | Gm29811 | 0 | 0.063873051 | 1 |
| ENSMUSG00000105957 | Gm43254 | 0 | 1.063873051 | 1 |
| ENSMUSG00000105952 | Gm40153 | 0 | 0.063873051 | 1 |
| ENSMUSG00000106069 | Gm6135 | 0 | 1.648835551 | 1 |
| ENSMUSG00000105765 | 4930539C22Rik | 0 | 1.648835551 | 1 |
| ENSMUSG00000106398 | Gm43357 | 0 | 1.648835551 | 1 |
| ENSMUSG00000106122 | Gm42610 | 0 | 0.326907457 | 1 |
| ENSMUSG00000105676 | Gm43403 | 0 | 1.648835551 | 1 |
| ENSMUSG00000106629 | Gm40155 | 0 | 1.648835551 | 1 |
| ENSMUSG00000104675 | Gm43689 | 0 | 0.648835551 | 1 |
| ENSMUSG00000105296 | Gm19708 | 0 | 1.063873051 | 1 |
| ENSMUSG00000104776 | Gm43691 | 0 | 0.063873051 | 1 |
| ENSMUSG00000099146 | 0610031O16Rik | 0 | 1.648835551 | 1 |
| ENSMUSG00000074207 | Adh1 | 0 | 1.063873051 | 1 |
| ENSMUSG00000053054 | Adh6a | 0 | 0.648835551 | 1 |
| ENSMUSG00000074206 | Adh6b | 0 | -0.52108945 | 1 |
| ENSMUSG00000104544 | Gm32754 | 0 | 1.063873051 | 1 |
| ENSMUSG00000089924 | Gm15689 | 0 | 1.648835551 | 1 |
| ENSMUSG00000040264 | Gbp2b | 0 | 0.326907457 | 1 |
| ENSMUSG00000028270 | Gbp2 | 0 | -0.52108945 | 1 |
| ENSMUSG00000040213 | Kyat3 | 0 | -1.158519371 | 0.930796482 |
| ENSMUSG00000105652 | 4930519L02Rik | 0 | 1.648835551 | 1 |
| ENSMUSG00000106052 | Gm43614 | 0 | 0.326907457 | 1 |
| ENSMUSG00000105923 | A830019L24Rik | 0 | 1.063873051 | 1 |
| ENSMUSG00000106162 | Gm43613 | 0 | 1.648835551 | 1 |
| ENSMUSG00000106210 | 1700001N15Rik | 0 | 1.063873051 | 1 |
| ENSMUSG00000104865 | Gm42708 | 0 | 1.648835551 | 1 |
| ENSMUSG00000106367 | Gm34078 | 0 | 0.063873051 | 1 |
| ENSMUSG00000104969 | Gm43445 | 0 | -1.051604167 | 0.858831093 |
| ENSMUSG00000104903 | Gm43707 | 0 | 1.648835551 | 1 |
| ENSMUSG00000106057 | Gm43560 | 0 | 1.648835551 | 1 |
| ENSMUSG00000106424 | Gm34866 | 0 | 1.063873051 | 1 |
| ENSMUSG00000105900 | Gm43240 | 0 | 0.648835551 | 1 |
| ENSMUSG00000074195 | Clca4b | 0 | -0.158519371 | 1 |
| ENSMUSG00000085773 | Gm16233 | 0 | 0.648835551 | 1 |
| ENSMUSG00000068547 | Clca4a | 0 | 0.648835551 | 1 |
| ENSMUSG00000106361 | Gm35066 | 0 | 0.648835551 | 1 |
| ENSMUSG00000036960 | Clca2 | 0 | 1.063873051 | 1 |
| ENSMUSG00000105619 | 9530034A14Rik | 0 | 1.648835551 | 1 |
| ENSMUSG00000028185 | Dnase2b | 0 | 0.326907457 | 1 |
| ENSMUSG00000085810 | Gm16325 | 0 | 1.648835551 | 1 |
| ENSMUSG00000090202 | 4930503B20Rik | 0 | 1.648835551 | 1 |
| ENSMUSG00000103334 | Gm29771 | 0 | -0.351164449 | 1 |
| ENSMUSG00000105876 | Gm43572 | 0 | 1.648835551 | 1 |
| ENSMUSG00000070342 | Gm10287 | 0 | 1.063873051 | 1 |
| ENSMUSG00000105190 | Gm43470 | 0 | 0.648835551 | 1 |
| ENSMUSG00000105125 | Gm43468 | 0 | 1.648835551 | 1 |
| ENSMUSG00000105698 | Gm31881 | 0 | 1.648835551 | 1 |
| ENSMUSG00000097745 | AI115009 | 0 | 1.063873051 | 1 |
| ENSMUSG00000105337 | 1700094M23Rik | 0 | 0.648835551 | 1 |
| ENSMUSG00000105023 | 1700012D16Rik | 0 | 1.648835551 | 1 |

| | | | | |
|---|---|---|---|---|
| ENSMUSG00000062046 | 5730460C07Rik | 0 | 1.648835551 | 1 |
| ENSMUSG00000086438 | Asb17os | 0 | 0.326907457 | 1 |
| ENSMUSG00000038997 | Asb17 | 0 | 1.648835551 | 1 |
| ENSMUSG00000096225 | Lhx8 | 0 | 0.063873051 | 1 |
| ENSMUSG00000093738 | AI606473 | 0 | 1.648835551 | 1 |
| ENSMUSG00000103204 | Gm38139 | 0 | 1.648835551 | 1 |
| ENSMUSG00000104553 | 9330178D15Rik | 0 | -0.52108945 | 1 |
| ENSMUSG00000106632 | Gm33466 | 0 | 1.063873051 | 1 |
| ENSMUSG00000028175 | Depdc1a | 0 | -1.52108945 | 0.544908989 |
| ENSMUSG00000028174 | Rpe65 | 0 | 0.063873051 | 1 |
| ENSMUSG00000085180 | AI838599 | 0 | 0.063873051 | 1 |
| ENSMUSG00000078365 | Mos | 0 | 1.648835551 | 1 |
| ENSMUSG00000028236 | Sdr16c5 | 0 | 0.648835551 | 1 |
| ENSMUSG00000071019 | Sdr16c6 | 0 | 1.648835551 | 1 |
| ENSMUSG00000028240 | Cyp7a1 | 0 | 1.648835551 | 1 |
| ENSMUSG00000086233 | Gm11816 | 0 | 1.648835551 | 1 |
| ENSMUSG00000051279 | Gdf6 | 0 | 0.326907457 | 1 |
| ENSMUSG00000086765 | Gm11827 | 0 | 1.648835551 | 1 |
| ENSMUSG00000028214 | Gem | 0 | -1.258055044 | 0.94561855 |
| ENSMUSG00000085614 | 1700123M08Rik | 0 | -1.258055044 | 0.673811773 |
| ENSMUSG00000049488 | Tmem67 | 0 | -2.599091962 | 0.14143852 |
| ENSMUSG00000046667 | Rbm12b1 | 0 | -2.106051951 | 0.279592421 |
| ENSMUSG00000085909 | Gm11846 | 0 | 0.063873051 | 1 |
| ENSMUSG00000086227 | Gm11820 | 0 | 1.063873051 | 1 |
| ENSMUSG00000086068 | 1700120G11Rik | 0 | 0.063873051 | 1 |
| ENSMUSG00000086587 | Gm11837 | 0 | 0.648835551 | 1 |
| ENSMUSG00000028222 | Calb1 | 0 | 1.648835551 | 1 |
| ENSMUSG00000041153 | Osgin2 | 0 | 0.326907457 | 1 |
| ENSMUSG00000086332 | 4930480G23Rik | 0 | 1.648835551 | 1 |
| ENSMUSG00000028244 | 1700025O08Rik | 0 | 1.063873051 | 1 |
| ENSMUSG00000087399 | Gm11899 | 0 | 1.063873051 | 1 |
| ENSMUSG00000087316 | Gm11917 | 0 | 0.648835551 | 1 |
| ENSMUSG00000051056 | Gja10 | 0 | 1.648835551 | 1 |
| ENSMUSG00000023267 | Gabrr2 | 0 | 0.063873051 | 1 |
| ENSMUSG00000086127 | Gm11934 | 0 | 1.648835551 | 1 |
| ENSMUSG00000044288 | Cnr1 | 0 | 1.063873051 | 1 |
| ENSMUSG00000028264 | Spaca1 | 0 | 0.063873051 | 1 |
| ENSMUSG00000087602 | Gm12753 | 0 | 1.648835551 | 1 |
| ENSMUSG00000028298 | Cga | 0 | 0.648835551 | 1 |
| ENSMUSG00000028300 | C9orf72 | 0 | -1.995020638 | 0.370945697 |
| ENSMUSG00000087137 | 4930509K18Rik | 0 | 1.648835551 | 1 |
| ENSMUSG00000046593 | Tmem215 | 0 | 1.648835551 | 1 |
| ENSMUSG00000086355 | 4933428C19Rik | 0 | 0.648835551 | 1 |
| ENSMUSG00000028415 | Spink4 | 0 | -0.52108945 | 1 |
| ENSMUSG00000028427 | Aqp7 | 0 | 1.648835551 | 1 |
| ENSMUSG00000028435 | Aqp3 | 0 | 1.063873051 | 1 |
| ENSMUSG00000046312 | Myorg | 0 | 1.063873051 | 1 |
| ENSMUSG00000078747 | Gm20878 | 0 | 1.648835551 | 1 |
| ENSMUSG00000054885 | 4930578G10Rik | 0 | -0.158519371 | 1 |
| ENSMUSG00000071005 | Ccl19 | 0 | 1.648835551 | 1 |
| ENSMUSG00000036052 | Dnajb5 | 0 | 0.326907457 | 1 |

| | | | | |
|---|---|---|---|---|
| ENSMUSG00000028453 | Fancg | 0 | -1.351164449 | 0.687112677 |
| ENSMUSG00000096974 | Gm26881 | 0 | -0.673092543 | 1 |
| ENSMUSG00000028459 | Cd72 | 0 | 1.063873051 | 1 |
| ENSMUSG00000028460 | Sit1 | 0 | 1.063873051 | 1 |
| ENSMUSG00000051517 | Arhgef39 | 0 | 0.063873051 | 1 |
| ENSMUSG00000028467 | Gba2 | 0 | -1.158519371 | 0.915416636 |
| ENSMUSG00000078719 | Msmp | 0 | 1.648835551 | 1 |
| ENSMUSG00000085351 | Gm12472 | 0 | 0.648835551 | 1 |
| ENSMUSG00000066196 | Spag8 | 0 | 1.063873051 | 1 |
| ENSMUSG00000043633 | Fam221b | 0 | 1.063873051 | 1 |
| ENSMUSG00000046450 | Olfr71 | 0 | 1.063873051 | 1 |
| ENSMUSG00000071001 | Hrct1 | 0 | -1.810596067 | 0.465580427 |
| ENSMUSG00000111611 | Olfr157 | 0 | 1.648835551 | 1 |
| ENSMUSG00000084854 | Gm12678 | 0 | 1.648835551 | 1 |
| ENSMUSG00000049657 | Zbtb5 | 0 | -2.051604167 | 0.274319469 |
| ENSMUSG00000073860 | Gm829 | 0 | -0.52108945 | 1 |
| ENSMUSG00000085876 | Gm12409 | 0 | 1.648835551 | 1 |
| ENSMUSG00000035561 | Aldh1b1 | 0 | 0.326907457 | 1 |
| ENSMUSG00000035551 | Igfbpl1 | 0 | -0.810596067 | 1 |
| ENSMUSG00000028328 | Tmod1 | 0 | -1.52108945 | 0.586600655 |
| ENSMUSG00000070990 | Foxe1 | 0 | 1.648835551 | 1 |
| ENSMUSG00000087500 | Gm12426 | 0 | 1.648835551 | 1 |
| ENSMUSG00000028341 | Nr4a3 | 0 | 0.063873051 | 1 |
| ENSMUSG00000084787 | Gm12436 | 0 | -0.52108945 | 1 |
| ENSMUSG00000039653 | Baat | 0 | 1.648835551 | 1 |
| ENSMUSG00000028307 | Aldob | 0 | 1.648835551 | 1 |
| ENSMUSG00000039555 | Cylc2 | 0 | -0.936126949 | 0.867531944 |
| ENSMUSG00000089717 | Olfr275 | 0 | 1.063873051 | 1 |
| ENSMUSG00000070983 | Olfr270 | 0 | 0.326907457 | 1 |
| ENSMUSG00000086712 | AI427809 | 0 | 1.648835551 | 1 |
| ENSMUSG00000028417 | Tal2 | 0 | 0.326907457 | 1 |
| ENSMUSG00000086987 | Gm12484 | 0 | 1.063873051 | 1 |
| ENSMUSG00000086371 | Gm12478 | 0 | 0.326907457 | 1 |
| ENSMUSG00000087070 | Gm12505 | 0 | 1.063873051 | 1 |
| ENSMUSG00000085643 | Gm12519 | 0 | 1.648835551 | 1 |
| ENSMUSG00000070980 | Actl7b | 0 | 1.063873051 | 1 |
| ENSMUSG00000070979 | Actl7a | 0 | 1.648835551 | 1 |
| ENSMUSG00000038827 | Abitram | 0 | -0.158519371 | 1 |
| ENSMUSG00000038709 | Txndc8 | 0 | 0.326907457 | 1 |
| ENSMUSG00000086470 | Gm12580 | 0 | 1.063873051 | 1 |
| ENSMUSG00000043385 | Olfr267 | 0 | 0.648835551 | 1 |
| ENSMUSG00000038630 | Zkscan16 | 0 | 1.063873051 | 1 |
| ENSMUSG00000073843 | Gm12542 | 0 | 1.648835551 | 1 |
| ENSMUSG00000028386 | Slc46a2 | 0 | 0.648835551 | 1 |
| ENSMUSG00000078689 | Mup6 | 0 | 0.648835551 | 1 |
| ENSMUSG00000081207 | Gm13775 | 0 | 1.063873051 | 1 |
| ENSMUSG00000086530 | Gm12910 | 0 | 1.648835551 | 1 |
| ENSMUSG00000063851 | Rnf183 | 0 | 1.063873051 | 1 |
| ENSMUSG00000058046 | 4933430I17Rik | 0 | 1.648835551 | 1 |
| ENSMUSG00000085177 | Gm11209 | 0 | 1.648835551 | 1 |
| ENSMUSG00000028357 | Kif12 | 0 | -3.023589791 | 0.034614 |

| | | | | |
|---|---|---|---|---|
| ENSMUSG00000061540 | Orm2 | 0 | 1.648835551 | 1 |
| ENSMUSG00000086953 | Aknaos | 0 | 1.063873051 | 1 |
| ENSMUSG00000045917 | Tmem268 | 0 | -1.051604167 | 0.889426255 |
| ENSMUSG00000086790 | Gm12911 | 0 | -1.051604167 | 0.899941735 |
| ENSMUSG00000095935 | Gm11238 | 0 | 0.648835551 | 1 |
| ENSMUSG00000085034 | Gm11240 | 0 | 1.648835551 | 1 |
| ENSMUSG00000085482 | A230083N12Rik | 0 | 1.648835551 | 1 |
| ENSMUSG00000110373 | Gm42303 | 0 | -0.52108945 | 1 |
| ENSMUSG00000005994 | Tyrp1 | 0 | 1.648835551 | 1 |
| ENSMUSG00000084851 | Gm11267 | 0 | 1.648835551 | 1 |
| ENSMUSG00000086321 | Gm11413 | 0 | 1.063873051 | 1 |
| ENSMUSG00000087592 | Gm11184 | 0 | 1.063873051 | 1 |
| ENSMUSG00000085217 | 6030471H07Rik | 0 | 1.648835551 | 1 |
| ENSMUSG00000087378 | Gm12414 | 0 | 0.648835551 | 1 |
| ENSMUSG00000084809 | Gm12631 | 0 | -0.351164449 | 1 |
| ENSMUSG00000095101 | Gm13285 | 0 | 1.648835551 | 1 |
| ENSMUSG00000070908 | Gm13288 | 0 | 1.648835551 | 1 |
| ENSMUSG00000043753 | Dmrta1 | 0 | 1.648835551 | 1 |
| ENSMUSG00000085690 | Gm12629 | 0 | 1.648835551 | 1 |
| ENSMUSG00000087128 | Gm12655 | 0 | 0.326907457 | 1 |
| ENSMUSG00000028575 | Eqtn | 0 | 0.326907457 | 1 |
| ENSMUSG00000052767 | Gm12703 | 0 | 1.063873051 | 1 |
| ENSMUSG00000084918 | Gm12708 | 0 | 1.063873051 | 1 |
| ENSMUSG00000085835 | Gm12707 | 0 | 1.063873051 | 1 |
| ENSMUSG00000081225 | Cyp2j12 | 0 | -1.158519371 | 0.886203497 |
| ENSMUSG00000066097 | Cyp2j11 | 0 | 0.648835551 | 1 |
| ENSMUSG00000015224 | Cyp2j9 | 0 | 1.063873051 | 1 |
| ENSMUSG00000046683 | 0610025J13Rik | 0 | 1.063873051 | 1 |
| ENSMUSG00000087166 | L1td1 | 0 | 0.648835551 | 1 |
| ENSMUSG00000073794 | I0C0044D17Rik | 0 | 1.648835551 | 1 |
| ENSMUSG00000028553 | Angptl3 | 0 | 0.326907457 | 1 |
| ENSMUSG00000087395 | Gm12705 | 0 | 1.063873051 | 1 |
| ENSMUSG00000086498 | Gm12681 | 0 | 1.063873051 | 1 |
| ENSMUSG00000087332 | Gm12690 | 0 | 1.063873051 | 1 |
| ENSMUSG00000085142 | Gm12702 | 0 | 1.063873051 | 1 |
| ENSMUSG00000086385 | Gm12701 | 0 | 1.648835551 | 1 |
| ENSMUSG00000084758 | Gm12798 | 0 | 1.648835551 | 1 |
| ENSMUSG00000085181 | Gm12709 | 0 | -1.158519371 | 0.886203497 |
| ENSMUSG00000107726 | Gm44037 | 0 | 1.648835551 | 1 |
| ENSMUSG00000086136 | Gm12718 | 0 | -0.673092543 | 1 |
| ENSMUSG00000029656 | C8b | 0 | 1.063873051 | 1 |
| ENSMUSG00000035031 | C8a | 0 | 0.063873051 | 1 |
| ENSMUSG00000085790 | Gm12729 | 0 | 1.648835551 | 1 |
| ENSMUSG00000086172 | 2700068H02Rik | 0 | 1.063873051 | 1 |
| ENSMUSG00000085023 | Gm12744 | 0 | 1.648835551 | 1 |
| ENSMUSG00000034919 | Ttc22 | 0 | -0.351164449 | 1 |
| ENSMUSG00000043572 | Pars2 | 0 | 0.326907457 | 1 |
| ENSMUSG00000025413 | Ttc4 | 0 | -3.636566667 | 0.005443828 |
| ENSMUSG00000034871 | Fam151a | 0 | 0.326907457 | 1 |
| ENSMUSG00000028621 | Cyb5rl | 0 | -2.258055044 | 0.222101622 |
| ENSMUSG00000070877 | Ldlrad1 | 0 | -0.158519371 | 1 |

| | | | | |
|---|---|---|---|---|
| ENSMUSG00000034785 | Dio1 | 0 | 1.063873051 | 1 |
| ENSMUSG00000028610 | Dmrtb1 | 0 | 1.063873051 | 1 |
| ENSMUSG00000087195 | Gm12907 | 0 | 1.063873051 | 1 |
| ENSMUSG00000073779 | Lrp8os2 | 0 | 0.326907457 | 1 |
| ENSMUSG00000028612 | Lrp8os1 | 0 | 1.648835551 | 1 |
| ENSMUSG00000086713 | 4930407G08Rik | 0 | 1.063873051 | 1 |
| ENSMUSG00000028587 | Orc1 | 0 | -2.560617814 | 0.116909584 |
| ENSMUSG00000091985 | Gm17354 | 0 | -0.158519371 | 1 |
| ENSMUSG00000028558 | Calr4 | 0 | 1.648835551 | 1 |
| ENSMUSG00000085873 | Ttc39aos1 | 0 | 1.063873051 | 1 |
| ENSMUSG00000059027 | 9630013D21Rik | 0 | 0.063873051 | 1 |
| ENSMUSG00000047143 | Dmrta2 | 0 | 1.063873051 | 1 |
| ENSMUSG00000028545 | Bend5 | 0 | 1.063873051 | 1 |
| ENSMUSG00000078599 | Skint8 | 0 | 1.063873051 | 1 |
| ENSMUSG00000055960 | Skint4 | 0 | 0.063873051 | 1 |
| ENSMUSG00000070868 | Skint3 | 0 | 0.648835551 | 1 |
| ENSMUSG00000049972 | Skint9 | 0 | -0.351164449 | 1 |
| ENSMUSG00000034359 | Skint2 | 0 | 0.648835551 | 1 |
| ENSMUSG00000048766 | Skint10 | 0 | 0.648835551 | 1 |
| ENSMUSG00000085878 | Gm12823 | 0 | 1.648835551 | 1 |
| ENSMUSG00000085323 | 9130410C08Rik | 0 | -0.351164449 | 1 |
| ENSMUSG00000028717 | Tal1 | 0 | 0.648835551 | 1 |
| ENSMUSG00000083138 | Cyp4a29 | 0 | 1.648835551 | 1 |
| ENSMUSG00000078597 | Cyp4a12b | 0 | 0.326907457 | 1 |
| ENSMUSG00000084346 | Cyp4a30b | 0 | 1.063873051 | 1 |
| ENSMUSG00000063929 | Cyp4a32 | 0 | 1.063873051 | 1 |
| ENSMUSG00000073774 | Kncn | 0 | 1.063873051 | 1 |
| ENSMUSG00000034185 | 6430628N08Rik | 0 | -0.52108945 | 1 |
| ENSMUSG00000028707 | Dmbx1 | 0 | 1.648835551 | 1 |
| ENSMUSG00000028701 | Lurap1 | 0 | 0.063873051 | 1 |
| ENSMUSG00000087299 | Gm12953 | 0 | 1.648835551 | 1 |
| ENSMUSG00000085408 | C530005A16Rik | 0 | 0.326907457 | 1 |
| ENSMUSG00000034035 | Ccdc17 | 0 | -0.158519371 | 1 |
| ENSMUSG00000028687 | Mutyh | 0 | -0.52108945 | 1 |
| ENSMUSG00000043155 | Hpdl | 0 | 1.063873051 | 1 |
| ENSMUSG00000073771 | Btbd19 | 0 | -0.351164449 | 1 |
| ENSMUSG00000047671 | Tctex1d4 | 0 | 1.648835551 | 1 |
| ENSMUSG00000085749 | Gm12843 | 0 | 1.648835551 | 1 |
| ENSMUSG00000028541 | B4galt2 | 0 | -0.936126949 | 0.847265357 |
| ENSMUSG00000028536 | 2610528J11Rik | 0 | 1.648835551 | 1 |
| ENSMUSG00000028730 | Cfap57 | 0 | -0.673092543 | 1 |
| ENSMUSG00000085292 | Gm12853 | 0 | 1.063873051 | 1 |
| ENSMUSG00000043698 | Olfr62 | 0 | 1.063873051 | 1 |
| ENSMUSG00000095218 | Olfr1338 | 0 | 1.648835551 | 1 |
| ENSMUSG00000110947 | Olfr1333 | 0 | -2.158519371 | 0.330002444 |
| ENSMUSG00000024903 | Lao1 | 0 | 0.648835551 | 1 |
| ENSMUSG00000085505 | Gm12868 | 0 | 1.648835551 | 1 |
| ENSMUSG00000028644 | Ermap | 0 | 0.648835551 | 1 |
| ENSMUSG00000028642 | Tmem269 | 0 | 1.648835551 | 1 |
| ENSMUSG00000028641 | P3h1 | 0 | -3.132524162 | 0.032604666 |
| ENSMUSG00000066058 | Cldn19 | 0 | 1.648835551 | 1 |

| | | | | |
|---|---|---|---|---|
| ENSMUSG00000048899 | Rimkla | 0 | -0.810596067 | 0.847265357 |
| ENSMUSG00000087385 | Frg2f1 | 0 | -0.351164449 | 1 |
| ENSMUSG00000052135 | Foxo6 | 0 | -0.52108945 | 1 |
| ENSMUSG00000084929 | Foxo6os | 0 | 1.063873051 | 1 |
| ENSMUSG00000110389 | Gm45579 | 0 | 1.648835551 | 1 |
| ENSMUSG00000023263 | 9530002B09Rik | 0 | 0.648835551 | 1 |
| ENSMUSG00000086443 | 4933421A08Rik | 0 | -0.351164449 | 1 |
| ENSMUSG00000028654 | Mycl | 0 | 0.063873051 | 1 |
| ENSMUSG00000054958 | Nt5c1a | 0 | -0.158519371 | 1 |
| ENSMUSG00000032744 | Heyl | 0 | 0.648835551 | 1 |
| ENSMUSG00000087307 | Gm12925 | 0 | 0.326907457 | 1 |
| ENSMUSG00000090125 | Pou3f1 | 0 | 1.648835551 | 1 |
| ENSMUSG00000028873 | Cdca8 | 0 | -1.874726405 | 0.352215149 |
| ENSMUSG00000028871 | Rspo1 | 0 | -0.351164449 | 1 |
| ENSMUSG00000042707 | Dnali1 | 0 | 1.063873051 | 1 |
| ENSMUSG00000086342 | Gm12932 | 0 | 1.648835551 | 1 |
| ENSMUSG00000087010 | 1700041M05Rik | 0 | 0.648835551 | 1 |
| ENSMUSG00000042558 | Adprhl2 | 0 | -0.936126949 | 0.847265357 |
| ENSMUSG00000028845 | Tekt2 | 0 | 0.648835551 | 1 |
| ENSMUSG00000042477 | Tfap2e | 0 | 1.648835551 | 1 |
| ENSMUSG00000050234 | Gja4 | 0 | 0.648835551 | 1 |
| ENSMUSG00000086771 | 1700080G11Rik | 0 | 1.648835551 | 1 |
| ENSMUSG00000028813 | CK137956 | 0 | 0.063873051 | 1 |
| ENSMUSG00000087459 | Csmd2os | 0 | -1.258055044 | 0.708502317 |
| ENSMUSG00000041000 | Trim62 | 0 | 0.063873051 | 1 |
| ENSMUSG00000085085 | 1700086P04Rik | 0 | -0.52108945 | 1 |
| ENSMUSG00000028786 | Tmem54 | 0 | 1.648835551 | 1 |
| ENSMUSG00000001334 | Fndc5 | 0 | -0.158519371 | 1 |
| ENSMUSG00000048485 | Zbtb8b | 0 | -0.158519371 | 1 |
| ENSMUSG00000000411 | Tssk3 | 0 | 1.648835551 | 1 |
| ENSMUSG00000050493 | Fam167b | 0 | 1.648835551 | 1 |
| ENSMUSG00000086369 | E330017L17Rik | 0 | 1.063873051 | 1 |
| ENSMUSG00000028778 | Hcrtr1 | 0 | 0.648835551 | 1 |
| ENSMUSG00000023120 | Ldc1 | 0 | -0.351164449 | 1 |
| ENSMUSG00000040533 | Matn1 | 0 | 1.648835551 | 1 |
| ENSMUSG00000086856 | Gm16080 | 0 | 1.648835551 | 1 |
| ENSMUSG00000085667 | Gm12992 | 0 | 0.648835551 | 1 |
| ENSMUSG00000070720 | Tmem200b | 0 | 1.648835551 | 1 |
| ENSMUSG00000066042 | Med18 | 0 | -1.258055044 | 0.673811773 |
| ENSMUSG00000086997 | Gm12981 | 0 | 1.648835551 | 1 |
| ENSMUSG00000037752 | Xkr8 | 0 | 0.326907457 | 1 |
| ENSMUSG00000028865 | Cd164l2 | 0 | 0.063873051 | 1 |
| ENSMUSG00000037600 | Kdf1 | 0 | -1.743481871 | 0.463009681 |
| ENSMUSG00000085009 | Gm12977 | 0 | 1.648835551 | 1 |
| ENSMUSG00000012126 | Ubxn11 | 0 | -0.158519371 | 1 |
| ENSMUSG00000073747 | Gm7534 | 0 | 0.326907457 | 1 |
| ENSMUSG00000050105 | Grrp1 | 0 | 1.648835551 | 1 |
| ENSMUSG00000050890 | Pdik1l | 0 | -2.395558568 | 0.174915761 |
| ENSMUSG00000028834 | Trim63 | 0 | 0.063873051 | 1 |
| ENSMUSG00000086884 | Gm16225 | 0 | 1.648835551 | 1 |
| ENSMUSG00000087016 | Gm15979 | 0 | 1.648835551 | 1 |

| | | | | |
|---|---|---|---|---|
| ENSMUSG00000085714 | Gm13008 | 0 | 1.063873051 | 1 |
| ENSMUSG00000018983 | E2f2 | 0 | -1.936126949 | 0.403518669 |
| ENSMUSG00000066178 | 6030445D17Rik | 0 | 1.063873051 | 1 |
| ENSMUSG00000036921 | Tex46 | 0 | 1.648835551 | 1 |
| ENSMUSG00000070683 | Lactbl1 | 0 | 0.326907457 | 1 |
| ENSMUSG00000028661 | Epha8 | 0 | 1.648835551 | 1 |
| ENSMUSG00000085833 | Gm13003 | 0 | 1.648835551 | 1 |
| ENSMUSG00000101493 | 2810405F17Rik | 0 | 0.063873051 | 1 |
| ENSMUSG00000078520 | Cela3a | 0 | 1.648835551 | 1 |
| ENSMUSG00000023433 | Cela3b | 0 | 1.648835551 | 1 |
| ENSMUSG00000043621 | Ubxn10 | 0 | -0.52108945 | 1 |
| ENSMUSG00000028750 | Pla2g2c | 0 | 1.648835551 | 1 |
| ENSMUSG00000041202 | Pla2g2d | 0 | 0.326907457 | 1 |
| ENSMUSG00000073739 | Gm16287 | 0 | 0.648835551 | 1 |
| ENSMUSG00000028738 | Tas1r2 | 0 | 1.648835551 | 1 |
| ENSMUSG00000087628 | Gm13028 | 0 | 0.648835551 | 1 |
| ENSMUSG00000087084 | Gm13016 | 0 | 1.063873051 | 1 |
| ENSMUSG00000086437 | Gm13021 | 0 | 0.648835551 | 1 |
| ENSMUSG00000085047 | Gm13026 | 0 | -0.673092543 | 1 |
| ENSMUSG00000025328 | Padi3 | 0 | 0.063873051 | 1 |
| ENSMUSG00000087045 | 4930515B02Rik | 0 | 1.648835551 | 1 |
| ENSMUSG00000087698 | Gm13031 | 0 | -0.810596067 | 0.848985747 |
| ENSMUSG00000045004 | Spata21 | 0 | -2.258055044 | 0.275000936 |
| ENSMUSG00000073733 | Cplane2 | 0 | -0.52108945 | 1 |
| ENSMUSG00000085774 | Gm13055 | 0 | 1.648835551 | 1 |
| ENSMUSG00000086061 | Gm13075 | 0 | -0.351164449 | 1 |
| ENSMUSG00000006221 | Hspb7 | 0 | -0.158519371 | 1 |
| ENSMUSG00000070637 | Srarp | 0 | 1.063873051 | 1 |
| ENSMUSG00000043085 | Tmem82 | 0 | 1.063873051 | 1 |
| ENSMUSG00000073731 | AI507597 | 0 | 1.648835551 | 1 |
| ENSMUSG00000040740 | Slc25a34 | 0 | 1.648835551 | 1 |
| ENSMUSG00000040706 | Agmat | 0 | 0.648835551 | 1 |
| ENSMUSG00000086818 | Ctrcos | 0 | 1.648835551 | 1 |
| ENSMUSG00000100396 | Gm29367 | 0 | 1.648835551 | 1 |
| ENSMUSG00000085761 | 4930455G09Rik | 0 | 1.648835551 | 1 |
| ENSMUSG00000073728 | Tmem51os1 | 0 | 0.326907457 | 1 |
| ENSMUSG00000028584 | Lrrc38 | 0 | -0.158519371 | 1 |
| ENSMUSG00000047799 | Oog4 | 0 | 1.648835551 | 1 |
| ENSMUSG00000070617 | Gm13089 | 0 | 1.648835551 | 1 |
| ENSMUSG00000066031 | Gm13023 | 0 | 1.648835551 | 1 |
| ENSMUSG00000073724 | Pramel4 | 0 | 1.648835551 | 1 |
| ENSMUSG00000046262 | C87977 | 0 | 1.648835551 | 1 |
| ENSMUSG00000036749 | Pramel5 | 0 | 1.063873051 | 1 |
| ENSMUSG00000070619 | Gm13119 | 0 | 1.648835551 | 1 |
| ENSMUSG00000078507 | Aadacl3 | 0 | 1.648835551 | 1 |
| ENSMUSG00000028593 | 9430007A20Rik | 0 | 1.648835551 | 1 |
| ENSMUSG00000078506 | Gm13124 | 0 | 0.648835551 | 1 |
| ENSMUSG00000070609 | Aadacl4 | 0 | 1.063873051 | 1 |
| ENSMUSG00000078503 | Zfp990 | 0 | 0.648835551 | 1 |
| ENSMUSG00000078502 | Gm13212 | 0 | 1.063873051 | 1 |
| ENSMUSG00000058186 | Zfp980 | 0 | 0.063873051 | 1 |

| | | | | |
|---|---|---|---|---|
| ENSMUSG00000087513 | Gm13236 | 0 | 1.063873051 | 1 |
| ENSMUSG00000066007 | Zfp600 | 0 | 1.648835551 | 1 |
| ENSMUSG00000086654 | Gm13165 | 0 | -0.52108945 | 1 |
| ENSMUSG00000063245 | Zfp993 | 0 | 1.648835551 | 1 |
| ENSMUSG00000067919 | Rex2 | 0 | 1.648835551 | 1 |
| ENSMUSG00000078497 | Zfp978 | 0 | 1.063873051 | 1 |
| ENSMUSG00000065999 | Zfp985 | 0 | 1.063873051 | 1 |
| ENSMUSG00000066000 | Zfp979 | 0 | 0.648835551 | 1 |
| ENSMUSG00000086130 | Gm16211 | 0 | 0.326907457 | 1 |
| ENSMUSG00000070583 | Fv1 | 0 | -0.52108945 | 1 |
| ENSMUSG00000078494 | Gm13201 | 0 | 0.648835551 | 1 |
| ENSMUSG00000029005 | Draxin | 0 | -0.52108945 | 1 |
| ENSMUSG00000047719 | Ubiad1 | 0 | -2.051604167 | 0.281836872 |
| ENSMUSG00000028979 | Masp2 | 0 | 0.326907457 | 1 |
| ENSMUSG00000070577 | Gm572 | 0 | -0.158519371 | 1 |
| ENSMUSG00000087273 | Gm13203 | 0 | 1.648835551 | 1 |
| ENSMUSG00000028996 | Rbp7 | 0 | 0.648835551 | 1 |
| ENSMUSG00000087198 | Tmem274 | 0 | 1.648835551 | 1 |
| ENSMUSG00000085368 | Gm16188 | 0 | 1.648835551 | 1 |
| ENSMUSG00000086665 | Gm13067 | 0 | 1.648835551 | 1 |
| ENSMUSG00000062064 | Slc2a7 | 0 | 0.648835551 | 1 |
| ENSMUSG00000085213 | Gm13091 | 0 | 0.063873051 | 1 |
| ENSMUSG00000039838 | Slc45a1 | 0 | 0.648835551 | 1 |
| ENSMUSG00000085869 | Gm16079 | 0 | 1.063873051 | 1 |
| ENSMUSG00000085097 | Gm13049 | 0 | 0.648835551 | 1 |
| ENSMUSG00000078492 | 1700045H11Rik | 0 | -0.673092543 | 1 |
| ENSMUSG00000028950 | Tas1r1 | 0 | 0.648835551 | 1 |
| ENSMUSG00000028940 | Hes2 | 0 | 1.648835551 | 1 |
| ENSMUSG00000090057 | Gm16333 | 0 | 1.648835551 | 1 |
| ENSMUSG00000039546 | Ajap1 | 0 | -1.051604167 | 0.858831093 |
| ENSMUSG00000047613 | A430005L14Rik | 0 | -1.874726405 | 0.352215149 |
| ENSMUSG00000039492 | Ccdc27 | 0 | 1.063873051 | 1 |
| ENSMUSG00000085069 | Prdm16os | 0 | -0.351164449 | 1 |
| ENSMUSG00000087437 | Gm13112 | 0 | 0.063873051 | 1 |
| ENSMUSG00000042333 | Tnfrsf14 | 0 | 0.648835551 | 1 |
| ENSMUSG00000073686 | Gm10564 | 0 | 1.063873051 | 1 |
| ENSMUSG00000029054 | Gabrd | 0 | -1.810596067 | 0.499921258 |
| ENSMUSG00000086682 | Gm16023 | 0 | -1.258055044 | 0.673811773 |
| ENSMUSG00000084845 | Tmem240 | 0 | 0.648835551 | 1 |
| ENSMUSG00000054514 | Atad3aos | 0 | 1.063873051 | 1 |
| ENSMUSG00000042116 | Vwa1 | 0 | -0.936126949 | 0.847265357 |
| ENSMUSG00000073680 | Tmem88b | 0 | 0.063873051 | 1 |
| ENSMUSG00000078487 | Ankrd65 | 0 | 1.648835551 | 1 |
| ENSMUSG00000029072 | Tas1r3 | 0 | 1.063873051 | 1 |
| ENSMUSG00000051557 | Pusl1 | 0 | -0.673092543 | 1 |
| ENSMUSG00000041954 | Tnfrsf18 | 0 | 1.063873051 | 1 |
| ENSMUSG00000086549 | Gm13648 | 0 | -0.158519371 | 1 |
| ENSMUSG00000096351 | Samd11 | 0 | 1.063873051 | 1 |
| ENSMUSG00000057021 | Vmn2r-ps159 | 0 | 1.648835551 | 1 |
| ENSMUSG00000106658 | Gm36548 | 0 | 1.063873051 | 1 |
| ENSMUSG00000105534 | Gm42435 | 0 | 1.648835551 | 1 |

| | | | | |
|---|---|---|---|---|
| ENSMUSG00000105656 | Gm43017 | 0 | 0.063873051 | 1 |
| ENSMUSG00000008307 | 1700109H08Rik | 0 | -2.965874293 | 0.044171611 |
| ENSMUSG00000104891 | 2510017J16Rik | 0 | 1.648835551 | 1 |
| ENSMUSG00000105130 | Gm43825 | 0 | 1.648835551 | 1 |
| ENSMUSG00000104950 | 4833413G10Rik | 0 | 1.063873051 | 1 |
| ENSMUSG00000053178 | Mterf1b | 0 | -0.810596067 | 0.844591606 |
| ENSMUSG00000105560 | Gm42744 | 0 | 1.648835551 | 1 |
| ENSMUSG00000105287 | Gm43577 | 0 | 1.063873051 | 1 |
| ENSMUSG00000079666 | 1700015F17Rik | 0 | -0.351164449 | 1 |
| ENSMUSG00000073234 | Gm8773 | 0 | 1.063873051 | 1 |
| ENSMUSG00000106064 | Gm40263 | 0 | 1.063873051 | 1 |
| ENSMUSG00000097279 | Gm5106 | 0 | 0.063873051 | 1 |
| ENSMUSG00000114430 | Gm40264 | 0 | 1.063873051 | 1 |
| ENSMUSG00000095710 | Gm8871 | 0 | 1.648835551 | 1 |
| ENSMUSG00000104811 | Gm33093 | 0 | 1.648835551 | 1 |
| ENSMUSG00000105688 | 4930437M23Rik | 0 | 1.063873051 | 1 |
| ENSMUSG00000099391 | 1700003C15Rik | 0 | 1.063873051 | 1 |
| ENSMUSG00000067848 | 4933402N22Rik | 0 | 1.648835551 | 1 |
| ENSMUSG00000104638 | 4930568A13Rik | 0 | 0.648835551 | 1 |
| ENSMUSG00000033219 | Gm9758 | 0 | 1.648835551 | 1 |
| ENSMUSG00000091255 | Speer4e | 0 | 1.648835551 | 1 |
| ENSMUSG00000105300 | Gm30613 | 0 | -2.599091962 | 0.119055904 |
| ENSMUSG00000070933 | Speer4d | 0 | 1.648835551 | 1 |
| ENSMUSG00000105185 | 4930519H02Rik | 0 | 1.648835551 | 1 |
| ENSMUSG00000096936 | Gm3510 | 0 | 1.063873051 | 1 |
| ENSMUSG00000028777 | Gnat3 | 0 | 1.063873051 | 1 |
| ENSMUSG00000104718 | Gm31752 | 0 | 1.063873051 | 1 |
| ENSMUSG00000097626 | 4921504A21Rik | 0 | 1.063873051 | 1 |
| ENSMUSG00000105076 | A930003O13Rik | 0 | 1.648835551 | 1 |
| ENSMUSG00000097180 | 2700038G22Rik | 0 | -2.351164449 | 0.169846604 |
| ENSMUSG00000048439 | Nupl2 | 0 | -0.936126949 | 0.847265357 |
| ENSMUSG00000028978 | Nos3 | 0 | -1.051604167 | 0.908885177 |
| ENSMUSG00000038276 | Asic3 | 0 | 0.648835551 | 1 |
| ENSMUSG00000028958 | Tmub1 | 0 | -1.810596067 | 0.465580427 |
| ENSMUSG00000038204 | Asb10 | 0 | 1.063873051 | 1 |
| ENSMUSG00000038199 | Iqca1l | 0 | 1.648835551 | 1 |
| ENSMUSG00000097275 | Gm26648 | 0 | 1.648835551 | 1 |
| ENSMUSG00000038135 | Crygn | 0 | 1.063873051 | 1 |
| ENSMUSG00000073117 | Gm7347 | 0 | 1.648835551 | 1 |
| ENSMUSG00000039106 | Htr5a | 0 | 1.648835551 | 1 |
| ENSMUSG00000097673 | Gm26608 | 0 | 0.063873051 | 1 |
| ENSMUSG00000039095 | En2 | 0 | 1.648835551 | 1 |
| ENSMUSG00000044681 | Cnpy1 | 0 | 1.063873051 | 1 |
| ENSMUSG00000097561 | 4632411P08Rik | 0 | 1.063873051 | 1 |
| ENSMUSG00000025746 | Il6 | 0 | -1.351164449 | 0.97950322 |
| ENSMUSG00000044576 | Garem2 | 0 | 0.648835551 | 1 |
| ENSMUSG00000029182 | 1700001C02Rik | 0 | 1.648835551 | 1 |
| ENSMUSG00000106300 | Gm42765 | 0 | 1.063873051 | 1 |
| ENSMUSG00000049265 | Kcnk3 | 0 | -2.209145444 | 0.217194106 |
| ENSMUSG00000106229 | Gm19409 | 0 | 0.063873051 | 1 |
| ENSMUSG00000029161 | Cgref1 | 0 | -0.52108945 | 1 |

| | | | | |
|---|---|---|---|---|
| ENSMUSG00000006641 | Slc5a6 | 0 | -0.936126949 | 0.847265357 |
| ENSMUSG00000038552 | Fndc4 | 0 | 0.063873051 | 1 |
| ENSMUSG00000029138 | 4930548H24Rik | 0 | 1.648835551 | 1 |
| ENSMUSG00000106894 | Gm17130 | 0 | 1.648835551 | 1 |
| ENSMUSG00000107036 | Gm7596 | 0 | 1.648835551 | 1 |
| ENSMUSG00000029112 | Nkx1-1 | 0 | 1.648835551 | 1 |
| ENSMUSG00000019295 | Tmem129 | 0 | 0.063873051 | 1 |
| ENSMUSG00000106688 | Gm42851 | 0 | 1.648835551 | 1 |
| ENSMUSG00000048142 | Nat8l | 0 | 1.648835551 | 1 |
| ENSMUSG00000086291 | Gm15513 | 0 | 1.063873051 | 1 |
| ENSMUSG00000054520 | Sh3bp2 | 0 | -3.209145444 | 0.025125353 |
| ENSMUSG00000107246 | Gm42560 | 0 | 0.063873051 | 1 |
| ENSMUSG00000051246 | Msantd1 | 0 | -0.351164449 | 1 |
| ENSMUSG00000044716 | Dok7 | 0 | -0.936126949 | 0.847265357 |
| ENSMUSG00000067438 | Hmx1 | 0 | 1.648835551 | 1 |
| ENSMUSG00000029096 | Htra3 | 0 | -0.673092543 | 1 |
| ENSMUSG00000043430 | Psapl1 | 0 | 1.063873051 | 1 |
| ENSMUSG00000050677 | Ccdc96 | 0 | 1.648835551 | 1 |
| ENSMUSG00000029120 | Ppp2r2c | 0 | -0.158519371 | 1 |
| ENSMUSG00000062329 | Cytl1 | 0 | 0.648835551 | 1 |
| ENSMUSG00000029127 | Zbtb49 | 0 | -1.258055044 | 0.673811773 |
| ENSMUSG00000051596 | Otop1 | 0 | 0.326907457 | 1 |
| ENSMUSG00000079528 | Clnkos | 0 | 1.063873051 | 1 |
| ENSMUSG00000106679 | Gm42524 | 0 | 1.648835551 | 1 |
| ENSMUSG00000107244 | Gm43699 | 0 | 1.648835551 | 1 |
| ENSMUSG00000033036 | Gm7879 | 0 | 0.326907457 | 1 |
| ENSMUSG00000105503 | Gm42982 | 0 | 1.063873051 | 1 |
| ENSMUSG00000104784 | Gm42984 | 0 | -0.158519371 | 1 |
| ENSMUSG00000106656 | Gm42750 | 0 | 0.648835551 | 1 |
| ENSMUSG00000062496 | 4930431F12Rik | 0 | 1.063873051 | 1 |
| ENSMUSG00000049530 | Clrn2 | 0 | 1.648835551 | 1 |
| ENSMUSG00000106464 | C130083M11Rik | 0 | 1.648835551 | 1 |
| ENSMUSG00000039252 | Lgi2 | 0 | -1.258055044 | 0.927814401 |
| ENSMUSG00000052295 | 8030423F21Rik | 0 | 0.648835551 | 1 |
| ENSMUSG00000106808 | Gm42745 | 0 | -0.158519371 | 1 |
| ENSMUSG00000029193 | Cckar | 0 | 0.063873051 | 1 |
| ENSMUSG00000105234 | 4930459L07Rik | 0 | 1.063873051 | 1 |
| ENSMUSG00000106634 | Gm43042 | 0 | 1.063873051 | 1 |
| ENSMUSG00000105903 | 1700014F14Rik | 0 | 1.648835551 | 1 |
| ENSMUSG00000100911 | 1700027F09Rik | 0 | 1.648835551 | 1 |
| ENSMUSG00000097668 | Gm26761 | 0 | 1.648835551 | 1 |
| ENSMUSG00000044827 | Tlr1 | 0 | -0.158519371 | 1 |
| ENSMUSG00000051498 | Tlr6 | 0 | 1.648835551 | 1 |
| ENSMUSG00000107102 | Gm42726 | 0 | -0.810596067 | 0.847265357 |
| ENSMUSG00000107014 | 1700126H18Rik | 0 | -0.673092543 | 1 |
| ENSMUSG00000107266 | Gm43698 | 0 | 1.648835551 | 1 |
| ENSMUSG00000092060 | Bend4 | 0 | 0.326907457 | 1 |
| ENSMUSG00000097840 | Gm26756 | 0 | 1.063873051 | 1 |
| ENSMUSG00000095346 | Gm21905 | 0 | 1.648835551 | 1 |
| ENSMUSG00000094719 | Gm5108 | 0 | 0.648835551 | 1 |
| ENSMUSG00000029158 | Yipf7 | 0 | 1.063873051 | 1 |

| | | | | |
|---|---|---|---|---|
| ENSMUSG00000029208 | Guf1 | 0 | -2.351164449 | 0.169846604 |
| ENSMUSG00000063935 | Zar1 | 0 | 1.648835551 | 1 |
| ENSMUSG00000060204 | Gm5868 | 0 | 0.648835551 | 1 |
| ENSMUSG00000067220 | Cnga1 | 0 | 0.648835551 | 1 |
| ENSMUSG00000029219 | Slc10a4 | 0 | 1.648835551 | 1 |
| ENSMUSG00000029156 | Sgcb | 0 | -1.051604167 | 1 |
| ENSMUSG00000029155 | Spata18 | 0 | 1.063873051 | 1 |
| ENSMUSG00000106968 | C78283 | 0 | 1.648835551 | 1 |
| ENSMUSG00000049907 | Rasl11b | 0 | -0.351164449 | 1 |
| ENSMUSG00000085657 | Gm15984 | 0 | 1.648835551 | 1 |
| ENSMUSG00000072874 | Gm6116 | 0 | 1.063873051 | 1 |
| ENSMUSG00000029235 | Pdcl2 | 0 | 0.326907457 | 1 |
| ENSMUSG00000029236 | Nmu | 0 | -0.52108945 | 1 |
| ENSMUSG00000091204 | Gm7271 | 0 | -0.810596067 | 1 |
| ENSMUSG00000086632 | 1700112J05Rik | 0 | 0.063873051 | 1 |
| ENSMUSG00000059325 | Hopx | 0 | -1.051604167 | 0.858831093 |
| ENSMUSG00000087541 | Hopxos | 0 | 1.063873051 | 1 |
| ENSMUSG00000053030 | Spink2 | 0 | 1.648835551 | 1 |
| ENSMUSG00000106828 | Gm34648 | 0 | 1.648835551 | 1 |
| ENSMUSG00000106906 | Gm34728 | 0 | 0.063873051 | 1 |
| ENSMUSG00000107232 | 1700017L05Rik | 0 | 1.648835551 | 1 |
| ENSMUSG00000087431 | Gm16054 | 0 | 1.063873051 | 1 |
| ENSMUSG00000049537 | Tecrl | 0 | -0.158519371 | 1 |
| ENSMUSG00000104878 | Gm43567 | 0 | 1.648835551 | 1 |
| ENSMUSG00000029255 | Gnrhr | 0 | 1.063873051 | 1 |
| ENSMUSG00000072845 | Tmprss11a | 0 | 0.063873051 | 1 |
| ENSMUSG00000079451 | Tmprss11g | 0 | 0.326907457 | 1 |
| ENSMUSG00000089851 | Gm16579 | 0 | 0.326907457 | 1 |
| ENSMUSG00000104937 | Gm43057 | 0 | -1.051604167 | 0.858831093 |
| ENSMUSG00000057425 | Ugt2b37 | 0 | 1.648835551 | 1 |
| ENSMUSG00000035780 | Ugt2a3 | 0 | -0.673092543 | 1 |
| ENSMUSG00000061906 | Ugt2b38 | 0 | 0.648835551 | 1 |
| ENSMUSG00000106677 | Ugt2a1 | 0 | 1.063873051 | 1 |
| ENSMUSG00000070702 | Csn1s1 | 0 | 1.648835551 | 1 |
| ENSMUSG00000061388 | Csn1s2b | 0 | 0.648835551 | 1 |
| ENSMUSG00000001622 | Csn3 | 0 | 1.648835551 | 1 |
| ENSMUSG00000090302 | Gm7714 | 0 | 0.063873051 | 1 |
| ENSMUSG00000029286 | Enam | 0 | 0.648835551 | 1 |
| ENSMUSG00000029366 | Dck | 0 | -1.810596067 | 0.549953572 |
| ENSMUSG00000105640 | Gm42912 | 0 | -0.158519371 | 1 |
| ENSMUSG00000035505 | Cox18 | 0 | -1.43862729 | 0.771569813 |
| ENSMUSG00000054932 | Afp | 0 | -0.936126949 | 0.847265357 |
| ENSMUSG00000029369 | Afm | 0 | 0.326907457 | 1 |
| ENSMUSG00000029372 | Ppbp | 0 | 0.326907457 | 1 |
| ENSMUSG00000029380 | Cxcl1 | 0 | -1.43862729 | 0.538110273 |
| ENSMUSG00000035020 | Epgn | 0 | 1.063873051 | 1 |
| ENSMUSG00000107350 | Gm19610 | 0 | -2.158519371 | 0.290284345 |
| ENSMUSG00000107198 | Gm19619 | 0 | 1.063873051 | 1 |
| ENSMUSG00000099468 | Gm28271 | 0 | 1.648835551 | 1 |
| ENSMUSG00000096035 | Odaph | 0 | 0.063873051 | 1 |
| ENSMUSG00000106698 | Gm43599 | 0 | 1.063873051 | 1 |

| | | | | |
|---|---|---|---|---|
| ENSMUSG00000029417 | Cxcl9 | 0 | 1.063873051 | 1 |
| ENSMUSG00000057068 | Fam47e | 0 | 0.063873051 | 1 |
| ENSMUSG00000092266 | Gm20500 | 0 | 1.648835551 | 1 |
| ENSMUSG00000070686 | C87414 | 0 | 1.648835551 | 1 |
| ENSMUSG00000095074 | Gm3139 | 0 | 1.648835551 | 1 |
| ENSMUSG00000092073 | Gm6205 | 0 | 1.063873051 | 1 |
| ENSMUSG00000096742 | Gm6367 | 0 | 1.648835551 | 1 |
| ENSMUSG00000070677 | D5Ertd577e | 0 | 1.063873051 | 1 |
| ENSMUSG00000072813 | E330014E10Rik | 0 | 1.648835551 | 1 |
| ENSMUSG00000106319 | Gm42601 | 0 | 1.648835551 | 1 |
| ENSMUSG00000105283 | Gm33370 | 0 | -0.52108945 | 1 |
| ENSMUSG00000105006 | Gm9484 | 0 | 1.063873051 | 1 |
| ENSMUSG00000055725 | Paqr3 | 0 | -0.673092543 | 1 |
| ENSMUSG00000104717 | Gm33969 | 0 | 0.063873051 | 1 |
| ENSMUSG00000035456 | Prdm8 | 0 | -1.351164449 | 0.687112677 |
| ENSMUSG00000029337 | Fgf5 | 0 | 1.648835551 | 1 |
| ENSMUSG00000091623 | Gm17092 | 0 | -2.43862729 | 0.208883511 |
| ENSMUSG00000103927 | Gm9932 | 0 | 1.063873051 | 1 |
| ENSMUSG00000100157 | 2310034O05Rik | 0 | 1.648835551 | 1 |
| ENSMUSG00000097772 | 5430416N02Rik | 0 | -3.636566667 | 0.005443828 |
| ENSMUSG00000106492 | Gm29707 | 0 | 1.648835551 | 1 |
| ENSMUSG00000104684 | 5430427N15Rik | 0 | 0.648835551 | 1 |
| ENSMUSG00000029298 | Gbp9 | 0 | -2.258055044 | 0.222101622 |
| ENSMUSG00000079363 | Gbp4 | 0 | 0.326907457 | 1 |
| ENSMUSG00000105096 | Gbp10 | 0 | -0.351164449 | 1 |
| ENSMUSG00000104713 | Gbp6 | 0 | 0.326907457 | 1 |
| ENSMUSG00000097237 | Gm26519 | 0 | 1.063873051 | 1 |
| ENSMUSG00000085771 | C230066G23Rik | 0 | 1.648835551 | 1 |
| ENSMUSG00000084878 | Lrrc8dos | 0 | 1.648835551 | 1 |
| ENSMUSG00000106483 | Gm32736 | 0 | 1.063873051 | 1 |
| ENSMUSG00000106594 | 4930432H08Rik | 0 | 1.648835551 | 1 |
| ENSMUSG00000034384 | Barhl2 | 0 | 1.648835551 | 1 |
| ENSMUSG00000043410 | Hfm1 | 0 | -3.023589791 | 0.061119858 |
| ENSMUSG00000106554 | Gm33474 | 0 | 1.648835551 | 1 |
| ENSMUSG00000106396 | Gm42902 | 0 | 0.648835551 | 1 |
| ENSMUSG00000085900 | A930041C12Rik | 0 | 0.648835551 | 1 |
| ENSMUSG00000029275 | Gfi1 | 0 | 1.648835551 | 1 |
| ENSMUSG00000104904 | 9330198I05Rik | 0 | 0.326907457 | 1 |
| ENSMUSG00000029490 | Mfsd7a | 0 | -0.673092543 | 1 |
| ENSMUSG00000033615 | Cplx1 | 0 | -0.158519371 | 1 |
| ENSMUSG00000033540 | Idua | 0 | -0.673092543 | 1 |
| ENSMUSG00000046959 | Slc26a1 | 0 | 1.648835551 | 1 |
| ENSMUSG00000055385 | Rnf212 | 0 | 0.326907457 | 1 |
| ENSMUSG00000004821 | Tmed11 | 0 | 1.648835551 | 1 |
| ENSMUSG00000091624 | Vmn2r9 | 0 | 1.648835551 | 1 |
| ENSMUSG00000093479 | Gm20629 | 0 | -1.051604167 | 0.871122629 |
| ENSMUSG00000067010 | Vmn2r10 | 0 | 1.648835551 | 1 |
| ENSMUSG00000091375 | Vmn2r15 | 0 | 1.648835551 | 1 |
| ENSMUSG00000091879 | Vmn2r17 | 0 | 1.648835551 | 1 |
| ENSMUSG00000072762 | 4930522L14Rik | 0 | -0.936126949 | 0.847265357 |
| ENSMUSG00000105471 | A430073D23Rik | 0 | 0.063873051 | 1 |

| | | | | |
|---|---|---|---|---|
| ENSMUSG00000097140 | Gm26779 | 0 | -0.52108945 | 1 |
| ENSMUSG00000090015 | Gm15446 | 0 | -0.810596067 | 1 |
| ENSMUSG00000064247 | Plcxd1 | 0 | 0.648835551 | 1 |
| ENSMUSG00000023284 | Zfp605 | 0 | -3.459688905 | 0.008890736 |
| ENSMUSG00000104720 | Gm43138 | 0 | 1.648835551 | 1 |
| ENSMUSG00000087439 | Gm15788 | 0 | 0.326907457 | 1 |
| ENSMUSG00000029503 | P2rx2 | 0 | 1.063873051 | 1 |
| ENSMUSG00000072754 | Lrcol1 | 0 | 1.648835551 | 1 |
| ENSMUSG00000105161 | Gm42595 | 0 | 1.648835551 | 1 |
| ENSMUSG00000104833 | Gm42489 | 0 | 1.648835551 | 1 |
| ENSMUSG00000104538 | 1700016B01Rik | 0 | 1.063873051 | 1 |
| ENSMUSG00000097767 | Miat | 0 | 0.326907457 | 1 |
| ENSMUSG00000097986 | Gm26953 | 0 | -0.158519371 | 1 |
| ENSMUSG00000066975 | Cryba4 | 0 | 1.063873051 | 1 |
| ENSMUSG00000097962 | 1700034G24Rik | 0 | 0.326907457 | 1 |
| ENSMUSG00000105152 | Gm42864 | 0 | 1.648835551 | 1 |
| ENSMUSG00000098230 | 1700095B10Rik | 0 | 1.063873051 | 1 |
| ENSMUSG00000042240 | Crybb2 | 0 | 1.648835551 | 1 |
| ENSMUSG00000029352 | Crybb3 | 0 | -0.158519371 | 1 |
| ENSMUSG00000042190 | Cmklr1 | 0 | -1.051604167 | 0.858831093 |
| ENSMUSG00000053334 | Ficd | 0 | 1.063873051 | 1 |
| ENSMUSG00000048163 | Selplg | 0 | -1.351164449 | 0.705975399 |
| ENSMUSG00000042002 | Foxn4 | 0 | 1.648835551 | 1 |
| ENSMUSG00000029575 | Mmab | 0 | -0.52108945 | 1 |
| ENSMUSG00000041939 | Mvk | 0 | -1.43862729 | 0.538110273 |
| ENSMUSG00000029556 | Hnf1a | 0 | -1.051604167 | 0.874796537 |
| ENSMUSG00000072693 | Gm10401 | 0 | 1.648835551 | 1 |
| ENSMUSG00000046562 | Unc119b | 0 | -2.305360759 | 0.545392675 |
| ENSMUSG00000029544 | Cabp1 | 0 | -0.158519371 | 1 |
| ENSMUSG00000107288 | Gm42903 | 0 | -0.158519371 | 1 |
| ENSMUSG00000107353 | 4930430O22Rik | 0 | 1.648835551 | 1 |
| ENSMUSG00000085376 | Gm14508 | 0 | 0.326907457 | 1 |
| ENSMUSG00000084992 | Gm13842 | 0 | 1.063873051 | 1 |
| ENSMUSG00000084921 | Gm13838 | 0 | 1.648835551 | 1 |
| ENSMUSG00000086219 | Srrm4os | 0 | 0.648835551 | 1 |
| ENSMUSG00000086047 | 9530046B11Rik | 0 | 0.648835551 | 1 |
| ENSMUSG00000097163 | BC051077 | 0 | 0.648835551 | 1 |
| ENSMUSG00000072688 | Gm10399 | 0 | 0.063873051 | 1 |
| ENSMUSG00000086075 | Gm15728 | 0 | 0.648835551 | 1 |
| ENSMUSG00000098072 | Gm26995 | 0 | 1.648835551 | 1 |
| ENSMUSG00000100573 | 1700081H04Rik | 0 | 1.648835551 | 1 |
| ENSMUSG00000107334 | Gm7538 | 0 | 1.063873051 | 1 |
| ENSMUSG00000106938 | Gm31314 | 0 | 1.063873051 | 1 |
| ENSMUSG00000106795 | Gm43050 | 0 | 1.648835551 | 1 |
| ENSMUSG00000107084 | Gm43269 | 0 | 1.063873051 | 1 |
| ENSMUSG00000097368 | Gm10390 | 0 | 1.648835551 | 1 |
| ENSMUSG00000029603 | Dtx1 | 0 | -0.810596067 | 0.844591606 |
| ENSMUSG00000032690 | Oas2 | 0 | 1.063873051 | 1 |
| ENSMUSG00000107179 | Gm43069 | 0 | 0.326907457 | 1 |
| ENSMUSG00000066867 | Oas1e | 0 | 1.063873051 | 1 |
| ENSMUSG00000053765 | Oas1f | 0 | 1.648835551 | 1 |

| | | | | |
|---|---|---|---|---|
| ENSMUSG00000066861 | Oas1g | 0 | 1.063873051 | 1 |
| ENSMUSG00000104985 | Gm42917 | 0 | -0.810596067 | 0.844591606 |
| ENSMUSG00000062438 | Adam1b | 0 | 1.063873051 | 1 |
| ENSMUSG00000072647 | Adam1a | 0 | -0.351164449 | 1 |
| ENSMUSG00000087386 | Gm15637 | 0 | 0.326907457 | 1 |
| ENSMUSG00000013936 | Myl2 | 0 | -0.351164449 | 1 |
| ENSMUSG00000105895 | Gm42829 | 0 | 1.063873051 | 1 |
| ENSMUSG00000056735 | A930024E05Rik | 0 | -1.43862729 | 0.538110273 |
| ENSMUSG00000029477 | Morn3 | 0 | -0.810596067 | 1 |
| ENSMUSG00000104156 | Gm38102 | 0 | 1.648835551 | 1 |
| ENSMUSG00000105881 | 4932422M17Rik | 0 | -1.051604167 | 0.912904617 |
| ENSMUSG00000029445 | Hpd | 0 | 1.063873051 | 1 |
| ENSMUSG00000087063 | Gm15857 | 0 | 0.326907457 | 1 |
| ENSMUSG00000104830 | 5830487J09Rik | 0 | 1.063873051 | 1 |
| ENSMUSG00000063409 | Lrrc43 | 0 | 0.326907457 | 1 |
| ENSMUSG00000114278 | Gm49027 | 0 | 1.648835551 | 1 |
| ENSMUSG00000045502 | Hcar2 | 0 | 0.326907457 | 1 |
| ENSMUSG00000085433 | Gm16001 | 0 | 0.648835551 | 1 |
| ENSMUSG00000089842 | Pitpnm2os2 | 0 | 1.063873051 | 1 |
| ENSMUSG00000090220 | Pitpnm2os1 | 0 | 1.648835551 | 1 |
| ENSMUSG00000037962 | Rflna | 0 | 0.648835551 | 1 |
| ENSMUSG00000106663 | Gm42839 | 0 | 1.648835551 | 1 |
| ENSMUSG00000072612 | Gm10382 | 0 | 1.648835551 | 1 |
| ENSMUSG00000105316 | Gm42953 | 0 | 1.648835551 | 1 |
| ENSMUSG00000087104 | Tmem132cos | 0 | -0.673092543 | 1 |
| ENSMUSG00000106015 | Gm42980 | 0 | 0.648835551 | 1 |
| ENSMUSG00000105857 | Gm42817 | 0 | 1.648835551 | 1 |
| ENSMUSG00000104727 | Gm42500 | 0 | 1.648835551 | 1 |
| ENSMUSG00000072591 | Fzd10os | 0 | 0.648835551 | 1 |
| ENSMUSG00000105271 | Gm42875 | 0 | 1.648835551 | 1 |
| ENSMUSG00000051034 | Zfp11 | 0 | -0.673092543 | 1 |
| ENSMUSG00000034219 | 14-Sep | 0 | -1.258055044 | 0.687112677 |
| ENSMUSG00000095789 | Nupr1l | 0 | -0.673092543 | 1 |
| ENSMUSG00000034110 | Kctd7 | 0 | -0.158519371 | 1 |
| ENSMUSG00000107143 | Gm6598 | 0 | 0.326907457 | 1 |
| ENSMUSG00000052014 | A330070K13Rik | 0 | 0.326907457 | 1 |
| ENSMUSG00000106710 | Gm43166 | 0 | 1.648835551 | 1 |
| ENSMUSG00000104987 | 2810432F15Rik | 0 | 1.063873051 | 1 |
| ENSMUSG00000100420 | Gm28616 | 0 | 1.648835551 | 1 |
| ENSMUSG00000040557 | Mettl27 | 0 | -1.874726405 | 0.352215149 |
| ENSMUSG00000054909 | Wbscr25 | 0 | -1.995020638 | 0.438745691 |
| ENSMUSG00000061118 | Dnajc30 | 0 | -1.051604167 | 0.886203497 |
| ENSMUSG00000043614 | Vps37d | 0 | -0.52108945 | 1 |
| ENSMUSG00000049551 | Fzd9 | 0 | 1.648835551 | 1 |
| ENSMUSG00000040013 | Fkbp6 | 0 | 0.063873051 | 1 |
| ENSMUSG00000000916 | Nsun5 | 0 | -0.351164449 | 1 |
| ENSMUSG00000070464 | Ccl26 | 0 | 1.648835551 | 1 |
| ENSMUSG00000105239 | Gm43091 | 0 | -0.673092543 | 1 |
| ENSMUSG00000019178 | Styxl1 | 0 | -0.673092543 | 1 |
| ENSMUSG00000086656 | Gm15701 | 0 | 1.648835551 | 1 |
| ENSMUSG00000005474 | Myl10 | 0 | 1.648835551 | 1 |

| | | | | |
|---|---|---|---|---|
| ENSMUSG00000001739 | Cldn15 | 0 | 0.063873051 | 1 |
| ENSMUSG00000079174 | Gm3054 | 0 | 1.648835551 | 1 |
| ENSMUSG00000118522 | CAAA01077340.1 | 0 | 0.326907457 | 1 |
| ENSMUSG00000105736 | Gm42456 | 0 | 1.648835551 | 1 |
| ENSMUSG00000023328 | Ache | 0 | 1.648835551 | 1 |
| ENSMUSG00000106237 | Gm8066 | 0 | -0.810596067 | 0.844591606 |
| ENSMUSG00000029712 | Actl6b | 0 | -0.158519371 | 1 |
| ENSMUSG00000029716 | Tfr2 | 0 | -1.52108945 | 0.544908989 |
| ENSMUSG00000106617 | Gm36266 | 0 | -0.673092543 | 1 |
| ENSMUSG00000079163 | Gm15498 | 0 | 1.063873051 | 1 |
| ENSMUSG00000066684 | Pilrb1 | 0 | -2.106051951 | 0.279592421 |
| ENSMUSG00000106478 | Gm36551 | 0 | 0.648835551 | 1 |
| ENSMUSG00000106155 | Gm43495 | 0 | 1.648835551 | 1 |
| ENSMUSG00000037053 | Azgp1 | 0 | 1.063873051 | 1 |
| ENSMUSG00000049285 | Mblac1 | 0 | 1.648835551 | 1 |
| ENSMUSG00000089783 | Gm454 | 0 | 1.648835551 | 1 |
| ENSMUSG00000036948 | Map11 | 0 | -3.373532262 | 0.013375059 |
| ENSMUSG00000075593 | Gal3st4 | 0 | 1.063873051 | 1 |
| ENSMUSG00000029510 | Gpc2 | 0 | -1.673092543 | 0.451785624 |
| ENSMUSG00000036928 | Stag3 | 0 | -1.599091962 | 0.559918909 |
| ENSMUSG00000075591 | Gm10874 | 0 | 1.063873051 | 1 |
| ENSMUSG00000036898 | Zfp157 | 0 | -1.351164449 | 0.687112677 |
| ENSMUSG00000075585 | 6330403L08Rik | 0 | -0.351164449 | 1 |
| ENSMUSG00000106127 | 4930520M14Rik | 0 | 1.063873051 | 1 |
| ENSMUSG00000053647 | Gper1 | 0 | 1.063873051 | 1 |
| ENSMUSG00000029546 | Uncx | 0 | 1.648835551 | 1 |
| ENSMUSG00000048988 | Elfn1 | 0 | -0.936126949 | 1 |
| ENSMUSG00000087606 | Gm16121 | 0 | 0.648835551 | 1 |
| ENSMUSG00000029557 | Mrm2 | 0 | -0.810596067 | 0.844591606 |
| ENSMUSG00000085540 | Gm16036 | 0 | 0.326907457 | 1 |
| ENSMUSG00000074817 | Papolb | 0 | 1.648835551 | 1 |
| ENSMUSG00000090981 | Olfr718-ps1 | 0 | 1.063873051 | 1 |
| ENSMUSG00000029586 | Spdye4b | 0 | -0.158519371 | 1 |
| ENSMUSG00000039296 | Spdye4a | 0 | -0.52108945 | 1 |
| ENSMUSG00000089879 | 4930448H16Rik | 0 | 0.648835551 | 1 |
| ENSMUSG00000105345 | BC030343 | 0 | 1.063873051 | 1 |
| ENSMUSG00000093910 | Zfp853 | 0 | 0.326907457 | 1 |
| ENSMUSG00000001844 | Zdhhc4 | 0 | -1.810596067 | 0.466464457 |
| ENSMUSG00000089889 | 0610040B10Rik | 0 | -0.52108945 | 1 |
| ENSMUSG00000083012 | Fam220a | 0 | 0.648835551 | 1 |
| ENSMUSG00000097287 | D130017N08Rik | 0 | 0.326907457 | 1 |
| ENSMUSG00000029607 | Ankrd61 | 0 | -1.874726405 | 0.521282131 |
| ENSMUSG00000029618 | Ocm | 0 | 1.063873051 | 1 |
| ENSMUSG00000086894 | Gm15708 | 0 | -0.351164449 | 1 |
| ENSMUSG00000029620 | 1700018F24Rik | 0 | 1.648835551 | 1 |
| ENSMUSG00000070420 | Zscan25 | 0 | -1.874726405 | 0.375105691 |
| ENSMUSG00000038656 | Cyp3a16 | 0 | 1.648835551 | 1 |
| ENSMUSG00000054417 | Cyp3a44 | 0 | 1.648835551 | 1 |
| ENSMUSG00000029630 | Cyp3a25 | 0 | 0.648835551 | 1 |
| ENSMUSG00000096527 | Gm6370 | 0 | 1.648835551 | 1 |
| ENSMUSG00000041468 | Gpr12 | 0 | 1.648835551 | 1 |

| | | | | |
|---|---|---|---|---|
| ENSMUSG00000105907 | Gm34333 | 0 | 1.648835551 | 1 |
| ENSMUSG00000104845 | 1700041I07Rik | 0 | 1.063873051 | 1 |
| ENSMUSG00000098053 | Gm27033 | 0 | 0.063873051 | 1 |
| ENSMUSG00000029646 | Cdx2 | 0 | 1.648835551 | 1 |
| ENSMUSG00000075543 | Urad | 0 | 1.063873051 | 1 |
| ENSMUSG00000029650 | Slc46a3 | 0 | -0.810596067 | 1 |
| ENSMUSG00000085565 | Gm15721 | 0 | 1.648835551 | 1 |
| ENSMUSG00000106733 | Gm36186 | 0 | 1.063873051 | 1 |
| ENSMUSG00000086494 | 2210417A02Rik | 0 | -1.051604167 | 0.858831093 |
| ENSMUSG00000085740 | 4930505K14Rik | 0 | 1.648835551 | 1 |
| ENSMUSG00000085237 | Gm15406 | 0 | 1.648835551 | 1 |
| ENSMUSG00000079069 | 8430423G03Rik | 0 | 0.326907457 | 1 |
| ENSMUSG00000106951 | 5930430L01Rik | 0 | -1.743481871 | 0.455131546 |
| ENSMUSG00000085971 | Gm15411 | 0 | 0.326907457 | 1 |
| ENSMUSG00000107388 | Gm42788 | 0 | -0.351164449 | 1 |
| ENSMUSG00000107313 | Gm43332 | 0 | 1.648835551 | 1 |
| ENSMUSG00000100783 | 2310047D07Rik | 0 | 1.648835551 | 1 |
| ENSMUSG00000107271 | BC028471 | 0 | 1.648835551 | 1 |
| ENSMUSG00000106777 | Gm43150 | 0 | 1.063873051 | 1 |
| ENSMUSG00000029660 | Tex26 | 0 | 1.063873051 | 1 |
| ENSMUSG00000106674 | Gm20005 | 0 | 1.648835551 | 1 |
| ENSMUSG00000106740 | Gm21221 | 0 | 1.648835551 | 1 |
| ENSMUSG00000107214 | 4930500F04Rik | 0 | 1.648835551 | 1 |
| ENSMUSG00000107112 | Gm36241 | 0 | 1.648835551 | 1 |
| ENSMUSG00000106927 | Gm43598 | 0 | 0.648835551 | 1 |
| ENSMUSG00000107048 | Gm43807 | 0 | 1.648835551 | 1 |
| ENSMUSG00000054622 | D730045B01Rik | 0 | 1.063873051 | 1 |
| ENSMUSG00000107241 | Gm43129 | 0 | 1.648835551 | 1 |
| ENSMUSG00000029664 | Tfpi2 | 0 | 0.326907457 | 1 |
| ENSMUSG00000107681 | 4930528H21Rik | 0 | 1.063873051 | 1 |
| ENSMUSG00000093570 | Gm20714 | 0 | 1.648835551 | 1 |
| ENSMUSG00000002588 | Pon1 | 0 | 1.063873051 | 1 |
| ENSMUSG00000093380 | Gm20685 | 0 | 0.326907457 | 1 |
| ENSMUSG00000093675 | Gm20618 | 0 | -0.936126949 | 0.874192437 |
| ENSMUSG00000090063 | Dlx6os1 | 0 | -0.673092543 | 1 |
| ENSMUSG00000029754 | Dlx6 | 0 | 0.063873051 | 1 |
| ENSMUSG00000029755 | Dlx5 | 0 | 1.648835551 | 1 |
| ENSMUSG00000097356 | Gm26774 | 0 | 0.648835551 | 1 |
| ENSMUSG00000090151 | Gm16043 | 0 | 0.648835551 | 1 |
| ENSMUSG00000107422 | Gm44129 | 0 | 0.063873051 | 1 |
| ENSMUSG00000101894 | 1700016P04Rik | 0 | 1.648835551 | 1 |
| ENSMUSG00000042717 | Ppp1r3a | 0 | 0.648835551 | 1 |
| ENSMUSG00000108295 | Gm36406 | 0 | 1.648835551 | 1 |
| ENSMUSG00000085171 | D830026I12Rik | 0 | 1.648835551 | 1 |
| ENSMUSG00000029517 | Ankrd7 | 0 | 0.648835551 | 1 |
| ENSMUSG00000107217 | 4930524B17Rik | 0 | 0.648835551 | 1 |
| ENSMUSG00000106871 | Gm3289 | 0 | 1.648835551 | 1 |
| ENSMUSG00000029683 | Lmod2 | 0 | 1.063873051 | 1 |
| ENSMUSG00000029680 | Hyal4 | 0 | 1.063873051 | 1 |
| ENSMUSG00000029682 | Spam1 | 0 | 1.648835551 | 1 |
| ENSMUSG00000029678 | Hyal5 | 0 | -0.52108945 | 1 |

| | | | | |
|---|---|---|---|---|
| ENSMUSG00000048022 | Tmem229a | 0 | 1.648835551 | 1 |
| ENSMUSG00000039904 | Gpr37 | 0 | 1.648835551 | 1 |
| ENSMUSG00000106799 | Gm20756 | 0 | 0.648835551 | 1 |
| ENSMUSG00000029707 | Fscn3 | 0 | 0.063873051 | 1 |
| ENSMUSG00000092090 | Gm3294 | 0 | -1.051604167 | 0.894729931 |
| ENSMUSG00000059201 | Lep | 0 | 1.648835551 | 1 |
| ENSMUSG00000039742 | Fam71f1 | 0 | -0.158519371 | 1 |
| ENSMUSG00000058831 | Opn1sw | 0 | 1.648835551 | 1 |
| ENSMUSG00000029769 | Ccdc136 | 0 | -1.52108945 | 0.544908989 |
| ENSMUSG00000090685 | Atp6v1fnb | 0 | 1.063873051 | 1 |
| ENSMUSG00000059022 | Kcp | 0 | -0.673092543 | 1 |
| ENSMUSG00000097137 | Gm26627 | 0 | 0.648835551 | 1 |
| ENSMUSG00000029771 | Irf5 | 0 | -0.351164449 | 1 |
| ENSMUSG00000001763 | Tspan33 | 0 | -1.051604167 | 0.886203497 |
| ENSMUSG00000103108 | Rncr4 | 0 | 1.648835551 | 1 |
| ENSMUSG00000029784 | Ssmem1 | 0 | 0.648835551 | 1 |
| ENSMUSG00000039070 | Cpa4 | 0 | 1.648835551 | 1 |
| ENSMUSG00000086542 | Gm13782 | 0 | 1.063873051 | 1 |
| ENSMUSG00000097998 | Gm27019 | 0 | 0.063873051 | 1 |
| ENSMUSG00000086931 | Gm14532 | 0 | -0.810596067 | 0.844591606 |
| ENSMUSG00000086732 | Gm6117 | 0 | 1.063873051 | 1 |
| ENSMUSG00000029766 | 1700012A03Rik | 0 | 1.648835551 | 1 |
| ENSMUSG00000086763 | Plxna4os1 | 0 | 1.648835551 | 1 |
| ENSMUSG00000106873 | BC049739 | 0 | 1.648835551 | 1 |
| ENSMUSG00000087441 | Gm13853 | 0 | 1.063873051 | 1 |
| ENSMUSG00000087356 | Gm13856 | 0 | 0.326907457 | 1 |
| ENSMUSG00000087613 | Gm13855 | 0 | 0.326907457 | 1 |
| ENSMUSG00000085187 | Gm13860 | 0 | 1.063873051 | 1 |
| ENSMUSG00000086834 | Gm14547 | 0 | 1.063873051 | 1 |
| ENSMUSG00000029848 | Stra8 | 0 | 1.063873051 | 1 |
| ENSMUSG00000047420 | Fam180a | 0 | 0.648835551 | 1 |
| ENSMUSG00000029837 | 1700111E14Rik | 0 | -0.52108945 | 1 |
| ENSMUSG00000079598 | Clec2l | 0 | -0.158519371 | 1 |
| ENSMUSG00000106737 | Gm42962 | 0 | -1.051604167 | 0.889311043 |
| ENSMUSG00000073144 | 4930599N23Rik | 0 | -0.936126949 | 0.847265357 |
| ENSMUSG00000068601 | Gm10244 | 0 | 1.648835551 | 1 |
| ENSMUSG00000086582 | Gm16272 | 0 | -0.158519371 | 1 |
| ENSMUSG00000097563 | Gm26638 | 0 | 1.063873051 | 1 |
| ENSMUSG00000097484 | Gm26807 | 0 | 0.326907457 | 1 |
| ENSMUSG00000029915 | Clec5a | 0 | 1.063873051 | 1 |
| ENSMUSG00000068587 | Mgam | 0 | -1.810596067 | 0.466464457 |
| ENSMUSG00000063252 | Gm4744 | 0 | 1.648835551 | 1 |
| ENSMUSG00000101629 | Gm29588 | 0 | 1.648835551 | 1 |
| ENSMUSG00000094808 | 1810009J06Rik | 0 | 1.648835551 | 1 |
| ENSMUSG00000029882 | 2210010C04Rik | 0 | 1.063873051 | 1 |
| ENSMUSG00000029868 | Trpv6 | 0 | 0.648835551 | 1 |
| ENSMUSG00000036899 | Trpv5 | 0 | 1.063873051 | 1 |
| ENSMUSG00000029866 | Kel | 0 | 0.648835551 | 1 |
| ENSMUSG00000045479 | Olfr459 | 0 | 1.648835551 | 1 |
| ENSMUSG00000014104 | Sval2 | 0 | 1.648835551 | 1 |
| ENSMUSG00000029865 | Sval1 | 0 | 1.648835551 | 1 |

| | | | | |
|---|---|---|---|---|
| ENSMUSG00000062833 | Sval3 | 0 | 1.648835551 | 1 |
| ENSMUSG00000071506 | Tmem139 | 0 | 0.326907457 | 1 |
| ENSMUSG00000029862 | Clcn1 | 0 | -0.936126949 | 0.847265357 |
| ENSMUSG00000029861 | Fam131b | 0 | 1.648835551 | 1 |
| ENSMUSG00000108033 | Gm44284 | 0 | 1.648835551 | 1 |
| ENSMUSG00000029859 | Epha1 | 0 | 0.063873051 | 1 |
| ENSMUSG00000101268 | 2010310C07Rik | 0 | 1.063873051 | 1 |
| ENSMUSG00000091983 | Olfr457 | 0 | 0.648835551 | 1 |
| ENSMUSG00000071494 | Olfr455 | 0 | 1.648835551 | 1 |
| ENSMUSG00000073110 | Olfr444 | 0 | 0.326907457 | 1 |
| ENSMUSG00000094200 | Olfr237-ps1 | 0 | 1.648835551 | 1 |
| ENSMUSG00000107875 | Gm44696 | 0 | -0.351164449 | 1 |
| ENSMUSG00000039347 | Atp6v0e2 | 0 | 1.063873051 | 1 |
| ENSMUSG00000073096 | Lrrc61 | 0 | 1.063873051 | 1 |
| ENSMUSG00000052730 | Gm5111 | 0 | 1.648835551 | 1 |
| ENSMUSG00000052751 | Repin1 | 0 | 0.326907457 | 1 |
| ENSMUSG00000053297 | AI854703 | 0 | 0.063873051 | 1 |
| ENSMUSG00000043931 | Gimap7 | 0 | 1.648835551 | 1 |
| ENSMUSG00000090019 | Gimap1 | 0 | 0.063873051 | 1 |
| ENSMUSG00000039264 | Gimap3 | 0 | 1.063873051 | 1 |
| ENSMUSG00000107597 | Gm44764 | 0 | 0.648835551 | 1 |
| ENSMUSG00000108084 | Gm7932 | 0 | 0.063873051 | 1 |
| ENSMUSG00000029811 | Aoc1 | 0 | 0.648835551 | 1 |
| ENSMUSG00000107656 | 4930563H07Rik | 0 | 1.648835551 | 1 |
| ENSMUSG00000039215 | Svs1 | 0 | 1.648835551 | 1 |
| ENSMUSG00000047115 | Fam221a | 0 | -0.673092543 | 1 |
| ENSMUSG00000029831 | Npvf | 0 | 1.648835551 | 1 |
| ENSMUSG00000107978 | Gm44109 | 0 | 1.648835551 | 1 |
| ENSMUSG00000107619 | Gm32479 | 0 | 1.648835551 | 1 |
| ENSMUSG00000108244 | Gm44077 | 0 | 0.063873051 | 1 |
| ENSMUSG00000087658 | Hotairm1 | 0 | 1.648835551 | 1 |
| ENSMUSG00000079560 | Hoxa3 | 0 | -1.43862729 | 0.538110273 |
| ENSMUSG00000101451 | Gm29430 | 0 | 0.326907457 | 1 |
| ENSMUSG00000038253 | Hoxa5 | 0 | 0.648835551 | 1 |
| ENSMUSG00000043219 | Hoxa6 | 0 | 1.063873051 | 1 |
| ENSMUSG00000038236 | Hoxa7 | 0 | 1.648835551 | 1 |
| ENSMUSG00000000938 | Hoxa10 | 0 | 0.648835551 | 1 |
| ENSMUSG00000038210 | Hoxa11 | 0 | 1.648835551 | 1 |
| ENSMUSG00000005503 | Evx1 | 0 | 1.648835551 | 1 |
| ENSMUSG00000052955 | Cpvl | 0 | -1.258055044 | 0.687112677 |
| ENSMUSG00000108060 | 4921529L05Rik | 0 | 0.326907457 | 1 |
| ENSMUSG00000108081 | Gm44026 | 0 | 1.063873051 | 1 |
| ENSMUSG00000019124 | Scrn1 | 0 | -0.810596067 | 1 |
| ENSMUSG00000107766 | Gm44073 | 0 | 0.648835551 | 1 |
| ENSMUSG00000029778 | Adcyap1r1 | 0 | 0.063873051 | 1 |
| ENSMUSG00000037984 | Neurod6 | 0 | 1.648835551 | 1 |
| ENSMUSG00000108063 | Gm3279 | 0 | 1.648835551 | 1 |
| ENSMUSG00000002930 | Ppp1r17 | 0 | 1.648835551 | 1 |
| ENSMUSG00000061208 | Vmn1r8 | 0 | 1.063873051 | 1 |
| ENSMUSG00000091541 | Vmn1r9 | 0 | 1.648835551 | 1 |
| ENSMUSG00000115236 | Vmn1r11 | 0 | 1.648835551 | 1 |

| | | | | |
|---|---|---|---|---|
| ENSMUSG00000114982 | Vmn1r14 | 0 | 1.063873051 | 1 |
| ENSMUSG00000115253 | Vmn1r20 | 0 | 1.063873051 | 1 |
| ENSMUSG00000115343 | Vmn1r21 | 0 | 1.063873051 | 1 |
| ENSMUSG00000115507 | Vmn1r24 | 0 | 1.648835551 | 1 |
| ENSMUSG00000115705 | Vmn1r28 | 0 | 1.063873051 | 1 |
| ENSMUSG00000091734 | Vmn1r29 | 0 | 1.648835551 | 1 |
| ENSMUSG00000095670 | Vmn1r30 | 0 | 1.063873051 | 1 |
| ENSMUSG00000055430 | Nap1l5 | 0 | 1.063873051 | 1 |
| ENSMUSG00000049232 | Tigd2 | 0 | -1.158519371 | 0.903712767 |
| ENSMUSG00000097924 | A730020E08Rik | 0 | 1.063873051 | 1 |
| ENSMUSG00000029919 | Hpgds | 0 | -1.051604167 | 0.878674585 |
| ENSMUSG00000085689 | Gm16838 | 0 | 1.063873051 | 1 |
| ENSMUSG00000107479 | 2610300M13Rik | 0 | -0.810596067 | 1 |
| ENSMUSG00000062905 | Vmn1r32 | 0 | 1.648835551 | 1 |
| ENSMUSG00000115482 | Vmn1r36 | 0 | 1.063873051 | 1 |
| ENSMUSG00000115170 | Vmn1r38 | 0 | 1.648835551 | 1 |
| ENSMUSG00000093755 | Vmn1r39 | 0 | 1.648835551 | 1 |
| ENSMUSG00000107634 | Gm36816 | 0 | 1.063873051 | 1 |
| ENSMUSG00000105650 | Gm42889 | 0 | 0.063873051 | 1 |
| ENSMUSG00000076514 | Igkv17-121 | 0 | 1.648835551 | 1 |
| ENSMUSG00000094335 | Igkv1-117 | 0 | 1.063873051 | 1 |
| ENSMUSG00000093861 | Igkv1-110 | 0 | 1.063873051 | 1 |
| ENSMUSG00000076522 | Igkv16-104 | 0 | 1.648835551 | 1 |
| ENSMUSG00000106403 | Igkv20-101-2 | 0 | 1.648835551 | 1 |
| ENSMUSG00000106124 | Gm42539 | 0 | 1.648835551 | 1 |
| ENSMUSG00000076564 | Igkv12-46 | 0 | 1.648835551 | 1 |
| ENSMUSG00000096422 | Igkv12-44 | 0 | 1.648835551 | 1 |
| ENSMUSG00000076577 | Igkv8-30 | 0 | 0.648835551 | 1 |
| ENSMUSG00000076583 | Igkv8-24 | 0 | 1.648835551 | 1 |
| ENSMUSG00000076586 | Igkv8-21 | 0 | 1.648835551 | 1 |
| ENSMUSG00000096594 | Igkv8-19 | 0 | 1.648835551 | 1 |
| ENSMUSG00000076591 | Igkv8-16 | 0 | 1.648835551 | 1 |
| ENSMUSG00000094117 | Igkv3-12 | 0 | 1.648835551 | 1 |
| ENSMUSG00000076598 | Igkv3-7 | 0 | 1.648835551 | 1 |
| ENSMUSG00000053044 | Cd8b1 | 0 | 0.063873051 | 1 |
| ENSMUSG00000108132 | Gm44175 | 0 | 1.063873051 | 1 |
| ENSMUSG00000053977 | Cd8a | 0 | 1.648835551 | 1 |
| ENSMUSG00000108371 | Gm38832 | 0 | -0.351164449 | 1 |
| ENSMUSG00000056666 | Retsat | 0 | 0.326907457 | 1 |
| ENSMUSG00000092386 | Gm20536 | 0 | 1.063873051 | 1 |
| ENSMUSG00000072999 | Gm15401 | 0 | 0.648835551 | 1 |
| ENSMUSG00000107546 | Gm20560 | 0 | -0.673092543 | 1 |
| ENSMUSG00000107441 | Gm31747 | 0 | 1.648835551 | 1 |
| ENSMUSG00000107880 | Gm44113 | 0 | 1.648835551 | 1 |
| ENSMUSG00000060780 | Lrrtm1 | 0 | 0.326907457 | 1 |
| ENSMUSG00000030017 | Reg3g | 0 | 1.648835551 | 1 |
| ENSMUSG00000101404 | 1700009C05Rik | 0 | 1.648835551 | 1 |
| ENSMUSG00000107412 | Gm20383 | 0 | 1.648835551 | 1 |
| ENSMUSG00000107453 | Gm44145 | 0 | -0.158519371 | 1 |
| ENSMUSG00000109556 | Gm38843 | 0 | 1.063873051 | 1 |
| ENSMUSG00000100291 | 2310069B03Rik | 0 | 0.063873051 | 1 |

| | | | | |
|---|---|---|---|---|
| ENSMUSG00000109076 | Gm45145 | 0 | 1.648835551 | 1 |
| ENSMUSG00000068329 | Htra2 | 0 | -2.743481871 | 0.119283157 |
| ENSMUSG00000069678 | Pcgf1 | 0 | -1.258055044 | 0.673811773 |
| ENSMUSG00000087578 | Ccdc142os | 0 | 0.648835551 | 1 |
| ENSMUSG00000030032 | Wdr54 | 0 | 1.063873051 | 1 |
| ENSMUSG00000085782 | Gm15624 | 0 | 0.063873051 | 1 |
| ENSMUSG00000051844 | B230319C09Rik | 0 | 1.648835551 | 1 |
| ENSMUSG00000014542 | Clec4f | 0 | 1.063873051 | 1 |
| ENSMUSG00000085794 | Vax2os | 0 | 0.326907457 | 1 |
| ENSMUSG00000006269 | Atp6v1b1 | 0 | 1.648835551 | 1 |
| ENSMUSG00000099354 | 1700124L16Rik | 0 | 1.648835551 | 1 |
| ENSMUSG00000107534 | Gm43920 | 0 | 1.648835551 | 1 |
| ENSMUSG00000107538 | Gm44127 | 0 | -0.810596067 | 0.848836955 |
| ENSMUSG00000033726 | Emx1 | 0 | 1.648835551 | 1 |
| ENSMUSG00000068302 | Noto | 0 | 1.648835551 | 1 |
| ENSMUSG00000089694 | Nat8f7 | 0 | -0.673092543 | 1 |
| ENSMUSG00000033634 | Nat8f2 | 0 | 1.648835551 | 1 |
| ENSMUSG00000068299 | Nat8f4 | 0 | -0.158519371 | 1 |
| ENSMUSG00000057103 | Nat8f1 | 0 | 1.063873051 | 1 |
| ENSMUSG00000107230 | Gm19265 | 0 | -0.673092543 | 1 |
| ENSMUSG00000085589 | A430078I02Rik | 0 | 1.063873051 | 1 |
| ENSMUSG00000101655 | 2310040G24Rik | 0 | -1.52108945 | 0.544908989 |
| ENSMUSG00000033508 | Asprv1 | 0 | 1.648835551 | 1 |
| ENSMUSG00000108197 | Gm44214 | 0 | 1.648835551 | 1 |
| ENSMUSG00000107961 | Gm44036 | 0 | 1.063873051 | 1 |
| ENSMUSG00000030046 | Bmp10 | 0 | 1.648835551 | 1 |
| ENSMUSG00000107593 | Gm44097 | 0 | 1.648835551 | 1 |
| ENSMUSG00000097763 | Gm26636 | 0 | 1.648835551 | 1 |
| ENSMUSG00000015053 | Gata2 | 0 | 1.063873051 | 1 |
| ENSMUSG00000097730 | Gm26588 | 0 | 1.648835551 | 1 |
| ENSMUSG00000033152 | Podxl2 | 0 | -0.936126949 | 0.847265357 |
| ENSMUSG00000060416 | Gm839 | 0 | 1.648835551 | 1 |
| ENSMUSG00000107622 | 4930512J16Rik | 0 | 1.063873051 | 1 |
| ENSMUSG00000097816 | Gm26811 | 0 | 1.648835551 | 1 |
| ENSMUSG00000094586 | Vmn1r41 | 0 | 1.648835551 | 1 |
| ENSMUSG00000068231 | Vmn1r43 | 0 | -1.351164449 | 0.687112677 |
| ENSMUSG00000044248 | Vmn1r45 | 0 | 1.063873051 | 1 |
| ENSMUSG00000060724 | Vmn1r47 | 0 | 0.063873051 | 1 |
| ENSMUSG00000095932 | Vmn1r49 | 0 | -0.158519371 | 1 |
| ENSMUSG00000060816 | Vmn1r52 | 0 | 1.648835551 | 1 |
| ENSMUSG00000048794 | Cfap100 | 0 | 1.063873051 | 1 |
| ENSMUSG00000030087 | Klf15 | 0 | 1.648835551 | 1 |
| ENSMUSG00000030089 | Slc41a3 | 0 | -0.673092543 | 1 |
| ENSMUSG00000108584 | Gm45216 | 0 | 0.648835551 | 1 |
| ENSMUSG00000108308 | Gm45218 | 0 | 1.648835551 | 1 |
| ENSMUSG00000079462 | Gm15737 | 0 | 0.648835551 | 1 |
| ENSMUSG00000085413 | Kbtbd8os | 0 | 1.648835551 | 1 |
| ENSMUSG00000030031 | Kbtbd8 | 0 | 1.648835551 | 1 |
| ENSMUSG00000044086 | Lmod3 | 0 | 0.326907457 | 1 |
| ENSMUSG00000107902 | Gm32592 | 0 | -0.673092543 | 1 |
| ENSMUSG00000093559 | Gm20705 | 0 | 1.063873051 | 1 |

| | | | | |
|---|---|---|---|---|
| ENSMUSG00000097548 | Gm26748 | 0 | 1.648835551 | 1 |
| ENSMUSG00000107204 | Gm43067 | 0 | 1.063873051 | 1 |
| ENSMUSG00000107887 | Gm43950 | 0 | 0.648835551 | 1 |
| ENSMUSG00000108274 | Gm44169 | 0 | 0.326907457 | 1 |
| ENSMUSG00000052236 | Gm9871 | 0 | 0.648835551 | 1 |
| ENSMUSG00000108237 | Gm19757 | 0 | 0.648835551 | 1 |
| ENSMUSG00000085067 | Gm15631 | 0 | 1.648835551 | 1 |
| ENSMUSG00000034648 | Lrrn1 | 0 | 1.648835551 | 1 |
| ENSMUSG00000107496 | 4933431M02Rik | 0 | 1.648835551 | 1 |
| ENSMUSG00000034639 | Setmar | 0 | -1.258055044 | 0.673811773 |
| ENSMUSG00000107458 | Gm44040 | 0 | 1.648835551 | 1 |
| ENSMUSG00000090576 | Gm17055 | 0 | 1.648835551 | 1 |
| ENSMUSG00000092448 | Gm20387 | 0 | 1.648835551 | 1 |
| ENSMUSG00000044574 | 5031434C07Rik | 0 | 1.648835551 | 1 |
| ENSMUSG00000030270 | Cpne9 | 0 | -0.673092543 | 1 |
| ENSMUSG00000085012 | Gm15492 | 0 | 1.648835551 | 1 |
| ENSMUSG00000051169 | Rpusd3 | 0 | 0.063873051 | 1 |
| ENSMUSG00000033963 | Fancd2os | 0 | 1.648835551 | 1 |
| ENSMUSG00000107937 | Gm44200 | 0 | 1.063873051 | 1 |
| ENSMUSG00000107560 | Gm44199 | 0 | 1.648835551 | 1 |
| ENSMUSG00000064177 | Ghrl | 0 | -1.158519371 | 0.909230539 |
| ENSMUSG00000108137 | Gm44053 | 0 | 1.063873051 | 1 |
| ENSMUSG00000030307 | Slc6a11 | 0 | -0.673092543 | 1 |
| ENSMUSG00000030310 | Slc6a1 | 0 | 0.648835551 | 1 |
| ENSMUSG00000097486 | Gm17733 | 0 | 1.648835551 | 1 |
| ENSMUSG00000108141 | Gm44079 | 0 | 0.648835551 | 1 |
| ENSMUSG00000107704 | Gm36355 | 0 | 1.063873051 | 1 |
| ENSMUSG00000107994 | D830050J10Rik | 0 | 0.326907457 | 1 |
| ENSMUSG00000107429 | Gm44206 | 0 | -0.936126949 | 0.886203497 |
| ENSMUSG00000030324 | Rho | 0 | 1.648835551 | 1 |
| ENSMUSG00000092625 | Gm20404 | 0 | 0.648835551 | 1 |
| ENSMUSG00000081649 | Olfr213 | 0 | 1.648835551 | 1 |
| ENSMUSG00000050654 | Olfr215 | 0 | 1.648835551 | 1 |
| ENSMUSG00000042129 | Rassf4 | 0 | -1.351164449 | 0.711791718 |
| ENSMUSG00000048108 | Tmem72 | 0 | -0.351164449 | 1 |
| ENSMUSG00000108038 | D030044L04Rik | 0 | 0.648835551 | 1 |
| ENSMUSG00000107785 | Gm45083 | 0 | 0.648835551 | 1 |
| ENSMUSG00000042097 | Zfp239 | 0 | -1.52108945 | 0.544908989 |
| ENSMUSG00000030110 | Ret | 0 | -0.673092543 | 1 |
| ENSMUSG00000107791 | Gm30557 | 0 | 1.648835551 | 1 |
| ENSMUSG00000055003 | Lrtm2 | 0 | 1.648835551 | 1 |
| ENSMUSG00000108282 | Gm44317 | 0 | 1.648835551 | 1 |
| ENSMUSG00000030166 | Rad52 | 0 | -2.743481871 | 0.109338528 |
| ENSMUSG00000107744 | 1700072O05Rik | 0 | 1.063873051 | 1 |
| ENSMUSG00000097011 | Gm4651 | 0 | -0.810596067 | 1 |
| ENSMUSG00000030137 | Tuba8 | 0 | -0.351164449 | 1 |
| ENSMUSG00000086527 | Gm15856 | 0 | 1.063873051 | 1 |
| ENSMUSG00000030107 | Usp18 | 0 | 0.326907457 | 1 |
| ENSMUSG00000030109 | Slc6a12 | 0 | 0.648835551 | 1 |
| ENSMUSG00000030131 | Mug2 | 0 | -0.158519371 | 1 |
| ENSMUSG00000030114 | Klrg1 | 0 | 1.648835551 | 1 |

| | | | | |
|---|---|---|---|---|
| ENSMUSG00000040627 | Aicda | 0 | -0.351164449 | 1 |
| ENSMUSG00000030117 | Gdf3 | 0 | 1.648835551 | 1 |
| ENSMUSG00000059639 | Clec4a4 | 0 | 0.063873051 | 1 |
| ENSMUSG00000030147 | Clec4b1 | 0 | 1.648835551 | 1 |
| ENSMUSG00000091260 | Vmn2r19 | 0 | 0.648835551 | 1 |
| ENSMUSG00000091620 | Vmn2r23 | 0 | 0.326907457 | 1 |
| ENSMUSG00000096630 | Vmn2r26 | 0 | -0.158519371 | 1 |
| ENSMUSG00000072778 | Vmn2r27 | 0 | 1.063873051 | 1 |
| ENSMUSG00000008845 | Cd163 | 0 | -0.52108945 | 1 |
| ENSMUSG00000107552 | Gm44096 | 0 | 1.063873051 | 1 |
| ENSMUSG00000089961 | Gm16567 | 0 | 1.648835551 | 1 |
| ENSMUSG00000087327 | Gm15884 | 0 | 0.063873051 | 1 |
| ENSMUSG00000004267 | Eno2 | 0 | -0.810596067 | 1 |
| ENSMUSG00000023439 | Gnb3 | 0 | 1.648835551 | 1 |
| ENSMUSG00000038390 | Gpr162 | 0 | 1.648835551 | 1 |
| ENSMUSG00000030124 | Lag3 | 0 | 0.326907457 | 1 |
| ENSMUSG00000067714 | Lpar5 | 0 | 0.326907457 | 1 |
| ENSMUSG00000099587 | Gm28967 | 0 | 1.063873051 | 1 |
| ENSMUSG00000067702 | Tuba3a | 0 | 0.648835551 | 1 |
| ENSMUSG00000101191 | Gm28809 | 0 | 1.648835551 | 1 |
| ENSMUSG00000097596 | Gm26673 | 0 | 1.648835551 | 1 |
| ENSMUSG00000107368 | 1700018A23Rik | 0 | 1.648835551 | 1 |
| ENSMUSG00000000183 | Fgf6 | 0 | 1.648835551 | 1 |
| ENSMUSG00000107379 | Gm43126 | 0 | 0.326907457 | 1 |
| ENSMUSG00000107879 | 9330102E08Rik | 0 | 1.063873051 | 1 |
| ENSMUSG00000056771 | Gm10010 | 0 | 1.063873051 | 1 |
| ENSMUSG00000108267 | Gm44190 | 0 | 0.648835551 | 1 |
| ENSMUSG00000097253 | Gm26770 | 0 | 0.648835551 | 1 |
| ENSMUSG00000001517 | Foxm1 | 0 | -1.051604167 | 0.874973657 |
| ENSMUSG00000085739 | Gm15862 | 0 | -0.351164449 | 1 |
| ENSMUSG00000001520 | Nrip2 | 0 | 0.063873051 | 1 |
| ENSMUSG00000030359 | Pzp | 0 | -1.258055044 | 0.73719709 |
| ENSMUSG00000047228 | A2ml1 | 0 | -0.810596067 | 1 |
| ENSMUSG00000030364 | Clec2h | 0 | 1.648835551 | 1 |
| ENSMUSG00000079298 | Klrb1b | 0 | 0.648835551 | 1 |
| ENSMUSG00000107946 | Gm44066 | 0 | 1.063873051 | 1 |
| ENSMUSG00000108095 | Gm44067 | 0 | 0.326907457 | 1 |
| ENSMUSG00000030365 | Clec2i | 0 | 1.648835551 | 1 |
| ENSMUSG00000085786 | Gm15987 | 0 | -1.810596067 | 0.465580427 |
| ENSMUSG00000089728 | Clec2f | 0 | -0.936126949 | 0.847265357 |
| ENSMUSG00000030154 | Klrb1f | 0 | 1.063873051 | 1 |
| ENSMUSG00000112144 | Gm47861 | 0 | 1.063873051 | 1 |
| ENSMUSG00000030156 | Cd69 | 0 | 0.648835551 | 1 |
| ENSMUSG00000053063 | Clec12a | 0 | 1.648835551 | 1 |
| ENSMUSG00000030158 | Clec12b | 0 | 1.063873051 | 1 |
| ENSMUSG00000030159 | Clec1b | 0 | 1.648835551 | 1 |
| ENSMUSG00000046080 | Clec9a | 0 | 1.648835551 | 1 |
| ENSMUSG00000033082 | Clec1a | 0 | 0.063873051 | 1 |
| ENSMUSG00000050241 | Klre1 | 0 | 1.648835551 | 1 |
| ENSMUSG00000030165 | Klrd1 | 0 | 0.326907457 | 1 |
| ENSMUSG00000033027 | Klrc3 | 0 | 1.063873051 | 1 |

| | | | | |
|---|---|---|---|---|
| ENSMUSG00000052736 | Klrc2 | 0 | -1.351164449 | 0.687112677 |
| ENSMUSG00000067610 | Klri1 | 0 | 0.326907457 | 1 |
| ENSMUSG00000043932 | Klri2 | 0 | 0.063873051 | 1 |
| ENSMUSG00000030173 | Klra5 | 0 | 1.063873051 | 1 |
| ENSMUSG00000061769 | Klra6 | 0 | 1.648835551 | 1 |
| ENSMUSG00000089727 | Klra8 | 0 | 0.648835551 | 1 |
| ENSMUSG00000033024 | Klra9 | 0 | 1.063873051 | 1 |
| ENSMUSG00000072718 | Klra10 | 0 | 1.648835551 | 1 |
| ENSMUSG00000055594 | 5530400C23Rik | 0 | 1.648835551 | 1 |
| ENSMUSG00000107767 | Gm44448 | 0 | 1.063873051 | 1 |
| ENSMUSG00000107728 | Gm38910 | 0 | 1.063873051 | 1 |
| ENSMUSG00000097496 | Gm26653 | 0 | 0.648835551 | 1 |
| ENSMUSG00000043298 | Smco3 | 0 | 1.063873051 | 1 |
| ENSMUSG00000030217 | Art4 | 0 | 1.648835551 | 1 |
| ENSMUSG00000100166 | B230110G15Rik | 0 | 1.063873051 | 1 |
| ENSMUSG00000108128 | 4922502N22Rik | 0 | 1.648835551 | 1 |
| ENSMUSG00000092164 | Rergl | 0 | 1.648835551 | 1 |
| ENSMUSG00000107638 | Gm30524 | 0 | -1.051604167 | 0.886493023 |
| ENSMUSG00000041698 | Slco1a1 | 0 | 0.326907457 | 1 |
| ENSMUSG00000079263 | Gm6614 | 0 | 0.648835551 | 1 |
| ENSMUSG00000041681 | Iapp | 0 | 0.326907457 | 1 |
| ENSMUSG00000030246 | Ldhb | 0 | -1.351164449 | 0.783214613 |
| ENSMUSG00000030247 | Kcnj8 | 0 | 0.326907457 | 1 |
| ENSMUSG00000104234 | 5330439B14Rik | 0 | 0.648835551 | 1 |
| ENSMUSG00000048473 | Sult6b2 | 0 | 0.648835551 | 1 |
| ENSMUSG00000072683 | Gm7457 | 0 | 0.648835551 | 1 |
| ENSMUSG00000085785 | Sox5os3 | 0 | 1.063873051 | 1 |
| ENSMUSG00000097419 | Gm26666 | 0 | 1.063873051 | 1 |
| ENSMUSG00000043541 | Casc1 | 0 | -3.184054463 | 0.023817286 |
| ENSMUSG00000087277 | 2010013B24Rik | 0 | 1.648835551 | 1 |
| ENSMUSG00000086722 | Gm15499 | 0 | 1.648835551 | 1 |
| ENSMUSG00000085658 | Gm15704 | 0 | 0.326907457 | 1 |
| ENSMUSG00000085238 | 4930479D17Rik | 0 | 1.063873051 | 1 |
| ENSMUSG00000030292 | Smco2 | 0 | -0.52108945 | 1 |
| ENSMUSG00000040163 | 1700034J05Rik | 0 | 1.063873051 | 1 |
| ENSMUSG00000072662 | Mansc4 | 0 | 1.063873051 | 1 |
| ENSMUSG00000048776 | Pthlh | 0 | -0.351164449 | 1 |
| ENSMUSG00000086777 | Far2os2 | 0 | 0.648835551 | 1 |
| ENSMUSG00000089948 | Far2os1 | 0 | 1.648835551 | 1 |
| ENSMUSG00000108148 | Gm44020 | 0 | 1.648835551 | 1 |
| ENSMUSG00000070995 | Gm6313 | 0 | 1.648835551 | 1 |
| ENSMUSG00000084909 | 4930528G23Rik | 0 | 0.063873051 | 1 |
| ENSMUSG00000107405 | Gm44115 | 0 | 0.326907457 | 1 |
| ENSMUSG00000067292 | Gm10203 | 0 | 1.648835551 | 1 |
| ENSMUSG00000107526 | 1700003I16Rik | 0 | -0.673092543 | 1 |
| ENSMUSG00000107956 | Speer9-ps1 | 0 | 0.648835551 | 1 |
| ENSMUSG00000069806 | Cacng7 | 0 | -2.258055044 | 0.222101622 |
| ENSMUSG00000107815 | Gm44257 | 0 | -0.158519371 | 1 |
| ENSMUSG00000107862 | Gm44151 | 0 | -0.158519371 | 1 |
| ENSMUSG00000078815 | Cacng6 | 0 | 0.648835551 | 1 |
| ENSMUSG00000108644 | Gm44920 | 0 | 1.648835551 | 1 |

| | | | | |
|---|---|---|---|---|
| ENSMUSG00000085039 | Gm15927 | 0 | -0.52108945 | 1 |
| ENSMUSG00000086032 | Gm15929 | 0 | -0.158519371 | 1 |
| ENSMUSG00000074419 | Gm15448 | 0 | 1.648835551 | 1 |
| ENSMUSG00000074417 | Gm14548 | 0 | 1.648835551 | 1 |
| ENSMUSG00000030427 | Lilra6 | 0 | 1.648835551 | 1 |
| ENSMUSG00000108593 | Gm44581 | 0 | 1.648835551 | 1 |
| ENSMUSG00000108505 | 1700095K22Rik | 0 | 1.648835551 | 1 |
| ENSMUSG00000055541 | Lair1 | 0 | -2.258055044 | 0.223176736 |
| ENSMUSG00000030428 | Ttyh1 | 0 | 0.326907457 | 1 |
| ENSMUSG00000085886 | D030047H15Rik | 0 | 0.326907457 | 1 |
| ENSMUSG00000070873 | Lilra5 | 0 | 1.063873051 | 1 |
| ENSMUSG00000062524 | Ncr1 | 0 | 1.648835551 | 1 |
| ENSMUSG00000035458 | Tnni3 | 0 | 0.326907457 | 1 |
| ENSMUSG00000085725 | Gm15873 | 0 | 1.648835551 | 1 |
| ENSMUSG00000004961 | Syt5 | 0 | 0.326907457 | 1 |
| ENSMUSG00000108320 | Gm44877 | 0 | 1.648835551 | 1 |
| ENSMUSG00000045282 | Tmem86b | 0 | -0.52108945 | 1 |
| ENSMUSG00000051811 | Cox6b2 | 0 | 0.648835551 | 1 |
| ENSMUSG00000004371 | Il11 | 0 | -0.52108945 | 1 |
| ENSMUSG00000052605 | Isoc2b | 0 | -0.936126949 | 0.886203497 |
| ENSMUSG00000110221 | Gm36210 | 0 | 1.648835551 | 1 |
| ENSMUSG00000035285 | Nat14 | 0 | 1.648835551 | 1 |
| ENSMUSG00000035279 | Ssc5d | 0 | 1.648835551 | 1 |
| ENSMUSG00000030433 | Sbk2 | 0 | 1.648835551 | 1 |
| ENSMUSG00000051550 | Zfp579 | 0 | -2.43862729 | 0.184141965 |
| ENSMUSG00000035228 | Ccdc106 | 0 | -0.351164449 | 1 |
| ENSMUSG00000043290 | Zfp784 | 0 | 1.648835551 | 1 |
| ENSMUSG00000086604 | Gm15510 | 0 | 0.326907457 | 1 |
| ENSMUSG00000118215 | Vmn1r55 | 0 | 1.063873051 | 1 |
| ENSMUSG00000066820 | Vmn2r28 | 0 | 1.648835551 | 1 |
| ENSMUSG00000058631 | Vmn1r63 | 0 | 1.063873051 | 1 |
| ENSMUSG00000109283 | Gm44733 | 0 | 1.063873051 | 1 |
| ENSMUSG00000109108 | Gm36371 | 0 | 1.063873051 | 1 |
| ENSMUSG00000034660 | Galp | 0 | 0.326907457 | 1 |
| ENSMUSG00000054893 | Zfp667 | 0 | -1.158519371 | 0.667778757 |
| ENSMUSG00000109398 | Gm3854 | 0 | -0.158519371 | 1 |
| ENSMUSG00000093536 | Smim17 | 0 | -0.158519371 | 1 |
| ENSMUSG00000093877 | Olfr1348 | 0 | 0.648835551 | 1 |
| ENSMUSG00000048067 | Olfr1349 | 0 | 1.648835551 | 1 |
| ENSMUSG00000063535 | Zfp773 | 0 | 0.648835551 | 1 |
| ENSMUSG00000066838 | Zfp772 | 0 | 1.063873051 | 1 |
| ENSMUSG00000096743 | Vmn2r32 | 0 | 0.648835551 | 1 |
| ENSMUSG00000070841 | Vmn2r34 | 0 | 1.648835551 | 1 |
| ENSMUSG00000070844 | Vmn2r42 | 0 | 1.648835551 | 1 |
| ENSMUSG00000090662 | Vmn2r45 | 0 | 1.063873051 | 1 |
| ENSMUSG00000095773 | Vmn2r38 | 0 | 0.648835551 | 1 |
| ENSMUSG00000096294 | Gm10302 | 0 | 1.063873051 | 1 |
| ENSMUSG00000095914 | Vmn2r48 | 0 | 1.648835551 | 1 |
| ENSMUSG00000058685 | Vmn2r51 | 0 | -0.52108945 | 1 |
| ENSMUSG00000091930 | Vmn2r52 | 0 | 0.063873051 | 1 |
| ENSMUSG00000043066 | Vmn1r66 | 0 | 1.063873051 | 1 |

| | | | | |
|---|---|---|---|---|
| ENSMUSG00000046716 | Vmn1r67 | 0 | 1.648835551 | 1 |
| ENSMUSG00000091662 | Vmn1r69 | 0 | 1.648835551 | 1 |
| ENSMUSG00000110034 | Gm45379 | 0 | 1.648835551 | 1 |
| ENSMUSG00000095430 | Vmn1r72 | 0 | 1.648835551 | 1 |
| ENSMUSG00000051687 | Vmn1r73 | 0 | 0.326907457 | 1 |
| ENSMUSG00000095864 | Vmn1r77 | 0 | 1.063873051 | 1 |
| ENSMUSG00000096735 | Vmn1r79 | 0 | 0.326907457 | 1 |
| ENSMUSG00000110042 | Gm4224 | 0 | 1.648835551 | 1 |
| ENSMUSG00000058132 | Vmn1r82 | 0 | 1.648835551 | 1 |
| ENSMUSG00000066804 | Vmn1r83 | 0 | 0.648835551 | 1 |
| ENSMUSG00000034071 | Zfp551 | 0 | -0.158519371 | 1 |
| ENSMUSG00000096002 | Vmn2r53 | 0 | 0.648835551 | 1 |
| ENSMUSG00000090762 | Vmn2r56 | 0 | 0.326907457 | 1 |
| ENSMUSG00000060397 | Zfp128 | 0 | -0.351164449 | 1 |
| ENSMUSG00000054715 | Zscan22 | 0 | -2.43862729 | 0.185472119 |
| ENSMUSG00000033967 | Rnf225 | 0 | 1.063873051 | 1 |
| ENSMUSG00000004500 | Zfp324 | 0 | -0.351164449 | 1 |
| ENSMUSG00000030382 | Slc27a5 | 0 | 1.648835551 | 1 |
| ENSMUSG00000030380 | Mzf1 | 0 | -1.051604167 | 0.858831093 |
| ENSMUSG00000110268 | Gm45436 | 0 | 1.648835551 | 1 |
| ENSMUSG00000070816 | Vmn1r86 | 0 | 0.063873051 | 1 |
| ENSMUSG00000070815 | Vmn1r87 | 0 | 1.063873051 | 1 |
| ENSMUSG00000117853 | Vmn1r88 | 0 | 0.063873051 | 1 |
| ENSMUSG00000005649 | Cabp5 | 0 | 1.063873051 | 1 |
| ENSMUSG00000078799 | Sult2a5 | 0 | -0.673092543 | 1 |
| ENSMUSG00000070810 | Sult2a6 | 0 | 1.063873051 | 1 |
| ENSMUSG00000030378 | Sult2a8 | 0 | -0.158519371 | 1 |
| ENSMUSG00000041583 | Obox6 | 0 | 1.648835551 | 1 |
| ENSMUSG00000041578 | Crx | 0 | 0.326907457 | 1 |
| ENSMUSG00000109984 | Gm38948 | 0 | 1.648835551 | 1 |
| ENSMUSG00000041420 | Meis3 | 0 | -1.43862729 | 0.748847221 |
| ENSMUSG00000109781 | Gm45509 | 0 | 0.648835551 | 1 |
| ENSMUSG00000109647 | Gm45510 | 0 | 1.063873051 | 1 |
| ENSMUSG00000078795 | Ceacam15 | 0 | 1.648835551 | 1 |
| ENSMUSG00000048920 | Fkrp | 0 | -0.52108945 | 1 |
| ENSMUSG00000063594 | Gng8 | 0 | 1.648835551 | 1 |
| ENSMUSG00000070802 | Pnmal2 | 0 | 1.648835551 | 1 |
| ENSMUSG00000041141 | Pnmal1 | 0 | 1.063873051 | 1 |
| ENSMUSG00000108348 | Gm42372 | 0 | 0.063873051 | 1 |
| ENSMUSG00000066760 | Psg16 | 0 | 0.326907457 | 1 |
| ENSMUSG00000108782 | Gm32772 | 0 | 1.063873051 | 1 |
| ENSMUSG00000008789 | Ceacam5 | 0 | 1.648835551 | 1 |
| ENSMUSG00000023185 | Ceacam14 | 0 | 0.648835551 | 1 |
| ENSMUSG00000108545 | Gm44613 | 0 | -2.158519371 | 0.290284345 |
| ENSMUSG00000030373 | Psg28 | 0 | 1.063873051 | 1 |
| ENSMUSG00000074359 | Psg23 | 0 | 1.648835551 | 1 |
| ENSMUSG00000070796 | Psg21 | 0 | 0.648835551 | 1 |
| ENSMUSG00000044903 | Psg22 | 0 | 1.648835551 | 1 |
| ENSMUSG00000004540 | Psg17 | 0 | 0.063873051 | 1 |
| ENSMUSG00000030411 | Nova2 | 0 | -0.810596067 | 0.844591606 |
| ENSMUSG00000086533 | Mypopos | 0 | 1.063873051 | 1 |

| | | | | |
|---|---|---|---|---|
| ENSMUSG00000085601 | Gm4969 | 0 | 1.063873051 | 1 |
| ENSMUSG00000050428 | Fbxo46 | 0 | -1.995020638 | 0.416543551 |
| ENSMUSG00000044317 | Gpr4 | 0 | 1.063873051 | 1 |
| ENSMUSG00000030402 | Ppm1n | 0 | 1.648835551 | 1 |
| ENSMUSG00000030401 | Rtn2 | 0 | -0.52108945 | 1 |
| ENSMUSG00000109366 | Gm44698 | 0 | 1.063873051 | 1 |
| ENSMUSG00000097231 | Gm26852 | 0 | 1.063873051 | 1 |
| ENSMUSG00000011267 | Zfp296 | 0 | 1.063873051 | 1 |
| ENSMUSG00000074336 | Apoc4 | 0 | 0.326907457 | 1 |
| ENSMUSG00000092381 | Gm20512 | 0 | 1.648835551 | 1 |
| ENSMUSG00000097226 | Gm26600 | 0 | 1.063873051 | 1 |
| ENSMUSG00000087593 | Gm16174 | 0 | 1.063873051 | 1 |
| ENSMUSG00000109424 | Gm44658 | 0 | 1.648835551 | 1 |
| ENSMUSG00000092216 | Gm19345 | 0 | 0.063873051 | 1 |
| ENSMUSG00000014686 | Ceacam16 | 0 | 0.063873051 | 1 |
| ENSMUSG00000109214 | Gm44659 | 0 | 0.326907457 | 1 |
| ENSMUSG00000090075 | Gm16251 | 0 | -0.673092543 | 1 |
| ENSMUSG00000074291 | Vmn1r168 | 0 | -0.810596067 | 0.844591606 |
| ENSMUSG00000046924 | Vmn1r179 | 0 | 1.063873051 | 1 |
| ENSMUSG00000092473 | Vmn1r180 | 0 | 0.648835551 | 1 |
| ENSMUSG00000118318 | Vmn1r183 | 0 | 0.648835551 | 1 |
| ENSMUSG00000057101 | Zfp180 | 0 | -2.395558568 | 0.174915761 |
| ENSMUSG00000068962 | Zfp114 | 0 | 1.063873051 | 1 |
| ENSMUSG00000087598 | Zfp111 | 0 | -1.258055044 | 0.673811773 |
| ENSMUSG00000074283 | Zfp109 | 0 | 0.063873051 | 1 |
| ENSMUSG00000055826 | Tescl | 0 | 1.648835551 | 1 |
| ENSMUSG00000030484 | Lypd5 | 0 | 1.648835551 | 1 |
| ENSMUSG00000011632 | Pinlyp | 0 | 0.648835551 | 1 |
| ENSMUSG00000057454 | Lypd3 | 0 | 1.063873051 | 1 |
| ENSMUSG00000058717 | Lypd11 | 0 | 1.648835551 | 1 |
| ENSMUSG00000091955 | Gm9844 | 0 | 0.326907457 | 1 |
| ENSMUSG00000074274 | D930028M14Rik | 0 | 0.648835551 | 1 |
| ENSMUSG00000046541 | Zfp526 | 0 | -1.051604167 | 0.858831093 |
| ENSMUSG00000090330 | 9130221H12Rik | 0 | 0.648835551 | 1 |
| ENSMUSG00000059479 | B3gnt8 | 0 | 0.648835551 | 1 |
| ENSMUSG00000097017 | Gm26707 | 0 | 1.648835551 | 1 |
| ENSMUSG00000108415 | Gm30146 | 0 | 1.063873051 | 1 |
| ENSMUSG00000030483 | Cyp2b10 | 0 | -0.673092543 | 1 |
| ENSMUSG00000108413 | BC026762 | 0 | 1.648835551 | 1 |
| ENSMUSG00000040583 | Cyp2b13 | 0 | 1.648835551 | 1 |
| ENSMUSG00000046130 | Vmn1r184 | 0 | 1.063873051 | 1 |
| ENSMUSG00000040650 | Cyp2b23 | 0 | 1.063873051 | 1 |
| ENSMUSG00000091867 | Cyp2a22 | 0 | 1.648835551 | 1 |
| ENSMUSG00000086868 | Gm15883 | 0 | 1.063873051 | 1 |
| ENSMUSG00000078787 | Cyp2t4 | 0 | 1.063873051 | 1 |
| ENSMUSG00000087371 | Gm15541 | 0 | 1.648835551 | 1 |
| ENSMUSG00000040424 | Hipk4 | 0 | 1.063873051 | 1 |
| ENSMUSG00000108592 | Gm38973 | 0 | -1.673092543 | 0.644599258 |
| ENSMUSG00000007944 | Ttc9b | 0 | 1.648835551 | 1 |
| ENSMUSG00000108520 | Gm44684 | 0 | 1.648835551 | 1 |
| ENSMUSG00000078779 | Zfp59 | 0 | -0.810596067 | 0.844591606 |

| | | | | |
|---|---|---|---|---|
| ENSMUSG00000047730 | Fcgbp | 0 | 0.063873051 | 1 |
| ENSMUSG00000078776 | 9530053A07Rik | 0 | 0.063873051 | 1 |
| ENSMUSG00000070705 | Eid2b | 0 | 0.326907457 | 1 |
| ENSMUSG00000003436 | Dll3 | 0 | 0.648835551 | 1 |
| ENSMUSG00000084174 | Sycn | 0 | 1.063873051 | 1 |
| ENSMUSG00000108865 | Gm44617 | 0 | 1.648835551 | 1 |
| ENSMUSG00000070699 | Sars2 | 0 | -0.52108945 | 1 |
| ENSMUSG00000109420 | Gm44702 | 0 | 0.063873051 | 1 |
| ENSMUSG00000108930 | Gm10648 | 0 | -0.351164449 | 1 |
| ENSMUSG00000054083 | Capn12 | 0 | 1.063873051 | 1 |
| ENSMUSG00000109314 | Gm44699 | 0 | 0.063873051 | 1 |
| ENSMUSG00000037337 | Map4k1 | 0 | -0.351164449 | 1 |
| ENSMUSG00000097079 | Gm26604 | 0 | 0.063873051 | 1 |
| ENSMUSG00000090009 | Gm16282 | 0 | 0.648835551 | 1 |
| ENSMUSG00000099098 | 1110035H17Rik | 0 | 1.648835551 | 1 |
| ENSMUSG00000031493 | Ggn | 0 | 1.648835551 | 1 |
| ENSMUSG00000037166 | Ppp1r14a | 0 | 0.326907457 | 1 |
| ENSMUSG00000109351 | Gm44632 | 0 | 1.063873051 | 1 |
| ENSMUSG00000100111 | A330087D11Rik | 0 | 1.648835551 | 1 |
| ENSMUSG00000049123 | Catsperg2 | 0 | 0.063873051 | 1 |
| ENSMUSG00000047473 | Zfp30 | 0 | -0.936126949 | 0.847265357 |
| ENSMUSG00000099689 | Zfp383 | 0 | -0.936126949 | 0.847265357 |
| ENSMUSG00000074220 | Zfp382 | 0 | -0.810596067 | 1 |
| ENSMUSG00000037029 | Zfp146 | 0 | -2.560617814 | 0.105255472 |
| ENSMUSG00000074218 | Cox7a1 | 0 | 0.326907457 | 1 |
| ENSMUSG00000097817 | Gm26810 | 0 | -0.52108945 | 1 |
| ENSMUSG00000074210 | E130208F15Rik | 0 | 1.648835551 | 1 |
| ENSMUSG00000062132 | Arhgap33os | 0 | 1.063873051 | 1 |
| ENSMUSG00000036882 | Arhgap33 | 0 | 0.648835551 | 1 |
| ENSMUSG00000036854 | Hspb6 | 0 | -0.158519371 | 1 |
| ENSMUSG00000036845 | Lin37 | 0 | -0.351164449 | 1 |
| ENSMUSG00000078765 | U2af1l4 | 0 | -0.52108945 | 1 |
| ENSMUSG00000036826 | Igflr1 | 0 | -1.43862729 | 0.545392675 |
| ENSMUSG00000006310 | Zbtb32 | 0 | 1.063873051 | 1 |
| ENSMUSG00000078762 | Haus5 | 0 | -0.810596067 | 0.852297462 |
| ENSMUSG00000049761 | Pmis2 | 0 | 1.648835551 | 1 |
| ENSMUSG00000097320 | Tmem147os | 0 | -0.351164449 | 1 |
| ENSMUSG00000061099 | Gapdhs | 0 | -0.158519371 | 1 |
| ENSMUSG00000098146 | Gm26935 | 0 | 1.063873051 | 1 |
| ENSMUSG00000044453 | Ffar1 | 0 | 1.063873051 | 1 |
| ENSMUSG00000108546 | Gm44662 | 0 | 0.326907457 | 1 |
| ENSMUSG00000056978 | Hamp2 | 0 | 1.063873051 | 1 |
| ENSMUSG00000050440 | Hamp | 0 | 1.063873051 | 1 |
| ENSMUSG00000090357 | Gm17077 | 0 | 1.648835551 | 1 |
| ENSMUSG00000097750 | Gm4673 | 0 | -0.52108945 | 1 |
| ENSMUSG00000036578 | Fxyd7 | 0 | 1.063873051 | 1 |
| ENSMUSG00000036570 | Fxyd1 | 0 | 0.648835551 | 1 |
| ENSMUSG00000036560 | Lgi4 | 0 | 0.063873051 | 1 |
| ENSMUSG00000097149 | G630030J09Rik | 0 | 1.063873051 | 1 |
| ENSMUSG00000100985 | Gm10640 | 0 | 1.648835551 | 1 |
| ENSMUSG00000099581 | Scgb1b10 | 0 | 1.648835551 | 1 |

| | | | | |
|---|---|---|---|---|
| ENSMUSG00000085312 | Gm12763 | 0 | 1.648835551 | 1 |
| ENSMUSG00000087127 | Gm12756 | 0 | 1.648835551 | 1 |
| ENSMUSG00000108486 | Gm44836 | 0 | 1.063873051 | 1 |
| ENSMUSG00000108739 | Gm45096 | 0 | 0.648835551 | 1 |
| ENSMUSG00000118454 | AC150683.1 | 0 | -0.810596067 | 1 |
| ENSMUSG00000056043 | Rgs9bp | 0 | 0.648835551 | 1 |
| ENSMUSG00000101965 | Gm29129 | 0 | 1.648835551 | 1 |
| ENSMUSG00000101342 | Gm28078 | 0 | 1.648835551 | 1 |
| ENSMUSG00000100616 | Gm28076 | 0 | 1.063873051 | 1 |
| ENSMUSG00000099715 | Gm28077 | 0 | 0.648835551 | 1 |
| ENSMUSG00000108726 | Gm30684 | 0 | 1.648835551 | 1 |
| ENSMUSG00000108334 | Gm30771 | 0 | 0.063873051 | 1 |
| ENSMUSG00000074170 | Plekhf1 | 0 | 0.648835551 | 1 |
| ENSMUSG00000030423 | Pop4 | 0 | 0.326907457 | 1 |
| ENSMUSG00000100169 | Gm29087 | 0 | 1.648835551 | 1 |
| ENSMUSG00000100431 | 4930558N11Rik | 0 | 1.063873051 | 1 |
| ENSMUSG00000109083 | Gm44992 | 0 | -1.158519371 | 0.890735403 |
| ENSMUSG00000100600 | A230077H06Rik | 0 | -0.351164449 | 1 |
| ENSMUSG00000039257 | Vstm2b | 0 | 1.063873051 | 1 |
| ENSMUSG00000091692 | 4930433I11Rik | 0 | 1.648835551 | 1 |
| ENSMUSG00000108364 | Gm38997 | 0 | 0.648835551 | 1 |
| ENSMUSG00000048312 | Gm4884 | 0 | 1.063873051 | 1 |
| ENSMUSG00000066537 | Vmn2r57 | 0 | -0.158519371 | 1 |
| ENSMUSG00000074166 | AW146154 | 0 | -3.673092543 | 0.00439237 |
| ENSMUSG00000090383 | Vmn2r58 | 0 | 1.063873051 | 1 |
| ENSMUSG00000030463 | 4933421I07Rik | 0 | 1.648835551 | 1 |
| ENSMUSG00000092335 | Zfp977 | 0 | 0.063873051 | 1 |
| ENSMUSG00000091926 | Vmn2r62 | 0 | 1.063873051 | 1 |
| ENSMUSG00000092225 | Gm2381 | 0 | -0.158519371 | 1 |
| ENSMUSG00000107181 | Gm17768 | 0 | 1.063873051 | 1 |
| ENSMUSG00000012640 | Zfp715 | 0 | -2.560617814 | 0.105255472 |
| ENSMUSG00000039013 | Siglecf | 0 | 0.326907457 | 1 |
| ENSMUSG00000118553 | AC149091.1 | 0 | 1.648835551 | 1 |
| ENSMUSG00000118560 | Lim2 | 0 | 1.063873051 | 1 |
| ENSMUSG00000038973 | Cldnd2 | 0 | 1.648835551 | 1 |
| ENSMUSG00000070604 | Vsig10l | 0 | 0.648835551 | 1 |
| ENSMUSG00000013367 | Iglon5 | 0 | 1.648835551 | 1 |
| ENSMUSG00000004609 | Cd33 | 0 | -1.158519371 | 0.886203497 |
| ENSMUSG00000055102 | Zfp819 | 0 | 1.648835551 | 1 |
| ENSMUSG00000030474 | Siglece | 0 | 1.063873051 | 1 |
| ENSMUSG00000108783 | Gm36546 | 0 | 1.063873051 | 1 |
| ENSMUSG00000044737 | Klk14 | 0 | 1.063873051 | 1 |
| ENSMUSG00000054046 | Klk13 | 0 | 1.063873051 | 1 |
| ENSMUSG00000044430 | Klk12 | 0 | 1.063873051 | 1 |
| ENSMUSG00000067616 | Klk11 | 0 | 1.648835551 | 1 |
| ENSMUSG00000064023 | Klk8 | 0 | 0.326907457 | 1 |
| ENSMUSG00000050063 | Klk6 | 0 | 0.648835551 | 1 |
| ENSMUSG00000074155 | Klk5 | 0 | 0.063873051 | 1 |
| ENSMUSG00000006948 | Klk4 | 0 | 1.648835551 | 1 |
| ENSMUSG00000038782 | 1700028J19Rik | 0 | 1.648835551 | 1 |
| ENSMUSG00000004473 | Clec11a | 0 | 1.063873051 | 1 |

| | | | | |
|---|---|---|---|---|
| ENSMUSG00000108322 | 5430431A17Rik | 0 | 0.063873051 | 1 |
| ENSMUSG00000051113 | Fam71e1 | 0 | 1.648835551 | 1 |
| ENSMUSG00000062785 | Kcnc3 | 0 | 0.326907457 | 1 |
| ENSMUSG00000070574 | 2310016G11Rik | 0 | 1.063873051 | 1 |
| ENSMUSG00000011658 | Fuz | 0 | -0.810596067 | 0.844591606 |
| ENSMUSG00000007783 | Cpt1c | 0 | -1.051604167 | 0.874192437 |
| ENSMUSG00000109111 | 6530437J22Rik | 0 | 0.326907457 | 1 |
| ENSMUSG00000110279 | Gm45552 | 0 | 1.648835551 | 1 |
| ENSMUSG00000110206 | Flt3l | 0 | -0.810596067 | 0.844591606 |
| ENSMUSG00000070570 | Slc17a7 | 0 | 0.648835551 | 1 |
| ENSMUSG00000094152 | Slc6a16 | 0 | 1.063873051 | 1 |
| ENSMUSG00000070568 | Slc6a21 | 0 | 1.063873051 | 1 |
| ENSMUSG00000038239 | Hrc | 0 | 1.648835551 | 1 |
| ENSMUSG00000003863 | Ppfia3 | 0 | -1.52108945 | 0.606122355 |
| ENSMUSG00000074121 | Ntf5 | 0 | 0.326907457 | 1 |
| ENSMUSG00000118462 | AC151602.1 | 0 | 1.648835551 | 1 |
| ENSMUSG00000040428 | Plekha4 | 0 | -1.874726405 | 0.353226088 |
| ENSMUSG00000109644 | 0610005C13Rik | 0 | 0.326907457 | 1 |
| ENSMUSG00000008461 | Fut1 | 0 | 1.648835551 | 1 |
| ENSMUSG00000064158 | Izumo1 | 0 | 0.648835551 | 1 |
| ENSMUSG00000109998 | Gm45437 | 0 | 1.063873051 | 1 |
| ENSMUSG00000040364 | Sec1 | 0 | 0.648835551 | 1 |
| ENSMUSG00000070564 | Ntn5 | 0 | 0.648835551 | 1 |
| ENSMUSG00000062044 | Lmtk3 | 0 | -0.351164449 | 1 |
| ENSMUSG00000058743 | Kcnj14 | 0 | 0.648835551 | 1 |
| ENSMUSG00000110020 | Gm45441 | 0 | 1.648835551 | 1 |
| ENSMUSG00000040231 | Syngr4 | 0 | 1.063873051 | 1 |
| ENSMUSG00000110208 | Gm45442 | 0 | 1.648835551 | 1 |
| ENSMUSG00000030838 | Ush1c | 0 | 0.063873051 | 1 |
| ENSMUSG00000087492 | Gm15700 | 0 | 0.326907457 | 1 |
| ENSMUSG00000040017 | Saa4 | 0 | 0.326907457 | 1 |
| ENSMUSG00000110001 | C86187 | 0 | 0.648835551 | 1 |
| ENSMUSG00000109118 | Gm32031 | 0 | -2.051604167 | 0.274319469 |
| ENSMUSG00000010307 | Tmem86a | 0 | 0.648835551 | 1 |
| ENSMUSG00000030854 | Ptpn5 | 0 | -0.52108945 | 1 |
| ENSMUSG00000109501 | Gm14377 | 0 | 1.063873051 | 1 |
| ENSMUSG00000074111 | Mrgpra9 | 0 | 1.063873051 | 1 |
| ENSMUSG00000050650 | Mrgpra1 | 0 | 1.063873051 | 1 |
| ENSMUSG00000093973 | Mrgpra2a | 0 | 1.648835551 | 1 |
| ENSMUSG00000078698 | Mrgpra3 | 0 | 0.648835551 | 1 |
| ENSMUSG00000067173 | Mrgpra4 | 0 | 1.648835551 | 1 |
| ENSMUSG00000070547 | Mrgprb1 | 0 | 1.648835551 | 1 |
| ENSMUSG00000074109 | Mrgprx2 | 0 | 0.648835551 | 1 |
| ENSMUSG00000050425 | Mrgprb2 | 0 | 1.648835551 | 1 |
| ENSMUSG00000109170 | Gm45086 | 0 | 1.063873051 | 1 |
| ENSMUSG00000030471 | Zdhhc13 | 0 | -2.965874293 | 0.044171611 |
| ENSMUSG00000030470 | Csrp3 | 0 | -0.351164449 | 1 |
| ENSMUSG00000046179 | E2f8 | 0 | -0.52108945 | 1 |
| ENSMUSG00000085995 | Gm2788 | 0 | -0.158519371 | 1 |
| ENSMUSG00000110093 | Gm45282 | 0 | 0.648835551 | 1 |
| ENSMUSG00000030507 | Dbx1 | 0 | 1.648835551 | 1 |

| | | | | |
|---|---|---|---|---|
| ENSMUSG00000030500 | Slc17a6 | 0 | -0.351164449 | 1 |
| ENSMUSG00000102124 | Gm29296 | 0 | 1.063873051 | 1 |
| ENSMUSG00000051504 | Siglech | 0 | -0.351164449 | 1 |
| ENSMUSG00000055078 | Gabra5 | 0 | -1.258055044 | 0.930100175 |
| ENSMUSG00000000948 | Gm38393 | 0 | 1.063873051 | 1 |
| ENSMUSG00000033585 | Ndn | 0 | 0.063873051 | 1 |
| ENSMUSG00000095061 | E030018B13Rik | 0 | 1.648835551 | 1 |
| ENSMUSG00000108313 | Gm45052 | 0 | 1.063873051 | 1 |
| ENSMUSG00000108469 | Gm45054 | 0 | 0.326907457 | 1 |
| ENSMUSG00000108620 | Gm20670 | 0 | 1.648835551 | 1 |
| ENSMUSG00000086662 | BC046251 | 0 | 0.326907457 | 1 |
| ENSMUSG00000100094 | 1810008I18Rik | 0 | 1.648835551 | 1 |
| ENSMUSG00000109119 | Gm44752 | 0 | 0.648835551 | 1 |
| ENSMUSG00000109039 | Gm33570 | 0 | 1.648835551 | 1 |
| ENSMUSG00000053091 | Lins1 | 0 | -1.673092543 | 0.451785624 |
| ENSMUSG00000097527 | 1700112J16Rik | 0 | -0.673092543 | 1 |
| ENSMUSG00000109329 | Gm33926 | 0 | -0.52108945 | 1 |
| ENSMUSG00000030554 | Synm | 0 | -1.258055044 | 0.692200558 |
| ENSMUSG00000097697 | 4833412C05Rik | 0 | 1.063873051 | 1 |
| ENSMUSG00000109464 | Gm34350 | 0 | 0.648835551 | 1 |
| ENSMUSG00000109458 | Gm44889 | 0 | 1.648835551 | 1 |
| ENSMUSG00000109524 | Gm44691 | 0 | 1.648835551 | 1 |
| ENSMUSG00000108885 | 4930402F11Rik | 0 | 0.326907457 | 1 |
| ENSMUSG00000109280 | Gm35325 | 0 | 1.648835551 | 1 |
| ENSMUSG00000108869 | Gm44532 | 0 | 1.063873051 | 1 |
| ENSMUSG00000108836 | Gm44813 | 0 | 1.063873051 | 1 |
| ENSMUSG00000109003 | Gm36011 | 0 | 1.063873051 | 1 |
| ENSMUSG00000100664 | 6030442E23Rik | 0 | 1.648835551 | 1 |
| ENSMUSG00000099558 | Gm29295 | 0 | 1.648835551 | 1 |
| ENSMUSG00000108978 | Gm44689 | 0 | 1.648835551 | 1 |
| ENSMUSG00000108427 | Gm36696 | 0 | 1.648835551 | 1 |
| ENSMUSG00000108329 | Gm44725 | 0 | 1.648835551 | 1 |
| ENSMUSG00000108345 | 4933435G04Rik | 0 | 1.648835551 | 1 |
| ENSMUSG00000108513 | Gm30075 | 0 | 1.063873051 | 1 |
| ENSMUSG00000108563 | Gm44686 | 0 | 0.648835551 | 1 |
| ENSMUSG00000108724 | Gm44734 | 0 | 0.648835551 | 1 |
| ENSMUSG00000108311 | Gm44737 | 0 | 0.063873051 | 1 |
| ENSMUSG00000108473 | Gm44739 | 0 | 1.648835551 | 1 |
| ENSMUSG00000074067 | Gm10619 | 0 | -0.351164449 | 1 |
| ENSMUSG00000108448 | Gm21269 | 0 | -1.673092543 | 0.612035282 |
| ENSMUSG00000085161 | Gm7580 | 0 | 1.063873051 | 1 |
| ENSMUSG00000108468 | Gm30459 | 0 | 1.648835551 | 1 |
| ENSMUSG00000108803 | 4930533N22Rik | 0 | 0.063873051 | 1 |
| ENSMUSG00000030610 | Det1 | 0 | -0.673092543 | 1 |
| ENSMUSG00000030606 | Hapln3 | 0 | 0.063873051 | 1 |
| ENSMUSG00000039194 | Rlbp1 | 0 | 1.648835551 | 1 |
| ENSMUSG00000108616 | Gm35040 | 0 | -0.936126949 | 0.847265357 |
| ENSMUSG00000108776 | Gm45169 | 0 | 1.648835551 | 1 |
| ENSMUSG00000030549 | Rhcg | 0 | -0.351164449 | 1 |
| ENSMUSG00000046591 | Ticrr | 0 | -1.743481871 | 0.472764596 |
| ENSMUSG00000050382 | Kif7 | 0 | 0.648835551 | 1 |

| | | | | |
|---|---|---|---|---|
| ENSMUSG00000030545 | Pex11a | 0 | -0.158519371 | 1 |
| ENSMUSG00000030544 | Mesp1 | 0 | 1.648835551 | 1 |
| ENSMUSG00000108714 | Gm21057 | 0 | 1.063873051 | 1 |
| ENSMUSG00000108757 | Gm45202 | 0 | 1.648835551 | 1 |
| ENSMUSG00000030532 | Hddc3 | 0 | -0.158519371 | 1 |
| ENSMUSG00000108436 | Gm44851 | 0 | 1.648835551 | 1 |
| ENSMUSG00000108321 | Gm44649 | 0 | 1.648835551 | 1 |
| ENSMUSG00000025723 | Nmb | 0 | 0.326907457 | 1 |
| ENSMUSG00000085194 | Platr32 | 0 | 1.648835551 | 1 |
| ENSMUSG00000097277 | 2900076A07Rik | 0 | -1.52108945 | 0.544908989 |
| ENSMUSG00000109485 | Gm45016 | 0 | 1.063873051 | 1 |
| ENSMUSG00000108868 | Gm32850 | 0 | 0.648835551 | 1 |
| ENSMUSG00000038570 | Saxo2 | 0 | 1.648835551 | 1 |
| ENSMUSG00000030636 | 1700010L04Rik | 0 | 1.648835551 | 1 |
| ENSMUSG00000109383 | Gm36584 | 0 | 0.648835551 | 1 |
| ENSMUSG00000108861 | 5930435M05Rik | 0 | -0.936126949 | 0.8730922 |
| ENSMUSG00000109506 | Gm44724 | 0 | 1.648835551 | 1 |
| ENSMUSG00000109450 | Gm39043 | 0 | -0.158519371 | 1 |
| ENSMUSG00000052353 | Cemip | 0 | -2.209145444 | 0.217194106 |
| ENSMUSG00000070459 | Olfr290 | 0 | 1.648835551 | 1 |
| ENSMUSG00000095664 | Vmn2r67 | 0 | 1.648835551 | 1 |
| ENSMUSG00000096861 | Vmn2r68 | 0 | 1.648835551 | 1 |
| ENSMUSG00000070458 | Vmn2r73 | 0 | 0.648835551 | 1 |
| ENSMUSG00000090774 | Vmn2r74 | 0 | 1.648835551 | 1 |
| ENSMUSG00000090436 | Vmn2r75 | 0 | 1.063873051 | 1 |
| ENSMUSG00000061549 | Olfr301 | 0 | 1.648835551 | 1 |
| ENSMUSG00000020168 | Olfr299 | 0 | 0.648835551 | 1 |
| ENSMUSG00000063394 | Olfr293 | 0 | 1.063873051 | 1 |
| ENSMUSG00000030559 | Rab38 | 0 | -1.43862729 | 0.538110273 |
| ENSMUSG00000109327 | Gm44995 | 0 | 0.063873051 | 1 |
| ENSMUSG00000039391 | Ccdc81 | 0 | 0.648835551 | 1 |
| ENSMUSG00000044362 | Ccdc89 | 0 | 1.648835551 | 1 |
| ENSMUSG00000098066 | Gm26944 | 0 | 0.648835551 | 1 |
| ENSMUSG00000074024 | 4632427E13Rik | 0 | -3.395558568 | 0.014189547 |
| ENSMUSG00000109315 | Gm45037 | 0 | 1.063873051 | 1 |
| ENSMUSG00000085751 | Gm15412 | 0 | 1.648835551 | 1 |
| ENSMUSG00000085792 | Gm15414 | 0 | 1.648835551 | 1 |
| ENSMUSG00000108635 | Gm44633 | 0 | 1.648835551 | 1 |
| ENSMUSG00000086993 | Rsf1os2 | 0 | -0.158519371 | 1 |
| ENSMUSG00000108449 | Gm44507 | 0 | -0.52108945 | 1 |
| ENSMUSG00000074003 | Gucy2d | 0 | 0.648835551 | 1 |
| ENSMUSG00000090958 | Lrrc32 | 0 | -0.673092543 | 1 |
| ENSMUSG00000055446 | A630091E08Rik | 0 | 0.063873051 | 1 |
| ENSMUSG00000015957 | Wnt11 | 0 | 1.648835551 | 1 |
| ENSMUSG00000109130 | Gm45187 | 0 | 1.648835551 | 1 |
| ENSMUSG00000109052 | Gm45012 | 0 | -0.52108945 | 1 |
| ENSMUSG00000097015 | Gm26705 | 0 | 0.063873051 | 1 |
| ENSMUSG00000109460 | Gm45591 | 0 | 1.648835551 | 1 |
| ENSMUSG00000035298 | Klhl35 | 0 | 1.063873051 | 1 |
| ENSMUSG00000109559 | Gm34280 | 0 | 1.648835551 | 1 |
| ENSMUSG00000073998 | Olfr520 | 0 | 1.648835551 | 1 |

| | | | | |
|---|---|---|---|---|
| ENSMUSG00000053360 | F730035P03Rik | 0 | 1.648835551 | 1 |
| ENSMUSG00000035239 | Neu3 | 0 | 1.648835551 | 1 |
| ENSMUSG00000109002 | Gm38405 | 0 | 0.648835551 | 1 |
| ENSMUSG00000032942 | Ucp3 | 0 | 1.648835551 | 1 |
| ENSMUSG00000108866 | Gm35082 | 0 | 0.326907457 | 1 |
| ENSMUSG00000032860 | P2ry2 | 0 | 0.326907457 | 1 |
| ENSMUSG00000030651 | Art2b | 0 | 1.063873051 | 1 |
| ENSMUSG00000032725 | Folr2 | 0 | 0.648835551 | 1 |
| ENSMUSG00000078630 | Tomt | 0 | 1.648835551 | 1 |
| ENSMUSG00000070427 | Il18bp | 0 | -0.52108945 | 1 |
| ENSMUSG00000070424 | Art5 | 0 | 1.063873051 | 1 |
| ENSMUSG00000044824 | Olfr545 | 0 | -0.351164449 | 1 |
| ENSMUSG00000030966 | Trim21 | 0 | -0.936126949 | 0.847265357 |
| ENSMUSG00000073974 | Olfr551 | 0 | 1.648835551 | 1 |
| ENSMUSG00000073973 | Olfr552 | 0 | 1.063873051 | 1 |
| ENSMUSG00000073972 | Olfr553 | 0 | 1.648835551 | 1 |
| ENSMUSG00000073968 | Trim68 | 0 | -1.673092543 | 0.451785624 |
| ENSMUSG00000070423 | Olfr558 | 0 | 1.063873051 | 1 |
| ENSMUSG00000043366 | Olfr78 | 0 | 1.063873051 | 1 |
| ENSMUSG00000073964 | Olfr570 | 0 | 1.648835551 | 1 |
| ENSMUSG00000073963 | Olfr572 | 0 | 1.648835551 | 1 |
| ENSMUSG00000073962 | Olfr576 | 0 | 1.648835551 | 1 |
| ENSMUSG00000045792 | Olfr578 | 0 | 1.063873051 | 1 |
| ENSMUSG00000073961 | Olfr582 | 0 | 1.063873051 | 1 |
| ENSMUSG00000066268 | Olfr586 | 0 | 1.648835551 | 1 |
| ENSMUSG00000073946 | Olfr617 | 0 | 1.648835551 | 1 |
| ENSMUSG00000045132 | Olfr620 | 0 | 1.648835551 | 1 |
| ENSMUSG00000045780 | Olfr624 | 0 | 1.063873051 | 1 |
| ENSMUSG00000094822 | Olfr243 | 0 | 1.063873051 | 1 |
| ENSMUSG00000061626 | Olfr68 | 0 | 1.648835551 | 1 |
| ENSMUSG00000047535 | Olfr67 | 0 | 1.648835551 | 1 |
| ENSMUSG00000073940 | Hbb-bt | 0 | -1.52108945 | 1 |
| ENSMUSG00000073938 | Olfr632 | 0 | 1.648835551 | 1 |
| ENSMUSG00000073937 | Olfr633 | 0 | 0.326907457 | 1 |
| ENSMUSG00000094063 | Olfr638 | 0 | 1.063873051 | 1 |
| ENSMUSG00000049797 | Olfr642 | 0 | -0.158519371 | 1 |
| ENSMUSG00000051437 | Ubqlnl | 0 | 1.648835551 | 1 |
| ENSMUSG00000044265 | Olfm5 | 0 | 1.648835551 | 1 |
| ENSMUSG00000056144 | Trim34a | 0 | 0.063873051 | 1 |
| ENSMUSG00000066258 | Trim12a | 0 | 1.648835551 | 1 |
| ENSMUSG00000057596 | Trim30d | 0 | 1.063873051 | 1 |
| ENSMUSG00000073928 | Olfr651 | 0 | 1.648835551 | 1 |
| ENSMUSG00000073926 | Olfr653 | 0 | 1.648835551 | 1 |
| ENSMUSG00000051182 | Olfr655 | 0 | 1.063873051 | 1 |
| ENSMUSG00000073924 | Olfr656 | 0 | 0.063873051 | 1 |
| ENSMUSG00000073923 | Olfr657 | 0 | 1.648835551 | 1 |
| ENSMUSG00000073922 | Olfr659 | 0 | 1.648835551 | 1 |
| ENSMUSG00000054568 | Usp17la | 0 | 1.648835551 | 1 |
| ENSMUSG00000073917 | Olfr665 | 0 | 1.648835551 | 1 |
| ENSMUSG00000057770 | Olfr668 | 0 | 1.063873051 | 1 |
| ENSMUSG00000073916 | Olfr669 | 0 | -0.673092543 | 1 |

| | | | | |
|---|---|---|---|---|
| ENSMUSG00000073913 | Olfr678 | 0 | 0.648835551 | 1 |
| ENSMUSG00000096029 | Olfr679 | 0 | 1.648835551 | 1 |
| ENSMUSG00000044120 | Olfr683 | 0 | 1.648835551 | 1 |
| ENSMUSG00000047225 | Olfr684 | 0 | 1.648835551 | 1 |
| ENSMUSG00000048425 | Olfr686 | 0 | 1.648835551 | 1 |
| ENSMUSG00000073907 | Olfr689 | 0 | 1.648835551 | 1 |
| ENSMUSG00000043948 | Olfr691 | 0 | 1.648835551 | 1 |
| ENSMUSG00000030898 | Cckbr | 0 | -0.810596067 | 0.844591606 |
| ENSMUSG00000037032 | Apbb1 | 0 | -0.52108945 | 1 |
| ENSMUSG00000030895 | Hpx | 0 | 1.648835551 | 1 |
| ENSMUSG00000064223 | Olfr694 | 0 | -1.258055044 | 0.936704655 |
| ENSMUSG00000094493 | Olfr700 | 0 | 1.648835551 | 1 |
| ENSMUSG00000069390 | Olfr707 | 0 | 1.063873051 | 1 |
| ENSMUSG00000062553 | Olfr709-ps1 | 0 | 0.326907457 | 1 |
| ENSMUSG00000045013 | Olfr711 | 0 | 1.648835551 | 1 |
| ENSMUSG00000073898 | Olfr713 | 0 | 1.648835551 | 1 |
| ENSMUSG00000110209 | Gm45868 | 0 | 1.648835551 | 1 |
| ENSMUSG00000073896 | Olfr716 | 0 | 0.648835551 | 1 |
| ENSMUSG00000109402 | Gm45040 | 0 | 1.648835551 | 1 |
| ENSMUSG00000073893 | Olfr472 | 0 | 0.648835551 | 1 |
| ENSMUSG00000109884 | Olfr490 | 0 | 1.648835551 | 1 |
| ENSMUSG00000094612 | Olfr491 | 0 | 1.063873051 | 1 |
| ENSMUSG00000058014 | Olfr502 | 0 | 1.648835551 | 1 |
| ENSMUSG00000096209 | Olfr510 | 0 | 1.063873051 | 1 |
| ENSMUSG00000051200 | Olfr513 | 0 | 0.648835551 | 1 |
| ENSMUSG00000062434 | Olfr516 | 0 | 1.648835551 | 1 |
| ENSMUSG00000048330 | Ric3 | 0 | -0.158519371 | 1 |
| ENSMUSG00000109001 | Gm45024 | 0 | 1.063873051 | 1 |
| ENSMUSG00000109499 | Gm44864 | 0 | 1.648835551 | 1 |
| ENSMUSG00000035951 | Ascl3 | 0 | 1.648835551 | 1 |
| ENSMUSG00000007279 | Scube2 | 0 | 0.326907457 | 1 |
| ENSMUSG00000109808 | 5430402P08Rik | 0 | 1.648835551 | 1 |
| ENSMUSG00000085049 | Parvaos | 0 | 1.063873051 | 1 |
| ENSMUSG00000110009 | 4930543E12Rik | 0 | -0.158519371 | 1 |
| ENSMUSG00000059077 | Pth | 0 | 1.648835551 | 1 |
| ENSMUSG00000101755 | Gm29507 | 0 | -0.158519371 | 1 |
| ENSMUSG00000110248 | Gm45615 | 0 | 1.648835551 | 1 |
| ENSMUSG00000030670 | Cyp2r1 | 0 | 1.063873051 | 1 |
| ENSMUSG00000030669 | Calca | 0 | 1.063873051 | 1 |
| ENSMUSG00000030666 | Calcb | 0 | 1.648835551 | 1 |
| ENSMUSG00000108650 | Gm44867 | 0 | 1.648835551 | 1 |
| ENSMUSG00000108738 | Gm44777 | 0 | 0.063873051 | 1 |
| ENSMUSG00000108736 | Gm45151 | 0 | 1.648835551 | 1 |
| ENSMUSG00000109284 | B230311B06Rik | 0 | 1.648835551 | 1 |
| ENSMUSG00000030954 | Gp2 | 0 | 1.648835551 | 1 |
| ENSMUSG00000030963 | Umod | 0 | 0.648835551 | 1 |
| ENSMUSG00000109254 | Gm44588 | 0 | 0.648835551 | 1 |
| ENSMUSG00000030972 | Acsm5 | 0 | 0.648835551 | 1 |
| ENSMUSG00000109478 | Gm39078 | 0 | 1.648835551 | 1 |
| ENSMUSG00000033533 | Acsm1 | 0 | -0.351164449 | 1 |
| ENSMUSG00000109093 | Gm19950 | 0 | -0.158519371 | 1 |

| | | | | |
|---|---|---|---|---|
| ENSMUSG00000109233 | Gm44866 | 0 | 1.063873051 | 1 |
| ENSMUSG00000030929 | Eri2 | 0 | -1.43862729 | 0.538110273 |
| ENSMUSG00000109392 | Gm5737 | 0 | 0.063873051 | 1 |
| ENSMUSG00000085118 | Gm15774 | 0 | 1.648835551 | 1 |
| ENSMUSG00000108627 | Gm45079 | 0 | 0.648835551 | 1 |
| ENSMUSG00000030873 | Scnn1b | 0 | -0.158519371 | 1 |
| ENSMUSG00000108511 | Gm44987 | 0 | -0.158519371 | 1 |
| ENSMUSG00000030757 | Zkscan2 | 0 | 1.063873051 | 1 |
| ENSMUSG00000098129 | Gm27040 | 0 | 0.648835551 | 1 |
| ENSMUSG00000085892 | Gm15339 | 0 | 1.063873051 | 1 |
| ENSMUSG00000108623 | Gm45092 | 0 | 0.326907457 | 1 |
| ENSMUSG00000108417 | Gm30928 | 0 | 1.648835551 | 1 |
| ENSMUSG00000030742 | Lat | 0 | 0.063873051 | 1 |
| ENSMUSG00000030724 | Cd19 | 0 | 0.648835551 | 1 |
| ENSMUSG00000030730 | Atp2a1 | 0 | 0.063873051 | 1 |
| ENSMUSG00000030714 | Sgf29 | 0 | -2.52108945 | 0.139351384 |
| ENSMUSG00000030711 | Sult1a1 | 0 | -0.52108945 | 1 |
| ENSMUSG00000108583 | Gm44854 | 0 | 1.063873051 | 1 |
| ENSMUSG00000042606 | Hirip3 | 0 | -0.810596067 | 0.844591606 |
| ENSMUSG00000046378 | Asphd1 | 0 | 1.648835551 | 1 |
| ENSMUSG00000097075 | Cdiptos | 0 | 0.063873051 | 1 |
| ENSMUSG00000030680 | Pagr1a | 0 | -0.158519371 | 1 |
| ENSMUSG00000107068 | Gm42742 | 0 | 1.648835551 | 1 |
| ENSMUSG00000045114 | Prrt2 | 0 | 1.648835551 | 1 |
| ENSMUSG00000108617 | Gm31749 | 0 | 1.648835551 | 1 |
| ENSMUSG00000108327 | Gm44939 | 0 | 1.648835551 | 1 |
| ENSMUSG00000030674 | Qprt | 0 | -0.673092543 | 1 |
| ENSMUSG00000051457 | Spn | 0 | 1.063873051 | 1 |
| ENSMUSG00000030672 | Mylpf | 0 | -0.158519371 | 1 |
| ENSMUSG00000108760 | Gm31897 | 0 | 1.063873051 | 1 |
| ENSMUSG00000030823 | 9130019O22Rik | 0 | -0.936126949 | 0.847265357 |
| ENSMUSG00000108483 | Gm45184 | 0 | 1.063873051 | 1 |
| ENSMUSG00000057176 | Ccdc189 | 0 | 1.063873051 | 1 |
| ENSMUSG00000108677 | Gm44759 | 0 | 1.063873051 | 1 |
| ENSMUSG00000108801 | Gm39090 | 0 | -1.52108945 | 0.61177833 |
| ENSMUSG00000030793 | Pycard | 0 | -1.673092543 | 0.458610057 |
| ENSMUSG00000042828 | Trim72 | 0 | 1.063873051 | 1 |
| ENSMUSG00000108604 | Gm44850 | 0 | 1.648835551 | 1 |
| ENSMUSG00000062944 | 9130023H24Rik | 0 | 0.648835551 | 1 |
| ENSMUSG00000084964 | Gm15503 | 0 | 1.648835551 | 1 |
| ENSMUSG00000108626 | Gm40457 | 0 | 1.648835551 | 1 |
| ENSMUSG00000108720 | Gm44672 | 0 | 1.648835551 | 1 |
| ENSMUSG00000108713 | Gm33027 | 0 | 1.648835551 | 1 |
| ENSMUSG00000108576 | Gm44778 | 0 | 0.648835551 | 1 |
| ENSMUSG00000109016 | Gm44647 | 0 | 1.648835551 | 1 |
| ENSMUSG00000050189 | Etos1 | 0 | 0.648835551 | 1 |
| ENSMUSG00000013668 | 4933402N03Rik | 0 | 1.648835551 | 1 |
| ENSMUSG00000040177 | 2310057M21Rik | 0 | -1.351164449 | 0.687112677 |
| ENSMUSG00000050100 | Hmx2 | 0 | 1.648835551 | 1 |
| ENSMUSG00000040125 | Gpr26 | 0 | 1.648835551 | 1 |
| ENSMUSG00000030862 | Cpxm2 | 0 | 1.063873051 | 1 |

| | | | | |
|---|---|---|---|---|
| ENSMUSG00000030930 | Chst15 | 0 | -0.351164449 | 1 |
| ENSMUSG00000109270 | Gm10584 | 0 | 1.648835551 | 1 |
| ENSMUSG00000109842 | Gm45502 | 0 | 0.063873051 | 1 |
| ENSMUSG00000030976 | Tex36 | 0 | 1.648835551 | 1 |
| ENSMUSG00000109635 | Gm45670 | 0 | 1.648835551 | 1 |
| ENSMUSG00000100717 | 1700120G07Rik | 0 | 1.648835551 | 1 |
| ENSMUSG00000097471 | 5830432E09Rik | 0 | 1.648835551 | 1 |
| ENSMUSG00000109633 | C230079O03Rik | 0 | 1.648835551 | 1 |
| ENSMUSG00000010476 | Ebf3 | 0 | -1.258055044 | 0.673811773 |
| ENSMUSG00000109973 | Gm45397 | 0 | 0.326907457 | 1 |
| ENSMUSG00000110090 | Gm45682 | 0 | 1.063873051 | 1 |
| ENSMUSG00000056856 | Jakmip3 | 0 | 0.063873051 | 1 |
| ENSMUSG00000015981 | Stk32c | 0 | -1.673092543 | 0.642927607 |
| ENSMUSG00000049571 | Cfap46 | 0 | -0.351164449 | 1 |
| ENSMUSG00000073795 | 6430531B16Rik | 0 | 0.326907457 | 1 |
| ENSMUSG00000025468 | Caly | 0 | -0.810596067 | 1 |
| ENSMUSG00000039018 | Mtg1 | 0 | 0.648835551 | 1 |
| ENSMUSG00000050366 | Olfr524 | 0 | 1.648835551 | 1 |
| ENSMUSG00000025461 | Cd163l1 | 0 | -0.936126949 | 1 |
| ENSMUSG00000061489 | Olfr525 | 0 | 1.648835551 | 1 |
| ENSMUSG00000062782 | Olfr527 | 0 | 1.648835551 | 1 |
| ENSMUSG00000060974 | Olfr530 | 0 | 1.648835551 | 1 |
| ENSMUSG00000063823 | Olfr532 | 0 | 1.648835551 | 1 |
| ENSMUSG00000063230 | Olfr535 | 0 | 0.648835551 | 1 |
| ENSMUSG00000096069 | Olfr61 | 0 | 1.648835551 | 1 |
| ENSMUSG00000057997 | Olfr541 | 0 | -0.936126949 | 0.847265357 |
| ENSMUSG00000025480 | Syce1 | 0 | 1.063873051 | 1 |
| ENSMUSG00000060314 | Zfp941 | 0 | -0.810596067 | 1 |
| ENSMUSG00000025482 | Odf3 | 0 | 1.648835551 | 1 |
| ENSMUSG00000109628 | BC024386 | 0 | 1.648835551 | 1 |
| ENSMUSG00000062031 | Pgghg | 0 | -2.305360759 | 0.231596497 |
| ENSMUSG00000025489 | Ifitm5 | 0 | 1.648835551 | 1 |
| ENSMUSG00000025491 | Ifitm1 | 0 | -0.52108945 | 1 |
| ENSMUSG00000025500 | Lmntd2 | 0 | -0.158519371 | 1 |
| ENSMUSG00000025496 | Drd4 | 0 | 1.648835551 | 1 |
| ENSMUSG00000019082 | Slc25a22 | 0 | 0.648835551 | 1 |
| ENSMUSG00000025507 | Pidd1 | 0 | -0.351164449 | 1 |
| ENSMUSG00000073787 | Gm10575 | 0 | -0.673092543 | 1 |
| ENSMUSG00000025515 | Muc2 | 0 | -0.52108945 | 1 |
| ENSMUSG00000066108 | Muc5b | 0 | 1.648835551 | 1 |
| ENSMUSG00000092536 | Gm20501 | 0 | 0.648835551 | 1 |
| ENSMUSG00000053046 | Brsk2 | 0 | 1.648835551 | 1 |
| ENSMUSG00000037887 | Dusp8 | 0 | -0.351164449 | 1 |
| ENSMUSG00000109958 | Gm32786 | 0 | 0.326907457 | 1 |
| ENSMUSG00000066101 | Gm10153 | 0 | 1.648835551 | 1 |
| ENSMUSG00000061723 | Tnnt3 | 0 | 0.063873051 | 1 |
| ENSMUSG00000000031 | H19 | 0 | 1.063873051 | 1 |
| ENSMUSG00000048583 | Igf2 | 0 | 0.648835551 | 1 |
| ENSMUSG00000000215 | Ins2 | 0 | -0.351164449 | 1 |
| ENSMUSG00000108952 | Gm39117 | 0 | 1.063873051 | 1 |
| ENSMUSG00000000214 | Th | 0 | 1.063873051 | 1 |

| | | | | |
|---|---|---|---|---|
| ENSMUSG00000009248 | Ascl2 | 0 | 1.648835551 | 1 |
| ENSMUSG00000052281 | Tspan32os | 0 | 1.648835551 | 1 |
| ENSMUSG00000059277 | R74862 | 0 | -0.351164449 | 1 |
| ENSMUSG00000045752 | Tssc4 | 0 | -1.599091962 | 0.559918909 |
| ENSMUSG00000037664 | Cdkn1c | 0 | -0.52108945 | 1 |
| ENSMUSG00000010760 | Phlda2 | 0 | 1.648835551 | 1 |
| ENSMUSG00000086828 | Gm15579 | 0 | 1.648835551 | 1 |
| ENSMUSG00000031090 | Nadsyn1 | 0 | -0.810596067 | 0.844591606 |
| ENSMUSG00000058454 | Dhcr7 | 0 | 0.063873051 | 1 |
| ENSMUSG00000108851 | Gm44930 | 0 | 1.648835551 | 1 |
| ENSMUSG00000050917 | Fgf4 | 0 | 1.648835551 | 1 |
| ENSMUSG00000031073 | Fgf15 | 0 | 1.063873051 | 1 |
| ENSMUSG00000069633 | Pex11g | 0 | -0.52108945 | 1 |
| ENSMUSG00000040340 | Tex45 | 0 | 0.063873051 | 1 |
| ENSMUSG00000108888 | Gm45231 | 0 | 1.063873051 | 1 |
| ENSMUSG00000074491 | Clec4g | 0 | 1.648835551 | 1 |
| ENSMUSG00000040197 | Cd209e | 0 | 1.648835551 | 1 |
| ENSMUSG00000040165 | Cd209c | 0 | 1.063873051 | 1 |
| ENSMUSG00000064125 | Prr36 | 0 | -0.52108945 | 1 |
| ENSMUSG00000097308 | Gm6410 | 0 | -0.351164449 | 1 |
| ENSMUSG00000109313 | Gm44693 | 0 | 1.063873051 | 1 |
| ENSMUSG00000109566 | Gm31401 | 0 | 1.063873051 | 1 |
| ENSMUSG00000109515 | Gm31343 | 0 | 1.648835551 | 1 |
| ENSMUSG00000108932 | Gm31463 | 0 | 1.648835551 | 1 |
| ENSMUSG00000040467 | 4921522P10Rik | 0 | 1.648835551 | 1 |
| ENSMUSG00000108848 | 4933430N04Rik | 0 | 1.063873051 | 1 |
| ENSMUSG00000109064 | Gm39128 | 0 | 1.648835551 | 1 |
| ENSMUSG00000109183 | Gm44761 | 0 | 1.648835551 | 1 |
| ENSMUSG00000095279 | Gm10067 | 0 | 0.648835551 | 1 |
| ENSMUSG00000049717 | Lig4 | 0 | -0.52108945 | 1 |
| ENSMUSG00000109258 | 5330413D20Rik | 0 | 1.648835551 | 1 |
| ENSMUSG00000109067 | Gm44784 | 0 | 1.063873051 | 1 |
| ENSMUSG00000109271 | Gm39129 | 0 | 1.063873051 | 1 |
| ENSMUSG00000109473 | B930025P03Rik | 0 | 1.648835551 | 1 |
| ENSMUSG00000108969 | Gm45044 | 0 | 1.648835551 | 1 |
| ENSMUSG00000109291 | Gm2814 | 0 | 1.648835551 | 1 |
| ENSMUSG00000109089 | 4833411C07Rik | 0 | 0.063873051 | 1 |
| ENSMUSG00000038917 | 3930402G23Rik | 0 | 0.063873051 | 1 |
| ENSMUSG00000109084 | Gm44955 | 0 | 0.648835551 | 1 |
| ENSMUSG00000109508 | Gm44956 | 0 | -0.158519371 | 1 |
| ENSMUSG00000108847 | B830042I05Rik | 0 | 1.063873051 | 1 |
| ENSMUSG00000084783 | Gm15419 | 0 | 1.648835551 | 1 |
| ENSMUSG00000096957 | E230013L22Rik | 0 | 1.648835551 | 1 |
| ENSMUSG00000031510 | 1700128E19Rik | 0 | -0.351164449 | 1 |
| ENSMUSG00000109772 | Gm39132 | 0 | 1.648835551 | 1 |
| ENSMUSG00000110002 | Gm45680 | 0 | 1.063873051 | 1 |
| ENSMUSG00000109995 | B020031H02Rik | 0 | -0.158519371 | 1 |
| ENSMUSG00000109921 | Gm33326 | 0 | 1.648835551 | 1 |
| ENSMUSG00000097421 | D630011A20Rik | 0 | -1.158519371 | 0.667778757 |
| ENSMUSG00000085465 | Gm15347 | 0 | 1.648835551 | 1 |
| ENSMUSG00000079146 | Gm15353 | 0 | 0.063873051 | 1 |

| | | | | |
|---|---|---|---|---|
| ENSMUSG00000031445 | Proz | 0 | 1.648835551 | 1 |
| ENSMUSG00000031448 | Adprhl1 | 0 | 1.648835551 | 1 |
| ENSMUSG00000031449 | Atp4b | 0 | 1.063873051 | 1 |
| ENSMUSG00000038457 | Tmem255b | 0 | 1.648835551 | 1 |
| ENSMUSG00000031452 | 1700029H14Rik | 0 | 1.648835551 | 1 |
| ENSMUSG00000057116 | AF366264 | 0 | 1.648835551 | 1 |
| ENSMUSG00000031458 | Coprs | 0 | 0.648835551 | 1 |
| ENSMUSG00000050052 | Tdrp | 0 | -0.810596067 | 0.844591606 |
| ENSMUSG00000049008 | BB014433 | 0 | 0.648835551 | 1 |
| ENSMUSG00000109886 | Gm45271 | 0 | 1.648835551 | 1 |
| ENSMUSG00000109748 | Gm35392 | 0 | 1.063873051 | 1 |
| ENSMUSG00000039814 | Xkr5 | 0 | 1.648835551 | 1 |
| ENSMUSG00000053678 | Defb40 | 0 | 1.063873051 | 1 |
| ENSMUSG00000053695 | Defb37 | 0 | 0.648835551 | 1 |
| ENSMUSG00000050756 | Defb6 | 0 | 0.326907457 | 1 |
| ENSMUSG00000071169 | Defb46 | 0 | 1.648835551 | 1 |
| ENSMUSG00000039785 | Defb5 | 0 | 1.648835551 | 1 |
| ENSMUSG00000095294 | Gm21119 | 0 | 1.648835551 | 1 |
| ENSMUSG00000061958 | Gm14851 | 0 | 1.063873051 | 1 |
| ENSMUSG00000074437 | Defa29 | 0 | 1.648835551 | 1 |
| ENSMUSG00000064213 | Defa24 | 0 | 1.648835551 | 1 |
| ENSMUSG00000044748 | Defb1 | 0 | -0.673092543 | 1 |
| ENSMUSG00000044222 | Defb13 | 0 | 0.648835551 | 1 |
| ENSMUSG00000079112 | Fam90a1a | 0 | 1.648835551 | 1 |
| ENSMUSG00000017049 | Ccdc70 | 0 | 1.648835551 | 1 |
| ENSMUSG00000006567 | Atp7b | 0 | -3.079084903 | 0.038177619 |
| ENSMUSG00000031480 | Thsd1 | 0 | -2.708716453 | 0.087122211 |
| ENSMUSG00000099825 | 1810012K16Rik | 0 | 1.648835551 | 1 |
| ENSMUSG00000086741 | Gm15816 | 0 | -0.158519371 | 1 |
| ENSMUSG00000063672 | Nkx6-3 | 0 | 0.326907457 | 1 |
| ENSMUSG00000109419 | Gm45163 | 0 | 1.648835551 | 1 |
| ENSMUSG00000031551 | Ido1 | 0 | 0.326907457 | 1 |
| ENSMUSG00000110140 | 5430421F17Rik | 0 | 1.648835551 | 1 |
| ENSMUSG00000110017 | Gm45237 | 0 | -0.351164449 | 1 |
| ENSMUSG00000110399 | Gm45238 | 0 | 0.648835551 | 1 |
| ENSMUSG00000086134 | Gm16159 | 0 | 1.648835551 | 1 |
| ENSMUSG00000037251 | Pomk | 0 | -1.995020638 | 0.370945697 |
| ENSMUSG00000085437 | 1700047A11Rik | 0 | 1.648835551 | 1 |
| ENSMUSG00000110317 | Gm45250 | 0 | 0.063873051 | 1 |
| ENSMUSG00000110229 | Gm31784 | 0 | -1.258055044 | 0.673811773 |
| ENSMUSG00000109996 | Gm31898 | 0 | 1.063873051 | 1 |
| ENSMUSG00000110222 | Gm39149 | 0 | 0.648835551 | 1 |
| ENSMUSG00000109680 | Gm32098 | 0 | 0.648835551 | 1 |
| ENSMUSG00000109689 | Gm45646 | 0 | 1.063873051 | 1 |
| ENSMUSG00000109603 | Gm32389 | 0 | 1.648835551 | 1 |
| ENSMUSG00000091154 | Proscos | 0 | 1.648835551 | 1 |
| ENSMUSG00000031487 | Brf2 | 0 | 0.326907457 | 1 |
| ENSMUSG00000097262 | 4933416M07Rik | 0 | 1.648835551 | 1 |
| ENSMUSG00000109589 | Gm45267 | 0 | 1.648835551 | 1 |
| ENSMUSG00000039720 | Got1l1 | 0 | 0.063873051 | 1 |
| ENSMUSG00000031489 | Adrb3 | 0 | 1.063873051 | 1 |

| | | | | |
|---|---|---|---|---|
| ENSMUSG00000109674 | Gm45470 | 0 | 1.648835551 | 1 |
| ENSMUSG00000071138 | Tex24 | 0 | 1.648835551 | 1 |
| ENSMUSG00000031492 | Chrnb3 | 0 | -0.52108945 | 1 |
| ENSMUSG00000074404 | Gm1698 | 0 | 0.648835551 | 1 |
| ENSMUSG00000110322 | Gm8110 | 0 | 1.648835551 | 1 |
| ENSMUSG00000110180 | Gm32567 | 0 | 0.648835551 | 1 |
| ENSMUSG00000110006 | Gm45490 | 0 | 1.063873051 | 1 |
| ENSMUSG00000031577 | Tti2 | 0 | -1.743481871 | 0.455131546 |
| ENSMUSG00000110329 | Gm45304 | 0 | 1.063873051 | 1 |
| ENSMUSG00000097442 | Gm26632 | 0 | -1.158519371 | 0.667778757 |
| ENSMUSG00000098166 | Gm26978 | 0 | 1.648835551 | 1 |
| ENSMUSG00000110663 | Gm39157 | 0 | -0.351164449 | 1 |
| ENSMUSG00000054618 | Gm9951 | 0 | 1.063873051 | 1 |
| ENSMUSG00000071113 | Mboat4 | 0 | 0.648835551 | 1 |
| ENSMUSG00000100605 | Gm29243 | 0 | 1.063873051 | 1 |
| ENSMUSG00000109819 | B930018H19Rik | 0 | -0.673092543 | 1 |
| ENSMUSG00000109717 | Gm34368 | 0 | 1.648835551 | 1 |
| ENSMUSG00000110367 | Gm34853 | 0 | 1.648835551 | 1 |
| ENSMUSG00000109714 | Gm35021 | 0 | 0.648835551 | 1 |
| ENSMUSG00000055976 | Cldn23 | 0 | 1.648835551 | 1 |
| ENSMUSG00000110321 | Gm39164 | 0 | 1.648835551 | 1 |
| ENSMUSG00000110242 | G630064G18Rik | 0 | 0.648835551 | 1 |
| ENSMUSG00000110312 | Gm45368 | 0 | 1.063873051 | 1 |
| ENSMUSG00000110236 | Gm40493 | 0 | 1.063873051 | 1 |
| ENSMUSG00000046723 | Adam24 | 0 | 0.648835551 | 1 |
| ENSMUSG00000071937 | Adam25 | 0 | 1.648835551 | 1 |
| ENSMUSG00000031594 | Fgl1 | 0 | -0.936126949 | 0.886203497 |
| ENSMUSG00000085445 | Gm16348 | 0 | 1.063873051 | 1 |
| ENSMUSG00000031651 | Triml1 | 0 | 0.648835551 | 1 |
| ENSMUSG00000091490 | Triml2 | 0 | 1.648835551 | 1 |
| ENSMUSG00000109993 | Gm45659 | 0 | 1.648835551 | 1 |
| ENSMUSG00000097491 | Gm2366 | 0 | 0.648835551 | 1 |
| ENSMUSG00000066158 | AY512931 | 0 | -0.158519371 | 1 |
| ENSMUSG00000031645 | F11 | 0 | 1.648835551 | 1 |
| ENSMUSG00000031636 | Pdlim3 | 0 | 1.648835551 | 1 |
| ENSMUSG00000086279 | Gm15634 | 0 | 1.648835551 | 1 |
| ENSMUSG00000109612 | Gm45253 | 0 | 1.648835551 | 1 |
| ENSMUSG00000031629 | Cenpu | 0 | -1.52108945 | 0.590972065 |
| ENSMUSG00000109888 | Gm45600 | 0 | 1.063873051 | 1 |
| ENSMUSG00000086190 | 4930579M01Rik | 0 | 1.648835551 | 1 |
| ENSMUSG00000097706 | E030037K01Rik | 0 | 1.648835551 | 1 |
| ENSMUSG00000110665 | Gm31786 | 0 | 1.648835551 | 1 |
| ENSMUSG00000110624 | Gm32052 | 0 | 0.648835551 | 1 |
| ENSMUSG00000110281 | Gm2516 | 0 | 0.063873051 | 1 |
| ENSMUSG00000089708 | 5330439A09Rik | 0 | -0.351164449 | 1 |
| ENSMUSG00000031518 | Spata4 | 0 | 1.648835551 | 1 |
| ENSMUSG00000046258 | Adam29 | 0 | 0.648835551 | 1 |
| ENSMUSG00000109697 | Gm34030 | 0 | 1.063873051 | 1 |
| ENSMUSG00000097910 | 5033428I22Rik | 0 | -0.351164449 | 1 |
| ENSMUSG00000031610 | Scrg1 | 0 | 0.326907457 | 1 |
| ENSMUSG00000110419 | AW046200 | 0 | -0.810596067 | 0.844591606 |

| | | | | |
|---|---|---|---|---|
| ENSMUSG00000109930 | Gm34730 | 0 | 1.648835551 | 1 |
| ENSMUSG00000045746 | B230317F23Rik | 0 | 1.063873051 | 1 |
| ENSMUSG00000086866 | 4930512H18Rik | 0 | -0.158519371 | 1 |
| ENSMUSG00000109767 | Gm45460 | 0 | 1.063873051 | 1 |
| ENSMUSG00000110364 | Gm45462 | 0 | 1.648835551 | 1 |
| ENSMUSG00000109825 | Gm45589 | 0 | -0.351164449 | 1 |
| ENSMUSG00000094443 | Sgo2b | 0 | -0.351164449 | 1 |
| ENSMUSG00000109579 | Gm35521 | 0 | 1.648835551 | 1 |
| ENSMUSG00000110157 | Gm32507 | 0 | 0.648835551 | 1 |
| ENSMUSG00000053490 | Trim60 | 0 | 1.648835551 | 1 |
| ENSMUSG00000086156 | Gm15354 | 0 | 1.063873051 | 1 |
| ENSMUSG00000085109 | Gm15356 | 0 | 1.648835551 | 1 |
| ENSMUSG00000025519 | Tktl2 | 0 | 1.648835551 | 1 |
| ENSMUSG00000044014 | Npy5r | 0 | 0.326907457 | 1 |
| ENSMUSG00000036437 | Npy1r | 0 | 1.648835551 | 1 |
| ENSMUSG00000110454 | Gm45911 | 0 | 1.648835551 | 1 |
| ENSMUSG00000074292 | Gm10660 | 0 | 1.063873051 | 1 |
| ENSMUSG00000025588 | Nat1 | 0 | 1.648835551 | 1 |
| ENSMUSG00000110594 | Gm33103 | 0 | -0.351164449 | 1 |
| ENSMUSG00000089995 | Gm15716 | 0 | -0.936126949 | 0.847265357 |
| ENSMUSG00000106824 | Gm7697 | 0 | 1.648835551 | 1 |
| ENSMUSG00000060427 | Zfp868 | 0 | -2.560617814 | 0.105255472 |
| ENSMUSG00000091764 | Zfp964 | 0 | -0.673092543 | 1 |
| ENSMUSG00000031861 | Lpar2 | 0 | -0.936126949 | 0.847265357 |
| ENSMUSG00000044006 | Cilp2 | 0 | 1.648835551 | 1 |
| ENSMUSG00000047654 | Tssk6 | 0 | 1.648835551 | 1 |
| ENSMUSG00000087408 | Cers1 | 0 | 1.648835551 | 1 |
| ENSMUSG00000031849 | Comp | 0 | -0.810596067 | 0.844591606 |
| ENSMUSG00000035964 | Tmem59l | 0 | 1.648835551 | 1 |
| ENSMUSG00000031840 | Rab3a | 0 | -1.351164449 | 0.687112677 |
| ENSMUSG00000100691 | 2010320M18Rik | 0 | -0.52108945 | 1 |
| ENSMUSG00000000791 | Il12rb1 | 0 | -0.351164449 | 1 |
| ENSMUSG00000000792 | Slc5a5 | 0 | 1.648835551 | 1 |
| ENSMUSG00000035439 | Haus8 | 0 | -2.480447466 | 0.163587195 |
| ENSMUSG00000043243 | Fam129c | 0 | -0.351164449 | 1 |
| ENSMUSG00000056019 | Zfp709 | 0 | -0.936126949 | 0.847265357 |
| ENSMUSG00000069998 | Olfr372 | 0 | 0.648835551 | 1 |
| ENSMUSG00000046881 | Olfr374 | 0 | -1.52108945 | 0.618397837 |
| ENSMUSG00000074240 | Cib3 | 0 | -0.52108945 | 1 |
| ENSMUSG00000070713 | Gm10282 | 0 | 0.648835551 | 1 |
| ENSMUSG00000110640 | Gm45903 | 0 | 1.648835551 | 1 |
| ENSMUSG00000034472 | Rasd2 | 0 | 1.063873051 | 1 |
| ENSMUSG00000110258 | Gm45406 | 0 | 1.063873051 | 1 |
| ENSMUSG00000109732 | Gm45285 | 0 | 1.063873051 | 1 |
| ENSMUSG00000097882 | 0610038B21Rik | 0 | -0.351164449 | 1 |
| ENSMUSG00000110388 | Gm30329 | 0 | 1.648835551 | 1 |
| ENSMUSG00000037022 | Mmaa | 0 | -2.810596067 | 0.070328284 |
| ENSMUSG00000109774 | Gm45602 | 0 | 0.326907457 | 1 |
| ENSMUSG00000110202 | Gm45435 | 0 | 1.648835551 | 1 |
| ENSMUSG00000109593 | Gm45431 | 0 | 1.063873051 | 1 |
| ENSMUSG00000109587 | Gm31105 | 0 | 0.648835551 | 1 |

| | | | | |
|---|---|---|---|---|
| ENSMUSG00000098001 | Gm27048 | 0 | 1.648835551 | 1 |
| ENSMUSG00000110045 | Gm38590 | 0 | 1.648835551 | 1 |
| ENSMUSG00000110471 | Gm45823 | 0 | 1.648835551 | 1 |
| ENSMUSG00000019464 | Ptger1 | 0 | 1.648835551 | 1 |
| ENSMUSG00000074219 | Gm10644 | 0 | -1.051604167 | 0.886203497 |
| ENSMUSG00000047986 | Palm3 | 0 | -0.158519371 | 1 |
| ENSMUSG00000005465 | Il27ra | 0 | 0.648835551 | 1 |
| ENSMUSG00000074215 | Gm10643 | 0 | 1.063873051 | 1 |
| ENSMUSG00000056753 | C330011M18Rik | 0 | 1.063873051 | 1 |
| ENSMUSG00000048617 | Rtbdn | 0 | 1.063873051 | 1 |
| ENSMUSG00000052819 | Best2 | 0 | 0.648835551 | 1 |
| ENSMUSG00000062687 | Cks1brt | 0 | 1.648835551 | 1 |
| ENSMUSG00000109731 | Gm45425 | 0 | 1.648835551 | 1 |
| ENSMUSG00000098439 | Hm629797 | 0 | 1.648835551 | 1 |
| ENSMUSG00000098434 | 2010110E17Rik | 0 | 1.648835551 | 1 |
| ENSMUSG00000098534 | Gm27167 | 0 | 0.648835551 | 1 |
| ENSMUSG00000098684 | Gm27246 | 0 | -0.810596067 | 1 |
| ENSMUSG00000099207 | Gm10637 | 0 | -0.52108945 | 1 |
| ENSMUSG00000098295 | Gm19872 | 0 | 1.648835551 | 1 |
| ENSMUSG00000097992 | Gm27016 | 0 | 1.648835551 | 1 |
| ENSMUSG00000062704 | 9430002A10Rik | 0 | 1.063873051 | 1 |
| ENSMUSG00000031659 | Adcy7 | 0 | -2.560617814 | 0.122576346 |
| ENSMUSG00000055994 | Nod2 | 0 | -0.351164449 | 1 |
| ENSMUSG00000109870 | Gm35850 | 0 | 0.063873051 | 1 |
| ENSMUSG00000110166 | Gm45641 | 0 | 1.648835551 | 1 |
| ENSMUSG00000109727 | Gm45464 | 0 | 1.648835551 | 1 |
| ENSMUSG00000090778 | Gm3235 | 0 | 1.063873051 | 1 |
| ENSMUSG00000110216 | Gm36325 | 0 | 1.063873051 | 1 |
| ENSMUSG00000110304 | Gm36380 | 0 | 1.648835551 | 1 |
| ENSMUSG00000099812 | Gm28515 | 0 | 1.063873051 | 1 |
| ENSMUSG00000110274 | Gm36843 | 0 | 1.648835551 | 1 |
| ENSMUSG00000110198 | Gm45335 | 0 | -1.810596067 | 0.47966612 |
| ENSMUSG00000031738 | Irx6 | 0 | 1.063873051 | 1 |
| ENSMUSG00000033192 | Lpcat2 | 0 | -1.810596067 | 0.503644124 |
| ENSMUSG00000109651 | Gm45463 | 0 | 0.326907457 | 1 |
| ENSMUSG00000071047 | Ces1a | 0 | -0.158519371 | 1 |
| ENSMUSG00000057400 | Ces1c | 0 | 0.648835551 | 1 |
| ENSMUSG00000056973 | Ces1d | 0 | -0.673092543 | 1 |
| ENSMUSG00000057074 | Ces1g | 0 | 0.326907457 | 1 |
| ENSMUSG00000110713 | Gm30606 | 0 | 1.063873051 | 1 |
| ENSMUSG00000097520 | 4930488L21Rik | 0 | -0.158519371 | 1 |
| ENSMUSG00000031760 | Mt3 | 0 | 1.648835551 | 1 |
| ENSMUSG00000110710 | C78859 | 0 | 0.063873051 | 1 |
| ENSMUSG00000031766 | Slc12a3 | 0 | 0.326907457 | 1 |
| ENSMUSG00000089712 | Gm15889 | 0 | -0.158519371 | 1 |
| ENSMUSG00000090105 | Gm15890 | 0 | 1.648835551 | 1 |
| ENSMUSG00000097194 | 9330175E14Rik | 0 | 1.063873051 | 1 |
| ENSMUSG00000110702 | Gm45767 | 0 | 0.326907457 | 1 |
| ENSMUSG00000031779 | Ccl22 | 0 | 1.063873051 | 1 |
| ENSMUSG00000031780 | Ccl17 | 0 | 1.648835551 | 1 |
| ENSMUSG00000061577 | Adgrg5 | 0 | 0.063873051 | 1 |

| | | | | |
|---|---|---|---|---|
| ENSMUSG00000031786 | Drc7 | 0 | 0.063873051 | 1 |
| ENSMUSG00000110428 | Gm45812 | 0 | 1.648835551 | 1 |
| ENSMUSG00000110475 | Gm31036 | 0 | 0.648835551 | 1 |
| ENSMUSG00000090206 | Tepp | 0 | 1.648835551 | 1 |
| ENSMUSG00000110427 | 4933406B17Rik | 0 | 1.063873051 | 1 |
| ENSMUSG00000036598 | Ccdc113 | 0 | 0.063873051 | 1 |
| ENSMUSG00000048400 | Prss54 | 0 | -0.158519371 | 1 |
| ENSMUSG00000074136 | 4930513N10Rik | 0 | -1.051604167 | 0.867271506 |
| ENSMUSG00000110661 | Gm31805 | 0 | -1.52108945 | 0.544908989 |
| ENSMUSG00000110717 | 1700047G07Rik | 0 | 1.648835551 | 1 |
| ENSMUSG00000110567 | Gm45703 | 0 | 1.648835551 | 1 |
| ENSMUSG00000110340 | 1600027J07Rik | 0 | 1.648835551 | 1 |
| ENSMUSG00000074128 | Gm10631 | 0 | 1.648835551 | 1 |
| ENSMUSG00000074127 | Cmtm2a | 0 | -0.158519371 | 1 |
| ENSMUSG00000110630 | K230015D01Rik | 0 | 0.648835551 | 1 |
| ENSMUSG00000031883 | Car7 | 0 | 1.063873051 | 1 |
| ENSMUSG00000048371 | Pdp2 | 0 | -0.158519371 | 1 |
| ENSMUSG00000031881 | Cdh16 | 0 | 1.648835551 | 1 |
| ENSMUSG00000031880 | Rrad | 0 | 1.063873051 | 1 |
| ENSMUSG00000069925 | 4932416K20Rik | 0 | 1.063873051 | 1 |
| ENSMUSG00000031886 | Ces2e | 0 | 1.648835551 | 1 |
| ENSMUSG00000062826 | Ces2f | 0 | 1.648835551 | 1 |
| ENSMUSG00000091813 | Ces2h | 0 | 1.648835551 | 1 |
| ENSMUSG00000069920 | B3gnt9 | 0 | 0.648835551 | 1 |
| ENSMUSG00000033313 | Fbxl8 | 0 | -0.936126949 | 0.879315575 |
| ENSMUSG00000033249 | Hsf4 | 0 | 1.648835551 | 1 |
| ENSMUSG00000014782 | Plekhg4 | 0 | 0.063873051 | 1 |
| ENSMUSG00000014846 | Tppp3 | 0 | 1.063873051 | 1 |
| ENSMUSG00000098120 | Gm5914 | 0 | 0.648835551 | 1 |
| ENSMUSG00000050357 | Carmil2 | 0 | 0.326907457 | 1 |
| ENSMUSG00000005699 | Pard6a | 0 | 1.063873051 | 1 |
| ENSMUSG00000013155 | Enkd1 | 0 | -0.158519371 | 1 |
| ENSMUSG00000110586 | 4930578M07Rik | 0 | 1.648835551 | 1 |
| ENSMUSG00000031896 | Ctrl | 0 | 1.063873051 | 1 |
| ENSMUSG00000035237 | Lcat | 0 | 1.063873051 | 1 |
| ENSMUSG00000031898 | Dpep3 | 0 | 1.648835551 | 1 |
| ENSMUSG00000045538 | Ddx28 | 0 | -0.673092543 | 1 |
| ENSMUSG00000054069 | 6030452D12Rik | 0 | 0.648835551 | 1 |
| ENSMUSG00000099139 | 1110028F18Rik | 0 | -0.52108945 | 1 |
| ENSMUSG00000031910 | Has3 | 0 | 0.063873051 | 1 |
| ENSMUSG00000046691 | Chtf8 | 0 | -0.673092543 | 1 |
| ENSMUSG00000085464 | Gm16208 | 0 | 1.648835551 | 1 |
| ENSMUSG00000078931 | Pdf | 0 | 1.063873051 | 1 |
| ENSMUSG00000031919 | Tmed6 | 0 | 0.326907457 | 1 |
| ENSMUSG00000110625 | 5033426E14Rik | 0 | 0.326907457 | 1 |
| ENSMUSG00000078143 | Gm17344 | 0 | 1.648835551 | 1 |
| ENSMUSG00000035930 | Chst4 | 0 | 1.063873051 | 1 |
| ENSMUSG00000085068 | Gm15895 | 0 | 1.648835551 | 1 |
| ENSMUSG00000033624 | Pdpr | 0 | -3.184054463 | 0.028368406 |
| ENSMUSG00000098075 | Gm26994 | 0 | 1.648835551 | 1 |
| ENSMUSG00000033409 | Syce1l | 0 | -0.810596067 | 0.844591606 |

| | | | | |
|---|---|---|---|---|
| ENSMUSG00000110582 | Gm30052 | 0 | 0.326907457 | 1 |
| ENSMUSG00000084848 | 4933408N05Rik | 0 | 1.063873051 | 1 |
| ENSMUSG00000110620 | Gm45733 | 0 | -0.158519371 | 1 |
| ENSMUSG00000085688 | Gm15395 | 0 | 1.063873051 | 1 |
| ENSMUSG00000034308 | Sdr42e1 | 0 | 0.063873051 | 1 |
| ENSMUSG00000110667 | Gm45720 | 0 | 1.648835551 | 1 |
| ENSMUSG00000110470 | Gm45723 | 0 | 1.648835551 | 1 |
| ENSMUSG00000074063 | Osgin1 | 0 | -1.743481871 | 0.466464457 |
| ENSMUSG00000034224 | Slc38a8 | 0 | 1.648835551 | 1 |
| ENSMUSG00000034189 | Hsdl1 | 0 | -1.874726405 | 0.514152672 |
| ENSMUSG00000045246 | Kcng4 | 0 | 1.063873051 | 1 |
| ENSMUSG00000089776 | Gm15684 | 0 | 0.648835551 | 1 |
| ENSMUSG00000097944 | A130014A01Rik | 0 | -0.351164449 | 1 |
| ENSMUSG00000031824 | 6430548M08Rik | 0 | -0.810596067 | 0.844591606 |
| ENSMUSG00000042269 | Fam92b | 0 | 0.648835551 | 1 |
| ENSMUSG00000097960 | A330074K22Rik | 0 | -0.810596067 | 0.844591606 |
| ENSMUSG00000097983 | Gm26971 | 0 | 1.648835551 | 1 |
| ENSMUSG00000097773 | Gm10614 | 0 | 0.063873051 | 1 |
| ENSMUSG00000097500 | Gm26537 | 0 | 0.326907457 | 1 |
| ENSMUSG00000042812 | Foxf1 | 0 | 1.063873051 | 1 |
| ENSMUSG00000046714 | Foxc2 | 0 | 1.648835551 | 1 |
| ENSMUSG00000097084 | Foxl1 | 0 | 1.648835551 | 1 |
| ENSMUSG00000097097 | 5033426O07Rik | 0 | 1.648835551 | 1 |
| ENSMUSG00000098211 | Gm27045 | 0 | 1.648835551 | 1 |
| ENSMUSG00000110673 | Gm45833 | 0 | -0.351164449 | 1 |
| ENSMUSG00000025318 | Jph3 | 0 | 1.063873051 | 1 |
| ENSMUSG00000092418 | Gm20406 | 0 | 1.648835551 | 1 |
| ENSMUSG00000039079 | Trhr2 | 0 | 1.648835551 | 1 |
| ENSMUSG00000093682 | Gm20681 | 0 | 1.063873051 | 1 |
| ENSMUSG00000046108 | Il17c | 0 | 0.063873051 | 1 |
| ENSMUSG00000110656 | Gm20735 | 0 | 1.648835551 | 1 |
| ENSMUSG00000006587 | Snai3 | 0 | -1.874726405 | 0.483128587 |
| ENSMUSG00000033594 | Spata2l | 0 | 1.648835551 | 1 |
| ENSMUSG00000097637 | 4933417D19Rik | 0 | 0.326907457 | 1 |
| ENSMUSG00000001062 | Vps9d1 | 0 | -1.43862729 | 0.538110273 |
| ENSMUSG00000110684 | Gm45866 | 0 | -0.351164449 | 1 |
| ENSMUSG00000085030 | 2810455O05Rik | 0 | 1.063873051 | 1 |
| ENSMUSG00000110547 | Gm29773 | 0 | -0.351164449 | 1 |
| ENSMUSG00000039509 | Nup133 | 0 | -3.328444372 | 0.016760227 |
| ENSMUSG00000110615 | Gm45890 | 0 | -0.810596067 | 0.844591606 |
| ENSMUSG00000110704 | Gm45732 | 0 | -0.158519371 | 1 |
| ENSMUSG00000089820 | Gm15775 | 0 | 0.648835551 | 1 |
| ENSMUSG00000031982 | Arv1 | 0 | -1.874726405 | 0.352215149 |
| ENSMUSG00000043068 | Fam89a | 0 | 0.326907457 | 1 |
| ENSMUSG00000050930 | Map10 | 0 | 1.648835551 | 1 |
| ENSMUSG00000110698 | Gm45875 | 0 | 0.063873051 | 1 |
| ENSMUSG00000078840 | Gm10999 | 0 | 1.648835551 | 1 |
| ENSMUSG00000096736 | Gm17535 | 0 | 0.326907457 | 1 |
| ENSMUSG00000025888 | Casp1 | 0 | 0.326907457 | 1 |
| ENSMUSG00000047619 | Ddi1 | 0 | 1.063873051 | 1 |
| ENSMUSG00000043613 | Mmp3 | 0 | 0.648835551 | 1 |

| | | | | |
|---|---|---|---|---|
| ENSMUSG00000047562 | Mmp10 | 0 | -0.158519371 | 1 |
| ENSMUSG00000005800 | Mmp8 | 0 | -0.52108945 | 1 |
| ENSMUSG00000086381 | AV064505 | 0 | -0.673092543 | 1 |
| ENSMUSG00000092185 | Gm20416 | 0 | 1.063873051 | 1 |
| ENSMUSG00000110161 | Gm45494 | 0 | -0.351164449 | 1 |
| ENSMUSG00000031870 | Pgr | 0 | -0.158519371 | 1 |
| ENSMUSG00000111284 | Gm47334 | 0 | 1.063873051 | 1 |
| ENSMUSG00000111503 | Gm47034 | 0 | 1.648835551 | 1 |
| ENSMUSG00000111830 | 4930568E12Rik | 0 | 1.063873051 | 1 |
| ENSMUSG00000111285 | Gm47051 | 0 | 1.648835551 | 1 |
| ENSMUSG00000111489 | 1700019J19Rik | 0 | 1.648835551 | 1 |
| ENSMUSG00000111803 | Gm47564 | 0 | 1.648835551 | 1 |
| ENSMUSG00000053914 | Kdm4d | 0 | -0.673092543 | 1 |
| ENSMUSG00000036912 | Piwil4 | 0 | 1.063873051 | 1 |
| ENSMUSG00000074508 | Gm10706 | 0 | 1.648835551 | 1 |
| ENSMUSG00000090023 | Gm16302 | 0 | 1.648835551 | 1 |
| ENSMUSG00000031933 | Izumo1r | 0 | 1.648835551 | 1 |
| ENSMUSG00000110766 | Gm47214 | 0 | 0.326907457 | 1 |
| ENSMUSG00000031936 | Hephl1 | 0 | -0.936126949 | 0.87893712 |
| ENSMUSG00000089702 | Gm16568 | 0 | 1.648835551 | 1 |
| ENSMUSG00000111299 | 4930540M03Rik | 0 | 1.063873051 | 1 |
| ENSMUSG00000047508 | Mbd3l2 | 0 | 0.648835551 | 1 |
| ENSMUSG00000074500 | Zfp558 | 0 | 0.063873051 | 1 |
| ENSMUSG00000095957 | Olfr832 | 0 | 1.648835551 | 1 |
| ENSMUSG00000059303 | Olfr836 | 0 | 1.648835551 | 1 |
| ENSMUSG00000110519 | Olfr839-ps1 | 0 | 0.326907457 | 1 |
| ENSMUSG00000048391 | Olfr843 | 0 | 1.063873051 | 1 |
| ENSMUSG00000061614 | Olfr845 | 0 | 1.648835551 | 1 |
| ENSMUSG00000058692 | Olfr846 | 0 | 0.648835551 | 1 |
| ENSMUSG00000052182 | Olfr849 | 0 | 1.648835551 | 1 |
| ENSMUSG00000051160 | Olfr853 | 0 | 1.648835551 | 1 |
| ENSMUSG00000094678 | Olfr857 | 0 | 1.648835551 | 1 |
| ENSMUSG00000095448 | Olfr859 | 0 | 1.648835551 | 1 |
| ENSMUSG00000110511 | Gm45699 | 0 | 1.063873051 | 1 |
| ENSMUSG00000063842 | Olfr862 | 0 | 1.648835551 | 1 |
| ENSMUSG00000066896 | Olfr18 | 0 | 1.648835551 | 1 |
| ENSMUSG00000053773 | Rdh8 | 0 | 0.063873051 | 1 |
| ENSMUSG00000001014 | Icam4 | 0 | 1.063873051 | 1 |
| ENSMUSG00000032174 | Icam5 | 0 | 1.648835551 | 1 |
| ENSMUSG00000079681 | Zglp1 | 0 | 1.648835551 | 1 |
| ENSMUSG00000097439 | Gm16754 | 0 | 0.063873051 | 1 |
| ENSMUSG00000111172 | Gm39302 | 0 | 1.648835551 | 1 |
| ENSMUSG00000110915 | Gm36198 | 0 | -0.351164449 | 1 |
| ENSMUSG00000032180 | Tmed1 | 0 | 0.326907457 | 1 |
| ENSMUSG00000048429 | Timm29 | 0 | -0.673092543 | 1 |
| ENSMUSG00000097155 | Gm26511 | 0 | 0.326907457 | 1 |
| ENSMUSG00000047822 | Angptl8 | 0 | 0.648835551 | 1 |
| ENSMUSG00000006241 | Ccdc159 | 0 | -0.936126949 | 1 |
| ENSMUSG00000006235 | Epor | 0 | 1.648835551 | 1 |
| ENSMUSG00000039632 | Ccdc151 | 0 | 0.326907457 | 1 |
| ENSMUSG00000111856 | 6530413G14Rik | 0 | -0.158519371 | 1 |

| | | | | |
|---|---|---|---|---|
| ENSMUSG00000074472 | Zfp872 | 0 | 1.648835551 | 1 |
| ENSMUSG00000062794 | Zfp599 | 0 | -0.52108945 | 1 |
| ENSMUSG00000085969 | Gm15600 | 0 | 1.648835551 | 1 |
| ENSMUSG00000111365 | Gm47772 | 0 | 1.648835551 | 1 |
| ENSMUSG00000111275 | Gm29824 | 0 | -0.351164449 | 1 |
| ENSMUSG00000100002 | Gm29642 | 0 | 1.648835551 | 1 |
| ENSMUSG00000111066 | Gm30313 | 0 | 0.648835551 | 1 |
| ENSMUSG00000110901 | Gm48393 | 0 | 1.648835551 | 1 |
| ENSMUSG00000036395 | Glb1l2 | 0 | 0.326907457 | 1 |
| ENSMUSG00000111390 | Gm48796 | 0 | 0.326907457 | 1 |
| ENSMUSG00000111413 | Gm47716 | 0 | 1.648835551 | 1 |
| ENSMUSG00000031994 | Adamts8 | 0 | -0.351164449 | 1 |
| ENSMUSG00000111246 | 4930517E14Rik | 0 | 1.063873051 | 1 |
| ENSMUSG00000042185 | Nfrkb | 0 | -3.395558568 | 0.014189547 |
| ENSMUSG00000111853 | Gm31408 | 0 | 1.063873051 | 1 |
| ENSMUSG00000032034 | Kcnj5 | 0 | 1.648835551 | 1 |
| ENSMUSG00000098843 | Gm27240 | 0 | 1.063873051 | 1 |
| ENSMUSG00000099148 | Gm3331 | 0 | 0.648835551 | 1 |
| ENSMUSG00000111878 | Gm47777 | 0 | 0.648835551 | 1 |
| ENSMUSG00000110771 | Gm47785 | 0 | 0.648835551 | 1 |
| ENSMUSG00000053889 | Kirrel3os | 0 | 1.648835551 | 1 |
| ENSMUSG00000050555 | Hyls1 | 0 | 0.063873051 | 1 |
| ENSMUSG00000032099 | Pate4 | 0 | 1.648835551 | 1 |
| ENSMUSG00000099201 | Gm27187 | 0 | 1.063873051 | 1 |
| ENSMUSG00000074448 | Pate9 | 0 | 1.648835551 | 1 |
| ENSMUSG00000032113 | Chek1 | 0 | -1.743481871 | 0.455131546 |
| ENSMUSG00000096993 | Gm26787 | 0 | -0.936126949 | 0.882653814 |
| ENSMUSG00000032125 | Robo4 | 0 | 0.063873051 | 1 |
| ENSMUSG00000032128 | Robo3 | 0 | 1.063873051 | 1 |
| ENSMUSG00000053310 | Nrgn | 0 | -0.158519371 | 1 |
| ENSMUSG00000001948 | Spa17 | 0 | 0.648835551 | 1 |
| ENSMUSG00000011118 | Panx3 | 0 | 1.648835551 | 1 |
| ENSMUSG00000061165 | Olfr160 | 0 | 1.648835551 | 1 |
| ENSMUSG00000063350 | Olfr874 | 0 | 1.648835551 | 1 |
| ENSMUSG00000066747 | Olfr878 | 0 | 1.648835551 | 1 |
| ENSMUSG00000060583 | Olfr881 | 0 | 1.648835551 | 1 |
| ENSMUSG00000096356 | Olfr889 | 0 | 1.648835551 | 1 |
| ENSMUSG00000094588 | Olfr898 | 0 | 1.648835551 | 1 |
| ENSMUSG00000049098 | Olfr147 | 0 | 1.648835551 | 1 |
| ENSMUSG00000052058 | Olfr901 | 0 | 1.648835551 | 1 |
| ENSMUSG00000094701 | Olfr916 | 0 | 1.648835551 | 1 |
| ENSMUSG00000061039 | Olfr920 | 0 | 1.648835551 | 1 |
| ENSMUSG00000049926 | Olfr921 | 0 | 1.648835551 | 1 |
| ENSMUSG00000043911 | Olfr922 | 0 | 1.648835551 | 1 |
| ENSMUSG00000062103 | Olfr924 | 0 | 0.648835551 | 1 |
| ENSMUSG00000096109 | Olfr1537 | 0 | 1.648835551 | 1 |
| ENSMUSG00000063380 | Olfr945 | 0 | 1.063873051 | 1 |
| ENSMUSG00000062649 | Olfr44 | 0 | -0.351164449 | 1 |
| ENSMUSG00000059366 | Olfr959 | 0 | 1.648835551 | 1 |
| ENSMUSG00000038112 | AW551984 | 0 | 0.063873051 | 1 |
| ENSMUSG00000095839 | Olfr965 | 0 | 1.648835551 | 1 |

| | | | | |
|---|---|---|---|---|
| ENSMUSG00000055820 | Olfr967 | 0 | 1.648835551 | 1 |
| ENSMUSG00000095774 | Olfr970 | 0 | 1.648835551 | 1 |
| ENSMUSG00000093934 | Olfr971 | 0 | 1.648835551 | 1 |
| ENSMUSG00000095390 | Olfr229 | 0 | 1.648835551 | 1 |
| ENSMUSG00000047352 | Olfr976 | 0 | 1.648835551 | 1 |
| ENSMUSG00000045812 | Olfr984 | 0 | 1.648835551 | 1 |
| ENSMUSG00000051095 | Olfr986 | 0 | -0.351164449 | 1 |
| ENSMUSG00000049281 | Scn3b | 0 | 0.063873051 | 1 |
| ENSMUSG00000111271 | Gm48127 | 0 | 1.648835551 | 1 |
| ENSMUSG00000110922 | Gm35177 | 0 | 0.326907457 | 1 |
| ENSMUSG00000111189 | Gm48293 | 0 | 0.063873051 | 1 |
| ENSMUSG00000087135 | Gm16096 | 0 | 1.063873051 | 1 |
| ENSMUSG00000100837 | 1700063D05Rik | 0 | -0.351164449 | 1 |
| ENSMUSG00000110981 | Gm35940 | 0 | 1.063873051 | 1 |
| ENSMUSG00000086360 | Gm16214 | 0 | 1.063873051 | 1 |
| ENSMUSG00000111167 | Gm39321 | 0 | 1.648835551 | 1 |
| ENSMUSG00000111074 | Gm36435 | 0 | 0.648835551 | 1 |
| ENSMUSG00000037705 | Tecta | 0 | -0.158519371 | 1 |
| ENSMUSG00000048503 | Tmem136 | 0 | -0.52108945 | 1 |
| ENSMUSG00000111400 | Gm29961 | 0 | 1.648835551 | 1 |
| ENSMUSG00000096322 | Gm3898 | 0 | 1.063873051 | 1 |
| ENSMUSG00000032109 | Nlrx1 | 0 | -1.599091962 | 0.602789838 |
| ENSMUSG00000032131 | Abcg4 | 0 | 0.063873051 | 1 |
| ENSMUSG00000111291 | Gm48604 | 0 | 0.648835551 | 1 |
| ENSMUSG00000111849 | Gm47186 | 0 | 1.648835551 | 1 |
| ENSMUSG00000074397 | Foxr1 | 0 | 1.648835551 | 1 |
| ENSMUSG00000110980 | Gm47204 | 0 | -0.936126949 | 0.847265357 |
| ENSMUSG00000110755 | BC049987 | 0 | 1.648835551 | 1 |
| ENSMUSG00000002032 | Tmem25 | 0 | -0.158519371 | 1 |
| ENSMUSG00000039438 | Ttc36 | 0 | 0.326907457 | 1 |
| ENSMUSG00000032094 | Cd3d | 0 | 0.648835551 | 1 |
| ENSMUSG00000074385 | Gm10684 | 0 | 1.648835551 | 1 |
| ENSMUSG00000046480 | Scn4b | 0 | -0.673092543 | 1 |
| ENSMUSG00000091659 | Gm17099 | 0 | 1.648835551 | 1 |
| ENSMUSG00000032089 | Il10ra | 0 | 0.063873051 | 1 |
| ENSMUSG00000085831 | Lncbate3 | 0 | 1.063873051 | 1 |
| ENSMUSG00000097074 | 4833428L15Rik | 0 | 1.063873051 | 1 |
| ENSMUSG00000089810 | Gm16536 | 0 | -0.158519371 | 1 |
| ENSMUSG00000111521 | Gm48529 | 0 | 0.648835551 | 1 |
| ENSMUSG00000111329 | A830035O19Rik | 0 | 1.063873051 | 1 |
| ENSMUSG00000056617 | 4931429L15Rik | 0 | -0.351164449 | 1 |
| ENSMUSG00000111126 | Gm39329 | 0 | 1.648835551 | 1 |
| ENSMUSG00000097463 | 4930448E22Rik | 0 | 0.648835551 | 1 |
| ENSMUSG00000111236 | Gm31614 | 0 | 0.063873051 | 1 |
| ENSMUSG00000099696 | 2900052N01Rik | 0 | 0.063873051 | 1 |
| ENSMUSG00000111117 | A730065G17Rik | 0 | 1.063873051 | 1 |
| ENSMUSG00000111570 | Gm32335 | 0 | 0.326907457 | 1 |
| ENSMUSG00000008590 | Htr3b | 0 | 1.648835551 | 1 |
| ENSMUSG00000105991 | 1700042D02Rik | 0 | 1.063873051 | 1 |
| ENSMUSG00000054156 | Gm4894 | 0 | 0.063873051 | 1 |
| ENSMUSG00000032259 | Drd2 | 0 | 0.063873051 | 1 |

| | | | | |
|---|---|---|---|---|
| ENSMUSG00000079564 | Gm11149 | 0 | -0.936126949 | 0.886203497 |
| ENSMUSG00000111654 | Gm47077 | 0 | 1.063873051 | 1 |
| ENSMUSG00000000167 | Pih1d2 | 0 | 0.063873051 | 1 |
| ENSMUSG00000032056 | Btg4 | 0 | 0.063873051 | 1 |
| ENSMUSG00000032053 | Pou2af1 | 0 | -0.936126949 | 1 |
| ENSMUSG00000079559 | Colca2 | 0 | 1.063873051 | 1 |
| ENSMUSG00000111357 | Gm6980 | 0 | 1.648835551 | 1 |
| ENSMUSG00000111561 | Gm47957 | 0 | -0.936126949 | 0.886203497 |
| ENSMUSG00000037493 | Cib2 | 0 | -0.52108945 | 1 |
| ENSMUSG00000032285 | Dnaja4 | 0 | -0.351164449 | 1 |
| ENSMUSG00000044820 | AY074887 | 0 | -1.258055044 | 0.727318386 |
| ENSMUSG00000032303 | Chrna3 | 0 | 1.648835551 | 1 |
| ENSMUSG00000035200 | Chrnb4 | 0 | 0.648835551 | 1 |
| ENSMUSG00000032318 | Isl2 | 0 | 1.063873051 | 1 |
| ENSMUSG00000050702 | 4930563M21Rik | 0 | 1.063873051 | 1 |
| ENSMUSG00000099543 | Gm29322 | 0 | 1.648835551 | 1 |
| ENSMUSG00000097111 | Peak1os | 0 | 0.326907457 | 1 |
| ENSMUSG00000097870 | Gm26868 | 0 | 1.648835551 | 1 |
| ENSMUSG00000045620 | Odf3l1 | 0 | 1.648835551 | 1 |
| ENSMUSG00000110888 | Gm47237 | 0 | 1.648835551 | 1 |
| ENSMUSG00000086728 | Man2c1os | 0 | -1.810596067 | 0.72408384 |
| ENSMUSG00000032298 | Neil1 | 0 | -1.158519371 | 0.667778757 |
| ENSMUSG00000097445 | Gm26631 | 0 | -0.52108945 | 1 |
| ENSMUSG00000032306 | Mpi | 0 | -1.43862729 | 0.747404803 |
| ENSMUSG00000039714 | Cplx3 | 0 | 1.648835551 | 1 |
| ENSMUSG00000032310 | Cyp1a2 | 0 | 1.648835551 | 1 |
| ENSMUSG00000038264 | Sema7a | 0 | -1.673092543 | 0.607652185 |
| ENSMUSG00000097604 | Gm17322 | 0 | 1.063873051 | 1 |
| ENSMUSG00000086624 | Gm16130 | 0 | 1.648835551 | 1 |
| ENSMUSG00000036244 | Tbc1d21 | 0 | 1.648835551 | 1 |
| ENSMUSG00000066607 | Insyn1 | 0 | 0.063873051 | 1 |
| ENSMUSG00000074269 | Rec114 | 0 | -0.351164449 | 1 |
| ENSMUSG00000110789 | Gm33053 | 0 | -0.52108945 | 1 |
| ENSMUSG00000032297 | Celf6 | 0 | 0.648835551 | 1 |
| ENSMUSG00000084913 | Gm7616 | 0 | 1.648835551 | 1 |
| ENSMUSG00000034839 | Larp6 | 0 | 0.326907457 | 1 |
| ENSMUSG00000095956 | 1700036A12Rik | 0 | 0.326907457 | 1 |
| ENSMUSG00000052143 | Gm9869 | 0 | 1.648835551 | 1 |
| ENSMUSG00000111444 | Gm47923 | 0 | 1.648835551 | 1 |
| ENSMUSG00000111169 | Gm34004 | 0 | 0.063873051 | 1 |
| ENSMUSG00000110998 | I730028E13Rik | 0 | 0.648835551 | 1 |
| ENSMUSG00000111449 | Gm34105 | 0 | 1.648835551 | 1 |
| ENSMUSG00000111658 | B930092H01Rik | 0 | -0.936126949 | 0.847265357 |
| ENSMUSG00000111033 | Gm34322 | 0 | 1.648835551 | 1 |
| ENSMUSG00000046846 | Spesp1 | 0 | 1.648835551 | 1 |
| ENSMUSG00000111058 | Gm36033 | 0 | -1.051604167 | 0.858831093 |
| ENSMUSG00000093616 | 1700055C04Rik | 0 | 1.648835551 | 1 |
| ENSMUSG00000093759 | Scarletltr | 0 | 1.063873051 | 1 |
| ENSMUSG00000032396 | Dis3l | 0 | -2.305360759 | 0.231596497 |
| ENSMUSG00000032387 | Rbpms2 | 0 | -0.936126949 | 0.847265357 |
| ENSMUSG00000039452 | Snx22 | 0 | 1.648835551 | 1 |

| | | | | |
|---|---|---|---|---|
| ENSMUSG00000100130 | Gm28379 | 0 | 0.648835551 | 1 |
| ENSMUSG00000092131 | BC050972 | 0 | 1.648835551 | 1 |
| ENSMUSG00000111825 | Gm10646 | 0 | 1.063873051 | 1 |
| ENSMUSG00000111412 | Gm47047 | 0 | 1.063873051 | 1 |
| ENSMUSG00000111841 | Gm20745 | 0 | 1.648835551 | 1 |
| ENSMUSG00000059246 | Foxb1 | 0 | 1.648835551 | 1 |
| ENSMUSG00000074213 | Gm10642 | 0 | 0.648835551 | 1 |
| ENSMUSG00000111429 | Gm32511 | 0 | -0.52108945 | 1 |
| ENSMUSG00000099190 | Gm27188 | 0 | -0.351164449 | 1 |
| ENSMUSG00000098340 | Gm27253 | 0 | 0.648835551 | 1 |
| ENSMUSG00000098959 | Gm27230 | 0 | 1.063873051 | 1 |
| ENSMUSG00000096986 | 4930509E16Rik | 0 | -0.351164449 | 1 |
| ENSMUSG00000098294 | Gm27204 | 0 | 1.648835551 | 1 |
| ENSMUSG00000086158 | Ccpg1os | 0 | 0.063873051 | 1 |
| ENSMUSG00000098875 | Gm27211 | 0 | -0.351164449 | 1 |
| ENSMUSG00000066477 | Gm16551 | 0 | -0.351164449 | 1 |
| ENSMUSG00000101596 | Gm28182 | 0 | 1.648835551 | 1 |
| ENSMUSG00000111348 | Gm19531 | 0 | 1.648835551 | 1 |
| ENSMUSG00000032191 | Bcl2l10 | 0 | 1.648835551 | 1 |
| ENSMUSG00000097021 | 4933433G15Rik | 0 | 1.063873051 | 1 |
| ENSMUSG00000032179 | Bmp5 | 0 | -1.051604167 | 0.893427308 |
| ENSMUSG00000059383 | Gfral | 0 | 0.326907457 | 1 |
| ENSMUSG00000111222 | Gm34158 | 0 | 0.648835551 | 1 |
| ENSMUSG00000111610 | Gm34829 | 0 | 1.648835551 | 1 |
| ENSMUSG00000023333 | Gcm1 | 0 | -0.810596067 | 1 |
| ENSMUSG00000057933 | Gsta2 | 0 | 0.063873051 | 1 |
| ENSMUSG00000090724 | 4930542C12Rik | 0 | 1.063873051 | 1 |
| ENSMUSG00000070291 | Ddx43 | 0 | -0.810596067 | 1 |
| ENSMUSG00000032344 | Cgas | 0 | 0.326907457 | 1 |
| ENSMUSG00000111417 | Gm47430 | 0 | 1.648835551 | 1 |
| ENSMUSG00000111633 | Gm39377 | 0 | 1.648835551 | 1 |
| ENSMUSG00000084917 | Gm17477 | 0 | 1.648835551 | 1 |
| ENSMUSG00000111510 | Gm48387 | 0 | -0.52108945 | 1 |
| ENSMUSG00000098747 | Gm27216 | 0 | 1.063873051 | 1 |
| ENSMUSG00000100155 | Gm2109 | 0 | 1.648835551 | 1 |
| ENSMUSG00000111757 | Gm46124 | 0 | 1.648835551 | 1 |
| ENSMUSG00000111245 | Gm39388 | 0 | 1.063873051 | 1 |
| ENSMUSG00000110783 | Gm48821 | 0 | 1.063873051 | 1 |
| ENSMUSG00000111840 | Gm48832 | 0 | 1.063873051 | 1 |
| ENSMUSG00000056031 | 9330154J02Rik | 0 | -1.258055044 | 0.736456705 |
| ENSMUSG00000087466 | A330041J22Rik | 0 | -0.158519371 | 1 |
| ENSMUSG00000033491 | Prss35 | 0 | -1.351164449 | 0.687112677 |
| ENSMUSG00000111874 | Gm36539 | 0 | 1.648835551 | 1 |
| ENSMUSG00000047897 | Ripply2 | 0 | 1.648835551 | 1 |
| ENSMUSG00000032419 | Tbx18 | 0 | 0.648835551 | 1 |
| ENSMUSG00000100782 | Gm28231 | 0 | 1.648835551 | 1 |
| ENSMUSG00000101335 | Gm28229 | 0 | 1.648835551 | 1 |
| ENSMUSG00000097073 | 9430037G07Rik | 0 | 0.648835551 | 1 |
| ENSMUSG00000074149 | Gm10634 | 0 | -0.351164449 | 1 |
| ENSMUSG00000099974 | Bcl2a1d | 0 | 1.648835551 | 1 |
| ENSMUSG00000067399 | Trim43c | 0 | 1.063873051 | 1 |

| | | | | |
|---|---|---|---|---|
| ENSMUSG00000097677 | Gm26611 | 0 | 1.063873051 | 1 |
| ENSMUSG00000102037 | Bcl2a1a | 0 | 1.063873051 | 1 |
| ENSMUSG00000079162 | Trim43b | 0 | 1.648835551 | 1 |
| ENSMUSG00000074146 | 4930579C12Rik | 0 | 1.648835551 | 1 |
| ENSMUSG00000039313 | Minar1 | 0 | -0.52108945 | 1 |
| ENSMUSG00000111274 | Gm47409 | 0 | -1.52108945 | 0.586479347 |
| ENSMUSG00000098727 | 1700017I07Rik | 0 | 1.648835551 | 1 |
| ENSMUSG00000111666 | Gm30849 | 0 | -0.158519371 | 1 |
| ENSMUSG00000101608 | Gm28652 | 0 | 1.648835551 | 1 |
| ENSMUSG00000109896 | Gm31409 | 0 | 0.648835551 | 1 |
| ENSMUSG00000095654 | Plscr5 | 0 | 1.648835551 | 1 |
| ENSMUSG00000099823 | Gm28054 | 0 | 1.648835551 | 1 |
| ENSMUSG00000100294 | 1700034K08Rik | 0 | 0.326907457 | 1 |
| ENSMUSG00000101037 | Gm28424 | 0 | 1.648835551 | 1 |
| ENSMUSG00000086585 | Gm16126 | 0 | 0.648835551 | 1 |
| ENSMUSG00000096655 | 1700065D16Rik | 0 | 1.648835551 | 1 |
| ENSMUSG00000097777 | Gm16794 | 0 | 1.648835551 | 1 |
| ENSMUSG00000056418 | BC043934 | 0 | 1.648835551 | 1 |
| ENSMUSG00000085541 | Gm16010 | 0 | 1.648835551 | 1 |
| ENSMUSG00000101365 | Gm19325 | 0 | -1.051604167 | 0.858831093 |
| ENSMUSG00000051074 | 4930579K19Rik | 0 | 0.326907457 | 1 |
| ENSMUSG00000102861 | Gm37637 | 0 | 1.063873051 | 1 |
| ENSMUSG00000100281 | 4930422M22Rik | 0 | 1.063873051 | 1 |
| ENSMUSG00000100123 | Gm28087 | 0 | 0.648835551 | 1 |
| ENSMUSG00000046242 | Nme9 | 0 | 0.648835551 | 1 |
| ENSMUSG00000110929 | Gm47757 | 0 | 0.326907457 | 1 |
| ENSMUSG00000101584 | 4930519F24Rik | 0 | 1.648835551 | 1 |
| ENSMUSG00000101287 | Gm28166 | 0 | 1.648835551 | 1 |
| ENSMUSG00000070287 | Slc35g2 | 0 | -0.52108945 | 1 |
| ENSMUSG00000101941 | Gm28979 | 0 | 1.648835551 | 1 |
| ENSMUSG00000093812 | Gm5627 | 0 | -0.158519371 | 1 |
| ENSMUSG00000110884 | Gm47416 | 0 | -1.810596067 | 0.580137963 |
| ENSMUSG00000099722 | Gm29154 | 0 | 0.648835551 | 1 |
| ENSMUSG00000111035 | Gm33054 | 0 | 1.648835551 | 1 |
| ENSMUSG00000099599 | Gm28548 | 0 | 0.063873051 | 1 |
| ENSMUSG00000111044 | Gm47106 | 0 | 1.648835551 | 1 |
| ENSMUSG00000032565 | Nudt16 | 0 | -0.936126949 | 0.847265357 |
| ENSMUSG00000032566 | Nudt16l2 | 0 | 1.648835551 | 1 |
| ENSMUSG00000085430 | Gm15619 | 0 | 0.648835551 | 1 |
| ENSMUSG00000032567 | Aste1 | 0 | -1.158519371 | 0.886746766 |
| ENSMUSG00000032571 | Pik3r4 | 0 | -2.051604167 | 0.274319469 |
| ENSMUSG00000032572 | Col6a4 | 0 | -1.158519371 | 0.667778757 |
| ENSMUSG00000045322 | Tlr9 | 0 | 1.648835551 | 1 |
| ENSMUSG00000099879 | Gm29123 | 0 | 0.063873051 | 1 |
| ENSMUSG00000023262 | Acy1 | 0 | -0.351164449 | 1 |
| ENSMUSG00000023249 | Parp3 | 0 | -2.051604167 | 0.274319469 |
| ENSMUSG00000041009 | Iqcf4 | 0 | 1.648835551 | 1 |
| ENSMUSG00000032579 | Hemk1 | 0 | -2.351164449 | 0.169846604 |
| ENSMUSG00000037977 | 6430571L13Rik | 0 | 0.648835551 | 1 |
| ENSMUSG00000102144 | 4930429P21Rik | 0 | 1.648835551 | 1 |
| ENSMUSG00000091816 | Gm17141 | 0 | 1.648835551 | 1 |

| | | | | |
|---|---|---|---|---|
| ENSMUSG00000010057 | Nprl2 | 0 | -0.810596067 | 0.844591606 |
| ENSMUSG00000102300 | Gm34106 | 0 | 1.648835551 | 1 |
| ENSMUSG00000097099 | Gm9917 | 0 | 1.063873051 | 1 |
| ENSMUSG00000089935 | Gm16343 | 0 | 1.648835551 | 1 |
| ENSMUSG00000010064 | Slc38a3 | 0 | 1.648835551 | 1 |
| ENSMUSG00000034837 | Gnat1 | 0 | 0.326907457 | 1 |
| ENSMUSG00000032936 | Camkv | 0 | -0.351164449 | 1 |
| ENSMUSG00000093619 | 4930535L15Rik | 0 | 1.648835551 | 1 |
| ENSMUSG00000102540 | Gm38134 | 0 | 0.063873051 | 1 |
| ENSMUSG00000041528 | Rnf123 | 0 | -3.258055044 | 0.02002031 |
| ENSMUSG00000032607 | Amt | 0 | -1.351164449 | 0.718318299 |
| ENSMUSG00000032611 | 1700102P08Rik | 0 | -1.936126949 | 0.432479223 |
| ENSMUSG00000070283 | Ndufaf3 | 0 | 1.648835551 | 1 |
| ENSMUSG00000066357 | Wdr6 | 0 | -0.936126949 | 0.847265357 |
| ENSMUSG00000102770 | Gm38163 | 0 | 1.648835551 | 1 |
| ENSMUSG00000023473 | Celsr3 | 0 | 1.648835551 | 1 |
| ENSMUSG00000023259 | Slc26a6 | 0 | -0.673092543 | 1 |
| ENSMUSG00000025644 | Gm7628 | 0 | 0.648835551 | 1 |
| ENSMUSG00000061701 | Fbxw20 | 0 | -0.158519371 | 1 |
| ENSMUSG00000054087 | Fbxw28 | 0 | 1.648835551 | 1 |
| ENSMUSG00000074061 | Fbxw19 | 0 | 1.063873051 | 1 |
| ENSMUSG00000104674 | Gm42756 | 0 | 1.648835551 | 1 |
| ENSMUSG00000074060 | Fbxw15 | 0 | 0.326907457 | 1 |
| ENSMUSG00000062275 | Fbxw24 | 0 | 1.648835551 | 1 |
| ENSMUSG00000094992 | Fbxw25 | 0 | 1.063873051 | 1 |
| ENSMUSG00000074059 | Fbxw18 | 0 | 0.063873051 | 1 |
| ENSMUSG00000050074 | Spink8 | 0 | 1.648835551 | 1 |
| ENSMUSG00000038357 | Camp | 0 | 1.648835551 | 1 |
| ENSMUSG00000105484 | Gm42775 | 0 | -1.52108945 | 0.806496094 |
| ENSMUSG00000104945 | Gm35715 | 0 | 0.648835551 | 1 |
| ENSMUSG00000044664 | Prss42 | 0 | 1.648835551 | 1 |
| ENSMUSG00000047257 | Prss45 | 0 | 1.063873051 | 1 |
| ENSMUSG00000049555 | Tmie | 0 | -1.673092543 | 0.451785624 |
| ENSMUSG00000096393 | Fam240a | 0 | 1.648835551 | 1 |
| ENSMUSG00000032494 | Tdgf1 | 0 | 1.648835551 | 1 |
| ENSMUSG00000032495 | Lrrc2 | 0 | -1.351164449 | 0.687112677 |
| ENSMUSG00000066319 | Rtp3 | 0 | 1.063873051 | 1 |
| ENSMUSG00000057802 | Gm10030 | 0 | 0.648835551 | 1 |
| ENSMUSG00000043953 | Ccrl2 | 0 | 1.648835551 | 1 |
| ENSMUSG00000111054 | 4930545L08Rik | 0 | 1.648835551 | 1 |
| ENSMUSG00000110996 | Gm36251 | 0 | -0.158519371 | 1 |
| ENSMUSG00000111235 | Gm4657 | 0 | 0.648835551 | 1 |
| ENSMUSG00000111532 | Gm36485 | 0 | 0.648835551 | 1 |
| ENSMUSG00000111199 | Gm47875 | 0 | 1.648835551 | 1 |
| ENSMUSG00000111364 | Gm36745 | 0 | 1.648835551 | 1 |
| ENSMUSG00000111734 | Gm29825 | 0 | 1.063873051 | 1 |
| ENSMUSG00000064299 | 4921528I07Rik | 0 | -0.52108945 | 1 |
| ENSMUSG00000074039 | 4930520O04Rik | 0 | 0.648835551 | 1 |
| ENSMUSG00000079260 | Tmppe | 0 | 1.063873051 | 1 |
| ENSMUSG00000047898 | Ccr4 | 0 | -0.52108945 | 1 |
| ENSMUSG00000079259 | Trim71 | 0 | -0.673092543 | 1 |

| | | | | |
|---|---|---|---|---|
| ENSMUSG00000052724 | Gm9888 | 0 | 1.648835551 | 1 |
| ENSMUSG00000098127 | Gm4665 | 0 | 1.648835551 | 1 |
| ENSMUSG00000111398 | D330037F02Rik | 0 | 1.648835551 | 1 |
| ENSMUSG00000087469 | Gm16142 | 0 | 1.648835551 | 1 |
| ENSMUSG00000111243 | Gm4668 | 0 | 1.648835551 | 1 |
| ENSMUSG00000111324 | Gm31410 | 0 | 0.648835551 | 1 |
| ENSMUSG00000110752 | D730003K21Rik | 0 | 0.648835551 | 1 |
| ENSMUSG00000097402 | Gm17399 | 0 | -1.158519371 | 0.899603262 |
| ENSMUSG00000110676 | Gm45897 | 0 | 1.063873051 | 1 |
| ENSMUSG00000074029 | Gm47289 | 0 | 1.063873051 | 1 |
| ENSMUSG00000111204 | Gm47320 | 0 | 0.063873051 | 1 |
| ENSMUSG00000097504 | 4930516B21Rik | 0 | 1.063873051 | 1 |
| ENSMUSG00000097223 | Gm2449 | 0 | -0.158519371 | 1 |
| ENSMUSG00000032512 | Wdr48 | 0 | -3.079084903 | 0.038177619 |
| ENSMUSG00000032513 | Gorasp1 | 0 | -0.673092543 | 1 |
| ENSMUSG00000111369 | Gm47043 | 0 | 1.648835551 | 1 |
| ENSMUSG00000079243 | Xirp1 | 0 | 1.648835551 | 1 |
| ENSMUSG00000032519 | Slc25a38 | 0 | -2.051604167 | 0.466464457 |
| ENSMUSG00000111028 | Gm5922 | 0 | 1.648835551 | 1 |
| ENSMUSG00000111828 | D830035M03Rik | 0 | 1.063873051 | 1 |
| ENSMUSG00000041608 | Entpd3 | 0 | -1.874726405 | 0.370945697 |
| ENSMUSG00000110760 | Gm39456 | 0 | 0.326907457 | 1 |
| ENSMUSG00000111233 | Gm47069 | 0 | 1.648835551 | 1 |
| ENSMUSG00000110730 | Gm19385 | 0 | 1.063873051 | 1 |
| ENSMUSG00000111094 | Gm34425 | 0 | -0.810596067 | 1 |
| ENSMUSG00000086365 | 4930593C16Rik | 0 | 0.648835551 | 1 |
| ENSMUSG00000110863 | Gm34655 | 0 | 1.063873051 | 1 |
| ENSMUSG00000111598 | Gm47095 | 0 | -0.936126949 | 1 |
| ENSMUSG00000097820 | E530011L22Rik | 0 | 1.063873051 | 1 |
| ENSMUSG00000074001 | Klhl40 | 0 | 1.648835551 | 1 |
| ENSMUSG00000032523 | Hhatl | 0 | 1.648835551 | 1 |
| ENSMUSG00000111465 | Gm47113 | 0 | 1.648835551 | 1 |
| ENSMUSG00000043773 | 1700048O20Rik | 0 | 1.648835551 | 1 |
| ENSMUSG00000066235 | Pomgnt2 | 0 | -0.158519371 | 1 |
| ENSMUSG00000110729 | Gm47117 | 0 | 1.648835551 | 1 |
| ENSMUSG00000111389 | Gm39465 | 0 | -0.351164449 | 1 |
| ENSMUSG00000111434 | Gm47121 | 0 | -1.051604167 | 0.858831093 |
| ENSMUSG00000111268 | Gm47126 | 0 | 1.648835551 | 1 |
| ENSMUSG00000107504 | Gm35549 | 0 | 1.648835551 | 1 |
| ENSMUSG00000097322 | A530083I20Rik | 0 | 1.648835551 | 1 |
| ENSMUSG00000110945 | Gm9856 | 0 | -1.995020638 | 0.370945697 |
| ENSMUSG00000025784 | Clec3b | 0 | 0.326907457 | 1 |
| ENSMUSG00000054871 | Tmem158 | 0 | -0.158519371 | 1 |
| ENSMUSG00000036814 | Slc6a20a | 0 | -0.158519371 | 1 |
| ENSMUSG00000091172 | Gm17120 | 0 | 1.648835551 | 1 |
| ENSMUSG00000035448 | Ccr3 | 0 | 1.648835551 | 1 |
| ENSMUSG00000049103 | Ccr2 | 0 | 0.648835551 | 1 |
| ENSMUSG00000074829 | 2010315B03Rik | 0 | -1.743481871 | 0.455131546 |
| ENSMUSG00000110909 | B020014A21Rik | 0 | 1.648835551 | 1 |
| ENSMUSG00000111266 | Gm30570 | 0 | 1.648835551 | 1 |
| ENSMUSG00000095123 | Gm21781 | 0 | -1.874726405 | 0.352215149 |

| | | | | |
|---|---|---|---|---|
| ENSMUSG00000096553 | Gm10097 | 0 | 1.648835551 | 1 |
| ENSMUSG00000019772 | Vip | 0 | 1.648835551 | 1 |
| ENSMUSG00000019773 | Fbxo5 | 0 | -1.743481871 | 0.455131546 |
| ENSMUSG00000078488 | Gm10945 | 0 | 1.063873051 | 1 |
| ENSMUSG00000019796 | Lrp11 | 0 | -0.52108945 | 1 |
| ENSMUSG00000040034 | Nup43 | 0 | -1.051604167 | 0.858831093 |
| ENSMUSG00000039981 | Zc3h12d | 0 | -0.673092543 | 1 |
| ENSMUSG00000112360 | Gm48614 | 0 | 0.648835551 | 1 |
| ENSMUSG00000097597 | Gm26674 | 0 | 1.648835551 | 1 |
| ENSMUSG00000111674 | Gm48728 | 0 | 1.648835551 | 1 |
| ENSMUSG00000112019 | Gm48748 | 0 | 1.648835551 | 1 |
| ENSMUSG00000112573 | Gm48324 | 0 | 0.648835551 | 1 |
| ENSMUSG00000111995 | Gm48697 | 0 | 0.648835551 | 1 |
| ENSMUSG00000112775 | Gm48722 | 0 | 1.063873051 | 1 |
| ENSMUSG00000112156 | Gm48723 | 0 | 1.063873051 | 1 |
| ENSMUSG00000112131 | Gm48724 | 0 | 1.063873051 | 1 |
| ENSMUSG00000097378 | B230208H11Rik | 0 | -1.673092543 | 0.451785624 |
| ENSMUSG00000112498 | Gm47691 | 0 | 1.648835551 | 1 |
| ENSMUSG00000112102 | Gm32105 | 0 | 1.063873051 | 1 |
| ENSMUSG00000112266 | Gm48843 | 0 | 1.648835551 | 1 |
| ENSMUSG00000112284 | Gm48845 | 0 | -0.351164449 | 1 |
| ENSMUSG00000112679 | Gm40603 | 0 | 0.648835551 | 1 |
| ENSMUSG00000112227 | C330004P14Rik | 0 | 1.648835551 | 1 |
| ENSMUSG00000039891 | Txlnb | 0 | 0.648835551 | 1 |
| ENSMUSG00000019853 | Hebp2 | 0 | -0.158519371 | 1 |
| ENSMUSG00000110737 | 4930444F02Rik | 0 | 1.063873051 | 1 |
| ENSMUSG00000111318 | Gm48483 | 0 | 1.063873051 | 1 |
| ENSMUSG00000110902 | Gm33104 | 0 | -0.810596067 | 1 |
| ENSMUSG00000042940 | 9230106D20Rik | 0 | 1.063873051 | 1 |
| ENSMUSG00000020007 | Il20ra | 0 | -1.158519371 | 0.886203497 |
| ENSMUSG00000110958 | Gm33332 | 0 | 1.648835551 | 1 |
| ENSMUSG00000050473 | Slc35d3 | 0 | 1.648835551 | 1 |
| ENSMUSG00000111227 | Gm20149 | 0 | 1.648835551 | 1 |
| ENSMUSG00000110967 | Gm40608 | 0 | 1.648835551 | 1 |
| ENSMUSG00000112654 | 4930455C13Rik | 0 | 0.648835551 | 1 |
| ENSMUSG00000037542 | Aldh8a1 | 0 | 0.326907457 | 1 |
| ENSMUSG00000037535 | 1700021A07Rik | 0 | 1.648835551 | 1 |
| ENSMUSG00000097108 | Gm26581 | 0 | 1.648835551 | 1 |
| ENSMUSG00000045680 | Tcf21 | 0 | 1.648835551 | 1 |
| ENSMUSG00000020010 | Vnn3 | 0 | 0.648835551 | 1 |
| ENSMUSG00000086148 | Gm15271 | 0 | 1.063873051 | 1 |
| ENSMUSG00000087400 | Gm15270 | 0 | -1.673092543 | 0.614552622 |
| ENSMUSG00000112352 | Gm40617 | 0 | 1.648835551 | 1 |
| ENSMUSG00000096546 | Smlr1 | 0 | 0.648835551 | 1 |
| ENSMUSG00000112803 | Gm36543 | 0 | 1.063873051 | 1 |
| ENSMUSG00000112797 | Gm30034 | 0 | 1.648835551 | 1 |
| ENSMUSG00000015519 | 2310057J18Rik | 0 | 0.648835551 | 1 |
| ENSMUSG00000110830 | Gm46218 | 0 | -0.351164449 | 1 |
| ENSMUSG00000019880 | Rspo3 | 0 | -0.158519371 | 1 |
| ENSMUSG00000111195 | Gm48215 | 0 | 1.648835551 | 1 |
| ENSMUSG00000019789 | Hey2 | 0 | 1.063873051 | 1 |

| | | | | |
|---|---|---|---|---|
| ENSMUSG00000111256 | Gm48094 | 0 | -0.673092543 | 1 |
| ENSMUSG00000110976 | Gm48093 | 0 | 1.648835551 | 1 |
| ENSMUSG00000112469 | Gm47823 | 0 | 0.648835551 | 1 |
| ENSMUSG00000039552 | Rsph4a | 0 | 0.648835551 | 1 |
| ENSMUSG00000049872 | Calhm5 | 0 | 1.063873051 | 1 |
| ENSMUSG00000046031 | Calhm6 | 0 | 1.648835551 | 1 |
| ENSMUSG00000097274 | Gm47512 | 0 | 0.648835551 | 1 |
| ENSMUSG00000039462 | Col10a1 | 0 | 0.326907457 | 1 |
| ENSMUSG00000111753 | 1700054O05Rik | 0 | 0.648835551 | 1 |
| ENSMUSG00000063953 | Amd2 | 0 | 0.326907457 | 1 |
| ENSMUSG00000111550 | Gm49446 | 0 | 1.648835551 | 1 |
| ENSMUSG00000046463 | 5930403N24Rik | 0 | -0.936126949 | 1 |
| ENSMUSG00000097742 | Gm26535 | 0 | 0.648835551 | 1 |
| ENSMUSG00000111488 | Gm31378 | 0 | 1.648835551 | 1 |
| ENSMUSG00000111208 | Gm48197 | 0 | 1.648835551 | 1 |
| ENSMUSG00000019845 | Tube1 | 0 | -1.158519371 | 0.885143138 |
| ENSMUSG00000062074 | Ccn6 | 0 | 0.648835551 | 1 |
| ENSMUSG00000111055 | D030034A15Rik | 0 | 1.648835551 | 1 |
| ENSMUSG00000111008 | Gm31562 | 0 | 1.648835551 | 1 |
| ENSMUSG00000111183 | Gm48486 | 0 | -0.351164449 | 1 |
| ENSMUSG00000097890 | 4930547M16Rik | 0 | 1.648835551 | 1 |
| ENSMUSG00000038528 | Mfsd4b5 | 0 | 0.063873051 | 1 |
| ENSMUSG00000111045 | Gm47598 | 0 | 1.648835551 | 1 |
| ENSMUSG00000111665 | A930033M14Rik | 0 | 1.648835551 | 1 |
| ENSMUSG00000019823 | Mical1 | 0 | -2.106051951 | 0.279592421 |
| ENSMUSG00000090708 | Gm17196 | 0 | 1.648835551 | 1 |
| ENSMUSG00000091370 | 5730435O14Rik | 0 | 1.648835551 | 1 |
| ENSMUSG00000019803 | Nr2e1 | 0 | 1.063873051 | 1 |
| ENSMUSG00000111171 | Gm47815 | 0 | 1.648835551 | 1 |
| ENSMUSG00000111565 | Gm34481 | 0 | 1.648835551 | 1 |
| ENSMUSG00000111422 | 1700027J07Rik | 0 | 0.648835551 | 1 |
| ENSMUSG00000111342 | 4933404K13Rik | 0 | -0.52108945 | 1 |
| ENSMUSG00000111463 | Gm48061 | 0 | 1.063873051 | 1 |
| ENSMUSG00000111362 | Gm47389 | 0 | 1.648835551 | 1 |
| ENSMUSG00000111494 | 4930431F10Rik | 0 | 1.648835551 | 1 |
| ENSMUSG00000110898 | Gm40638 | 0 | -0.351164449 | 1 |
| ENSMUSG00000112766 | Gm40645 | 0 | -0.673092543 | 1 |
| ENSMUSG00000112843 | Gm46224 | 0 | -0.673092543 | 1 |
| ENSMUSG00000019905 | Gprc6a | 0 | 0.063873051 | 1 |
| ENSMUSG00000019900 | Rfx6 | 0 | -2.106051951 | 0.49106711 |
| ENSMUSG00000112270 | Gm47613 | 0 | 1.648835551 | 1 |
| ENSMUSG00000112552 | Gm36065 | 0 | 1.648835551 | 1 |
| ENSMUSG00000112376 | Gm40649 | 0 | 0.326907457 | 1 |
| ENSMUSG00000112835 | Gm36173 | 0 | 0.648835551 | 1 |
| ENSMUSG00000038583 | Pln | 0 | 1.063873051 | 1 |
| ENSMUSG00000112734 | Gm47644 | 0 | 1.063873051 | 1 |
| ENSMUSG00000112132 | Gm36229 | 0 | 1.648835551 | 1 |
| ENSMUSG00000112733 | Gm40650 | 0 | 0.326907457 | 1 |
| ENSMUSG00000112739 | Gm20597 | 0 | 1.648835551 | 1 |
| ENSMUSG00000047669 | Msl3l2 | 0 | 1.648835551 | 1 |
| ENSMUSG00000112833 | Gm36595 | 0 | 0.326907457 | 1 |

| | | | | |
|---|---|---|---|---|
| ENSMUSG00000112327 | Gm36827 | 0 | 1.648835551 | 1 |
| ENSMUSG00000112345 | Gm40652 | 0 | 0.648835551 | 1 |
| ENSMUSG00000038010 | Ccdc138 | 0 | -3.132524162 | 0.034227446 |
| ENSMUSG00000003227 | Edar | 0 | -2.106051951 | 0.295840171 |
| ENSMUSG00000058297 | Spock2 | 0 | -1.258055044 | 0.673811773 |
| ENSMUSG00000057337 | Chst3 | 0 | -1.351164449 | 0.707681972 |
| ENSMUSG00000090439 | Gm17455 | 0 | 1.648835551 | 1 |
| ENSMUSG00000037247 | Pldi | 0 | 1.063873051 | 1 |
| ENSMUSG00000098055 | Gm26947 | 0 | 0.648835551 | 1 |
| ENSMUSG00000020096 | Tbata | 0 | 0.648835551 | 1 |
| ENSMUSG00000037171 | Nodal | 0 | 0.648835551 | 1 |
| ENSMUSG00000112229 | Gm48086 | 0 | 0.326907457 | 1 |
| ENSMUSG00000020087 | Tysnd1 | 0 | -0.673092543 | 1 |
| ENSMUSG00000044312 | Neurog3 | 0 | 1.648835551 | 1 |
| ENSMUSG00000020081 | Tacr2 | 0 | 1.648835551 | 1 |
| ENSMUSG00000020079 | Supv3l1 | 0 | -1.673092543 | 0.451785624 |
| ENSMUSG00000097129 | 4930507D05Rik | 0 | -0.158519371 | 1 |
| ENSMUSG00000112347 | Gm47257 | 0 | 1.648835551 | 1 |
| ENSMUSG00000096991 | Gm26789 | 0 | 1.648835551 | 1 |
| ENSMUSG00000036764 | Dnajc12 | 0 | -0.52108945 | 1 |
| ENSMUSG00000112913 | Gm30853 | 0 | 0.326907457 | 1 |
| ENSMUSG00000112202 | Gm31181 | 0 | 0.648835551 | 1 |
| ENSMUSG00000097134 | 1110002J07Rik | 0 | 1.063873051 | 1 |
| ENSMUSG00000112726 | Gm47531 | 0 | 0.326907457 | 1 |
| ENSMUSG00000112120 | Gm32255 | 0 | 1.648835551 | 1 |
| ENSMUSG00000112141 | 4930563J15Rik | 0 | 1.648835551 | 1 |
| ENSMUSG00000112558 | Gm32364 | 0 | 0.648835551 | 1 |
| ENSMUSG00000112619 | 1700030E10Rik | 0 | 1.063873051 | 1 |
| ENSMUSG00000037855 | Zfp365 | 0 | 0.326907457 | 1 |
| ENSMUSG00000112899 | 4930521O17Rik | 0 | 1.648835551 | 1 |
| ENSMUSG00000019945 | Cabcoco1 | 0 | -2.52108945 | 0.139351384 |
| ENSMUSG00000060044 | Tmem26 | 0 | 1.063873051 | 1 |
| ENSMUSG00000090622 | A930033H14Rik | 0 | 1.063873051 | 1 |
| ENSMUSG00000112884 | Gm47946 | 0 | 0.063873051 | 1 |
| ENSMUSG00000112607 | Gm33979 | 0 | 0.648835551 | 1 |
| ENSMUSG00000112752 | Gm48021 | 0 | 1.648835551 | 1 |
| ENSMUSG00000112745 | Gm48025 | 0 | 1.063873051 | 1 |
| ENSMUSG00000112028 | 1700113B09Rik | 0 | 0.648835551 | 1 |
| ENSMUSG00000093732 | Gm20609 | 0 | 1.648835551 | 1 |
| ENSMUSG00000112108 | 4930551I15Rik | 0 | 1.063873051 | 1 |
| ENSMUSG00000112168 | Gm34776 | 0 | 1.648835551 | 1 |
| ENSMUSG00000009070 | Rsph14 | 0 | 0.326907457 | 1 |
| ENSMUSG00000033427 | Upb1 | 0 | 0.063873051 | 1 |
| ENSMUSG00000046807 | Lrrc75b | 0 | 0.648835551 | 1 |
| ENSMUSG00000006342 | Susd2 | 0 | 1.063873051 | 1 |
| ENSMUSG00000050157 | Gm867 | 0 | 1.063873051 | 1 |
| ENSMUSG00000000903 | Vpreb3 | 0 | 1.648835551 | 1 |
| ENSMUSG00000009114 | 2610028H24Rik | 0 | -0.351164449 | 1 |
| ENSMUSG00000001155 | Ftcd | 0 | 1.648835551 | 1 |
| ENSMUSG00000112291 | Gm48276 | 0 | 1.648835551 | 1 |
| ENSMUSG00000112143 | Gm35721 | 0 | -1.43862729 | 0.770478499 |

| | | | | |
|---|---|---|---|---|
| ENSMUSG00000074919 | Gm10787 | 0 | 1.648835551 | 1 |
| ENSMUSG00000112662 | Gm47922 | 0 | 0.063873051 | 1 |
| ENSMUSG00000112652 | 4921516A02Rik | 0 | 0.648835551 | 1 |
| ENSMUSG00000112224 | Gm47944 | 0 | 1.648835551 | 1 |
| ENSMUSG00000032763 | Ilvbl | 0 | -2.158519371 | 0.290284345 |
| ENSMUSG00000071185 | Olfr1357 | 0 | 1.648835551 | 1 |
| ENSMUSG00000005355 | Casp14 | 0 | 1.063873051 | 1 |
| ENSMUSG00000078442 | Ccdc105 | 0 | 1.648835551 | 1 |
| ENSMUSG00000005357 | Slc1a6 | 0 | 1.063873051 | 1 |
| ENSMUSG00000062873 | Olfr1355 | 0 | 1.648835551 | 1 |
| ENSMUSG00000094673 | Olfr1354 | 0 | 1.648835551 | 1 |
| ENSMUSG00000042774 | Olfr1353 | 0 | 1.648835551 | 1 |
| ENSMUSG00000063216 | Olfr1351 | 0 | 1.648835551 | 1 |
| ENSMUSG00000060205 | Olfr57 | 0 | -0.673092543 | 1 |
| ENSMUSG00000091468 | Vmn2r82 | 0 | 0.326907457 | 1 |
| ENSMUSG00000112907 | Gm47015 | 0 | 1.648835551 | 1 |
| ENSMUSG00000020317 | Theg | 0 | 1.648835551 | 1 |
| ENSMUSG00000111977 | Gm47163 | 0 | 1.063873051 | 1 |
| ENSMUSG00000035963 | Odf3l2 | 0 | 0.326907457 | 1 |
| ENSMUSG00000054206 | Gzmm | 0 | 0.326907457 | 1 |
| ENSMUSG00000020331 | Hcn2 | 0 | -0.52108945 | 1 |
| ENSMUSG00000091994 | E130317F20Rik | 0 | 1.063873051 | 1 |
| ENSMUSG00000057729 | Prtn3 | 0 | -0.158519371 | 1 |
| ENSMUSG00000061780 | Cfd | 0 | 1.063873051 | 1 |
| ENSMUSG00000035773 | Kiss1r | 0 | 1.648835551 | 1 |
| ENSMUSG00000035754 | Wdr18 | 0 | -3.43862729 | 0.011284174 |
| ENSMUSG00000035745 | Grin3b | 0 | 1.648835551 | 1 |
| ENSMUSG00000020150 | Gamt | 0 | 0.648835551 | 1 |
| ENSMUSG00000020135 | Apc2 | 0 | -0.810596067 | 0.844591606 |
| ENSMUSG00000035486 | Plk5 | 0 | 1.063873051 | 1 |
| ENSMUSG00000085712 | Gm15124 | 0 | 1.063873051 | 1 |
| ENSMUSG00000003341 | Atp8b3 | 0 | 0.648835551 | 1 |
| ENSMUSG00000035370 | Gm49322 | 0 | 1.648835551 | 1 |
| ENSMUSG00000020216 | Jsrp1 | 0 | 0.326907457 | 1 |
| ENSMUSG00000051067 | Lingo3 | 0 | 1.648835551 | 1 |
| ENSMUSG00000035206 | Sppl2b | 0 | -0.52108945 | 1 |
| ENSMUSG00000087114 | Gm16099 | 0 | 1.648835551 | 1 |
| ENSMUSG00000004939 | Nmrk2 | 0 | 0.648835551 | 1 |
| ENSMUSG00000034872 | Gipc3 | 0 | 0.063873051 | 1 |
| ENSMUSG00000020234 | 4930404N11Rik | 0 | 0.063873051 | 1 |
| ENSMUSG00000114004 | Gm48552 | 0 | 1.648835551 | 1 |
| ENSMUSG00000078439 | Smim24 | 0 | 0.326907457 | 1 |
| ENSMUSG00000085327 | Gm16104 | 0 | 1.063873051 | 1 |
| ENSMUSG00000084849 | Gm16105 | 0 | 1.063873051 | 1 |
| ENSMUSG00000034771 | Tle2 | 0 | -1.810596067 | 0.465580427 |
| ENSMUSG00000034758 | Tle6 | 0 | -0.673092543 | 1 |
| ENSMUSG00000094622 | Gm3055 | 0 | 0.648835551 | 1 |
| ENSMUSG00000094076 | Gm4767 | 0 | 1.648835551 | 1 |
| ENSMUSG00000096718 | Zfp781 | 0 | -0.673092543 | 1 |
| ENSMUSG00000061371 | Zfp873 | 0 | -1.810596067 | 0.465580427 |
| ENSMUSG00000078435 | AU041133 | 0 | -2.106051951 | 0.279592421 |

| | | | | |
|---|---|---|---|---|
| ENSMUSG00000112110 | Gm15608 | 0 | 0.326907457 | 1 |
| ENSMUSG00000044581 | 4932415D10Rik | 0 | 1.063873051 | 1 |
| ENSMUSG00000073427 | Gm4924 | 0 | -2.636566667 | 0.112813335 |
| ENSMUSG00000020246 | Hcfc2 | 0 | -2.305360759 | 0.231596497 |
| ENSMUSG00000056821 | 1700028I16Rik | 0 | 0.326907457 | 1 |
| ENSMUSG00000112134 | Gm40723 | 0 | 0.648835551 | 1 |
| ENSMUSG00000111514 | E230014E18Rik | 0 | 1.063873051 | 1 |
| ENSMUSG00000097076 | Platr7 | 0 | 0.648835551 | 1 |
| ENSMUSG00000097692 | A230060F14Rik | 0 | 1.648835551 | 1 |
| ENSMUSG00000112493 | 4930486F22Rik | 0 | 1.063873051 | 1 |
| ENSMUSG00000087256 | Gm15990 | 0 | 1.648835551 | 1 |
| ENSMUSG00000044937 | Ttc41 | 0 | -2.305360759 | 0.281836872 |
| ENSMUSG00000020051 | Pah | 0 | -0.351164449 | 1 |
| ENSMUSG00000035351 | Nup37 | 0 | -1.351164449 | 0.687112677 |
| ENSMUSG00000004359 | Spic | 0 | 0.326907457 | 1 |
| ENSMUSG00000097383 | 1500026H17Rik | 0 | 0.063873051 | 1 |
| ENSMUSG00000111544 | 4930534H03Rik | 0 | 0.063873051 | 1 |
| ENSMUSG00000111525 | Gm47082 | 0 | 1.063873051 | 1 |
| ENSMUSG00000110794 | 4930401A07Rik | 0 | 1.063873051 | 1 |
| ENSMUSG00000111531 | Gm32688 | 0 | 1.648835551 | 1 |
| ENSMUSG00000097407 | 4933408J17Rik | 0 | 1.648835551 | 1 |
| ENSMUSG00000097689 | 4930471D02Rik | 0 | 1.648835551 | 1 |
| ENSMUSG00000097164 | Cep83os | 0 | 0.326907457 | 1 |
| ENSMUSG00000112481 | Gm29684 | 0 | 0.648835551 | 1 |
| ENSMUSG00000112593 | Gm48882 | 0 | 0.648835551 | 1 |
| ENSMUSG00000100550 | 2310039L15Rik | 0 | -1.351164449 | 0.687112677 |
| ENSMUSG00000097766 | 5730420D15Rik | 0 | 0.326907457 | 1 |
| ENSMUSG00000112519 | Gm47671 | 0 | 1.063873051 | 1 |
| ENSMUSG00000112188 | Gm47708 | 0 | 0.326907457 | 1 |
| ENSMUSG00000112771 | Gm47718 | 0 | 1.648835551 | 1 |
| ENSMUSG00000112599 | Gm47719 | 0 | -2.158519371 | 0.377049624 |
| ENSMUSG00000110353 | Gm33543 | 0 | 1.063873051 | 1 |
| ENSMUSG00000112234 | Gm48427 | 0 | 1.648835551 | 1 |
| ENSMUSG00000112264 | Gm48505 | 0 | 1.648835551 | 1 |
| ENSMUSG00000112484 | 4930473O22Rik | 0 | 1.063873051 | 1 |
| ENSMUSG00000112647 | Gm5427 | 0 | -0.351164449 | 1 |
| ENSMUSG00000112346 | Gm48768 | 0 | 0.648835551 | 1 |
| ENSMUSG00000089994 | Gm16239 | 0 | 1.063873051 | 1 |
| ENSMUSG00000111923 | Gm34777 | 0 | 1.648835551 | 1 |
| ENSMUSG00000112544 | Gm48089 | 0 | -0.52108945 | 1 |
| ENSMUSG00000112265 | Gm34983 | 0 | 0.648835551 | 1 |
| ENSMUSG00000112810 | Gm35101 | 0 | 1.648835551 | 1 |
| ENSMUSG00000112304 | Gm35206 | 0 | 1.648835551 | 1 |
| ENSMUSG00000112831 | Gm35533 | 0 | -0.673092543 | 1 |
| ENSMUSG00000046567 | 4930430F08Rik | 0 | -1.158519371 | 0.909230539 |
| ENSMUSG00000112342 | Gm47031 | 0 | 1.648835551 | 1 |
| ENSMUSG00000112073 | Gm47101 | 0 | 1.648835551 | 1 |
| ENSMUSG00000112845 | Gm47143 | 0 | 0.648835551 | 1 |
| ENSMUSG00000019894 | Slc6a15 | 0 | -0.673092543 | 1 |
| ENSMUSG00000085282 | Gm15663 | 0 | 0.326907457 | 1 |
| ENSMUSG00000112466 | Gm29674 | 0 | 1.648835551 | 1 |

| | | | | |
|---|---|---|---|---|
| ENSMUSG00000112532 | Gm36283 | 0 | -1.673092543 | 0.6095359 |
| ENSMUSG00000020185 | E2f7 | 0 | -2.810596067 | 0.070328284 |
| ENSMUSG00000112246 | Gm40761 | 0 | 0.063873051 | 1 |
| ENSMUSG00000112137 | Gm47865 | 0 | 1.063873051 | 1 |
| ENSMUSG00000112533 | Gm20758 | 0 | 0.648835551 | 1 |
| ENSMUSG00000020213 | Glipr1l1 | 0 | 0.648835551 | 1 |
| ENSMUSG00000112564 | Gm47419 | 0 | 0.063873051 | 1 |
| ENSMUSG00000112228 | Gm47524 | 0 | 1.063873051 | 1 |
| ENSMUSG00000089934 | 4930473D10Rik | 0 | 1.063873051 | 1 |
| ENSMUSG00000084865 | Gm15723 | 0 | 1.063873051 | 1 |
| ENSMUSG00000112016 | Gm31267 | 0 | 1.648835551 | 1 |
| ENSMUSG00000054934 | Kcnmb4 | 0 | -0.673092543 | 1 |
| ENSMUSG00000085837 | Kcnmb4os2 | 0 | 1.648835551 | 1 |
| ENSMUSG00000112627 | 4933412E12Rik | 0 | -1.743481871 | 0.455131546 |
| ENSMUSG00000031294 | D630029K05Rik | 0 | 1.648835551 | 1 |
| ENSMUSG00000020169 | Best3 | 0 | 0.326907457 | 1 |
| ENSMUSG00000060187 | Lrrc10 | 0 | 1.648835551 | 1 |
| ENSMUSG00000069515 | Lyz1 | 0 | 1.648835551 | 1 |
| ENSMUSG00000112344 | Gm32141 | 0 | -0.158519371 | 1 |
| ENSMUSG00000112711 | Gm32552 | 0 | 1.063873051 | 1 |
| ENSMUSG00000112500 | 4933411E08Rik | 0 | 1.063873051 | 1 |
| ENSMUSG00000112470 | 5330439M10Rik | 0 | 1.063873051 | 1 |
| ENSMUSG00000020212 | Mdm1 | 0 | -1.351164449 | 0.687112677 |
| ENSMUSG00000112684 | Gm47958 | 0 | 1.648835551 | 1 |
| ENSMUSG00000090461 | Iltifb | 0 | 1.648835551 | 1 |
| ENSMUSG00000112854 | Gm38403 | 0 | 0.648835551 | 1 |
| ENSMUSG00000100681 | 1700064J06Rik | 0 | 1.063873051 | 1 |
| ENSMUSG00000112161 | Gm34045 | 0 | 0.326907457 | 1 |
| ENSMUSG00000085454 | Grip1os3 | 0 | -0.52108945 | 1 |
| ENSMUSG00000084764 | Grip1os2 | 0 | 0.326907457 | 1 |
| ENSMUSG00000112894 | Gm47009 | 0 | 1.648835551 | 1 |
| ENSMUSG00000090191 | 9230105E05Rik | 0 | 1.648835551 | 1 |
| ENSMUSG00000111907 | Gm48202 | 0 | 1.648835551 | 1 |
| ENSMUSG00000112192 | Gm48341 | 0 | 0.648835551 | 1 |
| ENSMUSG00000112846 | 4930432O09Rik | 0 | 1.648835551 | 1 |
| ENSMUSG00000111952 | Gm4489 | 0 | 1.648835551 | 1 |
| ENSMUSG00000112774 | Gm36041 | 0 | -0.158519371 | 1 |
| ENSMUSG00000020123 | Avpr1a | 0 | 1.063873051 | 1 |
| ENSMUSG00000112130 | Gm47839 | 0 | 1.648835551 | 1 |
| ENSMUSG00000112283 | Gm36719 | 0 | 0.326907457 | 1 |
| ENSMUSG00000112208 | Gm40797 | 0 | 1.648835551 | 1 |
| ENSMUSG00000112832 | 4930484H19Rik | 0 | 1.063873051 | 1 |
| ENSMUSG00000006724 | Cyp27b1 | 0 | 1.648835551 | 1 |
| ENSMUSG00000040502 | 9-Mar | 0 | -0.351164449 | 1 |
| ENSMUSG00000112639 | A730063M14Rik | 0 | -0.52108945 | 1 |
| ENSMUSG00000097331 | F420014N23Rik | 0 | 1.063873051 | 1 |
| ENSMUSG00000074657 | Kif5a | 0 | -2.051604167 | 0.305378889 |
| ENSMUSG00000040345 | Arhgap9 | 0 | -0.351164449 | 1 |
| ENSMUSG00000112426 | Gm47200 | 0 | -1.52108945 | 0.584901683 |
| ENSMUSG00000047492 | Inhbe | 0 | 0.326907457 | 1 |
| ENSMUSG00000040287 | Stac3 | 0 | 0.326907457 | 1 |

| | | | | |
|---|---|---|---|---|
| ENSMUSG00000040258 | Nxph4 | 0 | 1.648835551 | 1 |
| ENSMUSG00000086499 | Gm16217 | 0 | 1.648835551 | 1 |
| ENSMUSG00000025401 | Myo1a | 0 | 1.648835551 | 1 |
| ENSMUSG00000025400 | Tac2 | 0 | 0.326907457 | 1 |
| ENSMUSG00000058396 | Gpr182 | 0 | 0.063873051 | 1 |
| ENSMUSG00000089789 | Rdh1 | 0 | 1.063873051 | 1 |
| ENSMUSG00000056148 | Rdh9 | 0 | 0.648835551 | 1 |
| ENSMUSG00000040134 | Rdh7 | 0 | 1.648835551 | 1 |
| ENSMUSG00000040127 | Sdr9c7 | 0 | 1.063873051 | 1 |
| ENSMUSG00000025396 | Hsd17b6 | 0 | -0.158519371 | 1 |
| ENSMUSG00000025395 | Prim1 | 0 | -3.599091962 | 0.006654031 |
| ENSMUSG00000111947 | Gm47438 | 0 | 0.063873051 | 1 |
| ENSMUSG00000118550 | AC135859.1 | 0 | 1.063873051 | 1 |
| ENSMUSG00000051716 | Apon | 0 | 1.648835551 | 1 |
| ENSMUSG00000047631 | Apof | 0 | 0.648835551 | 1 |
| ENSMUSG00000090761 | Gm17201 | 0 | 1.648835551 | 1 |
| ENSMUSG00000039824 | Myl6b | 0 | 1.648835551 | 1 |
| ENSMUSG00000100548 | Gm29585 | 0 | -1.743481871 | 0.515527066 |
| ENSMUSG00000111936 | Gm47540 | 0 | 0.648835551 | 1 |
| ENSMUSG00000097372 | Gm26876 | 0 | 1.063873051 | 1 |
| ENSMUSG00000025359 | Pmel | 0 | -0.351164449 | 1 |
| ENSMUSG00000025352 | Gdf11 | 0 | 0.326907457 | 1 |
| ENSMUSG00000025347 | Mettl7b | 0 | 1.648835551 | 1 |
| ENSMUSG00000107748 | Olfr766-ps1 | 0 | 1.648835551 | 1 |
| ENSMUSG00000061367 | Olfr771 | 0 | 1.063873051 | 1 |
| ENSMUSG00000107662 | Olfr772 | 0 | 1.648835551 | 1 |
| ENSMUSG00000096220 | Olfr775 | 0 | 1.648835551 | 1 |
| ENSMUSG00000062914 | Olfr777 | 0 | 1.648835551 | 1 |
| ENSMUSG00000095696 | Olfr786 | 0 | 1.648835551 | 1 |
| ENSMUSG00000096497 | Olfr787 | 0 | 1.063873051 | 1 |
| ENSMUSG00000047626 | Olfr791 | 0 | 1.648835551 | 1 |
| ENSMUSG00000095002 | Olfr798 | 0 | 1.648835551 | 1 |
| ENSMUSG00000093866 | Olfr802 | 0 | 1.648835551 | 1 |
| ENSMUSG00000046041 | Olfr803 | 0 | 1.648835551 | 1 |
| ENSMUSG00000095401 | Olfr804 | 0 | 1.648835551 | 1 |
| ENSMUSG00000096858 | Olfr805 | 0 | 1.648835551 | 1 |
| ENSMUSG00000071065 | Olfr806 | 0 | 1.648835551 | 1 |
| ENSMUSG00000069421 | Olfr810 | 0 | 1.063873051 | 1 |
| ENSMUSG00000052818 | Olfr813 | 0 | 1.648835551 | 1 |
| ENSMUSG00000063715 | Olfr816 | 0 | 0.648835551 | 1 |
| ENSMUSG00000048745 | Olfr820 | 0 | 1.648835551 | 1 |
| ENSMUSG00000044897 | Olfr821 | 0 | 1.648835551 | 1 |
| ENSMUSG00000058251 | Olfr822 | 0 | 1.648835551 | 1 |
| ENSMUSG00000059862 | Olfr826 | 0 | 1.063873051 | 1 |
| ENSMUSG00000045559 | Olfr827 | 0 | 1.648835551 | 1 |
| ENSMUSG00000048015 | Neurod4 | 0 | 1.648835551 | 1 |
| ENSMUSG00000092162 | Vmn2r86 | 0 | 0.326907457 | 1 |
| ENSMUSG00000034579 | Pla2g3 | 0 | 1.648835551 | 1 |
| ENSMUSG00000034570 | Inpp5j | 0 | -0.52108945 | 1 |
| ENSMUSG00000087291 | Gm11946 | 0 | 1.063873051 | 1 |
| ENSMUSG00000020434 | 4921536K21Rik | 0 | 0.648835551 | 1 |

| | | | | |
|---|---|---|---|---|
| ENSMUSG00000019368 | Sec14l4 | 0 | -0.351164449 | 1 |
| ENSMUSG00000054986 | Sec14l3 | 0 | 0.063873051 | 1 |
| ENSMUSG00000004748 | Mtfp1 | 0 | 0.648835551 | 1 |
| ENSMUSG00000003585 | Sec14l2 | 0 | -1.673092543 | 0.66416768 |
| ENSMUSG00000003581 | Rnf215 | 0 | -0.673092543 | 1 |
| ENSMUSG00000056288 | Gm11961 | 0 | -0.158519371 | 1 |
| ENSMUSG00000009075 | Cabp7 | 0 | 1.063873051 | 1 |
| ENSMUSG00000020396 | Nefh | 0 | 0.648835551 | 1 |
| ENSMUSG00000034164 | Emid1 | 0 | -1.051604167 | 0.884614256 |
| ENSMUSG00000020475 | Pgam2 | 0 | 1.648835551 | 1 |
| ENSMUSG00000020474 | Polm | 0 | -1.599091962 | 0.559918909 |
| ENSMUSG00000096923 | A730071L15Rik | 0 | 1.648835551 | 1 |
| ENSMUSG00000020447 | Npc1l1 | 0 | -0.351164449 | 1 |
| ENSMUSG00000000381 | Wap | 0 | 1.648835551 | 1 |
| ENSMUSG00000041046 | Ramp3 | 0 | -0.52108945 | 1 |
| ENSMUSG00000020413 | Hus1 | 0 | -1.051604167 | 0.858831093 |
| ENSMUSG00000040978 | Gm11992 | 0 | 1.063873051 | 1 |
| ENSMUSG00000086391 | 1700042O10Rik | 0 | 0.326907457 | 1 |
| ENSMUSG00000086584 | Gm12002 | 0 | -0.673092543 | 1 |
| ENSMUSG00000085561 | 1700046C09Rik | 0 | -0.52108945 | 1 |
| ENSMUSG00000085091 | Egfros | 0 | 0.648835551 | 1 |
| ENSMUSG00000087060 | Eldr | 0 | 0.648835551 | 1 |
| ENSMUSG00000086468 | Etaa1os | 0 | 1.063873051 | 1 |
| ENSMUSG00000099449 | Gm28401 | 0 | 1.648835551 | 1 |
| ENSMUSG00000087507 | Gm16141 | 0 | 1.063873051 | 1 |
| ENSMUSG00000086025 | 4933406G16Rik | 0 | 0.648835551 | 1 |
| ENSMUSG00000044066 | Cep68 | 0 | -1.673092543 | 0.451785624 |
| ENSMUSG00000005917 | Otx1 | 0 | 1.648835551 | 1 |
| ENSMUSG00000085856 | Gm12050 | 0 | 1.648835551 | 1 |
| ENSMUSG00000085021 | Gm12051 | 0 | 0.648835551 | 1 |
| ENSMUSG00000049904 | Tmem17 | 0 | -0.158519371 | 1 |
| ENSMUSG00000085803 | Gm12056 | 0 | 1.063873051 | 1 |
| ENSMUSG00000092455 | Gm20456 | 0 | 1.648835551 | 1 |
| ENSMUSG00000092626 | 9130230N09Rik | 0 | 1.063873051 | 1 |
| ENSMUSG00000086366 | 1700061J23Rik | 0 | 1.063873051 | 1 |
| ENSMUSG00000085415 | Selenok-ps1 | 0 | -1.158519371 | 0.667778757 |
| ENSMUSG00000055010 | A830031A19Rik | 0 | 0.326907457 | 1 |
| ENSMUSG00000086117 | Gm12065 | 0 | 1.063873051 | 1 |
| ENSMUSG00000085579 | Gm12066 | 0 | 1.063873051 | 1 |
| ENSMUSG00000073103 | Gm10466 | 0 | 0.326907457 | 1 |
| ENSMUSG00000086725 | A630052C17Rik | 0 | 1.648835551 | 1 |
| ENSMUSG00000020461 | Clhc1 | 0 | -0.158519371 | 1 |
| ENSMUSG00000086707 | Gm12088 | 0 | 0.326907457 | 1 |
| ENSMUSG00000084850 | Gm12092 | 0 | -0.810596067 | 0.844591606 |
| ENSMUSG00000078157 | 4931440F15Rik | 0 | 1.648835551 | 1 |
| ENSMUSG00000073073 | Gm8098 | 0 | 1.648835551 | 1 |
| ENSMUSG00000087474 | Gm12100 | 0 | 1.648835551 | 1 |
| ENSMUSG00000043999 | Gpr75 | 0 | 0.648835551 | 1 |
| ENSMUSG00000085772 | D630024D03Rik | 0 | -0.810596067 | 1 |
| ENSMUSG00000020299 | 4930524B15Rik | 0 | 1.648835551 | 1 |
| ENSMUSG00000086386 | Gm12108 | 0 | 1.648835551 | 1 |

| | | | | |
|---|---|---|---|---|
| ENSMUSG00000020279 | Il9r | 0 | 0.326907457 | 1 |
| ENSMUSG00000055609 | Hba-x | 0 | 1.648835551 | 1 |
| ENSMUSG00000087501 | Gm12116 | 0 | 0.326907457 | 1 |
| ENSMUSG00000073052 | D130052B06Rik | 0 | 0.326907457 | 1 |
| ENSMUSG00000087328 | Gm12120 | 0 | 1.648835551 | 1 |
| ENSMUSG00000020155 | Kcnmb1 | 0 | 0.648835551 | 1 |
| ENSMUSG00000086311 | Gm12122 | 0 | -2.209145444 | 0.217194106 |
| ENSMUSG00000087388 | Gm12128 | 0 | 0.648835551 | 1 |
| ENSMUSG00000020428 | Gabra6 | 0 | -0.52108945 | 1 |
| ENSMUSG00000085533 | Gm12146 | 0 | 1.063873051 | 1 |
| ENSMUSG00000046491 | C1qtnf2 | 0 | -0.936126949 | 0.847265357 |
| ENSMUSG00000020405 | Fabp6 | 0 | 0.648835551 | 1 |
| ENSMUSG00000004296 | Il12b | 0 | 1.063873051 | 1 |
| ENSMUSG00000086501 | 4930597A21Rik | 0 | 1.648835551 | 1 |
| ENSMUSG00000084773 | Gm12159 | 0 | 0.326907457 | 1 |
| ENSMUSG00000085476 | Gm12162 | 0 | 0.326907457 | 1 |
| ENSMUSG00000011254 | Thg1l | 0 | 0.063873051 | 1 |
| ENSMUSG00000020395 | Itk | 0 | -0.810596067 | 1 |
| ENSMUSG00000020401 | Fam71b | 0 | 1.063873051 | 1 |
| ENSMUSG00000078924 | Gm12169 | 0 | 1.063873051 | 1 |
| ENSMUSG00000086981 | Gm12171 | 0 | 1.648835551 | 1 |
| ENSMUSG00000040413 | Timd2 | 0 | -0.673092543 | 1 |
| ENSMUSG00000055546 | Timd4 | 0 | 1.648835551 | 1 |
| ENSMUSG00000058163 | Gm5431 | 0 | 1.063873051 | 1 |
| ENSMUSG00000069893 | 9930111J21Rik1 | 0 | 0.648835551 | 1 |
| ENSMUSG00000078922 | Tgtp1 | 0 | 1.648835551 | 1 |
| ENSMUSG00000078920 | Ifi47 | 0 | 1.648835551 | 1 |
| ENSMUSG00000040283 | Btnl9 | 0 | 1.063873051 | 1 |
| ENSMUSG00000100923 | Olfr10 | 0 | 1.648835551 | 1 |
| ENSMUSG00000044170 | Olfr1384 | 0 | 0.326907457 | 1 |
| ENSMUSG00000063827 | Olfr1382 | 0 | 1.648835551 | 1 |
| ENSMUSG00000020363 | Gfpt2 | 0 | -1.810596067 | 0.491653643 |
| ENSMUSG00000044296 | Zfp879 | 0 | 1.648835551 | 1 |
| ENSMUSG00000000617 | Grm6 | 0 | 1.648835551 | 1 |
| ENSMUSG00000048728 | Zfp454 | 0 | 1.063873051 | 1 |
| ENSMUSG00000061952 | Olfr1377 | 0 | 1.648835551 | 1 |
| ENSMUSG00000059397 | Olfr54 | 0 | 0.648835551 | 1 |
| ENSMUSG00000020364 | Zfp354a | 0 | 1.063873051 | 1 |
| ENSMUSG00000086646 | Platr20 | 0 | 1.063873051 | 1 |
| ENSMUSG00000061469 | Gm12569 | 0 | 0.063873051 | 1 |
| ENSMUSG00000045942 | BC049762 | 0 | 1.063873051 | 1 |
| ENSMUSG00000110618 | Gm39822 | 0 | 0.063873051 | 1 |
| ENSMUSG00000086920 | Gm12207 | 0 | 1.063873051 | 1 |
| ENSMUSG00000086395 | A630014C17Rik | 0 | 1.063873051 | 1 |
| ENSMUSG00000044352 | Sowaha | 0 | 1.648835551 | 1 |
| ENSMUSG00000000869 | Il4 | 0 | -0.351164449 | 1 |
| ENSMUSG00000036117 | Il5 | 0 | 1.063873051 | 1 |
| ENSMUSG00000018900 | Slc22a5 | 0 | -2.480447466 | 0.13388699 |
| ENSMUSG00000063652 | Slc22a21 | 0 | -1.936126949 | 0.66416768 |
| ENSMUSG00000085888 | Gm12224 | 0 | 0.648835551 | 1 |
| ENSMUSG00000086039 | Gm12227 | 0 | 0.648835551 | 1 |

| | | | | |
|---|---|---|---|---|
| ENSMUSG00000086789 | Lyrm7os | 0 | 1.648835551 | 1 |
| ENSMUSG00000020268 | Lyrm7 | 0 | 0.063873051 | 1 |
| ENSMUSG00000049588 | Ccdc69 | 0 | -1.051604167 | 0.858831093 |
| ENSMUSG00000020264 | Slc36a2 | 0 | -0.158519371 | 1 |
| ENSMUSG00000084980 | Slc36a1os | 0 | 0.648835551 | 1 |
| ENSMUSG00000020261 | Slc36a1 | 0 | -1.351164449 | 0.687112677 |
| ENSMUSG00000087342 | Gm12238 | 0 | 1.648835551 | 1 |
| ENSMUSG00000086460 | Gm12236 | 0 | 0.648835551 | 1 |
| ENSMUSG00000085719 | 4933424L21Rik | 0 | 0.063873051 | 1 |
| ENSMUSG00000078853 | Igtp | 0 | 1.648835551 | 1 |
| ENSMUSG00000037243 | Zfp692 | 0 | -1.743481871 | 0.455131546 |
| ENSMUSG00000013643 | Lypd8 | 0 | 1.648835551 | 1 |
| ENSMUSG00000058287 | Gm12253 | 0 | 0.648835551 | 1 |
| ENSMUSG00000043314 | Olfr30 | 0 | 0.648835551 | 1 |
| ENSMUSG00000064252 | Olfr329-ps | 0 | 1.648835551 | 1 |
| ENSMUSG00000059279 | Olfr224 | 0 | 1.648835551 | 1 |
| ENSMUSG00000043880 | Olfr323 | 0 | -0.351164449 | 1 |
| ENSMUSG00000063549 | Olfr322 | 0 | 1.648835551 | 1 |
| ENSMUSG00000108265 | Olfr318 | 0 | 1.648835551 | 1 |
| ENSMUSG00000059504 | Olfr314 | 0 | 1.063873051 | 1 |
| ENSMUSG00000049154 | Fam183b | 0 | 1.648835551 | 1 |
| ENSMUSG00000094805 | Olfr311 | 0 | 0.648835551 | 1 |
| ENSMUSG00000072915 | Gm12258 | 0 | -1.673092543 | 0.451785624 |
| ENSMUSG00000085744 | Gm12259 | 0 | 1.648835551 | 1 |
| ENSMUSG00000020490 | Btnl10 | 0 | 1.063873051 | 1 |
| ENSMUSG00000078851 | Hist3h2a | 0 | 0.063873051 | 1 |
| ENSMUSG00000036964 | Trim17 | 0 | 1.063873051 | 1 |
| ENSMUSG00000085882 | 2610507I01Rik | 0 | -1.158519371 | 0.667778757 |
| ENSMUSG00000009900 | Wnt3a | 0 | 0.063873051 | 1 |
| ENSMUSG00000086910 | Gm15755 | 0 | 1.648835551 | 1 |
| ENSMUSG00000054519 | Zfp867 | 0 | -0.810596067 | 1 |
| ENSMUSG00000032691 | Nlrp3 | 0 | 0.648835551 | 1 |
| ENSMUSG00000059610 | Olfr222 | 0 | 1.648835551 | 1 |
| ENSMUSG00000048919 | Olfr223 | 0 | 1.648835551 | 1 |
| ENSMUSG00000043648 | Pld6 | 0 | 1.648835551 | 1 |
| ENSMUSG00000087249 | Gm16062 | 0 | 0.063873051 | 1 |
| ENSMUSG00000042709 | Atpaf2 | 0 | -0.936126949 | 0.855220644 |
| ENSMUSG00000018599 | Mief2 | 0 | -0.936126949 | 0.856471739 |
| ENSMUSG00000010142 | Tnfrsf13b | 0 | -0.52108945 | 1 |
| ENSMUSG00000109829 | Gm45605 | 0 | 1.648835551 | 1 |
| ENSMUSG00000060808 | B9d1os | 0 | 1.648835551 | 1 |
| ENSMUSG00000001039 | B9d1 | 0 | -0.673092543 | 1 |
| ENSMUSG00000042371 | Slc5a10 | 0 | 1.648835551 | 1 |
| ENSMUSG00000087120 | Gm12279 | 0 | 1.648835551 | 1 |
| ENSMUSG00000085762 | Lrrc75aos2 | 0 | 0.326907457 | 1 |
| ENSMUSG00000085170 | Lrrc75aos1 | 0 | 1.648835551 | 1 |
| ENSMUSG00000005267 | Zfp287 | 0 | -0.52108945 | 1 |
| ENSMUSG00000086674 | Zfp286os | 0 | 1.648835551 | 1 |
| ENSMUSG00000090173 | Fbxw10 | 0 | -0.936126949 | 0.886203497 |
| ENSMUSG00000086677 | Tvp23bos | 0 | 1.648835551 | 1 |
| ENSMUSG00000087622 | Gm12290 | 0 | 1.648835551 | 1 |

| | | | | |
|---|---|---|---|---|
| ENSMUSG00000020549 | Elac2 | 0 | -0.936126949 | 0.847265357 |
| ENSMUSG00000072834 | Gm12298 | 0 | 0.063873051 | 1 |
| ENSMUSG00000085926 | Gm12299 | 0 | 0.326907457 | 1 |
| ENSMUSG00000020908 | Myh3 | 0 | 1.063873051 | 1 |
| ENSMUSG00000087410 | 2310065F04Rik | 0 | 1.648835551 | 1 |
| ENSMUSG00000055775 | Myh8 | 0 | -0.351164449 | 1 |
| ENSMUSG00000087058 | Gm12301 | 0 | 1.648835551 | 1 |
| ENSMUSG00000020907 | Rcvrn | 0 | 1.063873051 | 1 |
| ENSMUSG00000097886 | Gsg1l2 | 0 | 1.648835551 | 1 |
| ENSMUSG00000033044 | Dhrs7c | 0 | 1.648835551 | 1 |
| ENSMUSG00000048329 | Mfsd6l | 0 | 1.648835551 | 1 |
| ENSMUSG00000045915 | Ccdc42 | 0 | 1.648835551 | 1 |
| ENSMUSG00000032921 | Odf4 | 0 | 1.063873051 | 1 |
| ENSMUSG00000018740 | Slc25a35 | 0 | 1.063873051 | 1 |
| ENSMUSG00000020898 | Ctc1 | 0 | -1.995020638 | 0.370945697 |
| ENSMUSG00000020897 | Aurkb | 0 | -0.810596067 | 0.844591606 |
| ENSMUSG00000045176 | Borcs6 | 0 | -1.52108945 | 0.613899307 |
| ENSMUSG00000020895 | Tmem107 | 0 | 1.648835551 | 1 |
| ENSMUSG00000020894 | Vamp2 | 0 | -0.936126949 | 0.847265357 |
| ENSMUSG00000023781 | Hes7 | 0 | 1.063873051 | 1 |
| ENSMUSG00000020891 | Alox8 | 0 | 1.063873051 | 1 |
| ENSMUSG00000018470 | Kcnab3 | 0 | 1.648835551 | 1 |
| ENSMUSG00000045377 | Tmem88 | 0 | 1.648835551 | 1 |
| ENSMUSG00000003934 | Efnb3 | 0 | 0.326907457 | 1 |
| ENSMUSG00000041346 | Wrap53 | 0 | -1.52108945 | 0.544908989 |
| ENSMUSG00000041329 | Atp1b2 | 0 | 0.326907457 | 1 |
| ENSMUSG00000085890 | Tnfsf13os | 0 | -0.936126949 | 0.886203497 |
| ENSMUSG00000089669 | Tnfsf13 | 0 | -0.351164449 | 1 |
| ENSMUSG00000089876 | Tmem102 | 0 | -1.051604167 | 0.858831093 |
| ENSMUSG00000041165 | Spem1 | 0 | 1.648835551 | 1 |
| ENSMUSG00000094845 | Tmem95 | 0 | -0.52108945 | 1 |
| ENSMUSG00000018570 | 2810408A11Rik | 0 | -0.936126949 | 0.847265357 |
| ENSMUSG00000018554 | Ybx2 | 0 | 1.648835551 | 1 |
| ENSMUSG00000020888 | Dvl2 | 0 | -1.158519371 | 0.667778757 |
| ENSMUSG00000020884 | Asgr1 | 0 | 1.648835551 | 1 |
| ENSMUSG00000040963 | Asgr2 | 0 | 0.648835551 | 1 |
| ENSMUSG00000000318 | Clec10a | 0 | -0.673092543 | 1 |
| ENSMUSG00000000317 | Bcl6b | 0 | 0.326907457 | 1 |
| ENSMUSG00000087523 | Gm12319 | 0 | 1.063873051 | 1 |
| ENSMUSG00000093858 | Gm19967 | 0 | 1.063873051 | 1 |
| ENSMUSG00000069830 | Nlrp1a | 0 | 0.326907457 | 1 |
| ENSMUSG00000111450 | Gm33351 | 0 | 0.648835551 | 1 |
| ENSMUSG00000085528 | 4930401O10Rik | 0 | 1.063873051 | 1 |
| ENSMUSG00000040554 | Aipl1 | 0 | 0.648835551 | 1 |
| ENSMUSG00000020808 | Pimreg | 0 | -0.52108945 | 1 |
| ENSMUSG00000020807 | 4933427D14Rik | 0 | 1.648835551 | 1 |
| ENSMUSG00000107877 | Gm43951 | 0 | 0.326907457 | 1 |
| ENSMUSG00000053574 | 4930563E22Rik | 0 | 0.326907457 | 1 |
| ENSMUSG00000020805 | Slc13a5 | 0 | 0.326907457 | 1 |
| ENSMUSG00000020799 | Tekt1 | 0 | 0.648835551 | 1 |
| ENSMUSG00000045667 | Smtnl2 | 0 | -0.936126949 | 1 |

| | | | | |
|---|---|---|---|---|
| ENSMUSG00000040471 | Ggt6 | 0 | 1.648835551 | 1 |
| ENSMUSG00000057778 | Cyb5d2 | 0 | -1.874726405 | 0.370799436 |
| ENSMUSG00000020785 | Camkk1 | 0 | -0.158519371 | 1 |
| ENSMUSG00000050107 | Haspin | 0 | -1.599091962 | 0.559918909 |
| ENSMUSG00000005950 | P2rx5 | 0 | 1.648835551 | 1 |
| ENSMUSG00000005952 | Trpv1 | 0 | 1.648835551 | 1 |
| ENSMUSG00000043029 | Trpv3 | 0 | 0.063873051 | 1 |
| ENSMUSG00000062128 | Olfr20 | 0 | 1.063873051 | 1 |
| ENSMUSG00000060335 | Olfr384 | 0 | 0.648835551 | 1 |
| ENSMUSG00000070382 | Olfr391-ps | 0 | 1.063873051 | 1 |
| ENSMUSG00000056921 | Olfr394 | 0 | 0.326907457 | 1 |
| ENSMUSG00000062186 | Olfr395 | 0 | 1.648835551 | 1 |
| ENSMUSG00000069816 | Olfr23 | 0 | 1.648835551 | 1 |
| ENSMUSG00000057050 | Olfr397 | 0 | 1.648835551 | 1 |
| ENSMUSG00000047444 | Olfr139 | 0 | 1.063873051 | 1 |
| ENSMUSG00000069476 | Zfp616 | 0 | 1.648835551 | 1 |
| ENSMUSG00000070378 | Olfr403 | 0 | 1.648835551 | 1 |
| ENSMUSG00000070375 | Olfr406 | 0 | 1.648835551 | 1 |
| ENSMUSG00000054406 | Olfr411 | 0 | 1.648835551 | 1 |
| ENSMUSG00000043099 | Hic1 | 0 | 1.063873051 | 1 |
| ENSMUSG00000078789 | Dph1 | 0 | -1.874726405 | 0.360452168 |
| ENSMUSG00000038224 | Serpinf2 | 0 | -0.351164449 | 1 |
| ENSMUSG00000038195 | Rilp | 0 | 1.063873051 | 1 |
| ENSMUSG00000109715 | Gm45606 | 0 | 1.648835551 | 1 |
| ENSMUSG00000087322 | Gm16075 | 0 | 1.648835551 | 1 |
| ENSMUSG00000049396 | Gemin4 | 0 | 1.648835551 | 1 |
| ENSMUSG00000038046 | Mrm3 | 0 | -0.158519371 | 1 |
| ENSMUSG00000044243 | Bhlha9 | 0 | 1.648835551 | 1 |
| ENSMUSG00000046275 | Trarg1 | 0 | 1.648835551 | 1 |
| ENSMUSG00000020839 | Tmigd1 | 0 | 1.063873051 | 1 |
| ENSMUSG00000109010 | Gm45027 | 0 | 1.648835551 | 1 |
| ENSMUSG00000044328 | Trp53i13 | 0 | -0.351164449 | 1 |
| ENSMUSG00000072673 | Gm10392 | 0 | 1.063873051 | 1 |
| ENSMUSG00000000724 | Cryba1 | 0 | -0.158519371 | 1 |
| ENSMUSG00000055045 | Gm11190 | 0 | 1.648835551 | 1 |
| ENSMUSG00000020832 | Eral1 | 0 | -0.810596067 | 0.844591606 |
| ENSMUSG00000017390 | Aldoc | 0 | 0.326907457 | 1 |
| ENSMUSG00000050132 | Sarm1 | 0 | 1.063873051 | 1 |
| ENSMUSG00000037278 | Tmem97 | 0 | -1.52108945 | 0.544908989 |
| ENSMUSG00000072640 | Lyrm9 | 0 | -1.351164449 | 0.687112677 |
| ENSMUSG00000085941 | Gm11201 | 0 | 1.063873051 | 1 |
| ENSMUSG00000091523 | AU040972 | 0 | 1.648835551 | 1 |
| ENSMUSG00000070354 | Evi2 | 0 | 0.648835551 | 1 |
| ENSMUSG00000087434 | Rab11fip4os1 | 0 | 1.063873051 | 1 |
| ENSMUSG00000086474 | 9130204K15Rik | 0 | -0.936126949 | 0.868550295 |
| ENSMUSG00000046909 | Tefm | 0 | 0.063873051 | 1 |
| ENSMUSG00000048895 | Cdk5r1 | 0 | -0.351164449 | 1 |
| ENSMUSG00000085078 | C030013C21Rik | 0 | 1.648835551 | 1 |
| ENSMUSG00000086703 | 4930507D10Rik | 0 | 1.063873051 | 1 |
| ENSMUSG00000053184 | Spaca3 | 0 | 0.326907457 | 1 |
| ENSMUSG00000087534 | Gm11418 | 0 | 1.648835551 | 1 |

| | | | | |
|---|---|---|---|---|
| ENSMUSG00000020703 | 5530401A14Rik | 0 | 0.648835551 | 1 |
| ENSMUSG00000035373 | Ccl7 | 0 | 0.648835551 | 1 |
| ENSMUSG00000020676 | Ccl11 | 0 | 1.648835551 | 1 |
| ENSMUSG00000009185 | Ccl8 | 0 | 1.063873051 | 1 |
| ENSMUSG00000069796 | Gm11426 | 0 | 1.063873051 | 1 |
| ENSMUSG00000046010 | Zfp830 | 0 | -3.233807498 | 0.019039678 |
| ENSMUSG00000085175 | Gm11423 | 0 | 0.648835551 | 1 |
| ENSMUSG00000086058 | Unc45bos | 0 | 1.063873051 | 1 |
| ENSMUSG00000086327 | Slfn5os | 0 | 1.063873051 | 1 |
| ENSMUSG00000111857 | 1190001M18Rik | 0 | 1.648835551 | 1 |
| ENSMUSG00000035208 | Slfn8 | 0 | -1.158519371 | 0.667778757 |
| ENSMUSG00000111713 | Gm20234 | 0 | 1.648835551 | 1 |
| ENSMUSG00000078763 | Slfn1 | 0 | 1.648835551 | 1 |
| ENSMUSG00000000204 | Slfn4 | 0 | 0.648835551 | 1 |
| ENSMUSG00000018986 | Slfn3 | 0 | 0.063873051 | 1 |
| ENSMUSG00000087107 | AI662270 | 0 | -1.743481871 | 0.482996469 |
| ENSMUSG00000082101 | Slfn14 | 0 | 1.648835551 | 1 |
| ENSMUSG00000093483 | AA465934 | 0 | -0.351164449 | 1 |
| ENSMUSG00000018733 | Pex12 | 0 | -1.995020638 | 0.370945697 |
| ENSMUSG00000020686 | Gas2l2 | 0 | 1.648835551 | 1 |
| ENSMUSG00000035042 | Ccl5 | 0 | 1.063873051 | 1 |
| ENSMUSG00000051452 | Gm11437 | 0 | -1.351164449 | 0.71205721 |
| ENSMUSG00000087211 | Lhx1os | 0 | 1.063873051 | 1 |
| ENSMUSG00000087407 | Gm11431 | 0 | 1.648835551 | 1 |
| ENSMUSG00000018405 | Mrm1 | 0 | -0.936126949 | 1 |
| ENSMUSG00000086799 | 4930502E09Rik | 0 | 1.648835551 | 1 |
| ENSMUSG00000018479 | 1700125H20Rik | 0 | -0.936126949 | 0.886203497 |
| ENSMUSG00000087013 | 2610027K06Rik | 0 | -0.158519371 | 1 |
| ENSMUSG00000069785 | Gm11444 | 0 | -2.874726405 | 0.054186866 |
| ENSMUSG00000085991 | Gm11479 | 0 | 1.648835551 | 1 |
| ENSMUSG00000020495 | Smg8 | 0 | -1.351164449 | 0.687112677 |
| ENSMUSG00000020492 | Ska2 | 0 | -3.743481871 | 0.004068424 |
| ENSMUSG00000085972 | 1110028F11Rik | 0 | -0.52108945 | 1 |
| ENSMUSG00000084796 | Mir142hg | 0 | 1.063873051 | 1 |
| ENSMUSG00000034156 | Tspoap1 | 0 | 0.063873051 | 1 |
| ENSMUSG00000085872 | Gm11505 | 0 | 1.648835551 | 1 |
| ENSMUSG00000034121 | Mks1 | 0 | 1.063873051 | 1 |
| ENSMUSG00000052234 | Epx | 0 | 1.648835551 | 1 |
| ENSMUSG00000060787 | Olfr464 | 0 | 0.648835551 | 1 |
| ENSMUSG00000108634 | Gm38534 | 0 | 0.648835551 | 1 |
| ENSMUSG00000085646 | Gm11493 | 0 | 1.648835551 | 1 |
| ENSMUSG00000057534 | Elobl | 0 | 1.063873051 | 1 |
| ENSMUSG00000085152 | Gm11496 | 0 | 1.648835551 | 1 |
| ENSMUSG00000063109 | Dgkeos | 0 | 1.648835551 | 1 |
| ENSMUSG00000072553 | Gm525 | 0 | 0.063873051 | 1 |
| ENSMUSG00000020553 | Pctp | 0 | -0.673092543 | 1 |
| ENSMUSG00000069763 | Tmem100 | 0 | -0.52108945 | 1 |
| ENSMUSG00000020544 | Cox11 | 0 | -1.43862729 | 0.740075579 |
| ENSMUSG00000086003 | B230206L02Rik | 0 | -0.351164449 | 1 |
| ENSMUSG00000044177 | Wfikkn2 | 0 | 1.063873051 | 1 |
| ENSMUSG00000039055 | Eme1 | 0 | -0.810596067 | 1 |

| | | | | |
|---|---|---|---|---|
| ENSMUSG00000075610 | Tmem92 | 0 | 1.648835551 | 1 |
| ENSMUSG00000038994 | Hils1 | 0 | 1.648835551 | 1 |
| ENSMUSG00000001510 | Dlx3 | 0 | -0.936126949 | 0.847265357 |
| ENSMUSG00000020871 | Dlx4 | 0 | 0.326907457 | 1 |
| ENSMUSG00000085992 | Gm11515 | 0 | 1.648835551 | 1 |
| ENSMUSG00000020872 | Tac4 | 0 | 0.648835551 | 1 |
| ENSMUSG00000046719 | Nxph3 | 0 | 1.063873051 | 1 |
| ENSMUSG00000084856 | Gm11528 | 0 | 1.648835551 | 1 |
| ENSMUSG00000086015 | 4833417C18Rik | 0 | -0.158519371 | 1 |
| ENSMUSG00000014351 | Gip | 0 | 1.648835551 | 1 |
| ENSMUSG00000038756 | Ttll6 | 0 | -0.810596067 | 0.847265357 |
| ENSMUSG00000020875 | Hoxb9 | 0 | 1.063873051 | 1 |
| ENSMUSG00000056648 | Hoxb8 | 0 | 0.063873051 | 1 |
| ENSMUSG00000038700 | Hoxb5 | 0 | 0.063873051 | 1 |
| ENSMUSG00000048763 | Hoxb3 | 0 | -1.52108945 | 0.544908989 |
| ENSMUSG00000075589 | Gm11536 | 0 | 0.063873051 | 1 |
| ENSMUSG00000075588 | Hoxb2 | 0 | 0.648835551 | 1 |
| ENSMUSG00000075586 | Gm11529 | 0 | 1.063873051 | 1 |
| ENSMUSG00000018672 | Copz2 | 0 | -1.158519371 | 0.667778757 |
| ENSMUSG00000018678 | Sp2 | 0 | -2.52108945 | 0.179008189 |
| ENSMUSG00000020877 | Scrn2 | 0 | 0.063873051 | 1 |
| ENSMUSG00000020878 | Lrrc46 | 0 | -0.351164449 | 1 |
| ENSMUSG00000001444 | Tbx21 | 0 | -0.351164449 | 1 |
| ENSMUSG00000086680 | Gm11592 | 0 | 0.326907457 | 1 |
| ENSMUSG00000047988 | 4933428G20Rik | 0 | 0.063873051 | 1 |
| ENSMUSG00000043439 | Epop | 0 | 1.063873051 | 1 |
| ENSMUSG00000018537 | Pcgf2 | 0 | -2.874726405 | 0.054186866 |
| ENSMUSG00000018543 | 1700001P01Rik | 0 | 1.063873051 | 1 |
| ENSMUSG00000050538 | B230217C12Rik | 0 | 0.063873051 | 1 |
| ENSMUSG00000020882 | Cacnb1 | 0 | -1.599091962 | 0.559918909 |
| ENSMUSG00000086716 | Gm11629 | 0 | 1.063873051 | 1 |
| ENSMUSG00000017400 | Stac2 | 0 | -1.258055044 | 0.673811773 |
| ENSMUSG00000093673 | Gm20644 | 0 | -0.52108945 | 1 |
| ENSMUSG00000085944 | 1700003D09Rik | 0 | 0.648835551 | 1 |
| ENSMUSG00000038208 | Pgap3 | 0 | 0.063873051 | 1 |
| ENSMUSG00000018168 | Ikzf3 | 0 | -1.158519371 | 0.667778757 |
| ENSMUSG00000038150 | Ormdl3 | 0 | -1.52108945 | 0.544908989 |
| ENSMUSG00000064224 | Gsdma3 | 0 | 1.648835551 | 1 |
| ENSMUSG00000017211 | Gsdma2 | 0 | -0.158519371 | 1 |
| ENSMUSG00000075576 | Gm12359 | 0 | 0.063873051 | 1 |
| ENSMUSG00000037944 | Ccr7 | 0 | 1.648835551 | 1 |
| ENSMUSG00000075570 | Krt26 | 0 | 1.648835551 | 1 |
| ENSMUSG00000087692 | Gm11940 | 0 | 1.648835551 | 1 |
| ENSMUSG00000020912 | Krt12 | 0 | 0.648835551 | 1 |
| ENSMUSG00000064165 | Krt39 | 0 | 1.648835551 | 1 |
| ENSMUSG00000078131 | Krtap1-3 | 0 | 1.648835551 | 1 |
| ENSMUSG00000078130 | Gm11555 | 0 | 1.648835551 | 1 |
| ENSMUSG00000069717 | Gm11568 | 0 | 1.648835551 | 1 |
| ENSMUSG00000051481 | Krtap31-2 | 0 | 1.648835551 | 1 |
| ENSMUSG00000035592 | Krt33a | 0 | 1.648835551 | 1 |
| ENSMUSG00000057723 | Krt33b | 0 | 1.648835551 | 1 |

| | | | | |
|---|---|---|---|---|
| ENSMUSG00000087581 | Gm11571 | 0 | -0.673092543 | 1 |
| ENSMUSG00000020916 | Krt36 | 0 | 1.648835551 | 1 |
| ENSMUSG00000044041 | Krt13 | 0 | 1.648835551 | 1 |
| ENSMUSG00000051617 | Krt9 | 0 | 0.648835551 | 1 |
| ENSMUSG00000045545 | Krt14 | 0 | -1.673092543 | 0.451785624 |
| ENSMUSG00000053797 | Krt16 | 0 | 1.648835551 | 1 |
| ENSMUSG00000086163 | Gm14206 | 0 | 0.648835551 | 1 |
| ENSMUSG00000006930 | Hap1 | 0 | 0.326907457 | 1 |
| ENSMUSG00000001558 | Klhl10 | 0 | -1.258055044 | 0.736456705 |
| ENSMUSG00000048732 | Klhl11 | 0 | -0.52108945 | 1 |
| ENSMUSG00000014198 | Zfp385c | 0 | -1.599091962 | 0.596517007 |
| ENSMUSG00000017830 | Dhx58 | 0 | 1.648835551 | 1 |
| ENSMUSG00000004043 | Stat5a | 0 | -2.106051951 | 0.279592421 |
| ENSMUSG00000019303 | Psmc3ip | 0 | -0.810596067 | 0.844591606 |
| ENSMUSG00000035198 | Tubg1 | 0 | -0.936126949 | 0.874796537 |
| ENSMUSG00000045007 | Tubg2 | 0 | -0.810596067 | 0.844591606 |
| ENSMUSG00000035172 | Plekhh3 | 0 | -1.743481871 | 0.455131546 |
| ENSMUSG00000017167 | Cntnap1 | 0 | -1.258055044 | 0.673811773 |
| ENSMUSG00000001240 | Ramp2 | 0 | 0.326907457 | 1 |
| ENSMUSG00000085083 | Gm11615 | 0 | 1.648835551 | 1 |
| ENSMUSG00000078653 | Cntd1 | 0 | 1.648835551 | 1 |
| ENSMUSG00000078651 | Aoc2 | 0 | -0.158519371 | 1 |
| ENSMUSG00000101925 | Gm28156 | 0 | 0.648835551 | 1 |
| ENSMUSG00000078650 | G6pc | 0 | 1.648835551 | 1 |
| ENSMUSG00000097487 | Ptges3l | 0 | -0.351164449 | 1 |
| ENSMUSG00000086347 | Gm11626 | 0 | 1.648835551 | 1 |
| ENSMUSG00000001313 | Rnd2 | 0 | 0.648835551 | 1 |
| ENSMUSG00000017146 | Brca1 | 0 | -3.636566667 | 0.005443828 |
| ENSMUSG00000085486 | Gm11634 | 0 | 1.648835551 | 1 |
| ENSMUSG00000086617 | Gm11635 | 0 | 0.648835551 | 1 |
| ENSMUSG00000085594 | Gm11551 | 0 | 1.648835551 | 1 |
| ENSMUSG00000052373 | Mpp3 | 0 | -0.158519371 | 1 |
| ENSMUSG00000017314 | Mpp2 | 0 | -2.599091962 | 0.108201045 |
| ENSMUSG00000017316 | Ppy | 0 | 1.063873051 | 1 |
| ENSMUSG00000017311 | Pyy | 0 | 1.648835551 | 1 |
| ENSMUSG00000006574 | Slc4a1 | 0 | 1.648835551 | 1 |
| ENSMUSG00000086105 | Gm11636 | 0 | 1.063873051 | 1 |
| ENSMUSG00000086283 | 2810433D01Rik | 0 | 1.063873051 | 1 |
| ENSMUSG00000051455 | Meioc | 0 | 1.063873051 | 1 |
| ENSMUSG00000020928 | Higd1b | 0 | 1.648835551 | 1 |
| ENSMUSG00000085338 | 2410004I01Rik | 0 | 1.648835551 | 1 |
| ENSMUSG00000043372 | Hexim2 | 0 | 0.063873051 | 1 |
| ENSMUSG00000044787 | Spata32 | 0 | 1.063873051 | 1 |
| ENSMUSG00000078635 | 1700028N14Rik | 0 | 0.063873051 | 1 |
| ENSMUSG00000034239 | Gm884 | 0 | 0.326907457 | 1 |
| ENSMUSG00000085154 | C130046K22Rik | 0 | -1.258055044 | 0.734875632 |
| ENSMUSG00000086794 | Gm11642 | 0 | 1.648835551 | 1 |
| ENSMUSG00000085134 | Gm11659 | 0 | 1.063873051 | 1 |
| ENSMUSG00000086487 | Gm11638 | 0 | 1.063873051 | 1 |
| ENSMUSG00000085297 | Gm11651 | 0 | 0.648835551 | 1 |
| ENSMUSG00000001901 | Kcnh6 | 0 | 0.063873051 | 1 |

| | | | | |
|---|---|---|---|---|
| ENSMUSG00000085255 | Taco1os | 0 | 0.326907457 | 1 |
| ENSMUSG00000040592 | Cd79b | 0 | 0.326907457 | 1 |
| ENSMUSG00000001027 | Scn4a | 0 | 1.648835551 | 1 |
| ENSMUSG00000009210 | Prr29 | 0 | 1.063873051 | 1 |
| ENSMUSG00000001029 | Icam2 | 0 | 1.648835551 | 1 |
| ENSMUSG00000078607 | 1810010H24Rik | 0 | 0.648835551 | 1 |
| ENSMUSG00000087596 | Gm11715 | 0 | 0.648835551 | 1 |
| ENSMUSG00000085245 | Gm11713 | 0 | 1.063873051 | 1 |
| ENSMUSG00000059706 | A830035A12Rik | 0 | -0.52108945 | 1 |
| ENSMUSG00000084978 | Gm11655 | 0 | 1.648835551 | 1 |
| ENSMUSG00000078605 | E030025P04Rik | 0 | 1.063873051 | 1 |
| ENSMUSG00000020599 | Rgs9 | 0 | -0.673092543 | 1 |
| ENSMUSG00000099630 | 1700096J18Rik | 0 | 1.648835551 | 1 |
| ENSMUSG00000056687 | Gm11696 | 0 | -1.810596067 | 0.473330531 |
| ENSMUSG00000045065 | 9930022D16Rik | 0 | 0.326907457 | 1 |
| ENSMUSG00000085326 | Gm15642 | 0 | 1.648835551 | 1 |
| ENSMUSG00000041920 | Slc16a6 | 0 | -1.52108945 | 0.557051363 |
| ENSMUSG00000086733 | Gm11685 | 0 | 0.648835551 | 1 |
| ENSMUSG00000085734 | Gm11684 | 0 | 0.326907457 | 1 |
| ENSMUSG00000044749 | Abca6 | 0 | -0.810596067 | 0.851250572 |
| ENSMUSG00000041695 | Kcnj2 | 0 | 1.648835551 | 1 |
| ENSMUSG00000086752 | Gm11674 | 0 | 1.648835551 | 1 |
| ENSMUSG00000075437 | Gm11681 | 0 | 1.648835551 | 1 |
| ENSMUSG00000047904 | Sstr2 | 0 | -0.351164449 | 1 |
| ENSMUSG00000085844 | Gm11690 | 0 | 0.648835551 | 1 |
| ENSMUSG00000034677 | Gpr142 | 0 | 1.063873051 | 1 |
| ENSMUSG00000058728 | Cd300c | 0 | 1.648835551 | 1 |
| ENSMUSG00000044811 | Cd300c2 | 0 | -1.351164449 | 0.736456705 |
| ENSMUSG00000018861 | Fdxr | 0 | -1.351164449 | 0.687112677 |
| ENSMUSG00000050201 | Otop2 | 0 | -0.810596067 | 0.844591606 |
| ENSMUSG00000050910 | Cdr2l | 0 | -2.995020638 | 0.046343118 |
| ENSMUSG00000085072 | Ict1os | 0 | 1.063873051 | 1 |
| ENSMUSG00000070332 | Trim80 | 0 | 1.063873051 | 1 |
| ENSMUSG00000046756 | Mrps7 | 0 | -2.480447466 | 0.13388699 |
| ENSMUSG00000057948 | Unc13d | 0 | -2.480447466 | 0.13388699 |
| ENSMUSG00000020793 | Galr2 | 0 | 0.326907457 | 1 |
| ENSMUSG00000070331 | Qrich2 | 0 | 0.326907457 | 1 |
| ENSMUSG00000100779 | Gm29292 | 0 | -0.810596067 | 0.848260747 |
| ENSMUSG00000061878 | Sphk1 | 0 | -2.636566667 | 0.112813335 |
| ENSMUSG00000020804 | Aanat | 0 | 1.648835551 | 1 |
| ENSMUSG00000020812 | Snhg16 | 0 | -1.743481871 | 0.455131546 |
| ENSMUSG00000057286 | St6galnac2 | 0 | -0.351164449 | 1 |
| ENSMUSG00000086902 | Gm11728 | 0 | 1.063873051 | 1 |
| ENSMUSG00000086050 | Gm16045 | 0 | 0.648835551 | 1 |
| ENSMUSG00000085419 | Gm11734 | 0 | 1.648835551 | 1 |
| ENSMUSG00000054418 | 2900041M22Rik | 0 | 0.648835551 | 1 |
| ENSMUSG00000050106 | Tmc8 | 0 | -2.9057533 | 0.056537412 |
| ENSMUSG00000025573 | 6030468B19Rik | 0 | 1.063873051 | 1 |
| ENSMUSG00000017713 | Tha1 | 0 | -1.051604167 | 0.858831093 |
| ENSMUSG00000085434 | Gm11725 | 0 | 1.648835551 | 1 |
| ENSMUSG00000017446 | C1qtnf1 | 0 | -0.52108945 | 1 |

| | | | | |
|---|---|---|---|---|
| ENSMUSG00000086514 | Gm11747 | 0 | 1.648835551 | 1 |
| ENSMUSG00000046697 | Enpp7 | 0 | 1.063873051 | 1 |
| ENSMUSG00000025578 | Cbx8 | 0 | 0.063873051 | 1 |
| ENSMUSG00000005043 | Sgsh | 0 | -1.43862729 | 1 |
| ENSMUSG00000025582 | Nptx1 | 0 | 0.326907457 | 1 |
| ENSMUSG00000087675 | Gm11762 | 0 | 0.326907457 | 1 |
| ENSMUSG00000085207 | Gm11767 | 0 | -0.158519371 | 1 |
| ENSMUSG00000039781 | Cep131 | 0 | -1.995020638 | 0.377748229 |
| ENSMUSG00000025377 | Tepsin | 0 | -1.351164449 | 0.703945114 |
| ENSMUSG00000085636 | Gm11769 | 0 | -0.158519371 | 1 |
| ENSMUSG00000043993 | 2900052L18Rik | 0 | -0.52108945 | 1 |
| ENSMUSG00000085501 | Gm11772 | 0 | 1.648835551 | 1 |
| ENSMUSG00000025380 | Fscn2 | 0 | 0.648835551 | 1 |
| ENSMUSG00000039670 | Oxld1 | 0 | 0.063873051 | 1 |
| ENSMUSG00000057594 | Arl16 | 0 | -0.52108945 | 1 |
| ENSMUSG00000025127 | Gcgr | 0 | 0.648835551 | 1 |
| ENSMUSG00000084822 | Myadml2os | 0 | 1.648835551 | 1 |
| ENSMUSG00000042988 | Notum | 0 | 1.063873051 | 1 |
| ENSMUSG00000090758 | Gm17178 | 0 | -1.158519371 | 0.886203497 |
| ENSMUSG00000025145 | Lrrc45 | 0 | -2.158519371 | 0.290284345 |
| ENSMUSG00000025150 | Cbr2 | 0 | 1.648835551 | 1 |
| ENSMUSG00000025158 | Rfng | 0 | -0.673092543 | 1 |
| ENSMUSG00000087638 | Gm11773 | 0 | 0.648835551 | 1 |
| ENSMUSG00000100629 | Gm28192 | 0 | -0.158519371 | 1 |
| ENSMUSG00000025163 | Cd7 | 0 | 1.648835551 | 1 |
| ENSMUSG00000025165 | Sectm1a | 0 | 0.326907457 | 1 |
| ENSMUSG00000039253 | Fn3krp | 0 | 0.326907457 | 1 |
| ENSMUSG00000025175 | Fn3k | 0 | 0.063873051 | 1 |
| ENSMUSG00000039238 | Zfp750 | 0 | 1.648835551 | 1 |
| ENSMUSG00000039208 | Metrnl | 0 | 0.326907457 | 1 |
| ENSMUSG00000097429 | Gm26520 | 0 | -1.673092543 | 0.451785624 |
| ENSMUSG00000073242 | Dnmt3aos | 0 | 1.063873051 | 1 |
| ENSMUSG00000020660 | Pomc | 0 | 1.648835551 | 1 |
| ENSMUSG00000111938 | 2900045O20Rik | 0 | -0.351164449 | 1 |
| ENSMUSG00000112505 | Gm48610 | 0 | 1.648835551 | 1 |
| ENSMUSG00000050545 | Fam228b | 0 | -1.43862729 | 0.538110273 |
| ENSMUSG00000020635 | Fkbp1b | 0 | 0.063873051 | 1 |
| ENSMUSG00000037336 | Mfsd2b | 0 | -0.810596067 | 0.844591606 |
| ENSMUSG00000112404 | Gm32442 | 0 | -0.810596067 | 1 |
| ENSMUSG00000102096 | 1700101O22Rik | 0 | 1.648835551 | 1 |
| ENSMUSG00000112006 | Gm48633 | 0 | 1.063873051 | 1 |
| ENSMUSG00000020605 | Hs1bp3 | 0 | -2.599091962 | 0.108201045 |
| ENSMUSG00000112452 | Gm48075 | 0 | 0.326907457 | 1 |
| ENSMUSG00000042976 | 9930038B18Rik | 0 | -0.158519371 | 1 |
| ENSMUSG00000114000 | Gm48762 | 0 | 1.648835551 | 1 |
| ENSMUSG00000112496 | Gm48764 | 0 | 1.648835551 | 1 |
| ENSMUSG00000099360 | 1700022H16Rik | 0 | 0.648835551 | 1 |
| ENSMUSG00000048387 | Osr1 | 0 | -0.158519371 | 1 |
| ENSMUSG00000113620 | 7420701I03Rik | 0 | 0.326907457 | 1 |
| ENSMUSG00000043673 | Kcns3 | 0 | 0.063873051 | 1 |
| ENSMUSG00000113226 | Gm47808 | 0 | 1.063873051 | 1 |

| | | | | |
|---|---|---|---|---|
| ENSMUSG00000113330 | Gm34237 | 0 | -0.52108945 | 1 |
| ENSMUSG00000091430 | Platr19 | 0 | 0.648835551 | 1 |
| ENSMUSG00000113363 | Gm48187 | 0 | 1.648835551 | 1 |
| ENSMUSG00000037169 | Mycn | 0 | -3.395558568 | 0.014189547 |
| ENSMUSG00000112988 | Gm35298 | 0 | 1.648835551 | 1 |
| ENSMUSG00000113284 | Gm48213 | 0 | 1.648835551 | 1 |
| ENSMUSG00000113058 | Gm48311 | 0 | 1.648835551 | 1 |
| ENSMUSG00000113283 | Gm35890 | 0 | 1.648835551 | 1 |
| ENSMUSG00000113336 | Gm16497 | 0 | 0.326907457 | 1 |
| ENSMUSG00000113644 | 4930448C13Rik | 0 | 0.326907457 | 1 |
| ENSMUSG00000113734 | Gm48539 | 0 | 1.648835551 | 1 |
| ENSMUSG00000020601 | Trib2 | 0 | -2.258055044 | 0.276617983 |
| ENSMUSG00000113930 | Gm36235 | 0 | 0.326907457 | 1 |
| ENSMUSG00000113975 | Gm36372 | 0 | -0.158519371 | 1 |
| ENSMUSG00000113246 | Gm36752 | 0 | 1.063873051 | 1 |
| ENSMUSG00000113093 | Gm36862 | 0 | 1.648835551 | 1 |
| ENSMUSG00000114062 | Gm48313 | 0 | 0.326907457 | 1 |
| ENSMUSG00000114188 | Gm40849 | 0 | 0.063873051 | 1 |
| ENSMUSG00000093908 | Gm5784 | 0 | 1.648835551 | 1 |
| ENSMUSG00000116705 | Gm29737 | 0 | 1.648835551 | 1 |
| ENSMUSG00000116537 | Gm49698 | 0 | 0.648835551 | 1 |
| ENSMUSG00000116883 | Gm49700 | 0 | 1.648835551 | 1 |
| ENSMUSG00000116627 | Gm30124 | 0 | -0.158519371 | 1 |
| ENSMUSG00000116819 | Gm49586 | 0 | 1.648835551 | 1 |
| ENSMUSG00000097055 | Gm4419 | 0 | -1.673092543 | 0.451785624 |
| ENSMUSG00000114006 | Gm47997 | 0 | 1.648835551 | 1 |
| ENSMUSG00000112937 | Gm17746 | 0 | -0.158519371 | 1 |
| ENSMUSG00000114033 | Gm47705 | 0 | 0.063873051 | 1 |
| ENSMUSG00000114105 | Gm47713 | 0 | 1.648835551 | 1 |
| ENSMUSG00000113115 | 4930480M12Rik | 0 | 0.063873051 | 1 |
| ENSMUSG00000114048 | 4930549C15Rik | 0 | 1.648835551 | 1 |
| ENSMUSG00000020638 | Cmpk2 | 0 | 0.648835551 | 1 |
| ENSMUSG00000113524 | Gm47847 | 0 | -0.810596067 | 1 |
| ENSMUSG00000113218 | 4933409F18Rik | 0 | 0.648835551 | 1 |
| ENSMUSG00000113421 | Gm4166 | 0 | 1.063873051 | 1 |
| ENSMUSG00000113967 | Gm31025 | 0 | 1.063873051 | 1 |
| ENSMUSG00000113235 | Gm47872 | 0 | -0.936126949 | 1 |
| ENSMUSG00000036655 | Colec11 | 0 | 1.648835551 | 1 |
| ENSMUSG00000020628 | Trappc12 | 0 | -3.579983139 | 0.006320625 |
| ENSMUSG00000112390 | Gm20208 | 0 | 1.648835551 | 1 |
| ENSMUSG00000043061 | Tmem18 | 0 | -1.051604167 | 0.858831093 |
| ENSMUSG00000054204 | Alkal2 | 0 | 1.063873051 | 1 |
| ENSMUSG00000095953 | 6030469F06Rik | 0 | 0.648835551 | 1 |
| ENSMUSG00000020651 | Slc26a4 | 0 | -0.673092543 | 1 |
| ENSMUSG00000112071 | 5430401H09Rik | 0 | -0.936126949 | 0.859947725 |
| ENSMUSG00000112865 | Gm40375 | 0 | -2.051604167 | 0.440632126 |
| ENSMUSG00000112166 | Gm47948 | 0 | 1.648835551 | 1 |
| ENSMUSG00000084979 | Gm16267 | 0 | 0.326907457 | 1 |
| ENSMUSG00000020562 | Efcab10 | 0 | 1.648835551 | 1 |
| ENSMUSG00000113386 | Gm47357 | 0 | 1.648835551 | 1 |
| ENSMUSG00000036231 | Agr3 | 0 | 1.063873051 | 1 |

| | | | | |
|---|---|---|---|---|
| ENSMUSG00000020545 | Lrrc72 | 0 | 0.648835551 | 1 |
| ENSMUSG00000113832 | Gm40392 | 0 | 0.063873051 | 1 |
| ENSMUSG00000114141 | Gm34215 | 0 | 1.648835551 | 1 |
| ENSMUSG00000113064 | Gm40394 | 0 | 1.063873051 | 1 |
| ENSMUSG00000113764 | Gm48617 | 0 | -1.43862729 | 0.538110273 |
| ENSMUSG00000112967 | 4930428E07Rik | 0 | -1.351164449 | 0.73705235 |
| ENSMUSG00000113092 | Gm34662 | 0 | 1.648835551 | 1 |
| ENSMUSG00000113272 | Gm34809 | 0 | -0.673092543 | 1 |
| ENSMUSG00000113343 | Gm34923 | 0 | 0.326907457 | 1 |
| ENSMUSG00000113874 | Gm47868 | 0 | 1.063873051 | 1 |
| ENSMUSG00000002565 | Scin | 0 | -0.936126949 | 0.874192437 |
| ENSMUSG00000090582 | Gm17024 | 0 | 1.063873051 | 1 |
| ENSMUSG00000112904 | Gm47030 | 0 | -0.52108945 | 1 |
| ENSMUSG00000086012 | Gm15902 | 0 | -1.158519371 | 0.667778757 |
| ENSMUSG00000112112 | Gm48508 | 0 | -0.673092543 | 1 |
| ENSMUSG00000112859 | Gm48522 | 0 | 1.063873051 | 1 |
| ENSMUSG00000112174 | Gm48535 | 0 | 1.648835551 | 1 |
| ENSMUSG00000112410 | B230217J21Rik | 0 | 1.648835551 | 1 |
| ENSMUSG00000112353 | Gm40418 | 0 | 0.063873051 | 1 |
| ENSMUSG00000100315 | 1700031P21Rik | 0 | 1.063873051 | 1 |
| ENSMUSG00000112412 | Gm35239 | 0 | 1.063873051 | 1 |
| ENSMUSG00000099407 | 1700060O08Rik | 0 | 1.063873051 | 1 |
| ENSMUSG00000112053 | Gm46328 | 0 | 1.648835551 | 1 |
| ENSMUSG00000113947 | Gm10465 | 0 | 1.063873051 | 1 |
| ENSMUSG00000095595 | Fam177a | 0 | -0.351164449 | 1 |
| ENSMUSG00000094103 | 1700047I17Rik2 | 0 | 1.648835551 | 1 |
| ENSMUSG00000112348 | Gm36634 | 0 | 1.648835551 | 1 |
| ENSMUSG00000112164 | Gm19990 | 0 | 1.063873051 | 1 |
| ENSMUSG00000098172 | Gm26973 | 0 | -0.673092543 | 1 |
| ENSMUSG00000086372 | Gm16246 | 0 | 1.063873051 | 1 |
| ENSMUSG00000035451 | Foxa1 | 0 | 1.648835551 | 1 |
| ENSMUSG00000035431 | Sstr1 | 0 | 1.648835551 | 1 |
| ENSMUSG00000045930 | Clec14a | 0 | 0.648835551 | 1 |
| ENSMUSG00000113429 | Gm20063 | 0 | 1.063873051 | 1 |
| ENSMUSG00000113152 | Gm48422 | 0 | 1.063873051 | 1 |
| ENSMUSG00000047227 | Gm527 | 0 | -0.936126949 | 0.847265357 |
| ENSMUSG00000035560 | Wdr20rt | 0 | 0.648835551 | 1 |
| ENSMUSG00000113867 | Gm47456 | 0 | 0.063873051 | 1 |
| ENSMUSG00000034883 | Lrr1 | 0 | -1.158519371 | 0.667778757 |
| ENSMUSG00000020973 | Dnaaf2 | 0 | -1.43862729 | 0.538110273 |
| ENSMUSG00000097061 | 9330151L19Rik | 0 | -0.158519371 | 1 |
| ENSMUSG00000052673 | Gm9887 | 0 | 1.648835551 | 1 |
| ENSMUSG00000113035 | 5830428M24Rik | 0 | -1.258055044 | 0.673811773 |
| ENSMUSG00000114099 | Gm48695 | 0 | 1.063873051 | 1 |
| ENSMUSG00000113944 | Gm48847 | 0 | 1.648835551 | 1 |
| ENSMUSG00000112950 | Gm32369 | 0 | 1.063873051 | 1 |
| ENSMUSG00000113584 | Gm46355 | 0 | 1.648835551 | 1 |
| ENSMUSG00000113707 | Gm10457 | 0 | 1.648835551 | 1 |
| ENSMUSG00000021078 | Tomm20l | 0 | -0.158519371 | 1 |
| ENSMUSG00000044548 | Dact1 | 0 | -1.936126949 | 0.358945041 |
| ENSMUSG00000043398 | Gpr135 | 0 | 0.326907457 | 1 |

| | | | | |
|---|---|---|---|---|
| ENSMUSG00000021094 | Dhrs7 | 0 | -2.351164449 | 0.169846604 |
| ENSMUSG00000021099 | Six6 | 0 | 1.648835551 | 1 |
| ENSMUSG00000097010 | Gm26709 | 0 | 0.063873051 | 1 |
| ENSMUSG00000051367 | Six1 | 0 | 1.648835551 | 1 |
| ENSMUSG00000034460 | Six4 | 0 | -0.351164449 | 1 |
| ENSMUSG00000113425 | Gm48653 | 0 | 1.648835551 | 1 |
| ENSMUSG00000114178 | Gm48656 | 0 | 1.648835551 | 1 |
| ENSMUSG00000113321 | Gm8075 | 0 | 1.648835551 | 1 |
| ENSMUSG00000029878 | Dbpht2 | 0 | 1.063873051 | 1 |
| ENSMUSG00000113031 | Gm34552 | 0 | 1.648835551 | 1 |
| ENSMUSG00000113974 | 4930442G10Rik | 0 | 1.648835551 | 1 |
| ENSMUSG00000048982 | Gphb5 | 0 | 1.063873051 | 1 |
| ENSMUSG00000021055 | Esr2 | 0 | -1.351164449 | 0.775097267 |
| ENSMUSG00000097323 | 4930426I24Rik | 0 | -0.351164449 | 1 |
| ENSMUSG00000073000 | Gm10451 | 0 | 0.326907457 | 1 |
| ENSMUSG00000112030 | Gm35189 | 0 | 1.063873051 | 1 |
| ENSMUSG00000112150 | 9530018F02Rik | 0 | -0.158519371 | 1 |
| ENSMUSG00000111903 | 4930458K08Rik | 0 | 1.648835551 | 1 |
| ENSMUSG00000099418 | Gm6657 | 0 | 1.648835551 | 1 |
| ENSMUSG00000112302 | Gm48226 | 0 | 0.063873051 | 1 |
| ENSMUSG00000056987 | Fam71d | 0 | 0.326907457 | 1 |
| ENSMUSG00000111929 | Gm48780 | 0 | 1.648835551 | 1 |
| ENSMUSG00000112765 | 9430078K24Rik | 0 | 1.063873051 | 1 |
| ENSMUSG00000111933 | Gm36660 | 0 | -1.351164449 | 0.733465149 |
| ENSMUSG00000112386 | Gm47752 | 0 | -1.351164449 | 0.707847898 |
| ENSMUSG00000112545 | 1300014J16Rik | 0 | -0.673092543 | 1 |
| ENSMUSG00000112336 | Gm47766 | 0 | 1.063873051 | 1 |
| ENSMUSG00000112285 | Gm50278 | 0 | 1.648835551 | 1 |
| ENSMUSG00000112701 | 2310002D06Rik | 0 | 0.648835551 | 1 |
| ENSMUSG00000112421 | 4933406B15Rik | 0 | 1.648835551 | 1 |
| ENSMUSG00000112288 | Gm47879 | 0 | 1.648835551 | 1 |
| ENSMUSG00000021135 | Slc10a1 | 0 | 0.648835551 | 1 |
| ENSMUSG00000098128 | Gm3693 | 0 | 1.648835551 | 1 |
| ENSMUSG00000072974 | Gm4787 | 0 | -1.810596067 | 0.465580427 |
| ENSMUSG00000099478 | Gm28370 | 0 | 0.063873051 | 1 |
| ENSMUSG00000008438 | Adam21 | 0 | 1.648835551 | 1 |
| ENSMUSG00000113950 | Gm47080 | 0 | -0.158519371 | 1 |
| ENSMUSG00000114150 | Gm46367 | 0 | 1.063873051 | 1 |
| ENSMUSG00000052392 | Acot4 | 0 | 0.648835551 | 1 |
| ENSMUSG00000097762 | 4732463B04Rik | 0 | 1.648835551 | 1 |
| ENSMUSG00000021228 | Acot3 | 0 | 1.063873051 | 1 |
| ENSMUSG00000043487 | Acot6 | 0 | 1.063873051 | 1 |
| ENSMUSG00000054383 | Pnma1 | 0 | 0.648835551 | 1 |
| ENSMUSG00000113052 | Gm31513 | 0 | 1.063873051 | 1 |
| ENSMUSG00000113529 | Gm47484 | 0 | -0.673092543 | 1 |
| ENSMUSG00000021234 | Fam161b | 0 | 0.326907457 | 1 |
| ENSMUSG00000021235 | Coq6 | 0 | -1.051604167 | 0.858831093 |
| ENSMUSG00000098134 | Rnf113a2 | 0 | -1.351164449 | 0.716109464 |
| ENSMUSG00000021239 | Vsx2 | 0 | -0.52108945 | 1 |
| ENSMUSG00000071234 | Syndig1l | 0 | -1.158519371 | 0.899603262 |
| ENSMUSG00000092130 | D030025P21Rik | 0 | 1.063873051 | 1 |

| | | | | |
|---|---|---|---|---|
| ENSMUSG00000079645 | Gm17193 | 0 | 1.648835551 | 1 |
| ENSMUSG00000093629 | Prox2os | 0 | -1.43862729 | 0.538110273 |
| ENSMUSG00000045064 | Zc2hc1c | 0 | -1.158519371 | 0.667778757 |
| ENSMUSG00000113488 | Gm47819 | 0 | 0.648835551 | 1 |
| ENSMUSG00000114163 | Gm32296 | 0 | 1.648835551 | 1 |
| ENSMUSG00000113165 | Gm47863 | 0 | 1.648835551 | 1 |
| ENSMUSG00000034266 | Batf | 0 | -1.051604167 | 0.886203497 |
| ENSMUSG00000097902 | Gm26531 | 0 | -0.351164449 | 1 |
| ENSMUSG00000114051 | Gm48008 | 0 | 1.648835551 | 1 |
| ENSMUSG00000113918 | Gm6566 | 0 | 1.648835551 | 1 |
| ENSMUSG00000090812 | Samd15 | 0 | 1.063873051 | 1 |
| ENSMUSG00000100306 | Gm29362 | 0 | 1.063873051 | 1 |
| ENSMUSG00000095724 | Gm21319 | 0 | 1.648835551 | 1 |
| ENSMUSG00000072905 | Gm2016 | 0 | 1.648835551 | 1 |
| ENSMUSG00000066027 | Gm10436 | 0 | 1.648835551 | 1 |
| ENSMUSG00000113434 | 3200001D21Rik | 0 | -0.158519371 | 1 |
| ENSMUSG00000114001 | Gm48664 | 0 | 1.648835551 | 1 |
| ENSMUSG00000113539 | Gm47684 | 0 | 0.326907457 | 1 |
| ENSMUSG00000113943 | Gm47804 | 0 | 1.648835551 | 1 |
| ENSMUSG00000090722 | Gm8378 | 0 | -1.351164449 | 0.687112677 |
| ENSMUSG00000086689 | Gm16876 | 0 | 1.648835551 | 1 |
| ENSMUSG00000021886 | Gpr65 | 0 | 0.648835551 | 1 |
| ENSMUSG00000033854 | Kcnk10 | 0 | -1.52108945 | 0.586600655 |
| ENSMUSG00000114021 | Gm47109 | 0 | 1.063873051 | 1 |
| ENSMUSG00000114089 | Gm47116 | 0 | 1.648835551 | 1 |
| ENSMUSG00000033731 | 3300002A11Rik | 0 | 0.648835551 | 1 |
| ENSMUSG00000113147 | Gm47207 | 0 | 1.063873051 | 1 |
| ENSMUSG00000113268 | 4930477G07Rik | 0 | 0.063873051 | 1 |
| ENSMUSG00000021179 | Nrde2 | 0 | -3.636566667 | 0.005443828 |
| ENSMUSG00000072884 | Gm10433 | 0 | 0.063873051 | 1 |
| ENSMUSG00000113362 | Gm40557 | 0 | 1.063873051 | 1 |
| ENSMUSG00000113895 | Gm48383 | 0 | -1.351164449 | 0.687112677 |
| ENSMUSG00000047415 | Gpr68 | 0 | -1.52108945 | 0.587484274 |
| ENSMUSG00000097160 | A630072L19Rik | 0 | 1.648835551 | 1 |
| ENSMUSG00000097121 | D130020L05Rik | 0 | -0.158519371 | 1 |
| ENSMUSG00000113265 | Gm30198 | 0 | -0.52108945 | 1 |
| ENSMUSG00000021194 | Chga | 0 | 0.326907457 | 1 |
| ENSMUSG00000021200 | Asb2 | 0 | -0.936126949 | 0.869664742 |
| ENSMUSG00000090185 | Gm15523 | 0 | 0.648835551 | 1 |
| ENSMUSG00000021203 | Otub2 | 0 | -2.158519371 | 0.303134739 |
| ENSMUSG00000061947 | Serpina10 | 0 | 1.648835551 | 1 |
| ENSMUSG00000066366 | Serpina1a | 0 | 1.648835551 | 1 |
| ENSMUSG00000079015 | Serpina1c | 0 | -0.351164449 | 1 |
| ENSMUSG00000063232 | Serpina11 | 0 | 0.648835551 | 1 |
| ENSMUSG00000066363 | Serpina3f | 0 | 1.648835551 | 1 |
| ENSMUSG00000041481 | Serpina3g | 0 | 1.063873051 | 1 |
| ENSMUSG00000079014 | Serpina3i | 0 | 0.063873051 | 1 |
| ENSMUSG00000079012 | Serpina3m | 0 | 0.063873051 | 1 |
| ENSMUSG00000112954 | Gm49732 | 0 | 0.326907457 | 1 |
| ENSMUSG00000113995 | Gm47648 | 0 | 1.063873051 | 1 |
| ENSMUSG00000060863 | Tcl1b2 | 0 | 0.648835551 | 1 |

| | | | | |
|---|---|---|---|---|
| ENSMUSG00000068940 | Tcl1b3 | 0 | 1.063873051 | 1 |
| ENSMUSG00000079007 | Tcl1b4 | 0 | 0.326907457 | 1 |
| ENSMUSG00000097929 | Tunar | 0 | 0.063873051 | 1 |
| ENSMUSG00000113314 | Gm2721 | 0 | 1.648835551 | 1 |
| ENSMUSG00000094910 | D430019H16Rik | 0 | -0.158519371 | 1 |
| ENSMUSG00000113535 | AU015791 | 0 | 1.648835551 | 1 |
| ENSMUSG00000113537 | Gm48807 | 0 | 0.063873051 | 1 |
| ENSMUSG00000113390 | Gm32635 | 0 | 1.648835551 | 1 |
| ENSMUSG00000092066 | 1700121N20Rik | 0 | 1.648835551 | 1 |
| ENSMUSG00000091583 | Gm17033 | 0 | -0.158519371 | 1 |
| ENSMUSG00000085820 | 4933406K04Rik | 0 | -0.351164449 | 1 |
| ENSMUSG00000086319 | Gm16087 | 0 | 1.648835551 | 1 |
| ENSMUSG00000084871 | Gm2800 | 0 | 1.063873051 | 1 |
| ENSMUSG00000085641 | 4930465M20Rik | 0 | 1.648835551 | 1 |
| ENSMUSG00000086975 | Gm15208 | 0 | -0.351164449 | 1 |
| ENSMUSG00000048251 | Bcl11b | 0 | -4.065409966 | 0.000965898 |
| ENSMUSG00000097847 | 4930478K11Rik | 0 | 0.326907457 | 1 |
| ENSMUSG00000097788 | Gm16596 | 0 | 0.648835551 | 1 |
| ENSMUSG00000113396 | Gm33467 | 0 | 1.648835551 | 1 |
| ENSMUSG00000021265 | Slc25a29 | 0 | 0.063873051 | 1 |
| ENSMUSG00000040856 | Dlk1 | 0 | -0.52108945 | 1 |
| ENSMUSG00000085925 | Rtl1 | 0 | -2.051604167 | 0.45888423 |
| ENSMUSG00000113733 | Gm34785 | 0 | 0.648835551 | 1 |
| ENSMUSG00000075707 | Dio3 | 0 | 1.063873051 | 1 |
| ENSMUSG00000113361 | B930059L03Rik | 0 | 1.648835551 | 1 |
| ENSMUSG00000056508 | 1700001K19Rik | 0 | 0.063873051 | 1 |
| ENSMUSG00000021276 | Cinp | 0 | -2.636566667 | 0.112813335 |
| ENSMUSG00000037904 | Ankrd9 | 0 | 0.326907457 | 1 |
| ENSMUSG00000083193 | 4930595D18Rik | 0 | 0.326907457 | 1 |
| ENSMUSG00000021278 | Amn | 0 | 0.648835551 | 1 |
| ENSMUSG00000097081 | Gm10425 | 0 | 0.648835551 | 1 |
| ENSMUSG00000113029 | Gm40578 | 0 | -0.936126949 | 0.877701541 |
| ENSMUSG00000096995 | 2810029C07Rik | 0 | 1.648835551 | 1 |
| ENSMUSG00000112991 | Gm36635 | 0 | 0.648835551 | 1 |
| ENSMUSG00000090026 | Gm15996 | 0 | 1.648835551 | 1 |
| ENSMUSG00000021287 | Xrcc3 | 0 | 1.063873051 | 1 |
| ENSMUSG00000021286 | Zfyve21 | 0 | -1.810596067 | 0.465580427 |
| ENSMUSG00000113769 | 5033406O09Rik | 0 | -1.051604167 | 0.858831093 |
| ENSMUSG00000037686 | Aspg | 0 | 1.648835551 | 1 |
| ENSMUSG00000113189 | B020018J22Rik | 0 | 1.063873051 | 1 |
| ENSMUSG00000043122 | A530016L24Rik | 0 | 0.648835551 | 1 |
| ENSMUSG00000052160 | Pld4 | 0 | -0.158519371 | 1 |
| ENSMUSG00000021298 | Gpr132 | 0 | -1.810596067 | 0.470945303 |
| ENSMUSG00000097100 | 9230104M06Rik | 0 | 1.063873051 | 1 |
| ENSMUSG00000037466 | Tedc1 | 0 | -1.351164449 | 0.687112677 |
| ENSMUSG00000049036 | Tmem121 | 0 | 1.063873051 | 1 |
| ENSMUSG00000076612 | Ighg2c | 0 | -0.351164449 | 1 |
| ENSMUSG00000076613 | Ighg2b | 0 | -0.158519371 | 1 |
| ENSMUSG00000076615 | Ighg3 | 0 | 1.648835551 | 1 |
| ENSMUSG00000104213 | Ighd | 0 | -2.480447466 | 0.13388699 |
| ENSMUSG00000076633 | Ighv5-2 | 0 | 1.648835551 | 1 |

| | | | | |
|---|---|---|---|---|
| ENSMUSG00000095866 | Ighv2-4 | 0 | 1.648835551 | 1 |
| ENSMUSG00000103262 | Ighv5-8 | 0 | 1.648835551 | 1 |
| ENSMUSG00000103240 | Gm37976 | 0 | 1.648835551 | 1 |
| ENSMUSG00000096824 | Ighv2-7 | 0 | 1.648835551 | 1 |
| ENSMUSG00000102878 | Ighv2-8 | 0 | 1.648835551 | 1 |
| ENSMUSG00000076665 | Ighv7-1 | 0 | 1.063873051 | 1 |
| ENSMUSG00000076655 | Ighv4-1 | 0 | 1.648835551 | 1 |
| ENSMUSG00000095583 | Ighv14-2 | 0 | 1.648835551 | 1 |
| ENSMUSG00000103271 | Ighv4-2 | 0 | 1.648835551 | 1 |
| ENSMUSG00000076680 | Ighv6-6 | 0 | 0.326907457 | 1 |
| ENSMUSG00000076695 | Ighv1-18 | 0 | 1.648835551 | 1 |
| ENSMUSG00000102942 | Ighv1-33 | 0 | 1.648835551 | 1 |
| ENSMUSG00000095519 | Ighv1-66 | 0 | 1.063873051 | 1 |
| ENSMUSG00000094124 | Ighv1-74 | 0 | 1.648835551 | 1 |
| ENSMUSG00000094075 | Ighv1-80 | 0 | 1.648835551 | 1 |
| ENSMUSG00000095127 | Ighv1-82 | 0 | 0.648835551 | 1 |
| ENSMUSG00000105757 | Ighv1-83 | 0 | 1.648835551 | 1 |
| ENSMUSG00000011171 | Vipr2 | 0 | -0.158519371 | 1 |
| ENSMUSG00000093765 | Gm20658 | 0 | 0.648835551 | 1 |
| ENSMUSG00000112980 | D430020J02Rik | 0 | 1.063873051 | 1 |
| ENSMUSG00000099907 | Gm10421 | 0 | 1.063873051 | 1 |
| ENSMUSG00000113271 | Gm40653 | 0 | 1.648835551 | 1 |
| ENSMUSG00000112968 | Gm48530 | 0 | -0.673092543 | 1 |
| ENSMUSG00000113259 | Gm35190 | 0 | 1.648835551 | 1 |
| ENSMUSG00000021216 | Tubal3 | 0 | -0.52108945 | 1 |
| ENSMUSG00000044988 | Ucn3 | 0 | 1.648835551 | 1 |
| ENSMUSG00000113788 | Gm47904 | 0 | 1.648835551 | 1 |
| ENSMUSG00000021214 | Akr1c18 | 0 | 1.648835551 | 1 |
| ENSMUSG00000021210 | Akr1c6 | 0 | 0.063873051 | 1 |
| ENSMUSG00000113717 | Gm40658 | 0 | -1.995020638 | 0.416376397 |
| ENSMUSG00000053499 | Gm5444 | 0 | -0.158519371 | 1 |
| ENSMUSG00000113866 | Gm47450 | 0 | 1.648835551 | 1 |
| ENSMUSG00000100625 | 1700016G22Rik | 0 | 1.063873051 | 1 |
| ENSMUSG00000114097 | Gm35615 | 0 | 1.648835551 | 1 |
| ENSMUSG00000101540 | Gm28465 | 0 | 1.063873051 | 1 |
| ENSMUSG00000021193 | Pitrm1 | 0 | -3.82689788 | 0.002928655 |
| ENSMUSG00000113455 | Gm48825 | 0 | 1.063873051 | 1 |
| ENSMUSG00000113680 | Gm47405 | 0 | 1.648835551 | 1 |
| ENSMUSG00000114134 | Gm47408 | 0 | -0.158519371 | 1 |
| ENSMUSG00000114115 | Gm30239 | 0 | 0.326907457 | 1 |
| ENSMUSG00000079092 | Prl2c2 | 0 | 0.648835551 | 1 |
| ENSMUSG00000113049 | Gm48544 | 0 | 1.648835551 | 1 |
| ENSMUSG00000055360 | Prl2c5 | 0 | 1.063873051 | 1 |
| ENSMUSG00000021303 | Gng4 | 0 | 0.326907457 | 1 |
| ENSMUSG00000113775 | Gm48027 | 0 | -1.743481871 | 0.455131546 |
| ENSMUSG00000113410 | Gm20043 | 0 | 0.648835551 | 1 |
| ENSMUSG00000097806 | Gm6556 | 0 | -0.158519371 | 1 |
| ENSMUSG00000113960 | 4933412O06Rik | 0 | 0.648835551 | 1 |
| ENSMUSG00000113322 | Gm48492 | 0 | 1.648835551 | 1 |
| ENSMUSG00000113100 | Gm48754 | 0 | 0.648835551 | 1 |
| ENSMUSG00000113847 | Gm48799 | 0 | 1.063873051 | 1 |

| | | | | |
|---|---|---|---|---|
| ENSMUSG00000113470 | 4930448F12Rik | 0 | 1.648835551 | 1 |
| ENSMUSG00000114118 | Gm48829 | 0 | 1.063873051 | 1 |
| ENSMUSG00000114080 | Gm47661 | 0 | 0.648835551 | 1 |
| ENSMUSG00000105715 | Gm31887 | 0 | 0.648835551 | 1 |
| ENSMUSG00000076752 | Tcrg-C2 | 0 | 1.648835551 | 1 |
| ENSMUSG00000076754 | Trgv2 | 0 | 1.648835551 | 1 |
| ENSMUSG00000002808 | Epdr1 | 0 | -0.158519371 | 1 |
| ENSMUSG00000021319 | Sfrp4 | 0 | 0.648835551 | 1 |
| ENSMUSG00000113183 | Gm47664 | 0 | 0.326907457 | 1 |
| ENSMUSG00000004344 | Gpx5 | 0 | 0.648835551 | 1 |
| ENSMUSG00000004341 | Gpx6 | 0 | 1.063873051 | 1 |
| ENSMUSG00000055313 | Pgbd1 | 0 | -1.351164449 | 0.732525536 |
| ENSMUSG00000085788 | 4930470G03Rik | 0 | -0.158519371 | 1 |
| ENSMUSG00000114702 | Gm40828 | 0 | 1.648835551 | 1 |
| ENSMUSG00000054931 | Zkscan4 | 0 | 0.326907457 | 1 |
| ENSMUSG00000078139 | AK157302 | 0 | 1.063873051 | 1 |
| ENSMUSG00000049737 | Olfr1361 | 0 | 1.648835551 | 1 |
| ENSMUSG00000108674 | Olfr1359 | 0 | 1.648835551 | 1 |
| ENSMUSG00000094338 | Hist1h2bl | 0 | 0.648835551 | 1 |
| ENSMUSG00000101355 | Hist1h3h | 0 | 1.648835551 | 1 |
| ENSMUSG00000064288 | Hist1h4k | 0 | 1.648835551 | 1 |
| ENSMUSG00000069309 | Hist1h2an | 0 | 1.648835551 | 1 |
| ENSMUSG00000094777 | Hist1h2ap | 0 | -0.158519371 | 1 |
| ENSMUSG00000069303 | Hist1h2br | 0 | 1.648835551 | 1 |
| ENSMUSG00000016982 | Pom121l2 | 0 | 1.648835551 | 1 |
| ENSMUSG00000006179 | Prss16 | 0 | 1.648835551 | 1 |
| ENSMUSG00000085827 | Gm11290 | 0 | -2.480447466 | 0.882296038 |
| ENSMUSG00000043145 | Gm11292 | 0 | 0.648835551 | 1 |
| ENSMUSG00000062727 | Hist1h2bk | 0 | 1.648835551 | 1 |
| ENSMUSG00000060639 | Hist1h4i | 0 | -0.158519371 | 1 |
| ENSMUSG00000069299 | Vmn1r188 | 0 | 1.648835551 | 1 |
| ENSMUSG00000046932 | Vmn1r193 | 0 | 1.648835551 | 1 |
| ENSMUSG00000069296 | Vmn1r195 | 0 | 1.063873051 | 1 |
| ENSMUSG00000069292 | Vmn1r199 | 0 | 1.648835551 | 1 |
| ENSMUSG00000094898 | Vmn1r201 | 0 | 1.648835551 | 1 |
| ENSMUSG00000094379 | Vmn1r202 | 0 | 1.648835551 | 1 |
| ENSMUSG00000069289 | Vmn1r203 | 0 | 0.648835551 | 1 |
| ENSMUSG00000087688 | Gm11300 | 0 | 1.648835551 | 1 |
| ENSMUSG00000100296 | Vmn1r205 | 0 | 1.063873051 | 1 |
| ENSMUSG00000101578 | Vmn1r206 | 0 | 1.063873051 | 1 |
| ENSMUSG00000085221 | Gm11309 | 0 | 0.326907457 | 1 |
| ENSMUSG00000071493 | Vmn1r208 | 0 | 1.063873051 | 1 |
| ENSMUSG00000071491 | Vmn1r209 | 0 | 1.648835551 | 1 |
| ENSMUSG00000061296 | Vmn1r210 | 0 | 1.648835551 | 1 |
| ENSMUSG00000063998 | Vmn1r211 | 0 | 0.648835551 | 1 |
| ENSMUSG00000060024 | Vmn1r213 | 0 | 1.063873051 | 1 |
| ENSMUSG00000099917 | Vmn1r215 | 0 | 1.648835551 | 1 |
| ENSMUSG00000115791 | Vmn1r217 | 0 | 1.648835551 | 1 |
| ENSMUSG00000061376 | Vmn1r219 | 0 | 1.648835551 | 1 |
| ENSMUSG00000096099 | Vmn1r220 | 0 | 0.648835551 | 1 |
| ENSMUSG00000061022 | Vmn1r222 | 0 | 0.648835551 | 1 |

| | | | | |
|---|---|---|---|---|
| ENSMUSG00000069280 | Vmn1r223 | 0 | 1.648835551 | 1 |
| ENSMUSG00000085122 | 4933404K08Rik | 0 | 0.648835551 | 1 |
| ENSMUSG00000046351 | Zfp322a | 0 | -4.717486663 | 3.90E-05 |
| ENSMUSG00000085024 | C230035I16Rik | 0 | 0.326907457 | 1 |
| ENSMUSG00000113440 | Gm46404 | 0 | 0.063873051 | 1 |
| ENSMUSG00000000706 | Btn1a1 | 0 | 0.063873051 | 1 |
| ENSMUSG00000053216 | Btn2a2 | 0 | 0.648835551 | 1 |
| ENSMUSG00000061991 | Hist1h2af | 0 | 1.063873051 | 1 |
| ENSMUSG00000100210 | Hist1h3f | 0 | 1.063873051 | 1 |
| ENSMUSG00000069273 | Hist1h3e | 0 | 0.648835551 | 1 |
| ENSMUSG00000058385 | Hist1h2bg | 0 | 1.648835551 | 1 |
| ENSMUSG00000099583 | Hist1h3d | 0 | 1.648835551 | 1 |
| ENSMUSG00000069270 | Hist1h2ac | 0 | 1.063873051 | 1 |
| ENSMUSG00000060678 | Hist1h4c | 0 | 1.648835551 | 1 |
| ENSMUSG00000006611 | Hfe | 0 | 1.648835551 | 1 |
| ENSMUSG00000069310 | Hist1h3c | 0 | 1.648835551 | 1 |
| ENSMUSG00000075031 | Hist1h2bb | 0 | 0.648835551 | 1 |
| ENSMUSG00000069267 | Hist1h3b | 0 | 1.648835551 | 1 |
| ENSMUSG00000049539 | Hist1h1a | 0 | 0.648835551 | 1 |
| ENSMUSG00000036083 | Slc17a3 | 0 | 0.326907457 | 1 |
| ENSMUSG00000021335 | Slc17a1 | 0 | -0.52108945 | 1 |
| ENSMUSG00000021337 | Scgn | 0 | 0.063873051 | 1 |
| ENSMUSG00000085861 | Armh2 | 0 | 1.648835551 | 1 |
| ENSMUSG00000086448 | 9330162012Rik | 0 | 0.648835551 | 1 |
| ENSMUSG00000021340 | Gpld1 | 0 | 0.063873051 | 1 |
| ENSMUSG00000048978 | Nrsn1 | 0 | 1.648835551 | 1 |
| ENSMUSG00000084811 | 1700092E19Rik | 0 | 0.063873051 | 1 |
| ENSMUSG00000017922 | Prl3c1 | 0 | 0.648835551 | 1 |
| ENSMUSG00000038891 | Prl3b1 | 0 | 1.648835551 | 1 |
| ENSMUSG00000038883 | Prl3a1 | 0 | 0.326907457 | 1 |
| ENSMUSG00000018259 | Prl8a2 | 0 | 1.063873051 | 1 |
| ENSMUSG00000006490 | Prl8a9 | 0 | 0.648835551 | 1 |
| ENSMUSG00000019756 | Prl8a1 | 0 | 0.063873051 | 1 |
| ENSMUSG00000021347 | Prl7b1 | 0 | 1.648835551 | 1 |
| ENSMUSG00000062551 | Prl2c1 | 0 | 1.648835551 | 1 |
| ENSMUSG00000113352 | Gm34639 | 0 | -0.158519371 | 1 |
| ENSMUSG00000113424 | Gm48161 | 0 | 0.326907457 | 1 |
| ENSMUSG00000086263 | 4930519D14Rik | 0 | -0.351164449 | 1 |
| ENSMUSG00000021356 | Irf4 | 0 | 0.063873051 | 1 |
| ENSMUSG00000076430 | Hus1b | 0 | 0.648835551 | 1 |
| ENSMUSG00000087460 | 4930401O12Rik | 0 | 1.063873051 | 1 |
| ENSMUSG00000090863 | A530084C06Rik | 0 | 1.648835551 | 1 |
| ENSMUSG00000087220 | Gm11377 | 0 | 1.063873051 | 1 |
| ENSMUSG00000086144 | Gm11379 | 0 | 1.063873051 | 1 |
| ENSMUSG00000050295 | Foxc1 | 0 | -0.351164449 | 1 |
| ENSMUSG00000085770 | Gm11381 | 0 | 1.648835551 | 1 |
| ENSMUSG00000079049 | Serpinb1c | 0 | 1.648835551 | 1 |
| ENSMUSG00000114185 | Gm47662 | 0 | 0.063873051 | 1 |
| ENSMUSG00000021403 | Serpinb9b | 0 | 0.063873051 | 1 |
| ENSMUSG00000113726 | Gm36099 | 0 | 1.648835551 | 1 |
| ENSMUSG00000021404 | Serpinb9c | 0 | 1.648835551 | 1 |

| | | | | |
|---|---|---|---|---|
| ENSMUSG00000038327 | Serpinb9f | 0 | 1.648835551 | 1 |
| ENSMUSG00000047889 | Serpinb6d | 0 | 1.648835551 | 1 |
| ENSMUSG00000069248 | Serpinb6e | 0 | 1.648835551 | 1 |
| ENSMUSG00000113591 | 4930447K03Rik | 0 | 0.648835551 | 1 |
| ENSMUSG00000113998 | Gm40909 | 0 | 1.648835551 | 1 |
| ENSMUSG00000113185 | Gm47127 | 0 | 1.648835551 | 1 |
| ENSMUSG00000086786 | Gm15908 | 0 | 0.326907457 | 1 |
| ENSMUSG00000113170 | Gm36839 | 0 | 1.063873051 | 1 |
| ENSMUSG00000021414 | Fam217a | 0 | 0.648835551 | 1 |
| ENSMUSG00000021416 | Eci3 | 0 | 1.648835551 | 1 |
| ENSMUSG00000085962 | Gm16984 | 0 | 1.648835551 | 1 |
| ENSMUSG00000113738 | Gm40910 | 0 | 0.326907457 | 1 |
| ENSMUSG00000113906 | 4930529N20Rik | 0 | 1.648835551 | 1 |
| ENSMUSG00000102099 | 1700011B04Rik | 0 | 1.063873051 | 1 |
| ENSMUSG00000113770 | 1700019C18Rik | 0 | 1.063873051 | 1 |
| ENSMUSG00000114373 | Gm48708 | 0 | 1.648835551 | 1 |
| ENSMUSG00000021418 | Rpp40 | 0 | 1.063873051 | 1 |
| ENSMUSG00000114270 | Gm48746 | 0 | -0.52108945 | 1 |
| ENSMUSG00000114708 | Gm30177 | 0 | 0.326907457 | 1 |
| ENSMUSG00000114483 | Gm49144 | 0 | -1.743481871 | 0.455131546 |
| ENSMUSG00000114329 | Gm30489 | 0 | 1.063873051 | 1 |
| ENSMUSG00000100760 | Gm4035 | 0 | -2.051604167 | 0.274319469 |
| ENSMUSG00000114389 | Gm47734 | 0 | 0.648835551 | 1 |
| ENSMUSG00000114355 | Gm40916 | 0 | -1.673092543 | 0.646117176 |
| ENSMUSG00000114800 | Gm31600 | 0 | 1.063873051 | 1 |
| ENSMUSG00000114370 | Gm46411 | 0 | 1.648835551 | 1 |
| ENSMUSG00000044566 | Cage1 | 0 | -1.995020638 | 0.370945697 |
| ENSMUSG00000114525 | Gm47990 | 0 | 0.648835551 | 1 |
| ENSMUSG00000114638 | Gm31834 | 0 | 1.648835551 | 1 |
| ENSMUSG00000096932 | 5033403F01Rik | 0 | 0.648835551 | 1 |
| ENSMUSG00000114821 | Gm3509 | 0 | 1.648835551 | 1 |
| ENSMUSG00000114300 | Gm32243 | 0 | 1.648835551 | 1 |
| ENSMUSG00000114710 | Gm40923 | 0 | 1.648835551 | 1 |
| ENSMUSG00000114957 | Gm47316 | 0 | 1.648835551 | 1 |
| ENSMUSG00000097752 | Gm26688 | 0 | 0.063873051 | 1 |
| ENSMUSG00000114597 | Gm47067 | 0 | 1.648835551 | 1 |
| ENSMUSG00000021362 | Gcm2 | 0 | 1.648835551 | 1 |
| ENSMUSG00000114247 | Gm32063 | 0 | 1.063873051 | 1 |
| ENSMUSG00000097373 | Gm26877 | 0 | 0.648835551 | 1 |
| ENSMUSG00000114585 | Gm32401 | 0 | -1.158519371 | 0.920245613 |
| ENSMUSG00000114545 | Gm32184 | 0 | 1.648835551 | 1 |
| ENSMUSG00000087370 | Tmem170b | 0 | -2.777429203 | 0.067360743 |
| ENSMUSG00000054258 | Gm5082 | 0 | 0.648835551 | 1 |
| ENSMUSG00000099672 | Gm28564 | 0 | 1.648835551 | 1 |
| ENSMUSG00000087202 | Gm15813 | 0 | 1.063873051 | 1 |
| ENSMUSG00000021368 | Tbc1d7 | 0 | -0.673092543 | 1 |
| ENSMUSG00000114121 | Gm47728 | 0 | 1.063873051 | 1 |
| ENSMUSG00000015396 | Cd83 | 0 | -0.158519371 | 1 |
| ENSMUSG00000113252 | Gm33489 | 0 | 1.063873051 | 1 |
| ENSMUSG00000113689 | Gm29676 | 0 | 0.063873051 | 1 |
| ENSMUSG00000112936 | Gm33958 | 0 | -0.673092543 | 1 |

| | | | | |
|---|---|---|---|---|
| ENSMUSG00000113553 | Gm40932 | 0 | 1.648835551 | 1 |
| ENSMUSG00000113772 | Gm47448 | 0 | -1.351164449 | 0.72614554 |
| ENSMUSG00000000253 | Gmpr | 0 | -0.52108945 | 1 |
| ENSMUSG00000114131 | Gm47460 | 0 | 0.648835551 | 1 |
| ENSMUSG00000113860 | Gm45949 | 0 | 1.063873051 | 1 |
| ENSMUSG00000114752 | Gm35110 | 0 | -0.52108945 | 1 |
| ENSMUSG00000021376 | Tpmt | 0 | -1.258055044 | 0.673811773 |
| ENSMUSG00000114681 | 1700026N04Rik | 0 | 1.648835551 | 1 |
| ENSMUSG00000021379 | Id4 | 0 | -0.158519371 | 1 |
| ENSMUSG00000114695 | Gm48648 | 0 | 0.326907457 | 1 |
| ENSMUSG00000086889 | Phf2os1 | 0 | 1.063873051 | 1 |
| ENSMUSG00000021388 | Aspn | 0 | 0.326907457 | 1 |
| ENSMUSG00000021390 | Ogn | 0 | 1.063873051 | 1 |
| ENSMUSG00000114073 | Gm30302 | 0 | 0.648835551 | 1 |
| ENSMUSG00000113211 | 4921525O09Rik | 0 | 1.063873051 | 1 |
| ENSMUSG00000056749 | Nfil3 | 0 | -2.480447466 | 0.13388699 |
| ENSMUSG00000107008 | Gm2762 | 0 | -1.599091962 | 0.603006227 |
| ENSMUSG00000107305 | Gm43262 | 0 | 1.648835551 | 1 |
| ENSMUSG00000021469 | Msx2 | 0 | 1.648835551 | 1 |
| ENSMUSG00000113599 | 9530014B07Rik | 0 | 1.648835551 | 1 |
| ENSMUSG00000112965 | Gm34278 | 0 | 0.063873051 | 1 |
| ENSMUSG00000090016 | Gm16578 | 0 | 1.648835551 | 1 |
| ENSMUSG00000112941 | Gm48623 | 0 | -1.158519371 | 0.886203497 |
| ENSMUSG00000113404 | Gm48622 | 0 | 1.648835551 | 1 |
| ENSMUSG00000025870 | Arl10 | 0 | -1.051604167 | 0.858831093 |
| ENSMUSG00000084949 | Gm16249 | 0 | -2.599091962 | 0.143740392 |
| ENSMUSG00000025876 | Unc5a | 0 | -0.52108945 | 1 |
| ENSMUSG00000021485 | Mxd3 | 0 | -1.258055044 | 0.673811773 |
| ENSMUSG00000044444 | Pfn3 | 0 | 1.648835551 | 1 |
| ENSMUSG00000034686 | Prr7 | 0 | -1.051604167 | 0.858831093 |
| ENSMUSG00000086506 | Gm15911 | 0 | 1.648835551 | 1 |
| ENSMUSG00000069223 | 4930451E10Rik | 0 | 0.648835551 | 1 |
| ENSMUSG00000021506 | Pitx1 | 0 | 0.648835551 | 1 |
| ENSMUSG00000049625 | Tifab | 0 | 0.063873051 | 1 |
| ENSMUSG00000048904 | Neurog1 | 0 | 1.648835551 | 1 |
| ENSMUSG00000021509 | Slc25a48 | 0 | -1.995020638 | 0.39066019 |
| ENSMUSG00000035509 | Fbxl21 | 0 | -0.158519371 | 1 |
| ENSMUSG00000114524 | Gm48357 | 0 | 1.648835551 | 1 |
| ENSMUSG00000114929 | Gm48441 | 0 | 0.648835551 | 1 |
| ENSMUSG00000114393 | Gm47359 | 0 | 1.063873051 | 1 |
| ENSMUSG00000114240 | Gm47360 | 0 | -0.351164449 | 1 |
| ENSMUSG00000114787 | 4930455M05Rik | 0 | 1.063873051 | 1 |
| ENSMUSG00000050876 | Spata31d1a | 0 | 0.648835551 | 1 |
| ENSMUSG00000114903 | Gm46419 | 0 | 1.648835551 | 1 |
| ENSMUSG00000051054 | 1700014D04Rik | 0 | 1.648835551 | 1 |
| ENSMUSG00000089875 | Etohd2 | 0 | 1.648835551 | 1 |
| ENSMUSG00000114490 | Gm19866 | 0 | 1.063873051 | 1 |
| ENSMUSG00000114795 | Gm48396 | 0 | 1.648835551 | 1 |
| ENSMUSG00000052957 | Gas1 | 0 | -0.673092543 | 1 |
| ENSMUSG00000114433 | Gm48488 | 0 | 1.648835551 | 1 |
| ENSMUSG00000050765 | Gm5084 | 0 | 1.648835551 | 1 |

| | | | | |
|---|---|---|---|---|
| ENSMUSG00000114274 | Gm35333 | 0 | 1.648835551 | 1 |
| ENSMUSG00000050345 | 4930486L24Rik | 0 | -1.051604167 | 0.858831093 |
| ENSMUSG00000033834 | Tpbpa | 0 | 1.648835551 | 1 |
| ENSMUSG00000114502 | Gm40975 | 0 | 1.648835551 | 1 |
| ENSMUSG00000021439 | Ctsq | 0 | 1.648835551 | 1 |
| ENSMUSG00000057446 | Cts8 | 0 | 0.648835551 | 1 |
| ENSMUSG00000113495 | Gm19792 | 0 | 1.648835551 | 1 |
| ENSMUSG00000094942 | Gm3604 | 0 | -0.936126949 | 0.847265357 |
| ENSMUSG00000072066 | 6720489N17Rik | 0 | -2.52108945 | 0.139351384 |
| ENSMUSG00000091183 | Gm5141 | 0 | -1.43862729 | 0.538110273 |
| ENSMUSG00000113057 | Gm48812 | 0 | 1.648835551 | 1 |
| ENSMUSG00000089854 | Gm16133 | 0 | 1.648835551 | 1 |
| ENSMUSG00000113459 | Gm30655 | 0 | 0.648835551 | 1 |
| ENSMUSG00000114071 | Gm47390 | 0 | 1.648835551 | 1 |
| ENSMUSG00000114092 | Gm47418 | 0 | 1.648835551 | 1 |
| ENSMUSG00000033122 | Hsd17b3 | 0 | 0.326907457 | 1 |
| ENSMUSG00000113816 | Gm47003 | 0 | -2.051604167 | 0.279592421 |
| ENSMUSG00000096537 | Fam240b | 0 | 1.648835551 | 1 |
| ENSMUSG00000114168 | Gm47194 | 0 | 1.648835551 | 1 |
| ENSMUSG00000056223 | Spata31 | 0 | 1.063873051 | 1 |
| ENSMUSG00000074849 | Spata31d1c | 0 | 1.648835551 | 1 |
| ENSMUSG00000090658 | Prss47 | 0 | 0.648835551 | 1 |
| ENSMUSG00000049806 | Olfr466 | 0 | 1.063873051 | 1 |
| ENSMUSG00000110393 | Gm36445 | 0 | -0.158519371 | 1 |
| ENSMUSG00000021514 | Zfp369 | 0 | -3.579983139 | 0.006320625 |
| ENSMUSG00000112935 | Gm40983 | 0 | 1.648835551 | 1 |
| ENSMUSG00000095909 | Gm10324 | 0 | 1.648835551 | 1 |
| ENSMUSG00000118427 | AC192334.1 | 0 | 1.648835551 | 1 |
| ENSMUSG00000114327 | Gm40988 | 0 | 0.648835551 | 1 |
| ENSMUSG00000090641 | Zfp712 | 0 | 0.063873051 | 1 |
| ENSMUSG00000074824 | Rslcan18 | 0 | 1.648835551 | 1 |
| ENSMUSG00000057396 | Zfp759 | 0 | 0.063873051 | 1 |
| ENSMUSG00000051037 | Zfp455 | 0 | 0.648835551 | 1 |
| ENSMUSG00000114771 | Gm49064 | 0 | -0.52108945 | 1 |
| ENSMUSG00000055480 | Zfp458 | 0 | -2.480447466 | 0.13388699 |
| ENSMUSG00000114442 | F630042J09Rik | 0 | 0.648835551 | 1 |
| ENSMUSG00000092103 | Gm17039 | 0 | 1.063873051 | 1 |
| ENSMUSG00000114726 | Gm48900 | 0 | 1.648835551 | 1 |
| ENSMUSG00000055560 | Zfp459 | 0 | 0.326907457 | 1 |
| ENSMUSG00000044081 | Zfp85os | 0 | 0.063873051 | 1 |
| ENSMUSG00000058331 | Zfp85 | 0 | 1.648835551 | 1 |
| ENSMUSG00000090659 | Zfp493 | 0 | 0.326907457 | 1 |
| ENSMUSG00000030446 | Zfp273 | 0 | -1.158519371 | 0.677768314 |
| ENSMUSG00000058246 | Gm10037 | 0 | 1.063873051 | 1 |
| ENSMUSG00000112985 | Gm48556 | 0 | 1.648835551 | 1 |
| ENSMUSG00000034575 | Tent4a | 0 | -3.351164449 | 0.012879784 |
| ENSMUSG00000051805 | A530095I07Rik | 0 | 1.648835551 | 1 |
| ENSMUSG00000091133 | Gm17108 | 0 | 0.648835551 | 1 |
| ENSMUSG00000113299 | Gm35618 | 0 | 0.326907457 | 1 |
| ENSMUSG00000113324 | Gm48819 | 0 | 1.648835551 | 1 |
| ENSMUSG00000112957 | 4930547H16Rik | 0 | 0.648835551 | 1 |

| | | | | |
|---|---|---|---|---|
| ENSMUSG00000099639 | 1700084F23Rik | 0 | -0.158519371 | 1 |
| ENSMUSG00000113573 | AU017674 | 0 | -1.158519371 | 0.887847724 |
| ENSMUSG00000113334 | D030007L05Rik | 0 | 1.648835551 | 1 |
| ENSMUSG00000113796 | Gm36426 | 0 | 1.648835551 | 1 |
| ENSMUSG00000113771 | Gm36529 | 0 | 0.648835551 | 1 |
| ENSMUSG00000093587 | Gm20554 | 0 | 1.648835551 | 1 |
| ENSMUSG00000001504 | Irx2 | 0 | 1.648835551 | 1 |
| ENSMUSG00000113082 | D730050B12Rik | 0 | 1.648835551 | 1 |
| ENSMUSG00000021604 | Irx4 | 0 | 1.063873051 | 1 |
| ENSMUSG00000113346 | Eprn | 0 | 0.648835551 | 1 |
| ENSMUSG00000021609 | Slc6a3 | 0 | 0.326907457 | 1 |
| ENSMUSG00000021566 | Slc6a19os | 0 | 1.648835551 | 1 |
| ENSMUSG00000069189 | Zdhhc11 | 0 | 0.648835551 | 1 |
| ENSMUSG00000021573 | Tppp | 0 | 1.063873051 | 1 |
| ENSMUSG00000021572 | Cep72 | 0 | -0.351164449 | 1 |
| ENSMUSG00000021575 | Ahrr | 0 | 0.063873051 | 1 |
| ENSMUSG00000021579 | Lrrc14b | 0 | 1.063873051 | 1 |
| ENSMUSG00000113311 | Gm47428 | 0 | 1.648835551 | 1 |
| ENSMUSG00000116646 | Gm49763 | 0 | 1.648835551 | 1 |
| ENSMUSG00000113019 | Gm47467 | 0 | 1.063873051 | 1 |
| ENSMUSG00000113047 | Gm47469 | 0 | -2.636566667 | 0.112813335 |
| ENSMUSG00000113204 | Gm46430 | 0 | -3.52108945 | 0.007344049 |
| ENSMUSG00000114125 | 1700037F03Rik | 0 | 1.063873051 | 1 |
| ENSMUSG00000045509 | Gpr150 | 0 | 1.648835551 | 1 |
| ENSMUSG00000114649 | 3110006O06Rik | 0 | 1.648835551 | 1 |
| ENSMUSG00000069171 | Nr2f1 | 0 | -0.673092543 | 1 |
| ENSMUSG00000114632 | Gm48399 | 0 | 1.648835551 | 1 |
| ENSMUSG00000101416 | Gm29318 | 0 | 0.326907457 | 1 |
| ENSMUSG00000089827 | 1700023H06Rik | 0 | 1.063873051 | 1 |
| ENSMUSG00000097265 | Gm26803 | 0 | 1.648835551 | 1 |
| ENSMUSG00000100891 | 2810049E08Rik | 0 | -0.52108945 | 1 |
| ENSMUSG00000114379 | Gm4241 | 0 | 1.648835551 | 1 |
| ENSMUSG00000078958 | Atp6ap1l | 0 | 0.648835551 | 1 |
| ENSMUSG00000097182 | A830009L08Rik | 0 | -1.673092543 | 0.451785624 |
| ENSMUSG00000021620 | Acot12 | 0 | 0.063873051 | 1 |
| ENSMUSG00000034334 | Fam151b | 0 | -1.673092543 | 0.451785624 |
| ENSMUSG00000046957 | Spz1 | 0 | 1.648835551 | 1 |
| ENSMUSG00000114823 | Gm31630 | 0 | 1.648835551 | 1 |
| ENSMUSG00000114968 | A630019I02Rik | 0 | 1.648835551 | 1 |
| ENSMUSG00000085834 | Gm15622 | 0 | 0.648835551 | 1 |
| ENSMUSG00000042102 | Dmgdh | 0 | -0.810596067 | 1 |
| ENSMUSG00000087334 | AW495222 | 0 | 1.063873051 | 1 |
| ENSMUSG00000114055 | Gm32089 | 0 | 1.648835551 | 1 |
| ENSMUSG00000112934 | Gm32305 | 0 | -1.743481871 | 0.455131546 |
| ENSMUSG00000113070 | Gm48420 | 0 | 1.063873051 | 1 |
| ENSMUSG00000113359 | Gm48730 | 0 | -2.106051951 | 0.279592421 |
| ENSMUSG00000021675 | F2rl2 | 0 | 1.648835551 | 1 |
| ENSMUSG00000118445 | AC171003.1 | 0 | -0.52108945 | 1 |
| ENSMUSG00000097932 | C430039J16Rik | 0 | 1.648835551 | 1 |
| ENSMUSG00000054944 | 5330416C01Rik | 0 | 0.063873051 | 1 |
| ENSMUSG00000114348 | Gm41031 | 0 | -2.106051951 | 0.3498121 |

| | | | | |
|---|---|---|---|---|
| ENSMUSG00000114474 | Gm34388 | 0 | 1.063873051 | 1 |
| ENSMUSG00000048603 | Gm9828 | 0 | 0.648835551 | 1 |
| ENSMUSG00000099446 | 2310005E17Rik | 0 | 1.063873051 | 1 |
| ENSMUSG00000114521 | Gm48050 | 0 | 1.063873051 | 1 |
| ENSMUSG00000046082 | Tmem174 | 0 | 1.648835551 | 1 |
| ENSMUSG00000052485 | Tmem171 | 0 | -2.395558568 | 0.174915761 |
| ENSMUSG00000114689 | Gm9465 | 0 | 0.063873051 | 1 |
| ENSMUSG00000114569 | Gm35215 | 0 | 1.648835551 | 1 |
| ENSMUSG00000100410 | 2310020H05Rik | 0 | -0.158519371 | 1 |
| ENSMUSG00000097848 | Gm807 | 0 | 1.648835551 | 1 |
| ENSMUSG00000097027 | Gm26559 | 0 | 0.648835551 | 1 |
| ENSMUSG00000071203 | Naip5 | 0 | -0.351164449 | 1 |
| ENSMUSG00000078942 | Naip6 | 0 | -0.158519371 | 1 |
| ENSMUSG00000114360 | Gm48596 | 0 | 0.063873051 | 1 |
| ENSMUSG00000114400 | Gm47533 | 0 | 1.648835551 | 1 |
| ENSMUSG00000114846 | Gm29927 | 0 | 1.648835551 | 1 |
| ENSMUSG00000114276 | Gm47849 | 0 | 1.063873051 | 1 |
| ENSMUSG00000042743 | Sgtb | 0 | -1.599091962 | 0.625377538 |
| ENSMUSG00000084860 | 2610204G07Rik | 0 | 0.648835551 | 1 |
| ENSMUSG00000114422 | Gm30411 | 0 | 1.063873051 | 1 |
| ENSMUSG00000101308 | Gm28989 | 0 | 0.326907457 | 1 |
| ENSMUSG00000114627 | B230220B15Rik | 0 | -1.743481871 | 0.466464457 |
| ENSMUSG00000090358 | Gm2822 | 0 | 0.648835551 | 1 |
| ENSMUSG00000046006 | Gapt | 0 | 1.648835551 | 1 |
| ENSMUSG00000114307 | 4930526H09Rik | 0 | 1.063873051 | 1 |
| ENSMUSG00000114824 | 3110015C05Rik | 0 | 0.326907457 | 1 |
| ENSMUSG00000086201 | Gm6270 | 0 | 1.648835551 | 1 |
| ENSMUSG00000086480 | Gm15287 | 0 | 1.063873051 | 1 |
| ENSMUSG00000114536 | Gm48837 | 0 | 1.648835551 | 1 |
| ENSMUSG00000114869 | Gm48876 | 0 | 1.063873051 | 1 |
| ENSMUSG00000114438 | Gm47827 | 0 | 1.063873051 | 1 |
| ENSMUSG00000074651 | Mcidas | 0 | 1.063873051 | 1 |
| ENSMUSG00000078926 | Cdc20b | 0 | -0.810596067 | 0.844591606 |
| ENSMUSG00000114864 | Gm41071 | 0 | 1.648835551 | 1 |
| ENSMUSG00000042364 | Snx18 | 0 | -3.184054463 | 0.023817286 |
| ENSMUSG00000114417 | Gm34586 | 0 | 1.648835551 | 1 |
| ENSMUSG00000114213 | A430090L17Rik | 0 | 0.326907457 | 1 |
| ENSMUSG00000114879 | 4930467J12Rik | 0 | 1.063873051 | 1 |
| ENSMUSG00000114255 | Gm10734 | 0 | 0.648835551 | 1 |
| ENSMUSG00000042258 | Isl1 | 0 | 0.648835551 | 1 |
| ENSMUSG00000114290 | 4930435F18Rik | 0 | 0.063873051 | 1 |
| ENSMUSG00000079048 | 4933413L06Rik | 0 | 1.063873051 | 1 |
| ENSMUSG00000097411 | B430218F22Rik | 0 | 0.648835551 | 1 |
| ENSMUSG00000100727 | 1700003P14Rik | 0 | -0.158519371 | 1 |
| ENSMUSG00000025453 | Nnt | 0 | -1.051604167 | 0.858831093 |
| ENSMUSG00000074635 | 3110070M22Rik | 0 | 0.648835551 | 1 |
| ENSMUSG00000114945 | Gm48265 | 0 | 1.063873051 | 1 |
| ENSMUSG00000095514 | D13Ertd608e | 0 | 1.648835551 | 1 |
| ENSMUSG00000096175 | Gm21761 | 0 | 1.648835551 | 1 |
| ENSMUSG00000114608 | Gm36161 | 0 | 1.648835551 | 1 |
| ENSMUSG00000094796 | BC147527 | 0 | 1.063873051 | 1 |

| | | | | |
|---|---|---|---|---|
| ENSMUSG00000096775 | Gm5796 | 0 | 1.648835551 | 1 |
| ENSMUSG00000097822 | Gm26655 | 0 | 1.648835551 | 1 |
| ENSMUSG00000094132 | Gm3264 | 0 | 1.648835551 | 1 |
| ENSMUSG00000079371 | Gm3476 | 0 | 1.648835551 | 1 |
| ENSMUSG00000094460 | Gm21560 | 0 | 1.648835551 | 1 |
| ENSMUSG00000079396 | Gm3411 | 0 | 1.648835551 | 1 |
| ENSMUSG00000094811 | Gm21103 | 0 | 1.063873051 | 1 |
| ENSMUSG00000095360 | Gm3594 | 0 | 1.648835551 | 1 |
| ENSMUSG00000097901 | Gm26680 | 0 | 1.648835551 | 1 |
| ENSMUSG00000092167 | Gm3696 | 0 | 1.648835551 | 1 |
| ENSMUSG00000091472 | Gm3739 | 0 | 0.326907457 | 1 |
| ENSMUSG00000071591 | Gm10339 | 0 | 1.648835551 | 1 |
| ENSMUSG00000021752 | Kctd6 | 0 | -1.258055044 | 0.673811773 |
| ENSMUSG00000097898 | 4930455B14Rik | 0 | 0.648835551 | 1 |
| ENSMUSG00000114849 | Gm48595 | 0 | 1.648835551 | 1 |
| ENSMUSG00000114853 | D030051J21Rik | 0 | 0.648835551 | 1 |
| ENSMUSG00000079357 | Gm11100 | 0 | 0.326907457 | 1 |
| ENSMUSG00000072707 | Olfr31 | 0 | -0.158519371 | 1 |
| ENSMUSG00000091809 | Olfr721-ps1 | 0 | 1.063873051 | 1 |
| ENSMUSG00000114538 | Gm31517 | 0 | -1.52108945 | 0.588477963 |
| ENSMUSG00000114770 | 4921522A10Rik | 0 | 1.648835551 | 1 |
| ENSMUSG00000114789 | Gm31577 | 0 | 1.648835551 | 1 |
| ENSMUSG00000114586 | Gm31804 | 0 | 0.063873051 | 1 |
| ENSMUSG00000114907 | Gm47794 | 0 | 1.648835551 | 1 |
| ENSMUSG00000114419 | 5430414B19Rik | 0 | 1.063873051 | 1 |
| ENSMUSG00000114253 | Gm47798 | 0 | 1.648835551 | 1 |
| ENSMUSG00000114463 | Gm48162 | 0 | 1.648835551 | 1 |
| ENSMUSG00000021772 | Nkiras1 | 0 | -1.874726405 | 0.352215149 |
| ENSMUSG00000068776 | 4930469B13Rik | 0 | 1.648835551 | 1 |
| ENSMUSG00000114350 | D14Ertd670e | 0 | 1.063873051 | 1 |
| ENSMUSG00000023243 | Kcnk5 | 0 | -1.995020638 | 0.370945697 |
| ENSMUSG00000114439 | Gm48395 | 0 | 1.648835551 | 1 |
| ENSMUSG00000023387 | Kcnk16 | 0 | 1.648835551 | 1 |
| ENSMUSG00000114937 | Gm30054 | 0 | 0.326907457 | 1 |
| ENSMUSG00000021822 | Plau | 0 | -0.936126949 | 0.847265357 |
| ENSMUSG00000114412 | 4930405A10Rik | 0 | 0.648835551 | 1 |
| ENSMUSG00000114496 | Gm47601 | 0 | 1.648835551 | 1 |
| ENSMUSG00000114953 | 4930519K11Rik | 0 | 1.648835551 | 1 |
| ENSMUSG00000048628 | 4930542C16Rik | 0 | -1.158519371 | 0.675561045 |
| ENSMUSG00000114965 | Gm47906 | 0 | 1.063873051 | 1 |
| ENSMUSG00000114924 | Gm38408 | 0 | 1.063873051 | 1 |
| ENSMUSG00000114288 | Gm32123 | 0 | 1.648835551 | 1 |
| ENSMUSG00000097601 | Gm26660 | 0 | -0.52108945 | 1 |
| ENSMUSG00000114232 | Gm17747 | 0 | 1.648835551 | 1 |
| ENSMUSG00000114922 | Ppifos | 0 | 1.648835551 | 1 |
| ENSMUSG00000098609 | Anxa11os | 0 | 0.648835551 | 1 |
| ENSMUSG00000095304 | Plac9a | 0 | 1.648835551 | 1 |
| ENSMUSG00000072676 | Tmem254a | 0 | 1.063873051 | 1 |
| ENSMUSG00000098463 | Cphx2 | 0 | 1.063873051 | 1 |
| ENSMUSG00000072672 | Duxbl3 | 0 | 1.648835551 | 1 |
| ENSMUSG00000021874 | 4933413J09Rik | 0 | 1.063873051 | 1 |

| | | | | |
|---|---|---|---|---|
| ENSMUSG00000043702 | Pde12 | 0 | -1.52108945 | 0.544908989 |
| ENSMUSG00000096052 | 9930004E17Rik | 0 | 1.648835551 | 1 |
| ENSMUSG00000097002 | Gm2670 | 0 | 1.648835551 | 1 |
| ENSMUSG00000114818 | Gm35164 | 0 | -0.158519371 | 1 |
| ENSMUSG00000114577 | Gm35217 | 0 | 1.648835551 | 1 |
| ENSMUSG00000045776 | Lrtm1 | 0 | -0.52108945 | 1 |
| ENSMUSG00000114815 | Gm41118 | 0 | 1.063873051 | 1 |
| ENSMUSG00000114424 | Gm47414 | 0 | 1.063873051 | 1 |
| ENSMUSG00000015966 | Il17rb | 0 | 0.326907457 | 1 |
| ENSMUSG00000015970 | Chdh | 0 | -1.43862729 | 0.747121178 |
| ENSMUSG00000118437 | CT009536.1 | 0 | 1.063873051 | 1 |
| ENSMUSG00000115723 | 1700087M22Rik | 0 | -0.673092543 | 1 |
| ENSMUSG00000042485 | Mustn1 | 0 | 1.648835551 | 1 |
| ENSMUSG00000021922 | Itih4 | 0 | 0.326907457 | 1 |
| ENSMUSG00000006522 | Itih3 | 0 | 0.648835551 | 1 |
| ENSMUSG00000006529 | Itih1 | 0 | 0.648835551 | 1 |
| ENSMUSG00000071547 | Nt5dc2 | 0 | -1.051604167 | 0.858831093 |
| ENSMUSG00000046005 | D830044D21Rik | 0 | -0.52108945 | 1 |
| ENSMUSG00000090459 | Gm17210 | 0 | 1.648835551 | 1 |
| ENSMUSG00000115813 | Gm46447 | 0 | -1.051604167 | 0.858831093 |
| ENSMUSG00000071540 | 3425401B19Rik | 0 | 0.326907457 | 1 |
| ENSMUSG00000041673 | Lrrc18 | 0 | 0.326907457 | 1 |
| ENSMUSG00000087639 | Gm15512 | 0 | 0.063873051 | 1 |
| ENSMUSG00000115134 | Gm49108 | 0 | 1.648835551 | 1 |
| ENSMUSG00000072624 | Gm5460 | 0 | -0.52108945 | 1 |
| ENSMUSG00000047441 | Antxrl | 0 | 0.326907457 | 1 |
| ENSMUSG00000071531 | Gprin2 | 0 | 1.648835551 | 1 |
| ENSMUSG00000041479 | Syt15 | 0 | -0.158519371 | 1 |
| ENSMUSG00000023064 | Sncg | 0 | 1.063873051 | 1 |
| ENSMUSG00000115575 | Gm49024 | 0 | 1.648835551 | 1 |
| ENSMUSG00000115007 | 5830448L01Rik | 0 | -0.810596067 | 1 |
| ENSMUSG00000097310 | A930038B10Rik | 0 | 0.326907457 | 1 |
| ENSMUSG00000021804 | Rgr | 0 | 1.063873051 | 1 |
| ENSMUSG00000043418 | Lrit2 | 0 | 0.648835551 | 1 |
| ENSMUSG00000114219 | Gm31456 | 0 | 1.648835551 | 1 |
| ENSMUSG00000101261 | 1700109I08Rik | 0 | 0.326907457 | 1 |
| ENSMUSG00000021791 | Dydc2 | 0 | 1.063873051 | 1 |
| ENSMUSG00000037798 | Mat1a | 0 | 0.648835551 | 1 |
| ENSMUSG00000037780 | Mbl1 | 0 | 1.063873051 | 1 |
| ENSMUSG00000091110 | Gm2832 | 0 | 1.063873051 | 1 |
| ENSMUSG00000093863 | Gm7954 | 0 | 1.648835551 | 1 |
| ENSMUSG00000090505 | Gm3486 | 0 | 1.063873051 | 1 |
| ENSMUSG00000093833 | Gm7970 | 0 | 1.648835551 | 1 |
| ENSMUSG00000091657 | Gm3072 | 0 | -1.158519371 | 0.912226588 |
| ENSMUSG00000094157 | Gm7995 | 0 | 1.648835551 | 1 |
| ENSMUSG00000092148 | Gm8005 | 0 | -2.874726405 | 0.084875472 |
| ENSMUSG00000090634 | Gm8126 | 0 | 1.063873051 | 1 |
| ENSMUSG00000115483 | Gm9732 | 0 | 0.063873051 | 1 |
| ENSMUSG00000091477 | Gm5799 | 0 | 1.063873051 | 1 |
| ENSMUSG00000095743 | Gm17654 | 0 | 1.648835551 | 1 |
| ENSMUSG00000095528 | Gm10375 | 0 | 1.063873051 | 1 |

| | | | | |
|---|---|---|---|---|
| ENSMUSG00000115354 | Gm49083 | 0 | 1.648835551 | 1 |
| ENSMUSG00000072601 | Ear1 | 0 | 0.063873051 | 1 |
| ENSMUSG00000115724 | Gm49087 | 0 | 1.648835551 | 1 |
| ENSMUSG00000115794 | Gm49101 | 0 | -0.158519371 | 1 |
| ENSMUSG00000072595 | 4930503E14Rik | 0 | 1.063873051 | 1 |
| ENSMUSG00000115536 | Gm32857 | 0 | 1.648835551 | 1 |
| ENSMUSG00000072145 | BC061237 | 0 | 1.648835551 | 1 |
| ENSMUSG00000091429 | Gm17093 | 0 | -0.810596067 | 1 |
| ENSMUSG00000091923 | Gm8267 | 0 | 0.063873051 | 1 |
| ENSMUSG00000115747 | Gm49145 | 0 | 1.648835551 | 1 |
| ENSMUSG00000115846 | Gm41144 | 0 | 1.063873051 | 1 |
| ENSMUSG00000085913 | Gm15601 | 0 | 0.326907457 | 1 |
| ENSMUSG00000086124 | A530076I17Rik | 0 | 1.648835551 | 1 |
| ENSMUSG00000086746 | Gm15222 | 0 | -1.351164449 | 0.687112677 |
| ENSMUSG00000086168 | Gm15221 | 0 | 1.648835551 | 1 |
| ENSMUSG00000115083 | Gm34756 | 0 | 1.063873051 | 1 |
| ENSMUSG00000115158 | Gm48912 | 0 | 0.326907457 | 1 |
| ENSMUSG00000115003 | Gm48933 | 0 | 1.648835551 | 1 |
| ENSMUSG00000115100 | Gm34934 | 0 | 0.648835551 | 1 |
| ENSMUSG00000115410 | 2810457G06Rik | 0 | 0.326907457 | 1 |
| ENSMUSG00000115082 | Gm49189 | 0 | -0.673092543 | 1 |
| ENSMUSG00000115145 | Gm35166 | 0 | 0.648835551 | 1 |
| ENSMUSG00000115310 | Gm35360 | 0 | 0.648835551 | 1 |
| ENSMUSG00000115425 | Gm49305 | 0 | 1.648835551 | 1 |
| ENSMUSG00000115060 | Gm49306 | 0 | 1.063873051 | 1 |
| ENSMUSG00000115756 | Gm49519 | 0 | 0.063873051 | 1 |
| ENSMUSG00000115725 | 4930572G02Rik | 0 | 1.063873051 | 1 |
| ENSMUSG00000115636 | Gm49120 | 0 | 1.648835551 | 1 |
| ENSMUSG00000115579 | Gm49156 | 0 | 1.648835551 | 1 |
| ENSMUSG00000115008 | 4933429O19Rik | 0 | 0.063873051 | 1 |
| ENSMUSG00000068437 | Olfr725 | 0 | 1.648835551 | 1 |
| ENSMUSG00000049011 | Olfr729 | 0 | 1.063873051 | 1 |
| ENSMUSG00000048080 | Olfr731 | 0 | 1.648835551 | 1 |
| ENSMUSG00000090874 | Olfr733 | 0 | 1.648835551 | 1 |
| ENSMUSG00000051969 | Tlr11 | 0 | 1.648835551 | 1 |
| ENSMUSG00000047716 | Olfr736 | 0 | 0.648835551 | 1 |
| ENSMUSG00000094692 | Olfr738 | 0 | 1.648835551 | 1 |
| ENSMUSG00000057903 | Olfr739 | 0 | 1.063873051 | 1 |
| ENSMUSG00000094285 | Olfr743 | 0 | 1.648835551 | 1 |
| ENSMUSG00000057179 | Olfr747 | 0 | 1.648835551 | 1 |
| ENSMUSG00000006289 | Osgep | 0 | -3.351164449 | 0.01810333 |
| ENSMUSG00000052382 | Rnase9 | 0 | 1.648835551 | 1 |
| ENSMUSG00000035932 | Olfr750 | 0 | 0.326907457 | 1 |
| ENSMUSG00000021880 | Rnase6 | 0 | 1.648835551 | 1 |
| ENSMUSG00000114985 | 1810028F09Rik | 0 | 1.648835551 | 1 |
| ENSMUSG00000047222 | Rnase2a | 0 | 1.063873051 | 1 |
| ENSMUSG00000079244 | Gm5622 | 0 | 0.326907457 | 1 |
| ENSMUSG00000094715 | Gm17078 | 0 | 1.648835551 | 1 |
| ENSMUSG00000068506 | Gm5800 | 0 | 1.648835551 | 1 |
| ENSMUSG00000050961 | AY358078 | 0 | 0.326907457 | 1 |
| ENSMUSG00000004561 | Mettl17 | 0 | -2.965874293 | 0.044171611 |

| | | | | |
|---|---|---|---|---|
| ENSMUSG00000115698 | Gm49076 | 0 | 1.648835551 | 1 |
| ENSMUSG00000097633 | Gm16617 | 0 | 1.648835551 | 1 |
| ENSMUSG00000072571 | Tmem253 | 0 | 0.648835551 | 1 |
| ENSMUSG00000063106 | Olfr1510 | 0 | 1.648835551 | 1 |
| ENSMUSG00000104658 | Gm30275 | 0 | -0.52108945 | 1 |
| ENSMUSG00000104876 | Trdc | 0 | -0.673092543 | 1 |
| ENSMUSG00000115767 | Gm10366 | 0 | 1.063873051 | 1 |
| ENSMUSG00000059674 | Cdh24 | 0 | -1.351164449 | 0.687112677 |
| ENSMUSG00000091306 | Gm17606 | 0 | 1.063873051 | 1 |
| ENSMUSG00000040840 | 4930579G18Rik | 0 | 0.648835551 | 1 |
| ENSMUSG00000052435 | Cebpe | 0 | 1.648835551 | 1 |
| ENSMUSG00000112858 | Rnf212b | 0 | -0.673092543 | 1 |
| ENSMUSG00000057156 | Homez | 0 | -1.351164449 | 0.687112677 |
| ENSMUSG00000022199 | Slc22a17 | 0 | -0.810596067 | 0.844591606 |
| ENSMUSG00000022203 | Efs | 0 | 1.063873051 | 1 |
| ENSMUSG00000040770 | Il25 | 0 | 1.648835551 | 1 |
| ENSMUSG00000097652 | Mhrt | 0 | 1.648835551 | 1 |
| ENSMUSG00000053093 | Myh7 | 0 | 0.326907457 | 1 |
| ENSMUSG00000045691 | Thtpa | 0 | -0.158519371 | 1 |
| ENSMUSG00000022212 | Cpne6 | 0 | 1.648835551 | 1 |
| ENSMUSG00000040632 | Nrl | 0 | 0.648835551 | 1 |
| ENSMUSG00000086341 | Gm15932 | 0 | 0.648835551 | 1 |
| ENSMUSG00000022217 | Emc9 | 0 | 0.326907457 | 1 |
| ENSMUSG00000115222 | Gm49747 | 0 | 0.648835551 | 1 |
| ENSMUSG00000002326 | Gmpr2 | 0 | -0.936126949 | 0.847265357 |
| ENSMUSG00000007589 | Tinf2 | 0 | -3.132524162 | 0.030128441 |
| ENSMUSG00000022219 | Cideb | 0 | -0.351164449 | 1 |
| ENSMUSG00000040432 | Ltb4r2 | 0 | 0.648835551 | 1 |
| ENSMUSG00000022223 | Sdr39u1 | 0 | -0.351164449 | 1 |
| ENSMUSG00000022227 | Mcpt1 | 0 | -1.673092543 | 0.631617217 |
| ENSMUSG00000015441 | Gzmf | 0 | 0.648835551 | 1 |
| ENSMUSG00000114345 | Gm10873 | 0 | 0.326907457 | 1 |
| ENSMUSG00000090038 | Gm16573 | 0 | -0.351164449 | 1 |
| ENSMUSG00000097743 | Gm16973 | 0 | -2.106051951 | 0.279592421 |
| ENSMUSG00000100017 | 2410022M11Rik | 0 | 0.063873051 | 1 |
| ENSMUSG00000048582 | Gja3 | 0 | 0.063873051 | 1 |
| ENSMUSG00000046352 | Gjb2 | 0 | 1.648835551 | 1 |
| ENSMUSG00000086486 | Ift88os | 0 | 1.648835551 | 1 |
| ENSMUSG00000050222 | Il17d | 0 | 1.063873051 | 1 |
| ENSMUSG00000097303 | 3110083C13Rik | 0 | 0.326907457 | 1 |
| ENSMUSG00000114441 | Gm33472 | 0 | 0.648835551 | 1 |
| ENSMUSG00000114289 | Gm48453 | 0 | 1.648835551 | 1 |
| ENSMUSG00000053868 | Gm5142 | 0 | 0.648835551 | 1 |
| ENSMUSG00000044703 | Phf11a | 0 | 1.063873051 | 1 |
| ENSMUSG00000068245 | Phf11d | 0 | -0.158519371 | 1 |
| ENSMUSG00000091144 | Phf11c | 0 | -0.158519371 | 1 |
| ENSMUSG00000044461 | Shisa2 | 0 | 1.063873051 | 1 |
| ENSMUSG00000101505 | 1700109G14Rik | 0 | -0.52108945 | 1 |
| ENSMUSG00000043157 | Arl11 | 0 | 0.326907457 | 1 |
| ENSMUSG00000115656 | 4930556J02Rik | 0 | -0.52108945 | 1 |
| ENSMUSG00000035235 | Trim13 | 0 | -1.43862729 | 0.769129357 |

| | | | | |
|---|---|---|---|---|
| ENSMUSG00000046168 | Kcnrg | 0 | -0.810596067 | 0.847265357 |
| ENSMUSG00000098088 | Gm26916 | 0 | 1.648835551 | 1 |
| ENSMUSG00000021933 | Gucy1b2 | 0 | -1.258055044 | 0.718012086 |
| ENSMUSG00000075574 | Defb48 | 0 | 1.648835551 | 1 |
| ENSMUSG00000075573 | Defb47 | 0 | 1.648835551 | 1 |
| ENSMUSG00000075571 | Defb30 | 0 | 1.063873051 | 1 |
| ENSMUSG00000114898 | Gm49390 | 0 | 0.063873051 | 1 |
| ENSMUSG00000114892 | Gm5463 | 0 | 1.648835551 | 1 |
| ENSMUSG00000035095 | Fam167a | 0 | -0.52108945 | 1 |
| ENSMUSG00000021953 | Tdh | 0 | 0.648835551 | 1 |
| ENSMUSG00000090374 | Gm17232 | 0 | 1.648835551 | 1 |
| ENSMUSG00000087652 | Gm15918 | 0 | 1.648835551 | 1 |
| ENSMUSG00000063060 | Sox7 | 0 | 1.648835551 | 1 |
| ENSMUSG00000046049 | Rp1l1 | 0 | 0.063873051 | 1 |
| ENSMUSG00000052099 | Prss51 | 0 | 1.648835551 | 1 |
| ENSMUSG00000091686 | 4930471C04Rik | 0 | 1.648835551 | 1 |
| ENSMUSG00000090785 | Gm17116 | 0 | 0.326907457 | 1 |
| ENSMUSG00000114680 | Gm47202 | 0 | 1.063873051 | 1 |
| ENSMUSG00000097545 | Mir124a-1hg | 0 | 1.648835551 | 1 |
| ENSMUSG00000114658 | Gm30008 | 0 | 1.648835551 | 1 |
| ENSMUSG00000114357 | Gm48449 | 0 | 1.648835551 | 1 |
| ENSMUSG00000045731 | Pnoc | 0 | -0.52108945 | 1 |
| ENSMUSG00000022041 | Chrna2 | 0 | 0.648835551 | 1 |
| ENSMUSG00000085984 | 1700001G11Rik | 0 | 0.063873051 | 1 |
| ENSMUSG00000022044 | Stmn4 | 0 | 0.326907457 | 1 |
| ENSMUSG00000057913 | Gm10032 | 0 | 1.063873051 | 1 |
| ENSMUSG00000046204 | Pnma2 | 0 | 1.648835551 | 1 |
| ENSMUSG00000075549 | Gm6878 | 0 | 1.063873051 | 1 |
| ENSMUSG00000093634 | Gm10860 | 0 | 1.063873051 | 1 |
| ENSMUSG00000015812 | Gnrh1 | 0 | -1.351164449 | 0.703932614 |
| ENSMUSG00000114621 | Gm31107 | 0 | 1.648835551 | 1 |
| ENSMUSG00000022054 | Nefm | 0 | 0.326907457 | 1 |
| ENSMUSG00000022055 | Nefl | 0 | 0.326907457 | 1 |
| ENSMUSG00000114925 | Gm47212 | 0 | 1.648835551 | 1 |
| ENSMUSG00000022056 | Adam7 | 0 | -0.936126949 | 0.847265357 |
| ENSMUSG00000092499 | 1700092C10Rik | 0 | 1.648835551 | 1 |
| ENSMUSG00000044186 | Nkx2-6 | 0 | 1.063873051 | 1 |
| ENSMUSG00000047977 | Synb | 0 | 1.648835551 | 1 |
| ENSMUSG00000033730 | Egr3 | 0 | 0.063873051 | 1 |
| ENSMUSG00000022091 | Sorbs3 | 0 | -3.52108945 | 0.007344049 |
| ENSMUSG00000022097 | Sftpc | 0 | 1.648835551 | 1 |
| ENSMUSG00000033595 | Lgi3 | 0 | 0.648835551 | 1 |
| ENSMUSG00000045211 | Nudt18 | 0 | -0.351164449 | 1 |
| ENSMUSG00000022101 | Fgf17 | 0 | 1.063873051 | 1 |
| ENSMUSG00000115671 | Gm49289 | 0 | -1.051604167 | 0.858831093 |
| ENSMUSG00000115139 | Gm4251 | 0 | 1.648835551 | 1 |
| ENSMUSG00000109446 | Gm9195 | 0 | 0.326907457 | 1 |
| ENSMUSG00000115088 | Gm49165 | 0 | 0.648835551 | 1 |
| ENSMUSG00000115845 | 4933402J15Rik | 0 | 1.648835551 | 1 |
| ENSMUSG00000115002 | Gm10847 | 0 | 0.063873051 | 1 |
| ENSMUSG00000115019 | Gm4278 | 0 | -0.52108945 | 1 |

| | | | | |
|---|---|---|---|---|
| ENSMUSG00000085829 | Gm4285 | 0 | 1.648835551 | 1 |
| ENSMUSG00000046523 | Kctd4 | 0 | 1.063873051 | 1 |
| ENSMUSG00000115625 | 2900040C04Rik | 0 | 1.648835551 | 1 |
| ENSMUSG00000101009 | 1700108F19Rik | 0 | -0.351164449 | 1 |
| ENSMUSG00000115817 | Gm30246 | 0 | 1.648835551 | 1 |
| ENSMUSG00000115353 | Gm48968 | 0 | 1.648835551 | 1 |
| ENSMUSG00000115596 | Gm48969 | 0 | 1.063873051 | 1 |
| ENSMUSG00000115517 | Gm48954 | 0 | 0.648835551 | 1 |
| ENSMUSG00000045655 | Fam216b | 0 | 1.063873051 | 1 |
| ENSMUSG00000022022 | Mtrf1 | 0 | -1.43862729 | 0.538110273 |
| ENSMUSG00000043881 | Kbtbd7 | 0 | 0.326907457 | 1 |
| ENSMUSG00000115740 | Gm49042 | 0 | 0.648835551 | 1 |
| ENSMUSG00000036422 | Pcdh8 | 0 | 1.063873051 | 1 |
| ENSMUSG00000095959 | Gm10845 | 0 | 0.648835551 | 1 |
| ENSMUSG00000115190 | Gm49542 | 0 | 1.648835551 | 1 |
| ENSMUSG00000022026 | Olfm4 | 0 | -0.351164449 | 1 |
| ENSMUSG00000115006 | Gm49054 | 0 | 1.648835551 | 1 |
| ENSMUSG00000115043 | Gm32455 | 0 | 0.326907457 | 1 |
| ENSMUSG00000115217 | Gm32729 | 0 | 1.648835551 | 1 |
| ENSMUSG00000115777 | Gm32815 | 0 | 0.326907457 | 1 |
| ENSMUSG00000115313 | Gm48941 | 0 | 0.648835551 | 1 |
| ENSMUSG00000115292 | Gm33246 | 0 | 0.648835551 | 1 |
| ENSMUSG00000115822 | Gm33450 | 0 | -0.158519371 | 1 |
| ENSMUSG00000115580 | Gm41230 | 0 | -0.673092543 | 1 |
| ENSMUSG00000054391 | 4930517O19Rik | 0 | -0.673092543 | 1 |
| ENSMUSG00000115821 | 6330576A10Rik | 0 | 1.648835551 | 1 |
| ENSMUSG00000115816 | Gm34589 | 0 | 0.063873051 | 1 |
| ENSMUSG00000022126 | Acod1 | 0 | 1.648835551 | 1 |
| ENSMUSG00000022125 | Cln5 | 0 | -1.258055044 | 0.7014206 |
| ENSMUSG00000110080 | Gm6145 | 0 | -0.158519371 | 1 |
| ENSMUSG00000115796 | Gm48970 | 0 | 0.648835551 | 1 |
| ENSMUSG00000033214 | Slitrk5 | 0 | 1.063873051 | 1 |
| ENSMUSG00000115320 | 4930524C18Rik | 0 | -0.52108945 | 1 |
| ENSMUSG00000115689 | 4930435M08Rik | 0 | 1.648835551 | 1 |
| ENSMUSG00000061517 | Sox21 | 0 | -0.351164449 | 1 |
| ENSMUSG00000098630 | Gm27198 | 0 | 1.063873051 | 1 |
| ENSMUSG00000115220 | Gm49768 | 0 | 0.648835551 | 1 |
| ENSMUSG00000044819 | Oxgr1 | 0 | 0.648835551 | 1 |
| ENSMUSG00000097903 | Gm26679 | 0 | 1.648835551 | 1 |
| ENSMUSG00000115648 | Gm49008 | 0 | 0.326907457 | 1 |
| ENSMUSG00000085133 | B930095G15Rik | 0 | -3.459688905 | 0.008644519 |
| ENSMUSG00000115123 | Gm33299 | 0 | 1.648835551 | 1 |
| ENSMUSG00000025557 | Slc15a1 | 0 | 0.063873051 | 1 |
| ENSMUSG00000115390 | Gm49035 | 0 | 1.648835551 | 1 |
| ENSMUSG00000102112 | 1810041H14Rik | 0 | 1.063873051 | 1 |
| ENSMUSG00000051212 | Gpr183 | 0 | 0.063873051 | 1 |
| ENSMUSG00000071229 | Timm8a2 | 0 | 1.648835551 | 1 |
| ENSMUSG00000085615 | A330035P11Rik | 0 | 0.648835551 | 1 |
| ENSMUSG00000115263 | Gm5089 | 0 | -1.673092543 | 0.658010381 |
| ENSMUSG00000075465 | Gm10837 | 0 | 1.063873051 | 1 |
| ENSMUSG00000085423 | 4930594M22Rik | 0 | 1.063873051 | 1 |

| | | | | |
|---|---|---|---|---|
| ENSMUSG00000099994 | 1700024B18Rik | 0 | -0.158519371 | 1 |
| ENSMUSG00000115490 | 4930404O17Rik | 0 | 0.648835551 | 1 |
| ENSMUSG00000115013 | Gm49231 | 0 | 0.063873051 | 1 |
| ENSMUSG00000091119 | Ccdc152 | 0 | -0.351164449 | 1 |
| ENSMUSG00000087532 | Gm15938 | 0 | 1.063873051 | 1 |
| ENSMUSG00000090199 | Gm16029 | 0 | -0.52108945 | 1 |
| ENSMUSG00000115497 | Gm49207 | 0 | 1.648835551 | 1 |
| ENSMUSG00000095663 | Gm2310 | 0 | 1.648835551 | 1 |
| ENSMUSG00000115667 | Gm10389 | 0 | 1.648835551 | 1 |
| ENSMUSG00000058914 | C1qtnf3 | 0 | 0.326907457 | 1 |
| ENSMUSG00000115585 | Gm41271 | 0 | 0.648835551 | 1 |
| ENSMUSG00000097221 | 1810049J17Rik | 0 | 1.648835551 | 1 |
| ENSMUSG00000022195 | 6030458C11Rik | 0 | -3.540989007 | 0.007698034 |
| ENSMUSG00000115419 | Gm49113 | 0 | 1.648835551 | 1 |
| ENSMUSG00000115782 | Gm49116 | 0 | 0.063873051 | 1 |
| ENSMUSG00000115460 | Gm49127 | 0 | 1.648835551 | 1 |
| ENSMUSG00000115182 | Gm49128 | 0 | 0.648835551 | 1 |
| ENSMUSG00000115352 | Gm49160 | 0 | 1.063873051 | 1 |
| ENSMUSG00000047565 | Acot10 | 0 | 1.648835551 | 1 |
| ENSMUSG00000115174 | Gm35996 | 0 | -0.351164449 | 1 |
| ENSMUSG00000115850 | Gm48956 | 0 | 0.648835551 | 1 |
| ENSMUSG00000115335 | Gm41279 | 0 | 1.648835551 | 1 |
| ENSMUSG00000115839 | Gm49233 | 0 | 1.648835551 | 1 |
| ENSMUSG00000115012 | 9630009A06Rik | 0 | 1.648835551 | 1 |
| ENSMUSG00000022236 | Ropn1l | 0 | 0.063873051 | 1 |
| ENSMUSG00000115710 | Gm49208 | 0 | 1.063873051 | 1 |
| ENSMUSG00000038984 | Tspyl5 | 0 | 0.326907457 | 1 |
| ENSMUSG00000022325 | Pop1 | 0 | -2.965874293 | 0.044171611 |
| ENSMUSG00000050963 | Kcns2 | 0 | 1.648835551 | 1 |
| ENSMUSG00000115345 | Gm33497 | 0 | -1.52108945 | 0.598433943 |
| ENSMUSG00000022330 | Osr2 | 0 | 1.648835551 | 1 |
| ENSMUSG00000115453 | BC048602 | 0 | -1.258055044 | 0.719591951 |
| ENSMUSG00000115829 | Gm48960 | 0 | -0.351164449 | 1 |
| ENSMUSG00000048230 | Fbxo43 | 0 | 1.063873051 | 1 |
| ENSMUSG00000115811 | Gm41293 | 0 | 1.648835551 | 1 |
| ENSMUSG00000013611 | Snx31 | 0 | -0.158519371 | 1 |
| ENSMUSG00000115834 | Gm10385 | 0 | 1.648835551 | 1 |
| ENSMUSG00000072618 | Gm10384 | 0 | 1.063873051 | 1 |
| ENSMUSG00000115430 | Gm41300 | 0 | 0.063873051 | 1 |
| ENSMUSG00000115771 | Gm49312 | 0 | 1.648835551 | 1 |
| ENSMUSG00000115680 | Gm35167 | 0 | 0.063873051 | 1 |
| ENSMUSG00000115456 | Gm35248 | 0 | 0.063873051 | 1 |
| ENSMUSG00000115546 | Gm49077 | 0 | 1.063873051 | 1 |
| ENSMUSG00000097875 | Gm26704 | 0 | 0.648835551 | 1 |
| ENSMUSG00000087695 | Gm16291 | 0 | 0.648835551 | 1 |
| ENSMUSG00000115230 | AU022793 | 0 | 0.648835551 | 1 |
| ENSMUSG00000115142 | Gm33251 | 0 | -0.52108945 | 1 |
| ENSMUSG00000094112 | 9330182O14Rik | 0 | -0.158519371 | 1 |
| ENSMUSG00000097232 | Gm26854 | 0 | -0.158519371 | 1 |
| ENSMUSG00000097805 | Gm17473 | 0 | 1.648835551 | 1 |
| ENSMUSG00000054409 | Tmem74 | 0 | 1.648835551 | 1 |

| | | | | |
|---|---|---|---|---|
| ENSMUSG00000022342 | Kcnv1 | 0 | 1.648835551 | 1 |
| ENSMUSG00000115177 | 4930523O13Rik | 0 | 0.648835551 | 1 |
| ENSMUSG00000115775 | Gm41318 | 0 | 1.063873051 | 1 |
| ENSMUSG00000115339 | Gm48913 | 0 | -0.351164449 | 1 |
| ENSMUSG00000115563 | Gm34562 | 0 | 0.063873051 | 1 |
| ENSMUSG00000115284 | Gm34678 | 0 | 0.326907457 | 1 |
| ENSMUSG00000115616 | Gm48923 | 0 | 1.648835551 | 1 |
| ENSMUSG00000068522 | Aard | 0 | 1.648835551 | 1 |
| ENSMUSG00000022315 | Slc30a8 | 0 | 1.063873051 | 1 |
| ENSMUSG00000115241 | Gm41322 | 0 | 1.648835551 | 1 |
| ENSMUSG00000115137 | Gm34794 | 0 | -0.351164449 | 1 |
| ENSMUSG00000038591 | Colec10 | 0 | 0.648835551 | 1 |
| ENSMUSG00000097286 | Gm26684 | 0 | 1.648835551 | 1 |
| ENSMUSG00000022422 | Dscc1 | 0 | 0.063873051 | 1 |
| ENSMUSG00000053749 | Gm9920 | 0 | -0.52108945 | 1 |
| ENSMUSG00000115200 | Gm46516 | 0 | -1.351164449 | 0.72817309 |
| ENSMUSG00000022357 | Klhl38 | 0 | 0.326907457 | 1 |
| ENSMUSG00000115576 | Gm2675 | 0 | 1.063873051 | 1 |
| ENSMUSG00000115426 | Gm19510 | 0 | 1.648835551 | 1 |
| ENSMUSG00000022347 | A1bg | 0 | -0.52108945 | 1 |
| ENSMUSG00000115676 | 4930449C09Rik | 0 | 0.063873051 | 1 |
| ENSMUSG00000115464 | Gm49019 | 0 | 0.648835551 | 1 |
| ENSMUSG00000072511 | Hhla1 | 0 | 1.063873051 | 1 |
| ENSMUSG00000099013 | Gm27153 | 0 | 1.648835551 | 1 |
| ENSMUSG00000090464 | Gm17140 | 0 | 1.648835551 | 1 |
| ENSMUSG00000005124 | Ccn4 | 0 | -1.673092543 | 0.462591844 |
| ENSMUSG00000116289 | Gm31342 | 0 | 1.063873051 | 1 |
| ENSMUSG00000102069 | 1700012I11Rik | 0 | -1.52108945 | 0.6134821 |
| ENSMUSG00000096967 | Gm26621 | 0 | -1.351164449 | 0.687112677 |
| ENSMUSG00000090237 | Gm16308 | 0 | 1.648835551 | 1 |
| ENSMUSG00000116088 | Gm49460 | 0 | 1.648835551 | 1 |
| ENSMUSG00000115693 | 1700085D07Rik | 0 | 1.063873051 | 1 |
| ENSMUSG00000115810 | Gm49066 | 0 | -2.52108945 | 0.139351384 |
| ENSMUSG00000045281 | Gpr20 | 0 | 0.326907457 | 1 |
| ENSMUSG00000116256 | Gm32405 | 0 | 0.063873051 | 1 |
| ENSMUSG00000022602 | Arc | 0 | 1.648835551 | 1 |
| ENSMUSG00000044276 | 4933427E11Rik | 0 | 1.648835551 | 1 |
| ENSMUSG00000101026 | Ly6g6g | 0 | 1.648835551 | 1 |
| ENSMUSG00000044678 | Ly6k | 0 | 1.648835551 | 1 |
| ENSMUSG00000068349 | Gml | 0 | 1.648835551 | 1 |
| ENSMUSG00000068600 | Gml2 | 0 | -0.158519371 | 1 |
| ENSMUSG00000075604 | Cyp11b1 | 0 | 0.648835551 | 1 |
| ENSMUSG00000063522 | Ly6m | 0 | 0.648835551 | 1 |
| ENSMUSG00000022586 | Ly6i | 0 | 1.648835551 | 1 |
| ENSMUSG00000099452 | Gm28502 | 0 | 1.648835551 | 1 |
| ENSMUSG00000047003 | Zfp41 | 0 | -0.52108945 | 1 |
| ENSMUSG00000115969 | Gm46519 | 0 | 1.063873051 | 1 |
| ENSMUSG00000022580 | Rhpn1 | 0 | 1.063873051 | 1 |
| ENSMUSG00000063704 | Mapk15 | 0 | -0.158519371 | 1 |
| ENSMUSG00000115332 | Gm48952 | 0 | 0.063873051 | 1 |
| ENSMUSG00000049653 | Spatc1 | 0 | -0.158519371 | 1 |

| | | | | |
|---|---|---|---|---|
| ENSMUSG00000071724 | Smpd5 | 0 | 0.648835551 | 1 |
| ENSMUSG00000022554 | Hgh1 | 0 | -1.258055044 | 0.673811773 |
| ENSMUSG00000048385 | Scrt1 | 0 | 0.648835551 | 1 |
| ENSMUSG00000004187 | Kifc2 | 0 | -1.599091962 | 0.572887115 |
| ENSMUSG00000019080 | Mfsd3 | 0 | 0.648835551 | 1 |
| ENSMUSG00000033762 | Recql4 | 0 | 0.063873051 | 1 |
| ENSMUSG00000033707 | Lrrc24 | 0 | 1.648835551 | 1 |
| ENSMUSG00000022526 | Zfp251 | 0 | -2.305360759 | 0.231596497 |
| ENSMUSG00000055041 | Commd5 | 0 | 0.063873051 | 1 |
| ENSMUSG00000116339 | Gm35933 | 0 | 1.648835551 | 1 |
| ENSMUSG00000033576 | Apol6 | 0 | 1.063873051 | 1 |
| ENSMUSG00000057346 | Apol9a | 0 | -0.351164449 | 1 |
| ENSMUSG00000068252 | Apol7b | 0 | 1.063873051 | 1 |
| ENSMUSG00000086467 | 4930571N24Rik | 0 | 0.648835551 | 1 |
| ENSMUSG00000044309 | Apol7c | 0 | 1.648835551 | 1 |
| ENSMUSG00000101113 | 1700025B11Rik | 0 | 1.063873051 | 1 |
| ENSMUSG00000050014 | Apol10b | 0 | -0.158519371 | 1 |
| ENSMUSG00000116079 | Gm36245 | 0 | 1.648835551 | 1 |
| ENSMUSG00000091694 | Apol11b | 0 | 1.063873051 | 1 |
| ENSMUSG00000116162 | Gm36329 | 0 | 1.648835551 | 1 |
| ENSMUSG00000068246 | Apol9b | 0 | 0.063873051 | 1 |
| ENSMUSG00000056656 | Apol8 | 0 | 0.648835551 | 1 |
| ENSMUSG00000016552 | Foxred2 | 0 | -0.52108945 | 1 |
| ENSMUSG00000005716 | Pvalb | 0 | 0.326907457 | 1 |
| ENSMUSG00000071714 | Csf2rb2 | 0 | 0.648835551 | 1 |
| ENSMUSG00000062154 | Tex33 | 0 | 1.063873051 | 1 |
| ENSMUSG00000018169 | Mfng | 0 | 1.063873051 | 1 |
| ENSMUSG00000097085 | Gm26634 | 0 | 1.648835551 | 1 |
| ENSMUSG00000116505 | Gm10865 | 0 | 1.063873051 | 1 |
| ENSMUSG00000115867 | Gm17753 | 0 | -1.258055044 | 0.736456705 |
| ENSMUSG00000092494 | Gm20420 | 0 | 1.648835551 | 1 |
| ENSMUSG00000044216 | Kcnj4 | 0 | 1.063873051 | 1 |
| ENSMUSG00000042524 | Sun2 | 0 | -1.810596067 | 0.466464457 |
| ENSMUSG00000087543 | Gm16576 | 0 | -1.599091962 | 0.57709448 |
| ENSMUSG00000022421 | Nptxr | 0 | 1.063873051 | 1 |
| ENSMUSG00000115971 | Gm10856 | 0 | -0.351164449 | 1 |
| ENSMUSG00000115888 | Gm31369 | 0 | 1.063873051 | 1 |
| ENSMUSG00000116480 | Gm49512 | 0 | 0.326907457 | 1 |
| ENSMUSG00000047938 | 4930483J18Rik | 0 | 0.648835551 | 1 |
| ENSMUSG00000047108 | Dnajb7 | 0 | -0.351164449 | 1 |
| ENSMUSG00000091492 | Gm17025 | 0 | -0.351164449 | 1 |
| ENSMUSG00000097613 | Gm17597 | 0 | 1.063873051 | 1 |
| ENSMUSG00000096883 | Shisa8 | 0 | 1.648835551 | 1 |
| ENSMUSG00000068105 | Tnfrsf13c | 0 | 1.648835551 | 1 |
| ENSMUSG00000116009 | Gm49503 | 0 | 1.063873051 | 1 |
| ENSMUSG00000022456 | 3-Sep | 0 | 1.063873051 | 1 |
| ENSMUSG00000061740 | Cyp2d22 | 0 | 1.063873051 | 1 |
| ENSMUSG00000068086 | Cyp2d9 | 0 | 1.648835551 | 1 |
| ENSMUSG00000096852 | Cyp2d12 | 0 | 1.648835551 | 1 |
| ENSMUSG00000094559 | Cyp2d34 | 0 | 1.648835551 | 1 |
| ENSMUSG00000100199 | Gm20324 | 0 | 0.648835551 | 1 |

| | | | | |
|---|---|---|---|---|
| ENSMUSG00000075511 | 1700001L05Rik | 0 | 0.326907457 | 1 |
| ENSMUSG00000016758 | Bik | 0 | -0.351164449 | 1 |
| ENSMUSG00000018865 | Sult4a1 | 0 | 0.648835551 | 1 |
| ENSMUSG00000018868 | Pnpla5 | 0 | 1.648835551 | 1 |
| ENSMUSG00000041653 | Pnpla3 | 0 | 0.648835551 | 1 |
| ENSMUSG00000022439 | Parvg | 0 | -1.810596067 | 0.466464457 |
| ENSMUSG00000116409 | Gm34095 | 0 | 1.648835551 | 1 |
| ENSMUSG00000116396 | Gm20556 | 0 | 0.648835551 | 1 |
| ENSMUSG00000100967 | Gm29666 | 0 | 1.063873051 | 1 |
| ENSMUSG00000022435 | Upk3a | 0 | 1.648835551 | 1 |
| ENSMUSG00000022431 | Ribc2 | 0 | 0.648835551 | 1 |
| ENSMUSG00000116004 | Gm49539 | 0 | -1.43862729 | 0.538110273 |
| ENSMUSG00000116435 | Gm49548 | 0 | 1.648835551 | 1 |
| ENSMUSG00000089680 | Gm15722 | 0 | 1.648835551 | 1 |
| ENSMUSG00000116040 | Gm49464 | 0 | 1.063873051 | 1 |
| ENSMUSG00000116229 | Gm49470 | 0 | 0.648835551 | 1 |
| ENSMUSG00000115930 | A930027H12Rik | 0 | 1.063873051 | 1 |
| ENSMUSG00000116061 | Gm49472 | 0 | 0.326907457 | 1 |
| ENSMUSG00000116331 | B230214G05Rik | 0 | -1.43862729 | 0.73806297 |
| ENSMUSG00000058441 | Panx2 | 0 | 1.648835551 | 1 |
| ENSMUSG00000035757 | Selenoo | 0 | -2.158519371 | 0.300120509 |
| ENSMUSG00000062906 | Hdac10 | 0 | -0.158519371 | 1 |
| ENSMUSG00000054136 | Adm2 | 0 | 0.326907457 | 1 |
| ENSMUSG00000078938 | Syce3 | 0 | 1.063873051 | 1 |
| ENSMUSG00000078937 | Cpt1b | 0 | -1.051604167 | 0.858831093 |
| ENSMUSG00000079242 | C730034F03Rik | 0 | 0.326907457 | 1 |
| ENSMUSG00000022622 | Acr | 0 | 0.648835551 | 1 |
| ENSMUSG00000116204 | Gm36480 | 0 | 1.063873051 | 1 |
| ENSMUSG00000086160 | 4933438A12Rik | 0 | 1.648835551 | 1 |
| ENSMUSG00000116309 | Gm49418 | 0 | 0.648835551 | 1 |
| ENSMUSG00000116237 | Gm30339 | 0 | 0.326907457 | 1 |
| ENSMUSG00000116360 | 9430014N10Rik | 0 | 0.063873051 | 1 |
| ENSMUSG00000045608 | Dbx2 | 0 | 0.648835551 | 1 |
| ENSMUSG00000097643 | A130051J06Rik | 0 | 1.648835551 | 1 |
| ENSMUSG00000078648 | Gm17546 | 0 | 1.648835551 | 1 |
| ENSMUSG00000078925 | D030018L15Rik | 0 | 0.648835551 | 1 |
| ENSMUSG00000097619 | 4833422M21Rik | 0 | 1.648835551 | 1 |
| ENSMUSG00000116295 | Gm32885 | 0 | 1.063873051 | 1 |
| ENSMUSG00000022483 | Col2a1 | 0 | 0.648835551 | 1 |
| ENSMUSG00000090129 | Olfr287 | 0 | 1.648835551 | 1 |
| ENSMUSG00000059460 | Olfr286 | 0 | 1.063873051 | 1 |
| ENSMUSG00000022987 | Zfp641 | 0 | -1.743481871 | 0.455131546 |
| ENSMUSG00000062037 | Olfr285 | 0 | 1.648835551 | 1 |
| ENSMUSG00000108728 | OR5BS1P | 0 | 1.648835551 | 1 |
| ENSMUSG00000022991 | Lalba | 0 | 1.648835551 | 1 |
| ENSMUSG00000091050 | 9330020H09Rik | 0 | 0.063873051 | 1 |
| ENSMUSG00000022996 | Wnt10b | 0 | 1.648835551 | 1 |
| ENSMUSG00000097414 | B130046B21Rik | 0 | 1.063873051 | 1 |
| ENSMUSG00000023000 | Dhh | 0 | -1.051604167 | 0.858831093 |
| ENSMUSG00000097587 | 4930578M01Rik | 0 | -1.051604167 | 0.858831093 |
| ENSMUSG00000116371 | Gm34284 | 0 | 1.648835551 | 1 |

| | | | | |
|---|---|---|---|---|
| ENSMUSG00000089968 | Nckap5los | 0 | 0.326907457 | 1 |
| ENSMUSG00000116249 | Gm49484 | 0 | 0.326907457 | 1 |
| ENSMUSG00000089671 | Gm16537 | 0 | 1.063873051 | 1 |
| ENSMUSG00000096111 | Gm21917 | 0 | -0.936126949 | 0.847265357 |
| ENSMUSG00000116311 | 4930478M13Rik | 0 | 0.648835551 | 1 |
| ENSMUSG00000045631 | Tmprss12 | 0 | 1.063873051 | 1 |
| ENSMUSG00000058057 | Mettl7a3 | 0 | 1.648835551 | 1 |
| ENSMUSG00000056487 | Mettl7a2 | 0 | 0.648835551 | 1 |
| ENSMUSG00000093550 | Higd1c | 0 | 0.063873051 | 1 |
| ENSMUSG00000097827 | 5330439K02Rik | 0 | -0.351164449 | 1 |
| ENSMUSG00000116164 | 1110013H19Rik | 0 | 1.648835551 | 1 |
| ENSMUSG00000097093 | C330013E15Rik | 0 | 1.648835551 | 1 |
| ENSMUSG00000090062 | Galnt6os | 0 | -0.52108945 | 1 |
| ENSMUSG00000116504 | I730030J21Rik | 0 | 1.648835551 | 1 |
| ENSMUSG00000097915 | A330009N23Rik | 0 | 1.648835551 | 1 |
| ENSMUSG00000075408 | Smim41 | 0 | 0.063873051 | 1 |
| ENSMUSG00000116114 | Gm35853 | 0 | -1.673092543 | 0.451785624 |
| ENSMUSG00000067614 | Krt86 | 0 | 0.648835551 | 1 |
| ENSMUSG00000116336 | Gm49425 | 0 | 1.648835551 | 1 |
| ENSMUSG00000064232 | Gm5414 | 0 | 1.648835551 | 1 |
| ENSMUSG00000061527 | Krt5 | 0 | 1.648835551 | 1 |
| ENSMUSG00000056605 | Krt72 | 0 | 0.326907457 | 1 |
| ENSMUSG00000046834 | Krt1 | 0 | 1.648835551 | 1 |
| ENSMUSG00000067594 | Krt77 | 0 | 1.648835551 | 1 |
| ENSMUSG00000050463 | Krt78 | 0 | 1.063873051 | 1 |
| ENSMUSG00000116010 | Gm36026 | 0 | -0.158519371 | 1 |
| ENSMUSG00000001281 | Itgb7 | 0 | -1.874726405 | 0.495276006 |
| ENSMUSG00000116091 | Gm49479 | 0 | 1.648835551 | 1 |
| ENSMUSG00000060284 | Sp7 | 0 | 1.648835551 | 1 |
| ENSMUSG00000116500 | Gm36560 | 0 | 1.648835551 | 1 |
| ENSMUSG00000001656 | Hoxc11 | 0 | 1.648835551 | 1 |
| ENSMUSG00000022485 | Hoxc5 | 0 | -0.158519371 | 1 |
| ENSMUSG00000036139 | Hoxc9 | 0 | 1.648835551 | 1 |
| ENSMUSG00000001657 | Hoxc8 | 0 | -0.52108945 | 1 |
| ENSMUSG00000001661 | Hoxc6 | 0 | 1.063873051 | 1 |
| ENSMUSG00000075394 | Hoxc4 | 0 | -0.158519371 | 1 |
| ENSMUSG00000058794 | Nfe2 | 0 | 0.648835551 | 1 |
| ENSMUSG00000116308 | Gm49482 | 0 | 0.648835551 | 1 |
| ENSMUSG00000036925 | Mucl2 | 0 | 1.648835551 | 1 |
| ENSMUSG00000116389 | Gm49504 | 0 | 1.063873051 | 1 |
| ENSMUSG00000115871 | Gm49507 | 0 | 0.326907457 | 1 |
| ENSMUSG00000116455 | Gm35974 | 0 | 1.063873051 | 1 |
| ENSMUSG00000022534 | Mefv | 0 | 0.648835551 | 1 |
| ENSMUSG00000005983 | 1700037C18Rik | 0 | -0.52108945 | 1 |
| ENSMUSG00000085048 | Gm15537 | 0 | 1.648835551 | 1 |
| ENSMUSG00000084790 | Gm15879 | 0 | -0.673092543 | 1 |
| ENSMUSG00000005718 | Tfap4 | 0 | -1.43862729 | 0.538110273 |
| ENSMUSG00000063445 | Nmral1 | 0 | 0.648835551 | 1 |
| ENSMUSG00000051669 | AU021092 | 0 | -0.158519371 | 1 |
| ENSMUSG00000039427 | Alg1 | 0 | 0.326907457 | 1 |
| ENSMUSG00000116450 | Gm49534 | 0 | 1.648835551 | 1 |

| | | | | |
|---|---|---|---|---|
| ENSMUSG00000022715 | Tmem114 | 0 | 0.326907457 | 1 |
| ENSMUSG00000043140 | Tmem186 | 0 | -1.52108945 | 0.554265357 |
| ENSMUSG00000107252 | Gm5767 | 0 | 0.063873051 | 1 |
| ENSMUSG00000116426 | Gm30567 | 0 | 1.648835551 | 1 |
| ENSMUSG00000115943 | Gm49455 | 0 | -0.351164449 | 1 |
| ENSMUSG00000116270 | Gm1600 | 0 | -0.673092543 | 1 |
| ENSMUSG00000039179 | Tekt5 | 0 | -0.936126949 | 0.85551189 |
| ENSMUSG00000050908 | Tvp23a | 0 | -0.673092543 | 1 |
| ENSMUSG00000038037 | Socs1 | 0 | 0.063873051 | 1 |
| ENSMUSG00000038015 | Prm2 | 0 | 1.648835551 | 1 |
| ENSMUSG00000022501 | Prm1 | 0 | 0.326907457 | 1 |
| ENSMUSG00000116038 | Gm46563 | 0 | 0.063873051 | 1 |
| ENSMUSG00000115992 | Gm10832 | 0 | 0.326907457 | 1 |
| ENSMUSG00000037972 | Snn | 0 | -1.051604167 | 0.858831093 |
| ENSMUSG00000096959 | 4930509G22Rik | 0 | -1.258055044 | 0.726471441 |
| ENSMUSG00000022496 | Tnfrsf17 | 0 | 1.648835551 | 1 |
| ENSMUSG00000101565 | 1700003L19Rik | 0 | -0.351164449 | 1 |
| ENSMUSG00000109857 | Gm32817 | 0 | 0.648835551 | 1 |
| ENSMUSG00000022676 | Snai2 | 0 | 1.648835551 | 1 |
| ENSMUSG00000115923 | Gm49521 | 0 | 1.648835551 | 1 |
| ENSMUSG00000116096 | 4933404G15Rik | 0 | 0.648835551 | 1 |
| ENSMUSG00000075370 | Igll1 | 0 | 1.063873051 | 1 |
| ENSMUSG00000084792 | 1700056N10Rik | 0 | -0.936126949 | 0.847265357 |
| ENSMUSG00000049916 | 2610318N02Rik | 0 | -1.673092543 | 0.451785624 |
| ENSMUSG00000022769 | Sdf2l1 | 0 | 0.326907457 | 1 |
| ENSMUSG00000022766 | Serpind1 | 0 | 0.326907457 | 1 |
| ENSMUSG00000022759 | Lrrc74b | 0 | 1.648835551 | 1 |
| ENSMUSG00000022756 | Slc7a4 | 0 | -1.051604167 | 0.858831093 |
| ENSMUSG00000041617 | Ccdc74a | 0 | 1.648835551 | 1 |
| ENSMUSG00000022721 | Trmt2a | 0 | -2.258055044 | 0.222101622 |
| ENSMUSG00000086965 | Rtl10 | 0 | 1.648835551 | 1 |
| ENSMUSG00000009097 | Tbx1 | 0 | 1.063873051 | 1 |
| ENSMUSG00000050761 | Gp1bb | 0 | 1.063873051 | 1 |
| ENSMUSG00000072214 | 5-Sep | 0 | 0.648835551 | 1 |
| ENSMUSG00000105906 | Iglc1 | 0 | 1.648835551 | 1 |
| ENSMUSG00000105547 | Iglc3 | 0 | 1.063873051 | 1 |
| ENSMUSG00000076937 | Iglc2 | 0 | 1.063873051 | 1 |
| ENSMUSG00000104927 | Gm43388 | 0 | 1.063873051 | 1 |
| ENSMUSG00000076940 | Iglv2 | 0 | 1.648835551 | 1 |
| ENSMUSG00000050742 | Olfr164 | 0 | 1.648835551 | 1 |
| ENSMUSG00000022686 | B3gnt5 | 0 | 0.326907457 | 1 |
| ENSMUSG00000086837 | Gm16618 | 0 | 0.648835551 | 1 |
| ENSMUSG00000033809 | Alg3 | 0 | -0.52108945 | 1 |
| ENSMUSG00000022847 | Thpo | 0 | -0.158519371 | 1 |
| ENSMUSG00000006958 | Chrd | 0 | 0.326907457 | 1 |
| ENSMUSG00000116922 | 4833419O12Rik | 0 | 1.648835551 | 1 |
| ENSMUSG00000090356 | Teddm3 | 0 | 1.648835551 | 1 |
| ENSMUSG00000005958 | Ephb3 | 0 | -1.874726405 | 0.352215149 |
| ENSMUSG00000116843 | Gm49750 | 0 | 0.326907457 | 1 |
| ENSMUSG00000022853 | Ehhadh | 0 | -0.52108945 | 1 |
| ENSMUSG00000097401 | Gm26744 | 0 | 1.648835551 | 1 |

| | | | | |
|---|---|---|---|---|
| ENSMUSG00000089874 | 9230117E06Rik | 0 | 0.648835551 | 1 |
| ENSMUSG00000089663 | Gm15651 | 0 | 1.063873051 | 1 |
| ENSMUSG00000033501 | Crygs | 0 | 1.648835551 | 1 |
| ENSMUSG00000116831 | Gm30505 | 0 | 1.063873051 | 1 |
| ENSMUSG00000022868 | Ahsg | 0 | 1.063873051 | 1 |
| ENSMUSG00000022877 | Hrg | 0 | -2.051604167 | 0.451785624 |
| ENSMUSG00000022878 | Adipoq | 0 | 1.648835551 | 1 |
| ENSMUSG00000004366 | Sst | 0 | 1.648835551 | 1 |
| ENSMUSG00000115919 | Gm31583 | 0 | 1.648835551 | 1 |
| ENSMUSG00000096960 | A230028O05Rik | 0 | 1.648835551 | 1 |
| ENSMUSG00000091972 | Tmem207 | 0 | 0.648835551 | 1 |
| ENSMUSG00000116530 | Gm32312 | 0 | 1.648835551 | 1 |
| ENSMUSG00000056423 | Uts2b | 0 | 0.326907457 | 1 |
| ENSMUSG00000022525 | Plaat1 | 0 | 1.063873051 | 1 |
| ENSMUSG00000116814 | Gm49636 | 0 | 1.648835551 | 1 |
| ENSMUSG00000116891 | Gm32679 | 0 | 1.063873051 | 1 |
| ENSMUSG00000097184 | 4632428C04Rik | 0 | -1.743481871 | 0.455131546 |
| ENSMUSG00000101888 | 1700025H01Rik | 0 | 1.648835551 | 1 |
| ENSMUSG00000097621 | Gm26562 | 0 | 1.063873051 | 1 |
| ENSMUSG00000047953 | Gp5 | 0 | 0.648835551 | 1 |
| ENSMUSG00000116570 | Gm49754 | 0 | 1.648835551 | 1 |
| ENSMUSG00000116951 | Gm49756 | 0 | 0.326907457 | 1 |
| ENSMUSG00000116737 | Gm49757 | 0 | 1.648835551 | 1 |
| ENSMUSG00000116946 | Gm41442 | 0 | 0.648835551 | 1 |
| ENSMUSG00000022548 | Apod | 0 | 1.063873051 | 1 |
| ENSMUSG00000116690 | Gm49736 | 0 | 1.063873051 | 1 |
| ENSMUSG00000116793 | Gm34256 | 0 | 1.063873051 | 1 |
| ENSMUSG00000045625 | Pigz | 0 | -0.52108945 | 1 |
| ENSMUSG00000086146 | Gm15729 | 0 | 1.063873051 | 1 |
| ENSMUSG00000075269 | Bex6 | 0 | 1.063873051 | 1 |
| ENSMUSG00000046345 | Smco1 | 0 | 1.063873051 | 1 |
| ENSMUSG00000035638 | Muc20 | 0 | 0.648835551 | 1 |
| ENSMUSG00000094430 | Smbd1 | 0 | 0.648835551 | 1 |
| ENSMUSG00000035578 | Iqcg | 0 | 1.648835551 | 1 |
| ENSMUSG00000097318 | 1700007L15Rik | 0 | -0.351164449 | 1 |
| ENSMUSG00000100593 | 1700119H24Rik | 0 | 1.063873051 | 1 |
| ENSMUSG00000116840 | Gm21691 | 0 | -1.258055044 | 0.678664944 |
| ENSMUSG00000099429 | 1600019K03Rik | 0 | 1.063873051 | 1 |
| ENSMUSG00000097914 | Gm26838 | 0 | 1.648835551 | 1 |
| ENSMUSG00000034379 | Wdr5b | 0 | -0.52108945 | 1 |
| ENSMUSG00000034362 | Csta1 | 0 | 1.063873051 | 1 |
| ENSMUSG00000059657 | Stfa2l1 | 0 | 1.648835551 | 1 |
| ENSMUSG00000094733 | Csta3 | 0 | 0.063873051 | 1 |
| ENSMUSG00000095620 | Csta2 | 0 | 0.648835551 | 1 |
| ENSMUSG00000071561 | Cstdc5 | 0 | 1.063873051 | 1 |
| ENSMUSG00000087689 | Gm15845 | 0 | 0.648835551 | 1 |
| ENSMUSG00000054905 | Stfa3 | 0 | 1.063873051 | 1 |
| ENSMUSG00000047746 | Fbxo40 | 0 | 0.648835551 | 1 |
| ENSMUSG00000022827 | Rabl3 | 0 | -1.874726405 | 0.352215149 |
| ENSMUSG00000087568 | Maats1os | 0 | 1.063873051 | 1 |
| ENSMUSG00000022803 | Popdc2 | 0 | -2.158519371 | 0.290284345 |

| | | | | |
|---|---|---|---|---|
| ENSMUSG00000002844 | Adprh | 0 | -1.52108945 | 0.544908989 |
| ENSMUSG00000097576 | D930030I03Rik | 0 | 0.326907457 | 1 |
| ENSMUSG00000091238 | Gm17103 | 0 | -0.936126949 | 0.847265357 |
| ENSMUSG00000116903 | Gm19522 | 0 | -0.351164449 | 1 |
| ENSMUSG00000055370 | Gm9968 | 0 | 1.063873051 | 1 |
| ENSMUSG00000085499 | Gm15713 | 0 | 1.063873051 | 1 |
| ENSMUSG00000071552 | Tigit | 0 | -0.673092543 | 1 |
| ENSMUSG00000043065 | Spice1 | 0 | -3.417253639 | 0.010795678 |
| ENSMUSG00000062082 | Cd200r4 | 0 | 0.063873051 | 1 |
| ENSMUSG00000052013 | Btla | 0 | -0.158519371 | 1 |
| ENSMUSG00000053182 | Gm609 | 0 | -1.52108945 | 0.544908989 |
| ENSMUSG00000116542 | Gm17783 | 0 | 1.648835551 | 1 |
| ENSMUSG00000022659 | Gcsam | 0 | 1.063873051 | 1 |
| ENSMUSG00000033177 | Tmprss7 | 0 | -0.351164449 | 1 |
| ENSMUSG00000087141 | Plcxd2 | 0 | -1.936126949 | 0.421719645 |
| ENSMUSG00000058550 | Dppa4 | 0 | 1.648835551 | 1 |
| ENSMUSG00000022650 | Retnlb | 0 | 1.648835551 | 1 |
| ENSMUSG00000087066 | Gm15518 | 0 | -0.158519371 | 1 |
| ENSMUSG00000102101 | Zbtb11os1 | 0 | 0.063873051 | 1 |
| ENSMUSG00000109588 | Lnp1 | 0 | 1.648835551 | 1 |
| ENSMUSG00000046748 | Tmem45a2 | 0 | 1.648835551 | 1 |
| ENSMUSG00000086827 | 4930461C15Rik | 0 | 1.648835551 | 1 |
| ENSMUSG00000116579 | Gm49701 | 0 | 1.648835551 | 1 |
| ENSMUSG00000049362 | Olfr173 | 0 | 1.648835551 | 1 |
| ENSMUSG00000060663 | Olfr175-ps1 | 0 | 1.648835551 | 1 |
| ENSMUSG00000047960 | Olfr186 | 0 | 1.063873051 | 1 |
| ENSMUSG00000096695 | Olfr196 | 0 | 1.648835551 | 1 |
| ENSMUSG00000052537 | Olfr198 | 0 | 1.648835551 | 1 |
| ENSMUSG00000068182 | Olfr203 | 0 | 1.648835551 | 1 |
| ENSMUSG00000074991 | Gabrr3 | 0 | 0.326907457 | 1 |
| ENSMUSG00000099983 | 1700022E09Rik | 0 | 1.648835551 | 1 |
| ENSMUSG00000022722 | Arl6 | 0 | 0.326907457 | 1 |
| ENSMUSG00000116619 | Gm49615 | 0 | 1.648835551 | 1 |
| ENSMUSG00000047854 | Stx19 | 0 | 0.326907457 | 1 |
| ENSMUSG00000116752 | Gm49622 | 0 | 1.648835551 | 1 |
| ENSMUSG00000116699 | Gm49631 | 0 | 1.063873051 | 1 |
| ENSMUSG00000116750 | 4931420L22Rik | 0 | 1.648835551 | 1 |
| ENSMUSG00000116982 | 4930567J20Rik | 0 | 0.648835551 | 1 |
| ENSMUSG00000116769 | 8030451O07Rik | 0 | 1.648835551 | 1 |
| ENSMUSG00000032948 | Lipi | 0 | 0.648835551 | 1 |
| ENSMUSG00000032940 | Rbm11 | 0 | 1.063873051 | 1 |
| ENSMUSG00000116695 | 4930578N18Rik | 0 | 0.063873051 | 1 |
| ENSMUSG00000079546 | Gm11146 | 0 | 1.648835551 | 1 |
| ENSMUSG00000091193 | Gm17333 | 0 | -0.936126949 | 0.877701541 |
| ENSMUSG00000094030 | Gm21833 | 0 | 1.648835551 | 1 |
| ENSMUSG00000116745 | Gm41481 | 0 | 0.326907457 | 1 |
| ENSMUSG00000116907 | 4930553E22Rik | 0 | 1.063873051 | 1 |
| ENSMUSG00000097418 | Mir155hg | 0 | -0.673092543 | 1 |
| ENSMUSG00000115518 | Gm10791 | 0 | 0.648835551 | 1 |
| ENSMUSG00000115469 | Gm49227 | 0 | 0.648835551 | 1 |
| ENSMUSG00000114988 | Gm49226 | 0 | 0.326907457 | 1 |

| | | | | |
|---|---|---|---|---|
| ENSMUSG00000116586 | 4930556C24Rik | 0 | 1.648835551 | 1 |
| ENSMUSG00000116799 | Gm32357 | 0 | 0.648835551 | 1 |
| ENSMUSG00000116993 | Gm32461 | 0 | -0.158519371 | 1 |
| ENSMUSG00000041079 | Rwdd2b | 0 | -1.743481871 | 0.455131546 |
| ENSMUSG00000055972 | 2810407A14Rik | 0 | -0.673092543 | 1 |
| ENSMUSG00000116754 | Gm33255 | 0 | 1.648835551 | 1 |
| ENSMUSG00000050239 | Krtap24-1 | 0 | 1.648835551 | 1 |
| ENSMUSG00000022931 | Krtap15 | 0 | 0.648835551 | 1 |
| ENSMUSG00000116600 | Gm49707 | 0 | 0.326907457 | 1 |
| ENSMUSG00000116989 | Gm49709 | 0 | 0.648835551 | 1 |
| ENSMUSG00000116753 | Gm36001 | 0 | 1.648835551 | 1 |
| ENSMUSG00000116830 | Gm49714 | 0 | 1.648835551 | 1 |
| ENSMUSG00000116582 | Gm36169 | 0 | 1.648835551 | 1 |
| ENSMUSG00000039956 | Mrap | 0 | -0.673092543 | 1 |
| ENSMUSG00000085279 | Gm15965 | 0 | -0.52108945 | 1 |
| ENSMUSG00000039851 | 4932438H23Rik | 0 | -0.158519371 | 1 |
| ENSMUSG00000116763 | Gm41492 | 0 | 0.648835551 | 1 |
| ENSMUSG00000046160 | Olig1 | 0 | 1.648835551 | 1 |
| ENSMUSG00000084866 | A930006K02Rik | 0 | 1.063873051 | 1 |
| ENSMUSG00000085337 | Gm15964 | 0 | 1.063873051 | 1 |
| ENSMUSG00000022960 | Donson | 0 | -1.43862729 | 0.538110273 |
| ENSMUSG00000087240 | Gm15976 | 0 | 1.063873051 | 1 |
| ENSMUSG00000087190 | D430001F17Rik | 0 | -0.158519371 | 1 |
| ENSMUSG00000086331 | Gm16310 | 0 | 0.063873051 | 1 |
| ENSMUSG00000051728 | Fam243 | 0 | 1.648835551 | 1 |
| ENSMUSG00000039639 | Kcne1 | 0 | 1.063873051 | 1 |
| ENSMUSG00000051297 | 2410124H12Rik | 0 | 1.063873051 | 1 |
| ENSMUSG00000097139 | Gm26626 | 0 | 1.063873051 | 1 |
| ENSMUSG00000116805 | Gm49723 | 0 | -1.599091962 | 0.606655093 |
| ENSMUSG00000116595 | Gm30695 | 0 | -0.158519371 | 1 |
| ENSMUSG00000117317 | Gm49961 | 0 | 0.326907457 | 1 |
| ENSMUSG00000116927 | Gm30881 | 0 | -0.936126949 | 0.876018808 |
| ENSMUSG00000116569 | Gm49620 | 0 | -0.351164449 | 1 |
| ENSMUSG00000115246 | Gm48984 | 0 | 1.648835551 | 1 |
| ENSMUSG00000062609 | Kcnj15 | 0 | -0.52108945 | 1 |
| ENSMUSG00000117289 | 1700093J21Rik | 0 | 0.063873051 | 1 |
| ENSMUSG00000117129 | Gm49928 | 0 | 1.648835551 | 1 |
| ENSMUSG00000074892 | B3galt5 | 0 | 0.326907457 | 1 |
| ENSMUSG00000116874 | Gm34035 | 0 | 1.648835551 | 1 |
| ENSMUSG00000036983 | Tfb1m | 0 | -0.936126949 | 1 |
| ENSMUSG00000116975 | 4930470H14Rik | 0 | 1.063873051 | 1 |
| ENSMUSG00000116615 | 4930548J01Rik | 0 | 1.648835551 | 1 |
| ENSMUSG00000096965 | 3300005D01Rik | 0 | 0.326907457 | 1 |
| ENSMUSG00000092074 | Dynlt1a | 0 | 0.326907457 | 1 |
| ENSMUSG00000000579 | Dynlt1c | 0 | 1.063873051 | 1 |
| ENSMUSG00000116644 | 4933426B08Rik | 0 | 0.326907457 | 1 |
| ENSMUSG00000099735 | 1700122H20Rik | 0 | 1.648835551 | 1 |
| ENSMUSG00000094083 | Gm1604b | 0 | 0.063873051 | 1 |
| ENSMUSG00000071322 | Tcp10a | 0 | 1.063873051 | 1 |
| ENSMUSG00000079722 | Ttll2 | 0 | 1.648835551 | 1 |
| ENSMUSG00000056133 | Unc93a2 | 0 | 1.648835551 | 1 |

| | | | | |
|---|---|---|---|---|
| ENSMUSG00000033450 | Tagap | 0 | 1.648835551 | 1 |
| ENSMUSG00000116760 | Gm1604a | 0 | -1.43862729 | 0.741065009 |
| ENSMUSG00000040899 | Ccr6 | 0 | 0.326907457 | 1 |
| ENSMUSG00000055945 | Prr18 | 0 | 1.648835551 | 1 |
| ENSMUSG00000117264 | 4930452A19Rik | 0 | 0.326907457 | 1 |
| ENSMUSG00000117110 | Gm49808 | 0 | 0.648835551 | 1 |
| ENSMUSG00000099384 | 1700110C19Rik | 0 | 1.648835551 | 1 |
| ENSMUSG00000089941 | Gm16168 | 0 | -0.673092543 | 1 |
| ENSMUSG00000040966 | Slc22a2 | 0 | -0.158519371 | 1 |
| ENSMUSG00000023829 | Slc22a1 | 0 | 1.648835551 | 1 |
| ENSMUSG00000068037 | Mas1 | 0 | 0.063873051 | 1 |
| ENSMUSG00000055602 | Tcp10b | 0 | 1.648835551 | 1 |
| ENSMUSG00000117282 | Gm49898 | 0 | 0.648835551 | 1 |
| ENSMUSG00000073457 | Smok2b | 0 | 0.648835551 | 1 |
| ENSMUSG00000090211 | Gm16050 | 0 | 1.648835551 | 1 |
| ENSMUSG00000067941 | Gm7168 | 0 | 1.063873051 | 1 |
| ENSMUSG00000048826 | Dact2 | 0 | 0.648835551 | 1 |
| ENSMUSG00000117254 | Gm34567 | 0 | 1.063873051 | 1 |
| ENSMUSG00000116801 | Gm38551 | 0 | 0.326907457 | 1 |
| ENSMUSG00000116895 | Gm3435 | 0 | 1.063873051 | 1 |
| ENSMUSG00000116655 | Gm35455 | 0 | 1.063873051 | 1 |
| ENSMUSG00000097624 | Gm5091 | 0 | 1.063873051 | 1 |
| ENSMUSG00000116718 | Gm49668 | 0 | 0.648835551 | 1 |
| ENSMUSG00000051977 | Prdm9 | 0 | -1.258055044 | 0.673811773 |
| ENSMUSG00000095990 | Zfp97 | 0 | 0.648835551 | 1 |
| ENSMUSG00000072761 | Gm6712 | 0 | -2.673092543 | 0.084161134 |
| ENSMUSG00000052270 | Fpr2 | 0 | 1.648835551 | 1 |
| ENSMUSG00000094396 | Vmn2r124 | 0 | 1.063873051 | 1 |
| ENSMUSG00000091350 | Vmn2r92 | 0 | 1.648835551 | 1 |
| ENSMUSG00000079698 | Vmn2r93 | 0 | 0.648835551 | 1 |
| ENSMUSG00000091491 | Vmn2r97 | 0 | 0.326907457 | 1 |
| ENSMUSG00000091859 | Vmn2r100 | 0 | -0.52108945 | 1 |
| ENSMUSG00000094892 | Vmn2r101 | 0 | 0.063873051 | 1 |
| ENSMUSG00000091670 | Vmn2r105 | 0 | 1.063873051 | 1 |
| ENSMUSG00000091656 | Vmn2r106 | 0 | 1.648835551 | 1 |
| ENSMUSG00000091805 | Vmn2r108 | 0 | 1.063873051 | 1 |
| ENSMUSG00000090572 | Vmn2r109 | 0 | 1.063873051 | 1 |
| ENSMUSG00000091259 | Vmn2r110 | 0 | 1.648835551 | 1 |
| ENSMUSG00000067951 | Vmn1r227 | 0 | 1.063873051 | 1 |
| ENSMUSG00000117148 | Vmn1r229 | 0 | 1.648835551 | 1 |
| ENSMUSG00000054142 | Vmn1r236 | 0 | 1.648835551 | 1 |
| ENSMUSG00000062743 | Zfp677 | 0 | -1.599091962 | 0.559918909 |
| ENSMUSG00000035868 | Zfp983 | 0 | 0.648835551 | 1 |
| ENSMUSG00000069743 | Zfp820 | 0 | -2.051604167 | 0.274319469 |
| ENSMUSG00000063383 | Zfp947 | 0 | 0.063873051 | 1 |
| ENSMUSG00000033972 | Zfp944 | 0 | -2.43862729 | 0.184141965 |
| ENSMUSG00000094921 | Vmn2r112 | 0 | 1.063873051 | 1 |
| ENSMUSG00000098164 | Gm5493 | 0 | 1.648835551 | 1 |
| ENSMUSG00000092111 | Vmn2r113 | 0 | 0.648835551 | 1 |
| ENSMUSG00000091076 | Vmn2r115 | 0 | 1.648835551 | 1 |
| ENSMUSG00000090966 | Vmn2r116 | 0 | 1.648835551 | 1 |

| | | | | |
|---|---|---|---|---|
| ENSMUSG00000117322 | 6330415G19Rik | 0 | 0.648835551 | 1 |
| ENSMUSG00000071256 | Zfp213 | 0 | -0.158519371 | 1 |
| ENSMUSG00000062012 | Zfp13 | 0 | -2.599091962 | 0.108201045 |
| ENSMUSG00000023902 | Zscan10 | 0 | 0.326907457 | 1 |
| ENSMUSG00000043747 | 1520401A03Rik | 0 | -0.158519371 | 1 |
| ENSMUSG00000023908 | Pkmyt1 | 0 | 0.063873051 | 1 |
| ENSMUSG00000097006 | 9530082P21Rik | 0 | 1.063873051 | 1 |
| ENSMUSG00000115692 | Gm49163 | 0 | 1.648835551 | 1 |
| ENSMUSG00000086719 | Gm16275 | 0 | 1.648835551 | 1 |
| ENSMUSG00000024114 | Prss41 | 0 | 1.648835551 | 1 |
| ENSMUSG00000096278 | Dcpp2 | 0 | 1.648835551 | 1 |
| ENSMUSG00000024128 | Sbp | 0 | 1.648835551 | 1 |
| ENSMUSG00000117523 | Gm50071 | 0 | 1.648835551 | 1 |
| ENSMUSG00000117406 | Ntn3 | 0 | -0.158519371 | 1 |
| ENSMUSG00000072082 | Ccnf | 0 | -3.209145444 | 0.026203604 |
| ENSMUSG00000073437 | D330041H03Rik | 0 | -2.395558568 | 0.174915761 |
| ENSMUSG00000033597 | Caskin1 | 0 | -0.158519371 | 1 |
| ENSMUSG00000079657 | Rab26 | 0 | 0.326907457 | 1 |
| ENSMUSG00000019320 | Noxo1 | 0 | 0.063873051 | 1 |
| ENSMUSG00000008482 | Rnf151 | 0 | 1.648835551 | 1 |
| ENSMUSG00000002500 | Rpl3l | 0 | 0.063873051 | 1 |
| ENSMUSG00000039628 | Hs3st6 | 0 | 1.063873051 | 1 |
| ENSMUSG00000024168 | Tmem204 | 0 | -0.52108945 | 1 |
| ENSMUSG00000044172 | Ptx4 | 0 | 1.063873051 | 1 |
| ENSMUSG00000059562 | Ccdc154 | 0 | 0.326907457 | 1 |
| ENSMUSG00000015126 | Tsr3 | 0 | -1.936126949 | 0.358945041 |
| ENSMUSG00000047507 | Baiap3 | 0 | 1.648835551 | 1 |
| ENSMUSG00000033200 | Tpsg1 | 0 | 1.648835551 | 1 |
| ENSMUSG00000024175 | Tekt4 | 0 | 1.648835551 | 1 |
| ENSMUSG00000117437 | Gm33727 | 0 | 1.648835551 | 1 |
| ENSMUSG00000117445 | Gm50026 | 0 | 0.326907457 | 1 |
| ENSMUSG00000019214 | Chtf18 | 0 | -0.351164449 | 1 |
| ENSMUSG00000025738 | Fbxl16 | 0 | 1.063873051 | 1 |
| ENSMUSG00000025736 | Jmjd8 | 0 | -1.351164449 | 0.695372548 |
| ENSMUSG00000071192 | Wfikkn1 | 0 | -1.351164449 | 0.707032059 |
| ENSMUSG00000025727 | A930017K11Rik | 0 | 1.063873051 | 1 |
| ENSMUSG00000024177 | Nme4 | 0 | -0.673092543 | 1 |
| ENSMUSG00000024184 | Pdia2 | 0 | 1.648835551 | 1 |
| ENSMUSG00000034413 | Neurl1b | 0 | -1.258055044 | 0.673811773 |
| ENSMUSG00000118232 | Gm50269 | 0 | 1.648835551 | 1 |
| ENSMUSG00000092539 | Gm20468 | 0 | 0.326907457 | 1 |
| ENSMUSG00000117799 | Gm50295 | 0 | 0.063873051 | 1 |
| ENSMUSG00000086898 | Itpr3os | 0 | 1.648835551 | 1 |
| ENSMUSG00000024210 | Ip6k3 | 0 | 0.648835551 | 1 |
| ENSMUSG00000117794 | Gm50328 | 0 | 0.648835551 | 1 |
| ENSMUSG00000116970 | Gm49796 | 0 | 1.063873051 | 1 |
| ENSMUSG00000086485 | Gm15458 | 0 | 0.063873051 | 1 |
| ENSMUSG00000085846 | Gm15597 | 0 | 1.063873051 | 1 |
| ENSMUSG00000117303 | Gm49889 | 0 | 1.648835551 | 1 |
| ENSMUSG00000117313 | Gm49838 | 0 | 1.063873051 | 1 |
| ENSMUSG00000066170 | E230001N04Rik | 0 | -2.051604167 | 0.274319469 |

| | | | | |
|---|---|---|---|---|
| ENSMUSG00000117131 | Gm41506 | 0 | 1.648835551 | 1 |
| ENSMUSG00000024224 | Clpsl2 | 0 | 1.063873051 | 1 |
| ENSMUSG00000062252 | Lhfpl5 | 0 | 1.648835551 | 1 |
| ENSMUSG00000117255 | Gm49864 | 0 | 0.648835551 | 1 |
| ENSMUSG00000048905 | 4930539E08Rik | 0 | -0.158519371 | 1 |
| ENSMUSG00000045378 | Pxt1 | 0 | 0.648835551 | 1 |
| ENSMUSG00000117614 | Gm50068 | 0 | 1.648835551 | 1 |
| ENSMUSG00000091614 | Gm17657 | 0 | 1.648835551 | 1 |
| ENSMUSG00000089900 | Tbc1d22bos | 0 | 0.648835551 | 1 |
| ENSMUSG00000117534 | Gm20161 | 0 | 1.063873051 | 1 |
| ENSMUSG00000086534 | Gm16758 | 0 | 1.648835551 | 1 |
| ENSMUSG00000117528 | Gm50017 | 0 | 1.648835551 | 1 |
| ENSMUSG00000117371 | Gm50019 | 0 | 0.648835551 | 1 |
| ENSMUSG00000117725 | Gm50240 | 0 | -1.351164449 | 0.687112677 |
| ENSMUSG00000118284 | Gm41562 | 0 | 1.063873051 | 1 |
| ENSMUSG00000086010 | Gm15318 | 0 | -0.810596067 | 0.844591606 |
| ENSMUSG00000024034 | Tmprss3 | 0 | 0.648835551 | 1 |
| ENSMUSG00000118049 | Gm50103 | 0 | 0.648835551 | 1 |
| ENSMUSG00000013562 | 4833413E03Rik | 0 | 1.063873051 | 1 |
| ENSMUSG00000024041 | Cryaa | 0 | 1.648835551 | 1 |
| ENSMUSG00000037577 | Ephx3 | 0 | 1.063873051 | 1 |
| ENSMUSG00000052142 | Rasal3 | 0 | -0.158519371 | 1 |
| ENSMUSG00000061126 | Cyp4f39 | 0 | -0.52108945 | 1 |
| ENSMUSG00000048440 | Cyp4f16 | 0 | -1.52108945 | 0.544908989 |
| ENSMUSG00000055202 | Zfp811 | 0 | -1.874726405 | 0.352215149 |
| ENSMUSG00000097030 | Gm26693 | 0 | -0.673092543 | 1 |
| ENSMUSG00000053600 | Zfp472 | 0 | -1.743481871 | 0.455131546 |
| ENSMUSG00000053390 | Zfp952 | 0 | -2.258055044 | 0.233570669 |
| ENSMUSG00000093884 | Olfr239 | 0 | 1.648835551 | 1 |
| ENSMUSG00000094441 | Zfp955a | 0 | -2.560617814 | 0.105255472 |
| ENSMUSG00000096910 | Zfp955b | 0 | -2.708716453 | 0.112236795 |
| ENSMUSG00000092519 | Actl9 | 0 | 1.648835551 | 1 |
| ENSMUSG00000032739 | Pram1 | 0 | -0.810596067 | 0.844591606 |
| ENSMUSG00000092349 | Smim40 | 0 | 0.648835551 | 1 |
| ENSMUSG00000092564 | BC051226 | 0 | 1.063873051 | 1 |
| ENSMUSG00000067370 | B3galt4 | 0 | 1.063873051 | 1 |
| ENSMUSG00000024325 | Ring1 | 0 | -1.673092543 | 0.451785624 |
| ENSMUSG00000024330 | Col11a2 | 0 | -1.051604167 | 0.858831093 |
| ENSMUSG00000098197 | BC051537 | 0 | 0.648835551 | 1 |
| ENSMUSG00000037649 | H2-DMa | 0 | -1.52108945 | 0.544908989 |
| ENSMUSG00000118026 | Gm50335 | 0 | 1.648835551 | 1 |
| ENSMUSG00000037548 | H2-DMb2 | 0 | -0.673092543 | 1 |
| ENSMUSG00000037321 | Tap1 | 0 | -0.810596067 | 0.844591606 |
| ENSMUSG00000092550 | Gm20496 | 0 | 1.648835551 | 1 |
| ENSMUSG00000060586 | H2-Eb1 | 0 | 0.326907457 | 1 |
| ENSMUSG00000067341 | H2-Eb2 | 0 | 0.326907457 | 1 |
| ENSMUSG00000024340 | Btnl2 | 0 | 0.326907457 | 1 |
| ENSMUSG00000062638 | Btnl1 | 0 | 0.326907457 | 1 |
| ENSMUSG00000015452 | Ager | 0 | 0.326907457 | 1 |
| ENSMUSG00000015467 | Egfl8 | 0 | 0.648835551 | 1 |
| ENSMUSG00000015476 | Prrt1 | 0 | 0.326907457 | 1 |

| | | | | |
|---|---|---|---|---|
| ENSMUSG00000015461 | Atf6b | 0 | -1.258055044 | 0.717944159 |
| ENSMUSG00000015451 | C4a | 0 | 0.648835551 | 1 |
| ENSMUSG00000073415 | Gm10501 | 0 | 1.648835551 | 1 |
| ENSMUSG00000007030 | Vwa7 | 0 | 1.063873051 | 1 |
| ENSMUSG00000036185 | Sapcd1 | 0 | -0.810596067 | 0.844591606 |
| ENSMUSG00000073413 | Ly6g6d | 0 | 0.648835551 | 1 |
| ENSMUSG00000043807 | Ly6g5b | 0 | 1.648835551 | 1 |
| ENSMUSG00000092241 | Gm20522 | 0 | 1.063873051 | 1 |
| ENSMUSG00000043311 | D17H6S53E | 0 | -0.673092543 | 1 |
| ENSMUSG00000090936 | Gm17705 | 0 | 0.063873051 | 1 |
| ENSMUSG00000024402 | Lta | 0 | 1.648835551 | 1 |
| ENSMUSG00000024403 | Atp6v1g2 | 0 | -0.936126949 | 0.886203497 |
| ENSMUSG00000079507 | H2-Q1 | 0 | 1.648835551 | 1 |
| ENSMUSG00000091705 | H2-Q2 | 0 | 1.648835551 | 1 |
| ENSMUSG00000079505 | Gm11131 | 0 | 0.326907457 | 1 |
| ENSMUSG00000073409 | H2-Q6 | 0 | 0.648835551 | 1 |
| ENSMUSG00000060550 | H2-Q7 | 0 | 1.648835551 | 1 |
| ENSMUSG00000024406 | Pou5f1 | 0 | 1.063873051 | 1 |
| ENSMUSG00000024409 | Psors1c2 | 0 | 0.326907457 | 1 |
| ENSMUSG00000038838 | Vars2 | 0 | -2.843017545 | 0.074369691 |
| ENSMUSG00000086867 | Gm4577 | 0 | 0.648835551 | 1 |
| ENSMUSG00000092219 | Gm20443 | 0 | 1.648835551 | 1 |
| ENSMUSG00000059791 | Nrm | 0 | -0.351164449 | 1 |
| ENSMUSG00000084998 | Gm16279 | 0 | -1.936126949 | 0.358945041 |
| ENSMUSG00000053835 | H2-T24 | 0 | 0.326907457 | 1 |
| ENSMUSG00000079492 | Gm11127 | 0 | 1.648835551 | 1 |
| ENSMUSG00000092277 | Gm19684 | 0 | -0.351164449 | 1 |
| ENSMUSG00000073402 | Gm8909 | 0 | 0.326907457 | 1 |
| ENSMUSG00000054128 | H2-T3 | 0 | 0.063873051 | 1 |
| ENSMUSG00000092392 | Gm20546 | 0 | 1.648835551 | 1 |
| ENSMUSG00000024448 | H2-M10.1 | 0 | 1.648835551 | 1 |
| ENSMUSG00000067201 | H2-M9 | 0 | 1.063873051 | 1 |
| ENSMUSG00000050747 | Trim15 | 0 | 0.063873051 | 1 |
| ENSMUSG00000058063 | Trim31 | 0 | 1.648835551 | 1 |
| ENSMUSG00000092239 | 1700031A10Rik | 0 | 1.648835551 | 1 |
| ENSMUSG00000036492 | Rnf39 | 0 | -1.599091962 | 0.559918909 |
| ENSMUSG00000036036 | Zfp57 | 0 | -4.470105521 | 0.000143457 |
| ENSMUSG00000056600 | Olfr90 | 0 | 1.648835551 | 1 |
| ENSMUSG00000096477 | Olfr92 | 0 | 0.648835551 | 1 |
| ENSMUSG00000091601 | Olfr93 | 0 | 1.648835551 | 1 |
| ENSMUSG00000049561 | Olfr95 | 0 | 0.648835551 | 1 |
| ENSMUSG00000016206 | H2-M3 | 0 | 1.648835551 | 1 |
| ENSMUSG00000095858 | Olfr104-ps | 0 | 0.648835551 | 1 |
| ENSMUSG00000095603 | Olfr106-ps | 0 | 1.063873051 | 1 |
| ENSMUSG00000029184 | Olfr109 | 0 | 1.648835551 | 1 |
| ENSMUSG00000092292 | Olfr113 | 0 | 1.648835551 | 1 |
| ENSMUSG00000092413 | Olfr115 | 0 | 1.648835551 | 1 |
| ENSMUSG00000095286 | Olfr117 | 0 | 1.648835551 | 1 |
| ENSMUSG00000083947 | Olfr122 | 0 | 1.648835551 | 1 |
| ENSMUSG00000044985 | Olfr124 | 0 | 1.648835551 | 1 |
| ENSMUSG00000109212 | Olfr126 | 0 | 1.648835551 | 1 |

| | | | | |
|---|---|---|---|---|
| ENSMUSG00000043312 | Olfr131 | 0 | 1.648835551 | 1 |
| ENSMUSG00000057443 | Olfr138 | 0 | 1.648835551 | 1 |
| ENSMUSG00000092322 | Esp36 | 0 | 1.648835551 | 1 |
| ENSMUSG00000094024 | Esp18 | 0 | 1.648835551 | 1 |
| ENSMUSG00000096345 | Esp16 | 0 | 1.648835551 | 1 |
| ENSMUSG00000095104 | Esp15 | 0 | 0.648835551 | 1 |
| ENSMUSG00000095886 | Esp38 | 0 | 0.326907457 | 1 |
| ENSMUSG00000025433 | Crisp3 | 0 | 1.648835551 | 1 |
| ENSMUSG00000090897 | Gm44501 | 0 | 0.648835551 | 1 |
| ENSMUSG00000093957 | Esp3 | 0 | 1.648835551 | 1 |
| ENSMUSG00000091043 | Glyatl3 | 0 | 1.648835551 | 1 |
| ENSMUSG00000057899 | Adgrf2 | 0 | -0.936126949 | 0.877701541 |
| ENSMUSG00000023914 | Mep1a | 0 | 0.326907457 | 1 |
| ENSMUSG00000040140 | Tdrd6 | 0 | 1.648835551 | 1 |
| ENSMUSG00000089849 | Runx2os2 | 0 | 1.648835551 | 1 |
| ENSMUSG00000117044 | Gm35097 | 0 | 1.063873051 | 1 |
| ENSMUSG00000097119 | B230354K17Rik | 0 | -2.810596067 | 0.070328284 |
| ENSMUSG00000023938 | Aars2 | 0 | 0.063873051 | 1 |
| ENSMUSG00000023949 | Tcte1 | 0 | -0.351164449 | 1 |
| ENSMUSG00000067389 | Gm17080 | 0 | 1.648835551 | 1 |
| ENSMUSG00000094690 | 1600014C23Rik | 0 | 1.648835551 | 1 |
| ENSMUSG00000023966 | Rsph9 | 0 | -1.52108945 | 0.573149109 |
| ENSMUSG00000071073 | Lrrc73 | 0 | 0.326907457 | 1 |
| ENSMUSG00000047428 | Dlk2 | 0 | -0.158519371 | 1 |
| ENSMUSG00000040658 | Dnph1 | 0 | 1.063873051 | 1 |
| ENSMUSG00000002769 | Gnmt | 0 | 0.326907457 | 1 |
| ENSMUSG00000036858 | Ptcra | 0 | 0.063873051 | 1 |
| ENSMUSG00000023979 | Guca1b | 0 | 1.648835551 | 1 |
| ENSMUSG00000084880 | Tomm6os | 0 | 1.648835551 | 1 |
| ENSMUSG00000096549 | Prickle4 | 0 | 0.326907457 | 1 |
| ENSMUSG00000023266 | Frs3 | 0 | -0.52108945 | 1 |
| ENSMUSG00000117157 | Gm49946 | 0 | 1.648835551 | 1 |
| ENSMUSG00000073386 | 9830107B12Rik | 0 | 1.063873051 | 1 |
| ENSMUSG00000043939 | A530064D06Rik | 0 | 1.063873051 | 1 |
| ENSMUSG00000043740 | B430306N03Rik | 0 | -1.051604167 | 0.886203497 |
| ENSMUSG00000023993 | Treml1 | 0 | 1.063873051 | 1 |
| ENSMUSG00000117104 | Gm49893 | 0 | 1.063873051 | 1 |
| ENSMUSG00000040694 | Apobec2 | 0 | 0.648835551 | 1 |
| ENSMUSG00000023995 | Tspo2 | 0 | 1.063873051 | 1 |
| ENSMUSG00000117117 | Gm49901 | 0 | 0.648835551 | 1 |
| ENSMUSG00000104256 | Gm37173 | 0 | 1.063873051 | 1 |
| ENSMUSG00000067103 | AY702103 | 0 | 0.326907457 | 1 |
| ENSMUSG00000010592 | Dazl | 0 | 1.063873051 | 1 |
| ENSMUSG00000117204 | C330011F03Rik | 0 | 0.063873051 | 1 |
| ENSMUSG00000117175 | Gm20098 | 0 | -0.351164449 | 1 |
| ENSMUSG00000117076 | Gm41600 | 0 | 1.648835551 | 1 |
| ENSMUSG00000117033 | Gm49912 | 0 | -0.158519371 | 1 |
| ENSMUSG00000117035 | Gm5094 | 0 | 1.648835551 | 1 |
| ENSMUSG00000098552 | Gm27217 | 0 | 1.648835551 | 1 |
| ENSMUSG00000117128 | Gm49915 | 0 | 1.648835551 | 1 |
| ENSMUSG00000023943 | Sult1c1 | 0 | 1.063873051 | 1 |

| | | | | |
|---|---|---|---|---|
| ENSMUSG00000117132 | Gm32055 | 0 | 1.648835551 | 1 |
| ENSMUSG00000091504 | Vmn2r118 | 0 | 1.648835551 | 1 |
| ENSMUSG00000057835 | Zfp119a | 0 | -1.743481871 | 0.455131546 |
| ENSMUSG00000062101 | Zfp119b | 0 | 0.326907457 | 1 |
| ENSMUSG00000097077 | Gm16712 | 0 | 0.326907457 | 1 |
| ENSMUSG00000011589 | Fsd1 | 0 | 0.648835551 | 1 |
| ENSMUSG00000011305 | Plin5 | 0 | 1.063873051 | 1 |
| ENSMUSG00000044469 | Tnfaip8l1 | 0 | -2.777429203 | 0.067360743 |
| ENSMUSG00000043683 | Fem1a | 0 | -1.810596067 | 0.465580427 |
| ENSMUSG00000073380 | Arrdc5 | 0 | -0.158519371 | 1 |
| ENSMUSG00000040828 | Catsperd | 0 | -1.874726405 | 0.364014601 |
| ENSMUSG00000039481 | Nrtn | 0 | -0.158519371 | 1 |
| ENSMUSG00000090655 | Vmn2r120 | 0 | -1.673092543 | 0.611141945 |
| ENSMUSG00000086594 | Nudt12os | 0 | 0.648835551 | 1 |
| ENSMUSG00000024228 | Nudt12 | 0 | -1.258055044 | 0.737916568 |
| ENSMUSG00000117066 | Gm49860 | 0 | -0.351164449 | 1 |
| ENSMUSG00000090565 | 4930405O22Rik | 0 | 1.648835551 | 1 |
| ENSMUSG00000117250 | Gm49871 | 0 | 1.063873051 | 1 |
| ENSMUSG00000117229 | Gm35550 | 0 | 1.648835551 | 1 |
| ENSMUSG00000035842 | Ddx11 | 0 | -3.89032326 | 0.002005926 |
| ENSMUSG00000117003 | Gm49937 | 0 | 1.648835551 | 1 |
| ENSMUSG00000117235 | Gm49938 | 0 | 1.063873051 | 1 |
| ENSMUSG00000114563 | 1700016K05Rik | 0 | 0.648835551 | 1 |
| ENSMUSG00000114291 | Gm47471 | 0 | 1.063873051 | 1 |
| ENSMUSG00000117029 | Gm36201 | 0 | 0.326907457 | 1 |
| ENSMUSG00000095407 | Tmem200c | 0 | 1.648835551 | 1 |
| ENSMUSG00000100572 | 2410021H03Rik | 0 | 1.648835551 | 1 |
| ENSMUSG00000117323 | Gm36487 | 0 | -0.351164449 | 1 |
| ENSMUSG00000117069 | Gm49894 | 0 | 1.648835551 | 1 |
| ENSMUSG00000117233 | Gm36594 | 0 | 0.063873051 | 1 |
| ENSMUSG00000097282 | 5031415H12Rik | 0 | 0.063873051 | 1 |
| ENSMUSG00000117079 | Gm41611 | 0 | 1.648835551 | 1 |
| ENSMUSG00000117137 | 4930471L23Rik | 0 | 0.326907457 | 1 |
| ENSMUSG00000117199 | Gm4566 | 0 | 1.648835551 | 1 |
| ENSMUSG00000091831 | Gm4707 | 0 | 0.648835551 | 1 |
| ENSMUSG00000052525 | Spdya | 0 | -0.52108945 | 1 |
| ENSMUSG00000117637 | Gm30091 | 0 | 0.326907457 | 1 |
| ENSMUSG00000044375 | Pcare | 0 | -1.258055044 | 0.723846946 |
| ENSMUSG00000117096 | Gm30420 | 0 | 1.648835551 | 1 |
| ENSMUSG00000117292 | E330032C10Rik | 0 | 1.648835551 | 1 |
| ENSMUSG00000117118 | Gm49878 | 0 | 1.063873051 | 1 |
| ENSMUSG00000024065 | Ehd3 | 0 | -0.810596067 | 0.85004325 |
| ENSMUSG00000038541 | Srd5a2 | 0 | 1.648835551 | 1 |
| ENSMUSG00000117226 | Gm31084 | 0 | -0.158519371 | 1 |
| ENSMUSG00000117444 | Gm50020 | 0 | -0.158519371 | 1 |
| ENSMUSG00000117598 | Gm50028 | 0 | 1.648835551 | 1 |
| ENSMUSG00000117525 | Gm50034 | 0 | 1.648835551 | 1 |
| ENSMUSG00000050668 | Gpatch11 | 0 | -3.106051951 | 0.028853812 |
| ENSMUSG00000038045 | Sult6b1 | 0 | 0.326907457 | 1 |
| ENSMUSG00000036533 | Cdc42ep3 | 0 | -0.52108945 | 1 |
| ENSMUSG00000117426 | Gm33055 | 0 | 0.648835551 | 1 |

| | | | | |
|---|---|---|---|---|
| ENSMUSG00000045257 | Morn2 | 0 | -1.051604167 | 0.859517261 |
| ENSMUSG00000117375 | Gm33373 | 0 | 0.063873051 | 1 |
| ENSMUSG00000112121 | C230072F16Rik | 0 | 0.063873051 | 1 |
| ENSMUSG00000100154 | Gm29052 | 0 | 1.648835551 | 1 |
| ENSMUSG00000117442 | 1810073O08Rik | 0 | 1.063873051 | 1 |
| ENSMUSG00000117431 | 8430430B14Rik | 0 | 1.648835551 | 1 |
| ENSMUSG00000117432 | Gm49967 | 0 | 1.648835551 | 1 |
| ENSMUSG00000117571 | 4933433H22Rik | 0 | 1.648835551 | 1 |
| ENSMUSG00000117485 | Gm19696 | 0 | 0.063873051 | 1 |
| ENSMUSG00000117391 | 4930500A05Rik | 0 | 0.648835551 | 1 |
| ENSMUSG00000040505 | Abcg5 | 0 | 0.326907457 | 1 |
| ENSMUSG00000024131 | Slc3a1 | 0 | -3.209145444 | 0.025125353 |
| ENSMUSG00000100345 | Gm29418 | 0 | 0.648835551 | 1 |
| ENSMUSG00000117501 | Gm30117 | 0 | -0.810596067 | 0.844591606 |
| ENSMUSG00000117628 | Gm50012 | 0 | 0.326907457 | 1 |
| ENSMUSG00000100847 | Gm28676 | 0 | 1.648835551 | 1 |
| ENSMUSG00000117414 | Gm46587 | 0 | 1.063873051 | 1 |
| ENSMUSG00000050138 | Kcnk12 | 0 | 0.326907457 | 1 |
| ENSMUSG00000117466 | Gm50039 | 0 | 0.648835551 | 1 |
| ENSMUSG00000117497 | 1700116H05Rik | 0 | 1.648835551 | 1 |
| ENSMUSG00000117524 | Gm4719 | 0 | 1.648835551 | 1 |
| ENSMUSG00000066878 | Gm10184 | 0 | 1.648835551 | 1 |
| ENSMUSG00000117635 | Gm41653 | 0 | 0.648835551 | 1 |
| ENSMUSG00000090160 | 4930480K15Rik | 0 | 1.648835551 | 1 |
| ENSMUSG00000117417 | Gm49991 | 0 | -0.810596067 | 0.844591606 |
| ENSMUSG00000024232 | Bambi | 0 | 0.326907457 | 1 |
| ENSMUSG00000024235 | Map3k8 | 0 | -2.395558568 | 0.174915761 |
| ENSMUSG00000117401 | 4833419F23Rik | 0 | 0.063873051 | 1 |
| ENSMUSG00000117579 | Gm50023 | 0 | 0.648835551 | 1 |
| ENSMUSG00000097888 | Gm26682 | 0 | 0.326907457 | 1 |
| ENSMUSG00000117617 | Gm50065 | 0 | 1.648835551 | 1 |
| ENSMUSG00000091165 | Gm17036 | 0 | 1.063873051 | 1 |
| ENSMUSG00000039540 | 4921524L21Rik | 0 | 1.063873051 | 1 |
| ENSMUSG00000117356 | Gm46633 | 0 | 1.648835551 | 1 |
| ENSMUSG00000117490 | Gm35031 | 0 | 0.648835551 | 1 |
| ENSMUSG00000036904 | Fzd8 | 0 | 1.648835551 | 1 |
| ENSMUSG00000095438 | Mir133a-1hg | 0 | 1.648835551 | 1 |
| ENSMUSG00000024430 | Cabyr | 0 | 0.326907457 | 1 |
| ENSMUSG00000044252 | Osbpl1a | 0 | -3.373532262 | 0.013375059 |
| ENSMUSG00000117494 | Gm49970 | 0 | 1.648835551 | 1 |
| ENSMUSG00000117608 | AC103362.1 | 0 | -1.051604167 | 0.886203497 |
| ENSMUSG00000117613 | Gm2629 | 0 | 0.063873051 | 1 |
| ENSMUSG00000086095 | Gm15328 | 0 | 1.063873051 | 1 |
| ENSMUSG00000117615 | 4930545E07Rik | 0 | 1.063873051 | 1 |
| ENSMUSG00000059898 | Dsc3 | 0 | -1.810596067 | 0.466464457 |
| ENSMUSG00000061928 | Dsg1b | 0 | 0.648835551 | 1 |
| ENSMUSG00000033107 | Rnf125 | 0 | -2.209145444 | 0.37114869 |
| ENSMUSG00000024313 | Mep1b | 0 | -0.158519371 | 1 |
| ENSMUSG00000043168 | 4930426D05Rik | 0 | 1.648835551 | 1 |
| ENSMUSG00000114088 | Gm47898 | 0 | 1.063873051 | 1 |
| ENSMUSG00000086287 | Gm15972 | 0 | 1.648835551 | 1 |

| | | | | |
|---|---|---|---|---|
| ENSMUSG00000051469 | Zfp24 | 0 | -3.708716453 | 0.003607261 |
| ENSMUSG00000117467 | Gm33674 | 0 | 0.648835551 | 1 |
| ENSMUSG00000097748 | Gm26533 | 0 | -1.158519371 | 0.886203497 |
| ENSMUSG00000044982 | Sft2d3 | 0 | -0.673092543 | 1 |
| ENSMUSG00000117650 | 4933435E02Rik | 0 | 1.648835551 | 1 |
| ENSMUSG00000097363 | Gm26717 | 0 | 0.648835551 | 1 |
| ENSMUSG00000109238 | Gm35060 | 0 | 1.063873051 | 1 |
| ENSMUSG00000097710 | 2310026I22Rik | 0 | 1.063873051 | 1 |
| ENSMUSG00000012282 | Wnt8a | 0 | 1.063873051 | 1 |
| ENSMUSG00000044201 | Cdc25c | 0 | -0.810596067 | 1 |
| ENSMUSG00000038555 | Reep2 | 0 | 1.063873051 | 1 |
| ENSMUSG00000117727 | Gm3587 | 0 | -0.158519371 | 1 |
| ENSMUSG00000117876 | Gm50146 | 0 | 1.648835551 | 1 |
| ENSMUSG00000024354 | Slc23a1 | 0 | 1.648835551 | 1 |
| ENSMUSG00000024353 | Mzb1 | 0 | 1.063873051 | 1 |
| ENSMUSG00000073600 | Prob1 | 0 | 0.648835551 | 1 |
| ENSMUSG00000024352 | Spata24 | 0 | 1.648835551 | 1 |
| ENSMUSG00000024349 | Tmem173 | 0 | -1.158519371 | 0.886203497 |
| ENSMUSG00000100320 | Gm29417 | 0 | 0.063873051 | 1 |
| ENSMUSG00000118347 | C330008A17Rik | 0 | 1.648835551 | 1 |
| ENSMUSG00000118122 | Gm50405 | 0 | 0.648835551 | 1 |
| ENSMUSG00000117679 | Apbb3 | 0 | -2.52108945 | 0.139351384 |
| ENSMUSG00000051678 | Pcdhb6 | 0 | 1.648835551 | 1 |
| ENSMUSG00000103985 | Gm37446 | 0 | 1.648835551 | 1 |
| ENSMUSG00000043458 | Pcdhb12 | 0 | 1.648835551 | 1 |
| ENSMUSG00000044043 | Pcdhb14 | 0 | 1.063873051 | 1 |
| ENSMUSG00000046387 | Pcdhb17 | 0 | 1.063873051 | 1 |
| ENSMUSG00000048347 | Pcdhb18 | 0 | 1.648835551 | 1 |
| ENSMUSG00000046191 | Pcdhb20 | 0 | 1.648835551 | 1 |
| ENSMUSG00000073591 | Pcdhb22 | 0 | 1.063873051 | 1 |
| ENSMUSG00000050304 | Slc25a2 | 0 | 1.648835551 | 1 |
| ENSMUSG00000051316 | Taf7 | 0 | -1.599091962 | 0.559918909 |
| ENSMUSG00000103037 | Pcdhgb1 | 0 | 0.648835551 | 1 |
| ENSMUSG00000103677 | Pcdhga4 | 0 | 1.648835551 | 1 |
| ENSMUSG00000102748 | Pcdhgb2 | 0 | -1.351164449 | 0.687112677 |
| ENSMUSG00000103567 | Pcdhga5 | 0 | -1.743481871 | 0.455131546 |
| ENSMUSG00000104512 | Gm37165 | 0 | 1.648835551 | 1 |
| ENSMUSG00000103585 | Pcdhgb4 | 0 | -0.351164449 | 1 |
| ENSMUSG00000103897 | Pcdhga8 | 0 | -0.52108945 | 1 |
| ENSMUSG00000103749 | Pcdhgb5 | 0 | -1.673092543 | 0.451785624 |
| ENSMUSG00000102440 | Pcdhga9 | 0 | -1.43862729 | 0.538110273 |
| ENSMUSG00000103088 | Pcdhgb6 | 0 | -2.051604167 | 0.274319469 |
| ENSMUSG00000102222 | Pcdhga10 | 0 | -2.106051951 | 0.279592421 |
| ENSMUSG00000102742 | Pcdhga11 | 0 | -3.051604167 | 0.036135464 |
| ENSMUSG00000102428 | Pcdhga12 | 0 | -1.599091962 | 0.559918909 |
| ENSMUSG00000023036 | Pcdhgc4 | 0 | -1.43862729 | 0.545533791 |
| ENSMUSG00000102543 | Pcdhgc5 | 0 | 1.648835551 | 1 |
| ENSMUSG00000102713 | Gm29994 | 0 | 0.648835551 | 1 |
| ENSMUSG00000044024 | Rell2 | 0 | 1.063873051 | 1 |
| ENSMUSG00000038524 | Fchsd1 | 0 | -1.599091962 | 0.559918909 |
| ENSMUSG00000089983 | 2010320O07Rik | 0 | 1.063873051 | 1 |

| | | | | |
|---|---|---|---|---|
| ENSMUSG00000097080 | 1700086O06Rik | 0 | 1.063873051 | 1 |
| ENSMUSG00000024442 | Dele1 | 0 | -1.936126949 | 0.423267288 |
| ENSMUSG00000024440 | Pcdh12 | 0 | -0.673092543 | 1 |
| ENSMUSG00000086991 | Gm15334 | 0 | 1.648835551 | 1 |
| ENSMUSG00000085289 | Gm15337 | 0 | 1.063873051 | 1 |
| ENSMUSG00000118117 | Gm30961 | 0 | 1.063873051 | 1 |
| ENSMUSG00000051732 | Pabpc2 | 0 | 1.648835551 | 1 |
| ENSMUSG00000118015 | Gm41708 | 0 | 0.648835551 | 1 |
| ENSMUSG00000117674 | Gm31087 | 0 | 0.648835551 | 1 |
| ENSMUSG00000117847 | Gm31205 | 0 | 1.648835551 | 1 |
| ENSMUSG00000073574 | Grxcr2 | 0 | 1.063873051 | 1 |
| ENSMUSG00000117902 | Gm41715 | 0 | 0.648835551 | 1 |
| ENSMUSG00000055561 | Spink5 | 0 | 0.326907457 | 1 |
| ENSMUSG00000055095 | Spink6 | 0 | 1.063873051 | 1 |
| ENSMUSG00000069385 | Gm10267 | 0 | 1.648835551 | 1 |
| ENSMUSG00000038071 | Npy6r | 0 | 1.063873051 | 1 |
| ENSMUSG00000117919 | Gm41717 | 0 | 1.063873051 | 1 |
| ENSMUSG00000024471 | Myot | 0 | 1.063873051 | 1 |
| ENSMUSG00000097610 | A930012L18Rik | 0 | -0.52108945 | 1 |
| ENSMUSG00000118013 | Gm31706 | 0 | 1.063873051 | 1 |
| ENSMUSG00000033949 | Trim36 | 0 | -0.158519371 | 1 |
| ENSMUSG00000056130 | Ticam2 | 0 | 1.063873051 | 1 |
| ENSMUSG00000117780 | Gm3734 | 0 | -2.395558568 | 0.234011885 |
| ENSMUSG00000024481 | Lvrn | 0 | -2.051604167 | 0.455131546 |
| ENSMUSG00000073568 | Arl14epl | 0 | 1.648835551 | 1 |
| ENSMUSG00000117588 | Gm49974 | 0 | -0.351164449 | 1 |
| ENSMUSG00000117602 | 9130209A04Rik | 0 | -0.351164449 | 1 |
| ENSMUSG00000118180 | Gm50223 | 0 | -0.158519371 | 1 |
| ENSMUSG00000118139 | Gm41720 | 0 | 1.648835551 | 1 |
| ENSMUSG00000087020 | Gm16283 | 0 | 1.648835551 | 1 |
| ENSMUSG00000086600 | C030005K06Rik | 0 | 1.648835551 | 1 |
| ENSMUSG00000048875 | Pudp | 0 | 0.648835551 | 1 |
| ENSMUSG00000118334 | Gm50446 | 0 | 1.648835551 | 1 |
| ENSMUSG00000118154 | 9330117O12Rik | 0 | -1.258055044 | 0.693687193 |
| ENSMUSG00000032900 | Tex43 | 0 | 0.648835551 | 1 |
| ENSMUSG00000024592 | C330018D20Rik | 0 | -1.52108945 | 0.545513258 |
| ENSMUSG00000118361 | Gm50237 | 0 | 1.063873051 | 1 |
| ENSMUSG00000054072 | Iigp1 | 0 | -0.351164449 | 1 |
| ENSMUSG00000052026 | Slc6a7 | 0 | 0.326907457 | 1 |
| ENSMUSG00000118229 | Lncbate10 | 0 | 0.063873051 | 1 |
| ENSMUSG00000097324 | Carmn | 0 | -0.158519371 | 1 |
| ENSMUSG00000024578 | Il17b | 0 | 0.063873051 | 1 |
| ENSMUSG00000054589 | Gm9949 | 0 | 1.648835551 | 1 |
| ENSMUSG00000110576 | Gm36368 | 0 | 0.648835551 | 1 |
| ENSMUSG00000060201 | Spink7 | 0 | 1.648835551 | 1 |
| ENSMUSG00000041789 | 2700046A07Rik | 0 | 1.648835551 | 1 |
| ENSMUSG00000097980 | Gm26972 | 0 | 0.648835551 | 1 |
| ENSMUSG00000118356 | Gm50353 | 0 | 1.648835551 | 1 |
| ENSMUSG00000118045 | Gm29860 | 0 | 0.648835551 | 1 |
| ENSMUSG00000118017 | Gm29966 | 0 | 1.063873051 | 1 |
| ENSMUSG00000118163 | Gm41757 | 0 | 1.063873051 | 1 |

| | | | | |
|---|---|---|---|---|
| ENSMUSG00000117992 | Gm41760 | 0 | 0.648835551 | 1 |
| ENSMUSG00000117775 | AC132307.1 | 0 | 0.648835551 | 1 |
| ENSMUSG00000024517 | Grp | 0 | 1.063873051 | 1 |
| ENSMUSG00000117907 | Gm50142 | 0 | 1.063873051 | 1 |
| ENSMUSG00000117764 | Gm50143 | 0 | -1.52108945 | 0.544908989 |
| ENSMUSG00000047259 | Mc4r | 0 | 1.648835551 | 1 |
| ENSMUSG00000117868 | Gm50165 | 0 | 1.063873051 | 1 |
| ENSMUSG00000117815 | Gm50166 | 0 | 0.326907457 | 1 |
| ENSMUSG00000046415 | B430212C06Rik | 0 | 0.326907457 | 1 |
| ENSMUSG00000024530 | Prelid3a | 0 | 1.063873051 | 1 |
| ENSMUSG00000097835 | Gm26910 | 0 | -0.52108945 | 1 |
| ENSMUSG00000117891 | Gm41764 | 0 | 0.063873051 | 1 |
| ENSMUSG00000118102 | Gm50260 | 0 | 1.648835551 | 1 |
| ENSMUSG00000117970 | Gm4402 | 0 | 1.648835551 | 1 |
| ENSMUSG00000118028 | Gm50215 | 0 | -0.351164449 | 1 |
| ENSMUSG00000024512 | Dynap | 0 | 1.063873051 | 1 |
| ENSMUSG00000118316 | Gm31908 | 0 | 1.063873051 | 1 |
| ENSMUSG00000090263 | D730045A05Rik | 0 | 0.648835551 | 1 |
| ENSMUSG00000053861 | Gm9925 | 0 | 1.648835551 | 1 |
| ENSMUSG00000036223 | Ska1 | 0 | -1.158519371 | 0.944314987 |
| ENSMUSG00000053846 | Lipg | 0 | 0.063873051 | 1 |
| ENSMUSG00000118384 | 9030625G05Rik | 0 | 0.326907457 | 1 |
| ENSMUSG00000092569 | Gm20544 | 0 | 0.648835551 | 1 |
| ENSMUSG00000117960 | Gm31933 | 0 | 1.063873051 | 1 |
| ENSMUSG00000091055 | Siglec15 | 0 | 0.648835551 | 1 |
| ENSMUSG00000090451 | Gm6133 | 0 | -0.52108945 | 1 |
| ENSMUSG00000078963 | Hsbp1l1 | 0 | 0.648835551 | 1 |
| ENSMUSG00000118070 | Gm50209 | 0 | 0.648835551 | 1 |
| ENSMUSG00000118265 | Gm50211 | 0 | 1.648835551 | 1 |
| ENSMUSG00000118262 | Gm50423 | 0 | 1.648835551 | 1 |
| ENSMUSG00000118300 | Gm50412 | 0 | 0.326907457 | 1 |
| ENSMUSG00000117870 | Gm41792 | 0 | 0.648835551 | 1 |
| ENSMUSG00000118172 | Gm50414 | 0 | 1.648835551 | 1 |
| ENSMUSG00000118363 | Gm50290 | 0 | 1.648835551 | 1 |
| ENSMUSG00000117817 | Gm50292 | 0 | 1.648835551 | 1 |
| ENSMUSG00000117923 | Gm30593 | 0 | 0.326907457 | 1 |
| ENSMUSG00000024553 | Galr1 | 0 | 0.326907457 | 1 |
| ENSMUSG00000117842 | Gm41793 | 0 | 0.063873051 | 1 |
| ENSMUSG00000097738 | 4930445N18Rik | 0 | 0.063873051 | 1 |
| ENSMUSG00000117762 | Gm30889 | 0 | 0.648835551 | 1 |
| ENSMUSG00000117976 | Gm50437 | 0 | 1.648835551 | 1 |
| ENSMUSG00000118333 | 4921531P14Rik | 0 | 1.648835551 | 1 |
| ENSMUSG00000117783 | Gm31265 | 0 | 1.648835551 | 1 |
| ENSMUSG00000117993 | Gm31503 | 0 | 0.648835551 | 1 |
| ENSMUSG00000118175 | Gm50416 | 0 | 0.326907457 | 1 |
| ENSMUSG00000118200 | Gm31621 | 0 | 1.063873051 | 1 |
| ENSMUSG00000118024 | Gm50419 | 0 | 0.326907457 | 1 |
| ENSMUSG00000118225 | Gm31854 | 0 | 1.648835551 | 1 |
| ENSMUSG00000118075 | Gm38576 | 0 | 1.648835551 | 1 |
| ENSMUSG00000069324 | Gm5096 | 0 | 1.648835551 | 1 |
| ENSMUSG00000024907 | Gal | 0 | 1.648835551 | 1 |

| | | | | |
|---|---|---|---|---|
| ENSMUSG00000114315 | Gm48696 | 0 | 0.648835551 | 1 |
| ENSMUSG00000114302 | Gm48683 | 0 | 1.648835551 | 1 |
| ENSMUSG00000024885 | Aldh3b1 | 0 | -1.995020638 | 0.370945697 |
| ENSMUSG00000075296 | Aldh3b2 | 0 | 1.063873051 | 1 |
| ENSMUSG00000024866 | Acy3 | 0 | 0.648835551 | 1 |
| ENSMUSG00000038155 | Gstp2 | 0 | 1.648835551 | 1 |
| ENSMUSG00000024857 | Cabp2 | 0 | 1.648835551 | 1 |
| ENSMUSG00000024842 | Cabp4 | 0 | 0.648835551 | 1 |
| ENSMUSG00000044724 | Gpr152 | 0 | 1.648835551 | 1 |
| ENSMUSG00000045826 | Ptprcap | 0 | 0.648835551 | 1 |
| ENSMUSG00000075289 | Carns1 | 0 | -0.52108945 | 1 |
| ENSMUSG00000040247 | Tbc1d10c | 0 | -0.158519371 | 1 |
| ENSMUSG00000117718 | 1500032F14Rik | 0 | 0.648835551 | 1 |
| ENSMUSG00000118344 | Gm50152 | 0 | 1.063873051 | 1 |
| ENSMUSG00000071691 | Gm960 | 0 | -3.281901786 | 0.022820613 |
| ENSMUSG00000095913 | Gm21844 | 0 | 1.648835551 | 1 |
| ENSMUSG00000083282 | Ctsf | 0 | -0.936126949 | 1 |
| ENSMUSG00000006463 | Zdhhc24 | 0 | 0.326907457 | 1 |
| ENSMUSG00000117893 | Gm46658 | 0 | -0.52108945 | 1 |
| ENSMUSG00000024873 | Cnih2 | 0 | 1.648835551 | 1 |
| ENSMUSG00000118243 | Gm50203 | 0 | 1.063873051 | 1 |
| ENSMUSG00000038498 | Catsper1 | 0 | 1.648835551 | 1 |
| ENSMUSG00000117959 | D330050I16Rik | 0 | 0.063873051 | 1 |
| ENSMUSG00000039330 | Tsga10ip | 0 | 0.326907457 | 1 |
| ENSMUSG00000024922 | Ovol1 | 0 | 0.326907457 | 1 |
| ENSMUSG00000089766 | Gm16538 | 0 | 0.648835551 | 1 |
| ENSMUSG00000110156 | Gm42067 | 0 | 1.063873051 | 1 |
| ENSMUSG00000024833 | Pola2 | 0 | -3.233807498 | 0.019039678 |
| ENSMUSG00000037451 | Slc22a20 | 0 | -0.936126949 | 1 |
| ENSMUSG00000097404 | Gm10814 | 0 | -0.810596067 | 0.844591606 |
| ENSMUSG00000047733 | Tmem262 | 0 | 1.648835551 | 1 |
| ENSMUSG00000117666 | Gm550 | 0 | -0.351164449 | 1 |
| ENSMUSG00000054999 | Naaladl1 | 0 | -0.810596067 | 1 |
| ENSMUSG00000032946 | Rasgrp2 | 0 | -0.158519371 | 1 |
| ENSMUSG00000084876 | Gm14965 | 0 | 1.648835551 | 1 |
| ENSMUSG00000052188 | Gm14964 | 0 | 1.063873051 | 1 |
| ENSMUSG00000061742 | Slc22a12 | 0 | 1.648835551 | 1 |
| ENSMUSG00000024952 | Rps6ka4 | 0 | -1.673092543 | 0.451785624 |
| ENSMUSG00000050623 | Catsperz | 0 | 1.648835551 | 1 |
| ENSMUSG00000047656 | Trpt1 | 0 | 0.063873051 | 1 |
| ENSMUSG00000091558 | Gm17227 | 0 | -1.936126949 | 0.358945041 |
| ENSMUSG00000117743 | Gm50232 | 0 | 1.063873051 | 1 |
| ENSMUSG00000024973 | Plaat5 | 0 | 1.648835551 | 1 |
| ENSMUSG00000118280 | Gm41804 | 0 | -1.351164449 | 0.728062885 |
| ENSMUSG00000024650 | Slc22a6 | 0 | 1.648835551 | 1 |
| ENSMUSG00000117763 | 9830166K06Rik | 0 | 0.648835551 | 1 |
| ENSMUSG00000090840 | 1700092M07Rik | 0 | 1.063873051 | 1 |
| ENSMUSG00000071661 | Zbtb3 | 0 | 1.648835551 | 1 |
| ENSMUSG00000071658 | Gng3 | 0 | 1.063873051 | 1 |
| ENSMUSG00000071648 | Rom1 | 0 | 0.326907457 | 1 |
| ENSMUSG00000071645 | Tut1 | 0 | -3.132524162 | 0.030128441 |

| | | | | |
|---|---|---|---|---|
| ENSMUSG00000118383 | Gm50321 | 0 | -1.673092543 | 0.451785624 |
| ENSMUSG00000118138 | Gm50322 | 0 | 1.648835551 | 1 |
| ENSMUSG00000118100 | Gm50340 | 0 | 1.648835551 | 1 |
| ENSMUSG00000118330 | Gm30042 | 0 | -0.158519371 | 1 |
| ENSMUSG00000090291 | Lrrc10b | 0 | 1.063873051 | 1 |
| ENSMUSG00000043789 | Vwce | 0 | -1.52108945 | 0.587484274 |
| ENSMUSG00000024738 | Pga5 | 0 | 1.063873051 | 1 |
| ENSMUSG00000118027 | Gm10802 | 0 | 0.326907457 | 1 |
| ENSMUSG00000034117 | Ptgdr2 | 0 | 1.648835551 | 1 |
| ENSMUSG00000094584 | Ms4a18 | 0 | 1.648835551 | 1 |
| ENSMUSG00000099398 | Ms4a14 | 0 | 0.326907457 | 1 |
| ENSMUSG00000024672 | Ms4a7 | 0 | -0.158519371 | 1 |
| ENSMUSG00000056290 | Ms4a4b | 0 | -1.051604167 | 0.858831093 |
| ENSMUSG00000024680 | Ms4a2 | 0 | 1.648835551 | 1 |
| ENSMUSG00000024681 | Ms4a3 | 0 | 0.648835551 | 1 |
| ENSMUSG00000055895 | Oosp2 | 0 | 1.063873051 | 1 |
| ENSMUSG00000055933 | Oosp3 | 0 | 1.063873051 | 1 |
| ENSMUSG00000067529 | Olfr1423 | 0 | 1.063873051 | 1 |
| ENSMUSG00000067526 | Olfr1425 | 0 | 1.063873051 | 1 |
| ENSMUSG00000094133 | Olfr1431 | 0 | 0.648835551 | 1 |
| ENSMUSG00000109022 | Olfr1432 | 0 | 1.648835551 | 1 |
| ENSMUSG00000060049 | Olfr235 | 0 | 0.326907457 | 1 |
| ENSMUSG00000109328 | Olfr1438-ps1 | 0 | 1.063873051 | 1 |
| ENSMUSG00000046650 | Olfr1440 | 0 | 1.648835551 | 1 |
| ENSMUSG00000114644 | Gm47243 | 0 | 0.648835551 | 1 |
| ENSMUSG00000071633 | Gm4952 | 0 | -0.158519371 | 1 |
| ENSMUSG00000063683 | Glyat | 0 | 0.063873051 | 1 |
| ENSMUSG00000044441 | Olfr1442 | 0 | 1.648835551 | 1 |
| ENSMUSG00000091014 | Gm5244 | 0 | 1.063873051 | 1 |
| ENSMUSG00000046272 | Olfr1444 | 0 | 1.648835551 | 1 |
| ENSMUSG00000048456 | Olfr1448 | 0 | 0.648835551 | 1 |
| ENSMUSG00000062892 | Olfr1450 | 0 | 1.063873051 | 1 |
| ENSMUSG00000094986 | Olfr1454 | 0 | 1.063873051 | 1 |
| ENSMUSG00000062844 | Olfr1458 | 0 | 0.063873051 | 1 |
| ENSMUSG00000057503 | Olfr1459 | 0 | 0.648835551 | 1 |
| ENSMUSG00000094721 | Olfr1462 | 0 | 1.648835551 | 1 |
| ENSMUSG00000062199 | Olfr1465 | 0 | 1.648835551 | 1 |
| ENSMUSG00000071629 | Olfr1477 | 0 | 1.648835551 | 1 |
| ENSMUSG00000045678 | Olfr1489 | 0 | 1.648835551 | 1 |
| ENSMUSG00000050865 | Olfr1494 | 0 | 0.326907457 | 1 |
| ENSMUSG00000045395 | Olfr1499 | 0 | 1.648835551 | 1 |
| ENSMUSG00000057270 | Olfr1501 | 0 | 1.648835551 | 1 |
| ENSMUSG00000059105 | Olfr1504 | 0 | 1.648835551 | 1 |
| ENSMUSG00000097557 | C130060C02Rik | 0 | -1.258055044 | 0.730310091 |
| ENSMUSG00000118126 | Gm32341 | 0 | 1.063873051 | 1 |
| ENSMUSG00000117941 | Gm50276 | 0 | 1.648835551 | 1 |
| ENSMUSG00000118367 | Gm50277 | 0 | 1.063873051 | 1 |
| ENSMUSG00000117687 | Gm50182 | 0 | 0.063873051 | 1 |
| ENSMUSG00000118109 | Gm41815 | 0 | 0.326907457 | 1 |
| ENSMUSG00000118167 | Gm50216 | 0 | 1.063873051 | 1 |
| ENSMUSG00000118226 | Gm50134 | 0 | 1.648835551 | 1 |

| | | | | |
|---|---|---|---|---|
| ENSMUSG00000118155 | Gm50136 | 0 | -0.351164449 | 1 |
| ENSMUSG00000044387 | 2410080I02Rik | 0 | 0.648835551 | 1 |
| ENSMUSG00000117837 | Gm41821 | 0 | 1.648835551 | 1 |
| ENSMUSG00000118092 | Gm50445 | 0 | 1.063873051 | 1 |
| ENSMUSG00000118340 | Gm50301 | 0 | 1.648835551 | 1 |
| ENSMUSG00000048572 | Tmem252 | 0 | -0.351164449 | 1 |
| ENSMUSG00000058927 | Gm10053 | 0 | 1.648835551 | 1 |
| ENSMUSG00000042372 | Dmrt3 | 0 | 0.648835551 | 1 |
| ENSMUSG00000114682 | Gm48775 | 0 | 0.063873051 | 1 |
| ENSMUSG00000074913 | Gm815 | 0 | 1.648835551 | 1 |
| ENSMUSG00000118209 | Gm35438 | 0 | 1.648835551 | 1 |
| ENSMUSG00000047298 | Kcnv2 | 0 | 0.063873051 | 1 |
| ENSMUSG00000092182 | Gm20407 | 0 | 1.648835551 | 1 |
| ENSMUSG00000097735 | D930032P07Rik | 0 | -1.673092543 | 0.458009808 |
| ENSMUSG00000024935 | Slc1a1 | 0 | -1.158519371 | 0.901106111 |
| ENSMUSG00000039097 | Rln1 | 0 | 1.648835551 | 1 |
| ENSMUSG00000016496 | Cd274 | 0 | -0.52108945 | 1 |
| ENSMUSG00000117964 | Gm36043 | 0 | 1.648835551 | 1 |
| ENSMUSG00000024863 | Mbl2 | 0 | 0.326907457 | 1 |
| ENSMUSG00000024868 | Dkk1 | 0 | 1.063873051 | 1 |
| ENSMUSG00000052595 | A1cf | 0 | -1.43862729 | 0.538110273 |
| ENSMUSG00000118366 | 9130016M20Rik | 0 | 1.648835551 | 1 |
| ENSMUSG00000118064 | Gm50210 | 0 | 1.063873051 | 1 |
| ENSMUSG00000097787 | 2700046G09Rik | 0 | -0.351164449 | 1 |
| ENSMUSG00000118168 | Gm36419 | 0 | 0.648835551 | 1 |
| ENSMUSG00000117914 | Gm29863 | 0 | 0.063873051 | 1 |
| ENSMUSG00000118196 | Gm29946 | 0 | 1.063873051 | 1 |
| ENSMUSG00000079344 | Lipo4 | 0 | 0.326907457 | 1 |
| ENSMUSG00000024768 | Lipf | 0 | -0.158519371 | 1 |
| ENSMUSG00000024770 | Lipn | 0 | -0.158519371 | 1 |
| ENSMUSG00000056078 | Lipm | 0 | 0.326907457 | 1 |
| ENSMUSG00000024774 | Ankrd22 | 0 | -0.351164449 | 1 |
| ENSMUSG00000050370 | Ch25h | 0 | 1.648835551 | 1 |
| ENSMUSG00000097558 | Gm26902 | 0 | 1.063873051 | 1 |
| ENSMUSG00000045932 | Ifit2 | 0 | 0.063873051 | 1 |
| ENSMUSG00000079339 | Ifit1bl1 | 0 | 1.063873051 | 1 |
| ENSMUSG00000062488 | Ifit3b | 0 | 1.648835551 | 1 |
| ENSMUSG00000117973 | Gm50140 | 0 | 1.063873051 | 1 |
| ENSMUSG00000024798 | Htr7 | 0 | -1.52108945 | 0.571786485 |
| ENSMUSG00000117990 | Gm32027 | 0 | 0.063873051 | 1 |
| ENSMUSG00000114235 | F530104D19Rik | 0 | -0.158519371 | 1 |
| ENSMUSG00000041180 | Hectd2 | 0 | -2.051604167 | 0.31180435 |
| ENSMUSG00000067279 | Ppp1r3c | 0 | 0.063873051 | 1 |
| ENSMUSG00000047632 | Fgfbp3 | 0 | -0.158519371 | 1 |
| ENSMUSG00000097591 | A330032B11Rik | 0 | -0.810596067 | 1 |
| ENSMUSG00000062432 | Cyp26c1 | 0 | 0.063873051 | 1 |
| ENSMUSG00000024987 | Cyp26a1 | 0 | 1.648835551 | 1 |
| ENSMUSG00000044026 | Slc35g1 | 0 | -0.52108945 | 1 |
| ENSMUSG00000101079 | Gm28991 | 0 | 1.063873051 | 1 |
| ENSMUSG00000025002 | Cyp2c55 | 0 | 0.063873051 | 1 |
| ENSMUSG00000062624 | Cyp2c67 | 0 | 0.648835551 | 1 |

| | | | | |
|---|---|---|---|---|
| ENSMUSG00000074882 | Cyp2c68 | 0 | 0.648835551 | 1 |
| ENSMUSG00000067225 | Cyp2c54 | 0 | 0.648835551 | 1 |
| ENSMUSG00000097792 | Gm27042 | 0 | 1.063873051 | 1 |
| ENSMUSG00000097740 | E030044B06Rik | 0 | -0.158519371 | 1 |
| ENSMUSG00000050121 | Opalin | 0 | 1.648835551 | 1 |
| ENSMUSG00000067199 | Frat1 | 0 | 1.063873051 | 1 |
| ENSMUSG00000090369 | 4933411K16Rik | 0 | 0.648835551 | 1 |
| ENSMUSG00000025185 | Loxl4 | 0 | -1.051604167 | 0.858831093 |
| ENSMUSG00000060224 | Pyroxd2 | 0 | 0.326907457 | 1 |
| ENSMUSG00000114721 | Gm47938 | 0 | 0.063873051 | 1 |
| ENSMUSG00000117991 | Gm34636 | 0 | 0.326907457 | 1 |
| ENSMUSG00000118236 | Gm34922 | 0 | 0.648835551 | 1 |
| ENSMUSG00000117698 | BC037704 | 0 | 1.063873051 | 1 |
| ENSMUSG00000118000 | Gm50217 | 0 | 0.648835551 | 1 |
| ENSMUSG00000037578 | Pkd2l1 | 0 | -1.051604167 | 0.917930288 |
| ENSMUSG00000117900 | Gm50323 | 0 | -1.051604167 | 0.858831093 |
| ENSMUSG00000117957 | Gm50334 | 0 | 0.648835551 | 1 |
| ENSMUSG00000097683 | Gm26644 | 0 | 1.063873051 | 1 |
| ENSMUSG00000117823 | Gm35610 | 0 | 1.063873051 | 1 |
| ENSMUSG00000092469 | Gm20395 | 0 | 1.648835551 | 1 |
| ENSMUSG00000092369 | 1700039E22Rik | 0 | 0.648835551 | 1 |
| ENSMUSG00000025213 | Kazald1 | 0 | 0.326907457 | 1 |
| ENSMUSG00000025219 | Fgf8 | 0 | 1.648835551 | 1 |
| ENSMUSG00000117956 | Gm50196 | 0 | 1.063873051 | 1 |
| ENSMUSG00000074811 | Hps6 | 0 | 0.063873051 | 1 |
| ENSMUSG00000025229 | Pitx3 | 0 | 1.063873051 | 1 |
| ENSMUSG00000099655 | 2310034G01Rik | 0 | 1.648835551 | 1 |
| ENSMUSG00000118040 | Gm36602 | 0 | 0.648835551 | 1 |
| ENSMUSG00000117814 | Gm50431 | 0 | 1.063873051 | 1 |
| ENSMUSG00000034336 | Ina | 0 | 1.063873051 | 1 |
| ENSMUSG00000025049 | Taf5 | 0 | -1.673092543 | 0.66012174 |
| ENSMUSG00000006435 | Neurl1a | 0 | -2.051604167 | 0.274319469 |
| ENSMUSG00000117659 | Gm50339 | 0 | 1.063873051 | 1 |
| ENSMUSG00000097990 | Gm19557 | 0 | 0.063873051 | 1 |
| ENSMUSG00000118206 | Gm50358 | 0 | 1.648835551 | 1 |
| ENSMUSG00000086121 | Gm16068 | 0 | 1.648835551 | 1 |
| ENSMUSG00000117702 | Gm50281 | 0 | 1.648835551 | 1 |
| ENSMUSG00000117996 | Gm50441 | 0 | 1.063873051 | 1 |
| ENSMUSG00000110007 | Gm45352 | 0 | -0.936126949 | 0.865906102 |
| ENSMUSG00000117758 | Gm50302 | 0 | -0.158519371 | 1 |
| ENSMUSG00000117867 | Gm50330 | 0 | 0.648835551 | 1 |
| ENSMUSG00000118173 | Gm50329 | 0 | -1.051604167 | 0.920002616 |
| ENSMUSG00000118114 | Gm50332 | 0 | 1.648835551 | 1 |
| ENSMUSG00000118233 | Gm30381 | 0 | 1.063873051 | 1 |
| ENSMUSG00000118314 | 1700001K23Rik | 0 | -0.158519371 | 1 |
| ENSMUSG00000117768 | 8030456M14Rik | 0 | 1.063873051 | 1 |
| ENSMUSG00000117921 | Gm50186 | 0 | -2.209145444 | 0.229785766 |
| ENSMUSG00000118207 | Gm50191 | 0 | 1.648835551 | 1 |
| ENSMUSG00000117827 | D730002M21Rik | 0 | 1.063873051 | 1 |
| ENSMUSG00000090990 | Gm17197 | 0 | 1.648835551 | 1 |
| ENSMUSG00000025077 | Dclre1a | 0 | -2.106051951 | 0.279592421 |

| | | | | |
|---|---|---|---|---|
| ENSMUSG00000117859 | Gm32492 | 0 | 1.648835551 | 1 |
| ENSMUSG00000117696 | Gm50100 | 0 | 0.648835551 | 1 |
| ENSMUSG00000117670 | Gm50270 | 0 | 0.326907457 | 1 |
| ENSMUSG00000087002 | Gm16277 | 0 | 0.648835551 | 1 |
| ENSMUSG00000100001 | 1810007D17Rik | 0 | 0.063873051 | 1 |
| ENSMUSG00000040901 | Kcnk18 | 0 | 1.063873051 | 1 |
| ENSMUSG00000110210 | Gm33756 | 0 | 1.063873051 | 1 |
| ENSMUSG00000090965 | Gm17203 | 0 | 0.326907457 | 1 |
| ENSMUSG00000053117 | E330013P04Rik | 0 | 0.648835551 | 1 |
| ENSMUSG00000063698 | Sfxn4 | 0 | -0.158519371 | 1 |
| ENSMUSG00000051592 | Ccnb3 | 0 | 1.648835551 | 1 |
| ENSMUSG00000050089 | Akap4 | 0 | 1.648835551 | 1 |
| ENSMUSG00000046269 | Usp27x | 0 | 0.063873051 | 1 |
| ENSMUSG00000087252 | Gm14379 | 0 | 1.063873051 | 1 |
| ENSMUSG00000039556 | Ppp1r3f | 0 | -1.810596067 | 0.465580427 |
| ENSMUSG00000039521 | Foxp3 | 0 | 1.063873051 | 1 |
| ENSMUSG00000031142 | Cacna1f | 0 | 1.648835551 | 1 |
| ENSMUSG00000087644 | Gm14703 | 0 | 1.063873051 | 1 |
| ENSMUSG00000031147 | Magix | 0 | 0.063873051 | 1 |
| ENSMUSG00000039278 | Pcsk1n | 0 | 1.648835551 | 1 |
| ENSMUSG00000031165 | Was | 0 | 1.063873051 | 1 |
| ENSMUSG00000086404 | Gm6787 | 0 | 0.063873051 | 1 |
| ENSMUSG00000079704 | Gm14459 | 0 | 1.648835551 | 1 |
| ENSMUSG00000052364 | B630019K06Rik | 0 | 1.648835551 | 1 |
| ENSMUSG00000079694 | Gm14862 | 0 | 1.648835551 | 1 |
| ENSMUSG00000048994 | H2al3 | 0 | 1.648835551 | 1 |
| ENSMUSG00000090084 | Srpx | 0 | 0.326907457 | 1 |
| ENSMUSG00000100714 | 2010308F09Rik | 0 | 0.648835551 | 1 |
| ENSMUSG00000040229 | Gpr34 | 0 | 1.063873051 | 1 |
| ENSMUSG00000025037 | Maoa | 0 | -2.636566667 | 0.129391482 |
| ENSMUSG00000040138 | Ndp | 0 | 0.326907457 | 1 |
| ENSMUSG00000037358 | Dipk2b | 0 | -1.051604167 | 0.872990655 |
| ENSMUSG00000097497 | Gm26652 | 0 | -0.673092543 | 1 |
| ENSMUSG00000037347 | Chst7 | 0 | 0.648835551 | 1 |
| ENSMUSG00000023070 | Rgn | 0 | 1.063873051 | 1 |
| ENSMUSG00000031066 | Usp11 | 0 | -3.106051951 | 0.028853812 |
| ENSMUSG00000031079 | Zfp300 | 0 | 1.648835551 | 1 |
| ENSMUSG00000087586 | Gm6938 | 0 | 1.648835551 | 1 |
| ENSMUSG00000031089 | Slc6a14 | 0 | 0.326907457 | 1 |
| ENSMUSG00000084063 | Gm5934 | 0 | 0.648835551 | 1 |
| ENSMUSG00000095063 | Slx | 0 | 1.063873051 | 1 |
| ENSMUSG00000071788 | Gm14525 | 0 | 1.648835551 | 1 |
| ENSMUSG00000085534 | Gm2117 | 0 | 1.648835551 | 1 |
| ENSMUSG00000085489 | Gm3669 | 0 | 0.648835551 | 1 |
| ENSMUSG00000118405 | FQ976806.1 | 0 | 1.648835551 | 1 |
| ENSMUSG00000073247 | Gm10486 | 0 | -0.158519371 | 1 |
| ENSMUSG00000085000 | Gm14553 | 0 | 1.063873051 | 1 |
| ENSMUSG00000073245 | Gm14819 | 0 | 1.648835551 | 1 |
| ENSMUSG00000016239 | Lonrf3 | 0 | -1.43862729 | 0.538110273 |
| ENSMUSG00000091556 | Gm14569 | 0 | 1.063873051 | 1 |
| ENSMUSG00000037636 | Slc25a43 | 0 | -0.810596067 | 0.844591606 |

| | | | | |
|---|---|---|---|---|
| ENSMUSG00000044400 | Sowahd | 0 | 1.648835551 | 1 |
| ENSMUSG00000036551 | Akap14 | 0 | 0.063873051 | 1 |
| ENSMUSG00000036537 | Rnf113a1 | 0 | 1.648835551 | 1 |
| ENSMUSG00000079639 | Rhox4a | 0 | 0.648835551 | 1 |
| ENSMUSG00000071771 | Rhox4b | 0 | 1.648835551 | 1 |
| ENSMUSG00000079633 | Rhox4d | 0 | 1.063873051 | 1 |
| ENSMUSG00000079630 | Rhox4f | 0 | 1.063873051 | 1 |
| ENSMUSG00000071769 | Rhox3h | 0 | 1.648835551 | 1 |
| ENSMUSG00000053909 | Rhox10 | 0 | 1.648835551 | 1 |
| ENSMUSG00000071766 | Rhox12 | 0 | 1.063873051 | 1 |
| ENSMUSG00000087390 | Gm7598 | 0 | 1.063873051 | 1 |
| ENSMUSG00000085635 | Gm14565 | 0 | 0.326907457 | 1 |
| ENSMUSG00000051361 | 6030498E09Rik | 0 | -1.051604167 | 0.886203497 |
| ENSMUSG00000005696 | Sh2d1a | 0 | 0.648835551 | 1 |
| ENSMUSG00000045284 | Dcaf12l1 | 0 | 1.648835551 | 1 |
| ENSMUSG00000085157 | 4930515L19Rik | 0 | 1.648835551 | 1 |
| ENSMUSG00000031099 | Smarca1 | 0 | -1.52108945 | 0.552613787 |
| ENSMUSG00000037010 | Apln | 0 | -0.351164449 | 1 |
| ENSMUSG00000037005 | Xpnpep2 | 0 | 1.648835551 | 1 |
| ENSMUSG00000031101 | Sash3 | 0 | 0.326907457 | 1 |
| ENSMUSG00000031104 | Rab33a | 0 | 1.648835551 | 1 |
| ENSMUSG00000031105 | Slc25a14 | 0 | -1.599091962 | 0.559918909 |
| ENSMUSG00000051209 | Gpr119 | 0 | 1.648835551 | 1 |
| ENSMUSG00000031107 | Rbmx2 | 0 | -0.351164449 | 1 |
| ENSMUSG00000079606 | Gm595 | 0 | -1.258055044 | 0.729762905 |
| ENSMUSG00000087369 | Gm14696 | 0 | 1.063873051 | 1 |
| ENSMUSG00000036198 | Arhgap36 | 0 | 1.648835551 | 1 |
| ENSMUSG00000071764 | Olfr1320 | 0 | 1.648835551 | 1 |
| ENSMUSG00000067971 | Olfr1321 | 0 | 1.648835551 | 1 |
| ENSMUSG00000036109 | Mbnl3 | 0 | -1.351164449 | 0.687112677 |
| ENSMUSG00000055780 | Usp26 | 0 | 0.648835551 | 1 |
| ENSMUSG00000031118 | 1700080O16Rik | 0 | 0.648835551 | 1 |
| ENSMUSG00000073207 | Ccdc160 | 0 | 1.648835551 | 1 |
| ENSMUSG00000087365 | C430049B03Rik | 0 | -3.540989007 | 0.007698034 |
| ENSMUSG00000079593 | Gm14597 | 0 | 1.063873051 | 1 |
| ENSMUSG00000059334 | Zfp36l3 | 0 | 1.648835551 | 1 |
| ENSMUSG00000031125 | 3830403N18Rik | 0 | 1.648835551 | 1 |
| ENSMUSG00000073177 | Gm773 | 0 | 1.648835551 | 1 |
| ENSMUSG00000085114 | 1600025M17Rik | 0 | 0.326907457 | 1 |
| ENSMUSG00000023092 | Fhl1 | 0 | -1.158519371 | 0.886203497 |
| ENSMUSG00000031132 | Cd40lg | 0 | 1.648835551 | 1 |
| ENSMUSG00000067860 | Zic3 | 0 | 0.648835551 | 1 |
| ENSMUSG00000087401 | Gm14664 | 0 | 1.063873051 | 1 |
| ENSMUSG00000085757 | Tslrn1 | 0 | 1.648835551 | 1 |
| ENSMUSG00000062170 | Fmr1nb | 0 | 0.326907457 | 1 |
| ENSMUSG00000071748 | Gm14698 | 0 | 1.648835551 | 1 |
| ENSMUSG00000073139 | Tmem185a | 0 | 1.063873051 | 1 |
| ENSMUSG00000085284 | Gm16189 | 0 | -0.351164449 | 1 |
| ENSMUSG00000033361 | Prrg3 | 0 | 0.063873051 | 1 |
| ENSMUSG00000073125 | Xlr3b | 0 | 0.326907457 | 1 |
| ENSMUSG00000078317 | F8a | 0 | 0.648835551 | 1 |

| | | | | |
|---|---|---|---|---|
| ENSMUSG00000031362 | Xlr4c | 0 | 1.648835551 | 1 |
| ENSMUSG00000096966 | Gm18336 | 0 | 1.063873051 | 1 |
| ENSMUSG00000031374 | Zfp92 | 0 | 1.063873051 | 1 |
| ENSMUSG00000031385 | Plxnb3 | 0 | 1.648835551 | 1 |
| ENSMUSG00000002007 | Srpk3 | 0 | 1.648835551 | 1 |
| ENSMUSG00000085669 | Gm14817 | 0 | 1.648835551 | 1 |
| ENSMUSG00000031391 | L1cam | 0 | -0.810596067 | 1 |
| ENSMUSG00000031394 | Opn1mw | 0 | 0.648835551 | 1 |
| ENSMUSG00000031397 | Tktl1 | 0 | 0.648835551 | 1 |
| ENSMUSG00000032806 | Slc10a3 | 0 | 1.063873051 | 1 |
| ENSMUSG00000031399 | Fam3a | 0 | -0.52108945 | 1 |
| ENSMUSG00000073094 | Smim9 | 0 | 1.648835551 | 1 |
| ENSMUSG00000090110 | Cmc4 | 0 | -0.158519371 | 1 |
| ENSMUSG00000031200 | Mtcp1 | 0 | -0.673092543 | 1 |
| ENSMUSG00000085598 | Gm14715 | 0 | 1.648835551 | 1 |
| ENSMUSG00000025666 | Tmem47 | 0 | -0.52108945 | 1 |
| ENSMUSG00000078314 | Gm14762 | 0 | 1.648835551 | 1 |
| ENSMUSG00000025056 | Nr0b1 | 0 | 1.648835551 | 1 |
| ENSMUSG00000035427 | Mageb4 | 0 | -0.673092543 | 1 |
| ENSMUSG00000097961 | Gm27000 | 0 | 1.648835551 | 1 |
| ENSMUSG00000071726 | Gm5941 | 0 | 1.648835551 | 1 |
| ENSMUSG00000035277 | Arx | 0 | 1.648835551 | 1 |
| ENSMUSG00000081044 | AU015836 | 0 | 1.063873051 | 1 |
| ENSMUSG00000118554 | Fam90a1b | 0 | 1.648835551 | 1 |
| ENSMUSG00000043549 | Fam90a1b | 0 | 1.648835551 | 1 |
| ENSMUSG00000084904 | Gm14827 | 0 | 1.648835551 | 1 |
| ENSMUSG00000071723 | Gspt2 | 0 | -1.351164449 | 0.687112677 |
| ENSMUSG00000109459 | Gm39526 | 0 | -1.258055044 | 0.673811773 |
| ENSMUSG00000035062 | Zc4h2 | 0 | 1.063873051 | 1 |
| ENSMUSG00000078122 | F630028O10Rik | 0 | -1.051604167 | 0.858831093 |
| ENSMUSG00000031210 | Gpr165 | 0 | 1.648835551 | 1 |
| ENSMUSG00000034457 | Eda2r | 0 | -2.480447466 | 0.152741466 |
| ENSMUSG00000073051 | Gm14812 | 0 | -0.158519371 | 1 |
| ENSMUSG00000031220 | Awat2 | 0 | 0.648835551 | 1 |
| ENSMUSG00000019359 | Gdpd2 | 0 | 1.063873051 | 1 |
| ENSMUSG00000085314 | Gm14866 | 0 | -0.158519371 | 1 |
| ENSMUSG00000031297 | Slc7a3 | 0 | -1.158519371 | 0.667778757 |
| ENSMUSG00000090141 | Gm614 | 0 | 1.648835551 | 1 |
| ENSMUSG00000079487 | Med12 | 0 | -4.009375931 | 0.001341203 |
| ENSMUSG00000031302 | Nlgn3 | 0 | 0.326907457 | 1 |
| ENSMUSG00000051159 | Cited1 | 0 | 0.648835551 | 1 |
| ENSMUSG00000031329 | Tsx | 0 | 0.326907457 | 1 |
| ENSMUSG00000085715 | Tsix | 0 | -1.351164449 | 0.741406232 |
| ENSMUSG00000031227 | Magee1 | 0 | 1.648835551 | 1 |
| ENSMUSG00000087620 | 5330434G04Rik | 0 | 1.648835551 | 1 |
| ENSMUSG00000031230 | Fgf16 | 0 | 0.648835551 | 1 |
| ENSMUSG00000047242 | Taf9b | 0 | -0.52108945 | 1 |
| ENSMUSG00000073012 | Fnd3c2 | 0 | -0.351164449 | 1 |
| ENSMUSG00000033737 | Fndc3c1 | 0 | -0.351164449 | 1 |
| ENSMUSG00000052821 | Cysltr1 | 0 | 1.063873051 | 1 |
| ENSMUSG00000050921 | P2ry10 | 0 | 1.648835551 | 1 |

| | | | | |
|---|---|---|---|---|
| ENSMUSG00000054293 | P2ry10b | 0 | -0.158519371 | 1 |
| ENSMUSG00000073008 | Gpr174 | 0 | 0.648835551 | 1 |
| ENSMUSG00000048621 | Gm6377 | 0 | 1.648835551 | 1 |
| ENSMUSG00000034555 | Tex16 | 0 | 1.648835551 | 1 |
| ENSMUSG00000025528 | 2010106E10Rik | 0 | -0.673092543 | 1 |
| ENSMUSG00000025529 | Zfp711 | 0 | 1.063873051 | 1 |
| ENSMUSG00000084963 | Gm14936 | 0 | 0.648835551 | 1 |
| ENSMUSG00000025597 | Klhl4 | 0 | -0.158519371 | 1 |
| ENSMUSG00000072995 | Cpxcr1 | 0 | 0.326907457 | 1 |
| ENSMUSG00000086431 | Gm14929 | 0 | 1.648835551 | 1 |
| ENSMUSG00000095474 | Cldn34c2 | 0 | 1.648835551 | 1 |
| ENSMUSG00000043569 | Cldn34c4 | 0 | 1.648835551 | 1 |
| ENSMUSG00000086244 | 4930558G05Rik | 0 | 1.063873051 | 1 |
| ENSMUSG00000031250 | Tnmd | 0 | 1.648835551 | 1 |
| ENSMUSG00000067377 | Tspan6 | 0 | 0.326907457 | 1 |
| ENSMUSG00000084875 | 4930570D08Rik | 0 | 1.063873051 | 1 |
| ENSMUSG00000031255 | Sytl4 | 0 | 1.648835551 | 1 |
| ENSMUSG00000031262 | Cenpi | 0 | -1.599091962 | 0.559918909 |
| ENSMUSG00000000223 | Drp2 | 0 | -0.351164449 | 1 |
| ENSMUSG00000009596 | Taf7l | 0 | 1.063873051 | 1 |
| ENSMUSG00000049804 | Armcx4 | 0 | 1.063873051 | 1 |
| ENSMUSG00000033460 | Armcx1 | 0 | -0.158519371 | 1 |
| ENSMUSG00000050394 | Armcx6 | 0 | 1.063873051 | 1 |
| ENSMUSG00000033436 | Armcx2 | 0 | 1.063873051 | 1 |
| ENSMUSG00000009941 | Nxf2 | 0 | 1.063873051 | 1 |
| ENSMUSG00000031409 | Tceal6 | 0 | 1.648835551 | 1 |
| ENSMUSG00000031411 | Prame | 0 | 1.648835551 | 1 |
| ENSMUSG00000058252 | Tcp11x2 | 0 | 1.648835551 | 1 |
| ENSMUSG00000060726 | Tmsb15a | 0 | 1.648835551 | 1 |
| ENSMUSG00000048040 | Arxes2 | 0 | -0.158519371 | 1 |
| ENSMUSG00000042750 | Bex2 | 0 | 1.648835551 | 1 |
| ENSMUSG00000087553 | 5730412P04Rik | 0 | 1.648835551 | 1 |
| ENSMUSG00000050071 | Bex1 | 0 | 1.648835551 | 1 |
| ENSMUSG00000031424 | Kir3dl1 | 0 | 1.648835551 | 1 |
| ENSMUSG00000044550 | Tceal3 | 0 | 0.326907457 | 1 |
| ENSMUSG00000049536 | Tceal1 | 0 | 0.648835551 | 1 |
| ENSMUSG00000087368 | BC065397 | 0 | -1.258055044 | 0.673811773 |
| ENSMUSG00000043463 | Rab9b | 0 | 1.648835551 | 1 |
| ENSMUSG00000089996 | Tmsb15b2 | 0 | -0.810596067 | 1 |
| ENSMUSG00000089768 | Tmsb15b1 | 0 | 1.648835551 | 1 |
| ENSMUSG00000044348 | Slc25a53 | 0 | -2.051604167 | 0.455131546 |
| ENSMUSG00000042515 | Pwwp3b | 0 | -0.810596067 | 0.844591606 |
| ENSMUSG00000072944 | Nup62cl | 0 | 0.648835551 | 1 |
| ENSMUSG00000042433 | Pih1h3b | 0 | 1.063873051 | 1 |
| ENSMUSG00000086441 | Gm15046 | 0 | -0.52108945 | 1 |
| ENSMUSG00000085146 | Eif2c5 | 0 | 1.063873051 | 1 |
| ENSMUSG00000042386 | Tex13b | 0 | 1.648835551 | 1 |
| ENSMUSG00000031285 | Dcx | 0 | 0.063873051 | 1 |
| ENSMUSG00000085139 | A730046J19Rik | 0 | 1.648835551 | 1 |
| ENSMUSG00000041700 | Lhfpl1 | 0 | 1.648835551 | 1 |
| ENSMUSG00000094885 | Ott | 0 | -2.395558568 | 0.219901826 |

| | | | | |
|---|---|---|---|---|
| ENSMUSG00000094378 | Gm15097 | 0 | 1.063873051 | 1 |
| ENSMUSG00000087360 | Gm15104 | 0 | 1.063873051 | 1 |
| ENSMUSG00000025272 | Tro | 0 | 0.648835551 | 1 |
| ENSMUSG00000041245 | Wnk3 | 0 | -1.52108945 | 0.544908989 |
| ENSMUSG00000025257 | Ribc1 | 0 | -0.52108945 | 1 |
| ENSMUSG00000084885 | 3010001F23Rik | 0 | -0.52108945 | 1 |
| ENSMUSG00000067215 | Usp51 | 0 | 1.648835551 | 1 |
| ENSMUSG00000047238 | Mageh1 | 0 | -0.673092543 | 1 |
| ENSMUSG00000085982 | 9530051G07Rik | 0 | 0.326907457 | 1 |
| ENSMUSG00000041658 | Rragb | 0 | -1.43862729 | 0.538110273 |
| ENSMUSG00000041633 | Kctd12b | 0 | 0.648835551 | 1 |
| ENSMUSG00000063728 | Magea6 | 0 | 1.063873051 | 1 |
| ENSMUSG00000041552 | Ptchd1 | 0 | -0.673092543 | 1 |
| ENSMUSG00000041476 | Smpx | 0 | -0.158519371 | 1 |
| ENSMUSG00000087505 | Gm15241 | 0 | 0.648835551 | 1 |
| ENSMUSG00000031293 | Rs1 | 0 | -0.936126949 | 1 |
| ENSMUSG00000108981 | Gm15262 | 0 | 1.063873051 | 1 |
| ENSMUSG00000031364 | Grpr | 0 | 1.063873051 | 1 |
| ENSMUSG00000031367 | Ap1s2 | 0 | -2.258055044 | 0.222101622 |
| ENSMUSG00000031382 | Asb11 | 0 | -0.158519371 | 1 |
| ENSMUSG00000047757 | Fancb | 0 | -0.673092543 | 1 |
| ENSMUSG00000051224 | Tceanc | 0 | -0.673092543 | 1 |
| ENSMUSG00000086860 | Gm1720 | 0 | 1.648835551 | 1 |
| ENSMUSG00000084806 | Gm15232 | 0 | 0.648835551 | 1 |
| ENSMUSG00000040522 | Tlr8 | 0 | 0.648835551 | 1 |
| ENSMUSG00000044583 | Tlr7 | 0 | 1.648835551 | 1 |
| ENSMUSG00000031354 | Amelx | 0 | 1.648835551 | 1 |
| ENSMUSG00000087159 | Gm15246 | 0 | -0.351164449 | 1 |
| ENSMUSG00000087587 | Gm15245 | 0 | 1.648835551 | 1 |
| ENSMUSG00000087263 | Gm15726 | 0 | 0.326907457 | 1 |
| ENSMUSG00000069053 | Uba1y | 0 | 1.648835551 | 1 |
| ENSMUSG00000101036 | Gm28588 | 0 | 1.063873051 | 1 |
| ENSMUSG00000101107 | Gm28587 | 0 | -0.52108945 | 1 |
| ENSMUSG00000099876 | Gm29650 | 0 | 0.648835551 | 1 |
| ENSMUSG00000069044 | Usp9y | 0 | -1.599091962 | 0.603006227 |
| ENSMUSG00000101243 | Gm28919 | 0 | 1.648835551 | 1 |
| ENSMUSG00000100939 | Gm28998 | 0 | 1.063873051 | 1 |
| ENSMUSG00000100645 | Gm28576 | 0 | 1.648835551 | 1 |
| ENSMUSG00000101928 | Gm20824 | 0 | 1.063873051 | 1 |
| ENSMUSG00000091077 | Gm21292 | 0 | 1.648835551 | 1 |
| ENSMUSG00000101396 | Gm28510 | 0 | -1.43862729 | 0.81567408 |
| ENSMUSG00000099482 | Gm28955 | 0 | 1.648835551 | 1 |
| ENSMUSG00000093903 | Gm21454 | 0 | -1.874726405 | 0.503117077 |
| ENSMUSG00000094325 | Gm20831 | 0 | 1.648835551 | 1 |
| ENSMUSG00000102396 | Gm37574 | 0 | 1.648835551 | 1 |
| ENSMUSG00000099799 | Gm28260 | 0 | 1.648835551 | 1 |
| ENSMUSG00000099832 | Gm29025 | 0 | 1.648835551 | 1 |
| ENSMUSG00000100043 | Gm29433 | 0 | 0.648835551 | 1 |
| ENSMUSG00000102116 | Gm28887 | 0 | 1.648835551 | 1 |
| ENSMUSG00000101399 | Gm28961 | 0 | 0.063873051 | 1 |
| ENSMUSG00000101182 | Gm29209 | 0 | 1.648835551 | 1 |

| | | | | |
|---|---|---|---|---|
| ENSMUSG00000099716 | Gm29082 | 0 | 1.063873051 | 1 |
| ENSMUSG00000096016 | Gm20937 | 0 | 1.648835551 | 1 |
| ENSMUSG00000094616 | Gm20888 | 0 | 1.648835551 | 1 |
| ENSMUSG00000101528 | Gm29110 | 0 | 0.648835551 | 1 |
| ENSMUSG00000100902 | Gm20850 | 0 | 1.648835551 | 1 |
| ENSMUSG00000099792 | Gm29564 | 0 | 1.648835551 | 1 |
| ENSMUSG00000101115 | Gm28664 | 0 | 1.063873051 | 1 |
| ENSMUSG00000101321 | Gm28817 | 0 | 1.648835551 | 1 |
| ENSMUSG00000095578 | Gm21760 | 0 | 1.648835551 | 1 |
| ENSMUSG00000100925 | Gm28898 | 0 | 1.648835551 | 1 |
| ENSMUSG00000099740 | Gm28897 | 0 | 1.063873051 | 1 |
| ENSMUSG00000100035 | Gm29473 | 0 | 1.648835551 | 1 |
| ENSMUSG00000095153 | Gm21095 | 0 | 1.648835551 | 1 |
| ENSMUSG00000094660 | Gm21394 | 0 | 1.648835551 | 1 |
| ENSMUSG00000099840 | Gm21409 | 0 | 1.648835551 | 1 |
| ENSMUSG00000099341 | Gm29090 | 0 | 0.648835551 | 1 |
| ENSMUSG00000095250 | AC133103.4 | 0 | 1.648835551 | 1 |
| ENSMUSG00000095672 | AC133103.5 | 0 | 1.648835551 | 1 |
| ENSMUSG00000079190 | AC133103.1 | 0 | 0.326907457 | 1 |
| ENSMUSG00000079808 | AC168977.1 | 0 | 0.326907457 | 1 |
| ENSMUSG00000095500 | AC132444.5 | 0 | 1.648835551 | 1 |
| ENSMUSG00000062783 | Csprs | 0 | -0.52108945 | 1 |
| ENSMUSG00000096808 | AC132444.6 | 0 | 0.326907457 | 1 |
| ENSMUSG00000095742 | CAAA01147332.1 | 0 | 0.326907457 | 1 |
