## Supplementary Table S6 for "Molecular profiling of XPO1 inhibitor and gemcitabine-nab-paclitaxel combination in cellular and LSL-Kras G12D/+; Trp53 fl/+; Pdx1-Cre (KPC) pancreatic cancer model"

| ID | gene_id | gene_nam | Descriptior | FPKMMia_ | FPKMMia_ | FPKMMia_Comb |
|---|---|---|---|---|---|---|
| 5274 | ENSG0000 | SRGN | serglycin [S | 474.76 | 585.887 | 614.7266 |
| 16793 | ENSG0000 | PSAP | prosaposin | 172.6889 | 197.5416 | 242.1063 |
| 20007 | ENSG0000 | UCA1 | urothelial c | 39.64245 | 178.2402 | 228.6547 |
| 4643 | ENSG0000 | ID3 | inhibitor of | 144.4047 | 205.0972 | 216.5926 |
| 8077 | ENSG0000 | APP | amyloid be | 158.0224 | 212.972 | 214.0516 |
| 7355 | ENSG0000 | TUBB2A | tubulin bet | 146.149 | 156.0496 | 205.211 |
| 6195 | ENSG0000 | ACTN4 | actinin alpl | 155.014 | 172.7959 | 181.886 |
| 6791 | ENSG0000 | LDHA | lactate deh | 108.3862 | 116.293 | 173.078 |
| 36613 | ENSG0000 | TXNIP | thioredoxir | 38.34644 | 139.0628 | 172.9522 |
| 16218 | ENSG0000 | H1F0 | H1 histone | 71.26847 | 106.2914 | 165.6948 |
| 26422 | ENSG0000 | LINC02154 | long interg | 82.99883 | 127.1095 | 156.4384 |
| 5602 | ENSG0000 | PPDPF | pancreatic | 74.78201 | 128.7974 | 153.8175 |
| 6350 | ENSG0000 | CAP1 | cyclase ass | 85.39724 | 149.5687 | 150.1635 |
| 5801 | ENSG0000 | CANX | calnexin [S | 110.4737 | 127.0265 | 139.1132 |
| 19200 | ENSG0000 | MT-TP | mitochond | 70.79537 | 115.8941 | 138.6957 |
| 14555 | ENSG0000 | F2R | coagulatior | 123.9148 | 129.9387 | 138.6099 |
| 6261 | ENSG0000 | EIF2S3 | eukaryotic | 122.2125 | 127.2116 | 134.215 |
| 10156 | ENSG0000 | STC1 | stanniocalc | 38.91062 | 129.8527 | 132.6082 |
| 5518 | ENSG0000 | GLO1 | glyoxalase | 107.5711 | 118.9669 | 129.4928 |
| 5691 | ENSG0000 | ID1 | inhibitor of | 96.51645 | 99.11083 | 126.7361 |
| 10187 | ENSG0000 | PTMS | parathymo | 117.5934 | 117.7694 | 123.5494 |
| 5273 | ENSG0000 | PLAU | plasminoge | 71.29424 | 94.09332 | 118.7048 |
| 13488 | ENSG0000 | MARCKSL1 | MARCKS lil | 85.14923 | 104.6187 | 118.5929 |
| 1056 | ENSG0000 | LAPTM4A | lysosomal | 80.00257 | 99.85801 | 114.1902 |
| 3000 | ENSG0000 | LIPA | lipase A, ly | 57.05968 | 89.08414 | 112.8399 |
| 9420 | ENSG0000 | TMEM123 | transmeml | 70.49719 | 76.67655 | 99.6226 |
| 125 | ENSG0000 | AP2B1 | adaptor re | 84.66512 | 86.4364 | 94.60738 |
| 3184 | ENSG0000 | FKBP8 | FK506 binc | 65.08876 | 68.34401 | 94.55668 |
| 5353 | ENSG0000 | MORF4L2 | mortality f | 69.97568 | 69.99743 | 91.18569 |
| 13965 | ENSG0000 | JUN | Jun proto-c | 47.23763 | 85.74876 | 87.4923 |
| 6077 | ENSG0000 | SLC44A2 | solute carr | 19.77866 | 77.88256 | 86.27645 |
| 10163 | ENSG0000 | RCAN1 | regulator c | 30.7134 | 82.47747 | 84.34677 |
| 3217 | ENSG0000 | MTPN | myotrophi | 71.94399 | 77.99501 | 83.38615 |
| 36399 | ENSG0000 | TIMM23 | translocase | 77.27995 | 77.79654 | 83.32447 |
| 2612 | ENSG0000 | SMARCA1 | SWI/SNF re | 56.64354 | 78.65683 | 82.7006 |
| 8606 | ENSG0000 | RPS3A | ribosomal | 74.16539 | 77.82211 | 82.54292 |
| 2559 | ENSG0000 | MYL12A | myosin ligh | 79.9496 | 81.2212 | 82.17297 |
| 14858 | ENSG0000 | HMGN4 | high mobil | 47.33176 | 60.92614 | 79.26108 |
| 7061 | ENSG0000 | ITM2C | integral me | 31.378 | 36.9636 | 79.21964 |
| 2505 | ENSG0000 | MYL9 | myosin ligh | 22.50609 | 74.34827 | 78.71454 |
| 8068 | ENSG0000 | COL6A1 | collagen ty | 33.4235 | 56.80317 | 76.94417 |
| 10927 | ENSG0000 | PTX3 | pentraxin 3 | 38.46195 | 69.18942 | 76.78552 |
| 24966 | ENSG0000 | AL353751.1 | | 44.15173 | 67.69505 | 76.65804 |
| 264 | ENSG0000 | RAB27B | RAB27B, m | 50.10907 | 58.81994 | 76.14585 |
| 515 | ENSG0000 | GRN | granulin pr | 38.50289 | 62.13206 | 75.70344 |
| 6723 | ENSG0000 | LOXL2 | lysyl oxidas | 53.79608 | 59.84339 | 75.30533 |

| | | | | | | |
|---|---|---|---|---|---|---|
| 5575 | ENSG0000 | SOX9 | SRY-box 9 | 63.11799 | 67.67138 | 72.56739 |
| 924 | ENSG0000 | OAT | ornithine a | 42.62889 | 57.93566 | 71.3234 |
| 888 | ENSG0000 | NGFR | nerve grow | 2.738906 | 54.40052 | 68.61533 |
| 12836 | ENSG0000 | CLSTN1 | calsyntenir | 46.95585 | 50.55652 | 66.82817 |
| 3204 | ENSG0000 | NAMPT | nicotinami | 33.7571 | 42.75802 | 66.6048 |
| 18236 | ENSG0000 | HLA-E | major histo | 26.25477 | 52.51671 | 64.59561 |
| 6955 | ENSG0000 | PRRG4 | proline rich | 31.16876 | 55.54109 | 64.25345 |
| 4786 | ENSG0000 | CCNI | cyclin I [So | 46.991 | 56.62973 | 64.0074 |
| 547 | ENSG0000 | VCL | vinculin [So | 37.97531 | 50.76885 | 63.16134 |
| 4112 | ENSG0000 | DAP | death asso | 42.06145 | 61.84688 | 63.1181 |
| 3293 | ENSG0000 | SERPINE1 | serpin fam | 39.93315 | 61.83135 | 63.0579 |
| 15408 | ENSG0000 | TCEAL9 | transcriptio | 31.3471 | 45.59976 | 62.33864 |
| 9744 | ENSG0000 | RRAGA | Ras relatec | 33.53133 | 38.52314 | 59.73574 |
| 3 | ENSG0000 | DPM1 | dolichyl-ph | 51.97838 | 53.40411 | 58.88402 |
| 18131 | ENSG0000 | AGPAT1 | 1-acylglyce | 45.53241 | 47.57936 | 58.19842 |
| 3165 | ENSG0000 | ISYNA1 | inositol-3-p | 17.95506 | 50.41481 | 58.05577 |
| 4399 | ENSG0000 | SF3B1 | splicing fac | 48.41648 | 57.06252 | 57.11345 |
| 473 | ENSG0000 | EHD2 | EH domain | 30.40995 | 53.0706 | 56.29711 |
| 46 | ENSG0000 | ARF5 | ADP ribosy | 47.42022 | 48.84176 | 55.7079 |
| 5615 | ENSG0000 | GTF2F1 | general tra | 48.27484 | 53.2597 | 55.55279 |
| 32568 | ENSG0000 | AP002990.1 | | 10.64395 | 43.27947 | 55.08875 |
| 252 | ENSG0000 | CD9 | CD9 molec | 20.9077 | 36.43845 | 54.55812 |
| 23480 | ENSG0000 | DHFR | dihydrofola | 31.91559 | 38.68975 | 54.33041 |
| 9215 | ENSG0000 | UBC | ubiquitin C | 46.24479 | 51.19844 | 54.15022 |
| 5204 | ENSG0000 | COPA | coatomer p | 36.02831 | 46.80665 | 53.9044 |
| 3742 | ENSG0000 | PDHX | pyruvate d | 33.50884 | 47.60456 | 53.30227 |
| 5281 | ENSG0000 | P4HA1 | prolyl 4-hy | 13.58651 | 20.66804 | 52.81543 |
| 2938 | ENSG0000 | GYS1 | glycogen sy | 23.77382 | 33.617 | 52.79777 |
| 4178 | ENSG0000 | C5orf15 | chromosor | 33.50441 | 38.20507 | 52.75124 |
| 8932 | ENSG0000 | GSN | gelsolin [So | 23.75339 | 33.04688 | 52.22685 |
| 5599 | ENSG0000 | SLC2A4RG | SLC2A4 reg | 29.34942 | 48.86758 | 52.21809 |
| 8686 | ENSG0000 | ZMAT2 | zinc finger | 43.27178 | 50.77806 | 51.71327 |
| 9342 | ENSG0000 | BAG3 | BCL2 assoc | 45.829 | 48.37945 | 51.61452 |
| 11603 | ENSG0000 | ILK | integrin lin | 36.3422 | 41.72796 | 51.45753 |
| 2527 | ENSG0000 | CHMP4B | charged m | 39.01318 | 42.10203 | 50.71707 |
| 24000 | ENSG0000 | AL590666.2 | | 23.14671 | 29.04963 | 50.70784 |
| 1664 | ENSG0000 | RAB10 | RAB10, me | 45.90297 | 49.99834 | 50.57765 |
| 6330 | ENSG0000 | ATP6V1E1 | ATPase H+ | 35.21887 | 41.86754 | 50.49018 |
| 1715 | ENSG0000 | EIF2AK1 | eukaryotic | 43.12499 | 45.50466 | 49.82204 |
| 16537 | ENSG0000 | C6orf106 | chromosor | 36.49912 | 46.78213 | 49.33472 |
| 2608 | ENSG0000 | PLS3 | plastin 3 [S | 35.02394 | 47.72982 | 48.82714 |
| 3039 | ENSG0000 | FAM32A | family with | 40.33219 | 45.65709 | 48.82305 |
| 11896 | ENSG0000 | CLIC4 | chloride in | 35.82301 | 39.32229 | 48.27048 |
| 12914 | ENSG0000 | ZNF217 | zinc finger | 27.72785 | 39.22019 | 47.6583 |
| 31677 | ENSG0000 | PRKDC | protein kin | 43.85798 | 43.90713 | 47.47029 |
| 41511 | ENSG0000 | MRPL45 | mitochond | 42.36447 | 45.35726 | 47.34613 |
| 7316 | ENSG0000 | TLN1 | talin 1 [Sou | 37.27407 | 44.79058 | 46.66115 |

| | | | | | | |
|---|---|---|---|---|---|---|
| 24223 | ENSG0000 | AL512488.1 | | 44.94532 | 45.34649 | 46.39069 |
| 5622 | ENSG0000 | TRIP10 | thyroid hor | 40.31917 | 41.28042 | 46.28814 |
| 3995 | ENSG0000 | C6orf62 | chromosor | 39.02915 | 46.05593 | 46.18081 |
| 7762 | ENSG0000 | DENR | density reg | 37.72651 | 44.21222 | 46.03075 |
| 2453 | ENSG0000 | TPD52L2 | tumor prot | 30.42595 | 43.67791 | 46.02286 |
| 3566 | ENSG0000 | VAT1 | vesicle ami | 25.05975 | 36.17844 | 45.86054 |
| 6158 | ENSG0000 | NECTIN2 | nectin cell | 33.61197 | 35.47136 | 45.86017 |
| 5743 | ENSG0000 | RRAS | related RA! | 13.64681 | 22.99297 | 45.82997 |
| 11082 | ENSG0000 | ANXA5 | annexin A5 | 21.13911 | 28.89803 | 45.54204 |
| 2332 | ENSG0000 | PYGL | glycogen p | 38.61639 | 39.63872 | 45.37693 |
| 4315 | ENSG0000 | MOB1A | MOB kinas | 32.63578 | 40.29776 | 45.19074 |
| 33778 | ENSG0000 | ARHGAP5-. | ARHGAP5 a | 16.6008 | 37.24681 | 45.14259 |
| 7636 | ENSG0000 | USO1 | USO1 vesic | 30.31758 | 44.91113 | 45.09872 |
| 3037 | ENSG0000 | TNNT1 | troponin T | 13.32587 | 22.1327 | 45.04676 |
| 4474 | ENSG0000 | SUMO1 | small ubiqu | 34.99853 | 39.27492 | 44.88237 |
| 25405 | ENSG0000 | BX322234.2 | | 21.60984 | 26.40221 | 44.84044 |
| 3404 | ENSG0000 | PDLIM1 | PDZ and LI | 36.52958 | 38.09519 | 44.83024 |
| 21522 | ENSG0000 | AC104823.1 | | 24.64955 | 26.26267 | 44.66695 |
| 10044 | ENSG0000 | WASF2 | WAS prote | 35.4081 | 36.28064 | 43.20173 |
| 1279 | ENSG0000 | LMAN1 | lectin, mar | 34.42375 | 39.01014 | 43.15246 |
| 1994 | ENSG0000 | MYL6 | myosin ligh | 32.30507 | 40.11471 | 42.95291 |
| 1607 | ENSG0000 | NFE2L1 | nuclear fac | 28.41654 | 40.96495 | 42.88191 |
| 8907 | ENSG0000 | PLIN2 | perilipin 2 | 16.68288 | 21.18127 | 42.39924 |
| 6652 | ENSG0000 | RTN3 | reticulon 3 | 30.36207 | 31.81269 | 42.29944 |
| 3418 | ENSG0000 | CUBN | cubilin [Sou | 26.86227 | 34.6037 | 42.27682 |
| 12762 | ENSG0000 | JUNB | JunB proto | 15.63337 | 24.46974 | 41.77806 |
| 10254 | ENSG0000 | DFFA | DNA fragm | 36.62039 | 39.80101 | 41.42379 |
| 2022 | ENSG0000 | MSH2 | mutS homo | 33.8946 | 36.84343 | 40.79261 |
| 8868 | ENSG0000 | GINS4 | GINS comp | 22.14231 | 40.46456 | 40.76997 |
| 38550 | ENSG0000 | LIX1L | limb and C | 34.93827 | 37.12412 | 40.65996 |
| 102 | ENSG0000 | RHBDD2 | rhomboid d | 25.35525 | 31.81201 | 40.62686 |
| 1962 | ENSG0000 | PSME1 | proteasom | 28.13528 | 28.55517 | 40.58161 |
| 10190 | ENSG0000 | ADIPOR1 | adiponecti | 29.27911 | 38.13739 | 40.26458 |
| 8071 | ENSG0000 | COL6A2 | collagen ty | 13.19491 | 23.30206 | 40.02957 |
| 4153 | ENSG0000 | GOLPH3 | golgi phosp | 32.51649 | 38.64542 | 39.97036 |
| 2932 | ENSG0000 | BNIP3L | BCL2 intera | 17.10525 | 18.31027 | 39.96564 |
| 12159 | ENSG0000 | COL3A1 | collagen ty | 12.42663 | 28.68115 | 39.72045 |
| 4007 | ENSG0000 | PERP | PERP, TP53 | 27.07294 | 31.78548 | 39.40744 |
| 6229 | ENSG0000 | H19 | H19, imprin | 2.902007 | 6.940318 | 39.31038 |
| 2946 | ENSG0000 | TUBB4A | tubulin bet | 16.64821 | 18.06234 | 39.04951 |
| 3152 | ENSG0000 | DNASE2 | deoxyribor | 19.16022 | 21.23238 | 38.97709 |
| 9829 | ENSG0000 | HK1 | hexokinase | 24.66973 | 30.99786 | 38.80444 |
| 10629 | ENSG0000 | DDR2 | discoidin d | 16.21713 | 28.04282 | 38.4979 |
| 311 | ENSG0000 | MRC2 | mannose r | 15.18069 | 20.59991 | 38.4197 |
| 8443 | ENSG0000 | UBXN4 | UBX doma | 27.86801 | 36.08099 | 38.40522 |
| 5693 | ENSG0000 | DYNLRB1 | dynein ligh | 33.58136 | 37.2876 | 38.33311 |
| 12571 | ENSG0000 | SERPINB9 | serpin fam | 26.96891 | 31.69346 | 38.26389 |

| | | | | | | |
|---|---|---|---|---|---|---|
| 3950 | ENSG0000 | STK38 | serine/thre | 24.62295 | 35.05004 | 37.96503 |
| 10248 | ENSG0000 | GNE | glucosamir | 19.45142 | 33.36006 | 37.58904 |
| 1167 | ENSG0000 | TRIB2 | tribbles ps | 19.52728 | 35.57071 | 37.34858 |
| 36740 | ENSG0000 | NCOA4 | nuclear rec | 33.22132 | 35.84809 | 36.78914 |
| 1874 | ENSG0000 | SPTLC1 | serine paln | 33.71926 | 36.44762 | 36.51275 |
| 1172 | ENSG0000 | FAM50A | family with | 30.02647 | 34.49468 | 36.43948 |
| 18683 | ENSG0000 | HLA-A | major histo | 23.74113 | 24.66309 | 36.36178 |
| 16910 | ENSG0000 | TOR4A | torsin fami | 31.05417 | 33.28732 | 36.33199 |
| 5595 | ENSG0000 | MBOAT7 | membrane | 23.12186 | 28.57863 | 36.29867 |
| 512 | ENSG0000 | BAK1 | BCL2 antag | 22.23133 | 22.91177 | 36.01858 |
| 38933 | ENSG0000 | AL365181.3 | | 18.74537 | 21.54519 | 36.0057 |
| 7746 | ENSG0000 | ESYT1 | extended s | 32.5201 | 33.06935 | 35.82888 |
| 5294 | ENSG0000 | RNF11 | ring finger | 24.02659 | 30.44309 | 35.73394 |
| 3894 | ENSG0000 | CMAS | cytidine m | 22.91452 | 33.61969 | 35.72228 |
| 11654 | ENSG0000 | CCDC68 | coiled-coil | 13.16243 | 24.73141 | 35.71497 |
| 3602 | ENSG0000 | CDR2L | cerebellar | 22.19259 | 34.6798 | 35.68162 |
| 13925 | ENSG0000 | MAGEF1 | MAGE fam | 31.99428 | 33.15176 | 35.57547 |
| 16613 | ENSG0000 | DYNC1H1 | dynein cyto | 27.62702 | 34.05045 | 35.53842 |
| 160 | ENSG0000 | ADIPOR2 | adiponecti | 29.83254 | 34.75171 | 35.53544 |
| 16260 | ENSG0000 | TSPYL1 | TSPY like 1 | 26.99231 | 32.95353 | 35.20106 |
| 9130 | ENSG0000 | PLCB3 | phospholip | 27.14795 | 29.79841 | 35.13897 |
| 610 | ENSG0000 | CAPG | capping ac | 20.73353 | 21.54235 | 35.04804 |
| 8235 | ENSG0000 | MPC2 | mitochond | 26.21935 | 30.48559 | 34.97306 |
| 17118 | ENSG0000 | ZNF358 | zinc finger | 26.97722 | 28.16615 | 34.90849 |
| 14523 | ENSG0000 | TCEAL8 | transcriptio | 20.36199 | 25.69577 | 34.90338 |
| 11605 | ENSG0000 | TPP1 | tripeptidyl | 23.16049 | 28.83997 | 34.8729 |
| 3324 | ENSG0000 | RHEB | Ras homol | 29.59803 | 33.02318 | 34.84356 |
| 3528 | ENSG0000 | CCDC47 | coiled-coil | 29.19694 | 32.78182 | 34.69774 |
| 1709 | ENSG0000 | CHMP5 | charged m | 23.03217 | 28.47046 | 34.61453 |
| 7808 | ENSG0000 | WARS | tryptophar | 20.27955 | 25.89851 | 34.54527 |
| 10961 | ENSG0000 | TOPBP1 | topoisome | 32.56909 | 33.78798 | 34.1429 |
| 1909 | ENSG0000 | PDPR | pyruvate d | 21.26977 | 28.29108 | 34.00272 |
| 8382 | ENSG0000 | EPHX1 | epoxide hy | 20.37672 | 25.12659 | 33.83008 |
| 12457 | ENSG0000 | NLGN2 | neuroligin | 21.61534 | 26.61863 | 33.73833 |
| 2636 | ENSG0000 | USP11 | ubiquitin s | 25.35273 | 27.30112 | 33.65705 |
| 2030 | ENSG0000 | PSMD5 | proteasom | 30.84677 | 32.52972 | 33.62666 |
| 4470 | ENSG0000 | PCYOX1 | prenylcyste | 20.2776 | 25.65526 | 33.57013 |
| 16415 | ENSG0000 | EPHB4 | EPH recept | 26.40851 | 31.73862 | 33.26129 |
| 10976 | ENSG0000 | CDCP1 | CUB doma | 18.53251 | 30.92382 | 33.23827 |
| 6953 | ENSG0000 | EHF | ETS homol | 14.72135 | 30.10765 | 33.17316 |
| 10446 | ENSG0000 | LSM12 | LSM12 hor | 30.475 | 33.02111 | 33.11592 |
| 4245 | ENSG0000 | TFG | TRK-fused | 25.79261 | 30.91108 | 32.88447 |
| 15082 | ENSG0000 | NOG | noggin [So | 21.49365 | 24.16944 | 32.85813 |
| 4879 | ENSG0000 | PCMT1 | protein-L-i | 29.17144 | 31.00219 | 32.7303 |
| 5957 | ENSG0000 | MRPS12 | mitochond | 24.03356 | 27.53537 | 32.7115 |
| 18020 | ENSG0000 | GDI1 | GDP dissoc | 25.0195 | 28.74562 | 32.68139 |
| 8208 | ENSG0000 | PTPRF | protein tyr | 23.74404 | 24.16628 | 32.67759 |

| | | | | | | |
|---|---|---|---|---|---|---|
| 6424 | ENSG0000 | MCCC2 | methylcrot | 31.43335 | 31.94107 | 32.63743 |
| 7913 | ENSG0000 | MAP1LC3B | microtubul | 12.57655 | 28.01685 | 32.54501 |
| 817 | ENSG0000 | SLC2A3 | solute carr | 6.29787 | 16.08274 | 32.37511 |
| 56 | ENSG0000 | RECQL | RecQ like h | 28.28477 | 31.83303 | 32.16067 |
| 9316 | ENSG0000 | CENPU | centromer | 22.85672 | 28.09022 | 32.1286 |
| 1943 | ENSG0000 | CDV3 | CDV3 hom | 25.95095 | 31.15271 | 32.0789 |
| 28752 | ENSG0000 | GNG10 | G protein s | 30.43941 | 30.791 | 31.94702 |
| 16847 | ENSG0000 | ERO1A | endoplasm | 17.35148 | 18.49074 | 31.8213 |
| 26998 | ENSG0000 | LINC01010 | long interg | 8.631243 | 21.28637 | 31.6791 |
| 6418 | ENSG0000 | PIAS3 | protein inh | 23.72578 | 29.61898 | 31.64019 |
| 11417 | ENSG0000 | C10orf10 | chromosor | 10.16264 | 14.0327 | 31.6078 |
| 8018 | ENSG0000 | CBX4 | chromobo: | 22.41276 | 26.0306 | 31.41002 |
| 6778 | ENSG0000 | ARF3 | ADP ribosy | 24.04956 | 27.49211 | 31.15449 |
| 6095 | ENSG0000 | SNX6 | sorting nex | 26.05187 | 27.03721 | 31.11615 |
| 10416 | ENSG0000 | DVL3 | dishevellec | 23.82179 | 30.63406 | 30.98227 |
| 10550 | ENSG0000 | AK4 | adenylate l | 10.39122 | 22.58551 | 30.91806 |
| 5076 | ENSG0000 | TNFRSF10E | TNF recept | 19.78672 | 25.75295 | 30.91082 |
| 15243 | ENSG0000 | COPB2 | coatomer p | 27.24393 | 30.56978 | 30.80922 |
| 13699 | ENSG0000 | CDC26 | cell divisior | 22.51872 | 27.38404 | 30.78752 |
| 14435 | ENSG0000 | MCFD2 | multiple cc | 25.00447 | 28.02469 | 30.74415 |
| 8844 | ENSG0000 | CETN2 | centrin 2 [S | 24.09403 | 30.02509 | 30.74062 |
| 4675 | ENSG0000 | DARS2 | aspartyl-tR | 28.06661 | 28.14618 | 30.70737 |
| 11742 | ENSG0000 | YWHAB | tyrosine 3- | 12.57871 | 29.72143 | 30.69627 |
| 4831 | ENSG0000 | GLE1 | GLE1, RNA | 28.06022 | 28.78997 | 30.656 |
| 1690 | ENSG0000 | AKR1B1 | aldo-keto r | 4.104633 | 20.10473 | 30.45536 |
| 6661 | ENSG0000 | GSTT2B | glutathione | 18.03046 | 21.15525 | 29.95159 |
| 9389 | ENSG0000 | UHMK1 | U2AF hom | 19.19591 | 28.78721 | 29.94762 |
| 4649 | ENSG0000 | SLC2A1 | solute carr | 17.06167 | 19.06907 | 29.90533 |
| 17952 | ENSG0000 | PBX2 | PBX home | 22.00083 | 28.95242 | 29.83238 |
| 17075 | ENSG0000 | LDB1 | LIM domai | 19.8946 | 26.94921 | 29.7943 |
| 1945 | ENSG0000 | ALKBH5 | alkB homo | 19.67267 | 21.2362 | 29.74485 |
| 9832 | ENSG0000 | CD109 | CD109 mol | 19.38767 | 21.11589 | 29.67376 |
| 18679 | ENSG0000 | RAB12 | RAB12, me | 23.5942 | 29.38163 | 29.66832 |
| 14593 | ENSG0000 | RAP2B | RAP2B, me | 26.23102 | 28.85118 | 29.66452 |
| 5284 | ENSG0000 | VPS26A | VPS26, retr | 24.69707 | 26.13416 | 29.66379 |
| 4264 | ENSG0000 | SNX4 | sorting nex | 22.75502 | 28.46295 | 29.61113 |
| 2071 | ENSG0000 | ITPR3 | inositol 1,4 | 22.61565 | 23.8303 | 29.56388 |
| 11124 | ENSG0000 | F2RL1 | F2R like try | 10.34988 | 17.65476 | 29.41769 |
| 11797 | ENSG0000 | PRRX2 | paired rela | 14.98901 | 17.53483 | 29.38553 |
| 2988 | ENSG0000 | BUD31 | BUD31 hor | 24.29367 | 28.62012 | 29.32129 |
| 3828 | ENSG0000 | CDKN1B | cyclin depe | 19.16276 | 23.93257 | 29.26294 |
| 5976 | ENSG0000 | TWSG1 | twisted gas | 23.63551 | 25.22261 | 29.21835 |
| 5875 | ENSG0000 | PTPN12 | protein tyr | 25.74266 | 28.68166 | 29.08771 |
| 7261 | ENSG0000 | ST6GALNA | ST6 N-acet | 20.72734 | 21.47178 | 28.94906 |
| 8866 | ENSG0000 | GOLGA7 | golgin A7 [ | 25.49986 | 28.76578 | 28.9206 |
| 6574 | ENSG0000 | OSER1 | oxidative s | 19.60852 | 27.31535 | 28.87253 |
| 2433 | ENSG0000 | ZMYND8 | zinc finger | 25.2949 | 28.36429 | 28.79299 |

| | | | | | | |
|---|---|---|---|---|---|---|
| 624 | ENSG0000 | CTNNA1 | catenin alp | 28.2271 | 28.24818 | 28.75201 |
| 1184 | ENSG0000 | ACTN1 | actinin alpl | 28.58936 | 28.65521 | 28.71173 |
| 5401 | ENSG0000 | FAM210B | family with | 17.1735 | 23.90986 | 28.68728 |
| 469 | ENSG0000 | DEF6 | DEF6, guar | 19.50343 | 24.67813 | 28.67305 |
| 10052 | ENSG0000 | CUL4B | cullin 4B [S | 14.0122 | 21.06906 | 28.59389 |
| 11279 | ENSG0000 | SLC4A2 | solute carr | 21.44751 | 22.93922 | 28.5921 |
| 7970 | ENSG0000 | SLC39A6 | solute carr | 21.59638 | 27.65591 | 28.44046 |
| 112 | ENSG0000 | LAMP2 | lysosomal | 21.10513 | 24.91706 | 28.33329 |
| 2122 | ENSG0000 | CDC34 | cell divisio | 24.54751 | 25.37103 | 28.2937 |
| 17197 | ENSG0000 | PJA2 | praja ring f | 20.56256 | 26.52246 | 28.09095 |
| 5710 | ENSG0000 | UROD | uroporphy | 18.33225 | 22.06997 | 28.06364 |
| 10400 | ENSG0000 | MGAT4B | mannosyl ( | 23.52511 | 27.42228 | 27.97625 |
| 16470 | ENSG0000 | PLXNB2 | plexin B2 [ | 20.57903 | 23.07285 | 27.82423 |
| 5153 | ENSG0000 | ZCCHC17 | zinc finger | 21.37875 | 25.53257 | 27.71018 |
| 7814 | ENSG0000 | MFAP1 | microfibril | 21.03008 | 23.6096 | 27.70431 |
| 3382 | ENSG0000 | CREB3 | cAMP resp | 18.66629 | 24.7787 | 27.65516 |
| 14444 | ENSG0000 | GAS1 | growth arr | 19.58873 | 23.63457 | 27.64719 |
| 15719 | ENSG0000 | UBE2H | ubiquitin c | 14.53247 | 21.1997 | 27.55696 |
| 820 | ENSG0000 | YBX3 | Y-box bind | 25.09914 | 26.8358 | 27.44139 |
| 17000 | ENSG0000 | YTHDF2 | YTH N6-me | 6.140164 | 26.5748 | 27.41211 |
| 11747 | ENSG0000 | TSC22D4 | TSC22 dom | 13.50093 | 26.51947 | 27.27576 |
| 1041 | ENSG0000 | COASY | Coenzyme | 23.83709 | 26.75978 | 27.17586 |
| 2808 | ENSG0000 | BCKDK | branched c | 13.67928 | 18.41173 | 27.14976 |
| 3599 | ENSG0000 | TNFAIP1 | TNF alpha | 20.39228 | 26.78434 | 27.09992 |
| 12476 | ENSG0000 | UBE2E1 | ubiquitin c | 19.64956 | 21.77345 | 26.94851 |
| 11699 | ENSG0000 | C16orf45 | chromosor | 15.91685 | 26.36213 | 26.85759 |
| 4018 | ENSG0000 | FAF2 | Fas associa | 25.7701 | 26.04526 | 26.8373 |
| 2242 | ENSG0000 | LMF2 | lipase matu | 15.40077 | 18.96161 | 26.77995 |
| 11843 | ENSG0000 | VKORC1 | vitamin K e | 10.33078 | 16.12287 | 26.77875 |
| 24881 | ENSG0000 | TAPBP | TAP bindin | 22.73492 | 24.11468 | 26.71388 |
| 865 | ENSG0000 | CA11 | carbonic ai | 12.32266 | 14.94244 | 26.66865 |
| 6739 | ENSG0000 | BHLHE40 | basic helix- | 12.64197 | 17.64075 | 26.66572 |
| 1411 | ENSG0000 | TULP3 | tubby like | 22.36706 | 22.6091 | 26.65974 |
| 10420 | ENSG0000 | FBXO27 | F-box prot | 10.02942 | 15.22271 | 26.61775 |
| 3825 | ENSG0000 | CREBL2 | cAMP resp | 11.99117 | 20.85127 | 26.47572 |
| 9002 | ENSG0000 | EIF4EBP2 | eukaryotic | 21.9278 | 25.68931 | 26.45461 |
| 4545 | ENSG0000 | IVNS1ABP | influenza v | 11.60348 | 22.5234 | 26.30914 |
| 309 | ENSG0000 | LYPLA2 | lysophosph | 22.784 | 24.99986 | 26.21556 |
| 2106 | ENSG0000 | HSD3B7 | hydroxy-de | 18.11493 | 19.40374 | 26.13481 |
| 13969 | ENSG0000 | GBA | glucosylcei | 19.1073 | 20.294 | 26.1158 |
| 5140 | ENSG0000 | DESI2 | desumoyla | 18.56869 | 23.98653 | 26.06908 |
| 3410 | ENSG0000 | PHYH | phytanoyl- | 9.253982 | 17.44138 | 26.01571 |
| 5910 | ENSG0000 | CDC42EP1 | CDC42 effe | 21.19411 | 22.93447 | 26.01384 |
| 7119 | ENSG0000 | SCRN1 | secernin 1 | 17.21698 | 18.7003 | 26.0005 |
| 11448 | ENSG0000 | QSOX2 | quiescin su | 24.46975 | 25.17425 | 25.94845 |
| 4960 | ENSG0000 | IFIT2 | interferon | 8.421953 | 15.57463 | 25.84469 |
| 8413 | ENSG0000 | CIAO1 | cytosolic ir | 23.05114 | 24.6093 | 25.73535 |

| | | | | | | |
|---|---|---|---|---|---|---|
| 14349 | ENSG0000 | SEPHS2 | selenopho: | 18.22693 | 21.54509 | 25.65903 |
| 3459 | ENSG0000 | STN1 | STN1, CST | 18.81941 | 19.5889 | 25.64003 |
| 13713 | ENSG0000 | DIRAS1 | DIRAS fami | 11.8256 | 14.63276 | 25.56668 |
| 13344 | ENSG0000 | ZBTB4 | zinc finger | 18.83557 | 22.17423 | 25.45291 |
| 20567 | ENSG0000 | TUBB2BP1 | tubulin bet | 17.44736 | 20.23799 | 25.32693 |
| 6993 | ENSG0000 | CD164 | CD164 mol | 19.89733 | 20.48537 | 25.26716 |
| 10678 | ENSG0000 | WDR26 | WD repeat | 17.81827 | 23.74761 | 25.21583 |
| 5055 | ENSG0000 | FAM53C | family with | 15.36753 | 23.82123 | 25.12237 |
| 13569 | ENSG0000 | RAB6A | RAB6A, me | 15.71073 | 21.61021 | 25.11931 |
| 4823 | ENSG0000 | PTBP3 | polypyrimi | 15.87616 | 23.85226 | 25.11169 |
| 12261 | ENSG0000 | KLK7 | kallikrein r | 11.86817 | 15.40992 | 25.09814 |
| 29207 | ENSG0000 | DBNDD2 | dysbindin | 19.20364 | 20.57673 | 25.05585 |
| 10786 | ENSG0000 | PBXIP1 | PBX home | 7.214095 | 15.10541 | 24.76723 |
| 3912 | ENSG0000 | TDP2 | tyrosyl-DN | 16.92995 | 24.7087 | 24.73554 |
| 897 | ENSG0000 | CTSA | cathepsin | 21.71392 | 23.68092 | 24.70169 |
| 4538 | ENSG0000 | FBXO2 | F-box prot | 5.808291 | 13.94977 | 24.67385 |
| 13971 | ENSG0000 | ACAD9 | acyl-CoA d | 22.41478 | 24.00358 | 24.66869 |
| 1231 | ENSG0000 | TP63 | tumor prot | 10.2497 | 16.56572 | 24.64444 |
| 7024 | ENSG0000 | DYNC1LI2 | dynein cyt | 19.85149 | 23.95751 | 24.59033 |
| 2153 | ENSG0000 | BCL2L13 | BCL2 like 1 | 18.29177 | 20.48099 | 24.57334 |
| 8720 | ENSG0000 | PNRC1 | proline rich | 7.234459 | 17.38811 | 24.44822 |
| 2929 | ENSG0000 | KCTD9 | potassium | 16.40789 | 22.77555 | 24.34554 |
| 13295 | ENSG0000 | HOXB2 | homeobox | 15.99807 | 17.2976 | 24.32289 |
| 33553 | ENSG0000 | RPL10P13 | ribosomal | 8.292119 | 17.36114 | 24.07983 |
| 11394 | ENSG0000 | CFL2 | cofilin 2 [S | 18.63095 | 19.41868 | 24.07596 |
| 6706 | ENSG0000 | SARAF | store-oper | 18.71768 | 20.36535 | 24.04371 |
| 4868 | ENSG0000 | NPC2 | NPC intrac | 16.88821 | 17.27721 | 23.94262 |
| 3115 | ENSG0000 | PTPRS | protein tyr | 12.3331 | 20.10663 | 23.9336 |
| 15322 | ENSG0000 | ARMCX2 | armadillo r | 19.9732 | 23.2853 | 23.90773 |
| 7991 | ENSG0000 | MINK1 | misshapen | 18.49402 | 23.43108 | 23.89618 |
| 986 | ENSG0000 | LRRC40 | leucine rich | 19.59165 | 20.33752 | 23.85803 |
| 5002 | ENSG0000 | CYSTM1 | cysteine ric | 15.75973 | 17.67773 | 23.78219 |
| 4989 | ENSG0000 | DUSP1 | dual specif | 9.688015 | 19.183 | 23.7769 |
| 7474 | ENSG0000 | KYAT3 | kynurenine | 14.65758 | 20.24786 | 23.75082 |
| 387 | ENSG0000 | COX15 | COX15, cyt | 16.8939 | 18.06277 | 23.73933 |
| 15131 | ENSG0000 | TOB2 | transducer | 17.24469 | 23.31451 | 23.73769 |
| 9412 | ENSG0000 | USP12 | ubiquitin s | 13.03799 | 21.47243 | 23.71751 |
| 3412 | ENSG0000 | DNMBP | dynamin b | 16.6319 | 17.98323 | 23.67365 |
| 3399 | ENSG0000 | UBE2R2 | ubiquitin c | 20.6223 | 21.36202 | 23.66101 |
| 10891 | ENSG0000 | CMBL | carboxyme | 11.87642 | 15.17404 | 23.63361 |
| 14562 | ENSG0000 | HIST3H2A | histone clu | 13.49132 | 22.36303 | 23.63265 |
| 11220 | ENSG0000 | KCNK5 | potassium | 8.328003 | 20.14399 | 23.46569 |
| 3135 | ENSG0000 | TMEM205 | transmeml | 16.87162 | 18.97347 | 23.44029 |
| 4270 | ENSG0000 | ATP6V1A | ATPase H+ | 13.99382 | 19.03908 | 23.34708 |
| 2961 | ENSG0000 | ERCC2 | ERCC excis | 19.40195 | 20.23266 | 23.34306 |
| 2357 | ENSG0000 | LGMN | legumain [ | 10.47014 | 21.58501 | 23.32632 |
| 11879 | ENSG0000 | COPS6 | COP9 signa | 22.24885 | 22.3258 | 23.25775 |

| | | | | | | |
|---|---|---|---|---|---|---|
| 1197 | ENSG0000 | AFF4 | AF4/FMR2 | 15.51892 | 21.77144 | 23.21296 |
| 6096 | ENSG0000 | EAPP | E2F associa | 15.8767 | 18.69433 | 23.20915 |
| 8294 | ENSG0000 | RFX5 | regulatory | 11.29425 | 20.45124 | 23.19791 |
| 5232 | ENSG0000 | SEPT7 | septin 7 [S | 21.05191 | 21.97382 | 23.18734 |
| 12431 | ENSG0000 | SYAP1 | synapse as | 19.56242 | 21.82383 | 23.16728 |
| 117 | ENSG0000 | CRLF1 | cytokine re | 8.211861 | 9.168638 | 22.97972 |
| 2402 | ENSG0000 | SRP54 | signal reco | 17.82579 | 19.15874 | 22.97904 |
| 11313 | ENSG0000 | UBAP1 | ubiquitin a | 17.84033 | 19.90741 | 22.92897 |
| 7467 | ENSG0000 | RSL24D1 | ribosomal | 18.67981 | 21.76199 | 22.90069 |
| 4152 | ENSG0000 | ARRDC3 | arrestin do | 3.243713 | 21.30071 | 22.89665 |
| 518 | ENSG0000 | CENPQ | centromer | 14.15973 | 18.08593 | 22.85675 |
| 465 | ENSG0000 | SNAPC1 | small nucle | 11.55648 | 19.13251 | 22.8397 |
| 18146 | ENSG0000 | NELFE | negative el | 19.55972 | 19.77658 | 22.83698 |
| 6005 | ENSG0000 | MTSS1L | MTSS1L, I-I | 15.04499 | 20.64519 | 22.81293 |
| 6686 | ENSG0000 | KRAS | KRAS proto | 12.77436 | 17.24762 | 22.80904 |
| 8810 | ENSG0000 | ZMYM3 | zinc finger | 15.88691 | 18.77308 | 22.76919 |
| 5846 | ENSG0000 | KLF2 | Kruppel lik | 15.39213 | 18.02654 | 22.74007 |
| 16838 | ENSG0000 | ADH5 | alcohol del | 12.4461 | 18.53207 | 22.67758 |
| 7257 | ENSG0000 | TOR1A | torsin fami | 20.67916 | 21.95024 | 22.54623 |
| 926 | ENSG0000 | PKN2 | protein kin | 18.17091 | 21.19227 | 22.52551 |
| 5296 | ENSG0000 | BHLHE41 | basic helix- | 8.05863 | 13.03372 | 22.46029 |
| 10217 | ENSG0000 | TMEM69 | transmeml | 16.06367 | 20.21808 | 22.42391 |
| 16608 | ENSG0000 | IGF2R | insulin like | 12.94992 | 20.33543 | 22.41232 |
| 14671 | ENSG0000 | PNMA8A | paraneopl | 15.50001 | 20.07788 | 22.39112 |
| 9311 | ENSG0000 | ADAM17 | ADAM met | 10.83719 | 17.30215 | 22.2182 |
| 10571 | ENSG0000 | KIAA1522 | KIAA1522 \| | 10.25449 | 11.57858 | 22.20844 |
| 13450 | ENSG0000 | SEZ6L2 | seizure rel | 12.0432 | 14.70551 | 22.15713 |
| 3420 | ENSG0000 | DDX50 | DExD-box I | 18.44218 | 20.88728 | 22.14648 |
| 9929 | ENSG0000 | IER2 | immediate | 11.16623 | 21.0555 | 22.09462 |
| 5159 | ENSG0000 | POLR3GL | RNA polym | 14.02589 | 14.50732 | 22.0363 |
| 6284 | ENSG0000 | SLC6A8 | solute carr | 15.13025 | 20.24754 | 21.98132 |
| 7366 | ENSG0000 | IER3 | immediate | 5.030133 | 11.59847 | 21.90007 |
| 12029 | ENSG0000 | KCTD5 | potassium | 17.84496 | 19.00865 | 21.89974 |
| 9737 | ENSG0000 | RNF20 | ring finger | 17.7391 | 21.27326 | 21.85109 |
| 9693 | ENSG0000 | HSPA13 | heat shock | 16.82231 | 19.68767 | 21.67528 |
| 15528 | ENSG0000 | IFIT1 | interferon | 6.748545 | 9.046599 | 21.67409 |
| 15080 | ENSG0000 | RFLNB | refilin B [S | 15.16511 | 19.78874 | 21.63321 |
| 3701 | ENSG0000 | OSBP | oxysterol b | 19.88087 | 21.56703 | 21.60552 |
| 5737 | ENSG0000 | NR1D1 | nuclear re | 10.66005 | 21.5229 | 21.54138 |
| 31445 | ENSG0000 | CCDC71L | coiled-coil | 13.30816 | 20.91096 | 21.48642 |
| 5360 | ENSG0000 | RAB9A | RAB9A, me | 13.20918 | 17.01956 | 21.47646 |
| 30 | ENSG0000 | LASP1 | LIM and SH | 16.79755 | 20.01264 | 21.45096 |
| 8549 | ENSG0000 | NCEH1 | neutral ch | 17.22203 | 20.79139 | 21.41653 |
| 11731 | ENSG0000 | NEMP1 | nuclear en | 12.31364 | 15.47773 | 21.38837 |
| 11982 | ENSG0000 | ANGPTL4 | angiopoiet | 4.834411 | 5.979155 | 21.38548 |
| 8200 | ENSG0000 | BCL10 | B-cell CLL/I | 11.6551 | 19.30188 | 21.34984 |
| 15191 | ENSG0000 | TSPYL2 | TSPY like 2 | 9.743012 | 21.22848 | 21.33155 |

| | | | | | | |
|---|---|---|---|---|---|---|
| 7630 | ENSG0000 | G3BP2 | G3BP stres | 19.0506 | 19.08966 | 21.27488 |
| 759 | ENSG0000 | GINM1 | glycoprote | 18.16811 | 18.81 | 21.26723 |
| 12389 | ENSG0000 | CENPX | centromer | 14.82247 | 16.40023 | 21.26279 |
| 14240 | ENSG0000 | ZNF664 | zinc finger | 10.16656 | 19.97216 | 21.08688 |
| 3370 | ENSG0000 | TBC1D13 | TBC1 doma | 16.98382 | 20.49949 | 21.06303 |
| 9835 | ENSG0000 | UBE2L6 | ubiquitin c | 5.311117 | 7.95341 | 21.05989 |
| 18 | ENSG0000 | CYP51A1 | cytochrom | 15.06393 | 18.88691 | 21.01267 |
| 3517 | ENSG0000 | RNF167 | ring finger | 15.94438 | 19.25399 | 21.00897 |
| 37 | ENSG0000 | DBNDD1 | dysbindin ( | 12.59787 | 15.77391 | 21.0005 |
| 8133 | ENSG0000 | RASSF3 | Ras associa | 15.05422 | 16.3413 | 20.99377 |
| 14726 | ENSG0000 | B4GALNT4 | beta-1,4-N | 15.07395 | 16.22481 | 20.98822 |
| 49 | ENSG0000 | PLXND1 | plexin D1 [ | 15.97047 | 16.35054 | 20.94707 |
| 14763 | ENSG0000 | EXOC7 | exocyst co | 11.36586 | 14.53565 | 20.94296 |
| 3066 | ENSG0000 | PLD3 | phospholip | 14.64834 | 14.92883 | 20.93529 |
| 3986 | ENSG0000 | WASF1 | WAS prote | 12.32073 | 20.67266 | 20.93271 |
| 6887 | ENSG0000 | TMED7 | transmeml | 16.00282 | 17.15994 | 20.8574 |
| 6830 | ENSG0000 | STK26 | serine/thre | 9.361362 | 13.46224 | 20.78383 |
| 6200 | ENSG0000 | CACNG6 | calcium vo | 6.654066 | 9.616201 | 20.77304 |
| 2800 | ENSG0000 | RBL2 | RB transcri | 13.40798 | 18.59268 | 20.75513 |
| 17149 | ENSG0000 | DCAF12 | DDB1 and ( | 14.72383 | 17.0062 | 20.74151 |
| 1165 | ENSG0000 | ATP6AP1 | ATPase H+ | 17.20293 | 19.22254 | 20.68459 |
| 6747 | ENSG0000 | EMC7 | ER membra | 20.00881 | 20.43608 | 20.68116 |
| 5874 | ENSG0000 | HIP1 | huntingtin | 13.22811 | 13.66723 | 20.66458 |
| 3469 | ENSG0000 | TFAM | transcriptio | 16.44427 | 17.67117 | 20.62669 |
| 10038 | ENSG0000 | TPRG1L | tumor prot | 12.31985 | 19.86821 | 20.551 |
| 749 | ENSG0000 | FAM168A | family with | 11.3032 | 13.23953 | 20.5423 |
| 11885 | ENSG0000 | ARF4 | ADP ribosy | 15.78409 | 19.14673 | 20.51149 |
| 10124 | ENSG0000 | PINK1 | PTEN induc | 10.19633 | 19.69415 | 20.44771 |
| 11125 | ENSG0000 | AGGF1 | angiogenic | 14.54651 | 19.26942 | 20.38351 |
| 7237 | ENSG0000 | BIN1 | bridging in | 16.42307 | 17.8585 | 20.29256 |
| 17145 | ENSG0000 | OSTC | oligosacch | 14.96456 | 18.25276 | 20.2887 |
| 5507 | ENSG0000 | MAD2L1BP | MAD2L1 bi | 16.49829 | 16.84584 | 20.26148 |
| 6580 | ENSG0000 | AP3B1 | adaptor re | 12.03669 | 16.2922 | 20.23782 |
| 11216 | ENSG0000 | CAMLG | calcium mo | 12.54232 | 13.64965 | 20.23166 |
| 6186 | ENSG0000 | SNX9 | sorting nex | 19.19472 | 19.88264 | 20.19802 |
| 7783 | ENSG0000 | TMX1 | thioredoxi | 16.6797 | 19.17276 | 20.17678 |
| 19712 | ENSG0000 | SUPT4H1 | SPT4 homo | 17.76153 | 20.01766 | 20.1679 |
| 7016 | ENSG0000 | GNS | glucosamir | 13.8792 | 14.89216 | 20.11595 |
| 10333 | ENSG0000 | ZER1 | zyg-11 rela | 13.58878 | 17.23103 | 20.10446 |
| 5173 | ENSG0000 | LRIF1 | ligand depe | 9.605488 | 18.71554 | 20.04595 |
| 13703 | ENSG0000 | EID2 | EP300 inte | 19.78612 | 19.81777 | 20.015 |
| 7255 | ENSG0000 | SMC2 | structural r | 14.89653 | 16.07652 | 19.9952 |
| 5922 | ENSG0000 | LIF | LIF, interle | 10.53113 | 16.60292 | 19.98592 |
| 9807 | ENSG0000 | SFR1 | SWI5 depe | 11.38226 | 12.45797 | 19.97953 |
| 11265 | ENSG0000 | SUN1 | Sad1 and U | 13.5849 | 18.10426 | 19.9771 |
| 7909 | ENSG0000 | CMTM3 | CKLF like M | 15.22181 | 18.62378 | 19.9623 |
| 14110 | ENSG0000 | H3F3AP6 | H3 histone | 14.96522 | 17.48957 | 19.93849 |

| | | | | | | |
|---|---|---|---|---|---|---|
| 14964 | ENSG0000 | RIPK4 | receptor ir | 7.511603 | 15.95736 | 19.92966 |
| 10041 | ENSG0000 | XDH | xanthine d | 13.11852 | 19.27861 | 19.82805 |
| 2232 | ENSG0000 | POLDIP3 | DNA polym | 12.45552 | 13.22394 | 19.81268 |
| 3497 | ENSG0000 | DHX40 | DEAH-box | 9.355987 | 14.13135 | 19.80146 |
| 6255 | ENSG0000 | CHMP2A | charged m | 17.87576 | 18.44475 | 19.74304 |
| 16482 | ENSG0000 | MMP1 | matrix met | 6.416416 | 13.88578 | 19.73765 |
| 8957 | ENSG0000 | PTGES | prostaglan | 11.13458 | 11.60459 | 19.62168 |
| 330 | ENSG0000 | PLAUR | plasminoge | 12.49585 | 13.08128 | 19.56768 |
| 4117 | ENSG0000 | MRPS30 | mitochond | 14.7576 | 19.48415 | 19.5441 |
| 3440 | ENSG0000 | TNKS2 | tankyrase 2 | 11.06473 | 18.54088 | 19.53099 |
| 5346 | ENSG0000 | HJURP | Holliday ju | 9.058485 | 17.7566 | 19.47124 |
| 6919 | ENSG0000 | TMEM60 | transmeml | 14.34589 | 14.76968 | 19.455 |
| 8394 | ENSG0000 | RHOB | ras homolc | 9.311349 | 15.90631 | 19.45485 |
| 261 | ENSG0000 | PHF23 | PHD finger | 16.24842 | 16.37915 | 19.43275 |
| 8967 | ENSG0000 | NOTCH1 | notch 1 [Sc | 10.3001 | 13.52171 | 19.40745 |
| 9939 | ENSG0000 | PYGO2 | pygopus fa | 17.25065 | 17.59129 | 19.38209 |
| 476 | ENSG0000 | RRAGD | Ras relatec | 4.65592 | 11.05257 | 19.35938 |
| 5001 | ENSG0000 | NUP43 | nucleopori | 16.36893 | 16.48355 | 19.3342 |
| 542 | ENSG0000 | PEX3 | peroxisom | 11.32712 | 13.12663 | 19.32177 |
| 15091 | ENSG0000 | TMEM50A | transmeml | 16.60634 | 18.26514 | 19.27131 |
| 15776 | ENSG0000 | C1QL4 | compleme | 2.688606 | 3.902225 | 19.265 |
| 3137 | ENSG0000 | PLPPR2 | phospholip | 14.19025 | 18.91302 | 19.21162 |
| 10268 | ENSG0000 | CCDC58 | coiled-coil | 15.97786 | 18.98005 | 19.13663 |
| 13632 | ENSG0000 | UNC119B | unc-119 lip | 16.17131 | 17.33128 | 19.05394 |
| 14517 | ENSG0000 | FAM83H | family with | 13.74698 | 15.45176 | 19.05072 |
| 14830 | ENSG0000 | ACBD3 | acyl-CoA b | 14.04692 | 17.56666 | 19.04479 |
| 12374 | ENSG0000 | FAM103A1 | family with | 11.93249 | 14.98152 | 19.04307 |
| 9908 | ENSG0000 | DHRS4 | dehydroge | 14.98076 | 16.38543 | 19.03093 |
| 5950 | ENSG0000 | SMO | smoothene | 11.14989 | 12.38104 | 18.99784 |
| 10364 | ENSG0000 | FAM189B | family with | 12.27713 | 15.53239 | 18.98411 |
| 11646 | ENSG0000 | LEO1 | LEO1 homo | 18.25178 | 18.79376 | 18.96638 |
| 40559 | ENSG0000 | PIP4K2B | phosphatic | 17.79217 | 18.05353 | 18.92438 |
| 4412 | ENSG0000 | IL1R1 | interleukin | 10.52686 | 18.51702 | 18.91892 |
| 4958 | ENSG0000 | IFIT3 | interferon | 3.828448 | 5.911501 | 18.87843 |
| 2650 | ENSG0000 | MAGED2 | MAGE fam | 7.005054 | 14.3174 | 18.86944 |
| 12098 | ENSG0000 | COA6 | cytochrom | 15.9011 | 18.0662 | 18.83283 |
| 21499 | ENSG0000 | CPTP | ceramide-1 | 12.83263 | 15.80525 | 18.79974 |
| 11880 | ENSG0000 | PAFAH1B2 | platelet ac | 16.97685 | 18.72386 | 18.78655 |
| 11492 | ENSG0000 | TRUB1 | TruB pseuc | 13.75565 | 17.09795 | 18.77129 |
| 12713 | ENSG0000 | CTDSP2 | CTD small p | 13.99257 | 15.50533 | 18.75714 |
| 18200 | ENSG0000 | NFKBIL1 | NFKB inhib | 13.29076 | 16.60725 | 18.71899 |
| 11918 | ENSG0000 | SERTAD3 | SERTA dom | 11.32998 | 14.05688 | 18.69148 |
| 10998 | ENSG0000 | POLR2H | RNA polym | 10.65837 | 16.18768 | 18.66475 |
| 564 | ENSG0000 | TUBG2 | tubulin gar | 10.30096 | 15.18783 | 18.65781 |
| 35347 | ENSG0000 | AC092368.3 | | 10.95681 | 18.46115 | 18.64891 |
| 11162 | ENSG0000 | GFM2 | G elongatio | 14.13219 | 14.97025 | 18.57757 |
| 3816 | ENSG0000 | VPS29 | VPS29, retr | 16.14458 | 17.33009 | 18.56756 |

| | | | | | | |
|---|---|---|---|---|---|---|
| 5997 | ENSG0000 | RLIM | ring finger | 15.79447 | 17.09027 | 18.55688 |
| 11977 | ENSG0000 | KRT80 | keratin 80 | 11.39662 | 17.34574 | 18.50836 |
| 3029 | ENSG0000 | MED25 | mediator c | 12.93006 | 15.07531 | 18.46639 |
| 30995 | ENSG0000 | SNORA70 | Small nucle | 7.364775 | 14.10797 | 18.43774 |
| 26139 | ENSG0000 | HLA-B | major histo | 6.131668 | 10.65773 | 18.43247 |
| 11026 | ENSG0000 | UBXN7 | UBX doma | 10.7248 | 14.95094 | 18.42626 |
| 16023 | ENSG0000 | AGRN | agrin [Sour | 12.8689 | 14.25391 | 18.32376 |
| 6036 | ENSG0000 | ARPP19 | cAMP regu | 16.44972 | 17.27461 | 18.32337 |
| 20173 | ENSG0000 | AS3MT | arsenite m | 6.709396 | 8.130545 | 18.31816 |
| 5318 | ENSG0000 | OPTN | optineurin | 5.259575 | 17.68319 | 18.30261 |
| 10690 | ENSG0000 | DPY30 | dpy-30, his | 13.06133 | 13.44832 | 18.24083 |
| 6678 | ENSG0000 | BTG1 | BTG anti-p | 8.915883 | 15.2541 | 18.14599 |
| 7473 | ENSG0000 | FNBP1L | formin bin | 11.97848 | 16.04192 | 18.11254 |
| 11156 | ENSG0000 | UBLCP1 | ubiquitin li | 17.53949 | 18.05138 | 18.09374 |
| 13862 | ENSG0000 | FBXO46 | F-box prot | 7.320203 | 12.31592 | 18.08923 |
| 1869 | ENSG0000 | RCOR1 | REST corep | 15.60814 | 15.83856 | 18.06384 |
| 20940 | ENSG0000 | PPP3R1 | protein ph | 15.68884 | 17.52993 | 18.02642 |
| 1065 | ENSG0000 | SIRT2 | sirtuin 2 [S | 12.1012 | 13.03253 | 17.99315 |
| 15232 | ENSG0000 | COLEC10 | collectin su | 13.20624 | 16.81032 | 17.92238 |
| 2956 | ENSG0000 | PIH1D1 | PIH1 doma | 10.87757 | 12.63872 | 17.8353 |
| 14635 | ENSG0000 | SIAH2 | siah E3 ubi | 11.95081 | 13.22802 | 17.80917 |
| 8623 | ENSG0000 | OSMR | oncostatin | 11.76892 | 16.06389 | 17.76915 |
| 509 | ENSG0000 | RABEP1 | rabaptin, R | 16.79074 | 16.92785 | 17.76454 |
| 9677 | ENSG0000 | AZIN1 | antizyme in | 15.11989 | 15.64954 | 17.72765 |
| 22050 | ENSG0000 | LINC02557 | long interg | 5.924581 | 6.782393 | 17.69261 |
| 29214 | ENSG0000 | GATA2-AS1 | GATA2 ant | 11.7003 | 12.96006 | 17.63959 |
| 11039 | ENSG0000 | METAP1 | methionyl | 13.32026 | 17.17728 | 17.61687 |
| 8984 | ENSG0000 | NRBF2 | nuclear rec | 9.888694 | 13.70751 | 17.59032 |
| 9912 | ENSG0000 | CNNM4 | cyclin and | 8.182263 | 12.83045 | 17.57945 |
| 4583 | ENSG0000 | ADPRHL2 | ADP-ribosy | 14.59169 | 15.14266 | 17.56976 |
| 4759 | ENSG0000 | RAB32 | RAB32, me | 13.46706 | 14.09886 | 17.53696 |
| 18144 | ENSG0000 | SKIV2L | Ski2 like RN | 15.53241 | 17.17003 | 17.5273 |
| 29182 | ENSG0000 | TMEM141 | transmeml | 10.23473 | 12.97524 | 17.52178 |
| 8031 | ENSG0000 | NT5C3B | 5'-nucleoti | 16.30909 | 17.35334 | 17.51773 |
| 9766 | ENSG0000 | MCU | mitochond | 16.19964 | 16.46393 | 17.51555 |
| 11490 | ENSG0000 | SALL2 | spalt like tr | 6.64242 | 8.591949 | 17.51134 |
| 6514 | ENSG0000 | WBP2 | WW doma | 13.94067 | 14.80974 | 17.5012 |
| 6617 | ENSG0000 | SLC41A1 | solute carr | 9.706945 | 15.5597 | 17.48725 |
| 13019 | ENSG0000 | GTPBP2 | GTP bindin | 12.21945 | 17.23567 | 17.46246 |
| 3345 | ENSG0000 | TMEM245 | transmeml | 15.31718 | 16.89643 | 17.45832 |
| 2066 | ENSG0000 | TMEM14A | transmeml | 7.957375 | 12.27121 | 17.45386 |
| 2499 | ENSG0000 | SEC23B | Sec23 hom | 13.38491 | 16.29357 | 17.44095 |
| 2408 | ENSG0000 | PSMA6 | proteasom | 15.22468 | 15.76777 | 17.40465 |
| 12596 | ENSG0000 | SOCS6 | suppressor | 12.60094 | 16.79142 | 17.37723 |
| 5504 | ENSG0000 | TBCC | tubulin fol | 16.13281 | 16.39842 | 17.31464 |
| 13267 | ENSG0000 | STAT5B | signal trans | 11.70389 | 17.1135 | 17.30264 |
| 3156 | ENSG0000 | TFPT | TCF3 fusion | 12.87101 | 16.1086 | 17.28951 |

| | | | | | | |
|---|---|---|---|---|---|---|
| 9037 | ENSG0000 | APIP | APAF1 inte | 15.4693 | 16.18417 | 17.28886 |
| 134 | ENSG0000 | RALA | RAS like pr | 13.13448 | 15.21154 | 17.23693 |
| 14227 | ENSG0000 | SAMD4B | sterile alph | 14.29031 | 16.73702 | 17.2281 |
| 576 | ENSG0000 | MAT2B | methionine | 11.79684 | 15.09868 | 17.20692 |
| 758 | ENSG0000 | TAB2 | TGF-beta a | 9.009442 | 16.94665 | 17.19594 |
| 3049 | ENSG0000 | SYDE1 | synapse de | 9.753275 | 15.51124 | 17.17426 |
| 9911 | ENSG0000 | DDX19B | DEAD-box | 13.25014 | 15.90899 | 17.16785 |
| 4334 | ENSG0000 | TP53I3 | tumor prot | 10.20567 | 15.3442 | 17.14377 |
| 1669 | ENSG0000 | CD59 | CD59 mole | 14.15864 | 14.62602 | 17.13055 |
| 2268 | ENSG0000 | RBFOX2 | RNA bindir | 13.17817 | 15.57413 | 17.12108 |
| 15716 | ENSG0000 | NF2 | neurofibro | 11.75416 | 16.70453 | 17.06113 |
| 16620 | ENSG0000 | SRC | SRC proto- | 11.96939 | 14.50295 | 16.98531 |
| 3842 | ENSG0000 | GTF2H3 | general tra | 11.81228 | 13.86362 | 16.98044 |
| 33886 | ENSG0000 | AL133370.1 | | 3.467552 | 12.09102 | 16.95603 |
| 1589 | ENSG0000 | STRADB | STE20-rela | 12.48959 | 13.09217 | 16.90485 |
| 7508 | ENSG0000 | FBXO11 | F-box prote | 14.19358 | 15.95949 | 16.87617 |
| 4703 | ENSG0000 | OSBPL9 | oxysterol b | 10.66546 | 14.70118 | 16.80167 |
| 5594 | ENSG0000 | PPP1R12C | protein ph | 9.148168 | 14.80617 | 16.72878 |
| 1454 | ENSG0000 | SAR1A | secretion a | 13.07685 | 15.0501 | 16.71815 |
| 475 | ENSG0000 | CCDC28A | coiled-coil | 9.378999 | 14.78816 | 16.71308 |
| 3102 | ENSG0000 | ICAM5 | intercellula | 12.35356 | 16.19446 | 16.71185 |
| 5464 | ENSG0000 | SNRNP27 | small nucle | 13.35437 | 14.6316 | 16.68419 |
| 329 | ENSG0000 | PIK3C2A | phosphatic | 9.332099 | 14.49606 | 16.68371 |
| 7568 | ENSG0000 | IDH1 | isocitrate c | 13.70009 | 14.31577 | 16.6803 |
| 14888 | ENSG0000 | NKX2-5 | NK2 home | 11.67479 | 15.28067 | 16.65555 |
| 8296 | ENSG0000 | PIP5K1A | phosphatic | 12.01664 | 16.45063 | 16.64501 |
| 8069 | ENSG0000 | IFNAR1 | interferon | 13.09617 | 14.73999 | 16.61659 |
| 2133 | ENSG0000 | PALM | paralemmi | 9.83591 | 9.997849 | 16.61504 |
| 4207 | ENSG0000 | TIMMDC1 | translocase | 13.02481 | 14.91162 | 16.61086 |
| 8400 | ENSG0000 | EML4 | echinoderr | 15.42544 | 16.40584 | 16.57079 |
| 2404 | ENSG0000 | CHD8 | chromodor | 13.31928 | 14.94042 | 16.56087 |
| 8937 | ENSG0000 | ALAD | aminolevu | 7.722715 | 11.24941 | 16.55291 |
| 511 | ENSG0000 | NUP160 | nucleopori | 15.96362 | 16.26637 | 16.54156 |
| 1171 | ENSG0000 | HLTF | helicase lik | 11.17118 | 14.03988 | 16.52338 |
| 10197 | ENSG0000 | BTG2 | BTG anti-p | 4.774033 | 9.009189 | 16.47418 |
| 3003 | ENSG0000 | LZTS2 | leucine zip | 10.26852 | 10.83272 | 16.45484 |
| 5268 | ENSG0000 | CALD1 | caldesmon | 8.559883 | 15.02105 | 16.44546 |
| 6386 | ENSG0000 | NDFIP1 | Nedd4 fam | 12.88403 | 15.40613 | 16.44226 |
| 7487 | ENSG0000 | RTCA | RNA 3'-terı | 13.75971 | 15.58602 | 16.43686 |
| 3697 | ENSG0000 | LPXN | leupaxin [S | 9.687989 | 12.7228 | 16.43685 |
| 2439 | ENSG0000 | NDRG3 | NDRG fami | 7.977653 | 12.65686 | 16.43016 |
| 15995 | ENSG0000 | S100A3 | S100 calciu | 1.665285 | 6.477173 | 16.41992 |
| 16596 | ENSG0000 | GMFB | glia matura | 12.25278 | 14.49189 | 16.37327 |
| 7377 | ENSG0000 | FAM8A1 | family with | 9.385172 | 13.17339 | 16.34873 |
| 15157 | ENSG0000 | DENND5A | DENN dom | 11.74297 | 13.50609 | 16.32798 |
| 16714 | ENSG0000 | GSTK1 | glutathione | 11.22872 | 13.70782 | 16.32649 |
| 6634 | ENSG0000 | TCEAL4 | transcriptio | 12.61249 | 12.6279 | 16.32614 |

| | | | | | | |
|---|---|---|---|---|---|---|
| 9980 | ENSG0000 | CREB3L1 | cAMP resp | 5.224886 | 8.874191 | 16.32263 |
| 91 | ENSG0000 | PRKAR2B | protein kin | 10.30354 | 15.90767 | 16.24923 |
| 9825 | ENSG0000 | FAM122B | family with | 9.145726 | 13.65081 | 16.2446 |
| 13309 | ENSG0000 | CEP19 | centrosom | 7.728295 | 14.67712 | 16.22697 |
| 36578 | ENSG0000 | AC067852.2 | | 11.2432 | 11.49537 | 16.20535 |
| 14353 | ENSG0000 | C14orf119 | chromosor | 14.84888 | 15.15817 | 16.1973 |
| 623 | ENSG0000 | CUL7 | cullin 7 [So | 9.37515 | 14.69568 | 16.18963 |
| 6110 | ENSG0000 | SEC14L1 | SEC14 like | 14.65721 | 15.72565 | 16.18302 |
| 20772 | ENSG0000 | NBPF1 | NBPF mem | 10.25552 | 15.98063 | 16.16775 |
| 8236 | ENSG0000 | CREG1 | cellular rep | 8.571481 | 15.00724 | 16.15564 |
| 9675 | ENSG0000 | KLF10 | Kruppel lik | 8.268111 | 12.10683 | 16.15344 |
| 8033 | ENSG0000 | ERBB2 | erb-b2 rec | 13.90675 | 14.69416 | 16.11433 |
| 16900 | ENSG0000 | CD2AP | CD2 associ | 11.89885 | 15.70732 | 16.08021 |
| 2316 | ENSG0000 | ABHD4 | abhydrolas | 6.358602 | 8.824447 | 16.0752 |
| 8017 | ENSG0000 | WDR45B | WD repeat | 11.94502 | 15.16762 | 16.06949 |
| 9983 | ENSG0000 | SLC38A10 | solute carr | 13.4036 | 13.6878 | 15.98382 |
| 9831 | ENSG0000 | PHF6 | PHD finger | 12.0906 | 13.40326 | 15.97359 |
| 7408 | ENSG0000 | TGS1 | trimethylgu | 14.60381 | 15.55336 | 15.96735 |
| 9699 | ENSG0000 | MOV10 | Mov10 RIS | 7.874634 | 11.38943 | 15.92497 |
| 7756 | ENSG0000 | SBNO1 | strawberry | 14.42227 | 15.08033 | 15.88628 |
| 10743 | ENSG0000 | SCNM1 | sodium cha | 8.184211 | 14.21796 | 15.84075 |
| 6252 | ENSG0000 | UCK1 | uridine-cyt | 8.991388 | 14.45464 | 15.80955 |
| 16768 | ENSG0000 | CDC42SE1 | CDC42 sma | 11.02475 | 14.56625 | 15.80481 |
| 3843 | ENSG0000 | EIF2B1 | eukaryotic | 14.2212 | 14.65788 | 15.80433 |
| 365 | ENSG0000 | NUB1 | negative re | 11.5692 | 14.5514 | 15.79988 |
| 6671 | ENSG0000 | ADCK2 | aarF doma | 8.596503 | 11.52387 | 15.7752 |
| 6779 | ENSG0000 | TMEM106 | transmeml | 9.558586 | 9.874612 | 15.69767 |
| 3805 | ENSG0000 | SLC6A12 | solute carr | 2.501713 | 11.86497 | 15.69047 |
| 20500 | ENSG0000 | VAMP2 | vesicle ass | 8.181634 | 14.55183 | 15.68238 |
| 13860 | ENSG0000 | SIX5 | SIX homeo | 9.682945 | 13.71579 | 15.67818 |
| 8335 | ENSG0000 | SNAPIN | SNAP asso | 9.652317 | 14.697 | 15.66249 |
| 16785 | ENSG0000 | FAM114A1 | family with | 9.106816 | 12.94114 | 15.64332 |
| 7034 | ENSG0000 | EGLN1 | egl-9 famil | 9.928809 | 10.89445 | 15.63479 |
| 4253 | ENSG0000 | FXR1 | FMR1 auto | 15.15617 | 15.58333 | 15.625 |
| 3084 | ENSG0000 | CCDC9 | coiled-coil | 10.62332 | 12.12423 | 15.58222 |
| 1691 | ENSG0000 | CPNE3 | copine 3 [S | 12.18713 | 14.90546 | 15.57635 |
| 2551 | ENSG0000 | ADNP2 | ADNP hom | 11.26653 | 14.55391 | 15.57429 |
| 7299 | ENSG0000 | CTSV | cathepsin V | 4.014536 | 7.197346 | 15.56542 |
| 4648 | ENSG0000 | P3H1 | prolyl 3-hy | 12.82202 | 14.42157 | 15.55895 |
| 2035 | ENSG0000 | NANS | N-acetylne | 14.0953 | 15.51441 | 15.54248 |
| 18305 | ENSG0000 | FAM216A | family with | 11.90143 | 13.24592 | 15.51885 |
| 9493 | ENSG0000 | TMEM87B | transmeml | 6.30948 | 14.81322 | 15.49709 |
| 11736 | ENSG0000 | ATP23 | ATP23 met | 11.60071 | 12.82268 | 15.48335 |
| 19613 | ENSG0000 | COX20P1 | COX20, cyt | 8.020679 | 13.17877 | 15.47625 |
| 6573 | ENSG0000 | VSTM2L | V-set and t | 12.15351 | 12.58834 | 15.47056 |
| 8909 | ENSG0000 | HAUS6 | HAUS augr | 12.2053 | 13.24055 | 15.46823 |
| 3514 | ENSG0000 | HOXB6 | homeobox | 9.646773 | 11.76169 | 15.45497 |

| | | | | | | |
|---|---|---|---|---|---|---|
| 1365 | ENSG0000 | NFKB2 | nuclear fac | 6.583634 | 12.48486 | 15.45013 |
| 16589 | ENSG0000 | SERTAD1 | SERTA dom | 9.93984 | 13.70452 | 15.40327 |
| 9794 | ENSG0000 | RWDD2B | RWD doma | 9.577718 | 12.39316 | 15.40121 |
| 13786 | ENSG0000 | GRB2 | growth fac | 10.27901 | 10.54394 | 15.39409 |
| 9123 | ENSG0000 | CABLES2 | Cdk5 and A | 10.81634 | 12.00341 | 15.38897 |
| 11075 | ENSG0000 | C4orf3 | chromosor | 6.863722 | 8.385294 | 15.37282 |
| 4973 | ENSG0000 | AVPI1 | arginine va | 11.03991 | 12.49892 | 15.32224 |
| 9197 | ENSG0000 | DOCK1 | dedicator c | 12.28858 | 14.42451 | 15.3091 |
| 38835 | ENSG0000 | HIST2H2A/ | histone clu | 4.039967 | 15.09671 | 15.30315 |
| 7409 | ENSG0000 | SDCBP | syndecan b | 10.01154 | 13.15235 | 15.2922 |
| 14481 | ENSG0000 | CHRM4 | cholinergic | 2.921714 | 5.212884 | 15.28823 |
| 13163 | ENSG0000 | IQCB1 | IQ motif cc | 13.1417 | 15.18651 | 15.27638 |
| 4403 | ENSG0000 | MOB4 | MOB famil | 6.630195 | 6.760912 | 15.26735 |
| 17 | ENSG0000 | ANKIB1 | ankyrin rep | 10.30363 | 14.59951 | 15.26282 |
| 3012 | ENSG0000 | MADD | MAP kinas | 10.8539 | 14.4737 | 15.25582 |
| 6899 | ENSG0000 | AGTPBP1 | ATP/GTP b | 9.617755 | 12.59851 | 15.2486 |
| 3942 | ENSG0000 | PPARD | peroxisom | 12.72126 | 14.87219 | 15.2231 |
| 9650 | ENSG0000 | SKA1 | spindle and | 13.34002 | 13.57036 | 15.19037 |
| 11069 | ENSG0000 | DUSP7 | dual specif | 11.84401 | 14.14034 | 15.1854 |
| 10325 | ENSG0000 | GPSM1 | G protein s | 12.63635 | 14.25859 | 15.12339 |
| 3762 | ENSG0000 | TCIRG1 | T-cell immu | 6.939844 | 13.06261 | 15.10531 |
| 12434 | ENSG0000 | TPST1 | tyrosylprot | 11.43466 | 13.67622 | 15.10244 |
| 6672 | ENSG0000 | MKRN1 | makorin rii | 11.86633 | 14.60593 | 15.0494 |
| 1260 | ENSG0000 | NOTCH3 | notch 3 [Sc | 4.244947 | 7.891728 | 15.0491 |
| 2229 | ENSG0000 | JOSD1 | Josephin d | 13.00963 | 15.02659 | 15.03887 |
| 6928 | ENSG0000 | SRPK2 | SRSF prote | 12.30232 | 13.83396 | 15.02584 |
| 506 | ENSG0000 | ANK1 | ankyrin 1 [ | 12.11944 | 13.63441 | 15.01132 |
| 14306 | ENSG0000 | CDC42EP4 | CDC42 effe | 11.66306 | 12.02317 | 14.99734 |
| 4041 | ENSG0000 | EHBP1 | EH domain | 12.76591 | 14.98398 | 14.99546 |
| 14800 | ENSG0000 | TTC3 | tetratricop | 7.382064 | 8.373712 | 14.97621 |
| 16337 | ENSG0000 | ACADSB | acyl-CoA d | 9.218933 | 11.86346 | 14.97228 |
| 6925 | ENSG0000 | HILPDA | hypoxia inc | 4.118607 | 4.72644 | 14.96728 |
| 31797 | ENSG0000 | LYN | LYN proto- | 8.677524 | 11.78961 | 14.93797 |
| 1661 | ENSG0000 | AGBL5 | ATP/GTP b | 8.727432 | 10.30628 | 14.92543 |
| 11786 | ENSG0000 | FAM102A | family with | 5.037072 | 6.664805 | 14.92404 |
| 16450 | ENSG0000 | TCEAL3 | transcripti | 4.066987 | 8.961192 | 14.91674 |
| 5187 | ENSG0000 | RASL11A | RAS like fa | 4.992406 | 9.978611 | 14.88154 |
| 6027 | ENSG0000 | FLNB | filamin B [S | 8.507775 | 13.32916 | 14.8795 |
| 2556 | ENSG0000 | LPIN2 | lipin 2 [Sou | 8.509789 | 11.20331 | 14.83551 |
| 848 | ENSG0000 | GPBP1 | GC-rich pro | 9.740375 | 13.14181 | 14.81078 |
| 7505 | ENSG0000 | PREPL | prolyl endc | 6.889725 | 13.8713 | 14.81062 |
| 297 | ENSG0000 | IDS | iduronate 2 | 6.026244 | 13.69341 | 14.78181 |
| 13917 | ENSG0000 | CCDC71 | coiled-coil | 10.58914 | 11.26465 | 14.77745 |
| 3108 | ENSG0000 | TYK2 | tyrosine ki | 13.30633 | 13.61971 | 14.75799 |
| 14189 | ENSG0000 | SELENOW | selenoprot | 8.615567 | 13.02604 | 14.72615 |
| 9178 | ENSG0000 | PDCD4 | programm | 9.973427 | 14.0135 | 14.68844 |
| 406 | ENSG0000 | ATP2C1 | ATPase sec | 11.20635 | 13.05546 | 14.67127 |

| | | | | | | |
|---|---|---|---|---|---|---|
| 8029 | ENSG0000 | PMAIP1 | phorbol-12 | 6.737547 | 7.559636 | 14.65724 |
| 10258 | ENSG0000 | BSDC1 | BSD domai | 10.7128 | 13.81566 | 14.65718 |
| 8556 | ENSG0000 | TCTA | T-cell leuk | 8.732775 | 13.53633 | 14.64735 |
| 5738 | ENSG0000 | FRMD8 | FERM dom | 8.377185 | 13.20194 | 14.62483 |
| 16893 | ENSG0000 | MARCH5 | membrane | 9.696624 | 11.93454 | 14.6203 |
| 9385 | ENSG0000 | TCF7L1 | transcripti | 7.660797 | 10.24017 | 14.59365 |
| 9678 | ENSG0000 | ATP6V1C1 | ATPase H+ | 8.276788 | 14.41281 | 14.57662 |
| 5741 | ENSG0000 | IRF3 | interferon | 9.571603 | 13.25821 | 14.54925 |
| 10781 | ENSG0000 | CGGBP1 | CGG triplet | 11.55234 | 13.31653 | 14.5426 |
| 7755 | ENSG0000 | RB1 | RB transcri | 12.77911 | 14.30533 | 14.52568 |
| 1562 | ENSG0000 | ARG2 | arginase 2 | 6.468681 | 14.48673 | 14.52177 |
| 9711 | ENSG0000 | CNOT8 | CCR4-NOT | 8.936222 | 10.11958 | 14.5001 |
| 7263 | ENSG0000 | DAB2IP | DAB2 inter | 10.43145 | 12.54972 | 14.49288 |
| 5570 | ENSG0000 | RNF113A | ring finger | 8.944772 | 9.786824 | 14.44385 |
| 12986 | ENSG0000 | CERS6 | ceramide s | 9.934367 | 13.22734 | 14.425 |
| 8365 | ENSG0000 | SDE2 | SDE2 telon | 10.89553 | 12.97015 | 14.42448 |
| 3470 | ENSG0000 | CCDC6 | coiled-coil | 10.74352 | 14.37781 | 14.41945 |
| 11911 | ENSG0000 | B3GNT2 | UDP-GlcNA | 9.043552 | 11.48237 | 14.41747 |
| 2633 | ENSG0000 | RP2 | RP2, ARL3 | 7.395069 | 9.915183 | 14.41072 |
| 9681 | ENSG0000 | CDK19 | cyclin depe | 6.304293 | 11.22859 | 14.38714 |
| 4514 | ENSG0000 | SRSF4 | serine and | 12.21262 | 14.26212 | 14.37602 |
| 2104 | ENSG0000 | FBXL19 | F-box and l | 12.26602 | 12.66099 | 14.3491 |
| 8311 | ENSG0000 | GOLPH3L | golgi phosp | 9.463274 | 10.90874 | 14.29107 |
| 3925 | ENSG0000 | ASF1A | anti-silenci | 11.1148 | 12.59326 | 14.26335 |
| 9631 | ENSG0000 | BTG3 | BTG anti-p | 10.03841 | 11.07068 | 14.26063 |
| 7112 | ENSG0000 | ITM2B | integral me | 8.681173 | 10.44517 | 14.25587 |
| 13486 | ENSG0000 | PACS1 | phosphofu | 12.45984 | 12.66258 | 14.24505 |
| 5077 | ENSG0000 | SORBS3 | sorbin and | 10.45154 | 10.74543 | 14.22874 |
| 7987 | ENSG0000 | ARRB2 | arrestin be | 9.048171 | 11.17746 | 14.20301 |
| 33493 | ENSG0000 | AC078923.1 | | 4.187488 | 12.69504 | 14.20032 |
| 1663 | ENSG0000 | KIF3C | kinesin fam | 5.706251 | 8.197834 | 14.17584 |
| 34568 | ENSG0000 | HOXB7 | homeobox | 9.89704 | 10.12463 | 14.10884 |
| 28287 | ENSG0000 | ARHGEF25 | Rho guanir | 3.335534 | 9.635778 | 14.10404 |
| 13875 | ENSG0000 | ANO6 | anoctamin | 8.799016 | 12.91604 | 14.0651 |
| 13872 | ENSG0000 | EPS8L2 | EPS8 like 2 | 9.069956 | 9.380424 | 14.0622 |
| 5011 | ENSG0000 | MRPS14 | mitochond | 11.88848 | 12.62707 | 14.05994 |
| 4883 | ENSG0000 | LYPLA1 | lysophosph | 13.26858 | 13.60251 | 14.05851 |
| 13666 | ENSG0000 | MYPOP | Myb relate | 8.716661 | 9.233299 | 13.9931 |
| 4871 | ENSG0000 | IRF2BPL | interferon | 8.942623 | 10.18676 | 13.99306 |
| 28265 | ENSG0000 | PNMA2 | paraneopla | 5.693871 | 8.910883 | 13.9753 |
| 34613 | ENSG0000 | AC026464.2 | | 8.743711 | 9.714742 | 13.97358 |
| 23034 | ENSG0000 | AL390728.4 | | 7.315106 | 12.82097 | 13.96301 |
| 4573 | ENSG0000 | CRYZ | crystallin z | 8.808003 | 13.19303 | 13.96062 |
| 1112 | ENSG0000 | SLC44A1 | solute carr | 5.725318 | 8.200728 | 13.95024 |
| 129 | ENSG0000 | SPATA20 | spermatog | 8.844916 | 10.34009 | 13.94824 |
| 16786 | ENSG0000 | RPE | ribulose-5- | 12.23872 | 13.00017 | 13.946 |
| 543 | ENSG0000 | GABARAPL | GABA type | 10.39811 | 11.86788 | 13.93434 |

| | | | | | | |
|---|---|---|---|---|---|---|
| 5856 | ENSG0000 | TICAM1 | toll like rec | 8.593271 | 11.2036 | 13.93381 |
| 11278 | ENSG0000 | CDK5 | cyclin depe | 10.10132 | 10.51445 | 13.8907 |
| 2897 | ENSG0000 | ARMC1 | armadillo r | 12.86405 | 13.29313 | 13.88445 |
| 15231 | ENSG0000 | CSF1 | colony stin | 8.816102 | 12.53429 | 13.8747 |
| 11516 | ENSG0000 | MOAP1 | modulator | 6.926082 | 11.95222 | 13.8104 |
| 6488 | ENSG0000 | PRKAA1 | protein kin | 10.99271 | 12.67235 | 13.7994 |
| 4339 | ENSG0000 | GPD2 | glycerol-3- | 5.992062 | 6.533196 | 13.78198 |
| 4807 | ENSG0000 | GTF3C3 | general tra | 12.21887 | 13.09238 | 13.72166 |
| 1044 | ENSG0000 | OTUD5 | OTU deubi | 12.45526 | 13.26714 | 13.71417 |
| 16957 | ENSG0000 | ZKSCAN8 | zinc finger | 7.056598 | 12.19405 | 13.71358 |
| 1671 | ENSG0000 | BCORL1 | BCL6 corep | 5.951269 | 8.011968 | 13.68698 |
| 4166 | ENSG0000 | SLC12A7 | solute carr | 8.882914 | 11.88787 | 13.68257 |
| 1581 | ENSG0000 | HSPB11 | heat shock | 7.586403 | 9.107975 | 13.66025 |
| 20730 | ENSG0000 | ZNF579 | zinc finger | 8.971946 | 11.44499 | 13.65701 |
| 27392 | ENSG0000 | RGL2 | ral guanine | 8.965609 | 10.08442 | 13.65215 |
| 12459 | ENSG0000 | CHD3 | chromodor | 9.093854 | 11.39562 | 13.63526 |
| 9686 | ENSG0000 | PI4K2A | phosphatic | 11.41624 | 12.93669 | 13.62823 |
| 6508 | ENSG0000 | GRSF1 | G-rich RNA | 11.84335 | 12.29018 | 13.62648 |
| 27186 | ENSG0000 | B3GNT9 | UDP-GlcNA | 7.775806 | 9.791173 | 13.61085 |
| 12630 | ENSG0000 | TRIAP1 | TP53 regul | 11.62336 | 13.10509 | 13.57292 |
| 13024 | ENSG0000 | TCEAL1 | transcriptio | 8.461807 | 10.88051 | 13.55563 |
| 7242 | ENSG0000 | STAM | signal trans | 10.95013 | 12.41352 | 13.52828 |
| 3889 | ENSG0000 | SUDS3 | SDS3 homo | 8.220807 | 11.36556 | 13.52599 |
| 164 | ENSG0000 | MARK4 | microtubul | 8.14352 | 11.72516 | 13.51165 |
| 17161 | ENSG0000 | CAPZA2 | capping ac | 12.03985 | 12.4198 | 13.49905 |
| 13408 | ENSG0000 | TMEM167A | transmeml | 11.79574 | 13.21436 | 13.49154 |
| 11597 | ENSG0000 | APBB1 | amyloid be | 4.924977 | 7.516498 | 13.48736 |
| 15846 | ENSG0000 | BCAM | basal cell a | 5.150639 | 6.977317 | 13.45214 |
| 4439 | ENSG0000 | STRN | striatin [So | 9.568927 | 12.31012 | 13.4519 |
| 3397 | ENSG0000 | ABCA2 | ATP bindin | 10.35431 | 11.45186 | 13.41211 |
| 347 | ENSG0000 | MBTPS2 | membrane | 10.76793 | 11.77165 | 13.35881 |
| 37777 | ENSG0000 | FBXO17 | F-box prote | 10.01766 | 10.91913 | 13.33781 |
| 28443 | ENSG0000 | PDXP | pyridoxal p | 2.305301 | 11.00303 | 13.30874 |
| 35189 | ENSG0000 | ISY1-RAB43 | ISY1-RAB43 | 6.072599 | 7.906122 | 13.28872 |
| 12400 | ENSG0000 | ZNF32 | zinc finger | 8.714251 | 12.09111 | 13.28201 |
| 7570 | ENSG0000 | CIR1 | corepresso | 7.945306 | 12.85346 | 13.27176 |
| 8847 | ENSG0000 | ATP6V1B2 | ATPase H+ | 12.33056 | 12.80757 | 13.19619 |
| 28907 | ENSG0000 | PRAF2 | PRA1 doma | 7.737265 | 9.035011 | 13.18353 |
| 25048 | ENSG0000 | AL078590.3 | | 3.380456 | 8.967686 | 13.17937 |
| 3449 | ENSG0000 | ACBD5 | acyl-CoA b | 9.972325 | 10.65011 | 13.16103 |
| 16507 | ENSG0000 | AMZ2 | archaelysir | 10.70207 | 12.03053 | 13.15928 |
| 5421 | ENSG0000 | PLCG1 | phospholip | 7.979194 | 12.40639 | 13.12616 |
| 13106 | ENSG0000 | OXSR1 | oxidative s | 10.04341 | 11.99277 | 13.12245 |
| 9461 | ENSG0000 | LMNTD1 | lamin tail c | 5.835839 | 7.660435 | 13.12238 |
| 12546 | ENSG0000 | DENND5B | DENN dom | 10.96182 | 13.01107 | 13.11529 |
| 12806 | ENSG0000 | WIPF2 | WAS/WASI | 8.669452 | 10.74068 | 13.10772 |
| 13500 | ENSG0000 | PARL | presenilin a | 7.736313 | 8.144197 | 13.10482 |

| | | | | | | |
|---|---|---|---|---|---|---|
| 2340 | ENSG0000 | ATP6V1D | ATPase H+ | 8.160264 | 9.17294 | 13.09028 |
| 9436 | ENSG0000 | SLC30A6 | solute carr | 9.326753 | 12.01697 | 13.08177 |
| 7131 | ENSG0000 | ZDHHC4 | zinc finger | 8.195516 | 9.547451 | 13.06741 |
| 18184 | ENSG0000 | C6orf47 | chromosor | 10.27283 | 11.15216 | 13.04375 |
| 6824 | ENSG0000 | ACP2 | acid phosp | 8.583851 | 10.7529 | 13.00537 |
| 6668 | ENSG0000 | ZDHHC8P1 | zinc finger | 4.930657 | 11.12599 | 12.98475 |
| 1892 | ENSG0000 | SPG21 | SPG21, ma | 10.23449 | 10.79051 | 12.94488 |
| 2737 | ENSG0000 | SLC7A6 | solute carr | 6.444897 | 8.317319 | 12.93715 |
| 10959 | ENSG0000 | HPS3 | HPS3, biog | 7.627006 | 8.657605 | 12.93458 |
| 4962 | ENSG0000 | CUTC | cutC coppe | 9.086028 | 9.668987 | 12.9204 |
| 8728 | ENSG0000 | ARHGAP18 | Rho GTPas | 7.651909 | 10.87462 | 12.8644 |
| 1038 | ENSG0000 | FGFR3 | fibroblast g | 4.518105 | 8.999496 | 12.84768 |
| 11286 | ENSG0000 | FOXK1 | forkhead b | 8.343768 | 12.63475 | 12.82533 |
| 20713 | ENSG0000 | CEBPZOS | CEBPZ opp | 8.772979 | 9.742291 | 12.81918 |
| 6436 | ENSG0000 | RHPN2 | rhophilin R | 8.359176 | 11.06726 | 12.78615 |
| 4729 | ENSG0000 | ATF6 | activating t | 10.29386 | 12.02744 | 12.77898 |
| 12568 | ENSG0000 | PFKFB3 | 6-phospho | 6.296673 | 8.282919 | 12.76727 |
| 4498 | ENSG0000 | TMEM59 | transmeml | 8.201069 | 10.00286 | 12.75584 |
| 10717 | ENSG0000 | COQ8A | coenzyme | 8.270637 | 11.95916 | 12.7531 |
| 18415 | ENSG0000 | SNX2 | sorting nex | 11.27826 | 12.60804 | 12.74853 |
| 7246 | ENSG0000 | DNAJC1 | DnaJ heat s | 8.369333 | 11.914 | 12.74827 |
| 7346 | ENSG0000 | TMEM63B | transmeml | 10.5021 | 11.19486 | 12.7467 |
| 13744 | ENSG0000 | BOK | BOK, BCL2 | 11.12378 | 11.14218 | 12.74538 |
| 15460 | ENSG0000 | TMEM179I | transmeml | 9.329775 | 11.04115 | 12.67248 |
| 601 | ENSG0000 | PARP3 | poly(ADP-r | 7.604609 | 8.86365 | 12.65859 |
| 5131 | ENSG0000 | TRMT1L | tRNA meth | 7.837646 | 11.79288 | 12.64874 |
| 75 | ENSG0000 | SKAP2 | src kinase a | 7.402317 | 10.25806 | 12.64051 |
| 3835 | ENSG0000 | CDK2AP1 | cyclin depe | 10.61932 | 12.46404 | 12.60192 |
| 1336 | ENSG0000 | BAZ2A | bromodom | 10.02516 | 10.39019 | 12.58766 |
| 10016 | ENSG0000 | WIPI2 | WD repeat | 9.742598 | 12.16029 | 12.57584 |
| 17178 | ENSG0000 | ATG9A | autophagy | 8.652126 | 9.829437 | 12.57186 |
| 2622 | ENSG0000 | PIM2 | Pim-2 prot | 7.845283 | 7.985658 | 12.5665 |
| 5967 | ENSG0000 | ORMDL1 | ORMDL spl | 8.191476 | 10.7967 | 12.55635 |
| 8035 | ENSG0000 | MIEN1 | migration a | 10.78079 | 11.03576 | 12.54782 |
| 1868 | ENSG0000 | DHX32 | DEAH-box | 9.948285 | 11.8351 | 12.54008 |
| 9830 | ENSG0000 | TYSND1 | trypsin dor | 9.392237 | 9.919482 | 12.53417 |
| 878 | ENSG0000 | CASP8 | caspase 8 [ | 9.600789 | 9.710667 | 12.52167 |
| 20750 | ENSG0000 | AC138392.1 | | 10.0011 | 11.10355 | 12.51906 |
| 14098 | ENSG0000 | FAM220A | family with | 7.986565 | 9.909805 | 12.48325 |
| 9857 | ENSG0000 | PRR14 | proline rich | 9.678791 | 11.01531 | 12.47937 |
| 17945 | ENSG0000 | COX20 | COX20, cyt | 8.989477 | 10.70122 | 12.45252 |
| 3777 | ENSG0000 | COQ5 | coenzyme | 9.464388 | 9.831755 | 12.44235 |
| 5705 | ENSG0000 | KDM5C | lysine dem | 11.97771 | 12.32027 | 12.43577 |
| 28854 | ENSG0000 | EIF4EBP3 | eukaryotic | 4.333023 | 7.842114 | 12.43556 |
| 10746 | ENSG0000 | ERCC3 | ERCC excis | 10.28627 | 12.41988 | 12.43192 |
| 10992 | ENSG0000 | YEATS2 | YEATS dor | 7.13412 | 8.868888 | 12.42859 |
| 6310 | ENSG0000 | ULBP2 | UL16 bindi | 3.944295 | 9.780538 | 12.42221 |

| | | | | | | |
|---|---|---|---|---|---|---|
| 101 | ENSG0000 | KMT2E | lysine metl | 8.495963 | 10.22351 | 12.39573 |
| 6432 | ENSG0000 | LLGL1 | LLGL1, scri | 7.905729 | 8.274099 | 12.38926 |
| 8032 | ENSG0000 | RETREG3 | reticulopha | 6.273139 | 8.285125 | 12.38571 |
| 14697 | ENSG0000 | IST1 | IST1, ESCR | 11.4052 | 12.31482 | 12.38353 |
| 16516 | ENSG0000 | GM2A | GM2 gangl | 6.464305 | 7.600751 | 12.35023 |
| 8521 | ENSG0000 | TMF1 | TATA elem | 6.863408 | 10.0609 | 12.33689 |
| 1371 | ENSG0000 | USP33 | ubiquitin s | 7.681215 | 10.31575 | 12.31998 |
| 7370 | ENSG0000 | TPMT | thiopurine | 8.839309 | 9.898989 | 12.3063 |
| 10589 | ENSG0000 | MYSM1 | Myb like, S | 2.803268 | 4.733256 | 12.26399 |
| 12622 | ENSG0000 | USP32 | ubiquitin s | 8.708469 | 12.08835 | 12.25607 |
| 10331 | ENSG0000 | SHKBP1 | SH3KBP1 b | 7.886319 | 8.18265 | 12.22938 |
| 37689 | ENSG0000 | AL118506.1 | | 4.943516 | 9.933162 | 12.21381 |
| 13220 | ENSG0000 | TNFRSF10I | TNF recept | 11.11391 | 12.18576 | 12.21067 |
| 7706 | ENSG0000 | NEDD1 | neural pre | 10.47053 | 11.13473 | 12.2091 |
| 12976 | ENSG0000 | C5orf34 | chromosor | 5.547704 | 10.05677 | 12.2082 |
| 1681 | ENSG0000 | WDR47 | WD repeat | 5.85114 | 9.615933 | 12.19551 |
| 13951 | ENSG0000 | IRX3 | iroquois hc | 9.72817 | 10.72306 | 12.18078 |
| 15484 | ENSG0000 | OLFML2A | olfactome | 1.883567 | 3.54558 | 12.17505 |
| 5141 | ENSG0000 | MAPK8IP1 | mitogen-ac | 8.426711 | 10.52 | 12.17104 |
| 36905 | ENSG0000 | ILF3-AS1 | ILF3 antise | 9.793729 | 12.14966 | 12.16879 |
| 10603 | ENSG0000 | HENMT1 | HEN methy | 9.229958 | 10.60373 | 12.16793 |
| 4390 | ENSG0000 | UNC50 | unc-50 inn | 9.841013 | 11.18534 | 12.14207 |
| 18295 | ENSG0000 | MRPL53 | mitochond | 10.8936 | 11.01882 | 12.13553 |
| 7979 | ENSG0000 | ESCO1 | establishm | 4.957882 | 8.075163 | 12.13457 |
| 17119 | ENSG0000 | SFT2D1 | SFT2 doma | 9.563295 | 10.28394 | 12.11243 |
| 6063 | ENSG0000 | PLD2 | phospholip | 6.729951 | 11.04404 | 12.08668 |
| 8197 | ENSG0000 | HSPG2 | heparan su | 6.568007 | 7.759274 | 12.07778 |
| 8186 | ENSG0000 | EVA1B | eva-1 hom | 6.350989 | 10.14209 | 12.07724 |
| 13271 | ENSG0000 | JUP | junction pl | 4.095488 | 4.221643 | 12.07617 |
| 4383 | ENSG0000 | FANCL | Fanconi an | 7.904508 | 7.92513 | 12.06641 |
| 9549 | ENSG0000 | HS2ST1 | heparan su | 9.513313 | 10.12432 | 12.06014 |
| 4809 | ENSG0000 | UBE2B | ubiquitin c | 8.882932 | 10.15024 | 12.00298 |
| 26987 | ENSG0000 | MTX1P1 | metaxin 1 | 8.868185 | 9.138457 | 11.99767 |
| 2722 | ENSG0000 | USB1 | U6 snRNA | 7.872822 | 10.72481 | 11.98815 |
| 4228 | ENSG0000 | SLC25A36 | solute carr | 6.681045 | 9.365488 | 11.98333 |
| 4520 | ENSG0000 | RAP1A | RAP1A, me | 8.666304 | 10.29517 | 11.98307 |
| 9524 | ENSG0000 | RMND5A | required fc | 6.785756 | 11.7321 | 11.98161 |
| 3133 | ENSG0000 | RAB3D | RAB3D, me | 4.937307 | 5.817079 | 11.97449 |
| 4638 | ENSG0000 | RAB29 | RAB29, me | 5.685175 | 8.962152 | 11.92843 |
| 1797 | ENSG0000 | PDRG1 | p53 and DI | 8.195633 | 11.39238 | 11.91988 |
| 5332 | ENSG0000 | CDK2 | cyclin depe | 9.494147 | 9.533271 | 11.91945 |
| 32138 | ENSG0000 | MEX3A | mex-3 RNA | 6.305582 | 7.819556 | 11.91863 |
| 609 | ENSG0000 | RETSAT | retinol satu | 11.18208 | 11.54545 | 11.89657 |
| 1391 | ENSG0000 | UBE2A | ubiquitin c | 9.512027 | 10.4465 | 11.89075 |
| 8228 | ENSG0000 | ITGA10 | integrin su | 7.485374 | 8.581848 | 11.88725 |
| 7958 | ENSG0000 | ABCA8 | ATP bindin | 5.099986 | 8.106529 | 11.85988 |
| 8325 | ENSG0000 | TP53BP2 | tumor prot | 8.625854 | 8.849359 | 11.85662 |

| | | | | | | |
|---|---|---|---|---|---|---|
| 12386 | ENSG0000 | LRRC45 | leucine ricl | 8.061433 | 10.57246 | 11.8359 |
| 1821 | ENSG0000 | MAVS | mitochond | 6.782001 | 11.45394 | 11.8098 |
| 2563 | ENSG0000 | RNMT | RNA guani | 2.649859 | 5.530414 | 11.80558 |
| 8700 | ENSG0000 | MUT | methylmal | 8.315855 | 9.43801 | 11.79757 |
| 11466 | ENSG0000 | FAM69B | family witl | 6.506818 | 7.413932 | 11.78235 |
| 23883 | ENSG0000 | NAMPTP1 | nicotinami | 7.170531 | 8.657821 | 11.7803 |
| 6695 | ENSG0000 | SWAP70 | SWAP swit | 6.905141 | 10.54795 | 11.74099 |
| 4943 | ENSG0000 | YPEL5 | yippee like | 5.014997 | 9.480824 | 11.71375 |
| 7405 | ENSG0000 | GGH | gamma-glu | 7.014777 | 7.464231 | 11.71156 |
| 8732 | ENSG0000 | ABRACL | ABRA C-ter | 5.332045 | 7.418858 | 11.70352 |
| 3014 | ENSG0000 | NAA40 | N(alpha)-a | 8.353344 | 10.26258 | 11.69821 |
| 18087 | ENSG0000 | ZDHHC18 | zinc finger | 9.965553 | 10.73994 | 11.69478 |
| 10971 | ENSG0000 | KIAA1143 | KIAA1143 \| | 9.470531 | 10.4848 | 11.65072 |
| 28174 | ENSG0000 | ACAD11 | acyl-CoA d | 4.902269 | 7.773766 | 11.64714 |
| 1497 | ENSG0000 | CPOX | coproporp | 9.995124 | 10.60799 | 11.63656 |
| 14197 | ENSG0000 | C3orf38 | chromosor | 7.68144 | 11.10446 | 11.63234 |
| 3080 | ENSG0000 | PRKD2 | protein kin | 6.325134 | 6.365678 | 11.6102 |
| 15090 | ENSG0000 | CMTM4 | CKLF like M | 9.811478 | 10.31572 | 11.60052 |
| 15202 | ENSG0000 | OAF | out at first | 10.14376 | 11.28395 | 11.59983 |
| 10305 | ENSG0000 | RALGDS | ral guanine | 7.576914 | 11.09788 | 11.58214 |
| 5482 | ENSG0000 | TRERF1 | transcriptio | 9.813239 | 11.24642 | 11.57683 |
| 3934 | ENSG0000 | HINT3 | histidine tr | 8.361744 | 10.67653 | 11.56113 |
| 12790 | ENSG0000 | ZNF581 | zinc finger | 7.674037 | 7.936214 | 11.52722 |
| 18390 | ENSG0000 | PSENEN | presenilin | 8.237251 | 8.3703 | 11.49423 |
| 3069 | ENSG0000 | NUMBL | NUMB like | 10.15462 | 10.50498 | 11.48595 |
| 16748 | ENSG0000 | TOPORS | TOP1 bindi | 10.79313 | 11.25781 | 11.48131 |
| 8653 | ENSG0000 | YIPF5 | Yip1 doma | 9.834261 | 10.96901 | 11.41659 |
| 15875 | ENSG0000 | PRR13P5 | proline ricl | 8.506644 | 10.11715 | 11.40517 |
| 12399 | ENSG0000 | DCXR | dicarbonyl | 9.435218 | 9.689767 | 11.40511 |
| 11222 | ENSG0000 | ZNF12 | zinc finger | 8.256689 | 10.04348 | 11.3918 |
| 12197 | ENSG0000 | SHOX2 | short statu | 6.645261 | 7.890232 | 11.39043 |
| 12088 | ENSG0000 | UBTD2 | ubiquitin d | 8.775454 | 9.863725 | 11.38525 |
| 9578 | ENSG0000 | NRGN | neurograni | 6.054954 | 7.729462 | 11.38423 |
| 1978 | ENSG0000 | TOX4 | TOX high m | 6.400086 | 8.25818 | 11.35674 |
| 9595 | ENSG0000 | ABCA10 | ATP bindin | 5.824827 | 9.203933 | 11.35598 |
| 6537 | ENSG0000 | PCED1A | PC-esteras | 4.742584 | 9.25248 | 11.34273 |
| 11746 | ENSG0000 | NYAP1 | neuronal ty | 0.7139 | 4.682734 | 11.30419 |
| 7902 | ENSG0000 | CPNE2 | copine 2 [S | 8.125471 | 8.205653 | 11.29681 |
| 4661 | ENSG0000 | PIK3R3 | phosphoin | 6.993178 | 9.731085 | 11.26717 |
| 8699 | ENSG0000 | RNF44 | ring finger | 5.386543 | 11.11749 | 11.26629 |
| 5236 | ENSG0000 | HERPUD2 | HERPUD fa | 8.462606 | 11.07833 | 11.26426 |
| 6278 | ENSG0000 | PPAN | peter pan l | 7.06914 | 10.84508 | 11.26238 |
| 16143 | ENSG0000 | ZDHHC9 | zinc finger | 7.649952 | 8.741504 | 11.2526 |
| 6677 | ENSG0000 | NTS | neurotensi | 7.940264 | 9.037931 | 11.24666 |
| 13886 | ENSG0000 | RPS6KA3 | ribosomal | 9.844075 | 10.51121 | 11.24651 |
| 7344 | ENSG0000 | YIPF3 | Yip1 doma | 9.215778 | 9.95363 | 11.24053 |
| 9638 | ENSG0000 | RABGEF1 | RAB guanir | 7.660177 | 10.47022 | 11.21639 |

| | | | | | | |
|---|---|---|---|---|---|---|
| 6126 | ENSG0000 | TMEM8A | transmeml | 7.787136 | 9.512317 | 11.21607 |
| 6465 | ENSG0000 | HSD17B7 | hydroxyste | 5.136889 | 9.333818 | 11.19873 |
| 1434 | ENSG0000 | TP53INP2 | tumor prot | 7.0944 | 9.093319 | 11.19431 |
| 9416 | ENSG0000 | ZFP36L2 | ZFP36 ring | 9.718325 | 10.98805 | 11.19165 |
| 333 | ENSG0000 | RABGAP1 | RAB GTPas | 9.193315 | 10.57022 | 11.18584 |
| 19901 | ENSG0000 | ZNF134 | zinc finger | 5.307227 | 10.02929 | 11.16726 |
| 1908 | ENSG0000 | EFNB1 | ephrin B1 [ | 6.981288 | 9.068959 | 11.16474 |
| 13435 | ENSG0000 | GLMN | glomulin, F | 8.394722 | 10.78319 | 11.15494 |
| 1382 | ENSG0000 | SIRT6 | sirtuin 6 [S | 9.271117 | 11.11053 | 11.14996 |
| 5888 | ENSG0000 | LRFN1 | leucine ricl | 5.55742 | 5.870575 | 11.13675 |
| 7971 | ENSG0000 | RPRD1A | regulation | 7.772008 | 11.06775 | 11.11998 |
| 2646 | ENSG0000 | FGD1 | FYVE, Rho( | 10.04745 | 10.21902 | 11.10984 |
| 6526 | ENSG0000 | DLG4 | discs large | 5.214019 | 8.757635 | 11.07725 |
| 5589 | ENSG0000 | TTF1 | transcriptio | 9.78424 | 10.75739 | 11.07599 |
| 3386 | ENSG0000 | DDX58 | DExD/H-bo | 2.563574 | 7.622172 | 11.06121 |
| 12642 | ENSG0000 | GSTA4 | glutathione | 8.227808 | 8.686834 | 11.02212 |
| 20424 | ENSG0000 | BX322557.1 | | 7.449834 | 8.621508 | 10.98111 |
| 7152 | ENSG0000 | PAPSS1 | 3'-phospho | 8.138649 | 8.937941 | 10.97649 |
| 37056 | ENSG0000 | LINC02001 | long interg | 5.62569 | 7.430075 | 10.96115 |
| 3680 | ENSG0000 | CCDC34 | coiled-coil | 6.390881 | 9.555373 | 10.9542 |
| 6044 | ENSG0000 | MBD4 | methyl-Cp( | 8.423778 | 9.858888 | 10.95345 |
| 18434 | ENSG0000 | MT1M | metallothi( | 8.448682 | 8.563884 | 10.948 |
| 11408 | ENSG0000 | SKA3 | spindle and | 10.12745 | 10.36054 | 10.93312 |
| 6091 | ENSG0000 | DTD2 | D-tyrosyl-t | 7.422667 | 10.37755 | 10.90559 |
| 1475 | ENSG0000 | PTPRH | protein tyr | 5.274958 | 6.146285 | 10.88898 |
| 9898 | ENSG0000 | SSBP3 | single strar | 8.631653 | 8.706106 | 10.86377 |
| 11153 | ENSG0000 | PAPD4 | poly(A) RN | 6.89372 | 8.929795 | 10.85158 |
| 7760 | ENSG0000 | VPS37B | VPS37B, ES | 9.911656 | 10.10935 | 10.82546 |
| 8618 | ENSG0000 | SRD5A1 | steroid 5 a | 6.771258 | 8.911377 | 10.823 |
| 7272 | ENSG0000 | ZNF189 | zinc finger | 5.191779 | 9.172333 | 10.81927 |
| 6161 | ENSG0000 | APOC1 | apolipopro | 3.506525 | 4.777256 | 10.81448 |
| 25512 | ENSG0000 | TLR8-AS1 | TLR8 antise | 2.910511 | 9.697547 | 10.80198 |
| 3011 | ENSG0000 | AMBRA1 | autophagy | 9.625717 | 9.682494 | 10.79824 |
| 5368 | ENSG0000 | LPGAT1 | lysophospl | 8.419938 | 10.15748 | 10.77707 |
| 32743 | ENSG0000 | LBX2-AS1 | LBX2 antise | 7.154773 | 9.895151 | 10.7726 |
| 1346 | ENSG0000 | ANKRD13A | ankyrin rep | 3.962748 | 9.017007 | 10.76942 |
| 241 | ENSG0000 | VTA1 | vesicle traf | 8.753352 | 10.73107 | 10.76873 |
| 11662 | ENSG0000 | TK2 | thymidine | 4.845594 | 8.351445 | 10.76566 |
| 6958 | ENSG0000 | DNAJC14 | DnaJ heat s | 5.398297 | 6.332245 | 10.74064 |
| 15034 | ENSG0000 | LTN1 | listerin E3 ( | 9.233248 | 9.619681 | 10.72092 |
| 4467 | ENSG0000 | TRAK2 | trafficking | 7.013664 | 9.550182 | 10.71387 |
| 12624 | ENSG0000 | PPM1D | protein ph( | 6.692797 | 10.41238 | 10.71252 |
| 3765 | ENSG0000 | HPS5 | HPS5, biog | 5.9728 | 8.874859 | 10.67531 |
| 19211 | ENSG0000 | STK38L | serine/thre | 6.090334 | 10.2252 | 10.65665 |
| 10703 | ENSG0000 | CFAP36 | cilia and fla | 6.752285 | 8.698373 | 10.64186 |
| 29944 | ENSG0000 | HAUS5 | HAUS augr | 8.719571 | 8.988023 | 10.62522 |
| 5457 | ENSG0000 | VAMP7 | vesicle ass( | 8.38623 | 9.837592 | 10.61432 |

| | | | | | | |
|---|---|---|---|---|---|---|
| 258 | ENSG0000 | RTEL1-TNF | RTEL1-TNF | 6.741312 | 7.129451 | 10.5944 |
| 8594 | ENSG0000 | CISD2 | CDGSH iro | 7.640415 | 9.131897 | 10.59313 |
| 18112 | ENSG0000 | OXLD1 | oxidoredu | 9.123789 | 10.55095 | 10.59004 |
| 1244 | ENSG0000 | FERMT2 | fermitin fa | 6.888574 | 8.582682 | 10.58082 |
| 3870 | ENSG0000 | UHRF1BP1 | UHRF1 bin | 8.794568 | 10.45283 | 10.56777 |
| 13349 | ENSG0000 | SLC16A13 | solute carr | 2.101161 | 5.776899 | 10.56593 |
| 43113 | ENSG0000 | AC069544.2 | | 6.706504 | 8.836124 | 10.53586 |
| 9446 | ENSG0000 | IFIT5 | interferon | 7.227143 | 7.682719 | 10.52891 |
| 4075 | ENSG0000 | PTK7 | protein tyr | 3.183666 | 3.952414 | 10.52762 |
| 9250 | ENSG0000 | CSNK1G3 | casein kina | 8.011688 | 8.929303 | 10.5015 |
| 10859 | ENSG0000 | RYBP | RING1 and | 4.207256 | 9.888249 | 10.48345 |
| 12260 | ENSG0000 | MAP2K1 | mitogen-ac | 8.242773 | 9.335811 | 10.47254 |
| 2416 | ENSG0000 | GMPR2 | guanosine | 6.718521 | 8.114996 | 10.47201 |
| 5609 | ENSG0000 | POLR1B | RNA polym | 7.719382 | 7.723593 | 10.45875 |
| 293 | ENSG0000 | NISCH | nischarin [S | 5.897964 | 7.161844 | 10.45226 |
| 7213 | ENSG0000 | MARCH7 | membrane | 8.806695 | 9.883207 | 10.45167 |
| 21935 | ENSG0000 | SLC9A3-AS | SLC9A3 ant | 3.839228 | 9.082634 | 10.43305 |
| 3591 | ENSG0000 | WSB1 | WD repeat | 4.902952 | 7.639519 | 10.40829 |
| 22661 | ENSG0000 | PARG | poly(ADP-r | 8.463084 | 10.07777 | 10.40228 |
| 13149 | ENSG0000 | MTX1 | metaxin 1 | 9.360254 | 9.7396 | 10.39685 |
| 12685 | ENSG0000 | TRMT61B | tRNA meth | 8.328764 | 9.899275 | 10.39361 |
| 1829 | ENSG0000 | RBBP9 | RB binding | 5.005065 | 6.757074 | 10.38951 |
| 3699 | ENSG0000 | ATG2A | autophagy | 6.07498 | 10.3016 | 10.38834 |
| 4101 | ENSG0000 | ERBIN | erbb2 inter | 6.456258 | 9.738376 | 10.37276 |
| 10721 | ENSG0000 | SGCB | sarcoglycar | 6.171065 | 8.50143 | 10.36842 |
| 8489 | ENSG0000 | RETREG2 | reticulopha | 5.242968 | 8.059327 | 10.35372 |
| 12728 | ENSG0000 | DNAJC30 | DnaJ heat s | 9.927337 | 10.23673 | 10.34441 |
| 9022 | ENSG0000 | BTBD10 | BTB domai | 8.061756 | 9.845908 | 10.32663 |
| 10986 | ENSG0000 | DTX3L | deltex E3 u | 7.210468 | 9.967422 | 10.32629 |
| 15725 | ENSG0000 | ARAP1 | ArfGAP wit | 6.429316 | 7.505683 | 10.30976 |
| 1258 | ENSG0000 | CLASP1 | cytoplasmi | 8.805942 | 9.776405 | 10.30808 |
| 14569 | ENSG0000 | FRAT2 | FRAT2, WN | 6.005908 | 8.746942 | 10.29017 |
| 4966 | ENSG0000 | MXI1 | MAX intera | 2.043981 | 5.45073 | 10.28193 |
| 3543 | ENSG0000 | LGALS3BP | galectin 3 b | 3.195021 | 3.850873 | 10.27314 |
| 15404 | ENSG0000 | TNFAIP2 | TNF alpha | 2.632945 | 3.375618 | 10.25778 |
| 5345 | ENSG0000 | STIL | STIL, centri | 7.198023 | 7.568341 | 10.25669 |
| 16476 | ENSG0000 | HDAC2 | histone de | 9.247003 | 9.864349 | 10.23827 |
| 39229 | ENSG0000 | PAXIP1-AS | PAXIP1 ant | 5.556016 | 7.873027 | 10.23717 |
| 4700 | ENSG0000 | STX12 | syntaxin 12 | 6.060759 | 8.424797 | 10.22242 |
| 9463 | ENSG0000 | RAD17 | RAD17 che | 7.194197 | 9.042573 | 10.20949 |
| 13129 | ENSG0000 | FAM222B | family with | 7.733349 | 8.807336 | 10.20739 |
| 12054 | ENSG0000 | LTBP3 | latent tran | 7.995363 | 9.472426 | 10.18942 |
| 4621 | ENSG0000 | KDM5B | lysine dem | 5.821271 | 9.748357 | 10.18649 |
| 12063 | ENSG0000 | SCARA3 | scavenger | 4.265011 | 5.177312 | 10.17188 |
| 15975 | ENSG0000 | PMS2CL | PMS2 C-ter | 5.386535 | 8.957856 | 10.16972 |
| 7846 | ENSG0000 | DNAJA4 | DnaJ heat s | 5.515487 | 8.322875 | 10.16948 |
| 6311 | ENSG0000 | AKAP12 | A-kinase ar | 2.998846 | 5.830657 | 10.16883 |

| | | | | | | |
|---|---|---|---|---|---|---|
| 651 | ENSG0000 | USP28 | ubiquitin s | 7.051723 | 7.37135 | 10.16669 |
| 9531 | ENSG0000 | GTF2E1 | general tra | 8.458506 | 8.781056 | 10.15516 |
| 4405 | ENSG0000 | KDM3A | lysine dem | 4.208047 | 6.653771 | 10.14388 |
| 8326 | ENSG0000 | ATP8B2 | ATPase pho | 5.223068 | 5.83277 | 10.14038 |
| 3332 | ENSG0000 | TBL2 | transducin | 6.242486 | 8.356371 | 10.14009 |
| 17017 | ENSG0000 | ZNF627 | zinc finger | 1.863334 | 8.653059 | 10.12051 |
| 11179 | ENSG0000 | MB21D1 | Mab-21 do | 5.41331 | 6.455486 | 10.10857 |
| 29399 | ENSG0000 | CEBPA | CCAAT/enh | 5.26326 | 7.911145 | 10.10736 |
| 11177 | ENSG0000 | SLC35A1 | solute carr | 7.523165 | 8.364366 | 10.1066 |
| 2513 | ENSG0000 | KIF3B | kinesin fam | 8.845333 | 9.287277 | 10.09415 |
| 4936 | ENSG0000 | SUPT7L | SPT7 like, S | 7.235394 | 9.410591 | 10.09131 |
| 2255 | ENSG0000 | THOC5 | THO comp | 7.471065 | 9.20619 | 10.08214 |
| 28774 | ENSG0000 | RTL5 | retrotransp | 2.345513 | 8.534246 | 10.06813 |
| 16944 | ENSG0000 | STYX | serine/thre | 7.511964 | 9.835477 | 10.05852 |
| 6572 | ENSG0000 | RBM38 | RNA bindir | 9.465024 | 9.554054 | 10.05703 |
| 11570 | ENSG0000 | DPCD | deleted in | 6.317022 | 6.517124 | 10.04996 |
| 36938 | ENSG0000 | LINC01842 | long interg | 6.268668 | 9.208255 | 10.04259 |
| 2347 | ENSG0000 | TIMM9 | translocase | 8.559596 | 8.684245 | 10.03285 |
| 11821 | ENSG0000 | TBC1D16 | TBC1 doma | 7.682012 | 9.013677 | 10.03133 |
| 4582 | ENSG0000 | TMEM9 | transmeml | 6.360106 | 9.102725 | 10.02235 |
| 15748 | ENSG0000 | SPIN4 | spindlin far | 6.39732 | 8.319771 | 10.01575 |
| 2933 | ENSG0000 | MAN2B1 | mannosida | 8.736899 | 8.849204 | 10.01493 |
| 6215 | ENSG0000 | TRPM4 | transient r | 2.024487 | 8.136501 | 10.01003 |
| 10651 | ENSG0000 | BPNT1 | 3'(2'), 5'-bi | 5.325034 | 7.166364 | 10.00424 |
| 7714 | ENSG0000 | SDSL | serine deh | 3.459311 | 7.146671 | 9.997953 |
| 3757 | ENSG0000 | PITPNM1 | phosphatic | 8.104795 | 9.056382 | 9.99072 |
| 7529 | ENSG0000 | DUSP5 | dual specif | 6.213051 | 9.175471 | 9.983439 |
| 1998 | ENSG0000 | RFFL | ring finger | 9.159621 | 9.770936 | 9.90987 |
| 472 | ENSG0000 | UBR2 | ubiquitin p | 7.1729 | 9.55507 | 9.899183 |
| 19986 | ENSG0000 | NAXD | NAD(P)HX | 6.658355 | 9.3635 | 9.892238 |
| 6939 | ENSG0000 | SNX14 | sorting nex | 7.516475 | 8.418375 | 9.883401 |
| 3603 | ENSG0000 | PMP22 | peripheral | 4.491963 | 6.645351 | 9.813476 |
| 7264 | ENSG0000 | STXBP1 | syntaxin bi | 4.814708 | 8.034357 | 9.811541 |
| 2601 | ENSG0000 | ABCD1 | ATP bindin | 5.948363 | 6.480035 | 9.809453 |
| 5761 | ENSG0000 | IFI6 | interferon | 2.187655 | 2.709357 | 9.805777 |
| 4877 | ENSG0000 | LRP11 | LDL recept | 8.064172 | 8.140259 | 9.805068 |
| 16528 | ENSG0000 | STRN3 | striatin 3 [S | 7.975145 | 9.048596 | 9.804082 |
| 1108 | ENSG0000 | PTPN3 | protein tyr | 6.13173 | 8.560033 | 9.801028 |
| 7315 | ENSG0000 | RNF38 | ring finger | 5.991349 | 9.569078 | 9.80074 |
| 14429 | ENSG0000 | ZNF609 | zinc finger | 7.838599 | 9.03525 | 9.794261 |
| 1008 | ENSG0000 | PHKA1 | phosphory | 7.689121 | 8.736798 | 9.790845 |
| 72 | ENSG0000 | DVL2 | dishevellec | 6.996633 | 8.435539 | 9.77893 |
| 2310 | ENSG0000 | CERK | ceramide k | 8.671257 | 8.870891 | 9.771712 |
| 15027 | ENSG0000 | ZNF770 | zinc finger | 7.669081 | 9.649035 | 9.756928 |
| 20384 | ENSG0000 | DHRS4-AS1 | DHRS4 ant | 3.324461 | 4.932276 | 9.746599 |
| 2807 | ENSG0000 | CDIPT | CDP-diacyl | 7.406064 | 7.822729 | 9.739468 |
| 13279 | ENSG0000 | TIGD3 | tigger tran | 2.72466 | 8.782513 | 9.737307 |

| | | | | | | |
|---|---|---|---|---|---|---|
| 3776 | ENSG0000 | CLEC2B | C-type lect | 4.233455 | 4.955702 | 9.709683 |
| 12121 | ENSG0000 | TAP1 | transporte | 4.932071 | 8.452757 | 9.707732 |
| 3606 | ENSG0000 | ALDOC | aldolase, fr | 0.991911 | 1.294981 | 9.7058 |
| 207 | ENSG0000 | CYTH3 | cytohesin 3 | 6.203969 | 7.835648 | 9.697813 |
| 9665 | ENSG0000 | VOPP1 | vesicular, c | 9.238535 | 9.579478 | 9.679804 |
| 14253 | ENSG0000 | MEIS3P1 | Meis home | 3.0834 | 6.862707 | 9.675029 |
| 2073 | ENSG0000 | DSP | desmoplak | 3.406249 | 7.151399 | 9.672169 |
| 12186 | ENSG0000 | PKIG | cAMP-depe | 5.770399 | 6.712565 | 9.648333 |
| 3582 | ENSG0000 | EFNB3 | ephrin B3 [ | 2.103012 | 4.580558 | 9.648285 |
| 18108 | ENSG0000 | PFDN6 | prefoldin s | 8.833327 | 9.269731 | 9.644473 |
| 3532 | ENSG0000 | AKAP10 | A-kinase ar | 7.416903 | 8.261195 | 9.637761 |
| 34847 | ENSG0000 | AL049796.1 | | 2.441126 | 7.396166 | 9.6198 |
| 13625 | ENSG0000 | ARL4D | ADP ribosy | 2.396619 | 3.78805 | 9.61237 |
| 10337 | ENSG0000 | BRSK1 | BR serine/t | 2.409497 | 4.060086 | 9.60777 |
| 24352 | ENSG0000 | AC018638.2 | | 4.450418 | 8.134148 | 9.60717 |
| 643 | ENSG0000 | C12orf4 | chromosor | 7.631278 | 9.103512 | 9.601601 |
| 5814 | ENSG0000 | ABHD8 | abhydrolas | 3.650716 | 4.021607 | 9.591075 |
| 28645 | ENSG0000 | ARFGAP3 | ADP ribosy | 7.86945 | 8.285766 | 9.58986 |
| 295 | ENSG0000 | FUZ | fuzzy plana | 5.194126 | 5.615859 | 9.582358 |
| 28059 | ENSG0000 | C1orf226 | chromosor | 5.711784 | 6.622221 | 9.580117 |
| 6455 | ENSG0000 | TRIM21 | tripartite n | 4.990563 | 9.245935 | 9.575765 |
| 5260 | ENSG0000 | ACO1 | aconitase 1 | 8.177674 | 8.400286 | 9.575401 |
| 11525 | ENSG0000 | PTER | phosphotri | 7.772074 | 8.748913 | 9.574965 |
| 10133 | ENSG0000 | TOMM40L | translocase | 3.030077 | 7.656914 | 9.565935 |
| 11743 | ENSG0000 | C15orf48 | chromosor | 6.355849 | 7.995489 | 9.562834 |
| 10911 | ENSG0000 | WBP1L | WW doma | 6.278104 | 8.031875 | 9.551668 |
| 16686 | ENSG0000 | UAP1L1 | UDP-N-ace | 3.711751 | 4.461107 | 9.546638 |
| 5555 | ENSG0000 | DOK4 | docking pr | 1.898527 | 5.149701 | 9.544852 |
| 10213 | ENSG0000 | RSPRY1 | ring finger | 7.691131 | 7.937468 | 9.533153 |
| 14808 | ENSG0000 | CMC4 | C-X9-C mo | 6.917921 | 7.004792 | 9.522962 |
| 2325 | ENSG0000 | COCH | cochlin [So | 3.582102 | 4.723621 | 9.516658 |
| 4718 | ENSG0000 | IFT46 | intraflagell | 4.518417 | 8.790957 | 9.516592 |
| 2573 | ENSG0000 | RIOK3 | RIO kinase | 4.265215 | 8.32375 | 9.500099 |
| 12157 | ENSG0000 | TRAPPC11 | trafficking | 7.362923 | 8.828142 | 9.471789 |
| 4985 | ENSG0000 | HOXB5 | homeobox | 6.465029 | 6.637121 | 9.470827 |
| 5571 | ENSG0000 | SEPT6 | septin 6 [S | 3.576795 | 3.669149 | 9.459278 |
| 7992 | ENSG0000 | SAT2 | spermidine | 4.74335 | 6.827577 | 9.456142 |
| 1387 | ENSG0000 | GPR137B | G protein-c | 4.23777 | 6.200252 | 9.45524 |
| 6570 | ENSG0000 | LPIN3 | lipin 3 [Sou | 3.334415 | 8.149063 | 9.449706 |
| 9940 | ENSG0000 | HIPK1 | homeodon | 8.22584 | 9.022538 | 9.424401 |
| 7941 | ENSG0000 | PRPSAP2 | phosphorik | 7.689561 | 8.310334 | 9.418368 |
| 7980 | ENSG0000 | OSBPL1A | oxysterol b | 6.544648 | 7.347074 | 9.39642 |
| 11251 | ENSG0000 | MED30 | mediator c | 6.240828 | 6.521365 | 9.395729 |
| 251 | ENSG0000 | STARD3NL | STARD3 N- | 6.946821 | 8.488713 | 9.385159 |
| 6209 | ENSG0000 | GDF15 | growth diff | 1.388483 | 3.028072 | 9.38119 |
| 3574 | ENSG0000 | MPP2 | membrane | 3.954914 | 4.356319 | 9.37309 |
| 11291 | ENSG0000 | FZD6 | frizzled cla | 5.842719 | 8.004747 | 9.364389 |

| | | | | | | |
|---|---|---|---|---|---|---|
| 9318 | ENSG0000 | SLC25A4 | solute carr | 6.500365 | 8.449727 | 9.358871 |
| 2445 | ENSG0000 | STK4 | serine/thre | 2.446191 | 2.962096 | 9.354634 |
| 14996 | ENSG0000 | SETD3 | SET domai | 7.169242 | 8.081309 | 9.346283 |
| 8157 | ENSG0000 | PDE6D | phosphodi | 7.297433 | 7.735908 | 9.33594 |
| 13404 | ENSG0000 | SLC29A2 | solute carr | 4.717982 | 7.555063 | 9.335037 |
| 5668 | ENSG0000 | MGME1 | mitochond | 7.944952 | 8.464737 | 9.334275 |
| 5028 | ENSG0000 | NUDCD1 | NudC dom | 7.374426 | 9.264629 | 9.320553 |
| 15400 | ENSG0000 | NRBP2 | nuclear rec | 3.453142 | 7.731689 | 9.319487 |
| 20052 | ENSG0000 | SH3D21 | SH3 domai | 2.166315 | 7.628705 | 9.318797 |
| 10714 | ENSG0000 | CCDC74A | coiled-coil | 5.537688 | 6.648407 | 9.315414 |
| 7680 | ENSG0000 | CLSTN3 | calsytenir | 4.355681 | 8.418525 | 9.310447 |
| 14175 | ENSG0000 | APOLD1 | apolipopro | 4.296297 | 7.288227 | 9.307193 |
| 11214 | ENSG0000 | RP9 | retinitis pig | 6.882536 | 9.302774 | 9.305177 |
| 16191 | ENSG0000 | FAM120AC | family with | 5.424657 | 9.089389 | 9.303445 |
| 11166 | ENSG0000 | FOXQ1 | forkhead b | 2.533521 | 4.86443 | 9.286798 |
| 3258 | ENSG0000 | FKBP14 | FK506 binc | 5.318761 | 7.429188 | 9.285743 |
| 6269 | ENSG0000 | SESN2 | sestrin 2 [S | 8.298199 | 8.912402 | 9.266587 |
| 190 | ENSG0000 | TEAD3 | TEA domai | 8.358103 | 8.809318 | 9.254284 |
| 29024 | ENSG0000 | AC018638.5 | | 4.538082 | 8.532215 | 9.248397 |
| 42835 | ENSG0000 | AC116366.3 | | 0.034119 | 0.082712 | 9.237527 |
| 7912 | ENSG0000 | NOL3 | nucleolar p | 3.618987 | 5.091813 | 9.236903 |
| 13976 | ENSG0000 | AGTRAP | angiotensir | 6.756547 | 6.99214 | 9.230241 |
| 15537 | ENSG0000 | WDR53 | WD repeat | 8.36279 | 8.818419 | 9.226669 |
| 10896 | ENSG0000 | SHH | sonic hedg | 2.826166 | 3.733768 | 9.208631 |
| 2741 | ENSG0000 | WDR59 | WD repeat | 6.814364 | 7.889378 | 9.201445 |
| 531 | ENSG0000 | UBA6 | ubiquitin li | 6.294263 | 8.617846 | 9.195032 |
| 2683 | ENSG0000 | MRPS31 | mitochond | 7.721421 | 8.736646 | 9.18899 |
| 9848 | ENSG0000 | BAG4 | BCL2 assoc | 6.671511 | 7.254072 | 9.179897 |
| 8011 | ENSG0000 | TRIM65 | tripartite n | 7.986624 | 8.434704 | 9.179669 |
| 15194 | ENSG0000 | C22orf46 | chromosor | 4.301715 | 8.456473 | 9.161675 |
| 12081 | ENSG0000 | RBPJ | recombina | 5.26837 | 7.08632 | 9.155184 |
| 991 | ENSG0000 | MSANTD3 | Myb/SANT | 7.053457 | 9.022909 | 9.149761 |
| 7191 | ENSG0000 | CALCOCO2 | calcium bir | 7.333631 | 8.307713 | 9.143364 |
| 8013 | ENSG0000 | SECTM1 | secreted ai | 2.533221 | 3.55458 | 9.140686 |
| 4975 | ENSG0000 | C10orf76 | chromosor | 5.610324 | 7.965425 | 9.133153 |
| 6153 | ENSG0000 | DOCK6 | dedicator c | 2.924407 | 8.133477 | 9.128782 |
| 6398 | ENSG0000 | NINJ1 | ninjurin 1 [ | 5.687673 | 6.020209 | 9.119246 |
| 3383 | ENSG0000 | RGP1 | RGP1 hom | 7.348403 | 8.295633 | 9.117235 |
| 5404 | ENSG0000 | SNX21 | sorting nex | 6.709959 | 7.738535 | 9.112753 |
| 4605 | ENSG0000 | RLF | rearrangec | 4.861811 | 7.211699 | 9.083658 |
| 14224 | ENSG0000 | SPTY2D1 | SPT2 chror | 7.528662 | 8.901684 | 9.08042 |
| 929 | ENSG0000 | NTN1 | netrin 1 [S | 5.471875 | 6.523691 | 9.075666 |
| 28539 | ENSG0000 | TMEFF1 | transmeml | 1.378298 | 3.940423 | 9.072471 |
| 12191 | ENSG0000 | SEMA4C | semaphori | 6.817834 | 7.971548 | 9.067226 |
| 9141 | ENSG0000 | HIRIP3 | HIRA intera | 8.064893 | 8.270707 | 9.067172 |
| 3409 | ENSG0000 | HPS1 | HPS1, biog | 6.428459 | 8.826394 | 9.056715 |
| 19675 | ENSG0000 | KLHL23 | kelch like f | 6.459209 | 7.919532 | 9.056522 |

| | | | | | | |
|---|---|---|---|---|---|---|
| 2428 | ENSG0000 | PROCR | protein C r | 2.16052 | 2.966295 | 9.053453 |
| 7399 | ENSG0000 | PRCP | prolylcarbo | 3.723274 | 6.443313 | 9.051928 |
| 5889 | ENSG0000 | ZFP36 | ZFP36 ring | 5.181104 | 5.483575 | 9.044331 |
| 960 | ENSG0000 | SLC9A7 | solute carr | 4.744702 | 6.394134 | 9.041106 |
| 1287 | ENSG0000 | ACTR6 | ARP6 actin | 6.331342 | 7.418479 | 9.038192 |
| 7538 | ENSG0000 | DBR1 | debranchir | 7.900855 | 8.846853 | 9.032481 |
| 20094 | ENSG0000 | SDHAP2 | succinate d | 5.62677 | 7.677498 | 9.019538 |
| 4952 | ENSG0000 | SLC17A5 | solute carr | 6.007925 | 7.569823 | 9.014658 |
| 24276 | ENSG0000 | AC021078.1 | | 7.583615 | 8.32694 | 9.002895 |
| 34417 | ENSG0000 | RAB43P1 | RAB43 pse | 5.594397 | 8.801853 | 8.996743 |
| 2669 | ENSG0000 | FNDC3A | fibronectin | 6.922241 | 8.55957 | 8.991594 |
| 1971 | ENSG0000 | RNF31 | ring finger | 5.757417 | 7.606222 | 8.981428 |
| 7081 | ENSG0000 | C2orf49 | chromosor | 6.250529 | 7.597075 | 8.973215 |
| 6910 | ENSG0000 | MSI1 | musashi RI | 3.199098 | 3.272901 | 8.955207 |
| 24599 | ENSG0000 | AC125232.1 | | 5.450231 | 8.58245 | 8.950496 |
| 14039 | ENSG0000 | MLF1 | myeloid le | 3.498164 | 5.931139 | 8.944128 |
| 20110 | ENSG0000 | CROCCP2 | ciliary root | 4.328248 | 7.413831 | 8.944111 |
| 7905 | ENSG0000 | KIFC3 | kinesin fan | 4.691226 | 7.962159 | 8.939281 |
| 4241 | ENSG0000 | ACAP2 | ArfGAP wit | 7.521954 | 8.410915 | 8.933911 |
| 7671 | ENSG0000 | AEBP2 | AE binding | 8.194563 | 8.40027 | 8.927665 |
| 3993 | ENSG0000 | SMAP1 | small ArfG | 7.499876 | 7.536887 | 8.925882 |
| 4789 | ENSG0000 | RAB3GAP2 | RAB3 GTPa | 6.212937 | 8.600163 | 8.924887 |
| 6288 | ENSG0000 | DUSP9 | dual specif | 3.747689 | 4.620859 | 8.922235 |
| 5478 | ENSG0000 | USP9X | ubiquitin s | 5.951883 | 8.706931 | 8.9164 |
| 8988 | ENSG0000 | POLR3A | RNA polym | 7.804536 | 8.465657 | 8.916121 |
| 5095 | ENSG0000 | TBX2 | T-box 2 [Sc | 5.828002 | 7.887719 | 8.909834 |
| 2761 | ENSG0000 | ABCC1 | ATP bindin | 7.68102 | 7.933869 | 8.909738 |
| 6440 | ENSG0000 | ACTR10 | actin relate | 7.686218 | 8.189929 | 8.901624 |
| 16660 | ENSG0000 | KANK2 | KN motif a | 7.269414 | 8.051383 | 8.899896 |
| 3733 | ENSG0000 | BIRC2 | baculoviral | 5.005583 | 8.871326 | 8.890189 |
| 3510 | ENSG0000 | GALK1 | galactokina | 5.364459 | 6.052168 | 8.88564 |
| 8634 | ENSG0000 | IQGAP2 | IQ motif cc | 4.126262 | 5.923802 | 8.876525 |
| 1792 | ENSG0000 | EDEM2 | ER degrada | 5.940554 | 6.523455 | 8.86659 |
| 14755 | ENSG0000 | TRAPPC6B | trafficking | 4.268584 | 8.019595 | 8.858407 |
| 3739 | ENSG0000 | HIPK3 | homeodon | 6.352976 | 8.708424 | 8.84573 |
| 23477 | ENSG0000 | AP001065.1 | | 4.592177 | 8.637036 | 8.844477 |
| 4458 | ENSG0000 | ORC2 | origin reco | 7.289164 | 7.369087 | 8.838612 |
| 4322 | ENSG0000 | PIKFYVE | phosphoin | 6.754899 | 8.570374 | 8.834688 |
| 493 | ENSG0000 | IFNGR1 | interferon | 6.135966 | 7.788921 | 8.821421 |
| 12202 | ENSG0000 | CHTF8 | chromosor | 7.150204 | 7.497096 | 8.808471 |
| 10414 | ENSG0000 | YDJC | YdjC chitoc | 8.47427 | 8.569278 | 8.798433 |
| 11713 | ENSG0000 | TMEM170A | transmeml | 5.612721 | 8.561032 | 8.792927 |
| 14814 | ENSG0000 | PAQR7 | progestin a | 4.941328 | 6.498063 | 8.785751 |
| 41470 | ENSG0000 | AC005332.9 | | 4.971638 | 6.248984 | 8.784807 |
| 15095 | ENSG0000 | TBK1 | TANK bindi | 7.403386 | 7.805383 | 8.762717 |
| 8174 | ENSG0000 | ARHGEF19 | Rho guanir | 6.779425 | 7.80372 | 8.746552 |
| 7587 | ENSG0000 | TMOD3 | tropomodu | 6.07439 | 7.645663 | 8.739421 |

| | | | | | | |
|---|---|---|---|---|---|---|
| 18235 | ENSG0000 | GNL1 | G protein r | 6.187816 | 6.22887 | 8.739201 |
| 26292 | ENSG0000 | LINC00094 | long interg | 3.930031 | 5.194365 | 8.738928 |
| 12334 | ENSG0000 | SNUPN | snurportin | 5.966396 | 6.661754 | 8.736978 |
| 5003 | ENSG0000 | WDR55 | WD repeat | 7.678188 | 8.402379 | 8.73483 |
| 5773 | ENSG0000 | SIX1 | SIX homeo | 6.253349 | 8.194837 | 8.731961 |
| 4287 | ENSG0000 | COMMD2 | COMM do | 7.448979 | 7.554519 | 8.725688 |
| 9478 | ENSG0000 | CARHSP1 | calcium re | 5.654731 | 6.275851 | 8.714581 |
| 3400 | ENSG0000 | ABHD17B | abhydrolas | 7.794972 | 8.089594 | 8.709498 |
| 4369 | ENSG0000 | HTRA2 | HtrA serine | 6.913306 | 7.474131 | 8.709031 |
| 14629 | ENSG0000 | GPR3 | G protein-c | 5.898863 | 6.211634 | 8.695515 |
| 18275 | ENSG0000 | TRIM27 | tripartite n | 7.639184 | 7.87564 | 8.693686 |
| 5628 | ENSG0000 | RTN2 | reticulon 2 | 3.93932 | 5.619244 | 8.692359 |
| 6741 | ENSG0000 | EDEM1 | ER degrada | 5.95002 | 7.320784 | 8.675655 |
| 11529 | ENSG0000 | ARL5B | ADP ribosy | 5.032116 | 7.168485 | 8.667933 |
| 712 | ENSG0000 | SIKE1 | suppressor | 7.348419 | 7.863147 | 8.665168 |
| 8643 | ENSG0000 | SLC30A5 | solute carr | 8.187923 | 8.620448 | 8.662622 |
| 16970 | ENSG0000 | GFPT1 | glutamine- | 5.507264 | 7.432574 | 8.640685 |
| 11935 | ENSG0000 | ZNF526 | zinc finger | 7.090389 | 7.947323 | 8.639717 |
| 6735 | ENSG0000 | IRAK2 | interleukin | 2.574082 | 6.871431 | 8.627723 |
| 5834 | ENSG0000 | FGFRL1 | fibroblast g | 7.653347 | 8.263435 | 8.625016 |
| 9876 | ENSG0000 | NTAN1 | N-terminal | 4.603859 | 6.759912 | 8.621806 |
| 8584 | ENSG0000 | SCD5 | stearoyl-Co | 4.479572 | 5.861798 | 8.618853 |
| 10159 | ENSG0000 | HOXB13 | homeobox | 4.174166 | 5.681839 | 8.613832 |
| 314 | ENSG0000 | TSPAN9 | tetraspanin | 5.799752 | 6.011273 | 8.602766 |
| 6823 | ENSG0000 | DDB2 | damage sp | 4.053015 | 5.801193 | 8.598125 |
| 37265 | ENSG0000 | FDX2 | ferredoxin | 5.655268 | 6.251164 | 8.58166 |
| 1034 | ENSG0000 | PDK3 | pyruvate d | 3.379419 | 3.568823 | 8.578371 |
| 4950 | ENSG0000 | CRIPT | CXXC repea | 4.257588 | 6.702585 | 8.549709 |
| 10012 | ENSG0000 | PEX10 | peroxisom | 4.38503 | 4.808087 | 8.546128 |
| 6251 | ENSG0000 | POMT1 | protein O-r | 7.594011 | 7.850351 | 8.539211 |
| 26625 | ENSG0000 | LINC00941 | long interg | 4.216598 | 6.545744 | 8.536028 |
| 15777 | ENSG0000 | RTN4RL2 | reticulon 4 | 2.651048 | 3.701807 | 8.533799 |
| 4841 | ENSG0000 | HDHD3 | haloacid de | 3.246918 | 5.618327 | 8.521684 |
| 8615 | ENSG0000 | NKD2 | naked cuti | 5.095298 | 5.638684 | 8.512317 |
| 7339 | ENSG0000 | PIM1 | Pim-1 prot | 5.288432 | 6.22348 | 8.504789 |
| 591 | ENSG0000 | PNKP | polynucleo | 6.111526 | 8.274641 | 8.503717 |
| 9900 | ENSG0000 | CLDN12 | claudin 12 | 2.930518 | 7.609793 | 8.502013 |
| 39857 | ENSG0000 | AC244154.1 | | 6.239771 | 8.092919 | 8.496682 |
| 15247 | ENSG0000 | KNTC1 | kinetochor | 6.19166 | 6.604041 | 8.496195 |
| 5051 | ENSG0000 | ALG5 | ALG5, dolic | 6.767893 | 7.484019 | 8.493449 |
| 10628 | ENSG0000 | IGSF8 | immunoglo | 3.631262 | 3.999587 | 8.493161 |
| 8954 | ENSG0000 | SLC25A25 | solute carr | 3.831879 | 6.882101 | 8.484997 |
| 4592 | ENSG0000 | TSNAX | translin ass | 6.854461 | 8.0152 | 8.481509 |
| 2832 | ENSG0000 | RAB11A | RAB11A, m | 6.031987 | 6.900486 | 8.480963 |
| 8012 | ENSG0000 | CBX8 | chromobox | 5.61323 | 7.085612 | 8.470822 |
| 11055 | ENSG0000 | SPRY1 | sprouty RT | 7.646237 | 7.877976 | 8.469113 |
| 2634 | ENSG0000 | JADE3 | jade family | 7.648775 | 7.957383 | 8.44547 |

| | | | | | | |
|---|---|---|---|---|---|---|
| 7331 | ENSG0000 | DENND4C | DENN dom | 5.170886 | 6.92186 | 8.445421 |
| 2920 | ENSG0000 | TNFRSF10A | TNF recept | 6.853843 | 8.236527 | 8.435205 |
| 5917 | ENSG0000 | TST | thiosulfate | 7.253381 | 7.545228 | 8.427309 |
| 13093 | ENSG0000 | CES2 | carboxyles | 5.098853 | 6.747247 | 8.423034 |
| 1649 | ENSG0000 | SSH1 | slingshot p | 6.795455 | 7.672271 | 8.421334 |
| 4905 | ENSG0000 | ATG4C | autophagy | 5.108215 | 5.712181 | 8.411053 |
| 7776 | ENSG0000 | CDH24 | cadherin 2 | 5.246178 | 6.770838 | 8.405018 |
| 32086 | ENSG0000 | WAC-AS1 | WAC antise | 6.410376 | 7.945242 | 8.402055 |
| 5278 | ENSG0000 | CISD1 | CDGSH iror | 7.961794 | 8.029151 | 8.401945 |
| 28102 | ENSG0000 | LINC00888 | long interg | 3.092054 | 5.310886 | 8.401878 |
| 13094 | ENSG0000 | PDP2 | pyruvate d | 6.448047 | 7.418256 | 8.399632 |
| 11834 | ENSG0000 | FN3K | fructosami | 4.564222 | 4.793027 | 8.376287 |
| 7219 | ENSG0000 | GATA4 | GATA bind | 6.392345 | 6.702738 | 8.37128 |
| 9059 | ENSG0000 | CAPN5 | calpain 5 [S | 1.196376 | 7.810506 | 8.369631 |
| 437 | ENSG0000 | MRE11 | MRE11 hor | 6.283561 | 8.135255 | 8.352422 |
| 12201 | ENSG0000 | ZBTB5 | zinc finger | 6.081816 | 7.716652 | 8.331095 |
| 13812 | ENSG0000 | PJA1 | praja ring f | 4.646341 | 7.66547 | 8.324002 |
| 4663 | ENSG0000 | DR1 | down-regu | 6.428577 | 7.995358 | 8.313155 |
| 17015 | ENSG0000 | ZNF511 | zinc finger | 7.480481 | 7.703455 | 8.306048 |
| 10362 | ENSG0000 | RUSC1 | RUN and S | 5.988692 | 7.298465 | 8.299061 |
| 8502 | ENSG0000 | POMGNT2 | protein O-l | 5.498256 | 5.598484 | 8.273846 |
| 38011 | ENSG0000 | AC127502.2 | | 5.323314 | 7.244302 | 8.268334 |
| 996 | ENSG0000 | MTFR1 | mitochond | 6.834913 | 7.575856 | 8.254957 |
| 5516 | ENSG0000 | CDKN1A | cyclin depe | 2.393704 | 4.00079 | 8.249094 |
| 9878 | ENSG0000 | ZFYVE9 | zinc finger | 7.165126 | 8.006547 | 8.242479 |
| 14107 | ENSG0000 | GLDC | glycine dec | 2.213417 | 3.819725 | 8.241401 |
| 16586 | ENSG0000 | ZNF138 | zinc finger | 4.851598 | 7.393812 | 8.23755 |
| 26972 | ENSG0000 | PITPNA-AS | PITPNA ant | 4.838494 | 6.409363 | 8.234639 |
| 4248 | ENSG0000 | TUSC2 | tumor supp | 7.430128 | 7.505203 | 8.233313 |
| 8903 | ENSG0000 | AK3 | adenylate k | 5.622209 | 6.445439 | 8.231741 |
| 4375 | ENSG0000 | CCDC88A | coiled-coil | 6.801944 | 7.693625 | 8.225506 |
| 869 | ENSG0000 | MED29 | mediator c | 5.274906 | 6.708396 | 8.22207 |
| 8114 | ENSG0000 | PRIM2 | primase (D | 6.340008 | 7.851651 | 8.212914 |
| 12599 | ENSG0000 | ZNF296 | zinc finger | 5.724443 | 7.625089 | 8.203341 |
| 8194 | ENSG0000 | WDTC1 | WD and te | 5.262577 | 6.052974 | 8.2004 |
| 36490 | ENSG0000 | SEC22B | SEC22 hom | 7.476756 | 7.794208 | 8.188178 |
| 13307 | ENSG0000 | CCS | copper cha | 5.916862 | 7.211326 | 8.174832 |
| 15005 | ENSG0000 | ATL3 | atlastin GT | 5.687135 | 7.869983 | 8.166604 |
| 8306 | ENSG0000 | ARNT | aryl hydroc | 6.17035 | 8.054885 | 8.12753 |
| 2916 | ENSG0000 | LEPROTL1 | leptin rece | 6.286173 | 6.451743 | 8.105813 |
| 10888 | ENSG0000 | F2RL2 | coagulatio | 5.251716 | 7.183073 | 8.102553 |
| 535 | ENSG0000 | SLC4A7 | solute carr | 4.785276 | 7.394998 | 8.102017 |
| 12942 | ENSG0000 | ORMDL3 | ORMDL spl | 6.962499 | 7.716963 | 8.090641 |
| 13565 | ENSG0000 | UCP2 | uncoupling | 2.561136 | 3.873637 | 8.090195 |
| 39739 | ENSG0000 | NATD1 | N-acetyltra | 2.094671 | 5.730394 | 8.085258 |
| 10605 | ENSG0000 | C1orf52 | chromosor | 6.255936 | 7.772455 | 8.072406 |
| 15823 | ENSG0000 | AKR1C1 | aldo-keto r | 2.88557 | 6.998866 | 8.065116 |

| | | | | | | |
|---|---|---|---|---|---|---|
| 6116 | ENSG0000 | ASH2L | ASH2 like h | 6.408367 | 6.502752 | 8.061157 |
| 8931 | ENSG0000 | STOM | stomatin [S | 1.128904 | 2.666171 | 8.057708 |
| 16619 | ENSG0000 | PGAP1 | post-GPI at | 5.076327 | 5.833055 | 8.054959 |
| 12307 | ENSG0000 | THBS3 | thrombosp | 5.44472 | 5.487468 | 8.04964 |
| 11318 | ENSG0000 | NFIL3 | nuclear fac | 3.668849 | 4.786488 | 8.048385 |
| 3992 | ENSG0000 | ACOT13 | acyl-CoA th | 5.489779 | 5.807362 | 8.044238 |
| 9132 | ENSG0000 | CDC42EP2 | CDC42 effe | 4.674513 | 7.688051 | 8.034898 |
| 2350 | ENSG0000 | TMED8 | transmemb | 4.648339 | 6.639045 | 8.032614 |
| 1863 | ENSG0000 | GMIP | GEM intera | 4.106713 | 7.158444 | 8.032026 |
| 2142 | ENSG0000 | MED15 | mediator c | 6.958351 | 7.799901 | 8.030706 |
| 4350 | ENSG0000 | FNDC4 | fibronectin | 2.006735 | 3.749081 | 8.017659 |
| 35626 | ENSG0000 | AC139530.1 | | 4.948109 | 7.282209 | 8.00173 |
| 5566 | ENSG0000 | C17orf53 | chromosor | 4.747859 | 6.588816 | 7.988777 |
| 741 | ENSG0000 | KIF1B | kinesin fam | 6.494773 | 7.436758 | 7.988007 |
| 10451 | ENSG0000 | FAM171A2 | family with | 4.336099 | 5.008533 | 7.97961 |
| 37431 | ENSG0000 | PTOV1-AS1 | PTOV1 ant | 4.586075 | 7.170576 | 7.978976 |
| 11484 | ENSG0000 | CASP7 | caspase 7 [ | 4.843895 | 6.993727 | 7.97846 |
| 9338 | ENSG0000 | CACUL1 | CDK2 asso | 4.790517 | 7.82776 | 7.975425 |
| 13412 | ENSG0000 | KIAA1551 | KIAA1551 [ | 6.669172 | 7.10204 | 7.967017 |
| 9213 | ENSG0000 | RILPL2 | Rab intera | 6.485378 | 6.885986 | 7.958186 |
| 1610 | ENSG0000 | ITGB5 | integrin su | 4.786352 | 6.392018 | 7.955204 |
| 7540 | ENSG0000 | GPR87 | G protein-c | 5.282571 | 7.665828 | 7.949146 |
| 40557 | ENSG0000 | FRG1HP | FSHD regio | 2.547291 | 5.075736 | 7.934777 |
| 16454 | ENSG0000 | AFAP1 | actin filam | 6.185572 | 7.45587 | 7.934013 |
| 9779 | ENSG0000 | DCK | deoxycytid | 4.529676 | 5.165219 | 7.924084 |
| 8308 | ENSG0000 | C1orf56 | chromosor | 5.197918 | 6.698884 | 7.915389 |
| 13239 | ENSG0000 | PC | pyruvate c | 7.050358 | 7.158829 | 7.911809 |
| 10003 | ENSG0000 | DPYSL5 | dihydropyr | 3.350126 | 3.931783 | 7.907395 |
| 1622 | ENSG0000 | ULK2 | unc-51 like | 4.060292 | 7.380523 | 7.894106 |
| 3511 | ENSG0000 | INTS2 | integrator | 6.124387 | 7.882761 | 7.889434 |
| 1764 | ENSG0000 | L2HGDH | L-2-hydrox | 6.055512 | 6.997716 | 7.888031 |
| 13548 | ENSG0000 | POLD4 | DNA polym | 4.219506 | 5.320348 | 7.882555 |
| 3970 | ENSG0000 | BAG2 | BCL2 assoc | 6.515546 | 6.641314 | 7.878876 |
| 12852 | ENSG0000 | PLEKHG5 | pleckstrin l | 5.369533 | 6.280157 | 7.875255 |
| 10117 | ENSG0000 | DEDD | death effe | 6.098301 | 6.550196 | 7.865304 |
| 11868 | ENSG0000 | TP53I13 | tumor prot | 5.981702 | 6.515859 | 7.861554 |
| 5093 | ENSG0000 | SCPEP1 | serine carb | 4.619863 | 5.096042 | 7.861305 |
| 7592 | ENSG0000 | INTS14 | integrator | 7.028272 | 7.560503 | 7.859527 |
| 5629 | ENSG0000 | EML2 | echinoderr | 3.132566 | 5.522412 | 7.856173 |
| 8407 | ENSG0000 | ETAA1 | ETAA1, ATI | 5.433322 | 7.524593 | 7.856032 |
| 621 | ENSG0000 | TRIT1 | tRNA isope | 3.50173 | 7.102087 | 7.855481 |
| 2542 | ENSG0000 | MAP1LC3A | microtubul | 2.97147 | 5.32457 | 7.848331 |
| 6388 | ENSG0000 | EXOC4 | exocyst co | 6.254736 | 7.379072 | 7.843998 |
| 12082 | ENSG0000 | LMBRD1 | LMBR1 dor | 4.888528 | 7.444816 | 7.843471 |
| 8362 | ENSG0000 | SNAP47 | synaptosor | 6.075351 | 6.775765 | 7.840552 |
| 10885 | ENSG0000 | TIPARP | TCDD indu | 3.365776 | 5.812578 | 7.838258 |
| 1995 | ENSG0000 | AGO1 | argonaute | 2.465589 | 6.885344 | 7.817778 |

| | | | | | | |
|---|---|---|---|---|---|---|
| 14018 | ENSG0000 | CENPBD1 | CENPB DN/ | 4.649169 | 7.574479 | 7.816252 |
| 6021 | ENSG0000 | OASL | 2'-5'-oligoa | 1.439653 | 2.090214 | 7.811185 |
| 6016 | ENSG0000 | LRP4 | LDL recept | 2.342631 | 4.428773 | 7.801484 |
| 11367 | ENSG0000 | GAPVD1 | GTPase act | 6.895575 | 7.416446 | 7.795225 |
| 22150 | ENSG0000 | AC012512.1 | | 3.198873 | 6.262276 | 7.791314 |
| 3451 | ENSG0000 | LARP4B | La ribonucl | 7.092931 | 7.675506 | 7.782121 |
| 2893 | ENSG0000 | NDRG1 | N-myc dow | 1.354134 | 3.578087 | 7.779339 |
| 953 | ENSG0000 | TLE2 | transducin | 3.073599 | 3.640419 | 7.779317 |
| 12944 | ENSG0000 | KLF11 | Kruppel lik | 2.197763 | 4.777604 | 7.77794 |
| 4594 | ENSG0000 | RRAGC | Ras relatec | 4.146021 | 6.781076 | 7.776639 |
| 7721 | ENSG0000 | NUP58 | nucleopori | 6.289166 | 7.397183 | 7.774694 |
| 8283 | ENSG0000 | TUFT1 | tuftelin 1 [! | 3.107874 | 7.754389 | 7.75889 |
| 10594 | ENSG0000 | DNAJB4 | DnaJ heat : | 3.515125 | 4.99695 | 7.757268 |
| 2203 | ENSG0000 | ANKRD54 | ankyrin rep | 5.242118 | 6.345356 | 7.755683 |
| 13531 | ENSG0000 | NRIP3 | nuclear rec | 3.854092 | 4.869859 | 7.752923 |
| 15770 | ENSG0000 | POFUT2 | protein O-f | 5.823992 | 7.541488 | 7.743957 |
| 15345 | ENSG0000 | USP18 | ubiquitin s| | 3.28897 | 4.143517 | 7.742966 |
| 9688 | ENSG0000 | ZFYVE27 | zinc finger | 4.398004 | 7.303711 | 7.742348 |
| 2286 | ENSG0000 | SGSM3 | small G prc | 5.953214 | 7.328819 | 7.706035 |
| 10322 | ENSG0000 | C9orf116 | chromosor | 4.730974 | 5.220527 | 7.697248 |
| 20053 | ENSG0000 | SMIM30 | small integ | 3.969587 | 6.119701 | 7.692116 |
| 4972 | ENSG0000 | FAM45A | family with | 4.990452 | 6.368143 | 7.688627 |
| 28622 | ENSG0000 | AL365356.5 | | 1.338939 | 2.730062 | 7.685404 |
| 19494 | ENSG0000 | RNA5SP18 | RNA, 5S rib | 2.604542 | 5.468935 | 7.682042 |
| 11932 | ENSG0000 | CDC42EP5 | CDC42 effe | 0.195827 | 1.337215 | 7.678792 |
| 3161 | ENSG0000 | ARRDC2 | arrestin do | 4.335141 | 5.665766 | 7.678097 |
| 7917 | ENSG0000 | TLDC1 | TBC/LysM- | 3.71432 | 6.40509 | 7.676278 |
| 141 | ENSG0000 | PHTF2 | putative hc | 6.044347 | 6.633293 | 7.669642 |
| 2466 | ENSG0000 | TCFL5 | transcriptic | 6.593851 | 6.691464 | 7.661913 |
| 14369 | ENSG0000 | SOCS4 | suppressor | 5.408183 | 7.260758 | 7.655216 |
| 14077 | ENSG0000 | GALNT11 | polypeptid | 6.058196 | 6.803686 | 7.650341 |
| 501 | ENSG0000 | SNX1 | sorting nex | 6.273957 | 6.692392 | 7.648283 |
| 10002 | ENSG0000 | SPPL3 | signal pept | 5.717151 | 6.468186 | 7.640399 |
| 939 | ENSG0000 | TBC1D22B | TBC1 doma | 5.530118 | 6.294923 | 7.634748 |
| 17131 | ENSG0000 | DENND4B | DENN dom | 6.223407 | 6.91343 | 7.629018 |
| 3673 | ENSG0000 | UGDH | UDP-gluco: | 6.56033 | 7.309179 | 7.625799 |
| 10575 | ENSG0000 | CAMK2N1 | calcium/ca | 3.556908 | 6.124574 | 7.625121 |
| 1682 | ENSG0000 | WDFY1 | WD repeat | 5.039663 | 6.03212 | 7.623493 |
| 8736 | ENSG0000 | SLC2A12 | solute carr | 3.523432 | 5.620709 | 7.622292 |
| 6019 | ENSG0000 | FBXO21 | F-box prote | 6.691629 | 7.274315 | 7.609104 |
| 28204 | ENSG0000 | LRRFIP1P1 | LRR bindin | 1.707544 | 6.537942 | 7.608591 |
| 104 | ENSG0000 | IBTK | inhibitor of | 7.504229 | 7.520382 | 7.603625 |
| 215 | ENSG0000 | SS18L2 | SS18 like 2 | 6.070669 | 6.511833 | 7.591291 |
| 9009 | ENSG0000 | FUOM | fucose mut | 4.257631 | 5.427587 | 7.591071 |
| 6756 | ENSG0000 | GSTM3 | glutathione | 5.132105 | 5.513225 | 7.587491 |
| 7256 | ENSG0000 | KLF4 | Kruppel lik | 2.758908 | 4.460119 | 7.579576 |
| 22023 | ENSG0000 | PPP1R26-A | PPP1R26 a | 2.822384 | 5.315191 | 7.575815 |

| | | | | | | |
|---|---|---|---|---|---|---|
| 4473 | ENSG0000 | ARID3A | AT-rich int | 4.017137 | 4.155282 | 7.569909 |
| 4885 | ENSG0000 | COPS5 | COP9 signa | 6.916612 | 7.147514 | 7.564329 |
| 8496 | ENSG0000 | EAF1 | ELL associa | 4.34777 | 7.506767 | 7.563883 |
| 18311 | ENSG0000 | AC021218.1 | | 1.95562 | 5.640614 | 7.562387 |
| 372 | ENSG0000 | DNASE1L1 | deoxyribor | 2.993544 | 5.830465 | 7.558149 |
| 8121 | ENSG0000 | ZNF92 | zinc finger | 6.108047 | 6.756548 | 7.557476 |
| 41173 | ENSG0000 | AL445363.3 | | 5.811996 | 6.030978 | 7.535033 |
| 1802 | ENSG0000 | TGDS | TDP-glucos | 4.473906 | 6.564494 | 7.527637 |
| 9842 | ENSG0000 | RAB11FIP1 | RAB11 fam | 3.814382 | 5.243314 | 7.525143 |
| 7722 | ENSG0000 | MTMR6 | myotubula | 6.077261 | 6.788896 | 7.517923 |
| 1101 | ENSG0000 | LRP6 | LDL recept | 6.512417 | 7.276701 | 7.517307 |
| 15355 | ENSG0000 | DGAT1 | diacylglyce | 5.135563 | 5.619518 | 7.498807 |
| 13410 | ENSG0000 | SH3PXD2B | SH3 and PX | 4.464262 | 6.197664 | 7.495758 |
| 5850 | ENSG0000 | GFER | growth fac | 6.21059 | 6.482542 | 7.494716 |
| 22695 | ENSG0000 | HSBP1L1 | heat shock | 3.626468 | 6.187712 | 7.493586 |
| 3872 | ENSG0000 | ING4 | inhibitor of | 3.398009 | 5.893214 | 7.490668 |
| 2358 | ENSG0000 | ALKBH1 | alkB homo | 6.06228 | 6.992262 | 7.489056 |
| 11981 | ENSG0000 | RCOR2 | REST corep | 1.141377 | 2.477656 | 7.487107 |
| 14396 | ENSG0000 | MED14 | mediator c | 6.573727 | 7.356888 | 7.485895 |
| 4106 | ENSG0000 | MAN2A1 | mannosida | 5.039724 | 7.41402 | 7.46755 |
| 14261 | ENSG0000 | RAB39A | RAB39A, m | 1.943722 | 5.781771 | 7.464817 |
| 3016 | ENSG0000 | CSRNP2 | cysteine ar | 4.649813 | 7.424795 | 7.442251 |
| 12578 | ENSG0000 | STAT2 | signal trans | 4.448206 | 5.796215 | 7.439048 |
| 18674 | ENSG0000 | HLA-H | major histo | 4.577986 | 4.883338 | 7.420689 |
| 11027 | ENSG0000 | RNF168 | ring finger | 6.280036 | 6.562463 | 7.420311 |
| 1634 | ENSG0000 | ZNF416 | zinc finger | 2.889066 | 6.545407 | 7.413771 |
| 27381 | ENSG0000 | FOXD2-AS1 | FOXD2 ant | 5.7316 | 6.465889 | 7.411969 |
| 15466 | ENSG0000 | SDHAP1 | succinate c | 5.544426 | 6.465422 | 7.404989 |
| 5390 | ENSG0000 | DNPEP | aspartyl an | 5.303556 | 6.430396 | 7.400578 |
| 11641 | ENSG0000 | AKIP1 | A-kinase in | 6.215663 | 6.699251 | 7.390834 |
| 61 | ENSG0000 | PDK4 | pyruvate d | 0.922646 | 2.620456 | 7.389492 |
| 5728 | ENSG0000 | KIRREL2 | kirre like n | 1.173763 | 2.960407 | 7.375846 |
| 11567 | ENSG0000 | BTRC | beta-trans | 6.310593 | 6.936616 | 7.369492 |
| 2029 | ENSG0000 | TMEM38B | transmeml | 5.931778 | 6.303137 | 7.362709 |
| 199 | ENSG0000 | JARID2 | jumonji an | 4.182057 | 6.229035 | 7.340856 |
| 12245 | ENSG0000 | MFF | mitochond | 5.73897 | 7.336197 | 7.339662 |
| 11623 | ENSG0000 | TUB | tubby bipa | 3.18863 | 5.66553 | 7.337011 |
| 6951 | ENSG0000 | PHF21A | PHD finger | 3.875458 | 6.49565 | 7.335025 |
| 38825 | ENSG0000 | U47924.2 | | 3.076582 | 4.306068 | 7.330499 |
| 8976 | ENSG0000 | MINDY3 | MINDY lysi | 3.764537 | 6.181658 | 7.321124 |
| 18503 | ENSG0000 | ETFRF1 | electron tr | 5.115689 | 6.903563 | 7.317444 |
| 568 | ENSG0000 | FBXO42 | F-box prot | 5.500579 | 5.679858 | 7.313641 |
| 20507 | ENSG0000 | AL441992.1 | | 2.283729 | 4.322904 | 7.30754 |
| 6088 | ENSG0000 | RAB2B | RAB2B, me | 3.859176 | 6.052061 | 7.303622 |
| 5722 | ENSG0000 | LRFN3 | leucine ricl | 5.945279 | 6.383759 | 7.292999 |
| 14958 | ENSG0000 | FHL3 | four and a | 4.850543 | 5.520501 | 7.291512 |
| 3179 | ENSG0000 | TMEM59L | transmeml | 2.57098 | 2.932497 | 7.283838 |

| | | | | | | |
|---|---|---|---|---|---|---|
| 21450 | ENSG0000 | ROR1-AS1 | ROR1 antis | 1.953559 | 4.741169 | 7.282885 |
| 5484 | ENSG0000 | BTN2A2 | butyrophili | 1.358252 | 5.149211 | 7.278839 |
| 40130 | ENSG0000 | NUDT18 | nudix hydr | 2.322647 | 2.962721 | 7.267776 |
| 6030 | ENSG0000 | MINDY2 | MINDY lysi | 2.843754 | 6.91551 | 7.251032 |
| 15035 | ENSG0000 | RUNDC1 | RUN doma | 4.631797 | 5.785759 | 7.250647 |
| 18206 | ENSG0000 | MICB | MHC class | 3.324323 | 4.527423 | 7.240791 |
| 42 | ENSG0000 | NDUFAF7 | NADH:ubic | 4.584306 | 6.375424 | 7.240361 |
| 10330 | ENSG0000 | ST6GALNA | ST6 N-acet | 5.37453 | 6.470942 | 7.238992 |
| 4204 | ENSG0000 | SELENOK | selenoprot | 5.920813 | 6.061497 | 7.235997 |
| 16690 | ENSG0000 | ZNF786 | zinc finger | 4.961924 | 6.507938 | 7.231201 |
| 2394 | ENSG0000 | CCNB1IP1 | cyclin B1 ir | 5.217437 | 5.794269 | 7.229032 |
| 12424 | ENSG0000 | TRIM56 | tripartite n | 0.846649 | 5.491333 | 7.227722 |
| 10302 | ENSG0000 | LRRC3 | leucine ricl | 3.763304 | 6.014251 | 7.217308 |
| 4587 | ENSG0000 | AL591845.1 | | 1.276762 | 3.661946 | 7.214222 |
| 12233 | ENSG0000 | SLC35G2 | solute carr | 2.481852 | 5.77624 | 7.206421 |
| 12242 | ENSG0000 | CEP120 | centrosom | 4.959534 | 6.424139 | 7.201314 |
| 10256 | ENSG0000 | IQCC | IQ motif cc | 4.195167 | 6.32856 | 7.199088 |
| 24074 | ENSG0000 | AL158209.1 | | 1.897624 | 6.029296 | 7.193628 |
| 37002 | ENSG0000 | AC011591.1 | | 3.101754 | 3.820934 | 7.190566 |
| 11582 | ENSG0000 | SGPL1 | sphingosin | 5.387217 | 7.154734 | 7.190253 |
| 12071 | ENSG0000 | SETD5 | SET domair | 4.164115 | 5.469887 | 7.182363 |
| 8668 | ENSG0000 | ZNF300 | zinc finger | 3.256871 | 6.365211 | 7.17275 |
| 20468 | ENSG0000 | IFI30 | IFI30, lysos | 2.926095 | 4.474753 | 7.167047 |
| 10708 | ENSG0000 | FBXO41 | F-box prot | 4.631685 | 6.171388 | 7.159495 |
| 1765 | ENSG0000 | TXNDC16 | thioredoxi | 3.773525 | 5.143185 | 7.159337 |
| 8155 | ENSG0000 | MPV17L | MPV17 mi | 3.834848 | 6.404682 | 7.15905 |
| 10053 | ENSG0000 | GPR153 | G protein-c | 4.507841 | 5.533501 | 7.158079 |
| 20965 | ENSG0000 | FAM200A | family with | 4.783687 | 7.100892 | 7.157693 |
| 8291 | ENSG0000 | ADAMTSL4 | ADAMTS li | 1.55981 | 3.576638 | 7.144166 |
| 10230 | ENSG0000 | ZDHHC1 | zinc finger | 4.826959 | 6.216481 | 7.135608 |
| 15144 | ENSG0000 | HTR7P1 | 5-hydroxyt | 3.280836 | 7.036031 | 7.128551 |
| 3319 | ENSG0000 | COA1 | cytochrom | 3.351121 | 5.583483 | 7.127187 |
| 13174 | ENSG0000 | GPR137 | G protein-c | 5.880755 | 6.490139 | 7.126459 |
| 5287 | ENSG0000 | IFT81 | intraflagell | 3.574051 | 5.265137 | 7.117364 |
| 1641 | ENSG0000 | CHMP2B | charged m | 4.954352 | 5.976404 | 7.111346 |
| 945 | ENSG0000 | TMEM206 | transmeml | 4.45631 | 5.244426 | 7.090596 |
| 6264 | ENSG0000 | NPAS1 | neuronal P | 2.372334 | 3.423603 | 7.089138 |
| 3560 | ENSG0000 | DLX4 | distal-less | 1.059457 | 2.255454 | 7.082743 |
| 1914 | ENSG0000 | PLEKHG2 | pleckstrin | 3.747267 | 5.823046 | 7.071413 |
| 10767 | ENSG0000 | DCAF16 | DDB1 and | 5.737757 | 6.279837 | 7.05547 |
| 9733 | ENSG0000 | FZD7 | frizzled cla | 4.326999 | 6.75429 | 7.052826 |
| 1067 | ENSG0000 | PPP2R5B | protein ph | 2.57686 | 4.461318 | 7.036409 |
| 15269 | ENSG0000 | SOCS3 | suppressor | 6.024034 | 6.083651 | 7.036136 |
| 13993 | ENSG0000 | TMEM94 | transmeml | 5.138016 | 6.237202 | 7.031638 |
| 521 | ENSG0000 | ARID4A | AT-rich int | 4.954262 | 6.840788 | 7.027183 |
| 5945 | ENSG0000 | DNAJB9 | DnaJ heat | 3.634753 | 5.27244 | 7.02172 |
| 14357 | ENSG0000 | BBS10 | Bardet-Bie | 5.052262 | 6.045362 | 7.008245 |

| | | | | | | |
|---|---|---|---|---|---|---|
| 9309 | ENSG0000 | RNF144A | ring finger | 3.521008 | 5.025495 | 7.003895 |
| 4006 | ENSG0000 | FIG4 | FIG4 phosp | 4.513385 | 5.172752 | 6.995484 |
| 2364 | ENSG0000 | SIX4 | SIX homeo | 4.704836 | 5.65175 | 6.994094 |
| 5904 | ENSG0000 | GNAZ | G protein s | 3.776441 | 3.898932 | 6.982564 |
| 30146 | ENSG0000 | AC124854.1 | | 2.039543 | 6.825202 | 6.975515 |
| 12667 | ENSG0000 | LRRC8E | leucine rich | 4.62797 | 5.371391 | 6.974949 |
| 13398 | ENSG0000 | ANGEL2 | angel hom | 4.738668 | 5.848515 | 6.967439 |
| 5754 | ENSG0000 | BECN1 | beclin 1 [S | 6.653292 | 6.876764 | 6.965314 |
| 1030 | ENSG0000 | PDZD4 | PDZ domai | 4.993883 | 5.384102 | 6.965257 |
| 10737 | ENSG0000 | MSX1 | msh home | 3.886208 | 4.931397 | 6.964312 |
| 16603 | ENSG0000 | ZSCAN26 | zinc finger | 3.592901 | 5.573348 | 6.960118 |
| 1527 | ENSG0000 | MOSPD1 | motile spe | 3.501115 | 6.201925 | 6.956121 |
| 18460 | ENSG0000 | RGL3 | ral guanine | 2.289268 | 5.005242 | 6.954171 |
| 9617 | ENSG0000 | FAM69A | family with | 3.960739 | 3.991807 | 6.952598 |
| 14222 | ENSG0000 | HES7 | hes family | 0.77154 | 2.626767 | 6.939336 |
| 4443 | ENSG0000 | QPCT | glutaminyl- | 1.339579 | 3.408522 | 6.932265 |
| 4461 | ENSG0000 | ORC4 | origin reco | 4.356386 | 6.313162 | 6.93013 |
| 13278 | ENSG0000 | RNF213 | ring finger | 3.698509 | 5.463044 | 6.925547 |
| 4716 | ENSG0000 | KMT2A | lysine metl | 4.195499 | 5.981308 | 6.917782 |
| 3908 | ENSG0000 | FGFR1OP2 | FGFR1 onc | 5.548336 | 6.715942 | 6.90891 |
| 10661 | ENSG0000 | PPP1R21 | protein ph | 5.273217 | 5.99418 | 6.90434 |
| 11658 | ENSG0000 | ZSCAN21 | zinc finger | 3.256462 | 5.011165 | 6.893999 |
| 1684 | ENSG0000 | SLC25A24 | solute carr | 5.375132 | 6.819263 | 6.882086 |
| 13705 | ENSG0000 | EID2B | EP300 inte | 4.789099 | 5.198881 | 6.879874 |
| 5560 | ENSG0000 | RAP2A | RAP2A, me | 4.048341 | 6.055958 | 6.872987 |
| 4271 | ENSG0000 | ABTB1 | ankyrin rep | 2.326658 | 6.515762 | 6.859174 |
| 18211 | ENSG0000 | HLA-C | major histo | 3.628626 | 4.297693 | 6.855095 |
| 5550 | ENSG0000 | BBS2 | Bardet-Bie | 3.565801 | 4.795676 | 6.848799 |
| 14337 | ENSG0000 | CITED4 | Cbp/p300 i | 2.967715 | 3.707205 | 6.843843 |
| 7190 | ENSG0000 | CIB2 | calcium an | 3.326183 | 3.565463 | 6.830953 |
| 3163 | ENSG0000 | RAB3A | RAB3A, me | 1.49611 | 1.579198 | 6.829261 |
| 12016 | ENSG0000 | TMEM99 | transmeml | 2.431622 | 5.226954 | 6.828154 |
| 20563 | ENSG0000 | AL109918.1 | | 2.04024 | 4.38359 | 6.824307 |
| 9379 | ENSG0000 | C18orf25 | chromosor | 5.427261 | 6.441523 | 6.822994 |
| 37186 | ENSG0000 | S1PR2 | sphingosin | 1.35795 | 4.72381 | 6.820626 |
| 2856 | ENSG0000 | BLOC1S6 | biogenesis | 4.904993 | 6.180791 | 6.819902 |
| 919 | ENSG0000 | SLC9A3R2 | SLC9A3 reg | 5.578818 | 5.973255 | 6.811694 |
| 13707 | ENSG0000 | LPCAT4 | lysophosph | 6.212851 | 6.796719 | 6.806266 |
| 3447 | ENSG0000 | SUFU | SUFU nega | 4.358362 | 5.053183 | 6.805153 |
| 857 | ENSG0000 | APPBP2 | amyloid be | 4.312929 | 6.302402 | 6.800916 |
| 5335 | ENSG0000 | ATG101 | autophagy | 5.084348 | 6.431017 | 6.794122 |
| 19928 | ENSG0000 | KCTD11 | potassium | 1.787377 | 3.085861 | 6.787198 |
| 13303 | ENSG0000 | UBXN2A | UBX doma | 4.743295 | 5.493418 | 6.781273 |
| 7877 | ENSG0000 | MFGE8 | milk fat glc | 1.945947 | 4.51811 | 6.78068 |
| 6372 | ENSG0000 | GFPT2 | glutamine- | 2.018035 | 2.598447 | 6.771559 |
| 12200 | ENSG0000 | ABHD15 | abhydrolas | 3.387754 | 5.096188 | 6.771214 |
| 3148 | ENSG0000 | CACNG7 | calcium vo | 0.964966 | 1.374598 | 6.76805 |

| | | | | | | |
|---|---|---|---|---|---|---|
| 8869 | ENSG0000 | NSD3 | nuclear rec | 5.214473 | 6.220779 | 6.766848 |
| 4780 | ENSG0000 | TGIF2 | TGFB induc | 4.891301 | 5.177982 | 6.76095 |
| 1050 | ENSG0000 | GRIPAP1 | GRIP1 assc | 4.188471 | 6.14873 | 6.760017 |
| 1643 | ENSG0000 | PPIE | peptidylpro | 5.256329 | 6.696693 | 6.759128 |
| 31 | ENSG0000 | SNX11 | sorting nex | 3.898161 | 5.04999 | 6.757819 |
| 11574 | ENSG0000 | ZNF319 | zinc finger | 5.169884 | 6.523002 | 6.754129 |
| 26665 | ENSG0000 | PNMA6A | paraneopla | 6.106043 | 6.264634 | 6.75088 |
| 16317 | ENSG0000 | TDRD7 | tudor dom | 2.971119 | 5.497972 | 6.740417 |
| 6224 | ENSG0000 | HELZ2 | helicase wi | 3.990958 | 4.387351 | 6.734078 |
| 617 | ENSG0000 | DCUN1D1 | defective ir | 5.192823 | 6.472243 | 6.732515 |
| 21387 | ENSG0000 | CERS1 | ceramide s | 3.851705 | 4.698928 | 6.731249 |
| 6266 | ENSG0000 | MAP3K10 | mitogen-ac | 3.642813 | 6.607907 | 6.728282 |
| 11305 | ENSG0000 | FAM219A | family with | 0.779441 | 1.159395 | 6.726637 |
| 37665 | ENSG0000 | SRGAP2D | SLIT-ROBO | 4.975014 | 5.184117 | 6.722278 |
| 16521 | ENSG0000 | ZNF700 | zinc finger | 3.56231 | 6.518464 | 6.717714 |
| 16473 | ENSG0000 | MYO6 | myosin VI [ | 5.824035 | 6.625435 | 6.705138 |
| 5757 | ENSG0000 | GLIS2 | GLIS family | 2.992961 | 3.633923 | 6.701273 |
| 22437 | ENSG0000 | BAIAP2-AS | BAIAP2 ant | 3.846211 | 4.169083 | 6.700117 |
| 5420 | ENSG0000 | CHD6 | chromodor | 3.96857 | 6.180903 | 6.698482 |
| 35621 | ENSG0000 | DLGAP1-AS | DLGAP1 ar | 2.00925 | 5.155427 | 6.682032 |
| 6079 | ENSG0000 | CDKN2D | cyclin depe | 3.988029 | 4.667765 | 6.673917 |
| 14345 | ENSG0000 | R3HDM2 | R3H domai | 5.57745 | 6.204538 | 6.668288 |
| 6718 | ENSG0000 | COX16 | COX16, cyt | 5.79364 | 6.598079 | 6.665862 |
| 20282 | ENSG0000 | LINC01588 | long interg | 1.282242 | 4.579867 | 6.653037 |
| 2124 | ENSG0000 | MISP | mitotic spi | 1.269511 | 1.659796 | 6.646788 |
| 9506 | ENSG0000 | FAM49B | family with | 5.685133 | 6.355947 | 6.642896 |
| 5525 | ENSG0000 | SLC35B3 | solute carr | 5.156014 | 6.209653 | 6.627648 |
| 16600 | ENSG0000 | ZMYM1 | zinc finger | 5.50896 | 5.733655 | 6.597856 |
| 10071 | ENSG0000 | AHCYL2 | adenosylhc | 4.33261 | 6.288604 | 6.592343 |
| 12746 | ENSG0000 | ZNF672 | zinc finger | 1.716666 | 5.350334 | 6.580237 |
| 32824 | ENSG0000 | URB1-AS1 | URB1 antis | 4.585579 | 4.642444 | 6.580067 |
| 13089 | ENSG0000 | RARG | retinoic aci | 5.287679 | 5.295638 | 6.570704 |
| 4010 | ENSG0000 | HECA | hdc homol | 2.318483 | 6.359756 | 6.567696 |
| 7645 | ENSG0000 | CASP6 | caspase 6 [ | 2.196771 | 3.71885 | 6.566718 |
| 9782 | ENSG0000 | DPY19L4 | dpy-19 like | 4.284789 | 5.335342 | 6.566555 |
| 7204 | ENSG0000 | BRIP1 | BRCA1 inte | 4.459099 | 5.765946 | 6.564207 |
| 15239 | ENSG0000 | SS18L1 | SS18L1, nB | 5.1433 | 5.492552 | 6.562149 |
| 11126 | ENSG0000 | WDR41 | WD repeat | 4.659735 | 5.792897 | 6.557847 |
| 37480 | ENSG0000 | AC007192.1 | | 0.006011 | 0.178476 | 6.555109 |
| 18388 | ENSG0000 | SDHAF1 | succinate c | 3.686739 | 3.780311 | 6.546761 |
| 11376 | ENSG0000 | BRWD3 | bromodom | 3.362273 | 4.683495 | 6.546707 |
| 14943 | ENSG0000 | ZNF623 | zinc finger | 4.011252 | 5.768306 | 6.541538 |
| 16442 | ENSG0000 | C20orf96 | chromosor | 3.33634 | 3.916007 | 6.538411 |
| 9352 | ENSG0000 | CCDC74B | coiled-coil | 2.40627 | 4.391444 | 6.534458 |
| 6615 | ENSG0000 | DSTYK | dual serine | 4.444536 | 6.140473 | 6.528003 |
| 19599 | ENSG0000 | RTL8B | retrotransp | 4.57793 | 5.340989 | 6.522352 |
| 7009 | ENSG0000 | RAB11FIP5 | RAB11 fam | 3.366507 | 4.564267 | 6.522266 |

| | | | | | | |
|---|---|---|---|---|---|---|
| 1176 | ENSG0000 | CYBRD1 | cytochrom | 5.43191 | 6.293733 | 6.519038 |
| 15468 | ENSG0000 | MUC1 | mucin 1, ce | 1.342489 | 2.586145 | 6.504457 |
| 14719 | ENSG0000 | SYNM | synemin [S | 2.907187 | 4.761628 | 6.498958 |
| 4127 | ENSG0000 | TMCO6 | transmeml | 3.05133 | 5.676574 | 6.493435 |
| 1060 | ENSG0000 | SRBD1 | S1 RNA bin | 5.787269 | 6.390648 | 6.490416 |
| 8034 | ENSG0000 | GRB7 | growth fac | 2.252567 | 3.833698 | 6.48765 |
| 14648 | ENSG0000 | RNF41 | ring finger | 4.790974 | 5.477977 | 6.48687 |
| 2586 | ENSG0000 | NKAP | NFKB activ | 4.951618 | 6.231452 | 6.486715 |
| 5214 | ENSG0000 | FAM213A | family with | 2.935935 | 3.671325 | 6.485834 |
| 16036 | ENSG0000 | NCR3LG1 | natural kill | 2.918477 | 5.67288 | 6.482975 |
| 15543 | ENSG0000 | NAT8L | N-acetyltra | 3.929612 | 5.039271 | 6.482879 |
| 10006 | ENSG0000 | FAM213B | family with | 4.212358 | 6.021987 | 6.482183 |
| 2037 | ENSG0000 | WHRN | whirlin [So | 5.014311 | 5.93121 | 6.460188 |
| 1722 | ENSG0000 | ITPKC | inositol-tris | 5.0686 | 6.418396 | 6.459777 |
| 7634 | ENSG0000 | CCNG2 | cyclin G2 [S | 0.864733 | 3.419857 | 6.427731 |
| 2991 | ENSG0000 | ZKSCAN1 | zinc finger | 5.041022 | 6.317997 | 6.426454 |
| 14545 | ENSG0000 | METTL23 | methyltran | 3.490741 | 6.068504 | 6.42346 |
| 9061 | ENSG0000 | PAK1 | p21 (RAC1) | 5.043153 | 5.900327 | 6.419887 |
| 2072 | ENSG0000 | ZNF184 | zinc finger | 3.16348 | 4.878892 | 6.411323 |
| 15077 | ENSG0000 | LINC00518 | long interg | 1.188296 | 4.056683 | 6.40945 |
| 10764 | ENSG0000 | CCNYL1 | cyclin Y like | 4.460319 | 5.484588 | 6.399021 |
| 11046 | ENSG0000 | PGRMC2 | progestero | 4.202783 | 4.310331 | 6.392017 |
| 1017 | ENSG0000 | TP53BP1 | tumor prot | 4.20603 | 5.853842 | 6.374221 |
| 13701 | ENSG0000 | CRLF3 | cytokine re | 5.815432 | 6.253548 | 6.373951 |
| 13979 | ENSG0000 | THAP5 | THAP dom | 5.121964 | 5.957118 | 6.373819 |
| 5126 | ENSG0000 | ZNF211 | zinc finger | 2.479041 | 4.742416 | 6.373392 |
| 5584 | ENSG0000 | SLC25A19 | solute carr | 5.097422 | 5.403926 | 6.368531 |
| 6878 | ENSG0000 | KDELC1 | KDEL motif | 5.008503 | 5.308948 | 6.365565 |
| 2099 | ENSG0000 | OCEL1 | occludin/E | 1.694202 | 3.065604 | 6.36152 |
| 6194 | ENSG0000 | AFDN | afadin, adh | 3.992691 | 5.730014 | 6.35898 |
| 5882 | ENSG0000 | PEX1 | peroxisom | 4.839185 | 5.568761 | 6.357317 |
| 32874 | ENSG0000 | SMIM3 | small integ | 3.971557 | 5.010787 | 6.353605 |
| 4294 | ENSG0000 | EIF1B | eukaryotic | 5.501422 | 5.620351 | 6.350952 |
| 9944 | ENSG0000 | ANKZF1 | ankyrin rep | 3.282961 | 4.437329 | 6.346853 |
| 916 | ENSG0000 | ANKS1A | ankyrin rep | 5.173539 | 5.959951 | 6.34248 |
| 10413 | ENSG0000 | TUBA3FP | tubulin alp | 1.846191 | 4.60024 | 6.327756 |
| 12719 | ENSG0000 | LYSMD3 | LysM doma | 3.894603 | 6.005193 | 6.327256 |
| 14573 | ENSG0000 | ZNF322 | zinc finger | 4.619743 | 6.310375 | 6.321664 |
| 3824 | ENSG0000 | DUSP16 | dual specif | 5.236225 | 6.08994 | 6.317699 |
| 3215 | ENSG0000 | CBLL1 | Cbl proto-c | 3.561584 | 5.980949 | 6.311938 |
| 22494 | ENSG0000 | GTF2H2B | general tra | 4.72301 | 5.826546 | 6.306705 |
| 17923 | ENSG0000 | PDCD4-AS1 | PDCD4 ant | 1.892083 | 2.87869 | 6.304481 |
| 5376 | ENSG0000 | COQ8B | coenzyme | 3.964834 | 4.289666 | 6.304261 |
| 12494 | ENSG0000 | ADPRM | ADP-ribose | 3.45203 | 5.821994 | 6.303735 |
| 16947 | ENSG0000 | HELZ | helicase wi | 4.329991 | 4.615618 | 6.300591 |
| 1115 | ENSG0000 | EXOC5 | exocyst co | 6.249422 | 6.273407 | 6.299696 |
| 13890 | ENSG0000 | CHD9 | chromodo | 4.064281 | 5.908222 | 6.299278 |

| | | | | | | |
|---|---|---|---|---|---|---|
| 5436 | ENSG0000 | SNAI1 | snail family | 3.348013 | 5.268586 | 6.298255 |
| 35749 | ENSG0000 | OVCA2 | OVCA2, ser | 4.742037 | 5.976182 | 6.293936 |
| 15447 | ENSG0000 | MAPK11 | mitogen-ac | 4.653359 | 5.389577 | 6.292504 |
| 17183 | ENSG0000 | TBKBP1 | TBK1 bindi | 3.593519 | 6.121483 | 6.287406 |
| 2089 | ENSG0000 | RAB18 | RAB18, me | 5.784992 | 6.187399 | 6.26779 |
| 572 | ENSG0000 | HOXC8 | homeobox | 4.980636 | 6.0213 | 6.265395 |
| 31941 | ENSG0000 | RHPN1-AS1 | RHPN1 ant | 1.786494 | 3.883542 | 6.262384 |
| 3166 | ENSG0000 | ELL | elongation | 4.88973 | 5.818074 | 6.249131 |
| 4062 | ENSG0000 | PDE10A | phosphodi | 4.446171 | 4.657003 | 6.24808 |
| 18865 | ENSG0000 | SNORA14B | small nucle | 3.254606 | 5.173641 | 6.246309 |
| 3694 | ENSG0000 | DNAJC4 | DnaJ heat s | 3.665593 | 3.891128 | 6.240261 |
| 2263 | ENSG0000 | PDGFB | platelet de | 2.432052 | 2.457217 | 6.227843 |
| 9023 | ENSG0000 | ADM | adrenomed | 0.920271 | 4.254661 | 6.224176 |
| 11397 | ENSG0000 | ZCCHC24 | zinc finger | 3.673094 | 4.402135 | 6.21876 |
| 1187 | ENSG0000 | EPN2 | epsin 2 [So | 4.353389 | 4.743205 | 6.215441 |
| 27933 | ENSG0000 | C8orf58 | chromosor | 3.925395 | 5.175976 | 6.211375 |
| 4318 | ENSG0000 | RTKN | rhotekin [S | 4.538223 | 5.568934 | 6.211241 |
| 17997 | ENSG0000 | EEF1AKMT | EEF1A lysir | 5.274633 | 5.330999 | 6.20624 |
| 478 | ENSG0000 | PHF20 | PHD finger | 5.126924 | 5.674341 | 6.206177 |
| 40790 | ENSG0000 | STAG3L2 | stromal an | 4.928009 | 6.069777 | 6.204837 |
| 8109 | ENSG0000 | ZNF473 | zinc finger | 3.488898 | 5.322357 | 6.202078 |
| 9728 | ENSG0000 | KCTD18 | potassium | 1.567656 | 4.513467 | 6.201595 |
| 7965 | ENSG0000 | TAF4B | TATA-box b | 4.445791 | 5.731564 | 6.198286 |
| 6954 | ENSG0000 | ELF5 | E74 like ET | 0.264599 | 1.586666 | 6.197542 |
| 5582 | ENSG0000 | ARMC7 | armadillo r | 4.221515 | 4.97099 | 6.197323 |
| 11535 | ENSG0000 | R3HCC1L | R3H domai | 4.051671 | 5.931282 | 6.196244 |
| 12730 | ENSG0000 | VPS37D | VPS37D, ES | 1.931687 | 2.672018 | 6.192129 |
| 2097 | ENSG0000 | MAST3 | microtubul | 3.352735 | 4.009335 | 6.187995 |
| 2233 | ENSG0000 | RAB36 | RAB36, me | 2.174204 | 5.982305 | 6.186465 |
| 5674 | ENSG0000 | MCM8 | minichrom | 5.455728 | 5.798178 | 6.185282 |
| 13978 | ENSG0000 | SRRM3 | serine/argi | 2.536633 | 2.842313 | 6.177604 |
| 9363 | ENSG0000 | GPATCH11 | G-patch do | 5.765998 | 5.990008 | 6.177312 |
| 8185 | ENSG0000 | KIAA0319L | KIAA0319 l | 5.950316 | 6.009244 | 6.169722 |
| 9089 | ENSG0000 | EML3 | echinoderr | 5.081878 | 5.786708 | 6.142589 |
| 845 | ENSG0000 | TNK2 | tyrosine ki | 5.111754 | 6.000117 | 6.137917 |
| 10116 | ENSG0000 | NIT1 | nitrilase 1 | | 3.538818 | 4.042753 | 6.136419 |
| 12626 | ENSG0000 | AC093323.1 | | 4.232107 | 5.231484 | 6.130201 |
| 23217 | ENSG0000 | MELTF-AS1 | MELTF ant | 4.148851 | 4.16518 | 6.128093 |
| 27324 | ENSG0000 | GS1-124K5 | uncharacte | 2.909893 | 3.063139 | 6.125862 |
| 7162 | ENSG0000 | TMTC3 | transmeml | 3.705801 | 4.862783 | 6.114453 |
| 17123 | ENSG0000 | INPP5F | inositol po | 3.659042 | 4.585327 | 6.112251 |
| 2738 | ENSG0000 | PLA2G15 | phospholip | 5.889553 | 6.011294 | 6.109445 |
| 17002 | ENSG0000 | TMA16 | translation | 4.368572 | 4.816136 | 6.108376 |
| 9376 | ENSG0000 | PSTPIP2 | proline-ser | 1.576845 | 3.218707 | 6.106763 |
| 787 | ENSG0000 | ATG5 | autophagy | 4.875306 | 6.034155 | 6.106514 |
| 17127 | ENSG0000 | SELENOM | selenoprot | 4.018631 | 4.759078 | 6.105297 |
| 11218 | ENSG0000 | RELL2 | RELT like 2 | 2.452553 | 4.079105 | 6.104789 |

| | | | | | | |
|---|---|---|---|---|---|---|
| 38 | ENSG0000 | ALS2 | ALS2, alsin | 4.533066 | 5.859646 | 6.101051 |
| 23710 | ENSG0000 | BX664615.1 | | 1.829227 | 5.016324 | 6.099892 |
| 6090 | ENSG0000 | AJUBA | ajuba LIM | 3.116851 | 5.020071 | 6.099625 |
| 14985 | ENSG0000 | COA5 | cytochrom | 5.241604 | 5.354091 | 6.089444 |
| 9569 | ENSG0000 | TBCEL | tubulin fol | 3.419803 | 4.990883 | 6.082441 |
| 1570 | ENSG0000 | ZNF510 | zinc finger | 3.259715 | 5.296348 | 6.081915 |
| 7274 | ENSG0000 | STX17 | syntaxin 17 | 4.513321 | 6.051284 | 6.079509 |
| 20146 | ENSG0000 | AC009053.1 | | 3.673174 | 5.668815 | 6.071813 |
| 8348 | ENSG0000 | RIT1 | Ras like wi | 3.124013 | 3.86096 | 6.037096 |
| 12060 | ENSG0000 | MAP4K2 | mitogen-ac | 3.538232 | 3.981535 | 6.035591 |
| 13337 | ENSG0000 | ADCY6 | adenylate | 2.742041 | 3.458195 | 6.032338 |
| 1439 | ENSG0000 | TOLLIP | toll interac | 4.386878 | 5.887009 | 6.028425 |
| 89 | ENSG0000 | FAM214B | family with | 2.229243 | 3.215095 | 6.020356 |
| 18106 | ENSG0000 | BMPR2 | bone morp | 3.515696 | 5.621512 | 6.014031 |
| 6419 | ENSG0000 | PRKAB2 | protein kin | 3.121962 | 4.708213 | 6.005903 |
| 448 | ENSG0000 | FHL1 | four and a | 0.877365 | 2.364724 | 5.988583 |
| 5664 | ENSG0000 | BFSP1 | beaded fila | 3.186553 | 5.5713 | 5.985311 |
| 3792 | ENSG0000 | RSRC2 | arginine ar | 4.966233 | 5.302511 | 5.981774 |
| 17160 | ENSG0000 | CIPC | CLOCK inte | 3.80083 | 5.639074 | 5.980624 |
| 1511 | ENSG0000 | TESC | tescalcin [S | 3.086496 | 3.806535 | 5.975748 |
| 5536 | ENSG0000 | RAB17 | RAB17, me | 1.295727 | 4.405538 | 5.974495 |
| 27864 | ENSG0000 | ALKBH6 | alkB homo | 4.598735 | 5.163456 | 5.972536 |
| 290 | ENSG0000 | GIPR | gastric inhi | 1.189938 | 3.381199 | 5.966384 |
| 12385 | ENSG0000 | SPNS1 | sphingolipi | 4.415391 | 5.0735 | 5.959742 |
| 10062 | ENSG0000 | MITD1 | microtubul | 4.021241 | 4.863763 | 5.940959 |
| 8307 | ENSG0000 | POGZ | pogo trans | 3.157963 | 3.974835 | 5.936267 |
| 40818 | ENSG0000 | F8A3 | coagulatio | 4.902859 | 5.727373 | 5.935607 |
| 14778 | ENSG0000 | MXRA7 | matrix rem | 3.962843 | 4.727334 | 5.929257 |
| 12899 | ENSG0000 | PTEN | phosphata | 5.299391 | 5.912929 | 5.923564 |
| 4567 | ENSG0000 | CTH | cystathioni | 3.593216 | 5.034688 | 5.920004 |
| 1985 | ENSG0000 | TRPM7 | transient r | 3.576383 | 5.578115 | 5.917814 |
| 19792 | ENSG0000 | SYNJ2BP | synaptojan | 3.153548 | 4.506374 | 5.913938 |
| 5460 | ENSG0000 | STAMBP | STAM bind | 3.694435 | 5.329287 | 5.911635 |
| 15100 | ENSG0000 | KREMEN1 | kringle con | 2.801126 | 3.319687 | 5.9104 |
| 8281 | ENSG0000 | PRUNE1 | prune exop | 3.885486 | 4.377789 | 5.909893 |
| 37697 | ENSG0000 | PNMA6B | paraneopla | 4.579622 | 4.843129 | 5.90972 |
| 13839 | ENSG0000 | TOB2P1 | transducer | 1.097455 | 4.675572 | 5.90943 |
| 1879 | ENSG0000 | YPEL3 | yippee like | 2.11056 | 3.039128 | 5.902439 |
| 3043 | ENSG0000 | OLFM2 | olfactomed | 1.645014 | 2.201057 | 5.89508 |
| 2258 | ENSG0000 | ARSA | arylsulfata | 2.518422 | 4.763077 | 5.891845 |
| 3557 | ENSG0000 | CNTNAP1 | contactin a | 1.681894 | 2.330605 | 5.884526 |
| 19880 | ENSG0000 | PIGCP1 | phosphatic | 4.01992 | 4.782602 | 5.881095 |
| 29100 | ENSG0000 | ZNF512 | zinc finger | 4.218214 | 5.678227 | 5.878829 |
| 20992 | ENSG0000 | EXOC3-AS1 | EXOC3 ant | 4.07742 | 5.366 | 5.873215 |
| 6517 | ENSG0000 | TRIM47 | tripartite n | 2.957083 | 3.106523 | 5.873078 |
| 27480 | ENSG0000 | C2orf74 | chromosor | 3.01805 | 4.350931 | 5.869992 |
| 17894 | ENSG0000 | AL590705.1 | | 2.74503 | 3.825401 | 5.865712 |

| | | | | | | |
|---|---|---|---|---|---|---|
| 10681 | ENSG0000 | PEX13 | peroxisom | 4.511571 | 5.070316 | 5.865622 |
| 4872 | ENSG0000 | ACOT2 | acyl-CoA tł | 4.361241 | 5.651159 | 5.863066 |
| 8289 | ENSG0000 | SNX27 | sorting nex | 4.604701 | 4.869297 | 5.8626 |
| 12938 | ENSG0000 | EPHX4 | epoxide hy | 4.494197 | 5.806163 | 5.859613 |
| 7586 | ENSG0000 | SECISBP2L | SECIS bindi | 3.511541 | 5.233707 | 5.855217 |
| 6840 | ENSG0000 | AGO4 | argonaute | 3.585563 | 4.929308 | 5.847402 |
| 34468 | ENSG0000 | AL050341.2 | | 4.747664 | 5.761924 | 5.84256 |
| 18482 | ENSG0000 | LCMT1 | leucine car | 3.944322 | 4.488451 | 5.833617 |
| 10756 | ENSG0000 | DHX57 | DExH-box l | 5.610393 | 5.731262 | 5.833417 |
| 15712 | ENSG0000 | SMYD4 | SET and M | 3.928245 | 5.466767 | 5.822396 |
| 7394 | ENSG0000 | CCDC90B | coiled-coil | 3.722846 | 5.092493 | 5.819528 |
| 16750 | ENSG0000 | GPX1P1 | glutathione | 1.004687 | 3.160522 | 5.808985 |
| 5760 | ENSG0000 | AHDC1 | AT-hook D | 3.275417 | 3.357463 | 5.805491 |
| 8440 | ENSG0000 | FAHD2B | fumarylace | 4.50809 | 5.312576 | 5.804127 |
| 6976 | ENSG0000 | B4GALNT1 | beta-1,4-N | 3.989959 | 4.10058 | 5.803305 |
| 40525 | ENSG0000 | RN7SL1 | RNA, 7SL, c | 5.274338 | 5.65834 | 5.799775 |
| 5787 | ENSG0000 | WDR60 | WD repeat | 4.415739 | 4.657878 | 5.79949 |
| 16280 | ENSG0000 | SLC35E2B | solute carr | 4.067607 | 5.182649 | 5.796238 |
| 9658 | ENSG0000 | RAB6B | RAB6B, me | 1.910334 | 2.514289 | 5.787071 |
| 10313 | ENSG0000 | DIP2A | disco inter | 4.356859 | 5.283673 | 5.782364 |
| 12017 | ENSG0000 | GHDC | GH3 doma | 3.948279 | 4.644302 | 5.778556 |
| 5333 | ENSG0000 | LRP1 | LDL recept | 0.676412 | 1.875572 | 5.773397 |
| 8215 | ENSG0000 | MTF2 | metal resp | 4.660449 | 5.615109 | 5.768379 |
| 4297 | ENSG0000 | KLHL24 | kelch like f | 2.204817 | 4.185289 | 5.768072 |
| 2214 | ENSG0000 | SEPT3 | septin 3 [S | 1.525265 | 2.236931 | 5.766429 |
| 5144 | ENSG0000 | DEPDC7 | DEP domai | 3.328434 | 3.874389 | 5.763213 |
| 6924 | ENSG0000 | PNPLA8 | patatin like | 3.552584 | 5.222508 | 5.760817 |
| 17103 | ENSG0000 | CNOT7 | CCR4-NOT | 5.15866 | 5.524525 | 5.760097 |
| 10787 | ENSG0000 | COL6A3 | collagen ty | 3.280714 | 3.994579 | 5.752804 |
| 3609 | ENSG0000 | PHF12 | PHD finger | 4.294739 | 5.031237 | 5.750453 |
| 12072 | ENSG0000 | VASN | vasorin [Sc | 1.342864 | 5.388544 | 5.743044 |
| 14066 | ENSG0000 | SH2B1 | SH2B adap | 4.516238 | 5.305317 | 5.740559 |
| 6245 | ENSG0000 | LAMA5 | laminin sul | 2.860697 | 4.017097 | 5.733046 |
| 32655 | ENSG0000 | GOLGA2P1 | golgin A2 p | 4.538976 | 4.768031 | 5.730975 |
| 10075 | ENSG0000 | SPATA2 | spermatog | 5.21782 | 5.548371 | 5.725213 |
| 9981 | ENSG0000 | C2CD2 | C2 calcium | 3.257141 | 3.751955 | 5.718459 |
| 7772 | ENSG0000 | GRTP1 | growth hor | 2.346625 | 2.935428 | 5.718004 |
| 7247 | ENSG0000 | NIPSNAP3/ | nipsnap ho | 4.35777 | 4.940432 | 5.716232 |
| 10115 | ENSG0000 | SPATA2L | spermatog | 3.425667 | 4.677463 | 5.700836 |
| 186 | ENSG0000 | BAIAP3 | BAI1 assoc | 0.925821 | 3.676294 | 5.700716 |
| 5004 | ENSG0000 | ARAP3 | ArfGAP wit | 3.673016 | 4.187594 | 5.697742 |
| 39620 | ENSG0000 | LHX1 | LIM homec | 3.416296 | 4.19215 | 5.693066 |
| 7946 | ENSG0000 | C17orf80 | chromosor | 4.426059 | 5.632688 | 5.69117 |
| 13629 | ENSG0000 | ORAI3 | ORAI calciu | 1.926659 | 2.319405 | 5.690704 |
| 11817 | ENSG0000 | ENGASE | endo-beta- | 4.708114 | 5.552342 | 5.688152 |
| 989 | ENSG0000 | TRMT11 | tRNA meth | 4.626187 | 5.504839 | 5.685358 |
| 16460 | ENSG0000 | BORCS6 | BLOC-1 rel | 3.454387 | 3.45557 | 5.682602 |

| | | | | | | |
|---|---|---|---|---|---|---|
| 2094 | ENSG0000 | TSPAN15 | tetraspanin | 3.640008 | 3.831306 | 5.67464 |
| 6412 | ENSG0000 | RARA | retinoic aci | 4.088413 | 4.389433 | 5.673434 |
| 3436 | ENSG0000 | KAZALD1 | Kazal type | 3.185018 | 3.929978 | 5.66965 |
| 1496 | ENSG0000 | PSEN1 | presenilin : | 4.46159 | 5.412399 | 5.669041 |
| 3955 | ENSG0000 | RNF8 | ring finger | 4.034091 | 5.498913 | 5.667036 |
| 21296 | ENSG0000 | TINCR | tissue diffe | 0.908778 | 2.55885 | 5.666784 |
| 6225 | ENSG0000 | SAMD10 | sterile alph | 3.226724 | 3.648302 | 5.666262 |
| 3169 | ENSG0000 | UPK1A | uroplakin 1 | 0.155547 | 0.808163 | 5.665262 |
| 2614 | ENSG0000 | ASB9 | ankyrin rep | 2.233864 | 2.940719 | 5.663225 |
| 36450 | ENSG0000 | AC015922.3 | | 2.216318 | 4.69985 | 5.661397 |
| 10326 | ENSG0000 | C19orf47 | chromosor | 4.48709 | 4.716265 | 5.660223 |
| 14615 | ENSG0000 | ZFP41 | ZFP41 zinc | 2.877231 | 5.205053 | 5.660052 |
| 1058 | ENSG0000 | IP6K2 | inositol he: | 4.009323 | 4.973149 | 5.658691 |
| 13512 | ENSG0000 | TP53I11 | tumor prot | 1.004332 | 2.123013 | 5.657021 |
| 5150 | ENSG0000 | TBC1D15 | TBC1 doma | 3.488253 | 5.15913 | 5.655696 |
| 2890 | ENSG0000 | EMC2 | ER membra | 4.815688 | 5.648258 | 5.652038 |
| 12584 | ENSG0000 | COMMD5 | COMM dor | 4.388987 | 4.39365 | 5.651558 |
| 16400 | ENSG0000 | SRGAP2B | SLIT-ROBO | 3.830879 | 4.296519 | 5.649057 |
| 11840 | ENSG0000 | PPP2R3B | protein ph | 4.914315 | 5.458987 | 5.641895 |
| 14862 | ENSG0000 | CNOT10 | CCR4-NOT | 5.283572 | 5.440004 | 5.637629 |
| 3114 | ENSG0000 | MEIS3 | Meis home | 1.018392 | 3.090514 | 5.635892 |
| 14975 | ENSG0000 | GTF2H2C | GTF2H2 fa | 4.02854 | 5.123567 | 5.631261 |
| 5920 | ENSG0000 | APOL2 | apolipopro | 3.507981 | 4.571147 | 5.626775 |
| 32547 | ENSG0000 | AP001528.2 | | 1.91933 | 2.828381 | 5.62504 |
| 4624 | ENSG0000 | CTBS | chitobiase | 2.149358 | 3.622678 | 5.622649 |
| 1335 | ENSG0000 | RBMS2 | RNA bindir | 2.909243 | 5.384595 | 5.62126 |
| 3920 | ENSG0000 | GCNT2 | glucosamir | 1.093435 | 5.583576 | 5.619724 |
| 3703 | ENSG0000 | PUS3 | pseudouric | 4.51594 | 4.533435 | 5.61273 |
| 8445 | ENSG0000 | SPOPL | speckle typ | 3.942031 | 4.39405 | 5.610445 |
| 9697 | ENSG0000 | ZCCHC10 | zinc finger | 3.794543 | 4.518105 | 5.607348 |
| 13924 | ENSG0000 | PPFIA3 | PTPRF inte | 2.049839 | 2.728369 | 5.603122 |
| 5073 | ENSG0000 | APAF1 | apoptotic p | 3.620073 | 3.754952 | 5.602585 |
| 6673 | ENSG0000 | AGAP3 | ArfGAP wit | 5.474343 | 5.591162 | 5.601544 |
| 2223 | ENSG0000 | HSCB | HscB mitoc | 3.615449 | 4.412931 | 5.598743 |
| 4785 | ENSG0000 | STBD1 | starch binc | 1.373292 | 3.08888 | 5.597822 |
| 15989 | ENSG0000 | TPRG1 | tumor prot | 2.597728 | 4.777648 | 5.597228 |
| 4003 | ENSG0000 | TRIM38 | tripartite n | 2.686595 | 4.921426 | 5.593432 |
| 3960 | ENSG0000 | CD83 | CD83 mole | 2.192864 | 4.281555 | 5.591925 |
| 9055 | ENSG0000 | CCDC82 | coiled-coil | 3.219729 | 5.016588 | 5.591873 |
| 7936 | ENSG0000 | GID4 | GID compl | 4.552359 | 5.472788 | 5.583408 |
| 26076 | ENSG0000 | LINC01659 | long interg | 1.48008 | 2.127928 | 5.579886 |
| 11634 | ENSG0000 | RNF169 | ring finger | 4.494251 | 5.477891 | 5.577979 |
| 5779 | ENSG0000 | TRMT5 | tRNA meth | 4.838527 | 5.279949 | 5.577893 |
| 502 | ENSG0000 | TBPL1 | TATA-box I | 3.704063 | 5.064339 | 5.570472 |
| 3939 | ENSG0000 | ULBP1 | UL16 bindi | 2.175364 | 2.967294 | 5.56969 |
| 17174 | ENSG0000 | DZIP3 | DAZ intera | 3.571486 | 4.978331 | 5.56931 |
| 5327 | ENSG0000 | SPATS2 | spermatog | 3.775713 | 4.39238 | 5.56891 |

| | | | | | | |
|---|---|---|---|---|---|---|
| 790 | ENSG0000 | MTA3 | metastasis | 3.913076 | 4.854806 | 5.566929 |
| 15483 | ENSG0000 | AHNAK2 | AHNAK nu | 3.710147 | 4.112314 | 5.559218 |
| 4974 | ENSG0000 | WDR11 | WD repeat | 3.811198 | 5.553645 | 5.557036 |
| 2826 | ENSG0000 | TRIP4 | thyroid ho | 3.654164 | 5.181947 | 5.556022 |
| 20128 | ENSG0000 | ANG | angiogenin | 0.813574 | 1.388746 | 5.551277 |
| 17067 | ENSG0000 | GLMP | glycosylate | 3.584788 | 4.344267 | 5.545403 |
| 11297 | ENSG0000 | INTS8 | integrator | 5.222458 | 5.481165 | 5.531993 |
| 14293 | ENSG0000 | LBX2 | ladybird ho | 2.245369 | 3.174667 | 5.527858 |
| 1268 | ENSG0000 | NCBP3 | nuclear ca | 5.319905 | 5.439839 | 5.526045 |
| 6425 | ENSG0000 | ZNF304 | zinc finger | 2.978417 | 4.501399 | 5.518586 |
| 505 | ENSG0000 | SLC39A9 | solute carr | 5.082559 | 5.39773 | 5.515986 |
| 13960 | ENSG0000 | PIDD1 | p53-induce | 3.133257 | 4.960232 | 5.515791 |
| 6690 | ENSG0000 | IMPA1 | inositol mo | 3.554773 | 5.372405 | 5.514809 |
| 26629 | ENSG0000 | TSPEAR-AS | TSPEAR an | 1.323537 | 5.208294 | 5.512312 |
| 27950 | ENSG0000 | RASSF8-AS | RASSF8 an | 2.424157 | 5.151451 | 5.511741 |
| 9231 | ENSG0000 | UEVLD | UEV and la | 3.831212 | 4.962501 | 5.511462 |
| 4611 | ENSG0000 | ETV3 | ETS variant | 3.714923 | 5.480679 | 5.509902 |
| 6707 | ENSG0000 | RNF122 | ring finger | 1.263705 | 2.509422 | 5.505575 |
| 1907 | ENSG0000 | USP48 | ubiquitin s | 4.494789 | 5.088052 | 5.505452 |
| 2588 | ENSG0000 | NXT2 | nuclear tra | 1.209467 | 2.316963 | 5.502578 |
| 38076 | ENSG0000 | BIVM-ERC | BIVM-ERC | 0.388758 | 4.282967 | 5.502252 |
| 8970 | ENSG0000 | PROSER2 | proline an | 2.010261 | 3.266724 | 5.491526 |
| 11958 | ENSG0000 | NXN | nucleored | 2.592489 | 3.222444 | 5.49079 |
| 16709 | ENSG0000 | IPP | intraciste | 3.971492 | 4.020392 | 5.485061 |
| 10146 | ENSG0000 | SYNJ1 | synaptojan | 4.332552 | 5.235113 | 5.484406 |
| 14064 | ENSG0000 | PARD6G | par-6 fami | 2.445024 | 2.893308 | 5.482795 |
| 4232 | ENSG0000 | XRN1 | 5'-3' exorib | 3.294003 | 4.940323 | 5.47758 |
| 12436 | ENSG0000 | TOR1AIP2 | torsin 1A i | 3.691367 | 4.783035 | 5.475986 |
| 7919 | ENSG0000 | OSGIN1 | oxidative s | 2.917372 | 4.884057 | 5.469801 |
| 11258 | ENSG0000 | SPIDR | scaffolding | 4.609215 | 4.838692 | 5.466487 |
| 5965 | ENSG0000 | EIF2S2P4 | eukaryotic | 4.864035 | 5.318668 | 5.465581 |
| 5264 | ENSG0000 | CNTFR | ciliary neu | 0.943969 | 2.483902 | 5.455665 |
| 10242 | ENSG0000 | ZNF222 | zinc finger | 2.3587 | 5.266328 | 5.453442 |
| 17069 | ENSG0000 | DLL1 | delta like c | 1.158072 | 3.173373 | 5.44879 |
| 4701 | ENSG0000 | MARC2 | mitochond | 2.786012 | 3.698955 | 5.447124 |
| 11432 | ENSG0000 | AKR1E2 | aldo-keto r | 4.871981 | 5.021274 | 5.446775 |
| 4040 | ENSG0000 | GGCX | gamma-glu | 3.467503 | 4.031664 | 5.427952 |
| 1855 | ENSG0000 | ZNF302 | zinc finger | 2.324464 | 5.267447 | 5.422542 |
| 29229 | ENSG0000 | AL662791.4 | | 1.109539 | 3.117606 | 5.419833 |
| 3143 | ENSG0000 | MIER2 | MIER famil | 4.29741 | 5.188215 | 5.417901 |
| 9324 | ENSG0000 | NBAS | neuroblast | 4.777425 | 5.345283 | 5.40929 |
| 708 | ENSG0000 | HEBP2 | heme bind | 4.474966 | 4.970279 | 5.405121 |
| 9321 | ENSG0000 | SAV1 | salvador fa | 3.736225 | 4.135666 | 5.404642 |
| 4343 | ENSG0000 | TANC1 | tetratricop | 3.724119 | 4.973686 | 5.401873 |
| 9620 | ENSG0000 | FAM27C | family with | 3.561222 | 4.277635 | 5.399413 |
| 13763 | ENSG0000 | VSIG10 | V-set and i | 3.741114 | 5.24989 | 5.398035 |
| 26214 | ENSG0000 | SNHG20 | small nucle | 2.927479 | 4.88934 | 5.390848 |

| | | | | | | |
|---|---|---|---|---|---|---|
| 88 | ENSG0000 | SPPL2B | signal pept | 4.293715 | 5.340777 | 5.388064 |
| 13675 | ENSG0000 | RTTN | rotatin [So | 4.125781 | 4.410957 | 5.387492 |
| 6963 | ENSG0000 | GDF11 | growth diff | 2.760204 | 3.87857 | 5.385304 |
| 9961 | ENSG0000 | RNF111 | ring finger | 3.584779 | 4.416679 | 5.381202 |
| 3238 | ENSG0000 | HOXA3 | homeobox | 4.419741 | 5.093731 | 5.379789 |
| 12007 | ENSG0000 | EVPL | envoplakin | 2.68764 | 4.270526 | 5.378167 |
| 28627 | ENSG0000 | ARHGDIG | Rho GDP d | 2.327932 | 4.422662 | 5.376203 |
| 15880 | ENSG0000 | SBK2 | SH3 domai | 0.598174 | 1.181299 | 5.374744 |
| 20945 | ENSG0000 | AP4M1 | adaptor re | 3.818231 | 4.374352 | 5.365974 |
| 4288 | ENSG0000 | GORASP1 | golgi reass | 1.656912 | 3.199621 | 5.365241 |
| 1160 | ENSG0000 | VASH1 | vasohibin 1 | 3.444691 | 4.355677 | 5.361909 |
| 12305 | ENSG0000 | RAB24 | RAB24, me | 3.204264 | 5.261609 | 5.35723 |
| 5618 | ENSG0000 | TNFSF9 | TNF superf | 1.196902 | 2.906052 | 5.350108 |
| 27853 | ENSG0000 | ATP6V1B1- | ATP6V1B1 | 2.330929 | 3.775417 | 5.345614 |
| 11087 | ENSG0000 | FBXO8 | F-box prot | 3.049714 | 5.130093 | 5.342407 |
| 38178 | ENSG0000 | AL391422.3 | | 0.865178 | 4.935389 | 5.336912 |
| 8219 | ENSG0000 | CTTNBP2N | CTTNBP2 N | 2.640848 | 4.297132 | 5.336014 |
| 10532 | ENSG0000 | ZYG11B | zyg-11 fam | 4.081281 | 4.448684 | 5.334555 |
| 33095 | ENSG0000 | KIAA1147 | KIAA1147 \| | 2.816446 | 4.839922 | 5.326726 |
| 16469 | ENSG0000 | LAMA2 | laminin sub | 1.882597 | 3.375989 | 5.319932 |
| 3259 | ENSG0000 | PLEKHA8 | pleckstrin l | 3.01668 | 3.647736 | 5.319183 |
| 355 | ENSG0000 | CALCOCO1 | calcium bir | 1.651915 | 4.295386 | 5.317785 |
| 40864 | ENSG0000 | AC243773.2 | | 3.453902 | 4.250506 | 5.316666 |
| 10505 | ENSG0000 | NEU3 | neuraminic | 4.614628 | 4.848932 | 5.314122 |
| 17031 | ENSG0000 | LRBA | LPS respon | 2.691541 | 4.774144 | 5.31308 |
| 34368 | ENSG0000 | AC004943.2 | | 3.219668 | 4.039722 | 5.308784 |
| 21950 | ENSG0000 | AL590617.2 | | 2.730662 | 4.401305 | 5.307667 |
| 13483 | ENSG0000 | ZNF654 | zinc finger | 2.823129 | 4.261249 | 5.306263 |
| 14952 | ENSG0000 | JRKL | JRK like [Sc | 3.148719 | 3.933695 | 5.302385 |
| 14235 | ENSG0000 | FUCA1 | fucosidase, | 1.840533 | 2.81176 | 5.299812 |
| 14727 | ENSG0000 | AP1S2 | adaptor re | 3.162477 | 4.292309 | 5.295752 |
| 13175 | ENSG0000 | SNCG | synuclein g | 0.840178 | 0.917753 | 5.289809 |
| 6416 | ENSG0000 | PEX11B | peroxisom | 3.646153 | 3.986617 | 5.285056 |
| 20077 | ENSG0000 | TMEM167I | transmeml | 4.623746 | 4.778477 | 5.284754 |
| 854 | ENSG0000 | ELMO2 | engulfmen | 4.106415 | 5.232341 | 5.283222 |
| 19634 | ENSG0000 | SFT2D2 | SFT2 doma | 3.437514 | 5.109797 | 5.281647 |
| 27931 | ENSG0000 | PLEKHO2 | pleckstrin l | 3.125805 | 4.164867 | 5.279055 |
| 8096 | ENSG0000 | CACNG8 | calcium vo | 0.022769 | 5.134736 | 5.276137 |
| 3349 | ENSG0000 | CORO2A | coronin 2A | 1.132528 | 3.172046 | 5.275586 |
| 14695 | ENSG0000 | TDRKH | tudor and l | 3.054906 | 4.096627 | 5.271368 |
| 11774 | ENSG0000 | TEF | TEF, PAR b | 4.25914 | 5.009584 | 5.263948 |
| 12769 | ENSG0000 | NPTX1 | neuronal p | 1.939923 | 2.344428 | 5.263614 |
| 39024 | ENSG0000 | AL136370.1 | | 4.393769 | 4.607505 | 5.262559 |
| 3187 | ENSG0000 | ZNF14 | zinc finger | 1.856423 | 3.919975 | 5.25819 |
| 15738 | ENSG0000 | LYRM7 | LYR motif c | 4.185195 | 4.829168 | 5.254179 |
| 470 | ENSG0000 | PLEKHO1 | pleckstrin l | 2.282907 | 3.147611 | 5.253678 |
| 4093 | ENSG0000 | FAM46A | family with | 2.811341 | 3.457836 | 5.251507 |

| | | | | | | |
|---|---|---|---|---|---|---|
| 6429 | ENSG0000 | CHSY1 | chondroiti | 4.0962 | 4.11789 | 5.250849 |
| 11495 | ENSG0000 | ZFYVE1 | zinc finger | 2.571443 | 3.775781 | 5.249155 |
| 2289 | ENSG0000 | KIAA0930 | KIAA0930 | | 4.706535 | 4.927294 | 5.247115 |
| 9488 | ENSG0000 | SCOC | short coile | 3.145319 | 5.104847 | 5.238685 |
| 36674 | ENSG0000 | AC080112.2 | | 1.194461 | 3.048936 | 5.236462 |
| 14385 | ENSG0000 | TRNAU1AF | tRNA selen | 3.393226 | 3.71188 | 5.23395 |
| 12443 | ENSG0000 | KLF13 | Kruppel lik | 3.630742 | 4.331131 | 5.222171 |
| 11317 | ENSG0000 | ABCA1 | ATP bindin | 1.947075 | 3.949799 | 5.220495 |
| 36480 | ENSG0000 | MIR4664 | microRNA | 2.483489 | 3.825983 | 5.219299 |
| 5263 | ENSG0000 | DCAF10 | DDB1 and | 4.578734 | 4.802163 | 5.213742 |
| 1677 | ENSG0000 | SCAMP1 | secretory c | 3.170883 | 3.26121 | 5.213433 |
| 18518 | ENSG0000 | PLPP6 | phospholip | 3.756387 | 4.618855 | 5.209621 |
| 40010 | ENSG0000 | AL627230.2 | | 2.289375 | 5.193662 | 5.207765 |
| 12916 | ENSG0000 | SRGAP2C | SLIT-ROBO | 3.824083 | 4.489165 | 5.207611 |
| 20169 | ENSG0000 | LRRC37A4I | leucine ricl | 2.880722 | 4.104582 | 5.207191 |
| 4688 | ENSG0000 | MTFR1L | mitochond | 3.306598 | 4.637355 | 5.201113 |
| 6287 | ENSG0000 | PLXNA3 | plexin A3 [ | 1.893881 | 4.482805 | 5.200108 |
| 13127 | ENSG0000 | ZNF680 | zinc finger | 3.552674 | 4.113331 | 5.198052 |
| 4331 | ENSG0000 | EPB41L5 | erythrocyt | 2.996891 | 4.31758 | 5.19622 |
| 1064 | ENSG0000 | IFT80 | intraflagell | 2.421622 | 3.233196 | 5.186252 |
| 39054 | ENSG0000 | AC012306.2 | | 3.901711 | 4.297176 | 5.184096 |
| 316 | ENSG0000 | APBA3 | amyloid be | 3.575291 | 3.795798 | 5.181988 |
| 6520 | ENSG0000 | KDM6B | lysine dem | 2.259947 | 2.33887 | 5.17938 |
| 39303 | ENSG0000 | AL391121.1 | | 2.438146 | 4.417476 | 5.176553 |
| 6294 | ENSG0000 | C12orf65 | chromosor | 3.338752 | 4.117993 | 5.174553 |
| 4324 | ENSG0000 | FAHD2A | fumarylace | 4.210398 | 4.473862 | 5.174155 |
| 10191 | ENSG0000 | CYB5R1 | cytochrom | 3.153203 | 4.042351 | 5.171757 |
| 5951 | ENSG0000 | IRF5 | interferon | 2.444468 | 3.678563 | 5.163997 |
| 343 | ENSG0000 | BRCA1 | BRCA1, DN | 3.900716 | 4.76056 | 5.161667 |
| 1043 | ENSG0000 | MEF2A | myocyte er | 3.366204 | 4.064943 | 5.158968 |
| 15101 | ENSG0000 | TRAIP | TRAF inter | 4.100328 | 4.152657 | 5.157981 |
| 16420 | ENSG0000 | S100A5 | S100 calciu | 3.043158 | 3.881954 | 5.150683 |
| 24724 | ENSG0000 | SNORD62B | small nucle | 4.38602 | 4.599537 | 5.150445 |
| 35788 | ENSG0000 | AC068152.1 | | 3.134988 | 4.15633 | 5.148571 |
| 16991 | ENSG0000 | ZNF480 | zinc finger | 4.535987 | 5.120182 | 5.142073 |
| 1905 | ENSG0000 | CERS4 | ceramide s | 3.281906 | 3.843867 | 5.136354 |
| 6240 | ENSG0000 | ZNF337 | zinc finger | 2.297988 | 5.066596 | 5.131317 |
| 27427 | ENSG0000 | LINC00857 | long interg | 3.208317 | 4.522542 | 5.12597 |
| 4201 | ENSG0000 | EHHADH | enoyl-CoA | 2.641384 | 3.350767 | 5.122636 |
| 19040 | ENSG0000 | AC016747.1 | | 2.200916 | 4.659287 | 5.114582 |
| 7019 | ENSG0000 | KLHL36 | kelch like f | 3.728601 | 3.760531 | 5.114516 |
| 4233 | ENSG0000 | KAT2B | lysine acet | 2.994082 | 4.162148 | 5.109721 |
| 10123 | ENSG0000 | CDA | cytidine de | 0.64197 | 1.045041 | 5.103141 |
| 10464 | ENSG0000 | LEMD2 | LEM doma | 2.191977 | 2.997737 | 5.098575 |
| 14186 | ENSG0000 | CTC1 | CST telome | 3.785399 | 4.449757 | 5.095059 |
| 16848 | ENSG0000 | ZNF823 | zinc finger | 3.159362 | 3.625727 | 5.090932 |
| 9127 | ENSG0000 | TRPT1 | tRNA phos | 2.534789 | 2.695094 | 5.085378 |

| | | | | | | |
|---|---|---|---|---|---|---|
| 10045 | ENSG0000 | ABHD3 | abhydrolas | 3.319828 | 4.959408 | 5.08454 |
| 1029 | ENSG0000 | ROGDI | rogdi hom | 4.691863 | 4.796782 | 5.080466 |
| 11787 | ENSG0000 | ACSF2 | acyl-CoA sy | 2.114713 | 2.8215 | 5.079848 |
| 30730 | ENSG0000 | AC093297.2 | | 0.51292 | 3.911328 | 5.057855 |
| 10105 | ENSG0000 | SLC45A3 | solute carr | 2.080708 | 3.772707 | 5.055795 |
| 15992 | ENSG0000 | MORN2 | MORN rep | 3.129066 | 3.460092 | 5.053861 |
| 16575 | ENSG0000 | FUT11 | fucosyltrar | 2.205706 | 2.563704 | 5.053596 |
| 9198 | ENSG0000 | DIXDC1 | DIX domai | 1.41579 | 4.038334 | 5.047023 |
| 1308 | ENSG0000 | MARK3 | microtubul | 4.097177 | 4.707938 | 5.045054 |
| 3947 | ENSG0000 | MAPK14 | mitogen-ac | 4.481893 | 4.778266 | 5.041527 |
| 11710 | ENSG0000 | LDHD | lactate deh | 0.794295 | 1.404137 | 5.040847 |
| 6715 | ENSG0000 | NUMB | NUMB, enc | 3.212389 | 4.921906 | 5.034823 |
| 8351 | ENSG0000 | HCN3 | hyperpolar | 1.394476 | 2.470228 | 5.03093 |
| 6384 | ENSG0000 | ANKHD1 | ankyrin rep | 4.362989 | 4.595358 | 5.023201 |
| 1989 | ENSG0000 | SNAP23 | synaptosor | 3.930151 | 4.820333 | 5.023007 |
| 1970 | ENSG0000 | SLC22A17 | solute carr | 1.189961 | 2.188802 | 5.020451 |
| 2784 | ENSG0000 | CLUAP1 | clusterin as | 3.389773 | 4.613954 | 5.019851 |
| 10388 | ENSG0000 | MUM1 | melanoma | 2.967235 | 4.581974 | 5.01862 |
| 15733 | ENSG0000 | C17orf58 | chromosor | 3.683578 | 3.720634 | 5.013629 |
| 5151 | ENSG0000 | ADGRB2 | adhesion G | 1.562897 | 1.996992 | 4.99884 |
| 8376 | ENSG0000 | MBOAT2 | membrane | 3.281861 | 4.409752 | 4.99759 |
| 12924 | ENSG0000 | ZNF57 | zinc finger | 2.255745 | 3.892818 | 4.987259 |
| 7633 | ENSG0000 | SCARB2 | scavenger | 2.582216 | 2.657449 | 4.973873 |
| 38913 | ENSG0000 | AC008035.1 | | 1.568314 | 4.521276 | 4.97043 |
| 7967 | ENSG0000 | PRELID3A | PRELI dom | 2.120704 | 4.497233 | 4.968088 |
| 8279 | ENSG0000 | LYPLAL1 | lysophospl | 3.484228 | 4.284893 | 4.963565 |
| 15952 | ENSG0000 | PLSCR3 | phospholip | 3.677025 | 4.291427 | 4.962152 |
| 10363 | ENSG0000 | GBAP1 | glucosylcer | 3.143365 | 3.974851 | 4.956254 |
| 9433 | ENSG0000 | CCNO | cyclin O [S | 4.448076 | 4.908704 | 4.954962 |
| 42549 | ENSG0000 | ABALON | apoptotic I | 3.137515 | 4.679279 | 4.951739 |
| 18473 | ENSG0000 | STAG3L1 | stromal an | 3.402605 | 4.658915 | 4.94964 |
| 11531 | ENSG0000 | CEP295 | centrosom | 3.390021 | 4.382132 | 4.943009 |
| 4104 | ENSG0000 | NUDT12 | nudix hydr | 3.250766 | 3.362817 | 4.9392 |
| 4223 | ENSG0000 | UBE3A | ubiquitin p | 3.741944 | 4.679814 | 4.938835 |
| 12745 | ENSG0000 | MORN4 | MORN rep | 3.485475 | 3.769255 | 4.936044 |
| 13331 | ENSG0000 | ZDHHC24 | zinc finger | 3.057741 | 4.118743 | 4.935357 |
| 6276 | ENSG0000 | HIP1R | huntingtin | 3.898158 | 4.054192 | 4.934634 |
| 10616 | ENSG0000 | AGL | amylo-alph | 2.908637 | 4.352923 | 4.926276 |
| 22243 | ENSG0000 | SLC26A6 | solute carr | 3.558289 | 4.557825 | 4.92577 |
| 2801 | ENSG0000 | QPRT | quinolinate | 0.801614 | 1.050369 | 4.920056 |
| 30850 | ENSG0000 | FOXD1 | forkhead b | 3.62564 | 4.002101 | 4.919084 |
| 10780 | ENSG0000 | MRPS18C | mitochond | 4.50864 | 4.590391 | 4.914501 |
| 42812 | ENSG0000 | AC019117.3 | | 0.530885 | 1.034452 | 4.907874 |
| 12805 | ENSG0000 | ZNF561 | zinc finger | 3.08376 | 4.687764 | 4.90739 |
| 12023 | ENSG0000 | MLST8 | MTOR asso | 4.633197 | 4.847365 | 4.906949 |
| 16695 | ENSG0000 | ADARB1 | adenosine | 4.06118 | 4.772059 | 4.897338 |
| 11211 | ENSG0000 | BMT2 | base meth | 1.604878 | 2.605845 | 4.89632 |

| | | | | | | |
|---|---|---|---|---|---|---|
| 6642 | ENSG0000 | KMT5C | lysine metl | 2.894382 | 3.426633 | 4.896151 |
| 11722 | ENSG0000 | DCTN5 | dynactin su | 4.470437 | 4.557536 | 4.894938 |
| 3231 | ENSG0000 | MET | MET proto | 2.961963 | 4.596198 | 4.890504 |
| 2355 | ENSG0000 | SPTLC2 | serine paln | 3.616542 | 3.62746 | 4.890403 |
| 10511 | ENSG0000 | LBHD1 | LBH domai | 1.523002 | 3.762907 | 4.890322 |
| 1357 | ENSG0000 | CAMSAP3 | calmodulir | 1.314261 | 2.747262 | 4.884566 |
| 23303 | ENSG0000 | AL669831.1 | | 2.58251 | 4.815385 | 4.870474 |
| 32441 | ENSG0000 | AP006621.3 | | 2.382857 | 4.012453 | 4.870309 |
| 9047 | ENSG0000 | PTPRJ | protein tyr | 4.438727 | 4.448733 | 4.869371 |
| 12490 | ENSG0000 | SLC16A5 | solute carr | 2.638931 | 3.433015 | 4.864698 |
| 2493 | ENSG0000 | CDS2 | CDP-diacyl | 3.354563 | 4.193555 | 4.861383 |
| 18096 | ENSG0000 | TMEM57 | transmeml | 3.647857 | 4.508456 | 4.853565 |
| 7849 | ENSG0000 | IGF1R | insulin like | 3.664873 | 4.788336 | 4.850611 |
| 20398 | ENSG0000 | AC135983.1 | | 1.717328 | 2.217509 | 4.837719 |
| 25531 | ENSG0000 | AK4P3 | adenylate l | 4.028254 | 4.43412 | 4.832805 |
| 15532 | ENSG0000 | ADAMTSL5 | ADAMTS li | 3.36567 | 4.109093 | 4.831727 |
| 17167 | ENSG0000 | L1CAM | L1 cell adh | 1.284708 | 4.371173 | 4.831471 |
| 14976 | ENSG0000 | ASB7 | ankyrin rep | 2.637789 | 3.86119 | 4.829372 |
| 34624 | ENSG0000 | ULK4P2 | ULK4 pseu | 1.846994 | 2.809952 | 4.826748 |
| 35187 | ENSG0000 | GOLGA8H | golgin A8 f | 1.602718 | 3.189668 | 4.826594 |
| 1551 | ENSG0000 | RSBN1 | round sper | 3.030274 | 3.765187 | 4.826136 |
| 12264 | ENSG0000 | MECP2 | methyl-CpG | 4.255498 | 4.81047 | 4.824995 |
| 40723 | ENSG0000 | AC015922.4 | | 1.071256 | 3.046862 | 4.817876 |
| 14407 | ENSG0000 | HERC2P3 | hect doma | 2.108487 | 3.781202 | 4.80298 |
| 9060 | ENSG0000 | INTS4 | integrator | 4.474256 | 4.704253 | 4.802592 |
| 28749 | ENSG0000 | DECR2 | 2,4-dienoy | 3.466195 | 3.942362 | 4.794372 |
| 15867 | ENSG0000 | COL4A1 | collagen ty | 3.376695 | 3.825929 | 4.794157 |
| 23077 | ENSG0000 | AP000344.1 | | 0.832327 | 1.437987 | 4.791939 |
| 41572 | ENSG0000 | AL031587.5 | | 3.311386 | 3.92138 | 4.789563 |
| 6864 | ENSG0000 | TCN1 | transcobal | 2.896369 | 3.324331 | 4.788378 |
| 16818 | ENSG0000 | OCLN | occludin [S | 2.188871 | 3.073899 | 4.78714 |
| 2487 | ENSG0000 | RASSF2 | Ras associa | 1.483103 | 1.814064 | 4.783167 |
| 15383 | ENSG0000 | FAM43A | family witl | 1.679711 | 2.922303 | 4.780734 |
| 1741 | ENSG0000 | HSD17B14 | hydroxyste | 0.778353 | 1.836712 | 4.780145 |
| 5881 | ENSG0000 | PMS2P3 | PMS1 hom | 2.75385 | 4.204022 | 4.778638 |
| 15107 | ENSG0000 | ZNF703 | zinc finger | 2.487818 | 3.015004 | 4.776206 |
| 7139 | ENSG0000 | C7orf69 | chromosor | 0.705539 | 2.726042 | 4.774421 |
| 386 | ENSG0000 | MTMR11 | myotubula | 1.97861 | 2.864551 | 4.772129 |
| 18659 | ENSG0000 | HERC2P9 | hect doma | 2.925866 | 4.480459 | 4.764354 |
| 21265 | ENSG0000 | AC135983.2 | | 2.941959 | 3.997078 | 4.76314 |
| 35978 | ENSG0000 | AP005230.1 | | 1.156394 | 3.814166 | 4.75249 |
| 9471 | ENSG0000 | SREK1IP1 | SREK1 inte | 2.98387 | 4.662424 | 4.747925 |
| 7793 | ENSG0000 | DCAF5 | DDB1 and | 2.119994 | 2.946004 | 4.744778 |
| 7638 | ENSG0000 | SHROOM3 | shroom far | 2.698722 | 3.676471 | 4.739297 |
| 3801 | ENSG0000 | PPM1H | protein ph | 0.487425 | 3.272323 | 4.738842 |
| 7726 | ENSG0000 | LNX2 | ligand of n | 2.755312 | 4.051352 | 4.735199 |
| 36465 | ENSG0000 | AP000275.2 | | 2.371972 | 4.403235 | 4.734322 |

| | | | | | | |
|---|---|---|---|---|---|---|
| 16853 | ENSG0000 | FCHSD1 | FCH and do | 1.787001 | 4.525724 | 4.732257 |
| 1569 | ENSG0000 | CDC14B | cell divisio | 1.752445 | 4.329881 | 4.727131 |
| 7954 | ENSG0000 | SCRN2 | secernin 2 | 1.796373 | 2.48828 | 4.724855 |
| 11826 | ENSG0000 | STIM1 | stromal int | 3.04065 | 3.961186 | 4.719833 |
| 9572 | ENSG0000 | OTULIN | OTU deubi | 4.045483 | 4.288468 | 4.714504 |
| 13109 | ENSG0000 | LCLAT1 | lysocardiol | 3.060627 | 3.967762 | 4.712866 |
| 29767 | ENSG0000 | AC004997.1 | | 0.908473 | 2.991699 | 4.710616 |
| 13700 | ENSG0000 | HSD11B2 | hydroxyste | 0.618419 | 1.458128 | 4.708573 |
| 16668 | ENSG0000 | SYNGAP1 | synaptic Ra | 2.447078 | 3.38112 | 4.704843 |
| 3371 | ENSG0000 | RIC1 | RIC1 homo | 3.801135 | 4.200845 | 4.68693 |
| 16827 | ENSG0000 | SGTB | small gluta | 1.738201 | 3.142652 | 4.683208 |
| 12310 | ENSG0000 | EFNA1 | ephrin A1 \| | 1.669673 | 1.891492 | 4.682429 |
| 9201 | ENSG0000 | C11orf57 | chromosor | 2.206233 | 3.87369 | 4.681451 |
| 42207 | ENSG0000 | AL627171.2 | | 0.734281 | 4.376556 | 4.680304 |
| 12446 | ENSG0000 | ZNF764 | zinc finger | 2.689184 | 4.198752 | 4.672807 |
| 6704 | ENSG0000 | ZFC3H1 | zinc finger | 3.632002 | 4.340794 | 4.672323 |
| 16738 | ENSG0000 | TTC30A | tetratricop | 1.122584 | 3.053167 | 4.669716 |
| 39041 | ENSG0000 | GTF2IP20 | general tra | 2.068388 | 4.494674 | 4.667592 |
| 8298 | ENSG0000 | MINDY1 | MINDY lysi | 3.297994 | 3.52762 | 4.666946 |
| 16914 | ENSG0000 | HIBCH | 3-hydroxyi | 3.411285 | 3.74923 | 4.665109 |
| 5929 | ENSG0000 | EMC4 | ER membra | 4.102791 | 4.443573 | 4.662194 |
| 9935 | ENSG0000 | ACOX1 | acyl-CoA o | 3.929548 | 4.298395 | 4.661476 |
| 9111 | ENSG0000 | DUSP15 | dual specif | 0.103117 | 3.290628 | 4.660113 |
| 39902 | ENSG0000 | AL355338.1 | | 2.825542 | 3.758813 | 4.657326 |
| 11344 | ENSG0000 | ANKS6 | ankyrin rep | 3.952418 | 4.612641 | 4.655236 |
| 1667 | ENSG0000 | MAPRE3 | microtubul | 2.88427 | 3.367989 | 4.654939 |
| 6849 | ENSG0000 | ELP2 | elongator a | 4.053402 | 4.612115 | 4.650089 |
| 1037 | ENSG0000 | RASSF1 | Ras associa | 3.139406 | 3.67911 | 4.63798 |
| 35090 | ENSG0000 | CCPG1 | cell cycle p | 2.497866 | 4.339619 | 4.634308 |
| 11998 | ENSG0000 | ZNF232 | zinc finger | 1.70767 | 2.341809 | 4.633545 |
| 24602 | ENSG0000 | SBK3 | SH3 domai | 0.821261 | 1.183537 | 4.631616 |
| 18159 | ENSG0000 | NEU1 | neuraminic | 2.77968 | 2.860163 | 4.631391 |
| 3487 | ENSG0000 | RAPGEFL1 | Rap guanir | 2.464353 | 3.1391 | 4.630569 |
| 7486 | ENSG0000 | DBT | dihydrolipo | 3.070259 | 4.057432 | 4.626686 |
| 9600 | ENSG0000 | NEIL2 | nei like DN | 4.320655 | 4.560479 | 4.625521 |
| 11966 | ENSG0000 | CYB5D2 | cytochrom | 2.560245 | 3.611282 | 4.624015 |
| 26436 | ENSG0000 | AC068768.1 | | 2.11687 | 3.219231 | 4.623076 |
| 10379 | ENSG0000 | FGFR4 | fibroblast g | 1.618544 | 2.882184 | 4.621192 |
| 12447 | ENSG0000 | ZNF747 | zinc finger | 1.49654 | 3.142195 | 4.617652 |
| 11544 | ENSG0000 | JAM3 | junctional a | 3.053441 | 3.341531 | 4.613147 |
| 42316 | ENSG0000 | AC022107.1 | | 1.75197 | 4.377535 | 4.611688 |
| 6595 | ENSG0000 | ZMYM5 | zinc finger | 3.197475 | 4.041666 | 4.610598 |
| 5044 | ENSG0000 | PROSER1 | proline and | 1.727631 | 2.265292 | 4.60488 |
| 11700 | ENSG0000 | MARF1 | meiosis reg | 2.727435 | 4.474812 | 4.604238 |
| 840 | ENSG0000 | SPAG4 | sperm asso | 0.476366 | 0.75992 | 4.592472 |
| 1408 | ENSG0000 | PIK3C3 | phosphatic | 2.119198 | 4.058613 | 4.592069 |
| 7105 | ENSG0000 | RCBTB1 | RCC1 and E | 3.462053 | 3.927859 | 4.582523 |

| | | | | | | |
|---|---|---|---|---|---|---|
| 2101 | ENSG0000 | KCNK6 | potassium | 1.327473 | 3.077971 | 4.582235 |
| 12502 | ENSG0000 | FAM161A | family with | 2.175204 | 3.181222 | 4.580963 |
| 20428 | ENSG0000 | RPL17-C18 | RPL17-C18 | 3.561314 | 3.843744 | 4.577637 |
| 18078 | ENSG0000 | C2orf72 | chromosor | 0.996066 | 1.730353 | 4.57686 |
| 5934 | ENSG0000 | POT1 | protection | 2.164819 | 4.181227 | 4.573188 |
| 22962 | ENSG0000 | GEMIN8P4 | gem nucle | 2.454844 | 3.734468 | 4.568638 |
| 15188 | ENSG0000 | KCNJ12 | potassium | 2.749559 | 3.65873 | 4.562289 |
| 1022 | ENSG0000 | PMS2P4 | PMS1 hom | 3.016674 | 3.563798 | 4.561922 |
| 12380 | ENSG0000 | LUZP1 | leucine zip | 2.915933 | 4.478254 | 4.559094 |
| 3711 | ENSG0000 | ST3GAL4 | ST3 beta-g | 3.54519 | 4.077725 | 4.558578 |
| 18306 | ENSG0000 | ZBTB48 | zinc finger | 3.629594 | 4.536501 | 4.556325 |
| 20220 | ENSG0000 | AL161911. | Putative U | 2.626491 | 3.79714 | 4.554732 |
| 7018 | ENSG0000 | MDM2 | MDM2 pro | 2.01167 | 4.424754 | 4.554162 |
| 12802 | ENSG0000 | DLK2 | delta like n | 1.73094 | 2.865217 | 4.554155 |
| 35669 | ENSG0000 | AC004771.3 | | 1.409188 | 2.581578 | 4.552614 |
| 431 | ENSG0000 | ADGRA2 | adhesion G | 1.117628 | 1.270426 | 4.545721 |
| 11568 | ENSG0000 | POLL | DNA polym | 3.274267 | 3.895545 | 4.545655 |
| 15624 | ENSG0000 | ZBTB6 | zinc finger | 2.512448 | 4.344234 | 4.545026 |
| 11957 | ENSG0000 | ZNF444 | zinc finger | 3.310209 | 4.305968 | 4.544833 |
| 14983 | ENSG0000 | PI4KAP2 | phosphatic | 2.819216 | 4.532181 | 4.541105 |
| 5125 | ENSG0000 | ZSCAN18 | zinc finger | 1.418742 | 4.475819 | 4.539287 |
| 29159 | ENSG0000 | DNAJC25-C | DNAJC25-C | 3.640078 | 3.700454 | 4.538417 |
| 2682 | ENSG0000 | SUPT20H | SPT20 hom | 3.479091 | 4.42722 | 4.537019 |
| 1328 | ENSG0000 | PAX2 | paired box | 1.348899 | 2.055669 | 4.536387 |
| 1395 | ENSG0000 | FBLN1 | fibulin 1 [S | 2.664723 | 2.796038 | 4.535908 |
| 7300 | ENSG0000 | LMX1B | LIM homec | 3.800394 | 3.991288 | 4.535402 |
| 7939 | ENSG0000 | GFOD2 | glucose-fru | 4.008739 | 4.325071 | 4.532933 |
| 6246 | ENSG0000 | OSBPL2 | oxysterol b | 3.341244 | 4.369329 | 4.523748 |
| 14469 | ENSG0000 | ZNF594 | zinc finger | 2.160144 | 4.171086 | 4.523087 |
| 40887 | ENSG0000 | ZNF670 | zinc finger | 1.970083 | 2.813151 | 4.521417 |
| 28182 | ENSG0000 | RPL23AP7 | ribosomal | 3.883639 | 4.376099 | 4.520314 |
| 15900 | ENSG0000 | ZKSCAN4 | zinc finger | 2.318021 | 3.972541 | 4.514773 |
| 39986 | ENSG0000 | AC068473.5 | | 2.600933 | 3.403768 | 4.511526 |
| 34351 | ENSG0000 | ZNF444P1 | zinc finger | 2.382863 | 2.401276 | 4.511148 |
| 11361 | ENSG0000 | PIGA | phosphatic | 3.934642 | 4.041087 | 4.510115 |
| 17159 | ENSG0000 | SHISA4 | shisa famil | 1.155247 | 1.912669 | 4.509934 |
| 16440 | ENSG0000 | SIAH1 | siah E3 ubi | 3.273915 | 3.852888 | 4.506159 |
| 11469 | ENSG0000 | RET | ret proto-c | 1.026978 | 1.785435 | 4.505984 |
| 12947 | ENSG0000 | EIF2AK3 | eukaryotic | 2.862264 | 3.316073 | 4.505303 |
| 42610 | ENSG0000 | LINC02246 | long interg | 2.873556 | 3.398385 | 4.504694 |
| 12089 | ENSG0000 | POLR2J3 | RNA polym | 3.87701 | 4.324819 | 4.494655 |
| 12143 | ENSG0000 | BMP1 | bone morp | 3.955537 | 4.091792 | 4.493768 |
| 39292 | ENSG0000 | AC009237.14 | | 2.14842 | 3.604666 | 4.492232 |
| 16290 | ENSG0000 | C8orf76 | chromosor | 3.832398 | 4.309966 | 4.489526 |
| 14798 | ENSG0000 | LINC01006 | long interg | 0.313769 | 3.279777 | 4.488994 |
| 14889 | ENSG0000 | AFMID | arylformar | 2.835531 | 3.414682 | 4.487504 |
| 13438 | ENSG0000 | CNIH2 | cornichon | 1.788263 | 2.59944 | 4.486924 |

| | | | | | | |
|---|---|---|---|---|---|---|
| 17972 | ENSG0000 | TATDN3 | TatD DNas | 2.604235 | 3.259247 | 4.48563 |
| 34428 | ENSG0000 | AC009113.1 | | 2.224134 | 3.952229 | 4.483593 |
| 22284 | ENSG0000 | TRAM2-AS | TRAM2 ant | 2.07918 | 3.157839 | 4.480585 |
| 7790 | ENSG0000 | NAA30 | N(alpha)-a | 4.168635 | 4.33458 | 4.480207 |
| 1780 | ENSG0000 | AAMDC | adipogene | 2.968066 | 3.641144 | 4.476378 |
| 10206 | ENSG0000 | UBR1 | ubiquitin p | 2.549943 | 4.12056 | 4.469397 |
| 872 | ENSG0000 | ZNF275 | zinc finger | 2.643826 | 3.576728 | 4.465545 |
| 9391 | ENSG0000 | POC5 | POC5 centr | 2.825111 | 4.168497 | 4.461093 |
| 16062 | ENSG0000 | SBK1 | SH3 domai | 1.43314 | 2.058836 | 4.457628 |
| 10425 | ENSG0000 | THAP8 | THAP dom | 1.977462 | 2.843686 | 4.457612 |
| 1717 | ENSG0000 | EPDR1 | ependymir | 3.414071 | 4.043371 | 4.45644 |
| 11446 | ENSG0000 | ZNF503 | zinc finger | 2.837465 | 3.477836 | 4.456234 |
| 1217 | ENSG0000 | DERL2 | derlin 2 [Sc | 2.968335 | 4.222107 | 4.455046 |
| 15181 | ENSG0000 | ADRA2C | adrenocep | 2.295949 | 4.183216 | 4.453662 |
| 8998 | ENSG0000 | ADD3 | adducin 3 \| | 1.643364 | 2.811132 | 4.453131 |
| 1660 | ENSG0000 | NCOA1 | nuclear rec | 3.859572 | 3.903196 | 4.45312 |
| 32074 | ENSG0000 | AP003068.2 | | 3.285742 | 3.766189 | 4.45296 |
| 10320 | ENSG0000 | SLC2A6 | solute carr | 3.234458 | 3.507963 | 4.449925 |
| 11552 | ENSG0000 | RAB8B | RAB8B, me | 2.114038 | 2.591185 | 4.446831 |
| 6103 | ENSG0000 | TEP1 | telomerase | 1.999802 | 3.917585 | 4.445724 |
| 36273 | ENSG0000 | AC026271.4 | | 2.429624 | 3.981996 | 4.444674 |
| 40836 | ENSG0000 | AC005696.4 | | 0.720397 | 1.895251 | 4.443825 |
| 14711 | ENSG0000 | MOB2 | MOB kinas | 3.208899 | 3.658746 | 4.441256 |
| 7945 | ENSG0000 | TOM1L1 | target of m | 3.042794 | 4.242871 | 4.439242 |
| 5147 | ENSG0000 | ZMYM2 | zinc finger | 3.666109 | 3.696998 | 4.437649 |
| 11919 | ENSG0000 | NCKAP5L | NCK associ | 2.568002 | 3.703184 | 4.433164 |
| 321 | ENSG0000 | MBTD1 | mbt domai | 3.096874 | 3.775623 | 4.431842 |
| 42461 | ENSG0000 | PCAT14 | prostate ca | 0.500494 | 1.007761 | 4.43093 |
| 13156 | ENSG0000 | AHSA2 | activator o | 2.544077 | 4.242573 | 4.42528 |
| 13710 | ENSG0000 | WDR25 | WD repeat | 1.336533 | 2.545566 | 4.423927 |
| 16712 | ENSG0000 | OGDHL | oxoglutara | 0.835398 | 2.17357 | 4.420215 |
| 292 | ENSG0000 | SEMA3G | semaphori | 1.111902 | 2.352716 | 4.411358 |
| 39943 | ENSG0000 | AC005332.7 | | 1.300265 | 3.425872 | 4.408253 |
| 4160 | ENSG0000 | LNPEP | leucyl and | 3.51524 | 4.37935 | 4.397119 |
| 5632 | ENSG0000 | GPCPD1 | glyceropho | 1.125872 | 3.860303 | 4.394306 |
| 2847 | ENSG0000 | DMXL2 | Dmx like 2 | 2.220917 | 3.083168 | 4.390537 |
| 15023 | ENSG0000 | TRMT2B | tRNA meth | 3.771549 | 3.800263 | 4.390455 |
| 18441 | ENSG0000 | CNEP1R1 | CTD nuclea | 1.668562 | 1.70487 | 4.389734 |
| 8304 | ENSG0000 | SEMA6C | semaphori | 1.065295 | 2.150203 | 4.388719 |
| 19605 | ENSG0000 | RNF208 | ring finger | 3.191709 | 3.911184 | 4.385332 |
| 13927 | ENSG0000 | AC021054.1 | | 0.66428 | 2.691697 | 4.381744 |
| 8659 | ENSG0000 | RNF145 | ring finger | 3.694698 | 4.209599 | 4.377156 |
| 10360 | ENSG0000 | ANO10 | anoctamin | 2.57982 | 3.280349 | 4.371395 |
| 12839 | ENSG0000 | PIK3CD | phosphatic | 1.169821 | 2.236796 | 4.355464 |
| 7687 | ENSG0000 | PIANP | PILR alpha | 0.827427 | 1.684618 | 4.354414 |
| 16340 | ENSG0000 | TMEM63A | transmeml | 2.944164 | 3.781938 | 4.353382 |
| 4424 | ENSG0000 | STK16 | serine/thre | 3.03598 | 4.159763 | 4.34861 |

| | | | | | | |
|---|---|---|---|---|---|---|
| 13128 | ENSG0000 | HECTD4 | HECT dom: | 2.539662 | 3.745164 | 4.345733 |
| 12078 | ENSG0000 | HOOK3 | hook micr | 3.281853 | 4.020275 | 4.338428 |
| 4086 | ENSG0000 | VEGFA | vascular er | 2.167021 | 2.326664 | 4.336801 |
| 4215 | ENSG0000 | NPHP3 | nephrocyst | 2.99911 | 3.829894 | 4.336259 |
| 15300 | ENSG0000 | SEPT5 | septin 5 [S | 2.725019 | 2.977589 | 4.331571 |
| 18097 | ENSG0000 | PTPN20 | protein tyr | 3.241492 | 4.28552 | 4.331313 |
| 2482 | ENSG0000 | RNF24 | ring finger | 1.602353 | 3.261429 | 4.329473 |
| 5608 | ENSG0000 | INSIG2 | insulin indu | 1.595641 | 2.412635 | 4.325611 |
| 11155 | ENSG0000 | ANKRA2 | ankyrin rep | 2.00361 | 3.213916 | 4.321039 |
| 762 | ENSG0000 | KMT2C | lysine metl | 3.55133 | 4.292381 | 4.319586 |
| 10649 | ENSG0000 | IER5 | immediate | 3.429282 | 3.692691 | 4.314488 |
| 11751 | ENSG0000 | CCNDBP1 | cyclin D1 b | 2.294872 | 3.712405 | 4.308374 |
| 3975 | ENSG0000 | FBXL4 | F-box and l | 3.094267 | 4.058521 | 4.303729 |
| 30058 | ENSG0000 | AL731577.2 | | 2.50054 | 3.830614 | 4.296473 |
| 7543 | ENSG0000 | ASCC1 | activating s | 2.866744 | 3.051373 | 4.29616 |
| 1163 | ENSG0000 | SEL1L | SEL1L ERAI | 3.650886 | 4.249112 | 4.295913 |
| 5855 | ENSG0000 | KDM4B | lysine dem | 2.849753 | 2.959545 | 4.292196 |
| 25430 | ENSG0000 | HMGA1P8 | high mobil | 1.801437 | 3.006937 | 4.289654 |
| 4863 | ENSG0000 | IFI27L2 | interferon | 1.936455 | 2.338266 | 4.287077 |
| 9645 | ENSG0000 | FLCN | folliculin [S | 2.250327 | 3.802939 | 4.279557 |
| 7950 | ENSG0000 | SGSM2 | small G pro | 3.542775 | 4.224103 | 4.276904 |
| 33702 | ENSG0000 | AP001781.3 | | 0.038502 | 2.897407 | 4.276715 |
| 42329 | ENSG0000 | AC233723.2 | | 2.01292 | 3.196213 | 4.270186 |
| 8758 | ENSG0000 | LINC00525 | long interg | 1.49301 | 3.661683 | 4.268109 |
| 40485 | ENSG0000 | AC027682.6 | | 0.331891 | 1.322025 | 4.256885 |
| 36735 | ENSG0000 | AP002478.1 | | 1.030865 | 2.900825 | 4.255791 |
| 5181 | ENSG0000 | POLK | DNA polym | 3.057416 | 3.742534 | 4.255052 |
| 13227 | ENSG0000 | SNX33 | sorting nex | 1.994508 | 2.490308 | 4.251337 |
| 10967 | ENSG0000 | ZNF513 | zinc finger | 1.89717 | 3.769566 | 4.249631 |
| 2382 | ENSG0000 | PCNX1 | pecanex hc | 2.987189 | 3.817648 | 4.243724 |
| 6978 | ENSG0000 | COQ10A | coenzyme | 1.722495 | 3.180828 | 4.23929 |
| 11200 | ENSG0000 | DAGLB | diacylglyce | 3.850438 | 4.184487 | 4.236141 |
| 16190 | ENSG0000 | USP32P1 | ubiquitin sp | 1.725411 | 4.113969 | 4.229072 |
| 4531 | ENSG0000 | GON4L | gon-4 like | | 3.078775 | 3.731657 | 4.228535 |
| 23144 | ENSG0000 | AC007383.2 | | 3.316676 | 3.937959 | 4.227482 |
| 9767 | ENSG0000 | ELMSAN1 | ELM2 and | 2.458993 | 3.51546 | 4.220643 |
| 103 | ENSG0000 | SOX8 | SRY-box 8 | 2.353683 | 3.547243 | 4.220133 |
| 444 | ENSG0000 | OSBPL5 | oxysterol b | 3.742924 | 4.049745 | 4.217503 |
| 5521 | ENSG0000 | RREB1 | ras respons | 2.571834 | 3.33639 | 4.214684 |
| 7573 | ENSG0000 | FAM117B | family with | 3.04967 | 3.809243 | 4.214646 |
| 8008 | ENSG0000 | NARF | nuclear pre | 2.277686 | 3.204503 | 4.213956 |
| 16186 | ENSG0000 | FBF1 | Fas binding | 2.93402 | 3.245641 | 4.21202 |
| 4796 | ENSG0000 | HS1BP3 | HCLS1 binc | 2.461731 | 3.469326 | 4.211622 |
| 8140 | ENSG0000 | DDAH1 | dimethylar | 3.060108 | 3.955985 | 4.207279 |
| 19220 | ENSG0000 | MIR760 | microRNA | 1.651892 | 3.815655 | 4.205673 |
| 15505 | ENSG0000 | BRWD1 | bromodom | 3.336247 | 3.833172 | 4.20538 |
| 5362 | ENSG0000 | TTC21B | tetratricop | 3.210205 | 3.961658 | 4.204995 |

| | | | | | | |
|---|---|---|---|---|---|---|
| 11461 | ENSG0000 | SPACA9 | sperm acro | 1.887029 | 2.388869 | 4.204319 |
| 23086 | ENSG0000 | AL031282.1 | | 3.403713 | 4.152027 | 4.199172 |
| 14849 | ENSG0000 | TSPEAR-AS | TSPEAR an | 0.341771 | 2.996296 | 4.19537 |
| 280 | ENSG0000 | ANO8 | anoctamin | 3.146554 | 3.177794 | 4.195315 |
| 6628 | ENSG0000 | RFXAP | regulatory | 3.061264 | 3.446327 | 4.190702 |
| 2362 | ENSG0000 | DHRS7 | dehydroge | 1.941469 | 2.721331 | 4.190005 |
| 11351 | ENSG0000 | METTL27 | methyltrar | 1.412802 | 2.685676 | 4.18544 |
| 21454 | ENSG0000 | AC009948.1 | | 1.886504 | 3.748774 | 4.182128 |
| 1392 | ENSG0000 | FGFR1 | fibroblast g | 2.776724 | 3.580259 | 4.182081 |
| 14860 | ENSG0000 | GJC1 | gap junctic | 3.648994 | 3.714405 | 4.179223 |
| 6201 | ENSG0000 | ZSWIM6 | zinc finger | 2.429576 | 4.074699 | 4.176828 |
| 1438 | ENSG0000 | TP73 | tumor prot | 2.27574 | 2.854783 | 4.174218 |
| 5149 | ENSG0000 | GJA3 | gap junctic | 2.32209 | 3.050651 | 4.164457 |
| 6745 | ENSG0000 | DPH6 | diphthamir | 2.266271 | 2.536149 | 4.161747 |
| 7001 | ENSG0000 | MICAL1 | microtubul | 2.778416 | 4.043359 | 4.160458 |
| 9953 | ENSG0000 | DHRS1 | dehydroge | 1.587117 | 2.398822 | 4.159592 |
| 7235 | ENSG0000 | CCDC115 | coiled-coil | 2.285004 | 3.909998 | 4.158704 |
| 8196 | ENSG0000 | NBPF3 | NBPF mem | 2.419884 | 3.156555 | 4.158054 |
| 11874 | ENSG0000 | RILP | Rab intera | 2.003031 | 3.072034 | 4.152558 |
| 14645 | ENSG0000 | RFX7 | regulatory | 2.787922 | 3.779516 | 4.151665 |
| 14701 | ENSG0000 | CREB3L2 | cAMP resp | 2.893486 | 3.296104 | 4.15067 |
| 3966 | ENSG0000 | RBM24 | RNA bindir | 1.693334 | 2.426948 | 4.144944 |
| 10718 | ENSG0000 | SLC16A14 | solute carr | 0.721723 | 2.890074 | 4.143443 |
| 9397 | ENSG0000 | JMY | junction m | 2.080652 | 3.939947 | 4.138815 |
| 246 | ENSG0000 | SPRTN | SprT-like N | 3.260203 | 4.096237 | 4.138114 |
| 31750 | ENSG0000 | AC100810.1 | | 2.16547 | 2.644145 | 4.132879 |
| 9319 | ENSG0000 | AMN1 | antagonist | 2.077309 | 2.876516 | 4.130164 |
| 7203 | ENSG0000 | LIMD2 | LIM domai | 1.638416 | 2.514655 | 4.129999 |
| 23613 | ENSG0000 | AC091729.3 | | 2.995987 | 3.941186 | 4.123213 |
| 6369 | ENSG0000 | PDLIM4 | PDZ and LI | 2.957295 | 3.127813 | 4.117203 |
| 11410 | ENSG0000 | DDIAS | DNA dama | 3.110885 | 3.775683 | 4.106215 |
| 3067 | ENSG0000 | PRX | periaxin [S | 2.969536 | 3.474542 | 4.106033 |
| 19941 | ENSG0000 | PPM1N | protein ph | 0.624479 | 1.295653 | 4.094638 |
| 8806 | ENSG0000 | KRBOX4 | KRAB box c | 2.49246 | 3.286173 | 4.092088 |
| 37343 | ENSG0000 | AC024592.3 | | 3.758336 | 3.904193 | 4.091461 |
| 24657 | ENSG0000 | WASIR2 | WASH and | 1.047407 | 2.053531 | 4.080714 |
| 18121 | ENSG0000 | PSMB8 | proteasom | 0.503532 | 1.635911 | 4.080246 |
| 39212 | ENSG0000 | AC005229.4 | | 2.041407 | 2.639519 | 4.079909 |
| 39753 | ENSG0000 | AC009163.7 | | 0.520809 | 2.80289 | 4.077229 |
| 462 | ENSG0000 | BIRC3 | baculoviral | 1.667022 | 2.373266 | 4.068525 |
| 10264 | ENSG0000 | FNDC5 | fibronectir | 1.269837 | 3.853269 | 4.065531 |
| 16885 | ENSG0000 | ZNF84 | zinc finger | 1.597277 | 3.721848 | 4.060327 |
| 7092 | ENSG0000 | APPL2 | adaptor pr | 3.579278 | 3.907473 | 4.05825 |
| 8316 | ENSG0000 | DYRK3 | dual specif | 2.116111 | 3.113184 | 4.055885 |
| 16054 | ENSG0000 | ZNF669 | zinc finger | 1.345732 | 2.927422 | 4.054589 |
| 16523 | ENSG0000 | CD47 | CD47 mole | 2.335387 | 3.029585 | 4.04787 |
| 2239 | ENSG0000 | DNAL4 | dynein axo | 2.692956 | 3.021415 | 4.046928 |

| | | | | | | |
|---|---|---|---|---|---|---|
| 7496 | ENSG0000 | DYNC2LI1 | dynein cyto | 2.693833 | 3.426397 | 4.045986 |
| 18401 | ENSG0000 | PSMB10 | proteasom | 3.449787 | 3.861503 | 4.045198 |
| 11308 | ENSG0000 | KIAA1161 | KIAA1161 \| | 1.451641 | 2.580626 | 4.041148 |
| 9282 | ENSG0000 | ACAD8 | acyl-CoA d | 3.353493 | 3.916396 | 4.037617 |
| 10395 | ENSG0000 | ORAI2 | ORAI calciu | 2.125496 | 2.685981 | 4.036733 |
| 6084 | ENSG0000 | KLK8 | kallikrein r | 1.767542 | 1.951505 | 4.036042 |
| 11536 | ENSG0000 | AMOTL1 | angiomotir | 2.864126 | 3.230625 | 4.035131 |
| 19922 | ENSG0000 | AC098614.1 | | 1.701403 | 2.659666 | 4.033663 |
| 11419 | ENSG0000 | C10orf25 | chromosor | 0.770248 | 2.362171 | 4.031142 |
| 1334 | ENSG0000 | RBM7 | RNA bindir | 2.820015 | 3.444628 | 4.025179 |
| 2822 | ENSG0000 | LACTB | lactamase | 2.675664 | 3.046013 | 4.023744 |
| 8597 | ENSG0000 | TIFA | TRAF intera | 1.879555 | 2.202133 | 4.017202 |
| 32997 | ENSG0000 | PLBD1-AS1 | PLBD1 anti | 2.53341 | 3.31767 | 4.014918 |
| 626 | ENSG0000 | CNTLN | centlein [S | 2.896443 | 3.779125 | 4.005034 |
| 32299 | ENSG0000 | AC136475.3 | | 1.507021 | 2.346948 | 3.99892 |
| 19809 | ENSG0000 | RBMXL1 | RNA bindir | 3.586086 | 3.991931 | 3.997872 |
| 28965 | ENSG0000 | NAT6 | N-acetyltra | 2.205112 | 3.725567 | 3.996596 |
| 1822 | ENSG0000 | LZTS3 | leucine zip | 1.526317 | 2.718561 | 3.994576 |
| 16265 | ENSG0000 | GJB5 | gap junctic | 2.007791 | 2.932121 | 3.990393 |
| 8386 | ENSG0000 | PPFIA4 | PTPRF inte | 0.173362 | 0.537148 | 3.989601 |
| 14783 | ENSG0000 | EPHB3 | EPH recept | 0.22364 | 0.543093 | 3.986914 |
| 7688 | ENSG0000 | SLC38A4 | solute carr | 3.528727 | 3.876264 | 3.984896 |
| 16578 | ENSG0000 | ANXA4 | annexin A4 | 3.07023 | 3.295274 | 3.981241 |
| 2164 | ENSG0000 | SEC14L2 | SEC14 like | 2.157233 | 3.484146 | 3.972977 |
| 14439 | ENSG0000 | C11orf71 | chromosor | 3.045795 | 3.370662 | 3.971346 |
| 10185 | ENSG0000 | ARHGAP27 | Rho GTPas | 2.138075 | 2.883759 | 3.970212 |
| 13546 | ENSG0000 | PPP2R2D | protein ph | 3.781538 | 3.957671 | 3.969353 |
| 14218 | ENSG0000 | PER1 | period circ | 1.995735 | 2.923104 | 3.96732 |
| 23365 | ENSG0000 | SDAD1P1 | SDA1 doma | 0.40602 | 1.323196 | 3.966159 |
| 13224 | ENSG0000 | MOB1B | MOB kinas | 3.255884 | 3.835273 | 3.959895 |
| 717 | ENSG0000 | ALX4 | ALX homeo | 1.389565 | 1.989118 | 3.952763 |
| 744 | ENSG0000 | SYNE2 | spectrin re | 2.090493 | 3.648815 | 3.950022 |
| 1352 | ENSG0000 | ICAM3 | intercellula | 2.386262 | 2.407453 | 3.946623 |
| 35569 | ENSG0000 | MAFG-AS1 | MAFG anti | 1.927983 | 2.568524 | 3.946133 |
| 6674 | ENSG0000 | KRBA1 | KRAB-A do | 1.960637 | 3.358116 | 3.946035 |
| 10181 | ENSG0000 | SIM2 | single-mino | 1.509221 | 2.893818 | 3.943003 |
| 8702 | ENSG0000 | DOK3 | docking pro | 1.316821 | 2.476819 | 3.941217 |
| 5684 | ENSG0000 | ZNF436 | zinc finger | 2.159221 | 3.520632 | 3.937762 |
| 7583 | ENSG0000 | PARP9 | poly(ADP-r | 2.391883 | 2.7689 | 3.936117 |
| 20136 | ENSG0000 | APTR | Alu-mediat | 2.11073 | 3.892292 | 3.929655 |
| 7795 | ENSG0000 | WDR89 | WD repeat | 1.79352 | 3.014174 | 3.927076 |
| 9107 | ENSG0000 | SIDT2 | SID1 transr | 2.287052 | 3.348661 | 3.925459 |
| 6684 | ENSG0000 | TMEM254 | transmeml | 2.009697 | 3.363388 | 3.921001 |
| 11133 | ENSG0000 | GRPEL2 | GrpE like 2 | 2.462012 | 3.004301 | 3.91824 |
| 35741 | ENSG0000 | AC005670.2 | | 2.413841 | 3.619366 | 3.917101 |
| 4310 | ENSG0000 | SLC4A3 | solute carr | 1.368228 | 3.750908 | 3.915817 |
| 1039 | ENSG0000 | IFI35 | interferon | 1.037426 | 1.837085 | 3.91264 |

| | | | | | | |
|---|---|---|---|---|---|---|
| 10705 | ENSG0000 | CCDC138 | coiled-coil | 2.476985 | 3.600601 | 3.910134 |
| 3481 | ENSG0000 | FBXL20 | F-box and l | 2.693826 | 3.848672 | 3.906118 |
| 8320 | ENSG0000 | TAF1A | TATA-box b | 2.681506 | 3.675491 | 3.90573 |
| 2432 | ENSG0000 | UQCC1 | ubiquinol-c | 2.71949 | 3.759424 | 3.905507 |
| 3570 | ENSG0000 | HDAC5 | histone de | 1.576072 | 2.372674 | 3.904293 |
| 17184 | ENSG0000 | MAGEE1 | MAGE fam | 2.199719 | 2.403043 | 3.90222 |
| 7120 | ENSG0000 | C7orf25 | chromosor | 3.13712 | 3.562108 | 3.899297 |
| 18373 | ENSG0000 | TMEM231 | transmeml | 2.697535 | 3.487529 | 3.895177 |
| 208 | ENSG0000 | ADAM22 | ADAM met | 1.882888 | 2.596051 | 3.892454 |
| 14895 | ENSG0000 | ARHGEF37 | Rho guanir | 1.354833 | 2.23917 | 3.886818 |
| 33120 | ENSG0000 | TMPO-AS1 | TMPO anti | 1.951508 | 2.673222 | 3.88047 |
| 5400 | ENSG0000 | HMGB1P1 | high mobil | 0.545386 | 1.410693 | 3.877942 |
| 8711 | ENSG0000 | NFKBIE | NFKB inhib | 2.360077 | 3.26831 | 3.873759 |
| 8254 | ENSG0000 | PPOX | protoporpl | 1.86239 | 3.088411 | 3.866945 |
| 15670 | ENSG0000 | NUDT17 | nudix hydr | 2.210187 | 3.51375 | 3.864345 |
| 589 | ENSG0000 | RAI14 | retinoic aci | 3.005653 | 3.508659 | 3.861982 |
| 750 | ENSG0000 | RELT | RELT, TNF r | 3.47274 | 3.68019 | 3.860798 |
| 3529 | ENSG0000 | MED31 | mediator c | 2.983071 | 3.049108 | 3.849462 |
| 11395 | ENSG0000 | SUGT1 | SGT1 homo | 3.492464 | 3.670459 | 3.832643 |
| 36444 | ENSG0000 | RNF115 | ring finger | 3.687603 | 3.805475 | 3.83163 |
| 2269 | ENSG0000 | SYNGR1 | synaptogyr | 1.532855 | 1.786866 | 3.831457 |
| 10742 | ENSG0000 | LYSMD1 | LysM doma | 1.75685 | 2.993078 | 3.827546 |
| 34513 | ENSG0000 | AC026464.3 | | 0.022452 | 0.140391 | 3.82492 |
| 22610 | ENSG0000 | HCG15 | HLA compl | 0.512506 | 2.479978 | 3.820447 |
| 36494 | ENSG0000 | FSBP | fibrinogen | 3.395677 | 3.522797 | 3.813723 |
| 1842 | ENSG0000 | SIRT4 | sirtuin 4 [S | 1.808984 | 2.251171 | 3.813679 |
| 4105 | ENSG0000 | CEP72 | centrosom | 3.411303 | 3.4178 | 3.813243 |
| 11804 | ENSG0000 | GPRC5B | G protein-c | 0.39378 | 1.35028 | 3.81322 |
| 6347 | ENSG0000 | NFATC1 | nuclear fac | 2.041032 | 2.147585 | 3.812986 |
| 4039 | ENSG0000 | TTC31 | tetratricop | 2.030518 | 2.592272 | 3.810956 |
| 30889 | ENSG0000 | LINC02482 | long interg | 1.968998 | 2.491636 | 3.808482 |
| 4344 | ENSG0000 | SLC30A3 | solute carr | 1.355885 | 1.893343 | 3.807767 |
| 16854 | ENSG0000 | ZNF71 | zinc finger | 2.126081 | 3.162409 | 3.807154 |
| 14169 | ENSG0000 | TMEM52 | transmeml | 1.015973 | 1.970817 | 3.806808 |
| 15546 | ENSG0000 | ARL17A | ADP ribosy | 2.557628 | 3.511566 | 3.805332 |
| 5954 | ENSG0000 | NDUFA5 | NADH:ubic | 3.401341 | 3.483663 | 3.800283 |
| 11528 | ENSG0000 | HACD1 | 3-hydroxya | 2.364699 | 2.723388 | 3.79924 |
| 4641 | ENSG0000 | HMGCL | 3-hydroxyr | 1.414989 | 1.894259 | 3.799007 |
| 11892 | ENSG0000 | TMUB2 | transmeml | 2.984265 | 3.342455 | 3.795711 |
| 8642 | ENSG0000 | GTF2H2 | general tra | 3.054733 | 3.542942 | 3.794544 |
| 16837 | ENSG0000 | KIF13B | kinesin fan | 2.059112 | 3.104342 | 3.793018 |
| 6060 | ENSG0000 | RPAIN | RPA intera | 3.231923 | 3.59068 | 3.789141 |
| 14450 | ENSG0000 | MIGA1 | mitoguardi | 2.366224 | 2.765656 | 3.788243 |
| 2621 | ENSG0000 | FMR1 | fragile X m | 2.810565 | 3.368303 | 3.787707 |
| 9760 | ENSG0000 | VPS37A | VPS37A, ES | 2.702597 | 3.050552 | 3.787305 |
| 829 | ENSG0000 | PTPRU | protein tyr | 0.655184 | 1.13312 | 3.785542 |
| 4873 | ENSG0000 | LTBP2 | latent tran | 2.444863 | 2.618816 | 3.785345 |

| | | | | | | |
|---|---|---|---|---|---|---|
| 5043 | ENSG0000 | DNAJC15 | DnaJ heat | 3.288877 | 3.505346 | 3.7812 |
| 9238 | ENSG0000 | IPMK | inositol po | 2.103937 | 2.799753 | 3.779318 |
| 20019 | ENSG0000 | ARL16 | ADP ribosy | 3.096143 | 3.70321 | 3.778141 |
| 3038 | ENSG0000 | VRK3 | vaccinia re | 2.6688 | 3.490349 | 3.776829 |
| 18493 | ENSG0000 | BX890604.1 | | 3.384345 | 3.722348 | 3.771907 |
| 1603 | ENSG0000 | TRAF5 | TNF recept | 1.651281 | 3.133294 | 3.767486 |
| 6797 | ENSG0000 | NAV1 | neuron nav | 2.175588 | 3.570121 | 3.766644 |
| 4526 | ENSG0000 | TRIM62 | tripartite n | 2.599677 | 2.726489 | 3.76653 |
| 4126 | ENSG0000 | APBB3 | amyloid be | 1.817407 | 3.645523 | 3.766256 |
| 13141 | ENSG0000 | ADCK5 | aarF doma | 3.192824 | 3.323754 | 3.759504 |
| 11861 | ENSG0000 | SPATA33 | spermatog | 2.613953 | 3.134357 | 3.758331 |
| 9304 | ENSG0000 | KIN | Kin17 DNA | 2.120301 | 3.146298 | 3.75701 |
| 5687 | ENSG0000 | ARMCX5 | armadillo r | 2.744687 | 3.635398 | 3.747449 |
| 25210 | ENSG0000 | GOLGA8N | golgin A8 f | 1.382935 | 2.594929 | 3.742155 |
| 4769 | ENSG0000 | MED28 | mediator c | 3.051608 | 3.38386 | 3.740478 |
| 9679 | ENSG0000 | TMEM55A | transmeml | 2.739879 | 2.968729 | 3.740403 |
| 14546 | ENSG0000 | SLC26A11 | solute carr | 2.10352 | 3.16575 | 3.736451 |
| 4776 | ENSG0000 | FOXO3 | forkhead b | 1.977675 | 3.548634 | 3.734669 |
| 1963 | ENSG0000 | PPP2R3C | protein ph | 3.071483 | 3.517324 | 3.727755 |
| 7771 | ENSG0000 | RAB20 | RAB20, me | 0.093482 | 0.438103 | 3.725515 |
| 9382 | ENSG0000 | PDK1 | pyruvate d | 0.90953 | 1.208976 | 3.722261 |
| 9860 | ENSG0000 | PHKG2 | phosphory | 2.76786 | 3.63049 | 3.721076 |
| 13359 | ENSG0000 | PMS2P6 | PMS1 hom | 2.763613 | 3.542062 | 3.718445 |
| 16327 | ENSG0000 | WDSUB1 | WD repeat | 2.972506 | 3.135821 | 3.71746 |
| 12031 | ENSG0000 | ZNF597 | zinc finger | 1.719852 | 3.172774 | 3.716612 |
| 9234 | ENSG0000 | TMEM263 | transmeml | 1.766057 | 2.693008 | 3.712048 |
| 9729 | ENSG0000 | FAM126B | family with | 3.048429 | 3.347071 | 3.711959 |
| 4727 | ENSG0000 | DDX59 | DEAD-box | 2.884227 | 3.559721 | 3.71087 |
| 7805 | ENSG0000 | FBLN5 | fibulin 5 [S | 0.301987 | 1.251505 | 3.705628 |
| 27678 | ENSG0000 | BX470102.1 | | 1.588044 | 2.914117 | 3.70504 |
| 19940 | ENSG0000 | LINC01521 | long interg | 1.506868 | 1.96361 | 3.704982 |
| 4812 | ENSG0000 | KLF9 | Kruppel lik | 2.066024 | 2.598869 | 3.702206 |
| 22097 | ENSG0000 | LINC01703 | long interg | 1.475401 | 1.54603 | 3.698828 |
| 2016 | ENSG0000 | UPRT | uracil phos | 1.317847 | 3.039047 | 3.698658 |
| 10841 | ENSG0000 | CLASP2 | cytoplasmi | 3.131189 | 3.24989 | 3.697228 |
| 38658 | ENSG0000 | AL162258.1 | | 2.358434 | 3.261612 | 3.692618 |
| 11099 | ENSG0000 | ARFIP1 | ADP ribosy | 2.668823 | 3.054652 | 3.690071 |
| 619 | ENSG0000 | ZIC2 | Zic family r | 2.641929 | 3.577245 | 3.689664 |
| 6067 | ENSG0000 | ATP1B2 | ATPase Na | 0.086518 | 0.716917 | 3.689118 |
| 16808 | ENSG0000 | ZNF780A | zinc finger | 1.961701 | 3.086739 | 3.688166 |
| 1582 | ENSG0000 | PHLPP1 | PH domain | 2.647815 | 3.411244 | 3.687538 |
| 12691 | ENSG0000 | ZNF440 | zinc finger | 1.44701 | 2.582461 | 3.685255 |
| 8454 | ENSG0000 | SCRN3 | secernin 3 | 2.373278 | 2.465918 | 3.681658 |
| 6190 | ENSG0000 | RSPH3 | radial spok | 1.554294 | 2.317692 | 3.679417 |
| 7635 | ENSG0000 | CNOT6L | CCR4-NOT | 2.364246 | 3.177242 | 3.678573 |
| 4805 | ENSG0000 | CYP20A1 | cytochrom | 2.72384 | 3.310328 | 3.678264 |
| 6174 | ENSG0000 | GTPBP3 | GTP bindin | 3.51585 | 3.545427 | 3.675994 |

| | | | | | | |
|---|---|---|---|---|---|---|
| 40731 | ENSG0000 | AC023157.3 | | 0.523687 | 1.879288 | 3.67325 |
| 5804 | ENSG0000 | IPPK | inositol-pe | 3.14905 | 3.436763 | 3.673134 |
| 5867 | ENSG0000 | RNF6 | ring finger | 3.395711 | 3.666675 | 3.670421 |
| 12832 | ENSG0000 | ZNF584 | zinc finger | 2.544944 | 2.563086 | 3.668956 |
| 4920 | ENSG0000 | ABCD4 | ATP bindin | 1.544888 | 3.075067 | 3.668577 |
| 5474 | ENSG0000 | ZNF45 | zinc finger | 1.995708 | 2.994746 | 3.66461 |
| 5223 | ENSG0000 | RWDD3 | RWD doma | 2.14329 | 2.715445 | 3.663646 |
| 1277 | ENSG0000 | ZNF532 | zinc finger | 1.753758 | 2.801549 | 3.662802 |
| 41728 | ENSG0000 | AC135048.4 | | 2.36564 | 3.263325 | 3.659188 |
| 1124 | ENSG0000 | POLB | DNA polym | 2.136904 | 3.019003 | 3.65667 |
| 7585 | ENSG0000 | USP8 | ubiquitin s | 2.763872 | 3.632475 | 3.656121 |
| 8997 | ENSG0000 | FRA10AC1 | FRA10A as | 2.808393 | 3.302852 | 3.652547 |
| 5100 | ENSG0000 | FAM117A | family with | 0.94093 | 2.332631 | 3.649888 |
| 11443 | ENSG0000 | COMTD1 | catechol-O | 3.417241 | 3.447987 | 3.649703 |
| 30791 | ENSG0000 | ZNF550 | zinc finger | 1.898292 | 3.19523 | 3.649318 |
| 31749 | ENSG0000 | ALG1L13P | asparagine | 0.462885 | 1.419461 | 3.648555 |
| 1832 | ENSG0000 | SLC8B1 | solute carr | 1.417232 | 2.008784 | 3.646505 |
| 15259 | ENSG0000 | PROS1 | protein S (a | 1.063155 | 1.89973 | 3.646443 |
| 4354 | ENSG0000 | ASB3 | ankyrin rep | 2.946232 | 3.52247 | 3.64178 |
| 1011 | ENSG0000 | EVI5 | ecotropic v | 2.533358 | 3.167574 | 3.638269 |
| 7685 | ENSG0000 | RBP5 | retinol bin | 1.682679 | 3.419047 | 3.635131 |
| 36611 | ENSG0000 | GACAT2 | gastric can | 1.46922 | 2.066042 | 3.632388 |
| 7049 | ENSG0000 | PIGC | phosphatic | 2.795608 | 3.617819 | 3.631899 |
| 16924 | ENSG0000 | ZNF607 | zinc finger | 1.544959 | 2.847907 | 3.62907 |
| 7041 | ENSG0000 | STX6 | syntaxin 6 | 2.668315 | 3.463951 | 3.627024 |
| 1727 | ENSG0000 | ERO1B | endoplasm | 1.504854 | 3.173282 | 3.617105 |
| 15254 | ENSG0000 | FOXO4 | forkhead b | 0.819144 | 2.771584 | 3.616664 |
| 17088 | ENSG0000 | PLXNB3 | plexin B3 [ | 1.165628 | 1.955722 | 3.613186 |
| 2014 | ENSG0000 | HDAC6 | histone de | 2.085626 | 3.352221 | 3.612874 |
| 35817 | ENSG0000 | ZNF234 | zinc finger | 1.752991 | 3.359482 | 3.612099 |
| 6338 | ENSG0000 | GSE1 | Gse1 coile | 2.421399 | 2.758586 | 3.61109 |
| 17189 | ENSG0000 | L3MBTL3 | L3MBTL3, I | 2.891319 | 3.039679 | 3.611082 |
| 16494 | ENSG0000 | TTC30B | tetratricop | 1.426223 | 3.048255 | 3.607921 |
| 36640 | ENSG0000 | BAHCC1 | BAH doma | 2.22365 | 2.456174 | 3.603308 |
| 2272 | ENSG0000 | MTMR3 | myotubula | 2.536418 | 2.995299 | 3.601961 |
| 11045 | ENSG0000 | BDH2 | 3-hydroxyb | 0.887568 | 2.241312 | 3.600695 |
| 18234 | ENSG0000 | LINC01123 | long interg | 2.446956 | 2.938544 | 3.599492 |
| 12763 | ENSG0000 | C10orf35 | chromosor | 2.403251 | 2.866111 | 3.596507 |
| 37940 | ENSG0000 | AC091057.2 | | 1.696666 | 2.470559 | 3.594796 |
| 14365 | ENSG0000 | TSHZ1 | teashirt zir | 2.901873 | 2.954607 | 3.587281 |
| 8930 | ENSG0000 | SNX30 | sorting nex | 2.39637 | 3.195163 | 3.584899 |
| 37165 | ENSG0000 | ZNF887P | zinc finger | 0.922033 | 1.830683 | 3.584033 |
| 9140 | ENSG0000 | DOC2A | double C2 | 0.757135 | 2.124178 | 3.58146 |
| 9998 | ENSG0000 | SLC37A3 | solute carr | 2.792281 | 3.35526 | 3.578322 |
| 6931 | ENSG0000 | MDFIC | MyoD fam | 1.071637 | 1.112825 | 3.576175 |
| 37038 | ENSG0000 | CEBPA-AS1 | CEBPA anti | 1.784023 | 3.088227 | 3.574763 |
| 16163 | ENSG0000 | ZNF548 | zinc finger | 1.235142 | 3.161877 | 3.574433 |

| | | | | | | |
|---|---|---|---|---|---|---|
| 8299 | ENSG0000 | ANXA9 | annexin A9 | 1.036212 | 1.633366 | 3.572866 |
| 28897 | ENSG0000 | STON1 | stonin 1 [S | 0.325164 | 1.120361 | 3.572338 |
| 2168 | ENSG0000 | YPEL1 | yippee like | 0.727838 | 1.139544 | 3.570745 |
| 5517 | ENSG0000 | SOX4 | SRY-box 4 | 1.909577 | 2.356647 | 3.565049 |
| 10281 | ENSG0000 | PDE9A | phosphodi | 1.486741 | 1.805723 | 3.563984 |
| 16553 | ENSG0000 | HIST3H2BE | histone clu | 0.233083 | 3.148069 | 3.563906 |
| 16761 | ENSG0000 | C5orf42 | chromosor | 2.058985 | 3.088758 | 3.557547 |
| 8855 | ENSG0000 | SLC25A37 | solute carr | 2.790924 | 3.287353 | 3.556166 |
| 1487 | ENSG0000 | SESN1 | sestrin 1 [S | 1.935457 | 2.739912 | 3.555517 |
| 8867 | ENSG0000 | PLPP5 | phospholip | 2.059791 | 2.977279 | 3.55393 |
| 42846 | ENSG0000 | C2orf81 | chromosor | 1.292549 | 1.935865 | 3.552873 |
| 26643 | ENSG0000 | ASH1L-AS1 | ASH1L anti | 2.043732 | 2.986656 | 3.541573 |
| 903 | ENSG0000 | ABCA7 | ATP bindin | 2.344015 | 3.375793 | 3.540452 |
| 14104 | ENSG0000 | NT5DC1 | 5'-nucleoti | 2.926667 | 3.163422 | 3.539801 |
| 40686 | ENSG0000 | MIR6821 | microRNA | 2.404066 | 2.750022 | 3.537682 |
| 40817 | ENSG0000 | TYW1B | tRNA-yW s | 1.895722 | 3.152181 | 3.524111 |
| 12196 | ENSG0000 | TCTN2 | tectonic fa | 1.631062 | 2.795723 | 3.523333 |
| 15756 | ENSG0000 | ZSCAN30 | zinc finger | 1.458822 | 2.34912 | 3.518202 |
| 16618 | ENSG0000 | SLC25A29 | solute carr | 3.219849 | 3.30656 | 3.510849 |
| 12711 | ENSG0000 | PIGG | phosphatic | 2.98596 | 3.050878 | 3.510486 |
| 11888 | ENSG0000 | MFSD2A | major facil | 2.416018 | 2.97102 | 3.509484 |
| 11770 | ENSG0000 | NKX3-1 | NK3 home | 1.292213 | 2.904512 | 3.509484 |
| 20768 | ENSG0000 | FAM19A5 | family with | 2.152739 | 2.747406 | 3.503151 |
| 11421 | ENSG0000 | KLHDC2 | kelch dom | 2.521696 | 2.97443 | 3.501887 |
| 35220 | ENSG0000 | ULK4P1 | ULK4 pseu | 1.36812 | 2.200228 | 3.501777 |
| 40372 | ENSG0000 | AC091982.4 | | 0.623225 | 2.212907 | 3.500525 |
| 8229 | ENSG0000 | GPR161 | G protein-c | 1.36305 | 2.536252 | 3.499493 |
| 4945 | ENSG0000 | YIPF4 | Yip1 doma | 2.634331 | 3.010492 | 3.499465 |
| 13151 | ENSG0000 | PARP14 | poly(ADP-r | 0.879933 | 2.318194 | 3.495142 |
| 28152 | ENSG0000 | COX19 | COX19, cyt | 2.945251 | 3.26509 | 3.494765 |
| 33008 | ENSG0000 | C17orf100 | chromosor | 1.472757 | 1.760298 | 3.492248 |
| 11866 | ENSG0000 | CACNB3 | calcium vo | 1.578044 | 2.150566 | 3.491864 |
| 5437 | ENSG0000 | MOCS3 | molybdenu | 2.630949 | 3.43889 | 3.489493 |
| 37623 | ENSG0000 | IPO5P1 | importin 5 | 1.177401 | 2.25833 | 3.489466 |
| 10495 | ENSG0000 | AMDHD2 | amidohydr | 2.815318 | 3.255546 | 3.489194 |
| 19701 | ENSG0000 | AC093724.1 | | 1.937999 | 2.27277 | 3.486698 |
| 5374 | ENSG0000 | PLA2G12A | phospholip | 1.966428 | 2.622193 | 3.486581 |
| 5235 | ENSG0000 | KLHL7 | kelch like f | 2.963663 | 3.463345 | 3.485935 |
| 20190 | ENSG0000 | STARD10 | StAR relate | 1.72998 | 2.793039 | 3.481513 |
| 9048 | ENSG0000 | C11orf49 | chromosor | 1.888273 | 2.540145 | 3.480579 |
| 42902 | ENSG0000 | AC008505.1 | | 2.707819 | 3.067964 | 3.480118 |
| 3113 | ENSG0000 | ATP1A3 | ATPase Na | 1.962918 | 2.36931 | 3.479823 |
| 15455 | ENSG0000 | SH3BGR | SH3 domai | 1.591048 | 2.207966 | 3.477059 |
| 18330 | ENSG0000 | ATP6V0E2- | ATP6V0E2 | 1.825086 | 3.088075 | 3.476889 |
| 2764 | ENSG0000 | FOXF1 | forkhead b | 1.634763 | 2.811751 | 3.47679 |
| 7943 | ENSG0000 | PCTP | phosphatic | 2.713104 | 3.301029 | 3.475249 |
| 18354 | ENSG0000 | SPIRE2 | spire type | 2.436413 | 2.962282 | 3.474624 |

| | | | | | | |
|---|---|---|---|---|---|---|
| 8217 | ENSG0000 | IGSF3 | immunogl | 0.609836 | 1.682285 | 3.469299 |
| 15055 | ENSG0000 | SMG1P5 | SMG1P5, r | 2.027603 | 3.069982 | 3.466577 |
| 10579 | ENSG0000 | SCNN1D | sodium ch | 1.768997 | 3.351854 | 3.462647 |
| 9030 | ENSG0000 | PGAP2 | post-GPI a | 2.304803 | 2.329569 | 3.461986 |
| 9046 | ENSG0000 | SLC43A1 | solute carr | 1.055299 | 2.065997 | 3.460417 |
| 9575 | ENSG0000 | ROBO3 | roundabou | 0.811424 | 1.872361 | 3.457282 |
| 8804 | ENSG0000 | ZNF182 | zinc finger | 1.840403 | 2.893862 | 3.452673 |
| 2690 | ENSG0000 | KATNAL1 | katanin ca | 2.226011 | 3.064262 | 3.452084 |
| 10069 | ENSG0000 | TSPAN33 | tetraspani | 2.327811 | 2.554726 | 3.451812 |
| 3550 | ENSG0000 | DHX58 | DExH-box l | 1.328652 | 1.804388 | 3.448042 |
| 7368 | ENSG0000 | PGBD1 | piggyBac t | 2.093782 | 2.467928 | 3.446068 |
| 1330 | ENSG0000 | KIFAP3 | kinesin ass | 2.479994 | 3.263422 | 3.441649 |
| 137 | ENSG0000 | ETV1 | ETS variant | 2.143994 | 2.8418 | 3.44072 |
| 7094 | ENSG0000 | DRAM1 | DNA dama | 0.811847 | 2.605471 | 3.43983 |
| 20575 | ENSG0000 | AC009403.1 | | 2.39022 | 3.303692 | 3.439206 |
| 495 | ENSG0000 | SH2D2A | SH2 domai | 1.388834 | 1.888367 | 3.433865 |
| 10864 | ENSG0000 | SPICE1 | spindle and | 1.711132 | 2.984046 | 3.431868 |
| 13913 | ENSG0000 | DLGAP1-AS | DLGAP1 ar | 0.833746 | 2.879519 | 3.431281 |
| 5752 | ENSG0000 | STAT5A | signal trans | 2.198252 | 2.714491 | 3.424814 |
| 6223 | ENSG0000 | ZBTB46 | zinc finger | 2.082752 | 2.893218 | 3.424122 |
| 6937 | ENSG0000 | CEP162 | centrosom | 1.195442 | 2.508776 | 3.422491 |
| 26229 | ENSG0000 | LINC01849 | long interg | 0 | 0.775232 | 3.42133 |
| 24092 | ENSG0000 | HOXB-AS1 | HOXB clust | 1.196788 | 2.537507 | 3.420052 |
| 13316 | ENSG0000 | SLC25A30 | solute carr | 1.620216 | 2.535724 | 3.420007 |
| 32575 | ENSG0000 | AL356215.1 | | 0.797138 | 2.765751 | 3.417852 |
| 9698 | ENSG0000 | C16orf87 | chromosor | 1.980249 | 2.434879 | 3.417488 |
| 41940 | ENSG0000 | AC022400.8 | | 2.713616 | 2.782079 | 3.417208 |
| 7812 | ENSG0000 | SLC12A6 | solute carr | 1.504528 | 3.1512 | 3.416825 |
| 9375 | ENSG0000 | EPG5 | ectopic P-g | 2.382421 | 3.165116 | 3.412962 |
| 1922 | ENSG0000 | DTX2 | deltex E3 u | 2.387314 | 2.674293 | 3.410061 |
| 9740 | ENSG0000 | SLC26A2 | solute carr | 2.25826 | 2.847154 | 3.401529 |
| 16514 | ENSG0000 | COL27A1 | collagen ty | 0.967619 | 3.039484 | 3.401476 |
| 15990 | ENSG0000 | AC026412.1 | | 1.687231 | 3.02875 | 3.397654 |
| 2887 | ENSG0000 | GDAP1 | ganglioside | 1.556516 | 3.062626 | 3.395717 |
| 3772 | ENSG0000 | PPFIBP1 | PPFIA bind | 1.654775 | 3.095571 | 3.394102 |
| 37268 | ENSG0000 | ZNF224 | zinc finger | 1.420102 | 2.815329 | 3.393486 |
| 11116 | ENSG0000 | LMBRD2 | LMBR1 dor | 1.936077 | 2.477017 | 3.392454 |
| 27850 | ENSG0000 | RNF103 | ring finger | 2.282723 | 3.276161 | 3.388836 |
| 8267 | ENSG0000 | PIGM | phosphatic | 2.853627 | 3.100153 | 3.387417 |
| 40715 | ENSG0000 | PMS2P10 | PMS1 hom | 1.756623 | 2.772064 | 3.382544 |
| 15162 | ENSG0000 | VPS33B | VPS33B, la | 2.548635 | 3.219777 | 3.379249 |
| 4447 | ENSG0000 | DLX2 | distal-less | 0.916905 | 2.570987 | 3.378907 |
| 3662 | ENSG0000 | GLRB | glycine rec | 2.327556 | 3.236597 | 3.376831 |
| 1081 | ENSG0000 | KCNAB2 | potassium | 1.588274 | 2.056566 | 3.375806 |
| 17001 | ENSG0000 | NBR2 | neighbor o | 1.747503 | 2.897509 | 3.37444 |
| 4609 | ENSG0000 | RIMS3 | regulating | 1.800471 | 1.820878 | 3.373633 |
| 5485 | ENSG0000 | SIRT5 | sirtuin 5 [S | 2.487084 | 3.060295 | 3.371222 |

| | | | | | | |
|---|---|---|---|---|---|---|
| 15325 | ENSG0000 | RBM43 | RNA bindir | 1.404591 | 2.214369 | 3.367345 |
| 3969 | ENSG0000 | ZNF451 | zinc finger | 2.7487 | 3.031245 | 3.365381 |
| 7542 | ENSG0000 | FAM149B1 | family with | 2.456261 | 2.955466 | 3.364829 |
| 13794 | ENSG0000 | ZNF552 | zinc finger | 2.62317 | 3.197427 | 3.363529 |
| 18143 | ENSG0000 | DXO | decapping | 2.047069 | 2.891794 | 3.362843 |
| 18230 | ENSG0000 | DDR1 | discoidin d | 1.772969 | 2.278033 | 3.360631 |
| 9492 | ENSG0000 | MERTK | MER proto | 1.284276 | 1.770987 | 3.3594 |
| 16792 | ENSG0000 | PTMAP2 | prothymos | 2.004882 | 3.014545 | 3.357749 |
| 15233 | ENSG0000 | ACTRT3 | actin relate | 2.152148 | 3.211661 | 3.356378 |
| 33293 | ENSG0000 | TMEM179 | transmeml | 1.592264 | 2.16493 | 3.354593 |
| 9594 | ENSG0000 | ABCA6 | ATP bindin | 2.062807 | 2.864522 | 3.353765 |
| 850 | ENSG0000 | ZNF112 | zinc finger | 1.801781 | 2.837763 | 3.351959 |
| 34107 | ENSG0000 | AC090515.4 | | 1.291005 | 2.884715 | 3.351393 |
| 11900 | ENSG0000 | METTL15 | methyltrar | 2.580253 | 3.228879 | 3.3468 |
| 1568 | ENSG0000 | STK17B | serine/thre | 2.784766 | 3.278827 | 3.341608 |
| 4302 | ENSG0000 | ZBTB47 | zinc finger | 2.859493 | 3.118468 | 3.341252 |
| 5926 | ENSG0000 | RIBC2 | RIB43A dor | 1.612445 | 2.277431 | 3.332755 |
| 11165 | ENSG0000 | CCDC127 | coiled-coil | 2.816225 | 3.289645 | 3.332252 |
| 20397 | ENSG0000 | AC127502.1 | | 2.260342 | 2.78109 | 3.332236 |
| 20858 | ENSG0000 | SNORA11F | small nucle | 2.732505 | 2.805203 | 3.332124 |
| 5328 | ENSG0000 | ORMDL2 | ORMDL spl | 2.685102 | 3.025203 | 3.330591 |
| 573 | ENSG0000 | AGA | aspartylglu | 1.852067 | 2.232312 | 3.325713 |
| 11107 | ENSG0000 | PRMT9 | protein arg | 2.719499 | 3.172307 | 3.324324 |
| 22835 | ENSG0000 | LINC00511 | long interg | 1.247515 | 1.898334 | 3.320203 |
| 27070 | ENSG0000 | NIFK-AS1 | NIFK antise | 1.546183 | 2.131344 | 3.317772 |
| 18241 | ENSG0000 | ZNF616 | zinc finger | 1.627176 | 3.162339 | 3.316202 |
| 20323 | ENSG0000 | CD27-AS1 | CD27 antis | 1.413542 | 2.166646 | 3.316047 |
| 8505 | ENSG0000 | CSRNP1 | cysteine ar | 1.513055 | 2.460247 | 3.313625 |
| 16157 | ENSG0000 | SPRED3 | sprouty rel | 1.039909 | 1.796672 | 3.312627 |
| 5265 | ENSG0000 | KIAA1549 | KIAA1549 \| | 2.634662 | 3.184273 | 3.310142 |
| 40276 | ENSG0000 | CU633906.1 | | 2.688066 | 2.782222 | 3.309529 |
| 29499 | ENSG0000 | AC120114.1 | | 0.486603 | 1.543939 | 3.305377 |
| 4042 | ENSG0000 | OTX1 | orthodenti | 1.402643 | 1.779976 | 3.295227 |
| 7059 | ENSG0000 | USP37 | ubiquitin s | 2.587058 | 2.671065 | 3.293871 |
| 8734 | ENSG0000 | SLC18B1 | solute carr | 1.527887 | 2.142618 | 3.29365 |
| 7807 | ENSG0000 | C14orf79 | chromosor | 1.855105 | 3.160828 | 3.289911 |
| 15219 | ENSG0000 | RPL23AP82 | ribosomal | 2.306615 | 2.890872 | 3.286121 |
| 16774 | ENSG0000 | ZNF433 | zinc finger | 0.642828 | 2.874417 | 3.286062 |
| 11912 | ENSG0000 | FOS | Fos proto-c | 0.99206 | 1.80655 | 3.283029 |
| 2343 | ENSG0000 | PIGH | phosphatic | 2.535364 | 2.570384 | 3.278224 |
| 22140 | ENSG0000 | UPK1A-AS1 | UPK1A ant | 0.221796 | 0.427665 | 3.276031 |
| 7618 | ENSG0000 | KIAA1109 | KIAA1109 \| | 1.758603 | 3.245027 | 3.271626 |
| 13081 | ENSG0000 | HOXC5 | homeobox | 1.465885 | 1.720987 | 3.271533 |
| 6571 | ENSG0000 | ZSWIM3 | zinc finger | 1.718015 | 2.235677 | 3.268821 |
| 5064 | ENSG0000 | NR2C1 | nuclear rec | 2.360898 | 2.896295 | 3.266031 |
| 11721 | ENSG0000 | C18orf54 | chromosor | 2.403552 | 2.74948 | 3.265093 |
| 4866 | ENSG0000 | ACYP1 | acylphosph | 3.093437 | 3.174442 | 3.251286 |

| | | | | | | |
|---|---|---|---|---|---|---|
| 13181 | ENSG0000 | PPP1R3B | protein ph | 0.938229 | 1.984502 | 3.250749 |
| 8900 | ENSG0000 | SLC39A4 | solute carr | 2.024632 | 2.50761 | 3.240565 |
| 6563 | ENSG0000 | TESMIN | testis expr | 2.141399 | 2.574926 | 3.237193 |
| 11356 | ENSG0000 | KIAA1958 | KIAA1958 \| | 1.758854 | 2.199927 | 3.236194 |
| 8385 | ENSG0000 | ETNK2 | ethanolam | 0.90957 | 1.602874 | 3.235334 |
| 16905 | ENSG0000 | ZNF649 | zinc finger | 1.864221 | 2.901318 | 3.231988 |
| 11565 | ENSG0000 | CKB | creatine ki | 1.433214 | 1.769775 | 3.231624 |
| 30513 | ENSG0000 | GLI4 | GLI family | 2.372832 | 2.950199 | 3.231567 |
| 28236 | ENSG0000 | TM4SF1-AS | TM4SF1 ar | 0.32275 | 2.096664 | 3.230686 |
| 4574 | ENSG0000 | PHTF1 | putative hc | 1.76551 | 2.524213 | 3.229682 |
| 4468 | ENSG0000 | C2orf42 | chromosor | 1.365893 | 2.336571 | 3.22284 |
| 22700 | ENSG0000 | PSMD5-AS | PSMD5 ant | 1.873789 | 2.841848 | 3.212984 |
| 32041 | ENSG0000 | BBOX1-AS | BBOX1 ant | 1.305047 | 1.892675 | 3.211932 |
| 24188 | ENSG0000 | TPI1P2 | triosephos | 1.68221 | 2.784786 | 3.208554 |
| 13684 | ENSG0000 | SLC35G1 | solute carr | 2.293948 | 2.983978 | 3.20569 |
| 3749 | ENSG0000 | SLC22A18 | solute carr | 1.403247 | 2.334286 | 3.205674 |
| 11988 | ENSG0000 | ZNF558 | zinc finger | 1.312394 | 3.184014 | 3.203634 |
| 35918 | ENSG0000 | IKBKE | inhibitor of | 1.535295 | 2.606075 | 3.202587 |
| 12162 | ENSG0000 | CDKN2AIP | CDKN2A in | 1.571482 | 3.139703 | 3.200869 |
| 3285 | ENSG0000 | PCOLCE | procollage | 0.560861 | 1.23241 | 3.199749 |
| 7683 | ENSG0000 | TAPBPL | TAP bindin | 2.373616 | 2.584199 | 3.196669 |
| 13330 | ENSG0000 | CYB561D1 | cytochrom | 1.127658 | 2.969787 | 3.190246 |
| 26969 | ENSG0000 | ZNF853 | zinc finger | 1.556735 | 2.439758 | 3.18449 |
| 5805 | ENSG0000 | ZNF484 | zinc finger | 1.575139 | 2.441578 | 3.180539 |
| 15451 | ENSG0000 | SMYD3 | SET and M | 2.532478 | 2.891484 | 3.179357 |
| 20 | ENSG0000 | RAD52 | RAD52 hor | 1.459805 | 2.732725 | 3.171104 |
| 11934 | ENSG0000 | TMEM145 | transmeml | 1.585355 | 1.871213 | 3.163493 |
| 13243 | ENSG0000 | NMNAT1 | nicotinami | 2.147246 | 2.341169 | 3.163247 |
| 7855 | ENSG0000 | PML | promyeloc | 1.99296 | 2.251153 | 3.154191 |
| 10321 | ENSG0000 | ZNF761 | zinc finger | 0.946473 | 2.567055 | 3.153724 |
| 15644 | ENSG0000 | ZNF749 | zinc finger | 1.934107 | 2.421734 | 3.153387 |
| 13097 | ENSG0000 | DMXL1 | Dmx like 1 | 2.083609 | 3.029651 | 3.151839 |
| 16069 | ENSG0000 | PRR19 | proline rich | 2.578799 | 2.671196 | 3.149849 |
| 7347 | ENSG0000 | FRS3 | fibroblast g | 1.571787 | 1.590302 | 3.148074 |
| 14882 | ENSG0000 | CAMK1D | calcium/ca | 1.683737 | 2.110548 | 3.145642 |
| 2193 | ENSG0000 | HPS4 | HPS4, biog | 1.997939 | 2.797017 | 3.140837 |
| 1399 | ENSG0000 | PIAS2 | protein inh | 2.031162 | 2.608167 | 3.140353 |
| 3747 | ENSG0000 | ACCS | 1-aminocy | 0.982757 | 2.323544 | 3.14 |
| 5048 | ENSG0000 | SMAD9 | SMAD fam | 0.853607 | 2.001642 | 3.139239 |
| 15933 | ENSG0000 | SEMA4D | semaphori | 1.436039 | 2.063456 | 3.138585 |
| 11613 | ENSG0000 | C11orf74 | chromosor | 2.458557 | 3.110732 | 3.13817 |
| 20441 | ENSG0000 | LINC00189 | long interg | 0.878293 | 2.184091 | 3.138039 |
| 32674 | ENSG0000 | LINC02361 | long interg | 0.843777 | 1.685267 | 3.137984 |
| 9925 | ENSG0000 | TLCD1 | TLC domai | 1.688256 | 2.252563 | 3.134481 |
| 39692 | ENSG0000 | AC211476.3 | | 2.494142 | 2.603635 | 3.131725 |
| 9640 | ENSG0000 | JAM2 | junctional | 0.833439 | 1.732466 | 3.125769 |
| 846 | ENSG0000 | MON2 | MON2 hon | 2.223241 | 3.068056 | 3.124435 |

| | | | | | | |
|---|---|---|---|---|---|---|
| 1791 | ENSG0000 | ASAP3 | ArfGAP wit | 1.213479 | 1.591091 | 3.12433 |
| 2567 | ENSG0000 | RNF125 | ring finger | 1.388638 | 2.892804 | 3.122211 |
| 27898 | ENSG0000 | KMT2E-AS | KMT2E ant | 2.500667 | 2.543378 | 3.120493 |
| 7369 | ENSG0000 | ATAT1 | alpha tubu | 1.960864 | 2.710443 | 3.119546 |
| 2971 | ENSG0000 | LYL1 | LYL1, basic | 1.191769 | 1.67476 | 3.115481 |
| 11335 | ENSG0000 | GKAP1 | G kinase ar | 1.693194 | 2.954584 | 3.11285 |
| 16407 | ENSG0000 | INCA1 | inhibitor of | 0.969974 | 2.334081 | 3.110522 |
| 6444 | ENSG0000 | PODNL1 | podocan lil | 2.129235 | 2.606891 | 3.108984 |
| 18037 | ENSG0000 | EFCAB7 | EF-hand ca | 1.486453 | 2.424844 | 3.108192 |
| 20960 | ENSG0000 | ZBED8 | zinc finger | 1.848003 | 2.222266 | 3.108116 |
| 18133 | ENSG0000 | PRRT1 | proline rich | 2.444444 | 2.472803 | 3.107589 |
| 7594 | ENSG0000 | PARP16 | poly(ADP-r | 1.906341 | 2.213983 | 3.102612 |
| 38316 | ENSG0000 | LPP-AS2 | LPP antiser | 0.980131 | 2.641803 | 3.100359 |
| 1694 | ENSG0000 | WNT11 | Wnt family | 0.140107 | 0.492944 | 3.099378 |
| 20352 | ENSG0000 | AC141586.1 | | 1.813885 | 2.928425 | 3.098758 |
| 42486 | ENSG0000 | LINC00294 | long interg | 1.522928 | 2.289887 | 3.096557 |
| 12003 | ENSG0000 | HID1 | HID1 doma | 0.27237 | 1.646333 | 3.096514 |
| 42653 | ENSG0000 | AL358472.4 | | 0.623797 | 0.855074 | 3.094853 |
| 12797 | ENSG0000 | ZBTB26 | zinc finger | 2.658472 | 2.832351 | 3.092312 |
| 16306 | ENSG0000 | ZNF724 | zinc finger | 2.017075 | 2.47189 | 3.090415 |
| 4772 | ENSG0000 | ZNF430 | zinc finger | 2.34069 | 2.698017 | 3.087627 |
| 10682 | ENSG0000 | KIAA1841 | KIAA1841 | | 2.015579 | 2.305164 | 3.087102 |
| 17106 | ENSG0000 | SCAMP5 | secretory c | 1.48239 | 1.551616 | 3.087018 |
| 27874 | ENSG0000 | Z68871.1 | | 1.113623 | 2.751504 | 3.08685 |
| 15997 | ENSG0000 | RILPL1 | Rab interac | 2.42298 | 2.742905 | 3.086567 |
| 26231 | ENSG0000 | AC016745.1 | | 0.950256 | 2.890302 | 3.085941 |
| 7025 | ENSG0000 | FBXL8 | F-box and l | 2.045914 | 2.2967 | 3.084654 |
| 1738 | ENSG0000 | ACOX3 | acyl-CoA o | 2.139516 | 2.747289 | 3.084135 |
| 14971 | ENSG0000 | OR7E38P | olfactory re | 2.71653 | 2.722021 | 3.083988 |
| 36389 | ENSG0000 | MIR4263 | microRNA | 1.290428 | 2.962313 | 3.080727 |
| 2443 | ENSG0000 | RIMS4 | regulating | 0.802 | 1.260958 | 3.076841 |
| 1595 | ENSG0000 | CCNT2 | cyclin T2 [S | 0.9914 | 1.671328 | 3.076422 |
| 948 | ENSG0000 | CYB5R4 | cytochrom | 1.872175 | 2.974793 | 3.07161 |
| 6911 | ENSG0000 | P2RX4 | purinergic | 1.682861 | 1.792279 | 3.071545 |
| 27807 | ENSG0000 | NCK1-AS1 | NCK1 antis | 1.171148 | 2.301959 | 3.067978 |
| 33877 | ENSG0000 | RNASE4 | ribonuclea | 0.750796 | 0.929258 | 3.066561 |
| 16298 | ENSG0000 | ZFP92 | ZFP92 zinc | 1.239123 | 2.348415 | 3.060534 |
| 4186 | ENSG0000 | TTC33 | tetratricop | 1.505956 | 2.325508 | 3.059676 |
| 3081 | ENSG0000 | TJP3 | tight juncti | 0.453667 | 0.684194 | 3.058662 |
| 2431 | ENSG0000 | CD40 | CD40 mole | 0.75714 | 1.707989 | 3.05711 |
| 37016 | ENSG0000 | PARD6G-A | PARD6G ar | 1.433247 | 1.810805 | 3.048252 |
| 13452 | ENSG0000 | KCTD13 | potassium | 2.336741 | 3.045089 | 3.04776 |
| 1455 | ENSG0000 | CDC14A | cell divisio | 1.682188 | 2.365333 | 3.047601 |
| 1960 | ENSG0000 | CCDC80 | coiled-coil | 1.003752 | 1.70705 | 3.047228 |
| 14496 | ENSG0000 | ZFP3 | ZFP3 zinc f | 1.969228 | 2.829121 | 3.04639 |
| 12878 | ENSG0000 | KNDC1 | kinase non | 1.562638 | 1.654244 | 3.044074 |
| 21329 | ENSG0000 | C1GALT1C | C1GALT1-s | 1.939315 | 2.412699 | 3.043758 |

| | | | | | | |
|---|---|---|---|---|---|---|
| 41508 | ENSG0000 | AC073508.3 | | 1.4948 | 2.450041 | 3.041317 |
| 4466 | ENSG0000 | AAK1 | AP2 associ | 2.158134 | 2.727609 | 3.037963 |
| 32359 | ENSG0000 | AP003392.4 | | 1.137615 | 2.080304 | 3.036538 |
| 1871 | ENSG0000 | LTBP4 | latent tran | 1.739251 | 1.908347 | 3.035982 |
| 7727 | ENSG0000 | SUOX | sulfite oxid | 0.843837 | 0.947927 | 3.035798 |
| 40125 | ENSG0000 | SYNRG | synergin ga | 2.437407 | 2.851822 | 3.033647 |
| 6936 | ENSG0000 | KHDC1 | KH homolo | 2.141414 | 2.737608 | 3.031416 |
| 4669 | ENSG0000 | VAMP4 | vesicle ass | 2.272099 | 2.583728 | 3.028351 |
| 13178 | ENSG0000 | TNKS | tankyrase | | 2.205814 | 2.951167 | 3.027528 |
| 2749 | ENSG0000 | NECAB2 | N-terminal | 1.235548 | 2.561589 | 3.026838 |
| 4476 | ENSG0000 | GRIN3B | glutamate | 0.490943 | 1.321629 | 3.023728 |
| 40931 | ENSG0000 | AC018553.1 | | 2.382764 | 2.694012 | 3.02341 |
| 11961 | ENSG0000 | MFSD3 | major facil | 2.338196 | 2.461421 | 3.021501 |
| 13026 | ENSG0000 | MANEA | mannosida | 2.673349 | 2.993405 | 3.018342 |
| 13074 | ENSG0000 | TMCC1 | transmeml | 2.729735 | 2.911568 | 3.018226 |
| 36438 | ENSG0000 | DTX2P1-UI | DTX2P1-UI | 2.002278 | 2.701205 | 3.017937 |
| 10204 | ENSG0000 | THEM4 | thioesteras | 2.540581 | 2.771648 | 3.016722 |
| 39167 | ENSG0000 | AP000525.1 | | 2.535123 | 2.699033 | 3.014093 |
| 188 | ENSG0000 | PIGQ | phosphatic | 2.392838 | 2.92507 | 3.011358 |
| 15722 | ENSG0000 | KTN1-AS1 | KTN1 antis | 1.312193 | 2.57651 | 3.010479 |
| 2955 | ENSG0000 | FCGRT | Fc fragmer | 1.120069 | 1.436103 | 3.006907 |
| 10880 | ENSG0000 | PRICKLE2 | prickle plai | 1.806623 | 2.012122 | 3.00525 |
| 16754 | ENSG0000 | DMBX1 | diencephal | 0.839201 | 1.018665 | 3.001888 |
| 2113 | ENSG0000 | CBARP | CACN beta | 1.756305 | 2.190709 | 2.993121 |
| 39276 | ENSG0000 | LINC01578 | long interg | 2.00096 | 2.965595 | 2.992139 |
| 3087 | ENSG0000 | BBC3 | BCL2 bindi | 1.677816 | 1.942616 | 2.991872 |
| 20139 | ENSG0000 | MBLAC1 | metallo-be | 1.843253 | 2.240769 | 2.991113 |
| 9307 | ENSG0000 | INPP1 | inositol po | 2.355648 | 2.750932 | 2.986821 |
| 5563 | ENSG0000 | EFNB2 | ephrin B2 [ | 1.670918 | 2.80823 | 2.98623 |
| 11901 | ENSG0000 | ZNF280A | zinc finger | 0.546676 | 1.562899 | 2.98429 |
| 32870 | ENSG0000 | ZNF10 | zinc finger | 1.035601 | 2.439185 | 2.97846 |
| 4602 | ENSG0000 | MYCL | MYCL prot | 0.475463 | 1.926906 | 2.978437 |
| 10341 | ENSG0000 | PLPP7 | phospholip | 1.074355 | 1.14185 | 2.976454 |
| 13091 | ENSG0000 | CES3 | carboxyles | 1.749005 | 1.994038 | 2.973368 |
| 536 | ENSG0000 | APBA2 | amyloid be | 2.701799 | 2.765881 | 2.969467 |
| 7697 | ENSG0000 | MARCH9 | membrane | 1.104207 | 1.344572 | 2.968304 |
| 7742 | ENSG0000 | CERS5 | ceramide s | 2.156314 | 2.817675 | 2.966047 |
| 36105 | ENSG0000 | AC239868.3 | | 1.960903 | 2.679155 | 2.963671 |
| 14598 | ENSG0000 | ACBD4 | acyl-CoA b | 0.952741 | 1.210711 | 2.963185 |
| 12387 | ENSG0000 | CHRNA5 | cholinergic | 2.167561 | 2.395097 | 2.961782 |
| 13222 | ENSG0000 | TNFRSF10C | TNF recept | 1.354263 | 1.4695 | 2.961348 |
| 13716 | ENSG0000 | PHLDB3 | pleckstrin l | 1.614641 | 2.030251 | 2.96036 |
| 34985 | ENSG0000 | SLC7A5P1 | solute carr | 2.090141 | 2.862743 | 2.958274 |
| 6917 | ENSG0000 | TMEM243 | transmeml | 2.423325 | 2.557301 | 2.957785 |
| 5568 | ENSG0000 | IRF1 | interferon | 1.517935 | 2.37554 | 2.956125 |
| 1483 | ENSG0000 | RAB21 | RAB21, me | 2.007277 | 2.646089 | 2.955753 |
| 16745 | ENSG0000 | HHLA3 | HERV-H LT | 2.35874 | 2.764072 | 2.952651 |

| | | | | | | |
|---|---|---|---|---|---|---|
| 15481 | ENSG0000 | TLCD2 | TLC domai | 1.096513 | 1.301422 | 2.95246 |
| 16343 | ENSG0000 | HRCT1 | histidine ri | 0.102277 | 0.704057 | 2.950947 |
| 35037 | ENSG0000 | AC004656.1 | | 0.85391 | 1.107827 | 2.948915 |
| 21426 | ENSG0000 | OSER1-AS1 | OSER1 anti | 1.016748 | 1.597267 | 2.944415 |
| 15850 | ENSG0000 | EPOR | erythropoi | 0.954818 | 2.070194 | 2.941654 |
| 7185 | ENSG0000 | ADAMTS7 | ADAM met | 1.809094 | 1.815205 | 2.937251 |
| 3203 | ENSG0000 | BET1 | Bet1 golgi | 1.856833 | 2.273768 | 2.935979 |
| 18452 | ENSG0000 | AC004980.1 | | 1.453476 | 2.575487 | 2.935603 |
| 26066 | ENSG0000 | MAPKAPK5 | MAPKAPK5 | 1.91188 | 2.338786 | 2.93514 |
| 22411 | ENSG0000 | ZBTB45P2 | zinc finger | 1.778302 | 2.493644 | 2.928369 |
| 6438 | ENSG0000 | FAAP24 | Fanconi an | 2.038007 | 2.705084 | 2.92699 |
| 38428 | ENSG0000 | TMCC1-AS | TMCC1 ant | 2.238104 | 2.673373 | 2.926014 |
| 14052 | ENSG0000 | GOLGA8Q | golgin A8 f | 0.824115 | 1.744145 | 2.922864 |
| 39035 | ENSG0000 | ZNF595 | zinc finger | 2.359795 | 2.798455 | 2.921089 |
| 10979 | ENSG0000 | LZTFL1 | leucine zip | 1.095964 | 2.870053 | 2.920823 |
| 24620 | ENSG0000 | LINC01816 | long interg | 2.372108 | 2.750443 | 2.919291 |
| 15420 | ENSG0000 | NOTUM | NOTUM, p | 0.333588 | 1.12215 | 2.917879 |
| 34714 | ENSG0000 | AL133367.1 | | 1.271192 | 1.530887 | 2.916869 |
| 6835 | ENSG0000 | SPOCD1 | SPOC dom | 0.363498 | 1.698032 | 2.915168 |
| 13674 | ENSG0000 | ZNF404 | zinc finger | 1.180821 | 1.832565 | 2.911393 |
| 10110 | ENSG0000 | HTR6 | 5-hydroxyt | 1.383904 | 1.658237 | 2.910568 |
| 5781 | ENSG0000 | PLEKHG3 | pleckstrin l | 1.522692 | 2.018251 | 2.9081 |
| 20241 | ENSG0000 | SEPT7P2 | septin 7 ps | 1.977117 | 2.738371 | 2.907746 |
| 13215 | ENSG0000 | ZNF417 | zinc finger | 1.915953 | 2.645095 | 2.905063 |
| 12156 | ENSG0000 | SERINC2 | serine inco | 1.09072 | 1.606886 | 2.904409 |
| 38350 | ENSG0000 | AL390955.2 | | 1.296048 | 1.598111 | 2.901917 |
| 9441 | ENSG0000 | GPR180 | G protein-c | 1.995731 | 2.667979 | 2.901636 |
| 4855 | ENSG0000 | C19orf25 | chromosor | 2.420294 | 2.770408 | 2.899174 |
| 23103 | ENSG0000 | SLC9A7P1 | solute carr | 0.824407 | 1.878585 | 2.897975 |
| 35592 | ENSG0000 | ZNF488 | zinc finger | 2.033553 | 2.676414 | 2.897047 |
| 2476 | ENSG0000 | PTK6 | protein tyr | 0.803452 | 2.047879 | 2.896128 |
| 4423 | ENSG0000 | ABCB6 | ATP bindin | 2.259201 | 2.264693 | 2.893167 |
| 692 | ENSG0000 | LETMD1 | LETM1 dor | 2.001701 | 2.14593 | 2.891935 |
| 5851 | ENSG0000 | SYNGR3 | synaptogyr | 0.464997 | 0.874913 | 2.891825 |
| 15474 | ENSG0000 | FAM131C | family with | 0.895497 | 0.975071 | 2.891433 |
| 5498 | ENSG0000 | ZNF391 | zinc finger | 1.520912 | 2.787179 | 2.888165 |
| 14200 | ENSG0000 | TMEM107 | transmeml | 1.052952 | 1.619179 | 2.88786 |
| 39552 | ENSG0000 | AC026356.1 | | 1.502418 | 2.781102 | 2.882921 |
| 20082 | ENSG0000 | TNFRSF25 | TNF recept | 2.204511 | 2.414667 | 2.882915 |
| 16406 | ENSG0000 | ZNF140 | zinc finger | 0.736477 | 1.996434 | 2.882226 |
| 8609 | ENSG0000 | CBR4 | carbonyl re | 1.803433 | 2.224374 | 2.879894 |
| 5462 | ENSG0000 | MCEE | methylmal | 2.082961 | 2.617475 | 2.879724 |
| 3628 | ENSG0000 | ELF2 | E74 like ET | 1.230371 | 1.609887 | 2.879417 |
| 15121 | ENSG0000 | NUDT14 | nudix hydr | 1.857089 | 2.189738 | 2.876916 |
| 16741 | ENSG0000 | RAB40C | RAB40C, m | 1.094938 | 2.121605 | 2.874819 |
| 7642 | ENSG0000 | GSTCD | glutathione | 2.205856 | 2.533406 | 2.871383 |
| 29133 | ENSG0000 | LINC01011 | long interg | 1.201528 | 1.507625 | 2.868636 |

| | | | | | | |
|---|---|---|---|---|---|---|
| 3937 | ENSG0000 | SASH1 | SAM and S | 1.857375 | 2.100144 | 2.868102 |
| 19713 | ENSG0000 | RBMS2P1 | RNA bindir | 1.685795 | 2.812131 | 2.866473 |
| 31992 | ENSG0000 | AL354920.1 | | 1.363411 | 2.002911 | 2.866378 |
| 21021 | ENSG0000 | C10orf55 | chromosor | 1.226666 | 1.649169 | 2.865734 |
| 3901 | ENSG0000 | PHC1 | polyhomeo | 1.610956 | 2.108414 | 2.864819 |
| 6622 | ENSG0000 | COG6 | componen | 1.868855 | 2.549489 | 2.864036 |
| 35116 | ENSG0000 | AL137802.2 | | 0.655258 | 1.245543 | 2.863364 |
| 7658 | ENSG0000 | B4GALNT3 | beta-1,4-N | 0.55537 | 1.638572 | 2.859114 |
| 6210 | ENSG0000 | PGPEP1 | pyroglutan | 1.189014 | 1.824269 | 2.858373 |
| 17133 | ENSG0000 | ZNF277 | zinc finger | 2.317543 | 2.755882 | 2.856819 |
| 2560 | ENSG0000 | CEP76 | centrosom | 2.123561 | 2.407242 | 2.849504 |
| 42984 | ENSG0000 | AC018553.2 | | 1.90355 | 2.46217 | 2.848319 |
| 35 | ENSG0000 | CYP26B1 | cytochrom | 0.842972 | 1.788314 | 2.845569 |
| 14048 | ENSG0000 | TSSK6 | testis speci | 1.125293 | 2.231553 | 2.844508 |
| 3621 | ENSG0000 | LAMTOR3 | late endos | 2.244523 | 2.755134 | 2.844134 |
| 5063 | ENSG0000 | ZFP30 | ZFP30 zinc | 1.149861 | 2.001563 | 2.843425 |
| 452 | ENSG0000 | SLC45A4 | solute carr | 2.287886 | 2.702512 | 2.837366 |
| 37025 | ENSG0000 | TBX2-AS1 | TBX2 antis | 1.96072 | 2.024187 | 2.837283 |
| 11497 | ENSG0000 | HSPA12A | heat shock | 1.311867 | 2.219491 | 2.834264 |
| 6052 | ENSG0000 | SERGEF | secretion r | 2.026304 | 2.066805 | 2.832641 |
| 39834 | ENSG0000 | AL121772.1 | | 0.562833 | 2.599748 | 2.831776 |
| 15567 | ENSG0000 | POMK | protein-O- | 1.593458 | 1.958384 | 2.824777 |
| 26418 | ENSG0000 | SSR4P1 | signal sequ | 0.84182 | 2.048125 | 2.824272 |
| 2492 | ENSG0000 | RSPO4 | R-spondin | 1.455452 | 2.278616 | 2.823468 |
| 2732 | ENSG0000 | HAS3 | hyaluronar | 1.751478 | 2.168124 | 2.823243 |
| 225 | ENSG0000 | MAPK8IP2 | mitogen-ac | 1.061159 | 1.585673 | 2.821369 |
| 6007 | ENSG0000 | GPALPP1 | GPALPP mo | 1.731867 | 2.440854 | 2.818643 |
| 3041 | ENSG0000 | C19orf44 | chromosor | 1.318385 | 2.301838 | 2.816559 |
| 33611 | ENSG0000 | PPT2-EGFL | PPT2-EGFL | 1.986082 | 2.636146 | 2.813072 |
| 12087 | ENSG0000 | GNG4 | G protein s | 1.176161 | 2.345847 | 2.810876 |
| 13260 | ENSG0000 | HEG1 | heart deve | 0.887456 | 1.963086 | 2.807645 |
| 12379 | ENSG0000 | HIC2 | HIC ZBTB tr | 2.098966 | 2.564689 | 2.805505 |
| 4991 | ENSG0000 | MSX2 | msh home | 1.830103 | 2.350747 | 2.791389 |
| 16380 | ENSG0000 | MFSD14C | major facil | 2.094567 | 2.543755 | 2.791298 |
| 8152 | ENSG0000 | GALK2 | galactokina | 2.485983 | 2.7029 | 2.790072 |
| 15672 | ENSG0000 | ZNF75D | zinc finger | 1.593059 | 2.00195 | 2.787647 |
| 24629 | ENSG0000 | BEND3P1 | BEN domai | 1.301246 | 1.857855 | 2.786697 |
| 9167 | ENSG0000 | TIRAP | TIR domair | 2.114741 | 2.737328 | 2.779567 |
| 6304 | ENSG0000 | PRRG1 | proline rich | 1.568165 | 2.05928 | 2.775423 |
| 14654 | ENSG0000 | ZNF101 | zinc finger | 0.94388 | 2.151975 | 2.774039 |
| 13160 | ENSG0000 | MFSD4B | major facil | 1.674194 | 2.116514 | 2.77393 |
| 11021 | ENSG0000 | ARHGEF3 | Rho guanir | 2.314791 | 2.586206 | 2.773303 |
| 14340 | ENSG0000 | TIGD5 | tigger trans | 2.314555 | 2.763354 | 2.771007 |
| 18323 | ENSG0000 | ZNF155 | zinc finger | 0.920649 | 1.315757 | 2.770288 |
| 2430 | ENSG0000 | NINL | ninein like | 0.993413 | 1.788406 | 2.769985 |
| 1636 | ENSG0000 | ZNF446 | zinc finger | 1.858739 | 2.645532 | 2.767566 |
| 16202 | ENSG0000 | KCTD21 | potassium | 1.676438 | 2.060556 | 2.763209 |

| | | | | | | |
|---|---|---|---|---|---|---|
| 10644 | ENSG0000 | ATF3 | activating t | 0.775398 | 1.563528 | 2.762663 |
| 24255 | ENSG0000 | THAP7-AS1 | THAP7 ant | 1.087182 | 2.214098 | 2.762374 |
| 24747 | ENSG0000 | AL590428.1 | | 0.795178 | 1.850946 | 2.759199 |
| 9005 | ENSG0000 | TCF7L2 | transcriptio | 2.205842 | 2.537366 | 2.758933 |
| 20157 | ENSG0000 | HAUS3 | HAUS augr | 2.047307 | 2.197541 | 2.758185 |
| 3490 | ENSG0000 | WNT3 | Wnt family | 1.181912 | 2.215147 | 2.756146 |
| 979 | ENSG0000 | ZNRD1 | zinc ribbon | 2.27285 | 2.663696 | 2.755564 |
| 16208 | ENSG0000 | ZNF567 | zinc finger | 1.999485 | 2.613762 | 2.751471 |
| 10029 | ENSG0000 | DUSP2 | dual specif | 1.183734 | 1.734459 | 2.750757 |
| 9431 | ENSG0000 | GPD1L | glycerol-3- | 2.43645 | 2.572264 | 2.746378 |
| 6322 | ENSG0000 | GGT7 | gamma-glu | 0.852616 | 1.834719 | 2.74462 |
| 11916 | ENSG0000 | ZNF528 | zinc finger | 0.755251 | 2.395159 | 2.744421 |
| 9458 | ENSG0000 | ZNF117 | zinc finger | 0.410733 | 2.552445 | 2.743717 |
| 1010 | ENSG0000 | CACNB1 | calcium vo | 0.929233 | 1.817381 | 2.742557 |
| 30902 | ENSG0000 | AL928654.2 | | 0.944629 | 1.907199 | 2.742537 |
| 32978 | ENSG0000 | AC026369.2 | | 1.424224 | 2.407329 | 2.740991 |
| 12912 | ENSG0000 | TVP23B | trans-golgi | 1.907723 | 2.185308 | 2.740132 |
| 35342 | ENSG0000 | AC233699.1 | | 1.679668 | 2.330415 | 2.737415 |
| 33513 | ENSG0000 | AC044802.2 | | 0.502709 | 1.194098 | 2.731336 |
| 10806 | ENSG0000 | FSTL1 | follistatin l | 1.087698 | 1.391963 | 2.729894 |
| 10607 | ENSG0000 | GBP2 | guanylate l | 0.625047 | 2.649936 | 2.726615 |
| 1689 | ENSG0000 | ZNF213 | zinc finger | 2.000588 | 2.357115 | 2.725783 |
| 11003 | ENSG0000 | TMEM41A | transmeml | 2.357426 | 2.463344 | 2.725465 |
| 2549 | ENSG0000 | ZNF516 | zinc finger | 1.188882 | 1.496228 | 2.724651 |
| 15338 | ENSG0000 | WT1 | Wilms tum | 0.813192 | 1.247656 | 2.724424 |
| 9958 | ENSG0000 | AASDH | aminoadip | 2.286361 | 2.547664 | 2.722666 |
| 18334 | ENSG0000 | ZNF425 | zinc finger | 1.4976 | 2.126682 | 2.722525 |
| 2384 | ENSG0000 | GSKIP | GSK3B inte | 0.822725 | 2.125369 | 2.718724 |
| 3693 | ENSG0000 | VWA5A | von Willeb | 0.947814 | 1.156114 | 2.71866 |
| 11478 | ENSG0000 | NDRG2 | NDRG fami | 1.706132 | 1.75566 | 2.715304 |
| 1118 | ENSG0000 | FSTL3 | follistatin l | 1.484688 | 1.983712 | 2.714195 |
| 13384 | ENSG0000 | PELI3 | pellino E3 u | 1.205081 | 1.549023 | 2.712657 |
| 367 | ENSG0000 | RWDD2A | RWD doma | 0.897666 | 1.672351 | 2.710012 |
| 3189 | ENSG0000 | PBX4 | PBX home | 1.0369 | 1.556019 | 2.708795 |
| 23666 | ENSG0000 | AC006001.3 | | 1.726956 | 2.293321 | 2.704904 |
| 15923 | ENSG0000 | GABRD | gamma-am | 1.483149 | 1.81019 | 2.704881 |
| 2285 | ENSG0000 | TNRC6B | trinucleoti | 2.173376 | 2.637971 | 2.703188 |
| 1600 | ENSG0000 | DLG3 | discs large | 1.247607 | 2.059623 | 2.701709 |
| 9164 | ENSG0000 | DCUN1D2 | defective i | 2.10802 | 2.456185 | 2.700339 |
| 754 | ENSG0000 | KCNH2 | potassium | 0.707171 | 1.137913 | 2.697144 |
| 15726 | ENSG0000 | KIF24 | kinesin fan | 2.294048 | 2.603304 | 2.693597 |
| 27579 | ENSG0000 | AL355574.1 | | 1.320937 | 1.712611 | 2.692458 |
| 6589 | ENSG0000 | CASP9 | caspase 9 [ | 1.990784 | 2.621857 | 2.692364 |
| 341 | ENSG0000 | TMEM159 | transmeml | 1.096825 | 1.504023 | 2.689546 |
| 34390 | ENSG0000 | AC012640.2 | | 0.845109 | 1.825889 | 2.689468 |
| 42277 | ENSG0000 | AC026471.6 | | 0.793602 | 2.367954 | 2.689041 |
| 11300 | ENSG0000 | GEM | GTP bindin | 1.217261 | 2.599926 | 2.686651 |

| | | | | | | |
|---|---|---|---|---|---|---|
| 32956 | ENSG0000 | AC007406.3 | | 0.943321 | 1.546539 | 2.68324 |
| 2883 | ENSG0000 | PLAT | plasminog | 1.596512 | 2.158253 | 2.683023 |
| 3649 | ENSG0000 | SEPSECS | Sep (O-pho | 1.861741 | 2.31016 | 2.680392 |
| 14445 | ENSG0000 | ARHGAP45 | Rho GTPas | 1.212549 | 1.271078 | 2.677181 |
| 40806 | ENSG0000 | AC211476.4 | | 1.907534 | 2.140742 | 2.676625 |
| 18399 | ENSG0000 | LGR4 | leucine rich | 1.413238 | 1.704952 | 2.674666 |
| 16994 | ENSG0000 | FLVCR1-AS | FLVCR1 an | 1.592471 | 1.940836 | 2.673119 |
| 10726 | ENSG0000 | INHBB | inhibin bet | 0.544365 | 1.116385 | 2.671854 |
| 4061 | ENSG0000 | C6orf118 | chromosor | 1.233564 | 1.97677 | 2.670493 |
| 14266 | ENSG0000 | ARID3B | AT-rich inte | 1.669766 | 2.210713 | 2.667859 |
| 6576 | ENSG0000 | PPP1R3D | protein ph | 1.288159 | 1.977075 | 2.666532 |
| 29433 | ENSG0000 | SBF2-AS1 | SBF2 antise | 0.833244 | 1.633044 | 2.666167 |
| 20043 | ENSG0000 | AMZ2P1 | archaelysir | 1.090235 | 1.791552 | 2.659176 |
| 18079 | ENSG0000 | RUFY2 | RUN and F | 2.154869 | 2.6404 | 2.658396 |
| 1098 | ENSG0000 | RAB27A | RAB27A, m | 1.465223 | 2.115315 | 2.655794 |
| 42175 | ENSG0000 | MIR4697H | MIR4697 h | 1.470229 | 2.368074 | 2.65318 |
| 15713 | ENSG0000 | FOXD2 | forkhead b | 1.529846 | 1.909926 | 2.652794 |
| 22113 | ENSG0000 | BX322234.1 | | 1.652149 | 2.082831 | 2.650666 |
| 7509 | ENSG0000 | SIX3 | SIX homeo | 0.58724 | 0.863627 | 2.650453 |
| 8131 | ENSG0000 | BMP6 | bone morp | 1.330772 | 1.571934 | 2.648659 |
| 7962 | ENSG0000 | BCAS3 | BCAS3, mic | 1.873403 | 2.148365 | 2.646629 |
| 10994 | ENSG0000 | ZC3H12A | zinc finger | 1.199848 | 2.157108 | 2.646326 |
| 21010 | ENSG0000 | AC112721.2 | | 1.618352 | 2.190288 | 2.639778 |
| 14057 | ENSG0000 | DALRD3 | DALR antic | 1.856292 | 2.588015 | 2.63804 |
| 18654 | ENSG0000 | GOLGA8O | golgin A8 f | 0.956603 | 1.777931 | 2.637853 |
| 13347 | ENSG0000 | PHLDA3 | pleckstrin l | 1.208473 | 1.521397 | 2.637423 |
| 2087 | ENSG0000 | ABLIM1 | actin bindi | 1.563076 | 2.027382 | 2.636235 |
| 29430 | ENSG0000 | CASC8 | cancer sus | 0.940426 | 1.761643 | 2.635201 |
| 37032 | ENSG0000 | AC138150.2 | | 0.85196 | 2.497506 | 2.631478 |
| 5908 | ENSG0000 | A4GALT | alpha 1,4-g | 1.133025 | 1.13911 | 2.630587 |
| 38998 | ENSG0000 | LINC01023 | long interg | 1.248481 | 2.46378 | 2.629888 |
| 38999 | ENSG0000 | AC099522.2 | | 1.383935 | 2.370531 | 2.626568 |
| 15053 | ENSG0000 | TANGO2 | transport a | 2.179878 | 2.21586 | 2.626245 |
| 7492 | ENSG0000 | KHK | ketohexok | 1.492419 | 2.079215 | 2.626212 |
| 14124 | ENSG0000 | EPM2AIP1 | EPM2A inte | 1.842418 | 2.285216 | 2.624648 |
| 2672 | ENSG0000 | CAB39L | calcium bir | 1.459612 | 2.465242 | 2.622764 |
| 35737 | ENSG0000 | AC025283.2 | | 1.290871 | 2.478795 | 2.621574 |
| 27319 | ENSG0000 | SMG1P1 | SMG1P1, n | 1.316738 | 2.463618 | 2.621471 |
| 6891 | ENSG0000 | OSTF1 | osteoclast | 1.9801 | 2.487002 | 2.620932 |
| 14736 | ENSG0000 | GLTPD2 | glycolipid t | 1.724304 | 2.240397 | 2.620822 |
| 3008 | ENSG0000 | DNAJC12 | DnaJ heat s | 0.165556 | 0.940543 | 2.619452 |
| 16942 | ENSG0000 | SLC29A3 | solute carr | 1.529738 | 2.351753 | 2.615751 |
| 39524 | ENSG0000 | CASTOR2 | cytosolic a | 1.478629 | 2.493849 | 2.613367 |
| 10939 | ENSG0000 | IL17RE | interleukin | 1.415062 | 1.559678 | 2.609371 |
| 14851 | ENSG0000 | C11orf54 | chromosor | 1.78778 | 2.536175 | 2.607166 |
| 29540 | ENSG0000 | AC091849.1 | | 1.400045 | 1.772995 | 2.606882 |
| 11799 | ENSG0000 | C15orf39 | chromosor | 1.753145 | 2.067993 | 2.598528 |

| | | | | | | |
|---|---|---|---|---|---|---|
| 39666 | ENSG0000 | USP27X | ubiquitin s | 1.589383 | 2.00488 | 2.598086 |
| 9538 | ENSG0000 | JAZF1 | JAZF zinc fi | 1.350648 | 1.49092 | 2.597192 |
| 7183 | ENSG0000 | ZFHX2 | zinc finger | 0.861896 | 1.638428 | 2.595852 |
| 8553 | ENSG0000 | TMEM44 | transmeml | 1.347732 | 2.539827 | 2.594862 |
| 4255 | ENSG0000 | BBX | BBX, HMG- | 0.924986 | 1.967856 | 2.59465 |
| 9676 | ENSG0000 | PTPRN2 | protein tyr | 1.518426 | 1.852491 | 2.594068 |
| 20083 | ENSG0000 | SLC35E2 | solute carr | 1.341609 | 1.934867 | 2.590794 |
| 23125 | ENSG0000 | SNAP25-AS | SNAP25 an | 0.308343 | 1.471936 | 2.589564 |
| 12587 | ENSG0000 | DPY19L2P2 | DPY19L2 p | 1.227966 | 1.371517 | 2.589562 |
| 5489 | ENSG0000 | BTN2A3P | butyrophili | 0.808442 | 2.351231 | 2.588739 |
| 14656 | ENSG0000 | C5orf24 | chromosor | 1.856708 | 2.223754 | 2.588673 |
| 5913 | ENSG0000 | TPST2 | tyrosylprot | 1.483421 | 1.533522 | 2.588487 |
| 39582 | ENSG0000 | AC211486.3 | | 1.234723 | 1.902459 | 2.582223 |
| 1503 | ENSG0000 | IGSF9B | immunoglc | 1.043465 | 2.168063 | 2.580832 |
| 3813 | ENSG0000 | PARP11 | poly(ADP-r | 1.307244 | 2.426402 | 2.578577 |
| 30660 | ENSG0000 | AC025470.2 | | 0.618557 | 1.828745 | 2.573886 |
| 33744 | ENSG0000 | LINC01629 | long interg | 0.371479 | 1.235673 | 2.573006 |
| 4560 | ENSG0000 | RGS2 | regulator c | 1.28139 | 2.179078 | 2.572456 |
| 3625 | ENSG0000 | MANBA | mannosida | 1.615665 | 1.838083 | 2.571503 |
| 6037 | ENSG0000 | VPS13C | vacuolar pi | 1.392199 | 2.211461 | 2.570688 |
| 7730 | ENSG0000 | TARBP2 | TARBP2, RI | 2.027281 | 2.526027 | 2.56836 |
| 18470 | ENSG0000 | SERF1B | small EDRK | 2.417486 | 2.445079 | 2.563285 |
| 14486 | ENSG0000 | SMG1P3 | SMG1P3, n | 1.164996 | 2.41138 | 2.561475 |
| 24226 | ENSG0000 | LRRC37A6I | leucine ricl | 1.209584 | 1.25431 | 2.56037 |
| 16256 | ENSG0000 | PAX8-AS1 | PAX8 antis | 0.538418 | 1.630473 | 2.559259 |
| 2555 | ENSG0000 | METTL4 | methyltrar | 2.399124 | 2.458139 | 2.559127 |
| 12198 | ENSG0000 | PPIP5K1 | diphospho | 1.589522 | 2.193066 | 2.553775 |
| 11426 | ENSG0000 | TTC8 | tetratricop | 1.362759 | 2.235562 | 2.553713 |
| 3935 | ENSG0000 | NCOA7 | nuclear rec | 2.187645 | 2.543091 | 2.552773 |
| 28065 | ENSG0000 | PRKAG2-AS | PRKAG2 ar | 1.671872 | 1.996801 | 2.551804 |
| 20983 | ENSG0000 | FADS3 | fatty acid c | 2.050404 | 2.309017 | 2.549307 |
| 39399 | ENSG0000 | AC144652.1 | | 1.424203 | 1.721886 | 2.547225 |
| 15164 | ENSG0000 | ADAP2 | ArfGAP wit | 0.517826 | 1.374111 | 2.545732 |
| 11146 | ENSG0000 | ERAP2 | endoplasm | 1.169943 | 1.858828 | 2.545603 |
| 4108 | ENSG0000 | C7 | compleme | 0.783139 | 2.147536 | 2.545523 |
| 37974 | ENSG0000 | AC146944.4 | | 1.101174 | 1.232641 | 2.543456 |
| 30168 | ENSG0000 | AC011330.1 | | 1.248734 | 1.842378 | 2.543311 |
| 19952 | ENSG0000 | MDP1 | magnesiun | 1.775841 | 2.048159 | 2.538678 |
| 6489 | ENSG0000 | CARD6 | caspase re | 0.708372 | 2.363477 | 2.538522 |
| 42947 | ENSG0000 | AC002384.2 | | 0.544218 | 1.93875 | 2.536888 |
| 27532 | ENSG0000 | FAM200B | family with | 2.026762 | 2.408287 | 2.536272 |
| 15523 | ENSG0000 | ZNF696 | zinc finger | 0.338828 | 0.708151 | 2.535604 |
| 16766 | ENSG0000 | ZNF615 | zinc finger | 0.895276 | 2.410561 | 2.533257 |
| 42832 | ENSG0000 | AL137798.3 | | 0.143006 | 1.929262 | 2.533225 |
| 10640 | ENSG0000 | C1orf74 | chromosor | 1.443825 | 2.525511 | 2.527216 |
| 16940 | ENSG0000 | AC131392.1 | | 1.465568 | 2.412103 | 2.525273 |
| 11530 | ENSG0000 | SMCO4 | single-pass | 1.8011 | 2.185793 | 2.521016 |

| | | | | | | |
|---|---|---|---|---|---|---|
| 25459 | ENSG0000 | AC016876.1 | | 0.488288 | 1.306907 | 2.519899 |
| 3116 | ENSG0000 | CNFN | cornifelin [ | 0.377158 | 1.228234 | 2.519898 |
| 25463 | ENSG0000 | AC079807.2 | | 1.590433 | 1.825349 | 2.519019 |
| 9669 | ENSG0000 | CYP2U1 | cytochrom | 1.356622 | 1.580057 | 2.518045 |
| 17098 | ENSG0000 | FAM169A | family with | 1.410713 | 2.415781 | 2.517782 |
| 26803 | ENSG0000 | KDM4A-AS | KDM4A an | 1.274789 | 1.509029 | 2.515416 |
| 13821 | ENSG0000 | LIMK2 | LIM domai | 1.591495 | 1.753486 | 2.51113 |
| 12107 | ENSG0000 | PCMTD1 | protein-L-i | 1.116075 | 1.699148 | 2.510696 |
| 7148 | ENSG0000 | FXYD6 | FXYD doma | 0.381874 | 0.883721 | 2.510528 |
| 36882 | ENSG0000 | PTGES3L | prostaglan | 0.337006 | 1.868916 | 2.503707 |
| 4987 | ENSG0000 | HOXB3 | homeobox | 1.630649 | 1.950153 | 2.502379 |
| 776 | ENSG0000 | IL17RB | interleukin | 1.7164 | 2.113726 | 2.500475 |
| 4848 | ENSG0000 | GALNT12 | polypeptid | 0.446661 | 0.892242 | 2.497971 |
| 32816 | ENSG0000 | APOPT1 | apoptogen | 1.711774 | 2.003451 | 2.497938 |
| 10870 | ENSG0000 | WDFY3 | WD repeat | 1.479902 | 2.482728 | 2.496297 |
| 27226 | ENSG0000 | LINC00649 | long interg | 0.186524 | 1.901294 | 2.496062 |
| 10851 | ENSG0000 | EIF5A2 | eukaryotic | 1.310599 | 2.240702 | 2.495757 |
| 39253 | ENSG0000 | AC083880.1 | | 1.343509 | 2.193665 | 2.494931 |
| 16666 | ENSG0000 | RAD54B | RAD54 hor | 2.126196 | 2.217697 | 2.493937 |
| 15601 | ENSG0000 | ZNF284 | zinc finger | 0.732935 | 1.549801 | 2.493098 |
| 14156 | ENSG0000 | ERFE | erythroferi | 1.163597 | 1.509088 | 2.489325 |
| 9630 | ENSG0000 | CXADR | CXADR, Ig- | 1.100327 | 1.847673 | 2.484919 |
| 8689 | ENSG0000 | DCDC2 | doublecort | 1.32877 | 2.014856 | 2.483427 |
| 13735 | ENSG0000 | FOXL1 | forkhead b | 0.319353 | 1.131861 | 2.480692 |
| 5249 | ENSG0000 | POLM | DNA polym | 1.440832 | 2.023691 | 2.479156 |
| 12506 | ENSG0000 | FAXDC2 | fatty acid h | 0.430816 | 0.95613 | 2.479023 |
| 1798 | ENSG0000 | EPB41L1 | erythrocyt | 0.876233 | 1.246331 | 2.47696 |
| 16533 | ENSG0000 | CTBP1-AS2 | CTBP1 anti | 1.561046 | 2.270861 | 2.475075 |
| 6536 | ENSG0000 | ANKEF1 | ankyrin rep | 0.889244 | 1.852562 | 2.473628 |
| 26469 | ENSG0000 | AC073046.1 | | 1.736201 | 1.757834 | 2.472353 |
| 30301 | ENSG0000 | AC106791.1 | | 1.36408 | 1.959283 | 2.469737 |
| 4364 | ENSG0000 | CLIP4 | CAP-Gly do | 1.548769 | 2.422076 | 2.465753 |
| 12410 | ENSG0000 | LINGO1 | leucine rich | 1.896176 | 2.052639 | 2.462779 |
| 8577 | ENSG0000 | CENPC | centromer | 1.956211 | 2.29495 | 2.460592 |
| 29171 | ENSG0000 | AC010973.2 | | 1.747397 | 2.142807 | 2.458587 |
| 11808 | ENSG0000 | FBXO22 | F-box prote | 1.895354 | 2.411822 | 2.457782 |
| 11580 | ENSG0000 | GABRB3 | gamma-am | 1.496354 | 2.351162 | 2.454765 |
| 25433 | ENSG0000 | AC138356.2 | | 2.060342 | 2.442345 | 2.452567 |
| 41147 | ENSG0000 | ZNF8 | zinc finger | 1.538287 | 2.079668 | 2.451934 |
| 32707 | ENSG0000 | GALNT4 | polypeptid | 0.606636 | 1.126306 | 2.447957 |
| 1451 | ENSG0000 | MKNK1 | MAP kinase | 1.703471 | 2.030906 | 2.446871 |
| 11370 | ENSG0000 | ATP7A | ATPase cop | 1.233246 | 1.313812 | 2.446534 |
| 2899 | ENSG0000 | SPAG1 | sperm asso | 1.097109 | 2.151942 | 2.446463 |
| 14816 | ENSG0000 | MAFA | MAF bZIP t | 0.26182 | 0.743375 | 2.442683 |
| 7006 | ENSG0000 | SEMA4F | ssemaphor | 0.862664 | 0.999277 | 2.44222 |
| 11838 | ENSG0000 | ZNF226 | zinc finger | 1.510136 | 2.270016 | 2.440806 |
| 4682 | ENSG0000 | RCAN3 | RCAN fami | 1.246914 | 2.091857 | 2.440544 |

| | | | | | | |
|---|---|---|---|---|---|---|
| 10932 | ENSG0000 | SMIM14 | small integ | 0.982234 | 1.271386 | 2.440167 |
| 6364 | ENSG0000 | KCNC3 | potassium | 1.246333 | 1.252316 | 2.439205 |
| 1810 | ENSG0000 | PPP1R13B | protein ph | 2.058496 | 2.260107 | 2.437163 |
| 2025 | ENSG0000 | HOOK2 | hook micro | 0.743286 | 1.437132 | 2.435388 |
| 4834 | ENSG0000 | TRIM32 | tripartite n | 1.653685 | 2.149843 | 2.435226 |
| 12333 | ENSG0000 | SLC33A1 | solute carr | 1.658297 | 2.36333 | 2.430176 |
| 7053 | ENSG0000 | SP110 | SP110 nucl | 1.648875 | 1.976208 | 2.429192 |
| 7537 | ENSG0000 | RBP4 | retinol bin | 0.430733 | 1.13891 | 2.428677 |
| 7923 | ENSG0000 | ZSCAN32 | zinc finger | 1.751718 | 2.192993 | 2.426424 |
| 14770 | ENSG0000 | CEP97 | centrosom | 1.92355 | 2.126396 | 2.425002 |
| 16737 | ENSG0000 | SIPA1L1 | signal indu | 1.576748 | 2.093768 | 2.424607 |
| 1679 | ENSG0000 | HACE1 | HECT doma | 1.41886 | 2.115351 | 2.424415 |
| 21295 | ENSG0000 | CKMT1A | creatine ki | 0.943771 | 1.68497 | 2.421491 |
| 9510 | ENSG0000 | LINC00467 | long interg | 1.950952 | 2.126727 | 2.420041 |
| 1090 | ENSG0000 | KIAA1107 | KIAA1107 \| | 0.673116 | 1.384432 | 2.417146 |
| 16268 | ENSG0000 | ZKSCAN3 | zinc finger | 1.160161 | 1.871752 | 2.41704 |
| 12654 | ENSG0000 | ZNF160 | zinc finger | 0.835776 | 1.471287 | 2.41139 |
| 22083 | ENSG0000 | AL354707.1 | | 0.448947 | 0.97681 | 2.411187 |
| 10022 | ENSG0000 | PAFAH2 | platelet ac | 2.187944 | 2.303875 | 2.406535 |
| 11609 | ENSG0000 | CYB5A | cytochrom | 1.885773 | 2.07653 | 2.405767 |
| 5084 | ENSG0000 | RNF170 | ring finger | 1.796345 | 1.983603 | 2.405485 |
| 2339 | ENSG0000 | CGRRF1 | cell growth | 1.493049 | 1.973446 | 2.404192 |
| 18062 | ENSG0000 | SYS1 | SYS1, golgi | 2.042791 | 2.167599 | 2.403979 |
| 41842 | ENSG0000 | AC046185.3 | | 0.595206 | 1.259926 | 2.398161 |
| 21884 | ENSG0000 | LINC00863 | long interg | 1.224894 | 2.152758 | 2.395649 |
| 13930 | ENSG0000 | UBE2U | ubiquitin c | 1.353804 | 1.654278 | 2.394617 |
| 15475 | ENSG0000 | LMNTD2 | lamin tail c | 0.641503 | 1.34397 | 2.393021 |
| 14165 | ENSG0000 | MPI | mannose p | 1.428967 | 1.637623 | 2.39224 |
| 10055 | ENSG0000 | GPRASP2 | G protein-c | 1.62975 | 2.237785 | 2.391342 |
| 326 | ENSG0000 | DPF1 | double PHI | 1.353881 | 1.585177 | 2.386368 |
| 16607 | ENSG0000 | KIAA1671 | KIAA1671 \| | 1.77105 | 2.195204 | 2.38627 |
| 24354 | ENSG0000 | LINC01001 | long interg | 0.960691 | 2.363905 | 2.386252 |
| 2673 | ENSG0000 | KLF5 | Kruppel lik | 0.685223 | 2.030308 | 2.385785 |
| 28659 | ENSG0000 | AC108488.1 | | 1.864333 | 1.866427 | 2.384572 |
| 25599 | ENSG0000 | ZNF529-AS | ZNF529 an | 1.341253 | 2.322088 | 2.384514 |
| 35489 | ENSG0000 | AC099518.5 | | 0.062988 | 0.259535 | 2.384181 |
| 12862 | ENSG0000 | TMEM51 | transmeml | 1.642232 | 2.019945 | 2.383922 |
| 19050 | ENSG0000 | ZNF611 | zinc finger | 0.671593 | 1.812832 | 2.382546 |
| 15540 | ENSG0000 | PIGP | phosphatic | 1.150517 | 2.267982 | 2.380473 |
| 37972 | ENSG0000 | AL627309.8 | | 0.464435 | 0.820826 | 2.379586 |
| 6696 | ENSG0000 | ARNTL | aryl hydroc | 1.150075 | 2.135991 | 2.372108 |
| 14637 | ENSG0000 | ADGRB1 | adhesion G | 1.033123 | 1.50859 | 2.366665 |
| 32183 | ENSG0000 | AP006284.1 | | 0.234769 | 0.690282 | 2.363532 |
| 15739 | ENSG0000 | DTX2P1 | DTX2 pseu | 0.827013 | 1.426412 | 2.361449 |
| 10954 | ENSG0000 | RCHY1 | ring finger | 1.976175 | 1.993272 | 2.359417 |
| 5755 | ENSG0000 | PRKCG | protein kin | 0.384239 | 0.992945 | 2.358944 |
| 34969 | ENSG0000 | AC026150.1 | | 0.745532 | 1.930272 | 2.356431 |

| | | | | | | |
|---|---|---|---|---|---|---|
| 10765 | ENSG0000 | FZD5 | frizzled cla | 1.721773 | 2.059668 | 2.356065 |
| 3836 | ENSG0000 | OAS3 | 2'-5'-oligoa | 0.814091 | 1.321773 | 2.354322 |
| 39104 | ENSG0000 | AC093726.1 | | 1.135863 | 1.547221 | 2.354057 |
| 25795 | ENSG0000 | AC008443.1 | | 1.725572 | 1.835439 | 2.35404 |
| 35798 | ENSG0000 | TMPOP2 | thymopoie | 0.75777 | 1.549541 | 2.35268 |
| 40 | ENSG0000 | CFLAR | CASP8 and | 2.1933 | 2.207914 | 2.352155 |
| 1743 | ENSG0000 | ACHE | acetylcholi | 1.181546 | 2.162869 | 2.352133 |
| 10454 | ENSG0000 | FMNL3 | formin like | 1.412013 | 1.984066 | 2.350324 |
| 7161 | ENSG0000 | POC1B | POC1 centr | 1.485028 | 2.105539 | 2.348665 |
| 491 | ENSG0000 | PRKCH | protein kin | 0.667135 | 1.327844 | 2.347171 |
| 15519 | ENSG0000 | C16orf52 | chromosor | 1.561179 | 2.106273 | 2.344708 |
| 2141 | ENSG0000 | KLHL22 | kelch like f | 1.673835 | 1.898618 | 2.337029 |
| 7589 | ENSG0000 | GLCE | glucuronic | 1.287631 | 1.989197 | 2.335672 |
| 24794 | ENSG0000 | AL136985.1 | | 0.534181 | 0.932517 | 2.335494 |
| 12943 | ENSG0000 | SERF1A | small EDRK | 2.080942 | 2.201107 | 2.332024 |
| 216 | ENSG0000 | MPND | MPN doma | 0.82309 | 1.367622 | 2.331449 |
| 6663 | ENSG0000 | SLC2A11 | solute carr | 1.621518 | 1.892682 | 2.331259 |
| 16598 | ENSG0000 | ZNF420 | zinc finger | 1.335493 | 2.28224 | 2.329939 |
| 15752 | ENSG0000 | HYAL3 | hyalurono | 1.625843 | 1.839793 | 2.329695 |
| 40073 | ENSG0000 | AC244196.3 | | 0.821445 | 1.469047 | 2.32934 |
| 13864 | ENSG0000 | SLC38A9 | solute carr | 0.976471 | 1.821833 | 2.324343 |
| 16230 | ENSG0000 | SP6 | Sp6 transcr | 0.510506 | 0.901813 | 2.322598 |
| 21007 | ENSG0000 | AC112721.1 | | 0.673385 | 0.987254 | 2.317324 |
| 14924 | ENSG0000 | WT1-AS | WT1 antise | 1.568617 | 1.987615 | 2.316947 |
| 27424 | ENSG0000 | DGCR5 | DiGeorge s | 1.116662 | 1.81815 | 2.314031 |
| 1348 | ENSG0000 | ACACB | acetyl-CoA | 0.965701 | 1.571873 | 2.312517 |
| 6272 | ENSG0000 | MED18 | mediator c | 1.557356 | 2.278407 | 2.310116 |
| 3474 | ENSG0000 | TBC1D12 | TBC1 doma | 1.735422 | 2.271396 | 2.309404 |
| 39249 | ENSG0000 | AL356488.3 | | 0.552748 | 1.856699 | 2.30789 |
| 1189 | ENSG0000 | LIMS2 | LIM zinc fir | 0.795218 | 1.803589 | 2.306887 |
| 8341 | ENSG0000 | CREB3L4 | cAMP resp | 1.111802 | 1.517244 | 2.304222 |
| 22070 | ENSG0000 | JPX | JPX transcr | 1.725264 | 1.90227 | 2.303406 |
| 37489 | ENSG0000 | AC010503.2 | | 0.743111 | 1.013243 | 2.301212 |
| 4004 | ENSG0000 | PEX7 | peroxisom | 2.016787 | 2.248142 | 2.300919 |
| 14487 | ENSG0000 | GPR157 | G protein-c | 1.246801 | 1.680775 | 2.29713 |
| 109 | ENSG0000 | PDK2 | pyruvate d | 1.123037 | 1.532902 | 2.295771 |
| 33001 | ENSG0000 | ZNF253 | zinc finger | 1.234967 | 2.223119 | 2.295647 |
| 3905 | ENSG0000 | RIC8B | RIC8 guani | 1.589193 | 2.03081 | 2.287918 |
| 8414 | ENSG0000 | ZNF514 | zinc finger | 1.324349 | 2.132223 | 2.28627 |
| 14262 | ENSG0000 | CLK3 | CDC like ki | 1.752594 | 2.074504 | 2.283478 |
| 4358 | ENSG0000 | PCSK4 | proprotein | 0.737887 | 1.994543 | 2.283258 |
| 10944 | ENSG0000 | PCOLCE2 | procollage | 1.407881 | 1.781675 | 2.281399 |
| 3687 | ENSG0000 | SC5D | sterol-C5-c | 0.587218 | 1.16768 | 2.281022 |
| 11380 | ENSG0000 | OTUD1 | OTU deubi | 1.045616 | 1.805446 | 2.278559 |
| 18695 | ENSG0000 | METTL6 | methyltrar | 1.662037 | 1.736549 | 2.278306 |
| 8939 | ENSG0000 | WDR31 | WD repeat | 1.159008 | 1.48553 | 2.278187 |
| 20686 | ENSG0000 | RAET1K | retinoic aci | 0.155624 | 0.619909 | 2.270186 |

| | | | | | | |
|---|---|---|---|---|---|---|
| 12779 | ENSG0000 | LURAP1 | leucine rich | 1.320471 | 1.890806 | 2.269826 |
| 37522 | ENSG0000 | VN1R81P | vomeronas | 1.641235 | 1.813515 | 2.266827 |
| 36910 | ENSG0000 | AP001029.1 | | 0.794856 | 1.207024 | 2.266629 |
| 5607 | ENSG0000 | PAX8 | paired box | 1.427503 | 1.934899 | 2.266376 |
| 14045 | ENSG0000 | ULK4P3 | ULK4 pseu | 0.775552 | 1.444657 | 2.265551 |
| 4867 | ENSG0000 | IFT43 | intraflagell | 2.043141 | 2.044977 | 2.262908 |
| 7580 | ENSG0000 | SENP7 | SUMO1/se | 1.955593 | 2.176014 | 2.262737 |
| 9548 | ENSG0000 | DGKE | diacylglyce | 1.433608 | 2.091274 | 2.262473 |
| 6538 | ENSG0000 | SNAP25 | synaptosor | 0.522042 | 1.020377 | 2.261356 |
| 20413 | ENSG0000 | MYL5 | myosin ligh | 2.032852 | 2.093082 | 2.258005 |
| 12548 | ENSG0000 | DNAJC18 | DnaJ heat s | 0.875657 | 1.26152 | 2.256328 |
| 2390 | ENSG0000 | C14orf93 | chromosor | 1.167475 | 1.446896 | 2.255661 |
| 37357 | ENSG0000 | EML2-AS1 | EML2 antis | 0.556034 | 0.86008 | 2.255013 |
| 16588 | ENSG0000 | ZNF470 | zinc finger | 0.822911 | 1.746145 | 2.254674 |
| 36867 | ENSG0000 | AC027097.1 | | 1.093818 | 2.130545 | 2.253033 |
| 14411 | ENSG0000 | ZNF816 | zinc finger | 0.806083 | 2.21679 | 2.250615 |
| 8516 | ENSG0000 | IL17RD | interleukin | 1.104378 | 1.679241 | 2.247789 |
| 2150 | ENSG0000 | P2RX6 | purinergic | 0.55234 | 1.175599 | 2.244253 |
| 6273 | ENSG0000 | THEMIS2 | thymocyte | 1.394099 | 1.649911 | 2.242775 |
| 5500 | ENSG0000 | MOCS1 | molybdenu | 1.091741 | 1.639424 | 2.241498 |
| 34892 | ENSG0000 | AC132872.1 | | 1.000864 | 1.487812 | 2.232321 |
| 11636 | ENSG0000 | ST5 | suppressio | 1.250808 | 1.584122 | 2.231515 |
| 30659 | ENSG0000 | GLUD1P3 | glutamate | 0.834015 | 2.052494 | 2.230657 |
| 12 | ENSG0000 | NIPAL3 | NIPA like d | 0.88798 | 1.529241 | 2.228284 |
| 19682 | ENSG0000 | FAM24B | family with | 0.912668 | 1.569255 | 2.228157 |
| 4335 | ENSG0000 | DNAJC27 | DnaJ heat s | 1.219088 | 1.751193 | 2.227356 |
| 5111 | ENSG0000 | ECHDC2 | enoyl-CoA | 1.364333 | 1.927918 | 2.226225 |
| 39657 | ENSG0000 | BX322562.1 | | 0.902322 | 1.182147 | 2.225732 |
| 18384 | ENSG0000 | TMEM88B | transmeml | 0.040563 | 0.080461 | 2.220818 |
| 11617 | ENSG0000 | PPFIBP2 | PPFIA bind | 1.722297 | 1.765762 | 2.220734 |
| 2180 | ENSG0000 | CRYBB2P1 | crystallin b | 1.179803 | 1.653076 | 2.220221 |
| 43408 | ENSG0000 | AC007240.1 | | 0.335508 | 1.003732 | 2.219815 |
| 28010 | ENSG0000 | CDRT4 | CMT1A du | 0.02115 | 0.950193 | 2.21604 |
| 42657 | ENSG0000 | AC092299.1 | | 0.859742 | 1.74672 | 2.215949 |
| 5396 | ENSG0000 | ENKD1 | enkurin do | 1.631192 | 1.725952 | 2.21149 |
| 1471 | ENSG0000 | STX7 | syntaxin 7 | 1.102245 | 1.993086 | 2.210465 |
| 6908 | ENSG0000 | USP30 | ubiquitin s | 1.646449 | 1.890146 | 2.208814 |
| 30251 | ENSG0000 | AC069360.1 | | 0.403362 | 0.636944 | 2.20703 |
| 21540 | ENSG0000 | AC009506.1 | | 1.641187 | 2.170948 | 2.205794 |
| 36868 | ENSG0000 | ZNF850 | zinc finger | 0.898229 | 1.852919 | 2.203395 |
| 3748 | ENSG0000 | MDK | midkine (n | 0.772103 | 1.313001 | 2.201747 |
| 24417 | ENSG0000 | AC044797.1 | | 0.647596 | 0.758569 | 2.198515 |
| 15755 | ENSG0000 | ZNF397 | zinc finger | 1.542524 | 2.18024 | 2.198376 |
| 41028 | ENSG0000 | AC068870.2 | | 1.649101 | 2.106077 | 2.198179 |
| 2163 | ENSG0000 | RNF215 | ring finger | 1.185539 | 1.562501 | 2.198108 |
| 17958 | ENSG0000 | LINC01556 | long interg | 0.074813 | 0.656818 | 2.198057 |
| 16428 | ENSG0000 | ZNF569 | zinc finger | 1.601752 | 2.192815 | 2.19517 |

| | | | | | | |
|---|---|---|---|---|---|---|
| 15945 | ENSG0000 | ZFP69B | ZFP69 zinc | 1.720782 | 1.900958 | 2.192217 |
| 16691 | ENSG0000 | ZNF517 | zinc finger | 0.597457 | 1.460991 | 2.191668 |
| 36006 | ENSG0000 | AC022211.2 | | 1.520401 | 1.666898 | 2.18971 |
| 3445 | ENSG0000 | FBXL15 | F-box and l | 1.780162 | 1.899189 | 2.1852 |
| 15268 | ENSG0000 | DUSP8 | dual specif | 0.845345 | 1.295203 | 2.183386 |
| 15651 | ENSG0000 | KDM4D | lysine dem | 1.223762 | 1.843913 | 2.18318 |
| 11269 | ENSG0000 | GPR146 | G protein-c | 0.699117 | 1.270958 | 2.178836 |
| 3616 | ENSG0000 | USP46 | ubiquitin sp | 1.814975 | 2.071315 | 2.178129 |
| 8548 | ENSG0000 | TRPC1 | transient re | 0.841556 | 1.766316 | 2.172742 |
| 4758 | ENSG0000 | AKAP7 | A-kinase ar | 1.596522 | 2.013255 | 2.17251 |
| 2720 | ENSG0000 | ZNF821 | zinc finger | 0.837934 | 1.027125 | 2.169368 |
| 11915 | ENSG0000 | ZNF610 | zinc finger | 0.321911 | 1.161807 | 2.165536 |
| 27589 | ENSG0000 | LRRC37A2 | leucine rich | 1.342881 | 2.014364 | 2.164537 |
| 20088 | ENSG0000 | LINC01139 | long interg | 1.228614 | 1.321805 | 2.163545 |
| 34553 | ENSG0000 | AL133338.1 | | 0.734658 | 1.409026 | 2.16337 |
| 7778 | ENSG0000 | CBLN3 | cerebellin 3 | 0.557766 | 1.402675 | 2.162957 |
| 15 | ENSG0000 | SEMA3F | semaphori | 0.545742 | 0.808054 | 2.162667 |
| 26871 | ENSG0000 | AC005220.1 | | 0.910501 | 1.534988 | 2.162436 |
| 6588 | ENSG0000 | RSG1 | REM2 and | 1.59107 | 2.011543 | 2.161685 |
| 3473 | ENSG0000 | TSPAN14 | tetraspanir | 1.234032 | 1.780638 | 2.160472 |
| 35431 | ENSG0000 | C15orf65 | chromosor | 1.553001 | 1.672937 | 2.160317 |
| 11945 | ENSG0000 | DNAAF3 | dynein axo | 0.605886 | 1.094828 | 2.157056 |
| 28891 | ENSG0000 | AC007686.1 | | 0.735307 | 1.321702 | 2.155733 |
| 12960 | ENSG0000 | MTBP | MDM2 bin | 1.479074 | 1.793562 | 2.153989 |
| 39842 | ENSG0000 | AC114271.1 | | 1.591511 | 1.716124 | 2.153406 |
| 10613 | ENSG0000 | GFI1 | growth fac | 0.468937 | 0.933502 | 2.153336 |
| 3004 | ENSG0000 | SFXN3 | sideroflexir | 0.63741 | 0.718602 | 2.151622 |
| 9914 | ENSG0000 | DZIP1L | DAZ intera | 0.742574 | 1.531226 | 2.150689 |
| 4030 | ENSG0000 | UNC5A | unc-5 netri | 0.734431 | 0.817474 | 2.149817 |
| 23362 | ENSG0000 | TSTD3 | thiosulfate | 1.433828 | 1.481119 | 2.148444 |
| 5445 | ENSG0000 | BCAS4 | breast carc | 1.38296 | 1.459373 | 2.148346 |
| 14158 | ENSG0000 | ZHX2 | zinc fingers | 0.800013 | 1.597823 | 2.146807 |
| 9945 | ENSG0000 | HDAC11 | histone de | 0.878242 | 1.176637 | 2.145643 |
| 175 | ENSG0000 | MATK | megakaryo | 1.339998 | 1.68727 | 2.145502 |
| 4941 | ENSG0000 | FKBP1B | FK506 binc | 0.416214 | 0.637019 | 2.145077 |
| 4923 | ENSG0000 | TGFB3 | transformi | 1.390642 | 1.769189 | 2.144409 |
| 8887 | ENSG0000 | EBAG9 | estrogen re | 1.540255 | 2.028417 | 2.142797 |
| 12588 | ENSG0000 | ZNF16 | zinc finger | 1.62473 | 2.105355 | 2.137197 |
| 24181 | ENSG0000 | AL109811.2 | | 1.155078 | 1.363455 | 2.13538 |
| 11001 | ENSG0000 | CAMK2N2 | calcium/ca | 1.247726 | 1.302116 | 2.134295 |
| 13741 | ENSG0000 | CCDC121 | coiled-coil | 1.698937 | 1.928043 | 2.131085 |
| 4984 | ENSG0000 | KANSL1 | KAT8 regul | 1.569671 | 2.076069 | 2.130487 |
| 32377 | ENSG0000 | TOLLIP-AS1 | TOLLIP ant | 1.104579 | 1.6006 | 2.129558 |
| 33234 | ENSG0000 | AC010203.1 | | 0.365479 | 1.290783 | 2.127329 |
| 15813 | ENSG0000 | PLCD1 | phospholip | 0.361588 | 0.662138 | 2.124085 |
| 37867 | ENSG0000 | TMEM185A | transmeml | 1.491214 | 1.865273 | 2.122732 |
| 14269 | ENSG0000 | PACS2 | phosphofu | 1.995443 | 2.063386 | 2.12187 |

| | | | | | | |
|---|---|---|---|---|---|---|
| 10127 | ENSG0000 | ADAMTS4 | ADAM met | 0.607273 | 0.898799 | 2.121019 |
| 13583 | ENSG0000 | ZNF77 | zinc finger | 1.175953 | 1.452755 | 2.120167 |
| 13449 | ENSG0000 | C3orf33 | chromosor | 0.536384 | 1.66497 | 2.119843 |
| 7854 | ENSG0000 | BBS4 | Bardet-Bie | 1.275959 | 1.854811 | 2.119443 |
| 2105 | ENSG0000 | STX1B | syntaxin 1E | 0.951083 | 1.320072 | 2.116925 |
| 10720 | ENSG0000 | EN1 | engrailed h | 1.143934 | 1.856291 | 2.115374 |
| 8002 | ENSG0000 | RAB40B | RAB40B, m | 1.492312 | 1.808936 | 2.107658 |
| 16416 | ENSG0000 | PRTN3 | proteinase | 0.134393 | 0.16714 | 2.107551 |
| 12670 | ENSG0000 | TSNARE1 | t-SNARE dc | 1.430738 | 1.804489 | 2.10406 |
| 15973 | ENSG0000 | ARHGAP11 | Rho GTPas | 1.659906 | 2.094521 | 2.10344 |
| 37435 | ENSG0000 | AC005253.1 | | 1.095723 | 1.452096 | 2.102458 |
| 11686 | ENSG0000 | ZNF606 | zinc finger | 1.024194 | 1.761764 | 2.098481 |
| 4608 | ENSG0000 | KCNQ4 | potassium | 1.093406 | 1.743202 | 2.09771 |
| 1927 | ENSG0000 | LAMB1 | laminin sul | 0.677802 | 1.194883 | 2.094757 |
| 16874 | ENSG0000 | CCDC151 | coiled-coil | 0.42477 | 1.394405 | 2.091897 |
| 7458 | ENSG0000 | PAK6 | p21 (RAC1) | 0.135039 | 1.034245 | 2.088245 |
| 7856 | ENSG0000 | CYP1A1 | cytochrom | 0.408719 | 1.716782 | 2.087624 |
| 9554 | ENSG0000 | GDPD1 | glycerophc | 0.353846 | 0.93247 | 2.083683 |
| 11112 | ENSG0000 | TMEM161 | transmeml | 1.479381 | 1.742433 | 2.08161 |
| 6267 | ENSG0000 | ARHGEF16 | Rho guanir | 0.481277 | 0.930443 | 2.081394 |
| 9561 | ENSG0000 | CABYR | calcium bir | 0.959456 | 1.867388 | 2.080437 |
| 10664 | ENSG0000 | PKDCC | protein kin | 1.105347 | 1.495129 | 2.078927 |
| 7457 | ENSG0000 | TMEM62 | transmeml | 1.414992 | 1.735267 | 2.077589 |
| 39414 | ENSG0000 | BX537318.1 | | 0.690814 | 1.332069 | 2.076794 |
| 37046 | ENSG0000 | AC092295.2 | | 0.735472 | 1.309654 | 2.075592 |
| 3248 | ENSG0000 | HOXA13 | homeobox | 0.428453 | 0.735266 | 2.072559 |
| 16356 | ENSG0000 | LCOR | ligand depe | 1.603192 | 1.710555 | 2.072153 |
| 29500 | ENSG0000 | USP51 | ubiquitin sp | 1.203466 | 1.543441 | 2.069287 |
| 2814 | ENSG0000 | CCP110 | centriolar c | 1.489661 | 1.980176 | 2.068589 |
| 24606 | ENSG0000 | IGKV1OR2- | immunoglc | 0.572603 | 1.767149 | 2.068309 |
| 8214 | ENSG0000 | SYPL2 | synaptophy | 1.120022 | 1.44862 | 2.066817 |
| 38238 | ENSG0000 | AC079610.2 | | 0.634947 | 1.42654 | 2.066659 |
| 29246 | ENSG0000 | ENO1P1 | enolase 1 p | 0.835027 | 0.951054 | 2.061558 |
| 11499 | ENSG0000 | FRAT1 | FRAT1, WN | 0.517212 | 1.603495 | 2.060279 |
| 7004 | ENSG0000 | TEC | tec protein | 0.910161 | 1.14807 | 2.059745 |
| 15039 | ENSG0000 | GRK5 | G protein-c | 1.58276 | 1.698829 | 2.059592 |
| 9967 | ENSG0000 | AFAP1L1 | actin filame | 1.028953 | 1.627723 | 2.058573 |
| 29448 | ENSG0000 | AC138904.1 | | 0.208242 | 0.98246 | 2.057255 |
| 2277 | ENSG0000 | PNPLA3 | patatin like | 1.146294 | 1.418775 | 2.052097 |
| 14955 | ENSG0000 | KIAA2026 | KIAA2026 \| | 1.602641 | 1.692482 | 2.051366 |
| 28890 | ENSG0000 | AC006252.1 | | 0.4209 | 1.01237 | 2.046792 |
| 4761 | ENSG0000 | ALDH8A1 | aldehyde d | 0.151763 | 0.846112 | 2.042205 |
| 3128 | ENSG0000 | ZNF175 | zinc finger | 1.019698 | 1.824137 | 2.042156 |
| 29331 | ENSG0000 | N4BP2L2 | NEDD4 bin | 1.244201 | 1.397666 | 2.037513 |
| 10200 | ENSG0000 | C1R | compleme | 0.378002 | 0.506968 | 2.036956 |
| 14271 | ENSG0000 | ELMOD2 | ELMO dom | 1.498374 | 1.897133 | 2.036285 |
| 43278 | ENSG0000 | AL022238.4 | | 0.564796 | 1.209006 | 2.035245 |

| | | | | | | |
|---|---|---|---|---|---|---|
| 5254 | ENSG0000 | GLIPR2 | GLI pathog | 1.041698 | 1.185012 | 2.03357 |
| 14628 | ENSG0000 | AMIGO1 | adhesion n | 1.461025 | 1.564973 | 2.031547 |
| 22192 | ENSG0000 | AP000320.1 | | 0.712656 | 0.725369 | 2.03086 |
| 4744 | ENSG0000 | USP35 | ubiquitin s | 0.971947 | 1.669965 | 2.027211 |
| 9586 | ENSG0000 | PITPNC1 | phosphatic | 1.160004 | 1.379735 | 2.026398 |
| 10523 | ENSG0000 | TPCN2 | two pore s | 1.609617 | 1.687673 | 2.02426 |
| 41463 | ENSG0000 | NKILA | NF-kappaB | 0.453536 | 1.128128 | 2.023001 |
| 35597 | ENSG0000 | AC006441.3 | | 1.084278 | 1.747577 | 2.02023 |
| 38984 | ENSG0000 | AL662844.4 | | 0.948196 | 1.432013 | 2.019492 |
| 41841 | ENSG0000 | AL031123.3 | | 0.164705 | 1.312516 | 2.018633 |
| 20233 | ENSG0000 | AC005562.1 | | 0.623248 | 1.605294 | 2.017889 |
| 3025 | ENSG0000 | NOVA2 | NOVA alte | 0.232862 | 0.302411 | 2.01683 |
| 2053 | ENSG0000 | CREM | cAMP resp | 1.266726 | 1.845541 | 2.013278 |
| 774 | ENSG0000 | TRAF1 | TNF recept | 1.280677 | 1.79486 | 2.012075 |
| 16851 | ENSG0000 | ZNF347 | zinc finger | 0.527045 | 1.45802 | 2.004849 |
| 674 | ENSG0000 | PER3 | period circ | 1.941345 | 1.951339 | 2.004398 |
| 13865 | ENSG0000 | ACER2 | alkaline ce | 0.595494 | 0.961399 | 2.00432 |
| 1405 | ENSG0000 | NEBL | nebulette | | 0.542231 | 1.957754 | 2.00215 |
| 265 | ENSG0000 | FAM214A | family with | 1.013016 | 1.931979 | 2.001241 |
| 22522 | ENSG0000 | NUP50-AS: | NUP50 ant | 0.67967 | 1.264035 | 2.000206 |
| 2703 | ENSG0000 | CORO1A | coronin 1A | 1.191574 | 1.308572 | 1.998954 |
| 3296 | ENSG0000 | C1GALT1 | core 1 synt | 1.312376 | 1.498233 | 1.998932 |
| 23493 | ENSG0000 | AC002384.1 | | 0.338453 | 1.858345 | 1.997757 |
| 29704 | ENSG0000 | AC008517.1 | | 0.697492 | 1.499225 | 1.997067 |
| 487 | ENSG0000 | AGPAT4 | 1-acylglyce | 0.700095 | 1.60408 | 1.995069 |
| 5983 | ENSG0000 | TTBK2 | tau tubulin | 1.195808 | 1.68755 | 1.994302 |
| 14508 | ENSG0000 | ZNF792 | zinc finger | 0.693886 | 0.910946 | 1.994135 |
| 34042 | ENSG0000 | AC018904.1 | | 0.720759 | 1.507468 | 1.99242 |
| 3586 | ENSG0000 | MMD | monocyte | 0.998118 | 1.177474 | 1.990633 |
| 16503 | ENSG0000 | TRPV1 | transient r | 1.303081 | 1.34956 | 1.990335 |
| 20027 | ENSG0000 | LDHAP4 | lactate deh | 0.993084 | 1.176931 | 1.990248 |
| 3253 | ENSG0000 | CPVL | carboxype | 1.293208 | 1.388885 | 1.988835 |
| 7329 | ENSG0000 | ARHGEF39 | Rho guanir | 1.400483 | 1.561183 | 1.98792 |
| 16109 | ENSG0000 | RALGAPA2 | Ral GTPase | 0.563128 | 1.325302 | 1.987445 |
| 13496 | ENSG0000 | PPM1E | protein ph | 0.9737 | 1.493152 | 1.986764 |
| 13287 | ENSG0000 | ZNF791 | zinc finger | 0.981439 | 1.550287 | 1.984315 |
| 5170 | ENSG0000 | ZSCAN20 | zinc finger | 0.849409 | 1.934535 | 1.983488 |
| 19957 | ENSG0000 | GALT | galactose-1 | 1.413027 | 1.931468 | 1.982806 |
| 1637 | ENSG0000 | ZNF264 | zinc finger | 0.242142 | 0.506566 | 1.981274 |
| 4349 | ENSG0000 | ITGB6 | integrin su | 0.409956 | 1.014655 | 1.980196 |
| 9103 | ENSG0000 | ESAM | endothelia | 0.721423 | 1.021081 | 1.979313 |
| 3660 | ENSG0000 | NKX3-2 | NK3 home | 0.569531 | 0.776697 | 1.97846 |
| 15377 | ENSG0000 | RPS27L | ribosomal | 1.776148 | 1.825885 | 1.978148 |
| 16852 | ENSG0000 | PLCG2 | phospholip | 1.077506 | 1.472327 | 1.97401 |
| 562 | ENSG0000 | USP2 | ubiquitin s | 0.701574 | 0.98972 | 1.973924 |
| 3061 | ENSG0000 | DYRK1B | dual specif | 1.216954 | 1.561362 | 1.972445 |
| 9100 | ENSG0000 | CCDC15 | coiled-coil | 1.265056 | 1.534919 | 1.972151 |

| | | | | | | |
|---|---|---|---|---|---|---|
| 21495 | ENSG0000 | AC005076.1 | | 0.607324 | 1.046931 | 1.964947 |
| 14914 | ENSG0000 | SMDT1 | single-pass | 1.44159 | 1.499335 | 1.963484 |
| 27203 | ENSG0000 | NALT1 | NOTCH1 as | 0.286656 | 1.000305 | 1.962713 |
| 15729 | ENSG0000 | CARMIL3 | capping pr | 0.145757 | 1.01398 | 1.960711 |
| 9859 | ENSG0000 | FRRS1 | ferric chela | 0.844729 | 1.232107 | 1.960226 |
| 16884 | ENSG0000 | ZNF273 | zinc finger | 1.322796 | 1.683479 | 1.959911 |
| 18065 | ENSG0000 | INPP5B | inositol po | 1.434308 | 1.56872 | 1.959366 |
| 12782 | ENSG0000 | TPPP | tubulin pol | 1.111795 | 1.12266 | 1.959054 |
| 20024 | ENSG0000 | PAXIP1-AS1 | PAXIP1 ant | 1.080357 | 1.583285 | 1.958564 |
| 24304 | ENSG0000 | AC092683.1 | | 1.250674 | 1.87055 | 1.958371 |
| 11102 | ENSG0000 | ANAPC10 | anaphase p | 1.463798 | 1.896774 | 1.958195 |
| 12012 | ENSG0000 | TMEM68 | transmeml | 1.258627 | 1.92296 | 1.957626 |
| 18163 | ENSG0000 | HSPA1L | heat shock | 0.513081 | 0.980783 | 1.956942 |
| 39334 | ENSG0000 | AP000347.2 | | 0.696642 | 1.93042 | 1.956153 |
| 9028 | ENSG0000 | IMMP1L | inner mito | 1.454085 | 1.526608 | 1.955996 |
| 18687 | ENSG0000 | LNP1 | leukemia N | 0.984929 | 1.261046 | 1.954501 |
| 13923 | ENSG0000 | HIC1 | HIC ZBTB tr | 0.697238 | 1.870092 | 1.954432 |
| 10710 | ENSG0000 | ACTG2 | actin, gam | 0.101849 | 1.481037 | 1.951703 |
| 15275 | ENSG0000 | TMEM173 | transmeml | 0.445963 | 0.705622 | 1.948646 |
| 17056 | ENSG0000 | PAPSS2 | 3'-phospho | 0.562494 | 0.80261 | 1.947781 |
| 27391 | ENSG0000 | ZNF737 | zinc finger | 0.472637 | 1.413086 | 1.946423 |
| 30515 | ENSG0000 | AL732372.3 | | 0.821439 | 1.468013 | 1.945768 |
| 13491 | ENSG0000 | YPEL2 | yippee like | 0.059385 | 0.170594 | 1.944022 |
| 39280 | ENSG0000 | AL355987.3 | | 1.210117 | 1.864999 | 1.943683 |
| 13802 | ENSG0000 | PSMG4 | proteasom | 1.730353 | 1.939553 | 1.942916 |
| 10048 | ENSG0000 | FAM46B | family with | 0.789698 | 0.798476 | 1.942116 |
| 39809 | ENSG0000 | ZNF658 | zinc finger | 1.216046 | 1.852485 | 1.942066 |
| 327 | ENSG0000 | SYT7 | synaptotag | 0.958431 | 1.544507 | 1.940532 |
| 38073 | ENSG0000 | AC104109.2 | | 0.266628 | 1.638551 | 1.940232 |
| 3645 | ENSG0000 | AADAT | aminoadip | 0.924487 | 1.902675 | 1.937185 |
| 27406 | ENSG0000 | LINC01637 | long interg | 1.044036 | 1.253219 | 1.936498 |
| 299 | ENSG0000 | ZNF200 | zinc finger | 1.317947 | 1.402127 | 1.936377 |
| 7744 | ENSG0000 | LMBR1L | limb devel | 0.723159 | 1.443887 | 1.934747 |
| 41360 | ENSG0000 | MIR6781 | microRNA | 0.378559 | 1.78248 | 1.934391 |
| 12646 | ENSG0000 | PAQR8 | progestin a | 0.488999 | 1.43984 | 1.933989 |
| 11741 | ENSG0000 | MTMR10 | myotubula | 1.135225 | 1.86773 | 1.931696 |
| 2023 | ENSG0000 | MAP3K1 | mitogen-ac | 1.466203 | 1.821015 | 1.924934 |
| 32280 | ENSG0000 | AC124276.1 | | 0 | 1.304176 | 1.924422 |
| 17897 | ENSG0000 | TDRKH-AS1 | TDRKH ant | 0.899615 | 1.200671 | 1.922335 |
| 38592 | ENSG0000 | AC010501.1 | | 0.119723 | 1.852643 | 1.922109 |
| 41314 | ENSG0000 | AL161772.1 | | 0.962845 | 1.520308 | 1.92169 |
| 4637 | ENSG0000 | CDK18 | cyclin depe | 0.324332 | 0.743681 | 1.917473 |
| 43018 | ENSG0000 | MIR936 | microRNA | 0.63874 | 0.826931 | 1.915554 |
| 38737 | ENSG0000 | ANKRD34A | ankyrin rep | 0.653146 | 1.277961 | 1.915445 |
| 12933 | ENSG0000 | RAB33B | RAB33B, m | 1.194719 | 1.601092 | 1.913518 |
| 17082 | ENSG0000 | ZNF652 | zinc finger | 1.070658 | 1.07306 | 1.912695 |
| 14283 | ENSG0000 | KLHL28 | kelch like f | 0.884735 | 1.375799 | 1.912582 |

| | | | | | | |
|---|---|---|---|---|---|---|
| 2852 | ENSG0000 | RHOV | ras homolo | 0.592863 | 1.223639 | 1.912567 |
| 6582 | ENSG0000 | PATJ | PATJ, crum | 1.202692 | 1.716357 | 1.912223 |
| 38604 | ENSG0000 | AC114490.2 | | 0.480963 | 0.910018 | 1.911436 |
| 7667 | ENSG0000 | ALG10 | ALG10, alp | 0.985397 | 1.890267 | 1.9111 |
| 22526 | ENSG0000 | AL359182.1 | | 0.370108 | 1.327983 | 1.911028 |
| 14477 | ENSG0000 | TMEM64 | transmeml | 1.127394 | 1.445054 | 1.908551 |
| 12972 | ENSG0000 | AZU1 | azurocidin | 0.545861 | 0.84023 | 1.90839 |
| 13444 | ENSG0000 | PQLC2L | PQ loop re | 0.841836 | 1.538778 | 1.908269 |
| 4665 | ENSG0000 | CNN3 | calponin 3 | 0.540323 | 0.690576 | 1.906031 |
| 2835 | ENSG0000 | TTC23 | tetratricop | 1.46465 | 1.61304 | 1.903999 |
| 9406 | ENSG0000 | DCLRE1C | DNA cross- | 0.743409 | 1.357383 | 1.903922 |
| 28803 | ENSG0000 | ZNF709 | zinc finger | 0.3257 | 1.257428 | 1.903334 |
| 13957 | ENSG0000 | C18orf32 | chromosor | 1.407237 | 1.493697 | 1.900988 |
| 1457 | ENSG0000 | CEACAM1 | carcinoeml | 0.602723 | 1.317713 | 1.900988 |
| 4541 | ENSG0000 | SWT1 | SWT1, RNA | 0.901904 | 1.404586 | 1.899481 |
| 31733 | ENSG0000 | CLDN23 | claudin 23 | 0.292451 | 0.541026 | 1.898981 |
| 10259 | ENSG0000 | ZBTB8A | zinc finger | 1.541772 | 1.716413 | 1.898917 |
| 4493 | ENSG0000 | RALGPS2 | Ral GEF wi | 0.869925 | 1.614599 | 1.89859 |
| 19991 | ENSG0000 | GANC | glucosidase | 1.74179 | 1.770779 | 1.898207 |
| 15668 | ENSG0000 | ANKRD37 | ankyrin rep | 0.220418 | 0.484172 | 1.895763 |
| 5210 | ENSG0000 | SERAC1 | serine activ | 1.306635 | 1.496096 | 1.895374 |
| 1358 | ENSG0000 | RAP1GAP | RAP1 GTPa | 0.532178 | 0.985438 | 1.892185 |
| 34803 | ENSG0000 | LINC02256 | long interg | 0.669824 | 1.108706 | 1.891046 |
| 27133 | ENSG0000 | ZNRF2P1 | zinc and rir | 1.228661 | 1.245237 | 1.890556 |
| 15778 | ENSG0000 | ZDHHC17 | zinc finger | 1.287776 | 1.78923 | 1.889715 |
| 21547 | ENSG0000 | AL390729.1 | | 0.895684 | 1.793419 | 1.887124 |
| 19951 | ENSG0000 | DNASE1 | deoxyribor | 1.109353 | 1.615144 | 1.886259 |
| 12062 | ENSG0000 | CCDC88B | coiled-coil | 0.598633 | 0.708721 | 1.881471 |
| 14342 | ENSG0000 | PHC1P1 | polyhomeo | 1.323138 | 1.525731 | 1.881305 |
| 9276 | ENSG0000 | CCDC3 | coiled-coil | 1.174212 | 1.505411 | 1.881076 |
| 7488 | ENSG0000 | IFT172 | intraflagell | 1.576561 | 1.811655 | 1.880308 |
| 6765 | ENSG0000 | PTGFRN | prostaglan | 0.6008 | 1.324805 | 1.880108 |
| 8210 | ENSG0000 | MOB3C | MOB kinas | 0.778191 | 0.945099 | 1.879445 |
| 2370 | ENSG0000 | SUSD6 | sushi doma | 1.573997 | 1.716067 | 1.878683 |
| 42534 | ENSG0000 | GHET1 | gastric carc | 0.393555 | 0.876598 | 1.872512 |
| 5942 | ENSG0000 | STRIP2 | striatin inte | 1.072287 | 1.259578 | 1.87096 |
| 39535 | ENSG0000 | AL627230.1 | | 1.234128 | 1.387331 | 1.866574 |
| 9474 | ENSG0000 | MR1 | major histo | 0.919551 | 1.142998 | 1.866407 |
| 14094 | ENSG0000 | PLEKHM3 | pleckstrin l | 0.998968 | 1.674794 | 1.864955 |
| 11586 | ENSG0000 | ARIH1 | ariadne RB | 1.254696 | 1.656743 | 1.864447 |
| 13219 | ENSG0000 | PEAK1 | pseudopoc | 1.302007 | 1.788015 | 1.8638 |
| 11709 | ENSG0000 | KIF7 | kinesin fan | 1.163938 | 1.77387 | 1.860666 |
| 11296 | ENSG0000 | TP53INP1 | tumor prot | 0.294767 | 0.838595 | 1.860442 |
| 30110 | ENSG0000 | NOP14-AS: | NOP14 ant | 0.971291 | 1.49595 | 1.860372 |
| 7336 | ENSG0000 | KLC4 | kinesin ligh | 0.770845 | 1.206245 | 1.857962 |
| 1837 | ENSG0000 | TASP1 | taspase 1 [ | 1.2494 | 1.831497 | 1.857847 |
| 5439 | ENSG0000 | PPP4R1L | protein ph | 0.747251 | 1.479268 | 1.856616 |

| | | | | | | |
|---|---|---|---|---|---|---|
| 40769 | ENSG0000 | AC008556.1 | | 0.352589 | 0.75221 | 1.8565 |
| 18490 | ENSG0000 | LIN52 | lin-52 DRE/ | 1.134234 | 1.556301 | 1.855444 |
| 15889 | ENSG0000 | PLEKHN1 | pleckstrin l | 0.927037 | 1.512868 | 1.85523 |
| 6530 | ENSG0000 | MATN2 | matrilin 2 [ | 0.554463 | 0.918477 | 1.851739 |
| 7269 | ENSG0000 | SLC31A2 | solute carr | 1.099954 | 1.310225 | 1.850602 |
| 1489 | ENSG0000 | COL5A3 | collagen ty | 0.485935 | 1.785252 | 1.848931 |
| 4395 | ENSG0000 | EFHD1 | EF-hand dc | 0.099343 | 0.290053 | 1.848908 |
| 3315 | ENSG0000 | TSPAN13 | tetraspanir | 1.34716 | 1.559113 | 1.848455 |
| 25215 | ENSG0000 | AL391069.2 | | 0.770248 | 1.287359 | 1.847255 |
| 13117 | ENSG0000 | SH3RF3 | SH3 domai | 1.110799 | 1.790458 | 1.846311 |
| 9230 | ENSG0000 | OXSM | 3-oxoacyl-/ | 1.228764 | 1.604521 | 1.845752 |
| 17171 | ENSG0000 | RASGEF1A | RasGEF do | 0.297144 | 0.950151 | 1.845336 |
| 33488 | ENSG0000 | AC009779.2 | | 1.25309 | 1.669388 | 1.842428 |
| 11639 | ENSG0000 | PRTG | protogenir | 0.963345 | 1.839576 | 1.841534 |
| 12486 | ENSG0000 | CHRNB1 | cholinergic | 0.565706 | 1.11316 | 1.840904 |
| 7937 | ENSG0000 | ZNF287 | zinc finger | 1.344779 | 1.503396 | 1.839136 |
| 11006 | ENSG0000 | HEYL | hes relatec | 0.680946 | 0.776604 | 1.838169 |
| 18624 | ENSG0000 | PDPK2P | 3-phospho | 1.364062 | 1.816929 | 1.835055 |
| 15771 | ENSG0000 | MAPT | microtubul | 0.886882 | 1.271414 | 1.834753 |
| 37383 | ENSG0000 | LIN37 | lin-37 DRE/ | 1.273548 | 1.689932 | 1.833912 |
| 38809 | ENSG0000 | NBPF25P | NBPF mem | 1.383985 | 1.760014 | 1.830765 |
| 25006 | ENSG0000 | Z99943.1 | | 0.047504 | 1.121438 | 1.829599 |
| 16997 | ENSG0000 | ZNF808 | zinc finger | 0.724933 | 0.884949 | 1.829165 |
| 39034 | ENSG0000 | AC010655.3 | | 0.195664 | 0.870451 | 1.829015 |
| 6129 | ENSG0000 | SHC2 | SHC adapt | 0.388146 | 1.010764 | 1.827444 |
| 10288 | ENSG0000 | HSF2BP | heat shock | 0.314053 | 1.406579 | 1.82678 |
| 39870 | ENSG0000 | Six3os1_1 | Six3os1 co | 0.348642 | 0.746521 | 1.824715 |
| 7907 | ENSG0000 | NUDT7 | nudix hydr | 1.017996 | 1.211621 | 1.823812 |
| 13430 | ENSG0000 | BTC | betacellulir | 1.215901 | 1.799846 | 1.816869 |
| 7982 | ENSG0000 | GREB1L | growth reg | 1.332393 | 1.365831 | 1.81529 |
| 36839 | ENSG0000 | DHX40P1 | DEAH-box | 0.345301 | 1.119088 | 1.814601 |
| 5083 | ENSG0000 | EPHX2 | epoxide hy | 1.024845 | 1.365614 | 1.813487 |
| 11502 | ENSG0000 | E2F7 | E2F transc | 0.827195 | 1.35975 | 1.810061 |
| 6368 | ENSG0000 | LRRC4B | leucine ricl | 0.460849 | 1.05542 | 1.809228 |
| 32169 | ENSG0000 | CKLF-CMTI | CKLF-CMTI | 1.34636 | 1.372276 | 1.808878 |
| 1233 | ENSG0000 | LLGL2 | LLGL2, scri | 0.9429 | 1.035151 | 1.808732 |
| 8564 | ENSG0000 | EAF2 | ELL associa | 0.874366 | 1.537629 | 1.805406 |
| 10941 | ENSG0000 | CRELD1 | cysteine ric | 0.916312 | 1.495786 | 1.804858 |
| 30207 | ENSG0000 | GOLGA8K | golgin A8 f | 0.736526 | 1.187709 | 1.804698 |
| 14336 | ENSG0000 | AC104581.1 | | 0.650218 | 0.801551 | 1.802396 |
| 13698 | ENSG0000 | B3GNT4 | UDP-GlcNA | 0.594314 | 0.770965 | 1.801673 |
| 15574 | ENSG0000 | PTCH1 | patched 1 | 1.024223 | 1.694652 | 1.799802 |
| 28944 | ENSG0000 | MRPS31P5 | mitochond | 0.968693 | 1.720025 | 1.799311 |
| 41291 | ENSG0000 | PRICKLE4 | prickle plar | 0.518585 | 0.979815 | 1.797812 |
| 16336 | ENSG0000 | ZNF681 | zinc finger | 1.382544 | 1.734419 | 1.797776 |
| 3885 | ENSG0000 | NT5DC3 | 5'-nucleoti | 1.247629 | 1.720976 | 1.796984 |
| 10924 | ENSG0000 | KATNAL2 | katanin cat | 0.672072 | 1.165255 | 1.796547 |

| | | | | | | |
|---|---|---|---|---|---|---|
| 10656 | ENSG0000 | ACP6 | acid phosp | 0.882018 | 1.293683 | 1.796197 |
| 5887 | ENSG0000 | ZNF780B | zinc finger | 1.261172 | 1.516686 | 1.791434 |
| 37353 | ENSG0000 | RUNDC3A- | RUNDC3A | 0.346888 | 0.842251 | 1.790011 |
| 13983 | ENSG0000 | DNAJC28 | DnaJ heat | 0.412775 | 1.073782 | 1.789846 |
| 3858 | ENSG0000 | RAB5B | RAB5B, me | 1.45953 | 1.648233 | 1.78659 |
| 8233 | ENSG0000 | NME7 | NME/NM2 | 1.061917 | 1.287546 | 1.786307 |
| 27945 | ENSG0000 | AC100861.1 | | 0.779682 | 1.210531 | 1.7863 |
| 16314 | ENSG0000 | ZNF699 | zinc finger | 0.706117 | 1.541262 | 1.7847 |
| 11712 | ENSG0000 | PEX11A | peroxisom | 0.908475 | 0.931821 | 1.784196 |
| 9968 | ENSG0000 | TSC22D3 | TSC22 dom | 0.908512 | 1.443167 | 1.784191 |
| 38959 | ENSG0000 | AL139246.5 | | 1.158883 | 1.535704 | 1.784174 |
| 29407 | ENSG0000 | LINC01184 | long interg | 1.672451 | 1.751985 | 1.783244 |
| 16433 | ENSG0000 | ZNF775 | zinc finger | 1.261762 | 1.378501 | 1.781325 |
| 24925 | ENSG0000 | FAAHP1 | fatty acid a | 0.5118 | 0.713727 | 1.781102 |
| 480 | ENSG0000 | NR1H3 | nuclear rec | 1.147025 | 1.369003 | 1.779655 |
| 9536 | ENSG0000 | C7orf31 | chromosor | 0.378654 | 1.523552 | 1.776604 |
| 6202 | ENSG0000 | FCHO1 | FCH domai | 0.441303 | 0.507726 | 1.775321 |
| 2147 | ENSG0000 | MMP11 | matrix met | 1.044372 | 1.424618 | 1.772435 |
| 15750 | ENSG0000 | SPIN2B | spindlin fai | 0.883957 | 1.268525 | 1.771641 |
| 11302 | ENSG0000 | TMEM67 | transmeml | 0.508231 | 1.075649 | 1.770475 |
| 22121 | ENSG0000 | AC018641.1 | | 0.216141 | 0.834959 | 1.770237 |
| 17951 | ENSG0000 | NOTCH4 | notch 4 [Sc | 0.836966 | 1.570042 | 1.766806 |
| 14145 | ENSG0000 | DHFR2 | dihydrofol | 1.138958 | 1.282889 | 1.766541 |
| 16948 | ENSG0000 | TMEM116 | transmeml | 1.237947 | 1.322809 | 1.766253 |
| 3540 | ENSG0000 | C17orf75 | chromosor | 1.563213 | 1.70378 | 1.764736 |
| 12378 | ENSG0000 | RGPD8 | RANBP2-lik | 1.210652 | 1.727621 | 1.764385 |
| 14212 | ENSG0000 | CCDC89 | coiled-coil | 0.523935 | 0.99404 | 1.763236 |
| 33699 | ENSG0000 | AL049830.3 | | 0.731471 | 0.882123 | 1.762167 |
| 41927 | ENSG0000 | AP006621.5 | | 1.179688 | 1.535503 | 1.761673 |
| 4265 | ENSG0000 | C3orf52 | chromosor | 0.848686 | 1.292116 | 1.759935 |
| 14317 | ENSG0000 | FLJ37453 | uncharacte | 0.8279 | 1.200247 | 1.759259 |
| 13241 | ENSG0000 | SCAI | suppressor | 1.050358 | 1.365759 | 1.755861 |
| 9623 | ENSG0000 | PDLIM3 | PDZ and LI | 1.151439 | 1.194207 | 1.754123 |
| 37066 | ENSG0000 | AC060780.2 | | 0.660439 | 0.877315 | 1.750895 |
| 3462 | ENSG0000 | EBF3 | early B-cell | 0.66704 | 1.561547 | 1.750769 |
| 34785 | ENSG0000 | AC005730.2 | | 0.310075 | 0.708925 | 1.747497 |
| 26423 | ENSG0000 | SPAAR | small regul | 0.10514 | 0.376761 | 1.745369 |
| 10770 | ENSG0000 | GNPDA2 | glucosamir | 1.517288 | 1.59527 | 1.744209 |
| 42274 | ENSG0000 | AC023632.6 | | 0.647445 | 1.16194 | 1.743152 |
| 4658 | ENSG0000 | PTCH2 | patched 2 | 0.271536 | 1.173815 | 1.742167 |
| 29365 | ENSG0000 | AC008393.1 | | 0.875036 | 1.327598 | 1.740666 |
| 36256 | ENSG0000 | AC068025.1 | | 0.38919 | 0.750881 | 1.736798 |
| 15071 | ENSG0000 | KLHL25 | kelch like f | 1.085929 | 1.464966 | 1.735094 |
| 27202 | ENSG0000 | DGUOK-AS | DGUOK an | 1.158116 | 1.524575 | 1.734159 |
| 33094 | ENSG0000 | AP001453.3 | | 0.878733 | 1.467635 | 1.732982 |
| 1243 | ENSG0000 | PPP2R3A | protein ph | 0.55331 | 1.300904 | 1.732654 |
| 29916 | ENSG0000 | AC008771.1 | | 0.507726 | 1.219515 | 1.732385 |

| | | | | | | |
|---|---|---|---|---|---|---|
| 34419 | ENSG0000 | AL390728.6 | | 1.141918 | 1.520385 | 1.730331 |
| 43192 | ENSG0000 | SCO2 | SCO2, cyto | 0.338818 | 0.614883 | 1.729061 |
| 15876 | ENSG0000 | IFT140 | intraflagell | 1.13476 | 1.332128 | 1.728924 |
| 4008 | ENSG0000 | ARFGEF3 | ARFGEF fa | 0.403353 | 0.815256 | 1.7286 |
| 32760 | ENSG0000 | DENND5B- | DENND5B | 1.084086 | 1.595782 | 1.725363 |
| 8620 | ENSG0000 | FAM105A | family with | 0.323296 | 0.707204 | 1.725163 |
| 9949 | ENSG0000 | GLB1L | galactosida | 0.809694 | 1.320553 | 1.72444 |
| 8669 | ENSG0000 | N4BP3 | NEDD4 bin | 0.569178 | 0.778371 | 1.724333 |
| 410 | ENSG0000 | RUFY3 | RUN and F | 0.816076 | 1.524865 | 1.722719 |
| 36846 | ENSG0000 | AC060780.1 | | 0.73167 | 1.553046 | 1.721895 |
| 1552 | ENSG0000 | MAGI3 | membrane | 1.24412 | 1.254101 | 1.714893 |
| 10240 | ENSG0000 | ZNF230 | zinc finger | 0.751828 | 1.114141 | 1.714615 |
| 33111 | ENSG0000 | AC007249.2 | | 0.358592 | 1.096574 | 1.712761 |
| 15942 | ENSG0000 | ZNF70 | zinc finger | 0.766766 | 1.18239 | 1.711551 |
| 11599 | ENSG0000 | NUDT13 | nudix hydr | 0.797717 | 1.177426 | 1.711278 |
| 1617 | ENSG0000 | DOPEY1 | dopey fam | 1.379405 | 1.696004 | 1.710662 |
| 16321 | ENSG0000 | MYT1 | myelin trar | 0.459965 | 0.509538 | 1.709578 |
| 15675 | ENSG0000 | GOLGA8R | golgin A8 f | 0.705058 | 1.120908 | 1.709247 |
| 37893 | ENSG0000 | AC010422.6 | | 0.022874 | 0.051156 | 1.70645 |
| 15143 | ENSG0000 | DUSP5P1 | dual specif | 0.811174 | 1.661222 | 1.706105 |
| 10582 | ENSG0000 | MEGF6 | multiple E( | 0.30395 | 1.057031 | 1.705848 |
| 42448 | ENSG0000 | Metazoa_S | Metazoan | 0.904516 | 1.197628 | 1.705783 |
| 11668 | ENSG0000 | IQCD | IQ motif co | 0.495069 | 0.971246 | 1.704528 |
| 11473 | ENSG0000 | JCAD | junctional | 0.536271 | 1.242748 | 1.703998 |
| 9248 | ENSG0000 | MAGI1 | membrane | 1.275396 | 1.620563 | 1.702962 |
| 33931 | ENSG0000 | FOXN3-AS1 | FOXN3 ant | 0.832746 | 1.386129 | 1.701824 |
| 4359 | ENSG0000 | APC2 | APC2, WN | 1.047298 | 1.4218 | 1.701295 |
| 16552 | ENSG0000 | LAMB3 | laminin sul | 0.977772 | 1.168603 | 1.700043 |
| 28633 | ENSG0000 | SRRM1P2 | serine/argi | 0.660334 | 1.320291 | 1.699823 |
| 3651 | ENSG0000 | TRIM2 | tripartite n | 0.912585 | 1.277786 | 1.693993 |
| 4588 | ENSG0000 | OSCP1 | organic sol | 1.067813 | 1.295193 | 1.693701 |
| 33808 | ENSG0000 | LINC00638 | long interg | 0.512731 | 0.837379 | 1.692209 |
| 22010 | ENSG0000 | AC016027.1 | | 0.890476 | 1.57913 | 1.689756 |
| 11232 | ENSG0000 | SYTL3 | synaptotag | 0.563141 | 0.878108 | 1.689551 |
| 16902 | ENSG0000 | SFI1 | SFI1 centri | 1.166143 | 1.566524 | 1.689329 |
| 32792 | ENSG0000 | AC131009.1 | | 0.162198 | 0.831327 | 1.689073 |
| 16574 | ENSG0000 | ZNF585A | zinc finger | 0.703083 | 1.283247 | 1.688926 |
| 26495 | ENSG0000 | AC146949.1 | | 1.141068 | 1.424806 | 1.686662 |
| 12623 | ENSG0000 | CEL | carboxyl es | 1.25105 | 1.322952 | 1.684731 |
| 23849 | ENSG0000 | LINC01204 | long interg | 0.549365 | 1.640824 | 1.683741 |
| 13289 | ENSG0000 | GPR160 | G protein-c | 0.383486 | 0.588288 | 1.677774 |
| 1266 | ENSG0000 | SNCB | synuclein b | 0.37458 | 0.39011 | 1.677619 |
| 6334 | ENSG0000 | ZNF141 | zinc finger | 0.83663 | 1.182301 | 1.677236 |
| 8518 | ENSG0000 | SLC25A26 | solute carr | 0.840329 | 1.617897 | 1.675104 |
| 5461 | ENSG0000 | NAGK | N-acetylglu | 1.582325 | 1.618769 | 1.675062 |
| 8470 | ENSG0000 | NBEAL1 | neurobeac | 0.881903 | 1.342087 | 1.6739 |
| 9827 | ENSG0000 | FBXO43 | F-box prot | 0.831685 | 0.943792 | 1.66983 |

| | | | | | | |
|---|---|---|---|---|---|---|
| 1864 | ENSG0000 | ZBTB25 | zinc finger | 0.440938 | 0.906014 | 1.668989 |
| 32829 | ENSG0000 | ZNF432 | zinc finger | 0.714547 | 1.572428 | 1.66536 |
| 1116 | ENSG0000 | CLTCL1 | clathrin he | 1.32206 | 1.478586 | 1.664208 |
| 22250 | ENSG0000 | AL031846.1 | | 0.247588 | 0.937322 | 1.663819 |
| 13056 | ENSG0000 | TMEM134 | transmeml | 1.090606 | 1.208201 | 1.663621 |
| 22875 | ENSG0000 | AC073174.1 | | 0.285684 | 1.085312 | 1.661877 |
| 12308 | ENSG0000 | CA5B | carbonic ai | 0.689296 | 1.027565 | 1.66091 |
| 23287 | ENSG0000 | PCAT6 | prostate ca | 0.256242 | 1.001622 | 1.660552 |
| 5422 | ENSG0000 | TOX2 | TOX high m | 0.668647 | 1.082492 | 1.660123 |
| 31718 | ENSG0000 | TNFRSF10A | TNFRSF10A | 1.058921 | 1.580422 | 1.659856 |
| 870 | ENSG0000 | AHRR | aryl-hydro | 0.30275 | 1.42979 | 1.656977 |
| 3125 | ENSG0000 | CARD8 | caspase re | 1.166101 | 1.245886 | 1.653072 |
| 2879 | ENSG0000 | UBE2W | ubiquitin c | 1.293107 | 1.507782 | 1.650821 |
| 5450 | ENSG0000 | NEURL2 | neuralized | 0.058899 | 0.306479 | 1.650294 |
| 9741 | ENSG0000 | LSM11 | LSM11, U7 | 1.364409 | 1.540446 | 1.65019 |
| 1897 | ENSG0000 | CHRD | chordin [So | 0.353945 | 0.905984 | 1.649963 |
| 13884 | ENSG0000 | RIMKLA | ribosomal | 0.140419 | 0.15884 | 1.648528 |
| 4528 | ENSG0000 | DLGAP3 | DLG associ | 0.762018 | 1.316918 | 1.646594 |
| 21279 | ENSG0000 | LINC00630 | long interg | 0.690539 | 1.431588 | 1.644643 |
| 659 | ENSG0000 | SNX29 | sorting nex | 1.227488 | 1.568156 | 1.644602 |
| 12967 | ENSG0000 | MBOAT1 | membrane | 0.920261 | 1.037921 | 1.643395 |
| 7597 | ENSG0000 | SEMA7A | semaphori | 0.920293 | 1.154788 | 1.641636 |
| 2846 | ENSG0000 | BMF | Bcl2 modif | 0.214171 | 0.75959 | 1.640992 |
| 27421 | ENSG0000 | AC007384.1 | | 0.126171 | 0.279628 | 1.640617 |
| 7737 | ENSG0000 | N4BP2L1 | NEDD4 bin | 0.393727 | 0.621835 | 1.640554 |
| 38972 | ENSG0000 | AL024507.2 | | 0.449695 | 1.253603 | 1.640238 |
| 16694 | ENSG0000 | DACT3 | dishevelled | 0.435962 | 0.602142 | 1.637866 |
| 21905 | ENSG0000 | LINC00337 | long interg | 0.371894 | 0.93689 | 1.635663 |
| 15903 | ENSG0000 | SAMD11 | sterile alph | 0.263657 | 0.860245 | 1.634816 |
| 12027 | ENSG0000 | CASKIN1 | CASK intera | 0.63593 | 0.859567 | 1.633356 |
| 43293 | ENSG0000 | RABGEF1 | RAB guanir | 1.334336 | 1.409802 | 1.633229 |
| 3924 | ENSG0000 | ADTRP | androgen c | 0.464403 | 1.158782 | 1.630861 |
| 4427 | ENSG0000 | PASK | PAS domai | 1.499221 | 1.557195 | 1.626505 |
| 16961 | ENSG0000 | ZNF442 | zinc finger | 0.169314 | 1.2091 | 1.626373 |
| 16681 | ENSG0000 | PELI1 | pellino E3 u | 0.450985 | 1.047442 | 1.625397 |
| 36880 | ENSG0000 | AC006213.2 | | 0.878506 | 1.088319 | 1.625229 |
| 15287 | ENSG0000 | ZNF93 | zinc finger | 0.278218 | 0.971909 | 1.624834 |
| 6468 | ENSG0000 | EMILIN2 | elastin mic | 0.15193 | 0.70865 | 1.624092 |
| 421 | ENSG0000 | PHLDB1 | pleckstrin l | 1.413955 | 1.599948 | 1.621804 |
| 41091 | ENSG0000 | AC006449.6 | | 0.578752 | 0.643112 | 1.620786 |
| 13426 | ENSG0000 | THAP6 | THAP dom | 1.193993 | 1.411051 | 1.618796 |
| 12865 | ENSG0000 | LRRC34 | leucine rich | 1.297203 | 1.370302 | 1.616617 |
| 20421 | ENSG0000 | MCM3AP-A | MCM3AP a | 1.069205 | 1.497911 | 1.615396 |
| 9211 | ENSG0000 | SEC24D | SEC24 hom | 1.139938 | 1.182381 | 1.612731 |
| 37446 | ENSG0000 | NAPA-AS1 | NAPA antis | 0.344283 | 0.878465 | 1.611151 |
| 16037 | ENSG0000 | DCUN1D3 | defective ir | 1.317583 | 1.563014 | 1.610079 |
| 9708 | ENSG0000 | SLC7A7 | solute carr | 0.498525 | 0.85314 | 1.609949 |

| | | | | | | |
|---|---|---|---|---|---|---|
| 23237 | ENSG0000 | AL357140.2 | | 0.153318 | 0.496335 | 1.606873 |
| 22902 | ENSG0000 | DLG1-AS1 | DLG1 antis | 0.612795 | 1.143653 | 1.60628 |
| 3515 | ENSG0000 | ENO3 | enolase 3 [ | 0.51517 | 1.071268 | 1.605816 |
| 11062 | ENSG0000 | ABHD18 | abhydrolas | 0.975917 | 1.507314 | 1.604094 |
| 8630 | ENSG0000 | LHFPL2 | LHFPL tetra | 1.112859 | 1.488681 | 1.60375 |
| 16184 | ENSG0000 | ZNF563 | zinc finger | 0.181761 | 1.159604 | 1.6037 |
| 5218 | ENSG0000 | ODF2L | outer dens | 0.604379 | 1.310359 | 1.602937 |
| 16623 | ENSG0000 | ZNF772 | zinc finger | 0.78679 | 1.328952 | 1.602231 |
| 32571 | ENSG0000 | AP002748.3 | | 1.257484 | 1.41893 | 1.601618 |
| 38673 | ENSG0000 | AL136162.1 | | 0.404895 | 0.847201 | 1.601359 |
| 35612 | ENSG0000 | CCDC92B | coiled-coil | 0.397361 | 0.568631 | 1.601234 |
| 29610 | ENSG0000 | WHAMMP | WAS prote | 0.554065 | 1.526745 | 1.599153 |
| 12294 | ENSG0000 | CCDC126 | coiled-coil | 0.958202 | 1.155649 | 1.597212 |
| 14379 | ENSG0000 | C10orf91 | chromosor | 0.365113 | 0.772475 | 1.597142 |
| 40654 | ENSG0000 | AL161891.1 | | 0.740539 | 1.29677 | 1.595105 |
| 5825 | ENSG0000 | DYRK2 | dual specif | 1.242012 | 1.446399 | 1.594648 |
| 2568 | ENSG0000 | ANKRD12 | ankyrin rep | 1.068332 | 1.458216 | 1.594043 |
| 36325 | ENSG0000 | MIR3942 | microRNA | 0.929505 | 1.28911 | 1.593424 |
| 6231 | ENSG0000 | COL5A1 | collagen ty | 0.286436 | 0.486985 | 1.592834 |
| 10937 | ENSG0000 | RBM47 | RNA bindir | 1.074533 | 1.093958 | 1.586347 |
| 29383 | ENSG0000 | DDX11-AS1 | DDX11 ant | 0.747402 | 1.39002 | 1.584472 |
| 32207 | ENSG0000 | AC067930.4 | | 0.245205 | 1.457523 | 1.582088 |
| 22337 | ENSG0000 | RIMKLBP2 | ribosomal | 0.114889 | 0.942636 | 1.581832 |
| 15063 | ENSG0000 | DGCR6 | DiGeorge s | 1.059469 | 1.218536 | 1.58114 |
| 26090 | ENSG0000 | MED14OS | MED14 op | 0.65928 | 0.903067 | 1.580418 |
| 8472 | ENSG0000 | SPAG16 | sperm assc | 1.099236 | 1.210153 | 1.580243 |
| 1250 | ENSG0000 | MAP3K13 | mitogen-ac | 0.992386 | 1.342063 | 1.57997 |
| 26595 | ENSG0000 | OLMALINC | oligodendr | 0.676227 | 1.123909 | 1.579353 |
| 38315 | ENSG0000 | AC009948.4 | | 0.788899 | 1.477714 | 1.578567 |
| 118 | ENSG0000 | OSBPL7 | oxysterol b | 0.578289 | 1.240551 | 1.576536 |
| 16466 | ENSG0000 | SULF2 | sulfatase 2 | 0.204733 | 1.246297 | 1.575951 |
| 41750 | ENSG0000 | AC009303.4 | | 0.425485 | 1.042594 | 1.575019 |
| 28675 | ENSG0000 | BMS1P4 | BMS1, ribc | 0.316891 | 0.820678 | 1.573961 |
| 37823 | ENSG0000 | AC008655.2 | | 0.442575 | 1.058717 | 1.573793 |
| 891 | ENSG0000 | HIPK2 | homeodon | 0.745501 | 1.194501 | 1.572922 |
| 29665 | ENSG0000 | ABHD14A | abhydrolas | 0.964221 | 1.253304 | 1.57079 |
| 3637 | ENSG0000 | CPE | carboxypep | 1.087306 | 1.098888 | 1.570645 |
| 18875 | ENSG0000 | RNVU1-15 | RNA, variai | 0.215104 | 1.394723 | 1.569888 |
| 27315 | ENSG0000 | CKMT1B | creatine ki | 0.562814 | 1.1011 | 1.569291 |
| 16154 | ENSG0000 | TMEM198 | transmeml | 0.775972 | 1.432267 | 1.568246 |
| 9175 | ENSG0000 | HNMT | histamine l | 1.133426 | 1.133643 | 1.566318 |
| 15753 | ENSG0000 | VSIG10L | V-set and i | 0.758089 | 0.759915 | 1.565144 |
| 1520 | ENSG0000 | OTUB2 | OTU deubi | 0.390406 | 0.628 | 1.563761 |
| 973 | ENSG0000 | SLC9A3 | solute carr | 0.511334 | 1.337095 | 1.563657 |
| 3237 | ENSG0000 | HOXA2 | homeobox | 1.358892 | 1.560827 | 1.563145 |
| 1251 | ENSG0000 | ST6GAL1 | ST6 beta-g | 0.85564 | 1.121162 | 1.562137 |
| 11186 | ENSG0000 | CREBRF | CREB3 regu | 0.562212 | 1.416263 | 1.561047 |

| | | | | | | |
|---|---|---|---|---|---|---|
| 14333 | ENSG0000 | AKAP5 | A-kinase ar | 0.309033 | 1.162585 | 1.559604 |
| 24328 | ENSG0000 | KLHL7-AS1 | KLHL7 anti | 0.471208 | 0.905986 | 1.559211 |
| 5625 | ENSG0000 | FOSB | FosB proto | 0.704996 | 1.528106 | 1.558603 |
| 798 | ENSG0000 | CAMK2B | calcium/ca | 0.631756 | 1.036443 | 1.558197 |
| 14255 | ENSG0000 | TMEM151 | transmeml | 0.262672 | 0.449665 | 1.558096 |
| 31840 | ENSG0000 | AC008708.2 | | 0.646872 | 1.553519 | 1.557897 |
| 17854 | ENSG0000 | Y_RNA | Y RNA [Sou | 0 | 0.872168 | 1.557351 |
| 24850 | ENSG0000 | AL162377.1 | | 0.827383 | 1.536344 | 1.557069 |
| 3938 | ENSG0000 | UST | uronyl 2-su | 0.728465 | 1.416083 | 1.556219 |
| 11136 | ENSG0000 | RHOBTB3 | Rho relate | 0.60307 | 1.365939 | 1.556199 |
| 23025 | ENSG0000 | LINC01714 | long interg | 0.203139 | 0.689132 | 1.556063 |
| 23468 | ENSG0000 | ARL17B | ADP ribosy | 0.976715 | 1.282031 | 1.555069 |
| 5597 | ENSG0000 | OPRL1 | opioid rela | 0.450967 | 0.538488 | 1.552589 |
| 6074 | ENSG0000 | PUS7L | pseudouri | 0.903216 | 1.282384 | 1.552519 |
| 3509 | ENSG0000 | PIGL | phosphatic | 0.981517 | 1.266333 | 1.550849 |
| 11061 | ENSG0000 | MFSD8 | major facil | 1.088723 | 1.302882 | 1.550007 |
| 15894 | ENSG0000 | ZNF286A | zinc finger | 1.074185 | 1.42477 | 1.545702 |
| 13272 | ENSG0000 | HAP1 | huntingtin | 0.1296 | 0.905092 | 1.543822 |
| 4214 | ENSG0000 | ARL6 | ADP ribosy | 0.44533 | 1.144133 | 1.542774 |
| 11508 | ENSG0000 | TTC7B | tetratricop | 1.232187 | 1.380405 | 1.53927 |
| 8089 | ENSG0000 | ADAMTS1( | ADAM met | 0.897554 | 0.919505 | 1.538431 |
| 28425 | ENSG0000 | SUCLG2-AS | SUCLG2 an | 0.336298 | 1.139714 | 1.537602 |
| 39196 | ENSG0000 | AP001033.2 | | 0.752892 | 1.031218 | 1.537172 |
| 1941 | ENSG0000 | SEL1L3 | SEL1L fami | 0.577197 | 0.731454 | 1.536791 |
| 41811 | ENSG0000 | AC020910.5 | | 0.109988 | 1.532656 | 1.534575 |
| 4757 | ENSG0000 | TNFAIP3 | TNF alpha | 0.285449 | 1.513477 | 1.530271 |
| 9801 | ENSG0000 | TIAM1 | T-cell lymp | 1.067509 | 1.288686 | 1.529687 |
| 33993 | ENSG0000 | AL163636.2 | | 0.503488 | 0.561998 | 1.529202 |
| 15454 | ENSG0000 | IFNLR1 | interferon | 0.154952 | 0.449188 | 1.529173 |
| 10653 | ENSG0000 | C1orf115 | chromosor | 0.245005 | 0.871123 | 1.52609 |
| 38512 | ENSG0000 | CCL5 | C-C motif c | 0.379835 | 0.497577 | 1.525744 |
| 4947 | ENSG0000 | LGALSL | galectin lik | 0.601294 | 0.689822 | 1.525635 |
| 15165 | ENSG0000 | AL021453.1 | | 0.927086 | 1.213078 | 1.521641 |
| 22369 | ENSG0000 | HAR1A | highly acce | 0.220152 | 0.592418 | 1.521364 |
| 34492 | ENSG0000 | AC007541.1 | | 0.840741 | 1.44187 | 1.521182 |
| 31460 | ENSG0000 | TMEM200I | transmeml | 0.460858 | 1.300824 | 1.514411 |
| 8205 | ENSG0000 | AZIN2 | antizyme ir | 0.839577 | 1.001915 | 1.513568 |
| 39203 | ENSG0000 | AC009275.1 | | 0.088571 | 1.002747 | 1.513169 |
| 13561 | ENSG0000 | ALG10B | ALG10B, al | 0.920367 | 1.276341 | 1.512931 |
| 2759 | ENSG0000 | ZNF500 | zinc finger | 1.260931 | 1.371131 | 1.512602 |
| 5338 | ENSG0000 | SMUG1 | single-stra | 0.663566 | 1.259961 | 1.511811 |
| 29828 | ENSG0000 | AC034231.1 | | 0.55602 | 1.189143 | 1.509402 |
| 5643 | ENSG0000 | NAPB | NSF attach | 1.032988 | 1.067919 | 1.50861 |
| 14330 | ENSG0000 | MROH1 | maestro he | 1.050275 | 1.477325 | 1.505874 |
| 9511 | ENSG0000 | INO80C | INO80 con | 1.27326 | 1.371859 | 1.505853 |
| 41553 | ENSG0000 | CEP83-AS1 | CEP83 anti | 0.585835 | 0.730027 | 1.505818 |
| 10112 | ENSG0000 | F11R | F11 recept | 0.45122 | 0.743157 | 1.505522 |

| | | | | | | |
|---|---|---|---|---|---|---|
| 5949 | ENSG0000 | CCDC136 | coiled-coil | 0.376466 | 0.665308 | 1.505422 |
| 10001 | ENSG0000 | GAREM2 | GRB2 asso | 0.882605 | 1.312025 | 1.505353 |
| 8395 | ENSG0000 | ATP6V1C2 | ATPase H+ | 0.912473 | 1.042292 | 1.504358 |
| 24958 | ENSG0000 | DCST1-AS1 | DCST1 anti | 0.870291 | 0.942492 | 1.504348 |
| 34846 | ENSG0000 | AL133355.1 | | 0.548004 | 1.15809 | 1.502496 |
| 1205 | ENSG0000 | CHFR | checkpoint | 1.2516 | 1.472613 | 1.49794 |
| 40676 | ENSG0000 | AC123768.4 | | 0.595906 | 0.776406 | 1.496605 |
| 37705 | ENSG0000 | AL627309.7 | | 0.443158 | 0.947568 | 1.495668 |
| 15619 | ENSG0000 | ANKRD46 | ankyrin rep | 0.911973 | 0.980531 | 1.491559 |
| 28693 | ENSG0000 | PCDHGC4 | protocadh | 0.537611 | 0.721809 | 1.491394 |
| 6992 | ENSG0000 | MAP7 | microtubul | 0.669081 | 1.170729 | 1.489951 |
| 12297 | ENSG0000 | RAB3B | RAB3B, me | 0.103801 | 0.559576 | 1.489042 |
| 6817 | ENSG0000 | CABLES1 | Cdk5 and A | 0.589876 | 1.176141 | 1.487336 |
| 35654 | ENSG0000 | LINC02175 | long interg | 0.521826 | 0.69585 | 1.486018 |
| 10938 | ENSG0000 | APBB2 | amyloid be | 0.763731 | 1.362659 | 1.485529 |
| 5675 | ENSG0000 | TMEM74B | transmeml | 0.032119 | 0.716268 | 1.48414 |
| 8679 | ENSG0000 | TBC1D7 | TBC1 doma | 1.144611 | 1.1779 | 1.482562 |
| 7600 | ENSG0000 | FAM13A | family with | 0.749314 | 1.135642 | 1.481621 |
| 9655 | ENSG0000 | MPPE1 | metallopho | 0.972749 | 1.180929 | 1.481491 |
| 36856 | ENSG0000 | LRRC37A16 | leucine rich | 0.634059 | 1.036661 | 1.479668 |
| 36887 | ENSG0000 | AP005264.1 | | 0 | 0.178808 | 1.478734 |
| 21796 | ENSG0000 | AC025165.1 | | 0.498156 | 1.424912 | 1.478557 |
| 1493 | ENSG0000 | CHRNA3 | cholinergic | 0.177408 | 0.483426 | 1.478329 |
| 22367 | ENSG0000 | LINC01534 | long interg | 1.01852 | 1.207724 | 1.47718 |
| 34677 | ENSG0000 | JHDM1D-A | JHDM1D a | 0.462715 | 1.042822 | 1.47323 |
| 3519 | ENSG0000 | RASD1 | ras related | 0.27447 | 0.809689 | 1.471819 |
| 40106 | ENSG0000 | SRCIN1 | SRC kinase | 0.504505 | 0.788511 | 1.4715 |
| 7769 | ENSG0000 | ZIC5 | Zic family r | 0.926412 | 1.379606 | 1.469178 |
| 3121 | ENSG0000 | GRIN2D | glutamate | 0.100754 | 0.586109 | 1.467319 |
| 27832 | ENSG0000 | GATS | GATS, stro | 0.802396 | 1.055 | 1.466484 |
| 3460 | ENSG0000 | MAP3K8 | mitogen-ac | 0.233726 | 1.019887 | 1.465608 |
| 40474 | ENSG0000 | Six3os1_5 | Six3os1 co | 0 | 0.73343 | 1.465333 |
| 32974 | ENSG0000 | ZNF350 | zinc finger | 0.377675 | 1.432389 | 1.465037 |
| 18746 | ENSG0000 | SNORA51 | Small nucle | 0.580963 | 1.00122 | 1.464821 |
| 9735 | ENSG0000 | DEPTOR | DEP domai | 0.736586 | 1.433245 | 1.464648 |
| 7998 | ENSG0000 | TMC6 | transmeml | 0.281029 | 0.634761 | 1.463196 |
| 4 | ENSG0000 | SCYL3 | SCY1 like p | 1.157589 | 1.393227 | 1.46256 |
| 43276 | ENSG0000 | AC092329.4 | | 0.473402 | 1.014413 | 1.461552 |
| 34908 | ENSG0000 | AC027702.1 | | 0.572905 | 1.330165 | 1.460145 |
| 37941 | ENSG0000 | AL353593.2 | | 0.41884 | 1.024381 | 1.45993 |
| 678 | ENSG0000 | LTBP1 | latent tran | 0.428036 | 0.620742 | 1.45963 |
| 24301 | ENSG0000 | TEX48 | testis expr | 0.75604 | 0.77959 | 1.459562 |
| 16417 | ENSG0000 | ZNF765 | zinc finger | 0.595594 | 1.170149 | 1.458876 |
| 2246 | ENSG0000 | RASL10A | RAS like fa | 0.29084 | 0.311581 | 1.457516 |
| 7903 | ENSG0000 | NLRC5 | NLR family | 0.387678 | 1.158952 | 1.455771 |
| 668 | ENSG0000 | COL9A2 | collagen ty | 0.355131 | 0.47835 | 1.455567 |
| 6409 | ENSG0000 | TNS4 | tensin 4 [S | 0.087896 | 0.422116 | 1.454174 |

| | | | | | | |
|---|---|---|---|---|---|---|
| 37202 | ENSG0000 | AC011477.3 | | 1.081686 | 1.430377 | 1.452666 |
| 3429 | ENSG0000 | VSIR | V-set immu | 0.691633 | 1.383365 | 1.452403 |
| 2877 | ENSG0000 | SFRP1 | secreted fr | 0.314941 | 0.651827 | 1.452187 |
| 6768 | ENSG0000 | TRIM45 | tripartite n | 1.318055 | 1.346599 | 1.451872 |
| 20175 | ENSG0000 | FAM187A | family with | 0.529 | 0.837113 | 1.450385 |
| 27946 | ENSG0000 | AC139795.1 | | 1.114919 | 1.39885 | 1.449726 |
| 10212 | ENSG0000 | ISL2 | ISL LIM hor | 0.576365 | 0.862154 | 1.448667 |
| 2858 | ENSG0000 | SGK3 | serum/gluc | 0.873358 | 1.237724 | 1.448185 |
| 12287 | ENSG0000 | CPT1C | carnitine p | 0.548308 | 0.581089 | 1.447935 |
| 2970 | ENSG0000 | DKKL1 | dickkopf lik | 0.609398 | 0.728206 | 1.44715 |
| 10769 | ENSG0000 | NPPC | natriuretic | 0.049826 | 0.353647 | 1.445258 |
| 3653 | ENSG0000 | FBXW7 | F-box and ' | 1.157686 | 1.318038 | 1.443334 |
| 22363 | ENSG0000 | ABHD11-A: | ABHD11 ar | 0.275197 | 0.815151 | 1.439517 |
| 31494 | ENSG0000 | TRNP1 | TMF1-regu | 0.927633 | 1.10995 | 1.438781 |
| 2596 | ENSG0000 | XIAP | X-linked in | 1.091045 | 1.262653 | 1.438385 |
| 28730 | ENSG0000 | AP001432.1 | | 0.400448 | 0.719377 | 1.43745 |
| 27353 | ENSG0000 | BX088651.4 | | 0.720897 | 1.291811 | 1.436019 |
| 13052 | ENSG0000 | EFEMP2 | EGF contai | 0.133853 | 0.649879 | 1.435754 |
| 20102 | ENSG0000 | LINC01356 | long interg | 0.446256 | 0.994977 | 1.435072 |
| 37903 | ENSG0000 | AC010326.3 | | 0.933174 | 1.112843 | 1.433458 |
| 12920 | ENSG0000 | CYP4F22 | cytochrom | 0.127001 | 0.259287 | 1.430052 |
| 38736 | ENSG0000 | AL162258.2 | | 0.576811 | 0.824599 | 1.429579 |
| 16168 | ENSG0000 | NHLRC3 | NHL repeat | 0.800512 | 1.16003 | 1.42946 |
| 15441 | ENSG0000 | C14orf80 | chromosor | 1.39303 | 1.406829 | 1.428692 |
| 1656 | ENSG0000 | NKAIN1 | sodium/po | 0.594867 | 1.029685 | 1.428551 |
| 10813 | ENSG0000 | TRIM46 | tripartite n | 0.669409 | 0.835418 | 1.426534 |
| 14276 | ENSG0000 | LINC00174 | long interg | 0.588194 | 1.236491 | 1.425914 |
| 6906 | ENSG0000 | CCNJL | cyclin J like | 0.673983 | 0.793167 | 1.425025 |
| 154 | ENSG0000 | ARHGAP44 | Rho GTPas | 0.987896 | 1.420181 | 1.424612 |
| 42247 | ENSG0000 | AL353763.2 | | 0.994608 | 1.321222 | 1.424193 |
| 21986 | ENSG0000 | TAF1A-AS1 | TAF1A anti | 0.261265 | 0.902843 | 1.421504 |
| 2953 | ENSG0000 | LIN7B | lin-7 homo | 0.992854 | 1.159632 | 1.421241 |
| 10101 | ENSG0000 | ZSCAN12 | zinc finger | 0.554783 | 1.10859 | 1.42049 |
| 41415 | ENSG0000 | HOXA11-A. | HOXA11 ar | 0.458174 | 0.89938 | 1.419891 |
| 34767 | ENSG0000 | DOCK9-AS: | DOCK9 ant | 0.227075 | 0.943148 | 1.419575 |
| 40541 | ENSG0000 | AC006064.5 | | 1.058296 | 1.1727 | 1.417273 |
| 36149 | ENSG0000 | NOTCH2NI | notch 2 N- | 1.033191 | 1.114312 | 1.415295 |
| 4685 | ENSG0000 | SLC35A3 | solute carr | 0.78501 | 1.01641 | 1.41478 |
| 36698 | ENSG0000 | LINC01910 | long interg | 0.540232 | 0.849279 | 1.412329 |
| 35614 | ENSG0000 | MMP25-AS | MMP25 ar | 0.587065 | 1.251216 | 1.411521 |
| 15510 | ENSG0000 | SNAI3 | snail family | 0.220883 | 0.561296 | 1.410406 |
| 32178 | ENSG0000 | SYS1-DBNI | SYS1-DBNI | 1.103499 | 1.213216 | 1.409771 |
| 3996 | ENSG0000 | B3GAT2 | beta-1,3-gl | 0.025876 | 0.419515 | 1.408875 |
| 15518 | ENSG0000 | SMG1P4 | SMG1P4, n | 0.624663 | 1.345764 | 1.406877 |
| 17464 | ENSG0000 | Y_RNA | Y RNA [Sou | 0.630563 | 1.251487 | 1.406874 |
| 29508 | ENSG0000 | AC006064.2 | | 1.051905 | 1.403042 | 1.406488 |
| 41946 | ENSG0000 | AP001273.1 | | 1.199757 | 1.349711 | 1.405324 |

| | | | | | | |
|---|---|---|---|---|---|---|
| 10899 | ENSG0000 | AQP3 | aquaporin | 0.140363 | 0.34623 | 1.404917 |
| 8724 | ENSG0000 | TBC1D32 | TBC1 doma | 0.678006 | 1.321641 | 1.404595 |
| 6458 | ENSG0000 | CCT6B | chaperonir | 1.102931 | 1.234532 | 1.404432 |
| 32682 | ENSG0000 | AC010186.2 | | 0.571669 | 1.14414 | 1.401675 |
| 8162 | ENSG0000 | RCN3 | reticulocal | 0.555179 | 0.807747 | 1.401253 |
| 14762 | ENSG0000 | CAPN12 | calpain 12 | 0.100348 | 0.516276 | 1.401025 |
| 16125 | ENSG0000 | ZNF177 | zinc finger | 0.369202 | 0.460164 | 1.399936 |
| 26611 | ENSG0000 | AC005540.1 | | 1.174654 | 1.334583 | 1.39916 |
| 38791 | ENSG0000 | AC233992.3 | | 1.099769 | 1.192242 | 1.399012 |
| 23679 | ENSG0000 | AC044860.1 | | 1.038817 | 1.054014 | 1.398856 |
| 37376 | ENSG0000 | AC027097.2 | | 1.176875 | 1.362088 | 1.398288 |
| 11088 | ENSG0000 | CEP44 | centrosom | 1.167435 | 1.293617 | 1.397346 |
| 1544 | ENSG0000 | HOMER2 | homer scaf | 0.989644 | 1.100116 | 1.39665 |
| 4906 | ENSG0000 | CD70 | CD70 mole | 0.243502 | 0.652301 | 1.395398 |
| 6815 | ENSG0000 | TMEM241 | transmeml | 1.02292 | 1.25982 | 1.393901 |
| 15734 | ENSG0000 | BCDIN3D | BCDIN3 do | 0.702236 | 0.993059 | 1.393302 |
| 39214 | ENSG0000 | AC007032.1 | | 0.406325 | 1.097986 | 1.392262 |
| 15865 | ENSG0000 | KCNJ11 | potassium | 0.452251 | 0.572666 | 1.390019 |
| 12203 | ENSG0000 | ADAL | adenosine | 0.827498 | 0.95122 | 1.387303 |
| 14489 | ENSG0000 | CHST13 | carbohydra | 0.424334 | 0.797875 | 1.386401 |
| 3124 | ENSG0000 | CCDC114 | coiled-coil | 0.265719 | 0.535566 | 1.386111 |
| 7325 | ENSG0000 | TMEM8B | transmeml | 0.895062 | 1.266555 | 1.38562 |
| 14693 | ENSG0000 | FAM89A | family with | 0.763731 | 1.28531 | 1.384762 |
| 517 | ENSG0000 | ARHGAP31 | Rho GTPas | 0.561622 | 1.179493 | 1.383898 |
| 40262 | ENSG0000 | AL117332.1 | | 0.691121 | 1.174102 | 1.382843 |
| 9885 | ENSG0000 | TMEM171 | transmeml | 0.379297 | 0.739502 | 1.381282 |
| 35634 | ENSG0000 | AC040977.1 | | 0.576545 | 0.756203 | 1.380495 |
| 9073 | ENSG0000 | SLX4IP | SLX4 intera | 0.803971 | 0.930995 | 1.380067 |
| 20982 | ENSG0000 | APOL6 | apolipopro | 0.574427 | 0.61906 | 1.379335 |
| 37386 | ENSG0000 | AC087289.5 | | 0.819803 | 1.094205 | 1.378065 |
| 12194 | ENSG0000 | TET2 | tet methyl | 0.968842 | 1.336159 | 1.377398 |
| 16326 | ENSG0000 | ZNF250 | zinc finger | 0.714581 | 1.092294 | 1.376865 |
| 14568 | ENSG0000 | TMEM136 | transmeml | 0.524697 | 1.242497 | 1.376758 |
| 10428 | ENSG0000 | LRRC56 | leucine rich | 0.730725 | 0.997905 | 1.375514 |
| 18074 | ENSG0000 | CHIC1 | cysteine ric | 0.691767 | 1.319659 | 1.375008 |
| 16587 | ENSG0000 | ZNF429 | zinc finger | 0.616342 | 1.243811 | 1.37427 |
| 20430 | ENSG0000 | CES5AP1 | carboxyles | 0.328599 | 0.884209 | 1.374002 |
| 6039 | ENSG0000 | ISLR | immunoglo | 0.027056 | 0.271834 | 1.373774 |
| 40849 | ENSG0000 | GTSE1-AS1 | GTSE1 anti | 0.500132 | 0.854551 | 1.373747 |
| 3562 | ENSG0000 | COL1A1 | collagen ty | 0.028284 | 0.212732 | 1.372765 |
| 2964 | ENSG0000 | SLC17A7 | solute carr | 0.083664 | 0.764331 | 1.372546 |
| 2705 | ENSG0000 | GDPD3 | glycerophc | 0.592945 | 0.897925 | 1.371308 |
| 9102 | ENSG0000 | FEZ1 | fasciculatic | 0.246524 | 0.499987 | 1.369988 |
| 7082 | ENSG0000 | ANKRD36 | ankyrin rep | 0.661746 | 1.117387 | 1.369111 |
| 26655 | ENSG0000 | ZNF815P | zinc finger | 0.500842 | 0.966142 | 1.368946 |
| 5537 | ENSG0000 | CXCL6 | C-X-C moti | 0.443301 | 0.519705 | 1.368273 |
| 4251 | ENSG0000 | CYB561D2 | cytochrom | 1.142256 | 1.268903 | 1.368064 |

| | | | | | | |
|---|---|---|---|---|---|---|
| 15662 | ENSG0000 | RGS9BP | regulator c | 0.122249 | 0.412791 | 1.367285 |
| 1783 | ENSG0000 | SULT2B1 | sulfotransf | 0.314668 | 0.412254 | 1.363221 |
| 31815 | ENSG0000 | PCDHGB7 | protocadh | 1.075627 | 1.260423 | 1.359246 |
| 10555 | ENSG0000 | RBP7 | retinol bin | 0.310918 | 0.752122 | 1.357691 |
| 16557 | ENSG0000 | ANKRD36B | ankyrin rep | 0.633113 | 1.234611 | 1.356995 |
| 12551 | ENSG0000 | SPATA24 | spermatog | 0.851675 | 1.101391 | 1.355432 |
| 33028 | ENSG0000 | AC022509.3 | | 0.607394 | 0.898766 | 1.354246 |
| 24796 | ENSG0000 | LINC01273 | long interg | 0.485414 | 0.656196 | 1.35409 |
| 4712 | ENSG0000 | COLEC11 | collectin su | 0.556419 | 0.819959 | 1.353799 |
| 1113 | ENSG0000 | TMEM260 | transmeml | 0.941097 | 1.110258 | 1.353592 |
| 9823 | ENSG0000 | FAM122C | family with | 0.375565 | 0.639275 | 1.352624 |
| 4862 | ENSG0000 | PGF | placental g | 0.091408 | 0.532141 | 1.350565 |
| 37781 | ENSG0000 | LINC00528 | long interg | 0.378682 | 0.559507 | 1.350372 |
| 13484 | ENSG0000 | TVP23C | trans-golgi | 0.698792 | 0.994753 | 1.350038 |
| 40093 | ENSG0000 | Six3os1_2 | Six3os1 co | 0.223297 | 0.457496 | 1.349324 |
| 38623 | ENSG0000 | AL118558.3 | | 0.656353 | 0.812212 | 1.347459 |
| 18395 | ENSG0000 | ZBTB10 | zinc finger | 0.819205 | 1.323839 | 1.346878 |
| 25722 | ENSG0000 | AC131011.1 | | 0.12881 | 0.714273 | 1.34596 |
| 29247 | ENSG0000 | AC147067.1 | | 0.123896 | 0.458301 | 1.345704 |
| 36152 | ENSG0000 | MIR4258 | microRNA | 0.251717 | 0.46221 | 1.345619 |
| 11051 | ENSG0000 | PLXNB1 | plexin B1 [ | 0.706923 | 1.044334 | 1.345502 |
| 11881 | ENSG0000 | ANKS3 | ankyrin rep | 0.92381 | 1.178 | 1.345473 |
| 24164 | ENSG0000 | BANF1P2 | barrier to a | 0.740849 | 1.009042 | 1.342759 |
| 136 | ENSG0000 | KDM7A | lysine dem | 0.443628 | 1.195176 | 1.338574 |
| 29417 | ENSG0000 | AC016065.1 | | 0.360163 | 0.79483 | 1.337603 |
| 40829 | ENSG0000 | AC006449.5 | | 0.588678 | 0.800286 | 1.337031 |
| 11398 | ENSG0000 | PGM2L1 | phosphogl | 0.384343 | 1.006955 | 1.333729 |
| 8141 | ENSG0000 | ABCA5 | ATP bindin | 0.442836 | 1.276107 | 1.333329 |
| 7047 | ENSG0000 | NPL | N-acetylne | 0.7979 | 1.225753 | 1.332659 |
| 41036 | ENSG0000 | AC006213.4 | | 0.214396 | 0.633506 | 1.331512 |
| 23518 | ENSG0000 | AC064807.1 | | 0.624025 | 1.196816 | 1.330712 |
| 13912 | ENSG0000 | C8orf31 | chromosor | 0.759004 | 1.073235 | 1.329614 |
| 8557 | ENSG0000 | NICN1 | nicolin 1 [S | 0.776083 | 0.947241 | 1.328858 |
| 7801 | ENSG0000 | PTGR2 | prostaglan | 0.813556 | 0.933361 | 1.328147 |
| 21669 | ENSG0000 | PRKAR2A-/ | PRKAR2A a | 0.176117 | 0.686038 | 1.327507 |
| 42718 | ENSG0000 | GTF2IP12 | general tra | 0.54997 | 1.192861 | 1.326929 |
| 39852 | ENSG0000 | AC242376.2 | | 0.782275 | 1.271619 | 1.325594 |
| 18258 | ENSG0000 | LINC02210 | long interg | 0.259893 | 0.635677 | 1.324657 |
| 5391 | ENSG0000 | INHA | inhibin alpl | 0.220999 | 0.885801 | 1.324256 |
| 19782 | ENSG0000 | AC091982.1 | | 0.031305 | 0.642737 | 1.322992 |
| 39464 | ENSG0000 | AC064836.3 | | 0.219364 | 0.805121 | 1.322966 |
| 41640 | ENSG0000 | FP236315.1 | | 0.487736 | 0.951888 | 1.322852 |
| 43390 | ENSG0000 | AP000553.6 | | 0.008533 | 0.285684 | 1.321977 |
| 16005 | ENSG0000 | TMEM221 | transmeml | 0.160677 | 0.448759 | 1.319046 |
| 34705 | ENSG0000 | AC068338.2 | | 0.691536 | 1.136432 | 1.318209 |
| 21920 | ENSG0000 | ZBTB45P1 | zinc finger | 0.8328 | 0.985151 | 1.317937 |
| 14735 | ENSG0000 | C1S | compleme | 0.510554 | 0.573757 | 1.317206 |

| | | | | | | |
|---|---|---|---|---|---|---|
| 10245 | ENSG0000 | ZNF221 | zinc finger | 0.241205 | 0.828228 | 1.317206 |
| 12045 | ENSG0000 | ATG16L2 | autophagy | 0.749158 | 1.254886 | 1.316768 |
| 32616 | ENSG0000 | AC073585.1 | | 0.888074 | 1.061024 | 1.314631 |
| 7084 | ENSG0000 | ARHGEF4 | Rho guanir | 0.937721 | 1.029632 | 1.314045 |
| 10830 | ENSG0000 | EOMES | eomesode | 0.210303 | 1.202492 | 1.313131 |
| 37438 | ENSG0000 | AC010616.1 | | 0.764417 | 0.918 | 1.312712 |
| 30784 | ENSG0000 | WWC2-AS2 | WWC2 ant | 0.407671 | 0.950251 | 1.312242 |
| 38903 | ENSG0000 | AC116351.2 | | 0.175567 | 0.426153 | 1.308285 |
| 5178 | ENSG0000 | CXCR4 | C-X-C moti | 0.534559 | 0.68117 | 1.305437 |
| 18057 | ENSG0000 | LRRC73 | leucine ricl | 0.537317 | 1.028674 | 1.30481 |
| 15018 | ENSG0000 | WWOX | WW doma | 0.923363 | 1.085565 | 1.303475 |
| 1262 | ENSG0000 | PPP2R2C | protein ph | 0.632001 | 0.695043 | 1.302713 |
| 3221 | ENSG0000 | DFNA5 | DFNA5, de | 0.613285 | 0.794963 | 1.300129 |
| 15308 | ENSG0000 | OSBP2 | oxysterol b | 0.754136 | 0.789544 | 1.299333 |
| 8381 | ENSG0000 | WNT9A | Wnt family | 0.522118 | 1.284094 | 1.29912 |
| 12057 | ENSG0000 | BATF2 | basic leuci | 0.148079 | 0.816703 | 1.298402 |
| 5340 | ENSG0000 | EEF1AKMT | EEF1A lysir | 0.537818 | 0.891331 | 1.295791 |
| 36720 | ENSG0000 | AC004477.2 | | 0.714837 | 1.265012 | 1.295676 |
| 23888 | ENSG0000 | AL365203.1 | | 0.138661 | 0.750856 | 1.295326 |
| 6619 | ENSG0000 | TMCC2 | transmeml | 0.696728 | 0.834201 | 1.292152 |
| 35504 | ENSG0000 | LINC00662 | long interg | 0.722605 | 1.180345 | 1.29079 |
| 16481 | ENSG0000 | ZNF846 | zinc finger | 0.349555 | 0.736511 | 1.28868 |
| 12694 | ENSG0000 | KCNK3 | potassium | 0.267432 | 0.500292 | 1.288628 |
| 38082 | ENSG0000 | AC097359.2 | | 0.67075 | 1.074298 | 1.286504 |
| 35562 | ENSG0000 | SUZ12P1 | SUZ12 poly | 1.087444 | 1.104635 | 1.284731 |
| 35248 | ENSG0000 | AC005837.1 | | 0.577981 | 0.587381 | 1.284197 |
| 21847 | ENSG0000 | AL161908.1 | | 0.732431 | 0.870928 | 1.279493 |
| 301 | ENSG0000 | LRRC23 | leucine ricl | 0.633897 | 0.682712 | 1.278577 |
| 11486 | ENSG0000 | BTNL9 | butyrophili | 0.693097 | 1.028836 | 1.278118 |
| 37045 | ENSG0000 | LINC01764 | long interg | 0.406993 | 1.102901 | 1.277604 |
| 9088 | ENSG0000 | ROM1 | retinal out | 0.881128 | 0.944851 | 1.277023 |
| 19037 | ENSG0000 | BX005266.1 | | 1.251239 | 1.260159 | 1.273607 |
| 12735 | ENSG0000 | KCNG3 | potassium | 0.564929 | 0.681088 | 1.271752 |
| 23856 | ENSG0000 | ACVR2B-AS | ACVR2B ar | 0.340123 | 1.153906 | 1.270278 |
| 7470 | ENSG0000 | GCHFR | GTP cycloh | 0.202893 | 0.336621 | 1.26989 |
| 8218 | ENSG0000 | ZNF697 | zinc finger | 0.553228 | 1.135108 | 1.26726 |
| 35091 | ENSG0000 | AL158212.3 | | 1.118497 | 1.119475 | 1.266171 |
| 185 | ENSG0000 | CACNA2D2 | calcium vo | 0.306263 | 0.644993 | 1.264615 |
| 37070 | ENSG0000 | AC003070.1 | | 0.740784 | 1.241656 | 1.264589 |
| 15349 | ENSG0000 | TMEM121 | transmeml | 0.493278 | 0.669676 | 1.264112 |
| 7419 | ENSG0000 | C11orf70 | chromosor | 0.407396 | 0.64378 | 1.263478 |
| 20775 | ENSG0000 | AP000580.1 | | 0.698968 | 1.159177 | 1.260234 |
| 21809 | ENSG0000 | AC099850.1 | | 0.403343 | 0.820927 | 1.259825 |
| 1898 | ENSG0000 | FLT3LG | fms relatec | 0.426635 | 0.580355 | 1.258963 |
| 40075 | ENSG0000 | AC025287.2 | | 0.958069 | 1.081529 | 1.258926 |
| 2902 | ENSG0000 | SNX16 | sorting nex | 0.765712 | 0.892359 | 1.258183 |
| 29897 | ENSG0000 | AC145098.1 | | 0.193942 | 0.761781 | 1.257746 |

| | | | | | | |
|---|---|---|---|---|---|---|
| 5228 | ENSG0000 | BBS9 | Bardet-Bie | 0.614359 | 1.005584 | 1.25692 |
| 35502 | ENSG0000 | AC012184.3 | | 0.220169 | 0.887665 | 1.2554 |
| 14237 | ENSG0000 | GGN | gametoger | 0.394028 | 0.744279 | 1.253857 |
| 36825 | ENSG0000 | AC061992.1 | | 0.218842 | 0.329518 | 1.253671 |
| 14906 | ENSG0000 | GJD3 | gap junctic | 0.288594 | 0.698208 | 1.253136 |
| 3361 | ENSG0000 | AKNA | AT-hook tr | 0.61041 | 0.818574 | 1.250344 |
| 43387 | ENSG0000 | AP000553.5 | | 0.125858 | 0.164182 | 1.246551 |
| 9390 | ENSG0000 | ATG10 | autophagy | 0.872551 | 0.943644 | 1.244034 |
| 28575 | ENSG0000 | ELOCP19 | elongin C p | 0.880751 | 0.925042 | 1.244025 |
| 3859 | ENSG0000 | MDM1 | Mdm1 nuc | 0.955741 | 1.226623 | 1.241502 |
| 3895 | ENSG0000 | HCFC2 | host cell fa | 0.976781 | 1.062682 | 1.241277 |
| 7262 | ENSG0000 | TMOD1 | tropomodu | 0.086247 | 0.741633 | 1.241095 |
| 14948 | ENSG0000 | CCDC125 | coiled-coil | 0.734951 | 0.796552 | 1.240034 |
| 14994 | ENSG0000 | SERHL2 | serine hydı | 0.37545 | 0.849618 | 1.239804 |
| 3411 | ENSG0000 | RASSF4 | Ras associa | 0.873466 | 1.049373 | 1.238786 |
| 30702 | ENSG0000 | AC112184.1 | | 0.15426 | 0.910307 | 1.236692 |
| 18467 | ENSG0000 | CPT1B | carnitine p | 0.816946 | 1.028907 | 1.235901 |
| 18696 | ENSG0000 | AC022007.1 | | 0.339992 | 0.624783 | 1.234734 |
| 8536 | ENSG0000 | ADPRH | ADP-ribosy | 0.926841 | 1.108094 | 1.234515 |
| 35435 | ENSG0000 | Z92544.2 | | 0.738733 | 0.865044 | 1.232737 |
| 7340 | ENSG0000 | GMPR | guanosine | 0.27045 | 0.403888 | 1.231792 |
| 4259 | ENSG0000 | IQCG | IQ motif cc | 0.787329 | 1.040868 | 1.230203 |
| 10246 | ENSG0000 | ZNF233 | zinc finger | 0.425663 | 0.974611 | 1.229443 |
| 16438 | ENSG0000 | ZNF799 | zinc finger | 0.866903 | 0.900672 | 1.228154 |
| 12365 | ENSG0000 | CLIC3 | chloride in | 0.112195 | 0.260271 | 1.227565 |
| 18438 | ENSG0000 | SAMD9 | sterile alph | 0.353027 | 0.900644 | 1.226661 |
| 27270 | ENSG0000 | AC242426.3 | | 0.624892 | 1.091547 | 1.225842 |
| 14638 | ENSG0000 | LINC00471 | long interg | 0.178578 | 0.451687 | 1.224323 |
| 34678 | ENSG0000 | SSSCA1-AS | SSSCA1 ant | 0.620465 | 0.983558 | 1.223465 |
| 5985 | ENSG0000 | CCDC32 | coiled-coil | 0.694992 | 0.936938 | 1.222972 |
| 15248 | ENSG0000 | CCR10 | C-C motif c | 0.255513 | 0.265028 | 1.222252 |
| 6994 | ENSG0000 | AFG1L | AFG1 like A | 1.157671 | 1.157783 | 1.221429 |
| 14288 | ENSG0000 | C14orf28 | chromosor | 0.430668 | 0.955416 | 1.220632 |
| 42120 | ENSG0000 | AC011815.3 | | 0.553458 | 0.812741 | 1.220584 |
| 31616 | ENSG0000 | AC107959.3 | | 0.51294 | 0.766922 | 1.219107 |
| 37590 | ENSG0000 | AC010522.1 | | 0.647106 | 1.091987 | 1.218691 |
| 15843 | ENSG0000 | DYNC2H1 | dynein cyto | 0.463735 | 0.993574 | 1.216711 |
| 10691 | ENSG0000 | TYW5 | tRNA-yW s | 0.783638 | 0.969439 | 1.215428 |
| 2134 | ENSG0000 | MADCAM1 | mucosal va | 0.506339 | 0.983284 | 1.214453 |
| 36482 | ENSG0000 | RARA-AS1 | RARA antis | 0.8081 | 1.207804 | 1.21246 |
| 13863 | ENSG0000 | ZDHHC13 | zinc finger | 0.643203 | 1.16451 | 1.211493 |
| 42051 | ENSG0000 | AC006511.3 | | 0.651675 | 1.190259 | 1.209527 |
| 456 | ENSG0000 | GRAMD1B | GRAM don | 0.672433 | 0.786278 | 1.207862 |
| 11783 | ENSG0000 | PIP5KL1 | phosphatic | 0.225709 | 0.679647 | 1.207801 |
| 10090 | ENSG0000 | GDPD5 | glycerophc | 0.862347 | 1.128081 | 1.206366 |
| 15222 | ENSG0000 | GDF3 | growth dif | 0 | 0.586346 | 1.205948 |
| 16498 | ENSG0000 | LINC00173 | long interg | 0.110133 | 0.388702 | 1.205655 |

| | | | | | | |
|---|---|---|---|---|---|---|
| 12533 | ENSG0000 | GPRC5C | G protein-c | 0.383482 | 0.573072 | 1.204506 |
| 18058 | ENSG0000 | LINC00963 | long interg | 0.626005 | 0.717755 | 1.202957 |
| 10445 | ENSG0000 | NAGS | N-acetylglu | 0.447463 | 0.466834 | 1.20044 |
| 9654 | ENSG0000 | CCDC144B | coiled-coil | 0.392903 | 1.159201 | 1.199688 |
| 24107 | ENSG0000 | AL080317.1 | | 0.771871 | 0.970573 | 1.199431 |
| 5686 | ENSG0000 | CHURC1-FI | CHURC1-FI | 0.20013 | 1.083504 | 1.197075 |
| 23952 | ENSG0000 | AC139149.1 | | 0.132471 | 1.048814 | 1.196951 |
| 32770 | ENSG0000 | AP000786.1 | | 0.412406 | 0.694863 | 1.195691 |
| 14009 | ENSG0000 | ZNF620 | zinc finger | 0.43102 | 0.816579 | 1.195578 |
| 2558 | ENSG0000 | MYOM1 | myomesin | 0.189647 | 1.093838 | 1.195512 |
| 9720 | ENSG0000 | PIK3AP1 | phosphoin | 0.390077 | 0.529241 | 1.192297 |
| 23559 | ENSG0000 | UBAC2-AS1 | UBAC2 ant | 0.492632 | 1.116479 | 1.190707 |
| 22771 | ENSG0000 | AL451069.1 | | 0.154445 | 0.176751 | 1.190552 |
| 40741 | ENSG0000 | AC009041.4 | | 0.600885 | 1.144504 | 1.190109 |
| 14431 | ENSG0000 | CCDC66 | coiled-coil | 0.845793 | 1.148628 | 1.189995 |
| 35957 | ENSG0000 | AC007639.1 | | 0.450552 | 1.17744 | 1.189379 |
| 30681 | ENSG0000 | AC136604.3 | | 0.208357 | 0.390897 | 1.188669 |
| 42488 | ENSG0000 | AC090114.3 | | 0.80111 | 1.150489 | 1.188523 |
| 21241 | ENSG0000 | LINC00473 | long interg | 0.692675 | 1.026385 | 1.187688 |
| 8839 | ENSG0000 | FBXO25 | F-box prote | 0.866527 | 1.05867 | 1.187135 |
| 37621 | ENSG0000 | ZNF528-AS | ZNF528 an | 0.167083 | 0.93307 | 1.186041 |
| 40821 | ENSG0000 | AL589743.5 | | 0.616646 | 1.000768 | 1.185621 |
| 14868 | ENSG0000 | C12orf60 | chromosor | 0.599122 | 0.628198 | 1.184355 |
| 28632 | ENSG0000 | AL359075.1 | | 0.533724 | 0.724432 | 1.184106 |
| 29836 | ENSG0000 | AL589765.4 | | 0.7912 | 1.157312 | 1.183794 |
| 8085 | ENSG0000 | LMTK3 | lemur tyro | 0.043442 | 0.255882 | 1.182394 |
| 26753 | ENSG0000 | COX10-AS1 | COX10 ant | 0.915166 | 0.969904 | 1.181732 |
| 23458 | ENSG0000 | PROB1 | proline rich | 0.359073 | 0.50835 | 1.181487 |
| 41020 | ENSG0000 | Vax2os1_3 | Vax2os1 cc | 0 | 0.891188 | 1.180769 |
| 40719 | ENSG0000 | AC016590.3 | | 0.326931 | 0.514497 | 1.179819 |
| 6454 | ENSG0000 | MATN3 | matrilin 3 [ | 0.748008 | 0.968239 | 1.179587 |
| 34064 | ENSG0000 | ZNF710-AS | ZNF710 an | 0.500398 | 0.518119 | 1.179336 |
| 13136 | ENSG0000 | HSPA6 | heat shock | 0.398224 | 0.772791 | 1.17893 |
| 37406 | ENSG0000 | AC022144.1 | | 0.021826 | 0.527983 | 1.178289 |
| 3048 | ENSG0000 | ZNF419 | zinc finger | 0.399319 | 0.91757 | 1.178191 |
| 29067 | ENSG0000 | APOBEC3D | apolipopro | 0.656337 | 0.78758 | 1.178143 |
| 10569 | ENSG0000 | SYNC | syncoilin, ir | 0.493166 | 0.740063 | 1.177346 |
| 42456 | ENSG0000 | BTG3-AS1 | BTG3 antis | 0.443272 | 1.017024 | 1.176343 |
| 38868 | ENSG0000 | AC092687.3 | | 0.241604 | 1.137017 | 1.176012 |
| 14941 | ENSG0000 | TMEM121I | transmeml | 0.451566 | 1.124214 | 1.175918 |
| 15001 | ENSG0000 | APOBR | apolipopro | 0.412229 | 1.100622 | 1.175427 |
| 27686 | ENSG0000 | AL034417.2 | | 0.255113 | 0.82912 | 1.175168 |
| 12965 | ENSG0000 | ISG20 | interferon | 0.443637 | 0.478626 | 1.172317 |
| 11760 | ENSG0000 | MAPRE2 | microtubul | 0.212999 | 0.422571 | 1.172254 |
| 16393 | ENSG0000 | NTNG2 | netrin G2 [ | 0.547439 | 1.002427 | 1.169948 |
| 29525 | ENSG0000 | DPH6-AS1 | DPH6 antis | 0.317072 | 0.617603 | 1.168281 |
| 10347 | ENSG0000 | S100A1 | S100 calciu | 0.652004 | 0.758502 | 1.167702 |

| | | | | | | |
|---|---|---|---|---|---|---|
| 35627 | ENSG0000 | AC005696.1 | | 0.285992 | 0.607192 | 1.167219 |
| 15433 | ENSG0000 | C12orf56 | chromosor | 0.515042 | 0.703849 | 1.161562 |
| 38008 | ENSG0000 | AC009061.2 | | 0.405064 | 0.747513 | 1.161142 |
| 424 | ENSG0000 | PRDM11 | PR/SET dor | 0.827125 | 0.908615 | 1.15836 |
| 35158 | ENSG0000 | AL031123.2 | | 0.30119 | 0.828883 | 1.157854 |
| 38897 | ENSG0000 | AC067838.1 | | 0.889298 | 1.091121 | 1.15706 |
| 16550 | ENSG0000 | CBWD3 | COBW don | 0.962458 | 1.041063 | 1.153587 |
| 6011 | ENSG0000 | STARD13 | StAR relate | 0.719454 | 1.107361 | 1.152115 |
| 15058 | ENSG0000 | FAM167B | family with | 0.199948 | 0.535831 | 1.151644 |
| 28927 | ENSG0000 | RN7SL738I | RNA, 7SL, c | 0.508519 | 1.11005 | 1.151406 |
| 29279 | ENSG0000 | RPL30P11 | ribosomal | 0.355844 | 0.537833 | 1.150591 |
| 43039 | ENSG0000 | AL137002.2 | | 0.410939 | 0.78456 | 1.150223 |
| 11092 | ENSG0000 | TMEM144 | transmeml | 0.497631 | 0.643087 | 1.148861 |
| 11430 | ENSG0000 | CDX2 | caudal typ | 0.527433 | 0.747764 | 1.148064 |
| 3957 | ENSG0000 | MDGA1 | MAM dom | 0.540386 | 0.986419 | 1.146944 |
| 13328 | ENSG0000 | FAM53A | family with | 0.465816 | 0.87021 | 1.145592 |
| 12427 | ENSG0000 | WNT10B | Wnt family | 0.351041 | 0.660543 | 1.144507 |
| 2712 | ENSG0000 | TSNAXIP1 | translin as | 0.516387 | 0.967789 | 1.142657 |
| 39341 | ENSG0000 | AC010864.1 | | 0.418425 | 0.888384 | 1.138256 |
| 25733 | ENSG0000 | AP000695.2 | | 0.10152 | 0.171345 | 1.137645 |
| 27639 | ENSG0000 | PAXBP1-AS | PAXBP1 an | 0.298404 | 1.119822 | 1.136792 |
| 27526 | ENSG0000 | AC079922.2 | | 0.83501 | 0.970076 | 1.136376 |
| 8527 | ENSG0000 | NFKBIZ | NFKB inhib | 0.595956 | 0.842313 | 1.135746 |
| 29088 | ENSG0000 | AC108751.4 | | 0.262781 | 0.439553 | 1.135617 |
| 23798 | ENSG0000 | AC018638.1 | | 0.728072 | 0.827852 | 1.132687 |
| 10602 | ENSG0000 | FAM102B | family with | 0.290504 | 0.743556 | 1.131883 |
| 16508 | ENSG0000 | ZNF431 | zinc finger | 0.840381 | 1.100801 | 1.131696 |
| 33178 | ENSG0000 | AC012555.1 | | 0.194107 | 0.276962 | 1.130292 |
| 29980 | ENSG0000 | AP000311.1 | | 0 | 0.048444 | 1.129181 |
| 37636 | ENSG0000 | PDLIM1P1 | PDZ and LI | 0.129365 | 1.033995 | 1.128546 |
| 34995 | ENSG0000 | AC009962.1 | | 0.469997 | 0.944068 | 1.128424 |
| 36103 | ENSG0000 | MIR4701 | microRNA | 0.154826 | 0.742092 | 1.12773 |
| 8713 | ENSG0000 | TPBG | trophoblas | 0.156886 | 0.453875 | 1.12707 |
| 39192 | ENSG0000 | AC090114.2 | | 0.669109 | 0.700621 | 1.127041 |
| 34139 | ENSG0000 | TGIF2-C20 | TGIF2-C20 | 0.436749 | 0.62203 | 1.126124 |
| 18365 | ENSG0000 | AC118344.1 | | 0.41147 | 0.712188 | 1.126081 |
| 40789 | ENSG0000 | CU634019.1 | | 0.828433 | 0.940655 | 1.12478 |
| 34400 | ENSG0000 | AC083843.2 | | 0.39435 | 0.888487 | 1.124431 |
| 15660 | ENSG0000 | PRELID2 | PRELI dom | 0.510502 | 0.55558 | 1.123913 |
| 27983 | ENSG0000 | AC073610.1 | | 0.023586 | 0.315618 | 1.122715 |
| 26929 | ENSG0000 | AC093582.1 | | 0.385094 | 0.621491 | 1.122443 |
| 1583 | ENSG0000 | ATP8B1 | ATPase ph | 0.496612 | 0.899906 | 1.122254 |
| 14033 | ENSG0000 | LRRC75B | leucine ricl | 0.692238 | 0.758542 | 1.12176 |
| 4530 | ENSG0000 | RHOU | ras homolo | 0.386543 | 0.695581 | 1.121216 |
| 34342 | ENSG0000 | AL354993.2 | | 0.427504 | 1.006754 | 1.120511 |
| 33280 | ENSG0000 | ZNF625 | zinc finger | 0.251537 | 1.10288 | 1.118944 |
| 7840 | ENSG0000 | SCAPER | S-phase cy | 0.54731 | 0.727351 | 1.118818 |

| | | | | | | |
|---|---|---|---|---|---|---|
| 39775 | ENSG0000 | AL137060.1 | | 0.199719 | 0.456668 | 1.118767 |
| 37296 | ENSG0000 | AC008878.2 | | 0.329649 | 0.495449 | 1.11825 |
| 21999 | ENSG0000 | AC023283.1 | | 0.523088 | 1.045075 | 1.116915 |
| 2867 | ENSG0000 | FZD3 | frizzled cla | 0.746841 | 0.904106 | 1.116201 |
| 11772 | ENSG0000 | AL357033.1 | | 0.107366 | 0.772782 | 1.115361 |
| 3923 | ENSG0000 | CEP85L | centrosom | 0.702337 | 0.824947 | 1.114843 |
| 22751 | ENSG0000 | AC010894.2 | | 0.337956 | 0.454548 | 1.114665 |
| 9402 | ENSG0000 | ZNF773 | zinc finger | 0.343371 | 0.982096 | 1.114064 |
| 24498 | ENSG0000 | AL109917.1 | | 0.411638 | 0.575063 | 1.113418 |
| 1804 | ENSG0000 | C3orf18 | chromosor | 0.230021 | 0.446556 | 1.113266 |
| 29479 | ENSG0000 | DNAJC3-AS | DNAJC3 an | 0.498541 | 0.821047 | 1.111422 |
| 12722 | ENSG0000 | ZNF613 | zinc finger | 0.341405 | 1.024493 | 1.111022 |
| 38952 | ENSG0000 | AL645608.7 | | 0.177381 | 0.604249 | 1.109754 |
| 15731 | ENSG0000 | PRR5 | proline ricl | 0.584793 | 1.041478 | 1.109718 |
| 15656 | ENSG0000 | ZNF555 | zinc finger | 0.332579 | 0.885075 | 1.106324 |
| 32899 | ENSG0000 | AC138466.1 | | 0.140064 | 0.228857 | 1.102868 |
| 10318 | ENSG0000 | ADAMTS13 | ADAM met | 0.412327 | 0.878652 | 1.102788 |
| 39045 | ENSG0000 | AL358472.2 | | 0.827932 | 1.090889 | 1.099851 |
| 34820 | ENSG0000 | AC007066.2 | | 0.361196 | 0.470579 | 1.099366 |
| 24565 | ENSG0000 | SPRY4-AS1 | SPRY4 anti | 0.463032 | 0.719081 | 1.09933 |
| 4306 | ENSG0000 | FOXP1 | forkhead b | 0.460401 | 0.916729 | 1.098949 |
| 12171 | ENSG0000 | AXIN2 | axin 2 [Sou | 0.532357 | 0.543756 | 1.097812 |
| 4667 | ENSG0000 | F3 | coagulatio | 0.439462 | 0.519207 | 1.097757 |
| 6816 | ENSG0000 | KCTD1 | potassium | 0.817279 | 0.858627 | 1.097014 |
| 31585 | ENSG0000 | HOXA-AS2 | HOXA clust | 0.671521 | 1.079006 | 1.096469 |
| 7867 | ENSG0000 | HAPLN3 | hyaluronar | 0.090858 | 0.670374 | 1.093893 |
| 39886 | ENSG0000 | AC080038.1 | | 0.461663 | 0.614941 | 1.093641 |
| 5279 | ENSG0000 | EGR2 | early grow | 0.077105 | 0.594603 | 1.092648 |
| 8240 | ENSG0000 | TBX19 | T-box 19 [S | 0.309922 | 1.008932 | 1.092193 |
| 42652 | ENSG0000 | AC104073.4 | | 0 | 0.383763 | 1.091648 |
| 16279 | ENSG0000 | KAZN | kazrin, per | 0.590816 | 1.062317 | 1.09035 |
| 972 | ENSG0000 | ZMYND12 | zinc finger | 0.345339 | 0.551793 | 1.088982 |
| 24322 | ENSG0000 | USP12-AS2 | USP12 anti | 0 | 0.363959 | 1.086117 |
| 22701 | ENSG0000 | AL606760.1 | | 0.453249 | 0.543345 | 1.083772 |
| 32600 | ENSG0000 | BRWD1-AS | BRWD1 an | 0.312075 | 0.927017 | 1.083107 |
| 9200 | ENSG0000 | PIH1D2 | PIH1 doma | 0.296653 | 0.953609 | 1.082043 |
| 479 | ENSG0000 | HSD17B6 | hydroxyste | 0.274215 | 0.532801 | 1.081854 |
| 5456 | ENSG0000 | IQSEC2 | IQ motif ar | 0.825135 | 0.848582 | 1.081652 |
| 12932 | ENSG0000 | ZNF554 | zinc finger | 0.554693 | 0.963306 | 1.079835 |
| 34104 | ENSG0000 | AC011921.1 | | 0.223483 | 0.790156 | 1.079155 |
| 773 | ENSG0000 | PHF21B | PHD finger | 0.73978 | 1.074322 | 1.078909 |
| 7040 | ENSG0000 | GLUL | glutamate- | 0.524048 | 0.544523 | 1.076489 |
| 42738 | ENSG0000 | AC010642.2 | | 0.877045 | 0.907627 | 1.07534 |
| 4247 | ENSG0000 | HYAL1 | hyalurono | 0.050574 | 0.240024 | 1.07431 |
| 33754 | ENSG0000 | AL121820.1 | | 0.22322 | 0.480569 | 1.074195 |
| 28350 | ENSG0000 | TPM3P9 | tropomyos | 0.403623 | 0.992346 | 1.074049 |
| 15809 | ENSG0000 | TMEM262 | transmeml | 0.965914 | 0.981562 | 1.073038 |

| | | | | | | |
|---|---|---|---|---|---|---|
| 15452 | ENSG0000 | METTL7A | methyltrar | 0.185536 | 0.19982 | 1.071133 |
| 18675 | ENSG0000 | HCG27 | HLA compl | 0.548438 | 0.791063 | 1.07076 |
| 37014 | ENSG0000 | AC011476.3 | | 0.455712 | 0.946135 | 1.069971 |
| 9110 | ENSG0000 | JPH2 | junctophili | 0.167213 | 0.332674 | 1.06977 |
| 37269 | ENSG0000 | AC135721.2 | | 0.723889 | 1.020129 | 1.068268 |
| 6631 | ENSG0000 | BEX2 | brain expre | 0.272051 | 0.379938 | 1.065524 |
| 19589 | ENSG0000 | LINC01089 | long interg | 0.664769 | 1.048839 | 1.065408 |
| 29845 | ENSG0000 | AC024560.2 | | 0.550711 | 0.953316 | 1.065277 |
| 33859 | ENSG0000 | AL162171.1 | | 0.292739 | 0.334816 | 1.064573 |
| 3089 | ENSG0000 | DENND3 | DENN dom | 0.480413 | 0.867599 | 1.063877 |
| 9661 | ENSG0000 | ACSS1 | acyl-CoA sy | 0.433503 | 0.826768 | 1.062774 |
| 14463 | ENSG0000 | SKIDA1 | SKI/DACH ( | 0.390948 | 1.04469 | 1.062617 |
| 1819 | ENSG0000 | EBF4 | early B-cell | 0.345611 | 0.521991 | 1.062321 |
| 39873 | ENSG0000 | AC090970.3 | | 0.346765 | 0.96377 | 1.061154 |
| 33475 | ENSG0000 | AC144548.1 | | 0.457791 | 0.628728 | 1.059533 |
| 34796 | ENSG0000 | SULT1A3 | sulfotransf | 0.837398 | 0.962384 | 1.059346 |
| 11714 | ENSG0000 | MESP1 | mesoderm | 0.390645 | 0.65262 | 1.057976 |
| 16030 | ENSG0000 | SMTNL2 | smoothelir | 0.01442 | 0.186517 | 1.056161 |
| 7449 | ENSG0000 | PAQR5 | progestin a | 0.183471 | 0.332274 | 1.054739 |
| 37524 | ENSG0000 | AC092279.1 | | 0.576887 | 0.708407 | 1.054418 |
| 42375 | ENSG0000 | AP001528.3 | | 0.513893 | 0.741056 | 1.054229 |
| 30952 | ENSG0000 | SCARNA8 | small Cajal | 0.315888 | 0.679195 | 1.053997 |
| 32135 | ENSG0000 | AC080023.2 | | 0.585121 | 0.862891 | 1.05371 |
| 2959 | ENSG0000 | PPP1R13L | protein ph | 0.58599 | 0.688718 | 1.053025 |
| 34932 | ENSG0000 | SNAI3-AS1 | SNAI3 antis | 0.691429 | 0.802402 | 1.052258 |
| 34036 | ENSG0000 | MIR4713H | MIR4713 h | 0.180897 | 1.048184 | 1.052257 |
| 10516 | ENSG0000 | SLC25A45 | solute carr | 0.235847 | 0.976703 | 1.051492 |
| 9621 | ENSG0000 | MAGED4 | MAGE fam | 0.177147 | 0.527337 | 1.050387 |
| 15395 | ENSG0000 | LINC00482 | long interg | 0.128386 | 0.239331 | 1.050332 |
| 19074 | ENSG0000 | MIR641 | microRNA | 0.124587 | 0.608133 | 1.050073 |
| 42662 | ENSG0000 | AL356095.3 | | 0.004161 | 0.23688 | 1.049375 |
| 13049 | ENSG0000 | SMPDL3A | sphingomy | 0.272667 | 0.672044 | 1.046959 |
| 14832 | ENSG0000 | PLCXD3 | phosphatic | 0.344766 | 0.534637 | 1.046845 |
| 13833 | ENSG0000 | C8G | compleme | 0.30423 | 0.826078 | 1.044916 |
| 37485 | ENSG0000 | AC010503.1 | | 0.326214 | 0.54288 | 1.044862 |
| 28114 | ENSG0000 | PSMB9 | proteasom | 0.209723 | 0.303617 | 1.043887 |
| 37895 | ENSG0000 | AC005498.3 | | 0.151892 | 0.86662 | 1.043322 |
| 14532 | ENSG0000 | ZFP82 | ZFP82 zinc | 0.564351 | 0.841813 | 1.043011 |
| 12791 | ENSG0000 | NAT1 | N-acetyltra | 0.521312 | 0.748043 | 1.041625 |
| 21510 | ENSG0000 | SNHG14 | small nucle | 0.656766 | 0.965722 | 1.040887 |
| 14577 | ENSG0000 | LRRC75A | leucine rich | 0.285716 | 0.699079 | 1.040871 |
| 8718 | ENSG0000 | FAXC | failed axon | 0.303751 | 0.857782 | 1.040399 |
| 39 | ENSG0000 | CASP10 | caspase 10 | 0.435186 | 0.46155 | 1.040389 |
| 2137 | ENSG0000 | ARVCF | armadillo r | 0.835797 | 0.836207 | 1.039766 |
| 985 | ENSG0000 | FGFR2 | fibroblast g | 0.421724 | 0.697194 | 1.039318 |
| 12772 | ENSG0000 | CHST11 | carbohydra | 0.237926 | 0.687019 | 1.039308 |
| 40355 | ENSG0000 | HIST1H3A | histone clu | 0.209458 | 0.911013 | 1.036344 |

| | | | | | | |
|---|---|---|---|---|---|---|
| 38608 | ENSG0000 | AP003392.5 | | 0.40396 | 0.687108 | 1.034936 |
| 11520 | ENSG0000 | CLMN | calmin [Sou | 0.582513 | 0.720391 | 1.033178 |
| 33348 | ENSG0000 | AC074091.1 | | 0.397412 | 0.756676 | 1.033089 |
| 4332 | ENSG0000 | TFCP2L1 | transcriptio | 0.377119 | 0.646574 | 1.031746 |
| 25460 | ENSG0000 | AC004987.2 | | 0.833812 | 0.837046 | 1.030527 |
| 5123 | ENSG0000 | ZNF549 | zinc finger | 0.432552 | 0.818858 | 1.030113 |
| 10783 | ENSG0000 | GPR155 | G protein-c | 0.465337 | 0.518014 | 1.029997 |
| 10637 | ENSG0000 | FCGR3B | Fc fragmer | 0.730503 | 0.969548 | 1.029686 |
| 788 | ENSG0000 | TMCC3 | transmeml | 0.189087 | 0.460972 | 1.029603 |
| 26291 | ENSG0000 | AL356968.2 | | 0.75465 | 0.928871 | 1.029579 |
| 4113 | ENSG0000 | NME5 | NME/NM2 | 0.47915 | 0.795502 | 1.029445 |
| 22878 | ENSG0000 | LBX1-AS1 | LBX1 antise | 0.685239 | 0.822416 | 1.028588 |
| 10086 | ENSG0000 | PPP1R9A | protein ph | 0.661588 | 0.693325 | 1.026869 |
| 161 | ENSG0000 | CDKL3 | cyclin depe | 0.459402 | 0.670035 | 1.023311 |
| 489 | ENSG0000 | BTN3A1 | butyrophili | 0.582566 | 0.932587 | 1.02321 |
| 7751 | ENSG0000 | WDFY2 | WD repeat | 0.751145 | 0.881633 | 1.023085 |
| 9277 | ENSG0000 | C4orf33 | chromosor | 0.491937 | 0.625968 | 1.02213 |
| 20347 | ENSG0000 | CBWD6 | COBW don | 0.753385 | 0.983773 | 1.021587 |
| 15582 | ENSG0000 | C7orf61 | chromosor | 0.455562 | 0.773108 | 1.021203 |
| 16408 | ENSG0000 | ZNF774 | zinc finger | 0.353057 | 0.533841 | 1.02111 |
| 19208 | ENSG0000 | GNRHR2 | gonadotro | 0.615514 | 0.865441 | 1.020256 |
| 14296 | ENSG0000 | HTR1D | 5-hydroxyt | 0.025172 | 0.190218 | 1.020174 |
| 10336 | ENSG0000 | SPTBN4 | spectrin be | 0.197777 | 0.384799 | 1.019402 |
| 2855 | ENSG0000 | SLC30A4 | solute carr | 0.627981 | 0.791685 | 1.019287 |
| 19729 | ENSG0000 | ZNF625-ZN | ZNF625-ZN | 0.021296 | 0.137893 | 1.018852 |
| 10855 | ENSG0000 | ICA1L | islet cell au | 0.813368 | 0.908326 | 1.01849 |
| 24365 | ENSG0000 | AC021188.1 | | 0.441015 | 0.583997 | 1.018306 |
| 9984 | ENSG0000 | C9orf43 | chromosor | 0.566555 | 0.908714 | 1.015032 |
| 5842 | ENSG0000 | ADGRE2 | adhesion G | 0.416247 | 0.557683 | 1.014388 |
| 2028 | ENSG0000 | EPB41L4B | erythrocyt | 0.272267 | 0.872871 | 1.014257 |
| 935 | ENSG0000 | ANKRD44 | ankyrin rep | 0.649897 | 0.729992 | 1.013545 |
| 11719 | ENSG0000 | ANKDD1A | ankyrin rep | 0.292943 | 0.55445 | 1.012227 |
| 11310 | ENSG0000 | TMEM65 | transmeml | 0.350059 | 0.843579 | 1.01187 |
| 29424 | ENSG0000 | KCTD21-AS | KCTD21 an | 0.339892 | 0.517788 | 1.011369 |
| 16699 | ENSG0000 | C5AR1 | compleme | 0.165546 | 0.658087 | 1.009799 |
| 9747 | ENSG0000 | PXYLP1 | 2-phospho | 0.578742 | 0.711941 | 1.009704 |
| 10324 | ENSG0000 | ZNF714 | zinc finger | 0.732435 | 0.977803 | 1.009363 |
| 17935 | ENSG0000 | GP1BB | glycoprote | 0.42812 | 0.442591 | 1.008473 |
| 4577 | ENSG0000 | TFAP2E | transcriptio | 0.320244 | 0.654462 | 1.006673 |
| 32020 | ENSG0000 | FLJ20021 | uncharacte | 0.425529 | 0.805415 | 1.005911 |
| 14675 | ENSG0000 | USH1G | USH1 prote | 0.104811 | 0.207602 | 1.00545 |
| 11930 | ENSG0000 | TTYH1 | tweety fan | 0.456016 | 0.462011 | 1.002851 |
| 9520 | ENSG0000 | ADPRHL1 | ADP-ribosy | 0.546052 | 0.662455 | 1.000822 |
| 29352 | ENSG0000 | LINC01024 | long interg | 0.23062 | 0.689713 | 0.998599 |
| 11863 | ENSG0000 | PROCA1 | protein int | 0.287424 | 0.708882 | 0.99754 |
| 14905 | ENSG0000 | GPR19 | G protein-c | 0.270041 | 0.709942 | 0.995656 |
| 20297 | ENSG0000 | AP000317.1 | | 0.321783 | 0.727331 | 0.995584 |

| | | | | | | |
|---|---|---|---|---|---|---|
| 4598 | ENSG0000 | NT5C1A | 5'-nucleoti | 0.204946 | 0.490218 | 0.995331 |
| 4543 | ENSG0000 | DNAJC6 | DnaJ heat s | 0.320665 | 0.796672 | 0.994525 |
| 40968 | ENSG0000 | AC007906.2 | | 0.165336 | 0.433181 | 0.993315 |
| 39082 | ENSG0000 | AL121658.1 | | 0.646969 | 0.864835 | 0.991912 |
| 11268 | ENSG0000 | FAM86FP | family with | 0.839568 | 0.921364 | 0.99149 |
| 41773 | ENSG0000 | AL078621.3 | | 0.316865 | 0.764449 | 0.990401 |
| 20247 | ENSG0000 | POLR2J4 | RNA polym | 0.527146 | 0.735116 | 0.990167 |
| 15845 | ENSG0000 | MAGED4B | MAGE fam | 0.200901 | 0.536959 | 0.989617 |
| 15403 | ENSG0000 | WASIR1 | WASH and | 0 | 0.412682 | 0.988764 |
| 20827 | ENSG0000 | LINC01881 | long interg | 0.566328 | 0.97676 | 0.988193 |
| 21894 | ENSG0000 | PRRT4 | proline rich | 0.210771 | 0.279594 | 0.988108 |
| 39234 | ENSG0000 | AP000894.4 | | 0.514639 | 0.770088 | 0.985788 |
| 11633 | ENSG0000 | TRIM66 | tripartite n | 0.589994 | 0.82487 | 0.985306 |
| 39209 | ENSG0000 | DGCR11 | DiGeorge s | 0.572673 | 0.781005 | 0.984903 |
| 10635 | ENSG0000 | OLFML2B | olfactomed | 0.083144 | 0.711421 | 0.984046 |
| 12597 | ENSG0000 | CAVIN4 | caveolae a | 0.177309 | 0.585214 | 0.982697 |
| 41813 | ENSG0000 | RBM12B-A | RBM12B ar | 0.6742 | 0.768525 | 0.982463 |
| 34929 | ENSG0000 | AC026471.2 | | 0.205024 | 0.362303 | 0.982094 |
| 6365 | ENSG0000 | NAPSA | napsin A as | 0.206508 | 0.569657 | 0.981357 |
| 6472 | ENSG0000 | CNGA4 | cyclic nucle | 0.142474 | 0.80175 | 0.980213 |
| 40384 | ENSG0000 | RIMBP3 | RIMS bindi | 0.18262 | 0.223403 | 0.979781 |
| 30786 | ENSG0000 | AC012625.1 | | 0.416841 | 0.764358 | 0.978875 |
| 18386 | ENSG0000 | C4orf47 | chromosor | 0.015552 | 0.222682 | 0.97855 |
| 2564 | ENSG0000 | SMAD7 | SMAD fam | 0.235366 | 0.855695 | 0.978124 |
| 42985 | ENSG0000 | Y_RNA | Y RNA [Sou | 0.376957 | 0.432437 | 0.975962 |
| 42135 | ENSG0000 | CU639417.4 | | 0.514378 | 0.878017 | 0.975877 |
| 12473 | ENSG0000 | NIPA1 | non imprin | 0.75401 | 0.899355 | 0.975211 |
| 212 | ENSG0000 | CELSR3 | cadherin E | 0.50035 | 0.930822 | 0.975066 |
| 28925 | ENSG0000 | KCTD7 | potassium | 0.57868 | 0.756535 | 0.974243 |
| 1257 | ENSG0000 | GLI2 | GLI family z | 0.540897 | 0.805594 | 0.973994 |
| 16979 | ENSG0000 | ZNF658B | zinc finger | 0.817951 | 0.895638 | 0.973638 |
| 15648 | ENSG0000 | MKL2 | MKL1/myo | 0.48628 | 0.890379 | 0.972947 |
| 19000 | ENSG0000 | MIR635 | microRNA | 0.435367 | 0.439715 | 0.972878 |
| 14500 | ENSG0000 | MAP3K15 | mitogen-ac | 0.377012 | 0.697727 | 0.972726 |
| 4139 | ENSG0000 | THBS4 | thrombosp | 0.484089 | 0.918348 | 0.972404 |
| 34418 | ENSG0000 | SH3RF3-AS | SH3RF3 an | 0.649772 | 0.965803 | 0.972383 |
| 39335 | ENSG0000 | AL731533.2 | | 0.239305 | 0.356895 | 0.970631 |
| 2894 | ENSG0000 | ZC2HC1A | zinc finger | 0.502484 | 0.927215 | 0.970624 |
| 34897 | ENSG0000 | AC069224.1 | | 0.217372 | 0.594261 | 0.969797 |
| 13633 | ENSG0000 | DENND2C | DENN dom | 0.234323 | 0.728671 | 0.969058 |
| 6159 | ENSG0000 | APOE | apolipopro | 0.253926 | 0.481959 | 0.968106 |
| 37912 | ENSG0000 | AL353593.1 | | 0.447348 | 0.807002 | 0.967973 |
| 5984 | ENSG0000 | ELL3 | elongation | 0.384812 | 0.641955 | 0.967676 |
| 20644 | ENSG0000 | AL390719.1 | | 0.269981 | 0.408021 | 0.965966 |
| 15212 | ENSG0000 | TACSTD2 | tumor asso | 0.173771 | 0.510412 | 0.965231 |
| 1062 | ENSG0000 | RASGRP2 | RAS guanyl | 0.160099 | 0.24045 | 0.96264 |
| 21372 | ENSG0000 | AL645608.1 | | 0.264812 | 0.67245 | 0.961814 |

| | | | | | | |
|---|---|---|---|---|---|---|
| 15152 | ENSG0000 | NPB | neuropept | 0.332817 | 0.644411 | 0.961573 |
| 12647 | ENSG0000 | NUDT6 | nudix hydr | 0.509751 | 0.635198 | 0.961313 |
| 16401 | ENSG0000 | FUT4 | fucosyltran | 0.874281 | 0.935901 | 0.958943 |
| 31414 | ENSG0000 | GUSBP3 | glucuronid | 0.35177 | 0.713384 | 0.958665 |
| 25113 | ENSG0000 | AL133517.1 | | 0.57795 | 0.58478 | 0.95784 |
| 24539 | ENSG0000 | HAR1B | highly acce | 0.49655 | 0.726699 | 0.956907 |
| 17027 | ENSG0000 | ARC | activity reg | 0.065265 | 0.514834 | 0.955252 |
| 13372 | ENSG0000 | C12orf76 | chromosor | 0.633949 | 0.688364 | 0.955182 |
| 38268 | ENSG0000 | AL035413.2 | | 0.263197 | 0.497064 | 0.95517 |
| 10203 | ENSG0000 | STARD9 | StAR relate | 0.500652 | 0.922736 | 0.954511 |
| 7603 | ENSG0000 | HERC5 | HECT and I | 0.189976 | 0.318792 | 0.95422 |
| 39429 | ENSG0000 | SNURF | SNRPN ups | 0.47546 | 0.878208 | 0.95317 |
| 3488 | ENSG0000 | RGS9 | regulator c | 0.127781 | 0.341646 | 0.95195 |
| 36802 | ENSG0000 | AC024267.6 | | 0.11433 | 0.327594 | 0.95125 |
| 3257 | ENSG0000 | COBL | cordon-ble | 0.459675 | 0.830906 | 0.951063 |
| 5167 | ENSG0000 | LIAS | lipoic acid | 0.74709 | 0.818358 | 0.950253 |
| 34593 | ENSG0000 | AC104794.2 | | 0.229351 | 0.495299 | 0.950028 |
| 30675 | ENSG0000 | SNHG21 | small nucle | 0.639312 | 0.847568 | 0.949697 |
| 13602 | ENSG0000 | SLC35E3 | solute carr | 0.489437 | 0.64428 | 0.949177 |
| 37637 | ENSG0000 | AC135983.5 | | 0.371457 | 0.872694 | 0.948474 |
| 37098 | ENSG0000 | AC036176.1 | | 0.180831 | 0.833921 | 0.948079 |
| 22854 | ENSG0000 | AC104170.1 | | 0.341133 | 0.71601 | 0.947538 |
| 15608 | ENSG0000 | C15orf41 | chromosor | 0.656644 | 0.903373 | 0.947308 |
| 18118 | ENSG0000 | HLA-DMA | major histo | 0.292834 | 0.355708 | 0.946131 |
| 28131 | ENSG0000 | ERICH6-AS | ERICH6 ant | 0.29159 | 0.438526 | 0.945373 |
| 24949 | ENSG0000 | LINC01063 | long interg | 0.064339 | 0.323196 | 0.945036 |
| 9216 | ENSG0000 | ITPR1 | inositol 1,4 | 0.613118 | 0.905387 | 0.944304 |
| 23968 | ENSG0000 | LINC01315 | long interg | 0.53244 | 0.829693 | 0.944282 |
| 33495 | ENSG0000 | AL391832.3 | | 0.025403 | 0.357772 | 0.943685 |
| 40516 | ENSG0000 | WHAMMP | WAS prote | 0.362819 | 0.656711 | 0.941162 |
| 1097 | ENSG0000 | GNB5 | G protein s | 0.667685 | 0.763157 | 0.940851 |
| 932 | ENSG0000 | DGKA | diacylglyce | 0.447262 | 0.52361 | 0.940726 |
| 33943 | ENSG0000 | AL132639.2 | | 0.48839 | 0.846085 | 0.940275 |
| 34113 | ENSG0000 | AC090825.1 | | 0.403751 | 0.578179 | 0.939379 |
| 9701 | ENSG0000 | PPM1J | protein ph | 0.210023 | 0.688855 | 0.938495 |
| 42644 | ENSG0000 | AL157392.5 | | 0.225294 | 0.355127 | 0.937589 |
| 1006 | ENSG0000 | NEO1 | neogenin 1 | 0.321286 | 0.612775 | 0.93721 |
| 13830 | ENSG0000 | MAMSTR | MEF2 activ | 0.348277 | 0.373864 | 0.93713 |
| 7396 | ENSG0000 | RAB30 | RAB30, me | 0.30332 | 0.773989 | 0.935534 |
| 13381 | ENSG0000 | ANKRD36C | ankyrin rep | 0.355495 | 0.662483 | 0.934128 |
| 13050 | ENSG0000 | RND1 | Rho family | 0.265781 | 0.570775 | 0.933745 |
| 18476 | ENSG0000 | DENND6B | DENN dom | 0.305564 | 0.55106 | 0.933521 |
| 31418 | ENSG0000 | AC040970.1 | | 0.717253 | 0.864585 | 0.933478 |
| 14507 | ENSG0000 | CAPS2 | calcyphosi | 0.477378 | 0.562157 | 0.932854 |
| 14610 | ENSG0000 | GPR135 | G protein-c | 0.432011 | 0.755599 | 0.932434 |
| 9418 | ENSG0000 | PLEKHH2 | pleckstrin l | 0.125624 | 0.427164 | 0.932399 |
| 38370 | ENSG0000 | AC133552.5 | | 0.096958 | 0.572728 | 0.930233 |

| | | | | | | |
|---|---|---|---|---|---|---|
| 42987 | ENSG0000 | AL592295.4 | | 0.268191 | 0.630334 | 0.930135 |
| 13297 | ENSG0000 | MARCH3 | membrane | 0.655756 | 0.830944 | 0.929335 |
| 24015 | ENSG0000 | TOB1-AS1 | TOB1 antis | 0.438659 | 0.794257 | 0.929156 |
| 33082 | ENSG0000 | AP003721.4 | | 0.067124 | 0.880983 | 0.927184 |
| 13478 | ENSG0000 | DES | desmin [Sc | 0 | 0.065101 | 0.9262 |
| 29380 | ENSG0000 | AP001258.1 | | 0.242294 | 0.622167 | 0.925791 |
| 827 | ENSG0000 | CREB3L3 | cAMP resp | 0.087602 | 0.153453 | 0.924365 |
| 41407 | ENSG0000 | TMEM191 | transmeml | 0.231254 | 0.779527 | 0.923005 |
| 2604 | ENSG0000 | SYP | synaptoph | 0.50275 | 0.504394 | 0.922383 |
| 25584 | ENSG0000 | TMEM238 | transmeml | 0.408536 | 0.410133 | 0.919953 |
| 4558 | ENSG0000 | WLS | wntless W | 0.540803 | 0.546352 | 0.919534 |
| 6698 | ENSG0000 | AMPD3 | adenosine | 0.39218 | 0.541985 | 0.918873 |
| 20059 | ENSG0000 | C19orf38 | chromosor | 0.240119 | 0.383755 | 0.918539 |
| 33981 | ENSG0000 | AC005476.2 | | 0.398013 | 0.447725 | 0.918251 |
| 35797 | ENSG0000 | AC027796.4 | | 0.224039 | 0.87474 | 0.917776 |
| 8714 | ENSG0000 | IRAK1BP1 | interleukin | 0.562842 | 0.662574 | 0.917647 |
| 28023 | ENSG0000 | AC009120.1 | | 0.20763 | 0.568338 | 0.916632 |
| 40206 | ENSG0000 | AL136981.2 | | 0.435875 | 0.887369 | 0.916612 |
| 13748 | ENSG0000 | PFN4 | profilin fan | 0.482038 | 0.755805 | 0.914282 |
| 4924 | ENSG0000 | ZC2HC1C | zinc finger | 0.503227 | 0.665016 | 0.912605 |
| 13060 | ENSG0000 | ZNF738 | zinc finger | 0.38718 | 0.804018 | 0.912414 |
| 4540 | ENSG0000 | C1orf21 | chromosor | 0.333448 | 0.40378 | 0.912268 |
| 13498 | ENSG0000 | CSRP2 | cysteine ar | 0.237115 | 0.269183 | 0.90844 |
| 37401 | ENSG0000 | AC011483.2 | | 0.600329 | 0.7878 | 0.906529 |
| 20959 | ENSG0000 | ARIH2OS | ariadne RB | 0.483783 | 0.497185 | 0.905935 |
| 10280 | ENSG0000 | SLC37A1 | solute carr | 0.553738 | 0.82278 | 0.905888 |
| 16813 | ENSG0000 | ZNF461 | zinc finger | 0.47718 | 0.664117 | 0.90557 |
| 15994 | ENSG0000 | AC087499.1 | | 0.327817 | 0.558849 | 0.904159 |
| 37009 | ENSG0000 | AC020928.1 | | 0.042391 | 0.331802 | 0.903758 |
| 9571 | ENSG0000 | ANKH | ANKH inor | 0.707624 | 0.855785 | 0.902841 |
| 26230 | ENSG0000 | AC098934.2 | | 0.190283 | 0.480262 | 0.902589 |
| 41408 | ENSG0000 | PTPN20CP | protein tyr | 0.515706 | 0.73626 | 0.902375 |
| 1235 | ENSG0000 | CLCN4 | chloride vc | 0.368931 | 0.514168 | 0.902079 |
| 18247 | ENSG0000 | AC115618.1 | | 0.76455 | 0.860594 | 0.902011 |
| 11795 | ENSG0000 | CCDC103 | coiled-coil | 0.104976 | 0.392864 | 0.901921 |
| 14657 | ENSG0000 | AP003774.1 | | 0.155028 | 0.319534 | 0.900092 |
| 38962 | ENSG0000 | AC113194.1 | | 0.284188 | 0.637309 | 0.898282 |
| 7086 | ENSG0000 | ALDH1L2 | aldehyde d | 0.24638 | 0.391866 | 0.896605 |
| 1796 | ENSG0000 | FER1L4 | fer-1 like fa | 0.112924 | 0.269574 | 0.896571 |
| 27111 | ENSG0000 | AL139412.1 | | 0.071956 | 0.309787 | 0.896275 |
| 15179 | ENSG0000 | LRTOMT | leucine rich | 0.762198 | 0.880302 | 0.895667 |
| 25977 | ENSG0000 | USP27X-AS | USP27X an | 0.318963 | 0.733107 | 0.893767 |
| 3357 | ENSG0000 | LHX6 | LIM homeo | 0.099079 | 0.201791 | 0.891063 |
| 15666 | ENSG0000 | THBS2 | thrombosp | 0.427161 | 0.583493 | 0.890316 |
| 21364 | ENSG0000 | CCDC18-AS | CCDC18 an | 0.453709 | 0.831819 | 0.889751 |
| 13877 | ENSG0000 | LINC00982 | long interg | 0.359269 | 0.683516 | 0.889241 |
| 5377 | ENSG0000 | PFKFB2 | 6-phospho | 0.642499 | 0.860296 | 0.888849 |

| | | | | | | |
|---|---|---|---|---|---|---|
| 13761 | ENSG0000 | LRRC37A3 | leucine ricl | 0.504968 | 0.508378 | 0.888282 |
| 15615 | ENSG0000 | GSAP | gamma-se | 0.58442 | 0.73588 | 0.887706 |
| 2829 | ENSG0000 | AP3B2 | adaptor re | 0.193671 | 0.727144 | 0.885202 |
| 4024 | ENSG0000 | PDE8B | phosphodi | 0.313213 | 0.70517 | 0.884252 |
| 10677 | ENSG0000 | OBSCN-AS: | OBSCN ant | 0.036569 | 0.390622 | 0.884189 |
| 1481 | ENSG0000 | RFX3 | regulatory | 0.62611 | 0.810303 | 0.883412 |
| 16693 | ENSG0000 | SLC22A5 | solute carr | 0.601582 | 0.676943 | 0.882738 |
| 40037 | ENSG0000 | ZNF280B | zinc finger | 0.521819 | 0.81347 | 0.882581 |
| 32819 | ENSG0000 | DNAAF4 | dynein axo | 0.423818 | 0.764815 | 0.881835 |
| 42489 | ENSG0000 | GSEC | G-quadrup | 0.512639 | 0.723792 | 0.881626 |
| 9542 | ENSG0000 | PID1 | phosphoty | 0.569468 | 0.839346 | 0.88068 |
| 10843 | ENSG0000 | NUAK2 | NUAK fami | 0.267445 | 0.62268 | 0.880068 |
| 25471 | ENSG0000 | AC233728.1 | | 0.088553 | 0.264821 | 0.879967 |
| 29523 | ENSG0000 | UBA6-AS1 | UBA6 antis | 0.510487 | 0.831859 | 0.878559 |
| 15432 | ENSG0000 | ARL15 | ADP ribosy | 0.36223 | 0.623056 | 0.877783 |
| 1136 | ENSG0000 | TESK2 | testis-spec | 0.050907 | 0.260008 | 0.87565 |
| 42477 | ENSG0000 | LINC01232 | long interg | 0.491144 | 0.565584 | 0.875588 |
| 22636 | ENSG0000 | KLLN | killin, p53-r | 0.249569 | 0.763625 | 0.874102 |
| 17319 | ENSG0000 | RNY4P19 | RNA, Ro-as | 0.125858 | 0.206182 | 0.871096 |
| 25539 | ENSG0000 | LINC01719 | long interg | 0.149946 | 0.242264 | 0.870594 |
| 26685 | ENSG0000 | AC011290.2 | | 0.434158 | 0.733412 | 0.870527 |
| 13312 | ENSG0000 | SPERT | spermatid | 0.318466 | 0.479162 | 0.870291 |
| 37422 | ENSG0000 | AC008878.1 | | 0.246788 | 0.673429 | 0.8697 |
| 8799 | ENSG0000 | CCNB3 | cyclin B3 [S | 0.31816 | 0.609797 | 0.867705 |
| 41191 | ENSG0000 | Z99129.4 | | 0.453349 | 0.810714 | 0.86768 |
| 8515 | ENSG0000 | PTPRG | protein tyr | 0.765468 | 0.850012 | 0.867322 |
| 35828 | ENSG0000 | AC090617.6 | | 0.372691 | 0.711295 | 0.866709 |
| 9072 | ENSG0000 | GLB1L2 | galactosida | 0.248427 | 0.404463 | 0.866269 |
| 1596 | ENSG0000 | FAM135A | family with | 0.6439 | 0.764744 | 0.865831 |
| 1948 | ENSG0000 | PITPNM3 | PITPNM fa | 0.199542 | 0.451768 | 0.865192 |
| 35639 | ENSG0000 | AC127496.2 | | 0.070142 | 0.312611 | 0.864117 |
| 27007 | ENSG0000 | LINC01844 | long interg | 0.23737 | 0.617716 | 0.863149 |
| 11562 | ENSG0000 | DEPDC4 | DEP domai | 0.438439 | 0.657647 | 0.86229 |
| 9535 | ENSG0000 | FAM92B | family with | 0 | 0.172179 | 0.862272 |
| 21378 | ENSG0000 | LGALS8-AS | LGALS8 ant | 0.302541 | 0.469894 | 0.860724 |
| 36321 | ENSG0000 | AC093484.3 | | 0.093903 | 0.324886 | 0.860629 |
| 9278 | ENSG0000 | FRMD4A | FERM dom | 0.350334 | 0.691385 | 0.859081 |
| 39975 | ENSG0000 | NPHP3-AC | NPHP3-AC | 0.193552 | 0.668141 | 0.858525 |
| 26889 | ENSG0000 | DDX11L2 | DEAD/H-bo | 0.234251 | 0.661855 | 0.857614 |
| 1759 | ENSG0000 | GNAO1 | G protein s | 0.200245 | 0.207802 | 0.856793 |
| 32641 | ENSG0000 | TDGP1 | thymine-D | 0.116181 | 0.569936 | 0.85657 |
| 1631 | ENSG0000 | SLC27A5 | solute carr | 0.732011 | 0.789795 | 0.85654 |
| 1269 | ENSG0000 | ATP2A3 | ATPase sar | 0.198322 | 0.223142 | 0.856502 |
| 37116 | ENSG0000 | AC016582.2 | | 0.469797 | 0.747398 | 0.856219 |
| 27251 | ENSG0000 | AL391832.2 | | 0.217322 | 0.370354 | 0.855248 |
| 21639 | ENSG0000 | LINC01058 | long interg | 0.159336 | 0.370943 | 0.853832 |
| 9770 | ENSG0000 | FAM161B | family with | 0.473911 | 0.83213 | 0.85299 |

| | | | | | | |
|---|---|---|---|---|---|---|
| 10604 | ENSG0000 | AKNAD1 | AKNA dom | 0.255251 | 0.579447 | 0.852632 |
| 14781 | ENSG0000 | NXPH3 | neurexoph | 0.040913 | 0.257878 | 0.852604 |
| 16027 | ENSG0000 | TMPPE | transmeml | 0.391679 | 0.71599 | 0.850633 |
| 7173 | ENSG0000 | NKD1 | naked cuti | 0.269305 | 0.614318 | 0.849484 |
| 14140 | ENSG0000 | PARP10 | poly(ADP-r | 0.239286 | 0.359794 | 0.84916 |
| 5712 | ENSG0000 | TMEM53 | transmeml | 0.491608 | 0.775786 | 0.848974 |
| 26249 | ENSG0000 | MNX1-AS2 | MNX1 anti | 0.529117 | 0.609065 | 0.848931 |
| 16427 | ENSG0000 | NPIPB15 | nuclear po | 0.32112 | 0.840543 | 0.848545 |
| 14649 | ENSG0000 | SLC2A4 | solute carr | 0.166982 | 0.32439 | 0.8477 |
| 9010 | ENSG0000 | LRRC27 | leucine rich | 0.554644 | 0.636472 | 0.847107 |
| 22686 | ENSG0000 | FAM66C | family with | 0.291943 | 0.369285 | 0.847058 |
| 9114 | ENSG0000 | KIAA1755 | KIAA1755 \| | 0.150785 | 0.466578 | 0.845669 |
| 25838 | ENSG0000 | AC062029.1 | | 0.248925 | 0.485007 | 0.844836 |
| 8814 | ENSG0000 | CCDC120 | coiled-coil | 0.115869 | 0.578612 | 0.84415 |
| 23277 | ENSG0000 | RALY-AS1 | RALY antis | 0.181542 | 0.667663 | 0.841719 |
| 6934 | ENSG0000 | ANKRD6 | ankyrin rep | 0.158826 | 0.569351 | 0.841549 |
| 41004 | ENSG0000 | AC097641.3 | | 0.145659 | 0.602612 | 0.839491 |
| 23680 | ENSG0000 | LINC00242 | long interg | 0.185815 | 0.456777 | 0.838743 |
| 34168 | ENSG0000 | LINC01833 | long interg | 0.162106 | 0.392096 | 0.838135 |
| 29015 | ENSG0000 | ZNF487 | zinc finger | 0.551589 | 0.823521 | 0.838085 |
| 37959 | ENSG0000 | AC010359.1 | | 0.416301 | 0.495122 | 0.838074 |
| 15636 | ENSG0000 | EDARADD | EDAR asso | 0.151131 | 0.386437 | 0.835741 |
| 10702 | ENSG0000 | DUSP19 | dual specif | 0.258846 | 0.784222 | 0.83526 |
| 36249 | ENSG0000 | AC005722.3 | | 0.351495 | 0.601351 | 0.833903 |
| 3634 | ENSG0000 | GAB1 | GRB2 asso | 0.288121 | 0.737923 | 0.833837 |
| 18649 | ENSG0000 | ZDHHC11B | zinc finger | 0.215795 | 0.591616 | 0.833737 |
| 35589 | ENSG0000 | AC090772.3 | | 0.487225 | 0.5828 | 0.831704 |
| 1860 | ENSG0000 | CMTM1 | CKLF like M | 0.157272 | 0.831131 | 0.83135 |
| 7720 | ENSG0000 | FOXN4 | forkhead b | 0.03577 | 0.228528 | 0.829865 |
| 3504 | ENSG0000 | TRIM16L | tripartite n | 0.62447 | 0.709577 | 0.829013 |
| 9232 | ENSG0000 | TMEM86A | transmeml | 0.147607 | 0.548268 | 0.828496 |
| 42545 | ENSG0000 | AC120114.5 | | 0.22428 | 0.278423 | 0.826964 |
| 10507 | ENSG0000 | PPP1R32 | protein ph | 0.328635 | 0.671709 | 0.826671 |
| 35596 | ENSG0000 | AC018413.1 | | 0.371793 | 0.592422 | 0.826437 |
| 20886 | ENSG0000 | MIR1229 | microRNA | 0.298838 | 0.792605 | 0.826286 |
| 37616 | ENSG0000 | AC008403.3 | | 0.117819 | 0.373989 | 0.826102 |
| 41350 | ENSG0000 | PMS2P2 | PMS1 hom | 0.37293 | 0.642079 | 0.825531 |
| 3363 | ENSG0000 | DNM1 | dynamin 1 | 0.210448 | 0.470791 | 0.824153 |
| 19695 | ENSG0000 | CCDC183 | coiled-coil | 0.220968 | 0.587198 | 0.82328 |
| 13740 | ENSG0000 | SCAND2P | SCAN dom | 0.414581 | 0.592439 | 0.822087 |
| 23185 | ENSG0000 | AL160408.1 | | 0.49827 | 0.696445 | 0.821697 |
| 15905 | ENSG0000 | VMAC | vimentin ty | 0.618013 | 0.818645 | 0.821416 |
| 33269 | ENSG0000 | LINC02298 | long interg | 0.313724 | 0.497914 | 0.820554 |
| 28751 | ENSG0000 | AC022415.1 | | 0.30777 | 0.68399 | 0.819896 |
| 1662 | ENSG0000 | EFR3B | EFR3 homo | 0.283974 | 0.558814 | 0.81923 |
| 12760 | ENSG0000 | CDC42BPG | CDC42 bin | 0.118075 | 0.220436 | 0.817968 |
| 8465 | ENSG0000 | CCDC150 | coiled-coil | 0.792131 | 0.797732 | 0.817965 |

| | | | | | | |
|---|---|---|---|---|---|---|
| 13509 | ENSG0000 | CHST1 | carbohydra | 0.030722 | 0.141234 | 0.81702 |
| 42300 | ENSG0000 | CU638689.5 | | 0.531038 | 0.67391 | 0.816719 |
| 13762 | ENSG0000 | FKBP9P1 | FK506 binc | 0.748598 | 0.810174 | 0.815682 |
| 31591 | ENSG0000 | OSGEPL1-A | OSGEPL1 a | 0.059798 | 0.46265 | 0.8148 |
| 20251 | ENSG0000 | AC098934.1 | | 0.421934 | 0.571532 | 0.814257 |
| 8288 | ENSG0000 | CGN | cingulin [Sc | 0.317565 | 0.333665 | 0.814039 |
| 6450 | ENSG0000 | C19orf57 | chromosor | 0.421553 | 0.474868 | 0.813853 |
| 13573 | ENSG0000 | SUGCT | succinyl-Cc | 0.452775 | 0.518136 | 0.813236 |
| 38083 | ENSG0000 | AC016773.1 | | 0.5706 | 0.659551 | 0.812938 |
| 10762 | ENSG0000 | S100A12 | S100 calciu | 0.137278 | 0.47804 | 0.812072 |
| 889 | ENSG0000 | CDON | cell adhesi | 0.410307 | 0.561491 | 0.811212 |
| 30746 | ENSG0000 | AC004069.1 | | 0.2246 | 0.314343 | 0.811133 |
| 13395 | ENSG0000 | ZNF497 | zinc finger | 0.38378 | 0.721476 | 0.810856 |
| 21589 | ENSG0000 | AL451074.2 | | 0.266111 | 0.698142 | 0.810768 |
| 6285 | ENSG0000 | PNCK | pregnancy | 0.085392 | 0.422764 | 0.809357 |
| 12047 | ENSG0000 | TRANK1 | tetratricop | 0.1273 | 0.387123 | 0.80877 |
| 22478 | ENSG0000 | GAS1RR | GAS1 adjac | 0.182779 | 0.239135 | 0.808161 |
| 9525 | ENSG0000 | CD8A | CD8a mole | 0.135185 | 0.643567 | 0.807693 |
| 8633 | ENSG0000 | ANKRD31 | ankyrin rep | 0.201491 | 0.610497 | 0.807269 |
| 36976 | ENSG0000 | AC007773.1 | | 0.182831 | 0.368833 | 0.806671 |
| 15068 | ENSG0000 | ZNF530 | zinc finger | 0.665654 | 0.771041 | 0.806242 |
| 29674 | ENSG0000 | SRP14-AS1 | SRP14 anti | 0.220643 | 0.487925 | 0.805403 |
| 39756 | ENSG0000 | AC073575.2 | | 0.38953 | 0.767353 | 0.804909 |
| 7584 | ENSG0000 | MNS1 | meiosis spe | 0.226445 | 0.420207 | 0.804626 |
| 39336 | ENSG0000 | AL355388.2 | | 0.33371 | 0.468864 | 0.804501 |
| 35751 | ENSG0000 | AC004771.5 | | 0.260585 | 0.673878 | 0.804102 |
| 25959 | ENSG0000 | AP000346.2 | | 0.155265 | 0.347289 | 0.802928 |
| 14095 | ENSG0000 | ZNF223 | zinc finger | 0.350536 | 0.752892 | 0.802007 |
| 27051 | ENSG0000 | ANKRD20A | ankyrin rep | 0.264024 | 0.478454 | 0.801742 |
| 16051 | ENSG0000 | ZNF383 | zinc finger | 0.422699 | 0.613994 | 0.80102 |
| 4760 | ENSG0000 | MYB | MYB proto | 0.455065 | 0.789385 | 0.800818 |
| 7386 | ENSG0000 | MYO7A | myosin VII/ | 0.174669 | 0.380652 | 0.800795 |
| 36885 | ENSG0000 | UXT-AS1 | UXT antise | 0.571575 | 0.753973 | 0.800764 |
| 26711 | ENSG0000 | FAM157A | family with | 0.269764 | 0.65607 | 0.800008 |
| 37995 | ENSG0000 | AC026691.1 | | 0.407081 | 0.69047 | 0.799419 |
| 4900 | ENSG0000 | GRK4 | G protein-c | 0.59941 | 0.705356 | 0.79941 |
| 42517 | ENSG0000 | AL606763.1 | | 0.497168 | 0.758746 | 0.797486 |
| 3129 | ENSG0000 | PLA2G4C | phospholip | 0.193633 | 0.732547 | 0.797146 |
| 901 | ENSG0000 | EYA2 | EYA transc | 0.398517 | 0.531408 | 0.797096 |
| 30523 | ENSG0000 | AC093535.1 | | 0.216369 | 0.535841 | 0.796929 |
| 22910 | ENSG0000 | KIF9-AS1 | KIF9 antise | 0.242122 | 0.485629 | 0.796786 |
| 38318 | ENSG0000 | AC016355.1 | | 0.316822 | 0.563335 | 0.795997 |
| 8572 | ENSG0000 | VWA5B2 | von Willeb | 0.207315 | 0.212727 | 0.795965 |
| 15707 | ENSG0000 | ARHGAP30 | Rho GTPas | 0.226988 | 0.780605 | 0.795581 |
| 29409 | ENSG0000 | AC093752.1 | | 0.437761 | 0.750331 | 0.79552 |
| 41271 | ENSG0000 | GLIDR | glioblaston | 0.488277 | 0.729655 | 0.795508 |
| 36000 | ENSG0000 | AC009831.1 | | 0.552035 | 0.759616 | 0.794707 |

| | | | | | | |
|---|---|---|---|---|---|---|
| 8529 | ENSG0000 | NXPE3 | neurexoph | 0.393297 | 0.41111 | 0.794637 |
| 20072 | ENSG0000 | AC091305.1 | | 0.175921 | 0.401191 | 0.793822 |
| 1884 | ENSG0000 | ICAM1 | intercellula | 0.148612 | 0.318554 | 0.793806 |
| 39246 | ENSG0000 | AC069222.1 | | 0.160057 | 0.411965 | 0.793286 |
| 15956 | ENSG0000 | CCDC157 | coiled-coil | 0.271238 | 0.347931 | 0.793146 |
| 7175 | ENSG0000 | KSR1 | kinase sup | 0.357176 | 0.574108 | 0.792935 |
| 16299 | ENSG0000 | USP32P3 | ubiquitin s | 0.501823 | 0.638485 | 0.792226 |
| 10524 | ENSG0000 | FGF19 | fibroblast g | 0.058326 | 0.142314 | 0.792072 |
| 37367 | ENSG0000 | AC011498.6 | | 0.096952 | 0.310407 | 0.791701 |
| 9924 | ENSG0000 | NEK8 | NIMA relat | 0.579572 | 0.719838 | 0.791311 |
| 29458 | ENSG0000 | AC009126.1 | | 0.433211 | 0.706552 | 0.791165 |
| 12269 | ENSG0000 | ROR2 | receptor ty | 0.043016 | 0.228899 | 0.791093 |
| 13150 | ENSG0000 | ADCY5 | adenylate | 0.216535 | 0.412525 | 0.791033 |
| 16682 | ENSG0000 | AC008543.1 | | 0.185902 | 0.717883 | 0.790959 |
| 22353 | ENSG0000 | NTF4 | neurotropl | 0.013537 | 0.25189 | 0.79037 |
| 3018 | ENSG0000 | BIN2 | bridging in | 0.25274 | 0.455984 | 0.790202 |
| 39219 | ENSG0000 | AC078846.1 | | 0.56948 | 0.671409 | 0.788066 |
| 32349 | ENSG0000 | AP006621.1 | | 0.448302 | 0.492558 | 0.786851 |
| 8303 | ENSG0000 | AC116050.1 | | 0.466854 | 0.622019 | 0.78529 |
| 1133 | ENSG0000 | ST6GALNA | ST6 N-acet | 0.264851 | 0.286116 | 0.785153 |
| 2545 | ENSG0000 | TNNC2 | troponin C | 0.151 | 0.183631 | 0.782788 |
| 14822 | ENSG0000 | CCDC87 | coiled-coil | 0.393286 | 0.5324 | 0.781652 |
| 35002 | ENSG0000 | AC093520.1 | | 0.349478 | 0.365541 | 0.781467 |
| 10056 | ENSG0000 | RHBDL2 | rhomboid l | 0.260259 | 0.423072 | 0.781115 |
| 15266 | ENSG0000 | CEND1 | cell cycle e | 0.427186 | 0.556841 | 0.780962 |
| 13624 | ENSG0000 | A2M | alpha-2-ma | 0.201464 | 0.326705 | 0.780828 |
| 23793 | ENSG0000 | MIR5689H | MIR5689 h | 0.169689 | 0.525884 | 0.780247 |
| 24345 | ENSG0000 | AL645608.3 | | 0.193445 | 0.601623 | 0.779638 |
| 10488 | ENSG0000 | SSTR5 | somatosta | 0.026059 | 0.113309 | 0.778281 |
| 37311 | ENSG0000 | AC008687.4 | | 0.299501 | 0.597156 | 0.777493 |
| 10942 | ENSG0000 | PRRT3 | proline ricl | 0.553496 | 0.70397 | 0.777193 |
| 40109 | ENSG0000 | AC010654.1 | | 0.176638 | 0.524896 | 0.776793 |
| 38219 | ENSG0000 | AL353622.1 | | 0.192638 | 0.447374 | 0.776686 |
| 19648 | ENSG0000 | CFAP45 | cilia and fla | 0.413931 | 0.545965 | 0.77515 |
| 11468 | ENSG0000 | STOX1 | storkhead | 0.264797 | 0.429361 | 0.77514 |
| 24472 | ENSG0000 | RPL36AP39 | ribosomal | 0.056992 | 0.127405 | 0.77483 |
| 38649 | ENSG0000 | AL445222.1 | | 0.468562 | 0.70555 | 0.774274 |
| 200 | ENSG0000 | CDKL5 | cyclin depe | 0.093569 | 0.505739 | 0.774183 |
| 3139 | ENSG0000 | RASIP1 | Ras interac | 0.191947 | 0.25097 | 0.77405 |
| 389 | ENSG0000 | YAF2 | YY1 associa | 0.580095 | 0.613711 | 0.772922 |
| 23844 | ENSG0000 | AL353194.1 | | 0.523982 | 0.586606 | 0.772918 |
| 32472 | ENSG0000 | AP001830.1 | | 0.176656 | 0.24937 | 0.772631 |
| 35545 | ENSG0000 | AC138207.4 | | 0.265659 | 0.49336 | 0.772386 |
| 24506 | ENSG0000 | AC234582.1 | | 0.314153 | 0.508064 | 0.771574 |
| 12868 | ENSG0000 | GATM | glycine am | 0.349607 | 0.62837 | 0.770658 |
| 17978 | ENSG0000 | RAET1G | retinoic aci | 0.016267 | 0.399042 | 0.770652 |
| 2183 | ENSG0000 | LRP5L | LDL recept | 0.381736 | 0.720576 | 0.769753 |

| | | | | | | |
|---|---|---|---|---|---|---|
| 18014 | ENSG0000 | ATP1A1-AS | ATP1A1 an | 0.251154 | 0.548081 | 0.769648 |
| 9209 | ENSG0000 | FOXO1 | forkhead b | 0.24277 | 0.561341 | 0.769252 |
| 7527 | ENSG0000 | CUZD1 | CUB and zc | 0.457043 | 0.724891 | 0.769195 |
| 4406 | ENSG0000 | PLCD4 | phospholip | 0.379427 | 0.424642 | 0.768611 |
| 532 | ENSG0000 | GAB2 | GRB2 asso | 0.466667 | 0.739906 | 0.768547 |
| 5337 | ENSG0000 | IKZF4 | IKAROS far | 0.260967 | 0.450114 | 0.767515 |
| 25479 | ENSG0000 | HMGB1P3 | high mobil | 0.343672 | 0.594239 | 0.766965 |
| 42600 | ENSG0000 | AL732314.6 | | 0.221032 | 0.585631 | 0.766353 |
| 19084 | ENSG0000 | MIR659 | microRNA | 0.192031 | 0.201931 | 0.766194 |
| 32166 | ENSG0000 | AP003084.1 | | 0.119922 | 0.687762 | 0.766138 |
| 1548 | ENSG0000 | CROCCP3 | ciliary root | 0.465396 | 0.526565 | 0.765388 |
| 27139 | ENSG0000 | AC245052.4 | | 0.45295 | 0.71494 | 0.765376 |
| 20783 | ENSG0000 | PPP1R3G | protein ph | 0.07197 | 0.185845 | 0.76537 |
| 33337 | ENSG0000 | AC005520.2 | | 0.181105 | 0.217924 | 0.763206 |
| 8688 | ENSG0000 | KLHL3 | kelch like f | 0.437006 | 0.531598 | 0.762786 |
| 27048 | ENSG0000 | ELOA-AS1 | ELOA antis | 0.321118 | 0.419502 | 0.76222 |
| 30279 | ENSG0000 | AC107982.1 | | 0.214566 | 0.76126 | 0.761966 |
| 29780 | ENSG0000 | AC092903.2 | | 0.155485 | 0.592301 | 0.761593 |
| 31730 | ENSG0000 | HMGN1P3 | high mobil | 0.562832 | 0.696119 | 0.76125 |
| 861 | ENSG0000 | SLC6A16 | solute carr | 0.299037 | 0.620213 | 0.760804 |
| 23023 | ENSG0000 | LINC01132 | long interg | 0.107132 | 0.34214 | 0.759714 |
| 33073 | ENSG0000 | AC010186.3 | | 0.1549 | 0.619533 | 0.758745 |
| 15467 | ENSG0000 | AC138393.1 | | 0.258921 | 0.424612 | 0.753846 |
| 23797 | ENSG0000 | AL117328.1 | | 0 | 0.532637 | 0.753745 |
| 20008 | ENSG0000 | FBXO16 | F-box prot | 0.343982 | 0.448197 | 0.753728 |
| 13221 | ENSG0000 | MST1 | macrophag | 0.35108 | 0.634569 | 0.753712 |
| 41943 | ENSG0000 | AC110285.7 | | 0.314236 | 0.511697 | 0.753684 |
| 3444 | ENSG0000 | CPEB3 | cytoplasmi | 0.263806 | 0.343817 | 0.753328 |
| 9166 | ENSG0000 | TMEM218 | transmeml | 0.565073 | 0.727204 | 0.752197 |
| 43373 | ENSG0000 | AP000553.4 | | 0.044307 | 0.363511 | 0.751959 |
| 2496 | ENSG0000 | SNPH | syntaphilin | 0.456239 | 0.600555 | 0.751149 |
| 16347 | ENSG0000 | GREB1 | growth reg | 0.288395 | 0.553415 | 0.75084 |
| 35205 | ENSG0000 | GOLGA8T | golgin A8 f | 0.219693 | 0.558335 | 0.750172 |
| 4516 | ENSG0000 | KCNC4 | potassium | 0.393941 | 0.694614 | 0.749859 |
| 14026 | ENSG0000 | ODF3B | outer dens | 0.305121 | 0.326584 | 0.748816 |
| 9626 | ENSG0000 | LY96 | lymphocyt | 0.058886 | 0.144701 | 0.747884 |
| 23396 | ENSG0000 | AL845321.1 | | 0.101783 | 0.504808 | 0.747835 |
| 35766 | ENSG0000 | AC135050.5 | | 0.311925 | 0.660179 | 0.747395 |
| 42423 | ENSG0000 | AC096887.2 | | 0.234481 | 0.743869 | 0.745803 |
| 22967 | ENSG0000 | AL512353.1 | | 0.116003 | 0.24975 | 0.745462 |
| 34014 | ENSG0000 | ITGB3 | integrin su | 0.176577 | 0.263695 | 0.743805 |
| 8882 | ENSG0000 | ATP6V0D2 | ATPase H+ | 0.271578 | 0.406501 | 0.743334 |
| 2354 | ENSG0000 | ISM2 | isthmin 2 [ | 0.025836 | 0.060043 | 0.743311 |
| 2537 | ENSG0000 | PPP1R16B | protein ph | 0.022222 | 0.313412 | 0.742078 |
| 7354 | ENSG0000 | SLC22A23 | solute carr | 0.572345 | 0.729969 | 0.741204 |
| 1276 | ENSG0000 | SLC24A1 | solute carr | 0.454179 | 0.585853 | 0.740479 |
| 42280 | ENSG0000 | AC240565.1 | | 0.304566 | 0.571403 | 0.740299 |

| | | | | | | |
|---|---|---|---|---|---|---|
| 30266 | ENSG0000 | AP000866.2 | | 0.045476 | 0.117257 | 0.740178 |
| 14570 | ENSG0000 | TMEM102 | transmeml | 0.058407 | 0.266407 | 0.739254 |
| 8764 | ENSG0000 | SSC4D | scavenger | 0.59221 | 0.621439 | 0.738311 |
| 32202 | ENSG0000 | NPIPA2 | nuclear po | 0.354629 | 0.657769 | 0.735038 |
| 8127 | ENSG0000 | CLGN | calmegin [S | 0.119298 | 0.37956 | 0.73466 |
| 35725 | ENSG0000 | AC090617.4 | | 0.178439 | 0.578616 | 0.734243 |
| 3663 | ENSG0000 | BST1 | bone marr | 0.206122 | 0.373482 | 0.733895 |
| 10840 | ENSG0000 | SERPINI1 | serpin fam | 0.397939 | 0.720798 | 0.733835 |
| 9599 | ENSG0000 | FAM167A | family with | 0.141251 | 0.699145 | 0.733782 |
| 9785 | ENSG0000 | C8orf37 | chromosor | 0.581878 | 0.607414 | 0.733321 |
| 3232 | ENSG0000 | RNF32 | ring finger | 0.626043 | 0.672754 | 0.73328 |
| 14703 | ENSG0000 | TP53TG1 | TP53 targe | 0.452411 | 0.536803 | 0.731238 |
| 18831 | ENSG0000 | Y_RNA | Y RNA [Sou | 0.097479 | 0.396193 | 0.730888 |
| 34968 | ENSG0000 | ANKRD20A | ankyrin rep | 0.326317 | 0.536606 | 0.730371 |
| 29493 | ENSG0000 | LRP4-AS1 | LRP4 antis | 0.124929 | 0.250656 | 0.72997 |
| 5744 | ENSG0000 | PRRG2 | proline rich | 0.33329 | 0.672408 | 0.729488 |
| 35666 | ENSG0000 | AC144836.1 | | 0.297988 | 0.425099 | 0.729424 |
| 34734 | ENSG0000 | AC099518.1 | | 0.121847 | 0.342897 | 0.729103 |
| 18325 | ENSG0000 | FBXO48 | F-box prot | 0.297164 | 0.441366 | 0.728672 |
| 35437 | ENSG0000 | AC009065.8 | | 0.391872 | 0.60618 | 0.728601 |
| 14760 | ENSG0000 | TEX19 | testis expr | 0.156023 | 0.165446 | 0.728023 |
| 18705 | ENSG0000 | RNU1-27P | RNA, U1 sr | 0.190493 | 0.507844 | 0.727769 |
| 20015 | ENSG0000 | CPSF1P1 | cleavage ar | 0.204728 | 0.372176 | 0.727049 |
| 12576 | ENSG0000 | SIX2 | SIX homeo | 0.096748 | 0.238582 | 0.726155 |
| 19532 | ENSG0000 | RNA5SP32 | RNA, 5S rib | 0.338777 | 0.486776 | 0.725494 |
| 38000 | ENSG0000 | AC136475.7 | | 0.059953 | 0.227601 | 0.725116 |
| 24053 | ENSG0000 | LRRC37A1 | leucine rich | 0.12633 | 0.316157 | 0.723801 |
| 26396 | ENSG0000 | AC012360.1 | | 0.178594 | 0.316909 | 0.723586 |
| 15639 | ENSG0000 | MARC1 | mitochond | 0.385491 | 0.567547 | 0.723007 |
| 9592 | ENSG0000 | GAL3ST2 | galactose-3 | 0.182681 | 0.697765 | 0.722842 |
| 39283 | ENSG0000 | TBC1D8-AS | TBC1 doma | 0.144855 | 0.204934 | 0.72241 |
| 4098 | ENSG0000 | PRSS16 | protease, s | 0.25715 | 0.535983 | 0.722163 |
| 15389 | ENSG0000 | INPP5J | inositol po | 0.223238 | 0.395866 | 0.71972 |
| 25369 | ENSG0000 | FAM157B | family with | 0.240822 | 0.416218 | 0.719443 |
| 26805 | ENSG0000 | LINC01376 | long interg | 0.32432 | 0.632766 | 0.719165 |
| 43129 | ENSG0000 | AP001273.2 | | 0.027613 | 0.236307 | 0.718928 |
| 36755 | ENSG0000 | AC007448.3 | | 0.087686 | 0.188077 | 0.717895 |
| 26740 | ENSG0000 | GAPDHP14 | glyceraldeh | 0.402828 | 0.710745 | 0.71748 |
| 39142 | ENSG0000 | AC002470.1 | | 0.207746 | 0.420455 | 0.717428 |
| 15419 | ENSG0000 | CDNF | cerebral do | 0.260466 | 0.576895 | 0.717081 |
| 12784 | ENSG0000 | APLN | apelin [Sou | 0.127233 | 0.131751 | 0.716942 |
| 22726 | ENSG0000 | CACTIN-AS | CACTIN ant | 0.519517 | 0.599974 | 0.715861 |
| 12228 | ENSG0000 | LRRC28 | leucine rich | 0.595482 | 0.672529 | 0.715638 |
| 20785 | ENSG0000 | FAM228B | family with | 0.41743 | 0.50126 | 0.71523 |
| 5161 | ENSG0000 | TNFSF10 | TNF superf | 0.123035 | 0.259702 | 0.714902 |
| 13639 | ENSG0000 | RPL23P2 | ribosomal | 0.165918 | 0.531639 | 0.714321 |
| 40703 | ENSG0000 | AC092117.1 | | 0.354495 | 0.68002 | 0.713713 |

| | | | | | | |
|---|---|---|---|---|---|---|
| 23896 | ENSG0000 | MAST4-AS | MAST4 ant | 0.527569 | 0.658084 | 0.713317 |
| 33029 | ENSG0000 | AC018410.2 | | 0.177501 | 0.236811 | 0.713156 |
| 42107 | ENSG0000 | GVQW2 | GVQW mo | 0.261767 | 0.365146 | 0.712076 |
| 26986 | ENSG0000 | PRKG1-AS1 | PRKG1 ant | 0.331884 | 0.486342 | 0.71145 |
| 32906 | ENSG0000 | ABCC6P1 | ATP bindin | 0.103409 | 0.283799 | 0.710395 |
| 81 | ENSG0000 | MEOX1 | mesenchyr | 0.183139 | 0.209914 | 0.709804 |
| 11679 | ENSG0000 | ATF7IP2 | activating | 0.348662 | 0.592113 | 0.709445 |
| 28892 | ENSG0000 | AC011005.4 | | 0.239061 | 0.692279 | 0.708598 |
| 39687 | ENSG0000 | FRMD6-AS | FRMD6 ant | 0.200956 | 0.646943 | 0.708338 |
| 32208 | ENSG0000 | TMEM9B-A | TMEM9B a | 0.439621 | 0.663603 | 0.708262 |
| 38690 | ENSG0000 | AC016877.3 | | 0.092146 | 0.282273 | 0.706997 |
| 5329 | ENSG0000 | NR4A1 | nuclear rec | 0.15365 | 0.3837 | 0.70678 |
| 40761 | ENSG0000 | AL513314.2 | | 0.252332 | 0.697541 | 0.706628 |
| 11540 | ENSG0000 | TCP11L2 | t-complex | 0.062449 | 0.393467 | 0.705679 |
| 32103 | ENSG0000 | STT3A-AS1 | STT3A anti | 0.220605 | 0.580472 | 0.704732 |
| 42343 | ENSG0000 | FLJ30679 | uncharacte | 0.232838 | 0.679463 | 0.704104 |
| 8501 | ENSG0000 | OSBPL10 | oxysterol b | 0.596395 | 0.602014 | 0.703616 |
| 27599 | ENSG0000 | GOLGA2P5 | golgin A2 p | 0.241605 | 0.608587 | 0.702691 |
| 17993 | ENSG0000 | FAM229B | family with | 0.290717 | 0.628843 | 0.702003 |
| 25132 | ENSG0000 | AC098820.1 | | 0.147613 | 0.60279 | 0.701919 |
| 2047 | ENSG0000 | SORBS1 | sorbin and | 0.293005 | 0.565197 | 0.701677 |
| 27162 | ENSG0000 | AL732372.2 | | 0.383792 | 0.612812 | 0.700893 |
| 21948 | ENSG0000 | AL662890.2 | | 0.1129 | 0.393898 | 0.700885 |
| 26088 | ENSG0000 | AC245060.2 | | 0.288895 | 0.573742 | 0.700668 |
| 30742 | ENSG0000 | ZNF345 | zinc finger | 0.323526 | 0.668493 | 0.700261 |
| 35869 | ENSG0000 | AC133552.4 | | 0.342858 | 0.444066 | 0.700242 |
| 24534 | ENSG0000 | AF129075.1 | | 0.260543 | 0.547676 | 0.70011 |
| 23071 | ENSG0000 | CALR4P | calreticulin | 0.086247 | 0.568357 | 0.699799 |
| 20046 | ENSG0000 | GCC2-AS1 | GCC2 antis | 0.242574 | 0.355332 | 0.699426 |
| 37848 | ENSG0000 | AC010319.4 | | 0.50391 | 0.657294 | 0.699306 |
| 38930 | ENSG0000 | CD24 | CD24 mole | 0.333908 | 0.464625 | 0.699253 |
| 24222 | ENSG0000 | SERPINB9P | serpin fam | 0.243404 | 0.501034 | 0.699166 |
| 11316 | ENSG0000 | NIPSNAP3I | nipsnap ho | 0.154028 | 0.315757 | 0.698793 |
| 38340 | ENSG0000 | LRRC37A9I | leucine rich | 0.203045 | 0.378192 | 0.698108 |
| 30415 | ENSG0000 | AC122718.1 | | 0.396804 | 0.627183 | 0.697737 |
| 18271 | ENSG0000 | AL133464.1 | | 0.342336 | 0.520383 | 0.697725 |
| 37632 | ENSG0000 | AC137695.2 | | 0.057087 | 0.094204 | 0.696989 |
| 12291 | ENSG0000 | MN1 | MN1 protc | 0.184706 | 0.275758 | 0.696223 |
| 20000 | ENSG0000 | ZNF891 | zinc finger | 0.295915 | 0.589002 | 0.695506 |
| 13138 | ENSG0000 | LRRN3 | leucine rich | 0.440327 | 0.694633 | 0.695355 |
| 5007 | ENSG0000 | PCDHB14 | protocadh | 0.125363 | 0.505697 | 0.695298 |
| 12300 | ENSG0000 | RSPO1 | R-spondin | 0.125899 | 0.161194 | 0.694634 |
| 9193 | ENSG0000 | PPP1R1C | protein ph | 0.258604 | 0.368352 | 0.692788 |
| 9115 | ENSG0000 | SPATA25 | spermatog | 0.231766 | 0.552166 | 0.692753 |
| 22869 | ENSG0000 | AC104667.1 | | 0.049536 | 0.50178 | 0.692286 |
| 3077 | ENSG0000 | CLIP3 | CAP-Gly dc | 0.335188 | 0.48396 | 0.692224 |
| 25457 | ENSG0000 | AC133785.1 | | 0.035498 | 0.170801 | 0.692149 |

| | | | | | | |
|---|---|---|---|---|---|---|
| 28733 | ENSG0000 | HLA-DMB | major histo | 0.050474 | 0.266917 | 0.692128 |
| 15820 | ENSG0000 | CMC1 | C-X9-C mo | 0.520565 | 0.625404 | 0.690644 |
| 20160 | ENSG0000 | AC068781.1 | | 0.22372 | 0.520318 | 0.689982 |
| 13002 | ENSG0000 | CFAP53 | cilia and fla | 0.189502 | 0.569312 | 0.689002 |
| 24373 | ENSG0000 | SNAP23P1 | synaptosor | 0.216927 | 0.637823 | 0.688752 |
| 22837 | ENSG0000 | ITGB2-AS1 | ITGB2 antis | 0.019474 | 0.121733 | 0.686937 |
| 35016 | ENSG0000 | AC007225.1 | | 0.116193 | 0.510885 | 0.686459 |
| 21795 | ENSG0000 | NPIPA3 | nuclear po | 0.344572 | 0.616115 | 0.68602 |
| 26320 | ENSG0000 | AC131097.4 | | 0.062599 | 0.24943 | 0.686001 |
| 5937 | ENSG0000 | CDHR3 | cadherin re | 0.388301 | 0.6708 | 0.685518 |
| 18761 | ENSG0000 | SNORA7 | Small nucle | 0.421943 | 0.474664 | 0.684261 |
| 2173 | ENSG0000 | PRODH | proline deh | 0.244741 | 0.416114 | 0.684163 |
| 28768 | ENSG0000 | AC004893.2 | | 0.368314 | 0.679902 | 0.684156 |
| 11927 | ENSG0000 | TMC4 | transmeml | 0.194661 | 0.439703 | 0.683785 |
| 20332 | ENSG0000 | AC025171.2 | | 0.423507 | 0.541086 | 0.683495 |
| 37283 | ENSG0000 | AC011815.1 | | 0.297596 | 0.597318 | 0.683397 |
| 14982 | ENSG0000 | MEX3B | mex-3 RNA | 0.206755 | 0.316833 | 0.682988 |
| 15217 | ENSG0000 | ZDHHC23 | zinc finger | 0.39558 | 0.514427 | 0.681812 |
| 14605 | ENSG0000 | TMIE | transmeml | 0.019803 | 0.214547 | 0.681221 |
| 18292 | ENSG0000 | TTC25 | tetratricop | 0.256214 | 0.504677 | 0.680607 |
| 20978 | ENSG0000 | C1orf229 | chromosor | 0.11053 | 0.230791 | 0.68052 |
| 23385 | ENSG0000 | AC106875.1 | | 0.045571 | 0.118497 | 0.68008 |
| 29886 | ENSG0000 | KRT8P46 | keratin 8 p | 0.100528 | 0.373904 | 0.679311 |
| 18070 | ENSG0000 | MAFB | MAF bZIP t | 0.120851 | 0.302467 | 0.679266 |
| 18515 | ENSG0000 | LOH12CR2 | loss of hete | 0.145146 | 0.632269 | 0.67818 |
| 362 | ENSG0000 | CLDN11 | claudin 11 | 0.524236 | 0.526574 | 0.677707 |
| 23435 | ENSG0000 | AC245100.3 | | 0 | 0.398869 | 0.677227 |
| 8934 | ENSG0000 | NR6A1 | nuclear rec | 0.426071 | 0.545958 | 0.675391 |
| 1232 | ENSG0000 | ALPK1 | alpha kinas | 0.329692 | 0.494384 | 0.674365 |
| 13523 | ENSG0000 | ZNF519 | zinc finger | 0.401935 | 0.511556 | 0.673137 |
| 29411 | ENSG0000 | AC090515.2 | | 0.169907 | 0.61244 | 0.672927 |
| 32933 | ENSG0000 | TAS2R31 | taste 2 rec | 0.106875 | 0.237281 | 0.672681 |
| 28097 | ENSG0000 | LINC02004 | long interg | 0 | 0.198468 | 0.672028 |
| 6784 | ENSG0000 | GRHL1 | grainyhead | 0.264142 | 0.345021 | 0.670898 |
| 16863 | ENSG0000 | ELOVL2 | ELOVL fatty | 0.10621 | 0.353679 | 0.670252 |
| 31275 | ENSG0000 | RNA5SP49 | RNA, 5S rib | 0.056451 | 0.331256 | 0.669793 |
| 33167 | ENSG0000 | AL132780.1 | | 0.222165 | 0.284558 | 0.668809 |
| 65 | ENSG0000 | ARX | aristaless r | 0.108506 | 0.584236 | 0.667389 |
| 22428 | ENSG0000 | AC004383.1 | | 0.229507 | 0.490485 | 0.667358 |
| 22606 | ENSG0000 | AC079145.1 | | 0.172387 | 0.304029 | 0.667243 |
| 132 | ENSG0000 | DLX6 | distal-less l | 0.046417 | 0.203964 | 0.667197 |
| 15946 | ENSG0000 | SOWAHD | sosondowa | 0.011434 | 0.11166 | 0.666941 |
| 22903 | ENSG0000 | FTH1P16 | ferritin hea | 0.209308 | 0.365301 | 0.66612 |
| 19174 | ENSG0000 | SNORD83B | small nucle | 0.001256 | 0.167209 | 0.665128 |
| 5807 | ENSG0000 | FGD3 | FYVE, RhoG | 0.197404 | 0.337167 | 0.665016 |
| 36815 | ENSG0000 | AL135905.2 | | 0.165032 | 0.484635 | 0.664944 |
| 30379 | ENSG0000 | LINC01021 | long interg | 0.091284 | 0.20837 | 0.664058 |

| | | | | | | |
|---|---|---|---|---|---|---|
| 1299 | ENSG0000 | WNT8B | Wnt family | 0 | 0.280927 | 0.663873 |
| 3837 | ENSG0000 | OAS2 | 2'-5'-oligoa | 0.173819 | 0.231908 | 0.663849 |
| 12206 | ENSG0000 | IL12A | interleukin | 0.188094 | 0.560876 | 0.663586 |
| 35487 | ENSG0000 | AC005592.1 | | 0.094471 | 0.414438 | 0.662743 |
| 7558 | ENSG0000 | CARF | calcium res | 0.270443 | 0.373649 | 0.662721 |
| 29450 | ENSG0000 | AP001528.1 | | 0.459794 | 0.529442 | 0.662254 |
| 16805 | ENSG0000 | KLHDC1 | kelch doma | 0.292802 | 0.308803 | 0.662038 |
| 16080 | ENSG0000 | TCTEX1D4 | Tctex1 dor | 0.165117 | 0.388409 | 0.661598 |
| 33151 | ENSG0000 | AC124947.1 | | 0.281399 | 0.315828 | 0.660668 |
| 7602 | ENSG0000 | HERC6 | HECT and F | 0.343792 | 0.57088 | 0.660645 |
| 17961 | ENSG0000 | SLCO1B7 | solute carr | 0.47372 | 0.604234 | 0.660017 |
| 15599 | ENSG0000 | AC021092.1 | | 0.098227 | 0.270125 | 0.659 |
| 2167 | ENSG0000 | UPB1 | beta-ureid | 0.043905 | 0.312064 | 0.658693 |
| 14470 | ENSG0000 | PCGF5 | polycomb g | 0.567841 | 0.602503 | 0.658157 |
| 20127 | ENSG0000 | AGGF1P1 | angiogenic | 0.304203 | 0.632147 | 0.657994 |
| 345 | ENSG0000 | CD22 | CD22 mole | 0.403453 | 0.482286 | 0.657634 |
| 7591 | ENSG0000 | APH1B | aph-1 hom | 0.171479 | 0.363386 | 0.656754 |
| 24166 | ENSG0000 | AC010422.1 | | 0.052278 | 0.162629 | 0.656737 |
| 28664 | ENSG0000 | DEFB109A | defensin be | 0 | 0.163955 | 0.656106 |
| 14550 | ENSG0000 | MAPK15 | mitogen-ac | 0.173875 | 0.37467 | 0.655875 |
| 35029 | ENSG0000 | AC092803.2 | | 0.176427 | 0.60568 | 0.655829 |
| 22043 | ENSG0000 | RAB28P5 | RAB28, me | 0.080812 | 0.477607 | 0.655731 |
| 16489 | ENSG0000 | ZNF136 | zinc finger | 0.421892 | 0.471763 | 0.655694 |
| 11485 | ENSG0000 | PPP1R36 | protein ph | 0.15323 | 0.50493 | 0.655052 |
| 18502 | ENSG0000 | LINC00634 | long interg | 0.116342 | 0.375237 | 0.654887 |
| 34999 | ENSG0000 | AC092720.1 | | 0.224598 | 0.331934 | 0.654102 |
| 14166 | ENSG0000 | ADORA2A- | ADORA2A | 0.183983 | 0.640591 | 0.653631 |
| 14718 | ENSG0000 | UBE2E2 | ubiquitin c | 0.301652 | 0.558287 | 0.652551 |
| 22297 | ENSG0000 | UBXN7-AS1 | UBXN7 ant | 0.023476 | 0.380849 | 0.652516 |
| 22505 | ENSG0000 | TMEM191A | transmeml | 0.234551 | 0.41143 | 0.652251 |
| 33934 | ENSG0000 | AC002094.1 | | 0 | 0.43429 | 0.652127 |
| 30257 | ENSG0000 | RARRES2P4 | retinoic aci | 0.453246 | 0.513219 | 0.651958 |
| 39997 | ENSG0000 | HOTAIRM1 | HOX antise | 0.006642 | 0.027385 | 0.651571 |
| 23452 | ENSG0000 | AC090587.1 | | 0 | 0.442193 | 0.650619 |
| 24828 | ENSG0000 | PCAT7 | prostate ca | 0.0306 | 0.45771 | 0.649252 |
| 40822 | ENSG0000 | HIST1H4D | histone clu | 0.420738 | 0.462918 | 0.647984 |
| 33006 | ENSG0000 | AC022613.1 | | 0.097476 | 0.222648 | 0.646208 |
| 40862 | ENSG0000 | MIR6833 | microRNA | 0.172282 | 0.267868 | 0.644966 |
| 33501 | ENSG0000 | AC025031.2 | | 0.372566 | 0.637619 | 0.644608 |
| 12528 | ENSG0000 | LRRN2 | leucine rich | 0.062608 | 0.105214 | 0.643783 |
| 20510 | ENSG0000 | LINC01615 | long interg | 0.155285 | 0.193187 | 0.643537 |
| 23447 | ENSG0000 | HNRNPCP7 | heterogene | 0.379383 | 0.510215 | 0.643525 |
| 40370 | ENSG0000 | C18orf65 | chromosom | 0.168539 | 0.233345 | 0.643515 |
| 4116 | ENSG0000 | NNT | nicotinami | 0.312352 | 0.354204 | 0.643357 |
| 26219 | ENSG0000 | AC104667.2 | | 0.059461 | 0.305607 | 0.642732 |
| 9306 | ENSG0000 | ANKAR | ankyrin an | 0.260367 | 0.435548 | 0.642439 |
| 39271 | ENSG0000 | AL645608.9 | | 0.259612 | 0.381182 | 0.640908 |

| | | | | | | |
|---|---|---|---|---|---|---|
| 8886 | ENSG0000 | LRP12 | LDL recept | 0.420745 | 0.610512 | 0.640261 |
| 15359 | ENSG0000 | F8 | coagulatio | 0.365866 | 0.472027 | 0.639507 |
| 39119 | ENSG0000 | AC253536.6 | | 0.376057 | 0.38593 | 0.638628 |
| 37566 | ENSG0000 | AC005523.1 | | 0.097555 | 0.234298 | 0.63728 |
| 35452 | ENSG0000 | AC134682.1 | | 0.062687 | 0.365443 | 0.636938 |
| 39789 | ENSG0000 | MIR6864 | microRNA | 0.152828 | 0.237846 | 0.636707 |
| 23359 | ENSG0000 | AL139260.1 | | 0.363547 | 0.468759 | 0.63661 |
| 35486 | ENSG0000 | AC008555.1 | | 0.122389 | 0.199414 | 0.636555 |
| 36104 | ENSG0000 | AGAP7P | ArfGAP wit | 0.229577 | 0.482653 | 0.635852 |
| 17004 | ENSG0000 | ATL1 | atlastin GT | 0.101239 | 0.269333 | 0.634987 |
| 40799 | ENSG0000 | AL445670.1 | | 0.432255 | 0.555224 | 0.634724 |
| 38730 | ENSG0000 | AL121944.1 | | 0.067983 | 0.581427 | 0.634631 |
| 43132 | ENSG0000 | MIR4469 | microRNA | 0.116292 | 0.296816 | 0.634287 |
| 27952 | ENSG0000 | MGC1627 | uncharacte | 0.1117 | 0.283675 | 0.633155 |
| 21494 | ENSG0000 | AL354761.1 | | 0 | 0.187431 | 0.633127 |
| 42811 | ENSG0000 | AL831711.1 | | 0.275739 | 0.307642 | 0.633094 |
| 36556 | ENSG0000 | AC006213.1 | | 0.404131 | 0.588617 | 0.632932 |
| 16074 | ENSG0000 | JAKMIP3 | Janus kina | 0.376249 | 0.55507 | 0.632771 |
| 37934 | ENSG0000 | AC068620.2 | | 0.184862 | 0.475115 | 0.632327 |
| 16367 | ENSG0000 | ZNF836 | zinc finger | 0.216479 | 0.371281 | 0.63226 |
| 34539 | ENSG0000 | AC009120.2 | | 0.526938 | 0.613957 | 0.632246 |
| 37028 | ENSG0000 | AC005306.1 | | 0.227327 | 0.283435 | 0.63191 |
| 22842 | ENSG0000 | C14orf132 | chromosor | 0.070998 | 0.209603 | 0.630019 |
| 35522 | ENSG0000 | AC087500.1 | | 0.426768 | 0.486027 | 0.629359 |
| 36957 | ENSG0000 | AC067852.4 | | 0.193001 | 0.480085 | 0.629141 |
| 11695 | ENSG0000 | SLFN5 | schlafen fa | 0.305485 | 0.50643 | 0.628379 |
| 34837 | ENSG0000 | LCMT1-AS | LCMT1 ant | 0.202904 | 0.292414 | 0.627707 |
| 1340 | ENSG0000 | FMO4 | flavin cont | 0.208191 | 0.346696 | 0.627452 |
| 8423 | ENSG0000 | DQX1 | DEAQ-box | 0.057566 | 0.086261 | 0.627214 |
| 37887 | ENSG0000 | NOP53-AS | NOP53 ant | 0.300676 | 0.538346 | 0.626797 |
| 23301 | ENSG0000 | AC008440.1 | | 0.073138 | 0.219779 | 0.626632 |
| 41259 | ENSG0000 | AL845472.1 | | 0.259579 | 0.399295 | 0.626072 |
| 29996 | ENSG0000 | AC010226.1 | | 0.078588 | 0.205513 | 0.625971 |
| 13373 | ENSG0000 | ZCCHC12 | zinc finger | 0.048121 | 0.293008 | 0.625168 |
| 42944 | ENSG0000 | AC098484.3 | | 0.296707 | 0.580545 | 0.623903 |
| 11614 | ENSG0000 | WDR88 | WD repeat | 0.313612 | 0.480424 | 0.623283 |
| 339 | ENSG0000 | ZBTB32 | zinc finger | 0.063148 | 0.392588 | 0.623189 |
| 43300 | ENSG0000 | AC009477.2 | | 0.019382 | 0.342064 | 0.622865 |
| 11994 | ENSG0000 | AC011511.1 | | 0.31408 | 0.612732 | 0.622089 |
| 28345 | ENSG0000 | AL157400.4 | | 0.121647 | 0.411591 | 0.621882 |
| 13655 | ENSG0000 | AL445665.1 | | 0.07891 | 0.400918 | 0.621617 |
| 22381 | ENSG0000 | BX005266.2 | | 0.36859 | 0.590899 | 0.621567 |
| 10866 | ENSG0000 | CCDC191 | coiled-coil | 0.335949 | 0.454804 | 0.621421 |
| 41426 | ENSG0000 | AC008567.3 | | 0.448726 | 0.504848 | 0.620696 |
| 16320 | ENSG0000 | HLA-DRB1 | major histo | 0.071992 | 0.257266 | 0.618746 |
| 11319 | ENSG0000 | LETM2 | leucine zip | 0.314102 | 0.484947 | 0.618331 |
| 9919 | ENSG0000 | ZFYVE28 | zinc finger | 0.380132 | 0.396152 | 0.617081 |

| | | | | | | |
|---|---|---|---|---|---|---|
| 33782 | ENSG0000 | TRIM34 | tripartite m | 0.020717 | 0.231553 | 0.616779 |
| 40013 | ENSG0000 | AL139384.1 | | 0.239119 | 0.343276 | 0.61621 |
| 7222 | ENSG0000 | HLX | H2.0 like h | 0.172733 | 0.454465 | 0.614813 |
| 18132 | ENSG0000 | DFNB59 | deafness, a | 0.395514 | 0.433437 | 0.614787 |
| 41759 | ENSG0000 | AP003108.3 | | 0.305133 | 0.362829 | 0.614563 |
| 20788 | ENSG0000 | AC008770.1 | | 0.355499 | 0.474769 | 0.614239 |
| 14079 | ENSG0000 | LINC01658 | long interg | 0.096388 | 0.323966 | 0.614035 |
| 3883 | ENSG0000 | PTPN6 | protein tyr | 0.230752 | 0.382866 | 0.613863 |
| 10064 | ENSG0000 | RIBC1 | RIB43A doi | 0.092106 | 0.256776 | 0.61136 |
| 6025 | ENSG0000 | SLC41A2 | solute carr | 0.45558 | 0.581887 | 0.610615 |
| 8762 | ENSG0000 | AC004453.1 | | 0.156398 | 0.458098 | 0.610246 |
| 27761 | ENSG0000 | RNU7-59P | RNA, U7 sr | 0.125858 | 0.211252 | 0.610128 |
| 11965 | ENSG0000 | HSD11B1L | hydroxyste | 0.538152 | 0.550106 | 0.609993 |
| 34321 | ENSG0000 | LINC01220 | long interg | 0.102571 | 0.330954 | 0.609964 |
| 4282 | ENSG0000 | HEMK1 | HemK met | 0.433684 | 0.581817 | 0.609928 |
| 13324 | ENSG0000 | TLR1 | toll like rec | 0.291313 | 0.361078 | 0.609569 |
| 10306 | ENSG0000 | FTCD | formimido | 0.072792 | 0.314945 | 0.609232 |
| 20340 | ENSG0000 | TTC3P1 | tetratricop | 0.268326 | 0.590473 | 0.609031 |
| 9203 | ENSG0000 | IL18 | interleukin | 0.162734 | 0.355146 | 0.608557 |
| 30113 | ENSG0000 | AC106897.1 | | 0.116944 | 0.25014 | 0.608461 |
| 25053 | ENSG0000 | AC008440.3 | | 0 | 0.379742 | 0.607887 |
| 5895 | ENSG0000 | ASPHD2 | aspartate b | 0.254704 | 0.430526 | 0.607862 |
| 4360 | ENSG0000 | IFIH1 | interferon | 0.228174 | 0.239229 | 0.607519 |
| 8137 | ENSG0000 | ZNF599 | zinc finger | 0.397756 | 0.455818 | 0.606917 |
| 32097 | ENSG0000 | RTL1 | retrotransp | 0.078193 | 0.112163 | 0.606868 |
| 3676 | ENSG0000 | CRYAB | crystallin a | 0 | 0.204174 | 0.606676 |
| 37879 | ENSG0000 | RPARP-AS1 | RPARP ant | 0.461843 | 0.470229 | 0.606597 |
| 18678 | ENSG0000 | H1FX-AS1 | H1FX antis | 0.278237 | 0.359757 | 0.606585 |
| 1867 | ENSG0000 | ANKRD24 | ankyrin rep | 0.266106 | 0.392263 | 0.606476 |
| 6941 | ENSG0000 | MRAP2 | melanocor | 0.197527 | 0.318897 | 0.604174 |
| 37113 | ENSG0000 | AC011477.1 | | 0.330366 | 0.556095 | 0.603515 |
| 1478 | ENSG0000 | CRYBG3 | crystallin b | 0.178509 | 0.342851 | 0.603031 |
| 41130 | ENSG0000 | AC114341.1 | | 0.098451 | 0.586348 | 0.602985 |
| 35079 | ENSG0000 | AC106886.2 | | 0.304746 | 0.557795 | 0.602273 |
| 7723 | ENSG0000 | SLC46A3 | solute carr | 0.329886 | 0.431308 | 0.601272 |
| 42520 | ENSG0000 | TRG-AS1 | T-cell recep | 0.29496 | 0.37255 | 0.600383 |
| 23343 | ENSG0000 | AL022068.1 | | 0.292511 | 0.576333 | 0.599912 |
| 39785 | ENSG0000 | AC084018.2 | | 0.476023 | 0.526539 | 0.599538 |
| 5239 | ENSG0000 | WIPF3 | WAS/WASI | 0.259434 | 0.278576 | 0.598586 |
| 24094 | ENSG0000 | AC018463.1 | | 0.466456 | 0.535447 | 0.59744 |
| 5078 | ENSG0000 | PTK2B | protein tyr | 0.430976 | 0.446534 | 0.597148 |
| 16622 | ENSG0000 | ZNF682 | zinc finger | 0.163738 | 0.456063 | 0.596665 |
| 16121 | ENSG0000 | NANOS1 | nanos C2H | 0.175619 | 0.46228 | 0.596311 |
| 5342 | ENSG0000 | SARDH | sarcosine d | 0.071556 | 0.164131 | 0.595558 |
| 27494 | ENSG0000 | AL139039.3 | | 0.074227 | 0.172862 | 0.595449 |
| 36986 | ENSG0000 | AC104971.2 | | 0.075438 | 0.340593 | 0.595379 |
| 19504 | ENSG0000 | SNORA16 | Small nucle | 0 | 0.023053 | 0.593415 |

| | | | | | | |
|---|---|---|---|---|---|---|
| 15913 | ENSG0000 | TRPV2 | transient r | 0.125934 | 0.127535 | 0.592614 |
| 6786 | ENSG0000 | RSAD2 | radical S-a | 0.035413 | 0.201607 | 0.592243 |
| 13402 | ENSG0000 | SLCO2A1 | solute carr | 0.062707 | 0.132466 | 0.592098 |
| 41564 | ENSG0000 | CR381670.1 | | 0.16613 | 0.331191 | 0.591877 |
| 13915 | ENSG0000 | FLJ13224 | uncharacte | 0.105741 | 0.411275 | 0.591421 |
| 26340 | ENSG0000 | OR6E1P | olfactory r | 0.018136 | 0.33817 | 0.591223 |
| 5549 | ENSG0000 | LRRC29 | leucine rich | 0.355665 | 0.404062 | 0.591115 |
| 15372 | ENSG0000 | AC000068.1 | | 0.251955 | 0.586499 | 0.591001 |
| 19915 | ENSG0000 | ZNF321P | zinc finger | 0.198275 | 0.457348 | 0.58975 |
| 37714 | ENSG0000 | LINC01711 | long interg | 0 | 0.1855 | 0.589554 |
| 41380 | ENSG0000 | AC006213.5 | | 0 | 0.146761 | 0.588708 |
| 37910 | ENSG0000 | AC112491.1 | | 0.45965 | 0.51135 | 0.588616 |
| 1866 | ENSG0000 | ARHGAP4 | Rho GTPas | 0.27093 | 0.400074 | 0.588445 |
| 22309 | ENSG0000 | RUSC1-AS1 | RUSC1 anti | 0.408642 | 0.415157 | 0.588409 |
| 16823 | ENSG0000 | FAM212B | family with | 0.21253 | 0.226872 | 0.587836 |
| 10788 | ENSG0000 | INAVA | innate imm | 0.036155 | 0.165332 | 0.587792 |
| 17076 | ENSG0000 | PPP1R14C | protein ph | 0.181711 | 0.392148 | 0.587721 |
| 36851 | ENSG0000 | AC011498.1 | | 0.044338 | 0.306838 | 0.587662 |
| 38016 | ENSG0000 | AL606834.3 | | 0.449893 | 0.482823 | 0.58755 |
| 10650 | ENSG0000 | SNED1 | sushi, nido | 0.241095 | 0.313423 | 0.587183 |
| 35131 | ENSG0000 | AL592424.1 | | 0.061805 | 0.254124 | 0.586163 |
| 42056 | ENSG0000 | AC091152.4 | | 0.036882 | 0.478085 | 0.585992 |
| 21637 | ENSG0000 | AL162586.1 | | 0.452909 | 0.498016 | 0.58585 |
| 36517 | ENSG0000 | AP005136.2 | | 0.201925 | 0.375904 | 0.585081 |
| 6048 | ENSG0000 | PALLD | palladin, cy | 0.335034 | 0.380536 | 0.584868 |
| 20455 | ENSG0000 | AC114730.1 | | 0.401134 | 0.558669 | 0.58424 |
| 27320 | ENSG0000 | TTN-AS1 | TTN antise | 0.315551 | 0.463514 | 0.583778 |
| 8843 | ENSG0000 | ZNF185 | zinc finger | 0.446429 | 0.534436 | 0.583763 |
| 9875 | ENSG0000 | EXOG | exo/endon | 0.270096 | 0.499233 | 0.582975 |
| 27968 | ENSG0000 | Z84485.1 | | 0.181184 | 0.511868 | 0.582973 |
| 40598 | ENSG0000 | AC025287.3 | | 0.422543 | 0.426768 | 0.582374 |
| 39630 | ENSG0000 | AC007686.3 | | 0.351638 | 0.433835 | 0.58119 |
| 38976 | ENSG0000 | AC254633.1 | | 0.059828 | 0.213652 | 0.580343 |
| 4457 | ENSG0000 | WIPF1 | WAS/WASL | 0.266874 | 0.380495 | 0.579492 |
| 28349 | ENSG0000 | AC114490.1 | | 0.427981 | 0.528666 | 0.578468 |
| 26874 | ENSG0000 | AL445183.3 | | 0.427604 | 0.498512 | 0.577464 |
| 33626 | ENSG0000 | AL158801.2 | | 0 | 0.34869 | 0.577005 |
| 15782 | ENSG0000 | ZACN | zinc activat | 0.240619 | 0.399654 | 0.576998 |
| 38447 | ENSG0000 | AC112220.3 | | 0.249962 | 0.420989 | 0.57618 |
| 36840 | ENSG0000 | AL050343.1 | | 0.1926 | 0.379111 | 0.576009 |
| 20326 | ENSG0000 | GRID2IP | Grid2 inter | 0.04789 | 0.266962 | 0.575957 |
| 26068 | ENSG0000 | AL450992.2 | | 0.127939 | 0.271174 | 0.575711 |
| 33657 | ENSG0000 | AL132780.2 | | 0.209737 | 0.2835 | 0.575457 |
| 4919 | ENSG0000 | FLVCR2 | feline leuk | 0.224702 | 0.447183 | 0.574696 |
| 12085 | ENSG0000 | TTC39C | tetratricop | 0.115735 | 0.505978 | 0.574338 |
| 13725 | ENSG0000 | B3GNT5 | UDP-GlcNA | 0.096682 | 0.268357 | 0.573743 |
| 3157 | ENSG0000 | JAK3 | Janus kinas | 0.091056 | 0.255047 | 0.572983 |

| | | | | | | |
|---|---|---|---|---|---|---|
| 16174 | ENSG0000 | LINC00910 | long interg | 0.138378 | 0.14036 | 0.572591 |
| 15798 | ENSG0000 | C17orf82 | chromosor | 0.119113 | 0.14864 | 0.572247 |
| 39108 | ENSG0000 | JMJD1C-AS | JMJD1C an | 0.379558 | 0.488945 | 0.571678 |
| 9603 | ENSG0000 | OBSCN | obscurin, c | 0.07659 | 0.254798 | 0.571438 |
| 20394 | ENSG0000 | TMCO5B | transmeml | 0.133927 | 0.251319 | 0.571303 |
| 13162 | ENSG0000 | GLRX | glutaredox | 0.114762 | 0.338643 | 0.571268 |
| 4475 | ENSG0000 | CD207 | CD207 mol | 0.155426 | 0.249791 | 0.570785 |
| 607 | ENSG0000 | SPATA7 | spermatog | 0.356565 | 0.538732 | 0.570706 |
| 6453 | ENSG0000 | RTBDN | retbindin [ | 0.082914 | 0.293943 | 0.569892 |
| 41219 | ENSG0000 | AL928654.5 | | 0.440733 | 0.466156 | 0.569251 |
| 17924 | ENSG0000 | IQANK1 | IQ motif ar | 0.169244 | 0.426368 | 0.568338 |
| 41542 | ENSG0000 | AL358852.1 | | 0.134986 | 0.465968 | 0.567633 |
| 1191 | ENSG0000 | SPEG | SPEG comp | 0.057586 | 0.165475 | 0.567404 |
| 30815 | ENSG0000 | AC138956.2 | | 0.20166 | 0.24019 | 0.566853 |
| 35977 | ENSG0000 | RN7SL4P | RNA, 7SL, c | 0.273689 | 0.555119 | 0.566756 |
| 39724 | ENSG0000 | AC121761.2 | | 0.446076 | 0.495256 | 0.56668 |
| 221 | ENSG0000 | ST3GAL1 | ST3 beta-g | 0.186616 | 0.275367 | 0.566365 |
| 23108 | ENSG0000 | SUCLA2-AS | SUCLA2 an | 0.21474 | 0.468765 | 0.565921 |
| 6135 | ENSG0000 | TNNI3 | troponin I3 | 0.087588 | 0.167485 | 0.565918 |
| 34220 | ENSG0000 | AC022613.2 | | 0.24391 | 0.50359 | 0.56572 |
| 8084 | ENSG0000 | NTN5 | netrin 5 [S | 0.087775 | 0.423211 | 0.565535 |
| 10273 | ENSG0000 | ABCG1 | ATP bindin | 0.113832 | 0.324767 | 0.564865 |
| 10408 | ENSG0000 | CELF5 | CUGBP Ela | 0.046872 | 0.222938 | 0.563003 |
| 14221 | ENSG0000 | TMTC2 | transmeml | 0.507241 | 0.549853 | 0.562757 |
| 15285 | ENSG0000 | KRBA2 | KRAB-A do | 0.217082 | 0.449143 | 0.561133 |
| 7356 | ENSG0000 | LRRC1 | leucine ricl | 0.192424 | 0.435981 | 0.561062 |
| 18568 | ENSG0000 | RNU1-73P | RNA, U1 sr | 0 | 0.057832 | 0.55964 |
| 10546 | ENSG0000 | ZSWIM5 | zinc finger | 0.14324 | 0.281949 | 0.558989 |
| 25190 | ENSG0000 | AL683813.1 | | 0.164535 | 0.539595 | 0.558663 |
| 9579 | ENSG0000 | RETREG1 | reticulopha | 0.248066 | 0.428947 | 0.557605 |
| 24143 | ENSG0000 | HERC2P4 | hect doma | 0.31215 | 0.530347 | 0.55757 |
| 7045 | ENSG0000 | KIAA1614 | KIAA1614 \| | 0.260171 | 0.293596 | 0.557287 |
| 11025 | ENSG0000 | SLC51A | solute carr | 0.204985 | 0.262293 | 0.556763 |
| 37825 | ENSG0000 | AC008764.6 | | 0.421738 | 0.523826 | 0.556583 |
| 5418 | ENSG0000 | PARD6B | par-6 famil | 0.213315 | 0.337195 | 0.556071 |
| 13637 | ENSG0000 | NUPR1 | nuclear pro | 0.094959 | 0.138539 | 0.556051 |
| 14058 | ENSG0000 | ZNF114 | zinc finger | 0.005046 | 0.053063 | 0.555808 |
| 16778 | ENSG0000 | AL157838.1 | | 0.19347 | 0.540224 | 0.555457 |
| 42344 | ENSG0000 | AC240565.2 | | 0.231221 | 0.388952 | 0.555373 |
| 16124 | ENSG0000 | GOLGA8M | golgin A8 f | 0.017557 | 0.371614 | 0.555235 |
| 25519 | ENSG0000 | AL356124.2 | | 0.136803 | 0.218488 | 0.55496 |
| 10249 | ENSG0000 | TNFRSF13C | TNF recept | 0.396063 | 0.514445 | 0.554578 |
| 30026 | ENSG0000 | UGDH-AS1 | UGDH anti | 0.195847 | 0.307314 | 0.554347 |
| 30607 | ENSG0000 | ZBED3-AS1 | ZBED3 anti | 0.22369 | 0.269936 | 0.553634 |
| 20993 | ENSG0000 | ZNF630 | zinc finger | 0.30177 | 0.492346 | 0.553472 |
| 21543 | ENSG0000 | HCG14 | HLA compl | 0 | 0.256355 | 0.55182 |
| 26919 | ENSG0000 | AL135791.1 | | 0.331397 | 0.530086 | 0.551414 |

| | | | | | | |
|---|---|---|---|---|---|---|
| 8426 | ENSG0000 | NPHP1 | nephrocyst | 0.302251 | 0.476322 | 0.550304 |
| 41918 | ENSG0000 | AC005703.6 | | 0.361192 | 0.489003 | 0.549802 |
| 15673 | ENSG0000 | CYP4X1 | cytochrom | 0.155717 | 0.290762 | 0.5489 |
| 14753 | ENSG0000 | IFNL1 | interferon | 0 | 0.169613 | 0.548413 |
| 32093 | ENSG0000 | AP000911.3 | | 0.024114 | 0.18976 | 0.548326 |
| 41094 | ENSG0000 | AC010332.1 | | 0.208899 | 0.363476 | 0.548296 |
| 11917 | ENSG0000 | ZNF701 | zinc finger | 0.187258 | 0.409269 | 0.547687 |
| 5243 | ENSG0000 | HOXA7 | homeobox | 0.152729 | 0.351414 | 0.54753 |
| 42497 | ENSG0000 | FOXCUT | FOXC1 ups | 0.446729 | 0.529777 | 0.547443 |
| 6218 | ENSG0000 | CRB3 | crumbs 3, | 0.033362 | 0.458222 | 0.547366 |
| 21001 | ENSG0000 | AC005481.1 | | 0.206875 | 0.35781 | 0.547343 |
| 9975 | ENSG0000 | TSPAN18 | tetraspanin | 0.017943 | 0.049772 | 0.547019 |
| 29048 | ENSG0000 | HLA-L | major histo | 0.032033 | 0.182039 | 0.546675 |
| 34651 | ENSG0000 | AC141586.2 | | 0.289893 | 0.401529 | 0.546636 |
| 6969 | ENSG0000 | AGAP2 | ArfGAP wit | 0.029454 | 0.169244 | 0.546119 |
| 21850 | ENSG0000 | AL133351.1 | | 0.091797 | 0.159904 | 0.545315 |
| 20632 | ENSG0000 | HIST1H2BF | histone clu | 0.06515 | 0.473006 | 0.544956 |
| 36644 | ENSG0000 | SNORA59B | small nucle | 0.110151 | 0.187064 | 0.544117 |
| 39204 | ENSG0000 | AC016717.2 | | 0.376779 | 0.515744 | 0.543578 |
| 7197 | ENSG0000 | CHAD | chondroad | 0.164297 | 0.286622 | 0.543009 |
| 15821 | ENSG0000 | SLIT1 | slit guidanc | 0.173527 | 0.28587 | 0.542445 |
| 177 | ENSG0000 | CD79B | CD79b mol | 0 | 0.169604 | 0.542326 |
| 25175 | ENSG0000 | KIAA1614- | KIAA1614 a | 0.193318 | 0.330247 | 0.541851 |
| 23771 | ENSG0000 | AC046143.1 | | 0 | 0.48141 | 0.54185 |
| 12052 | ENSG0000 | ULK4 | unc-51 like | 0.327642 | 0.423601 | 0.541687 |
| 40545 | ENSG0000 | AL442125.1 | | 0.307983 | 0.437263 | 0.541481 |
| 41739 | ENSG0000 | CR381653.2 | | 0.311011 | 0.399665 | 0.541272 |
| 7779 | ENSG0000 | TSSK4 | testis speci | 0.346677 | 0.507955 | 0.54046 |
| 23846 | ENSG0000 | NKX1-2 | NK1 home | 0.031652 | 0.220524 | 0.539768 |
| 5074 | ENSG0000 | DUSP4 | dual specif | 0.133182 | 0.258033 | 0.538932 |
| 25612 | ENSG0000 | B4GALT1-A | B4GALT1 a | 0.152842 | 0.280372 | 0.538739 |
| 13133 | ENSG0000 | HPSE | heparanas | 0.090455 | 0.171309 | 0.538668 |
| 40304 | ENSG0000 | AL031670.1 | | 0.285909 | 0.46176 | 0.538535 |
| 35177 | ENSG0000 | AL136038.3 | | 0.371238 | 0.377561 | 0.538295 |
| 36823 | ENSG0000 | MYO15B | myosin XV | 0.134845 | 0.317086 | 0.537938 |
| 33430 | ENSG0000 | AC125603.2 | | 0 | 0.127061 | 0.537633 |
| 23908 | ENSG0000 | AC009237.3 | | 0.2111 | 0.4058 | 0.537107 |
| 35843 | ENSG0000 | LRRC37A1 | leucine ricl | 0.383528 | 0.396975 | 0.536679 |
| 11661 | ENSG0000 | BEAN1 | brain expre | 0.191583 | 0.296026 | 0.536325 |
| 4100 | ENSG0000 | TBX18 | T-box 18 [S | 0.245642 | 0.295402 | 0.536179 |
| 4487 | ENSG0000 | MORN1 | MORN rep | 0.397009 | 0.523859 | 0.535778 |
| 35754 | ENSG0000 | AC040160.1 | | 0.178033 | 0.498685 | 0.535764 |
| 17549 | ENSG0000 | Y_RNA | Y RNA [Sou | 0.108038 | 0.486261 | 0.535721 |
| 38072 | ENSG0000 | AC023509.4 | | 0.279812 | 0.39015 | 0.534478 |
| 997 | ENSG0000 | STAG3 | stromal an | 0.29469 | 0.331746 | 0.533953 |
| 14088 | ENSG0000 | KCNG2 | potassium | 0.266279 | 0.397773 | 0.532808 |
| 23937 | ENSG0000 | MRPS18AF | mitochond | 0.166031 | 0.240137 | 0.532695 |

| | | | | | | |
|---|---|---|---|---|---|---|
| 42429 | ENSG0000 | AC084876.2 | | 0.139114 | 0.144955 | 0.532004 |
| 33220 | ENSG0000 | ZNF878 | zinc finger | 0.313449 | 0.42401 | 0.530803 |
| 36553 | ENSG0000 | ZNF793-AS | ZNF793 an | 0.294324 | 0.447532 | 0.530244 |
| 39436 | ENSG0000 | AC010655.4 | | 0.151977 | 0.251804 | 0.529018 |
| 10636 | ENSG0000 | FCRLB | Fc recepto | 0.134841 | 0.282198 | 0.528041 |
| 42494 | ENSG0000 | AC009133.6 | | 0 | 0.173137 | 0.52781 |
| 3848 | ENSG0000 | VDR | vitamin D r | 0.309215 | 0.34287 | 0.527644 |
| 7091 | ENSG0000 | PLXNC1 | plexin C1 [ | 0.121094 | 0.14142 | 0.527565 |
| 8596 | ENSG0000 | ANK2 | ankyrin 2 [ | 0.277898 | 0.427857 | 0.526884 |
| 24375 | ENSG0000 | VN1R108P | vomeronas | 0.029721 | 0.236612 | 0.526824 |
| 14790 | ENSG0000 | TSPAN10 | tetraspanir | 0.107623 | 0.269308 | 0.526589 |
| 26913 | ENSG0000 | SIX3-AS1 | SIX3 antise | 0.046364 | 0.124229 | 0.526545 |
| 34142 | ENSG0000 | AC020661.2 | | 0.313707 | 0.500068 | 0.526344 |
| 5099 | ENSG0000 | TEX14 | testis expr | 0 | 0.389015 | 0.525392 |
| 34877 | ENSG0000 | FAM157C | family with | 0.14424 | 0.221624 | 0.525036 |
| 35255 | ENSG0000 | AL031432.2 | | 0.163325 | 0.371767 | 0.525035 |
| 6249 | ENSG0000 | PRDM12 | PR/SET dor | 0.179638 | 0.469671 | 0.525026 |
| 11685 | ENSG0000 | PLEKHA7 | pleckstrin l | 0.359386 | 0.359801 | 0.523629 |
| 32269 | ENSG0000 | AC105219.4 | | 0.029569 | 0.324476 | 0.523169 |
| 26951 | ENSG0000 | AC116049.1 | | 0.046094 | 0.245276 | 0.523031 |
| 17847 | ENSG0000 | SNORA70 | Small nucle | 0.200998 | 0.206182 | 0.522915 |
| 39130 | ENSG0000 | AC015712.6 | | 0.009565 | 0.387988 | 0.522561 |
| 37023 | ENSG0000 | MAP3K14- | MAP3K14 a | 0.168142 | 0.339011 | 0.522549 |
| 1397 | ENSG0000 | CST7 | cystatin F [ | 0.15306 | 0.226357 | 0.522473 |
| 31989 | ENSG0000 | AP002495. | Putative sh | 0.293382 | 0.311121 | 0.522005 |
| 13058 | ENSG0000 | ZFAND4 | zinc finger | 0.354812 | 0.419865 | 0.52188 |
| 39182 | ENSG0000 | AL022328.4 | | 0.39646 | 0.416243 | 0.520372 |
| 3831 | ENSG0000 | GSG1 | germ cell a | 0.097342 | 0.350382 | 0.520138 |
| 30530 | ENSG0000 | AC012645.1 | | 0.22123 | 0.356695 | 0.519908 |
| 9921 | ENSG0000 | C16orf86 | chromosor | 0.136548 | 0.202091 | 0.51887 |
| 3173 | ENSG0000 | ATP4A | ATPase H+, | 0.025802 | 0.140245 | 0.518472 |
| 11460 | ENSG0000 | AK8 | adenylate l | 0.218475 | 0.290889 | 0.516073 |
| 21326 | ENSG0000 | AC106786.1 | | 0.151247 | 0.360778 | 0.515992 |
| 38782 | ENSG0000 | AL024498.1 | | 0.23684 | 0.403603 | 0.515313 |
| 16385 | ENSG0000 | CGB7 | chorionic g | 0.094457 | 0.300093 | 0.515157 |
| 33912 | ENSG0000 | AC007040.2 | | 0.183031 | 0.40343 | 0.514877 |
| 39323 | ENSG0000 | AL008721.2 | | 0.094846 | 0.360736 | 0.514772 |
| 7425 | ENSG0000 | POU2F3 | POU class 2 | 0.005341 | 0.287933 | 0.51469 |
| 1834 | ENSG0000 | DZANK1 | double zin | 0.135504 | 0.434145 | 0.513697 |
| 727 | ENSG0000 | LAMA3 | laminin sul | 0.072425 | 0.274143 | 0.513428 |
| 20478 | ENSG0000 | RPL23AP47 | ribosomal | 0.419819 | 0.461996 | 0.513401 |
| 25507 | ENSG0000 | AL078581.1 | | 0.104759 | 0.128531 | 0.51332 |
| 9401 | ENSG0000 | ZNF547 | zinc finger | 0.376221 | 0.38579 | 0.512821 |
| 12455 | ENSG0000 | TIGD4 | tigger tran: | 0.226871 | 0.405233 | 0.512546 |
| 11208 | ENSG0000 | HCN1 | hyperpolar | 0.221795 | 0.453239 | 0.511772 |
| 33413 | ENSG0000 | AC011611.4 | | 0.022781 | 0.348885 | 0.51167 |
| 20461 | ENSG0000 | HIST1H1PS | histone clu | 0.055157 | 0.374242 | 0.510961 |

| | | | | | | |
|---|---|---|---|---|---|---|
| 1214 | ENSG0000 | ACAP1 | ArfGAP wit | 0.040171 | 0.26081 | 0.510765 |
| 18840 | ENSG0000 | RNA5SP18 | RNA, 5S rik | 0.234795 | 0.279818 | 0.51064 |
| 10303 | ENSG0000 | ITGB2 | integrin su | 0.427769 | 0.446986 | 0.510598 |
| 23787 | ENSG0000 | LINC01715 | long interg | 0.193806 | 0.274909 | 0.510032 |
| 4377 | ENSG0000 | EVA1A | eva-1 hom | 0.090954 | 0.2561 | 0.510014 |
| 40976 | ENSG0000 | AL354696.1 | | 0 | 0.166771 | 0.509403 |
| 14834 | ENSG0000 | VMO1 | vitelline m | 0.139837 | 0.462691 | 0.50908 |
| 5862 | ENSG0000 | TUBA4A | tubulin alp | 0.132173 | 0.262673 | 0.509077 |
| 10773 | ENSG0000 | ALPPL2 | alkaline ph | 0.04998 | 0.110638 | 0.508901 |
| 27168 | ENSG0000 | CYP2F2P | cytochrom | 0.248488 | 0.404828 | 0.508703 |
| 17201 | ENSG0000 | MIR219A1 | microRNA | 0 | 0.325205 | 0.50856 |
| 39532 | ENSG0000 | AC009032.1 | | 0.019565 | 0.227948 | 0.508519 |
| 29374 | ENSG0000 | AC026250.1 | | 0.115577 | 0.227621 | 0.50823 |
| 34167 | ENSG0000 | AC090970.2 | | 0.159751 | 0.417388 | 0.508099 |
| 6812 | ENSG0000 | IL15RA | interleukin | 0.445671 | 0.47597 | 0.508052 |
| 35332 | ENSG0000 | AC026471.4 | | 0.276945 | 0.487339 | 0.507584 |
| 4864 | ENSG0000 | BBOF1 | basal body | 0.252598 | 0.4104 | 0.507457 |
| 17129 | ENSG0000 | GJC2 | gap junctic | 0.28428 | 0.423027 | 0.506851 |
| 36927 | ENSG0000 | AC008738.2 | | 0 | 0.411383 | 0.506628 |
| 40567 | ENSG0000 | AL133243.2 | | 0.336891 | 0.411283 | 0.506108 |
| 2939 | ENSG0000 | MAP4K1 | mitogen-ac | 0.231164 | 0.316446 | 0.504784 |
| 19726 | ENSG0000 | RPL23AP45 | ribosomal | 0.430459 | 0.486609 | 0.504522 |
| 4854 | ENSG0000 | ONECUT2 | one cut ho | 0.350611 | 0.437594 | 0.503859 |
| 43368 | ENSG0000 | AP000553.3 | | 0.125858 | 0.155362 | 0.503162 |
| 1686 | ENSG0000 | PILRA | paired imn | 0.304204 | 0.394372 | 0.502944 |
| 30236 | ENSG0000 | AC079848.1 | | 0.234155 | 0.261643 | 0.502934 |
| 17698 | ENSG0000 | RNVU1-6 | RNA, variar | 0 | 0.385334 | 0.502906 |
| 2841 | ENSG0000 | DTWD1 | DTW doma | 0.415243 | 0.494275 | 0.502894 |
| 5402 | ENSG0000 | PI3 | peptidase i | 0 | 0.049669 | 0.502879 |
| 3569 | ENSG0000 | ALOX12 | arachidona | 0.262619 | 0.408997 | 0.502793 |
| 31678 | ENSG0000 | AC015909.2 | | 0 | 0.132282 | 0.502517 |
| 42608 | ENSG0000 | AC018638.8 | | 0 | 0.014335 | 0.502464 |
| 18410 | ENSG0000 | TMEM170I | transmeml | 0.279441 | 0.34796 | 0.502367 |
| 763 | ENSG0000 | MCOLN3 | mucolipin : | 0.174955 | 0.372893 | 0.50236 |
| 43238 | ENSG0000 | AL807752.7 | | 0 | 0.173347 | 0.502025 |
| 36428 | ENSG0000 | AC012447.1 | | 0.104821 | 0.266041 | 0.501503 |
| 23209 | ENSG0000 | AC118553.1 | | 0 | 0.0981 | 0.501334 |
| 33694 | ENSG0000 | LINC00239 | long interg | 0.081443 | 0.209731 | 0.501275 |
| 4472 | ENSG0000 | EPAS1 | endothelia | 0.292045 | 0.486265 | 0.501268 |
| 18729 | ENSG0000 | Y_RNA | Y RNA [Sou | 0.074034 | 0.110166 | 0.501148 |
| 13268 | ENSG0000 | CD7 | CD7 molec | 0.038709 | 0.138598 | 0.500978 |
| 234 | ENSG0000 | TENM1 | teneurin tr | 0.162599 | 0.489313 | 0.500859 |
| 26039 | ENSG0000 | AL354751.1 | | 0.046133 | 0.33129 | 0.500727 |
| 31472 | ENSG0000 | AL606534.5 | | 0.093558 | 0.370309 | 0.500316 |
| 40291 | ENSG0000 | STPG3-AS1 | STPG3 anti | 0.041583 | 0.298801 | 0.500196 |
| 12928 | ENSG0000 | SYNPO | synaptopo | 0.119332 | 0.146414 | 0.499312 |
| 28493 | ENSG0000 | ACTG1P20 | actin gamr | 0.15957 | 0.409941 | 0.498751 |

| | | | | | | |
|---|---|---|---|---|---|---|
| 33270 | ENSG0000 | PPP1R12A- | PPP1R12A | 0.4004 | 0.459122 | 0.498598 |
| 22660 | ENSG0000 | HAUS6P1 | HAUS augr | 0.3312 | 0.373281 | 0.498214 |
| 29297 | ENSG0000 | Z98749.2 | | 0.235553 | 0.334553 | 0.498165 |
| 27225 | ENSG0000 | PRKCQ-AS1 | PRKCQ ant | 0.206272 | 0.24281 | 0.497962 |
| 21424 | ENSG0000 | AC073073.1 | | 0.154945 | 0.327531 | 0.497935 |
| 24798 | ENSG0000 | AC079922.1 | | 0.099915 | 0.105388 | 0.497556 |
| 30272 | ENSG0000 | AL390198.2 | | 0.109134 | 0.421187 | 0.497252 |
| 2194 | ENSG0000 | PIK3IP1 | phosphoin | 0.141223 | 0.183749 | 0.496316 |
| 40173 | ENSG0000 | AL512791.2 | | 0.135116 | 0.475194 | 0.496066 |
| 23321 | ENSG0000 | AL662791.2 | | 0 | 0.063654 | 0.495328 |
| 13584 | ENSG0000 | GPR156 | G protein-c | 0.076268 | 0.21625 | 0.494872 |
| 22970 | ENSG0000 | UGT1A12P | UDP glucur | 0.041756 | 0.224794 | 0.494627 |
| 20795 | ENSG0000 | KRT8P42 | keratin 8 p | 0.087742 | 0.362476 | 0.494159 |
| 40199 | ENSG0000 | LINC02340 | long interg | 0.162194 | 0.205821 | 0.493898 |
| 33098 | ENSG0000 | CLIP1-AS1 | CLIP1 antis | 0.041218 | 0.452854 | 0.493706 |
| 4915 | ENSG0000 | RASL11B | RAS like fa | 0.085026 | 0.485491 | 0.493494 |
| 7282 | ENSG0000 | GARNL3 | GTPase act | 0.272923 | 0.443136 | 0.493025 |
| 13837 | ENSG0000 | EFCAB5 | EF-hand ca | 0.174794 | 0.433844 | 0.492876 |
| 25851 | ENSG0000 | AL390879.2 | | 0.215785 | 0.29766 | 0.492549 |
| 41389 | ENSG0000 | MIR6865 | microRNA | 0.230418 | 0.490079 | 0.492173 |
| 40394 | ENSG0000 | AC073107.2 | | 0.219677 | 0.491223 | 0.492086 |
| 11427 | ENSG0000 | TMEM63C | transmeml | 0.134147 | 0.228448 | 0.491884 |
| 37725 | ENSG0000 | AC005253.2 | | 0.14378 | 0.240523 | 0.491154 |
| 15086 | ENSG0000 | IFNL2 | interferon | 0 | 0.077493 | 0.490808 |
| 35735 | ENSG0000 | AC133552.2 | | 0.142849 | 0.274573 | 0.490085 |
| 32037 | ENSG0000 | AC080023.1 | | 0 | 0.363203 | 0.490029 |
| 34260 | ENSG0000 | PRELID1P4 | PRELI dom | 0.112584 | 0.354375 | 0.489992 |
| 19177 | ENSG0000 | SNORD105 | small nucle | 0.118455 | 0.214672 | 0.48921 |
| 28893 | ENSG0000 | PCDHAC2 | protocadh | 0.086545 | 0.102013 | 0.488818 |
| 16615 | ENSG0000 | IFNL3 | interferon | 0 | 0.02574 | 0.487862 |
| 5964 | ENSG0000 | GAD1 | glutamate | 0.08674 | 0.222666 | 0.487184 |
| 21376 | ENSG0000 | CD99P1 | CD99 mole | 0.203767 | 0.288171 | 0.486901 |
| 25336 | ENSG0000 | AL157400.2 | | 0 | 0.401885 | 0.486757 |
| 35142 | ENSG0000 | AL031058.1 | | 0 | 0.212103 | 0.486515 |
| 14139 | ENSG0000 | ZNF713 | zinc finger | 0.331052 | 0.483692 | 0.486502 |
| 7993 | ENSG0000 | ASGR1 | asialoglyco | 0.140343 | 0.247781 | 0.485547 |
| 22377 | ENSG0000 | GTF2IP14 | general tra | 0 | 0.37393 | 0.485018 |
| 37153 | ENSG0000 | AP001542.3 | | 0.077142 | 0.094679 | 0.484982 |
| 43207 | ENSG0000 | AL356275.1 | | 0.050363 | 0.27272 | 0.484461 |
| 14528 | ENSG0000 | GPR137C | G protein-c | 0.167561 | 0.367767 | 0.484445 |
| 6945 | ENSG0000 | LCA5 | LCA5, lebe | 0.11001 | 0.391864 | 0.484082 |
| 23279 | ENSG0000 | AL021707.2 | | 0.249323 | 0.287449 | 0.484004 |
| 33662 | ENSG0000 | AL139011.2 | | 0.016299 | 0.344928 | 0.483722 |
| 30353 | ENSG0000 | AC021491.2 | | 0.127832 | 0.15546 | 0.483215 |
| 21871 | ENSG0000 | AC138028.2 | | 0.052655 | 0.201728 | 0.483087 |
| 26617 | ENSG0000 | B3GALT4 | beta-1,3-ga | 0.155258 | 0.412962 | 0.482429 |
| 32782 | ENSG0000 | AC006064.3 | | 0.218999 | 0.301492 | 0.482357 |

| | | | | | | |
|---|---|---|---|---|---|---|
| 27425 | ENSG0000 | AL391832.1 | | 0 | 0.107549 | 0.482307 |
| 41853 | ENSG0000 | AC021106.3 | | 0.197913 | 0.234527 | 0.481978 |
| 35123 | ENSG0000 | EPPK1 | epiplakin 1 | 0.013347 | 0.167043 | 0.481936 |
| 4689 | ENSG0000 | MAN1C1 | mannosida | 0.125278 | 0.212789 | 0.48114 |
| 14231 | ENSG0000 | ALOXE3 | arachidona | 0.16385 | 0.422209 | 0.480433 |
| 34709 | ENSG0000 | AC137932.1 | | 0.084491 | 0.478909 | 0.479862 |
| 42565 | ENSG0000 | AC104662.3 | | 0.196806 | 0.389124 | 0.479732 |
| 41802 | ENSG0000 | CU634019.4 | | 0.209913 | 0.215138 | 0.479641 |
| 8371 | ENSG0000 | ITPKB | inositol-tri | 0.060318 | 0.233654 | 0.479632 |
| 31156 | ENSG0000 | RNA5SP82 | RNA, 5S rib | 0.251717 | 0.453824 | 0.479457 |
| 3070 | ENSG0000 | EBI3 | Epstein-Ba | 0 | 0.165114 | 0.479341 |
| 35802 | ENSG0000 | AC118754.2 | | 0 | 0.064488 | 0.478304 |
| 14592 | ENSG0000 | TMEM45A | transmeml | 0.020729 | 0.02194 | 0.477799 |
| 36397 | ENSG0000 | MIR5188 | microRNA | 0.062372 | 0.407802 | 0.477777 |
| 25315 | ENSG0000 | AC127070.1 | | 0.249637 | 0.457733 | 0.477754 |
| 28555 | ENSG0000 | AC079944.2 | | 0.121743 | 0.453797 | 0.476694 |
| 32306 | ENSG0000 | AP006287.2 | | 0 | 0.10246 | 0.476641 |
| 43376 | ENSG0000 | AC092821.3 | | 0.216458 | 0.350207 | 0.476118 |
| 35181 | ENSG0000 | AC009054.2 | | 0.187105 | 0.272474 | 0.475604 |
| 32509 | ENSG0000 | AC124276.2 | | 0 | 0.425672 | 0.475352 |
| 41546 | ENSG0000 | CU633906.2 | | 0.320922 | 0.34418 | 0.47513 |
| 14074 | ENSG0000 | PRSS36 | protease, s | 0.048973 | 0.173904 | 0.474687 |
| 2353 | ENSG0000 | DAAM1 | dishevellec | 0.318379 | 0.440755 | 0.47444 |
| 21826 | ENSG0000 | CETN4P | centrin 4, p | 0.063754 | 0.451113 | 0.473589 |
| 39330 | ENSG0000 | LINC02012 | long interg | 0.305344 | 0.470864 | 0.473464 |
| 4617 | ENSG0000 | PADI2 | peptidyl ar | 0.050415 | 0.208577 | 0.472572 |
| 16234 | ENSG0000 | UBE2Q2P1 | ubiquitin c | 0.241542 | 0.46505 | 0.472387 |
| 42879 | ENSG0000 | U6 | U6 spliceos | 0.125858 | 0.435794 | 0.471228 |
| 7656 | ENSG0000 | KIAA1644 | KIAA1644 \| | 0.200057 | 0.365128 | 0.471203 |
| 25276 | ENSG0000 | TNF | tumor necr | 0 | 0.104075 | 0.470994 |
| 6918 | ENSG0000 | CCDC146 | coiled-coil | 0.189706 | 0.41158 | 0.4706 |
| 37771 | ENSG0000 | LYPLA2P2 | lysophosph | 0.092253 | 0.272215 | 0.470279 |
| 15793 | ENSG0000 | KANK3 | KN motif a | 0.111908 | 0.331456 | 0.470198 |
| 20362 | ENSG0000 | MSMP | microsemi | 0.168463 | 0.296736 | 0.46984 |
| 16629 | ENSG0000 | AC107956.1 | | 0 | 0.087385 | 0.469611 |
| 8706 | ENSG0000 | ANO7 | anoctamin | 0.056013 | 0.097027 | 0.468649 |
| 3708 | ENSG0000 | NRXN2 | neurexin 2 | 0.030275 | 0.109872 | 0.467996 |
| 35785 | ENSG0000 | AC127496.5 | | 0.128396 | 0.227862 | 0.467714 |
| 14043 | ENSG0000 | GRAMD1C | GRAM don | 0.157685 | 0.215921 | 0.4673 |
| 14614 | ENSG0000 | TNFSF15 | TNF superf | 0.136748 | 0.279274 | 0.467104 |
| 32181 | ENSG0000 | AC067930.3 | | 0.120387 | 0.236183 | 0.467017 |
| 7822 | ENSG0000 | SLC27A2 | solute carr | 0.063513 | 0.257504 | 0.466797 |
| 29977 | ENSG0000 | AC106822.1 | | 0.10217 | 0.244988 | 0.466468 |
| 14320 | ENSG0000 | PIPOX | pipecolic a | 0.029954 | 0.209555 | 0.466279 |
| 22214 | ENSG0000 | ZNF469 | zinc finger | 0.22134 | 0.238035 | 0.466206 |
| 11592 | ENSG0000 | FAM227B | family with | 0.132087 | 0.275845 | 0.465832 |
| 1896 | ENSG0000 | THPO | thrombopo | 0.018599 | 0.112938 | 0.46554 |

| | | | | | | |
|---|---|---|---|---|---|---|
| 37948 | ENSG0000 | AL158152.2 | | 0 | 0.164946 | 0.464856 |
| 38258 | ENSG0000 | AC124312.4 | | 0.138852 | 0.308005 | 0.464838 |
| 37183 | ENSG0000 | LINC01836 | long interg | 0 | 0.204115 | 0.464629 |
| 12648 | ENSG0000 | TPT1-AS1 | TPT1 antis | 0.238107 | 0.398386 | 0.464419 |
| 31402 | ENSG0000 | AC009630.2 | | 0.03505 | 0.329646 | 0.463927 |
| 15679 | ENSG0000 | GLDN | gliomedin | 0.116698 | 0.363488 | 0.463876 |
| 39286 | ENSG0000 | AL353708.3 | | 0.10537 | 0.366914 | 0.46364 |
| 38676 | ENSG0000 | AC064834.1 | | 0.078782 | 0.14412 | 0.463592 |
| 15769 | ENSG0000 | PDZD7 | PDZ domai | 0.09576 | 0.164793 | 0.463458 |
| 8025 | ENSG0000 | TNFRSF11A | TNF recept | 0.130031 | 0.203847 | 0.463257 |
| 10963 | ENSG0000 | SNRK | SNF relate | 0.42286 | 0.44694 | 0.46295 |
| 12345 | ENSG0000 | NPR1 | natriuretic | 0.077381 | 0.15647 | 0.462717 |
| 31298 | ENSG0000 | MIR2116 | microRNA | 0.047197 | 0.265459 | 0.462351 |
| 19947 | ENSG0000 | LIPE-AS1 | LIPE antise | 0.157647 | 0.254176 | 0.462291 |
| 18167 | ENSG0000 | VWA7 | von Willeb | 0.162852 | 0.199217 | 0.462265 |
| 16936 | ENSG0000 | AFDN-AS1 | AFDN antis | 0.045292 | 0.185839 | 0.46222 |
| 23932 | ENSG0000 | AC096649.1 | | 0.111925 | 0.452368 | 0.461975 |
| 17726 | ENSG0000 | RNU1-59P | RNA, U1 sr | 0 | 0.064909 | 0.461961 |
| 43239 | ENSG0000 | MIR940 | microRNA | 0 | 0.007129 | 0.461432 |
| 41251 | ENSG0000 | AC005261.6 | | 0.123297 | 0.418075 | 0.46088 |
| 37301 | ENSG0000 | AL121761.2 | | 0.043226 | 0.153721 | 0.460664 |
| 26118 | ENSG0000 | AF015262.1 | | 0.263978 | 0.372043 | 0.460302 |
| 37690 | ENSG0000 | AC008878.3 | | 0.082332 | 0.119561 | 0.460246 |
| 4739 | ENSG0000 | CA14 | carbonic ai | 0.105567 | 0.117218 | 0.459143 |
| 35891 | ENSG0000 | AC004477.1 | | 0.21181 | 0.437179 | 0.458818 |
| 1152 | ENSG0000 | MGAT4A | mannosyl ( | 0.31494 | 0.332623 | 0.458047 |
| 42632 | ENSG0000 | AL160408.6 | | 0 | 0.125991 | 0.457292 |
| 17615 | ENSG0000 | Y_RNA | Y RNA [Sou | 0.106193 | 0.293165 | 0.457086 |
| 19875 | ENSG0000 | RPL17P50 | ribosomal | 0 | 0.132312 | 0.456907 |
| 2535 | ENSG0000 | WFDC2 | WAP four-c | 0.101026 | 0.138231 | 0.45657 |
| 5475 | ENSG0000 | LYPD3 | LY6/PLAUR | 0.09167 | 0.147215 | 0.455669 |
| 22234 | ENSG0000 | LINC01771 | long interg | 0.062596 | 0.289273 | 0.454415 |
| 9922 | ENSG0000 | MPZL3 | myelin pro | 0.147459 | 0.373565 | 0.454153 |
| 39189 | ENSG0000 | AC092636.1 | | 0.114803 | 0.329891 | 0.453944 |
| 6226 | ENSG0000 | LSP1 | lymphocyt | 0.313272 | 0.368747 | 0.453927 |
| 41214 | ENSG0000 | AC004623.1 | | 0.193652 | 0.42762 | 0.453609 |
| 18899 | ENSG0000 | SNORD116 | small nucle | 0.261293 | 0.32384 | 0.453237 |
| 10021 | ENSG0000 | ANKRD61 | ankyrin rep | 0.261229 | 0.323912 | 0.452306 |
| 31646 | ENSG0000 | AP003356.1 | | 0.248869 | 0.396621 | 0.452089 |
| 25548 | ENSG0000 | AL359962.1 | | 0.187671 | 0.365316 | 0.449754 |
| 8912 | ENSG0000 | C9orf72 | chromosor | 0.368726 | 0.378221 | 0.449584 |
| 13817 | ENSG0000 | UBA7 | ubiquitin li | 0.152866 | 0.34651 | 0.449554 |
| 17435 | ENSG0000 | RNY1P16 | RNA, Ro-as | 0 | 0.077086 | 0.449131 |
| 28081 | ENSG0000 | RN7SL368P | RNA, 7SL, c | 0.109501 | 0.432833 | 0.449112 |
| 3160 | ENSG0000 | KCNN1 | potassium | 0.048017 | 0.155845 | 0.448436 |
| 7043 | ENSG0000 | RNASEL | ribonuclea | 0.073953 | 0.258965 | 0.448133 |
| 9435 | ENSG0000 | CLEC4F | C-type lect | 0.107806 | 0.217159 | 0.448109 |

| | | | | | | |
|---|---|---|---|---|---|---|
| 2831 | ENSG0000 | IGDCC4 | immunogl | 0.050528 | 0.143481 | 0.44779 |
| 36902 | ENSG0000 | AC025048.1 | | 0 | 0.103852 | 0.447505 |
| 27781 | ENSG0000 | Y_RNA | Y RNA [Sou | 0 | 0.300265 | 0.447212 |
| 25762 | ENSG0000 | DLG5-AS1 | DLG5 antis | 0.120968 | 0.248698 | 0.446607 |
| 3198 | ENSG0000 | RASA4 | RAS p21 pr | 0.142626 | 0.210569 | 0.4465 |
| 27965 | ENSG0000 | AP000487.1 | | 0.251985 | 0.416767 | 0.446081 |
| 6214 | ENSG0000 | HRC | histidine ri | 0.013172 | 0.244917 | 0.446046 |
| 20209 | ENSG0000 | EML6 | echinoderr | 0.16932 | 0.30904 | 0.444545 |
| 13017 | ENSG0000 | RSPH9 | radial spok | 0.300208 | 0.426507 | 0.443946 |
| 42197 | ENSG0000 | BISPR | BST2 interf | 0.024874 | 0.102718 | 0.443637 |
| 39422 | ENSG0000 | AL359962.2 | | 0.058568 | 0.192573 | 0.443504 |
| 19281 | ENSG0000 | TRGV5 | T-cell recep | 0 | 0.105519 | 0.443456 |
| 41955 | ENSG0000 | AC080038.3 | | 0.318421 | 0.34986 | 0.443221 |
| 31253 | ENSG0000 | RNA5SP31 | RNA, 5S rib | 0.145931 | 0.172855 | 0.44314 |
| 12484 | ENSG0000 | CR848007.1 | | 0.094005 | 0.23288 | 0.443084 |
| 11645 | ENSG0000 | PKD1L2 | polycystin | 0.058517 | 0.306303 | 0.44305 |
| 36908 | ENSG0000 | AC011511.3 | | 0.066575 | 0.111265 | 0.442911 |
| 27175 | ENSG0000 | ZNF503-AS | ZNF503 an | 0.199361 | 0.315074 | 0.442458 |
| 18702 | ENSG0000 | RNU1-28P | RNA, U1 sr | 0.190493 | 0.377485 | 0.442359 |
| 11757 | ENSG0000 | MAP1A | microtubul | 0.242481 | 0.264134 | 0.441999 |
| 24842 | ENSG0000 | FAM192BF | family with | 0.090843 | 0.273015 | 0.441842 |
| 31252 | ENSG0000 | Y_RNA | Y RNA [Sou | 0.019578 | 0.0504 | 0.441611 |
| 25777 | ENSG0000 | LINC01503 | long interg | 0.022266 | 0.05304 | 0.441318 |
| 35383 | ENSG0000 | AP003119.3 | | 0.335603 | 0.369352 | 0.44112 |
| 39554 | ENSG0000 | Y_RNA | Y RNA [Sou | 0.125858 | 0.130705 | 0.441018 |
| 39026 | ENSG0000 | AL592494.3 | | 0.240325 | 0.262615 | 0.440912 |
| 14356 | ENSG0000 | GOLGA8J | golgin A8 f | 0.13773 | 0.337051 | 0.440855 |
| 6965 | ENSG0000 | ITGA7 | integrin su | 0.173544 | 0.193292 | 0.440512 |
| 17280 | ENSG0000 | Y_RNA | Y RNA [Sou | 0 | 0.148909 | 0.440334 |
| 2034 | ENSG0000 | SH2D3C | SH2 domai | 0.068481 | 0.139899 | 0.440237 |
| 26844 | ENSG0000 | GUSBP5 | glucuronid | 0.070554 | 0.339155 | 0.440164 |
| 38292 | ENSG0000 | AL645939.4 | | 0 | 0.053455 | 0.439909 |
| 39522 | ENSG0000 | MIR6514 | microRNA | 0.334424 | 0.363764 | 0.439788 |
| 11501 | ENSG0000 | ANKRD2 | ankyrin rep | 0.188287 | 0.343591 | 0.439785 |
| 41242 | ENSG0000 | AC008764.8 | | 0.241855 | 0.318882 | 0.439128 |
| 2091 | ENSG0000 | HSD17B7P | hydroxyste | 0.116847 | 0.320965 | 0.438618 |
| 10098 | ENSG0000 | PAGE5 | PAGE fami | 0 | 0.274031 | 0.438608 |
| 27668 | ENSG0000 | AL513320.1 | | 0.200522 | 0.210651 | 0.438426 |
| 42666 | ENSG0000 | LINC01002 | long interg | 0.205856 | 0.368564 | 0.437792 |
| 5336 | ENSG0000 | NFE2 | nuclear fac | 0.129707 | 0.153697 | 0.43766 |
| 1102 | ENSG0000 | GUCY2C | guanylate | 0.200071 | 0.382219 | 0.437518 |
| 42703 | ENSG0000 | FRG1EP | FSHD regio | 0.110605 | 0.264999 | 0.437326 |
| 32419 | ENSG0000 | AP002992.1 | | 0 | 0.132517 | 0.43722 |
| 5370 | ENSG0000 | G0S2 | G0/G1 swi | 0.151461 | 0.189635 | 0.436665 |
| 13118 | ENSG0000 | GXYLT2 | glucoside x | 0.371672 | 0.383387 | 0.436551 |
| 29342 | ENSG0000 | AC100803.1 | | 0.054003 | 0.401457 | 0.436256 |
| 10232 | ENSG0000 | FAM131B | family with | 0.146508 | 0.191038 | 0.435857 |

| | | | | | | |
|---|---|---|---|---|---|---|
| 11600 | ENSG0000 | C11orf65 | chromosor | 0.235434 | 0.247365 | 0.435103 |
| 16376 | ENSG0000 | AC005154.1 | | 0.233098 | 0.364945 | 0.435032 |
| 37400 | ENSG0000 | AC005261.1 | | 0.133285 | 0.202878 | 0.434576 |
| 18278 | ENSG0000 | AC008429.1 | | 0.04918 | 0.301246 | 0.434478 |
| 9207 | ENSG0000 | C2orf50 | chromosor | 0.035213 | 0.164397 | 0.434352 |
| 19847 | ENSG0000 | LBH | limb bud a | 0.003941 | 0.035318 | 0.434223 |
| 12697 | ENSG0000 | NEUROD2 | neuronal d | 0.070813 | 0.135049 | 0.433716 |
| 41233 | ENSG0000 | AC007382.1 | | 0.259072 | 0.30758 | 0.433548 |
| 36053 | ENSG0000 | AC104564.3 | | 0 | 0.203597 | 0.433373 |
| 9415 | ENSG0000 | TRIM36 | tripartite n | 0.223686 | 0.342137 | 0.432223 |
| 30674 | ENSG0000 | AC141928.1 | | 0.133285 | 0.321717 | 0.432063 |
| 2982 | ENSG0000 | GRIK5 | glutamate | 0.266377 | 0.327917 | 0.432021 |
| 34990 | ENSG0000 | AC139256.1 | | 0.119367 | 0.408995 | 0.431499 |
| 39557 | ENSG0000 | MIR6735 | microRNA | 0 | 0.240075 | 0.431155 |
| 26252 | ENSG0000 | C19orf81 | chromosor | 0.079654 | 0.194224 | 0.430892 |
| 12136 | ENSG0000 | HR | HR, lysine ( | 0.097747 | 0.099724 | 0.430775 |
| 7482 | ENSG0000 | SLC44A5 | solute carr | 0.271887 | 0.379706 | 0.430693 |
| 24262 | ENSG0000 | COL6A4P1 | collagen ty | 0.113808 | 0.427005 | 0.430472 |
| 17517 | ENSG0000 | SNORA71D | small nucle | 0.197241 | 0.24064 | 0.430316 |
| 23281 | ENSG0000 | AC112518.1 | | 0.107451 | 0.282774 | 0.430001 |
| 36907 | ENSG0000 | TBC1D3P1- | TBC1D3P1- | 0.171163 | 0.251955 | 0.429925 |
| 6967 | ENSG0000 | FAM186B | family with | 0.115905 | 0.133995 | 0.428956 |
| 7279 | ENSG0000 | KIF12 | kinesin fam | 0.011768 | 0.118675 | 0.428926 |
| 27700 | ENSG0000 | RNU7-57P | RNA, U7 sr | 0 | 0.103091 | 0.428847 |
| 5993 | ENSG0000 | CNN1 | calponin 1 | 0.036968 | 0.295275 | 0.428421 |
| 16274 | ENSG0000 | FAM180A | family with | 0.0283 | 0.222612 | 0.428373 |
| 34913 | ENSG0000 | AC012531.1 | | 0.073308 | 0.407594 | 0.42819 |
| 28545 | ENSG0000 | AL627309.5 | | 0.198156 | 0.352231 | 0.427998 |
| 7733 | ENSG0000 | ACVRL1 | activin A re | 0 | 0.045348 | 0.427661 |
| 2388 | ENSG0000 | RPS6KA5 | ribosomal | 0.248606 | 0.412996 | 0.427193 |
| 3745 | ENSG0000 | SLC15A3 | solute carr | 0.153303 | 0.207352 | 0.427098 |
| 43131 | ENSG0000 | AC002472.2 | | 0.256996 | 0.367452 | 0.426754 |
| 23779 | ENSG0000 | DPY19L1P1 | DPY19L1 p | 0.184353 | 0.334233 | 0.42658 |
| 22938 | ENSG0000 | STK4-AS1 | STK4 antis | 0.112023 | 0.133943 | 0.42538 |
| 25781 | ENSG0000 | AC026202.2 | | 0 | 0.055997 | 0.424865 |
| 3310 | ENSG0000 | MEST | mesoderm | 0.04277 | 0.15512 | 0.424573 |
| 20263 | ENSG0000 | LINC01347 | long interg | 0.103803 | 0.326851 | 0.424562 |
| 31959 | ENSG0000 | AL139349.1 | | 0.23681 | 0.358403 | 0.424102 |
| 27137 | ENSG0000 | AL031186.1 | | 0.227034 | 0.317252 | 0.424089 |
| 25435 | ENSG0000 | AC008105.1 | | 0.087706 | 0.188435 | 0.422932 |
| 32975 | ENSG0000 | AC019209.3 | | 0.194086 | 0.269612 | 0.42243 |
| 15183 | ENSG0000 | C1QTNF12 | C1q and TN | 0.103275 | 0.198647 | 0.4223 |
| 8612 | ENSG0000 | ROPN1L | rhophilin a | 0.039136 | 0.288589 | 0.4222 |
| 11266 | ENSG0000 | OXR1 | oxidation r | 0.332169 | 0.391923 | 0.421635 |
| 29497 | ENSG0000 | AC091057.1 | | 0.187709 | 0.33178 | 0.421545 |
| 31697 | ENSG0000 | AC099548.2 | | 0.003522 | 0.046967 | 0.42128 |
| 39747 | ENSG0000 | U1 | U1 spliceos | 0 | 0.395849 | 0.42119 |

| | | | | | | |
|---|---|---|---|---|---|---|
| 11113 | ENSG0000 | ELOVL7 | ELOVL fatt | 0.085647 | 0.25766 | 0.420997 |
| 15703 | ENSG0000 | ZNF396 | zinc finger | 0.095776 | 0.201507 | 0.420122 |
| 11559 | ENSG0000 | SPINT1 | serine pep | 0.163708 | 0.201865 | 0.419953 |
| 24238 | ENSG0000 | AP000695.1 | | 0 | 0.080491 | 0.419873 |
| 26797 | ENSG0000 | TCEA1P4 | transcripti | 0.262454 | 0.306952 | 0.419706 |
| 5325 | ENSG0000 | MMP19 | matrix met | 0.037946 | 0.328372 | 0.418698 |
| 8924 | ENSG0000 | C9orf3 | chromosor | 0.222044 | 0.392866 | 0.418628 |
| 13869 | ENSG0000 | SCN4B | sodium vol | 0.022061 | 0.052831 | 0.418345 |
| 28066 | ENSG0000 | RPL39P36 | ribosomal | 0 | 0.241207 | 0.418244 |
| 24511 | ENSG0000 | AL590133.1 | | 0.121317 | 0.292551 | 0.418212 |
| 42182 | ENSG0000 | AL157888.1 | | 0.126495 | 0.264942 | 0.418026 |
| 42925 | ENSG0000 | AC005622.1 | | 0.04183 | 0.051394 | 0.417592 |
| 39438 | ENSG0000 | AL022328.3 | | 0.276555 | 0.303352 | 0.417355 |
| 38180 | ENSG0000 | AP000811.1 | | 0.070372 | 0.25749 | 0.417308 |
| 29692 | ENSG0000 | AC109454.2 | | 0.282232 | 0.402763 | 0.417168 |
| 39328 | ENSG0000 | AC009570.1 | | 0.3546 | 0.360473 | 0.417078 |
| 26284 | ENSG0000 | ID2-AS1 | ID2 antiser | 0.166394 | 0.197792 | 0.416439 |
| 21505 | ENSG0000 | AL049795.1 | | 0.065204 | 0.289093 | 0.416308 |
| 37365 | ENSG0000 | LINC01801 | long interg | 0.100821 | 0.205583 | 0.416224 |
| 9113 | ENSG0000 | C20orf144 | chromosor | 0.128847 | 0.379471 | 0.415734 |
| 36429 | ENSG0000 | MIR3682 | microRNA | 0.251717 | 0.405 | 0.415719 |
| 12526 | ENSG0000 | TCAF2 | TRPM8 cha | 0.03758 | 0.075105 | 0.415585 |
| 23369 | ENSG0000 | AP006222.1 | | 0.224262 | 0.307886 | 0.415169 |
| 17060 | ENSG0000 | ABCA4 | ATP bindin | 0.062392 | 0.332283 | 0.4134 |
| 1564 | ENSG0000 | PTPRC | protein tyr | 0.315557 | 0.354107 | 0.413344 |
| 11560 | ENSG0000 | FBN1 | fibrillin 1 [S | 0.06639 | 0.213771 | 0.413226 |
| 29691 | ENSG0000 | AC024558.1 | | 0.179539 | 0.199833 | 0.412995 |
| 37923 | ENSG0000 | AC010531.5 | | 0.14646 | 0.33452 | 0.412728 |
| 14925 | ENSG0000 | RIMBP3C | RIMS bindi | 0.115455 | 0.12005 | 0.41257 |
| 23678 | ENSG0000 | SERPINH1F | serpin fam | 0.057135 | 0.140068 | 0.412483 |
| 20677 | ENSG0000 | HIST1H2AF | histone clu | 0 | 0.391592 | 0.412331 |
| 16040 | ENSG0000 | ZNF793 | zinc finger | 0.247096 | 0.317437 | 0.412228 |
| 14813 | ENSG0000 | SLC35D3 | solute carr | 0.052766 | 0.292273 | 0.4121 |
| 34697 | ENSG0000 | LINC02562 | long interg | 0.093693 | 0.16776 | 0.411774 |
| 34557 | ENSG0000 | AC107871.1 | | 0.246886 | 0.393916 | 0.411667 |
| 9889 | ENSG0000 | SYN2 | synapsin II | 0.150286 | 0.261283 | 0.411201 |
| 36569 | ENSG0000 | AC022916.1 | | 0.106823 | 0.301247 | 0.41104 |
| 16988 | ENSG0000 | ZNF568 | zinc finger | 0.151646 | 0.334793 | 0.410886 |
| 28883 | ENSG0000 | AC092944.1 | | 0.199675 | 0.275267 | 0.410657 |
| 11236 | ENSG0000 | SLC13A4 | solute carr | 0.032151 | 0.266568 | 0.409983 |
| 33084 | ENSG0000 | AP001363.2 | | 0 | 0.342795 | 0.40949 |
| 19282 | ENSG0000 | TRGV4 | T-cell rece | 0.152671 | 0.30406 | 0.409391 |
| 10811 | ENSG0000 | TMEM169 | transmeml | 0.159102 | 0.196 | 0.40939 |
| 5996 | ENSG0000 | RAB11FIP4 | RAB11 fam | 0.083748 | 0.158959 | 0.409032 |
| 22787 | ENSG0000 | IL9RP3 | interleukin | 0.326335 | 0.376488 | 0.408708 |
| 10886 | ENSG0000 | STARD4 | StAR relate | 0.172166 | 0.35405 | 0.408189 |
| 31695 | ENSG0000 | AC087752.3 | | 0 | 0.148489 | 0.407812 |

| | | | | | | |
|---|---|---|---|---|---|---|
| 11159 | ENSG0000 | TLR3 | toll like rec | 0.079457 | 0.235956 | 0.407565 |
| 7673 | ENSG0000 | ETFBKMT | electron tr | 0.083914 | 0.180388 | 0.407341 |
| 6062 | ENSG0000 | SHBG | sex hormo | 0.162085 | 0.343479 | 0.407211 |
| 33897 | ENSG0000 | AL359397.1 | | 0.138771 | 0.172616 | 0.406745 |
| 12577 | ENSG0000 | DLGAP1 | DLG associ | 0.134262 | 0.241376 | 0.406232 |
| 14245 | ENSG0000 | AP002360.1 | | 0.245038 | 0.337458 | 0.406013 |
| 39696 | ENSG0000 | AC012313.9 | | 0.262056 | 0.30683 | 0.405765 |
| 11785 | ENSG0000 | TMEM92 | transmeml | 0 | 0.04683 | 0.40554 |
| 33669 | ENSG0000 | LINC02313 | long interg | 0.138215 | 0.365317 | 0.405386 |
| 22268 | ENSG0000 | DBH-AS1 | DBH antise | 0 | 0.054654 | 0.40524 |
| 2503 | ENSG0000 | CCM2L | CCM2 like | 0.185756 | 0.258767 | 0.404799 |
| 23706 | ENSG0000 | AC016708.1 | | 0.107409 | 0.249007 | 0.404465 |
| 8544 | ENSG0000 | MED12L | mediator c | 0.128043 | 0.261236 | 0.404317 |
| 8067 | ENSG0000 | HUNK | hormonally | 0.102854 | 0.372588 | 0.40391 |
| 37861 | ENSG0000 | AC003002.4 | | 0.098399 | 0.385864 | 0.403283 |
| 300 | ENSG0000 | CD4 | CD4 molec | 0.117361 | 0.263329 | 0.402879 |
| 20257 | ENSG0000 | MPRIPP1 | myosin pho | 0.25946 | 0.266295 | 0.402625 |
| 37931 | ENSG0000 | AP006621.4 | | 0.148806 | 0.38887 | 0.402476 |
| 545 | ENSG0000 | SH3YL1 | SH3 and SY | 0.356507 | 0.390826 | 0.402165 |
| 6950 | ENSG0000 | LMO2 | LIM domai | 0.137234 | 0.315052 | 0.401754 |
| 20381 | ENSG0000 | AC116351.1 | | 0 | 0.071116 | 0.401433 |
| 32728 | ENSG0000 | AC117498.2 | | 0 | 0.084695 | 0.401078 |
| 26580 | ENSG0000 | AL662800.1 | | 0 | 0.231049 | 0.400762 |
| 39321 | ENSG0000 | AP000350.5 | | 0.080542 | 0.091375 | 0.400744 |
| 22274 | ENSG0000 | DHRS4L1 | dehydroge | 0.107719 | 0.253028 | 0.400712 |
| 34765 | ENSG0000 | AL133297.2 | | 0.03659 | 0.207103 | 0.400415 |
| 28483 | ENSG0000 | AC098820.3 | | 0.051762 | 0.086736 | 0.400237 |
| 36552 | ENSG0000 | LINC01841 | long interg | 0.13711 | 0.167249 | 0.400114 |
| 41528 | ENSG0000 | AC145207.9 | | 0.17504 | 0.322913 | 0.399394 |
| 2067 | ENSG0000 | EFHC1 | EF-hand dc | 0.196668 | 0.288079 | 0.399393 |
| 37975 | ENSG0000 | AC078795.1 | | 0.330943 | 0.357042 | 0.399228 |
| 8170 | ENSG0000 | PADI3 | peptidyl ar | 0.005644 | 0.045678 | 0.399127 |
| 23766 | ENSG0000 | ACAP2-IT1 | ACAP2 intr | 0.187382 | 0.241287 | 0.398469 |
| 27582 | ENSG0000 | SPATA3-AS | SPATA3 an | 0.096004 | 0.249793 | 0.398442 |
| 21384 | ENSG0000 | UBE2E1-AS | UBE2E1 an | 0.054781 | 0.21665 | 0.397763 |
| 18662 | ENSG0000 | Z69890.1 | | 0.097271 | 0.347286 | 0.39747 |
| 10394 | ENSG0000 | FOXH1 | forkhead b | 0.115048 | 0.392168 | 0.397299 |
| 37488 | ENSG0000 | AC020915.3 | | 0.083307 | 0.103443 | 0.397275 |
| 42490 | ENSG0000 | AL355480.1 | | 0.283487 | 0.331292 | 0.397026 |
| 32029 | ENSG0000 | AL354919.2 | | 0 | 0.186936 | 0.396943 |
| 2460 | ENSG0000 | HRH3 | histamine r | 0.130571 | 0.226388 | 0.396583 |
| 10706 | ENSG0000 | C2orf48 | chromosor | 0.0458 | 0.242427 | 0.396418 |
| 39091 | ENSG0000 | AC007881.3 | | 0.208382 | 0.328974 | 0.396273 |
| 22055 | ENSG0000 | RBMS1P1 | RNA bindir | 0.261026 | 0.318949 | 0.395784 |
| 40520 | ENSG0000 | TTC28-AS1 | TTC28 anti | 0.311577 | 0.373474 | 0.393957 |
| 8495 | ENSG0000 | GRIP2 | glutamate | 0.290549 | 0.391285 | 0.393917 |
| 15343 | ENSG0000 | RPL7AP49 | ribosomal | 0.001405 | 0.09276 | 0.393873 |

| | | | | | | |
|---|---|---|---|---|---|---|
| 2045 | ENSG0000 | CYP26A1 | cytochrom | 0.046998 | 0.078044 | 0.393675 |
| 22235 | ENSG0000 | AC002378.1 | | 0.110654 | 0.340754 | 0.393564 |
| 41405 | ENSG0000 | MIR6736 | microRNA | 0.031998 | 0.128427 | 0.393283 |
| 908 | ENSG0000 | BCAS1 | breast carc | 0.034607 | 0.370196 | 0.393232 |
| 29993 | ENSG0000 | TMEM150 | transmeml | 0.11899 | 0.165856 | 0.393187 |
| 34816 | ENSG0000 | CDC37P2 | cell divisio | 0 | 0.092121 | 0.392646 |
| 14672 | ENSG0000 | AL591379.1 | | 0.035918 | 0.330238 | 0.391451 |
| 10327 | ENSG0000 | HIPK4 | homeodon | 0.144799 | 0.18844 | 0.390942 |
| 39441 | ENSG0000 | AC062037.2 | | 0.036206 | 0.290914 | 0.3905 |
| 13040 | ENSG0000 | NIPAL4 | NIPA like d | 0.220162 | 0.291046 | 0.390006 |
| 18903 | ENSG0000 | RNU6-30P | RNA, U6 sr | 0 | 0.263027 | 0.389699 |
| 4722 | ENSG0000 | SLC8A2 | solute carr | 0.077294 | 0.228253 | 0.389493 |
| 37739 | ENSG0000 | AP001462.1 | | 0.220252 | 0.313606 | 0.389338 |
| 2260 | ENSG0000 | RASD2 | RASD famil | 0.010439 | 0.217512 | 0.389036 |
| 36739 | ENSG0000 | AC005972.1 | | 0.119754 | 0.253262 | 0.388756 |
| 676 | ENSG0000 | TNFRSF9 | TNF recept | 0.072889 | 0.163194 | 0.388645 |
| 18127 | ENSG0000 | TNRC6C-AS | TNRC6C an | 0.106816 | 0.209678 | 0.388574 |
| 37260 | ENSG0000 | AC004156.1 | | 0.099706 | 0.192888 | 0.388218 |
| 7265 | ENSG0000 | SLC2A8 | solute carr | 0.248182 | 0.385182 | 0.388034 |
| 11098 | ENSG0000 | FAM160A1 | family with | 0.088247 | 0.194581 | 0.387637 |
| 39319 | ENSG0000 | AC107294.2 | | 0 | 0.089142 | 0.387461 |
| 42172 | ENSG0000 | AC015813.7 | | 0.144055 | 0.374038 | 0.386722 |
| 26632 | ENSG0000 | TM4SF19-A | TM4SF19 a | 0.206346 | 0.271929 | 0.386352 |
| 22529 | ENSG0000 | NMD3P1 | NMD3 ribo | 0.208483 | 0.285648 | 0.386296 |
| 32618 | ENSG0000 | AC025576.1 | | 0.153754 | 0.363692 | 0.385366 |
| 26835 | ENSG0000 | ENO1-IT1 | ENO1 intro | 0.090435 | 0.163198 | 0.385046 |
| 3690 | ENSG0000 | B3GAT1 | beta-1,3-gl | 0.054474 | 0.114879 | 0.384823 |
| 23971 | ENSG0000 | AL157373.2 | | 0.043621 | 0.159621 | 0.384544 |
| 42546 | ENSG0000 | ARRDC3-AS | ARRDC3 ar | 0.268698 | 0.324835 | 0.384414 |
| 32331 | ENSG0000 | GRM5-AS1 | GRM5 anti | 0.091137 | 0.365402 | 0.384114 |
| 31224 | ENSG0000 | Y_RNA | Y RNA [Sou | 0 | 0.050472 | 0.384096 |
| 27146 | ENSG0000 | NDUFA6-A | NDUFA6 ar | 0.321355 | 0.329216 | 0.383532 |
| 6559 | ENSG0000 | SYT11 | synaptotag | 0.225801 | 0.307384 | 0.383477 |
| 22518 | ENSG0000 | ZNF192P1 | zinc finger | 0.223724 | 0.245727 | 0.38347 |
| 35834 | ENSG0000 | AC009121.2 | | 0.212048 | 0.283909 | 0.382494 |
| 24594 | ENSG0000 | ITPR1-AS1 | ITPR1 antis | 0.062049 | 0.25953 | 0.382364 |
| 18536 | ENSG0000 | SNORA80B | small nucle | 0.286883 | 0.338836 | 0.381773 |
| 23482 | ENSG0000 | AL022313.2 | | 0 | 0.166553 | 0.381664 |
| 38767 | ENSG0000 | AC005618.1 | | 0.142408 | 0.241877 | 0.380881 |
| 15507 | ENSG0000 | PMEL | premelano | 0.081367 | 0.214093 | 0.380706 |
| 34 | ENSG0000 | KLHL13 | kelch like f | 0.236704 | 0.282161 | 0.380602 |
| 42682 | ENSG0000 | AL021920.3 | | 0.124593 | 0.277205 | 0.380344 |
| 33970 | ENSG0000 | UBE2F-SCL | UBE2F-SCL | 0.000447 | 0.104144 | 0.380011 |
| 18672 | ENSG0000 | FOXL2NB | FOXL2 neig | 0.048155 | 0.083646 | 0.38001 |
| 33306 | ENSG0000 | AC005479.2 | | 0.090601 | 0.351927 | 0.379714 |
| 33101 | ENSG0000 | NHLRC4 | NHL repea | 0.016861 | 0.119233 | 0.378614 |
| 24150 | ENSG0000 | HNRNPA1F | heterogen | 0.171787 | 0.275134 | 0.378395 |

| | | | | | | |
|---|---|---|---|---|---|---|
| 42474 | ENSG0000 | AL139035.1 | | 0.280602 | 0.349667 | 0.378307 |
| 38947 | ENSG0000 | AL356056.3 | | 0 | 0.095408 | 0.378126 |
| 40528 | ENSG0000 | AC087683.2 | | 0.184541 | 0.230215 | 0.378084 |
| 1286 | ENSG0000 | TACR2 | tachykinin | 0.077705 | 0.245313 | 0.377538 |
| 34878 | ENSG0000 | AL031598.1 | | 0.023212 | 0.072158 | 0.376939 |
| 7506 | ENSG0000 | SLC3A1 | solute carr | 0.107009 | 0.240508 | 0.376731 |
| 24477 | ENSG0000 | AL049649.1 | | 0.116735 | 0.208302 | 0.376402 |
| 21987 | ENSG0000 | LINC00705 | long interg | 0.196993 | 0.366496 | 0.376243 |
| 39118 | ENSG0000 | AC019193.2 | | 0.04546 | 0.161628 | 0.376135 |
| 30183 | ENSG0000 | AC021106.1 | | 0.190614 | 0.282648 | 0.376117 |
| 33510 | ENSG0000 | AC008083.2 | | 0.054733 | 0.179326 | 0.375982 |
| 7803 | ENSG0000 | AK7 | adenylate l | 0.092488 | 0.19259 | 0.375958 |
| 18597 | ENSG0000 | TPTE2P6 | transmeml | 0.058391 | 0.240589 | 0.375924 |
| 25118 | ENSG0000 | AL355994.2 | | 0.056475 | 0.107366 | 0.375922 |
| 1838 | ENSG0000 | OAS1 | 2'-5'-oligoa | 0.074346 | 0.242979 | 0.375716 |
| 28 | ENSG0000 | HECW1 | HECT, C2 a | 0.206542 | 0.277301 | 0.375609 |
| 25896 | ENSG0000 | AL355355.2 | | 0.105751 | 0.280298 | 0.375536 |
| 34599 | ENSG0000 | AC007611.1 | | 0.083272 | 0.126972 | 0.375509 |
| 36909 | ENSG0000 | ZNF561-AS | ZNF561 an | 0.284169 | 0.288986 | 0.375324 |
| 5212 | ENSG0000 | LDB3 | LIM domai | 0.069239 | 0.128863 | 0.375243 |
| 7080 | ENSG0000 | GPR45 | G protein-c | 0.010304 | 0.056396 | 0.375195 |
| 2135 | ENSG0000 | IGF2-AS | IGF2 antise | 0.093091 | 0.124136 | 0.374988 |
| 29452 | ENSG0000 | AC079089.1 | | 0.126434 | 0.152985 | 0.374752 |
| 42098 | ENSG0000 | AC129778.1 | | 0.024976 | 0.301434 | 0.37462 |
| 4635 | ENSG0000 | KIF17 | kinesin fam | 0.059117 | 0.134739 | 0.374463 |
| 12420 | ENSG0000 | ONECUT1 | one cut ho | 0.1051 | 0.195659 | 0.373673 |
| 29484 | ENSG0000 | MIR4458H | MIR4458 h | 0.247168 | 0.271615 | 0.373014 |
| 31576 | ENSG0000 | PCDHGA7 | protocadh | 0.013065 | 0.032123 | 0.372495 |
| 22383 | ENSG0000 | KCNIP2-AS | KCNIP2 ant | 0.213811 | 0.217825 | 0.371638 |
| 5547 | ENSG0000 | SH3TC1 | SH3 domai | 0.071911 | 0.178046 | 0.371411 |
| 30425 | ENSG0000 | FAM218A | family with | 0.179558 | 0.260358 | 0.371122 |
| 24597 | ENSG0000 | C17orf105 | chromosor | 0.018981 | 0.090986 | 0.370995 |
| 12482 | ENSG0000 | AL512625.1 | | 0.158846 | 0.356357 | 0.370163 |
| 38910 | ENSG0000 | AL158211.3 | | 0.159561 | 0.36923 | 0.370083 |
| 32371 | ENSG0000 | STX17-AS1 | STX17 anti | 0 | 0.239856 | 0.369344 |
| 30196 | ENSG0000 | AC006487.1 | | 0.0856 | 0.157998 | 0.368921 |
| 41633 | ENSG0000 | AC080080.1 | | 0.154151 | 0.240347 | 0.368865 |
| 27288 | ENSG0000 | BMS1P10 | BMS1, ribo | 0.162605 | 0.336464 | 0.368801 |
| 38233 | ENSG0000 | AC104695.3 | | 0.099921 | 0.195734 | 0.36867 |
| 305 | ENSG0000 | FYN | FYN proto- | 0.142333 | 0.277314 | 0.368201 |
| 10229 | ENSG0000 | TPPP3 | tubulin pol | 0.074612 | 0.363152 | 0.367947 |
| 34075 | ENSG0000 | AC020891.2 | | 0.037141 | 0.213026 | 0.367674 |
| 25984 | ENSG0000 | CCDC188 | coiled-coil | 0.17909 | 0.354367 | 0.366906 |
| 10771 | ENSG0000 | ALPP | alkaline ph | 0.101769 | 0.181075 | 0.366746 |
| 13815 | ENSG0000 | RGMA | repulsive g | 0.132758 | 0.158017 | 0.366702 |
| 13737 | ENSG0000 | FOXC2 | forkhead b | 0.014069 | 0.16433 | 0.365933 |
| 7725 | ENSG0000 | PDX1 | pancreatic | 0.097225 | 0.3352 | 0.365921 |

| | | | | | | |
|---|---|---|---|---|---|---|
| 32940 | ENSG0000 | AC080075.1 | | 0.080762 | 0.241914 | 0.365324 |
| 29750 | ENSG0000 | CCDC153 | coiled-coil | 0.04732 | 0.070139 | 0.36507 |
| 2446 | ENSG0000 | SALL4 | spalt like tr | 0.07573 | 0.221814 | 0.36477 |
| 36128 | ENSG0000 | MIR4786 | microRNA | 0 | 0.043814 | 0.364712 |
| 31561 | ENSG0000 | AC004080.2 | | 0.23741 | 0.32254 | 0.364358 |
| 41915 | ENSG0000 | AL132780.4 | | 0.326678 | 0.328165 | 0.364244 |
| 13563 | ENSG0000 | LONRF3 | LON peptic | 0.09165 | 0.138754 | 0.36397 |
| 24089 | ENSG0000 | AC016738.2 | | 0 | 0.092664 | 0.363938 |
| 26296 | ENSG0000 | AL138895.1 | | 0.124403 | 0.162085 | 0.363865 |
| 21622 | ENSG0000 | VTA1P1 | vesicle traf | 0.083679 | 0.20856 | 0.363704 |
| 39531 | ENSG0000 | AC106739.1 | | 0.124098 | 0.182968 | 0.363246 |
| 34248 | ENSG0000 | CERNA1 | competing | 0.034106 | 0.112141 | 0.362531 |
| 9161 | ENSG0000 | ARID5B | AT-rich inte | 0.202 | 0.271618 | 0.361339 |
| 15887 | ENSG0000 | DPPA3 | developme | 0 | 0.154302 | 0.361324 |
| 40538 | ENSG0000 | AL118522.1 | | 0.101136 | 0.146352 | 0.36102 |
| 23457 | ENSG0000 | TRGV5P | T-cell recep | 0 | 0.322378 | 0.360185 |
| 3482 | ENSG0000 | RUNDC3A | RUN doma | 0.115036 | 0.205286 | 0.35976 |
| 38689 | ENSG0000 | AC008972.1 | | 0.055456 | 0.1999 | 0.358968 |
| 29124 | ENSG0000 | RPL35P6 | ribosomal | 0 | 0.341022 | 0.358814 |
| 25607 | ENSG0000 | AL391845.2 | | 0.018228 | 0.099662 | 0.35866 |
| 26568 | ENSG0000 | SEPT14P12 | septin 14 p | 0.025846 | 0.156293 | 0.358558 |
| 11761 | ENSG0000 | EVA1C | eva-1 hom | 0.05258 | 0.238678 | 0.358182 |
| 6109 | ENSG0000 | FOXJ1 | forkhead b | 0.087591 | 0.255678 | 0.358156 |
| 17966 | ENSG0000 | CATSPER2P | cation char | 0.240814 | 0.306199 | 0.357807 |
| 11702 | ENSG0000 | YPEL4 | yippee like | 0.079442 | 0.296644 | 0.357548 |
| 40226 | ENSG0000 | AC012645.4 | | 0.05077 | 0.221145 | 0.357493 |
| 22226 | ENSG0000 | AC064875.1 | | 0.049172 | 0.165409 | 0.35729 |
| 15991 | ENSG0000 | SNHG28 | small nucle | 0.067266 | 0.173385 | 0.356988 |
| 10237 | ENSG0000 | TCAF2P1 | TRPM8 cha | 0.047665 | 0.064242 | 0.356766 |
| 14384 | ENSG0000 | SEPT1 | septin 1 [S | 0.203184 | 0.307735 | 0.355993 |
| 826 | ENSG0000 | GNA15 | G protein s | 0.013582 | 0.224634 | 0.355819 |
| 11875 | ENSG0000 | SERPINF2 | serpin fam | 0.118517 | 0.155602 | 0.355693 |
| 42296 | ENSG0000 | AC135050.7 | | 0.263563 | 0.294283 | 0.355687 |
| 4416 | ENSG0000 | IL18R1 | interleukin | 0.116923 | 0.190028 | 0.355463 |
| 39364 | ENSG0000 | AC012531.3 | | 0 | 0.131505 | 0.355242 |
| 8278 | ENSG0000 | RGL1 | ral guanine | 0.164292 | 0.271865 | 0.354526 |
| 6134 | ENSG0000 | SYT5 | synaptotag | 0.029288 | 0.235589 | 0.353905 |
| 38150 | ENSG0000 | AC145285.6 | | 0.066881 | 0.347305 | 0.353536 |
| 22033 | ENSG0000 | IGKV1OR9- | immunoglo | 0 | 0.303877 | 0.353525 |
| 42896 | ENSG0000 | AL136985.3 | | 0.196555 | 0.254024 | 0.353362 |
| 14377 | ENSG0000 | C3orf80 | chromosor | 0.058993 | 0.272831 | 0.35264 |
| 16998 | ENSG0000 | ANKRD35 | ankyrin rep | 0.066503 | 0.228283 | 0.352247 |
| 14327 | ENSG0000 | PCBP1-AS1 | PCBP1 anti | 0.237847 | 0.33136 | 0.351363 |
| 25935 | ENSG0000 | AC123595.1 | | 0.135845 | 0.218797 | 0.351308 |
| 9513 | ENSG0000 | PLEKHG4B | pleckstrin l | 0.062242 | 0.224938 | 0.351246 |
| 37338 | ENSG0000 | AP005264.6 | | 0.056582 | 0.151179 | 0.351204 |
| 29296 | ENSG0000 | MIATNB | MIAT neigh | 0.166224 | 0.302432 | 0.350666 |

| | | | | | | |
|---|---|---|---|---|---|---|
| 24921 | ENSG0000 | AC005537.1 | | 0.124288 | 0.253399 | 0.350599 |
| 25729 | ENSG0000 | LINC01237 | long interg | 0.241109 | 0.260035 | 0.350505 |
| 26314 | ENSG0000 | AC003984.1 | | 0.090509 | 0.307296 | 0.350466 |
| 12246 | ENSG0000 | GRM5 | glutamate | 0.076841 | 0.132842 | 0.350358 |
| 10928 | ENSG0000 | HESX1 | HESX home | 0.252423 | 0.344545 | 0.349285 |
| 37239 | ENSG0000 | AC040904.1 | | 0.012417 | 0.045305 | 0.349246 |
| 39974 | ENSG0000 | TBC1D3B | TBC1 doma | 0.169858 | 0.337394 | 0.349027 |
| 28136 | ENSG0000 | RN7SL263F | RNA, 7SL, c | 0.185316 | 0.196228 | 0.348967 |
| 27762 | ENSG0000 | SNORD121 | small nucle | 0.165765 | 0.184809 | 0.348812 |
| 11782 | ENSG0000 | SAMD14 | sterile alph | 0.048853 | 0.067232 | 0.348565 |
| 28243 | ENSG0000 | AQP1 | aquaporin | 0.037763 | 0.140244 | 0.34805 |
| 38661 | ENSG0000 | AL096865.1 | | 0.180646 | 0.265679 | 0.34802 |
| 8593 | ENSG0000 | CAMK2D | calcium/ca | 0.253062 | 0.339125 | 0.347823 |
| 5369 | ENSG0000 | BATF3 | basic leuci | 0.085452 | 0.300951 | 0.346162 |
| 9979 | ENSG0000 | MX1 | MX dynam | 0.137092 | 0.302232 | 0.345631 |
| 14204 | ENSG0000 | EXOC3L1 | exocyst co | 0.11716 | 0.11944 | 0.345488 |
| 32796 | ENSG0000 | AC084117.1 | | 0.042498 | 0.272557 | 0.345464 |
| 25712 | ENSG0000 | AC092835.1 | | 0 | 0.114501 | 0.345158 |
| 18015 | ENSG0000 | RBM20 | RNA bindir | 0.107139 | 0.151656 | 0.345108 |
| 39825 | ENSG0000 | AC103691.1 | | 0.159599 | 0.222592 | 0.344838 |
| 17672 | ENSG0000 | SNORA63C | small nucle | 0 | 0.130262 | 0.344665 |
| 36346 | ENSG0000 | MIR4262 | microRNA | 0 | 0.101182 | 0.34461 |
| 34212 | ENSG0000 | AC027243.1 | | 0.067013 | 0.231337 | 0.343527 |
| 7151 | ENSG0000 | MMP13 | matrix met | 0.044011 | 0.240958 | 0.343516 |
| 30093 | ENSG0000 | LINC01234 | long interg | 0.039605 | 0.089644 | 0.343478 |
| 14722 | ENSG0000 | PRR34 | proline rich | 0.094628 | 0.102652 | 0.343305 |
| 31047 | ENSG0000 | RNA5SP31 | RNA, 5S rib | 0.084086 | 0.090205 | 0.343124 |
| 6912 | ENSG0000 | BICDL1 | BICD family | 0.211966 | 0.330326 | 0.343055 |
| 22371 | ENSG0000 | AC102953.1 | | 0.059358 | 0.079998 | 0.342995 |
| 12480 | ENSG0000 | RNF150 | ring finger | 0 | 0.330305 | 0.342939 |
| 15132 | ENSG0000 | SCN5A | sodium vol | 0.033657 | 0.190894 | 0.342754 |
| 38552 | ENSG0000 | MILR1 | mast cell ir | 0.039227 | 0.291596 | 0.342698 |
| 26500 | ENSG0000 | NFAM1 | NFAT activ | 0.145028 | 0.154379 | 0.342575 |
| 20308 | ENSG0000 | AC129492.1 | | 0.019674 | 0.11332 | 0.341935 |
| 5718 | ENSG0000 | F10 | coagulatio | 0.009563 | 0.171454 | 0.341338 |
| 17073 | ENSG0000 | TEX45 | testis expr | 0.039549 | 0.144746 | 0.341324 |
| 13069 | ENSG0000 | PURG | purine rich | 0.052398 | 0.096503 | 0.340802 |
| 33821 | ENSG0000 | AC105001.2 | | 0.194745 | 0.31944 | 0.340496 |
| 13251 | ENSG0000 | TAS1R1 | taste 1 rec | 0.068171 | 0.209384 | 0.340142 |
| 35348 | ENSG0000 | AC009097.4 | | 0.077714 | 0.231668 | 0.340027 |
| 21748 | ENSG0000 | TNK2-AS1 | TNK2 antis | 0.072198 | 0.265835 | 0.339898 |
| 21964 | ENSG0000 | AC078883.1 | | 0.044233 | 0.046412 | 0.339656 |
| 30115 | ENSG0000 | AC106795.2 | | 0.260252 | 0.336726 | 0.339583 |
| 22858 | ENSG0000 | AL158166.1 | | 0 | 0.115211 | 0.339561 |
| 9299 | ENSG0000 | AKR1C2 | aldo-keto r | 0.191228 | 0.308046 | 0.339547 |
| 41447 | ENSG0000 | GOLGA6L1 | golgin A6 f | 0.160888 | 0.226931 | 0.339256 |
| 23098 | ENSG0000 | FAM212B-. | FAM212B a | 0.041756 | 0.07513 | 0.339166 |

| | | | | | | |
|---|---|---|---|---|---|---|
| 5519 | ENSG0000 | CPNE5 | copine 5 [S | 0.041724 | 0.071959 | 0.339027 |
| 6113 | ENSG0000 | ARHGEF6 | Rac/Cdc42 | 0.055339 | 0.227564 | 0.338771 |
| 40959 | ENSG0000 | AC008406.3 | | 0.207016 | 0.242147 | 0.338617 |
| 26033 | ENSG0000 | PTGES3P1 | prostaglan | 0.165205 | 0.265305 | 0.338545 |
| 18656 | ENSG0000 | TMEM191 | transmeml | 0.159027 | 0.300804 | 0.338377 |
| 16351 | ENSG0000 | SRGAP3 | SLIT-ROBO | 0.11553 | 0.217502 | 0.338268 |
| 5731 | ENSG0000 | HCST | hematopoi | 0 | 0.071139 | 0.338239 |
| 34940 | ENSG0000 | HERC2P5 | hect doma | 0.119475 | 0.23537 | 0.337617 |
| 622 | ENSG0000 | ADRB1 | adrenocep | 0.143284 | 0.15292 | 0.337322 |
| 33789 | ENSG0000 | HIF1A-AS2 | HIF1A antis | 0 | 0.027559 | 0.337282 |
| 38277 | ENSG0000 | AL136141.1 | | 0.100277 | 0.142763 | 0.337138 |
| 3019 | ENSG0000 | IL23A | interleukin | 0.153639 | 0.319695 | 0.336643 |
| 25933 | ENSG0000 | H2BFS | H2B histon | 0.191706 | 0.270054 | 0.336536 |
| 10339 | ENSG0000 | TMEM190 | transmeml | 0.204402 | 0.28779 | 0.336438 |
| 1270 | ENSG0000 | CA12 | carbonic ai | 0.091352 | 0.198278 | 0.336198 |
| 33489 | ENSG0000 | BCDIN3D-A | BCDIN3D a | 0.120363 | 0.188825 | 0.335911 |
| 16087 | ENSG0000 | CERKL | ceramide k | 0.197676 | 0.264361 | 0.335747 |
| 38098 | ENSG0000 | AC096720.1 | | 0 | 0.113231 | 0.335194 |
| 16278 | ENSG0000 | S100A14 | S100 calciu | 0.069779 | 0.227656 | 0.334625 |
| 31315 | ENSG0000 | AC004707.1 | | 0.075814 | 0.234075 | 0.334506 |
| 29070 | ENSG0000 | RPL12P6 | ribosomal | 0.317553 | 0.324626 | 0.334499 |
| 8830 | ENSG0000 | FRMPD3 | FERM and | 0.048499 | 0.085846 | 0.33445 |
| 27357 | ENSG0000 | AL355803.1 | | 0 | 0.175384 | 0.334223 |
| 9173 | ENSG0000 | FAM124A | family with | 0.243524 | 0.32496 | 0.333988 |
| 32861 | ENSG0000 | AP003721.1 | | 0.035157 | 0.209738 | 0.333827 |
| 13226 | ENSG0000 | CSPG4 | chondroitir | 0.070868 | 0.121496 | 0.333667 |
| 14904 | ENSG0000 | ANKRD20A | ankyrin rep | 0.208054 | 0.253048 | 0.333422 |
| 40186 | ENSG0000 | AC010503.4 | | 0.002561 | 0.155685 | 0.332922 |
| 2594 | ENSG0000 | SRPX | sushi repea | 0.231828 | 0.240108 | 0.332719 |
| 6 | ENSG0000 | FGR | FGR proto- | 0.258594 | 0.305841 | 0.332673 |
| 9139 | ENSG0000 | FAM57B | family with | 0.086062 | 0.104352 | 0.332534 |
| 24384 | ENSG0000 | AC012456.2 | | 0 | 0.137113 | 0.331682 |
| 41261 | ENSG0000 | AC093525.9 | | 0.067579 | 0.252768 | 0.331432 |
| 35731 | ENSG0000 | PCDHGA4 | protocadh | 0.112621 | 0.322563 | 0.33143 |
| 15334 | ENSG0000 | NUTM2A | NUT family | 0.268963 | 0.284193 | 0.331174 |
| 9347 | ENSG0000 | SCHIP1 | schwannor | 0.160019 | 0.319086 | 0.331013 |
| 23395 | ENSG0000 | AC099552.3 | | 0 | 0.230194 | 0.330908 |
| 3430 | ENSG0000 | SPOCK2 | SPARC/ost | 0.210612 | 0.245329 | 0.330795 |
| 23241 | ENSG0000 | TLE1P1 | transducin | 0.094127 | 0.147373 | 0.330665 |
| 35232 | ENSG0000 | FIGNL2 | fidgetin lik | 0 | 0.039605 | 0.330611 |
| 16931 | ENSG0000 | RPS6KL1 | ribosomal | 0.261312 | 0.323089 | 0.330302 |
| 15038 | ENSG0000 | STKLD1 | serine/thre | 0.290354 | 0.320164 | 0.329608 |
| 21202 | ENSG0000 | RNU6-387I | RNA, U6 sr | 0 | 0.094255 | 0.329294 |
| 35472 | ENSG0000 | AL034346.1 | | 0.027793 | 0.0687 | 0.328868 |
| 39275 | ENSG0000 | AC006480.2 | | 0.136119 | 0.278199 | 0.328801 |
| 13383 | ENSG0000 | MFSD4A | major facil | 0.04304 | 0.090414 | 0.328335 |
| 4191 | ENSG0000 | DPYSL3 | dihydropyr | 0.230292 | 0.231538 | 0.327429 |

| | | | | | | |
|---|---|---|---|---|---|---|
| 23046 | ENSG0000 | MED28P3 | mediator c | 0.099312 | 0.298401 | 0.327303 |
| 12595 | ENSG0000 | RASA4B | RAS p21 pr | 0.089623 | 0.262813 | 0.327218 |
| 16096 | ENSG0000 | LCTL | lactase like | 0.111063 | 0.130564 | 0.326878 |
| 8132 | ENSG0000 | RGPD3 | RANBP2-lik | 0.216439 | 0.273866 | 0.326529 |
| 24451 | ENSG0000 | LINC02541 | long interg | 0 | 0.082397 | 0.325868 |
| 14069 | ENSG0000 | VN1R1 | vomeronas | 0.122978 | 0.263398 | 0.325674 |
| 6904 | ENSG0000 | ADAM19 | ADAM met | 0.153582 | 0.178644 | 0.3251 |
| 28963 | ENSG0000 | INGX | inhibitor of | 0.156541 | 0.190916 | 0.325081 |
| 38515 | ENSG0000 | AL161756.2 | | 0 | 0.099319 | 0.324998 |
| 17202 | ENSG0000 | MIR210 | microRNA | 0 | 0.027178 | 0.324769 |
| 20580 | ENSG0000 | CCDC7 | coiled-coil | 0.198991 | 0.288737 | 0.324562 |
| 24064 | ENSG0000 | TMEM254- | TMEM254 | 0.147325 | 0.315198 | 0.324457 |
| 31167 | ENSG0000 | RNA5SP73 | RNA, 5S rib | 0 | 0.294955 | 0.324401 |
| 26072 | ENSG0000 | AC098484.2 | | 0.074761 | 0.139673 | 0.323967 |
| 29476 | ENSG0000 | AC090061.1 | | 0.093786 | 0.256971 | 0.323524 |
| 40378 | ENSG0000 | RN7SL185P | RNA, 7SL, c | 0 | 0.218031 | 0.323158 |
| 35014 | ENSG0000 | AC009065.4 | | 0.089514 | 0.207373 | 0.323089 |
| 3378 | ENSG0000 | KCNT1 | potassium | 0.019678 | 0.091151 | 0.322664 |
| 22848 | ENSG0000 | LINC00629 | long interg | 0.071306 | 0.205292 | 0.322622 |
| 39289 | ENSG0000 | AC090425.2 | | 0.298945 | 0.301659 | 0.322599 |
| 36831 | ENSG0000 | AC011511.2 | | 0.258111 | 0.321038 | 0.322168 |
| 15605 | ENSG0000 | TAL2 | TAL bHLH t | 0.027244 | 0.176795 | 0.322021 |
| 7194 | ENSG0000 | MYCBPAP | MYCBP ass | 0.115718 | 0.182663 | 0.321827 |
| 15625 | ENSG0000 | CYP4Z1 | cytochrom | 0.137538 | 0.17571 | 0.321064 |
| 23041 | ENSG0000 | CROCCP4 | ciliary root | 0.044431 | 0.21205 | 0.321006 |
| 27996 | ENSG0000 | AL157392.3 | | 0.109973 | 0.270705 | 0.320728 |
| 39083 | ENSG0000 | AC112236.2 | | 0.041853 | 0.237414 | 0.320412 |
| 38015 | ENSG0000 | AC068790.5 | | 0 | 0.312467 | 0.320353 |
| 18119 | ENSG0000 | PSMB8-AS1 | PSMB8 ant | 0.061667 | 0.224023 | 0.320242 |
| 2360 | ENSG0000 | CHGA | chromogra | 0.175916 | 0.313119 | 0.32008 |
| 18685 | ENSG0000 | CFAP44 | cilia and fla | 0.199419 | 0.315223 | 0.319961 |
| 11835 | ENSG0000 | PRRT2 | proline rich | 0.098123 | 0.123342 | 0.319745 |
| 43090 | ENSG0000 | AP002851.1 | | 0.050094 | 0.084264 | 0.31916 |
| 34353 | ENSG0000 | AC009269.5 | | 0.047176 | 0.187451 | 0.318904 |
| 114 | ENSG0000 | ASB4 | ankyrin rep | 0.019431 | 0.101434 | 0.318031 |
| 35379 | ENSG0000 | AC097639.1 | | 0.018969 | 0.129804 | 0.317699 |
| 22898 | ENSG0000 | AC254562.1 | | 0.060077 | 0.073048 | 0.31731 |
| 37723 | ENSG0000 | AC022150.2 | | 0.102071 | 0.199057 | 0.31707 |
| 20825 | ENSG0000 | MTMR9LP | myotubula | 0.115751 | 0.267538 | 0.316945 |
| 37209 | ENSG0000 | AC010327.3 | | 0.029842 | 0.195309 | 0.316851 |
| 25403 | ENSG0000 | AL353764.1 | | 0 | 0.236413 | 0.31674 |
| 10918 | ENSG0000 | FAM86GP | family with | 0.202233 | 0.263904 | 0.316632 |
| 13010 | ENSG0000 | PRSS27 | protease, s | 0.140393 | 0.160886 | 0.316581 |
| 11002 | ENSG0000 | LIPH | lipase H [So | 0.008579 | 0.240921 | 0.316524 |
| 39071 | ENSG0000 | AC073107.1 | | 0.222488 | 0.291328 | 0.31652 |
| 37345 | ENSG0000 | FAM60CP | family with | 0.067384 | 0.120868 | 0.316498 |
| 24698 | ENSG0000 | AL157932.1 | | 0.174715 | 0.20673 | 0.316446 |

| | | | | | | |
|---|---|---|---|---|---|---|
| 14847 | ENSG0000 | SLC25A18 | solute carr | 0.172554 | 0.186029 | 0.316414 |
| 19497 | ENSG0000 | RNA5SP21 | RNA, 5S rib | 0 | 0.20527 | 0.316146 |
| 15891 | ENSG0000 | ZNF385C | zinc finger | 0.080907 | 0.167078 | 0.316071 |
| 11173 | ENSG0000 | GDF9 | growth diff | 0.173255 | 0.255457 | 0.31583 |
| 24423 | ENSG0000 | AGAP10P | ArfGAP wit | 0.175773 | 0.298901 | 0.315823 |
| 35389 | ENSG0000 | AC093249.4 | | 0 | 0.131749 | 0.315781 |
| 30079 | ENSG0000 | AC006160.1 | | 0.027881 | 0.193903 | 0.315226 |
| 5898 | ENSG0000 | GAL3ST1 | galactose-3 | 0 | 0.038544 | 0.314576 |
| 33042 | ENSG0000 | AC148477.3 | | 0 | 0.058297 | 0.314392 |
| 14871 | ENSG0000 | SPNS2 | sphingolipi | 0.186584 | 0.295512 | 0.314006 |
| 2993 | ENSG0000 | AHR | aryl hydroc | 0.095585 | 0.278927 | 0.313679 |
| 20544 | ENSG0000 | AP000697.1 | | 0 | 0.302689 | 0.313546 |
| 19670 | ENSG0000 | CRIP1 | cysteine ric | 0.08505 | 0.199594 | 0.313473 |
| 24428 | ENSG0000 | AC005077.4 | | 0.035255 | 0.11837 | 0.31332 |
| 37712 | ENSG0000 | AC005387.1 | | 0 | 0.09641 | 0.313309 |
| 16865 | ENSG0000 | LEKR1 | leucine, glu | 0.098057 | 0.258459 | 0.312964 |
| 16963 | ENSG0000 | HOXC4 | homeobox | 0.134246 | 0.167835 | 0.31293 |
| 19899 | ENSG0000 | ATP6V1G2 | ATPase H+ | 0.041386 | 0.20596 | 0.3124 |
| 12430 | ENSG0000 | MUC3A | mucin 3A, | 0.037401 | 0.112141 | 0.312292 |
| 43335 | ENSG0000 | AL022318.4 | | 0.101069 | 0.146453 | 0.312225 |
| 36400 | ENSG0000 | MIR3136 | microRNA | 0 | 0.103091 | 0.31221 |
| 27966 | ENSG0000 | LINC00920 | long interg | 0.066417 | 0.178381 | 0.312187 |
| 27948 | ENSG0000 | PICART1 | p53-induci | 0.128519 | 0.22843 | 0.31213 |
| 15922 | ENSG0000 | DNAJB13 | DnaJ heat s | 0.014377 | 0.30786 | 0.312117 |
| 22767 | ENSG0000 | AL359541.1 | | 0 | 0.172296 | 0.311447 |
| 7920 | ENSG0000 | IRF8 | interferon | 0.044404 | 0.086042 | 0.311002 |
| 39783 | ENSG0000 | ADRA2B | adrenocep | 0.015756 | 0.120247 | 0.310943 |
| 34511 | ENSG0000 | AC027373.1 | | 0.062378 | 0.153553 | 0.310913 |
| 1252 | ENSG0000 | TBX21 | T-box 21 [S | 0.054118 | 0.100074 | 0.310905 |
| 15904 | ENSG0000 | PERM1 | PPARGC1 a | 0.218051 | 0.28133 | 0.310767 |
| 2297 | ENSG0000 | IL2RB | interleukin | 0.026564 | 0.238096 | 0.310735 |
| 11962 | ENSG0000 | GPT | glutamic--p | 0.164038 | 0.272159 | 0.31047 |
| 18723 | ENSG0000 | Y_RNA | Y RNA [Sou | 0 | 0.090089 | 0.310459 |
| 10542 | ENSG0000 | PLPP3 | phospholip | 0.045848 | 0.280838 | 0.310028 |
| 24557 | ENSG0000 | AC002451.1 | | 0 | 0.06647 | 0.309807 |
| 17187 | ENSG0000 | ZFP2 | ZFP2 zinc f | 0.035862 | 0.294042 | 0.309538 |
| 35395 | ENSG0000 | HERC2P8 | hect doma | 0.106427 | 0.149991 | 0.309273 |
| 18858 | ENSG0000 | SNORA15 | small nucle | 0 | 0.292995 | 0.309199 |
| 7125 | ENSG0000 | GPNMB | glycoprote | 0.0913 | 0.116159 | 0.309169 |
| 21680 | ENSG0000 | AC005020.1 | | 0.05934 | 0.084276 | 0.309013 |
| 42369 | ENSG0000 | AC061999.1 | | 0 | 0.253359 | 0.308916 |
| 23474 | ENSG0000 | LINC00659 | long interg | 0 | 0.252 | 0.30866 |
| 15686 | ENSG0000 | SPATA12 | spermatog | 0.088774 | 0.14496 | 0.308621 |
| 12885 | ENSG0000 | ZNF540 | zinc finger | 0.061877 | 0.178014 | 0.308597 |
| 8209 | ENSG0000 | BEST4 | bestrophin | 0.017339 | 0.125552 | 0.307966 |
| 19283 | ENSG0000 | TRGV3 | T-cell recep | 0 | 0.229603 | 0.307817 |
| 18925 | ENSG0000 | RNU6-658I | RNA, U6 sr | 0.078365 | 0.158527 | 0.307765 |

| | | | | | | |
|---|---|---|---|---|---|---|
| 10921 | ENSG0000 | NOD2 | nucleotide | 0.036982 | 0.099696 | 0.307666 |
| 8300 | ENSG0000 | SELENBP1 | selenium b | 0.075896 | 0.09109 | 0.307497 |
| 37707 | ENSG0000 | AC012313.5 | | 0.207514 | 0.235672 | 0.30705 |
| 24631 | ENSG0000 | HLA-DPA1 | major histo | 0.042498 | 0.151561 | 0.306358 |
| 30260 | ENSG0000 | AC091976.1 | | 0.046713 | 0.147942 | 0.306277 |
| 29106 | ENSG0000 | AL390195.1 | | 0.043482 | 0.156181 | 0.305801 |
| 42961 | ENSG0000 | EXOC3L2 | exocyst co | 0.048281 | 0.191504 | 0.305796 |
| 13167 | ENSG0000 | C11orf86 | chromosor | 0.047275 | 0.105697 | 0.305774 |
| 41303 | ENSG0000 | MIR6803 | microRNA | 0.077451 | 0.275967 | 0.305731 |
| 12465 | ENSG0000 | GPR37L1 | G protein-c | 0.005387 | 0.037352 | 0.30557 |
| 34067 | ENSG0000 | CSPG4P12 | chondroitin | 0.140879 | 0.20297 | 0.305329 |
| 16633 | ENSG0000 | ZNF785 | zinc finger | 0.109948 | 0.121893 | 0.304968 |
| 3698 | ENSG0000 | DTX4 | deltex E3 u | 0.011774 | 0.019236 | 0.304722 |
| 3806 | ENSG0000 | WNT5B | Wnt family | 0.142856 | 0.262648 | 0.304486 |
| 26928 | ENSG0000 | AL033528.2 | | 0 | 0.066375 | 0.30405 |
| 37657 | ENSG0000 | AL121912.1 | | 0 | 0.027289 | 0.303585 |
| 13033 | ENSG0000 | CARNS1 | carnosine s | 0.107114 | 0.215352 | 0.303019 |
| 22776 | ENSG0000 | BSN-AS2 | BSN antise | 0.116288 | 0.220402 | 0.302113 |
| 40084 | ENSG0000 | AL109976.1 | | 0.050006 | 0.187663 | 0.301882 |
| 35185 | ENSG0000 | AC005606.2 | | 0.109036 | 0.109528 | 0.301765 |
| 23625 | ENSG0000 | LINC01165 | long interg | 0.005351 | 0.189032 | 0.301507 |
| 11678 | ENSG0000 | CHRFAM7A | CHRNA7 (e | 0.16911 | 0.228376 | 0.301471 |
| 18377 | ENSG0000 | TMEM240 | transmeml | 0.110492 | 0.2357 | 0.301242 |
| 12996 | ENSG0000 | STARD5 | StAR relate | 0.174354 | 0.218909 | 0.30099 |
| 39001 | ENSG0000 | SNORA28 | small nucle | 0.194781 | 0.245455 | 0.30085 |
| 6280 | ENSG0000 | ANGPTL6 | angiopoiet | 0.044458 | 0.143884 | 0.300715 |
| 6022 | ENSG0000 | HRK | harakiri, BC | 0.090634 | 0.133741 | 0.300713 |
| 40530 | ENSG0000 | Y_RNA | Y RNA [Sou | 0.057377 | 0.176872 | 0.300689 |
| 33517 | ENSG0000 | AC106782.1 | | 0.013111 | 0.031993 | 0.300624 |
| 40556 | ENSG0000 | KCNE1B | potassium | 0.119305 | 0.255608 | 0.30016 |
| 42892 | ENSG0000 | AL139352.1 | | 0 | 0.030801 | 0.300085 |
| 25615 | ENSG0000 | AL050331.1 | | 0 | 0.232809 | 0.299905 |
| 34531 | ENSG0000 | AC012414.5 | | 0.127665 | 0.168086 | 0.29983 |
| 11231 | ENSG0000 | INTS4P1 | integrator | 0.139422 | 0.296281 | 0.299794 |
| 29299 | ENSG0000 | LINC02086 | long interg | 0.009351 | 0.13728 | 0.299732 |
| 22193 | ENSG0000 | C2CD4D | C2 calcium | 0.151646 | 0.156729 | 0.299529 |
| 6719 | ENSG0000 | TTC9 | tetratricop | 0.108923 | 0.116667 | 0.298481 |
| 26975 | ENSG0000 | AL354928.1 | | 0.029149 | 0.194784 | 0.29819 |
| 26298 | ENSG0000 | AC128709.2 | | 0.060497 | 0.1727 | 0.297468 |
| 42262 | ENSG0000 | AC008555.8 | | 0.024672 | 0.121113 | 0.297454 |
| 22009 | ENSG0000 | AL391811.1 | | 0.175102 | 0.23127 | 0.297178 |
| 10833 | ENSG0000 | TGFBR2 | transformi | 0.181617 | 0.216658 | 0.296747 |
| 17994 | ENSG0000 | FANK1 | fibronectin | 0.094213 | 0.12869 | 0.29665 |
| 14620 | ENSG0000 | PLAG1 | PLAG1 zinc | 0.090426 | 0.120768 | 0.296583 |
| 35933 | ENSG0000 | MIR3936 | microRNA | 0 | 0.095593 | 0.296573 |
| 29013 | ENSG0000 | RN7SL834F | RNA, 7SL, c | 0 | 0.081103 | 0.296413 |
| 39371 | ENSG0000 | AL449106.1 | | 0 | 0.290019 | 0.296329 |

| | | | | | | |
|---|---|---|---|---|---|---|
| 34204 | ENSG0000 | AL132640.1 | | 0 | 0.19818 | 0.296266 |
| 14617 | ENSG0000 | ATG9B | autophagy | 0.071154 | 0.160791 | 0.295828 |
| 25972 | ENSG0000 | AL513174.1 | | 0.069936 | 0.225689 | 0.295667 |
| 12268 | ENSG0000 | ACTBL2 | actin, beta | 0.095452 | 0.286581 | 0.295397 |
| 34728 | ENSG0000 | LINC02182 | long interg | 0 | 0.103919 | 0.294325 |
| 40937 | ENSG0000 | pRNA | NoRC asso | 0 | 0.100774 | 0.294273 |
| 3101 | ENSG0000 | NKG7 | natural kill | 0.118183 | 0.257328 | 0.294159 |
| 28761 | ENSG0000 | AC108693.2 | | 0 | 0.117451 | 0.293902 |
| 29174 | ENSG0000 | RPS27AP1 | ribosomal | 0.026552 | 0.173993 | 0.293718 |
| 9125 | ENSG0000 | GPHA2 | glycoprote | 0 | 0.183582 | 0.293291 |
| 35554 | ENSG0000 | AC090616.4 | | 0.041953 | 0.079933 | 0.293001 |
| 818 | ENSG0000 | PSD | pleckstrin | 0.217859 | 0.266656 | 0.292608 |
| 6456 | ENSG0000 | SPATA6 | spermatog | 0.103519 | 0.169447 | 0.292431 |
| 34777 | ENSG0000 | AL136982.4 | | 0.115537 | 0.141098 | 0.292158 |
| 27057 | ENSG0000 | CBR3-AS1 | CBR3 antis | 0.149584 | 0.233105 | 0.291865 |
| 10500 | ENSG0000 | ZG16B | zymogen g | 0.053366 | 0.28124 | 0.291811 |
| 36945 | ENSG0000 | AC005379.1 | | 0.13391 | 0.170021 | 0.291469 |
| 30572 | ENSG0000 | AC100861.2 | | 0.204208 | 0.247668 | 0.291214 |
| 30682 | ENSG0000 | AC008592.3 | | 0 | 0.180265 | 0.291159 |
| 16722 | ENSG0000 | PCDHB11 | protocadh | 0.050787 | 0.166448 | 0.290992 |
| 42518 | ENSG0000 | LINC01395 | long interg | 0 | 0.249447 | 0.290684 |
| 27426 | ENSG0000 | NONOP2 | non-POU d | 0.091379 | 0.187139 | 0.290674 |
| 38887 | ENSG0000 | AC026801.3 | | 0.022679 | 0.102498 | 0.290588 |
| 11671 | ENSG0000 | CDH16 | cadherin 1 | 0.039161 | 0.067675 | 0.290246 |
| 6228 | ENSG0000 | TNNI2 | troponin I2 | 0 | 0.215203 | 0.290201 |
| 8463 | ENSG0000 | FAM171B | family with | 0.097515 | 0.250331 | 0.289876 |
| 36376 | ENSG0000 | AC124066.1 | | 0.230456 | 0.254317 | 0.28944 |
| 41578 | ENSG0000 | AC092645.1 | | 0.04559 | 0.057943 | 0.289146 |
| 14880 | ENSG0000 | ABAT | 4-aminobu | 0.100936 | 0.208086 | 0.289093 |
| 18492 | ENSG0000 | FAM239B | family with | 0.111297 | 0.251501 | 0.288868 |
| 28595 | ENSG0000 | RPS4XP19 | ribosomal | 0.07165 | 0.137128 | 0.288735 |
| 36091 | ENSG0000 | MIR3127 | microRNA | 0 | 0.10716 | 0.288686 |
| 40347 | ENSG0000 | FAS-AS1 | FAS antiser | 0 | 0.08126 | 0.288526 |
| 8319 | ENSG0000 | VASH2 | vasohibin 2 | 0.105321 | 0.171227 | 0.28804 |
| 34282 | ENSG0000 | AC104046.1 | | 0.02542 | 0.176981 | 0.287741 |
| 28723 | ENSG0000 | AC083798.1 | | 0 | 0.204608 | 0.287613 |
| 18313 | ENSG0000 | GPR20 | G protein-c | 0 | 0.084242 | 0.287353 |
| 39073 | ENSG0000 | GAS6-AS2 | GAS6 antis | 0.018182 | 0.148616 | 0.287234 |
| 22442 | ENSG0000 | TUBBP10 | tubulin bet | 0.011648 | 0.202661 | 0.286933 |
| 41506 | ENSG0000 | AC099811.5 | | 0.031871 | 0.248602 | 0.286804 |
| 29528 | ENSG0000 | INSL3 | insulin like | 0 | 0.047246 | 0.286744 |
| 18289 | ENSG0000 | AL590399.1 | | 0.185923 | 0.274249 | 0.286552 |
| 36777 | ENSG0000 | AC005838.2 | | 0.057172 | 0.25041 | 0.286269 |
| 37303 | ENSG0000 | AP003680.1 | | 0.02414 | 0.154613 | 0.286264 |
| 29051 | ENSG0000 | HOTTIP | HOXA dista | 0.024765 | 0.149913 | 0.286046 |
| 13618 | ENSG0000 | TMEM270 | transmeml | 0 | 0.129681 | 0.285822 |
| 40506 | ENSG0000 | AC026356.2 | | 0.086335 | 0.220991 | 0.28532 |

| | | | | | | |
|---|---|---|---|---|---|---|
| 2291 | ENSG0000 | CSF2RB | colony stin | 0.181755 | 0.19526 | 0.285061 |
| 35191 | ENSG0000 | AL033527.3 | | 0.177317 | 0.193856 | 0.284966 |
| 13429 | ENSG0000 | CD248 | CD248 mol | 0.006937 | 0.073872 | 0.28475 |
| 37967 | ENSG0000 | AC010969.2 | | 0.235902 | 0.238504 | 0.284704 |
| 32463 | ENSG0000 | AP001922.5 | | 0.081972 | 0.171185 | 0.284616 |
| 34974 | ENSG0000 | AL031713.1 | | 0 | 0.044137 | 0.284358 |
| 5199 | ENSG0000 | FMOD | fibromodu | 0.182324 | 0.22026 | 0.284294 |
| 37394 | ENSG0000 | AC245884.8 | | 0.157895 | 0.196224 | 0.284042 |
| 5446 | ENSG0000 | KCNK15 | potassium | 0.072396 | 0.106106 | 0.283661 |
| 39171 | ENSG0000 | AC092574.1 | | 0.042252 | 0.124936 | 0.283619 |
| 847 | ENSG0000 | CDH3 | cadherin 3 | 0.070364 | 0.08544 | 0.283549 |
| 4400 | ENSG0000 | ST3GAL5 | ST3 beta-g | 0.126407 | 0.151976 | 0.283441 |
| 19755 | ENSG0000 | GSTM2 | glutathione | 0.078569 | 0.180663 | 0.283324 |
| 30101 | ENSG0000 | AC106772.1 | | 0.07724 | 0.154001 | 0.283306 |
| 9388 | ENSG0000 | KCNK13 | potassium | 0.028246 | 0.080936 | 0.283303 |
| 27887 | ENSG0000 | AC007405.3 | | 0 | 0.120315 | 0.283258 |
| 6885 | ENSG0000 | SLC37A2 | solute carr | 0.037946 | 0.23609 | 0.282698 |
| 27236 | ENSG0000 | EIF4HP2 | eukaryotic | 0.250599 | 0.254523 | 0.282584 |
| 9777 | ENSG0000 | KCNMA1 | potassium | 0.067996 | 0.189512 | 0.282452 |
| 40684 | ENSG0000 | AC105137.3 | | 0.228054 | 0.265899 | 0.282272 |
| 23878 | ENSG0000 | AL137159.1 | | 0 | 0.082553 | 0.282239 |
| 16795 | ENSG0000 | CFAP43 | cilia and fla | 0.081829 | 0.221069 | 0.281946 |
| 20488 | ENSG0000 | AL391422.1 | | 0 | 0.213117 | 0.281899 |
| 28298 | ENSG0000 | AC055855.1 | | 0.092166 | 0.220426 | 0.281872 |
| 29506 | ENSG0000 | NR2F2-AS1 | NR2F2 anti | 0.126946 | 0.250103 | 0.281473 |
| 34345 | ENSG0000 | CSPG4P11 | chondroitin | 0.091384 | 0.222952 | 0.28147 |
| 130 | ENSG0000 | CACNA1G | calcium vo | 0.010979 | 0.068769 | 0.281462 |
| 41069 | ENSG0000 | AC135279.3 | | 0.074026 | 0.230402 | 0.28126 |
| 4804 | ENSG0000 | DNAH7 | dynein axo | 0.114072 | 0.279697 | 0.281214 |
| 1732 | ENSG0000 | LAT2 | linker for a | 0.19946 | 0.269603 | 0.280984 |
| 1354 | ENSG0000 | MCAM | melanoma | 0.174796 | 0.184108 | 0.280904 |
| 8435 | ENSG0000 | FBLN7 | fibulin 7 [S | 0.146561 | 0.174796 | 0.280818 |
| 18673 | ENSG0000 | HCP5 | HLA compl | 0.03268 | 0.278491 | 0.280572 |
| 22775 | ENSG0000 | ISCA2P1 | iron-sulfur | 0.118835 | 0.205492 | 0.280508 |
| 2698 | ENSG0000 | MSLN | mesothelir | 0.057177 | 0.082513 | 0.280261 |
| 20954 | ENSG0000 | MAGEA3 | MAGE fam | 0.109212 | 0.187213 | 0.279979 |
| 24742 | ENSG0000 | LUARIS | lncRNA upr | 0.1095 | 0.270181 | 0.279847 |
| 34466 | ENSG0000 | AC109449.1 | | 0.031577 | 0.162709 | 0.2797 |
| 12258 | ENSG0000 | MFSD7 | major facil | 0.210034 | 0.224623 | 0.2797 |
| 39094 | ENSG0000 | AC135457.1 | | 0.102809 | 0.120953 | 0.279542 |
| 41279 | ENSG0000 | LINC01971 | long interg | 0 | 0.055677 | 0.279173 |
| 13082 | ENSG0000 | RAB37 | RAB37, me | 0.025038 | 0.054614 | 0.279148 |
| 25375 | ENSG0000 | CRYZP1 | crystallin z | 0.105126 | 0.228063 | 0.279108 |
| 35973 | ENSG0000 | AP001178.1 | | 0.130525 | 0.136424 | 0.278981 |
| 14275 | ENSG0000 | VWA1 | von Willeb | 0.048472 | 0.052236 | 0.278979 |
| 7838 | ENSG0000 | BCL2A1 | BCL2 relate | 0.179792 | 0.200123 | 0.278723 |
| 19119 | ENSG0000 | MIR27A | microRNA | 0 | 0.103091 | 0.278339 |

| | | | | | | |
|---|---|---|---|---|---|---|
| 786 | ENSG0000 | PRDM1 | PR/SET doı | 0.104496 | 0.276657 | 0.278144 |
| 5903 | ENSG0000 | POM121L9 | POM121 tr | 0.055322 | 0.126108 | 0.278105 |
| 710 | ENSG0000 | PLEKHA5 | pleckstrin l | 0.162292 | 0.181027 | 0.277976 |
| 38729 | ENSG0000 | AL139274.2 | | 0.15175 | 0.195865 | 0.27792 |
| 13838 | ENSG0000 | GCNT4 | glucosamir | 0.083392 | 0.23553 | 0.277723 |
| 10807 | ENSG0000 | LMOD1 | leiomodin | 0.062882 | 0.115838 | 0.277701 |
| 35166 | ENSG0000 | AC092384.3 | | 0.160304 | 0.257673 | 0.277398 |
| 9602 | ENSG0000 | WNT3A | Wnt family | 0.059195 | 0.099258 | 0.277106 |
| 35229 | ENSG0000 | AC106779.1 | | 0.090279 | 0.233461 | 0.276954 |
| 34000 | ENSG0000 | AC090971.1 | | 0 | 0.1312 | 0.276952 |
| 15816 | ENSG0000 | MITF | melanoger | 0.075166 | 0.238401 | 0.276891 |
| 34314 | ENSG0000 | AC027176.2 | | 0.122316 | 0.154637 | 0.27655 |
| 24566 | ENSG0000 | AL356056.2 | | 0.096917 | 0.215492 | 0.276453 |
| 12167 | ENSG0000 | GDNF | glial cell de | 0.033283 | 0.102102 | 0.276382 |
| 33409 | ENSG0000 | AC121757.1 | | 0.029039 | 0.185735 | 0.276242 |
| 16496 | ENSG0000 | TLR7 | toll like rec | 0.184475 | 0.230666 | 0.275626 |
| 16776 | ENSG0000 | DNAH10 | dynein axo | 0.041862 | 0.130674 | 0.275582 |
| 33237 | ENSG0000 | AC025031.1 | | 0.187464 | 0.256075 | 0.275411 |
| 23745 | ENSG0000 | AL391825.1 | | 0.098056 | 0.173153 | 0.275274 |
| 31174 | ENSG0000 | RNU6-1098 | RNA, U6 sr | 0 | 0.256654 | 0.275209 |
| 9443 | ENSG0000 | WDR78 | WD repeat | 0.140609 | 0.241368 | 0.275198 |
| 18449 | ENSG0000 | ATP6AP1L | ATPase H+ | 0.080688 | 0.211719 | 0.275109 |
| 5785 | ENSG0000 | EVI2A | ecotropic v | 0.086235 | 0.239035 | 0.274778 |
| 22952 | ENSG0000 | LTB | lymphotox | 0.027057 | 0.268101 | 0.27464 |
| 739 | ENSG0000 | PTPRN | protein tyr | 0.079002 | 0.118092 | 0.274449 |
| 12000 | ENSG0000 | CD300C | CD300c mc | 0.046792 | 0.076928 | 0.273886 |
| 22534 | ENSG0000 | ACTG1P24 | actin gamn | 0.020235 | 0.200069 | 0.273831 |
| 20816 | ENSG0000 | AL139094.1 | | 0.099797 | 0.182232 | 0.273444 |
| 19872 | ENSG0000 | S1PR3 | sphingosin | 0.0651 | 0.26961 | 0.273357 |
| 34504 | ENSG0000 | AC092287.1 | | 0.089705 | 0.101755 | 0.273157 |
| 13197 | ENSG0000 | GLIPR1L1 | GLI pathog | 0.172952 | 0.177935 | 0.273106 |
| 8558 | ENSG0000 | UCN2 | urocortin 2 | 0.047781 | 0.127987 | 0.273097 |
| 25841 | ENSG0000 | TXNP6 | thioredoxiı | 0 | 0.039574 | 0.272723 |
| 22896 | ENSG0000 | AC017074.1 | | 0.127204 | 0.188628 | 0.272514 |
| 9952 | ENSG0000 | IL34 | interleukin | 0 | 0.095723 | 0.272503 |
| 18179 | ENSG0000 | LY6G5C | lymphocyt | 0.07315 | 0.090987 | 0.272362 |
| 16238 | ENSG0000 | CRYM-AS1 | CRYM antis | 0.110813 | 0.246427 | 0.272336 |
| 5006 | ENSG0000 | PCDHB10 | protocadh | 0.075783 | 0.107927 | 0.271521 |
| 27257 | ENSG0000 | AC093110.1 | | 0.107623 | 0.260772 | 0.271444 |
| 13338 | ENSG0000 | REP15 | RAB15 effe | 0.066081 | 0.107873 | 0.271191 |
| 25163 | ENSG0000 | AC027644.1 | | 0 | 0.134641 | 0.270874 |
| 26287 | ENSG0000 | ANKRD65 | ankyrin rep | 0.023098 | 0.096774 | 0.270004 |
| 38858 | ENSG0000 | AC113361.1 | | 0 | 0.035503 | 0.269722 |
| 13540 | ENSG0000 | PCSK1 | proprotein | 0.199954 | 0.262657 | 0.269501 |
| 37859 | ENSG0000 | AC245052.7 | | 0.049432 | 0.210346 | 0.269416 |
| 39178 | ENSG0000 | AC097376.2 | | 0.178258 | 0.238556 | 0.26936 |
| 9652 | ENSG0000 | APCDD1 | APC down- | 0.038925 | 0.139935 | 0.269131 |

| | | | | | | |
|---|---|---|---|---|---|---|
| 27552 | ENSG0000 | AC002454.1 | | 0.15605 | 0.223368 | 0.269128 |
| 6705 | ENSG0000 | TEX15 | testis expr | 0.121845 | 0.244898 | 0.268752 |
| 9768 | ENSG0000 | CFAP70 | cilia and fla | 0.078293 | 0.266567 | 0.268531 |
| 42787 | ENSG0000 | AC074141.1 | | 0.068553 | 0.24833 | 0.267852 |
| 30216 | ENSG0000 | CCT7P2 | chaperonir | 0.142997 | 0.157685 | 0.267819 |
| 40975 | ENSG0000 | AL662791.5 | | 0 | 0.033893 | 0.267347 |
| 37894 | ENSG0000 | AC005197.1 | | 0.050843 | 0.148921 | 0.267019 |
| 31752 | ENSG0000 | AC104561.3 | | 0.143026 | 0.190742 | 0.266425 |
| 34876 | ENSG0000 | AC109347.1 | | 0.140818 | 0.260715 | 0.266412 |
| 22208 | ENSG0000 | LINC01623 | long interg | 0 | 0.064981 | 0.266285 |
| 40484 | ENSG0000 | AL034549.2 | | 0.172053 | 0.266148 | 0.266213 |
| 29139 | ENSG0000 | RN7SL417I | RNA, 7SL, c | 0.07694 | 0.128377 | 0.266151 |
| 31475 | ENSG0000 | AC024451.2 | | 0.080649 | 0.25689 | 0.266025 |
| 27395 | ENSG0000 | BHMG1 | basic helix- | 0.019412 | 0.125929 | 0.265571 |
| 42573 | ENSG0000 | AC241585.3 | | 0 | 0.16036 | 0.265425 |
| 19778 | ENSG0000 | GPC2 | glypican 2 | 0.070429 | 0.094959 | 0.264953 |
| 5290 | ENSG0000 | NME2P1 | NME/NM2 | 0 | 0.073459 | 0.264423 |
| 24696 | ENSG0000 | CR769775.1 | | 0.051641 | 0.159269 | 0.264329 |
| 18876 | ENSG0000 | Y_RNA | Y RNA [Sou | 0 | 0.092066 | 0.264302 |
| 40160 | ENSG0000 | AC027348.1 | | 0.086771 | 0.141813 | 0.264294 |
| 2937 | ENSG0000 | DHDH | dihydrodio | 0 | 0.172981 | 0.264167 |
| 17858 | ENSG0000 | RN7SKP26! | RNA, 7SK s | 0.054447 | 0.071744 | 0.263589 |
| 38740 | ENSG0000 | AP002907.1 | | 0.092942 | 0.157953 | 0.263237 |
| 15156 | ENSG0000 | TMPRSS2 | transmeml | 0 | 0.077144 | 0.263229 |
| 32675 | ENSG0000 | AC007406.4 | | 0.019266 | 0.161045 | 0.262675 |
| 29552 | ENSG0000 | AC098487.1 | | 0.107335 | 0.230811 | 0.262403 |
| 17910 | ENSG0000 | AC105020.1 | | 0.069636 | 0.230681 | 0.262351 |
| 37133 | ENSG0000 | ZNF582-AS | ZNF582 an | 0.093003 | 0.164717 | 0.262303 |
| 17318 | ENSG0000 | RNU5F-1 | RNA, U5F s | 0 | 0.046699 | 0.262139 |
| 544 | ENSG0000 | MYOC | myocilin [S | 0 | 0.020923 | 0.262083 |
| 38306 | ENSG0000 | AC010719.1 | | 0.006234 | 0.019459 | 0.261725 |
| 6486 | ENSG0000 | PTPRE | protein tyr | 0.082039 | 0.214216 | 0.261354 |
| 2162 | ENSG0000 | GGT5 | gamma-glu | 0.038273 | 0.107672 | 0.260921 |
| 12137 | ENSG0000 | TXNDC2 | thioredoxi | 0.120218 | 0.20451 | 0.260861 |
| 25295 | ENSG0000 | PTGES2-AS | PTGES2 an | 0.109437 | 0.207017 | 0.2607 |
| 24559 | ENSG0000 | AC007099.1 | | 0.128939 | 0.259685 | 0.260566 |
| 14582 | ENSG0000 | SYNE4 | spectrin re | 0.16943 | 0.18253 | 0.260324 |
| 40438 | ENSG0000 | TBC1D3F | TBC1 doma | 0 | 0.159365 | 0.260187 |
| 38795 | ENSG0000 | AL359715.4 | | 0.11946 | 0.212069 | 0.259897 |
| 24517 | ENSG0000 | TSSK1A | testis speci | 0.128051 | 0.189537 | 0.259583 |
| 10338 | ENSG0000 | COX6B2 | cytochrom | 0.024637 | 0.088165 | 0.259544 |
| 39085 | ENSG0000 | AL022322.1 | | 0.073073 | 0.188269 | 0.259479 |
| 12534 | ENSG0000 | TMEM182 | transmeml | 0.155234 | 0.193782 | 0.259436 |
| 43437 | ENSG0000 | AC116562.4 | | 0.070781 | 0.198391 | 0.259416 |
| 18519 | ENSG0000 | KLRC2 | killer cell le | 0 | 0.111135 | 0.259402 |
| 3417 | ENSG0000 | PKD2L1 | polycystin | 0.037162 | 0.067199 | 0.259329 |
| 10252 | ENSG0000 | PTGIR | prostaglan | 0.008611 | 0.105957 | 0.259061 |

| | | | | | | |
|---|---|---|---|---|---|---|
| 13301 | ENSG0000 | PIFO | primary cil | 0.044867 | 0.226706 | 0.25906 |
| 38311 | ENSG0000 | MTDHP1 | metadheri | 0.222981 | 0.234979 | 0.259043 |
| 16399 | ENSG0000 | NUDT11 | nudix hydr | 0.152165 | 0.253902 | 0.258927 |
| 30173 | ENSG0000 | AC138819.1 | | 0 | 0.23632 | 0.25879 |
| 27964 | ENSG0000 | DNM1P35 | dynamin 1 | 0.027354 | 0.029039 | 0.258637 |
| 4976 | ENSG0000 | KCNIP2 | potassium | 0.193131 | 0.206744 | 0.258486 |
| 12479 | ENSG0000 | AL391987.1 | | 0 | 0.05442 | 0.258341 |
| 35349 | ENSG0000 | LINC01976 | long interg | 0 | 0.033968 | 0.258325 |
| 40209 | ENSG0000 | PDCD6IPP1 | PDCD6IP p | 0 | 0.233997 | 0.258278 |
| 7324 | ENSG0000 | CD72 | CD72 mole | 0.043872 | 0.145724 | 0.258053 |
| 8555 | ENSG0000 | AMT | aminometl | 0.067238 | 0.161521 | 0.257963 |
| 32918 | ENSG0000 | AC069234.2 | | 0.179202 | 0.187673 | 0.257878 |
| 32567 | ENSG0000 | AP003031.2 | | 0.035135 | 0.125112 | 0.257592 |
| 3339 | ENSG0000 | LHX2 | LIM home | 0.033468 | 0.197309 | 0.257451 |
| 43372 | ENSG0000 | AC079781.4 | | 0.062642 | 0.229582 | 0.257423 |
| 36964 | ENSG0000 | AC011461.1 | | 0.12244 | 0.152263 | 0.257405 |
| 11946 | ENSG0000 | PSCA | prostate st | 0.050329 | 0.067533 | 0.256909 |
| 21409 | ENSG0000 | AL360093.1 | | 0.063153 | 0.084859 | 0.256667 |
| 18791 | ENSG0000 | RNU6-574I | RNA, U6 sr | 0.017644 | 0.231232 | 0.256579 |
| 34741 | ENSG0000 | FOXC2-AS1 | FOXC2 anti | 0 | 0.151156 | 0.256396 |
| 5988 | ENSG0000 | DLL4 | delta like c | 0.010195 | 0.171396 | 0.256359 |
| 17918 | ENSG0000 | LINC00449 | long interg | 0 | 0.100135 | 0.256294 |
| 2670 | ENSG0000 | MLNR | motilin rec | 0.066868 | 0.137836 | 0.256163 |
| 40999 | ENSG0000 | AC079174.2 | | 0.078941 | 0.229871 | 0.256078 |
| 1084 | ENSG0000 | CLEC2D | C-type lect | 0.111083 | 0.192717 | 0.255997 |
| 15954 | ENSG0000 | P2RX2 | purinergic | 0.106063 | 0.255448 | 0.255608 |
| 4948 | ENSG0000 | CNRIP1 | cannabinoi | 0.040894 | 0.202768 | 0.254826 |
| 37576 | ENSG0000 | AL158151.4 | | 0 | 0.193372 | 0.254662 |
| 17180 | ENSG0000 | CSAG1 | chondrosa | 0 | 0.089609 | 0.254565 |
| 38611 | ENSG0000 | AL031118.1 | | 0.048299 | 0.050272 | 0.254554 |
| 30133 | ENSG0000 | OSMR-AS1 | OSMR anti | 0.083272 | 0.142635 | 0.254513 |
| 9948 | ENSG0000 | FBLN2 | fibulin 2 [S | 0.051734 | 0.060564 | 0.254232 |
| 33884 | ENSG0000 | AC137834.1 | | 0 | 0.220671 | 0.253964 |
| 13724 | ENSG0000 | KBTBD11 | kelch repea | 0.068012 | 0.184319 | 0.253895 |
| 29706 | ENSG0000 | TMEM110- | TMEM110- | 0.088976 | 0.089538 | 0.253743 |
| 5163 | ENSG0000 | SLITRK3 | SLIT and N | 0.065425 | 0.152086 | 0.253667 |
| 10412 | ENSG0000 | USP41 | ubiquitin s | 0.028539 | 0.150331 | 0.253642 |
| 23838 | ENSG0000 | AC053503.2 | | 0.02221 | 0.053158 | 0.253461 |
| 42217 | ENSG0000 | RHOXF1P3 | Rhox home | 0.033245 | 0.206593 | 0.253322 |
| 38697 | ENSG0000 | AC098614.4 | | 0.032963 | 0.189223 | 0.253112 |
| 33022 | ENSG0000 | AC048338.1 | | 0.012643 | 0.083279 | 0.25298 |
| 29377 | ENSG0000 | RORA-AS1 | RORA antis | 0.246579 | 0.246883 | 0.252625 |
| 15341 | ENSG0000 | FAM227A | family with | 0.054924 | 0.126331 | 0.252442 |
| 30605 | ENSG0000 | AC112484.3 | | 0.103811 | 0.107631 | 0.25235 |
| 17211 | ENSG0000 | MIR30C2 | microRNA | 0 | 0.234818 | 0.252076 |
| 11538 | ENSG0000 | LIPC | lipase C, he | 0.025212 | 0.171496 | 0.252071 |
| 13164 | ENSG0000 | SYT12 | synaptotag | 0.042445 | 0.098308 | 0.251983 |

| | | | | | | |
|---|---|---|---|---|---|---|
| 27893 | ENSG0000 | HOXB-AS2 | HOXB clust | 0.107019 | 0.109205 | 0.251893 |
| 30931 | ENSG0000 | BRCC3P1 | BRCA1/BR( | 0.076097 | 0.200111 | 0.251602 |
| 28178 | ENSG0000 | AC009078.2 | | 0.022391 | 0.25115 | 0.251448 |
| 12010 | ENSG0000 | TMC8 | transmeml | 0.042256 | 0.100919 | 0.251269 |
| 34198 | ENSG0000 | AC023355.1 | | 0.173945 | 0.235437 | 0.251015 |
| 19977 | ENSG0000 | AC010615.1 | | 0.147834 | 0.222384 | 0.251001 |
| 15022 | ENSG0000 | FAM196A | family witl | 0.034142 | 0.234906 | 0.250957 |
| 17959 | ENSG0000 | SPDYC | speedy/RIN | 0 | 0.060648 | 0.250372 |
| 18358 | ENSG0000 | AC092384.1 | | 0.022014 | 0.106695 | 0.250279 |
| 13980 | ENSG0000 | CRACR2B | calcium rel | 0.128405 | 0.168819 | 0.25027 |
| 10274 | ENSG0000 | TFF3 | trefoil fact | 0.091484 | 0.124275 | 0.250196 |
| 35921 | ENSG0000 | MIR5582 | microRNA | 0.031465 | 0.168281 | 0.250011 |
| 34873 | ENSG0000 | BMS1P8 | BMS1, ribc | 0.104847 | 0.148621 | 0.249785 |
| 12290 | ENSG0000 | GSG1L | GSG1 like [ | 0.010068 | 0.050406 | 0.249369 |
| 35178 | ENSG0000 | AC022165.1 | | 0.09831 | 0.237411 | 0.249323 |
| 17920 | ENSG0000 | AL354956.1 | | 0.146191 | 0.194411 | 0.249056 |
| 41006 | ENSG0000 | TRAC | T-cell rece | 0.036623 | 0.078955 | 0.248453 |
| 27823 | ENSG0000 | RN7SL333I | RNA, 7SL, c | 0.051699 | 0.143355 | 0.247918 |
| 38941 | ENSG0000 | AC090607.4 | | 0.153684 | 0.167594 | 0.246763 |
| 36981 | ENSG0000 | AC010504.1 | | 0.091005 | 0.091194 | 0.246471 |
| 25391 | ENSG0000 | LINC00884 | long interg | 0.024336 | 0.047847 | 0.24645 |
| 23472 | ENSG0000 | AL645998.1 | | 0.086824 | 0.237259 | 0.246388 |
| 37788 | ENSG0000 | AC011445.2 | | 0 | 0.095344 | 0.246339 |
| 16640 | ENSG0000 | ADGRA1 | adhesion G | 0.063937 | 0.163078 | 0.246142 |
| 35681 | ENSG0000 | AC027796.3 | | 0 | 0.010532 | 0.245959 |
| 11865 | ENSG0000 | ZNF641 | zinc finger | 0.049298 | 0.145411 | 0.245798 |
| 29453 | ENSG0000 | LINC02288 | long interg | 0.048639 | 0.109226 | 0.245677 |
| 4462 | ENSG0000 | PLEK | pleckstrin | | 0.085906 | 0.14523 | 0.245303 |
| 26213 | ENSG0000 | TEX21P | testis expr | 0.105493 | 0.240998 | 0.244778 |
| 29425 | ENSG0000 | AL353807.3 | | 0.096743 | 0.143706 | 0.244705 |
| 34065 | ENSG0000 | LIPC-AS1 | LIPC antise | 0.047105 | 0.178531 | 0.244606 |
| 38720 | ENSG0000 | AL139424.1 | | 0 | 0.030352 | 0.244585 |
| 25656 | ENSG0000 | LINC01137 | long interg | 0.045026 | 0.055177 | 0.244479 |
| 13380 | ENSG0000 | GCSAM | germinal c | 0.166344 | 0.200741 | 0.244243 |
| 25677 | ENSG0000 | ANAPC1P1 | anaphase p | 0.179347 | 0.196483 | 0.244027 |
| 38784 | ENSG0000 | AL133406.3 | | 0.081192 | 0.124433 | 0.243966 |
| 14921 | ENSG0000 | GDPGP1 | GDP-D-glu | 0.085174 | 0.097478 | 0.24391 |
| 33663 | ENSG0000 | AL049779.1 | | 0.11875 | 0.156081 | 0.243626 |
| 11907 | ENSG0000 | KCNAB3 | potassium | 0.19193 | 0.237851 | 0.242768 |
| 4303 | ENSG0000 | TNNC1 | troponin C | 0.086541 | 0.167845 | 0.242741 |
| 18102 | ENSG0000 | RPL12P1 | ribosomal | 0.098058 | 0.112406 | 0.242635 |
| 30189 | ENSG0000 | RNF103-CH | RNF103-CH | 0.059633 | 0.210499 | 0.242412 |
| 6401 | ENSG0000 | MAP1B | microtubul | 0.160267 | 0.164051 | 0.242096 |
| 33979 | ENSG0000 | AL049780.2 | | 0 | 0.177067 | 0.241947 |
| 10010 | ENSG0000 | MEGF11 | multiple E( | 0.188935 | 0.225783 | 0.241895 |
| 17007 | ENSG0000 | C1orf228 | chromosor | 0.029897 | 0.173828 | 0.241751 |
| 28373 | ENSG0000 | HMGN1P8 | high mobil | 0.032776 | 0.190074 | 0.241306 |

| | | | | | | |
|---|---|---|---|---|---|---|
| 10801 | ENSG0000 | SLC15A2 | solute carr | 0.029673 | 0.076296 | 0.240799 |
| 39642 | ENSG0000 | AC007318.2 | | 0.041953 | 0.056848 | 0.240612 |
| 17528 | ENSG0000 | RNU6-509I | RNA, U6 sr | 0 | 0.012525 | 0.240475 |
| 34517 | ENSG0000 | AC007610.2 | | 0.049723 | 0.139622 | 0.240367 |
| 20389 | ENSG0000 | AP000523.1 | | 0.113441 | 0.197141 | 0.240362 |
| 7287 | ENSG0000 | GABBR2 | gamma-am | 0.14916 | 0.236237 | 0.240222 |
| 29478 | ENSG0000 | AC055713.1 | | 0.12387 | 0.225873 | 0.239834 |
| 1992 | ENSG0000 | COL9A3 | collagen ty | 0.037162 | 0.068896 | 0.239791 |
| 21891 | ENSG0000 | AL391684.1 | | 0.100199 | 0.149191 | 0.239531 |
| 23861 | ENSG0000 | PRDX3P1 | peroxiredo | 0.158519 | 0.159568 | 0.239526 |
| 29199 | ENSG0000 | RN7SL130I | RNA, 7SL, c | 0.042916 | 0.047658 | 0.239155 |
| 34657 | ENSG0000 | LINC02137 | long interg | 0.086319 | 0.238561 | 0.238869 |
| 14977 | ENSG0000 | SH2D7 | SH2 domai | 0.140715 | 0.149079 | 0.238779 |
| 7708 | ENSG0000 | ASCL1 | achaete-sc | 0.135277 | 0.227117 | 0.238708 |
| 27340 | ENSG0000 | FTCD-AS1 | FTCD antis | 0 | 0.117477 | 0.23824 |
| 26449 | ENSG0000 | AC098934.3 | | 0 | 0.056429 | 0.238204 |
| 42465 | ENSG0000 | AC245100.8 | | 0.016773 | 0.137097 | 0.237649 |
| 12189 | ENSG0000 | CA7 | carbonic ar | 0.086047 | 0.177752 | 0.237591 |
| 11283 | ENSG0000 | GBX1 | gastrulatio | 0 | 0.099226 | 0.237257 |
| 7022 | ENSG0000 | CHST5 | carbohydra | 0.112998 | 0.159391 | 0.236965 |
| 31709 | ENSG0000 | PCDHGB2 | protocadhe | 0.115927 | 0.23156 | 0.236617 |
| 37229 | ENSG0000 | AC004223.3 | | 0.089886 | 0.153389 | 0.236439 |
| 25402 | ENSG0000 | LINC01399 | long interg | 0 | 0.140434 | 0.236176 |
| 42286 | ENSG0000 | CU638689.4 | | 0.0611 | 0.186244 | 0.235985 |
| 40646 | ENSG0000 | AL117335.1 | | 0.146148 | 0.191835 | 0.235655 |
| 28597 | ENSG0000 | AL160314.1 | | 0 | 0.06708 | 0.235509 |
| 5857 | ENSG0000 | METTL25 | methyltrar | 0.079027 | 0.139046 | 0.23534 |
| 38102 | ENSG0000 | AC104819.3 | | 0 | 0.050381 | 0.234674 |
| 40616 | ENSG0000 | AC130650.2 | | 0.048186 | 0.159806 | 0.234663 |
| 19881 | ENSG0000 | FAM209B | family with | 0 | 0.084226 | 0.234043 |
| 34375 | ENSG0000 | AL138976.2 | | 0.086868 | 0.14201 | 0.233878 |
| 22673 | ENSG0000 | LINC01535 | long interg | 0.065087 | 0.178705 | 0.233721 |
| 29257 | ENSG0000 | AL021707.5 | | 0.149734 | 0.218652 | 0.233355 |
| 13733 | ENSG0000 | MYO1D | myosin ID | 0.139348 | 0.228115 | 0.233338 |
| 12429 | ENSG0000 | REPS2 | RALBP1 ass | 0.010993 | 0.053264 | 0.233157 |
| 35526 | ENSG0000 | AC108134.2 | | 0.047449 | 0.07718 | 0.233118 |
| 5660 | ENSG0000 | OVOL2 | ovo like zir | 0.033505 | 0.056831 | 0.233067 |
| 2226 | ENSG0000 | RSPH14 | radial spok | 0.102402 | 0.125813 | 0.232882 |
| 22956 | ENSG0000 | AL592301.1 | | 0 | 0.045192 | 0.232714 |
| 31630 | ENSG0000 | LINCR-000 | uncharacte | 0.101295 | 0.165222 | 0.2326 |
| 35311 | ENSG0000 | AC124068.2 | | 0.096352 | 0.152452 | 0.232536 |
| 42912 | ENSG0000 | AC017104.5 | | 0.062594 | 0.101656 | 0.232258 |
| 26176 | ENSG0000 | AL136309.2 | | 0.062404 | 0.137061 | 0.232189 |
| 35419 | ENSG0000 | AC026992.2 | | 0.007416 | 0.012063 | 0.231909 |
| 2995 | ENSG0000 | GIMAP2 | GTPase, IM | 0.076103 | 0.221225 | 0.231384 |
| 6857 | ENSG0000 | SLC43A3 | solute carr | 0.133643 | 0.147379 | 0.231285 |
| 15912 | ENSG0000 | KRT18P59 | keratin 18 | 0.045541 | 0.227801 | 0.230454 |

| | | | | | | |
|---|---|---|---|---|---|---|
| 14008 | ENSG0000 | PCDHB9 | protocadh | 0.048604 | 0.063323 | 0.23004 |
| 26245 | ENSG0000 | AP001626.1 | | 0 | 0.066102 | 0.229893 |
| 16518 | ENSG0000 | CLPSL2 | colipase lik | 0 | 0.065971 | 0.229843 |
| 40924 | ENSG0000 | HOXB13-A | HOXB13 ar | 0 | 0.039476 | 0.22974 |
| 27754 | ENSG0000 | RNU7-45P | RNA, U7 sr | 0 | 0.050727 | 0.22974 |
| 26043 | ENSG0000 | LINC01759 | long interg | 0.085723 | 0.132058 | 0.229705 |
| 14076 | ENSG0000 | TMEM151 | transmeml | 0.040234 | 0.193022 | 0.229655 |
| 40016 | ENSG0000 | AL157813.1 | | 0.062384 | 0.121707 | 0.229348 |
| 39297 | ENSG0000 | AC005070.3 | | 0.086474 | 0.185338 | 0.229219 |
| 29190 | ENSG0000 | TMEM225 | transmeml | 0.097337 | 0.158956 | 0.229192 |
| 22128 | ENSG0000 | UPF3AP1 | UPF3A pse | 0.188825 | 0.220541 | 0.228941 |
| 38757 | ENSG0000 | DOC2B | double C2 | 0.112976 | 0.113052 | 0.228457 |
| 14581 | ENSG0000 | DDX60L | DEAD-box | 0.126102 | 0.169403 | 0.228148 |
| 35660 | ENSG0000 | OCLM | oculomedi | 0.046129 | 0.200001 | 0.22805 |
| 41838 | ENSG0000 | AP000695.3 | | 0 | 0.021119 | 0.22786 |
| 26373 | ENSG0000 | MCF2L-AS | MCF2L ant | 0.07542 | 0.103005 | 0.227664 |
| 42809 | ENSG0000 | AC017104.4 | | 0.047987 | 0.166681 | 0.227647 |
| 5633 | ENSG0000 | SDCBP2 | syndecan b | 0.021849 | 0.206046 | 0.227547 |
| 32387 | ENSG0000 | AC134775.1 | | 0 | 0.210797 | 0.227398 |
| 29607 | ENSG0000 | LUCAT1 | lung cance | 0.128456 | 0.197711 | 0.227389 |
| 23120 | ENSG0000 | FAM66A | family with | 0.101861 | 0.161107 | 0.226513 |
| 22885 | ENSG0000 | THEM7P | thioestera | 0.059767 | 0.165551 | 0.226226 |
| 4838 | ENSG0000 | BSPRY | B-box and | 0.030456 | 0.080966 | 0.226204 |
| 23867 | ENSG0000 | AL354707.2 | | 0.12363 | 0.20951 | 0.226201 |
| 12432 | ENSG0000 | ITGAM | integrin su | 0.166697 | 0.21112 | 0.226098 |
| 7785 | ENSG0000 | PELI2 | pellino E3 | 0.033921 | 0.074322 | 0.225953 |
| 7122 | ENSG0000 | SPDYE1 | speedy/RI | 0.158377 | 0.171897 | 0.225746 |
| 37635 | ENSG0000 | AC091133.5 | | 0 | 0.005137 | 0.225614 |
| 21806 | ENSG0000 | PCOLCE-AS | PCOLCE an | 0 | 0.028955 | 0.225194 |
| 2891 | ENSG0000 | ESRP1 | epithelial s | 0.146744 | 0.147001 | 0.224996 |
| 687 | ENSG0000 | DKK3 | dickkopf W | 0.055358 | 0.140627 | 0.224478 |
| 41797 | ENSG0000 | AL139288.1 | | 0.075947 | 0.108947 | 0.224446 |
| 34399 | ENSG0000 | AL606760.3 | | 0.009884 | 0.133723 | 0.224206 |
| 43401 | ENSG0000 | AC009093.9 | | 0 | 0.039115 | 0.224169 |
| 34506 | ENSG0000 | AL353708.1 | | 0.046559 | 0.182562 | 0.223983 |
| 4740 | ENSG0000 | CASC1 | cancer sus | 0.102537 | 0.130432 | 0.223924 |
| 30639 | ENSG0000 | GMDS-AS1 | GMDS anti | 0.12526 | 0.204303 | 0.2239 |
| 3869 | ENSG0000 | ACRBP | acrosin bin | 0.063221 | 0.174217 | 0.223757 |
| 23380 | ENSG0000 | LINC01125 | long interg | 0.062436 | 0.196875 | 0.223617 |
| 25974 | ENSG0000 | LINC01049 | long interg | 0.173188 | 0.183266 | 0.223558 |
| 24309 | ENSG0000 | HM13-AS1 | HM13 anti | 0 | 0.081979 | 0.223457 |
| 41207 | ENSG0000 | AC138466.4 | | 0.026025 | 0.081553 | 0.223268 |
| 15192 | ENSG0000 | GOLGA6L4 | golgin A6 f | 0.097709 | 0.19449 | 0.223095 |
| 21193 | ENSG0000 | SNORD71 | small nucle | 0.076944 | 0.17022 | 0.223061 |
| 21342 | ENSG0000 | FO393418.1 | | 0 | 0.093995 | 0.222983 |
| 8540 | ENSG0000 | BOC | BOC cell ac | 0.062571 | 0.122635 | 0.222912 |
| 31282 | ENSG0000 | SNORD116 | Small nucle | 0 | 0.183713 | 0.222376 |

| | | | | | | |
|---|---|---|---|---|---|---|
| 5601 | ENSG0000 | FNDC11 | fibronectin | 0.009791 | 0.025282 | 0.222265 |
| 33072 | ENSG0000 | AC087241.3 | | 0.105398 | 0.123875 | 0.22189 |
| 23718 | ENSG0000 | AL139161.1 | | 0.047616 | 0.12588 | 0.22188 |
| 22988 | ENSG0000 | SLC2A1-AS | SLC2A1 ant | 0.031745 | 0.13015 | 0.221818 |
| 8654 | ENSG0000 | ARHGAP26 | Rho GTPas | 0.160452 | 0.216049 | 0.221798 |
| 31612 | ENSG0000 | CERNA3 | competing | 0 | 0.060522 | 0.221766 |
| 14483 | ENSG0000 | SHISA2 | shisa famil | 0.01219 | 0.132576 | 0.221747 |
| 10882 | ENSG0000 | PPM1K | protein ph | 0.07051 | 0.185756 | 0.221485 |
| 223 | ENSG0000 | IL32 | interleukin | 0.064329 | 0.172423 | 0.221427 |
| 17018 | ENSG0000 | KCNRG | potassium | 0.001106 | 0.002934 | 0.221096 |
| 39697 | ENSG0000 | WT1-AS_7 | WT1 antise | 0 | 0.025364 | 0.221037 |
| 41486 | ENSG0000 | Metazoa_S | Metazoan | 0 | 0.177566 | 0.220746 |
| 39277 | ENSG0000 | AL133551.1 | | 0.188376 | 0.214365 | 0.220474 |
| 2264 | ENSG0000 | ACR | acrosin [So | 0.040346 | 0.105019 | 0.220446 |
| 14038 | ENSG0000 | ALS2CL | ALS2 C-ter | 0.068284 | 0.178223 | 0.220266 |
| 14451 | ENSG0000 | KCNE1 | potassium | 0.099195 | 0.176463 | 0.220165 |
| 40722 | ENSG0000 | AC026124.2 | | 0.014691 | 0.079212 | 0.220092 |
| 3598 | ENSG0000 | VTN | vitronectin | 0.043923 | 0.166305 | 0.219928 |
| 34857 | ENSG0000 | AC008870.1 | | 0 | 0.129071 | 0.219775 |
| 10121 | ENSG0000 | FGF17 | fibroblast g | 0.010798 | 0.069346 | 0.219605 |
| 21820 | ENSG0000 | MUC20P1 | mucin 20, | 0.062854 | 0.180102 | 0.219605 |
| 21869 | ENSG0000 | EIPR1-IT1 | EIPR1 intro | 0.013497 | 0.030439 | 0.219275 |
| 30417 | ENSG0000 | AC253536.5 | | 0.018135 | 0.112072 | 0.219234 |
| 26140 | ENSG0000 | AC012618.2 | | 0.079903 | 0.176675 | 0.219186 |
| 41588 | ENSG0000 | AC092611.3 | | 0.049607 | 0.060304 | 0.219181 |
| 42971 | ENSG0000 | AC138904.3 | | 0.072342 | 0.119216 | 0.218797 |
| 16549 | ENSG0000 | KIAA1211L | KIAA1211 l | 0.086761 | 0.1392 | 0.218793 |
| 15950 | ENSG0000 | C2orf78 | chromosor | 0.058445 | 0.105461 | 0.218649 |
| 32486 | ENSG0000 | AP000640.2 | | 0 | 0.209503 | 0.218611 |
| 35859 | ENSG0000 | AC003965.1 | | 0 | 0.138949 | 0.218546 |
| 29693 | ENSG0000 | AC079140.1 | | 0.1057 | 0.205335 | 0.218418 |
| 32297 | ENSG0000 | AF131216.3 | | 0 | 0.052718 | 0.218008 |
| 34815 | ENSG0000 | AP003469.4 | | 0.062907 | 0.170748 | 0.217785 |
| 29325 | ENSG0000 | C4A | compleme | 0.140899 | 0.149779 | 0.217636 |
| 37930 | ENSG0000 | FAM226B | family with | 0 | 0.195077 | 0.217559 |
| 37703 | ENSG0000 | AC009955.3 | | 0 | 0.166602 | 0.217176 |
| 19205 | ENSG0000 | GPX3 | glutathione | 0.152986 | 0.153431 | 0.217112 |
| 28845 | ENSG0000 | AC005486.1 | | 0 | 0.140958 | 0.217099 |
| 7156 | ENSG0000 | SLC39A8 | solute carr | 0.038398 | 0.062978 | 0.216976 |
| 15622 | ENSG0000 | TEX38 | testis expr | 0.051211 | 0.194689 | 0.216897 |
| 10004 | ENSG0000 | DRC1 | dynein reg | 0.034975 | 0.082797 | 0.216662 |
| 5717 | ENSG0000 | MCF2L | MCF.2 cell | 0.073126 | 0.121683 | 0.216636 |
| 30255 | ENSG0000 | AC114316.2 | | 0.047775 | 0.100274 | 0.216493 |
| 14360 | ENSG0000 | SSC5D | scavenger | 0.077375 | 0.101223 | 0.216399 |
| 2276 | ENSG0000 | APOL1 | apolipopro | 0.060587 | 0.198434 | 0.216193 |
| 2178 | ENSG0000 | CYTH4 | cytohesin 4 | 0.117458 | 0.152468 | 0.216175 |
| 41585 | ENSG0000 | AC091305.2 | | 0.01142 | 0.18368 | 0.215673 |

| | | | | | | |
|---|---|---|---|---|---|---|
| 36546 | ENSG0000 | LINC00663 | long interg | 0.114852 | 0.143805 | 0.21535 |
| 21765 | ENSG0000 | AF124730.1 | | 0 | 0.164706 | 0.215301 |
| 39236 | ENSG0000 | AC021016.3 | | 0.056729 | 0.079966 | 0.215291 |
| 35273 | ENSG0000 | AC027682.4 | | 0.009992 | 0.19286 | 0.215284 |
| 37478 | ENSG0000 | AC004076.1 | | 0.050625 | 0.065192 | 0.215174 |
| 4128 | ENSG0000 | SPARC | secreted p | 0.132516 | 0.197625 | 0.215045 |
| 16368 | ENSG0000 | ZNF493 | zinc finger | 0.016177 | 0.196731 | 0.215003 |
| 41972 | ENSG0000 | AP000974.1 | | 0.119057 | 0.1341 | 0.214886 |
| 42690 | ENSG0000 | AC063952.4 | | 0.051635 | 0.071726 | 0.214772 |
| 41340 | ENSG0000 | AL354696.2 | | 0.113784 | 0.1254 | 0.214736 |
| 36975 | ENSG0000 | AC018761.3 | | 0 | 0.034807 | 0.213982 |
| 42585 | ENSG0000 | LINC01012 | long interg | 0.060222 | 0.104333 | 0.213526 |
| 28083 | ENSG0000 | RN7SL273I | RNA, 7SL, c | 0 | 0.076373 | 0.213314 |
| 12892 | ENSG0000 | FAM90A1 | family with | 0.026981 | 0.045981 | 0.213171 |
| 21220 | ENSG0000 | AC137630.1 | | 0.066315 | 0.095101 | 0.212595 |
| 43321 | ENSG0000 | AC015802.6 | | 0.055268 | 0.134585 | 0.212556 |
| 7820 | ENSG0000 | DUOX2 | dual oxidas | 0.010892 | 0.037723 | 0.21229 |
| 26854 | ENSG0000 | AC104333.2 | | 0.027532 | 0.063895 | 0.211791 |
| 12422 | ENSG0000 | P2RY1 | purinergic | 0.052642 | 0.075557 | 0.211659 |
| 10154 | ENSG0000 | SV2A | synaptic ve | 0.106943 | 0.157222 | 0.211499 |
| 22463 | ENSG0000 | SGMS1-AS | SGMS1 ant | 0.139599 | 0.172729 | 0.211288 |
| 33487 | ENSG0000 | AC073569.2 | | 0.089019 | 0.159224 | 0.210985 |
| 24615 | ENSG0000 | AC007388.1 | | 0.154471 | 0.20484 | 0.210863 |
| 20242 | ENSG0000 | AL161756.1 | | 0.069867 | 0.149854 | 0.210567 |
| 29304 | ENSG0000 | SNRPCP3 | small nucle | 0.128716 | 0.199834 | 0.210483 |
| 12342 | ENSG0000 | PTAFR | platelet ac | 0.121098 | 0.126903 | 0.210261 |
| 39049 | ENSG0000 | AC090425.1 | | 0 | 0.174518 | 0.210048 |
| 16689 | ENSG0000 | FBXL22 | F-box and l | 0.117241 | 0.176261 | 0.209926 |
| 35985 | ENSG0000 | AL645941.3 | | 0.04943 | 0.170725 | 0.209877 |
| 41121 | ENSG0000 | uc_338 | TUC338 [So | 0.043451 | 0.153409 | 0.20966 |
| 10100 | ENSG0000 | PKD1L1 | polycystin | 0.041844 | 0.126946 | 0.209404 |
| 18101 | ENSG0000 | GGNBP1 | gametoger | 0.01831 | 0.103749 | 0.209301 |
| 37871 | ENSG0000 | AP001350.1 | | 0.075069 | 0.158621 | 0.209087 |
| 39424 | ENSG0000 | DGCR10 | DiGeorge s | 0.097636 | 0.168486 | 0.208854 |
| 358 | ENSG0000 | EHD3 | EH domain | 0.042769 | 0.067112 | 0.208654 |
| 8154 | ENSG0000 | B3GNT7 | UDP-GlcNA | 0.088888 | 0.175618 | 0.208311 |
| 6072 | ENSG0000 | LRRC6 | leucine rich | 0.037547 | 0.17308 | 0.208153 |
| 4266 | ENSG0000 | FRMD4B | FERM dom | 0.096408 | 0.145163 | 0.208079 |
| 7757 | ENSG0000 | MORN3 | MORN rep | 0.027646 | 0.056572 | 0.207993 |
| 4897 | ENSG0000 | PTGER2 | prostaglan | 0.057102 | 0.089313 | 0.207955 |
| 37525 | ENSG0000 | SMC5-AS1 | SMC5 antis | 0.029492 | 0.202242 | 0.207941 |
| 9261 | ENSG0000 | KCTD14 | potassium | 0.178119 | 0.180662 | 0.207898 |
| 21370 | ENSG0000 | TSSC2 | tumor sup | 0 | 0.108693 | 0.207727 |
| 26537 | ENSG0000 | AC064850.1 | | 0.048345 | 0.200651 | 0.207637 |
| 12935 | ENSG0000 | ANKRD20A | ankyrin rep | 0.081169 | 0.126866 | 0.207394 |
| 16577 | ENSG0000 | SMIM10L2 | small integ | 0.012922 | 0.042638 | 0.207328 |
| 6949 | ENSG0000 | PRR5L | proline rich | 0.059239 | 0.121571 | 0.206827 |

| | | | | | | |
|---|---|---|---|---|---|---|
| 11683 | ENSG0000 | TMPRSS5 | transmeml | 0 | 0.059394 | 0.206639 |
| 22946 | ENSG0000 | AL138921.1 | | 0 | 0.176231 | 0.206531 |
| 15198 | ENSG0000 | OLIG1 | oligodendr | 0.078047 | 0.089554 | 0.206029 |
| 41942 | ENSG0000 | AC113410.3 | | 0 | 0.059417 | 0.205974 |
| 17028 | ENSG0000 | AC073343.1 | | 0.095811 | 0.12875 | 0.205571 |
| 23029 | ENSG0000 | ISCA1P6 | iron-sulfur | 0.060885 | 0.099919 | 0.205293 |
| 34253 | ENSG0000 | AC021755.3 | | 0.056957 | 0.066401 | 0.205277 |
| 14643 | ENSG0000 | KCTD9P2 | potassium | 0.066789 | 0.168794 | 0.205131 |
| 31469 | ENSG0000 | AZIN1-AS1 | AZIN1 anti: | 0.160291 | 0.184923 | 0.205082 |
| 20195 | ENSG0000 | AC087491.1 | | 0.038039 | 0.110607 | 0.205028 |
| 29672 | ENSG0000 | AL356235.1 | | 0.040889 | 0.088028 | 0.205017 |
| 14836 | ENSG0000 | LCK | LCK proto- | 0.031926 | 0.037933 | 0.204943 |
| 35392 | ENSG0000 | TPBGL | trophoblas | 0.002614 | 0.052155 | 0.204857 |
| 32724 | ENSG0000 | AC124312.2 | | 0 | 0.175255 | 0.204852 |
| 34035 | ENSG0000 | AC092755.2 | | 0 | 0.00436 | 0.204835 |
| 15864 | ENSG0000 | C11orf96 | chromosor | 0.015009 | 0.03671 | 0.204665 |
| 1537 | ENSG0000 | ZNF423 | zinc finger | 0.077121 | 0.116772 | 0.203777 |
| 41009 | ENSG0000 | Metazoa_S | Metazoan | 0 | 0.066706 | 0.203506 |
| 14133 | ENSG0000 | GPR35 | G protein-c | 0.091769 | 0.196466 | 0.203466 |
| 5981 | ENSG0000 | CGNL1 | cingulin lik | 0.061644 | 0.087276 | 0.203442 |
| 20840 | ENSG0000 | MIR1250 | microRNA | 0.015593 | 0.090319 | 0.20331 |
| 12662 | ENSG0000 | PLAC1 | placenta sp | 0 | 0.068548 | 0.2033 |
| 29439 | ENSG0000 | EXTL3-AS1 | EXTL3 anti: | 0.075432 | 0.098575 | 0.203227 |
| 8216 | ENSG0000 | SLC44A3 | solute carr | 0.049747 | 0.081322 | 0.202957 |
| 35260 | ENSG0000 | AC010491.1 | | 0.076406 | 0.104234 | 0.202881 |
| 15244 | ENSG0000 | LRRC19 | leucine rich | 0 | 0.164927 | 0.202817 |
| 16836 | ENSG0000 | MEIG1 | meiosis/sp | 0.174059 | 0.196125 | 0.202815 |
| 32957 | ENSG0000 | AC148477.2 | | 0 | 0.039583 | 0.202714 |
| 21602 | ENSG0000 | C4B | compleme | 0.152797 | 0.168371 | 0.202439 |
| 34788 | ENSG0000 | LINC02544 | long interg | 0.02952 | 0.083864 | 0.202234 |
| 7379 | ENSG0000 | FGFBP1 | fibroblast g | 0.026209 | 0.064555 | 0.2021 |
| 21640 | ENSG0000 | RPL18AP2 | ribosomal | 0 | 0.121057 | 0.202095 |
| 26272 | ENSG0000 | GAPDHP52 | glyceraldel | 0 | 0.108125 | 0.202059 |
| 8507 | ENSG0000 | ITGA9 | integrin su | 0.02102 | 0.109917 | 0.201948 |
| 39398 | ENSG0000 | AC133528.1 | | 0.022045 | 0.172886 | 0.201914 |
| 39704 | ENSG0000 | MIR6853 | microRNA | 0 | 0.00209 | 0.201798 |
| 38967 | ENSG0000 | AL357054.4 | | 0 | 0.139689 | 0.201777 |
| 3887 | ENSG0000 | APOBEC1 | apolipopro | 0 | 0.171827 | 0.201751 |
| 43440 | ENSG0000 | AC083906.6 | | 0 | 0.132747 | 0.201655 |
| 37340 | ENSG0000 | AC020934.2 | | 0 | 0.142499 | 0.201574 |
| 2129 | ENSG0000 | CDHR5 | cadherin re | 0.05537 | 0.161784 | 0.201544 |
| 33114 | ENSG0000 | AC084032.1 | | 0 | 0.172323 | 0.201532 |
| 20627 | ENSG0000 | AL158066.1 | | 0 | 0.11229 | 0.201514 |
| 18255 | ENSG0000 | HLA-F | major histo | 0.139938 | 0.153177 | 0.201345 |
| 5958 | ENSG0000 | MYO1B | myosin IB \| | 0.056731 | 0.09724 | 0.201133 |
| 35858 | ENSG0000 | AC020978.8 | | 0.149026 | 0.15042 | 0.201111 |
| 5747 | ENSG0000 | TSKS | testis speci | 0.012086 | 0.079075 | 0.201106 |

| | | | | | | |
|---|---|---|---|---|---|---|
| 39644 | ENSG0000 | ZNRD1-AS1 | ZNRD1 ant | 0.077833 | 0.169558 | 0.201022 |
| 39390 | ENSG0000 | AP000569.1 | | 0.060054 | 0.098381 | 0.200682 |
| 22464 | ENSG0000 | LINC01733 | long interg | 0.079887 | 0.121478 | 0.200665 |
| 6961 | ENSG0000 | AVIL | advillin [So | 0.111773 | 0.117546 | 0.200399 |
| 22095 | ENSG0000 | AL158824.1 | | 0 | 0.03276 | 0.200129 |
| 41143 | ENSG0000 | MIR6757 | microRNA | 0 | 0.086656 | 0.199774 |
| 25563 | ENSG0000 | PMS2P9 | PMS1 hom | 0.045392 | 0.145607 | 0.199681 |
| 14134 | ENSG0000 | ACTG1P1 | actin gamn | 0.085058 | 0.103456 | 0.199408 |
| 12179 | ENSG0000 | SLC16A4 | solute carr | 0.054644 | 0.150696 | 0.199161 |
| 26288 | ENSG0000 | AL157400.3 | | 0 | 0.179835 | 0.198943 |
| 24054 | ENSG0000 | RPL4P6 | ribosomal | 0.12248 | 0.152897 | 0.198908 |
| 34103 | ENSG0000 | AC087482.1 | | 0.042432 | 0.135893 | 0.198857 |
| 41989 | ENSG0000 | CU639417.3 | | 0 | 0.121154 | 0.198772 |
| 37876 | ENSG0000 | AC010422.5 | | 0.140705 | 0.14236 | 0.198728 |
| 23166 | ENSG0000 | AC073343.2 | | 0.06061 | 0.160757 | 0.198246 |
| 9771 | ENSG0000 | GNAQ | G protein s | 0.150066 | 0.173493 | 0.198168 |
| 27183 | ENSG0000 | AC128709.3 | | 0.140172 | 0.161648 | 0.197887 |
| 25088 | ENSG0000 | AL583856.1 | | 0.160495 | 0.170976 | 0.197866 |
| 24711 | ENSG0000 | AC245100.4 | | 0.006811 | 0.19358 | 0.197825 |
| 41218 | ENSG0000 | AL117328.2 | | 0.051317 | 0.098946 | 0.197706 |
| 35013 | ENSG0000 | AC061975.1 | | 0.045403 | 0.136513 | 0.197496 |
| 23494 | ENSG0000 | AL450306.1 | | 0.051079 | 0.083302 | 0.197487 |
| 40942 | ENSG0000 | AL023803.3 | | 0 | 0.052533 | 0.197418 |
| 27200 | ENSG0000 | LINC00398 | long interg | 0 | 0.12801 | 0.197409 |
| 6762 | ENSG0000 | PTPN22 | protein tyr | 0.00791 | 0.069426 | 0.197238 |
| 40587 | ENSG0000 | MIR6829 | microRNA | 0.054476 | 0.107707 | 0.197165 |
| 38922 | ENSG0000 | AC016405.3 | | 0 | 0.054902 | 0.197149 |
| 35831 | ENSG0000 | AC124319.2 | | 0.067459 | 0.147419 | 0.1971 |
| 17471 | ENSG0000 | Y_RNA | Y RNA [Sou | 0 | 0.192032 | 0.197081 |
| 27596 | ENSG0000 | ABCA17P | ATP bindin | 0.066141 | 0.116861 | 0.196721 |
| 36930 | ENSG0000 | AC005954.1 | | 0.027263 | 0.141627 | 0.196657 |
| 19779 | ENSG0000 | KRT222 | keratin 222 | 0 | 0.002408 | 0.19643 |
| 41722 | ENSG0000 | CR559946.2 | | 0.053849 | 0.170018 | 0.196125 |
| 40932 | ENSG0000 | GOLGA8IP | golgin A8 f | 0.01371 | 0.09721 | 0.196104 |
| 15310 | ENSG0000 | B3GALT5-A | B3GALT5 a | 0.100208 | 0.187846 | 0.195954 |
| 42233 | ENSG0000 | AC004241.5 | | 0.02838 | 0.161014 | 0.195825 |
| 41158 | ENSG0000 | AL359183.1 | | 0.098172 | 0.115595 | 0.195796 |
| 28904 | ENSG0000 | RN7SL614P | RNA, 7SL, c | 0 | 0.103827 | 0.195689 |
| 5193 | ENSG0000 | SASH3 | SAM and S | 0.077809 | 0.114975 | 0.195565 |
| 21922 | ENSG0000 | AC083900.1 | | 0 | 0.062206 | 0.195206 |
| 37872 | ENSG0000 | AC008764.7 | | 0.065665 | 0.0782 | 0.195204 |
| 38916 | ENSG0000 | AC009102.2 | | 0.021655 | 0.045413 | 0.194977 |
| 34266 | ENSG0000 | AC015712.4 | | 0 | 0.061552 | 0.194836 |
| 36371 | ENSG0000 | AC015917.2 | | 0.075611 | 0.13757 | 0.194789 |
| 27083 | ENSG0000 | AP001615.1 | | 0.119068 | 0.142419 | 0.194665 |
| 9923 | ENSG0000 | JAML | junction ac | 0.105412 | 0.142654 | 0.194625 |
| 31825 | ENSG0000 | AL359075.2 | | 0.03854 | 0.181058 | 0.194542 |

| | | | | | | |
|---|---|---|---|---|---|---|
| 26748 | ENSG0000 | LINC01447 | long interg | 0.015221 | 0.173421 | 0.194418 |
| 23269 | ENSG0000 | AC012442.1 | | 0 | 0.094906 | 0.193911 |
| 34170 | ENSG0000 | AC079075.1 | | 0 | 0.031211 | 0.193863 |
| 11824 | ENSG0000 | ART5 | ADP-ribosy | 0 | 0.072301 | 0.193725 |
| 1968 | ENSG0000 | SLC7A8 | solute carr | 0.083044 | 0.143961 | 0.193702 |
| 40428 | ENSG0000 | Y_RNA | Y RNA [Sou | 0.057377 | 0.068727 | 0.193702 |
| 36789 | ENSG0000 | MIR4254 | microRNA | 0 | 0.075962 | 0.193465 |
| 26239 | ENSG0000 | AL512622.1 | | 0 | 0.075561 | 0.193465 |
| 24378 | ENSG0000 | ANKRD63 | ankyrin rep | 0.004955 | 0.13998 | 0.19346 |
| 38646 | ENSG0000 | PLS3-AS1 | PLS3 antise | 0.084007 | 0.174634 | 0.193369 |
| 43267 | ENSG0000 | BTBD8 | BTB domai | 0.09198 | 0.098854 | 0.193365 |
| 38840 | ENSG0000 | AL023583.1 | | 0 | 0.091185 | 0.193259 |
| 9387 | ENSG0000 | SH2D6 | SH2 domai | 0.050573 | 0.061859 | 0.192904 |
| 21893 | ENSG0000 | LINC00184 | long interg | 0.094033 | 0.155367 | 0.1929 |
| 19836 | ENSG0000 | U73169.1 | | 0.168726 | 0.172193 | 0.192806 |
| 21212 | ENSG0000 | RN7SKP30 | RNA, 7SK s | 0.053509 | 0.086115 | 0.192785 |
| 9884 | ENSG0000 | RBPMS | RNA bindir | 0.05822 | 0.105509 | 0.192708 |
| 10023 | ENSG0000 | EXTL1 | exostosin l | 0.025628 | 0.074394 | 0.192479 |
| 18378 | ENSG0000 | FRG2 | FSHD regio | 0 | 0.131989 | 0.192324 |
| 9414 | ENSG0000 | CAMK4 | calcium/ca | 0.123012 | 0.141333 | 0.192294 |
| 20961 | ENSG0000 | HMSD | histocomp | 0.057234 | 0.09112 | 0.192123 |
| 34578 | ENSG0000 | AC023813.3 | | 0 | 0.033089 | 0.192018 |
| 1161 | ENSG0000 | LMCD1 | LIM and cy | 0.10496 | 0.120417 | 0.191926 |
| 10693 | ENSG0000 | KCNF1 | potassium | 0.031231 | 0.038282 | 0.191751 |
| 15849 | ENSG0000 | WDR86 | WD repeat | 0.136026 | 0.179674 | 0.191516 |
| 40713 | ENSG0000 | AC020922.4 | | 0 | 0.061429 | 0.19145 |
| 11899 | ENSG0000 | CCDC8 | coiled-coil | 0.015904 | 0.036289 | 0.19145 |
| 25261 | ENSG0000 | FLJ22447 | uncharacte | 0.072882 | 0.159241 | 0.19103 |
| 35250 | ENSG0000 | AC021016.2 | | 0.079768 | 0.185242 | 0.190926 |
| 8687 | ENSG0000 | GFRA3 | GDNF fami | 0.034752 | 0.145415 | 0.190811 |
| 37971 | ENSG0000 | AC068768.2 | | 0 | 0.121425 | 0.190557 |
| 31109 | ENSG0000 | RNU6-118I | RNA, U6 sr | 0.090075 | 0.164743 | 0.190324 |
| 11513 | ENSG0000 | TC2N | tandem C2 | 0.043304 | 0.129908 | 0.189668 |
| 14933 | ENSG0000 | PLGLB1 | plasminoge | 0.067569 | 0.18647 | 0.18956 |
| 12176 | ENSG0000 | FAM84B | family with | 0.120688 | 0.131554 | 0.189331 |
| 7728 | ENSG0000 | CCDC65 | coiled-coil | 0.132182 | 0.184701 | 0.189179 |
| 28505 | ENSG0000 | RPL23AP8 | ribosomal | 0 | 0.054692 | 0.189112 |
| 27993 | ENSG0000 | GUSBP4 | glucuronid | 0.075683 | 0.127541 | 0.189102 |
| 42315 | ENSG0000 | AL359258.3 | | 0.002791 | 0.07395 | 0.188953 |
| 10265 | ENSG0000 | CPAMD8 | C3 and PZP | 0.015164 | 0.075291 | 0.188788 |
| 24070 | ENSG0000 | AC018712.1 | | 0.142163 | 0.149009 | 0.188774 |
| 15986 | ENSG0000 | RINL | Ras and Ra | 0.029531 | 0.109104 | 0.188698 |
| 40156 | ENSG0000 | SCARNA16 | small Cajal | 0.062336 | 0.11006 | 0.187839 |
| 4581 | ENSG0000 | KIF21B | kinesin fan | 0.09935 | 0.112096 | 0.187822 |
| 12241 | ENSG0000 | SPRY3 | sprouty RT | 0.08278 | 0.14288 | 0.187595 |
| 41349 | ENSG0000 | uc_338 | TUC338 [So | 0 | 0.004147 | 0.187489 |
| 20762 | ENSG0000 | HSPA8P15 | heat shock | 0.128904 | 0.153712 | 0.187349 |

| | | | | | | |
|---|---|---|---|---|---|---|
| 10165 | ENSG0000 | CIART | circadian a | 0.069277 | 0.14696 | 0.187241 |
| 9312 | ENSG0000 | FLI1 | Fli-1 proto- | 0.115227 | 0.139006 | 0.187032 |
| 499 | ENSG0000 | POU2F2 | POU class 2 | 0.061683 | 0.181517 | 0.187008 |
| 26308 | ENSG0000 | AC010148.1 | | 0.114675 | 0.118293 | 0.186794 |
| 40293 | ENSG0000 | AC243965.2 | | 0.031352 | 0.114263 | 0.186785 |
| 24693 | ENSG0000 | AC007285.2 | | 0.06122 | 0.16282 | 0.186691 |
| 5930 | ENSG0000 | RNF112 | ring finger | 0.047852 | 0.12645 | 0.186616 |
| 15221 | ENSG0000 | SRPK3 | SRSF prote | 0.16939 | 0.176102 | 0.186526 |
| 14984 | ENSG0000 | FAM46C | family with | 0.024795 | 0.111878 | 0.186376 |
| 28991 | ENSG0000 | HNRNPKP4 | heterogen | 0.139661 | 0.177843 | 0.186311 |
| 15094 | ENSG0000 | ASCL2 | achaete-sc | 0.0193 | 0.025547 | 0.186293 |
| 10606 | ENSG0000 | WDR63 | WD repeat | 0.020084 | 0.179326 | 0.186169 |
| 6112 | ENSG0000 | AANAT | aralkylami | 0.033211 | 0.144073 | 0.185998 |
| 3914 | ENSG0000 | DSE | dermatan s | 0.097321 | 0.185145 | 0.185925 |
| 25446 | ENSG0000 | AC096536.1 | | 0.077406 | 0.151119 | 0.185916 |
| 18623 | ENSG0000 | SRRM2-AS | SRRM2 ant | 0.069783 | 0.150742 | 0.185886 |
| 10683 | ENSG0000 | TRIM17 | tripartite n | 0.123492 | 0.180583 | 0.185778 |
| 5343 | ENSG0000 | DBH | dopamine | 0.025034 | 0.042147 | 0.18555 |
| 26776 | ENSG0000 | AC245014.1 | | 0.069778 | 0.075731 | 0.185179 |
| 22351 | ENSG0000 | AL121574.1 | | 0 | 0.024951 | 0.185129 |
| 43426 | ENSG0000 | AP003071.5 | | 0.012239 | 0.060691 | 0.185044 |
| 30709 | ENSG0000 | AC137794.1 | | 0.085204 | 0.09847 | 0.185019 |
| 37194 | ENSG0000 | AC006116.8 | | 0.06092 | 0.109105 | 0.184824 |
| 2365 | ENSG0000 | GALNT16 | polypeptid | 0.030452 | 0.089483 | 0.184651 |
| 29389 | ENSG0000 | AF250324.1 | | 0.084555 | 0.161027 | 0.184543 |
| 35543 | ENSG0000 | AC036222.1 | | 0 | 0.182358 | 0.184513 |
| 21214 | ENSG0000 | RNU6-516I | RNA, U6 sr | 0 | 0.174462 | 0.184455 |
| 19971 | ENSG0000 | AC019080.1 | | 0.16209 | 0.17941 | 0.184256 |
| 40534 | ENSG0000 | ANKRD20A | ankyrin rep | 0.129678 | 0.141095 | 0.184145 |
| 11076 | ENSG0000 | PRSS12 | protease, s | 0.092017 | 0.144386 | 0.184093 |
| 17769 | ENSG0000 | RNY1P13 | RNA, Ro-as | 0 | 0.092291 | 0.183792 |
| 41104 | ENSG0000 | AC009093.6 | | 0.011152 | 0.106819 | 0.183626 |
| 18932 | ENSG0000 | RNU6-2 | RNA, U6 sr | 0 | 0.146447 | 0.183577 |
| 18529 | ENSG0000 | SNORD116 | small nucle | 0.056089 | 0.123506 | 0.18341 |
| 5982 | ENSG0000 | TMOD2 | tropomodu | 0.112259 | 0.176356 | 0.183137 |
| 39127 | ENSG0000 | AC021851.1 | | 0.139468 | 0.143013 | 0.182497 |
| 5811 | ENSG0000 | BCL11B | B-cell CLL/l | 0.013588 | 0.057847 | 0.182494 |
| 34716 | ENSG0000 | TBC1D3G | TBC1 doma | 0 | 0.172935 | 0.182179 |
| 3897 | ENSG0000 | CLEC4A | C-type lect | 0 | 0.100536 | 0.18201 |
| 37102 | ENSG0000 | AC090229.1 | | 0.11084 | 0.177862 | 0.181978 |
| 31440 | ENSG0000 | AC100860.1 | | 0 | 0.097894 | 0.181825 |
| 23665 | ENSG0000 | AC233280.1 | | 0.04133 | 0.156821 | 0.181597 |
| 12022 | ENSG0000 | RAB26 | RAB26, me | 0.078155 | 0.082114 | 0.181514 |
| 4393 | ENSG0000 | IGFBP5 | insulin like | 0.049663 | 0.07691 | 0.18131 |
| 29496 | ENSG0000 | STX18-AS1 | STX18 anti | 0.098307 | 0.101617 | 0.181274 |
| 33312 | ENSG0000 | AL442663.4 | | 0.120228 | 0.178505 | 0.180901 |
| 28246 | ENSG0000 | AL096701.3 | | 0.120797 | 0.166903 | 0.180418 |

| | | | | | | |
|---|---|---|---|---|---|---|
| 10496 | ENSG0000 | NTN3 | netrin 3 [S | 0 | 0.148951 | 0.180376 |
| 25224 | ENSG0000 | AP001596.1 | | 0.031647 | 0.069914 | 0.180138 |
| 10405 | ENSG0000 | FBXL13 | F-box and l | 0.012892 | 0.163052 | 0.179983 |
| 12661 | ENSG0000 | PDGFD | platelet de | 0.132601 | 0.149949 | 0.179837 |
| 33472 | ENSG0000 | AC126614.1 | | 0 | 0.032973 | 0.179738 |
| 39451 | ENSG0000 | AC002059.1 | | 0.063853 | 0.09142 | 0.17965 |
| 3631 | ENSG0000 | TBC1D9 | TBC1 doma | 0.06357 | 0.151425 | 0.179203 |
| 12376 | ENSG0000 | APLF | aprataxin a | 0.125334 | 0.17111 | 0.179027 |
| 15329 | ENSG0000 | AMYH02020865.1 | | 0.043206 | 0.121375 | 0.178918 |
| 42225 | ENSG0000 | AC095031.1 | | 0 | 0.108767 | 0.1788 |
| 2739 | ENSG0000 | ESRP2 | epithelial s | 0.052335 | 0.129534 | 0.178766 |
| 18629 | ENSG0000 | C21orf62-A | C21orf62 a | 0.041836 | 0.046279 | 0.178721 |
| 22292 | ENSG0000 | DEFB131B | defensin be | 0.068178 | 0.113086 | 0.178707 |
| 6473 | ENSG0000 | TRIM22 | tripartite n | 0.078371 | 0.084707 | 0.178638 |
| 22691 | ENSG0000 | TMEM256 | transmeml | 0 | 0.047162 | 0.178537 |
| 9184 | ENSG0000 | CCDC102B | coiled-coil | 0.05793 | 0.171135 | 0.178463 |
| 42478 | ENSG0000 | EIF1B-AS1 | EIF1B antis | 0.139529 | 0.143947 | 0.178433 |
| 23999 | ENSG0000 | FLJ31356 | uncharacte | 0.065299 | 0.102405 | 0.17841 |
| 18194 | ENSG0000 | AIF1 | allograft in | 0.052211 | 0.135869 | 0.178336 |
| 12602 | ENSG0000 | TTLL6 | tubulin tyr | 0.046774 | 0.092754 | 0.17832 |
| 21609 | ENSG0000 | AL645608.2 | | 0.108064 | 0.167126 | 0.17799 |
| 8095 | ENSG0000 | NLRP12 | NLR family | 0.041953 | 0.082619 | 0.177967 |
| 7132 | ENSG0000 | AOAH | acyloxyacy | 0.050056 | 0.15183 | 0.177919 |
| 40119 | ENSG0000 | MIR6891 | microRNA | 0.01218 | 0.012194 | 0.177863 |
| 24950 | ENSG0000 | TMEM253 | transmeml | 0.07689 | 0.125839 | 0.177863 |
| 42028 | ENSG0000 | AC136469.2 | | 0.008687 | 0.120563 | 0.177756 |
| 19068 | ENSG0000 | MIR193A | microRNA | 0.055655 | 0.16518 | 0.177526 |
| 24155 | ENSG0000 | AL445928.1 | | 0 | 0.058047 | 0.177479 |
| 32399 | ENSG0000 | AC090559.1 | | 0.126989 | 0.145908 | 0.177414 |
| 10450 | ENSG0000 | SHANK1 | SH3 and m | 0 | 0.032765 | 0.177356 |
| 34610 | ENSG0000 | AC012435.1 | | 0.063947 | 0.113787 | 0.177344 |
| 15099 | ENSG0000 | ACP7 | acid phosp | 0 | 0.126616 | 0.17724 |
| 28502 | ENSG0000 | ATP5A1P7 | ATP syntha | 0.04388 | 0.073416 | 0.177207 |
| 5732 | ENSG0000 | FFAR1 | free fatty a | 0.015412 | 0.032877 | 0.177002 |
| 8191 | ENSG0000 | FCN3 | ficolin 3 [S | 0.01333 | 0.041057 | 0.176924 |
| 36358 | ENSG0000 | AC010761.3 | | 0.115396 | 0.163301 | 0.176765 |
| 24622 | ENSG0000 | AC103563.3 | | 0 | 0.082997 | 0.176685 |
| 26182 | ENSG0000 | AL096677.1 | | 0.043602 | 0.123911 | 0.17668 |
| 23048 | ENSG0000 | DNMBP-AS | DNMBP an | 0.056572 | 0.08081 | 0.176644 |
| 16544 | ENSG0000 | TOMM20L | translocase | 0 | 0.168604 | 0.176598 |
| 30152 | ENSG0000 | AC092490.1 | | 0.027957 | 0.153105 | 0.176508 |
| 28851 | ENSG0000 | AL117692.1 | | 0 | 0.05798 | 0.176413 |
| 15705 | ENSG0000 | CLCNKA | chloride vo | 0.109855 | 0.13868 | 0.176246 |
| 15741 | ENSG0000 | CCDC73 | coiled-coil | 0.06786 | 0.08923 | 0.176198 |
| 35478 | ENSG0000 | GOLGA8S | golgin A8 f | 0.100478 | 0.175698 | 0.176161 |
| 18214 | ENSG0000 | POU5F1 | POU class 5 | 0.07915 | 0.088501 | 0.175893 |
| 42320 | ENSG0000 | AD000864.1 | | 0.013957 | 0.076049 | 0.175842 |

| | | | | | | |
|---|---|---|---|---|---|---|
| 29333 | ENSG0000 | AC087667.1 | | 0 | 0.122593 | 0.175822 |
| 29099 | ENSG0000 | AP002884.2 | | 0 | 0.014459 | 0.175814 |
| 1540 | ENSG0000 | CCL17 | C-C motif c | 0 | 0.079731 | 0.175753 |
| 16056 | ENSG0000 | LRRIQ4 | leucine rich | 0.086361 | 0.126846 | 0.175629 |
| 2479 | ENSG0000 | SPEF1 | sperm flag | 0.04642 | 0.156056 | 0.175469 |
| 23334 | ENSG0000 | LINC01635 | long interg | 0.03152 | 0.035227 | 0.175217 |
| 18113 | ENSG0000 | AP000911.1 | | 0.056474 | 0.056592 | 0.174991 |
| 12234 | ENSG0000 | INPP5D | inositol po | 0.076816 | 0.077764 | 0.174903 |
| 10672 | ENSG0000 | FCMR | Fc fragmer | 0.030071 | 0.14442 | 0.174901 |
| 24484 | ENSG0000 | AC013275.1 | | 0 | 0.04175 | 0.174862 |
| 15008 | ENSG0000 | SPDYE16 | speedy/RIN | 0.009518 | 0.078264 | 0.1748 |
| 21396 | ENSG0000 | CFL1P1 | cofilin 1 ps | 0.027183 | 0.115051 | 0.174538 |
| 4283 | ENSG0000 | CISH | cytokine in | 0.041855 | 0.099542 | 0.174462 |
| 30654 | ENSG0000 | AC091180.4 | | 0 | 0.107431 | 0.174415 |
| 27003 | ENSG0000 | AC108448.2 | | 0.047932 | 0.091822 | 0.174291 |
| 6854 | ENSG0000 | FHOD3 | formin hor | 0.077519 | 0.144104 | 0.174199 |
| 36463 | ENSG0000 | WDR45BP1 | WD repeat | 0.052349 | 0.084957 | 0.174147 |
| 37573 | ENSG0000 | AC010643.1 | | 0.065301 | 0.071592 | 0.17403 |
| 18453 | ENSG0000 | CALML3-AS | CALML3 ar | 0.022489 | 0.090369 | 0.174007 |
| 19427 | ENSG0000 | IGHJ6 | immunoglo | 0 | 0.103091 | 0.173246 |
| 40087 | ENSG0000 | ATP5A1P10 | ATP syntha | 0.039604 | 0.095352 | 0.172944 |
| 7981 | ENSG0000 | GATA6 | GATA bind | 0.008996 | 0.059137 | 0.172785 |
| 2687 | ENSG0000 | RGCC | regulator c | 0 | 0.084511 | 0.17275 |
| 5792 | ENSG0000 | AVPR2 | arginine va | 0.015909 | 0.028249 | 0.172669 |
| 42707 | ENSG0000 | AL139353.3 | | 0 | 0.012291 | 0.171886 |
| 31589 | ENSG0000 | AC100849.1 | | 0.039563 | 0.04117 | 0.171839 |
| 8478 | ENSG0000 | TRPM8 | transient re | 0.053207 | 0.124334 | 0.171719 |
| 7994 | ENSG0000 | PIK3R5 | phosphoin | 0.032999 | 0.060846 | 0.17159 |
| 9877 | ENSG0000 | NMNAT2 | nicotinami | 0.064554 | 0.104819 | 0.17156 |
| 9656 | ENSG0000 | CCDC144C | coiled-coil | 0.08816 | 0.166048 | 0.171419 |
| 23822 | ENSG0000 | PATL2 | PAT1 homo | 0.022502 | 0.080005 | 0.171041 |
| 9937 | ENSG0000 | CYGB | cytoglobin | 0.047672 | 0.085424 | 0.170763 |
| 33614 | ENSG0000 | AL442163.1 | | 0 | 0.133221 | 0.170678 |
| 1394 | ENSG0000 | SMC1B | structural r | 0.031221 | 0.066346 | 0.170655 |
| 21738 | ENSG0000 | ZMIZ1-AS1 | ZMIZ1 anti | 0.031756 | 0.058939 | 0.1706 |
| 28164 | ENSG0000 | RPL12P37 | ribosomal | 0.118485 | 0.146202 | 0.170564 |
| 22008 | ENSG0000 | LINC01678 | long interg | 0 | 0.065042 | 0.170363 |
| 15470 | ENSG0000 | IRF7 | interferon | 0.079247 | 0.115982 | 0.169984 |
| 28868 | ENSG0000 | MTND4LP1 | mitochond | 0 | 0.089489 | 0.169912 |
| 14170 | ENSG0000 | TMEM139 | transmeml | 0.076515 | 0.078347 | 0.169824 |
| 7150 | ENSG0000 | FXYD2 | FXYD doma | 0.012732 | 0.073872 | 0.169759 |
| 14772 | ENSG0000 | FES | FES proto-c | 0.055434 | 0.168034 | 0.169734 |
| 15857 | ENSG0000 | MAGI2 | membrane | 0.064065 | 0.150211 | 0.169607 |
| 14273 | ENSG0000 | CATSPERE | catsper cha | 0.021378 | 0.164485 | 0.169433 |
| 35304 | ENSG0000 | AC021087.4 | | 0.031241 | 0.04789 | 0.16925 |
| 23852 | ENSG0000 | AL358074.1 | | 0.011796 | 0.107922 | 0.169181 |
| 42223 | ENSG0000 | DNM1P41 | dynamin 1 | 0.031897 | 0.130048 | 0.168979 |

| | | | | | | |
|---|---|---|---|---|---|---|
| 39788 | ENSG0000 | AC009269.6 | | 0 | 0.126409 | 0.168593 |
| 22916 | ENSG0000 | AL022238.1 | | 0 | 0.035719 | 0.16833 |
| 40273 | ENSG0000 | LINC01730 | long interg | 0 | 0.140356 | 0.168183 |
| 35632 | ENSG0000 | DKFZP434A | uncharacte | 0.02825 | 0.063649 | 0.168038 |
| 24424 | ENSG0000 | FBXW11P1 | F-box and ' | 0.048499 | 0.050986 | 0.167863 |
| 27849 | ENSG0000 | AC080162.1 | | 0 | 0.144017 | 0.167858 |
| 23464 | ENSG0000 | AL590723.1 | | 0 | 0.138166 | 0.167799 |
| 8260 | ENSG0000 | NR1I3 | nuclear rec | 0.082224 | 0.106054 | 0.167322 |
| 16720 | ENSG0000 | SPN | sialophorir | 0.024074 | 0.104184 | 0.167295 |
| 38691 | ENSG0000 | AL353135.1 | | 0.062929 | 0.090614 | 0.167291 |
| 7382 | ENSG0000 | FHDC1 | FH2 domai | 0.045758 | 0.105968 | 0.167191 |
| 30574 | ENSG0000 | SELENOP | selenoprot | 0.080944 | 0.102276 | 0.166941 |
| 43066 | ENSG0000 | Z98749.3 | | 0.008797 | 0.042436 | 0.16692 |
| 711 | ENSG0000 | PRSS8 | protease, s | 0.011875 | 0.130336 | 0.166752 |
| 34530 | ENSG0000 | AC110079.1 | | 0.103083 | 0.133222 | 0.166587 |
| 13230 | ENSG0000 | ADGRF3 | adhesion G | 0.066986 | 0.161624 | 0.166536 |
| 8151 | ENSG0000 | FGD5 | FYVE, RhoG | 0.008786 | 0.061144 | 0.166499 |
| 35763 | ENSG0000 | AC123768.3 | | 0.127308 | 0.141578 | 0.166346 |
| 33701 | ENSG0000 | ASB9P1 | ankyrin rep | 0 | 0.071501 | 0.166148 |
| 376 | ENSG0000 | MAMLD1 | mastermin | 0.12675 | 0.129547 | 0.166096 |
| 14759 | ENSG0000 | KCNK4 | potassium | 0.082233 | 0.110004 | 0.16595 |
| 38882 | ENSG0000 | AL133255.1 | | 0.041382 | 0.118987 | 0.165923 |
| 27274 | ENSG0000 | ZBTB40-IT1 | ZBTB40 int | 0 | 0.161868 | 0.16579 |
| 13751 | ENSG0000 | C15orf56 | chromosor | 0 | 0.090685 | 0.165729 |
| 14769 | ENSG0000 | BGN | biglycan [S | 0 | 0.083423 | 0.165632 |
| 42958 | ENSG0000 | AL137785.1 | | 0 | 0.059961 | 0.165506 |
| 26154 | ENSG0000 | AC012618.3 | | 0.062288 | 0.110311 | 0.165391 |
| 16465 | ENSG0000 | CACNA1H | calcium vo | 0.030595 | 0.053421 | 0.165382 |
| 9558 | ENSG0000 | GRAP | GRB2-relat | 0.011504 | 0.035898 | 0.165261 |
| 41013 | ENSG0000 | H2AFB3 | H2A histon | 0.021935 | 0.036457 | 0.16521 |
| 31549 | ENSG0000 | PCDHGA5 | protocadh | 0.05811 | 0.075344 | 0.165182 |
| 23812 | ENSG0000 | AC114495.2 | | 0.122402 | 0.134702 | 0.165076 |
| 35445 | ENSG0000 | SCRT1 | scratch fan | 0.04253 | 0.155928 | 0.164874 |
| 40139 | ENSG0000 | MIR6782 | microRNA | 0.038305 | 0.147913 | 0.164813 |
| 20487 | ENSG0000 | BTF3P7 | basic trans | 0 | 0.047954 | 0.164772 |
| 34862 | ENSG0000 | AC011773.4 | | 0.034652 | 0.085285 | 0.164771 |
| 5578 | ENSG0000 | HS3ST3B1 | heparan su | 0.039795 | 0.148194 | 0.164707 |
| 41330 | ENSG0000 | MIR6754 | microRNA | 0 | 0.103091 | 0.164473 |
| 7910 | ENSG0000 | CMTM2 | CKLF like M | 0.062218 | 0.114018 | 0.164163 |
| 16919 | ENSG0000 | SVIP | small VCP i | 0.04614 | 0.083141 | 0.164103 |
| 16142 | ENSG0000 | CYP51A1-A | CYP51A1 a | 0.028211 | 0.066152 | 0.164048 |
| 9648 | ENSG0000 | PLCL2 | phospholip | 0.059261 | 0.096099 | 0.164034 |
| 7639 | ENSG0000 | ANXA3 | annexin A3 | 0.083532 | 0.158571 | 0.163981 |
| 24266 | ENSG0000 | LIMD1-AS1 | LIMD1 anti | 0.136589 | 0.151706 | 0.163925 |
| 12326 | ENSG0000 | IL1RAPL1 | interleukin | 0.067802 | 0.077704 | 0.163662 |
| 27512 | ENSG0000 | Z97987.1 | | 0 | 0.013579 | 0.16346 |
| 15336 | ENSG0000 | LCN12 | lipocalin 12 | 0.01201 | 0.04725 | 0.163237 |

| | | | | | | |
|---|---|---|---|---|---|---|
| 14025 | ENSG0000 | LCN15 | lipocalin 1! | 0.012337 | 0.029924 | 0.162974 |
| 40677 | ENSG0000 | AC137834.2 | | 0.050631 | 0.069855 | 0.162528 |
| 8409 | ENSG0000 | ABHD1 | abhydrolas | 0.042784 | 0.103891 | 0.162524 |
| 25432 | ENSG0000 | RPS12P17 | ribosomal | 0 | 0.060159 | 0.162402 |
| 31701 | ENSG0000 | FAM85B | family witl | 0.021461 | 0.116254 | 0.162272 |
| 13350 | ENSG0000 | GLIS1 | GLIS family | 0.011401 | 0.023413 | 0.162231 |
| 18514 | ENSG0000 | DPP9-AS1 | DPP9 antis | 0.066066 | 0.120776 | 0.162105 |
| 18625 | ENSG0000 | ONECUT3 | one cut ho | 0.006416 | 0.027626 | 0.162016 |
| 6641 | ENSG0000 | PRAM1 | PML-RARA | 0.07084 | 0.118999 | 0.161785 |
| 1342 | ENSG0000 | RGS11 | regulator c | 0.029476 | 0.137617 | 0.161561 |
| 21812 | ENSG0000 | AC015987.1 | | 0.022072 | 0.104758 | 0.161475 |
| 36925 | ENSG0000 | RNF157-AS | RNF157 an | 0.061776 | 0.089252 | 0.1612 |
| 37906 | ENSG0000 | AC004754.1 | | 0.056484 | 0.136426 | 0.161149 |
| 26109 | ENSG0000 | NUS1P2 | NUS1 dehy | 0.04844 | 0.097777 | 0.161055 |
| 27184 | ENSG0000 | RPL12P27 | ribosomal | 0 | 0.073144 | 0.160983 |
| 35160 | ENSG0000 | AL162231.4 | | 0.033376 | 0.091651 | 0.160578 |
| 14049 | ENSG0000 | BOLA1 | bolA family | 0.024293 | 0.156006 | 0.160363 |
| 33184 | ENSG0000 | AL928654.4 | | 0 | 0.038019 | 0.160311 |
| 23702 | ENSG0000 | AL596223.1 | | 0 | 0.071909 | 0.160093 |
| 18219 | ENSG0000 | PSORS1C1 | psoriasis su | 0.009032 | 0.029038 | 0.159984 |
| 38647 | ENSG0000 | AC008937.3 | | 0.021026 | 0.081139 | 0.159737 |
| 2102 | ENSG0000 | CATSPERG | cation char | 0.063919 | 0.097967 | 0.15972 |
| 4977 | ENSG0000 | CFAP58 | cilia and fla | 0 | 0.104808 | 0.159646 |
| 28954 | ENSG0000 | RPL5P30 | ribosomal | 0 | 0.066413 | 0.159348 |
| 36828 | ENSG0000 | FARSA-AS1 | FARSA anti | 0 | 0.034171 | 0.159253 |
| 10296 | ENSG0000 | GAB3 | GRB2 asso | 0.023586 | 0.074619 | 0.159184 |
| 38224 | ENSG0000 | AC011451.2 | | 0 | 0.038961 | 0.159051 |
| 36683 | ENSG0000 | NARF-IT1 | NARF intro | 0.046645 | 0.110668 | 0.158982 |
| 4791 | ENSG0000 | PPL | periplakin | 0.023394 | 0.086033 | 0.158942 |
| 20274 | ENSG0000 | MED28P1 | mediator c | 0 | 0.124395 | 0.158927 |
| 16201 | ENSG0000 | HUS1B | HUS1 chec | 0 | 0.119384 | 0.158801 |
| 41829 | ENSG0000 | AP001324.3 | | 0.024854 | 0.06057 | 0.158791 |
| 7547 | ENSG0000 | ADAMTS14 | ADAM met | 0.040607 | 0.052543 | 0.158704 |
| 33610 | ENSG0000 | AL352984.1 | | 0 | 0.083206 | 0.15857 |
| 6708 | ENSG0000 | DUSP26 | dual specif | 0.034654 | 0.120022 | 0.158533 |
| 35333 | ENSG0000 | AC009163.3 | | 0 | 0.130986 | 0.158515 |
| 21238 | ENSG0000 | AC072022.1 | | 0 | 0.087071 | 0.158349 |
| 15784 | ENSG0000 | CHCHD2P9 | coiled-coil- | 0.066517 | 0.110551 | 0.158198 |
| 14339 | ENSG0000 | NLRP11 | NLR family | 0.033041 | 0.091184 | 0.158107 |
| 24207 | ENSG0000 | LINC01747 | long interg | 0.040804 | 0.087151 | 0.158032 |
| 39205 | ENSG0000 | AC016747.3 | | 0.090058 | 0.110259 | 0.158001 |
| 7013 | ENSG0000 | EMX1 | empty spir | 0.00648 | 0.047635 | 0.157993 |
| 21429 | ENSG0000 | AP001469.1 | | 0 | 0.066447 | 0.157953 |
| 34975 | ENSG0000 | LINC00543 | long interg | 0 | 0.031402 | 0.157705 |
| 38172 | ENSG0000 | AC110048.1 | | 0.016454 | 0.054055 | 0.157686 |
| 26720 | ENSG0000 | AC067956.1 | | 0.094913 | 0.105541 | 0.157477 |
| 37708 | ENSG0000 | CSAG3 | CSAG famil | 0.043832 | 0.045549 | 0.157418 |

| | | | | | | |
|---|---|---|---|---|---|---|
| 35578 | ENSG0000 | MIR4284 | microRNA | 0.041953 | 0.080182 | 0.157414 |
| 41614 | ENSG0000 | AC009090.4 | | 0.038088 | 0.078736 | 0.157403 |
| 43369 | ENSG0000 | AC092902.5 | | 0.033604 | 0.081778 | 0.157224 |
| 122 | ENSG0000 | ABCC8 | ATP bindin | 0.101528 | 0.103477 | 0.157173 |
| 21947 | ENSG0000 | DUTP6 | deoxyuridi | 0.104337 | 0.155083 | 0.156889 |
| 1711 | ENSG0000 | AQP6 | aquaporin | 0.062289 | 0.111794 | 0.15677 |
| 26882 | ENSG0000 | LINC00479 | long interg | 0.031087 | 0.104937 | 0.156615 |
| 16912 | ENSG0000 | MB | myoglobin | 0 | 0.026866 | 0.156356 |
| 33309 | ENSG0000 | TPRX2P | tetrapeptic | 0 | 0.032999 | 0.156333 |
| 35701 | ENSG0000 | AC007613.1 | | 0.076398 | 0.12599 | 0.156301 |
| 12470 | ENSG0000 | SPDYE5 | speedy/RII | 0.046091 | 0.090445 | 0.156043 |
| 35301 | ENSG0000 | AL023803.1 | | 0 | 0.134514 | 0.156026 |
| 38794 | ENSG0000 | AP006545.2 | | 0.038495 | 0.070442 | 0.155984 |
| 4298 | ENSG0000 | PLCH1 | phospholip | 0.069255 | 0.126593 | 0.155825 |
| 28877 | ENSG0000 | AL162431.2 | | 0.041167 | 0.07699 | 0.15572 |
| 24820 | ENSG0000 | RPL31P50 | ribosomal | 0 | 0.094732 | 0.155618 |
| 8388 | ENSG0000 | PTPN7 | protein tyr | 0.006031 | 0.091047 | 0.155522 |
| 37429 | ENSG0000 | AC010605.1 | | 0 | 0.137311 | 0.155446 |
| 16305 | ENSG0000 | SYCP2 | synaptone | 0.030228 | 0.096809 | 0.155445 |
| 26708 | ENSG0000 | AC009237.8 | | 0 | 0.038353 | 0.15518 |
| 6827 | ENSG0000 | RAB33A | RAB33A, m | 0.04242 | 0.070477 | 0.154956 |
| 16160 | ENSG0000 | SKOR1 | SKI family t | 0.057379 | 0.110433 | 0.154878 |
| 41089 | ENSG0000 | HOXA11-A | HOXA11 ar | 0.006702 | 0.016373 | 0.154829 |
| 23160 | ENSG0000 | MTND5P1 | mitochond | 0 | 0.042511 | 0.154636 |
| 38812 | ENSG0000 | AC055822.1 | | 0 | 0.094751 | 0.154106 |
| 1683 | ENSG0000 | OVGP1 | oviductal g | 0.108532 | 0.138138 | 0.15407 |
| 29794 | ENSG0000 | ZNF807 | zinc finger | 0.092416 | 0.134918 | 0.154066 |
| 27533 | ENSG0000 | GGTA2P | glycoprote | 0 | 0.039129 | 0.153568 |
| 17838 | ENSG0000 | RN7SKP20 | RNA, 7SK s | 0 | 0.060255 | 0.153054 |
| 42176 | ENSG0000 | AC022400.9 | | 0.033945 | 0.116067 | 0.152766 |
| 25465 | ENSG0000 | NPIPB7 | nuclear po | 0.011953 | 0.032358 | 0.152689 |
| 3063 | ENSG0000 | CNTD2 | cyclin N-te | 0.103494 | 0.133078 | 0.152606 |
| 42483 | ENSG0000 | AC243547.3 | | 0.044121 | 0.068213 | 0.152363 |
| 17182 | ENSG0000 | GPRASP1 | G protein-c | 0.013796 | 0.079948 | 0.15232 |
| 17305 | ENSG0000 | RNU4-39P | RNA, U4 sr | 0.112975 | 0.150984 | 0.151954 |
| 14249 | ENSG0000 | SMCO3 | single-pass | 0.025782 | 0.097162 | 0.151845 |
| 2056 | ENSG0000 | SMIM24 | small integ | 0 | 0.117164 | 0.15183 |
| 5058 | ENSG0000 | MYOT | myotilin [S | 0.109686 | 0.126452 | 0.151754 |
| 26020 | ENSG0000 | RHOXF1P1 | Rhox home | 0 | 0.037791 | 0.151626 |
| 24390 | ENSG0000 | AC007279.1 | | 0.026916 | 0.034733 | 0.151513 |
| 12481 | ENSG0000 | CCDC144A | coiled-coil | 0.011593 | 0.014061 | 0.151265 |
| 42476 | ENSG0000 | LINC01943 | long interg | 0.09212 | 0.14123 | 0.15114 |
| 38124 | ENSG0000 | POC1B-AS1 | POC1B ant | 0.053374 | 0.134784 | 0.151027 |
| 32414 | ENSG0000 | AC069287.3 | | 0 | 0.093634 | 0.150977 |
| 29526 | ENSG0000 | NNT-AS1 | NNT antise | 0.042846 | 0.098548 | 0.150807 |
| 33839 | ENSG0000 | AL358334.3 | | 0 | 0.059708 | 0.150727 |
| 7975 | ENSG0000 | ADCYAP1 | adenylate | 0.138955 | 0.149779 | 0.150699 |

| | | | | | | |
|---|---|---|---|---|---|---|
| 31719 | ENSG0000 | AC105118.1 | | 0 | 0.024444 | 0.150584 |
| 42406 | ENSG0000 | AL109806.1 | | 0 | 0.023295 | 0.150583 |
| 9253 | ENSG0000 | AKAP6 | A-kinase ar | 0.103571 | 0.117541 | 0.150573 |
| 34478 | ENSG0000 | AC009119.2 | | 0.018519 | 0.044697 | 0.150491 |
| 13499 | ENSG0000 | INHBC | inhibin bet | 0.066278 | 0.102211 | 0.150385 |
| 29107 | ENSG0000 | AL109754.1 | | 0.026379 | 0.071237 | 0.150333 |
| 18069 | ENSG0000 | NEU4 | neuraminic | 0.018019 | 0.038803 | 0.150012 |
| 13318 | ENSG0000 | CD34 | CD34 mole | 0 | 0.14394 | 0.149958 |
| 18879 | ENSG0000 | SNORA80D | small nucle | 0.038224 | 0.142037 | 0.149756 |
| 37977 | ENSG0000 | AC040174.2 | | 0.038434 | 0.063407 | 0.149701 |
| 10685 | ENSG0000 | DISC1 | disrupted i | 0.019271 | 0.077558 | 0.149685 |
| 40365 | ENSG0000 | AC244472.2 | | 0 | 0.071859 | 0.14967 |
| 17456 | ENSG0000 | Y_RNA | Y RNA [Sou | 0 | 0.074226 | 0.149331 |
| 12956 | ENSG0000 | SLC9C1 | solute carr | 0.022385 | 0.116832 | 0.149326 |
| 12698 | ENSG0000 | MAP6 | microtubul | 0.062674 | 0.129598 | 0.149231 |
| 29275 | ENSG0000 | KCNK15-AS | KCNK15 an | 0.012855 | 0.080051 | 0.149174 |
| 13892 | ENSG0000 | SPACA4 | sperm acro | 0.036256 | 0.051227 | 0.149142 |
| 3854 | ENSG0000 | TBC1D30 | TBC1 doma | 0.047342 | 0.048633 | 0.149017 |
| 34306 | ENSG0000 | AC021231.3 | | 0 | 0.138476 | 0.14871 |
| 7067 | ENSG0000 | CYP27A1 | cytochrom | 0.022213 | 0.120863 | 0.148698 |
| 36941 | ENSG0000 | AC006557.1 | | 0.062987 | 0.0991 | 0.148643 |
| 32485 | ENSG0000 | AC022239.2 | | 0 | 0.028538 | 0.148582 |
| 10897 | ENSG0000 | TAGAP | T-cell activ | 0.115958 | 0.122063 | 0.148403 |
| 9077 | ENSG0000 | ST14 | suppressio | 0.040969 | 0.072428 | 0.148397 |
| 20462 | ENSG0000 | AL139039.1 | | 0 | 0.104134 | 0.148143 |
| 9243 | ENSG0000 | SLC2A13 | solute carr | 0.110764 | 0.124812 | 0.147892 |
| 12575 | ENSG0000 | EMB | embigin [S | 0.097728 | 0.105542 | 0.147755 |
| 39487 | ENSG0000 | TBC1D3K | TBC1 doma | 0.089047 | 0.143313 | 0.147104 |
| 34236 | ENSG0000 | AC048382.4 | | 0.043004 | 0.045241 | 0.147059 |
| 15108 | ENSG0000 | SLC35F3 | solute carr | 0.022217 | 0.081957 | 0.146868 |
| 34979 | ENSG0000 | AC133552.1 | | 0 | 0.066742 | 0.14678 |
| 18510 | ENSG0000 | DENND1C | DENN dom | 0.045826 | 0.056724 | 0.146647 |
| 37036 | ENSG0000 | AC012569.1 | | 0.036502 | 0.074274 | 0.146602 |
| 25998 | ENSG0000 | KBTBD13 | kelch repea | 0 | 0.033914 | 0.14657 |
| 31333 | ENSG0000 | TPTE2P1 | transmeml | 0 | 0.0991 | 0.146466 |
| 40161 | ENSG0000 | AC244196.2 | | 0 | 0.077432 | 0.146367 |
| 12065 | ENSG0000 | SCARA5 | scavenger | 0.020092 | 0.082148 | 0.146054 |
| 2528 | ENSG0000 | BPI | bactericida | 0.025391 | 0.056914 | 0.146051 |
| 35680 | ENSG0000 | AC003688.1 | | 0 | 0.095147 | 0.146022 |
| 10739 | ENSG0000 | BNIPL | BCL2 intera | 0.060415 | 0.130455 | 0.145971 |
| 2740 | ENSG0000 | FA2H | fatty acid 2 | 0.021648 | 0.085734 | 0.145628 |
| 1158 | ENSG0000 | RPS6KA2 | ribosomal | 0.1163 | 0.119631 | 0.145429 |
| 31473 | ENSG0000 | RAD21-AS1 | RAD21 ant | 0.071285 | 0.09826 | 0.145225 |
| 13248 | ENSG0000 | HSPB7 | heat shock | 0.02664 | 0.069737 | 0.145071 |
| 14626 | ENSG0000 | C3orf58 | chromosor | 0.074478 | 0.083539 | 0.144974 |
| 26957 | ENSG0000 | SNRPGP14 | small nucle | 0 | 0.081908 | 0.144934 |
| 32465 | ENSG0000 | AC136475.5 | | 0 | 0.067583 | 0.144833 |

| | | | | | | |
|---|---|---|---|---|---|---|
| 16627 | ENSG0000 | ACSL5 | acyl-CoA sy | 0.014294 | 0.073011 | 0.144627 |
| 25622 | ENSG0000 | EIF4EP1 | eukaryotic | 0 | 0.046407 | 0.144571 |
| 21373 | ENSG0000 | LINC01626 | long interg | 0 | 0.039046 | 0.14435 |
| 30139 | ENSG0000 | AC084357.2 | | 0 | 0.141869 | 0.144236 |
| 30821 | ENSG0000 | AC108062.1 | | 0.107309 | 0.129438 | 0.144096 |
| 7563 | ENSG0000 | CDK15 | cyclin depe | 0.048088 | 0.092573 | 0.144046 |
| 34111 | ENSG0000 | AC012409.1 | | 0.031004 | 0.077696 | 0.143585 |
| 602 | ENSG0000 | TNC | tenascin C | 0.111208 | 0.122605 | 0.143441 |
| 22503 | ENSG0000 | AL031123.1 | | 0.058803 | 0.06425 | 0.143245 |
| 11987 | ENSG0000 | SOAT2 | sterol O-ac | 0.030744 | 0.051412 | 0.143138 |
| 5512 | ENSG0000 | TREM1 | triggering r | 0.055333 | 0.08183 | 0.143065 |
| 17901 | ENSG0000 | ZNF525 | zinc finger | 0.02964 | 0.1015 | 0.142878 |
| 29436 | ENSG0000 | AC113382.1 | | 0 | 0.117772 | 0.142556 |
| 28848 | ENSG0000 | AC012158.1 | | 0.03326 | 0.07957 | 0.142533 |
| 13942 | ENSG0000 | GPR4 | G protein-c | 0.011285 | 0.12502 | 0.142483 |
| 6603 | ENSG0000 | WASF3 | WAS prote | 0.0345 | 0.062989 | 0.142419 |
| 23736 | ENSG0000 | SNRPFP2 | small nucle | 0 | 0.08543 | 0.142329 |
| 20546 | ENSG0000 | AC131097.3 | | 0.028797 | 0.119082 | 0.142185 |
| 1721 | ENSG0000 | HBQ1 | hemoglobi | 0 | 0.057514 | 0.142166 |
| 4537 | ENSG0000 | DLEU2L | deleted in | 0.04521 | 0.110399 | 0.142077 |
| 15163 | ENSG0000 | TBX1 | T-box 1 [Sc | 0.033612 | 0.080671 | 0.141892 |
| 5447 | ENSG0000 | TP53TG5 | TP53 targe | 0.077862 | 0.120459 | 0.141758 |
| 27627 | ENSG0000 | TNFRSF14- | TNFRSF14 | 0.023112 | 0.079693 | 0.141693 |
| 13419 | ENSG0000 | C4orf32 | chromosor | 0.095529 | 0.127691 | 0.141648 |
| 38152 | ENSG0000 | NRBF2P5 | nuclear rec | 0.02252 | 0.035716 | 0.141547 |
| 15588 | ENSG0000 | GRK1 | G protein-c | 0.043074 | 0.046361 | 0.141405 |
| 27737 | ENSG0000 | RNY4P25 | RNA, Ro-as | 0 | 0.089131 | 0.141194 |
| 39508 | ENSG0000 | AL590282.1 | | 0 | 0.066451 | 0.140946 |
| 8660 | ENSG0000 | C1QTNF2 | C1q and TN | 0 | 0.114579 | 0.140946 |
| 2059 | ENSG0000 | KCNK16 | potassium | 0.051538 | 0.093798 | 0.140772 |
| 23879 | ENSG0000 | PRAC2 | prostate ca | 0.015108 | 0.085828 | 0.140763 |
| 36302 | ENSG0000 | AC011825.4 | | 0.075568 | 0.094276 | 0.140427 |
| 1572 | ENSG0000 | ZNF506 | zinc finger | 0.08268 | 0.132187 | 0.14037 |
| 36808 | ENSG0000 | AC129510.2 | | 0 | 0.058269 | 0.140353 |
| 21361 | ENSG0000 | AC098828.2 | | 0.056641 | 0.07424 | 0.14033 |
| 8684 | ENSG0000 | PSD2 | pleckstrin a | 0.027498 | 0.077484 | 0.140183 |
| 35529 | ENSG0000 | MYZAP | myocardia | 0.013982 | 0.035407 | 0.140134 |
| 40138 | ENSG0000 | AC025594.3 | | 0 | 0.067857 | 0.139734 |
| 40108 | ENSG0000 | AL138966.2 | | 0 | 0.09176 | 0.139626 |
| 40555 | ENSG0000 | AC022960.2 | | 0 | 0.043806 | 0.139534 |
| 10565 | ENSG0000 | MATN1 | matrilin 1, | 0.014183 | 0.08557 | 0.139407 |
| 16097 | ENSG0000 | NCCRP1 | non-specifi | 0.071965 | 0.099504 | 0.139319 |
| 39258 | ENSG0000 | AC107294.3 | | 0.032781 | 0.065657 | 0.139111 |
| 15841 | ENSG0000 | SESTD1 | SEC14 and | 0.083041 | 0.111467 | 0.139044 |
| 15382 | ENSG0000 | MYADML2 | myeloid as | 0.079149 | 0.13639 | 0.139027 |
| 5088 | ENSG0000 | TNFRSF8 | TNF recept | 0.009495 | 0.031137 | 0.139023 |
| 19738 | ENSG0000 | LTC4S | leukotriene | 0.034843 | 0.073334 | 0.138931 |

| | | | | | | |
|---|---|---|---|---|---|---|
| 40952 | ENSG0000 | AC130343.2 | | 0 | 0.034122 | 0.138868 |
| 36320 | ENSG0000 | AC007922.2 | | 0 | 0.001834 | 0.138789 |
| 3456 | ENSG0000 | NEURL1 | neuralized | 0.010732 | 0.096857 | 0.138626 |
| 41706 | ENSG0000 | AC093503.3 | | 0.015251 | 0.074162 | 0.138313 |
| 29427 | ENSG0000 | AC022113.1 | | 0 | 0.057894 | 0.138313 |
| 3428 | ENSG0000 | CDH23 | cadherin re | 0.024023 | 0.027682 | 0.138151 |
| 36283 | ENSG0000 | AC115989.2 | | 0 | 0.074306 | 0.138089 |
| 10411 | ENSG0000 | AC007663.1 | | 0.0706 | 0.083084 | 0.138064 |
| 2181 | ENSG0000 | MFNG | MFNG O-fu | 0.043111 | 0.06117 | 0.138055 |
| 7407 | ENSG0000 | SULF1 | sulfatase 1 | 0.072085 | 0.125064 | 0.138034 |
| 42943 | ENSG0000 | AL160035.1 | | 0 | 0.101301 | 0.138003 |
| 24740 | ENSG0000 | AGBL5-AS1 | AGBL5 anti | 0 | 0.109316 | 0.137986 |
| 13495 | ENSG0000 | FAM182B | family with | 0.086201 | 0.115897 | 0.137908 |
| 9567 | ENSG0000 | DNAAF1 | dynein axo | 0.091195 | 0.135236 | 0.137886 |
| 9614 | ENSG0000 | CCDC173 | coiled-coil | 0 | 0.110191 | 0.137665 |
| 37273 | ENSG0000 | KF456478.1 | | 0 | 0.050616 | 0.137638 |
| 21270 | ENSG0000 | AC093690.1 | | 0.082861 | 0.120345 | 0.13756 |
| 38478 | ENSG0000 | AC091167.5 | | 0.019709 | 0.029718 | 0.137314 |
| 26271 | ENSG0000 | AC012074.1 | | 0.10443 | 0.132934 | 0.136586 |
| 16232 | ENSG0000 | PLAC9 | placenta sp | 0.027608 | 0.051179 | 0.136407 |
| 34497 | ENSG0000 | HEXA-AS1 | HEXA antis | 0.106153 | 0.131258 | 0.136387 |
| 25490 | ENSG0000 | RASA4DP | RAS p21 pr | 0.038839 | 0.059262 | 0.136314 |
| 42004 | ENSG0000 | AC018892.3 | | 0.057062 | 0.122109 | 0.136198 |
| 35448 | ENSG0000 | AC018362.1 | | 0.052545 | 0.053411 | 0.136098 |
| 28306 | ENSG0000 | AC093627.4 | | 0.089804 | 0.09914 | 0.135439 |
| 13132 | ENSG0000 | RXFP4 | relaxin/ins | 0.023243 | 0.105585 | 0.135435 |
| 35625 | ENSG0000 | AC091180.5 | | 0.077274 | 0.08261 | 0.135258 |
| 14488 | ENSG0000 | PIPSL | PIP5K1A ar | 0.067231 | 0.074441 | 0.135032 |
| 35312 | ENSG0000 | AC023830.1 | | 0.032101 | 0.046834 | 0.135005 |
| 38221 | ENSG0000 | AC009549.1 | | 0.021051 | 0.060534 | 0.134832 |
| 27141 | ENSG0000 | AL050331.3 | | 0.025311 | 0.094178 | 0.134788 |
| 42391 | ENSG0000 | AP002364.1 | | 0.031446 | 0.129122 | 0.134682 |
| 31666 | ENSG0000 | AC023632.2 | | 0.039748 | 0.114097 | 0.134646 |
| 16952 | ENSG0000 | CARD11 | caspase re | 0 | 0.043474 | 0.134467 |
| 2777 | ENSG0000 | MEFV | MEFV, pyri | 0.058532 | 0.05855 | 0.134464 |
| 29773 | ENSG0000 | AC139495.2 | | 0 | 0.036893 | 0.134394 |
| 34493 | ENSG0000 | AC079148.1 | | 0.013062 | 0.09214 | 0.134365 |
| 27640 | ENSG0000 | AL357055.3 | | 0.018682 | 0.053285 | 0.134303 |
| 21864 | ENSG0000 | AC083949.1 | | 0 | 0.076669 | 0.134279 |
| 12117 | ENSG0000 | INSM2 | INSM trans | 0.035381 | 0.07719 | 0.134275 |
| 30568 | ENSG0000 | AP000892.1 | | 0 | 0.134154 | 0.134257 |
| 31512 | ENSG0000 | AC083973.1 | | 0.016072 | 0.047968 | 0.134061 |
| 16502 | ENSG0000 | HSH2D | hematopoi | 0.076127 | 0.09257 | 0.134003 |
| 24353 | ENSG0000 | AC114878.1 | | 0 | 0.112306 | 0.133801 |
| 36477 | ENSG0000 | AC106037.2 | | 0.013742 | 0.020538 | 0.13378 |
| 6555 | ENSG0000 | HAPLN2 | hyaluronar | 0.021813 | 0.042484 | 0.13368 |
| 24869 | ENSG0000 | DARS-AS1 | DARS antis | 0.026783 | 0.034692 | 0.13364 |

| | | | | | | |
|---|---|---|---|---|---|---|
| 5490 | ENSG0000 | BTN1A1 | butyrophili | 0 | 0.02116 | 0.133598 |
| 235 | ENSG0000 | PAX7 | paired box | 0.111714 | 0.131325 | 0.133466 |
| 12945 | ENSG0000 | LRRC15 | leucine rich | 0.02388 | 0.063729 | 0.13326 |
| 4623 | ENSG0000 | ACTL8 | actin like 8 | 0.076095 | 0.125159 | 0.133093 |
| 11444 | ENSG0000 | SLC18A2 | solute carr | 0.088411 | 0.104892 | 0.132991 |
| 21343 | ENSG0000 | AF230666.1 | | 0.052897 | 0.111875 | 0.132872 |
| 38907 | ENSG0000 | AC116036.2 | | 0 | 0.031463 | 0.132772 |
| 29928 | ENSG0000 | C5orf66-AS | C5orf66 an | 0 | 0.036242 | 0.132433 |
| 7513 | ENSG0000 | TRIM54 | tripartite n | 0 | 0.054791 | 0.132334 |
| 27246 | ENSG0000 | RPL35P1 | ribosomal | 0.048302 | 0.115629 | 0.132256 |
| 9290 | ENSG0000 | TEX9 | testis expr | 0.064156 | 0.121842 | 0.132068 |
| 43157 | ENSG0000 | AC002472.3 | | 0 | 0.106976 | 0.131867 |
| 34084 | ENSG0000 | AF111167.2 | | 0.017612 | 0.096437 | 0.131822 |
| 41831 | ENSG0000 | AGPAT4-IT | AGPAT4 in | 0.018899 | 0.02344 | 0.131813 |
| 35391 | ENSG0000 | AC010531.3 | | 0 | 0.089883 | 0.131771 |
| 12140 | ENSG0000 | TNXB | tenascin XI | 0.043794 | 0.076481 | 0.131631 |
| 34128 | ENSG0000 | AC090695.1 | | 0.028632 | 0.094908 | 0.131602 |
| 15459 | ENSG0000 | KPNA7 | karyopheri | 0.051803 | 0.066994 | 0.131551 |
| 15480 | ENSG0000 | DLK1 | delta like n | 0.016289 | 0.057422 | 0.131483 |
| 14452 | ENSG0000 | PRR26 | proline rich | 0.007676 | 0.097784 | 0.13123 |
| 28613 | ENSG0000 | AC092978.1 | | 0 | 0.114356 | 0.131209 |
| 25688 | ENSG0000 | GAS6-AS1 | GAS6 antis | 0.004951 | 0.017723 | 0.131182 |
| 5415 | ENSG0000 | MATN4 | matrilin 4 [ | 0.028671 | 0.040676 | 0.131013 |
| 28881 | ENSG0000 | RN7SL861P | RNA, 7SL, c | 0.061825 | 0.092239 | 0.130992 |
| 36685 | ENSG0000 | MIR5091 | microRNA | 0 | 0.103091 | 0.130927 |
| 42384 | ENSG0000 | AC011466.4 | | 0.044058 | 0.054037 | 0.130599 |
| 38543 | ENSG0000 | AL671883.3 | | 0 | 0.044756 | 0.130483 |
| 42596 | ENSG0000 | AL499606.1 | | 0 | 0.060194 | 0.130432 |
| 28056 | ENSG0000 | AC068760.1 | | 0 | 0.014991 | 0.130398 |
| 23784 | ENSG0000 | CICP3 | capicua tra | 0.029719 | 0.100589 | 0.130208 |
| 6270 | ENSG0000 | SMPDL3B | sphingomy | 0.026492 | 0.033016 | 0.130175 |
| 6078 | ENSG0000 | AP1M2 | adaptor re | 0.016031 | 0.101374 | 0.130141 |
| 35730 | ENSG0000 | AC018512.1 | | 0 | 0.029627 | 0.130116 |
| 39641 | ENSG0000 | AC245041.1 | | 0 | 0.035445 | 0.130049 |
| 37192 | ENSG0000 | AC015802.4 | | 0.078127 | 0.099997 | 0.129479 |
| 32808 | ENSG0000 | CBX3P4 | chromobox | 0.056983 | 0.101684 | 0.129426 |
| 35377 | ENSG0000 | AC007216.3 | | 0.070043 | 0.073095 | 0.129238 |
| 12170 | ENSG0000 | WFDC13 | WAP four-c | 0.013263 | 0.041885 | 0.128984 |
| 42918 | ENSG0000 | AC137936.2 | | 0.033338 | 0.048983 | 0.128958 |
| 11771 | ENSG0000 | SGSM1 | small G pro | 0.030605 | 0.057577 | 0.12865 |
| 27962 | ENSG0000 | AC012377.1 | | 0.020651 | 0.045208 | 0.128577 |
| 34722 | ENSG0000 | AC040169.2 | | 0 | 0.077868 | 0.128499 |
| 29625 | ENSG0000 | PCDHAC1 | protocadhe | 0.012 | 0.081918 | 0.128238 |
| 9033 | ENSG0000 | SYT8 | synaptotag | 0.026348 | 0.113017 | 0.128232 |
| 13184 | ENSG0000 | GPR148 | G protein-c | 0 | 0.031082 | 0.128107 |
| 14468 | ENSG0000 | SSTR2 | somatosta | 0.042957 | 0.115578 | 0.128022 |
| 26183 | ENSG0000 | PHBP13 | prohibitin p | 0.055146 | 0.095902 | 0.12802 |

| | | | | | | |
|---|---|---|---|---|---|---|
| 14236 | ENSG0000 | PXT1 | peroxisom | 0.016837 | 0.043876 | 0.127584 |
| 24841 | ENSG0000 | AC073342.1 | | 0 | 0.062741 | 0.127318 |
| 23285 | ENSG0000 | LYPLA2P1 | lysophospł | 0.112929 | 0.119606 | 0.126918 |
| 39081 | ENSG0000 | AL157904.1 | | 0.046334 | 0.075905 | 0.126867 |
| 26286 | ENSG0000 | LINC00330 | long interg | 0.017494 | 0.035798 | 0.126831 |
| 18217 | ENSG0000 | PSORS1C2 | psoriasis sı | 0 | 0.03792 | 0.126756 |
| 3769 | ENSG0000 | VWF | von Willeb | 0.003428 | 0.096613 | 0.126684 |
| 16960 | ENSG0000 | MYL4 | myosin ligł | 0 | 0.080154 | 0.1265 |
| 3929 | ENSG0000 | MAN1A1 | mannosida | 0.05334 | 0.102054 | 0.12621 |
| 25009 | ENSG0000 | AC098824.1 | | 0 | 0.094344 | 0.126157 |
| 39170 | ENSG0000 | AC113189.4 | | 0 | 0.022983 | 0.126113 |
| 36352 | ENSG0000 | ANXA8 | annexin A8 | 0.038296 | 0.041195 | 0.126102 |
| 10130 | ENSG0000 | SLC5A11 | solute carr | 0 | 0.033869 | 0.125663 |
| 43319 | ENSG0000 | AC004834.1 | | 0.011588 | 0.114766 | 0.125633 |
| 3164 | ENSG0000 | PDE4C | phosphodi | 0.024046 | 0.067014 | 0.125587 |
| 16206 | ENSG0000 | FAM35DP | family witł | 0.077316 | 0.090664 | 0.125544 |
| 29281 | ENSG0000 | LINC01391 | long interg | 0.027379 | 0.029542 | 0.125474 |
| 23192 | ENSG0000 | CFL1P3 | cofilin 1 ps | 0 | 0.072795 | 0.125419 |
| 20912 | ENSG0000 | MIR1249 | microRNA | 0.102975 | 0.124959 | 0.125313 |
| 26158 | ENSG0000 | TPT1P9 | tumor prot | 0.033056 | 0.057161 | 0.12515 |
| 35434 | ENSG0000 | BEAN1-AS1 | BEAN1 ant | 0.053888 | 0.101165 | 0.125093 |
| 41391 | ENSG0000 | MIR7851 | microRNA | 0.020452 | 0.112111 | 0.124921 |
| 15944 | ENSG0000 | PEAR1 | platelet en | 0.048272 | 0.088889 | 0.124658 |
| 38559 | ENSG0000 | AC138894.3 | | 0.010521 | 0.093497 | 0.124654 |
| 879 | ENSG0000 | LIMCH1 | LIM and ca | 0.061155 | 0.110665 | 0.124636 |
| 23555 | ENSG0000 | CICP9 | capicua tra | 0.101976 | 0.106486 | 0.124623 |
| 34864 | ENSG0000 | AC009148.1 | | 0.052904 | 0.1078 | 0.12451 |
| 6789 | ENSG0000 | CMPK2 | cytidine/ur | 0.024809 | 0.094189 | 0.124424 |
| 11219 | ENSG0000 | SMAD5-AS | SMAD5 an | 0 | 0.022322 | 0.12434 |
| 16178 | ENSG0000 | ENTPD8 | ectonucleo | 0.011905 | 0.116405 | 0.124229 |
| 8958 | ENSG0000 | LCN2 | lipocalin 2 | 0 | 0.018193 | 0.124163 |
| 38091 | ENSG0000 | AC015849.1 | | 0 | 0.066528 | 0.124117 |
| 42930 | ENSG0000 | AC073264.4 | | 0 | 0.046829 | 0.123997 |
| 31686 | ENSG0000 | AC025442.1 | | 0 | 0.123172 | 0.123991 |
| 36 | ENSG0000 | ICA1 | islet cell au | 0.058233 | 0.087325 | 0.123979 |
| 33208 | ENSG0000 | AC008740.1 | | 0 | 0.055386 | 0.123879 |
| 31804 | ENSG0000 | AC005740.3 | | 0 | 0.05926 | 0.12382 |
| 35580 | ENSG0000 | AP001099.1 | | 0 | 0.076806 | 0.123603 |
| 13870 | ENSG0000 | DSCAML1 | DS cell adh | 0 | 0.052139 | 0.123377 |
| 37342 | ENSG0000 | AC087645.2 | | 0.010771 | 0.017281 | 0.123344 |
| 14900 | ENSG0000 | PTGDR2 | prostaglan | 0.003138 | 0.070402 | 0.123167 |
| 16413 | ENSG0000 | THEM5 | thioesteras | 0.05131 | 0.082569 | 0.123145 |
| 24411 | ENSG0000 | LINC01293 | long interg | 0 | 0.022963 | 0.123121 |
| 39472 | ENSG0000 | AL133215.3 | | 0 | 0.075247 | 0.123075 |
| 20847 | ENSG0000 | RNU6ATAC | RNA, U6ata | 0 | 0.117 | 0.123075 |
| 20542 | ENSG0000 | RPSAP18 | ribosomal | 0.06023 | 0.090001 | 0.122973 |
| 12948 | ENSG0000 | TEX37 | testis expr | 0 | 0.036568 | 0.122961 |

| | | | | | | |
|---|---|---|---|---|---|---|
| 10429 | ENSG0000 | PLXDC1 | plexin dom | 0.013697 | 0.064594 | 0.122885 |
| 10744 | ENSG0000 | TMOD4 | tropomodu | 0.031098 | 0.05006 | 0.122883 |
| 24920 | ENSG0000 | CAP1P2 | CAP1 pseu | 0.01322 | 0.095043 | 0.122696 |
| 39819 | ENSG0000 | AC130371.2 | | 0 | 0.092461 | 0.122578 |
| 32012 | ENSG0000 | AP001107.5 | | 0 | 0.042191 | 0.122436 |
| 40995 | ENSG0000 | AC239859.6 | | 0 | 0.021296 | 0.12242 |
| 40385 | ENSG0000 | AP001059.2 | | 0.079327 | 0.104239 | 0.122289 |
| 34108 | ENSG0000 | AC025580.2 | | 0.057664 | 0.082489 | 0.122233 |
| 14747 | ENSG0000 | CLN8 | CLN8, tran | 0.056254 | 0.085949 | 0.121986 |
| 31381 | ENSG0000 | AC009630.1 | | 0 | 0.087685 | 0.121931 |
| 9119 | ENSG0000 | CDH22 | cadherin 2 | 0.007303 | 0.038379 | 0.121844 |
| 26631 | ENSG0000 | BX284632.1 | | 0.0057 | 0.061169 | 0.121824 |
| 36096 | ENSG0000 | MIR3189 | microRNA | 0 | 0.103091 | 0.121772 |
| 14322 | ENSG0000 | FOXS1 | forkhead b | 0 | 0.022041 | 0.121489 |
| 6658 | ENSG0000 | PDZD2 | PDZ domai | 0.038579 | 0.062732 | 0.121489 |
| 29708 | ENSG0000 | AC139491.2 | | 0.034526 | 0.109416 | 0.121339 |
| 3206 | ENSG0000 | PIK3CG | phosphatic | 0.060021 | 0.069048 | 0.121232 |
| 677 | ENSG0000 | EPN3 | epsin 3 [So | 0.047802 | 0.092262 | 0.121045 |
| 25971 | ENSG0000 | LINC01426 | long interg | 0.016578 | 0.017306 | 0.120776 |
| 3408 | ENSG0000 | ATRNL1 | attractin lil | 0.014464 | 0.054673 | 0.120774 |
| 42069 | ENSG0000 | AP001767.4 | | 0 | 0.039001 | 0.120715 |
| 29461 | ENSG0000 | AC090204.1 | | 0.023197 | 0.057273 | 0.120418 |
| 25579 | ENSG0000 | AC008105.2 | | 0.042814 | 0.04409 | 0.120319 |
| 26448 | ENSG0000 | LURAP1L-A | LURAP1L a | 0.013159 | 0.054926 | 0.120289 |
| 32936 | ENSG0000 | AP000763.3 | | 0.104272 | 0.115978 | 0.120213 |
| 24964 | ENSG0000 | RFX3-AS1 | RFX3 antis | 0.00963 | 0.104645 | 0.120067 |
| 14457 | ENSG0000 | C9orf139 | chromosor | 0.042134 | 0.065462 | 0.120026 |
| 11225 | ENSG0000 | STEAP1 | STEAP fam | 0 | 0.017954 | 0.119818 |
| 42849 | ENSG0000 | MIR6511B | microRNA | 0 | 0.000363 | 0.119723 |
| 38019 | ENSG0000 | AC125494.3 | | 0.052351 | 0.084102 | 0.119667 |
| 40339 | ENSG0000 | AC091167.7 | | 0.000972 | 0.076622 | 0.11966 |
| 725 | ENSG0000 | MCF2L2 | MCF.2 cell | 0.048507 | 0.080421 | 0.119653 |
| 27865 | ENSG0000 | ASB14 | ankyrin rep | 0.11208 | 0.114174 | 0.11964 |
| 7922 | ENSG0000 | RPL3L | ribosomal | 0.016151 | 0.073875 | 0.119531 |
| 9653 | ENSG0000 | PIEZO2 | piezo type | 0.035023 | 0.076116 | 0.119528 |
| 5505 | ENSG0000 | SPDEF | SAM point | 0.027815 | 0.037486 | 0.119491 |
| 11065 | ENSG0000 | MST1R | macrophag | 0.047377 | 0.056099 | 0.119265 |
| 43073 | ENSG0000 | MIR6132 | microRNA | 0 | 0.000005 | 0.119085 |
| 34566 | ENSG0000 | AL031716.1 | | 0 | 0.095845 | 0.118974 |
| 20032 | ENSG0000 | AC005005.1 | | 0 | 0.042778 | 0.118912 |
| 37797 | ENSG0000 | AC005614.1 | | 0.056822 | 0.059798 | 0.118891 |
| 22728 | ENSG0000 | AL136311.1 | | 0.032288 | 0.101415 | 0.118716 |
| 24110 | ENSG0000 | CNOT6LP1 | CCR4-NOT | 0.028839 | 0.072206 | 0.118563 |
| 11983 | ENSG0000 | AC010323.1 | | 0.003436 | 0.073707 | 0.118464 |
| 35830 | ENSG0000 | AF001548.1 | | 0 | 0.026429 | 0.11842 |
| 34239 | ENSG0000 | AC090971.3 | | 0.033852 | 0.035571 | 0.118019 |
| 34188 | ENSG0000 | KRT8P9 | keratin 8 p | 0 | 0.007399 | 0.117962 |

| | | | | | | |
|---|---|---|---|---|---|---|
| 27177 | ENSG0000 | MCFD2P1 | multiple cc | 0.076354 | 0.099998 | 0.117933 |
| 38662 | ENSG0000 | CYB561D2 | cytochrom | 0 | 0.035288 | 0.117811 |
| 34769 | ENSG0000 | BX255925.1 | | 0.003869 | 0.1136 | 0.117658 |
| 34488 | ENSG0000 | COX6CP1 | cytochrom | 0 | 0.056101 | 0.117541 |
| 28040 | ENSG0000 | SUGT1P3 | SGT1 hom | 0.104775 | 0.115391 | 0.117535 |
| 32176 | ENSG0000 | AC022182.2 | | 0 | 0.077599 | 0.117377 |
| 8763 | ENSG0000 | IGFBP1 | insulin like | 0.026598 | 0.063979 | 0.117369 |
| 26416 | ENSG0000 | DNM1P51 | dynamin 1 | 0.023597 | 0.043273 | 0.117257 |
| 38587 | ENSG0000 | SNX29P2 | sorting ne | 0 | 0.041373 | 0.117234 |
| 436 | ENSG0000 | RUNX3 | runt relate | 0.047137 | 0.064321 | 0.117037 |
| 7787 | ENSG0000 | C14orf37 | chromosor | 0.039746 | 0.048397 | 0.116925 |
| 35645 | ENSG0000 | MTCO1P28 | mitochond | 0.0668 | 0.08845 | 0.1169 |
| 5933 | ENSG0000 | DOCK4 | dedicator c | 0.062632 | 0.065009 | 0.116741 |
| 5355 | ENSG0000 | RAB9B | RAB9B, me | 0 | 0.077621 | 0.116709 |
| 40326 | ENSG0000 | MIR6793 | microRNA | 0.01798 | 0.018 | 0.116693 |
| 5524 | ENSG0000 | NRN1 | neuritin 1 \| | 0 | 0.062527 | 0.116652 |
| 14926 | ENSG0000 | PRR36 | proline rich | 0.027878 | 0.100222 | 0.116367 |
| 18080 | ENSG0000 | NHSL2 | NHS like 2 | 0.016029 | 0.099263 | 0.116211 |
| 14752 | ENSG0000 | CACNB4 | calcium vo | 0.022245 | 0.030584 | 0.116095 |
| 38526 | ENSG0000 | AC117498.3 | | 0 | 0.066777 | 0.115994 |
| 11887 | ENSG0000 | FILIP1L | filamin A ir | 0.01194 | 0.062844 | 0.115984 |
| 39815 | ENSG0000 | AC087742.2 | | 0.068372 | 0.089438 | 0.115956 |
| 23875 | ENSG0000 | AC073115.1 | | 0 | 0.039746 | 0.115915 |
| 10837 | ENSG0000 | NFASC | neurofasci | 0.061315 | 0.063226 | 0.115893 |
| 19284 | ENSG0000 | TRGV1 | T-cell recep | 0 | 0.017071 | 0.115686 |
| 39883 | ENSG0000 | AC010205.1 | | 0 | 0.065436 | 0.115637 |
| 14768 | ENSG0000 | XKRX | XK related, | 0 | 0.030236 | 0.115619 |
| 29201 | ENSG0000 | AC082651.4 | | 0 | 0.029043 | 0.115448 |
| 5991 | ENSG0000 | ANGPTL8 | angiopoiet | 0.007718 | 0.0821 | 0.115401 |
| 29091 | ENSG0000 | ELFN2 | extracellul | 0 | 0.010166 | 0.115174 |
| 1436 | ENSG0000 | MYH7B | myosin hea | 0.064111 | 0.068586 | 0.114902 |
| 36636 | ENSG0000 | MIR4771-2 | microRNA | 0 | 0.023683 | 0.11487 |
| 17544 | ENSG0000 | RNU6-236I | RNA, U6 sr | 0 | 0.103091 | 0.11487 |
| 17705 | ENSG0000 | RNU6-343I | RNA, U6 sr | 0 | 0.103091 | 0.11487 |
| 28614 | ENSG0000 | NPM1P23 | nucleopho | 0 | 0.098477 | 0.11487 |
| 31230 | ENSG0000 | SCARNA24 | Small Cajal | 0 | 0.027755 | 0.11487 |
| 17252 | ENSG0000 | RNU6-137I | RNA, U6 sr | 0 | 0.069682 | 0.11487 |
| 18867 | ENSG0000 | RNA5SP12 | RNA, 5S rib | 0.051276 | 0.103091 | 0.11487 |
| 18871 | ENSG0000 | SNORD116 | small nucle | 0 | 0.056155 | 0.11487 |
| 15044 | ENSG0000 | MIR429 | microRNA | 0 | 0.103091 | 0.11487 |
| 33034 | ENSG0000 | AC090023.2 | | 0.047644 | 0.104056 | 0.114819 |
| 549 | ENSG0000 | DAPK2 | death asso | 0.046964 | 0.057932 | 0.114777 |
| 36669 | ENSG0000 | AP001020.3 | | 0.029232 | 0.039442 | 0.114607 |
| 28353 | ENSG0000 | AC063943.1 | | 0.038731 | 0.050137 | 0.114569 |
| 29429 | ENSG0000 | AC006299.1 | | 0.011089 | 0.05065 | 0.114343 |
| 20791 | ENSG0000 | HNRNPH1F | heterogen | 0.014976 | 0.070119 | 0.114288 |
| 22393 | ENSG0000 | AC092802.1 | | 0.00211 | 0.032184 | 0.114105 |

| | | | | | | |
|---|---|---|---|---|---|---|
| 12904 | ENSG0000( | KLF17 | Kruppel lik | 0 | 0.065193 | 0.114026 |
| 6122 | ENSG0000( | RPS4Y1 | ribosomal | 0.078198 | 0.094139 | 0.113835 |
| 40514 | ENSG0000( | AC016957.2 | | 0 | 0.076224 | 0.113786 |
| 13529 | ENSG0000( | CHRNA7 | cholinergic | 0.058314 | 0.068391 | 0.113636 |
| 15273 | ENSG0000( | LPAR5 | lysophosph | 0 | 0.035787 | 0.1136 |
| 3612 | ENSG0000( | GNRHR | gonadotro | 0 | 0.107789 | 0.113534 |
| 16339 | ENSG0000( | RPS2P4 | ribosomal | 0.009763 | 0.065757 | 0.113528 |
| 42342 | ENSG0000( | AL079342.3 | | 0.060273 | 0.069027 | 0.113309 |
| 43136 | ENSG0000( | AP000356.2 | | 0.057254 | 0.07396 | 0.113187 |
| 7398 | ENSG0000( | LRRC32 | leucine ricl | 0 | 0.075301 | 0.113144 |
| 27066 | ENSG0000( | LINC01474 | long interg | 0 | 0.107219 | 0.113108 |
| 26574 | ENSG0000( | AP001625.2 | | 0 | 0.087774 | 0.112672 |
| 22753 | ENSG0000( | AC093901.1 | | 0.070641 | 0.085069 | 0.112495 |
| 25347 | ENSG0000( | AP001062.2 | | 0 | 0.02703 | 0.112156 |
| 25732 | ENSG0000( | AL162727.1 | | 0.020015 | 0.052409 | 0.112111 |
| 4652 | ENSG0000( | MPL | MPL proto | 0.015773 | 0.04893 | 0.112089 |
| 40200 | ENSG0000( | AL162731.1 | | 0.019065 | 0.075377 | 0.111766 |
| 28214 | ENSG0000( | RPS19P3 | ribosomal | 0 | 0.08439 | 0.111484 |
| 41338 | ENSG0000( | AC121338.2 | | 0.020423 | 0.033635 | 0.111445 |
| 5035 | ENSG0000( | IQSEC3 | IQ motif ar | 0.011833 | 0.042088 | 0.111417 |
| 3001 | ENSG0000( | TLX1 | T-cell leuke | 0.014628 | 0.035028 | 0.111172 |
| 40303 | ENSG0000( | AC022306.2 | | 0 | 0.088273 | 0.111142 |
| 27182 | ENSG0000( | LINC01792 | long interg | 0 | 0.028463 | 0.11106 |
| 2489 | ENSG0000( | SLC52A3 | solute carr | 0 | 0.056567 | 0.110853 |
| 30465 | ENSG0000( | LINC01337 | long interg | 0.03973 | 0.058809 | 0.110693 |
| 31679 | ENSG0000( | PCDHGA6 | protocadh( | 0.04059 | 0.0975 | 0.110593 |
| 10913 | ENSG0000( | C2 | compleme | 0.038262 | 0.08784 | 0.11039 |
| 15677 | ENSG0000( | CCDC30 | coiled-coil | 0.017587 | 0.092846 | 0.110208 |
| 25076 | ENSG0000( | AL136380.2 | | 0 | 0.059089 | 0.1102 |
| 20095 | ENSG0000( | AL451074.1 | | 0.019144 | 0.07436 | 0.110123 |
| 30788 | ENSG0000( | AC107884.1 | | 0.033408 | 0.03713 | 0.1099 |
| 15258 | ENSG0000( | TMEM255I | transmeml | 0.038793 | 0.059485 | 0.109845 |
| 13135 | ENSG0000( | CCDC63 | coiled-coil | 0.031568 | 0.080434 | 0.109812 |
| 13947 | ENSG0000( | RBM44 | RNA bindir | 0.038143 | 0.055441 | 0.109699 |
| 16976 | ENSG0000( | NTRK1 | neurotropl | 0 | 0.088202 | 0.109677 |
| 3098 | ENSG0000( | ICAM4 | intercellula | 0 | 0.022195 | 0.109627 |
| 2157 | ENSG0000( | OSM | oncostatin | 0.025124 | 0.043259 | 0.109621 |
| 20621 | ENSG0000( | AL356776.1 | | 0 | 0.01822 | 0.109473 |
| 34972 | ENSG0000( | AL591848.3 | | 0.012043 | 0.04578 | 0.109413 |
| 18115 | ENSG0000( | HLA-DOA | major histo | 0 | 0.069616 | 0.10924 |
| 38711 | ENSG0000( | AC103810.6 | | 0.03856 | 0.100185 | 0.109098 |
| 39791 | ENSG0000( | MIR5088 | microRNA | 0.015931 | 0.066552 | 0.109054 |
| 13199 | ENSG0000( | INSM1 | INSM trans | 0.037682 | 0.066869 | 0.108982 |
| 40333 | ENSG0000( | RN7SL313I | RNA, 7SL, c | 0 | 0.036511 | 0.108887 |
| 28698 | ENSG0000( | AC127024.1 | | 0 | 0.070043 | 0.108824 |
| 7138 | ENSG0000( | NACAD | NAC alpha | 0.028847 | 0.047342 | 0.108749 |
| 43205 | ENSG0000( | AP000356.4 | | 0 | 0.045102 | 0.108724 |

| | | | | | | |
|---|---|---|---|---|---|---|
| 22533 | ENSG0000 | AL353803.1 | | 0 | 0.030513 | 0.108696 |
| 42500 | ENSG0000 | AL353658.2 | | 0 | 0.091443 | 0.108656 |
| 40720 | ENSG0000 | AC245041.2 | | 0 | 0.075741 | 0.108631 |
| 22421 | ENSG0000 | EIF3LP3 | eukaryotic | 0 | 0.053051 | 0.108581 |
| 29335 | ENSG0000 | MTCYBP18 | mitochond | 0 | 0.092464 | 0.108565 |
| 28398 | ENSG0000 | IQCJ-SCHIP | IQCJ-SCHIP | 0 | 0.012364 | 0.108517 |
| 6020 | ENSG0000 | TBX3 | T-box 3 [Sc | 0.06209 | 0.089244 | 0.108278 |
| 38700 | ENSG0000 | AC093423.3 | | 0 | 0.100822 | 0.108273 |
| 34883 | ENSG0000 | AL031722.1 | | 0 | 0.083412 | 0.108243 |
| 28525 | ENSG0000 | AC099340.1 | | 0 | 0.010498 | 0.108158 |
| 9689 | ENSG0000 | GOLGA7B | golgin A7 f | 0.019157 | 0.07897 | 0.108129 |
| 37280 | ENSG0000 | AC003956.1 | | 0.072136 | 0.099836 | 0.10796 |
| 35114 | ENSG0000 | AC140504.1 | | 0.01541 | 0.07181 | 0.107946 |
| 41890 | ENSG0000 | AC124283.5 | | 0.086619 | 0.096138 | 0.107695 |
| 24951 | ENSG0000 | MRPL35P2 | mitochond | 0 | 0.057093 | 0.107676 |
| 16739 | ENSG0000 | SSPO | SCO-spond | 0.041513 | 0.084006 | 0.107672 |
| 11247 | ENSG0000 | HNF4G | hepatocyte | 0.014823 | 0.042022 | 0.10758 |
| 16908 | ENSG0000 | AC025279.1 | | 0.02636 | 0.056572 | 0.107553 |
| 28501 | ENSG0000 | PAQR9-AS1 | PAQR9 ant | 0.016183 | 0.05968 | 0.107365 |
| 37641 | ENSG0000 | AC005776.1 | | 0.089403 | 0.101314 | 0.107344 |
| 37226 | ENSG0000 | AC011511.5 | | 0.021973 | 0.070597 | 0.107245 |
| 3729 | ENSG0000 | CEP126 | centrosom | 0.049208 | 0.06196 | 0.107091 |
| 16551 | ENSG0000 | SCN8A | sodium vol | 0.022669 | 0.090641 | 0.107036 |
| 42781 | ENSG0000 | AC068205.1 | | 0.02317 | 0.040203 | 0.10698 |
| 1149 | ENSG0000 | TRPM5 | transient r | 0.037706 | 0.10246 | 0.106812 |
| 1109 | ENSG0000 | SPTB | spectrin be | 0.021337 | 0.026444 | 0.106656 |
| 11246 | ENSG0000 | C7orf57 | chromosor | 0 | 0.0191 | 0.106589 |
| 4381 | ENSG0000 | EFEMP1 | EGF contai | 0.064381 | 0.105735 | 0.106578 |
| 17902 | ENSG0000 | AC012358.1 | | 0.024359 | 0.078782 | 0.106468 |
| 30951 | ENSG0000 | SNORA4 | Small nucle | 0 | 0.102327 | 0.106361 |
| 39110 | ENSG0000 | AC097532.2 | | 0.021101 | 0.085378 | 0.106265 |
| 23316 | ENSG0000 | CELSR3-AS | CELSR3 ant | 0 | 0.021924 | 0.106162 |
| 22604 | ENSG0000 | CNN2P1 | calponin 2 | 0.033315 | 0.062888 | 0.105861 |
| 12699 | ENSG0000 | OTP | orthopedia | 0.016033 | 0.049813 | 0.105708 |
| 28314 | ENSG0000 | LINC00886 | long interg | 0.031086 | 0.060959 | 0.105692 |
| 34386 | ENSG0000 | AC013726.1 | | 0.023464 | 0.078367 | 0.10557 |
| 1538 | ENSG0000 | CCL22 | C-C motif c | 0 | 0.019464 | 0.105497 |
| 33322 | ENSG0000 | AL121820.2 | | 0 | 0.099396 | 0.105443 |
| 16511 | ENSG0000 | ZNF418 | zinc finger | 0.031597 | 0.051463 | 0.105351 |
| 24526 | ENSG0000 | AL031728.1 | | 0.028634 | 0.058844 | 0.105338 |
| 39964 | ENSG0000 | AC016747.4 | | 0.058191 | 0.061061 | 0.105295 |
| 14144 | ENSG0000 | KCTD12 | potassium | 0.045895 | 0.060927 | 0.105237 |
| 35049 | ENSG0000 | AC010542.3 | | 0 | 0.032961 | 0.105226 |
| 15591 | ENSG0000 | PLK5 | polo like ki | 0.014345 | 0.082264 | 0.105132 |
| 40945 | ENSG0000 | NEFL | neurofilam | 0.049148 | 0.100614 | 0.105078 |
| 37761 | ENSG0000 | TRABD2B | TraB doma | 0.02743 | 0.060302 | 0.105068 |
| 15398 | ENSG0000 | LINC00313 | long interg | 0.015466 | 0.045875 | 0.104857 |

| | | | | | | |
|---|---|---|---|---|---|---|
| 12058 | ENSG0000 | SLC22A11 | solute carr | 0 | 0.031147 | 0.10485 |
| 42319 | ENSG0000 | AC132219.2 | | 0.041105 | 0.085104 | 0.104849 |
| 29986 | ENSG0000 | CDH12P2 | cadherin 1 | 0.010559 | 0.040764 | 0.104848 |
| 42074 | ENSG0000 | AC026748.1 | | 0.043824 | 0.099666 | 0.1048 |
| 33363 | ENSG0000 | AL157791.1 | | 0.020964 | 0.025485 | 0.104661 |
| 10465 | ENSG0000 | ALOX15 | arachidona | 0.010464 | 0.046297 | 0.104544 |
| 35161 | ENSG0000 | AC007216.2 | | 0.044502 | 0.069584 | 0.104538 |
| 25786 | ENSG0000 | LINC01694 | long interg | 0.010164 | 0.053332 | 0.104513 |
| 38529 | ENSG0000 | AC018638.7 | | 0.010733 | 0.053766 | 0.104352 |
| 42590 | ENSG0000 | AC241377.3 | | 0.005437 | 0.067037 | 0.104312 |
| 10668 | ENSG0000 | C1orf147 | chromosor | 0.02534 | 0.048742 | 0.104257 |
| 22973 | ENSG0000 | SATB1-AS1 | SATB1 anti | 0.065402 | 0.080344 | 0.104029 |
| 15509 | ENSG0000 | POU3F1 | POU class 3 | 0 | 0.040075 | 0.103983 |
| 31917 | ENSG0000 | CDH12P1 | cadherin 1 | 0.009599 | 0.041178 | 0.103967 |
| 26196 | ENSG0000 | ATP5A1P1 | ATP syntha | 0.016856 | 0.053184 | 0.10395 |
| 23475 | ENSG0000 | AL691426.1 | | 0 | 0.083045 | 0.103896 |
| 16833 | ENSG0000 | OR7E13P | olfactory r | 0 | 0.027982 | 0.103888 |
| 5704 | ENSG0000 | GRPR | gastrin rele | 0.018595 | 0.02282 | 0.103859 |
| 27377 | ENSG0000 | PRSS56 | protease, s | 0 | 0.015846 | 0.103781 |
| 1542 | ENSG0000 | CEMIP | cell migrat | 0.05306 | 0.094836 | 0.103685 |
| 29404 | ENSG0000 | AC025164.2 | | 0.03501 | 0.092716 | 0.10365 |
| 5977 | ENSG0000 | ARHGAP22 | Rho GTPas | 0.010428 | 0.052596 | 0.103559 |
| 13423 | ENSG0000 | PCP2 | Purkinje ce | 0.037763 | 0.061992 | 0.103544 |
| 26752 | ENSG0000 | HMGN2P2 | high mobil | 0 | 0.065317 | 0.103471 |
| 12068 | ENSG0000 | OR1F1 | olfactory r | 0 | 0.061723 | 0.103407 |
| 9444 | ENSG0000 | ANKRD22 | ankyrin rep | 0.041251 | 0.067518 | 0.103204 |
| 31924 | ENSG0000 | AC120036.4 | | 0 | 0.083359 | 0.103203 |
| 30511 | ENSG0000 | AC098591.2 | | 0.076913 | 0.098182 | 0.103201 |
| 21689 | ENSG0000 | AL354953.1 | | 0.054743 | 0.088733 | 0.103082 |
| 18763 | ENSG0000 | RNU6-481P | RNA, U6 sr | 0 | 0.090566 | 0.103061 |
| 11325 | ENSG0000 | TMEM71 | transmeml | 0.038674 | 0.071127 | 0.102896 |
| 38557 | ENSG0000 | ATP2B1-AS | ATP2B1 an | 0.06837 | 0.076637 | 0.102725 |
| 22412 | ENSG0000 | LINC00623 | long interg | 0.008552 | 0.038745 | 0.102691 |
| 8166 | ENSG0000 | MMEL1 | membrane | 0 | 0.004234 | 0.102626 |
| 9105 | ENSG0000 | MPZL2 | myelin pro | 0.022799 | 0.040674 | 0.102602 |
| 8167 | ENSG0000 | CFAP74 | cilia and fla | 0.008939 | 0.034944 | 0.102569 |
| 34348 | ENSG0000 | AC092757.2 | | 0 | 0.040116 | 0.10253 |
| 27322 | ENSG0000 | AL121992.1 | | 0.018447 | 0.035982 | 0.102474 |
| 13076 | ENSG0000 | EFCAB12 | EF-hand ca | 0.010195 | 0.067301 | 0.102434 |
| 34112 | ENSG0000 | AC046168.1 | | 0.004881 | 0.007595 | 0.102311 |
| 19674 | ENSG0000 | AC004217.1 | | 0 | 0.079331 | 0.102283 |
| 36778 | ENSG0000 | AP001381.1 | | 0 | 0.073667 | 0.102162 |
| 7469 | ENSG0000 | GCOM1 | GRINL1A c | 0.042322 | 0.045287 | 0.102126 |
| 32363 | ENSG0000 | AP000880.1 | | 0.013575 | 0.060907 | 0.10204 |
| 27352 | ENSG0000 | AL365295.1 | | 0.032538 | 0.062508 | 0.101824 |
| 2342 | ENSG0000 | PLEK2 | pleckstrin 2 | 0.012784 | 0.053507 | 0.101541 |
| 42200 | ENSG0000 | AC084035.1 | | 0 | 0.016644 | 0.101055 |

| | | | | | | |
|---|---|---|---|---|---|---|
| 11979 | ENSG0000 | ACER1 | alkaline ce | 0 | 0.052062 | 0.100958 |
| 23907 | ENSG0000 | ISPD-AS1 | ISPD antise | 0.038353 | 0.065743 | 0.100921 |
| 26433 | ENSG0000 | AC009269.1 | | 0.037518 | 0.062074 | 0.100801 |
| 29006 | ENSG0000 | LINC00880 | long interg | 0 | 0.014559 | 0.100696 |
| 28277 | ENSG0000 | RN7SL559I | RNA, 7SL, c | 0 | 0.005554 | 0.10056 |
| 12882 | ENSG0000 | COL8A2 | collagen ty | 0.037865 | 0.077594 | 0.100555 |
| 21490 | ENSG0000 | HMGN1P4 | high mobil | 0 | 0.043369 | 0.100214 |
| 33781 | ENSG0000 | AL139021.2 | | 0 | 0.026589 | 0.100088 |
| 23790 | ENSG0000 | AL139246.4 | | 0 | 0.082995 | 0.100077 |
| 15150 | ENSG0000 | PP2D1 | protein ph | 0.021251 | 0.023045 | 0.099929 |
| 20786 | ENSG0000 | BMPR1APS | bone morp | 0.044473 | 0.089672 | 0.099846 |
| 6227 | ENSG0000 | TNNT3 | troponin T | 0 | 0.071741 | 0.099839 |
| 33748 | ENSG0000 | AF111169.2 | | 0.082506 | 0.085341 | 0.099801 |
| 7796 | ENSG0000 | ESR2 | estrogen re | 0.050406 | 0.076625 | 0.09961 |
| 38872 | ENSG0000 | LINC02084 | long interg | 0.016192 | 0.090863 | 0.099358 |
| 32375 | ENSG0000 | GLYATL1P3 | glycine-N-a | 0 | 0.040135 | 0.099328 |
| 35963 | ENSG0000 | AC005277.1 | | 0.031208 | 0.063086 | 0.099312 |
| 29684 | ENSG0000 | AC010931.2 | | 0.006692 | 0.089719 | 0.099214 |
| 13516 | ENSG0000 | CATSPER1 | cation char | 0 | 0.007402 | 0.09907 |
| 33774 | ENSG0000 | SEC23A-AS | SEC23A ant | 0 | 0.058893 | 0.098788 |
| 19985 | ENSG0000 | AL157395.1 | | 0.016092 | 0.086902 | 0.098653 |
| 967 | ENSG0000 | TIE1 | tyrosine ki | 0.010679 | 0.036832 | 0.098471 |
| 6362 | ENSG0000 | GALNT15 | polypeptid | 0.006228 | 0.070176 | 0.098294 |
| 8087 | ENSG0000 | CBLC | Cbl proto-c | 0.044676 | 0.054777 | 0.098211 |
| 30032 | ENSG0000 | AC011411.1 | | 0 | 0.033187 | 0.098037 |
| 24086 | ENSG0000 | LINC01721 | long interg | 0 | 0.028995 | 0.097912 |
| 34080 | ENSG0000 | ACTG1P17 | actin gamn | 0.042453 | 0.062519 | 0.097881 |
| 39771 | ENSG0000 | AC245297.3 | | 0 | 0.050257 | 0.097872 |
| 15586 | ENSG0000 | CCIN | calicin [Sou | 0 | 0.030221 | 0.097826 |
| 18866 | ENSG0000 | RNU6-115 | RNA, U6 sr | 0 | 0.040847 | 0.097823 |
| 28237 | ENSG0000 | THOC7-AS | THOC7 ant | 0.053298 | 0.069762 | 0.097789 |
| 24258 | ENSG0000 | HMGB1P4 | high mobil | 0 | 0.030076 | 0.09771 |
| 24550 | ENSG0000 | KLF3-AS1 | KLF3 antise | 0.07591 | 0.083526 | 0.09757 |
| 25995 | ENSG0000 | AC092171.3 | | 0 | 0.055065 | 0.097538 |
| 13937 | ENSG0000 | NIM1K | NIM1 serin | 0.024449 | 0.055571 | 0.097537 |
| 22890 | ENSG0000 | AC011290.1 | | 0.033116 | 0.054713 | 0.09748 |
| 5796 | ENSG0000 | ARMCX1 | armadillo r | 0.033213 | 0.034332 | 0.097437 |
| 7681 | ENSG0000 | KLRG1 | killer cell le | 0.039482 | 0.056684 | 0.097402 |
| 21882 | ENSG0000 | AP001347.1 | | 0.020407 | 0.060382 | 0.09733 |
| 10733 | ENSG0000 | NEURL3 | neuralized | 0.014214 | 0.058337 | 0.097113 |
| 10174 | ENSG0000 | AC005041.1 | | 0.018476 | 0.053758 | 0.097051 |
| 36176 | ENSG0000 | AC104982.3 | | 0 | 0.015644 | 0.096991 |
| 35519 | ENSG0000 | MFSD1P1 | major facil | 0.026936 | 0.036358 | 0.096951 |
| 34372 | ENSG0000 | AC012236.1 | | 0 | 0.022469 | 0.09695 |
| 11350 | ENSG0000 | DYNLT3 | dynein ligh | 0.01671 | 0.027374 | 0.096806 |
| 40694 | ENSG0000 | MIR6780B | microRNA | 0.000797 | 0.001305 | 0.096694 |
| 25887 | ENSG0000 | LINC02535 | long interg | 0.020777 | 0.05212 | 0.096494 |

| | | | | | | |
|---|---|---|---|---|---|---|
| 39134 | ENSG0000 | AC098850.5 | | 0.076135 | 0.080376 | 0.096442 |
| 23112 | ENSG0000 | AL358075.2 | | 0 | 0.075418 | 0.096233 |
| 8038 | ENSG0000 | STAC2 | SH3 and cy | 0.010557 | 0.025972 | 0.096215 |
| 30587 | ENSG0000 | AC025419.1 | | 0.044423 | 0.046594 | 0.096131 |
| 42160 | ENSG0000 | AC027682.7 | | 0 | 0.054729 | 0.096077 |
| 30584 | ENSG0000 | LINC02381 | long interg | 0.061513 | 0.062658 | 0.096073 |
| 15791 | ENSG0000 | EFCAB6 | EF-hand ca | 0.022425 | 0.089554 | 0.095987 |
| 31200 | ENSG0000 | SNORA81 | Small nucle | 0 | 0.083079 | 0.09595 |
| 395 | ENSG0000 | NPC1L1 | NPC1 like i | 0.006806 | 0.011169 | 0.095934 |
| 7819 | ENSG0000 | DUOXA2 | dual oxidas | 0 | 0.038277 | 0.095818 |
| 42524 | ENSG0000 | SCHLAP1 | SWI/SNF co | 0 | 0.030511 | 0.095439 |
| 18032 | ENSG0000 | C10orf62 | chromosor | 0.029693 | 0.069808 | 0.095376 |
| 19832 | ENSG0000 | AP000662.1 | | 0 | 0.004234 | 0.095292 |
| 40870 | ENSG0000 | AL121890.4 | | 0 | 0.047948 | 0.095184 |
| 12525 | ENSG0000 | SP7 | Sp7 transcı | 0.020865 | 0.055562 | 0.095153 |
| 19096 | ENSG0000 | MIR222 | microRNA | 0 | 0.005623 | 0.095029 |
| 34401 | ENSG0000 | AC018845.1 | | 0.010996 | 0.029007 | 0.094995 |
| 21293 | ENSG0000 | AL592045.1 | | 0 | 0.061796 | 0.094963 |
| 24310 | ENSG0000 | AL139220.2 | | 0.041798 | 0.053406 | 0.094946 |
| 20036 | ENSG0000 | PRCD | photorece | 0.028381 | 0.029812 | 0.094858 |
| 23723 | ENSG0000 | AL358876.1 | | 0 | 0.054408 | 0.094805 |
| 9076 | ENSG0000 | GRIK4 | glutamate | 0.018784 | 0.0449 | 0.094729 |
| 5133 | ENSG0000 | CSTA | cystatin A | | 0.077734 | 0.081241 | 0.094456 |
| 18136 | ENSG0000 | SMIM5 | small integ | 0.00701 | 0.057887 | 0.094297 |
| 40038 | ENSG0000 | AL354950.1 | | 0 | 0.006171 | 0.094242 |
| 42155 | ENSG0000 | AC106795.5 | | 0 | 0.011127 | 0.094122 |
| 33958 | ENSG0000 | AL117694.1 | | 0.049011 | 0.070445 | 0.093783 |
| 163 | ENSG0000 | PRSS21 | protease, s | 0.012753 | 0.016232 | 0.093719 |
| 38806 | ENSG0000 | ARF4-AS1 | ARF4 antis | 0.047478 | 0.04893 | 0.093494 |
| 25389 | ENSG0000 | ERVH48-1 | endogenou | 0.012533 | 0.040954 | 0.093461 |
| 34101 | ENSG0000 | AC013652.1 | | 0.034591 | 0.063986 | 0.093247 |
| 5371 | ENSG0000 | KCNJ2 | potassium | 0.054139 | 0.054758 | 0.093181 |
| 13553 | ENSG0000 | AL078621.2 | | 0 | 0.041502 | 0.093147 |
| 20318 | ENSG0000 | AC131097.1 | | 0.036208 | 0.050494 | 0.093115 |
| 12846 | ENSG0000 | S100Z | S100 calciu | 0 | 0.04321 | 0.093104 |
| 10724 | ENSG0000 | CFAP221 | cilia and fla | 0.019579 | 0.074128 | 0.093015 |
| 271 | ENSG0000 | FOXP3 | forkhead b | 0.063792 | 0.080993 | 0.092998 |
| 14837 | ENSG0000 | GALNT9 | polypeptid | 0.0079 | 0.05182 | 0.092951 |
| 35668 | ENSG0000 | AC110285.1 | | 0.021949 | 0.026662 | 0.092798 |
| 3973 | ENSG0000 | GPR63 | G protein-c | 0.050538 | 0.073421 | 0.092772 |
| 23554 | ENSG0000 | AL591848.1 | | 0 | 0.035538 | 0.092752 |
| 36408 | ENSG0000 | AC104984.2 | | 0 | 0.027848 | 0.092675 |
| 31978 | ENSG0000 | NAV2-AS2 | NAV2 antis | 0 | 0.018588 | 0.092609 |
| 10850 | ENSG0000 | EFHB | EF-hand do | 0.0169 | 0.030912 | 0.092357 |
| 17141 | ENSG0000 | CD3E | CD3e mole | 0 | 0.025547 | 0.092163 |
| 10757 | ENSG0000 | SPRR2D | small proli | 0 | 0.06837 | 0.092048 |
| 37908 | ENSG0000 | AC022382.1 | | 0 | 0.044489 | 0.091906 |

| | | | | | | |
|---|---|---|---|---|---|---|
| 40544 | ENSG0000 | AC243829.2 | | 0.024955 | 0.036752 | 0.09176 |
| 35438 | ENSG0000 | TTC39A-AS | TTC39A an | 0 | 0.023706 | 0.091745 |
| 27297 | ENSG0000 | AC073069.1 | | 0 | 0.047221 | 0.091688 |
| 6292 | ENSG0000 | SLC7A10 | solute carr | 0 | 0.006992 | 0.091675 |
| 1433 | ENSG0000 | PKD2L2 | polycystin | 0.048456 | 0.064707 | 0.091454 |
| 43046 | ENSG0000 | MIR6872 | microRNA | 0.064959 | 0.080379 | 0.091424 |
| 1456 | ENSG0000 | RAPGEF3 | Rap guanir | 0.010336 | 0.041686 | 0.0914 |
| 9294 | ENSG0000 | ZNF827 | zinc finger | 0.037397 | 0.053562 | 0.091295 |
| 24998 | ENSG0000 | AL161909.1 | | 0 | 0.032144 | 0.091154 |
| 35238 | ENSG0000 | AC055855.3 | | 0 | 0.047488 | 0.091021 |
| 41620 | ENSG0000 | AC010223.1 | | 0 | 0.014041 | 0.09099 |
| 29001 | ENSG0000 | AP001318.1 | | 0.055284 | 0.090887 | 0.090894 |
| 24929 | ENSG0000 | AC007391.1 | | 0 | 0.045277 | 0.090849 |
| 30003 | ENSG0000 | UNC93B8 | unc-93 hor | 0.012497 | 0.081303 | 0.090837 |
| 42130 | ENSG0000 | AC008079.2 | | 0.015288 | 0.080372 | 0.090806 |
| 23827 | ENSG0000 | AC005189.1 | | 0 | 0.030262 | 0.090784 |
| 37382 | ENSG0000 | SMIM22 | small integ | 0.020143 | 0.055907 | 0.090706 |
| 29648 | ENSG0000 | PCDHGB8F | protocadh | 0.012397 | 0.04044 | 0.09044 |
| 30061 | ENSG0000 | LINC01091 | long interg | 0.022686 | 0.056376 | 0.090383 |
| 38379 | ENSG0000 | AC004130.1 | | 0 | 0.064892 | 0.090383 |
| 5364 | ENSG0000 | TNFAIP6 | TNF alpha | 0.022558 | 0.043874 | 0.090281 |
| 25112 | ENSG0000 | AL354824.2 | | 0 | 0.089057 | 0.09028 |
| 37200 | ENSG0000 | AC027514.2 | | 0 | 0.027016 | 0.090272 |
| 1452 | ENSG0000 | TNS1 | tensin 1 [S | 0.027198 | 0.034523 | 0.090037 |
| 40012 | ENSG0000 | AC116447.1 | | 0 | 0.029357 | 0.090019 |
| 3718 | ENSG0000 | HPX | hemopexir | 0.034541 | 0.082208 | 0.08982 |
| 12757 | ENSG0000 | NETO2 | neuropilin | 0.054401 | 0.060682 | 0.089761 |
| 7691 | ENSG0000 | COL2A1 | collagen ty | 0.025017 | 0.045408 | 0.089682 |
| 22504 | ENSG0000 | ARPC3P1 | actin relate | 0.026919 | 0.027272 | 0.089655 |
| 22900 | ENSG0000 | AL121983.2 | | 0 | 0.081102 | 0.089545 |
| 38706 | ENSG0000 | AL583856.2 | | 0.009271 | 0.042717 | 0.089484 |
| 27527 | ENSG0000 | AL592435.1 | | 0.042864 | 0.046397 | 0.089285 |
| 37946 | ENSG0000 | AP001267.4 | | 0 | 0.053342 | 0.089156 |
| 10424 | ENSG0000 | NPHS1 | NPHS1, ne | 0.006271 | 0.037392 | 0.089011 |
| 42445 | ENSG0000 | AC005329.3 | | 0.050353 | 0.077005 | 0.088939 |
| 5152 | ENSG0000 | HCRTR1 | hypocretin | 0.017226 | 0.020404 | 0.088911 |
| 10634 | ENSG0000 | SLAMF6 | SLAM fami | 0.013063 | 0.031198 | 0.088851 |
| 11111 | ENSG0000 | EDIL3 | EGF like re | 0.043495 | 0.083849 | 0.088665 |
| 1937 | ENSG0000 | RAPGEF4 | Rap guanir | 0.007468 | 0.068151 | 0.08859 |
| 34430 | ENSG0000 | AC092384.2 | | 0.047674 | 0.067686 | 0.088564 |
| 10497 | ENSG0000 | BICDL2 | BICD famil | 0 | 0.062063 | 0.088454 |
| 15478 | ENSG0000 | PRKG1 | protein kin | 0.009456 | 0.032863 | 0.088288 |
| 20126 | ENSG0000 | AL136309.1 | | 0 | 0.041774 | 0.088272 |
| 22627 | ENSG0000 | AL158834.2 | | 0 | 0.055666 | 0.088227 |
| 28674 | ENSG0000 | TFP1 | transferrin | 0.047802 | 0.047881 | 0.088209 |
| 1271 | ENSG0000 | MGLL | monoglyce | 0.043967 | 0.074669 | 0.088193 |
| 17011 | ENSG0000 | PABPC1P2 | poly(A) bin | 0.049367 | 0.080759 | 0.088141 |

| | | | | | | |
|---|---|---|---|---|---|---|
| 32115 | ENSG0000 | AC233992.1 | | 0.050901 | 0.062624 | 0.087858 |
| 21671 | ENSG0000 | LINC00539 | long interg | 0 | 0.045098 | 0.087824 |
| 34736 | ENSG0000 | AC130456.5 | | 0.017658 | 0.059241 | 0.08776 |
| 21937 | ENSG0000 | AC073957.1 | | 0 | 0.036034 | 0.08774 |
| 15435 | ENSG0000 | TMEM105 | transmeml | 0 | 0.030677 | 0.087654 |
| 26393 | ENSG0000 | LINC00957 | long interg | 0.021605 | 0.031961 | 0.087566 |
| 21402 | ENSG0000 | AC005006.1 | | 0 | 0.077853 | 0.087557 |
| 9251 | ENSG0000 | AL136982.1 | | 0 | 0.000025 | 0.087551 |
| 9664 | ENSG0000 | CA10 | carbonic ar | 0.027971 | 0.036001 | 0.087482 |
| 1736 | ENSG0000 | MYBPC2 | myosin bin | 0 | 0.007868 | 0.087423 |
| 40615 | ENSG0000 | AL359075.3 | | 0 | 0.036581 | 0.087343 |
| 9179 | ENSG0000 | ADRA2A | adrenocep | 0.009511 | 0.027252 | 0.087295 |
| 22200 | ENSG0000 | RPL35P5 | ribosomal | 0.047962 | 0.078851 | 0.087239 |
| 30362 | ENSG0000 | SALL4P1 | spalt like tr | 0 | 0.054926 | 0.087141 |
| 37413 | ENSG0000 | AC009955.2 | | 0 | 0.052687 | 0.086894 |
| 4341 | ENSG0000 | CYTIP | cytohesin 1 | 0.025774 | 0.042434 | 0.08689 |
| 37213 | ENSG0000 | AC007998.3 | | 0.031809 | 0.085716 | 0.086663 |
| 15255 | ENSG0000 | POU3F2 | POU class 3 | 0.007274 | 0.071439 | 0.086547 |
| 17139 | ENSG0000 | TOX | thymocyte | 0.02154 | 0.047222 | 0.086396 |
| 12927 | ENSG0000 | LDHAL6B | lactate deh | 0 | 0.034343 | 0.086373 |
| 23704 | ENSG0000 | AC004691.1 | | 0 | 0.030299 | 0.086303 |
| 22398 | ENSG0000 | AC005740.1 | | 0 | 0.028241 | 0.086262 |
| 20521 | ENSG0000 | AC234781.1 | | 0 | 0.045234 | 0.086203 |
| 36936 | ENSG0000 | AC067968.2 | | 0 | 0.03176 | 0.086051 |
| 18260 | ENSG0000 | ASPDH | aspartate c | 0 | 0.077185 | 0.085997 |
| 32392 | ENSG0000 | CCDC166 | coiled-coil | 0.037565 | 0.07407 | 0.085978 |
| 2151 | ENSG0000 | DERL3 | derlin 3 [Sc | 0.035103 | 0.080943 | 0.085971 |
| 21447 | ENSG0000 | AC244033.1 | | 0 | 0.060462 | 0.085884 |
| 15711 | ENSG0000 | CYP4F3 | cytochrom | 0.048413 | 0.057342 | 0.085766 |
| 22905 | ENSG0000 | EIF4A2P2 | eukaryotic | 0 | 0.070865 | 0.085597 |
| 18787 | ENSG0000 | U8 | U8 small n | 0 | 0.081388 | 0.085505 |
| 21394 | ENSG0000 | AC007255.1 | | 0 | 0.030426 | 0.085499 |
| 11412 | ENSG0000 | PKNOX2 | PBX/knotte | 0.016247 | 0.068354 | 0.08538 |
| 2754 | ENSG0000 | SEC14L5 | SEC14 like | 0 | 0.026995 | 0.085128 |
| 26838 | ENSG0000 | AC114737.2 | | 0 | 0.036151 | 0.084844 |
| 566 | ENSG0000 | FLT4 | fms relatec | 0 | 0.019421 | 0.084734 |
| 39409 | ENSG0000 | AC005021.1 | | 0 | 0.02178 | 0.084699 |
| 34016 | ENSG0000 | AC013356.2 | | 0.062551 | 0.076239 | 0.084683 |
| 33946 | ENSG0000 | AC004846.2 | | 0.042763 | 0.083245 | 0.084563 |
| 2873 | ENSG0000 | CPQ | carboxypep | 0.066952 | 0.067336 | 0.084499 |
| 6061 | ENSG0000 | USP6 | ubiquitin s | 0.019518 | 0.070368 | 0.084495 |
| 41882 | ENSG0000 | AC005052.1 | | 0 | 0.046829 | 0.084493 |
| 37626 | ENSG0000 | AC092364.2 | | 0 | 0.010799 | 0.084488 |
| 39505 | ENSG0000 | LINC01138 | long interg | 0.012613 | 0.022664 | 0.084476 |
| 41555 | ENSG0000 | AC080112.4 | | 0.00733 | 0.043771 | 0.084342 |
| 14173 | ENSG0000 | EFCAB13 | EF-hand ca | 0.038455 | 0.069924 | 0.08414 |
| 8676 | ENSG0000 | FAM50B | family with | 0 | 0.021717 | 0.084131 |

| | | | | | | |
|---|---|---|---|---|---|---|
| 9343 | ENSG0000 | GLT1D1 | glycosyltra | 0.049829 | 0.050331 | 0.083857 |
| 11386 | ENSG0000 | INTS6L | integrator | 0.027191 | 0.061894 | 0.083821 |
| 9188 | ENSG0000 | DLG2 | discs large | 0.026931 | 0.060002 | 0.083791 |
| 4695 | ENSG0000 | PROX1 | prospero h | 0.034075 | 0.060037 | 0.083707 |
| 7285 | ENSG0000 | WDR38 | WD repeat | 0.02577 | 0.049463 | 0.083563 |
| 6082 | ENSG0000 | SIGLEC9 | sialic acid b | 0.025881 | 0.06666 | 0.083555 |
| 11716 | ENSG0000 | SCNN1G | sodium cha | 0.050781 | 0.062187 | 0.083327 |
| 3743 | ENSG0000 | SLC1A2 | solute carr | 0.035938 | 0.055646 | 0.083286 |
| 38904 | ENSG0000 | AC092354.1 | | 0 | 0.037793 | 0.083129 |
| 30081 | ENSG0000 | PCDHA14 | protocadh | 0.023167 | 0.037728 | 0.082918 |
| 35724 | ENSG0000 | AL022341.2 | | 0 | 0.010439 | 0.082903 |
| 3266 | ENSG0000 | GHRHR | growth hor | 0 | 0.058645 | 0.082873 |
| 3281 | ENSG0000 | TFR2 | transferrin | 0.023475 | 0.030278 | 0.082702 |
| 41324 | ENSG0000 | uc_338 | TUC338 [So | 0.022995 | 0.067331 | 0.082701 |
| 19486 | ENSG0000 | U3 | Small nucle | 0 | 0.058383 | 0.082636 |
| 24147 | ENSG0000 | AC087163.1 | | 0 | 0.038152 | 0.08261 |
| 40167 | ENSG0000 | AC106028.5 | | 0 | 0.07386 | 0.08259 |
| 9034 | ENSG0000 | ZNF214 | zinc finger | 0.018138 | 0.029889 | 0.082578 |
| 1344 | ENSG0000 | PLXNA2 | plexin A2 [ | 0.05676 | 0.064491 | 0.082529 |
| 10490 | ENSG0000 | MEIOB | meiosis spe | 0.016522 | 0.021946 | 0.082435 |
| 5166 | ENSG0000 | TMEM156 | transmeml | 0.041412 | 0.073315 | 0.082364 |
| 1608 | ENSG0000 | SEMA5B | semaphori | 0 | 0.022731 | 0.082221 |
| 24439 | ENSG0000 | AC004840.1 | | 0 | 0.028031 | 0.08218 |
| 882 | ENSG0000 | DLX3 | distal-less l | 0 | 0.032047 | 0.082121 |
| 5596 | ENSG0000 | SRMS | src-related | 0.035866 | 0.07561 | 0.082069 |
| 20074 | ENSG0000 | KCTD9P4 | potassium | 0 | 0.067259 | 0.082034 |
| 1786 | ENSG0000 | GP6 | glycoprote | 0.032966 | 0.052525 | 0.082032 |
| 20673 | ENSG0000 | AC136632.1 | | 0 | 0.048842 | 0.081828 |
| 15797 | ENSG0000 | RHD | Rh blood g | 0.02909 | 0.03862 | 0.081679 |
| 38329 | ENSG0000 | AC139493.1 | | 0 | 0.064399 | 0.081595 |
| 39452 | ENSG0000 | AC005776.2 | | 0 | 0.034177 | 0.081585 |
| 22582 | ENSG0000 | AP006216.1 | | 0 | 0.047628 | 0.081564 |
| 32132 | ENSG0000 | OR7E154P | olfactory re | 0 | 0.003429 | 0.081541 |
| 17079 | ENSG0000 | F5 | coagulatio | 0.039773 | 0.040499 | 0.081523 |
| 25870 | ENSG0000 | RHBDF1P1 | RHBDF1 ps | 0 | 0.01195 | 0.081492 |
| 13636 | ENSG0000 | AC007160.1 | | 0 | 0.069805 | 0.081381 |
| 9160 | ENSG0000 | FCGR1A | Fc fragmer | 0 | 0.067914 | 0.081375 |
| 33291 | ENSG0000 | AC089983.1 | | 0 | 0.062649 | 0.08127 |
| 22122 | ENSG0000 | ADAM1B | ADAM met | 0.021648 | 0.050205 | 0.08127 |
| 20068 | ENSG0000 | AC010336.1 | | 0.039165 | 0.062474 | 0.081236 |
| 12582 | ENSG0000 | FOXA3 | forkhead b | 0.048231 | 0.078134 | 0.081016 |
| 13874 | ENSG0000 | MRVI1-AS1 | MRVI1 ant | 0 | 0.051876 | 0.080974 |
| 6998 | ENSG0000 | PKIB | cAMP-depe | 0.004321 | 0.039728 | 0.08096 |
| 5829 | ENSG0000 | TAS2R5 | taste 2 rec | 0.057517 | 0.073892 | 0.080738 |
| 19785 | ENSG0000 | YBX2P1 | Y-box bind | 0 | 0.024823 | 0.080711 |
| 25715 | ENSG0000 | AL591479.1 | | 0 | 0.062408 | 0.080638 |
| 33056 | ENSG0000 | KHDC1L | KH domain | 0 | 0.075279 | 0.080432 |

| | | | | | | |
|---|---|---|---|---|---|---|
| 4922 | ENSG0000 | PPP4R4 | protein ph | 0.04294 | 0.064364 | 0.080182 |
| 26 | ENSG0000 | AOC1 | amine oxid | 0.037093 | 0.075178 | 0.080115 |
| 38216 | ENSG0000 | AL049840.6 | | 0 | 0.021052 | 0.079926 |
| 20641 | ENSG0000 | AL157777.1 | | 0 | 0.072401 | 0.079918 |
| 4122 | ENSG0000 | SLC4A9 | solute carr | 0 | 0.024101 | 0.079829 |
| 21860 | ENSG0000 | LINC01398 | long interg | 0.042334 | 0.063411 | 0.079822 |
| 33279 | ENSG0000 | AC025154.2 | | 0 | 0.042362 | 0.079722 |
| 14458 | ENSG0000 | TSPYL5 | TSPY like 5 | 0.047502 | 0.064618 | 0.07947 |
| 6433 | ENSG0000 | NR0B2 | nuclear rec | 0.030811 | 0.050297 | 0.079395 |
| 7169 | ENSG0000 | IGSF6 | immunoglo | 0.03264 | 0.070021 | 0.079388 |
| 27517 | ENSG0000 | RNF216-IT1 | RNF216 int | 0 | 0.033038 | 0.07925 |
| 14303 | ENSG0000 | ALOX15B | arachidona | 0 | 0.012961 | 0.078994 |
| 20934 | ENSG0000 | MIR1289-1 | microRNA | 0 | 0.053693 | 0.078973 |
| 35485 | ENSG0000 | AC007603.1 | | 0 | 0.020915 | 0.078913 |
| 31912 | ENSG0000 | AC008429.4 | | 0 | 0.03172 | 0.078778 |
| 39235 | ENSG0000 | LINC02019 | long interg | 0 | 0.020233 | 0.078736 |
| 23006 | ENSG0000 | RPS3AP37 | ribosomal | 0 | 0.077768 | 0.078327 |
| 4430 | ENSG0000 | PROC | protein C, i | 0.028551 | 0.06392 | 0.078314 |
| 20834 | ENSG0000 | LL22NC03- | uncharacte | 0.011449 | 0.056036 | 0.078278 |
| 19797 | ENSG0000 | TTLL13P | tubulin tyr | 0.010322 | 0.011063 | 0.078253 |
| 23553 | ENSG0000 | SEPT7-AS1 | SEPT7 antis | 0.072967 | 0.078133 | 0.078249 |
| 38313 | ENSG0000 | AC007938.3 | | 0 | 0.046664 | 0.077994 |
| 1956 | ENSG0000 | RGS17 | regulator o | 0.049569 | 0.064734 | 0.077952 |
| 7705 | ENSG0000 | AMDHD1 | amidohydr | 0.036789 | 0.054907 | 0.077939 |
| 4880 | ENSG0000 | PLEKHG1 | pleckstrin l | 0.004862 | 0.021769 | 0.077798 |
| 10868 | ENSG0000 | NKX6-1 | NK6 home | 0.011501 | 0.013458 | 0.077758 |
| 6909 | ENSG0000 | SDS | serine deh | 0.011933 | 0.067882 | 0.077618 |
| 4639 | ENSG0000 | CD160 | CD160 mol | 0.017908 | 0.029362 | 0.077505 |
| 43235 | ENSG0000 | MIR4651 | microRNA | 0.002776 | 0.009548 | 0.077497 |
| 43096 | ENSG0000 | AC084337.2 | | 0 | 0.057705 | 0.077467 |
| 1074 | ENSG0000 | SDK2 | sidekick ce | 0.046949 | 0.053895 | 0.077425 |
| 5736 | ENSG0000 | CCR7 | C-C motif c | 0.014252 | 0.057979 | 0.077175 |
| 15810 | ENSG0000 | C3orf70 | chromosor | 0 | 0.004961 | 0.077067 |
| 37700 | ENSG0000 | AC008750.7 | | 0 | 0.034221 | 0.077059 |
| 32449 | ENSG0000 | AP002893.1 | | 0.026547 | 0.04254 | 0.076863 |
| 42367 | ENSG0000 | Z97633.1 | | 0.054645 | 0.065801 | 0.076788 |
| 25016 | ENSG0000 | AC008440.2 | | 0 | 0.031975 | 0.076697 |
| 10869 | ENSG0000 | CDS1 | CDP-diacyl | 0.011991 | 0.022948 | 0.076554 |
| 9673 | ENSG0000 | PROM2 | prominin 2 | 0 | 0.0694 | 0.076533 |
| 21953 | ENSG0000 | AATK-AS1 | AATK antis | 0 | 0.01593 | 0.076348 |
| 42604 | ENSG0000 | PAUPAR | PAX6 upstr | 0.017998 | 0.047677 | 0.076167 |
| 7836 | ENSG0000 | PSTPIP1 | proline-ser | 0.027929 | 0.059597 | 0.076126 |
| 3583 | ENSG0000 | FAM20A | FAM20A, g | 0.031935 | 0.049497 | 0.076123 |
| 13967 | ENSG0000 | PGBD5 | piggyBac tr | 0.034593 | 0.058923 | 0.076082 |
| 14394 | ENSG0000 | TH | tyrosine hy | 0 | 0.015274 | 0.075982 |
| 9943 | ENSG0000 | DCST1 | DC-STAMP | 0.054517 | 0.057588 | 0.07598 |
| 41017 | ENSG0000 | FO681492.1 | | 0.016866 | 0.033717 | 0.075859 |

| | | | | | | |
|---|---|---|---|---|---|---|
| 20770 | ENSG0000 | RPL23P8 | ribosomal | 0 | 0.067924 | 0.075685 |
| 35357 | ENSG0000 | AC002400.1 | | 0 | 0.033654 | 0.075631 |
| 29788 | ENSG0000 | LINC02432 | long interg | 0.029559 | 0.045196 | 0.07563 |
| 35218 | ENSG0000 | AC138512.1 | | 0 | 0.035712 | 0.075528 |
| 20935 | ENSG0000 | SNORD23 | small nucle | 0.064936 | 0.067844 | 0.075374 |
| 30174 | ENSG0000 | KRT18P31 | keratin 18 | 0.054941 | 0.055376 | 0.075306 |
| 10471 | ENSG0000 | SCIMP | SLP adapto | 0 | 0.056323 | 0.075204 |
| 26757 | ENSG0000 | AL807757.2 | | 0 | 0.007808 | 0.075159 |
| 25994 | ENSG0000 | AC007283.1 | | 0 | 0.065821 | 0.075151 |
| 24675 | ENSG0000 | LINC00441 | long interg | 0 | 0.069473 | 0.075127 |
| 2044 | ENSG0000 | TLL2 | tolloid like | 0.023945 | 0.036436 | 0.0751 |
| 26312 | ENSG0000 | AL136528.2 | | 0 | 0.041046 | 0.075031 |
| 41388 | ENSG0000 | AC068831.8 | | 0 | 0.070623 | 0.074996 |
| 11169 | ENSG0000 | ACSL6 | acyl-CoA sy | 0.007192 | 0.010426 | 0.074697 |
| 26130 | ENSG0000 | LINC01287 | long interg | 0.036841 | 0.039824 | 0.074679 |
| 3244 | ENSG0000 | ANKRD7 | ankyrin rep | 0 | 0.066283 | 0.07446 |
| 11548 | ENSG0000 | ADAMTS15 | ADAM met | 0 | 0.017954 | 0.074434 |
| 7525 | ENSG0000 | BTBD16 | BTB domai | 0.014751 | 0.018253 | 0.074431 |
| 1803 | ENSG0000 | DOCK3 | dedicator c | 0.030419 | 0.040076 | 0.074364 |
| 22289 | ENSG0000 | AC025918.1 | | 0 | 0.066721 | 0.074315 |
| 17922 | ENSG0000 | AL133216.1 | | 0 | 0.025479 | 0.074248 |
| 38945 | ENSG0000 | BX284668.6 | | 0 | 0.064865 | 0.07422 |
| 41641 | ENSG0000 | AC132938.5 | | 0.019646 | 0.03207 | 0.07413 |
| 11630 | ENSG0000 | PLD4 | phospholip | 0.020164 | 0.034666 | 0.074094 |
| 30144 | ENSG0000 | AC026427.1 | | 0 | 0.007802 | 0.07404 |
| 1446 | ENSG0000 | OSBPL6 | oxysterol b | 0.060479 | 0.060934 | 0.074034 |
| 14267 | ENSG0000 | HMGN2P4 | high mobil | 0.029095 | 0.058871 | 0.07401 |
| 24669 | ENSG0000 | AC024937.2 | | 0.047369 | 0.048797 | 0.073984 |
| 22331 | ENSG0000 | RPS27P16 | ribosomal | 0 | 0.008719 | 0.073722 |
| 2653 | ENSG0000 | SRPX2 | sushi repea | 0.036381 | 0.041178 | 0.073717 |
| 21262 | ENSG0000 | AL353807.1 | | 0 | 0.048587 | 0.073572 |
| 36707 | ENSG0000 | MIR5093 | microRNA | 0 | 0.026804 | 0.073517 |
| 30035 | ENSG0000 | CASC11 | cancer susc | 0.0226 | 0.047777 | 0.07342 |
| 40397 | ENSG0000 | AC243654.2 | | 0 | 0.032812 | 0.073382 |
| 28587 | ENSG0000 | RN7SL68P | RNA, 7SL, c | 0.002842 | 0.037246 | 0.073369 |
| 43195 | ENSG0000 | MIR658 | microRNA | 0.000012 | 0.044843 | 0.073297 |
| 8417 | ENSG0000 | ANKRD53 | ankyrin rep | 0 | 0.047995 | 0.073197 |
| 18345 | ENSG0000 | PCDHA5 | protocadhe | 0.005007 | 0.059371 | 0.073097 |
| 39843 | ENSG0000 | AC145423.2 | | 0 | 0.066427 | 0.073055 |
| 9834 | ENSG0000 | NODAL | nodal grow | 0.031446 | 0.032076 | 0.073052 |
| 37956 | ENSG0000 | MKNK1-AS | MKNK1 ant | 0 | 0.023696 | 0.073009 |
| 37548 | ENSG0000 | AC008403.1 | | 0 | 0.066057 | 0.072874 |
| 25672 | ENSG0000 | SPIN4-AS1 | SPIN4 antis | 0.02659 | 0.03388 | 0.072805 |
| 15833 | ENSG0000 | AC092118.1 | | 0 | 0.059567 | 0.072796 |
| 4196 | ENSG0000 | PDGFRB | platelet de | 0.00499 | 0.04329 | 0.072685 |
| 8021 | ENSG0000 | MAPK4 | mitogen-ac | 0.041892 | 0.051566 | 0.072585 |
| 37997 | ENSG0000 | C8orf44-SG | C8orf44-SG | 0.007226 | 0.011952 | 0.072514 |

| | | | | | | |
|---|---|---|---|---|---|---|
| 29012 | ENSG0000 | CFB | compleme | 0 | 0.034714 | 0.072501 |
| 6683 | ENSG0000 | DYDC2 | DPY30 don | 0 | 0.061429 | 0.072463 |
| 7624 | ENSG0000 | PRDM5 | PR/SET dor | 0.052555 | 0.069498 | 0.072118 |
| 425 | ENSG0000 | SYT13 | synaptotag | 0.045898 | 0.048176 | 0.072069 |
| 43030 | ENSG0000 | MIR636 | microRNA | 0.008112 | 0.04804 | 0.072006 |
| 29310 | ENSG0000 | FCGR2C | Fc fragmer | 0.014836 | 0.061929 | 0.071861 |
| 43085 | ENSG0000 | MIR4709 | microRNA | 0.001497 | 0.064682 | 0.071768 |
| 4714 | ENSG0000 | A4GNT | alpha-1,4-I | 0.020041 | 0.053957 | 0.071737 |
| 14585 | ENSG0000 | UTS2R | urotensin 2 | 0 | 0.024766 | 0.071614 |
| 34254 | ENSG0000 | RBM17P4 | RNA bindir | 0 | 0.048951 | 0.071601 |
| 28865 | ENSG0000 | AC079203.1 | | 0.043926 | 0.069918 | 0.071367 |
| 36060 | ENSG0000 | LINC02003 | long interg | 0 | 0.015916 | 0.071353 |
| 29418 | ENSG0000 | AP002026.1 | | 0.033038 | 0.046712 | 0.071293 |
| 9622 | ENSG0000 | SRSF12 | serine and | 0.007859 | 0.060696 | 0.071251 |
| 9185 | ENSG0000 | CD226 | CD226 mol | 0.040384 | 0.040603 | 0.07124 |
| 35133 | ENSG0000 | AC133919.2 | | 0.042837 | 0.05255 | 0.071215 |
| 43047 | ENSG0000 | AC092053.2 | | 0.039045 | 0.056286 | 0.071209 |
| 38462 | ENSG0000 | AL512631.2 | | 0.061096 | 0.061795 | 0.071039 |
| 12114 | ENSG0000 | CX3CR1 | C-X3-C mo | 0.038877 | 0.068088 | 0.070964 |
| 5822 | ENSG0000 | BEST3 | bestrophin | 0.012305 | 0.035059 | 0.070949 |
| 13257 | ENSG0000 | ADGRG2 | adhesion G | 0.017844 | 0.059488 | 0.070903 |
| 29952 | ENSG0000 | PRDX2P3 | peroxiredo | 0.008161 | 0.049786 | 0.070765 |
| 31365 | ENSG0000 | AC010834.2 | | 0 | 0.055988 | 0.070714 |
| 9204 | ENSG0000 | TEX12 | testis expr | 0 | 0.013001 | 0.070692 |
| 24392 | ENSG0000 | C2orf27AP | chromosor | 0.030649 | 0.043088 | 0.070595 |
| 4289 | ENSG0000 | PEX5L | peroxisom | 0.043367 | 0.047074 | 0.070552 |
| 120 | ENSG0000 | YBX2 | Y-box bind | 0 | 0.023728 | 0.07042 |
| 14289 | ENSG0000 | ALOX12B | arachidona | 0.013411 | 0.01593 | 0.070403 |
| 1317 | ENSG0000 | MOCOS | molybdenu | 0.028393 | 0.046367 | 0.070347 |
| 32629 | ENSG0000 | AC092471.1 | | 0.002215 | 0.021364 | 0.070268 |
| 2590 | ENSG0000 | CHRDL1 | chordin lik | 0.040626 | 0.051282 | 0.070153 |
| 18841 | ENSG0000 | SNORD116 | small nucle | 0 | 0.058599 | 0.070131 |
| 37730 | ENSG0000 | VN1R82P | vomeronas | 0.043791 | 0.050173 | 0.069965 |
| 12796 | ENSG0000 | MCC | mutated in | 0.047938 | 0.048437 | 0.069611 |
| 20048 | ENSG0000 | ST7-OT4 | ST7 overla | 0.026895 | 0.032135 | 0.069607 |
| 22584 | ENSG0000 | PLCG1-AS1 | PLCG1 anti | 0 | 0.005476 | 0.069568 |
| 1085 | ENSG0000 | FUNDC1 | FUN14 dor | 0 | 0.041781 | 0.069546 |
| 35020 | ENSG0000 | LINC02280 | long interg | 0.018602 | 0.05168 | 0.069236 |
| 24009 | ENSG0000 | AL136982.2 | | 0 | 0.00921 | 0.069216 |
| 23543 | ENSG0000 | AC104653.1 | | 0 | 0.063247 | 0.069161 |
| 37925 | ENSG0000 | AC025165.4 | | 0 | 0.067616 | 0.068975 |
| 41902 | ENSG0000 | AP003108.4 | | 0.020212 | 0.041095 | 0.068817 |
| 6097 | ENSG0000 | EGLN3 | egl-9 famil | 0.017564 | 0.029685 | 0.06851 |
| 14376 | ENSG0000 | FAM71E2 | family with | 0 | 0.057686 | 0.068416 |
| 22124 | ENSG0000 | AC008013.1 | | 0.012054 | 0.034055 | 0.068296 |
| 32394 | ENSG0000 | PDXDC2P | pyridoxal c | 0.03282 | 0.059756 | 0.068258 |
| 34269 | ENSG0000 | AC023034.2 | | 0 | 0.035526 | 0.068053 |

| | | | | | | |
|---|---|---|---|---|---|---|
| 2313 | ENSG0000 | MLC1 | megalence | 0.00826 | 0.013195 | 0.067927 |
| 2434 | ENSG0000 | SGK2 | SGK2, serir | 0.016229 | 0.040501 | 0.067907 |
| 17808 | ENSG0000 | RNU6-128: | RNA, U6 sr | 0 | 0.060873 | 0.067828 |
| 9849 | ENSG0000 | MS4A1 | membrane | 0.007337 | 0.034944 | 0.067775 |
| 14499 | ENSG0000 | HOXC9 | homeobox | 0 | 0.03463 | 0.067663 |
| 38510 | ENSG0000 | AC011477.6 | | 0 | 0.015086 | 0.06742 |
| 6929 | ENSG0000 | KCP | kielin/chor | 0.029272 | 0.034625 | 0.067394 |
| 40361 | ENSG0000 | AL049794.2 | | 0 | 0.038694 | 0.067348 |
| 1087 | ENSG0000 | RORA | RAR relate | 0.041509 | 0.048656 | 0.067303 |
| 27604 | ENSG0000 | LINC01410 | long interg | 0.011683 | 0.018907 | 0.067217 |
| 28076 | ENSG0000 | AC108718.1 | | 0.04036 | 0.052848 | 0.067064 |
| 33863 | ENSG0000 | DUX4L27 | double hor | 0 | 0.034663 | 0.067045 |
| 27291 | ENSG0000 | LINC00987 | long interg | 0 | 0.023562 | 0.066899 |
| 21599 | ENSG0000 | PRR29 | proline rich | 0.003768 | 0.017254 | 0.066881 |
| 9643 | ENSG0000 | ADAMTS1 | ADAM met | 0.032752 | 0.066017 | 0.06673 |
| 14802 | ENSG0000 | PPP1R27 | protein ph | 0 | 0.057344 | 0.06667 |
| 17927 | ENSG0000 | AL139353.1 | | 0.0352 | 0.041582 | 0.066646 |
| 19696 | ENSG0000 | ARHGEF35 | Rho guanir | 0 | 0.024363 | 0.066578 |
| 3945 | ENSG0000 | TULP1 | tubby like | 0.027496 | 0.038695 | 0.066475 |
| 22569 | ENSG0000 | AL390067.1 | | 0.024122 | 0.061123 | 0.066421 |
| 37575 | ENSG0000 | BNIP3P26 | BCL2 intera | 0 | 0.029588 | 0.066351 |
| 18026 | ENSG0000 | OOEP | oocyte exp | 0 | 0.019817 | 0.066244 |
| 3263 | ENSG0000 | CRHR2 | corticotrop | 0.043539 | 0.048005 | 0.066238 |
| 24626 | ENSG0000 | ACTG1P10 | actin gamn | 0.032919 | 0.064218 | 0.065965 |
| 23583 | ENSG0000 | AC113554.1 | | 0 | 0.029875 | 0.065874 |
| 33185 | ENSG0000 | AC025165.2 | | 0.012753 | 0.021483 | 0.065607 |
| 14301 | ENSG0000 | CIITA | class II maj | 0.036561 | 0.059562 | 0.065597 |
| 1272 | ENSG0000 | NTN4 | netrin 4 [S | 0.057512 | 0.059291 | 0.065569 |
| 41919 | ENSG0000 | AC131649.2 | | 0.007996 | 0.044938 | 0.065548 |
| 12638 | ENSG0000 | PLA2G1B | phospholip | 0.03474 | 0.065119 | 0.065537 |
| 25144 | ENSG0000 | ST3GAL5-A | ST3GAL5 a | 0 | 0.03784 | 0.065514 |
| 4155 | ENSG0000 | NPR3 | natriuretic | 0.01439 | 0.0368 | 0.065474 |
| 35517 | ENSG0000 | AC130462.1 | | 0 | 0.037719 | 0.065387 |
| 25990 | ENSG0000 | AL353743.4 | | 0 | 0.031229 | 0.065318 |
| 14702 | ENSG0000 | P2RY8 | purinergic | 0.022919 | 0.062007 | 0.065305 |
| 14298 | ENSG0000 | LSMEM2 | leucine rich | 0.037375 | 0.050508 | 0.065277 |
| 29766 | ENSG0000 | ACTN3 | actinin alpl | 0 | 0.041345 | 0.065253 |
| 42309 | ENSG0000 | CU639417.6 | | 0.031191 | 0.047183 | 0.065123 |
| 33300 | ENSG0000 | AL136040.1 | | 0 | 0.035417 | 0.065117 |
| 31747 | ENSG0000 | CTB-178M. | uncharacte | 0 | 0.057797 | 0.065085 |
| 14350 | ENSG0000 | GPBAR1 | G protein-c | 0.013138 | 0.037913 | 0.064986 |
| 1944 | ENSG0000 | MYO15A | myosin XV. | 0.012963 | 0.043266 | 0.06494 |
| 15804 | ENSG0000 | TMPRSS6 | transmeml | 0.043476 | 0.049783 | 0.064829 |
| 41417 | ENSG0000 | AL133396.2 | | 0 | 0.0483 | 0.064813 |
| 10623 | ENSG0000 | NLRP3 | NLR family | 0.036013 | 0.064278 | 0.06474 |
| 37913 | ENSG0000 | ARHGAP19 | ARHGAP19 | 0 | 0.00711 | 0.064691 |
| 19859 | ENSG0000 | RPL31P44 | ribosomal | 0 | 0.010938 | 0.064595 |

| | | | | | | |
|---|---|---|---|---|---|---|
| 15282 | ENSG0000( | RPL7P26 | ribosomal | 0 | 0.025021 | 0.064595 |
| 16231 | ENSG0000( | ANKRD34B | ankyrin rep | 0 | 0.014475 | 0.064572 |
| 36597 | ENSG0000( | AP000919.2 | | 0.010416 | 0.017139 | 0.064567 |
| 40245 | ENSG0000( | MIR6882 | microRNA | 0 | 0.010153 | 0.064397 |
| 8268 | ENSG0000( | CASQ1 | calsequest | 0 | 0.023063 | 0.064372 |
| 24956 | ENSG0000( | AL606534.3 | | 0 | 0.038474 | 0.064369 |
| 28721 | ENSG0000( | OR7E100P | olfactory re | 0 | 0.057487 | 0.064055 |
| 25521 | ENSG0000( | Z83844.2 | | 0 | 0.029387 | 0.064001 |
| 42308 | ENSG0000( | AC004408.2 | | 0 | 0.021086 | 0.063991 |
| 36460 | ENSG0000( | MIR3143 | microRNA | 0 | 0.009818 | 0.063817 |
| 35224 | ENSG0000( | AC007906.1 | | 0 | 0.050131 | 0.063742 |
| 6413 | ENSG0000( | PPP1R1B | protein pho | 0.007794 | 0.014217 | 0.063741 |
| 28690 | ENSG0000( | AC007537.2 | | 0 | 0.029455 | 0.063617 |
| 10376 | ENSG0000( | LRRC71 | leucine rich | 0.034596 | 0.042426 | 0.063616 |
| 33315 | ENSG0000( | DNAJC8P1 | DnaJ heat s | 0 | 0.039075 | 0.063611 |
| 40118 | ENSG0000( | GPR179 | G protein-c | 0.034512 | 0.046304 | 0.06353 |
| 29271 | ENSG0000( | AC098583.1 | | 0 | 0.053365 | 0.063516 |
| 14877 | ENSG0000( | OTOP2 | otopetrin 2 | 0 | 0.031395 | 0.06342 |
| 9879 | ENSG0000( | ATP2B2 | ATPase pla | 0.003516 | 0.057418 | 0.063339 |
| 16850 | ENSG0000( | ZNF311 | zinc finger | 0 | 0.023838 | 0.063269 |
| 3326 | ENSG0000( | AEBP1 | AE binding | 0.008266 | 0.049619 | 0.063076 |
| 30278 | ENSG0000( | CACNA1G- | CACNA1G a | 0 | 0.057786 | 0.063062 |
| 7792 | ENSG0000( | RDH12 | retinol deh | 0.013382 | 0.029 | 0.063001 |
| 38192 | ENSG0000( | AC093915.1 | | 0.034503 | 0.037749 | 0.062981 |
| 8871 | ENSG0000( | CRH | corticotrop | 0 | 0.018908 | 0.062907 |
| 26963 | ENSG0000( | MED15P5 | mediator c | 0 | 0.012367 | 0.062646 |
| 5946 | ENSG0000( | FLNC | filamin C [S | 0.007595 | 0.028703 | 0.062624 |
| 38841 | ENSG0000( | AC114760.2 | | 0.017187 | 0.038719 | 0.06258 |
| 30143 | ENSG0000( | ENPP7P10 | ectonucleo | 0 | 0.03694 | 0.062575 |
| 35213 | ENSG0000( | AC026470.3 | | 0 | 0.03618 | 0.062546 |
| 39254 | ENSG0000( | AC128687.2 | | 0 | 0.030599 | 0.062509 |
| 28018 | ENSG0000( | TLR9 | toll like rec | 0.014096 | 0.026266 | 0.062481 |
| 41576 | ENSG0000( | AC130686.1 | | 0 | 0.021153 | 0.062466 |
| 3388 | ENSG0000( | PIP5K1B | phosphatic | 0.007217 | 0.034337 | 0.062456 |
| 36439 | ENSG0000( | KRT18P55 | keratin 18 | 0 | 0.020411 | 0.062281 |
| 38215 | ENSG0000( | AL158163.1 | | 0.010762 | 0.053077 | 0.062207 |
| 31908 | ENSG0000( | AC022973.4 | | 0 | 0.022452 | 0.062198 |
| 34604 | ENSG0000( | AC034111.1 | | 0.009548 | 0.050424 | 0.062188 |
| 4059 | ENSG0000( | PACRG | parkin core | 0.02043 | 0.039034 | 0.062184 |
| 7218 | ENSG0000( | BLK | BLK proto- | 0 | 0.035357 | 0.062171 |
| 13805 | ENSG0000( | CSNK1G2-/ | CSNK1G2 a | 0 | 0.031478 | 0.062167 |
| 17093 | ENSG0000( | EGFL6 | EGF like do | 0 | 0.011191 | 0.062127 |
| 14419 | ENSG0000( | PNPLA1 | patatin like | 0 | 0.019232 | 0.062072 |
| 41431 | ENSG0000( | AC037198.3 | | 0 | 0.041158 | 0.062021 |
| 27514 | ENSG0000( | RNGTTP1 | RNA guany | 0.010892 | 0.017559 | 0.062011 |
| 5506 | ENSG0000( | TCP11 | t-complex | 0.032436 | 0.053758 | 0.061991 |
| 30203 | ENSG0000( | AC007663.2 | | 0 | 0.032621 | 0.061981 |

| | | | | | | |
|---|---|---|---|---|---|---|
| 2896 | ENSG0000 | STMN2 | stathmin 2 | 0 | 0.012439 | 0.061861 |
| 6868 | ENSG0000 | PDGFRA | platelet de | 0.018325 | 0.04467 | 0.061706 |
| 40003 | ENSG0000 | AC011487.4 | | 0 | 0.024545 | 0.061702 |
| 43198 | ENSG0000 | MIR29B2 | microRNA | 0 | 0.034364 | 0.061689 |
| 26016 | ENSG0000 | AP1B1P1 | adaptor re | 0 | 0.017577 | 0.061255 |
| 5688 | ENSG0000 | GDF5 | growth diff | 0 | 0.055013 | 0.061211 |
| 18413 | ENSG0000 | LINC01602 | long interg | 0 | 0.032262 | 0.061111 |
| 38361 | ENSG0000 | AL139317.4 | | 0 | 0.03147 | 0.061062 |
| 41326 | ENSG0000 | VWA8-AS1 | VWA8 anti | 0.033308 | 0.037886 | 0.061025 |
| 18347 | ENSG0000 | PCDHA2 | protocadh | 0.021933 | 0.036946 | 0.060961 |
| 34381 | ENSG0000 | AC093525.1 | | 0.003069 | 0.05432 | 0.060948 |
| 36842 | ENSG0000 | AP001542.1 | | 0 | 0.054638 | 0.060881 |
| 3030 | ENSG0000 | LILRA1 | leukocyte i | 0.026233 | 0.048998 | 0.060797 |
| 8870 | ENSG0000 | DNAJC5B | DnaJ heat s | 0.01217 | 0.025355 | 0.060767 |
| 41903 | ENSG0000 | AL137918.1 | | 0 | 0.038726 | 0.060722 |
| 33031 | ENSG0000 | IQSEC3P1 | IQ motif ar | 0 | 0.053084 | 0.06065 |
| 41145 | ENSG0000 | SNORD116 | small nucle | 0 | 0.010453 | 0.060591 |
| 18952 | ENSG0000 | Y_RNA | Y RNA [Sou | 0.036709 | 0.054859 | 0.06058 |
| 24312 | ENSG0000 | UQCRHP1 | ubiquinol-c | 0 | 0.053427 | 0.06037 |
| 38051 | ENSG0000 | AC027020.2 | | 0.029924 | 0.036593 | 0.060248 |
| 20493 | ENSG0000 | AL079342.1 | | 0 | 0.032047 | 0.060231 |
| 18067 | ENSG0000 | TDRG1 | testis deve | 0 | 0.03583 | 0.060099 |
| 41888 | ENSG0000 | AL158070.2 | | 0 | 0.015518 | 0.060092 |
| 8037 | ENSG0000 | ARL5C | ADP ribosy | 0 | 0.017132 | 0.059973 |
| 40725 | ENSG0000 | AC074050.4 | | 0 | 0.02176 | 0.059913 |
| 7171 | ENSG0000 | MYLK3 | myosin ligh | 0.006698 | 0.032068 | 0.059862 |
| 8280 | ENSG0000 | LHX9 | LIM homeo | 0 | 0.003525 | 0.05982 |
| 5947 | ENSG0000 | LRRC4 | leucine rich | 0.052794 | 0.054151 | 0.05974 |
| 31084 | ENSG0000 | RNA5SP61 | RNA, 5S rib | 0 | 0.017525 | 0.059732 |
| 16796 | ENSG0000 | LHFPL5 | LHFPL tetra | 0 | 0.010635 | 0.059641 |
| 6382 | ENSG0000 | AC091132.1 | | 0 | 0.003832 | 0.05943 |
| 39736 | ENSG0000 | SNX18P1Y | sorting nex | 0 | 0.028446 | 0.059346 |
| 8503 | ENSG0000 | ACKR2 | atypical ch | 0.009583 | 0.035975 | 0.059201 |
| 15230 | ENSG0000 | MAP7D2 | MAP7 dom | 0.020277 | 0.023205 | 0.059193 |
| 35425 | ENSG0000 | AC007728.2 | | 0.010374 | 0.042254 | 0.059174 |
| 37509 | ENSG0000 | AC006547.3 | | 0.033581 | 0.053097 | 0.059127 |
| 26583 | ENSG0000 | AC114488.2 | | 0 | 0.048397 | 0.059124 |
| 16350 | ENSG0000 | RYR1 | ryanodine | 0 | 0.019541 | 0.059065 |
| 9996 | ENSG0000 | CABP1 | calcium bir | 0.018176 | 0.021188 | 0.059043 |
| 41814 | ENSG0000 | AC090772.4 | | 0 | 0.052115 | 0.059022 |
| 34225 | ENSG0000 | AC091045.1 | | 0 | 0.021468 | 0.059008 |
| 981 | ENSG0000 | CLDN18 | claudin 18 | 0.009782 | 0.033536 | 0.058952 |
| 16292 | ENSG0000 | OR7E12P | olfactory re | 0 | 0.045347 | 0.058911 |
| 12440 | ENSG0000 | OTUD7A | OTU deubi | 0.029724 | 0.044544 | 0.058881 |
| 1633 | ENSG0000 | ZNF671 | zinc finger | 0.058587 | 0.05879 | 0.058836 |
| 10014 | ENSG0000 | RADIL | Rap associa | 0.016127 | 0.053728 | 0.058812 |
| 26954 | ENSG0000 | TCF3P1 | transcriptio | 0 | 0.015115 | 0.058769 |

| | | | | | | |
|---|---|---|---|---|---|---|
| 42208 | ENSG0000 | AL691447.3 | | 0 | 0.022024 | 0.058711 |
| 4090 | ENSG0000 | WISP3 | WNT1 indu | 0.011163 | 0.019575 | 0.058613 |
| 30073 | ENSG0000 | GAPDHP70 | glyceraldeh | 0 | 0.02782 | 0.058589 |
| 28109 | ENSG0000 | AL590542.1 | | 0 | 0.004522 | 0.058454 |
| 36657 | ENSG0000 | FBXO36P1 | F-box prot | 0 | 0.039651 | 0.058337 |
| 37740 | ENSG0000 | BNIP3P24 | BCL2 intera | 0.039317 | 0.044975 | 0.058157 |
| 13616 | ENSG0000 | CALCB | calcitonin r | 0.013523 | 0.037078 | 0.05777 |
| 2366 | ENSG0000 | ASB2 | ankyrin rep | 0.006917 | 0.023114 | 0.057754 |
| 6623 | ENSG0000 | SPART | spartin [So | 0.026818 | 0.036429 | 0.057684 |
| 1488 | ENSG0000 | MID2 | midline 2 [ | 0 | 0.018741 | 0.057676 |
| 34422 | ENSG0000 | AL163952.1 | | 0.023412 | 0.02545 | 0.057674 |
| 31770 | ENSG0000 | AC083843.1 | | 0.023477 | 0.051849 | 0.057638 |
| 39525 | ENSG0000 | mascRNA-r | MALAT1-as | 0.02208 | 0.039461 | 0.057579 |
| 19813 | ENSG0000 | AL359073.1 | | 0 | 0.024558 | 0.057577 |
| 784 | ENSG0000 | F7 | coagulation | 0.018771 | 0.054336 | 0.057541 |
| 15140 | ENSG0000 | DNAH2 | dynein axo | 0.008765 | 0.045321 | 0.057403 |
| 35140 | ENSG0000 | AC079322.1 | | 0 | 0.038602 | 0.057384 |
| 40809 | ENSG0000 | AC073569.3 | | 0 | 0.042929 | 0.057267 |
| 27415 | ENSG0000 | AL357060.2 | | 0.017426 | 0.023573 | 0.057235 |
| 39897 | ENSG0000 | AL391669.1 | | 0 | 0.004702 | 0.057233 |
| 1450 | ENSG0000 | SP140 | SP140 nucl | 0.025478 | 0.029051 | 0.057229 |
| 31378 | ENSG0000 | AC055854.1 | | 0.027219 | 0.037489 | 0.057158 |
| 423 | ENSG0000 | CYP24A1 | cytochrom | 0.047118 | 0.052915 | 0.057101 |
| 14490 | ENSG0000 | WDFY3-AS | WDFY3 ant | 0.014709 | 0.042315 | 0.057094 |
| 15168 | ENSG0000 | AC091868.1 | | 0 | 0.032555 | 0.057057 |
| 27503 | ENSG0000 | AL031726.1 | | 0 | 0.024321 | 0.05703 |
| 25239 | ENSG0000 | TRIM80P | tripartite n | 0 | 0.037145 | 0.056991 |
| 19708 | ENSG0000 | YWHAZP2 | tyrosine 3- | 0.0306 | 0.031012 | 0.056804 |
| 12199 | ENSG0000 | TSPAN5 | tetraspanin | 0.01638 | 0.020826 | 0.056758 |
| 21956 | ENSG0000 | AC073283.1 | | 0 | 0.020019 | 0.056633 |
| 38434 | ENSG0000 | U73023.1 | | 0 | 0.041531 | 0.056614 |
| 19619 | ENSG0000 | AL445305.1 | | 0 | 0.009413 | 0.056536 |
| 12131 | ENSG0000 | KLHL30 | kelch like f | 0 | 0.02349 | 0.05651 |
| 41375 | ENSG0000 | ST7-OT3_3 | ST7 overla | 0 | 0.020021 | 0.056482 |
| 2686 | ENSG0000 | FLT1 | fms relatec | 0 | 0.008146 | 0.056395 |
| 10104 | ENSG0000 | SLAMF8 | SLAM fami | 0.011888 | 0.03902 | 0.056171 |
| 15512 | ENSG0000 | LYG2 | lysozyme g | 0 | 0.041354 | 0.056113 |
| 40249 | ENSG0000 | AC253576.2 | | 0 | 0.011731 | 0.055851 |
| 41674 | ENSG0000 | Z99129.3 | | 0.015657 | 0.020301 | 0.05583 |
| 29466 | ENSG0000 | AC091948.1 | | 0 | 0.049267 | 0.055673 |
| 35951 | ENSG0000 | AC023389.1 | | 0 | 0.047224 | 0.05564 |
| 42575 | ENSG0000 | AC006978.2 | | 0.025701 | 0.049533 | 0.055622 |
| 40332 | ENSG0000 | MIR6796 | microRNA | 0.015225 | 0.015796 | 0.055582 |
| 41054 | ENSG0000 | SCARNA15 | small Cajal | 0.041534 | 0.048393 | 0.055576 |
| 19873 | ENSG0000 | RPS7P14 | ribosomal | 0.013816 | 0.038195 | 0.055563 |
| 26037 | ENSG0000 | CSNK1G2P | casein kina | 0.01845 | 0.019224 | 0.055532 |
| 18137 | ENSG0000 | ERICH2 | glutamate | 0 | 0.043832 | 0.055513 |

| | | | | | | |
|---|---|---|---|---|---|---|
| 25378 | ENSG0000 | AC243772.2 | | 0 | 0.03512 | 0.055489 |
| 9262 | ENSG0000 | THRSP | thyroid hor | 0 | 0.021255 | 0.055282 |
| 23943 | ENSG0000 | PDE4DIPP1 | phosphodi | 0 | 0.02373 | 0.05485 |
| 20738 | ENSG0000 | AL033519.2 | | 0 | 0.023406 | 0.054798 |
| 30245 | ENSG0000 | AC009927.1 | | 0 | 0.016457 | 0.054692 |
| 12954 | ENSG0000 | SLFN12 | schlafen fa | 0 | 0.036893 | 0.05466 |
| 41153 | ENSG0000 | AL358075.4 | | 0 | 0.012229 | 0.054647 |
| 10537 | ENSG0000 | FAM151A | family with | 0.017757 | 0.022969 | 0.054532 |
| 15799 | ENSG0000 | ESPN | espin [Sour | 0.020071 | 0.025082 | 0.054531 |
| 8929 | ENSG0000 | ACTL7B | actin like 7 | 0 | 0.048664 | 0.054465 |
| 6655 | ENSG0000 | MYH11 | myosin hea | 0.016713 | 0.050884 | 0.054448 |
| 8726 | ENSG0000 | RNF217 | ring finger | 0.033447 | 0.048468 | 0.054427 |
| 22861 | ENSG0000 | AC012441.1 | | 0 | 0.04244 | 0.054417 |
| 3104 | ENSG0000 | CD33 | CD33 mole | 0.01849 | 0.018962 | 0.05435 |
| 13379 | ENSG0000 | IGDCC3 | immunoglo | 0.006438 | 0.018419 | 0.054323 |
| 40929 | ENSG0000 | HOTTIP_4 | HOXA tran | 0 | 0.032739 | 0.054302 |
| 33443 | ENSG0000 | DDN-AS1 | DDN and P | 0.011708 | 0.04783 | 0.054219 |
| 29890 | ENSG0000 | LAMTOR3P | late endos | 0 | 0.006665 | 0.053969 |
| 21349 | ENSG0000 | AC069213.1 | | 0 | 0.01232 | 0.053914 |
| 21553 | ENSG0000 | MIR3681H | MIR3681 h | 0.018379 | 0.02487 | 0.053912 |
| 29765 | ENSG0000 | LINC02428 | long interg | 0.028992 | 0.050156 | 0.053909 |
| 8826 | ENSG0000 | NXF3 | nuclear RN | 0.030114 | 0.034577 | 0.053794 |
| 35407 | ENSG0000 | AC036108.3 | | 0.036121 | 0.053599 | 0.05371 |
| 32666 | ENSG0000 | AC025576.2 | | 0 | 0.017446 | 0.053699 |
| 7422 | ENSG0000 | TRIM29 | tripartite m | 0.013956 | 0.014859 | 0.053694 |
| 11072 | ENSG0000 | ETNPPL | ethanolam | 0 | 0.014433 | 0.053687 |
| 9056 | ENSG0000 | KLHL35 | kelch like f | 0 | 0.048713 | 0.053686 |
| 27949 | ENSG0000 | LINC00535 | long interg | 0 | 0.041188 | 0.053599 |
| 41080 | ENSG0000 | MIAT_exon | Myocardia | 0 | 0.018796 | 0.053516 |
| 28195 | ENSG0000 | AC012358.3 | | 0.024178 | 0.024997 | 0.053486 |
| 13389 | ENSG0000 | KLK15 | kallikrein r | 0.006653 | 0.022106 | 0.053458 |
| 22515 | ENSG0000 | SMARCE1P | SWI/SNF re | 0 | 0.023764 | 0.053336 |
| 11798 | ENSG0000 | UGT1A6 | UDP glucur | 0.013728 | 0.019703 | 0.053255 |
| 1657 | ENSG0000 | COL16A1 | collagen ty | 0 | 0.04939 | 0.053212 |
| 5880 | ENSG0000 | GNAI1 | G protein s | 0.035153 | 0.0414 | 0.053178 |
| 42783 | ENSG0000 | AC068547.1 | | 0 | 0.050443 | 0.053178 |
| 5157 | ENSG0000 | CCRL2 | C-C motif c | 0 | 0.014448 | 0.053101 |
| 40088 | ENSG0000 | AC084876.1 | | 0 | 0.015922 | 0.053033 |
| 25633 | ENSG0000 | ERI3-IT1 | ERI3 intror | 0 | 0.00483 | 0.053031 |
| 5331 | ENSG0000 | HOXC13 | homeobox | 0.026907 | 0.048206 | 0.053002 |
| 32786 | ENSG0000 | CYP2A6 | cytochrom | 0 | 0.018329 | 0.052961 |
| 24731 | ENSG0000 | AC104462.1 | | 0.005913 | 0.040475 | 0.052938 |
| 38928 | ENSG0000 | IFNL4 | interferon | 0 | 0.015778 | 0.052931 |
| 42148 | ENSG0000 | AC007431.3 | | 0 | 0.020372 | 0.052836 |
| 19639 | ENSG0000 | LPAL2 | lipoprotein | 0 | 0.02656 | 0.052815 |
| 19921 | ENSG0000 | SUGT1P2 | SGT1 homo | 0 | 0.038778 | 0.05269 |
| 40967 | ENSG0000 | RN7SL469P | RNA, 7SL, c | 0 | 0.005365 | 0.052672 |

| | | | | | | |
|---|---|---|---|---|---|---|
| 37054 | ENSG0000 | AC005702.2 | | 0 | 0.048118 | 0.052643 |
| 33230 | ENSG0000 | AC068888.2 | | 0 | 0.010386 | 0.052632 |
| 12471 | ENSG0000 | SERPINA6 | serpin fam | 0 | 0.015809 | 0.052535 |
| 6366 | ENSG0000 | NAPSB | napsin B as | 0 | 0.031977 | 0.052486 |
| 42777 | ENSG0000 | AC001226.2 | | 0.005796 | 0.021372 | 0.052308 |
| 39540 | ENSG0000 | AC004706.3 | | 0.021268 | 0.03096 | 0.052217 |
| 26113 | ENSG0000 | AL353653.1 | | 0 | 0.018579 | 0.052037 |
| 40497 | ENSG0000 | Metazoa_S | Metazoan | 0 | 0.042839 | 0.052002 |
| 39003 | ENSG0000 | AC005014.3 | | 0 | 0.023331 | 0.051994 |
| 1967 | ENSG0000 | CEBPE | CCAAT/enh | 0 | 0.037681 | 0.051965 |
| 34202 | ENSG0000 | AC079328.2 | | 0.005291 | 0.029263 | 0.051962 |
| 26639 | ENSG0000 | APOA1-AS | APOA1 ant | 0 | 0.048128 | 0.051919 |
| 27348 | ENSG0000 | AL513366.1 | | 0 | 0.047085 | 0.051912 |
| 36980 | ENSG0000 | AC005336.2 | | 0 | 0.03107 | 0.051869 |
| 34717 | ENSG0000 | AC019294.2 | | 0.007928 | 0.028768 | 0.051612 |
| 3264 | ENSG0000 | EPHB6 | EPH recept | 0.006688 | 0.018997 | 0.051599 |
| 15621 | ENSG0000 | CYP4F2 | cytochrom | 0 | 0.029758 | 0.051586 |
| 16046 | ENSG0000 | IL17REL | interleukin | 0.019342 | 0.044852 | 0.05156 |
| 17380 | ENSG0000 | RNU6-127. | RNA, U6 sr | 0 | 0.006744 | 0.05153 |
| 11463 | ENSG0000 | GFI1B | growth fac | 0.018219 | 0.022024 | 0.051343 |
| 38419 | ENSG0000 | HMGB1P5( | high mobil | 0 | 0.033711 | 0.051333 |
| 26895 | ENSG0000 | NRADDP | neurotropl | 0 | 0.031399 | 0.051314 |
| 17052 | ENSG0000 | FAM19A2 | family with | 0.013154 | 0.03317 | 0.051302 |
| 14355 | ENSG0000 | LINC00652 | long interg | 0.006057 | 0.026456 | 0.051272 |
| 13909 | ENSG0000 | OR7E125P | olfactory r | 0 | 0.004143 | 0.051268 |
| 41669 | ENSG0000 | AC138470.1 | | 0 | 0.016847 | 0.051228 |
| 24273 | ENSG0000 | HMGB1P1: | high mobil | 0.027859 | 0.035119 | 0.051214 |
| 13635 | ENSG0000 | ASCL3 | achaete-sc | 0 | 0.009446 | 0.051174 |
| 35012 | ENSG0000 | AC104758.3 | | 0 | 0.041302 | 0.051013 |
| 31725 | ENSG0000 | KRT18P37 | keratin 18 | 0 | 0.022355 | 0.050984 |
| 41765 | ENSG0000 | AC020604.1 | | 0 | 0.023491 | 0.050966 |
| 5014 | ENSG0000 | SEC16B | SEC16 hom | 0 | 0.006084 | 0.050928 |
| 5481 | ENSG0000 | GRM4 | glutamate | 0.010547 | 0.030548 | 0.05079 |
| 13044 | ENSG0000 | PDE3A | phosphodi | 0.027057 | 0.045264 | 0.050771 |
| 13791 | ENSG0000 | ZNF354C | zinc finger | 0.006583 | 0.03235 | 0.050695 |
| 41818 | ENSG0000 | AC100788.2 | | 0 | 0.02976 | 0.050457 |
| 20310 | ENSG0000 | CHCHD2P2 | coiled-coil- | 0 | 0.018627 | 0.050457 |
| 30718 | ENSG0000 | CTBP2P4 | C-terminal | 0.038852 | 0.050257 | 0.050443 |
| 38846 | ENSG0000 | AL645933.2 | | 0 | 0.048086 | 0.05044 |
| 17941 | ENSG0000 | LINC01285 | long interg | 0 | 0.00485 | 0.050339 |
| 35104 | ENSG0000 | AL049555.1 | | 0.004883 | 0.024319 | 0.050274 |
| 41608 | ENSG0000 | AL031587.6 | | 0.03099 | 0.047687 | 0.050186 |
| 39962 | ENSG0000 | AC018926.1 | | 0 | 0.037208 | 0.050173 |
| 7055 | ENSG0000 | CHRND | cholinergic | 0 | 0.009348 | 0.050019 |
| 33015 | ENSG0000 | AP000721.2 | | 0 | 0.036735 | 0.04997 |
| 19656 | ENSG0000 | AL365223.1 | | 0 | 0.044829 | 0.049951 |
| 22908 | ENSG0000 | AC007386.1 | | 0 | 0.004783 | 0.049935 |

| | | | | | | |
|---|---|---|---|---|---|---|
| 2069 | ENSG0000 | MLN | motilin [So | 0 | 0.025383 | 0.049933 |
| 1579 | ENSG0000 | PCDHA6 | protocadh | 0 | 0.030778 | 0.049878 |
| 42368 | ENSG0000 | CU634019.7 | | 0 | 0.006133 | 0.049874 |
| 19769 | ENSG0000 | RPL12P18 | ribosomal | 0 | 0.044726 | 0.049836 |
| 33523 | ENSG0000 | AC133555.3 | | 0 | 0.033572 | 0.049827 |
| 15684 | ENSG0000 | ZNF501 | zinc finger | 0.009174 | 0.029794 | 0.049767 |
| 9563 | ENSG0000 | ANKRD29 | ankyrin rep | 0.021918 | 0.043424 | 0.049673 |
| 28368 | ENSG0000 | CEACAMP1 | carcinoeml | 0 | 0.029214 | 0.049648 |
| 36341 | ENSG0000 | TSPOAP1-A | TSPOAP1 a | 0 | 0.044846 | 0.049576 |
| 39287 | ENSG0000 | AC006033.2 | | 0 | 0.014877 | 0.049497 |
| 24170 | ENSG0000 | ELOVL2-AS | ELOVL2 an | 0 | 0.029348 | 0.049437 |
| 9053 | ENSG0000 | SESN3 | sestrin 3 [S | 0.026927 | 0.037217 | 0.049226 |
| 32493 | ENSG0000 | AC127526.4 | | 0 | 0.041689 | 0.049218 |
| 35836 | ENSG0000 | AC007114.2 | | 0.008261 | 0.0308 | 0.049157 |
| 9598 | ENSG0000 | TDH | L-threonin | 0 | 0.013053 | 0.049134 |
| 43000 | ENSG0000 | MIR6884 | microRNA | 0.003518 | 0.008843 | 0.049013 |
| 5902 | ENSG0000 | C22orf24 | chromosor | 0 | 0.037229 | 0.048912 |
| 31738 | ENSG0000 | AC008514.2 | | 0 | 0.031342 | 0.0488 |
| 884 | ENSG0000 | TSPAN32 | tetraspanir | 0.022963 | 0.023593 | 0.048749 |
| 21344 | ENSG0000 | RAET1E-AS | RAET1E an | 0.000631 | 0.007157 | 0.048606 |
| 17097 | ENSG0000 | RASSF9 | Ras associa | 0.012493 | 0.024439 | 0.04856 |
| 22178 | ENSG0000 | AL807761.2 | | 0.024861 | 0.035778 | 0.048532 |
| 43031 | ENSG0000 | AC007326.4 | | 0.012938 | 0.036906 | 0.048522 |
| 13646 | ENSG0000 | CRYBG2 | crystallin b | 0.005866 | 0.0192 | 0.048504 |
| 18607 | ENSG0000 | NANOGNB | NANOG ne | 0 | 0.043679 | 0.048439 |
| 12190 | ENSG0000 | FAM178B | family with | 0 | 0.008201 | 0.048386 |
| 3249 | ENSG0000 | CPED1 | cadherin lil | 0.025466 | 0.033601 | 0.048358 |
| 29877 | ENSG0000 | LDHBP3 | lactate deh | 0 | 0.035492 | 0.048352 |
| 5262 | ENSG0000 | DNAI1 | dynein axo | 0.025207 | 0.046427 | 0.048309 |
| 30890 | ENSG0000 | AC008427.1 | | 0 | 0.044384 | 0.048228 |
| 17899 | ENSG0000 | C9orf41-AS | C9orf41 an | 0 | 0.021651 | 0.048156 |
| 9301 | ENSG0000 | VENTX | VENT hom | 0 | 0.020044 | 0.048142 |
| 22603 | ENSG0000 | AL162724.1 | | 0 | 0.016583 | 0.048046 |
| 40229 | ENSG0000 | AC113385.3 | | 0 | 0.019402 | 0.047936 |
| 13731 | ENSG0000 | RNF152 | ring finger | 0.006724 | 0.042257 | 0.047787 |
| 42181 | ENSG0000 | AC005578.1 | | 0 | 0.042955 | 0.047637 |
| 18344 | ENSG0000 | PCDHA7 | protocadh | 0.015352 | 0.046348 | 0.047631 |
| 32053 | ENSG0000 | PSMA2P1 | proteasom | 0 | 0.03447 | 0.047477 |
| 20047 | ENSG0000 | XPOTP1 | exportin fc | 0 | 0.018185 | 0.047471 |
| 21916 | ENSG0000 | AL445250.1 | | 0 | 0.021473 | 0.047423 |
| 5399 | ENSG0000 | CTCFL | CCCTC-bin | 0 | 0.00978 | 0.047373 |
| 20547 | ENSG0000 | AC005077.2 | | 0 | 0.04034 | 0.04728 |
| 9566 | ENSG0000 | THY1 | Thy-1 cell s | 0.012044 | 0.04409 | 0.047225 |
| 42282 | ENSG0000 | AC027290.2 | | 0.029419 | 0.044079 | 0.047146 |
| 13103 | ENSG0000 | MRGPRF | MAS relate | 0 | 0.010662 | 0.047103 |
| 30202 | ENSG0000 | AC034207.1 | | 0 | 0.025685 | 0.047071 |
| 13814 | ENSG0000 | PENK | proenkeph | 0.010236 | 0.03175 | 0.046979 |

| | | | | | | |
|---|---|---|---|---|---|---|
| 38849 | ENSG0000 | AL590438.1 | | 0 | 0.008459 | 0.04693 |
| 36437 | ENSG0000 | AC107982.3 | | 0.001789 | 0.033858 | 0.046881 |
| 30752 | ENSG0000 | LINC02272 | long interg | 0 | 0.020635 | 0.046742 |
| 14896 | ENSG0000 | FAM43B | family with | 0 | 0.011343 | 0.046716 |
| 40449 | ENSG0000 | Y_RNA | Y RNA [Sou | 0 | 0.033121 | 0.046681 |
| 18636 | ENSG0000 | NYNRIN | NYN doma | 0.015599 | 0.021888 | 0.04667 |
| 5564 | ENSG0000 | SOX21 | SRY-box 21 | 0 | 0.010465 | 0.046643 |
| 1388 | ENSG0000 | NAALAD2 | N-acetylate | 0.02462 | 0.03699 | 0.046592 |
| 23990 | ENSG0000 | AL137003.1 | | 0 | 0.041502 | 0.046573 |
| 28149 | ENSG0000 | CPHL1P | ceruloplas | 0.009022 | 0.015602 | 0.046415 |
| 6644 | ENSG0000 | PDE6B | phosphodi | 0.024548 | 0.033858 | 0.046414 |
| 18610 | ENSG0000 | KRTAP5-6 | keratin ass | 0.007852 | 0.02297 | 0.046276 |
| 42090 | ENSG0000 | FLJ38576 | uncharacte | 0 | 0.005911 | 0.046153 |
| 9065 | ENSG0000 | NCAM1 | neural cell | 0.025934 | 0.035168 | 0.04613 |
| 20429 | ENSG0000 | SIAH3 | siah E3 ubi | 0 | 0.041225 | 0.046114 |
| 17026 | ENSG0000 | SH2D1B | SH2 domai | 0 | 0.043087 | 0.04608 |
| 302 | ENSG0000 | BTK | Bruton tyr | 0.007758 | 0.022907 | 0.045998 |
| 20660 | ENSG0000 | ADAMTS7F | ADAMTS7 | 0.021071 | 0.021599 | 0.045979 |
| 5567 | ENSG0000 | KIF25 | kinesin fan | 0 | 0.016202 | 0.045748 |
| 19704 | ENSG0000 | RPL12P38 | ribosomal | 0.002551 | 0.03672 | 0.045658 |
| 39665 | ENSG0000 | RN7SL673F | RNA, 7SL, c | 0 | 0.005365 | 0.04556 |
| 2992 | ENSG0000 | AGR2 | anterior gr | 0.035369 | 0.041019 | 0.045509 |
| 22581 | ENSG0000 | AL121899.1 | | 0 | 0.037564 | 0.04548 |
| 14989 | ENSG0000 | COL18A1-A | COL18A1 a | 0 | 0.036166 | 0.045471 |
| 25188 | ENSG0000 | TIMM9P2 | TIMM9 pse | 0 | 0.042308 | 0.045343 |
| 42684 | ENSG0000 | DUXB | double hor | 0 | 0.026943 | 0.045245 |
| 35691 | ENSG0000 | AC004233.1 | | 0 | 0.042844 | 0.045045 |
| 5359 | ENSG0000 | ATXN3L | ataxin 3 lik | 0.008259 | 0.026964 | 0.045014 |
| 38749 | ENSG0000 | AP000919.4 | | 0.018879 | 0.040377 | 0.044991 |
| 15700 | ENSG0000 | ANKRD20A | ankyrin rep | 0.008108 | 0.031456 | 0.044989 |
| 313 | ENSG0000 | SLC6A7 | solute carr | 0 | 0.006621 | 0.044656 |
| 14352 | ENSG0000 | ZNF648 | zinc finger | 0 | 0.012007 | 0.044512 |
| 6105 | ENSG0000 | CDO1 | cysteine di | 0 | 0.026562 | 0.0445 |
| 21428 | ENSG0000 | SEC13P1 | SEC13 hom | 0 | 0.015333 | 0.044488 |
| 41791 | ENSG0000 | PRAL | p53 regula | 0.021755 | 0.035451 | 0.044431 |
| 7632 | ENSG0000 | FRAS1 | Fraser extr | 0.014969 | 0.035304 | 0.044363 |
| 28580 | ENSG0000 | WWTR1-IT | WWTR1 in | 0 | 0.015544 | 0.044318 |
| 7020 | ENSG0000 | BCO1 | beta-carot | 0 | 0.018085 | 0.044301 |
| 32995 | ENSG0000 | AC009511.2 | | 0 | 0.023239 | 0.044279 |
| 8534 | ENSG0000 | PLA1A | phospholip | 0 | 0.034574 | 0.044263 |
| 34449 | ENSG0000 | AC017071.1 | | 0.024384 | 0.026568 | 0.044197 |
| 2802 | ENSG0000 | XYLT1 | xylosyltran | 0.022635 | 0.031806 | 0.044152 |
| 6318 | ENSG0000 | TTLL9 | tubulin tyr | 0.021879 | 0.02803 | 0.044071 |
| 12734 | ENSG0000 | FUT3 | fucosyltrar | 0.022913 | 0.040608 | 0.044004 |
| 42754 | ENSG0000 | AC068620.4 | | 0 | 0.006944 | 0.043926 |
| 42470 | ENSG0000 | SH3PXD2A | SH3PXD2A | 0 | 0.013113 | 0.043885 |
| 22152 | ENSG0000 | RPL7P22 | ribosomal | 0 | 0.019409 | 0.043879 |

| | | | | | | |
|---|---|---|---|---|---|---|
| 26343 | ENSG0000 | TSPY26P | testis speci | 0.013111 | 0.017916 | 0.043867 |
| 33243 | ENSG0000 | AC073487.1 | | 0 | 0.006041 | 0.043863 |
| 28859 | ENSG0000 | ARHGEF26 | ARHGEF26 | 0.031714 | 0.04004 | 0.043834 |
| 12145 | ENSG0000 | PHYHIP | phytanoyl- | 0.016561 | 0.028109 | 0.043827 |
| 14571 | ENSG0000 | TMEM132I | transmeml | 0.002861 | 0.03327 | 0.043818 |
| 12886 | ENSG0000 | ANGPTL7 | angiopoiet | 0 | 0.013036 | 0.043782 |
| 16217 | ENSG0000 | APOD | apolipopro | 0 | 0.013279 | 0.043762 |
| 33150 | ENSG0000 | PHBP19 | prohibitin | 0 | 0.006973 | 0.043724 |
| 27010 | ENSG0000 | OVAAL | ovarian ad | 0 | 0.019663 | 0.043664 |
| 33519 | ENSG0000 | AC016954.1 | | 0 | 0.012592 | 0.043635 |
| 9763 | ENSG0000 | MAGEA8 | MAGE fam | 0.01502 | 0.024736 | 0.043604 |
| 6243 | ENSG0000 | GATA5 | GATA bind | 0 | 0.028087 | 0.043602 |
| 29855 | ENSG0000 | AC097460.2 | | 0 | 0.038259 | 0.043561 |
| 39661 | ENSG0000 | AC083805.3 | | 0 | 0.011054 | 0.043525 |
| 28073 | ENSG0000 | PRDX3P4 | peroxiredo | 0 | 0.03257 | 0.043365 |
| 23161 | ENSG0000 | AC127537.1 | | 0 | 0.020811 | 0.043307 |
| 1813 | ENSG0000 | SIGLEC1 | sialic acid b | 0 | 0.010218 | 0.043288 |
| 13508 | ENSG0000 | C1orf127 | chromosor | 0.009934 | 0.016187 | 0.043236 |
| 11611 | ENSG0000 | RAG1 | recombina | 0.009083 | 0.018991 | 0.043176 |
| 21460 | ENSG0000 | DDX11L1 | DEAD/H-bo | 0 | 0.023837 | 0.043036 |
| 13008 | ENSG0000 | ARNT2 | aryl hydroc | 0.004618 | 0.015102 | 0.042982 |
| 3344 | ENSG0000 | NMRK1 | nicotinami | 0.016738 | 0.030615 | 0.042913 |
| 21793 | ENSG0000 | E2F3-IT1 | E2F3 intror | 0 | 0.012371 | 0.042902 |
| 18288 | ENSG0000 | LINC01291 | long interg | 0.008439 | 0.018403 | 0.042828 |
| 40467 | ENSG0000 | AC073534.2 | | 0 | 0.001623 | 0.042812 |
| 2663 | ENSG0000 | RUBCNL | RUN and cy | 0.023234 | 0.040122 | 0.042806 |
| 19762 | ENSG0000 | AC022217.1 | | 0 | 0.01802 | 0.042804 |
| 40626 | ENSG0000 | GXYLT1P5 | glucoside x | 0 | 0.008412 | 0.042756 |
| 41007 | ENSG0000 | AC126175.1 | | 0 | 0.038353 | 0.042735 |
| 12124 | ENSG0000 | BDKRB2 | bradykinin | 0 | 0.004277 | 0.042676 |
| 36534 | ENSG0000 | AC138207.8 | | 0.021359 | 0.033217 | 0.042666 |
| 29503 | ENSG0000 | PCED1B-AS | PCED1B an | 0.0116 | 0.026733 | 0.042649 |
| 7142 | ENSG0000 | MYO1G | myosin IG | 0.036969 | 0.037444 | 0.042642 |
| 26934 | ENSG0000 | RC3H1-IT1 | RC3H1 intr | 0 | 0.021861 | 0.042563 |
| 30738 | ENSG0000 | AC131392.2 | | 0.011265 | 0.03623 | 0.042546 |
| 27904 | ENSG0000 | FABP5P3 | fatty acid b | 0 | 0.021281 | 0.042534 |
| 40608 | ENSG0000 | AC005618.2 | | 0 | 0.006373 | 0.042407 |
| 25883 | ENSG0000 | NAP1L4P3 | nucleosom | 0 | 0.012403 | 0.042363 |
| 10633 | ENSG0000 | VANGL2 | VANGL pla | 0.006694 | 0.021796 | 0.042356 |
| 8480 | ENSG0000 | ESPNL | espin like [ | 0 | 0.027128 | 0.04231 |
| 40645 | ENSG0000 | Metazoa_S | Metazoan | 0 | 0.005976 | 0.042285 |
| 12061 | ENSG0000 | MAJIN | membrane | 0 | 0.025313 | 0.042109 |
| 33174 | ENSG0000 | AL589743.1 | | 0.006079 | 0.03079 | 0.042102 |
| 29521 | ENSG0000 | FAM13A-A | FAM13A ar | 0.016952 | 0.028596 | 0.042054 |
| 11929 | ENSG0000 | LAIR1 | leukocyte a | 0 | 0.010844 | 0.042012 |
| 43048 | ENSG0000 | MIR4738 | microRNA | 0.00434 | 0.034561 | 0.041921 |
| 22249 | ENSG0000 | AL512504.1 | | 0 | 0.018684 | 0.041784 |

| | | | | | | |
|---|---|---|---|---|---|---|
| 1318 | ENSG0000 | PLD1 | phospholip | 0.024139 | 0.030207 | 0.04169 |
| 12545 | ENSG0000 | KRT75 | keratin 75 | 0 | 0.012509 | 0.041668 |
| 2491 | ENSG0000 | ANGPT4 | angiopoiet | 0.007617 | 0.032907 | 0.041639 |
| 25195 | ENSG0000 | AC099336.1 | | 0 | 0.024491 | 0.041624 |
| 34454 | ENSG0000 | AC022819.1 | | 0.00911 | 0.022543 | 0.041512 |
| 37230 | ENSG0000 | AC005357.2 | | 0 | 0.019284 | 0.041511 |
| 15979 | ENSG0000 | CPSF4L | cleavage ar | 0 | 0.019228 | 0.041498 |
| 35643 | ENSG0000 | AC027601.2 | | 0 | 0.035295 | 0.041378 |
| 25296 | ENSG0000 | CICP4 | capicua tra | 0.006535 | 0.016199 | 0.041242 |
| 35608 | ENSG0000 | PCDHGA9 | protocadh | 0.010018 | 0.018821 | 0.041241 |
| 38322 | ENSG0000 | AC015849.5 | | 0.015584 | 0.023598 | 0.041228 |
| 10646 | ENSG0000 | DENND2D | DENN dom | 0 | 0.011929 | 0.041158 |
| 1665 | ENSG0000 | GCKR | glucokinas | 0 | 0.032572 | 0.04115 |
| 12418 | ENSG0000 | PCDH7 | protocadh | 0.029017 | 0.037562 | 0.041047 |
| 10875 | ENSG0000 | C3orf49 | chromosor | 0 | 0.009625 | 0.041014 |
| 20519 | ENSG0000 | LINC00466 | long interg | 0.033159 | 0.038566 | 0.041014 |
| 2387 | ENSG0000 | PAPLN | papilin, pro | 0.017471 | 0.04003 | 0.040989 |
| 1441 | ENSG0000 | CLUL1 | clusterin lil | 0 | 0.018978 | 0.040718 |
| 28681 | ENSG0000 | IGKV1-39 | immunoglo | 0 | 0.036652 | 0.040695 |
| 34416 | ENSG0000 | AP005233.1 | | 0 | 0.017524 | 0.040659 |
| 28150 | ENSG0000 | AL512306.2 | | 0 | 0.03949 | 0.040566 |
| 43242 | ENSG0000 | MIR34A | microRNA | 0.017163 | 0.019681 | 0.040553 |
| 43127 | ENSG0000 | MIR7112 | microRNA | 0.003626 | 0.021783 | 0.040522 |
| 10299 | ENSG0000 | AIRE | autoimmu | 0.005438 | 0.008605 | 0.040488 |
| 43400 | ENSG0000 | AC092053.3 | | 0.007134 | 0.017551 | 0.040453 |
| 22311 | ENSG0000 | AL645939.2 | | 0 | 0.035962 | 0.040445 |
| 21269 | ENSG0000 | KIF28P | kinesin fan | 0.026588 | 0.033871 | 0.040364 |
| 37356 | ENSG0000 | AC006557.4 | | 0.028583 | 0.030648 | 0.040359 |
| 5424 | ENSG0000 | GDAP1L1 | ganglioside | 0 | 0.036112 | 0.040308 |
| 4783 | ENSG0000 | ABCG2 | ATP bindin | 0 | 0.029941 | 0.040304 |
| 42910 | ENSG0000 | AC068205.4 | | 0 | 0.025388 | 0.04029 |
| 37753 | ENSG0000 | AC008555.2 | | 0.005143 | 0.010592 | 0.040225 |
| 12423 | ENSG0000 | CTNND2 | catenin del | 0.010361 | 0.025536 | 0.040186 |
| 9314 | ENSG0000 | TMEM45B | transmeml | 0.022702 | 0.037731 | 0.040071 |
| 24385 | ENSG0000 | SMARCE1P | SWI/SNF re | 0.014487 | 0.039133 | 0.039947 |
| 34365 | ENSG0000 | LINC02157 | long interg | 0 | 0.019603 | 0.039905 |
| 22280 | ENSG0000 | PROSER2-A | PROSER2 a | 0 | 0.035014 | 0.039892 |
| 34090 | ENSG0000 | AC023906.3 | | 0.027894 | 0.035136 | 0.039715 |
| 35759 | ENSG0000 | AC027455.1 | | 0 | 0.028984 | 0.039644 |
| 13296 | ENSG0000 | C1QTNF1 | C1q and TN | 0.007338 | 0.007619 | 0.039548 |
| 21360 | ENSG0000 | AL359706.1 | | 0.000799 | 0.002618 | 0.039384 |
| 14100 | ENSG0000 | NEUROG2 | neurogenir | 0 | 0.012712 | 0.03934 |
| 39939 | ENSG0000 | AC099521.2 | | 0 | 0.035697 | 0.039268 |
| 43049 | ENSG0000 | MIR7703 | microRNA | 0.02706 | 0.031653 | 0.039233 |
| 21280 | ENSG0000 | ZNF844 | zinc finger | 0.008023 | 0.030694 | 0.039181 |
| 2490 | ENSG0000 | RPS10P5 | ribosomal | 0 | 0.009513 | 0.039155 |
| 3176 | ENSG0000 | GAPDHS | glyceraldeh | 0 | 0.011514 | 0.039143 |

| | | | | | | |
|---|---|---|---|---|---|---|
| 4085 | ENSG0000 | IMPG1 | interphoto | 0 | 0.034192 | 0.039046 |
| 27349 | ENSG0000 | LINC01358 | long interg | 0.010408 | 0.038819 | 0.038996 |
| 18158 | ENSG0000 | SLC44A4 | solute carr | 0 | 0.025194 | 0.038963 |
| 5259 | ENSG0000 | TAF1L | TATA-box l | 0.014214 | 0.019852 | 0.038937 |
| 12688 | ENSG0000 | GIMAP8 | GTPase, IM | 0.008513 | 0.017519 | 0.038848 |
| 28564 | ENSG0000 | AC114480.1 | | 0 | 0.009995 | 0.038802 |
| 4843 | ENSG0000 | SLC46A2 | solute carr | 0.014286 | 0.023074 | 0.038795 |
| 1127 | ENSG0000 | FRMPD1 | FERM and | 0.012099 | 0.029678 | 0.038688 |
| 10410 | ENSG0000 | AC008132.1 | | 0.008747 | 0.024033 | 0.038682 |
| 14979 | ENSG0000 | GPR132 | G protein-c | 0.008421 | 0.013942 | 0.038618 |
| 4963 | ENSG0000 | PPP1R3C | protein ph | 0 | 0.011498 | 0.038593 |
| 24172 | ENSG0000 | AL022476.1 | | 0 | 0.007124 | 0.038537 |
| 29358 | ENSG0000 | AC105285.1 | | 0 | 0.012982 | 0.038519 |
| 29757 | ENSG0000 | NPHP3-AS1 | NPHP3 ant | 0.030003 | 0.033186 | 0.038472 |
| 36591 | ENSG0000 | MIR3174 | microRNA | 0 | 0.024884 | 0.03829 |
| 6865 | ENSG0000 | C5AR2 | compleme | 0.020927 | 0.021157 | 0.038183 |
| 23837 | ENSG0000 | MYCBP2-A | MYCBP2 ar | 0 | 0.022882 | 0.038111 |
| 27197 | ENSG0000 | AC073130.1 | | 0.013864 | 0.017115 | 0.038095 |
| 29847 | ENSG0000 | AC005699.1 | | 0 | 0.018858 | 0.038066 |
| 15141 | ENSG0000 | SH2D1A | SH2 domai | 0 | 0.009757 | 0.038034 |
| 42778 | ENSG0000 | AC106858.1 | | 0 | 0.034364 | 0.038022 |
| 23633 | ENSG0000 | YY1P2 | YY1 transc | 0 | 0.030666 | 0.037926 |
| 29994 | ENSG0000 | AC110373.1 | | 0.010126 | 0.024912 | 0.037919 |
| 16016 | ENSG0000 | C6orf132 | chromosor | 0 | 0.004243 | 0.037904 |
| 37117 | ENSG0000 | AC020934.1 | | 0 | 0.035712 | 0.03788 |
| 23492 | ENSG0000 | SPATA13 | spermatog | 0 | 0.025518 | 0.037778 |
| 34095 | ENSG0000 | AC021231.1 | | 0 | 0.032126 | 0.037766 |
| 27378 | ENSG0000 | NRBF2P3 | nuclear rec | 0.010331 | 0.016566 | 0.037715 |
| 9987 | ENSG0000 | DGKI | diacylglyce | 0.017645 | 0.034614 | 0.037598 |
| 13237 | ENSG0000 | SULT1B1 | sulfotransf | 0.021729 | 0.021834 | 0.037483 |
| 23843 | ENSG0000 | AC079776.2 | | 0 | 0.018821 | 0.037452 |
| 23232 | ENSG0000 | AL031283.1 | | 0 | 0.036474 | 0.03738 |
| 42185 | ENSG0000 | AC008993.3 | | 0 | 0.010946 | 0.037303 |
| 21941 | ENSG0000 | AC012354.1 | | 0 | 0.010789 | 0.03728 |
| 4793 | ENSG0000 | KLF12 | Kruppel lik | 0.008146 | 0.02938 | 0.037143 |
| 8252 | ENSG0000 | NECTIN4 | nectin cell | 0.010135 | 0.024934 | 0.037098 |
| 36838 | ENSG0000 | AC004528.1 | | 0 | 0.019309 | 0.037093 |
| 6041 | ENSG0000 | LOXL1 | lysyl oxidas | 0 | 0.016213 | 0.037079 |
| 37638 | ENSG0000 | AL050403.2 | | 0 | 0.027943 | 0.037072 |
| 16576 | ENSG0000 | NXF4 | nuclear RN | 0 | 0.008009 | 0.037062 |
| 942 | ENSG0000 | MYLK | myosin ligh | 0.011742 | 0.018928 | 0.036932 |
| 20998 | ENSG0000 | LINC01118 | long interg | 0 | 0.020233 | 0.036898 |
| 29285 | ENSG0000 | RAD21L1 | RAD21 coh | 0.005655 | 0.007439 | 0.036814 |
| 13939 | ENSG0000 | ERICH5 | glutamate | 0 | 0.016567 | 0.03679 |
| 41995 | ENSG0000 | AL445437.1 | | 0 | 0.02351 | 0.03669 |
| 14578 | ENSG0000 | OFCC1 | orofacial cl | 0 | 0.029929 | 0.03657 |
| 6179 | ENSG0000 | USHBP1 | USH1 prote | 0.003789 | 0.007429 | 0.036496 |

| | | | | | | |
|---|---|---|---|---|---|---|
| 18516 | ENSG0000 | AC098591.1 | | 0 | 0.004154 | 0.036454 |
| 1091 | ENSG0000 | PLA2G10 | phospholip | 0 | 0.035006 | 0.036411 |
| 35470 | ENSG0000 | AL138690.1 | | 0 | 0.025968 | 0.03631 |
| 24616 | ENSG0000 | AC078875.1 | | 0 | 0.019792 | 0.036174 |
| 6620 | ENSG0000 | DCLK1 | doublecort | 0.006199 | 0.032466 | 0.036128 |
| 40022 | ENSG0000 | AL139260.3 | | 0.012358 | 0.035098 | 0.036092 |
| 2279 | ENSG0000 | CACNA1I | calcium vo | 0 | 0.012142 | 0.036019 |
| 3852 | ENSG0000 | ADGRD1 | adhesion G | 0.006369 | 0.013519 | 0.036007 |
| 40511 | ENSG0000 | AL591441.1 | | 0 | 0.012984 | 0.035938 |
| 33132 | ENSG0000 | HNRNPA1P | heterogen | 0 | 0.031549 | 0.035773 |
| 8238 | ENSG0000 | GPA33 | glycoprote | 0 | 0.009117 | 0.035754 |
| 3535 | ENSG0000 | ICAM2 | intercellula | 0.026643 | 0.026989 | 0.035718 |
| 734 | ENSG0000 | ENTPD2 | ectonuclec | 0 | 0.016601 | 0.035713 |
| 15541 | ENSG0000 | IKZF1 | IKAROS far | 0.019544 | 0.022196 | 0.035665 |
| 42112 | ENSG0000 | AC008764.10 | | 0 | 0.015943 | 0.035654 |
| 16707 | ENSG0000 | GGT3P | gamma-glu | 0.012889 | 0.035287 | 0.035638 |
| 13734 | ENSG0000 | C20orf197 | chromosor | 0 | 0.006909 | 0.035613 |
| 25684 | ENSG0000 | OR6L1P | olfactory r | 0 | 0.016087 | 0.035597 |
| 6029 | ENSG0000 | ALDH1A2 | aldehyde d | 0.012167 | 0.029636 | 0.035597 |
| 14193 | ENSG0000 | EXD1 | exonucleas | 0.00471 | 0.007997 | 0.035581 |
| 15430 | ENSG0000 | SFTPA2 | surfactant | 0 | 0.010884 | 0.035571 |
| 39217 | ENSG0000 | OR6L2P | olfactory r | 0 | 0.015826 | 0.035517 |
| 1228 | ENSG0000 | MOV10L1 | Mov10 RIS | 0 | 0.034582 | 0.03542 |
| 4717 | ENSG0000 | TREH | trehalase [ | 0.020589 | 0.027445 | 0.035418 |
| 23165 | ENSG0000 | AC105935.1 | | 0 | 0.024233 | 0.035296 |
| 18327 | ENSG0000 | AC007389.1 | | 0.00947 | 0.027109 | 0.035267 |
| 9472 | ENSG0000 | LGI2 | leucine rich | 0.019482 | 0.024907 | 0.035258 |
| 9080 | ENSG0000 | ADAM33 | ADAM met | 0 | 0.018366 | 0.035209 |
| 40328 | ENSG0000 | SMAD5-AS | SMAD5 an | 0 | 0.019648 | 0.035182 |
| 36294 | ENSG0000 | AC090844.2 | | 0.017273 | 0.023647 | 0.035098 |
| 29562 | ENSG0000 | AC092658.1 | | 0 | 0.031008 | 0.035067 |
| 4882 | ENSG0000 | CXorf21 | chromosor | 0 | 0.015783 | 0.034987 |
| 10065 | ENSG0000 | TMSB15B | thymosin b | 0 | 0.009036 | 0.034804 |
| 12953 | ENSG0000 | CD8B | CD8b mole | 0 | 0.023339 | 0.034757 |
| 32512 | ENSG0000 | PCDHA3 | protocadh | 0.02146 | 0.023396 | 0.034661 |
| 33389 | ENSG0000 | OR5BK1P | olfactory r | 0 | 0.017537 | 0.034615 |
| 28113 | ENSG0000 | AC069431.1 | | 0 | 0.019665 | 0.034516 |
| 12069 | ENSG0000 | OR2W6P | olfactory r | 0 | 0.03107 | 0.034498 |
| 37980 | ENSG0000 | KC877982.1 | | 0 | 0.030927 | 0.034461 |
| 28101 | ENSG0000 | AL133163.1 | | 0.018965 | 0.030851 | 0.034376 |
| 14480 | ENSG0000 | LINC02363 | long interg | 0.015137 | 0.018532 | 0.034368 |
| 17971 | ENSG0000 | CAPN8 | calpain 8 [S | 0.019708 | 0.028166 | 0.034315 |
| 25413 | ENSG0000 | AC093797.1 | | 0 | 0.020923 | 0.034158 |
| 8587 | ENSG0000 | GC | GC, vitami | 0.029918 | 0.030741 | 0.034085 |
| 30726 | ENSG0000 | AC025171.3 | | 0 | 0.017292 | 0.034075 |
| 22573 | ENSG0000 | AC092155.1 | | 0 | 0.024287 | 0.034012 |
| 9468 | ENSG0000 | JAKMIP1 | janus kinas | 0 | 0.007573 | 0.033961 |

| | | | | | | |
|---|---|---|---|---|---|---|
| 37075 | ENSG0000 | AC020928.2 | | 0 | 0.000938 | 0.033948 |
| 8858 | ENSG0000 | STAR | steroidoge | 0 | 0.007744 | 0.03394 |
| 13612 | ENSG0000 | FAM172BP | family with | 0 | 0.026696 | 0.03386 |
| 43194 | ENSG0000 | AL158064.2 | | 0.008366 | 0.024069 | 0.033822 |
| 69 | ENSG0000 | SLC4A1 | solute carr | 0 | 0.011036 | 0.033763 |
| 28217 | ENSG0000 | LINC00973 | long interg | 0 | 0.030327 | 0.033673 |
| 13873 | ENSG0000 | ZDHHC22 | zinc finger | 0.013786 | 0.022636 | 0.033581 |
| 12152 | ENSG0000 | HFE2 | hemochroı | 0 | 0.024046 | 0.03354 |
| 26991 | ENSG0000 | AC068282.1 | | 0.013334 | 0.015486 | 0.033522 |
| 13730 | ENSG0000 | HORMAD2 | HORMA dc | 0 | 0.01133 | 0.033512 |
| 42168 | ENSG0000 | AC079753.1 | | 0.01256 | 0.027025 | 0.033469 |
| 27632 | ENSG0000 | AC017079.2 | | 0.001557 | 0.022744 | 0.033395 |
| 11781 | ENSG0000 | TTC16 | tetratricop | 0 | 0.027422 | 0.033343 |
| 28253 | ENSG0000 | AC063952.1 | | 0.016644 | 0.023094 | 0.033335 |
| 6685 | ENSG0000 | TMTC1 | transmeml | 0.024232 | 0.02964 | 0.033293 |
| 37337 | ENSG0000 | AC005332.5 | | 0.006079 | 0.008971 | 0.033292 |
| 1356 | ENSG0000 | MBNL3 | musclebliı | 0.005839 | 0.01396 | 0.033277 |
| 35544 | ENSG0000 | snoZ6 | Z6 small nu | 0 | 0.015799 | 0.033252 |
| 5911 | ENSG0000 | APOL3 | apolipopro | 0.009043 | 0.019759 | 0.033034 |
| 20156 | ENSG0000 | EIF4EP5 | eukaryotic | 0 | 0.022873 | 0.033025 |
| 43357 | ENSG0000 | AL121936.1 | | 0.01235 | 0.025272 | 0.032979 |
| 24436 | ENSG0000 | AL391807.1 | | 0 | 0.019723 | 0.032965 |
| 41996 | ENSG0000 | AC138907.9 | | 0 | 0.026661 | 0.032853 |
| 41166 | ENSG0000 | AC099494.2 | | 0 | 0.029831 | 0.03277 |
| 24712 | ENSG0000 | RSL24D1P3 | ribosomal | 0 | 0.005207 | 0.03272 |
| 42065 | ENSG0000 | AC012183.1 | | 0 | 0.02913 | 0.032688 |
| 6064 | ENSG0000 | AIPL1 | aryl hydroc | 0 | 0.010995 | 0.032659 |
| 40734 | ENSG0000 | MIR7106 | microRNA | 0 | 0.004641 | 0.032641 |
| 25571 | ENSG0000 | ST6GALNA | ST6 N-acet | 0.003531 | 0.031609 | 0.032458 |
| 14923 | ENSG0000 | AL109615.1 | | 0 | 0.002719 | 0.032315 |
| 7030 | ENSG0000 | ZNF670-ZN | ZNF670-ZN | 0.007073 | 0.01188 | 0.032209 |
| 41136 | ENSG0000 | AC036108.4 | | 0 | 0.017414 | 0.032187 |
| 27612 | ENSG0000 | SLC9A3P2 | solute carr | 0 | 0.007294 | 0.03217 |
| 13511 | ENSG0000 | VWA3A | von Willeb | 0.002612 | 0.020274 | 0.032165 |
| 39726 | ENSG0000 | SDR42E1P1 | short chair | 0 | 0.005295 | 0.032149 |
| 25062 | ENSG0000 | AC022540.1 | | 0.009551 | 0.019733 | 0.031841 |
| 5455 | ENSG0000 | CHST8 | carbohydra | 0.023341 | 0.025275 | 0.031841 |
| 6725 | ENSG0000 | PEBP4 | phosphatic | 0 | 0.026995 | 0.031835 |
| 10609 | ENSG0000 | GBP4 | guanylate l | 0.00871 | 0.026181 | 0.031816 |
| 25477 | ENSG0000 | AC006042.3 | | 0 | 0.019176 | 0.031748 |
| 7610 | ENSG0000 | PRKG2 | protein kin | 0.006962 | 0.028402 | 0.031727 |
| 41626 | ENSG0000 | AC015656.1 | | 0 | 0.018518 | 0.031654 |
| 36531 | ENSG0000 | AC090912.2 | | 0 | 0.008902 | 0.031573 |
| 1374 | ENSG0000 | DCX | doublecort | 0.010448 | 0.017107 | 0.031504 |
| 13280 | ENSG0000 | KCNH6 | potassium | 0.008458 | 0.010779 | 0.031477 |
| 23235 | ENSG0000 | CASP16P | caspase 16 | 0.015531 | 0.031331 | 0.031471 |
| 40432 | ENSG0000 | MIAT_exor | Myocardia | 0 | 0.018952 | 0.03142 |

| | | | | | | |
|---|---|---|---|---|---|---|
| 15534 | ENSG0000 | SPATA31A | SPATA31 s | 0 | 0.0135 | 0.031338 |
| 15037 | ENSG0000 | MTND4LP3 | mitochond | 0.02366 | 0.030545 | 0.031239 |
| 12348 | ENSG0000 | SCN9A | sodium vol | 0.013722 | 0.023371 | 0.0312 |
| 40708 | ENSG0000 | AL138955.1 | | 0 | 0.028053 | 0.031147 |
| 169 | ENSG0000 | NOS2 | nitric oxide | 0 | 0.013997 | 0.031028 |
| 7898 | ENSG0000 | MARVELD3 | MARVEL d | 0.005565 | 0.019377 | 0.031027 |
| 12358 | ENSG0000 | CT55 | cancer/tes | 0 | 0.030142 | 0.031022 |
| 18393 | ENSG0000 | LINC00654 | long interg | 0.011318 | 0.023144 | 0.031008 |
| 24965 | ENSG0000 | VN1R54P | vomeronas | 0 | 0.027714 | 0.03099 |
| 38156 | ENSG0000 | AL049829.3 | | 0 | 0.015938 | 0.030975 |
| 14335 | ENSG0000 | GIPC3 | GIPC PDZ c | 0 | 0.019193 | 0.030964 |
| 11924 | ENSG0000 | CYP2S1 | cytochrom | 0 | 0.02041 | 0.030919 |
| 18043 | ENSG0000 | ZYG11A | zyg-11 fam | 0 | 0.006193 | 0.030916 |
| 13360 | ENSG0000 | C20orf166 | C20orf166 | 0 | 0.00868 | 0.030895 |
| 15365 | ENSG0000 | LRRC14B | leucine ricl | 0 | 0.018547 | 0.03089 |
| 7476 | ENSG0000 | BRDT | bromodom | 0 | 0.013487 | 0.030865 |
| 10561 | ENSG0000 | DRAXIN | dorsal inhil | 0 | 0.011915 | 0.030821 |
| 20631 | ENSG0000 | AL355864.1 | | 0 | 0.004116 | 0.030792 |
| 11928 | ENSG0000 | ANKRD33 | ankyrin rep | 0 | 0.023596 | 0.030787 |
| 16067 | ENSG0000 | FOCAD | focadhesin | 0.01414 | 0.019313 | 0.030772 |
| 11234 | ENSG0000 | FNDC1 | fibronectin | 0 | 0.006287 | 0.030737 |
| 10183 | ENSG0000 | GOLGA6A | golgin A6 f | 0 | 0.017256 | 0.030541 |
| 7212 | ENSG0000 | TBR1 | T-box, brai | 0 | 0.0074 | 0.030524 |
| 29295 | ENSG0000 | OR2AE1 | olfactory r | 0 | 0.023968 | 0.03046 |
| 13356 | ENSG0000 | C11orf45 | chromosor | 0 | 0.007835 | 0.030326 |
| 10894 | ENSG0000 | ZNF704 | zinc finger | 0.014223 | 0.0165 | 0.030256 |
| 14761 | ENSG0000 | TSHZ2 | teashirt zir | 0.009904 | 0.026631 | 0.030252 |
| 25926 | ENSG0000 | SDR42E1P3 | short chair | 0 | 0.004723 | 0.030205 |
| 12160 | ENSG0000 | GFRA2 | GDNF fami | 0.007428 | 0.02079 | 0.030171 |
| 12605 | ENSG0000 | KCNS3 | potassium | 0 | 0.018307 | 0.030157 |
| 23246 | ENSG0000 | RNF144A-A | RNF144A a | 0 | 0.021703 | 0.030108 |
| 21818 | ENSG0000 | FDPSP8 | farnesyl di | 0 | 0.02679 | 0.030062 |
| 10578 | ENSG0000 | TTLL10 | tubulin tyr | 0 | 0.007373 | 0.029953 |
| 17903 | ENSG0000 | AP003027.1 | | 0 | 0.020965 | 0.029932 |
| 26837 | ENSG0000 | CLEC2L | C-type lect | 0 | 0.001876 | 0.029651 |
| 3819 | ENSG0000 | CUX2 | cut like hor | 0 | 0.001901 | 0.029603 |
| 2649 | ENSG0000 | ITIH6 | inter-alpha | 0.01937 | 0.022472 | 0.029548 |
| 28719 | ENSG0000 | LINC00960 | long interg | 0 | 0.009191 | 0.029428 |
| 24002 | ENSG0000 | Z98749.1 | | 0 | 0.026868 | 0.029089 |
| 62 | ENSG0000 | SLC22A16 | solute carr | 0 | 0.011293 | 0.029078 |
| 3250 | ENSG0000 | EVX1 | even-skipp | 0 | 0.012936 | 0.029051 |
| 2798 | ENSG0000 | TOX3 | TOX high n | 0.00708 | 0.011639 | 0.029019 |
| 21002 | ENSG0000 | RAB6C | RAB6C, me | 0.002886 | 0.014198 | 0.028988 |
| 16411 | ENSG0000 | EVL | Enah/Vasp | 0.005558 | 0.024965 | 0.028975 |
| 17024 | ENSG0000 | RD3 | retinal deg | 0 | 0.006815 | 0.028968 |
| 4388 | ENSG0000 | DNAH6 | dynein axo | 0.027642 | 0.027844 | 0.028785 |
| 24888 | ENSG0000 | PGM5P4-A | PGM5P4 a | 0 | 0.020331 | 0.028768 |

| | | | | | | |
|---|---|---|---|---|---|---|
| 43427 | ENSG0000 | AC080128.2 | | 0.010524 | 0.019666 | 0.028768 |
| 9583 | ENSG0000 | ABI3BP | ABI family | 0.006501 | 0.025763 | 0.028763 |
| 13600 | ENSG0000 | AP002986.1 | | 0.013553 | 0.022035 | 0.028735 |
| 6753 | ENSG0000 | TSPAN2 | tetraspanin | 0.010445 | 0.025606 | 0.028566 |
| 25301 | ENSG0000 | AL138787.1 | | 0 | 0.021554 | 0.028546 |
| 35803 | ENSG0000 | ALOX12P2 | arachidona | 0.006276 | 0.010431 | 0.028517 |
| 914 | ENSG0000 | CALCRL | calcitonin r | 0 | 0.022758 | 0.028512 |
| 19642 | ENSG0000 | FAM106A | family with | 0 | 0.025581 | 0.028403 |
| 17033 | ENSG0000 | ZNF536 | zinc finger | 0.006241 | 0.018525 | 0.028255 |
| 492 | ENSG0000 | INSRR | insulin rece | 0 | 0.002462 | 0.028212 |
| 22269 | ENSG0000 | LINC00431 | long interg | 0.019106 | 0.027225 | 0.02821 |
| 37516 | ENSG0000 | LRRC2-AS1 | LRRC2 anti | 0 | 0.00763 | 0.028205 |
| 17785 | ENSG0000 | RNU6-124( | RNA, U6 sr | 0 | 0.024534 | 0.028165 |
| 40917 | ENSG0000 | SMAD5-AS | SMAD5 an | 0 | 0.017459 | 0.02814 |
| 23026 | ENSG0000 | HNRNPA1F | heterogen | 0 | 0.015767 | 0.028134 |
| 5588 | ENSG0000 | C1orf61 | chromosor | 0.002882 | 0.007911 | 0.028056 |
| 1476 | ENSG0000 | DCT | dopachron | 0 | 0.024419 | 0.02799 |
| 10270 | ENSG0000 | KALRN | kalirin Rho | 0.005848 | 0.018467 | 0.027885 |
| 43273 | ENSG0000 | MIR4724 | microRNA | 0 | 0.025097 | 0.027842 |
| 40795 | ENSG0000 | RNU2-4P | RNA, U2 sr | 0 | 0.00179 | 0.02782 |
| 24812 | ENSG0000 | LINC01354 | long interg | 0 | 0.010201 | 0.027817 |
| 11399 | ENSG0000 | PHYHIPL | phytanoyl- | 0 | 0.007153 | 0.027756 |
| 25756 | ENSG0000 | ADH5P4 | alcohol del | 0 | 0.026284 | 0.027734 |
| 13173 | ENSG0000 | SLC2A14 | solute carr | 0.008836 | 0.022052 | 0.027527 |
| 15019 | ENSG0000 | AC099489.1 | | 0.015821 | 0.022379 | 0.027487 |
| 35245 | ENSG0000 | AC016597.1 | | 0 | 0.025519 | 0.027335 |
| 25582 | ENSG0000 | AC008163.1 | | 0.00083 | 0.010577 | 0.027308 |
| 8171 | ENSG0000 | FHAD1 | forkhead a | 0 | 0.027085 | 0.027277 |
| 11493 | ENSG0000 | ERICH6B | glutamate | 0.019056 | 0.024791 | 0.027271 |
| 16239 | ENSG0000 | GRAPL | GRB2 relat | 0 | 0.014335 | 0.027236 |
| 16878 | ENSG0000 | FCGR1B | Fc fragmer | 0 | 0.016214 | 0.027235 |
| 13999 | ENSG0000 | YIPF7 | Yip1 doma | 0 | 0.016035 | 0.027227 |
| 25349 | ENSG0000 | CYP1B1-AS | CYP1B1 an | 0.019772 | 0.020285 | 0.027197 |
| 11505 | ENSG0000 | OTOGL | otogelin lik | 0 | 0.006889 | 0.026975 |
| 41837 | ENSG0000 | AC079331.1 | | 0 | 0.012199 | 0.026899 |
| 18308 | ENSG0000 | IGFL2 | IGF like far | 0 | 0.011983 | 0.0268 |
| 37896 | ENSG0000 | ZIM2 | zinc finger | 0 | 0.005385 | 0.026794 |
| 14474 | ENSG0000 | MAB21L1 | mab-21 lik | 0 | 0.009601 | 0.026745 |
| 529 | ENSG0000 | LRRC7 | leucine ricl | 0.010187 | 0.021275 | 0.02668 |
| 15206 | ENSG0000 | KCNK12 | potassium | 0.013311 | 0.017426 | 0.026669 |
| 23920 | ENSG0000 | EEF1DP3 | eukaryotic | 0 | 0.004599 | 0.026661 |
| 34672 | ENSG0000 | CCDC187 | coiled-coil | 0 | 0.008995 | 0.026534 |
| 34641 | ENSG0000 | AC011468.1 | | 0.013876 | 0.01823 | 0.02652 |
| 31461 | ENSG0000 | PCDHGB6 | protocadh | 0 | 0.011826 | 0.026456 |
| 3563 | ENSG0000 | SGCA | sarcoglycar | 0 | 0.023713 | 0.026422 |
| 41060 | ENSG0000 | AC108359.1 | | 0 | 0.023121 | 0.026418 |
| 43375 | ENSG0000 | AL031590.1 | | 0 | 0.00643 | 0.026414 |

| | | | | | | |
|---|---|---|---|---|---|---|
| 16849 | ENSG0000 | CYYR1-AS1 | cysteine ar | 0 | 0.011243 | 0.026235 |
| 38036 | ENSG0000 | AL365273.2 | | 0 | 0.007155 | 0.026151 |
| 13205 | ENSG0000 | SAA1 | serum amy | 0 | 0.006604 | 0.026127 |
| 16286 | ENSG0000 | KIAA0408 | KIAA0408 \| | 0 | 0.011913 | 0.026037 |
| 4750 | ENSG0000 | CNR1 | cannabinoi | 0.005583 | 0.018373 | 0.026028 |
| 28312 | ENSG0000 | RPS29P22 | ribosomal | 0 | 0.014727 | 0.025982 |
| 1942 | ENSG0000 | TF | transferrin | 0.01662 | 0.020985 | 0.025763 |
| 8099 | ENSG0000 | FBN3 | fibrillin 3 [S | 0 | 0.002316 | 0.025738 |
| 34357 | ENSG0000 | AC018868.2 | | 0 | 0.010072 | 0.025729 |
| 2778 | ENSG0000 | CRYM | crystallin m | 0.00381 | 0.014497 | 0.025695 |
| 34334 | ENSG0000 | AC105339.3 | | 0 | 0.023179 | 0.025466 |
| 5514 | ENSG0000 | KLHL31 | kelch like f | 0 | 0.019979 | 0.025402 |
| 33527 | ENSG0000 | AC025569.1 | | 0 | 0.009146 | 0.025381 |
| 9501 | ENSG0000 | CD96 | CD96 mole | 0.009284 | 0.019144 | 0.025338 |
| 5527 | ENSG0000 | ATXN1 | ataxin 1 [S | 0.008619 | 0.019434 | 0.025181 |
| 6327 | ENSG0000 | GFAP | glial fibrilla | 0 | 0.013118 | 0.025151 |
| 39966 | ENSG0000 | AC090241.3 | | 0 | 0.011186 | 0.025148 |
| 23984 | ENSG0000 | KIF25-AS1 | KIF25 antis | 0 | 0.009315 | 0.025107 |
| 35162 | ENSG0000 | AC099524.1 | | 0 | 0.015094 | 0.025005 |
| 5207 | ENSG0000 | HS3ST2 | heparan su | 0 | 0.022359 | 0.024958 |
| 36896 | ENSG0000 | KRT18P61 | keratin 18 | 0 | 0.022477 | 0.024956 |
| 15993 | ENSG0000 | RTP5 | receptor tr | 0 | 0.011156 | 0.024949 |
| 5638 | ENSG0000 | FOXA2 | forkhead b | 0 | 0.016792 | 0.024934 |
| 40057 | ENSG0000 | BSNDP1 | barttin CLC | 0 | 0.002996 | 0.024883 |
| 27614 | ENSG0000 | CASC4P1 | cancer sus | 0 | 0.022158 | 0.024865 |
| 4243 | ENSG0000 | GNAT1 | G protein s | 0 | 0.010556 | 0.024859 |
| 8369 | ENSG0000 | LEFTY2 | left-right d | 0 | 0.024351 | 0.024841 |
| 6464 | ENSG0000 | FCRLA | Fc recepto | 0 | 0.024495 | 0.024798 |
| 40005 | ENSG0000 | AC011700.1 | | 0 | 0.018915 | 0.024749 |
| 14612 | ENSG0000 | ANKRD62 | ankyrin rep | 0 | 0.016013 | 0.024663 |
| 17138 | ENSG0000 | ARHGEF15 | Rho guanir | 0 | 0.005374 | 0.024652 |
| 144 | ENSG0000 | USH1C | USH1 prote | 0 | 0.005949 | 0.024614 |
| 6682 | ENSG0000 | SFTPD | surfactant | 0 | 0.022085 | 0.024609 |
| 13261 | ENSG0000 | WFIKKN2 | WAP, follis | 0 | 0.007321 | 0.024559 |
| 40133 | ENSG0000 | AC026469.1 | | 0 | 0.008794 | 0.024496 |
| 11624 | ENSG0000 | RIC3 | RIC3 acetyl | 0.010652 | 0.01491 | 0.024479 |
| 10892 | ENSG0000 | IQUB | IQ motif ar | 0.017488 | 0.021019 | 0.024405 |
| 6872 | ENSG0000 | DZIP1 | DAZ intera | 0.004191 | 0.010315 | 0.024396 |
| 7798 | ENSG0000 | STON2 | stonin 2 [S | 0 | 0.019599 | 0.024368 |
| 26739 | ENSG0000 | RRM2P2 | ribonucleo | 0 | 0.022294 | 0.024313 |
| 29260 | ENSG0000 | AC108751.5 | | 0 | 0.021915 | 0.024247 |
| 36476 | ENSG0000 | AC069061.4 | | 0.006073 | 0.021506 | 0.024219 |
| 5470 | ENSG0000 | POF1B | premature | 0.008841 | 0.010849 | 0.024177 |
| 10477 | ENSG0000 | FGF11 | fibroblast g | 0.001409 | 0.015081 | 0.024173 |
| 3962 | ENSG0000 | GLP1R | glucagon li | 0 | 0.018726 | 0.024142 |
| 14694 | ENSG0000 | KCNIP1 | potassium | 0 | 0.008675 | 0.024132 |
| 28267 | ENSG0000 | LINC01168 | long interg | 0.006779 | 0.01355 | 0.02411 |

| | | | | | | |
|---|---|---|---|---|---|---|
| 8074 | ENSG0000 | TRPM2 | transient r | 0 | 0.020983 | 0.024079 |
| 22981 | ENSG0000 | LINC01483 | long interg | 0 | 0.013415 | 0.024045 |
| 28524 | ENSG0000 | INMT | indolethyla | 0.010551 | 0.017131 | 0.023988 |
| 28475 | ENSG0000 | ARHGAP8 | Rho GTPas | 0.002805 | 0.023277 | 0.023895 |
| 7 | ENSG0000 | CFH | compleme | 0.004357 | 0.023304 | 0.023877 |
| 19600 | ENSG0000 | EWSAT1 | Ewing sarc | 0 | 0.009229 | 0.023868 |
| 27232 | ENSG0000 | TUBG1P | tubulin gar | 0 | 0.013892 | 0.023854 |
| 41629 | ENSG0000 | AC102945.2 | | 0 | 0.020517 | 0.02383 |
| 22512 | ENSG0000 | NPY6R | neuropept | 0 | 0.015417 | 0.023792 |
| 6997 | ENSG0000 | HEY2 | hes relatec | 0 | 0.021325 | 0.023762 |
| 8178 | ENSG0000 | MYOM3 | myomesin | 0 | 0.012388 | 0.023699 |
| 20184 | ENSG0000 | NOTO | notochord | 0 | 0.019801 | 0.023693 |
| 18309 | ENSG0000 | IGFL4 | IGF like far | 0 | 0.006793 | 0.023671 |
| 42949 | ENSG0000 | FAM169B | family with | 0 | 0.010302 | 0.023626 |
| 18088 | ENSG0000 | C10orf128 | chromosor | 0 | 0.013977 | 0.023598 |
| 488 | ENSG0000 | SLAMF7 | SLAM fami | 0.007395 | 0.009794 | 0.02353 |
| 36521 | ENSG0000 | AC127540.1 | | 0 | 0.021247 | 0.023507 |
| 43078 | ENSG0000 | AL117339.5 | | 0.003198 | 0.02213 | 0.023503 |
| 24525 | ENSG0000 | AC022395.1 | | 0 | 0.007169 | 0.02347 |
| 39647 | ENSG0000 | AC024337.2 | | 0 | 0.00903 | 0.023327 |
| 11379 | ENSG0000 | ARMC3 | armadillo r | 0 | 0.008337 | 0.023305 |
| 15261 | ENSG0000 | NUTM1 | NUT midlir | 0.017037 | 0.02081 | 0.02327 |
| 13919 | ENSG0000 | AC024940.1 | | 0.012594 | 0.015738 | 0.023235 |
| 8497 | ENSG0000 | CNTN4 | contactin 4 | 0 | 0.011195 | 0.023163 |
| 20199 | ENSG0000 | LRRC37A1 | leucine rich | 0 | 0.005149 | 0.023072 |
| 7556 | ENSG0000 | STAT4 | signal trans | 0 | 0.013753 | 0.023028 |
| 2543 | ENSG0000 | SYNDIG1 | synapse di | 0 | 0.010247 | 0.022998 |
| 14917 | ENSG0000 | CHST6 | carbohydra | 0 | 0.006877 | 0.02299 |
| 40080 | ENSG0000 | AC092111.2 | | 0 | 0.01028 | 0.02299 |
| 16103 | ENSG0000 | CFAP77 | cilia and fla | 0.002814 | 0.016751 | 0.022952 |
| 39469 | ENSG0000 | AC096733.2 | | 0.013543 | 0.020707 | 0.022826 |
| 18541 | ENSG0000 | Y_RNA | Y RNA [Sou | 0.005195 | 0.020214 | 0.022805 |
| 13522 | ENSG0000 | GRAMD2A | GRAM don | 0 | 0.005892 | 0.022758 |
| 14284 | ENSG0000 | MKRN3 | makorin rii | 0.006193 | 0.020219 | 0.022669 |
| 42224 | ENSG0000 | RN7SKP23 | RNA, 7SK s | 0 | 0.010163 | 0.022595 |
| 21289 | ENSG0000 | AC073136.1 | | 0 | 0.01008 | 0.022544 |
| 41719 | ENSG0000 | FP236241.2 | | 0 | 0.006133 | 0.022441 |
| 42173 | ENSG0000 | AC124303.2 | | 0 | 0.019494 | 0.022356 |
| 41357 | ENSG0000 | AC023830.3 | | 0 | 0.01998 | 0.022342 |
| 23568 | ENSG0000 | AC097381.1 | | 0 | 0.013682 | 0.022325 |
| 7189 | ENSG0000 | TM6SF1 | transmeml | 0 | 0.017563 | 0.022291 |
| 8115 | ENSG0000 | MLIP | muscular L | 0 | 0.016001 | 0.02224 |
| 16144 | ENSG0000 | ZBED6CL | ZBED6 C-te | 0.012101 | 0.019964 | 0.022089 |
| 3159 | ENSG0000 | SLC5A5 | solute carr | 0 | 0.015921 | 0.022081 |
| 21565 | ENSG0000 | OLFM5P | olfactomec | 0 | 0.002999 | 0.022072 |
| 23981 | ENSG0000 | RPL7P46 | ribosomal | 0 | 0.019643 | 0.022043 |
| 26729 | ENSG0000 | KCNMA1-A | KCNMA1 a | 0 | 0.010491 | 0.022018 |

| | | | | | | |
|---|---|---|---|---|---|---|
| 15016 | ENSG0000 | UBL4B | ubiquitin li | 0 | 0.01967 | 0.021917 |
| 14028 | ENSG0000 | SPATA31E1 | SPATA31 s | 0 | 0.013053 | 0.021702 |
| 23198 | ENSG0000 | LINC01515 | long interg | 0.012342 | 0.016737 | 0.021645 |
| 32683 | ENSG0000 | AC026358.1 | | 0 | 0.012886 | 0.02164 |
| 18244 | ENSG0000 | TRIM31 | tripartite m | 0.01004 | 0.016448 | 0.021638 |
| 10219 | ENSG0000 | DRC7 | dynein reg | 0.00689 | 0.011088 | 0.021632 |
| 15524 | ENSG0000 | ADARB2 | adenosine | 0.005088 | 0.017281 | 0.02146 |
| 15766 | ENSG0000 | KRT14 | keratin 14 | 0 | 0.01279 | 0.021352 |
| 9822 | ENSG0000 | KCNS2 | potassium | 0.00666 | 0.016308 | 0.021264 |
| 5200 | ENSG0000 | MYOG | myogenin | 0 | 0.019026 | 0.0212 |
| 41952 | ENSG0000 | AC133637.1 | | 0 | 0.018915 | 0.021076 |
| 13593 | ENSG0000 | C15orf54 | chromosor | 0 | 0.01899 | 0.020974 |
| 23434 | ENSG0000 | HDAC2-AS2 | HDAC2 anc | 0.007883 | 0.009078 | 0.02085 |
| 3573 | ENSG0000 | PPY | pancreatic | 0 | 0.014832 | 0.020833 |
| 30928 | ENSG0000 | PCDHA12 | protocadhe | 0 | 0.013383 | 0.02083 |
| 26893 | ENSG0000 | VLDLR-AS1 | VLDLR anti | 0 | 0.012425 | 0.020754 |
| 16634 | ENSG0000 | SULT1A2 | sulfotransf | 0 | 0.007885 | 0.020702 |
| 17968 | ENSG0000 | STUM | stum, mecl | 0.009109 | 0.011245 | 0.020695 |
| 33023 | ENSG0000 | AC005842.1 | | 0 | 0.017552 | 0.020688 |
| 38113 | ENSG0000 | PHACTR2P | phosphata | 0 | 0.013423 | 0.020666 |
| 831 | ENSG0000 | RIMBP2 | RIMS bindi | 0.008316 | 0.015002 | 0.020658 |
| 14068 | ENSG0000 | ZC3H12D | zinc finger | 0.007907 | 0.015189 | 0.020536 |
| 35061 | ENSG0000 | AC002310.4 | | 0.005561 | 0.015655 | 0.020467 |
| 14176 | ENSG0000 | RFLNA | refilin A [Sc | 0 | 0.006498 | 0.020418 |
| 30126 | ENSG0000 | FER1L5 | fer-1 like fa | 0 | 0.001504 | 0.020386 |
| 9235 | ENSG0000 | BTBD11 | BTB domai | 0.00675 | 0.007377 | 0.020353 |
| 7431 | ENSG0000 | CASP5 | caspase 5 [ | 0 | 0.018261 | 0.020348 |
| 8344 | ENSG0000 | KCNN3 | potassium | 0 | 0.006414 | 0.020316 |
| 42043 | ENSG0000 | AC004453.2 | | 0 | 0.016472 | 0.020274 |
| 23914 | ENSG0000 | AL391987.3 | | 0 | 0.019977 | 0.020269 |
| 11198 | ENSG0000 | PI16 | peptidase i | 0 | 0.012467 | 0.020248 |
| 10803 | ENSG0000 | PROK2 | prokinetici | 0 | 0.019857 | 0.020238 |
| 9460 | ENSG0000 | RAB3C | RAB3C, me | 0.003662 | 0.012096 | 0.020236 |
| 18671 | ENSG0000 | PRR23A | proline rich | 0 | 0.018276 | 0.020221 |
| 33303 | ENSG0000 | AL445363.1 | | 0 | 0.012296 | 0.020201 |
| 8683 | ENSG0000 | PCDHB18P | protocadhe | 0 | 0.009094 | 0.020193 |
| 3746 | ENSG0000 | CD5 | CD5 molec | 0 | 0.009058 | 0.020185 |
| 8462 | ENSG0000 | GULP1 | GULP, engu | 0.014609 | 0.016246 | 0.02018 |
| 21695 | ENSG0000 | AC245100.1 | | 0 | 0.010766 | 0.020175 |
| 29602 | ENSG0000 | LINC00616 | long interg | 0.012537 | 0.019529 | 0.020109 |
| 22794 | ENSG0000 | AC010890.1 | | 0 | 0.0042 | 0.019996 |
| 23540 | ENSG0000 | AL133410.2 | | 0.003872 | 0.007838 | 0.019858 |
| 9371 | ENSG0000 | ARL11 | ADP ribosy | 0 | 0.015304 | 0.019815 |
| 12704 | ENSG0000 | LVRN | laeverin [Sc | 0 | 0.011015 | 0.019811 |
| 1284 | ENSG0000 | WSCD2 | WSC doma | 0.005432 | 0.011116 | 0.019796 |
| 29402 | ENSG0000 | AP004609.1 | | 0 | 0.012305 | 0.01975 |
| 23037 | ENSG0000 | AL355581.1 | | 0 | 0.006563 | 0.019663 |

| | | | | | | |
|---|---|---|---|---|---|---|
| 23143 | ENSG0000 | FRMD8P1 | FERM dom | 0 | 0.009267 | 0.019597 |
| 9181 | ENSG0000 | WDR17 | WD repeat | 0.004562 | 0.014986 | 0.019373 |
| 16289 | ENSG0000 | TBC1D28 | TBC1 doma | 0.003644 | 0.013847 | 0.019357 |
| 10993 | ENSG0000 | GRIK3 | glutamate | 0.006996 | 0.008626 | 0.019211 |
| 16875 | ENSG0000 | DLGAP2 | DLG associ | 0 | 0.017566 | 0.019179 |
| 4482 | ENSG0000 | PRRX1 | paired rela | 0.004129 | 0.01414 | 0.019176 |
| 9637 | ENSG0000 | PDE1C | phosphodi | 0.015426 | 0.015836 | 0.019144 |
| 30777 | ENSG0000 | HSPD1P11 | heat shock | 0 | 0.017081 | 0.0191 |
| 2115 | ENSG0000 | PRKY | protein kin | 0 | 0.005682 | 0.018936 |
| 2895 | ENSG0000 | IL7 | interleukin | 0 | 0.008571 | 0.018901 |
| 22109 | ENSG0000 | ANKRD18D | ankyrin rep | 0 | 0.010192 | 0.018817 |
| 6172 | ENSG0000 | NCAN | neurocan [ | 0 | 0.012645 | 0.018799 |
| 14181 | ENSG0000 | FOXE1 | forkhead b | 0.010252 | 0.016854 | 0.018747 |
| 19926 | ENSG0000 | VDAC1P2 | voltage de | 0 | 0.017101 | 0.018649 |
| 28742 | ENSG0000 | MBL1P | mannose b | 0 | 0.016495 | 0.018628 |
| 5848 | ENSG0000 | F2RL3 | F2R like th | 0 | 0.0084 | 0.018621 |
| 14473 | ENSG0000 | OR2A4 | olfactory r | 0 | 0.00821 | 0.018524 |
| 15178 | ENSG0000 | CNTN2 | contactin 2 | 0 | 0.000788 | 0.018507 |
| 18382 | ENSG0000 | FAM205A | family with | 0 | 0.00793 | 0.018502 |
| 22058 | ENSG0000 | LINC01671 | long interg | 0 | 0.016262 | 0.018315 |
| 12111 | ENSG0000 | FAM107A | family with | 0 | 0.005464 | 0.018311 |
| 36122 | ENSG0000 | AC016866.1 | | 0 | 0.011566 | 0.018302 |
| 28227 | ENSG0000 | TNFRSF13B | TNF recept | 0 | 0.008189 | 0.018282 |
| 9479 | ENSG0000 | TEKT5 | tektin 5 [Sc | 0 | 0.016494 | 0.01825 |
| 33068 | ENSG0000 | GPR142 | G protein-c | 0 | 0.018144 | 0.018173 |
| 37984 | ENSG0000 | AC005479.3 | | 0 | 0.014591 | 0.018123 |
| 1422 | ENSG0000 | ADCYAP1R | ADCYAP re | 0.005195 | 0.014893 | 0.018018 |
| 16869 | ENSG0000 | AL592490.1 | | 0.008248 | 0.01379 | 0.01799 |
| 33037 | ENSG0000 | ADGRA1-A | adhesion G | 0 | 0.010525 | 0.017933 |
| 30120 | ENSG0000 | AC026369.1 | | 0 | 0.005345 | 0.01793 |
| 8392 | ENSG0000 | GDF7 | growth dif | 0.007294 | 0.008914 | 0.017872 |
| 10082 | ENSG0000 | CPA2 | carboxype | 0 | 0.015153 | 0.017774 |
| 20226 | ENSG0000 | ARHGEF33 | Rho guanir | 0 | 0.003037 | 0.017639 |
| 16024 | ENSG0000 | NHS | NHS actin r | 0.010509 | 0.012907 | 0.017568 |
| 27493 | ENSG0000 | AC234775.2 | | 0 | 0.002244 | 0.017564 |
| 15754 | ENSG0000 | CXCR3 | C-X-C moti | 0 | 0.015121 | 0.017341 |
| 41643 | ENSG0000 | AC244517.6 | | 0 | 0.005343 | 0.017341 |
| 1400 | ENSG0000 | AMPH | amphiphys | 0.012595 | 0.014106 | 0.017324 |
| 11775 | ENSG0000 | MEI1 | meiotic do | 0 | 0.010858 | 0.017289 |
| 3774 | ENSG0000 | CD69 | CD69 mole | 0 | 0.010338 | 0.017198 |
| 32983 | ENSG0000 | PGA5 | pepsinoger | 0 | 0.016969 | 0.017133 |
| 11334 | ENSG0000 | RASEF | RAS and EF | 0 | 0.013148 | 0.017052 |
| 8118 | ENSG0000 | FGD2 | FYVE, RhoG | 0.005224 | 0.012896 | 0.016969 |
| 41887 | ENSG0000 | AL353795.3 | | 0 | 0.015314 | 0.016962 |
| 15936 | ENSG0000 | FAM69C | family with | 0 | 0.007646 | 0.016829 |
| 266 | ENSG0000 | ARAP2 | ArfGAP wit | 0 | 0.011729 | 0.016794 |
| 9369 | ENSG0000 | CYSLTR2 | cysteinyl le | 0 | 0.010901 | 0.016733 |

| | | | | | | |
|---|---|---|---|---|---|---|
| 10611 | ENSG0000 | HFM1 | HFM1, ATF | 0.006112 | 0.01001 | 0.016678 |
| 16185 | ENSG0000 | TMC3 | transmeml | 0 | 0.011993 | 0.016633 |
| 3245 | ENSG0000 | VIPR2 | vasoactive | 0.004943 | 0.005567 | 0.016621 |
| 16662 | ENSG0000 | C6orf141 | chromosor | 0 | 0.009748 | 0.016566 |
| 1934 | ENSG0000 | ABCC6 | ATP bindin | 0.0059 | 0.011969 | 0.016524 |
| 1755 | ENSG0000 | CETP | cholesteryl | 0 | 0.005426 | 0.016488 |
| 42018 | ENSG0000 | AC093330.2 | | 0.004458 | 0.014595 | 0.016363 |
| 8313 | ENSG0000 | SYT14 | synaptotag | 0.002549 | 0.01257 | 0.016332 |
| 40244 | ENSG0000 | HNF1B | HNF1 hom | 0 | 0.00977 | 0.016242 |
| 12665 | ENSG0000 | HS6ST2 | heparan su | 0.007096 | 0.014489 | 0.016236 |
| 6434 | ENSG0000 | LIN28A | lin-28 hom | 0 | 0.014507 | 0.016164 |
| 26801 | ENSG0000 | AC002429.2 | | 0 | 0.003484 | 0.016096 |
| 12817 | ENSG0000 | OR1L8 | olfactory re | 0.00597 | 0.007182 | 0.016005 |
| 40201 | ENSG0000 | CCL4 | C-C motif c | 0 | 0.014307 | 0.015998 |
| 15010 | ENSG0000 | ANKS1B | ankyrin rep | 0.003222 | 0.014576 | 0.015992 |
| 7216 | ENSG0000 | SCN7A | sodium vol | 0 | 0.007104 | 0.015901 |
| 27449 | ENSG0000 | AL590666.3 | | 0 | 0.006465 | 0.015829 |
| 19597 | ENSG0000 | KRTAP2-1 | keratin ass | 0 | 0.011596 | 0.015765 |
| 3946 | ENSG0000 | SLC26A8 | solute carr | 0 | 0.006579 | 0.015753 |
| 2283 | ENSG0000 | GRAP2 | GRB2-relat | 0 | 0.003781 | 0.015718 |
| 1146 | ENSG0000 | SLC12A3 | solute carr | 0 | 0.004737 | 0.015707 |
| 41253 | ENSG0000 | AP000437.1 | | 0 | 0.013993 | 0.015702 |
| 36816 | ENSG0000 | snoZ6 | Z6 small nu | 0 | 0.009344 | 0.015664 |
| 5021 | ENSG0000 | KCNJ5 | potassium | 0 | 0.014293 | 0.015654 |
| 14208 | ENSG0000 | IGSF22 | immunoglo | 0 | 0.013676 | 0.015648 |
| 24808 | ENSG0000 | DLX6-AS1 | DLX6 antise | 0 | 0.006787 | 0.015642 |
| 43065 | ENSG0000 | MIR761 | microRNA | 0.010866 | 0.012104 | 0.015637 |
| 6973 | ENSG0000 | PPP1R1A | protein ph | 0 | 0.006968 | 0.015607 |
| 16962 | ENSG0000 | ZNF813 | zinc finger | 0 | 0.002345 | 0.015527 |
| 6013 | ENSG0000 | KLRC1 | killer cell le | 0 | 0.010775 | 0.015455 |
| 14559 | ENSG0000 | HIST3H2BA | histone clu | 0 | 0.002361 | 0.015455 |
| 2419 | ENSG0000 | NFATC4 | nuclear fac | 0 | 0.003446 | 0.015412 |
| 4281 | ENSG0000 | PLSCR4 | phospholip | 0 | 0.006926 | 0.015409 |
| 1965 | ENSG0000 | JPH4 | junctophili | 0.00085 | 0.001991 | 0.015385 |
| 1469 | ENSG0000 | RIMS1 | regulating | 0 | 0.004563 | 0.015342 |
| 513 | ENSG0000 | MUSK | muscle ass | 0 | 0.013694 | 0.015339 |
| 41692 | ENSG0000 | LINC01451 | long interg | 0.010421 | 0.014951 | 0.01521 |
| 16133 | ENSG0000 | LINC00319 | long interg | 0 | 0.009779 | 0.015157 |
| 15612 | ENSG0000 | KRT5 | keratin 5 [S | 0 | 0.013379 | 0.015122 |
| 35408 | ENSG0000 | LINC02177 | long interg | 0 | 0.005507 | 0.015115 |
| 34501 | ENSG0000 | MOCS1P1 | molybdenu | 0 | 0.011319 | 0.015107 |
| 30595 | ENSG0000 | AC005865.2 | | 0.005612 | 0.005988 | 0.015091 |
| 10732 | ENSG0000 | STPG2 | sperm tail | 0 | 0.002438 | 0.015024 |
| 14061 | ENSG0000 | AMER3 | APC memb | 0 | 0.004428 | 0.014905 |
| 4752 | ENSG0000 | SGIP1 | SH3 domai | 0.006751 | 0.006956 | 0.014845 |
| 3767 | ENSG0000 | POU2AF1 | POU class 2 | 0.004786 | 0.013456 | 0.014826 |
| 29733 | ENSG0000 | LY75-CD30 | LY75-CD30 | 0.001302 | 0.011337 | 0.014734 |

| | | | | | | |
|---|---|---|---|---|---|---|
| 41319 | ENSG0000 | ZNF229 | zinc finger | 0 | 0.010524 | 0.014711 |
| 1687 | ENSG0000 | IGSF9 | immunoglo | 0.004899 | 0.008262 | 0.014662 |
| 15854 | ENSG0000 | DCC | DCC netrin | 0.012898 | 0.014389 | 0.014631 |
| 682 | ENSG0000 | ARID1B | AT-rich inte | 0.007667 | 0.008401 | 0.0146 |
| 22425 | ENSG0000 | LINC00937 | long interg | 0.001934 | 0.006007 | 0.014575 |
| 34643 | ENSG0000 | AC012508.1 | | 0 | 0.013082 | 0.014551 |
| 23747 | ENSG0000 | CCNB1IP1F | cyclin B1 ir | 0 | 0.010813 | 0.014513 |
| 26269 | ENSG0000 | AC009303.2 | | 0 | 0.009105 | 0.014482 |
| 8567 | ENSG0000 | MUC4 | mucin 4, ce | 0 | 0.009688 | 0.014425 |
| 9954 | ENSG0000 | CACNA1D | calcium vo | 0.007134 | 0.013414 | 0.01437 |
| 5409 | ENSG0000 | KCNS1 | potassium | 0.004129 | 0.006429 | 0.014327 |
| 9731 | ENSG0000 | C2CD6 | C2 calcium | 0 | 0.004266 | 0.01431 |
| 5024 | ENSG0000 | TEX11 | testis expr | 0 | 0.008685 | 0.014244 |
| 22014 | ENSG0000 | AC079630.1 | | 0 | 0.012751 | 0.014208 |
| 13023 | ENSG0000 | FUT9 | fucosyltrar | 0.009082 | 0.010603 | 0.014126 |
| 38320 | ENSG0000 | AC136475.9 | | 0 | 0.011823 | 0.014099 |
| 37907 | ENSG0000 | AC007375.3 | | 0 | 0.005034 | 0.014098 |
| 11831 | ENSG0000 | AC087380.1 | | 0 | 0.004811 | 0.014095 |
| 13949 | ENSG0000 | ZBED2 | zinc finger | 0 | 0.012624 | 0.014067 |
| 30330 | ENSG0000 | AC253576.1 | | 0 | 0.00214 | 0.014011 |
| 14 | ENSG0000 | ENPP4 | ectonuclec | 0.007658 | 0.012435 | 0.013954 |
| 11422 | ENSG0000 | EML5 | echinoderr | 0.004817 | 0.006704 | 0.013851 |
| 38938 | ENSG0000 | FAM47E-S | FAM47E-S | 6.26E-05 | 0.004117 | 0.013818 |
| 10089 | ENSG0000 | POM121L2 | POM121 tr | 0 | 0.006625 | 0.013714 |
| 7546 | ENSG0000 | OIT3 | oncoprotei | 0 | 0.012336 | 0.013697 |
| 5828 | ENSG0000 | TAS2R4 | taste 2 rec | 0 | 0.012312 | 0.01367 |
| 15105 | ENSG0000 | KCTD16 | potassium | 0 | 0.011237 | 0.013646 |
| 15879 | ENSG0000 | AGMO | alkylglycer | 0 | 0.008076 | 0.013642 |
| 28592 | ENSG0000 | AC093583.1 | | 0 | 0.012293 | 0.013553 |
| 32478 | ENSG0000 | NOX5 | NADPH oxi | 0.008511 | 0.010922 | 0.01352 |
| 10184 | ENSG0000 | SCUBE1 | signal pept | 0 | 0.009595 | 0.013418 |
| 5257 | ENSG0000 | RECK | reversion i | 0.005913 | 0.012074 | 0.013416 |
| 11167 | ENSG0000 | LINC01600 | long interg | 0 | 0.011986 | 0.013356 |
| 3355 | ENSG0000 | ECM2 | extracellul | 0.006499 | 0.012065 | 0.013249 |
| 9881 | ENSG0000 | SLC6A1 | solute carr | 0 | 0.005949 | 0.013187 |
| 11212 | ENSG0000 | GPR85 | G protein-c | 0 | 0.003818 | 0.01314 |
| 1555 | ENSG0000 | COL4A4 | collagen ty | 0.003625 | 0.008845 | 0.013106 |
| 33177 | ENSG0000 | AC138123.2 | | 0 | 0.00948 | 0.013024 |
| 9811 | ENSG0000 | FUT6 | fucosyltrar | 0 | 0.010051 | 0.013022 |
| 15890 | ENSG0000 | TERF1P2 | telomeric r | 0 | 0.011908 | 0.012984 |
| 38695 | ENSG0000 | AC103724.4 | | 0 | 0.010592 | 0.012982 |
| 31827 | ENSG0000 | AC009812.3 | | 0.009458 | 0.011689 | 0.012932 |
| 14865 | ENSG0000 | ZNF662 | zinc finger | 0 | 0.005775 | 0.012846 |
| 9260 | ENSG0000 | ALLC | allantoicas | 0 | 0.011315 | 0.012743 |
| 8850 | ENSG0000 | CHRNB3 | cholinergic | 0 | 0.011238 | 0.012566 |
| 14229 | ENSG0000 | MTUS2-AS | MTUS2 ant | 0 | 0.011445 | 0.01244 |
| 18599 | ENSG0000 | C16orf96 | chromosor | 0 | 0.010237 | 0.012409 |

| | | | | | | |
|---|---|---|---|---|---|---|
| 20196 | ENSG0000 | MEG3 | maternally | 0 | 0.00555 | 0.012333 |
| 10030 | ENSG0000 | GRHL3 | grainyheac | 0 | 0.004975 | 0.012333 |
| 24127 | ENSG0000 | AL157384.1 | | 0 | 0.011881 | 0.012309 |
| 94 | ENSG0000 | TSPOAP1 | TSPO assoc | 0.006252 | 0.007682 | 0.012275 |
| 10544 | ENSG0000 | PRKAA2 | protein kin | 0.005705 | 0.009324 | 0.01217 |
| 42330 | ENSG0000 | AC004947.2 | | 0 | 0.010961 | 0.012128 |
| 34009 | ENSG0000 | AC020658.2 | | 0 | 0.006873 | 0.012087 |
| 13826 | ENSG0000 | SPNS3 | sphingolipi | 0 | 0.010832 | 0.012069 |
| 2926 | ENSG0000 | ARHGEF10 | Rho guanir | 0 | 0.001238 | 0.012006 |
| 43351 | ENSG0000 | MIR4467 | microRNA | 0.002991 | 0.008821 | 0.012002 |
| 22969 | ENSG0000 | SLC25A6P5 | solute carr | 0 | 0.010667 | 0.011971 |
| 7215 | ENSG0000 | GALNT5 | polypeptid | 0 | 0.004294 | 0.011962 |
| 43308 | ENSG0000 | MIR564 | microRNA | 0.003507 | 0.006686 | 0.011928 |
| 30724 | ENSG0000 | AC091173.1 | | 0 | 0.010609 | 0.011821 |
| 14091 | ENSG0000 | CALML3 | calmodulir | 0 | 0.010596 | 0.011807 |
| 38135 | ENSG0000 | HEATR9 | HEAT repe | 0.006228 | 0.008359 | 0.011738 |
| 10943 | ENSG0000 | FANCD2OS | FANCD2 op | 0.00704 | 0.010475 | 0.011679 |
| 5433 | ENSG0000 | PTGIS | prostaglan | 0 | 0.005248 | 0.011654 |
| 16333 | ENSG0000 | COLCA1 | colorectal | 0 | 0.003352 | 0.011654 |
| 3384 | ENSG0000 | MPDZ | multiple PI | 0.009445 | 0.011504 | 0.011634 |
| 6308 | ENSG0000 | TXLNGY | taxilin gam | 0 | 0.010783 | 0.011506 |
| 16881 | ENSG0000 | ZNF560 | zinc finger | 0 | 0.010251 | 0.011503 |
| 16166 | ENSG0000 | SHISA6 | shisa famil | 0 | 0.010253 | 0.011478 |
| 29569 | ENSG0000 | CICP16 | capicua tra | 0 | 0.000517 | 0.011434 |
| 12169 | ENSG0000 | DPCR1 | diffuse par | 0 | 0.005135 | 0.011423 |
| 25097 | ENSG0000 | AL121895.1 | | 0 | 0.010204 | 0.01133 |
| 40031 | ENSG0000 | AC254629.1 | | 0 | 0.005881 | 0.011258 |
| 33587 | ENSG0000 | AC125611.4 | | 0.002762 | 0.008889 | 0.01125 |
| 1086 | ENSG0000 | MAOB | monoamin | 0 | 0.010092 | 0.011245 |
| 13203 | ENSG0000 | CCDC36 | coiled-coil | 0.005684 | 0.006714 | 0.011142 |
| 8579 | ENSG0000 | CORIN | corin, serir | 0 | 0.006366 | 0.011122 |
| 15527 | ENSG0000 | C11orf87 | chromosor | 0 | 0.004941 | 0.01103 |
| 15858 | ENSG0000 | LUZP2 | leucine zip | 0.004053 | 0.009877 | 0.011005 |
| 24855 | ENSG0000 | AL807752.3 | | 0 | 0.009858 | 0.011004 |
| 14295 | ENSG0000 | SLITRK4 | SLIT and N | 0 | 0.009552 | 0.010902 |
| 41607 | ENSG0000 | AL022323.2 | | 0 | 0.004399 | 0.010798 |
| 6306 | ENSG0000 | RGN | regucalcin | 0 | 0.010415 | 0.010706 |
| 40674 | ENSG0000 | AC092375.3 | | 0 | 0.005106 | 0.010656 |
| 28824 | ENSG0000 | ERVW-1 | endogenou | 0 | 0.007097 | 0.010606 |
| 9329 | ENSG0000 | GABRA2 | gamma-am | 0.002986 | 0.007354 | 0.010582 |
| 15045 | ENSG0000 | MIR345 | microRNA | 0.007706 | 0.01052 | 0.010549 |
| 1565 | ENSG0000 | CACNA1S | calcium vo | 0 | 0.002892 | 0.010504 |
| 42969 | ENSG0000 | AL117258.1 | | 0 | 0.003907 | 0.010501 |
| 41132 | ENSG0000 | AC025279.3 | | 0 | 0.004708 | 0.010493 |
| 6312 | ENSG0000 | SYNE1 | spectrin re | 0.004014 | 0.009324 | 0.010483 |
| 42353 | ENSG0000 | AL137058.3 | | 0 | 0.009381 | 0.010453 |
| 42421 | ENSG0000 | AC018470.1 | | 0 | 0.004711 | 0.010426 |

| | | | | | | |
|---|---|---|---|---|---|---|
| 41654 | ENSG0000 | AL022322.2 | | 0.006121 | 0.009429 | 0.010334 |
| 11197 | ENSG0000 | RAET1E | retinoic aci | 0 | 0.009677 | 0.010325 |
| 16842 | ENSG0000 | HRNR | hornerin [S | 0 | 0.004467 | 0.010087 |
| 6087 | ENSG0000 | ADCY4 | adenylate c | 0 | 0.006837 | 0.010053 |
| 28544 | ENSG0000 | ANOS2P | anosmin 2, | 0 | 0.009266 | 0.009922 |
| 27455 | ENSG0000 | AL138828.1 | | 0 | 0.004415 | 0.00988 |
| 2038 | ENSG0000 | PDE6C | phosphodi | 0 | 0.004995 | 0.009821 |
| 2840 | ENSG0000 | OCA2 | OCA2 mela | 0 | 0.008843 | 0.009791 |
| 1200 | ENSG0000 | RHOBTB1 | Rho relatec | 0 | 0.006026 | 0.009766 |
| 9734 | ENSG0000 | SPAG17 | sperm asso | 0.004426 | 0.009015 | 0.00969 |
| 12616 | ENSG0000 | HTRA3 | HtrA serine | 0 | 0.008754 | 0.009685 |
| 30901 | ENSG0000 | AC139713.2 | | 0.002774 | 0.004315 | 0.009682 |
| 36508 | ENSG0000 | MIR3661 | microRNA | 0 | 0.001195 | 0.00968 |
| 3186 | ENSG0000 | HPN | hepsin [Sou | 0 | 0.00857 | 0.009584 |
| 11488 | ENSG0000 | VWA2 | von Willeb | 0.008505 | 0.008651 | 0.009558 |
| 43356 | ENSG0000 | AC105233.5 | | 0 | 0.007947 | 0.009308 |
| 6851 | ENSG0000 | DSC3 | desmocolli | 0 | 0.00827 | 0.009198 |
| 10910 | ENSG0000 | MYRFL | myelin reg | 0 | 0.008279 | 0.009193 |
| 9180 | ENSG0000 | GPM6A | glycoprote | 0 | 0.008332 | 0.00917 |
| 4995 | ENSG0000 | MOB3B | MOB kinas | 0 | 0.008219 | 0.009158 |
| 8723 | ENSG0000 | SCML4 | sex comb c | 0 | 0.0041 | 0.009137 |
| 9584 | ENSG0000 | ANGPT1 | angiopoiet | 0 | 0.004079 | 0.009089 |
| 9043 | ENSG0000 | SERPING1 | serpin fam | 0 | 0.008097 | 0.009023 |
| 12585 | ENSG0000 | SGCD | sarcoglycar | 0 | 0.002766 | 0.009008 |
| 13332 | ENSG0000 | AC020659.1 | | 0 | 0.002822 | 0.008931 |
| 904 | ENSG0000 | SNCAIP | synuclein a | 0 | 0.003983 | 0.008907 |
| 2438 | ENSG0000 | HNF4A | hepatocyte | 0.004871 | 0.008009 | 0.008893 |
| 10867 | ENSG0000 | CADPS | calcium de | 0.002425 | 0.007958 | 0.008868 |
| 1396 | ENSG0000 | ITGA8 | integrin su | 0 | 0.007751 | 0.008698 |
| 2474 | ENSG0000 | CHRNA4 | cholinergic | 0.003224 | 0.007912 | 0.008666 |
| 9025 | ENSG0000 | SLC5A12 | solute carr | 0 | 0.003853 | 0.008619 |
| 13789 | ENSG0000 | SPATA31C2 | SPATA31 su | 0 | 0.007724 | 0.008576 |
| 8028 | ENSG0000 | CBLN2 | cerebellin 2 | 0 | 0.002599 | 0.008441 |
| 3389 | ENSG0000 | GLIS3 | GLIS family | 0.003071 | 0.00504 | 0.008408 |
| 26004 | ENSG0000 | MAGI2-AS3 | MAGI2 ant | 0 | 0.000756 | 0.008369 |
| 28783 | ENSG0000 | LINC00882 | long interg | 0 | 0.003771 | 0.008344 |
| 31870 | ENSG0000 | PCDHGA3 | protocadhe | 0 | 0.007484 | 0.008304 |
| 3544 | ENSG0000 | ASIC2 | acid sensin | 0 | 0.005194 | 0.008297 |
| 4227 | ENSG0000 | RBP1 | retinol binc | 0 | 0.003643 | 0.008295 |
| 35631 | ENSG0000 | SNORD3B- | small nucle | 0 | 0.004236 | 0.008278 |
| 11749 | ENSG0000 | MS4A14 | membrane | 0 | 0.007309 | 0.008172 |
| 22675 | ENSG0000 | AC013470.2 | | 0 | 0.007335 | 0.00813 |
| 23628 | ENSG0000 | ANKRD20A | ankyrin rep | 0.003759 | 0.006675 | 0.008026 |
| 15249 | ENSG0000 | NCMAP | non-compa | 0 | 0.003527 | 0.007914 |
| 39076 | ENSG0000 | AC007938.4 | | 0 | 0.007085 | 0.007894 |
| 37320 | ENSG0000 | AP005131.6 | | 0 | 0.006946 | 0.007821 |
| 37776 | ENSG0000 | AC011452.1 | | 0.003003 | 0.006922 | 0.007784 |

| | | | | | | |
|---|---|---|---|---|---|---|
| 26718 | ENSG0000 | AC010894.3 | | 0 | 0.004474 | 0.00776 |
| 10696 | ENSG0000 | FAM84A | family with | 0 | 0.006239 | 0.007728 |
| 12329 | ENSG0000 | OR5AU1 | olfactory r | 0.004236 | 0.006914 | 0.007718 |
| 10873 | ENSG0000 | SYNPR | synaptopo | 0 | 0.006907 | 0.007669 |
| 4613 | ENSG0000 | ST6GALNA | ST6 N-acet | 0 | 0.003483 | 0.007652 |
| 12987 | ENSG0000 | SPTLC3 | serine palm | 0 | 0.003436 | 0.007631 |
| 26674 | ENSG0000 | AC004448.4 | | 0 | 0.0046 | 0.007616 |
| 42659 | ENSG0000 | TEX13D | TEX13 fam | 0 | 0.006797 | 0.007575 |
| 15977 | ENSG0000 | COL14A1 | collagen ty | 0 | 0.005067 | 0.007532 |
| 34119 | ENSG0000 | AC103740.1 | | 0 | 0.006813 | 0.007524 |
| 18301 | ENSG0000 | SPATA31A1 | SPATA31 s | 0 | 0.006677 | 0.00744 |
| 41879 | ENSG0000 | AC009948.5 | | 0 | 0.006645 | 0.007404 |
| 1878 | ENSG0000 | RGS1 | regulator c | 0 | 0.007137 | 0.007396 |
| 10349 | ENSG0000 | CXCR5 | C-X-C moti | 0 | 0.006561 | 0.007337 |
| 11243 | ENSG0000 | ADCY1 | adenylate c | 0.003963 | 0.004343 | 0.007216 |
| 41277 | ENSG0000 | SSTR3 | somatosta | 0 | 0.006431 | 0.007191 |
| 2652 | ENSG0000 | KLF8 | Kruppel lik | 0 | 0.003234 | 0.00718 |
| 37259 | ENSG0000 | AC021683.3 | | 0 | 0.006302 | 0.007047 |
| 20483 | ENSG0000 | AL445238.1 | | 0 | 0.006267 | 0.006983 |
| 22769 | ENSG0000 | LINC01359 | long interg | 0 | 0.004409 | 0.006958 |
| 13399 | ENSG0000 | UGT8 | UDP glycos | 0 | 0.006209 | 0.006894 |
| 6743 | ENSG0000 | CHL1 | cell adhesi | 0 | 0.004954 | 0.006837 |
| 3840 | ENSG0000 | RASAL1 | RAS protei | 0 | 0.006061 | 0.006754 |
| 41569 | ENSG0000 | AL137141.1 | | 0 | 0.006012 | 0.006698 |
| 20224 | ENSG0000 | C10orf105 | chromosor | 0 | 0.005894 | 0.006683 |
| 22388 | ENSG0000 | PRICKLE2-A | PRICKLE2 a | 0 | 0.00619 | 0.006554 |
| 34872 | ENSG0000 | AC009093.2 | | 0 | 0.005097 | 0.00651 |
| 10697 | ENSG0000 | KCNJ3 | potassium | 0 | 0.005737 | 0.006438 |
| 42876 | ENSG0000 | AC087280.2 | | 0 | 0.003877 | 0.006406 |
| 29251 | ENSG0000 | OR2A1-AS1 | OR2A1 ant | 0 | 0.005741 | 0.006397 |
| 24189 | ENSG0000 | DDX10P2 | DEAD-box | 0 | 0.005521 | 0.006196 |
| 23345 | ENSG0000 | LINC01185 | long interg | 0 | 0.003811 | 0.005963 |
| 2665 | ENSG0000 | FGF14 | fibroblast g | 0.002579 | 0.004249 | 0.005961 |
| 7443 | ENSG0000 | ITGA11 | integrin su | 0 | 0.005371 | 0.005942 |
| 11512 | ENSG0000 | AGBL2 | ATP/GTP b | 0 | 0.004977 | 0.005929 |
| 15397 | ENSG0000 | GOLGA8DP | golgin A8 f | 0 | 0.004537 | 0.005912 |
| 32451 | ENSG0000 | OR5BA1P | olfactory r | 0.000321 | 0.0053 | 0.005906 |
| 614 | ENSG0000 | TG | thyroglobu | 0 | 0.005223 | 0.00582 |
| 42457 | ENSG0000 | AJ239328.1 | | 0 | 0.004584 | 0.005742 |
| 11086 | ENSG0000 | GUCY1A3 | guanylate c | 0 | 0.005042 | 0.005658 |
| 33071 | ENSG0000 | HP | haptoglobi | 0 | 0.005398 | 0.005621 |
| 14494 | ENSG0000 | ANKRD30B | ankyrin rep | 0 | 0.004877 | 0.005435 |
| 20434 | ENSG0000 | LINC00598 | long interg | 0.000158 | 0.000309 | 0.005402 |
| 5182 | ENSG0000 | SV2C | synaptic ve | 0 | 0.002451 | 0.005386 |
| 14665 | ENSG0000 | GOLGA8CP | golgin A8 f | 0.000697 | 0.001529 | 0.005285 |
| 11190 | ENSG0000 | TMEM200A | transmemb | 0 | 0.004737 | 0.005279 |
| 20111 | ENSG0000 | C1orf167 | chromosor | 0.000659 | 0.000951 | 0.005276 |

| | | | | | | |
|---|---|---|---|---|---|---|
| 22669 | ENSG0000 | TEX41 | testis expr | 0.00366 | 0.003795 | 0.005226 |
| 1566 | ENSG0000 | PKP1 | plakophilin | 0 | 0.004697 | 0.005215 |
| 11250 | ENSG0000 | SLC30A8 | solute carr | 0 | 0.00463 | 0.005159 |
| 25233 | ENSG0000 | KIZ-AS1 | KIZ antisen | 0 | 0.001286 | 0.005068 |
| 6141 | ENSG0000 | PRRG3 | proline rich | 0 | 0.004568 | 0.005054 |
| 33110 | ENSG0000 | CLLU1 | chronic lyn | 0 | 0.004284 | 0.004858 |
| 8802 | ENSG0000 | CXorf36 | chromosor | 0 | 0.002965 | 0.004855 |
| 7860 | ENSG0000 | GOLGA6D | golgin A6 f | 0 | 0.004207 | 0.004771 |
| 8247 | ENSG0000 | ILDR2 | immunoglc | 0 | 0.00418 | 0.004707 |
| 9757 | ENSG0000 | AFF2 | AF4/FMR2 | 0 | 0.004166 | 0.004633 |
| 23083 | ENSG0000 | AC072062.1 | | 0 | 0.003812 | 0.004609 |
| 9780 | ENSG0000 | ADAMTS3 | ADAM met | 0 | 0.004053 | 0.004581 |
| 30109 | ENSG0000 | CARMN | cardiac me | 0 | 0.002281 | 0.004572 |
| 2875 | ENSG0000 | CALB1 | calbindin 1 | 0 | 0.003708 | 0.00449 |
| 10317 | ENSG0000 | ZNF208 | zinc finger | 0 | 0.004219 | 0.004434 |
| 42293 | ENSG0000 | AC006548.3 | | 0.000427 | 0.001732 | 0.004432 |
| 22646 | ENSG0000 | AC067942.1 | | 0.004327 | 0.004419 | 0.004429 |
| 6793 | ENSG0000 | ANO3 | anoctamin | 0 | 0.002543 | 0.004222 |
| 650 | ENSG0000 | ROS1 | ROS proto- | 0 | 7.84E-05 | 0.004182 |
| 42075 | ENSG0000 | AC068448.1 | | 0 | 0.003703 | 0.004156 |
| 15374 | ENSG0000 | FLRT2 | fibronectin | 0.001021 | 0.002502 | 0.00374 |
| 7766 | ENSG0000 | SRRM4 | serine/argi | 0 | 0.00325 | 0.003595 |
| 16846 | ENSG0000 | ZNF677 | zinc finger | 0 | 0.003055 | 0.003478 |
| 7478 | ENSG0000 | IFI44L | interferon | 0 | 0.002477 | 0.003174 |
| 43221 | ENSG0000 | MIR7855 | microRNA | 0 | 0.00169 | 0.003138 |
| 9843 | ENSG0000 | UNC5D | unc-5 netri | 0 | 0.002688 | 0.003026 |
| 16443 | ENSG0000 | ESRRG | estrogen re | 0 | 0.002664 | 0.002969 |
| 40392 | ENSG0000 | MIR6818 | microRNA | 0 | 0.002379 | 0.002651 |
| 43322 | ENSG0000 | MIR4723 | microRNA | 0.000039 | 0.001294 | 0.002553 |
| 22872 | ENSG0000 | LINC01630 | long interg | 0 | 0.002169 | 0.002417 |
| 5319 | ENSG0000 | ITIH5 | inter-alpha | 0 | 0.001987 | 0.002222 |
| 40755 | ENSG0000 | HYDIN2 | HYDIN2, ax | 0 | 0.001981 | 0.002214 |
| 16933 | ENSG0000 | CACNA1E | calcium vo | 0 | 0.001636 | 0.001823 |
| 22618 | ENSG0000 | CICP17 | capicua tra | 0 | 0.00155 | 0.001704 |
| 15064 | ENSG0000 | GOLGA8G | golgin A8 f | 0 | 0.000745 | 0.000839 |
| 42979 | ENSG0000 | MIR6830 | microRNA | 0 | 0.000736 | 0.000821 |
| 36414 | ENSG0000 | AC009084.1 | | 0 | 0.000662 | 0.000725 |
| 18560 | ENSG0000 | SNORD21 | small nucle | 0.000418 | 0.000619 | 0.000637 |
| 43083 | ENSG0000 | MIR3917 | microRNA | 0.000298 | 0.000382 | 0.000552 |
| 36622 | ENSG0000 | AC087393.3 | | 0 | 0.000373 | 0.000461 |
| 38029 | ENSG0000 | AC010997.3 | | 0 | 6.5E-05 | 7.21E-05 |
| 19176 | ENSG0000 | SNORA48 | small nucle | 0.00001 | 0.000016 | 0.00002 |
| 36179 | ENSG0000 | SNORD95 | small nucle | 0.000004 | 0.00001 | 0.000017 |
