## Supplementary Table S7 for "Molecular profiling of XPO1 inhibitor and gemcitabine-nab-paclitaxel combination in cellular and LSL-Kras G12D/+; Trp53 fl/+; Pdx1-Cre (KPC) pancreatic cancer model"

| ID | gene_id | gene_nam | Descriptior | FPKMMia_ | FPKMMia_ | FPKMMia_Comb |
|---|---|---|---|---|---|---|
| 2068 | ENSG0000 | HSP90AB1 | heat shock | 1475.553 | 970.2953 | 828.2642 |
| 257 | ENSG0000 | VIM | vimentin [S | 932.0973 | 751.9406 | 714.522 |
| 962 | ENSG0000 | YBX1 | Y-box bind | 819.4009 | 707.7249 | 623.254 |
| 1500 | ENSG0000 | HSP90AA1 | heat shock | 541.8988 | 407.955 | 363.5016 |
| 42459 | ENSG0000 | FP236383.2 | | 413.9112 | 382.9158 | 344.1888 |
| 42487 | ENSG0000 | FP671120.3 | | 405.2669 | 372.6671 | 335.5116 |
| 3516 | ENSG0000 | PFN1 | profilin 1 [S | 400.2773 | 276.4836 | 232.8374 |
| 7363 | ENSG0000 | HMGA1 | high mobil | 394.443 | 172.9014 | 172.472 |
| 13810 | ENSG0000 | NPM1 | nucleopho | 386.2254 | 292.8586 | 213.6961 |
| 16364 | ENSG0000 | PPIA | peptidylpro | 343.9865 | 280.8191 | 270.0926 |
| 29680 | ENSG0000 | MTATP6P1 | mitochond | 322.1021 | 140.2634 | 41.89375 |
| 42531 | ENSG0000 | FP236383.3 | | 319.4685 | 287.2165 | 259.1218 |
| 3691 | ENSG0000 | HSPA8 | heat shock | 314.0257 | 246.777 | 201.008 |
| 7296 | ENSG0000 | ANP32B | acidic nucl | 242.6993 | 167.2287 | 144.4196 |
| 8181 | ENSG0000 | RPL11 | ribosomal | 239.2966 | 195.9959 | 193.4852 |
| 8464 | ENSG0000 | HSPD1 | heat shock | 234.9916 | 151.6276 | 123.6987 |
| 12955 | ENSG0000 | CALB2 | calbindin 2 | 220.2751 | 72.16252 | 58.42411 |
| 13368 | ENSG0000 | RPL4 | ribosomal | 219.6774 | 169.0994 | 164.1791 |
| 11338 | ENSG0000 | HNRNPK | heterogen | 211.5305 | 162.2323 | 151.1101 |
| 1655 | ENSG0000 | EIF3I | eukaryotic | 204.6872 | 156.6341 | 154.7508 |
| 6519 | ENSG0000 | EIF5A | eukaryotic | 203.9598 | 126.8945 | 99.99256 |
| 13798 | ENSG0000 | TUFM | Tu translat | 200.8322 | 143.7463 | 137.9745 |
| 5238 | ENSG0000 | HNRNPA2B | heterogen | 198.6255 | 131.0495 | 130.4619 |
| 628 | ENSG0000 | HSPA5 | heat shock | 198.3531 | 137.5439 | 114.0699 |
| 2426 | ENSG0000 | PYGB | glycogen p | 195.2495 | 135.4793 | 118.0112 |
| 4829 | ENSG0000 | SET | SET nuclea | 188.0319 | 162.3331 | 130.986 |
| 16419 | ENSG0000 | XRCC6 | X-ray repai | 184.2881 | 134.0772 | 131.0545 |
| 7007 | ENSG0000 | CCT7 | chaperonir | 184.2447 | 149.9557 | 131.227 |
| 16715 | ENSG0000 | HNRNPAB | heterogen | 177.2525 | 99.36469 | 77.5255 |
| 8347 | ENSG0000 | ILF2 | interleukin | 174.206 | 145.0648 | 121.6566 |
| 8879 | ENSG0000 | RPL7 | ribosomal | 173.046 | 137.1191 | 135.9606 |
| 4651 | ENSG0000 | CDC20 | cell divisio | 166.974 | 89.07318 | 81.20977 |
| 36485 | ENSG0000 | RPL17 | ribosomal | 163.2363 | 112.4225 | 102.4506 |
| 4325 | ENSG0000 | NCL | nucleolin [S | 160.9018 | 144.7288 | 91.95365 |
| 1847 | ENSG0000 | PEBP1 | phosphatic | 153.2224 | 148.5587 | 142.9899 |
| 3584 | ENSG0000 | YWHAE | tyrosine 3- | 153.0656 | 119.4245 | 115.7071 |
| 18477 | ENSG0000 | EIF3CL | eukaryotic | 150.0183 | 108.28 | 100.1821 |
| 8769 | ENSG0000 | CCT6A | chaperonir | 148.0245 | 110.6912 | 82.88884 |
| 10818 | ENSG0000 | CCT3 | chaperonir | 145.1415 | 109.5064 | 102.6456 |
| 4118 | ENSG0000 | HSPA9 | heat shock | 135.393 | 123.4827 | 109.6634 |
| 4312 | ENSG0000 | EEF1B2 | eukaryotic | 134.729 | 88.6056 | 86.75938 |
| 48 | ENSG0000 | POLDIP2 | DNA polym | 132.1228 | 113.5807 | 109.4221 |
| 11652 | ENSG0000 | MCM7 | minichrom | 128.5122 | 71.47008 | 64.13769 |
| 7289 | ENSG0000 | PSMB7 | proteasom | 127.196 | 82.23739 | 74.97201 |
| 8368 | ENSG0000 | ARF1 | ADP ribosy | 126.6993 | 94.52652 | 89.60404 |
| 6985 | ENSG0000 | HNRNPA1 | heterogen | 126.5492 | 63.36398 | 58.55693 |

| | | | | | | |
|---|---|---|---|---|---|---|
| 11675 | ENSG0000 | HSP90B1 | heat shock | 125.0857 | 76.49761 | 72.63431 |
| 6257 | ENSG0000 | TRIM28 | tripartite n | 124.0838 | 79.65654 | 67.52219 |
| 9796 | ENSG0000 | CCT8 | chaperonir | 123.4612 | 123.0053 | 107.8797 |
| 12453 | ENSG0000 | SF3B5 | splicing fac | 123.1518 | 80.3476 | 75.72411 |
| 33324 | ENSG0000 | BLOC1S5-T | BLOC1S5-T | 122.93 | 70.30203 | 56.36565 |
| 11703 | ENSG0000 | PPIB | peptidylpr | 122.8112 | 79.91131 | 79.42536 |
| 1077 | ENSG0000 | NUCKS1 | nuclear cas | 120.4492 | 90.94545 | 89.65653 |
| 12134 | ENSG0000 | STIP1 | stress indu | 119.7968 | 85.12825 | 82.17828 |
| 5430 | ENSG0000 | CSE1L | chromosor | 119.3462 | 86.84439 | 82.49909 |
| 4307 | ENSG0000 | EIF4G1 | eukaryotic | 116.073 | 90.32769 | 79.56601 |
| 13562 | ENSG0000 | DRAP1 | DR1 associ | 113.6242 | 62.48908 | 58.15966 |
| 17163 | ENSG0000 | TOP1 | topoisome | 111.4281 | 109.8928 | 91.97653 |
| 3523 | ENSG0000 | C1QBP | compleme | 111.1778 | 90.49941 | 77.8569 |
| 4355 | ENSG0000 | PPM1G | protein ph | 109.0911 | 75.74908 | 68.80701 |
| 15079 | ENSG0000 | ALYREF | Aly/REF ex | 107.2962 | 55.45862 | 48.72279 |
| 1789 | ENSG0000 | KHSRP | KH-type sp | 107.0802 | 90.65306 | 83.91229 |
| 9195 | ENSG0000 | CCT5 | chaperonir | 106.3588 | 85.49926 | 69.50395 |
| 867 | ENSG0000 | U2AF2 | U2 small n | 106.3478 | 77.68987 | 60.47884 |
| 8199 | ENSG0000 | SERBP1 | SERPINE1 r | 106.2464 | 79.27835 | 64.91293 |
| 8798 | ENSG0000 | MSN | moesin [Sc | 105.8651 | 105.7827 | 98.52478 |
| 7250 | ENSG0000 | TXN | thioredoxi | 105.3342 | 79.76066 | 79.47936 |
| 4536 | ENSG0000 | SRM | spermidine | 105.3302 | 57.56432 | 34.02852 |
| 35776 | ENSG0000 | MRPL12 | mitochond | 103.6307 | 55.44695 | 53.36125 |
| 5554 | ENSG0000 | GOT2 | glutamic-o | 103.5001 | 74.70201 | 68.28673 |
| 37818 | ENSG0000 | AC022149.1 | | 100.4915 | 85.09444 | 79.70648 |
| 12413 | ENSG0000 | HNRNPF | heterogen | 100.4098 | 68.85263 | 63.19055 |
| 12536 | ENSG0000 | KRT8 | keratin 8 [S | 100.0216 | 87.16081 | 67.35816 |
| 467 | ENSG0000 | STRAP | serine/thre | 99.43346 | 87.96394 | 79.76669 |
| 15503 | ENSG0000 | UBE2L3 | ubiquitin c | 98.01224 | 64.09664 | 61.85828 |
| 5742 | ENSG0000 | PRMT1 | protein arg | 97.26182 | 68.48871 | 55.59313 |
| 6378 | ENSG0000 | ACLY | ATP citrate | 97.01415 | 88.87579 | 85.84921 |
| 11470 | ENSG0000 | DDX21 | DExD-box l | 96.99461 | 80.90355 | 51.01286 |
| 10903 | ENSG0000 | STOML2 | stomatin li | 96.09746 | 86.52023 | 84.73795 |
| 6602 | ENSG0000 | HMGB1P5 | high mobil | 95.16289 | 71.63344 | 67.91337 |
| 10901 | ENSG0000 | VCP | valosin cor | 94.13085 | 61.85503 | 53.92674 |
| 6247 | ENSG0000 | ADRM1 | adhesion r | 93.68288 | 73.76706 | 64.60221 |
| 12477 | ENSG0000 | HNRNPA3 | heterogen | 93.30827 | 73.7055 | 69.94309 |
| 8259 | ENSG0000 | PFDN2 | prefoldin s | 92.11097 | 66.32465 | 54.04823 |
| 12564 | ENSG0000 | PA2G4 | proliferatic | 91.43484 | 74.12469 | 52.67883 |
| 3416 | ENSG0000 | EIF3A | eukaryotic | 90.79463 | 89.80486 | 66.83257 |
| 7483 | ENSG0000 | RPL7P9 | ribosomal | 90.38126 | 70.87605 | 64.20156 |
| 3954 | ENSG0000 | MCM3 | minichrom | 90.12896 | 68.94202 | 61.6888 |
| 3714 | ENSG0000 | CCDC86 | coiled-coil | 89.99973 | 76.55608 | 50.6796 |
| 10800 | ENSG0000 | ATP1A1 | ATPase Na | 89.89946 | 83.62625 | 82.43611 |
| 3715 | ENSG0000 | PRPF19 | pre-mRNA | 89.79121 | 67.86096 | 67.85762 |
| 7195 | ENSG0000 | SRSF1 | serine and | 89.34314 | 74.03672 | 54.90246 |
| 10382 | ENSG0000 | NACC1 | nucleus ac | 88.93138 | 71.49063 | 68.32253 |

| | | | | | | |
|---|---|---|---|---|---|---|
| 3500 | ENSG0000 | KPNB1 | karyopheri | 86.19378 | 60.46195 | 45.98421 |
| 7997 | ENSG0000 | ARHGDIA | Rho GDP d | 85.63368 | 78.27354 | 69.1739 |
| 2770 | ENSG0000 | SLC7A5 | solute carr | 85.26761 | 55.78341 | 48.12307 |
| 8813 | ENSG0000 | NONO | non-POU d | 84.39926 | 65.01636 | 64.39252 |
| 6940 | ENSG0000 | NT5E | 5'-nucleoti | 84.0686 | 67.69463 | 63.61455 |
| 11577 | ENSG0000 | NOLC1 | nucleolar a | 83.76434 | 69.77825 | 50.49578 |
| 15326 | ENSG0000 | SUMO3 | small ubiqu | 83.50653 | 72.40498 | 67.17954 |
| 13571 | ENSG0000 | FOSL1 | FOS like 1, | 82.78459 | 71.48494 | 63.09127 |
| 9045 | ENSG0000 | SSRP1 | structure s | 82.76899 | 64.84215 | 60.64361 |
| 10675 | ENSG0000 | CAPN2 | calpain 2 [S | 82.32399 | 73.87095 | 56.86104 |
| 936 | ENSG0000 | KARS | lysyl-tRNA | 82.31622 | 56.70198 | 49.01547 |
| 35710 | ENSG0000 | AC021224.1 | | 79.50916 | 33.88595 | 31.68424 |
| 4529 | ENSG0000 | SFPQ | splicing fac | 79.32103 | 50.68222 | 33.96461 |
| 13211 | ENSG0000 | PPP1R14B | protein ph | 78.3682 | 37.27016 | 34.1854 |
| 483 | ENSG0000 | TOMM34 | translocase | 77.67248 | 64.92062 | 62.31958 |
| 1977 | ENSG0000 | SUPT16H | SPT16 hom | 77.6611 | 57.6913 | 53.49476 |
| 9682 | ENSG0000 | GTF3C6 | general tra | 77.04021 | 71.47468 | 64.36558 |
| 12229 | ENSG0000 | MAT2A | methionine | 76.96736 | 74.11768 | 61.30016 |
| 15196 | ENSG0000 | IRAK1 | interleukin | 76.51196 | 54.96539 | 53.21669 |
| 732 | ENSG0000 | THRAP3 | thyroid hor | 75.20608 | 41.54446 | 36.87089 |
| 8670 | ENSG0000 | NHP2 | NHP2 ribor | 74.86492 | 55.26717 | 49.70821 |
| 12581 | ENSG0000 | HSPA4 | heat shock | 74.57292 | 66.33248 | 61.12924 |
| 6956 | ENSG0000 | CAPRIN1 | cell cycle a | 74.33413 | 58.19876 | 52.00839 |
| 325 | ENSG0000 | PTBP1 | polypyrimi | 73.98295 | 55.18553 | 48.05399 |
| 3567 | ENSG0000 | LRRC59 | leucine ricl | 73.88209 | 59.13801 | 50.18324 |
| 352 | ENSG0000 | ELOVL5 | ELOVL fatty | 72.21899 | 46.10899 | 45.97858 |
| 13494 | ENSG0000 | PSMD2 | proteasom | 72.09048 | 59.28843 | 56.20718 |
| 5434 | ENSG0000 | STAU1 | staufen do | 71.31232 | 61.56903 | 61.19978 |
| 14809 | ENSG0000 | ANXA2 | annexin A2 | 71.10531 | 56.63633 | 46.13568 |
| 42840 | ENSG0000 | AC004922.1 | | 70.99205 | 35.67643 | 19.69989 |
| 6160 | ENSG0000 | TOMM40 | translocase | 70.66381 | 49.5782 | 39.02675 |
| 14206 | ENSG0000 | RCC2 | regulator c | 70.6454 | 60.38714 | 58.30519 |
| 12607 | ENSG0000 | KIF5B | kinesin fan | 69.95082 | 63.69929 | 59.76733 |
| 359 | ENSG0000 | PSMC4 | proteasom | 69.8538 | 57.01404 | 49.37498 |
| 12393 | ENSG0000 | FASN | fatty acid s | 69.77043 | 58.26077 | 47.9321 |
| 12712 | ENSG0000 | PRPF8 | pre-mRNA | 68.91279 | 64.00533 | 59.85183 |
| 1975 | ENSG0000 | HNRNPC | heterogene | 68.56422 | 49.76936 | 49.08961 |
| 1448 | ENSG0000 | XRCC5 | X-ray repai | 68.08056 | 56.28289 | 52.52344 |
| 7402 | ENSG0000 | MRPL15 | mitochond | 68.0354 | 54.71826 | 50.17246 |
| 4824 | ENSG0000 | RAD23B | RAD23 hor | 67.34169 | 54.82667 | 51.02551 |
| 16966 | ENSG0000 | ASNA1 | arsA arseni | 66.59394 | 43.29643 | 39.69705 |
| 3876 | ENSG0000 | USP5 | ubiquitin sp | 66.35928 | 43.33565 | 37.31191 |
| 8284 | ENSG0000 | SF3B4 | splicing fac | 66.21201 | 62.91997 | 53.42433 |
| 6938 | ENSG0000 | SYNCRIP | synaptotag | 66.14557 | 48.7935 | 39.59941 |
| 753 | ENSG0000 | SZRD1 | SUZ RNA b | 65.61074 | 63.06728 | 57.40277 |
| 8940 | ENSG0000 | POLE3 | DNA polym | 65.28371 | 53.11649 | 52.76343 |
| 7940 | ENSG0000 | NOB1 | NIN1/PSM | 64.96198 | 64.59427 | 59.04843 |

| | | | | | | |
|---|---|---|---|---|---|---|
| 3485 | ENSG0000 | PSMD3 | proteasom | 64.95095 | 47.21618 | 43.4852 |
| 3439 | ENSG0000 | NPM3 | nucleopho | 64.90887 | 18.74452 | 18.10093 |
| 3145 | ENSG0000 | PPP2R1A | protein ph | 64.88529 | 47.33368 | 46.0778 |
| 12714 | ENSG0000 | CKAP5 | cytoskelet | 64.20207 | 56.43181 | 54.96792 |
| 14819 | ENSG0000 | RPS17 | ribosomal | 64.03579 | 49.05214 | 48.07589 |
| 6804 | ENSG0000 | NARS | asparaginy | 63.64907 | 62.65673 | 57.12611 |
| 34378 | ENSG0000 | HMGB1P6 | high mobil | 62.79012 | 42.68718 | 37.62356 |
| 16977 | ENSG0000 | BZW1P2 | basic leuci | 62.55259 | 56.04928 | 52.5569 |
| 2942 | ENSG0000 | HNRNPL | heterogen | 62.25622 | 41.98719 | 36.33497 |
| 6422 | ENSG0000 | PDHA1 | pyruvate d | 62.14276 | 54.4239 | 51.28331 |
| 8401 | ENSG0000 | CALM2 | calmodulir | 61.77148 | 44.46832 | 42.69345 |
| 19730 | ENSG0000 | HNRNPA3F | heterogen | 61.54736 | 48.63825 | 42.6831 |
| 5727 | ENSG0000 | RBM42 | RNA bindir | 61.39627 | 55.17436 | 51.90308 |
| 6487 | ENSG0000 | RAN | RAN, mem | 61.25415 | 43.18681 | 36.62173 |
| 14967 | ENSG0000 | SF3A3 | splicing fac | 61.15839 | 52.7429 | 48.40564 |
| 4547 | ENSG0000 | MFN2 | mitofusin 2 | 61.10621 | 49.84278 | 44.19707 |
| 2518 | ENSG0000 | MAPRE1 | microtubul | 60.78191 | 53.54227 | 47.93391 |
| 1852 | ENSG0000 | FUS | FUS RNA b | 60.0833 | 52.0054 | 46.28145 |
| 2075 | ENSG0000 | HNRNPH3 | heterogen | 59.90657 | 37.85385 | 37.31861 |
| 14037 | ENSG0000 | IMPDH2 | inosine mc | 59.7893 | 50.61652 | 48.08009 |
| 5469 | ENSG0000 | USP22 | ubiquitin s | 59.20645 | 55.35548 | 50.20205 |
| 41427 | ENSG0000 | C11orf98 | chromosor | 58.8906 | 42.01062 | 36.92181 |
| 7259 | ENSG0000 | FAM129B | family with | 58.72104 | 37.82053 | 29.03585 |
| 243 | ENSG0000 | BAZ1B | bromodom | 58.61208 | 47.45561 | 41.74227 |
| 35172 | ENSG0000 | BOP1 | block of pr | 57.93867 | 33.51373 | 25.76259 |
| 1777 | ENSG0000 | AURKA | aurora kina | 57.65487 | 39.48939 | 35.31118 |
| 3026 | ENSG0000 | SGTA | small gluta | 57.56165 | 53.17015 | 52.17484 |
| 12306 | ENSG0000 | PRELID1 | PRELI dom | 57.53436 | 35.53167 | 34.636 |
| 14131 | ENSG0000 | GTPBP6 | GTP bindin | 57.2976 | 33.7919 | 19.12068 |
| 15172 | ENSG0000 | EIF3C | eukaryotic | 56.94016 | 42.68653 | 39.48674 |
| 166 | ENSG0000 | CCDC124 | coiled-coil | 56.93585 | 35.92405 | 34.86387 |
| 6076 | ENSG0000 | ILF3 | interleukin | 56.87626 | 46.72877 | 40.73403 |
| 13314 | ENSG0000 | GNG5 | G protein s | 56.57621 | 45.27164 | 40.31118 |
| 40472 | ENSG0000 | UHRF1 | ubiquitin li | 56.15461 | 31.05217 | 28.35087 |
| 12059 | ENSG0000 | SF1 | splicing fac | 56.15091 | 50.80575 | 40.02311 |
| 8916 | ENSG0000 | SIGMAR1 | sigma non- | 56.03405 | 34.8894 | 27.58774 |
| 16379 | ENSG0000 | IARS | isoleucyl-tl | 55.97308 | 40.67566 | 39.89814 |
| 14223 | ENSG0000 | FARSA | phenylalan | 55.35845 | 45.34136 | 42.73518 |
| 2968 | ENSG0000 | SF3A2 | splicing fac | 55.34093 | 44.57305 | 43.04251 |
| 842 | ENSG0000 | MRPS35 | mitochond | 55.33743 | 54.63802 | 47.0609 |
| 8179 | ENSG0000 | SH3BGRL3 | SH3 domai | 55.00347 | 49.2295 | 48.35707 |
| 2417 | ENSG0000 | PNN | pinin, desn | 54.88885 | 51.16191 | 41.39915 |
| 16194 | ENSG0000 | NOC2L | NOC2 like r | 54.67969 | 39.11415 | 32.71346 |
| 8828 | ENSG0000 | PRPS1 | phosphorib | 54.62156 | 43.03203 | 39.59753 |
| 1654 | ENSG0000 | WBP11 | WW doma | 54.25425 | 47.41238 | 43.2448 |
| 666 | ENSG0000 | FAM120A | family with | 53.6262 | 46.95403 | 36.89827 |
| 8100 | ENSG0000 | CARM1 | coactivator | 53.44061 | 39.88124 | 36.5591 |

| | | | | | | |
|---|---|---|---|---|---|---|
| 12893 | ENSG0000 | RRM2 | ribonucleo | 53.22127 | 31.50127 | 31.41496 |
| 8057 | ENSG0000 | SH3GL1 | SH3 domai | 52.64055 | 37.51895 | 37.49847 |
| 5018 | ENSG0000 | TCP1 | t-complex | 52.59875 | 39.82147 | 37.00587 |
| 2117 | ENSG0000 | HNRNPM | heterogen | 52.43961 | 46.41329 | 45.29794 |
| 9710 | ENSG0000 | LARP1 | La ribonucl | 52.24385 | 43.7158 | 35.22912 |
| 1249 | ENSG0000 | IGF2BP2 | insulin like | 52.20715 | 48.45924 | 40.35277 |
| 4519 | ENSG0000 | ATP5F1 | ATP syntha | 51.9318 | 39.05492 | 37.46261 |
| 26492 | ENSG0000 | AP005202.1 | | 51.8996 | 43.13385 | 41.29981 |
| 21732 | ENSG0000 | HNRNPA1F | heterogen | 51.37401 | 25.0182 | 22.90378 |
| 673 | ENSG0000 | VAMP3 | vesicle ass | 51.20477 | 45.88686 | 45.7384 |
| 11584 | ENSG0000 | CCT2 | chaperonir | 51.07666 | 50.38098 | 38.41755 |
| 13253 | ENSG0000 | EIF1AX | eukaryotic | 51.03323 | 42.51891 | 37.73208 |
| 5795 | ENSG0000 | HNRNPH2 | heterogen | 50.53189 | 29.34925 | 27.98805 |
| 5053 | ENSG0000 | ETF1 | eukaryotic | 50.33456 | 41.30706 | 36.95263 |
| 10373 | ENSG0000 | UBQLN4 | ubiquilin 4 | 50.22744 | 41.07965 | 37.03078 |
| 8301 | ENSG0000 | CERS2 | ceramide s | 50.1733 | 21.26068 | 20.95054 |
| 16983 | ENSG0000 | TXNRD1 | thioredoxi | 49.95941 | 29.29481 | 25.77318 |
| 5756 | ENSG0000 | TRAP1 | TNF recept | 49.95197 | 26.62932 | 25.7895 |
| 16395 | ENSG0000 | WDR5 | WD repeat | 49.88169 | 43.78537 | 37.30375 |
| 3716 | ENSG0000 | TMEM109 | transmeml | 49.82623 | 40.88944 | 37.96736 |
| 20180 | ENSG0000 | RPL7P1 | ribosomal | 49.67465 | 35.62361 | 34.58285 |
| 9084 | ENSG0000 | MTA2 | metastasis | 49.4419 | 43.069 | 41.28003 |
| 6491 | ENSG0000 | CLUH | clustered r | 48.754 | 45.06061 | 31.19816 |
| 3539 | ENSG0000 | DDX5 | DEAD-box | 48.72586 | 40.21854 | 29.69525 |
| 16294 | ENSG0000 | HMGB1 | high mobil | 48.56942 | 35.35036 | 34.69605 |
| 7553 | ENSG0000 | ATIC | 5-aminoim | 48.55873 | 38.94705 | 30.0006 |
| 2396 | ENSG0000 | APEX1 | apurinic/ap | 48.20761 | 46.70518 | 41.38221 |
| 2457 | ENSG0000 | PRPF6 | pre-mRNA | 47.89364 | 41.47186 | 34.66376 |
| 9846 | ENSG0000 | AIFM1 | apoptosis i | 47.61696 | 35.75524 | 34.73102 |
| 10407 | ENSG0000 | PSMC2 | proteasom | 47.54385 | 33.79925 | 32.63724 |
| 8338 | ENSG0000 | UBAP2L | ubiquitin a | 47.43002 | 45.51328 | 36.24203 |
| 8415 | ENSG0000 | SNRNP200 | small nucle | 47.39873 | 46.71103 | 46.55357 |
| 5729 | ENSG0000 | UBA2 | ubiquitin li | 47.14779 | 47.00218 | 44.43437 |
| 6374 | ENSG0000 | PSME3 | proteasom | 47.1245 | 40.14597 | 10.72088 |
| 5197 | ENSG0000 | MRPS2 | mitochond | 46.96465 | 42.18951 | 38.01826 |
| 29320 | ENSG0000 | BX679664.3 | | 46.92724 | 30.56472 | 29.21885 |
| 2324 | ENSG0000 | PRMT5 | protein arg | 46.85393 | 40.81254 | 35.08112 |
| 3225 | ENSG0000 | OGDH | oxoglutara | 46.78574 | 35.9093 | 34.66426 |
| 3953 | ENSG0000 | MRPL18 | mitochond | 46.77611 | 40.61487 | 31.49916 |
| 9451 | ENSG0000 | HNRNPDL | heterogen | 46.66639 | 37.04811 | 31.4522 |
| 11572 | ENSG0000 | API5 | apoptosis i | 46.55572 | 43.43836 | 42.68463 |
| 11779 | ENSG0000 | PHB | prohibitin | 46.52088 | 44.06958 | 39.88344 |
| 1645 | ENSG0000 | STARD7 | StAR relate | 46.42918 | 40.7902 | 35.82078 |
| 14400 | ENSG0000 | RCC1 | regulator c | 46.41104 | 32.66552 | 26.74629 |
| 13527 | ENSG0000 | BANF1 | barrier to a | 46.32107 | 31.52761 | 30.28274 |
| 2149 | ENSG0000 | SMARCB1 | SWI/SNF re | 46.31431 | 39.11327 | 26.98296 |
| 2994 | ENSG0000 | CHCHD3 | coiled-coil- | 46.24302 | 42.77261 | 37.75451 |

| | | | | | | |
|---|---|---|---|---|---|---|
| 18224 | ENSG0000 | MRPS18B | mitochond | 45.91495 | 45.04576 | 42.59699 |
| 4705 | ENSG0000 | ESYT2 | extended s | 45.872 | 43.89097 | 39.83019 |
| 16809 | ENSG0000 | ATAD3A | ATPase fan | 45.14046 | 29.837 | 22.60392 |
| 6286 | ENSG0000 | DKC1 | dyskerin ps | 45.06301 | 37.38471 | 25.1245 |
| 6004 | ENSG0000 | VPS4A | vacuolar pr | 44.93815 | 32.40957 | 28.19157 |
| 19833 | ENSG0000 | TMX2 | thioredoxi | 44.92157 | 33.60752 | 29.46211 |
| 1940 | ENSG0000 | FH | fumarate h | 44.67483 | 32.23069 | 30.61149 |
| 12952 | ENSG0000 | CYCS | cytochrom | 44.60747 | 34.89322 | 29.99891 |
| 519 | ENSG0000 | SARS | seryl-tRNA | 44.4213 | 33.96511 | 33.86754 |
| 4014 | ENSG0000 | SLC39A7 | solute carr | 44.04074 | 41.46893 | 40.44504 |
| 4175 | ENSG0000 | PPP2CA | protein ph | 44.02424 | 37.48819 | 35.6501 |
| 28481 | ENSG0000 | PSMC1P1 | proteasom | 43.59159 | 35.22961 | 31.17924 |
| 18166 | ENSG0000 | VARS | valyl-tRNA | 43.11881 | 29.85119 | 21.75335 |
| 7609 | ENSG0000 | HNRNPD | heterogen | 43.00272 | 29.63913 | 29.19158 |
| 13987 | ENSG0000 | POLR2L | RNA polym | 42.59741 | 27.76358 | 25.88185 |
| 2728 | ENSG0000 | PSMD7 | proteasom | 42.57131 | 21.56347 | 19.57336 |
| 3273 | ENSG0000 | EIF3B | eukaryotic | 42.51753 | 38.43948 | 29.16301 |
| 527 | ENSG0000 | ABCF2 | ATP bindin | 42.24644 | 35.21817 | 26.03973 |
| 4348 | ENSG0000 | NRBP1 | nuclear rec | 42.11542 | 39.4231 | 35.78705 |
| 2547 | ENSG0000 | APMAP | adipocyte | 41.95133 | 39.33143 | 38.34243 |
| 11013 | ENSG0000 | TKT | transketola | 41.94139 | 21.31561 | 19.83847 |
| 4048 | ENSG0000 | FARSB | phenylalan | 41.73128 | 33.48999 | 21.94922 |
| 16616 | ENSG0000 | PCBP2 | poly(rC) bi | 41.56104 | 33.98478 | 33.27618 |
| 16179 | ENSG0000 | RPL14 | ribosomal | 40.70534 | 33.66296 | 33.49989 |
| 3110 | ENSG0000 | CDC37 | cell divisio | 40.60675 | 38.65833 | 34.15127 |
| 6165 | ENSG0000 | XPO7 | exportin 7 | 40.60399 | 33.91722 | 33.25904 |
| 4028 | ENSG0000 | DBN1 | drebrin 1 [ | 40.31362 | 35.12191 | 33.34912 |
| 4502 | ENSG0000 | MRPL37 | mitochond | 40.27863 | 38.08701 | 34.49692 |
| 2161 | ENSG0000 | SF3A1 | splicing fac | 40.13532 | 28.53215 | 25.23326 |
| 1912 | ENSG0000 | KIF4A | kinesin fan | 39.8361 | 24.68697 | 23.63361 |
| 7569 | ENSG0000 | OLA1 | Obg like AT | 39.71014 | 39.24163 | 37.94494 |
| 16922 | ENSG0000 | TFDP1 | transcripti | 39.45929 | 33.63585 | 32.79958 |
| 14250 | ENSG0000 | RAD23A | RAD23 hor | 39.29843 | 32.63431 | 29.60292 |
| 2261 | ENSG0000 | TTLL12 | tubulin tyr | 39.25541 | 29.51381 | 26.18562 |
| 6212 | ENSG0000 | LSM4 | LSM4 hom | 39.25344 | 24.51208 | 24.16974 |
| 7883 | ENSG0000 | IQGAP1 | IQ motif cc | 39.1795 | 37.26823 | 32.48541 |
| 14839 | ENSG0000 | RBM10 | RNA bindir | 38.99129 | 29.31203 | 26.8329 |
| 14987 | ENSG0000 | PSMG1 | proteasom | 38.92817 | 31.15736 | 27.38368 |
| 1339 | ENSG0000 | UNG | uracil DNA | 38.66526 | 21.93694 | 20.21112 |
| 6279 | ENSG0000 | EIF3G | eukaryotic | 38.63189 | 32.38083 | 26.27072 |
| 4290 | ENSG0000 | RRP9 | ribosomal | 38.43383 | 26.76342 | 19.14593 |
| 4425 | ENSG0000 | HDLBP | high densit | 38.22077 | 38.02482 | 34.06547 |
| 22338 | ENSG0000 | NOL7 | nucleolar p | 38.14216 | 36.57728 | 33.09249 |
| 3458 | ENSG0000 | PITRM1 | pitrilysin m | 37.99797 | 32.3427 | 28.54467 |
| 966 | ENSG0000 | ELAVL1 | ELAV like R | 37.98772 | 27.53227 | 27.41355 |
| 9489 | ENSG0000 | HNRNPU | heterogen | 37.94853 | 28.68826 | 23.78264 |
| 4280 | ENSG0000 | MRPL3 | mitochond | 37.85405 | 34.79021 | 23.70243 |

| | | | | | | |
|---|---|---|---|---|---|---|
| 2515 | ENSG0000 | NOP56 | NOP56 ribo | 37.84663 | 37.22866 | 25.63958 |
| 16632 | ENSG0000 | SND1 | staphyloco | 37.83397 | 24.31532 | 23.33762 |
| 10144 | ENSG0000 | FBXW5 | F-box and ' | 37.80994 | 36.76434 | 28.87068 |
| 8422 | ENSG0000 | TEX261 | testis expr | 37.79226 | 36.36293 | 31.94931 |
| 4368 | ENSG0000 | RTN4 | reticulon 4 | 37.78078 | 37.71788 | 36.45148 |
| 15409 | ENSG0000 | RAB11B | RAB11B, m | 37.60484 | 29.67026 | 28.08388 |
| 546 | ENSG0000 | FAM136A | family with | 37.59379 | 32.67989 | 27.02927 |
| 30029 | ENSG0000 | NPM1P27 | nucleopho | 37.55001 | 26.47575 | 19.03553 |
| 13605 | ENSG0000 | RUVBL1 | RuvB like A | 37.49732 | 30.31508 | 23.14245 |
| 8923 | ENSG0000 | MFSD14B | major facil | 37.39219 | 35.78611 | 31.03661 |
| 4489 | ENSG0000 | CACYBP | calcyclin bi | 37.28477 | 28.83374 | 23.35696 |
| 2032 | ENSG0000 | NUP188 | nucleopori | 37.14516 | 22.89733 | 22.41801 |
| 851 | ENSG0000 | CS | citrate syn | 37.09422 | 24.24451 | 6.396501 |
| 10478 | ENSG0000 | EIF4A1 | eukaryotic | 37.04842 | 20.71407 | 17.11631 |
| 10151 | ENSG0000 | GART | phosphorib | 37.03443 | 29.62509 | 27.69394 |
| 2302 | ENSG0000 | RANGAP1 | Ran GTPas | 36.85608 | 31.21621 | 23.93181 |
| 152 | ENSG0000 | PAF1 | PAF1 homo | 36.84485 | 36.78442 | 31.70908 |
| 5488 | ENSG0000 | RRP36 | ribosomal | 36.60735 | 26.32053 | 24.44329 |
| 3951 | ENSG0000 | SRSF3 | serine and | 36.49306 | 31.50314 | 28.49189 |
| 2889 | ENSG0000 | EIF3E | eukaryotic | 36.47697 | 34.37414 | 31.76274 |
| 8836 | ENSG0000 | RBMX | RNA bindir | 36.45279 | 24.47646 | 21.06141 |
| 1261 | ENSG0000 | CLNS1A | chloride nu | 35.79946 | 31.88139 | 18.61433 |
| 17022 | ENSG0000 | DDX39B | DExD-box l | 35.72794 | 26.06792 | 20.65113 |
| 12257 | ENSG0000 | UQCRFS1 | ubiquinol-c | 35.65709 | 27.45122 | 25.74692 |
| 2378 | ENSG0000 | MTHFD1 | methylene | 35.6202 | 23.31505 | 23.06052 |
| 2928 | ENSG0000 | MCM4 | minichrom | 35.61517 | 24.48994 | 22.92209 |
| 16444 | ENSG0000 | IPO4 | importin 4 | 35.49479 | 17.84539 | 10.08379 |
| 5681 | ENSG0000 | NCLN | nicalin [Sou | 35.36619 | 19.78967 | 13.26257 |
| 1769 | ENSG0000 | SF3B2 | splicing fac | 35.30631 | 27.9386 | 26.96808 |
| 13422 | ENSG0000 | SRP72 | signal reco | 35.22528 | 32.65495 | 28.42578 |
| 13214 | ENSG0000 | SMARCC1 | SWI/SNF re | 35.20898 | 30.42627 | 26.64896 |
| 722 | ENSG0000 | MRTO4 | MRT4 hom | 35.2013 | 28.32099 | 20.45468 |
| 19978 | ENSG0000 | TAX1BP3 | Tax1 bindi | 35.19574 | 35.0148 | 31.97221 |
| 18266 | ENSG0000 | AKT1S1 | AKT1 subst | 35.10552 | 25.15145 | 21.58662 |
| 8266 | ENSG0000 | MRPL24 | mitochond | 35.04367 | 18.35588 | 17.3627 |
| 13256 | ENSG0000 | PSMD1 | proteasom | 35.00653 | 25.83279 | 22.13236 |
| 14203 | ENSG0000 | RRS1 | ribosome b | 34.93147 | 25.86315 | 15.61234 |
| 29609 | ENSG0000 | CDK11B | cyclin depe | 34.49703 | 27.54385 | 23.31693 |
| 13545 | ENSG0000 | SART1 | SART1, U4/ | 34.47174 | 26.95152 | 21.95518 |
| 363 | ENSG0000 | SLC25A39 | solute carr | 34.32674 | 32.1081 | 28.52166 |
| 6282 | ENSG0000 | DNMT1 | DNA methy | 34.27613 | 26.0456 | 22.46111 |
| 41087 | ENSG0000 | SENP3-EIF4 | SENP3-EIF4 | 34.25004 | 21.48762 | 16.43742 |
| 4832 | ENSG0000 | RAB14 | RAB14, me | 34.23105 | 28.00307 | 27.64128 |
| 1719 | ENSG0000 | SEPHS1 | selenophos | 34.1209 | 23.29104 | 21.11505 |
| 3280 | ENSG0000 | AIMP2 | aminoacyl | 34.08922 | 24.67759 | 20.11954 |
| 36366 | ENSG0000 | RBM8A | RNA bindir | 34.00172 | 27.56263 | 24.95971 |
| 3327 | ENSG0000 | POLD2 | DNA polym | 33.71087 | 18.51181 | 18.11531 |

| | | | | | | |
|---|---|---|---|---|---|---|
| 29277 | ENSG0000 | AC006011.1 | | 33.51141 | 32.08088 | 31.85972 |
| 13800 | ENSG0000 | PPA1 | pyrophosp | 33.30512 | 25.75769 | 25.30413 |
| 18227 | ENSG0000 | ABCF1 | ATP bindin | 33.12084 | 31.92536 | 22.99576 |
| 892 | ENSG0000 | TNPO3 | transportir | 33.07982 | 29.13063 | 26.37792 |
| 2228 | ENSG0000 | RTCB | RNA 2',3'-c | 32.9198 | 28.28117 | 23.76344 |
| 13841 | ENSG0000 | THAP4 | THAP dom | 32.8851 | 25.43842 | 25.39014 |
| 2305 | ENSG0000 | ACO2 | aconitase 2 | 32.77006 | 16.98561 | 15.73272 |
| 7614 | ENSG0000 | GPAT3 | glycerol-3- | 32.62206 | 30.05146 | 28.77741 |
| 21859 | ENSG0000 | YBX1P1 | Y-box bind | 32.46593 | 25.88876 | 20.67 |
| 322 | ENSG0000 | UTP18 | UTP18, sm | 32.38473 | 29.96698 | 23.65729 |
| 6479 | ENSG0000 | IMMT | inner mem | 32.30869 | 25.45792 | 24.24001 |
| 2756 | ENSG0000 | USP10 | ubiquitin s | 32.27649 | 28.00446 | 26.32605 |
| 917 | ENSG0000 | AP3D1 | adaptor re | 32.24427 | 26.86572 | 25.80253 |
| 8672 | ENSG0000 | BOD1 | biorientati | 32.23725 | 23.9308 | 18.86526 |
| 1174 | ENSG0000 | CPSF1 | cleavage a | 32.23377 | 29.7348 | 27.53101 |
| 1164 | ENSG0000 | TRIP13 | thyroid ho | 32.18365 | 21.95535 | 21.21367 |
| 4525 | ENSG0000 | SCAMP3 | secretory c | 32.14763 | 24.33237 | 22.50597 |
| 2126 | ENSG0000 | POLR2E | RNA polym | 32.06266 | 17.92538 | 16.01997 |
| 38438 | ENSG0000 | SRXN1 | sulfiredoxi | 32.00322 | 27.45482 | 24.87159 |
| 27973 | ENSG0000 | PGAM5 | PGAM fam | 31.93462 | 19.96548 | 19.26356 |
| 1426 | ENSG0000 | NRDC | nardilysin c | 31.91755 | 28.25486 | 26.96876 |
| 5644 | ENSG0000 | CENPB | centromer | 31.89745 | 23.66036 | 22.73307 |
| 3191 | ENSG0000 | GSK3A | glycogen s | 31.76797 | 26.47278 | 25.96741 |
| 40349 | ENSG0000 | AATF | apoptosis a | 31.74673 | 28.35346 | 20.80014 |
| 2170 | ENSG0000 | PES1 | pescadillo | 31.67775 | 24.42403 | 20.23928 |
| 14196 | ENSG0000 | MRFAP1 | Morf4 fam | 31.64029 | 27.53465 | 24.54455 |
| 2681 | ENSG0000 | PARP4 | poly(ADP-r | 31.43924 | 24.37663 | 23.37494 |
| 10136 | ENSG0000 | CCAR2 | cell cycle a | 31.3459 | 28.08234 | 23.21627 |
| 7541 | ENSG0000 | ANXA7 | annexin A7 | 31.30696 | 24.31215 | 19.2541 |
| 5291 | ENSG0000 | DDX54 | DEAD-box | 31.27716 | 21.44867 | 14.13978 |
| 1375 | ENSG0000 | SNRPA | small nucle | 31.09391 | 7.49768 | 7.074593 |
| 10433 | ENSG0000 | SRSF2 | serine and | 31.02671 | 14.59884 | 13.02697 |
| 2062 | ENSG0000 | SRPK1 | SRSF prote | 30.99034 | 30.92193 | 28.92623 |
| 4158 | ENSG0000 | TARS | threonyl-tF | 30.96418 | 28.00311 | 23.25406 |
| 565 | ENSG0000 | RPL26L1 | ribosomal | 30.87751 | 25.55158 | 23.09396 |
| 15564 | ENSG0000 | LAMP1 | lysosomal | 30.82595 | 22.93724 | 22.85575 |
| 8003 | ENSG0000 | EIF4A3 | eukaryotic | 30.70569 | 19.96403 | 19.23642 |
| 8183 | ENSG0000 | ZNF593 | zinc finger | 30.633 | 18.10551 | 14.86821 |
| 2635 | ENSG0000 | CDK16 | cyclin depe | 30.49067 | 28.69606 | 27.75976 |
| 7133 | ENSG0000 | BZW2 | basic leuci | 30.45309 | 25.44581 | 21.30336 |
| 3452 | ENSG0000 | GTPBP4 | GTP bindin | 30.41309 | 30.32004 | 23.17888 |
| 7935 | ENSG0000 | COPS3 | COP9 signa | 30.33398 | 21.82493 | 21.60868 |
| 1535 | ENSG0000 | ARL2BP | ADP ribosy | 30.29662 | 28.19254 | 26.7637 |
| 12034 | ENSG0000 | DDB1 | damage sp | 30.28965 | 27.21427 | 25.64417 |
| 11780 | ENSG0000 | SNRPD1 | small nucle | 30.26311 | 20.66193 | 20.41906 |
| 3392 | ENSG0000 | NPDC1 | neural prol | 30.10491 | 29.35506 | 27.79235 |
| 10476 | ENSG0000 | SENP3 | SUMO1/se | 30.07708 | 26.58798 | 25.91349 |

| | | | | | | |
|---|---|---|---|---|---|---|
| 7335 | ENSG0000 | PPIL1 | peptidylpr | 30.06486 | 22.91697 | 19.27736 |
| 14659 | ENSG0000 | COA4 | cytochrom | 30.01849 | 22.5089 | 19.84825 |
| 9582 | ENSG0000 | TOMM70 | translocase | 29.76382 | 24.87263 | 21.52448 |
| 9377 | ENSG0000 | ATP5A1 | ATP syntha | 29.70212 | 22.14816 | 21.43663 |
| 16488 | ENSG0000 | RABL6 | RAB, meml | 29.57814 | 19.68803 | 19.23839 |
| 145 | ENSG0000 | GGCT | gamma-glu | 29.55965 | 26.2219 | 24.52659 |
| 6646 | ENSG0000 | CSNK1G2 | casein kina | 29.48044 | 27.74553 | 27.16859 |
| 1903 | ENSG0000 | PABPC4 | poly(A) bin | 29.42768 | 21.56419 | 18.8379 |
| 16840 | ENSG0000 | TEAD4 | TEA domai | 29.35384 | 27.67906 | 20.29131 |
| 2810 | ENSG0000 | NOMO1 | NODAL mc | 29.21903 | 23.20435 | 22.2086 |
| 10884 | ENSG0000 | GMPS | guanine m | 29.21081 | 27.80668 | 25.45538 |
| 9927 | ENSG0000 | SAFB | scaffold at | 29.20805 | 25.0082 | 18.57212 |
| 3297 | ENSG0000 | PLOD3 | procollage | 29.11506 | 22.05166 | 20.62716 |
| 1790 | ENSG0000 | GNA11 | G protein s | 29.0601 | 25.61944 | 16.87205 |
| 34569 | ENSG0000 | NORAD | non-coding | 28.96345 | 27.78901 | 25.90382 |
| 5683 | ENSG0000 | HNRNPR | heterogen | 28.94267 | 22.50715 | 19.3108 |
| 6721 | ENSG0000 | EIF2S1 | eukaryotic | 28.93207 | 26.85852 | 24.16232 |
| 4518 | ENSG0000 | WDR77 | WD repeat | 28.92387 | 26.21712 | 18.2896 |
| 5854 | ENSG0000 | SMARCA4 | SWI/SNF re | 28.9032 | 19.21052 | 17.42529 |
| 2256 | ENSG0000 | MCM5 | minichrom | 28.80703 | 13.12761 | 11.04972 |
| 3403 | ENSG0000 | DVL1 | dishevelled | 28.77244 | 16.15365 | 14.69264 |
| 15855 | ENSG0000 | TAF9B | TATA-box b | 28.65195 | 22.12379 | 21.24177 |
| 2079 | ENSG0000 | ABL1 | ABL proto- | 28.47546 | 26.40187 | 25.04738 |
| 13645 | ENSG0000 | SLC35A4 | solute carr | 28.41446 | 22.37656 | 13.77927 |
| 1028 | ENSG0000 | IDH3G | isocitrate d | 28.40107 | 20.28323 | 19.79737 |
| 10129 | ENSG0000 | NDUFS2 | NADH:ubic | 28.30759 | 22.46954 | 20.23545 |
| 14856 | ENSG0000 | EWSR1 | EWS RNA b | 28.08473 | 20.4254 | 16.95111 |
| 13994 | ENSG0000 | FLII | FLII, actin r | 28.03419 | 26.86303 | 21.68894 |
| 8263 | ENSG0000 | PRCC | papillary re | 27.847 | 25.75944 | 23.3525 |
| 10816 | ENSG0000 | ARPC2 | actin relate | 27.71195 | 24.08963 | 23.97738 |
| 16803 | ENSG0000 | MCMBP | minichrom | 27.68494 | 22.16931 | 20.88141 |
| 19214 | ENSG0000 | TSN | translin [Sc | 27.62305 | 25.89747 | 25.06989 |
| 192 | ENSG0000 | DNAJC11 | DnaJ heat s | 27.58606 | 18.2648 | 16.54091 |
| 7295 | ENSG0000 | NCBP1 | nuclear cap | 27.54326 | 22.53443 | 20.56264 |
| 1541 | ENSG0000 | CTCF | CCCTC-bind | 27.49258 | 26.67414 | 22.24187 |
| 6858 | ENSG0000 | TIMM10 | translocase | 27.44788 | 24.88486 | 22.37745 |
| 15918 | ENSG0000 | TMEM203 | transmeml | 27.35404 | 19.36001 | 18.67598 |
| 12004 | ENSG0000 | MRPL58 | mitochond | 27.35378 | 17.66491 | 14.91382 |
| 13340 | ENSG0000 | DDX23 | DEAD-box | 27.33856 | 22.96417 | 21.25977 |
| 2805 | ENSG0000 | MAZ | MYC assoc | 27.32483 | 18.19491 | 17.91809 |
| 2553 | ENSG0000 | USP14 | ubiquitin s | 27.27647 | 27.02605 | 24.38677 |
| 12876 | ENSG0000 | CTPS1 | CTP syntha | 27.25029 | 19.38439 | 12.17192 |
| 16182 | ENSG0000 | RPSAP47 | ribosomal | 27.18949 | 17.65285 | 16.34 |
| 20883 | ENSG0000 | SNORD88B | small nucle | 27.08856 | 26.63826 | 13.57472 |
| 2304 | ENSG0000 | PHF5A | PHD finger | 27.07429 | 23.488 | 19.8662 |
| 27851 | ENSG0000 | RBM14 | RNA bindir | 27.05111 | 10.6977 | 10.37905 |
| 6396 | ENSG0000 | TRAF7 | TNF recept | 26.88761 | 20.59976 | 20.12239 |

| | | | | | | |
|---|---|---|---|---|---|---|
| 11277 | ENSG0000 | INTS1 | integrator | 26.85883 | 20.14961 | 18.34776 |
| 6952 | ENSG0000 | NAT10 | N-acetyltra | 26.83988 | 26.38228 | 23.99945 |
| 3919 | ENSG0000 | PAK1IP1 | PAK1 inter | 26.74202 | 23.20332 | 15.17612 |
| 25482 | ENSG0000 | GPX1 | glutathione | 26.55657 | 17.4727 | 16.60141 |
| 9078 | ENSG0000 | HYOU1 | hypoxia up | 26.48334 | 20.64623 | 15.4012 |
| 13 | ENSG0000 | LAS1L | LAS1 like, r | 26.47717 | 20.54436 | 17.05902 |
| 10293 | ENSG0000 | RRP1 | ribosomal | 26.3898 | 23.93353 | 17.75824 |
| 8667 | ENSG0000 | G3BP1 | G3BP stres | 26.25709 | 20.72544 | 17.61156 |
| 3120 | ENSG0000 | GRWD1 | glutamate | 26.18338 | 18.92941 | 13.89824 |
| 3542 | ENSG0000 | PSMD11 | proteasom | 25.97909 | 19.02625 | 17.01276 |
| 3247 | ENSG0000 | SSBP1 | single strar | 25.83127 | 23.63644 | 21.91622 |
| 8330 | ENSG0000 | JTB | jumping tra | 25.80728 | 23.60539 | 23.19506 |
| 14510 | ENSG0000 | SCRIB | scribbled p | 25.79198 | 19.38034 | 16.85821 |
| 8059 | ENSG0000 | DPP9 | dipeptidyl | 25.51783 | 19.29642 | 17.85285 |
| 2782 | ENSG0000 | GSPT1 | G1 to S pha | 25.47451 | 23.37822 | 22.45413 |
| 648 | ENSG0000 | MAP4 | microtubul | 25.37872 | 19.34846 | 19.08783 |
| 10067 | ENSG0000 | CNOT11 | CCR4-NOT | 25.18629 | 23.39966 | 23.03573 |
| 38069 | ENSG0000 | NCBP2-AS2 | NCBP2 ant | 25.1795 | 19.9809 | 19.31128 |
| 2393 | ENSG0000 | ACIN1 | apoptotic c | 25.14139 | 19.80511 | 15.29801 |
| 7966 | ENSG0000 | AFG3L2 | AFG3 like r | 25.13342 | 20.80653 | 18.40176 |
| 10592 | ENSG0000 | FUBP1 | far upstrea | 25.07967 | 23.60227 | 16.15466 |
| 1762 | ENSG0000 | NOP14 | NOP14 nuc | 25.07079 | 20.528 | 14.6005 |
| 4836 | ENSG0000 | PHF19 | PHD finger | 24.94634 | 13.15088 | 11.61283 |
| 3647 | ENSG0000 | DHX15 | DEAH-box | 24.94068 | 21.44977 | 16.46816 |
| 39871 | ENSG0000 | RCC1L | RCC1 like [ | 24.90883 | 23.48036 | 22.58833 |
| 35816 | ENSG0000 | GTF2I | general tra | 24.88474 | 23.39719 | 21.7376 |
| 16409 | ENSG0000 | PTPN1 | protein tyr | 24.83003 | 22.66078 | 21.4672 |
| 6451 | ENSG0000 | DCAF15 | DDB1 and c | 24.82437 | 20.47278 | 19.26555 |
| 14359 | ENSG0000 | PUF60 | poly(U) bir | 24.8123 | 20.72416 | 16.15995 |
| 805 | ENSG0000 | NDC1 | NDC1 trans | 24.74041 | 21.00207 | 20.95097 |
| 12221 | ENSG0000 | USP39 | ubiquitin s | 24.64271 | 20.87127 | 19.20314 |
| 913 | ENSG0000 | HMG20B | high mobil | 24.55051 | 23.52664 | 22.88539 |
| 8788 | ENSG0000 | LUC7L2 | LUC7 like 2 | 24.54423 | 20.85417 | 15.75408 |
| 16726 | ENSG0000 | RPF2 | ribosome p | 24.48038 | 23.7601 | 20.41351 |
| 16636 | ENSG0000 | PSMD12 | proteasom | 24.35849 | 22.92449 | 21.73954 |
| 11103 | ENSG0000 | ABCE1 | ATP bindin | 24.29633 | 21.40644 | 14.62694 |
| 11161 | ENSG0000 | NSA2 | NSA2, ribo | 24.2845 | 21.83752 | 18.76385 |
| 19867 | ENSG0000 | ATF6B | activating t | 24.19463 | 23.02023 | 18.49833 |
| 4704 | ENSG0000 | TXNDC12 | thioredoxin | 24.14904 | 23.67573 | 20.38889 |
| 16128 | ENSG0000 | S100A16 | S100 calciu | 24.10157 | 20.35026 | 15.93361 |
| 1753 | ENSG0000 | PSMC5 | proteasom | 23.93669 | 19.51489 | 18.64888 |
| 187 | ENSG0000 | TSR3 | TSR3, acp t | 23.87376 | 19.13138 | 14.75099 |
| 654 | ENSG0000 | NOP16 | NOP16 nuc | 23.83328 | 18.54282 | 12.04289 |
| 1195 | ENSG0000 | SREBF1 | sterol regu | 23.71466 | 14.72574 | 13.6785 |
| 1853 | ENSG0000 | IGBP1 | immunoglo | 23.71188 | 23.01531 | 20.60552 |
| 16573 | ENSG0000 | AP2A1 | adaptor re | 23.62932 | 16.79785 | 16.01145 |
| 4450 | ENSG0000 | SRSF7 | serine and | 23.59305 | 16.94054 | 15.82481 |

| | | | | | | |
|---|---|---|---|---|---|---|
| 74 | ENSG0000 | UPF1 | UPF1, RNA | 23.5214 | 19.91185 | 17.15167 |
| 6567 | ENSG0000 | NASP | nuclear au | 23.46659 | 15.65125 | 13.7897 |
| 7665 | ENSG0000 | YARS2 | tyrosyl-tRN | 23.31256 | 19.63579 | 18.67277 |
| 22846 | ENSG0000 | WDR46 | WD repeat | 23.30134 | 20.99875 | 16.24922 |
| 1142 | ENSG0000 | TCOF1 | treacle ribo | 23.23361 | 18.28753 | 15.9379 |
| 4839 | ENSG0000 | PPP6C | protein ph | 23.2306 | 21.07272 | 20.64784 |
| 5237 | ENSG0000 | CBX3 | chromobox | 23.05696 | 21.12821 | 21.02901 |
| 4269 | ENSG0000 | PLXNA1 | plexin A1 [ | 23.044 | 20.70215 | 19.48839 |
| 10261 | ENSG0000 | SSU72 | SSU72 hom | 22.95009 | 17.10301 | 15.29149 |
| 37042 | ENSG0000 | AC011511.4 | | 22.93659 | 0.095419 | 0.079073 |
| 3479 | ENSG0000 | GIT1 | GIT ArfGAP | 22.86925 | 22.67737 | 20.8939 |
| 20071 | ENSG0000 | FIS1 | fission, mit | 22.71287 | 17.73369 | 17.64007 |
| 12715 | ENSG0000 | ARHGAP1 | Rho GTPas | 22.71118 | 20.05433 | 19.92534 |
| 2121 | ENSG0000 | TIMM13 | translocase | 22.68955 | 16.41008 | 12.32761 |
| 6837 | ENSG0000 | PHC2 | polyhomeo | 22.60389 | 17.95506 | 17.06864 |
| 4460 | ENSG0000 | PNO1 | partner of | 22.38944 | 21.01243 | 14.74521 |
| 16338 | ENSG0000 | STK40 | serine/thre | 22.37823 | 21.23726 | 18.54531 |
| 1168 | ENSG0000 | DAZAP1 | DAZ associ | 22.3492 | 16.6812 | 14.68522 |
| 4451 | ENSG0000 | SDC1 | syndecan 1 | 22.34583 | 18.36514 | 12.78455 |
| 13610 | ENSG0000 | CTDNEP1 | CTD nuclea | 22.3376 | 21.06184 | 19.61632 |
| 923 | ENSG0000 | IPO5 | importin 5 | 22.32671 | 18.77157 | 14.60175 |
| 3175 | ENSG0000 | TMEM147 | transmeml | 22.3253 | 16.72742 | 15.91566 |
| 10343 | ENSG0000 | MED27 | mediator c | 22.32225 | 18.33914 | 15.3588 |
| 13333 | ENSG0000 | TRMT10C | tRNA meth | 22.24071 | 21.58391 | 19.18817 |
| 41099 | ENSG0000 | SNORA67 | small nucle | 22.15157 | 16.77198 | 8.715496 |
| 11457 | ENSG0000 | PMPCA | peptidase, | 22.10833 | 16.09474 | 15.48462 |
| 14070 | ENSG0000 | KDELC2 | KDEL motif | 22.07573 | 17.10091 | 14.12453 |
| 13315 | ENSG0000 | FAM3C2 | family with | 22.06569 | 21.65913 | 16.74753 |
| 2524 | ENSG0000 | E2F1 | E2F transcı | 22.04959 | 19.24795 | 17.72888 |
| 8965 | ENSG0000 | SEC16A | SEC16 hom | 22.03476 | 19.55148 | 17.43024 |
| 7198 | ENSG0000 | TACO1 | translation | 21.95315 | 20.39128 | 20.22674 |
| 2817 | ENSG0000 | RNF40 | ring finger | 21.93783 | 21.13384 | 17.042 |
| 2345 | ENSG0000 | PSMA3 | proteasom | 21.84321 | 15.32829 | 14.42795 |
| 155 | ENSG0000 | ELAC2 | elaC ribonu | 21.8005 | 18.32983 | 12.78823 |
| 2517 | ENSG0000 | IDH3B | isocitrate c | 21.79517 | 17.89346 | 17.16746 |
| 5416 | ENSG0000 | NCOA5 | nuclear rec | 21.77248 | 19.04476 | 11.56646 |
| 1930 | ENSG0000 | DLD | dihydrolipo | 21.70385 | 18.80143 | 18.4693 |
| 1785 | ENSG0000 | CNOT3 | CCR4-NOT | 21.6714 | 19.2939 | 16.34795 |
| 7310 | ENSG0000 | POLR1E | RNA polym | 21.59777 | 20.83943 | 16.49949 |
| 1701 | ENSG0000 | CHERP | calcium ho | 21.51094 | 19.37665 | 14.78651 |
| 8944 | ENSG0000 | SURF2 | surfeit 2 [S | 21.47408 | 16.82733 | 13.28951 |
| 5230 | ENSG0000 | ZMIZ2 | zinc finger | 21.41211 | 19.76349 | 19.21262 |
| 10142 | ENSG0000 | MIS18A | MIS18 kine | 21.31204 | 15.71599 | 15.27703 |
| 2225 | ENSG0000 | TOMM22 | translocase | 21.28332 | 17.14352 | 15.79288 |
| 3760 | ENSG0000 | NUP98 | nucleopori | 21.17427 | 19.49286 | 18.29616 |
| 11564 | ENSG0000 | BRD7 | bromodom | 21.17095 | 20.15779 | 18.39674 |
| 14986 | ENSG0000 | UTP11 | UTP11, sm | 21.15843 | 18.863 | 16.32792 |

| | | | | | | |
|---|---|---|---|---|---|---|
| 13366 | ENSG0000 | ATP2A2 | ATPase sar | 21.07888 | 20.33213 | 15.0383 |
| 11989 | ENSG0000 | NDUFV1 | NADH:ubic | 21.07531 | 17.49857 | 16.66158 |
| 584 | ENSG0000 | SKIV2L2 | Ski2 like RI | 21.06351 | 17.34407 | 15.55846 |
| 4436 | ENSG0000 | NOL10 | nucleolar p | 21.05853 | 18.35853 | 13.0888 |
| 14757 | ENSG0000 | NPLOC4 | NPL4 homo | 20.99004 | 17.14483 | 14.9053 |
| 6512 | ENSG0000 | UTP3 | UTP3, sma | 20.92577 | 20.15004 | 19.03271 |
| 8253 | ENSG0000 | UFC1 | ubiquitin-fo | 20.89087 | 20.45957 | 16.44139 |
| 830 | ENSG0000 | SNRNP40 | small nucle | 20.85247 | 14.16503 | 12.76067 |
| 11467 | ENSG0000 | ZMYND19 | zinc finger | 20.84544 | 16.72569 | 12.51205 |
| 10419 | ENSG0000 | PCYT1A | phosphate | 20.80917 | 17.21766 | 16.93054 |
| 3171 | ENSG0000 | DDX49 | DEAD-box | 20.77251 | 15.09175 | 14.03703 |
| 13437 | ENSG0000 | YIF1A | Yip1 intera | 20.76096 | 11.68473 | 11.37238 |
| 2189 | ENSG0000 | HIRA | histone cel | 20.68582 | 15.9235 | 14.03422 |
| 10310 | ENSG0000 | MCM3AP | minichrom | 20.63938 | 19.02043 | 16.03992 |
| 2171 | ENSG0000 | MAPK1 | mitogen-ac | 20.55224 | 18.36132 | 17.64356 |
| 16753 | ENSG0000 | ENTPD6 | ectonucleo | 20.55146 | 16.362 | 14.70226 |
| 1364 | ENSG0000 | TM9SF3 | transmeml | 20.52472 | 19.89971 | 18.95779 |
| 7695 | ENSG0000 | AC078814.1 | | 20.44119 | 16.98619 | 16.49765 |
| 17172 | ENSG0000 | SPOUT1 | SPOUT dor | 20.34162 | 16.09804 | 13.16943 |
| 7754 | ENSG0000 | ESD | esterase D | 20.33697 | 16.67034 | 16.4706 |
| 7327 | ENSG0000 | ALDH1B1 | aldehyde d | 20.26353 | 18.16218 | 17.91854 |
| 13339 | ENSG0000 | PITPNA | phosphatic | 20.21633 | 19.28211 | 18.10696 |
| 1788 | ENSG0000 | DDX18 | DEAD-box | 20.17392 | 19.86309 | 16.4422 |
| 27839 | ENSG0000 | TXNDC5 | thioredoxi | 20.16534 | 10.34691 | 9.515434 |
| 10087 | ENSG0000 | ZC3H18 | zinc finger | 20.16214 | 16.91654 | 14.18 |
| 390 | ENSG0000 | ZMYND11 | zinc finger | 20.13797 | 19.76871 | 13.71897 |
| 6106 | ENSG0000 | REEP5 | receptor ac | 20.05059 | 19.47671 | 18.73052 |
| 689 | ENSG0000 | NFE2L3 | nuclear fac | 20.02779 | 19.37293 | 15.39543 |
| 2685 | ENSG0000 | KPNA3 | karyopheri | 19.967 | 19.77869 | 17.16817 |
| 1104 | ENSG0000 | PHRF1 | PHD and ri | 19.95306 | 15.60326 | 13.73455 |
| 9612 | ENSG0000 | BUB3 | BUB3, mito | 19.9249 | 17.24504 | 16.30267 |
| 11791 | ENSG0000 | SLC27A4 | solute carr | 19.90063 | 17.93703 | 14.18475 |
| 9706 | ENSG0000 | NIFK | nucleolar p | 19.8797 | 19.64058 | 14.38994 |
| 16891 | ENSG0000 | GRK6 | G protein-c | 19.82307 | 16.59218 | 16.12057 |
| 7210 | ENSG0000 | TRA2B | transforme | 19.79262 | 12.35388 | 10.80627 |
| 906 | ENSG0000 | BTBD1 | BTB domai | 19.78519 | 18.57693 | 16.95418 |
| 16445 | ENSG0000 | NCOR2 | nuclear rec | 19.76176 | 17.52382 | 17.44257 |
| 13524 | ENSG0000 | LSM1 | LSM1 hom | 19.68266 | 15.12055 | 12.22443 |
| 2584 | ENSG0000 | POLA1 | DNA polym | 19.58332 | 17.90522 | 17.08683 |
| 10748 | ENSG0000 | IWS1 | IWS1, SUP | 19.55555 | 16.52159 | 15.96738 |
| 2609 | ENSG0000 | NAA10 | N(alpha)-ac | 19.55294 | 10.9382 | 9.342878 |
| 7135 | ENSG0000 | TBRG4 | transformi | 19.46952 | 16.52403 | 14.75832 |
| 32703 | ENSG0000 | AC112777.1 | | 19.44879 | 10.10282 | 9.953546 |
| 10085 | ENSG0000 | TSR2 | TSR2, ribos | 19.42081 | 15.15213 | 14.53707 |
| 10568 | ENSG0000 | PEF1 | penta-EF-h | 19.38303 | 13.52337 | 9.987631 |
| 3146 | ENSG0000 | TNPO2 | transportir | 19.37649 | 18.50694 | 16.34405 |
| 10521 | ENSG0000 | RPS6KA4 | ribosomal | 19.34804 | 15.61627 | 13.71668 |

| | | | | | | |
|---|---|---|---|---|---|---|
| 9096 | ENSG0000 | CPSF7 | cleavage a | 19.34726 | 19.28889 | 13.98508 |
| 5304 | ENSG0000 | DDX39A | DExD-box l | 19.3468 | 13.03133 | 11.21089 |
| 10269 | ENSG0000 | VMA21 | VMA21, va | 19.21424 | 18.68616 | 17.04402 |
| 4893 | ENSG0000 | XPO5 | exportin 5 | 19.18486 | 18.64037 | 14.52181 |
| 15860 | ENSG0000 | CHP1 | calcineurin | 19.17581 | 18.09143 | 16.32076 |
| 8859 | ENSG0000 | PLPBP | pyridoxal p | 19.17455 | 16.5247 | 15.44304 |
| 11805 | ENSG0000 | CRK | CRK proto- | 19.11716 | 17.80351 | 16.50388 |
| 15344 | ENSG0000 | NOC4L | nucleolar c | 19.08294 | 14.21876 | 12.10847 |
| 10402 | ENSG0000 | MAML1 | mastermin | 19.03852 | 17.24278 | 16.45001 |
| 13417 | ENSG0000 | BRMS1 | breast canc | 18.90363 | 14.90799 | 12.45458 |
| 7515 | ENSG0000 | ACTR1A | ARP1 actin | 18.80928 | 13.89061 | 13.53579 |
| 12902 | ENSG0000 | RNASEH1 | ribonuclea | 18.78653 | 15.48441 | 12.33714 |
| 12008 | ENSG0000 | SRP68 | signal reco | 18.7693 | 13.83766 | 13.78883 |
| 1692 | ENSG0000 | RRN3 | RRN3 hom | 18.71536 | 18.32622 | 15.76275 |
| 7415 | ENSG0000 | BUD13 | BUD13 hor | 18.62903 | 16.58812 | 14.95262 |
| 15494 | ENSG0000 | PSMD13 | proteasom | 18.62562 | 14.8229 | 13.94174 |
| 4352 | ENSG0000 | PSMD14 | proteasom | 18.52292 | 14.91813 | 14.38386 |
| 2005 | ENSG0000 | CDC45 | cell divisio | 18.35533 | 13.00101 | 12.70619 |
| 16939 | ENSG0000 | DDX42 | DEAD-box | 18.34529 | 18.27044 | 16.64306 |
| 6546 | ENSG0000 | DAP3 | death asso | 18.29235 | 15.51524 | 12.6295 |
| 6168 | ENSG0000 | SAFB2 | scaffold at | 18.25885 | 15.47637 | 11.05684 |
| 2244 | ENSG0000 | PACSIN2 | protein kin | 18.08236 | 14.52494 | 13.44043 |
| 5438 | ENSG0000 | STX16 | syntaxin 16 | 18.05292 | 8.098707 | 8.009174 |
| 18630 | ENSG0000 | RNPS1 | RNA bindir | 18.03727 | 13.25561 | 12.91489 |
| 13276 | ENSG0000 | EIF1 | eukaryotic | 17.98412 | 16.50472 | 14.71009 |
| 16457 | ENSG0000 | MYO18A | myosin XV | 17.9205 | 16.11536 | 13.06597 |
| 14864 | ENSG0000 | MTA1 | metastasis | 17.7916 | 14.17205 | 13.85567 |
| 13797 | ENSG0000 | ZBTB7A | zinc finger | 17.78476 | 15.27803 | 14.27274 |
| 1129 | ENSG0000 | NDST1 | N-deacetyl | 17.76665 | 15.58051 | 13.63386 |
| 5186 | ENSG0000 | GTF3A | general tra | 17.72286 | 11.09634 | 10.43819 |
| 14873 | ENSG0000 | AP2A2 | adaptor re | 17.63949 | 17.09086 | 16.59673 |
| 1076 | ENSG0000 | NUP133 | nucleopori | 17.61817 | 17.24313 | 16.75148 |
| 8885 | ENSG0000 | MTDH | metadheri | 17.60804 | 16.6125 | 15.16968 |
| 10308 | ENSG0000 | LSS | lanosterol | 17.5592 | 13.03926 | 11.70728 |
| 17105 | ENSG0000 | MTOR | mechanisti | 17.55892 | 12.42501 | 11.09439 |
| 7010 | ENSG0000 | SMYD5 | SMYD fami | 17.55615 | 16.15968 | 12.05386 |
| 6845 | ENSG0000 | PRPF38A | pre-mRNA | 17.53424 | 17.11088 | 15.29074 |
| 1014 | ENSG0000 | DHX29 | DExH-box l | 17.52316 | 16.70829 | 13.84181 |
| 17009 | ENSG0000 | GPN1 | GPN-loop ( | 17.5016 | 17.27708 | 15.88494 |
| 6415 | ENSG0000 | CHD1L | chromodoi | 17.50046 | 17.34877 | 16.15453 |
| 6459 | ENSG0000 | DHX30 | DExH-box l | 17.48744 | 13.1458 | 10.93939 |
| 1225 | ENSG0000 | SCARB1 | scavenger | 17.43044 | 11.85388 | 11.01841 |
| 15681 | ENSG0000 | KPNA4 | karyopheri | 17.40356 | 16.18058 | 12.56946 |
| 1169 | ENSG0000 | MBD3 | methyl-Cp | 17.39692 | 14.9342 | 12.1474 |
| 19954 | ENSG0000 | CSNK1E | casein kina | 17.3653 | 17.04651 | 16.46809 |
| 3454 | ENSG0000 | BCCIP | BRCA2 and | 17.35388 | 16.09462 | 14.66749 |
| 8541 | ENSG0000 | SRPRB | SRP recept | 17.30717 | 15.22368 | 11.6554 |

| | | | | | | |
|---|---|---|---|---|---|---|
| 25740 | ENSG0000 | EIF4HP1 | eukaryotic | 17.30254 | 14.69439 | 13.09266 |
| 18135 | ENSG0000 | MRPL38 | mitochond | 17.22903 | 13.00819 | 10.5112 |
| 17050 | ENSG0000 | CALM1 | calmodulir | 17.22217 | 16.22314 | 15.56658 |
| 8241 | ENSG0000 | UCK2 | uridine-cyt | 17.22154 | 13.17752 | 11.66181 |
| 4088 | ENSG0000 | TTK | TTK proteir | 17.16913 | 14.58843 | 11.244 |
| 2651 | ENSG0000 | RBM3 | RNA bindir | 17.12098 | 12.78425 | 8.457757 |
| 11458 | ENSG0000 | SDCCAG3 | serological | 17.08341 | 15.82169 | 12.61911 |
| 6155 | ENSG0000 | LDLR | low density | 17.02132 | 13.54086 | 12.50369 |
| 7254 | ENSG0000 | C9orf78 | chromosor | 17.01304 | 14.7224 | 11.11769 |
| 5631 | ENSG0000 | SYMPK | symplekin | 16.98863 | 13.4751 | 10.50722 |
| 449 | ENSG0000 | RTFDC1 | replication | 16.974 | 16.44276 | 15.52664 |
| 7748 | ENSG0000 | ANKRD52 | ankyrin rep | 16.89836 | 16.28275 | 14.18367 |
| 5692 | ENSG0000 | RALY | RALY heter | 16.89691 | 10.85399 | 10.56469 |
| 6874 | ENSG0000 | ARGLU1 | arginine ar | 16.89598 | 16.67177 | 14.77217 |
| 679 | ENSG0000 | RCN1 | reticulocall | 16.88853 | 16.87306 | 16.50549 |
| 10162 | ENSG0000 | ATP5G1 | ATP syntha | 16.82382 | 11.23372 | 10.67612 |
| 7238 | ENSG0000 | IMP4 | IMP4, U3 s | 16.8008 | 14.98247 | 12.08718 |
| 336 | ENSG0000 | PPP5C | protein ph | 16.76932 | 13.61142 | 12.39317 |
| 4067 | ENSG0000 | BYSL | bystin like | 16.76353 | 15.29812 | 12.17575 |
| 10375 | ENSG0000 | GPATCH4 | G-patch dc | 16.75243 | 15.73718 | 8.073428 |
| 1359 | ENSG0000 | XAB2 | XPA bindin | 16.46672 | 15.49061 | 13.61866 |
| 11371 | ENSG0000 | ZNF367 | zinc finger | 16.31869 | 11.93623 | 8.708969 |
| 8355 | ENSG0000 | GALNT2 | polypeptid | 16.2684 | 13.33009 | 12.69245 |
| 255 | ENSG0000 | SEC63 | SEC63 hor | 16.23671 | 16.04713 | 15.72487 |
| 2510 | ENSG0000 | POFUT1 | protein O-f | 16.20574 | 15.04738 | 14.35483 |
| 2346 | ENSG0000 | VTI1B | vesicle trar | 16.19857 | 14.42275 | 14.38591 |
| 3522 | ENSG0000 | NUP88 | nucleopori | 16.17426 | 11.60076 | 10.82857 |
| 20042 | ENSG0000 | ALG3 | ALG3, alph | 16.14098 | 14.15668 | 14.13619 |
| 4263 | ENSG0000 | NCBP2 | nuclear car | 16.12133 | 15.03197 | 14.45909 |
| 3949 | ENSG0000 | KCTD20 | potassium | 16.08212 | 14.20452 | 11.88206 |
| 457 | ENSG0000 | RNH1 | ribonuclea | 16.06889 | 12.81026 | 12.18121 |
| 13077 | ENSG0000 | FAM192A | family with | 16.03883 | 15.98612 | 12.85023 |
| 9121 | ENSG0000 | LSM14B | LSM family | 15.96209 | 15.37167 | 14.87514 |
| 3486 | ENSG0000 | CASC3 | cancer sus | 15.94174 | 15.28482 | 13.24444 |
| 7239 | ENSG0000 | HS6ST1 | heparan su | 15.93614 | 15.19651 | 14.48731 |
| 2455 | ENSG0000 | NELFCD | negative el | 15.93077 | 14.74837 | 12.84026 |
| 8315 | ENSG0000 | DTL | denticleles | 15.91936 | 12.81015 | 12.60595 |
| 7442 | ENSG0000 | KIF23 | kinesin fan | 15.84587 | 11.8315 | 10.93824 |
| 15059 | ENSG0000 | MRPL54 | mitochond | 15.82114 | 11.58918 | 8.706231 |
| 7773 | ENSG0000 | CUL4A | cullin 4A [S | 15.8142 | 15.66392 | 13.81288 |
| 15306 | ENSG0000 | UBE2G2 | ubiquitin c | 15.76126 | 13.16538 | 13.02569 |
| 7811 | ENSG0000 | NIPA2 | non imprin | 15.74697 | 13.57905 | 12.75483 |
| 16789 | ENSG0000 | PHF2 | PHD finger | 15.74076 | 13.87704 | 12.19097 |
| 15883 | ENSG0000 | USP7 | ubiquitin s | 15.68061 | 14.10758 | 12.09063 |
| 9917 | ENSG0000 | ATP6V0D1 | ATPase H+ | 15.68049 | 11.87798 | 10.69427 |
| 41403 | ENSG0000 | ACACA | acetyl-CoA | 15.68038 | 13.26806 | 11.48626 |
| 6639 | ENSG0000 | SRRM1 | serine and | 15.64436 | 14.37735 | 13.23007 |

| | | | | | | |
|---|---|---|---|---|---|---|
| 10898 | ENSG0000 | NOL6 | nucleolar p | 15.6176 | 14.91305 | 8.080855 |
| 20229 | ENSG0000 | IFRD2 | interferon | 15.60904 | 10.44759 | 7.973887 |
| 11640 | ENSG0000 | CENPN | centromer | 15.60467 | 11.20557 | 9.851992 |
| 8721 | ENSG0000 | PM20D2 | peptidase | 15.57401 | 13.29308 | 12.04208 |
| 3871 | ENSG0000 | COPS7A | COP9 signa | 15.5739 | 9.966217 | 9.82135 |
| 4438 | ENSG0000 | GORASP2 | golgi reass | 15.54533 | 14.05079 | 13.53019 |
| 4656 | ENSG0000 | B4GALT2 | beta-1,4-ga | 15.54482 | 9.829388 | 8.734272 |
| 7301 | ENSG0000 | ARPC5L | actin relate | 15.52876 | 14.17617 | 13.14129 |
| 6569 | ENSG0000 | CTNNBL1 | catenin be | 15.48191 | 14.11072 | 13.4654 |
| 40463 | ENSG0000 | DUSP14 | dual specif | 15.47057 | 15.08588 | 14.00408 |
| 7334 | ENSG0000 | FOXP4 | forkhead b | 15.45938 | 14.96693 | 12.80454 |
| 10145 | ENSG0000 | C21orf59 | chromosor | 15.45186 | 13.4953 | 13.45997 |
| 66 | ENSG0000 | SLC25A13 | solute carr | 15.44883 | 14.96185 | 13.29465 |
| 2386 | ENSG0000 | PSMC1 | proteasom | 15.43056 | 13.02723 | 11.57014 |
| 37786 | ENSG0000 | AC010422.3 | | 15.41001 | 12.05149 | 9.63839 |
| 3330 | ENSG0000 | BCL7B | BCL tumor | 15.40003 | 10.49439 | 8.916034 |
| 5493 | ENSG0000 | AL365205.1 | | 15.38913 | 11.48842 | 10.29261 |
| 3033 | ENSG0000 | TIMM44 | translocase | 15.37167 | 13.17695 | 10.01106 |
| 1066 | ENSG0000 | ERLEC1 | endoplasm | 15.29397 | 15.15233 | 14.27846 |
| 332 | ENSG0000 | WIZ | widely inte | 15.24632 | 14.79405 | 13.36426 |
| 4221 | ENSG0000 | KPNA1 | karyopheri | 15.24084 | 14.2051 | 11.86452 |
| 1704 | ENSG0000 | POMGNT1 | protein O-l | 15.22068 | 12.94883 | 12.64206 |
| 3940 | ENSG0000 | FBXO5 | F-box prote | 15.21826 | 12.87831 | 12.3183 |
| 13466 | ENSG0000 | KLC2 | kinesin ligh | 15.19502 | 13.74575 | 8.684455 |
| 16506 | ENSG0000 | ZNF512B | zinc finger | 15.19457 | 14.97544 | 12.63376 |
| 7054 | ENSG0000 | MRPL44 | mitochond | 15.1712 | 12.26264 | 11.45682 |
| 8402 | ENSG0000 | CHAC2 | ChaC catio | 15.14679 | 14.4702 | 12.76464 |
| 4190 | ENSG0000 | TCERG1 | transcriptio | 15.09259 | 13.5948 | 9.486902 |
| 11763 | ENSG0000 | MARS | methionyl- | 15.08006 | 7.942515 | 7.577867 |
| 6303 | ENSG0000 | SLC35D2 | solute carr | 15.04587 | 13.15417 | 10.96271 |
| 13649 | ENSG0000 | CSTF3 | cleavage st | 14.97257 | 12.33078 | 11.60504 |
| 3181 | ENSG0000 | USF2 | upstream t | 14.92825 | 13.04924 | 12.24282 |
| 14686 | ENSG0000 | TNRC18 | trinucleoti | 14.91761 | 14.45564 | 13.86383 |
| 855 | ENSG0000 | WAPL | WAPL cohe | 14.89914 | 14.77561 | 13.54862 |
| 1162 | ENSG0000 | BUD23 | BUD23, rRI | 14.85018 | 10.3807 | 9.923153 |
| 969 | ENSG0000 | SMARCD1 | SWI/SNF re | 14.82579 | 10.85464 | 8.368578 |
| 683 | ENSG0000 | CLPTM1L | CLPTM1 lik | 14.74236 | 13.67185 | 10.97938 |
| 34476 | ENSG0000 | RBM15B | RNA bindir | 14.7014 | 13.95927 | 10.70354 |
| 3471 | ENSG0000 | CUL2 | cullin 2 [So | 14.64794 | 14.47428 | 13.25421 |
| 6430 | ENSG0000 | SNRPA1 | small nucle | 14.62133 | 14.06933 | 13.57127 |
| 2399 | ENSG0000 | ARHGAP5 | Rho GTPas | 14.58821 | 13.72213 | 11.1636 |
| 4706 | ENSG0000 | CD3EAP | CD3e mole | 14.55329 | 13.69623 | 11.27577 |
| 8097 | ENSG0000 | ZNF787 | zinc finger | 14.51287 | 13.45955 | 11.45103 |
| 13202 | ENSG0000 | NAA20 | N(alpha)-a | 14.50979 | 12.00343 | 11.78895 |
| 3183 | ENSG0000 | KXD1 | KxDL motif | 14.5051 | 13.9488 | 13.8934 |
| 249 | ENSG0000 | ZNF207 | zinc finger | 14.50216 | 13.95075 | 10.19937 |
| 1107 | ENSG0000 | PFN2 | profilin 2 [S | 14.46197 | 9.184132 | 6.08348 |

| | | | | | | |
|---|---|---|---|---|---|---|
| 5839 | ENSG0000 | EMC1 | ER membr | 14.45369 | 11.62784 | 10.63251 |
| 3111 | ENSG0000 | NAPA | NSF attach | 14.40333 | 11.91885 | 10.51675 |
| 287 | ENSG0000 | DNM2 | dynamin 2 | 14.4 | 11.34302 | 10.66001 |
| 7276 | ENSG0000 | FPGS | folylpolygl | 14.3934 | 11.36178 | 11.31418 |
| 1949 | ENSG0000 | SPAG7 | sperm assc | 14.37762 | 13.36812 | 11.84664 |
| 12322 | ENSG0000 | MRPL1 | mitochond | 14.36097 | 14.2407 | 13.47499 |
| 23995 | ENSG0000 | EIF4EP2 | eukaryotic | 14.35964 | 8.49132 | 8.072391 |
| 3055 | ENSG0000 | URI1 | URI1, prefc | 14.35351 | 13.73241 | 11.45209 |
| 12543 | ENSG0000 | HARS | histidyl-tRI | 14.35219 | 13.67604 | 12.83949 |
| 5248 | ENSG0000 | CCZ1 | CCZ1 homc | 14.28274 | 14.14994 | 13.80641 |
| 1325 | ENSG0000 | SART3 | squamous | 14.27738 | 12.62282 | 11.62989 |
| 11096 | ENSG0000 | NAA15 | N(alpha)-a | 14.25524 | 12.74676 | 8.993635 |
| 7502 | ENSG0000 | PREB | prolactin r | 14.13215 | 13.38262 | 11.32145 |
| 17166 | ENSG0000 | MAP3K3 | mitogen-a | 14.13107 | 12.50889 | 10.74548 |
| 10835 | ENSG0000 | CHCHD4 | coiled-coil- | 14.12855 | 13.6667 | 10.35037 |
| 9874 | ENSG0000 | SEC13 | SEC13 hom | 14.0775 | 10.8237 | 10.15096 |
| 598 | ENSG0000 | PSMA4 | proteasom | 14.0746 | 12.46809 | 10.96786 |
| 17185 | ENSG0000 | CCDC167 | coiled-coil | 14.06717 | 7.461356 | 5.805476 |
| 2309 | ENSG0000 | DESI1 | desumoyla | 14.0061 | 10.29677 | 9.26122 |
| 5530 | ENSG0000 | EEF1E1 | eukaryotic | 13.96165 | 12.99469 | 9.927638 |
| 13078 | ENSG0000 | RAB43 | RAB43, me | 13.94126 | 4.946095 | 3.233796 |
| 7704 | ENSG0000 | SNRPF | small nucle | 13.81253 | 9.259428 | 9.096483 |
| 12235 | ENSG0000 | LETM1 | leucine zip | 13.79856 | 10.05445 | 6.775822 |
| 3071 | ENSG0000 | CCDC94 | coiled-coil | 13.7969 | 12.09746 | 8.809407 |
| 39637 | ENSG0000 | CYFIP1 | cytoplasmi | 13.7934 | 11.77643 | 10.96613 |
| 667 | ENSG0000 | R3HDM1 | R3H domai | 13.78804 | 11.595 | 9.727117 |
| 14182 | ENSG0000 | PFAS | phosphoril | 13.74355 | 11.79849 | 9.355816 |
| 16949 | ENSG0000 | UCKL1 | uridine-cyt | 13.68334 | 13.33363 | 11.38997 |
| 381 | ENSG0000 | ZC3H3 | zinc finger | 13.67549 | 11.74581 | 11.63924 |
| 12397 | ENSG0000 | GPS1 | G protein p | 13.6393 | 10.50134 | 9.860771 |
| 713 | ENSG0000 | RRP12 | ribosomal | 13.60118 | 9.798311 | 6.708273 |
| 40492 | ENSG0000 | DLEU1_1 | Deleted in | 13.59937 | 8.540375 | 8.514193 |
| 7973 | ENSG0000 | GALNT1 | polypeptid | 13.47426 | 11.8865 | 11.28256 |
| 12789 | ENSG0000 | MRPL36 | mitochond | 13.45755 | 9.729096 | 8.628703 |
| 33639 | ENSG0000 | PDF | peptide de | 13.4443 | 10.19358 | 7.049363 |
| 12216 | ENSG0000 | DDX19A | DEAD-box | 13.42203 | 13.06928 | 11.42383 |
| 6895 | ENSG0000 | C9orf40 | chromosor | 13.41112 | 11.09748 | 10.91723 |
| 8547 | ENSG0000 | OSBPL11 | oxysterol k | 13.40515 | 11.98441 | 11.91497 |
| 2235 | ENSG0000 | PPP6R2 | protein ph | 13.34514 | 11.47402 | 10.99904 |
| 28889 | ENSG0000 | PPAN-P2R\ | PPAN-P2RY | 13.34071 | 5.143364 | 5.067168 |
| 4496 | ENSG0000 | FAM20B | FAM20B, g | 13.30686 | 12.55921 | 9.556894 |
| 12396 | ENSG0000 | DUS1L | dihydrouri | 13.29663 | 12.94355 | 9.179179 |
| 7106 | ENSG0000 | MED4 | mediator c | 13.2451 | 12.77328 | 11.91351 |
| 2140 | ENSG0000 | ZDHHC8 | zinc finger | 13.17671 | 13.12012 | 11.73037 |
| 17196 | ENSG0000 | ARMCX6 | armadillo r | 13.11431 | 11.94338 | 9.159315 |
| 2710 | ENSG0000 | NUP93 | nucleopori | 13.06558 | 8.078468 | 8.022559 |
| 8079 | ENSG0000 | URB1 | URB1 ribos | 13.01053 | 9.620778 | 6.870069 |

| | | | | | | |
|---|---|---|---|---|---|---|
| 6178 | ENSG0000 | NSUN5 | NOP2/Sun | 13.00417 | 11.45837 | 10.52552 |
| 32278 | ENSG0000 | DPP3 | dipeptidyl | 12.9993 | 11.11618 | 10.16598 |
| 3147 | ENSG0000 | WDR83OS | WD repeat | 12.99122 | 11.60067 | 10.14269 |
| 3422 | ENSG0000 | SEC23IP | SEC23 inte | 12.96967 | 12.41203 | 11.90747 |
| 10095 | ENSG0000 | PPP1R15B | protein ph | 12.95388 | 12.93981 | 12.18374 |
| 19183 | ENSG0000 | MT-TQ | mitochond | 12.94313 | 12.15758 | 1.582652 |
| 5671 | ENSG0000 | ITPA | inosine trip | 12.92719 | 12.3303 | 12.11253 |
| 5764 | ENSG0000 | EMG1 | EMG1, N1- | 12.89623 | 11.45995 | 9.49938 |
| 7367 | ENSG0000 | MDC1 | mediator c | 12.82954 | 10.64334 | 10.1626 |
| 6807 | ENSG0000 | FBXO18 | F-box prot | 12.79374 | 12.47081 | 12.11655 |
| 1002 | ENSG0000 | IDI1 | isopenteny | 12.74331 | 10.13989 | 10.01097 |
| 4971 | ENSG0000 | TCTN3 | tectonic fa | 12.72536 | 10.98705 | 10.26473 |
| 1534 | ENSG0000 | CD99L2 | CD99 mole | 12.68406 | 8.179602 | 4.008215 |
| 13536 | ENSG0000 | EIF3F | eukaryotic | 12.68246 | 8.45559 | 8.27731 |
| 7889 | ENSG0000 | GLYR1 | glyoxylate | 12.6799 | 9.975369 | 8.879056 |
| 21648 | ENSG0000 | RPL23AP2 | ribosomal | 12.63304 | 8.004908 | 7.939621 |
| 3735 | ENSG0000 | DDX6 | DEAD-box | 12.62527 | 11.85077 | 11.62022 |
| 8952 | ENSG0000 | NTMT1 | N-terminal | 12.52319 | 9.561048 | 9.458804 |
| 10294 | ENSG0000 | AGPAT3 | 1-acylglyce | 12.52002 | 8.7661 | 5.481633 |
| 4668 | ENSG0000 | ABCD3 | ATP bindin | 12.50608 | 11.82459 | 11.625 |
| 12285 | ENSG0000 | UBE2V2 | ubiquitin c | 12.50397 | 10.88067 | 9.878419 |
| 3162 | ENSG0000 | PIK3R2 | phosphoin | 12.48806 | 12.23263 | 9.203948 |
| 2334 | ENSG0000 | PSMC6 | proteasom | 12.4869 | 11.47287 | 10.40774 |
| 9977 | ENSG0000 | SLC35B2 | solute carr | 12.45448 | 11.40901 | 10.28209 |
| 15381 | ENSG0000 | FAF1 | Fas associa | 12.45298 | 11.18843 | 10.91944 |
| 7308 | ENSG0000 | DMAC1 | distal mem | 12.42806 | 8.597987 | 7.316236 |
| 7495 | ENSG0000 | PNPT1 | polyribonu | 12.42646 | 11.86188 | 8.322703 |
| 4114 | ENSG0000 | BRD8 | bromodom | 12.37635 | 12.27621 | 11.20142 |
| 6478 | ENSG0000 | PTCD3 | pentatrico | 12.37319 | 11.70829 | 11.13022 |
| 9284 | ENSG0000 | VPS26B | VPS26, retr | 12.34336 | 11.1483 | 10.15402 |
| 28027 | ENSG0000 | MRPS17 | mitochond | 12.32283 | 11.51311 | 10.80487 |
| 8704 | ENSG0000 | PPP1R18 | protein ph | 12.28106 | 3.637191 | 3.63682 |
| 21865 | ENSG0000 | NDUFAF8 | NADH:ubic | 12.26134 | 8.948186 | 8.884308 |
| 3556 | ENSG0000 | MLX | MLX, MAX | 12.25623 | 10.67305 | 10.53635 |
| 10282 | ENSG0000 | WDR4 | WD repeat | 12.22801 | 9.957887 | 6.575565 |
| 2125 | ENSG0000 | CEP170B | centrosom | 12.16495 | 11.80528 | 10.20617 |
| 1193 | ENSG0000 | ALDH3A2 | aldehyde d | 12.13448 | 6.679646 | 6.60426 |
| 4876 | ENSG0000 | MTHFD1L | methylene | 12.13241 | 5.039521 | 3.738666 |
| 14280 | ENSG0000 | FJX1 | four jointe | 12.13158 | 11.97667 | 6.993465 |
| 2917 | ENSG0000 | DCTN6 | dynactin su | 11.98874 | 11.59716 | 7.637448 |
| 6457 | ENSG0000 | LRRC41 | leucine rich | 11.94048 | 9.190827 | 8.665821 |
| 5647 | ENSG0000 | DTD1 | D-tyrosyl-t | 11.92126 | 7.874531 | 6.291351 |
| 1055 | ENSG0000 | POLR1A | RNA polym | 11.89848 | 10.94097 | 7.039722 |
| 1894 | ENSG0000 | DNAJB11 | DnaJ heat s | 11.86962 | 9.928019 | 8.856378 |
| 9704 | ENSG0000 | HEATR3 | HEAT repe | 11.81532 | 11.27689 | 9.389766 |
| 458 | ENSG0000 | NDUFS1 | NADH:ubic | 11.81494 | 10.37932 | 10.17192 |
| 10260 | ENSG0000 | ATAD3B | ATPase fan | 11.76303 | 8.979627 | 5.60573 |

| | | | | | | |
|---|---|---|---|---|---|---|
| 16625 | ENSG0000 | PCNX3 | pecanex ho | 11.75804 | 9.681745 | 8.730495 |
| 5388 | ENSG0000 | ACSL3 | acyl-CoA sy | 11.75515 | 11.25941 | 10.6882 |
| 2824 | ENSG0000 | CSK | C-terminal | 11.74049 | 10.95371 | 10.22228 |
| 1950 | ENSG0000 | ORC6 | origin reco | 11.68611 | 10.51651 | 10.30739 |
| 12743 | ENSG0000 | C1GALT1C1 | C1GALT1 s | 11.66732 | 10.07696 | 9.506718 |
| 8329 | ENSG0000 | ADAM15 | ADAM met | 11.63921 | 7.144787 | 6.967011 |
| 15629 | ENSG0000 | BCL9L | B-cell CLL/l | 11.63839 | 7.228583 | 6.604235 |
| 14539 | ENSG0000 | AEN | apoptosis e | 11.60808 | 9.890089 | 8.606939 |
| 4238 | ENSG0000 | PRKAR2A | protein kin | 11.60323 | 9.441982 | 8.480316 |
| 10125 | ENSG0000 | B4GALT3 | beta-1,4-ga | 11.58653 | 9.797139 | 8.051053 |
| 28619 | ENSG0000 | SNHG3 | small nucle | 11.54863 | 11.29873 | 7.438931 |
| 1140 | ENSG0000 | EIF2B3 | eukaryotic | 11.51805 | 8.568497 | 8.452391 |
| 13679 | ENSG0000 | ANAPC2 | anaphase p | 11.51032 | 9.784938 | 7.949562 |
| 9202 | ENSG0000 | TIMM8B | translocase | 11.494 | 8.751387 | 7.37787 |
| 2714 | ENSG0000 | LONP2 | lon peptida | 11.47881 | 9.765234 | 8.672917 |
| 13647 | ENSG0000 | IP6K1 | inositol he | 11.41573 | 10.52598 | 10.39497 |
| 18111 | ENSG0000 | RXRB | retinoid X r | 11.39632 | 9.462228 | 7.676331 |
| 4346 | ENSG0000 | GTF3C2 | general tra | 11.38491 | 10.78681 | 9.56366 |
| 2507 | ENSG0000 | TM9SF4 | transmeml | 11.37263 | 8.930163 | 8.659673 |
| 32639 | ENSG0000 | SNHG1 | small nucle | 11.36122 | 11.24584 | 7.647684 |
| 15653 | ENSG0000 | TOR3A | torsin fami | 11.33595 | 7.049702 | 6.044463 |
| 2589 | ENSG0000 | AMMECR1 | Alport sync | 11.32455 | 9.58631 | 8.228295 |
| 6206 | ENSG0000 | SCO2 | SCO2, cyto | 11.30582 | 8.118981 | 5.213175 |
| 1320 | ENSG0000 | WDR62 | WD repeat | 11.29944 | 6.820946 | 5.288884 |
| 6056 | ENSG0000 | E2F8 | E2F transcr | 11.29938 | 8.859406 | 8.185749 |
| 4942 | ENSG0000 | ATL2 | atlastin GT | 11.29662 | 8.999127 | 8.732166 |
| 4654 | ENSG0000 | IPO13 | importin 1 | 11.27213 | 10.3638 | 8.529566 |
| 2773 | ENSG0000 | STUB1 | STIP1 hom | 11.25891 | 6.743358 | 6.73634 |
| 16104 | ENSG0000 | SRSF10 | serine and | 11.25423 | 9.513754 | 9.404534 |
| 4631 | ENSG0000 | RBBP5 | RB binding | 11.18808 | 10.91343 | 9.859283 |
| 5746 | ENSG0000 | PRR12 | proline rich | 11.1649 | 10.1466 | 9.826169 |
| 13322 | ENSG0000 | C16orf91 | chromosor | 11.1491 | 9.046414 | 8.740505 |
| 3200 | ENSG0000 | PMPCB | peptidase, | 11.12917 | 7.584756 | 7.464706 |
| 6232 | ENSG0000 | ATXN10 | ataxin 10 [ | 11.12753 | 8.613546 | 7.92568 |
| 14817 | ENSG0000 | NGRN | neugrin, ne | 11.12181 | 9.806784 | 9.233466 |
| 10096 | ENSG0000 | COPG2 | coatomer p | 11.10292 | 7.537073 | 5.876103 |
| 10226 | ENSG0000 | CTBP1 | C-terminal | 11.08207 | 8.141599 | 7.802797 |
| 3495 | ENSG0000 | TRIM37 | tripartite n | 11.07729 | 10.64319 | 9.155938 |
| 16344 | ENSG0000 | MPHOSPH | M-phase p | 11.07315 | 10.22832 | 9.404909 |
| 2385 | ENSG0000 | VRK1 | vaccinia re | 11.02142 | 10.10018 | 8.655966 |
| 5465 | ENSG0000 | MPHOSPH | M-phase p | 11.01389 | 10.94665 | 9.155223 |
| 2943 | ENSG0000 | NFKBIB | NFKB inhib | 10.9972 | 10.18021 | 8.260978 |
| 226 | ENSG0000 | MED24 | mediator c | 10.98879 | 10.08273 | 9.185963 |
| 4479 | ENSG0000 | NFE2L2 | nuclear fac | 10.94749 | 6.903279 | 5.57947 |
| 13922 | ENSG0000 | TIMM22 | translocase | 10.94486 | 9.407328 | 8.685605 |
| 3587 | ENSG0000 | RANGRF | RAN guanii | 10.93274 | 8.300343 | 8.05294 |
| 5979 | ENSG0000 | EIF2AK4 | eukaryotic | 10.9226 | 10.46538 | 9.504775 |

| | | | | | | |
|---|---|---|---|---|---|---|
| 1916 | ENSG0000 | NAT14 | N-acetyltra | 10.87453 | 10.32237 | 9.787282 |
| 6541 | ENSG0000 | NXT1 | nuclear tra | 10.86613 | 10.82433 | 9.085613 |
| 10348 | ENSG0000 | CHTOP | chromatin | 10.84007 | 8.919769 | 8.74471 |
| 17721 | ENSG0000 | SNORD32A | small nucle | 10.78852 | 9.838426 | 7.621309 |
| 25554 | ENSG0000 | EIF3FP3 | eukaryotic | 10.78348 | 7.236384 | 7.202003 |
| 13867 | ENSG0000 | POLE | DNA polym | 10.74976 | 10.72887 | 9.919142 |
| 7641 | ENSG0000 | CENPE | centromer | 10.73794 | 8.9864 | 7.880678 |
| 4080 | ENSG0000 | EXOC2 | exocyst co | 10.674 | 8.788133 | 7.058491 |
| 5086 | ENSG0000 | UBIAD1 | UbiA preny | 10.6627 | 9.399008 | 5.744027 |
| 6395 | ENSG0000 | THOC6 | THO comp | 10.65896 | 9.945724 | 8.311307 |
| 10166 | ENSG0000 | SNF8 | SNF8, ESCF | 10.63588 | 9.132784 | 8.788898 |
| 399 | ENSG0000 | NUDCD3 | NudC dom | 10.62441 | 8.277659 | 7.745642 |
| 12091 | ENSG0000 | DNAJC7 | DnaJ heat : | 10.59563 | 10.42972 | 9.366477 |
| 8974 | ENSG0000 | PDSS1 | decapreny | 10.54993 | 6.465676 | 5.259617 |
| 4149 | ENSG0000 | DROSHA | drosha ribo | 10.52591 | 10.32473 | 9.179303 |
| 3642 | ENSG0000 | GAR1 | GAR1 ribor | 10.51075 | 8.77745 | 6.716299 |
| 6659 | ENSG0000 | MORC2 | MORC fam | 10.48215 | 9.915971 | 8.27614 |
| 23292 | ENSG0000 | C19orf24 | chromosor | 10.45913 | 10.23112 | 9.923554 |
| 13598 | ENSG0000 | TOMM5 | translocase | 10.45288 | 7.628587 | 7.579574 |
| 10208 | ENSG0000 | MED8 | mediator c | 10.42502 | 10.33998 | 10.05709 |
| 1593 | ENSG0000 | ME2 | malic enzy | 10.37806 | 8.733327 | 8.454326 |
| 9310 | ENSG0000 | ASAP2 | ArfGAP wit | 10.36628 | 9.80441 | 7.410347 |
| 5020 | ENSG0000 | SNX19 | sorting nex | 10.34926 | 7.700214 | 4.981924 |
| 13946 | ENSG0000 | ARIH2 | ariadne RB | 10.33161 | 8.998468 | 8.971765 |
| 5213 | ENSG0000 | FAM35A | family with | 10.30549 | 9.514958 | 6.423674 |
| 11303 | ENSG0000 | WASHC5 | WASH com | 10.3011 | 7.384181 | 7.259301 |
| 16567 | ENSG0000 | NOP9 | NOP9 nucl | 10.30067 | 10.04481 | 8.149533 |
| 9408 | ENSG0000 | RPP38 | ribonuclea | 10.29316 | 8.60552 | 7.088516 |
| 3174 | ENSG0000 | ARMC6 | armadillo r | 10.25199 | 7.32026 | 4.350003 |
| 3054 | ENSG0000 | CCNE1 | cyclin E1 [S | 10.24745 | 9.273823 | 9.011338 |
| 8490 | ENSG0000 | CTDSP1 | CTD small | 10.24276 | 8.748641 | 8.592838 |
| 2018 | ENSG0000 | AAAS | aladin WD | 10.19139 | 8.830209 | 8.537313 |
| 11489 | ENSG0000 | METTL3 | methyltrar | 10.17613 | 8.708148 | 7.328437 |
| 10013 | ENSG0000 | RER1 | retention i | 10.16477 | 9.29137 | 9.177712 |
| 10879 | ENSG0000 | PSMD6 | proteasom | 10.15837 | 8.335893 | 7.447548 |
| 8135 | ENSG0000 | CEBPG | CCAAT/enh | 10.15545 | 9.415021 | 9.019631 |
| 4480 | ENSG0000 | MSH6 | mutS hom | 10.14436 | 7.819013 | 7.437489 |
| 385 | ENSG0000 | SLC30A9 | solute carr | 10.13692 | 8.314429 | 7.689321 |
| 10409 | ENSG0000 | MFSD12 | major facil | 10.11709 | 9.128155 | 8.915621 |
| 520 | ENSG0000 | RANBP3 | RAN bindir | 10.11578 | 9.777705 | 8.169106 |
| 3472 | ENSG0000 | CCNY | cyclin Y [Sc | 10.11105 | 9.293322 | 9.003388 |
| 28574 | ENSG0000 | PI4KA | phosphatic | 10.09026 | 8.368799 | 8.242111 |
| 2684 | ENSG0000 | SLC25A15 | solute carr | 10.07272 | 7.263007 | 7.222983 |
| 3928 | ENSG0000 | RNGTT | RNA guany | 10.04732 | 9.61697 | 8.955565 |
| 2912 | ENSG0000 | INTS10 | integrator | 10.04468 | 8.734763 | 8.203423 |
| 2271 | ENSG0000 | ASCC2 | activating : | 10.04211 | 7.435282 | 6.470017 |
| 21260 | ENSG0000 | VPS52 | VPS52, GA | 10.01098 | 9.316594 | 7.574327 |

| | | | | | | |
|---|---|---|---|---|---|---|
| 4428 | ENSG0000 | STK25 | serine/thre | 10.00833 | 9.348163 | 8.268376 |
| 21257 | ENSG0000 | EXOSC6 | exosome c | 9.99445 | 9.939908 | 7.274628 |
| 7686 | ENSG0000 | PEX5 | peroxisom | 9.906281 | 8.819931 | 8.796271 |
| 3976 | ENSG0000 | CCNC | cyclin C [Sc | 9.903785 | 9.660874 | 7.02707 |
| 5350 | ENSG0000 | NDUFAF4 | NADH:ubic | 9.901931 | 8.722854 | 5.706028 |
| 10386 | ENSG0000 | TONSL | tonsoku lik | 9.897355 | 9.55402 | 9.293925 |
| 6957 | ENSG0000 | ATP5G2 | ATP syntha | 9.880371 | 8.18771 | 6.674259 |
| 15472 | ENSG0000 | BRCC3 | BRCA1/BR( | 9.86495 | 8.052597 | 7.581665 |
| 10995 | ENSG0000 | MEAF6 | MYST/Esa1 | 9.863044 | 8.945966 | 8.059137 |
| 11406 | ENSG0000 | REEP3 | receptor ac | 9.816437 | 8.460997 | 8.362246 |
| 34802 | ENSG0000 | AC092718.4 | | 9.803774 | 4.971981 | 4.560462 |
| 12421 | ENSG0000 | AVEN | apoptosis a | 9.795914 | 9.774052 | 6.041557 |
| 7390 | ENSG0000 | THAP12 | THAP dom | 9.756425 | 9.352377 | 6.866814 |
| 10143 | ENSG0000 | ALG8 | ALG8, alph | 9.741848 | 8.977944 | 8.633951 |
| 7864 | ENSG0000 | SCAMP2 | secretory c | 9.728609 | 9.273008 | 6.464719 |
| 394 | ENSG0000 | MATR3 | matrin 3 [S | 9.728228 | 8.11584 | 7.446114 |
| 6197 | ENSG0000 | NDUFA10 | NADH:ubic | 9.71856 | 6.716323 | 6.150398 |
| 6070 | ENSG0000 | MPDU1 | mannose-F | 9.707186 | 9.43391 | 7.899196 |
| 10481 | ENSG0000 | POLR3K | RNA polym | 9.682933 | 7.440772 | 5.639552 |
| 13783 | ENSG0000 | SLC25A22 | solute carr | 9.614138 | 7.542437 | 5.107058 |
| 20382 | ENSG0000 | FASTKD5 | FAST kinas | 9.587683 | 8.2973 | 8.098774 |
| 8782 | ENSG0000 | NOM1 | nucleolar p | 9.572533 | 8.173755 | 7.0949 |
| 1512 | ENSG0000 | SNX5 | sorting nex | 9.5548 | 7.879192 | 6.930326 |
| 7234 | ENSG0000 | WDR33 | WD repeat | 9.553056 | 8.285851 | 7.671002 |
| 9049 | ENSG0000 | ARFGAP2 | ADP ribosy | 9.523931 | 6.746917 | 6.417107 |
| 5766 | ENSG0000 | UXT | ubiquitous | 9.488899 | 7.467801 | 6.391869 |
| 8295 | ENSG0000 | PI4KB | phosphatic | 9.483056 | 8.024214 | 8.001454 |
| 9523 | ENSG0000 | UBP1 | upstream b | 9.455594 | 6.358212 | 5.441427 |
| 4276 | ENSG0000 | SCAP | SREBF chap | 9.426533 | 7.21001 | 7.132209 |
| 11960 | ENSG0000 | GLOD4 | glyoxalase | 9.426414 | 8.096734 | 7.729248 |
| 4109 | ENSG0000 | PAPD7 | poly(A) RN | 9.381234 | 9.296951 | 8.76449 |
| 731 | ENSG0000 | TRAPPC3 | trafficking | 9.369439 | 8.132455 | 8.110899 |
| 36412 | ENSG0000 | AC093484.4 | | 9.35732 | 8.595233 | 7.697592 |
| 33087 | ENSG0000 | AL049844.1 | | 9.324617 | 7.246032 | 5.235715 |
| 12211 | ENSG0000 | GFM1 | G elongatic | 9.301325 | 7.544041 | 6.198838 |
| 5208 | ENSG0000 | RBBP6 | RB binding | 9.29623 | 9.288048 | 6.468832 |
| 15056 | ENSG0000 | SFXN4 | sideroflexi | 9.289209 | 8.137269 | 6.936261 |
| 10689 | ENSG0000 | MEMO1 | mediator c | 9.285704 | 8.404976 | 7.067341 |
| 1238 | ENSG0000 | SMARCE1 | SWI/SNF re | 9.278719 | 8.525697 | 7.515623 |
| 1035 | ENSG0000 | HYAL2 | hyalurono | 9.246101 | 5.995676 | 5.981175 |
| 20029 | ENSG0000 | MYCBP | MYC bindir | 9.226014 | 7.048538 | 6.634435 |
| 5185 | ENSG0000 | MTIF3 | mitochond | 9.225045 | 8.829593 | 7.560717 |
| 30242 | ENSG0000 | C15orf38-A | C15orf38-A | 9.216645 | 7.995933 | 0.504396 |
| 8569 | ENSG0000 | EIF2B5 | eukaryotic | 9.204866 | 8.092118 | 6.327327 |
| 7193 | ENSG0000 | NMT1 | N-myristoy | 9.194857 | 7.032845 | 6.543411 |
| 31801 | ENSG0000 | PINX1 | PIN2/TERF | 9.173453 | 8.525684 | 7.2557 |
| 9142 | ENSG0000 | TAOK2 | TAO kinase | 9.169683 | 7.232283 | 6.940502 |

| | | | | | | |
|---|---|---|---|---|---|---|
| 373 | ENSG0000 | DDX11 | DEAD/H-bo | 9.153231 | 7.511886 | 7.452939 |
| 1432 | ENSG0000 | ITCH | itchy E3 ub | 9.146575 | 8.533188 | 7.806489 |
| 1680 | ENSG0000 | SEH1L | SEH1 like n | 9.115478 | 7.710175 | 6.88963 |
| 3683 | ENSG0000 | ZPR1 | ZPR1 zinc f | 9.087782 | 8.984687 | 6.764091 |
| 29722 | ENSG0000 | RBM14-RB | RBM14-RB | 9.065267 | 5.948453 | 4.556448 |
| 2236 | ENSG0000 | SBF1 | SET binding | 9.058635 | 8.927323 | 7.998462 |
| 7427 | ENSG0000 | PPP2R1B | protein ph | 9.051011 | 8.995127 | 7.831947 |
| 25889 | ENSG0000 | AL513165.1 | | 9.046348 | 7.165597 | 6.20496 |
| 4189 | ENSG0000 | H2AFY | H2A histon | 9.020362 | 7.628587 | 6.581438 |
| 16841 | ENSG0000 | SPG7 | SPG7, para | 9.000011 | 8.558242 | 8.41071 |
| 7745 | ENSG0000 | C12orf10 | chromosor | 8.996541 | 7.306556 | 7.014899 |
| 29491 | ENSG0000 | MARS2 | methionyl- | 8.946663 | 7.345173 | 7.090732 |
| 3256 | ENSG0000 | ABHD11 | abhydrolas | 8.909522 | 7.888203 | 5.695118 |
| 4955 | ENSG0000 | IDE | insulin deg | 8.898403 | 7.320346 | 7.056671 |
| 10404 | ENSG0000 | LRWD1 | leucine rich | 8.886779 | 8.444208 | 8.270918 |
| 4732 | ENSG0000 | FASTKD2 | FAST kinas | 8.872723 | 5.909459 | 5.361554 |
| 12755 | ENSG0000 | TMEM126I | transmeml | 8.844174 | 8.763195 | 8.373013 |
| 8451 | ENSG0000 | PKP4 | plakophilin | 8.829644 | 7.787978 | 5.48101 |
| 10485 | ENSG0000 | JMJD8 | jumonji do | 8.817064 | 7.056263 | 6.253615 |
| 7033 | ENSG0000 | URB2 | URB2 ribos | 8.805948 | 7.333161 | 4.708419 |
| 1384 | ENSG0000 | POLD3 | DNA polym | 8.802365 | 8.260916 | 7.843968 |
| 3597 | ENSG0000 | TMEM104 | transmeml | 8.741087 | 4.447272 | 3.757696 |
| 28567 | ENSG0000 | PWP2 | PWP2, sma | 8.625533 | 7.158429 | 5.504746 |
| 10958 | ENSG0000 | GYG1 | glycogenin | 8.610938 | 6.676612 | 6.655158 |
| 1620 | ENSG0000 | KAT6A | lysine acet | 8.600026 | 8.497467 | 3.910436 |
| 6873 | ENSG0000 | UBAC2 | UBA doma | 8.590283 | 6.255289 | 5.695586 |
| 2280 | ENSG0000 | SAMM50 | SAMM50 s | 8.586439 | 6.785892 | 6.504513 |
| 9039 | ENSG0000 | DGKZ | diacylglyce | 8.583393 | 7.541589 | 6.442582 |
| 3944 | ENSG0000 | FANCE | Fanconi an | 8.548 | 5.865289 | 5.835314 |
| 2300 | ENSG0000 | L3MBTL2 | L3MBTL2, p | 8.542132 | 5.404734 | 4.535903 |
| 2500 | ENSG0000 | FERMT1 | fermitin fa | 8.537574 | 6.753781 | 6.412248 |
| 6320 | ENSG0000 | RBM39 | RNA bindir | 8.531394 | 8.358386 | 6.147841 |
| 34499 | ENSG0000 | AC138811.2 | | 8.50181 | 6.763629 | 6.144462 |
| 13212 | ENSG0000 | SSSCA1 | Sjogren syr | 8.491331 | 7.483851 | 5.415517 |
| 6565 | ENSG0000 | DPH2 | DPH2 hom | 8.468084 | 8.341358 | 4.780741 |
| 40408 | ENSG0000 | IKBKGP1 | inhibitor of | 8.460176 | 6.301657 | 6.014976 |
| 10286 | ENSG0000 | U2AF1 | U2 small nu | 8.426784 | 7.199976 | 6.552566 |
| 7303 | ENSG0000 | DSCC1 | DNA replic | 8.42531 | 7.204708 | 6.755103 |
| 2793 | ENSG0000 | HMOX2 | heme oxyg | 8.413159 | 5.95714 | 5.892117 |
| 2197 | ENSG0000 | PATZ1 | POZ/BTB a | 8.41086 | 7.904053 | 6.975184 |
| 15522 | ENSG0000 | YTHDF3 | YTH N6-me | 8.399126 | 7.999017 | 4.571077 |
| 5645 | ENSG0000 | PSMF1 | proteasom | 8.397897 | 6.440797 | 5.582177 |
| 4774 | ENSG0000 | DCLRE1B | DNA cross- | 8.373776 | 7.490703 | 7.223162 |
| 7908 | ENSG0000 | GCSH | glycine clea | 8.363344 | 6.014386 | 5.710154 |
| 11047 | ENSG0000 | CDC25A | cell divisior | 8.358787 | 6.049488 | 4.43748 |
| 38889 | ENSG0000 | NUDT3 | nudix hydr | 8.35855 | 7.129074 | 6.248735 |
| 10300 | ENSG0000 | C21orf2 | chromosor | 8.334491 | 7.374202 | 6.540086 |

| | | | | | | |
|---|---|---|---|---|---|---|
| 7841 | ENSG0000 | TSPAN3 | tetraspanii | 8.328496 | 7.295377 | 5.363375 |
| 4407 | ENSG0000 | CHMP3 | charged m | 8.319394 | 6.607552 | 5.379664 |
| 28062 | ENSG0000 | ADSL | adenylosu | 8.295219 | 4.681795 | 4.235517 |
| 20987 | ENSG0000 | CCNL2 | cyclin L2 [S | 8.294653 | 6.903026 | 6.49938 |
| 4319 | ENSG0000 | TTL | tubulin tyr | 8.292836 | 7.764052 | 6.934944 |
| 6089 | ENSG0000 | BCL2L2 | BCL2 like 2 | 8.259896 | 4.560494 | 1.461276 |
| 13116 | ENSG0000 | KAT5 | lysine acet | 8.25498 | 7.931692 | 6.135072 |
| 14661 | ENSG0000 | GINS3 | GINS comp | 8.244622 | 8.036868 | 7.417781 |
| 9819 | ENSG0000 | PTDSS1 | phosphatic | 8.234533 | 6.280263 | 6.223547 |
| 20236 | ENSG0000 | TOMM6 | translocase | 8.210056 | 1.236434 | 1.062361 |
| 3981 | ENSG0000 | ASCC3 | activating s | 8.18994 | 8.171029 | 6.69769 |
| 2298 | ENSG0000 | RBX1 | ring-box 1 | 8.15892 | 6.28535 | 6.191306 |
| 15539 | ENSG0000 | SLC52A2 | solute carr | 8.148011 | 6.912997 | 6.120893 |
| 13335 | ENSG0000 | CTU2 | cytosolic th | 8.143389 | 2.949094 | 2.239957 |
| 1815 | ENSG0000 | NSFL1C | NSFL1 cofa | 8.126411 | 5.659761 | 4.629152 |
| 812 | ENSG0000 | TARBP1 | TAR (HIV-1 | 8.095022 | 7.80357 | 7.450447 |
| 5266 | ENSG0000 | TRIM24 | tripartite n | 8.035129 | 7.90797 | 6.355142 |
| 11514 | ENSG0000 | CPSF2 | cleavage ar | 8.017974 | 6.927559 | 6.243005 |
| 7620 | ENSG0000 | RAP1GDS1 | Rap1 GTPa | 7.992668 | 6.689042 | 5.807271 |
| 1222 | ENSG0000 | AP1M1 | adaptor re | 7.987803 | 6.044289 | 5.361143 |
| 2780 | ENSG0000 | CAPN15 | calpain 15 | 7.983488 | 7.969197 | 6.188996 |
| 2733 | ENSG0000 | TANGO6 | transport a | 7.9797 | 5.844864 | 5.127166 |
| 10856 | ENSG0000 | SNHG16 | small nucle | 7.965902 | 7.102767 | 6.927815 |
| 13442 | ENSG0000 | RSRC1 | arginine ar | 7.953102 | 7.193267 | 7.152529 |
| 15033 | ENSG0000 | TSEN15 | tRNA splici | 7.884409 | 4.222195 | 3.54326 |
| 15161 | ENSG0000 | DIABLO | diablo IAP- | 7.884054 | 7.435105 | 5.778881 |
| 2820 | ENSG0000 | AAGAB | alpha and g | 7.851003 | 6.996966 | 5.859477 |
| 20399 | ENSG0000 | VPS16 | VPS16, COI | 7.782236 | 6.200971 | 6.06309 |
| 11411 | ENSG0000 | PCF11 | PCF11 clea | 7.76702 | 6.396957 | 5.18688 |
| 6611 | ENSG0000 | MPRIP | myosin pho | 7.757428 | 7.025232 | 6.057091 |
| 2845 | ENSG0000 | TJP1 | tight juncti | 7.743931 | 6.72082 | 6.120538 |
| 17104 | ENSG0000 | TMEM184I | transmeml | 7.737094 | 5.20977 | 5.123468 |
| 10225 | ENSG0000 | CHCHD6 | coiled-coil- | 7.722597 | 4.951732 | 4.575455 |
| 2082 | ENSG0000 | CDC7 | cell divisior | 7.70772 | 7.53856 | 7.292995 |
| 14951 | ENSG0000 | BCOR | BCL6 corep | 7.690971 | 7.539103 | 6.488517 |
| 40411 | ENSG0000 | U2AF1L5 | U2 small nu | 7.678132 | 6.479262 | 5.912246 |
| 5087 | ENSG0000 | TARDBP | TAR DNA b | 7.675783 | 5.992464 | 5.208795 |
| 2674 | ENSG0000 | STK24 | serine/thre | 7.668555 | 7.622517 | 6.04278 |
| 13367 | ENSG0000 | ZWILCH | zwilch kine | 7.662556 | 6.683064 | 6.32813 |
| 32206 | ENSG0000 | MPV17L2 | MPV17 mit | 7.660579 | 5.840677 | 5.092953 |
| 32886 | ENSG0000 | HMBS | hydroxyme | 7.657816 | 5.239277 | 5.035284 |
| 3704 | ENSG0000 | DCPS | decapping | 7.656712 | 5.774767 | 5.289417 |
| 7612 | ENSG0000 | SEC31A | SEC31 hom | 7.656521 | 6.636598 | 6.064699 |
| 16472 | ENSG0000 | XRCC2 | X-ray repai | 7.63748 | 6.721907 | 5.992733 |
| 5587 | ENSG0000 | MSTO1 | misato 1, r | 7.634116 | 6.568103 | 5.033449 |
| 18995 | ENSG0000 | MIR25 | microRNA | 7.611438 | 7.575961 | 4.563343 |
| 1236 | ENSG0000 | NLE1 | notchless h | 7.602152 | 7.481042 | 6.507733 |

| | | | | | | |
|---|---|---|---|---|---|---|
| 15438 | ENSG0000 | GAS2L1 | growth arr | 7.579675 | 6.747352 | 5.570168 |
| 8636 | ENSG0000 | RASA1 | RAS p21 pr | 7.554484 | 6.974187 | 6.044137 |
| 4262 | ENSG0000 | UMPS | uridine mo | 7.547073 | 6.648535 | 5.411408 |
| 19746 | ENSG0000 | QTRT1 | queuine tR | 7.542755 | 6.873725 | 6.237172 |
| 9317 | ENSG0000 | ACSL1 | acyl-CoA sy | 7.486164 | 5.56809 | 5.209711 |
| 1273 | ENSG0000 | BCS1L | BCS1 homo | 7.430348 | 5.125093 | 3.694977 |
| 3359 | ENSG0000 | SUSD1 | sushi doma | 7.415343 | 7.364752 | 6.381788 |
| 13485 | ENSG0000 | MRPS22 | mitochond | 7.376991 | 6.749553 | 6.245345 |
| 11445 | ENSG0000 | PDZD8 | PDZ domai | 7.371519 | 7.025108 | 5.755921 |
| 1685 | ENSG0000 | MAP3K4 | mitogen-ac | 7.368469 | 6.374961 | 5.244355 |
| 3083 | ENSG0000 | CACTIN | cactin, spli | 7.346793 | 6.327783 | 4.873601 |
| 35800 | ENSG0000 | FAM58A | family with | 7.3141 | 6.735687 | 4.300014 |
| 15098 | ENSG0000 | TBL3 | transducin | 7.299543 | 5.269053 | 3.84424 |
| 42438 | ENSG0000 | FP236383.1 | | 7.297156 | 4.767543 | 4.256746 |
| 8498 | ENSG0000 | DYNC1LI1 | dynein cyto | 7.275766 | 6.568816 | 5.401376 |
| 12989 | ENSG0000 | TP53RK | TP53 regul | 7.25967 | 6.896627 | 6.404745 |
| 556 | ENSG0000 | CUL3 | cullin 3 [So | 7.251912 | 5.99577 | 4.418097 |
| 9101 | ENSG0000 | CHEK1 | checkpoint | 7.220088 | 6.949594 | 5.817353 |
| 2000 | ENSG0000 | MFSD11 | major facil | 7.204458 | 4.991221 | 4.052378 |
| 8189 | ENSG0000 | PLK4 | polo like ki | 7.201859 | 5.878519 | 5.596874 |
| 10513 | ENSG0000 | TAF6L | TATA-box b | 7.158055 | 6.301182 | 5.119861 |
| 6392 | ENSG0000 | PPFIA1 | PTPRF inte | 7.152625 | 6.999887 | 6.418054 |
| 12356 | ENSG0000 | TM2D2 | TM2 doma | 7.121272 | 5.699328 | 5.639732 |
| 14540 | ENSG0000 | FKRP | fukutin rela | 7.10814 | 5.598403 | 4.60665 |
| 15011 | ENSG0000 | NELFA | negative el | 7.10527 | 5.947824 | 5.685606 |
| 4222 | ENSG0000 | PCCB | propionyl-C | 7.09951 | 4.561657 | 3.903639 |
| 9817 | ENSG0000 | UQCRB | ubiquinol-c | 7.078947 | 6.199265 | 5.978307 |
| 6371 | ENSG0000 | MGAT1 | mannosyl ( | 7.07586 | 6.331751 | 5.748421 |
| 26996 | ENSG0000 | EIF1AXP1 | eukaryotic | 7.040864 | 5.901663 | 4.553337 |
| 328 | ENSG0000 | LARS2 | leucyl-tRNA | 7.027723 | 6.16391 | 5.457008 |
| 8865 | ENSG0000 | TACC1 | transformi | 7.021204 | 3.554134 | 3.242195 |
| 4715 | ENSG0000 | STK11 | serine/thre | 7.012387 | 6.72528 | 6.449911 |
| 12770 | ENSG0000 | FAM98B | family with | 6.992233 | 6.833113 | 5.91134 |
| 6609 | ENSG0000 | PEMT | phosphatid | 6.987673 | 5.008889 | 4.992207 |
| 10704 | ENSG0000 | NUP35 | nucleopori | 6.983962 | 6.102747 | 4.882257 |
| 11876 | ENSG0000 | WDR81 | WD repeat | 6.96082 | 5.811789 | 5.712916 |
| 15699 | ENSG0000 | INSIG1 | insulin indu | 6.958233 | 6.33576 | 5.222735 |
| 35915 | ENSG0000 | FAM72C | family with | 6.954453 | 5.992784 | 5.655994 |
| 7869 | ENSG0000 | POLG | DNA polym | 6.933625 | 5.195216 | 4.340462 |
| 20080 | ENSG0000 | FAM72D | family with | 6.93292 | 5.956487 | 5.590465 |
| 13447 | ENSG0000 | PTDSS2 | phosphatid | 6.915291 | 6.759537 | 6.016929 |
| 3274 | ENSG0000 | SNX8 | sorting nex | 6.904828 | 6.854821 | 6.834707 |
| 3915 | ENSG0000 | RWDD1 | RWD doma | 6.888759 | 6.793311 | 6.501355 |
| 17991 | ENSG0000 | MSTO2P | misato fam | 6.862811 | 6.300545 | 4.137292 |
| 42504 | ENSG0000 | MATR3 | matrin 3 [S | 6.850093 | 4.818399 | 4.776335 |
| 9001 | ENSG0000 | DNAJB12 | DnaJ heat s | 6.845819 | 4.87479 | 3.664747 |
| 10814 | ENSG0000 | KRTCAP2 | keratinocy | 6.84307 | 5.512477 | 5.104826 |

| | | | | | | |
|---|---|---|---|---|---|---|
| 20939 | ENSG0000 | C6orf226 | chromosor | 6.841044 | 3.477827 | 3.402812 |
| 8927 | ENSG0000 | INIP | INTS3 and | 6.826295 | 6.669529 | 6.62415 |
| 1410 | ENSG0000 | TIGAR | TP53 induc | 6.820669 | 5.765949 | 4.549657 |
| 17046 | ENSG0000 | STK39 | serine/thre | 6.803102 | 6.49978 | 5.646043 |
| 4579 | ENSG0000 | TTF2 | transcriptio | 6.796582 | 5.945746 | 5.016215 |
| 1288 | ENSG0000 | TIPIN | TIMELESS i | 6.786118 | 6.118845 | 6.097302 |
| 13247 | ENSG0000 | SLC19A1 | solute carr | 6.782359 | 5.0985 | 4.185842 |
| 19685 | ENSG0000 | BTF3L4P2 | basic trans | 6.781351 | 5.242445 | 5.033442 |
| 30219 | ENSG0000 | AL157400.5 | | 6.711695 | 4.516106 | 3.85733 |
| 40264 | ENSG0000 | FP565260.1 | | 6.709509 | 5.186905 | 4.336941 |
| 15826 | ENSG0000 | RNF220 | ring finger | 6.705455 | 6.346459 | 5.940943 |
| 11857 | ENSG0000 | MVD | mevalonat | 6.668791 | 5.34602 | 4.313724 |
| 36245 | ENSG0000 | AC016876.2 | | 6.642748 | 5.778657 | 4.221135 |
| 24914 | ENSG0000 | ANXA2P2 | annexin A2 | 6.63457 | 5.019907 | 3.744174 |
| 13530 | ENSG0000 | TMEM9B | TMEM9 dc | 6.631679 | 6.179509 | 3.698375 |
| 10390 | ENSG0000 | LRRC14 | leucine rich | 6.591194 | 5.327183 | 3.843736 |
| 16649 | ENSG0000 | ENTPD4 | ectonuclec | 6.558537 | 6.127062 | 5.659961 |
| 41601 | ENSG0000 | FP671120.1 | | 6.551807 | 4.285828 | 3.732527 |
| 14660 | ENSG0000 | PRKAG1 | protein kin | 6.551374 | 6.082129 | 5.315717 |
| 283 | ENSG0000 | DGKD | diacylglyce | 6.523818 | 5.488072 | 5.340297 |
| 7444 | ENSG0000 | KNL1 | kinetochor | 6.509364 | 4.531965 | 3.221258 |
| 13134 | ENSG0000 | COQ2 | coenzyme | 6.50585 | 3.209601 | 2.646845 |
| 2306 | ENSG0000 | POLR3H | RNA polym | 6.451982 | 6.316895 | 6.186534 |
| 11309 | ENSG0000 | NUDT2 | nudix hydr | 6.441376 | 4.993679 | 4.956502 |
| 42011 | ENSG0000 | BCLAF1P2 | BCL2 assoc | 6.412813 | 6.102315 | 4.277534 |
| 12192 | ENSG0000 | CNNM3 | cyclin and | 6.398804 | 4.437934 | 4.435025 |
| 19663 | ENSG0000 | TCTEX1D2 | Tctex1 dor | 6.387573 | 3.62631 | 2.669077 |
| 21649 | ENSG0000 | AC004552.1 | | 6.372096 | 3.857707 | 2.768913 |
| 26223 | ENSG0000 | FABP5P7 | fatty acid b | 6.364655 | 6.052524 | 5.967041 |
| 15521 | ENSG0000 | ANKFY1 | ankyrin rep | 6.347983 | 5.986451 | 5.663699 |
| 6505 | ENSG0000 | FIGNL1 | fidgetin like | 6.340331 | 5.691534 | 4.995361 |
| 14370 | ENSG0000 | ZADH2 | zinc bindin | 6.334963 | 5.063597 | 4.926524 |
| 8431 | ENSG0000 | TMEM177 | transmeml | 6.334168 | 5.057415 | 4.722258 |
| 8779 | ENSG0000 | TMEM140 | transmeml | 6.319578 | 5.814843 | 5.101949 |
| 11424 | ENSG0000 | RPUSD4 | RNA pseud | 6.311903 | 5.783061 | 4.696092 |
| 10295 | ENSG0000 | TRAPPC10 | trafficking | 6.292981 | 6.150488 | 6.017862 |
| 13177 | ENSG0000 | MZT2A | mitotic spi | 6.287716 | 5.368119 | 5.091711 |
| 4794 | ENSG0000 | UCHL3 | ubiquitin C | 6.279801 | 5.61859 | 4.050629 |
| 13641 | ENSG0000 | TPRN | taperin [So | 6.278987 | 5.38079 | 4.496599 |
| 1782 | ENSG0000 | METTL2A | methyltrar | 6.253889 | 5.422744 | 5.322843 |
| 12867 | ENSG0000 | SPATA5L1 | spermatog | 6.249833 | 3.619413 | 3.355234 |
| 22792 | ENSG0000 | DANCR | differentia | 6.238338 | 5.79427 | 3.323899 |
| 9394 | ENSG0000 | TADA1 | transcriptio | 6.232582 | 5.864898 | 5.115715 |
| 6357 | ENSG0000 | TBC1D5 | TBC1 doma | 6.186288 | 5.622271 | 4.87338 |
| 2254 | ENSG0000 | MCAT | malonyl-Co | 6.177552 | 5.058369 | 4.809226 |
| 844 | ENSG0000 | SFSWAP | splicing fac | 6.172583 | 5.908661 | 4.234056 |
| 3401 | ENSG0000 | EXOSC3 | exosome c | 6.134439 | 5.514108 | 4.839525 |

| | | | | | | |
|---|---|---|---|---|---|---|
| 9396 | ENSG0000 | CWF19L2 | CWF19 like | 6.076875 | 5.624583 | 4.402078 |
| 11414 | ENSG0000 | LRR1 | leucine rich | 6.047224 | 4.598668 | 4.27188 |
| 9866 | ENSG0000 | MALSU1 | mitochond | 6.006693 | 5.784858 | 5.368435 |
| 8650 | ENSG0000 | COMMD1C | COMM dor | 5.952386 | 5.284932 | 4.134609 |
| 12841 | ENSG0000 | SLC25A33 | solute carr | 5.948792 | 5.934718 | 5.69663 |
| 2407 | ENSG0000 | DCAF11 | DDB1 and | 5.933115 | 4.81372 | 4.576848 |
| 19588 | ENSG0000 | AC064799.1 | | 5.914703 | 4.253324 | 4.198339 |
| 2880 | ENSG0000 | POP1 | POP1 hom | 5.913683 | 5.579688 | 4.870246 |
| 4144 | ENSG0000 | MSH3 | mutS hom | 5.903284 | 5.105882 | 4.421234 |
| 5316 | ENSG0000 | NLN | neurolysin | 5.886964 | 4.907699 | 3.084765 |
| 7758 | ENSG0000 | SETD1B | SET domai | 5.866088 | 5.595341 | 3.915859 |
| 369 | ENSG0000 | POLR3B | RNA polym | 5.853717 | 5.417529 | 3.490453 |
| 4768 | ENSG0000 | FBXL5 | F-box and l | 5.827509 | 5.563697 | 5.334471 |
| 5716 | ENSG0000 | TUBGCP3 | tubulin gar | 5.82096 | 5.009154 | 3.719839 |
| 16542 | ENSG0000 | PPTC7 | PTC7 prote | 5.81683 | 5.638853 | 5.526065 |
| 1178 | ENSG0000 | PDCD2 | programm | 5.813871 | 5.097418 | 4.083261 |
| 10895 | ENSG0000 | FABP5 | fatty acid b | 5.805845 | 5.430727 | 4.782442 |
| 42555 | ENSG0000 | SNHG4 | small nucle | 5.786795 | 4.204543 | 2.390385 |
| 4111 | ENSG0000 | HMGCS1 | 3-hydroxy- | 5.775037 | 5.76042 | 3.098665 |
| 19747 | ENSG0000 | CHUK | conserved | 5.773218 | 5.674601 | 5.133292 |
| 4670 | ENSG0000 | DPH5 | diphthamid | 5.769405 | 4.039472 | 3.409795 |
| 4553 | ENSG0000 | SLC35D1 | solute carr | 5.758339 | 4.817187 | 4.051354 |
| 4330 | ENSG0000 | STEAP3 | STEAP3 me | 5.741435 | 4.19205 | 3.34712 |
| 14978 | ENSG0000 | TREX2 | three prim | 5.732505 | 2.999512 | 1.18776 |
| 38278 | ENSG0000 | HSPE1-MO | HSPE1-MO | 5.72809 | 4.614507 | 1.851598 |
| 41103 | ENSG0000 | AC107075.1 | | 5.722751 | 3.946186 | 3.702725 |
| 10262 | ENSG0000 | UBE2J2 | ubiquitin c | 5.720055 | 5.305919 | 5.032195 |
| 5502 | ENSG0000 | MED20 | mediator c | 5.716644 | 4.330554 | 3.726945 |
| 2451 | ENSG0000 | BMP7 | bone morp | 5.715756 | 5.068627 | 4.927605 |
| 22420 | ENSG0000 | UQCRFS1P | ubiquinol-c | 5.697396 | 3.829453 | 3.465638 |
| 11320 | ENSG0000 | METTL2B | methyltran | 5.694072 | 4.724545 | 4.446478 |
| 7293 | ENSG0000 | GOLGA1 | golgin A1 [ | 5.693509 | 5.296996 | 5.100606 |
| 2207 | ENSG0000 | POLR2F | RNA polym | 5.688864 | 3.672849 | 3.614642 |
| 1414 | ENSG0000 | PPP2R5C | protein ph | 5.685486 | 5.322232 | 4.575549 |
| 1329 | ENSG0000 | EXOSC7 | exosome c | 5.664106 | 4.522365 | 4.359769 |
| 7831 | ENSG0000 | CDAN1 | codanin 1 | | 5.661292 | 4.633522 | 4.164245 |
| 7963 | ENSG0000 | PTRH2 | peptidyl-tR | 5.655579 | 5.057104 | 4.391697 |
| 34681 | ENSG0000 | PMF1-BGL | PMF1-BGL | 5.64907 | 5.23419 | 4.387576 |
| 15743 | ENSG0000 | BCR | BCR, RhoG | 5.627285 | 5.336674 | 5.245213 |
| 3053 | ENSG0000 | POP4 | POP4 hom | 5.609863 | 5.427457 | 5.007831 |
| 2592 | ENSG0000 | SUV39H1 | suppressor | 5.58358 | 4.207565 | 2.328941 |
| 5894 | ENSG0000 | DGCR8 | DGCR8, mi | 5.569175 | 4.521459 | 3.282211 |
| 9481 | ENSG0000 | TXNDC11 | thioredoxi | 5.561773 | 5.358656 | 5.332547 |
| 6187 | ENSG0000 | RTN4IP1 | reticulon 4 | 5.53471 | 3.814054 | 3.469926 |
| 9840 | ENSG0000 | KAT6B | lysine acet | 5.514005 | 5.403837 | 5.288766 |
| 5861 | ENSG0000 | METTL16 | methyltran | 5.501244 | 2.741195 | 2.491977 |
| 16273 | ENSG0000 | FAM53B | family with | 5.495018 | 4.867406 | 4.775021 |

| | | | | | | |
|---|---|---|---|---|---|---|
| 2544 | ENSG0000 | PIGU | phosphatic | 5.472323 | 4.32876 | 2.868732 |
| 83 | ENSG0000 | LIG3 | DNA ligase | 5.454673 | 4.339067 | 4.061282 |
| 4037 | ENSG0000 | INO80B | INO80 com | 5.437811 | 5.15583 | 5.11758 |
| 2248 | ENSG0000 | HMGXB4 | HMG-box c | 5.435306 | 4.748295 | 4.368909 |
| 28878 | ENSG0000 | MICAL3 | microtubul | 5.397287 | 5.175759 | 4.948865 |
| 6001 | ENSG0000 | PRMT7 | protein arg | 5.377566 | 3.865504 | 3.528736 |
| 15384 | ENSG0000 | NSMCE3 | NSE3 homo | 5.374972 | 5.27409 | 5.146708 |
| 146 | ENSG0000 | DBF4 | DBF4 zinc f | 5.368781 | 3.775337 | 3.510551 |
| 2868 | ENSG0000 | INTS9 | integrator | 5.326029 | 5.00283 | 4.535659 |
| 12589 | ENSG0000 | ARMC10 | armadillo r | 5.325955 | 4.42783 | 4.209662 |
| 5393 | ENSG0000 | OBSL1 | obscurin lil | 5.294963 | 4.352203 | 4.257244 |
| 10385 | ENSG0000 | VPS28 | VPS28, ESC | 5.288621 | 5.012477 | 4.257182 |
| 10712 | ENSG0000 | SMC6 | structural r | 5.282131 | 4.745655 | 4.350147 |
| 18151 | ENSG0000 | ZBTB12 | zinc finger | 5.270045 | 5.219126 | 4.898879 |
| 2199 | ENSG0000 | TFIP11 | tuftelin int | 5.252053 | 5.038165 | 4.549871 |
| 4820 | ENSG0000 | C1orf198 | chromosor | 5.246862 | 5.241258 | 4.990447 |
| 7494 | ENSG0000 | PPM1B | protein ph | 5.231483 | 5.183876 | 4.887573 |
| 13434 | ENSG0000 | PDE12 | phosphodi | 5.22601 | 4.184103 | 3.85692 |
| 37806 | ENSG0000 | IKBKG | inhibitor of | 5.208982 | 4.951035 | 4.229335 |
| 29 | ENSG0000 | MAD1L1 | MAD1 mito | 5.185974 | 5.183794 | 4.55897 |
| 10504 | ENSG0000 | CLPB | ClpB homo | 5.180772 | 4.529509 | 4.433555 |
| 18009 | ENSG0000 | HIST2H2BC | histone clu | 5.172565 | 4.684597 | 4.036785 |
| 4790 | ENSG0000 | EEF2KMT | eukaryotic | 5.165313 | 2.629602 | 1.747499 |
| 12992 | ENSG0000 | BPGM | bisphosph | 5.157216 | 4.597331 | 4.025079 |
| 10371 | ENSG0000 | CCDC12 | coiled-coil | 5.131691 | 3.444158 | 2.518908 |
| 8874 | ENSG0000 | MRPS28 | mitochond | 5.115763 | 4.299886 | 4.132835 |
| 5836 | ENSG0000 | AUNIP | aurora kina | 5.114926 | 4.378766 | 4.194701 |
| 2540 | ENSG0000 | DHX35 | DEAH-box | 5.102273 | 3.455618 | 2.997801 |
| 20232 | ENSG0000 | ZBED1 | zinc finger | 5.093019 | 4.942993 | 2.293821 |
| 15542 | ENSG0000 | PCYT2 | phosphate | 5.087779 | 2.317664 | 1.905661 |
| 10482 | ENSG0000 | SNRNP25 | small nucle | 5.066679 | 4.504304 | 4.350875 |
| 4083 | ENSG0000 | GMDS | GDP-mann | 5.050212 | 3.04023 | 2.705069 |
| 24059 | ENSG0000 | PRRT3-AS1 | PRRT3 anti | 5.003181 | 4.980686 | 4.643156 |
| 2736 | ENSG0000 | SLC7A6OS | solute carr | 4.99704 | 4.995591 | 3.556196 |
| 6740 | ENSG0000 | ARL8B | ADP ribosy | 4.996417 | 4.016718 | 3.769361 |
| 11294 | ENSG0000 | DCAF13 | DDB1 and | 4.982423 | 4.571295 | 3.512846 |
| 7655 | ENSG0000 | RNF185 | ring finger | 4.979249 | 2.249909 | 2.21873 |
| 4309 | ENSG0000 | NEK4 | NIMA relat | 4.960703 | 4.565395 | 4.466094 |
| 3149 | ENSG0000 | GCDH | glutaryl-Co | 4.953018 | 4.793406 | 2.925249 |
| 13471 | ENSG0000 | ZDHHC14 | zinc finger | 4.902027 | 4.034066 | 3.945139 |
| 490 | ENSG0000 | MIPEP | mitochond | 4.885595 | 4.574912 | 4.14779 |
| 9452 | ENSG0000 | HHEX | hematopoi | 4.866394 | 4.850608 | 3.911405 |
| 18178 | ENSG0000 | ABHD16A | abhydrolas | 4.855365 | 4.578947 | 4.506682 |
| 1883 | ENSG0000 | MAEA | macrophag | 4.837235 | 4.454474 | 4.441221 |
| 10182 | ENSG0000 | HLCS | holocarbox | 4.83553 | 4.744149 | 4.248395 |
| 17086 | ENSG0000 | GPATCH3 | G-patch dc | 4.829777 | 4.811946 | 4.0703 |
| 3304 | ENSG0000 | NRF1 | nuclear res | 4.823943 | 3.580136 | 3.46646 |

| | | | | | | |
|---|---|---|---|---|---|---|
| 275 | ENSG0000 | ZZEF1 | zinc finger | 4.823219 | 4.762039 | 3.857852 |
| 20792 | ENSG0000 | AC239868.1 | | 4.80381 | 4.204356 | 3.866548 |
| 39509 | ENSG0000 | FAM27E3 | family with | 4.803351 | 4.778049 | 4.747342 |
| 6154 | ENSG0000 | ECSIT | ECSIT signa | 4.766491 | 4.109024 | 3.903183 |
| 6482 | ENSG0000 | ILKAP | ILK associa | 4.747546 | 4.596468 | 3.738818 |
| 9732 | ENSG0000 | TMEM237 | transmeml | 4.738062 | 4.229616 | 3.905466 |
| 10645 | ENSG0000 | RBM15 | RNA bindir | 4.726724 | 3.471066 | 3.265577 |
| 2699 | ENSG0000 | MGRN1 | mahogunir | 4.713356 | 4.482608 | 4.103829 |
| 6564 | ENSG0000 | MMACHC | methylmal | 4.711766 | 3.727292 | 3.694506 |
| 10423 | ENSG0000 | BDH1 | 3-hydroxyk | 4.704729 | 4.456809 | 3.553212 |
| 7719 | ENSG0000 | FAM222A | family with | 4.698489 | 4.67693 | 3.438267 |
| 6035 | ENSG0000 | CLN6 | CLN6, tran: | 4.658906 | 2.268604 | 2.198388 |
| 7452 | ENSG0000 | RMDN3 | regulator c | 4.656443 | 4.26866 | 4.261204 |
| 21667 | ENSG0000 | ADM5 | adrenomed | 4.655072 | 3.957926 | 2.286739 |
| 17177 | ENSG0000 | DCLRE1A | DNA cross- | 4.615435 | 4.348079 | 3.755404 |
| 34694 | ENSG0000 | SNHG19 | small nucle | 4.608186 | 2.967936 | 2.950342 |
| 2688 | ENSG0000 | VWA8 | von Willeb | 4.60554 | 4.433963 | 3.619329 |
| 10370 | ENSG0000 | NBEAL2 | neurobeac | 4.605152 | 4.472962 | 3.807106 |
| 7184 | ENSG0000 | MTHFS | methenylt | 4.6008 | 4.424108 | 3.566274 |
| 9112 | ENSG0000 | COMMD7 | COMM dor | 4.593527 | 4.205761 | 3.864109 |
| 7141 | ENSG0000 | CCM2 | CCM2 scaf | 4.591767 | 3.472653 | 2.988973 |
| 39591 | ENSG0000 | AL121832.2 | | 4.589688 | 4.176112 | 4.127061 |
| 14386 | ENSG0000 | EXOC3 | exocyst cor | 4.586626 | 4.540565 | 4.294636 |
| 2787 | ENSG0000 | EARS2 | glutamyl-tI | 4.586545 | 4.185282 | 3.598084 |
| 18223 | ENSG0000 | C6orf136 | chromosor | 4.581016 | 4.053905 | 3.952248 |
| 4930 | ENSG0000 | EIF2B2 | eukaryotic | 4.564579 | 3.49978 | 3.085963 |
| 15925 | ENSG0000 | NHEJ1 | non-homo | 4.545495 | 4.517356 | 3.747618 |
| 15469 | ENSG0000 | FAAP100 | Fanconi an | 4.54255 | 4.423826 | 3.458121 |
| 29591 | ENSG0000 | TRIM52-AS | TRIM52 an | 4.54243 | 4.261785 | 3.529187 |
| 10311 | ENSG0000 | C21orf58 | chromosor | 4.538451 | 4.090994 | 3.980876 |
| 2520 | ENSG0000 | CDK5RAP1 | CDK5 regul | 4.511507 | 3.4338 | 3.316618 |
| 2534 | ENSG0000 | ACTR5 | ARP5 actin | 4.50053 | 3.556901 | 3.186643 |
| 9082 | ENSG0000 | KAT14 | lysine acet | 4.494629 | 2.918396 | 2.771647 |
| 9332 | ENSG0000 | CENPJ | centromer | 4.474617 | 4.332217 | 3.759355 |
| 19079 | ENSG0000 | MIR570 | microRNA | 4.472516 | 3.914269 | 2.511742 |
| 11750 | ENSG0000 | DIS3L | DIS3 like e: | 4.470502 | 4.255717 | 4.220529 |
| 34976 | ENSG0000 | AL118516.1 | | 4.45877 | 4.148259 | 2.734075 |
| 2166 | ENSG0000 | PPIL2 | peptidylpr | 4.446863 | 4.339142 | 4.13722 |
| 7451 | ENSG0000 | TUBGCP4 | tubulin gar | 4.434422 | 3.756627 | 3.49468 |
| 143 | ENSG0000 | FARP2 | FERM, ARH | 4.427897 | 3.82333 | 3.412514 |
| 12150 | ENSG0000 | MTCL1 | microtubul | 4.40331 | 4.069009 | 4.056274 |
| 635 | ENSG0000 | WDR37 | WD repeat | 4.402142 | 2.463422 | 2.453358 |
| 13620 | ENSG0000 | RPL7AP66 | ribosomal | 4.400124 | 2.343027 | 1.997794 |
| 9247 | ENSG0000 | EIF4E | eukaryotic | 4.367029 | 3.264121 | 3.169905 |
| 30367 | ENSG0000 | SMIM20 | small integ | 4.349467 | 3.558971 | 3.019027 |
| 14551 | ENSG0000 | EHMT1 | euchromat | 4.346417 | 3.147129 | 2.824164 |
| 9894 | ENSG0000 | LRP8 | LDL recept | 4.343439 | 2.778985 | 2.562853 |

| | | | | | | |
|---|---|---|---|---|---|---|
| 12776 | ENSG0000 | ESCO2 | establishm | 4.341004 | 3.233043 | 3.192047 |
| 630 | ENSG0000 | GEMIN8 | gem nucle | 4.336676 | 4.311759 | 3.555965 |
| 16492 | ENSG0000 | TRAPPC4 | trafficking | 4.326501 | 2.703833 | 2.386547 |
| 500 | ENSG0000 | BRD9 | bromodom | 4.318836 | 3.617218 | 3.106863 |
| 2307 | ENSG0000 | TRMU | tRNA 5-me | 4.314001 | 3.645277 | 3.515189 |
| 6242 | ENSG0000 | TAF4 | TATA-box b | 4.305752 | 3.606796 | 3.018525 |
| 6363 | ENSG0000 | SLC6A6 | solute carr | 4.294587 | 1.536258 | 1.505443 |
| 2724 | ENSG0000 | CCDC113 | coiled-coil | 4.284363 | 3.482732 | 3.382914 |
| 597 | ENSG0000 | CTNS | cystinosin, | 4.271978 | 3.556195 | 3.334773 |
| 13463 | ENSG0000 | FBXW8 | F-box and W | 4.253036 | 3.796179 | 3.423363 |
| 9605 | ENSG0000 | TRIM11 | tripartite m | 4.249751 | 4.014653 | 3.956258 |
| 7890 | ENSG0000 | PMM2 | phosphom | 4.241889 | 3.022122 | 2.71186 |
| 2643 | ENSG0000 | KLHL4 | kelch like f | 4.238929 | 2.999314 | 2.24782 |
| 1298 | ENSG0000 | CELSR1 | cadherin E | 4.21835 | 3.863823 | 3.50184 |
| 13711 | ENSG0000 | SGF29 | SAGA comp | 4.216193 | 3.842896 | 3.520048 |
| 6066 | ENSG0000 | TXNDC17 | thioredoxi | 4.213965 | 3.713419 | 3.523801 |
| 10534 | ENSG0000 | C1orf123 | chromosor | 4.206745 | 2.868803 | 2.841415 |
| 3243 | ENSG0000 | IQCE | IQ motif cc | 4.200051 | 3.16703 | 3.137882 |
| 883 | ENSG0000 | SPA17 | sperm auto | 4.195581 | 2.336876 | 2.130054 |
| 15807 | ENSG0000 | RPS19BP1 | ribosomal | 4.186401 | 3.10458 | 2.897013 |
| 15073 | ENSG0000 | TRMT12 | tRNA meth | 4.185833 | 3.812922 | 3.249283 |
| 19472 | ENSG0000 | SNORD67 | small nucle | 4.181675 | 2.830823 | 2.472435 |
| 1302 | ENSG0000 | TIMM21 | translocase | 4.180913 | 3.704607 | 3.529735 |
| 4199 | ENSG0000 | BNIP1 | BCL2 intera | 4.174509 | 3.884568 | 3.023767 |
| 1440 | ENSG0000 | UBE2D4 | ubiquitin c | 4.165907 | 2.568588 | 2.049349 |
| 1150 | ENSG0000 | NCK2 | NCK adapt | 4.147126 | 3.734191 | 3.335227 |
| 2323 | ENSG0000 | RBM23 | RNA bindir | 4.129746 | 3.436297 | 3.330421 |
| 15184 | ENSG0000 | CRELD2 | cysteine ric | 4.128074 | 2.199203 | 2.091617 |
| 4029 | ENSG0000 | ZNF346 | zinc finger | 4.116558 | 3.867094 | 3.820173 |
| 2730 | ENSG0000 | SLC38A7 | solute carr | 4.105544 | 4.048318 | 3.075433 |
| 6542 | ENSG0000 | POLR3F | RNA polym | 4.093097 | 3.631321 | 3.164369 |
| 22405 | ENSG0000 | MEMO1P1 | mediator c | 4.087175 | 3.550196 | 3.345068 |
| 5225 | ENSG0000 | CCDC18 | coiled-coil | 4.059148 | 2.956863 | 2.741432 |
| 15501 | ENSG0000 | AC034236.1 | | 4.027297 | 2.956979 | 2.610236 |
| 5719 | ENSG0000 | PCID2 | PCI domair | 4.024189 | 3.974474 | 3.487741 |
| 3873 | ENSG0000 | GNB3 | G protein s | 4.01951 | 3.010789 | 2.528467 |
| 2708 | ENSG0000 | LYRM1 | LYR motif c | 4.008805 | 3.220897 | 2.6577 |
| 18277 | ENSG0000 | AC083899.1 | | 3.989114 | 3.844636 | 3.371128 |
| 10241 | ENSG0000 | CCDC107 | coiled-coil | 3.988574 | 2.85039 | 2.671639 |
| 13988 | ENSG0000 | FAM20C | FAM20C, g | 3.978897 | 2.769638 | 2.602632 |
| 1969 | ENSG0000 | OSGEP | O-sialoglyc | 3.972161 | 3.949749 | 3.751696 |
| 12313 | ENSG0000 | SH3TC2 | SH3 domai | 3.959735 | 3.534674 | 3.444069 |
| 26777 | ENSG0000 | TMEM147- | TMEM147 | 3.959304 | 3.941638 | 1.524824 |
| 11016 | ENSG0000 | SFMBT1 | Scm-like w | 3.935025 | 3.900358 | 3.133857 |
| 4847 | ENSG0000 | INVS | inversin [S | 3.896869 | 3.600443 | 2.901523 |
| 43015 | ENSG0000 | AC118553.2 | | 3.891091 | 1.096829 | 0.390628 |
| 21942 | ENSG0000 | NPM1P39 | nucleopho | 3.887095 | 2.84197 | 1.936425 |

| | | | | | | |
|---|---|---|---|---|---|---|
| 2963 | ENSG0000 | PLEKHJ1 | pleckstrin l | 3.879971 | 3.542987 | 3.315118 |
| 12679 | ENSG0000 | FAM86JP | family with | 3.870038 | 2.631929 | 2.144923 |
| 15587 | ENSG0000 | TMLHE | trimethylly | 3.865387 | 3.457242 | 3.296414 |
| 13888 | ENSG0000 | PUS1 | pseudouric | 3.852075 | 3.411416 | 2.719736 |
| 2461 | ENSG0000 | MTG2 | mitochond | 3.828576 | 2.997379 | 2.71502 |
| 12466 | ENSG0000 | SIMC1 | SUMO inte | 3.821328 | 3.715178 | 3.213886 |
| 2818 | ENSG0000 | KNOP1 | lysine rich | 3.79384 | 3.341692 | 3.222678 |
| 17116 | ENSG0000 | GK | glycerol kir | 3.782928 | 3.535213 | 3.264541 |
| 17406 | ENSG0000 | SNORD104 | small nucle | 3.755975 | 3.428512 | 2.279346 |
| 32111 | ENSG0000 | AP002387.1 | | 3.753543 | 3.061196 | 2.521394 |
| 16630 | ENSG0000 | ABCB8 | ATP bindin | 3.728235 | 1.922624 | 1.871619 |
| 38254 | ENSG0000 | AC005034.3 | | 3.716534 | 3.38247 | 3.347467 |
| 20651 | ENSG0000 | PAM16 | presequen | 3.711218 | 2.879194 | 2.744354 |
| 15630 | ENSG0000 | POLR1D | RNA polym | 3.70644 | 3.536135 | 2.913976 |
| 15976 | ENSG0000 | CYHR1 | cysteine ar | 3.702134 | 3.481943 | 3.141953 |
| 9085 | ENSG0000 | TMEM138 | transmeml | 3.676937 | 3.495657 | 2.498946 |
| 442 | ENSG0000 | SPAST | spastin [So | 3.643788 | 3.606789 | 3.455972 |
| 6776 | ENSG0000 | PPHLN1 | periphilin 1 | 3.627691 | 3.626192 | 3.559005 |
| 35904 | ENSG0000 | SRSF8 | serine and | 3.62415 | 3.258244 | 2.350946 |
| 4707 | ENSG0000 | MESD | mesoderm | 3.61488 | 3.270835 | 3.020394 |
| 14307 | ENSG0000 | DGKZP1 | diacylglyce | 3.577377 | 2.944722 | 2.437585 |
| 9855 | ENSG0000 | NSMCE2 | NSE2/MMS | 3.56729 | 3.558295 | 3.438644 |
| 29456 | ENSG0000 | SNHG10 | small nucle | 3.563358 | 2.362524 | 1.446172 |
| 2376 | ENSG0000 | DICER1 | dicer 1, rib | 3.543555 | 3.385639 | 2.93718 |
| 12860 | ENSG0000 | GPHN | gephyrin [S | 3.50492 | 3.368385 | 2.73913 |
| 310 | ENSG0000 | CLCN6 | chloride vo | 3.459212 | 3.293734 | 3.267623 |
| 38866 | ENSG0000 | AL138724.1 | | 3.451536 | 3.04699 | 2.970784 |
| 9205 | ENSG0000 | PTS | 6-pyruvoyl | 3.441859 | 3.354465 | 3.236907 |
| 34193 | ENSG0000 | SORD2P | sorbitol de | 3.44062 | 3.251206 | 2.779875 |
| 8354 | ENSG0000 | C1orf131 | chromosor | 3.424232 | 3.250231 | 2.704778 |
| 39866 | ENSG0000 | TBC1D3L | TBC1 doma | 3.415648 | 2.746879 | 2.296708 |
| 10853 | ENSG0000 | RPL22L1 | ribosomal | 3.407924 | 2.790057 | 2.22443 |
| 40802 | ENSG0000 | FP565260.3 | | 3.375658 | 3.256561 | 1.27008 |
| 14024 | ENSG0000 | ASB8 | ankyrin rep | 3.364993 | 2.932482 | 2.821364 |
| 1539 | ENSG0000 | DHODH | dihydroorc | 3.336429 | 2.089459 | 1.814668 |
| 12404 | ENSG0000 | LIMS1 | LIM zinc fir | 3.332683 | 3.027543 | 2.437758 |
| 17072 | ENSG0000 | UNC13B | unc-13 hor | 3.325839 | 2.876127 | 2.20659 |
| 3042 | ENSG0000 | MED26 | mediator c | 3.324333 | 2.567282 | 1.813638 |
| 3305 | ENSG0000 | TMEM106I | transmeml | 3.309898 | 3.247749 | 3.126204 |
| 13236 | ENSG0000 | CEP83 | centrosom | 3.29185 | 3.147938 | 2.5289 |
| 7575 | ENSG0000 | ABI2 | abl interac | 3.291791 | 3.12957 | 3.080219 |
| 25292 | ENSG0000 | PET117 | PET117 ho | 3.278698 | 3.161745 | 2.932846 |
| 10078 | ENSG0000 | FAM86C1 | family with | 3.269858 | 2.786126 | 2.109058 |
| 7072 | ENSG0000 | REV1 | REV1, DNA | 3.267862 | 3.197411 | 2.942564 |
| 1890 | ENSG0000 | TFAP4 | transcriptio | 3.253825 | 2.959578 | 2.498761 |
| 12491 | ENSG0000 | NANP | N-acetylne | 3.214693 | 2.894818 | 2.623374 |
| 6389 | ENSG0000 | ACAP3 | ArfGAP wit | 3.168083 | 2.977165 | 2.545168 |

| | | | | | | |
|---|---|---|---|---|---|---|
| 33579 | ENSG0000 | C17orf49 | chromosor | 3.155896 | 2.901688 | 2.38436 |
| 7555 | ENSG0000 | BARD1 | BRCA1 ass | 3.141104 | 2.5984 | 2.394784 |
| 5348 | ENSG0000 | COL10A1 | collagen ty | 3.123934 | 2.224793 | 1.303527 |
| 35671 | ENSG0000 | CORO7 | coronin 7 [ | 3.106254 | 2.416733 | 1.808497 |
| 2579 | ENSG0000 | ATG4A | autophagy | 3.085614 | 3.079065 | 2.910792 |
| 8535 | ENSG0000 | RABL3 | RAB, meml | 3.077542 | 2.349933 | 1.916466 |
| 18520 | ENSG0000 | SNORA57 | small nucle | 3.073874 | 2.250805 | 2.031254 |
| 11064 | ENSG0000 | MON1A | MON1 hon | 3.035314 | 2.226349 | 2.128646 |
| 12609 | ENSG0000 | AKAP13 | A-kinase ar | 3.016934 | 2.187665 | 2.079276 |
| 11032 | ENSG0000 | EXO5 | exonucleas | 3.008313 | 2.876099 | 2.771946 |
| 6862 | ENSG0000 | FADS2 | fatty acid c | 2.977069 | 2.320876 | 2.281192 |
| 16044 | ENSG0000 | COMMD6 | COMM dor | 2.965289 | 2.413288 | 1.933874 |
| 25960 | ENSG0000 | RPS26P47 | ribosomal | 2.950151 | 1.774612 | 1.671034 |
| 17248 | ENSG0000 | SNORA33 | small nucle | 2.924757 | 2.71843 | 1.759276 |
| 13462 | ENSG0000 | AC026271.1 | | 2.903224 | 2.643862 | 2.292314 |
| 918 | ENSG0000 | ZNF76 | zinc finger | 2.901801 | 2.815645 | 2.213782 |
| 10862 | ENSG0000 | GTPBP8 | GTP bindin | 2.89169 | 2.682659 | 2.534891 |
| 18237 | ENSG0000 | TRIM39 | tripartite n | 2.878421 | 2.583802 | 2.468604 |
| 15319 | ENSG0000 | TMEM186 | transmeml | 2.868402 | 2.819575 | 2.61477 |
| 1368 | ENSG0000 | DNAJC10 | DnaJ heat s | 2.864935 | 2.860842 | 2.700333 |
| 1761 | ENSG0000 | SH3BP2 | SH3 domai | 2.822645 | 1.93106 | 1.893621 |
| 6676 | ENSG0000 | ACTR3B | ARP3 actin | 2.819076 | 2.492332 | 2.133803 |
| 7535 | ENSG0000 | EXOC6 | exocyst coi | 2.800406 | 1.927964 | 1.779351 |
| 800 | ENSG0000 | POLR3E | RNA polym | 2.772819 | 2.363457 | 1.45323 |
| 11035 | ENSG0000 | ERMAP | erythrobla: | 2.761648 | 2.036008 | 1.410077 |
| 18836 | ENSG0000 | SNORD14D | small nucle | 2.759223 | 1.463093 | 0.696077 |
| 22823 | ENSG0000 | AL009174.1 | | 2.743774 | 2.039764 | 1.777584 |
| 13784 | ENSG0000 | RABEP2 | rabaptin, R | 2.742412 | 2.489065 | 2.069009 |
| 640 | ENSG0000 | DTNBP1 | dystrobrev | 2.738286 | 2.373781 | 1.9652 |
| 3390 | ENSG0000 | BAG1 | BCL2 assoc | 2.728597 | 2.533522 | 2.329857 |
| 28651 | ENSG0000 | C22orf39 | chromosor | 2.728164 | 1.639836 | 1.173533 |
| 8476 | ENSG0000 | RHBDD1 | rhomboid ( | 2.724561 | 2.418957 | 2.321764 |
| 3770 | ENSG0000 | PSMD9 | proteasom | 2.714824 | 2.127655 | 1.770569 |
| 39539 | ENSG0000 | MIR6777 | microRNA | 2.712631 | 1.301133 | 0.663983 |
| 37929 | ENSG0000 | AL049840.2 | | 2.701184 | 1.773832 | 1.69423 |
| 22498 | ENSG0000 | AC135977.1 | | 2.69771 | 2.647489 | 2.395858 |
| 4442 | ENSG0000 | DCAF17 | DDB1 and ( | 2.697682 | 1.900843 | 1.446028 |
| 12894 | ENSG0000 | TRAPPC12 | trafficking | 2.691799 | 2.082788 | 1.991197 |
| 6875 | ENSG0000 | BIVM | basic, imm | 2.678257 | 1.655319 | 1.493353 |
| 17534 | ENSG0000 | SNORA63 | Small nucle | 2.632725 | 2.310323 | 0.718545 |
| 26351 | ENSG0000 | SUMO2P1 | SUMO2 ps | 2.626497 | 1.315753 | 1.247019 |
| 9714 | ENSG0000 | SETD9 | SET domair | 2.621344 | 2.564546 | 2.082263 |
| 41150 | ENSG0000 | AC135050.6 | | 2.615175 | 2.465137 | 1.773732 |
| 10430 | ENSG0000 | PGAP3 | post-GPI at | 2.61331 | 2.025057 | 1.87411 |
| 19814 | ENSG0000 | TMEM110 | transmeml | 2.613022 | 2.080048 | 1.869922 |
| 35987 | ENSG0000 | SNORD43 | small nucle | 2.608721 | 1.653685 | 0.625238 |
| 18259 | ENSG0000 | RPS26P8 | ribosomal | 2.591717 | 1.329133 | 1.112439 |

| | | | | | | |
|---|---|---|---|---|---|---|
| 8894 | ENSG0000 | TATDN1 | TatD DNas | 2.578375 | 2.009464 | 1.587636 |
| 20005 | ENSG0000 | SMIM7 | small integ | 2.563638 | 2.377593 | 2.211816 |
| 28519 | ENSG0000 | AC069499.1 | | 2.535466 | 0.907565 | 0.532373 |
| 42825 | ENSG0000 | MIR6850 | microRNA | 2.519571 | 2.272992 | 2.023776 |
| 11803 | ENSG0000 | COQ7 | coenzyme | 2.518077 | 2.513417 | 2.042856 |
| 8717 | ENSG0000 | MMS22L | MMS22 lik | 2.496024 | 2.149538 | 2.030942 |
| 39419 | ENSG0000 | AL121845.3 | | 2.493068 | 2.079064 | 1.251677 |
| 8398 | ENSG0000 | CAMKMT | calmodulir | 2.485966 | 2.127935 | 1.814416 |
| 12391 | ENSG0000 | ASPSCR1 | ASPSCR1, U | 2.468 | 2.372729 | 2.151132 |
| 9275 | ENSG0000 | SCLT1 | sodium cha | 2.457288 | 2.402604 | 2.08214 |
| 28232 | ENSG0000 | AL365273.1 | | 2.455551 | 2.165447 | 1.547826 |
| 29176 | ENSG0000 | P2RY11 | purinergic | 2.43453 | 2.433387 | 2.006413 |
| 33204 | ENSG0000 | AC023055.1 | | 2.418998 | 1.298363 | 0.045796 |
| 30635 | ENSG0000 | RNPS1P1 | RNA bindir | 2.418456 | 2.073499 | 1.704425 |
| 16358 | ENSG0000 | XPNPEP3 | X-prolyl am | 2.417405 | 1.941703 | 1.915047 |
| 35605 | ENSG0000 | AC026954.2 | | 2.405597 | 1.649897 | 1.282708 |
| 16495 | ENSG0000 | TECPR2 | tectonin be | 2.404637 | 2.350869 | 2.20787 |
| 19845 | ENSG0000 | RPSAP54 | ribosomal | 2.394338 | 1.711724 | 1.373711 |
| 59 | ENSG0000 | ARHGAP33 | Rho GTPas | 2.385046 | 1.903554 | 1.721615 |
| 12775 | ENSG0000 | CHD7 | chromodor | 2.384931 | 2.047674 | 1.534101 |
| 16373 | ENSG0000 | SUPT3H | SPT3 homc | 2.377587 | 2.366485 | 2.070692 |
| 4858 | ENSG0000 | DCAF4 | DDB1 and | 2.363337 | 2.30042 | 2.104698 |
| 2249 | ENSG0000 | TOM1 | target of m | 2.345947 | 2.228271 | 1.911489 |
| 25804 | ENSG0000 | UBE2SP1 | ubiquitin c | 2.305335 | 1.115895 | 1.088001 |
| 6502 | ENSG0000 | POPDC3 | popeye do | 2.290883 | 1.28128 | 1.207211 |
| 40060 | ENSG0000 | HIST1H2AJ | histone clu | 2.287241 | 1.990029 | 1.073009 |
| 42321 | ENSG0000 | AC245140.2 | | 2.279643 | 2.076242 | 1.881112 |
| 7582 | ENSG0000 | COX17 | COX17, cyt | 2.263658 | 1.930336 | 1.547779 |
| 2677 | ENSG0000 | UGGT2 | UDP-gluco | 2.250259 | 2.124337 | 1.816578 |
| 10088 | ENSG0000 | ZFAND2B | zinc finger | 2.234492 | 1.939846 | 1.821037 |
| 37549 | ENSG0000 | LINC02486 | long interg | 2.228552 | 1.411042 | 1.397761 |
| 20028 | ENSG0000 | LYRM4 | LYR motif c | 2.199786 | 2.189003 | 1.471174 |
| 41092 | ENSG0000 | AC007731.5 | | 2.194973 | 0.138804 | 0.084843 |
| 9718 | ENSG0000 | C9orf85 | chromosor | 2.131944 | 2.097767 | 2.002213 |
| 25469 | ENSG0000 | AC002524.1 | | 2.11914 | 1.926313 | 1.670128 |
| 5142 | ENSG0000 | CRY2 | cryptochro | 2.106836 | 1.873713 | 1.84288 |
| 40162 | ENSG0000 | AL354718.1 | | 2.100329 | 1.478133 | 1.30595 |
| 37807 | ENSG0000 | ZNF587B | zinc finger | 2.088653 | 1.217037 | 1.205573 |
| 17992 | ENSG0000 | SPRN | shadow of | 2.088527 | 1.908044 | 1.646175 |
| 19760 | ENSG0000 | COG8 | componen | 2.085054 | 1.846746 | 1.567373 |
| 29826 | ENSG0000 | SERBP1P5 | SERPINE1 r | 2.078422 | 1.07473 | 0.913853 |
| 6700 | ENSG0000 | SBF2 | SET binding | 2.072402 | 1.7462 | 1.417128 |
| 42526 | ENSG0000 | MIR3651 | microRNA | 2.061265 | 0.266315 | 0.160615 |
| 3307 | ENSG0000 | CEP41 | centrosom | 2.056954 | 1.874106 | 1.61286 |
| 20302 | ENSG0000 | PPIAP29 | peptidylpro | 2.054998 | 1.374302 | 1.350243 |
| 2769 | ENSG0000 | FAM173A | family with | 2.04036 | 1.030122 | 0.857914 |
| 16653 | ENSG0000 | HIST1H4J | histone clu | 2.024701 | 0.920633 | 0.844894 |

| | | | | | | |
|---|---|---|---|---|---|---|
| 34955 | ENSG0000 | AL096870.2 | | 2.019904 | 1.281626 | 0.892077 |
| 6856 | ENSG0000 | DAGLA | diacylglyce | 2.019725 | 1.978248 | 1.93083 |
| 13223 | ENSG0000 | GMPPB | GDP-mann | 2.01411 | 1.947488 | 1.944557 |
| 39917 | ENSG0000 | HIST1H2BC | histone clu | 1.982023 | 1.849685 | 1.666669 |
| 670 | ENSG0000 | ERCC8 | ERCC excis | 1.981239 | 1.891625 | 1.709612 |
| 13479 | ENSG0000 | PDIK1L | PDLIM1 int | 1.966114 | 1.109297 | 0.83564 |
| 24179 | ENSG0000 | HMGN2P3 | high mobil | 1.963568 | 1.807397 | 1.759571 |
| 36881 | ENSG0000 | AC005943.1 | | 1.962003 | 1.858748 | 1.484936 |
| 38629 | ENSG0000 | AL589666.1 | | 1.947902 | 1.546117 | 1.082727 |
| 38787 | ENSG0000 | AL691432.2 | | 1.935799 | 1.422412 | 1.211645 |
| 23648 | ENSG0000 | EIF4A1P10 | eukaryotic | 1.934918 | 1.10772 | 0.639767 |
| 8048 | ENSG0000 | TPGS1 | tubulin pol | 1.925202 | 1.204829 | 1.164261 |
| 16039 | ENSG0000 | AD000671.1 | | 1.91833 | 0.550669 | 0.502942 |
| 33455 | ENSG0000 | P2RX5-TAX | P2RX5-TAX | 1.897133 | 1.752533 | 1.59346 |
| 36601 | ENSG0000 | MIR3685 | microRNA | 1.895997 | 1.524749 | 1.224661 |
| 17953 | ENSG0000 | AGER | advanced g | 1.886522 | 1.784247 | 1.618453 |
| 19578 | ENSG0000 | SNORA26 | small nucle | 1.881687 | 1.679876 | 0.732531 |
| 7012 | ENSG0000 | CCDC142 | coiled-coil | 1.876233 | 1.694606 | 1.437498 |
| 17565 | ENSG0000 | SNORD14E | small nucle | 1.867147 | 1.653095 | 1.308165 |
| 11040 | ENSG0000 | DNAJB14 | DnaJ heat s | 1.856246 | 1.577375 | 1.528772 |
| 37957 | ENSG0000 | AL049840.4 | | 1.851097 | 1.299887 | 0.676047 |
| 21532 | ENSG0000 | AC026271.2 | | 1.8428 | 0.847234 | 0.593995 |
| 5987 | ENSG0000 | ICE2 | interactor | 1.835328 | 1.824107 | 1.359106 |
| 11053 | ENSG0000 | ATRIP | ATR intera | 1.823823 | 1.367577 | 0.706172 |
| 41062 | ENSG0000 | AL391988.1 | | 1.823286 | 1.69433 | 1.097958 |
| 28517 | ENSG0000 | UBQLN4P1 | ubiquilin 4 | 1.80982 | 1.293198 | 1.059249 |
| 15549 | ENSG0000 | GNB1L | G protein s | 1.78082 | 1.384086 | 1.192319 |
| 18428 | ENSG0000 | TECPR1 | tectonin be | 1.779003 | 1.689539 | 1.335787 |
| 5966 | ENSG0000 | OSGEPL1 | O-sialoglyc | 1.77662 | 1.536658 | 1.289699 |
| 580 | ENSG0000 | VCAN | versican [S | 1.773676 | 1.479824 | 1.085 |
| 8737 | ENSG0000 | SHPRH | SNF2 histo | 1.76142 | 1.676767 | 1.400898 |
| 27706 | ENSG0000 | SNORA81 | Small nucle | 1.759174 | 0.6267 | 0.144723 |
| 3952 | ENSG0000 | SOD2 | superoxide | 1.746427 | 1.460964 | 1.43262 |
| 15364 | ENSG0000 | BRF1 | BRF1, RNA | 1.742303 | 1.531743 | 1.419797 |
| 15262 | ENSG0000 | HDDC3 | HD domair | 1.740851 | 1.650601 | 1.352954 |
| 36191 | ENSG0000 | AC010761.1 | | 1.734637 | 1.715938 | 0.935938 |
| 875 | ENSG0000 | ADCK1 | aarF doma | 1.721951 | 1.43452 | 0.963908 |
| 540 | ENSG0000 | ASTE1 | asteroid hc | 1.700558 | 1.461457 | 1.239819 |
| 22317 | ENSG0000 | LINC00115 | long interg | 1.699376 | 1.246837 | 1.148001 |
| 24752 | ENSG0000 | AL355472.1 | | 1.675207 | 0.967263 | 0.959313 |
| 4911 | ENSG0000 | GNG13 | G protein s | 1.664271 | 1.242406 | 0.832884 |
| 39776 | ENSG0000 | AC069281.2 | | 1.660518 | 1.566335 | 1.195179 |
| 19700 | ENSG0000 | DNLZ | DNL-type z | 1.650771 | 1.560793 | 1.21297 |
| 3501 | ENSG0000 | GOSR2 | golgi SNAP | 1.64473 | 1.625068 | 1.165032 |
| 14583 | ENSG0000 | OGFOD3 | 2-oxogluta | 1.637542 | 1.617361 | 1.30673 |
| 18140 | ENSG0000 | STK19 | serine/thre | 1.610316 | 1.54088 | 1.495567 |
| 6637 | ENSG0000 | SLC39A11 | solute carr | 1.603343 | 1.231964 | 0.857897 |

| | | | | | | |
|---|---|---|---|---|---|---|
| 26304 | ENSG0000 | GABPAP | GA binding | 1.599644 | 1.4721 | 0.772971 |
| 13369 | ENSG0000 | SNAPC5 | small nucle | 1.59361 | 1.325057 | 1.070685 |
| 41048 | ENSG0000 | SNORD30 | small nucle | 1.593532 | 1.423855 | 0.745664 |
| 35028 | ENSG0000 | LINC01963 | long interg | 1.587104 | 0.7554 | 0.573839 |
| 8901 | ENSG0000 | NAPRT | nicotinate | 1.585438 | 1.491375 | 0.700531 |
| 22936 | ENSG0000 | TSSK5P | testis speci | 1.584196 | 0.90221 | 0.648116 |
| 18980 | ENSG0000 | SNORA46 | small nucle | 1.582087 | 0.99731 | 0.962354 |
| 14363 | ENSG0000 | PPP1R14BI | protein ph | 1.568281 | 1.334275 | 1.172834 |
| 8905 | ENSG0000 | NFIB | nuclear fac | 1.566049 | 1.430558 | 1.090791 |
| 14046 | ENSG0000 | AC005412.1 | | 1.550507 | 1.058417 | 1.021488 |
| 40774 | ENSG0000 | RMRP | RNA comp | 1.54987 | 0.38581 | 0.367236 |
| 24482 | ENSG0000 | CDC20P1 | cell divisio | 1.545835 | 0.934677 | 0.767369 |
| 35857 | ENSG0000 | AC004148.2 | | 1.541563 | 1.419575 | 1.189783 |
| 15514 | ENSG0000 | EP400NL | EP400 N-te | 1.526872 | 1.520018 | 0.700522 |
| 11105 | ENSG0000 | LSM6 | LSM6 hom | 1.507196 | 1.345915 | 1.27501 |
| 35375 | ENSG0000 | SMG1P7 | SMG1P7, n | 1.489347 | 1.19564 | 1.083155 |
| 32597 | ENSG0000 | FDXACB1 | ferredoxin- | 1.486725 | 0.677844 | 0.386301 |
| 15227 | ENSG0000 | HIST1H1B | histone clu | 1.479992 | 1.064819 | 0.975972 |
| 10952 | ENSG0000 | MTHFD2L | methylene | 1.479151 | 1.256098 | 1.012174 |
| 41223 | ENSG0000 | CR392039.3 | | 1.467093 | 1.226863 | 0.997524 |
| 25229 | ENSG0000 | AL158050.1 | | 1.466926 | 0.874018 | 0.656166 |
| 9305 | ENSG0000 | PIGF | phosphatic | 1.461212 | 1.007444 | 0.958523 |
| 13325 | ENSG0000 | TLR6 | toll like rec | 1.446422 | 1.431819 | 1.387976 |
| 24658 | ENSG0000 | AL512422.2 | | 1.438759 | 1.15795 | 0.851622 |
| 629 | ENSG0000 | DSG2 | desmoglei | 1.434186 | 1.343585 | 1.06545 |
| 32756 | ENSG0000 | PXN-AS1 | PXN antise | 1.424063 | 1.089624 | 1.003171 |
| 34805 | ENSG0000 | AC109460.3 | | 1.421895 | 1.303858 | 0.670567 |
| 26374 | ENSG0000 | SNORD62A | small nucle | 1.396151 | 0.95539 | 0.817446 |
| 10233 | ENSG0000 | RGS12 | regulator c | 1.39518 | 1.23786 | 1.211565 |
| 7595 | ENSG0000 | PPCDC | phosphopa | 1.385238 | 1.254703 | 1.244932 |
| 8638 | ENSG0000 | GIN1 | gypsy retro | 1.381724 | 1.37024 | 1.340753 |
| 16917 | ENSG0000 | AC007537.1 | | 1.37867 | 1.16332 | 1.068606 |
| 5247 | ENSG0000 | ARL4A | ADP ribosy | 1.372197 | 1.303988 | 0.879729 |
| 39028 | ENSG0000 | ZSWIM8-A | ZSWIM8 ar | 1.369498 | 0.188643 | 0.096692 |
| 16058 | ENSG0000 | PLSCR1 | phospholip | 1.367964 | 1.252295 | 1.198208 |
| 39565 | ENSG0000 | AC131009.3 | | 1.358131 | 1.128972 | 1.01902 |
| 24911 | ENSG0000 | AC112907.2 | | 1.353736 | 0.90119 | 0.878355 |
| 19580 | ENSG0000 | SNORA3B | small nucle | 1.352738 | 0.807415 | 0.555935 |
| 7606 | ENSG0000 | ZGRF1 | zinc finger | 1.350608 | 1.315143 | 1.297915 |
| 24592 | ENSG0000 | C1DP1 | C1D nuclea | 1.347812 | 1.096004 | 0.938104 |
| 38865 | ENSG0000 | AC021242.3 | | 1.343949 | 0.638669 | 0.318978 |
| 2706 | ENSG0000 | ELMO3 | engulfmen | 1.338153 | 0.903188 | 0.636306 |
| 22934 | ENSG0000 | AC073415.1 | | 1.324393 | 1.28291 | 0.927968 |
| 35097 | ENSG0000 | LINC01311 | long interg | 1.303789 | 1.283251 | 0.925001 |
| 25002 | ENSG0000 | FTH1P7 | ferritin hea | 1.302627 | 0.791765 | 0.671011 |
| 13502 | ENSG0000 | PCCA | propionyl-( | 1.290709 | 1.193338 | 1.185352 |
| 15598 | ENSG0000 | ZNF566 | zinc finger | 1.290305 | 1.1466 | 0.692909 |

| | | | | | | |
|---|---|---|---|---|---|---|
| 38810 | ENSG0000 | AL365330.1 | | 1.285368 | 1.208261 | 0.855791 |
| 35975 | ENSG0000 | AC145207.5 | | 1.284995 | 1.201534 | 0.622272 |
| 39004 | ENSG0000 | AL662797.3 | | 1.279233 | 0.911263 | 0.582761 |
| 15511 | ENSG0000 | ZBTB3 | zinc finger | 1.275609 | 1.085055 | 0.434212 |
| 16572 | ENSG0000 | CASP4 | caspase 4 [ | 1.249268 | 0.985894 | 0.972472 |
| 20367 | ENSG0000 | PEX26 | peroxisom | 1.244367 | 1.16257 | 1.079783 |
| 38632 | ENSG0000 | SNORA51 | small nucle | 1.244115 | 0.670783 | 0.39775 |
| 28702 | ENSG0000 | RPL7AP11 | ribosomal | 1.239498 | 0.774795 | 0.689902 |
| 9054 | ENSG0000 | ENDOD1 | endonucle | 1.237371 | 0.952036 | 0.840337 |
| 35030 | ENSG0000 | AL163051.1 | | 1.227702 | 0.694364 | 0.633532 |
| 37945 | ENSG0000 | AL049840.3 | | 1.212571 | 1.030822 | 0.387711 |
| 26781 | ENSG0000 | AC104076.1 | | 1.207291 | 1.200913 | 1.060271 |
| 4932 | ENSG0000 | COQ6 | coenzyme | 1.206764 | 1.144237 | 1.134757 |
| 21769 | ENSG0000 | SH3BP5-AS | SH3BP5 an | 1.200452 | 0.802034 | 0.710968 |
| 14143 | ENSG0000 | NSUN3 | NOP2/Sun | 1.199569 | 1.110728 | 1.07036 |
| 4392 | ENSG0000 | ELMOD3 | ELMO dom | 1.181293 | 1.18072 | 1.104858 |
| 31621 | ENSG0000 | EIF5AL1 | eukaryotic | 1.175329 | 0.631428 | 0.540008 |
| 42271 | ENSG0000 | AC073857.1 | | 1.168987 | 1.168732 | 1.131246 |
| 18834 | ENSG0000 | SNORA25 | small nucle | 1.167343 | 0.783856 | 0.005603 |
| 18991 | ENSG0000 | SNORA66 | small nucle | 1.166793 | 0.934795 | 0.537211 |
| 27529 | ENSG0000 | BANF1P3 | barrier to a | 1.162141 | 0.351536 | 0.349891 |
| 26138 | ENSG0000 | EIF5AP4 | eukaryotic | 1.159702 | 0.44318 | 0.217265 |
| 26614 | ENSG0000 | AC006978.1 | | 1.153328 | 1.139431 | 0.823172 |
| 32932 | ENSG0000 | AC005840.3 | | 1.153074 | 1.029501 | 0.613828 |
| 14803 | ENSG0000 | BRICD5 | BRICHOS d | 1.151984 | 0.612533 | 0.557013 |
| 28581 | ENSG0000 | PRR34-AS1 | PRR34 anti | 1.148352 | 0.573562 | 0.525086 |
| 40532 | ENSG0000 | AL163051.2 | | 1.145933 | 0.754738 | 0.629705 |
| 7861 | ENSG0000 | PCSK6 | proprotein | 1.137609 | 0.987243 | 0.881942 |
| 41241 | ENSG0000 | AC115618.3 | | 1.126803 | 1.046677 | 0.841658 |
| 9219 | ENSG0000 | SLC7A11 | solute carr | 1.125966 | 0.746136 | 0.629746 |
| 18949 | ENSG0000 | SNORA20 | small nucle | 1.122401 | 0.360051 | 0.281251 |
| 15327 | ENSG0000 | IMMP2L | inner mito | 1.120217 | 1.064229 | 1.010859 |
| 38678 | ENSG0000 | AL109811.3 | | 1.115583 | 1.099422 | 0.835909 |
| 30933 | ENSG0000 | FAM86EP | family with | 1.108374 | 0.858289 | 0.850985 |
| 2671 | ENSG0000 | CDADC1 | cytidine an | 1.10657 | 1.09086 | 0.989019 |
| 25052 | ENSG0000 | AC009299.2 | | 1.097998 | 0.849341 | 0.708289 |
| 21073 | ENSG0000 | SNORD12B | small nucle | 1.096766 | 1.044505 | 0.564251 |
| 41435 | ENSG0000 | AC141557.2 | | 1.094146 | 1.023255 | 0.825708 |
| 23880 | ENSG0000 | RPL4P4 | ribosomal | 1.092472 | 0.878401 | 0.661709 |
| 31977 | ENSG0000 | AP001107.2 | | 1.092402 | 0.977668 | 0.75838 |
| 18145 | ENSG0000 | C9orf129 | chromosor | 1.090566 | 0.986511 | 0.687278 |
| 25569 | ENSG0000 | AL445524.1 | | 1.084943 | 0.57087 | 0.519264 |
| 29390 | ENSG0000 | CRNDE | colorectal | 1.070626 | 0.759471 | 0.56477 |
| 16081 | ENSG0000 | SELL | selectin L [ | 1.061882 | 0.878946 | 0.870262 |
| 36430 | ENSG0000 | AC132938.3 | | 1.058481 | 0.867909 | 0.776631 |
| 38773 | ENSG0000 | AC004233.3 | | 1.055069 | 0.825865 | 0.812779 |
| 19989 | ENSG0000 | MEF2B | myocyte er | 1.053682 | 0.913812 | 0.393669 |

| | | | | | | |
|---|---|---|---|---|---|---|
| 23366 | ENSG0000 | AC098484.1 | | 1.051912 | 0.79401 | 0.310146 |
| 20774 | ENSG0000 | FTH1P8 | ferritin hea | 1.04727 | 0.520068 | 0.26058 |
| 23401 | ENSG0000 | AC005000.1 | | 1.01788 | 0.365544 | 0.321177 |
| 26742 | ENSG0000 | HSPB1P1 | heat shock | 1.017192 | 0.473172 | 0.308713 |
| 9738 | ENSG0000 | PPARGC1B | PPARG coa | 1.014516 | 0.655883 | 0.61946 |
| 16723 | ENSG0000 | ZNF628 | zinc finger | 1.004388 | 0.892405 | 0.643849 |
| 39408 | ENSG0000 | AL022328.2 | | 1.003598 | 0.721116 | 0.409964 |
| 41067 | ENSG0000 | MIR6746 | microRNA | 0.988888 | 0.618546 | 0.37196 |
| 40331 | ENSG0000 | MIR6851 | microRNA | 0.986204 | 0.91551 | 0.159446 |
| 32376 | ENSG0000 | MSH5-SAP | MSH5-SAP | 0.981441 | 0.853925 | 0.388834 |
| 11718 | ENSG0000 | NAV2 | neuron nav | 0.968676 | 0.919635 | 0.775779 |
| 23910 | ENSG0000 | SOS1-IT1 | SOS1 intro | 0.96686 | 0.599302 | 0.571876 |
| 3811 | ENSG0000 | PRR4 | proline rich | 0.96279 | 0.54843 | 0.363882 |
| 4886 | ENSG0000 | RDH10 | retinol deh | 0.961169 | 0.919892 | 0.804565 |
| 38665 | ENSG0000 | AC114810.1 | | 0.960036 | 0.789856 | 0.415766 |
| 11920 | ENSG0000 | RAB4B | RAB4B, me | 0.952883 | 0.6255 | 0.578246 |
| 17144 | ENSG0000 | FICD | FIC domair | 0.947579 | 0.851668 | 0.787676 |
| 29438 | ENSG0000 | PRR7-AS1 | PRR7 antis | 0.94603 | 0.559474 | 0.366382 |
| 28029 | ENSG0000 | RPL21P39 | ribosomal | 0.939969 | 0.466896 | 0.373936 |
| 24507 | ENSG0000 | NPM1P9 | nucleopho | 0.928131 | 0.659953 | 0.624806 |
| 5655 | ENSG0000 | AP5S1 | adaptor re | 0.926753 | 0.823728 | 0.740265 |
| 29181 | ENSG0000 | AC091153.3 | | 0.914874 | 0.739328 | 0.683135 |
| 40874 | ENSG0000 | MIR3687-1 | microRNA | 0.911958 | 0.673472 | 0.402045 |
| 39447 | ENSG0000 | AP001437.1 | | 0.910818 | 0.535576 | 0.106557 |
| 11862 | ENSG0000 | SGK494 | uncharacte | 0.905169 | 0.897948 | 0.786024 |
| 38493 | ENSG0000 | AL358072.1 | | 0.900052 | 0.367542 | 0.293068 |
| 34062 | ENSG0000 | AC010809.1 | | 0.896909 | 0.779603 | 0.653041 |
| 43174 | ENSG0000 | MIR3940 | microRNA | 0.8957 | 0.784029 | 0.292474 |
| 2815 | ENSG0000 | C16orf62 | chromosor | 0.892738 | 0.847966 | 0.778182 |
| 8588 | ENSG0000 | TRMT10A | tRNA meth | 0.892418 | 0.747547 | 0.51663 |
| 37470 | ENSG0000 | AC026304.1 | | 0.890187 | 0.352075 | 0.211504 |
| 20866 | ENSG0000 | SNORD88A | small nucle | 0.886541 | 0.357986 | 0.316839 |
| 37084 | ENSG0000 | UPK3BL1 | uroplakin 3 | 0.882532 | 0.13718 | 0.099237 |
| 25948 | ENSG0000 | AC012146.1 | | 0.881152 | 0.538024 | 0.424263 |
| 35700 | ENSG0000 | AC145207.2 | | 0.875771 | 0.561997 | 0.340024 |
| 27347 | ENSG0000 | CDC42P6 | cell divisio | 0.870521 | 0.74213 | 0.649035 |
| 18946 | ENSG0000 | Y_RNA | Y RNA [Sou | 0.869794 | 0.706326 | 0.491325 |
| 40211 | ENSG0000 | AL138781.2 | | 0.860317 | 0.334668 | 0.302476 |
| 25800 | ENSG0000 | UQCRHL | ubiquinol-c | 0.854307 | 0.590054 | 0.576352 |
| 38667 | ENSG0000 | AC024060.1 | | 0.851965 | 0.692421 | 0.431576 |
| 30308 | ENSG0000 | EEF1A1P13 | eukaryotic | 0.850111 | 0.61221 | 0.559422 |
| 36132 | ENSG0000 | RN7SL657F | RNA, 7SL, c | 0.843591 | 0.829523 | 0.708131 |
| 34289 | ENSG0000 | ST20-AS1 | ST20 antise | 0.841159 | 0.78694 | 0.745532 |
| 22715 | ENSG0000 | TBCAP1 | tubulin folc | 0.839826 | 0.736139 | 0.727158 |
| 7891 | ENSG0000 | SLC5A2 | solute carr | 0.839022 | 0.510133 | 0.362358 |
| 26953 | ENSG0000 | HNRNPA3F | heterogen | 0.838586 | 0.719639 | 0.63086 |
| 26676 | ENSG0000 | MORC2-AS | MORC2 an | 0.837923 | 0.021819 | 0 |

| | | | | | | |
|---|---|---|---|---|---|---|
| 8846 | ENSG0000 | CSGALNAC | chondroiti | 0.826721 | 0.585402 | 0.517091 |
| 25669 | ENSG0000 | AC108047.1 | | 0.823834 | 0.36653 | 0.289803 |
| 38202 | ENSG0000 | AC025449.1 | | 0.821768 | 0.758469 | 0.643402 |
| 2348 | ENSG0000 | GSTZ1 | glutathione | 0.816521 | 0.452389 | 0.425212 |
| 13390 | ENSG0000 | IL20RB | interleukin | 0.815527 | 0.51608 | 0.429288 |
| 41691 | ENSG0000 | AL590326.2 | | 0.811147 | 0.494161 | 0.273721 |
| 39690 | ENSG0000 | AL133520.1 | | 0.810528 | 0.666852 | 0.646389 |
| 23485 | ENSG0000 | SAPCD1 | suppressor | 0.80706 | 0.614603 | 0.418143 |
| 23123 | ENSG0000 | AL731569.1 | | 0.790604 | 0.652829 | 0.570706 |
| 26784 | ENSG0000 | CHCHD4P3 | coiled-coil- | 0.787995 | 0.508696 | 0.0816 |
| 27033 | ENSG0000 | AL391650.1 | | 0.786269 | 0.724579 | 0.528121 |
| 3182 | ENSG0000 | LSR | lipolysis sti | 0.782277 | 0.601124 | 0.601068 |
| 21984 | ENSG0000 | ST13P6 | ST13, Hsp7 | 0.777806 | 0.671651 | 0.644123 |
| 15169 | ENSG0000 | BRD7P2 | bromodom | 0.77699 | 0.775436 | 0.700889 |
| 37868 | ENSG0000 | AC093677.2 | | 0.776161 | 0.54766 | 0.431731 |
| 34998 | ENSG0000 | AC022968.1 | | 0.775937 | 0.589741 | 0.491056 |
| 16602 | ENSG0000 | HIST1H4C | histone clu | 0.77251 | 0.391509 | 0.386332 |
| 14366 | ENSG0000 | PSTK | phosphose | 0.769025 | 0.615725 | 0.50367 |
| 29850 | ENSG0000 | AC008438.1 | | 0.76753 | 0.503435 | 0.434438 |
| 12248 | ENSG0000 | JMJD7-PLA | JMJD7-PLA | 0.762709 | 0.631431 | 0.508254 |
| 40452 | ENSG0000 | SNORA24 | small nucle | 0.760976 | 0.740679 | 0.459402 |
| 18819 | ENSG0000 | SNORD116 | small nucle | 0.7578 | 0.32121 | 0.236995 |
| 6869 | ENSG0000 | GGACT | gamma-glu | 0.753413 | 0.360844 | 0.328213 |
| 41826 | ENSG0000 | AC012513.3 | | 0.745337 | 0.464007 | 0.362144 |
| 8852 | ENSG0000 | GNRH1 | gonadotro | 0.744079 | 0.697766 | 0.398634 |
| 28024 | ENSG0000 | AC017116.1 | | 0.742317 | 0.377184 | 0 |
| 14733 | ENSG0000 | KCNJ14 | potassium | 0.740647 | 0.658632 | 0.280706 |
| 39467 | ENSG0000 | AC012510.1 | | 0.738829 | 0.423662 | 0.31341 |
| 32370 | ENSG0000 | AP006621.2 | | 0.737574 | 0.721487 | 0.211622 |
| 33370 | ENSG0000 | NDUFC2-K | NDUFC2-K | 0.736705 | 0.551163 | 0.013474 |
| 24859 | ENSG0000 | IPO9-AS1 | IPO9 antise | 0.736317 | 0.24577 | 0.230505 |
| 38807 | ENSG0000 | AC013403.2 | | 0.729979 | 0.278131 | 0.251996 |
| 1367 | ENSG0000 | PPP1R12B | protein ph | 0.728517 | 0.651063 | 0.501593 |
| 33238 | ENSG0000 | AC121761.1 | | 0.725087 | 0.642521 | 0.370701 |
| 30071 | ENSG0000 | RBBP4P1 | RB binding | 0.721763 | 0.606144 | 0.590824 |
| 41146 | ENSG0000 | AC139768.1 | | 0.719411 | 0.376431 | 0.3661 |
| 29359 | ENSG0000 | USP2-AS1 | USP2 antis | 0.718452 | 0.645991 | 0.382141 |
| 40254 | ENSG0000 | AL121832.3 | | 0.708481 | 0.430391 | 0.305057 |
| 40784 | ENSG0000 | MIR6763 | microRNA | 0.706744 | 0.188736 | 0.001767 |
| 9233 | ENSG0000 | C12orf45 | chromosor | 0.70519 | 0.636661 | 0.498707 |
| 39619 | ENSG0000 | HIST1H2BN | histone clu | 0.703948 | 0.333084 | 0.085323 |
| 39560 | ENSG0000 | MIR6812 | microRNA | 0.703037 | 0.144972 | 0.115767 |
| 10328 | ENSG0000 | CFAP157 | cilia and fla | 0.701137 | 0.451197 | 0.363665 |
| 18537 | ENSG0000 | SNORA22 | small nucle | 0.690343 | 0.667783 | 0.582065 |
| 39459 | ENSG0000 | AC093788.1 | | 0.680477 | 0.182272 | 0.182032 |
| 3341 | ENSG0000 | FSD1L | fibronectin | 0.675526 | 0.612712 | 0.509639 |
| 37389 | ENSG0000 | AP001160.2 | | 0.674002 | 0.450627 | 0.437855 |

| | | | | | | |
|---|---|---|---|---|---|---|
| 39420 | ENSG0000 | AC092587.1 | | 0.671907 | 0.367928 | 0.202029 |
| 42454 | ENSG0000 | AC099329.3 | | 0.671008 | 0.556387 | 0.255161 |
| 38802 | ENSG0000 | AL359643.3 | | 0.668681 | 0.521894 | 0.212613 |
| 37254 | ENSG0000 | AC125437.1 | | 0.668601 | 0.61638 | 0.43229 |
| 36083 | ENSG0000 | AC015813.1 | | 0.667374 | 0.655459 | 0.253335 |
| 17335 | ENSG0000 | SNORA31 | small nucle | 0.665046 | 0.47324 | 0.18488 |
| 43393 | ENSG0000 | AC092902.6 | | 0.664104 | 0.55367 | 0.023489 |
| 32099 | ENSG0000 | AP000781.1 | | 0.656317 | 0.485316 | 0.321571 |
| 35582 | ENSG0000 | RN7SL5P | RNA, 7SL, c | 0.651314 | 0.615388 | 0.387194 |
| 33695 | ENSG0000 | AL355075.2 | | 0.648213 | 0.489018 | 0.25638 |
| 29844 | ENSG0000 | AP000295.1 | | 0.643782 | 0.641197 | 0.63876 |
| 29833 | ENSG0000 | AC139887.2 | | 0.639564 | 0.606519 | 0.55469 |
| 41190 | ENSG0000 | AC006277.1 | | 0.629761 | 0.561622 | 0.452273 |
| 16744 | ENSG0000 | ZNF624 | zinc finger | 0.628082 | 0.572223 | 0.530783 |
| 36801 | ENSG0000 | MIR4754 | microRNA | 0.623636 | 0.436689 | 0.32525 |
| 37173 | ENSG0000 | AC011451.1 | | 0.623486 | 0.244512 | 0.215746 |
| 39920 | ENSG0000 | AC106782.6 | | 0.622113 | 0.583228 | 0.559516 |
| 32898 | ENSG0000 | CCDC150P | coiled-coil | 0.621792 | 0.574377 | 0.27467 |
| 30854 | ENSG0000 | CENPS-COI | CENPS-COI | 0.620157 | 0.490877 | 0.253355 |
| 40146 | ENSG0000 | HIST1H4L | histone clu | 0.61598 | 0.418972 | 0.208754 |
| 9635 | ENSG0000 | NCAM2 | neural cell | 0.612351 | 0.458724 | 0.257018 |
| 39388 | ENSG0000 | AL021707.8 | | 0.610723 | 0.422504 | 0.268657 |
| 25638 | ENSG0000 | AC019068.1 | | 0.608887 | 0.607487 | 0.369852 |
| 43317 | ENSG0000 | MIRLET7B | microRNA | 0.604559 | 0.00207 | 0 |
| 39640 | ENSG0000 | AL117379.1 | | 0.604009 | 0.501598 | 0.369721 |
| 31277 | ENSG0000 | Y_RNA | Y RNA [Sou | 0.603171 | 0.578671 | 0.369534 |
| 8797 | ENSG0000 | SPIN2A | spindlin fa | 0.601658 | 0.486474 | 0.446095 |
| 29410 | ENSG0000 | AP003352.1 | | 0.599034 | 0.46587 | 0.288624 |
| 35339 | ENSG0000 | AC128688.2 | | 0.591801 | 0.502682 | 0.388019 |
| 39799 | ENSG0000 | AL136531.2 | | 0.590806 | 0.354329 | 0.030484 |
| 27782 | ENSG0000 | SNORD127 | small nucle | 0.589779 | 0.370408 | 0.332588 |
| 36515 | ENSG0000 | MIR4444-2 | microRNA | 0.587299 | 0.476253 | 0.143619 |
| 40610 | ENSG0000 | PCDHGB5 | protocadh | 0.584766 | 0.514092 | 0.009993 |
| 30688 | ENSG0000 | GAPDHP62 | glyceraldel | 0.584502 | 0.45472 | 0.26837 |
| 36195 | ENSG0000 | MIR4449 | microRNA | 0.581619 | 0.367066 | 0 |
| 37436 | ENSG0000 | AC243960.2 | | 0.581274 | 0.252483 | 0.240888 |
| 29447 | ENSG0000 | AL049840.1 | | 0.576946 | 0.555989 | 0.253507 |
| 2330 | ENSG0000 | CDKL1 | cyclin depe | 0.576418 | 0.525129 | 0.493605 |
| 40622 | ENSG0000 | MIR6895 | microRNA | 0.576044 | 0.359497 | 0.11487 |
| 5031 | ENSG0000 | KIAA1217 | KIAA1217 \| | 0.574272 | 0.41488 | 0.405627 |
| 21751 | ENSG0000 | AL451042.1 | | 0.571106 | 0.144707 | 0.091725 |
| 16353 | ENSG0000 | FAM217B | family with | 0.570039 | 0.51871 | 0.508227 |
| 19818 | ENSG0000 | RARRES2P1 | retinoic aci | 0.565195 | 0.536073 | 0.420992 |
| 41856 | ENSG0000 | AL158801.5 | | 0.562303 | 0.472888 | 0.256913 |
| 42613 | ENSG0000 | SNORA50A | small nucle | 0.559885 | 0.00371 | 0.002268 |
| 11593 | ENSG0000 | STXBP4 | syntaxin bi | 0.55929 | 0.449592 | 0.411424 |
| 39437 | ENSG0000 | AL359091.4 | | 0.558516 | 0.432528 | 0.158122 |

| | | | | | | |
|---|---|---|---|---|---|---|
| 19780 | ENSG0000 | HSPD1P1 | heat shock | 0.555462 | 0.230074 | 0.228722 |
| 30644 | ENSG0000 | AL035458.2 | | 0.555134 | 0.224194 | 0.147392 |
| 5873 | ENSG0000 | GNGT1 | G protein s | 0.553117 | 0.503785 | 0.448574 |
| 38148 | ENSG0000 | CAHM | colon aden | 0.551193 | 0.500728 | 0.484992 |
| 33454 | ENSG0000 | TEN1 | TEN1, CST | 0.545245 | 0.2699 | 0.200746 |
| 31995 | ENSG0000 | AC015689.1 | | 0.545114 | 0.187661 | 0 |
| 21255 | ENSG0000 | AC004471.1 | | 0.544482 | 0.371139 | 0.202414 |
| 34993 | ENSG0000 | AC026471.3 | | 0.544272 | 0.504268 | 0.4473 |
| 37676 | ENSG0000 | AC027307.3 | | 0.541535 | 0.391652 | 0.381577 |
| 43149 | ENSG0000 | AL772307.1 | | 0.541224 | 0.109742 | 0.031339 |
| 8436 | ENSG0000 | AL078621.1 | | 0.541208 | 0.44986 | 0.426948 |
| 39473 | ENSG0000 | AC099676.1 | | 0.537856 | 0.467768 | 0.462267 |
| 31393 | ENSG0000 | PCDHGA12 | protocadh | 0.536495 | 0.497468 | 0.127187 |
| 7677 | ENSG0000 | PRICKLE1 | prickle plar | 0.53459 | 0.443689 | 0.409428 |
| 910 | ENSG0000 | CHI3L2 | chitinase 3 | 0.530205 | 0.154874 | 0.112778 |
| 39689 | ENSG0000 | AL731566.1 | | 0.528385 | 0.367658 | 0.324276 |
| 18123 | ENSG0000 | SPIN3 | spindlin far | 0.518937 | 0.441031 | 0.328983 |
| 34719 | ENSG0000 | AC092115.1 | | 0.515687 | 0.266037 | 0.225643 |
| 35664 | ENSG0000 | PCDHGB3 | protocadh | 0.515524 | 0.143055 | 0.130397 |
| 6427 | ENSG0000 | ZNF132 | zinc finger | 0.514392 | 0.391295 | 0.358231 |
| 10214 | ENSG0000 | CCDC17 | coiled-coil | 0.512385 | 0.405969 | 0.177021 |
| 37987 | ENSG0000 | AC010531.6 | | 0.511635 | 0.268012 | 0.225013 |
| 24866 | ENSG0000 | PRH1 | proline rich | 0.51006 | 0.426999 | 0.368897 |
| 5963 | ENSG0000 | CHN1 | chimerin 1 | 0.507193 | 0.40391 | 0.358955 |
| 16643 | ENSG0000 | MIRLET7BH | MIRLET7B | 0.506439 | 0.270074 | 0.22155 |
| 28283 | ENSG0000 | PCDHGC5 | protocadh | 0.504517 | 0.49896 | 0.199423 |
| 42887 | ENSG0000 | MIR1244-3 | microRNA | 0.503434 | 0.423279 | 0.11487 |
| 39517 | ENSG0000 | uc_338 | TUC338 [So | 0.501921 | 0.451941 | 0.38112 |
| 35130 | ENSG0000 | AC107027.3 | | 0.500498 | 0.320464 | 0.248262 |
| 28635 | ENSG0000 | AC008897.1 | | 0.499807 | 0.12567 | 0.080111 |
| 13777 | ENSG0000 | TCEANC | transcriptio | 0.498431 | 0.448483 | 0.389338 |
| 15464 | ENSG0000 | STAC3 | SH3 and cy | 0.493095 | 0.421395 | 0.345487 |
| 21636 | ENSG0000 | EIF4BP7 | eukaryotic | 0.491852 | 0.391875 | 0.375429 |
| 28794 | ENSG0000 | AC073842.2 | | 0.491788 | 0.44696 | 0.133863 |
| 16145 | ENSG0000 | QRFP | pyroglutam | 0.491047 | 0.420534 | 0.270929 |
| 39164 | ENSG0000 | AC008575.2 | | 0.485474 | 0.267893 | 0.168372 |
| 19565 | ENSG0000 | SNORA75 | Small nucle | 0.484974 | 0.388997 | 0.293301 |
| 23417 | ENSG0000 | NUTM2E | NUT family | 0.483719 | 0.385195 | 0.265955 |
| 27796 | ENSG0000 | SNORA84 | small nucle | 0.479787 | 0.405117 | 0.128463 |
| 37750 | ENSG0000 | AC007842.1 | | 0.478262 | 0.30853 | 0.260807 |
| 37737 | ENSG0000 | AC016629.2 | | 0.478086 | 0.418793 | 0.104519 |
| 33875 | ENSG0000 | AL442663.3 | | 0.477884 | 0.379948 | 0.268336 |
| 29205 | ENSG0000 | AL139099.1 | | 0.476576 | 0.424592 | 0.328363 |
| 26407 | ENSG0000 | AC114730.3 | | 0.474365 | 0.454495 | 0.119442 |
| 40442 | ENSG0000 | AC110285.6 | | 0.471136 | 0.309903 | 0.262961 |
| 36560 | ENSG0000 | AC125232.2 | | 0.469109 | 0.387853 | 0.220806 |
| 38054 | ENSG0000 | AC025766.1 | | 0.468344 | 0.211863 | 0.145045 |

| | | | | | | |
|---|---|---|---|---|---|---|
| 7996 | ENSG0000 | CCDC40 | coiled-coil | 0.466337 | 0.364972 | 0.279795 |
| 31553 | ENSG0000 | CDH12P3 | cadherin 1 | 0.464396 | 0.405142 | 0.37654 |
| 30368 | ENSG0000 | AC003072.1 | | 0.463755 | 0.29612 | 0.228167 |
| 37086 | ENSG0000 | AC008752.3 | | 0.462131 | 0.433905 | 0.22861 |
| 33852 | ENSG0000 | HIF1A-AS1 | HIF1A antis | 0.461352 | 0.429915 | 0.352714 |
| 26156 | ENSG0000 | C11orf94 | chromosor | 0.460068 | 0.362827 | 0.095874 |
| 13020 | ENSG0000 | FGGY | FGGY carb | 0.460063 | 0.271186 | 0.232647 |
| 15971 | ENSG0000 | C2orf66 | chromosor | 0.458867 | 0.294634 | 0.220724 |
| 39151 | ENSG0000 | AC211476.2 | | 0.451171 | 0.328202 | 0.160461 |
| 25511 | ENSG0000 | UBE2FP3 | ubiquitin c | 0.450112 | 0.304551 | 0.098867 |
| 38906 | ENSG0000 | RNU4-89P | RNA, U4 sr | 0.440978 | 0.281369 | 0.240968 |
| 30418 | ENSG0000 | GMPSP1 | guanine m | 0.439526 | 0.227192 | 0.197699 |
| 32627 | ENSG0000 | AP002373.1 | | 0.438034 | 0.294903 | 0.003252 |
| 4653 | ENSG0000 | ARTN | artemin [S | 0.437358 | 0.283391 | 0.174168 |
| 39814 | ENSG0000 | AL355987.5 | | 0.434119 | 0.371079 | 0.09465 |
| 37449 | ENSG0000 | AC004466.1 | | 0.42836 | 0.317971 | 0.240615 |
| 25737 | ENSG0000 | AL391839.2 | | 0.427627 | 0.082593 | 0 |
| 25938 | ENSG0000 | AL592293.2 | | 0.42692 | 0.240649 | 0.22824 |
| 17979 | ENSG0000 | C1orf53 | chromosor | 0.426601 | 0.310285 | 0 |
| 25805 | ENSG0000 | AL359715.2 | | 0.423294 | 0.24292 | 0.220395 |
| 17987 | ENSG0000 | FCGR3A | Fc fragmer | 0.42288 | 0.308907 | 0.300445 |
| 35782 | ENSG0000 | AC136624.3 | | 0.421101 | 0.19282 | 0.034761 |
| 20918 | ENSG0000 | SNORA77 | small nucle | 0.418857 | 0.170719 | 0.100167 |
| 33768 | ENSG0000 | SYNJ2BP-C | SYNJ2BP-C | 0.418783 | 0.2674 | 0.264895 |
| 8611 | ENSG0000 | CYP4V2 | cytochrom | 0.418289 | 0.341461 | 0.328733 |
| 24445 | ENSG0000 | AC099513.1 | | 0.417716 | 0.389759 | 0.385825 |
| 27811 | ENSG0000 | AC008026.1 | | 0.415399 | 0.153714 | 0.045657 |
| 36645 | ENSG0000 | AC015813.2 | | 0.414425 | 0.042249 | 0.015574 |
| 39281 | ENSG0000 | AL109827.1 | | 0.413243 | 0.331339 | 0.210879 |
| 7802 | ENSG0000 | JDP2 | Jun dimeri: | 0.413142 | 0.314158 | 0.309919 |
| 40954 | ENSG0000 | AC005363.2 | | 0.40982 | 0.201496 | 0.089712 |
| 36391 | ENSG0000 | MIR4524B | microRNA | 0.408219 | 0.304791 | 0.094893 |
| 38631 | ENSG0000 | AC008494.3 | | 0.408148 | 0.363801 | 0.246068 |
| 21873 | ENSG0000 | RPS4XP16 | ribosomal | 0.403819 | 0.081074 | 0.047354 |
| 38721 | ENSG0000 | AC013400.1 | | 0.401592 | 0.225854 | 0.19138 |
| 1253 | ENSG0000 | VDAC1P1 | voltage de | 0.396993 | 0.295171 | 0.23631 |
| 29783 | ENSG0000 | AC026436.1 | | 0.395755 | 0.323018 | 0.145502 |
| 37650 | ENSG0000 | AC016727.1 | | 0.394045 | 0.300361 | 0.271752 |
| 32236 | ENSG0000 | SCARNA9 | small Cajal | 0.393016 | 0.288334 | 0.101543 |
| 28572 | ENSG0000 | AC079447.1 | | 0.392402 | 0.32835 | 0.206403 |
| 28605 | ENSG0000 | AC084864.1 | | 0.392317 | 0.24111 | 0.204479 |
| 27623 | ENSG0000 | MLLT10P1 | myeloid/ly | 0.390545 | 0.323832 | 0.166605 |
| 37188 | ENSG0000 | MTCO2P2 | mitochond | 0.388203 | 0.381666 | 0.118444 |
| 35991 | ENSG0000 | AC022211.1 | | 0.385444 | 0.324431 | 0.263427 |
| 21266 | ENSG0000 | CDRT15 | CMT1A du | 0.385269 | 0.292298 | 0.245767 |
| 27986 | ENSG0000 | RPSAP41 | ribosomal | 0.384058 | 0.116215 | 0.020214 |
| 26336 | ENSG0000 | AL162724.2 | | 0.383924 | 0.369486 | 0.284763 |

| | | | | | | |
|---|---|---|---|---|---|---|
| 18762 | ENSG0000 | Y_RNA | Y RNA [Sou | 0.382511 | 0.044471 | 0 |
| 36296 | ENSG0000 | MIR4436B | microRNA | 0.378267 | 0.158035 | 0 |
| 36016 | ENSG0000 | MIR4436B | microRNA | 0.378267 | 0.158035 | 0 |
| 32534 | ENSG0000 | AP003072.2 | | 0.377017 | 0.26825 | 0.246053 |
| 38664 | ENSG0000 | AC104118.1 | | 0.375278 | 0.263145 | 0.081688 |
| 20051 | ENSG0000 | UBE2V1P2 | ubiquitin c | 0.374858 | 0.308101 | 0.251408 |
| 33379 | ENSG0000 | AC073896.3 | | 0.374624 | 0.349299 | 0.123656 |
| 20481 | ENSG0000 | ZSCAN12P | zinc finger | 0.374347 | 0.186551 | 0.151681 |
| 32524 | ENSG0000 | AP001458.2 | | 0.373837 | 0.294559 | 0.263558 |
| 36932 | ENSG0000 | AC007998.2 | | 0.370766 | 0.247336 | 0.126764 |
| 31890 | ENSG0000 | KRT8P3 | keratin 8 p | 0.370192 | 0.308628 | 0.251699 |
| 6983 | ENSG0000 | KRT7 | keratin 7 [S | 0.369924 | 0.308821 | 0.292284 |
| 30348 | ENSG0000 | PSMC1P5 | proteasom | 0.367857 | 0.162888 | 0.060788 |
| 21558 | ENSG0000 | AC022400.1 | | 0.364871 | 0.345014 | 0.226428 |
| 37172 | ENSG0000 | ZNF285 | zinc finger | 0.364554 | 0.345072 | 0.238021 |
| 23195 | ENSG0000 | AL365277.2 | | 0.363931 | 0.197404 | 0.153104 |
| 35733 | ENSG0000 | AC009163.5 | | 0.362648 | 0.335202 | 0.25038 |
| 42502 | ENSG0000 | MIR3653 | microRNA | 0.359996 | 0.344831 | 0.324326 |
| 37667 | ENSG0000 | AL136038.4 | | 0.356616 | 0.350722 | 0.108949 |
| 35571 | ENSG0000 | LINC01970 | long interg | 0.356089 | 0.166221 | 0.157241 |
| 30116 | ENSG0000 | AC079921.2 | | 0.349361 | 0.248098 | 0.099975 |
| 30877 | ENSG0000 | AC091887.1 | | 0.347723 | 0.211942 | 0.201121 |
| 40375 | ENSG0000 | MIR6505 | microRNA | 0.343895 | 0.079859 | 0 |
| 24326 | ENSG0000 | AC140479.2 | | 0.343859 | 0.22927 | 0.173167 |
| 10020 | ENSG0000 | KRTCAP3 | keratinocy | 0.343701 | 0.234824 | 0.191149 |
| 25444 | ENSG0000 | RNF224 | ring finger | 0.342625 | 0.3254 | 0.247287 |
| 41366 | ENSG0000 | HIST1H2AE | histone clu | 0.341707 | 0.281992 | 0.27826 |
| 34605 | ENSG0000 | AC106820.3 | | 0.339532 | 0.206442 | 0.157693 |
| 34706 | ENSG0000 | AC022167.2 | | 0.339338 | 0.226324 | 0.211447 |
| 40961 | ENSG0000 | AL391987.6 | | 0.338996 | 0.286738 | 0.202321 |
| 40869 | ENSG0000 | AC069234.5 | | 0.335219 | 0.271357 | 0.167335 |
| 24563 | ENSG0000 | AL954705.1 | | 0.334122 | 0.301158 | 0.205587 |
| 37724 | ENSG0000 | MIA-RAB4I | MIA-RAB4I | 0.333773 | 0.271026 | 0.11395 |
| 32907 | ENSG0000 | AP006333.1 | | 0.333361 | 0.244975 | 0.169308 |
| 6601 | ENSG0000 | ALOX5AP | arachidona | 0.332075 | 0.316607 | 0.227194 |
| 24121 | ENSG0000 | FTH1P5 | ferritin hea | 0.330734 | 0.137991 | 0.091896 |
| 17682 | ENSG0000 | RNA5SP78 | RNA, 5S rib | 0.330645 | 0.26559 | 0.027257 |
| 22995 | ENSG0000 | SPAG5-AS1 | SPAG5 anti | 0.329949 | 0.311043 | 0.303503 |
| 20936 | ENSG0000 | PPP3CB-AS | PPP3CB an | 0.327008 | 0.287497 | 0.160502 |
| 24984 | ENSG0000 | ST13P4 | ST13, Hsp7 | 0.326238 | 0.257256 | 0.205673 |
| 27221 | ENSG0000 | AL450998.3 | | 0.325501 | 0.303857 | 0.187131 |
| 27213 | ENSG0000 | AC011753.4 | | 0.32537 | 0.297008 | 0.265567 |
| 26463 | ENSG0000 | DBF4P1 | DBF4 zinc f | 0.325323 | 0.130716 | 0.091263 |
| 11175 | ENSG0000 | LEAP2 | liver enrich | 0.322836 | 0.207433 | 0.079874 |
| 32272 | ENSG0000 | ALG1L9P | asparagine | 0.322418 | 0.107491 | 0.098574 |
| 37982 | ENSG0000 | AC068790.3 | | 0.32227 | 0.149883 | 0.035728 |
| 2997 | ENSG0000 | GLI3 | GLI family | 0.321068 | 0.245319 | 0.199938 |

| | | | | | | |
|---|---|---|---|---|---|---|
| 35086 | ENSG0000 | LINC00565 | long interg | 0.320666 | 0.261509 | 0.17982 |
| 39610 | ENSG0000 | AC009318.2 | | 0.320156 | 0.296604 | 0.178922 |
| 38027 | ENSG0000 | AL021878.2 | | 0.319535 | 0.282661 | 0.185404 |
| 39239 | ENSG0000 | Z83851.2 | | 0.31837 | 0.280645 | 0.232703 |
| 41419 | ENSG0000 | MIR6775 | microRNA | 0.318294 | 0.134467 | 0.015815 |
| 24868 | ENSG0000 | TRAF3IP2-/ | TRAF3IP2 a | 0.318272 | 0.308604 | 0.199299 |
| 10560 | ENSG0000 | AKR7A3 | aldo-keto r | 0.317148 | 0.254517 | 0.208244 |
| 19668 | ENSG0000 | CRYGS | crystallin g | 0.316664 | 0.104539 | 0.073466 |
| 27054 | ENSG0000 | BCYRN1 | brain cytop | 0.316534 | 0.285562 | 0.184366 |
| 43121 | ENSG0000 | AC073111.3 | | 0.313917 | 0.203982 | 0.128498 |
| 340 | ENSG0000 | TYROBP | TYRO prote | 0.313187 | 0.303811 | 0.24114 |
| 19672 | ENSG0000 | CNN2P9 | calponin 2 | 0.312362 | 0.29487 | 0.230372 |
| 37026 | ENSG0000 | AC023509.3 | | 0.311817 | 0.288425 | 0.143822 |
| 38002 | ENSG0000 | AL359881.2 | | 0.308905 | 0.168201 | 0.070131 |
| 29059 | ENSG0000 | JMJD7 | jumonji do | 0.307732 | 0.089539 | 0.079723 |
| 34575 | ENSG0000 | AC009090.1 | | 0.307609 | 0.254901 | 0.085497 |
| 8604 | ENSG0000 | MARCH1 | membrane | 0.306933 | 0.190433 | 0.121044 |
| 21535 | ENSG0000 | AC000123.1 | | 0.304671 | 0.213515 | 0.164881 |
| 24816 | ENSG0000 | LINC01611 | long interg | 0.30403 | 0.251397 | 0.229563 |
| 34747 | ENSG0000 | AL133297.1 | | 0.303964 | 0.179644 | 0.174357 |
| 25015 | ENSG0000 | AL021937.2 | | 0.303062 | 0.105655 | 0 |
| 35299 | ENSG0000 | AC097461.1 | | 0.302756 | 0.255444 | 0.117476 |
| 38771 | ENSG0000 | AC124045.1 | | 0.30179 | 0.186968 | 0.157748 |
| 3671 | ENSG0000 | FAM149A | family with | 0.29869 | 0.282251 | 0.262014 |
| 40620 | ENSG0000 | AC002550.2 | | 0.297716 | 0.068553 | 0 |
| 39893 | ENSG0000 | AP001282.2 | | 0.296521 | 0.269571 | 0.166921 |
| 19114 | ENSG0000 | MIR26A2 | microRNA | 0.290944 | 0.284804 | 0.002835 |
| 19049 | ENSG0000 | AL590762.1 | | 0.289008 | 0.189 | 0.185866 |
| 23387 | ENSG0000 | EEF1A1P11 | eukaryotic | 0.288935 | 0.229999 | 0.202082 |
| 3903 | ENSG0000 | AL021546.1 | | 0.288887 | 0.263066 | 0.23109 |
| 42170 | ENSG0000 | AL031719.2 | | 0.287683 | 0.264309 | 0.082601 |
| 16001 | ENSG0000 | NRN1L | neuritin 1 l | 0.28683 | 0.193282 | 0.146514 |
| 35496 | ENSG0000 | AC006504.1 | | 0.286185 | 0.25843 | 0.11089 |
| 27151 | ENSG0000 | PRMT5-AS | PRMT5 ant | 0.285299 | 0.267816 | 0.228999 |
| 25953 | ENSG0000 | AC026462.1 | | 0.283905 | 0.186582 | 0.163756 |
| 38862 | ENSG0000 | AC034198.2 | | 0.283552 | 0.171212 | 0.06712 |
| 26389 | ENSG0000 | AL445991.1 | | 0.281658 | 0.178836 | 0.146066 |
| 17632 | ENSG0000 | Y_RNA | Y RNA [Sou | 0.280675 | 0.153268 | 0 |
| 20629 | ENSG0000 | AL627402.1 | | 0.273549 | 0.107229 | 0.099085 |
| 41185 | ENSG0000 | AL513497.1 | | 0.273469 | 0.216328 | 0.202718 |
| 34502 | ENSG0000 | AC012173.1 | | 0.272991 | 0.185652 | 0.131781 |
| 31802 | ENSG0000 | AC078852.2 | | 0.272734 | 0.145301 | 0.0945 |
| 39373 | ENSG0000 | AC017083.2 | | 0.272415 | 0.128364 | 0.065355 |
| 33171 | ENSG0000 | AC073896.2 | | 0.271333 | 0.042904 | 0 |
| 36393 | ENSG0000 | AC130324.2 | | 0.271216 | 0.081878 | 0.026202 |
| 39916 | ENSG0000 | SNORD109 | small nucle | 0.269613 | 0.050481 | 0.046539 |
| 28576 | ENSG0000 | AKAP2 | A-kinase ar | 0.269025 | 0.211475 | 0.100253 |

| | | | | | | |
|---|---|---|---|---|---|---|
| 3936 | ENSG0000 | RIPOR2 | RHO family | 0.268044 | 0.176599 | 0.171408 |
| 22707 | ENSG0000 | SRRM5 | serine/argi | 0.266967 | 0.242662 | 0.230291 |
| 16421 | ENSG0000 | C20orf204 | chromosor | 0.266893 | 0.20798 | 0.155966 |
| 42396 | ENSG0000 | AC019080.5 | | 0.266181 | 0.254799 | 0.194687 |
| 40845 | ENSG0000 | AL139123.1 | | 0.265181 | 0.048189 | 0 |
| 32950 | ENSG0000 | AP003419.2 | | 0.263932 | 0.145135 | 0.101652 |
| 23722 | ENSG0000 | AL121845.1 | | 0.263495 | 0.260344 | 0.250961 |
| 20889 | ENSG0000 | MIR1238 | microRNA | 0.26309 | 0.214255 | 0.149469 |
| 33567 | ENSG0000 | POLR2KP1 | RNA polym | 0.262954 | 0.223977 | 0.182561 |
| 5927 | ENSG0000 | KRT17 | keratin 17 | 0.262529 | 0.148683 | 0.095899 |
| 4356 | ENSG0000 | PDE1A | phosphodi | 0.262506 | 0.180839 | 0.176848 |
| 18290 | ENSG0000 | FAM27E4 | family with | 0.260065 | 0.179616 | 0.136547 |
| 24643 | ENSG0000 | AC018868.1 | | 0.259708 | 0.074291 | 0 |
| 26234 | ENSG0000 | EIF4A1P2 | eukaryotic | 0.259145 | 0.177198 | 0.170328 |
| 39221 | ENSG0000 | AL662884.1 | | 0.258798 | 0.108265 | 0.091877 |
| 10391 | ENSG0000 | ZNF333 | zinc finger | 0.25876 | 0.25788 | 0.2166 |
| 42569 | ENSG0000 | AC254562.3 | | 0.258715 | 0.149107 | 0.128485 |
| 32294 | ENSG0000 | AP001767.2 | | 0.258306 | 0.034813 | 0 |
| 17310 | ENSG0000 | RNU5E-1 | RNA, U5E s | 0.255912 | 0.065291 | 0 |
| 12469 | ENSG0000 | HGNC:249 | Neuron-sp | 0.252196 | 0.232804 | 0.210266 |
| 34301 | ENSG0000 | AC009996.1 | | 0.251913 | 0.130437 | 0.014521 |
| 41096 | ENSG0000 | AL591926.5 | | 0.251717 | 0.054808 | 0 |
| 29974 | ENSG0000 | MTND3P2 | mitochond | 0.251717 | 0.217637 | 0.11487 |
| 37423 | ENSG0000 | MIR3188 | microRNA | 0.251717 | 0.138263 | 0 |
| 18588 | ENSG0000 | SNORA30 | small nucle | 0.251056 | 0.086883 | 0.069283 |
| 32749 | ENSG0000 | AP000593.3 | | 0.250846 | 0.119568 | 0 |
| 42556 | ENSG0000 | LINC01176 | long interg | 0.250606 | 0.234949 | 0.218646 |
| 35503 | ENSG0000 | AC113208.4 | | 0.250189 | 0.173653 | 0.151801 |
| 39079 | ENSG0000 | AC046143.2 | | 0.250144 | 0.156569 | 0.011487 |
| 23329 | ENSG0000 | AC007040.1 | | 0.25004 | 0.227683 | 0.067346 |
| 40611 | ENSG0000 | AC008740.2 | | 0.249429 | 0.103091 | 0.057435 |
| 19092 | ENSG0000 | MIR584 | microRNA | 0.249122 | 0.01347 | 0 |
| 39224 | ENSG0000 | AC103591.3 | | 0.248706 | 0.157843 | 0.053862 |
| 23215 | ENSG0000 | AP000692.1 | | 0.248683 | 0.234898 | 0.101669 |
| 28194 | ENSG0000 | AC004801.2 | | 0.247945 | 0.187788 | 0.022709 |
| 24865 | ENSG0000 | NDUFB1P1 | NADH:ubic | 0.245424 | 0.169527 | 0.054882 |
| 33600 | ENSG0000 | AC073655.2 | | 0.245352 | 0.195158 | 0.095313 |
| 21928 | ENSG0000 | RANP4 | RAN, mem | 0.244031 | 0.224911 | 0.19862 |
| 41404 | ENSG0000 | AC003101.2 | | 0.241466 | 0.211437 | 0.047819 |
| 37921 | ENSG0000 | AC025857.2 | | 0.24114 | 0.109155 | 0 |
| 26604 | ENSG0000 | AC073333.1 | | 0.240554 | 0.188137 | 0.100458 |
| 40589 | ENSG0000 | HOXA11-A | HOXA11 ar | 0.240499 | 0.082271 | 0.062606 |
| 21036 | ENSG0000 | RN7SKP26 | RNA, 7SK s | 0.239689 | 0.103443 | 0 |
| 40320 | ENSG0000 | AC004816.2 | | 0.236649 | 0.133946 | 0 |
| 30309 | ENSG0000 | AC091180.3 | | 0.236571 | 0.138929 | 0.07113 |
| 18521 | ENSG0000 | RNU6-431 | RNA, U6 sr | 0.235249 | 0.082858 | 0 |
| 17550 | ENSG0000 | SNORA80A | small nucle | 0.235059 | 0.204666 | 0.097133 |

| | | | | | | |
|---|---|---|---|---|---|---|
| 39077 | ENSG0000 | AC010913.1 | | 0.234167 | 0.146175 | 0.097093 |
| 40317 | ENSG0000 | U2 | U2 spliceo: | 0.233925 | 0.155392 | 0.149932 |
| 25821 | ENSG0000 | GDI2P2 | GDP dissoc | 0.233708 | 0.180325 | 0.104762 |
| 19839 | ENSG0000 | RPL7AP50 | ribosomal | 0.233432 | 0.142815 | 0.08537 |
| 34688 | ENSG0000 | AC009087.1 | | 0.233304 | 0.212941 | 0.177005 |
| 26451 | ENSG0000 | SMIM27 | small integ | 0.232886 | 0.194089 | 0.158761 |
| 11922 | ENSG0000 | GPD1 | glycerol-3- | 0.232035 | 0.125328 | 0.120388 |
| 32336 | ENSG0000 | AC136475.4 | | 0.231702 | 0.066457 | 0.035623 |
| 35658 | ENSG0000 | AC116025.2 | | 0.231261 | 0.16313 | 0.146245 |
| 36946 | ENSG0000 | AC091152.2 | | 0.22745 | 0.208389 | 0.088533 |
| 26350 | ENSG0000 | Z82188.2 | | 0.224618 | 0.046284 | 0.025698 |
| 29762 | ENSG0000 | AC008906.1 | | 0.224326 | 0.195735 | 0.096968 |
| 36659 | ENSG0000 | AC005730.3 | | 0.223863 | 0.107316 | 0.044253 |
| 31958 | ENSG0000 | SPON1-AS1 | SPON1 ant | 0.22235 | 0.092782 | 0 |
| 36540 | ENSG0000 | AP000902.1 | | 0.22222 | 0.077534 | 0 |
| 21092 | ENSG0000 | RN7SKP15( | RNA, 7SK s | 0.221134 | 0.101164 | 0.054393 |
| 23539 | ENSG0000 | PIK3IP1-AS | PIK3IP1 an | 0.220851 | 0.117418 | 0.057983 |
| 34411 | ENSG0000 | AC007610.1 | | 0.220244 | 0.058377 | 0 |
| 36750 | ENSG0000 | AP001178.3 | | 0.22016 | 0.193598 | 0.071573 |
| 40292 | ENSG0000 | AL096712.1 | | 0.220054 | 0.164038 | 0 |
| 16183 | ENSG0000 | FAM78B | family with | 0.219627 | 0.21096 | 0.129424 |
| 20848 | ENSG0000 | MIR1302-3 | microRNA | 0.217972 | 0.203194 | 0 |
| 18828 | ENSG0000 | SNORD116 | small nucle | 0.217271 | 0.185564 | 0.082223 |
| 36822 | ENSG0000 | MIR3149 | microRNA | 0.216841 | 0.074524 | 0.012456 |
| 38041 | ENSG0000 | TSNAX-DIS | TSNAX-DIS | 0.21442 | 0.1646 | 0.11801 |
| 22354 | ENSG0000 | ODF2-AS1 | ODF2 antis | 0.214183 | 0.178752 | 0 |
| 39013 | ENSG0000 | AL512343.2 | | 0.213446 | 0.149166 | 0 |
| 40430 | ENSG0000 | uc_338 | TUC338 [So | 0.212544 | 0.080219 | 0.056227 |
| 17931 | ENSG0000 | IGBP1-AS1 | IGBP1 anti: | 0.212504 | 0.175872 | 0.042172 |
| 28440 | ENSG0000 | AC092849.2 | | 0.21195 | 0.202101 | 0 |
| 32499 | ENSG0000 | AP001258.2 | | 0.20985 | 0.033242 | 0 |
| 25287 | ENSG0000 | AC231533.1 | | 0.209165 | 0.153962 | 0.042805 |
| 27768 | ENSG0000 | ACA64 | Small nucle | 0.208788 | 0.143848 | 0 |
| 33698 | ENSG0000 | NT5CP1 | 5',3'-nucle | 0.208693 | 0.057115 | 0 |
| 17259 | ENSG0000 | Y_RNA | Y RNA [Sou | 0.20853 | 0.164743 | 0.11487 |
| 31483 | ENSG0000 | AC040934.1 | | 0.207403 | 0.084239 | 0.055353 |
| 5767 | ENSG0000 | CFP | compleme | 0.207184 | 0.14149 | 0.101602 |
| 39198 | ENSG0000 | AL121672.3 | | 0.206806 | 0.117409 | 0.046881 |
| 25823 | ENSG0000 | LINC01425 | long interg | 0.206721 | 0.186152 | 0.083474 |
| 5381 | ENSG0000 | RAB38 | RAB38, me | 0.206442 | 0.099432 | 0.088485 |
| 42598 | ENSG0000 | LINC00602 | long interg | 0.206186 | 0.177279 | 0.163531 |
| 19135 | ENSG0000 | MIR574 | microRNA | 0.205831 | 0.136381 | 0.015792 |
| 38096 | ENSG0000 | AC093668.1 | | 0.20561 | 0.177448 | 0.136956 |
| 10970 | ENSG0000 | SPDYA | speedy/RII | 0.204197 | 0.174416 | 0.130861 |
| 26877 | ENSG0000 | AL359313.1 | | 0.203811 | 0.184422 | 0.083505 |
| 40782 | ENSG0000 | AL122125.1 | | 0.202632 | 0.073195 | 0 |
| 31194 | ENSG0000 | Y_RNA | Y RNA [Sou | 0.201898 | 0.103091 | 0 |

| | | | | | | |
|---|---|---|---|---|---|---|
| 17679 | ENSG0000 | RNA5SP37 | RNA, 5S rib | 0.201596 | 0.082108 | 0.009149 |
| 34941 | ENSG0000 | AL359715.3 | | 0.201385 | 0.081531 | 0.076346 |
| 28017 | ENSG0000 | RN7SL688I | RNA, 7SL, c | 0.201374 | 0.169069 | 0.068086 |
| 4013 | ENSG0000 | EPM2A | EPM2A, laf | 0.199569 | 0.159051 | 0.124039 |
| 28064 | ENSG0000 | RN7SL75P | RNA, 7SL, c | 0.199012 | 0.146329 | 0 |
| 14461 | ENSG0000 | AC068775. | Putative eu | 0.198364 | 0.170402 | 0.140324 |
| 42599 | ENSG0000 | MIR1291 | microRNA | 0.197626 | 0.046817 | 0 |
| 26080 | ENSG0000 | AC010883.1 | | 0.197361 | 0.07952 | 0 |
| 41483 | ENSG0000 | AC092683.2 | | 0.197151 | 0.166515 | 0.131097 |
| 26988 | ENSG0000 | AP000688.4 | | 0.19621 | 0.118004 | 0.065623 |
| 17716 | ENSG0000 | RNU6-7 | RNA, U6 sr | 0.195257 | 0.032758 | 0 |
| 29832 | ENSG0000 | AC004832.3 | | 0.194581 | 0.192138 | 0.028197 |
| 40383 | ENSG0000 | TRBV10-3 | T-cell recep | 0.19426 | 0.128023 | 0.096609 |
| 28436 | ENSG0000 | AC007688.1 | | 0.194011 | 0.057454 | 0.043133 |
| 39538 | ENSG0000 | AC084824.3 | | 0.192757 | 0.169932 | 0.07582 |
| 16068 | ENSG0000 | AC092171.1 | | 0.192402 | 0.148758 | 0.093739 |
| 26825 | ENSG0000 | AC009404.1 | | 0.191784 | 0.120346 | 0.069012 |
| 17554 | ENSG0000 | RNU6-109 | RNA, U6 sr | 0.191728 | 0.130068 | 0.016103 |
| 26832 | ENSG0000 | RPL26P30 | ribosomal | 0.191324 | 0.182227 | 0.076402 |
| 8820 | ENSG0000 | ITGB1BP2 | integrin su | 0.190996 | 0.147314 | 0.107241 |
| 24902 | ENSG0000 | AC007364.1 | | 0.190635 | 0.140912 | 0.078706 |
| 34013 | ENSG0000 | PRKXP1 | protein kin | 0.190425 | 0.158747 | 0.041424 |
| 20610 | ENSG0000 | PRELID1P1 | PRELI dom | 0.189554 | 0.082465 | 0.053216 |
| 22314 | ENSG0000 | LINC01529 | long interg | 0.189362 | 0.184242 | 0.139128 |
| 21303 | ENSG0000 | AL513523.1 | | 0.189317 | 0.064816 | 0.049669 |
| 18416 | ENSG0000 | SAP25 | Sin3A asso | 0.189239 | 0.15436 | 0.06835 |
| 29049 | ENSG0000 | ST13P15 | ST13, Hsp7 | 0.189129 | 0.168 | 0.153108 |
| 23947 | ENSG0000 | KHSRPP1 | KH-type sp | 0.188859 | 0.15452 | 0.146186 |
| 26456 | ENSG0000 | LINC01372 | long interg | 0.187422 | 0.1208 | 0 |
| 30989 | ENSG0000 | RNU7-75P | RNA, U7 sr | 0.186758 | 0.088126 | 0 |
| 41416 | ENSG0000 | AL589182.2 | | 0.186568 | 0.178272 | 0.085413 |
| 28293 | ENSG0000 | AC023818.1 | | 0.186493 | 0.066848 | 0 |
| 15411 | ENSG0000 | GP1BA | glycoprote | 0.186375 | 0.182174 | 0.127527 |
| 24721 | ENSG0000 | RPL7P21 | ribosomal | 0.186013 | 0.091808 | 0.070188 |
| 4384 | ENSG0000 | FN1 | fibronectin | 0.185455 | 0.18246 | 0.176779 |
| 5172 | ENSG0000 | HPCA | hippocalcin | 0.184953 | 0.131829 | 0.108984 |
| 34780 | ENSG0000 | AC010132.4 | | 0.182347 | 0.11545 | 0.071122 |
| 29025 | ENSG0000 | RPL37P23 | ribosomal | 0.181938 | 0.098883 | 0.055481 |
| 11655 | ENSG0000 | CLEC4E | C-type lect | 0.181848 | 0.066192 | 0.033531 |
| 16035 | ENSG0000 | NUTM2B | NUT family | 0.181623 | 0.178072 | 0.150047 |
| 3839 | ENSG0000 | MGP | matrix Gla | 0.181515 | 0.10477 | 0.098547 |
| 39166 | ENSG0000 | AL159169.2 | | 0.181498 | 0.078476 | 0.025708 |
| 38943 | ENSG0000 | AL513477.2 | | 0.181254 | 0.101821 | 0.045705 |
| 7130 | ENSG0000 | IL6 | interleukin | 0.180898 | 0.083454 | 0 |
| 24481 | ENSG0000 | RPL7P32 | ribosomal | 0.180262 | 0.104627 | 0.075592 |
| 37828 | ENSG0000 | SPIB | Spi-B trans | 0.178785 | 0.12605 | 0.094239 |
| 39511 | ENSG0000 | AC092140.2 | | 0.178279 | 0.056196 | 0.031088 |

| | | | | | | |
|---|---|---|---|---|---|---|
| 30977 | ENSG0000 | RNU6-322I | RNA, U6 sr | 0.177179 | 0.06806 | 0 |
| 23309 | ENSG0000 | AC073063.1 | | 0.175829 | 0.152853 | 0.053366 |
| 34784 | ENSG0000 | AC079171.1 | | 0.175293 | 0.134108 | 0.039086 |
| 32024 | ENSG0000 | AL360181.3 | | 0.174899 | 0.148531 | 0.024204 |
| 40891 | ENSG0000 | RN7SL628I | RNA, 7SL, c | 0.174715 | 0.143722 | 0.068825 |
| 18199 | ENSG0000 | LST1 | leukocyte s | 0.174349 | 0.138685 | 0.104116 |
| 39348 | ENSG0000 | AP000240.1 | | 0.174308 | 0.074809 | 0.050573 |
| 23234 | ENSG0000 | AC078927.1 | | 0.172919 | 0.037195 | 0 |
| 36550 | ENSG0000 | SLC25A6P4 | solute carr | 0.172287 | 0.018269 | 0 |
| 5027 | ENSG0000 | SLC10A7 | solute carr | 0.172084 | 0.165732 | 0.150251 |
| 31712 | ENSG0000 | THAP12P7 | THAP dom | 0.170733 | 0.148086 | 0.042361 |
| 19937 | ENSG0000 | NPM1P6 | nucleopho | 0.170436 | 0.087775 | 0.04728 |
| 19841 | ENSG0000 | RPL11P3 | ribosomal | 0.170137 | 0.06623 | 0 |
| 37385 | ENSG0000 | AC012254.4 | | 0.170116 | 0.106867 | 0 |
| 23047 | ENSG0000 | RPL23AP74 | ribosomal | 0.169548 | 0.041878 | 0 |
| 35705 | ENSG0000 | MIR193BH | MIR193B h | 0.169233 | 0.099067 | 0.080958 |
| 41623 | ENSG0000 | AC004232.3 | | 0.16877 | 0.148647 | 0.141563 |
| 38064 | ENSG0000 | AC018695.3 | | 0.168142 | 0.160547 | 0.073225 |
| 14613 | ENSG0000 | P2RY13 | purinergic | 0.167659 | 0.114653 | 0.08183 |
| 19210 | ENSG0000 | AKR7L | aldo-keto r | 0.167411 | 0.106958 | 0.075962 |
| 33064 | ENSG0000 | AC137590.1 | | 0.166992 | 0.075292 | 0 |
| 1265 | ENSG0000 | CDHR2 | cadherin re | 0.166895 | 0.132783 | 0.123327 |
| 23180 | ENSG0000 | VN1R51P | vomeronas | 0.165947 | 0.11777 | 0.078646 |
| 24372 | ENSG0000 | RHOQP3 | ras homolc | 0.165278 | 0.164555 | 0.085228 |
| 35291 | ENSG0000 | AC012645.3 | | 0.165207 | 0.108653 | 0.042272 |
| 3076 | ENSG0000 | OVOL3 | ovo like zir | 0.164947 | 0.160957 | 0.044099 |
| 34885 | ENSG0000 | AC026771.1 | | 0.164742 | 0.090219 | 0 |
| 30238 | ENSG0000 | AC018752.1 | | 0.164598 | 0.077532 | 0 |
| 35175 | ENSG0000 | AC009065.6 | | 0.163531 | 0.122003 | 0 |
| 37003 | ENSG0000 | AC139100.1 | | 0.163039 | 0.107492 | 0.082037 |
| 33978 | ENSG0000 | AL109766.1 | | 0.163023 | 0.094313 | 0 |
| 41821 | ENSG0000 | AP000866.6 | | 0.162502 | 0.126711 | 0.021889 |
| 17740 | ENSG0000 | RNU6-384I | RNA, U6 sr | 0.162164 | 0.089213 | 0 |
| 37027 | ENSG0000 | AC011481.2 | | 0.161869 | 0.152959 | 0 |
| 25059 | ENSG0000 | AL683842.1 | | 0.161818 | 0.110553 | 0.032164 |
| 39909 | ENSG0000 | MIR378J | microRNA | 0.161653 | 0.015133 | 0 |
| 6316 | ENSG0000 | LILRB2 | leukocyte i | 0.161131 | 0.124464 | 0.12276 |
| 32352 | ENSG0000 | AP003392.3 | | 0.161072 | 0.106739 | 0.088762 |
| 26481 | ENSG0000 | AGAP1-IT1 | AGAP1 intr | 0.160545 | 0.110078 | 0 |
| 28828 | ENSG0000 | FTH1P23 | ferritin hea | 0.160412 | 0.15876 | 0.143483 |
| 32815 | ENSG0000 | AL583722.1 | | 0.160016 | 0.104547 | 0.09801 |
| 42079 | ENSG0000 | AC022417.1 | | 0.159086 | 0.156891 | 0.087188 |
| 39601 | ENSG0000 | MIR6792 | microRNA | 0.157793 | 0.063086 | 0.003429 |
| 6535 | ENSG0000 | HSPA12B | heat shock | 0.157573 | 0.139095 | 0.138365 |
| 35087 | ENSG0000 | AC135050.3 | | 0.156973 | 0.084264 | 0.066773 |
| 37904 | ENSG0000 | AC245884.10 | | 0.155736 | 0.145612 | 0.089987 |
| 41370 | ENSG0000 | AC009268.2 | | 0.154776 | 0.127679 | 0.070883 |

| | | | | | | |
|---|---|---|---|---|---|---|
| 40081 | ENSG0000 | AC036214.3 | | 0.154073 | 0.126386 | 0.042872 |
| 26392 | ENSG0000 | HM13-IT1 | HM13 intr | 0.153658 | 0.056212 | 0.005485 |
| 25149 | ENSG0000 | AL353807.2 | | 0.152746 | 0.149316 | 0 |
| 36883 | ENSG0000 | AC018761.2 | | 0.152025 | 0.068276 | 0 |
| 20662 | ENSG0000 | AL021407.3 | | 0.151755 | 0.097782 | 0.052471 |
| 38263 | ENSG0000 | AC104785.1 | | 0.15133 | 0.019987 | 0 |
| 31078 | ENSG0000 | SNORA26 | Small nucle | 0.149959 | 0.045331 | 0 |
| 8794 | ENSG0000 | SYTL5 | synaptotag | 0.149305 | 0.118412 | 0.049094 |
| 38127 | ENSG0000 | AL451085.1 | | 0.149206 | 0.085162 | 0 |
| 25865 | ENSG0000 | RPS15AP1 | ribosomal | 0.148742 | 0.103359 | 0.04744 |
| 23928 | ENSG0000 | AL136531.1 | | 0.148437 | 0.046422 | 0 |
| 40009 | ENSG0000 | AC093512.1 | | 0.147891 | 0.056543 | 0.04906 |
| 36845 | ENSG0000 | AC006538.2 | | 0.147813 | 0.098938 | 0.097415 |
| 17832 | ENSG0000 | Y_RNA | Y RNA [Sou | 0.147402 | 0.121666 | 0 |
| 27994 | ENSG0000 | PSMD6-AS | PSMD6 ant | 0.147049 | 0.122511 | 0.100196 |
| 8353 | ENSG0000 | ACTA1 | actin, alpha | 0.146653 | 0.146051 | 0.021668 |
| 40123 | ENSG0000 | AC020663.3 | | 0.146047 | 0.106776 | 0 |
| 41464 | ENSG0000 | SNORD116 | small nucle | 0.145524 | 0.039915 | 0.032694 |
| 29072 | ENSG0000 | AC011495.1 | | 0.145039 | 0.118978 | 0.106412 |
| 37833 | ENSG0000 | AC024075.3 | | 0.144721 | 0.08654 | 0.06938 |
| 9346 | ENSG0000 | ZNF37CP | zinc finger | 0.144659 | 0.060315 | 0.050047 |
| 24833 | ENSG0000 | AC019097.1 | | 0.143581 | 0.039764 | 0.033367 |
| 26974 | ENSG0000 | Z99916.1 | | 0.143304 | 0.120809 | 0 |
| 37460 | ENSG0000 | ZNF649-AS | ZNF649 an | 0.143145 | 0.087778 | 0.071587 |
| 37849 | ENSG0000 | Z69720.1 | | 0.143021 | 0.124723 | 0.085179 |
| 12873 | ENSG0000 | SLFNL1 | schlafen lik | 0.142634 | 0.110615 | 0.104566 |
| 38813 | ENSG0000 | AC008280.3 | | 0.142536 | 0.141738 | 0.127895 |
| 24887 | ENSG0000 | HNRNPA1F | heterogen | 0.142069 | 0.015329 | 0 |
| 34209 | ENSG0000 | UBE2Q2L | ubiquitin c | 0.141756 | 0.092011 | 0.047997 |
| 30000 | ENSG0000 | EEF1A1P9 | eukaryotic | 0.141329 | 0.125863 | 0.086838 |
| 16203 | ENSG0000 | SBSN | suprabasin | 0.141258 | 0.124841 | 0.081728 |
| 39466 | ENSG0000 | AP000238.1 | | 0.141221 | 0.048642 | 0 |
| 42398 | ENSG0000 | AC012186.3 | | 0.14065 | 0.07057 | 0.033395 |
| 31234 | ENSG0000 | SNORD19 | Small nucle | 0.140001 | 0.060212 | 0.055909 |
| 42844 | ENSG0000 | MIR3179-1 | microRNA | 0.139539 | 0.028964 | 0 |
| 36857 | ENSG0000 | AC008747.1 | | 0.1388 | 0.088006 | 0.074597 |
| 21389 | ENSG0000 | AC244669.1 | | 0.138137 | 0.033892 | 0.030836 |
| 10189 | ENSG0000 | PADI4 | peptidyl ar | 0.137465 | 0.113614 | 0.072451 |
| 40650 | ENSG0000 | AC009090.3 | | 0.137442 | 0.09561 | 0.029073 |
| 30546 | ENSG0000 | ST3GAL1P1 | ST3 beta-g | 0.136905 | 0.088654 | 0.032173 |
| 32746 | ENSG0000 | TAS2R20 | taste 2 rec | 0.13643 | 0.082861 | 0 |
| 39728 | ENSG0000 | AL049794.1 | | 0.135615 | 0.113214 | 0.062926 |
| 39098 | ENSG0000 | AC063962.1 | | 0.134555 | 0.110214 | 0.052915 |
| 3374 | ENSG0000 | KANK1 | KN motif a | 0.134367 | 0.094557 | 0.07244 |
| 16954 | ENSG0000 | PEG3 | paternally | 0.133984 | 0.109529 | 0.105494 |
| 22959 | ENSG0000 | AL031963.1 | | 0.132974 | 0.01723 | 0 |
| 39357 | ENSG0000 | AL390726.5 | | 0.132395 | 0.088799 | 0.073111 |

| | | | | | | |
|---|---|---|---|---|---|---|
| 32914 | ENSG0000 | HSPA8P5 | heat shock | 0.132337 | 0.101771 | 0.026511 |
| 27068 | ENSG0000 | RAP1BP1 | RAP1B, me | 0.132231 | 0.063756 | 0.014748 |
| 33123 | ENSG0000 | AC009318.1 | | 0.13033 | 0.114085 | 0.057659 |
| 28221 | ENSG0000 | SETP14 | SET pseudo | 0.129751 | 0.082367 | 0.017106 |
| 28094 | ENSG0000 | LILRA2 | leukocyte i | 0.129469 | 0.090512 | 0.084075 |
| 41355 | ENSG0000 | RN7SL331P | RNA, 7SL, c | 0.129425 | 0.098917 | 0.076735 |
| 25647 | ENSG0000 | ETV5-AS1 | ETV5 antis | 0.129235 | 0.043874 | 0 |
| 7430 | ENSG0000 | CASP1 | caspase 1 [ | 0.128764 | 0.107298 | 0.048272 |
| 767 | ENSG0000 | ITIH4 | inter-alpha | 0.12871 | 0.110358 | 0.044009 |
| 38628 | ENSG0000 | AL080317.2 | | 0.128403 | 0.126687 | 0.099508 |
| 28387 | ENSG0000 | AP001992.1 | | 0.128184 | 0.107116 | 0.072356 |
| 42145 | ENSG0000 | AD000813.1 | | 0.127371 | 0.070255 | 0 |
| 36988 | ENSG0000 | AC012254.2 | | 0.127088 | 0.103415 | 0.082681 |
| 21481 | ENSG0000 | FLJ37035 | uncharacte | 0.126937 | 0.115611 | 0.078015 |
| 26074 | ENSG0000 | ATP5HP4 | ATP syntha | 0.126918 | 0.064676 | 0 |
| 35873 | ENSG0000 | AC027796.5 | | 0.12686 | 0.017214 | 0 |
| 18919 | ENSG0000 | RNU6-883P | RNA, U6 sr | 0.125858 | 0.071297 | 0 |
| 19057 | ENSG0000 | MIR593 | microRNA | 0.125858 | 0.029896 | 0 |
| 40529 | ENSG0000 | MIR6090 | microRNA | 0.125858 | 0.070445 | 0 |
| 17793 | ENSG0000 | Y_RNA | Y RNA [Sou | 0.125858 | 0.098274 | 0 |
| 34559 | ENSG0000 | AC132938.1 | | 0.125584 | 0.09938 | 0.052654 |
| 29437 | ENSG0000 | AC010198.1 | | 0.125516 | 0.069083 | 0.063237 |
| 7060 | ENSG0000 | HTR2B | 5-hydroxyt | 0.124241 | 0.065335 | 0.0377 |
| 5458 | ENSG0000 | IL9R | interleukin | 0.123633 | 0.09298 | 0.083803 |
| 40335 | ENSG0000 | AC006449.3 | | 0.122188 | 0.09833 | 0.084489 |
| 22356 | ENSG0000 | SATB2-AS1 | SATB2 anti | 0.121706 | 0.057893 | 0.051549 |
| 20799 | ENSG0000 | AL139095.2 | | 0.121203 | 0.119614 | 0.089063 |
| 11619 | ENSG0000 | CYB5R2 | cytochrom | 0.121202 | 0.056729 | 0.031962 |
| 28181 | ENSG0000 | AC017002.3 | | 0.121134 | 0.111785 | 0.001745 |
| 35699 | ENSG0000 | AL160291.1 | | 0.120844 | 0.11879 | 0.088633 |
| 4998 | ENSG0000 | MLANA | melan-A [S | 0.120843 | 0.113135 | 0.050341 |
| 4328 | ENSG0000 | ZAP70 | zeta chain | 0.119442 | 0.110659 | 0.070198 |
| 32 | ENSG0000 | TMEM176A | transmeml | 0.119064 | 0.10881 | 0.093334 |
| 31098 | ENSG0000 | SNORD116 | small nucle | 0.119018 | 0.087403 | 0 |
| 26903 | ENSG0000 | AC133473.1 | | 0.118282 | 0.027583 | 0.013062 |
| 13458 | ENSG0000 | FUT1 | fucosyltrar | 0.117311 | 0.077099 | 0.05803 |
| 18642 | ENSG0000 | DEFA1 | defensin al | 0.117157 | 0.102525 | 0.026802 |
| 28990 | ENSG0000 | RN7SL172P | RNA, 7SL, c | 0.117136 | 0.047973 | 0 |
| 5114 | ENSG0000 | TAS2R10 | taste 2 rec | 0.116975 | 0.0279 | 0 |
| 30872 | ENSG0000 | AC106760.2 | | 0.116709 | 0.098428 | 0.078683 |
| 19150 | ENSG0000 | MIR613 | microRNA | 0.116585 | 0.09658 | 0 |
| 4764 | ENSG0000 | ARG1 | arginase 1 | 0.116577 | 0.011676 | 0 |
| 27567 | ENSG0000 | NFYAP1 | nuclear tra | 0.116406 | 0.075197 | 0.067007 |
| 14443 | ENSG0000 | SERTM1 | serine rich | 0.116283 | 0.021827 | 0.020417 |
| 23572 | ENSG0000 | RPS13P2 | ribosomal | 0.116177 | 0.031947 | 0 |
| 26876 | ENSG0000 | AL358115.1 | | 0.116153 | 0.106324 | 0.040543 |
| 26002 | ENSG0000 | AC005534.2 | | 0.115976 | 0.057617 | 0 |

| | | | | | | |
|---|---|---|---|---|---|---|
| 28159 | ENSG0000 | DEFA1B | defensin al | 0.115861 | 0.101089 | 0.026483 |
| 25771 | ENSG0000 | AL354685.1 | | 0.115747 | 0.072369 | 0.055308 |
| 27690 | ENSG0000 | SNORD121 | small nucle | 0.115613 | 0.112139 | 0.000039 |
| 26474 | ENSG0000 | AL035681.1 | | 0.115601 | 0.035568 | 0.013836 |
| 21908 | ENSG0000 | PFN1P4 | profilin 1 p | 0.115527 | 0.111554 | 0.075437 |
| 30748 | ENSG0000 | ALG1L7P | asparagine | 0.115509 | 0.017364 | 0 |
| 29510 | ENSG0000 | AC021086.1 | | 0.11447 | 0.091596 | 0.037955 |
| 39105 | ENSG0000 | AC115284.1 | | 0.114404 | 0.093987 | 0.093065 |
| 693 | ENSG0000 | SLC4A8 | solute carr | 0.113916 | 0.090619 | 0.065105 |
| 38723 | ENSG0000 | AC126118.1 | | 0.113834 | 0.032011 | 0 |
| 43386 | ENSG0000 | AC009093.8 | | 0.113748 | 0.101451 | 0.015249 |
| 35695 | ENSG0000 | AC090617.3 | | 0.113405 | 0.111555 | 0 |
| 8854 | ENSG0000 | DOK2 | docking pr | 0.112368 | 0.039492 | 0 |
| 28496 | ENSG0000 | AC018475.1 | | 0.1121 | 0.110407 | 0.060317 |
| 29120 | ENSG0000 | RPSAP39 | ribosomal | 0.112033 | 0.081025 | 0.042585 |
| 35827 | ENSG0000 | AC090286.3 | | 0.111725 | 0.020398 | 0 |
| 18669 | ENSG0000 | HNRNPA1P | heterogen | 0.111708 | 0.073159 | 0.003707 |
| 24564 | ENSG0000 | FAM58DP | family with | 0.111422 | 0.094098 | 0.025609 |
| 17796 | ENSG0000 | SNORD58C | small nucle | 0.111273 | 0.087069 | 0.084806 |
| 23831 | ENSG0000 | AC092155.2 | | 0.111136 | 0.057777 | 0.041386 |
| 36849 | ENSG0000 | AC012615.2 | | 0.110761 | 0.088932 | 0.028659 |
| 33900 | ENSG0000 | AC008575.1 | | 0.110738 | 0.033235 | 0.000309 |
| 35359 | ENSG0000 | AC009065.7 | | 0.110578 | 0.041024 | 0.014508 |
| 40772 | ENSG0000 | AL590096.1 | | 0.110435 | 0.080454 | 0.02917 |
| 41574 | ENSG0000 | AC138907.8 | | 0.110349 | 0.058126 | 0.057791 |
| 24021 | ENSG0000 | AKR1B1P1 | aldehyde r | 0.110192 | 0.021351 | 0 |
| 40286 | ENSG0000 | AC006449.2 | | 0.109754 | 0.10352 | 0.081285 |
| 19770 | ENSG0000 | MAGEA12 | MAGE fam | 0.109665 | 0.081974 | 0.033284 |
| 37148 | ENSG0000 | AC008569.2 | | 0.10942 | 0.034131 | 0.012097 |
| 20928 | ENSG0000 | SNORA11 | small nucle | 0.109143 | 0.082151 | 0 |
| 2458 | ENSG0000 | TUBB1 | tubulin bet | 0.108795 | 0.037802 | 0.02783 |
| 28843 | ENSG0000 | RN7SL737P | RNA, 7SL, c | 0.108787 | 0.061964 | 0.034318 |
| 25022 | ENSG0000 | LINC02043 | long interg | 0.108664 | 0.08483 | 0.020944 |
| 34608 | ENSG0000 | AL512604.3 | | 0.108364 | 0.030157 | 0.019423 |
| 42554 | ENSG0000 | SCARNA4 | small Cajal | 0.108297 | 0.09861 | 0.048078 |
| 43021 | ENSG0000 | MIR4453 | microRNA | 0.10825 | 0.06713 | 0.050806 |
| 42241 | ENSG0000 | AC138965.3 | | 0.107662 | 0.058377 | 0 |
| 22605 | ENSG0000 | SUB1P1 | SUB1 hom | 0.107408 | 0.068547 | 0.050819 |
| 30274 | ENSG0000 | AC106795.3 | | 0.106543 | 0.098159 | 0.088363 |
| 43327 | ENSG0000 | MIR4517 | microRNA | 0.106459 | 0.061885 | 0.046114 |
| 36056 | ENSG0000 | AC015908.3 | | 0.106319 | 0.062495 | 0.058327 |
| 19480 | ENSG0000 | SNORD91A | small nucle | 0.10609 | 0.043179 | 0.015035 |
| 15760 | ENSG0000 | TNFRSF4 | TNF recept | 0.105079 | 0.047922 | 0.017458 |
| 42512 | ENSG0000 | N4BP2L2-I | N4BPL2 int | 0.104599 | 0.103766 | 0.04327 |
| 21542 | ENSG0000 | AC073136.2 | | 0.104586 | 0.077166 | 0.028887 |
| 43107 | ENSG0000 | MIR675 | microRNA | 0.104518 | 0.005568 | 0.000065 |
| 10387 | ENSG0000 | PTGER1 | prostaglan | 0.103632 | 0.063803 | 0.033222 |

| | | | | | | |
|---|---|---|---|---|---|---|
| 40239 | ENSG0000 | FCGBP | Fc fragmer | 0.103568 | 0.089808 | 0.063599 |
| 28193 | ENSG0000 | RPL5P23 | ribosomal | 0.103529 | 0.064016 | 0.037187 |
| 34025 | ENSG0000 | SLC35G6 | solute carr | 0.103386 | 0.094055 | 0.043258 |
| 38151 | ENSG0000 | AC099343.2 | | 0.103292 | 0.083705 | 0 |
| 26839 | ENSG0000 | NDUFB9P2 | NADH:ubic | 0.102524 | 0.045344 | 0 |
| 32654 | ENSG0000 | AC108488.2 | | 0.102404 | 0.032483 | 0.000352 |
| 25829 | ENSG0000 | DDX39B-AS | DDX39B ar | 0.101259 | 0.040282 | 0 |
| 30512 | ENSG0000 | NTAN1P2 | N-terminal | 0.100285 | 0.044571 | 0.038767 |
| 13533 | ENSG0000 | SCUBE2 | signal pept | 0.100104 | 0.098417 | 0.057833 |
| 17996 | ENSG0000 | KPRP | keratinocy | 0.099949 | 0.070161 | 0.025998 |
| 11856 | ENSG0000 | NOS2P2 | nitric oxide | 0.099271 | 0.044312 | 0.041287 |
| 35905 | ENSG0000 | AC006441.1 | | 0.099182 | 0.040837 | 0 |
| 21789 | ENSG0000 | LAMTOR5- | LAMTOR5 | 0.099081 | 0.084369 | 0.070051 |
| 39300 | ENSG0000 | AC092139.2 | | 0.099006 | 0.079209 | 0 |
| 42759 | ENSG0000 | hsa-mir-54 | hsa-mir-54 | 0.098611 | 0.069082 | 0.023933 |
| 24836 | ENSG0000 | AL513523.2 | | 0.098057 | 0.029281 | 0.015529 |
| 40955 | ENSG0000 | Y_RNA | Y RNA [Sou | 0.098037 | 0.000271 | 0 |
| 23107 | ENSG0000 | AL627223.1 | | 0.097668 | 0.032887 | 0 |
| 30691 | ENSG0000 | HMMR-AS | HMMR ant | 0.097343 | 0.016552 | 0 |
| 35343 | ENSG0000 | AC040174.1 | | 0.097104 | 0.045738 | 0.016878 |
| 28110 | ENSG0000 | RPL23AP1( | ribosomal | 0.09696 | 0.053758 | 0 |
| 41163 | ENSG0000 | AC020763.3 | | 0.096951 | 0.091909 | 0.090108 |
| 38168 | ENSG0000 | AC073413.1 | | 0.096457 | 0.034364 | 0.027776 |
| 38606 | ENSG0000 | AF287957.1 | | 0.096347 | 0.08913 | 0.020381 |
| 39502 | ENSG0000 | AL136038.5 | | 0.096325 | 0.090065 | 0.087915 |
| 9941 | ENSG0000 | LENEP | lens epithe | 0.096032 | 0.012569 | 0 |
| 43209 | ENSG0000 | MIR6716 | microRNA | 0.095967 | 0.028971 | 0.01864 |
| 38189 | ENSG0000 | PEBP1P2 | phosphatic | 0.09586 | 0.074639 | 0 |
| 20456 | ENSG0000 | RSC1A1 | regulator c | 0.095758 | 0.090474 | 0.079931 |
| 27633 | ENSG0000 | AHCYP2 | adenosylhc | 0.095556 | 0.042505 | 0 |
| 31684 | ENSG0000 | RNU6-323I | RNA, U6 sr | 0.095276 | 0.034685 | 0 |
| 21226 | ENSG0000 | FTH1P10 | ferritin hea | 0.095274 | 0.079719 | 0.046318 |
| 26453 | ENSG0000 | AC006463.2 | | 0.094756 | 0.04177 | 0.005894 |
| 27757 | ENSG0000 | SCARNA18 | small Cajal | 0.093924 | 0.045391 | 0 |
| 43163 | ENSG0000 | MIR6845 | microRNA | 0.093673 | 0.082648 | 0.079099 |
| 18528 | ENSG0000 | SNORD45C | small nucle | 0.093114 | 0.049293 | 0.049276 |
| 42151 | ENSG0000 | AC116407.4 | | 0.092737 | 0.073232 | 0.042257 |
| 30934 | ENSG0000 | AC010260.1 | | 0.092547 | 0.010333 | 0 |
| 34986 | ENSG0000 | AC009690.1 | | 0.09233 | 0.030637 | 0 |
| 26216 | ENSG0000 | GTF3C2-AS | GTF3C2 an | 0.09208 | 0.056274 | 0.039523 |
| 1923 | ENSG0000 | NLRC4 | NLR family | 0.091801 | 0.077181 | 0.039238 |
| 33955 | ENSG0000 | AL118558.1 | | 0.091458 | 0.085791 | 0.026713 |
| 38562 | ENSG0000 | AL139041.1 | | 0.091196 | 0.032279 | 0 |
| 32651 | ENSG0000 | AC009533.2 | | 0.091082 | 0.051455 | 0 |
| 22689 | ENSG0000 | AL513175.1 | | 0.090683 | 0.037536 | 0 |
| 18565 | ENSG0000 | SNORD116 | small nucle | 0.09029 | 0.07915 | 0.054505 |
| 43246 | ENSG0000 | MIR6824 | microRNA | 0.089717 | 0.074023 | 0.070518 |

| | | | | | | |
|---|---|---|---|---|---|---|
| 8161 | ENSG0000 | IGLON5 | IgLON fami | 0.089194 | 0.055699 | 0.043506 |
| 19495 | ENSG0000 | RNA5SP24 | RNA, 5S rib | 0.088948 | 0.084439 | 0.082655 |
| 14449 | ENSG0000 | GLIPR1L2 | GLI pathog | 0.088931 | 0.060358 | 0.046584 |
| 43418 | ENSG0000 | AC006511.5 | | 0.088637 | 0.08102 | 0.059946 |
| 40168 | ENSG0000 | AC018529.1 | | 0.088554 | 0.080531 | 0 |
| 33636 | ENSG0000 | AC013451.1 | | 0.088096 | 0.088017 | 0.045043 |
| 33395 | ENSG0000 | AC011595.2 | | 0.087817 | 0.078201 | 0.043775 |
| 34743 | ENSG0000 | AL390195.2 | | 0.087535 | 0.085918 | 0.069217 |
| 39488 | ENSG0000 | FOXD4L6 | forkhead b | 0.087309 | 0.021465 | 0.015644 |
| 37237 | ENSG0000 | AC008686.1 | | 0.087036 | 0.065042 | 0.047409 |
| 35228 | ENSG0000 | AC122134.1 | | 0.087014 | 0.008606 | 0 |
| 40353 | ENSG0000 | AC007485.1 | | 0.086757 | 0.085576 | 0.036803 |
| 23019 | ENSG0000 | ALG1L8P | asparagine | 0.086673 | 0.066339 | 0.028168 |
| 32172 | ENSG0000 | FAR1-IT1 | FAR1 intro | 0.086492 | 0.065925 | 0 |
| 17496 | ENSG0000 | RNU6-529I | RNA, U6 sr | 0.085923 | 0.037668 | 0 |
| 38325 | ENSG0000 | AL589674.2 | | 0.085524 | 0.076171 | 0.04772 |
| 31633 | ENSG0000 | PRSS51 | protease, s | 0.085231 | 0.049204 | 0.047777 |
| 29123 | ENSG0000 | B3GAT3P1 | beta-1,3-gl | 0.085103 | 0.053976 | 0.000855 |
| 22553 | ENSG0000 | RBMX2P1 | RNA bindir | 0.08506 | 0.056108 | 0.03174 |
| 1778 | ENSG0000 | CASS4 | Cas scaffol | 0.084444 | 0.045846 | 0.043579 |
| 27969 | ENSG0000 | SOCS2-AS1 | SOCS2 anti | 0.083971 | 0.026664 | 0.025436 |
| 25685 | ENSG0000 | EBAG9P1 | estrogen re | 0.083674 | 0.083156 | 0.029828 |
| 43295 | ENSG0000 | MIR1306 | microRNA | 0.083229 | 0.07412 | 0.074091 |
| 18016 | ENSG0000 | C6orf163 | chromosor | 0.083048 | 0.081286 | 0.073209 |
| 32264 | ENSG0000 | AP001972.2 | | 0.08274 | 0.071924 | 0 |
| 39020 | ENSG0000 | MUSTN1 | musculosk | 0.082688 | 0.025782 | 0.020242 |
| 22310 | ENSG0000 | AL162431.1 | | 0.082545 | 0.072129 | 0.058067 |
| 21617 | ENSG0000 | AL590714.1 | | 0.082429 | 0.0785 | 0.051981 |
| 1215 | ENSG0000 | CRMP1 | collapsin re | 0.082393 | 0.064438 | 0.058925 |
| 33310 | ENSG0000 | AL049869.1 | | 0.08205 | 0.040589 | 0.018107 |
| 24627 | ENSG0000 | VDAC1P6 | voltage de | 0.081992 | 0.034484 | 0.006852 |
| 37983 | ENSG0000 | LINC01930 | long interg | 0.08188 | 0.0378 | 0 |
| 15380 | ENSG0000 | ANO9 | anoctamin | 0.081623 | 0.076357 | 0.06767 |
| 22233 | ENSG0000 | IPO7P2 | importin 7 | 0.081556 | 0.05673 | 0.027766 |
| 31571 | ENSG0000 | AC087752.2 | | 0.081517 | 0.057232 | 0 |
| 19694 | ENSG0000 | NCLP1 | nucleolin p | 0.080911 | 0.057401 | 0.047731 |
| 26363 | ENSG0000 | KDM5C-IT1 | KDM5C int | 0.080876 | 0.001502 | 0 |
| 30556 | ENSG0000 | AC114781.3 | | 0.080656 | 0.062435 | 0 |
| 43405 | ENSG0000 | AL031985.4 | | 0.080318 | 0.018153 | 0 |
| 30637 | ENSG0000 | AC008443.6 | | 0.079985 | 0.041629 | 0.038545 |
| 34653 | ENSG0000 | AL162741.1 | | 0.079974 | 0.030702 | 0 |
| 17114 | ENSG0000 | PAX9 | paired box | 0.079894 | 0.079432 | 0.066436 |
| 7384 | ENSG0000 | MGARP | mitochond | 0.079519 | 0.055191 | 0.036546 |
| 27417 | ENSG0000 | PKN2-AS1 | PKN2 antis | 0.079235 | 0.036382 | 0.01816 |
| 25942 | ENSG0000 | AC099794.1 | | 0.079119 | 0.065037 | 0.062345 |
| 41433 | ENSG0000 | AC005962.2 | | 0.078565 | 0.047435 | 0 |
| 34367 | ENSG0000 | AC022558.1 | | 0.078454 | 0.027249 | 0 |

| | | | | | | |
|---|---|---|---|---|---|---|
| 6307 | ENSG0000 | POLN | DNA polym | 0.078224 | 0.057418 | 0.051951 |
| 20766 | ENSG0000 | BTBD10P2 | BTB domai | 0.078196 | 0.072516 | 0 |
| 35308 | ENSG0000 | AL157394.1 | | 0.077526 | 0.01675 | 0 |
| 21310 | ENSG0000 | LINC00370 | long interg | 0.077475 | 0.070361 | 0.047316 |
| 33401 | ENSG0000 | AC010200.1 | | 0.077275 | 0.020107 | 0.002334 |
| 43155 | ENSG0000 | AP000356.3 | | 0.07725 | 0.060036 | 0.042341 |
| 39634 | ENSG0000 | MIR6075 | microRNA | 0.07684 | 0.016278 | 0 |
| 33017 | ENSG0000 | AC068790.1 | | 0.076579 | 0.041236 | 0 |
| 20683 | ENSG0000 | LYPLA1P3 | lysophospl | 0.076239 | 0.058443 | 0.042973 |
| 23542 | ENSG0000 | AC092802.2 | | 0.076177 | 0.061701 | 0.025838 |
| 20245 | ENSG0000 | AC092821.1 | | 0.075847 | 0.045126 | 0.041875 |
| 10442 | ENSG0000 | SIGLEC16 | sialic acid b | 0.075743 | 0.041822 | 0.038763 |
| 42192 | ENSG0000 | AC004554.2 | | 0.075395 | 0.034625 | 0.026138 |
| 32689 | ENSG0000 | AP002414.2 | | 0.075228 | 0.049151 | 0.046226 |
| 37588 | ENSG0000 | AC010519.1 | | 0.074932 | 0.07466 | 0.062684 |
| 18605 | ENSG0000 | RFPL3S | RFPL3 anti | 0.074298 | 0.06527 | 0.061499 |
| 33376 | ENSG0000 | AL109628.2 | | 0.073887 | 0.021029 | 0 |
| 38944 | ENSG0000 | AC009902.3 | | 0.073765 | 0.024871 | 0 |
| 35789 | ENSG0000 | AC113189.2 | | 0.073671 | 0.043683 | 0.037369 |
| 25242 | ENSG0000 | AC211433.1 | | 0.073141 | 0.038853 | 0.034134 |
| 41723 | ENSG0000 | FP236315.2 | | 0.072948 | 0.026822 | 0.0181 |
| 42687 | ENSG0000 | AC140479.6 | | 0.072512 | 0.059828 | 0.059787 |
| 29468 | ENSG0000 | AC009060.1 | | 0.07242 | 0.0392 | 0.030735 |
| 29367 | ENSG0000 | AP005717.1 | | 0.072265 | 0.056806 | 0.012302 |
| 21719 | ENSG0000 | AC114491.1 | | 0.072138 | 0.059508 | 0 |
| 37324 | ENSG0000 | AC015961.2 | | 0.072114 | 0.041472 | 0.023269 |
| 42378 | ENSG0000 | FP236240.4 | | 0.071365 | 0.021005 | 0 |
| 13258 | ENSG0000 | MUC13 | mucin 13, c | 0.071113 | 0.045904 | 0.037196 |
| 39686 | ENSG0000 | snoU2-30 | Small nucle | 0.071032 | 0.041586 | 0.009786 |
| 31765 | ENSG0000 | IGHEP2 | immunoglo | 0.070845 | 0.042024 | 0 |
| 25263 | ENSG0000 | MRPL51P2 | mitochond | 0.070795 | 0.021249 | 0 |
| 30039 | ENSG0000 | CASC9 | cancer susc | 0.070792 | 0.019806 | 0 |
| 19305 | ENSG0000 | AC245427.1 | | 0.070326 | 0.048811 | 0.041639 |
| 43169 | ENSG0000 | MIR4784 | microRNA | 0.069782 | 0.061721 | 0.058461 |
| 31744 | ENSG0000 | NRG1-IT1 | NRG1 intro | 0.069385 | 0.061763 | 0 |
| 38876 | ENSG0000 | AC018709.1 | | 0.069222 | 0.000031 | 0 |
| 19955 | ENSG0000 | CCL27 | C-C motif c | 0.068967 | 0.041951 | 0 |
| 29143 | ENSG0000 | AC104653.2 | | 0.068737 | 0.034553 | 0 |
| 12723 | ENSG0000 | C11orf16 | chromosor | 0.068718 | 0.019541 | 0.018921 |
| 33136 | ENSG0000 | AC112229.3 | | 0.068656 | 0.028919 | 0 |
| 18814 | ENSG0000 | SNORA27 | small nucle | 0.068534 | 0.066985 | 0.050689 |
| 30586 | ENSG0000 | AC105250.1 | | 0.068497 | 0.044253 | 0.031691 |
| 35075 | ENSG0000 | AC027682.3 | | 0.068147 | 0.056387 | 0 |
| 20151 | ENSG0000 | VAC14-AS1 | VAC14 anti | 0.067945 | 0.032044 | 0.00348 |
| 25956 | ENSG0000 | AC234782.4 | | 0.067853 | 0.027838 | 0 |
| 16733 | ENSG0000 | GZMM | granzyme I | 0.067426 | 0.045809 | 0 |
| 39075 | ENSG0000 | AC007620.3 | | 0.067361 | 0.047086 | 0 |

| | | | | | | |
|---|---|---|---|---|---|---|
| 20517 | ENSG0000 | AC018797.1 | | 0.066137 | 0.056195 | 0 |
| 37071 | ENSG0000 | AC010632.1 | | 0.06601 | 0.061638 | 0.04099 |
| 37115 | ENSG0000 | AC005498.2 | | 0.065693 | 0.018504 | 0 |
| 30460 | ENSG0000 | AC136632.2 | | 0.065487 | 0.001317 | 0 |
| 32784 | ENSG0000 | AC024145.1 | | 0.065132 | 0.025154 | 0 |
| 21783 | ENSG0000 | RPL36AP29 | ribosomal | 0.065086 | 0.053312 | 0.050457 |
| 20432 | ENSG0000 | BCRP3 | breakpoint | 0.065031 | 0.048144 | 0.027254 |
| 17379 | ENSG0000 | RNU6-133( | RNA, U6 sr | 0.064965 | 0.05982 | 0.053642 |
| 27208 | ENSG0000 | AC004000.1 | | 0.064688 | 0.056365 | 0 |
| 1592 | ENSG0000 | C1QTNF3 | C1q and TN | 0.064618 | 0.048287 | 0.008834 |
| 13505 | ENSG0000 | NPPA | natriuretic | 0.063644 | 0.061922 | 0.021769 |
| 14401 | ENSG0000 | MYLPF | myosin ligh | 0.063435 | 0.041779 | 0 |
| 19516 | ENSG0000 | RNU6-244I | RNA, U6 sr | 0.062929 | 0.050785 | 0.01469 |
| 19203 | ENSG0000 | SNORA40 | small nucle | 0.06265 | 0.041114 | 0.040485 |
| 633 | ENSG0000 | MAGEC2 | MAGE fam | 0.062455 | 0.051364 | 0.040732 |
| 38067 | ENSG0000 | AC010530.1 | | 0.062376 | 0.03533 | 0.034656 |
| 25152 | ENSG0000 | AC245595.1 | | 0.062104 | 0.017325 | 0 |
| 14459 | ENSG0000 | FUT7 | fucosyltrar | 0.061979 | 0.050787 | 0.037831 |
| 25924 | ENSG0000 | RPL30P2 | ribosomal | 0.061415 | 0.048534 | 0 |
| 25743 | ENSG0000 | EIF3LP2 | eukaryotic | 0.061292 | 0.053142 | 0 |
| 20170 | ENSG0000 | CYCSP6 | cytochrom | 0.061262 | 0.01502 | 0 |
| 32561 | ENSG0000 | AC068385.1 | | 0.061194 | 0.02113 | 0 |
| 3709 | ENSG0000 | MS4A6A | membrane | 0.061145 | 0.029539 | 0.022355 |
| 16060 | ENSG0000 | ENO4 | enolase fa | 0.061023 | 0.030953 | 0.02491 |
| 11692 | ENSG0000 | NNMT | nicotinami | 0.060795 | 0.059686 | 0.044381 |
| 12032 | ENSG0000 | NLRC3 | NLR family | 0.060713 | 0.047892 | 0.031544 |
| 29375 | ENSG0000 | LINC00461 | long interg | 0.060524 | 0.044733 | 0.044579 |
| 41074 | ENSG0000 | uc_338 | TUC338 [S | 0.060429 | 0.040947 | 0.029794 |
| 34377 | ENSG0000 | AC009065.1 | | 0.06039 | 0.028902 | 0 |
| 27268 | ENSG0000 | AP004245.1 | | 0.060256 | 0.009951 | 0.003548 |
| 31650 | ENSG0000 | TAGLN2P1 | transgelin | 0.060249 | 0.024763 | 0 |
| 42313 | ENSG0000 | AC067852.7 | | 0.060224 | 0.052027 | 0 |
| 37306 | ENSG0000 | AL132655.2 | | 0.059963 | 0.048769 | 0 |
| 28294 | ENSG0000 | AC132217.1 | | 0.059705 | 0.057587 | 0.052626 |
| 39881 | ENSG0000 | AC083806.2 | | 0.059484 | 0.040891 | 0.038761 |
| 40773 | ENSG0000 | AL031663.3 | | 0.059421 | 0.013808 | 0 |
| 43076 | ENSG0000 | MIR637 | microRNA | 0.059262 | 0.044807 | 0.040095 |
| 24844 | ENSG0000 | HMGB3P14 | high mobil | 0.059252 | 0.048362 | 0 |
| 14918 | ENSG0000 | AC093768.1 | | 0.059049 | 0.048217 | 0.036184 |
| 25133 | ENSG0000 | AL592437.2 | | 0.058586 | 0.043097 | 0 |
| 37857 | ENSG0000 | AC024257.3 | | 0.058389 | 0.055915 | 0.044309 |
| 29732 | ENSG0000 | ALG1L9P | asparagine | 0.058353 | 0.056 | 0.052147 |
| 41169 | ENSG0000 | LINC00244 | long interg | 0.05831 | 0.043557 | 0.028296 |
| 8562 | ENSG0000 | CCDC39 | coiled-coil | 0.058286 | 0.034479 | 0.019778 |
| 11306 | ENSG0000 | C9orf24 | chromosor | 0.058036 | 0.020203 | 0 |
| 34758 | ENSG0000 | AC016722.3 | | 0.057743 | 0.053115 | 0.019775 |
| 41889 | ENSG0000 | AC012313.10 | | 0.057221 | 0.037316 | 0.034982 |

| | | | | | | |
|---|---|---|---|---|---|---|
| 36924 | ENSG0000 | AC090360.1 | | 0.057175 | 0.047095 | 0.031635 |
| 40240 | ENSG0000 | AC006001.4 | | 0.057095 | 0.042974 | 0.041646 |
| 38056 | ENSG0000 | AF230666.2 | | 0.05694 | 0.008816 | 0 |
| 39523 | ENSG0000 | AL449266.1 | | 0.056652 | 0.021085 | 0.019635 |
| 26225 | ENSG0000 | AL627389.1 | | 0.055691 | 0.054441 | 0 |
| 41521 | ENSG0000 | AC090616.7 | | 0.055363 | 0.052492 | 0.050107 |
| 30037 | ENSG0000 | AL033397.2 | | 0.055251 | 0.052669 | 0 |
| 21763 | ENSG0000 | AC026954.1 | | 0.055094 | 0.022564 | 0 |
| 23446 | ENSG0000 | AC074327.1 | | 0.05499 | 0.022363 | 0 |
| 18387 | ENSG0000 | TRIQK | triple QxxK | 0.054749 | 0.049629 | 0.027531 |
| 39708 | ENSG0000 | AL031667.3 | | 0.054379 | 0.044699 | 0 |
| 29509 | ENSG0000 | SPCS2P3 | signal pept | 0.054296 | 0.018858 | 0 |
| 13614 | ENSG0000 | GAPT | GRB2 bindi | 0.054258 | 0.031119 | 0.029691 |
| 25217 | ENSG0000 | LINC00665 | long interg | 0.054182 | 0.037121 | 0.023032 |
| 22447 | ENSG0000 | CLCP2 | Charcot-Le | 0.053811 | 0.03869 | 0 |
| 8204 | ENSG0000 | TINAGL1 | tubulointer | 0.05364 | 0.044561 | 0.036458 |
| 31758 | ENSG0000 | AC003991.2 | | 0.053402 | 0.038096 | 0 |
| 27680 | ENSG0000 | TBC1D3P1 | TBC1 doma | 0.053358 | 0.034922 | 0.010921 |
| 28112 | ENSG0000 | AC078785.1 | | 0.053248 | 0.052294 | 0.03282 |
| 40776 | ENSG0000 | SNORA71 | Small nucle | 0.053219 | 0.04733 | 0.026873 |
| 35441 | ENSG0000 | AC008731.1 | | 0.053061 | 0.041295 | 0.033824 |
| 34290 | ENSG0000 | AC022613.3 | | 0.052831 | 0.043503 | 0.012033 |
| 17081 | ENSG0000 | LRRTM3 | leucine rich | 0.052247 | 0.006163 | 0 |
| 20216 | ENSG0000 | AC073592.1 | | 0.052141 | 0.03997 | 0.027239 |
| 25716 | ENSG0000 | AC098617.1 | | 0.05209 | 0.042634 | 0.037343 |
| 821 | ENSG0000 | STYK1 | serine/thre | 0.051791 | 0.025502 | 0.017242 |
| 29614 | ENSG0000 | AC093864.1 | | 0.051576 | 0.036284 | 0 |
| 33045 | ENSG0000 | AC069503.2 | | 0.051524 | 0.027461 | 0.020991 |
| 4619 | ENSG0000 | MFAP2 | microfibril | 0.051482 | 0.020477 | 0 |
| 8815 | ENSG0000 | LPAR4 | lysophosph | 0.051427 | 0.046056 | 0.023974 |
| 39401 | ENSG0000 | AP005229.2 | | 0.051282 | 0.037272 | 0.023238 |
| 17398 | ENSG0000 | RN7SKP74 | RNA, 7SK s | 0.050914 | 0.036365 | 0 |
| 25517 | ENSG0000 | ERP29P1 | endoplasm | 0.05045 | 0.041324 | 0.022941 |
| 24358 | ENSG0000 | AC016949.1 | | 0.050326 | 0.041441 | 0.040441 |
| 32537 | ENSG0000 | AP002812.5 | | 0.049848 | 0.040831 | 0 |
| 37909 | ENSG0000 | AL391001.1 | | 0.049815 | 0.033595 | 0 |
| 28987 | ENSG0000 | RPS26P28 | ribosomal | 0.049548 | 0.029239 | 0 |
| 32507 | ENSG0000 | AC022182.3 | | 0.049484 | 0.025642 | 0.003711 |
| 30497 | ENSG0000 | AP002784.1 | | 0.049264 | 0.032974 | 0 |
| 29434 | ENSG0000 | AC090510.1 | | 0.049238 | 0.011316 | 0 |
| 764 | ENSG0000 | CCDC85A | coiled-coil | 0.049149 | 0.0364 | 0.016299 |
| 33163 | ENSG0000 | AC127164.1 | | 0.049072 | 0.01579 | 0.014523 |
| 9483 | ENSG0000 | ACMSD | aminocarb | 0.048988 | 0.043622 | 0.002269 |
| 19716 | ENSG0000 | YWHAZP5 | tyrosine 3- | 0.048792 | 0.033328 | 0 |
| 38672 | ENSG0000 | MIR98 | microRNA | 0.048651 | 0.020791 | 0 |
| 29647 | ENSG0000 | SEMA6A-A | SEMA6A ai | 0.048419 | 0.040538 | 0 |
| 38088 | ENSG0000 | AC079880.1 | | 0.048298 | 0.045308 | 0.013893 |

| | | | | | | |
|---|---|---|---|---|---|---|
| 38327 | ENSG0000 | AL133338.2 | | 0.048223 | 0.025913 | 0 |
| 41386 | ENSG0000 | C17orf78 | chromosor | 0.048146 | 0.017226 | 0.006925 |
| 38702 | ENSG0000 | AC100803.3 | | 0.048074 | 0.019671 | 0.010824 |
| 21655 | ENSG0000 | RPL7P57 | ribosomal | 0.047962 | 0.039286 | 0 |
| 20415 | ENSG0000 | ACTG1P3 | actin gamr | 0.04779 | 0.026097 | 0 |
| 27307 | ENSG0000 | ANO7L1 | anoctamin | 0.047345 | 0.042567 | 0.041245 |
| 20564 | ENSG0000 | AL590617.1 | | 0.047281 | 0.038866 | 0 |
| 18692 | ENSG0000 | ZCWPW2 | zinc finger | 0.047152 | 0.045123 | 0.015827 |
| 12658 | ENSG0000 | CEACAM3 | carcinoeml | 0.04715 | 0.020147 | 0.014963 |
| 28162 | ENSG0000 | B4GALT4-A | B4GALT4 a | 0.046773 | 0.027872 | 0 |
| 35793 | ENSG0000 | AC027281.1 | | 0.046657 | 0.019177 | 0 |
| 43260 | ENSG0000 | MIR4750 | microRNA | 0.04635 | 0.040342 | 0.033638 |
| 43104 | ENSG0000 | MIR4458 | microRNA | 0.046148 | 0.035051 | 0.031908 |
| 32876 | ENSG0000 | SUPT16HP | SPT16 hom | 0.046144 | 0.018904 | 0.00517 |
| 33048 | ENSG0000 | AC131009.2 | | 0.046094 | 0.037755 | 0 |
| 27030 | ENSG0000 | INTS6-AS1 | INTS6 antis | 0.045677 | 0.045112 | 0.030048 |
| 35877 | ENSG0000 | ABHD17AP | abhydrolas | 0.045646 | 0.015748 | 0.005849 |
| 30066 | ENSG0000 | AC008467.1 | | 0.045365 | 0.034464 | 0.016369 |
| 28825 | ENSG0000 | AC007182.2 | | 0.04477 | 0.03462 | 0 |
| 10201 | ENSG0000 | CELF3 | CUGBP Ela | 0.044746 | 0.0425 | 0.014647 |
| 11346 | ENSG0000 | TMEM246 | transmeml | 0.044643 | 0.03822 | 0.026757 |
| 26378 | ENSG0000 | HLA-W | major histo | 0.04451 | 0.007097 | 0 |
| 24694 | ENSG0000 | AL162385.2 | | 0.044354 | 0.041734 | 0 |
| 32636 | ENSG0000 | AC002563.1 | | 0.044022 | 0.024444 | 0 |
| 25084 | ENSG0000 | RPS3AP25 | ribosomal | 0.04391 | 0.03055 | 0.01702 |
| 26051 | ENSG0000 | RAD1P1 | RAD1 pseu | 0.043656 | 0.024727 | 0 |
| 23304 | ENSG0000 | AL158201.1 | | 0.043264 | 0.024534 | 0.012978 |
| 28151 | ENSG0000 | ZNF542P | zinc finger | 0.043142 | 0.009364 | 0.00533 |
| 38630 | ENSG0000 | AC011337.1 | | 0.043066 | 0.026387 | 0.009766 |
| 5639 | ENSG0000 | FAM182A | family witl | 0.04304 | 0.025681 | 0.013316 |
| 42595 | ENSG0000 | SNORA17 | Small nucle | 0.042677 | 0.032016 | 0.023864 |
| 24686 | ENSG0000 | LINC01353 | long interg | 0.042555 | 0.033805 | 0 |
| 38204 | ENSG0000 | AC097634.1 | | 0.042548 | 0.025741 | 0 |
| 14855 | ENSG0000 | OTOP3 | otopetrin 3 | 0.04242 | 0.034623 | 0.025796 |
| 29685 | ENSG0000 | AC008676.1 | | 0.042334 | 0.025689 | 0 |
| 36891 | ENSG0000 | AC015911.3 | | 0.041805 | 0.034242 | 0 |
| 2481 | ENSG0000 | ISM1 | isthmin 1 [ | 0.041548 | 0.023964 | 0 |
| 41570 | ENSG0000 | AC079228.1 | | 0.041453 | 0.034642 | 0.02327 |
| 8142 | ENSG0000 | ENPP3 | ectonucleo | 0.041175 | 0.00751 | 0 |
| 12751 | ENSG0000 | GRIK1 | glutamate | 0.040956 | 0.032784 | 0.024864 |
| 39877 | ENSG0000 | SNORD28 | small nucle | 0.040609 | 0.021762 | 0.021732 |
| 31583 | ENSG0000 | CA3-AS1 | CA3 antise | 0.040551 | 0.019109 | 0 |
| 28826 | ENSG0000 | RPSAP3 | ribosomal | 0.040439 | 0.011041 | 0.003393 |
| 19765 | ENSG0000 | RPSAP46 | ribosomal | 0.040383 | 0.022442 | 0 |
| 15856 | ENSG0000 | PCDHB13 | protocadh | 0.040349 | 0.024433 | 0.0048 |
| 27553 | ENSG0000 | AC083873.1 | | 0.040311 | 0.024671 | 0.022106 |
| 42696 | ENSG0000 | PCBP2-OT1 | PCBP2 ove | 0.040276 | 0.032308 | 0.03014 |

| | | | | | | |
|---|---|---|---|---|---|---|
| 28106 | ENSG0000 | AMY2B | amylase, a | 0.04018 | 0.019048 | 0.01832 |
| 32765 | ENSG0000 | AC008813.1 | | 0.040104 | 0.032849 | 0.018301 |
| 950 | ENSG0000 | GSTO2 | glutathione | 0.040084 | 0.039779 | 0.03289 |
| 43225 | ENSG0000 | MIR6843 | microRNA | 0.040008 | 0.012061 | 0 |
| 10789 | ENSG0000 | LINC01116 | long interg | 0.039953 | 0.016276 | 0.014525 |
| 13216 | ENSG0000 | PTPRM | protein tyr | 0.039512 | 0.023461 | 0.006254 |
| 41243 | ENSG0000 | AC092881.1 | | 0.039052 | 0.035231 | 0.029716 |
| 12321 | ENSG0000 | KCNAB1 | potassium | 0.039049 | 0.036911 | 0.007091 |
| 10662 | ENSG0000 | KLHDC8A | kelch doma | 0.038988 | 0.024001 | 0.022213 |
| 31930 | ENSG0000 | AC011379.2 | | 0.038864 | 0.027505 | 0.015354 |
| 4237 | ENSG0000 | FGF12 | fibroblast g | 0.038793 | 0.013876 | 0.002108 |
| 20496 | ENSG0000 | HNRNPA1P | heterogene | 0.038766 | 0.003218 | 0 |
| 24767 | ENSG0000 | LINC00574 | long interg | 0.03855 | 0.034779 | 0.027999 |
| 11820 | ENSG0000 | CD3D | CD3d mole | 0.038475 | 0.031405 | 0 |
| 32846 | ENSG0000 | AC027290.1 | | 0.03844 | 0.031216 | 0 |
| 43105 | ENSG0000 | MIR6084 | microRNA | 0.038333 | 0.03734 | 0.037014 |
| 20440 | ENSG0000 | EFCAB8 | EF-hand ca | 0.038322 | 0.012932 | 0 |
| 40110 | ENSG0000 | SNORD49A | small nucle | 0.037621 | 0.028719 | 0.020854 |
| 17043 | ENSG0000 | KLHL9 | kelch like f | 0.037449 | 0.030548 | 0.01712 |
| 29362 | ENSG0000 | AC124242.1 | | 0.037162 | 0.024236 | 0 |
| 27004 | ENSG0000 | SMAD9-IT1 | SMAD9 int | 0.037067 | 0.012886 | 0.001368 |
| 43038 | ENSG0000 | MIR22 | microRNA | 0.037016 | 0.026724 | 0.021377 |
| 1000 | ENSG0000 | DDX3Y | DEAD-box | 0.03695 | 0.020338 | 0.014173 |
| 35329 | ENSG0000 | AC096921.2 | | 0.036917 | 0.0304 | 0.016967 |
| 23593 | ENSG0000 | AC091167.2 | | 0.036797 | 0.035437 | 0.003936 |
| 16135 | ENSG0000 | RHCE | Rh blood g | 0.036625 | 0.01029 | 0 |
| 13538 | ENSG0000 | ARL10 | ADP ribosy | 0.036462 | 0.023613 | 0.022914 |
| 6814 | ENSG0000 | HRH4 | histamine r | 0.036406 | 0.019786 | 0.008706 |
| 14089 | ENSG0000 | SHISA3 | shisa famil | 0.036334 | 0.014802 | 0 |
| 7088 | ENSG0000 | USP44 | ubiquitin s | 0.036307 | 0.026796 | 0.020926 |
| 28840 | ENSG0000 | RPL12P35 | ribosomal | 0.036286 | 0.02993 | 0 |
| 28644 | ENSG0000 | NECTIN3-A | NECTIN3 a | 0.03593 | 0.031685 | 0.024117 |
| 29464 | ENSG0000 | LINC01252 | long interg | 0.035625 | 0.023048 | 0.012901 |
| 28390 | ENSG0000 | AC005410.1 | | 0.035618 | 0.029278 | 0 |
| 33122 | ENSG0000 | CR383656.2 | | 0.035401 | 0.012829 | 0.003618 |
| 20387 | ENSG0000 | AP000533.2 | | 0.035307 | 0.019227 | 0.010877 |
| 3375 | ENSG0000 | ELAVL2 | ELAV like R | 0.03503 | 0.033936 | 0.028516 |
| 17120 | ENSG0000 | CD247 | CD247 mol | 0.035005 | 0.021569 | 0.00803 |
| 31761 | ENSG0000 | LINC00051 | long interg | 0.034873 | 0.013592 | 0 |
| 25472 | ENSG0000 | AC007743.1 | | 0.03464 | 0.007858 | 0 |
| 24158 | ENSG0000 | AC078817.1 | | 0.03443 | 0.028202 | 0 |
| 10861 | ENSG0000 | CD200R1 | CD200 rece | 0.034421 | 0.02808 | 0 |
| 3313 | ENSG0000 | ACTR3C | ARP3 actin | 0.034153 | 0.032603 | 0.019631 |
| 19201 | ENSG0000 | RNU6ATAC | RNA, U6ata | 0.033962 | 0.024545 | 0 |
| 13697 | ENSG0000 | PRR18 | proline rich | 0.033812 | 0.018529 | 0 |
| 39781 | ENSG0000 | AC022929.2 | | 0.03361 | 0.006882 | 0 |
| 40819 | ENSG0000 | AL138820.1 | | 0.03357 | 0.022115 | 0.012256 |

| | | | | | | |
|---|---|---|---|---|---|---|
| 25271 | ENSG0000 | SCAND3P1 | SCAN dom | 0.033483 | 0.008072 | 0 |
| 5201 | ENSG0000 | LAX1 | lymphocyt | 0.033413 | 0.026527 | 0.020617 |
| 40591 | ENSG0000 | AP000892.2 | | 0.033018 | 0.018368 | 0 |
| 10658 | ENSG0000 | KIF26B | kinesin fan | 0.033005 | 0.017936 | 0.017665 |
| 4133 | ENSG0000 | HAVCR1 | hepatitis A | 0.033001 | 0.027031 | 0 |
| 38266 | ENSG0000 | AJ271736.1 | | 0.032918 | 0.003307 | 0.000211 |
| 38828 | ENSG0000 | AC012557.2 | | 0.03282 | 0.030829 | 0.030779 |
| 42993 | ENSG0000 | MIR3656 | microRNA | 0.032717 | 0.025314 | 0.001583 |
| 39312 | ENSG0000 | AC093799.2 | | 0.032444 | 0.026932 | 0 |
| 17206 | ENSG0000 | MIR135B | microRNA | 0.032438 | 0.011691 | 0 |
| 26741 | ENSG0000 | RPL17P36 | ribosomal | 0.032324 | 0.026101 | 0 |
| 18138 | ENSG0000 | SP5 | Sp5 transc | 0.032015 | 0.028124 | 0.01578 |
| 23591 | ENSG0000 | MTND1P9 | mitochond | 0.031998 | 0.015617 | 0 |
| 26137 | ENSG0000 | AC144530.1 | | 0.031975 | 0.018018 | 0 |
| 6669 | ENSG0000 | GIMAP6 | GTPase, IM | 0.031907 | 0.022341 | 0.016527 |
| 20709 | ENSG0000 | AC064847.1 | | 0.031838 | 0.014465 | 0 |
| 14476 | ENSG0000 | EXOC5P1 | exocyst co | 0.031704 | 0.009599 | 0 |
| 15160 | ENSG0000 | CTAG1B | cancer/tes | 0.031647 | 0.028045 | 0 |
| 14214 | ENSG0000 | FAM133A | family with | 0.031642 | 0.017319 | 0 |
| 33436 | ENSG0000 | AC027288.3 | | 0.031619 | 0.025898 | 0.021392 |
| 19994 | ENSG0000 | AL034370.1 | | 0.031425 | 0.028139 | 0 |
| 5624 | ENSG0000 | TNFSF14 | TNF superf | 0.03121 | 0.028209 | 0.013608 |
| 9144 | ENSG0000 | HMGA2 | high mobil | 0.031205 | 0.031129 | 0.015673 |
| 28530 | ENSG0000 | AC117394.1 | | 0.031076 | 0.025273 | 0 |
| 13935 | ENSG0000 | NAP1L5 | nucleosom | 0.030909 | 0.02514 | 0 |
| 2504 | ENSG0000 | PLCB4 | phospholip | 0.030899 | 0.027461 | 0.023341 |
| 26770 | ENSG0000 | USP17L4 | ubiquitin s | 0.03078 | 0.006844 | 0 |
| 34708 | ENSG0000 | AC098818.2 | | 0.030759 | 0.018796 | 0.013963 |
| 20106 | ENSG0000 | MARCKSL1 | MARCKS lil | 0.030433 | 0.021865 | 0 |
| 7650 | ENSG0000 | FBN2 | fibrillin 2 [S | 0.030303 | 0.026522 | 0.02097 |
| 14919 | ENSG0000 | POTEC | POTE anky | 0.030077 | 0.016076 | 0.015961 |
| 37110 | ENSG0000 | AC011471.1 | | 0.030062 | 0.029081 | 0 |
| 5619 | ENSG0000 | GRIA3 | glutamate | 0.029682 | 0.018904 | 0.004996 |
| 13708 | ENSG0000 | SLCO3A1 | solute carr | 0.029587 | 0.024339 | 0.017747 |
| 13323 | ENSG0000 | TLR10 | toll like rec | 0.029577 | 0.024154 | 0.023045 |
| 14775 | ENSG0000 | TBPL2 | TATA-box b | 0.029548 | 0.018331 | 0.013506 |
| 42753 | ENSG0000 | AC116353.4 | | 0.029534 | 0.010793 | 0 |
| 7911 | ENSG0000 | CDH11 | cadherin 1 | 0.029468 | 0.024497 | 0.014525 |
| 40260 | ENSG0000 | PTGER4P2- | PTGER4P2- | 0.02946 | 0.029363 | 0.019471 |
| 37722 | ENSG0000 | ERVV-2 | endogenou | 0.0292 | 0.02396 | 0.016798 |
| 12621 | ENSG0000 | CELP | carboxyl es | 0.029179 | 0.01624 | 0.008504 |
| 19045 | ENSG0000 | PTTG3P | pituitary tu | 0.029076 | 0.023646 | 0 |
| 41197 | ENSG0000 | AC096720.2 | | 0.028981 | 0.028562 | 0.007991 |
| 21745 | ENSG0000 | HSD52 | uncharacte | 0.028951 | 0.027192 | 0.008008 |
| 25489 | ENSG0000 | AC092159.2 | | 0.028902 | 0.016564 | 0 |
| 18296 | ENSG0000 | ST8SIA6-AS | ST8SIA6 an | 0.028881 | 0.019425 | 0 |
| 26282 | ENSG0000 | RPL7L1P3 | ribosomal | 0.028862 | 0.024688 | 0.016339 |

| | | | | | | |
|---|---|---|---|---|---|---|
| 1758 | ENSG0000 | LPCAT2 | lysophosph | 0.028621 | 0.007844 | 0.00437 |
| 6054 | ENSG0000 | TPH1 | tryptophar | 0.028501 | 0.028033 | 0.020573 |
| 38121 | ENSG0000 | SMC2-AS1 | SMC2 antis | 0.028446 | 0.021 | 0.0158 |
| 7380 | ENSG0000 | FGFBP2 | fibroblast g | 0.028394 | 0.023339 | 0.022147 |
| 16090 | ENSG0000 | SLC24A5 | solute carr | 0.028317 | 0.018644 | 0.012858 |
| 33647 | ENSG0000 | AC005225.1 | | 0.028146 | 0.023985 | 0 |
| 28625 | ENSG0000 | ABCF2P1 | ATP bindin | 0.028111 | 0.01535 | 0.008429 |
| 5468 | ENSG0000 | ATP8A1 | ATPase pho | 0.027984 | 0.018162 | 0.011631 |
| 37247 | ENSG0000 | AC008543.5 | | 0.027924 | 0.011272 | 0 |
| 37718 | ENSG0000 | MRPS17P1 | mitochond | 0.027906 | 0.004018 | 0 |
| 30809 | ENSG0000 | AC008676.2 | | 0.026946 | 0.010958 | 0 |
| 12593 | ENSG0000 | TMEM133 | transmeml | 0.026937 | 0.020837 | 0.004852 |
| 16614 | ENSG0000 | SLC6A17 | solute carr | 0.026748 | 0.010939 | 0.004807 |
| 10802 | ENSG0000 | EIF4E3 | eukaryotic | 0.026353 | 0.017493 | 0.016529 |
| 25503 | ENSG0000 | EEF1A1P34 | eukaryotic | 0.026125 | 0.022945 | 0 |
| 8786 | ENSG0000 | SHROOM2 | shroom far | 0.026075 | 0.003549 | 0 |
| 9153 | ENSG0000 | FRMPD2B | FERM and | 0.026069 | 0.020511 | 0 |
| 22766 | ENSG0000 | ERVMER34 | endogenou | 0.026033 | 0.014312 | 0 |
| 2440 | ENSG0000 | SLA2 | Src like ada | 0.025747 | 0.010582 | 0 |
| 647 | ENSG0000 | FAM184B | family with | 0.025644 | 0.02268 | 0 |
| 8568 | ENSG0000 | SLIT2 | slit guidanc | 0.02562 | 0.023894 | 0.015068 |
| 41038 | ENSG0000 | MIR6738 | microRNA | 0.025565 | 0.011276 | 0.009872 |
| 3140 | ENSG0000 | THEG | theg sperm | 0.025337 | 0.021298 | 0 |
| 5205 | ENSG0000 | CD244 | CD244 mol | 0.024889 | 0.010175 | 0 |
| 30819 | ENSG0000 | AC098679.3 | | 0.024538 | 0.011974 | 0.00224 |
| 16015 | ENSG0000 | EYS | eyes shut h | 0.024499 | 0.013792 | 0.006567 |
| 16348 | ENSG0000 | SIRPB2 | signal regu | 0.024395 | 0.023059 | 0.022545 |
| 20592 | ENSG0000 | Z98755.1 | | 0.024099 | 0.019292 | 0 |
| 43443 | ENSG0000 | AL139424.3 | | 0.024084 | 0.023567 | 0 |
| 27798 | ENSG0000 | SNORD5 | small nucle | 0.024029 | 0.017312 | 0.010935 |
| 31628 | ENSG0000 | AC022893.1 | | 0.023967 | 0.016441 | 0.008205 |
| 10224 | ENSG0000 | SPON2 | spondin 2 \| | 0.023736 | 0.014302 | 0.007928 |
| 22912 | ENSG0000 | LINC01806 | long interg | 0.023104 | 0.01223 | 0 |
| 16830 | ENSG0000 | FAM49A | family with | 0.022987 | 0.021458 | 0.014865 |
| 41179 | ENSG0000 | AC015799.1 | | 0.022979 | 0.009312 | 0 |
| 26635 | ENSG0000 | CPB2-AS1 | CPB2 antis | 0.02295 | 0.022136 | 0.01929 |
| 26094 | ENSG0000 | YWHAEP5 | tyrosine 3- | 0.022868 | 0.019001 | 0 |
| 11314 | ENSG0000 | DIRAS2 | DIRAS fami | 0.022776 | 0.01243 | 0.006986 |
| 21603 | ENSG0000 | PPP4R3CP | protein pho | 0.022752 | 0.018603 | 0.010419 |
| 1577 | ENSG0000 | CADPS2 | calcium de | 0.022433 | 0.020615 | 0.005119 |
| 1767 | ENSG0000 | NID2 | nidogen 2 | 0.022408 | 0.008487 | 0.007799 |
| 4169 | ENSG0000 | IL5 | interleukin | 0.022364 | 0.009354 | 0.008179 |
| 29633 | ENSG0000 | PRR5-ARH( | PRR5-ARH( | 0.022356 | 0.019529 | 0 |
| 38059 | ENSG0000 | TERC | telomerase | 0.022332 | 0.019478 | 0 |
| 33262 | ENSG0000 | OR7E47P | olfactory re | 0.022178 | 0.018166 | 0 |
| 10440 | ENSG0000 | SIGLEC11 | sialic acid b | 0.021895 | 0.014727 | 0.010743 |
| 7087 | ENSG0000 | STAB2 | stabilin 2 [S | 0.021854 | 0.013286 | 0.012421 |

| | | | | | | |
|---|---|---|---|---|---|---|
| 14935 | ENSG0000 | CCBE1 | collagen ar | 0.021738 | 0.004421 | 0 |
| 41524 | ENSG0000 | AC148477.5 | | 0.021652 | 0.008836 | 0 |
| 43285 | ENSG0000 | MIR197 | microRNA | 0.021396 | 0.013745 | 0 |
| 4292 | ENSG0000 | AADAC | arylacetam | 0.021354 | 0.017367 | 0 |
| 27803 | ENSG0000 | AC093484.1 | | 0.020593 | 0.006952 | 0.000082 |
| 23172 | ENSG0000 | AL158835.1 | | 0.020549 | 0.009567 | 0.007458 |
| 7648 | ENSG0000 | EGF | epidermal | 0.020336 | 0.011845 | 0.002662 |
| 34611 | ENSG0000 | AOC4P | amine oxid | 0.02031 | 0.007139 | 0 |
| 22873 | ENSG0000 | HORMAD2 | HORMAD2 | 0.020204 | 0.002712 | 0 |
| 25965 | ENSG0000 | TATDN1P1 | TatD DNas | 0.019917 | 0.016314 | 0 |
| 156 | ENSG0000 | SCIN | scinderin [S | 0.019881 | 0.00753 | 0.002787 |
| 1216 | ENSG0000 | EVC | EvC ciliary | 0.019879 | 0.015974 | 0.006813 |
| 5033 | ENSG0000 | PLXDC2 | plexin dom | 0.019729 | 0.017416 | 0.015475 |
| 6104 | ENSG0000 | EPB41L4A | erythrocyt | 0.019701 | 0.013715 | 0.000107 |
| 33721 | ENSG0000 | AL359232.1 | | 0.019672 | 0.015719 | 0 |
| 31667 | ENSG0000 | AC011632.1 | | 0.019658 | 0.008108 | 0.004549 |
| 43114 | ENSG0000 | MIR6805 | microRNA | 0.019556 | 0.013089 | 0.009916 |
| 11418 | ENSG0000 | MAGEC3 | MAGE fam | 0.01936 | 0.010355 | 0.00884 |
| 1559 | ENSG0000 | IMPG2 | interphoto | 0.019203 | 0.015766 | 0.011698 |
| 3675 | ENSG0000 | DDX25 | DEAD-box | 0.019115 | 0.01033 | 0.009092 |
| 43161 | ENSG0000 | MIR1234 | microRNA | 0.019106 | 0.000773 | 0 |
| 19508 | ENSG0000 | SNORD12 | small nucle | 0.019076 | 0.00851 | 0.004915 |
| 6905 | ENSG0000 | HAVCR2 | hepatitis A | 0.019058 | 0.011277 | 0 |
| 29384 | ENSG0000 | AC022075.1 | | 0.019034 | 0.010309 | 0 |
| 14422 | ENSG0000 | KCTD4 | potassium | 0.019031 | 0.015643 | 0 |
| 43281 | ENSG0000 | MIR6869 | microRNA | 0.018624 | 0.01527 | 0.007027 |
| 20348 | ENSG0000 | C16orf90 | chromosor | 0.018559 | 0.009724 | 0 |
| 41863 | ENSG0000 | AC008739.5 | | 0.018314 | 0.012607 | 0 |
| 34612 | ENSG0000 | AC005606.1 | | 0.018226 | 0.014576 | 0 |
| 25791 | ENSG0000 | MT1XP1 | metallothi | 0.018175 | 0.014411 | 0 |
| 8968 | ENSG0000 | CACNA1B | calcium vo | 0.017969 | 0.014667 | 0.009831 |
| 7479 | ENSG0000 | GIPC2 | GIPC PDZ c | 0.017899 | 0.014712 | 0 |
| 20414 | ENSG0000 | MIR99AHG | mir-99a-let | 0.01777 | 0.014183 | 0.004741 |
| 11657 | ENSG0000 | CLEC4D | C-type lect | 0.017757 | 0.014545 | 0 |
| 70 | ENSG0000 | CALCR | calcitonin r | 0.017655 | 0.007218 | 0 |
| 28247 | ENSG0000 | AADACP1 | arylacetam | 0.01764 | 0.014551 | 0 |
| 42830 | ENSG0000 | MIR6849 | microRNA | 0.017535 | 0.008196 | 0.00697 |
| 15498 | ENSG0000 | SHC4 | SHC adapt | 0.017499 | 0.004829 | 0 |
| 15751 | ENSG0000 | FOXE3 | forkhead b | 0.017472 | 0.016809 | 0 |
| 41789 | ENSG0000 | AC092135.1 | | 0.017238 | 0.013669 | 0.008982 |
| 37323 | ENSG0000 | AC020922.1 | | 0.016998 | 0.010425 | 0 |
| 12271 | ENSG0000 | AR | androgen r | 0.016699 | 0.015954 | 0.007634 |
| 12435 | ENSG0000 | TM4SF4 | transmeml | 0.016585 | 0.013585 | 0 |
| 29394 | ENSG0000 | AC097382.2 | | 0.016566 | 0.013522 | 0.01301 |
| 20797 | ENSG0000 | AL356234.1 | | 0.016555 | 0.007881 | 0 |
| 23699 | ENSG0000 | ST13P13 | ST13, Hsp7 | 0.016251 | 0.013406 | 0 |
| 33795 | ENSG0000 | AC091544.2 | | 0.016177 | 0.006625 | 0 |

| | | | | | | |
|---|---|---|---|---|---|---|
| 30264 | ENSG0000 | AC093214.1 | | 0.016031 | 0.009778 | 0 |
| 24654 | ENSG0000 | AL365440.1 | | 0.01603 | 0.01118 | 0.007421 |
| 16250 | ENSG0000 | DCAF8L2 | DDB1 and | 0.016012 | 0.012374 | 0.007281 |
| 6920 | ENSG0000 | CD36 | CD36 mole | 0.015815 | 0.01433 | 0.009631 |
| 12015 | ENSG0000 | GSDMA | gasdermin | 0.015725 | 0.01279 | 0 |
| 40354 | ENSG0000 | HIST1H2BH | histone clu | 0.015702 | 0.014112 | 0 |
| 40582 | ENSG0000 | NPPA-AS1_ | NPPA antis | 0.015485 | 0.005489 | 0.004326 |
| 39794 | ENSG0000 | SNORA71E | small nucle | 0.015377 | 0.013672 | 0.005438 |
| 10643 | ENSG0000 | FAM71A | family with | 0.015234 | 0.008006 | 0 |
| 5783 | ENSG0000 | PRDM7 | PR/SET dor | 0.015225 | 0.010625 | 0.00324 |
| 28767 | ENSG0000 | AC107464.1 | | 0.015137 | 0.006177 | 0.004589 |
| 42963 | ENSG0000 | AC012485.3 | | 0.014694 | 0.01288 | 0 |
| 38725 | ENSG0000 | AC034229.3 | | 0.014673 | 0.011892 | 0 |
| 8829 | ENSG0000 | CXorf57 | chromosor | 0.014627 | 0.005991 | 0 |
| 40783 | ENSG0000 | Metazoa_S | Metazoan | 0.014569 | 0.006298 | 0 |
| 42012 | ENSG0000 | AC111170.3 | | 0.014455 | 0.011799 | 0 |
| 7418 | ENSG0000 | MMP20 | matrix met | 0.014325 | 0.011437 | 0 |
| 25225 | ENSG0000 | AC012370.1 | | 0.014248 | 0.009337 | 0 |
| 30357 | ENSG0000 | RDH10-AS1 | RDH10 ant | 0.013973 | 0.009588 | 0 |
| 28474 | ENSG0000 | HSPA8P9 | heat shock | 0.013765 | 0.011222 | 0.004168 |
| 21268 | ENSG0000 | CSNK1A1P | casein kina | 0.013548 | 0.011056 | 0 |
| 176 | ENSG0000 | CEACAM7 | carcinoeml | 0.013446 | 0.010975 | 0 |
| 27323 | ENSG0000 | SRRM1P3 | serine/argi | 0.013347 | 0.002541 | 0 |
| 559 | ENSG0000 | CYP46A1 | cytochrom | 0.013265 | 0.012821 | 0.003476 |
| 3674 | ENSG0000 | PPARGC1A | PPARG coa | 0.013107 | 0.005368 | 0.00298 |
| 32710 | ENSG0000 | OVCH1-AS1 | OVCH1 ant | 0.012946 | 0.010415 | 0.00632 |
| 38470 | ENSG0000 | AC012464.3 | | 0.012799 | 0.010484 | 0.005841 |
| 26661 | ENSG0000 | CICP20 | capicua tra | 0.012716 | 0.001509 | 0 |
| 14321 | ENSG0000 | ATP8B5P | ATPase pho | 0.012684 | 0.005604 | 0 |
| 39729 | ENSG0000 | RBM17P2 | RNA bindir | 0.012631 | 0.00532 | 0 |
| 9528 | ENSG0000 | PTPRD | protein tyr | 0.012578 | 0.007838 | 0.005018 |
| 253 | ENSG0000 | HHATL | hedgehog a | 0.012451 | 0.010127 | 0 |
| 14122 | ENSG0000 | CA8 | carbonic ar | 0.01234 | 0.004991 | 0 |
| 32216 | ENSG0000 | ATP6V1G2 | ATP6V1G2 | 0.012142 | 0.00732 | 0 |
| 8334 | ENSG0000 | NUP210L | nucleopori | 0.011968 | 0.006195 | 0.00236 |
| 20729 | ENSG0000 | AL049842.1 | | 0.011914 | 0.004949 | 0 |
| 3998 | ENSG0000 | EYA4 | EYA transc | 0.011887 | 0.011369 | 0 |
| 8685 | ENSG0000 | LRRTM2 | leucine ricl | 0.011687 | 0.009539 | 0.008029 |
| 42962 | ENSG0000 | AP000547.3 | | 0.011557 | 0.0095 | 0.004938 |
| 16341 | ENSG0000 | CTSE | cathepsin E | 0.011497 | 0.010923 | 0 |
| 28872 | ENSG0000 | PSG4 | pregnancy | 0.011354 | 0.004634 | 0 |
| 8528 | ENSG0000 | COL8A1 | collagen ty | 0.011345 | 0.007736 | 0.006898 |
| 17198 | ENSG0000 | RORB | RAR relate | 0.01122 | 0.003031 | 0 |
| 37944 | ENSG0000 | PCF11-AS1 | PCF11 anti | 0.010915 | 0.007264 | 0 |
| 43316 | ENSG0000 | MIR6741 | microRNA | 0.010769 | 0.009852 | 0.007355 |
| 15081 | ENSG0000 | EFHC2 | EF-hand dc | 0.010731 | 0.008635 | 0.004776 |
| 1599 | ENSG0000 | COBLL1 | cordon-ble | 0.010338 | 0.00914 | 0.003765 |

| | | | | | | |
|---|---|---|---|---|---|---|
| 31720 | ENSG0000 | AC019270.1 | | 0.010212 | 0.00749 | 0.001449 |
| 11375 | ENSG0000 | AQP7 | aquaporin | 0.010159 | 0.008753 | 0.006167 |
| 16240 | ENSG0000 | FAM47E | family with | 0.010124 | 0.009309 | 0.002505 |
| 9344 | ENSG0000 | TMEM132I | transmeml | 0.010033 | 0.008203 | 0.004546 |
| 22953 | ENSG0000 | LINC01624 | long interg | 0.009899 | 0.008051 | 0 |
| 25945 | ENSG0000 | LINC01505 | long interg | 0.009741 | 0.007969 | 0.00296 |
| 19709 | ENSG0000 | NPM1P32 | nucleopho | 0.009689 | 0.008295 | 0 |
| 10369 | ENSG0000 | CCR5 | C-C motif c | 0.009666 | 0.007945 | 0 |
| 27562 | ENSG0000 | AL157713.1 | | 0.009536 | 0.009158 | 0 |
| 100 | ENSG0000 | ABCB4 | ATP bindin | 0.009489 | 0.007772 | 0.004323 |
| 27769 | ENSG0000 | SNORD2 | small nucle | 0.009018 | 0.006628 | 0.003863 |
| 42544 | ENSG0000 | HELLPAR | HELLP assc | 0.008557 | 0.004246 | 0.003963 |
| 41537 | ENSG0000 | AC010533.1 | | 0.00849 | 0.003575 | 0.000273 |
| 11383 | ENSG0000 | DEUP1 | deuterosoi | 0.008456 | 0.006926 | 0 |
| 20134 | ENSG0000 | RPL39P5 | ribosomal | 0.008409 | 0.006888 | 0 |
| 22004 | ENSG0000 | USP17L3 | ubiquitin s| | 0.008375 | 0.004562 | 0 |
| 11377 | ENSG0000 | SLITRK5 | SLIT and N | 0.008218 | 0.002066 | 0.00154 |
| 41409 | ENSG0000 | NR2E3 | nuclear rec | 0.007691 | 0.006459 | 0 |
| 18856 | ENSG0000 | SNORA70 | small nucle | 0.007667 | 0.006487 | 0.00407 |
| 30728 | ENSG0000 | AC090502.1 | | 0.007577 | 0.003103 | 0 |
| 43056 | ENSG0000 | MIR4260 | microRNA | 0.007514 | 0.007176 | 0.006308 |
| 37458 | ENSG0000 | GABRQ | gamma-am | 0.007478 | 0.004727 | 0 |
| 43119 | ENSG0000 | MIR10A | microRNA | 0.007355 | 0.004284 | 0.002669 |
| 18366 | ENSG0000 | SLFN12L | schlafen fa | 0.007299 | 0.006128 | 0 |
| 42848 | ENSG0000 | MIR939 | microRNA | 0.007286 | 0.007013 | 0.004857 |
| 17385 | ENSG0000 | SNORD16 | small nucle | 0.006985 | 0.002111 | 0.001989 |
| 11752 | ENSG0000 | EPB42 | erythrocyt | 0.006923 | 0.001478 | 0 |
| 1372 | ENSG0000 | PAK3 | p21 (RAC1) | 0.006896 | 0.005628 | 0.001582 |
| 15482 | ENSG0000 | LSAMP | limbic syst | 0.006884 | 0.005638 | 0.00476 |
| 43291 | ENSG0000 | MIR1304 | microRNA | 0.006253 | 0.004624 | 0.002217 |
| 4146 | ENSG0000 | GABRG2 | gamma-am | 0.006207 | 0.004436 | 0.002121 |
| 41579 | ENSG0000 | AC005921.3 | | 0.006204 | 0.000059 | 0.000013 |
| 34807 | ENSG0000 | AC124312.3 | | 0.006125 | 0.000242 | 0 |
| 43135 | ENSG0000 | MIR6837 | microRNA | 0.006118 | 0.00102 | 0.000564 |
| 685 | ENSG0000 | NEXMIF | neurite ext | 0.006059 | 0.002402 | 0 |
| 9498 | ENSG0000 | PLA2R1 | phospholip | 0.005794 | 0.001893 | 0 |
| 23162 | ENSG0000 | DHX9P1 | DEAH-box | 0.005769 | 0.002279 | 0 |
| 14721 | ENSG0000 | GABRG3 | gamma-am | 0.005655 | 0.004666 | 0.002572 |
| 41281 | ENSG0000 | MIR6857 | microRNA | 0.005413 | 0.002771 | 0.001853 |
| 39952 | ENSG0000 | ELOA3D | elongin A3 | 0.005407 | 0.00445 | 0 |
| 19170 | ENSG0000 | SNORD73A | small nucle | 0.005346 | 0.004665 | 0.00315 |
| 3627 | ENSG0000 | MAPK10 | mitogen-ac | 0.005332 | 0.004773 | 0.001502 |
| 17681 | ENSG0000 | SNORD45B | small nucle | 0.005289 | 0.004507 | 0.003191 |
| 43041 | ENSG0000 | MIR6734 | microRNA | 0.005172 | 0.003427 | 0.000492 |
| 11189 | ENSG0000 | SAMD3 | sterile alph | 0.005123 | 0.002658 | 0.000378 |
| 16538 | ENSG0000 | POTEI | POTE anky | 0.005072 | 0.002066 | 0 |
| 11217 | ENSG0000 | BMPER | BMP bindi | 0.004963 | 0.004587 | 0 |

| | | | | | | |
|---|---|---|---|---|---|---|
| 13885 | ENSG0000 | CLVS1 | clavesin 1 \| | 0.004801 | 0.003974 | 0 |
| 29129 | ENSG0000 | AC073320.1 | | 0.004762 | 0.003766 | 0.003597 |
| 8453 | ENSG0000 | SLC4A10 | solute carr | 0.004734 | 0.003864 | 0 |
| 1591 | ENSG0000 | PGR | progesterc | 0.0046 | 0.003782 | 0 |
| 13386 | ENSG0000 | MYO1H | myosin IH | 0.004548 | 0.003397 | 0 |
| 582 | ENSG0000 | MSR1 | macrophag | 0.004543 | 0.003484 | 0 |
| 17845 | ENSG0000 | SNORA62 | small nucle | 0.00453 | 0.00259 | 0.001615 |
| 4150 | ENSG0000 | CDH6 | cadherin 6 | 0.004321 | 0.001174 | 0 |
| 5244 | ENSG0000 | INHBA | inhibin bet | 0.004306 | 0.003502 | 0 |
| 39555 | ENSG0000 | MIR6771 | microRNA | 0.004195 | 0.003436 | 0.002872 |
| 16359 | ENSG0000 | OR2C3 | olfactory r | 0.004165 | 0.0034 | 0 |
| 6964 | ENSG0000 | GLS2 | glutaminas | 0.004103 | 0.003825 | 0 |
| 4455 | ENSG0000 | KYNU | kynurenina | 0.004054 | 0.001669 | 0 |
| 10923 | ENSG0000 | LOXHD1 | lipoxygena | 0.004032 | 0.003246 | 0 |
| 17404 | ENSG0000 | SNORD36A | small nucle | 0.003851 | 0.00349 | 0.00304 |
| 41362 | ENSG0000 | AL606534.6 | | 0.003525 | 0.002636 | 0 |
| 43342 | ENSG0000 | MIR6787 | microRNA | 0.0035 | 0.003205 | 0.001048 |
| 612 | ENSG0000 | USH2A | usherin [Sc | 0.003461 | 0.002825 | 0 |
| 12488 | ENSG0000 | GYPA | glycophori | 0.003253 | 0.000733 | 0 |
| 4246 | ENSG0000 | USP9Y | ubiquitin s | 0.00317 | 0.002587 | 0 |
| 8019 | ENSG0000 | RNF165 | ring finger | 0.003108 | 0.001282 | 0 |
| 16151 | ENSG0000 | FSIP2 | fibrous she | 0.003032 | 0.001865 | 0.001379 |
| 43229 | ENSG0000 | MIR7114 | microRNA | 0.002891 | 0.000877 | 0.000835 |
| 32661 | ENSG0000 | RMST | rhabdomy | 0.002764 | 0.001052 | 0 |
| 7788 | ENSG0000 | SYT16 | synaptotag | 0.002338 | 0.001901 | 0 |
| 15357 | ENSG0000 | ROBO2 | roundabou | 0.00213 | 0.001751 | 0 |
| 17820 | ENSG0000 | SNORD14C | small nucle | 0.001635 | 0.001155 | 0.000792 |
| 19719 | ENSG0000 | TSGA13 | testis speci | 0.001501 | 0.000689 | 0 |
| 41647 | ENSG0000 | AC118758.3 | | 0.00147 | 0.000305 | 0 |
| 40825 | ENSG0000 | U85056.1 | | 0.001384 | 0.000983 | 0 |
| 18759 | ENSG0000 | SNORA29 | small nucle | 0.00122 | 0.001089 | 0.000458 |
| 17358 | ENSG0000 | SNORD18C | small nucle | 0.001189 | 0.000944 | 0.000851 |
| 17509 | ENSG0000 | SNORA63 | small nucle | 0.001127 | 0.001018 | 0.000393 |
| 33342 | ENSG0000 | AL359397.2 | | 0.001102 | 0.000669 | 0 |
| 10977 | ENSG0000 | CLEC3B | C-type lect | 0.001025 | 0.000746 | 0.000416 |
| 4183 | ENSG0000 | C9 | compleme | 0.000682 | 0.000157 | 0 |
| 35966 | ENSG0000 | MIR4639 | microRNA | 0.000608 | 0.000498 | 0 |
| 20891 | ENSG0000 | SNORA81 | small nucle | 0.000565 | 0.000337 | 0.000026 |
| 40651 | ENSG0000 | TUG1_1 | Taurine up | 0.000116 | 0.000073 | 0.000058 |
| 22166 | ENSG0000 | SNORD57 | small nucle | 0.000014 | 0.000012 | 0.000005 |
| 43089 | ENSG0000 | MIR1248 | microRNA | 0.000008 | 0.000006 | 0.000003 |
